# The co-evolution of the genome and epigenome in colorectal cancer

**DOI:** 10.1101/2021.07.12.451121

**Authors:** Timon Heide, Jacob Househam, George D Cresswell, Inmaculada Spiteri, Claire Lynn, Max Mossner, Chris Kimberley, Javier Fernandez-Mateos, Bingjie Chen, Luis Zapata, Chela James, Iros Barozzi, Ketevan Chkhaidze, Daniel Nichol, Alison Berner, Melissa Schmidt, Eszter Lakatos, Ann-Marie Baker, Helena Costa, Miriam Mitchinson, Marnix Jansen, Giulio Caravagna, Daniele Ramazzotti, Darryl Shibata, John Bridgewater, Manuel Rodriguez-Justo, Luca Magnani, Trevor A Graham, Andrea Sottoriva

## Abstract

Colorectal malignancies are a leading cause of cancer death. Despite large-scale genomic efforts, DNA mutations do not fully explain malignant evolution. Here we study the co-evolution of the genome and epigenome of colorectal tumours at single-clone resolution using spatial multi-omic profiling of individual glands. We collected 1,373 samples from 30 primary cancers and 9 concomitant adenomas and generated 1,212 chromatin accessibility profiles, 527 whole-genomes and 297 whole-transcriptomes. We found positive selection for DNA mutations in chromatin modifier genes and recurrent chromatin changes in regulatory regions of cancer drivers with otherwise no mutation. Genome-wide alterations in transcription factor binding accessibility involved *CTCF*, downregulation of interferon, and increased accessibility for *SOX* and *HOX*, indicating developmental genes reactivation. Epigenetic aberrations were heritable, distinguishing adenomas from cancers. Mutational signature analysis showed the epigenome influencing DNA mutation accumulation. This study provides a map of (epi)genetic tumour heterogeneity, with fundamental implications for understanding colorectal cancer biology.

## Introduction

Clonal evolution, fuelled by intra-tumour heterogeneity, drives tumour initiation, progression and treatment resistance (Greaves and Maley, 2012; Turajlic et al., 2019). Much is known about the genetic evolution and intra-tumour heterogeneity of colorectal malignancies (Atlas, 2012; Cross et al., 2018; Sottoriva et al., 2015). Although genetic heterogeneity is widespread (McGranahan and Swanton, 2017), epigenetic changes are also responsible for phenotypic variation between cancer cells (Black and McGranahan, 2021; Mazor et al., 2016). Epigenetic profiling of chromatin accessibility in colon cancer has been performed in seminal studies in cell lines (Akhtar-Zaidi et al., 2012) and human samples (Johnstone et al., 2020). However, current investigations are limited to single bulk samples and some also lack normal controls (Corces et al., 2018). Moreover, how cancer genomes and epigenomes concomitantly evolve and shape intra-tumour genetic and epigenetic heterogeneity remains unexplored. Measuring this complex co-evolution in a quantitative manner requires multi-omics profiling at single clone resolution and accurate spatial sampling of human neoplasms, as well as matched normal tissue.

Colorectal cancers are organised into glandular structures, reminiscent of the crypts of the normal intestinal epithelium (Humphries and Wright, 2008). Normal crypts are tube-like invaginations where cell proliferation is driven by a relatively small number of stem cells at the base (Baker et al., 2014; Barker et al., 2009; Lopez-Garcia et al., 2010; Snippert et al., 2010) and cancer glands are thought to have the same architecture (Merlos-Suárez et al., 2011). This implies that all cells within a gland share a recent common ancestor and are a few cell divisions apart: thus glands are largely clonal populations that, through cell proliferation, copy DNA with relatively high fidelity. Ultimately, the gland can be thought of as a natural “whole-genome amplification machine” that can be exploited to perform multi-omics analysis at single clone resolution. Indeed, single crypt and gland genomic profiling has been long used to study clonal dynamics in both normal (Nicolas et al., 2007; Shibata, 2009; Yatabe et al., 2001) and cancer cells (Cross et al., 2020; Humphries et al., 2013; Kang et al., 2015; Siegmund et al., 2009a; Sottoriva et al., 2015; Tsao et al., 1999, 2000). We have developed a new method to concomitantly profile single nucleotide variants (SNVs), copy number alterations (CNAs), chromatin accessibility with ATAC-seq (Buenrostro et al., 2001) and full transcriptomes with RNA-seq from the same individual gland or crypt (see associated Protocol manuscript).

Here we present the results of multi-region single gland multi-omics profiling of 1,373 samples from 39 lesions arising in 30 patients, with 23-57 tumour samples per patient (median=42.5).

## Results

### Single gland multi-omics

We collected fresh resection specimens from 30 stage I-III primary colorectal cancers and 9 concomitant adenomas belonging to 30 patients referred for surgery at the University College London Hospital (Figure 1A and Table S1 for clinical information). Single gland isolation was performed from normal and neoplastic tissue (Figure 1B), as done in previous work (Martinez et al., 2018) and similar to other methods pioneered by Shibata and colleagues (Siegmund et al., 2009b; Tsao et al., 2000; Yatabe et al., 2001). After gland isolation we also collected small bulk samples (a.k.a. ‘minibulks’), representing an agglomerate of a few dozen glands to verify that picked glands were representative of the sample (Figure 1B). Cell lysing and nuclei pelleting separated nuclei from cytosol (Figure 1C). We used nuclei to perform whole-genome sequencing and chromatin accessibility profiling with ATAC-seq. We use the cytosol to perform full transcriptome RNA-seq (Figure 1D). Full details of the methodology are available in an associated Protocol manuscript.

**Figure 1.**
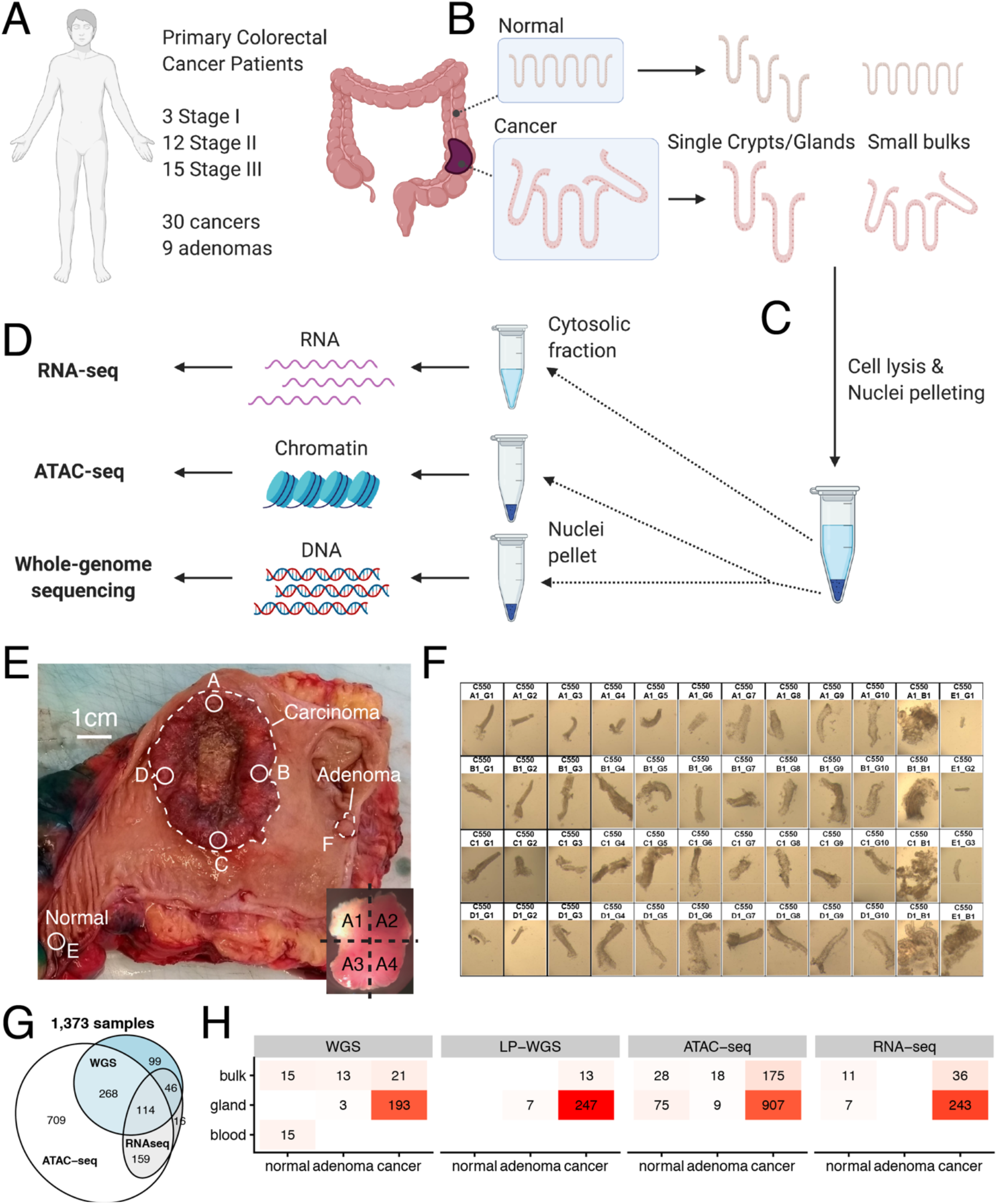
Spatial single gland collection and multi-omic data generation. **(A)** Fresh colectomy specimens from 30 stage I-III colorectal cancer patients were used to collect tissue from 30 cancers and 9 concomitant adenomas. **(B)** Single glands and small bulks were isolated from normal and neoplastic samples. **(C)** From each sample we performed cell lysis followed by nuclei pelleting. **(D)** Cytosolic fractions were used for RNA-seq whereas nuclei were used for whole-genome sequencing and ATAC-seq. **(E)** From each colectomy specimen we identified separate regions of the cancer (A, B, C, D), a distant normal sample (E) and adenomas if present (F-H). Each sample was split into 4 blocks (inlet square). **(F)** From each block (A-E) we collected individual glands (marked as _G) as well as small bulks, agglomerates of a few dozen crypts (marked as _B). **(G)** We performed multi-omic profiling using whole-genome sequencing, ATAC-seq and RNA-seq on the same sample, achieving a good level of overlap between assays. **(H)** For each assay we had representative samples from normal, adenoma and cancer.

We applied this method to a spatial sampling strategy designed to measure clonal evolution at multiple scales. We first sampled four spatially distant regions of a given cancer (A,B,C,D) located close to the tumour edge, one distant normal epithelium region (E), and concomitant adenomas if present (F,G,H). A bulk sample was collected from each region and was spatially annotated in the original resection specimen (Figure 1E and S1). Each piece was cut into 4 subregions (e.g., A1-A4, B1-B4,…) as shown in Figure 1E (bottom-right). We then collected and profiled 12-40 (median=37) individual tumour glands and 3-18 (median=4) minibulks per patient, a few healthy crypts and a minibulk from the matched normal, and blood when available (Figure 1F and S2). We note that C542 sample F was originally labelled as adenoma but confirmed to be part of the cancer upon histopathology revaluation (see Figure S1). We performed multi-omics profiling using deep whole-genome sequencing (WGS, median depth 35x – see Table S2) in 3-15 samples per patient (median=8.5), low-pass whole genome sequencing (lpWGS, median depth 1x – see Table S2) in 1-22 samples per patient (median=8), and chromatin characterisation with ATAC-seq in 18-61 samples per patient (median=42), see Table S3. For a proportion of samples (n=382/1,373) both WGS and ATAC-seq data were available (Figure 1G). We also generated a total of 623 whole-transcriptomes, of which 297 were of high quality to be used for analysis (1-40 samples per patient, median=7) with many also overlapping the WGS dataset, the ATAC-seq dataset or both (Figure 1H). We identified somatic single nucleotide variants (SNVs) and indels, ATAC peaks and copy number alterations (CNAs) from all samples (Methods).

### Genetic mutations affecting the epigenome

Five cases in the cohort were characterised by microsatellite instability (MSI), as reported in Figure 2A, leading to significantly higher SNV and indel burdens (Figure 2B). Copy number alterations recapitulated previous datasets (Atlas, 2012; Cross et al., 2018), with microsatellite stable (MSS) cases displaying high aneuploidy and MSI cases instead showing a largely flat copy number profile (Figure S3). Recurrent loss of chromosomal copies was confirmed for canonical tumour suppressor genes, such as *APC*, *PTEN*, *TP53* and *SMAD4*. Focal amplifications were found in *FGFR1* (2 cases) and *MYC* (1 case). Recurrent cancer driver events in colorectal cancers were recapitulated in this dataset, with stereotypical mutations in *APC*, *KRAS* and *TP53* (Figure 2C and S4). With the exception of a single case (C539), mutations in these genes were invariably clonal.

**Figure 2.**
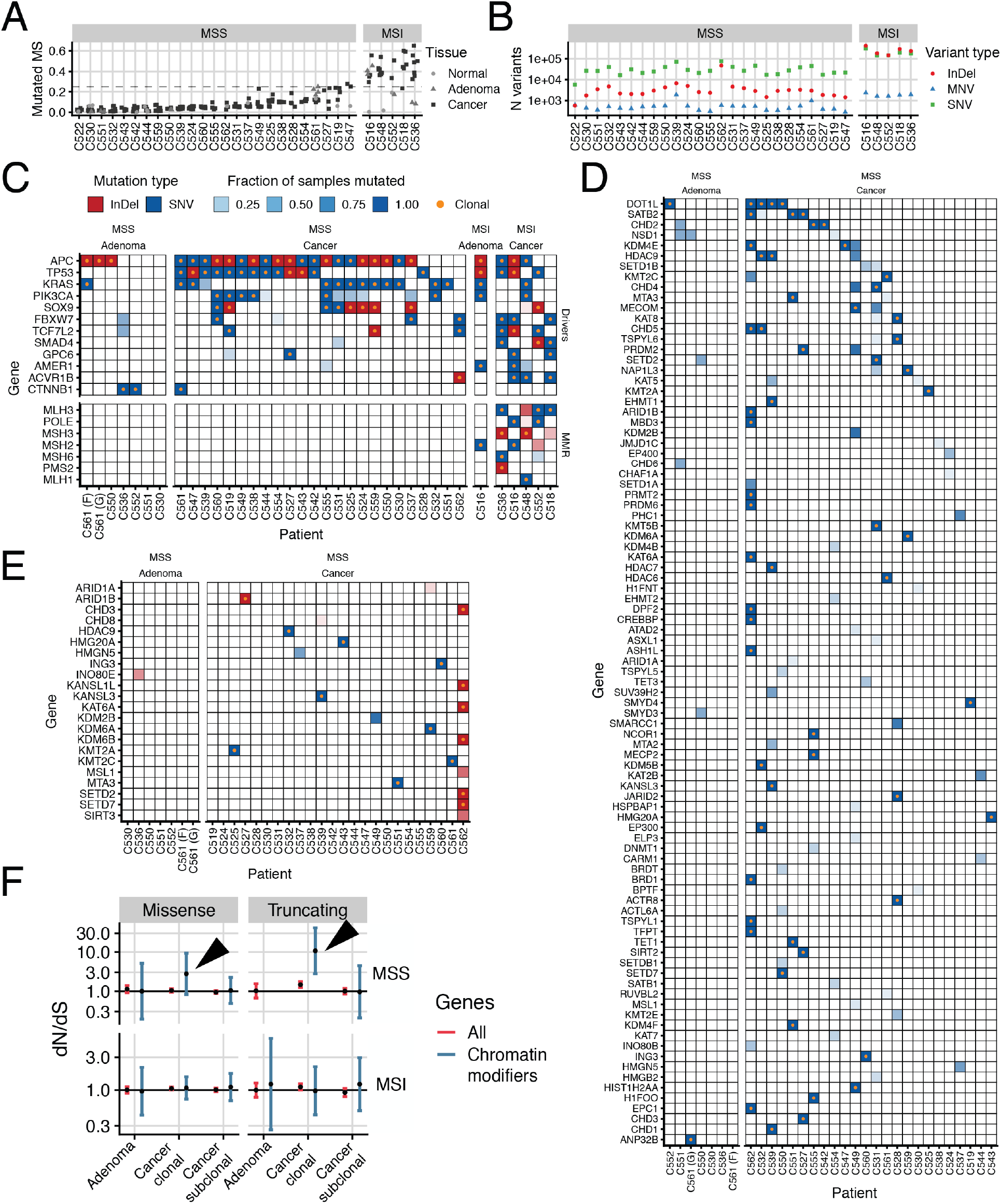
DNA alterations in canonical cancer drivers and chromatin modifier genes. **(A)** Microsatellite instability per case. **(B)** Mutational burden by type of mutation (InDel: small deletion or insertion, MNV: multiple nucleotide variant, SNV: single nucleotide variant). **(C)** Recurrently mutated colorectal cancer driver genes, with orange dot indicating whether the mutation is clonal or subclonal. **(D)** Missense mutations in chromatin modifier genes in MSS cases. **(E)** Truncating mutations and indels in MSS cases. **(F)** dN/dS analysis of clonal and subclonal chromatin modifier mutations in MSS and MSI cancers and adenomas.

We also identified recurrent somatic mutations in chromatin modifier genes, particularly in the lysine demethylases (*KDM*), lysine acetyltransferases (*KAT*), lysine methyltransferases (*KMT*) and SWI/SNF (*ARID1A*) families. Missense mutations in chromatin modifier genes for MSS cancers are reported in Figure 2D whereas truncating and indel variants are illustrated in Figure 2E (see Figure S5 for all cases). Selection on chromatin modifier genes was assessed by dN/dS (Martincorena et al., 2017; Zapata et al., 2018): clonal (occurring in all cancer cells) truncating mutations in chromatin modifier genes in MSS cases showed clear signs of positive selection, with dN/dS significantly >1 (Figure 2F, arrows). dN/dS was above 1 (positive selection) for missense mutations in chromatin modifiers but this was not significant, indicating that whereas a subset of missense mutations could be selected, a large proportion are likely neutral (Figure 2F, arrows). Subclonal chromatin modifier mutations were present but did not show signs of being positively selected, with dN/dS ≈ 1 (Figure 2F). No evidence of positive selection for chromatin modifier gene mutations was detected in MSI cancers, although their high mutational burden may limit the power of detection.

### Focal chromatin alterations are recurrent, hit known driver genes, and distinguish adenomas from cancers

We identified peaks in the ATAC-seq data in each region of a cancer using MACS2 (Feng et al., 2012) and compared each peak Counts per Million (CPM) versus normals (see Figure S6) to identify significant somatic chromatin changes (Figure 3A, see Methods). We found highly recurrent somatic chromatin accessibility alterations (SCAAs) in both promoters (Figure 3B) and putative enhancers (Figure 3C). Amongst recurrent events, gain of a peak (opening of chromatin in tumour vs normal) was more common than loss (closing of chromatin in tumour vs normal) both in promoters (68 gained vs 5 lost in >10 patients) and enhancers (6 gained vs 0 lost in >10 patients). This suggests an overall pattern of increased accessibility of chromatin in cancer versus normal tissue. We used matched RNA-seq (see associated TRANSCRIPTOME paper) to verify that SCAAs corresponded to changes in gene expression (Figure 3D). Indeed, 15.6% of promoters (92/586) and 11.9% of enhancers (29/244) with recurrent SCAAs (>5 patients) showed signs of altering the expression of associated genes (Figure S7, FDR<0.01 and Table S4). We note that our power to detect expression changes was limited by the recurrence of a given SCAA in the cohort and incomplete matched RNA data. Moreover, chromatin accessibility is necessary but not sufficient to induce changes in expression as it does not inform on whether the transcription factor is actually bound to the regulatory region. Therefore, more chromatin changes than those we report may actually cause a change in expression. ATAC peaks called in our dataset showed strong overlap with peaks from the TCGA dataset lacking normals (Corces et al., 2018) and the ENCODE normal colon tissue dataset (Dunham et al., 2012), both reanalysed with our pipeline (Figure S8). Due to unmatched normal controls however, in these orthogonal single bulk sample datasets it is not possible to distinguish chromatin changes occurred in the cancer versus those already present in the normal colon (e.g., to determine somatically-changed status of the peak), and indeed most of the signal of chromatin accessibility comes from the tissue of origin of the sample (Corces et al., 2018).

**Figure 3.**
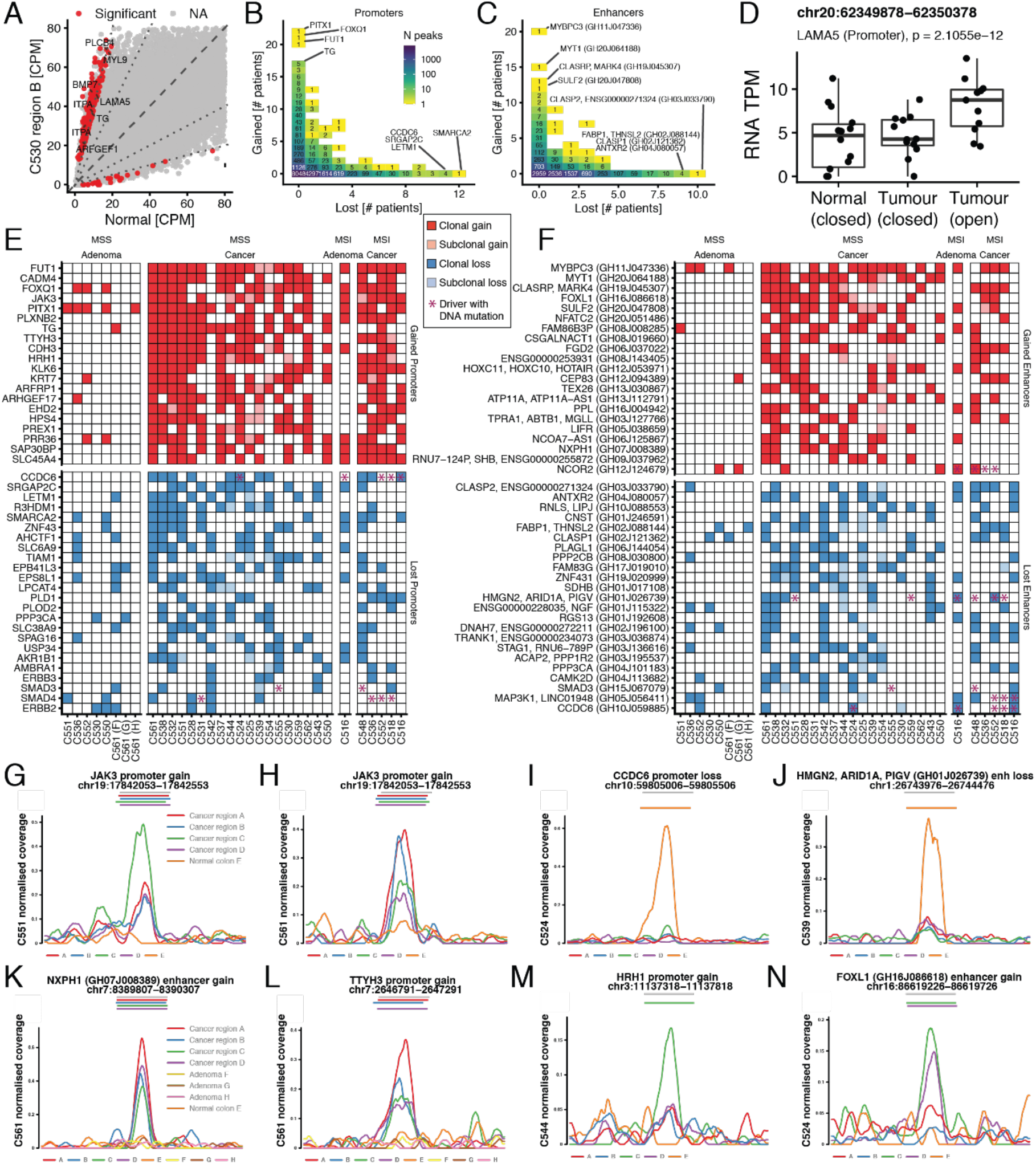
Somatic alterations in chromatin accessibility in cancers and adenomas. **(A)** SCAA detection of cancer C530 region B versus normal. Significantly altered peaks in red. **(B)** Recurrence of promoter SCAAs and **(C)** of enhancer SCAAs. **(D)** For a proportion of loci, we were able to confirm changes in gene expression, for this case of LAMA5 promoter opening. **(E)** Summary of the 20 most recurrent openings (gain) and closures (loss) of promoter and **(F)** of putative enhancers. Clonal changes are marked in solid squares, subclonal changes in shaded squares. Stars indicate DNA mutation in reported colorectal cancer driver gene. **(G)** Clonal somatic peak gained at the JAK3 promoter in cancer C551 and **(H)** in case C561. **(I)** Recurrent promoter loss of accessibility of colorectal cancer driver CCDC6, example from C524. **(J)** Example of recurrent enhancer loss of ARID1A in case C539. **(K)** Example of somatic peak in NXPH1 enhancer gain and **(L)** TTYH3 promoter gain found in the cancer but not in the concomitant adenomas of C561. **(M)** HRH1 promoter gain of accessibility was found in region C of C544 but not in other regions. **(N)** FOXL1 enhancer gain of accessibility was found in regions C and D of C524 but not in other regions. All heterogeneous peaks were identified accounting for purity differences.

We then leveraged our spatial multi-region profiling strategy to assess intra-tumour epigenetic heterogeneity at the level of chromatin accessibility and identify clonal versus subclonal SCAAs. The signal from ATAC peaks is notoriously difficult to compare between samples because it is confounded by variability in purity and transcription start site enrichment (TSSe). We used our matched WGS to identify clonal (truncal) DNA mutations present in all cancer samples and assessed the frequency of these variants in the reads from ATAC-seq to obtain an accurate estimate of sample purity (see Methods). To statistically test the clonality of ATAC peaks we used samples from each region as pseudo-“biological replicates”, and compared the signal between different cancer regions and the corresponding normal while accounting for sample purity (see Methods). This analysis was possible in 24/30 cancers and 9/9 adenomas due to the limited number of samples with sufficient purity in some cases.

Given the large number of SCAAs identified, we focused on the 20 most recurrently altered loci per category (promoter/enhancer, gained/lost), as well as those associated to colorectal cancer driver genes from the IntoGen list (Martínez-Jiménez et al., 2020). We found that a significant proportion of these events (782/854, 91.5%) were found altered in all distant regions of the same cancer and they were therefore ‘clonal’ epigenetic changes in the malignancy. A summary of clonal and subclonal SCAAs for the list of alteration described above, with assessment of clonality and comparison with adenomas, is reported in Figure 3E for promoters and Figure 3F for enhancers.

Amongst the recurrently altered and almost invariably clonal epigenetic changes, we report highly recurrent *JAK3* promoter gain of accessibility in 16/24 cancers (Figure 3G,H). We also found recurrent accessibility loss of the CRC driver gene *CCDC6* in 11/24 cancers (Figure 3I). Notably, this tumour suppressor gene appears rare in colorectal cancer based on DNA mutations alone (e.g., 3/30 cases in our cohort, annotated as purple star in Figure 3E,F). We also report *ARID1A* enhancer closure in 7 cancers and 1 adenoma, with only 2 of these cases reporting also a mutation in this gene (Figure 3J). Alterations in multiple other putative CRC drivers were also found, such as *SMAD3* and *SMAD4* promoter loss (Figure 3E), *NCOR2* enhancer gain, and *SMAD3*, *MAP3K1* and *CCDC6* enhancer loss (Figure 3F). *NFATC2* and *LIFR* cancer driver genes that were not reported in colorectal cancer were found epigenetically altered in our cohort again in the absence of DNA mutations. Of interest, we found typically-clonal promoter opening of *FOXQ1* (Figure 3E), a known oncogene reported to be involved in colorectal cancer tumourigenicity (Kaneda et al., 2010), angiogenesis and macrophage recruitment during progression (Tang et al., 2020). We also note that 11/24 cancers showed gain of *LAMA5* promoter (Figure 3A and D), a gene reported to be involved in colorectal cancer progression (Bartolini et al., 2016; Galatenko et al., 2018; Gordon-Weeks et al., 2019). We also found *MMP9*, a gene involved in EMT (Chae et al., 2018), promoter opening in 6 cases. This finding further highlights how cancer driver lists based on DNA mutations provide an incomplete picture of carcinogenesis, tumour progression and clonal evolution. We confirmed that the signal was not driven by copy number alterations, with 95.4% of loci in Figure 3E and F showing no difference in copy number alterations between altered and non-altered region accessibility (Figure S9). In supplementary figures we report all the results for each patient for promoter gains (Figure S10), promoter loss (Figure S11), enhancer gain (Figure S12) and enhancer loss (Figure S13).

When we compared cancers with adenomas, we found that out of the 235 SCAAs found recurrent in the cancers in Figure 3E and F, only 32 (13.6%) were also found in the matched adenoma, indicating that they likely occurred at the onset of malignant transformation rather than the initiation of neoplastic growth (Figure S14). This was exemplified by the opening of *NXPH1* enhancer and *TTYH3* promoter, which were present in each region of the cancer but not in the concomitant three adenomas (Figure 3K,L). It was previously noted that there were limited differences between adenomas and carcinomas in colorectal cancer at the level of point mutations in driver genes, and instead major differences at the level of chromosomal instability (Cross et al., 2018). Similarly, here we find major differences in epigenetic rewiring between adenomas and CRCs.

Although the majority of recurrent SCAAs were clonal in the cancer, a proportion of SCAAs were found to be subclonal and confined to one or more regions. This is exemplified by *HRH1* promoter gain in Figure 3M occurring only in region C of cancer C544, and *FOXL1* enhancer gain in Figure 3N occurring only in regions C and D of C524. All the per-region data for subclonal promoters and enhancers SCAAs is reported in Figures S15-S18.

### Transcription factor accessibility analysis reveals global epigenetic reprogramming

We then analysed the accessibility to transcription factor (TF) binding sites for 870 transcription factors (Dunham et al., 2012) using publicly available TF motif and ChIP-seq data (see Methods). To measure this, we piled the ATAC reads for all the TF sites of interest in the genome (Methods). When the number of reads (in CPM) is plotted versus the distance from the centre of the TF motif and the length of each read, this analysis produces a characteristic signature of TF accessibility for a given sample, which can also encode the footprint of the TF complex itself in the cancer (Figure 4A and S19) compared to the normal (Figure 4B). We divide the genomic loci analysed for each TF into three major subsets depending on their distance *d* from the closest Transcription Starting Site (TSS): proximal to TSS (pTSS, *d* ≤ 2,000 bp), close to TSS (cTSS, 2,000 < *d* ≤ 10,000 bp) and distal to TSS (dTSS, *d* > 10,000 bp). Each of these sets was further divided into those overlapping a called ATAC peak (oPeak) with non-overlapping a called ATAC peak (nPeak). Following normalisation and subtraction of the signal from the normal samples (Figure 4C), we use linear regression to account for confounding factors such as transcription start site enrichment (TSSe) per sample and identify signal differences driven by cancer cell purity (see Methods). We considered significant changes in TF chromatin accessibility if the signal difference of cancer-normal correlated with the purity of a sample, hence the more the amount of cancer cells in the sample, the stronger the signal difference. This allowed controlling for contamination from normal colon and stromal cells. As many transcription factors bind to similar loci in the genome, we considered only largely non-overlapping TF annotations to ensure the signal was not driven by the same loci in the genome for multiple TFs (Figure S20 and Methods). The signal was consistent when we looked at regions unique to each annotation (Figure S21), demonstrating it was not a small subset of common regions of the genome driving the signal.

**Figure 4.**
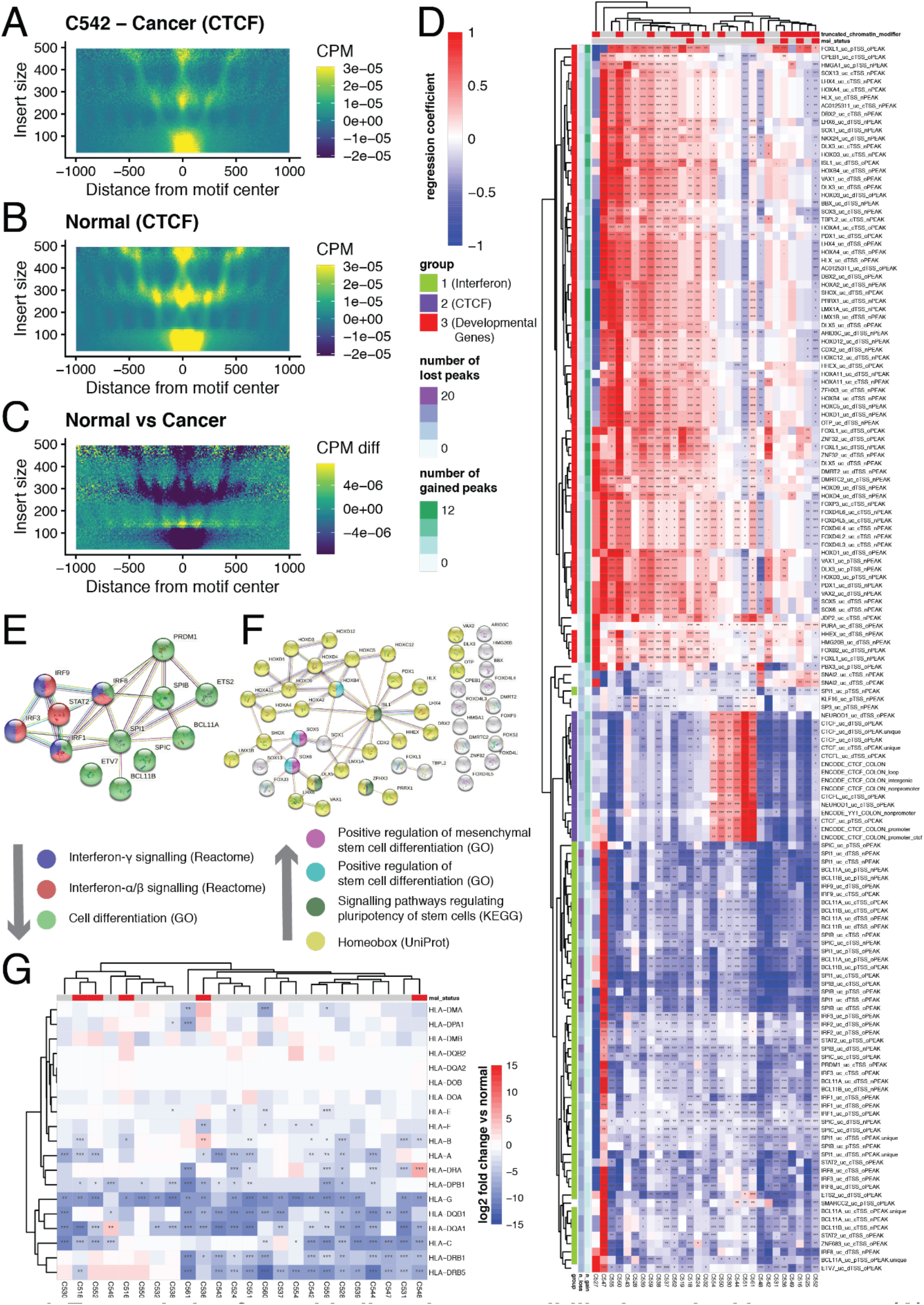
Transcription factor binding site accessibility is rewired in tumours. **(A)** TF binding site accessibility (in this example CTCF) is computed by summing the signal of ATAC-seq reads centred at binding site, plotted against read length. **(B)** The same is done for the normal controls. **(C)** Signal from the normal is subtracted to the signal from the cancer to assess differential accessibility. TF accessibility for CTCF is reduced in this example as demonstrated by fewer ATAC cuts at the binding site in the cancer (lower read count at TF binding site) which leads to fewer long reads initiating from the TF binding site. **(D)** The differential signal is then regressed against TSSe and purity to identify TF binding accessibility altered in tumours. Results here for the three major clusters of differentially accessible TF loci (colour is regression coefficient, star indicates significance). **(E)** String-db analysis of the green TF cluster highlights downregulation of interferon signalling and cell differentiation signalling. **(F)** String-db analysis of the red cluster indicates upregulation of stem cell differentiation and activity of developmental genes such as the homeobox family. **(G)** Differential gene expression of HLA genes.

The results are illustrated in Figure 4D where the colour scale indicates the regression coefficient of the model, and stars indicate significance. In red the signal from TF accessibility correlates positively with the purity, indicating increased accessibility in tumour. In blue we have loci that have decreased accessibility in tumour. The results support pervasive rewiring of TF chromatin accessibility in cancer, with three large clusters of altered TF binding sites. A first cluster (green in the left dendrogram) revealed downregulation of interferon signalling through closing of chromatin loci normally bound by transcription factors such of the IRF (interferon-regulatory factor) family, indicating suppression of immune signalling. As shown in Figure 4E, Reactome and GO analysis indicated that the signal was significantly enriched for downregulation of interferon-γ (FDR=4.2e-6) and interferon α/β (FDR=3e-8), as well as downregulation of cell differentiation (FDR=5e-5). The signal was particularly strong in MSI cancers (p=0.012, Fisher’s Exact Test). By analysing RNA-seq data from the same patients, we found that consistent with the downregulation of interferon signalling, gene expression of HLA genes was significantly reduced in the majority of patients (Figure 4G).

A second cluster (blue in the left dendrogram) identified two distinct subgroups of patients with differential chromatin accessibility for *CTCF*. CCCTC-Binding Factor (CTCF) is a key player in chromatin insulation, determining looping and TAD (Topological Associating Domain) formation. We report a larger group of cases characterised by loss of CTCF binding site accessibility, which was also enriched for MSI cancers. We also found a smaller but significant group showing increased *CTCF* accessibility. *CTCF* chromatin accessibility alterations were previously noted in single bulk cancer sample (Fang et al., 2020), *CTCF* somatic mutations are frequent in CRC (Katainen et al., 2015), and indeed a mouse model of chronic CTCF hemizygosity led to higher cancer incidence and dysregulation of oncogenic pathways (Aitken et al., 2018).

A third cluster (red in the left dendrogram) showed increased chromatin accessibility for TFs involved in stem cell differentiation and pluripotency (GO: ‘*positive regulation of stem cell differentiation’* – FDR=2.5e-4, and *‘mesenchymal stem cell differentiation’* – FDR=9e-4; KEGG: *signalling pathways regulating pluripotency of stem cells* – FDR=0.047), as well as TFs involved in development, such as the *HOX*, *FOX* and *SOX* families (UniProt: ‘*homeobox’* – FDR=2.7e-40, *‘developmental protein’* – FDR=1.7e-21). The chromatin accessibility of this cluster of TFs was higher in cancer in the majority of cases, pointing at a key role of reactivation of developmental genes in colorectal cancer tumorigenesis (Figure 4F). The expression of the TFs involved in this cluster is reported in Figure S22.

We also note a small cluster characterised by increased accessibility of *SNAI1* and *SNAI2* transcription factor binding sites, two genes involved in Epithelial to Mesenchymal Transition - EMT (Chae et al., 2018). This small cluster was significantly enriched with cases showing truncating mutations in chromatin modifier genes (p=0.047, Fisher’s Exact Test), consistently with previously reported regulation of EMT by chromatin modulators (Serresi et al., 2021)

### Binding sites of developmental TFs with increased accessibility are demethylated

Changes in chromatin accessibility can be accompanied by changes in DNA methylation, with heterochromatin regions often being methylated and vice-versa for open chromatin regions. This is particularly the case for regions that are permanently silenced after development (Smith and Meissner, 2013). We asked whether some of the SCAAs we identified at TF binding sites (Figure 4D) reflect in the methylation of the same loci.

We performed methylation profiling on a subset of 8 samples using Illumina EPIC 850k methylation arrays (1x sample from C516, 2x samples from C518, 2x samples from C560 and 3 samples from C561). Comparing the methylation of TF binding annotations in cluster 3 (Figure 4F), we found that methylation in these regions was significantly lower than in normal tissue, supporting the finding that these sites were accessible (Figure 5A). This was particularly clear for TF binding sites of DLX5, HOXA4, HOXB4, ISL1, SOX5 and SOX6 (Figure 5B). This suggests stable reactivation of regulatory regions involved in development and pluripotency.

**Figure 5.**
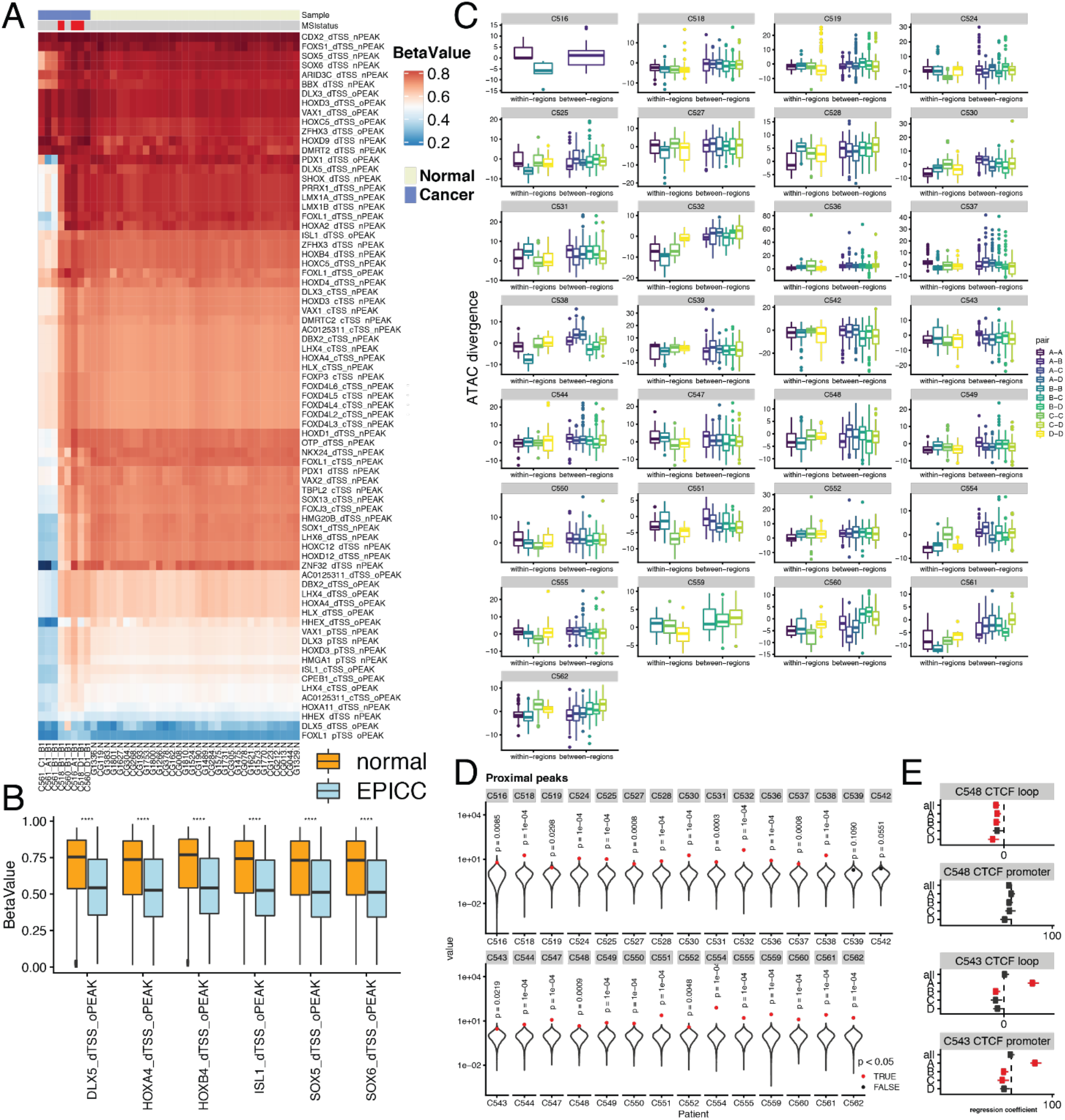
Demethylation in reactivated TF binding sites and heritability of chromatin. **(A)** We selected genomic regions in cluster 3 (enriched in developmental genes like SOX and HOX families) and verified their methylation status with CpG methylation arrays in EPICC samples versus normal. **(B)** In particular regions corresponding to binding sites of DLX5, HOXA4, HOXB4, ISL1, SOX5 and SOX6 showed decreased methylation in cancer vs normal. **(C)** We compared ATAC distance (euclidean on promoter peaks) between glands from the same region (within-region) and glands of different regions (between-regions) to evaluate divergence of chromatin against space and genetic distance. **(D)** For the large majority of patients within-region ATAC distance is significantly lower than between region, indicating heritability of the chromatin that follows the spatial and phylogenetic structure of the tumour. Here we plot the F statistics of the ANOVA model on TSSe, number of reads, and region. **(E)** A significant proportion of TF binding site accessibility changes were ‘clonal’ within the tumour, with distant regions showing the same pattern, again testimony of the heritability of chromatin accessibility. In this example CTCF loop and promoter loci in C548. However, there were some exceptions, as in this example of C543.

### Chromatin changes are stable and heritable, and can be a substrate for Darwinian clonal selection

Epigenetic alterations, and in particular chromatin modifications, are responsible for cell identity in all tissues, hence it remains unclear whether epigenetic changes in cancer are stable during tumour progression and growth. If they were stable, a key question would be whether they comprise a heritable trait passed on from mother to daughter cell, potentially providing the heritable substrate for Darwinian selection to operate. The overall picture in Figures 3E and F, with the majority of the most recurrent SCAAs being clonal, shows that chromatin alterations in cancer are stable during the course of tumour expansion, copied over thousands of cell generations as a heritable trait found several cm apart in different tumour regions.

To further test the heritability of epigenetic alterations we compared epigenetic divergence of samples within the same region versus samples in different regions (Figure 5C). In a majority of patients (23/29), ANOVA controlling for TSSe and number of reads, showed that samples from the same region are significantly less divergent in terms of chromatin accessibility with respect to samples from different regions (see Methods), indicating that chromatin profiles are heritable and follow, at least in part, genetic divergence (Figure 5D). We also report the ANOVA coefficients for each region in Figure S23.

Overall, the results support the finding that somatic chromatin modifications in cancer are in part heritable and stable over time. Moreover, many of the alteration that characterise cancers occur just before or at the onset of malignancy, with benign adenoma not sharing the widely altered epigenetic landscape observed in carcinomas.

We asked the same question of heritability of the chromatin in regards to the global rewiring for TF activity we report in Figure 4. When examining these patterns in each distinct region of a tumour, we observed the same overall trend of increased or decreased accessibility in all regions in a large proportion of cases (Figure S24), suggesting that such rewiring of the chromatin existed in a common ancestor of all the samples and was inherited during tumour expansion. There were however some interesting exceptions where different regions showed distinct profiles. For example, C548 showed homogeneous loss of accessibility to *CTCF* binding sites at loop loci. In C543 both promoter and loop binding sites of *CTCF* were altered and in a heterogeneous manner, with region displaying differential organisation of the chromatin (Figure 5E).

### Mutational signatures affecting the epigenome

With a median of 9 deep WGS samples per patient, we were powered to perform high resolution clonal and subclonal mutational signature analysis (Alexandrov et al., 2013, 2015). Moreover, we could exploit the matched chromatin accessibility and transcriptomic profiles to accurately measure clonal and subclonal per-signature mutation rates in regions of the genome with different chromatin accessibility and gene expression. We used a method based on sparse signatures identification, which is more robust to overfitting compared to previous methods (Lal et al., 2021), to perform mutational signature discovery in our cohort. However, we note that signature discovery with other methods, such as SigProfiler (Alexandrov et al., 2020) identified the same signatures (Figure S25).

We identified six signatures (Figure 6A):

- SparseSignature1, corresponding to COSMIC signature 1 of C>T deamination at methylated CpG sites
- SparseSignature2, corresponding to COSMIC signatures 2+13 caused by APOBEC enzymes
- SparseSignature3, corresponding to COSMIC clock-like signature 5
- SparseSignature4, corresponding to COSMIC signature 17a+b of unknown aetiology
- SparseSignature5, corresponding to COSMIC signature 9+41 also of unknown aetiology
- SparseSignature6, corresponding to COSMIC signature 44 caused by mismatch repair deficiency.

**Figure 6.**
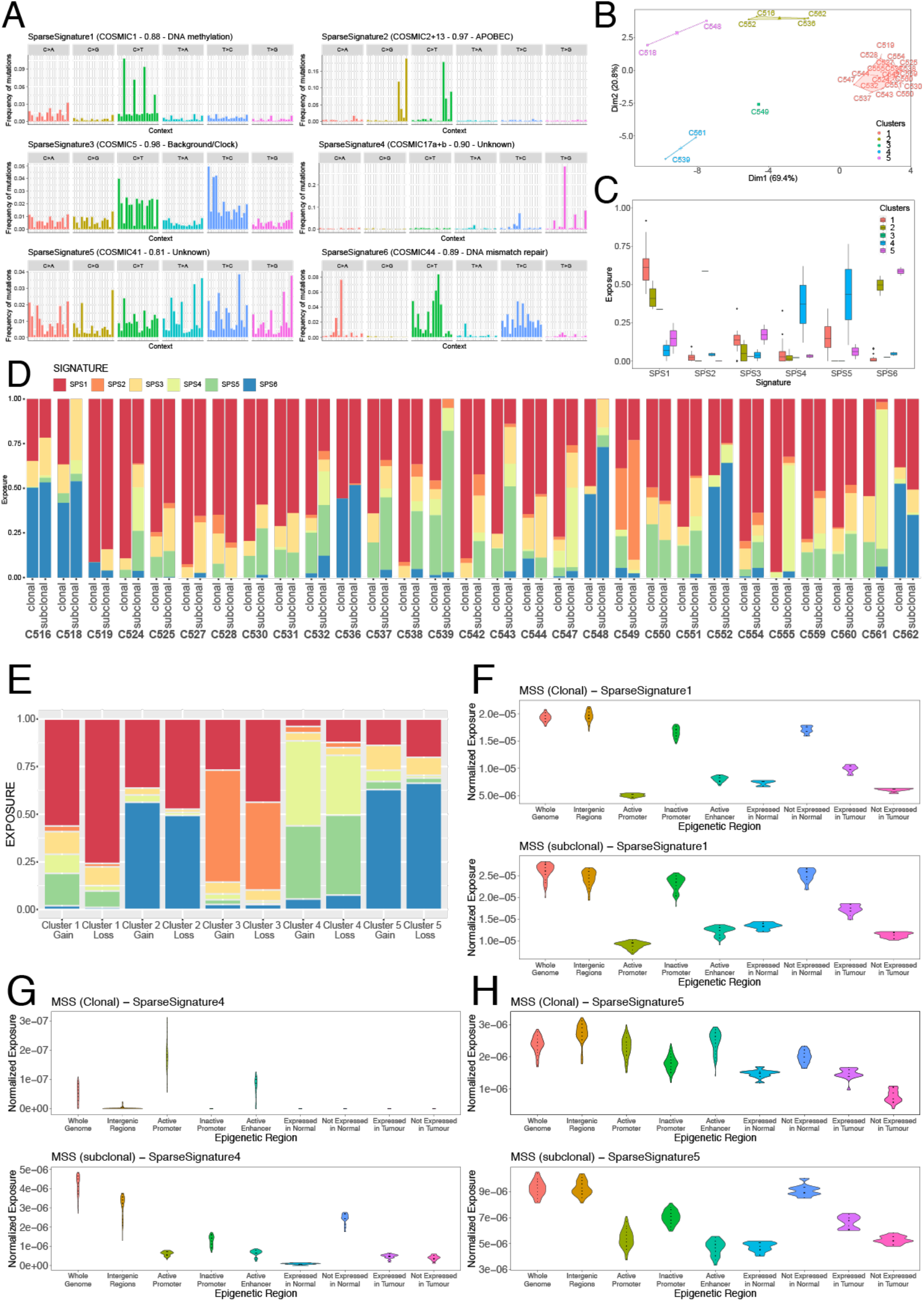
Mutational signatures and the epigenome. **(A)** Mutational signature discovery with sparse signatures identified 6 signatures in our cohort. **(B)** Principal Component Analysis divided the patients into 5 clusters depending on contribution from each signature. **(C)** Signature activity varied between clusters. **(D)** Clonal and subclonal mutational signature composition for each patient. **(E)** Proportion of each signature for every cluster responsible for generating loss or gain of CTCF binding affinity in our cohort. **(F)** The epigenome influences accumulation of deamination signature 1 in distinct regions, both for clonal and subclonal mutations. **(G)** Signature SparseSignature4, mostly present subclonally, is also influenced by the epigenome status. **(H)** Signature SparseSignature5, particularly at the subclonal level, is again depleted in active regions as SparseSignature1.

This analysis splits the cohort into 5 distinct clusters of patients with significantly distinct signature composition (Figure 6B), with the two major clusters enriched for MSS (cluster 1) and MSI cases (cluster 2). Cluster 3 contains only patient 549, which is the only case enriched with the APOBEC signature. Cluster 4 with patients 561 and 539 have higher SparseSignatures 4 and 5 of unknown aetiology. Cluster 5 with patients 518 and 548 have higher SparseSignature 3 (clock-like). The activity of each signature in each cluster of patients is reported in Figure 6C.

In Figure 6D we show the proportion of each signature between clonal and subclonal mutations of the same patient, highlighting the significant change in signature composition over time. As expected, SPS1 (deamination) is dominant in MSS cases whereas MSI cancers are dominated by both SPS1 (deamination) and SPS6 (mismatch repair). Interestingly, SPS2 (APOBEC), SPS4 and SPS5 (unknown) increase at the subclonal level, possibly due to their link to the altered tumour microenvironment.

We reasoned that different mutational processes may differentially alter TF binding site affinity, thus directly influencing the epigenome. It has been previously documented that point mutations can disrupt *CTCF* binding sites (Katainen et al., 2015) We examined all the somatic mutations in *CTCF* sites that were predicted to cause significant loss or gain of binding using deltaSVM (Lee et al., 2015) and assign this subset of mutations to the six signatures, thus constructing an ‘observed signature’ of gain or loss of *CTCF*. For different patient clusters, we then considered only *CTCF* binding sites in the genome and sampled the mutational signature composition to build the ‘predicted signature’ of mutations causing *CTCF* loss or gain based again on deltaSVM. We asked whether the most active mutational signatures could simply explain the accumulation of mutations in *CTCF* binding sites that change its binding affinity. Indeed, mutations predicted to cause loss of binding had a signature highly similar to the predicted signature (cosine similarity = 0.977; Figure S26A), and the same was true for gains (cosine similarity = 0.919; Figure S26B). Interestingly, SparseSignature6 in MSI cluster 2 (mismatch repair, Figure S26C) and SparseSignature4 (COSMIC signature 17, Figure S26D) in cluster 4, were also consistent with causing gain of *CTCF* binding affinity (cosine similarity = 0.925 and 0.977 respectively). These results indicate that signature 1 deamination is predominantly responsible for mutations altering *CTCF* binding in MSS cancers, with a higher tendency of generating loss of binding (Figure 6E). In MSI cases, signature 6 mismatch repair is also a dominant factor in causing altered binding of *CTCF*, with preference for generating increased affinity (Figure 6E). This analysis illustrates how mutational processes acting on the cancer genome can directly influence the cancer epigenome.

### The epigenome influences the mutational processes acting on the genome

Mutations in chromatin modifier genes, or in transcription factor binding sites, can determine the characteristics of the epigenome and the chromatin structure. At the same time, the epigenome also can determine how the cancer genome accumulates mutations due to its effect on different mutational processes and activity of DNA repair genes.

In this cohort, we have the unique opportunity of having matched chromatin accessibility and transcriptome profiles. We leverage on this by annotating each cancer’s genome by epigenetic regulatory regions: active/inactive promoter, active/inactive enhancer, intergenic and coding (expressed vs not expressed). For this analysis we considered the (in)activity of a promoter or an enhancer if the chromatin accessibility status was the same in normal and cancer (i.e., the epigenetic region did not change activity during tumourigenesis). Although we have information also on epigenetic regions that were changed from active to inactive in the cancer and vice versa, not enough mutations had the time to accumulate in the relatively brief final period between change of chromatin and sampling, hence analysis on those could not be done. We also used the RNA-seq data to split the coding genome into: genes expressed in the normal, genes not expressed in the normal, genes expressed in the tumour (were not expressed in the normal), genes not expressed in the tumour (were expressed in the normal). We then examined the mutational signatures activity in each of these genomic annotations per patient, using a Jackknife analysis to verify differences in mutation accumulation from distinct mutational processes.

We found that both clonal and subclonal mutational signature SparseSignature1 (deamination) was 2-4 folds higher in closed chromatin regions of the genome, consistent with the need for methyl-cytosine (enriched in inactive regions) to be present in order for it to become deaminated and produce the associated mutational signature (Figure 6F). Demethylated regions of the genome with high accessibility, such as active promoters and enhancers, suffer up to 4 times less from the accumulation of mutations from C>T deamination with respect to regions in closed chromatin. This could also be due to the contribution of transcription-coupled DNA repair. The same was true for expressed versus not expressed genes in the normal. Genes not expressed in cancer (they were in normal) did not have enough time to accumulate deamination. Genes expressed in tumour instead still carried the ageing mutations accumulated in the normal tissue before carcinogenesis, and hence showed an intermediate mutational load (Figure 6F). A very similar pattern was observed at the subclonal level. The same was true for SparseSignature4 (Figure 6G) and SparseSignature5 (Figure 6H), which were more active subclonally, hence the pattern visible more clearly for subclonal mutations. These effects could also be due to differences in replication timing of active and inactive regions of the genome (Tomkova et al., 2018). Whereas the signal for SparseSignature1-5 was consistent in MSI cases as well, the same magnitude of differences was not observed in mismatch repair signature, which seemed to be more uniformly distributed (Figure S27).

## Discussion

Seminal studies in clonal evolution have profiled intra-tumour heterogeneity using multi-region bulk sampling (reviewed in (McGranahan and Swanton, 2017). Although extremely useful, bulk sequencing data contain distinct cancer and normal cell populations, making it very difficult to deconvolute the signal (Tarabichi et al., 2021). Single-cell sequencing helps (Gawad et al., 2016) but remains extremely noisy, especially for single nucleotide variant (SNV) calling, and multi-omics profiling of the same cell is at an early stage. Moreover it is populations that evolve, not individual cells, and due to extensive cell death and lineage extinction in cancers it is likely that the large majority of profiled cells would have never contributed to the long-term evolution of a malignancy. Generally, we are interested in profiling single populations or ‘clones’ instead, with the aim of predicting their future evolution and impact on the clinical course of the disease.

Although we know a lot about the genetic lesions that lead to colorectal malignancies, epigenetic events in colorectal cancer and many other tumour types, despite being recognised as highly significant (Suvà et al., 2013), are severely understudied (Black and McGranahan, 2021). Recently, a pan-cancer analysis revealed the chromatin accessibility profile of multiple cancer types, but the lack of appropriate matched normal control precluded proper identification of cancerous events, as opposed to tissue specific and ‘cell of origin’ chromatin profiles which remained the dominant signal in the data (Corces et al., 2018). Seminal contributions by Scacheri and colleagues have used proper normal controls (single normal human colon crypts) in established colon cancer cell line models (Akhtar-Zaidi et al., 2012). Single bulk sample analyses added to this complex picture by discovering distinct chromatin compartmentalisation in colorectal cancers (Johnstone et al., 2020). All these studies point at the importance of studying the epigenome to fully comprehend carcinogenesis and malignant progression.

Using single-gland multi-omic profiling, our study highlights that:

- There are significant epigenetic events, also in cancer driver genes with otherwise no DNA mutation, which are highly recurrent in carcinogenesis.
- Chromatin alterations are stable and heritable, providing a substrate for Darwinian selection to act.
- Profiling the epigenome unveils a layer of cancer evolution that is largely invisible with genomic alterations alone, from immune escape to developmental genes reactivation.
- The cancer epigenome influences the accumulation of somatic mutations in the genome.

The fact that there are recurrent epigenetic changes in the promoters and enhancers of known cancer driver genes that are otherwise devoid of somatic mutations indicates that genomic medicine, which is based on the reliable identification of driver alterations to be targeted, needs to incorporate epigenetic assays. Furthermore, this could help explaining the many cases with limited or no driver mutations, thus facilitating tumour subtyping.

The results on TF accessibility again point at immune remodulation as a factor contributing to carcinogenesis of the colon, even in the context of MSS tumours, which are generally considered immunologically cold due to the limited number of infiltrating lymphocytes. Evidently, some level of immunogenicity must have been retained by MSS cancers in order for the emergence of interferon signalling suppression to be needed for carcinogenesis.

One of the most intriguing results has also been the evidence of reactivation of developmental genes during tumourigenesis. Those genes are usually silenced in somatic tissues as they almost only needed at the time of morphogenesis, where pluripotency and growth potential are at the highest. The reactivation of these gene families and their involvement in tumourigenesis has been postulated before in the context of glioblastoma tumourigenesis (Liau et al., 2017), and these results confirm that the signal of this process is there in the epigenome of colorectal malignancies. This is important also because besides copy number alterations, mostly non-focal chromosomal arm gains or losses, there is little difference in driver alterations between benign adenomas and malignant carcinomas (Cross et al., 2018). Moreover, there is no solid prognostic genetic alteration that predicts recurrence in colorectal cancer, and differentiate stage II/III that remain localise in the colon, versus those that infiltrate beyond the muscle wall and will re-present as metastatic months or years later. Instead, the epigenome of adenomas and carcinomas seem to be radically different, and we hypothesise that these differences may be prognostic in differentiating stage II/III with occult metastatic disease from the rest. We will follow these patients clinically to test this.

Finally, seminal papers have shown how the activity of mismatch repair genes (Polak et al., 2015) and nucleotide excision repair genes (Pich et al., 2018; Sabarinathan et al., 2016), is directly impacted by the 3D structure of the genome and its chromatin confirmation. The results we present here quantify the activity of the most common mutational signatures, such deamination, across the genome with matched epigenetic annotations, demonstrating how the epigenome of colorectal cancers directly influence the processes of accumulation of mutations, and vice-versa.

Necessarily, follow-up work is required to explore the significance of epigenetic alterations in cancer driver genes and other loci. And the involvement of developmental gene reactivation will need to be tested in different models of carcinogenesis. However, patient studies like this one are fundamental to unveiling the extent of non-genetic determinants of cancer clonal evolution that explain a substantial part of unknown tumour biology.

## Methods

### Sample preparation and sequencing

The method of sample collection and molecular processing is described in detail an accompany Protocol manuscript. Processing of RNAseq data is described in the associated manuscript TRANSCRIPTOME.

### Whole-genome sequencing – alignment

Contaminating adapter sequences were removed using Skewer v0.2.2 (Jiang et al., 2014). Adapter sequences were ‘AGATCGGAAGAGC’ and ‘ACGCTCTTCCGATCT’, with a maximum error rate of 0.1, minimum mean quality value of 10 and a minimum read length of 35 after trimming using options “-l 35 -r 0.1 -Q 10 -n”. The trimmed and filtered reads from each sequencing run and library where separately aligned to the GRCh38 reference assembly of the human genome (Schneider et al., 2017) using the BWA-MEM algorithm v0.7.17 (Li and Durbin, 2009) Following the GATK best practices and the associated set of tools v4.1.4.1,(Auwera et al., 2013; DePristo et al., 2011; McKenna et al., 2010) reads were sorted by coordinates (GATK SortSam), merged independent sequencing runs or libraries generated from the same tissue sample and marked duplicated reads using GATKs MarkDuplicates. The structure of the final bam files was verified using GATKs ValidateSamFile.

### Alignment – ATAC-seq

Adapter sequences were removed with Skewer v0.2.2 (Jiang et al., 2014) using the full-length adapter sequences below with the option “-m any.

Adapter sequences: CTGTCTCTTATACACATCTCCGAGCCCACGAGACNNNNNNNNATCTCGTATGCCGTCTTCTGCTTG

CTGTCTCTTATACACATCTGACGCTGCCGACGANNNNGTGTAGATCTCGGTGGTCGCCGTATCATT

The reads of each sequencing run and library were aligned to the GRCh38 reference genome using Bowtie2 v2.3.4.3 (Langmead and Salzberg, 2012) with the options “--very-sensitive -X 2000” set. After sorting the reads with samtools v1.9 (Li et al., 2009) reads mapping to non-canonical chromosomes and mitochondria (chrM) were removed (GATK PrintReads followed by RevertSam and SortSam). After of independent libraries of each sample, we removed duplicated reads using GATKs MarkDuplicates and removed all reads mapping to multiple-locations (multi-mappers). The final bam files were validated with GATK’s ValidateSamFile.

### Detection of germline variants

HaplotypeCaller v4.1.4.1 with the GATK package (Poplin et al., 2018) was used to identify germline variants from the reference normal samples in each patient (buffy coats or adjacent normal tissue) using know germline variant annotations from the build 146 of the dbSNP database (Sherry et al., 2001) separately for each chromosome. Resulting VCF files were then merged with GATK MergeVcfs. Variant recalibration was performed with gatk VariantRecalibrator with options set according to GATK best practices (Auton et al., 2015; Frazer et al., 2007; Mills et al., 2006; Sherry et al., 2001) and applied to VCF files using gatk ApplyVQSR with the options “-mode SNP -ts-filter-level 99.0” and “-mode INDEL -ts-filter-level 99.0” respectively. All germline variant calls marked as “PASS” were retained.

### Verification of sample-patient matches

For all samples we excluded the possibly of sample mismatch by comparing germline variants identified in normal tissue to neoplasia samples of a given patient. The reads of each read-group were extracted with samtools view using options ‘-bh {input_bam} -r {read_group_id}’ and GATK’s CheckFingerprint tool was applied to extract statistics on sample-patient matches (Javed et al., 2020). For virtually all high-purity samples without extensive loss of heterozygosity, we were able to confirm that the samples were obtained from the expected patient, for the latter group we inspected copy-number profiles (see below) to confirm that these matched the remaining samples.

### Copy number analysis

### Deep whole genome sequencing

Coverage of genomic loci relative to matched normal tissue samples (buffycoats or adjacent normals) were extracted with methods provided in the sequenza v2.1.2 package for R (Favero et al., 2015) and binned in non-overlapping windows of 10^6^ bp. B-allele frequencies (BAF) of germline mutations determined with the GATK HaplotypeCaller (see above) for each patient were added to these binned files. Joint segmentation on BAFs and read depth counts across all samples from a given tumour were used to determine a set of breakpoints to use for the subsequent analysis. specifically, GC content bias correction from was applied using the ‘gc.norm’ method from sequenza v2.1.2 and positions with non-unique mapability (i.e., < 1) determined by QDNAseq v.<u>3.8</u> (Scheinin et al., 2014) in windows of 50 bp were removed. Piecewise constant curves were fitted for each chromosome arm using the multipcf function (gamma = 80) from the copynumber v1.22.0 package for R (Nilsen et al., 2012). The per-patient set of break points, binned depth-ratio and BAF data were then inputted into the sequenza algorithm (version 2.1.2) to determine allele specific copy-numbers, ploidy Ψ and purity ρ estimates (Favero et al., 2015). The initial parameter space searched was restricted to {ρ │ 0.1 ≤ ρ ≤1} and {Ψ │ 1 ≤ Ψ ≤ 7}. Upon manuel review of the results, we identified several samples with unreasonable fits (cases where calls suggested extremely variable ploidy values across samples). For these samples, we manually identified alternative solutions consistent with the other samples and somatic variant calls.

### Low-pass whole genome sequencing

Low-pass WGS bam files were processed using QDNAseq (Scheinin et al., 2014) to convert read counts in 500kb bins across the autosomes of hg38 into log2ratio data. Data normalisation was performed in accordance with the QDNAseq workflow, except for outlier smoothing (smoothOutlierBins function) which was seen to artificially depress signal from highly amplified bins. Bins for hg38 were also generated according to QDNAseq instructions. Log2 ratio values in each bin were normalised by subtracting the median log2 ratio from all log2 ratios per sample. Samples in a patient were segmented jointly using the multipcf function in the R package copynumber (gamma = 10) (Nilsen et al., 2012) and the mean segment log2ratio was calculated across the bins.

Absolute copy number status was calculated using the approach taken by ASCAT (Loo et al., 2010). Using the ASCAT equation to describe LogR ratios, we took an integer ploidy value Ψ_t_ in the tumour *t* as determined by paired deep WGS in each case, and searched a range of purities from 0.1 to 1 (and assumed gamma was 1 as is the case in sequencing data). For each purity (ρ) value we calculated the continuous copy number status of each bin and calculated the sum of squared differences of these values to the nearest positive integer of the modulus. Purity estimates were given by local minima (goodness of fit to integer copy number values, measured as the sum of square distances) across the purity range considered. The absolute copy number state for each bin was taken as the closest integer value calculated using this purity. If no local minimum is found the purity is assumed to be 1. If the best solution produced negative copy number states at some loci, these were set to copy number zero to avoid impossible copy number states. In two patients per sample ploidies were determined by manual adjustment due to integer ploidy values producing poor fits.

### SNV detection

Somatic mutations were first called for each tumour sample separately against matched blood derived or adjacent normal tissue samples with Mutect2 (version 4.1.4.1) using options ”–af-of-alleles-not-in resource 0.0000025 –germline-resource af-onlygnomad.hg38.vcf.gz” (Cibulskis et al., 2013; Poplin et al., 2018) Variants detected in any tumour sample (marked PASS, coverage AD 10 in both normal and tumour, at least 3 variant reads in the tumour, 0 variant reads in the normal, reference genotype in normal and non-reference genotype in cancer) were merged into a single list of “candidate mutations”. The multi-sample caller Platypus v0.8.1.1 (Rimmer et al., 2014) was then used to recall variants at each candidate mutation position in all samples of the patient. In practice, this meant that the pipeline leverage information across samples to improve the sensitivity of variant calling. The platypus output of joint variant calls was then filtered to only keep high quality variants with flags ”PASS”, ”alleleBias”, ”QD” or ”Q20”, in canonical chromosomes (i.e., not in decoy), a minimum number of reads NR>5 in all samples, a genotyping quality GQ>10 in all samples, a reference genotype (i.e., 0/0) in the normal reference and a non-reference genotype (i.e., 0/1 or 1/1) in at least one tumour sample.

To alleviate concerns of false-negative calls of mutations in important driver alterations, we generated a second set of variant calls for the identification of known driver mutations and dNdS analysis (see details below) to which we did not apply the second step of filtering.

### SNV annotation

Somatic variants were annotated and candidate driver genes of colorectal cancers reported by (Cross et al., 2018) and IntOGen (Martínez-Jiménez et al., 2020) as well as pan-cancer driver genes reported (Martincorena et al., 2017) and (Tarabichi et al., 2021) filtered with the Variant Effect Predictor v93.2 (McLaren et al., 2016).

### MSI status detection

The identification of microsatellite instability (MSI) colorectal cancers was performed with the MSIsensor v0.2 (Niu et al., 2014). We first determined the position of microsatellites sites by applying the msisensor scan method to the GRCh38 reference assembly and subset these to the first chromosome. In a second step we identified the fraction of mutated microsatellites in each sample using the msisensor msi method with default options. Generally, in known MSI cases (e.g., those identified by mutation burden and mutational signature) more than 30% of microsatellites were mutated and we used this as a critical value to classify cases as MSS and MSI.

### Extraction of reads supporting variants

Using the VCF files from both somatic and germline variant calling, we extracted the number of reads supporting the reference and alternate alleles as well as the total number of reads covering the sites from WGS, LP-WGS and ATAC-seq samples using python and the pysam library (Li et al., 2009), pysam version 0.15.2, samtools version 1.9.

### dN/dS analysis

dndscv package for R (Martincorena et al., 2017) was used for dN/dS analysis. Per-patient variant calls were obtained from the vcf files (Obenchain et al., 2014)and lifted over to the hg19 reference genome using the rtracklayer package for R (Lawrence et al., 2009) Variants were divided into clonal mutations (i.e., present in all samples) and subclonal mutations (i.e., present in a subset of samples) present in the cancer and a set of mutation present in any of the adenoma samples. MSI and MSS patients were treated separately. dndscv was applied separately to each of the four sets (MSI/MSS & clonal/subclonal) (using default parameters apart from deactivated removal of cases due to number of variants). Further, dN/dS values for a set of 167 chromatin modifier genes were extracted.

### ATAC-seq

### ATAC peak calling analysis

### Extraction of cut-sites

For the detection of cut-sites (hereafter “peaks” where read density was high) bed-files of ATAC-seq cut-sites were produced. Aligned reads were sorted by read name using “samtools sort -n{bam}”, all proper reads pairs (i.e., reads mapped to the same chromosome and with correct read orientation) were isolated using ”samtools view -bf 0×2” and finally converted to the bed format using ”bedtools bamtobed -bedpe -mate1 -i{bam}”. Equivalent to (Buenrostro et al., 2013) the start site of reads was shifted to obtain the cut sites: specifically, forward reads were shifted by −4 bases and reverse reads by +5 bases. ATAC-seq reads spanning nucleosomes have an insertion size periodicity of multiples of 200 bp and reads in regions of open-chromatin have insertion sizes smaller than 100 bp (Buenrostro et al., 2013). For this reason, in line with previous studies, ATAC-seq reads were divided into a set of nucleosome-free reads (insertion size ≤ 100) and a set of nucleosome associated reads (180 ≤ insertion size ≤ 620).

### Peak detection

Peaks were called separately for each tumour region using MACS2 v2.21 (Zhang et al., 2008) using ”macs2 callpeak -f BED -g hs –shift −75 –extsize 150 –nomodel –call-summits–keep-dup all -p 0.01” with the concatenated and sorted bed read files of nucleosome-free cut-sites of all samples as input. A set of normal peaks (across patients) were also called using the concatenated normal sample bed files (i.e., region “E” samples) and per adenoma peak calls using all adenoma bulk samples as input.

### Filtering and concatenation of peaks

Strict filtering of per-region peak calls was applied (extended by 250 bp, q-value of 0.1%, enrichment of 4.0, maximum number of peaks 20,000). Iterative merging was then applied, using a method equivalent to that used by (Corces et al., 2018) on per-region peak calls of individual patients (per-tumour peaks set) as well as across all cancer samples and pan-patient normal peak calls (pan-patient peak set). This procedure resulted in a total of N = 343,240 peaks, of which filtered N = 67,215 peaks called in >2 tumour regions or the panel of normal. The ChIPseeker v2.14.0 package for R (Yu et al., 2015) was used to annotate peaks based on their genomic location. For peaks that were not proximal to known promotor regions (1000 bp), overlaps with known Enhancer elements reported in the double-elite annotations of the GeneHancer database was examined (Fishilevich et al., 2017). The general distribution of these features in the genome and overlaps of peaks with those reported by (Corces et al., 2018).

### Extraction of cut-sites in peaks

Read counts for each peak in the final set were collated using bedtools (Quinlan and Hall, 2010) as follows: ”bedtools coverage -a bed peaks -b bed cut sites -split -counts -sorted”.

### Purity estimation for ATAC-seq samples

Clonal variants identified by paired WGS sequencing (clonal variants were those present in all samples from the cancer) were used to estimate sample-specific ATACseq purity. First, variants in intervals with identical (clonal) copy-number states (i.e., A/B states) and regions of closed chromatin, were identified from WGS data. Copy-number values *c*_*i*_ and mutation multiplicity *m*_*i*_ of each variant site *i* were obtained from the WGS data. For a mutation at site *i* covered by *n*_*S,i*_ reads in sample *S* the number of reads *k*_*i*_ containing the alternate allele is expected to follow a binomial distribution with the likelihood:

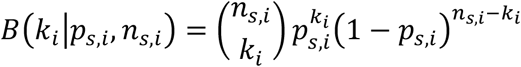

where the expected success probability *p*_*s,i*_ is a function of the sample purity as, the number of mutated alleles in the tumour cells *m*_*S,i*_, the total copy-number of the mutated site in the tumour cells *C*_*S,i*_ and the copy-number in contaminating normal cells CN=2

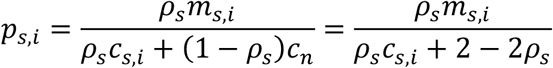

A maximum-likelihood estimate of the sample purity *ρ*_*S*_ was then obtained by minimising the negative-log-likelihood across all *N* mutated sites:

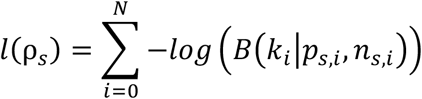

### Identification of recurrently altered peaks across patients

Analysis was restricted to samples with purity *ρ* > 0.4. Peaks proximal (≤ 1000 bp) to a transcription start site (i.e., promotors) and those more distant to a TSS (i.e., putative enhancers) were considered separately to account for the possibility of differential dispersion. An overdispersed Poisson model was fitted to each peak *edgeR* v3.30.3 (McCarthy et al., 2012; Robinson et al., 2010), and per sample set normalisation factors were calculated using the TMMwsp method (Robinson and Oshlack, 2010), estimated a global dispersion estimate across sets from all cancers and compared each set of pure glands (per-patient) against a large pool-of-normal tissue ATAC-seq samples. Independent filtering of events (CPM of at least 15 in tumour or normal, minimum fold change of 2) was applied, recurrently altered peaks called as those that were significantly altered at a level of p ≤ 0.01 in at least 5/25 (i.e., 20%) of cases.

### Identification of associated changes in gene expression

The basic processing of matched RNAseq data is described in the associated manuscript TRANSCRIPTOME. A subset of 27,731 peaks that where either adjacent to a known transcription start site (TSS) of a gene (Haeussler et al., 2018) or overlapped a previously characterised enhancer element described in the GenHancer database (Fishilevich et al., 2017) were identified. Of these 944/27731(≅ 3.40%) were recurrently altered. Changes in gene-expression of genes associated with these sites were tested for using *DESeq2* (Love et al., 2014) to compare coefficients of the fitted beta-binomial regression model (design: *~Patient*, with all normal samples as ‘Normal’) with the contrast argument being a list of vectors containing the significant and non-significant patient sets. For promotors, a one-tailed hypothesis test was applied by setting the *altHypothesis* argument to ‘less’ (for closed peaks) or ‘greater’ (for opened peaks). For enhancers a two-tailed hypothesis test on all associated genes was applied by setting the *altHypothesis* argument to ‘greaterAbs’. P-values were from all tests were adjusted for multiple hypothesis testing using FDR method (Benjamini and Hochberg, 1995) assoications at FDR<0.1% were reported. For the visualisation of gene expression values, the average gene expression values across samples from a given cancer and all normal samples on variance stabilised (log-transformed) FPM values (counts per million reads in gene) were calculated.

### Identification of subclonal changed is recurrently altered peaks

Subclonality was assessed only for a set of recurrent somatic accessibility changes, comprising recurrent events affecting drivers from CRC and the top 20 most recurrent in each of the of the 4 categories: gained promoter, lost promoter, gained enhancer, lost enhancer (total of 88 sites assessed).

Our previous analyses recognised that sample purity was highly correlated with tumour piece (regions A-D). To distinguish subclonal chromatin accessibility alterations from variability in ploidy, regression to account for purity was performed. Specifically, a log ratio test from DESeq2 (Love et al., 2014) was used to compare a “full model” *~purity + region* to a reduced model *~purity*. Samples from the same region were used as biological replicates. Events were considered putatively subclonal when the adjusted p-value was below 0.05 and if the direction of log fold change from analysis of matched bulk tissues was correlated with that observed in individual samples. In the case of gained events, subclonal events were filtered out if MACS peak-calling (see above) had not called a peak within 500 bp of the location of the putative gain event (this removed 33 sites). For losses, 5/45 subclonal events were removed as the log fold change was in the wrong direction.

For visualisation of peaks, coverage per region was calculated 1 kb upstream and 1kb downstream from the centre of the peak. Coverage was normalised per million reads in peaks and was plotted using functions from *GenomicRanges* (Lawrence et al., 2013) and *Gviz* (Hahne and Ivanek, 2016).

### TF Binding site prediction

The *motifmatchr* package for R (Schep, 2021), a reimplementation of the C++ library MOODS (Korhonen et al., 2009; Pizzi et al., 2011), was used to identify binding sites for all human TF motifs defined in a curated version of the CIS-BP database (Weirauch et al., 2014). The list of predicted binding sites was filtered using a minimum significance value of p ≤ 10^−6^, followed by removal of binding sites in centromeric regions and non-autosomal (i.e., sex and non-canonical) chromosome. After this initial filtering predicted binding sites were split into six distinct groups based on i) there distance to the next TSS (proximal: d ≤ 2000 bp, close: 2000 bp < d ≤ 10,000 bp, distal d > 10,000 bp) and ii) whether they overlapped with a peak observed in the ATAC-seq data. For a number of TF homotypic clustering of binding sites in specific intervals was observed; to account for this binding sites that where closer than d ≤ 1000 bp to the next predicted binding site of the same TF were removed.

### Extraction of signal values

For each of the TF sets described above, the counts of insertions around the centre of the TF binding site (±1000 bp) as well as the insertion size of the read pair (i.e., the distance to the second nick) for each sample (Lawrence et al., 2013) were tabulated. The insertion-sizes (rows) were binned into intervals of 5 bp and divided by total count of reads with an equivalent size in the entire genome. After this the background signal was estimated to be the average number of insertions 1000 bp – 750 bp from the centre of TF binding site per insertion size and subtracted from the counts. The difference between these “normalised and background corrected TF signals” in each sample and a pool of normal samples was calculated and integrated across the central region of the TF binding sites (insertion size [25;120], distances [-100 bp;100 bp]) as a summary statistic. Regression analysis linear regression was used to identify associations with purity estimates and in this context signals were found to correlated with TSSe (for both nucleosome-free and all reads). For this reason, an additional term was added to the regression model of each TF to correct for this effect: *signal ~ tsse*tsse_nf_ + purity:patient* where tsse and *tsse_nf_* are the TSSe differences of the sample and the pooled-normal samples) and weighted each observation by the square root of the number of reads in the sample. A second linear model in which a region-specific effect of the purity: *signal ~ tsse’tsse_nf_ + purity:region* was considered was also fitted to the data. For both models, the statistical significance of the ‘purity’ coefficient was determined. The estimates of the coefficients were also used as a patient specific summary for subsequent analysis.

### Cluster analysis

The analysis was focused on the 150 TF for which a significant association with the tumour cell content (i.e., the purity) and TF signal was most frequently observed. With the aim to identify general patterns in these data, a clustering analysis was conducted (hierarchical clustering with Euclidean distance and complete linkage). This method identified three major groups of TFs, and to each of these, analysis with String-DB (Szklarczyk et al., 2018) was applied to identify significantly overrepresented pathways.

### Methylation arrays analysis

A reference normal dataset methylation array dataset was downloaded from (Fennell et al., 2019) that including normal tissue sampled adjacent to colorectal cancers that was profiled using the HumanMethylation450 BeadChip array (Illumina).

Here, 8 bulk samples from 4 patients (C516, C518, C560 and C561) were profiled MethylationEPIC BeadChip (Infinium) microarray according to manufactorer’s instructions.

The ChAMP R package pipeline (Tian et al., 2017) was used to analyse the methylation beadarray data. Probes that had a detection of P > 0.01 and probes with <3 beads in at least 5% of samples per probe, probes that were on the X or Y chromosome, all SNP-related probe as well as all multi-hit probes were all removed. Subset-within-array normalization to was used to correct for biases resulting from type 1 and type 2 probes on the array. After QC and normalization, beta values were calculated for further comparison.

To compare the methylation patterns between our samples and the reference normal dataset, the overlapped probes of all samples located in the region of distal to TSS (dTSS), close to TSS (cTSS) and proximal to TSS (pTSS) in both on ATAC peak (oPEAK) and not on ATAC peak (nPEAK) were compared.

### Mutational signatures analysis

Mutational signatures analysis was performed with SparseSignatures (Lal et al., 2021). This method uses LASSO regularization (Tibshirani, 1996) to reduce noise in the signatures, controlled by a regularization parameter lambda (λ). It implements a procedure based on bi-cross-validation (Owen and Perry, 2009) to select the best values for both the regularization parameter λ and the number of signatures. Deconvolution using a maximum of 10 signatures was performed and values of λ of 0.000, 0.025, 0.050 and 0.100 were tested. Optimal parameters were selected based on the median bi-cross-validation error estimated over 1000 iterations, resulting in an optimal estimate with minimum cross-validation median error when 6 signatures were fitted and λ=0.025. A second analysis with SigProfiler (Alexandrov et al., 2020), with default parameters and a total of 1000 iterations, confirmed the existence of these signatures. Signatures based clustering was performed considering the 6 signatures solution by SparseSignatures; the low-rank signatures exposure matrix given as an output by the tool was used to compute the pairwise similarity matrix for each patient as 1 - cosine similarity of their exposures. Clustering was then performed on the similarity matrix by k-means with 6 clusters explaining all the variance.

Mutational signatures exposures were also analyzed across epigenetic regions. Mutations were first grouped in clonal or subclonal across whole genome and then in different genomic regions (as described above). Signatures activities in each region was estimated by Jackknife sampling (Efron and Stein, 1981). Specifically, data from each patient were partitioned based on their clusters as defined above, and repeated Jackknife sampling performed 100 times independently for each of the 3 clusters (including a random sample of 90% of the tissue samples each time). For each iteration the mutations within each genomic region were used to computed a data matrix normalised against trinucleotide count (across the 96 channels) in the whole genome versus region specific counts, and signatures assignments then performed on the normalized data by LASSO (Lal et al., 2021; Tibshirani, 1996). Finally, relative signature activities estimated over the 100 Jackknife samples were normalized based on total size of each region.

## Acknowledgments

This study was principally supported by funding from the Medical Research Council (MR/P000789/1 to A.S.) and the Wellcome Trust (202778/Z/16/Z to T.A.G. and 202778/B/16/Z to A.S.). A.S. and T.A.G. were also supported by Cancer Research UK (A22909 and A19771) and the National Institute of Health (NCI U54 CA217376 to D.S., T.A.G. and A.S). This work was also supported by a Wellcome Trust award to the Centre for Evolution and Cancer at the ICR (105104/Z/14/Z). D.R. was partially supported by a Bicocca 2020 Starting Grant and by a Premio Giovani Talenti dell’Università degli Studi di Milano-Bicocca. L.M. is supported by Cancer Research UK (A23110).

## Data availability

Analysed data are available on Mendeley: https://data.mendeley.com/datasets/dvv6kf856g/2. Sequence data have been deposited at the European Genome-phenome Archive (EGA), which is hosted by the EBI and the CRG, under accession number EGAS00001005230. Further information about EGA can be found on https://ega-archive.org.

## Code availability

Complete scripts to replicate all bioinformatic analysis and perform simulations and inference are available at: https://github.com/sottorivalab/EPICC2021_data_analysis.

## Author contributions

T.H. analysed and interpreted the data, with focus on ATAC and WGS data. J.H. analysed and interpreted the data, with focus on RNA data. GDC performed copy number analysis. I.S. devise multi-omics protocol, collected the samples and generated the data. C.K. collected the samples and contributed to data generation. C.L. contributed to ATAC data analysis. M.M., J.F.M., A.M.B., H.C., M.M., contributed to data generation. B.J. analysed methylation array data. L.Z.O. contributed to dN/dS data analysis. C.J., E.L., G.Car. contributed to data analysis. D.N. and K.C. contributed to signature analysis. A.B. generated methylation array data. I.B. contributed to analysis of CTCF binding site mutations. M.J. contributed to tissue collection. D.R. performed mutational signature analysis. D.S. contributed to experimental design and data interpretation. J.B. contributed to sample collection coordination. M.R.J. supervised sample collection. L.M. contributed to result interpretation. T.A.G. and A.S. conceived and supervised the study and wrote the manuscript.

## Supplementary Figure Legends

**Figure S1. Colectomy specimen collection images.** Resection specimens were collected from UCLH and sampled with the supervision of a pathologist. Spatial information on different regional samples was retained and indicated in the images. A, B, C, D are cancer regions. E is distant normal epithelium. Eventual concomitant adenomas are reported as F, G, H, etc.

**Figure S2. Gland and bulk collection from each tumour region.** We collected individual glands from cancer and normal samples from different regions of each tumour. We also collected ‘minibulks’, composed by agglomerate of a few dozen glands. Each sample was imaged individually.

**Figure S3. Copy number alteration profiles.** We estimated absolute copy number alterations for each sample in each patient, both for deep WGS and low-pass WGS.

**Figure S4. Single nucleotide variant profiles.** We called point mutations and indels in each sample and identified clusters of mutations found at the same frequency in the same samples. Values in Cancer Cell Fraction (CCF) are represented.

Figure S5. Mutations in chromatin modifier genes for all samples.

Figure S6. Images of all the normal samples used for ATAC-seq reference.

Figure S7. Gene expression differences for all the recurrent peaks that correlated with gene expression. EMT genes at the end.

Figure S8. Comparison of peak calling in our cohort from reanalysed TCGA ATAC-seq data.

Figure S9. Copy number differences for all the peaks in Figure 3E and F.

Figure S10. Peak densities for promoter gained loci in Figure 3E.

Figure S11. Peak densities for promoter lost loci in Figure 3E.

Figure S12. Peak densities for enhancer gained loci in Figure 3F.

Figure S13. Peak densities for enhancer lost loci in Figure 3F.

Figure S14. Peak densities for peaks found in cancers but not in concomitant adenomas in Figure 3E,F.

Figure S15. Peak densities for promoter gained peaks found subclonal from Figure 3E,F.

Figure S16. Peak densities for promoter lost peaks found subclonal from Figure 3E,F.

Figure S17. Peak densities for enhancer gained peaks found subclonal from Figure 3E,F.

Figure S18. Peak densities for enhancer lost peaks found subclonal from Figure 3E,F.

Figure S19. Transcription Factor binding sites density plots for annotations in Figure 4D.

Figure S20. Overlapping of TF annotations in Figure 4D.

Figure S21. Correlation of the TF signal in Figure 4D between all versus only unique loci.

Figure S22. Gene expression of TFs from cluster 1 of heatmap in Figure 4A.

Figure S23. Coefficients of the ANOVA model for the correlation between genetic and epigenetic distance for each region.

Figure S24. Mutational signature deconvolution with SigProfiler.

Figure S25. Coefficient of the purity model for TF loci in Figure 4D for each region.

Figure S26. Predicted versus observed mutational signatures that cause gain and loss of CTCF.

Figure S27. Accumulation of different mutational signatures in distinct epigenetic regions.

