## Supplementary Figures for "The co-evolution of the genome and epigenome in colorectal cancer": figS2.pdf

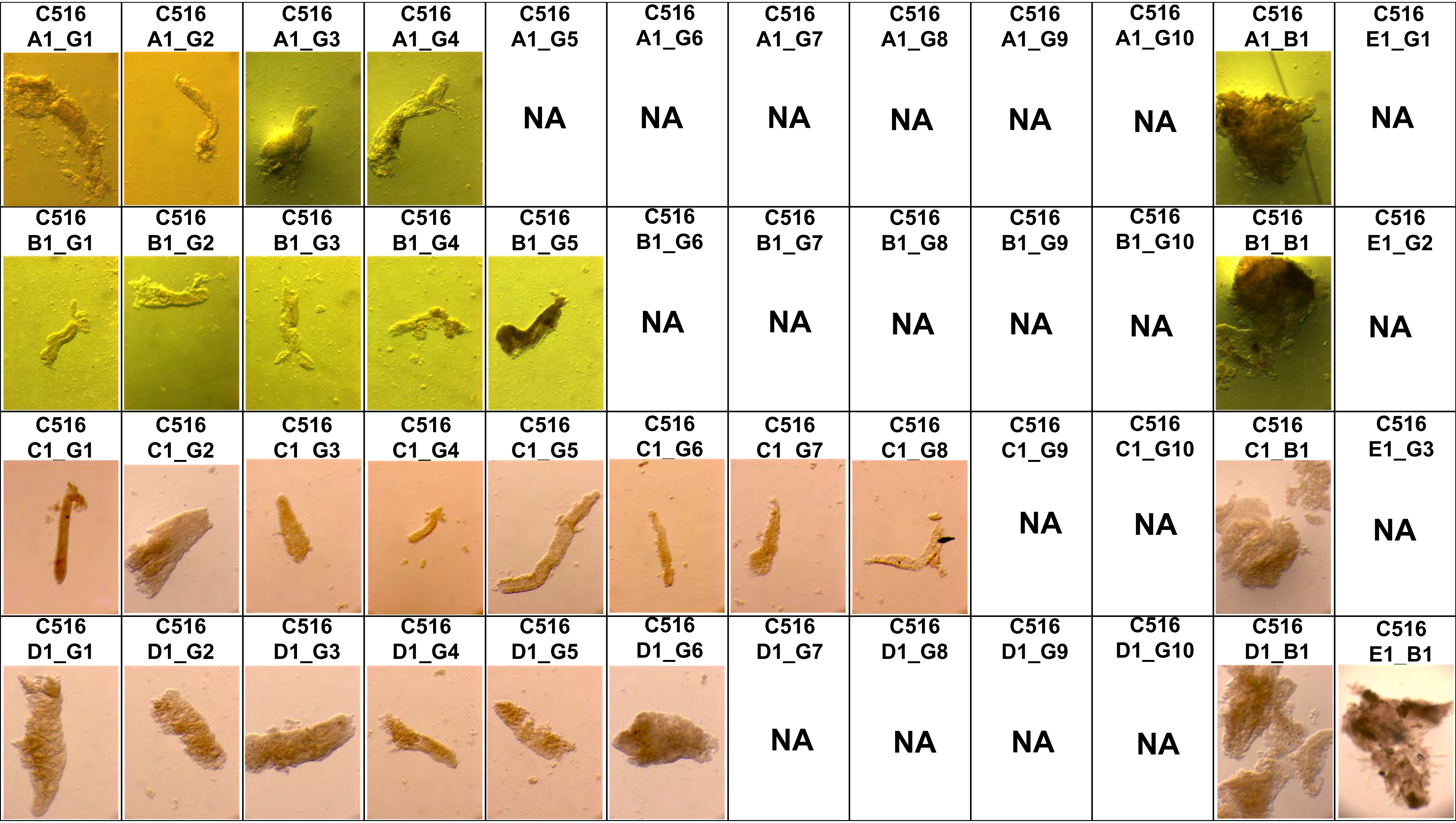
C516 A1_G1
C516 A1_G2
C516 A1_G3
C516 A1_G4
C516 A1_G5
NA
C516 A1_G6
NA
C516 A1_G7
NA
C516 A1_G8
NA
C516 A1_G9
NA
C516 A1_G10
NA
C516 A1_B1
C516 E1_G1
NA
C516 B1_G1
C516 B1_G2
C516 B1_G3
C516 B1_G4
C516 B1_G5
C516 B1_G6
NA
C516 B1_G7
NA
C516 B1_G8
NA
C516 B1_G9
NA
C516 B1_G10
NA
C516 B1_B1
C516 E1_G2
NA
C516 C1_G1
C516 C1_G2
C516 C1_G3
C516 C1_G4
C516 C1_G5
C516 C1_G6
C516 C1_G7
C516 C1_G8
C516 C1_G9
NA
C516 C1_G10
NA
C516 C1_B1
C516 E1_G3
NA
C516 D1_G1
C516 D1_G2
C516 D1_G3
C516 D1_G4
C516 D1_G5
C516 D1_G6
C516 D1_G7
NA
C516 D1_G8
NA
C516 D1_G9
NA
C516 D1_G10
NA
C516 D1_B1
C516 E1_B1

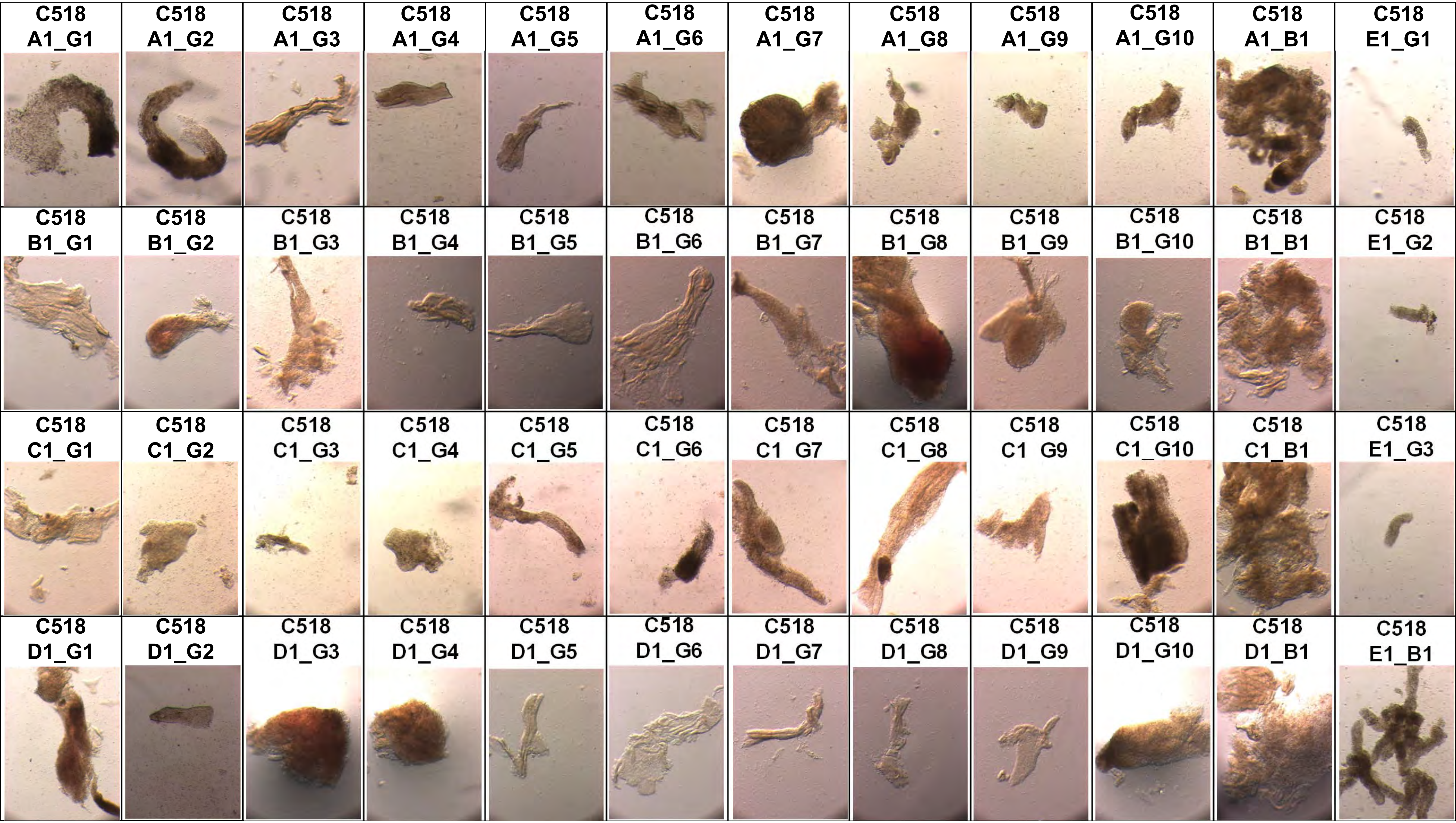
C518 A1_G1
C518 A1_G2
C518 A1_G3
C518 A1_G4
C518 A1_G5
C518 A1_G6
C518 A1_G7
C518 A1_G8
C518 A1_G9
C518 A1_G10
C518 A1_B1
C518 E1_G1
C518 B1_G1
C518 B1_G2
C518 B1_G3
C518 B1_G4
C518 B1_G5
C518 B1_G6
C518 B1_G7
C518 B1_G8
C518 B1_G9
C518 B1_G10
C518 B1_B1
C518 E1_G2
C518 C1_G1
C518 C1_G2
C518 C1_G3
C518 C1_G4
C518 C1_G5
C518 C1_G6
C518 C1 G7
C518 C1_G8
C518 C1 G9
C518 C1_G10
C518 C1_B1
C518 E1_G3
C518 D1_G1
C518 D1_G2
C518 D1_G3
C518 D1_G4
C518 D1_G5
C518 D1_G6
C518 D1_G7
C518 D1_G8
C518 D1_G9
C518 D1_G10
C518 D1_B1
C518 E1_B1

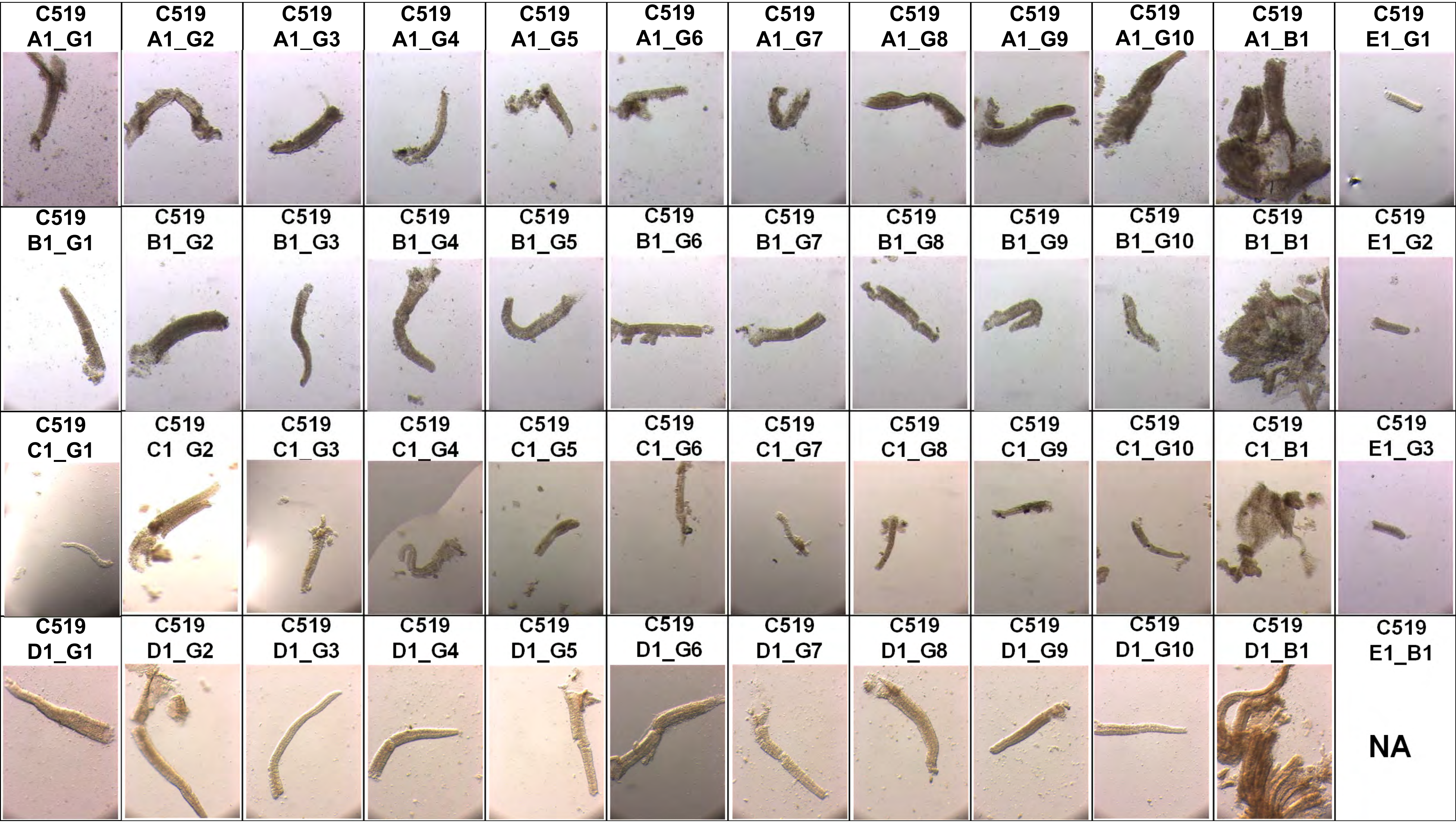
C519 A1_G1
C519 A1_G2
C519 A1_G3
C519 A1_G4
C519 A1_G5
C519 A1_G6
C519 A1_G7
C519 A1_G8
C519 A1_G9
C519 A1_G10
C519 A1_B1
C519 E1_G1
C519 B1_G1
C519 B1_G2
C519 B1_G3
C519 B1_G4
C519 B1_G5
C519 B1_G6
C519 B1_G7
C519 B1_G8
C519 B1_G9
C519 B1_G10
C519 B1_B1
C519 E1_G2
C519 C1_G1
C519 C1 G2
C519 C1_G3
C519 C1_G4
C519 C1_G5
C519 C1_G6
C519 C1_G7
C519 C1_G8
C519 C1_G9
C519 C1_G10
C519 C1_B1
C519 E1_G3
C519 D1_G1
C519 D1_G2
C519 D1_G3
C519 D1_G4
C519 D1_G5
C519 D1_G6
C519 D1_G7
C519 D1_G8
C519 D1_G9
C519 D1_G10
C519 D1_B1
C519 E1_B1
NA

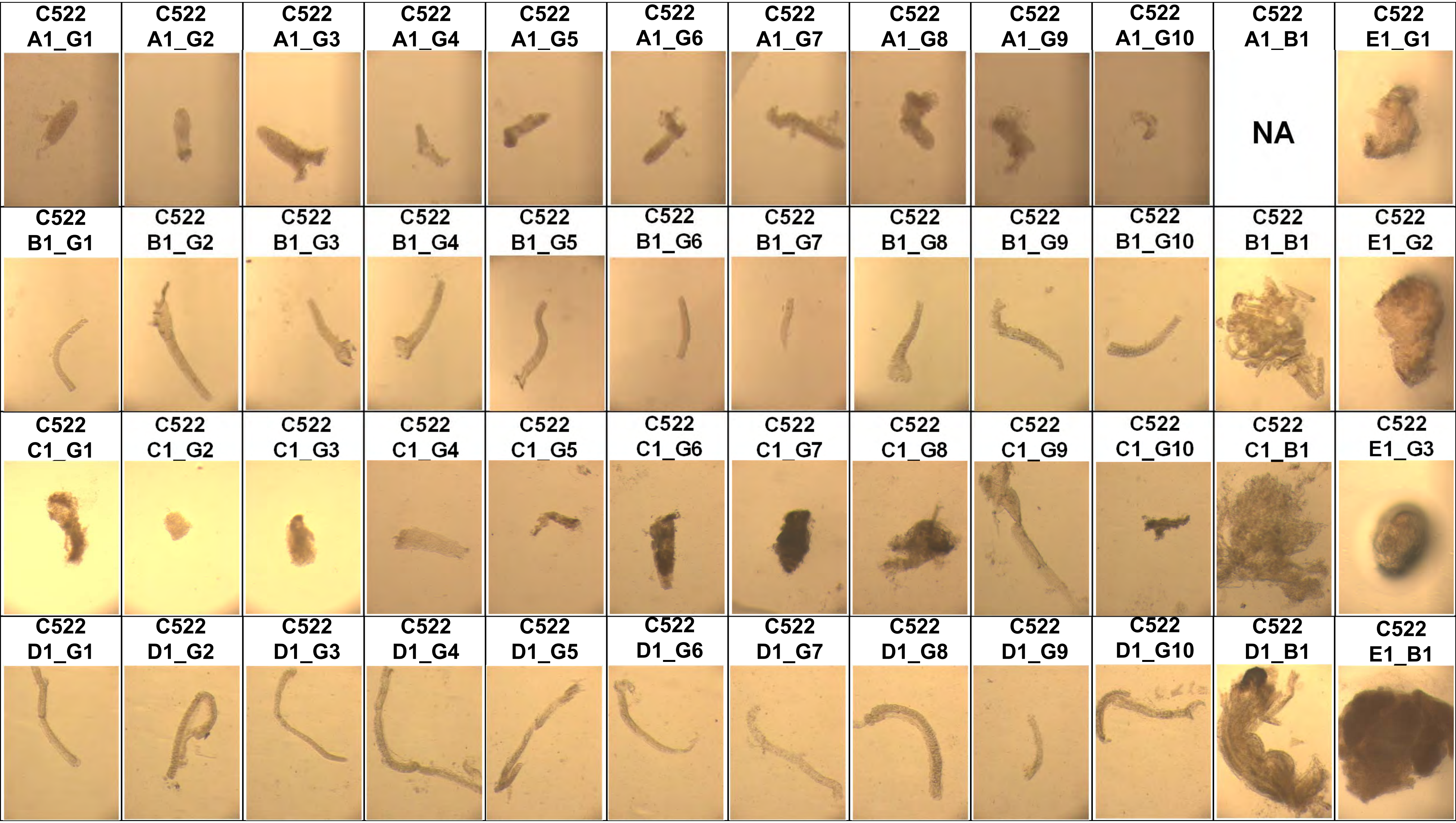
C522 A1_G1
C522 A1_G2
C522 A1_G3
C522 A1_G4
C522 A1_G5
C522 A1_G6
C522 A1_G7
C522 A1_G8
C522 A1_G9
C522 A1_G10
C522 A1_B1
NA
C522 E1_G1
C522 B1_G1
C522 B1_G2
C522 B1_G3
C522 B1_G4
C522 B1_G5
C522 B1_G6
C522 B1_G7
C522 B1_G8
C522 B1_G9
C522 B1_G10
C522 B1_B1
C522 E1_G2
C522 C1_G1
C522 C1_G2
C522 C1_G3
C522 C1_G4
C522 C1_G5
C522 C1_G6
C522 C1_G7
C522 C1_G8
C522 C1_G9
C522 C1_G10
C522 C1_B1
C522 E1_G3
C522 D1_G1
C522 D1_G2
C522 D1_G3
C522 D1_G4
C522 D1_G5
C522 D1_G6
C522 D1_G7
C522 D1_G8
C522 D1_G9
C522 D1_G10
C522 D1_B1
C522 E1_B1

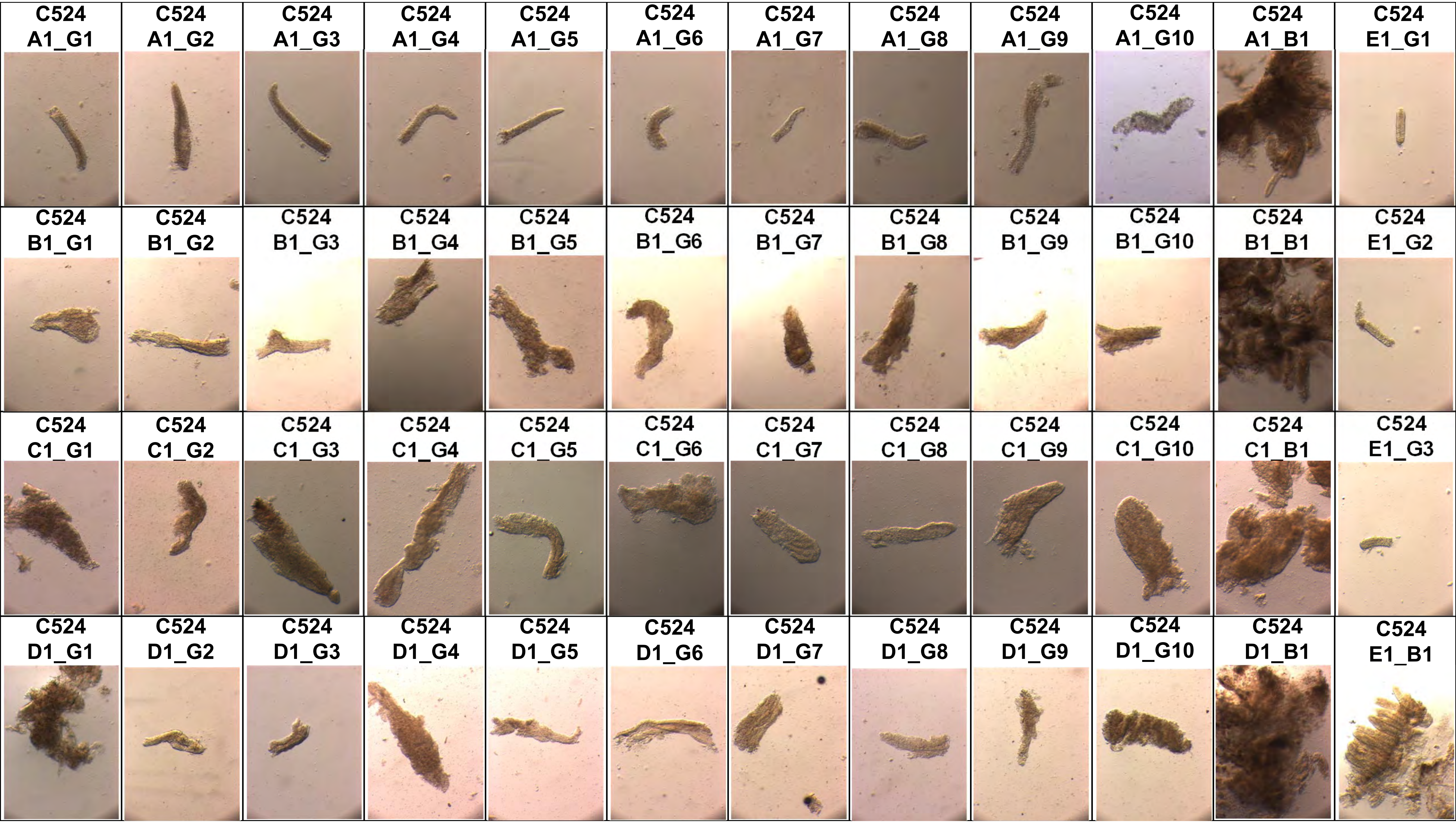
C524 A1_G1
C524 A1_G2
C524 A1_G3
C524 A1_G4
C524 A1_G5
C524 A1_G6
C524 A1_G7
C524 A1_G8
C524 A1_G9
C524 A1_G10
C524 A1_B1
C524 E1_G1
C524 B1_G1
C524 B1_G2
C524 B1_G3
C524 B1_G4
C524 B1_G5
C524 B1_G6
C524 B1_G7
C524 B1_G8
C524 B1_G9
C524 B1_G10
C524 B1_B1
C524 E1_G2
C524 C1_G1
C524 C1_G2
C524 C1_G3
C524 C1_G4
C524 C1_G5
C524 C1_G6
C524 C1_G7
C524 C1_G8
C524 C1_G9
C524 C1_G10
C524 C1_B1
C524 E1_G3
C524 D1_G1
C524 D1_G2
C524 D1_G3
C524 D1_G4
C524 D1_G5
C524 D1_G6
C524 D1_G7
C524 D1_G8
C524 D1_G9
C524 D1_G10
C524 D1_B1
C524 E1_B1

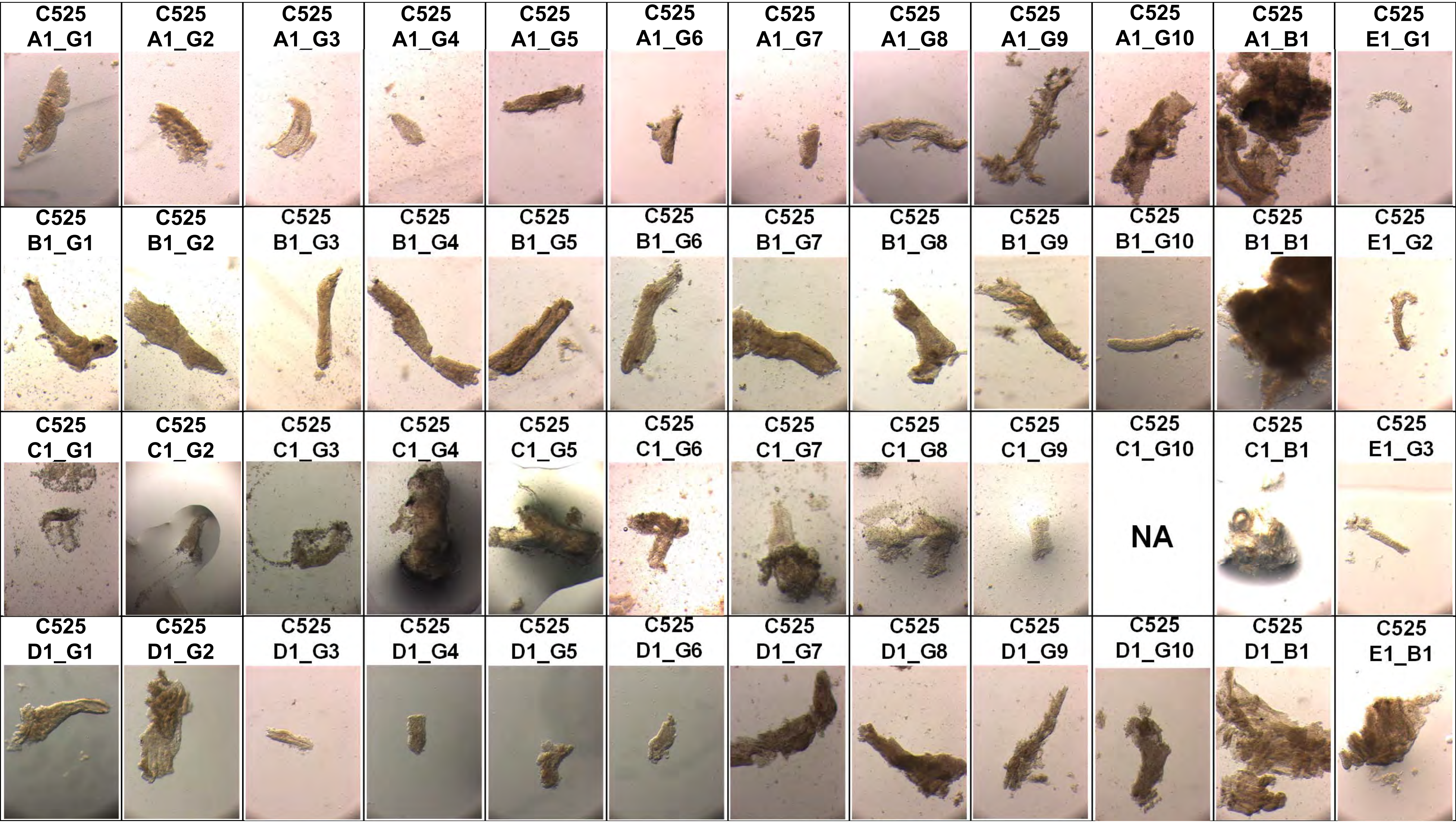
C525 A1_G1
C525 A1_G2
C525 A1_G3
C525 A1_G4
C525 A1_G5
C525 A1_G6
C525 A1_G7
C525 A1_G8
C525 A1_G9
C525 A1_G10
C525 A1_B1
C525 E1_G1
C525 B1_G1
C525 B1_G2
C525 B1_G3
C525 B1_G4
C525 B1_G5
C525 B1_G6
C525 B1_G7
C525 B1_G8
C525 B1_G9
C525 B1_G10
C525 B1_B1
C525 E1_G2
C525 C1_G1
C525 C1_G2
C525 C1_G3
C525 C1_G4
C525 C1_G5
C525 C1_G6
C525 C1_G7
C525 C1_G8
C525 C1_G9
C525 C1_G10
NA
C525 C1_B1
C525 E1_G3
C525 D1_G1
C525 D1_G2
C525 D1_G3
C525 D1_G4
C525 D1_G5
C525 D1_G6
C525 D1_G7
C525 D1_G8
C525 D1_G9
C525 D1_G10
C525 D1_B1
C525 E1_B1

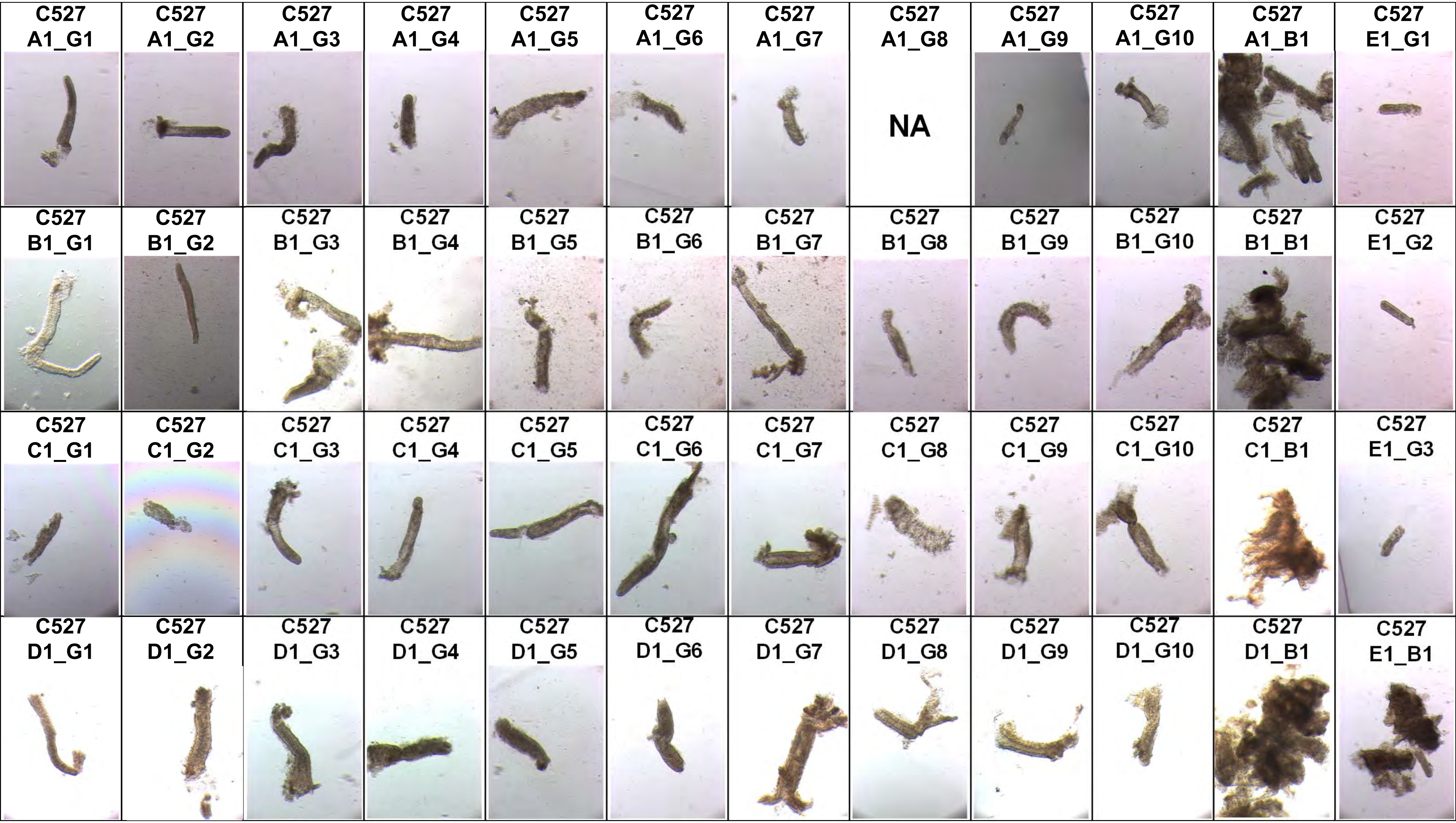
C527 A1_G1
C527 A1_G2
C527 A1_G3
C527 A1_G4
C527 A1_G5
C527 A1_G6
C527 A1_G7
C527 A1_G8
NA
C527 A1_G9
C527 A1_G10
C527 A1_B1
C527 E1_G1
C527 B1_G1
C527 B1_G2
C527 B1_G3
C527 B1_G4
C527 B1_G5
C527 B1_G6
C527 B1_G7
C527 B1_G8
C527 B1_G9
C527 B1_G10
C527 B1_B1
C527 E1_G2
C527 C1_G1
C527 C1_G2
C527 C1_G3
C527 C1_G4
C527 C1_G5
C527 C1_G6
C527 C1_G7
C527 C1_G8
C527 C1_G9
C527 C1_G10
C527 C1_B1
C527 E1_G3
C527 D1_G1
C527 D1_G2
C527 D1_G3
C527 D1_G4
C527 D1_G5
C527 D1_G6
C527 D1_G7
C527 D1_G8
C527 D1_G9
C527 D1_G10
C527 D1_B1
C527 E1_B1

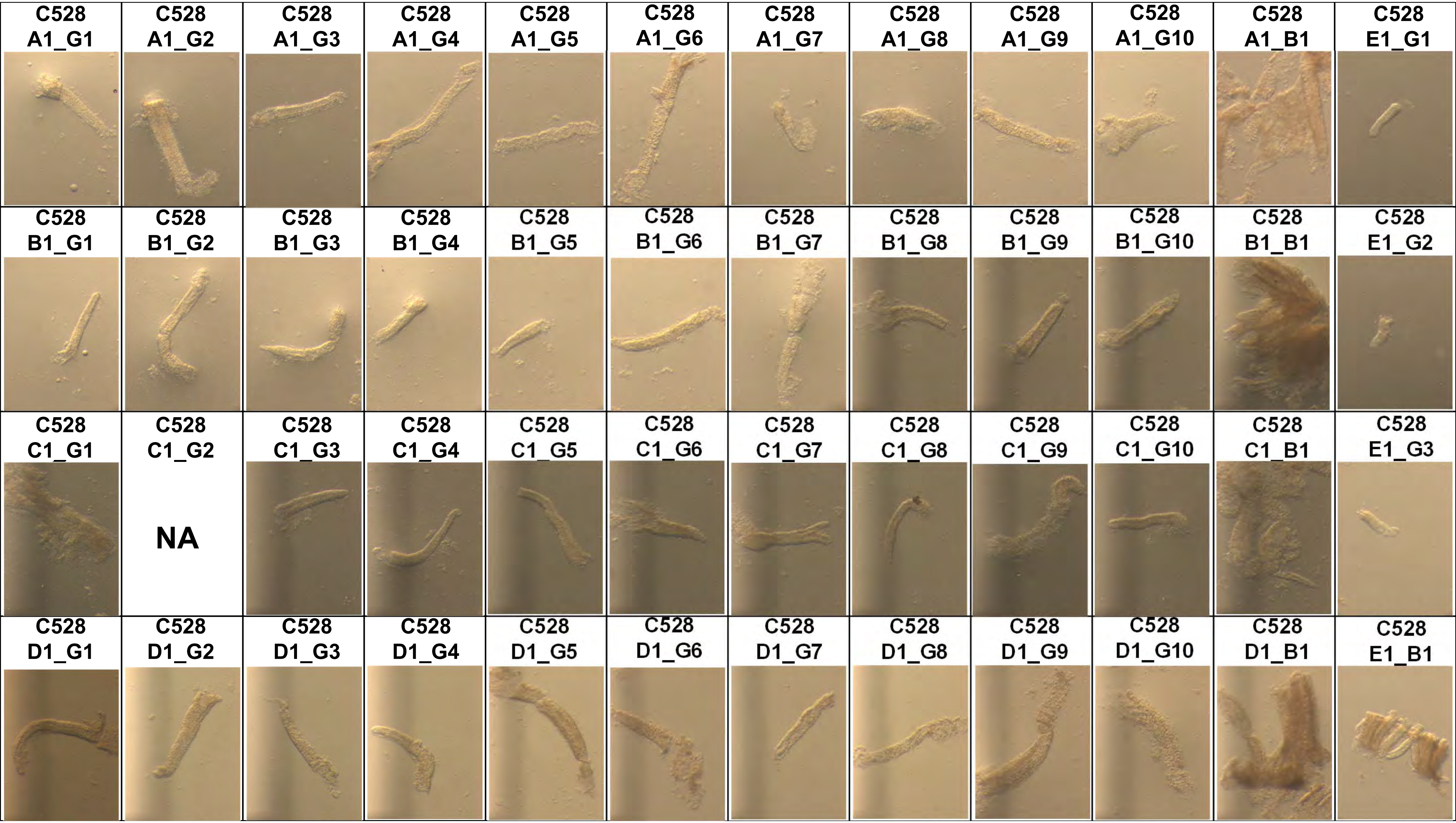
C528 A1_G1
C528 A1_G2
C528 A1_G3
C528 A1_G4
C528 A1_G5
C528 A1_G6
C528 A1_G7
C528 A1_G8
C528 A1_G9
C528 A1_G10
C528 A1_B1
C528 E1_G1
C528 B1_G1
C528 B1_G2
C528 B1_G3
C528 B1_G4
C528 B1_G5
C528 B1_G6
C528 B1_G7
C528 B1_G8
C528 B1_G9
C528 B1_G10
C528 B1_B1
C528 E1_G2
C528 C1_G1
C528 C1_G2
NA
C528 C1_G3
C528 C1_G4
C528 C1_G5
C528 C1_G6
C528 C1_G7
C528 C1_G8
C528 C1_G9
C528 C1_G10
C528 C1_B1
C528 E1_G3
C528 D1_G1
C528 D1_G2
C528 D1_G3
C528 D1_G4
C528 D1_G5
C528 D1_G6
C528 D1_G7
C528 D1_G8
C528 D1_G9
C528 D1_G10
C528 D1_B1
C528 E1_B1

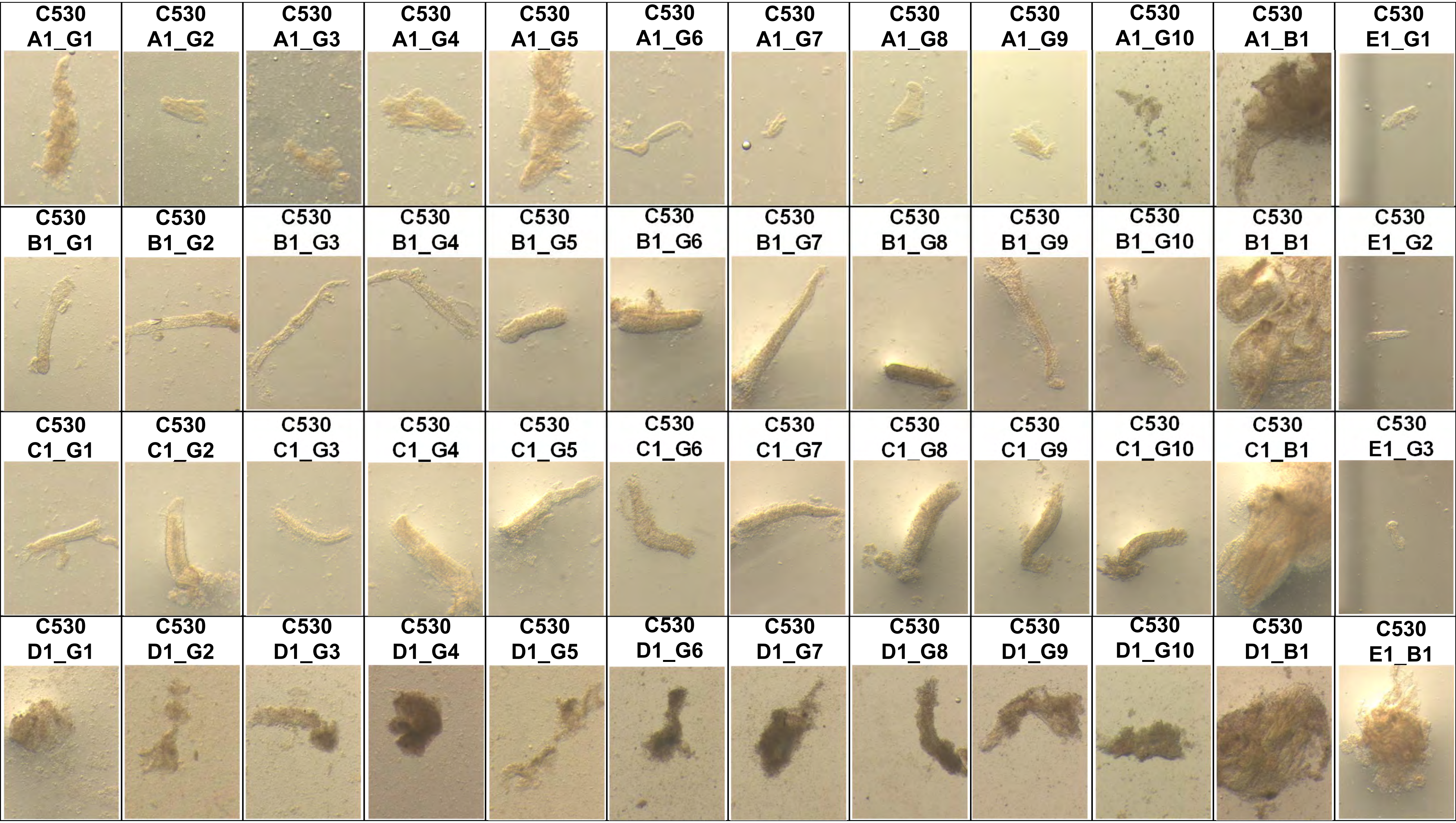
C530 A1_G1
C530 A1_G2
C530 A1_G3
C530 A1_G4
C530 A1_G5
C530 A1_G6
C530 A1_G7
C530 A1_G8
C530 A1_G9
C530 A1_G10
C530 A1_B1
C530 E1_G1
C530 B1_G1
C530 B1_G2
C530 B1_G3
C530 B1_G4
C530 B1_G5
C530 B1_G6
C530 B1_G7
C530 B1_G8
C530 B1_G9
C530 B1_G10
C530 B1_B1
C530 E1_G2
C530 C1_G1
C530 C1_G2
C530 C1_G3
C530 C1_G4
C530 C1_G5
C530 C1_G6
C530 C1_G7
C530 C1_G8
C530 C1_G9
C530 C1_G10
C530 C1_B1
C530 E1_G3
C530 D1_G1
C530 D1_G2
C530 D1_G3
C530 D1_G4
C530 D1_G5
C530 D1_G6
C530 D1_G7
C530 D1_G8
C530 D1_G9
C530 D1_G10
C530 D1_B1
C530 E1_B1

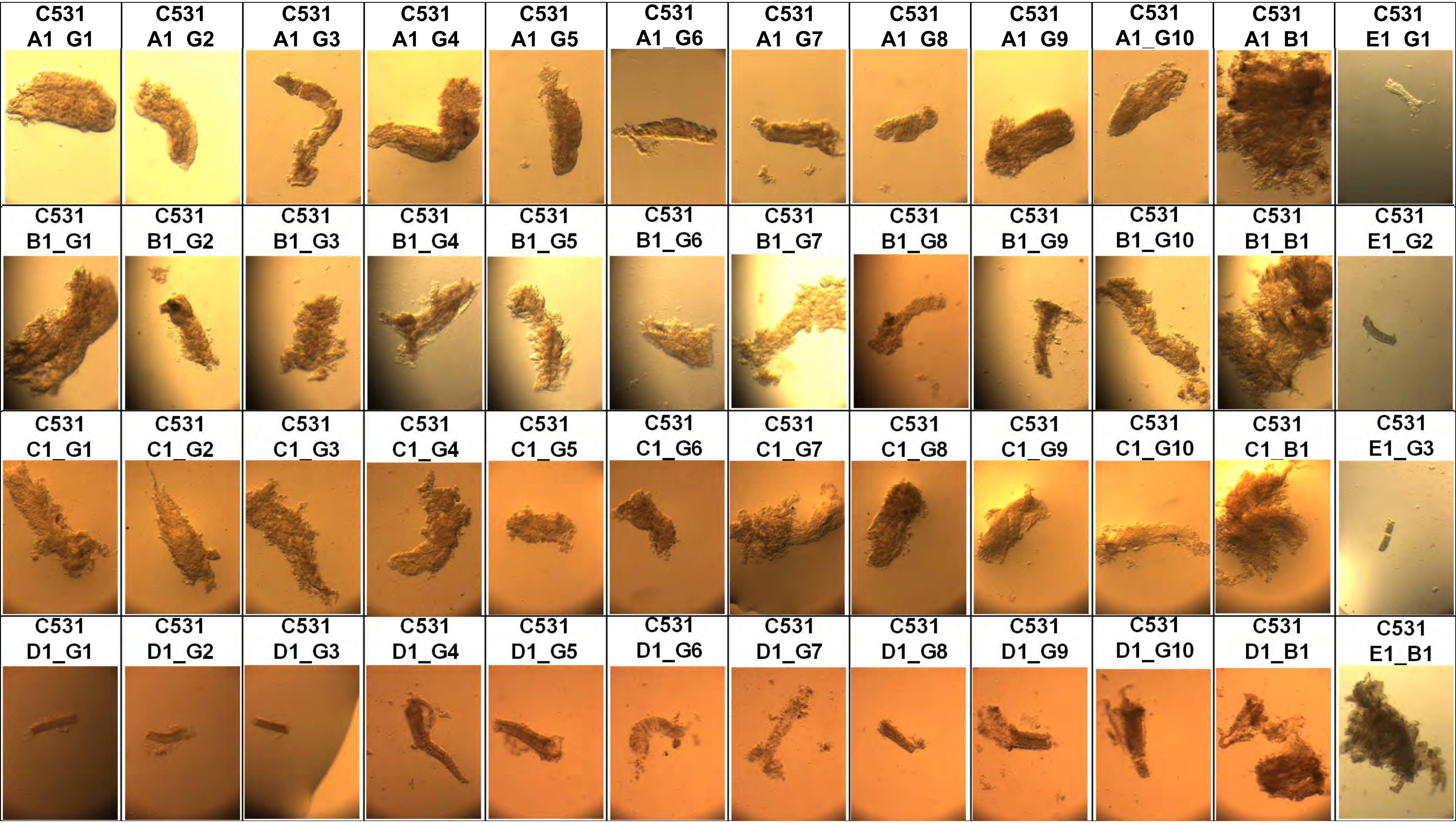
C531 A1 G1
C531 A1 G2
C531 A1 G3
C531 A1 G4
C531 A1 G5
C531 A1_G6
C531 A1 G7
C531 A1 G8
C531 A1 G9
C531 A1_G10
C531 A1 B1
C531 E1 G1
C531 B1_G1
C531 B1_G2
C531 B1_G3
C531 B1_G4
C531 B1_G5
C531 B1_G6
C531 B1_G7
C531 B1_G8
C531 B1_G9
C531 B1_G10
C531 B1_B1
C531 E1_G2
C531 C1_G1
C531 C1_G2
C531 C1_G3
C531 C1_G4
C531 C1_G5
C531 C1_G6
C531 C1_G7
C531 C1_G8
C531 C1_G9
C531 C1_G10
C531 C1_B1
C531 E1_G3
C531 D1_G1
C531 D1_G2
C531 D1_G3
C531 D1_G4
C531 D1_G5
C531 D1_G6
C531 D1_G7
C531 D1_G8
C531 D1_G9
C531 D1_G10
C531 D1_B1
C531 E1_B1

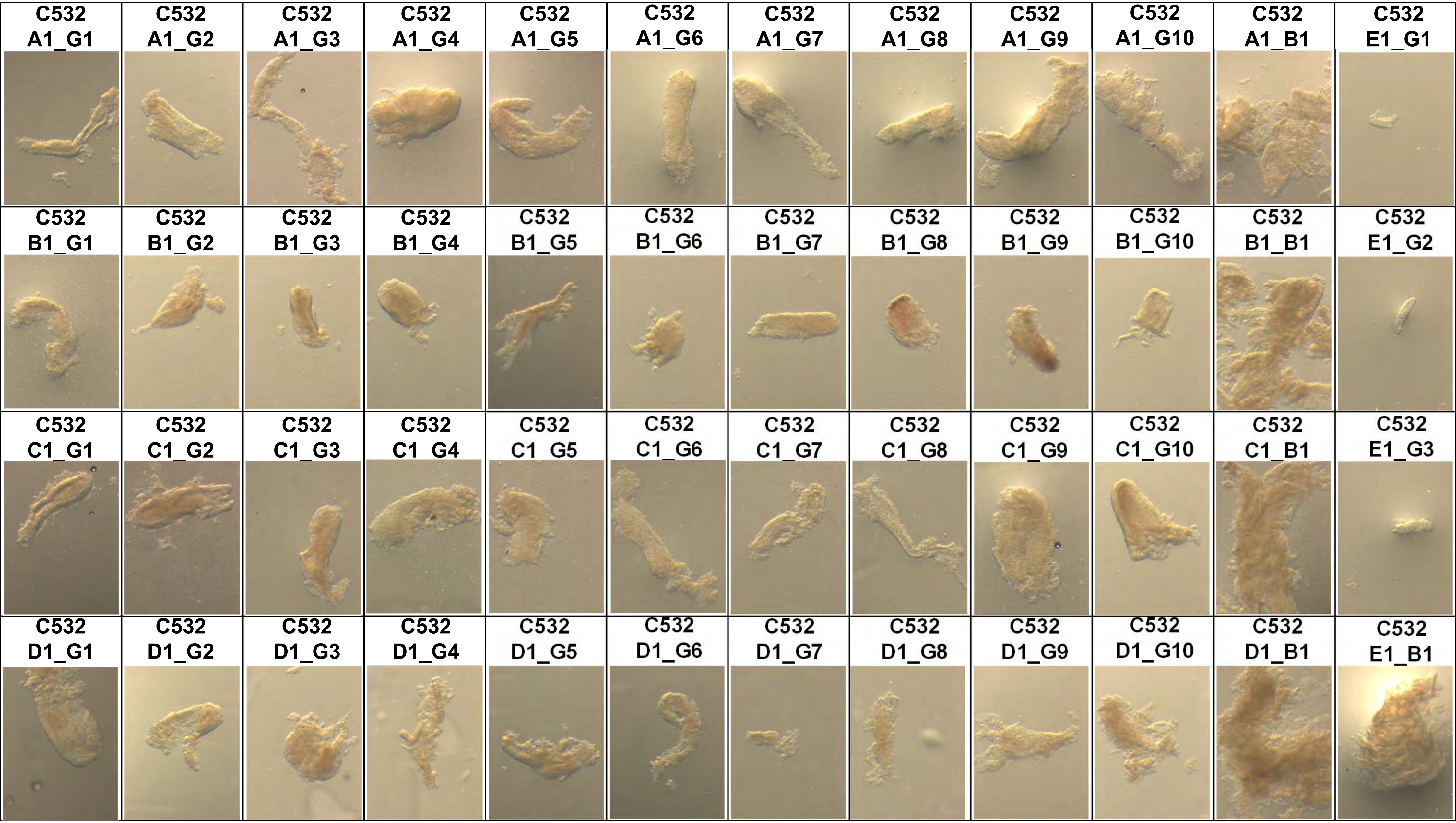
C532 A1_G1
C532 A1_G2
C532 A1_G3
C532 A1_G4
C532 A1_G5
C532 A1_G6
C532 A1_G7
C532 A1_G8
C532 A1_G9
C532 A1_G10
C532 A1_B1
C532 E1_G1
C532 B1_G1
C532 B1_G2
C532 B1_G3
C532 B1_G4
C532 B1_G5
C532 B1_G6
C532 B1_G7
C532 B1_G8
C532 B1_G9
C532 B1_G10
C532 B1_B1
C532 E1_G2
C532 C1_G1
C532 C1_G2
C532 C1_G3
C532 C1 G4
C532 C1 G5
C532 C1_G6
C532 C1_G7
C532 C1_G8
C532 C1_G9
C532 C1_G10
C532 C1_B1
C532 E1_G3
C532 D1_G1
C532 D1_G2
C532 D1_G3
C532 D1_G4
C532 D1_G5
C532 D1_G6
C532 D1_G7
C532 D1_G8
C532 D1_G9
C532 D1_G10
C532 D1_B1
C532 E1_B1

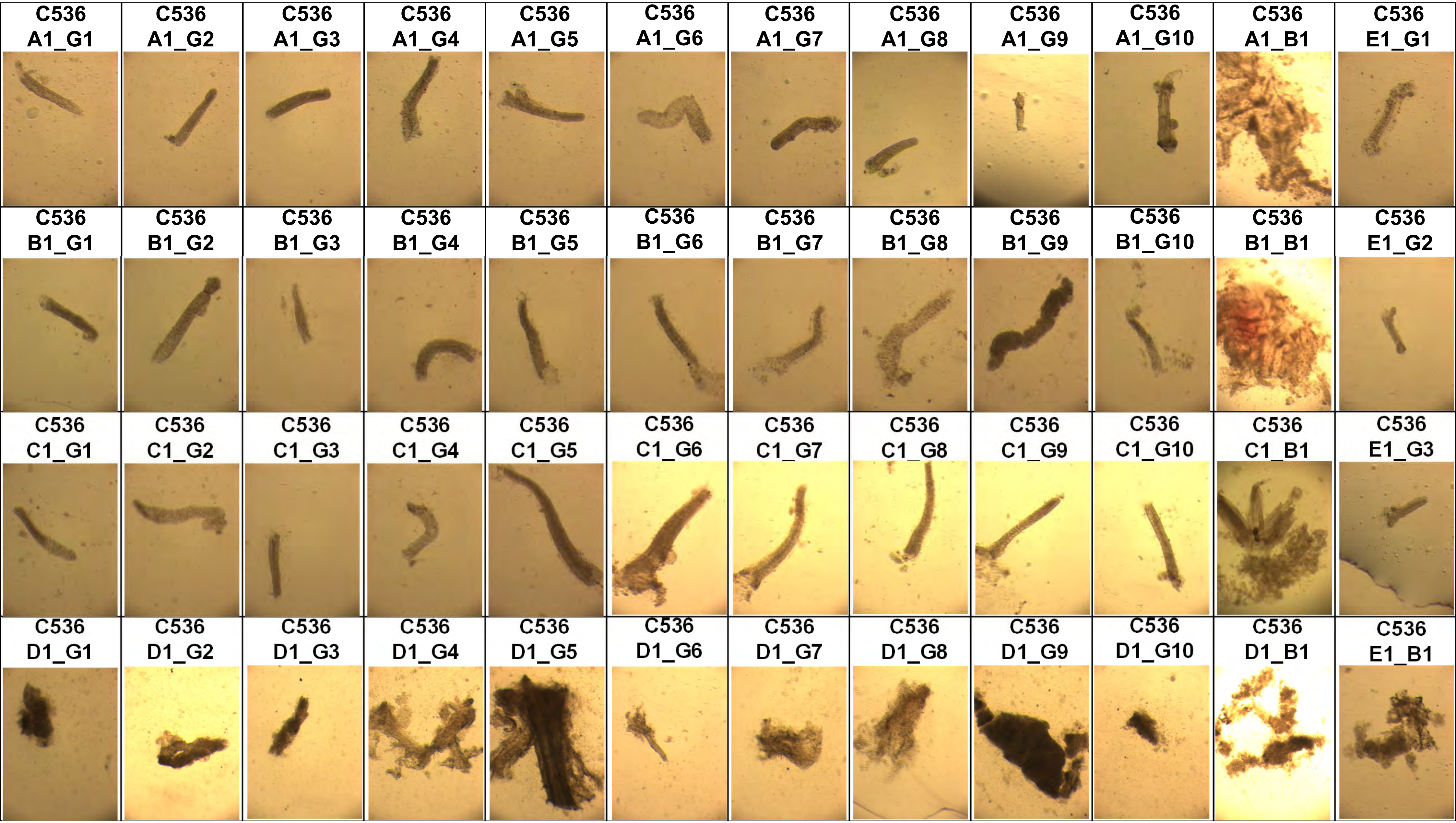
C536 A1_G1
C536 A1_G2
C536 A1_G3
C536 A1_G4
C536 A1_G5
C536 A1_G6
C536 A1_G7
C536 A1_G8
C536 A1_G9
C536 A1_G10
C536 A1_B1
C536 E1_G1
C536 B1_G1
C536 B1_G2
C536 B1_G3
C536 B1_G4
C536 B1_G5
C536 B1_G6
C536 B1_G7
C536 B1_G8
C536 B1_G9
C536 B1_G10
C536 B1_B1
C536 E1_G2
C536 C1_G1
C536 C1_G2
C536 C1_G3
C536 C1_G4
C536 C1_G5
C536 C1_G6
C536 C1_G7
C536 C1_G8
C536 C1_G9
C536 C1_G10
C536 C1_B1
C536 E1_G3
C536 D1_G1
C536 D1_G2
C536 D1_G3
C536 D1_G4
C536 D1_G5
C536 D1_G6
C536 D1_G7
C536 D1_G8
C536 D1_G9
C536 D1_G10
C536 D1_B1
C536 E1_B1

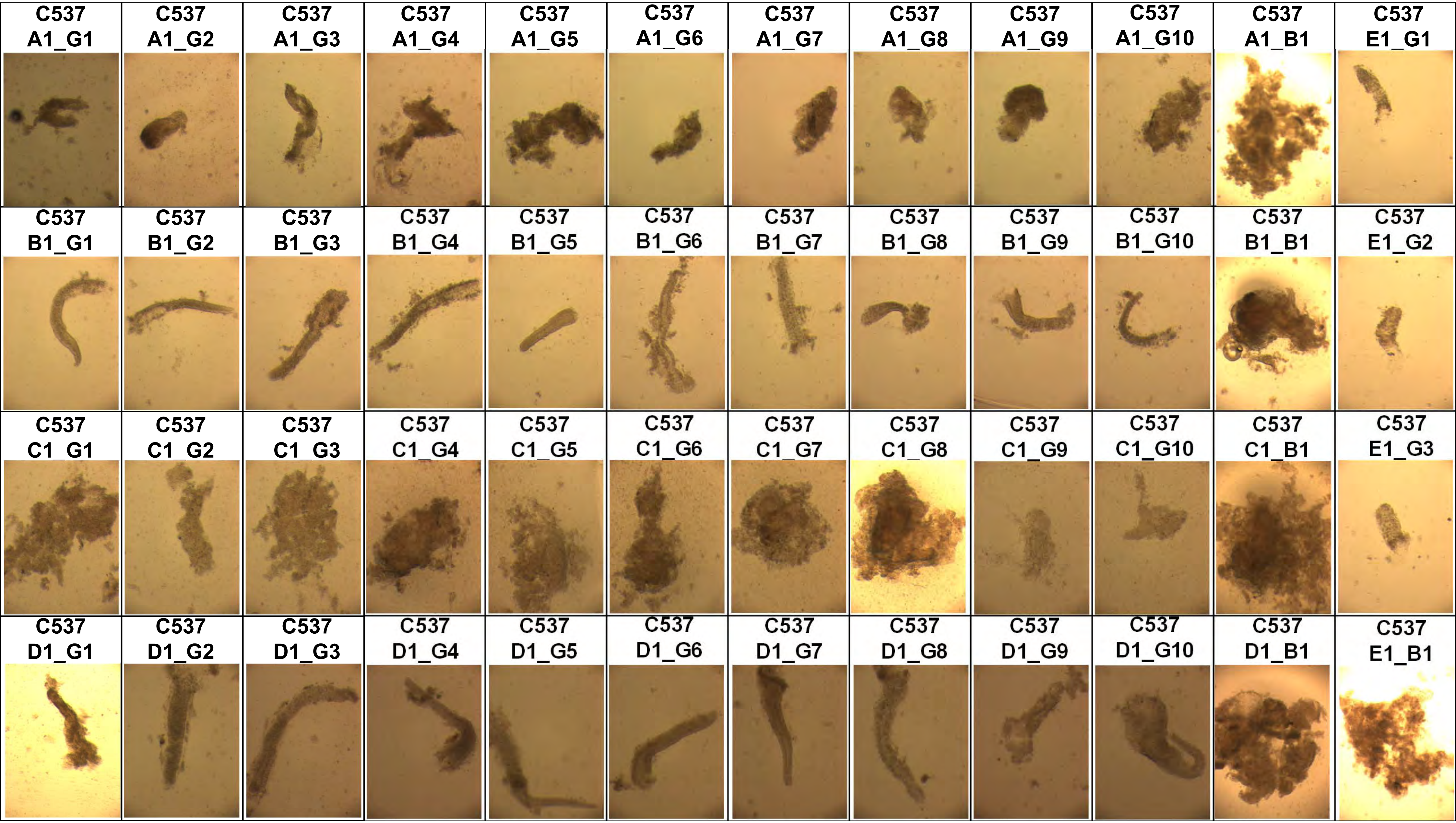
C537 A1_G1
C537 A1_G2
C537 A1_G3
C537 A1_G4
C537 A1_G5
C537 A1_G6
C537 A1_G7
C537 A1_G8
C537 A1_G9
C537 A1_G10
C537 A1_B1
C537 E1_G1
C537 B1_G1
C537 B1_G2
C537 B1_G3
C537 B1_G4
C537 B1_G5
C537 B1_G6
C537 B1_G7
C537 B1_G8
C537 B1_G9
C537 B1_G10
C537 B1_B1
C537 E1_G2
C537 C1_G1
C537 C1_G2
C537 C1_G3
C537 C1_G4
C537 C1_G5
C537 C1_G6
C537 C1_G7
C537 C1_G8
C537 C1_G9
C537 C1_G10
C537 C1_B1
C537 E1_G3
C537 D1_G1
C537 D1_G2
C537 D1_G3
C537 D1_G4
C537 D1_G5
C537 D1_G6
C537 D1_G7
C537 D1_G8
C537 D1_G9
C537 D1_G10
C537 D1_B1
C537 E1_B1

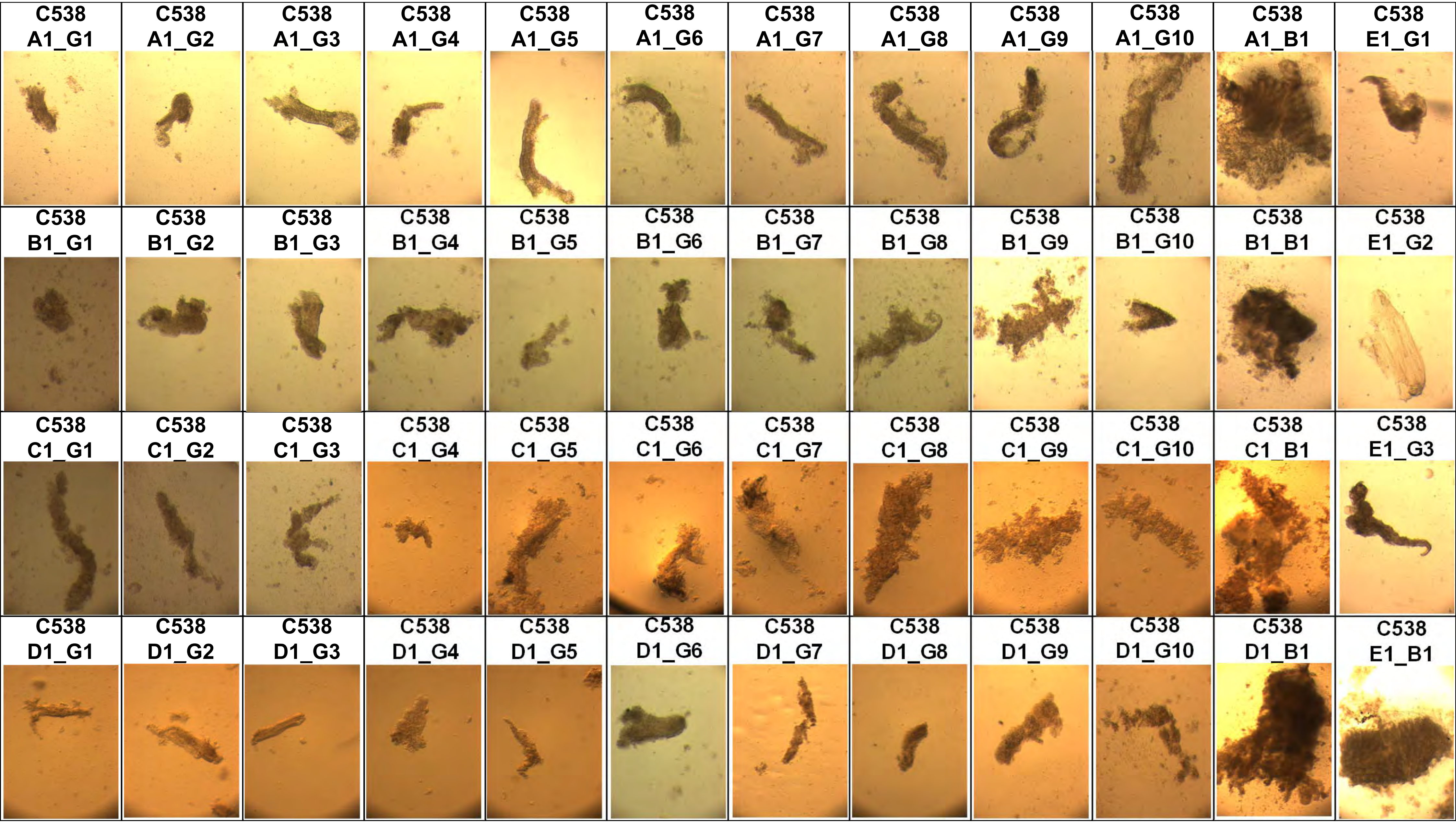
C538 A1_G1
C538 A1_G2
C538 A1_G3
C538 A1_G4
C538 A1_G5
C538 A1_G6
C538 A1_G7
C538 A1_G8
C538 A1_G9
C538 A1_G10
C538 A1_B1
C538 E1_G1
C538 B1_G1
C538 B1_G2
C538 B1_G3
C538 B1_G4
C538 B1_G5
C538 B1_G6
C538 B1_G7
C538 B1_G8
C538 B1_G9
C538 B1_G10
C538 B1_B1
C538 E1_G2
C538 C1_G1
C538 C1_G2
C538 C1_G3
C538 C1_G4
C538 C1_G5
C538 C1_G6
C538 C1_G7
C538 C1_G8
C538 C1_G9
C538 C1_G10
C538 C1_B1
C538 E1_G3
C538 D1_G1
C538 D1_G2
C538 D1_G3
C538 D1_G4
C538 D1_G5
C538 D1_G6
C538 D1_G7
C538 D1_G8
C538 D1_G9
C538 D1_G10
C538 D1_B1
C538 E1_B1

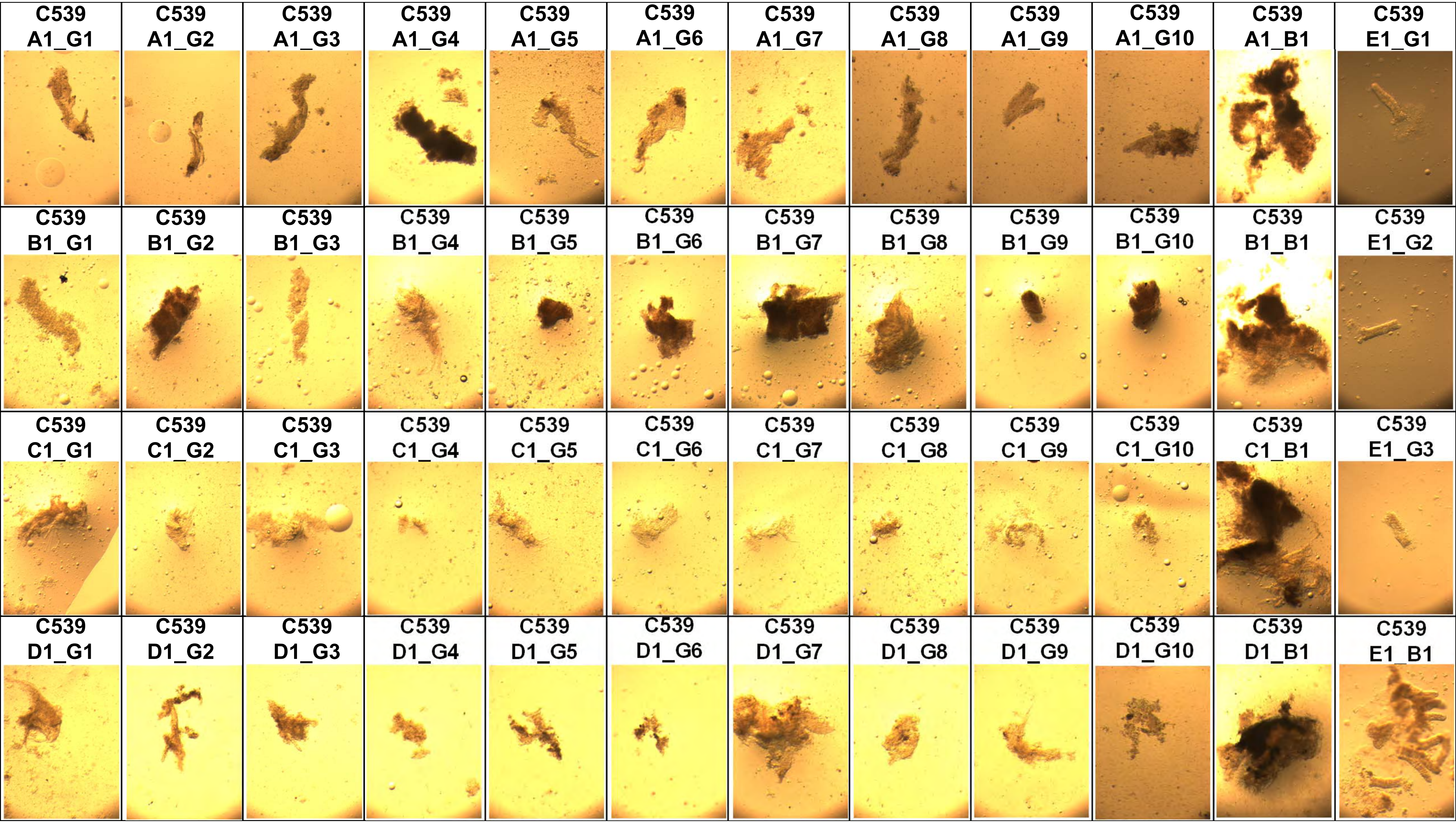
C539 A1_G1
C539 A1_G2
C539 A1_G3
C539 A1_G4
C539 A1_G5
C539 A1_G6
C539 A1_G7
C539 A1_G8
C539 A1_G9
C539 A1_G10
C539 A1_B1
C539 E1_G1
C539 B1_G1
C539 B1_G2
C539 B1_G3
C539 B1_G4
C539 B1_G5
C539 B1_G6
C539 B1_G7
C539 B1_G8
C539 B1_G9
C539 B1_G10
C539 B1_B1
C539 E1_G2
C539 C1_G1
C539 C1_G2
C539 C1_G3
C539 C1_G4
C539 C1_G5
C539 C1_G6
C539 C1_G7
C539 C1_G8
C539 C1_G9
C539 C1_G10
C539 C1_B1
C539 E1_G3
C539 D1_G1
C539 D1_G2
C539 D1_G3
C539 D1_G4
C539 D1_G5
C539 D1_G6
C539 D1_G7
C539 D1_G8
C539 D1_G9
C539 D1_G10
C539 D1_B1
C539 E1_B1

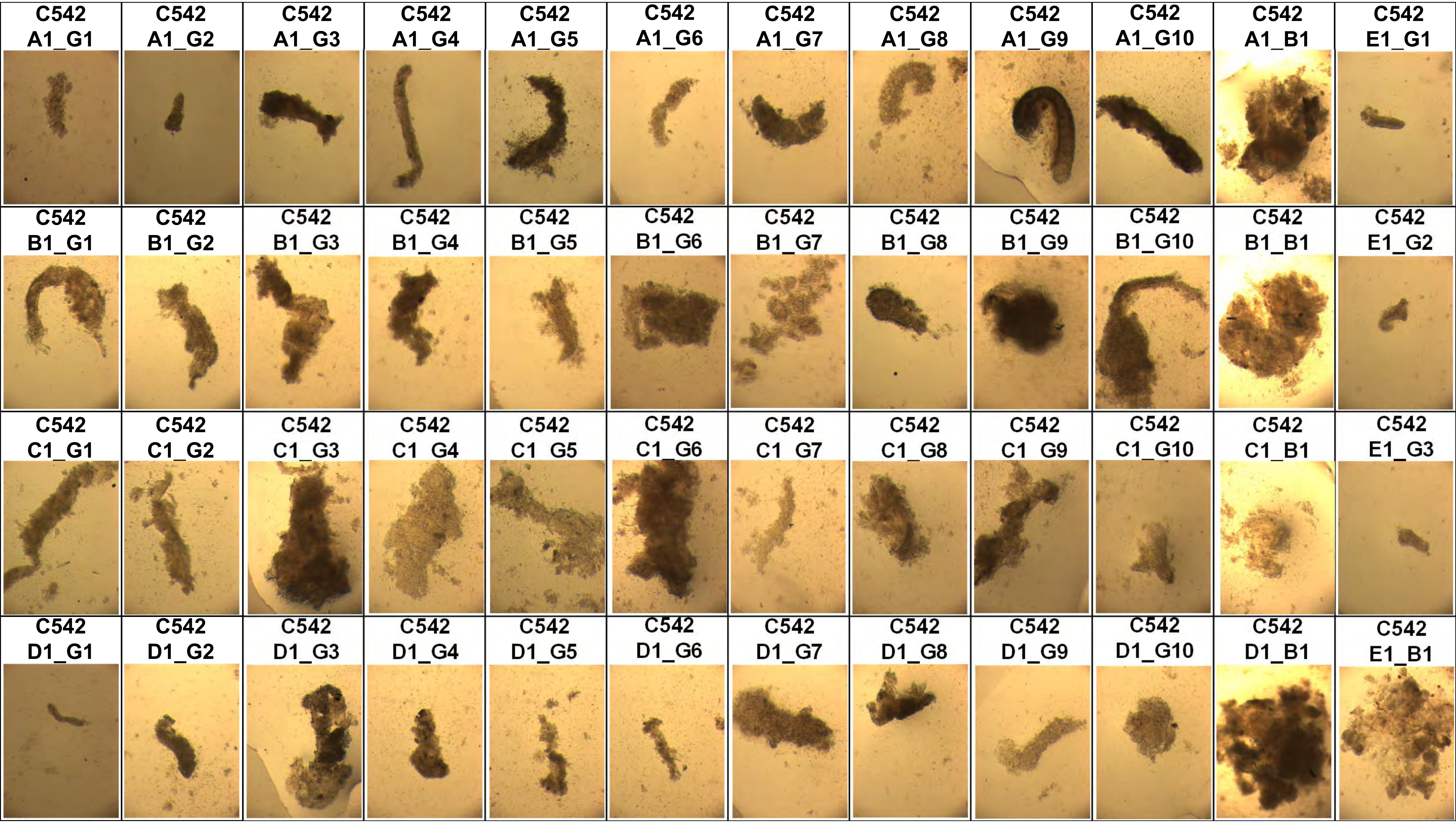
C542 A1_G1
C542 A1_G2
C542 A1_G3
C542 A1_G4
C542 A1_G5
C542 A1_G6
C542 A1_G7
C542 A1_G8
C542 A1_G9
C542 A1_G10
C542 A1_B1
C542 E1_G1
C542 B1_G1
C542 B1_G2
C542 B1_G3
C542 B1_G4
C542 B1_G5
C542 B1_G6
C542 B1_G7
C542 B1_G8
C542 B1_G9
C542 B1_G10
C542 B1_B1
C542 E1_G2
C542 C1_G1
C542 C1_G2
C542 C1_G3
C542 C1 G4
C542 C1 G5
C542 C1_G6
C542 C1 G7
C542 C1_G8
C542 C1 G9
C542 C1_G10
C542 C1_B1
C542 E1_G3
C542 D1_G1
C542 D1_G2
C542 D1_G3
C542 D1_G4
C542 D1_G5
C542 D1_G6
C542 D1_G7
C542 D1_G8
C542 D1_G9
C542 D1_G10
C542 D1_B1
C542 E1_B1

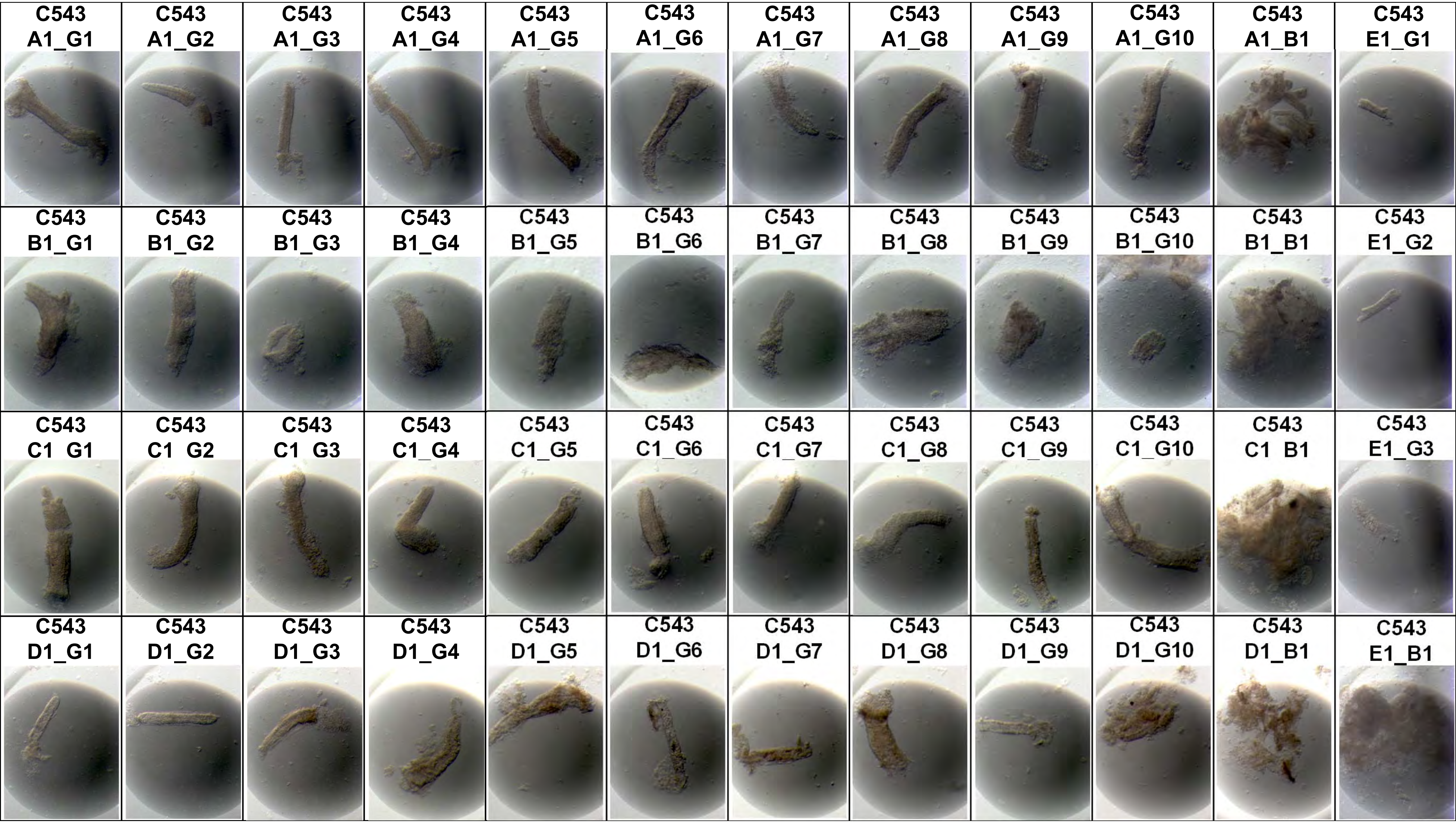
C543 A1_G1
C543 A1_G2
C543 A1_G3
C543 A1_G4
C543 A1_G5
C543 A1_G6
C543 A1_G7
C543 A1_G8
C543 A1_G9
C543 A1_G10
C543 A1_B1
C543 E1_G1
C543 B1_G1
C543 B1_G2
C543 B1_G3
C543 B1_G4
C543 B1_G5
C543 B1_G6
C543 B1_G7
C543 B1_G8
C543 B1_G9
C543 B1_G10
C543 B1_B1
C543 E1_G2
C543 C1 G1
C543 C1 G2
C543 C1 G3
C543 C1_G4
C543 C1_G5
C543 C1_G6
C543 C1_G7
C543 C1_G8
C543 C1_G9
C543 C1_G10
C543 C1 B1
C543 E1_G3
C543 D1_G1
C543 D1_G2
C543 D1_G3
C543 D1_G4
C543 D1_G5
C543 D1_G6
C543 D1_G7
C543 D1_G8
C543 D1_G9
C543 D1_G10
C543 D1_B1
C543 E1_B1

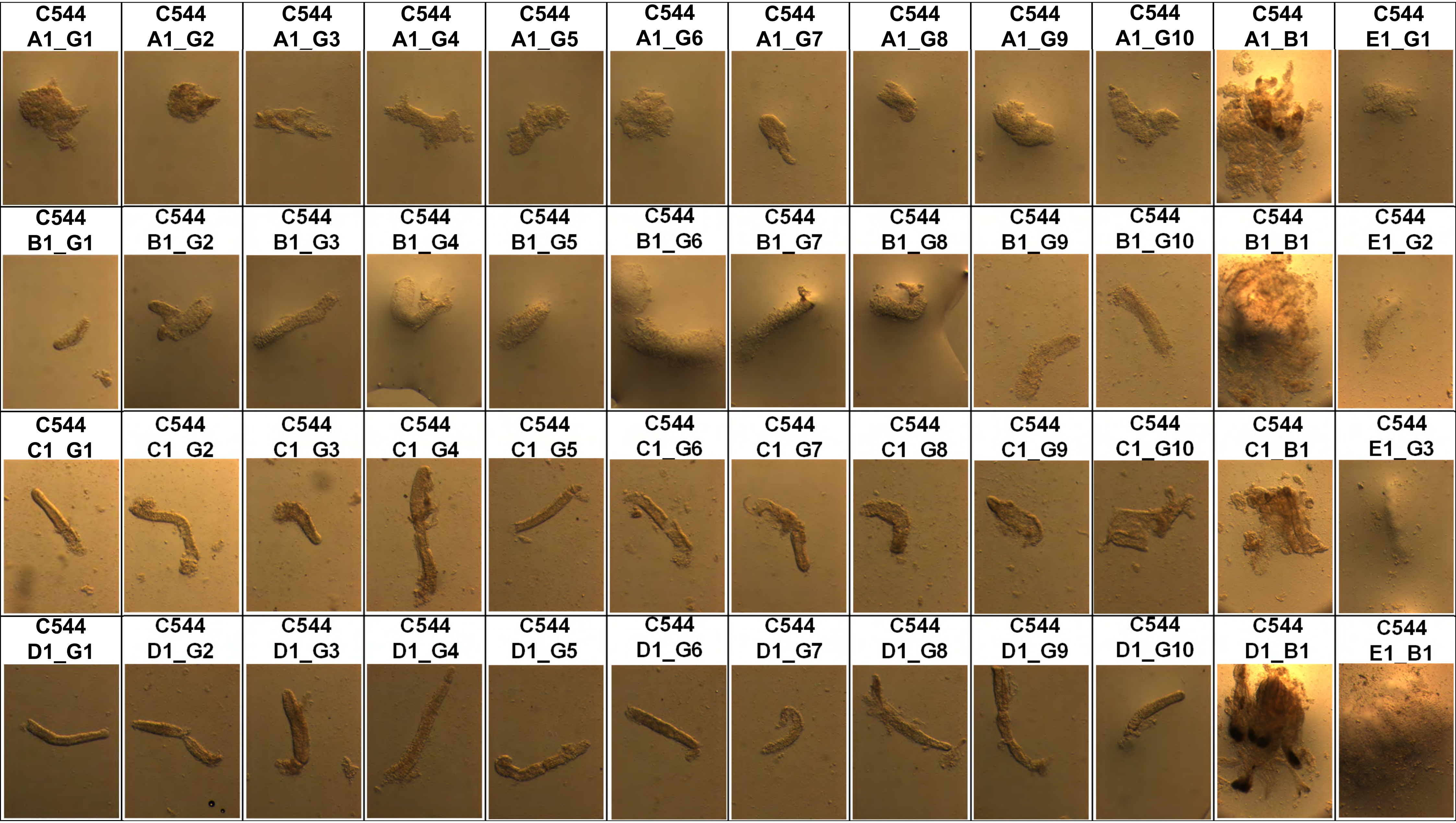
C544 A1_G1
C544 A1_G2
C544 A1_G3
C544 A1_G4
C544 A1_G5
C544 A1_G6
C544 A1_G7
C544 A1_G8
C544 A1_G9
C544 A1_G10
C544 A1_B1
C544 E1_G1
C544 B1_G1
C544 B1_G2
C544 B1_G3
C544 B1_G4
C544 B1_G5
C544 B1_G6
C544 B1_G7
C544 B1_G8
C544 B1_G9
C544 B1_G10
C544 B1_B1
C544 E1_G2
C544 C1_G1
C544 C1_G2
C544 C1_G3
C544 C1_G4
C544 C1_G5
C544 C1_G6
C544 C1_G7
C544 C1_G8
C544 C1_G9
C544 C1_G10
C544 C1_B1
C544 E1_G3
C544 D1_G1
C544 D1_G2
C544 D1_G3
C544 D1_G4
C544 D1_G5
C544 D1_G6
C544 D1_G7
C544 D1_G8
C544 D1_G9
C544 D1_G10
C544 D1_B1
C544 E1_B1

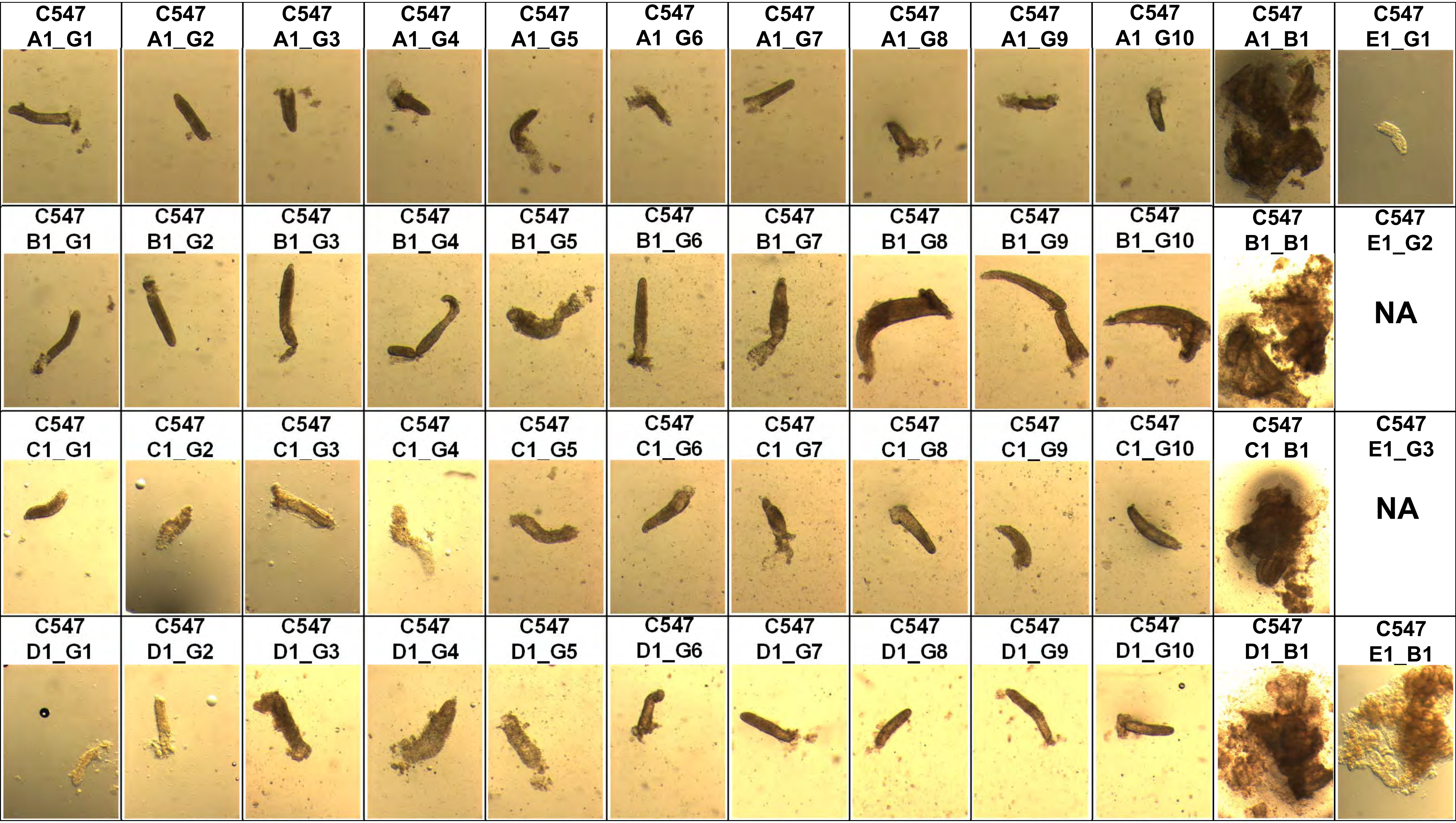
C547 A1_G1
C547 A1_G2
C547 A1_G3
C547 A1_G4
C547 A1_G5
C547 A1  G6
C547 A1_G7
C547 A1_G8
C547 A1_G9
C547 A1  G10
C547 A1_B1
C547 E1_G1
C547 B1_G1
C547 B1_G2
C547 B1_G3
C547 B1_G4
C547 B1_G5
C547 B1_G6
C547 B1_G7
C547 B1_G8
C547 B1_G9
C547 B1_G10
C547 B1_B1
C547 E1_G2
NA
C547 C1_G1
C547 C1_G2
C547 C1_G3
C547 C1_G4
C547 C1_G5
C547 C1_G6
C547 C1  G7
C547 C1_G8
C547 C1_G9
C547 C1_G10
C547 C1  B1
C547 E1_G3
NA
C547 D1_G1
C547 D1_G2
C547 D1_G3
C547 D1_G4
C547 D1_G5
C547 D1_G6
C547 D1_G7
C547 D1_G8
C547 D1_G9
C547 D1_G10
C547 D1_B1
C547 E1_B1

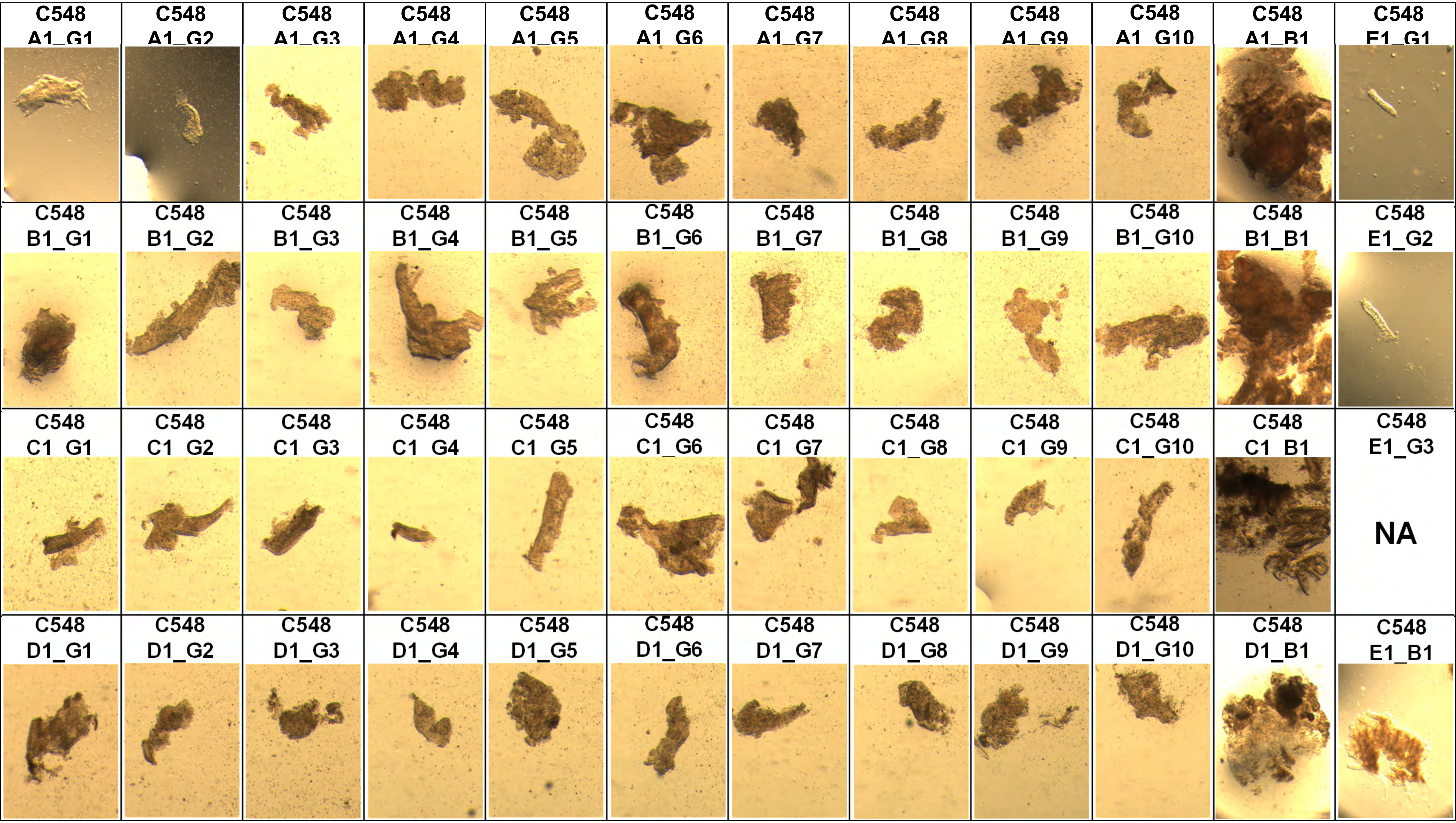
C548 A1_G1
C548 A1_G2
C548 A1_G3
C548 A1_G4
C548 A1_G5
C548 A1_G6
C548 A1_G7
C548 A1_G8
C548 A1_G9
C548 A1_G10
C548 A1_B1
C548 E1_G1
C548 B1_G1
C548 B1_G2
C548 B1_G3
C548 B1_G4
C548 B1_G5
C548 B1_G6
C548 B1_G7
C548 B1_G8
C548 B1_G9
C548 B1_G10
C548 B1_B1
C548 E1_G2
C548 C1_G1
C548 C1_G2
C548 C1_G3
C548 C1_G4
C548 C1_G5
C548 C1_G6
C548 C1_G7
C548 C1_G8
C548 C1_G9
C548 C1_G10
C548 C1_B1
C548 E1_G3
NA
C548 D1_G1
C548 D1_G2
C548 D1_G3
C548 D1_G4
C548 D1_G5
C548 D1_G6
C548 D1_G7
C548 D1_G8
C548 D1_G9
C548 D1_G10
C548 D1_B1
C548 E1_B1

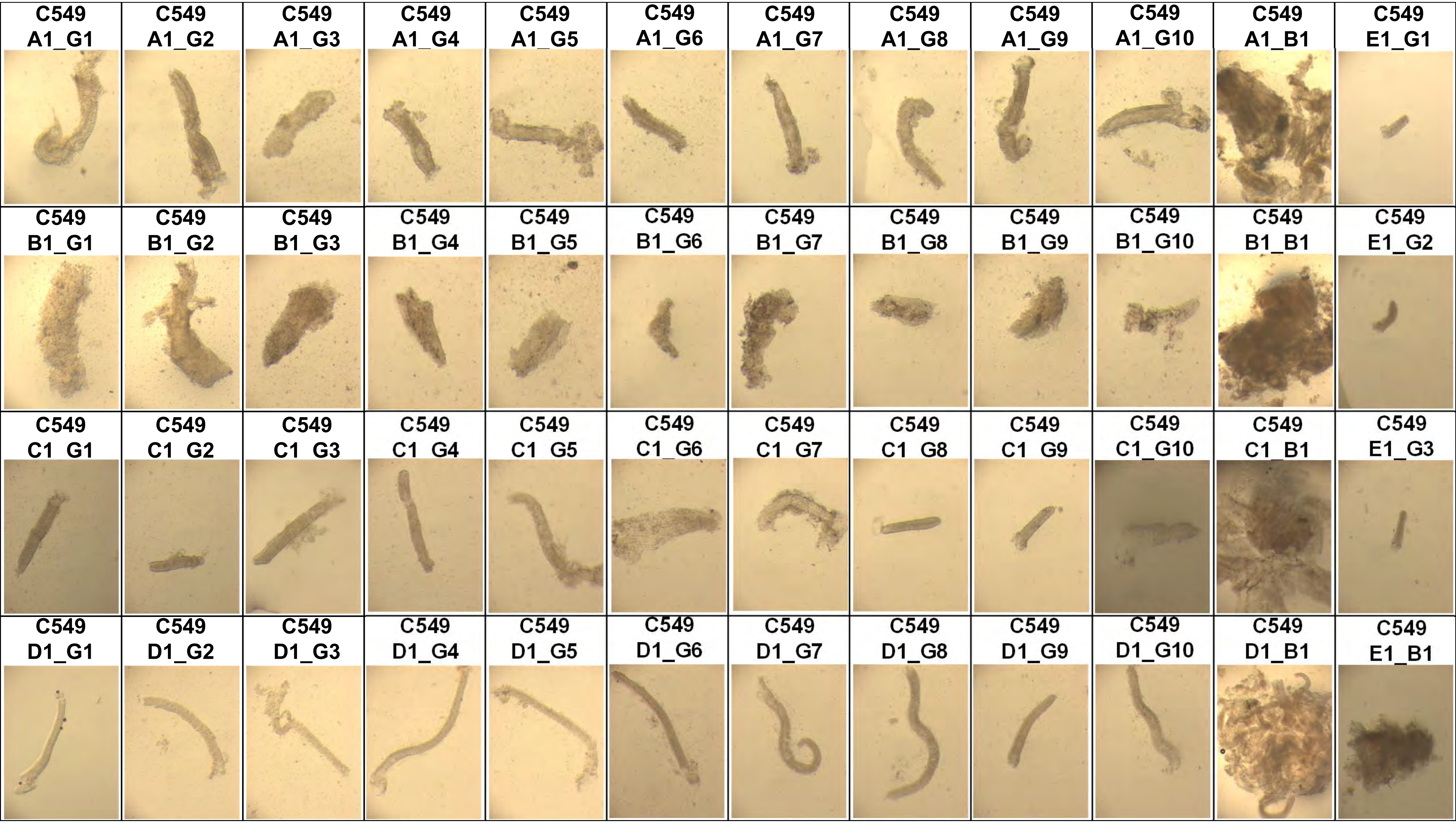
C549 A1_G1
C549 A1_G2
C549 A1_G3
C549 A1_G4
C549 A1_G5
C549 A1_G6
C549 A1_G7
C549 A1_G8
C549 A1_G9
C549 A1_G10
C549 A1_B1
C549 E1_G1
C549 B1_G1
C549 B1_G2
C549 B1_G3
C549 B1_G4
C549 B1_G5
C549 B1_G6
C549 B1_G7
C549 B1_G8
C549 B1_G9
C549 B1_G10
C549 B1_B1
C549 E1_G2
C549 C1 G1
C549 C1 G2
C549 C1 G3
C549 C1 G4
C549 C1 G5
C549 C1_G6
C549 C1 G7
C549 C1 G8
C549 C1 G9
C549 C1_G10
C549 C1_B1
C549 E1_G3
C549 D1_G1
C549 D1_G2
C549 D1_G3
C549 D1_G4
C549 D1_G5
C549 D1_G6
C549 D1_G7
C549 D1_G8
C549 D1_G9
C549 D1_G10
C549 D1_B1
C549 E1_B1

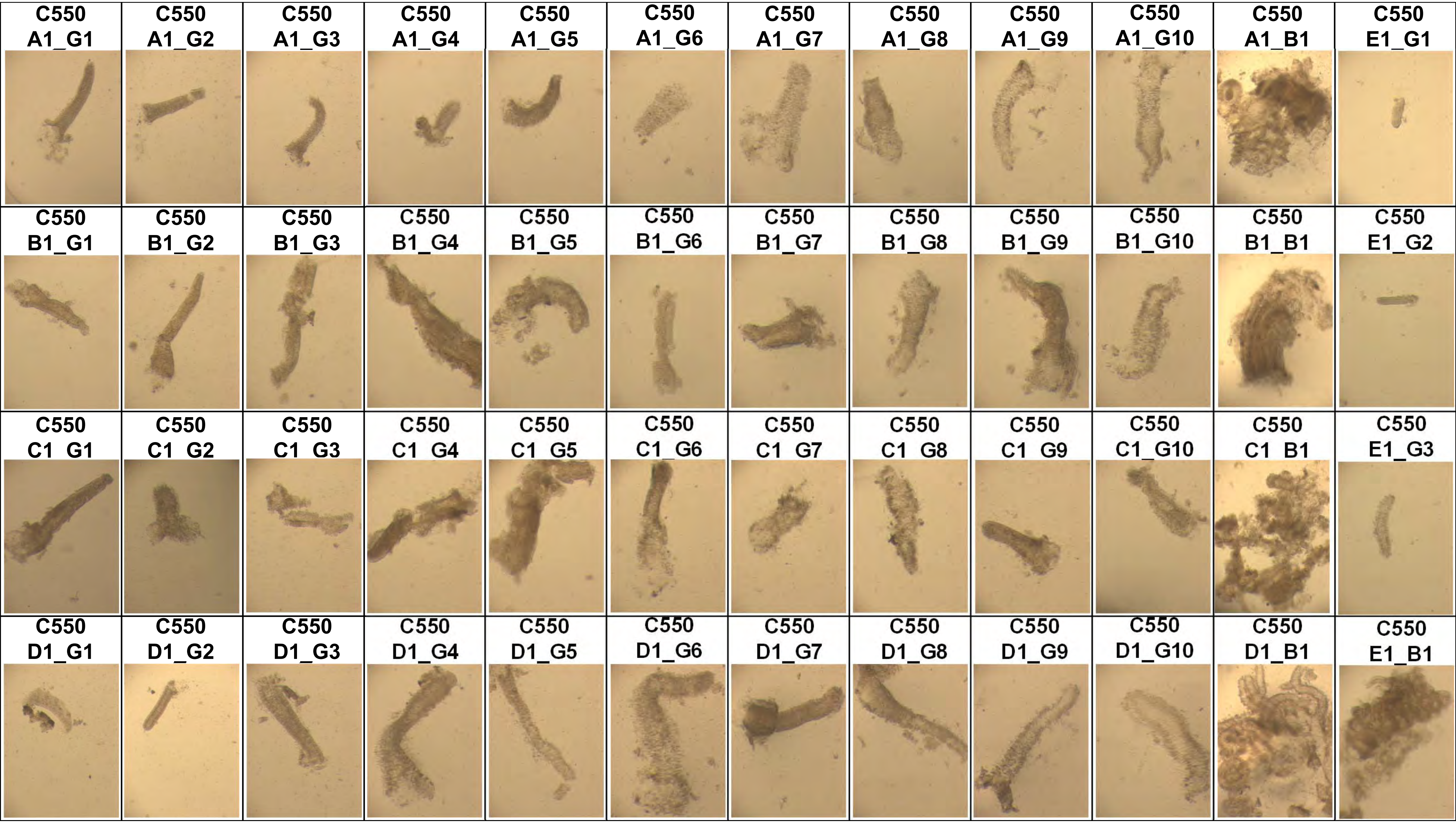
C550 A1_G1
C550 A1_G2
C550 A1_G3
C550 A1_G4
C550 A1_G5
C550 A1_G6
C550 A1_G7
C550 A1_G8
C550 A1_G9
C550 A1_G10
C550 A1_B1
C550 E1_G1
C550 B1_G1
C550 B1_G2
C550 B1_G3
C550 B1_G4
C550 B1_G5
C550 B1_G6
C550 B1_G7
C550 B1_G8
C550 B1_G9
C550 B1_G10
C550 B1_B1
C550 E1_G2
C550 C1 G1
C550 C1 G2
C550 C1 G3
C550 C1 G4
C550 C1 G5
C550 C1_G6
C550 C1 G7
C550 C1 G8
C550 C1 G9
C550 C1_G10
C550 C1 B1
C550 E1_G3
C550 D1_G1
C550 D1_G2
C550 D1_G3
C550 D1_G4
C550 D1_G5
C550 D1_G6
C550 D1_G7
C550 D1_G8
C550 D1_G9
C550 D1_G10
C550 D1_B1
C550 E1_B1

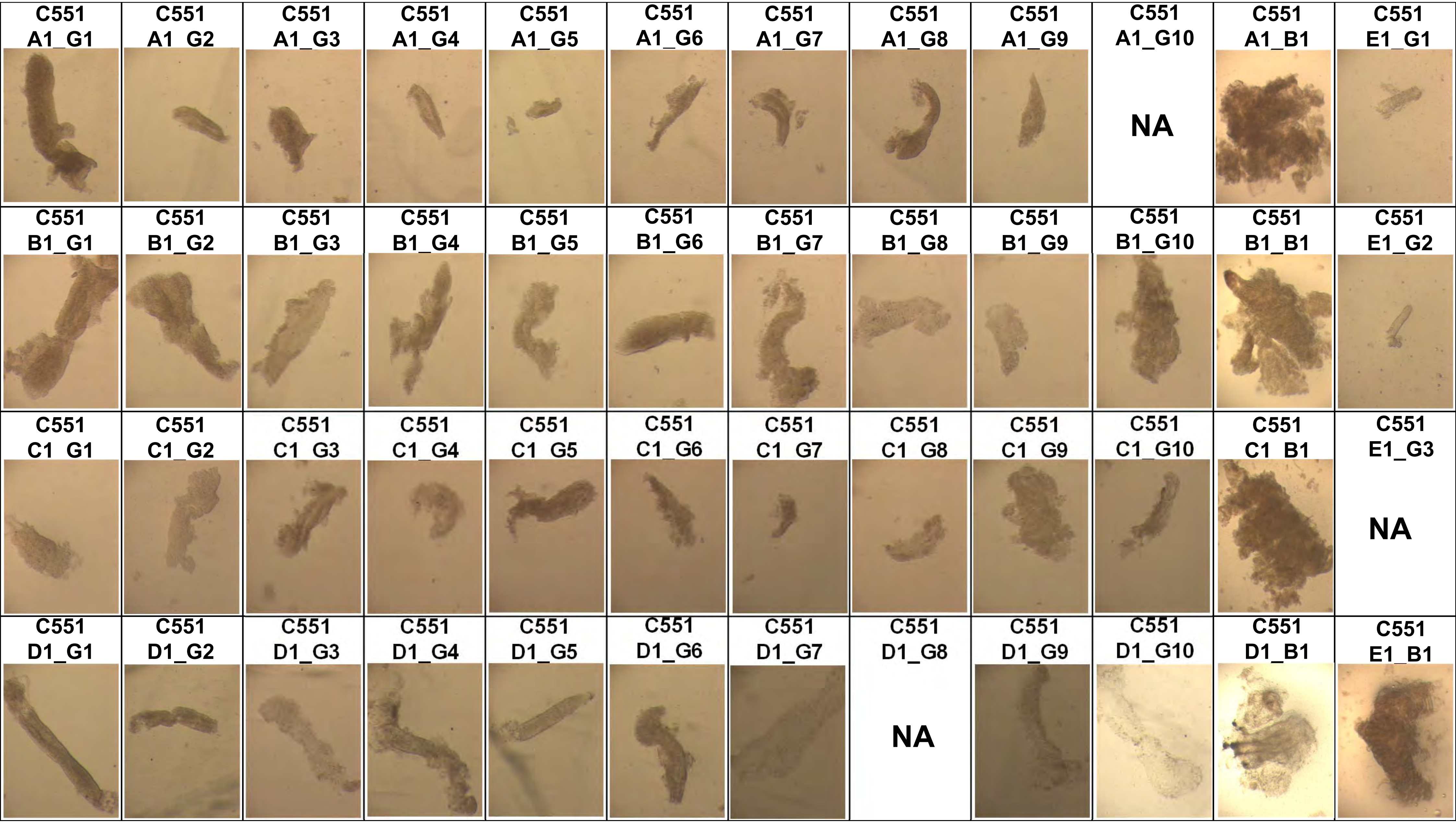
C551 A1_G1
C551 A1_G2
C551 A1_G3
C551 A1_G4
C551 A1_G5
C551 A1_G6
C551 A1_G7
C551 A1_G8
C551 A1_G9
C551 A1_G10
NA
C551 A1_B1
C551 E1_G1
C551 B1_G1
C551 B1_G2
C551 B1_G3
C551 B1_G4
C551 B1_G5
C551 B1_G6
C551 B1_G7
C551 B1_G8
C551 B1_G9
C551 B1_G10
C551 B1_B1
C551 E1_G2
C551 C1_G1
C551 C1_G2
C551 C1_G3
C551 C1_G4
C551 C1_G5
C551 C1_G6
C551 C1_G7
C551 C1_G8
C551 C1_G9
C551 C1_G10
C551 C1_B1
C551 E1_G3
NA
C551 D1_G1
C551 D1_G2
C551 D1_G3
C551 D1_G4
C551 D1_G5
C551 D1_G6
C551 D1_G7
C551 D1_G8
NA
C551 D1_G9
C551 D1_G10
C551 D1_B1
C551 E1_B1

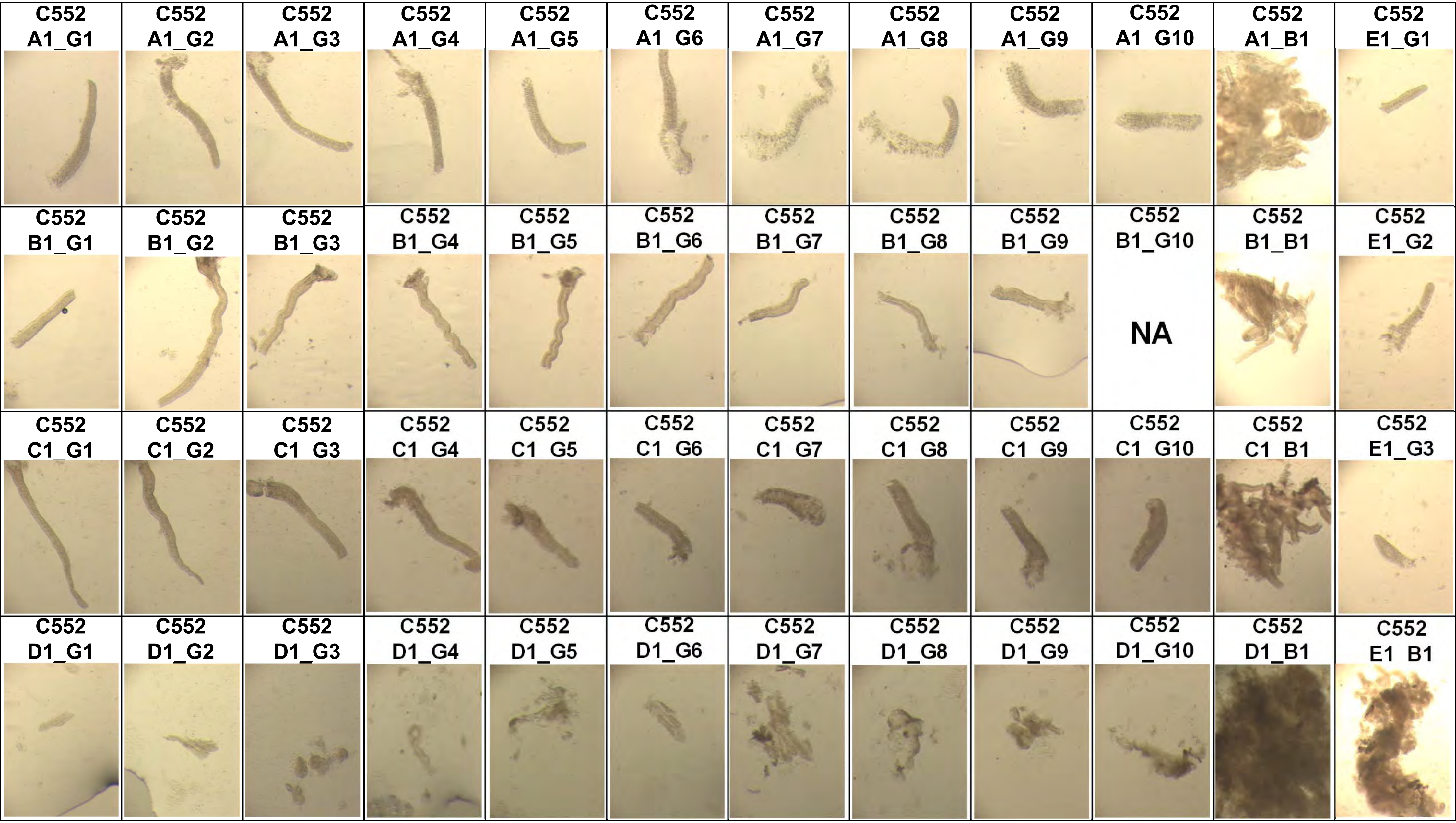
C552 A1_G1
C552 A1_G2
C552 A1_G3
C552 A1_G4
C552 A1_G5
C552 A1 G6
C552 A1_G7
C552 A1_G8
C552 A1_G9
C552 A1 G10
C552 A1_B1
C552 E1_G1
C552 B1_G1
C552 B1_G2
C552 B1_G3
C552 B1_G4
C552 B1_G5
C552 B1_G6
C552 B1_G7
C552 B1_G8
C552 B1_G9
C552 B1_G10
NA
C552 B1_B1
C552 E1_G2
C552 C1_G1
C552 C1_G2
C552 C1 G3
C552 C1 G4
C552 C1 G5
C552 C1 G6
C552 C1 G7
C552 C1 G8
C552 C1 G9
C552 C1 G10
C552 C1 B1
C552 E1_G3
C552 D1_G1
C552 D1_G2
C552 D1_G3
C552 D1_G4
C552 D1_G5
C552 D1_G6
C552 D1_G7
C552 D1_G8
C552 D1_G9
C552 D1_G10
C552 D1_B1
C552 E1 B1

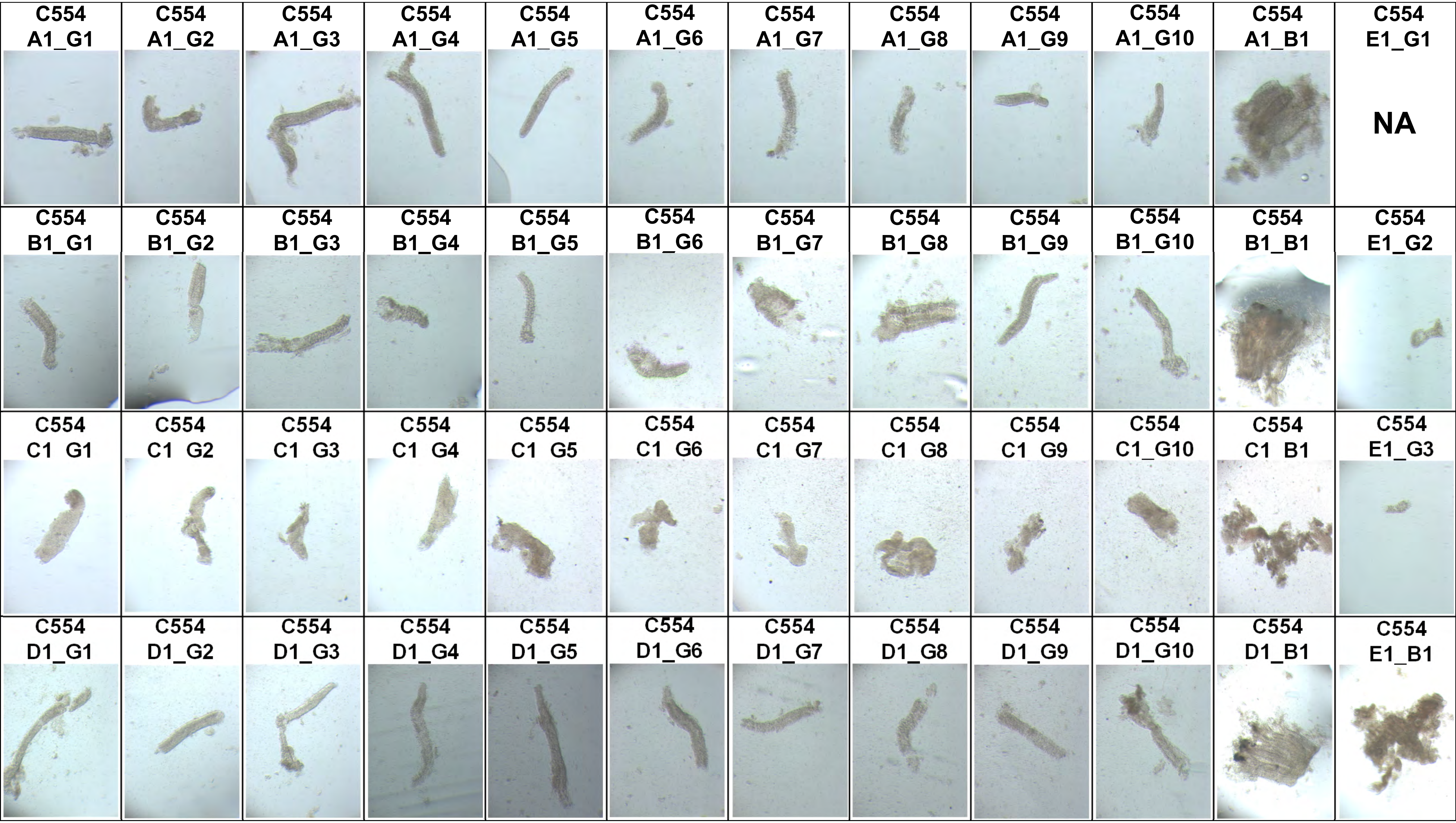
C554 A1_G1
C554 A1_G2
C554 A1_G3
C554 A1_G4
C554 A1_G5
C554 A1_G6
C554 A1_G7
C554 A1_G8
C554 A1_G9
C554 A1_G10
C554 A1_B1
C554 E1_G1
NA
C554 B1_G1
C554 B1_G2
C554 B1_G3
C554 B1_G4
C554 B1_G5
C554 B1_G6
C554 B1_G7
C554 B1_G8
C554 B1_G9
C554 B1_G10
C554 B1_B1
C554 E1_G2
C554 C1 G1
C554 C1 G2
C554 C1 G3
C554 C1 G4
C554 C1 G5
C554 C1 G6
C554 C1 G7
C554 C1 G8
C554 C1 G9
C554 C1_G10
C554 C1 B1
C554 E1_G3
C554 D1_G1
C554 D1_G2
C554 D1_G3
C554 D1_G4
C554 D1_G5
C554 D1_G6
C554 D1_G7
C554 D1_G8
C554 D1_G9
C554 D1_G10
C554 D1_B1
C554 E1_B1

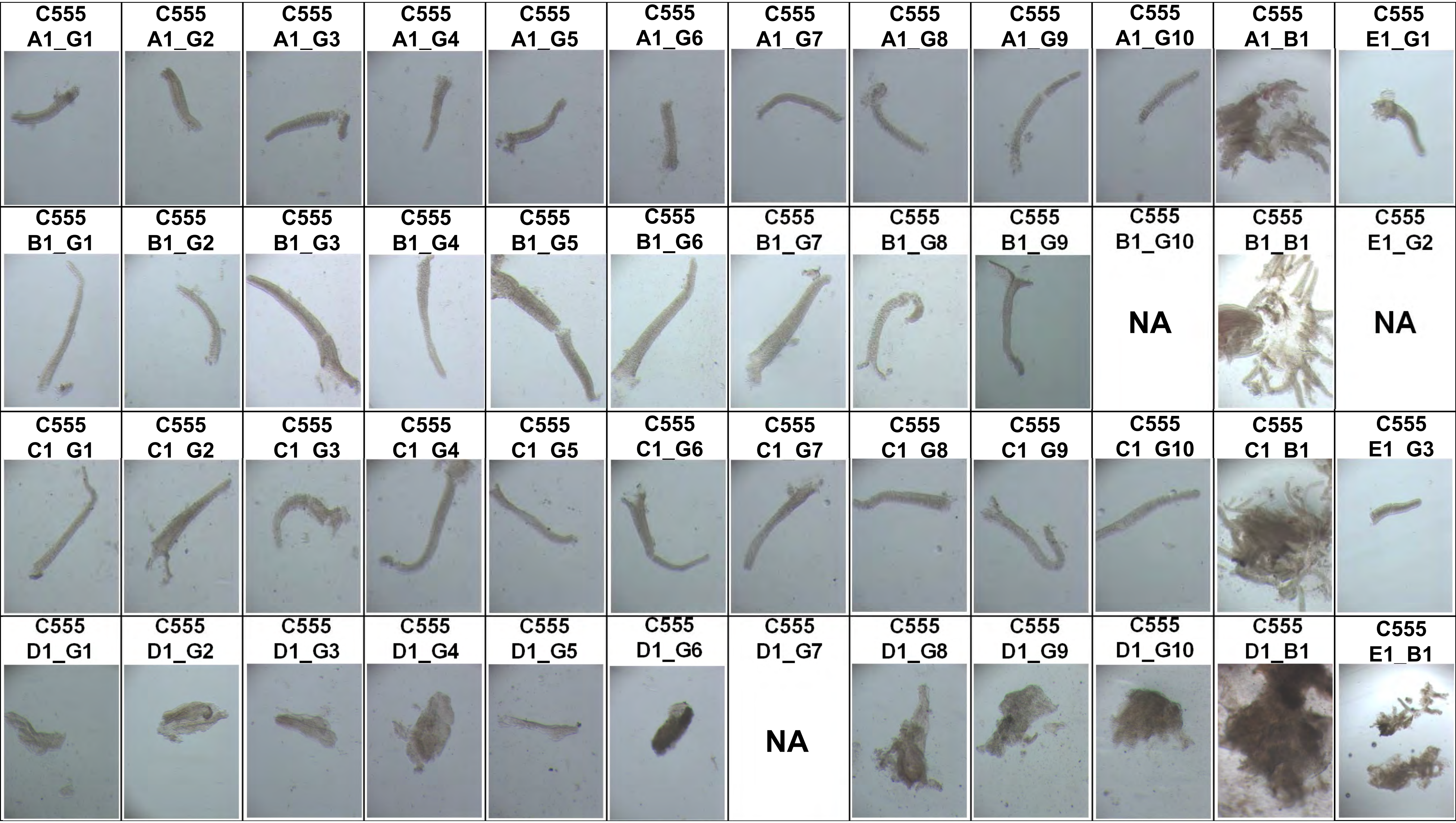
C555 A1_G1
C555 A1_G2
C555 A1_G3
C555 A1_G4
C555 A1_G5
C555 A1_G6
C555 A1_G7
C555 A1_G8
C555 A1_G9
C555 A1_G10
C555 A1_B1
C555 E1_G1
C555 B1_G1
C555 B1_G2
C555 B1_G3
C555 B1_G4
C555 B1_G5
C555 B1_G6
C555 B1_G7
C555 B1_G8
C555 B1_G9
C555 B1_G10
NA
C555 B1_B1
C555 E1_G2
NA
C555 C1_G1
C555 C1_G2
C555 C1_G3
C555 C1_G4
C555 C1_G5
C555 C1_G6
C555 C1_G7
C555 C1_G8
C555 C1_G9
C555 C1_G10
C555 C1_B1
C555 E1_G3
C555 D1_G1
C555 D1_G2
C555 D1_G3
C555 D1_G4
C555 D1_G5
C555 D1_G6
C555 D1_G7
NA
C555 D1_G8
C555 D1_G9
C555 D1_G10
C555 D1_B1
C555 E1_B1

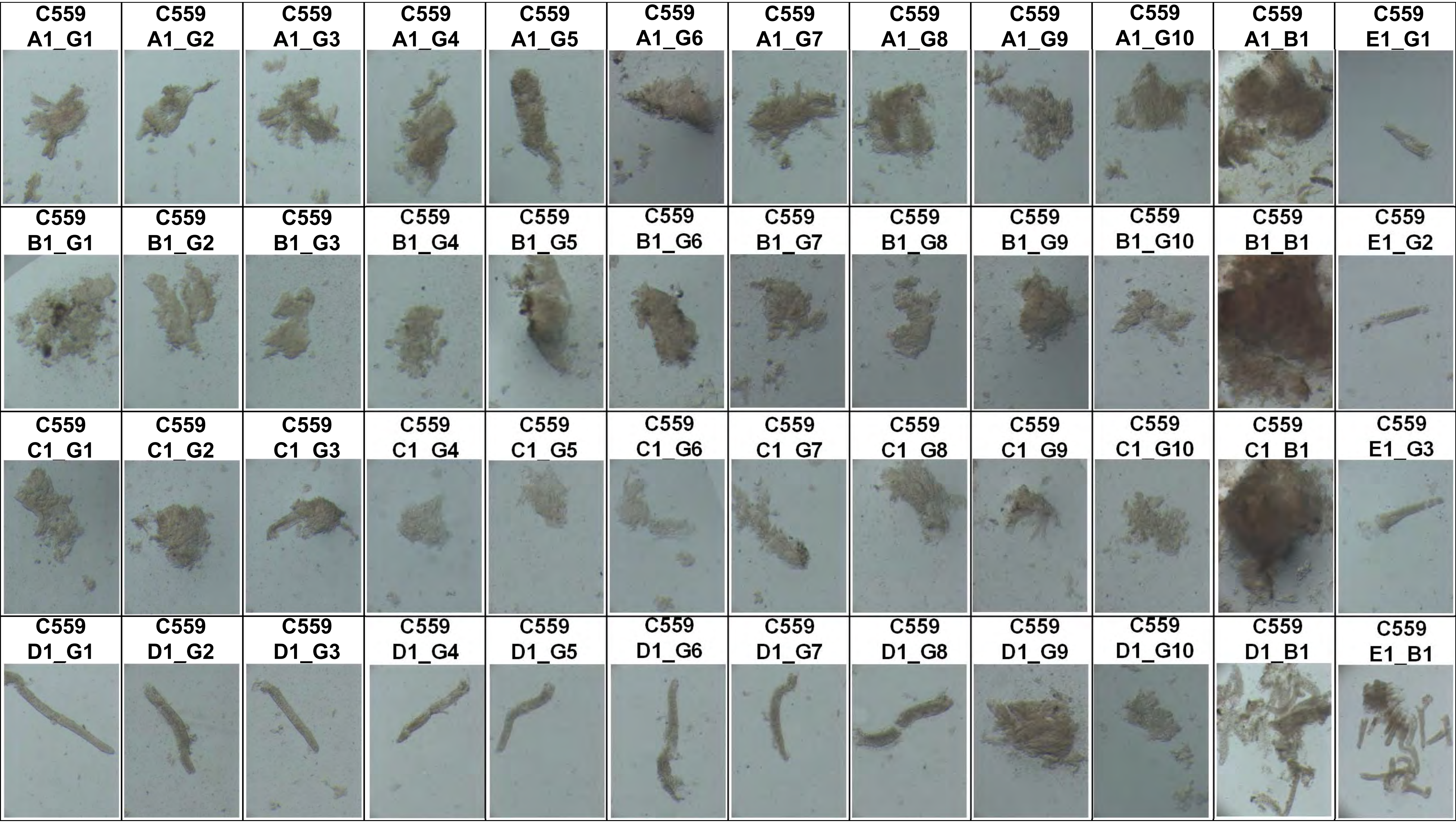
C559 A1_G1
C559 A1_G2
C559 A1_G3
C559 A1_G4
C559 A1_G5
C559 A1_G6
C559 A1_G7
C559 A1_G8
C559 A1_G9
C559 A1_G10
C559 A1_B1
C559 E1_G1
C559 B1_G1
C559 B1_G2
C559 B1_G3
C559 B1_G4
C559 B1_G5
C559 B1_G6
C559 B1_G7
C559 B1_G8
C559 B1_G9
C559 B1_G10
C559 B1_B1
C559 E1_G2
C559 C1_G1
C559 C1_G2
C559 C1_G3
C559 C1_G4
C559 C1_G5
C559 C1_G6
C559 C1_G7
C559 C1_G8
C559 C1_G9
C559 C1_G10
C559 C1_B1
C559 E1_G3
C559 D1_G1
C559 D1_G2
C559 D1_G3
C559 D1_G4
C559 D1_G5
C559 D1_G6
C559 D1_G7
C559 D1_G8
C559 D1_G9
C559 D1_G10
C559 D1_B1
C559 E1_B1

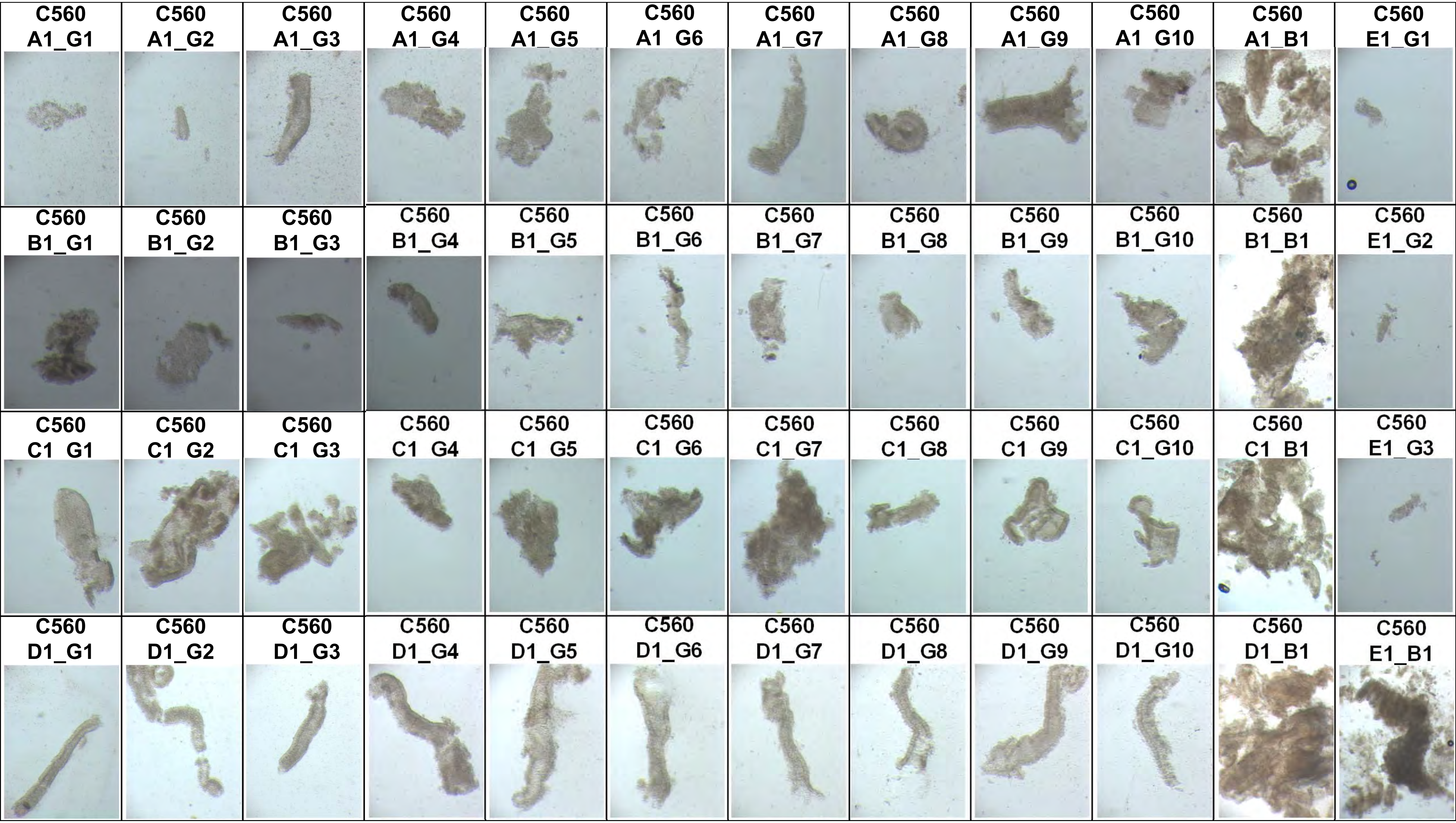
C560 A1_G1
C560 A1_G2
C560 A1_G3
C560 A1_G4
C560 A1_G5
C560 A1_G6
C560 A1_G7
C560 A1_G8
C560 A1_G9
C560 A1_G10
C560 A1_B1
C560 E1_G1
C560 B1_G1
C560 B1_G2
C560 B1_G3
C560 B1_G4
C560 B1_G5
C560 B1_G6
C560 B1_G7
C560 B1_G8
C560 B1_G9
C560 B1_G10
C560 B1_B1
C560 E1_G2
C560 C1_G1
C560 C1_G2
C560 C1_G3
C560 C1_G4
C560 C1_G5
C560 C1_G6
C560 C1_G7
C560 C1_G8
C560 C1_G9
C560 C1_G10
C560 C1_B1
C560 E1_G3
C560 D1_G1
C560 D1_G2
C560 D1_G3
C560 D1_G4
C560 D1_G5
C560 D1_G6
C560 D1_G7
C560 D1_G8
C560 D1_G9
C560 D1_G10
C560 D1_B1
C560 E1_B1

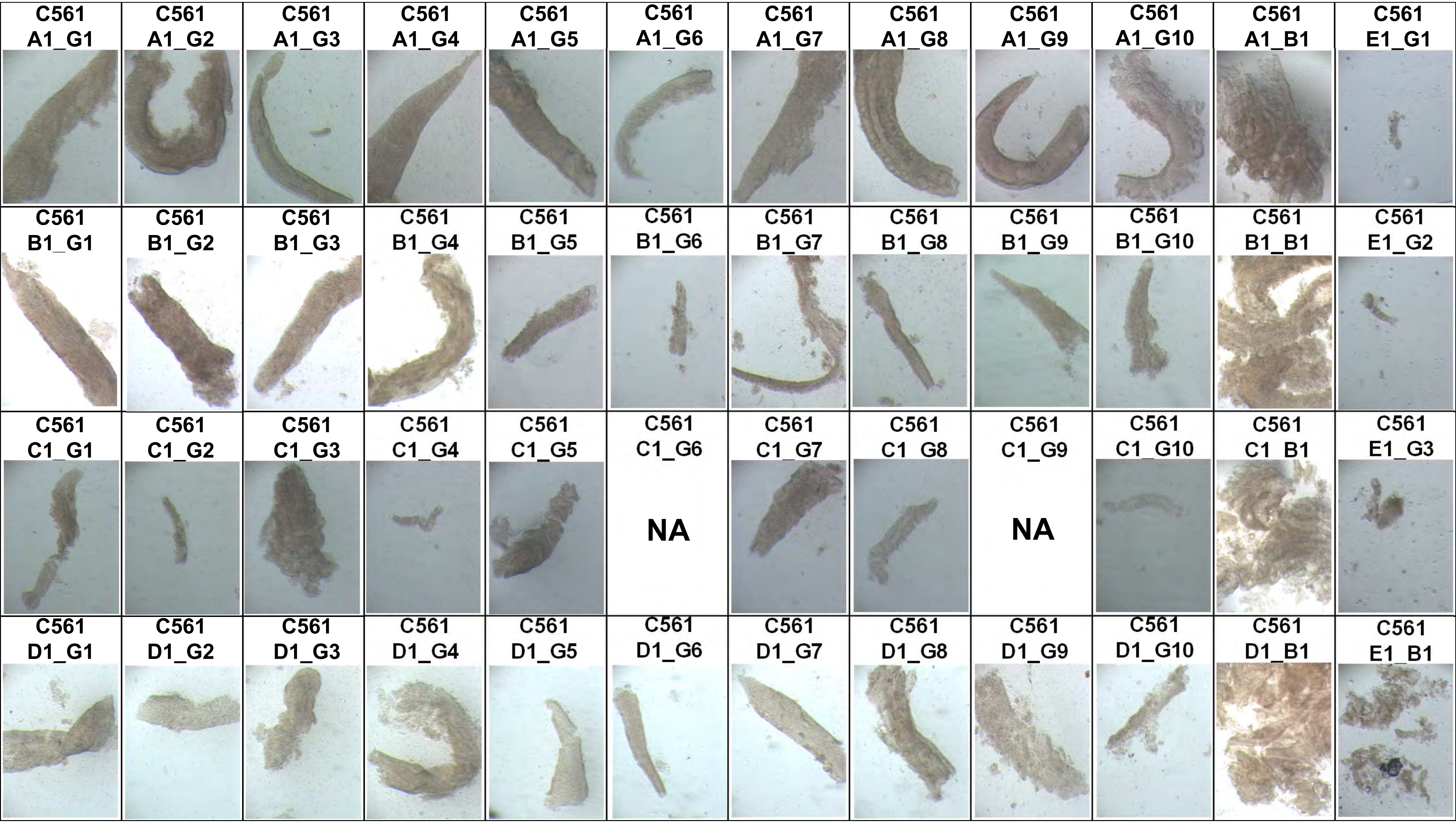
C561 A1_G1
C561 A1_G2
C561 A1_G3
C561 A1_G4
C561 A1_G5
C561 A1_G6
C561 A1_G7
C561 A1_G8
C561 A1_G9
C561 A1_G10
C561 A1_B1
C561 E1_G1
C561 B1_G1
C561 B1_G2
C561 B1_G3
C561 B1_G4
C561 B1_G5
C561 B1_G6
C561 B1_G7
C561 B1_G8
C561 B1_G9
C561 B1_G10
C561 B1_B1
C561 E1_G2
C561 C1_G1
C561 C1_G2
C561 C1_G3
C561 C1_G4
C561 C1_G5
C561 C1_G6
NA
C561 C1_G7
C561 C1_G8
C561 C1_G9
NA
C561 C1_G10
C561 C1_B1
C561 E1_G3
C561 D1_G1
C561 D1_G2
C561 D1_G3
C561 D1_G4
C561 D1_G5
C561 D1_G6
C561 D1_G7
C561 D1_G8
C561 D1_G9
C561 D1_G10
C561 D1_B1
C561 E1_B1

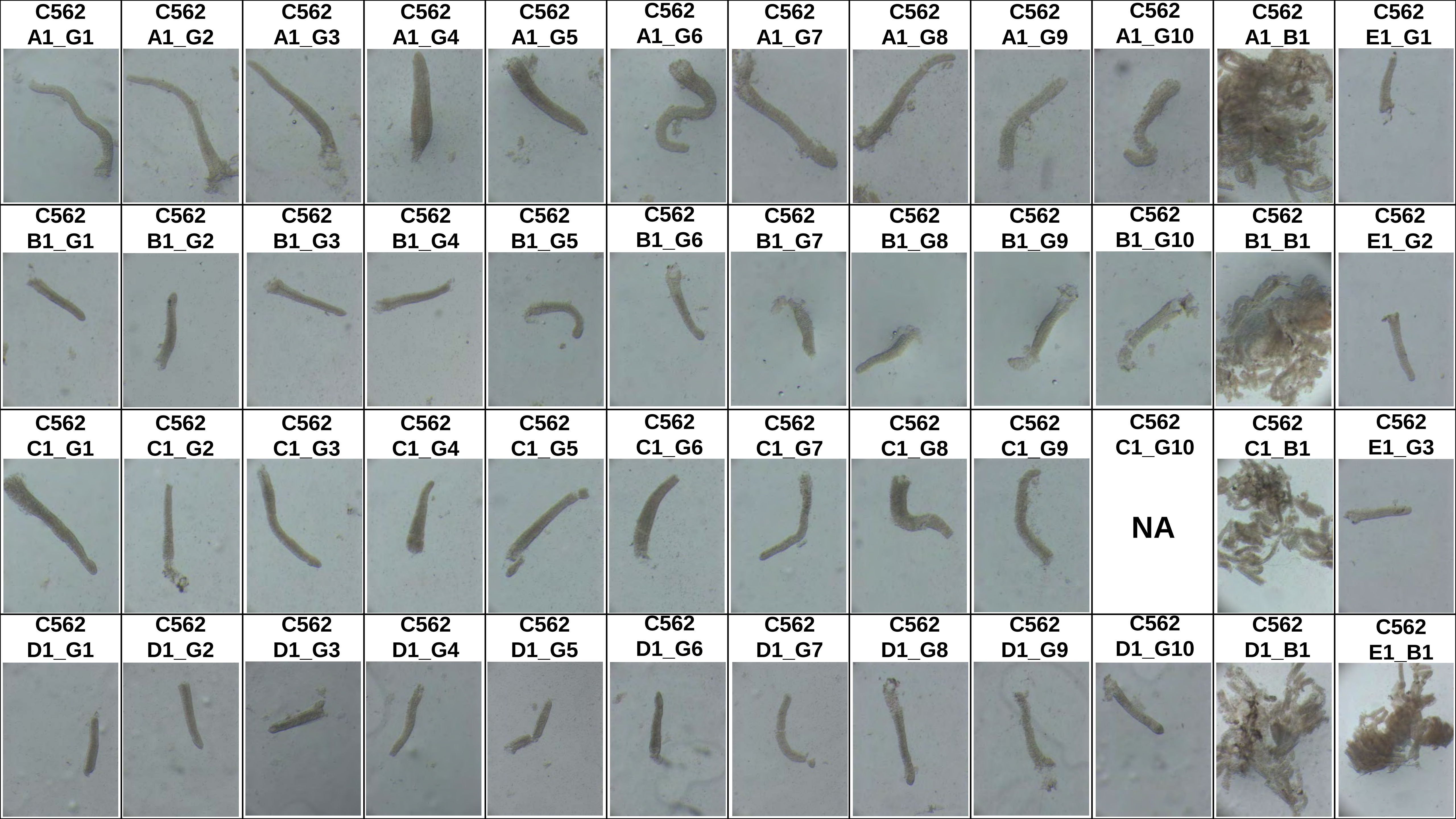
C562 A1_G1
C562 A1_G2
C562 A1_G3
C562 A1_G4
C562 A1_G5
C562 A1_G6
C562 A1_G7
C562 A1_G8
C562 A1_G9
C562 A1_G10
C562 A1_B1
C562 E1_G1
C562 B1_G1
C562 B1_G2
C562 B1_G3
C562 B1_G4
C562 B1_G5
C562 B1_G6
C562 B1_G7
C562 B1_G8
C562 B1_G9
C562 B1_G10
C562 B1_B1
C562 E1_G2
C562 C1_G1
C562 C1_G2
C562 C1_G3
C562 C1_G4
C562 C1_G5
C562 C1_G6
C562 C1_G7
C562 C1_G8
C562 C1_G9
C562 C1_G10
NA
C562 C1_B1
C562 E1_G3
C562 D1_G1
C562 D1_G2
C562 D1_G3
C562 D1_G4
C562 D1_G5
C562 D1_G6
C562 D1_G7
C562 D1_G8
C562 D1_G9
C562 D1_G10
C562 D1_B1
C562 E1_B1
