## Supplementary Figures for "The co-evolution of the genome and epigenome in colorectal cancer": figS4.pdf

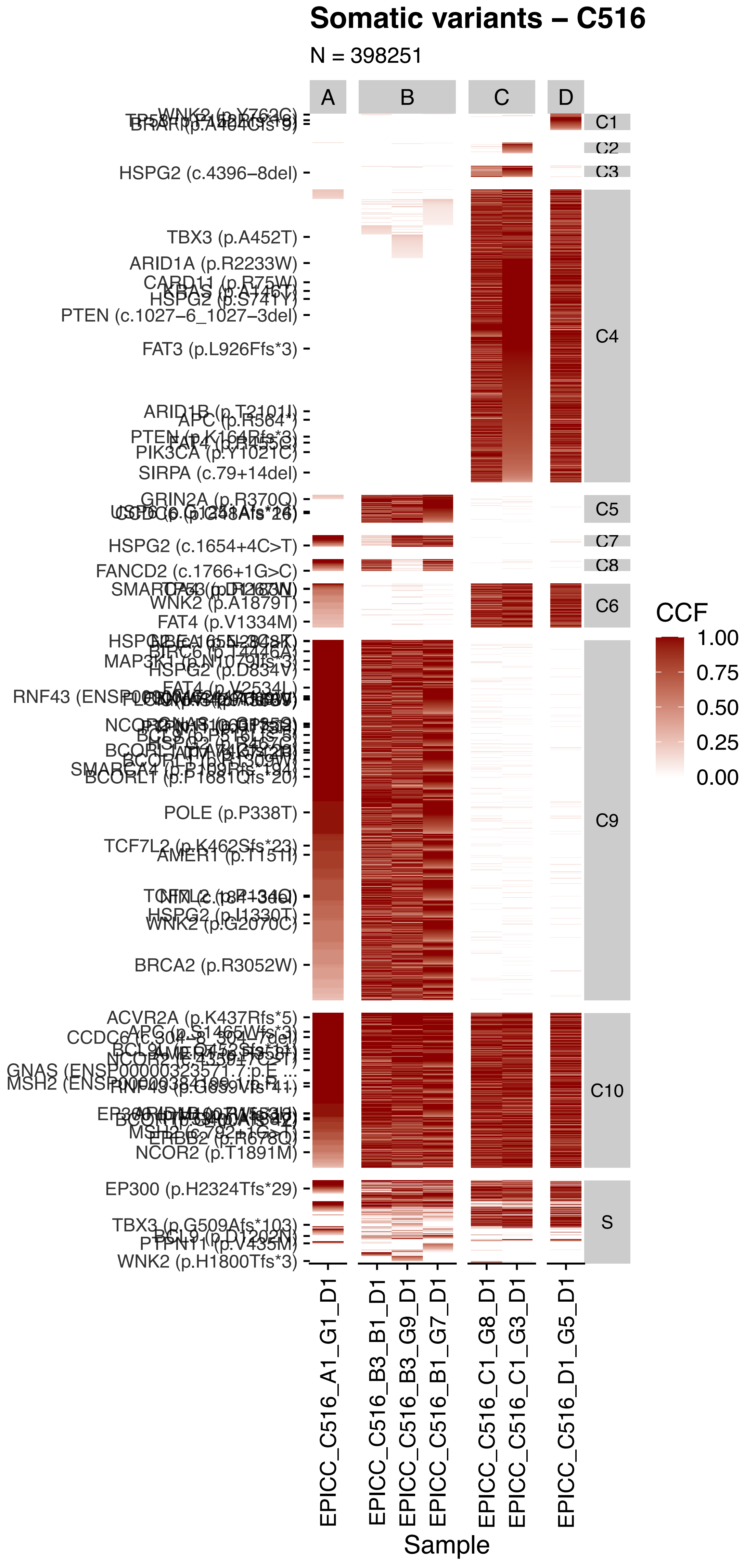

Somatic variants – C516
N = 398251
A
B
C
D
C1
C2
C3
C4
C5
C7
C8
C6
C9
C10
S
HSPG2 (c.4396–8del)
TBX3 (p.A452T)
ARID1A (p.R2233W)
PTEN (c.1027–6_1027–3del)
FAT3 (p.L926Ffs*3)
PIK3CA (p.Y1021C)
SIRPA (c.79+14del)
GRIN2A (p.R370Q)
HSPG2 (c.1654+4C>T)
FANCD2 (c.1766+1G>C)
WNK2 (p.A1879T)
FAT4 (p.V1334M)
HSPG2 (p.D834V)
POLE (p.P338T)
AMER1 (p.T151I)
HSPG2 (p.I1330T)
WNK2 (p.G2070C)
BRCA2 (p.R3052W)
ACVR2A (p.K437Rfs*5)
ERBB2 (p.R678Q)
NCOR2 (p.T1891M)
EP300 (p.H2324Tfs*29)
PTPN11 (p.V435M)
WNK2 (p.H1800Tfs*3)
CCF
1.00
0.75
0.50
0.25
0.00
EPICC_C516_A1_G1_D1
EPICC_C516_B3_B1_D1
EPICC_C516_B3_G9_D1
EPICC_C516_B1_G7_D1
EPICC_C516_C1_G8_D1
EPICC_C516_C1_G3_D1
EPICC_C516_D1_G5_D1
Sample

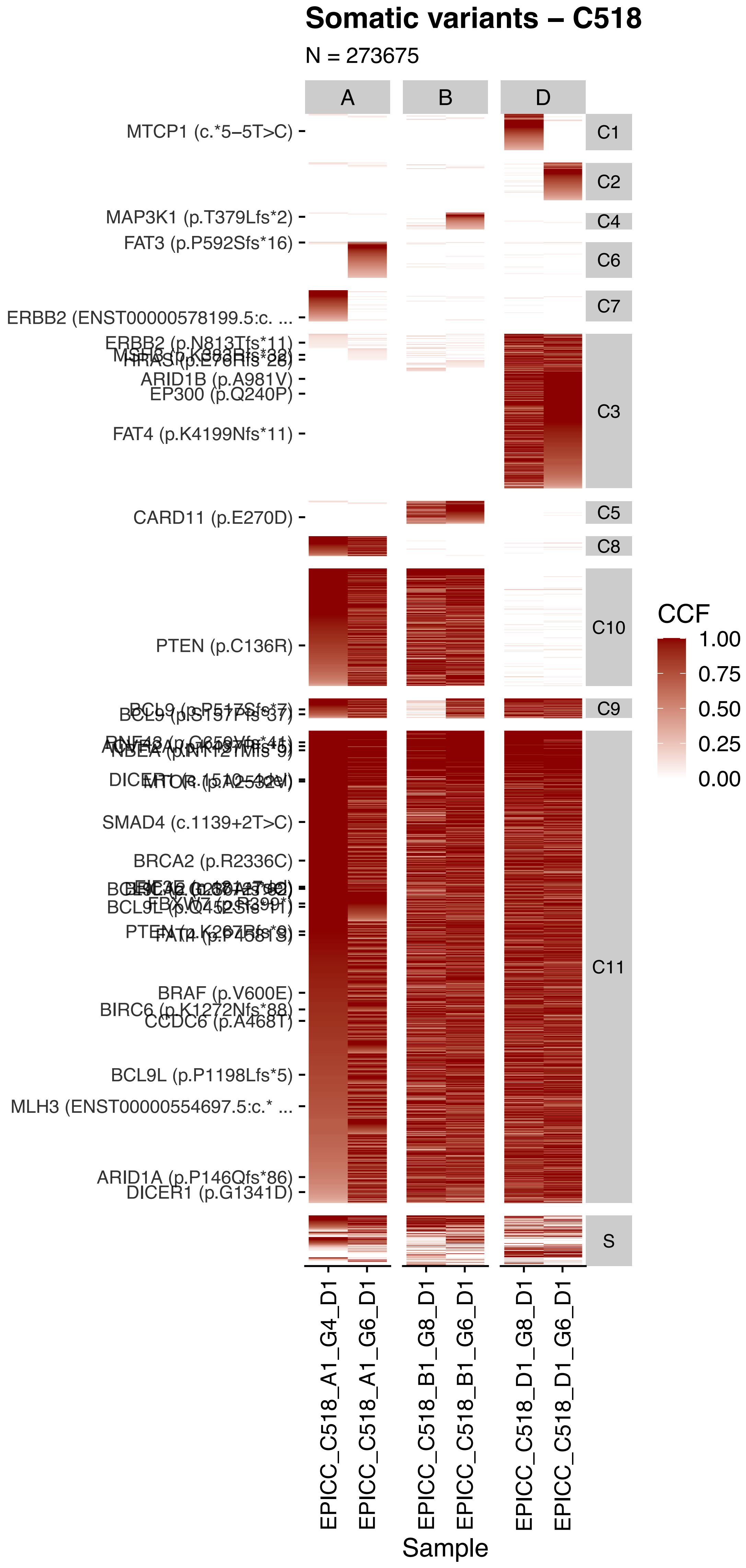

Somatic variants – C518
N = 273675
A
B
D
MTCP1 (c.*5–5T>C)
MAP3K1 (p.T379Lfs*2)
FAT3 (p.P592Sfs*16)
ERBB2 (ENST00000578199.5:c. ...
ARID1B (p.A981V)
EP300 (p.Q240P)
FAT4 (p.K4199Nfs*11)
CARD11 (p.E270D)
PTEN (p.C136R)
SMAD4 (c.1139+2T>C)
BRCA2 (p.R2336C)
BRAF (p.V600E)
CCDC6 (p.A468T)
BCL9L (p.P1198Lfs*5)
MLH3 (ENST00000554697.5:c.* ...
DICER1 (p.G1341D)
C1
C2
C4
C6
C7
C3
C5
C8
C10
C9
C11
S
CCF
1.00
0.75
0.50
0.25
0.00
EPICC_C518_A1_G4_D1
EPICC_C518_A1_G6_D1
EPICC_C518_B1_G8_D1
EPICC_C518_B1_G6_D1
EPICC_C518_D1_G8_D1
EPICC_C518_D1_G6_D1
Sample

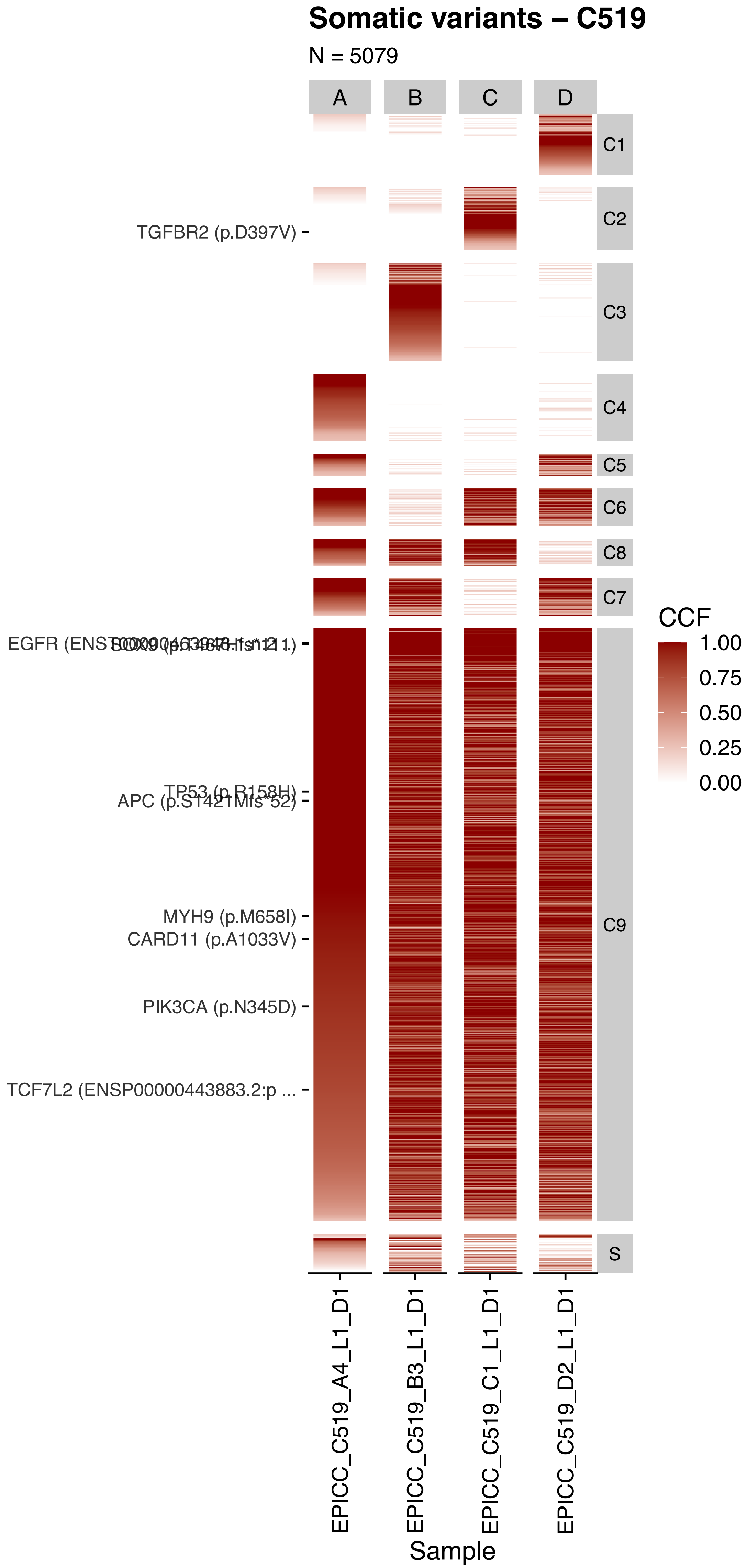

Somatic variants – C519
N = 5079
A
B
C
D
C1
C2
C3
C4
C5
C6
C8
C7
C9
S
TGFBR2 (p.D397V)
EGFR (ENST00000463948 fs*11)
SOX9 (p.T467Hfs*21)
TP53 (p.R158H)
APC (p.S1421Mfs*52)
MYH9 (p.M658I)
CARD11 (p.A1033V)
PIK3CA (p.N345D)
TCF7L2 (ENSP00000443883.2:p ...
CCF
1.00
0.75
0.50
0.25
0.00
EPICC_C519_A4_L1_D1
EPICC_C519_B3_L1_D1
EPICC_C519_C1_L1_D1
EPICC_C519_D2_L1_D1
Sample

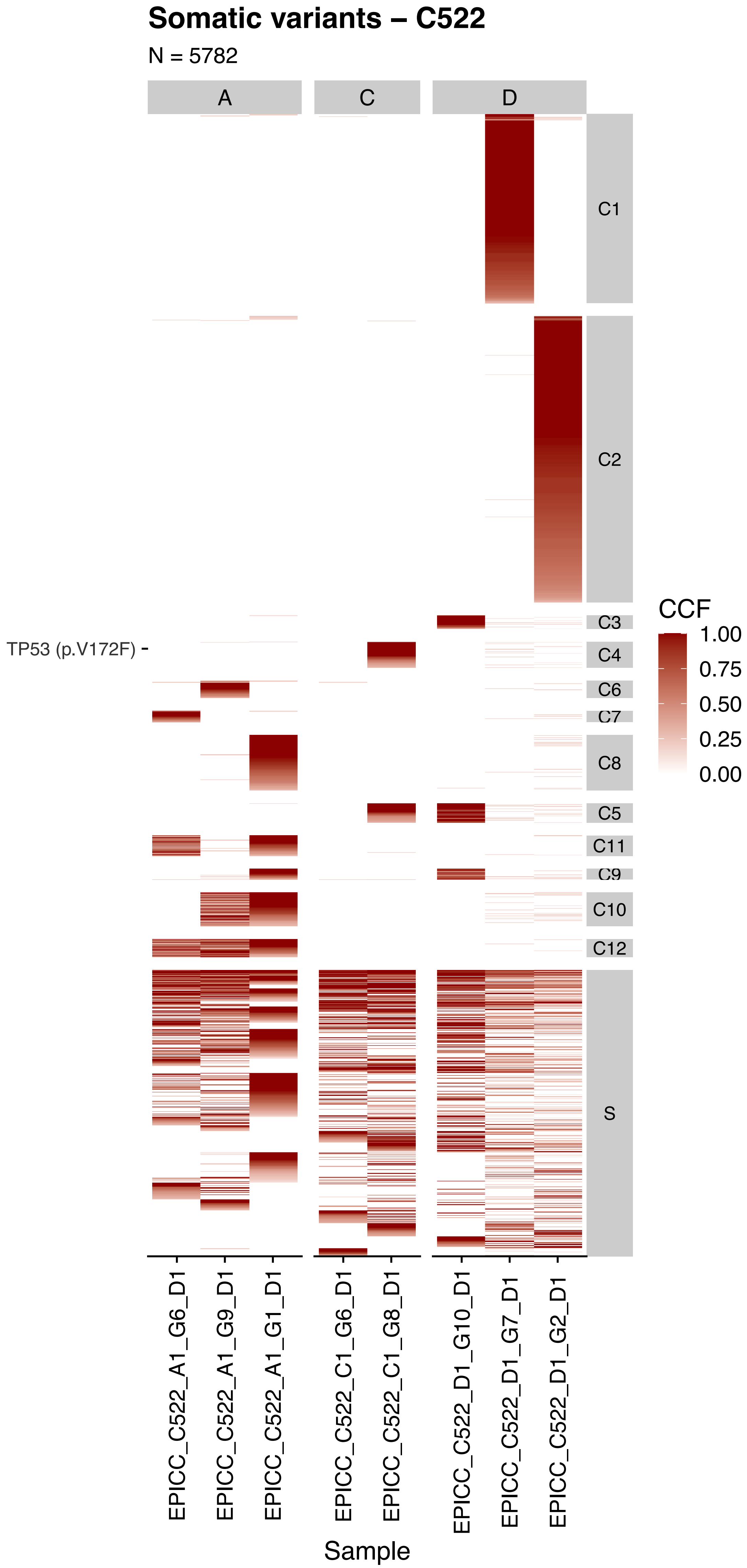

Somatic variants – C522
N = 5782
A
C
D
C1
C2
C3
C4
C6
C7
C8
C5
C11
C9
C10
C12
S
TP53 (p.V172F)
CCF
1.00
0.75
0.50
0.25
0.00
EPICC_C522_A1_G6_D1
EPICC_C522_A1_G9_D1
EPICC_C522_A1_G1_D1
EPICC_C522_C1_G6_D1
EPICC_C522_C1_G8_D1
EPICC_C522_D1_G10_D1
EPICC_C522_D1_G7_D1
EPICC_C522_D1_G2_D1
Sample

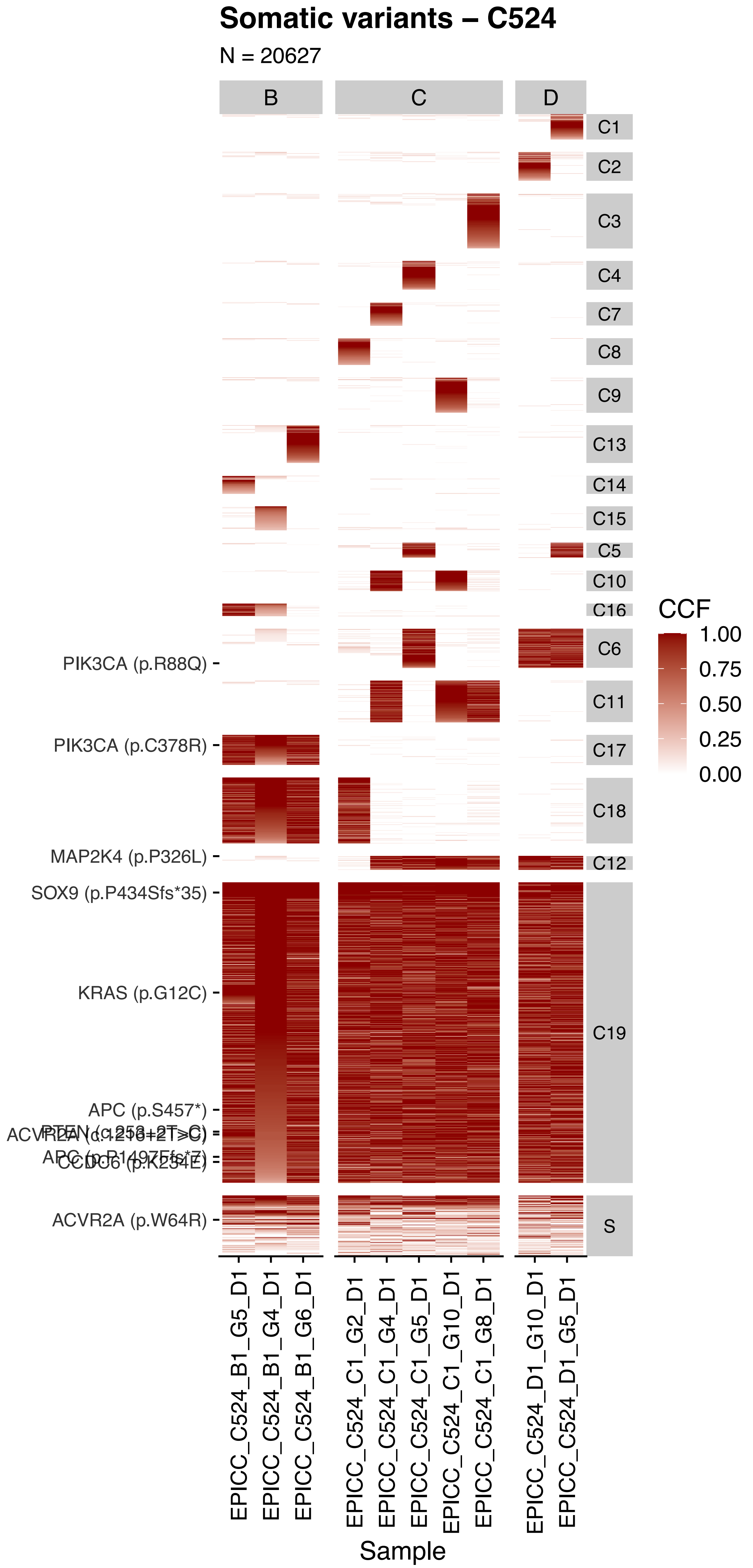

Somatic variants – C524
N = 20627
B
C
D
C1
C2
C3
C4
C7
C8
C9
C13
C14
C15
C5
C10
C16
C6
C11
C17
C18
C12
C19
S
PIK3CA (p.R88Q)
PIK3CA (p.C378R)
MAP2K4 (p.P326L)
SOX9 (p.P434Sfs*35)
KRAS (p.G12C)
APC (p.S457*)
ACVR2A (p.W64R)
CCF
1.00
0.75
0.50
0.25
0.00
EPICC_C524_B1_G5_D1
EPICC_C524_B1_G4_D1
EPICC_C524_B1_G6_D1
EPICC_C524_C1_G2_D1
EPICC_C524_C1_G4_D1
EPICC_C524_C1_G5_D1
EPICC_C524_C1_G10_D1
EPICC_C524_C1_G8_D1
EPICC_C524_D1_G10_D1
EPICC_C524_D1_G5_D1
Sample

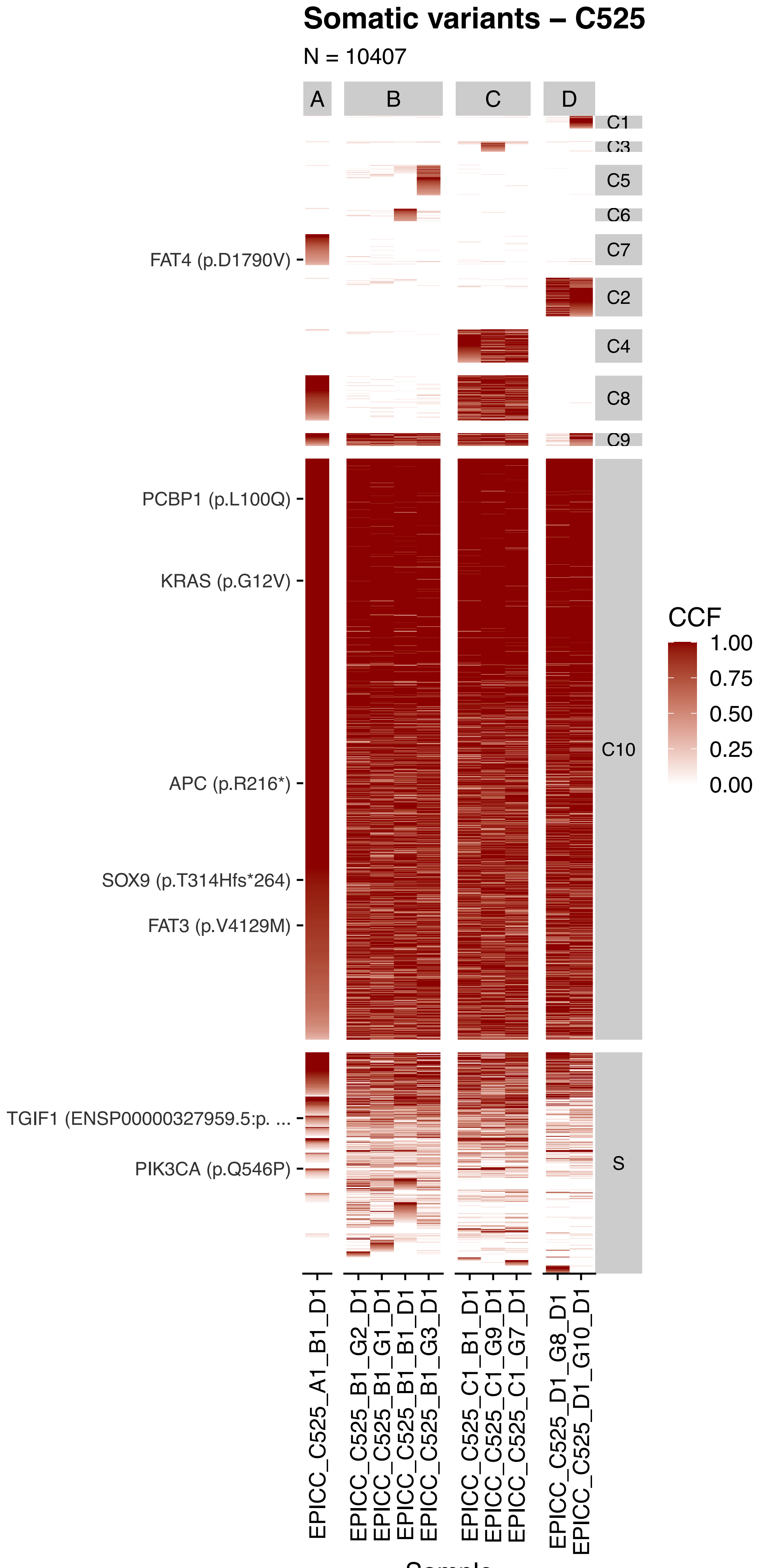

Somatic variants – C525
N = 10407
A
B
C
D
C1
C3
C5
C6
C7
C2
C4
C8
C9
C10
S
FAT4 (p.D1790V)
PCBP1 (p.L100Q)
KRAS (p.G12V)
APC (p.R216*)
SOX9 (p.T314Hfs*264)
FAT3 (p.V4129M)
TGIF1 (ENSP00000327959.5:p. ...
PIK3CA (p.Q546P)
CCF
1.00
0.75
0.50
0.25
0.00
EPICC_C525_A1_B1_D1
EPICC_C525_B1_G2_D1
EPICC_C525_B1_G1_D1
EPICC_C525_B1_B1_D1
EPICC_C525_B1_G3_D1
EPICC_C525_C1_B1_D1
EPICC_C525_C1_G9_D1
EPICC_C525_C1_G7_D1
EPICC_C525_D1_G8_D1
EPICC_C525_D1_G10_D1
Sample

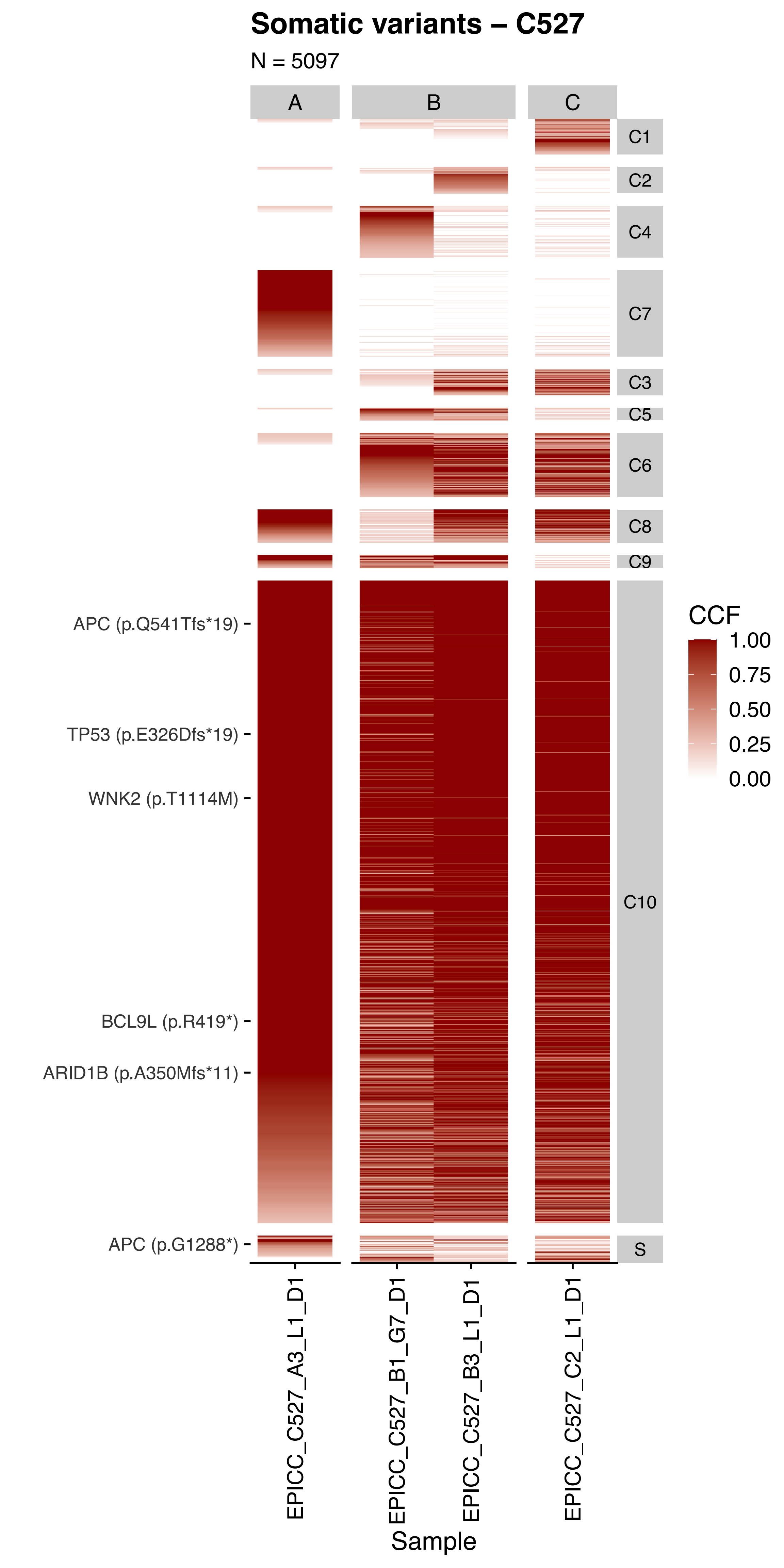
Somatic variants – C527
N = 5097
A
B
C
C1
C2
C4
C7
C3
C5
C6
C8
C9
C10
S
APC (p.Q541Tfs*19)
TP53 (p.E326Dfs*19)
WNK2 (p.T1114M)
BCL9L (p.R419*)
ARID1B (p.A350Mfs*11)
APC (p.G1288*)
CCF
1.00
0.75
0.50
0.25
0.00
EPICC_C527_A3_L1_D1
EPICC_C527_B1_G7_D1
EPICC_C527_B3_L1_D1
EPICC_C527_C2_L1_D1
Sample

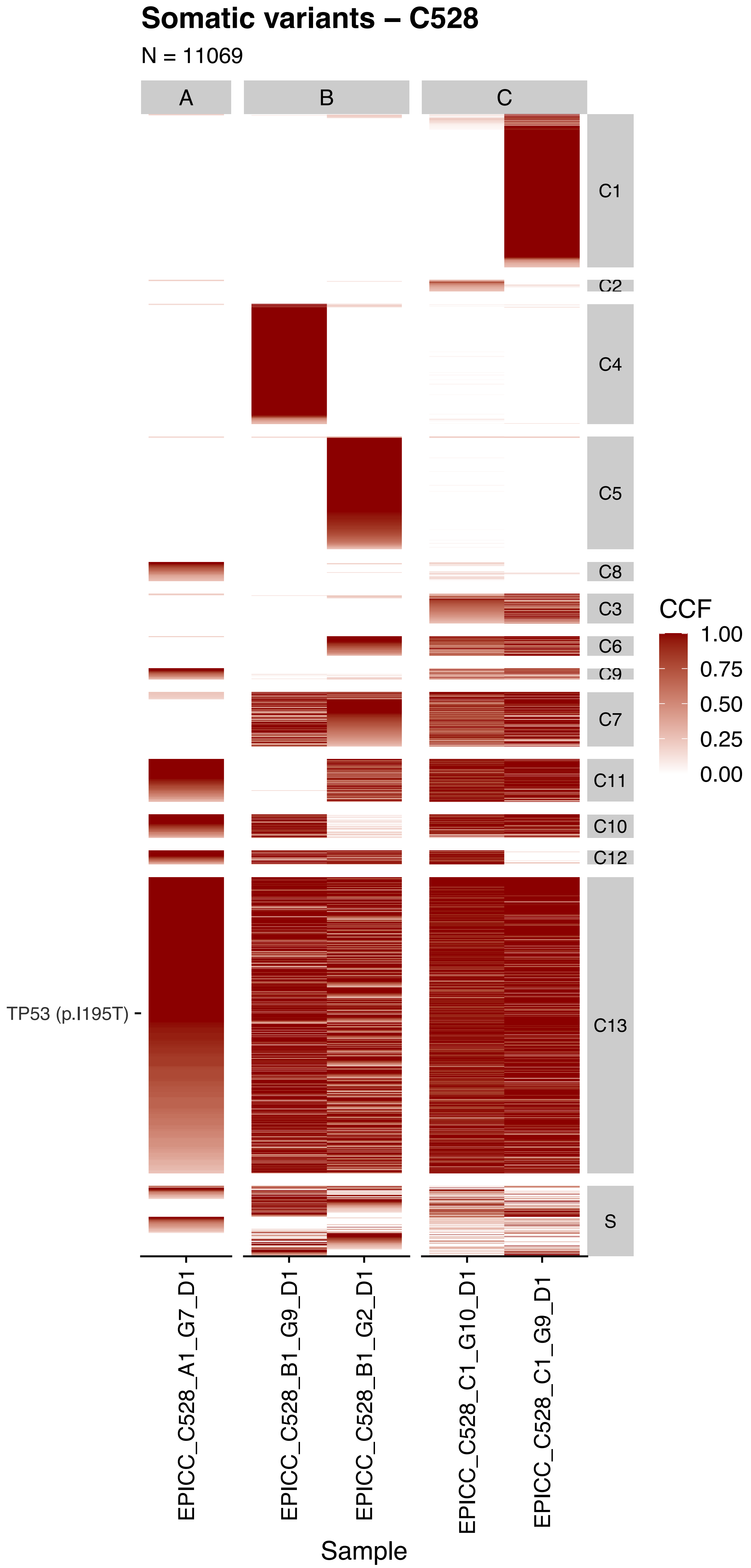

Somatic variants – C528
N = 11069
A
B
C
C1
C2
C4
C5
C8
C3
C6
C9
C7
C11
C10
C12
C13
S
TP53 (p.I195T)
CCF
1.00
0.75
0.50
0.25
0.00
EPICC_C528_A1_G7_D1
EPICC_C528_B1_G9_D1
EPICC_C528_B1_G2_D1
EPICC_C528_C1_G10_D1
EPICC_C528_C1_G9_D1
Sample

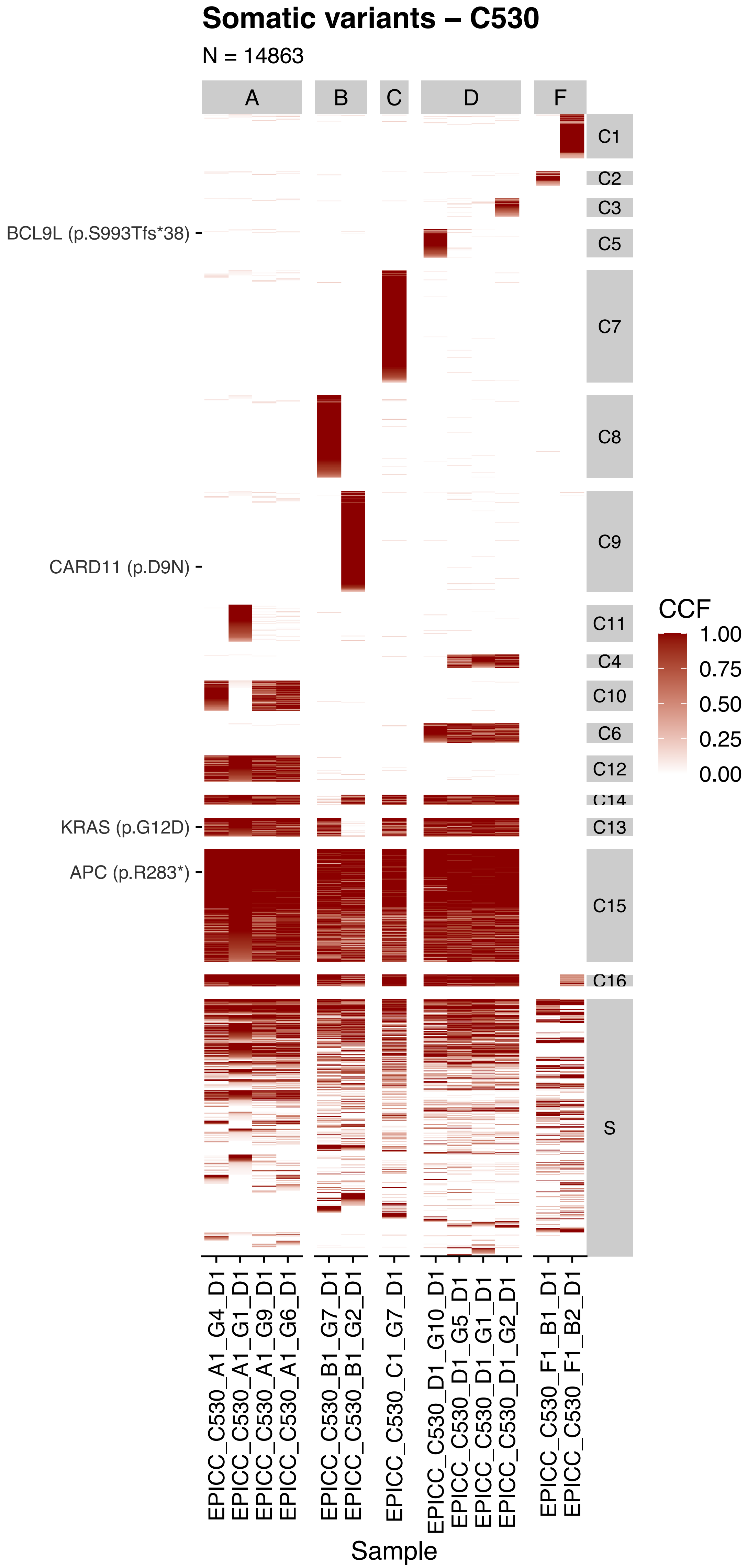

Somatic variants – C530
N = 14863
A
B
C
D
F
C1
C2
C3
C5
C7
C8
C9
C11
C4
C10
C6
C12
C14
C13
C15
C16
S
BCL9L (p.S993Tfs*38)
CARD11 (p.D9N)
KRAS (p.G12D)
APC (p.R283*)
CCF
1.00
0.75
0.50
0.25
0.00
EPICC_C530_A1_G4_D1
EPICC_C530_A1_G1_D1
EPICC_C530_A1_G9_D1
EPICC_C530_A1_G6_D1
EPICC_C530_B1_G7_D1
EPICC_C530_B1_G2_D1
EPICC_C530_C1_G7_D1
EPICC_C530_D1_G10_D1
EPICC_C530_D1_G5_D1
EPICC_C530_D1_G1_D1
EPICC_C530_D1_G2_D1
EPICC_C530_F1_B1_D1
EPICC_C530_F1_B2_D1
Sample

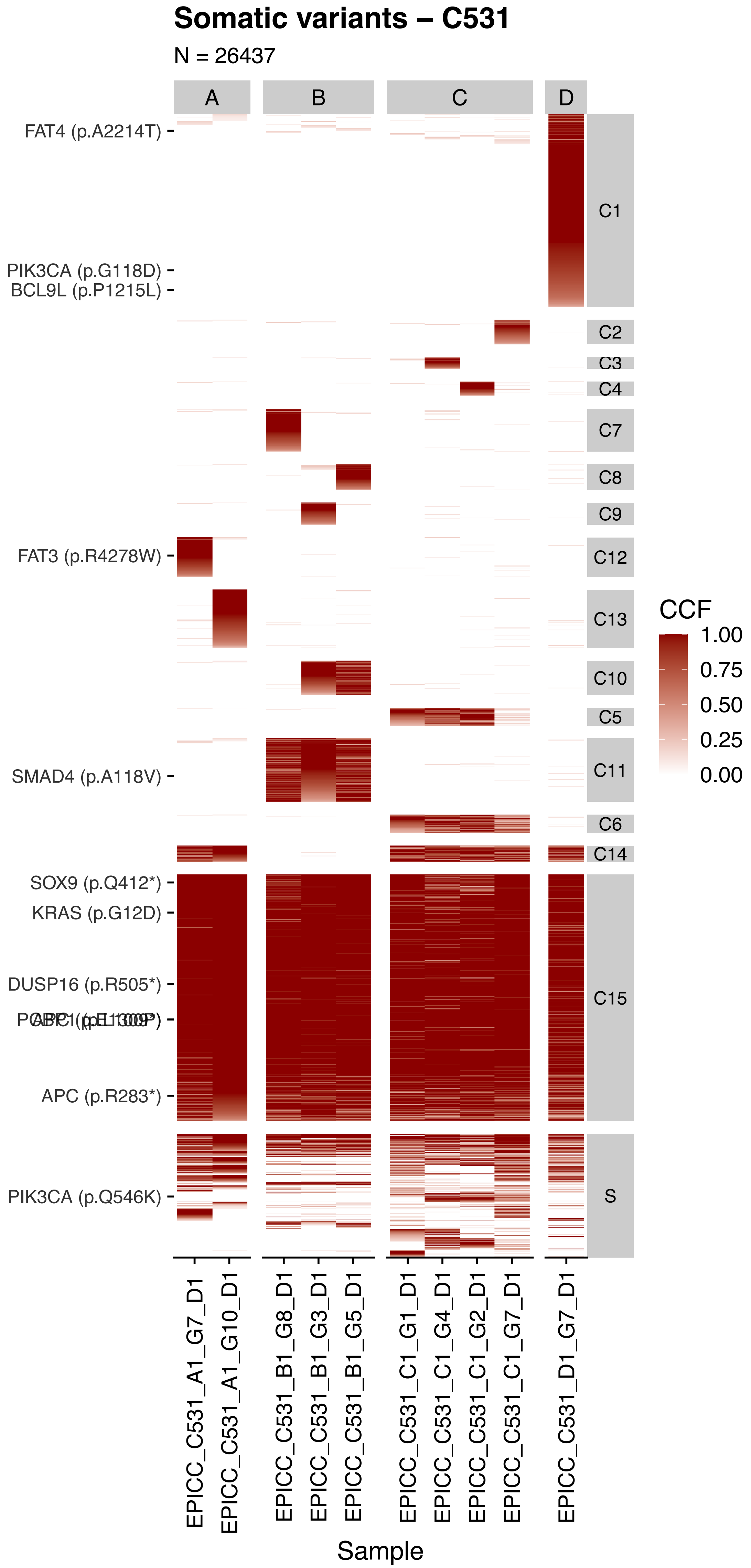
Somatic variants – C531
N = 26437
A
B
C
D
FAT4 (p.A2214T)
PIK3CA (p.G118D)
BCL9L (p.P1215L)
FAT3 (p.R4278W)
SMAD4 (p.A118V)
SOX9 (p.Q412*)
KRAS (p.G12D)
DUSP16 (p.R505*)
APC (p.R283*)
PIK3CA (p.Q546K)
C1
C2
C3
C4
C7
C8
C9
C12
C13
C10
C5
C11
C6
C14
C15
S
CCF
1.00
0.75
0.50
0.25
0.00
EPICC_C531_A1_G7_D1
EPICC_C531_A1_G10_D1
EPICC_C531_B1_G8_D1
EPICC_C531_B1_G3_D1
EPICC_C531_B1_G5_D1
EPICC_C531_C1_G1_D1
EPICC_C531_C1_G4_D1
EPICC_C531_C1_G2_D1
EPICC_C531_C1_G7_D1
EPICC_C531_D1_G7_D1
Sample

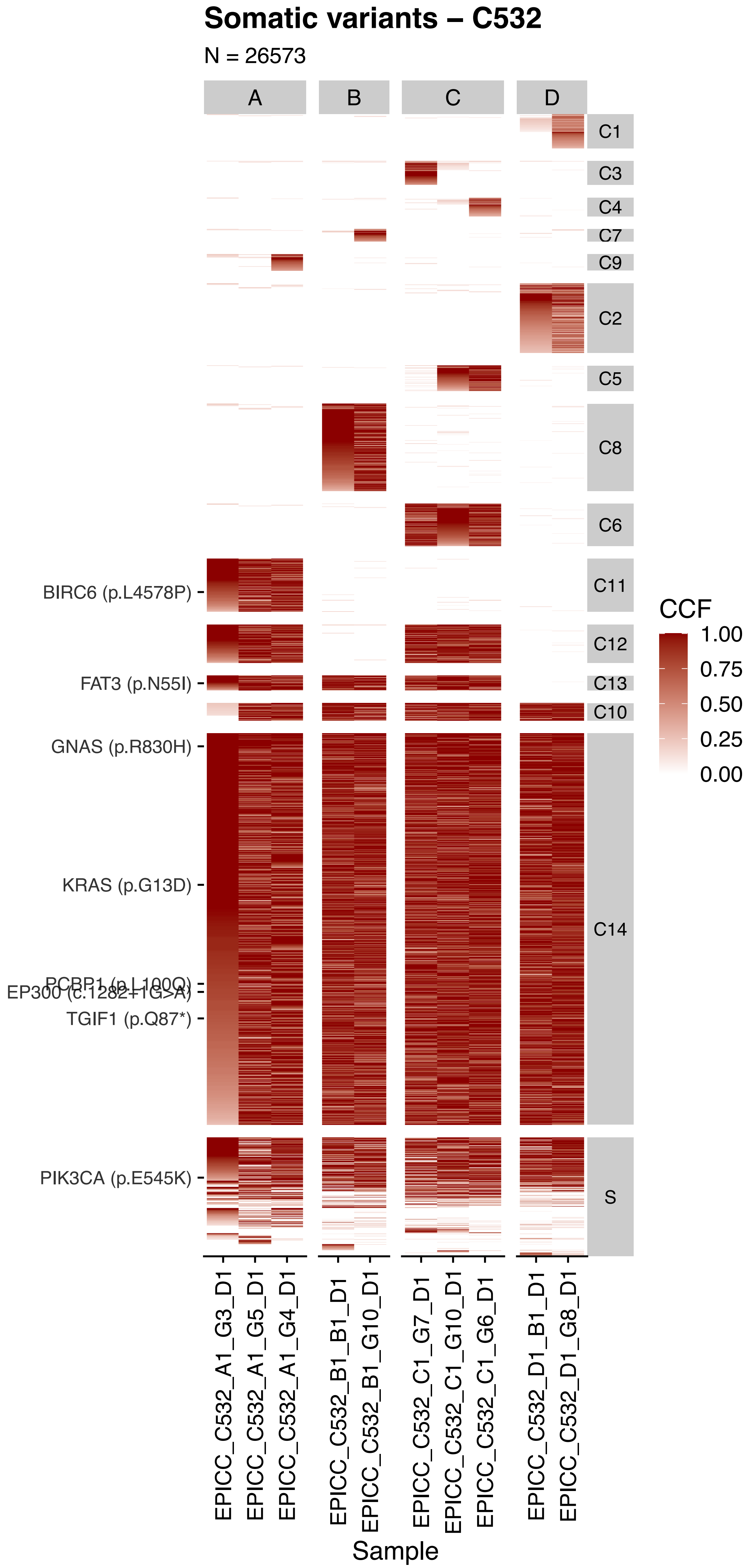

Somatic variants – C532
N = 26573
A
B
C
D
C1
C3
C4
C7
C9
C2
C5
C8
C6
C11
C12
C13
C10
C14
S
BIRC6 (p.L4578P)
FAT3 (p.N55I)
GNAS (p.R830H)
KRAS (p.G13D)
PCBP1 (p.L100Q)
EP300 (c.1282+1G>A)
TGIF1 (p.Q87*)
PIK3CA (p.E545K)
CCF
1.00
0.75
0.50
0.25
0.00
EPICC_C532_A1_G3_D1
EPICC_C532_A1_G5_D1
EPICC_C532_A1_G4_D1
EPICC_C532_B1_B1_D1
EPICC_C532_B1_G10_D1
EPICC_C532_C1_G7_D1
EPICC_C532_C1_G10_D1
EPICC_C532_C1_G6_D1
EPICC_C532_D1_B1_D1
EPICC_C532_D1_G8_D1
Sample

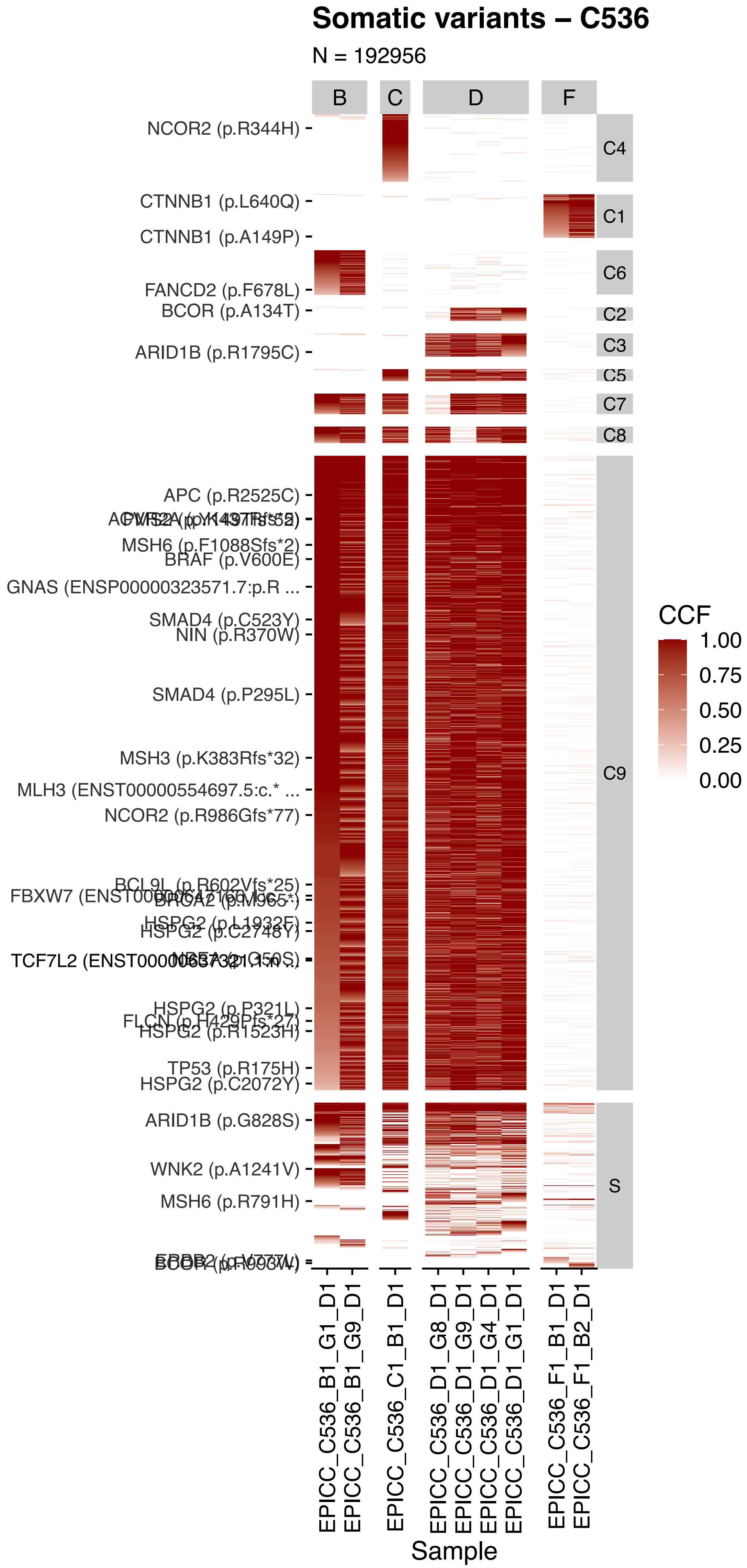

Somatic variants – C536
N = 192956
B
C
D
F
C4
C1
C6
C2
C3
C5
C7
C8
C9
S
NCOR2 (p.R344H)
CTNNB1 (p.L640Q)
CTNNB1 (p.A149P)
FANCD2 (p.F678L)
BCOR (p.A134T)
ARID1B (p.R1795C)
APC (p.R2525C)
MSH6 (p.F1088Sfs*2)
BRAF (p.V600E)
GNAS (ENSP00000323571.7:p.R ...
SMAD4 (p.C523Y)
NIN (p.R370W)
SMAD4 (p.P295L)
MSH3 (p.K383Rfs*32)
MLH3 (ENST00000554697.5:c.* ...
NCOR2 (p.R986Gfs*77)
HSPG2 (p.P321L)
HSPG2 (p.R1523H)
TP53 (p.R175H)
HSPG2 (p.C2072Y)
ARID1B (p.G828S)
WNK2 (p.A1241V)
MSH6 (p.R791H)
CCF
1.00
0.75
0.50
0.25
0.00
EPICC_C536_B1_G1_D1
EPICC_C536_B1_G9_D1
EPICC_C536_C1_B1_D1
EPICC_C536_D1_G8_D1
EPICC_C536_D1_G9_D1
EPICC_C536_D1_G4_D1
EPICC_C536_D1_G1_D1
EPICC_C536_F1_B1_D1
EPICC_C536_F1_B2_D1
Sample

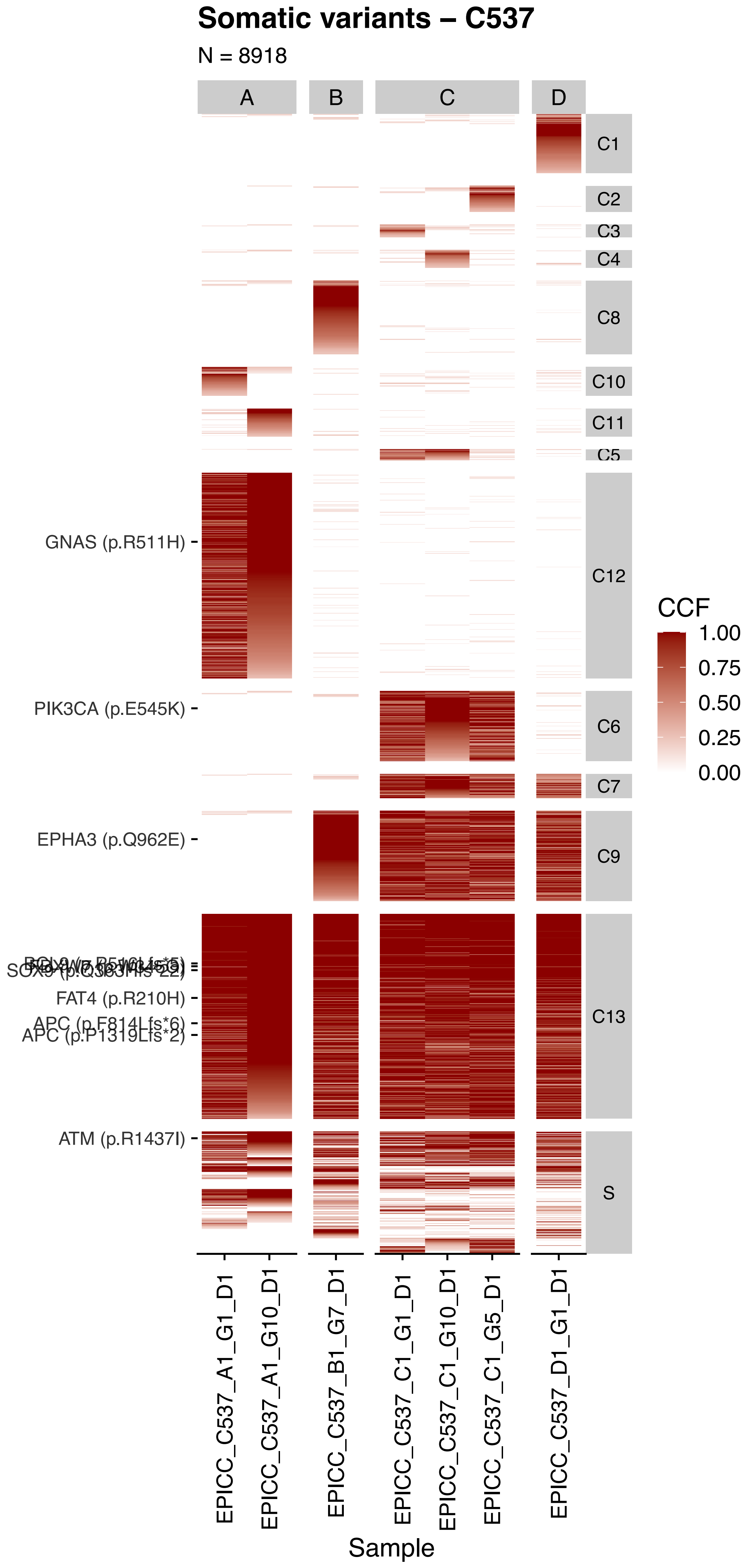
Somatic variants – C537
N = 8918
A
B
C
D
C1
C2
C3
C4
C8
C10
C11
C5
C12
C6
C7
C9
C13
S
GNAS (p.R511H)
PIK3CA (p.E545K)
EPHA3 (p.Q962E)
FAT4 (p.R210H)
APC (p.F814Lfs*6)
APC (p.P1319Lfs*2)
ATM (p.R1437I)
CCF
1.00
0.75
0.50
0.25
0.00
EPICC_C537_A1_G1_D1
EPICC_C537_A1_G10_D1
EPICC_C537_B1_G7_D1
EPICC_C537_C1_G1_D1
EPICC_C537_C1_G10_D1
EPICC_C537_C1_G5_D1
EPICC_C537_D1_G1_D1
Sample

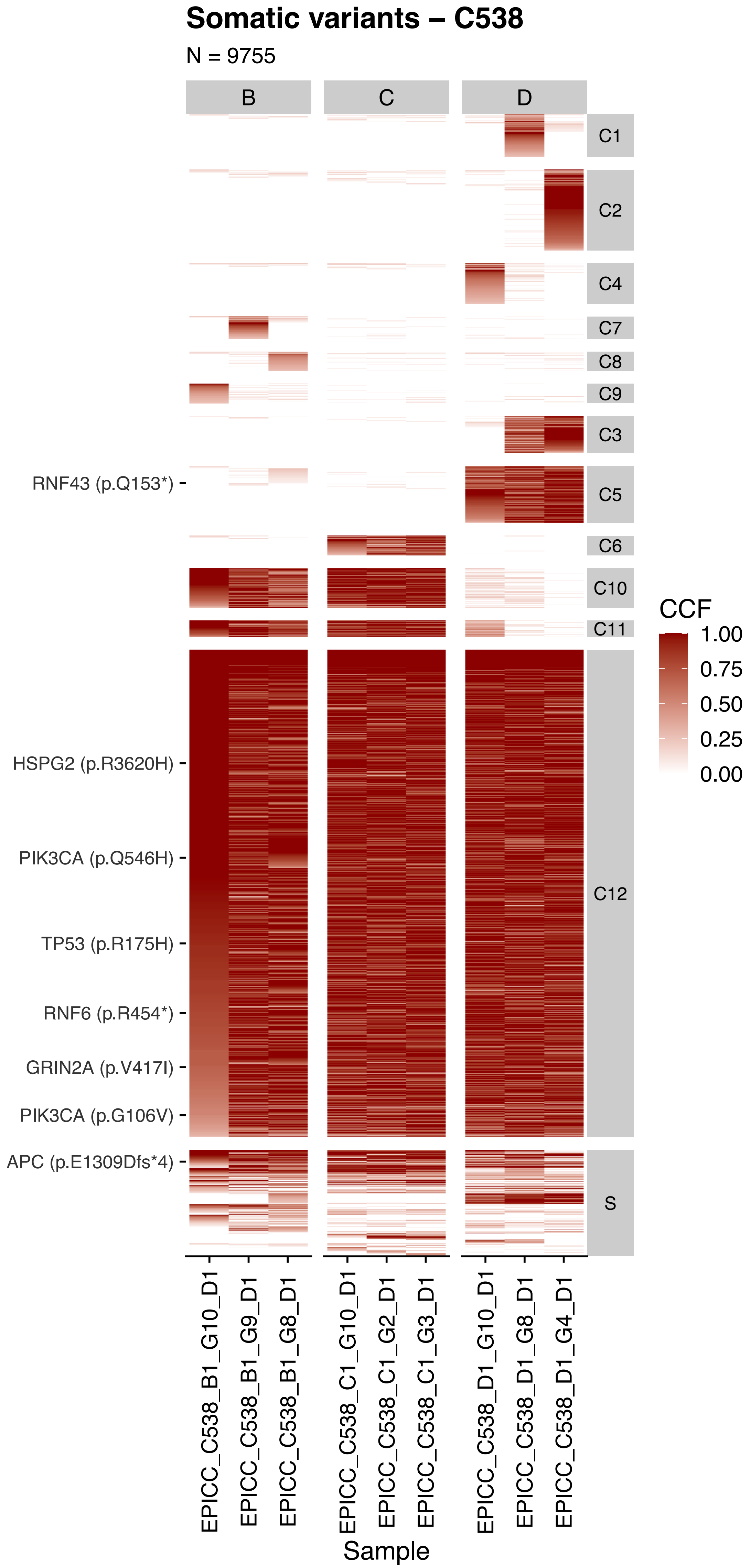
Somatic variants – C538
N = 9755
B
C
D
C1
C2
C4
C7
C8
C9
C3
C5
C6
C10
C11
C12
S
RNF43 (p.Q153*)
HSPG2 (p.R3620H)
PIK3CA (p.Q546H)
TP53 (p.R175H)
RNF6 (p.R454*)
GRIN2A (p.V417I)
PIK3CA (p.G106V)
APC (p.E1309Dfs*4)
CCF
1.00
0.75
0.50
0.25
0.00
EPICC_C538_B1_G10_D1
EPICC_C538_B1_G9_D1
EPICC_C538_B1_G8_D1
EPICC_C538_C1_G10_D1
EPICC_C538_C1_G2_D1
EPICC_C538_C1_G3_D1
EPICC_C538_D1_G10_D1
EPICC_C538_D1_G8_D1
EPICC_C538_D1_G4_D1
Sample

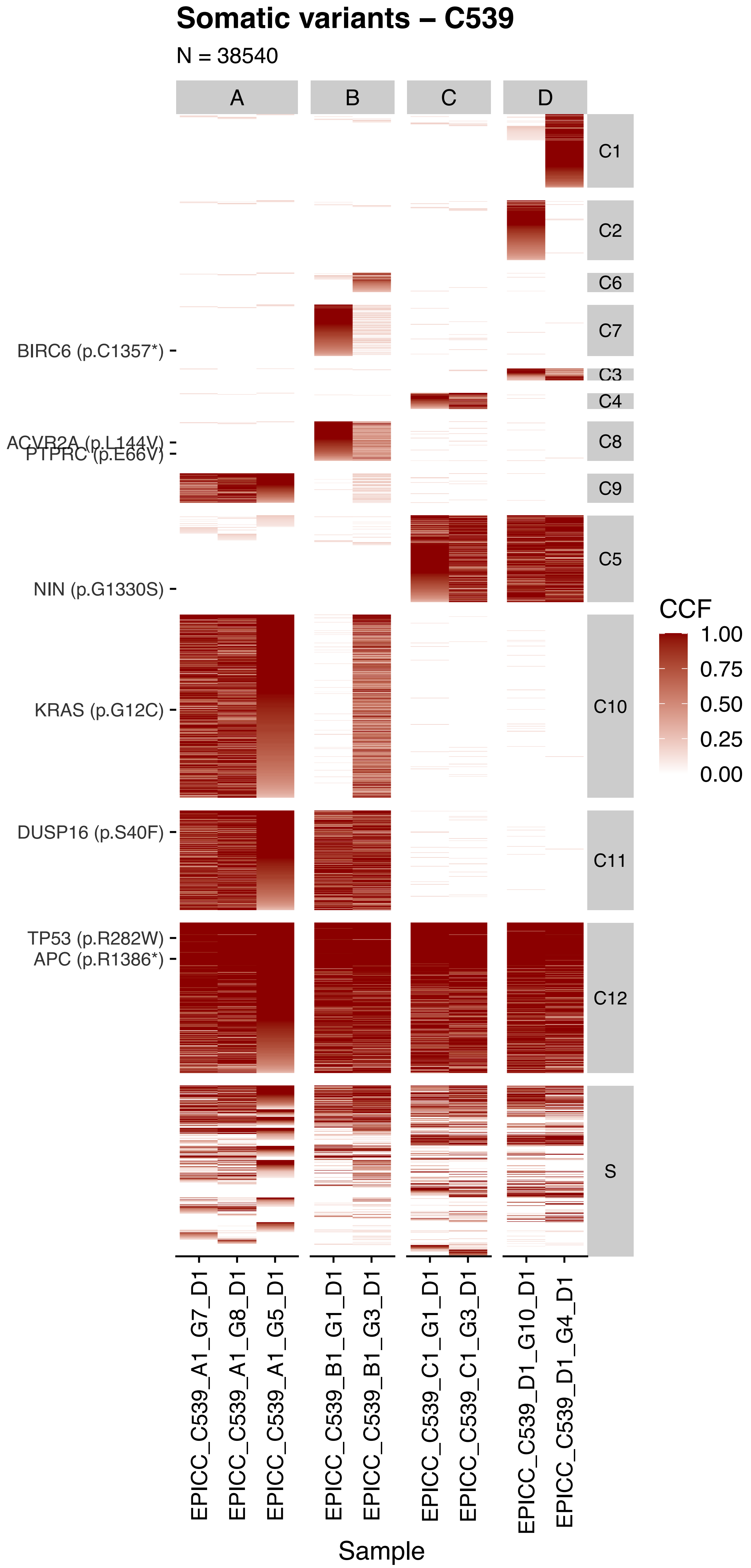

Somatic variants – C539
N = 38540
A
B
C
D
C1
C2
C6
C7
C3
C4
C8
C9
C5
C10
C11
C12
S
BIRC6 (p.C1357*)
ACVR2A (p.L144V)
PTPRC (p.E66V)
NIN (p.G1330S)
KRAS (p.G12C)
DUSP16 (p.S40F)
TP53 (p.R282W)
APC (p.R1386*)
CCF
1.00
0.75
0.50
0.25
0.00
EPICC_C539_A1_G7_D1
EPICC_C539_A1_G8_D1
EPICC_C539_A1_G5_D1
EPICC_C539_B1_G1_D1
EPICC_C539_B1_G3_D1
EPICC_C539_C1_G1_D1
EPICC_C539_C1_G3_D1
EPICC_C539_D1_G10_D1
EPICC_C539_D1_G4_D1
Sample

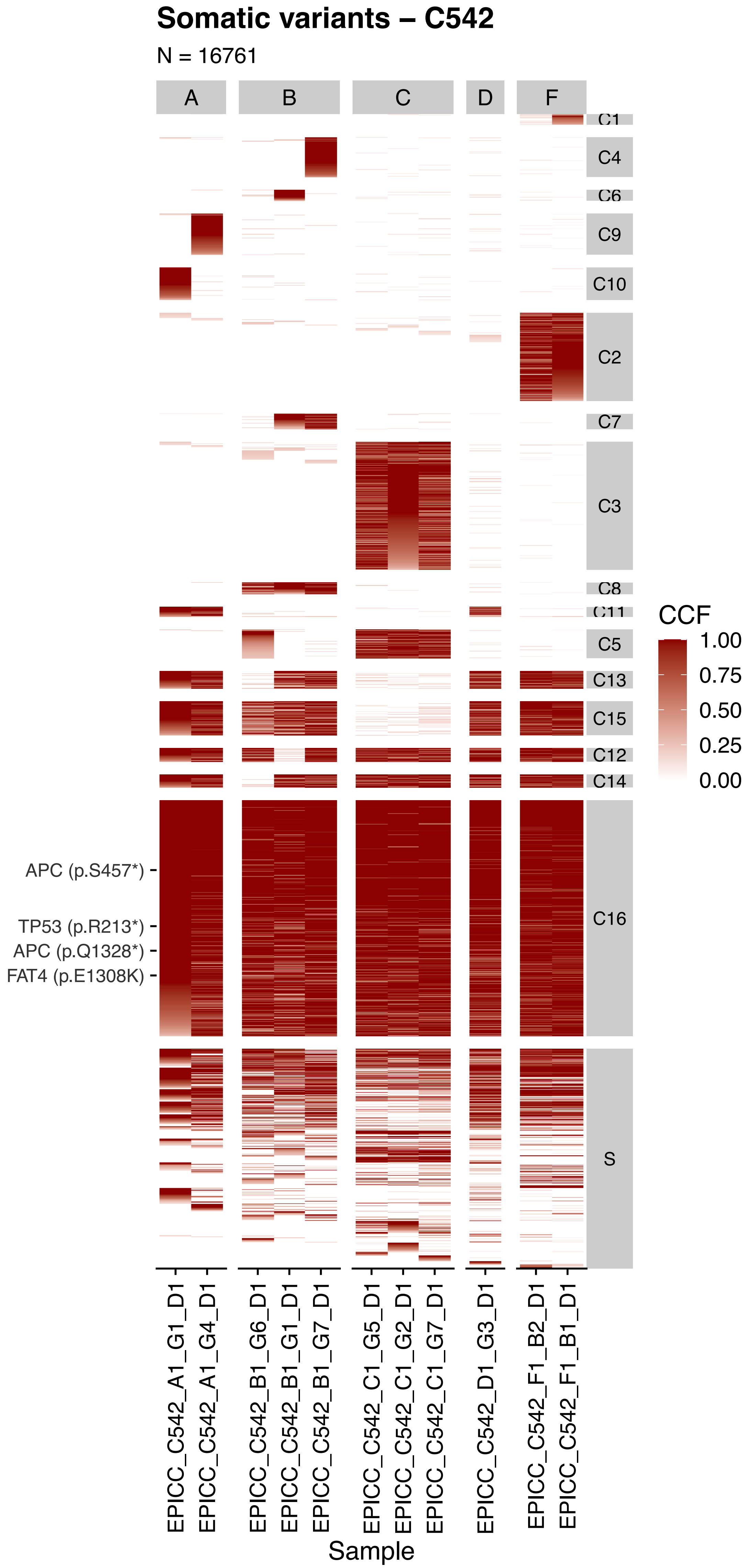

Somatic variants – C542
N = 16761
A
B
C
D
F
C1
C4
C6
C9
C10
C2
C7
C3
C8
C11
C5
C13
C15
C12
C14
C16
S
CCF
1.00
0.75
0.50
0.25
0.00
APC (p.S457*)
TP53 (p.R213*)
APC (p.Q1328*)
FAT4 (p.E1308K)
EPICC_C542_A1_G1_D1
EPICC_C542_A1_G4_D1
EPICC_C542_B1_G6_D1
EPICC_C542_B1_G1_D1
EPICC_C542_B1_G7_D1
EPICC_C542_C1_G5_D1
EPICC_C542_C1_G2_D1
EPICC_C542_C1_G7_D1
EPICC_C542_D1_G3_D1
EPICC_C542_F1_B2_D1
EPICC_C542_F1_B1_D1
Sample

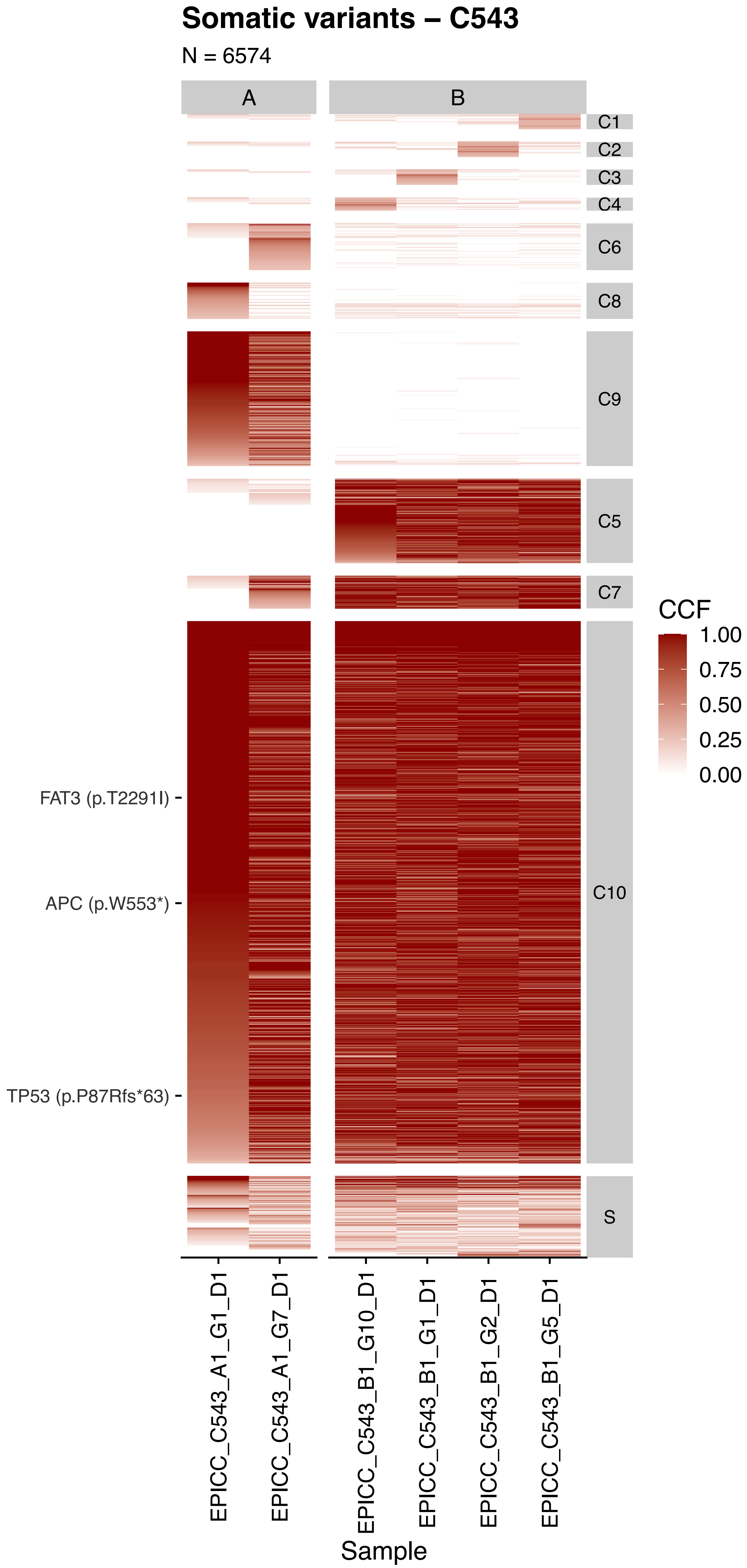

Somatic variants – C543
N = 6574
A
B
C1
C2
C3
C4
C6
C8
C9
C5
C7
C10
S
CCF
1.00
0.75
0.50
0.25
0.00
FAT3 (p.T2291I)
APC (p.W553*)
TP53 (p.P87Rfs*63)
EPICC_C543_A1_G1_D1
EPICC_C543_A1_G7_D1
EPICC_C543_B1_G10_D1
EPICC_C543_B1_G1_D1
EPICC_C543_B1_G2_D1
EPICC_C543_B1_G5_D1
Sample

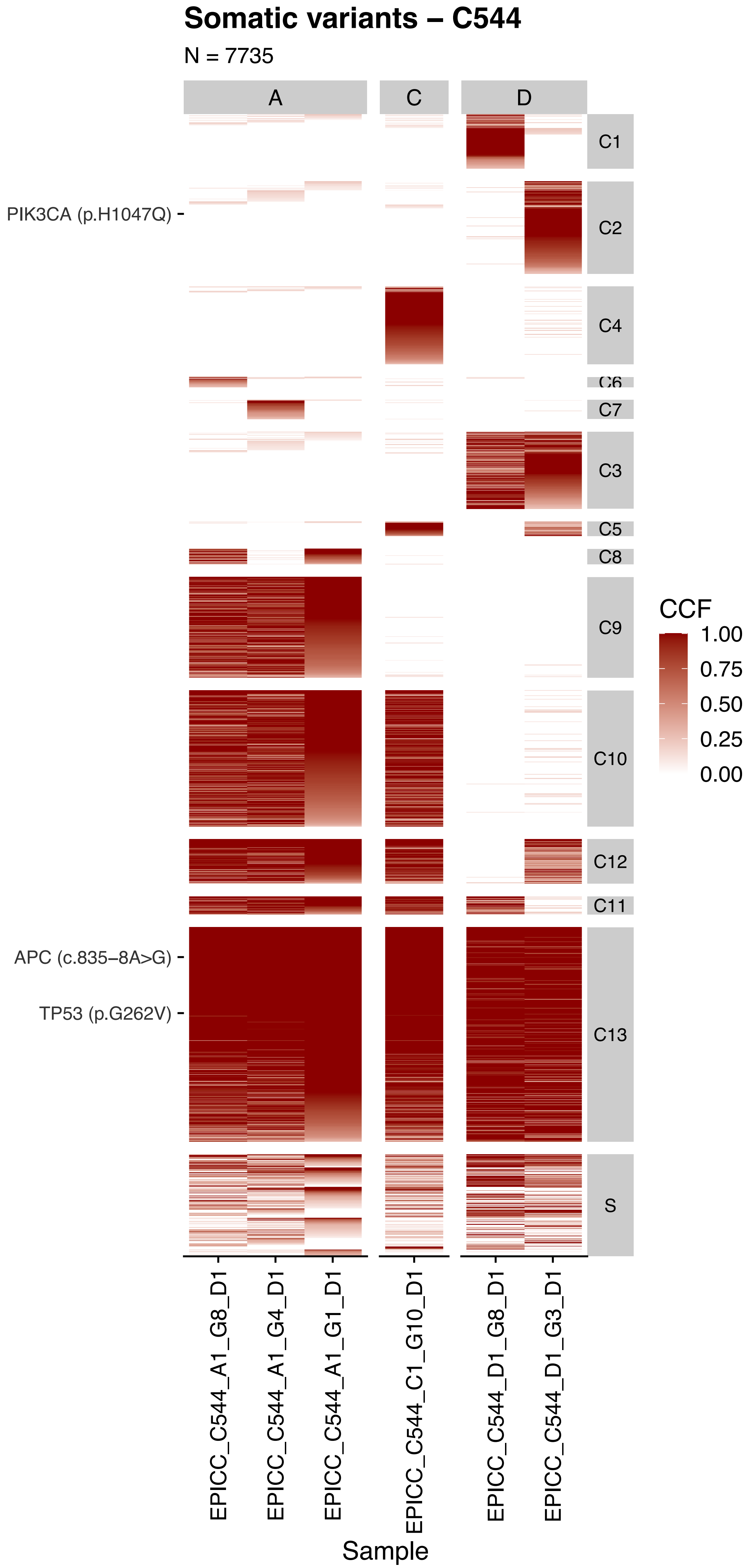
Somatic variants – C544
N = 7735
A
C
D
C1
C2
C4
C6
C7
C3
C5
C8
C9
C10
C12
C11
C13
S
PIK3CA (p.H1047Q)
APC (c.835–8A>G)
TP53 (p.G262V)
CCF
1.00
0.75
0.50
0.25
0.00
EPICC_C544_A1_G8_D1
EPICC_C544_A1_G4_D1
EPICC_C544_A1_G1_D1
EPICC_C544_C1_G10_D1
EPICC_C544_D1_G8_D1
EPICC_C544_D1_G3_D1
Sample

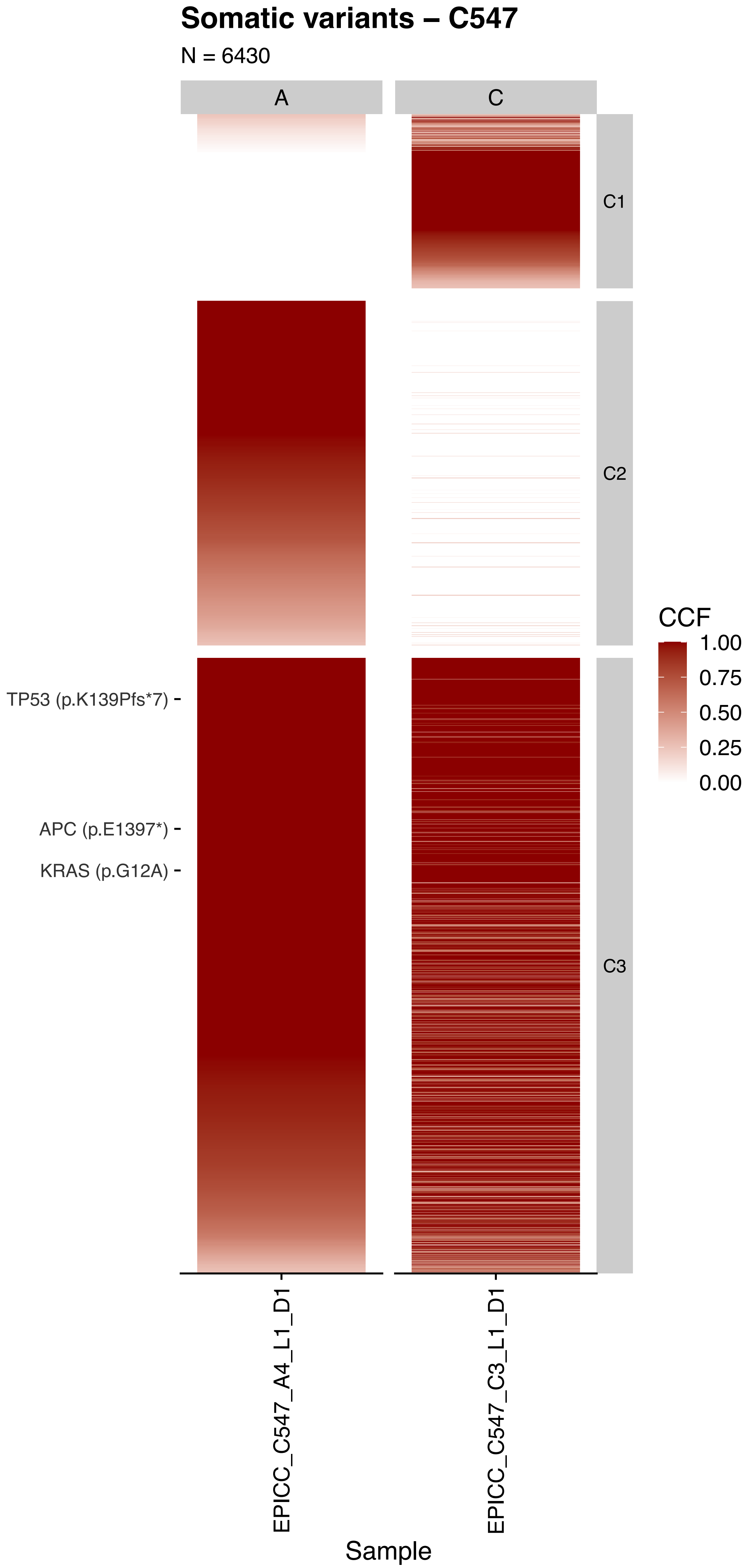

Somatic variants – C547
N = 6430
A
C
C1
C2
C3
TP53 (p.K139Pfs*7)
APC (p.E1397*)
KRAS (p.G12A)
CCF
1.00
0.75
0.50
0.25
0.00
EPICC_C547_A4_L1_D1
EPICC_C547_C3_L1_D1
Sample

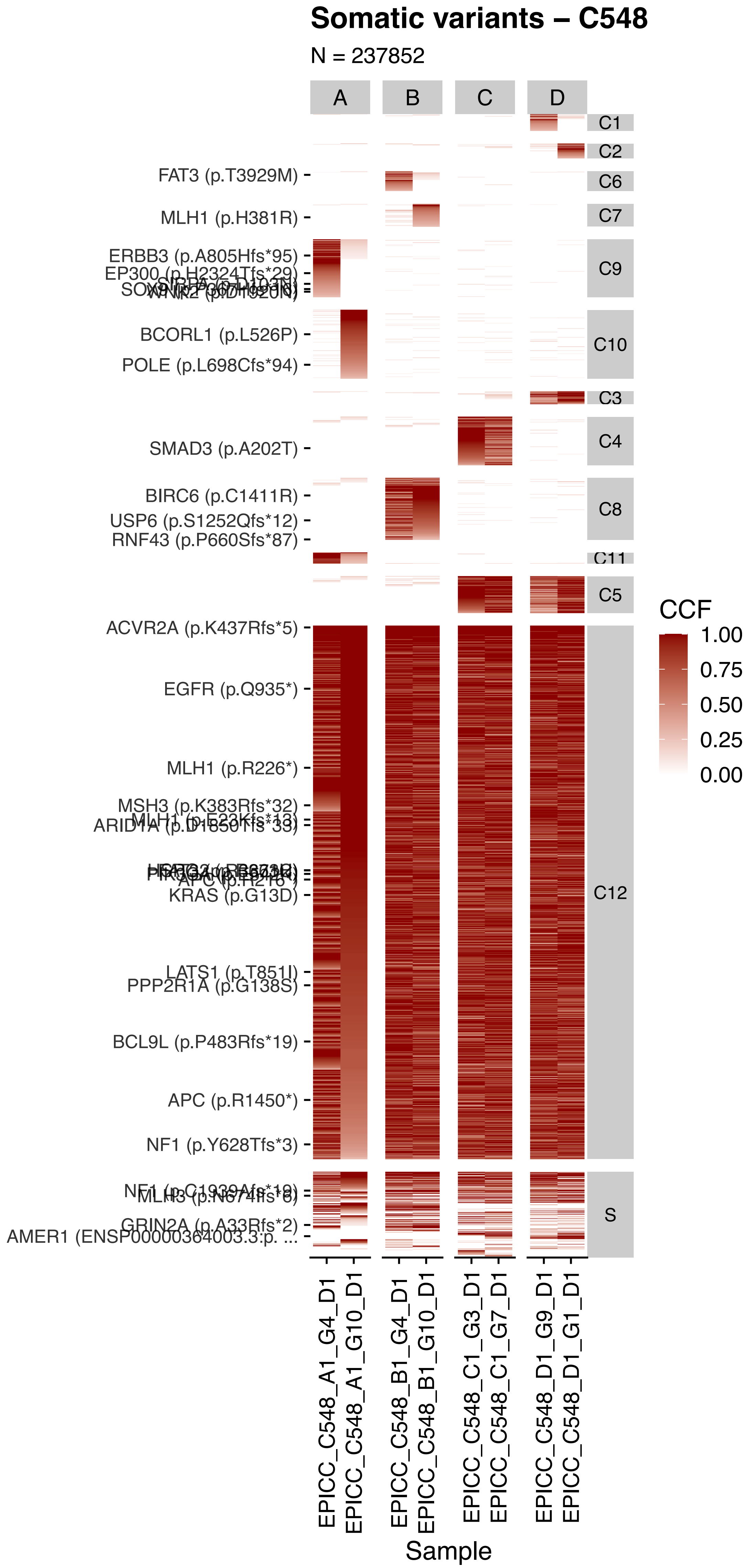

Somatic variants – C548
N = 237852
A
B
C
D
C1
C2
C6
C7
C9
C10
C3
C4
C8
C11
C5
C12
S
FAT3 (p.T3929M)
MLH1 (p.H381R)
ERBB3 (p.A805Hfs*95)
BCORL1 (p.L526P)
POLE (p.L698Cfs*94)
SMAD3 (p.A202T)
BIRC6 (p.C1411R)
USP6 (p.S1252Qfs*12)
RNF43 (p.P660Sfs*87)
ACVR2A (p.K437Rfs*5)
EGFR (p.Q935*)
MLH1 (p.R226*)
KRAS (p.G13D)
PPP2R1A (p.G138S)
BCL9L (p.P483Rfs*19)
APC (p.R1450*)
NF1 (p.Y628Tfs*3)
AMER1 (ENSP00000364003.3:p. ...
CCF
1.00
0.75
0.50
0.25
0.00
EPICC_C548_A1_G4_D1
EPICC_C548_A1_G10_D1
EPICC_C548_B1_G4_D1
EPICC_C548_B1_G10_D1
EPICC_C548_C1_G3_D1
EPICC_C548_C1_G7_D1
EPICC_C548_D1_G9_D1
EPICC_C548_D1_G1_D1
Sample

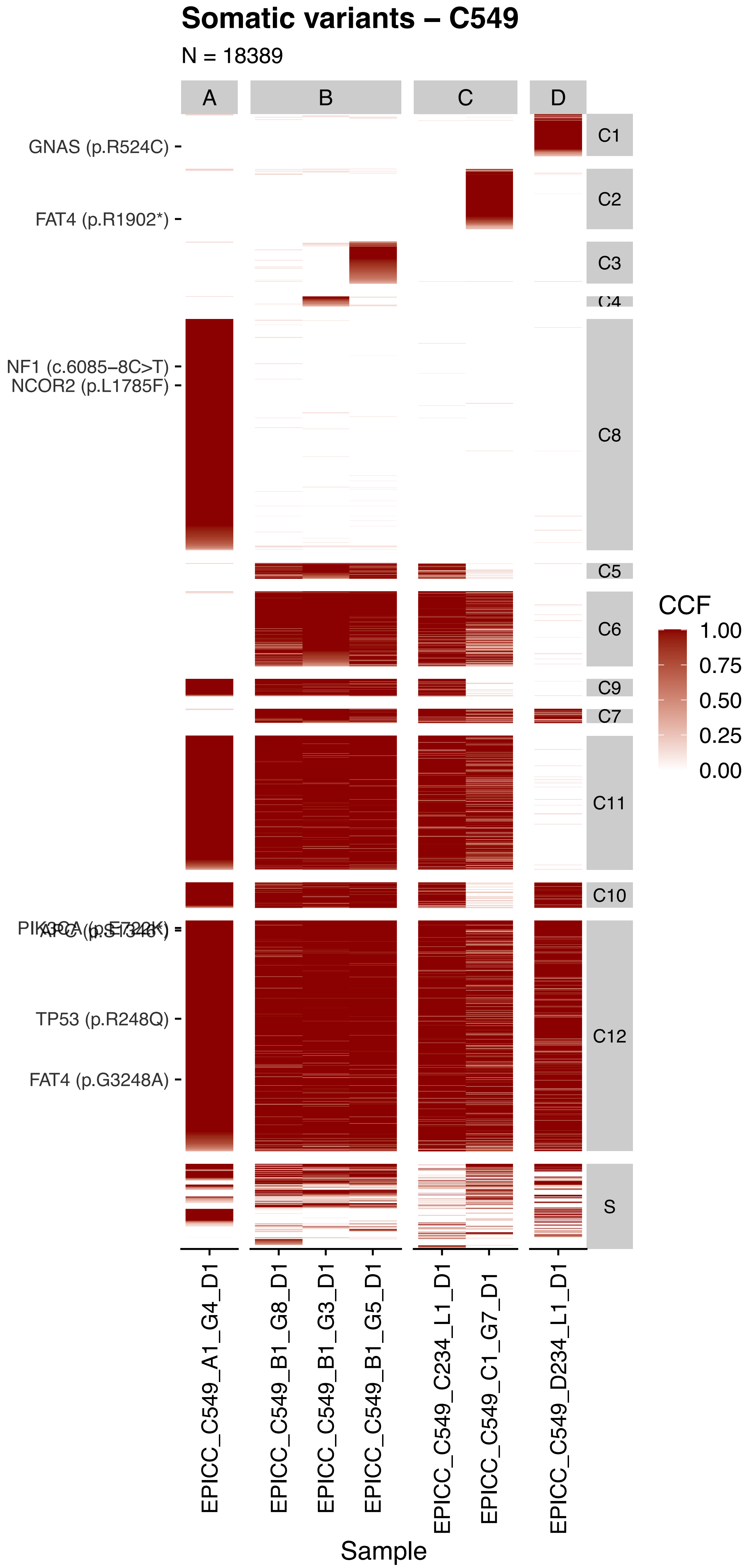

Somatic variants – C549
N = 18389
A
B
C
D
GNAS (p.R524C)
FAT4 (p.R1902*)
NF1 (c.6085−8C>T)
NCOR2 (p.L1785F)
PIK3CA (p.E722K)
APC (p.S1346*)
TP53 (p.R248Q)
FAT4 (p.G3248A)
C1
C2
C3
C4
C8
C5
C6
C9
C7
C11
C10
C12
S
CCF
1.00
0.75
0.50
0.25
0.00
EPICC_C549_A1_G4_D1
EPICC_C549_B1_G8_D1
EPICC_C549_B1_G3_D1
EPICC_C549_B1_G5_D1
EPICC_C549_C234_L1_D1
EPICC_C549_C1_G7_D1
EPICC_C549_D234_L1_D1
Sample

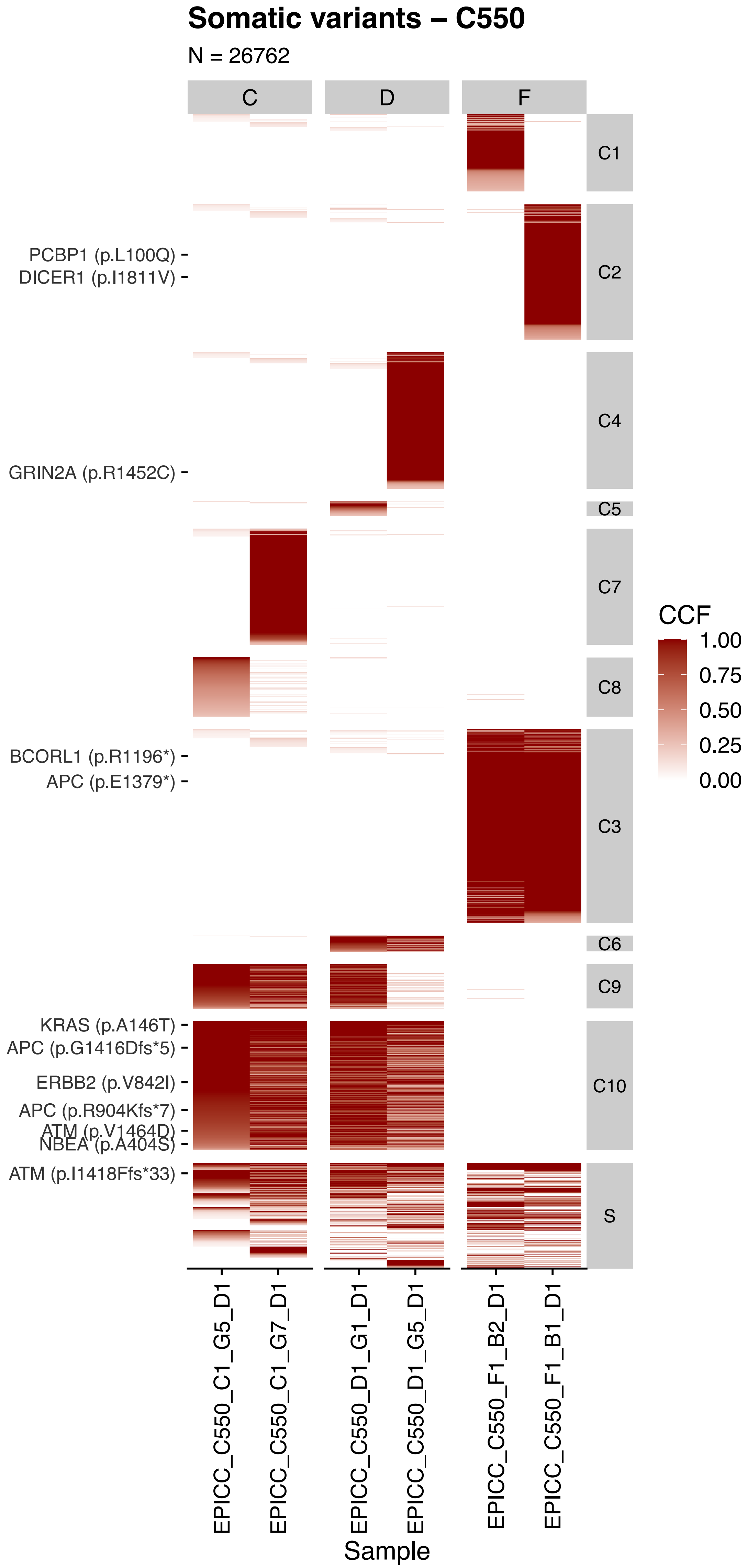

Somatic variants – C550
N = 26762
C
D
F
C1
C2
C4
C5
C7
C8
C3
C6
C9
C10
S
PCBP1 (p.L100Q)
DICER1 (p.I1811V)
GRIN2A (p.R1452C)
BCORL1 (p.R1196*)
APC (p.E1379*)
KRAS (p.A146T)
APC (p.G1416Dfs*5)
ERBB2 (p.V842I)
APC (p.R904Kfs*7)
ATM (p.V1464D)
NBEA (p.A404S)
ATM (p.I1418Ffs*33)
CCF
1.00
0.75
0.50
0.25
0.00
EPICC_C550_C1_G5_D1
EPICC_C550_C1_G7_D1
EPICC_C550_D1_G1_D1
EPICC_C550_D1_G5_D1
EPICC_C550_F1_B2_D1
EPICC_C550_F1_B1_D1
Sample

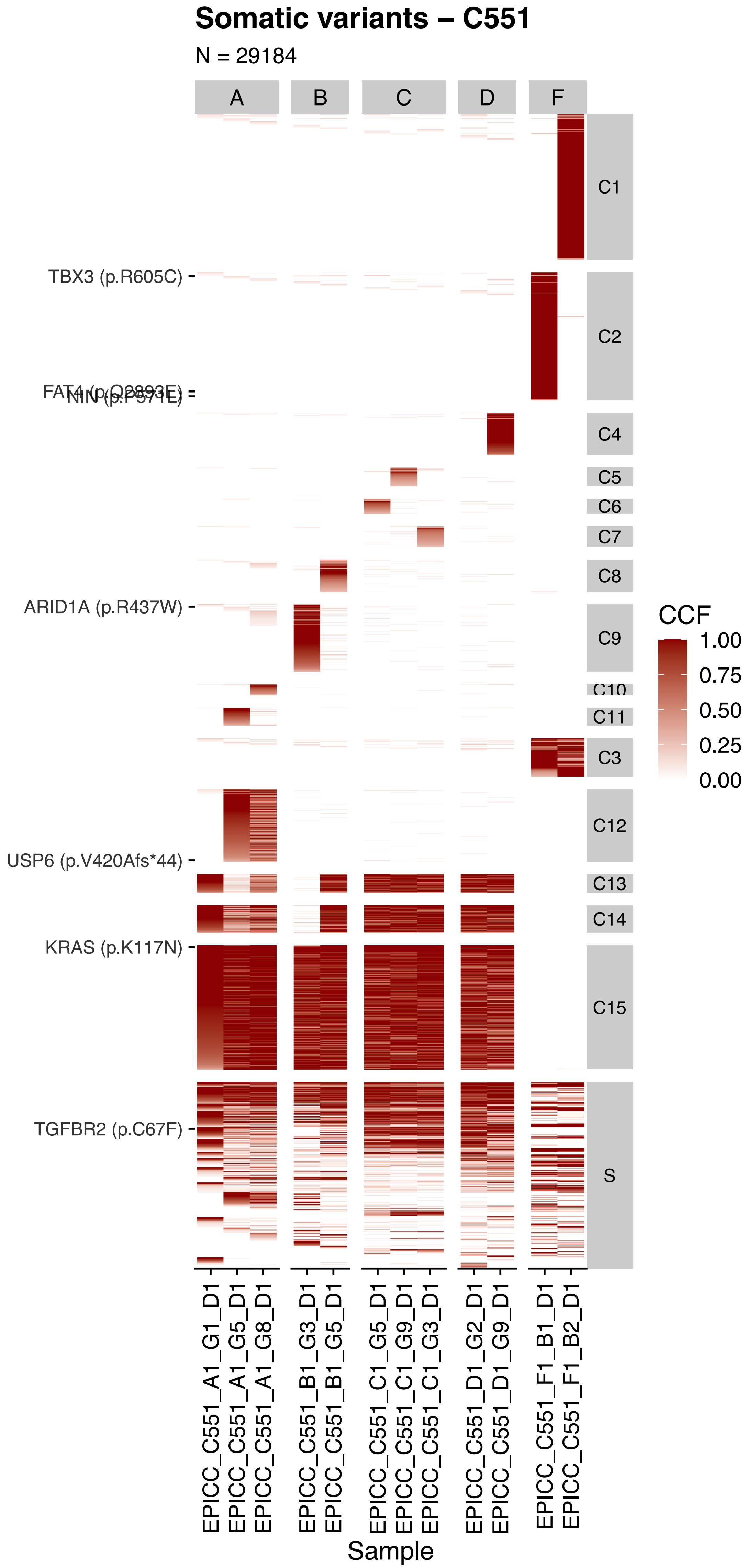

Somatic variants – C551
N = 29184
A
B
C
D
F
C1
C2
C4
C5
C6
C7
C8
C9
C10
C11
C3
C12
C13
C14
C15
S
TBX3 (p.R605C)
FAT4 (p.Q2893E)
NIN (p.P571E)
ARID1A (p.R437W)
USP6 (p.V420Afs*44)
KRAS (p.K117N)
TGFBR2 (p.C67F)
CCF
1.00
0.75
0.50
0.25
0.00
EPICC_C551_A1_G1_D1
EPICC_C551_A1_G5_D1
EPICC_C551_A1_G8_D1
EPICC_C551_B1_G3_D1
EPICC_C551_B1_G5_D1
EPICC_C551_C1_G5_D1
EPICC_C551_C1_G9_D1
EPICC_C551_C1_G3_D1
EPICC_C551_D1_G2_D1
EPICC_C551_D1_G9_D1
EPICC_C551_F1_B1_D1
EPICC_C551_F1_B2_D1
Sample

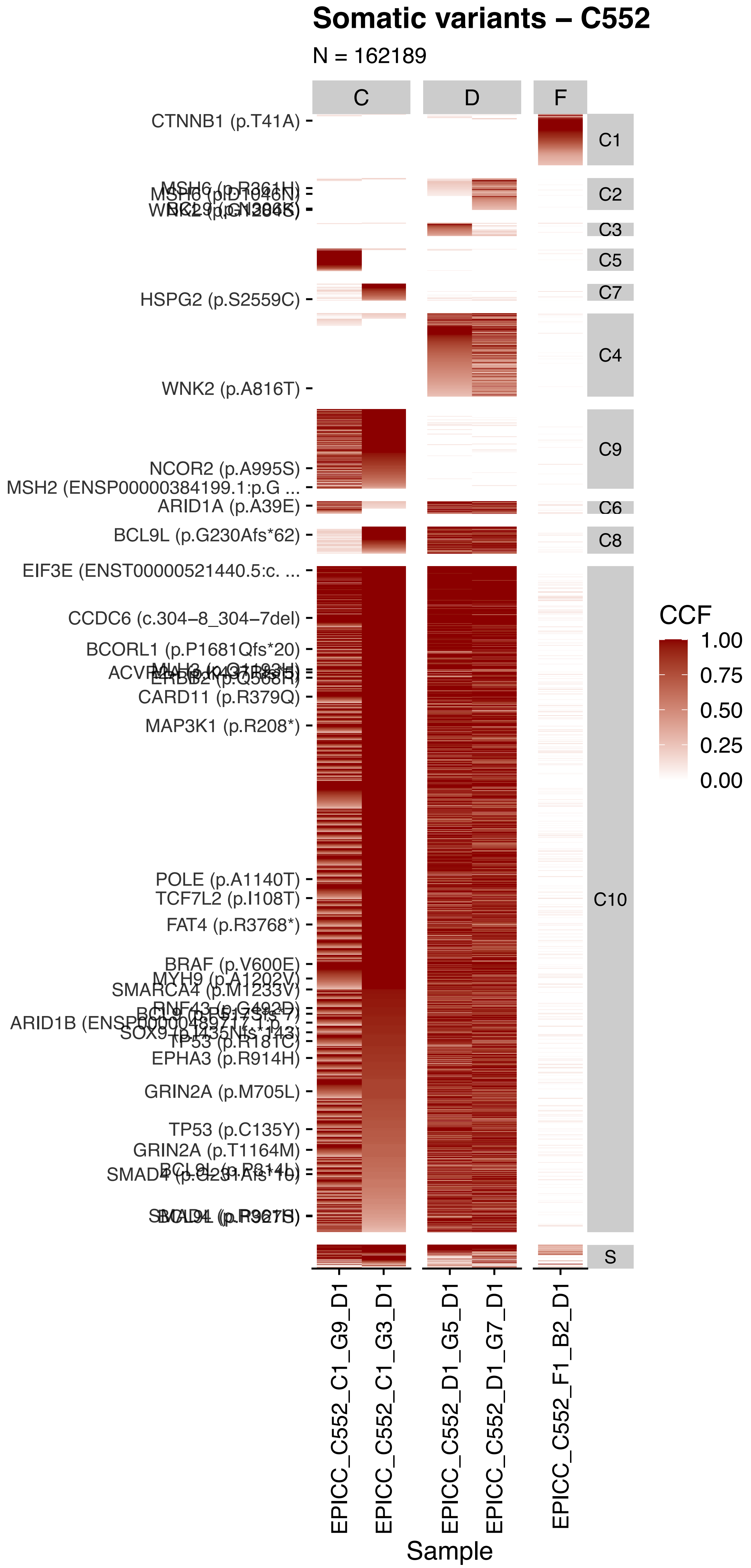

Somatic variants – C552
N = 162189
C
D
F
CTNNB1 (p.T41A)
HSPG2 (p.S2559C)
WNK2 (p.A816T)
NCOR2 (p.A995S)
MSH2 (ENSP00000384199.1:p.G ...
ARID1A (p.A39E)
BCL9L (p.G230Afs*62)
EIF3E (ENST00000521440.5:c. ...
CCDC6 (c.304−8_304−7del)
BCORL1 (p.P1681Qfs*20)
CARD11 (p.R379Q)
MAP3K1 (p.R208*)
POLE (p.A1140T)
TCF7L2 (p.I108T)
FAT4 (p.R3768*)
BRAF (p.V600E)
MYH9 (p.A1202V)
SMARCA4 (p.M1233V)
TP53 (p.R181C)
EPHA3 (p.R914H)
GRIN2A (p.M705L)
TP53 (p.C135Y)
GRIN2A (p.T1164M)
C1
C2
C3
C5
C7
C4
C9
C6
C8
C10
S
CCF
1.00
0.75
0.50
0.25
0.00
EPICC_C552_C1_G9_D1
EPICC_C552_C1_G3_D1
EPICC_C552_D1_G5_D1
EPICC_C552_D1_G7_D1
EPICC_C552_F1_B2_D1
Sample

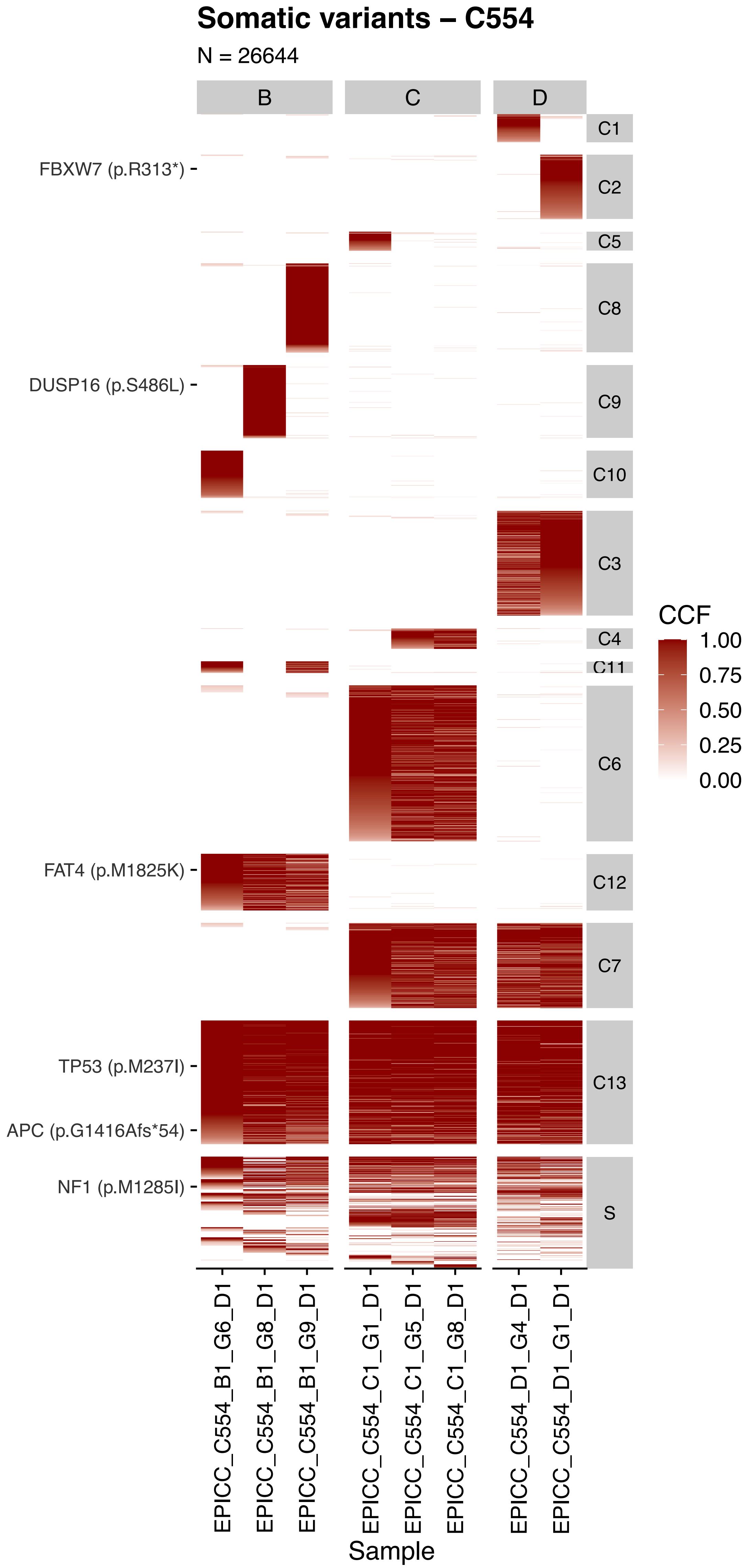

Somatic variants – C554
N = 26644
B
C
D
C1
C2
C5
C8
C9
C10
C3
C4
C11
C6
C12
C7
C13
S
FBXW7 (p.R313*)
DUSP16 (p.S486L)
FAT4 (p.M1825K)
TP53 (p.M237I)
APC (p.G1416Afs*54)
NF1 (p.M1285I)
CCF
1.00
0.75
0.50
0.25
0.00
EPICC_C554_B1_G6_D1
EPICC_C554_B1_G8_D1
EPICC_C554_B1_G9_D1
EPICC_C554_C1_G1_D1
EPICC_C554_C1_G5_D1
EPICC_C554_C1_G8_D1
EPICC_C554_D1_G4_D1
EPICC_C554_D1_G1_D1
Sample

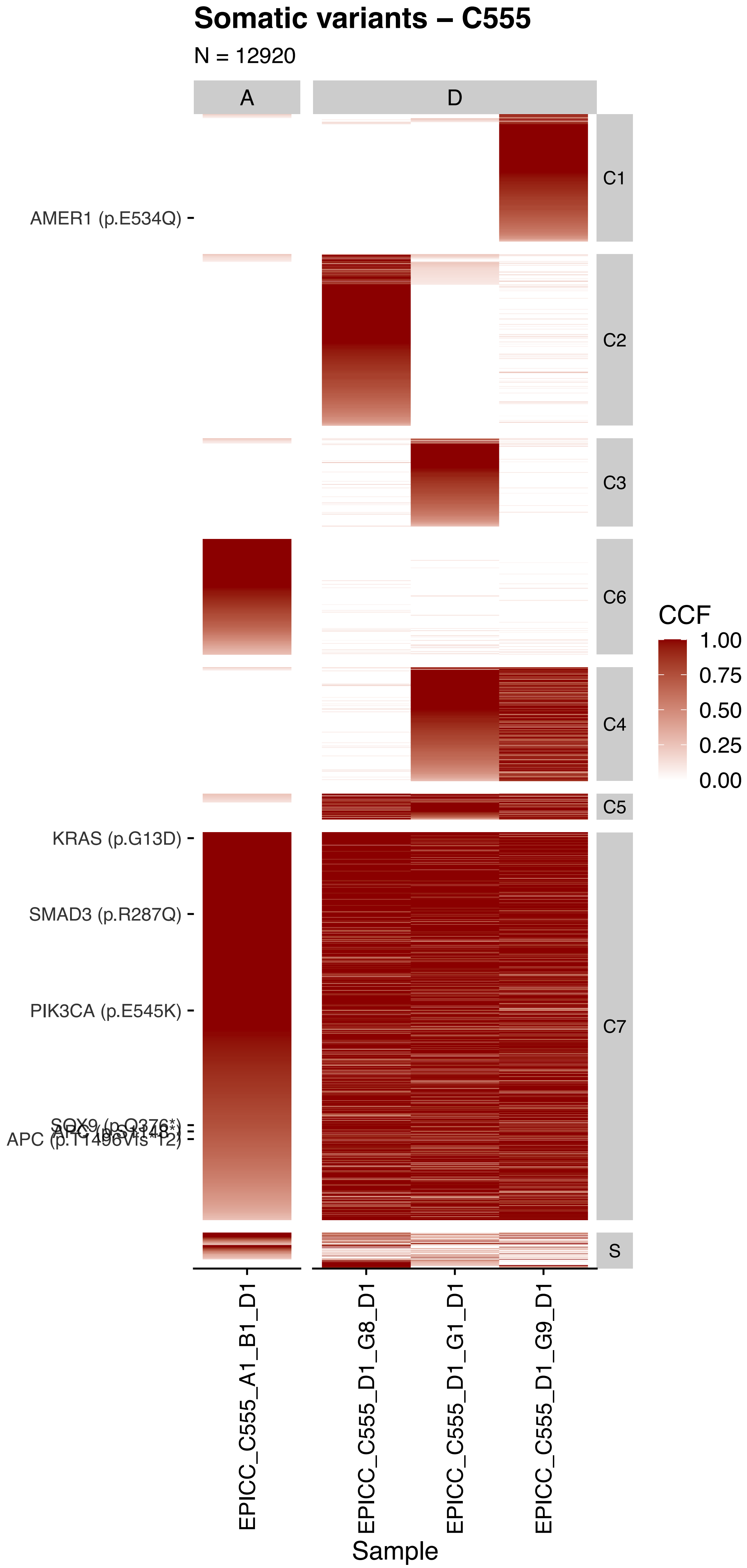

Somatic variants – C555
N = 12920
A
D
AMER1 (p.E534Q)
KRAS (p.G13D)
SMAD3 (p.R287Q)
PIK3CA (p.E545K)
SOX9 (p.Q376*)
APC (p.S1148*)
APC (p.T1496Vfs*12)
C1
C2
C3
C6
C4
C5
C7
S
CCF
1.00
0.75
0.50
0.25
0.00
EPICC_C555_A1_B1_D1
EPICC_C555_D1_G8_D1
EPICC_C555_D1_G1_D1
EPICC_C555_D1_G9_D1
Sample

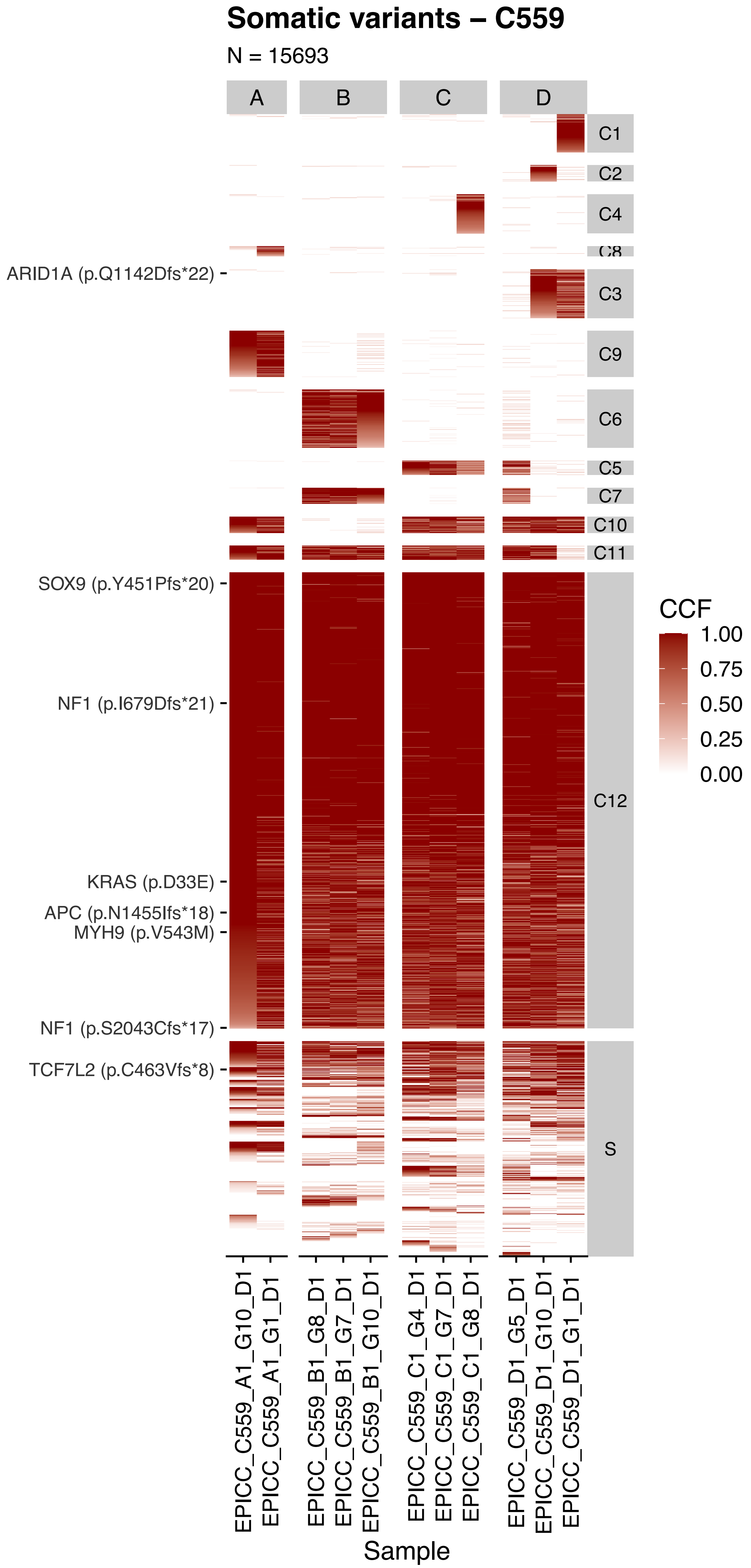

Somatic variants – C559
N = 15693
A
B
C
D
C1
C2
C4
C8
C3
C9
C6
C5
C7
C10
C11
C12
S
ARID1A (p.Q1142Dfs*22)
SOX9 (p.Y451Pfs*20)
NF1 (p.I679Dfs*21)
KRAS (p.D33E)
APC (p.N1455Ifs*18)
MYH9 (p.V543M)
NF1 (p.S2043Cfs*17)
TCF7L2 (p.C463Vfs*8)
CCF
1.00
0.75
0.50
0.25
0.00
EPICC_C559_A1_G10_D1
EPICC_C559_A1_G1_D1
EPICC_C559_B1_G8_D1
EPICC_C559_B1_G7_D1
EPICC_C559_B1_G10_D1
EPICC_C559_C1_G4_D1
EPICC_C559_C1_G7_D1
EPICC_C559_C1_G8_D1
EPICC_C559_D1_G5_D1
EPICC_C559_D1_G10_D1
EPICC_C559_D1_G1_D1
Sample

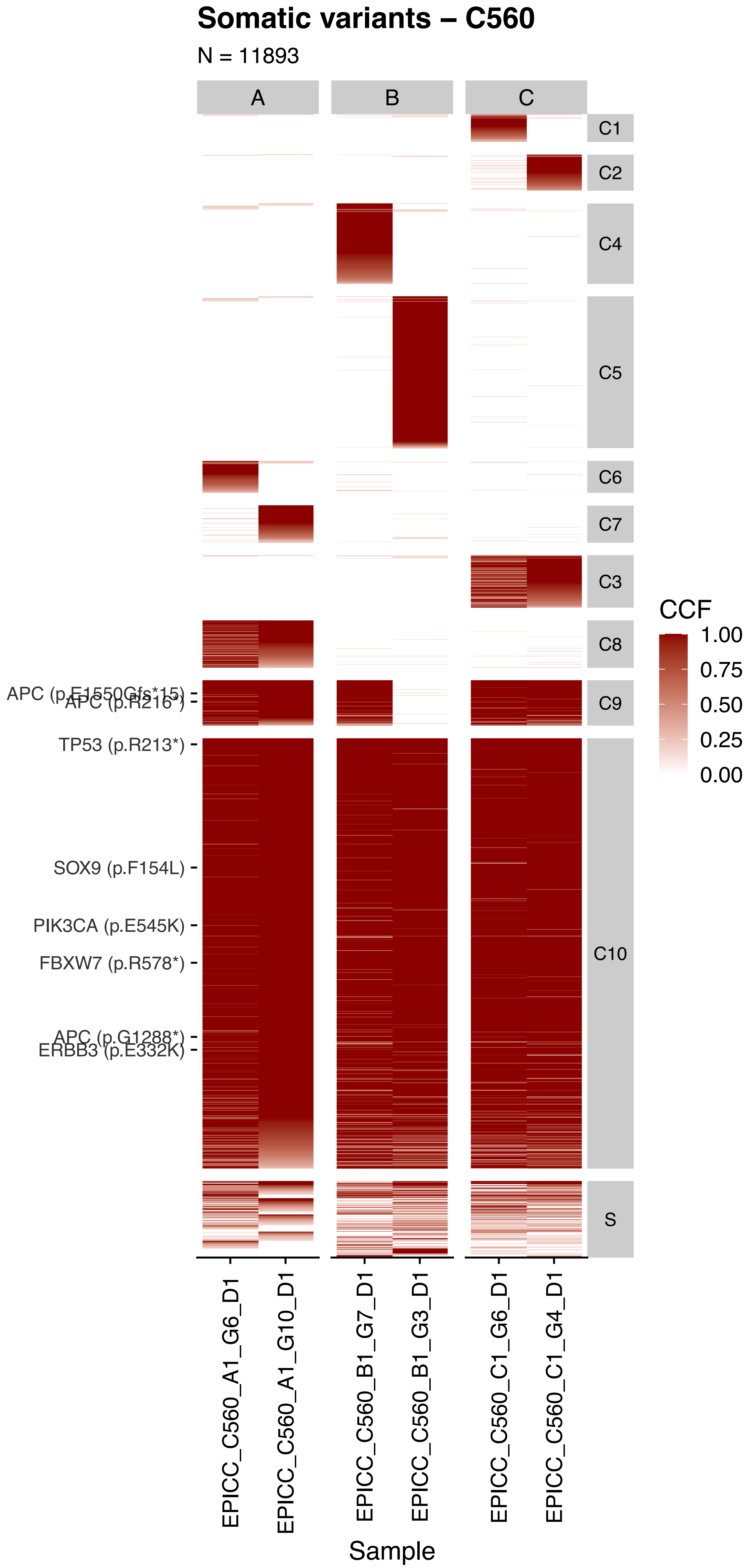

Somatic variants – C560
N = 11893
A
B
C
C1
C2
C4
C5
C6
C7
C3
C8
C9
C10
S
CCF
1.00
0.75
0.50
0.25
0.00
APC (p.E1550Gfs*15)
APC (p.R216*)
TP53 (p.R213*)
SOX9 (p.F154L)
PIK3CA (p.E545K)
FBXW7 (p.R578*)
APC (p.G1288*)
ERBB3 (p.E332K)
EPICC_C560_A1_G6_D1
EPICC_C560_A1_G10_D1
EPICC_C560_B1_G7_D1
EPICC_C560_B1_G3_D1
EPICC_C560_C1_G6_D1
EPICC_C560_C1_G4_D1
Sample

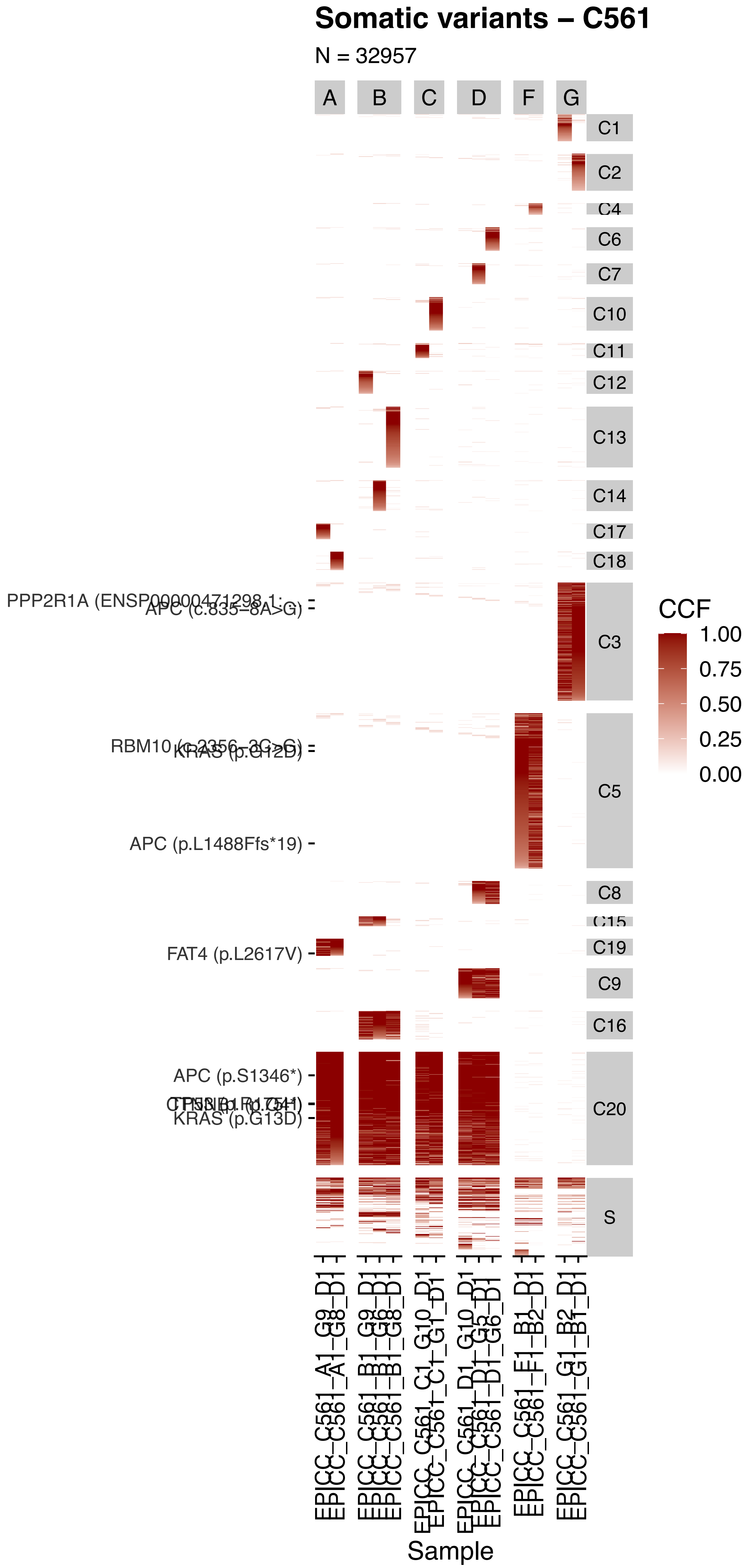

Somatic variants – C561
N = 32957
A
B
C
D
F
G
C1
C2
C4
C6
C7
C10
C11
C12
C13
C14
C17
C18
C3
C5
C8
C15
C19
C9
C16
C20
S
APC (c.835-8A>G)
APC (p.L1488Ffs*19)
FAT4 (p.L2617V)
APC (p.S1346*)
KRAS (p.G13D)
CCF
1.00
0.75
0.50
0.25
0.00
EPICC_C561_A1_G9_D1
EPICC_C561_A1_G8_D1
EPICC_C561_B1_G9_D1
EPICC_C561_B1_G6_D1
EPICC_C561_B1_G8_D1
EPICC_C561_C1_G10_D1
EPICC_C561_C1_G1_D1
EPICC_C561_D1_G10_D1
EPICC_C561_D1_G3_D1
EPICC_C561_D1_G6_D1
EPICC_C561_F1_B1_D1
EPICC_C561_F1_B2_D1
EPICC_C561_G1_B2_D1
EPICC_C561_G1_B1_D1
Sample

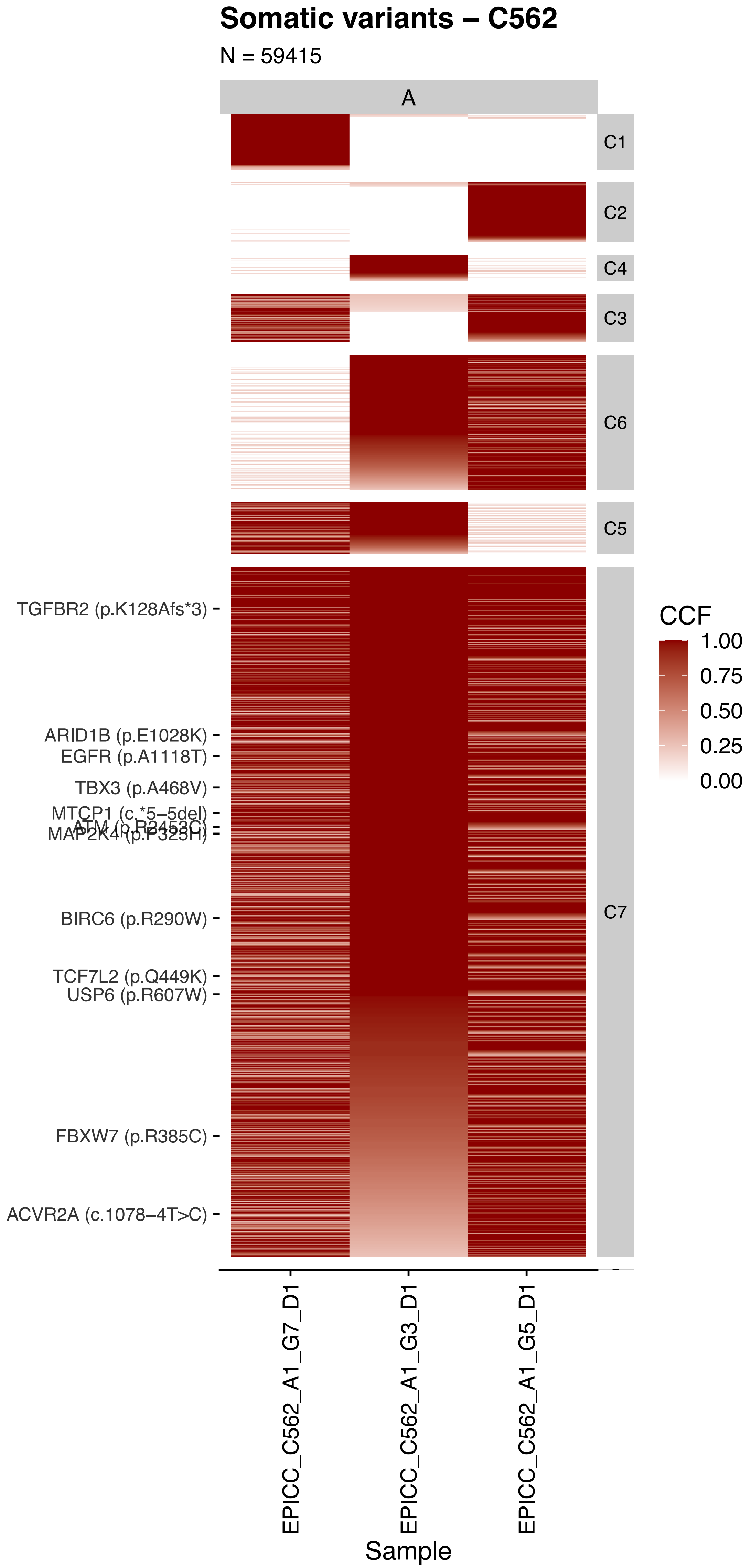

Somatic variants – C562
N = 59415
A
C1
C2
C4
C3
C6
C5
C7
TGFBR2 (p.K128Afs*3)
ARID1B (p.E1028K)
EGFR (p.A1118T)
TBX3 (p.A468V)
MTCP1 (c.*5–5del)
ATM (p.R2453C)
MAP2K4 (p.P325H)
BIRC6 (p.R290W)
TCF7L2 (p.Q449K)
USP6 (p.R607W)
FBXW7 (p.R385C)
ACVR2A (c.1078–4T>C)
CCF
1.00
0.75
0.50
0.25
0.00
EPICC_C562_A1_G7_D1
EPICC_C562_A1_G3_D1
EPICC_C562_A1_G5_D1
Sample
