## Supplementary Figures for "The co-evolution of the genome and epigenome in colorectal cancer": figS9.pdf

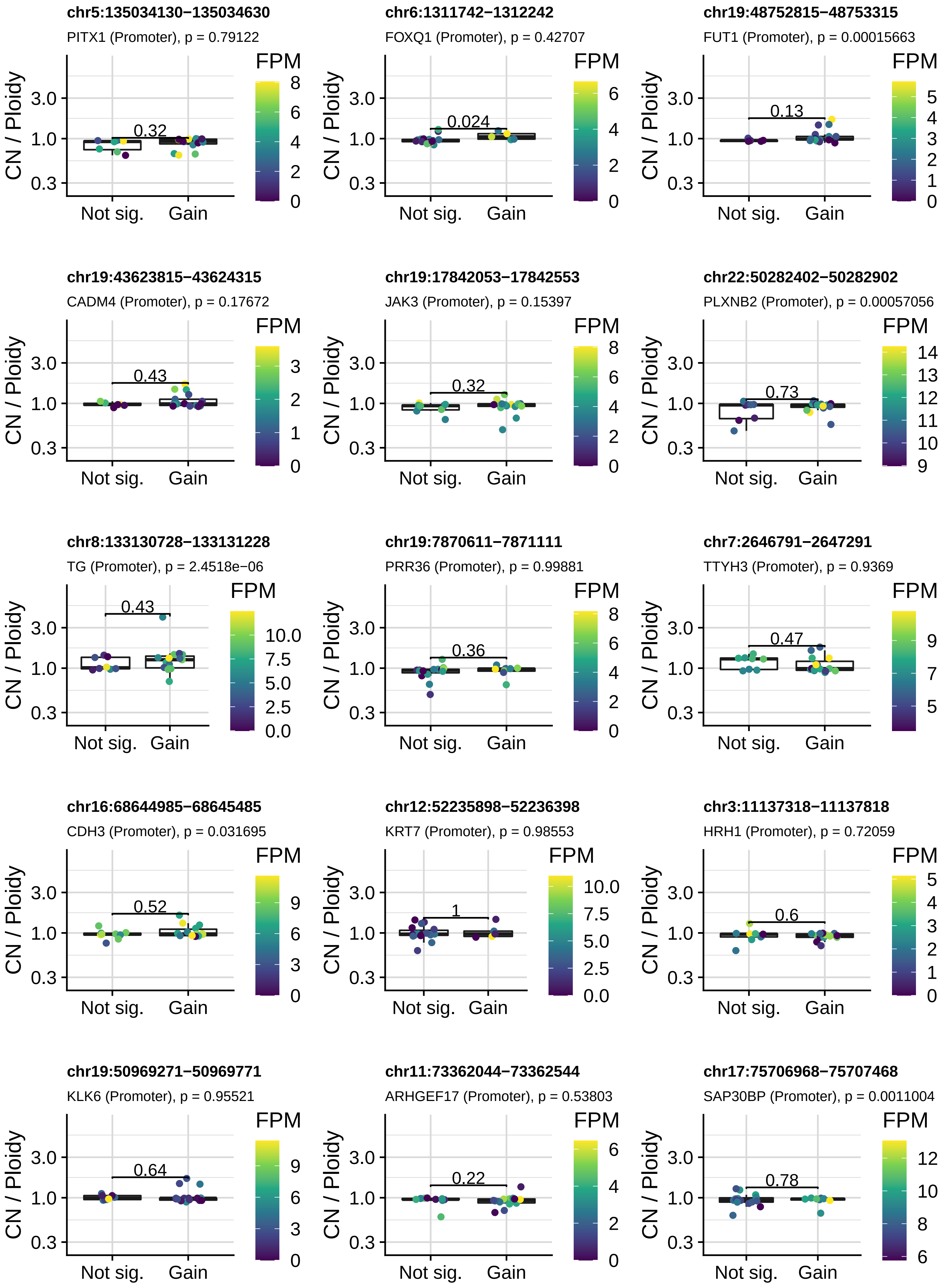
chr5:135034130−135034630
PITX1 (Promoter), p = 0.79122
0.32
chr6:1311742−1312242
FOXQ1 (Promoter), p = 0.42707
0.024
chr19:48752815−48753315
FUT1 (Promoter), p = 0.00015663
0.13
chr19:43623815−43624315
CADM4 (Promoter), p = 0.17672
0.43
chr19:17842053−17842553
JAK3 (Promoter), p = 0.15397
0.32
chr22:50282402−50282902
PLXNB2 (Promoter), p = 0.00057056
0.73
chr8:133130728−133131228
TG (Promoter), p = 2.4518e−06
0.43
chr19:7870611−7871111
PRR36 (Promoter), p = 0.99881
0.36
chr7:2646791−2647291
TTYH3 (Promoter), p = 0.9369
0.47
chr16:68644985−68645485
CDH3 (Promoter), p = 0.031695
0.52
chr12:52235898−52236398
KRT7 (Promoter), p = 0.98553
1
chr3:11137318−11137818
HRH1 (Promoter), p = 0.72059
0.6
chr19:50969271−50969771
KLK6 (Promoter), p = 0.95521
0.64
chr11:73362044−73362544
ARHGEF17 (Promoter), p = 0.53803
0.22
chr17:75706968−75707468
SAP30BP (Promoter), p = 0.0011004
0.78
CN / Ploidy
3.0
1.0
0.3
Not sig.
Gain
FPM

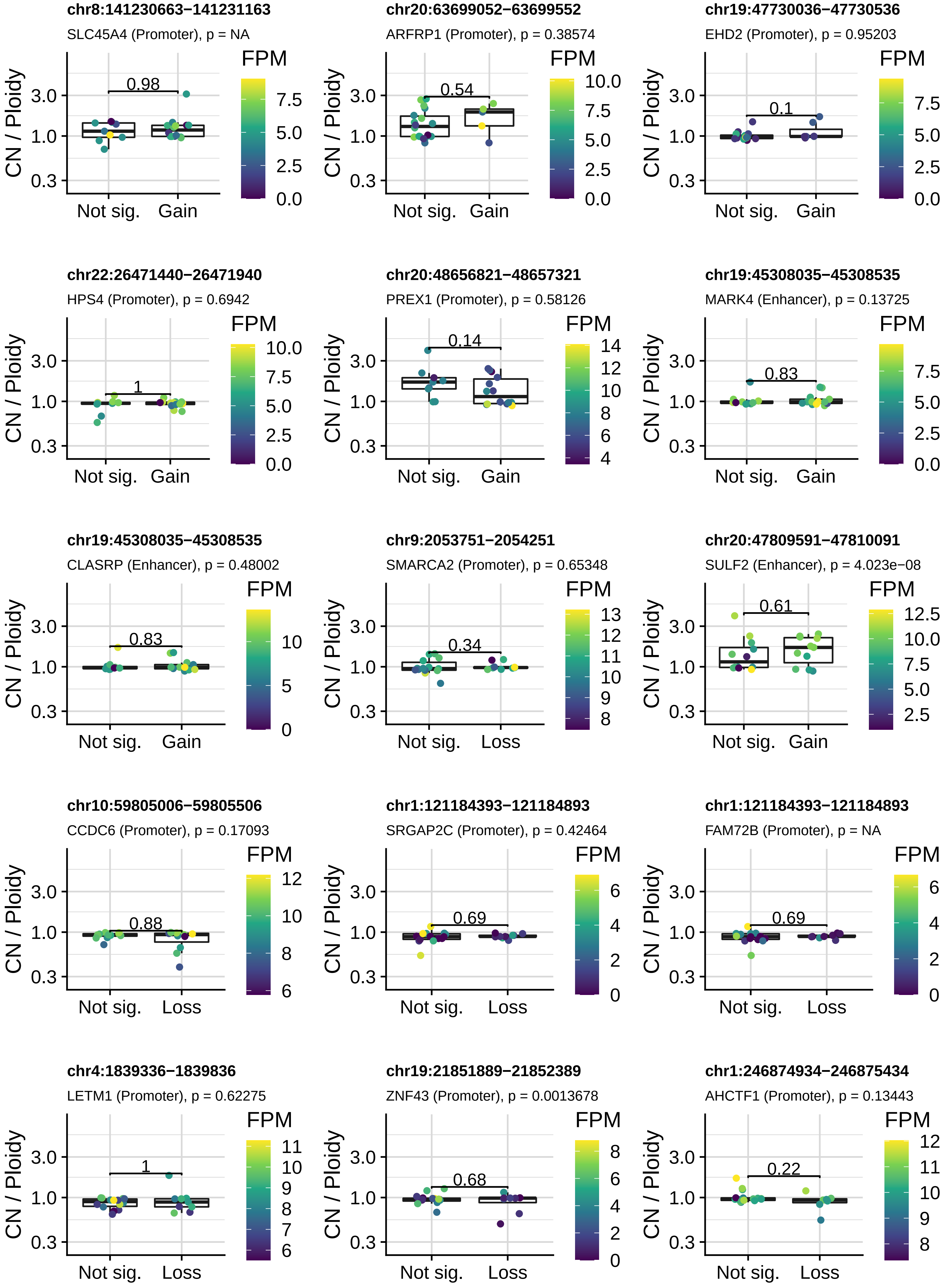
chr8:141230663−141231163
SLC45A4 (Promoter), p = NA
0.98
chr20:63699052−63699552
ARFRP1 (Promoter), p = 0.38574
0.54
chr19:47730036−47730536
EHD2 (Promoter), p = 0.95203
0.1
chr22:26471440−26471940
HPS4 (Promoter), p = 0.6942
1
chr20:48656821−48657321
PREX1 (Promoter), p = 0.58126
0.14
chr19:45308035−45308535
MARK4 (Enhancer), p = 0.13725
0.83
chr19:45308035−45308535
CLASRP (Enhancer), p = 0.48002
0.83
chr9:2053751−2054251
SMARCA2 (Promoter), p = 0.65348
0.34
chr20:47809591−47810091
SULF2 (Enhancer), p = 4.023e−08
0.61
chr10:59805006−59805506
CCDC6 (Promoter), p = 0.17093
0.88
chr1:121184393−121184893
SRGAP2C (Promoter), p = 0.42464
0.69
chr1:121184393−121184893
FAM72B (Promoter), p = NA
0.69
chr4:1839336−1839836
LETM1 (Promoter), p = 0.62275
1
chr19:21851889−21852389
ZNF43 (Promoter), p = 0.0013678
0.68
chr1:246874934−246875434
AHCTF1 (Promoter), p = 0.13443
0.22
CN / Ploidy
3.0
1.0
0.3
Not sig.
Gain
Loss
FPM

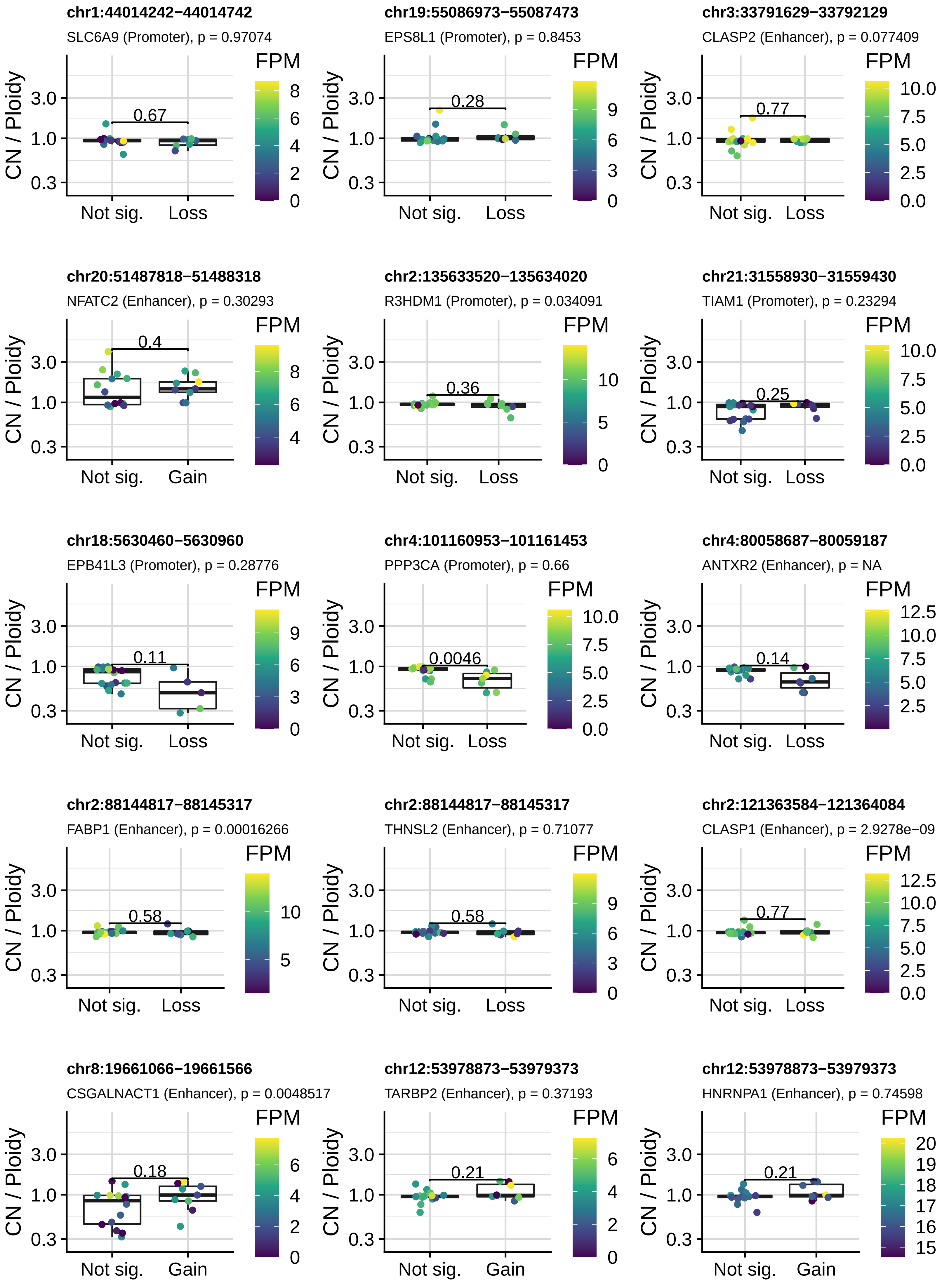
chr1:44014242−44014742
SLC6A9 (Promoter), p = 0.97074
0.67
Not sig.
Loss
chr19:55086973−55087473
EPS8L1 (Promoter), p = 0.8453
0.28
Not sig.
Loss
chr3:33791629−33792129
CLASP2 (Enhancer), p = 0.077409
0.77
Not sig.
Loss
chr20:51487818−51488318
NFATC2 (Enhancer), p = 0.30293
0.4
Not sig.
Gain
chr2:135633520−135634020
R3HDM1 (Promoter), p = 0.034091
0.36
Not sig.
Loss
chr21:31558930−31559430
TIAM1 (Promoter), p = 0.23294
0.25
Not sig.
Loss
chr18:5630460−5630960
EPB41L3 (Promoter), p = 0.28776
0.11
Not sig.
Loss
chr4:101160953−101161453
PPP3CA (Promoter), p = 0.66
0.0046
Not sig.
Loss
chr4:80058687−80059187
ANTXR2 (Enhancer), p = NA
0.14
Not sig.
Loss
chr2:88144817−88145317
FABP1 (Enhancer), p = 0.00016266
0.58
Not sig.
Loss
chr2:88144817−88145317
THNSL2 (Enhancer), p = 0.71077
0.58
Not sig.
Loss
chr2:121363584−121364084
CLASP1 (Enhancer), p = 2.9278e−09
0.77
Not sig.
Loss
chr8:19661066−19661566
CSGALNACT1 (Enhancer), p = 0.0048517
0.18
Not sig.
Gain
chr12:53978873−53979373
TARBP2 (Enhancer), p = 0.37193
0.21
Not sig.
Gain
chr12:53978873−53979373
HNRNPA1 (Enhancer), p = 0.74598
0.21
Not sig.
Gain
CN / Ploidy
FPM

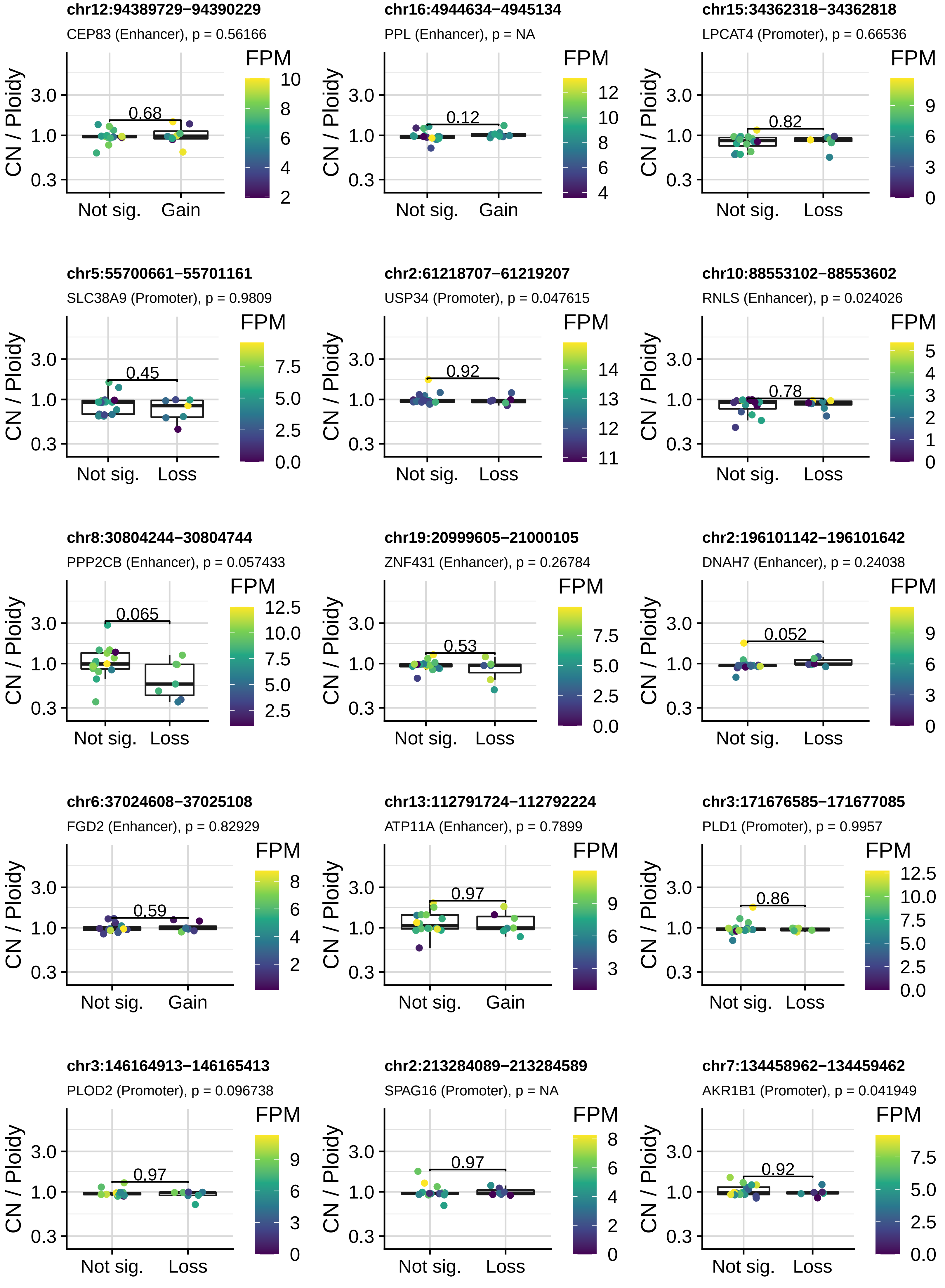
chr12:94389729−94390229
CEP83 (Enhancer), p = 0.56166
0.68
Not sig.
Gain
chr16:4944634−4945134
PPL (Enhancer), p = NA
0.12
Not sig.
Gain
chr15:34362318−34362818
LPCAT4 (Promoter), p = 0.66536
0.82
Not sig.
Loss
chr5:55700661−55701161
SLC38A9 (Promoter), p = 0.9809
0.45
Not sig.
Loss
chr2:61218707−61219207
USP34 (Promoter), p = 0.047615
0.92
Not sig.
Loss
chr10:88553102−88553602
RNLS (Enhancer), p = 0.024026
0.78
Not sig.
Loss
chr8:30804244−30804744
PPP2CB (Enhancer), p = 0.057433
0.065
Not sig.
Loss
chr19:20999605−21000105
ZNF431 (Enhancer), p = 0.26784
0.53
Not sig.
Loss
chr2:196101142−196101642
DNAH7 (Enhancer), p = 0.24038
0.052
Not sig.
Loss
chr6:37024608−37025108
FGD2 (Enhancer), p = 0.82929
0.59
Not sig.
Gain
chr13:112791724−112792224
ATP11A (Enhancer), p = 0.7899
0.97
Not sig.
Gain
chr3:171676585−171677085
PLD1 (Promoter), p = 0.9957
0.86
Not sig.
Loss
chr3:146164913−146165413
PLOD2 (Promoter), p = 0.096738
0.97
Not sig.
Loss
chr2:213284089−213284589
SPAG16 (Promoter), p = NA
0.97
Not sig.
Loss
chr7:134458962−134459462
AKR1B1 (Promoter), p = 0.041949
0.92
Not sig.
Loss
CN / Ploidy
3.0
1.0
0.3
FPM

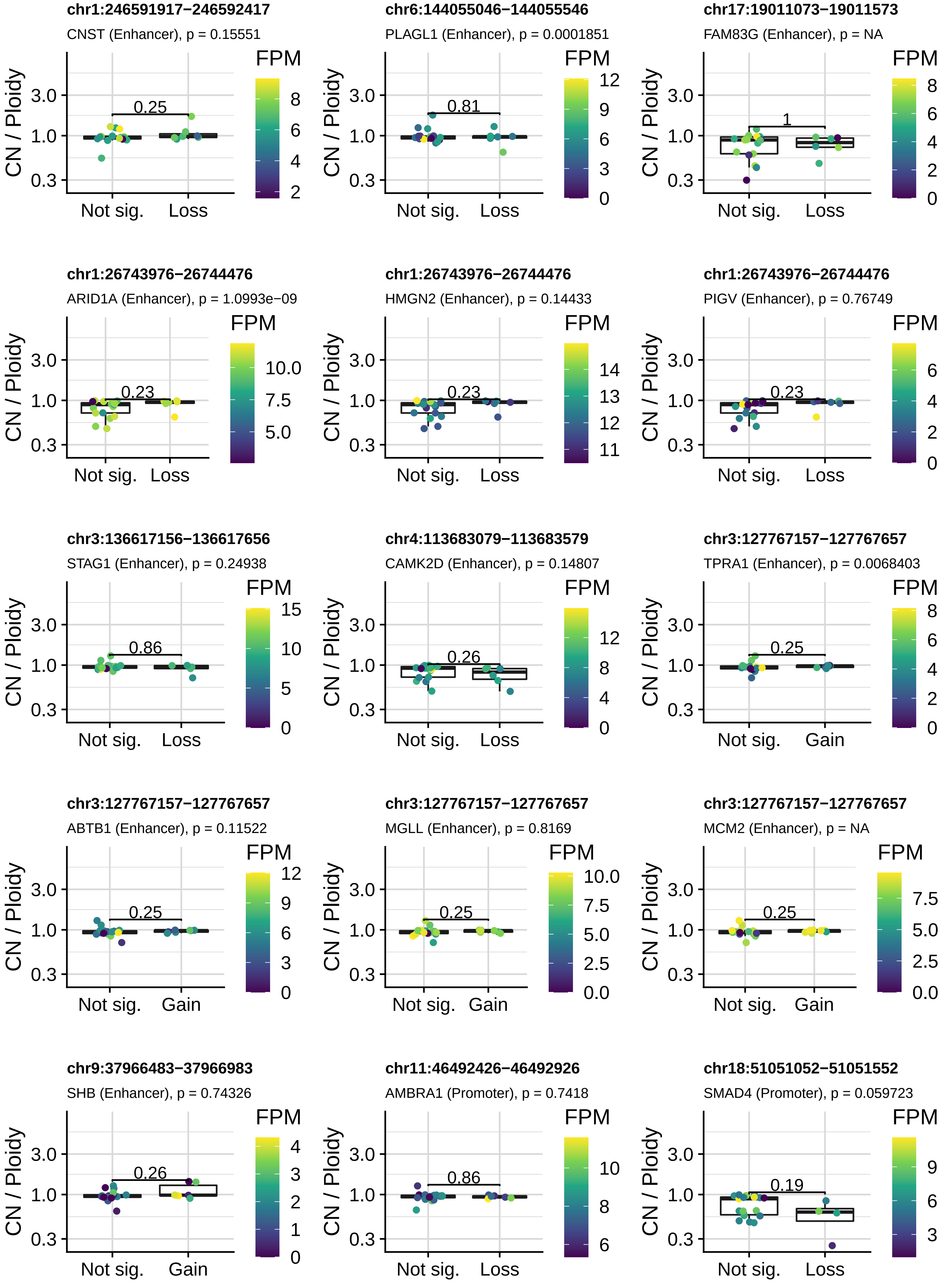
chr1:246591917−246592417
CNST (Enhancer), p = 0.15551
0.25
chr6:144055046−144055546
PLAGL1 (Enhancer), p = 0.0001851
0.81
chr17:19011073−19011573
FAM83G (Enhancer), p = NA
1
chr1:26743976−26744476
ARID1A (Enhancer), p = 1.0993e−09
0.23
chr1:26743976−26744476
HMGN2 (Enhancer), p = 0.14433
0.23
chr1:26743976−26744476
PIGV (Enhancer), p = 0.76749
0.23
chr3:136617156−136617656
STAG1 (Enhancer), p = 0.24938
0.86
chr4:113683079−113683579
CAMK2D (Enhancer), p = 0.14807
0.26
chr3:127767157−127767657
TPRA1 (Enhancer), p = 0.0068403
0.25
chr3:127767157−127767657
ABTB1 (Enhancer), p = 0.11522
0.25
chr3:127767157−127767657
MGLL (Enhancer), p = 0.8169
0.25
chr3:127767157−127767657
MCM2 (Enhancer), p = NA
0.25
chr9:37966483−37966983
SHB (Enhancer), p = 0.74326
0.26
chr11:46492426−46492926
AMBRA1 (Promoter), p = 0.7418
0.86
chr18:51051052−51051552
SMAD4 (Promoter), p = 0.059723
0.19
CN / Ploidy
3.0
1.0
0.3
Not sig.
Loss
Gain
FPM

**chr1:17108338−17108838**

SDHB (Enhancer), p = 0.46878

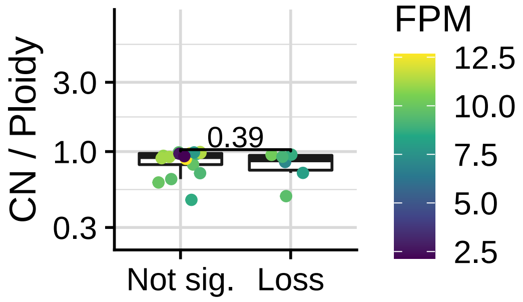


**chr3:36876260−36876760**

TRANK1 (Enhancer), p = 0.86871

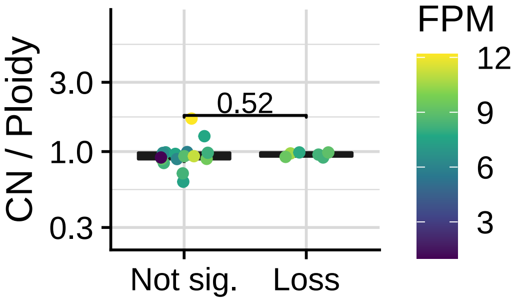


**chr3:195537917−195538417**

PPP1R2 (Enhancer), p = 0.068096

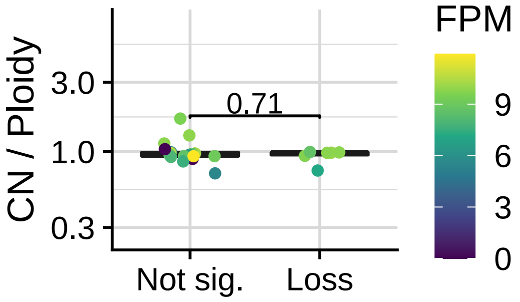


**chr3:195537917−195538417**

ACAP2 (Enhancer), p = NA

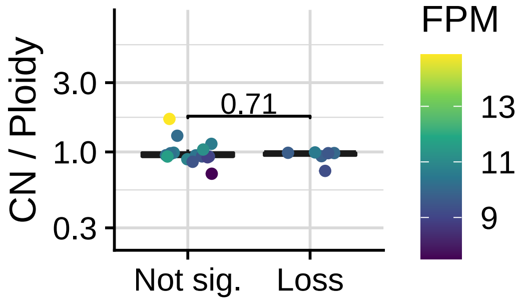


**chr4:101183634−101184134**

PPP3CA (Enhancer), p = 0.31102

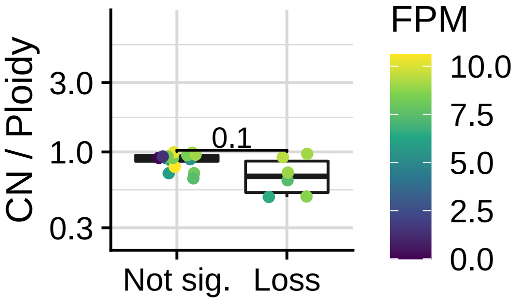


**chr5:56412924−56413424**

MAP3K1 (Enhancer), p = 0.33988

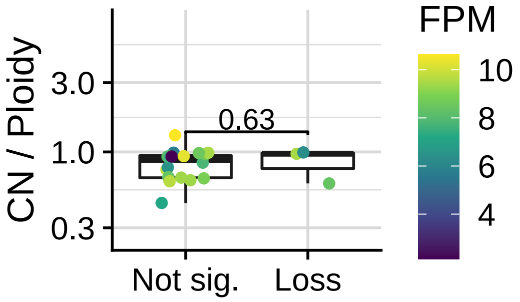


**chr12:124683111−124683611**

NCOR2 (Enhancer), p = 1.1628e−07

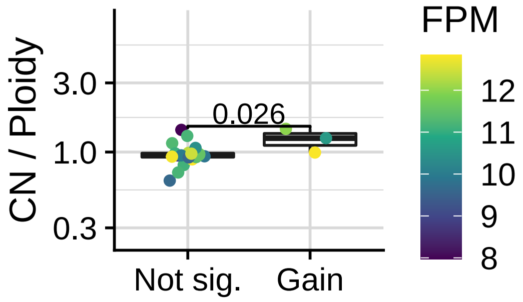


**chr12:56087145−56087645**

ERBB3 (Promoter), p = 0.84143

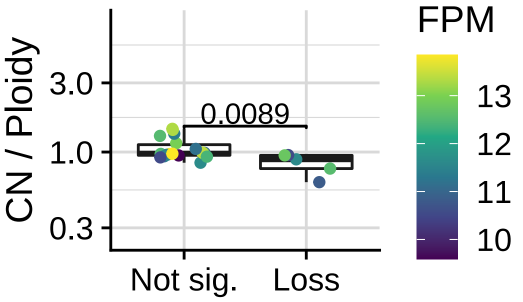


**chr15:67127194−67127694**

SMAD3 (Promoter), p = 0.60688

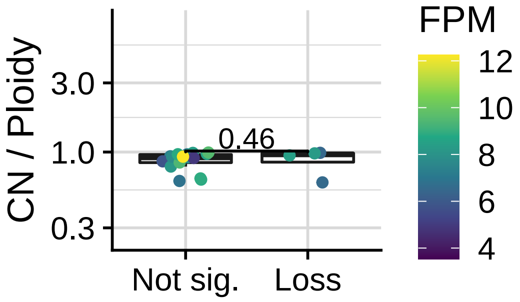


**chr17:39701512−39702012**

ERBB2 (Promoter), p = 1

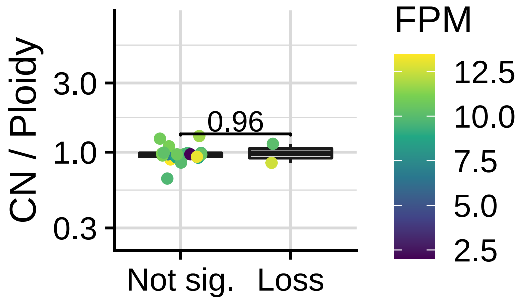


**chr15:67081050−67081550**

SMAD3 (Enhancer), p = 0.93416

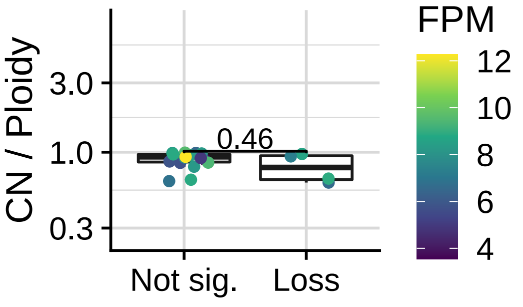


**chr10:59885378−59885878**

CCDC6 (Enhancer), p = 0.65672

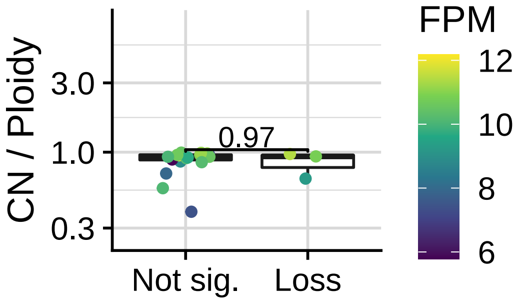
