## Supplementary Figures for "The co-evolution of the genome and epigenome in colorectal cancer": figS10_promo_gain.pdf

**ARFRP1 promoter gain**
**chr20:63699052−63699552**

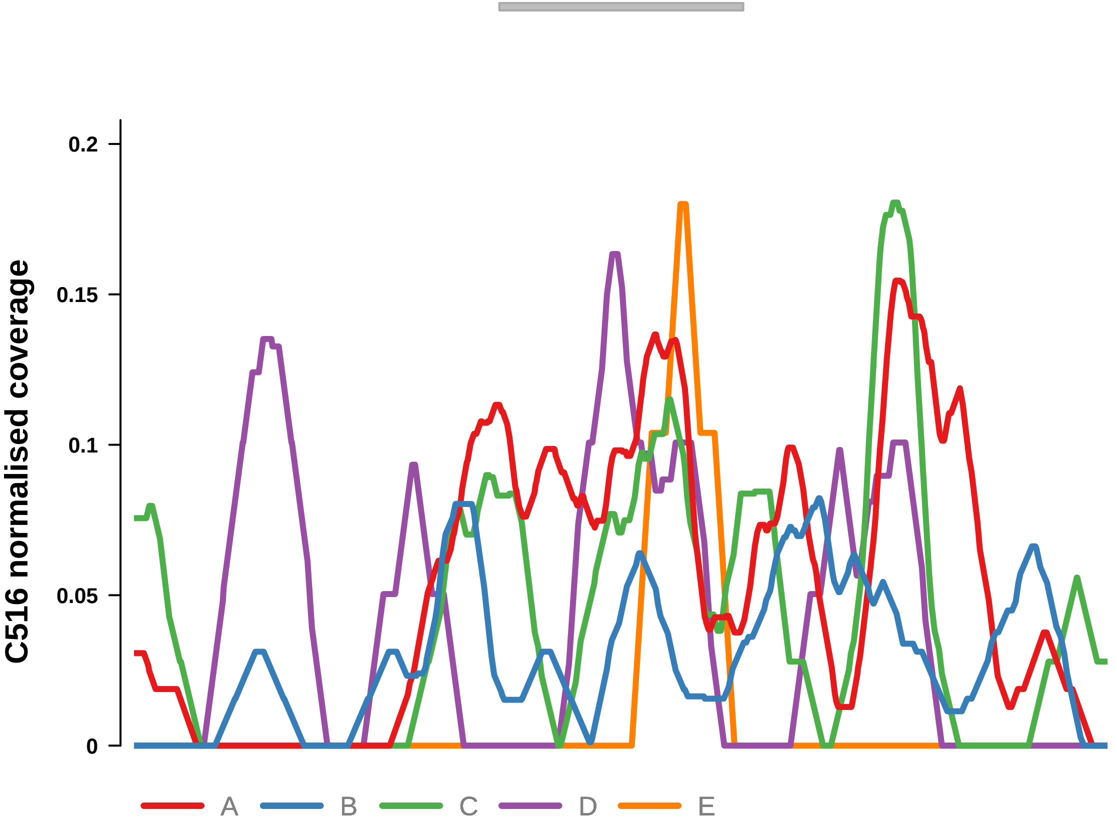


Putatively Clonal (logFC = 2.39)

| | **P.** | **Adjusted P.** |
|---|---|---|
| *~region vs. ~* | 5.86e−02 | 7.04e−01 |
| *~purity vs ~* | 1.63e−01 | 7.73e−01 |
| *~purity+region vs ~purity* | 2.05e−01 | 1e+00 |
| *Carcinoma vs. normal* | 9.01e−03 ** | 9.01e−03 ** |

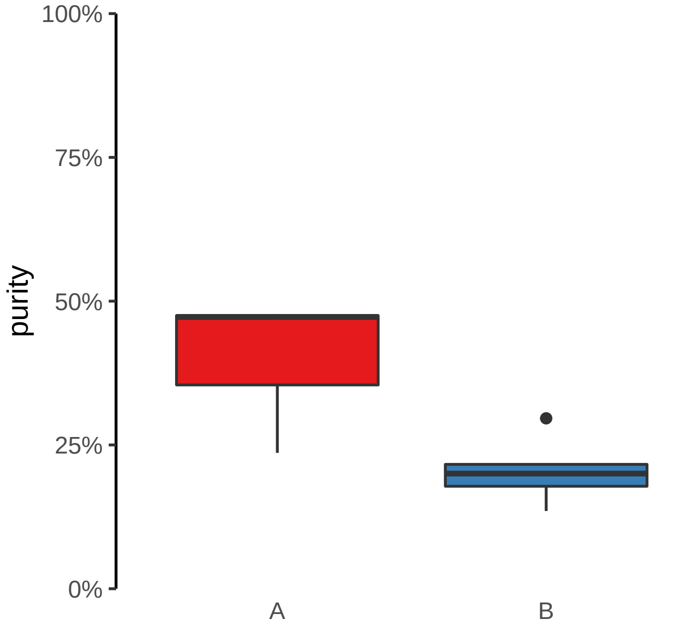

**ARHGEF17 promoter gain**
**chr11:73362044−73362544**

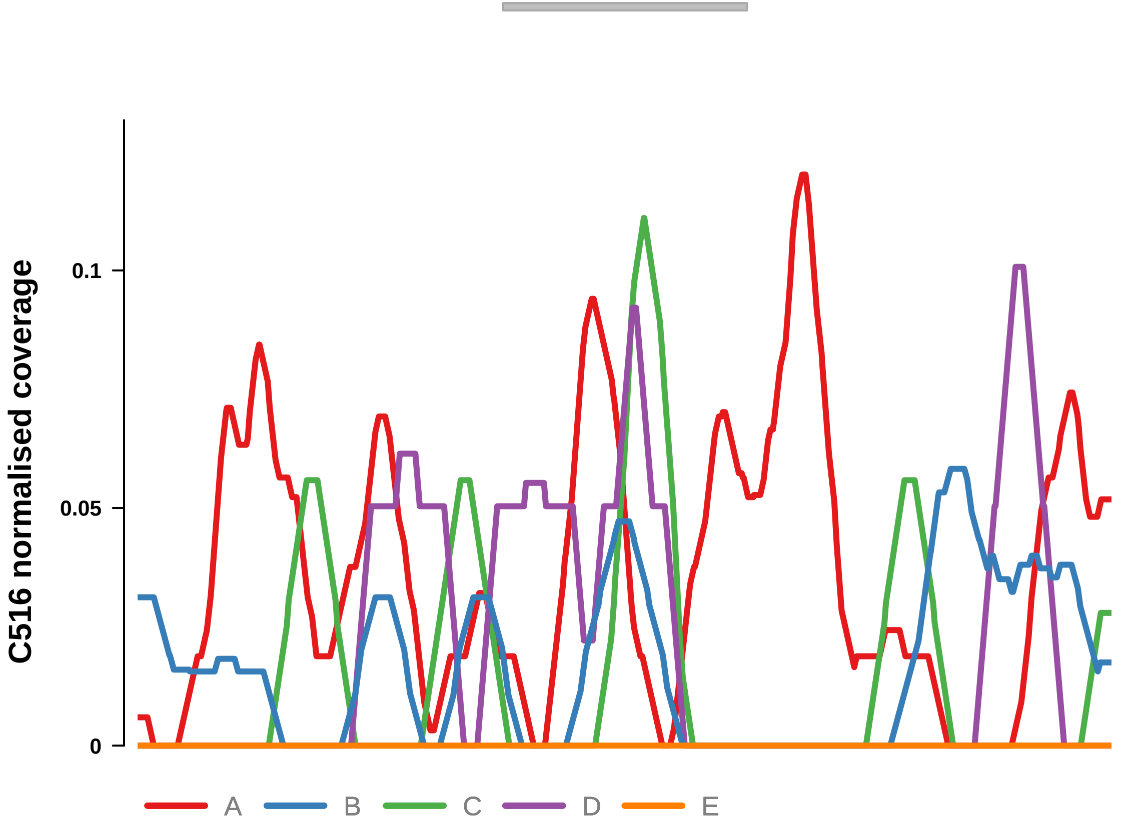


Putatively Clonal (logFC = 3.09)

| | P. | Adjusted P. |
|---|---|---|
| *~region vs. ~* | 4.71e−01 | 8.67e−01 |
| *~purity vs ~* | 5.23e−01 | 8.86e−01 |
| *~purity+region vs ~purity* | 1e+00 | 1e+00 |
| *Carcinoma vs. normal* | 4.95e−04 *** | 4.95e−04 *** |

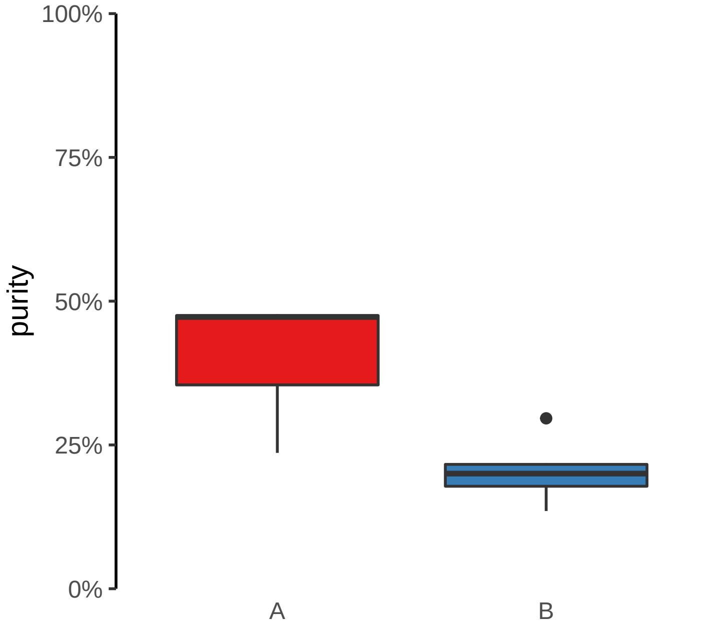

**EHD2 promoter gain**
**chr19:47730036−47730536**

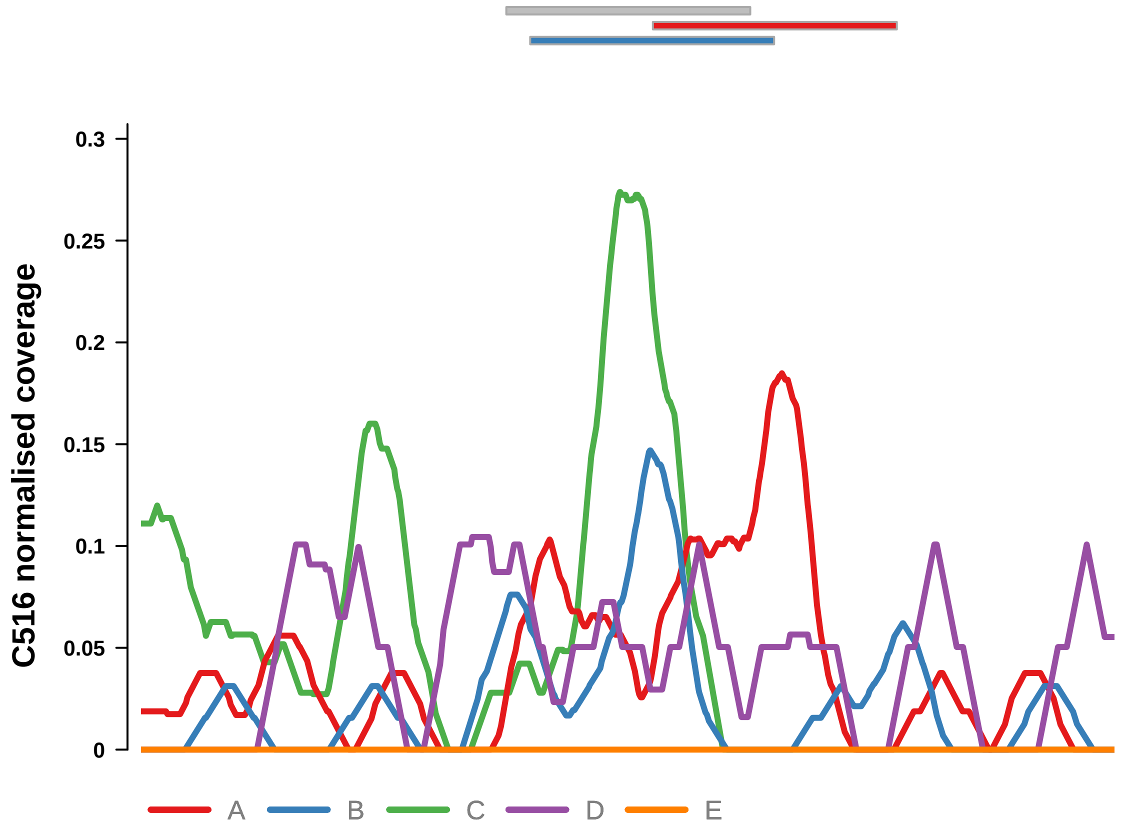


Putatively Clonal (logFC = 2.45)

| | P. | Adjusted P. |
|---|---|---|
| *~region vs. ~* | 6.8e−02 | 7.04e−01 |
| *~purity vs ~* | 4.52e−02 * | 5.76e−01 |
| *~purity+region vs ~purity* | 6.77e−01 | 1e+00 |
| *Carcinoma vs. normal* | 3.09e−03 ** | 3.09e−03 ** |

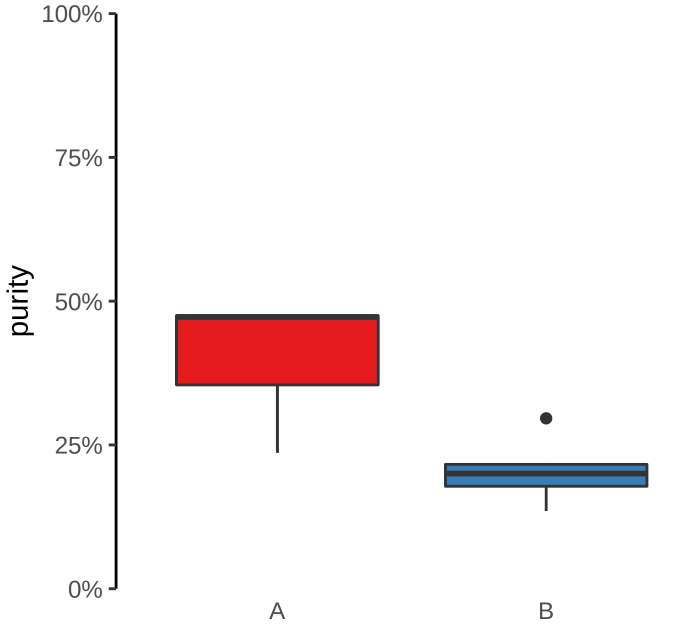

**FUT1 promoter gain**
**chr19:48752815−48753315**

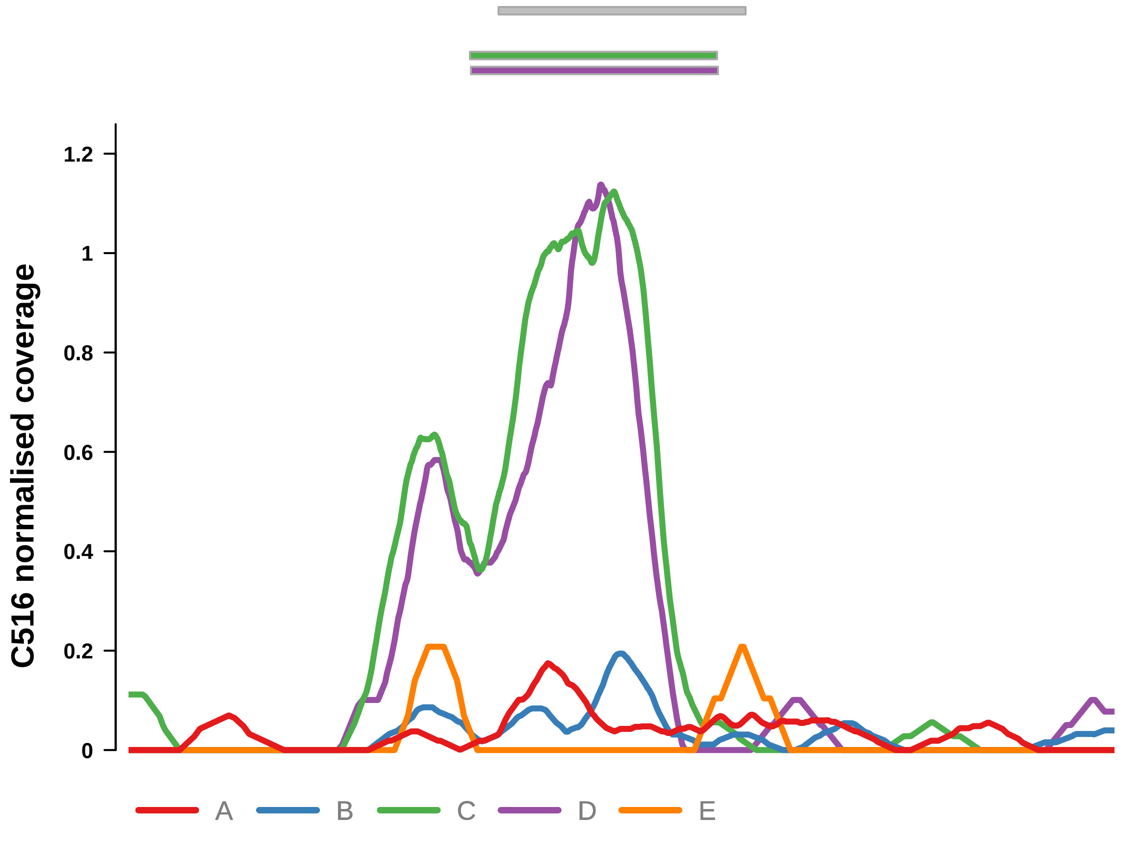


Putatively Clonal (logFC = 2.74)

| | P. | Adjusted P. |
|---|---|---|
| *~region vs. ~* | 3.17e−02 * | 7.04e−01 |
| *~purity vs ~* | 1.18e−02 * | 5.21e−01 |
| *~purity+region vs ~purity* | 3.81e−01 | 1e+00 |
| *Carcinoma vs. normal* | 3.11e−04 *** | 3.11e−04 *** |

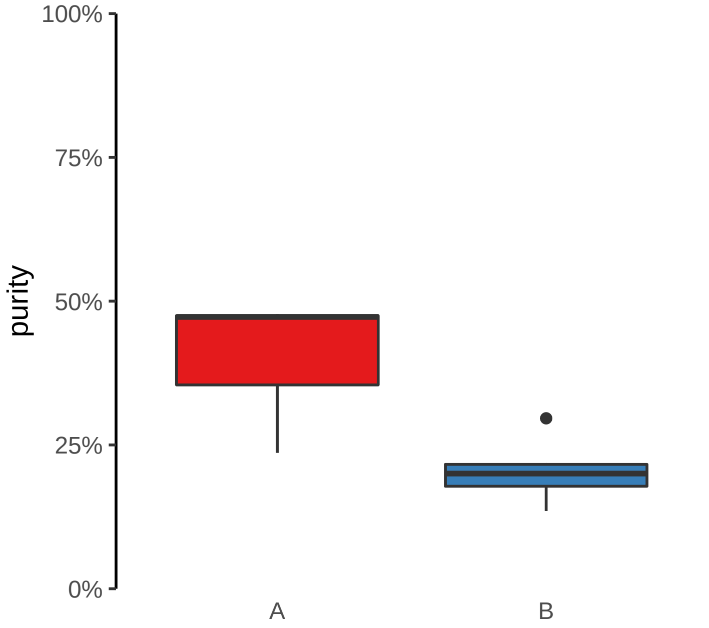

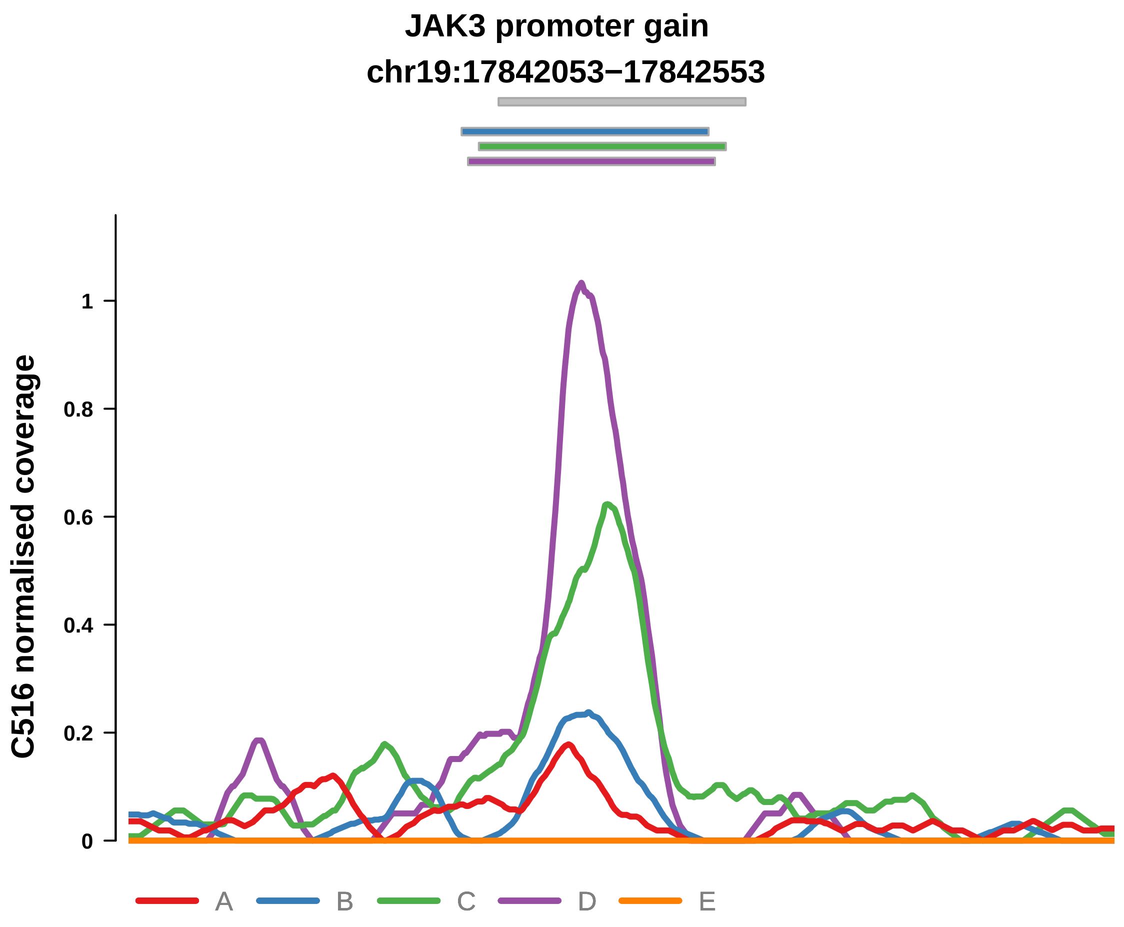


Putatively Clonal (logFC = 2.3)

| | P. | Adjusted P. |
|---|---|---|
| *~region vs. ~* | 2.91e−01 | 8.67e−01 |
| *~purity vs ~* | 3.04e−01 | 8.53e−01 |
| *~purity+region vs ~purity* | 9.06e−01 | 1e+00 |
| *Carcinoma vs. normal* | 5.97e−03 ** | 5.97e−03 ** |

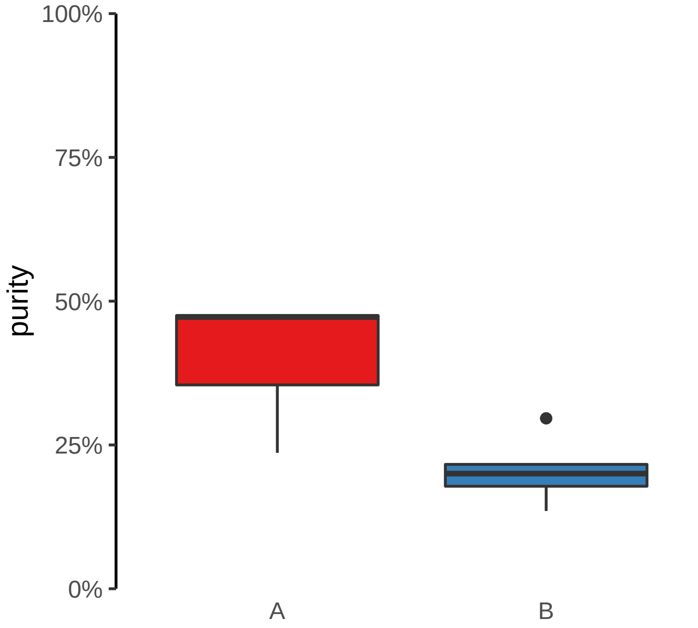

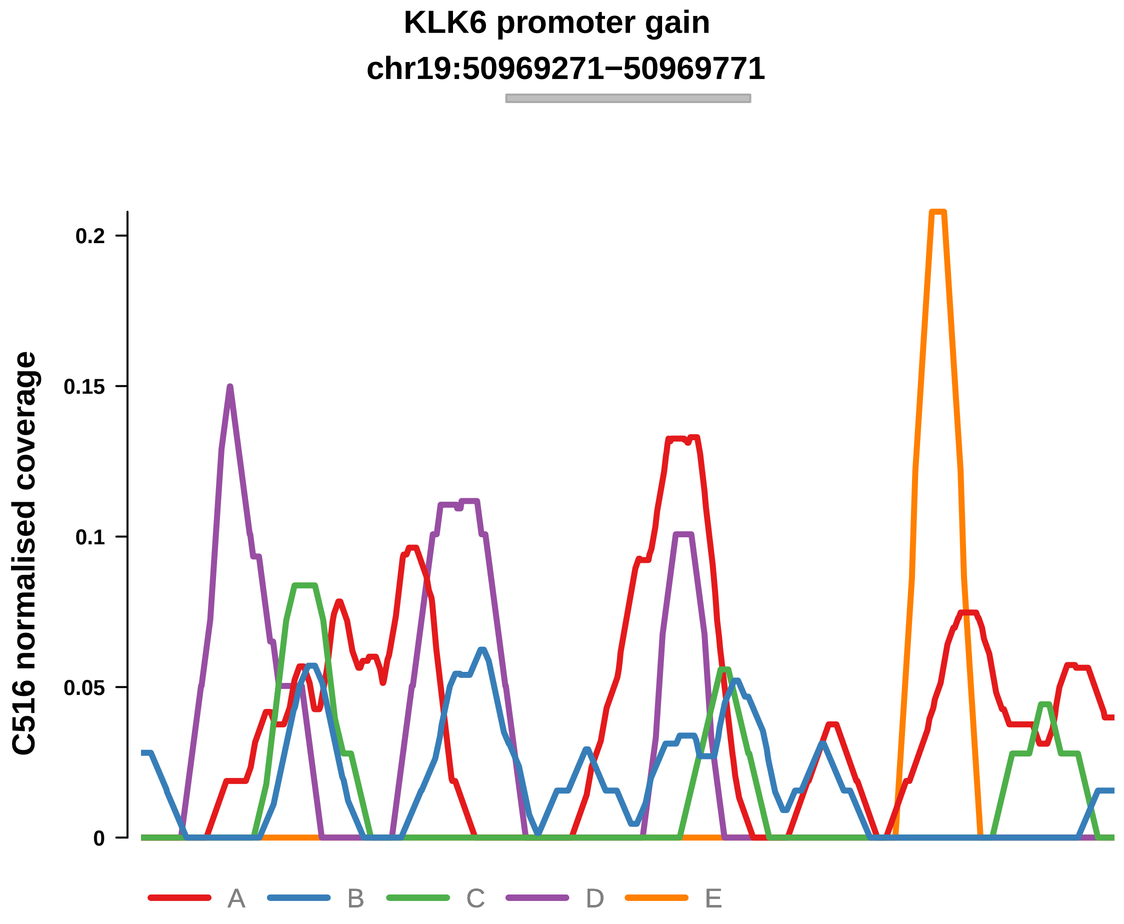


Putatively Clonal (logFC = 2.73)

| | P. | Adjusted P. |
|---|---|---|
| *~region vs. ~* | 1.73e−01 | 7.59e−01 |
| *~purity vs ~* | 2.81e−02 * | 5.76e−01 |
| *~purity+region vs ~purity* | 2.98e−01 | 1e+00 |
| *Carcinoma vs. normal* | 4.06e−03 ** | 4.06e−03 ** |

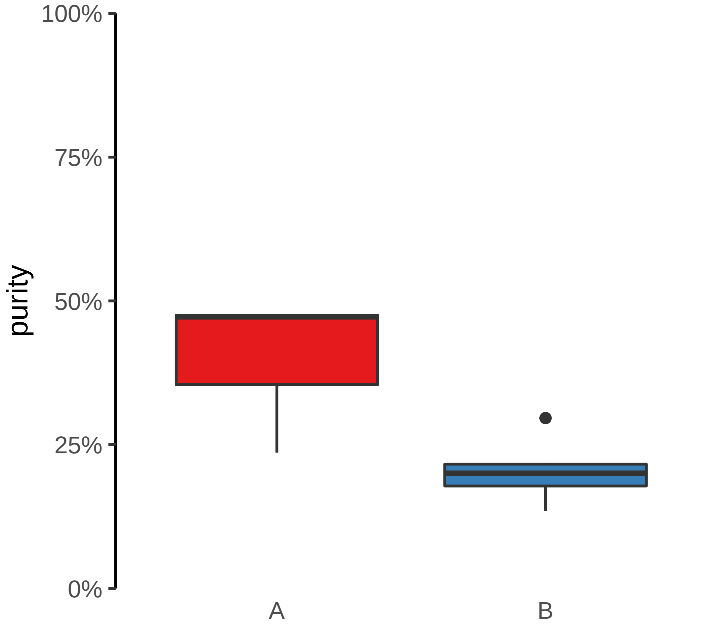

**PITX1 promoter gain**

**chr5:135034130−135034630**

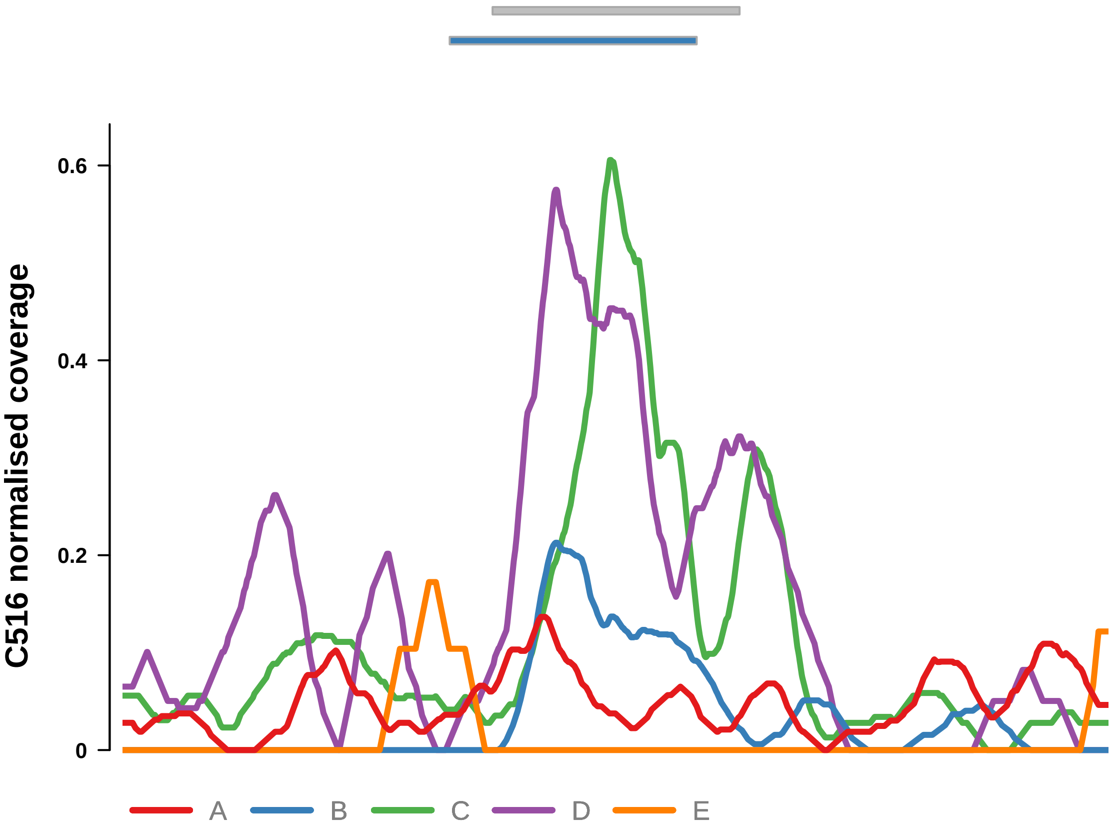


Putatively Clonal (logFC = 2.24)

| | **P.** | **Adjusted P.** |
|---|---|---|
| *~region vs. ~* | 8.45e−01 | 9.59e−01 |
| *~purity vs ~* | 5.16e−01 | 8.86e−01 |
| *~purity+region vs ~purity* | 3.21e−01 | 1e+00 |
| *Carcinoma vs. normal* | 7.03e−03 ** | 7.03e−03 ** |

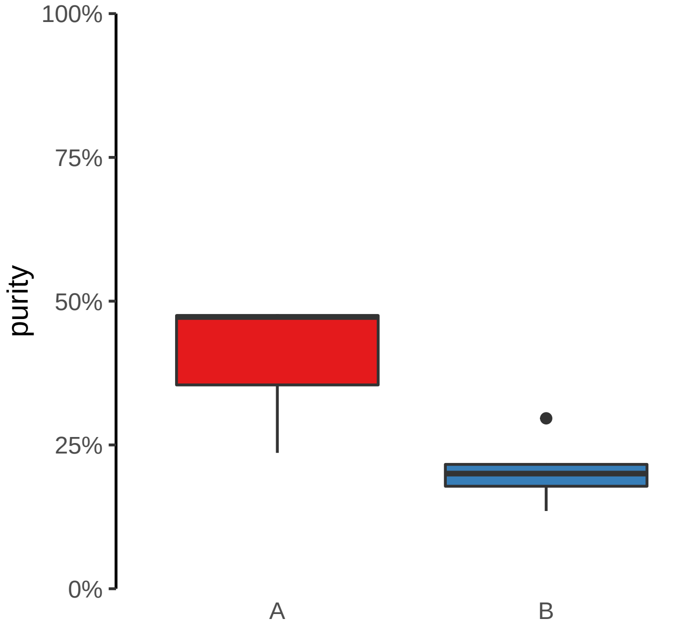

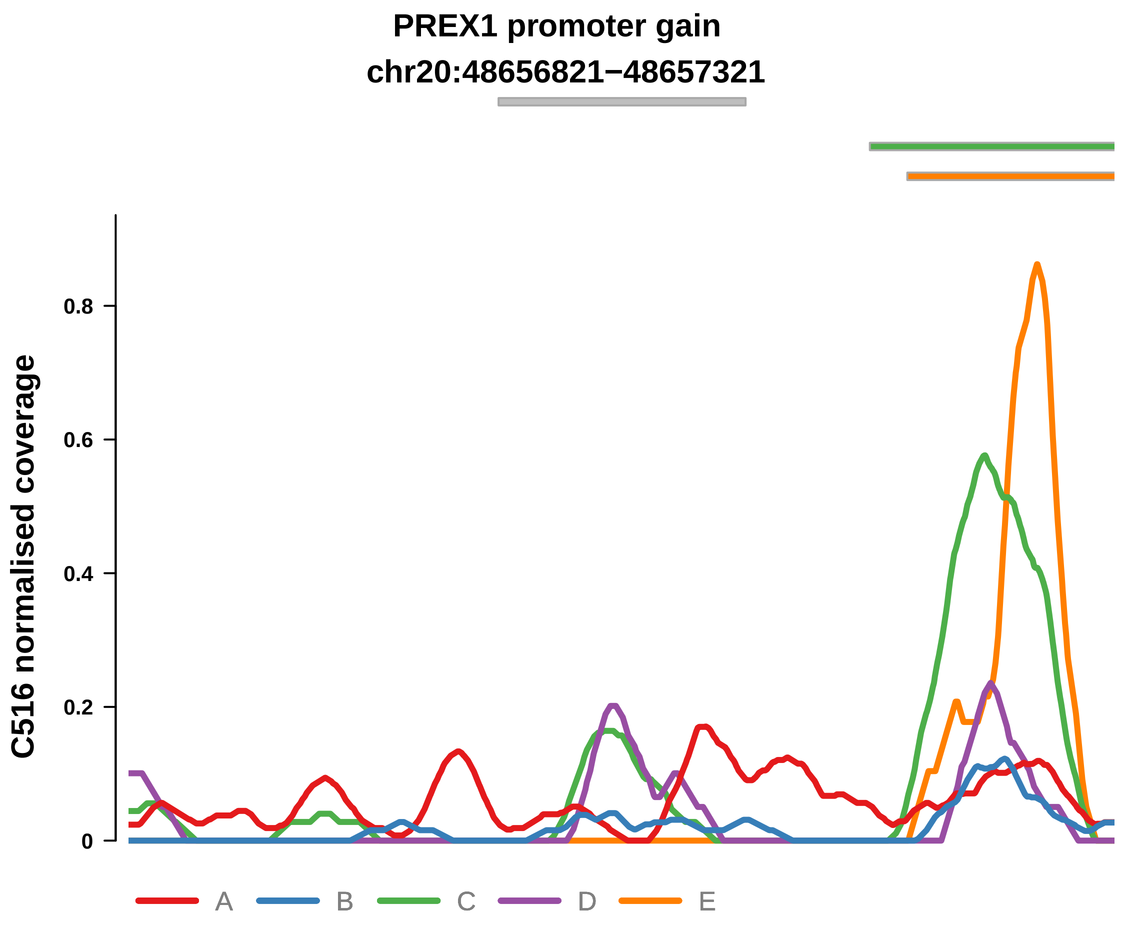


Putatively Clonal (logFC = 3.1)

| | P. | Adjusted P. |
|---|---|---|
| *~region vs. ~* | 8.68e−02 | 7.04e−01 |
| *~purity vs ~* | 5.89e−02 | 5.76e−01 |
| *~purity+region vs ~purity* | 8.25e−01 | 1e+00 |
| *Carcinoma vs. normal* | 1.06e−03 ** | 1.06e−03 ** |

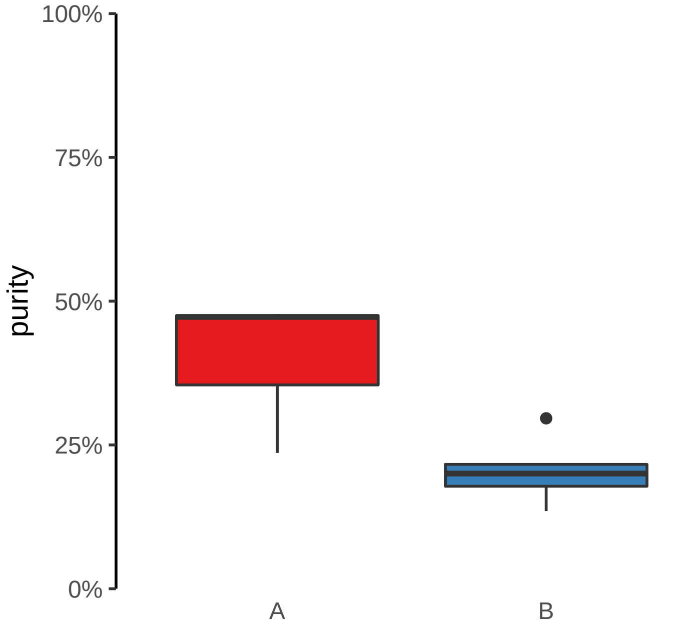

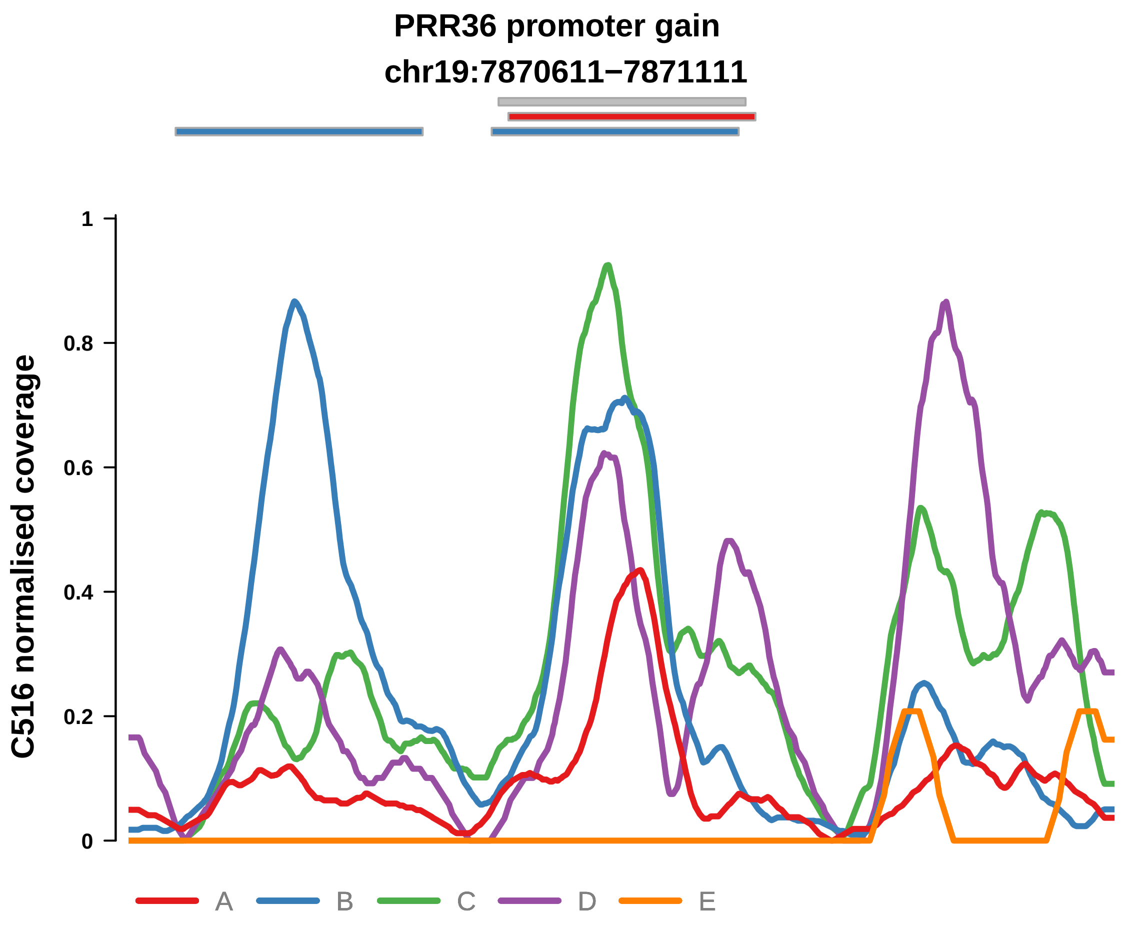


Putatively Clonal (logFC = 2.13)

| | P. | Adjusted P. |
|---|---|---|
| *~region vs. ~* | 4.1e−02 * | 7.04e−01 |
| *~purity vs ~* | 1.6e−01 | 7.73e−01 |
| *~purity+region vs ~purity* | 1.78e−01 | 1e+00 |
| *Carcinoma vs. normal* | 8.68e−03 ** | 8.68e−03 ** |

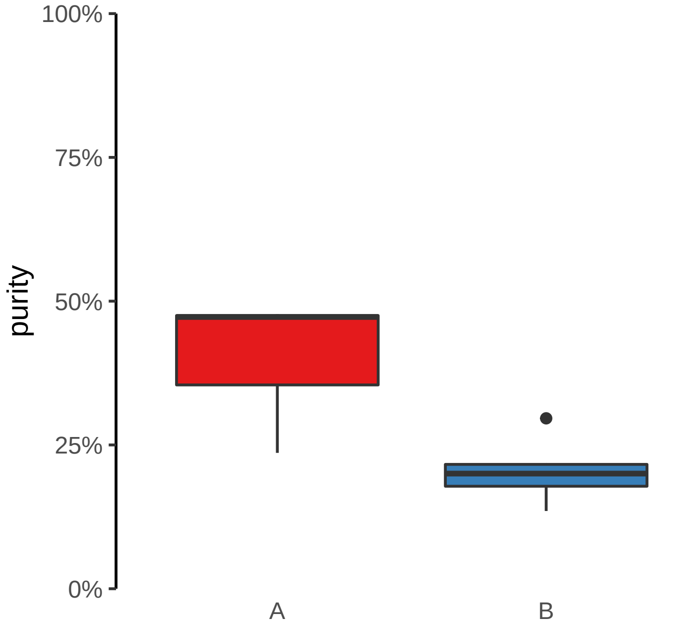

**SLC45A4 promoter gain**
**chr8:141230663−141231163**

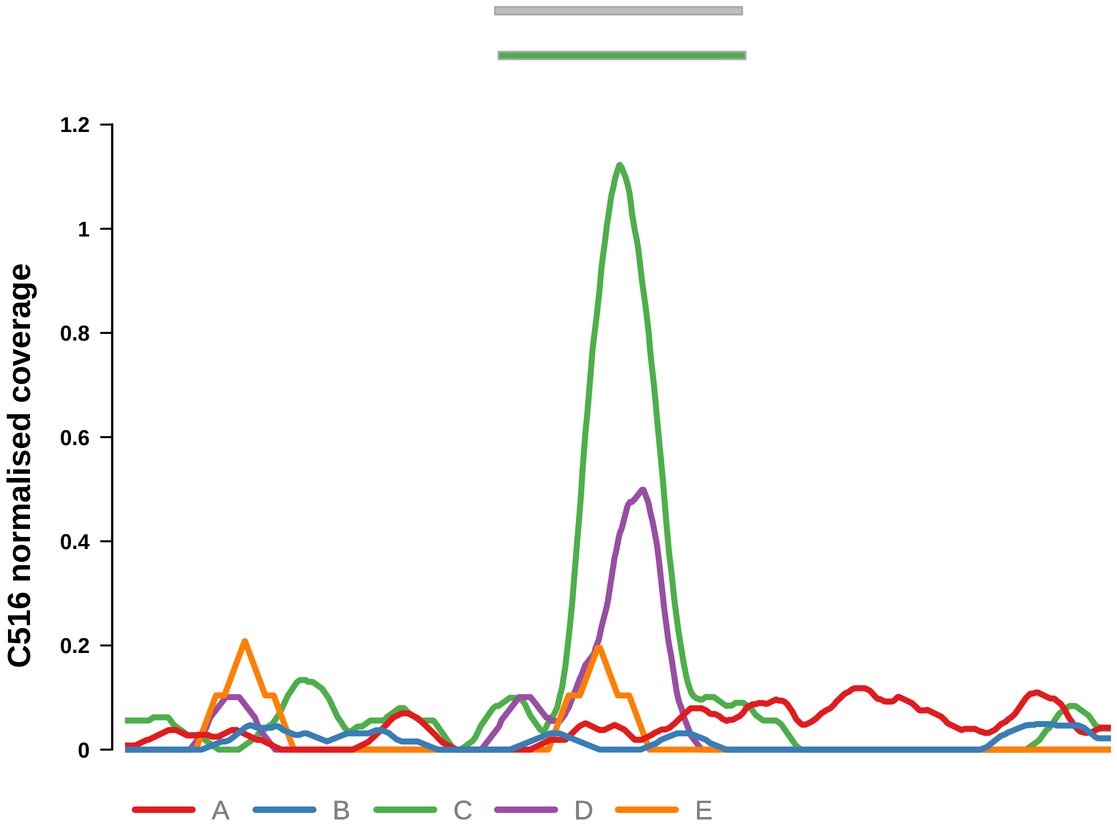


Putatively Clonal (logFC = 2.7)

| | P. | Adjusted P. |
|---|---|---|
| *~region vs. ~* | 2.24e−02 * | 7.04e−01 |
| *~purity vs ~* | 1.82e−02 * | 5.33e−01 |
| *~purity+region vs ~purity* | 2.65e−01 | 1e+00 |
| *Carcinoma vs. normal* | 5.66e−03 ** | 5.66e−03 ** |

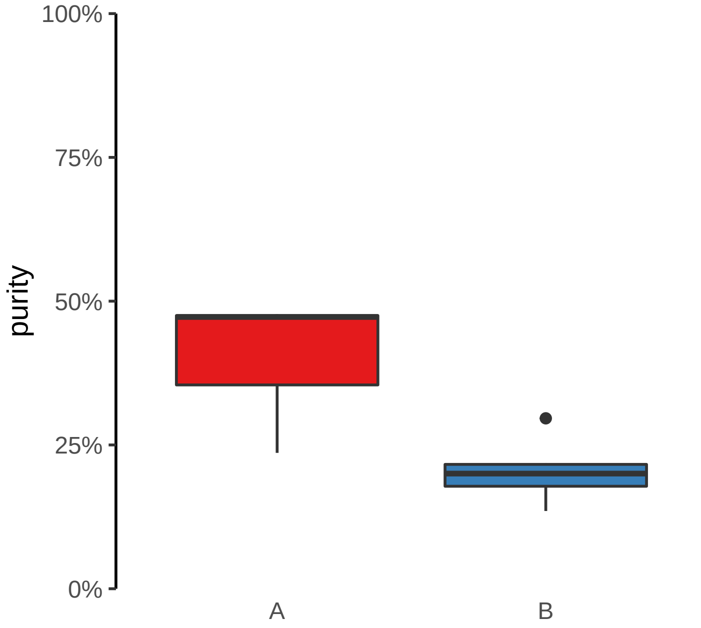

**CADM4 promoter gain**

**chr19:43623815−43624315**

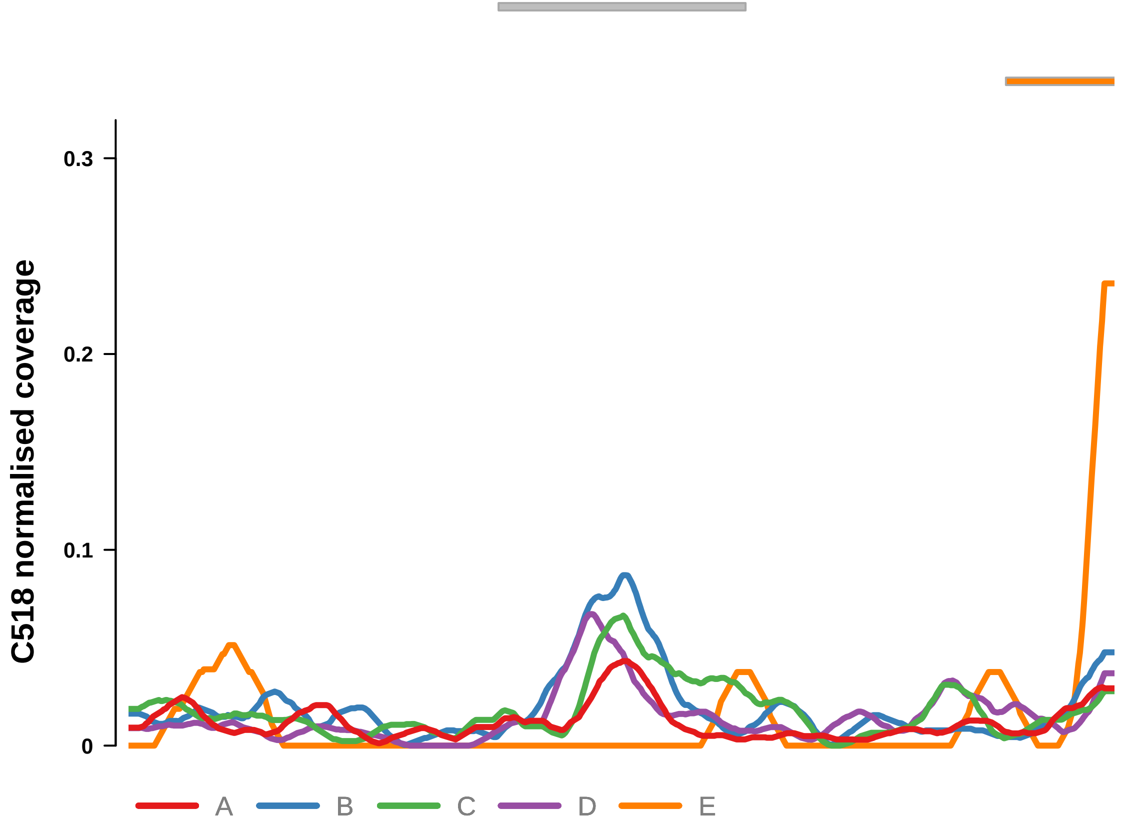


Putatively Clonal (logFC = 2.39)

| | **P.** | **Adjusted P.** |
|---|---|---|
| *~region vs. ~* | 4.9e−01 | 7.07e−01 |
| *~purity vs ~* | 3.24e−01 | 7.38e−01 |
| *~purity+region vs ~purity* | 7e−01 | 9.19e−01 |
| *Carcinoma vs. normal* | 5.16e−05 **** | 5.16e−05 **** |

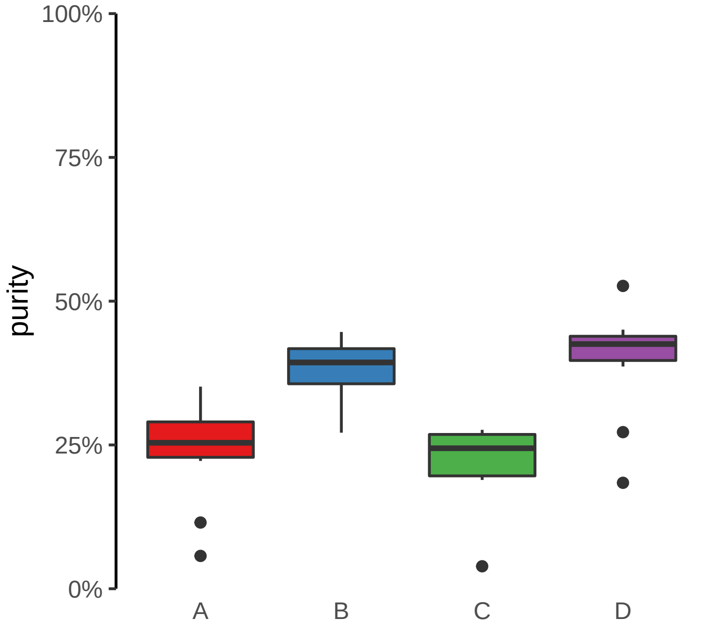

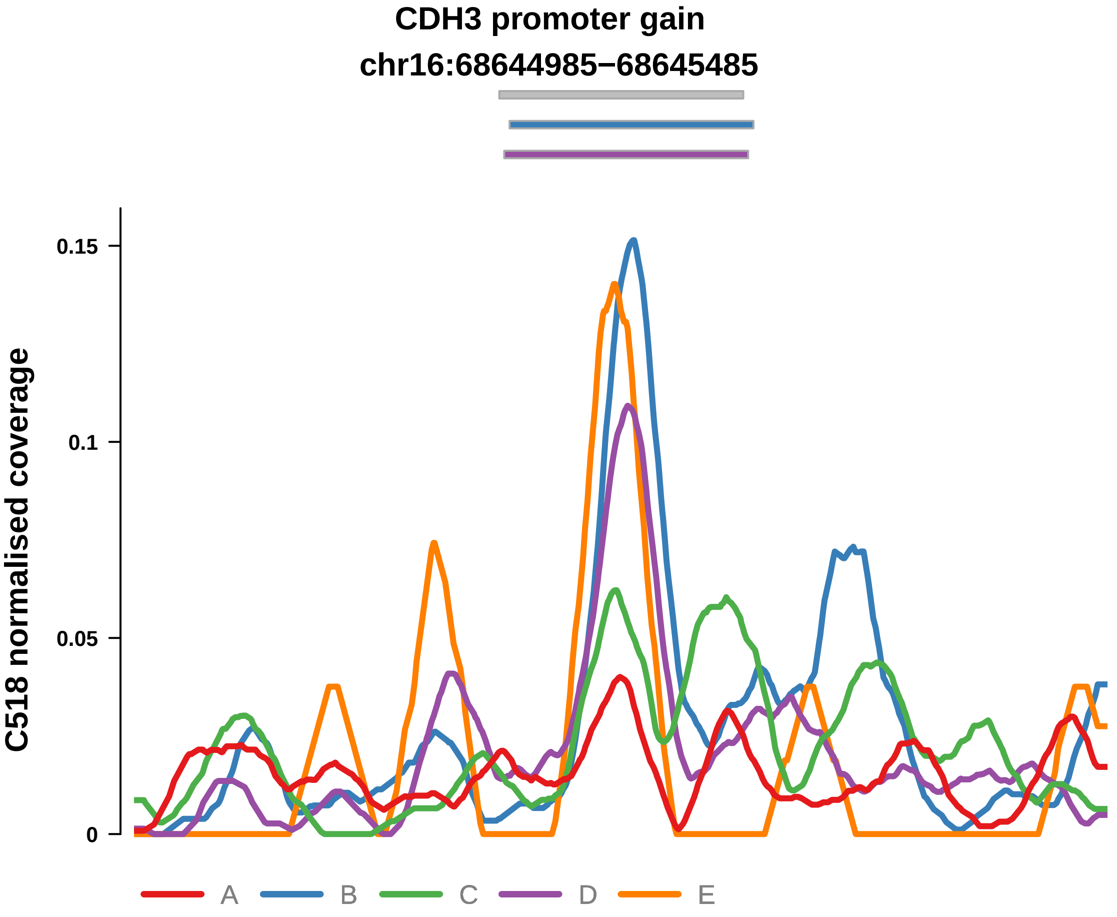


Putatively Clonal (logFC = 1.76)

| | P. | Adjusted P. |
|---|---|---|
| *~region vs. ~* | 9.27e−01 | 9.49e−01 |
| *~purity vs ~* | 8.78e−01 | 9.54e−01 |
| *~purity+region vs ~purity* | 9.34e−01 | 9.67e−01 |
| *Carcinoma vs. normal* | 1.7e−03 ** | 1.7e−03 ** |

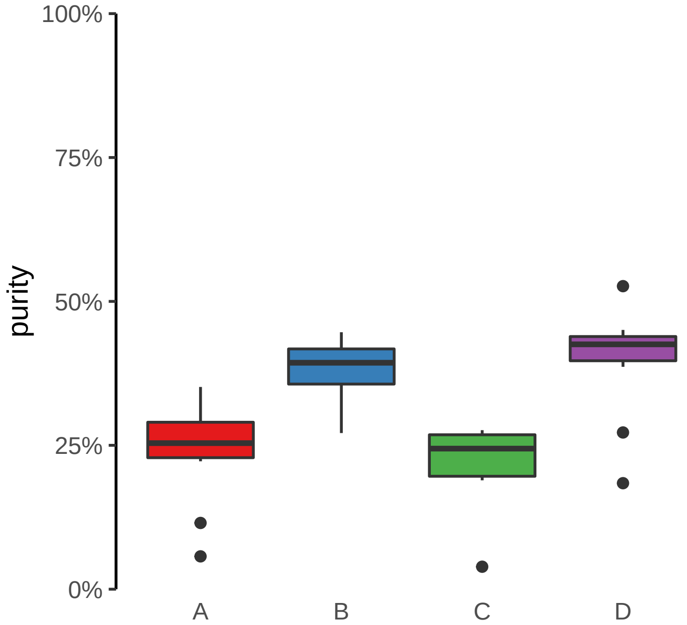

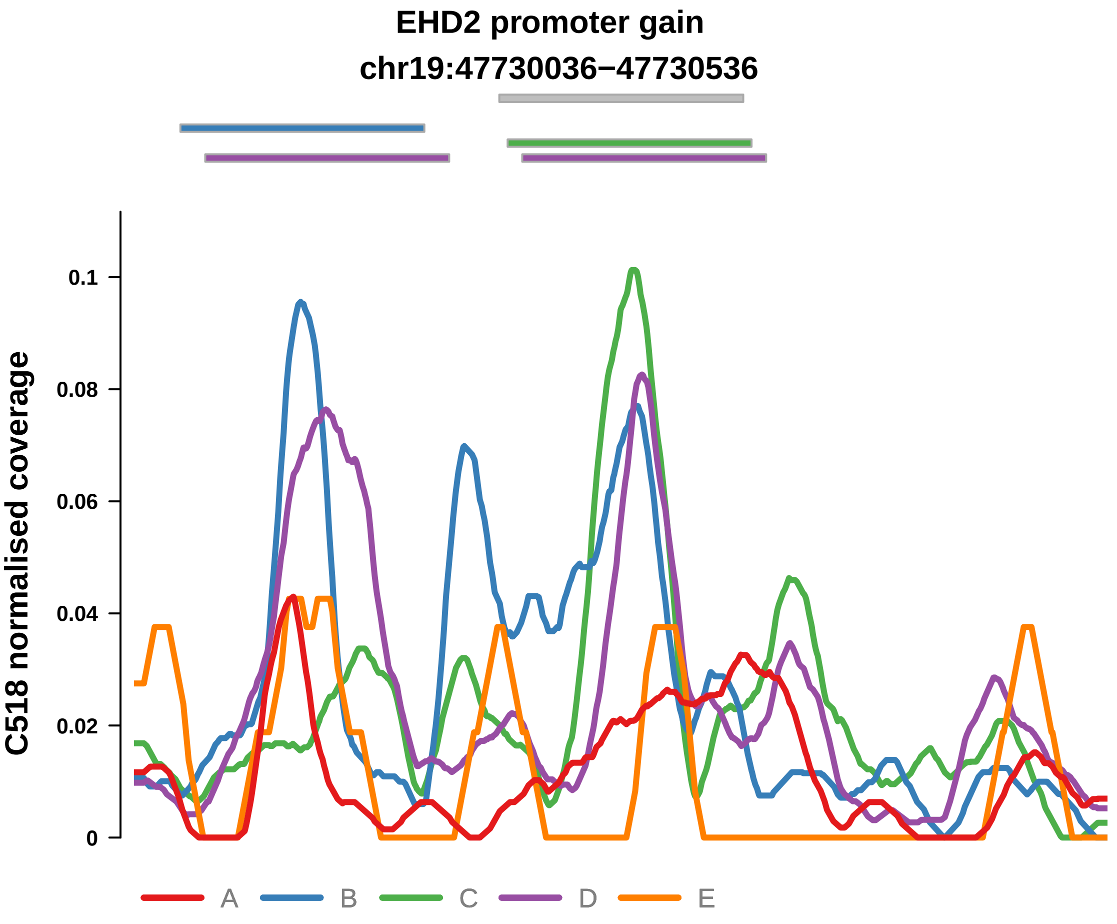


Putatively Clonal (logFC = 1.61)

| | P. | Adjusted P. |
|---|---|---|
| *~region vs. ~* | 4.61e−01 | 6.88e−01 |
| *~purity vs ~* | 4.79e−01 | 8.48e−01 |
| *~purity+region vs ~purity* | 5.64e−01 | 8.42e−01 |
| *Carcinoma vs. normal* | 5.13e−03 ** | 5.13e−03 ** |

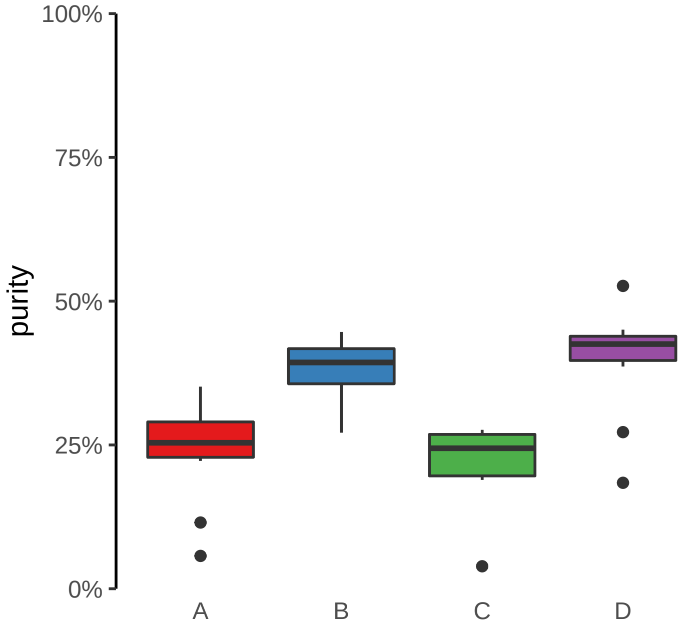

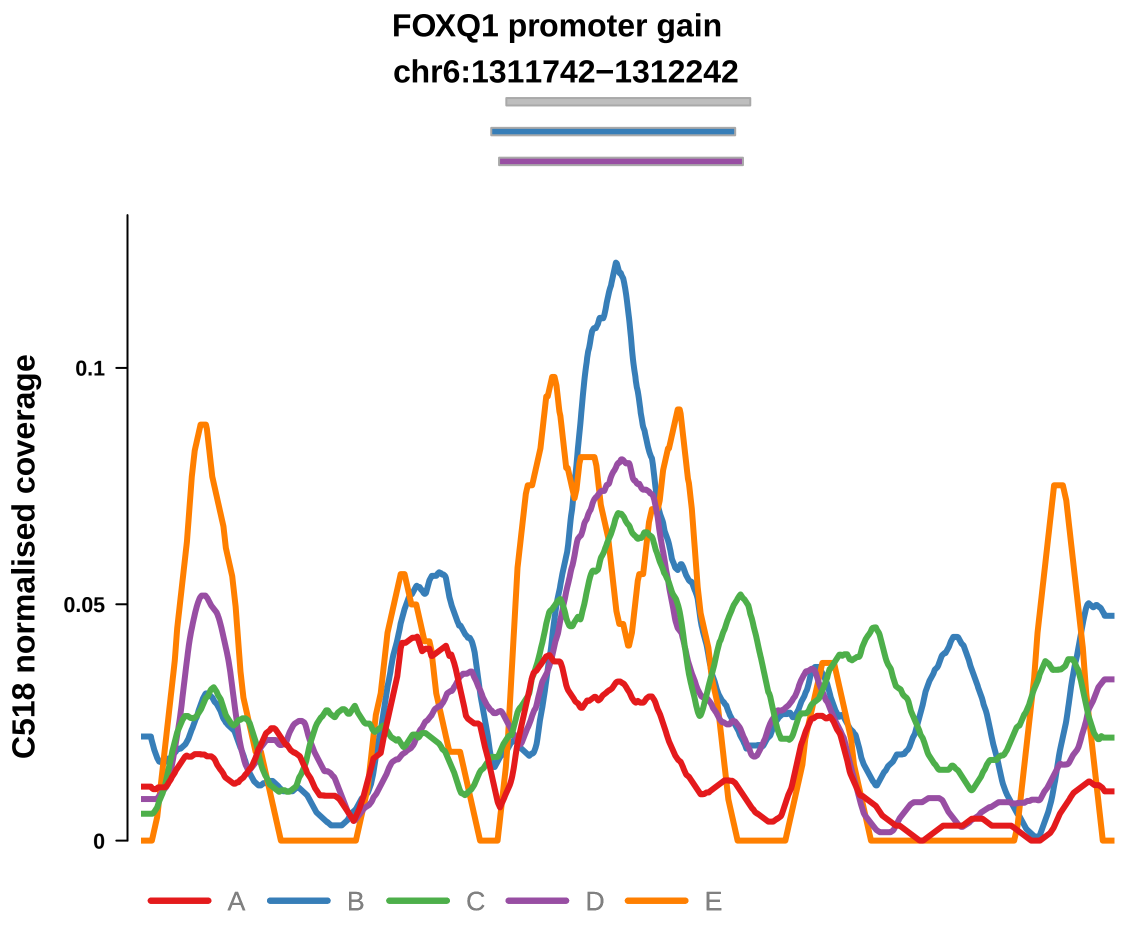


Putatively Clonal (logFC = 1.87)

| | P. | Adjusted P. |
|---|---|---|
| *~region vs. ~* | 4.15e−01 | 6.64e−01 |
| *~purity vs ~* | 4.26e−01 | 8.14e−01 |
| *~purity+region vs ~purity* | 2.54e−01 | 6.98e−01 |
| *Carcinoma vs. normal* | 8.05e−04 *** | 8.05e−04 *** |

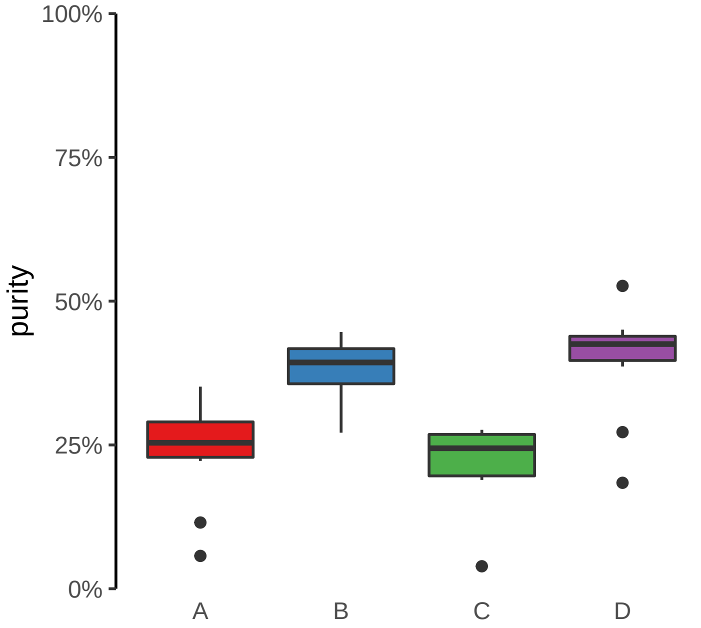

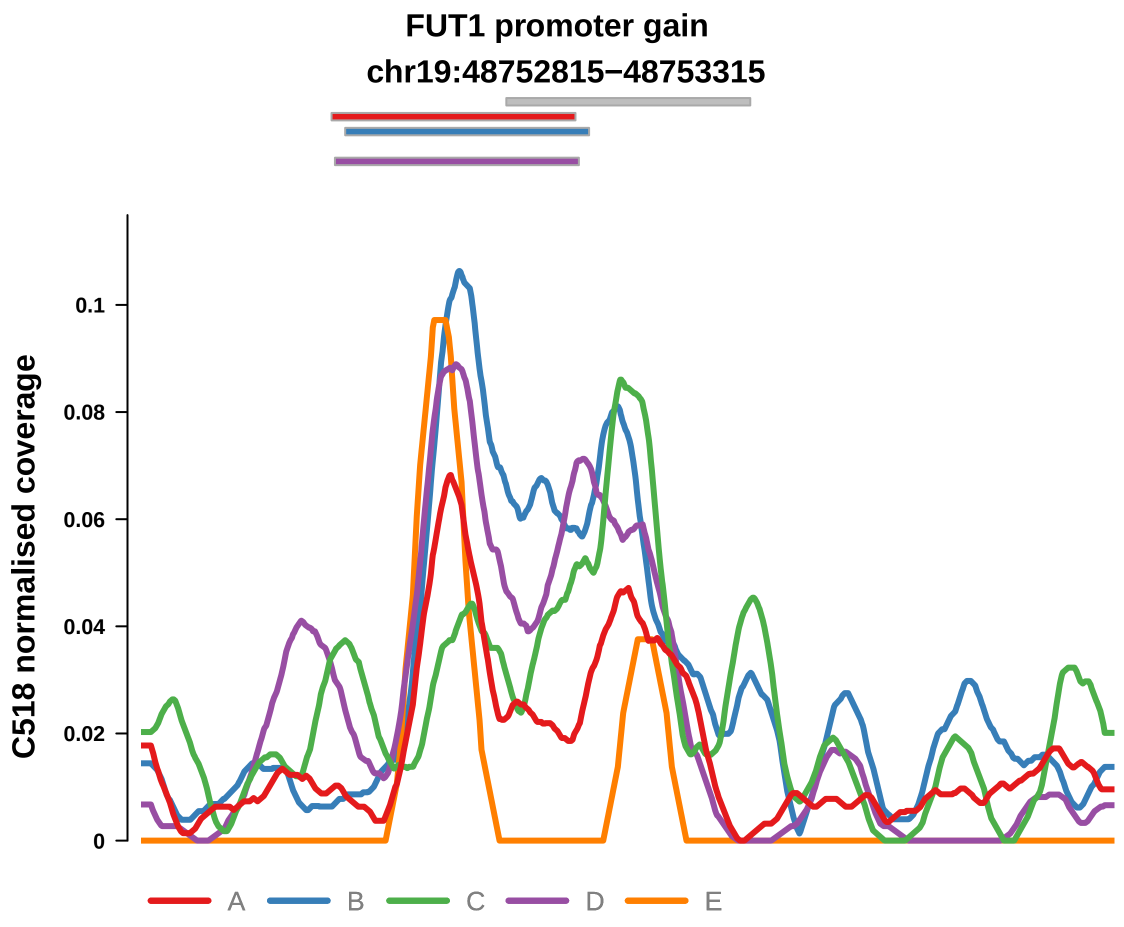


Putatively Clonal (logFC = 2.01)

| | P. | Adjusted P. |
|---|---|---|
| *~region vs. ~* | 7.43e−01 | 8.28e−01 |
| *~purity vs ~* | 3.8e−01 | 7.92e−01 |
| *~purity+region vs ~purity* | 8.81e−01 | 9.5e−01 |
| *Carcinoma vs. normal* | 3.61e−04 *** | 3.61e−04 *** |

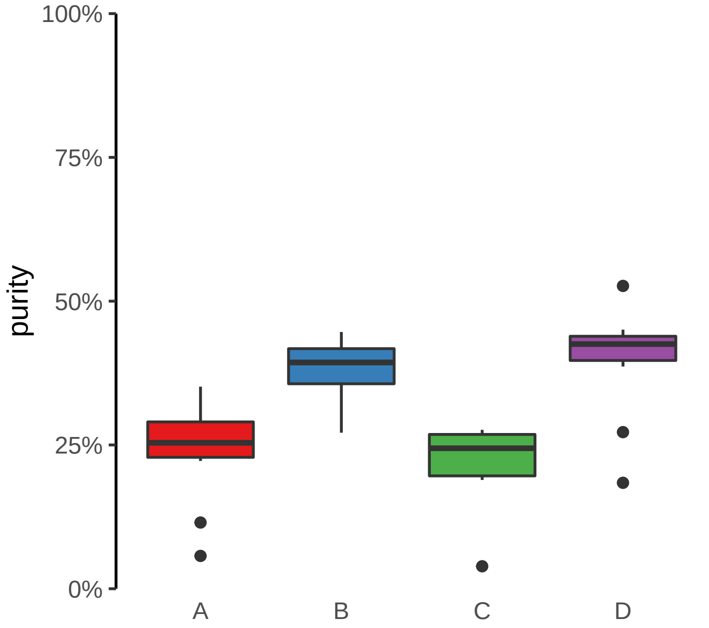

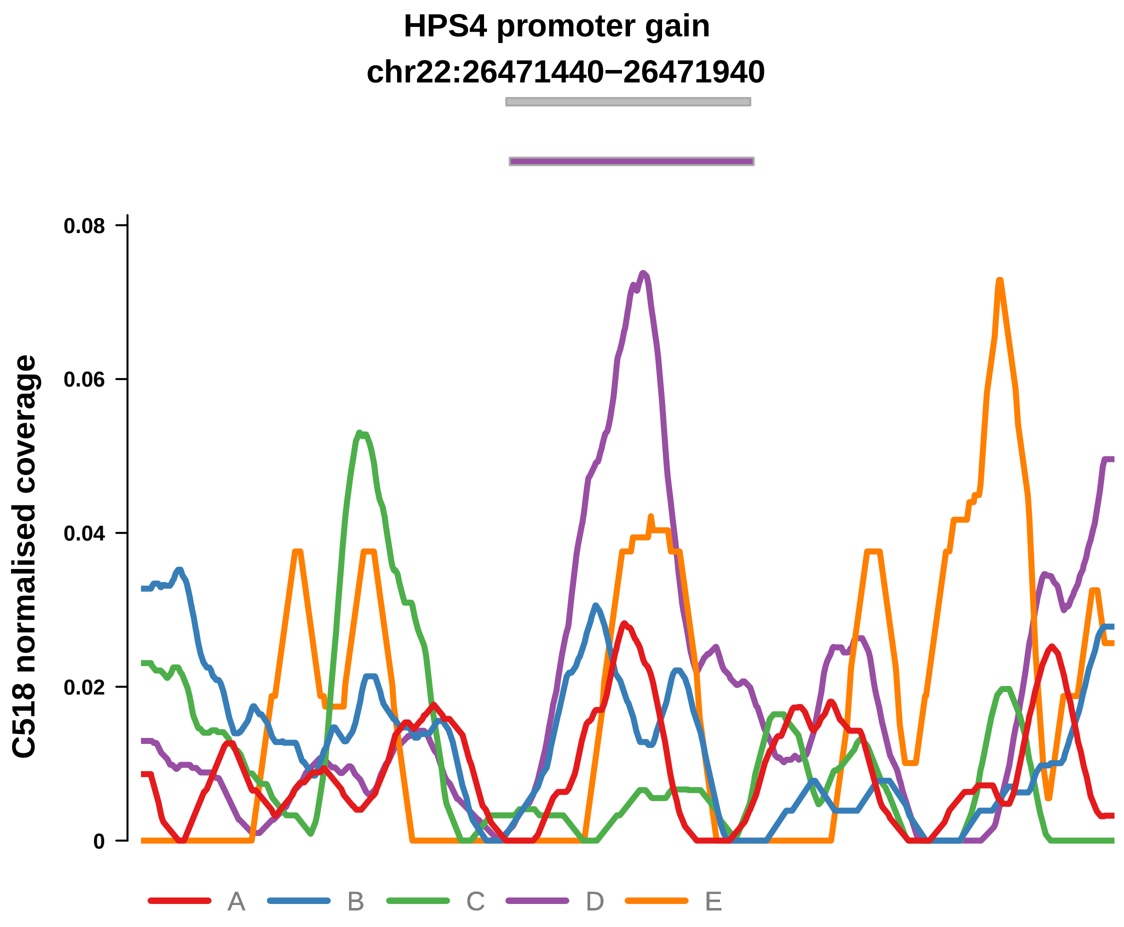


Putatively Clonal (logFC = 1.81)

| | P. | Adjusted P. |
|---|---|---|
| *~region vs. ~* | 3.96e−02 * | 2.62e−01 |
| *~purity vs ~* | 7.28e−01 | 9.22e−01 |
| *~purity+region vs ~purity* | 3.68e−02 * | 2.7e−01 |
| *Carcinoma vs. normal* | 3.23e−03 ** | 3.23e−03 ** |

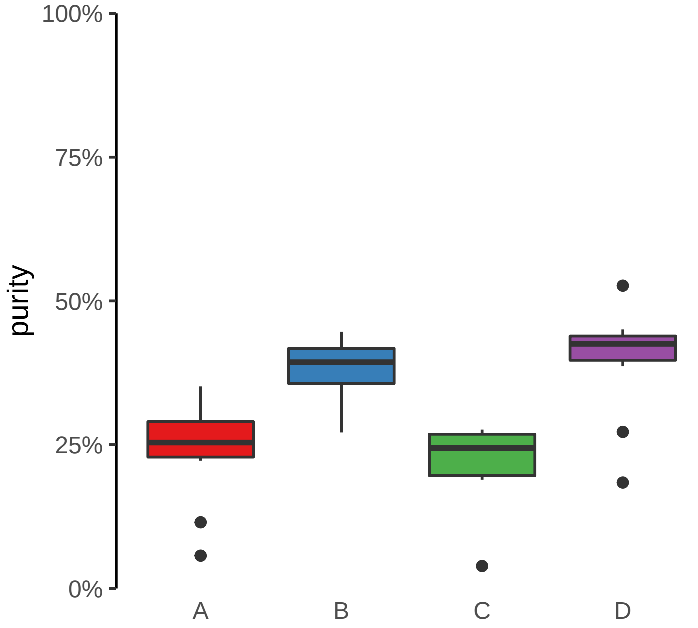

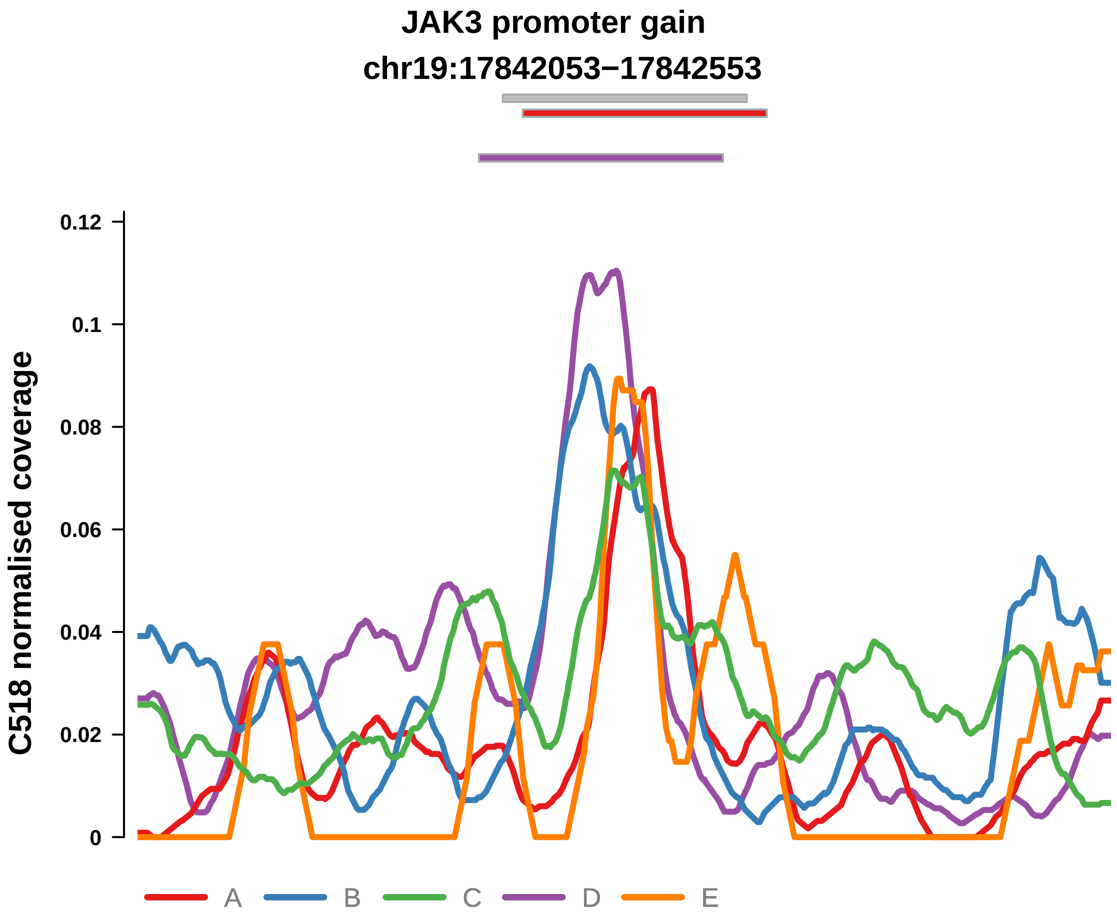


Putatively Clonal (logFC = 2.07)

| | P. | Adjusted P. |
|---|---|---|
| *~region vs. ~* | 3.75e−01 | 6.53e−01 |
| *~purity vs ~* | 3.05e−01 | 7.38e−01 |
| *~purity+region vs ~purity* | 5.57e−01 | 8.42e−01 |
| *Carcinoma vs. normal* | 2.65e−04 *** | 2.65e−04 *** |

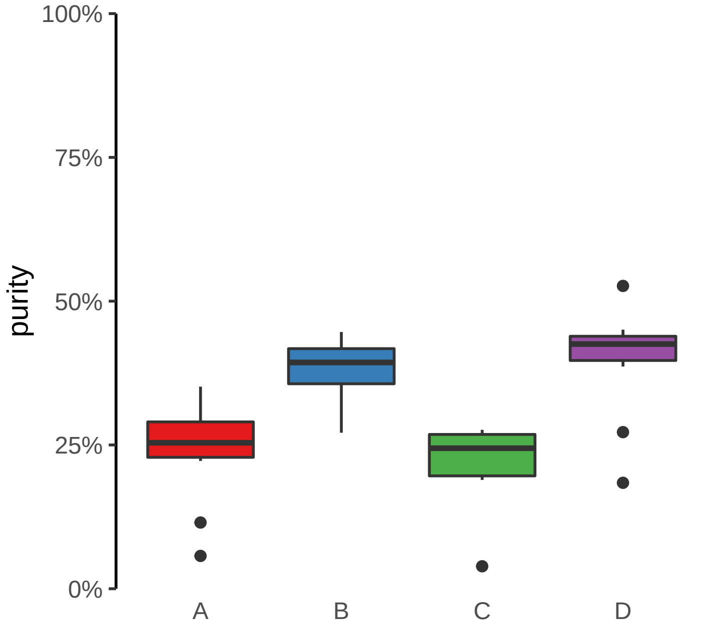

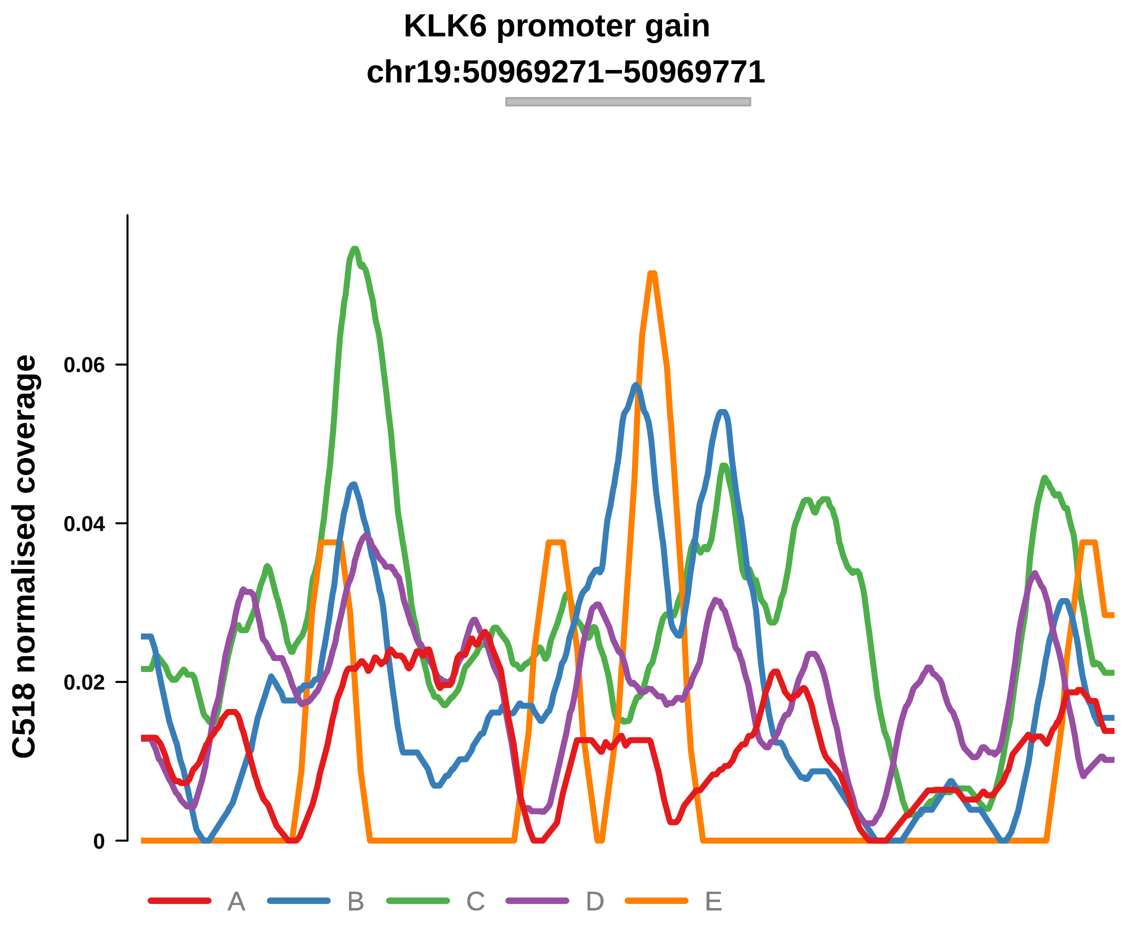


Putatively Clonal (logFC = 1.69)

| | P. | Adjusted P. |
|---|---|---|
| *~region vs. ~* | 1.38e−01 | 4.84e−01 |
| *~purity vs ~* | 6.42e−01 | 9.13e−01 |
| *~purity+region vs ~purity* | 1.6e−01 | 5.63e−01 |
| *Carcinoma vs. normal* | 4.75e−03 ** | 4.75e−03 ** |

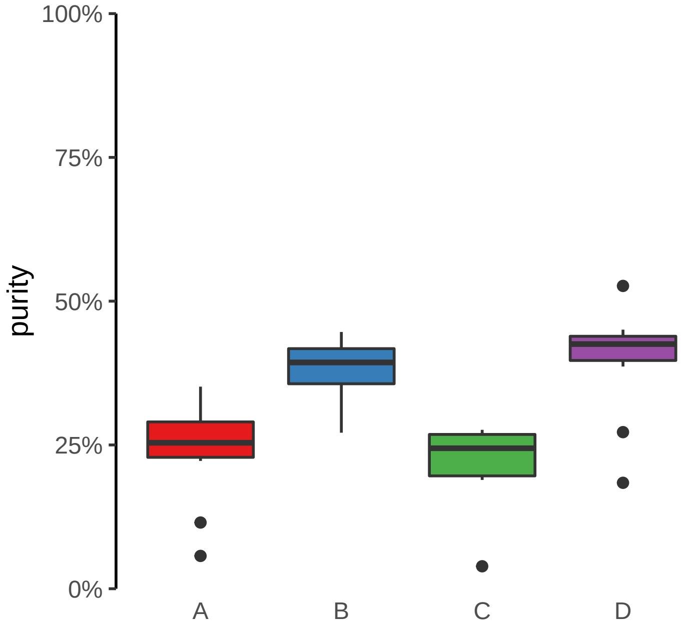

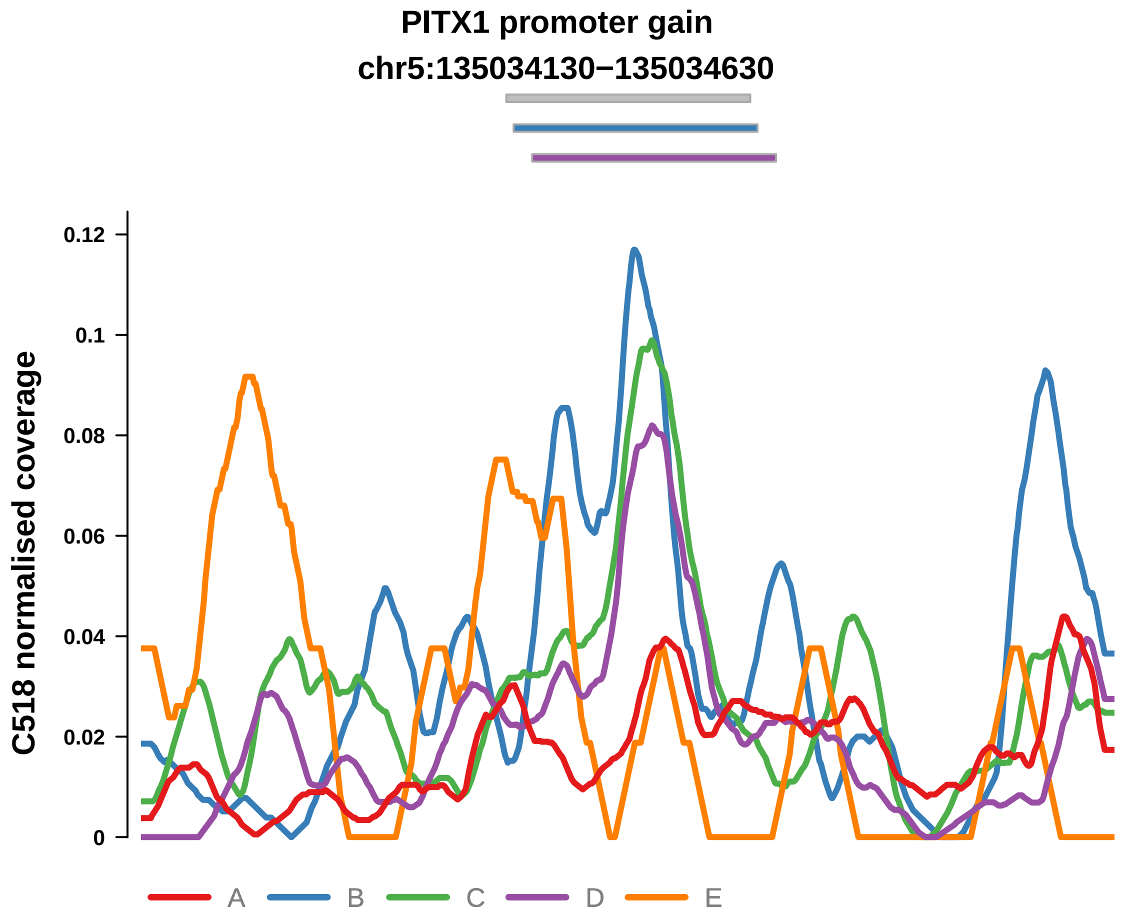


Putatively Clonal (logFC = 1.68)

| | P. | Adjusted P. |
|---|---|---|
| *~region vs. ~* | 7.26e−01 | 8.28e−01 |
| *~purity vs ~* | 2.82e−01 | 7.38e−01 |
| *~purity+region vs ~purity* | 8.19e−01 | 9.5e−01 |
| *Carcinoma vs. normal* | 2.93e−03 ** | 2.93e−03 ** |

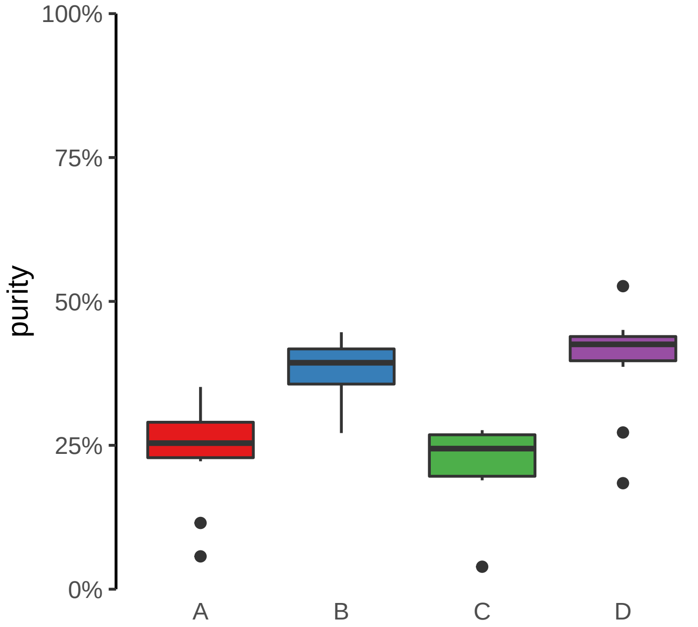

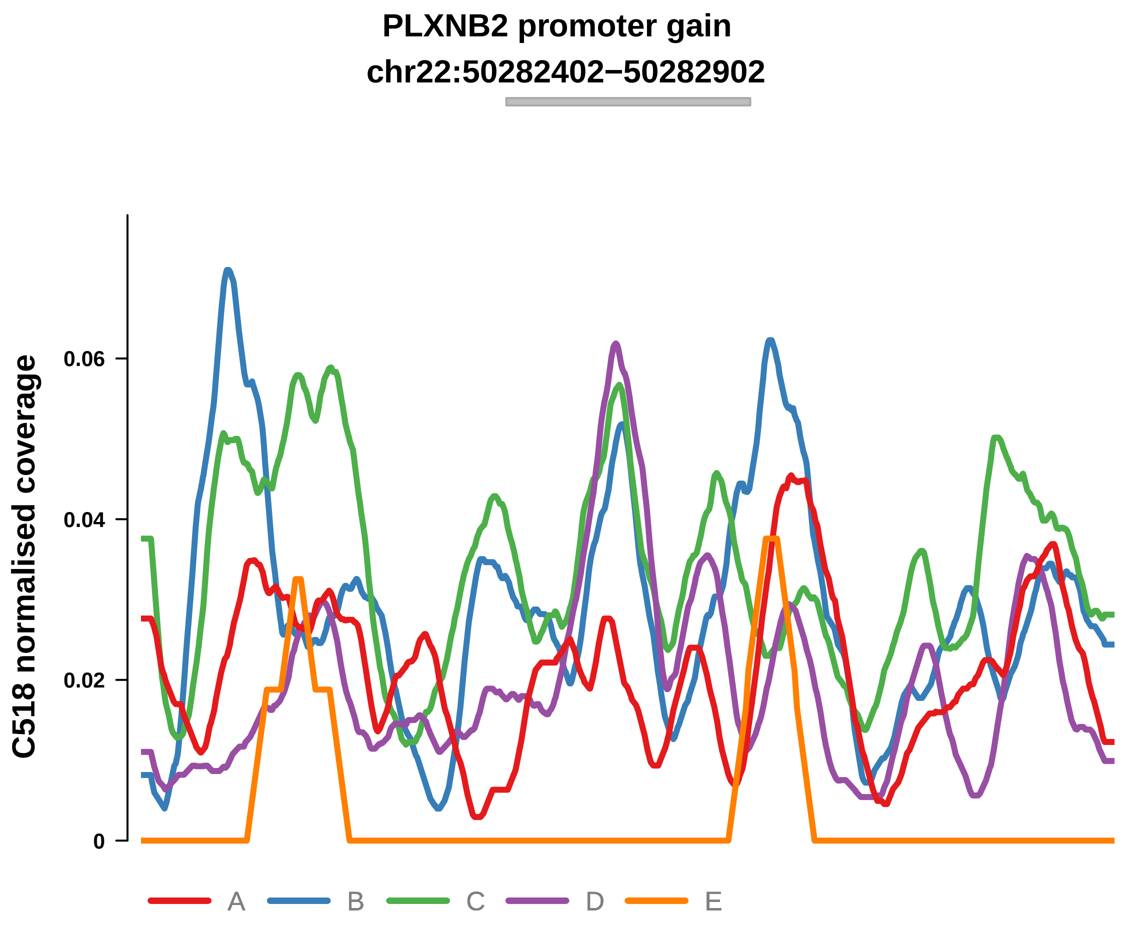


Putatively Clonal (logFC = 1.82)

| | P. | Adjusted P. |
|---|---|---|
| *~region vs. ~* | 3.07e−01 | 6.53e−01 |
| *~purity vs ~* | 3.23e−01 | 7.38e−01 |
| *~purity+region vs ~purity* | 4.66e−01 | 7.51e−01 |
| *Carcinoma vs. normal* | 1.55e−03 ** | 1.55e−03 ** |

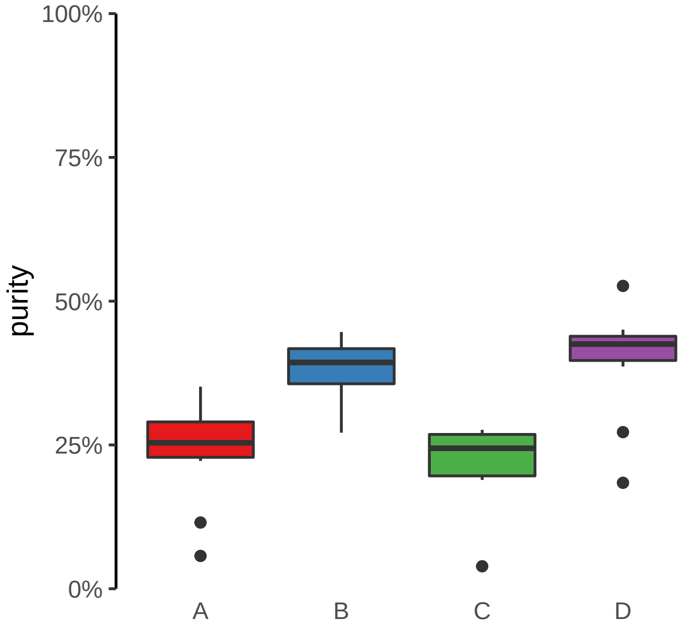

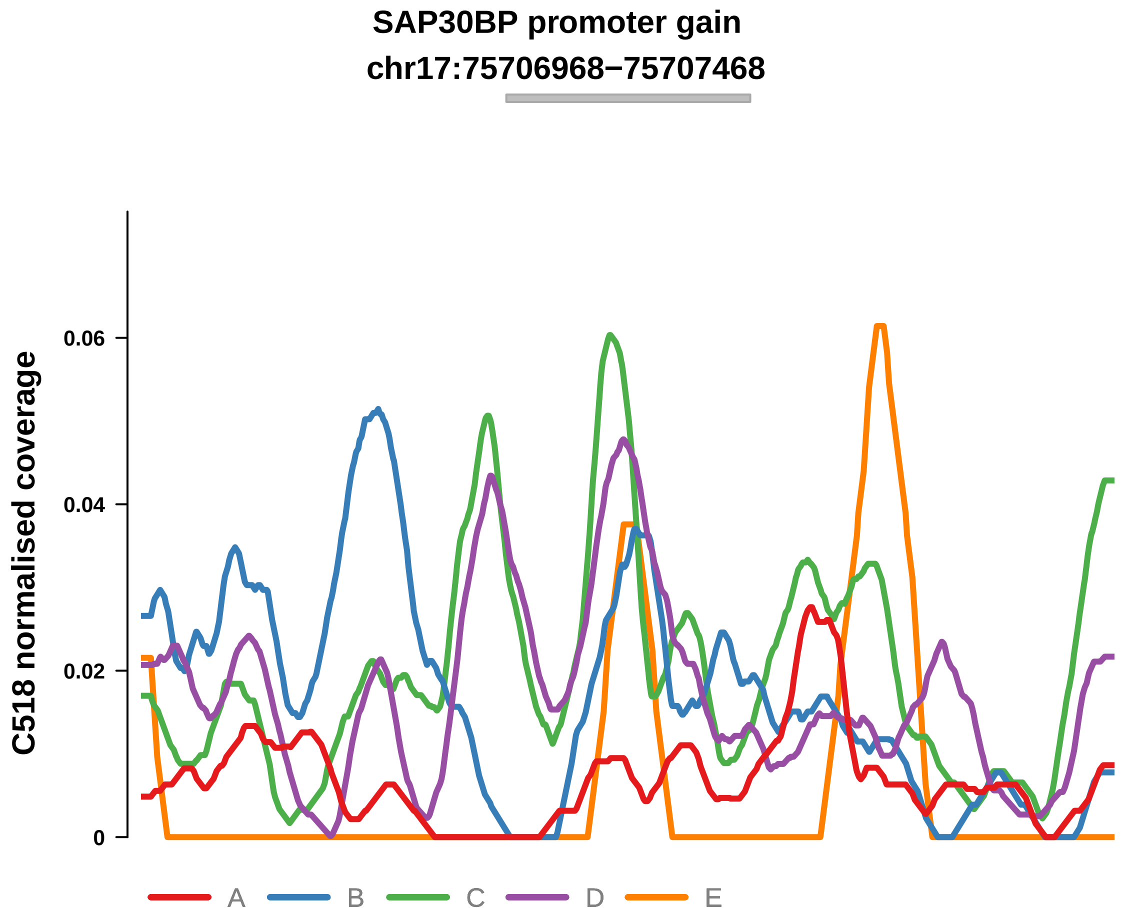


Putatively Clonal (logFC = 2.03)

| | P. | Adjusted P. |
|---|---|---|
| *~region vs. ~* | 1.96e−02 * | 1.57e−01 |
| *~purity vs ~* | 4.93e−01 | 8.48e−01 |
| *~purity+region vs ~purity* | 7.65e−03 ** | 1.12e−01 |
| *Carcinoma vs. normal* | 6.83e−04 *** | 6.83e−04 *** |

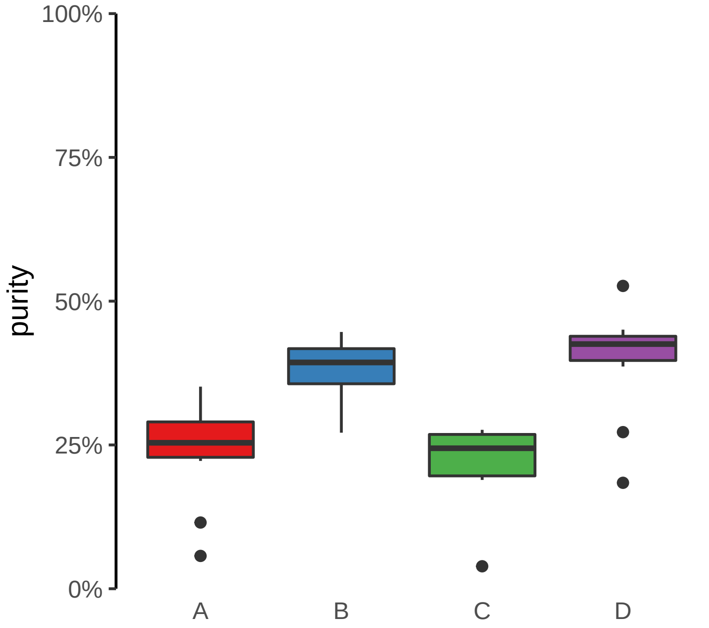

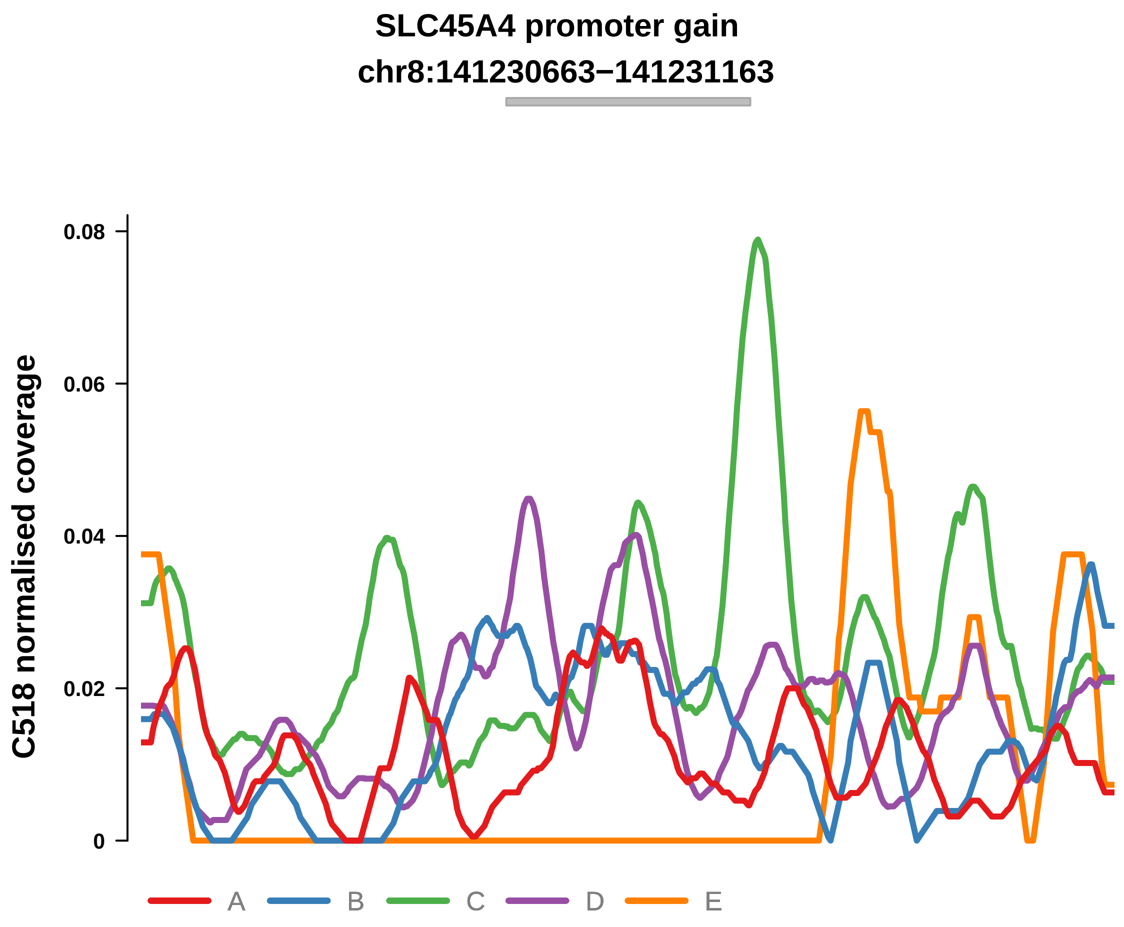


Putatively Clonal (logFC = 1.74)

| | P. | Adjusted P. |
|---|---|---|
| *~region vs. ~* | 1.25e−01 | 4.58e−01 |
| *~purity vs ~* | 4.19e−01 | 8.14e−01 |
| *~purity+region vs ~purity* | 1.6e−01 | 5.63e−01 |
| *Carcinoma vs. normal* | 4.21e−03 ** | 4.21e−03 ** |

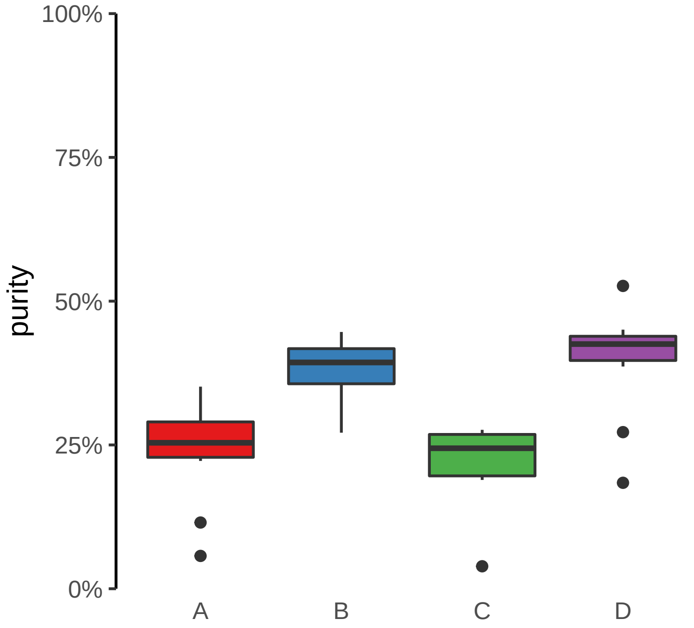

**ARHGEF17 promoter gain**

**chr11:73362044−73362544**

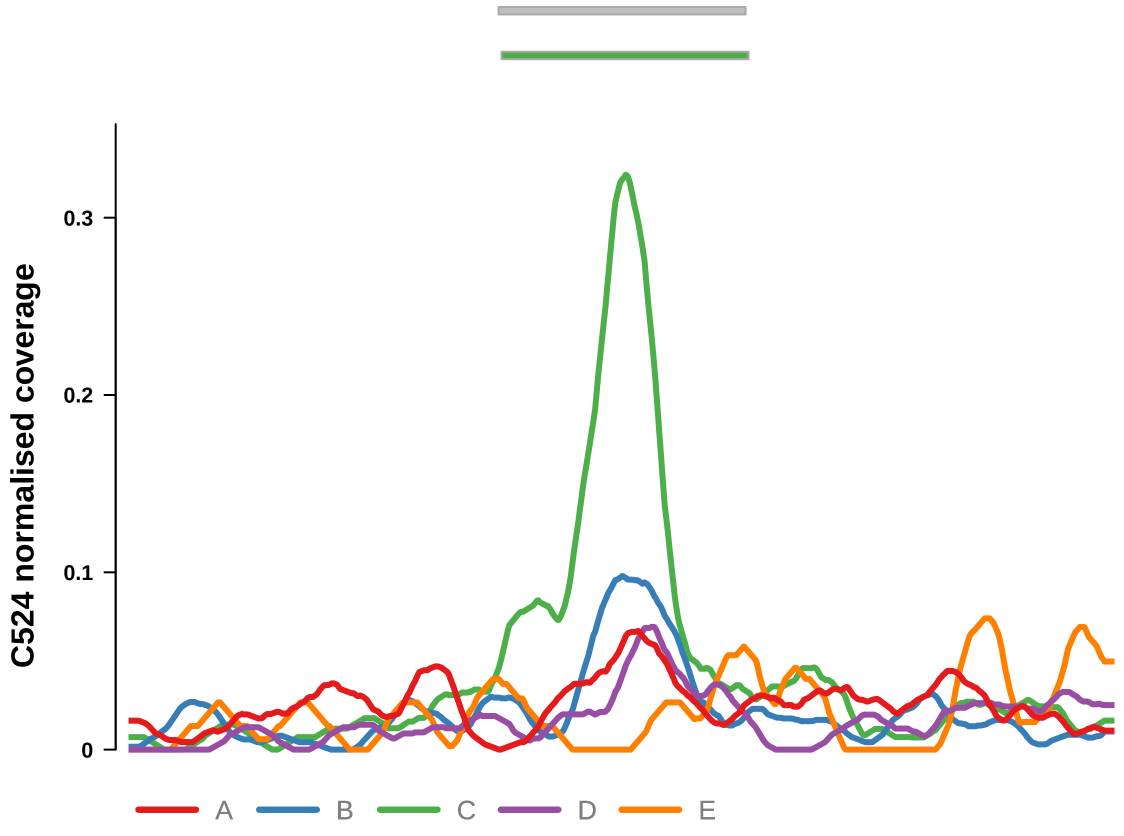


Putatively Clonal (logFC = 2.7)

| | P. | Adjusted P. |
|---|---|---|
| *~region vs. ~* | 3.21e−01 | 6.83e−01 |
| *~purity vs ~* | 9.49e−03 ** | 2.07e−01 |
| *~purity+region vs ~purity* | 6.2e−01 | 8.8e−01 |
| *Carcinoma vs. normal* | 1.62e−05 **** | 1.62e−05 **** |

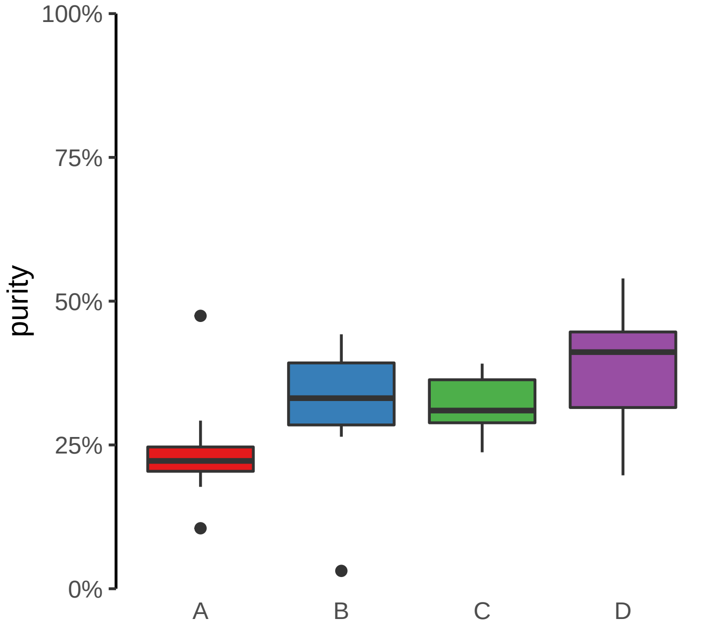

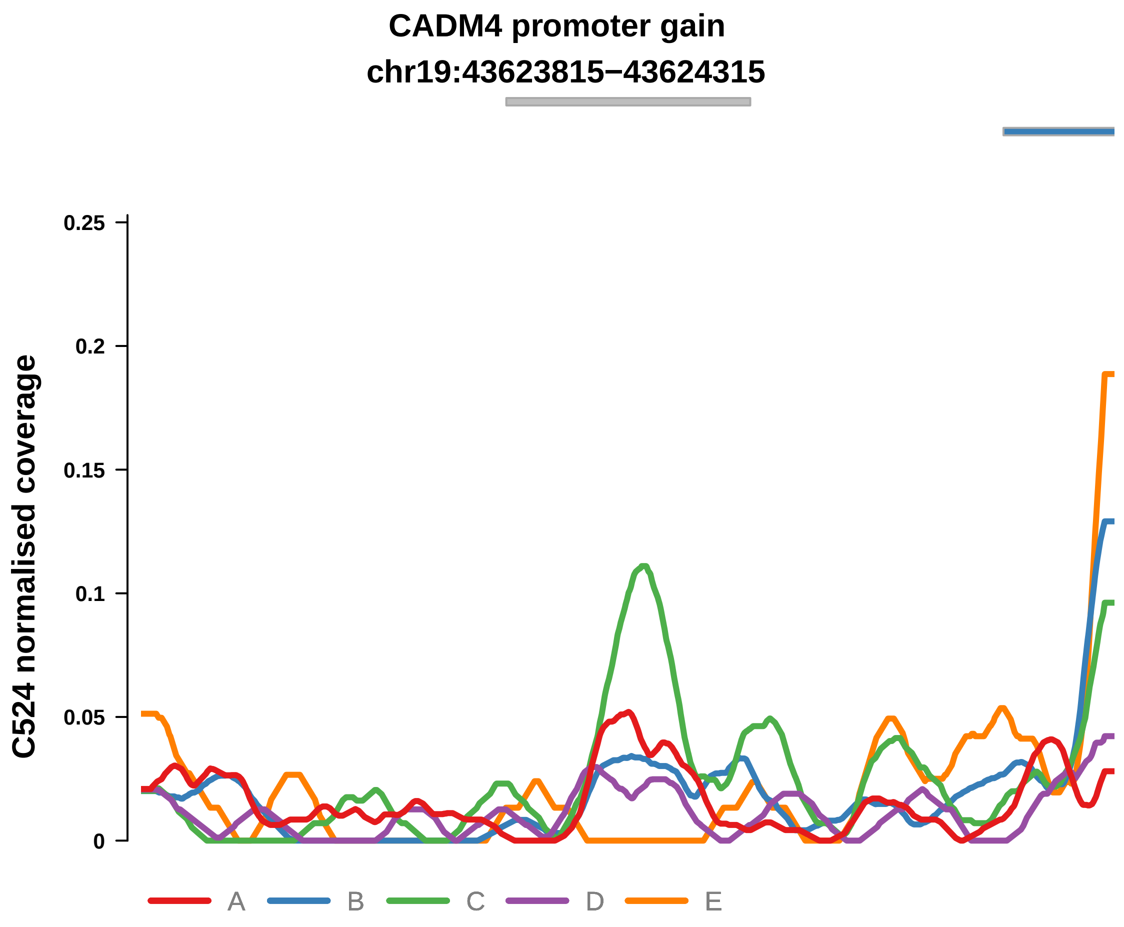


Putatively Clonal (logFC = 2.03)

| | P. | Adjusted P. |
|---|---|---|
| *~region vs. ~* | 7.52e−01 | 8.94e−01 |
| *~purity vs ~* | 5.82e−02 | 3.66e−01 |
| *~purity+region vs ~purity* | 9.83e−01 | 9.87e−01 |
| *Carcinoma vs. normal* | 2.19e−03 ** | 2.19e−03 ** |

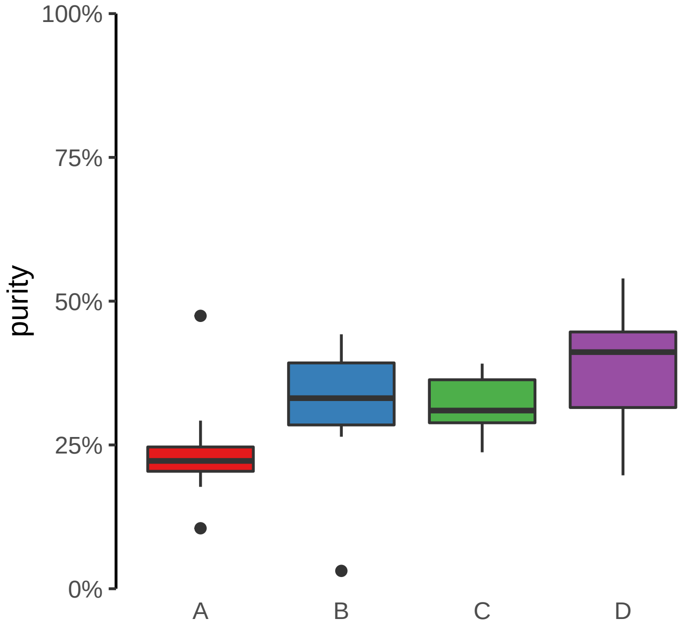

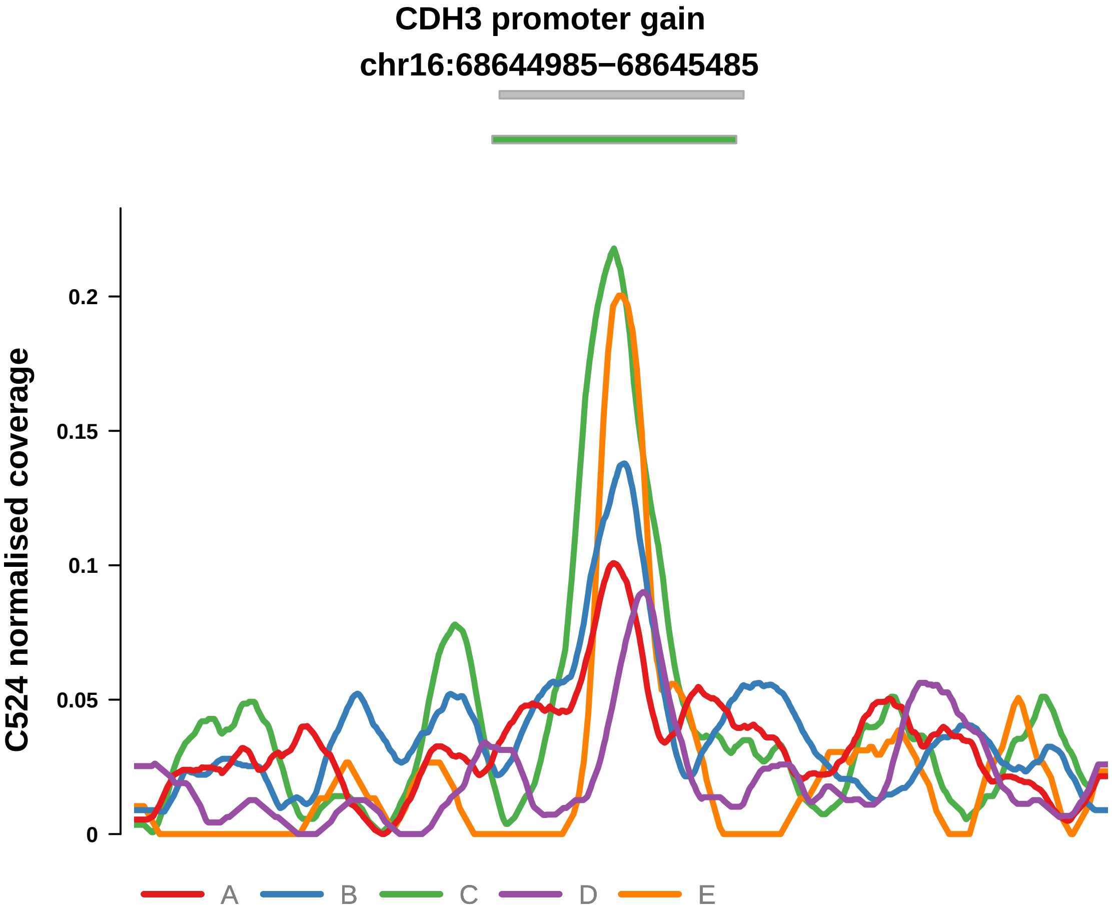


Putatively Clonal (logFC = 2.3)

| | P. | Adjusted P. |
|---|---|---|
| *~region vs. ~* | 4.05e−01 | 7.84e−01 |
| *~purity vs ~* | 1.41e−02 * | 2.07e−01 |
| *~purity+region vs ~purity* | 3.18e−01 | 6.8e−01 |
| *Carcinoma vs. normal* | 8.97e−05 **** | 8.97e−05 **** |

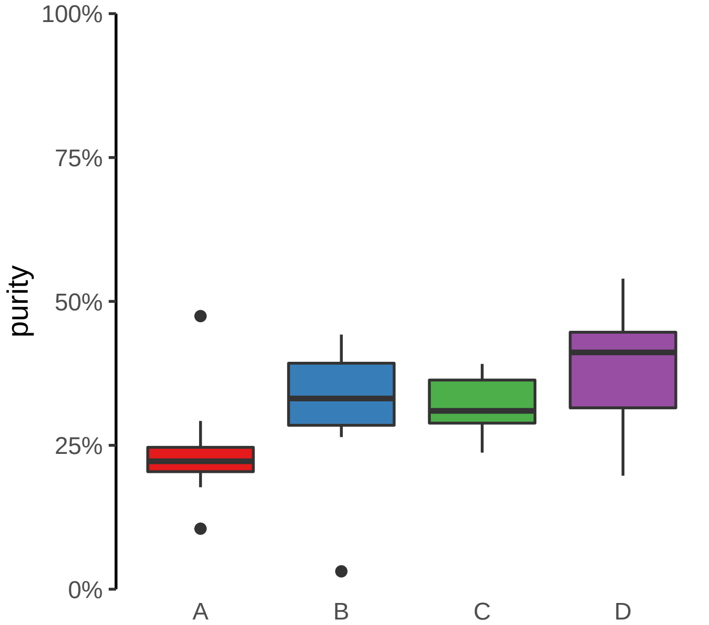

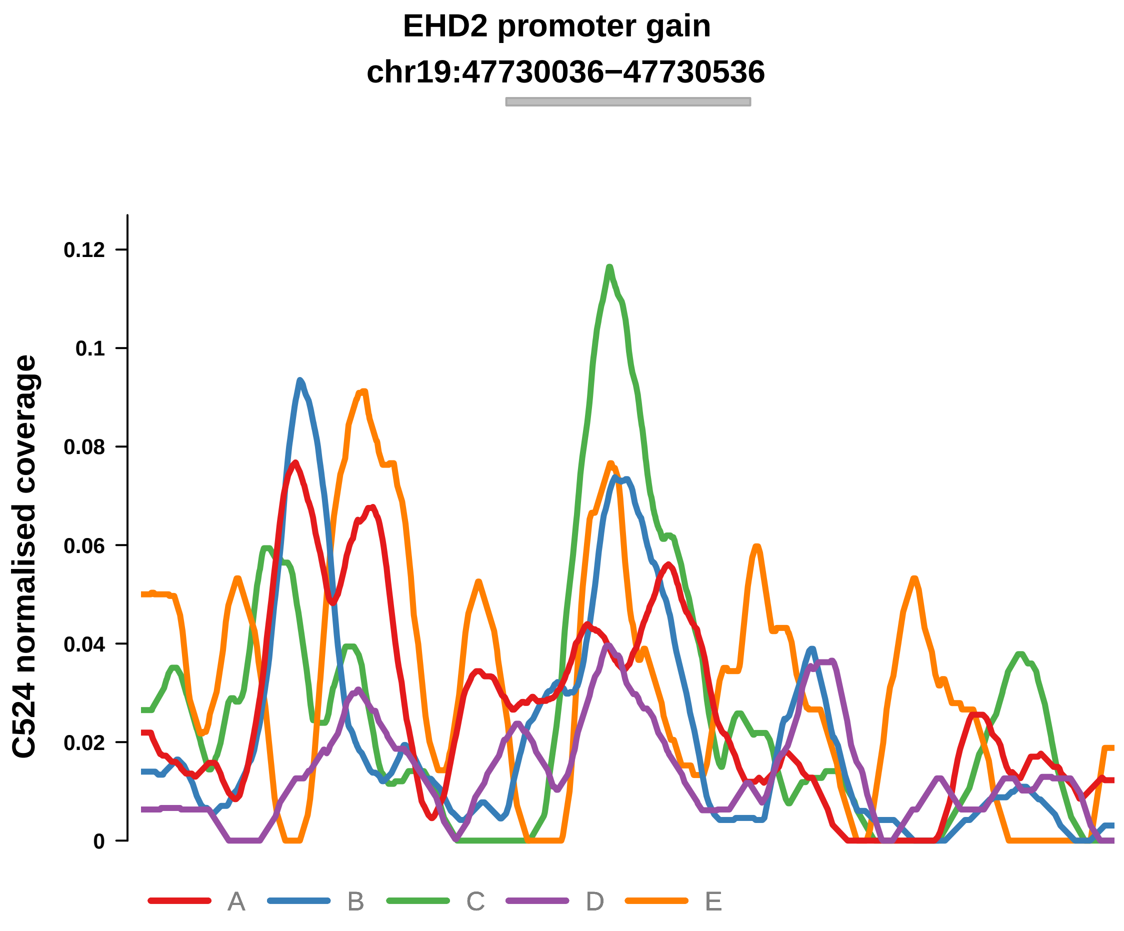


Putatively Clonal (logFC = 1.85)

| | P. | Adjusted P. |
|---|---|---|
| *~region vs. ~* | 7.94e−01 | 9.07e−01 |
| *~purity vs ~* | 9.14e−02 | 4.3e−01 |
| *~purity+region vs ~purity* | 6.96e−01 | 8.8e−01 |
| *Carcinoma vs. normal* | 2.53e−03 ** | 2.53e−03 ** |

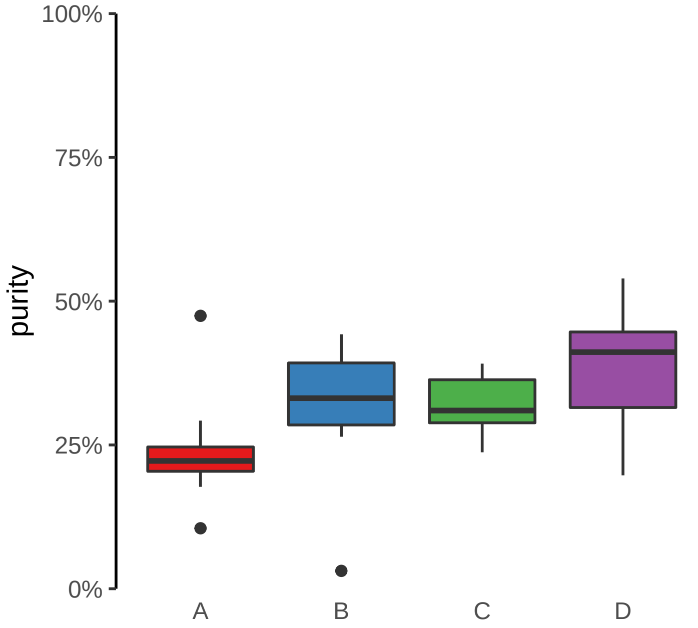

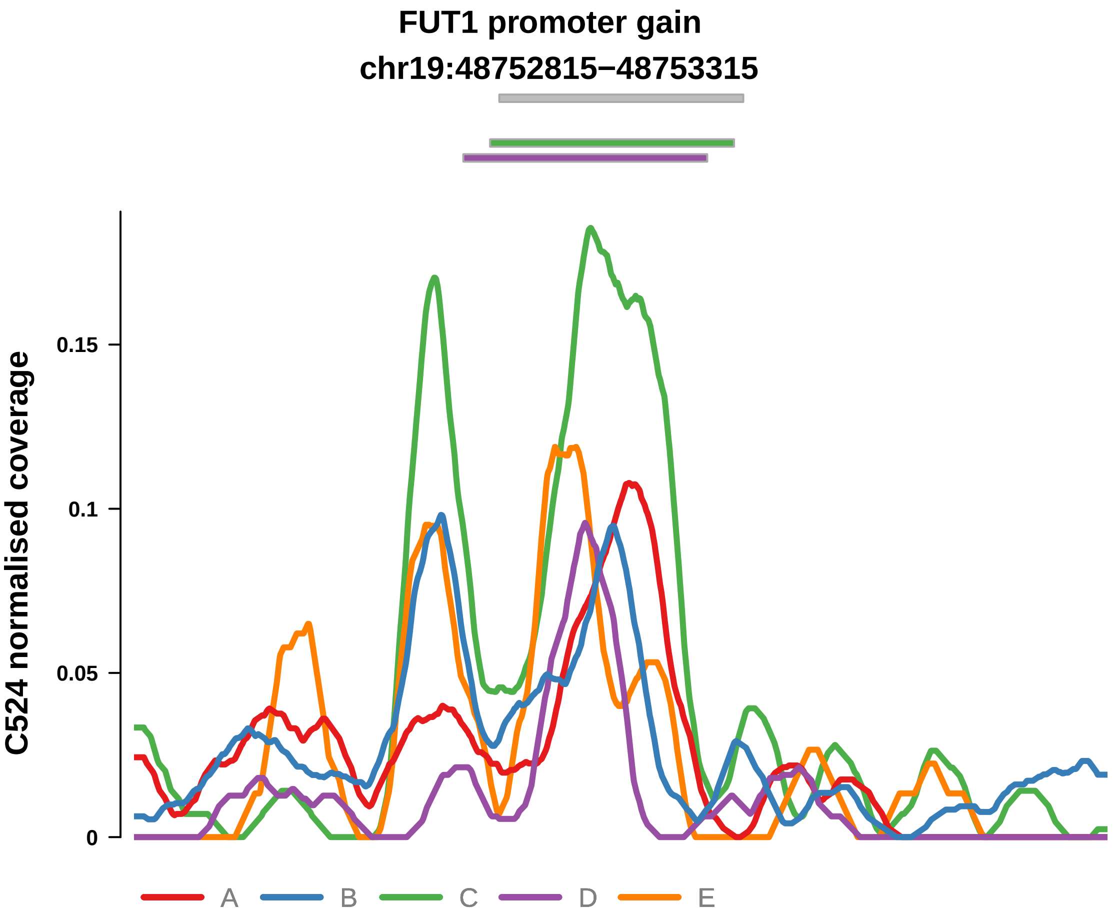


Putatively Clonal (logFC = 2.27)

| | P. | Adjusted P. |
|---|---|---|
| *~region vs. ~* | 6.02e−01 | 8.46e−01 |
| *~purity vs ~* | 3.64e−02 * | 2.8e−01 |
| *~purity+region vs ~purity* | 8.47e−01 | 9.68e−01 |
| *Carcinoma vs. normal* | 1.28e−04 *** | 1.28e−04 *** |

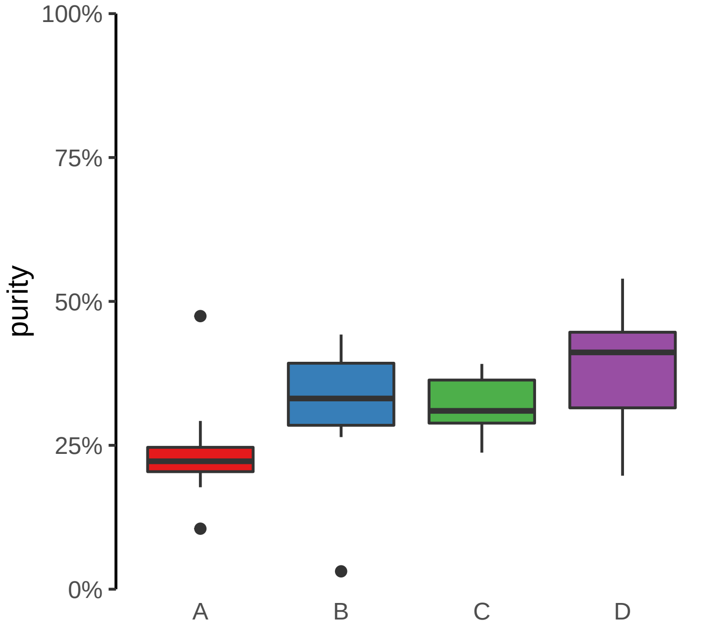

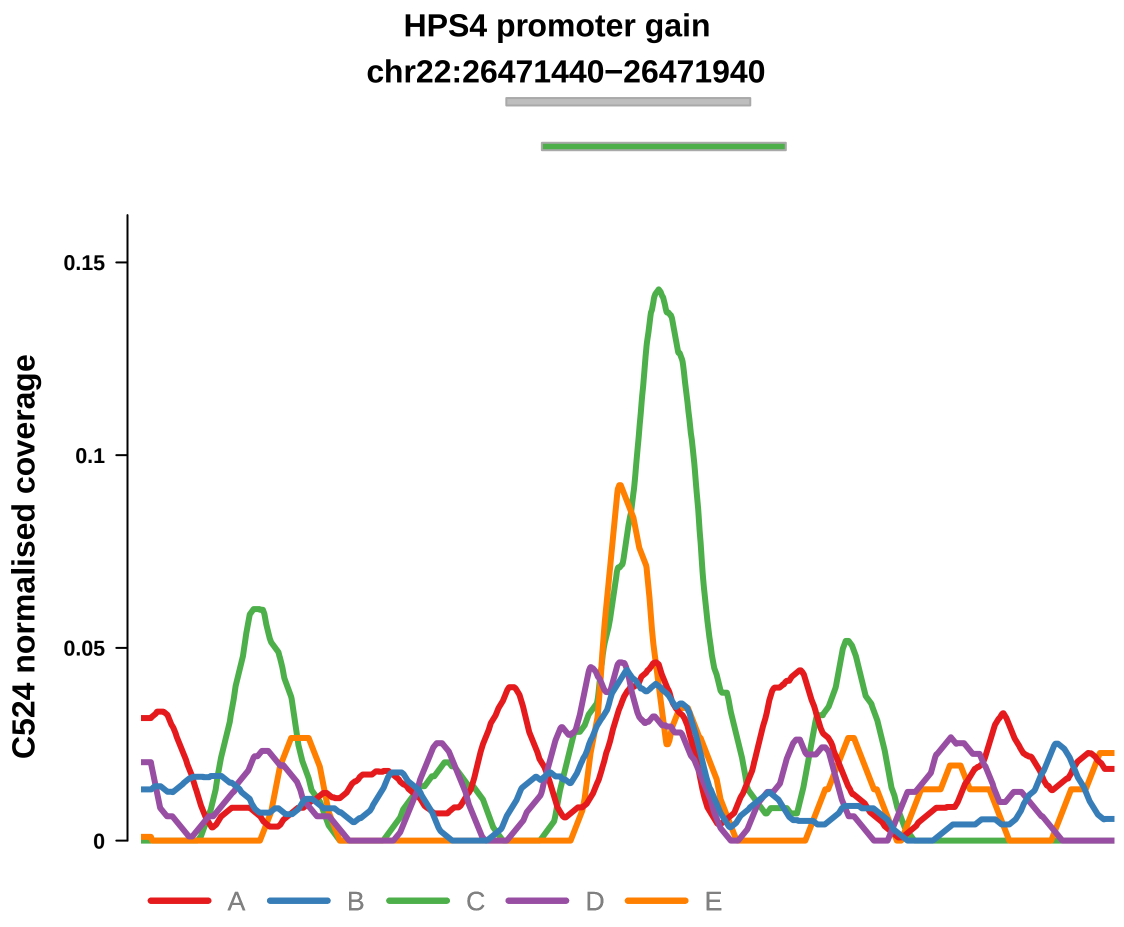


Putatively Clonal (logFC = 1.97)

| | P. | Adjusted P. |
|---|---|---|
| *~region vs. ~* | 6.66e−01 | 8.5e−01 |
| *~purity vs ~* | 1.91e−01 | 5e−01 |
| *~purity+region vs ~purity* | 5.28e−01 | 8.46e−01 |
| *Carcinoma vs. normal* | 4.55e−03 ** | 4.55e−03 ** |

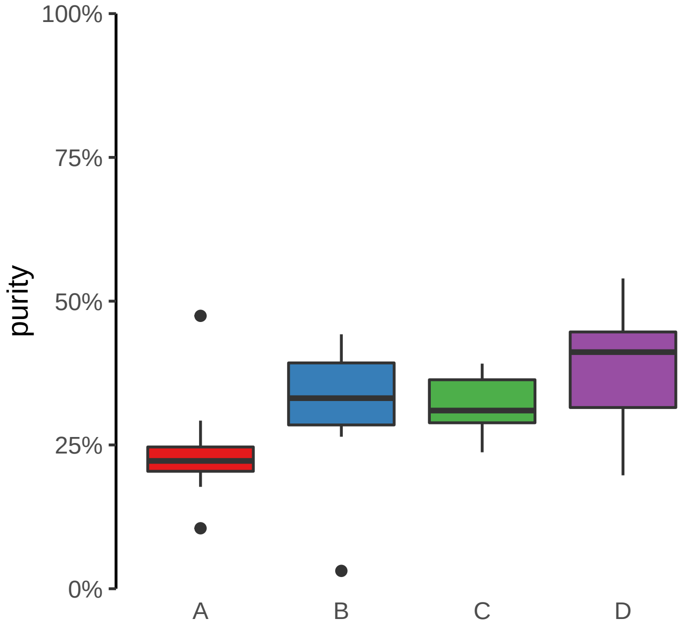

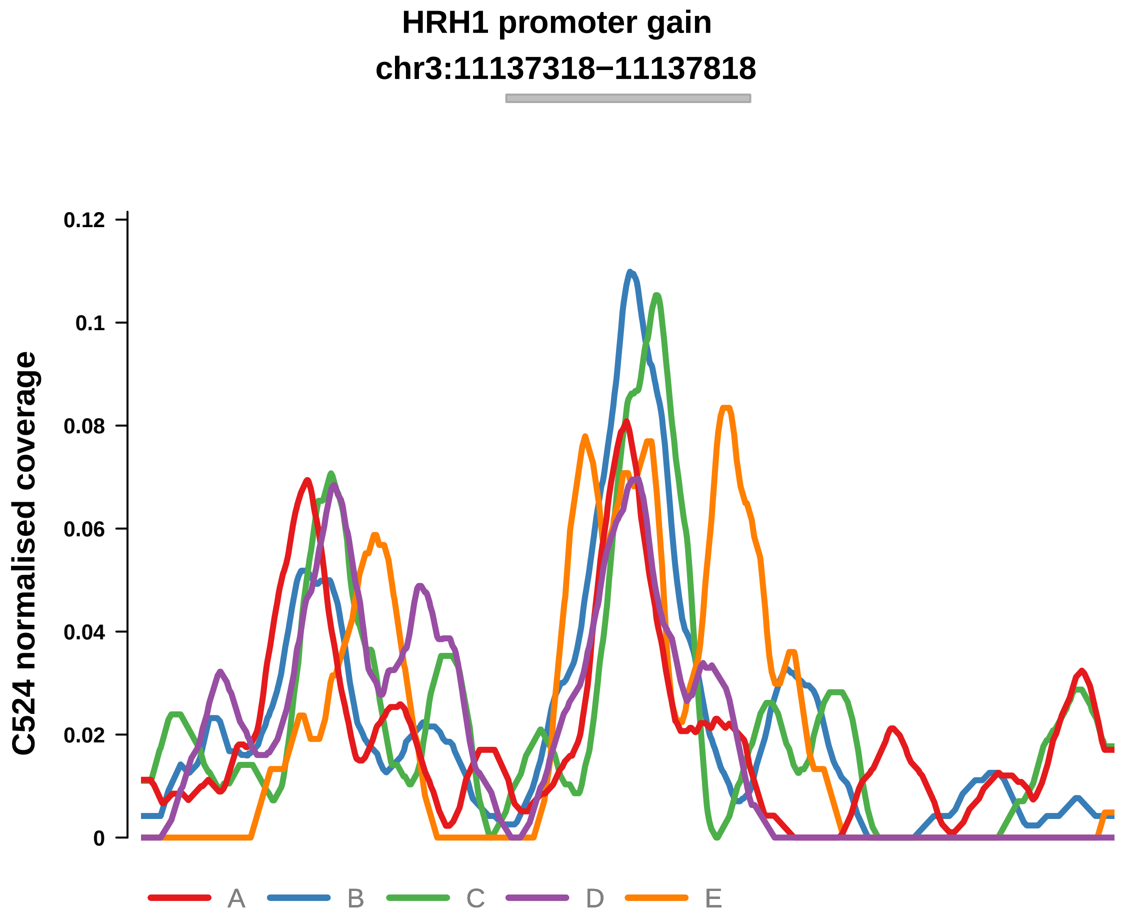


Putatively Clonal (logFC = 2.52)

| | P. | Adjusted P. |
|---|---|---|
| *~region vs. ~* | 5.93e−03 ** | 1.74e−01 |
| *~purity vs ~* | 8.92e−02 | 4.3e−01 |
| *~purity+region vs ~purity* | 1.49e−02 * | 1.87e−01 |
| *Carcinoma vs. normal* | 4.31e−05 **** | 4.31e−05 **** |

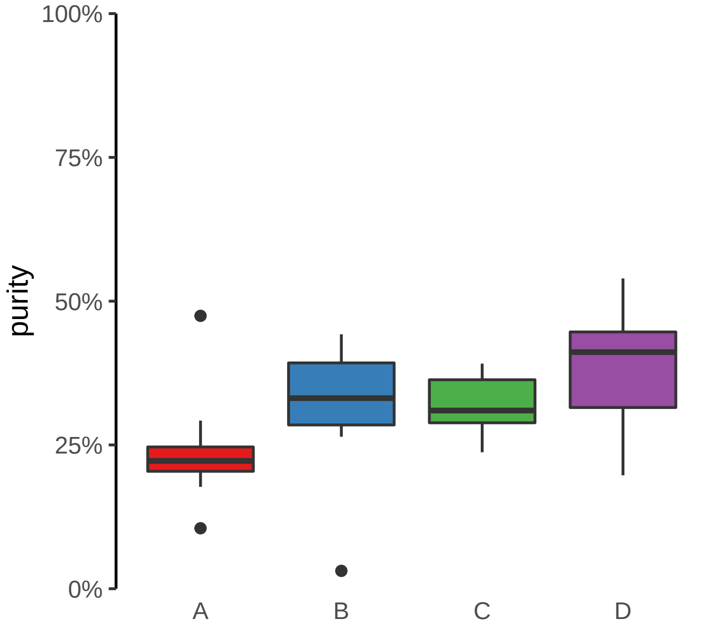

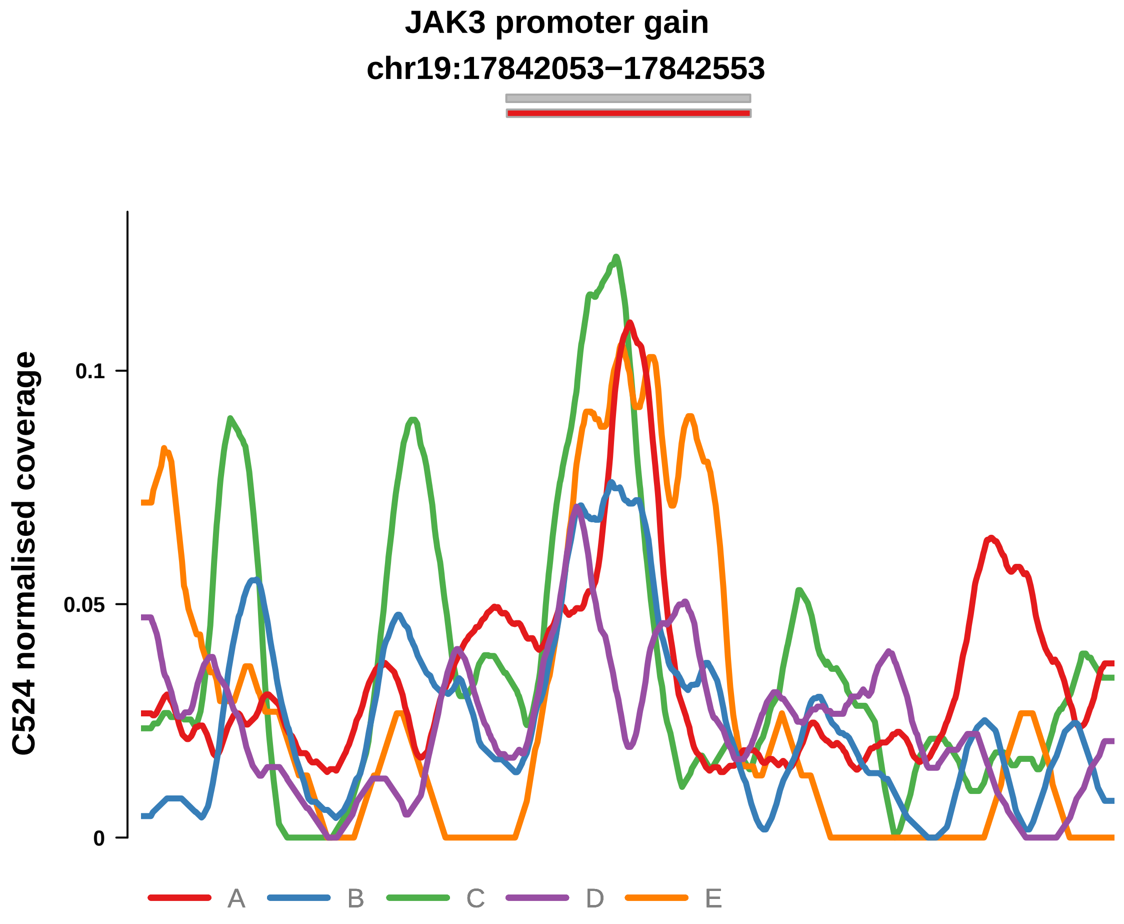


Putatively Clonal (logFC = 2.05)

| | P. | Adjusted P. |
|---|---|---|
| *~region vs. ~* | 1.81e−02 * | 3.17e−01 |
| *~purity vs ~* | 1.31e−01 | 5e−01 |
| *~purity+region vs ~purity* | 1.6e−02 * | 1.87e−01 |
| *Carcinoma vs. normal* | 5.59e−04 *** | 5.59e−04 *** |

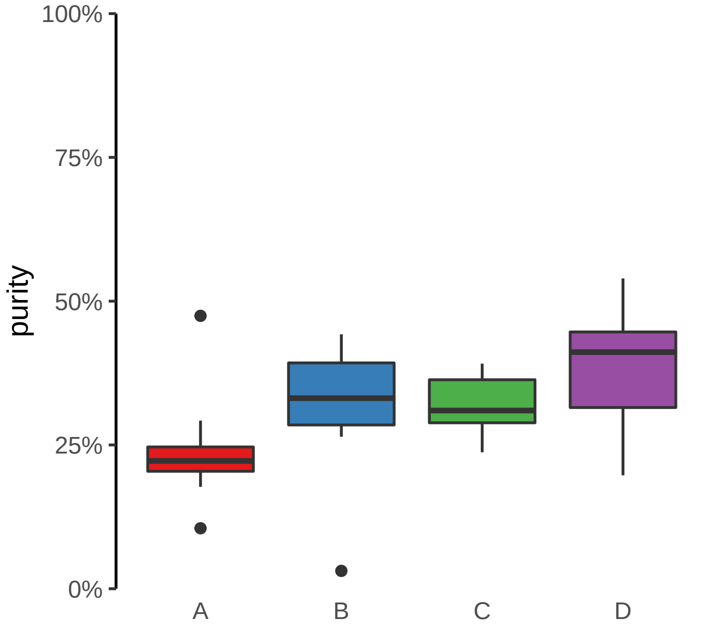

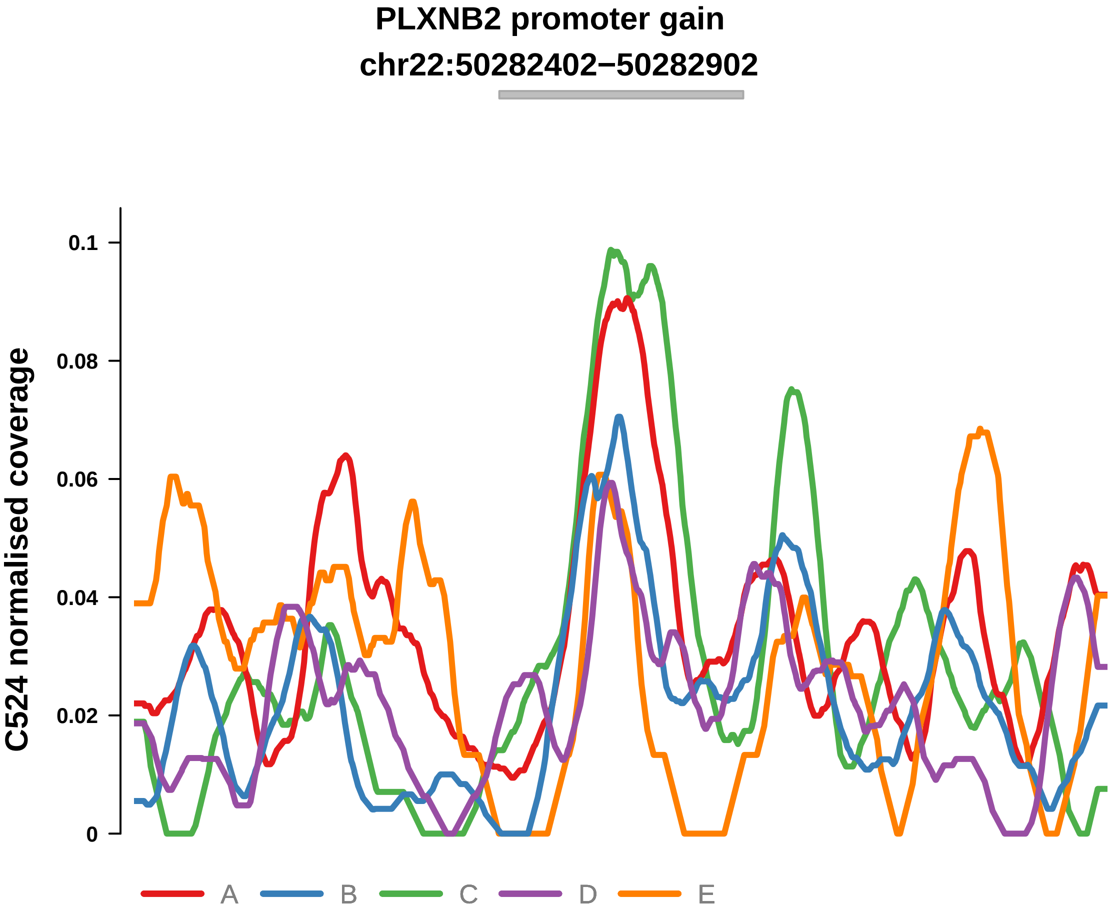


Putatively Clonal (logFC = 1.67)

| | P. | Adjusted P. |
|---|---|---|
| *~region vs. ~* | 4.92e−01 | 8.01e−01 |
| *~purity vs ~* | 3.28e−01 | 6.23e−01 |
| *~purity+region vs ~purity* | 5.75e−01 | 8.8e−01 |
| *Carcinoma vs. normal* | 9e−03 ** | 9e−03 ** |

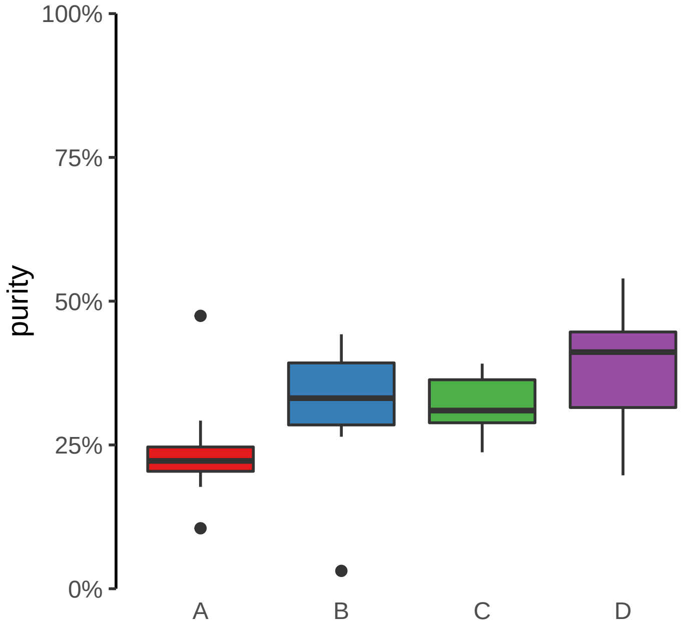

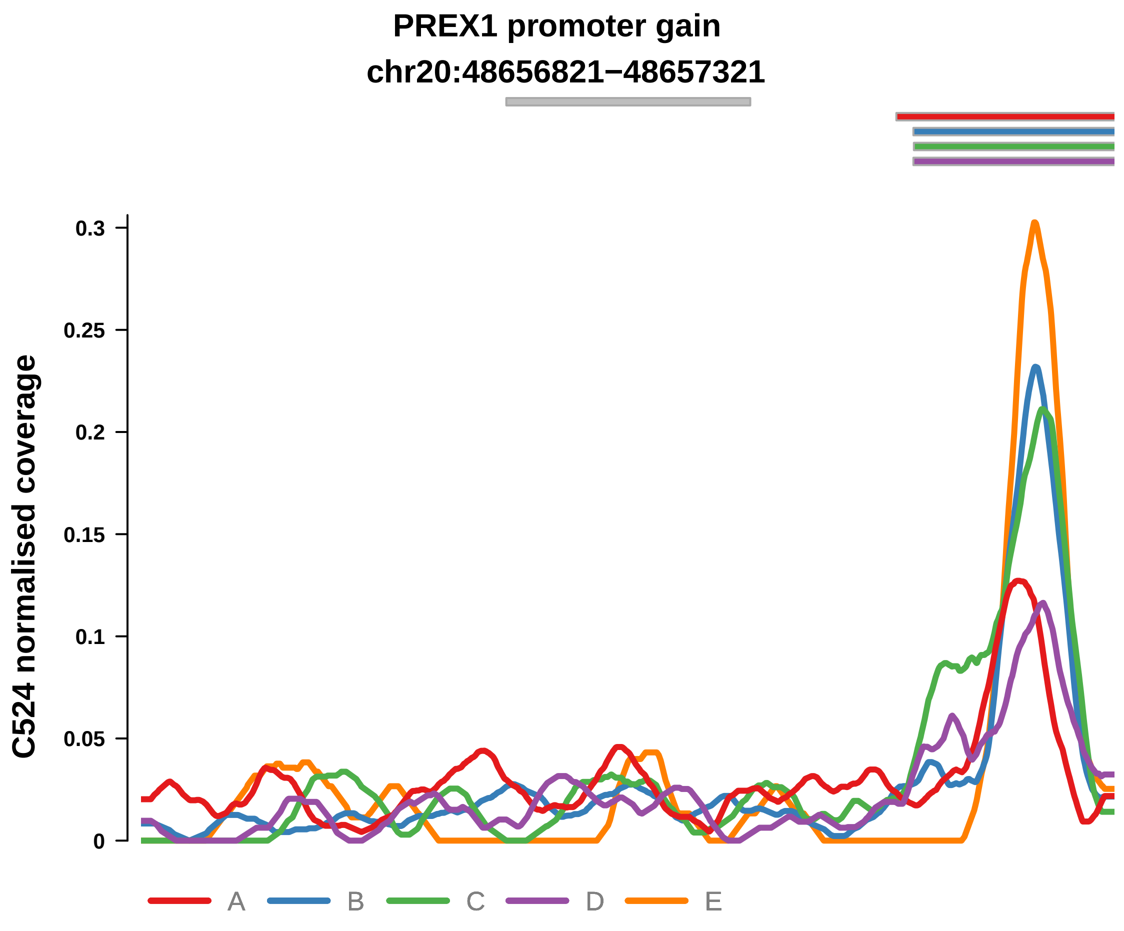


Putatively Clonal (logFC = 2.11)

| | P. | Adjusted P. |
|---|---|---|
| *~region vs. ~* | 8e−02 | 4.97e−01 |
| *~purity vs ~* | 5.37e−01 | 7.12e−01 |
| *~purity+region vs ~purity* | 7.34e−02 | 3.98e−01 |
| *Carcinoma vs. normal* | 1.45e−03 ** | 1.45e−03 ** |

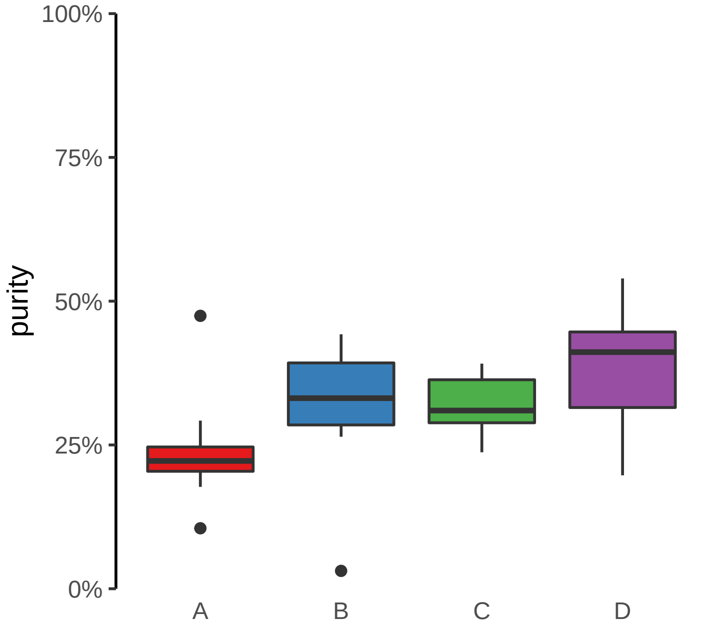

**PRR36 promoter gain**
**chr19:7870611−7871111**

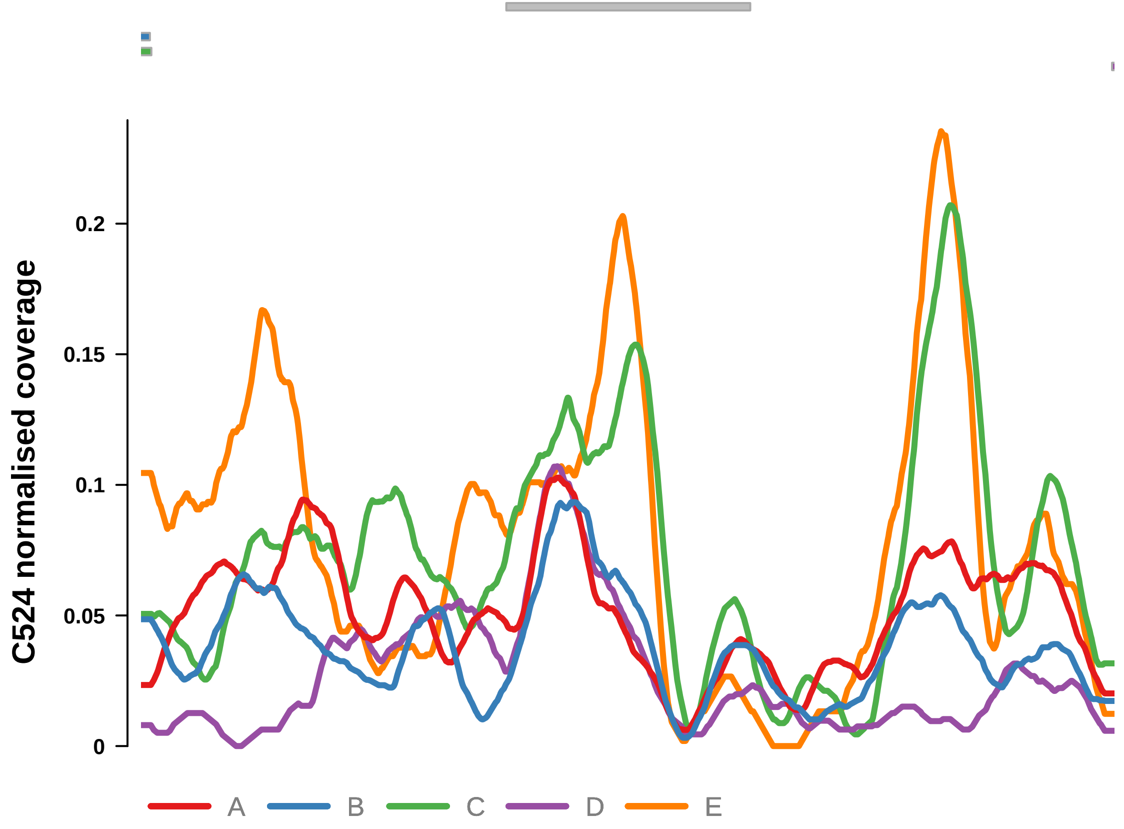


Putatively Clonal (logFC = 1.81)

| | P. | Adjusted P. |
|---|---|---|
| *~region vs. ~* | 3.27e−01 | 6.83e−01 |
| *~purity vs ~* | 1.41e−01 | 5e−01 |
| *~purity+region vs ~purity* | 4.04e−01 | 7.48e−01 |
| *Carcinoma vs. normal* | 2.45e−03 ** | 2.45e−03 ** |

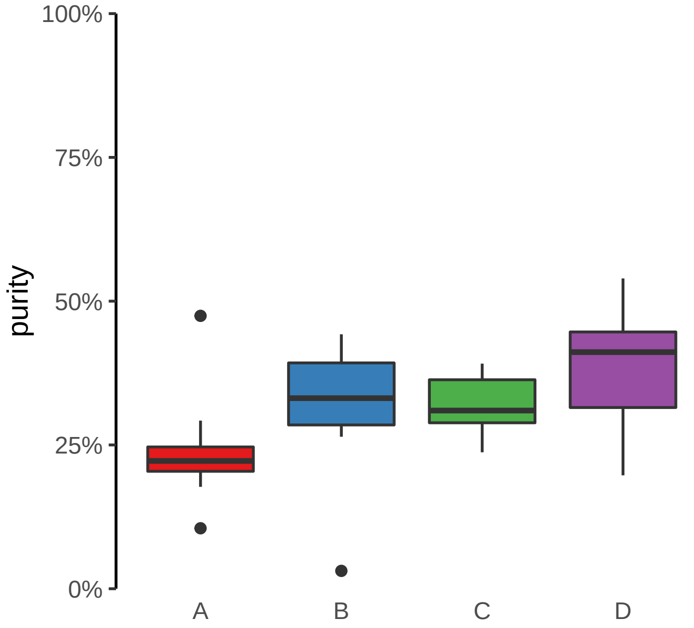

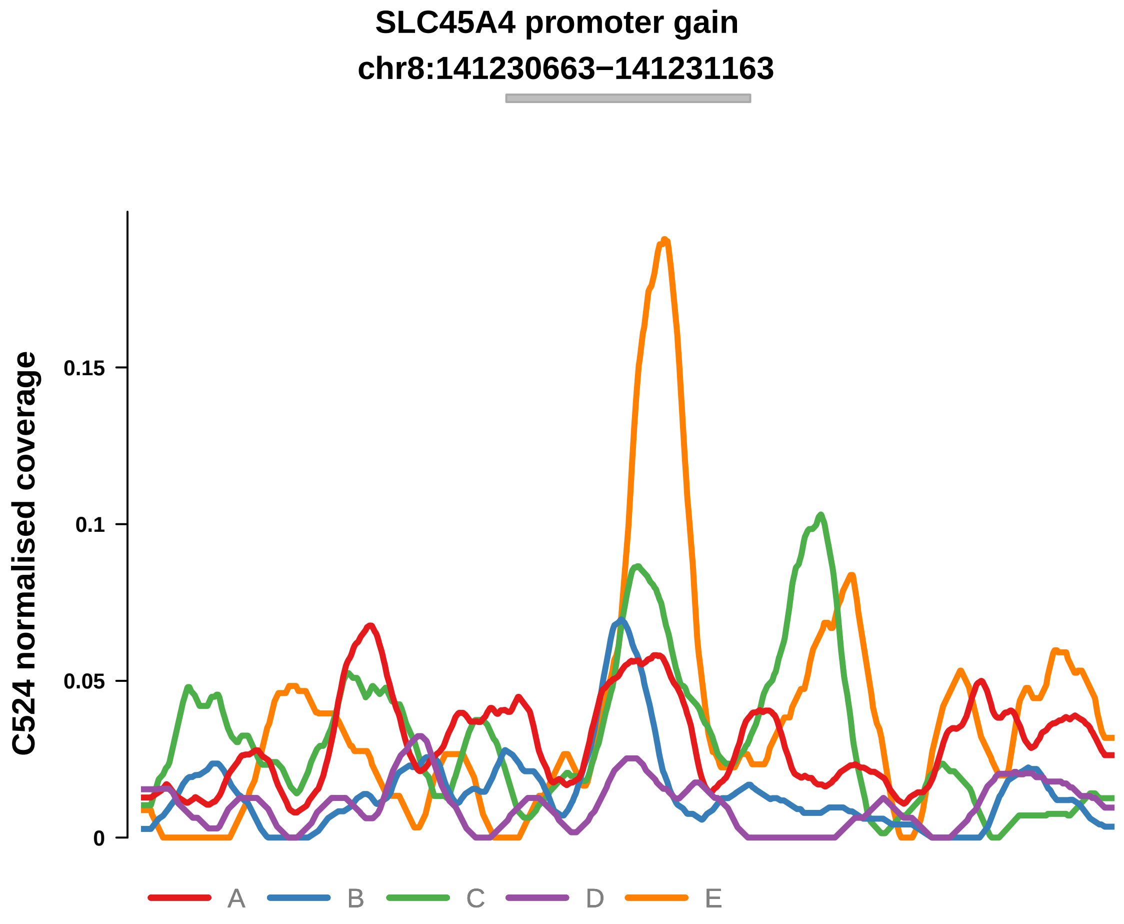


Putatively Clonal (logFC = 1.93)

| | P. | Adjusted P. |
|---|---|---|
| *~region vs. ~* | 5.3e−01 | 8.11e−01 |
| *~purity vs ~* | 8.33e−01 | 9.04e−01 |
| *~purity+region vs ~purity* | 4.08e−01 | 7.48e−01 |
| *Carcinoma vs. normal* | 3.92e−03 ** | 3.92e−03 ** |

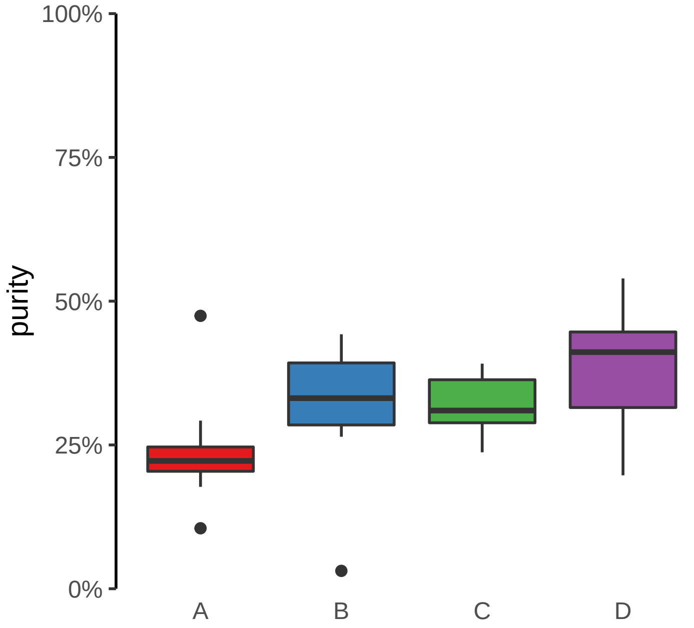

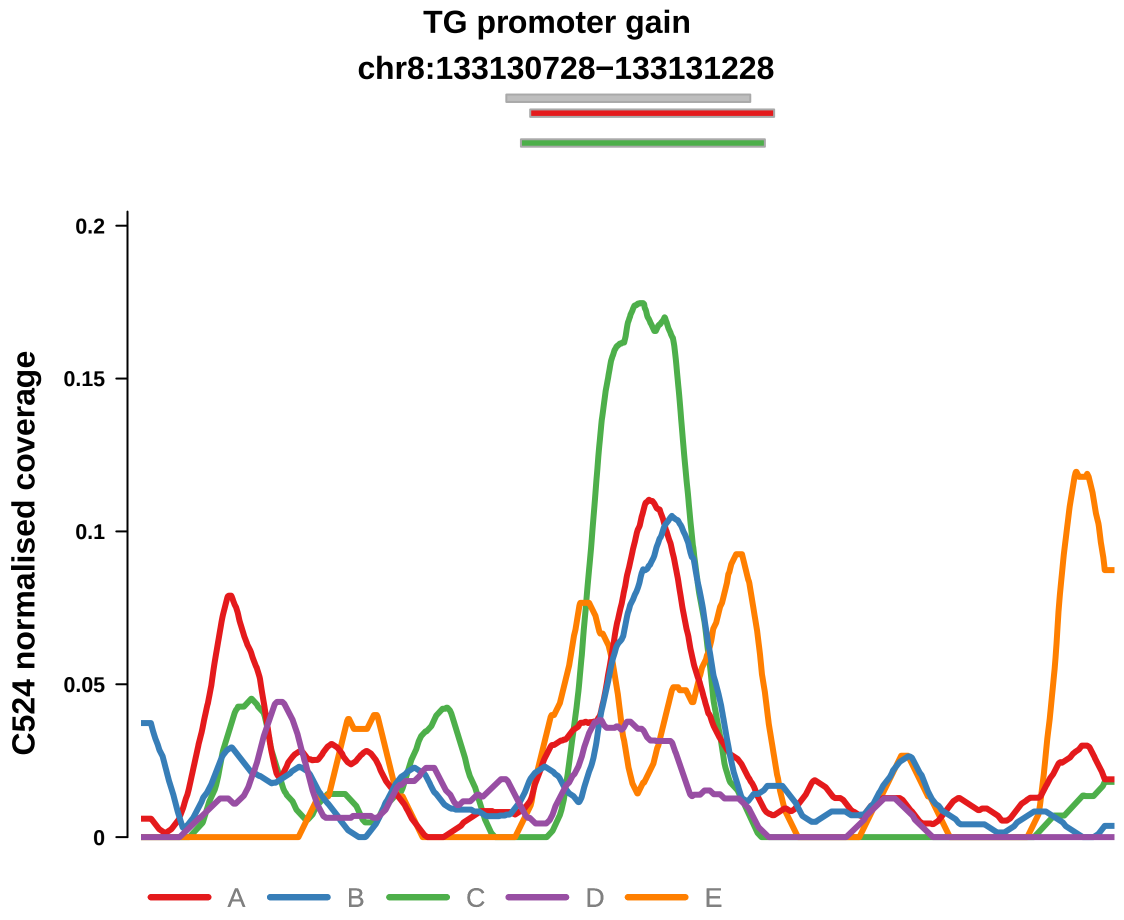


Putatively Clonal (logFC = 2.67)

| | P. | Adjusted P. |
|---|---|---|
| *~region vs. ~* | 6.6e−01 | 8.5e−01 |
| *~purity vs ~* | 3.82e−02 * | 2.8e−01 |
| *~purity+region vs ~purity* | 2.19e−01 | 6e−01 |
| *Carcinoma vs. normal* | 9.41e−06 **** | 9.41e−06 **** |

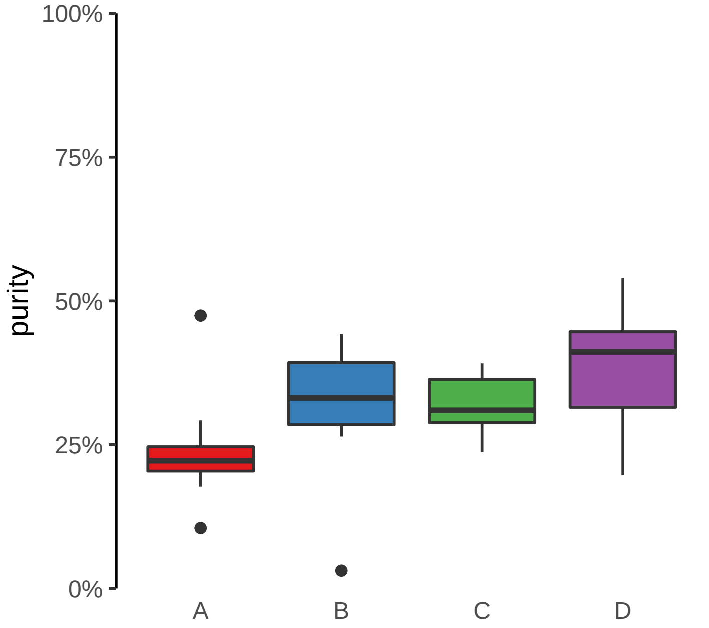

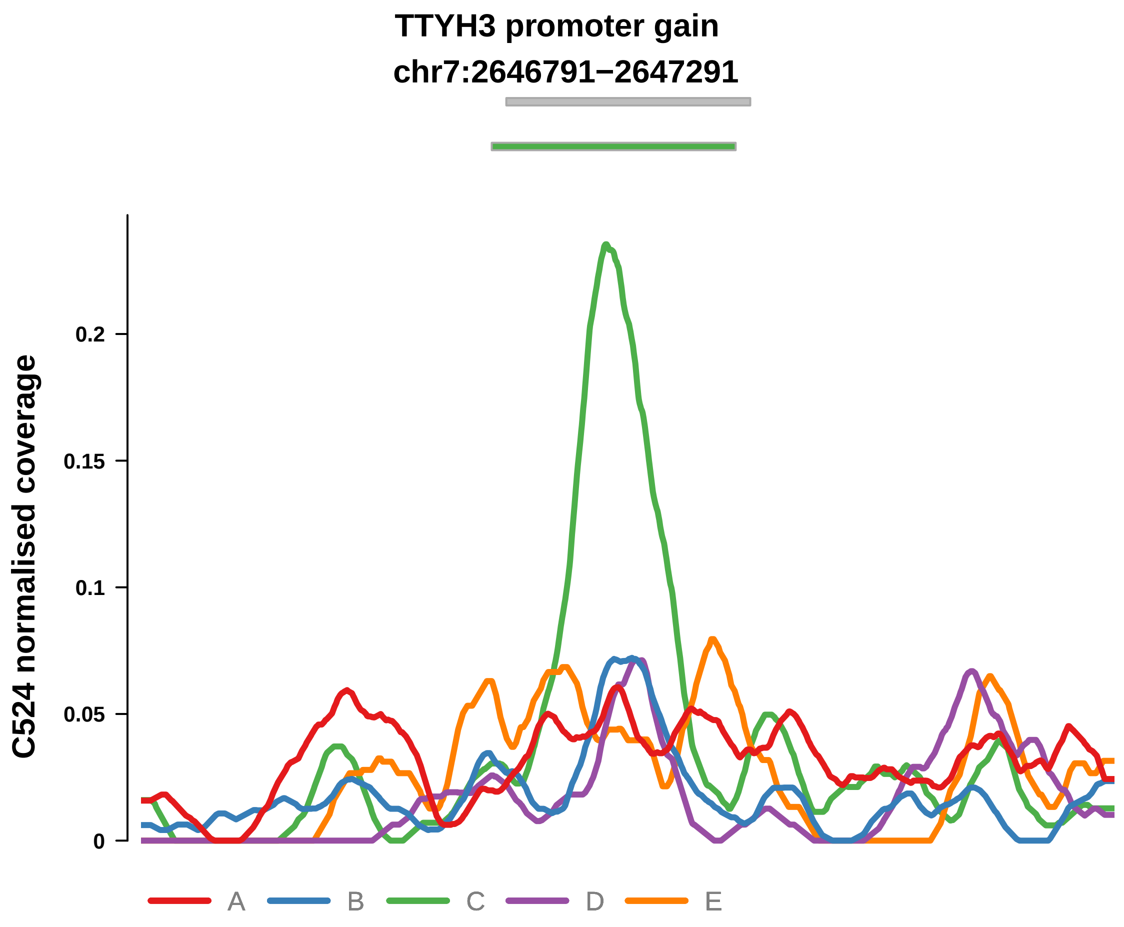


Putatively Clonal (logFC = 1.84)

| | P. | Adjusted P. |
|---|---|---|
| *~region vs. ~* | 7.24e−01 | 8.89e−01 |
| *~purity vs ~* | 7.11e−01 | 8.03e−01 |
| *~purity+region vs ~purity* | 5.94e−01 | 8.8e−01 |
| *Carcinoma vs. normal* | 4.69e−03 ** | 4.69e−03 ** |

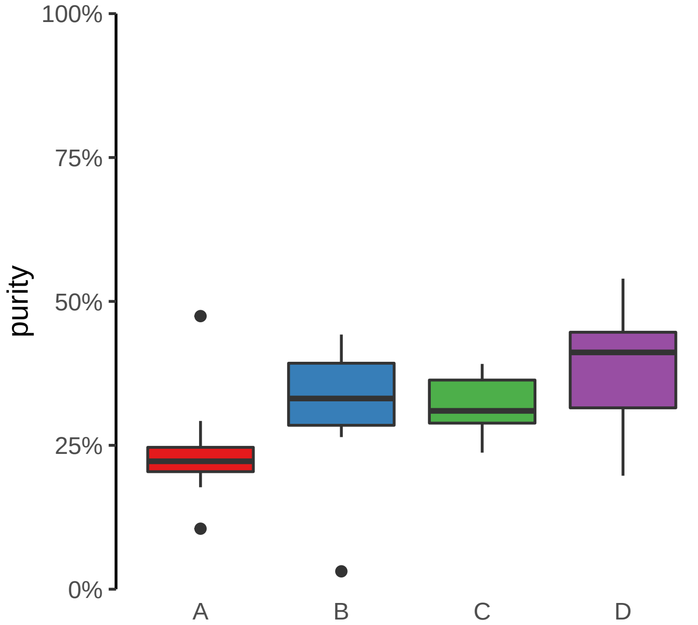

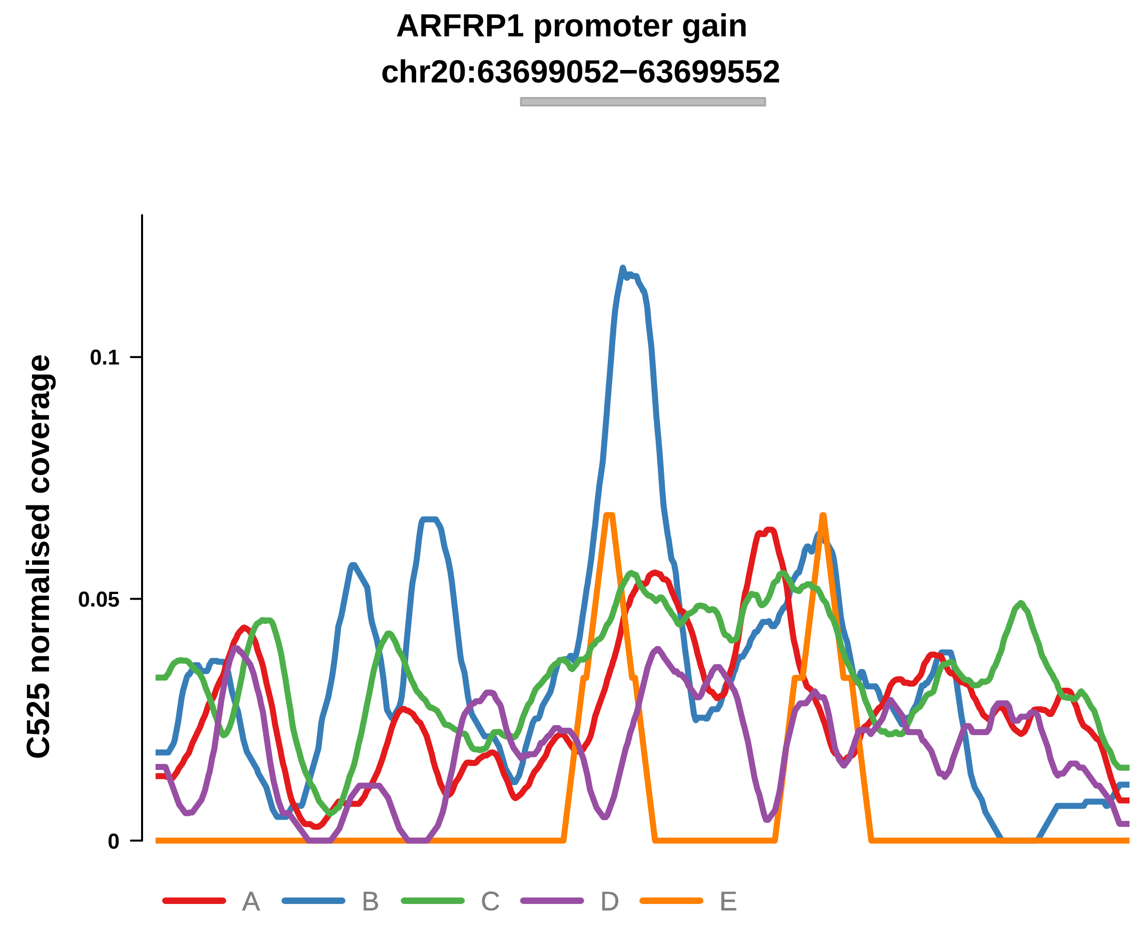


Putatively Clonal (logFC = 2.14)

| | P. | Adjusted P. |
|---|---|---|
| *~region vs. ~* | 2.83e−02 * | 6.41e−02 |
| *~purity vs ~* | 2.25e−01 | 7.37e−01 |
| *~purity+region vs ~purity* | 4.58e−02 * | 9.58e−02 |
| *Carcinoma vs. normal* | 3.45e−04 *** | 3.45e−04 *** |

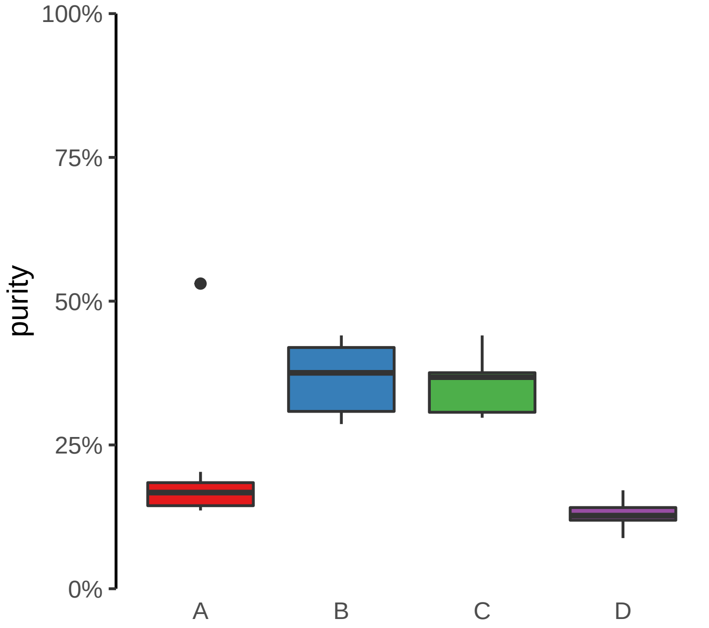

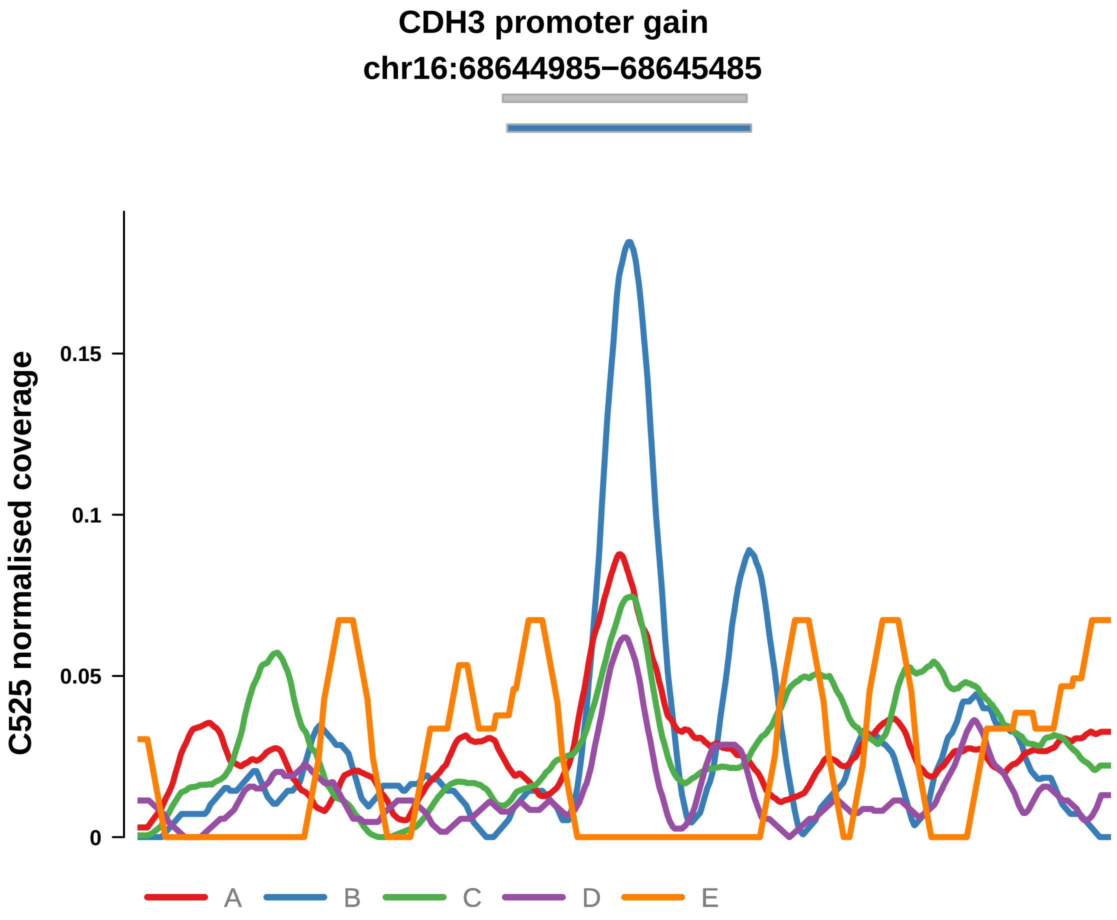


Putatively Clonal (logFC = 1.94)

| | P. | Adjusted P. |
|---|---|---|
| *~region vs. ~* | 5.58e−01 | 6.16e−01 |
| *~purity vs ~* | 2.4e−01 | 7.37e−01 |
| *~purity+region vs ~purity* | 5.26e−01 | 5.79e−01 |
| *Carcinoma vs. normal* | 8.51e−04 *** | 8.51e−04 *** |

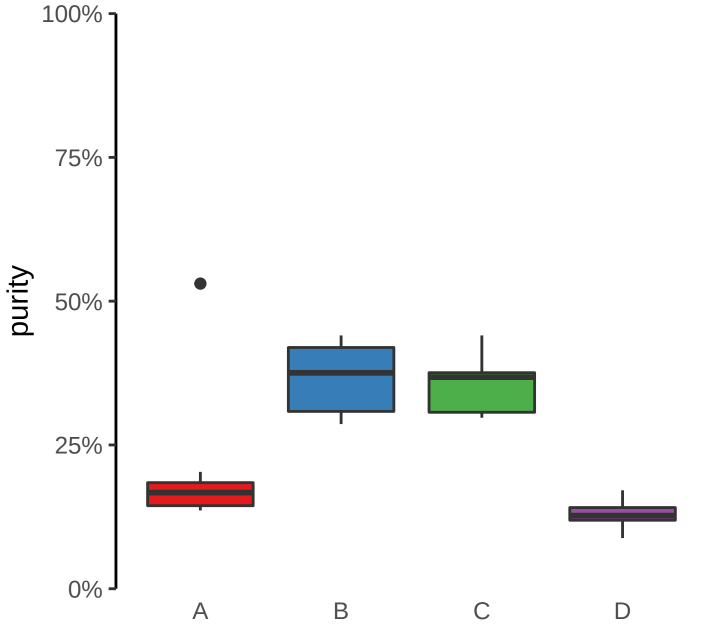

**FOXQ1 promoter gain**
**chr6:1311742−1312242**

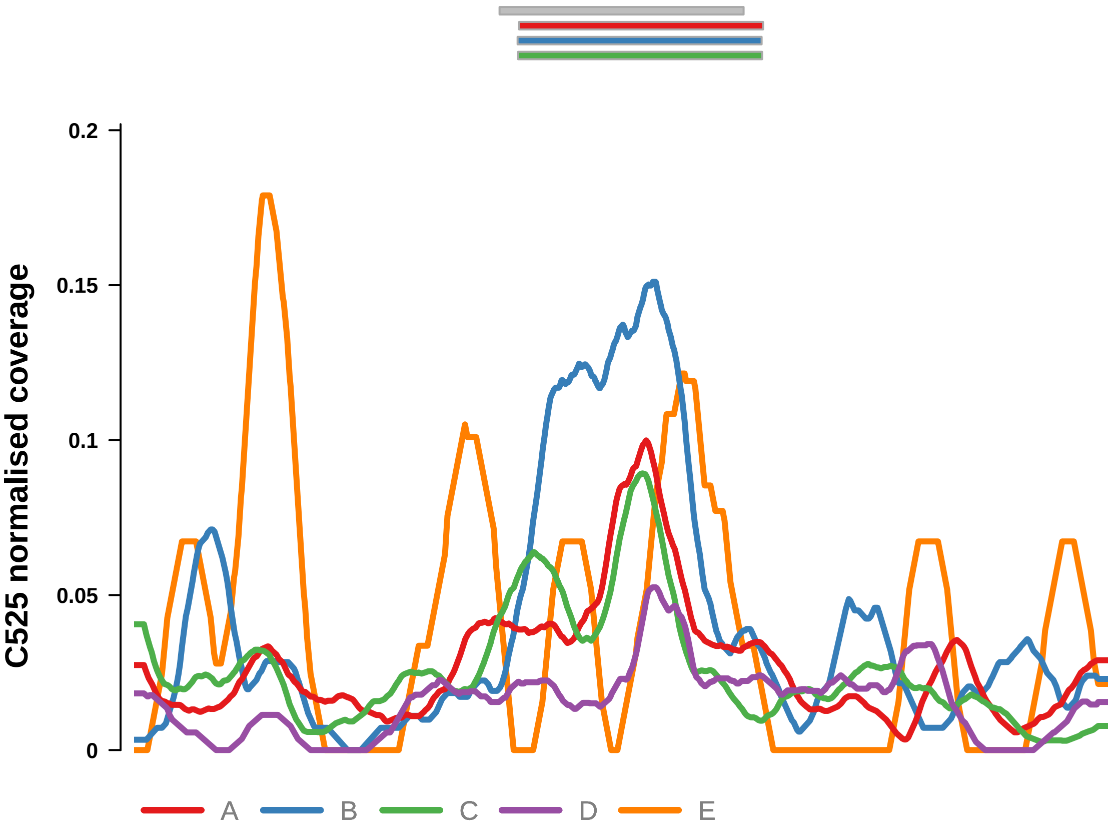


Putatively Clonal (logFC = 1.99)

| | **P.** | **Adjusted P.** |
|---|---|---|
| *~region vs. ~* | 6.61e−02 | 1.21e−01 |
| *~purity vs ~* | 1.4e−01 | 7.37e−01 |
| *~purity+region vs ~purity* | 1.65e−01 | 2.64e−01 |
| *Carcinoma vs. normal* | 5.15e−04 *** | 5.15e−04 *** |

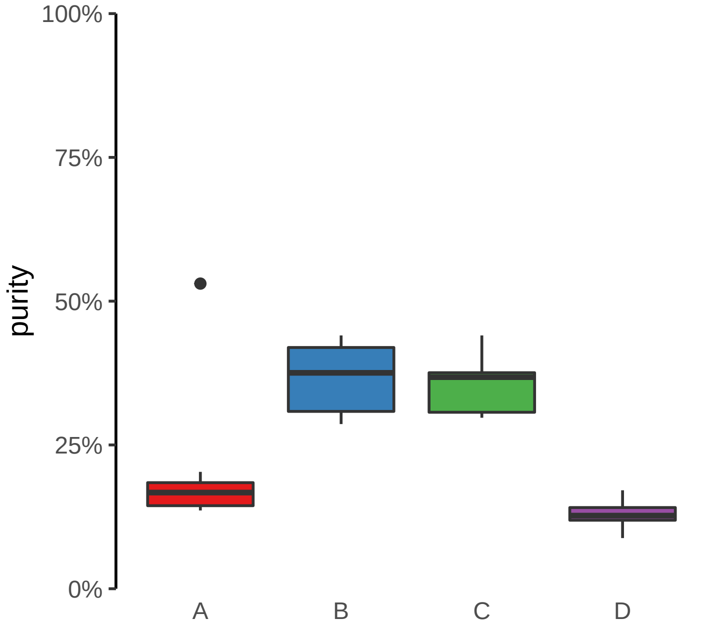

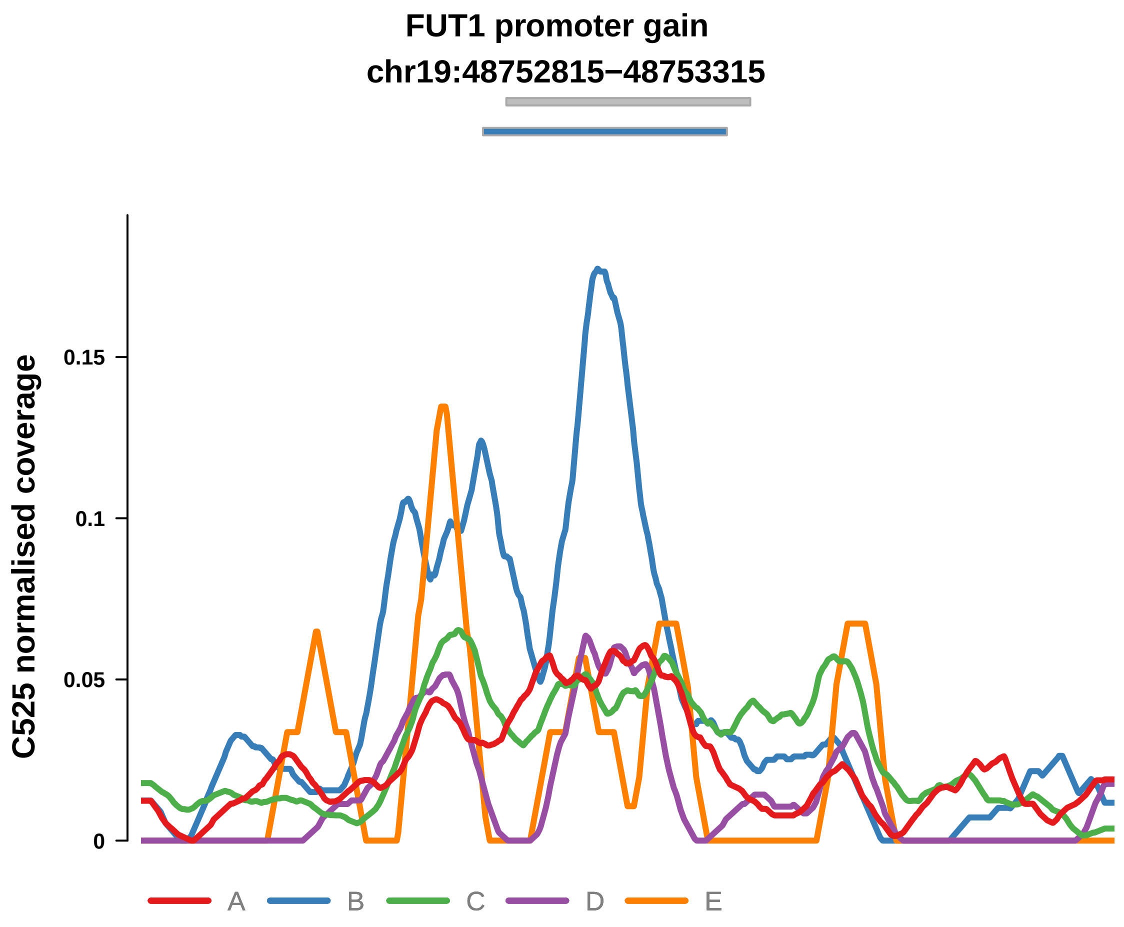


Putatively Clonal (logFC = 1.82)

| | P. | Adjusted P. |
|---|---|---|
| *~region vs. ~* | 5.4e−02 | 1.05e−01 |
| *~purity vs ~* | 3.22e−02 * | 5.67e−01 |
| *~purity+region vs ~purity* | 4.38e−01 | 5.5e−01 |
| *Carcinoma vs. normal* | 1.93e−03 ** | 1.93e−03 ** |

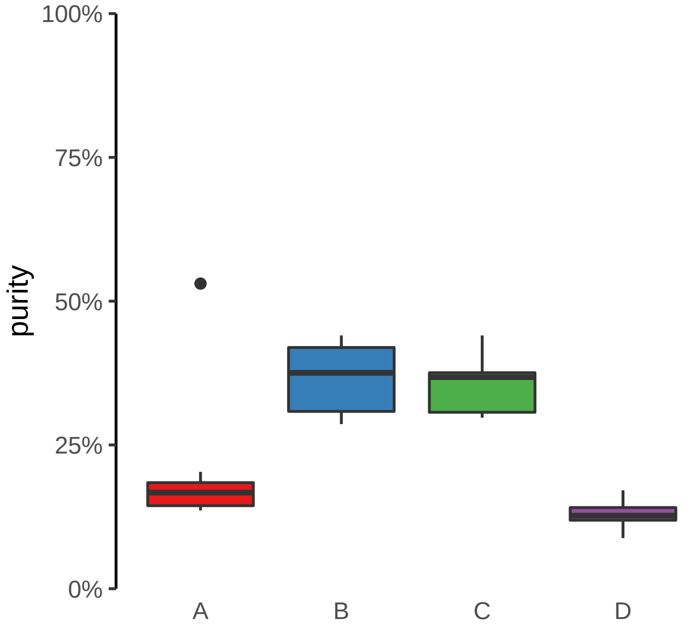

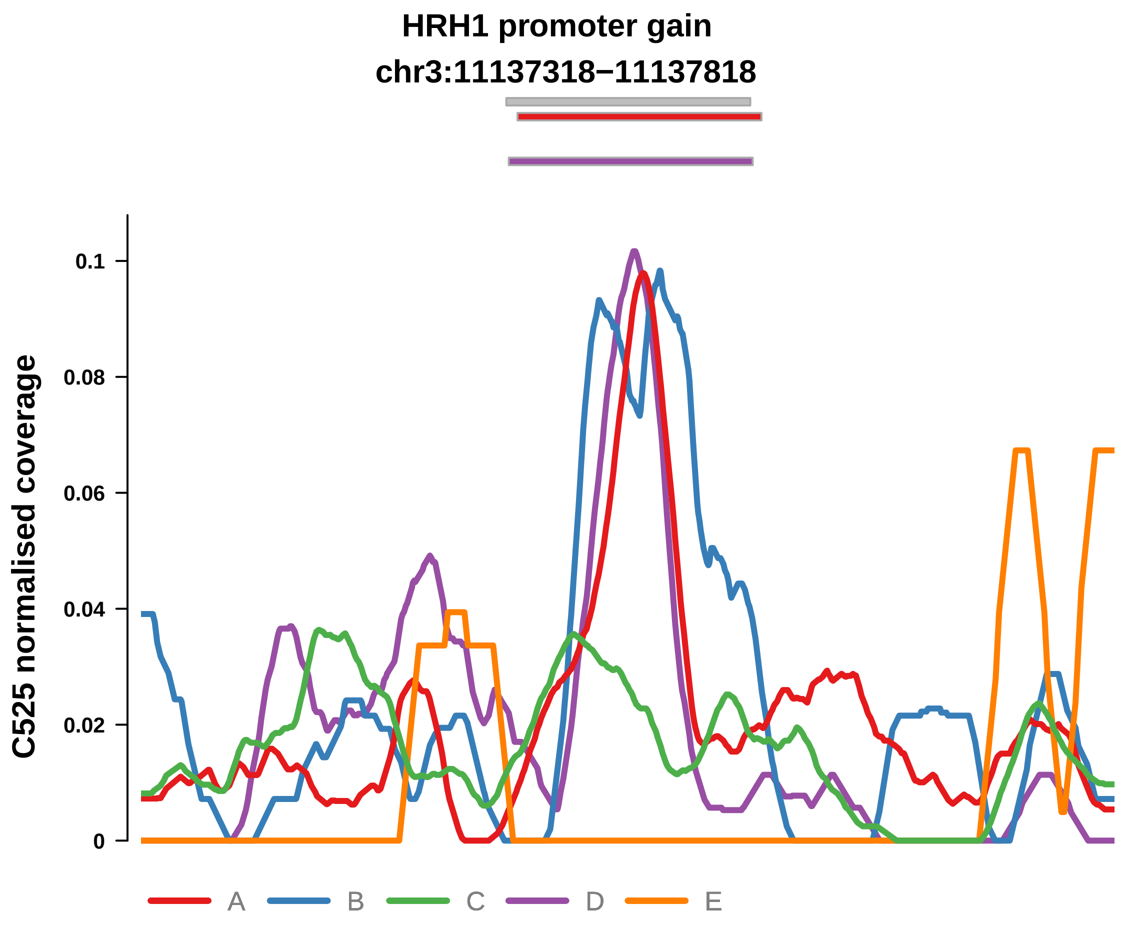


Putatively Clonal (logFC = 2.34)

| | P. | Adjusted P. |
|---|---|---|
| *~region vs. ~* | 6.74e−01 | 7.06e−01 |
| *~purity vs ~* | 9.77e−01 | 9.77e−01 |
| *~purity+region vs ~purity* | 4.21e−01 | 5.5e−01 |
| *Carcinoma vs. normal* | 1.07e−04 *** | 1.07e−04 *** |

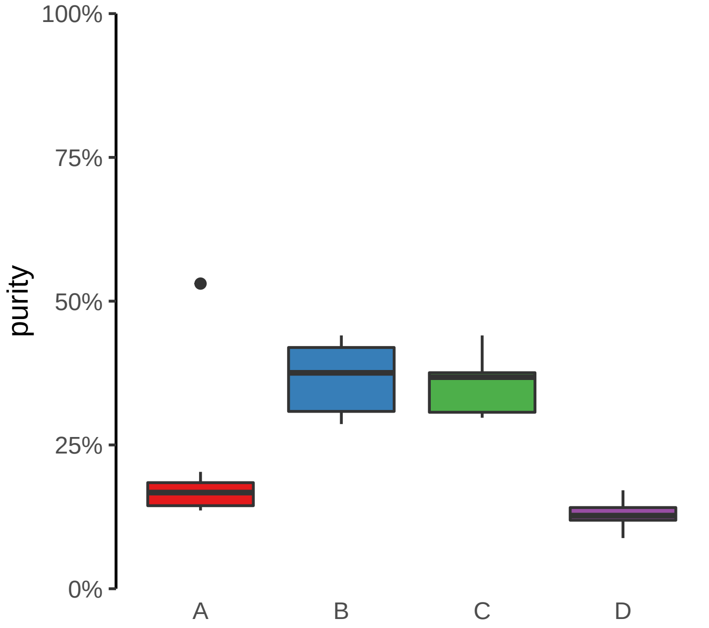

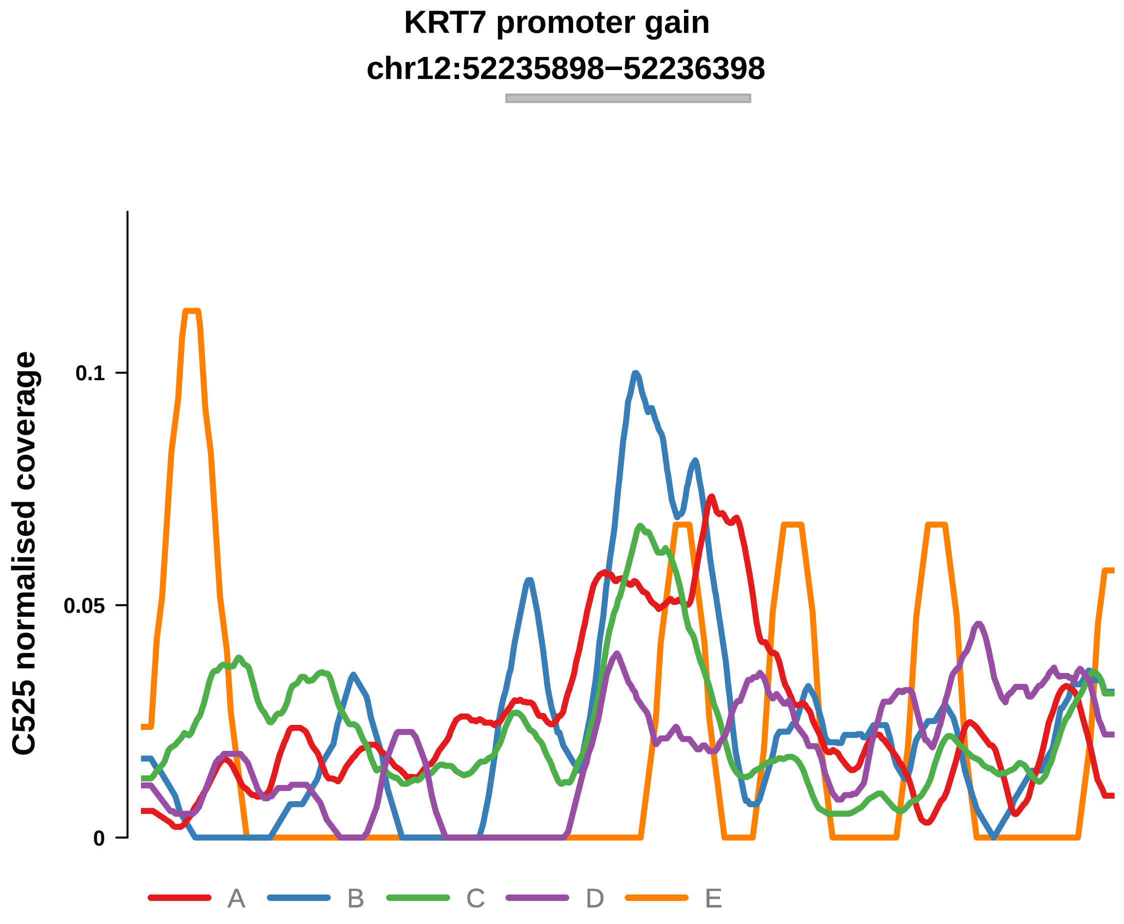


Putatively Clonal (logFC = 1.71)

| | P. | Adjusted P. |
|---|---|---|
| *~region vs. ~* | 2.65e−02 * | 6.41e−02 |
| *~purity vs ~* | 6.63e−01 | 8.63e−01 |
| *~purity+region vs ~purity* | 2.78e−02 * | 7.2e−02 |
| *Carcinoma vs. normal* | 3.95e−03 ** | 3.95e−03 ** |

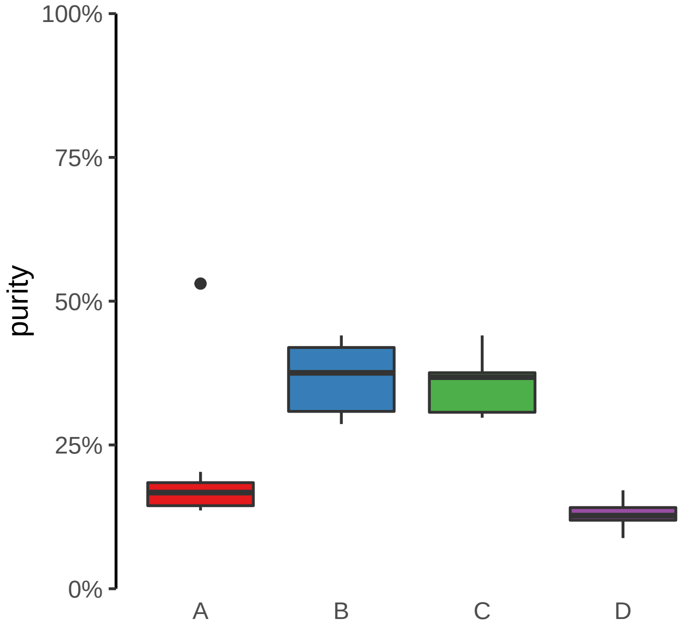

**PITX1 promoter gain**
**chr5:135034130−135034630**

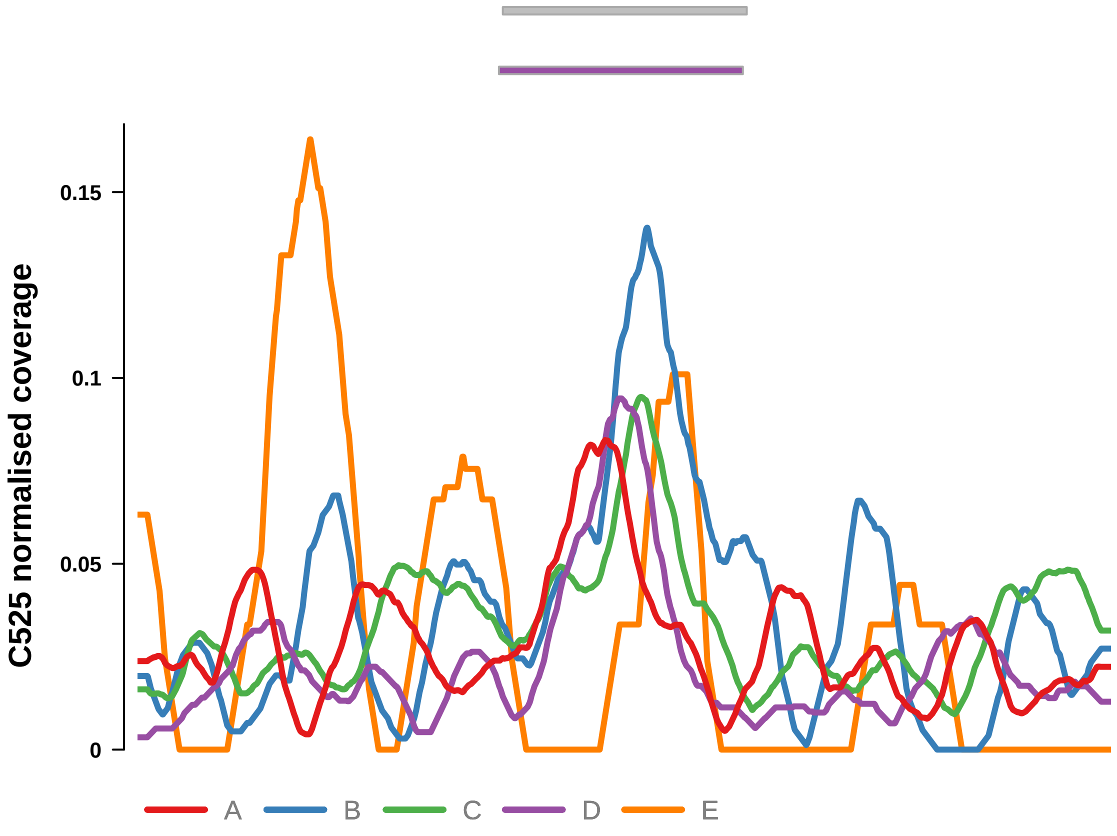


Putatively Clonal (logFC = 1.78)

| | P. | Adjusted P. |
|---|---|---|
| *~region vs. ~* | 2.75e−01 | 3.48e−01 |
| *~purity vs ~* | 2.74e−01 | 7.48e−01 |
| *~purity+region vs ~purity* | 4.31e−01 | 5.5e−01 |
| *Carcinoma vs. normal* | 2.36e−03 ** | 2.36e−03 ** |

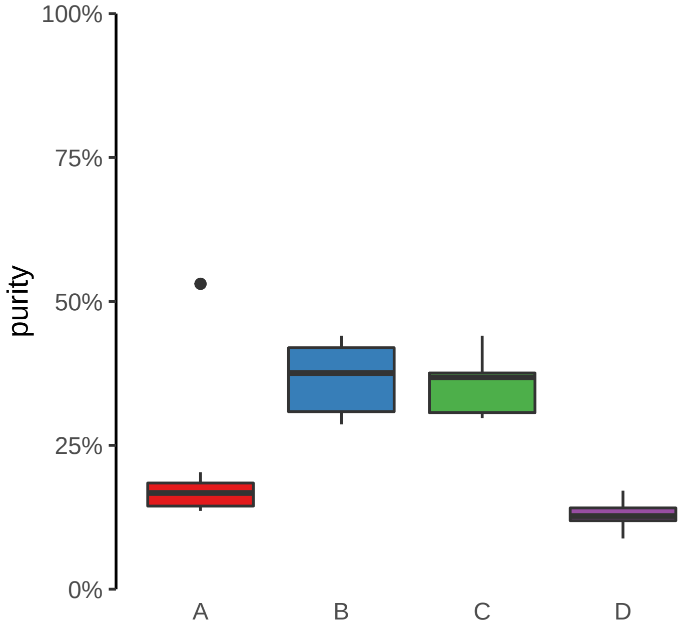

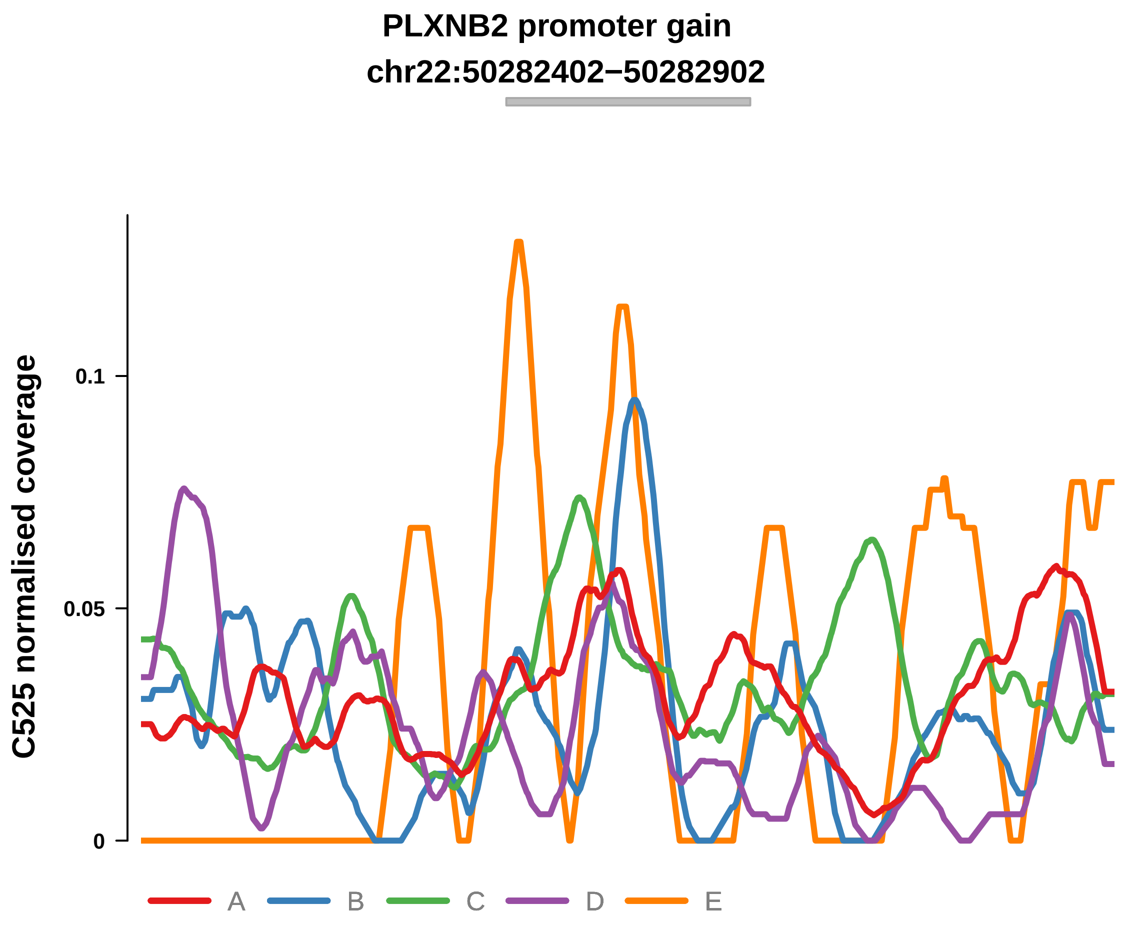


Putatively Clonal (logFC = 1.86)

| | P. | Adjusted P. |
|---|---|---|
| *~region vs. ~* | 7.11e−03 ** | 2.35e−02 * |
| *~purity vs ~* | 3e−01 | 7.48e−01 |
| *~purity+region vs ~purity* | 1.14e−02 * | 3.71e−02 * |
| *Carcinoma vs. normal* | 2.14e−03 ** | 2.14e−03 ** |

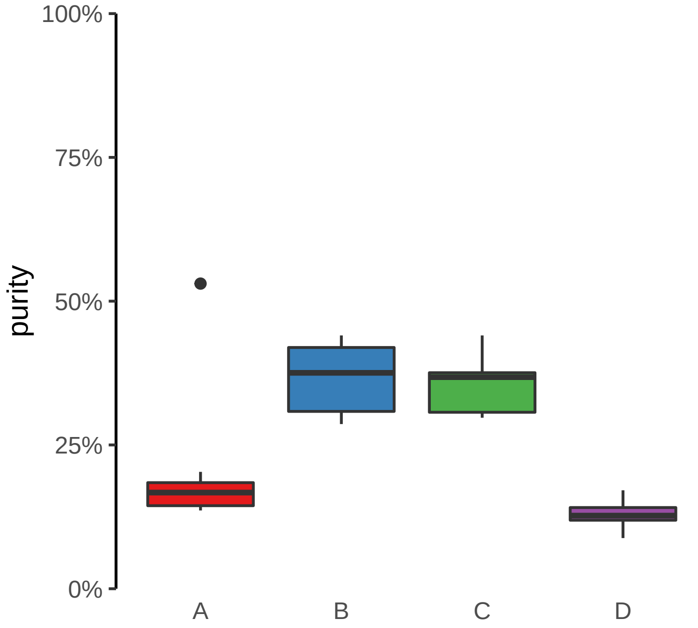

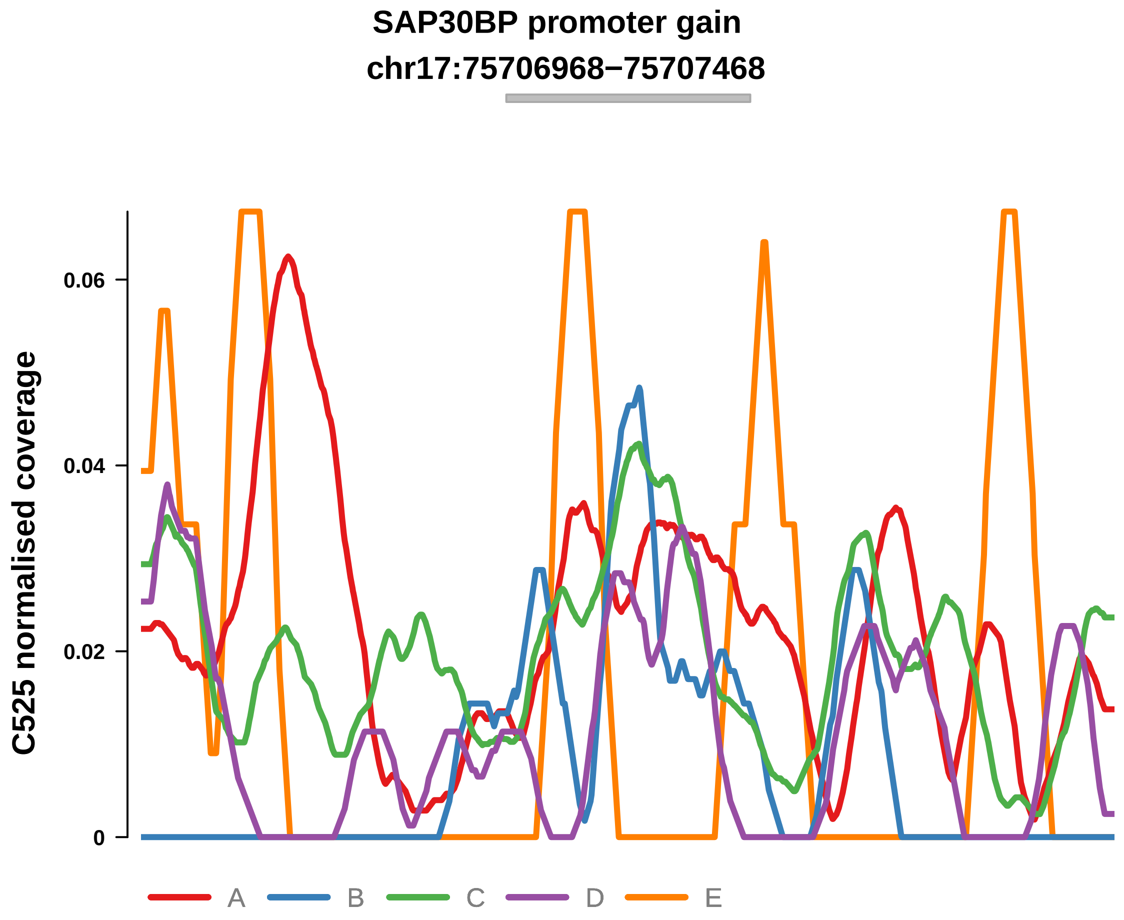


Putatively Clonal (logFC = 1.85)

| | P. | Adjusted P. |
|---|---|---|
| *~region vs. ~* | 9.42e−02 | 1.59e−01 |
| *~purity vs ~* | 6.29e−01 | 8.63e−01 |
| *~purity+region vs ~purity* | 1.27e−01 | 2.12e−01 |
| *Carcinoma vs. normal* | 4.41e−03 ** | 4.41e−03 ** |

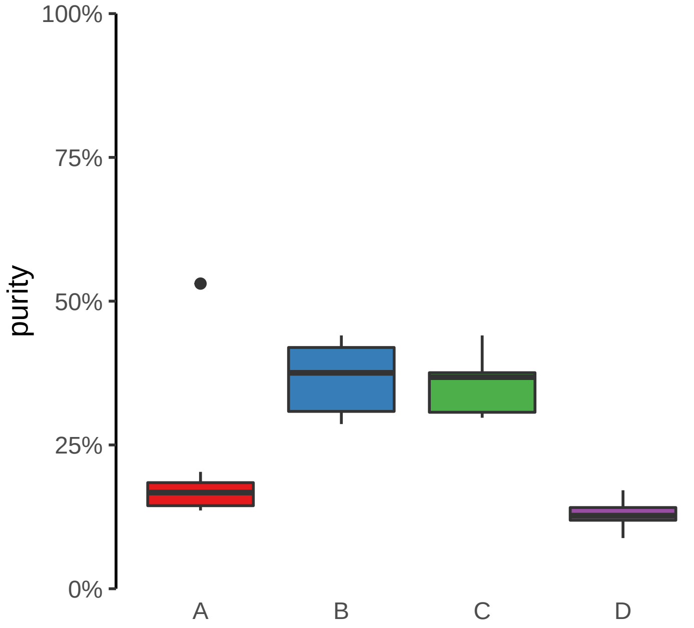

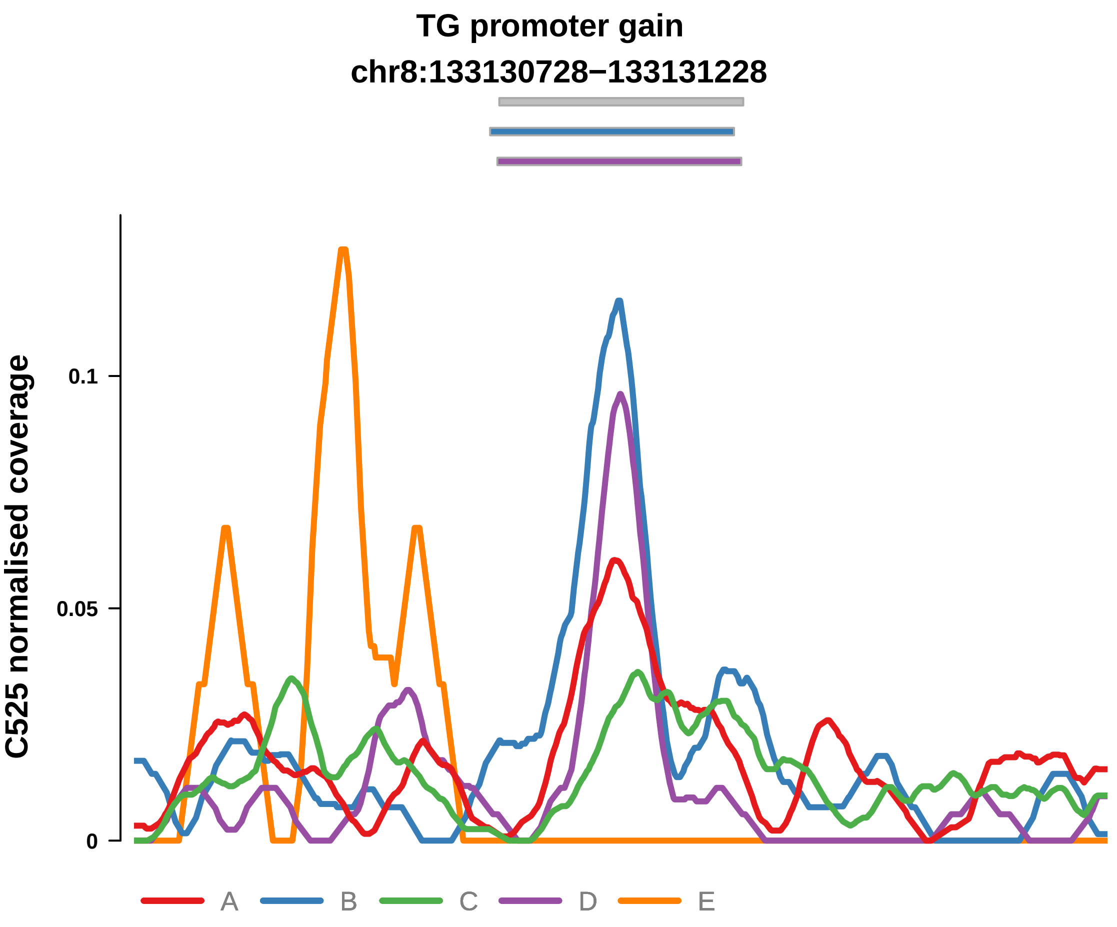


Putatively Clonal (logFC = 1.95)

| | P. | Adjusted P. |
|---|---|---|
| *~region vs. ~* | 6.36e−01 | 6.84e−01 |
| *~purity vs ~* | 3.3e−01 | 7.48e−01 |
| *~purity+region vs ~purity* | 8.46e−01 | 8.66e−01 |
| *Carcinoma vs. normal* | 1.12e−03 ** | 1.12e−03 ** |

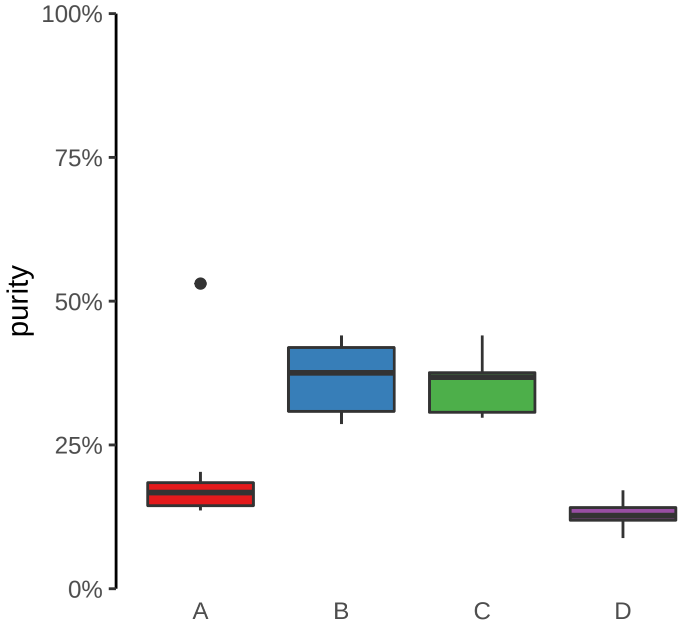

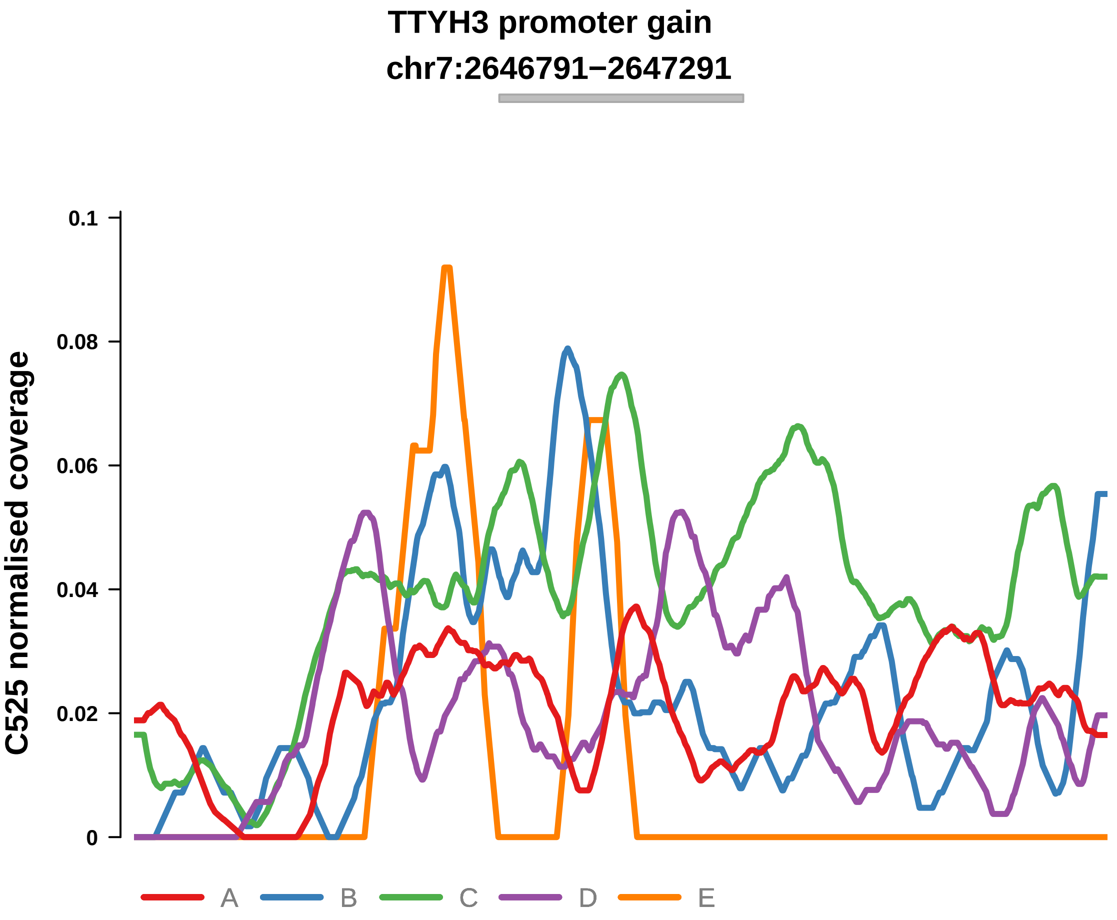


Putatively Clonal (logFC = 2.31)

| | P. | Adjusted P. |
|---|---|---|
| *~region vs. ~* | 5.21e−05 **** | 7.46e−04 *** |
| *~purity vs ~* | 1.15e−01 | 7.37e−01 |
| *~purity+region vs ~purity* | 3.02e−04 *** | 2.95e−03 ** |
| *Carcinoma vs. normal* | 1.53e−04 *** | 1.53e−04 *** |

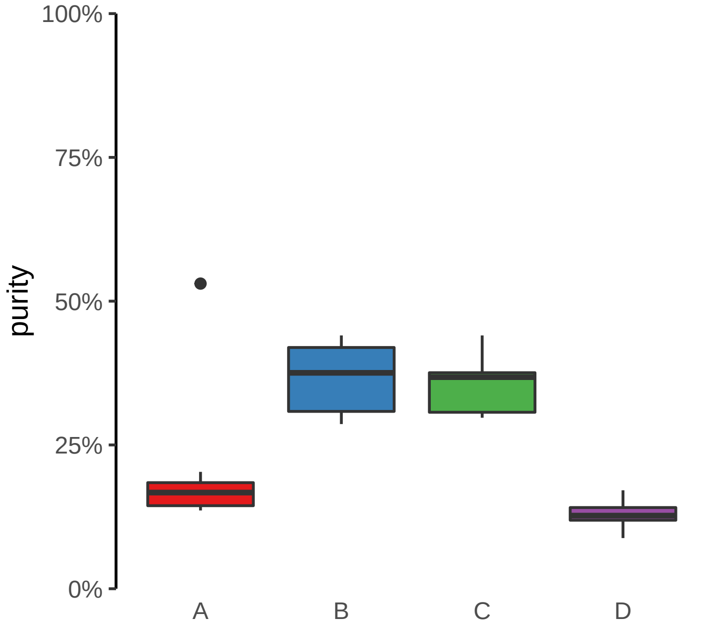

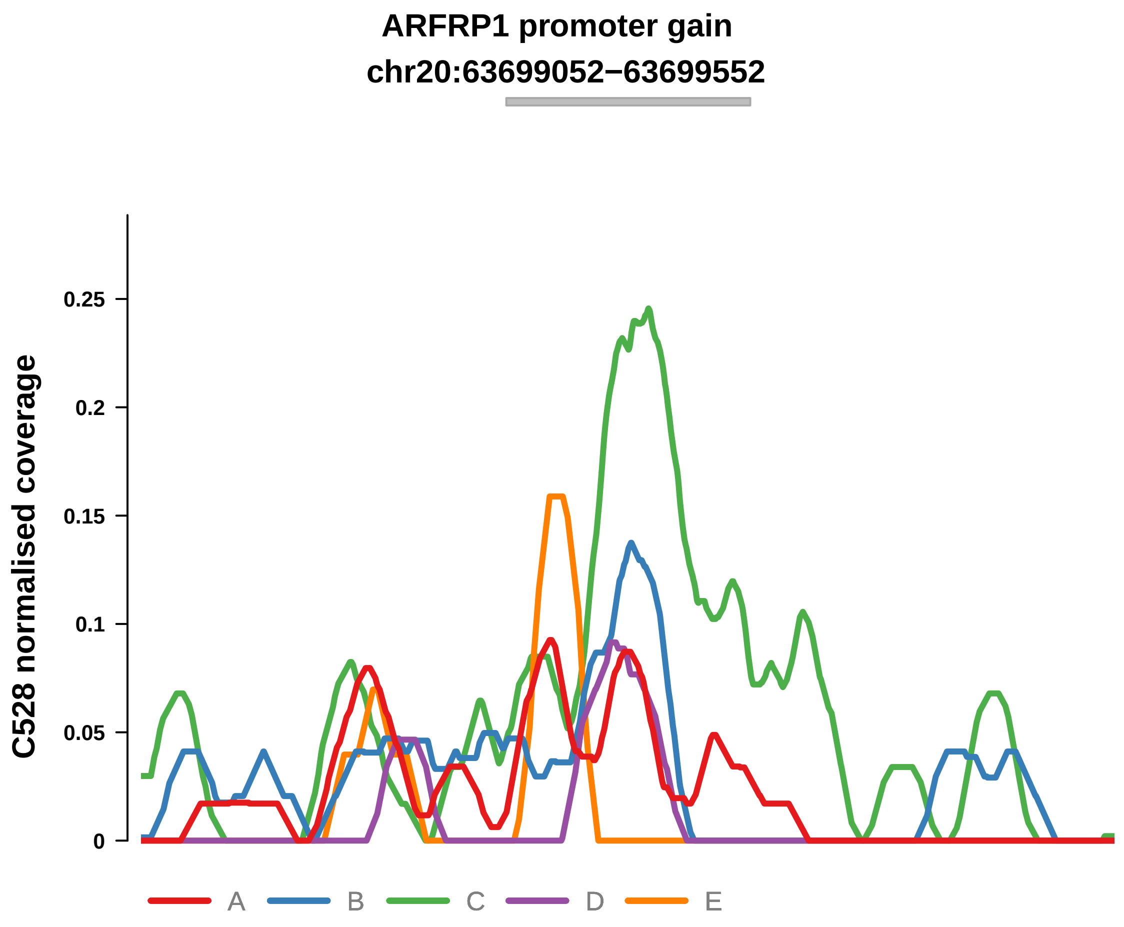


Putatively Clonal (logFC = 2.54)

| | P. | Adjusted P. |
|---|---|---|
| *~region vs. ~* | 7.75e−02 | 2.96e−01 |
| *~purity vs ~* | 1.86e−02 * | 9.87e−02 |
| *~purity+region vs ~purity* | 7.25e−01 | 9.17e−01 |
| *Carcinoma vs. normal* | 3.19e−05 **** | 3.19e−05 **** |

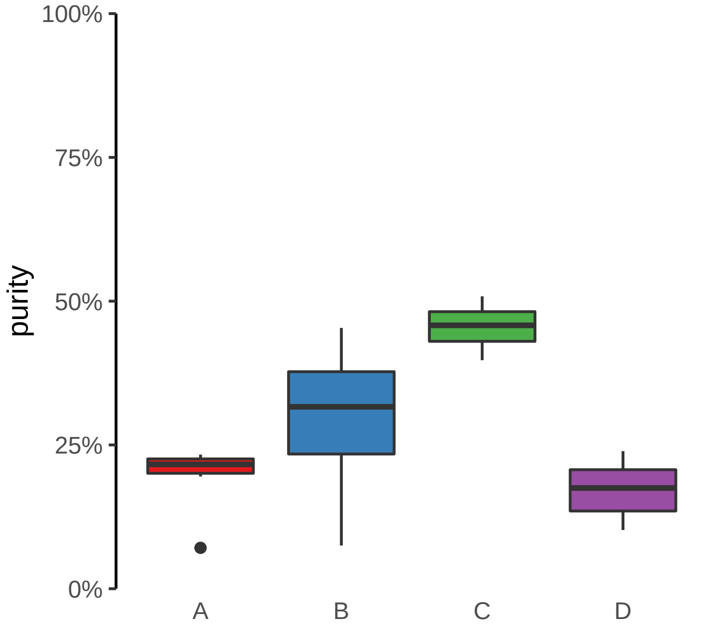

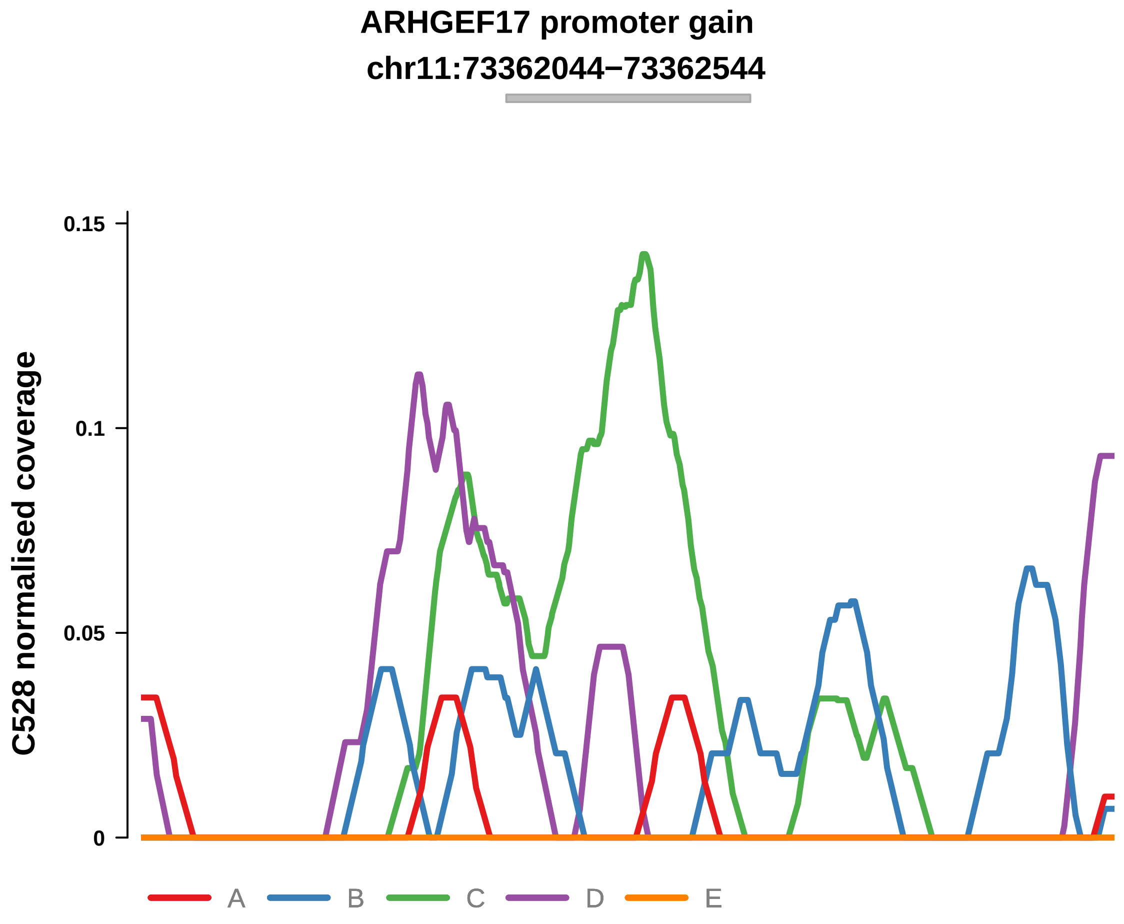


Putatively Clonal (logFC = 1.96)

| | P. | Adjusted P. |
|---|---|---|
| *~region vs. ~* | 3.94e−02 * | 2.31e−01 |
| *~purity vs ~* | 1.51e−03 ** | 2.6e−02 * |
| *~purity+region vs ~purity* | 5.43e−01 | 8.94e−01 |
| *Carcinoma vs. normal* | 3.59e−03 ** | 3.59e−03 ** |

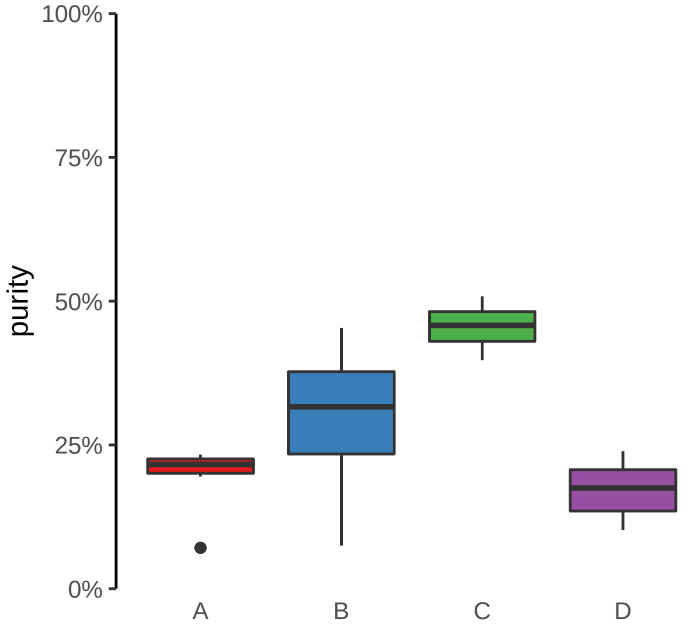

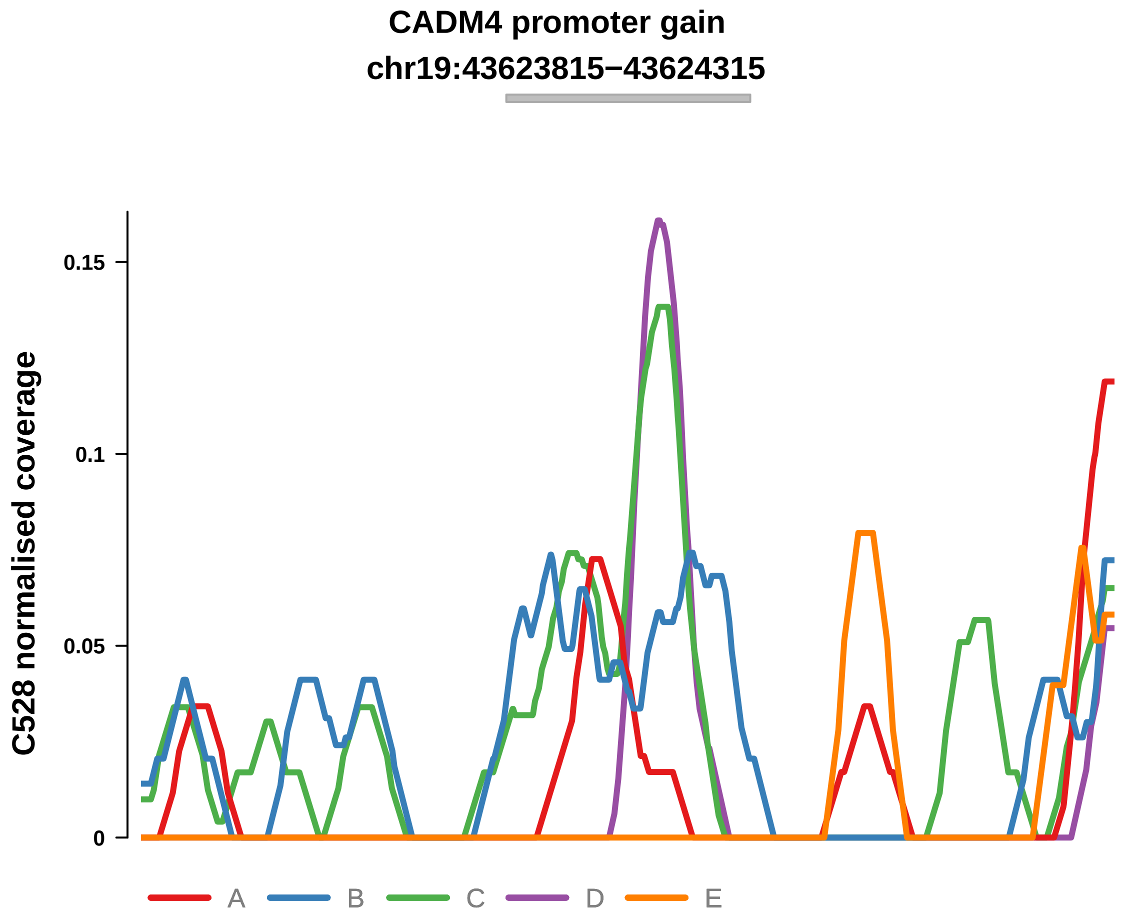


Putatively Clonal (logFC = 2.48)

| | P. | Adjusted P. |
|---|---|---|
| *~region vs. ~* | 2.52e−01 | 5.27e−01 |
| *~purity vs ~* | 1.48e−01 | 2.84e−01 |
| *~purity+region vs ~purity* | 5.64e−01 | 8.94e−01 |
| *Carcinoma vs. normal* | 1.2e−04 *** | 1.2e−04 *** |

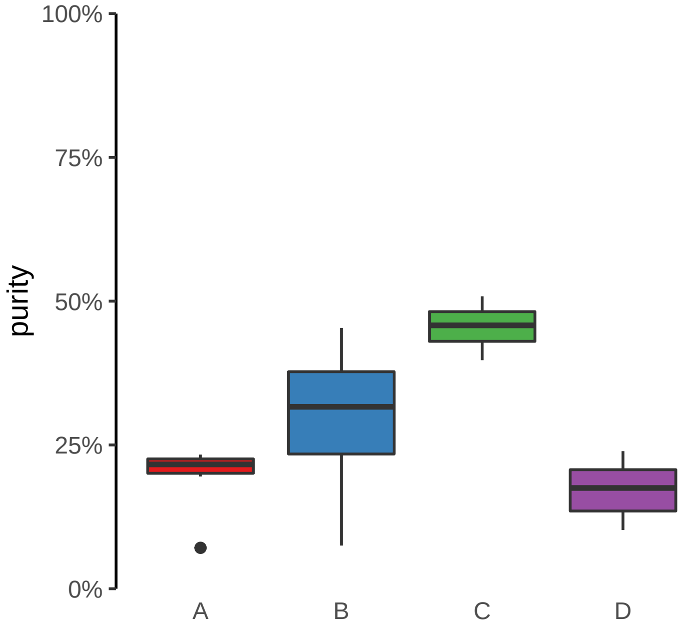

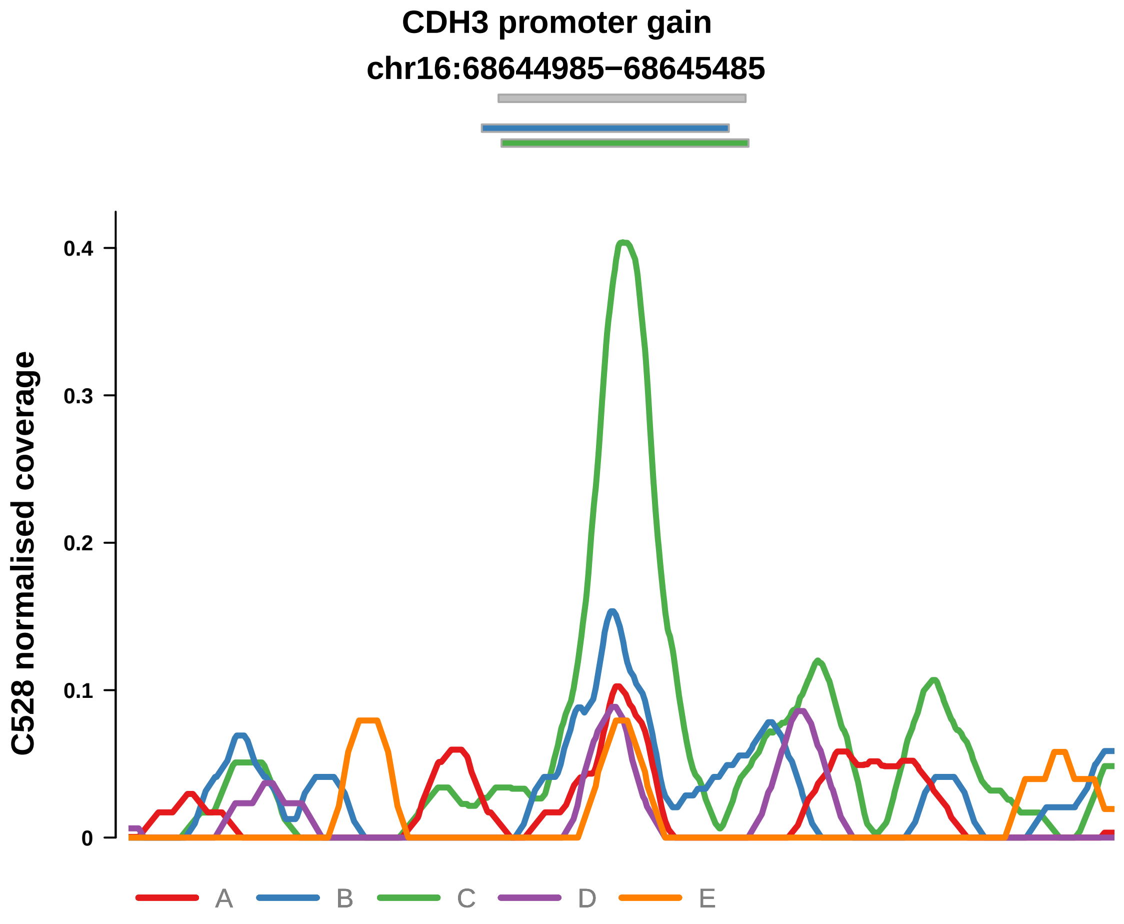


Putatively Clonal (logFC = 1.95)

| | P. | Adjusted P. |
|---|---|---|
| *~region vs. ~* | 8.69e−04 *** | 2.64e−02 * |
| *~purity vs ~* | 3.97e−04 *** | 9.13e−03 ** |
| *~purity+region vs ~purity* | 3.12e−01 | 7.26e−01 |
| *Carcinoma vs. normal* | 1.05e−03 ** | 1.05e−03 ** |

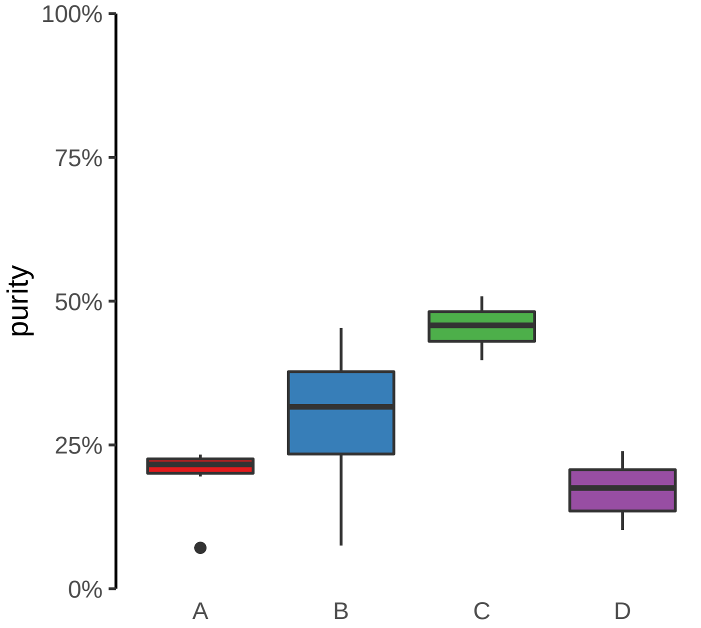

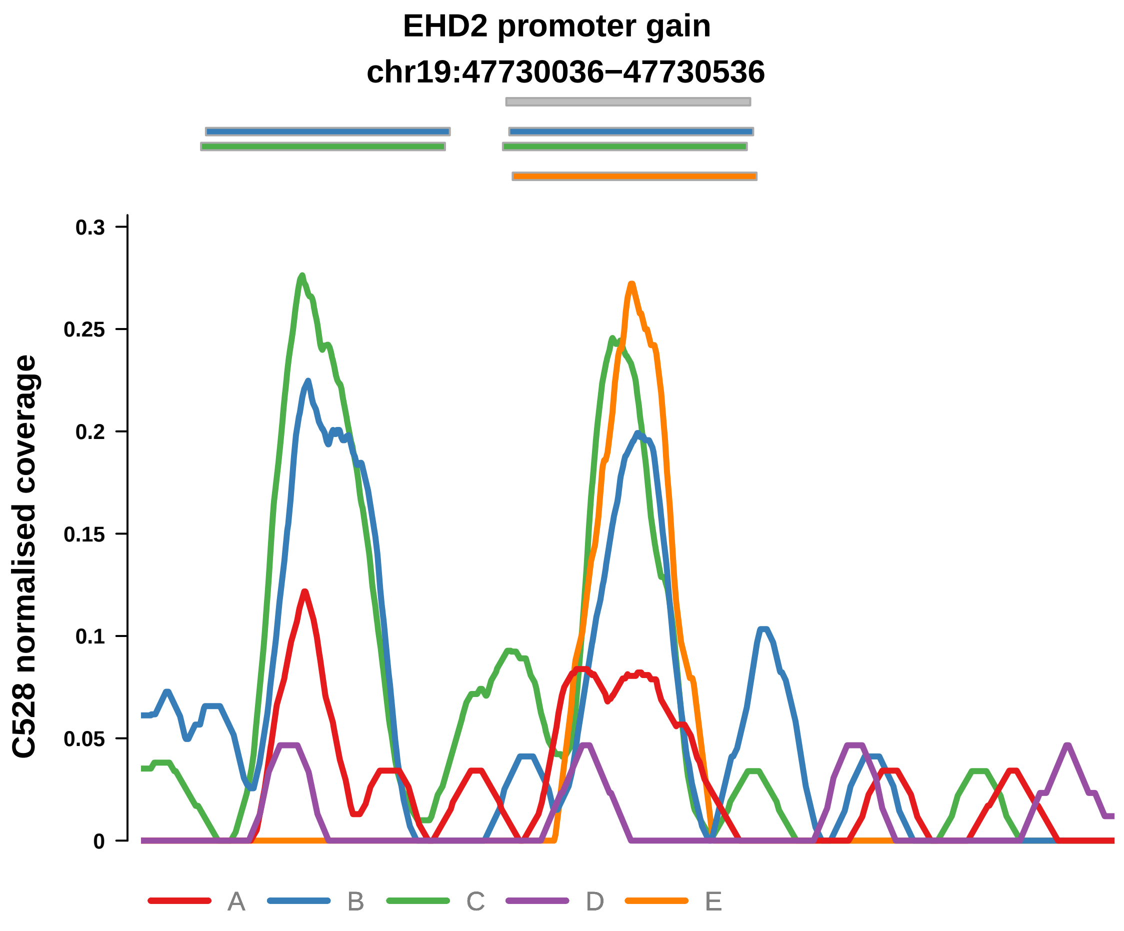


Putatively Clonal (logFC = 1.95)

| | P. | Adjusted P. |
|---|---|---|
| *~region vs. ~* | 2.92e−03 ** | 5.14e−02 |
| *~purity vs ~* | 6.53e−02 | 1.75e−01 |
| *~purity+region vs ~purity* | 1.44e−02 * | 4.21e−01 |
| *Carcinoma vs. normal* | 1.73e−03 ** | 1.73e−03 ** |

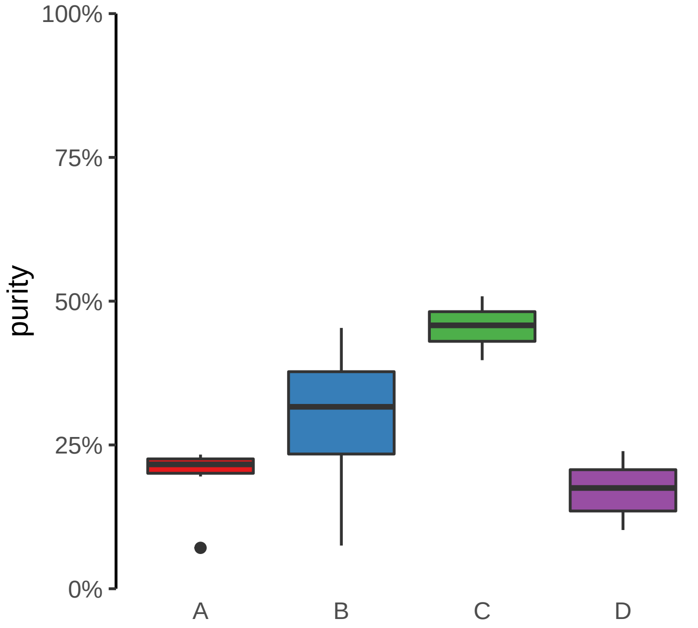

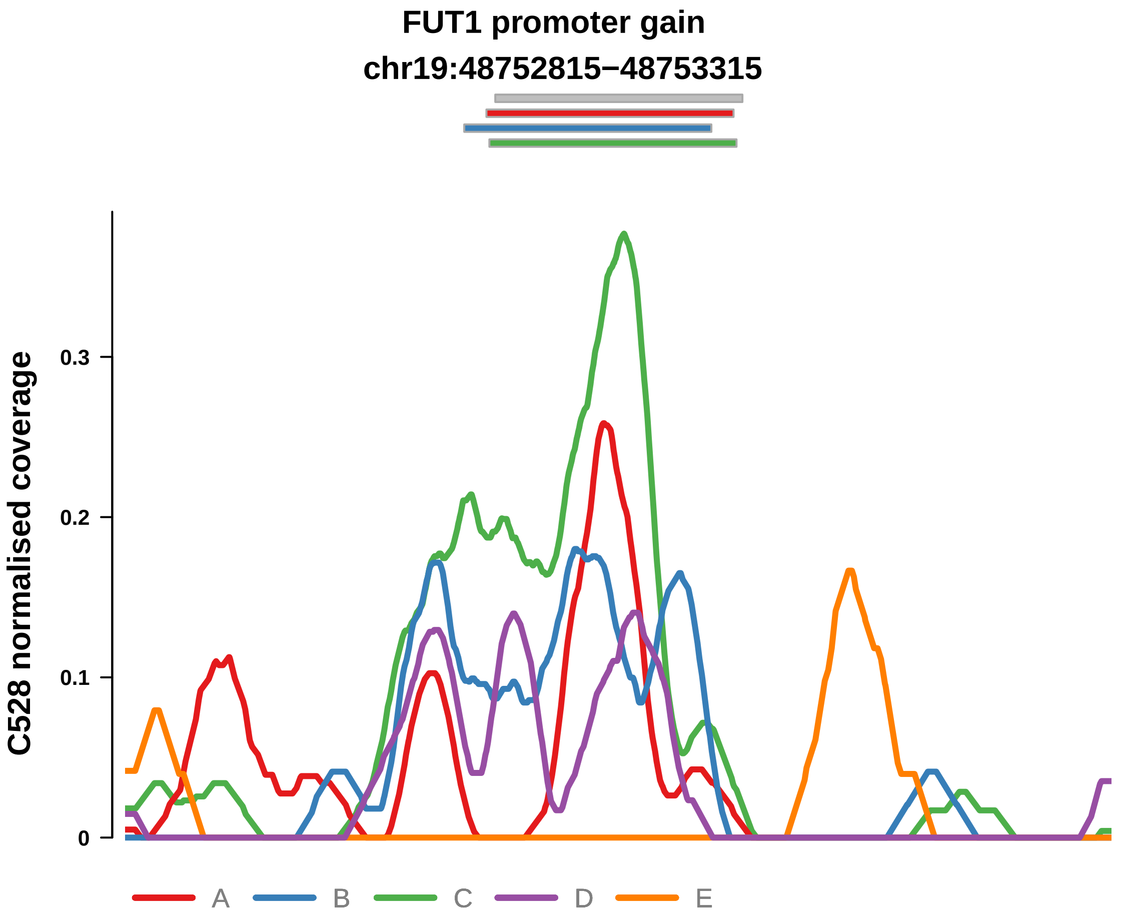


Putatively Clonal (logFC = 2.72)

| | P. | Adjusted P. |
|---|---|---|
| *~region vs. ~* | 4.27e−02 * | 2.35e−01 |
| *~purity vs ~* | 7.52e−03 ** | 7.41e−02 |
| *~purity+region vs ~purity* | 7.27e−01 | 9.17e−01 |
| *Carcinoma vs. normal* | 4.05e−06 **** | 4.05e−06 **** |

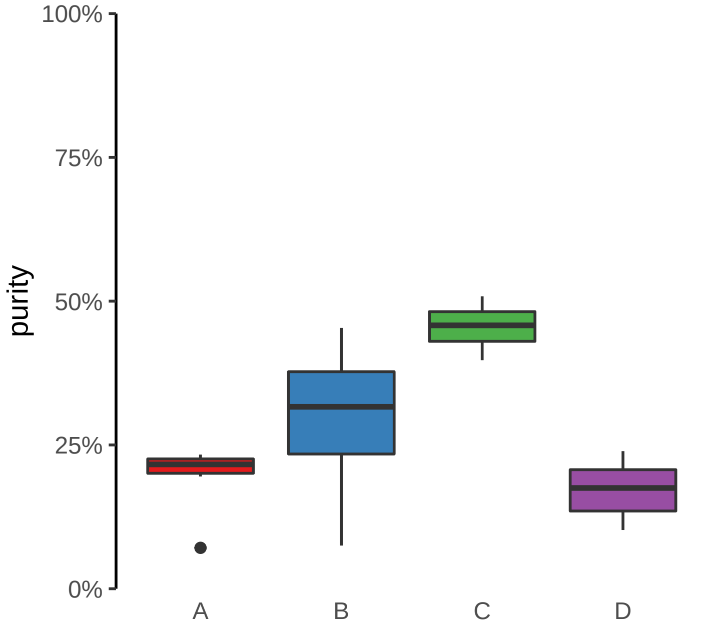

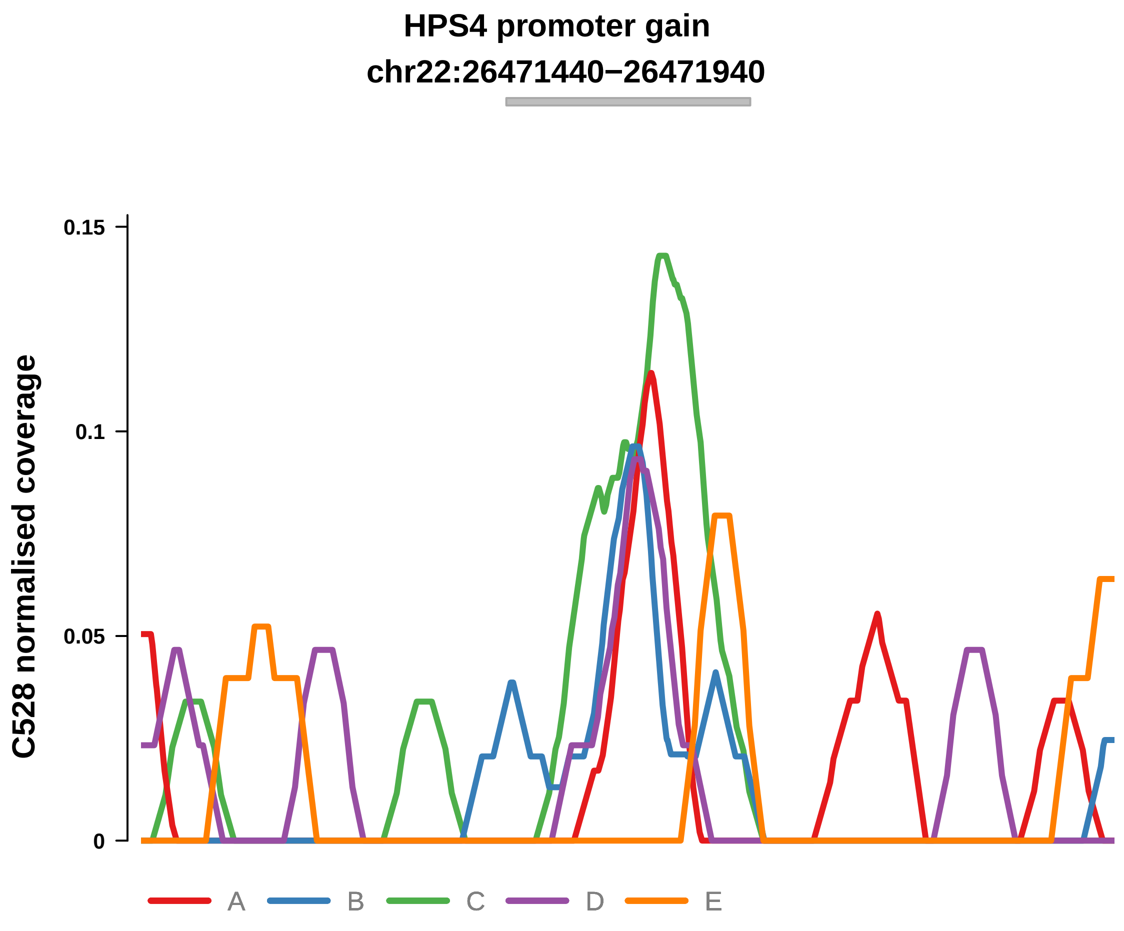


Putatively Clonal (logFC = 2.34)

| | P. | Adjusted P. |
|---|---|---|
| *~region vs. ~* | 5.58e−01 | 7.5e−01 |
| *~purity vs ~* | 9.79e−02 | 2.18e−01 |
| *~purity+region vs ~purity* | 9.25e−01 | 1e+00 |
| *Carcinoma vs. normal* | 7.28e−04 *** | 7.28e−04 *** |

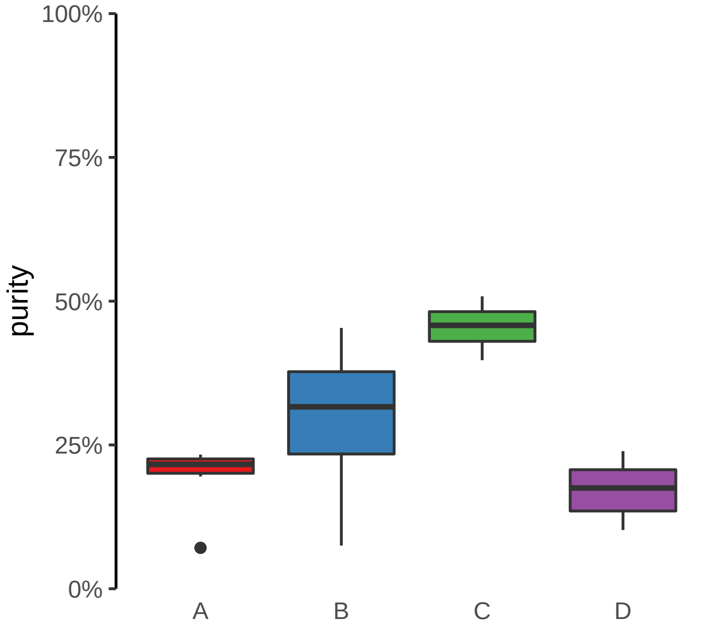

**HRH1 promoter gain**
**chr3:11137318−11137818**

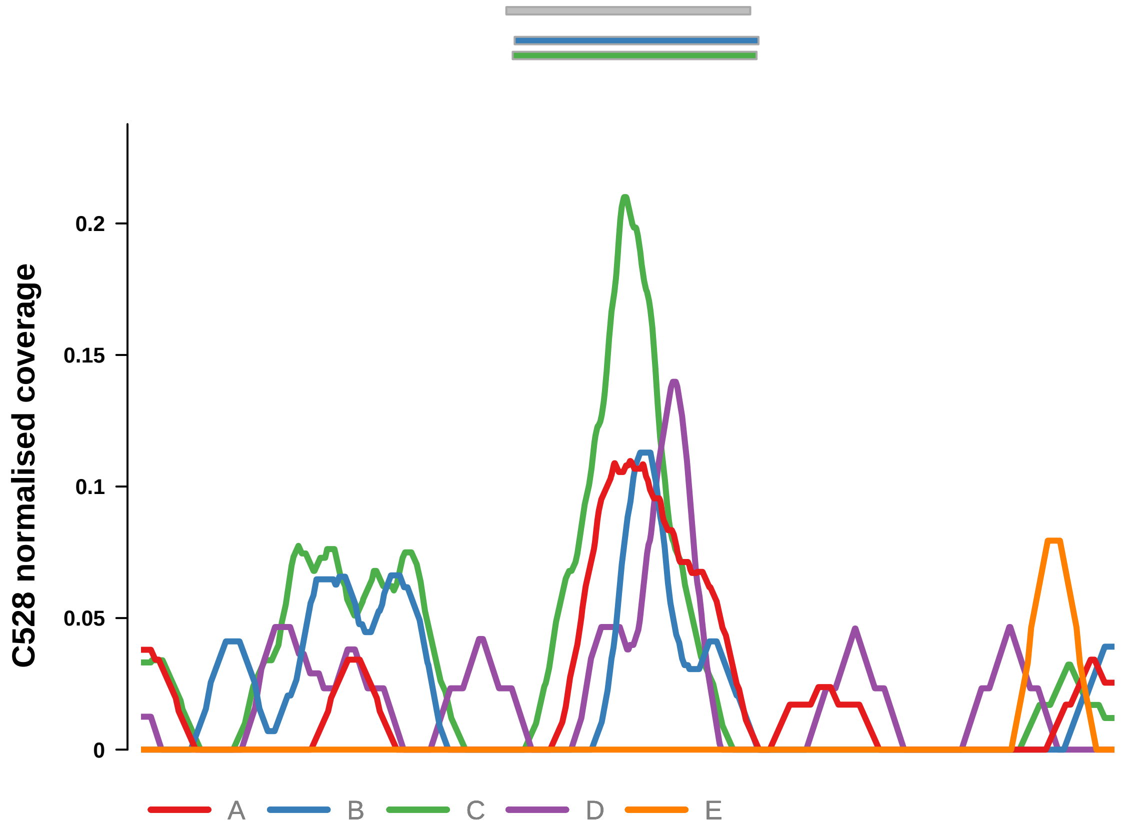


Putatively Clonal (logFC = 1.73)

| | **P.** | **Adjusted P.** |
|---|---|---|
| *~region vs. ~* | 6.73e−01 | 8.59e−01 |
| *~purity vs ~* | 8.82e−01 | 9.22e−01 |
| *~purity+region vs ~purity* | 1.79e−01 | 6.24e−01 |
| *Carcinoma vs. normal* | 8.88e−03 ** | 8.88e−03 ** |

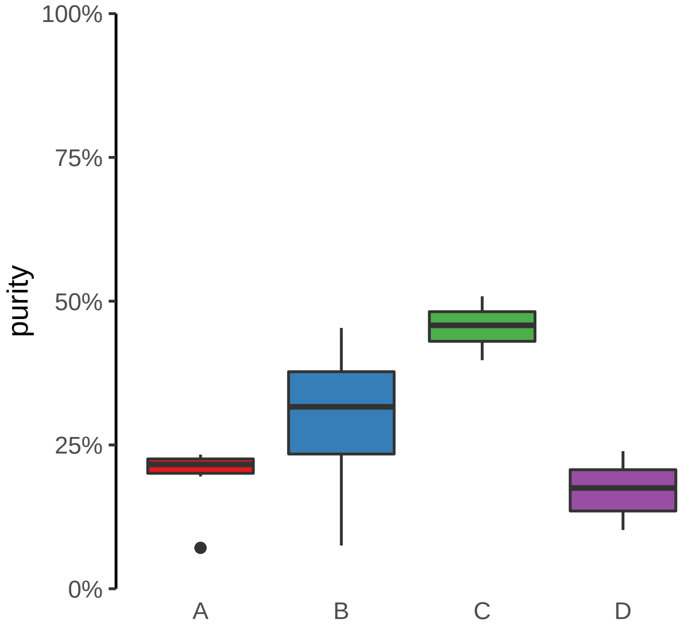

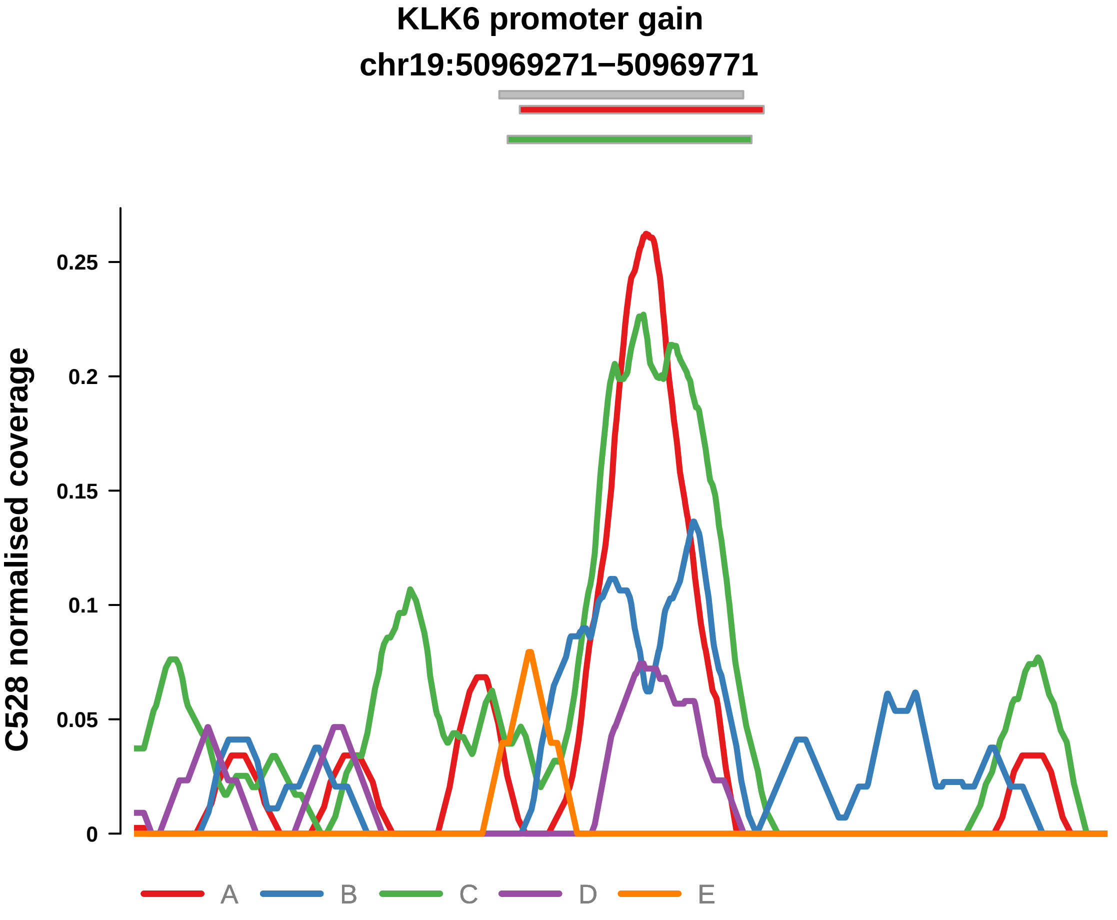


Putatively Clonal (logFC = 3.12)

| | P. | Adjusted P. |
|---|---|---|
| *~region vs. ~* | 1.54e−01 | 4.29e−01 |
| *~purity vs ~* | 2.63e−02 * | 1.21e−01 |
| *~purity+region vs ~purity* | 1.47e−01 | 6.24e−01 |
| *Carcinoma vs. normal* | 5.08e−07 **** | 5.08e−07 **** |

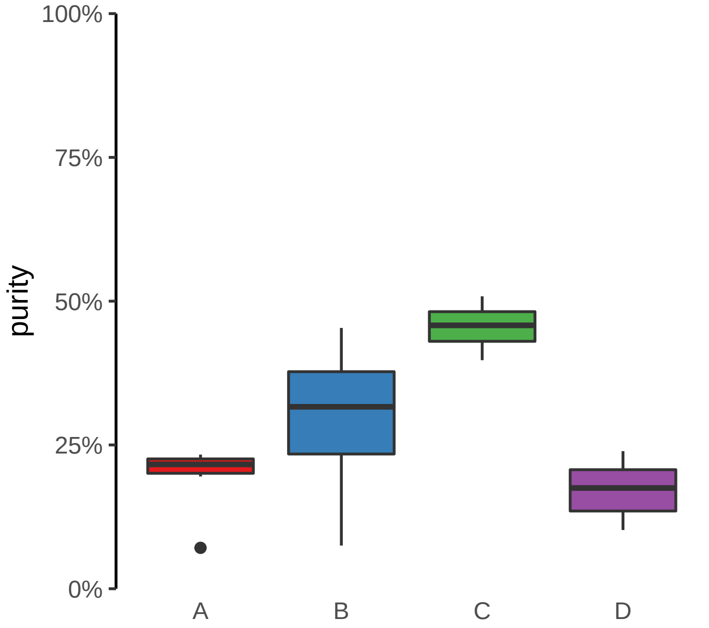

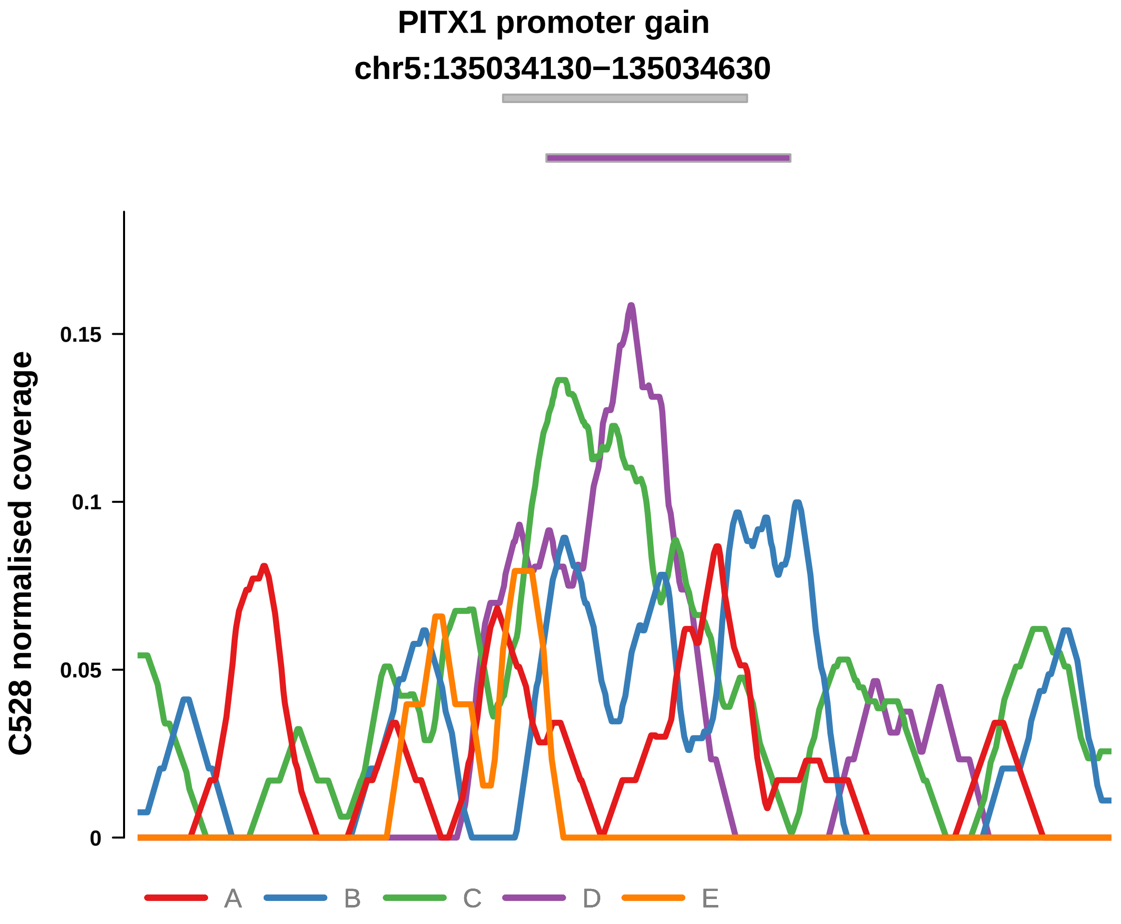


Putatively Clonal (logFC = 1.79)

| | P. | Adjusted P. |
|---|---|---|
| *~region vs. ~* | 3.39e−01 | 5.95e−01 |
| *~purity vs ~* | 3.5e−01 | 5.14e−01 |
| *~purity+region vs ~purity* | 1.56e−01 | 6.24e−01 |
| *Carcinoma vs. normal* | 3.39e−03 ** | 3.39e−03 ** |

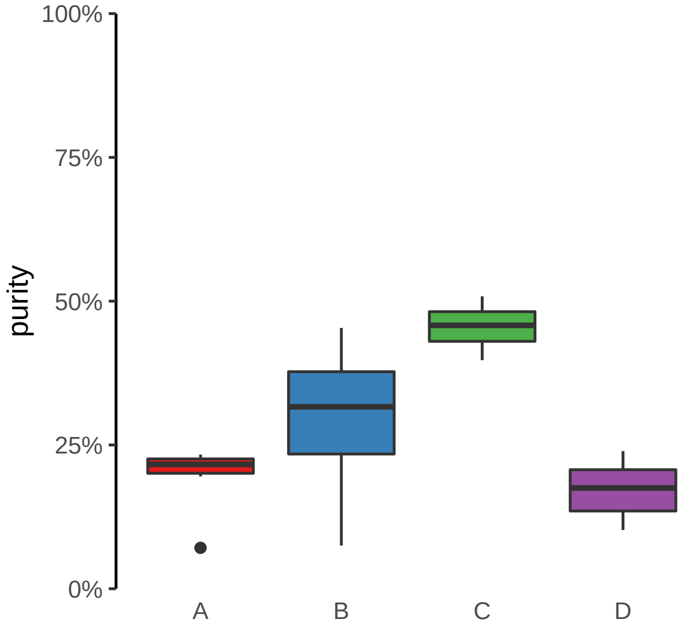

**PLXNB2 promoter gain**
**chr22:50282402−50282902**

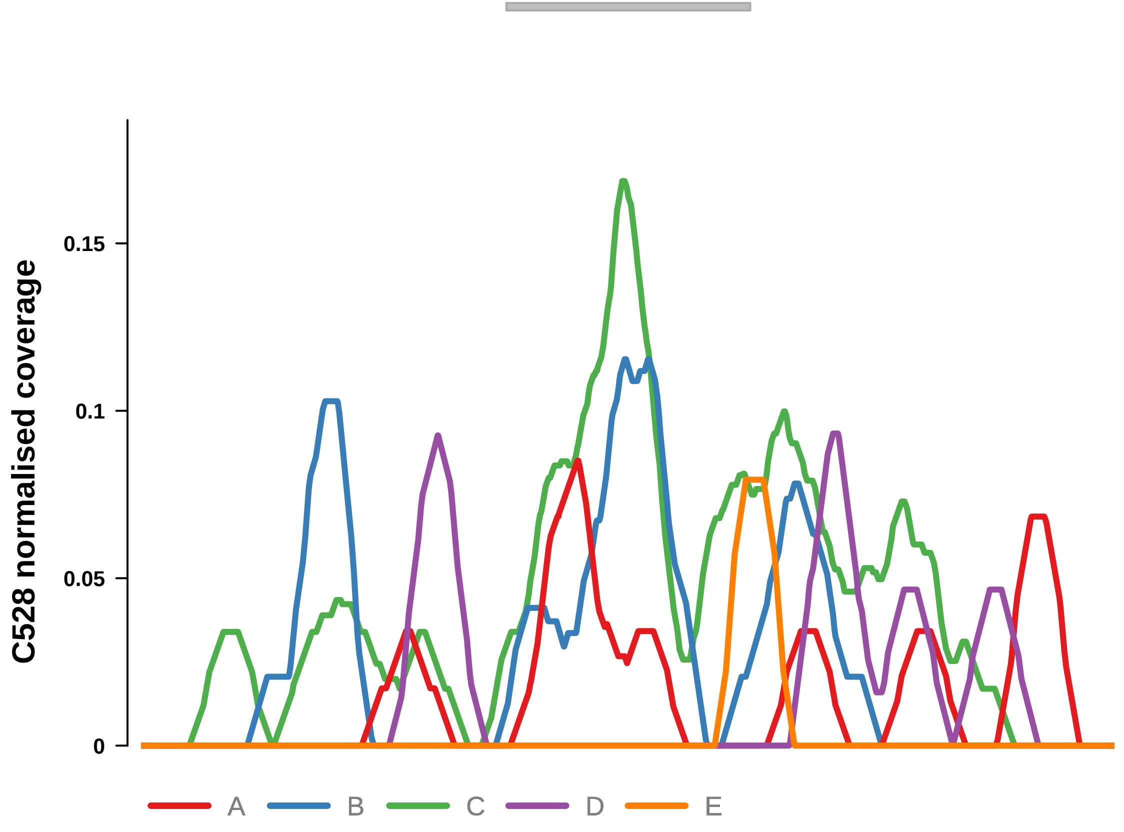


Putatively Clonal (logFC = 1.93)

| | P. | Adjusted P. |
|---|---|---|
| *~region vs. ~* | 4.61e−03 ** | 5.8e−02 |
| *~purity vs ~* | 1.38e−02 * | 9.7e−02 |
| *~purity+region vs ~purity* | 9.39e−02 | 6.11e−01 |
| *Carcinoma vs. normal* | 2.49e−03 ** | 2.49e−03 ** |

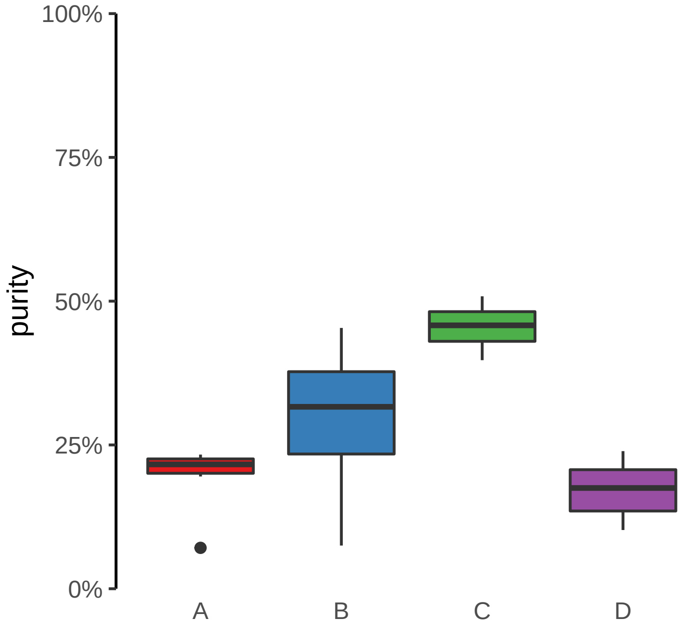

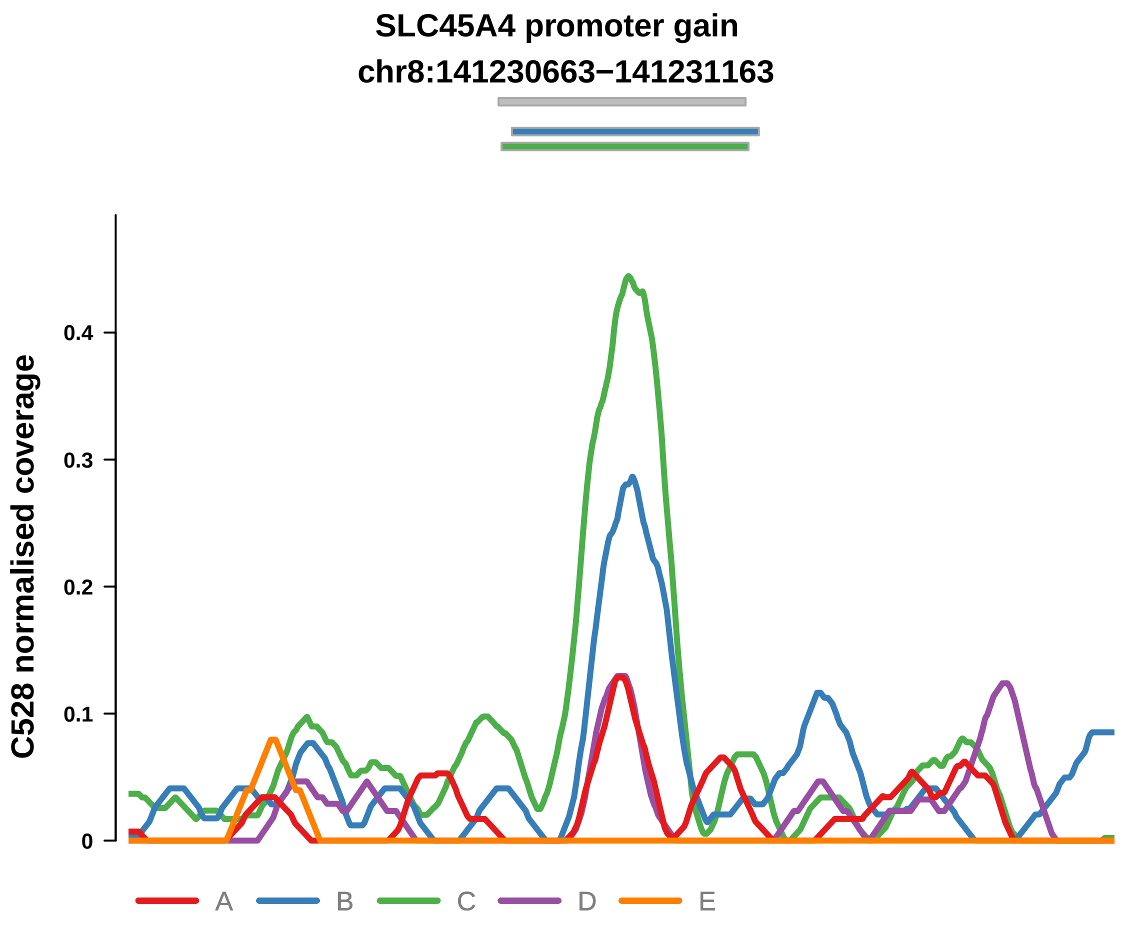


Putatively Clonal (logFC = 3.51)

| | P. | Adjusted P. |
|---|---|---|
| *~region vs. ~* | 9.02e−04 *** | 2.64e−02 * |
| *~purity vs ~* | 9.11e−06 **** | 6.28e−04 *** |
| *~purity+region vs ~purity* | 4.5e−01 | 7.92e−01 |
| *Carcinoma vs. normal* | 1.37e−08 **** | 1.37e−08 **** |

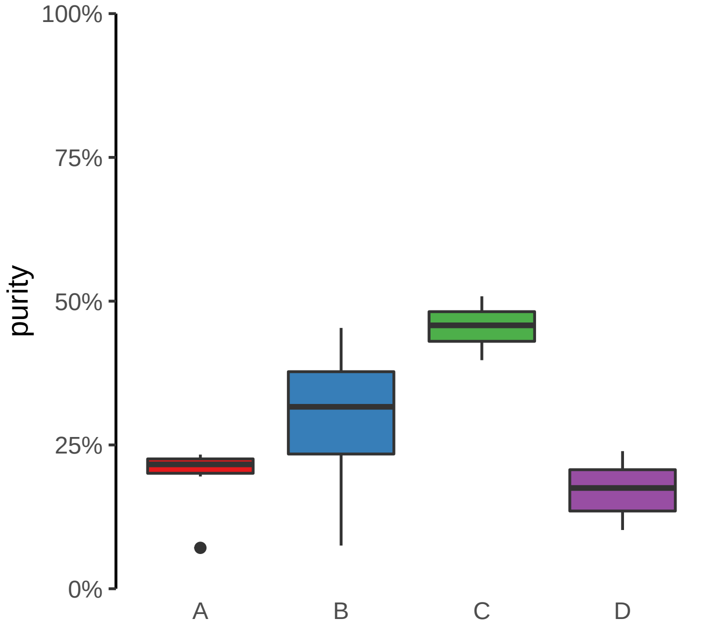

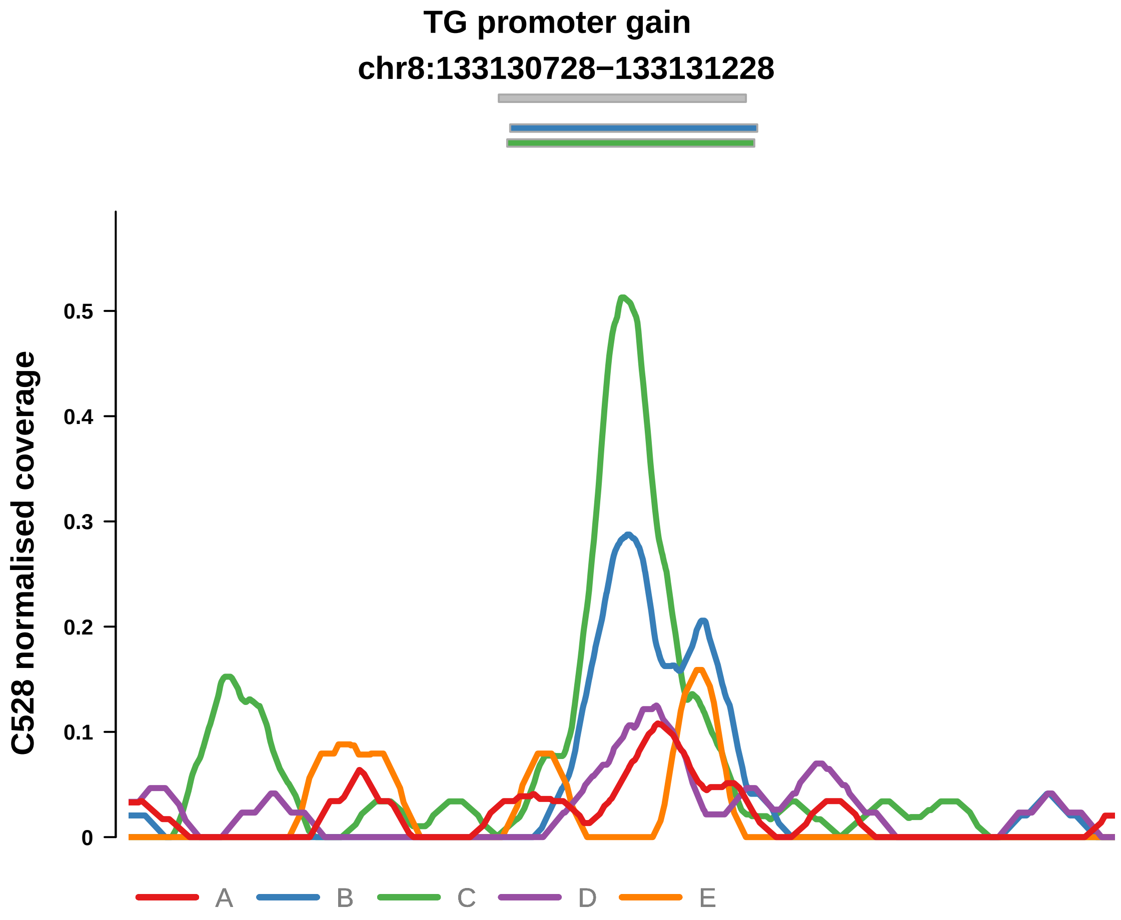


Putatively Clonal (logFC = 3.07)

| | P. | Adjusted P. |
|---|---|---|
| *~region vs. ~* | 7.57e−03 ** | 7.67e−02 |
| *~purity vs ~* | 1.91e−03 ** | 2.64e−02 * |
| *~purity+region vs ~purity* | 2.15e−01 | 6.38e−01 |
| *Carcinoma vs. normal* | 2.89e−07 **** | 2.89e−07 **** |

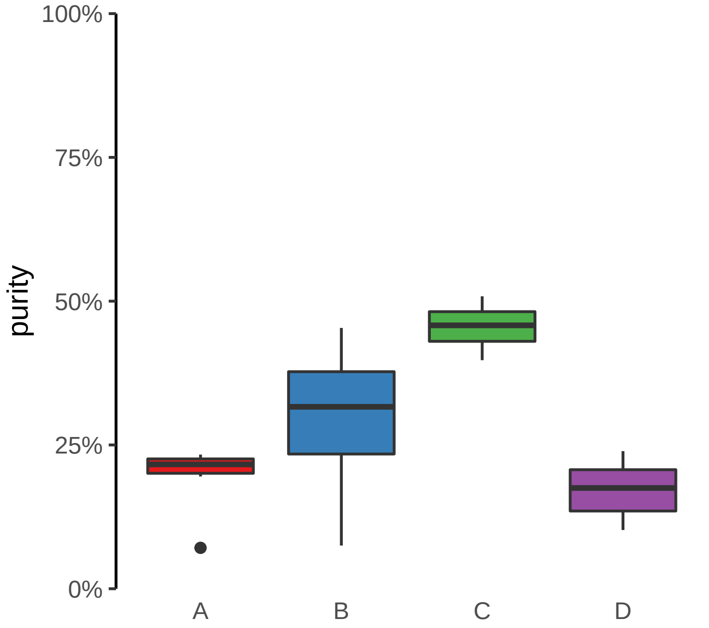

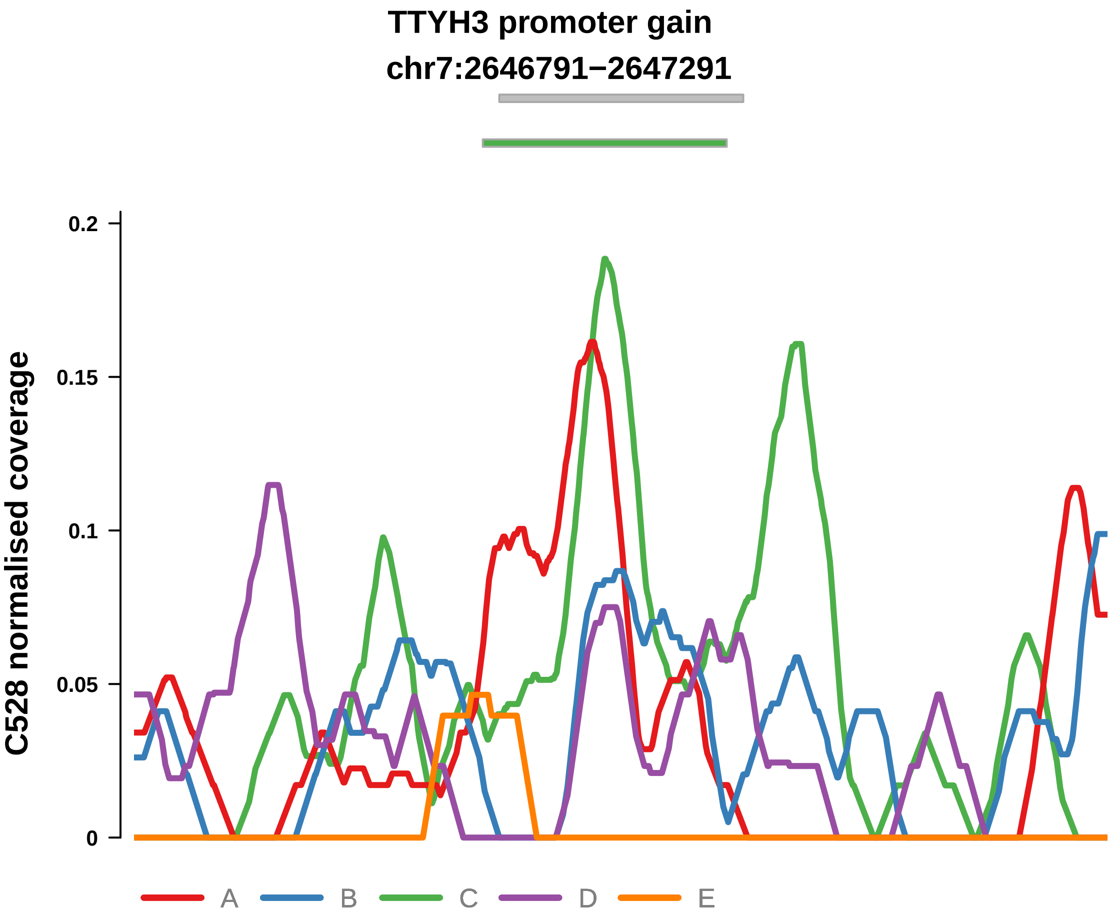


Putatively Clonal (logFC = 2.48)

| | P. | Adjusted P. |
|---|---|---|
| *~region vs. ~* | 4.44e−01 | 6.34e−01 |
| *~purity vs ~* | 1.68e−01 | 3.13e−01 |
| *~purity+region vs ~purity* | 3.94e−01 | 7.7e−01 |
| *Carcinoma vs. normal* | 9.39e−05 **** | 9.39e−05 **** |

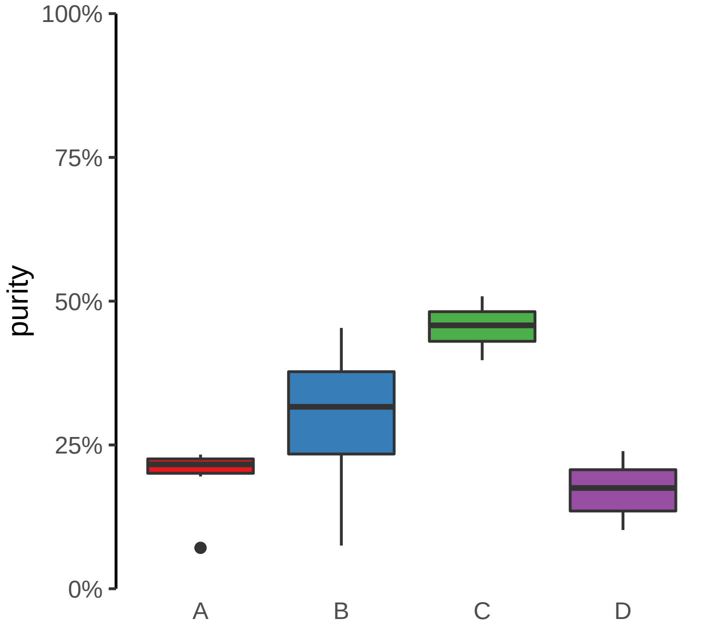

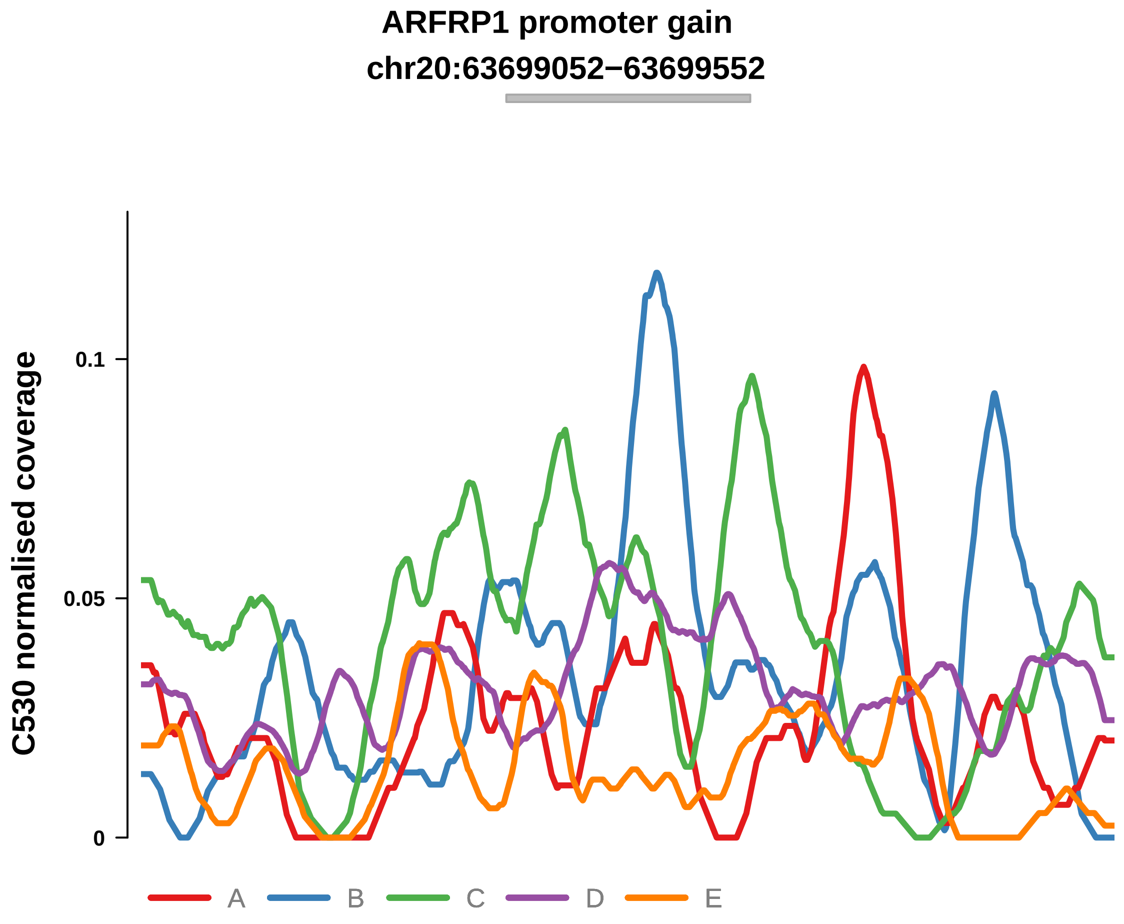


Putatively Clonal (logFC = 2.33)

| | P. | Adjusted P. |
|---|---|---|
| *~region vs. ~* | 1.16e−02 * | 1.34e−01 |
| *~purity vs ~* | 9.72e−02 | 4.33e−01 |
| *~purity+region vs ~purity* | 1.12e−02 * | 1.34e−01 |
| *Carcinoma vs. normal* | 4.99e−05 **** | 4.99e−05 **** |

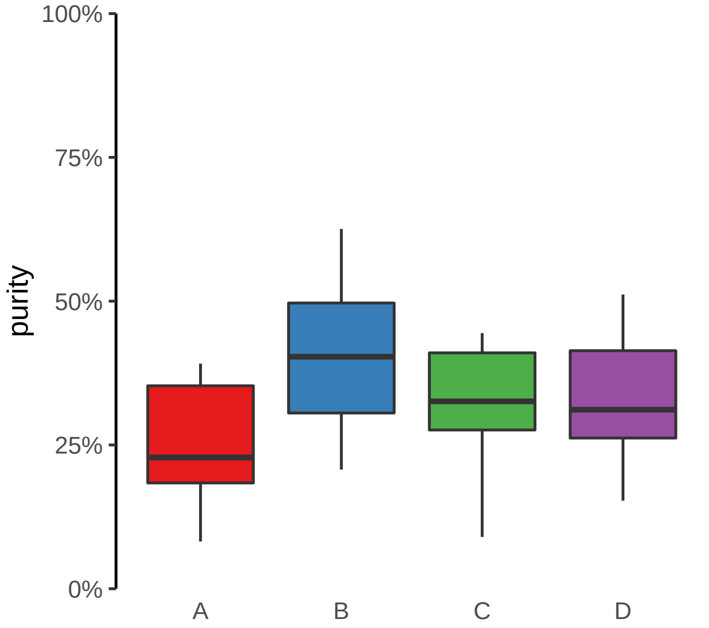

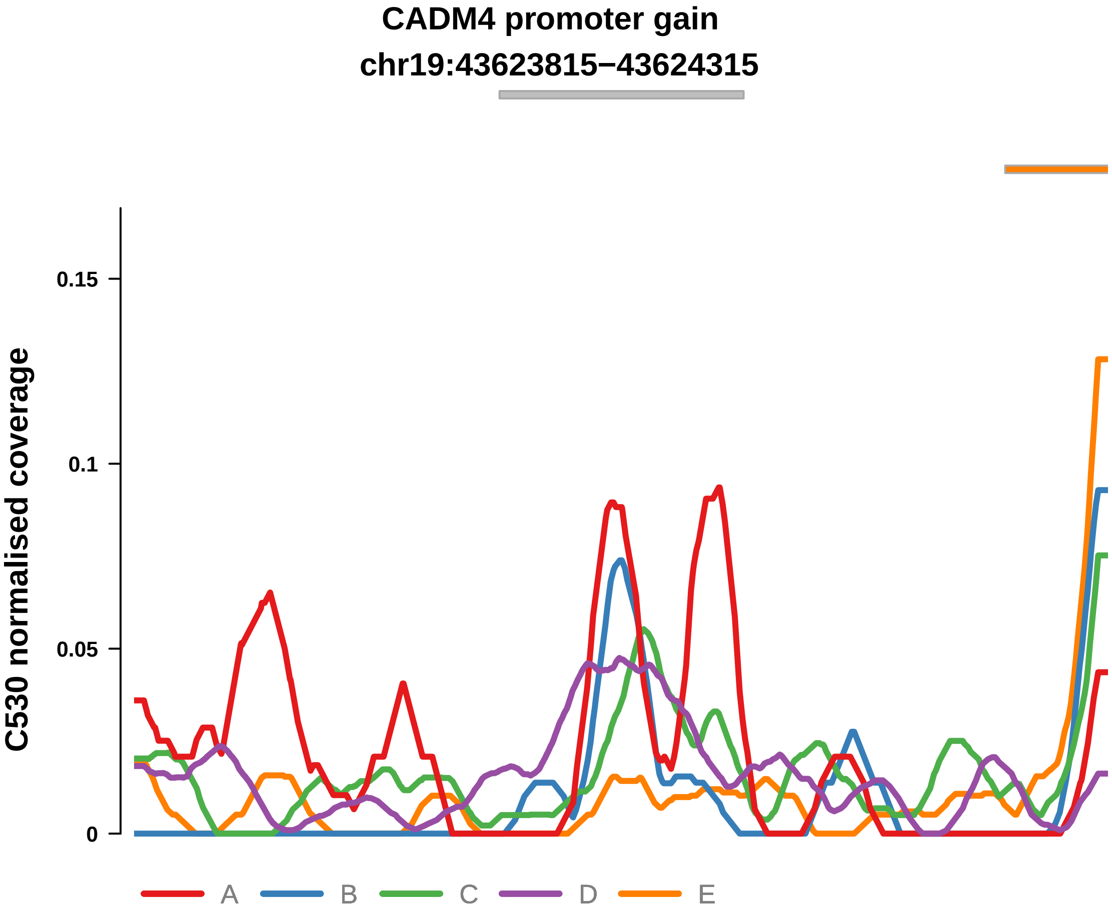


Putatively Clonal (logFC = 2.07)

| | P. | Adjusted P. |
|---|---|---|
| *~region vs. ~* | 1.98e−01 | 5.29e−01 |
| *~purity vs ~* | 5.81e−01 | 9.32e−01 |
| *~purity+region vs ~purity* | 9.93e−02 | 3.5e−01 |
| *Carcinoma vs. normal* | 4.76e−04 *** | 4.76e−04 *** |

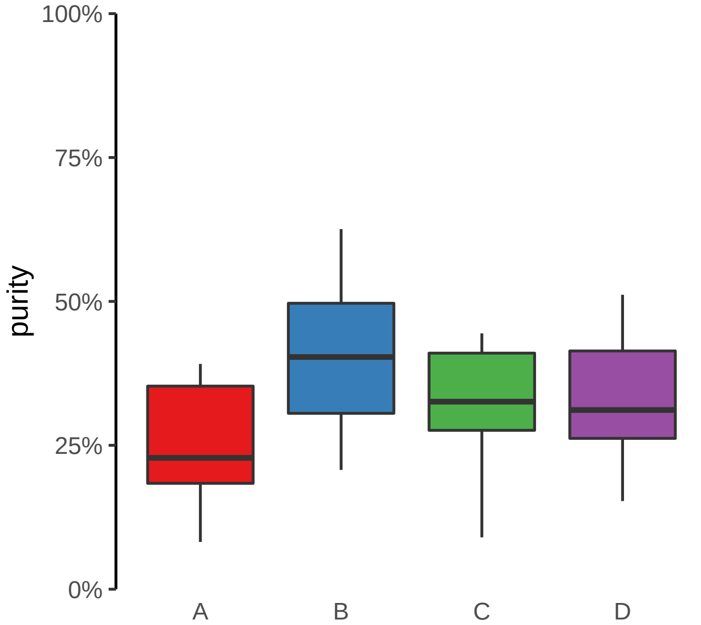

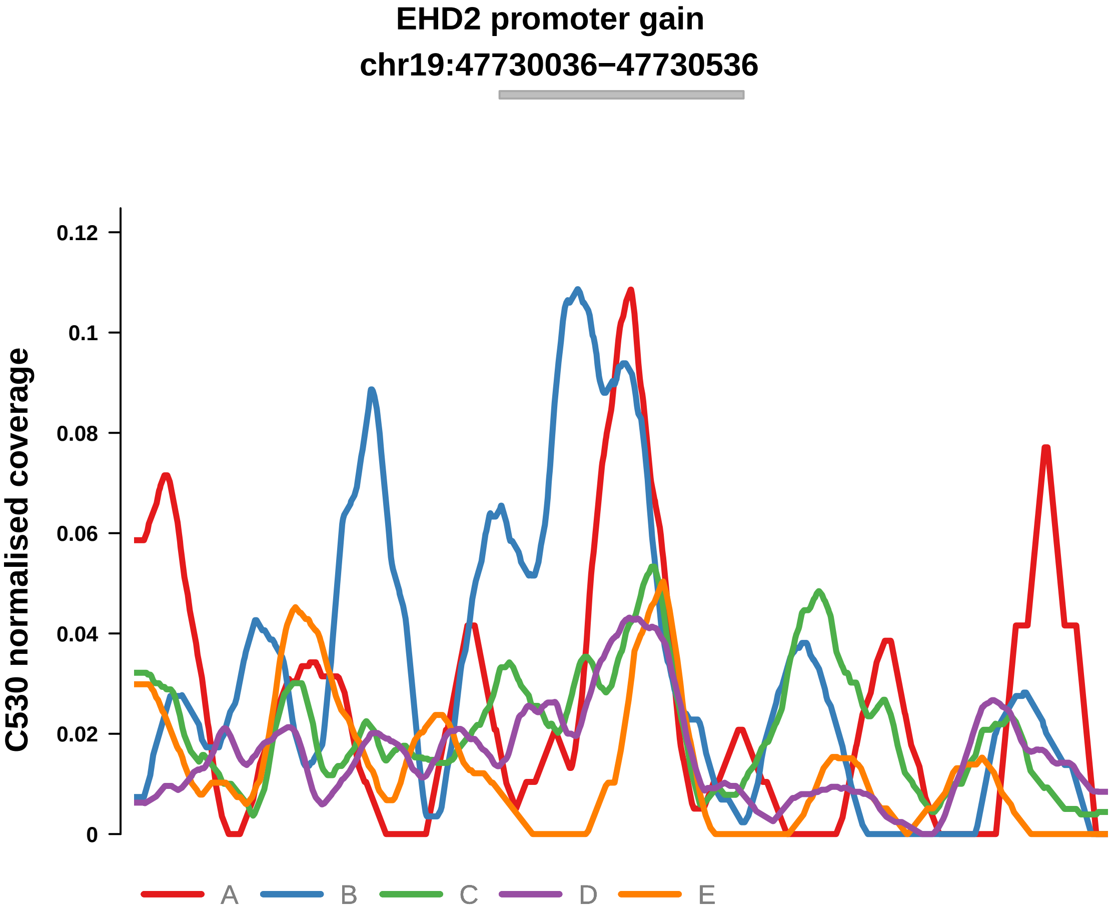


Putatively Clonal (logFC = 1.77)

| | P. | Adjusted P. |
|---|---|---|
| *~region vs. ~* | 4.42e−01 | 7.81e−01 |
| *~purity vs ~* | 1.45e−02 * | 2.55e−01 |
| *~purity+region vs ~purity* | 6.67e−01 | 8.58e−01 |
| *Carcinoma vs. normal* | 1.86e−03 ** | 1.86e−03 ** |

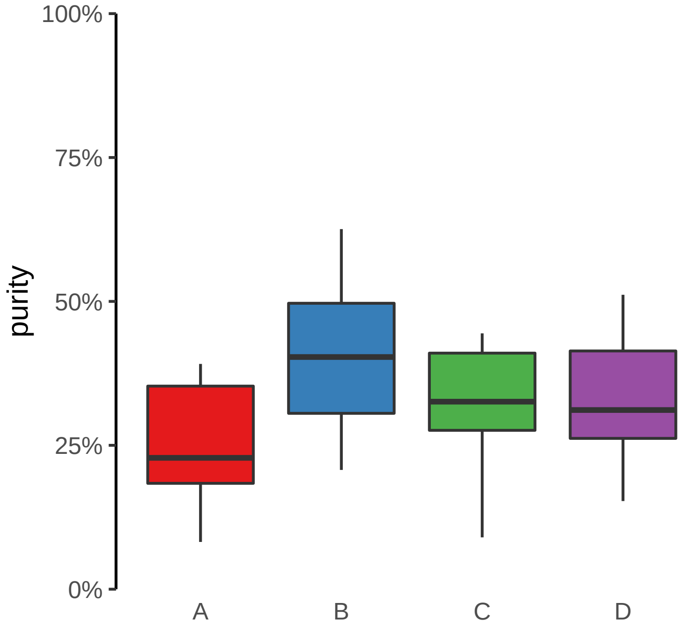

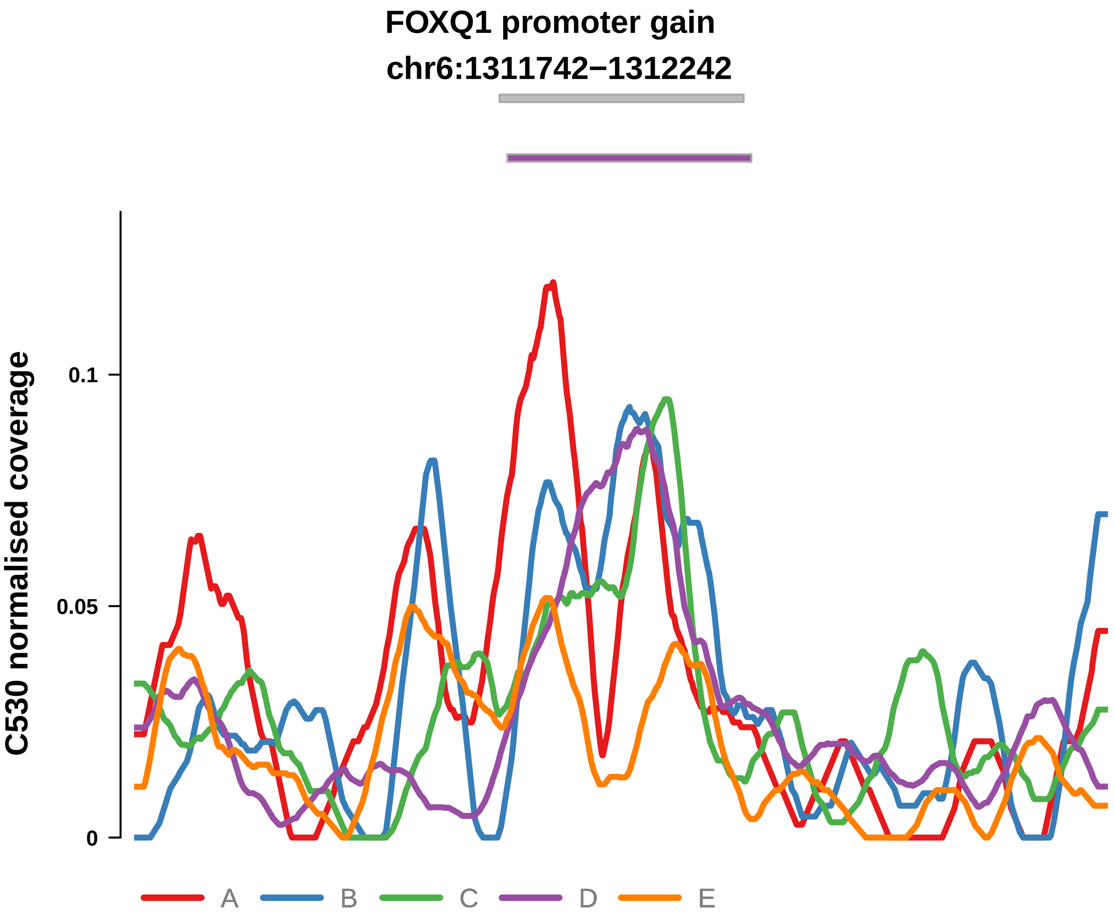


Putatively Clonal (logFC = 1.88)

| | P. | Adjusted P. |
|---|---|---|
| *~region vs. ~* | 8.06e−01 | 8.86e−01 |
| *~purity vs ~* | 5.08e−02 | 3.49e−01 |
| *~purity+region vs ~purity* | 3.88e−01 | 6.82e−01 |
| *Carcinoma vs. normal* | 8.05e−04 *** | 8.05e−04 *** |

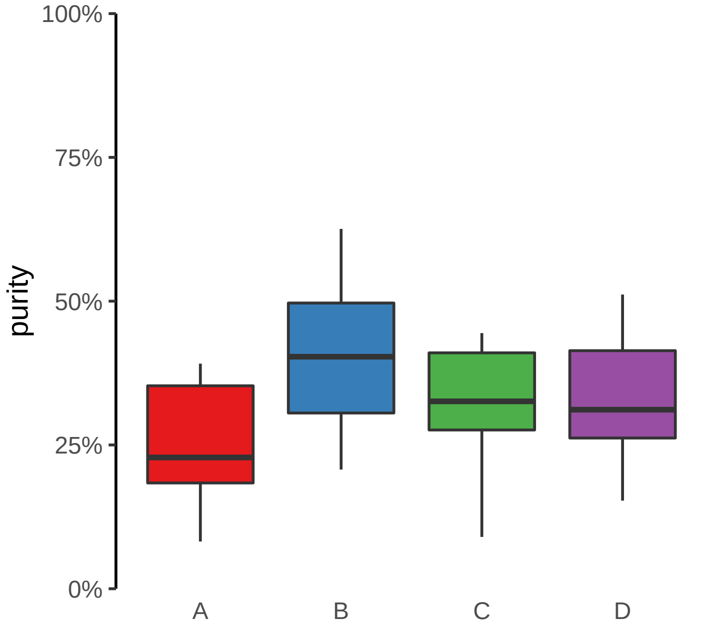

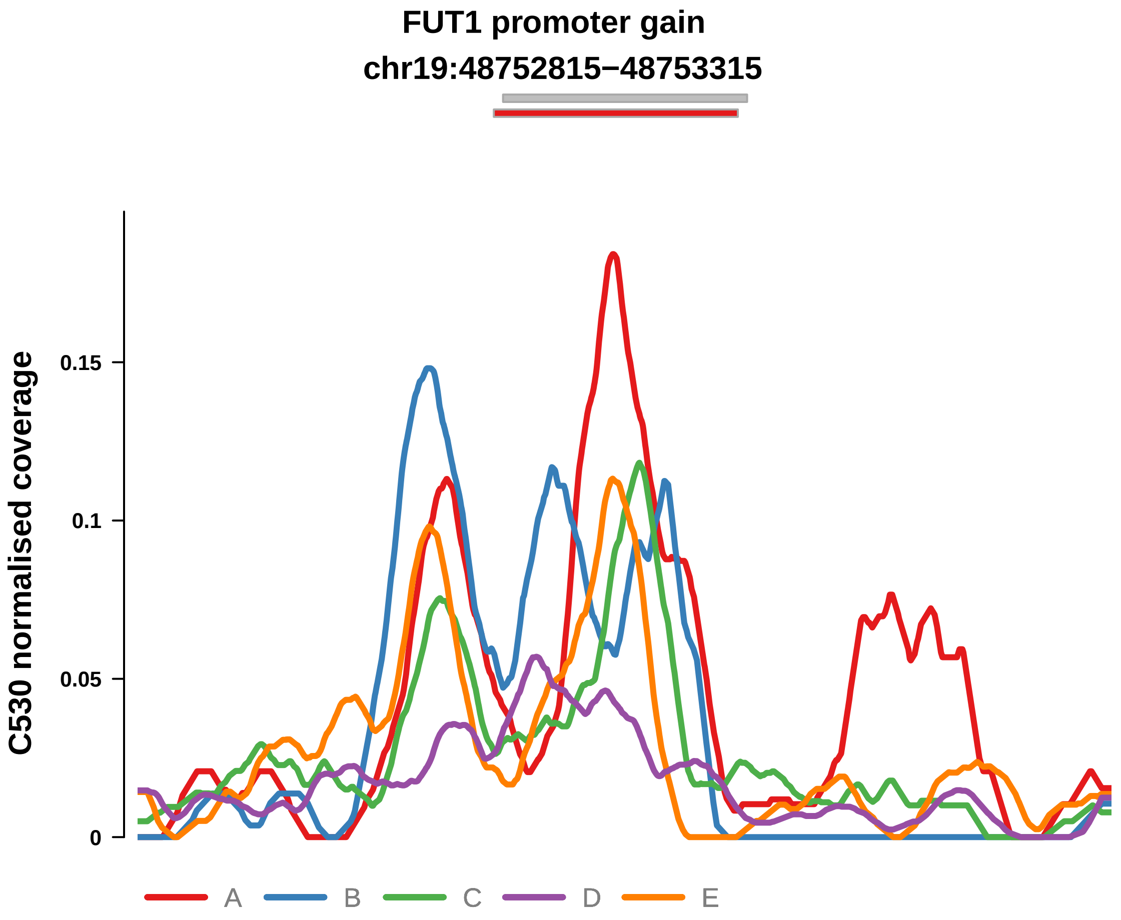


Putatively Clonal (logFC = 1.48)

| | P. | Adjusted P. |
|---|---|---|
| *~region vs. ~* | 4.75e−01 | 8.03e−01 |
| *~purity vs ~* | 7.89e−01 | 9.62e−01 |
| *~purity+region vs ~purity* | 4.23e−01 | 7.16e−01 |
| *Carcinoma vs. normal* | 9.39e−03 ** | 9.39e−03 ** |

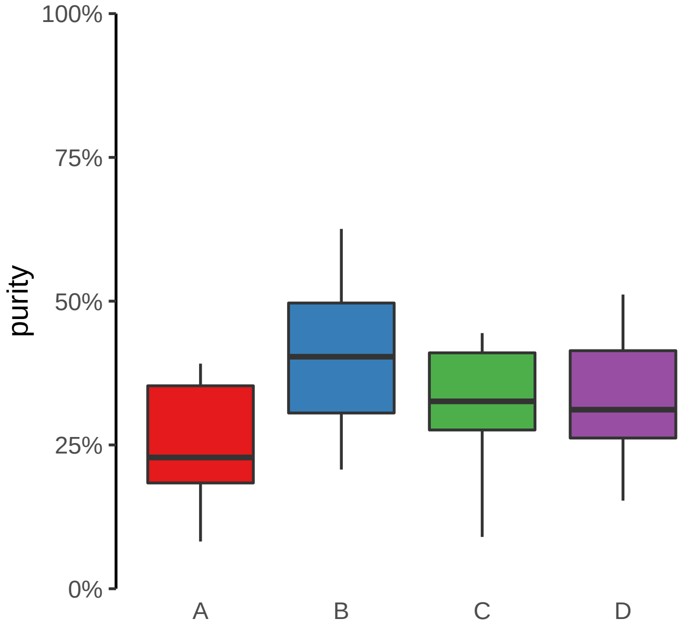

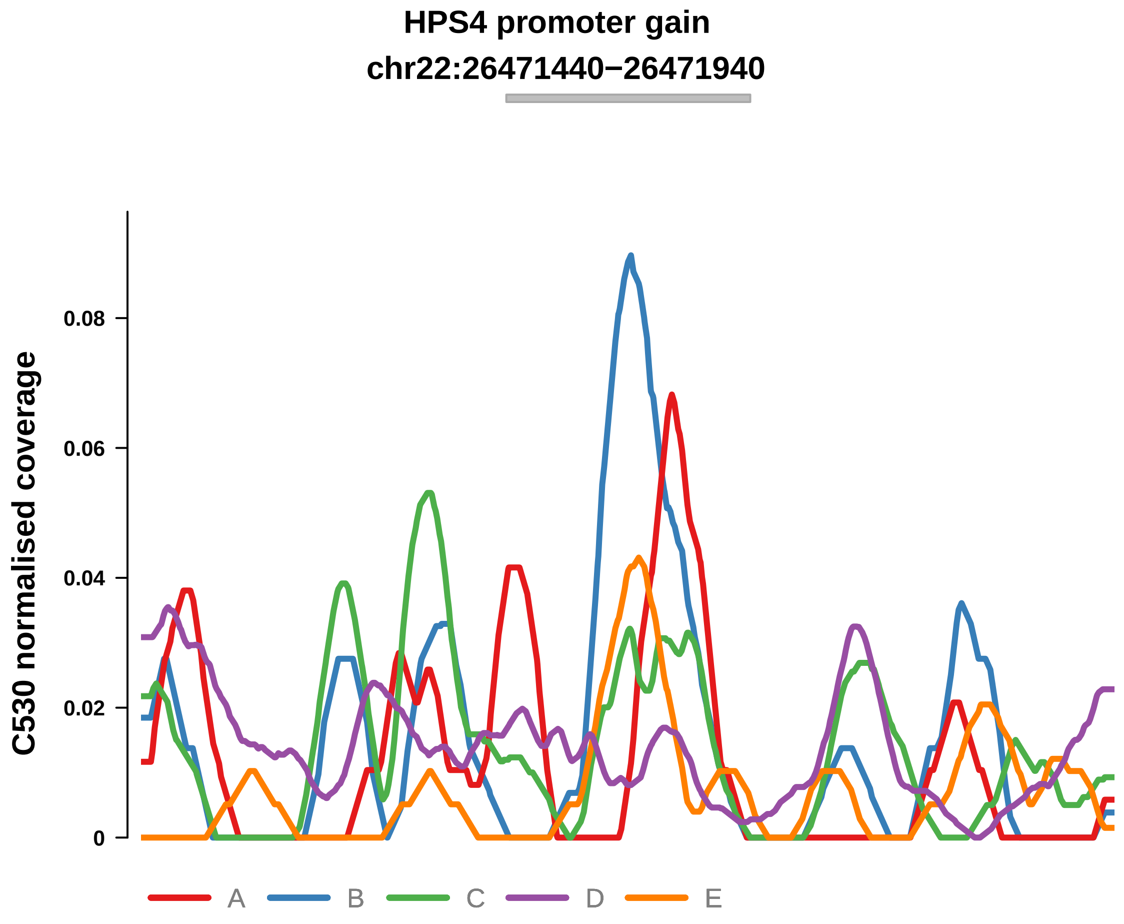


Putatively Clonal (logFC = 1.68)

| | P. | Adjusted P. |
|---|---|---|
| *~region vs. ~* | 4.91e−01 | 8.15e−01 |
| *~purity vs ~* | 5.44e−01 | 9.32e−01 |
| *~purity+region vs ~purity* | 5.35e−01 | 7.8e−01 |
| *Carcinoma vs. normal* | 6.49e−03 ** | 6.49e−03 ** |

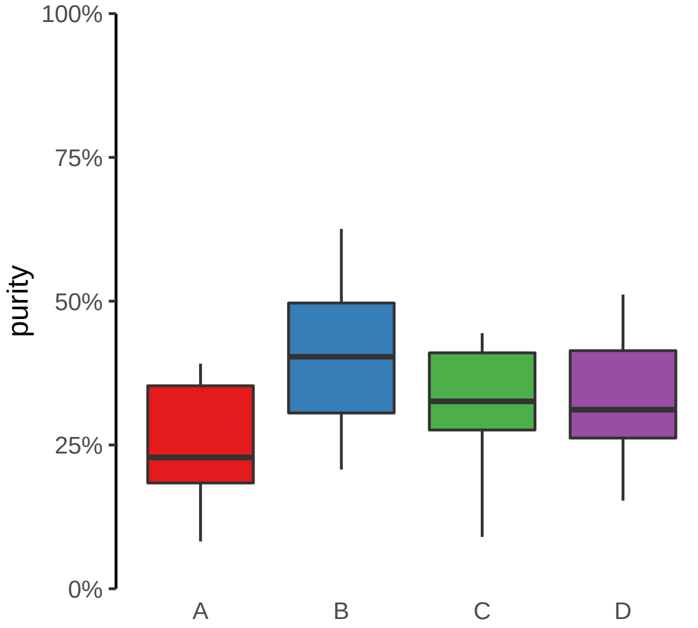

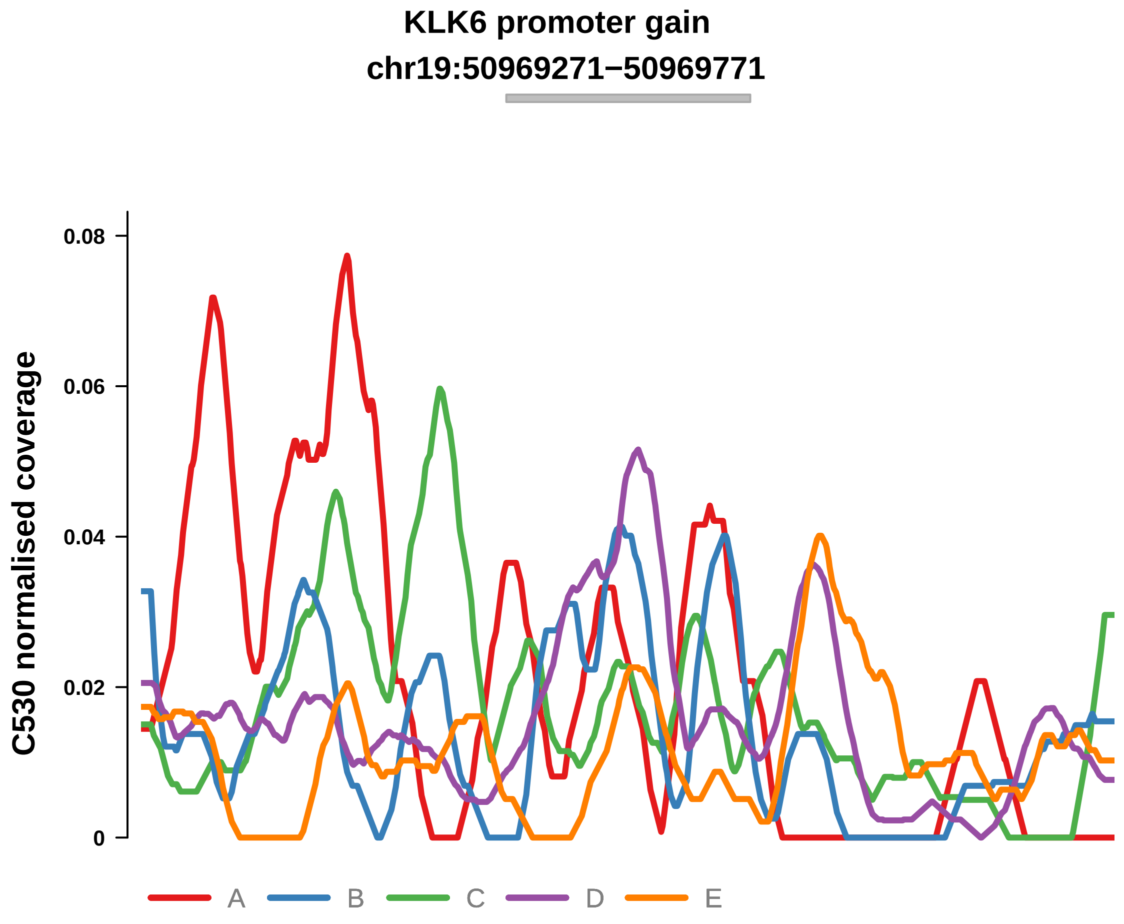


Putatively Clonal (logFC = 1.95)

| | P. | Adjusted P. |
|---|---|---|
| *~region vs. ~* | 9.22e−01 | 9.54e−01 |
| *~purity vs ~* | 5.81e−02 | 3.49e−01 |
| *~purity+region vs ~purity* | 3.37e−01 | 6.45e−01 |
| *Carcinoma vs. normal* | 8.99e−04 *** | 8.99e−04 *** |

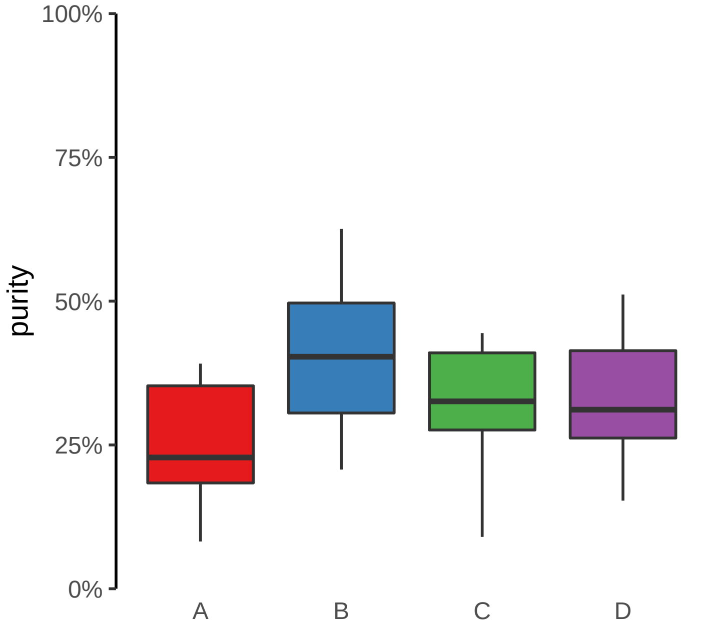

**KRT7 promoter gain**

**chr12:52235898−52236398**

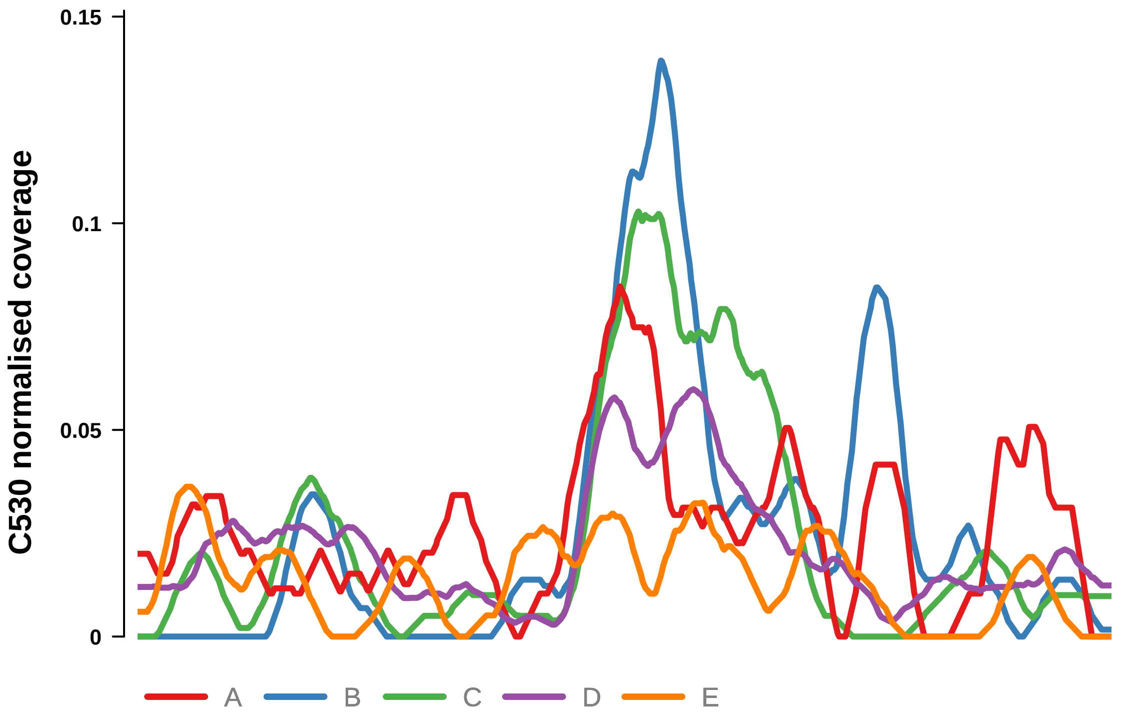


Putatively Clonal (logFC = 1.66)

| | **P.** | **Adjusted P.** |
|---|---|---|
| *~region vs. ~* | 5.55e−01 | 8.43e−01 |
| *~purity vs ~* | 7.73e−02 | 4e−01 |
| *~purity+region vs ~purity* | 4.89e−01 | 7.59e−01 |
| *Carcinoma vs. normal* | 3.38e−03 ** | 3.38e−03 ** |

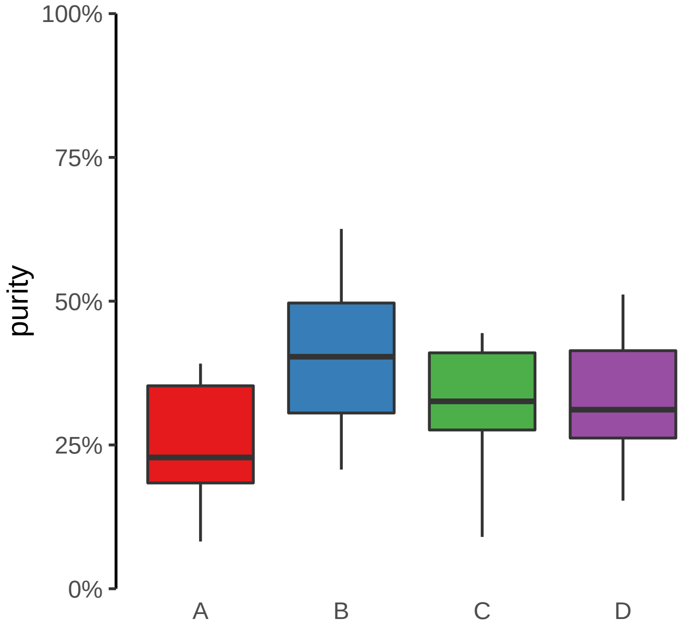

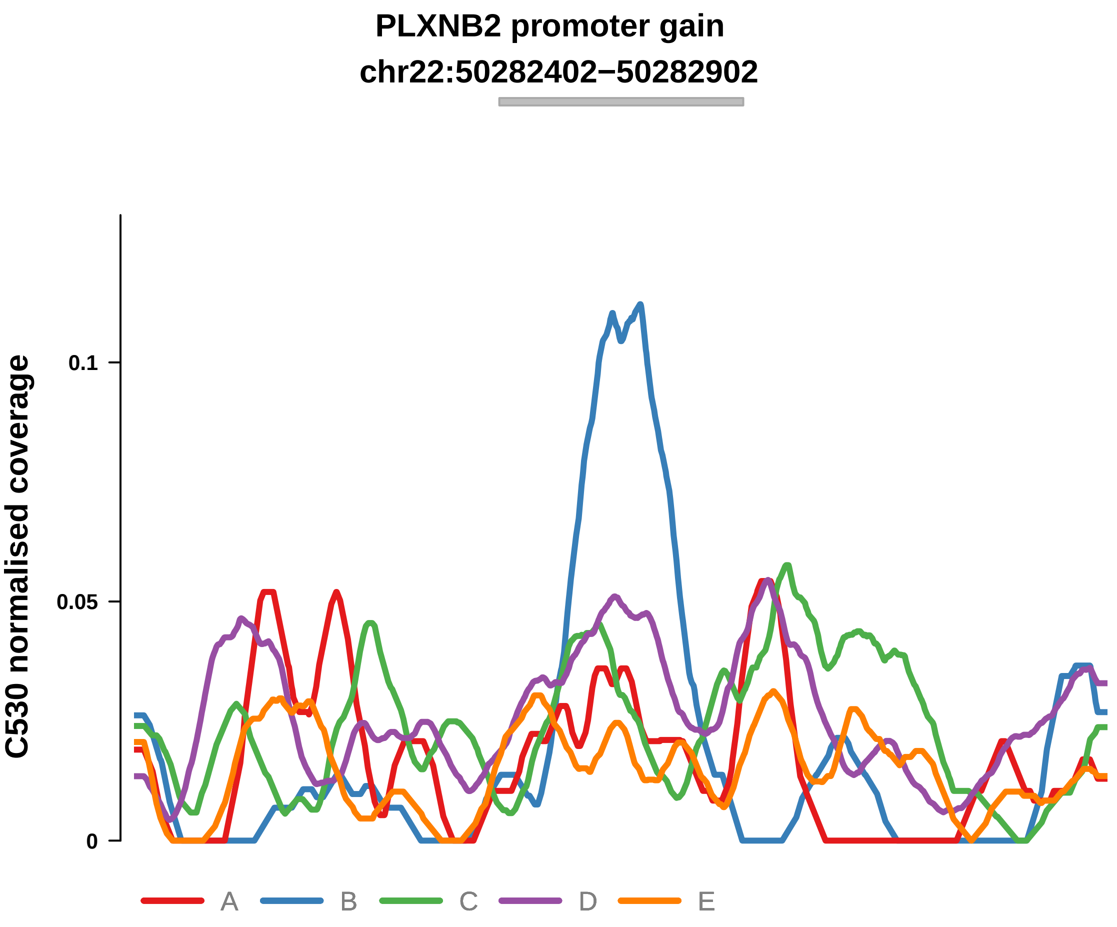


Putatively Clonal (logFC = 1.89)

| | P. | Adjusted P. |
|---|---|---|
| *~region vs. ~* | 6.77e−01 | 8.86e−01 |
| *~purity vs ~* | 1.26e−02 * | 2.55e−01 |
| *~purity+region vs ~purity* | 7.17e−01 | 8.58e−01 |
| *Carcinoma vs. normal* | 9.94e−04 *** | 9.94e−04 *** |

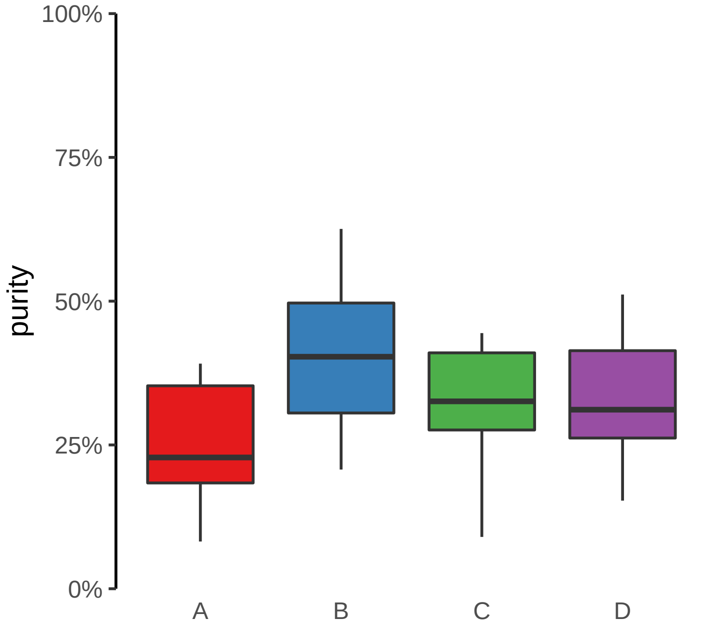

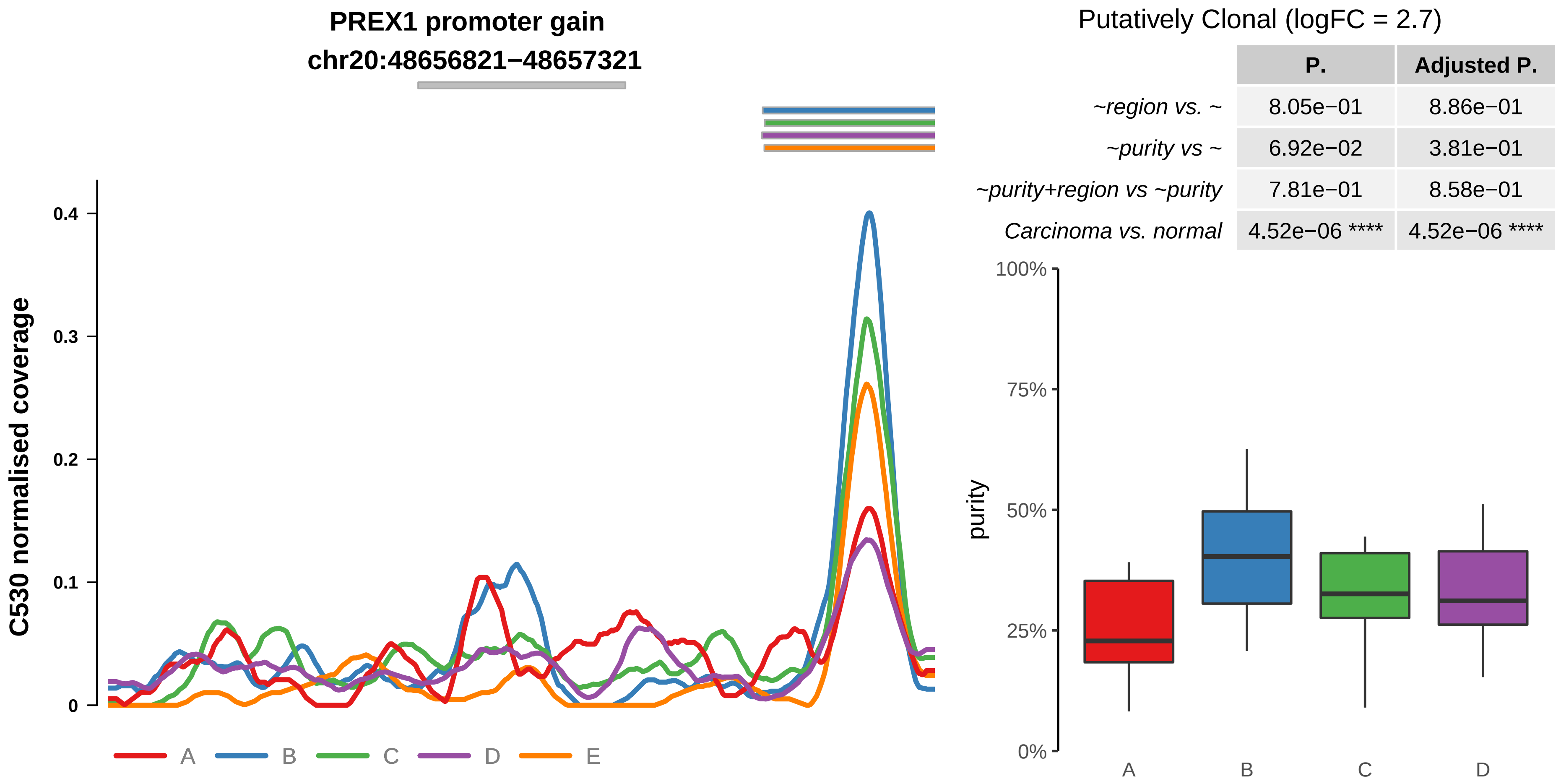
PREX1 promoter gain
chr20:48656821−48657321
C530 normalised coverage
0.4
0.3
0.2
0.1
0
A
B
C
D
E
Putatively Clonal (logFC = 2.7)
P.
Adjusted P.
~region vs. ~ 8.05e−01 8.86e−01
~purity vs ~ 6.92e−02 3.81e−01
~purity+region vs ~purity 7.81e−01 8.58e−01
Carcinoma vs. normal 4.52e−06 **** 4.52e−06 ****
100%
75%
50%
25%
0%
purity
A
B
C
D

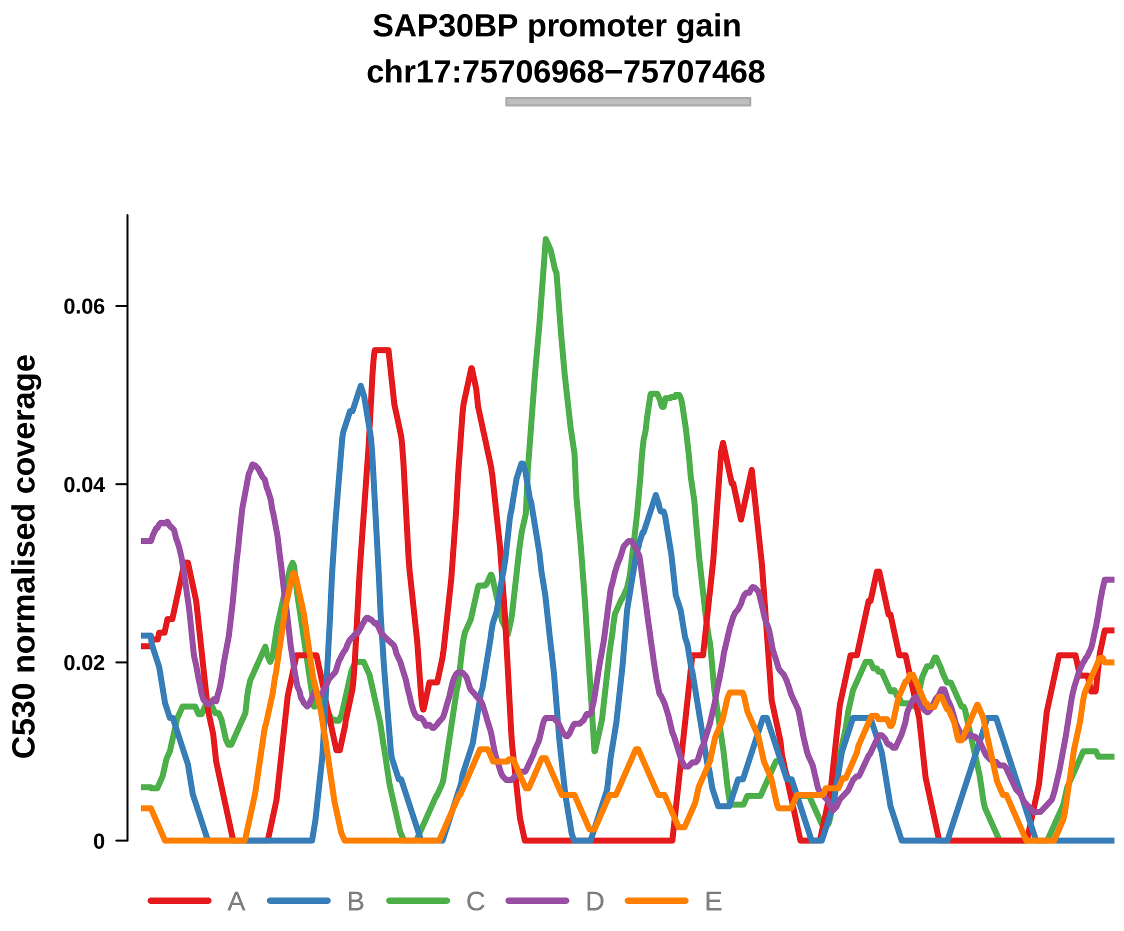


Putatively Clonal (logFC = 1.65)

| | P. | Adjusted P. |
|---|---|---|
| *~region vs. ~* | 1.37e−01 | 4.58e−01 |
| *~purity vs ~* | 3.07e−01 | 7.94e−01 |
| *~purity+region vs ~purity* | 1.13e−01 | 3.56e−01 |
| *Carcinoma vs. normal* | 6.47e−03 ** | 6.47e−03 ** |

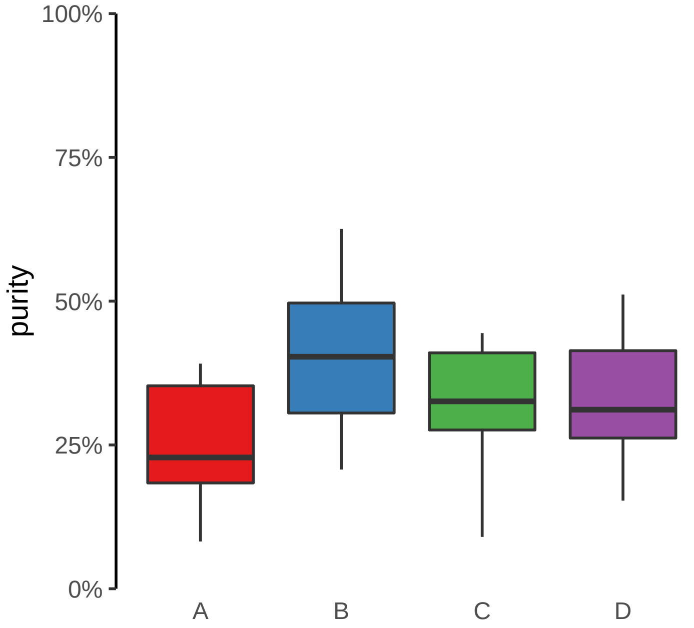

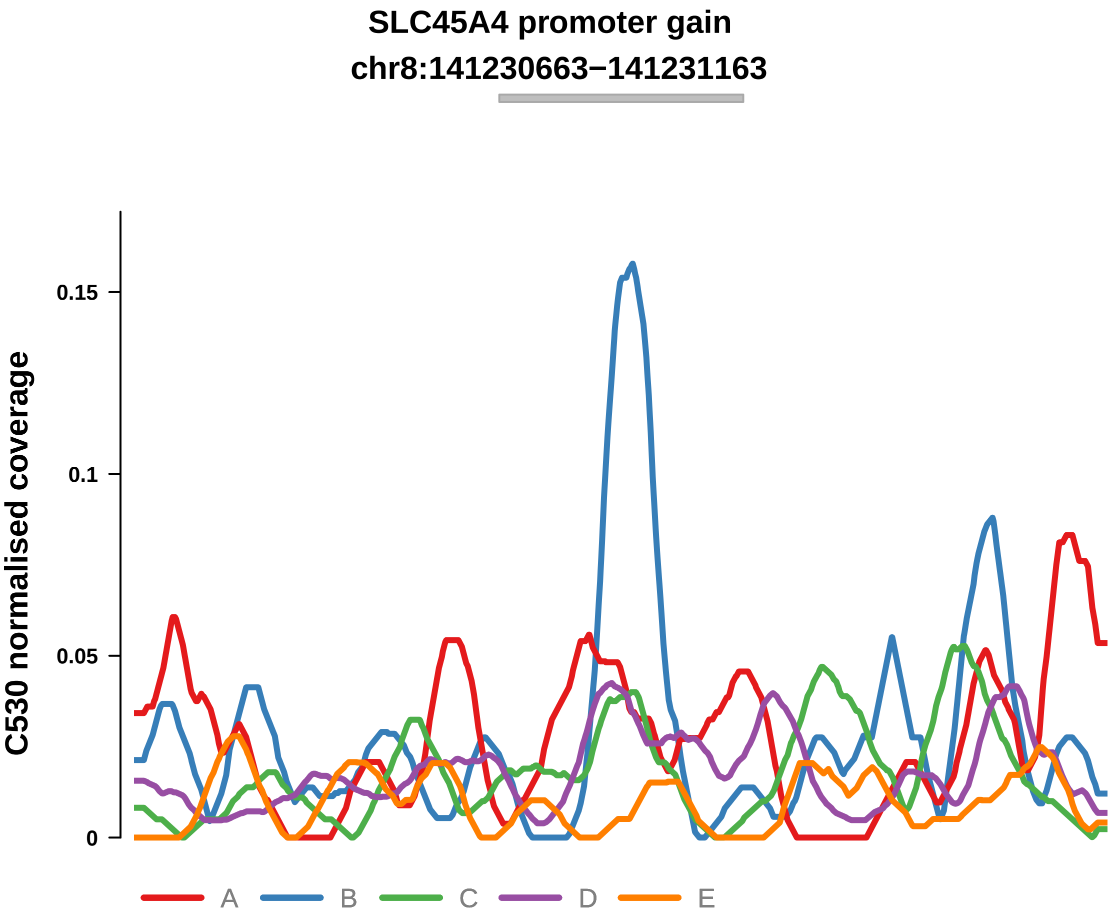


Putatively Clonal (logFC = 2.05)

| | P. | Adjusted P. |
|---|---|---|
| *~region vs. ~* | 6.21e−01 | 8.86e−01 |
| *~purity vs ~* | 4.28e−02 * | 3.43e−01 |
| *~purity+region vs ~purity* | 7.15e−01 | 8.58e−01 |
| *Carcinoma vs. normal* | 5.8e−04 *** | 5.8e−04 *** |

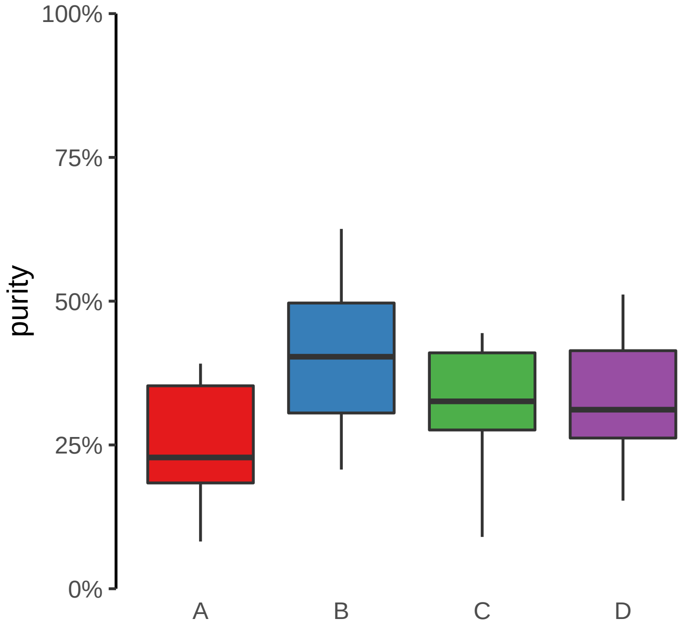

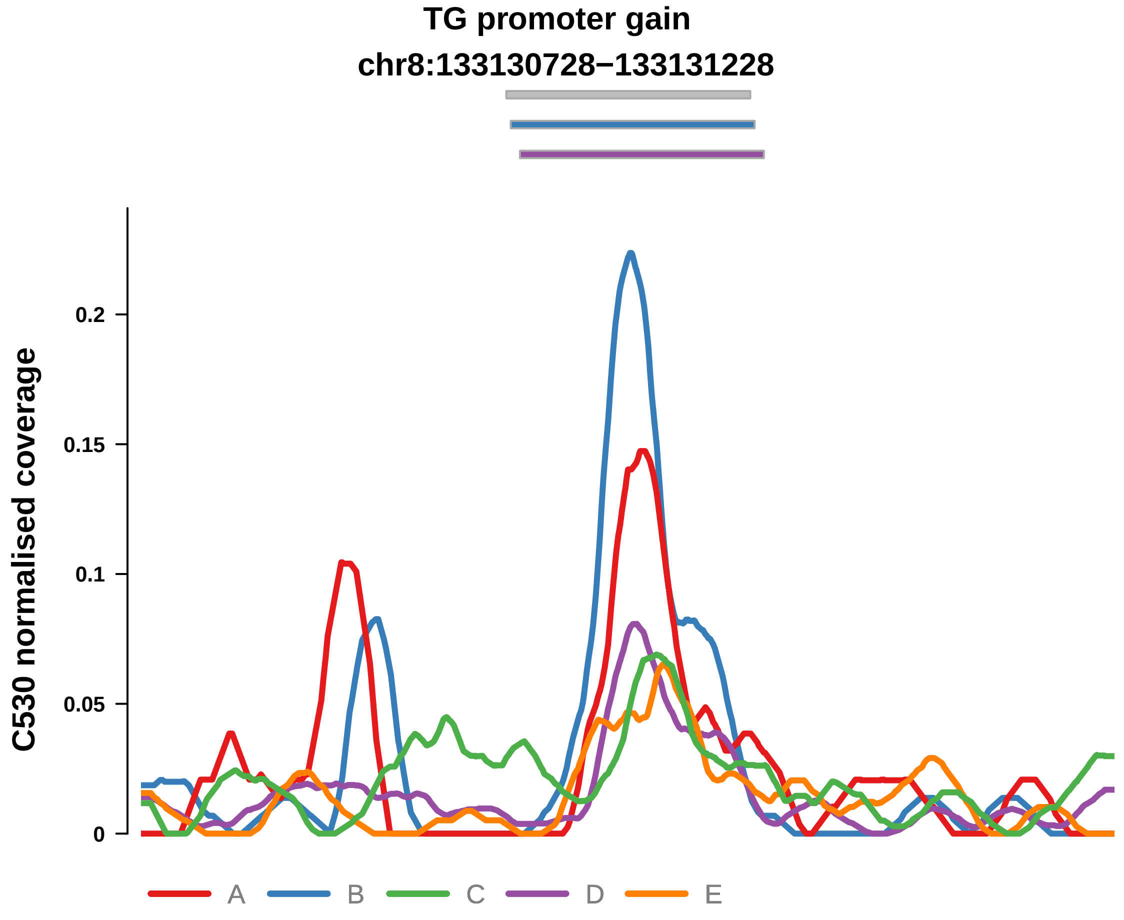


Putatively Clonal (logFC = 1.94)

| | P. | Adjusted P. |
|---|---|---|
| *~region vs. ~* | 7.66e−02 | 3.55e−01 |
| *~purity vs ~* | 1.39e−01 | 4.71e−01 |
| *~purity+region vs ~purity* | 1.52e−01 | 4.01e−01 |
| *Carcinoma vs. normal* | 7.34e−04 *** | 7.34e−04 *** |

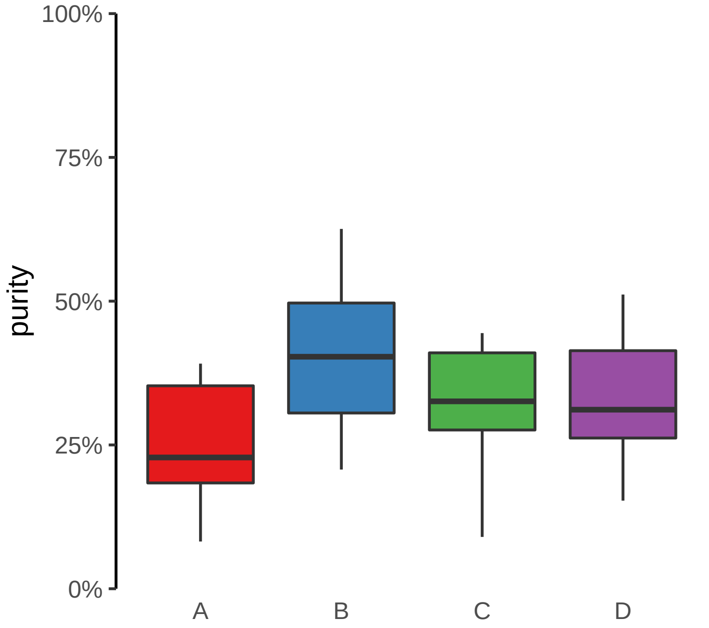

**ARHGEF17 promoter gain**
**chr11:73362044−73362544**

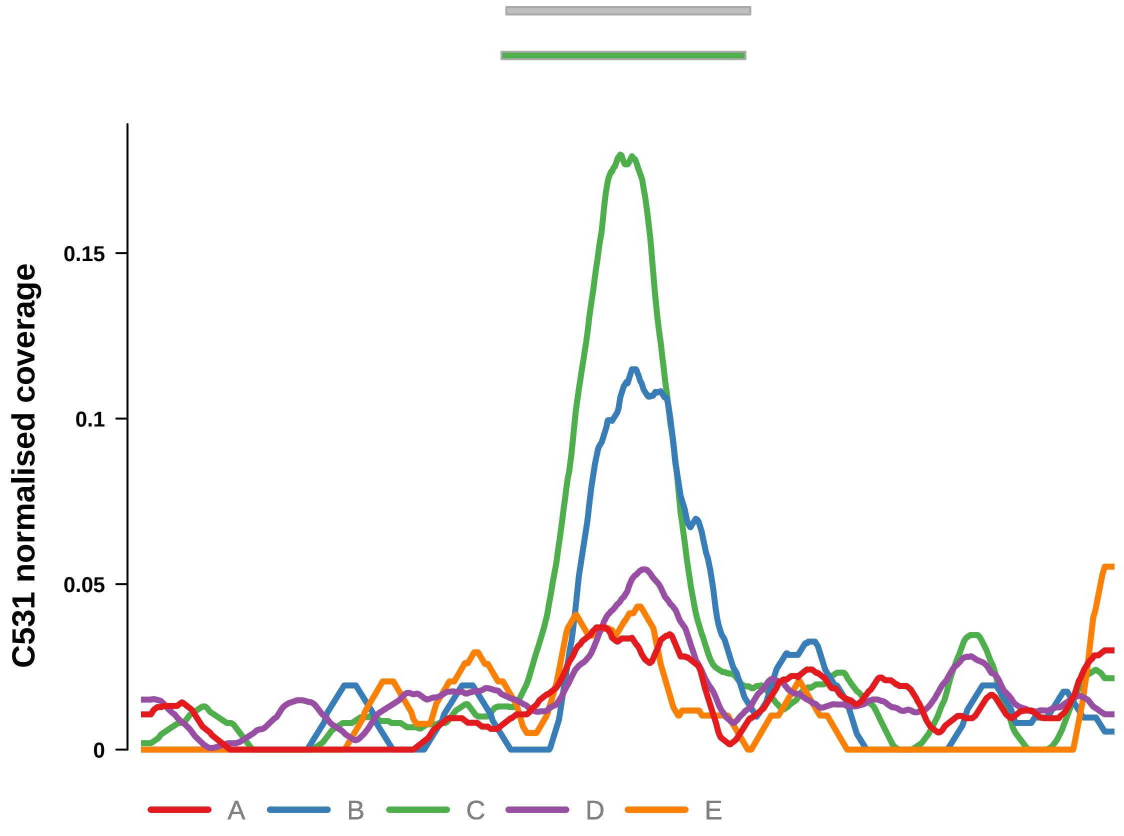


Putatively Clonal (logFC = 3.18)

| | **P.** | **Adjusted P.** |
|---|---|---|
| *~region vs. ~* | 3.08e−05 **** | 3.21e−04 *** |
| *~purity vs ~* | 1.2e−02 * | 5.87e−02 |
| *~purity+region vs ~purity* | 2.3e−03 ** | 6.55e−02 |
| *Carcinoma vs. normal* | 4.48e−07 **** | 4.48e−07 **** |

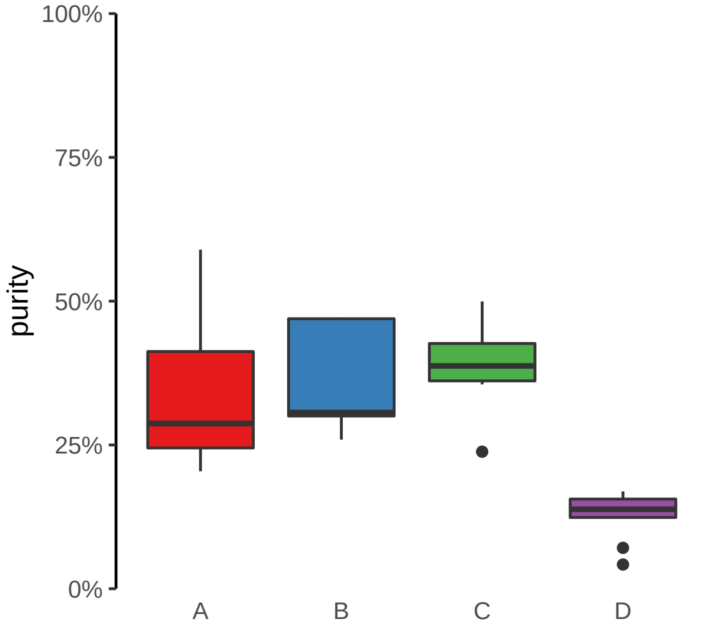

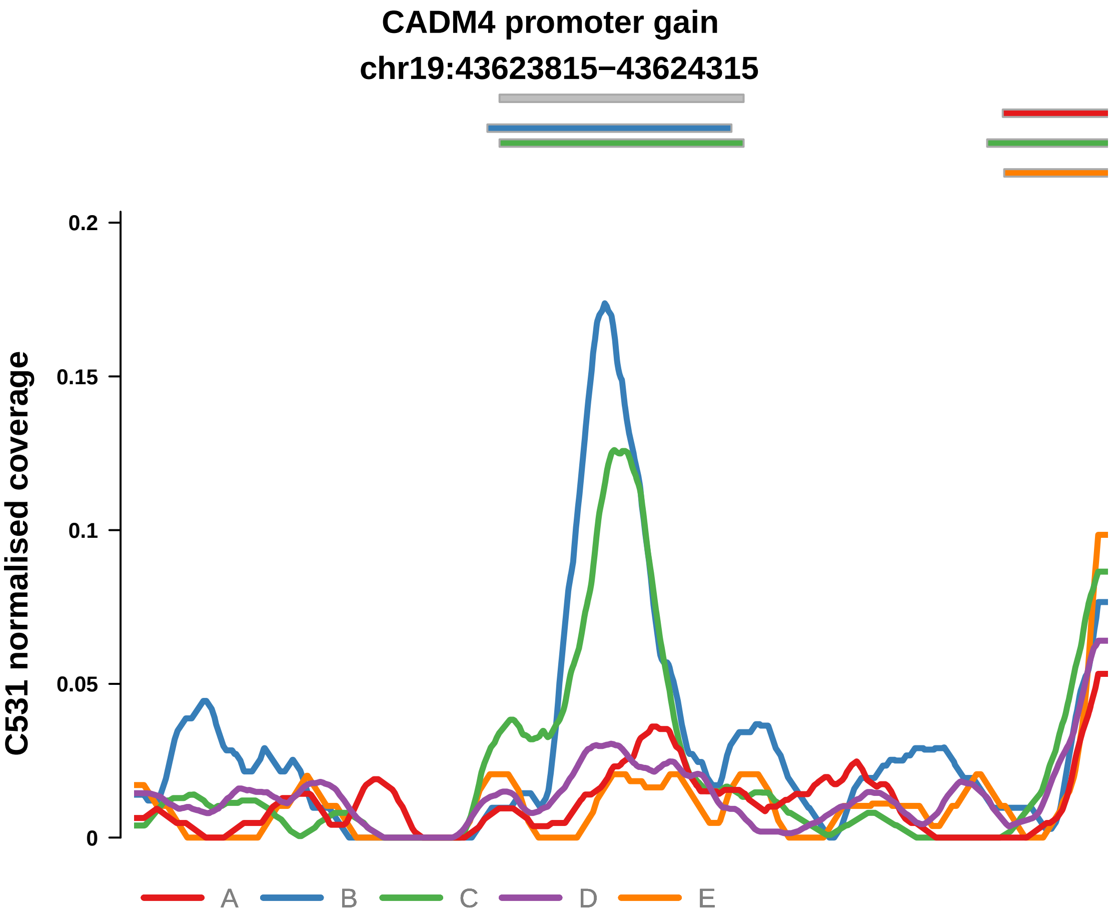


Putatively Clonal (logFC = 2.7)

| | P. | Adjusted P. |
|---|---|---|
| *~region vs. ~* | 1.61e−03 ** | 9.3e−03 ** |
| *~purity vs ~* | 1.27e−02 * | 5.89e−02 |
| *~purity+region vs ~purity* | 4.43e−02 * | 3.9e−01 |
| *Carcinoma vs. normal* | 4.58e−05 **** | 4.58e−05 **** |

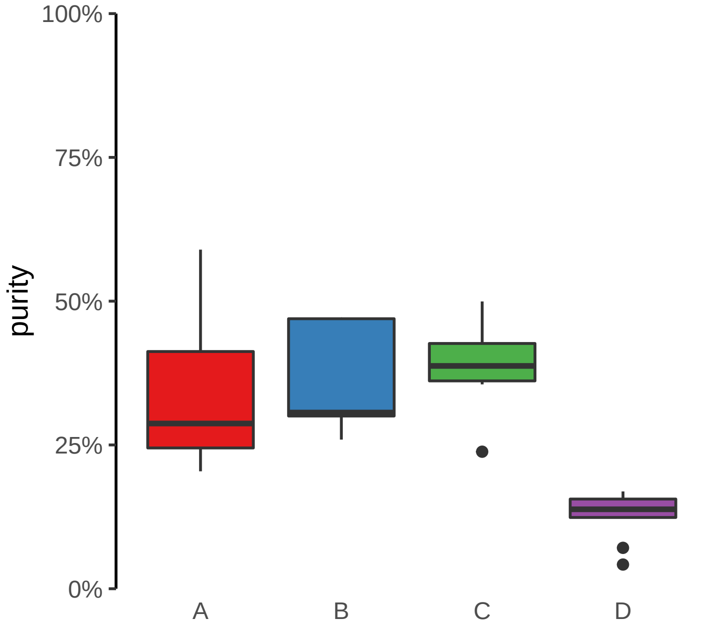

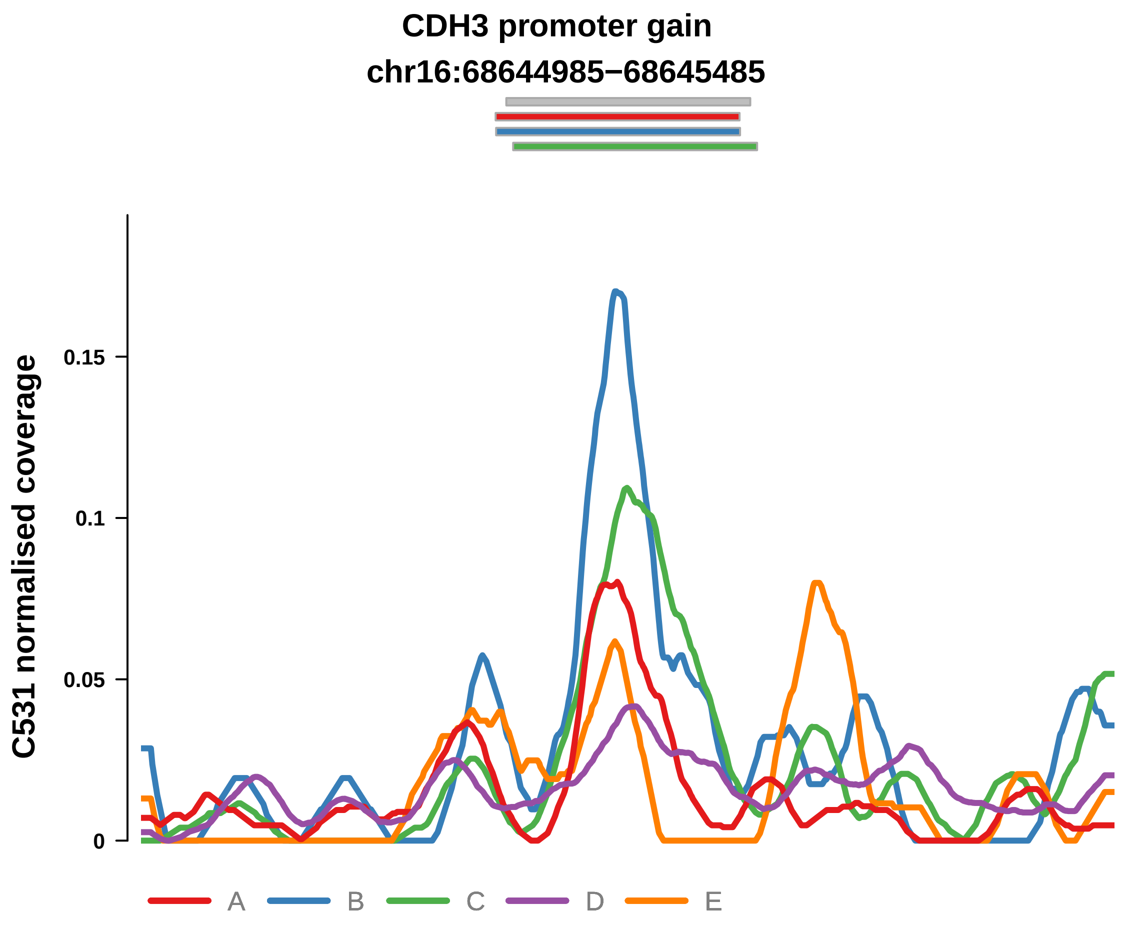


Putatively Clonal (logFC = 1.97)

| | P. | Adjusted P. |
|---|---|---|
| *~region vs. ~* | 8.24e−02 | 1.65e−01 |
| *~purity vs ~* | 9.68e−02 | 2.43e−01 |
| *~purity+region vs ~purity* | 2.66e−01 | 6.13e−01 |
| *Carcinoma vs. normal* | 1.24e−03 ** | 1.24e−03 ** |

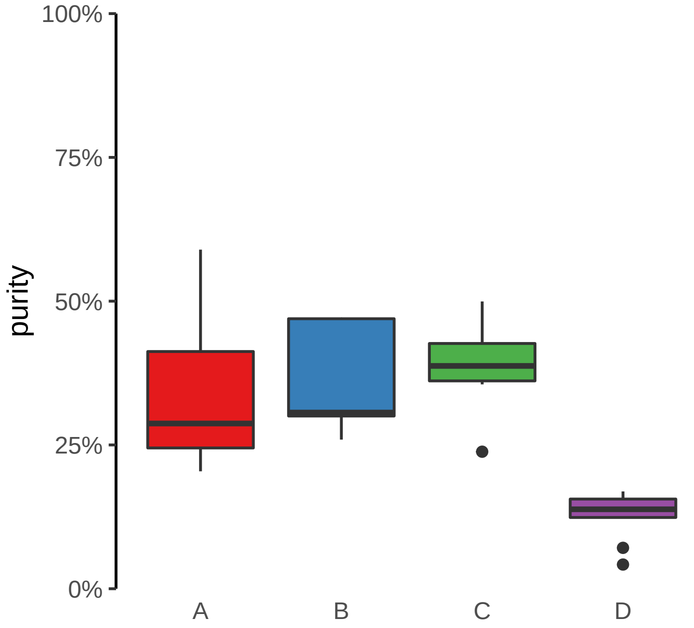

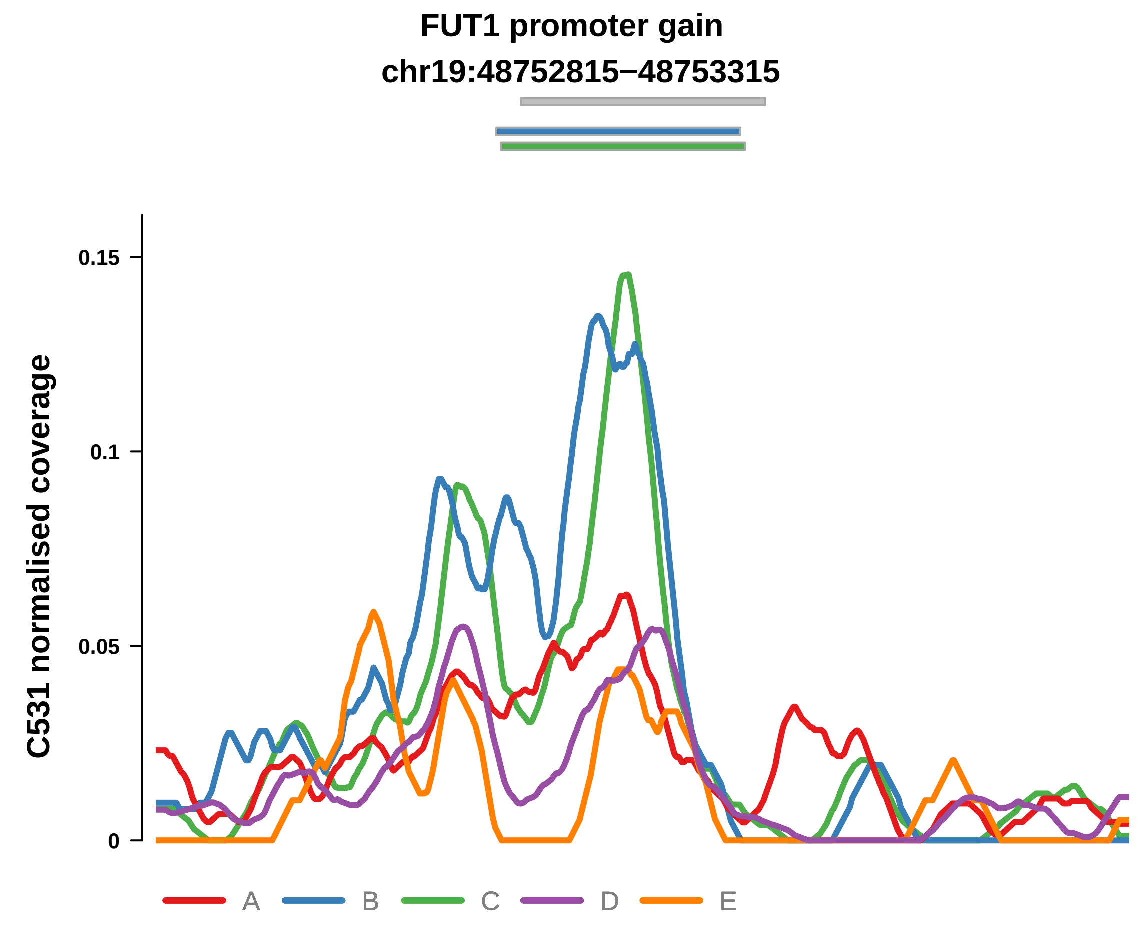


Putatively Clonal (logFC = 2.38)

| | P. | Adjusted P. |
|---|---|---|
| *~region vs. ~* | 1.54e−03 ** | 9.3e−03 ** |
| *~purity vs ~* | 3.31e−03 ** | 2.95e−02 * |
| *~purity+region vs ~purity* | 1.36e−01 | 5.51e−01 |
| *Carcinoma vs. normal* | 9.1e−05 **** | 9.1e−05 **** |

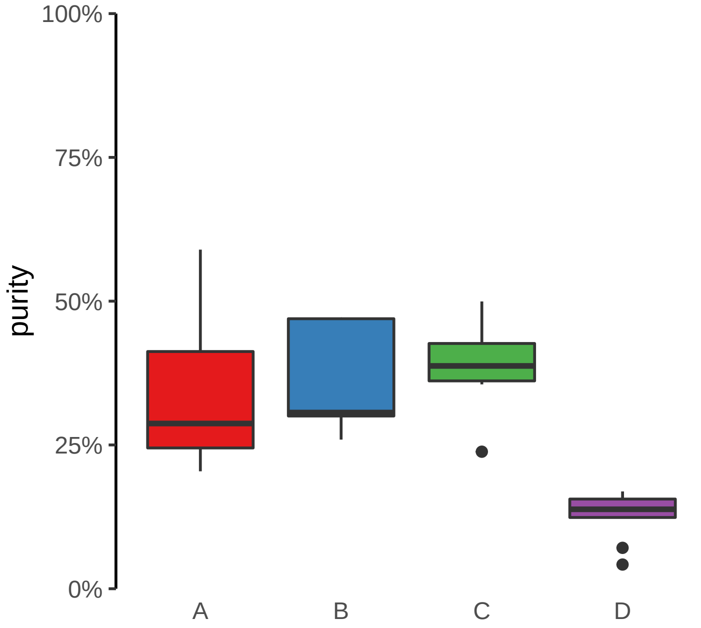

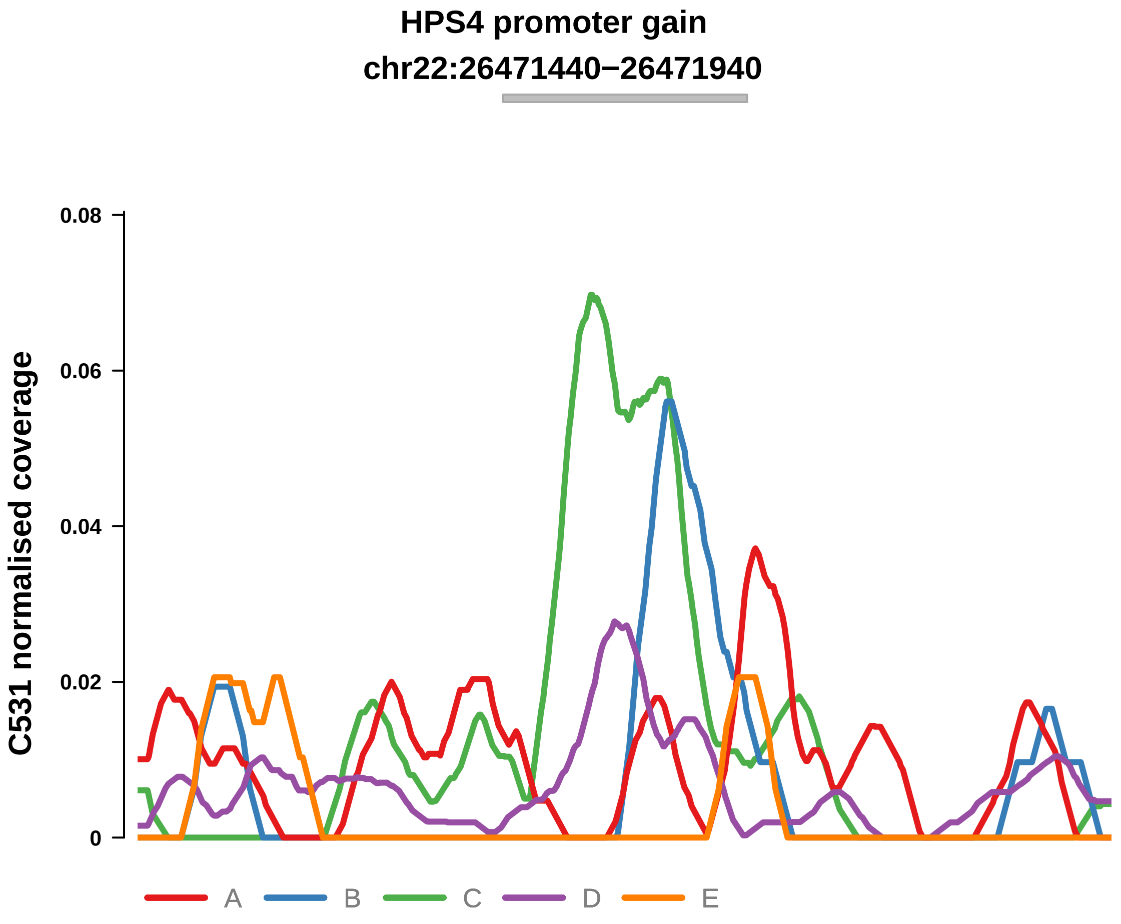


Putatively Clonal (logFC = 2.53)

| | P. | Adjusted P. |
|---|---|---|
| *~region vs. ~* | 3.22e−04 *** | 2.39e−03 ** |
| *~purity vs ~* | 1.67e−02 * | 7e−02 |
| *~purity+region vs ~purity* | 3.31e−03 ** | 6.55e−02 |
| *Carcinoma vs. normal* | 3.72e−04 *** | 3.72e−04 *** |

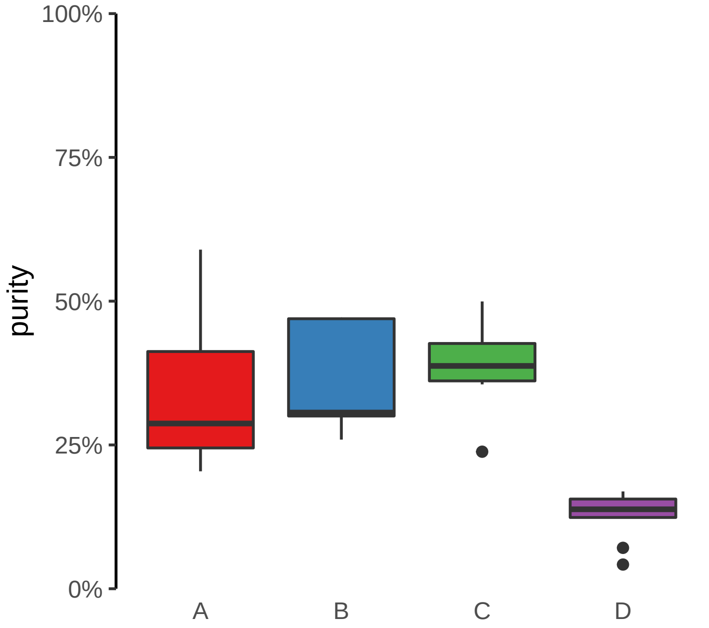

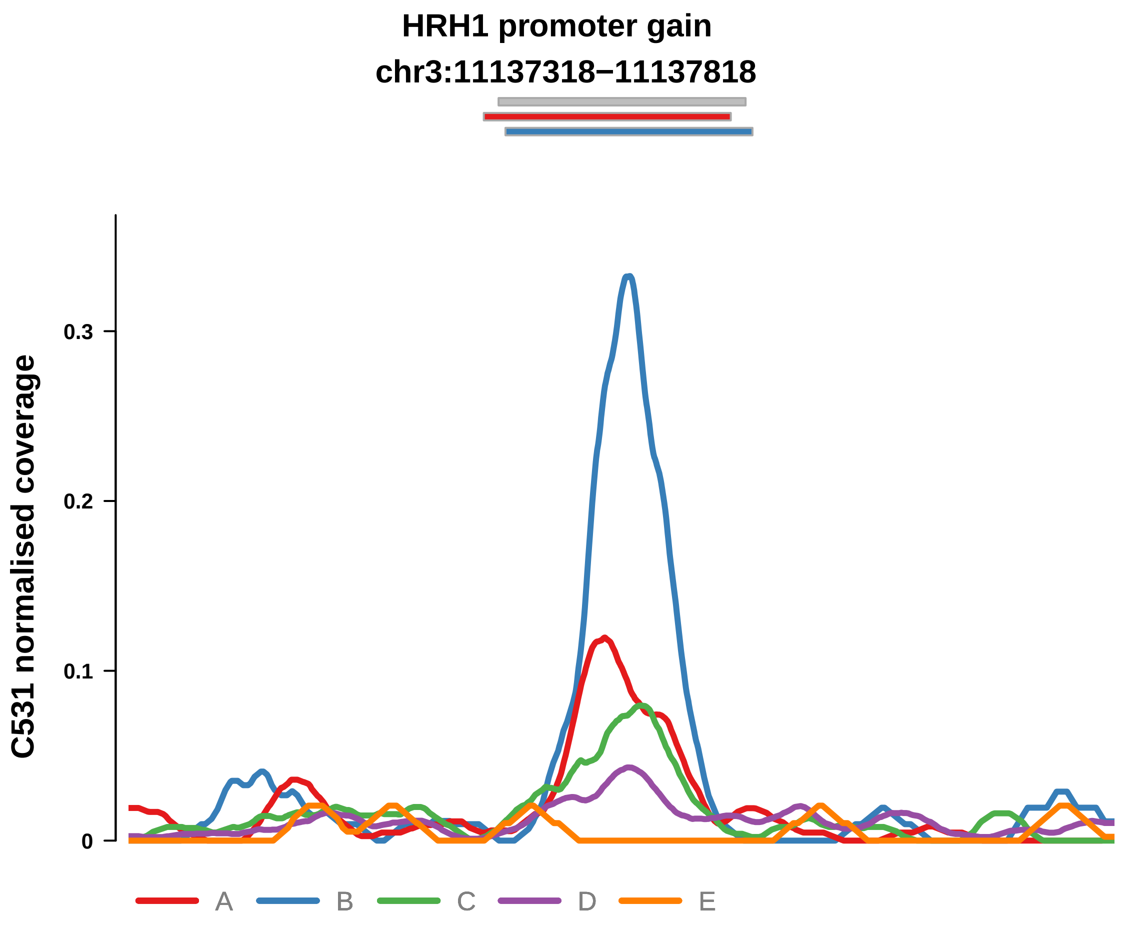


Putatively Clonal (logFC = 2.84)

| | P. | Adjusted P. |
|---|---|---|
| *~region vs. ~* | 2.51e−03 ** | 1.19e−02 * |
| *~purity vs ~* | 3.68e−03 ** | 2.95e−02 * |
| *~purity+region vs ~purity* | 1.38e−01 | 5.51e−01 |
| *Carcinoma vs. normal* | 6.62e−06 **** | 6.62e−06 **** |

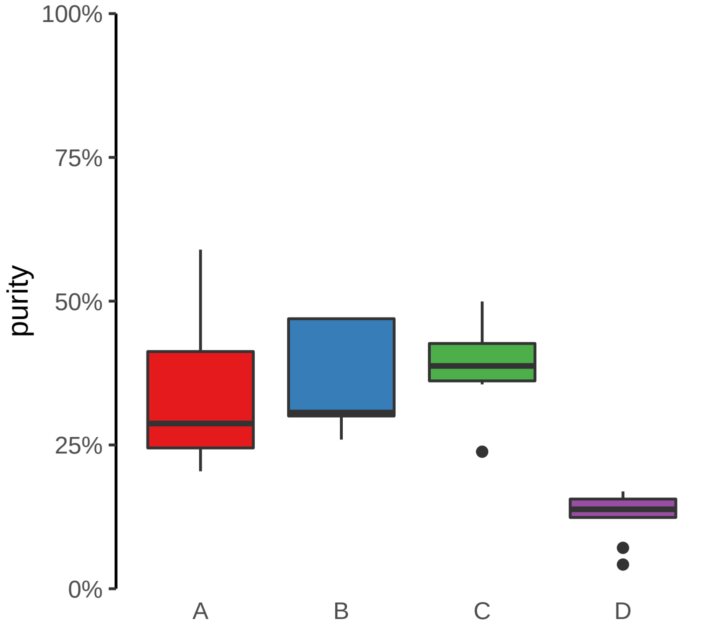

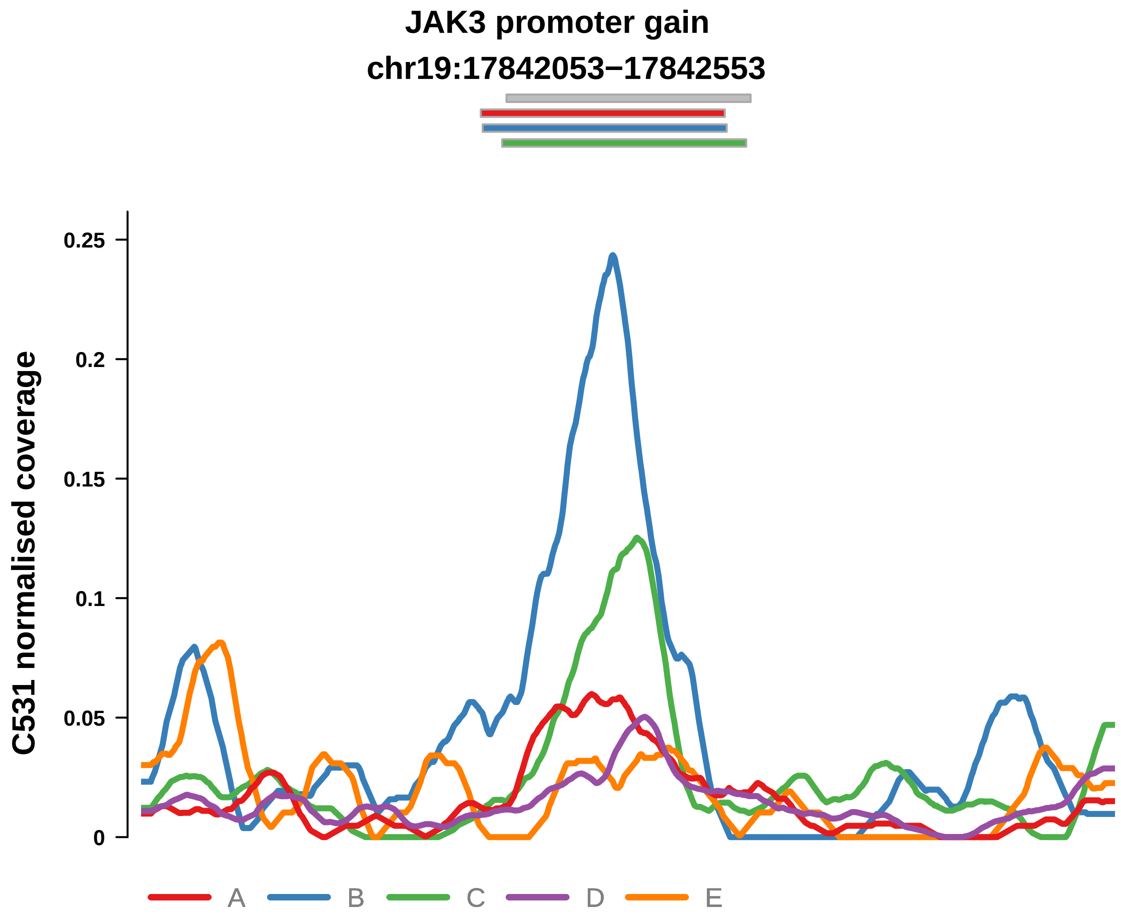


Putatively Clonal (logFC = 2.27)

| | P. | Adjusted P. |
|---|---|---|
| *~region vs. ~* | 1.85e−03 ** | 9.6e−03 ** |
| *~purity vs ~* | 7.98e−03 ** | 4.83e−02 * |
| *~purity+region vs ~purity* | 8.97e−02 | 5.23e−01 |
| *Carcinoma vs. normal* | 2.44e−04 *** | 2.44e−04 *** |

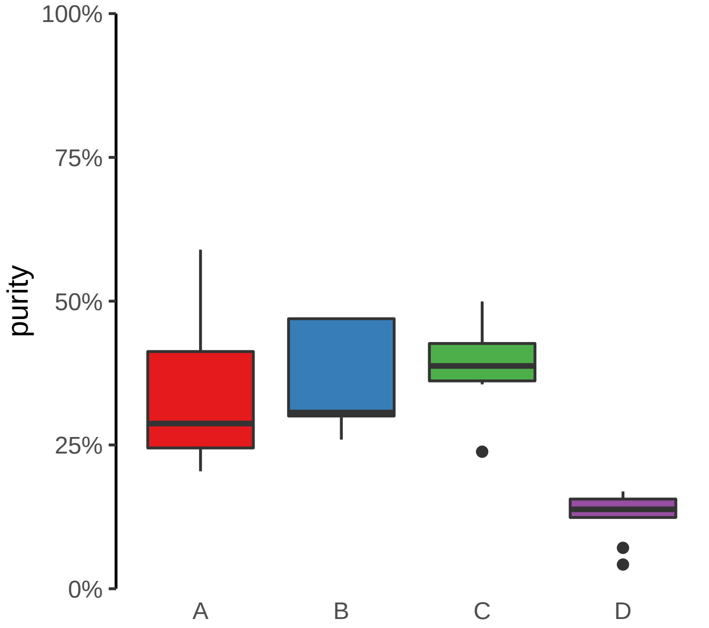

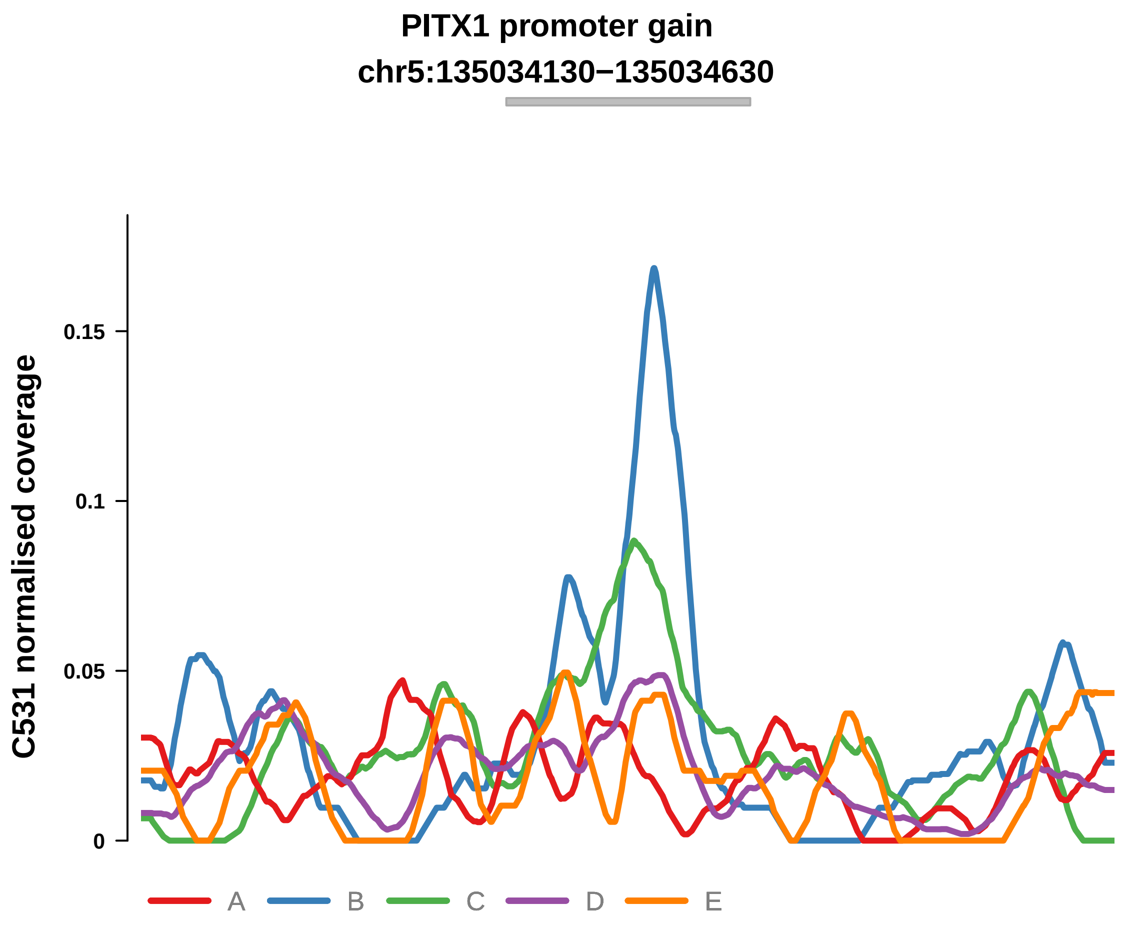


Putatively Clonal (logFC = 1.77)

| | P. | Adjusted P. |
|---|---|---|
| *~region vs. ~* | 3.17e−01 | 4.78e−01 |
| *~purity vs ~* | 4.49e−01 | 5.9e−01 |
| *~purity+region vs ~purity* | 3.24e−01 | 6.13e−01 |
| *Carcinoma vs. normal* | 5.07e−03 ** | 5.07e−03 ** |

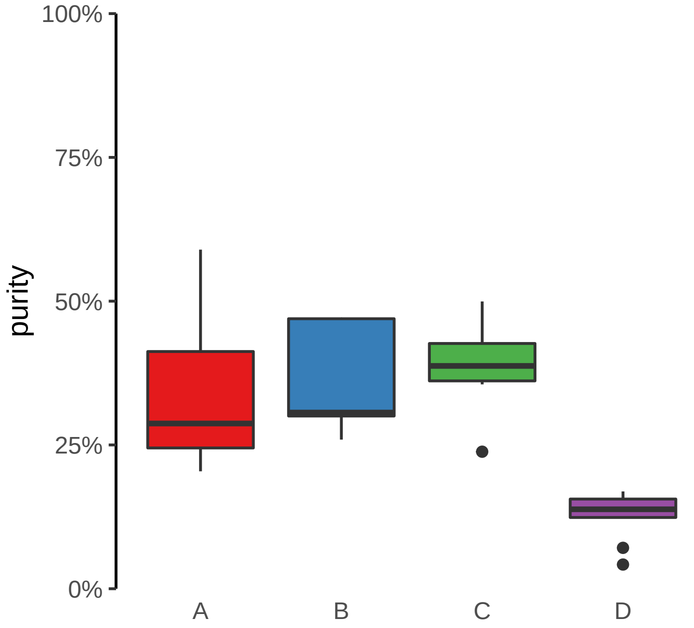

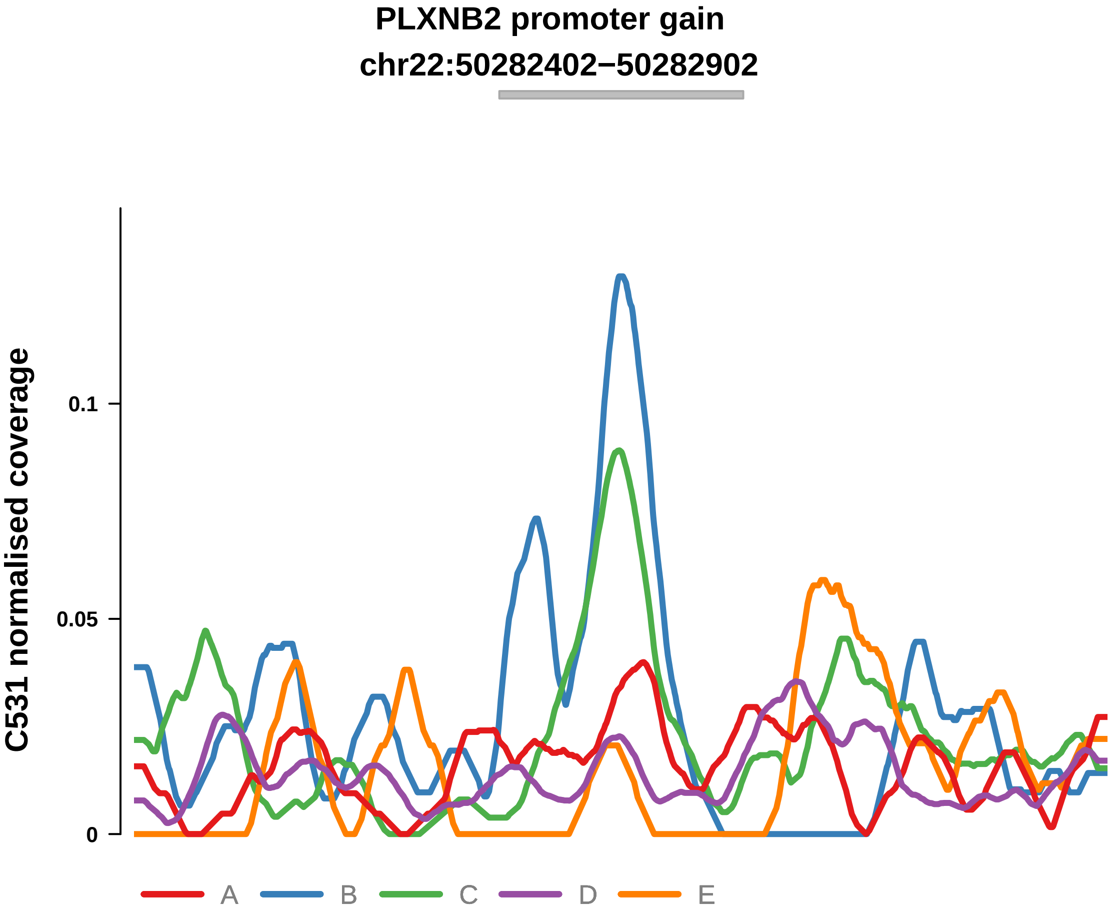


Putatively Clonal (logFC = 2.03)

| | P. | Adjusted P. |
|---|---|---|
| *~region vs. ~* | 1.72e−02 * | 5.59e−02 |
| *~purity vs ~* | 1.09e−02 * | 5.87e−02 |
| *~purity+region vs ~purity* | 3.42e−01 | 6.14e−01 |
| *Carcinoma vs. normal* | 1.99e−03 ** | 1.99e−03 ** |

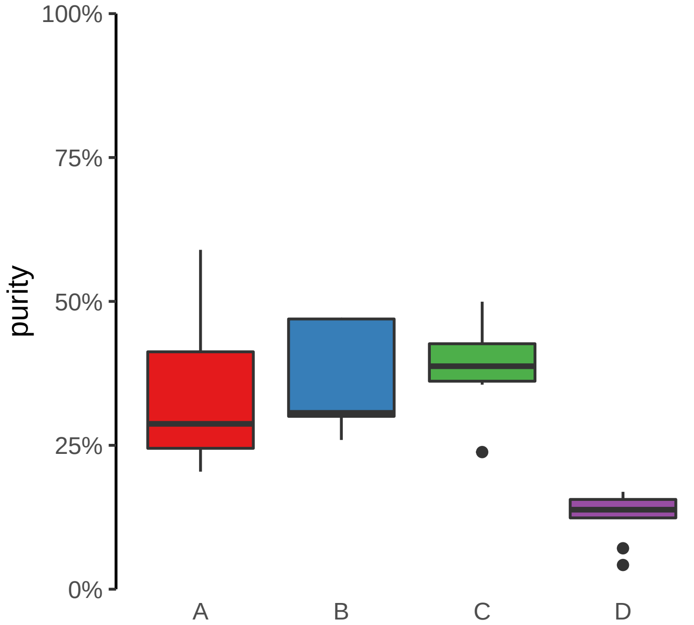

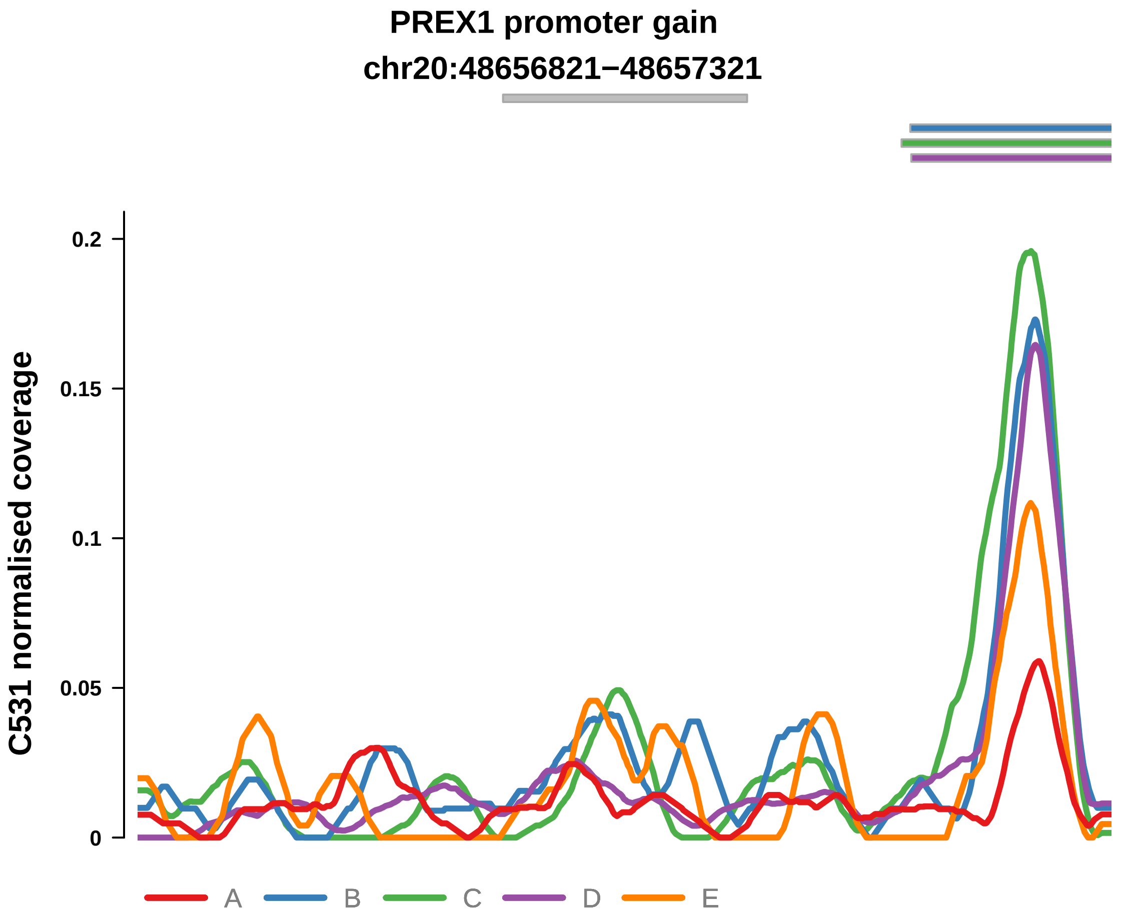


Putatively Clonal (logFC = 2.22)

| | P. | Adjusted P. |
|---|---|---|
| *~region vs. ~* | 4.77e−01 | 6.2e−01 |
| *~purity vs ~* | 5.13e−02 | 1.51e−01 |
| *~purity+region vs ~purity* | 8.88e−01 | 9.51e−01 |
| *Carcinoma vs. normal* | 1.74e−03 ** | 1.74e−03 ** |

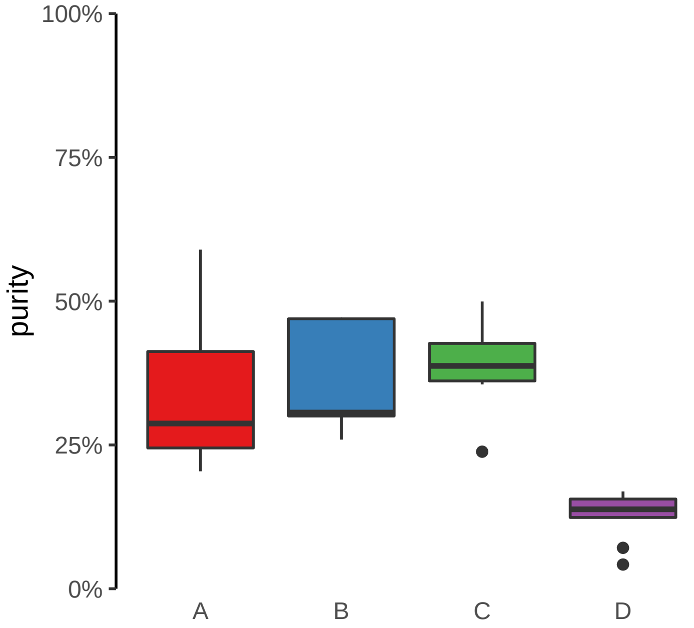

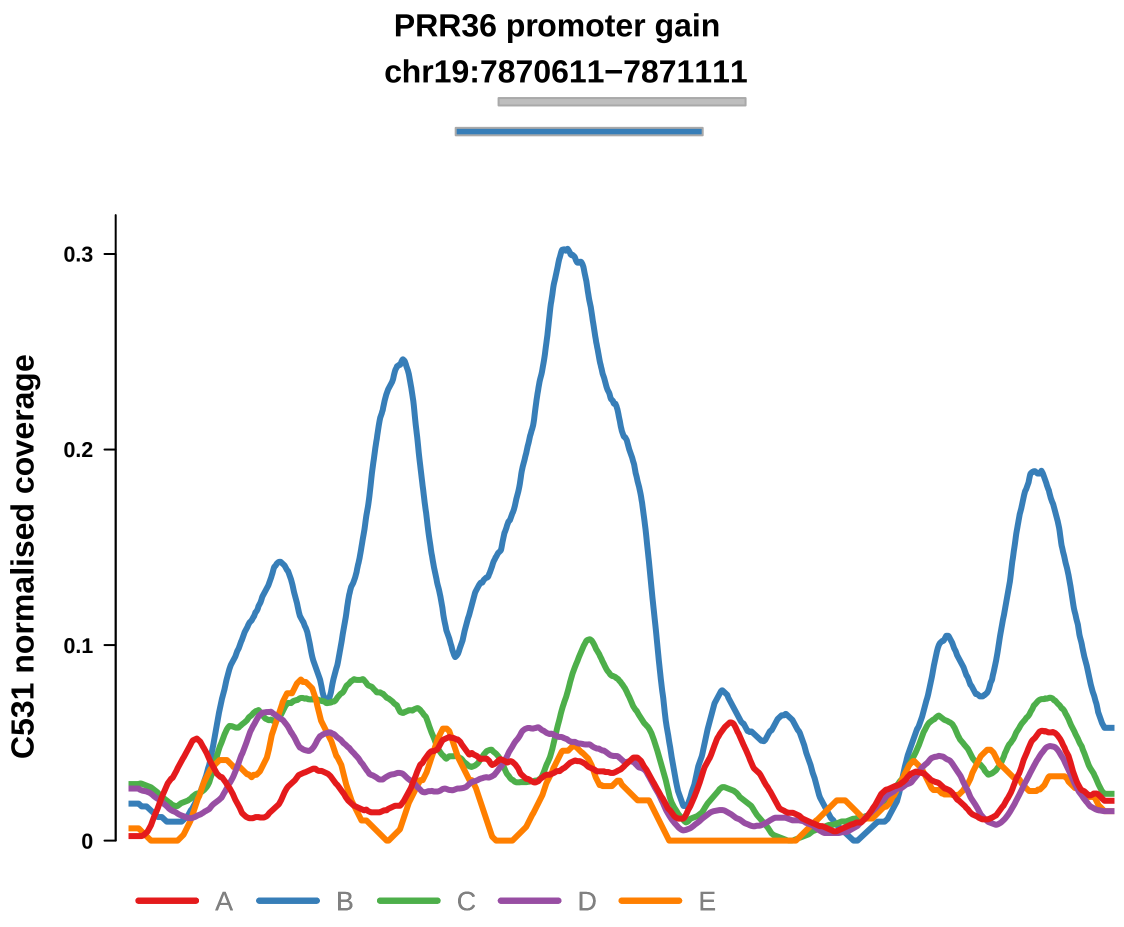


Putatively Clonal (logFC = 1.87)

| | P. | Adjusted P. |
|---|---|---|
| *~region vs. ~* | 2.74e−02 * | 7.13e−02 |
| *~purity vs ~* | 1.64e−01 | 3.27e−01 |
| *~purity+region vs ~purity* | 6.15e−02 | 4.16e−01 |
| *Carcinoma vs. normal* | 2.54e−03 ** | 2.54e−03 ** |

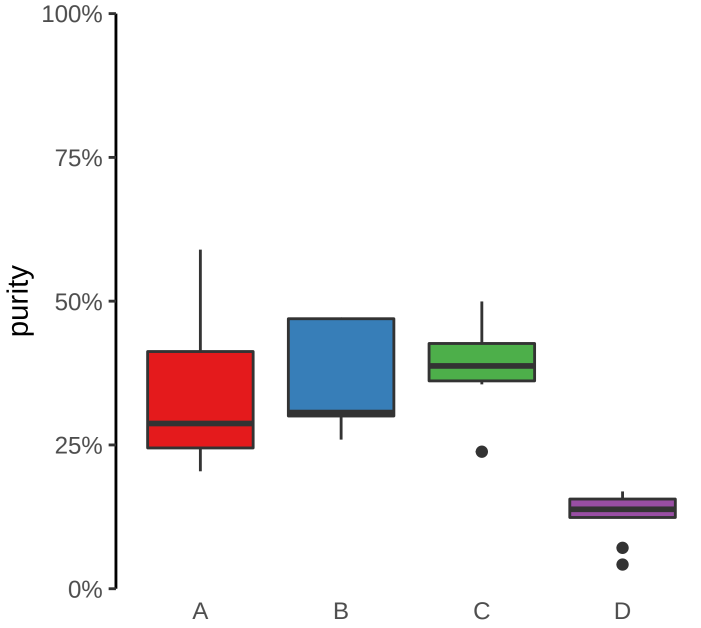

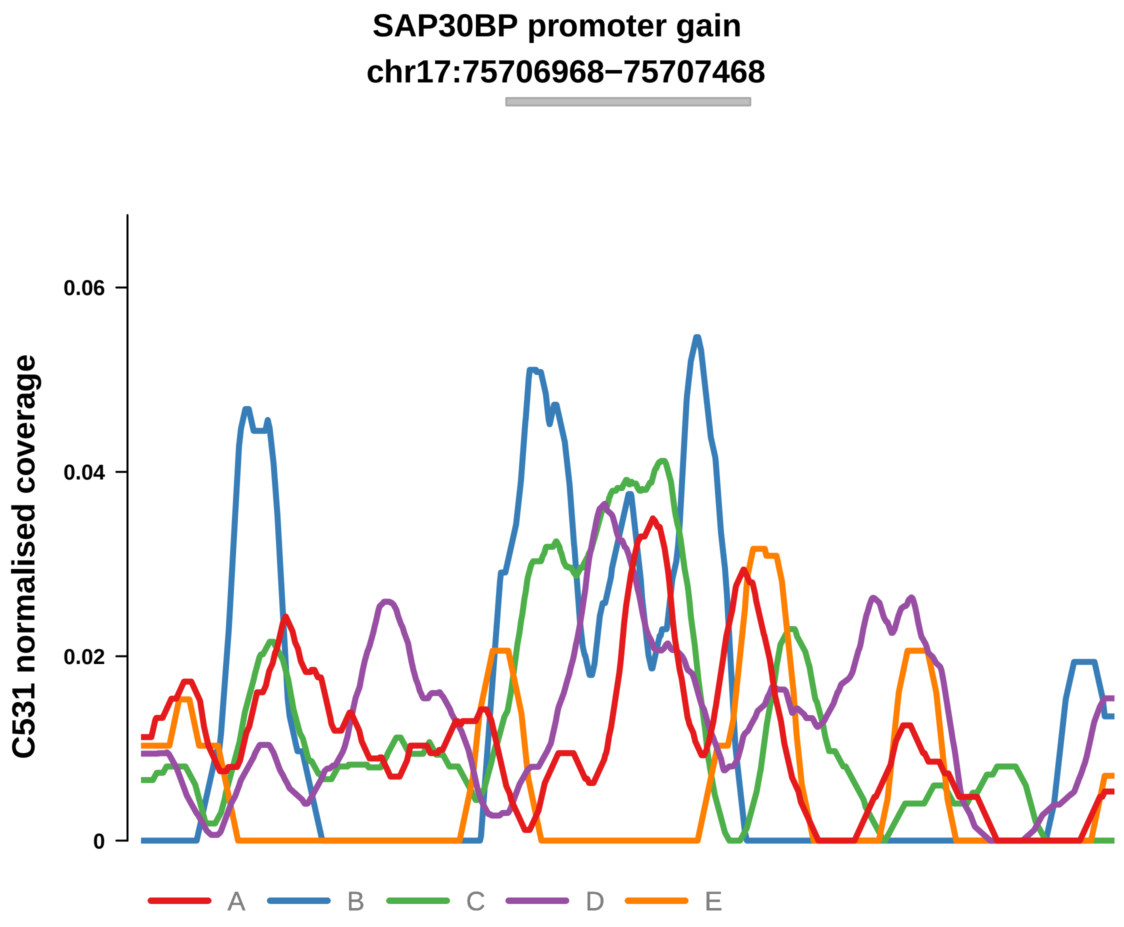


Putatively Clonal (logFC = 2.63)

| | P. | Adjusted P. |
|---|---|---|
| *~region vs. ~* | 3.31e−01 | 4.78e−01 |
| *~purity vs ~* | 7.18e−03 ** | 4.83e−02 * |
| *~purity+region vs ~purity* | 6.49e−01 | 8.66e−01 |
| *Carcinoma vs. normal* | 6.87e−05 **** | 6.87e−05 **** |

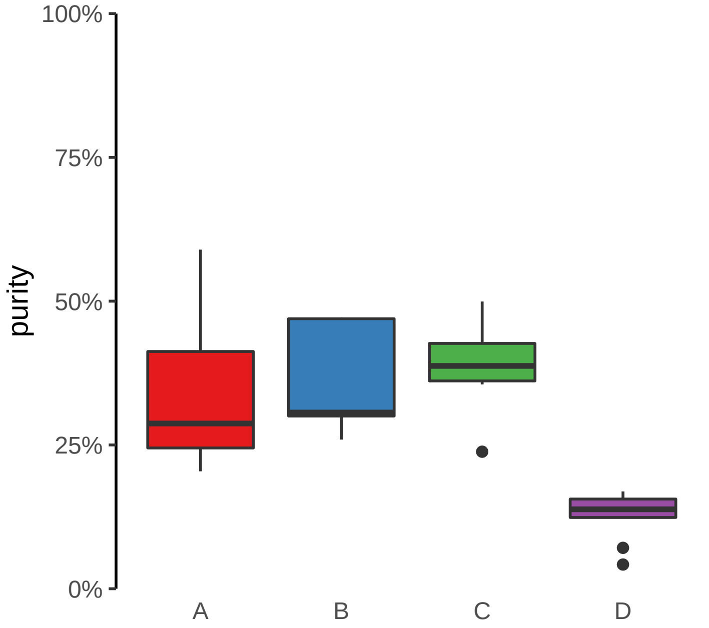

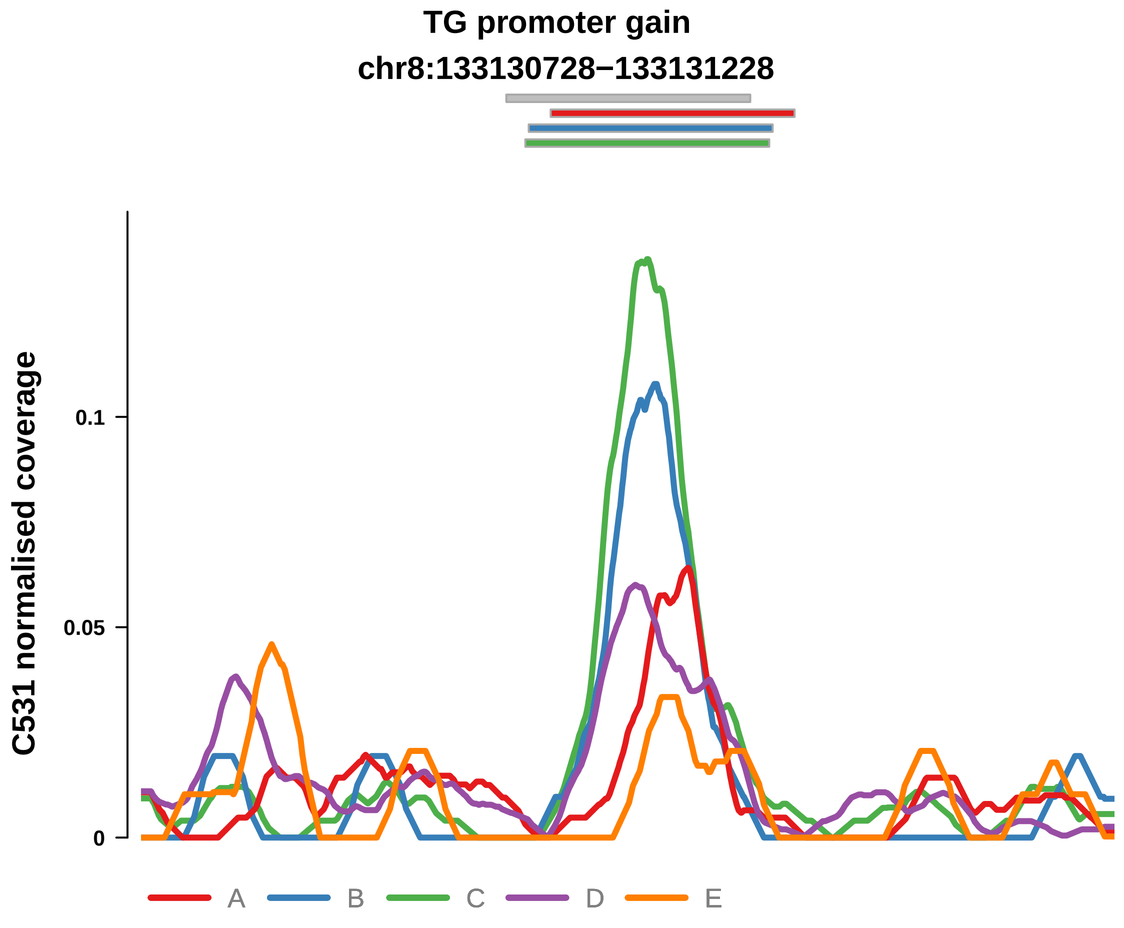


Putatively Clonal (logFC = 2.31)

| | P. | Adjusted P. |
|---|---|---|
| *~region vs. ~* | 2.44e−02 * | 7.04e−02 |
| *~purity vs ~* | 2.35e−01 | 3.89e−01 |
| *~purity+region vs ~purity* | 2.85e−02 * | 2.93e−01 |
| *Carcinoma vs. normal* | 2.44e−04 *** | 2.44e−04 *** |

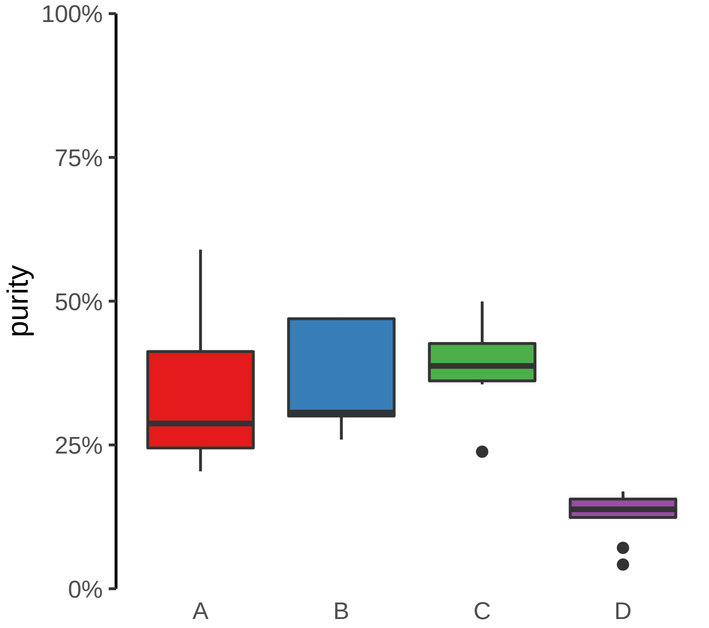

**TTYH3 promoter gain**
**chr7:2646791−2647291**

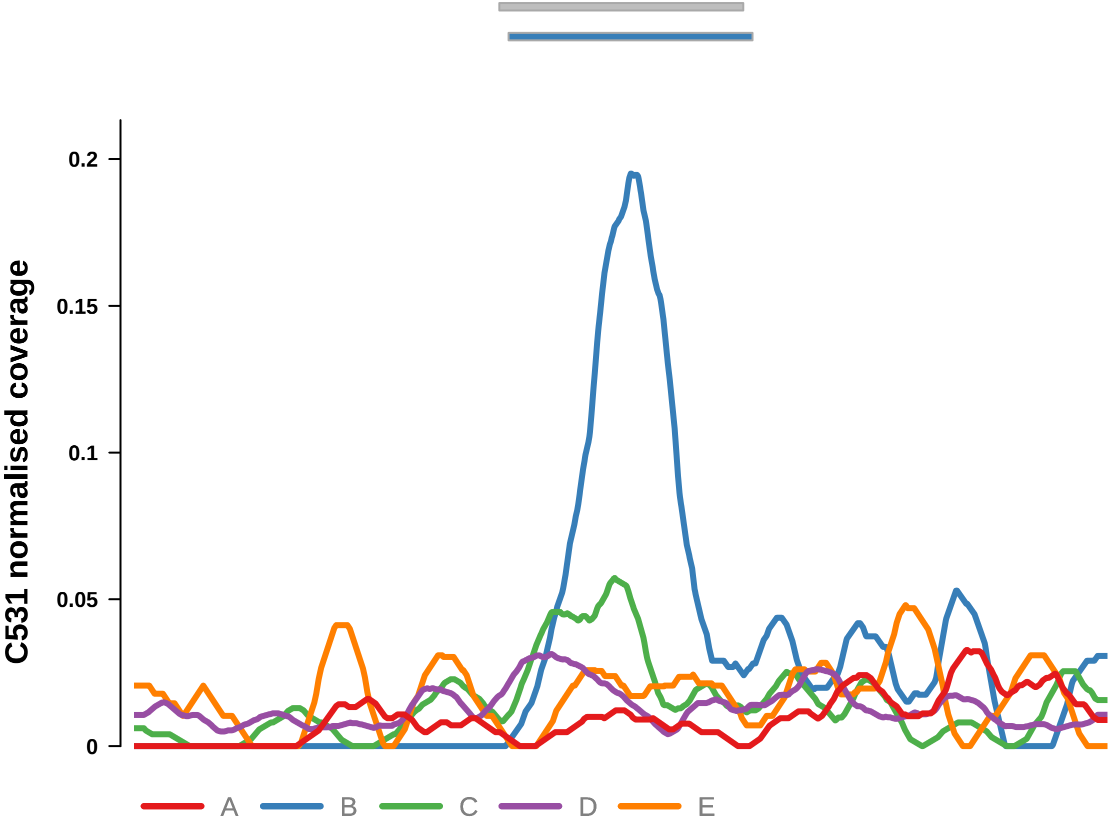


Putatively Clonal (logFC = 2.77)

| | P. | Adjusted P. |
|---|---|---|
| *~region vs. ~* | 1.06e−01 | 1.97e−01 |
| *~purity vs ~* | 1.86e−01 | 3.41e−01 |
| *~purity+region vs ~purity* | 1.73e−01 | 6.13e−01 |
| *Carcinoma vs. normal* | 1.6e−05 **** | 1.6e−05 **** |

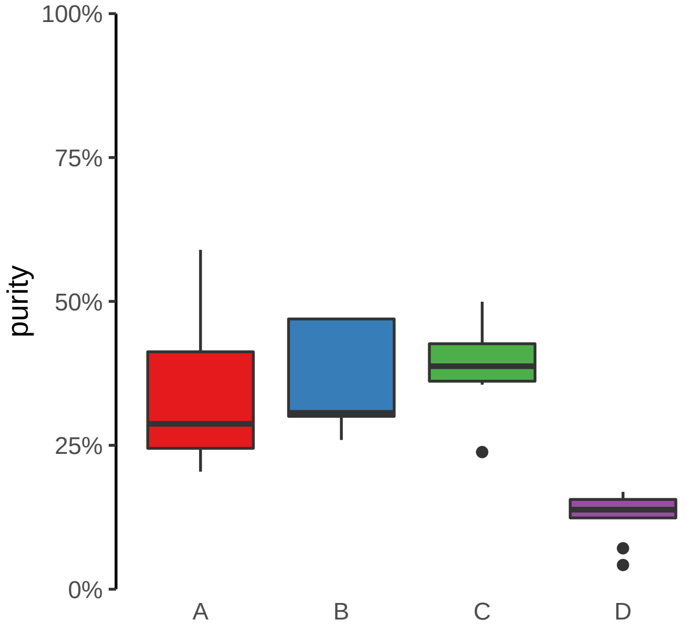

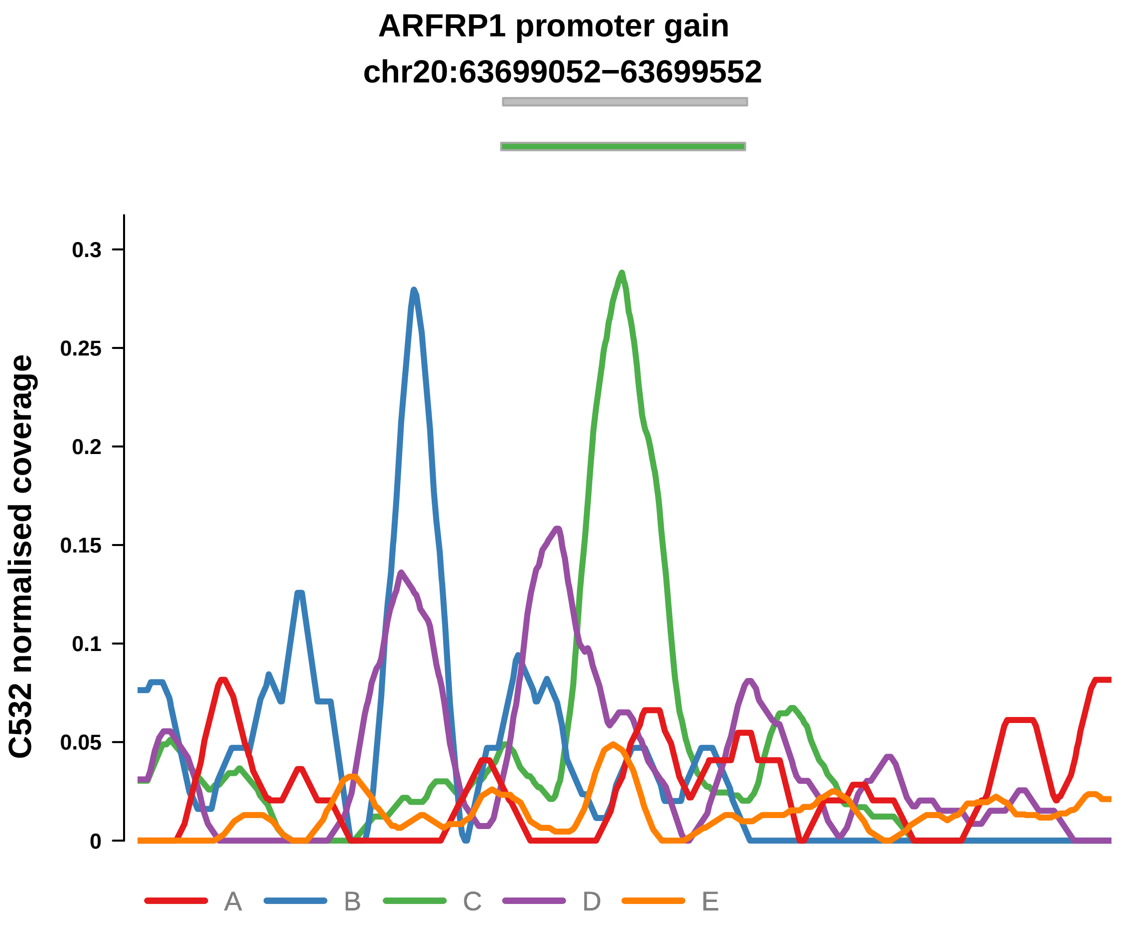


Putatively Clonal (logFC = 2.08)

| | P. | Adjusted P. |
|---|---|---|
| *~region vs. ~* | 2.1e−02 * | 2.06e−01 |
| *~purity vs ~* | 2.25e−01 | 8.56e−01 |
| *~purity+region vs ~purity* | 7.44e−02 | 5.15e−01 |
| *Carcinoma vs. normal* | 3.46e−04 *** | 3.46e−04 *** |

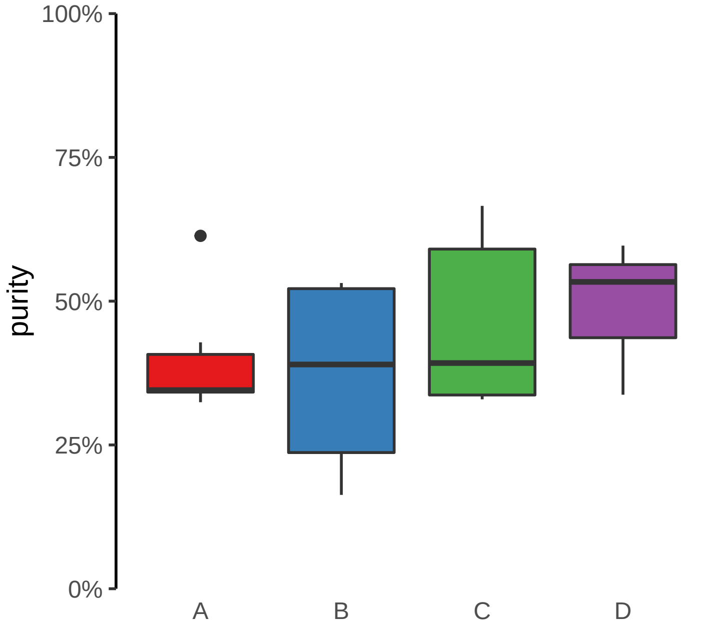

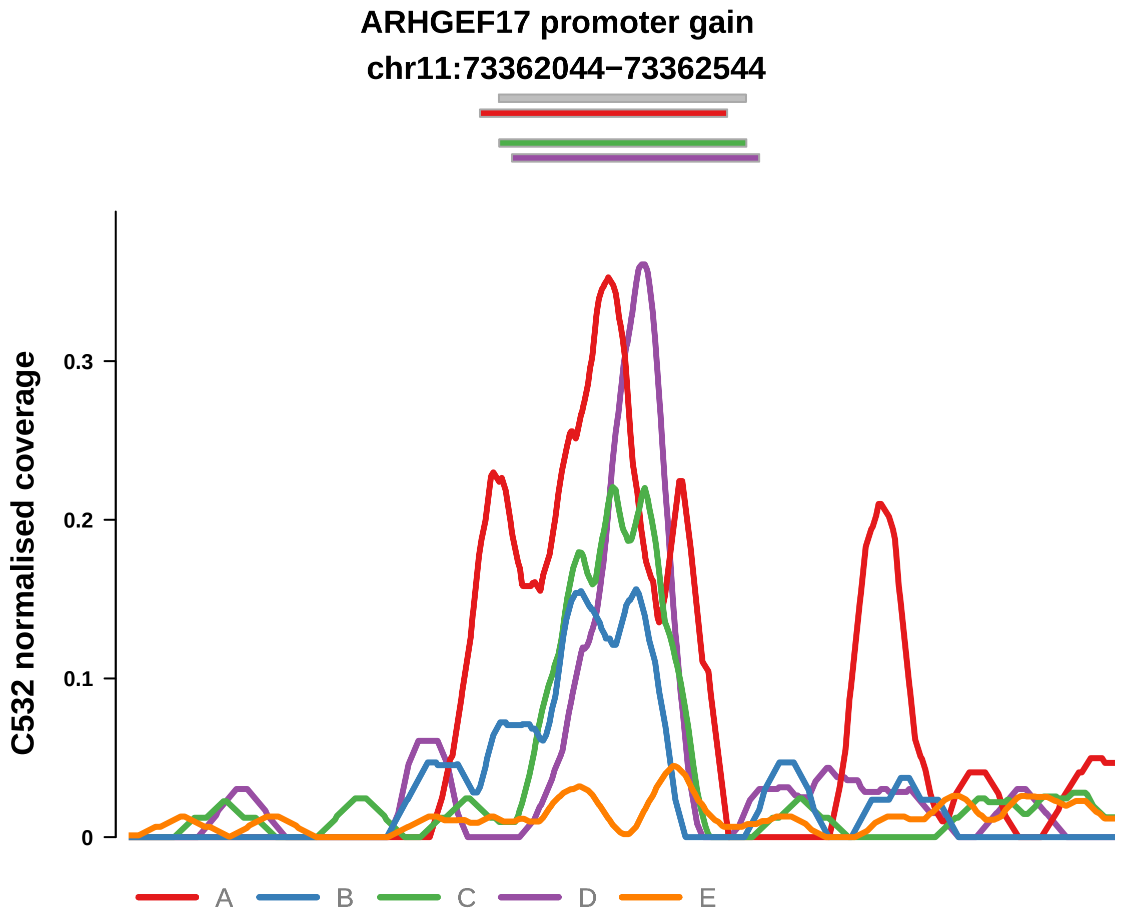


Putatively Clonal (logFC = 3.04)

| | P. | Adjusted P. |
|---|---|---|
| *~region vs. ~* | 2.93e−01 | 7.36e−01 |
| *~purity vs ~* | 8.46e−01 | 9.86e−01 |
| *~purity+region vs ~purity* | 2.58e−01 | 6.89e−01 |
| *Carcinoma vs. normal* | 2.29e−07 **** | 2.29e−07 **** |

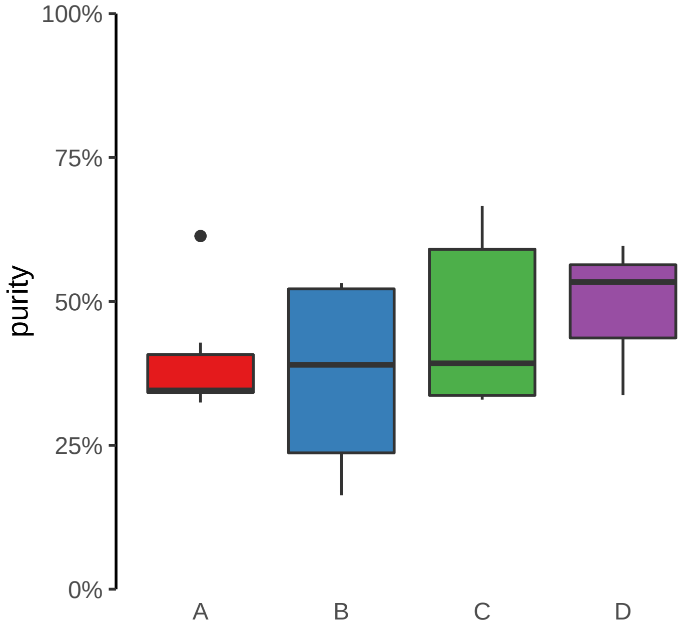

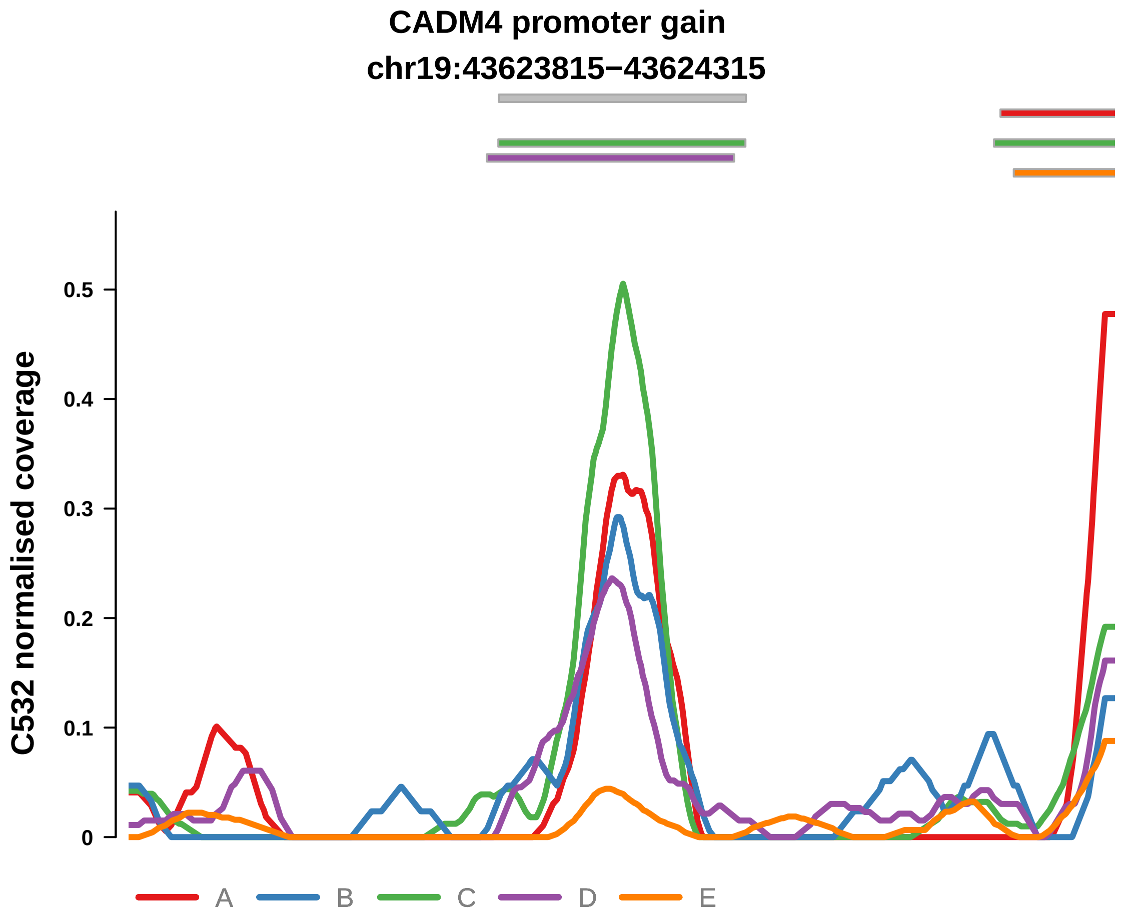


Putatively Clonal (logFC = 3.53)

| | P. | Adjusted P. |
|---|---|---|
| *~region vs. ~* | 1.66e−01 | 5.42e−01 |
| *~purity vs ~* | 1.29e−02 * | 2.84e−01 |
| *~purity+region vs ~purity* | 3.25e−01 | 7.99e−01 |
| *Carcinoma vs. normal* | 3.39e−09 **** | 3.39e−09 **** |

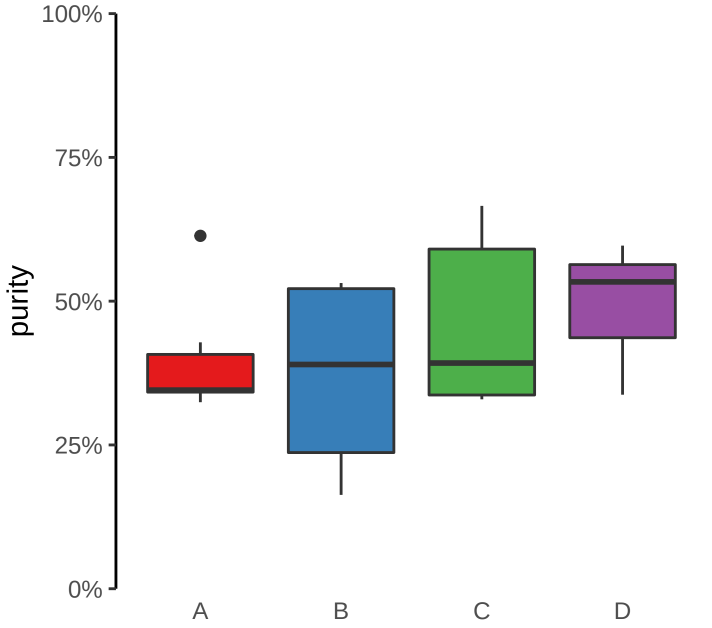

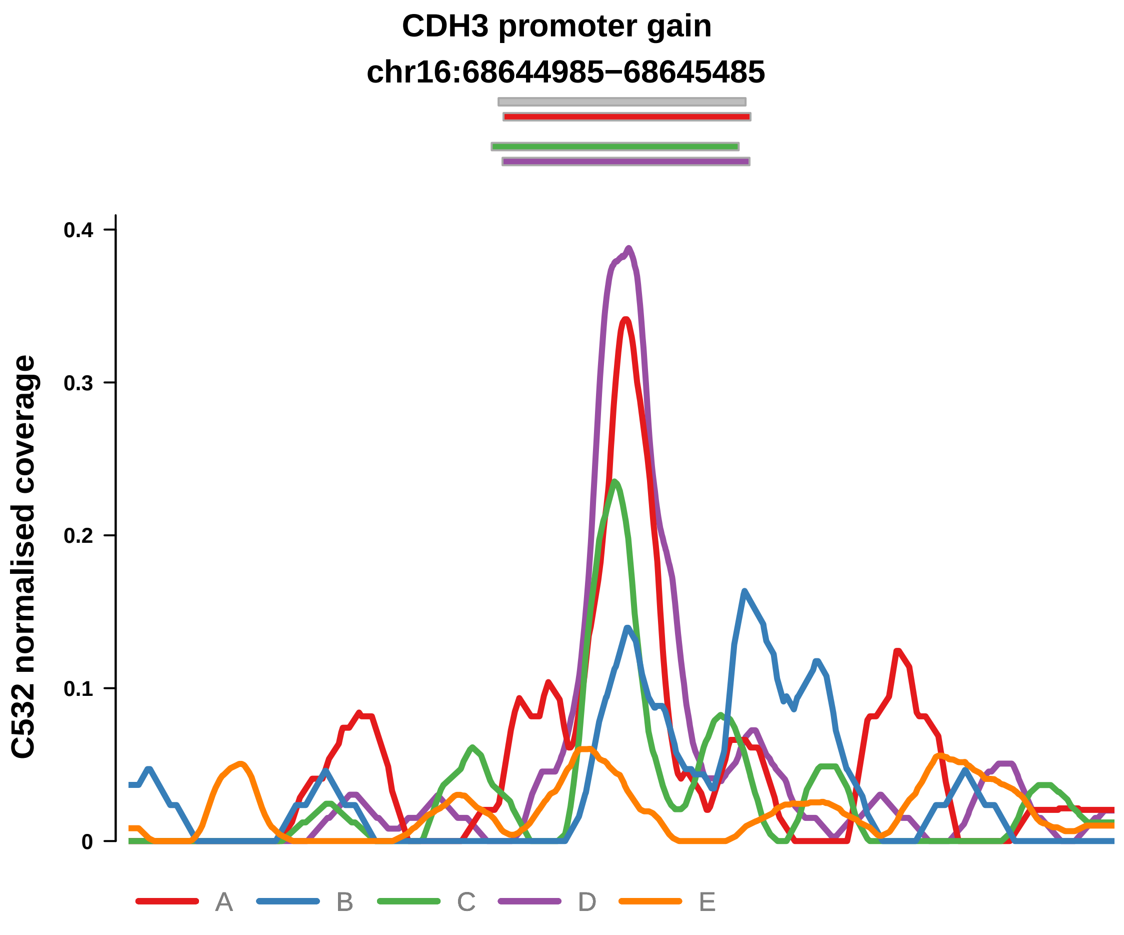


Putatively Clonal (logFC = 1.94)

| | P. | Adjusted P. |
|---|---|---|
| *~region vs. ~* | 3.52e−01 | 7.41e−01 |
| *~purity vs ~* | 1.02e−01 | 7.63e−01 |
| *~purity+region vs ~purity* | 4.98e−01 | 9.42e−01 |
| *Carcinoma vs. normal* | 5.88e−04 *** | 5.88e−04 *** |

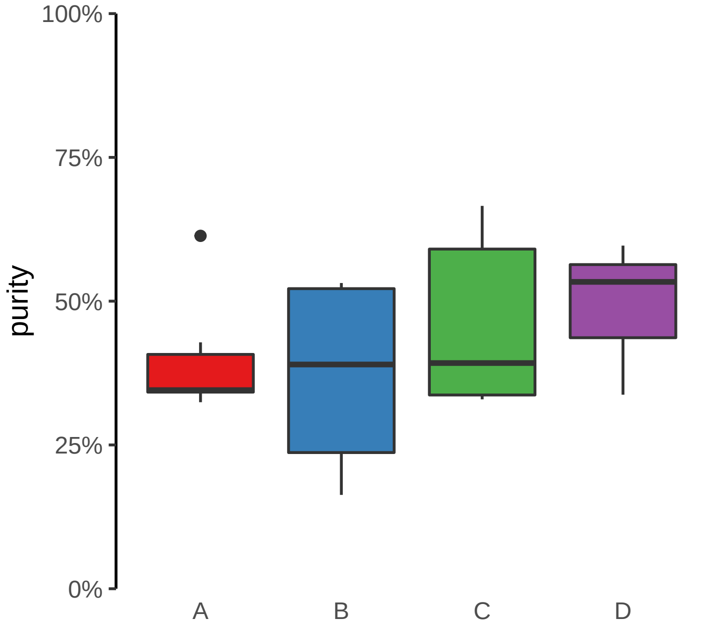

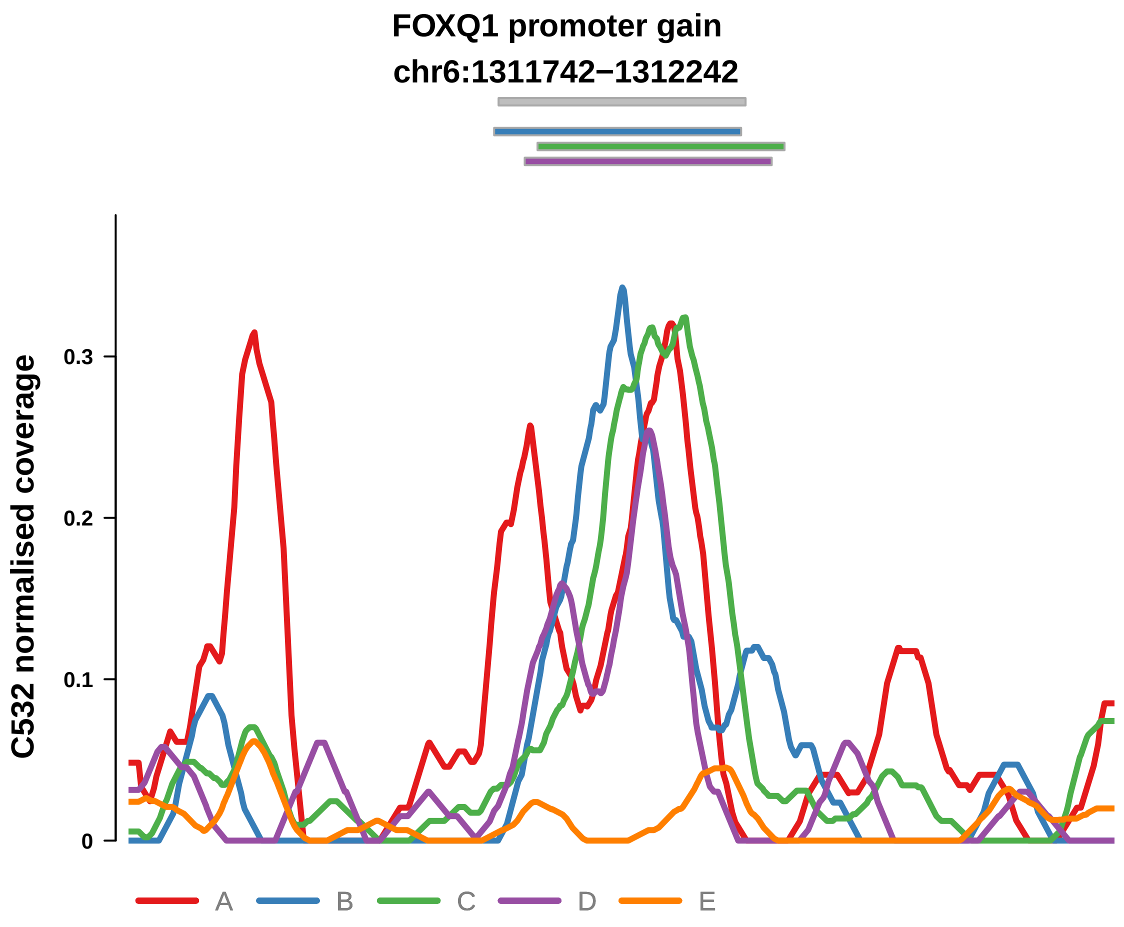


Putatively Clonal (logFC = 2.54)

| | P. | Adjusted P. |
|---|---|---|
| *~region vs. ~* | 1.93e−01 | 5.85e−01 |
| *~purity vs ~* | 1.82e−01 | 8.56e−01 |
| *~purity+region vs ~purity* | 1.78e−01 | 6.56e−01 |
| *Carcinoma vs. normal* | 6.66e−06 **** | 6.66e−06 **** |

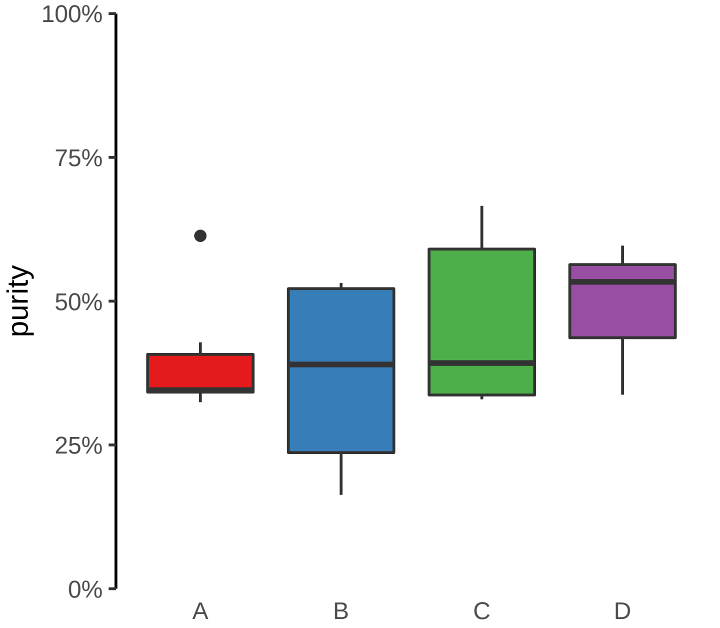

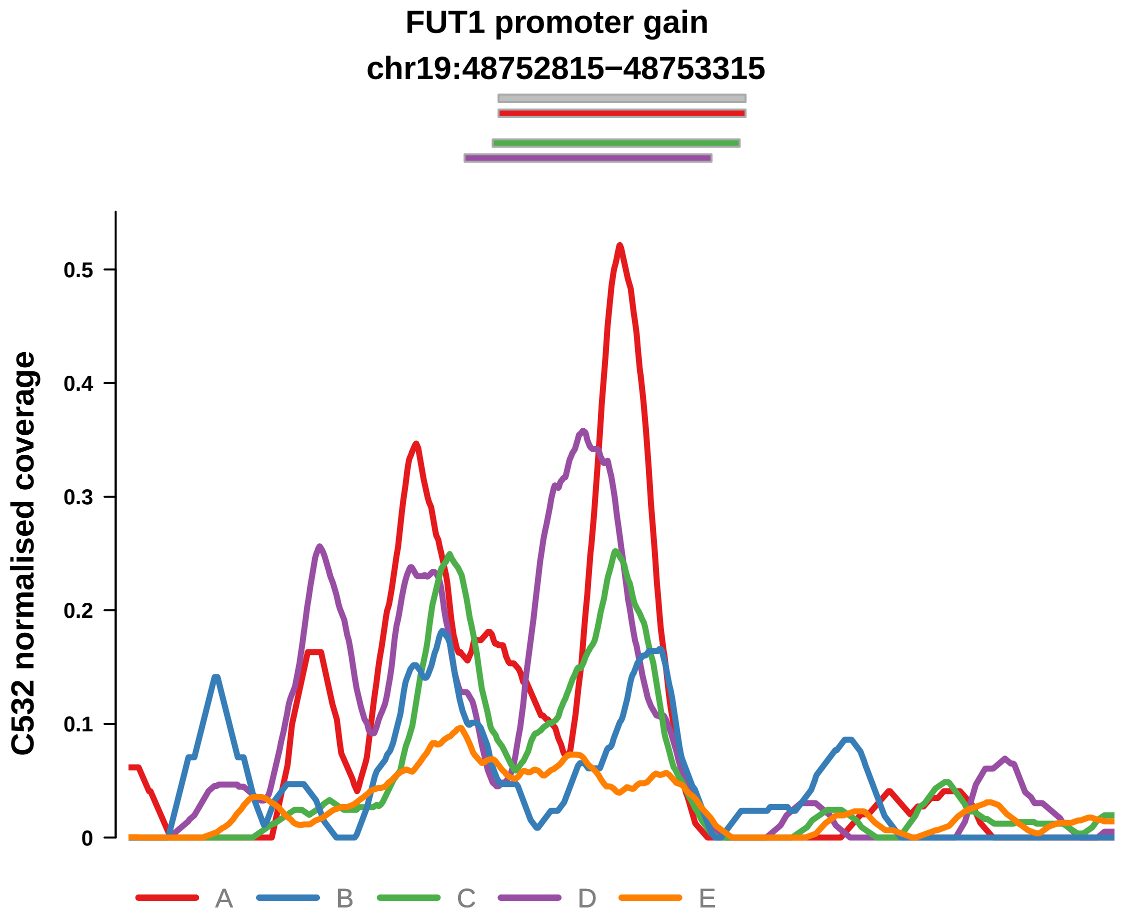


Putatively Clonal (logFC = 2.67)

| | P. | Adjusted P. |
|---|---|---|
| *~region vs. ~* | 1.39e−01 | 4.99e−01 |
| *~purity vs ~* | 6.66e−03 ** | 2.21e−01 |
| *~purity+region vs ~purity* | 2.24e−01 | 6.56e−01 |
| *Carcinoma vs. normal* | 2.78e−06 **** | 2.78e−06 **** |

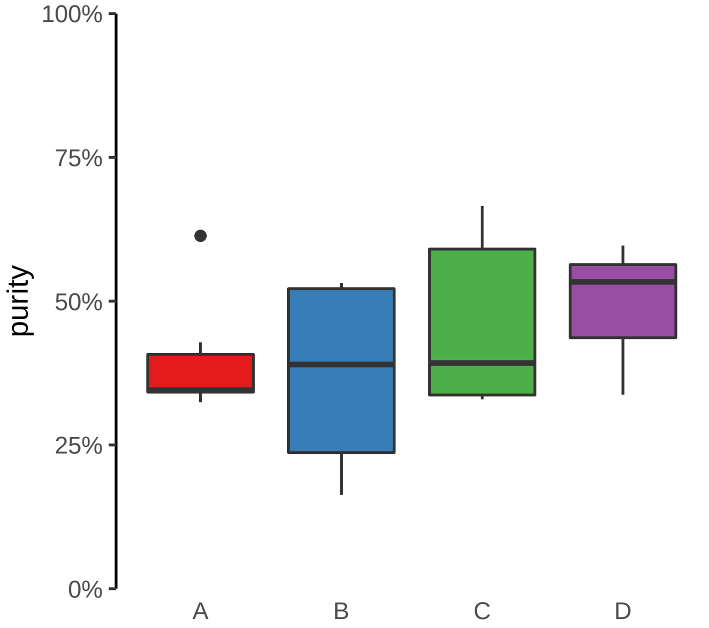

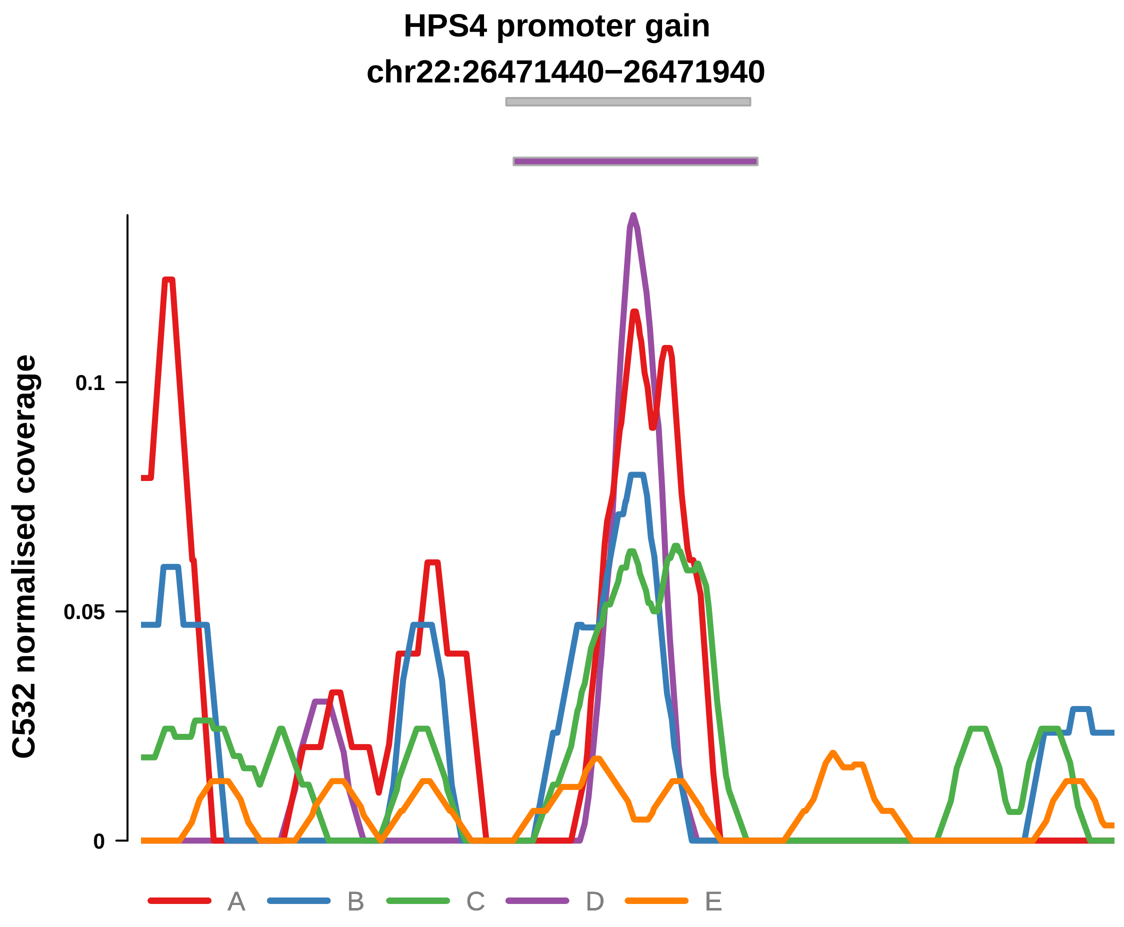


Putatively Clonal (logFC = 1.81)

| | P. | Adjusted P. |
|---|---|---|
| *~region vs. ~* | 9.01e−01 | 9.8e−01 |
| *~purity vs ~* | 6.71e−01 | 9.86e−01 |
| *~purity+region vs ~purity* | 9.42e−01 | 1e+00 |
| *Carcinoma vs. normal* | 3.77e−03 ** | 3.77e−03 ** |

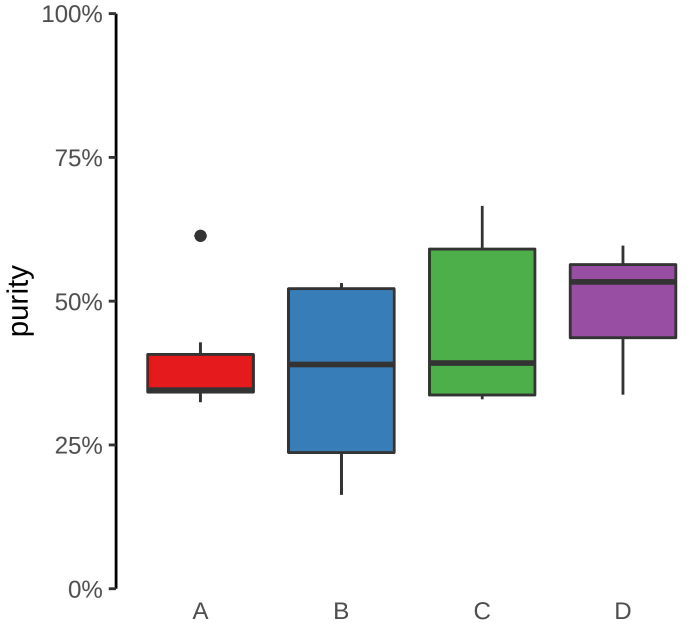

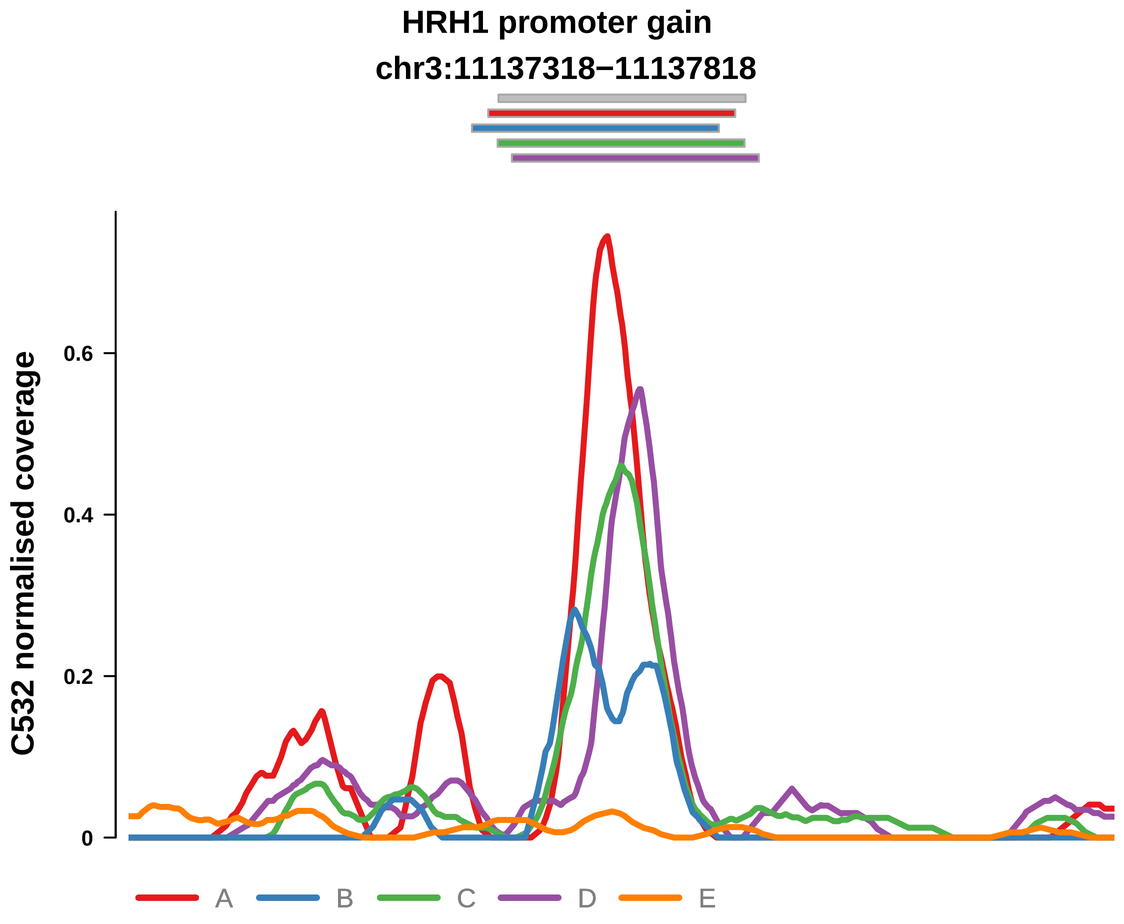


Putatively Clonal (logFC = 3.54)

| | P. | Adjusted P. |
|---|---|---|
| *~region vs. ~* | 4.48e−01 | 8.04e−01 |
| *~purity vs ~* | 5.36e−02 | 5.24e−01 |
| *~purity+region vs ~purity* | 7.24e−01 | 9.42e−01 |
| *Carcinoma vs. normal* | 2.19e−09 **** | 2.19e−09 **** |

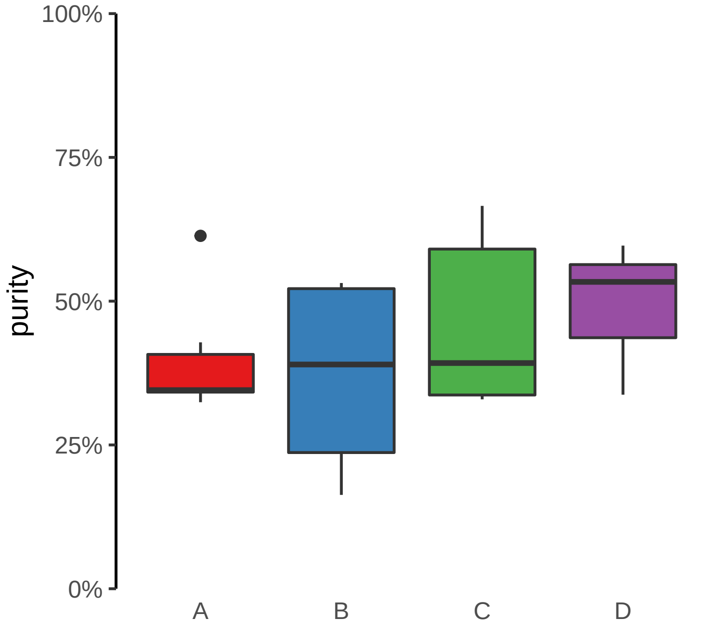

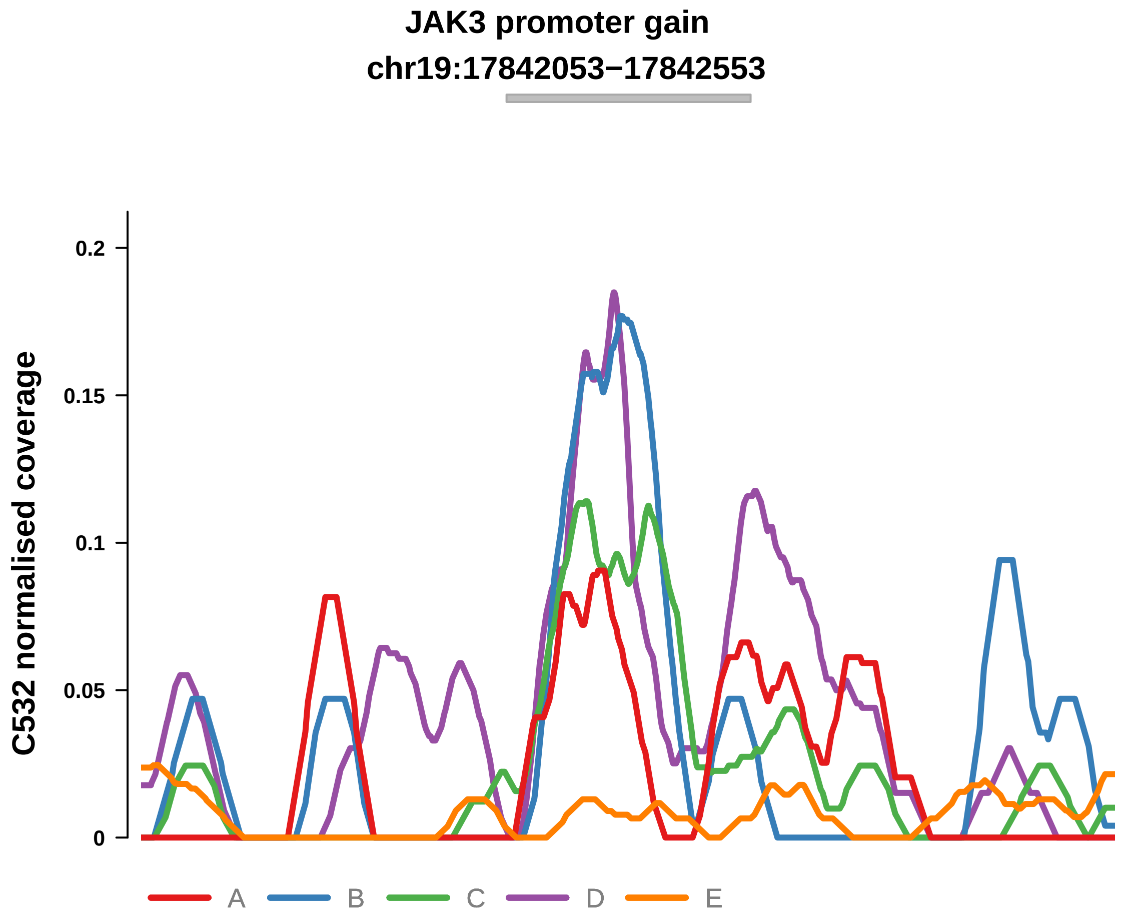


Putatively Clonal (logFC = 1.67)

| | P. | Adjusted P. |
|---|---|---|
| *~region vs. ~* | 3.45e−01 | 7.41e−01 |
| *~purity vs ~* | 4.76e−01 | 9.05e−01 |
| *~purity+region vs ~purity* | 4.08e−01 | 9.12e−01 |
| *Carcinoma vs. normal* | 3.28e−03 ** | 3.28e−03 ** |

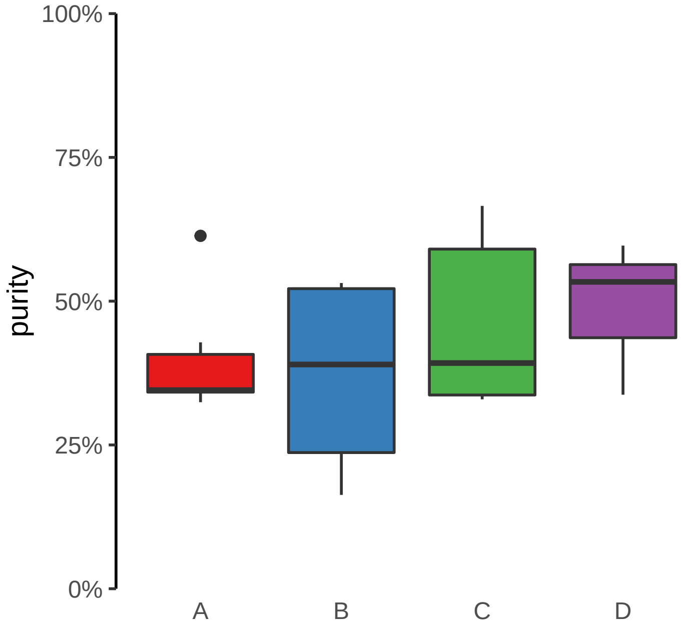

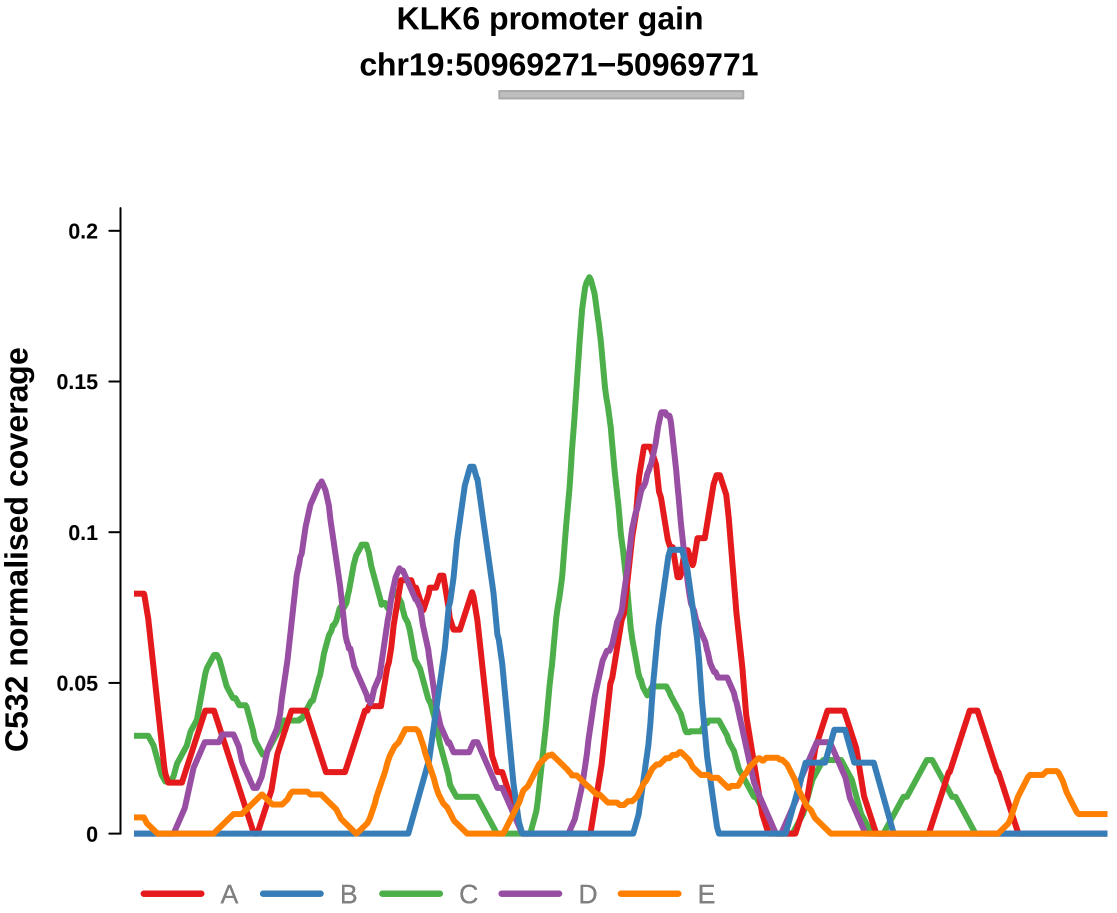


Putatively Clonal (logFC = 1.93)

| | P. | Adjusted P. |
|---|---|---|
| *~region vs. ~* | 3.58e−01 | 7.41e−01 |
| *~purity vs ~* | 5.02e−01 | 9.05e−01 |
| *~purity+region vs ~purity* | 4.51e−01 | 9.41e−01 |
| *Carcinoma vs. normal* | 1.1e−03 ** | 1.1e−03 ** |

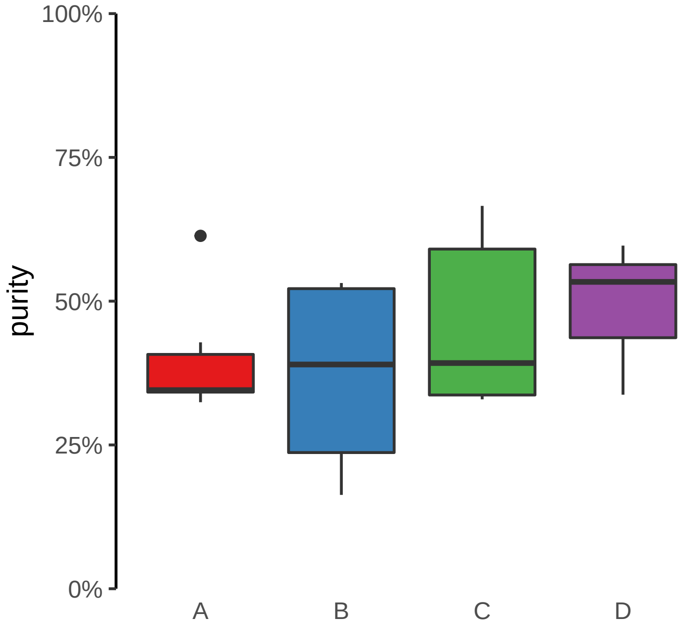

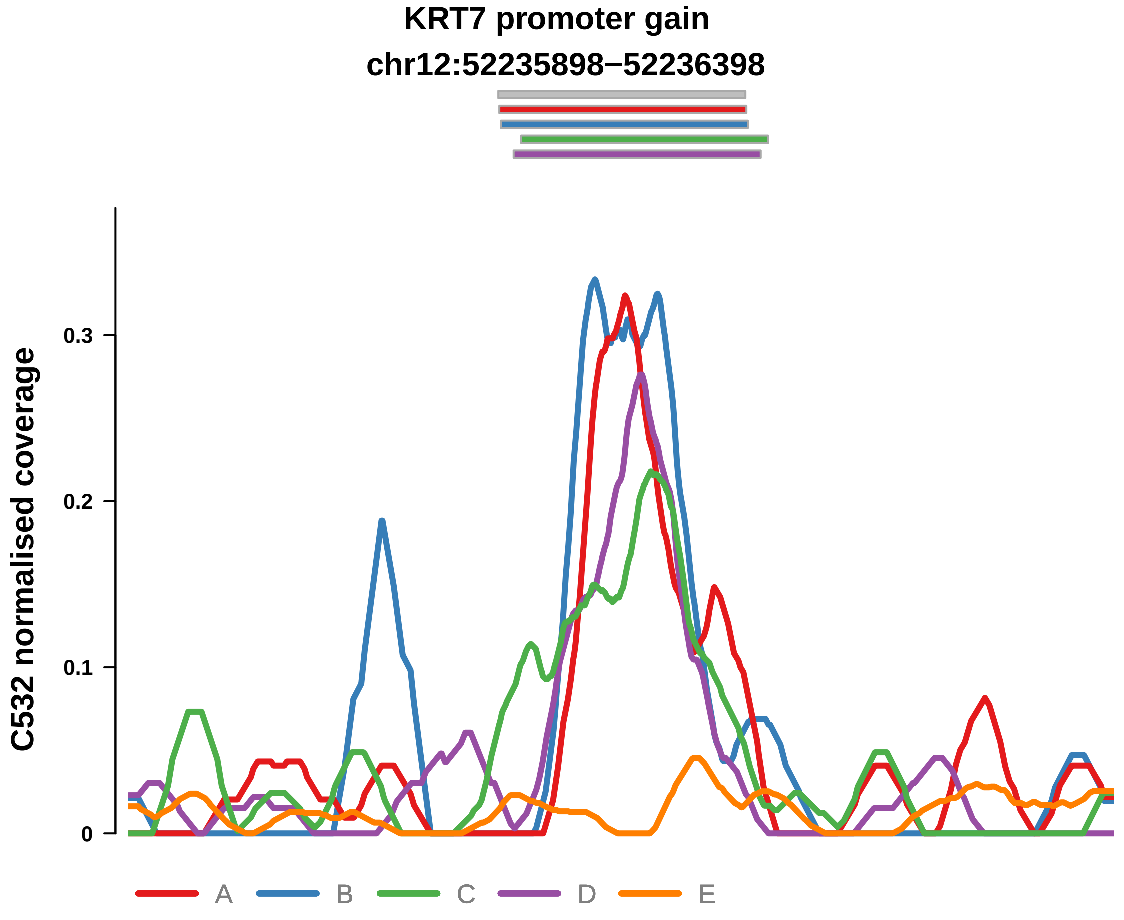


Putatively Clonal (logFC = 2.7)

| | P. | Adjusted P. |
|---|---|---|
| *~region vs. ~* | 5.37e−01 | 8.54e−01 |
| *~purity vs ~* | 3.11e−01 | 8.56e−01 |
| *~purity+region vs ~purity* | 5.21e−01 | 9.42e−01 |
| *Carcinoma vs. normal* | 2.43e−06 **** | 2.43e−06 **** |

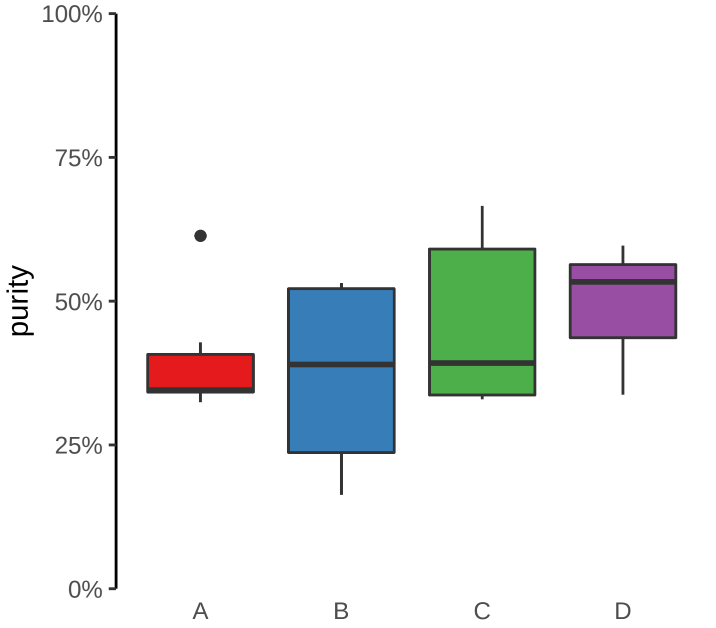

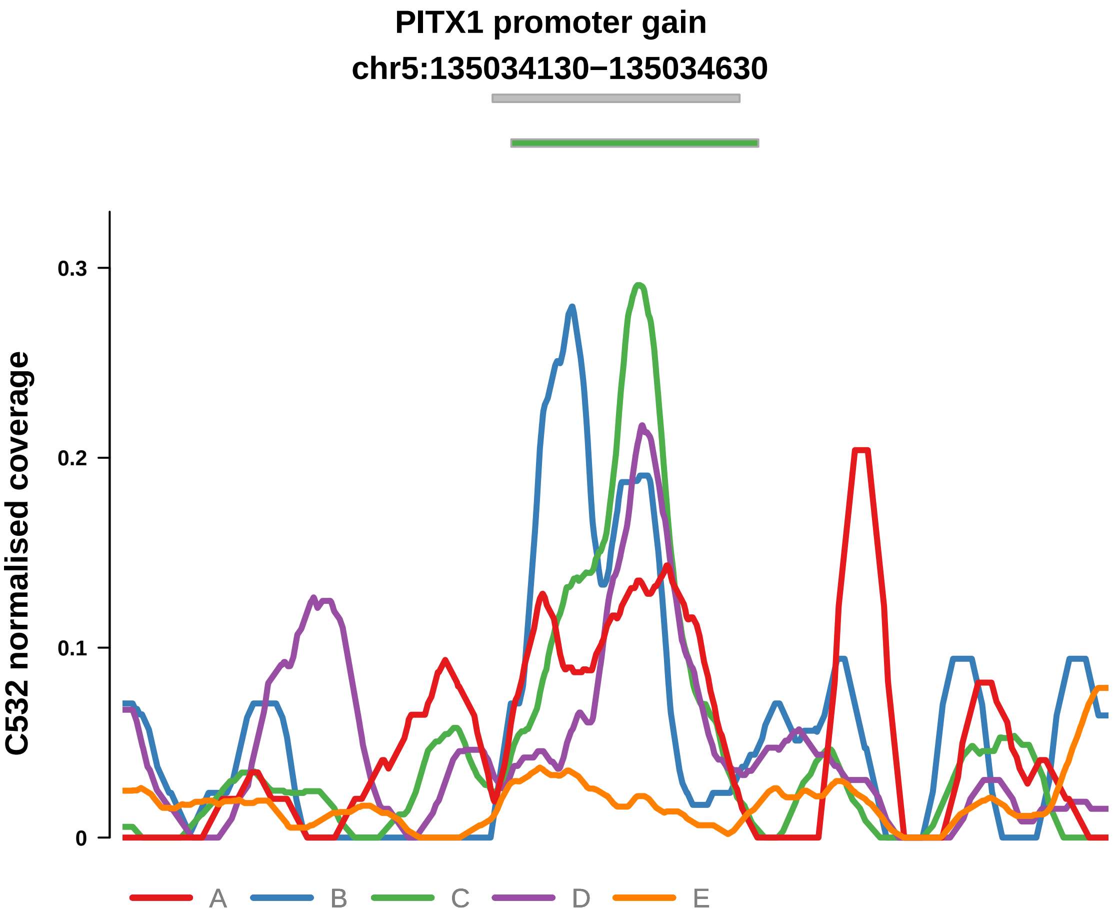


Putatively Clonal (logFC = 2.31)

| | P. | Adjusted P. |
|---|---|---|
| *~region vs. ~* | 8.7e−02 | 4.5e−01 |
| *~purity vs ~* | 3.67e−01 | 8.63e−01 |
| *~purity+region vs ~purity* | 1.07e−01 | 5.89e−01 |
| *Carcinoma vs. normal* | 5.01e−05 **** | 5.01e−05 **** |

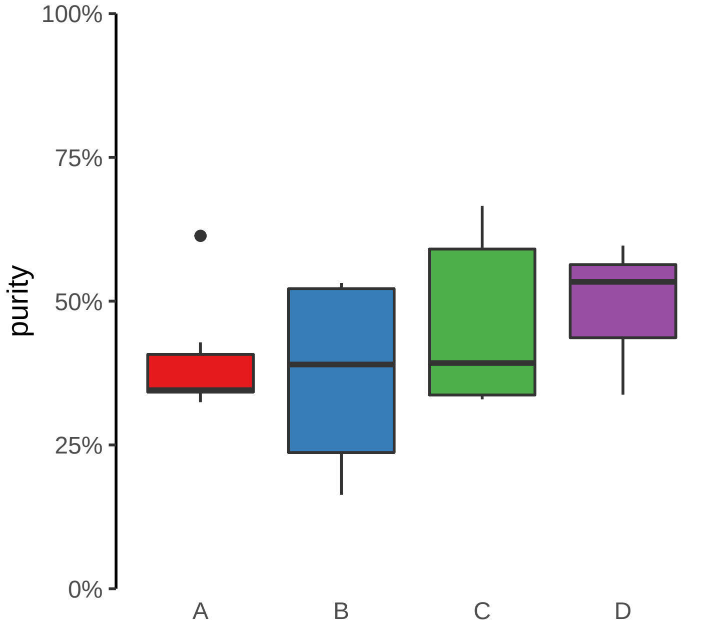

**PLXNB2 promoter gain**
**chr22:50282402−50282902**

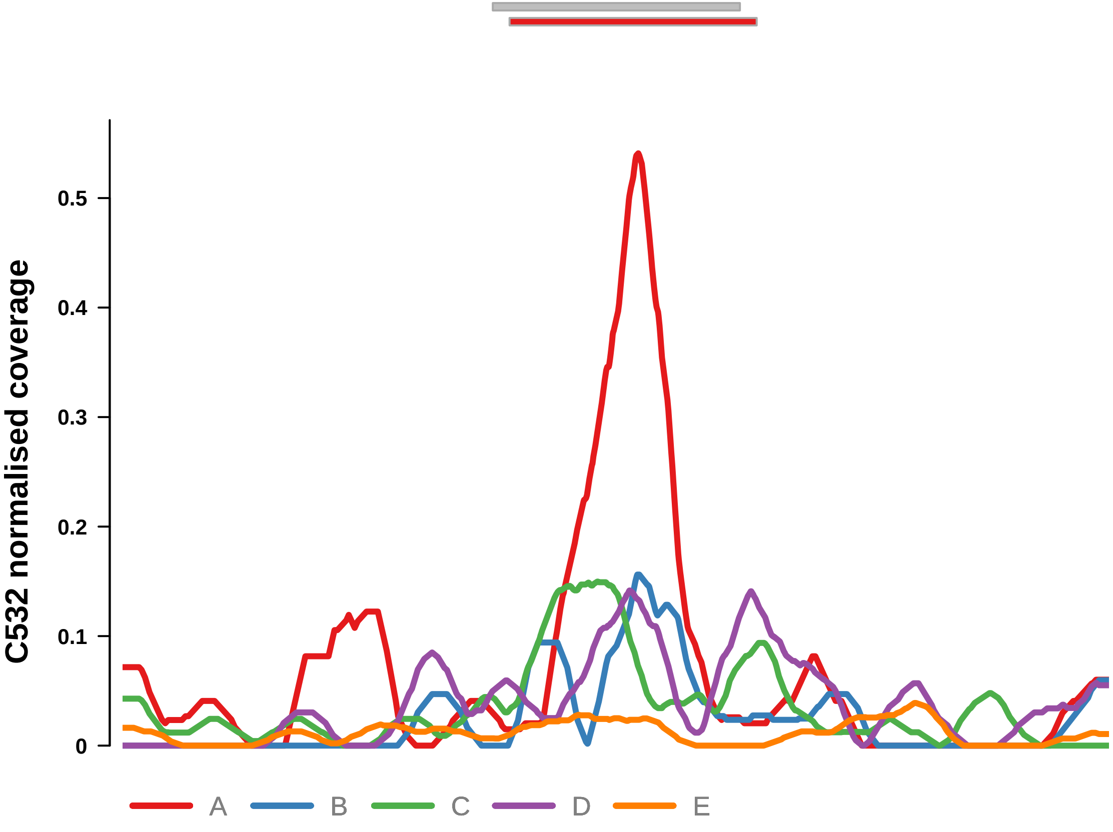


Putatively Clonal (logFC = 2.24)

| | P. | Adjusted P. |
|---|---|---|
| *~region vs. ~* | 2.11e−01 | 5.98e−01 |
| *~purity vs ~* | 4.68e−01 | 9.05e−01 |
| *~purity+region vs ~purity* | 1.98e−01 | 6.56e−01 |
| *Carcinoma vs. normal* | 1.03e−04 *** | 1.03e−04 *** |

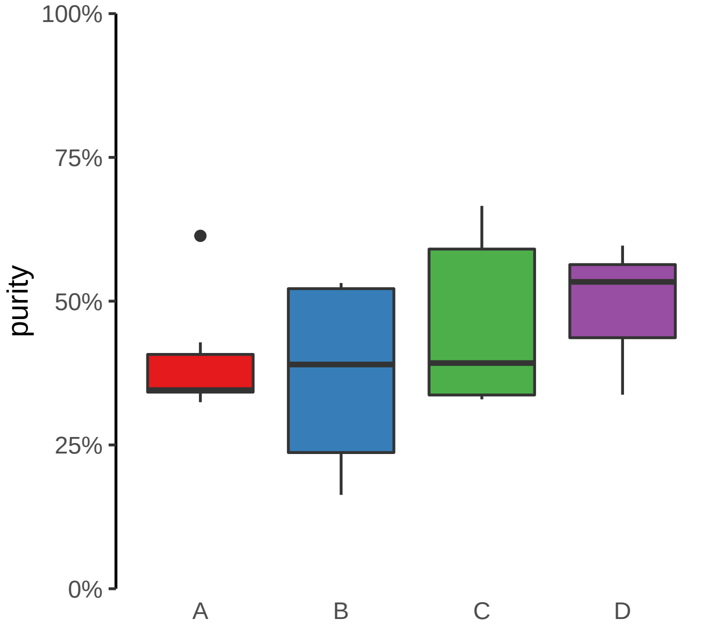

**PRR36 promoter gain**
**chr19:7870611−7871111**

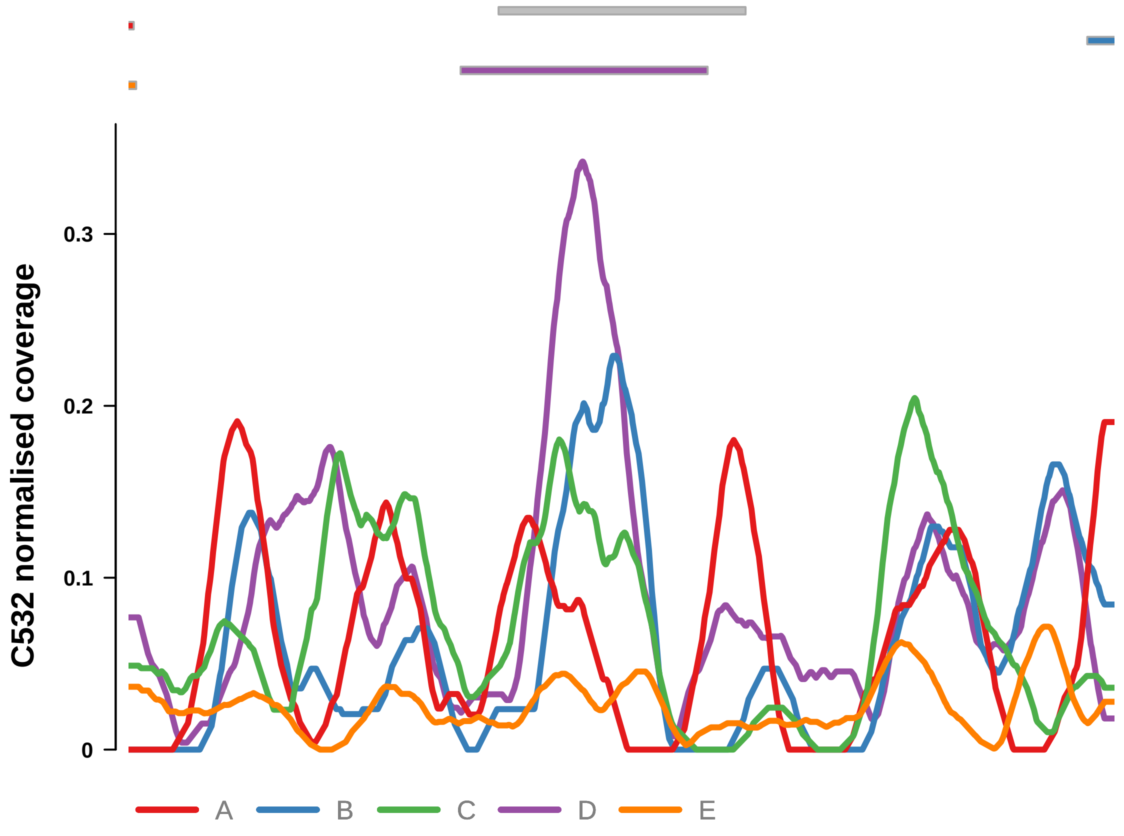


Putatively Clonal (logFC = 1.94)

| | P. | Adjusted P. |
|---|---|---|
| *~region vs. ~* | 3.96e−01 | 7.41e−01 |
| *~purity vs ~* | 9.79e−01 | 1e+00 |
| *~purity+region vs ~purity* | 5.03e−01 | 9.42e−01 |
| *Carcinoma vs. normal* | 6.03e−04 *** | 6.03e−04 *** |

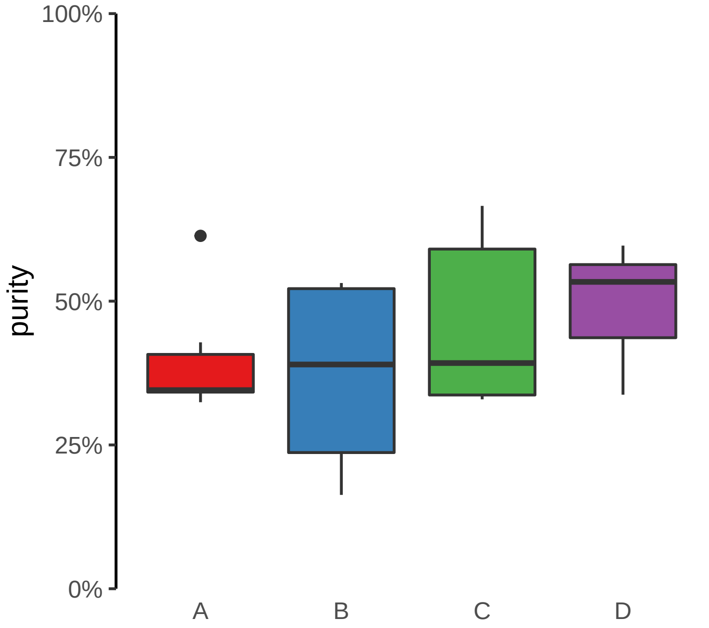

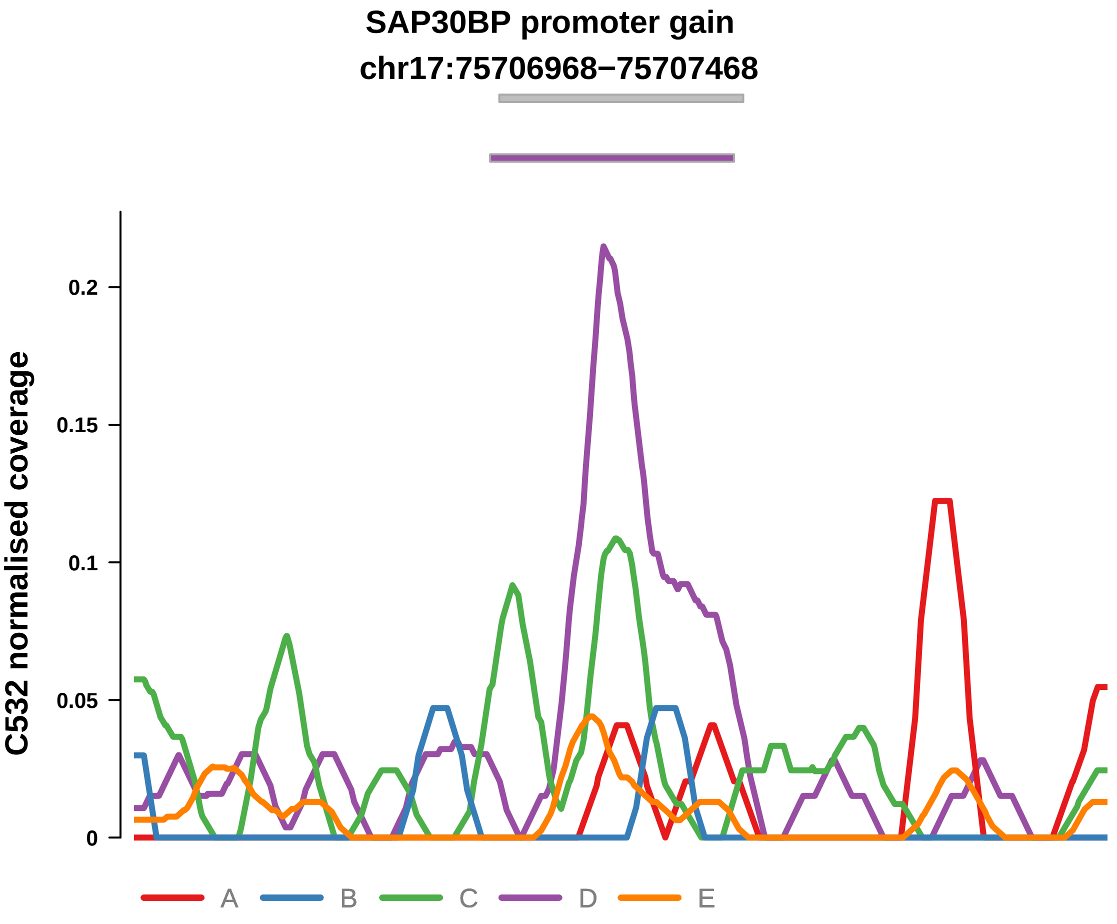


Putatively Clonal (logFC = 2.26)

| | P. | Adjusted P. |
|---|---|---|
| *~region vs. ~* | 3.94e−01 | 7.41e−01 |
| *~purity vs ~* | 3.62e−01 | 8.63e−01 |
| *~purity+region vs ~purity* | 7.54e−01 | 9.42e−01 |
| *Carcinoma vs. normal* | 1.54e−04 *** | 1.54e−04 *** |

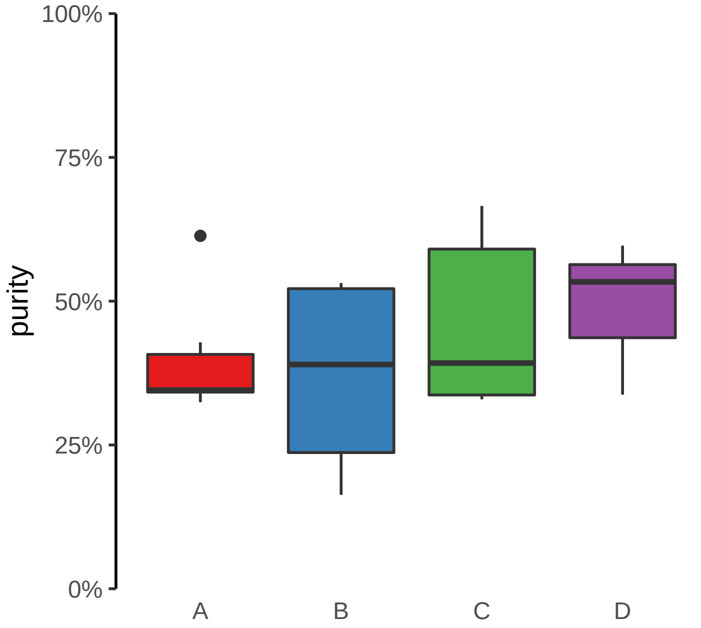

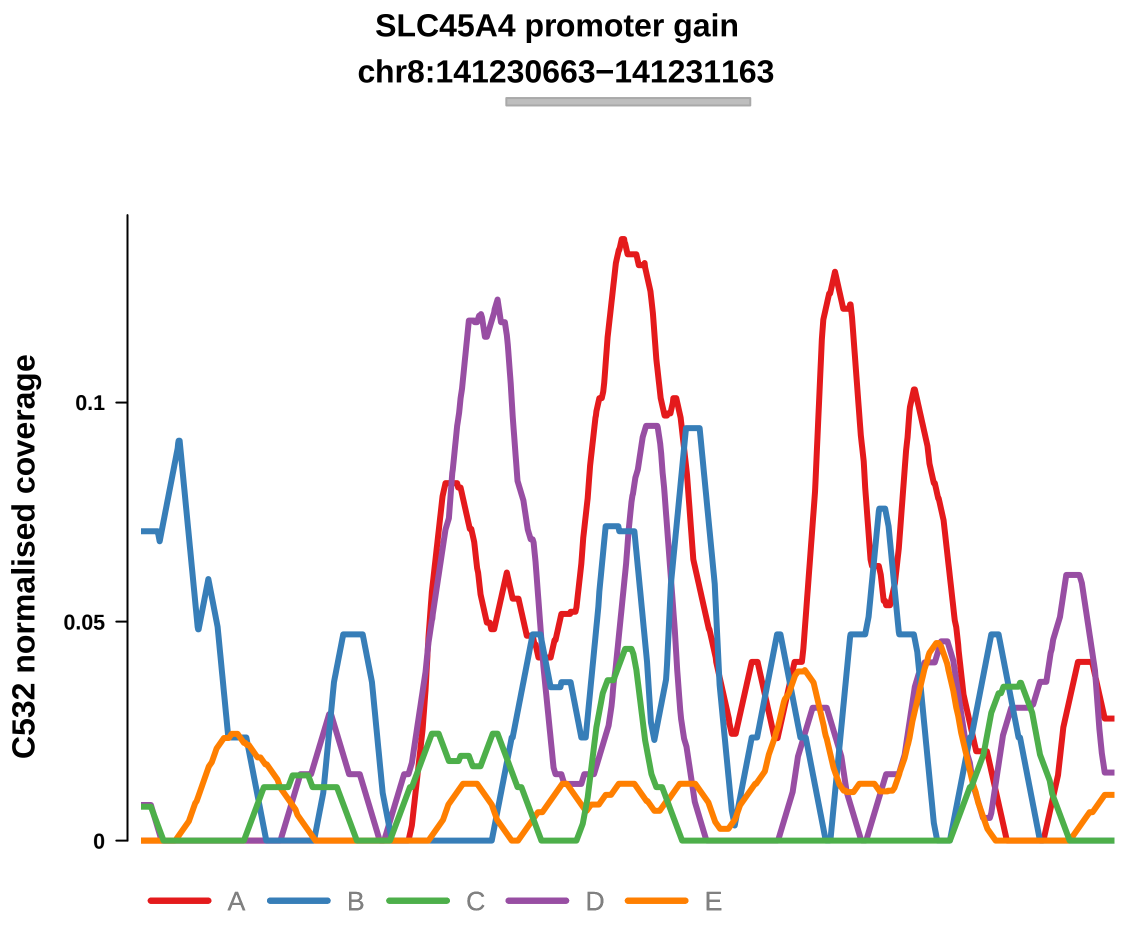


Putatively Clonal (logFC = 2.02)

| | P. | Adjusted P. |
|---|---|---|
| *~region vs. ~* | 6.06e−01 | 8.54e−01 |
| *~purity vs ~* | 4.95e−01 | 9.05e−01 |
| *~purity+region vs ~purity* | 5.34e−01 | 9.42e−01 |
| *Carcinoma vs. normal* | 8.1e−04 *** | 8.1e−04 *** |

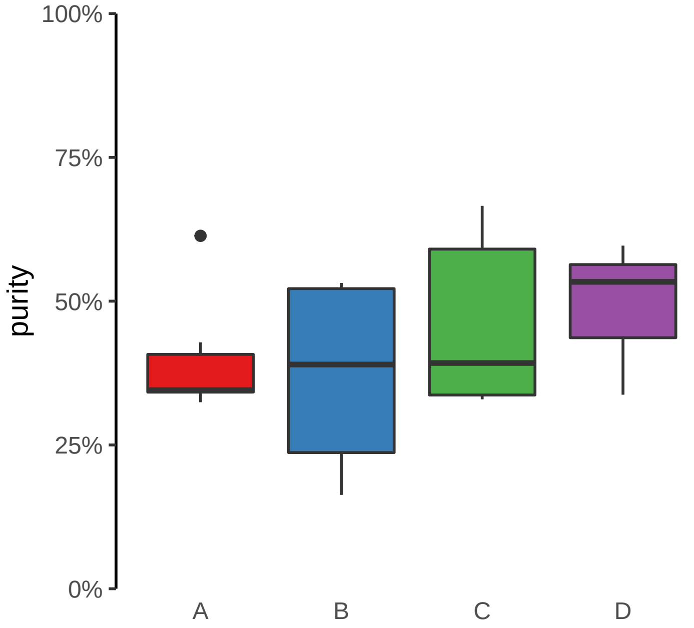

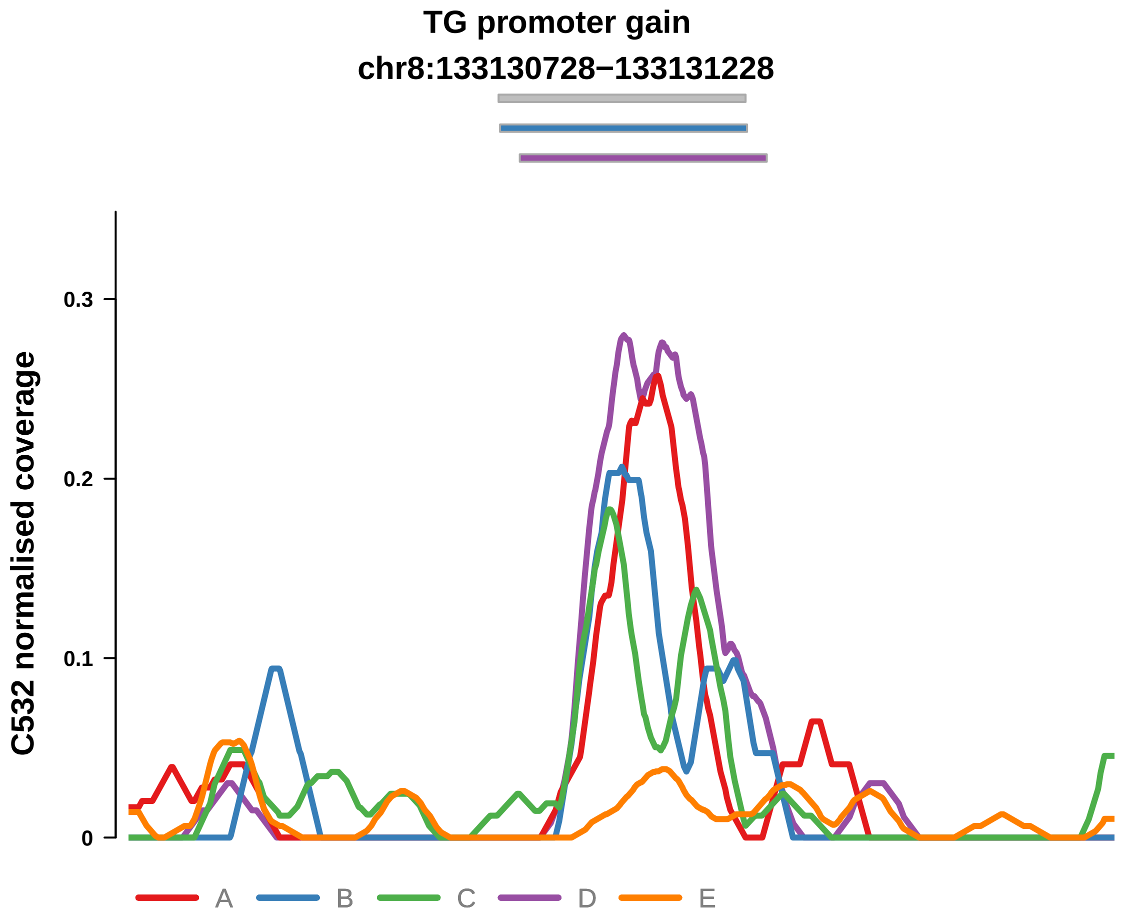


Putatively Clonal (logFC = 2.52)

| | P. | Adjusted P. |
|---|---|---|
| *~region vs. ~* | 1.39e−01 | 4.99e−01 |
| *~purity vs ~* | 5.32e−03 ** | 2.21e−01 |
| *~purity+region vs ~purity* | 4.37e−01 | 9.38e−01 |
| *Carcinoma vs. normal* | 1.26e−05 **** | 1.26e−05 **** |

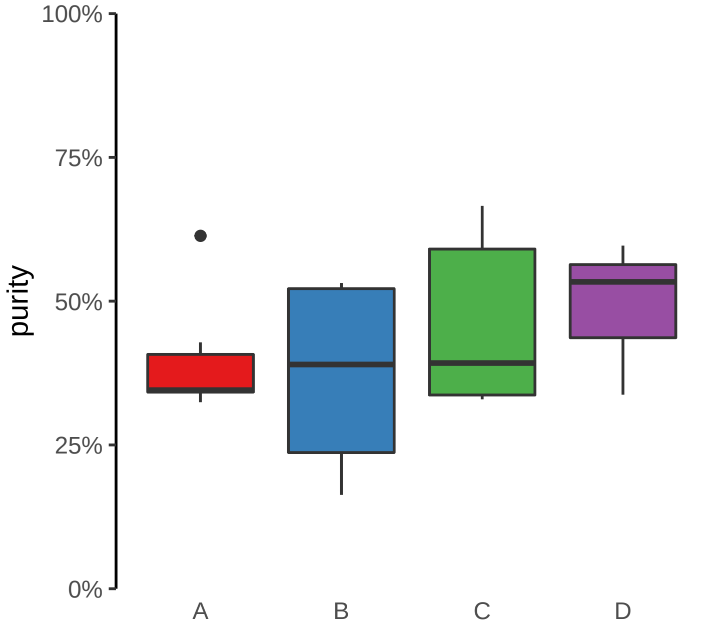

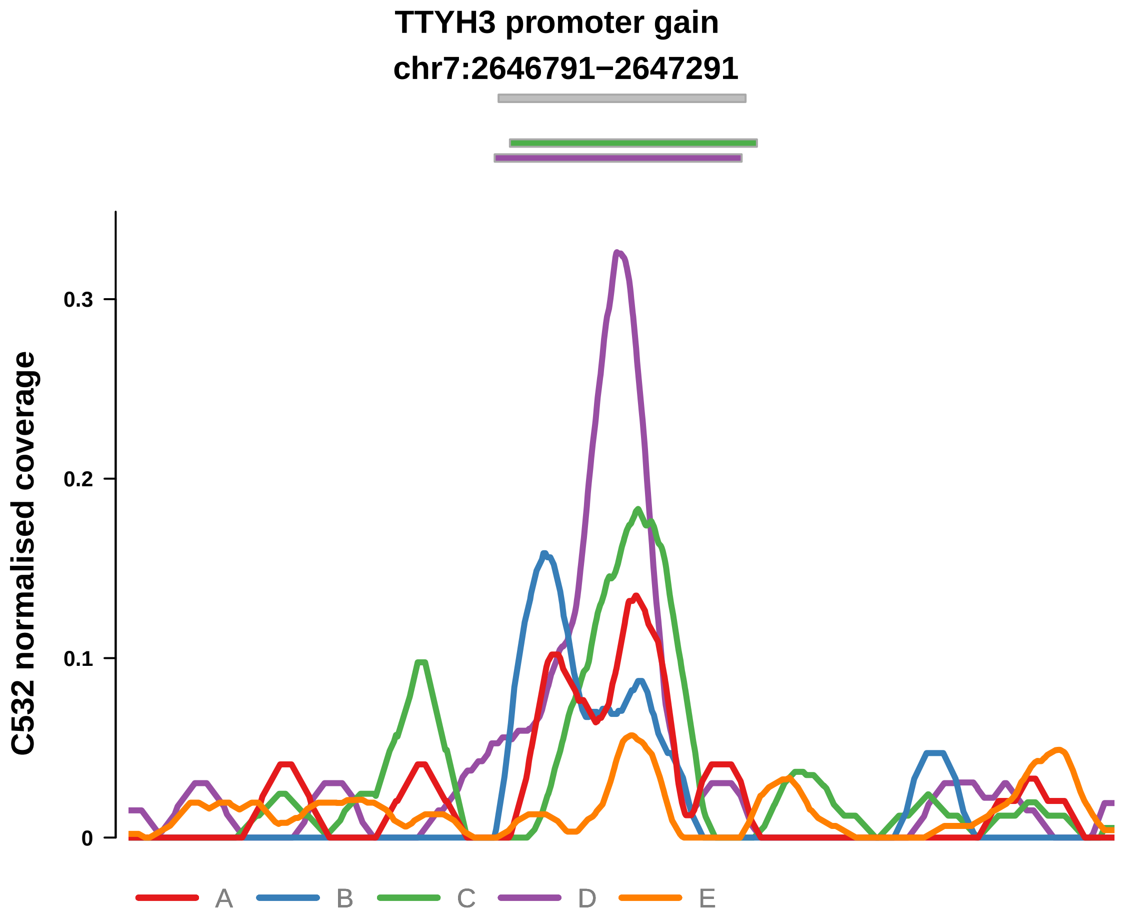


Putatively Clonal (logFC = 2.69)

| | P. | Adjusted P. |
|---|---|---|
| *~region vs. ~* | 5.73e−02 | 3.62e−01 |
| *~purity vs ~* | 2.54e−01 | 8.56e−01 |
| *~purity+region vs ~purity* | 4.48e−02 * | 4.93e−01 |
| *Carcinoma vs. normal* | 4.34e−06 **** | 4.34e−06 **** |

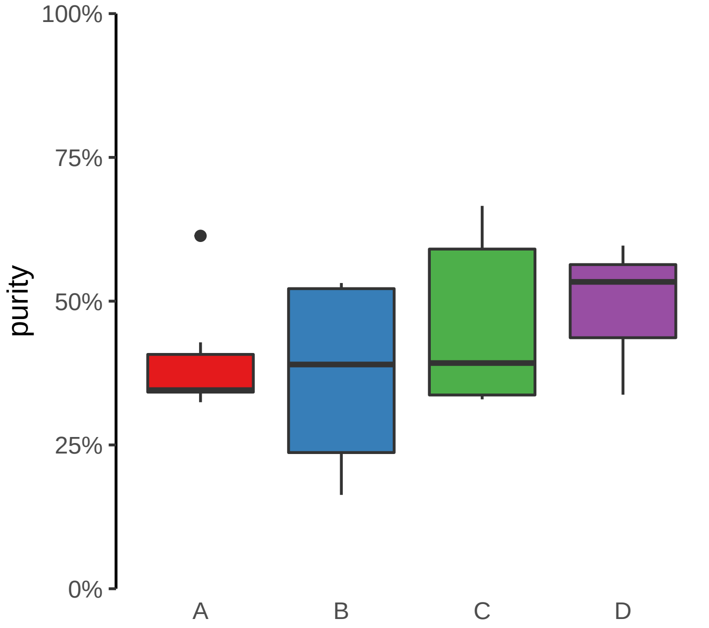

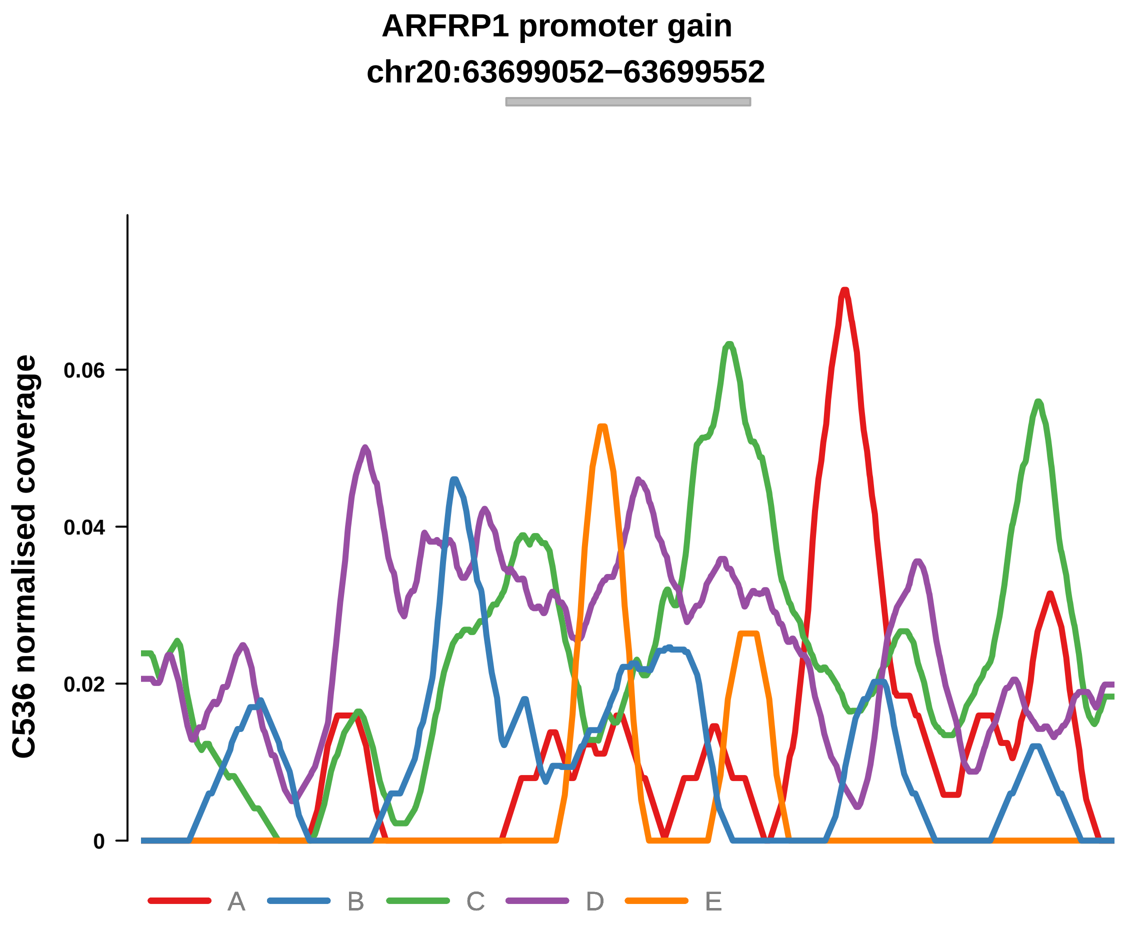


Putatively Clonal (logFC = 1.84)

| | P. | Adjusted P. |
|---|---|---|
| *~region vs. ~* | 4.25e−02 * | 1.32e−01 |
| *~purity vs ~* | 5.44e−01 | 7.21e−01 |
| *~purity+region vs ~purity* | 2.45e−02 * | 9.93e−02 |
| *Carcinoma vs. normal* | 1.97e−03 ** | 1.97e−03 ** |

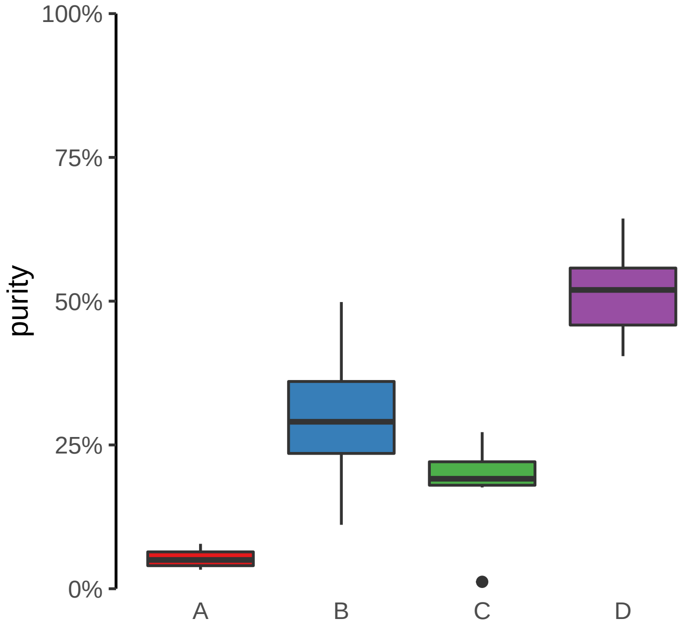

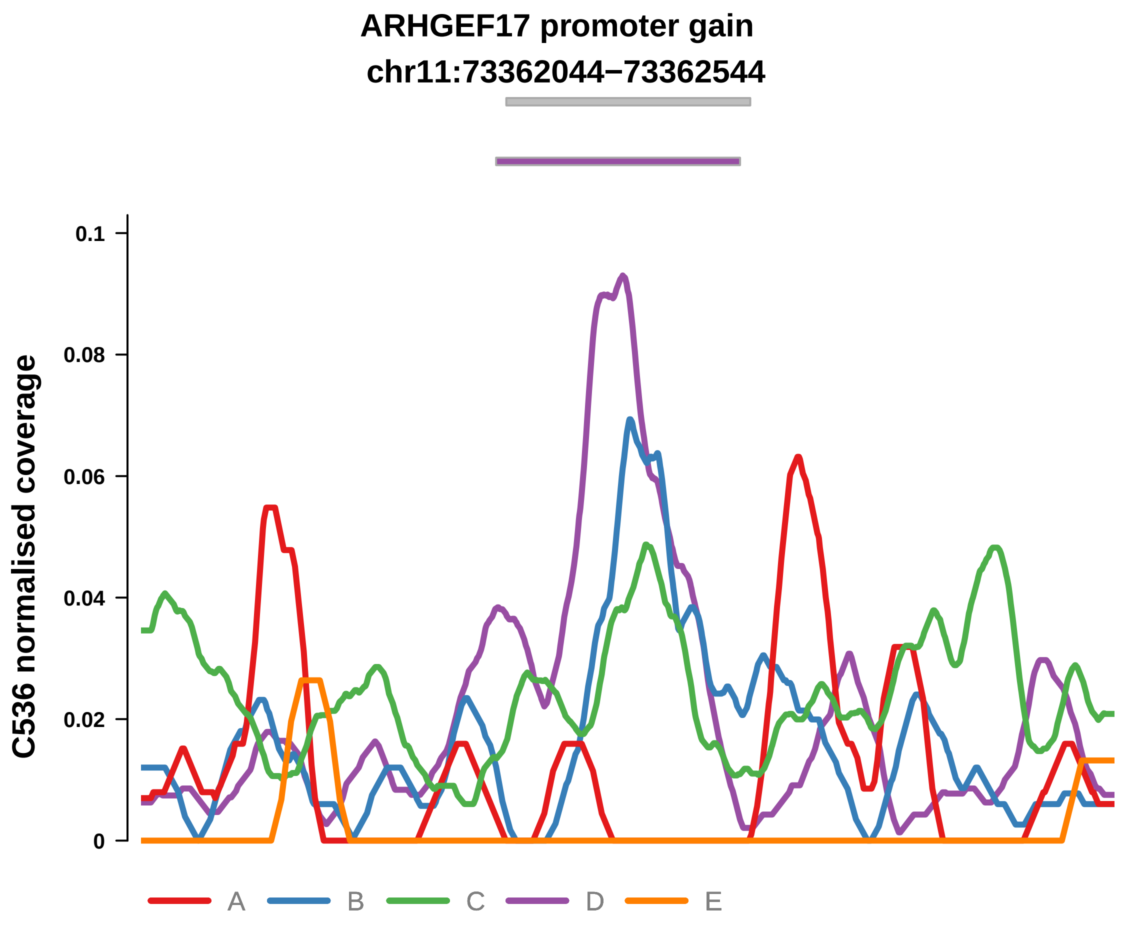


Putatively Clonal (logFC = 2.72)

| | P. | Adjusted P. |
|---|---|---|
| *~region vs. ~* | 6.25e−05 **** | 3.69e−03 ** |
| *~purity vs ~* | 5.51e−03 ** | 6.92e−02 |
| *~purity+region vs ~purity* | 6.6e−03 ** | 5.22e−02 |
| *Carcinoma vs. normal* | 4.74e−06 **** | 4.74e−06 **** |

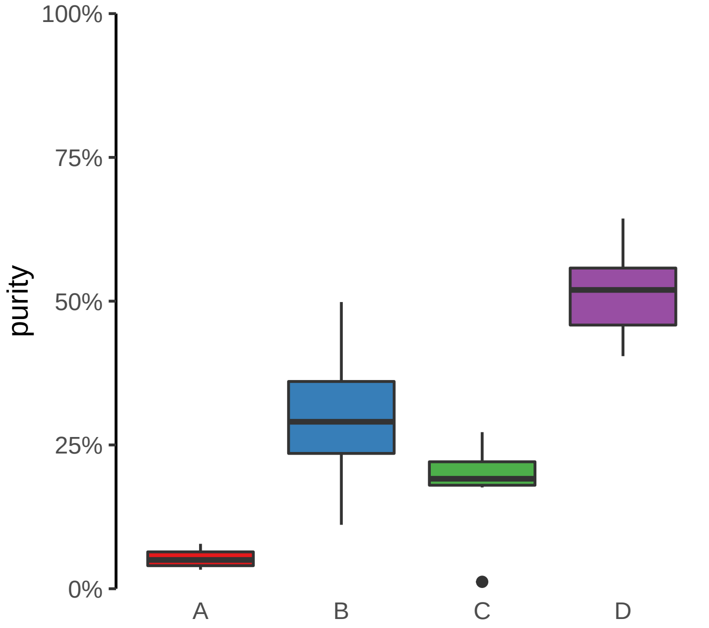

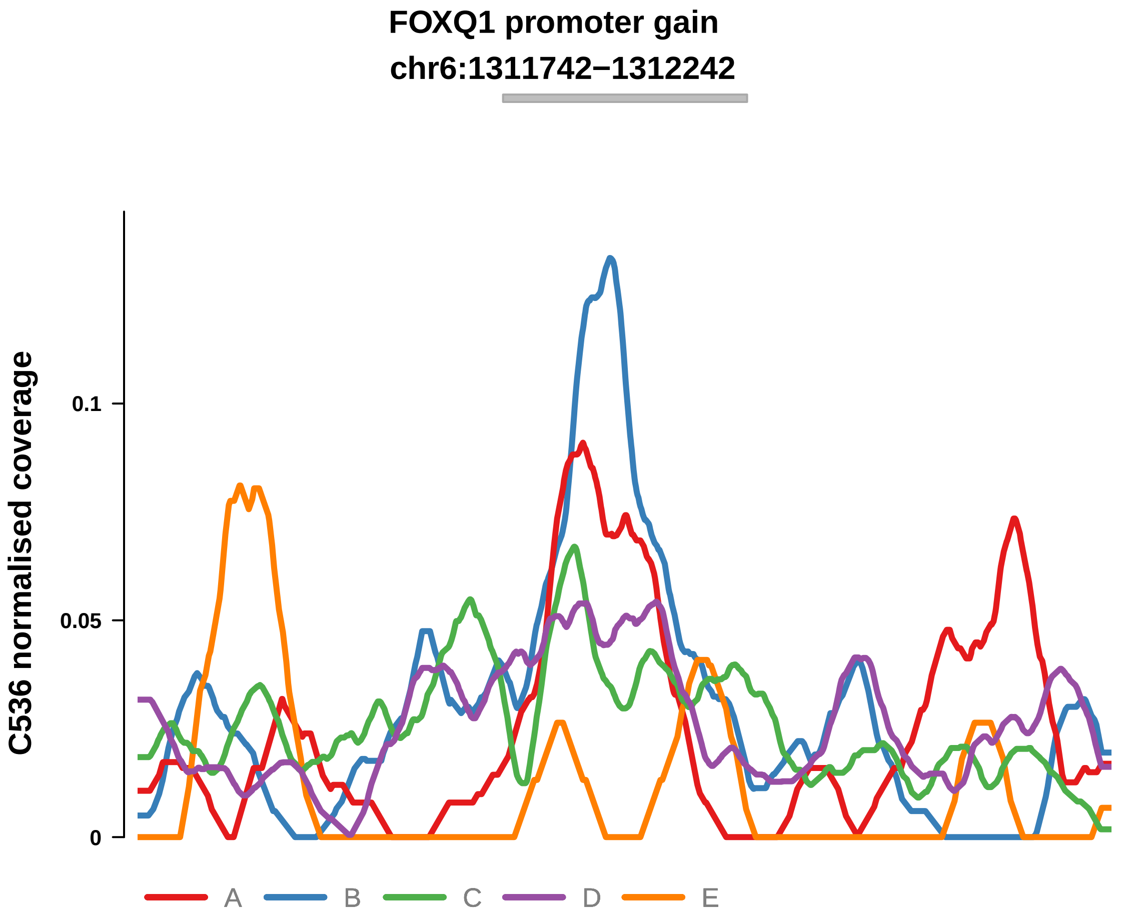


Putatively Clonal (logFC = 1.5)

| | P. | Adjusted P. |
|---|---|---|
| *~region vs. ~* | 5.41e−02 | 1.6e−01 |
| *~purity vs ~* | 5.1e−02 | 2.04e−01 |
| *~purity+region vs ~purity* | 2.75e−01 | 4.42e−01 |
| *Carcinoma vs. normal* | 8.91e−03 ** | 8.91e−03 ** |

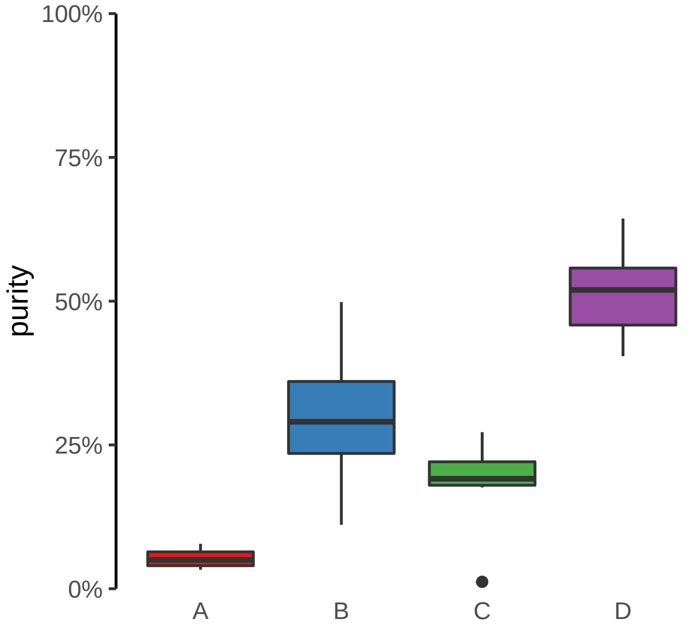

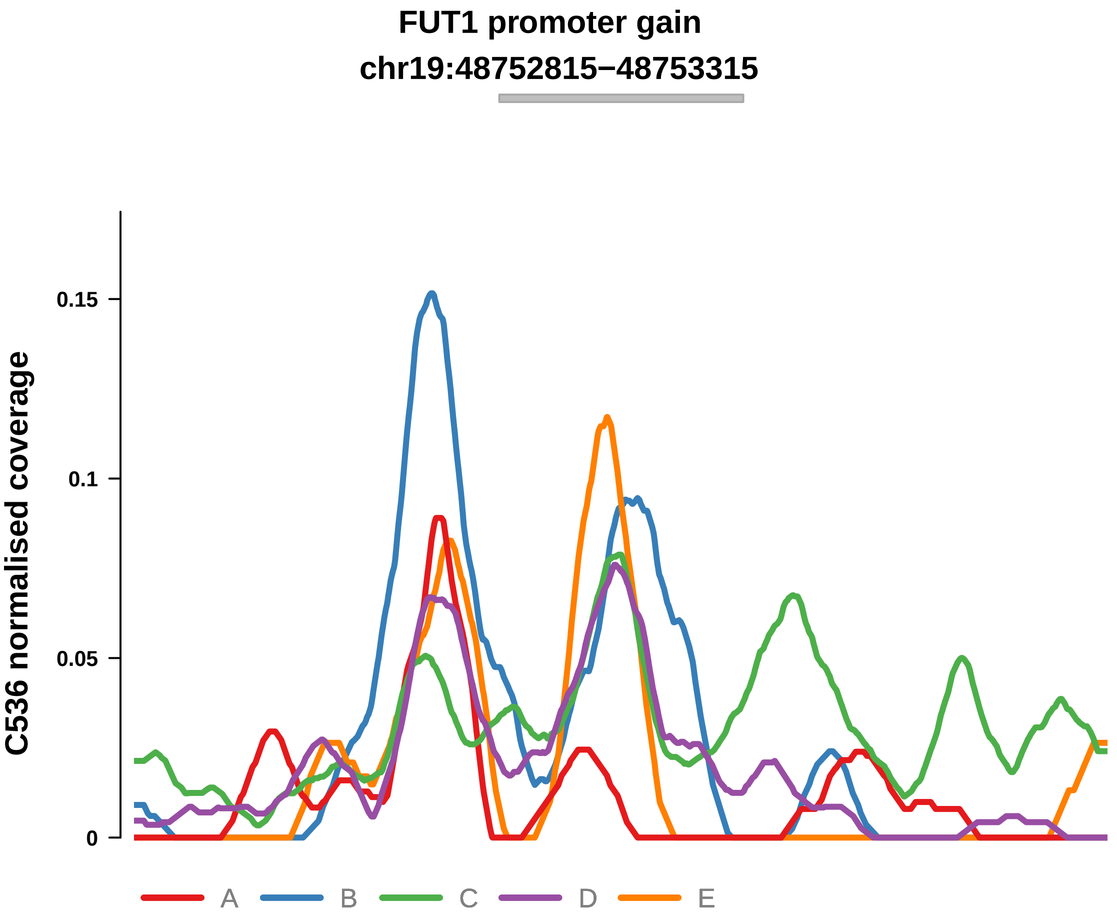


Putatively Clonal (logFC = 1.64)

| | P. | Adjusted P. |
|---|---|---|
| *~region vs. ~* | 1.58e−02 * | 7.76e−02 |
| *~purity vs ~* | 3.19e−01 | 5.93e−01 |
| *~purity+region vs ~purity* | 9.59e−03 ** | 6.02e−02 |
| *Carcinoma vs. normal* | 5.06e−03 ** | 5.06e−03 ** |

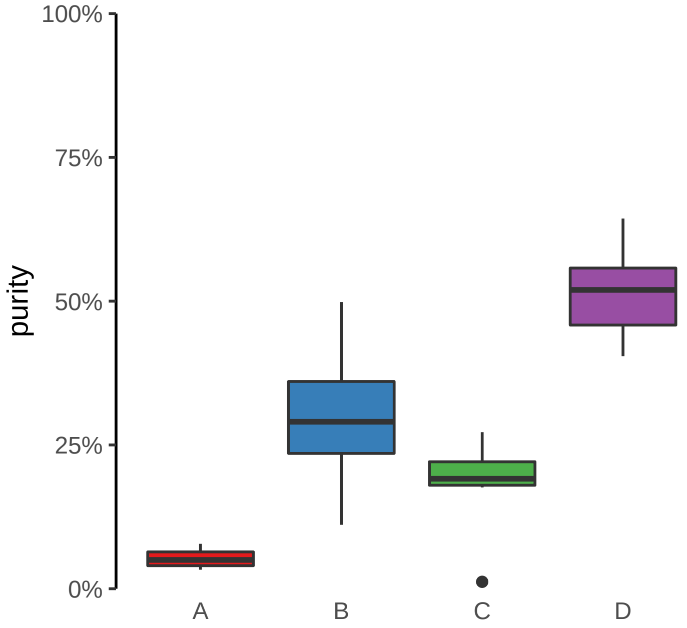

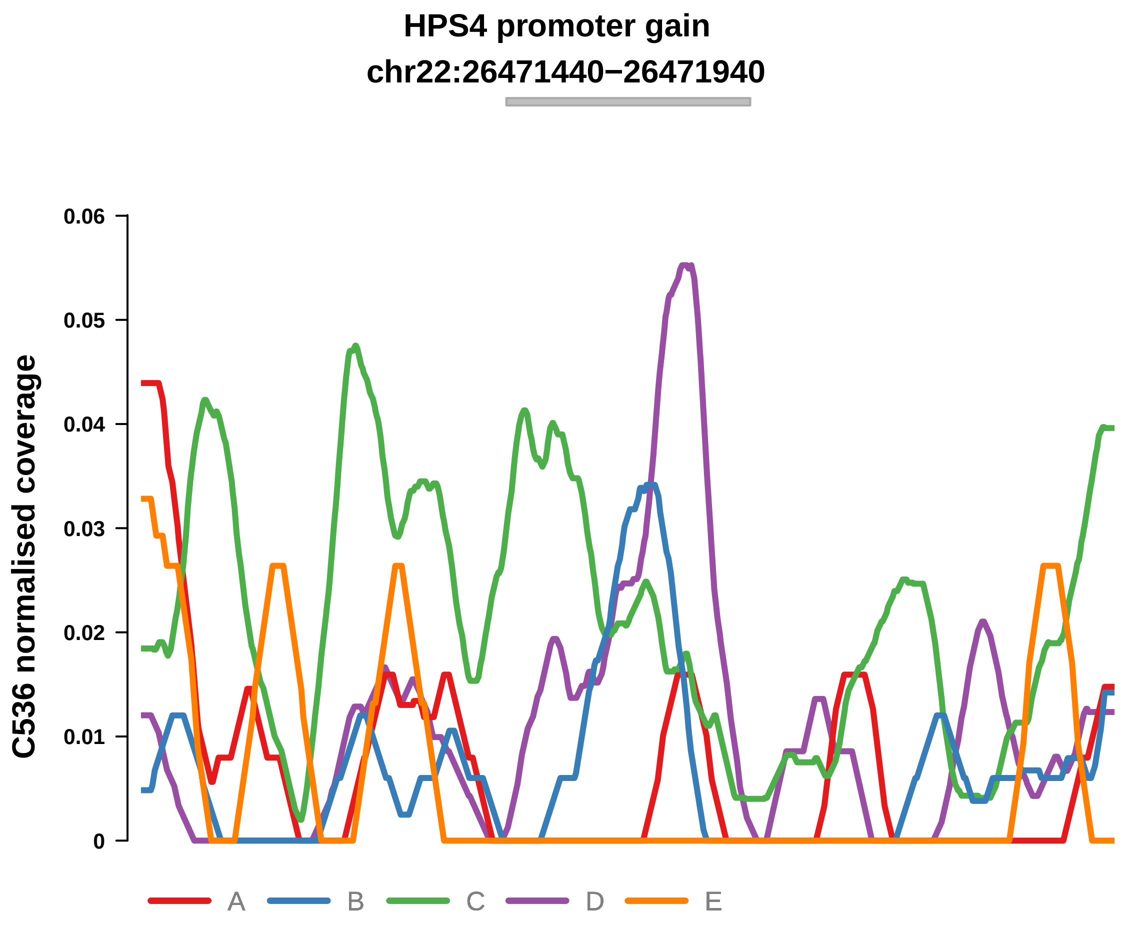


Putatively Clonal (logFC = 2.06)

| | P. | Adjusted P. |
|---|---|---|
| *~region vs. ~* | 1.15e−02 * | NA |
| *~purity vs ~* | 3.31e−01 | 5.93e−01 |
| *~purity+region vs ~purity* | 5.67e−03 ** | 5.22e−02 |
| *Carcinoma vs. normal* | 1.29e−03 ** | 1.29e−03 ** |

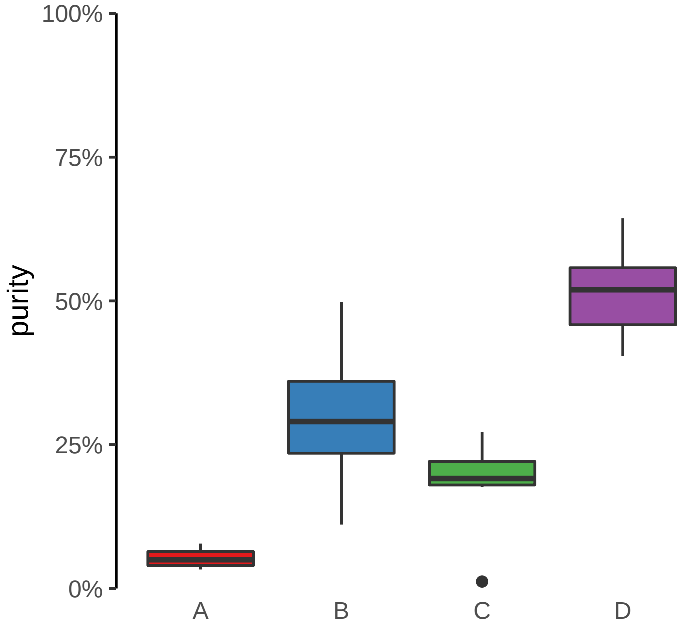

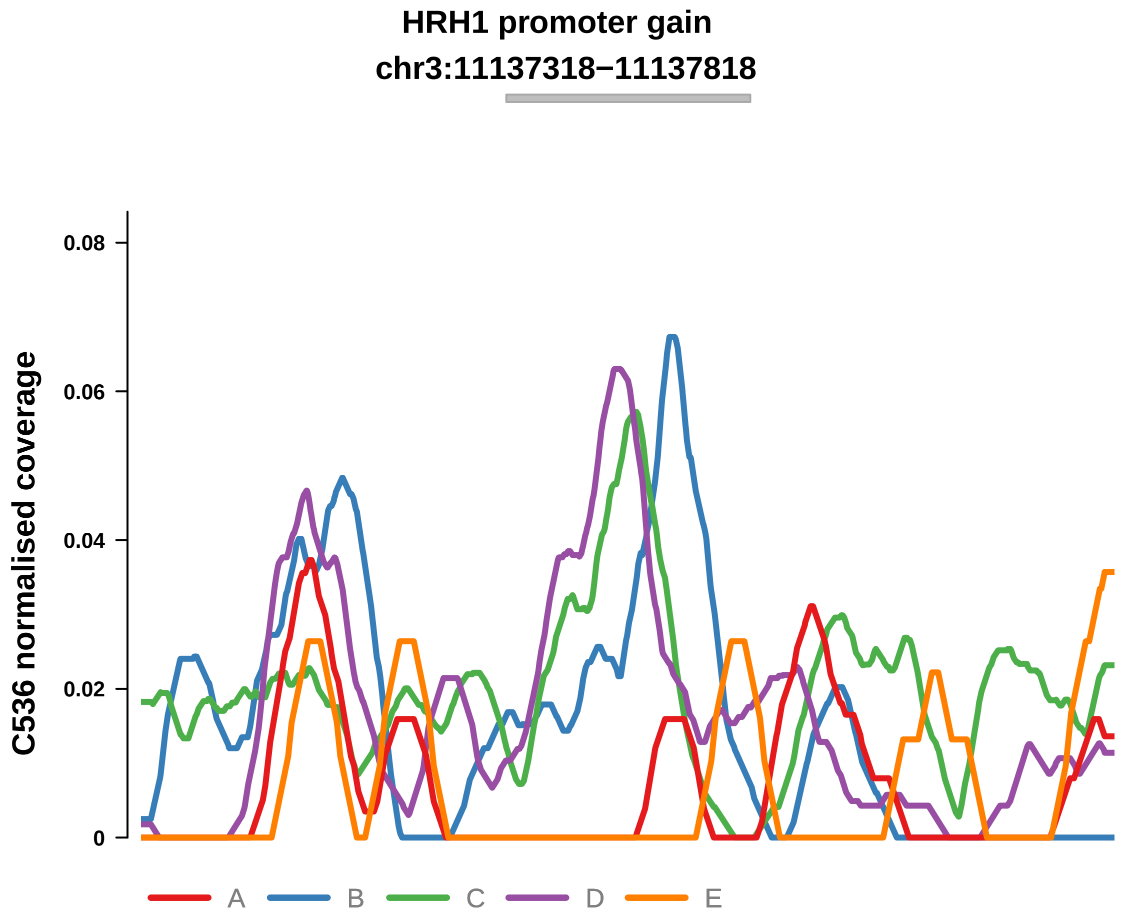


Putatively Clonal (logFC = 2.03)

| | P. | Adjusted P. |
|---|---|---|
| *~region vs. ~* | 1.63e−03 ** | 2.75e−02 * |
| *~purity vs ~* | 1.06e−02 * | 1.03e−01 |
| *~purity+region vs ~purity* | 5.1e−02 | 1.68e−01 |
| *Carcinoma vs. normal* | 6.15e−04 *** | 6.15e−04 *** |

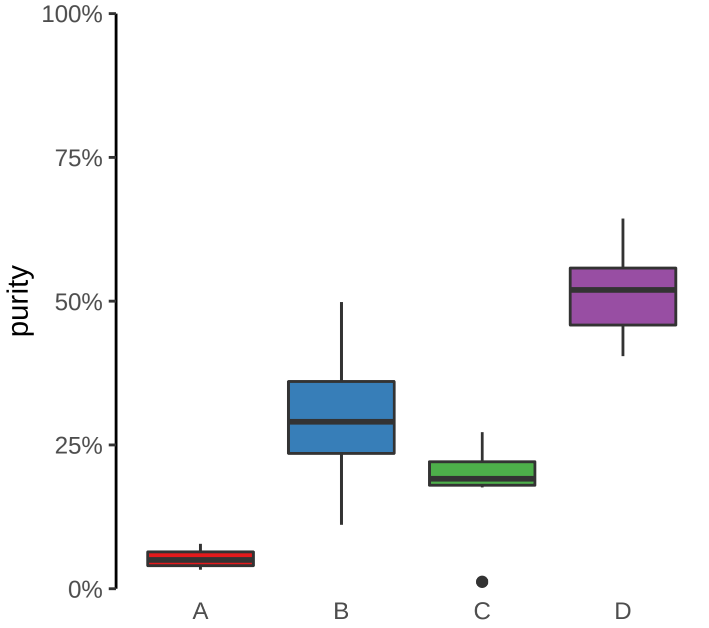

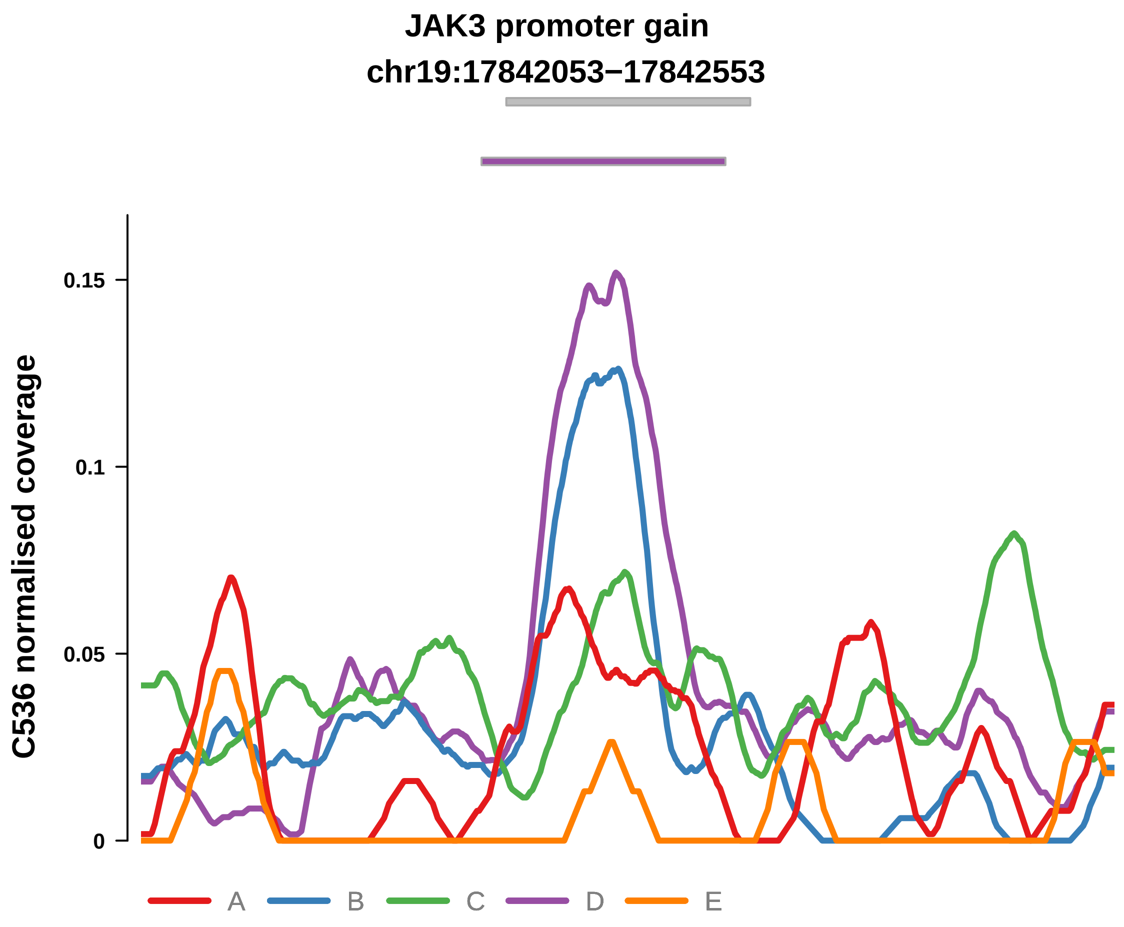


Putatively Clonal (logFC = 2.8)

| | P. | Adjusted P. |
|---|---|---|
| *~region vs. ~* | 1.26e−02 * | 7.43e−02 |
| *~purity vs ~* | 4.98e−02 * | 2.04e−01 |
| *~purity+region vs ~purity* | 3.94e−02 * | 1.43e−01 |
| *Carcinoma vs. normal* | 1.15e−06 **** | 1.15e−06 **** |

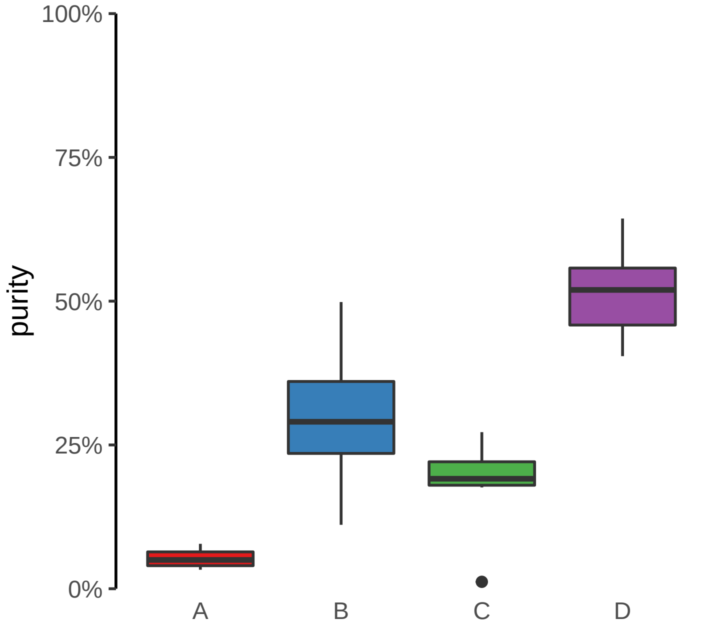

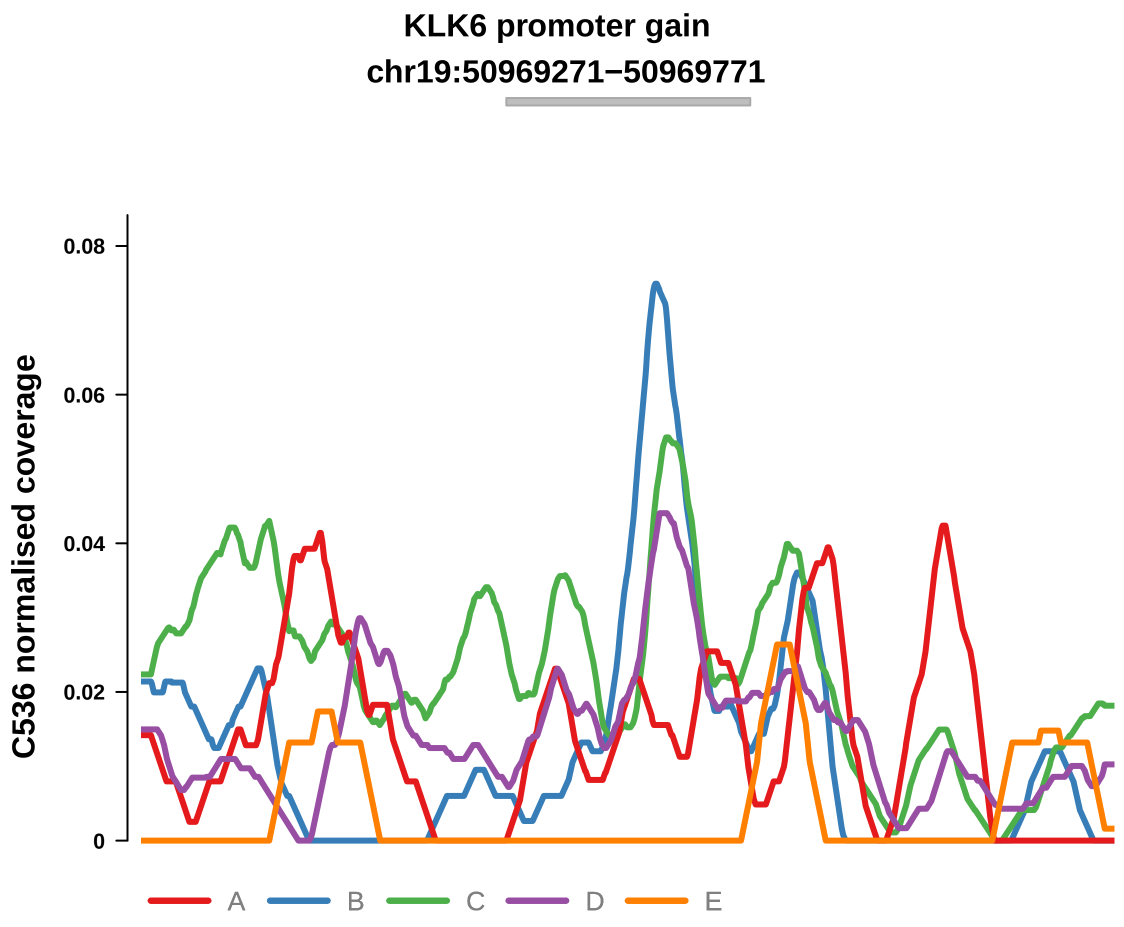


Putatively Clonal (logFC = 1.63)

| | P. | Adjusted P. |
|---|---|---|
| *~region vs. ~* | 8.87e−02 | 2.09e−01 |
| *~purity vs ~* | 5.39e−01 | 7.21e−01 |
| *~purity+region vs ~purity* | 1.08e−01 | 2.65e−01 |
| *Carcinoma vs. normal* | 9.84e−03 ** | 9.84e−03 ** |

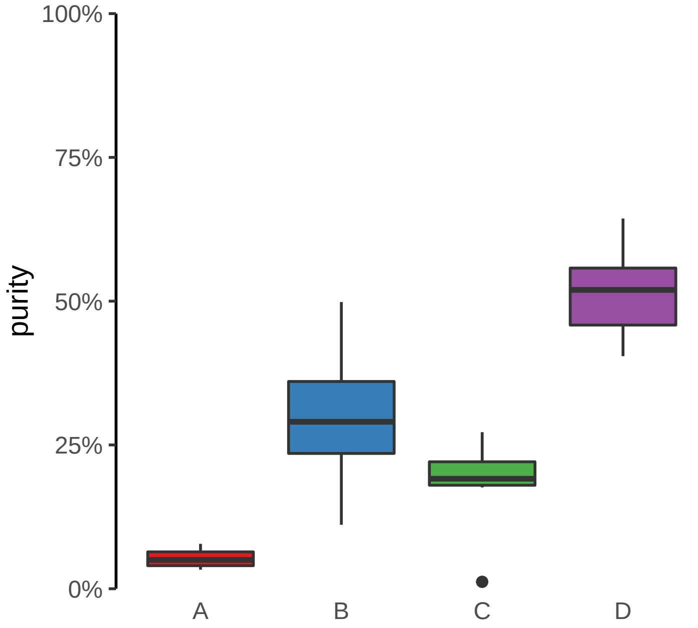

**PITX1 promoter gain**
**chr5:135034130−135034630**

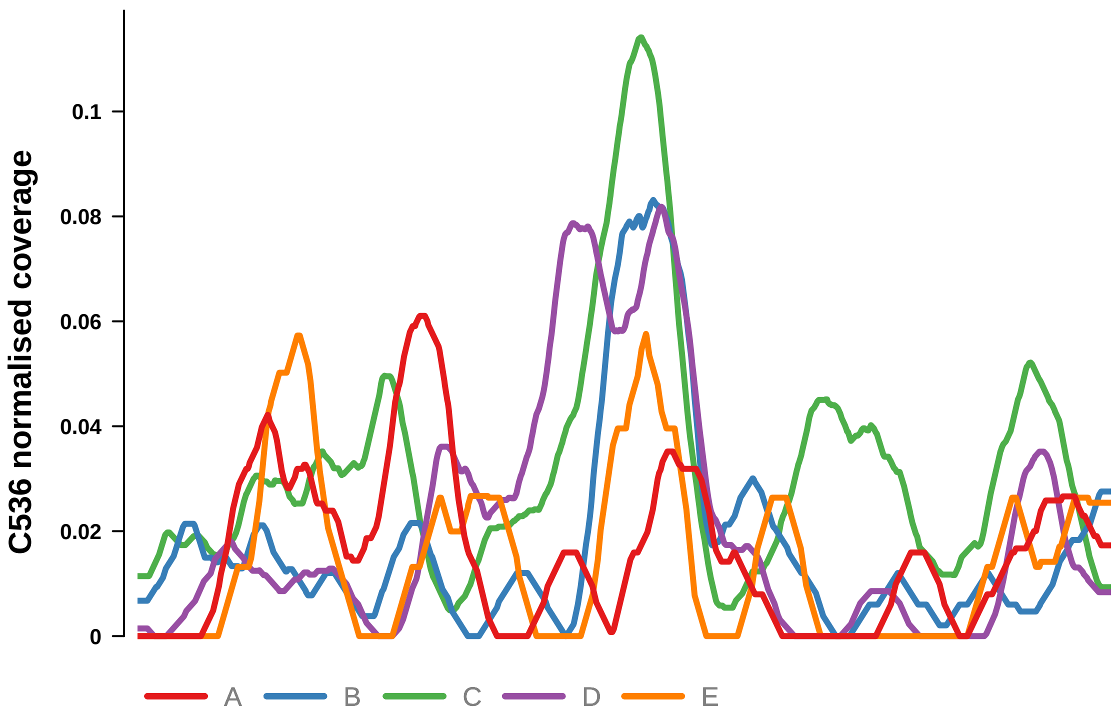


Putatively Clonal (logFC = 2.14)

| | **P.** | **Adjusted P.** |
|---|---|---|
| *~region vs. ~* | 2.36e−02 * | 9.95e−02 |
| *~purity vs ~* | 8.68e−02 | 2.55e−01 |
| *~purity+region vs ~purity* | 1.02e−01 | 2.6e−01 |
| *Carcinoma vs. normal* | 1.77e−04 *** | 1.77e−04 *** |

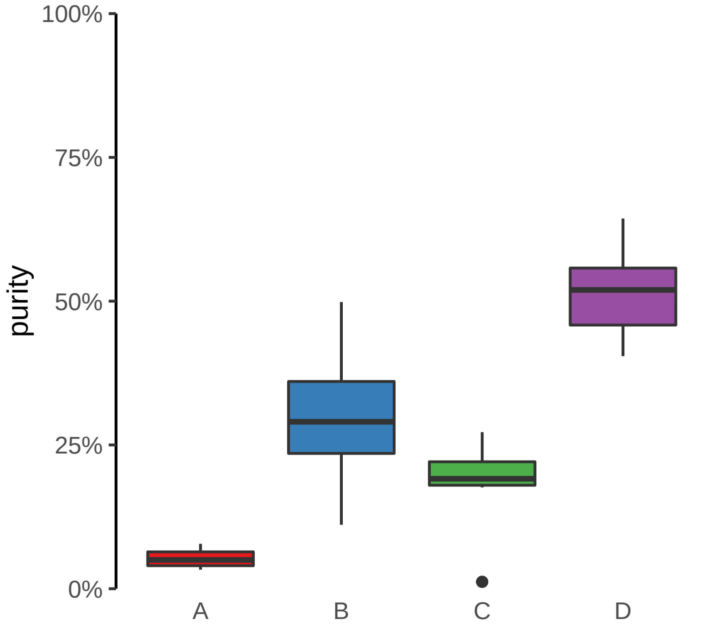

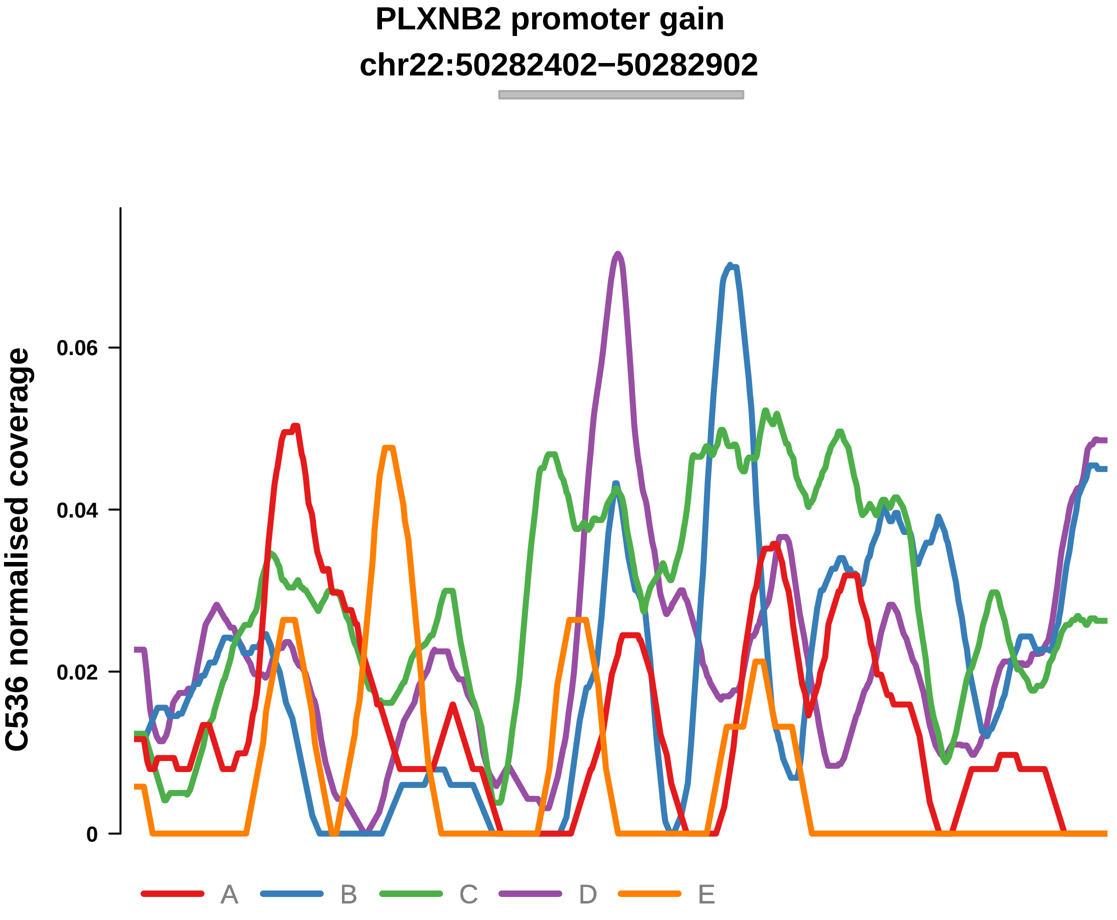


Putatively Clonal (logFC = 1.63)

| | P. | Adjusted P. |
|---|---|---|
| *~region vs. ~* | 9.32e−04 *** | 2.75e−02 * |
| *~purity vs ~* | 2.92e−01 | 5.85e−01 |
| *~purity+region vs ~purity* | 1.39e−03 ** | 2.4e−02 * |
| *Carcinoma vs. normal* | 6.58e−03 ** | 6.58e−03 ** |

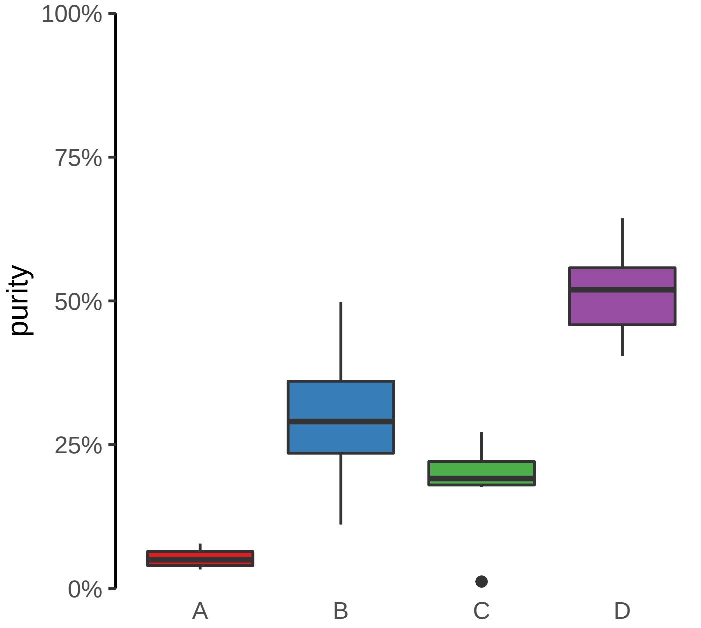

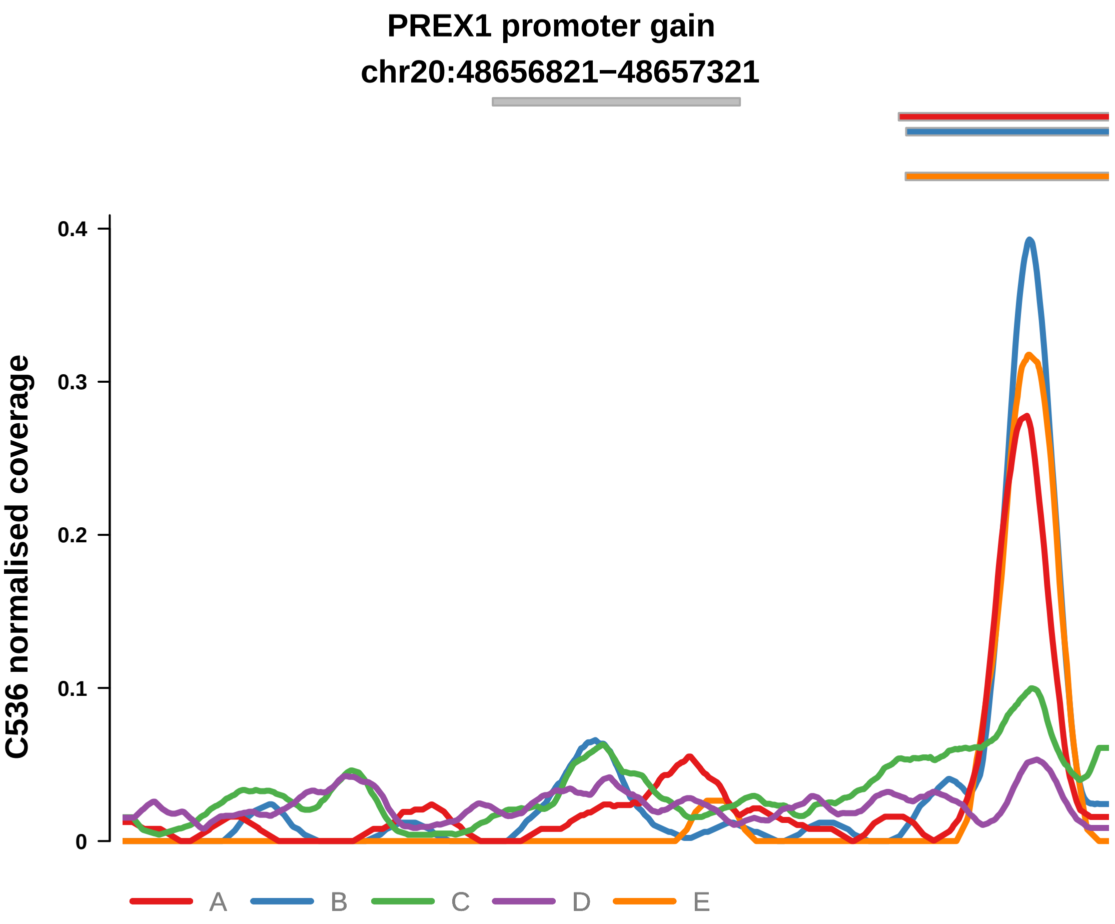


Putatively Clonal (logFC = 1.93)

| | P. | Adjusted P. |
|---|---|---|
| *~region vs. ~* | 2.56e−01 | 3.87e−01 |
| *~purity vs ~* | 9.42e−01 | 9.68e−01 |
| *~purity+region vs ~purity* | 3.97e−03 ** | 4.57e−02 * |
| *Carcinoma vs. normal* | 1.99e−03 ** | 1.99e−03 ** |

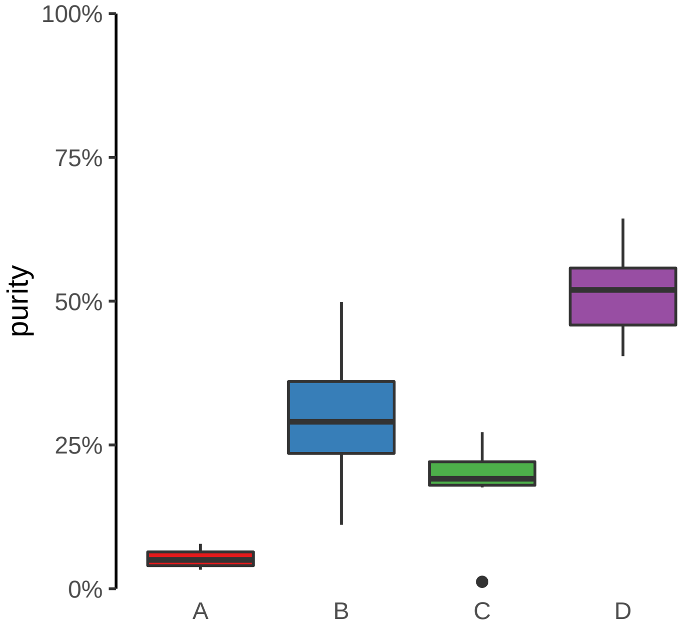

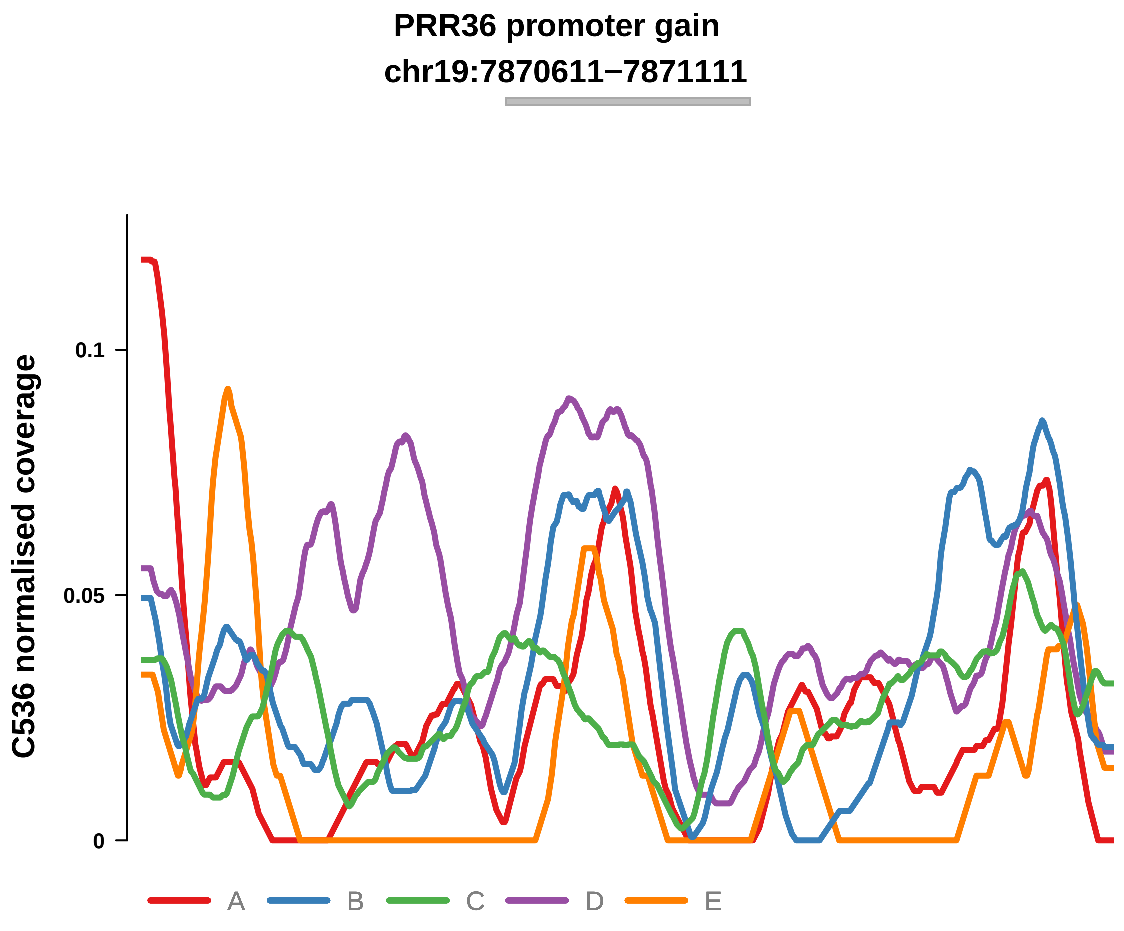


Putatively Clonal (logFC = 1.97)

| | P. | Adjusted P. |
|---|---|---|
| *~region vs. ~* | 3.63e−02 * | 1.19e−01 |
| *~purity vs ~* | 1.53e−01 | 3.63e−01 |
| *~purity+region vs ~purity* | 3.89e−02 * | 1.43e−01 |
| *Carcinoma vs. normal* | 6.16e−04 *** | 6.16e−04 *** |

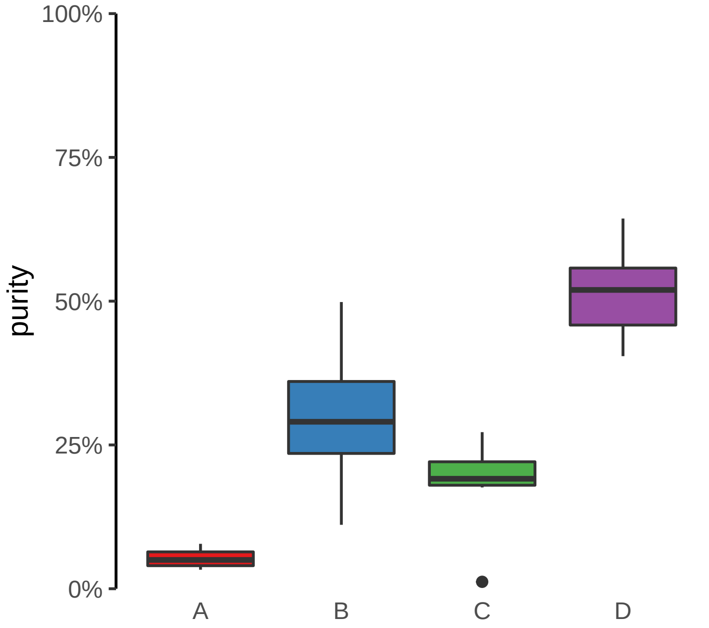

**SAP30BP promoter gain**
**chr17:75706968−75707468**

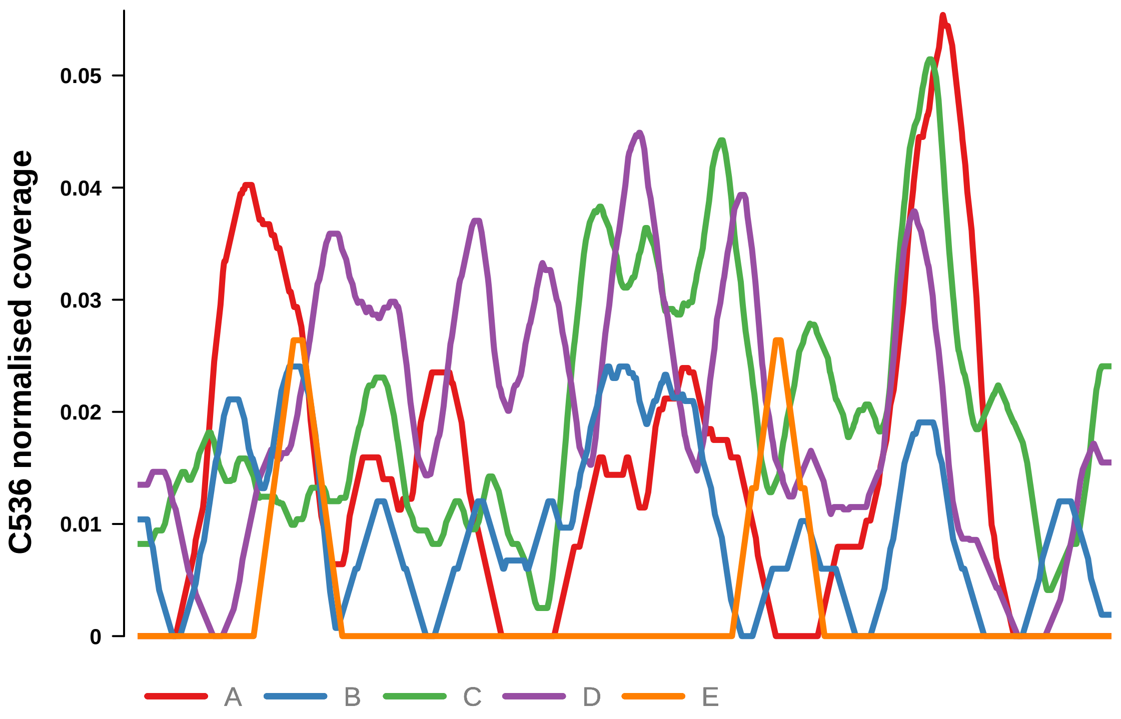


Putatively Clonal (logFC = 2.17)

| | **P.** | **Adjusted P.** |
|---|---|---|
| *~region vs. ~* | 5.51e−03 ** | 3.61e−02 * |
| *~purity vs ~* | 2.52e−01 | 5.28e−01 |
| *~purity+region vs ~purity* | 8.84e−03 ** | 6.02e−02 |
| *Carcinoma vs. normal* | 3.81e−04 *** | 3.81e−04 *** |

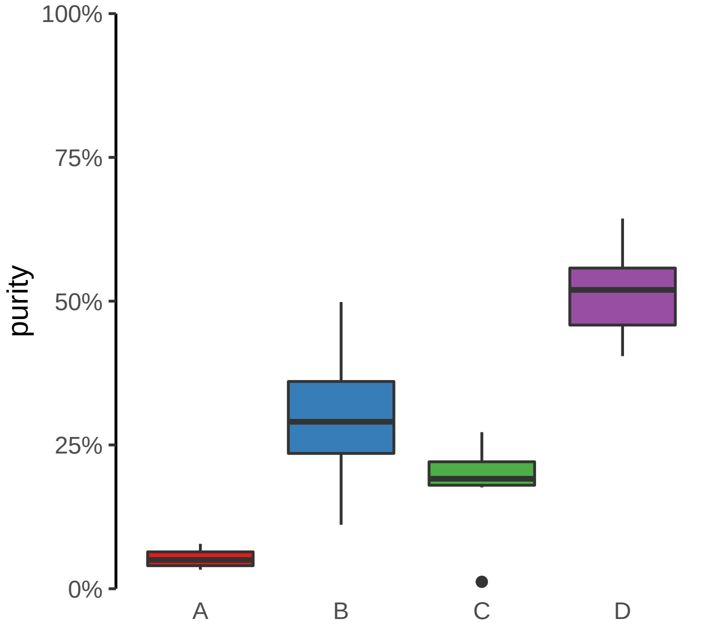

**SLC45A4 promoter gain**
**chr8:141230663−141231163**

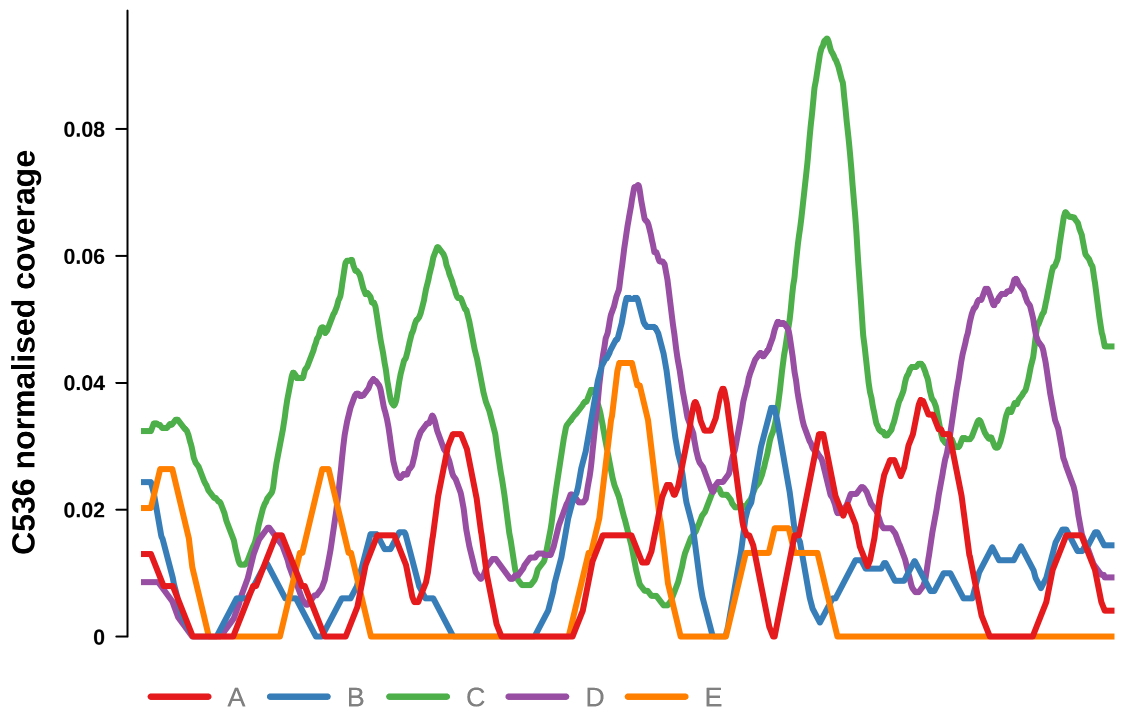


Putatively Clonal (logFC = 2.3)

| | P. | Adjusted P. |
|---|---|---|
| *~region vs. ~* | 1.57e−02 * | 7.76e−02 |
| *~purity vs ~* | 8.46e−02 | 2.55e−01 |
| *~purity+region vs ~purity* | 2.14e−02 * | 9.84e−02 |
| *Carcinoma vs. normal* | 1.7e−04 *** | 1.7e−04 *** |

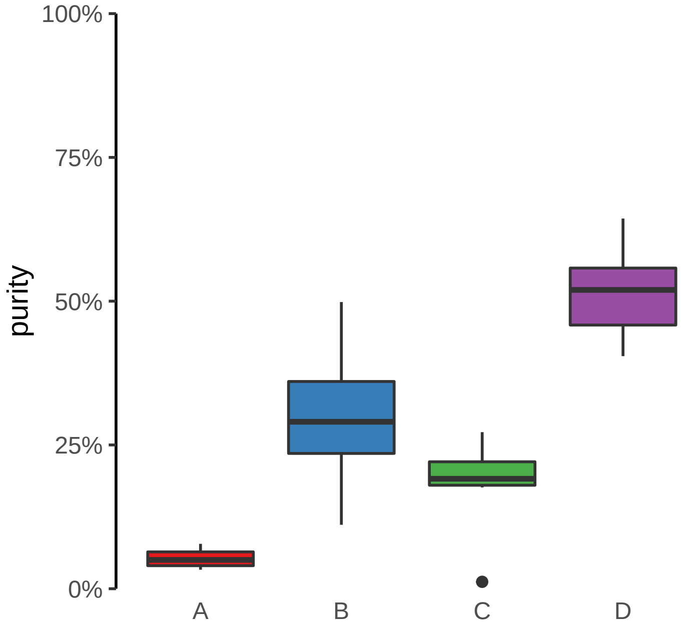

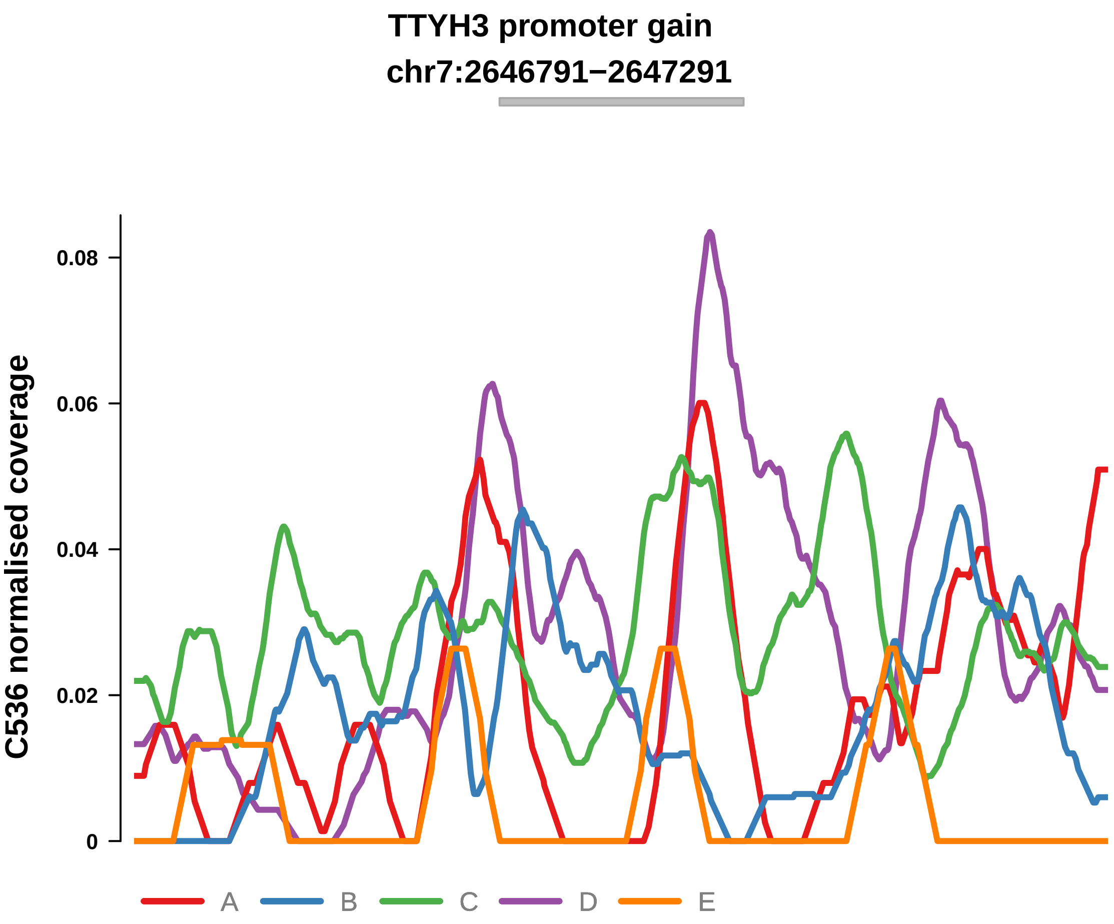


Putatively Clonal (logFC = 2.46)

| | P. | Adjusted P. |
|---|---|---|
| *~region vs. ~* | 4.45e−03 ** | 3.61e−02 * |
| *~purity vs ~* | 1.03e−01 | 2.7e−01 |
| *~purity+region vs ~purity* | 1.13e−02 * | 6.52e−02 |
| *Carcinoma vs. normal* | 4.05e−05 **** | 4.05e−05 **** |

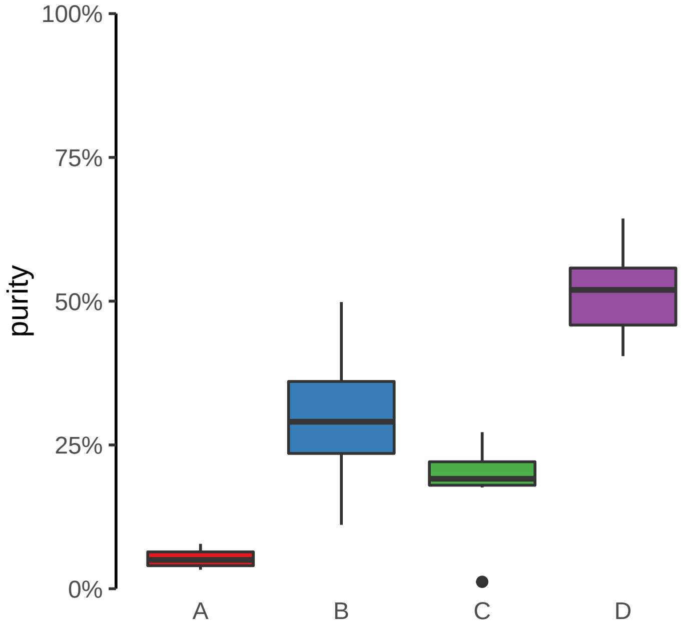

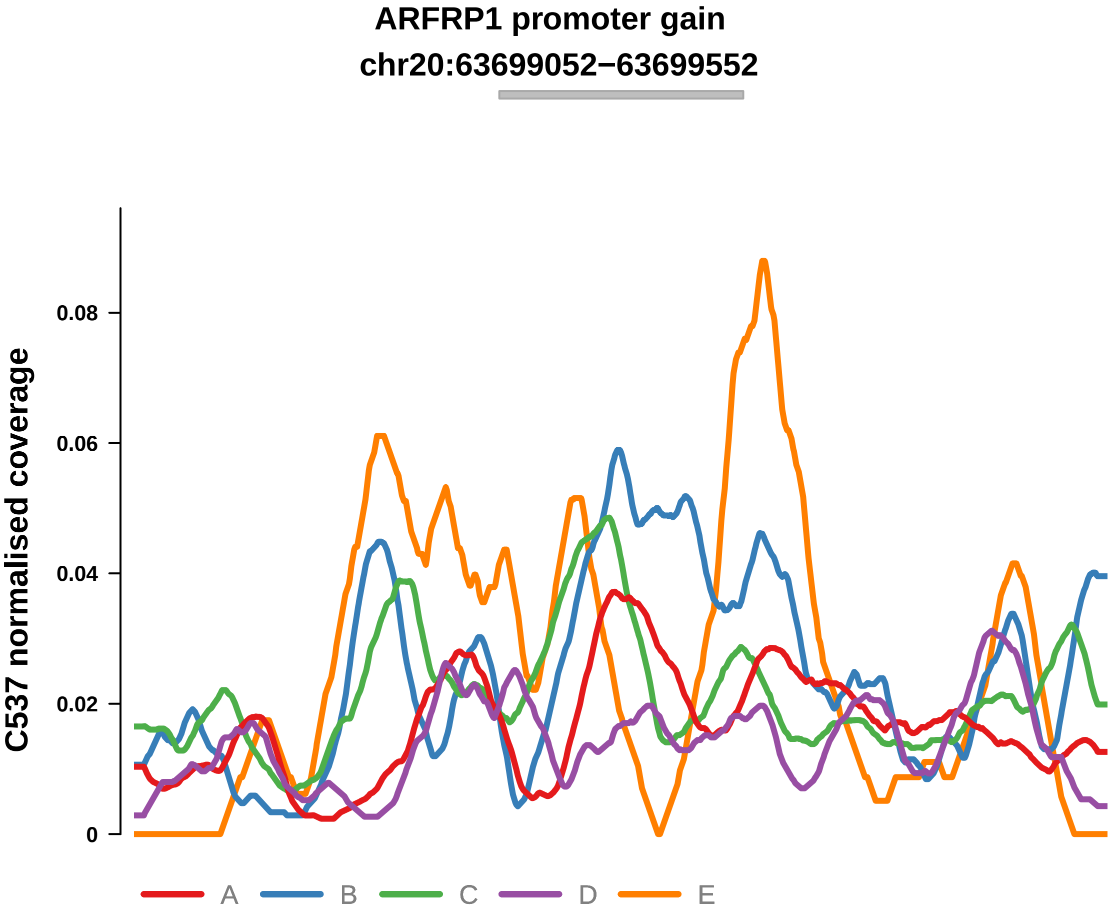


Putatively Clonal (logFC = 1.8)

| | P. | Adjusted P. |
|---|---|---|
| *~region vs. ~* | 1.29e−01 | 2.07e−01 |
| *~purity vs ~* | 2.81e−02 * | 9.23e−02 |
| *~purity+region vs ~purity* | 7.19e−01 | 8.6e−01 |
| *Carcinoma vs. normal* | 1.84e−03 ** | 1.84e−03 ** |

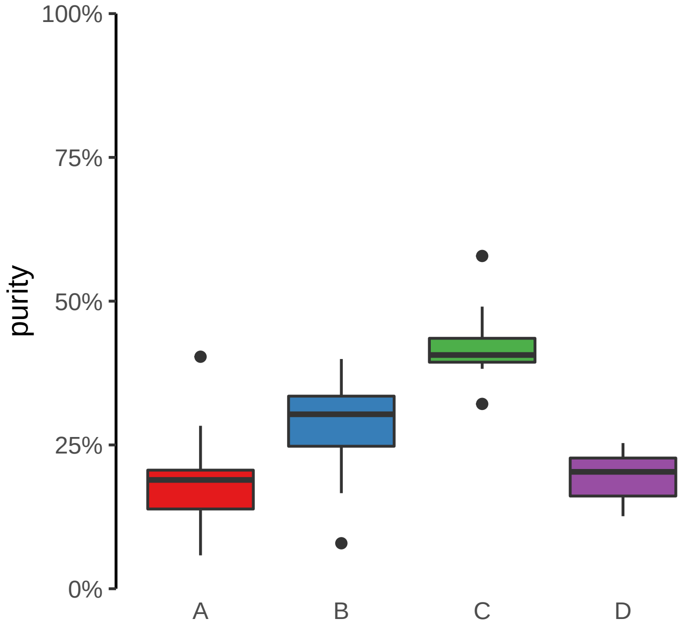

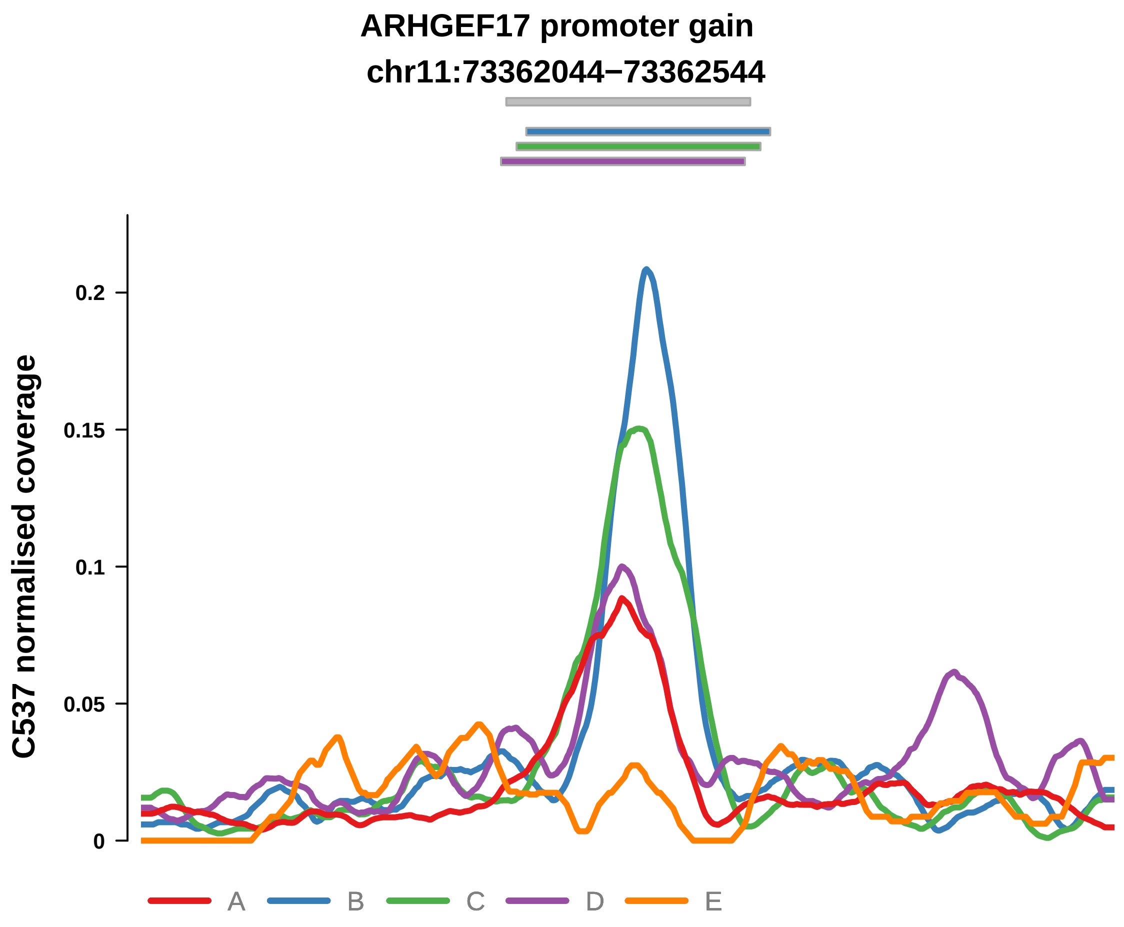


Putatively Clonal (logFC = 3.54)

| | P. | Adjusted P. |
|---|---|---|
| *~region vs. ~* | 9.34e−03 ** | 4.57e−02 * |
| *~purity vs ~* | 2.51e−03 ** | 1.73e−02 * |
| *~purity+region vs ~purity* | 1.62e−01 | 3.04e−01 |
| *Carcinoma vs. normal* | 2.21e−09 **** | 2.21e−09 **** |

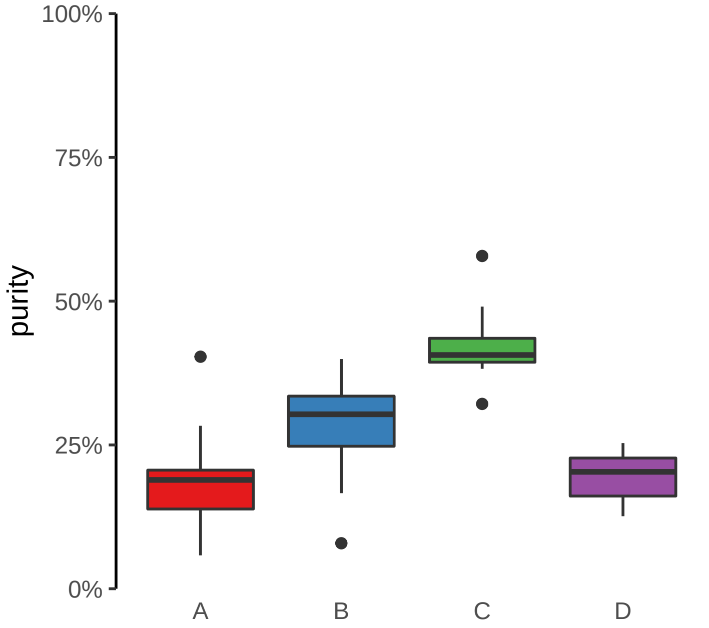

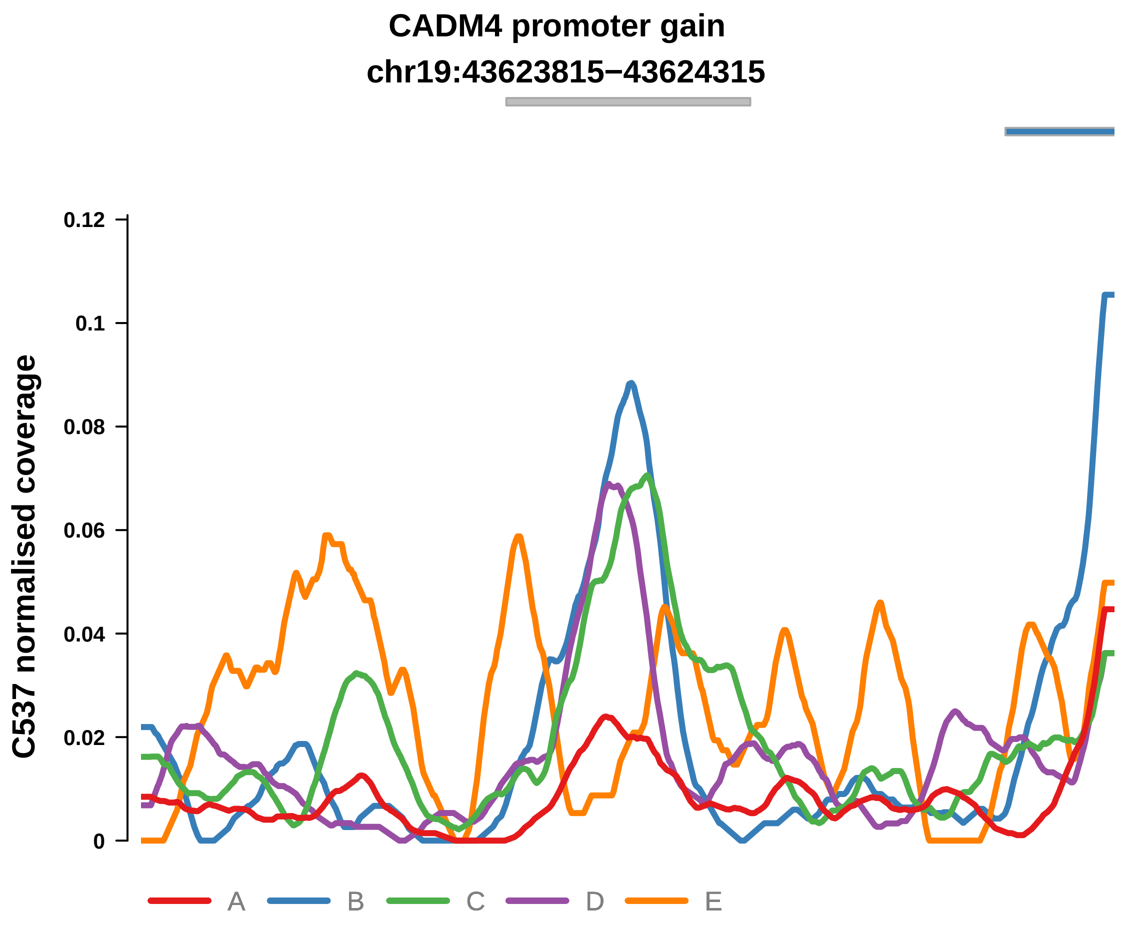


Putatively Clonal (logFC = 2.76)

| | P. | Adjusted P. |
|---|---|---|
| *~region vs. ~* | 3.13e−04 *** | 9.19e−03 ** |
| *~purity vs ~* | 8.04e−03 ** | 3.96e−02 * |
| *~purity+region vs ~purity* | 2.42e−03 ** | 3.88e−02 * |
| *Carcinoma vs. normal* | 2.69e−06 **** | 2.69e−06 **** |

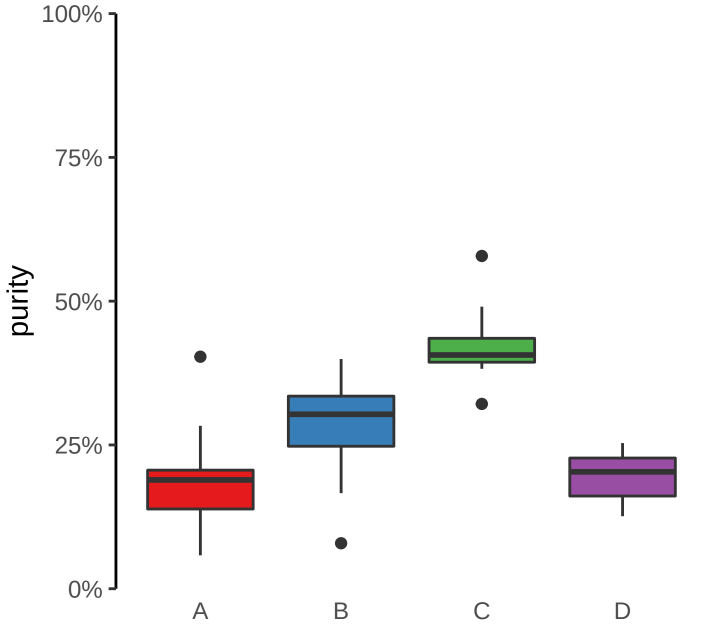

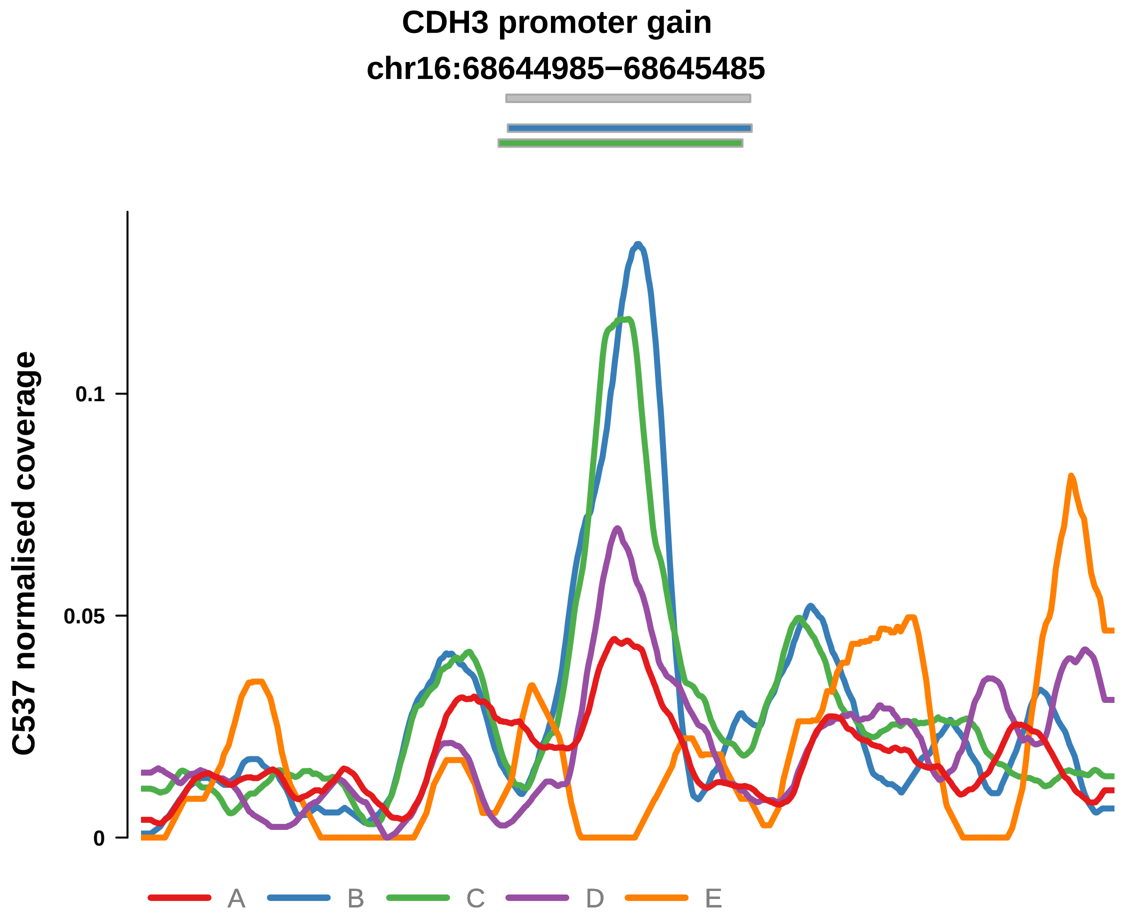


Putatively Clonal (logFC = 2.03)

| | P. | Adjusted P. |
|---|---|---|
| *~region vs. ~* | 2.77e−02 * | 7.46e−02 |
| *~purity vs ~* | 5.55e−04 *** | 7.12e−03 ** |
| *~purity+region vs ~purity* | 4.96e−01 | 6.42e−01 |
| *Carcinoma vs. normal* | 2.99e−04 *** | 2.99e−04 *** |

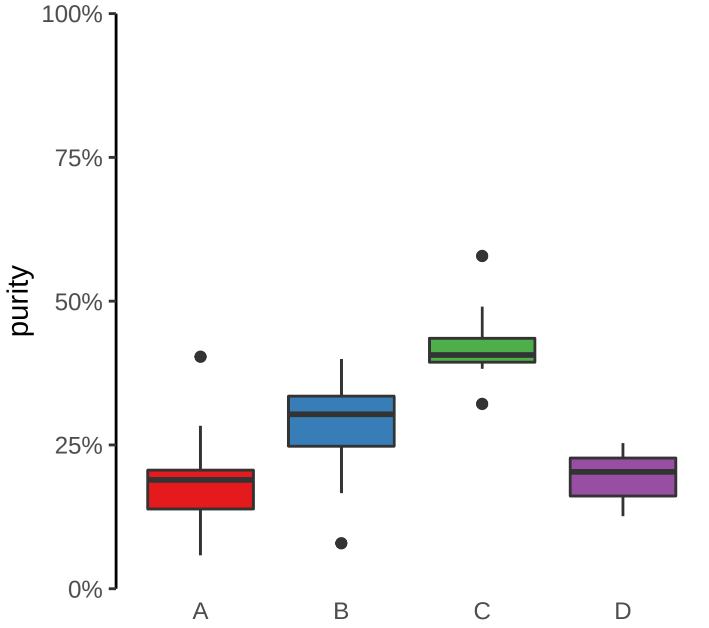

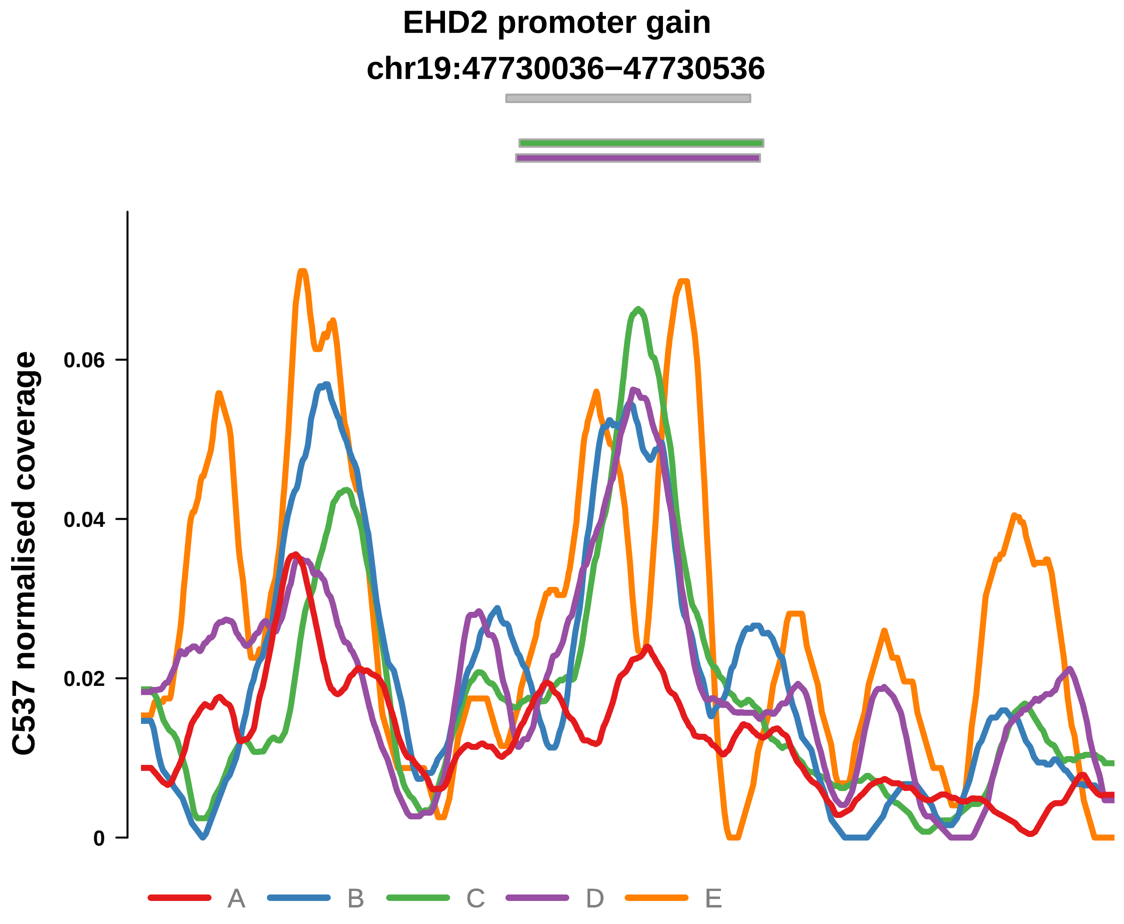


Putatively Clonal (logFC = 1.71)

| | P. | Adjusted P. |
|---|---|---|
| *~region vs. ~* | 4.91e−03 ** | 4.57e−02 * |
| *~purity vs ~* | 8.4e−02 | 1.81e−01 |
| *~purity+region vs ~purity* | 1.17e−02 * | 8.18e−02 |
| *Carcinoma vs. normal* | 2.67e−03 ** | 2.67e−03 ** |

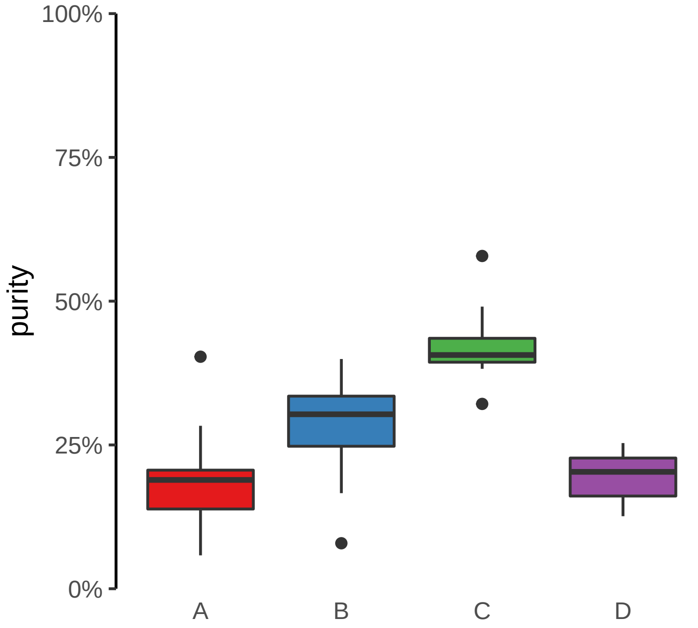

**FOXQ1 promoter gain**

**chr6:1311742−1312242**

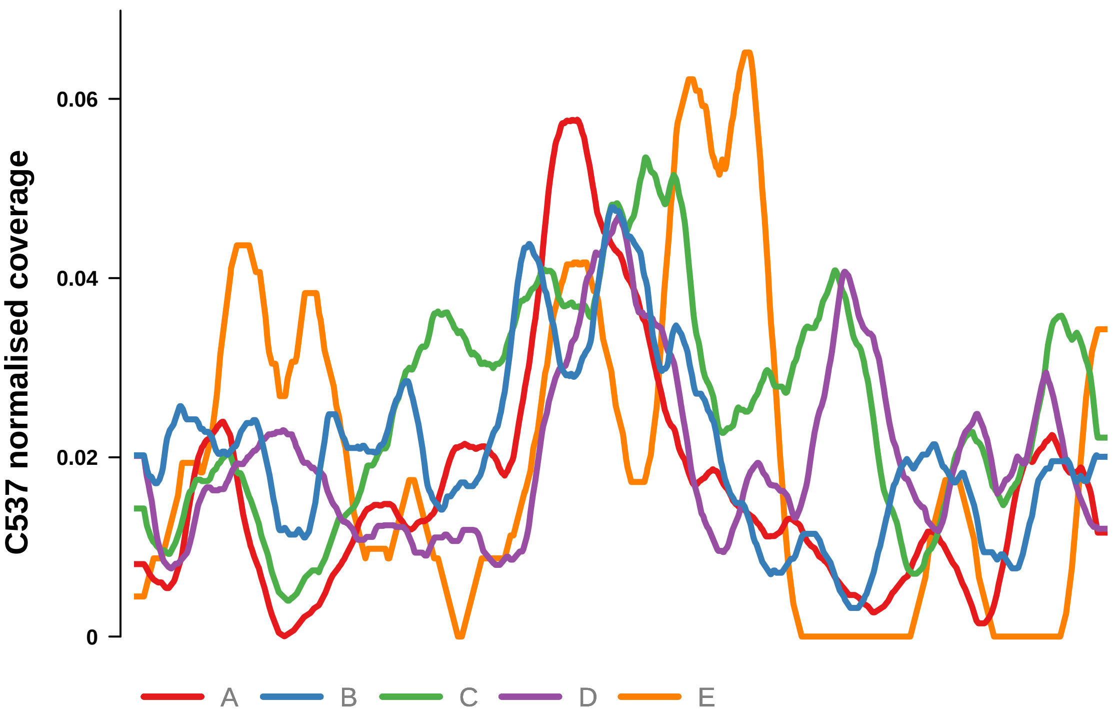


Putatively Clonal (logFC = 1.63)

| | **P.** | **Adjusted P.** |
|---|---|---|
| *~region vs. ~* | 3.05e−02 * | 7.46e−02 |
| *~purity vs ~* | 1.29e−01 | 2.41e−01 |
| *~purity+region vs ~purity* | 3.74e−02 * | 1.34e−01 |
| *Carcinoma vs. normal* | 3.47e−03 ** | 3.47e−03 ** |

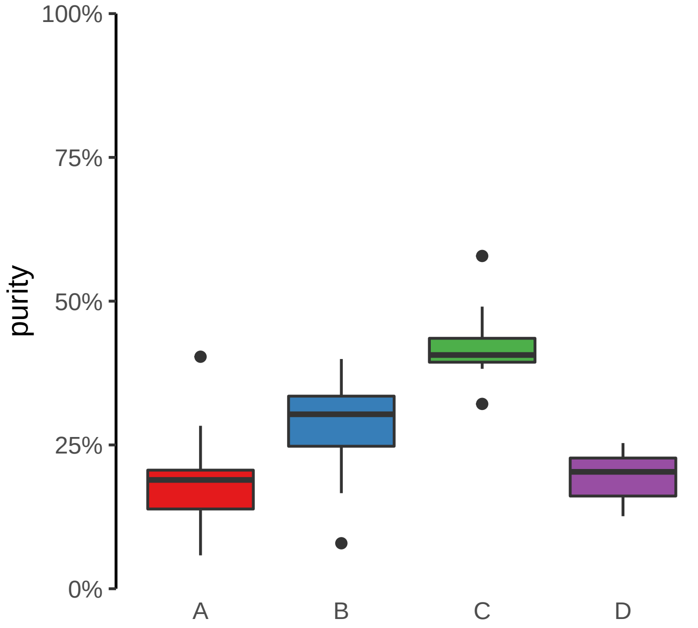

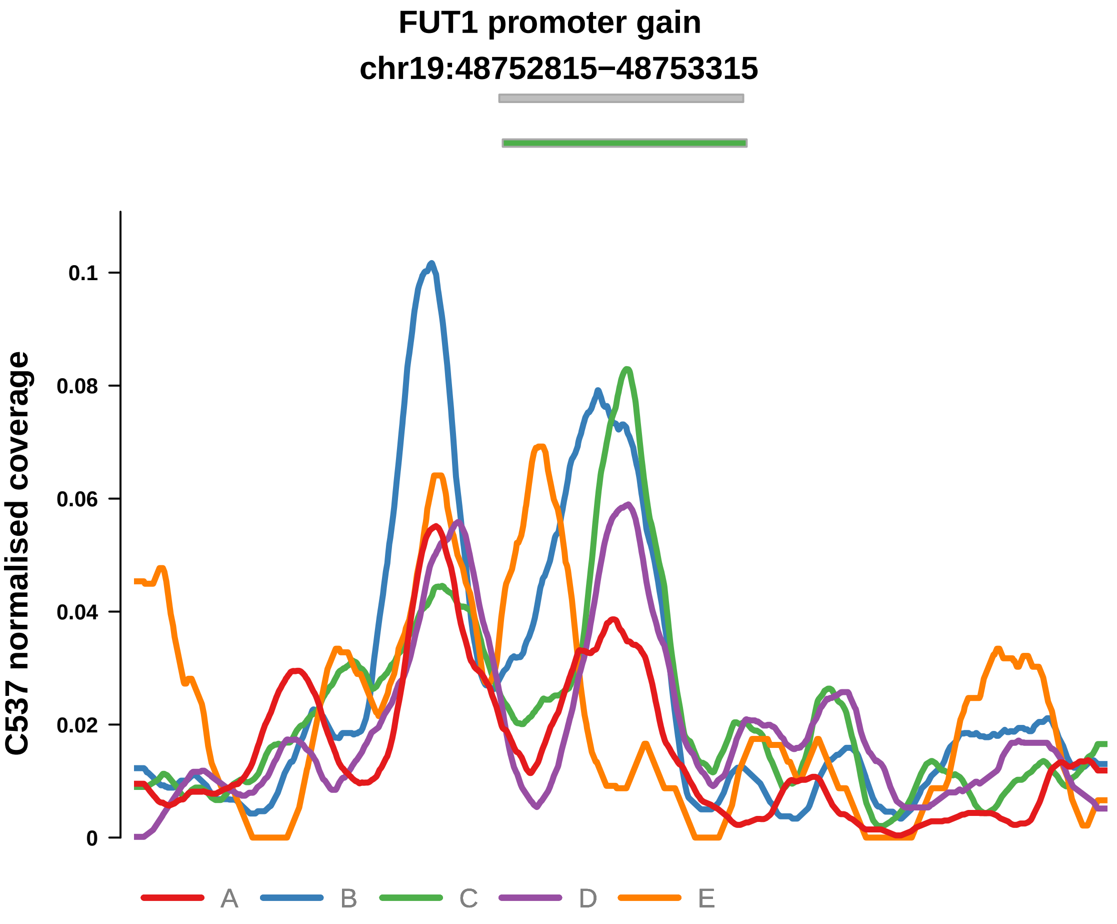


Putatively Clonal (logFC = 1.57)

| | P. | Adjusted P. |
|---|---|---|
| *~region vs. ~* | 4.49e−02 * | 1.01e−01 |
| *~purity vs ~* | 1.94e−02 * | 6.69e−02 |
| *~purity+region vs ~purity* | 2.7e−01 | 4.18e−01 |
| *Carcinoma vs. normal* | 5.65e−03 ** | 5.65e−03 ** |

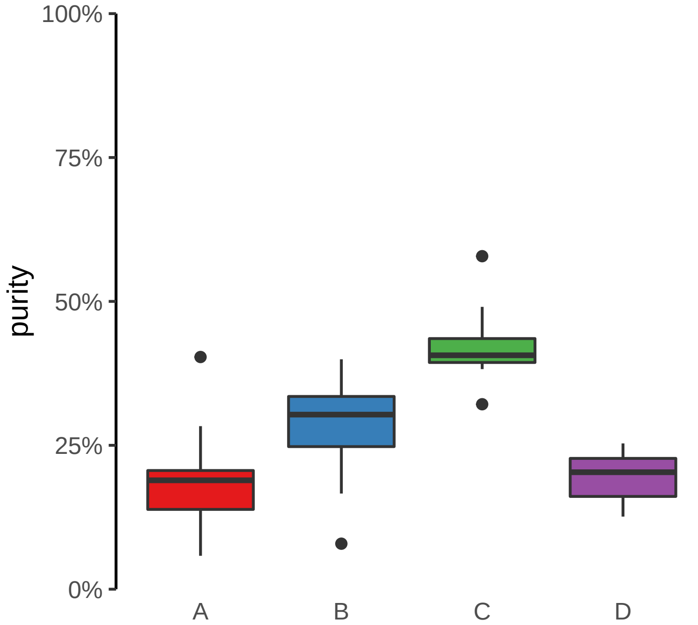

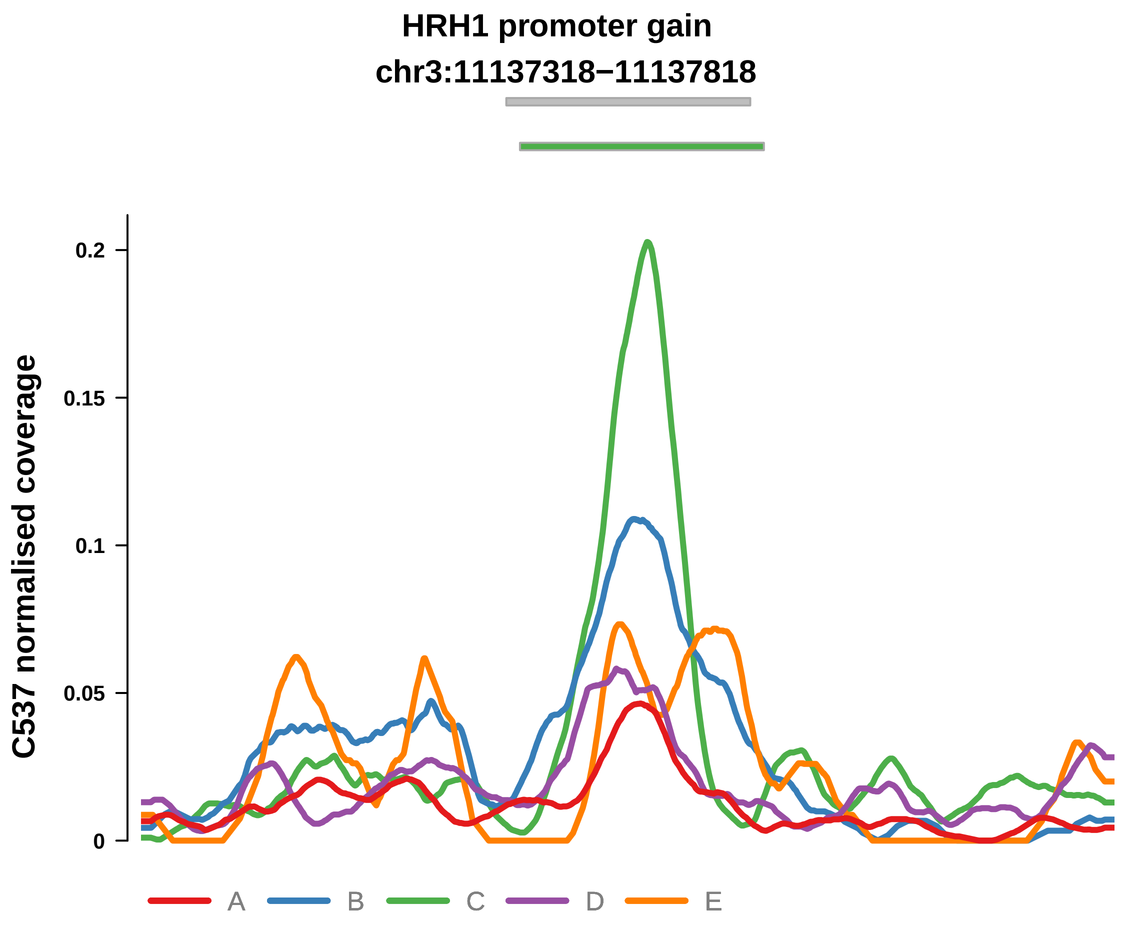


Putatively Clonal (logFC = 3.3)

| | P. | Adjusted P. |
|---|---|---|
| *~region vs. ~* | 1.22e−06 **** | 1.08e−04 *** |
| *~purity vs ~* | 1.11e−04 *** | 3.82e−03 ** |
| *~purity+region vs ~purity* | 5.88e−03 ** | 6.63e−02 |
| *Carcinoma vs. normal* | 1.62e−08 **** | 1.62e−08 **** |

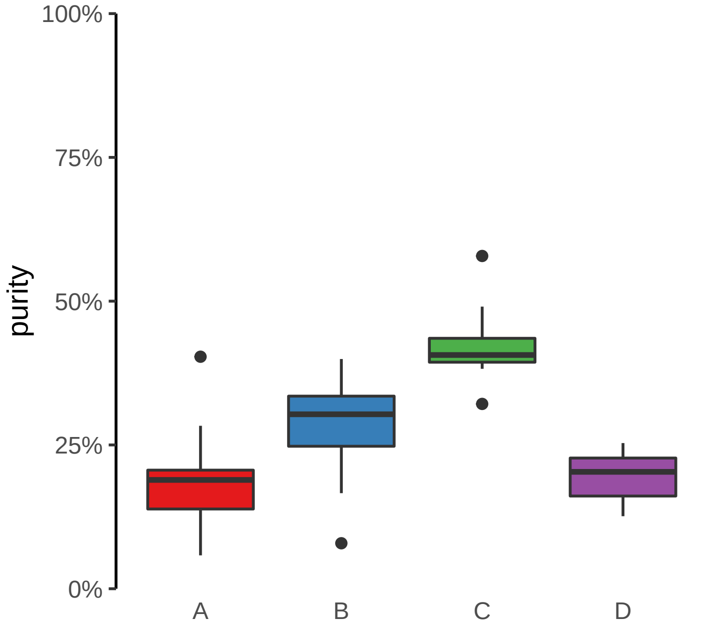

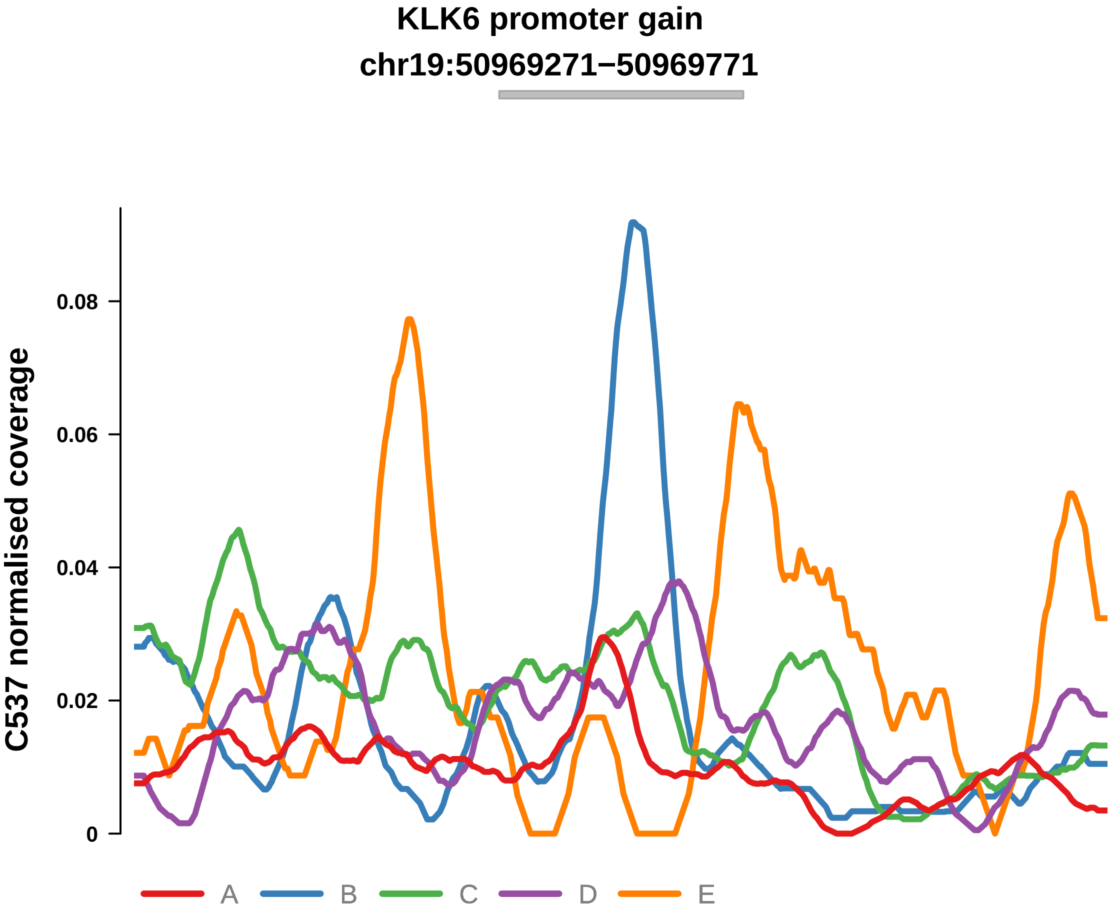


Putatively Clonal (logFC = 1.88)

| | P. | Adjusted P. |
|---|---|---|
| *~region vs. ~* | 6.67e−02 | 1.36e−01 |
| *~purity vs ~* | 6.66e−01 | 7.92e−01 |
| *~purity+region vs ~purity* | 6.03e−02 | 1.77e−01 |
| *Carcinoma vs. normal* | 1.3e−03 ** | 1.3e−03 ** |

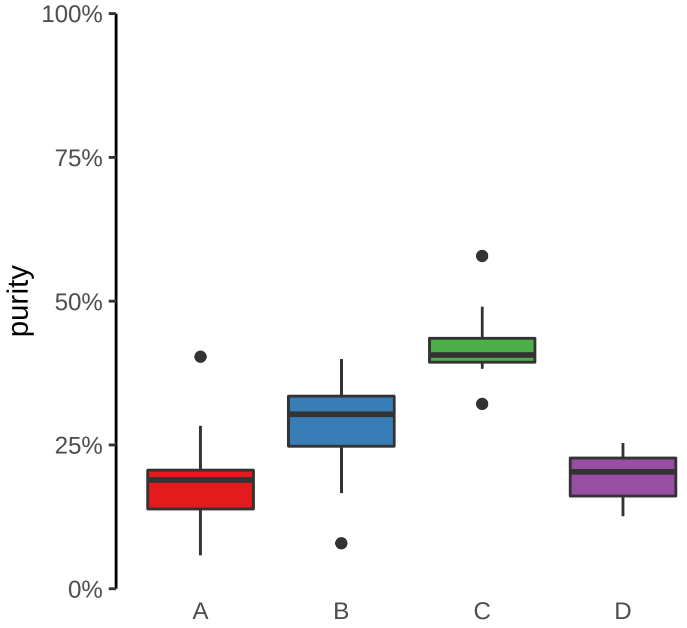

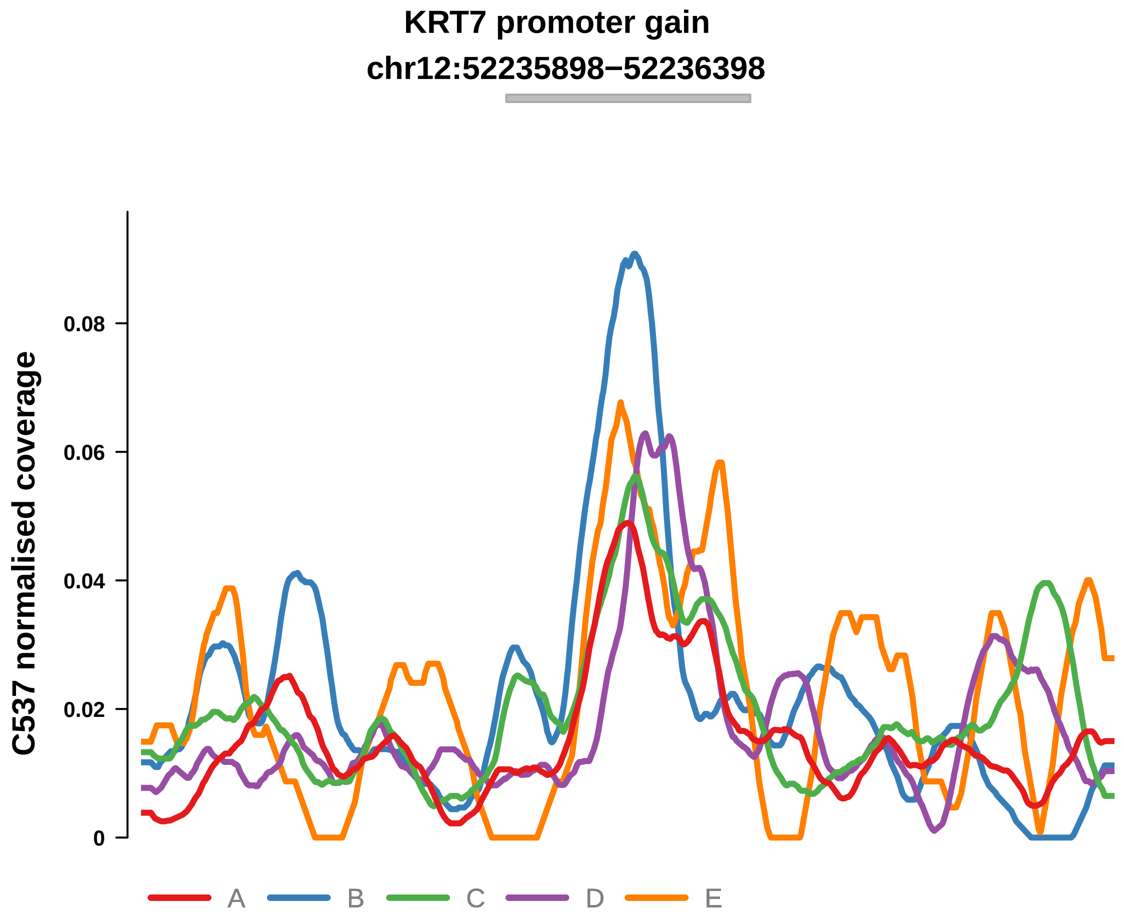


Putatively Clonal (logFC = 1.72)

| | P. | Adjusted P. |
|---|---|---|
| *~region vs. ~* | 4.34e−01 | 5.47e−01 |
| *~purity vs ~* | 3.32e−01 | 4.68e−01 |
| *~purity+region vs ~purity* | 4.24e−01 | 5.99e−01 |
| *Carcinoma vs. normal* | 2.34e−03 ** | 2.34e−03 ** |

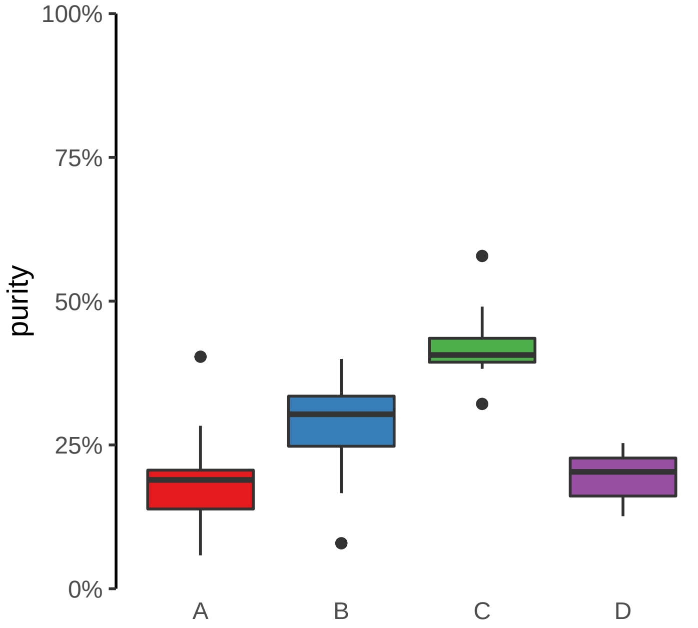

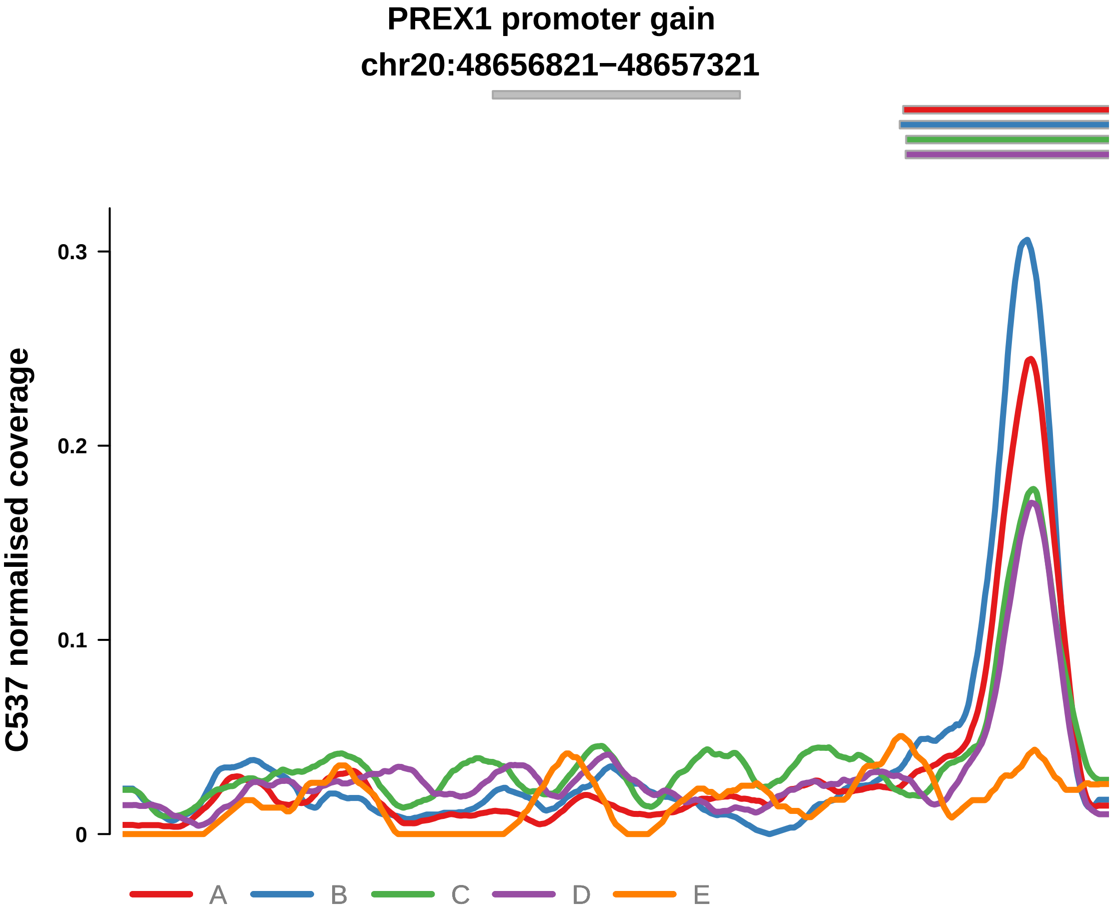


Putatively Clonal (logFC = 2.61)

| | P. | Adjusted P. |
|---|---|---|
| *~region vs. ~* | 6.2e−02 | 1.33e−01 |
| *~purity vs ~* | 6.96e−02 | 1.66e−01 |
| *~purity+region vs ~purity* | 7.93e−02 | 2.1e−01 |
| *Carcinoma vs. normal* | 8.5e−06 **** | 8.5e−06 **** |

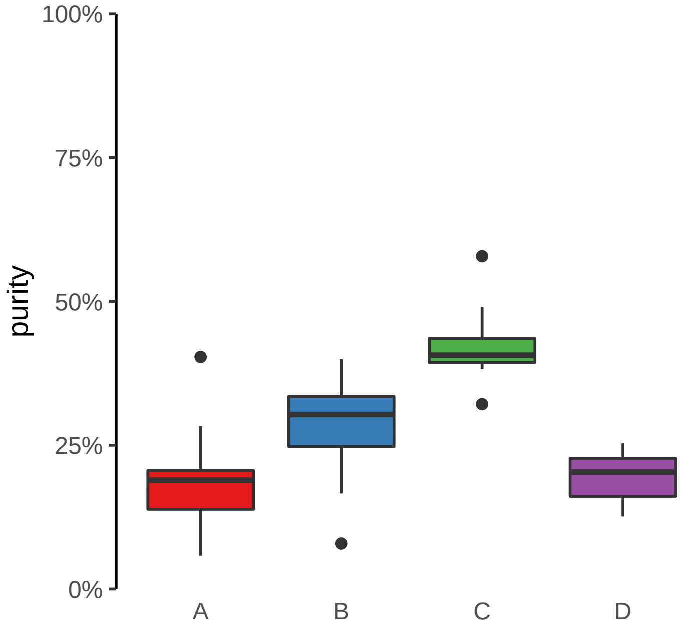

**PRR36 promoter gain**
**chr19:7870611−7871111**

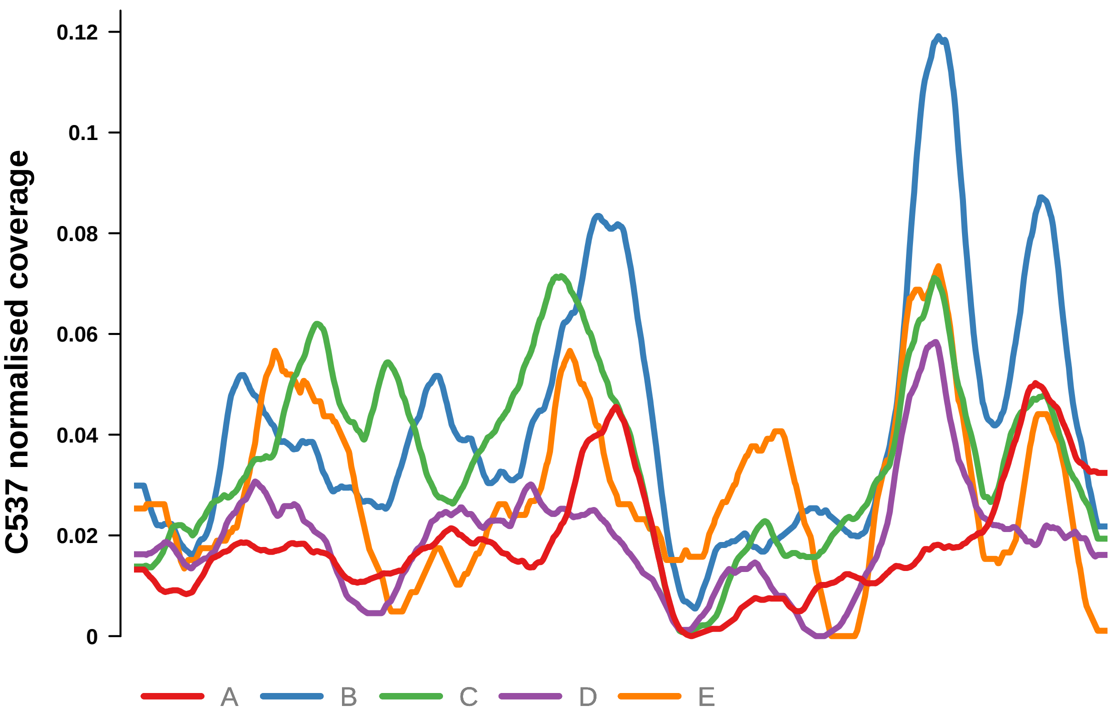


Putatively Clonal (logFC = 1.52)

| | P. | Adjusted P. |
|---|---|---|
| *~region vs. ~* | 1.73e−02 * | 6.61e−02 |
| *~purity vs ~* | 4.64e−04 *** | 7.12e−03 ** |
| *~purity+region vs ~purity* | 8.86e−01 | 9.09e−01 |
| *Carcinoma vs. normal* | 6.92e−03 ** | 6.92e−03 ** |

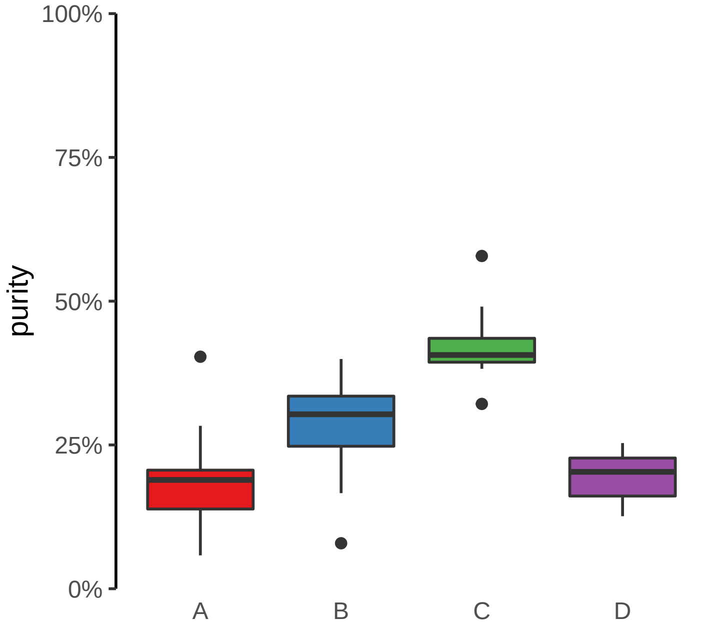

**SAP30BP promoter gain**
**chr17:75706968−75707468**

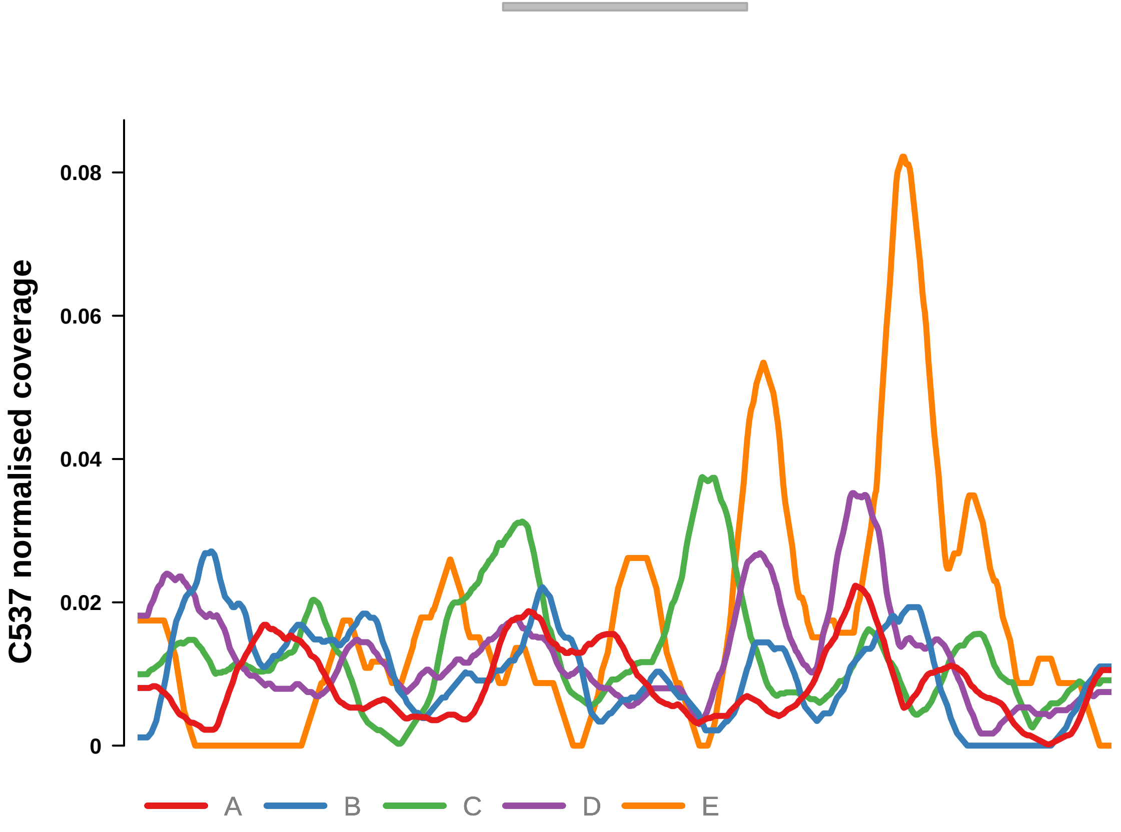


Putatively Clonal (logFC = 1.87)

| | **P.** | **Adjusted P.** |
|---|---|---|
| *~region vs. ~* | 1.82e−02 * | 6.67e−02 |
| *~purity vs ~* | 5.79e−01 | 7.26e−01 |
| *~purity+region vs ~purity* | 1.89e−02 * | 9.96e−02 |
| *Carcinoma vs. normal* | 1.78e−03 ** | 1.78e−03 ** |

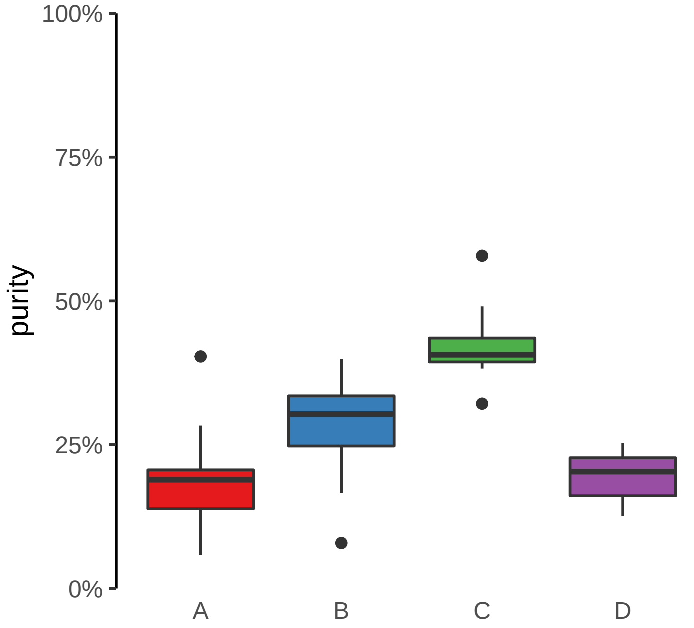

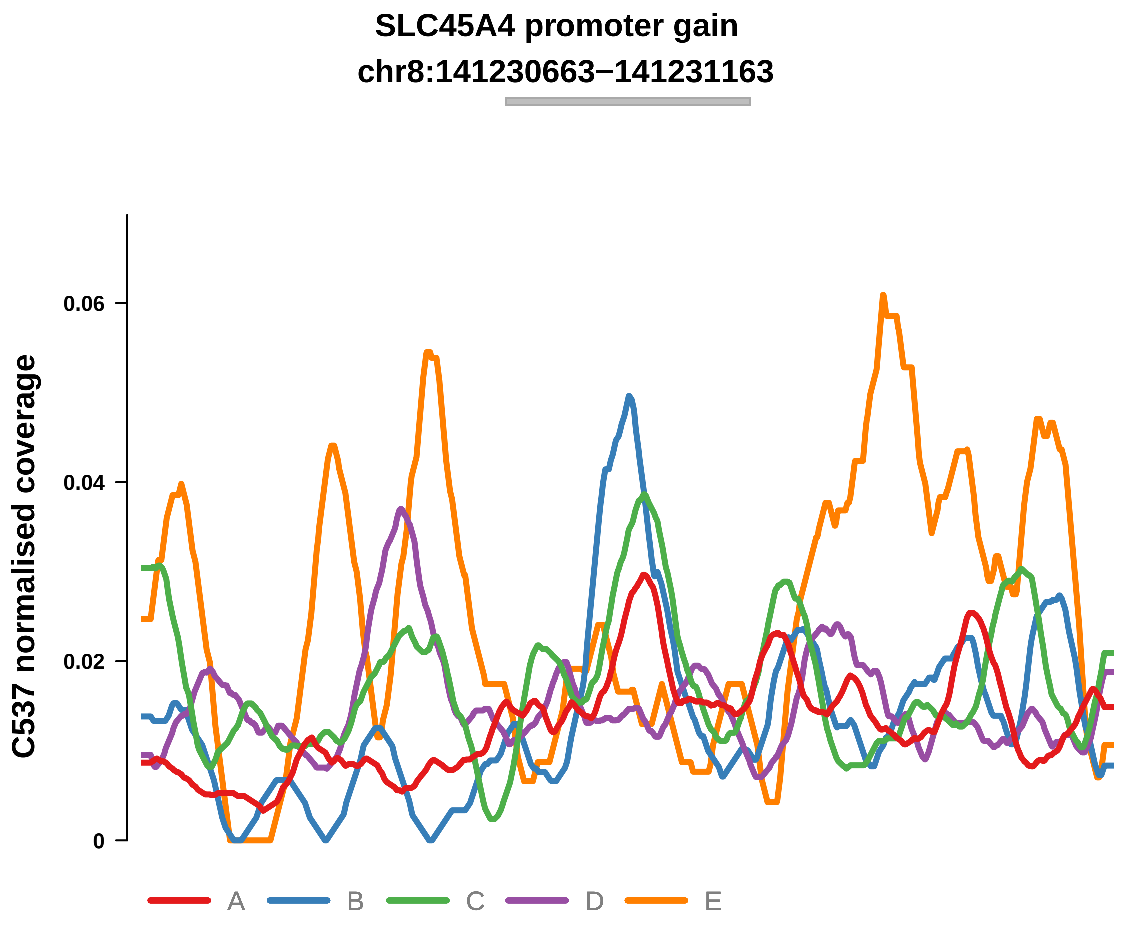


Putatively Clonal (logFC = 1.92)

| | P. | Adjusted P. |
|---|---|---|
| *~region vs. ~* | 8.9e−02 | 1.53e−01 |
| *~purity vs ~* | 8.41e−01 | 8.78e−01 |
| *~purity+region vs ~purity* | 9.39e−02 | 2.24e−01 |
| *Carcinoma vs. normal* | 1.21e−03 ** | 1.21e−03 ** |

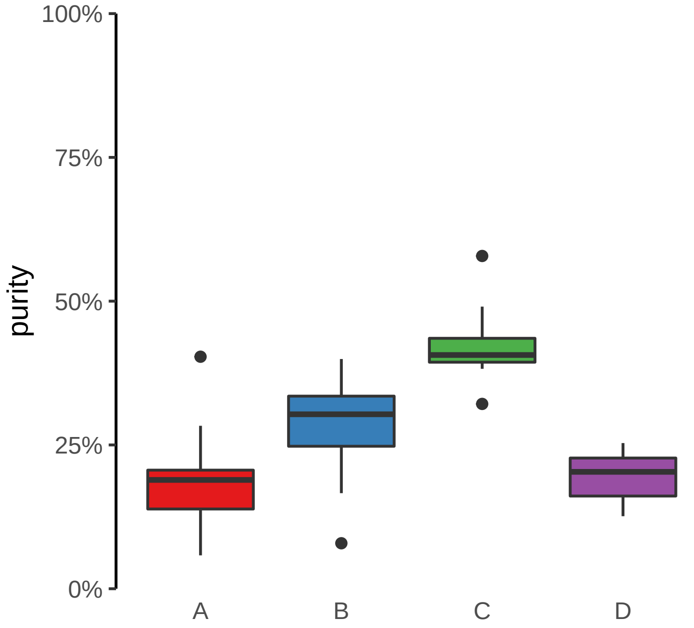

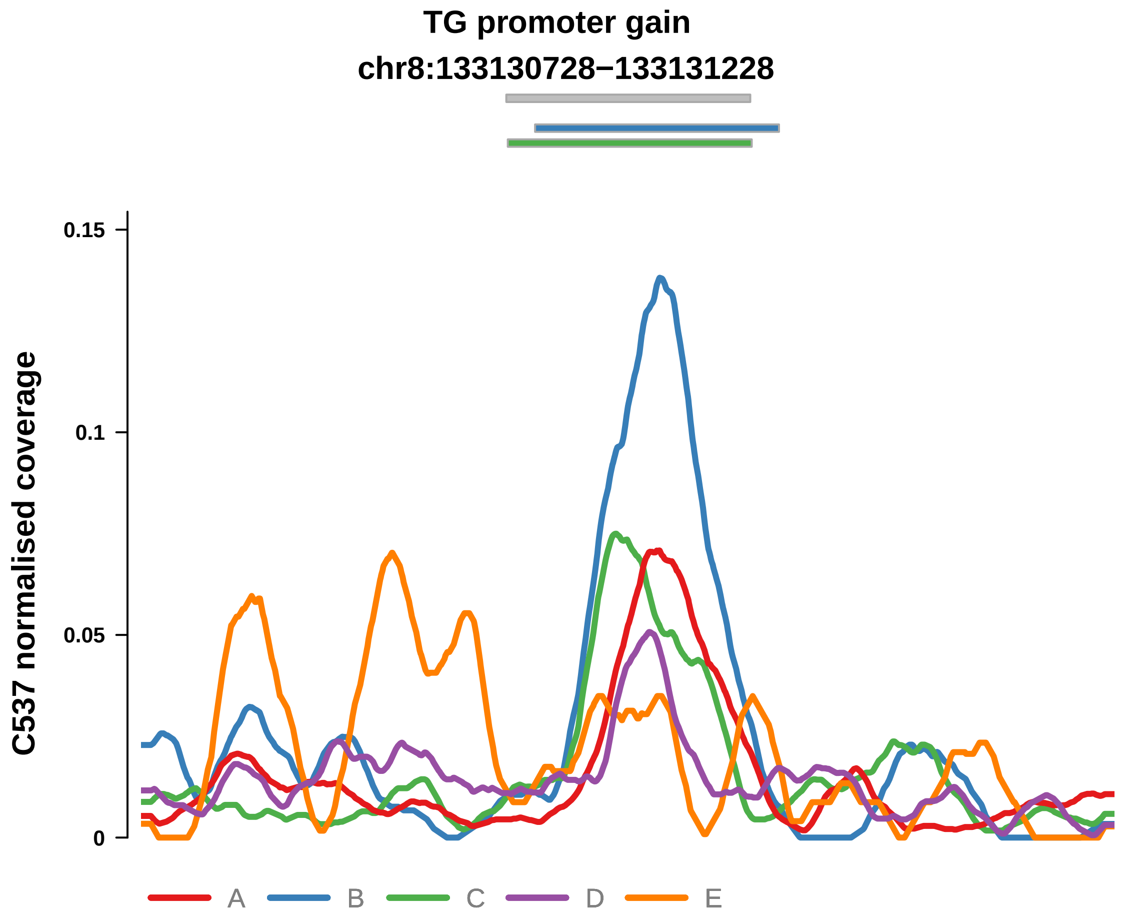


Putatively Clonal (logFC = 2.25)

| | P. | Adjusted P. |
|---|---|---|
| *~region vs. ~* | 6.8e−01 | 7.21e−01 |
| *~purity vs ~* | 7.47e−02 | 1.72e−01 |
| *~purity+region vs ~purity* | 5.63e−01 | 7e−01 |
| *Carcinoma vs. normal* | 7.82e−05 **** | 7.82e−05 **** |

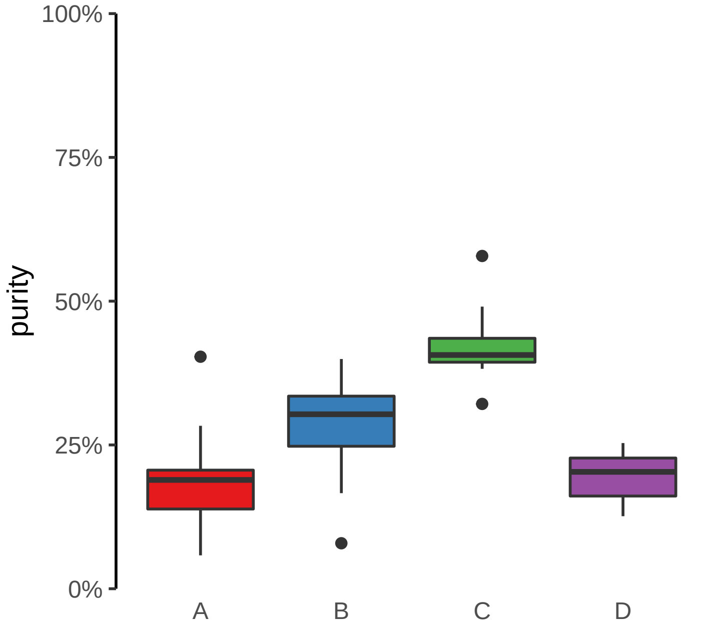

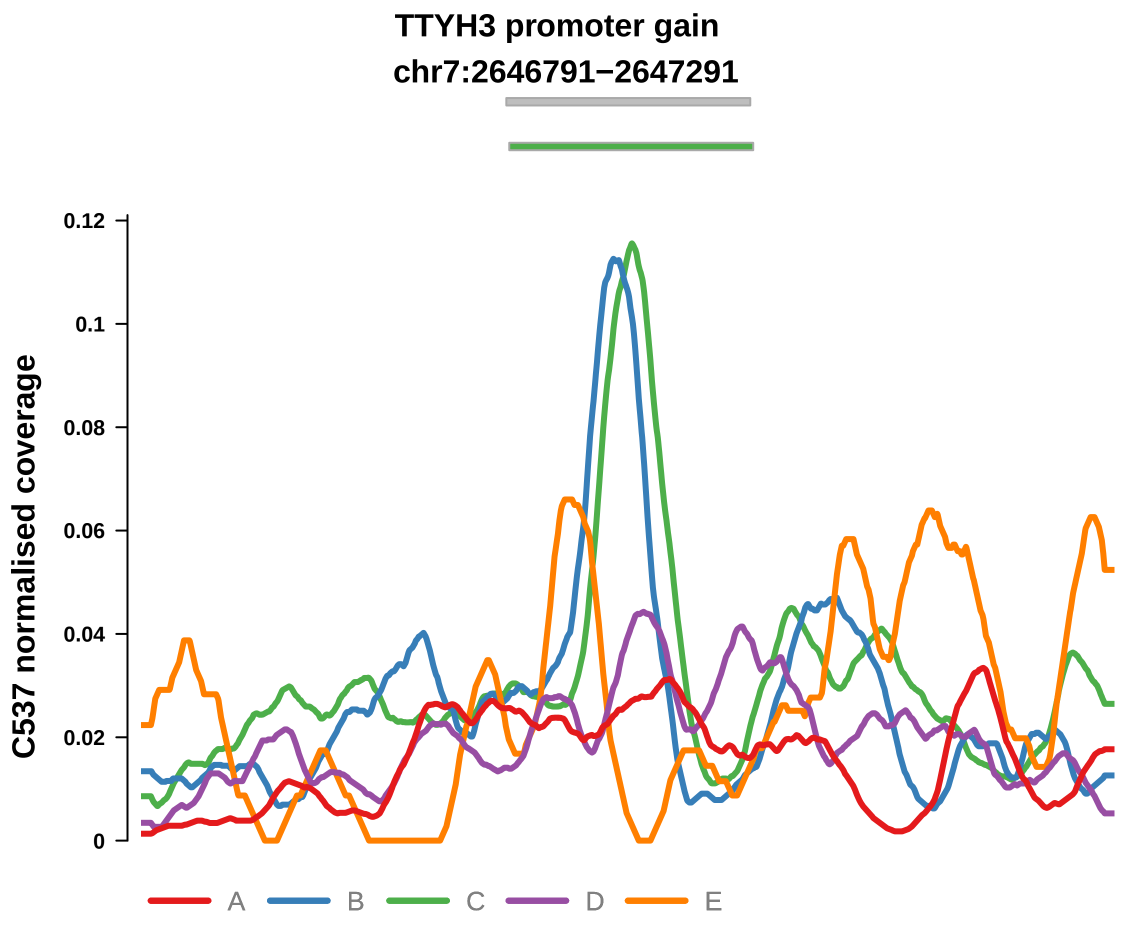


Putatively Clonal (logFC = 2.89)

| | P. | Adjusted P. |
|---|---|---|
| *~region vs. ~* | 2.57e−02 * | 7.46e−02 |
| *~purity vs ~* | 2.14e−03 ** | 1.73e−02 * |
| *~purity+region vs ~purity* | 1.94e−01 | 3.36e−01 |
| *Carcinoma vs. normal* | 6.96e−07 **** | 6.96e−07 **** |

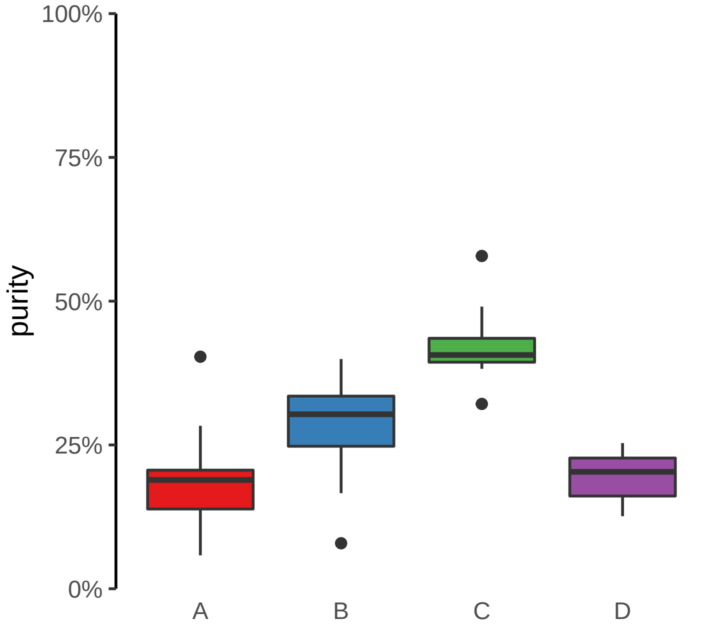

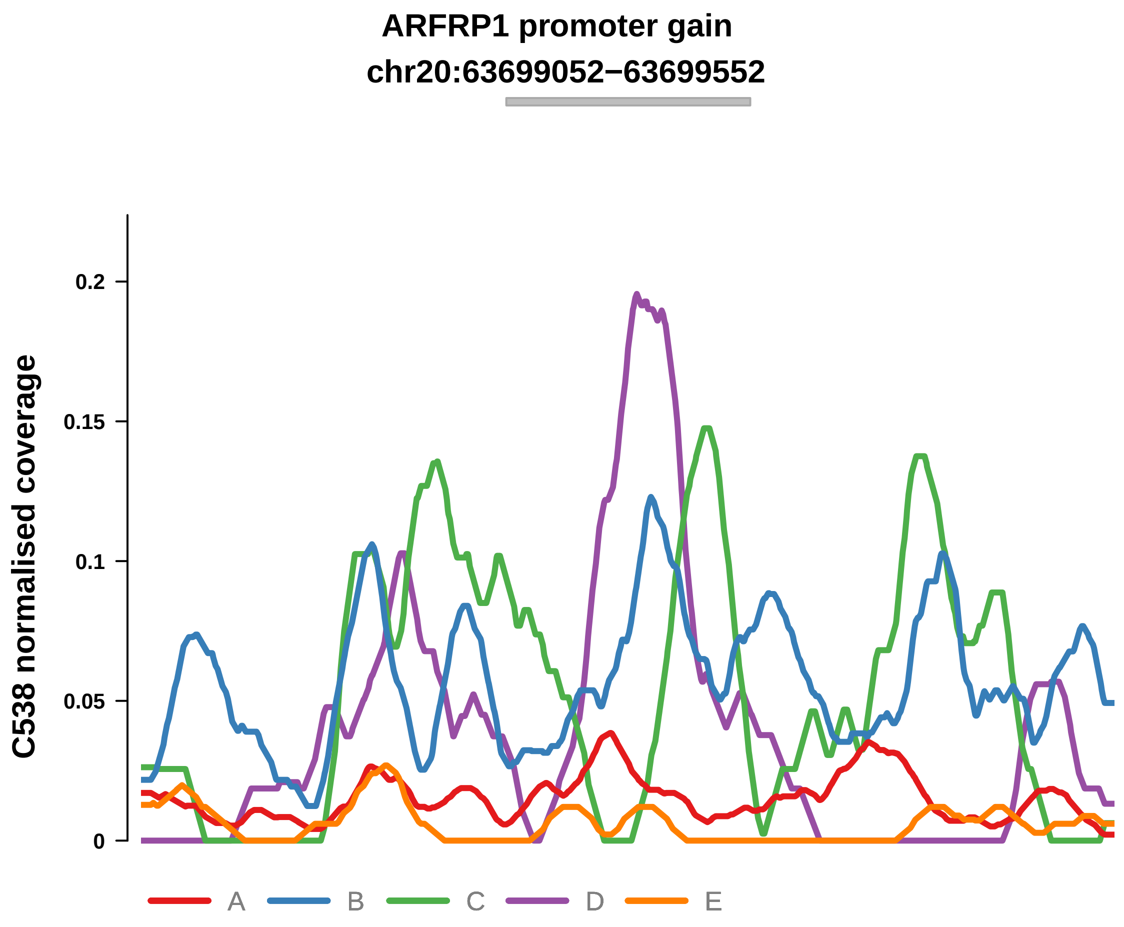


Putatively Clonal (logFC = 2.05)

| | P. | Adjusted P. |
|---|---|---|
| *~region vs. ~* | 1.86e−02 * | 7.78e−02 |
| *~purity vs ~* | 1.18e−01 | 2.31e−01 |
| *~purity+region vs ~purity* | 1.4e−02 * | 4.11e−01 |
| *Carcinoma vs. normal* | 3.75e−04 *** | 3.75e−04 *** |

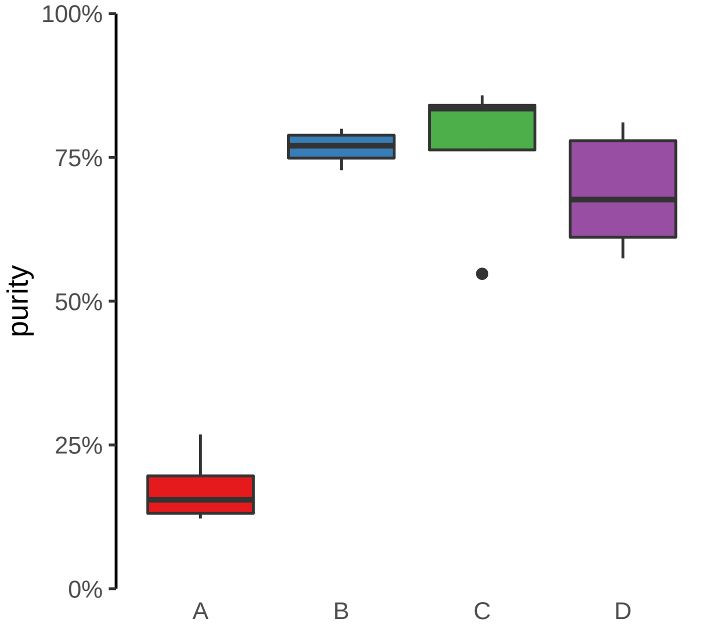

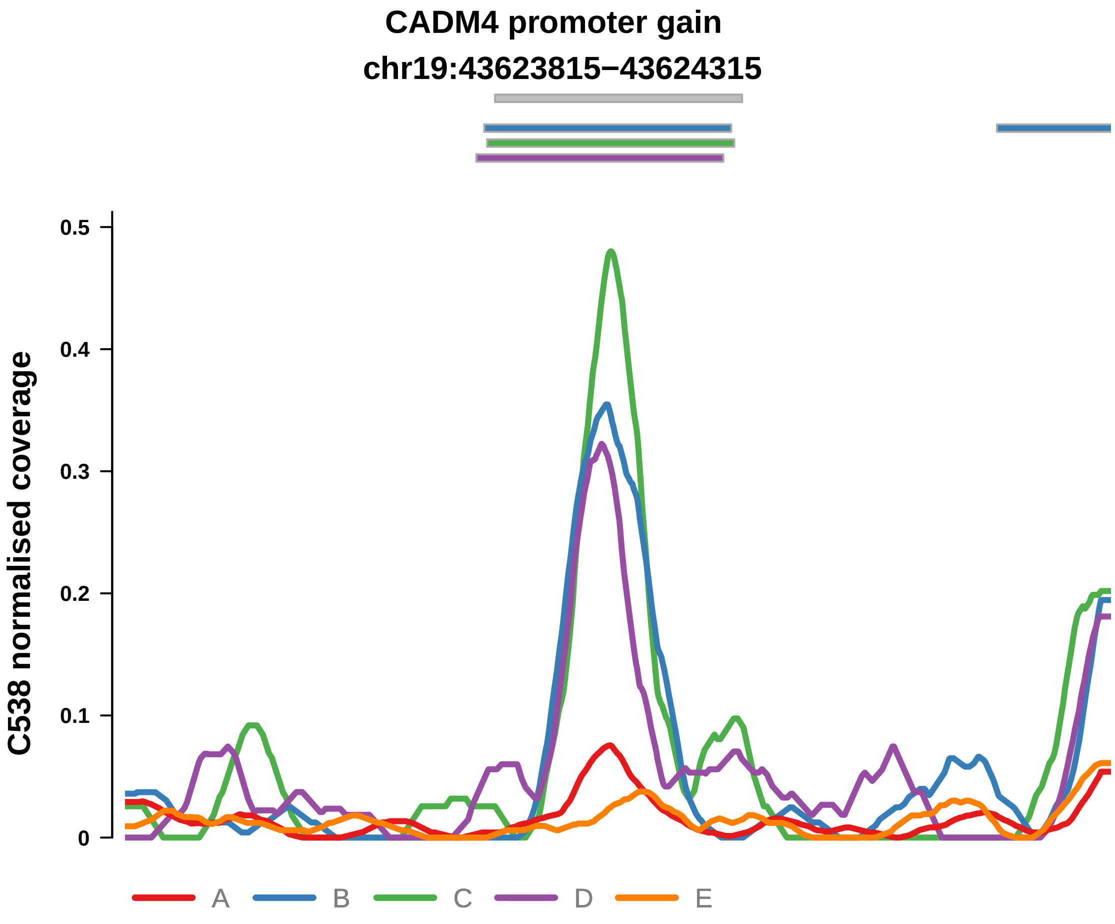


Putatively Clonal (logFC = 3.5)

| | P. | Adjusted P. |
|---|---|---|
| *~region vs. ~* | 7.65e−03 ** | 3.96e−02 * |
| *~purity vs ~* | 6.92e−03 ** | 2.54e−02 * |
| *~purity+region vs ~purity* | 1.55e−01 | 8.28e−01 |
| *Carcinoma vs. normal* | 3.92e−09 **** | 3.92e−09 **** |

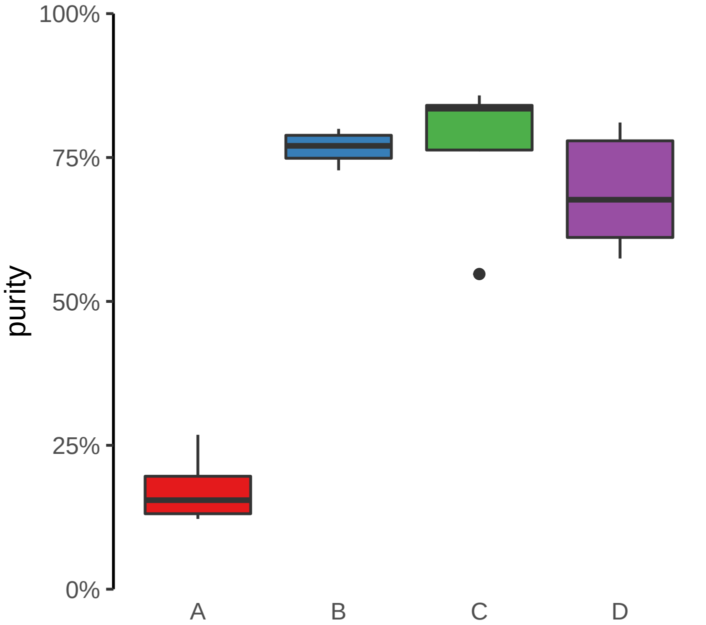

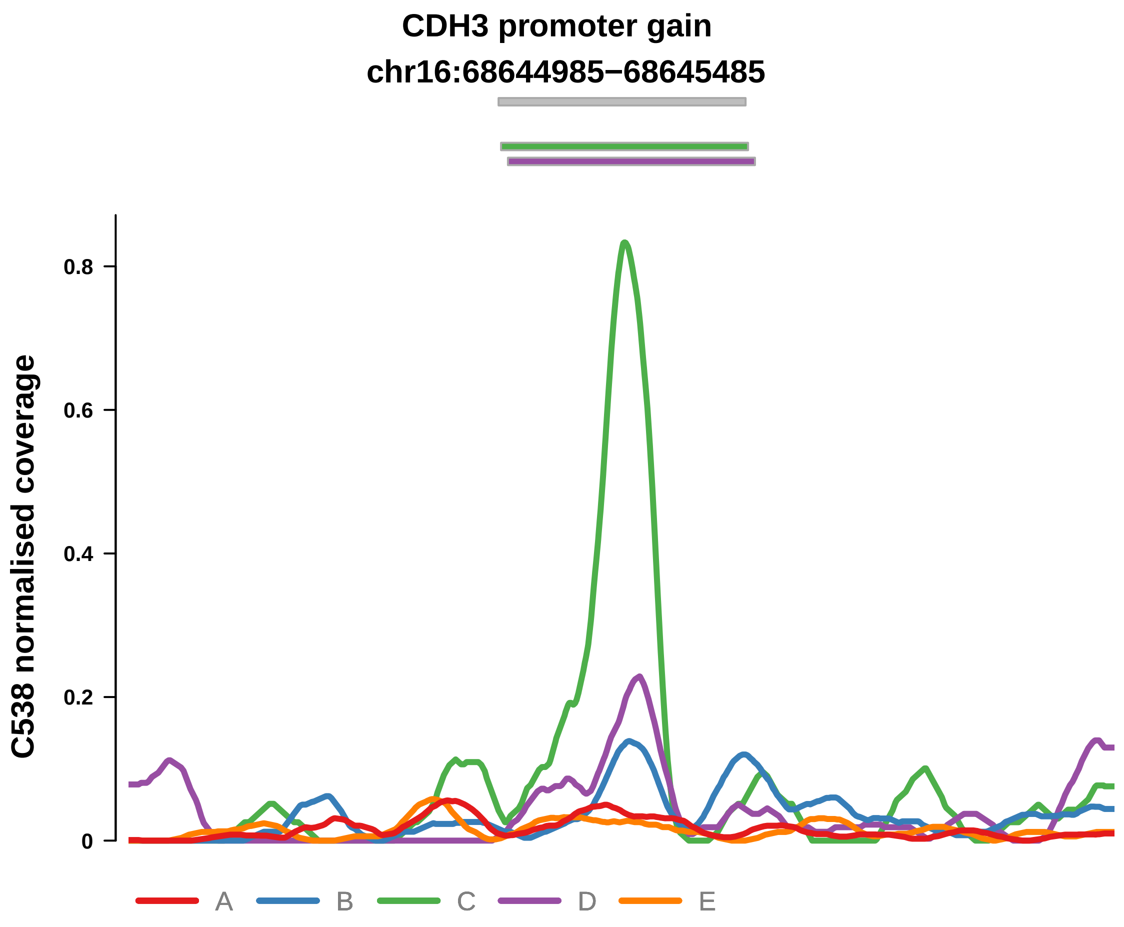


Putatively Clonal (logFC = 1.96)

| | P. | Adjusted P. |
|---|---|---|
| *~region vs. ~* | 3.15e−01 | 5.17e−01 |
| *~purity vs ~* | 3.47e−01 | 5.09e−01 |
| *~purity+region vs ~purity* | 4.68e−01 | 8.7e−01 |
| *Carcinoma vs. normal* | 5.2e−04 *** | 5.2e−04 *** |

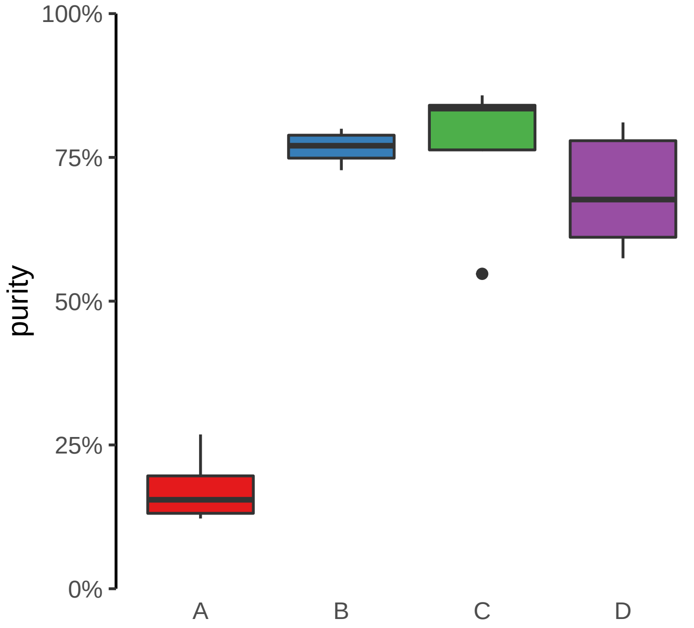

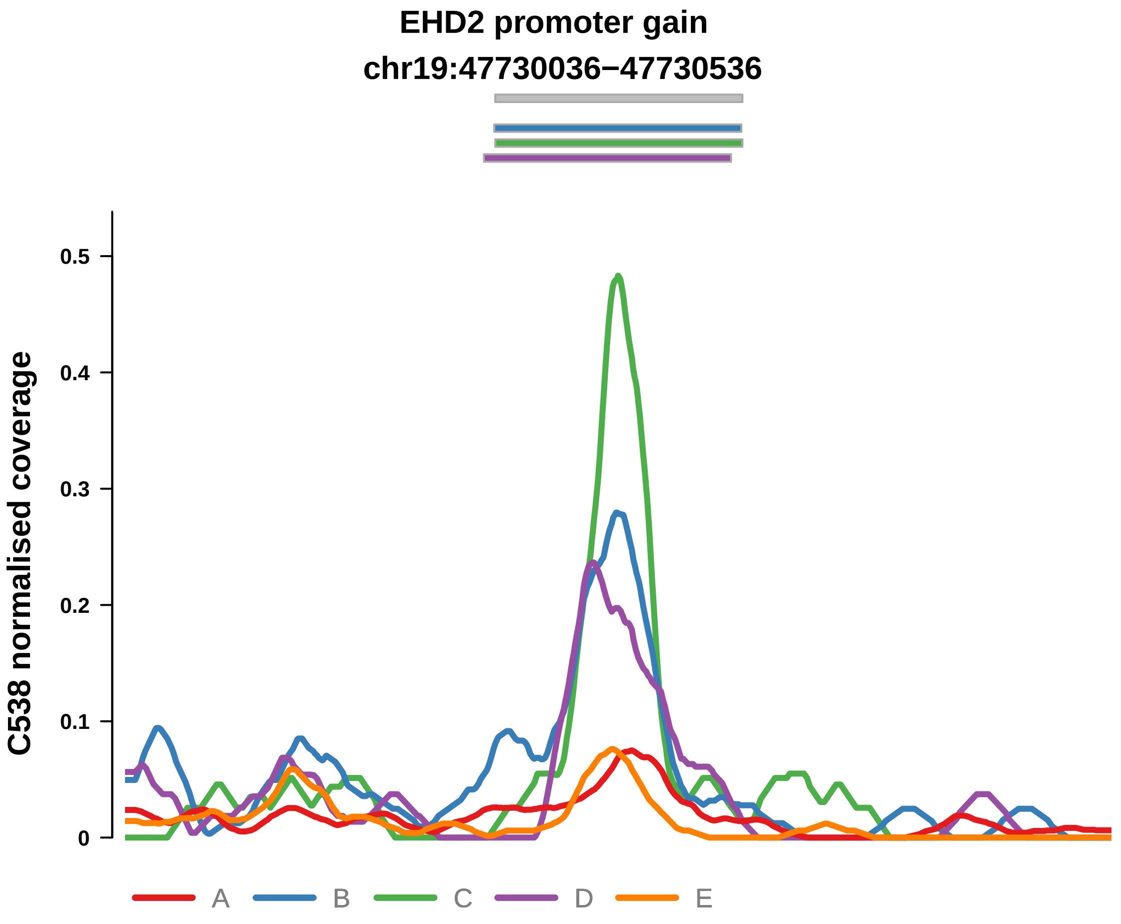


Putatively Clonal (logFC = 2.6)

| | P. | Adjusted P. |
|---|---|---|
| *~region vs. ~* | 6.86e−01 | 7.98e−01 |
| *~purity vs ~* | 5.03e−01 | 6.35e−01 |
| *~purity+region vs ~purity* | 8.04e−01 | 9.56e−01 |
| *Carcinoma vs. normal* | 5.47e−06 **** | 5.47e−06 **** |

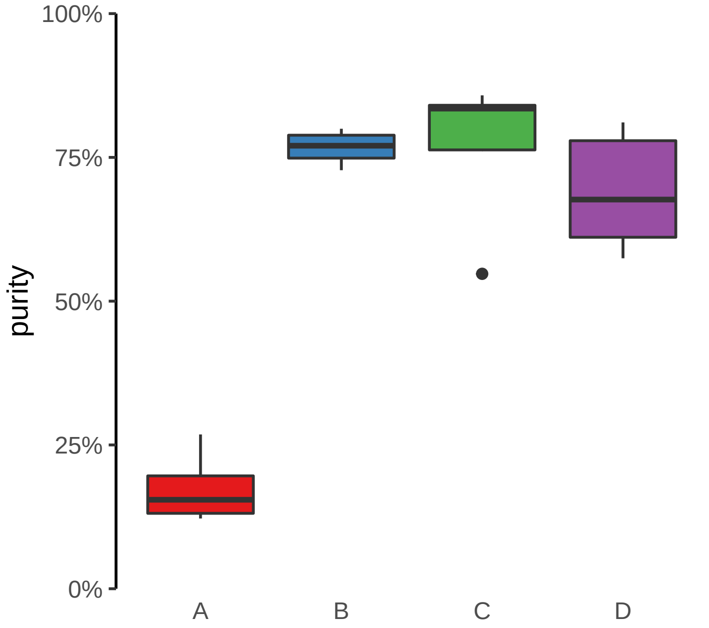

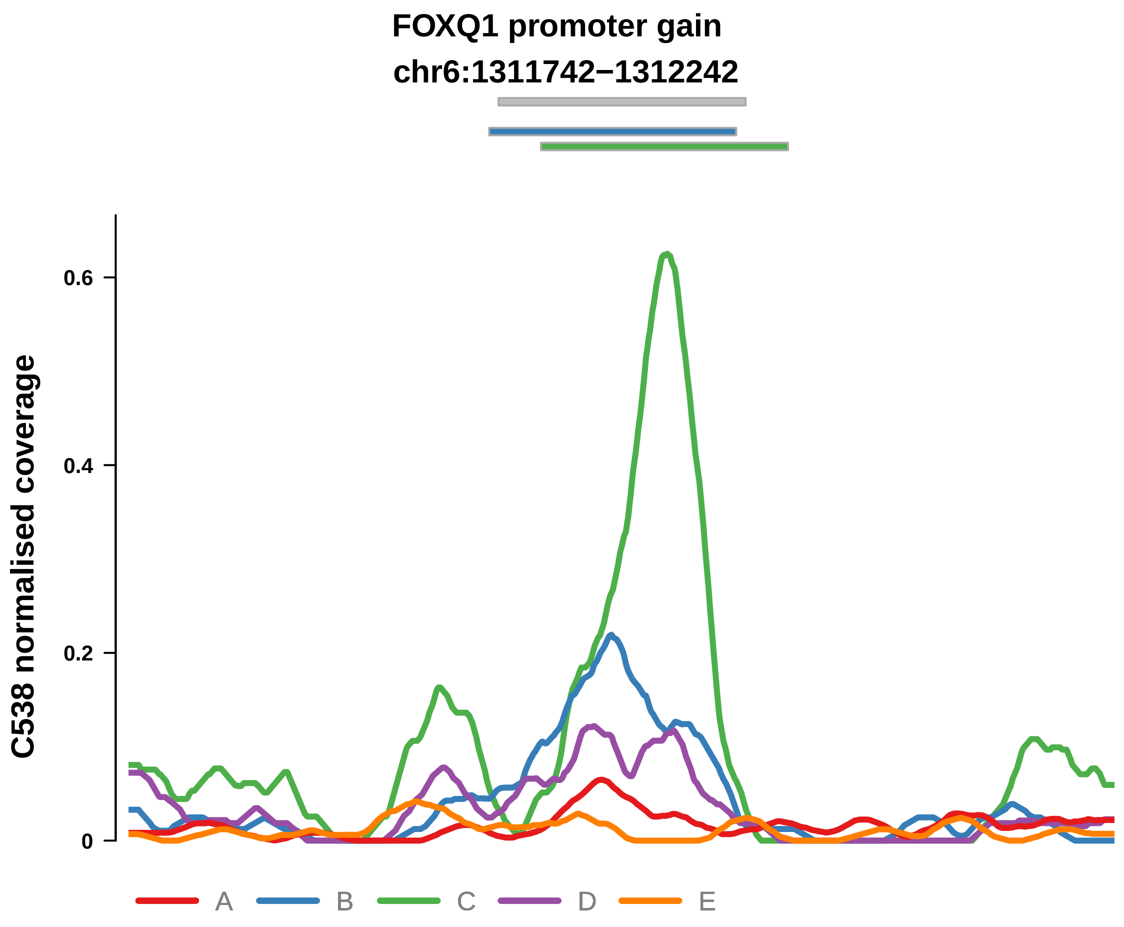


Putatively Clonal (logFC = 2.29)

| | P. | Adjusted P. |
|---|---|---|
| *~region vs. ~* | 2.69e−01 | 4.75e−01 |
| *~purity vs ~* | 8.03e−03 ** | 2.74e−02 * |
| *~purity+region vs ~purity* | 4.71e−01 | 8.7e−01 |
| *Carcinoma vs. normal* | 4.73e−05 **** | 4.73e−05 **** |

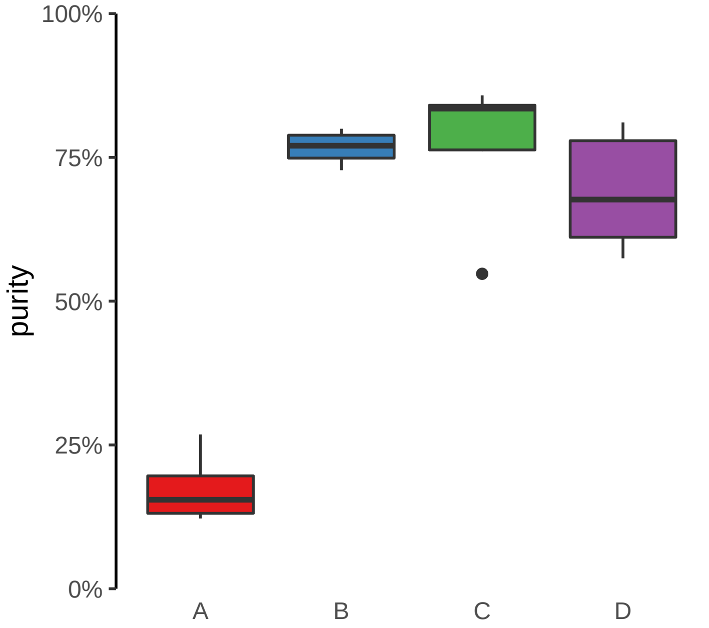

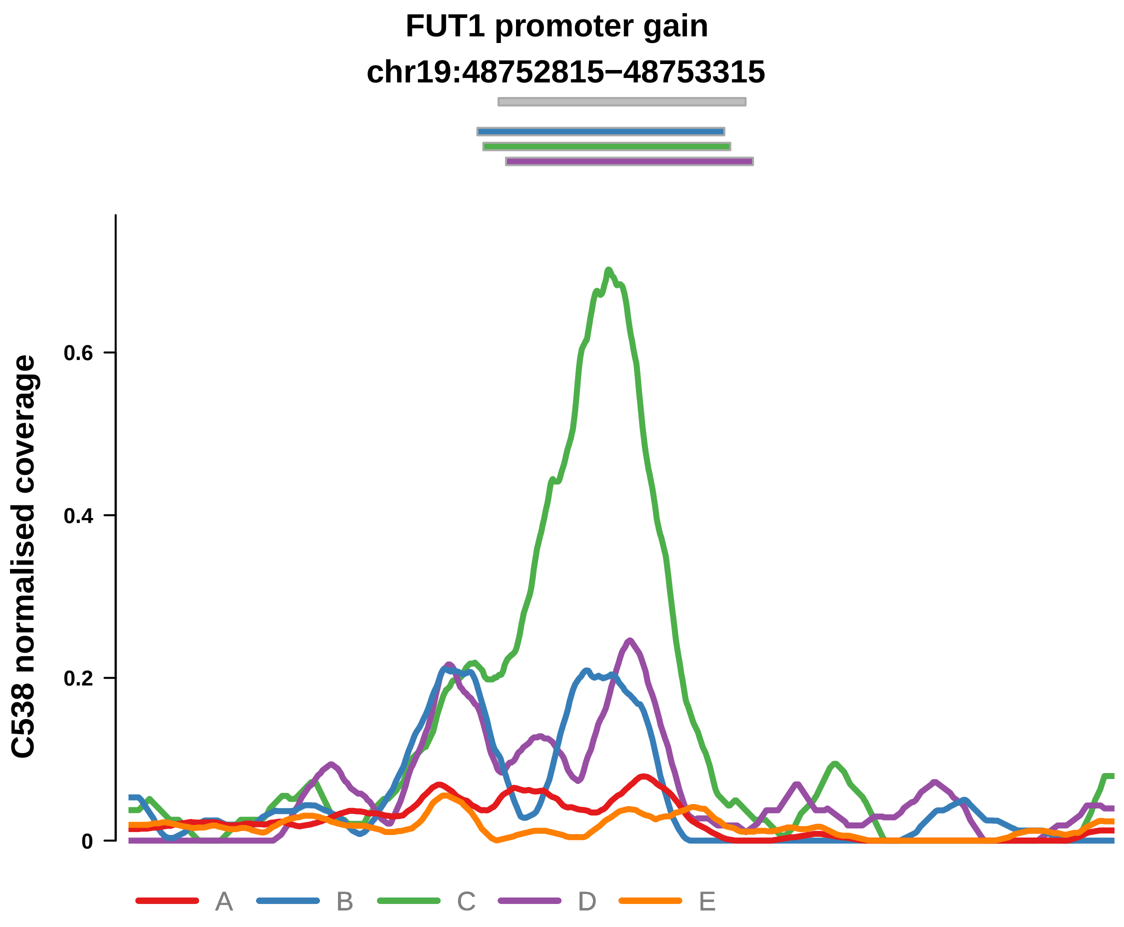


Putatively Clonal (logFC = 2.75)

| | P. | Adjusted P. |
|---|---|---|
| *~region vs. ~* | 7.37e−02 | 1.8e−01 |
| *~purity vs ~* | 1.18e−01 | 2.31e−01 |
| *~purity+region vs ~purity* | 1.85e−01 | 8.7e−01 |
| *Carcinoma vs. normal* | 1.47e−06 **** | 1.47e−06 **** |

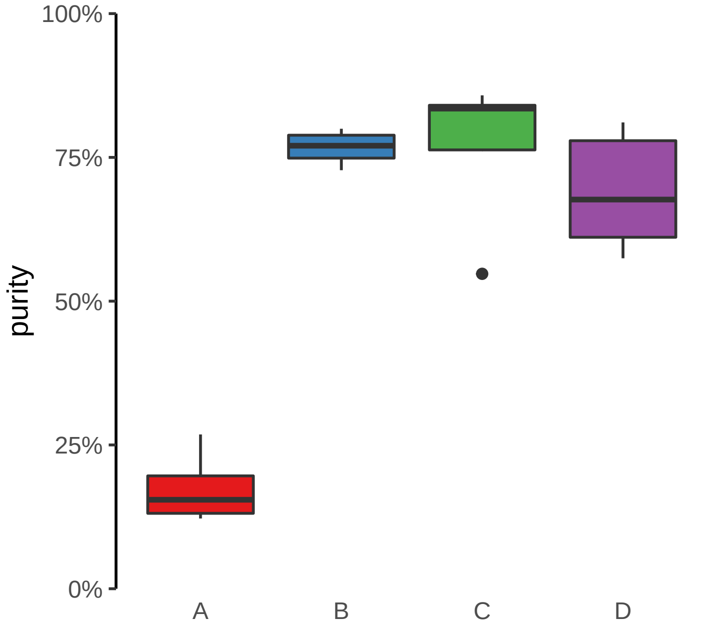

**HPS4 promoter gain**
**chr22:26471440−26471940**

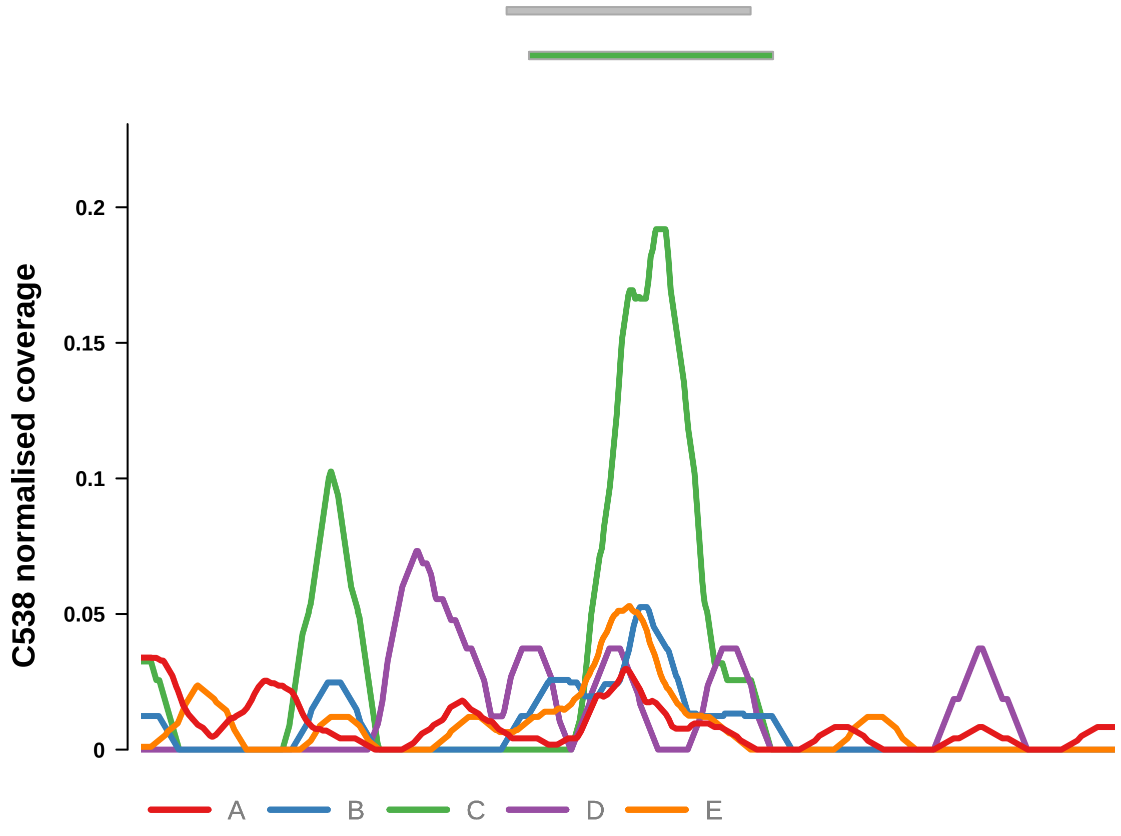


Putatively Clonal (logFC = 1.73)

| | P. | Adjusted P. |
|---|---|---|
| *~region vs. ~* | 7.82e−01 | 8.39e−01 |
| *~purity vs ~* | 3.59e−01 | 5.1e−01 |
| *~purity+region vs ~purity* | 9.49e−01 | 9.79e−01 |
| *Carcinoma vs. normal* | 5.43e−03 ** | 5.43e−03 ** |

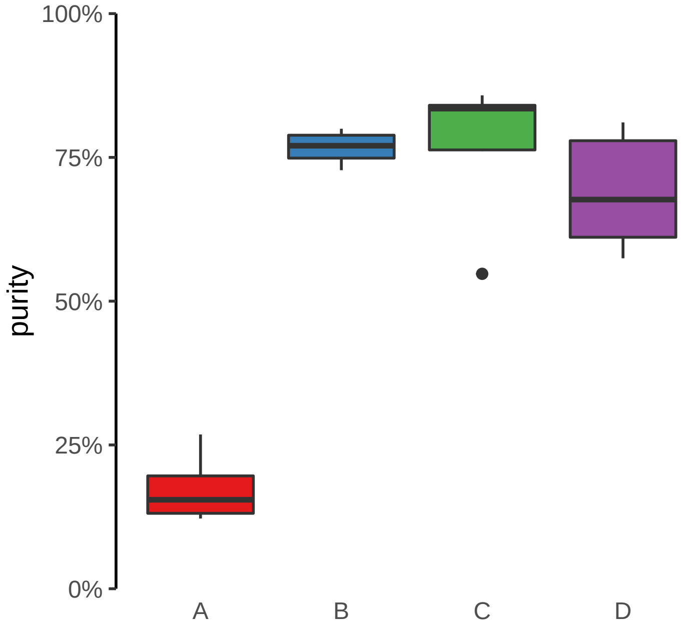

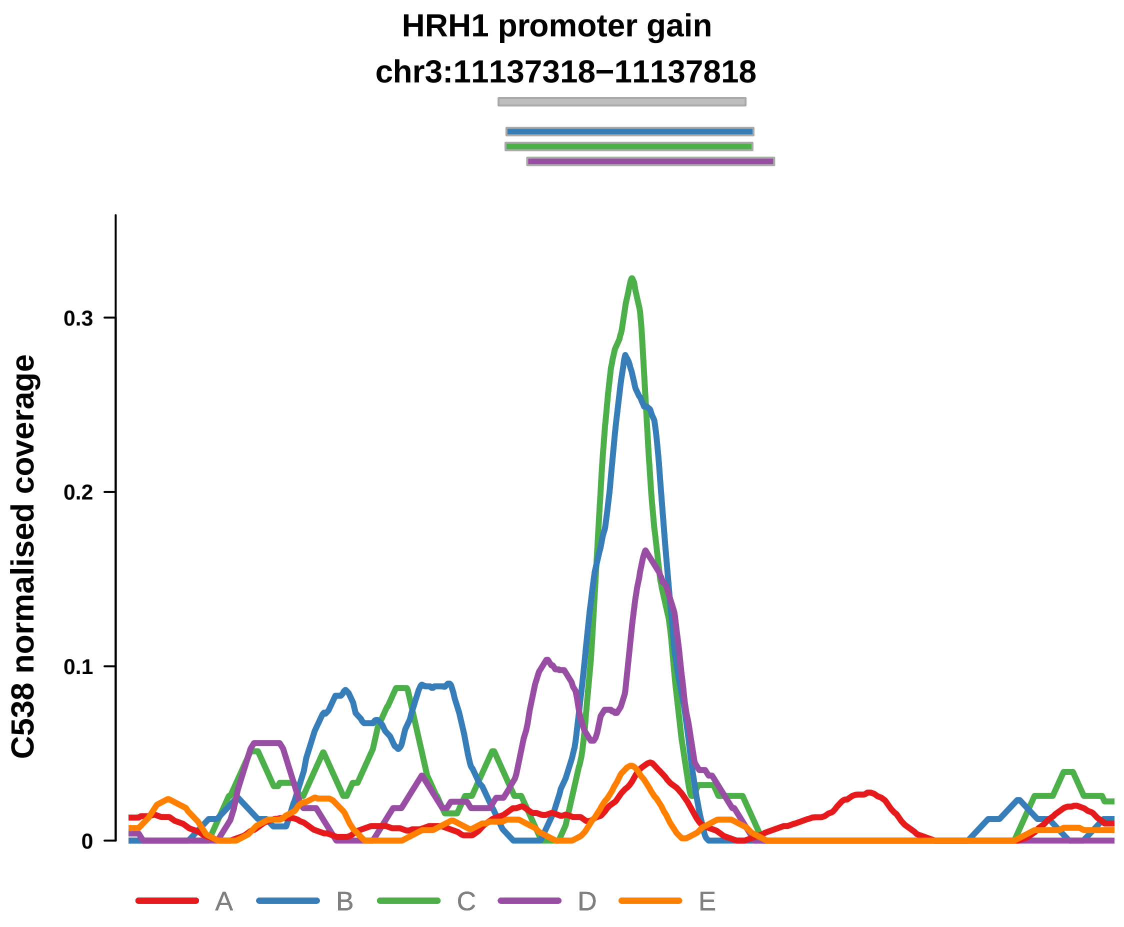


Putatively Clonal (logFC = 2.58)

| | P. | Adjusted P. |
|---|---|---|
| *~region vs. ~* | 1.35e−01 | 2.82e−01 |
| *~purity vs ~* | 5.72e−02 | 1.53e−01 |
| *~purity+region vs ~purity* | 1.47e−01 | 8.28e−01 |
| *Carcinoma vs. normal* | 9.64e−06 **** | 9.64e−06 **** |

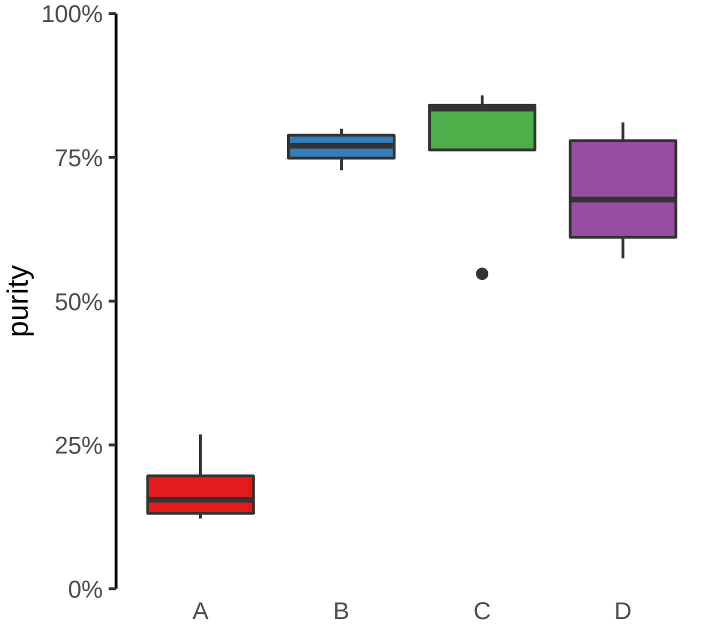

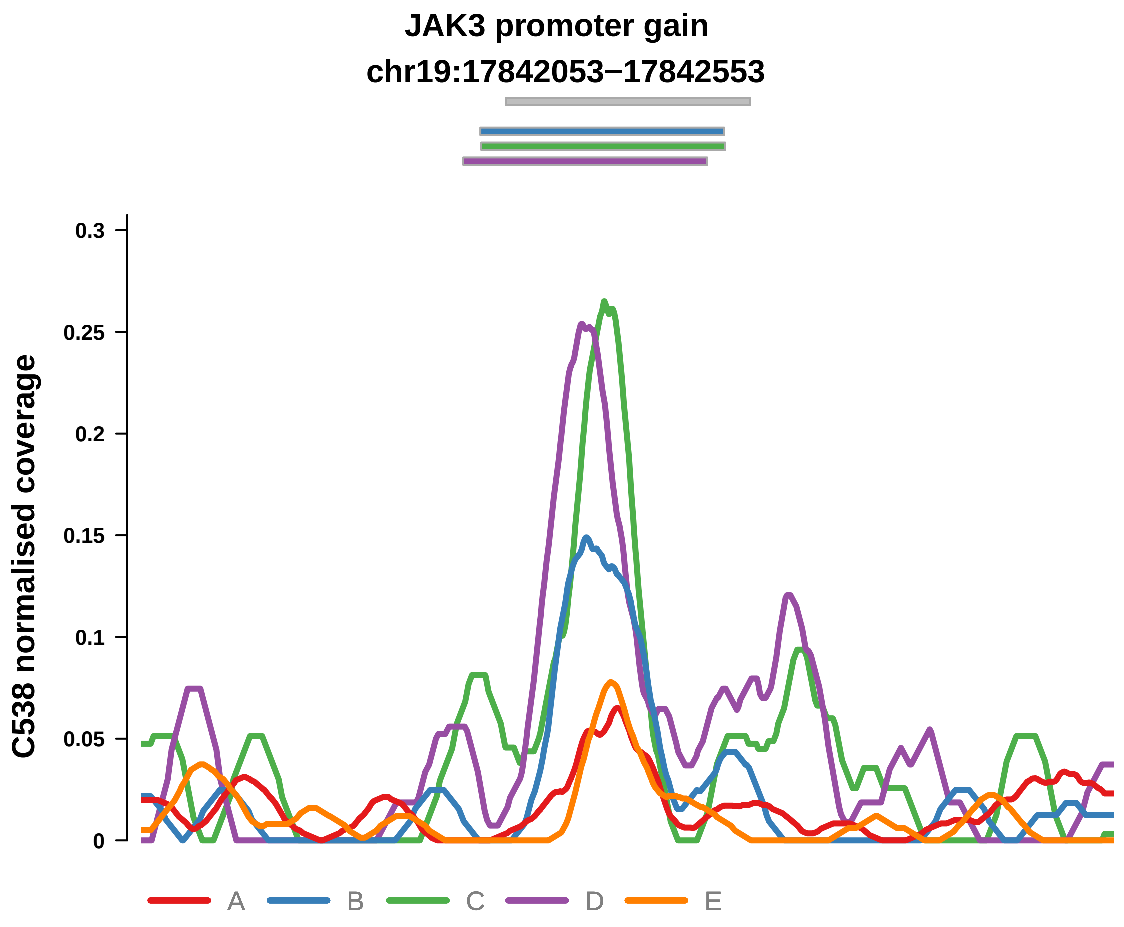


Putatively Clonal (logFC = 1.95)

| | P. | Adjusted P. |
|---|---|---|
| *~region vs. ~* | 1.84e−02 * | 7.78e−02 |
| *~purity vs ~* | 7.24e−02 | 1.77e−01 |
| *~purity+region vs ~purity* | 8.35e−02 | 7.35e−01 |
| *Carcinoma vs. normal* | 6.07e−04 *** | 6.07e−04 *** |

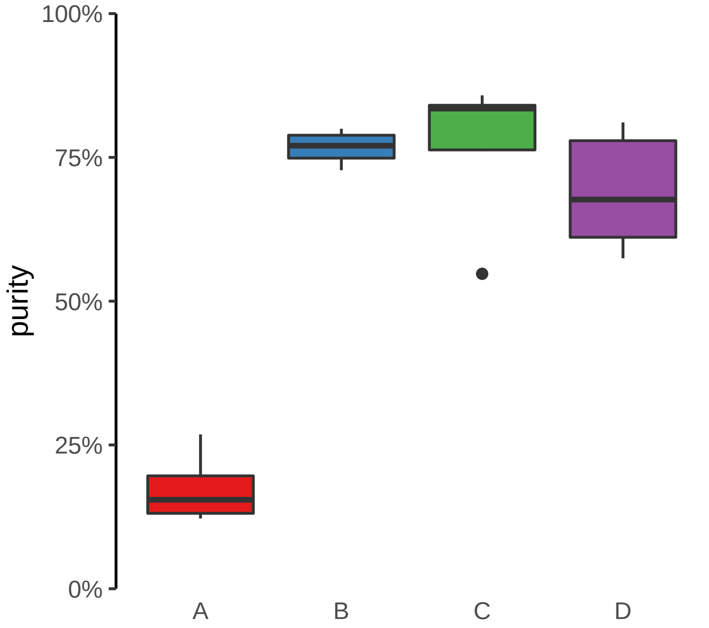

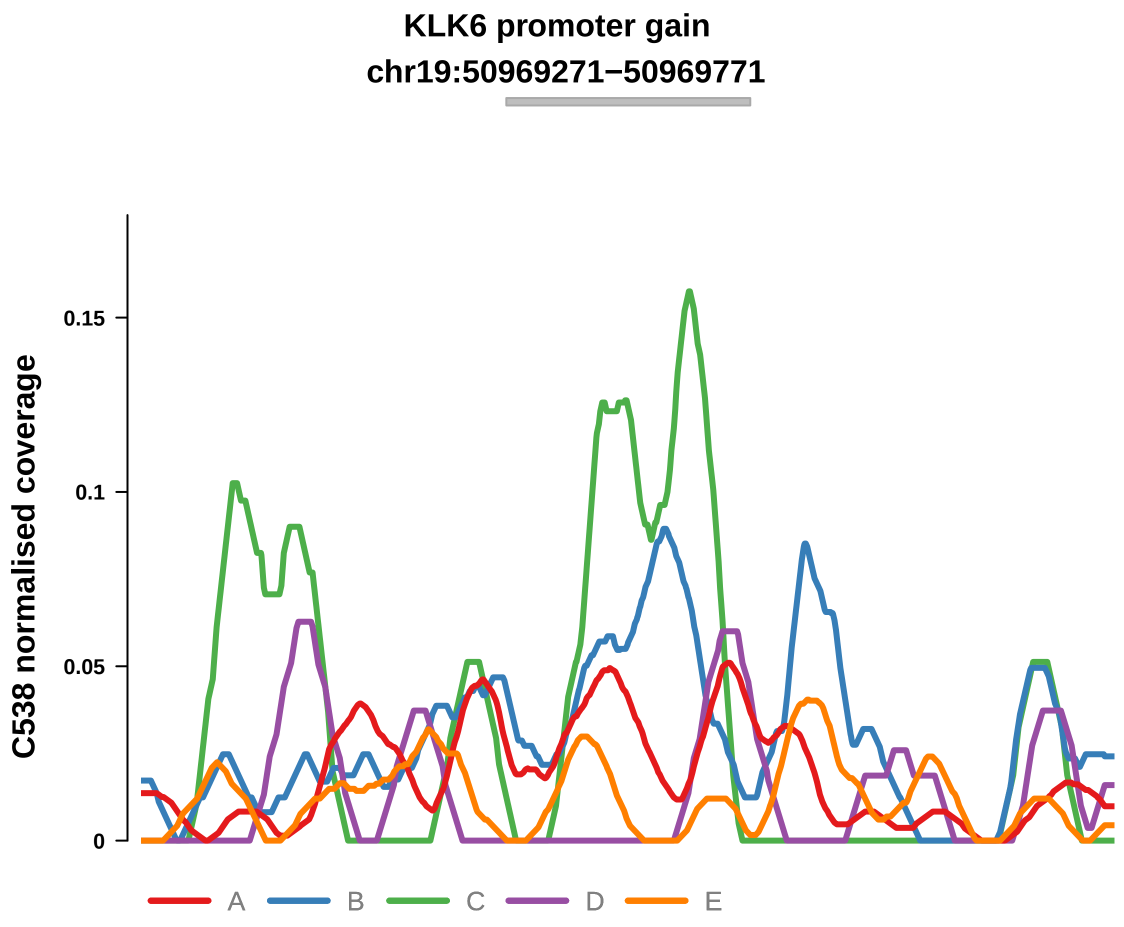


Putatively Clonal (logFC = 1.65)

| | P. | Adjusted P. |
|---|---|---|
| *~region vs. ~* | 7.61e−02 | 1.81e−01 |
| *~purity vs ~* | 1.12e−01 | 2.31e−01 |
| *~purity+region vs ~purity* | 2.33e−01 | 8.7e−01 |
| *Carcinoma vs. normal* | 5.81e−03 ** | 5.81e−03 ** |

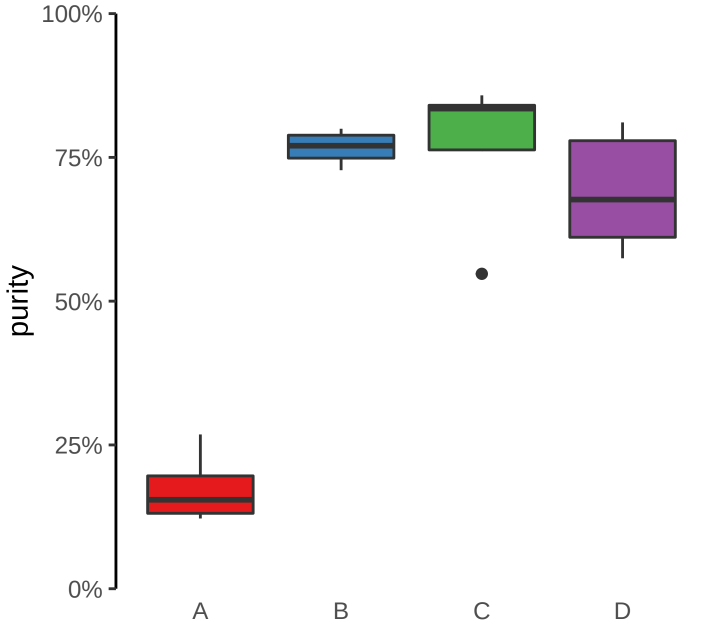

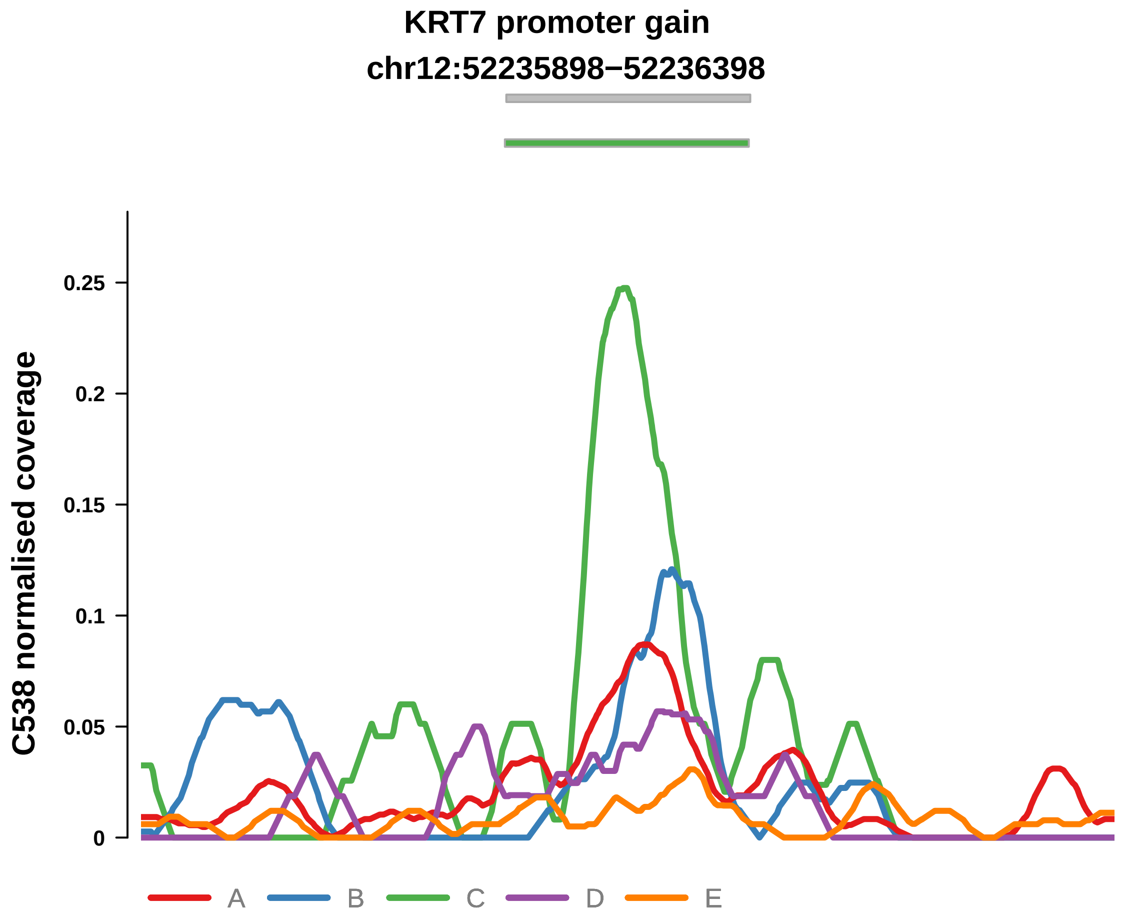


Putatively Clonal (logFC = 1.49)

| | P. | Adjusted P. |
|---|---|---|
| *~region vs. ~* | 5.87e−01 | 7.49e−01 |
| *~purity vs ~* | 4.62e−01 | 6.07e−01 |
| *~purity+region vs ~purity* | 7.15e−01 | 9.36e−01 |
| *Carcinoma vs. normal* | 9.69e−03 ** | 9.69e−03 ** |

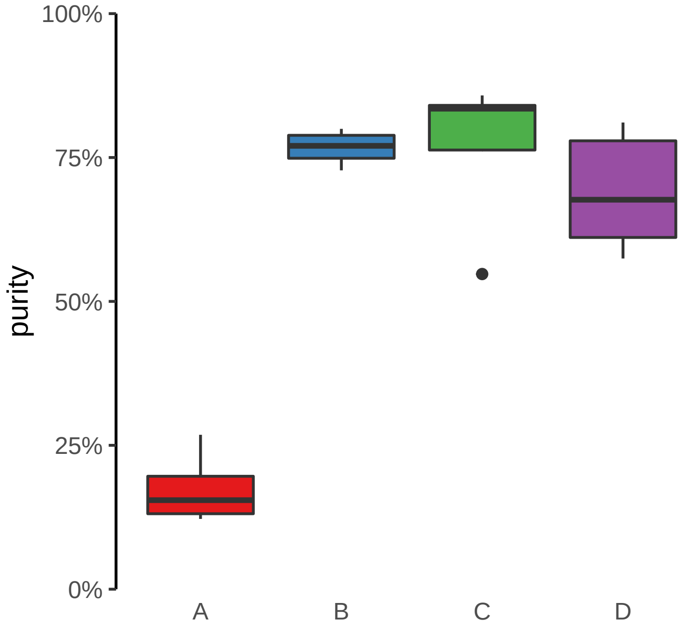

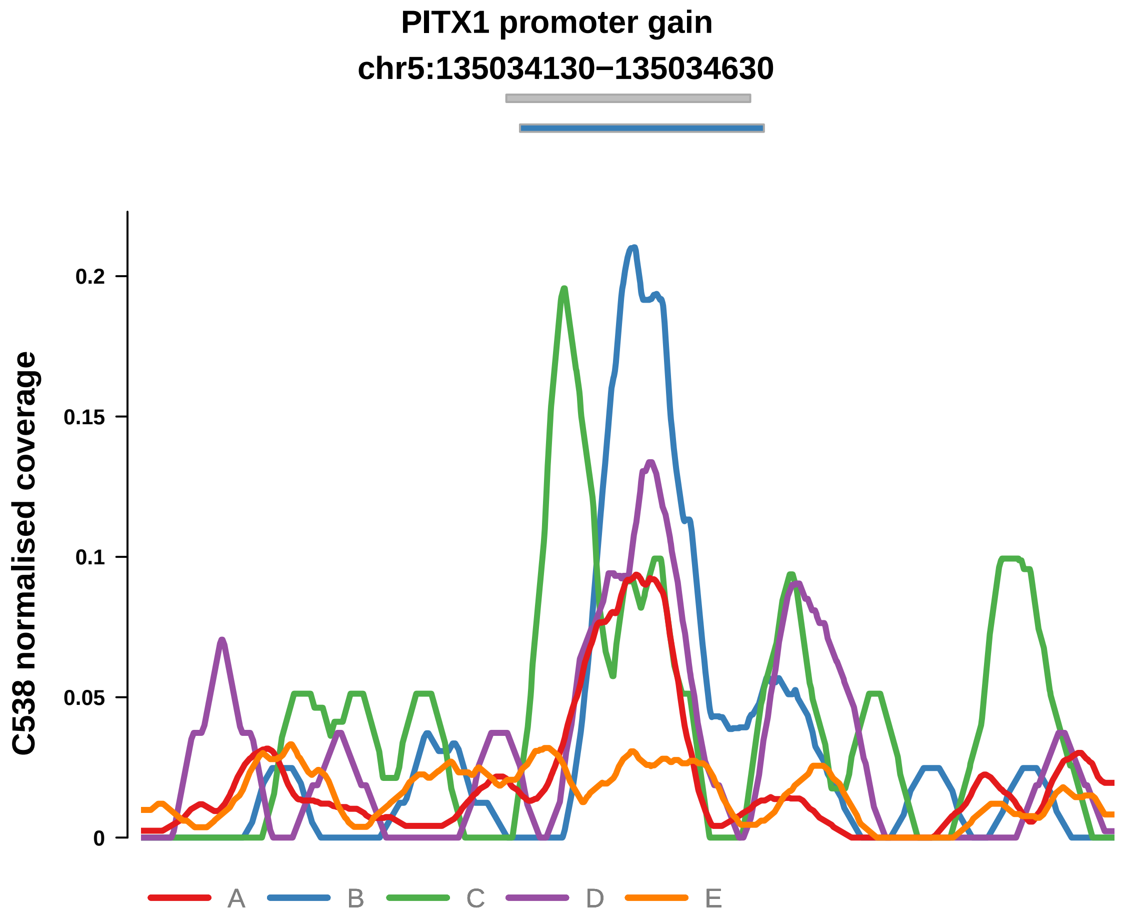


Putatively Clonal (logFC = 1.56)

| | P. | Adjusted P. |
|---|---|---|
| *~region vs. ~* | 1.03e−01 | 2.32e−01 |
| *~purity vs ~* | 8.76e−01 | 9.29e−01 |
| *~purity+region vs ~purity* | 6.95e−02 | 6.8e−01 |
| *Carcinoma vs. normal* | 5.96e−03 ** | 5.96e−03 ** |

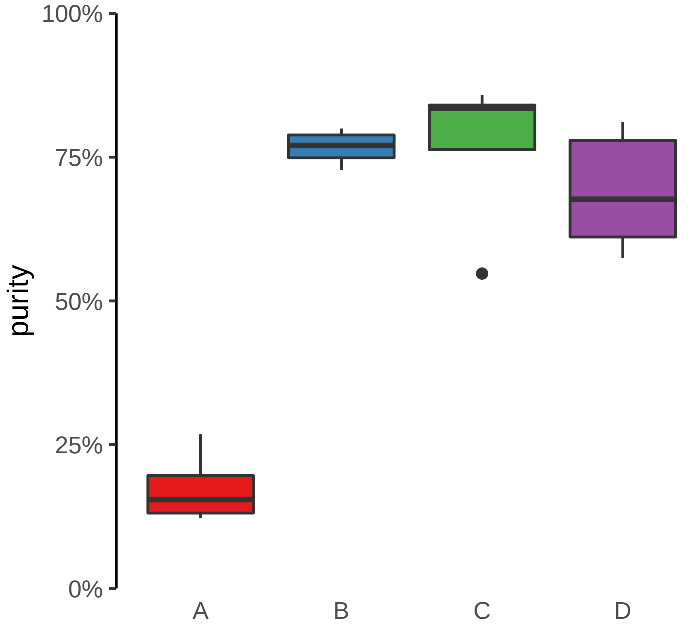

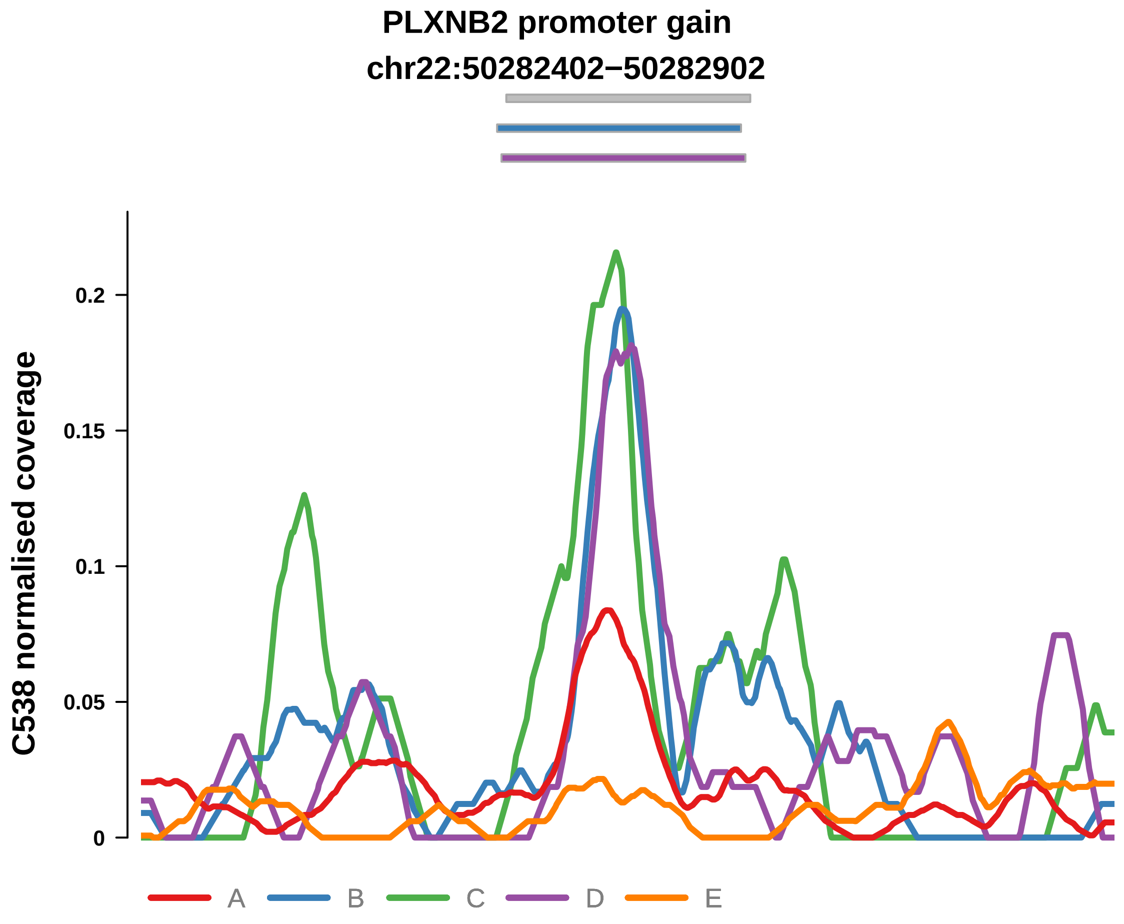


Putatively Clonal (logFC = 2.19)

| | P. | Adjusted P. |
|---|---|---|
| *~region vs. ~* | 7.27e−01 | 8e−01 |
| *~purity vs ~* | 8.1e−01 | 9.02e−01 |
| *~purity+region vs ~purity* | 5.81e−01 | 8.7e−01 |
| *Carcinoma vs. normal* | 1.57e−04 *** | 1.57e−04 *** |

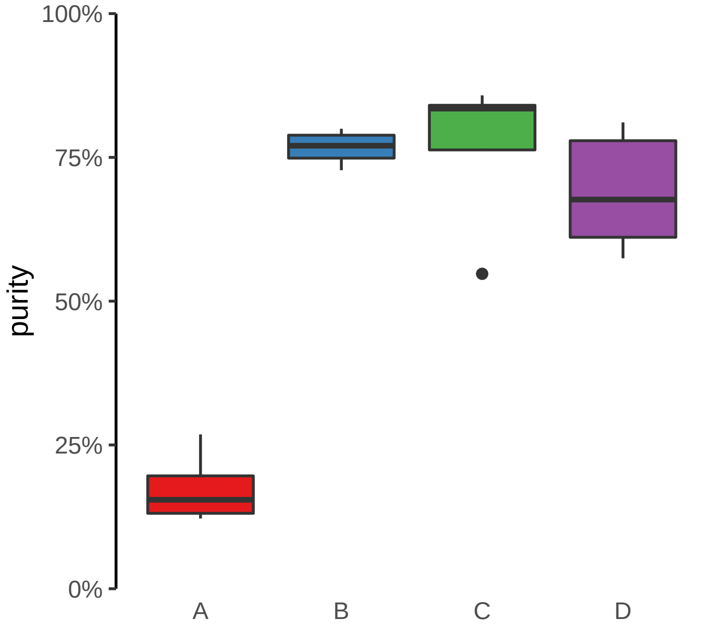

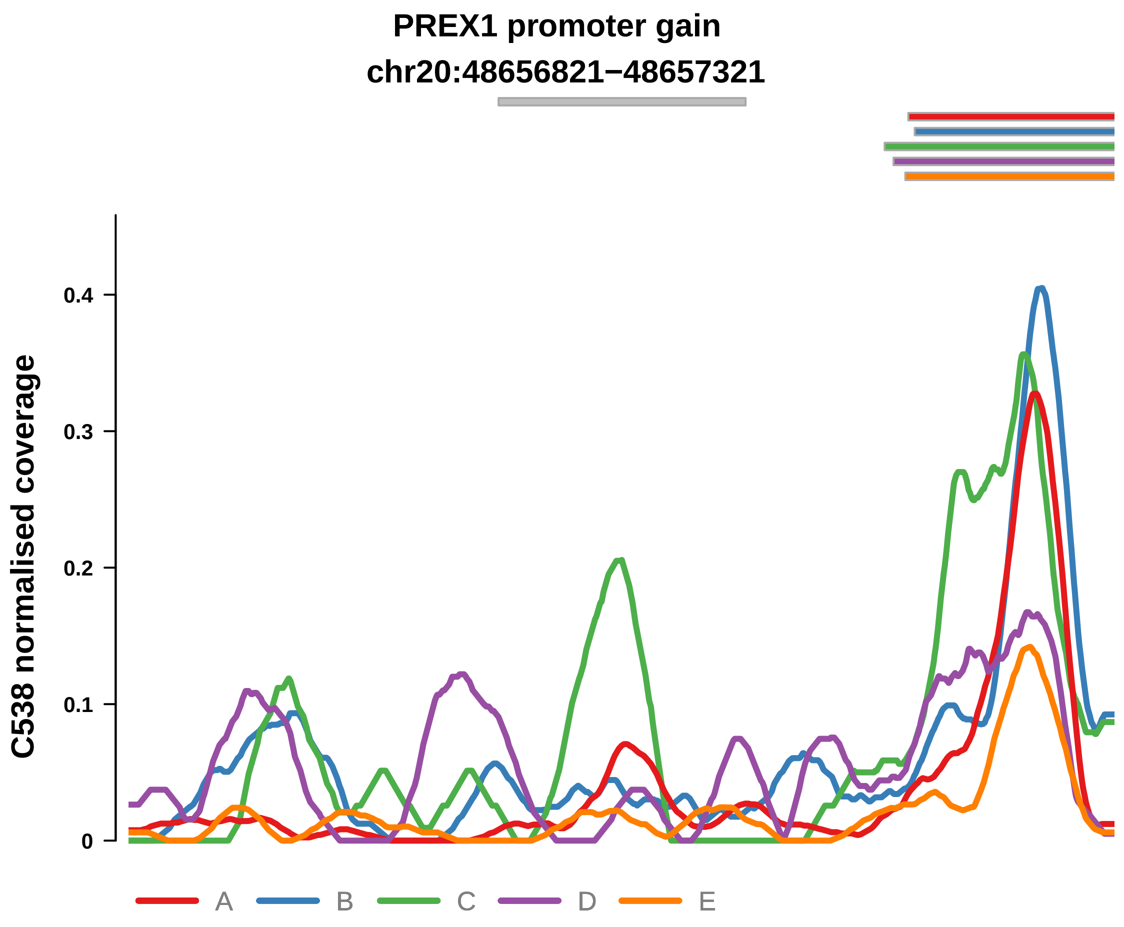


Putatively Clonal (logFC = 1.6)

| | P. | Adjusted P. |
|---|---|---|
| *~region vs. ~* | 6.77e−01 | 7.98e−01 |
| *~purity vs ~* | 6.89e−01 | 8.08e−01 |
| *~purity+region vs ~purity* | 6.41e−01 | 9.12e−01 |
| *Carcinoma vs. normal* | 8.89e−03 ** | 8.89e−03 ** |

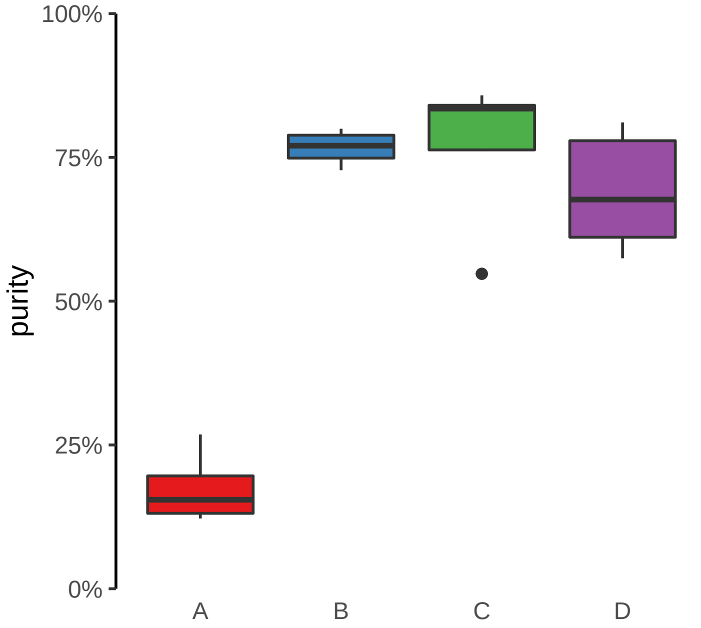

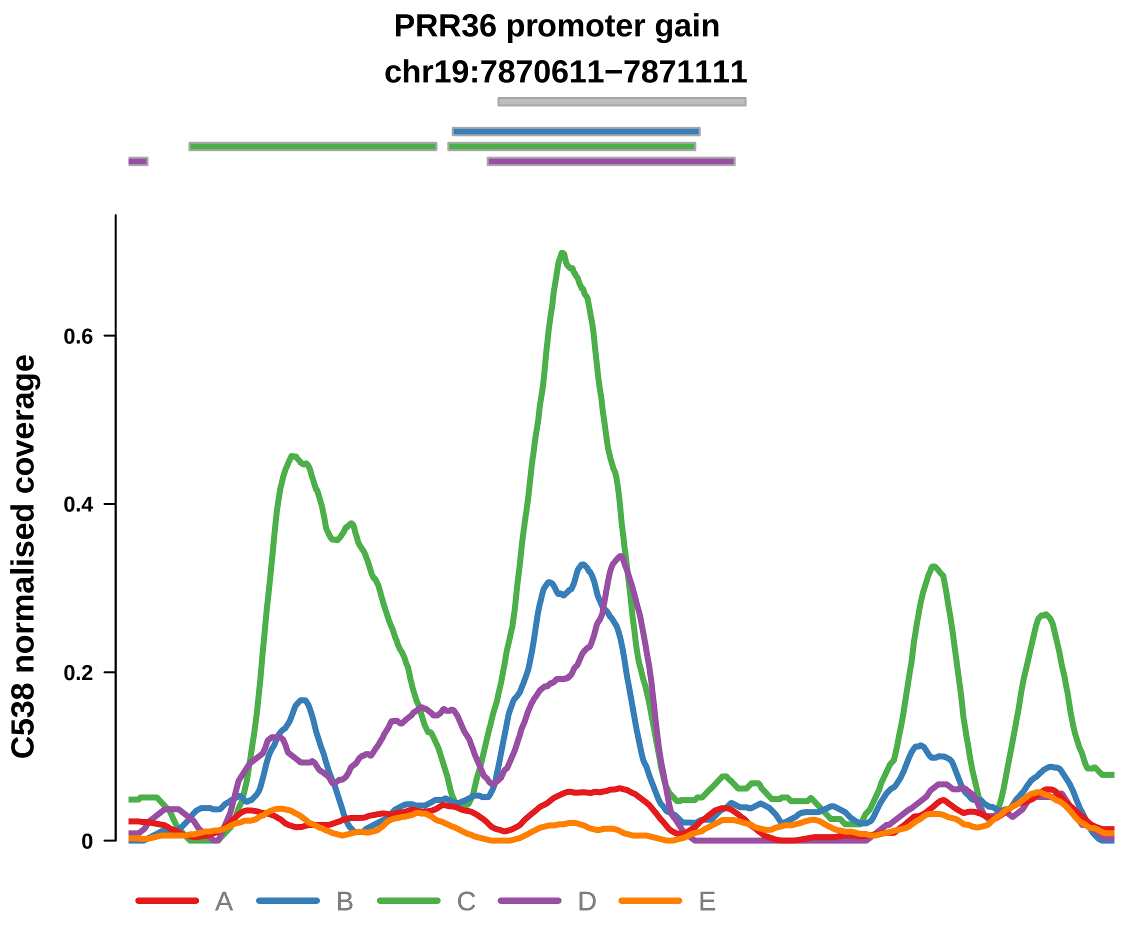


Putatively Clonal (logFC = 2.87)

| | P. | Adjusted P. |
|---|---|---|
| *~region vs. ~* | 2.13e−04 *** | 2.68e−03 ** |
| *~purity vs ~* | 3.85e−06 **** | 8.47e−05 **** |
| *~purity+region vs ~purity* | 9.36e−01 | 9.79e−01 |
| *Carcinoma vs. normal* | 4.62e−07 **** | 4.62e−07 **** |

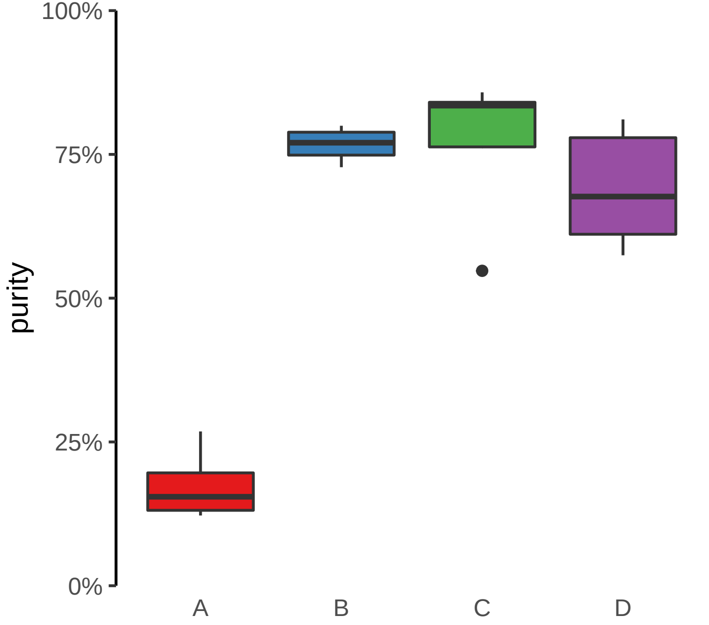

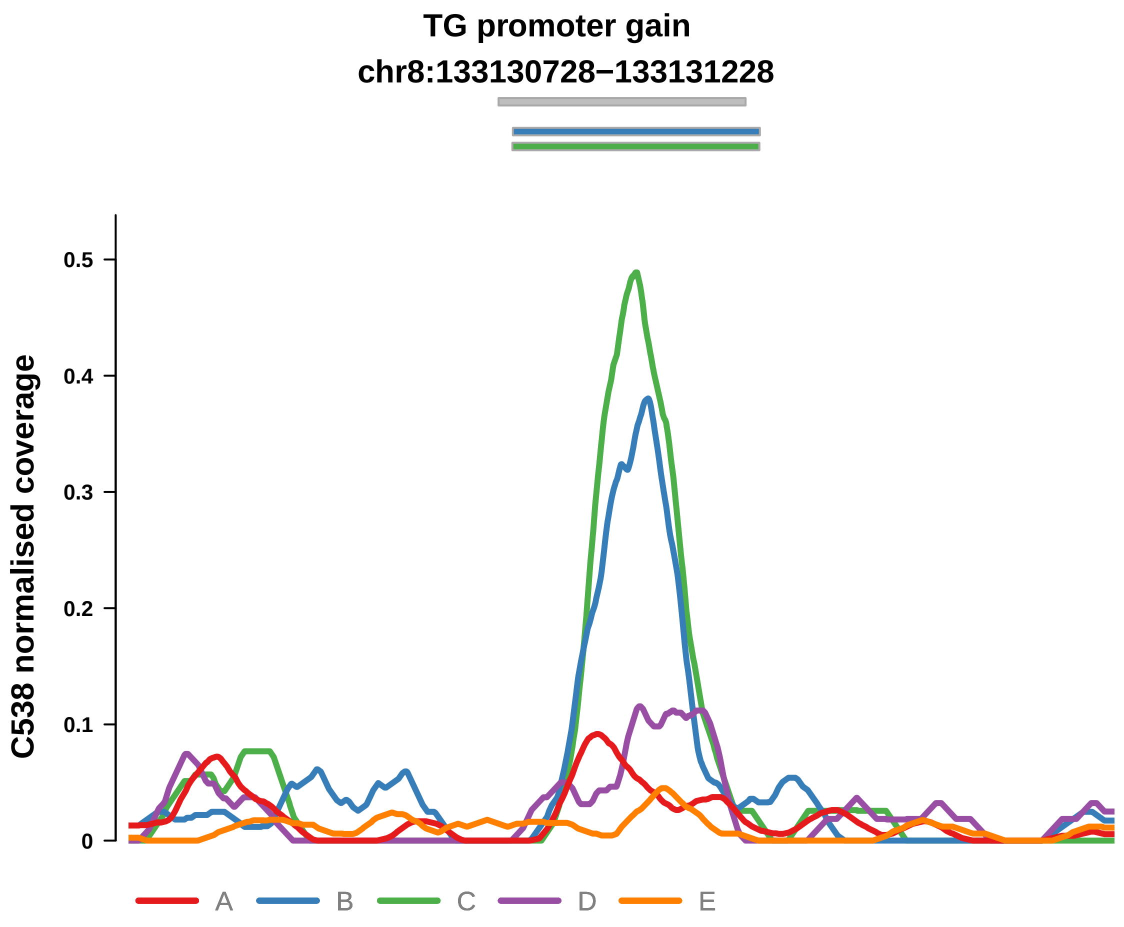


Putatively Clonal (logFC = 2.83)

| | P. | Adjusted P. |
|---|---|---|
| *~region vs. ~* | 5.52e−01 | 7.15e−01 |
| *~purity vs ~* | 2.4e−01 | 3.91e−01 |
| *~purity+region vs ~purity* | 5.83e−01 | 8.7e−01 |
| *Carcinoma vs. normal* | 8.69e−07 **** | 8.69e−07 **** |

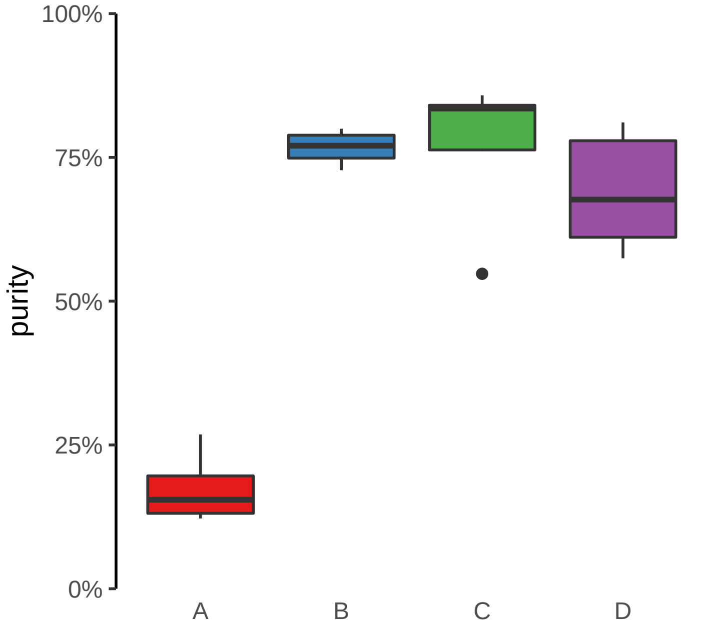

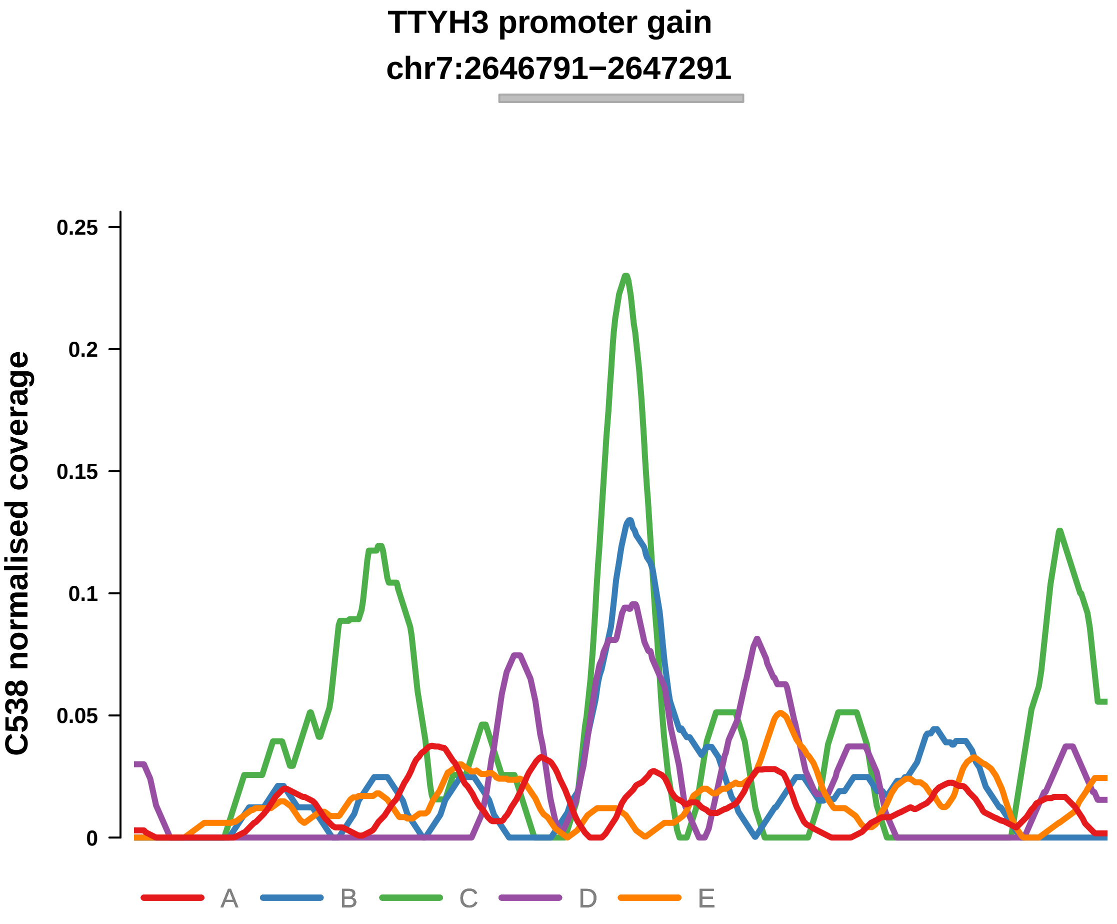


Putatively Clonal (logFC = 1.9)

| | P. | Adjusted P. |
|---|---|---|
| *~region vs. ~* | 3.81e−01 | 5.78e−01 |
| *~purity vs ~* | 2.1e−01 | 3.62e−01 |
| *~purity+region vs ~purity* | 7.01e−01 | 9.36e−01 |
| *Carcinoma vs. normal* | 1.28e−03 ** | 1.28e−03 ** |

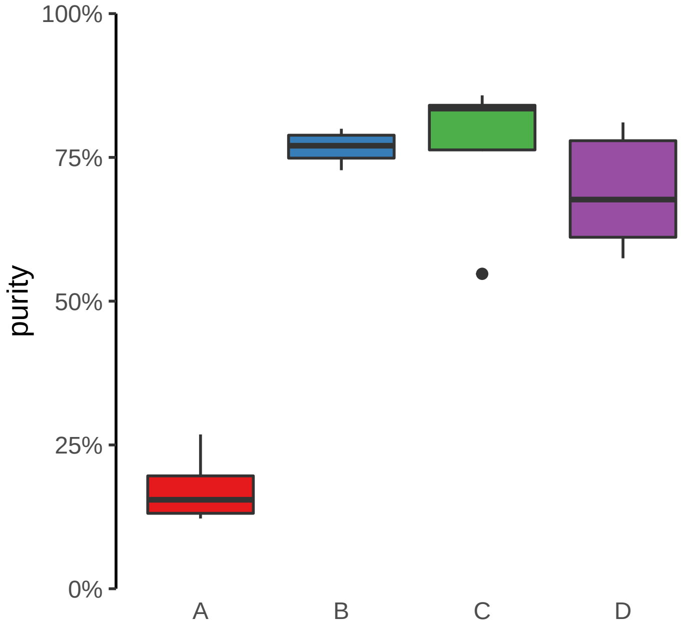

**ARHGEF17 promoter gain**
**chr11:73362044−73362544**

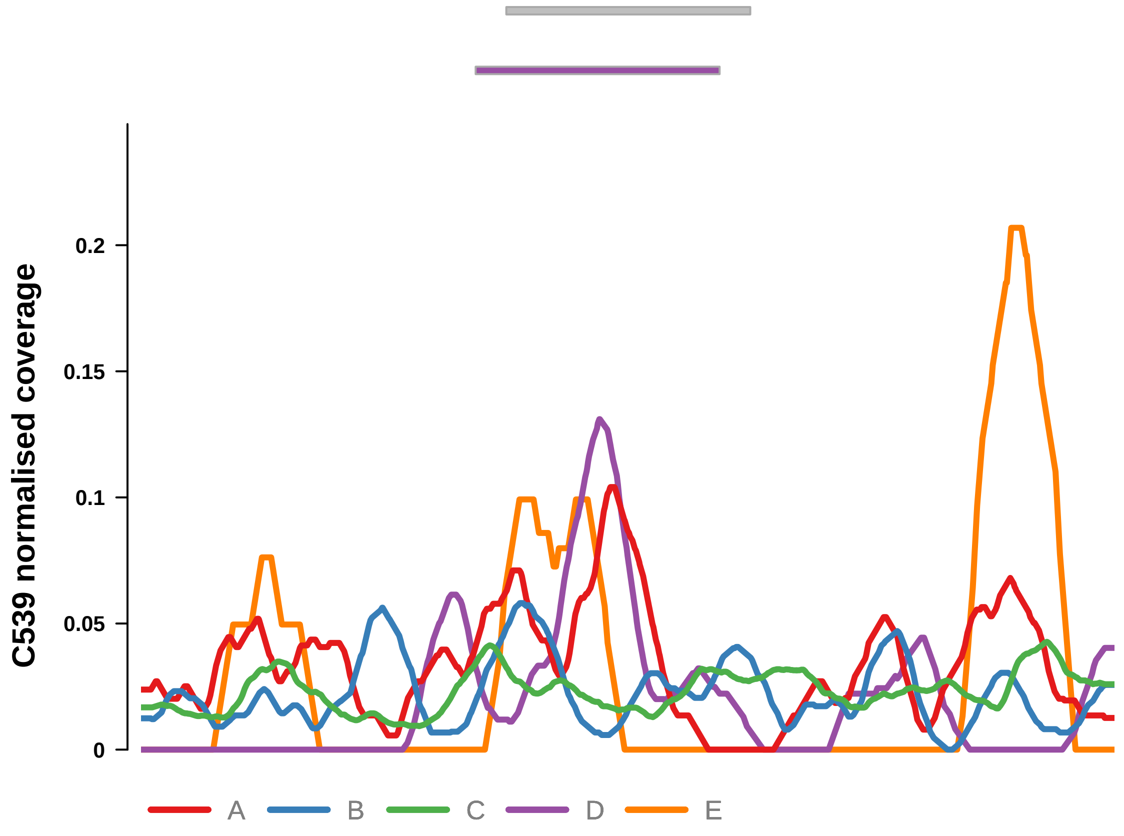


Putatively Clonal (logFC = 2.91)

| | P. | Adjusted P. |
|---|---|---|
| *~region vs. ~* | 3.59e−02 * | 4.26e−02 * |
| *~purity vs ~* | 6.34e−02 | 7.85e−02 |
| *~purity+region vs ~purity* | 5.13e−02 | 1.23e−01 |
| *Carcinoma vs. normal* | 3.46e−05 **** | 3.46e−05 **** |

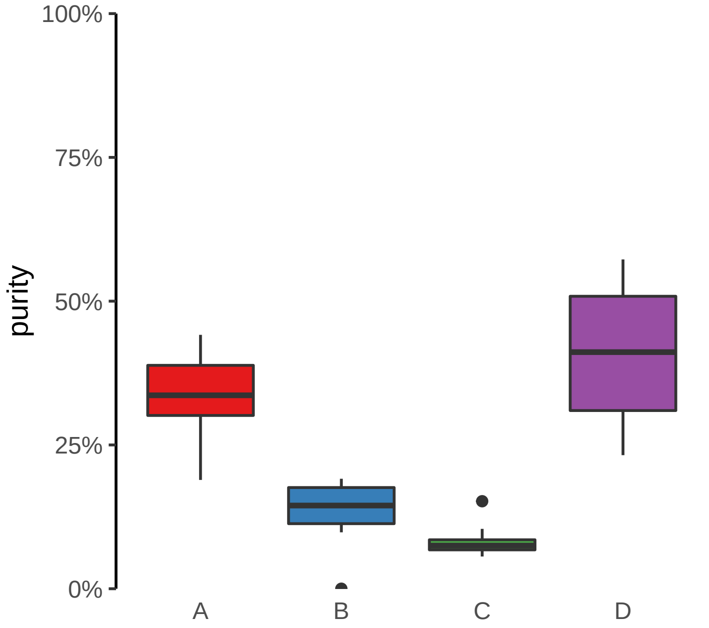

**CADM4 promoter gain**
**chr19:43623815−43624315**

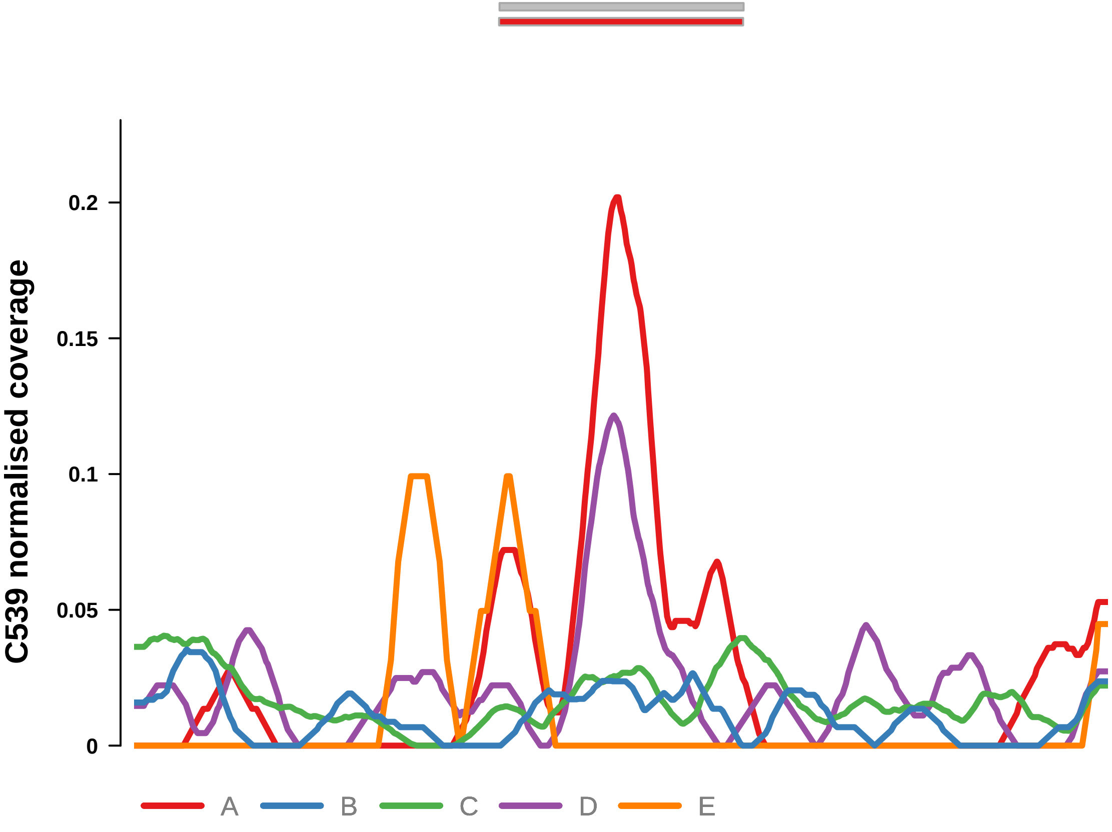


Putatively Clonal (logFC = 2.32)

| | P. | Adjusted P. |
|---|---|---|
| *~region vs. ~* | 2.09e−02 * | 2.79e−02 * |
| *~purity vs ~* | 3.62e−02 * | 4.98e−02 * |
| *~purity+region vs ~purity* | 1.99e−01 | 3.08e−01 |
| *Carcinoma vs. normal* | 2.76e−03 ** | 2.76e−03 ** |

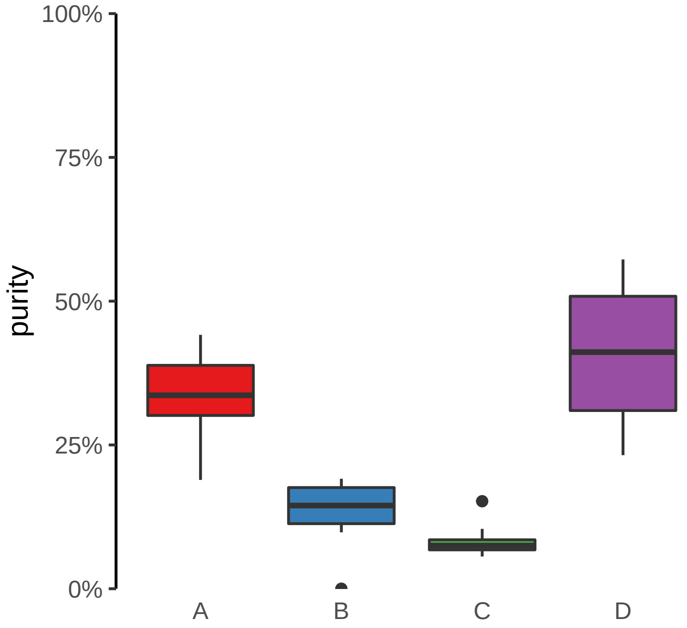

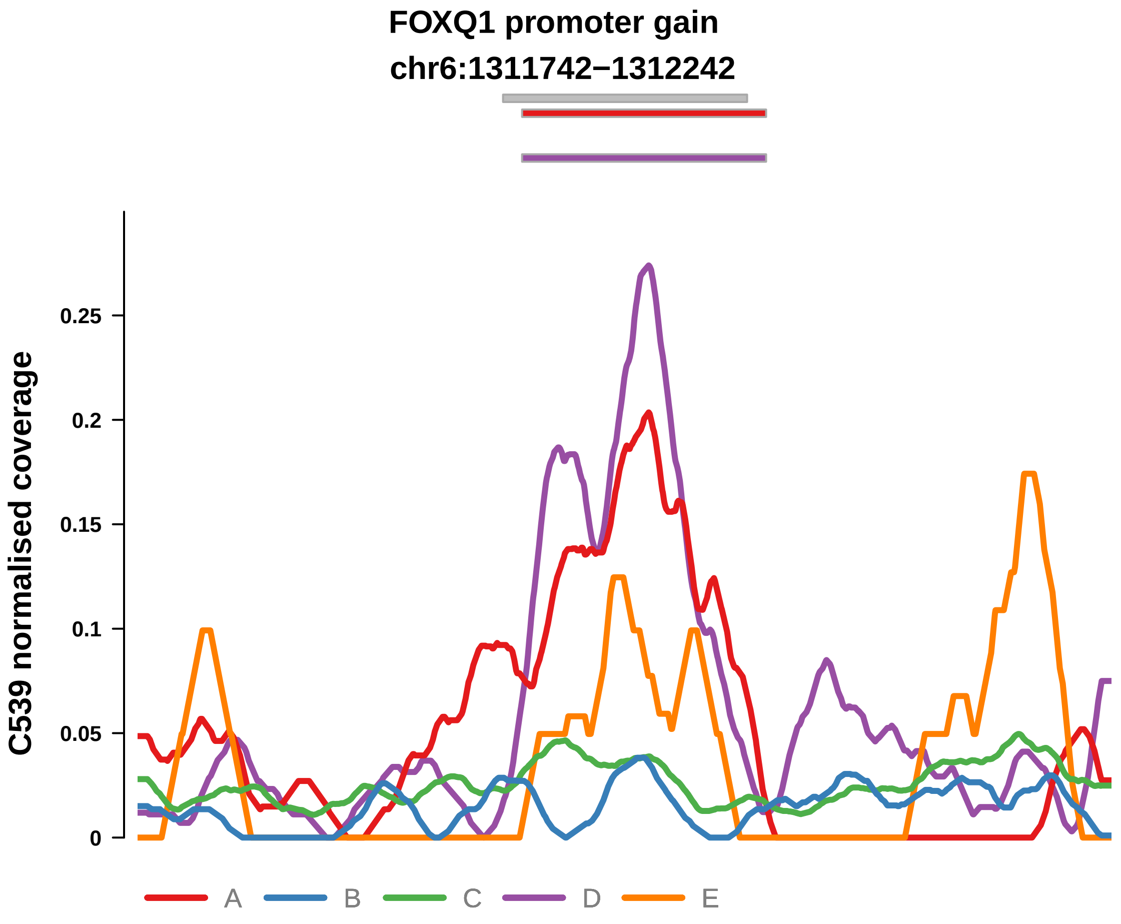


Putatively Clonal (logFC = 3.08)

| | P. | Adjusted P. |
|---|---|---|
| *~region vs. ~* | 2.26e−01 | 2.29e−01 |
| *~purity vs ~* | 6.83e−01 | 6.83e−01 |
| *~purity+region vs ~purity* | 1.56e−01 | 2.78e−01 |
| *Carcinoma vs. normal* | 4.3e−07 **** | 4.3e−07 **** |

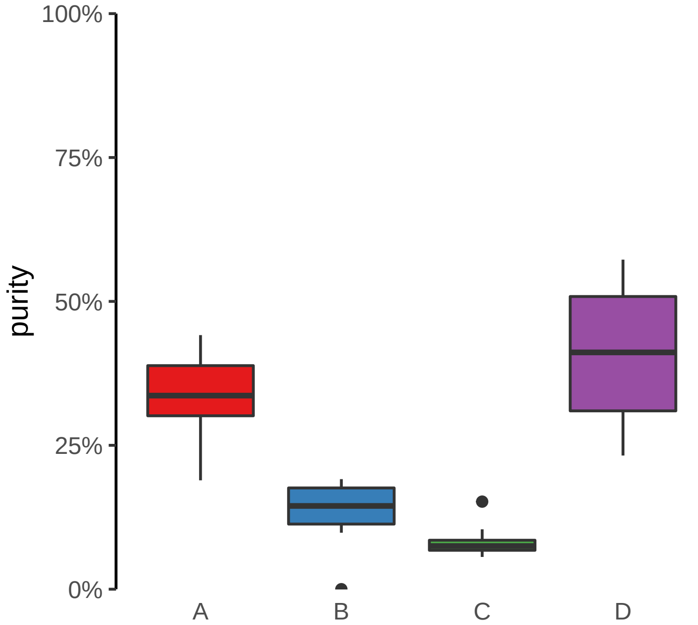

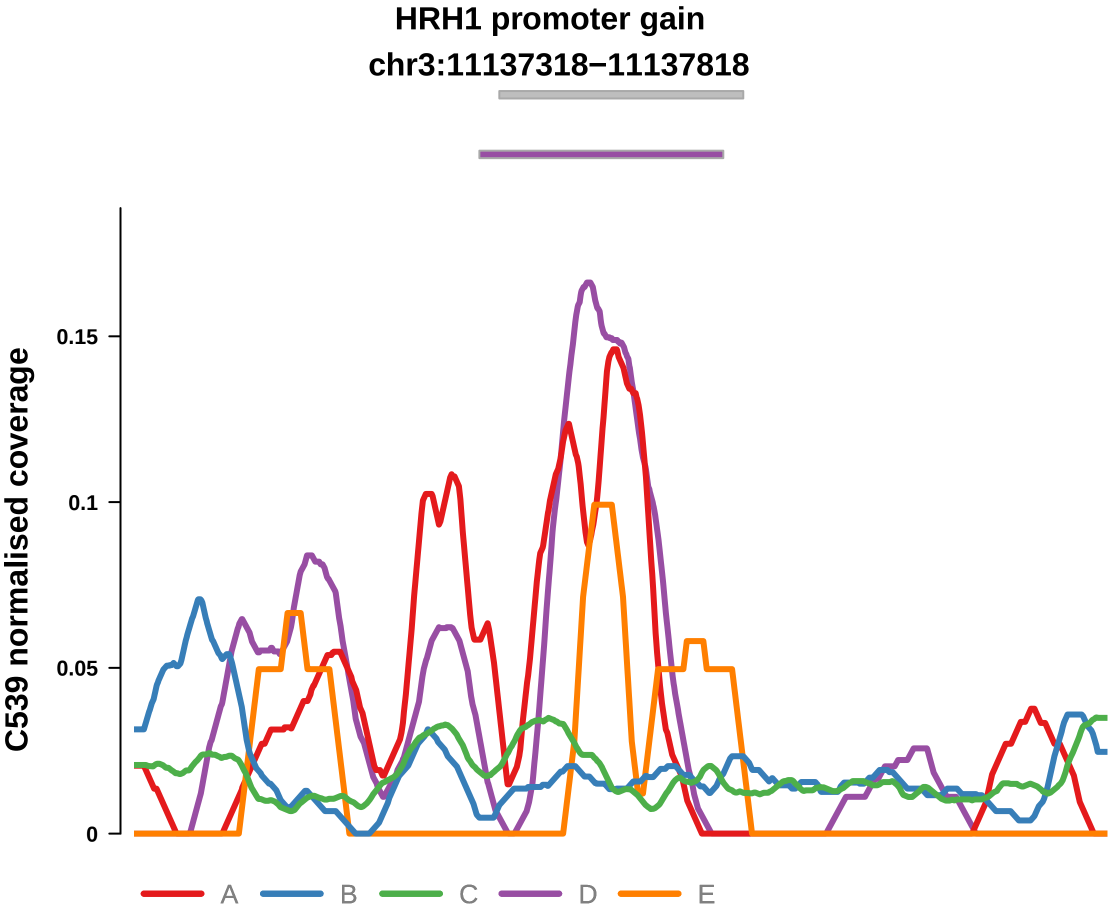


Putatively Clonal (logFC = 2.37)

| | P. | Adjusted P. |
|---|---|---|
| *~region vs. ~* | 1.81e−02 * | 2.48e−02 * |
| *~purity vs ~* | 6.75e−02 | 8.25e−02 |
| *~purity+region vs ~purity* | 9.01e−02 | 1.98e−01 |
| *Carcinoma vs. normal* | 1.04e−03 ** | 1.04e−03 ** |

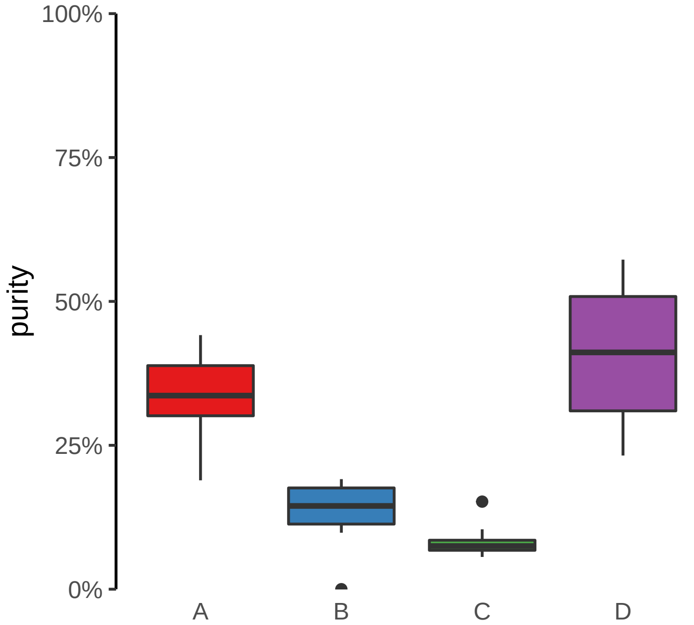

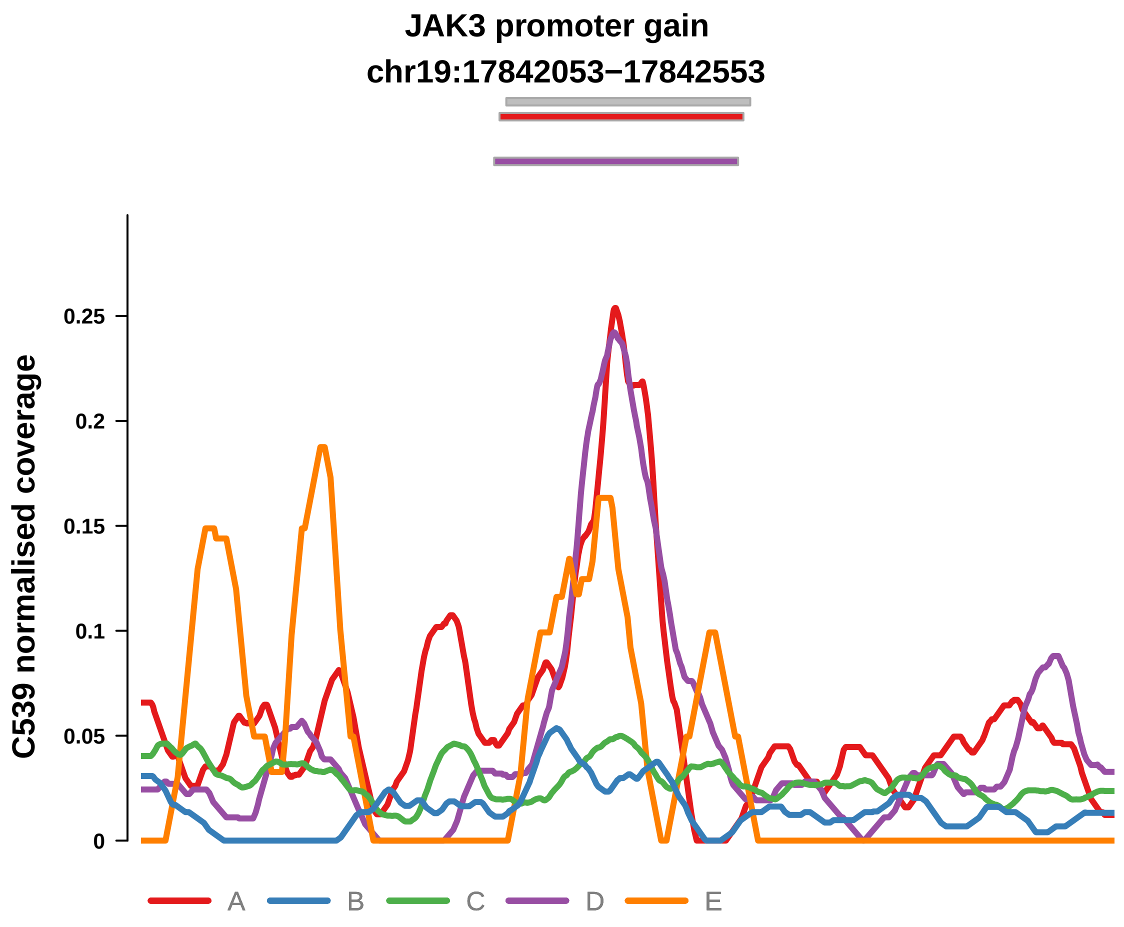


Putatively Clonal (logFC = 2.5)

| | P. | Adjusted P. |
|---|---|---|
| *~region vs. ~* | 5.47e−02 | 6.25e−02 |
| *~purity vs ~* | 8e−02 | 9.15e−02 |
| *~purity+region vs ~purity* | 2.34e−01 | 3.39e−01 |
| *Carcinoma vs. normal* | 1.21e−04 *** | 1.21e−04 *** |

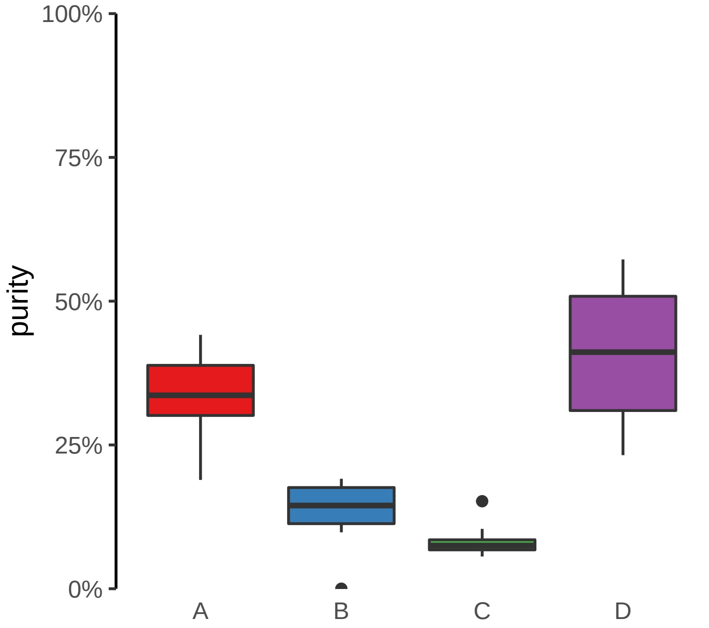

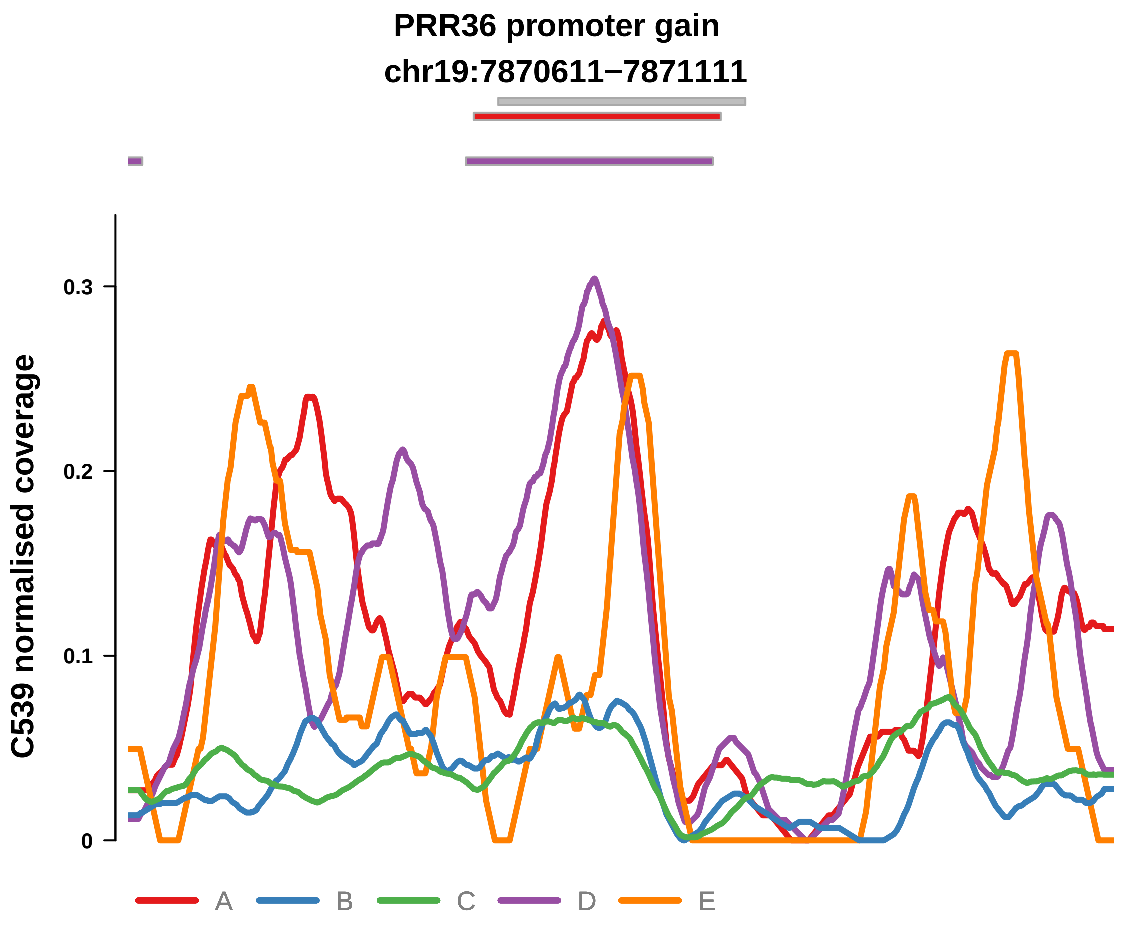


Putatively Clonal (logFC = 3.35)

| | P. | Adjusted P. |
|---|---|---|
| *~region vs. ~* | 2.59e−02 * | 3.25e−02 * |
| *~purity vs ~* | 9.08e−02 | 1.02e−01 |
| *~purity+region vs ~purity* | 2.85e−02 * | 9.39e−02 |
| *Carcinoma vs. normal* | 4.32e−08 **** | 4.32e−08 **** |

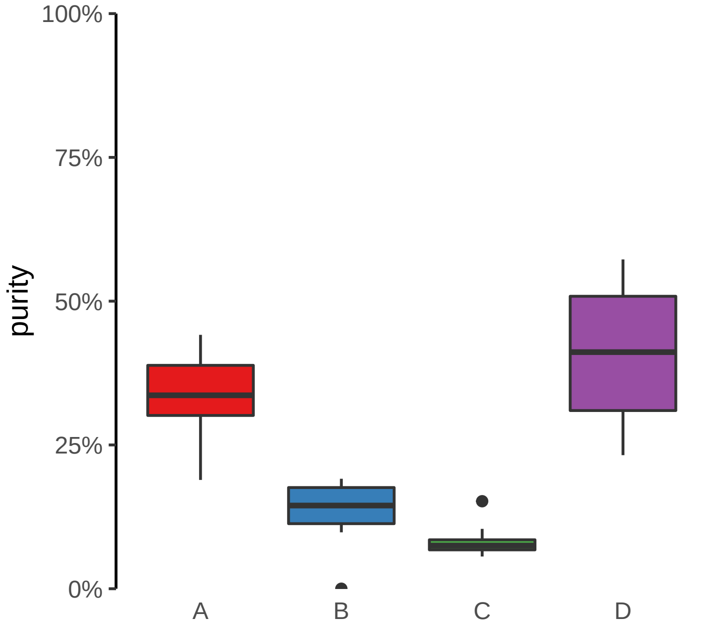

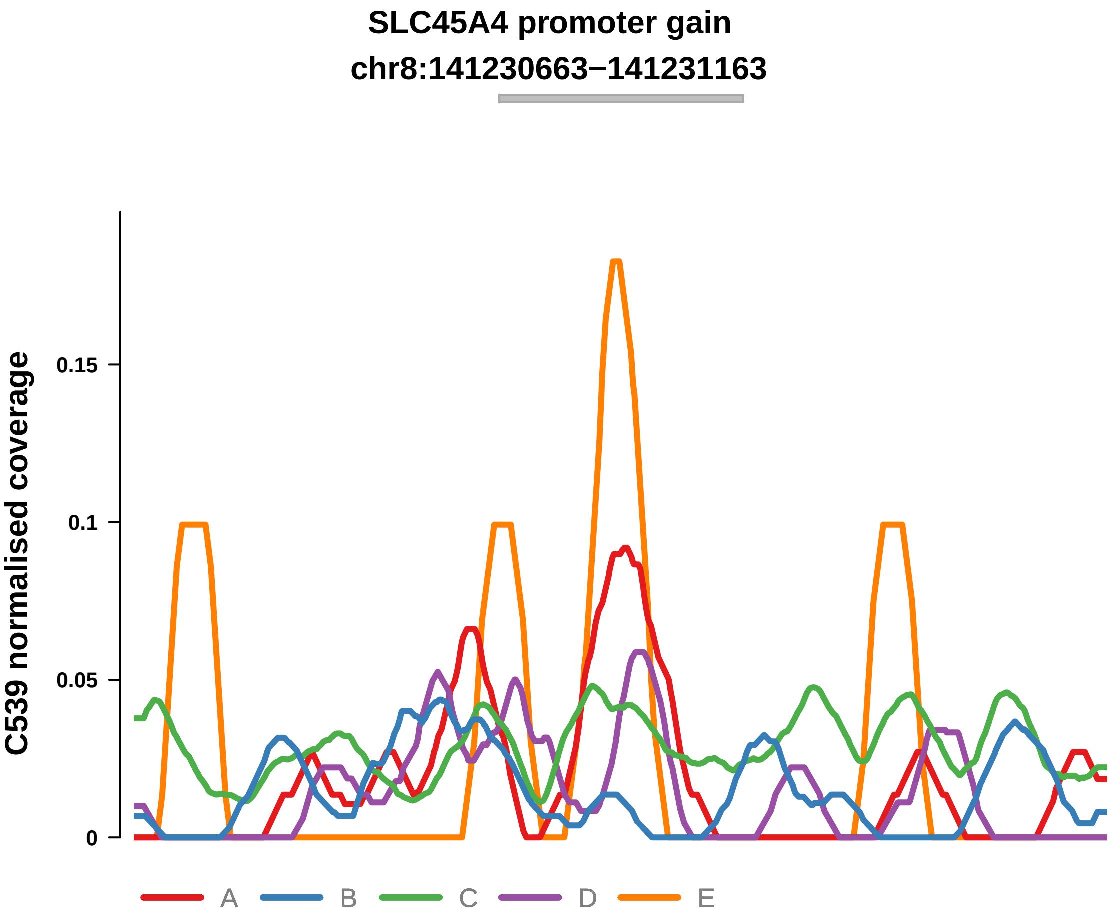


Putatively Clonal (logFC = 2.17)

| | P. | Adjusted P. |
|---|---|---|
| *~region vs. ~* | 1.38e−08 **** | 6.09e−07 **** |
| *~purity vs ~* | 9.78e−05 **** | 6.48e−04 *** |
| *~purity+region vs ~purity* | 4.96e−05 **** | 2.32e−03 ** |
| *Carcinoma vs. normal* | 6.85e−03 ** | 6.85e−03 ** |

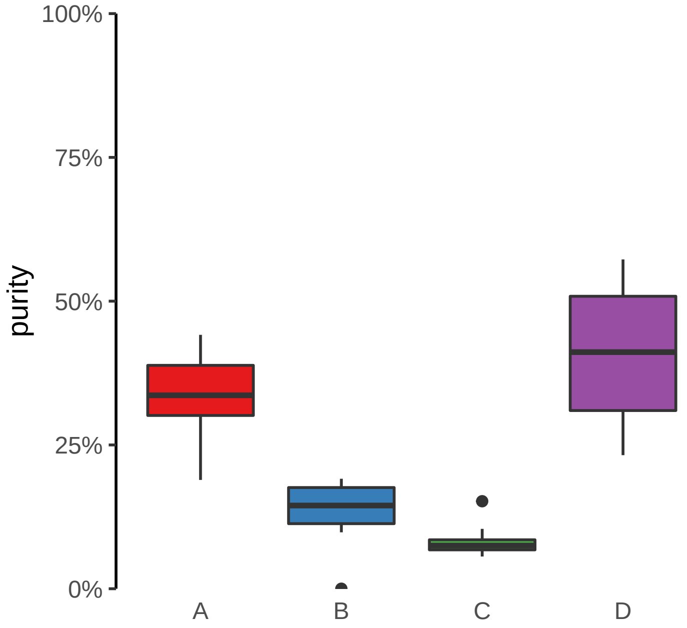

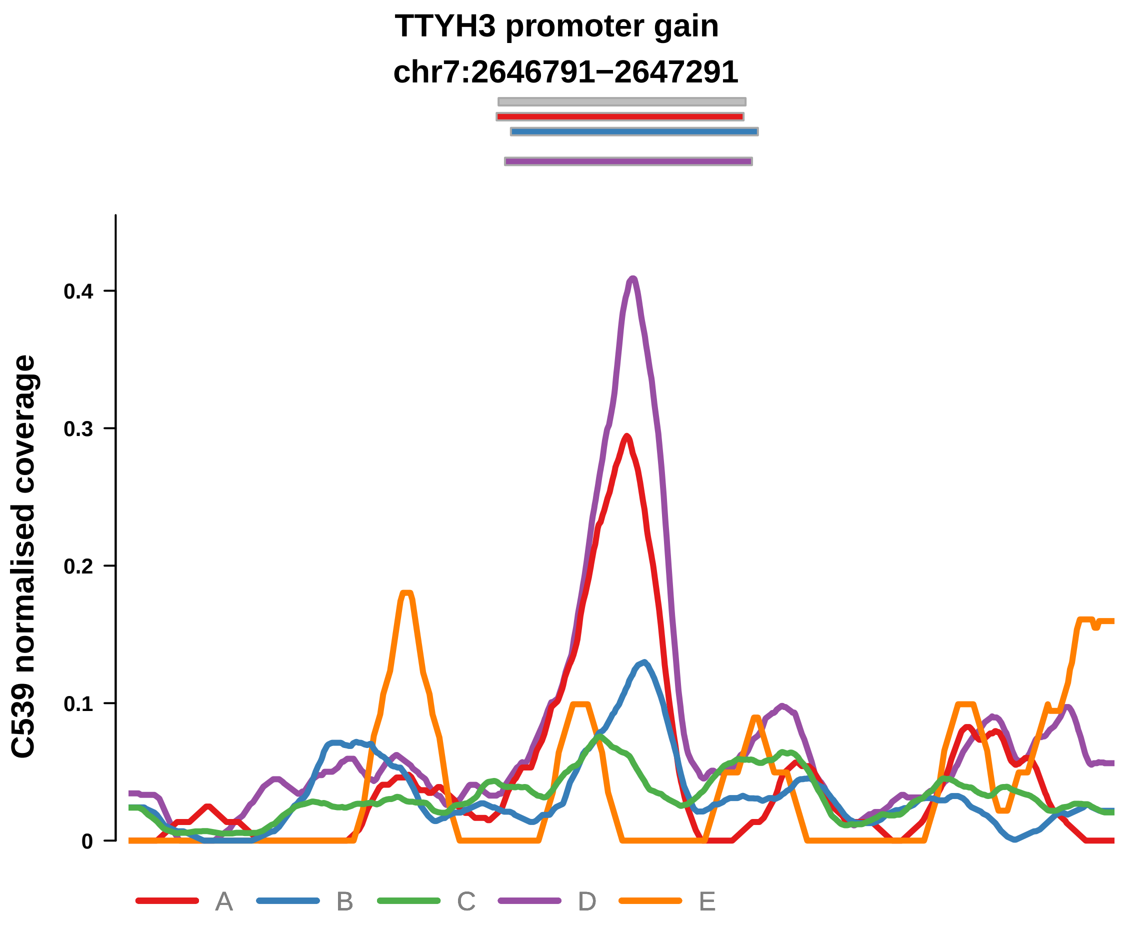


Putatively Clonal (logFC = 3.57)

| | P. | Adjusted P. |
|---|---|---|
| *~region vs. ~* | 2.62e−02 * | 3.25e−02 * |
| *~purity vs ~* | 2.48e−02 * | 3.7e−02 * |
| *~purity+region vs ~purity* | 2.76e−01 | 3.64e−01 |
| *Carcinoma vs. normal* | 4.52e−08 **** | 4.52e−08 **** |

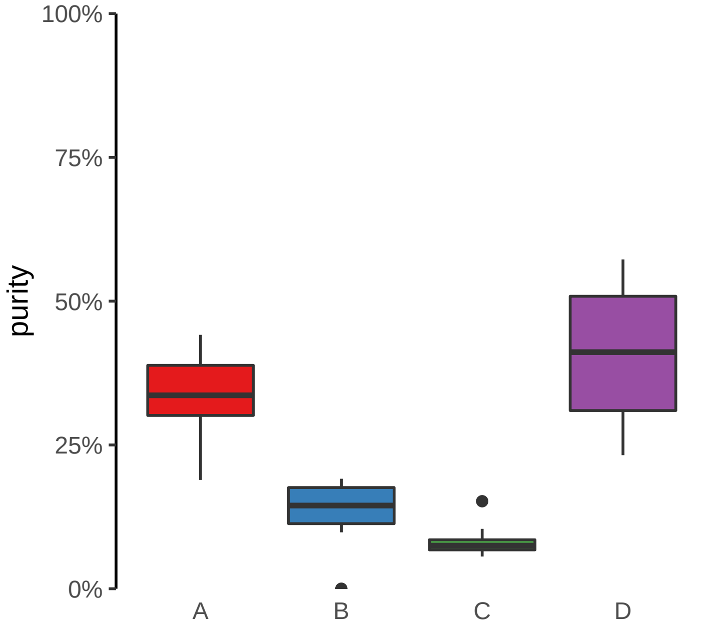

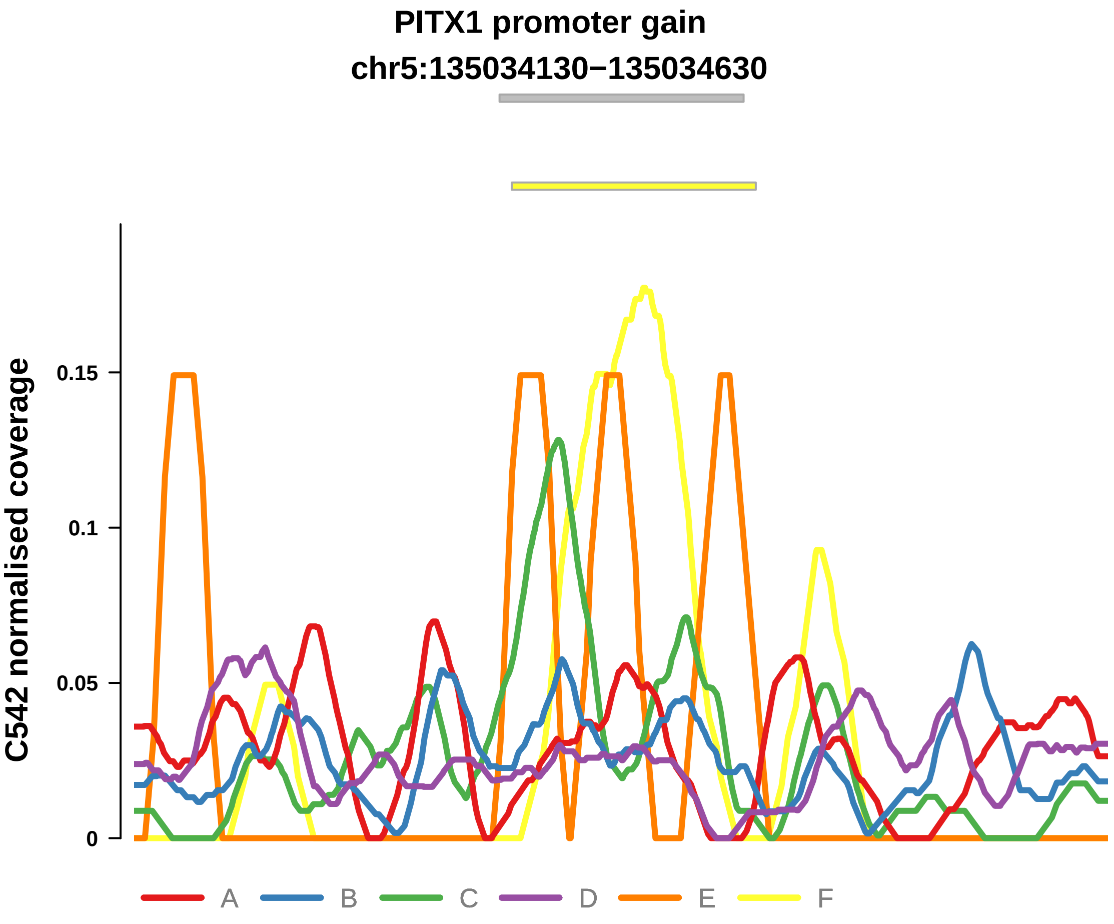


Putatively Clonal (logFC = 2.41)

| | P. | Adjusted P. |
|---|---|---|
| *~region vs. ~* | 4.63e−01 | 7.33e−01 |
| *~purity vs ~* | 1.48e−01 | 6.19e−01 |
| *~purity+region vs ~purity* | 7.18e−01 | 8.56e−01 |
| *Carcinoma vs. normal* | 3.75e−03 ** | 3.75e−03 ** |

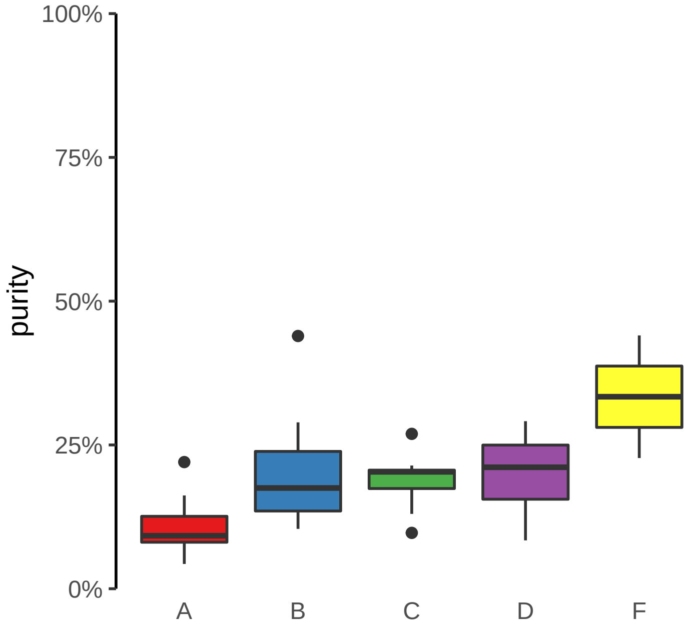

**TG promoter gain**
**chr8:133130728−133131228**

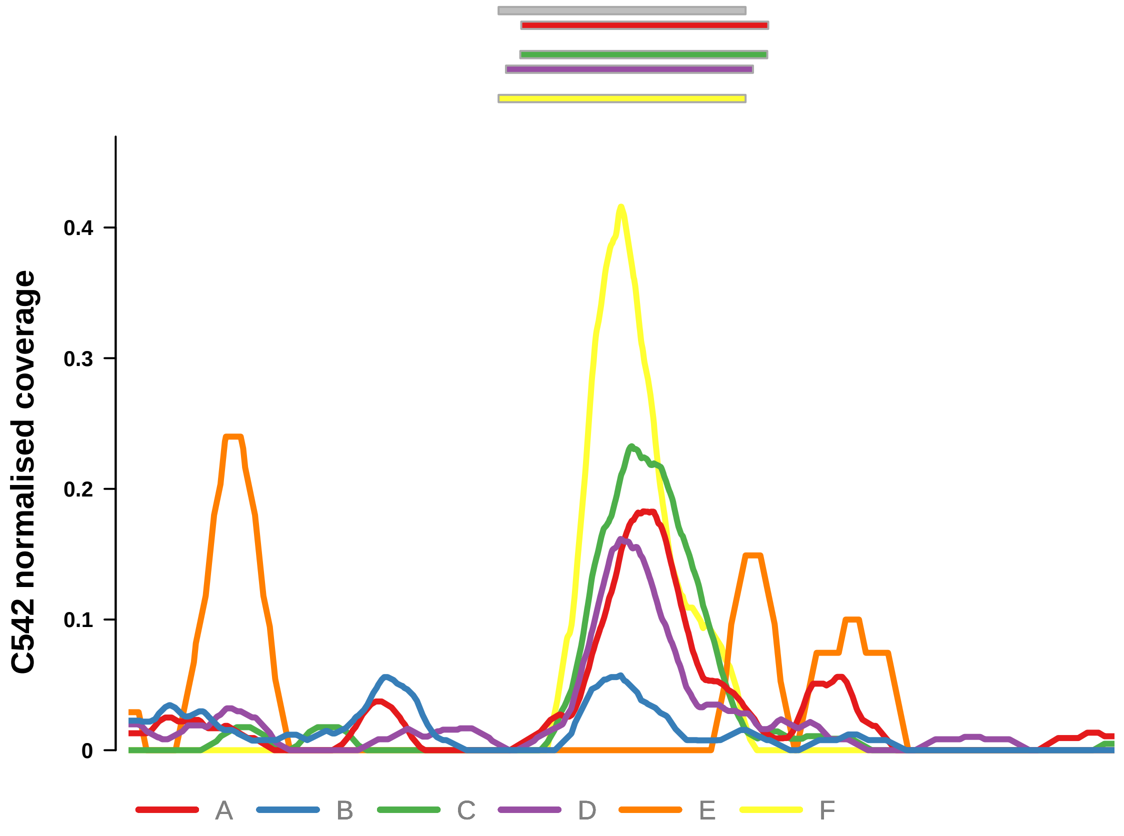


Putatively Clonal (logFC = 2.46)

| | P. | Adjusted P. |
|---|---|---|
| *~region vs. ~* | 1.34e−01 | 5.9e−01 |
| *~purity vs ~* | 7.82e−01 | 9.67e−01 |
| *~purity+region vs ~purity* | 6.63e−02 | 5.5e−01 |
| *Carcinoma vs. normal* | 7.08e−03 ** | 7.08e−03 ** |

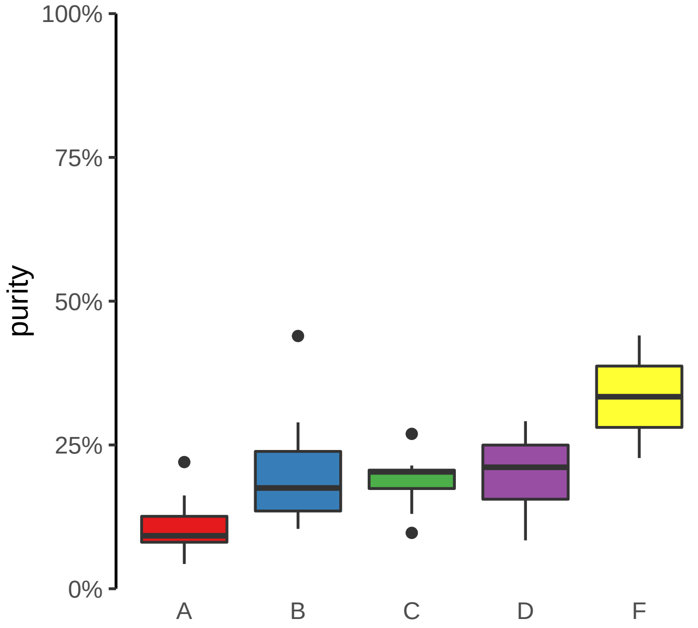

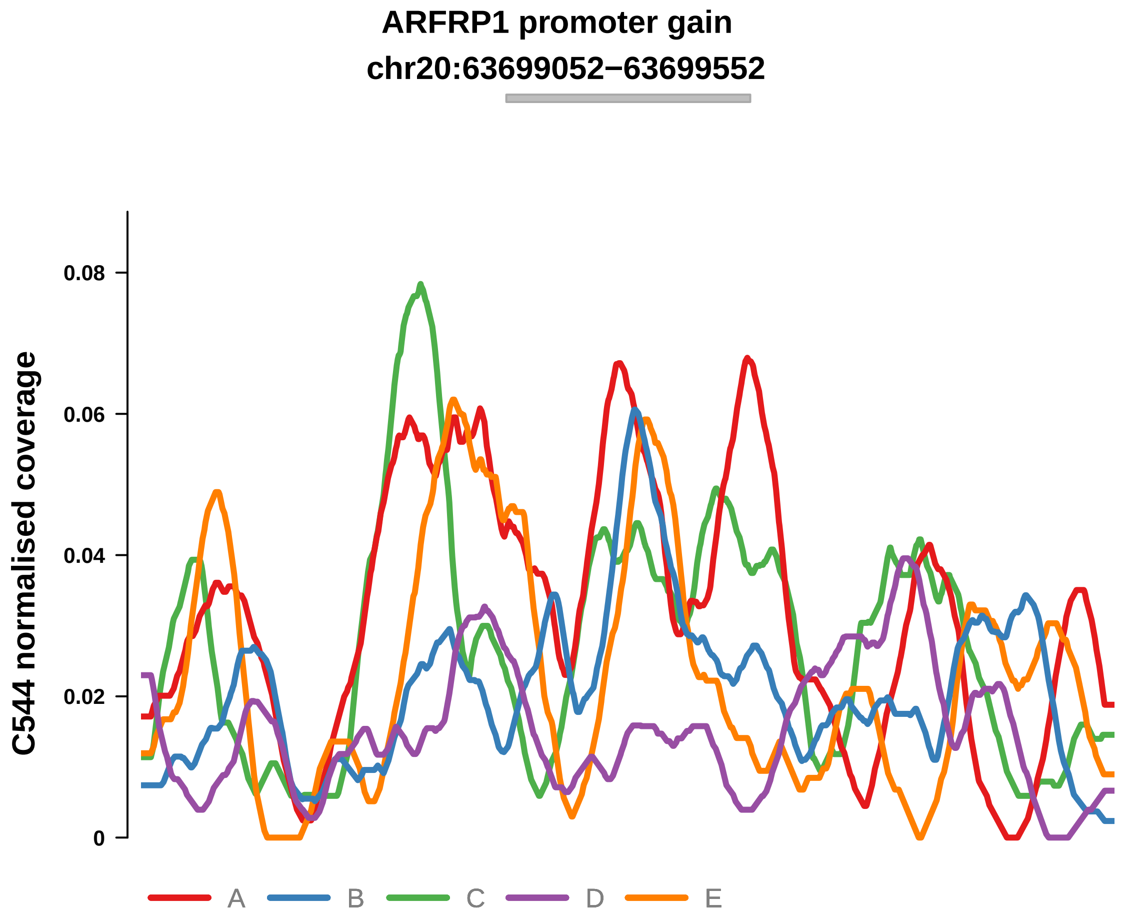


Putatively Clonal (logFC = 1.77)

| | P. | Adjusted P. |
|---|---|---|
| *~region vs. ~* | 3.04e−05 **** | 3.48e−04 *** |
| *~purity vs ~* | 1.42e−02 * | 6.75e−02 |
| *~purity+region vs ~purity* | 4.27e−04 *** | 9.1e−03 ** |
| *Carcinoma vs. normal* | 2.44e−03 ** | 2.44e−03 ** |

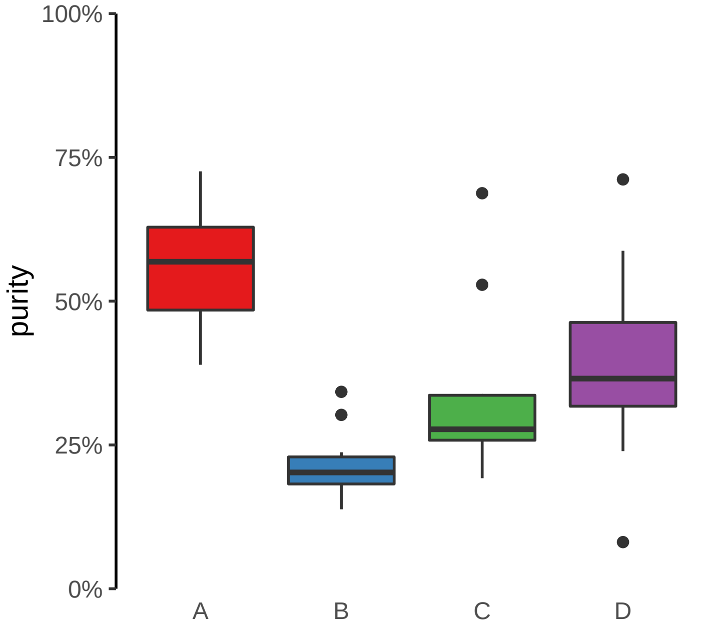

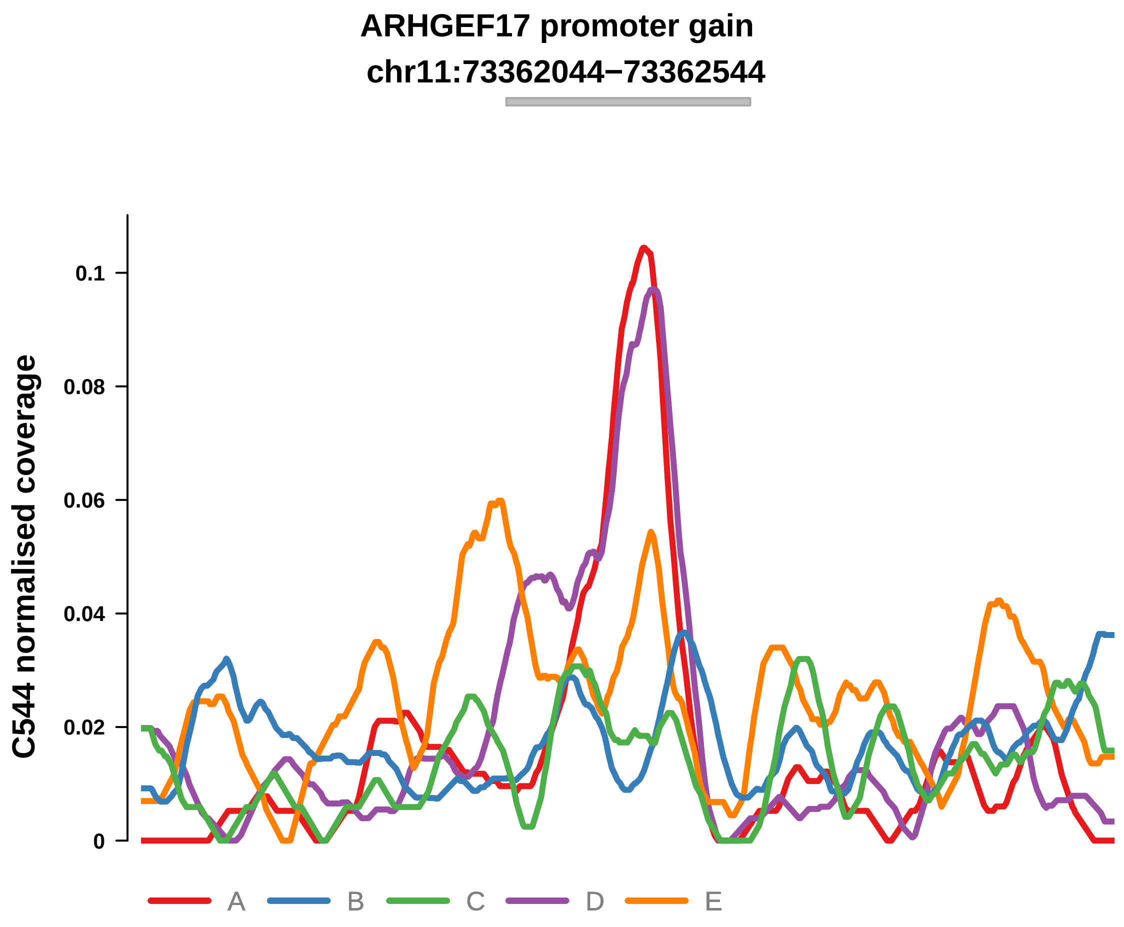


Putatively Clonal (logFC = 2.13)

| | P. | Adjusted P. |
|---|---|---|
| *~region vs. ~* | 3.18e−02 * | 7.77e−02 |
| *~purity vs ~* | 1.68e−01 | 3.24e−01 |
| *~purity+region vs ~purity* | 9.44e−02 | 1.83e−01 |
| *Carcinoma vs. normal* | 3.35e−04 *** | 3.35e−04 *** |

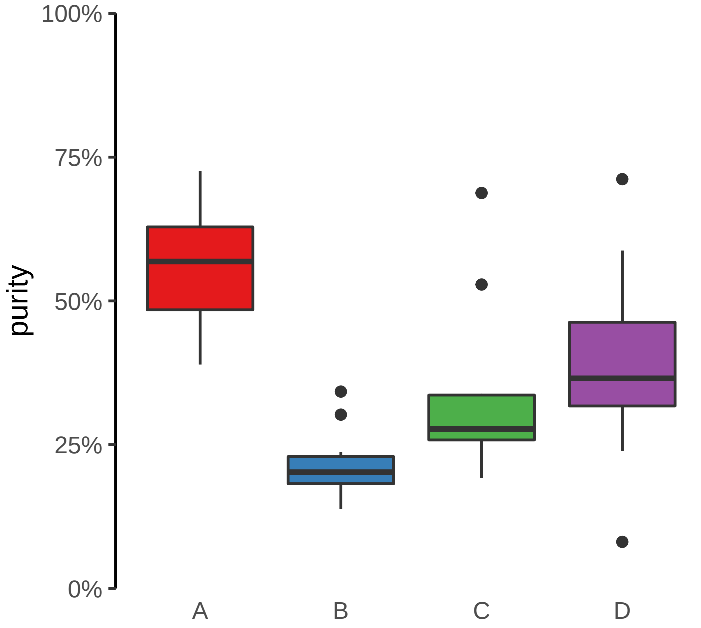

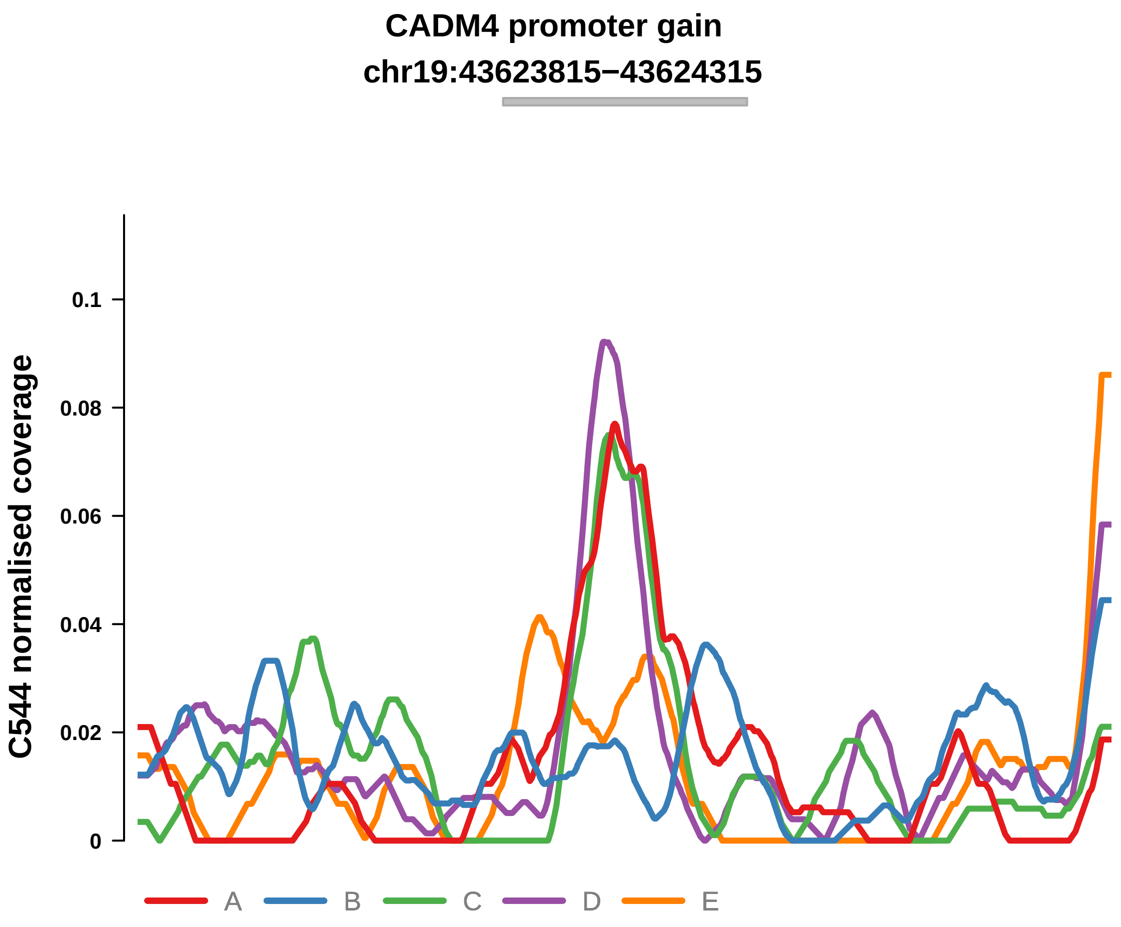


Putatively Clonal (logFC = 2.12)

| | P. | Adjusted P. |
|---|---|---|
| *~region vs. ~* | 1.78e−01 | 2.71e−01 |
| *~purity vs ~* | 2.99e−01 | 4.94e−01 |
| *~purity+region vs ~purity* | 2.47e−01 | 4.27e−01 |
| *Carcinoma vs. normal* | 4.43e−04 *** | 4.43e−04 *** |

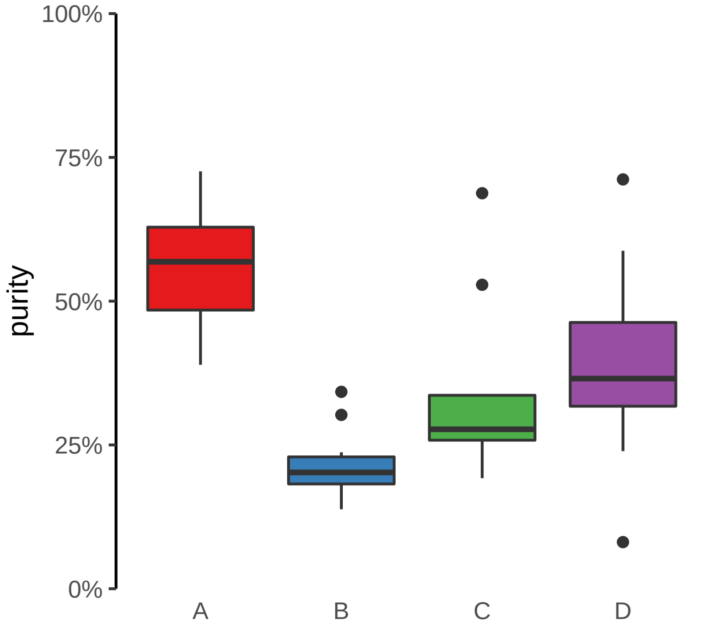

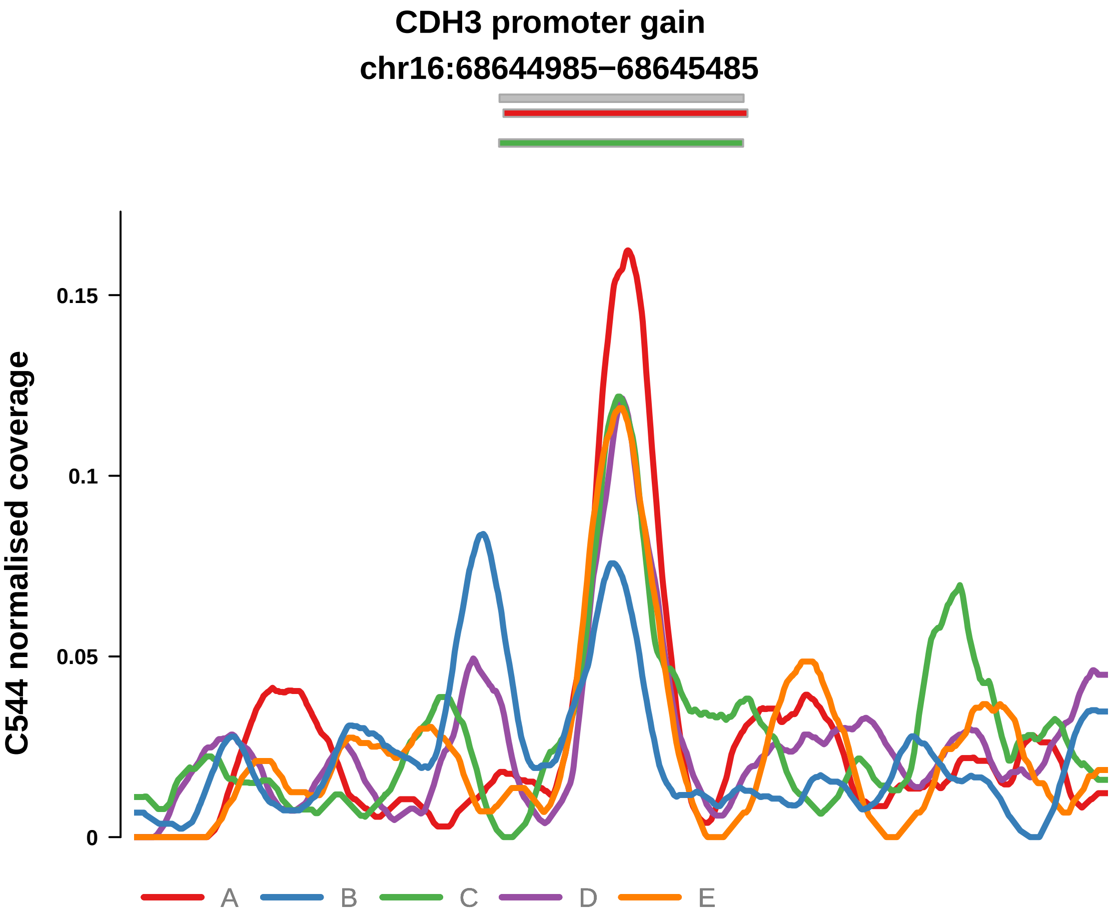


Putatively Clonal (logFC = 1.6)

| | P. | Adjusted P. |
|---|---|---|
| *~region vs. ~* | 3.04e−01 | 4.12e−01 |
| *~purity vs ~* | 1.86e−01 | 3.42e−01 |
| *~purity+region vs ~purity* | 6.11e−01 | 7e−01 |
| *Carcinoma vs. normal* | 4.62e−03 ** | 4.62e−03 ** |

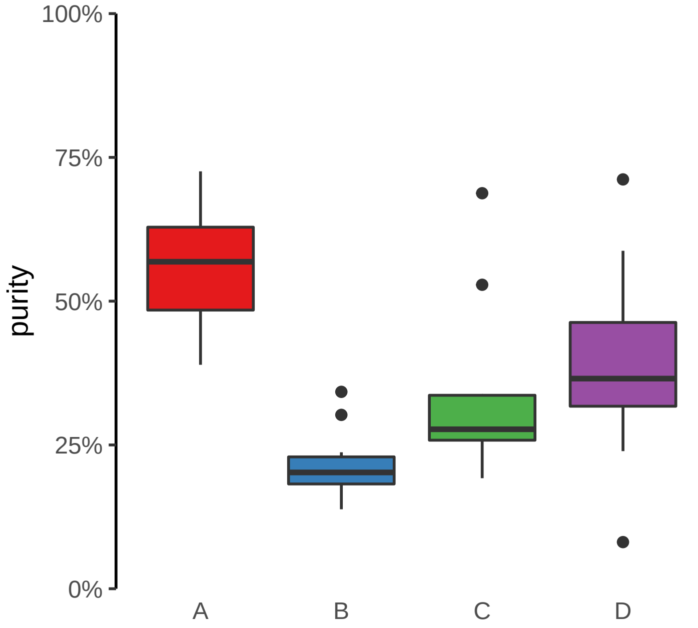

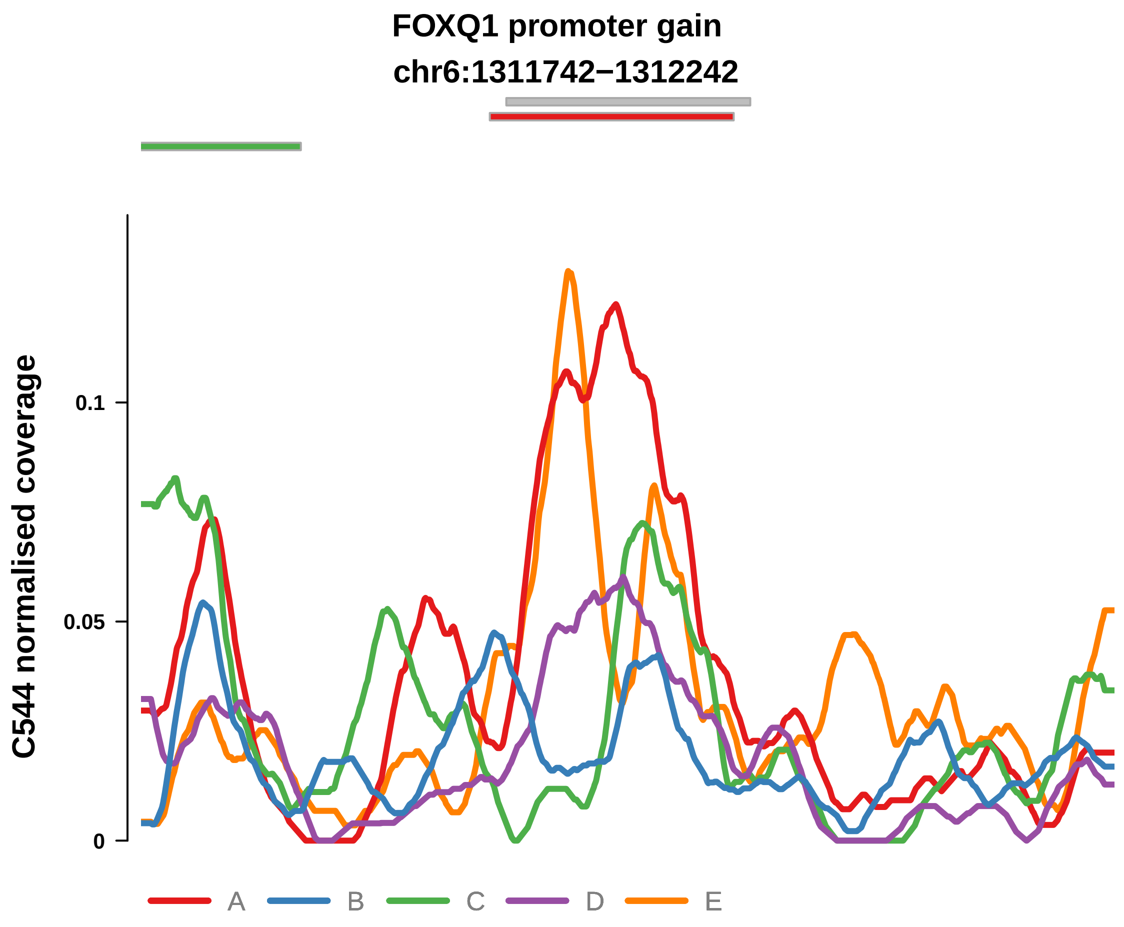


Putatively Clonal (logFC = 1.92)

| | P. | Adjusted P. |
|---|---|---|
| *~region vs. ~* | 3.16e−05 **** | 3.48e−04 *** |
| *~purity vs ~* | 1.47e−06 **** | 1.19e−04 *** |
| *~purity+region vs ~purity* | 2.67e−01 | 4.5e−01 |
| *Carcinoma vs. normal* | 6.24e−04 *** | 6.24e−04 *** |

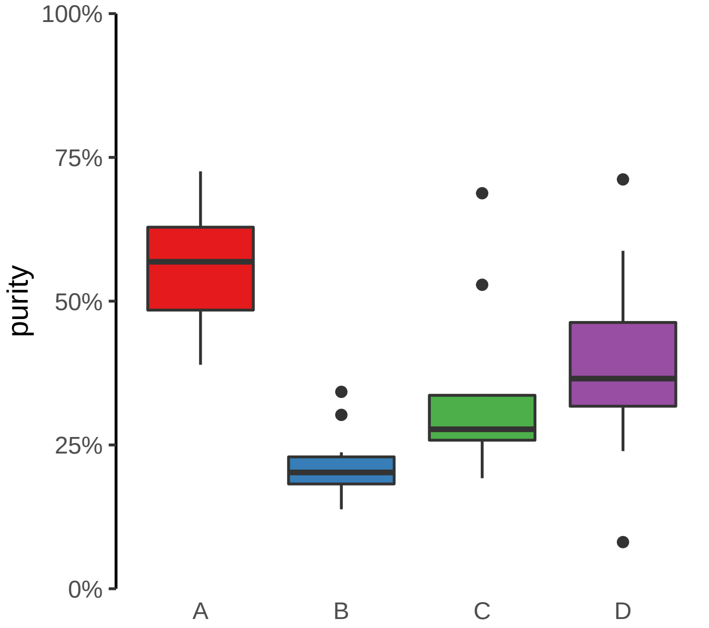

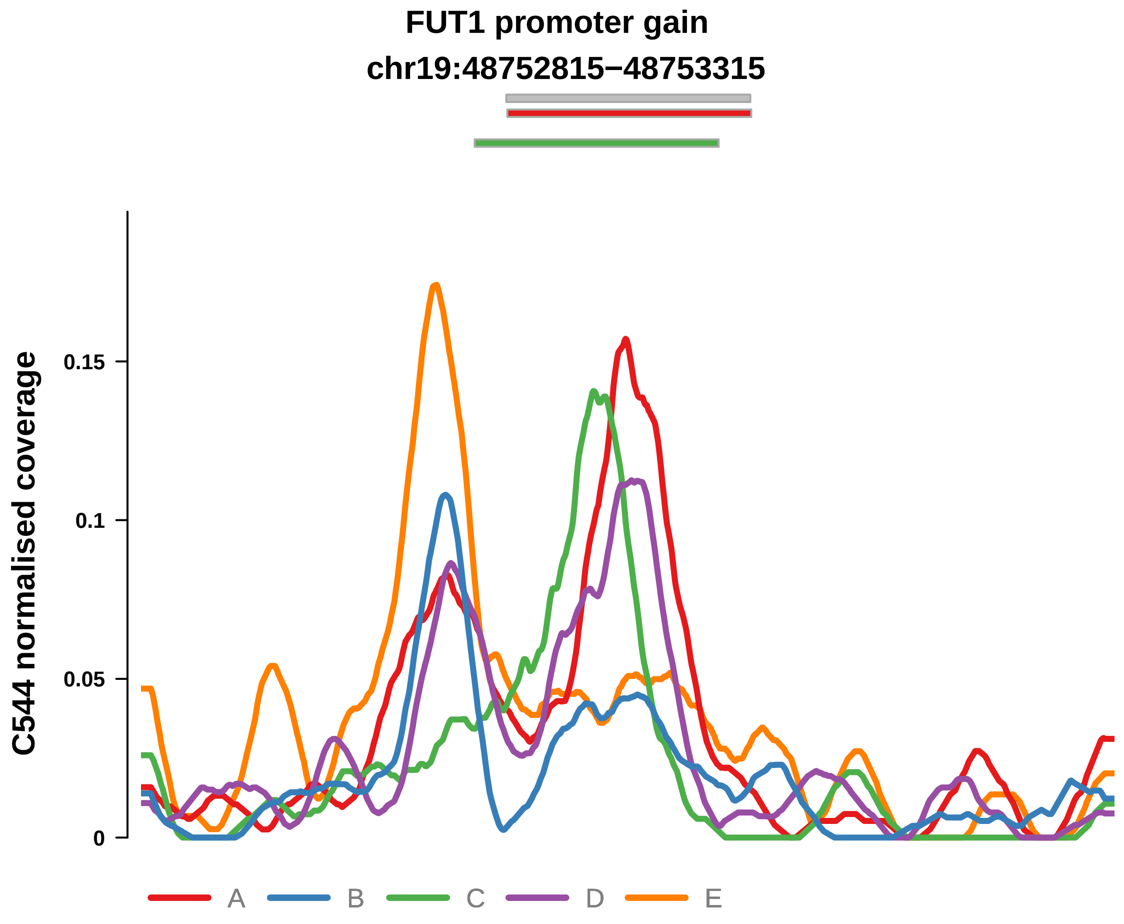


Putatively Clonal (logFC = 2.22)

| | P. | Adjusted P. |
|---|---|---|
| *~region vs. ~* | 5.54e−04 *** | 3.75e−03 ** |
| *~purity vs ~* | 2.24e−03 ** | 2.02e−02 * |
| *~purity+region vs ~purity* | 6.3e−02 | 1.36e−01 |
| *Carcinoma vs. normal* | 8.62e−05 **** | 8.62e−05 **** |

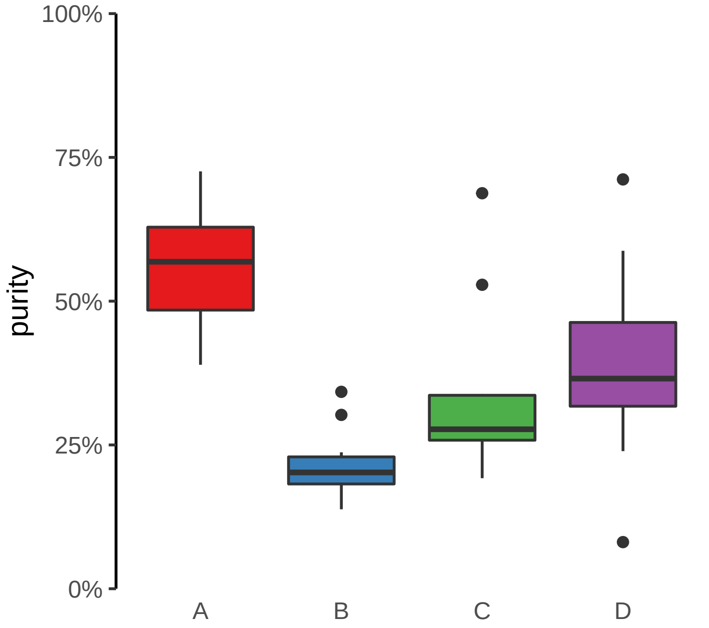

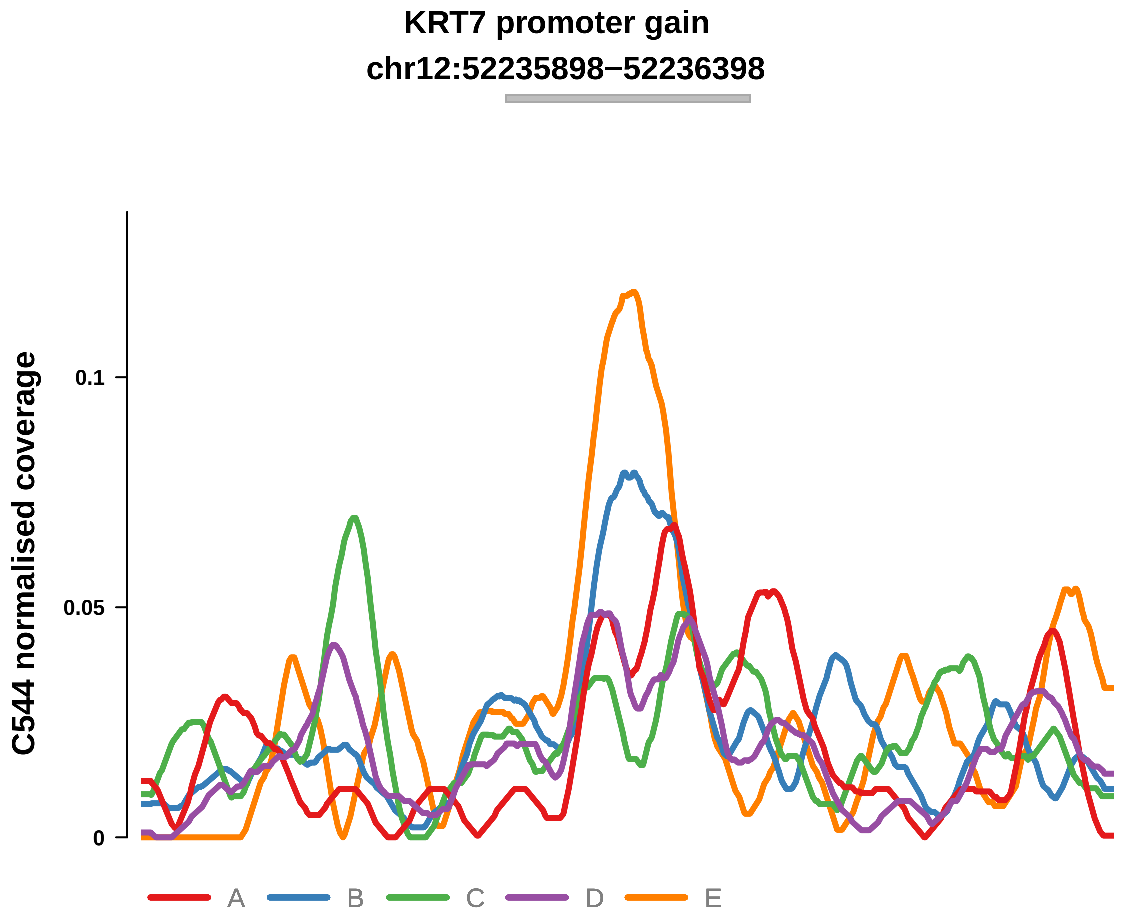


Putatively Clonal (logFC = 1.52)

| | P. | Adjusted P. |
|---|---|---|
| *~region vs. ~* | 1.28e−01 | 2.08e−01 |
| *~purity vs ~* | 1.38e−01 | 2.79e−01 |
| *~purity+region vs ~purity* | 2.35e−02 * | 8.64e−02 |
| *Carcinoma vs. normal* | 8.19e−03 ** | 8.19e−03 ** |

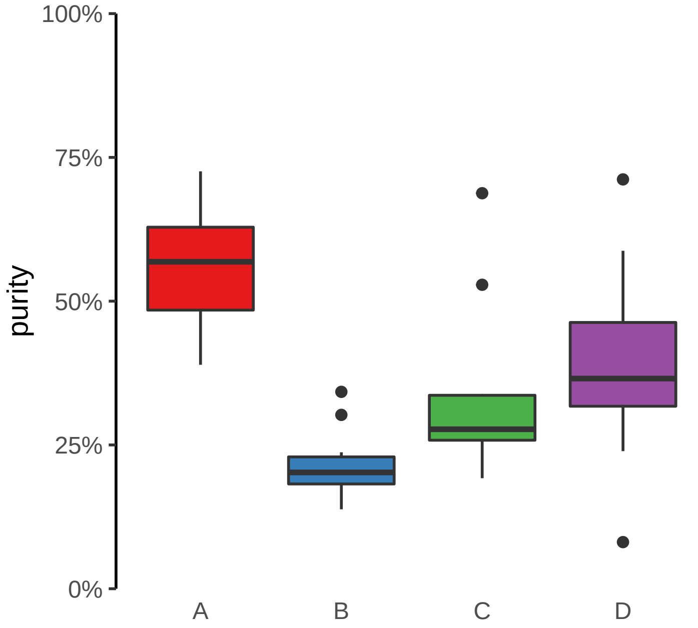

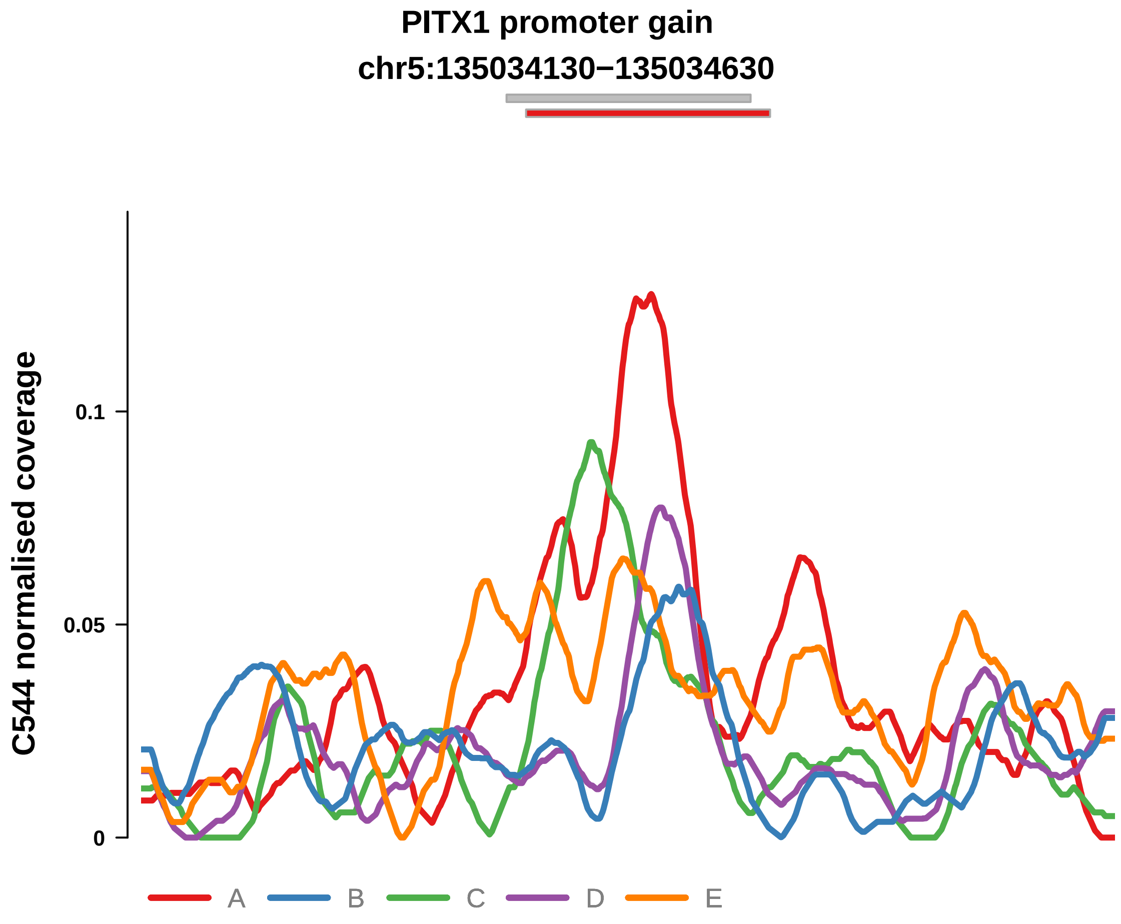


Putatively Clonal (logFC = 2.09)

| | P. | Adjusted P. |
|---|---|---|
| *~region vs. ~* | 5.57e−03 ** | 2.04e−02 * |
| *~purity vs ~* | 3.21e−02 * | 9.75e−02 |
| *~purity+region vs ~purity* | 6.37e−02 | 1.36e−01 |
| *Carcinoma vs. normal* | 2.38e−04 *** | 2.38e−04 *** |

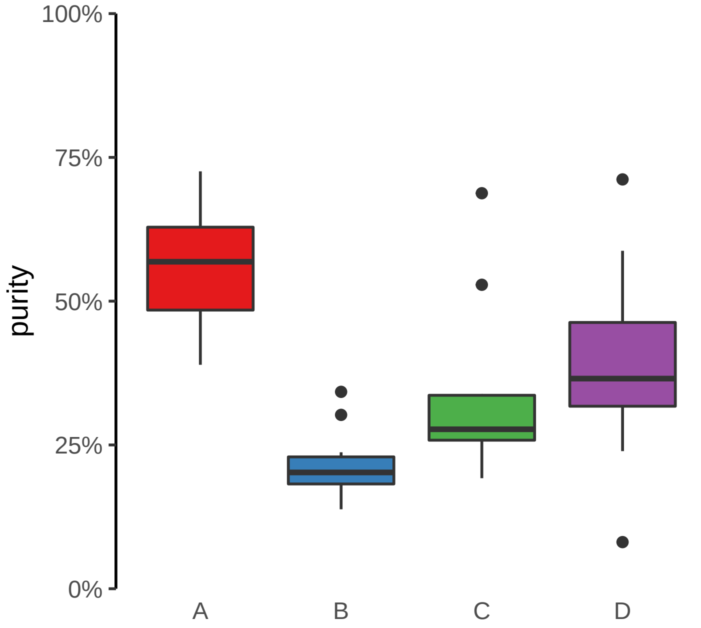

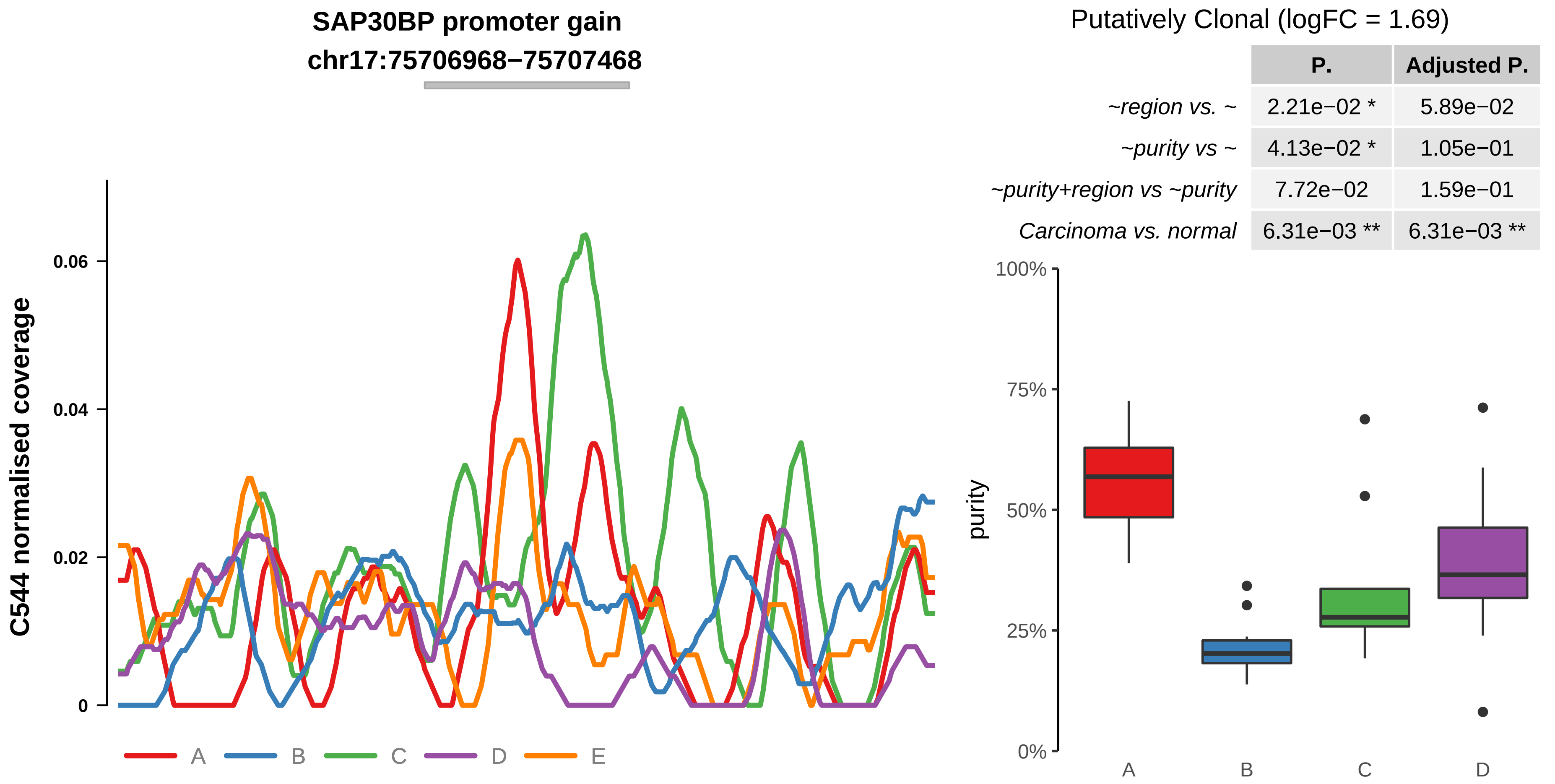


| | P. | Adjusted P. |
|---|---|---|
| *~region vs. ~* | 2.21e−02 * | 5.89e−02 |
| *~purity vs ~* | 4.13e−02 * | 1.05e−01 |
| *~purity+region vs ~purity* | 7.72e−02 | 1.59e−01 |
| *Carcinoma vs. normal* | 6.31e−03 ** | 6.31e−03 ** |

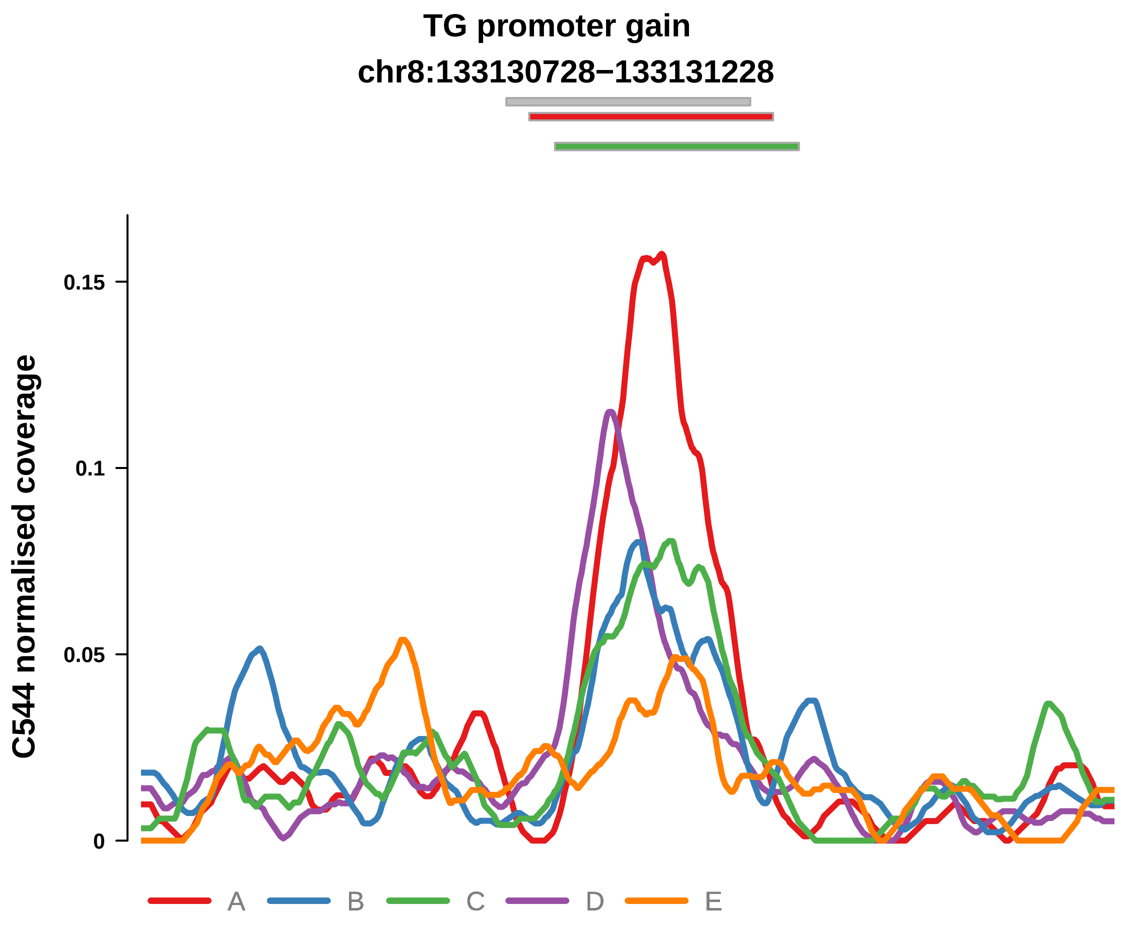


Putatively Clonal (logFC = 2.55)

| | P. | Adjusted P. |
|---|---|---|
| *~region vs. ~* | 8.7e−02 | 1.63e−01 |
| *~purity vs ~* | 6.76e−02 | 1.61e−01 |
| *~purity+region vs ~purity* | 4.06e−01 | 5.62e−01 |
| *Carcinoma vs. normal* | 9.79e−06 **** | 9.79e−06 **** |

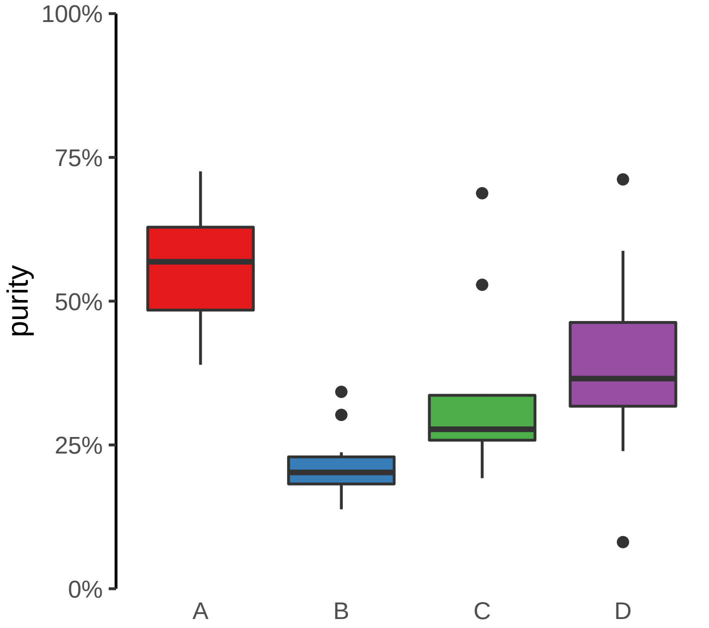

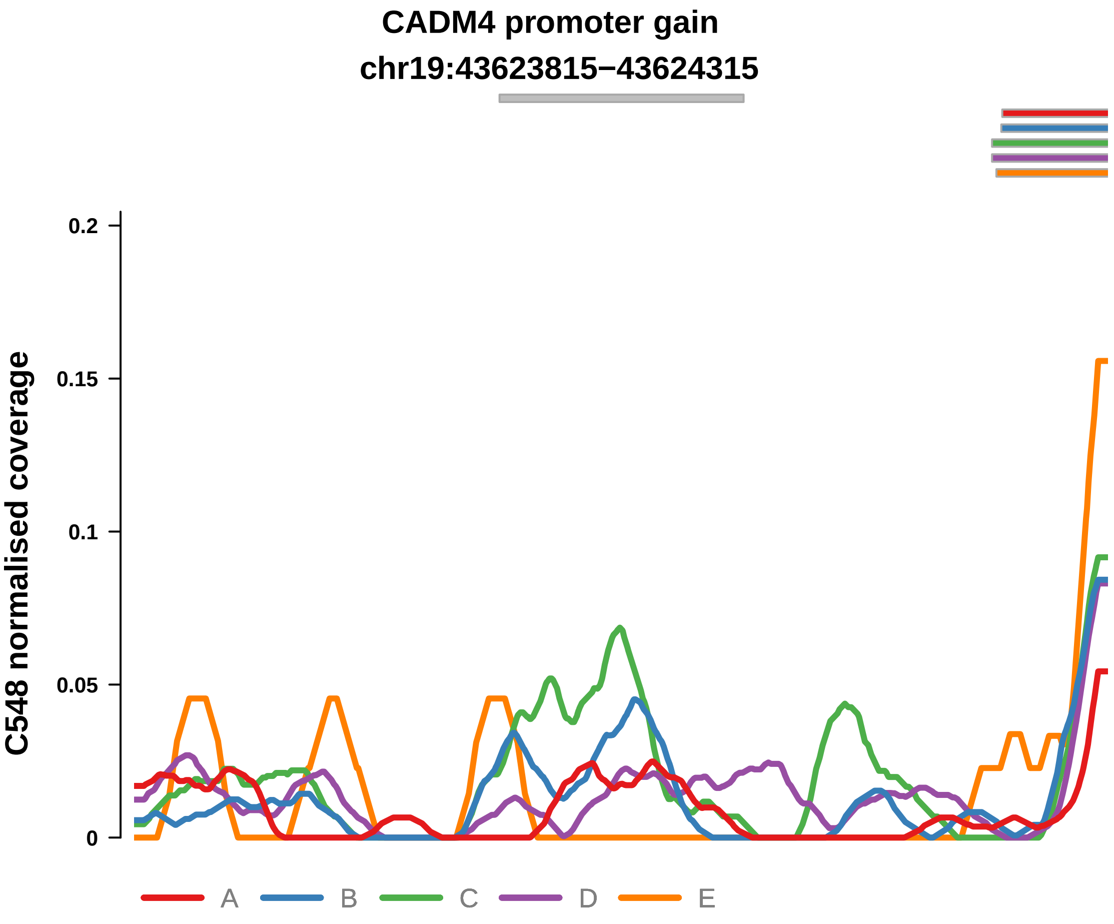


Putatively Clonal (logFC = 1.85)

| | P. | Adjusted P. |
|---|---|---|
| *~region vs. ~* | 8.86e−02 | 2e−01 |
| *~purity vs ~* | 1.78e−01 | 3.82e−01 |
| *~purity+region vs ~purity* | 1.18e−01 | 3.85e−01 |
| *Carcinoma vs. normal* | 6.66e−03 ** | 6.66e−03 ** |

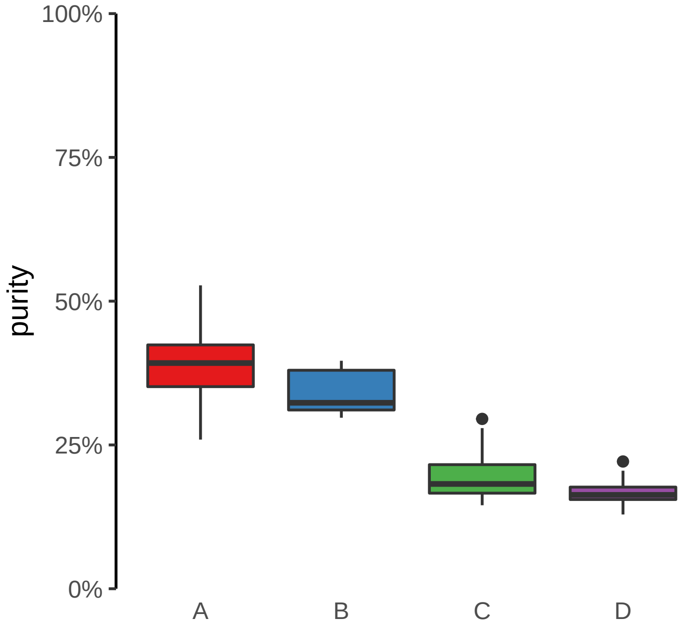

**EHD2 promoter gain**
**chr19:47730036−47730536**

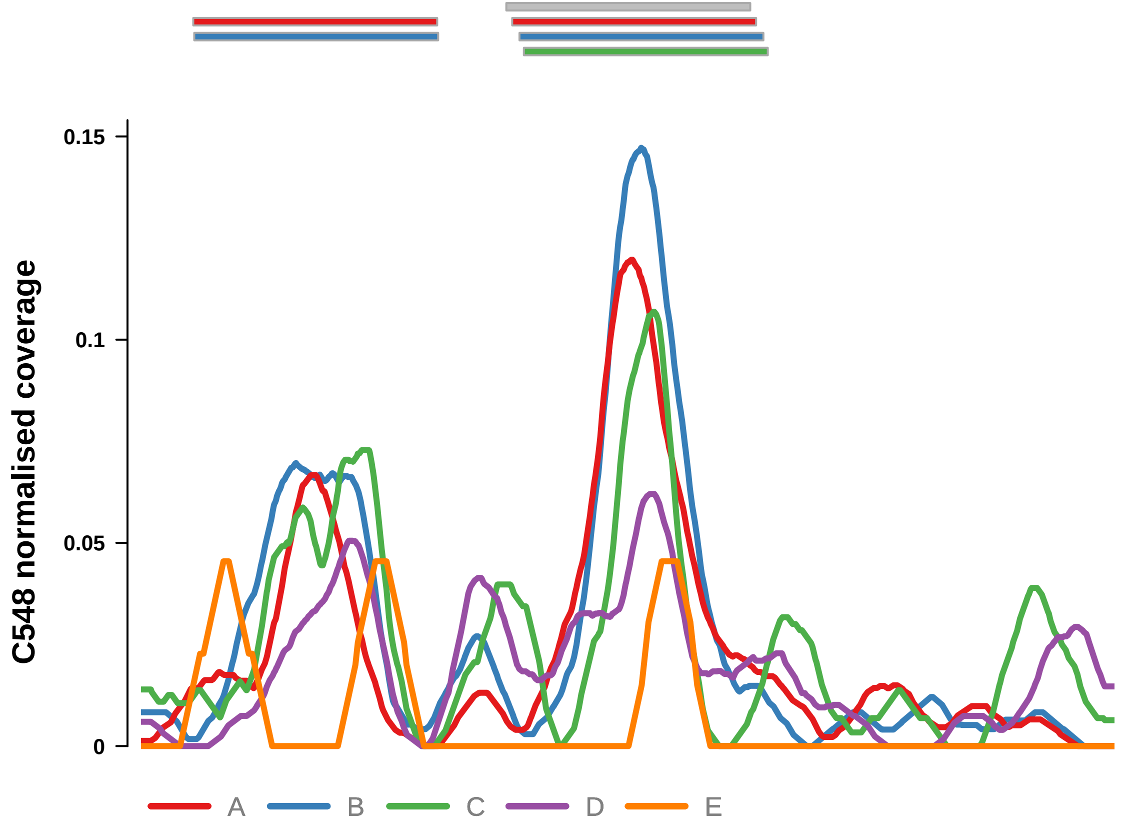


Putatively Clonal (logFC = 2.62)

| | **P.** | **Adjusted P.** |
|---|---|---|
| *~region vs. ~* | 8.83e−01 | 9.21e−01 |
| *~purity vs ~* | 8.54e−01 | 8.95e−01 |
| *~purity+region vs ~purity* | 8.88e−01 | 9.52e−01 |
| *Carcinoma vs. normal* | 1.17e−05 **** | 1.17e−05 **** |

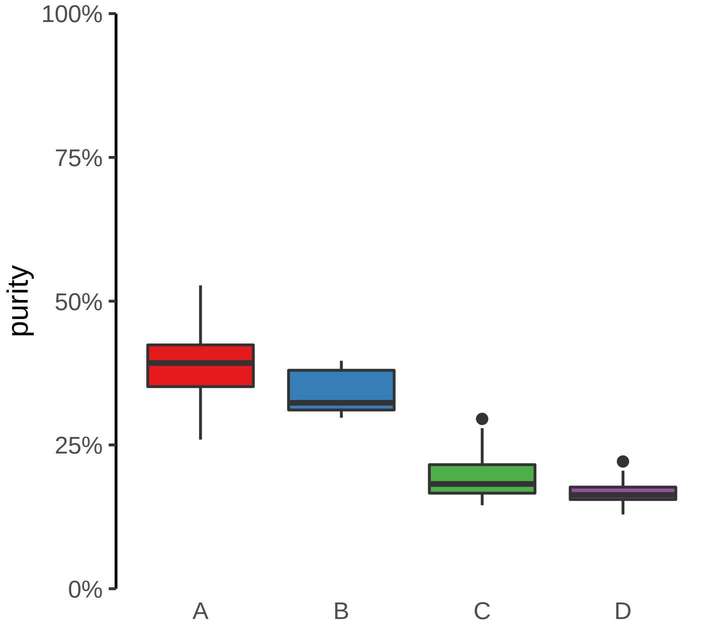

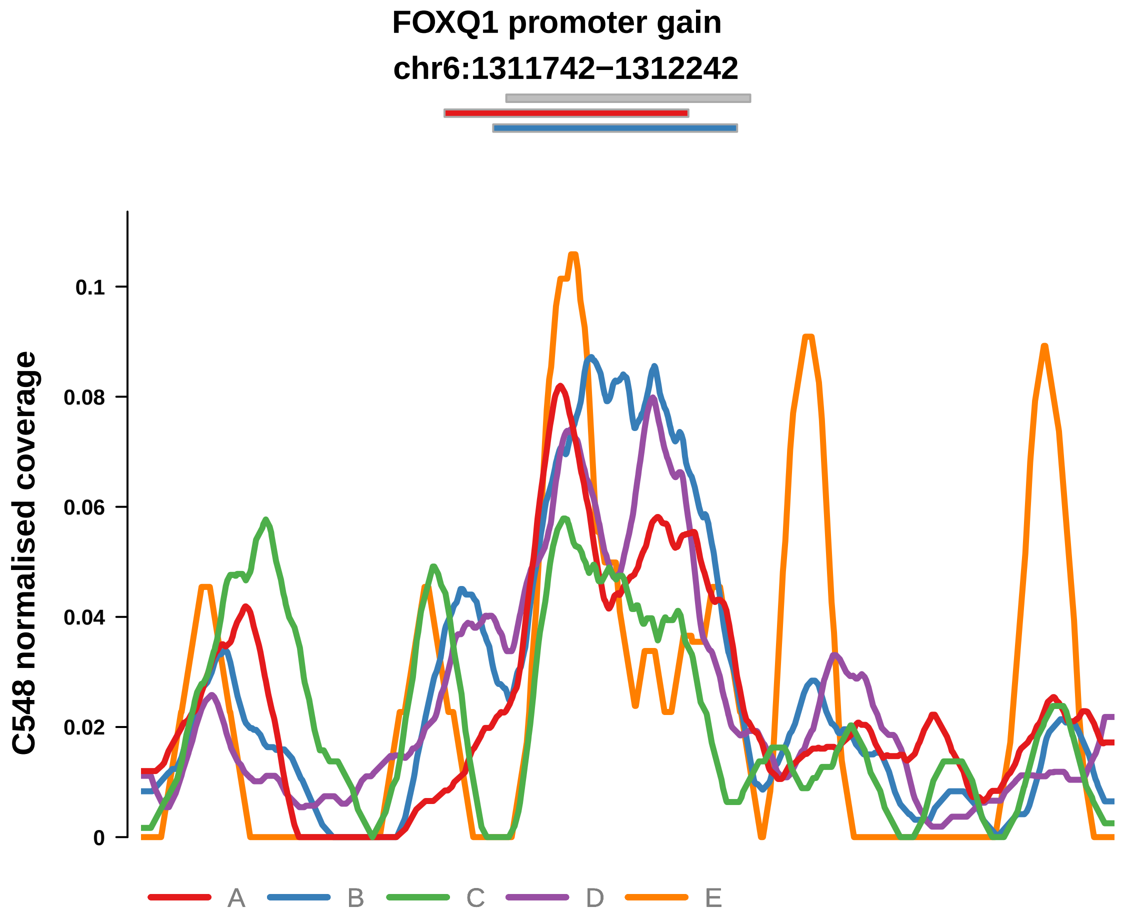


Putatively Clonal (logFC = 2.08)

| | P. | Adjusted P. |
|---|---|---|
| *~region vs. ~* | 4.08e−02 * | 1.35e−01 |
| *~purity vs ~* | 6.97e−01 | 8.07e−01 |
| *~purity+region vs ~purity* | 3.13e−02 * | 2.12e−01 |
| *Carcinoma vs. normal* | 3.72e−04 *** | 3.72e−04 *** |

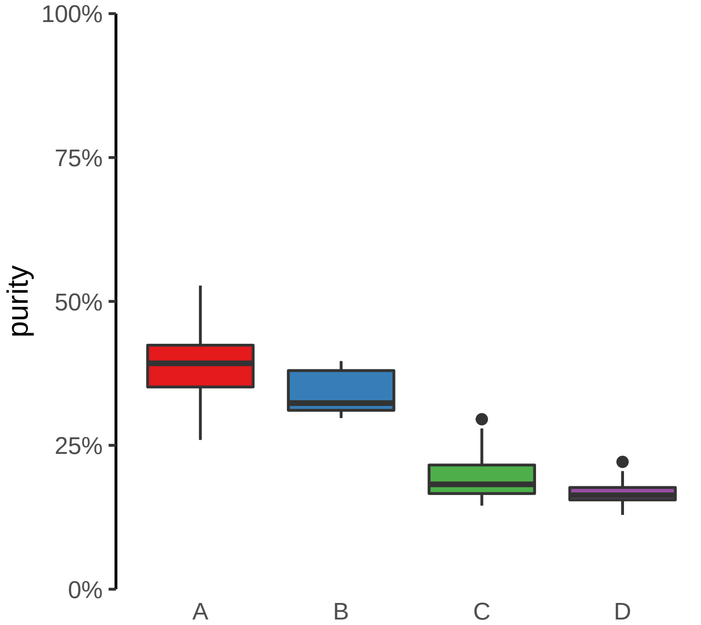

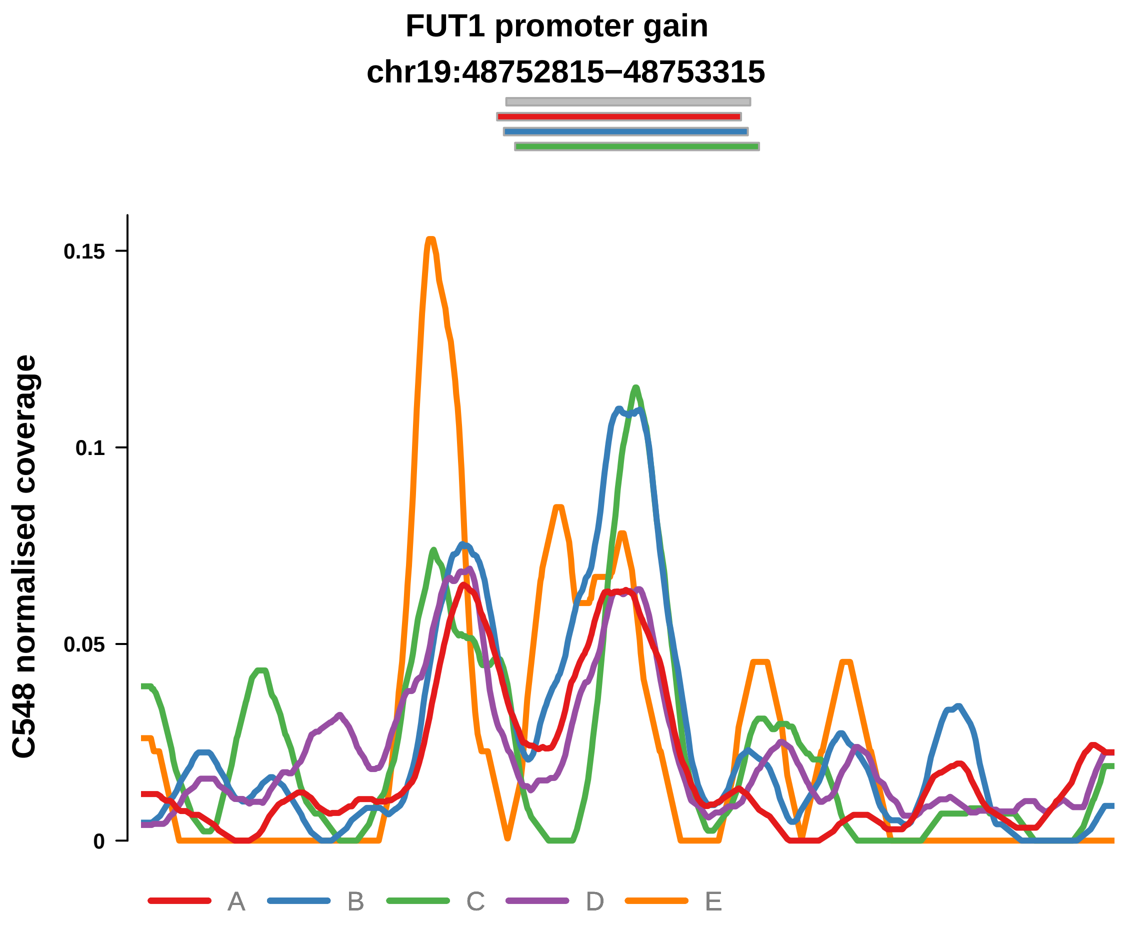


Putatively Clonal (logFC = 1.86)

| | P. | Adjusted P. |
|---|---|---|
| *~region vs. ~* | 8.92e−01 | 9.21e−01 |
| *~purity vs ~* | 8.3e−01 | 8.81e−01 |
| *~purity+region vs ~purity* | 7.73e−01 | 9.15e−01 |
| *Carcinoma vs. normal* | 2.12e−03 ** | 2.12e−03 ** |

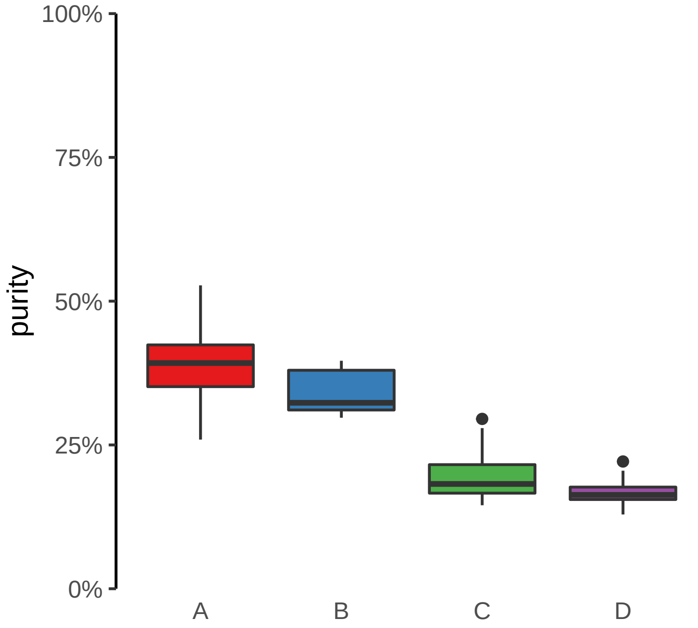

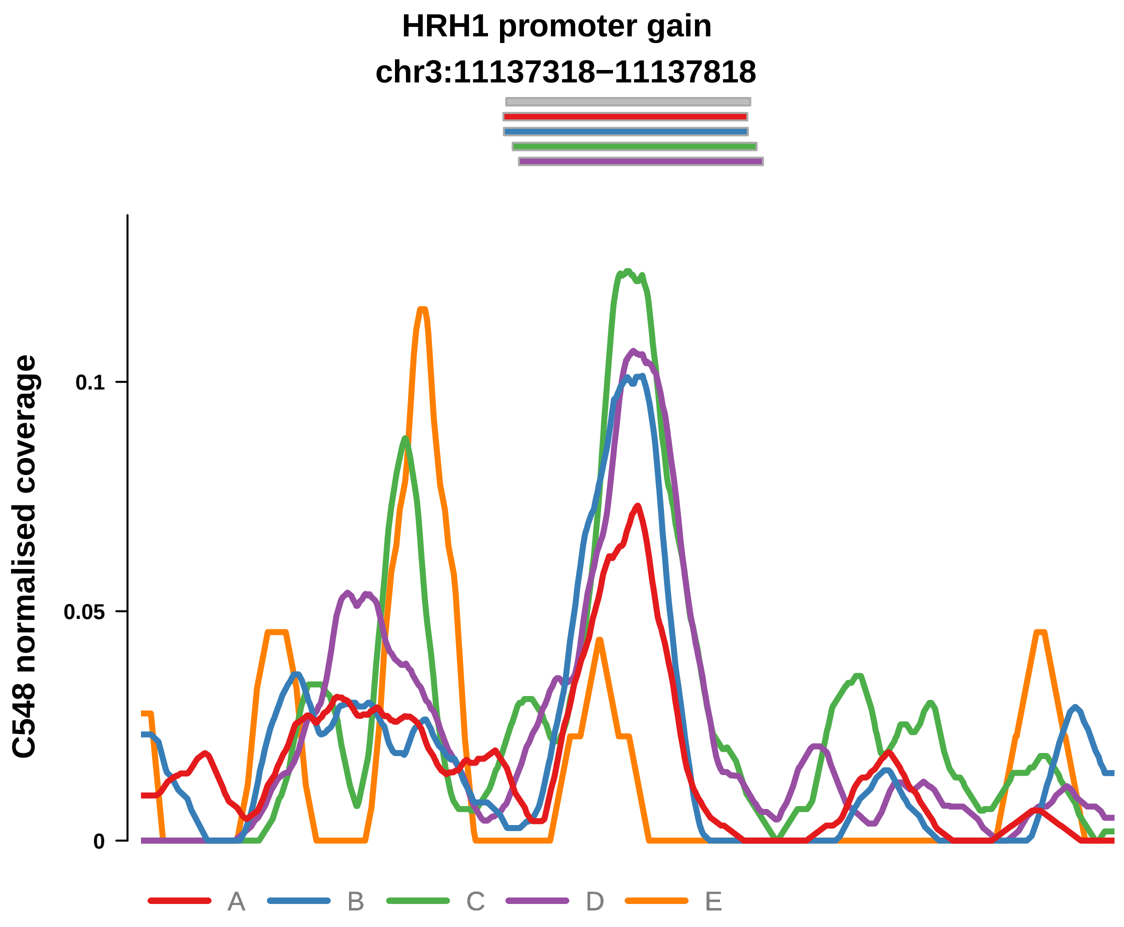


Putatively Clonal (logFC = 1.77)

| | P. | Adjusted P. |
|---|---|---|
| *~region vs. ~* | 1.68e−03 ** | 1.85e−02 * |
| *~purity vs ~* | 1.06e−03 ** | 1.55e−02 * |
| *~purity+region vs ~purity* | 3.49e−01 | 5.86e−01 |
| *Carcinoma vs. normal* | 6.05e−03 ** | 6.05e−03 ** |

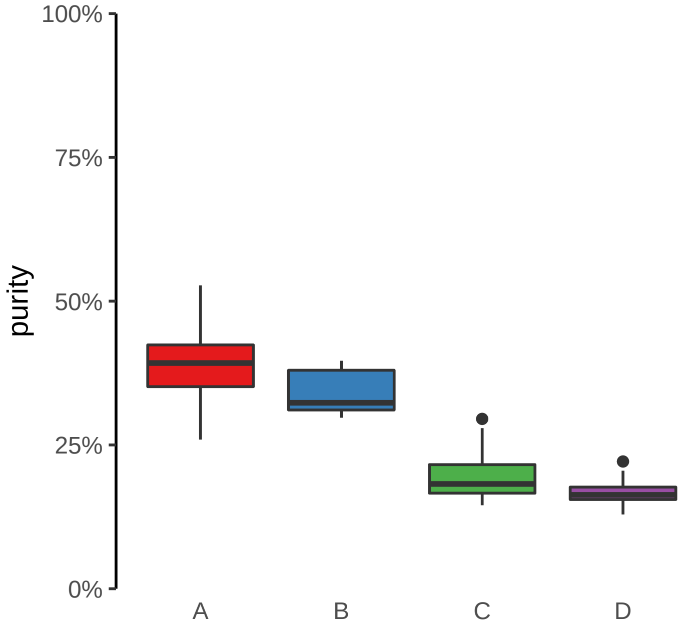

**KLK6 promoter gain**
**chr19:50969271−50969771**

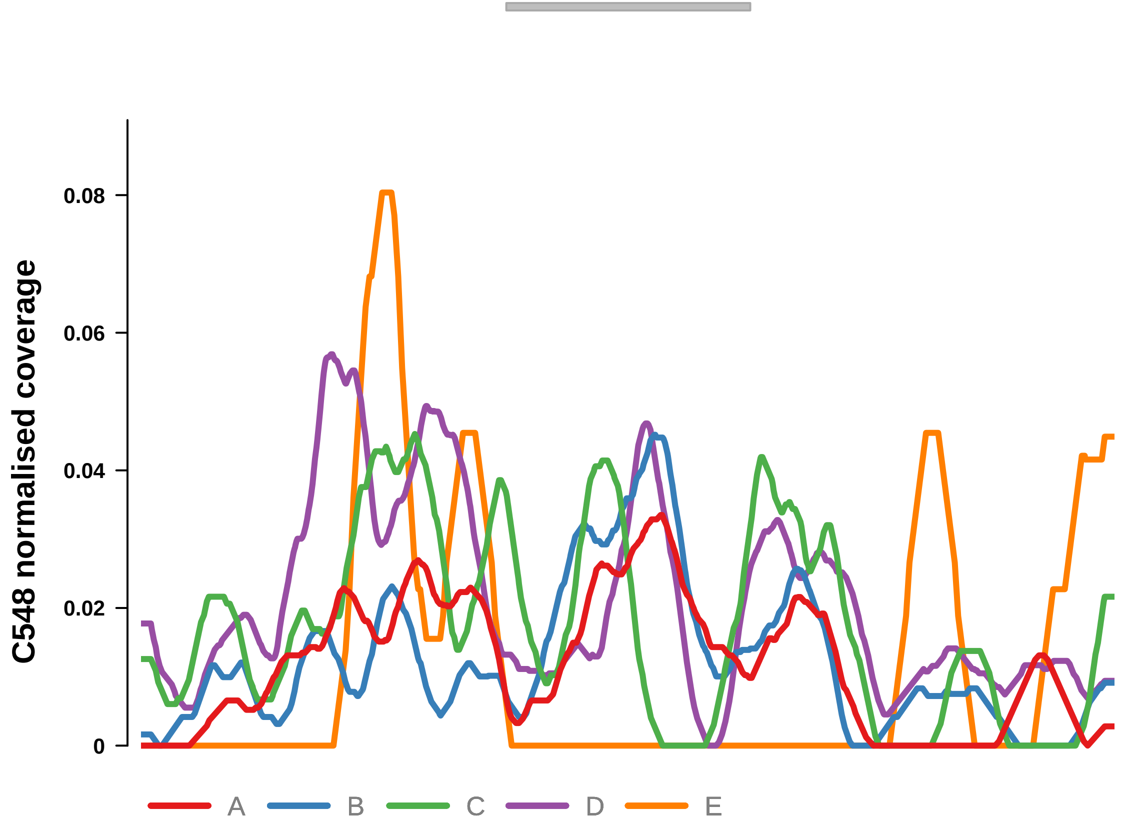


Putatively Clonal (logFC = 1.74)

| | P. | Adjusted P. |
|---|---|---|
| *~region vs. ~* | 9e−01 | 9.21e−01 |
| *~purity vs ~* | 6.32e−01 | 7.58e−01 |
| *~purity+region vs ~purity* | 8.95e−01 | 9.52e−01 |
| *Carcinoma vs. normal* | 9.01e−03 ** | 9.01e−03 ** |

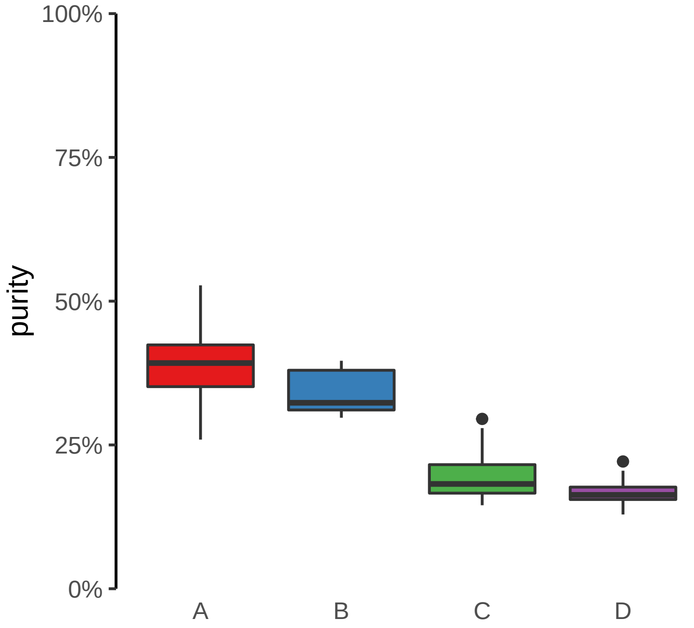

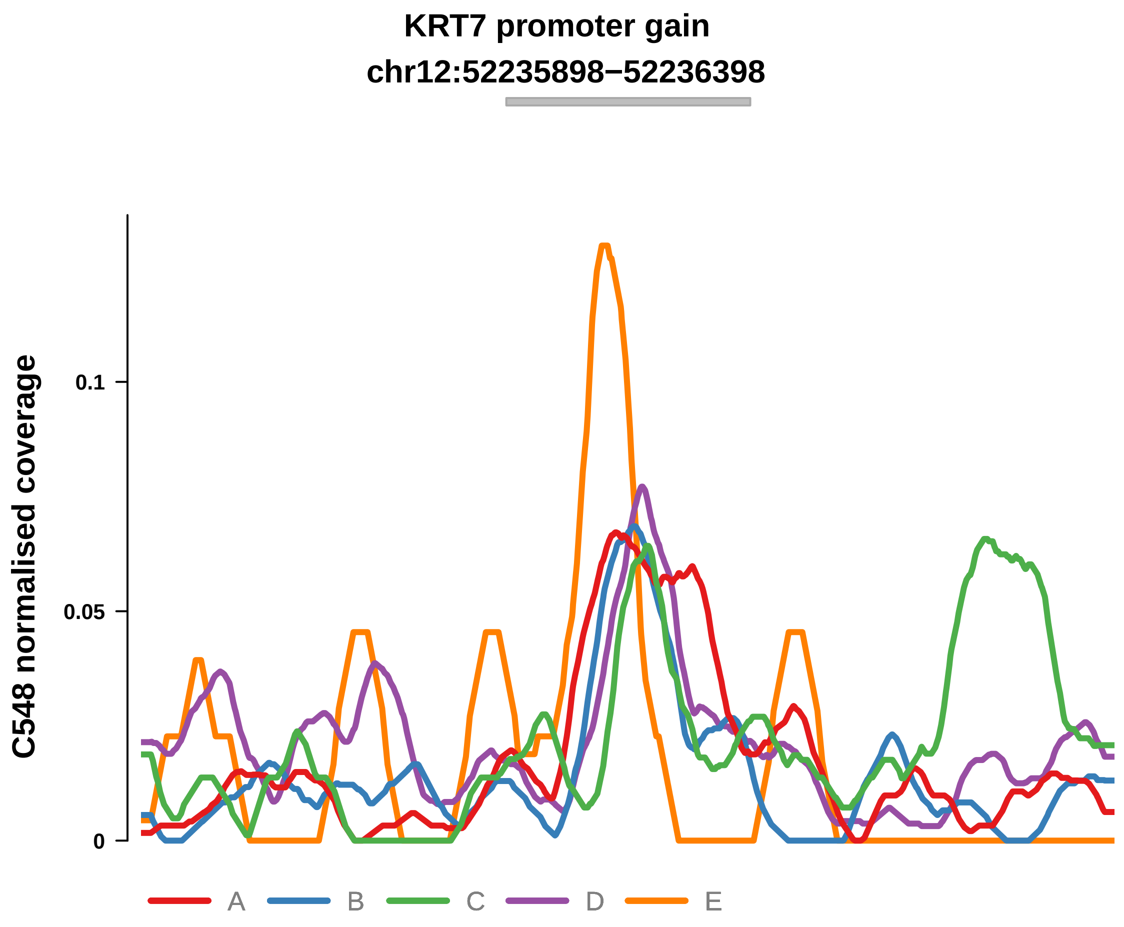


Putatively Clonal (logFC = 1.91)

| | P. | Adjusted P. |
|---|---|---|
| *~region vs. ~* | 9.55e−02 | 2.1e−01 |
| *~purity vs ~* | 4.26e−01 | 6.24e−01 |
| *~purity+region vs ~purity* | 1.31e−01 | 4.01e−01 |
| *Carcinoma vs. normal* | 1.56e−03 ** | 1.56e−03 ** |

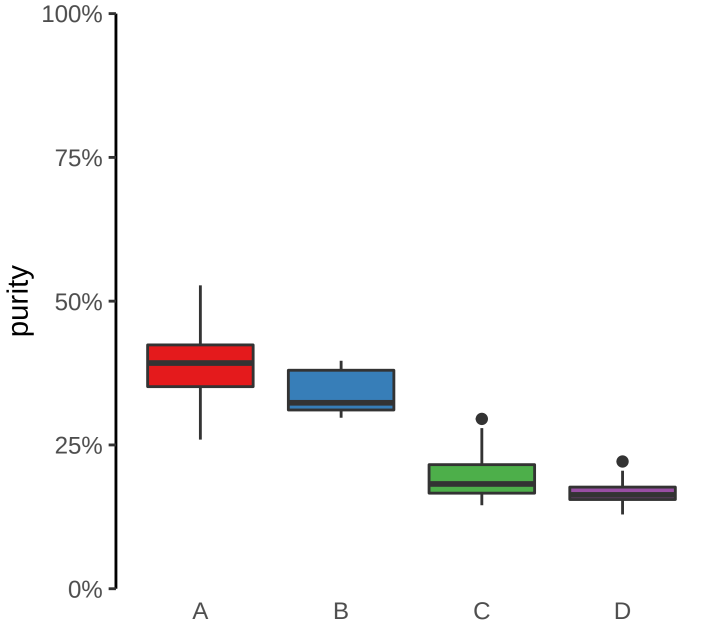

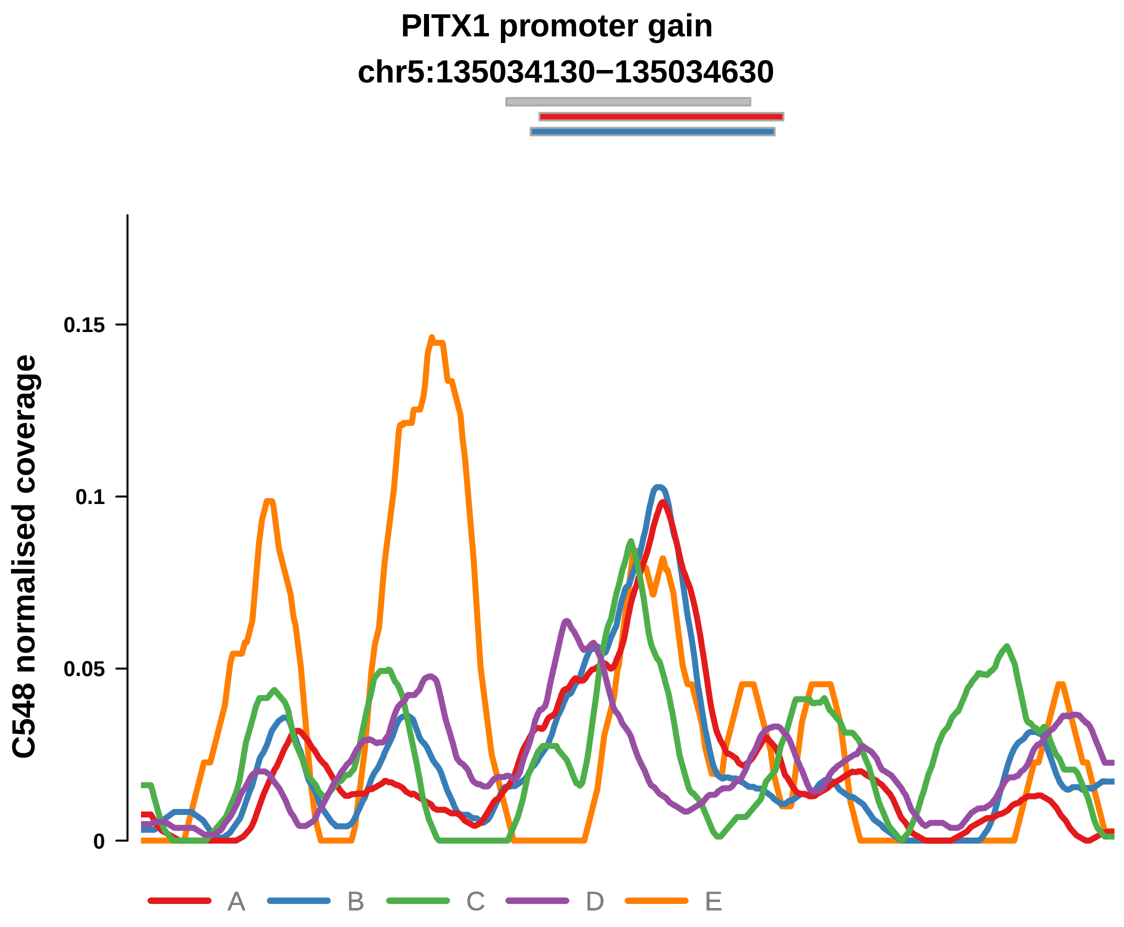


Putatively Clonal (logFC = 2.41)

| | P. | Adjusted P. |
|---|---|---|
| *~region vs. ~* | 1.12e−01 | 2.26e−01 |
| *~purity vs ~* | 6.38e−01 | 7.58e−01 |
| *~purity+region vs ~purity* | 1.07e−01 | 3.66e−01 |
| *Carcinoma vs. normal* | 4.09e−05 **** | 4.09e−05 **** |

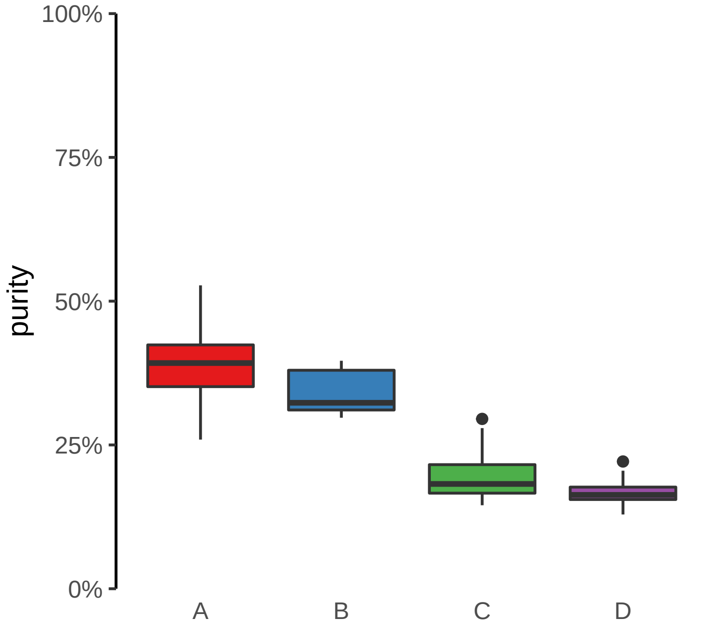

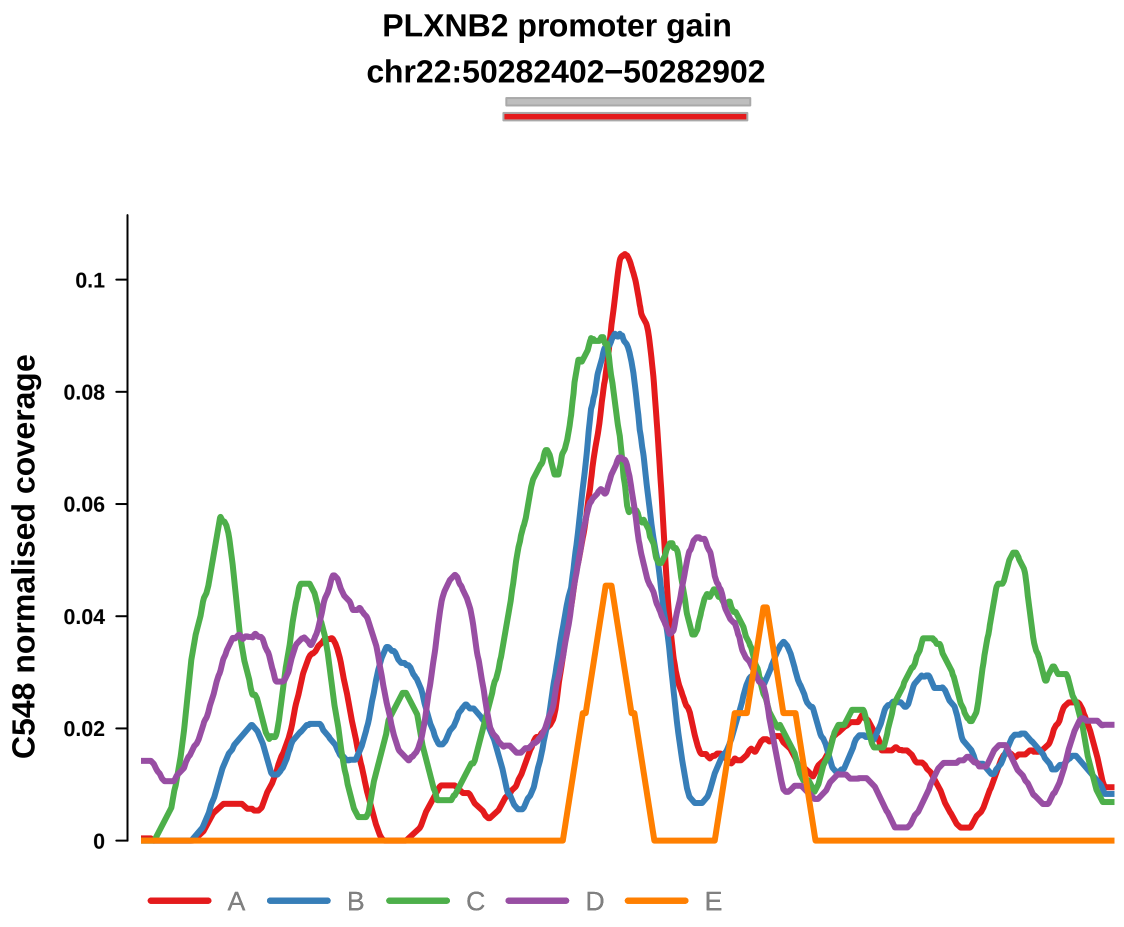


Putatively Clonal (logFC = 2.41)

| | P. | Adjusted P. |
|---|---|---|
| *~region vs. ~* | 8.71e−02 | 2e−01 |
| *~purity vs ~* | 3.94e−02 * | 1.93e−01 |
| *~purity+region vs ~purity* | 5.55e−01 | 7.44e−01 |
| *Carcinoma vs. normal* | 6.99e−05 **** | 6.99e−05 **** |

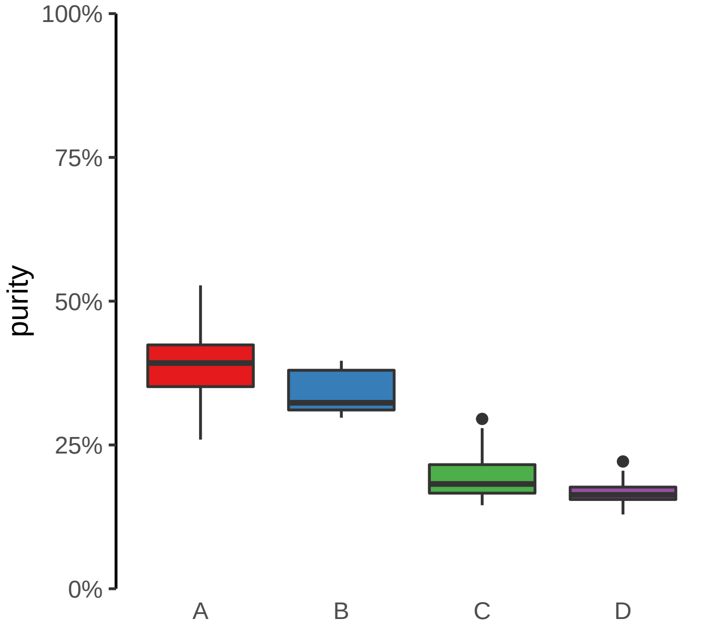

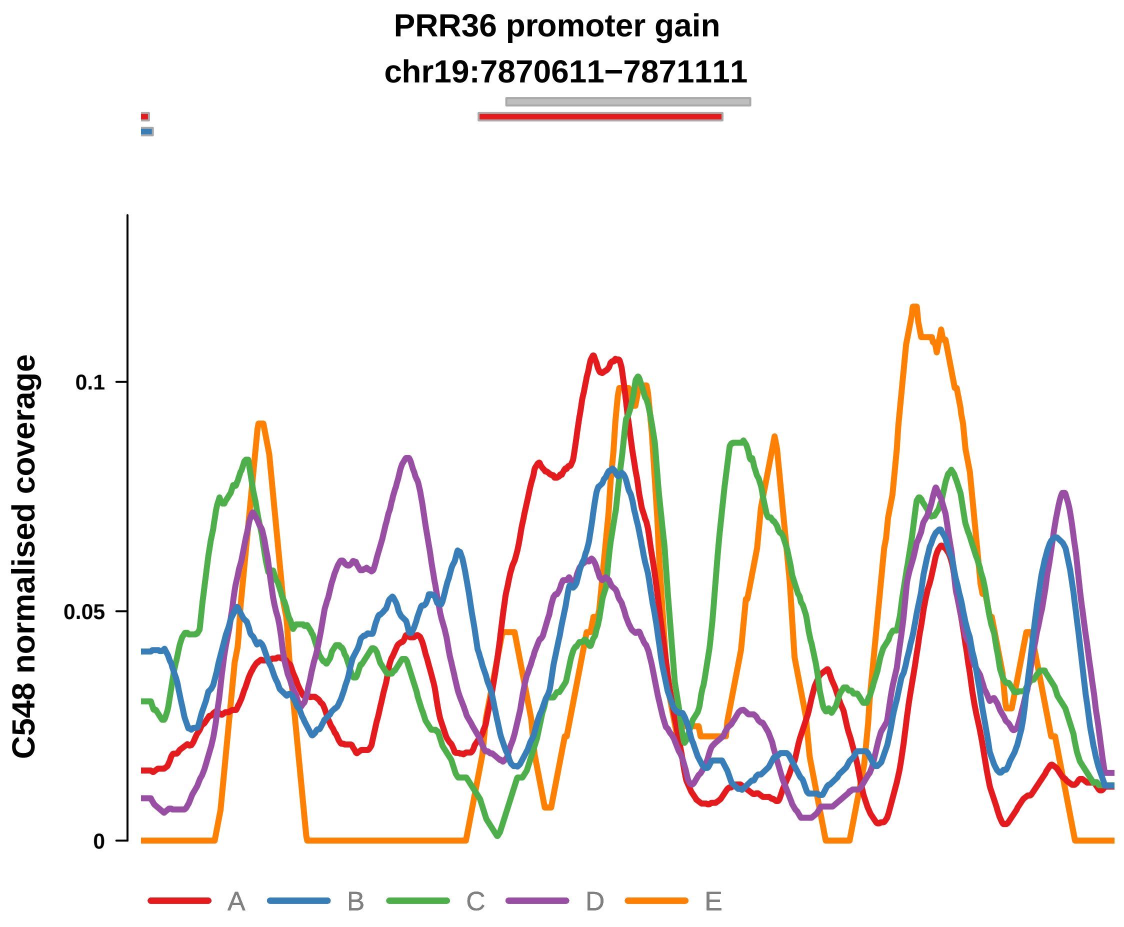


Putatively Clonal (logFC = 2.35)

| | P. | Adjusted P. |
|---|---|---|
| *~region vs. ~* | 6.39e−02 | 1.76e−01 |
| *~purity vs ~* | 6.66e−01 | 7.82e−01 |
| *~purity+region vs ~purity* | 7.69e−02 | 3.56e−01 |
| *Carcinoma vs. normal* | 6.28e−05 **** | 6.28e−05 **** |

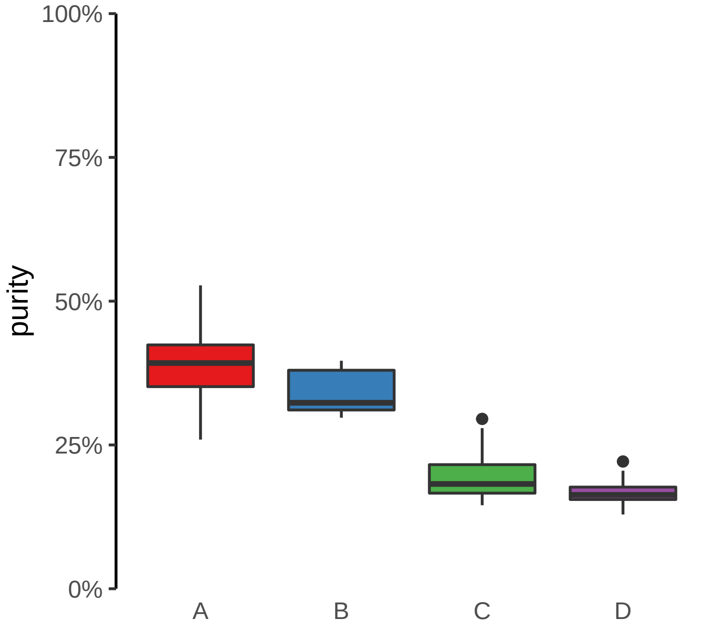

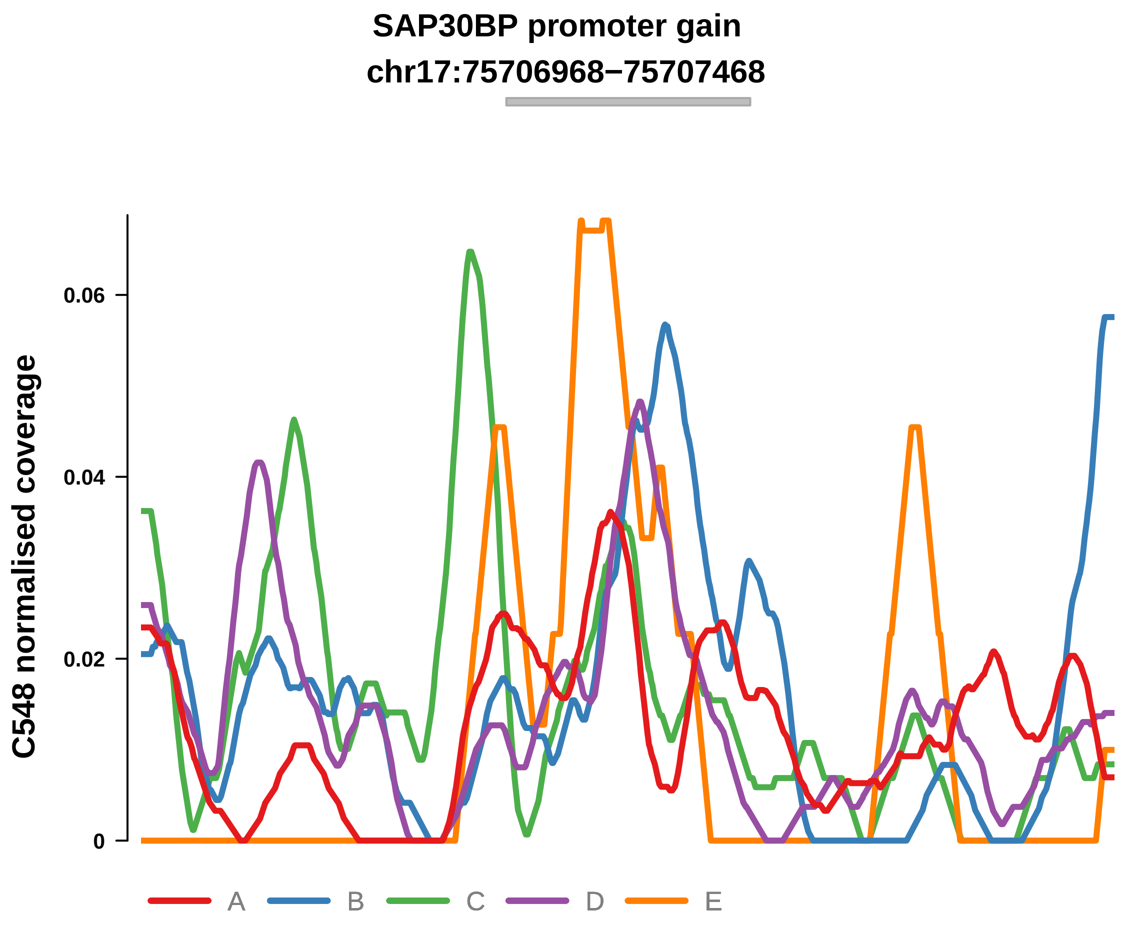


Putatively Clonal (logFC = 2.31)

| | P. | Adjusted P. |
|---|---|---|
| *~region vs. ~* | 9.24e−01 | 9.34e−01 |
| *~purity vs ~* | 9.23e−01 | 9.55e−01 |
| *~purity+region vs ~purity* | 4.36e−01 | 6.4e−01 |
| *Carcinoma vs. normal* | 4.06e−04 *** | 4.06e−04 *** |

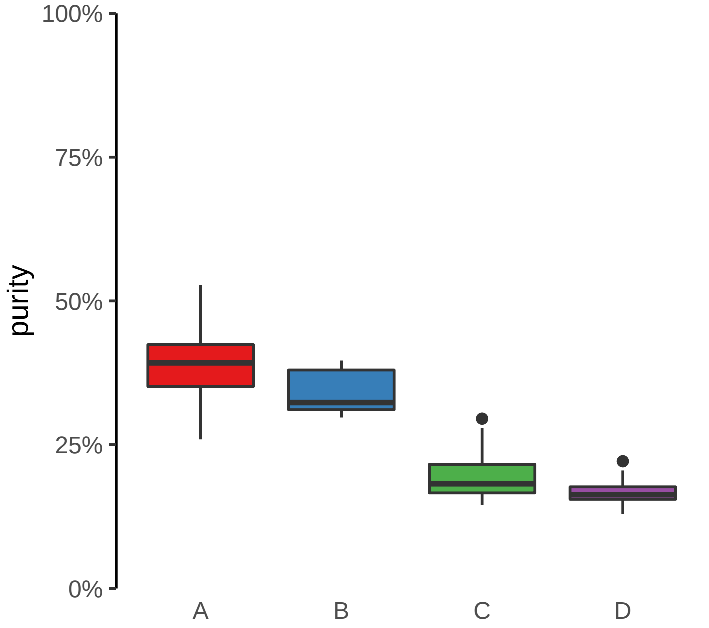

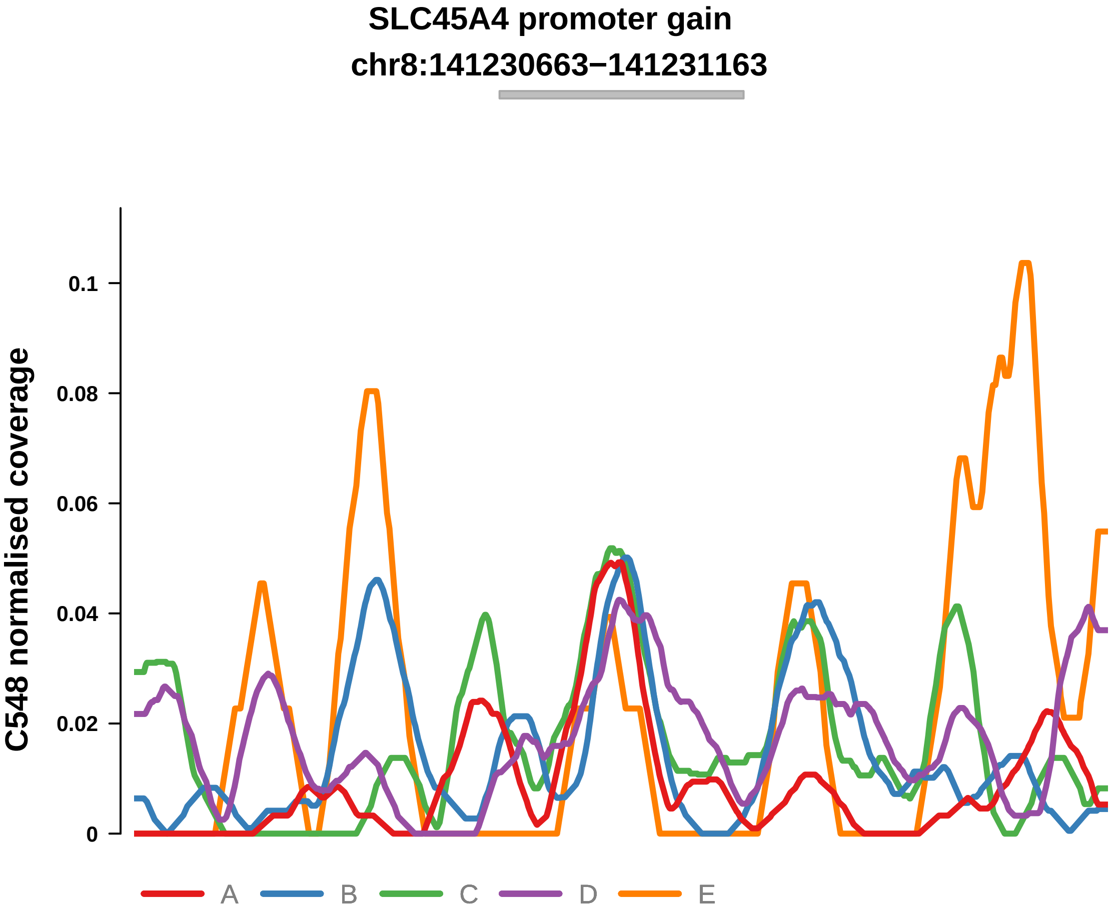


Putatively Clonal (logFC = 1.9)

| | P. | Adjusted P. |
|---|---|---|
| *~region vs. ~* | 2.61e−01 | 3.63e−01 |
| *~purity vs ~* | 2.75e−01 | 4.33e−01 |
| *~purity+region vs ~purity* | 1.59e−01 | 4.18e−01 |
| *Carcinoma vs. normal* | 5.49e−03 ** | 5.49e−03 ** |

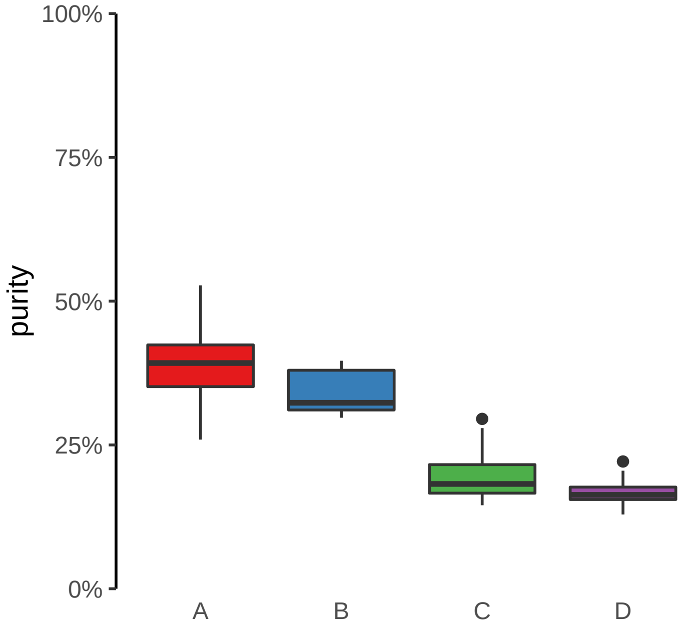

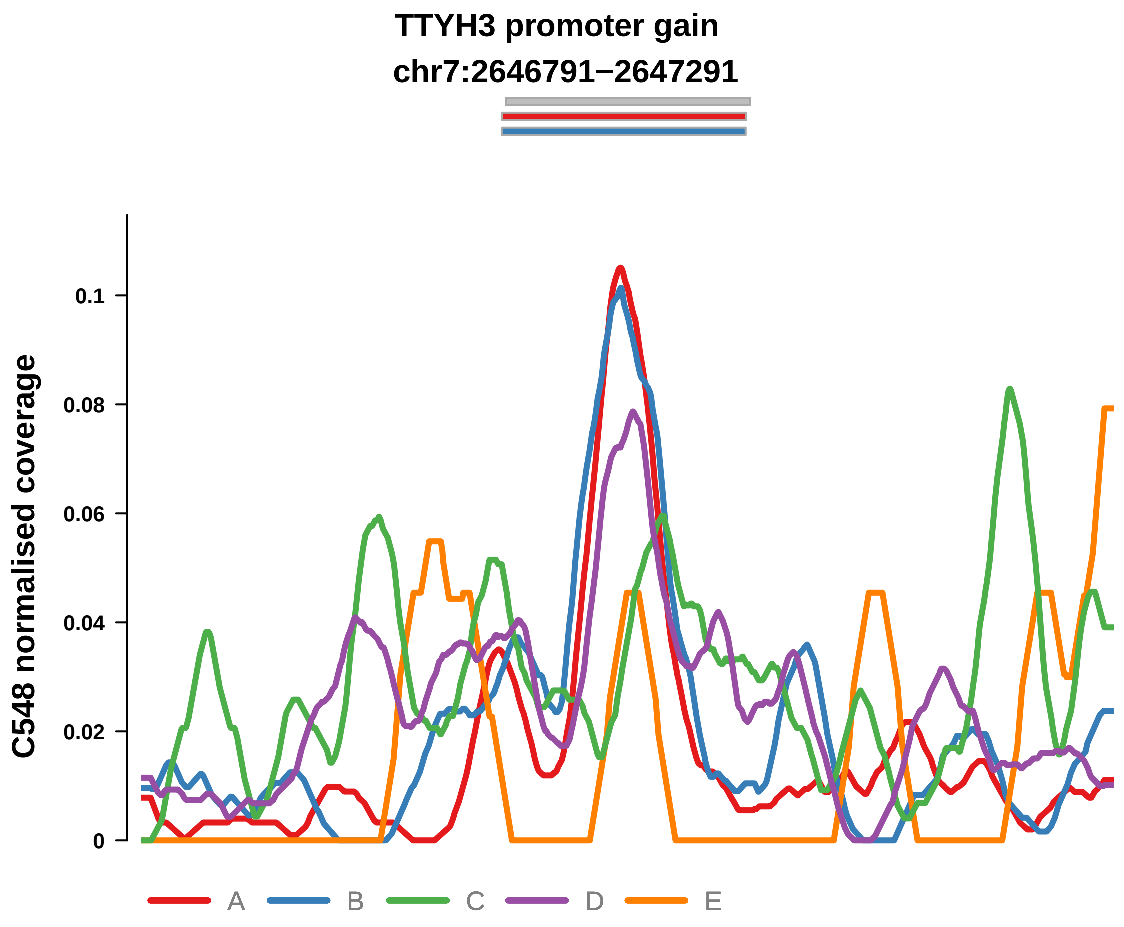


Putatively Clonal (logFC = 2.33)

| | P. | Adjusted P. |
|---|---|---|
| *~region vs. ~* | 6.84e−01 | 8.02e−01 |
| *~purity vs ~* | 6.29e−01 | 7.58e−01 |
| *~purity+region vs ~purity* | 7.03e−01 | 8.47e−01 |
| *Carcinoma vs. normal* | 2.33e−04 *** | 2.33e−04 *** |

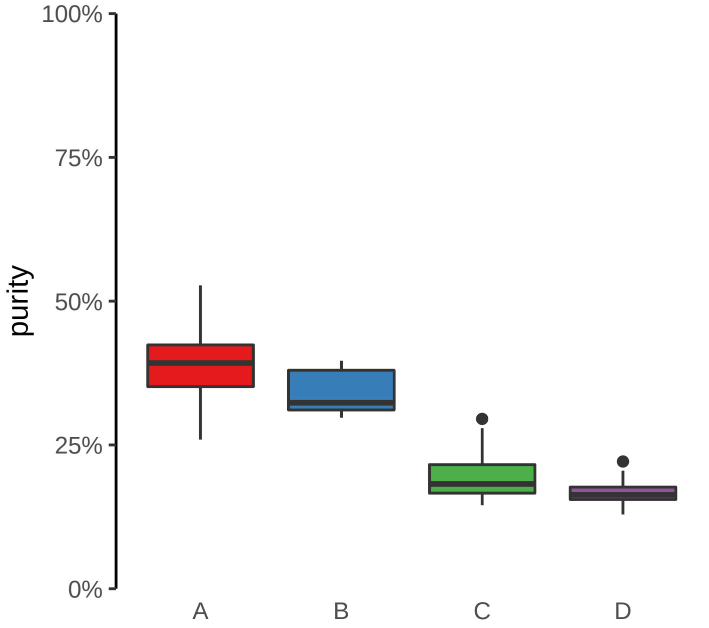

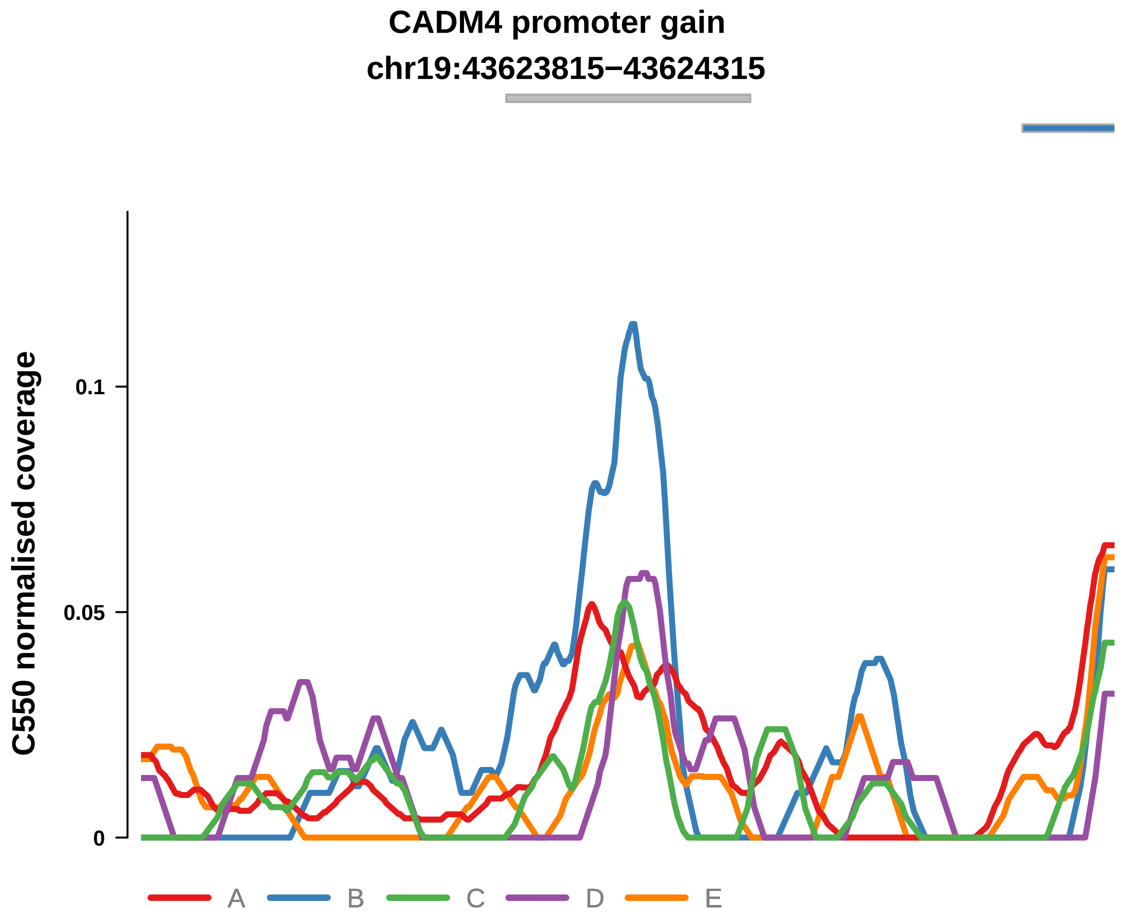


Putatively Clonal (logFC = 2.66)

| | P. | Adjusted P. |
|---|---|---|
| *~region vs. ~* | 3.06e−01 | 8.27e−01 |
| *~purity vs ~* | 1.91e−01 | 5.8e−01 |
| *~purity+region vs ~purity* | 4.15e−01 | 9.87e−01 |
| *Carcinoma vs. normal* | 9.56e−05 **** | 9.56e−05 **** |

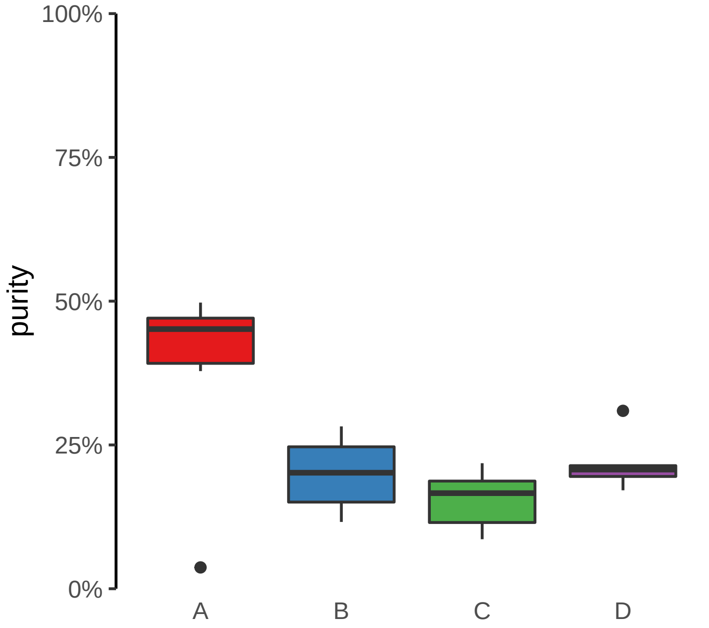

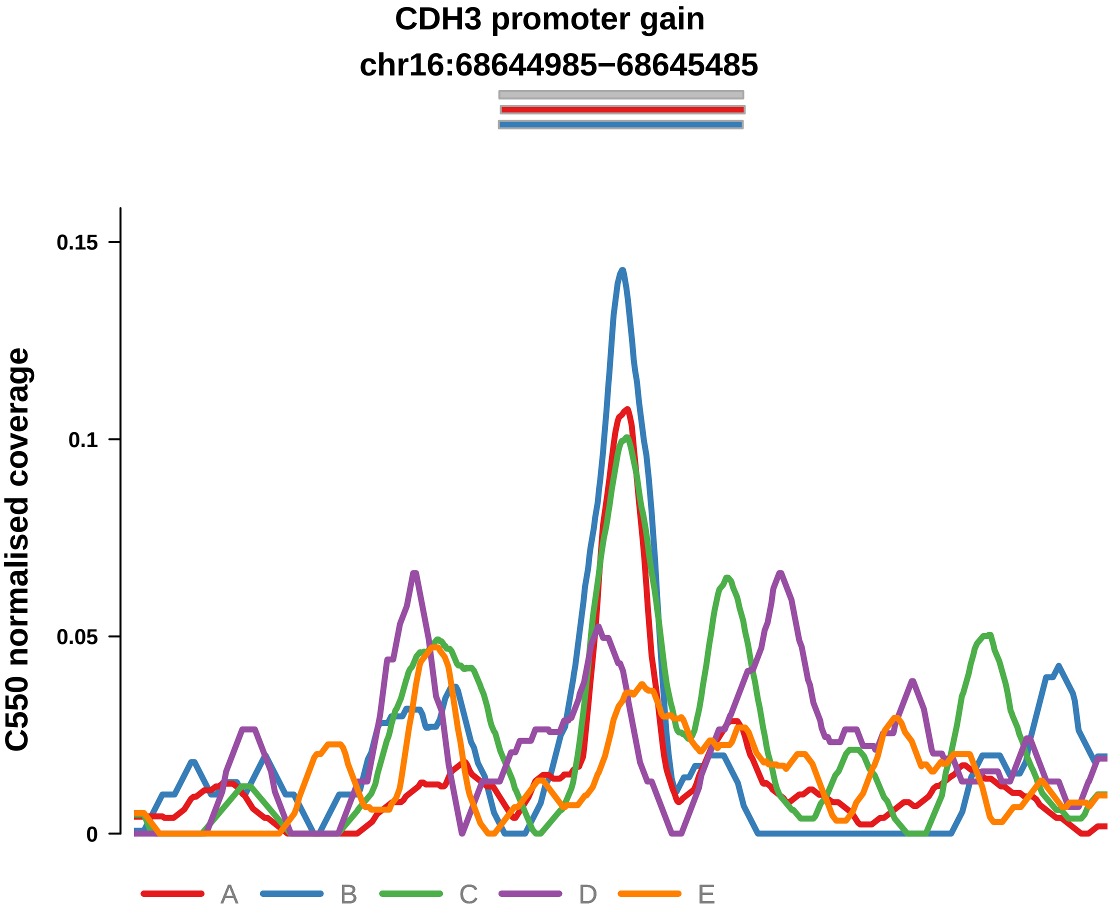


Putatively Clonal (logFC = 1.69)

| | P. | Adjusted P. |
|---|---|---|
| *~region vs. ~* | 6.58e−01 | 9.11e−01 |
| *~purity vs ~* | 9.08e−02 | 4.59e−01 |
| *~purity+region vs ~purity* | 9.91e−01 | 1e+00 |
| *Carcinoma vs. normal* | 6.93e−03 ** | 6.93e−03 ** |

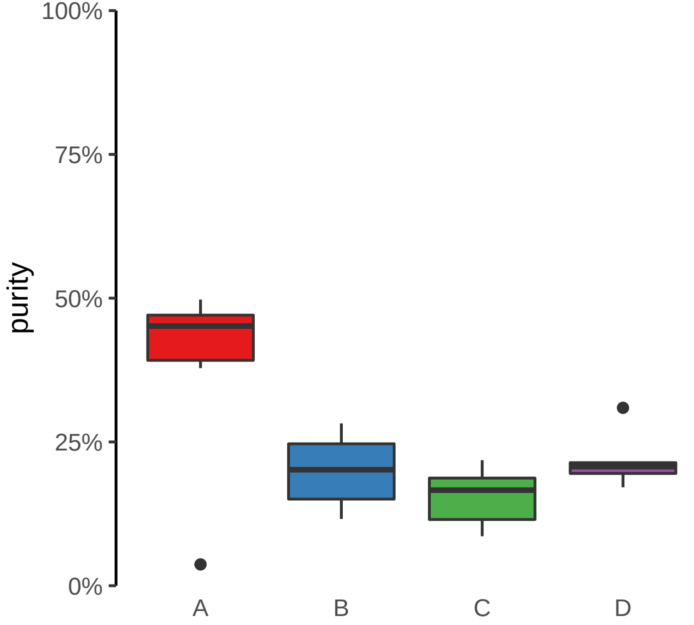

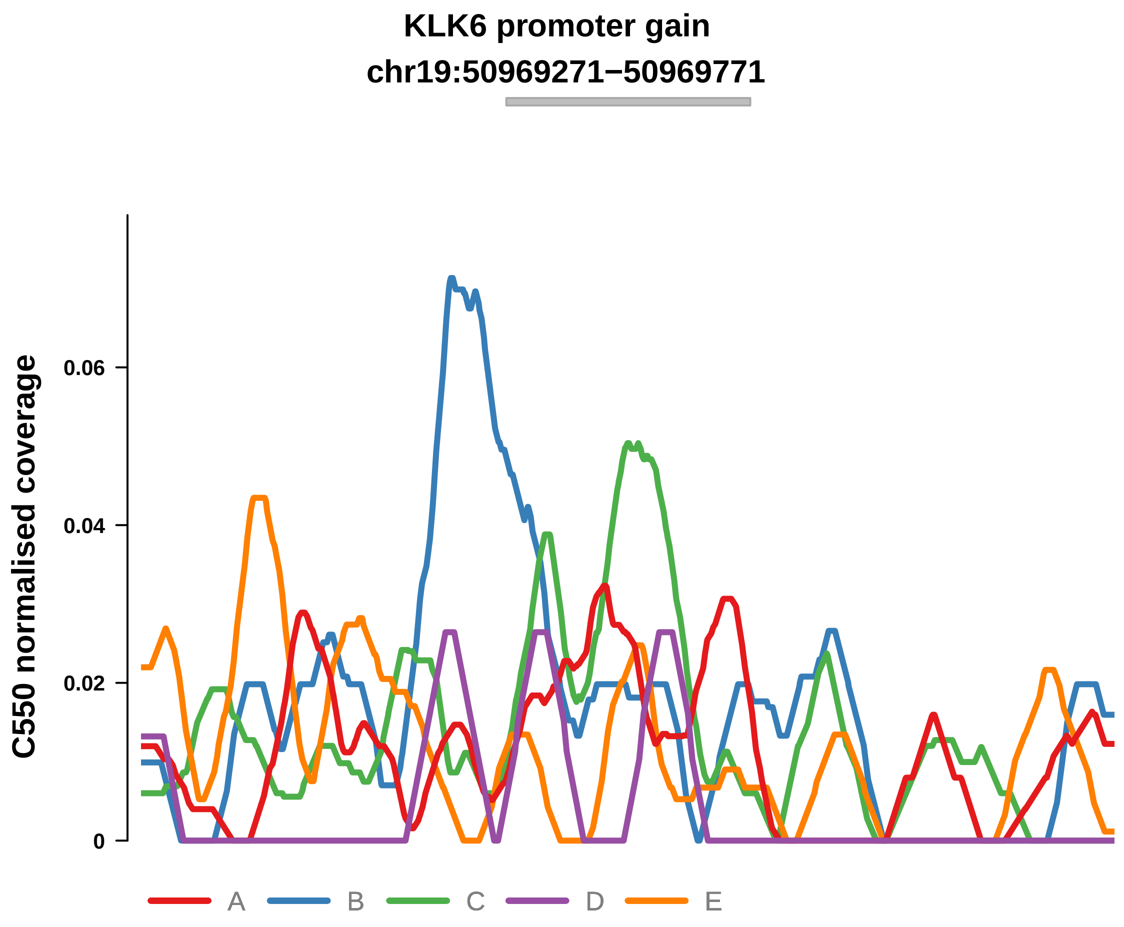


Putatively Clonal (logFC = 2.18)

| | P. | Adjusted P. |
|---|---|---|
| *~region vs. ~* | 6.73e−01 | 9.11e−01 |
| *~purity vs ~* | 1.99e−01 | 5.84e−01 |
| *~purity+region vs ~purity* | 5.15e−01 | 9.87e−01 |
| *Carcinoma vs. normal* | 1.39e−03 ** | 1.39e−03 ** |

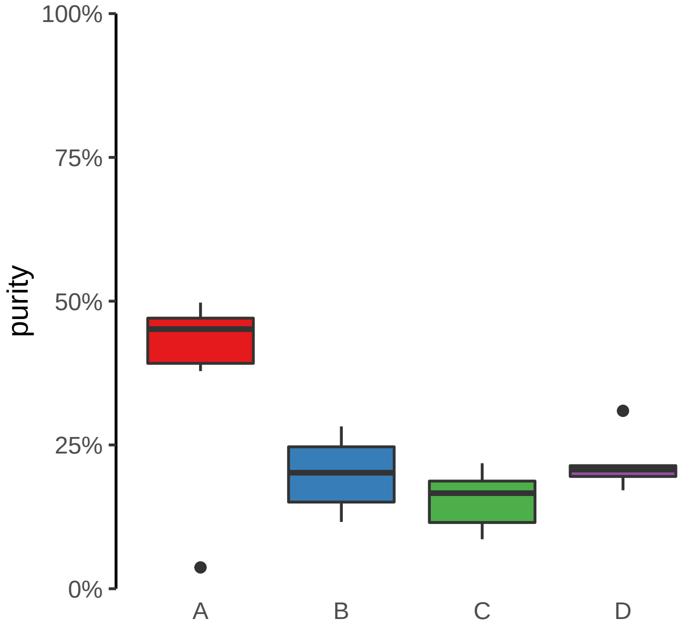

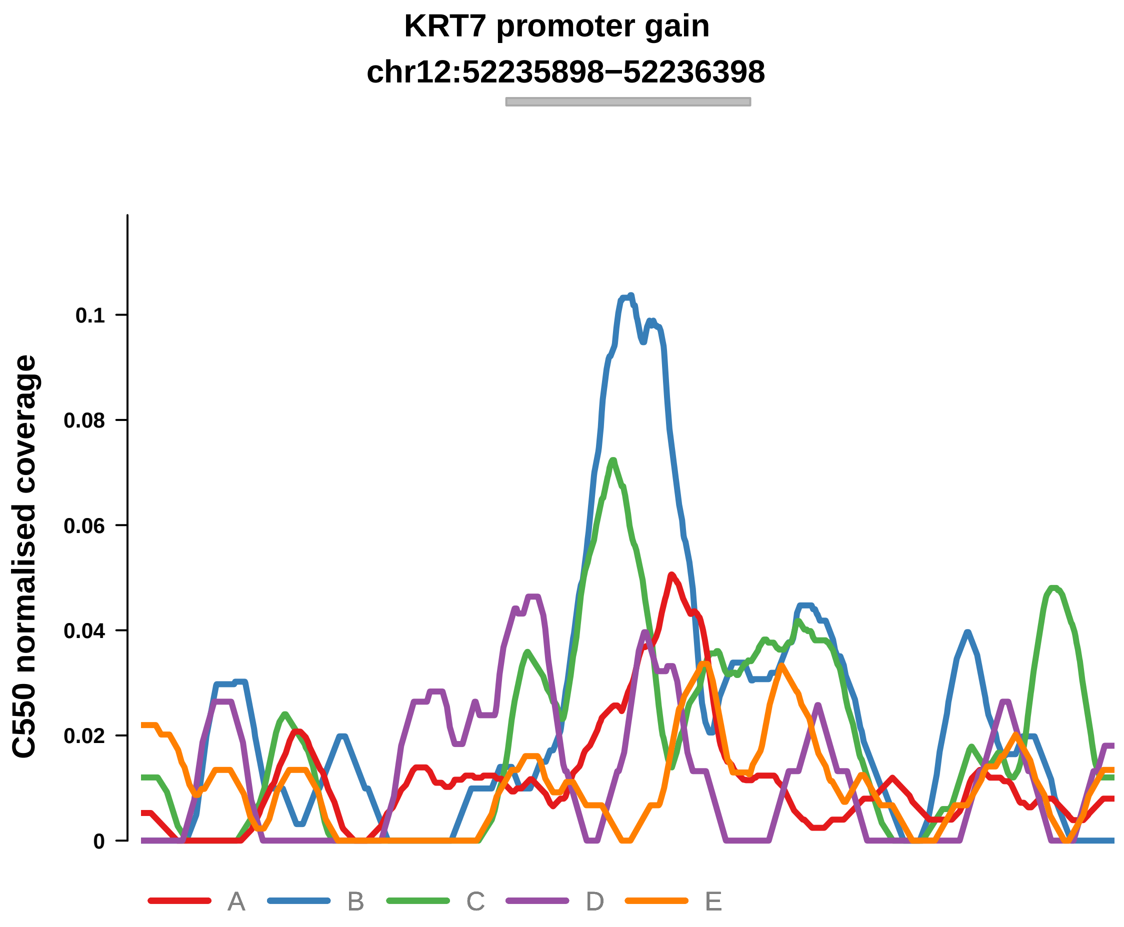


Putatively Clonal (logFC = 1.7)

| | P. | Adjusted P. |
|---|---|---|
| *~region vs. ~* | 8.01e−01 | 9.39e−01 |
| *~purity vs ~* | 8.47e−01 | 9.53e−01 |
| *~purity+region vs ~purity* | 5.95e−01 | 9.87e−01 |
| *Carcinoma vs. normal* | 9.66e−03 ** | 9.66e−03 ** |

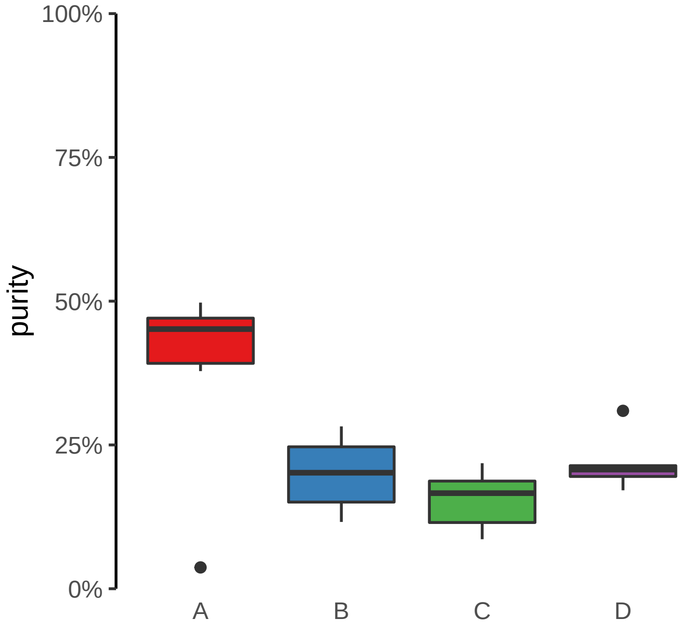

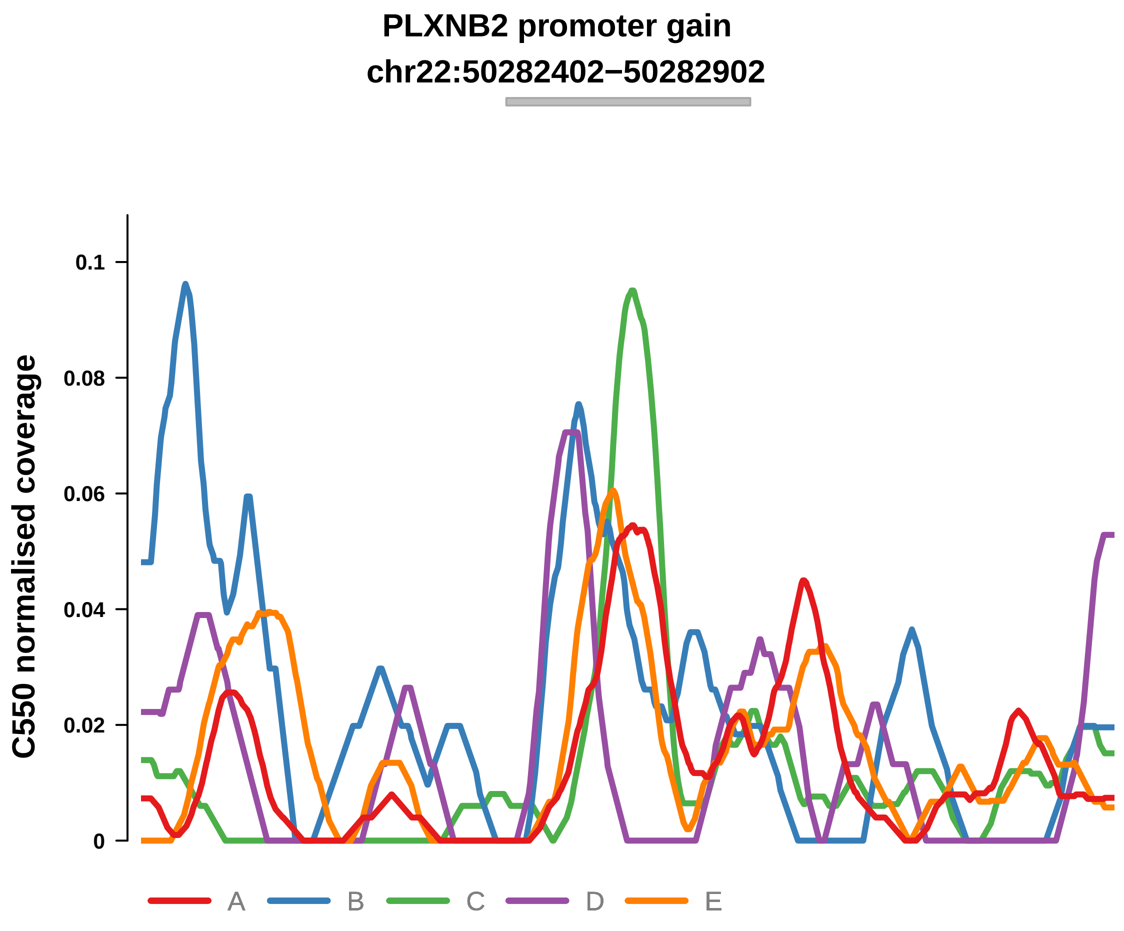


Putatively Clonal (logFC = 2.04)

| | P. | Adjusted P. |
|---|---|---|
| *~region vs. ~* | 9.02e−01 | 9.68e−01 |
| *~purity vs ~* | 9.41e−01 | 9.66e−01 |
| *~purity+region vs ~purity* | 8.6e−01 | 1e+00 |
| *Carcinoma vs. normal* | 1.98e−03 ** | 1.98e−03 ** |

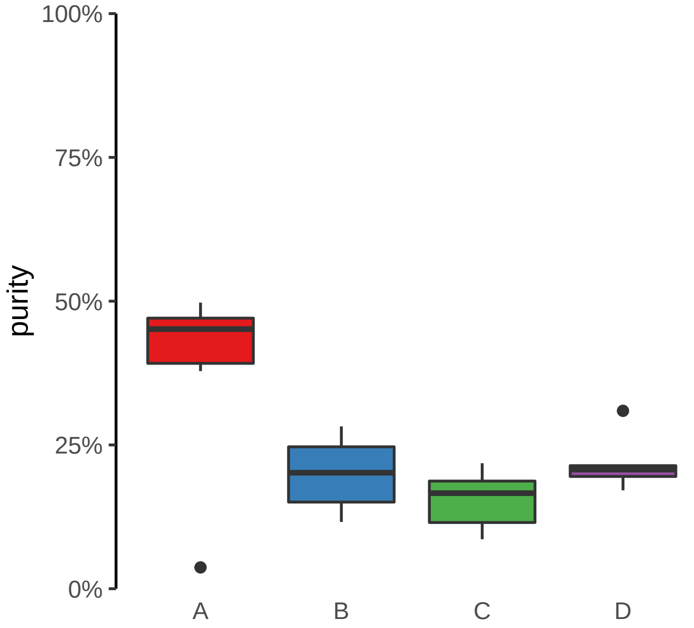

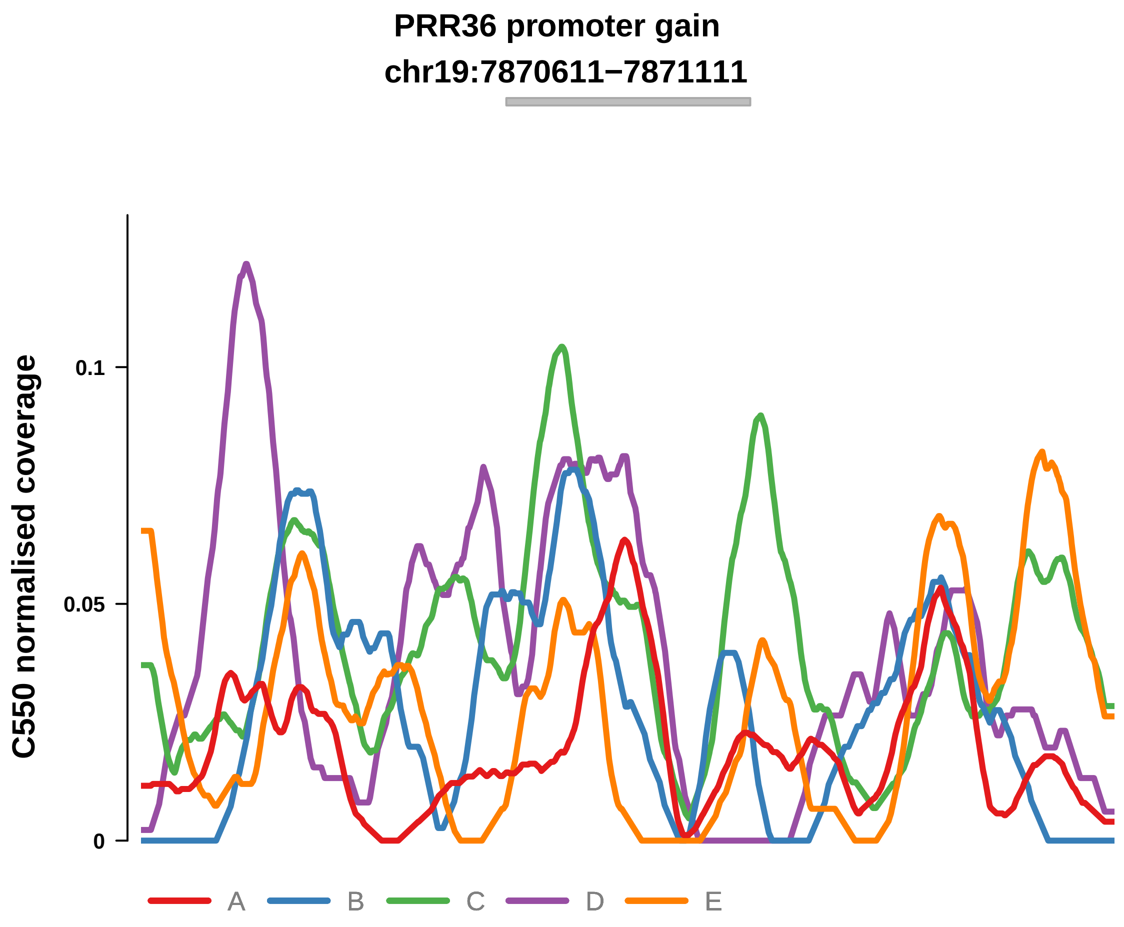


Putatively Clonal (logFC = 1.67)

| | P. | Adjusted P. |
|---|---|---|
| *~region vs. ~* | 2.17e−01 | 7.94e−01 |
| *~purity vs ~* | 6.71e−01 | 8.71e−01 |
| *~purity+region vs ~purity* | 1.04e−01 | 7.48e−01 |
| *Carcinoma vs. normal* | 8.69e−03 ** | 8.69e−03 ** |

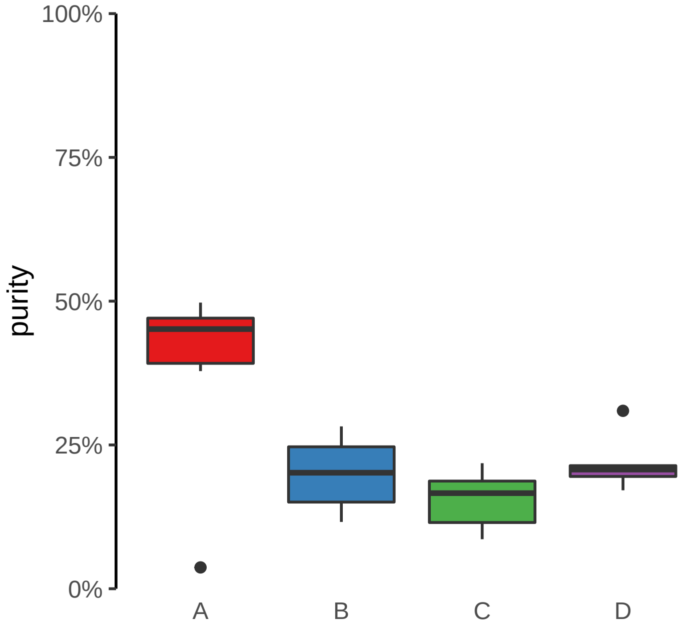

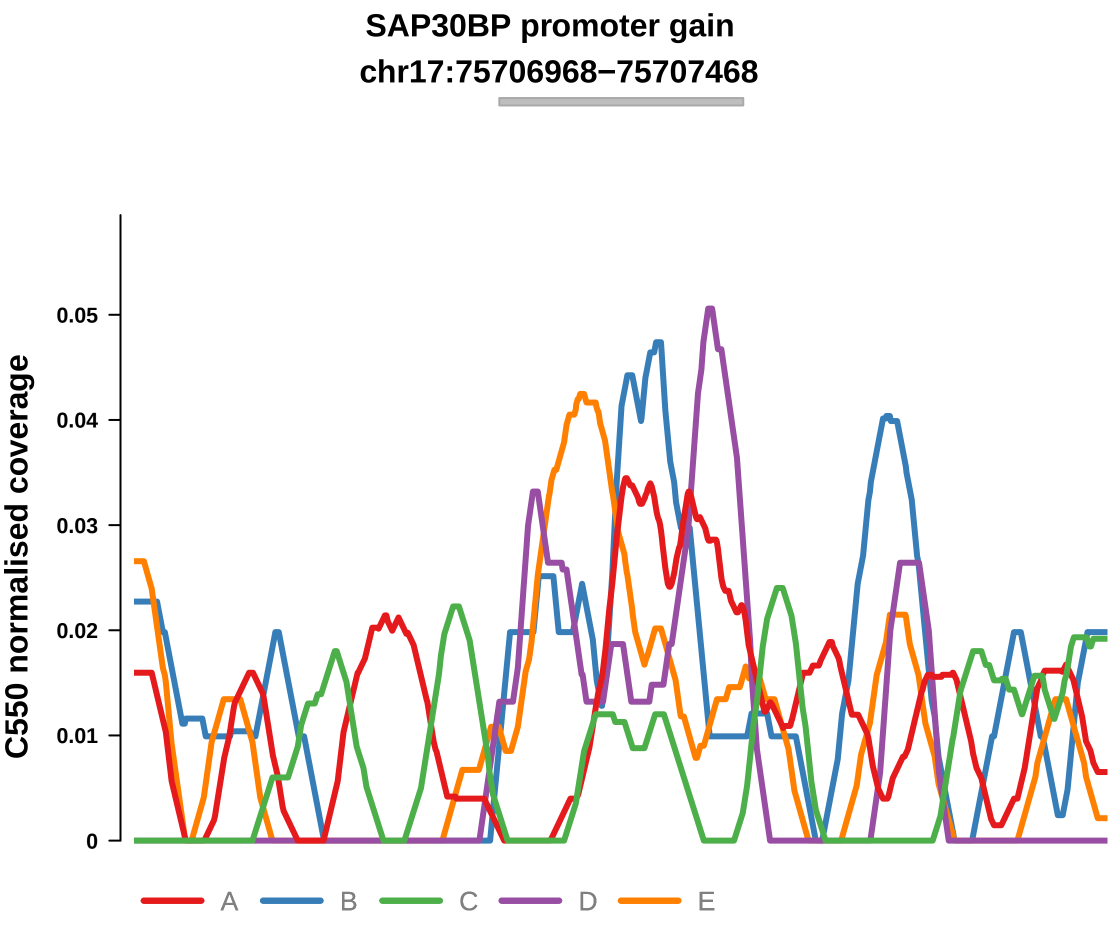


Putatively Clonal (logFC = 2)

| | P. | Adjusted P. |
|---|---|---|
| *~region vs. ~* | 2.1e−01 | 7.94e−01 |
| *~purity vs ~* | 3.93e−01 | 7.48e−01 |
| *~purity+region vs ~purity* | 2.35e−01 | 8.41e−01 |
| *Carcinoma vs. normal* | 7.46e−03 ** | 7.46e−03 ** |

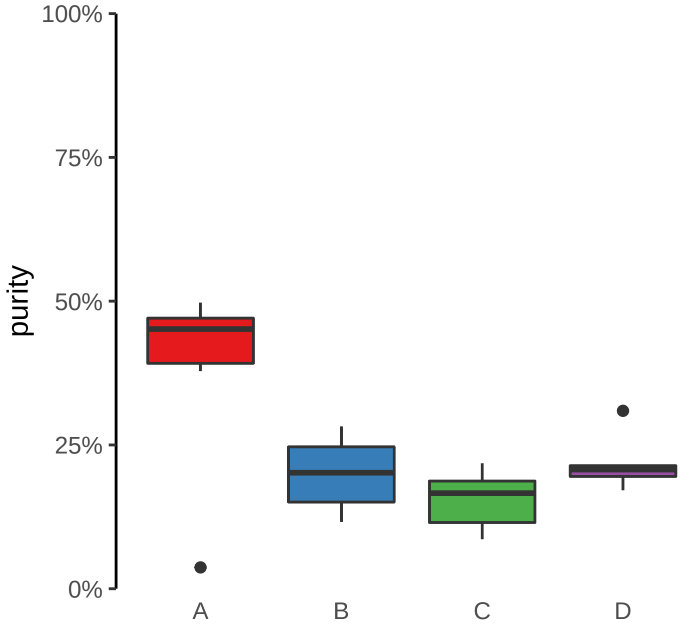

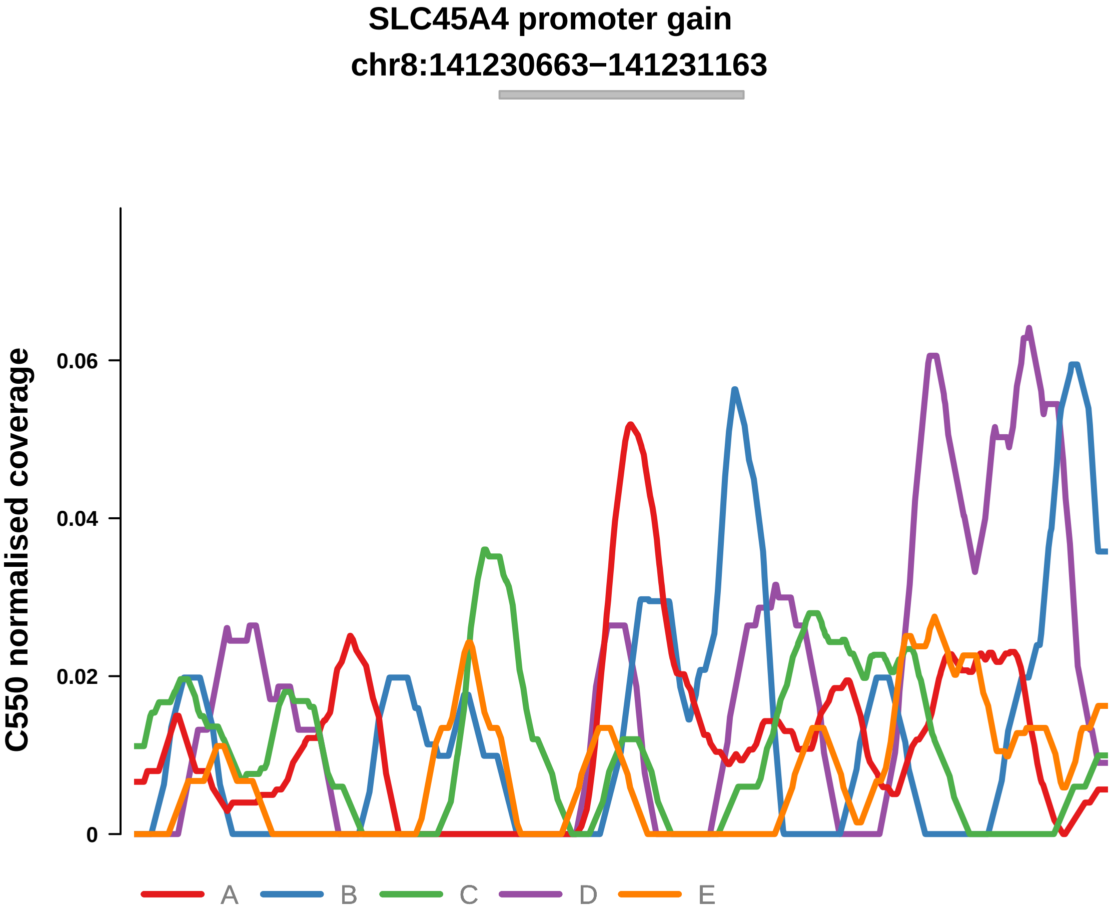


Putatively Clonal (logFC = 2.17)

| | P. | Adjusted P. |
|---|---|---|
| *~region vs. ~* | 5.36e−01 | 9.11e−01 |
| *~purity vs ~* | 2.99e−01 | 7.42e−01 |
| *~purity+region vs ~purity* | 7.63e−01 | 1e+00 |
| *Carcinoma vs. normal* | 2.55e−03 ** | 2.55e−03 ** |

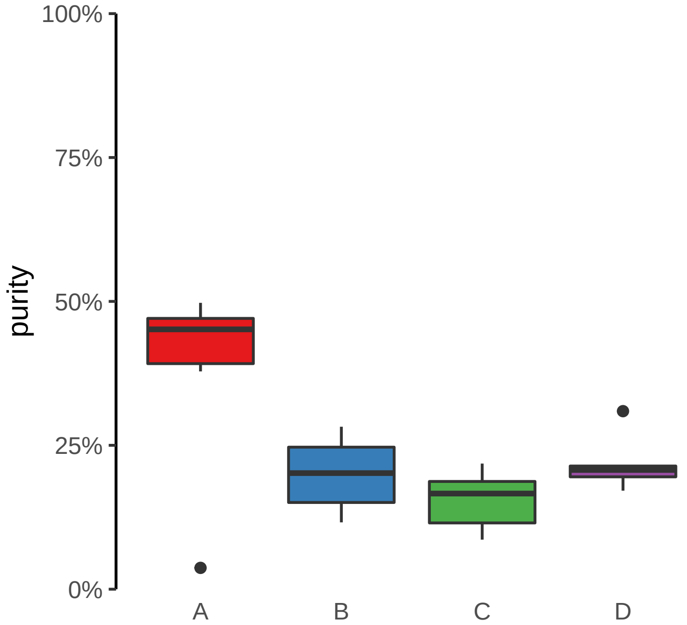

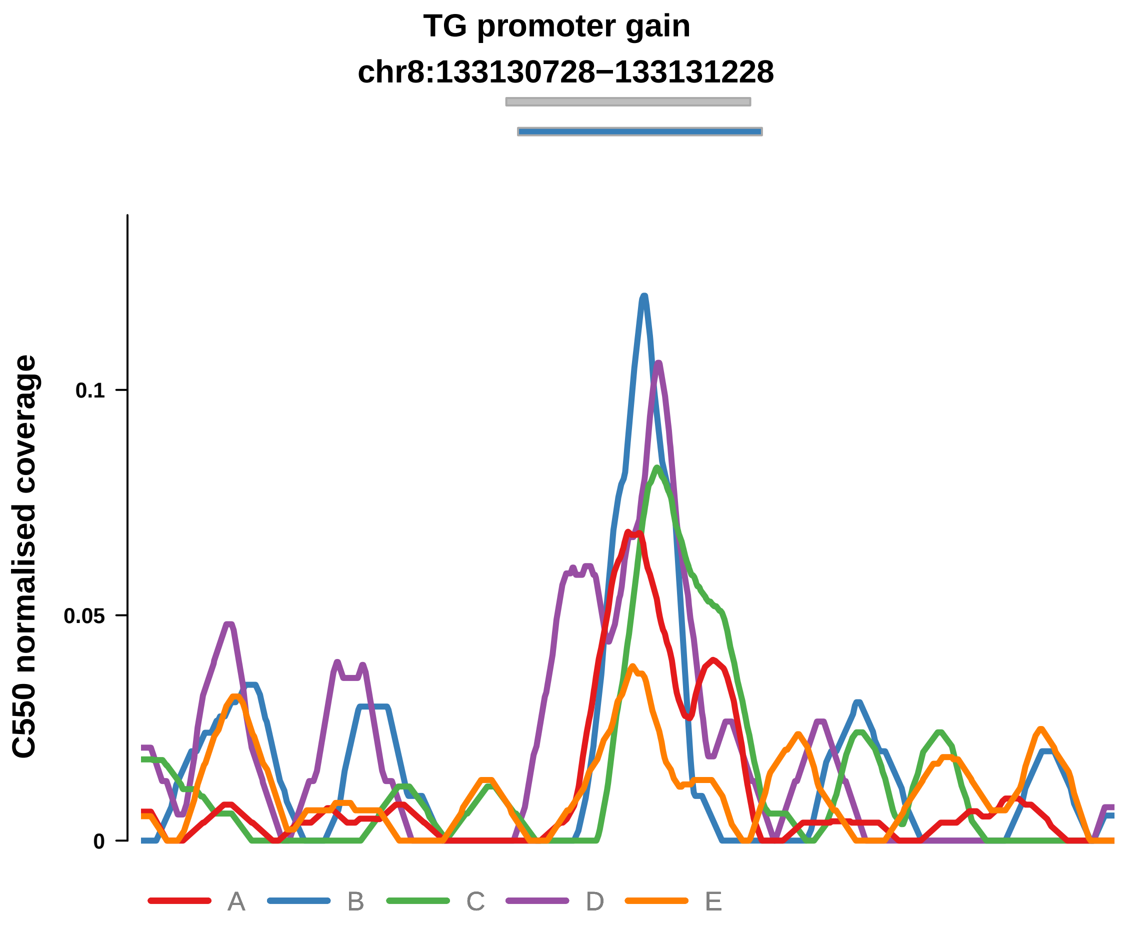


Putatively Clonal (logFC = 2.39)

| | P. | Adjusted P. |
|---|---|---|
| *~region vs. ~* | 3.97e−01 | 8.74e−01 |
| *~purity vs ~* | 1.16e−01 | 4.87e−01 |
| *~purity+region vs ~purity* | 5.52e−01 | 9.87e−01 |
| *Carcinoma vs. normal* | 1.6e−04 *** | 1.6e−04 *** |

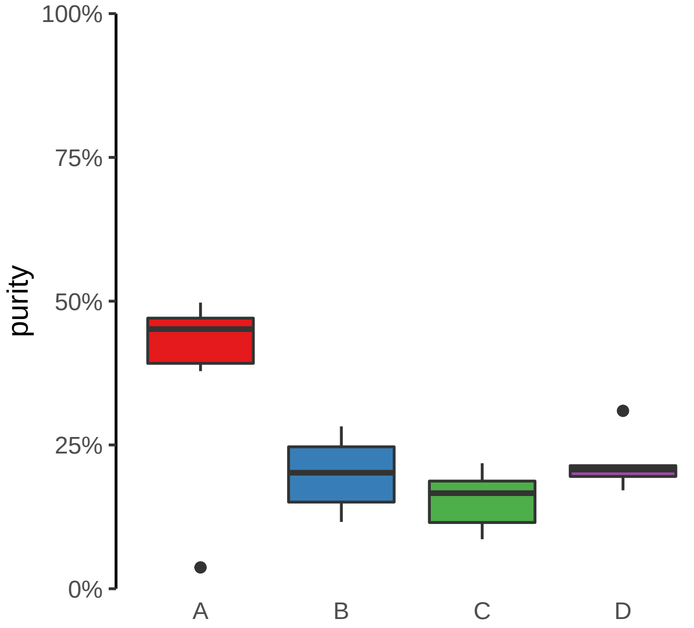

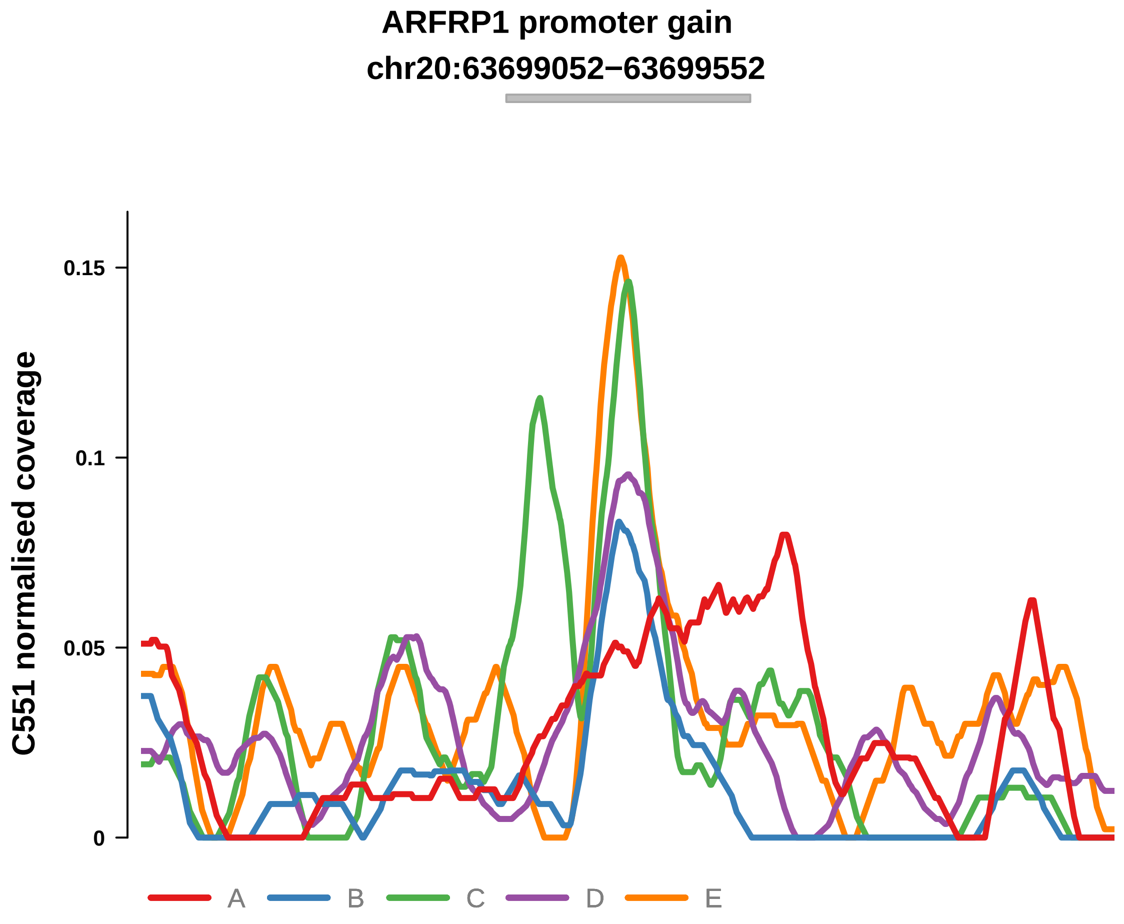


Putatively Clonal (logFC = 1.43)

| | P. | Adjusted P. |
|---|---|---|
| *~region vs. ~* | 6.76e−01 | 9.65e−01 |
| *~purity vs ~* | 8.3e−01 | 9.44e−01 |
| *~purity+region vs ~purity* | 6.25e−01 | 9.48e−01 |
| *Carcinoma vs. normal* | 9.82e−03 ** | 9.82e−03 ** |

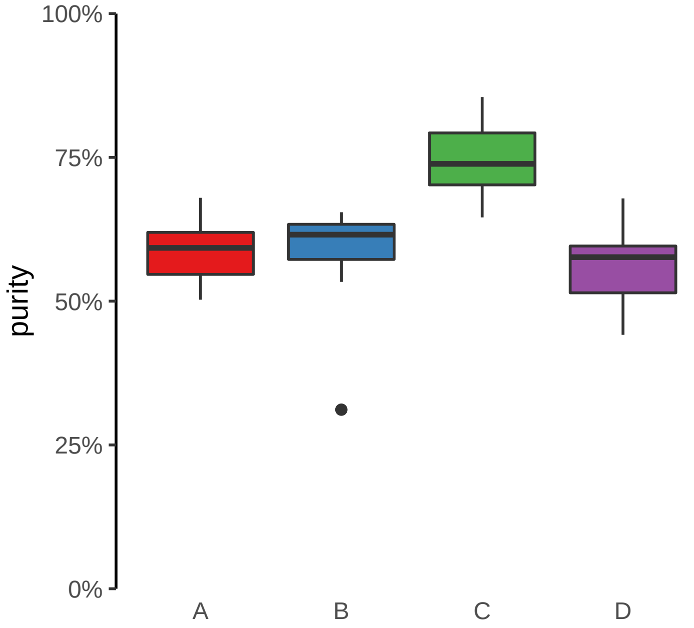

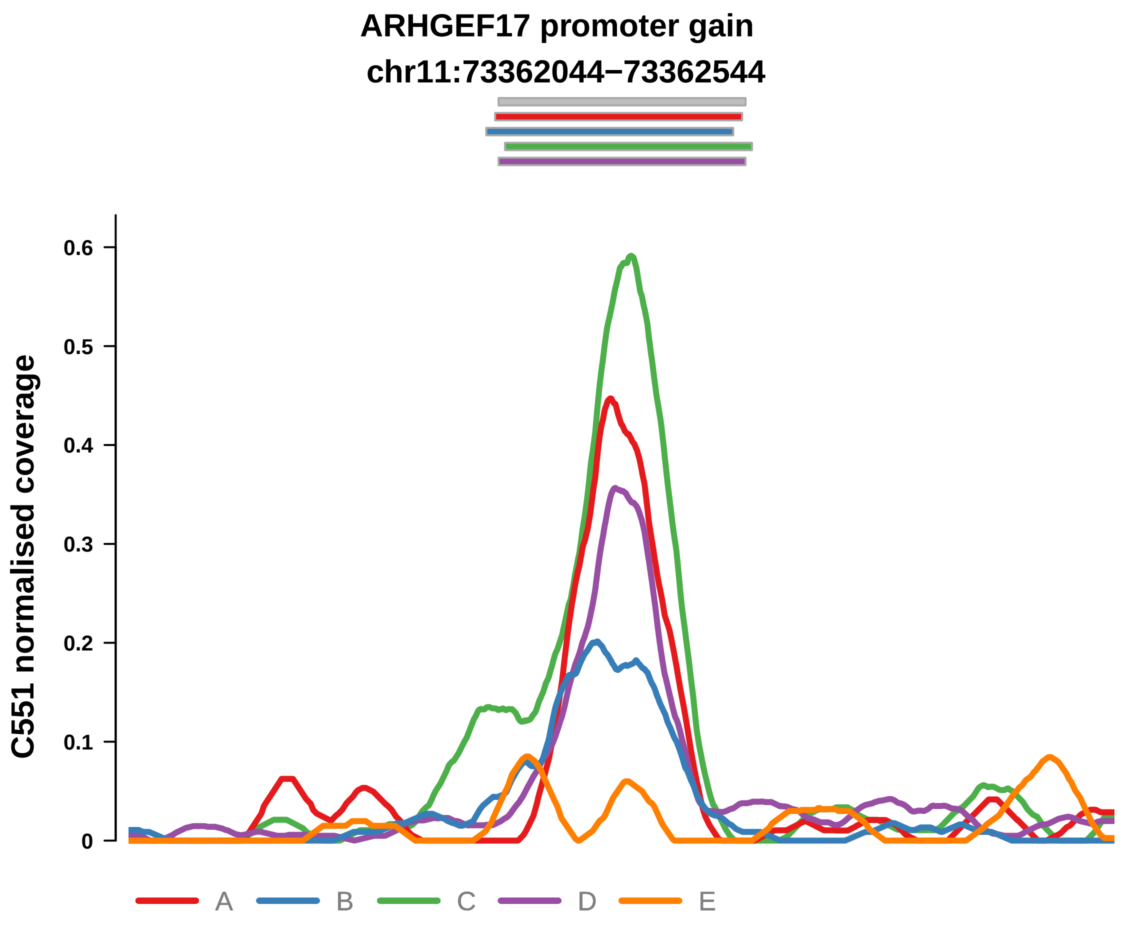


Putatively Clonal (logFC = 3.67)

| | P. | Adjusted P. |
|---|---|---|
| *~region vs. ~* | 4.24e−01 | 8.39e−01 |
| *~purity vs ~* | 2.99e−01 | 9.06e−01 |
| *~purity+region vs ~purity* | 5.06e−01 | 9.48e−01 |
| *Carcinoma vs. normal* | 3.6e−10 **** | 3.6e−10 **** |

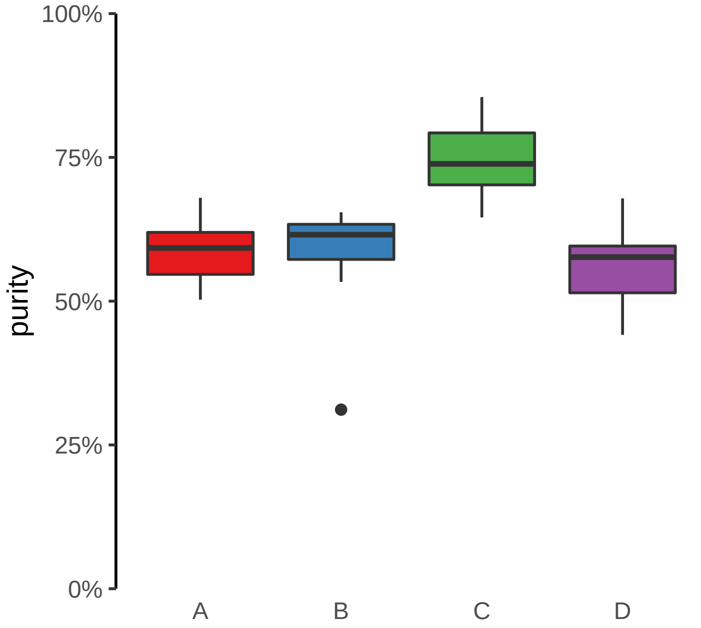

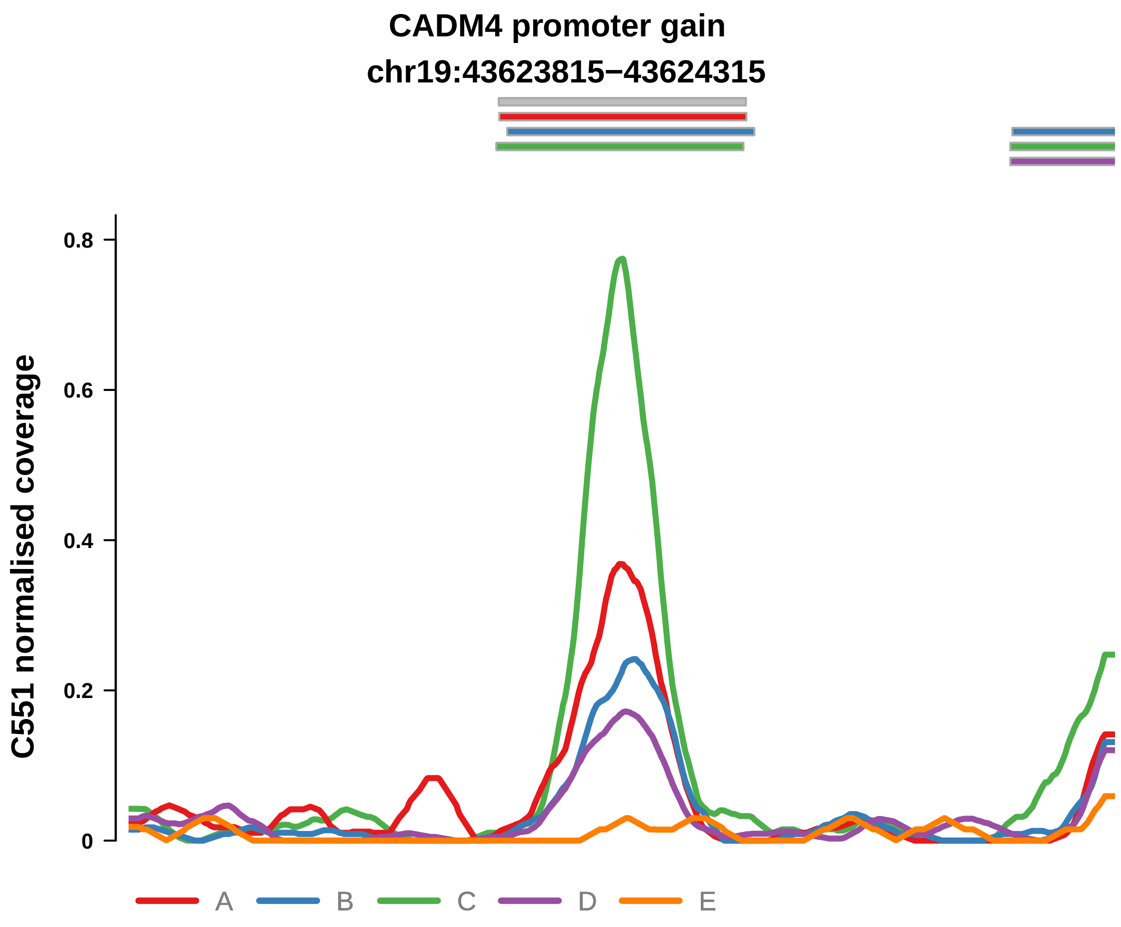


Putatively Clonal (logFC = 3.42)

| | P. | Adjusted P. |
|---|---|---|
| *~region vs. ~* | 2.67e−02 * | 2.61e−01 |
| *~purity vs ~* | 3.56e−02 * | 5.38e−01 |
| *~purity+region vs ~purity* | 1.6e−01 | 5.7e−01 |
| *Carcinoma vs. normal* | 4.33e−09 **** | 4.33e−09 **** |

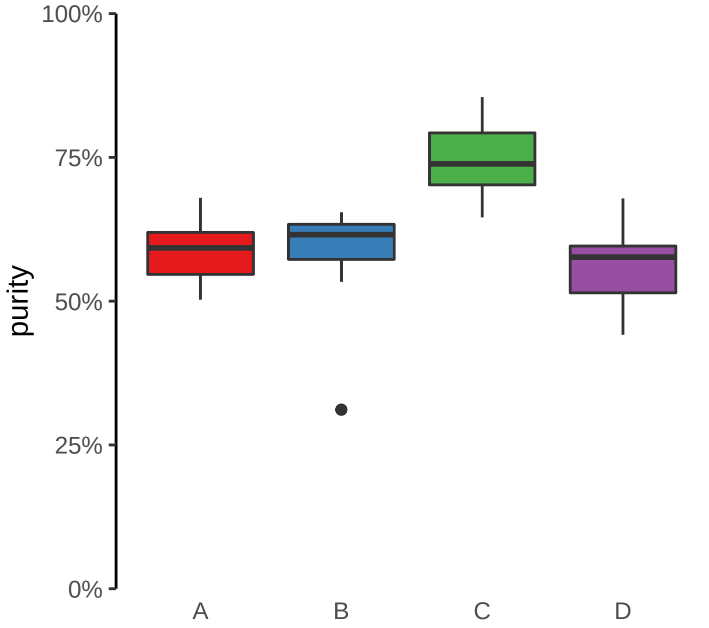

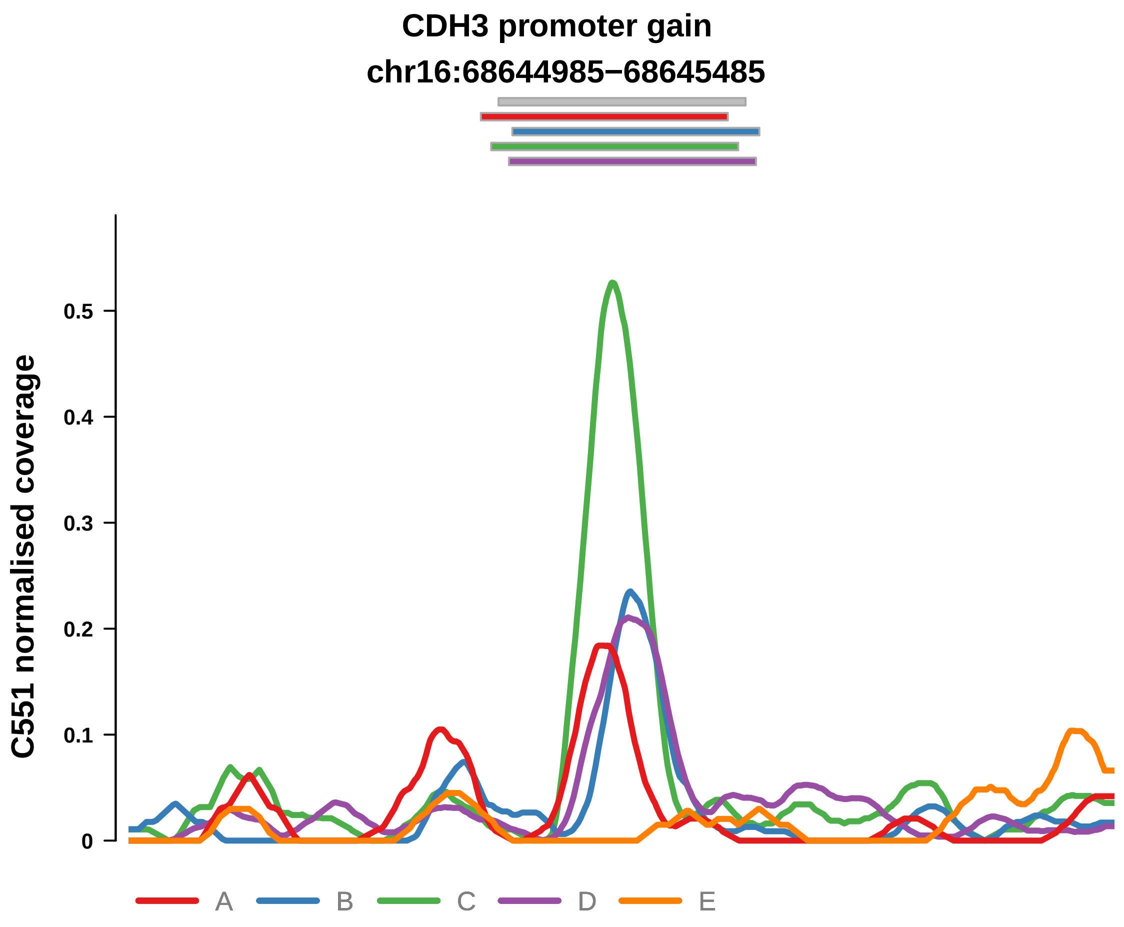


Putatively Clonal (logFC = 1.7)

| | P. | Adjusted P. |
|---|---|---|
| *~region vs. ~* | 3.17e−01 | 7.34e−01 |
| *~purity vs ~* | 5.17e−01 | 9.06e−01 |
| *~purity+region vs ~purity* | 3.85e−01 | 8.69e−01 |
| *Carcinoma vs. normal* | 1.7e−03 ** | 1.7e−03 ** |

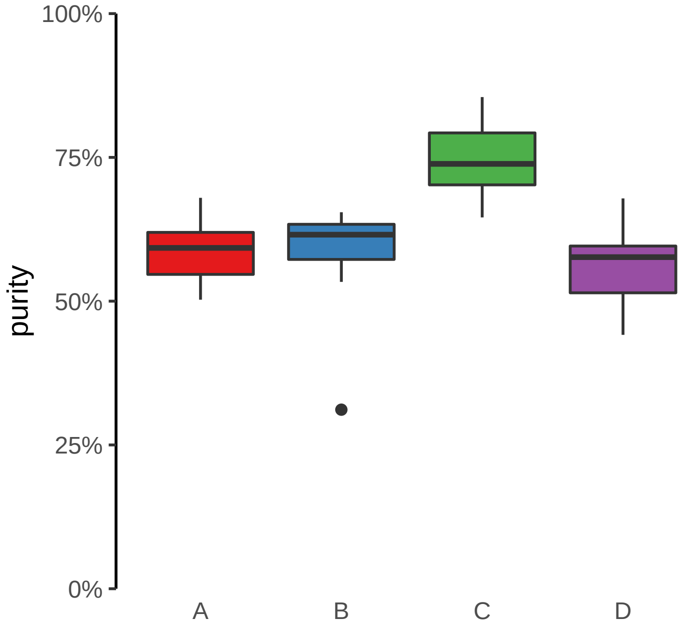

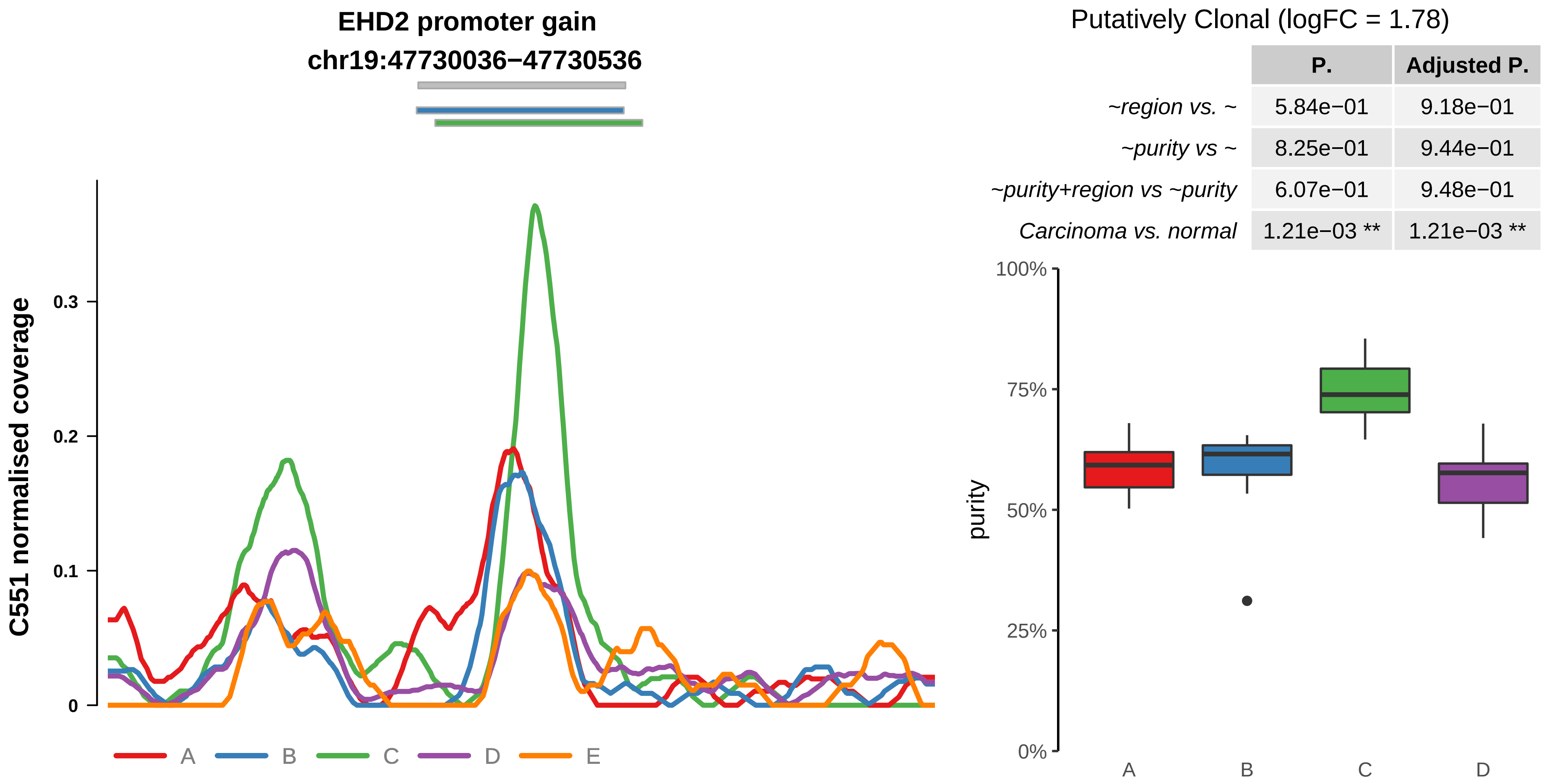


| | P. | Adjusted P. |
|---|---|---|
| *~region vs. ~* | 5.84e−01 | 9.18e−01 |
| *~purity vs ~* | 8.25e−01 | 9.44e−01 |
| *~purity+region vs ~purity* | 6.07e−01 | 9.48e−01 |
| *Carcinoma vs. normal* | 1.21e−03 ** | 1.21e−03 ** |

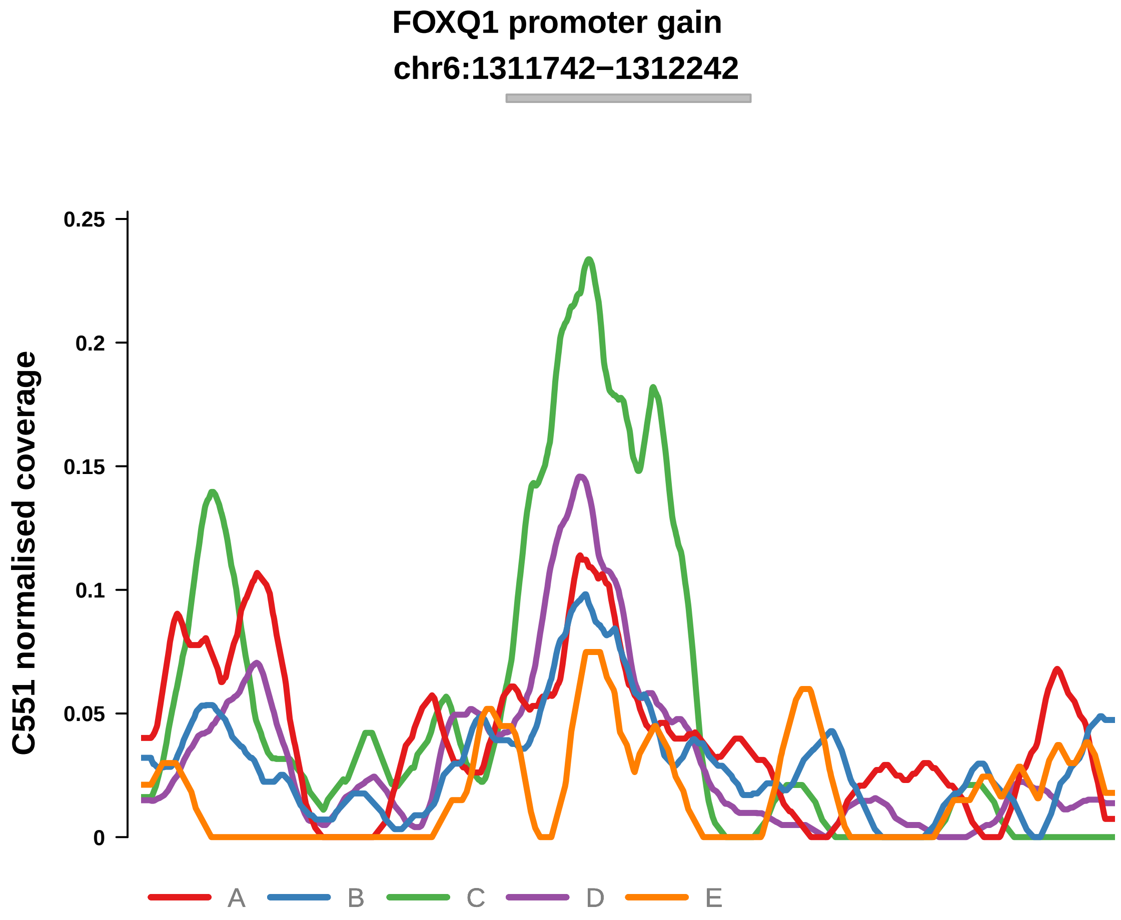


Putatively Clonal (logFC = 1.45)

| | P. | Adjusted P. |
|---|---|---|
| *~region vs. ~* | 8.44e−01 | 9.65e−01 |
| *~purity vs ~* | 8.37e−01 | 9.44e−01 |
| *~purity+region vs ~purity* | 8.26e−01 | 9.99e−01 |
| *Carcinoma vs. normal* | 7.59e−03 ** | 7.59e−03 ** |

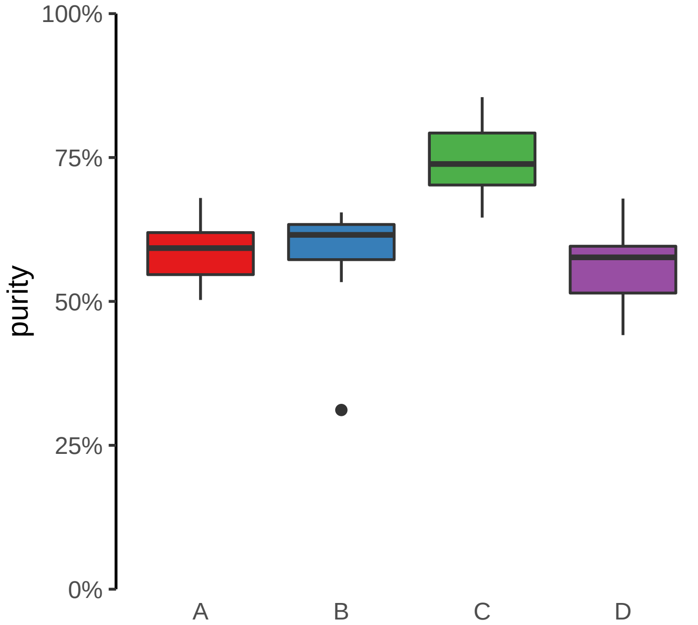

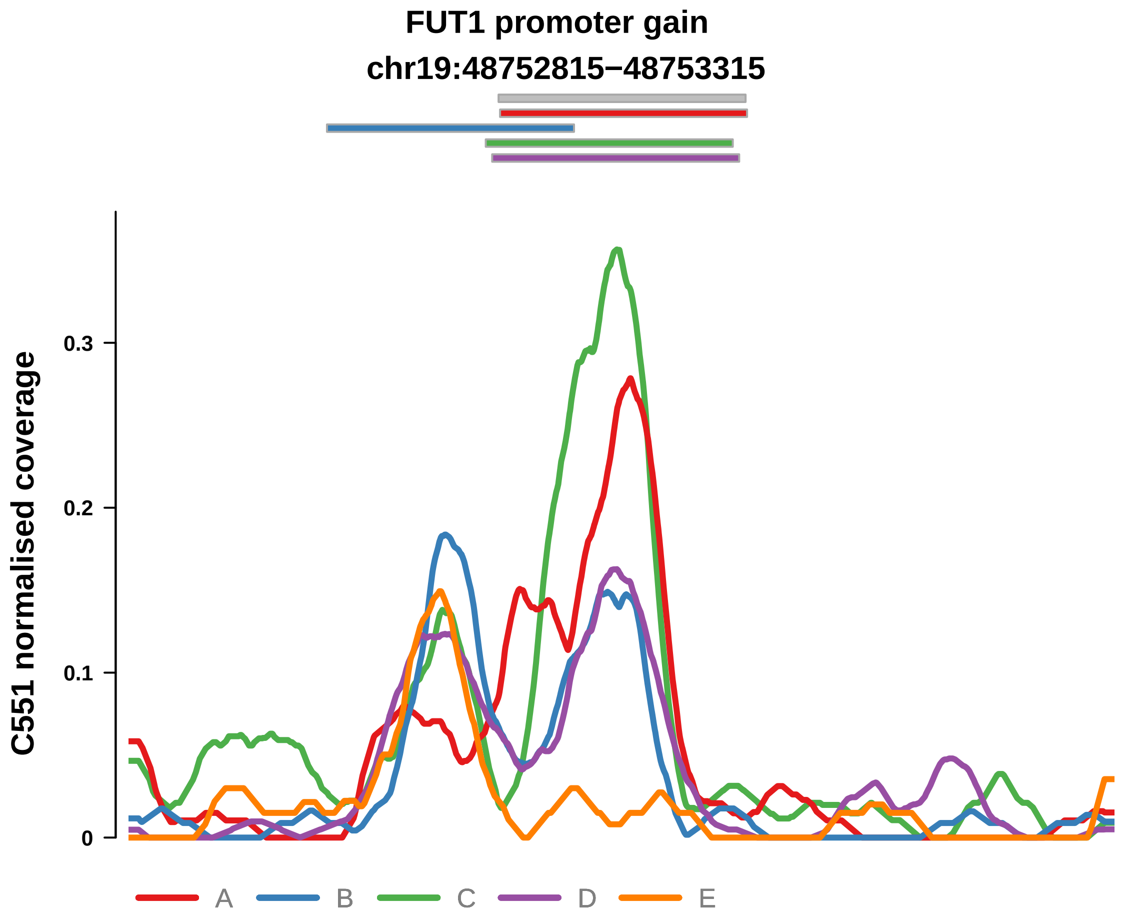


Putatively Clonal (logFC = 2.08)

| | P. | Adjusted P. |
|---|---|---|
| *~region vs. ~* | 8.02e−02 | 3.92e−01 |
| *~purity vs ~* | 3.59e−01 | 9.06e−01 |
| *~purity+region vs ~purity* | 5.56e−02 | 4.93e−01 |
| *Carcinoma vs. normal* | 1.5e−04 *** | 1.5e−04 *** |

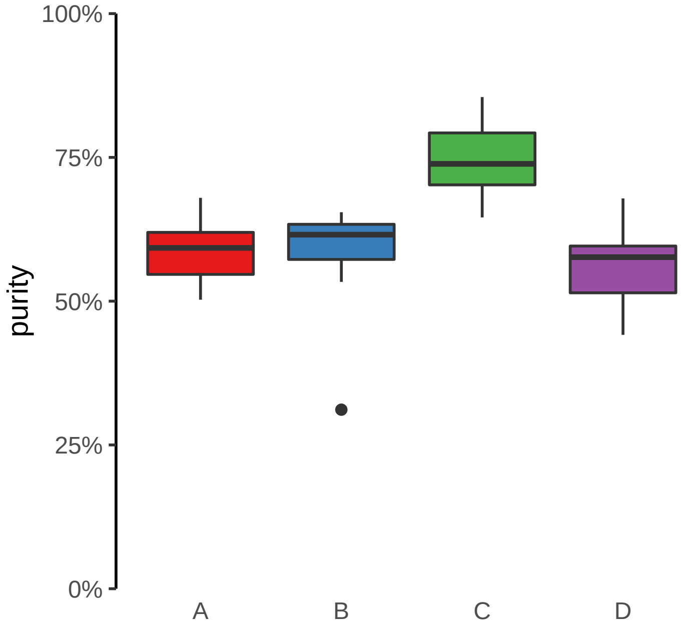

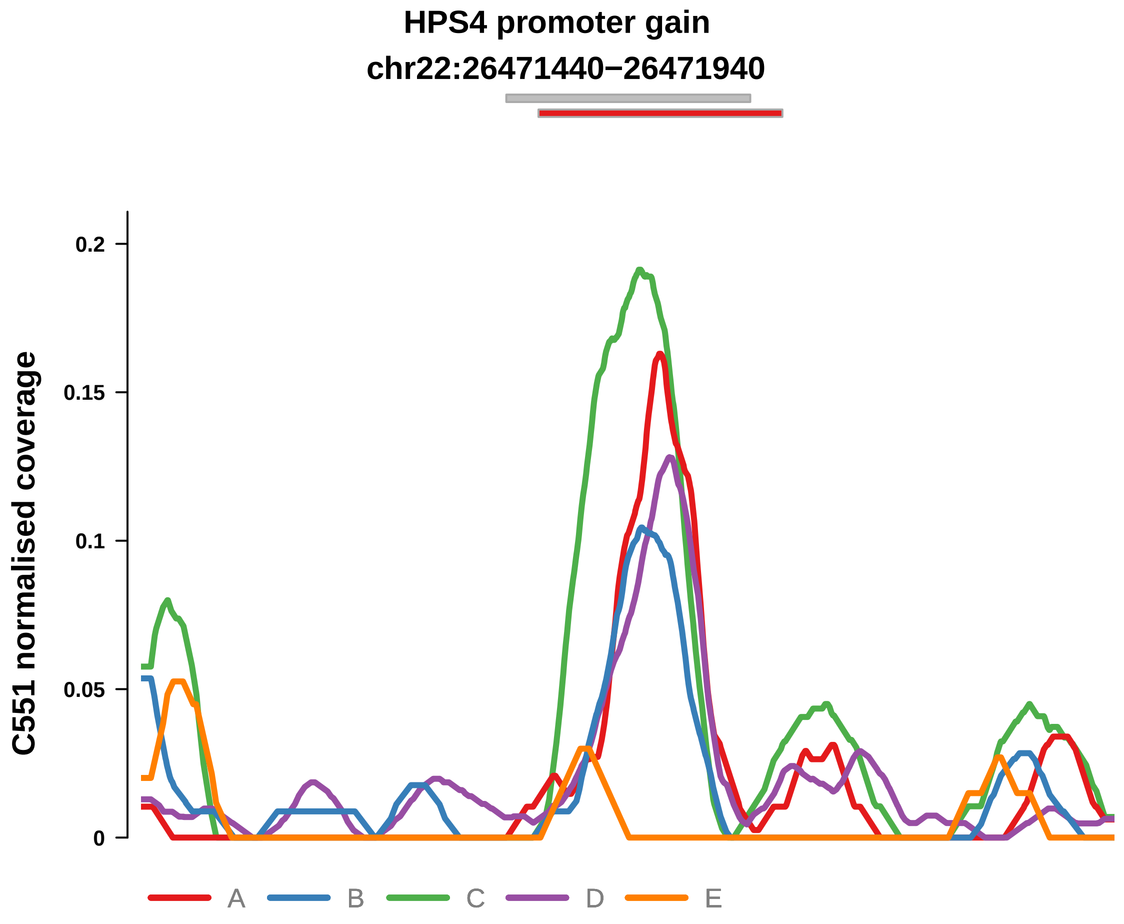


Putatively Clonal (logFC = 2.49)

| | P. | Adjusted P. |
|---|---|---|
| *~region vs. ~* | 9.51e−01 | 9.74e−01 |
| *~purity vs ~* | 1.98e−01 | 8.37e−01 |
| *~purity+region vs ~purity* | 8.95e−01 | 9.99e−01 |
| *Carcinoma vs. normal* | 1.29e−05 **** | 1.29e−05 **** |

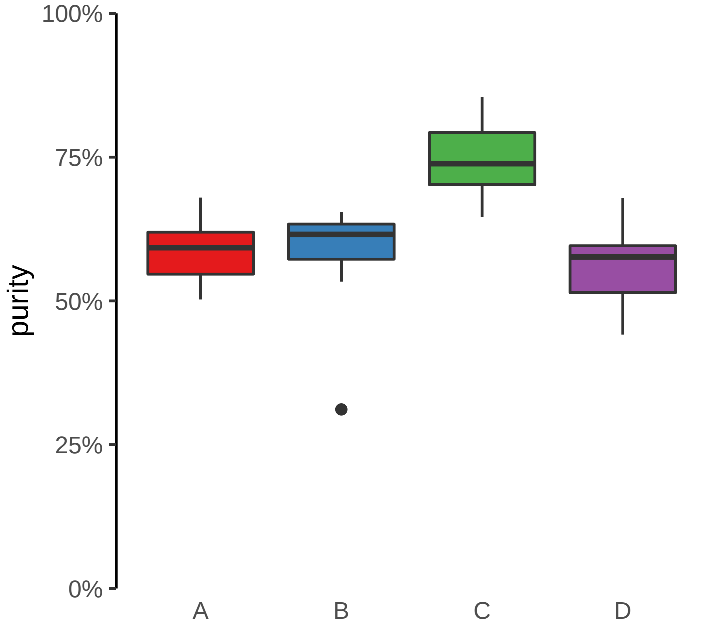

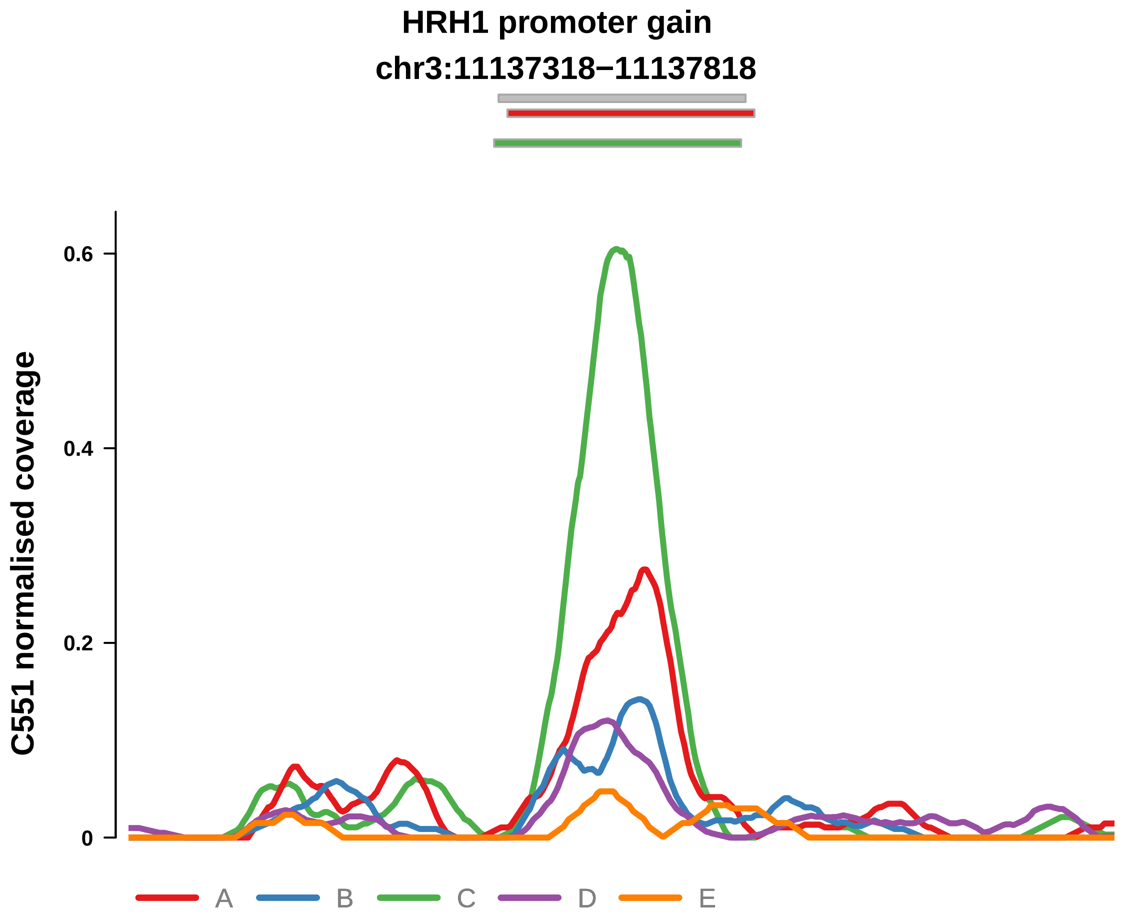


Putatively Clonal (logFC = 2.73)

| | P. | Adjusted P. |
|---|---|---|
| *~region vs. ~* | 2.47e−05 **** | 2.17e−03 ** |
| *~purity vs ~* | 1.11e−02 * | 4.89e−01 |
| *~purity+region vs ~purity* | 1.63e−03 ** | 7.18e−02 |
| *Carcinoma vs. normal* | 1.38e−06 **** | 1.38e−06 **** |

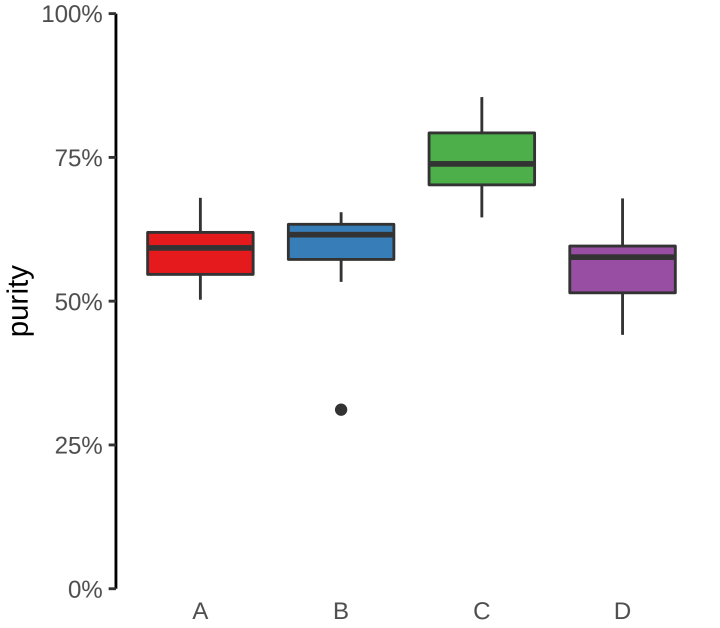

**JAK3 promoter gain**
**chr19:17842053−17842553**

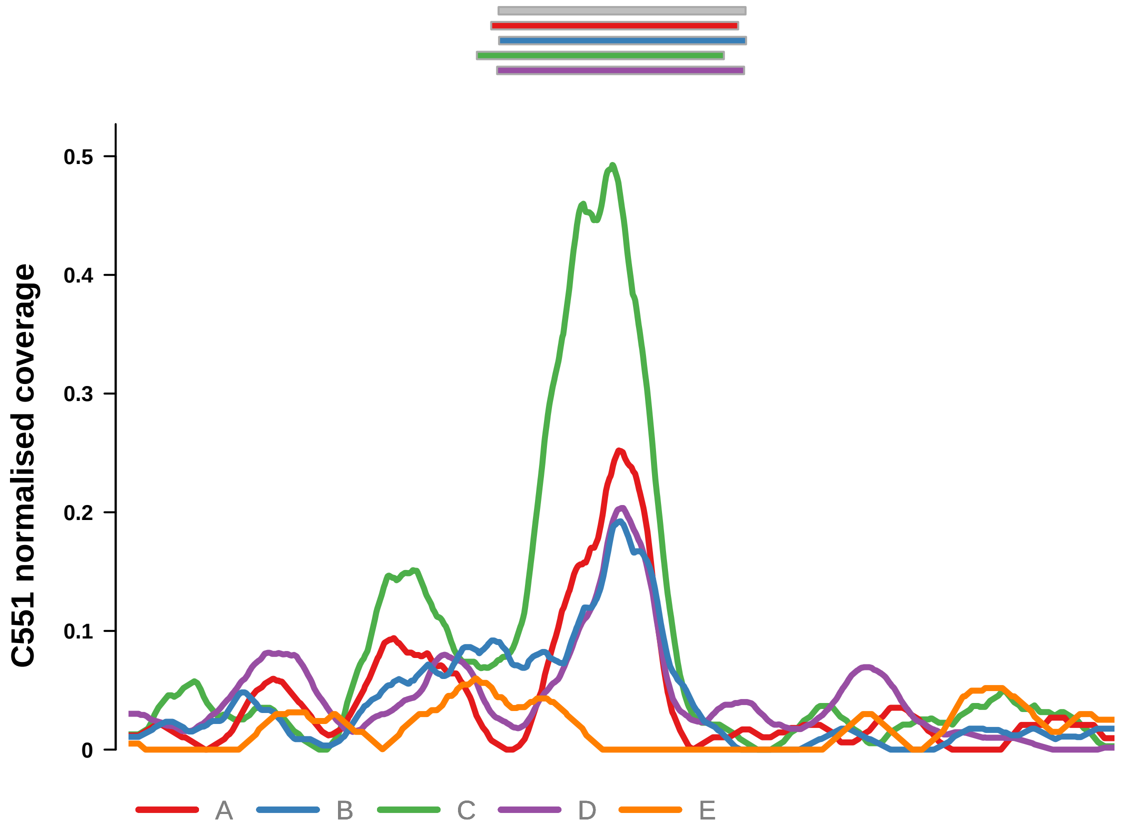


Putatively Clonal (logFC = 2.26)

| | P. | Adjusted P. |
|---|---|---|
| *~region vs. ~* | 4.38e−01 | 8.39e−01 |
| *~purity vs ~* | 1.48e−01 | 7.97e−01 |
| *~purity+region vs ~purity* | 5.95e−01 | 9.48e−01 |
| *Carcinoma vs. normal* | 4.53e−05 **** | 4.53e−05 **** |

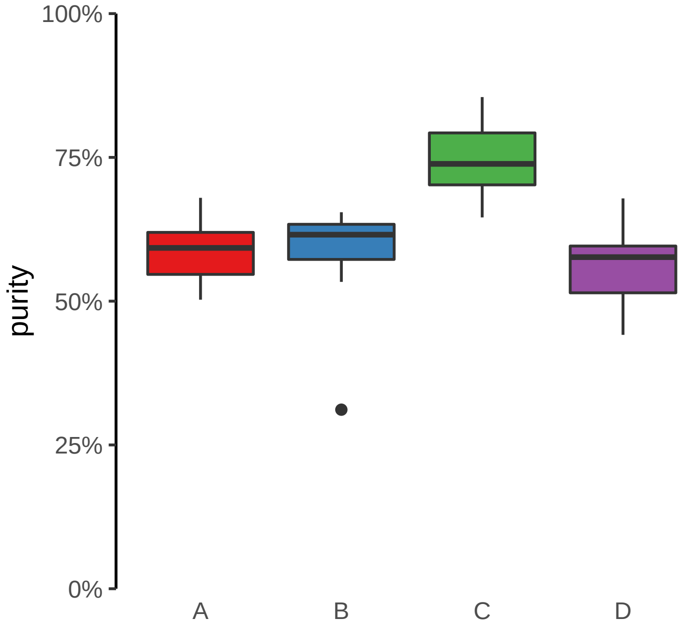

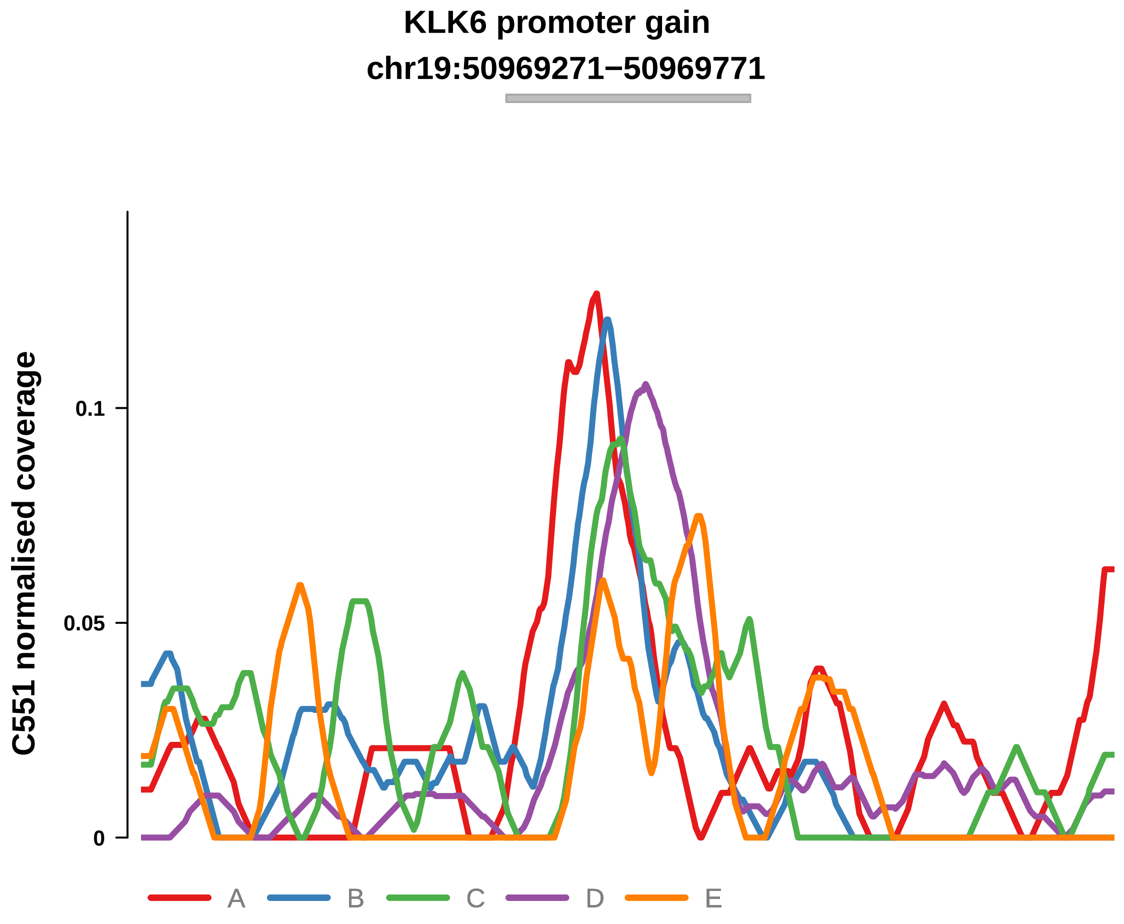


Putatively Clonal (logFC = 1.82)

| | P. | Adjusted P. |
|---|---|---|
| *~region vs. ~* | 5.39e−01 | 8.69e−01 |
| *~purity vs ~* | 9.38e−01 | 9.69e−01 |
| *~purity+region vs ~purity* | 3.2e−01 | 7.87e−01 |
| *Carcinoma vs. normal* | 1.21e−03 ** | 1.21e−03 ** |

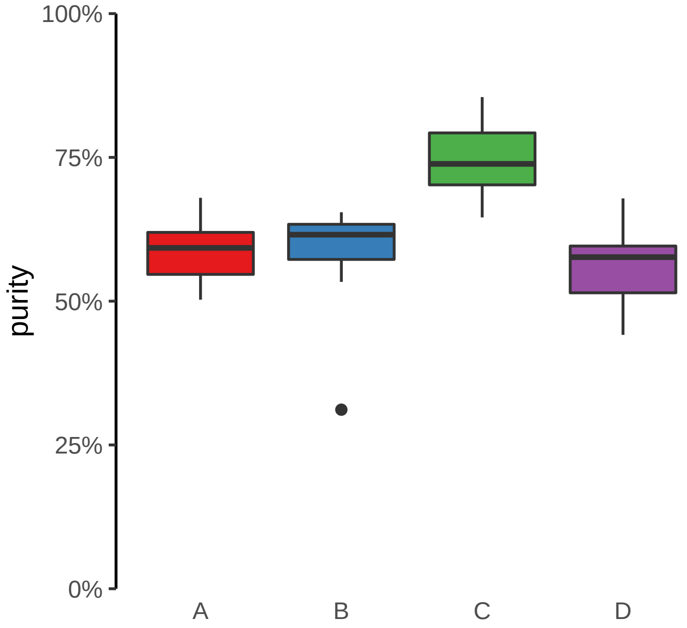

**KRT7 promoter gain**
**chr12:52235898−52236398**

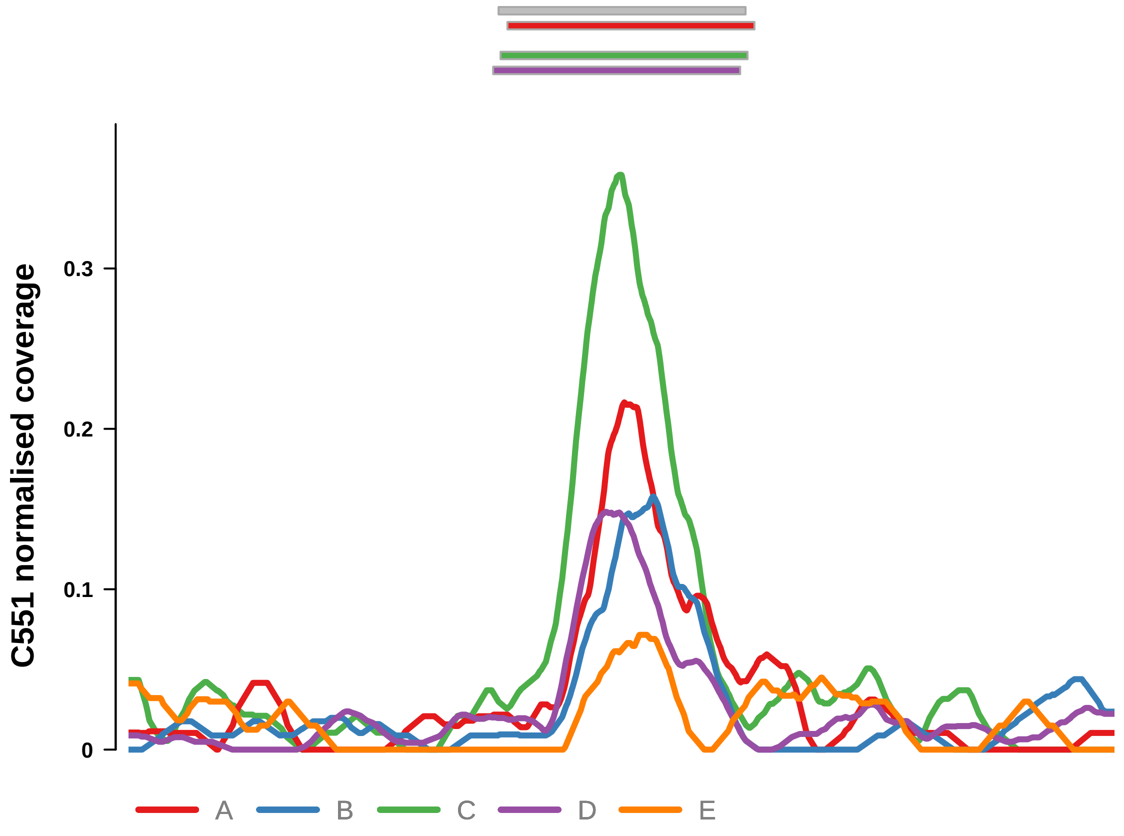


Putatively Clonal (logFC = 2.05)

| | P. | Adjusted P. |
|---|---|---|
| *~region vs. ~* | 4.67e−01 | 8.39e−01 |
| *~purity vs ~* | 1.36e−01 | 7.97e−01 |
| *~purity+region vs ~purity* | 7.62e−01 | 9.87e−01 |
| *Carcinoma vs. normal* | 1.89e−04 *** | 1.89e−04 *** |

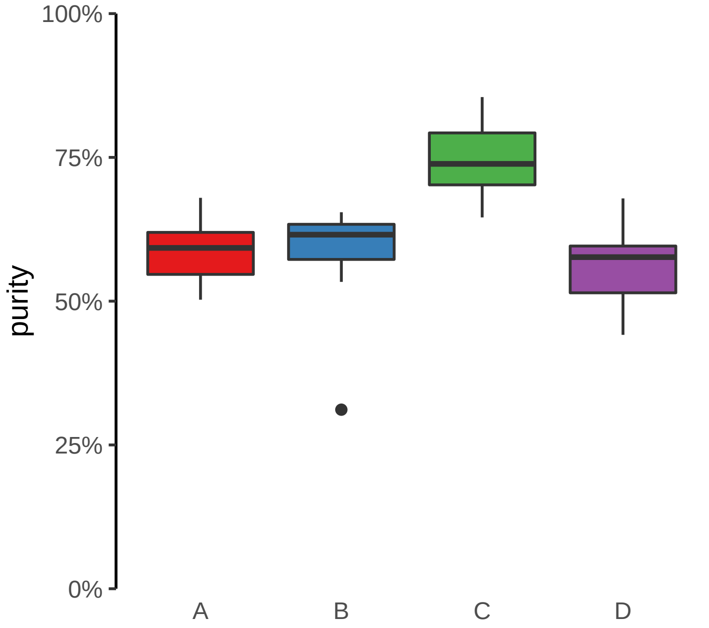

**PITX1 promoter gain**
**chr5:135034130−135034630**

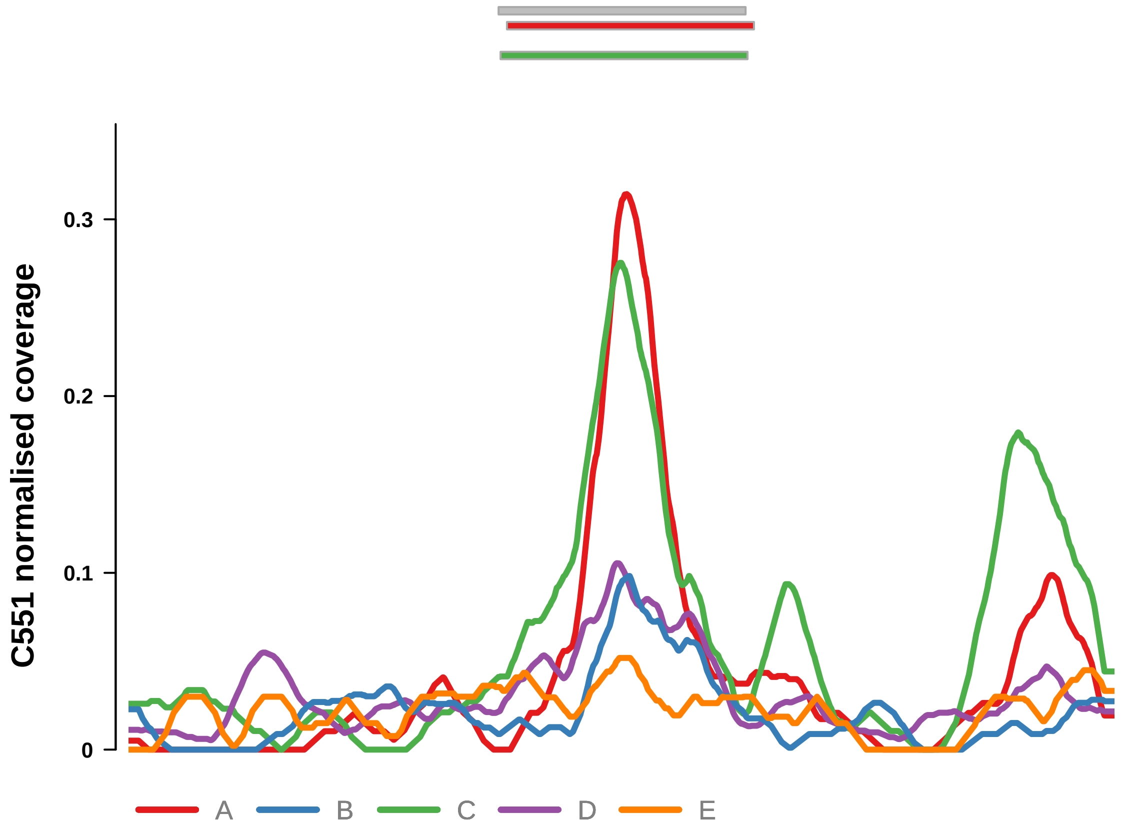


Putatively Clonal (logFC = 1.72)

| | P. | Adjusted P. |
|---|---|---|
| *~region vs. ~* | 9.92e−02 | 4.16e−01 |
| *~purity vs ~* | 7.79e−01 | 9.44e−01 |
| *~purity+region vs ~purity* | 8.56e−02 | 5.7e−01 |
| *Carcinoma vs. normal* | 1.69e−03 ** | 1.69e−03 ** |

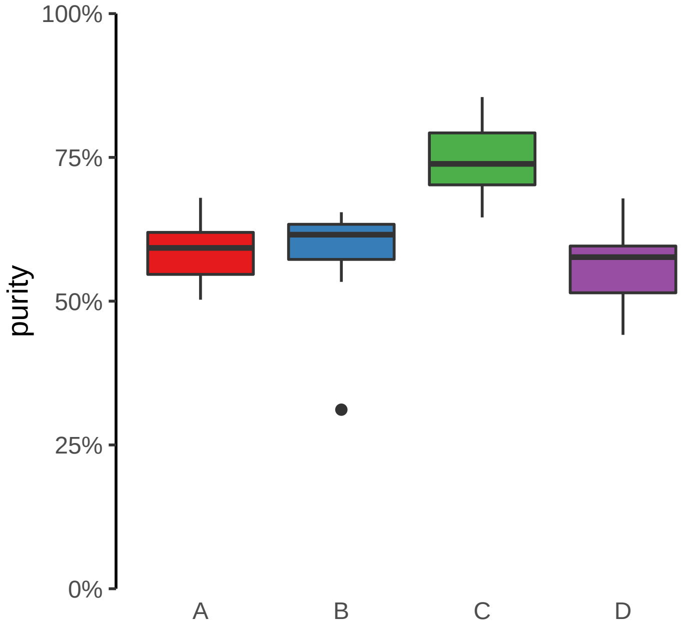

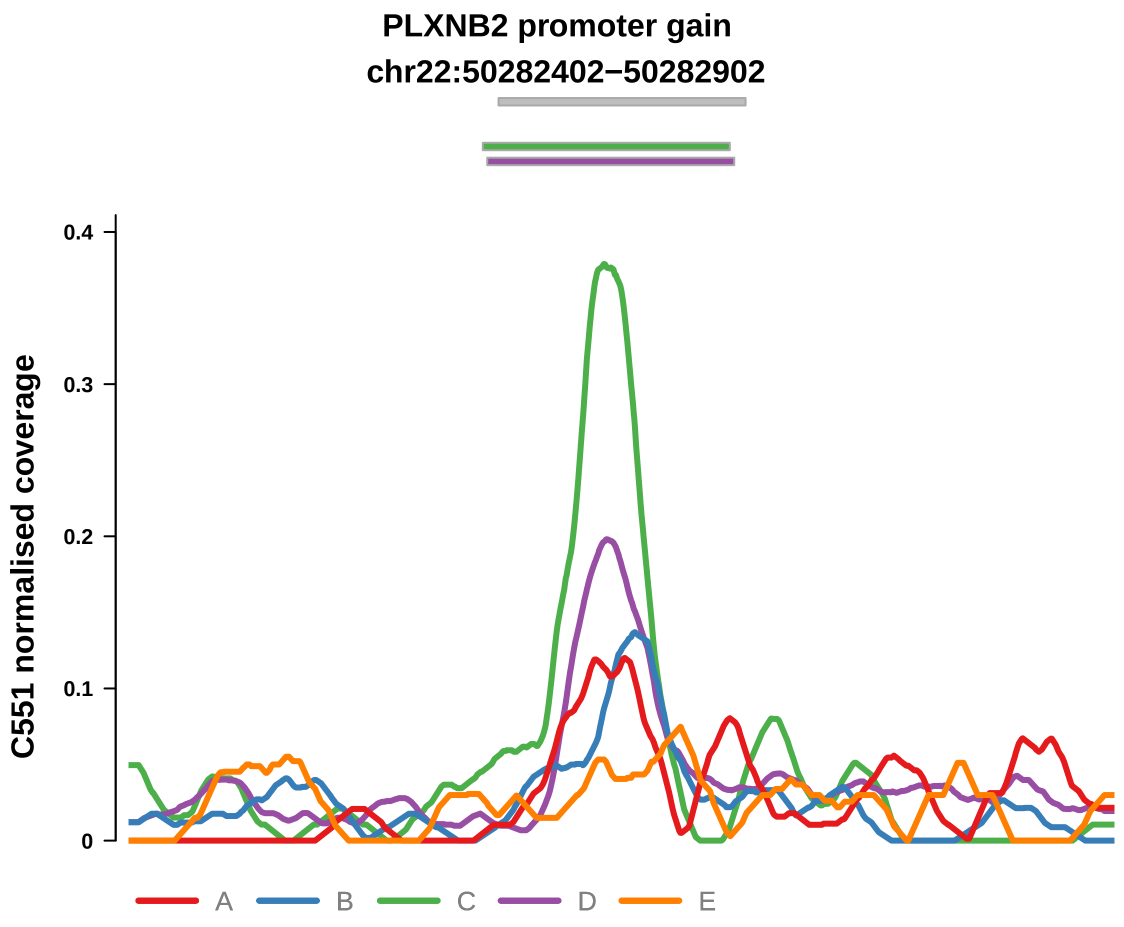


Putatively Clonal (logFC = 2.28)

| | P. | Adjusted P. |
|---|---|---|
| *~region vs. ~* | 3.43e−01 | 7.4e−01 |
| *~purity vs ~* | 5.87e−01 | 9.06e−01 |
| *~purity+region vs ~purity* | 3.57e−01 | 8.26e−01 |
| *Carcinoma vs. normal* | 4.41e−05 **** | 4.41e−05 **** |

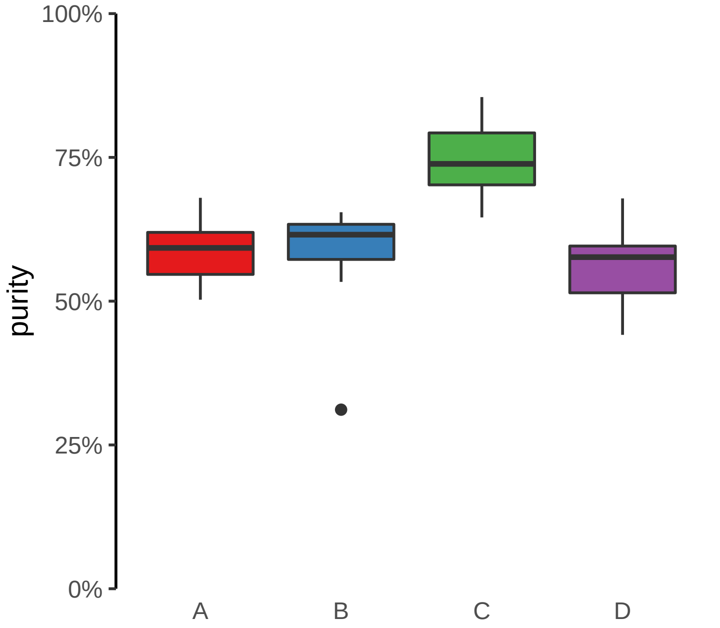

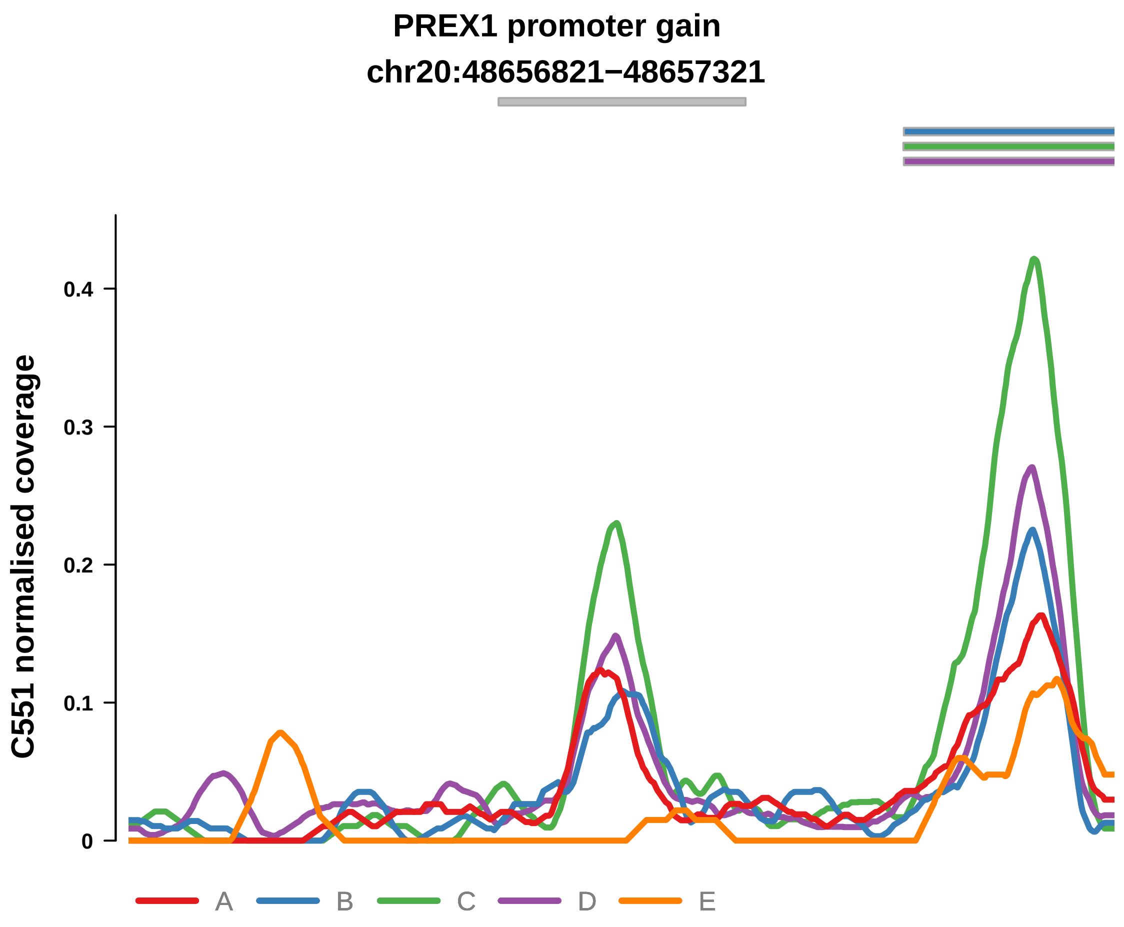


Putatively Clonal (logFC = 2.3)

| | P. | Adjusted P. |
|---|---|---|
| *~region vs. ~* | 7.83e−01 | 9.65e−01 |
| *~purity vs ~* | 8.84e−01 | 9.69e−01 |
| *~purity+region vs ~purity* | 5.94e−01 | 9.48e−01 |
| *Carcinoma vs. normal* | 4.8e−05 **** | 4.8e−05 **** |

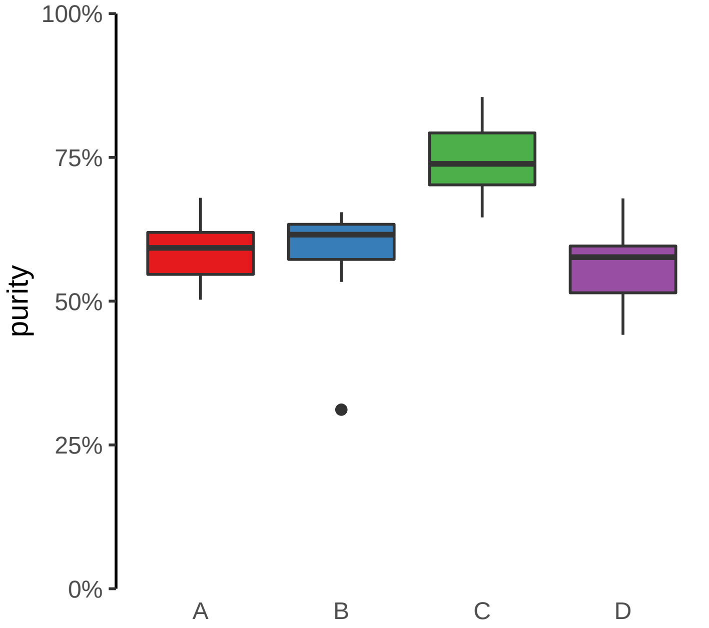

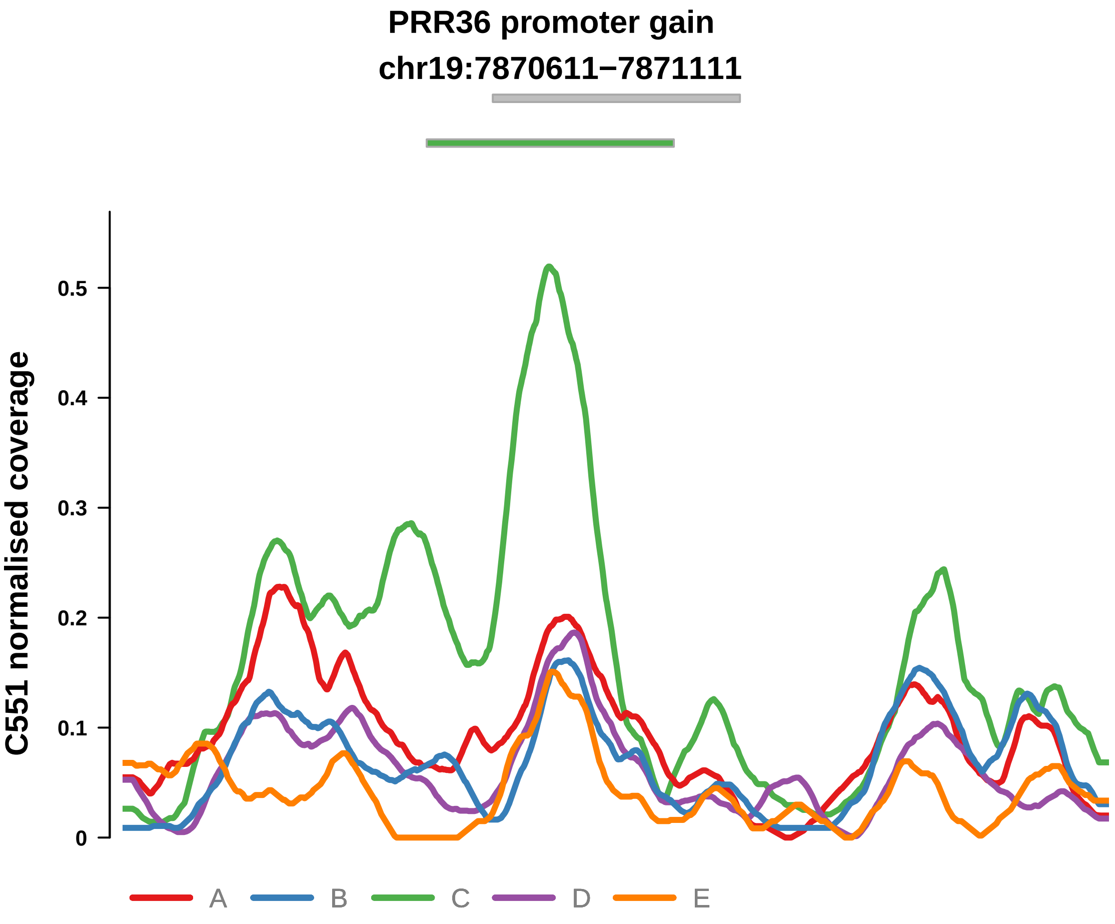


Putatively Clonal (logFC = 2.17)

| | P. | Adjusted P. |
|---|---|---|
| *~region vs. ~* | 4.23e−02 * | 2.67e−01 |
| *~purity vs ~* | 2.63e−01 | 9.06e−01 |
| *~purity+region vs ~purity* | 4.37e−02 * | 4.93e−01 |
| *Carcinoma vs. normal* | 7.73e−05 **** | 7.73e−05 **** |

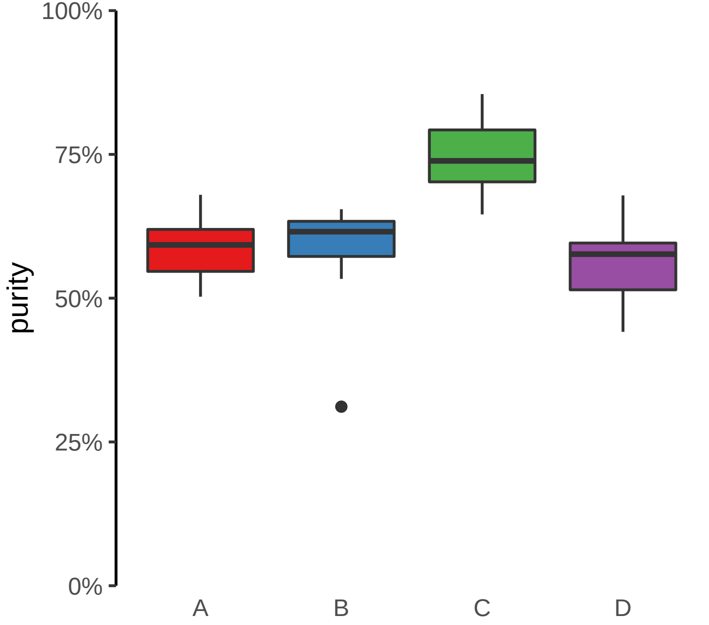

**SAP30BP promoter gain**
**chr17:75706968−75707468**

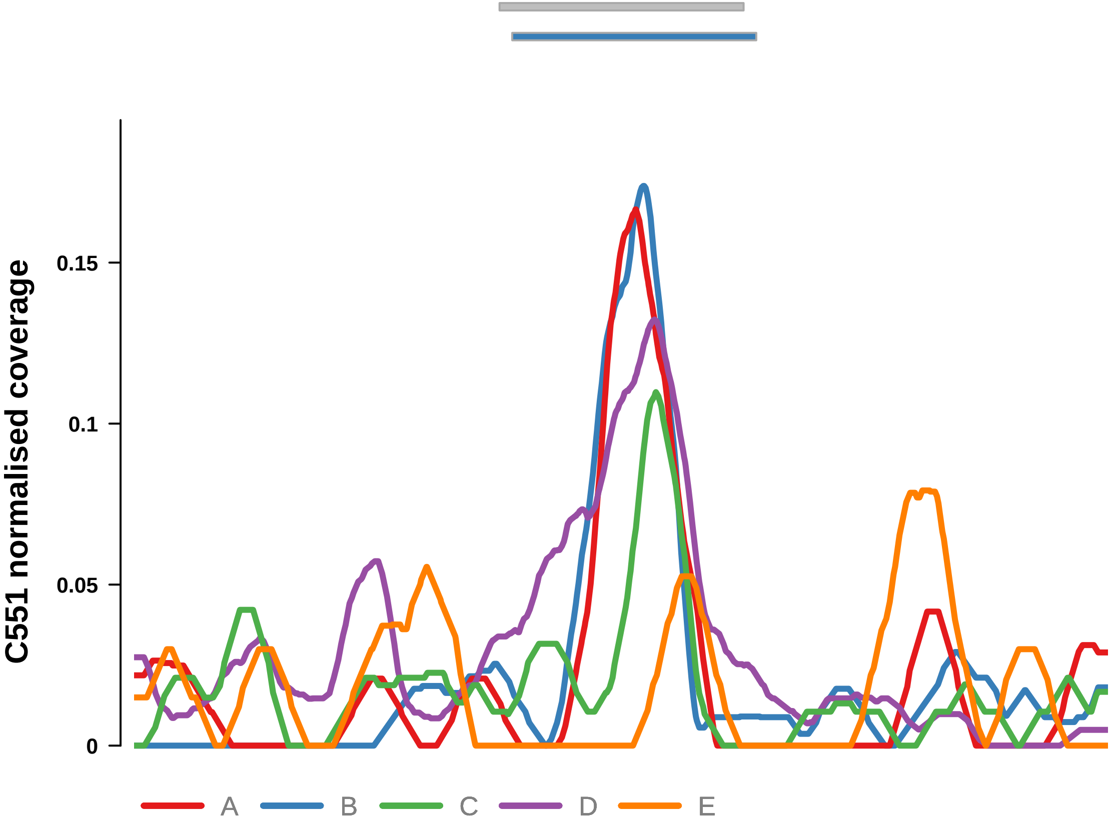


Putatively Clonal (logFC = 2.31)

| | **P.** | **Adjusted P.** |
|---|---|---|
| *~region vs. ~* | 8.71e−03 ** | 1.75e−01 |
| *~purity vs ~* | 1.38e−01 | 7.97e−01 |
| *~purity+region vs ~purity* | 2.41e−02 * | 4.25e−01 |
| *Carcinoma vs. normal* | 4.18e−05 **** | 4.18e−05 **** |

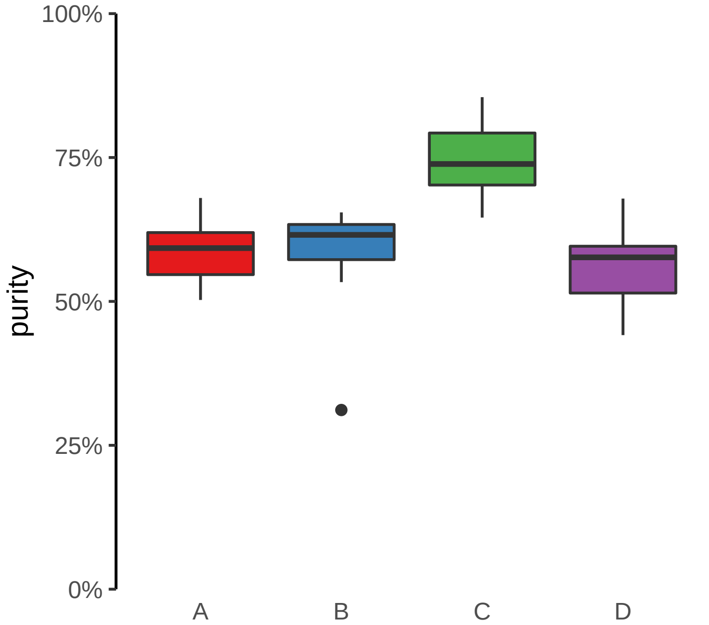

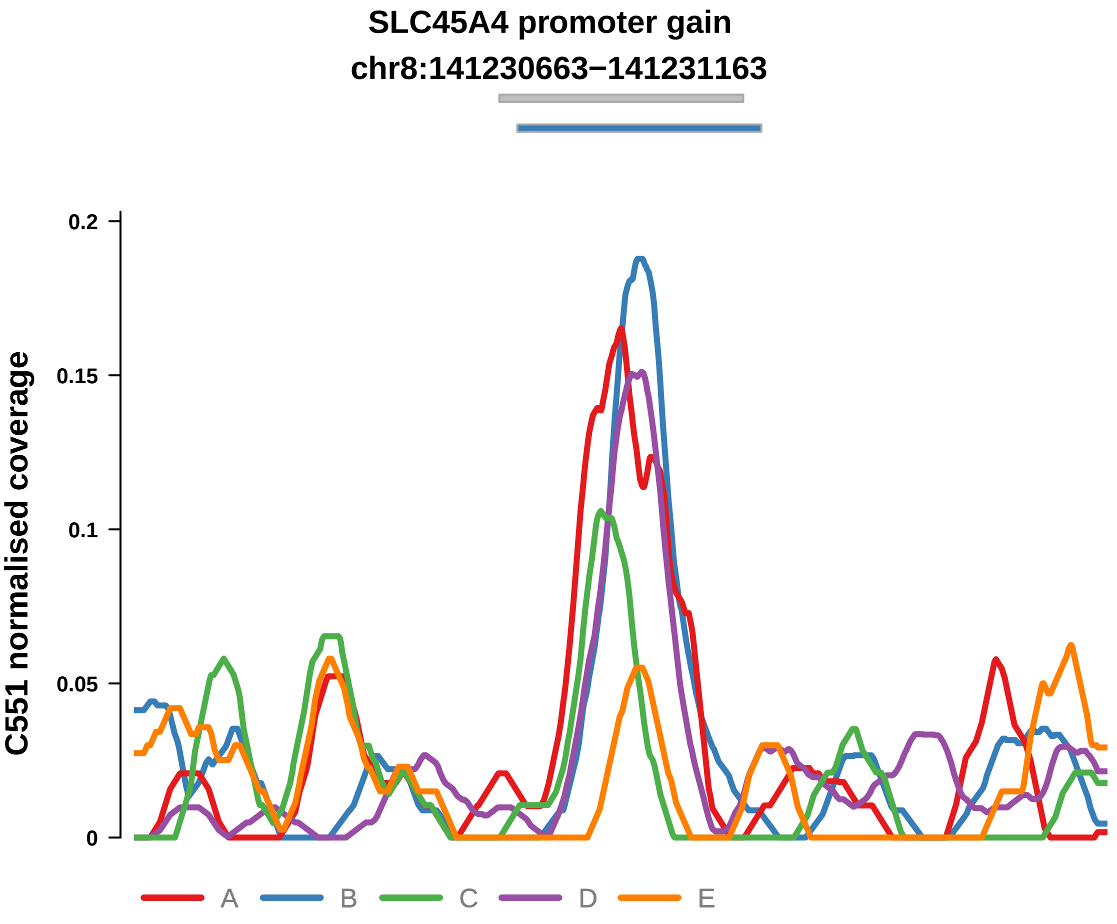


Putatively Clonal (logFC = 2.12)

| | P. | Adjusted P. |
|---|---|---|
| *~region vs. ~* | 9.61e−03 ** | 1.75e−01 |
| *~purity vs ~* | 3.25e−01 | 9.06e−01 |
| *~purity+region vs ~purity* | 7.33e−03 ** | 2.15e−01 |
| *Carcinoma vs. normal* | 1.66e−04 *** | 1.66e−04 *** |

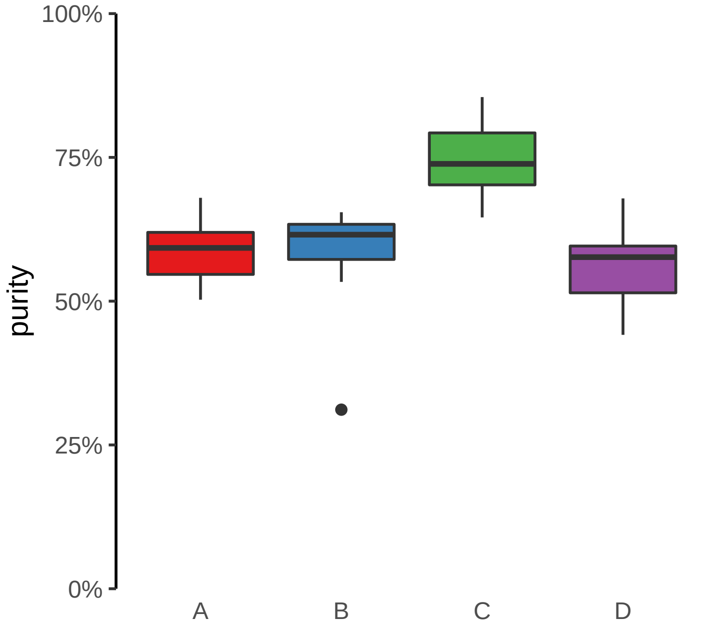

**TG promoter gain**
**chr8:133130728−133131228**

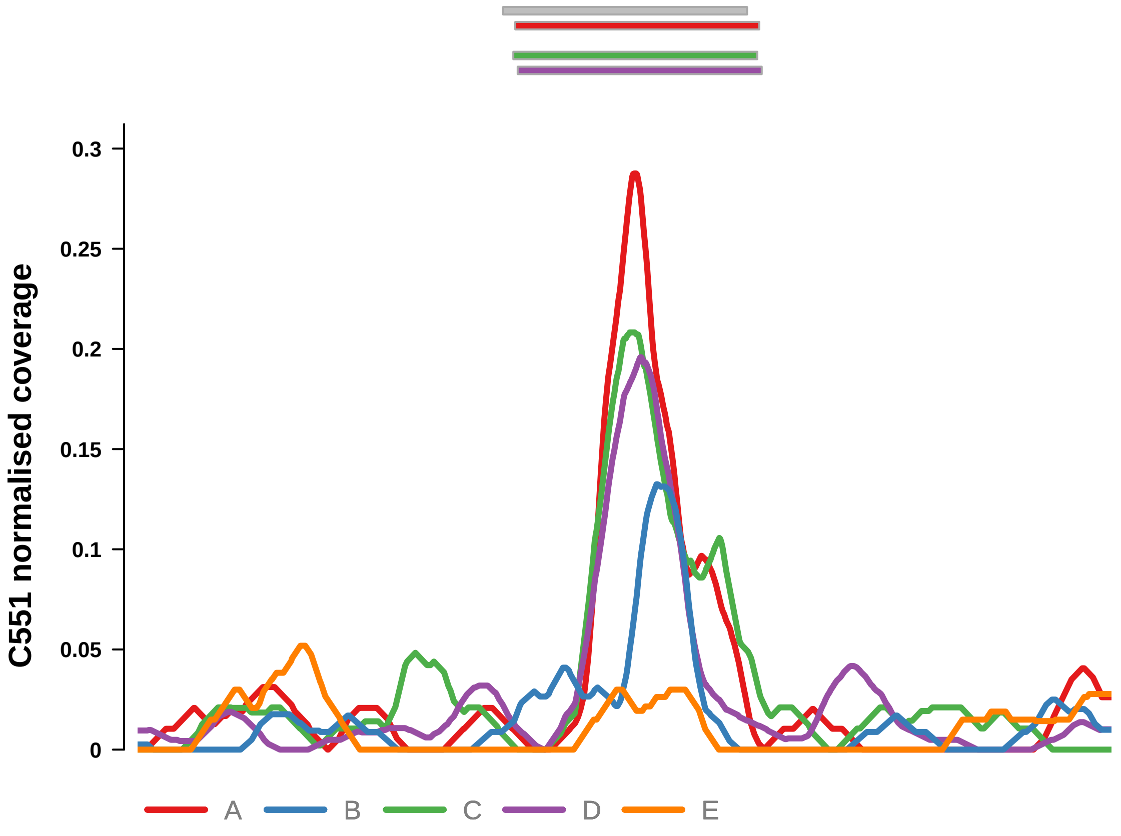


Putatively Clonal (logFC = 2.05)

| | P. | Adjusted P. |
|---|---|---|
| *~region vs. ~* | 5.43e−01 | 8.69e−01 |
| *~purity vs ~* | 2e−01 | 8.37e−01 |
| *~purity+region vs ~purity* | 1.68e−01 | 5.7e−01 |
| *Carcinoma vs. normal* | 2.18e−04 *** | 2.18e−04 *** |

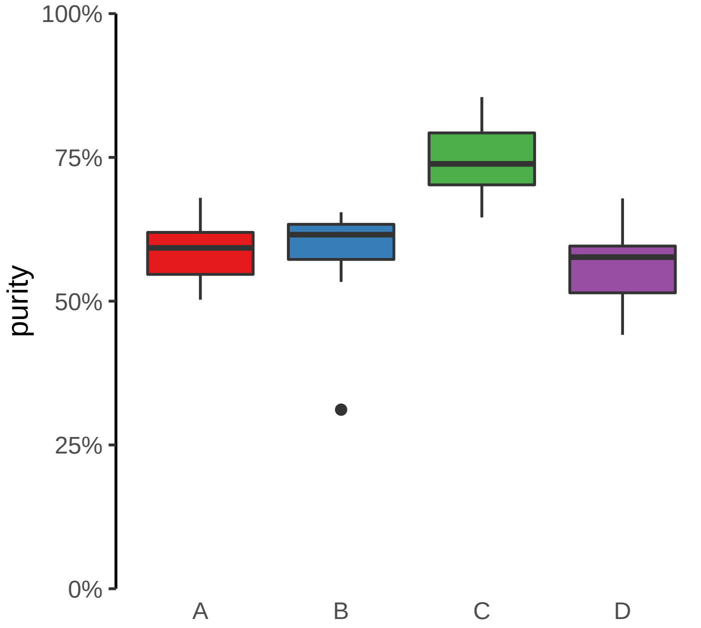

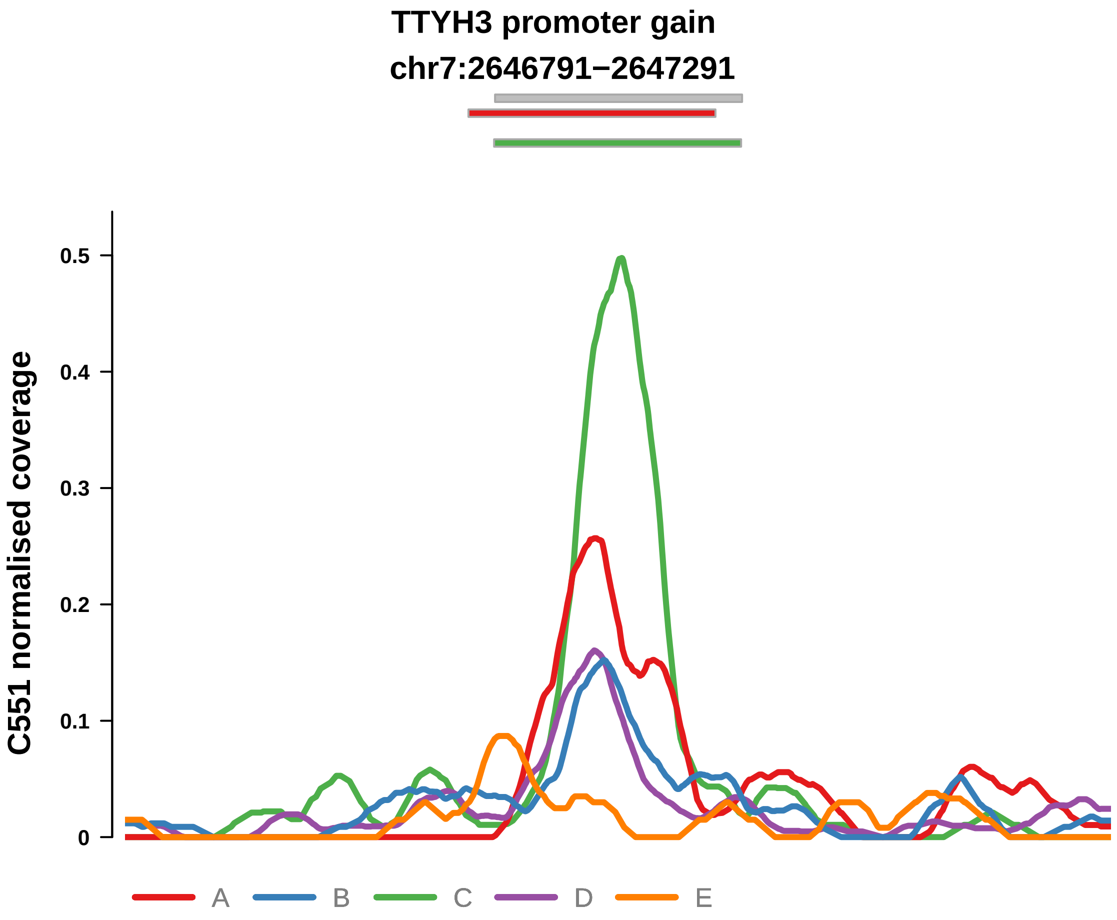


Putatively Clonal (logFC = 2.81)

| | P. | Adjusted P. |
|---|---|---|
| *~region vs. ~* | 2.37e−01 | 6.08e−01 |
| *~purity vs ~* | 9.77e−01 | 9.77e−01 |
| *~purity+region vs ~purity* | 1.18e−01 | 5.7e−01 |
| *Carcinoma vs. normal* | 7.38e−07 **** | 7.38e−07 **** |

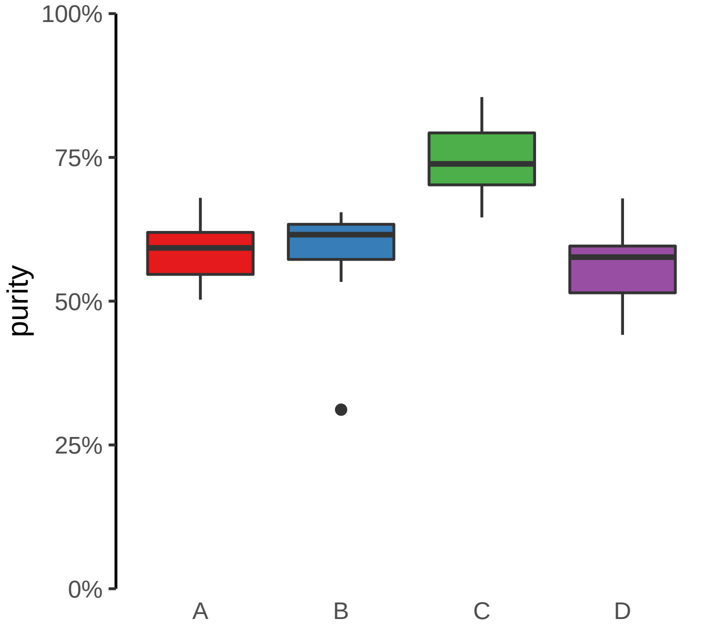

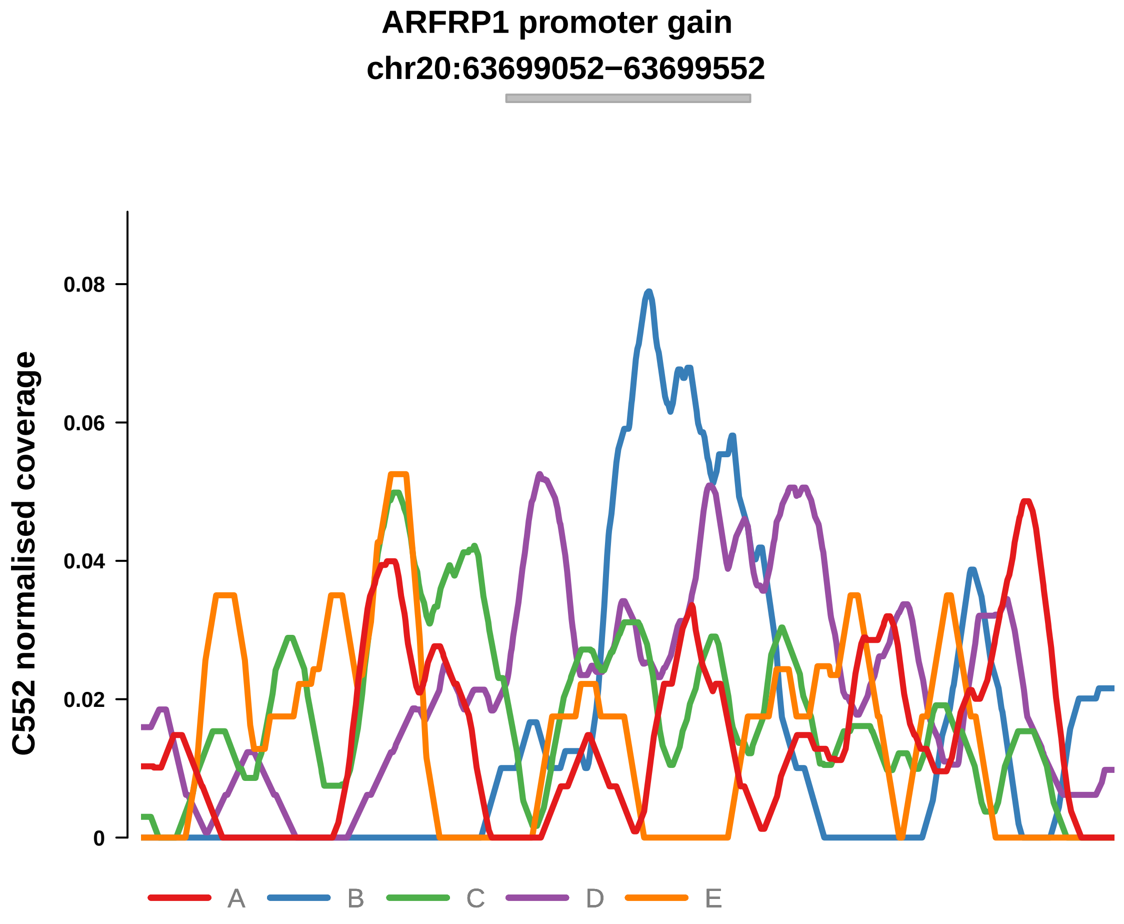


Putatively Clonal (logFC = 2.41)

| | P. | Adjusted P. |
|---|---|---|
| *~region vs. ~* | 1.34e−04 *** | 6.29e−03 ** |
| *~purity vs ~* | 5.73e−03 ** | 2.39e−02 * |
| *~purity+region vs ~purity* | 9.73e−03 ** | 3.6e−01 |
| *Carcinoma vs. normal* | 1.6e−04 *** | 1.6e−04 *** |

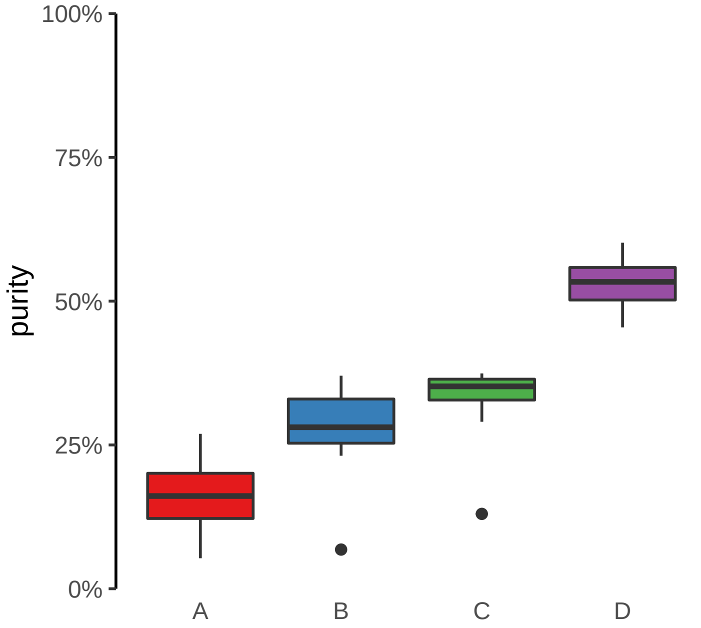

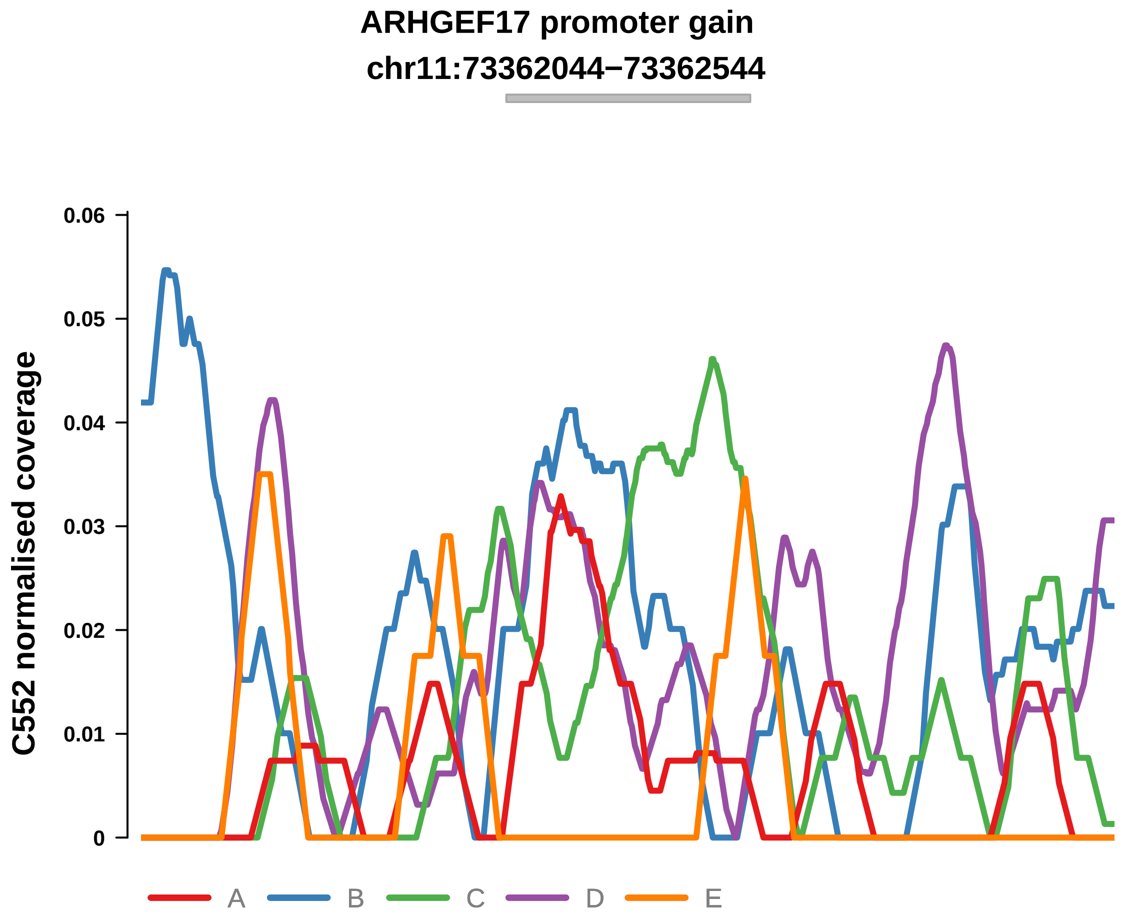


Putatively Clonal (logFC = 1.84)

| | P. | Adjusted P. |
|---|---|---|
| *~region vs. ~* | 1.41e−01 | 2.82e−01 |
| *~purity vs ~* | 4.02e−02 * | 8.93e−02 |
| *~purity+region vs ~purity* | 7.32e−01 | 9.85e−01 |
| *Carcinoma vs. normal* | 9.84e−03 ** | 9.84e−03 ** |

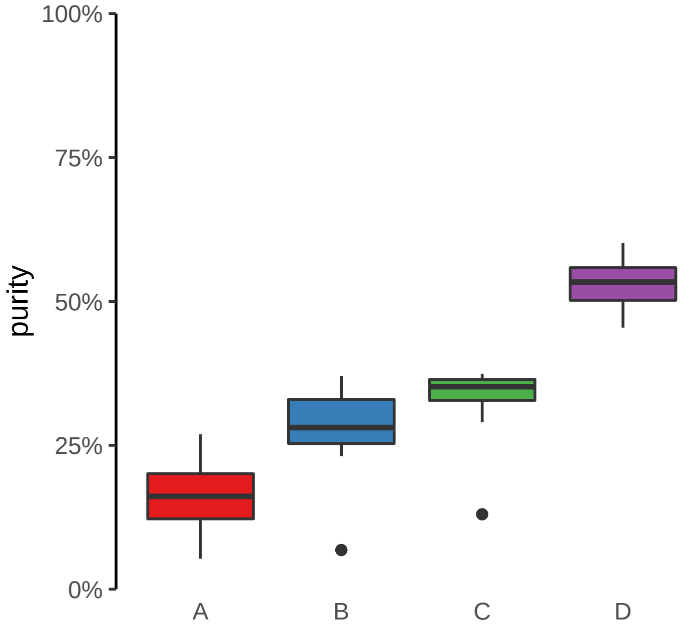

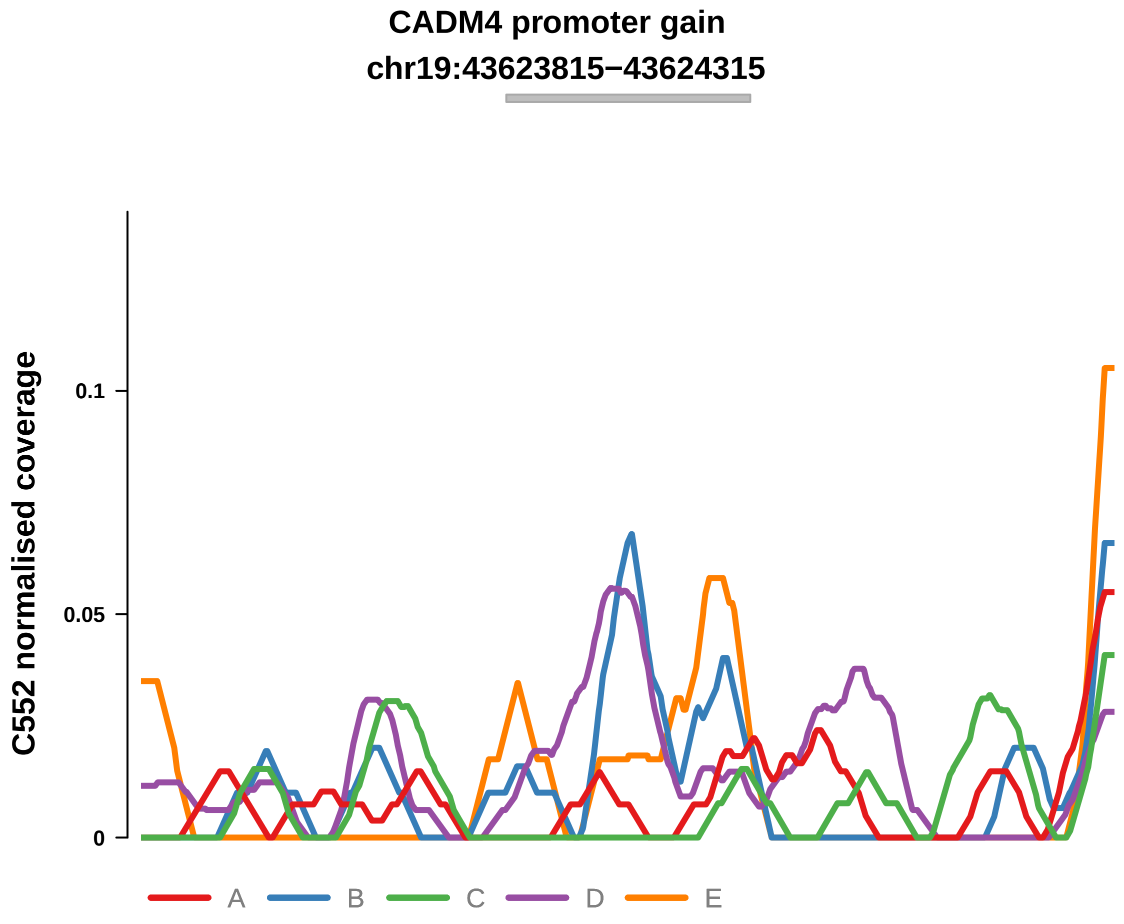


Putatively Clonal (logFC = 2.47)

| | P. | Adjusted P. |
|---|---|---|
| *~region vs. ~* | 5.77e−04 *** | 9.05e−03 ** |
| *~purity vs ~* | 2.1e−02 * | 6.46e−02 |
| *~purity+region vs ~purity* | 1.18e−02 * | 3.6e−01 |
| *Carcinoma vs. normal* | 2.9e−04 *** | 2.9e−04 *** |

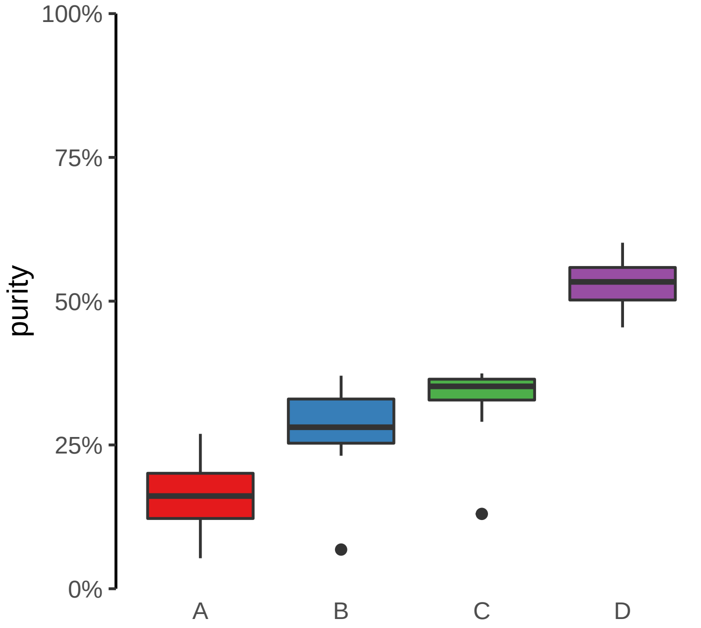

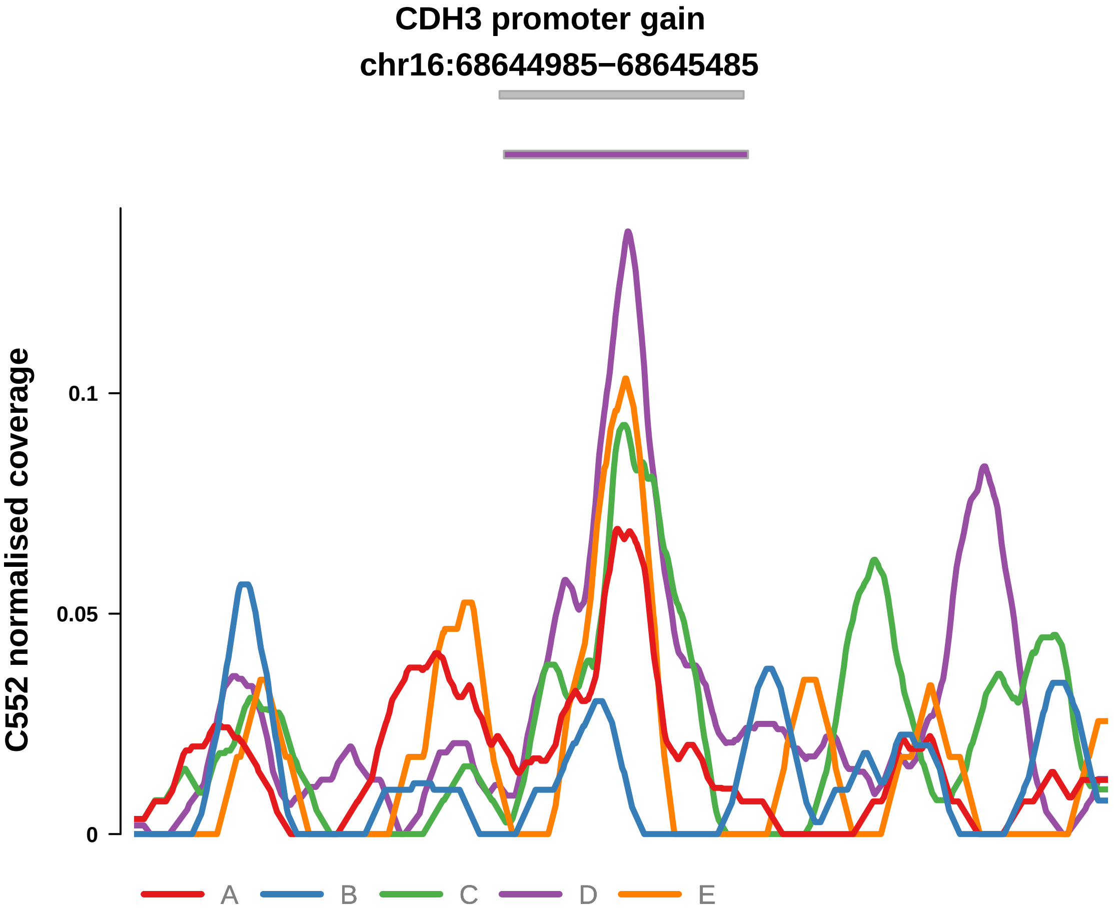


Putatively Clonal (logFC = 2.42)

| | P. | Adjusted P. |
|---|---|---|
| *~region vs. ~* | 3.77e−04 *** | 8.87e−03 ** |
| *~purity vs ~* | 1.76e−03 ** | 1.1e−02 * |
| *~purity+region vs ~purity* | 4.38e−02 * | 4.82e−01 |
| *Carcinoma vs. normal* | 4.74e−05 **** | 4.74e−05 **** |

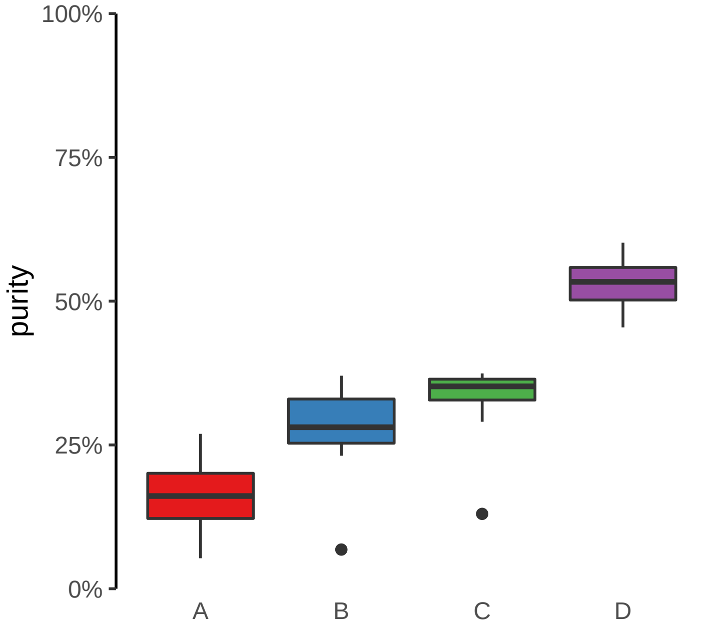

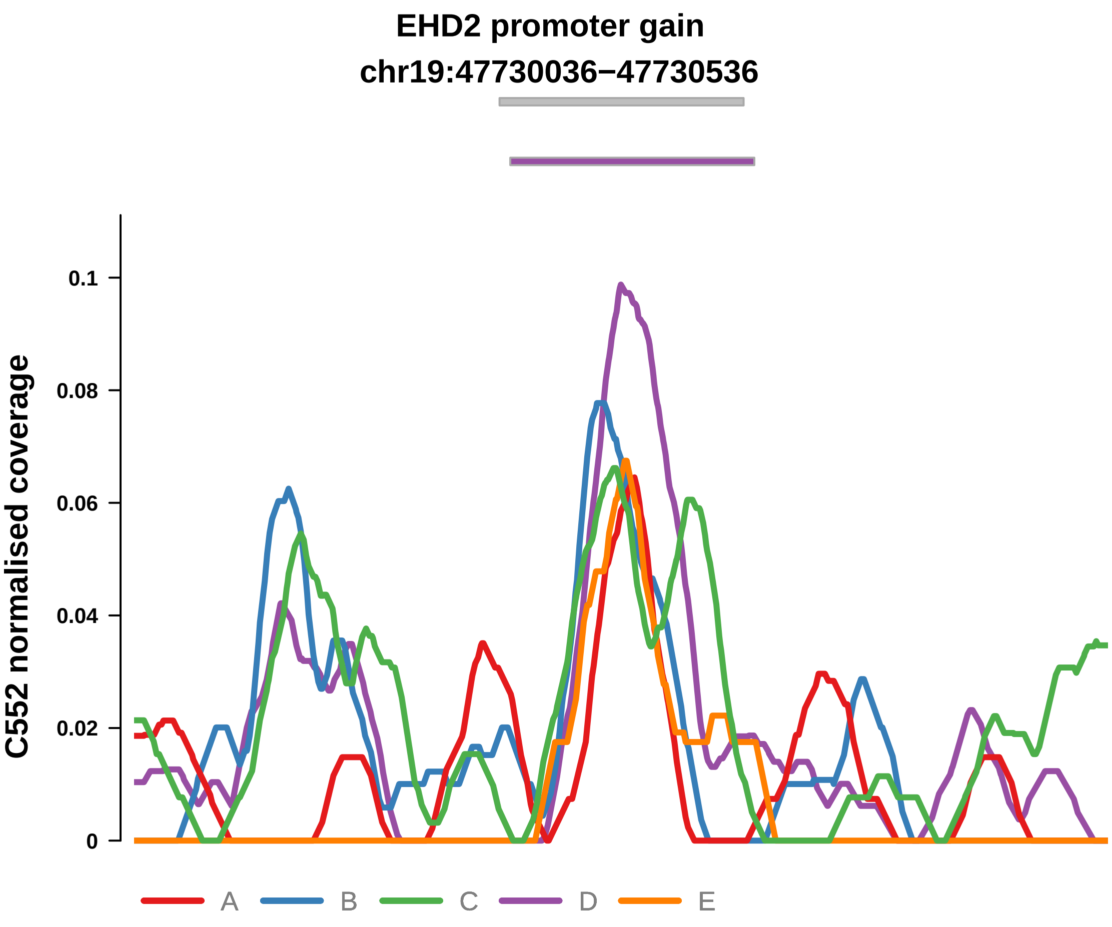


Putatively Clonal (logFC = 2.34)

| | P. | Adjusted P. |
|---|---|---|
| *~region vs. ~* | 1.47e−01 | 2.82e−01 |
| *~purity vs ~* | 2.15e−02 * | 6.46e−02 |
| *~purity+region vs ~purity* | 7.85e−01 | 9.85e−01 |
| *Carcinoma vs. normal* | 1.47e−04 *** | 1.47e−04 *** |

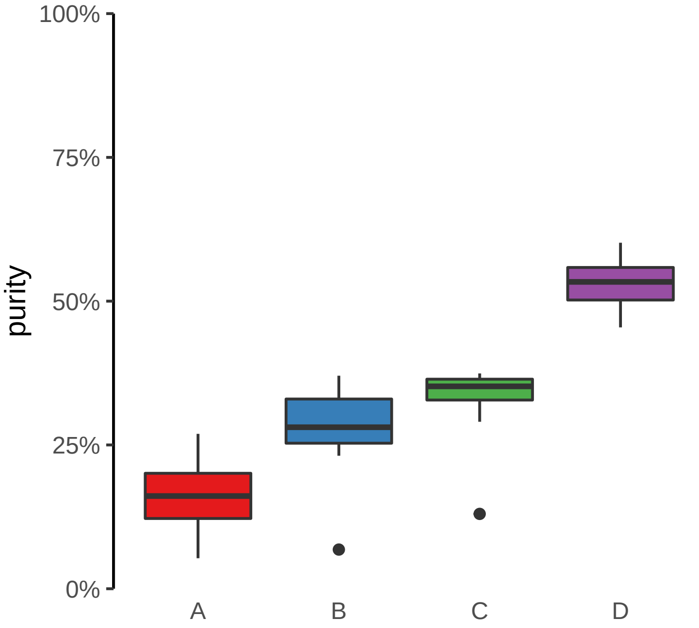

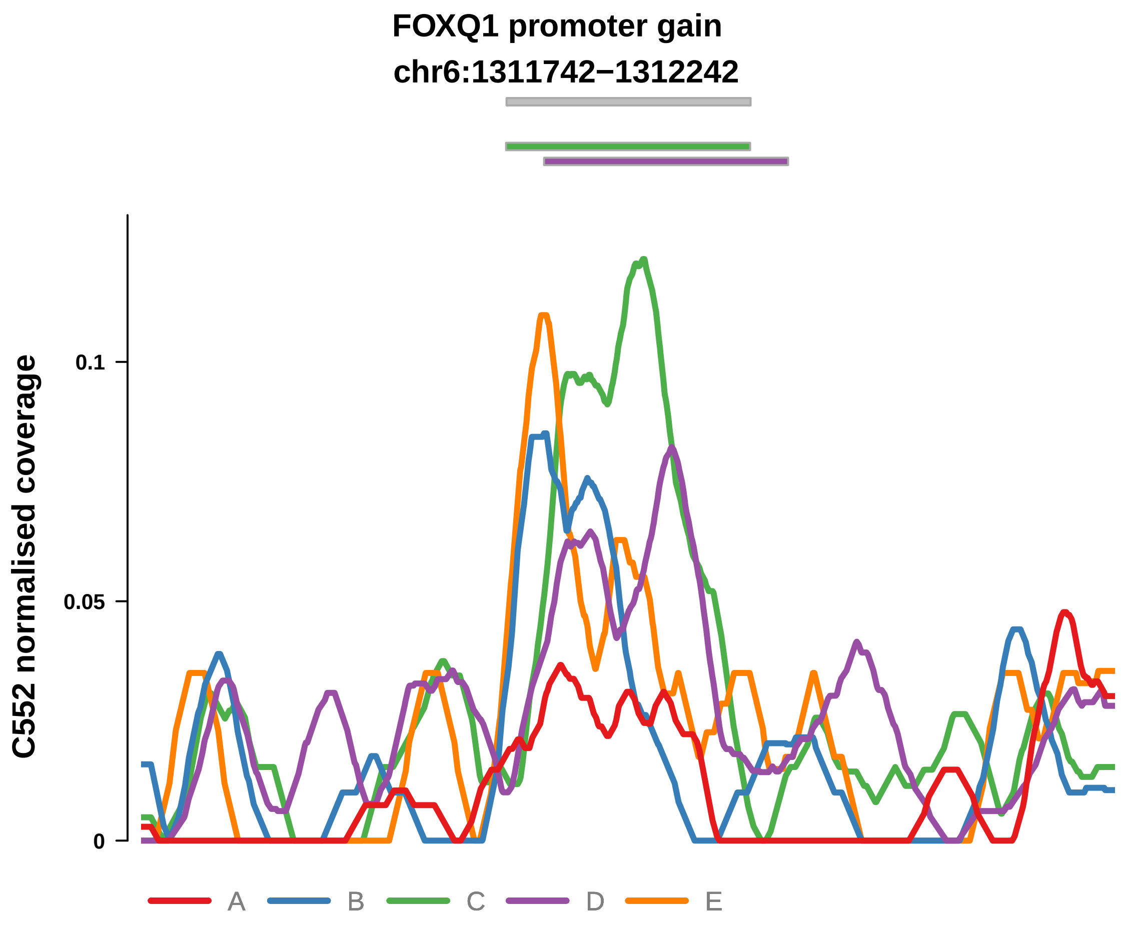


Putatively Clonal (logFC = 2.12)

| | P. | Adjusted P. |
|---|---|---|
| *~region vs. ~* | 3.38e−02 * | 9.92e−02 |
| *~purity vs ~* | 1.24e−04 *** | 2.06e−03 ** |
| *~purity+region vs ~purity* | 2.94e−01 | 9.1e−01 |
| *Carcinoma vs. normal* | 3.79e−04 *** | 3.79e−04 *** |

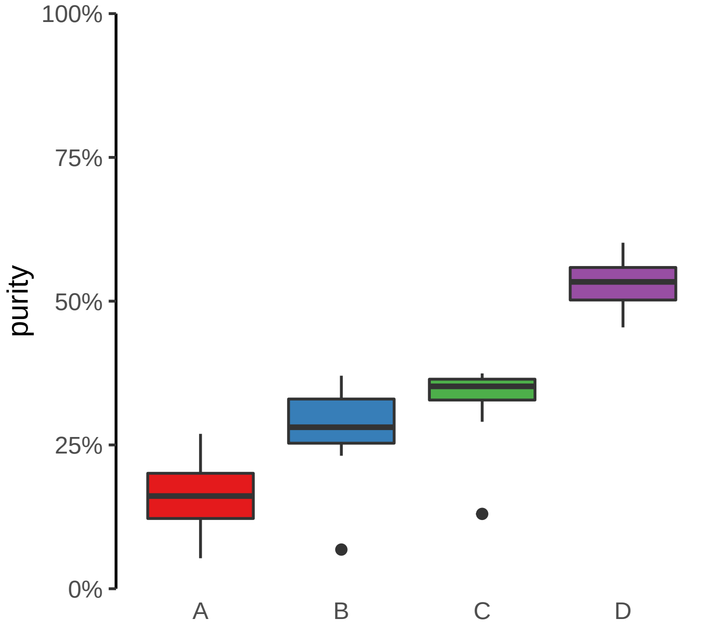

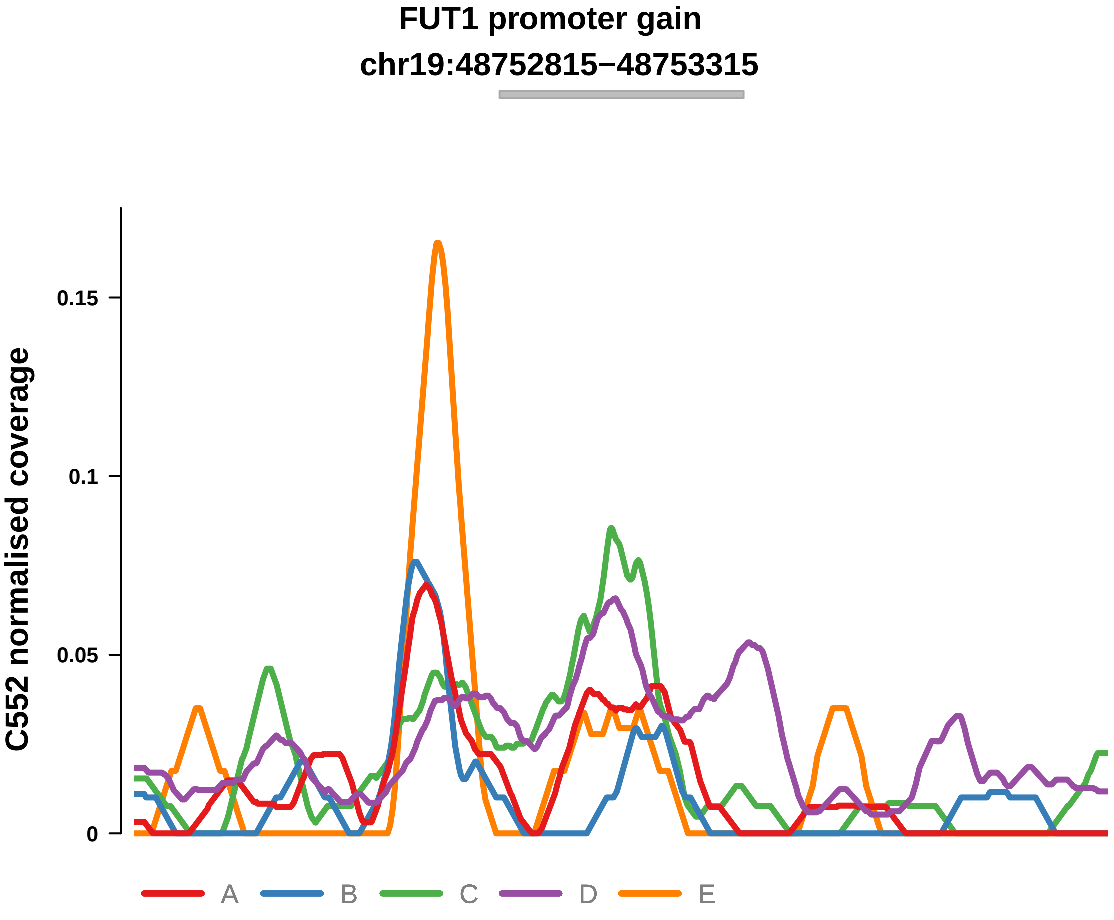


Putatively Clonal (logFC = 2.14)

| | P. | Adjusted P. |
|---|---|---|
| *~region vs. ~* | 8.53e−04 *** | 1e−02 * |
| *~purity vs ~* | 4.93e−06 **** | 2.46e−04 *** |
| *~purity+region vs ~purity* | 2.3e−01 | 9.1e−01 |
| *Carcinoma vs. normal* | 4.96e−04 *** | 4.96e−04 *** |

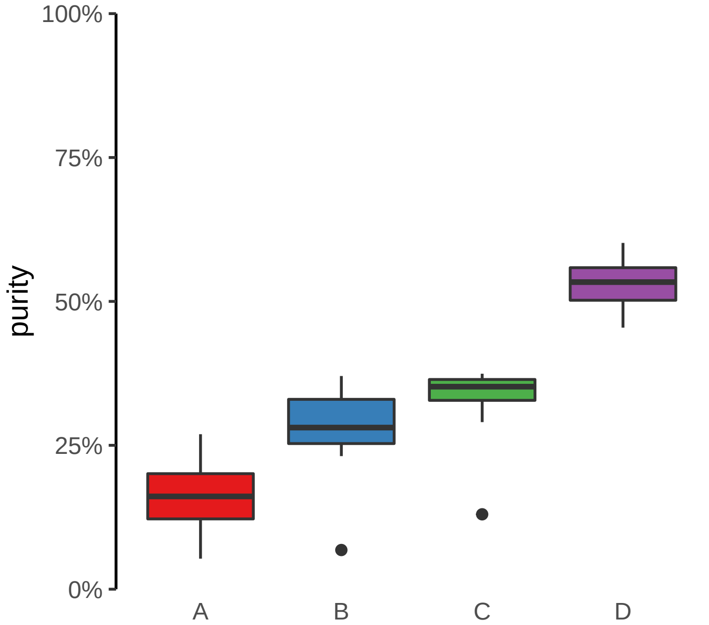

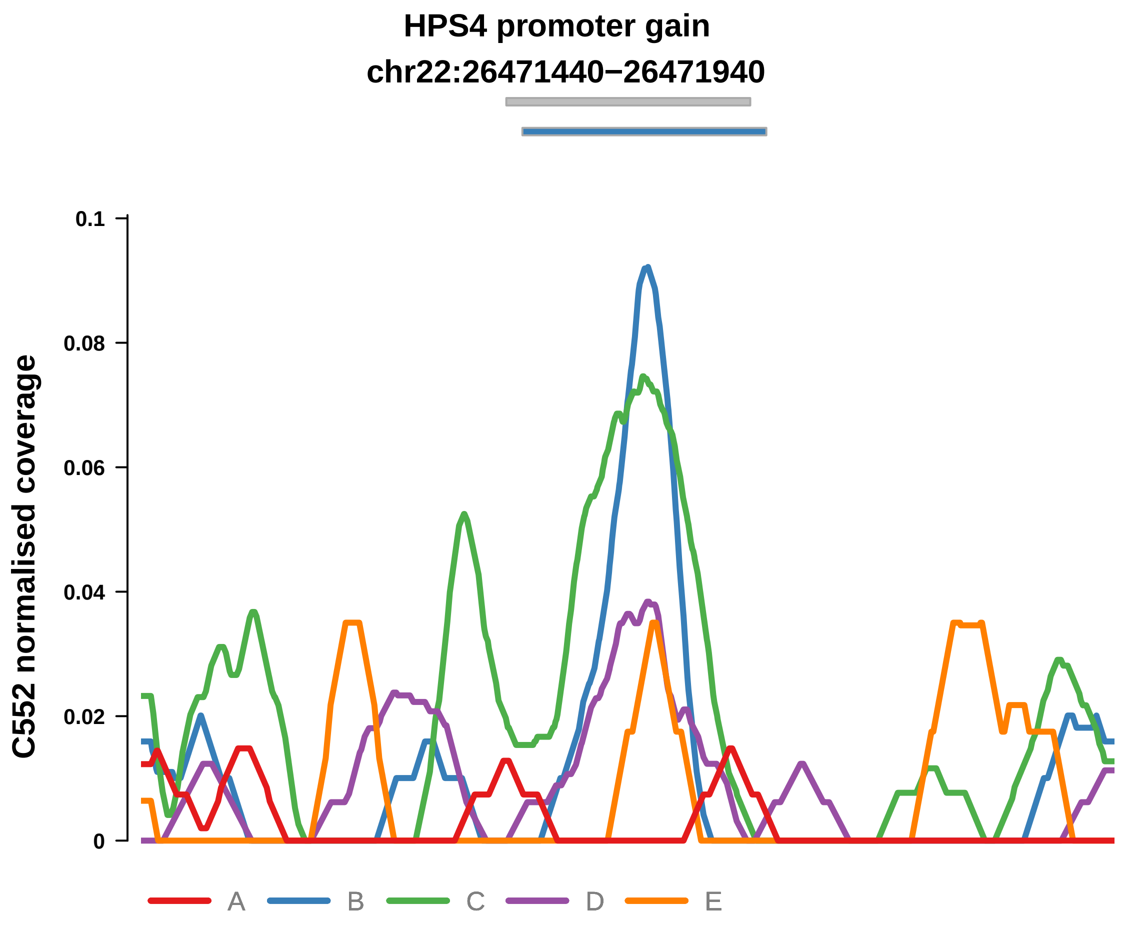


Putatively Clonal (logFC = 2.13)

| | P. | Adjusted P. |
|---|---|---|
| *~region vs. ~* | 1.63e−01 | 2.82e−01 |
| *~purity vs ~* | 4.99e−02 * | 9.98e−02 |
| *~purity+region vs ~purity* | 7.3e−01 | 9.85e−01 |
| *Carcinoma vs. normal* | 4.77e−03 ** | 4.77e−03 ** |

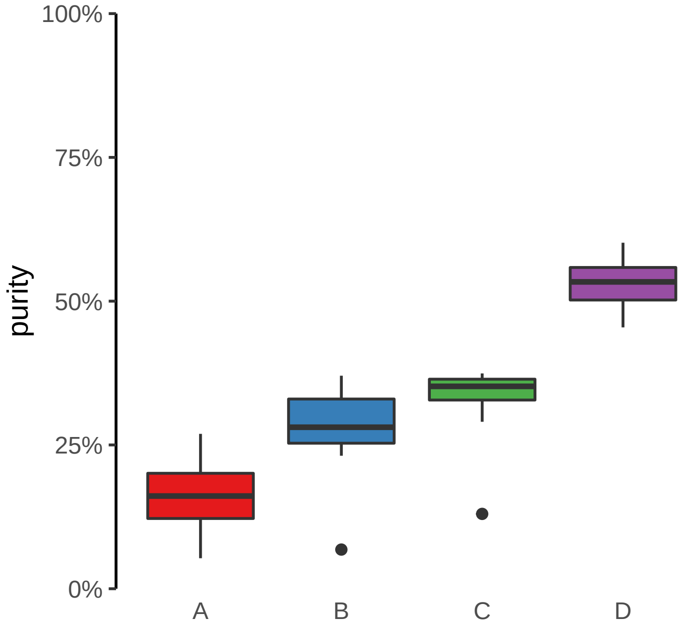

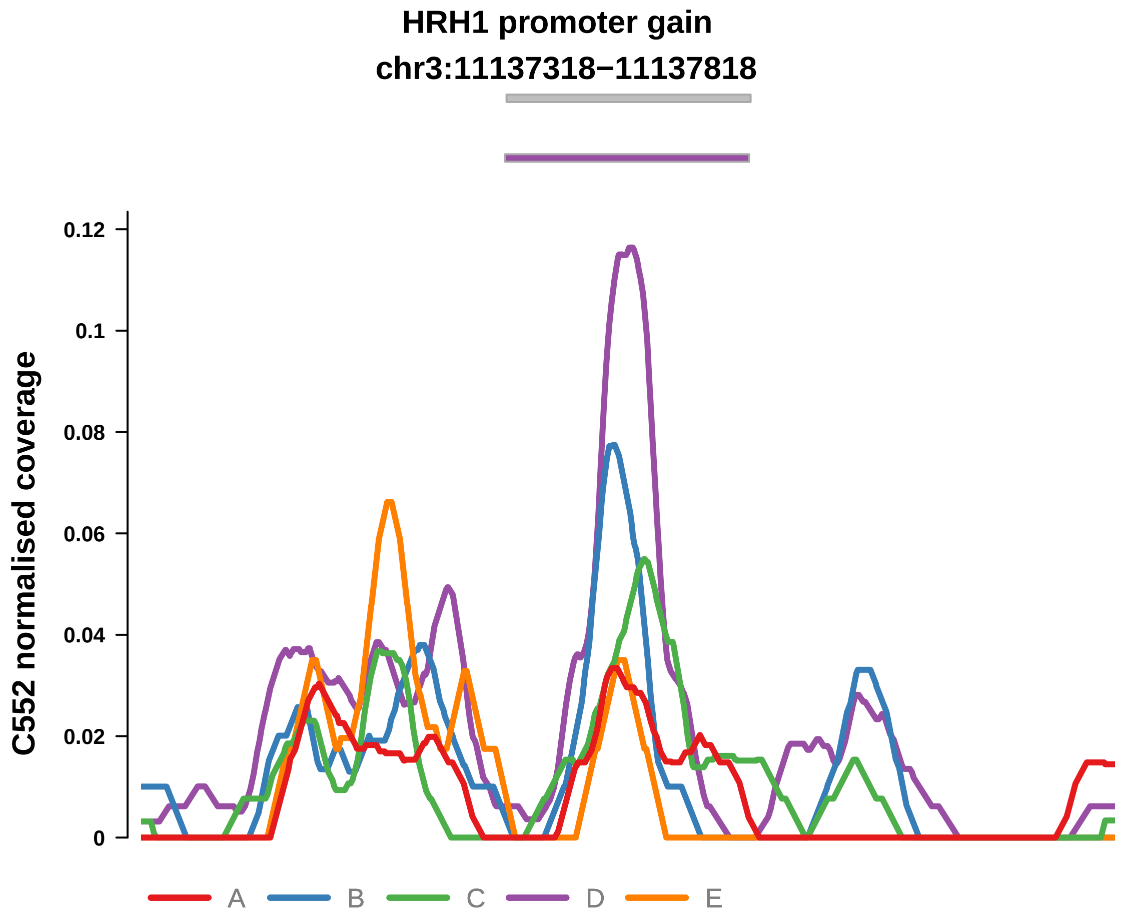


Putatively Clonal (logFC = 2.62)

| | P. | Adjusted P. |
|---|---|---|
| *~region vs. ~* | 6.08e−02 | 1.43e−01 |
| *~purity vs ~* | 1.72e−03 ** | 1.1e−02 * |
| *~purity+region vs ~purity* | 8.31e−01 | 9.85e−01 |
| *Carcinoma vs. normal* | 4.28e−05 **** | 4.28e−05 **** |

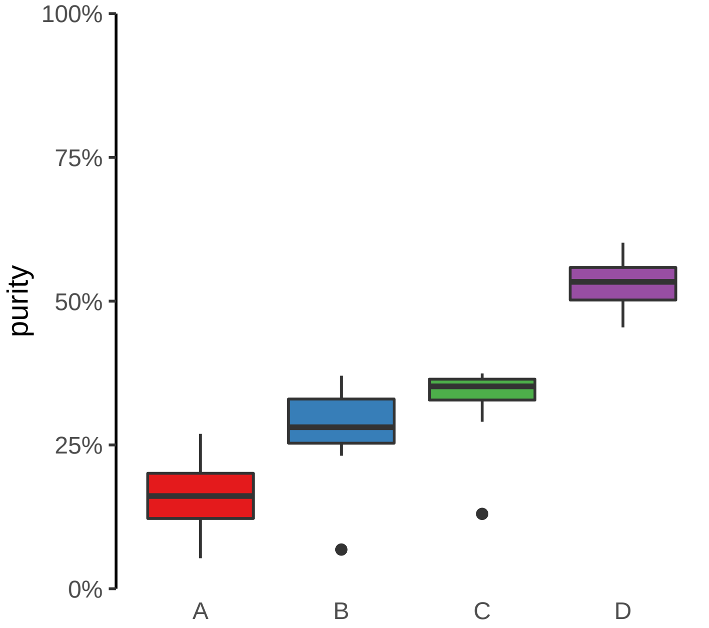

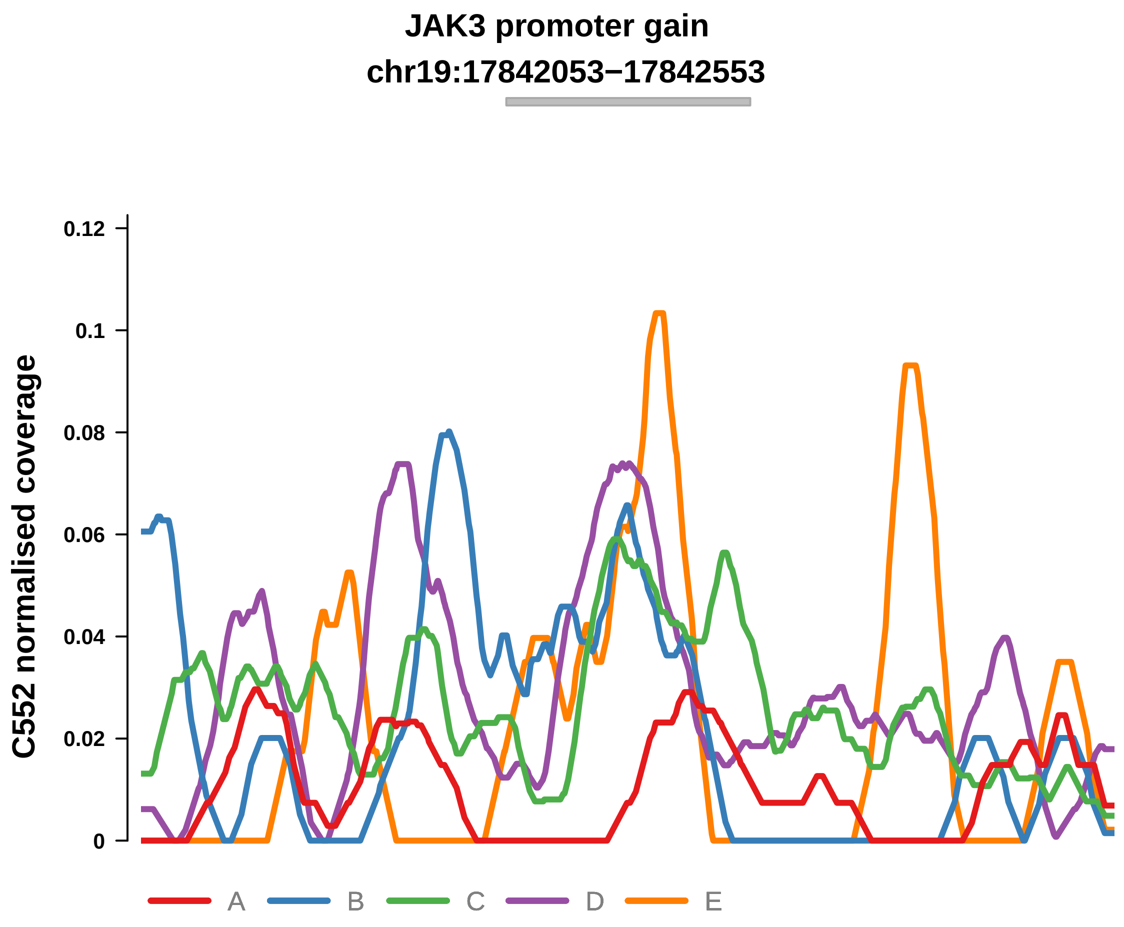


Putatively Clonal (logFC = 2.09)

| | P. | Adjusted P. |
|---|---|---|
| *~region vs. ~* | 2.73e−03 ** | 1.28e−02 * |
| *~purity vs ~* | 7.1e−04 *** | 7.77e−03 ** |
| *~purity+region vs ~purity* | 3e−01 | 9.1e−01 |
| *Carcinoma vs. normal* | 7.95e−04 *** | 7.95e−04 *** |

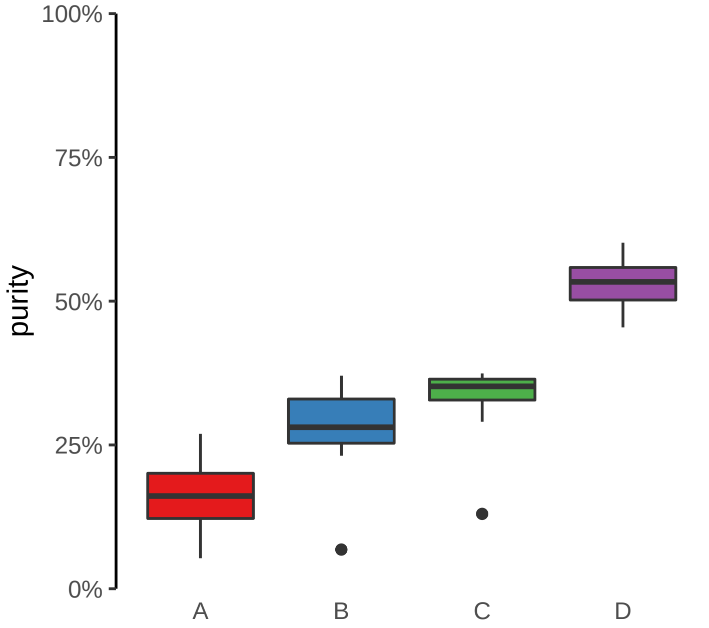

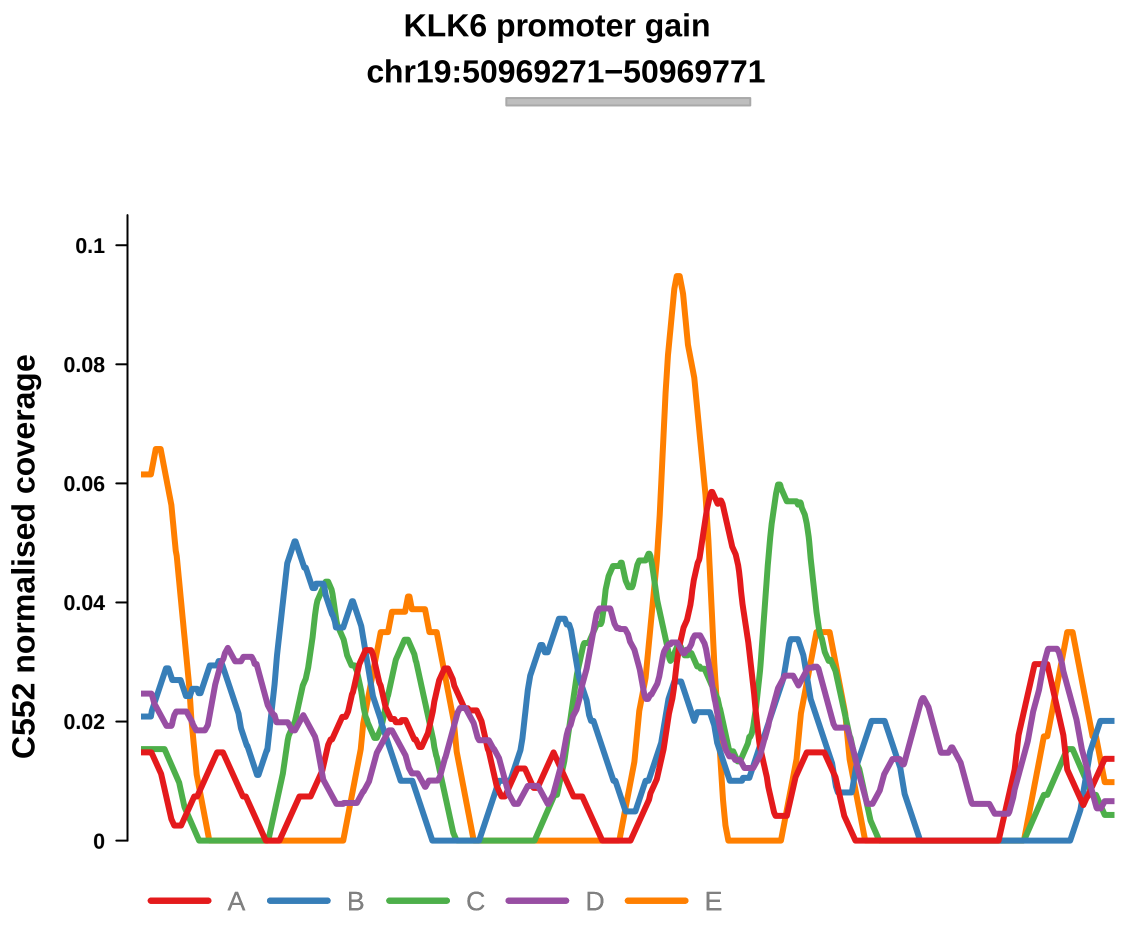


Putatively Clonal (logFC = 2.14)

| | P. | Adjusted P. |
|---|---|---|
| *~region vs. ~* | 8.22e−01 | 8.4e−01 |
| *~purity vs ~* | 1.35e−01 | 2.04e−01 |
| *~purity+region vs ~purity* | 3.79e−01 | 9.85e−01 |
| *Carcinoma vs. normal* | 1.85e−03 ** | 1.85e−03 ** |

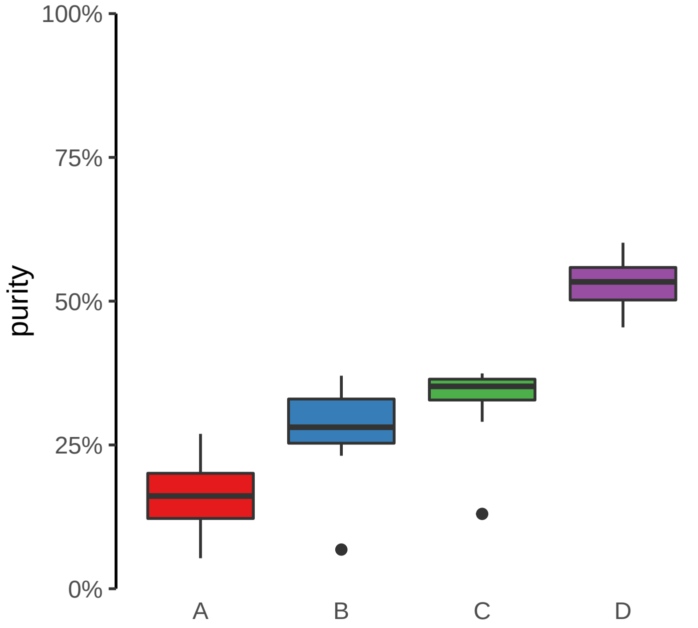

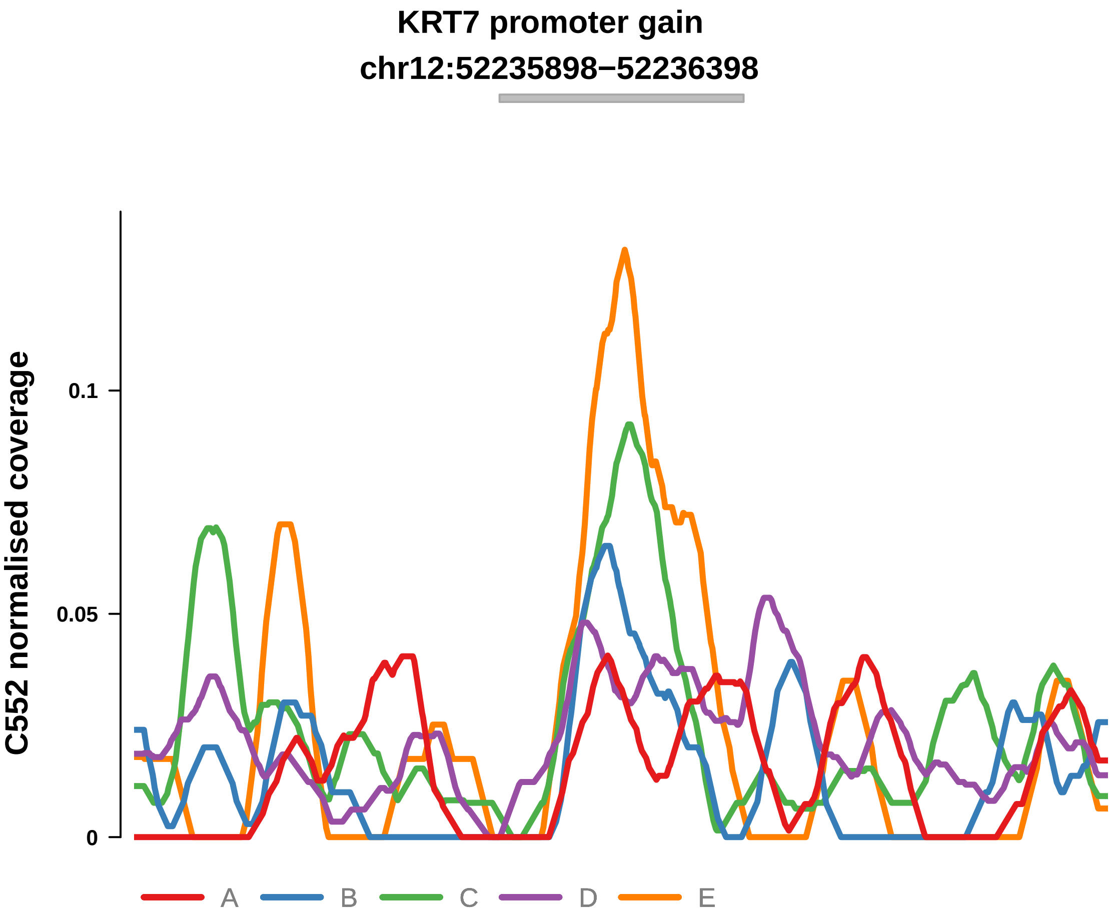


Putatively Clonal (logFC = 1.77)

| | P. | Adjusted P. |
|---|---|---|
| *~region vs. ~* | 2.33e−01 | 3.43e−01 |
| *~purity vs ~* | 6.47e−02 | 1.16e−01 |
| *~purity+region vs ~purity* | 8.25e−01 | 9.85e−01 |
| *Carcinoma vs. normal* | 4.96e−03 ** | 4.96e−03 ** |

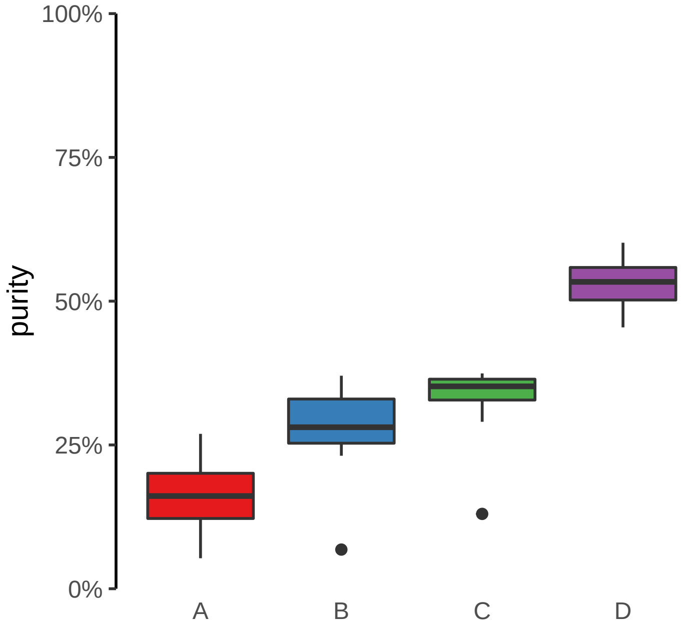

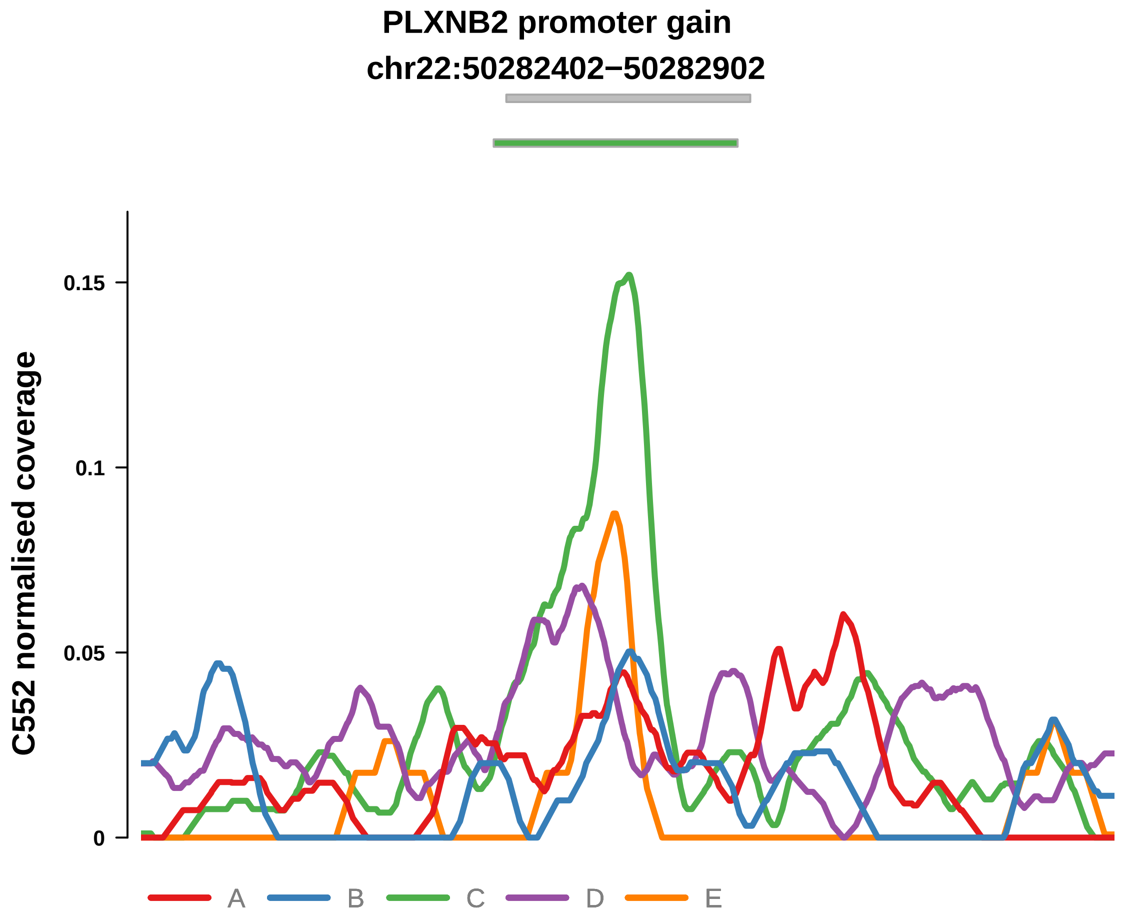


Putatively Clonal (logFC = 2.62)

| | P. | Adjusted P. |
|---|---|---|
| *~region vs. ~* | 3.26e−02 * | 9.92e−02 |
| *~purity vs ~* | 8.19e−04 *** | 7.77e−03 ** |
| *~purity+region vs ~purity* | 9.07e−01 | 9.85e−01 |
| *Carcinoma vs. normal* | 2.82e−05 **** | 2.82e−05 **** |

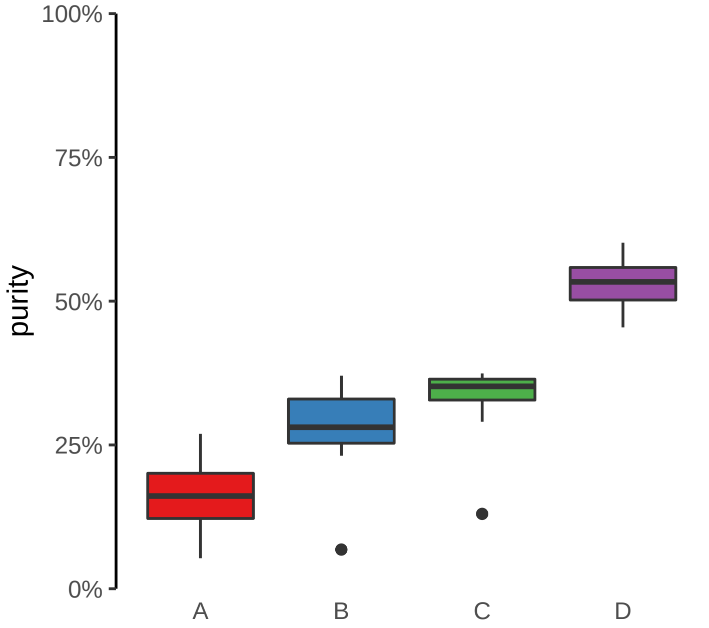

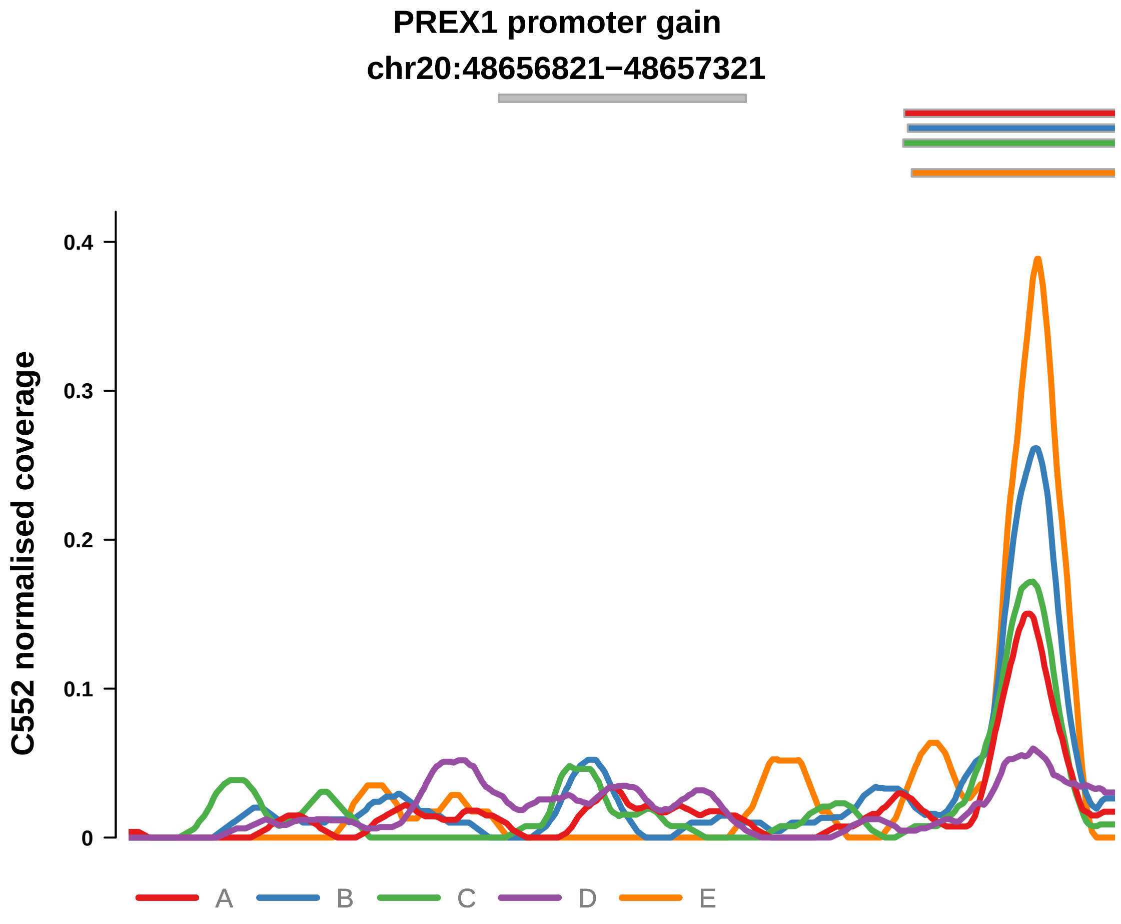


Putatively Clonal (logFC = 2.42)

| | P. | Adjusted P. |
|---|---|---|
| *~region vs. ~* | 4.95e−01 | NA |
| *~purity vs ~* | 5.51e−02 | NA |
| *~purity+region vs ~purity* | 9.5e−01 | 9.85e−01 |
| *Carcinoma vs. normal* | 4.47e−04 *** | 4.47e−04 *** |

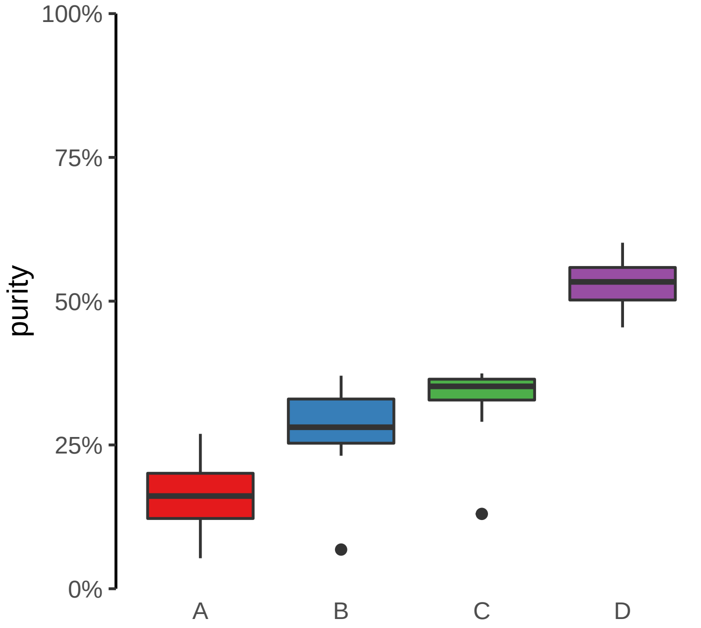

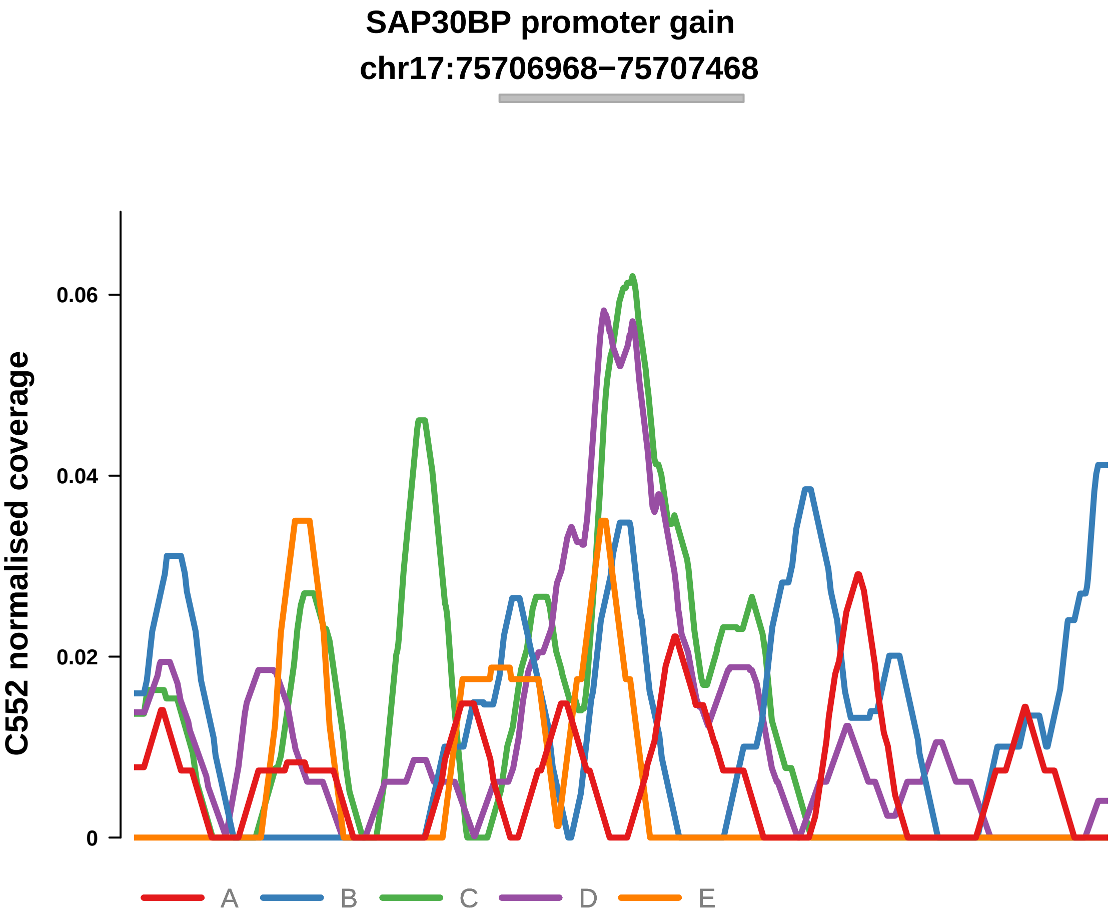


Putatively Clonal (logFC = 2.71)

| | P. | Adjusted P. |
|---|---|---|
| *~region vs. ~* | 6.55e−02 | NA |
| *~purity vs ~* | 1.51e−02 * | NA |
| *~purity+region vs ~purity* | 6.83e−01 | 9.85e−01 |
| *Carcinoma vs. normal* | 4.45e−05 **** | 4.45e−05 **** |

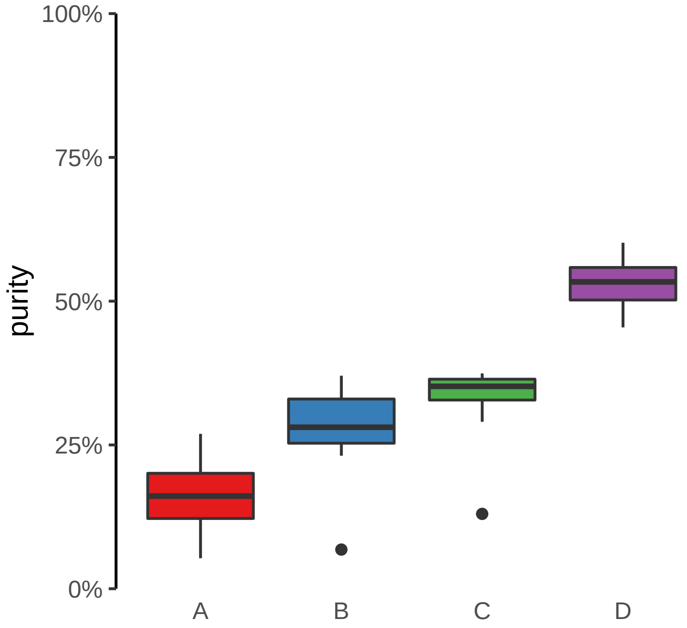

SLC45A4 promoter gain
chr8:141230663−141231163

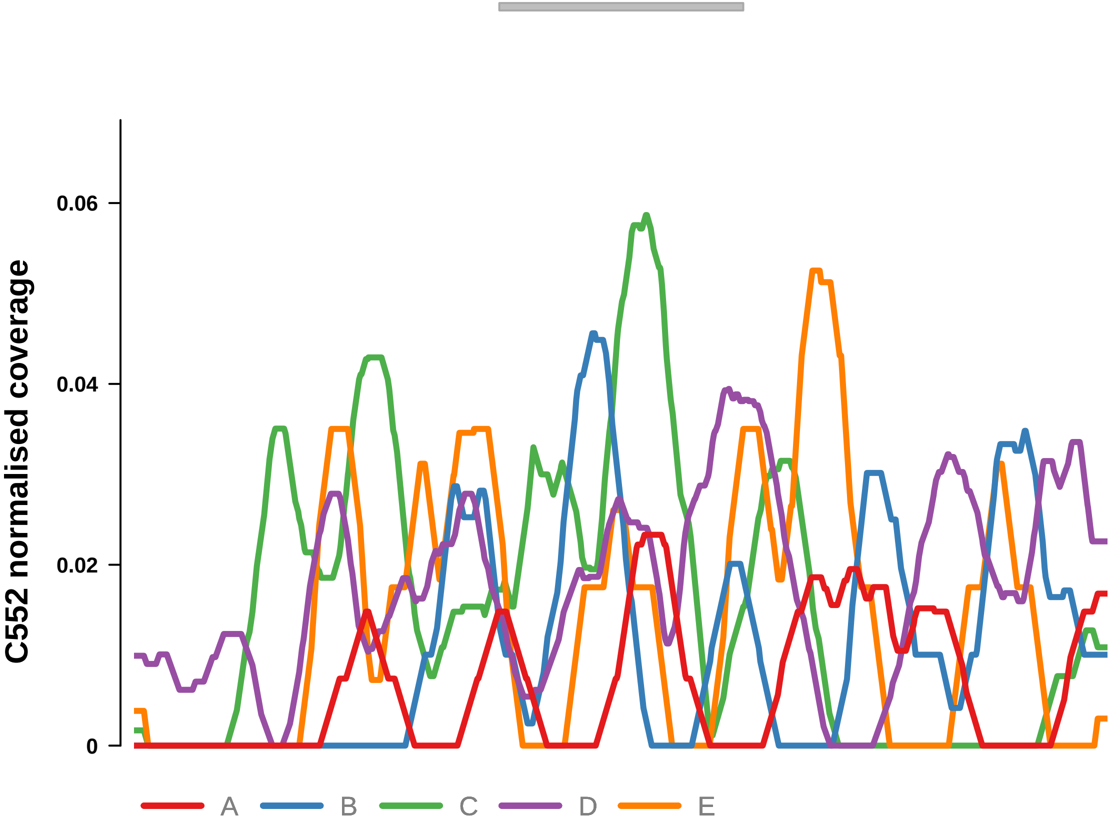


Putatively Clonal (logFC = 1.96)

| | P. | Adjusted P. |
|---|---|---|
| *~region vs. ~* | 1.88e−01 | 2.86e−01 |
| *~purity vs ~* | 2.5e−02 * | 6.94e−02 |
| *~purity+region vs ~purity* | 7.32e−01 | 9.85e−01 |
| *Carcinoma vs. normal* | 6.57e−03 ** | 6.57e−03 ** |

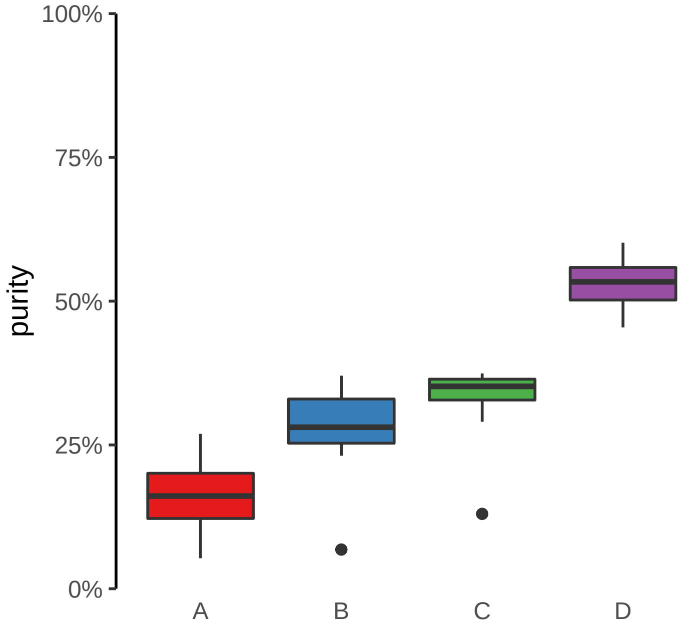

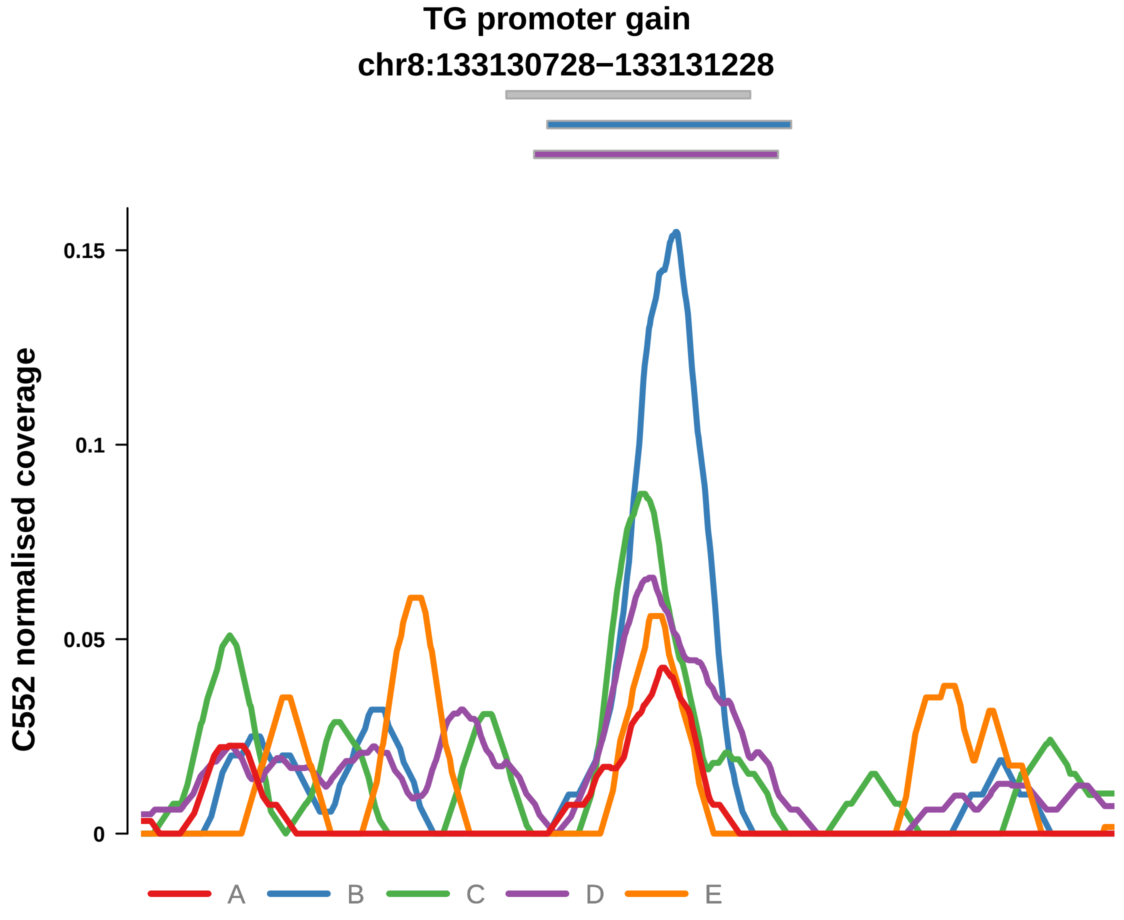


Putatively Clonal (logFC = 2.11)

| | P. | Adjusted P. |
|---|---|---|
| *~region vs. ~* | 1.76e−03 ** | 1.28e−02 * |
| *~purity vs ~* | 2.07e−02 * | 6.46e−02 |
| *~purity+region vs ~purity* | 2.55e−02 * | 3.73e−01 |
| *Carcinoma vs. normal* | 9.94e−04 *** | 9.94e−04 *** |

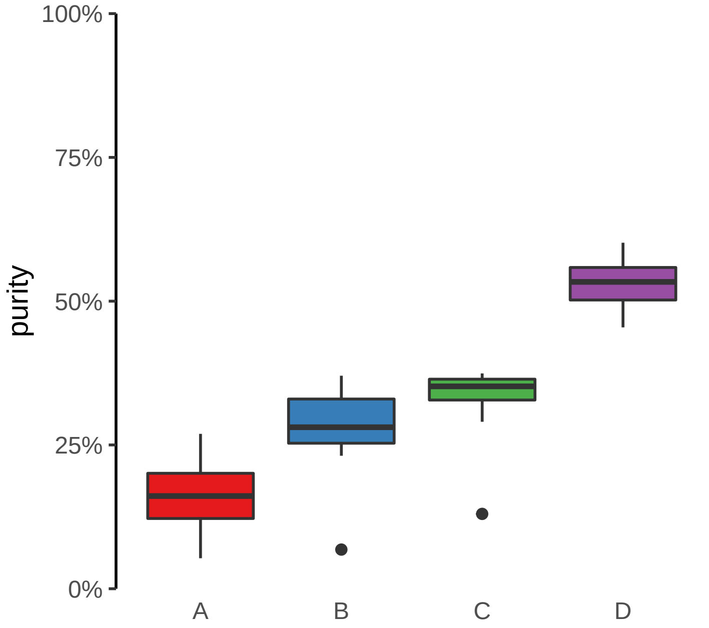

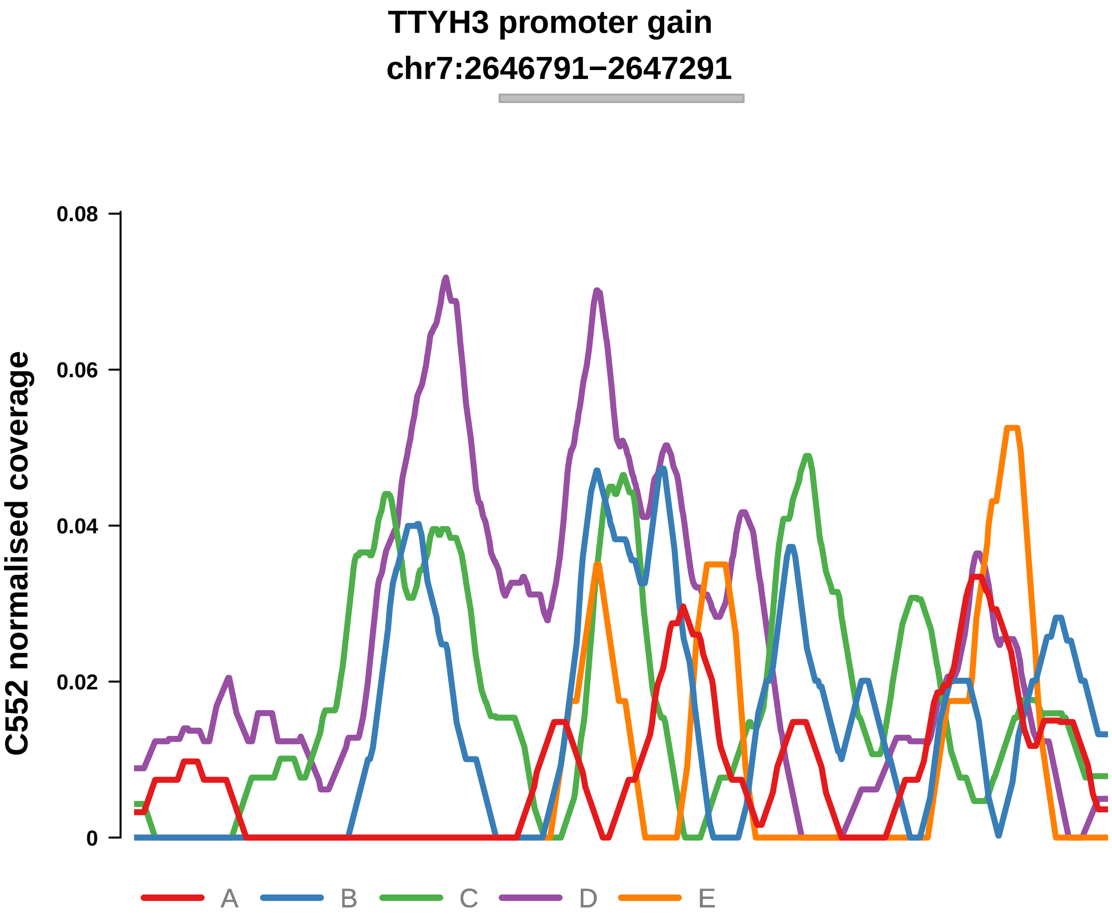


Putatively Clonal (logFC = 2.87)

| | P. | Adjusted P. |
|---|---|---|
| *~region vs. ~* | 2.21e−03 ** | 1.28e−02 * |
| *~purity vs ~* | 8.41e−05 **** | 2.06e−03 ** |
| *~purity+region vs ~purity* | 2.98e−01 | 9.1e−01 |
| *Carcinoma vs. normal* | 5.87e−06 **** | 5.87e−06 **** |

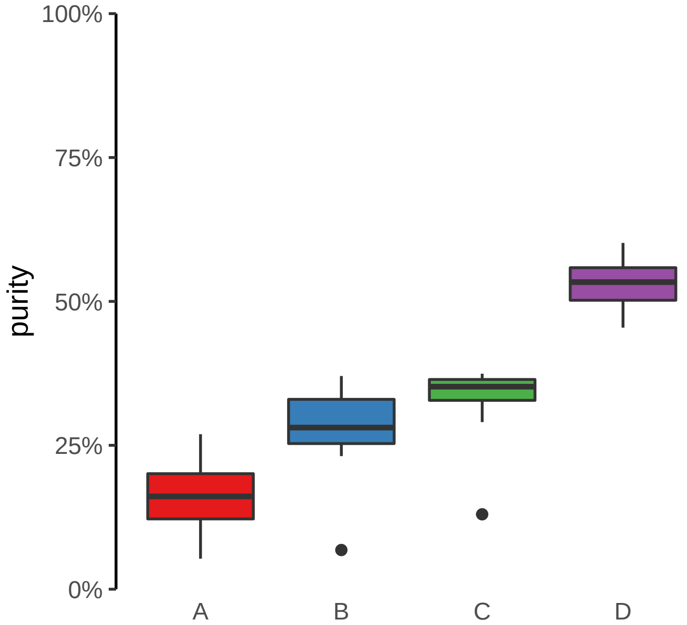

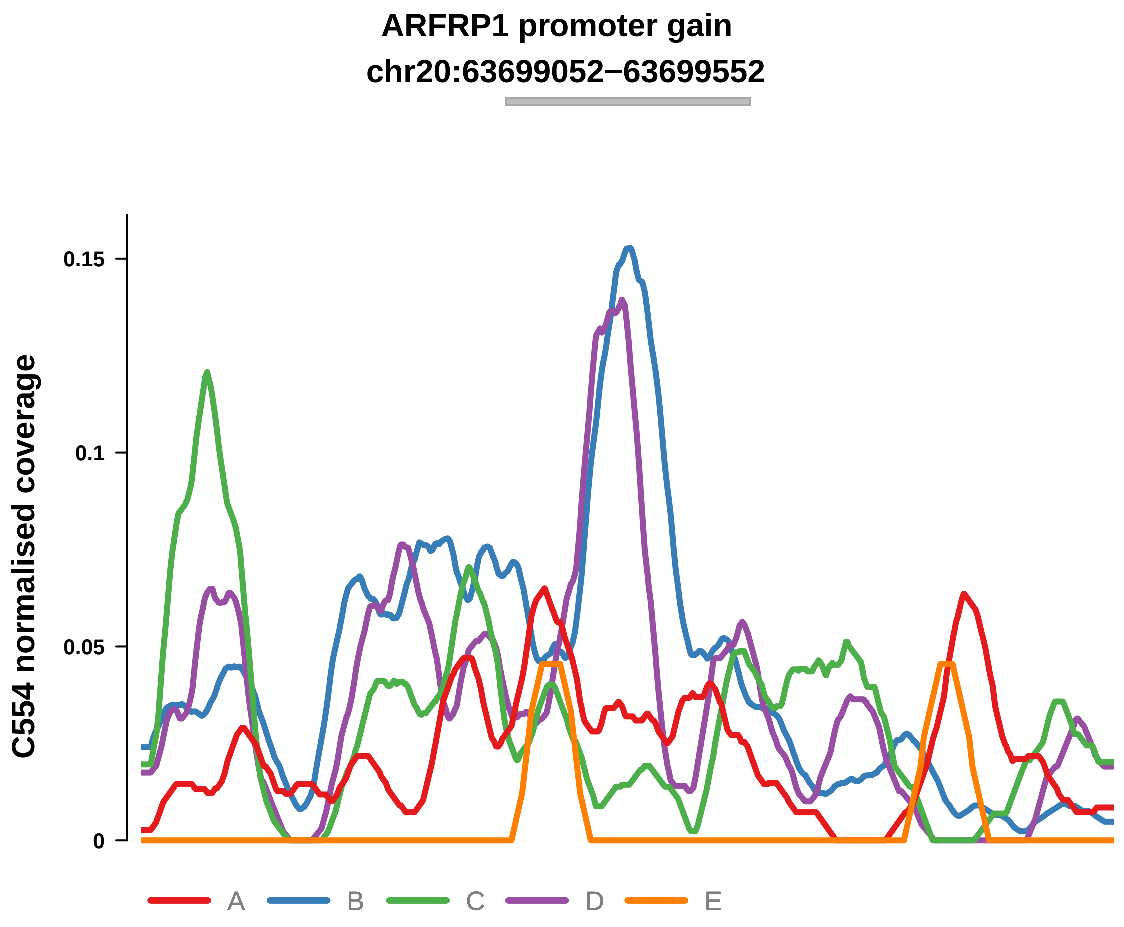


Putatively Clonal (logFC = 1.78)

| | P. | Adjusted P. |
|---|---|---|
| *~region vs. ~* | 1.88e−03 ** | 7.35e−03 ** |
| *~purity vs ~* | 5.64e−03 ** | 4.21e−02 * |
| *~purity+region vs ~purity* | 1.29e−01 | 2.65e−01 |
| *Carcinoma vs. normal* | 1.41e−03 ** | 1.41e−03 ** |

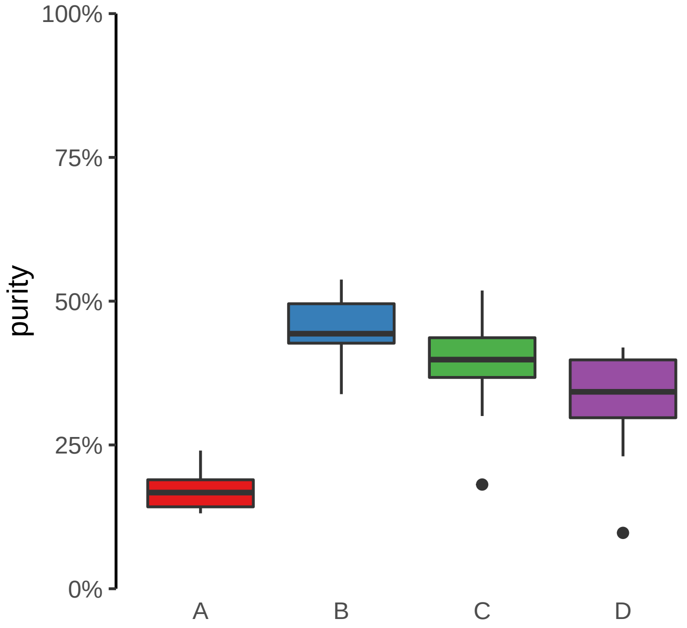

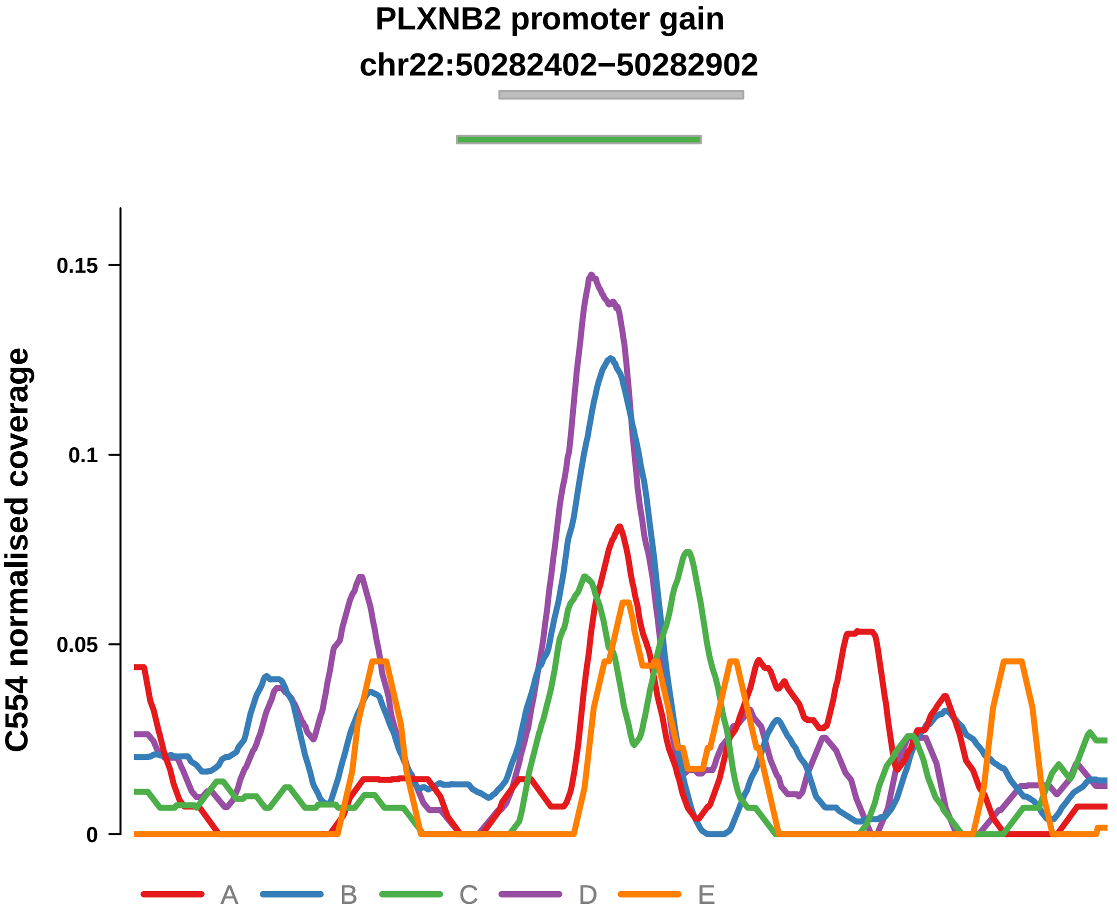


Putatively Clonal (logFC = 1.68)

| | P. | Adjusted P. |
|---|---|---|
| *~region vs. ~* | 2.59e−02 * | 6.86e−02 |
| *~purity vs ~* | 1.85e−02 * | 8.43e−02 |
| *~purity+region vs ~purity* | 1.39e−01 | 2.65e−01 |
| *Carcinoma vs. normal* | 2.51e−03 ** | 2.51e−03 ** |

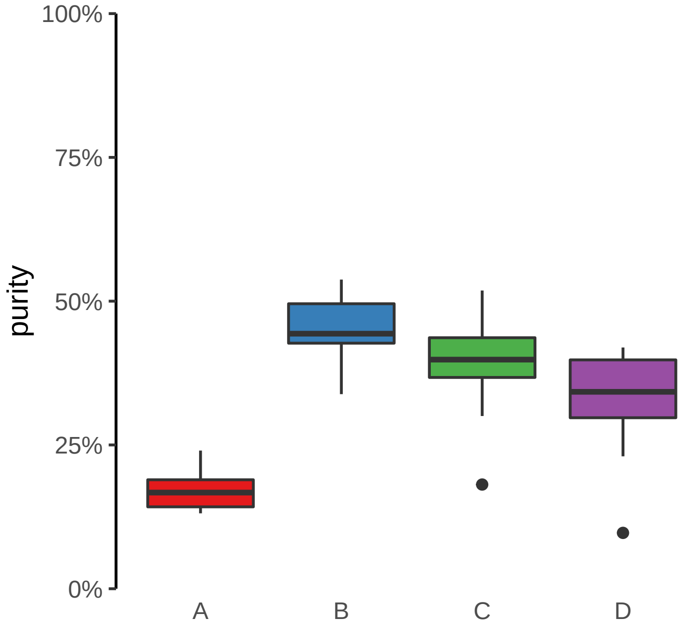

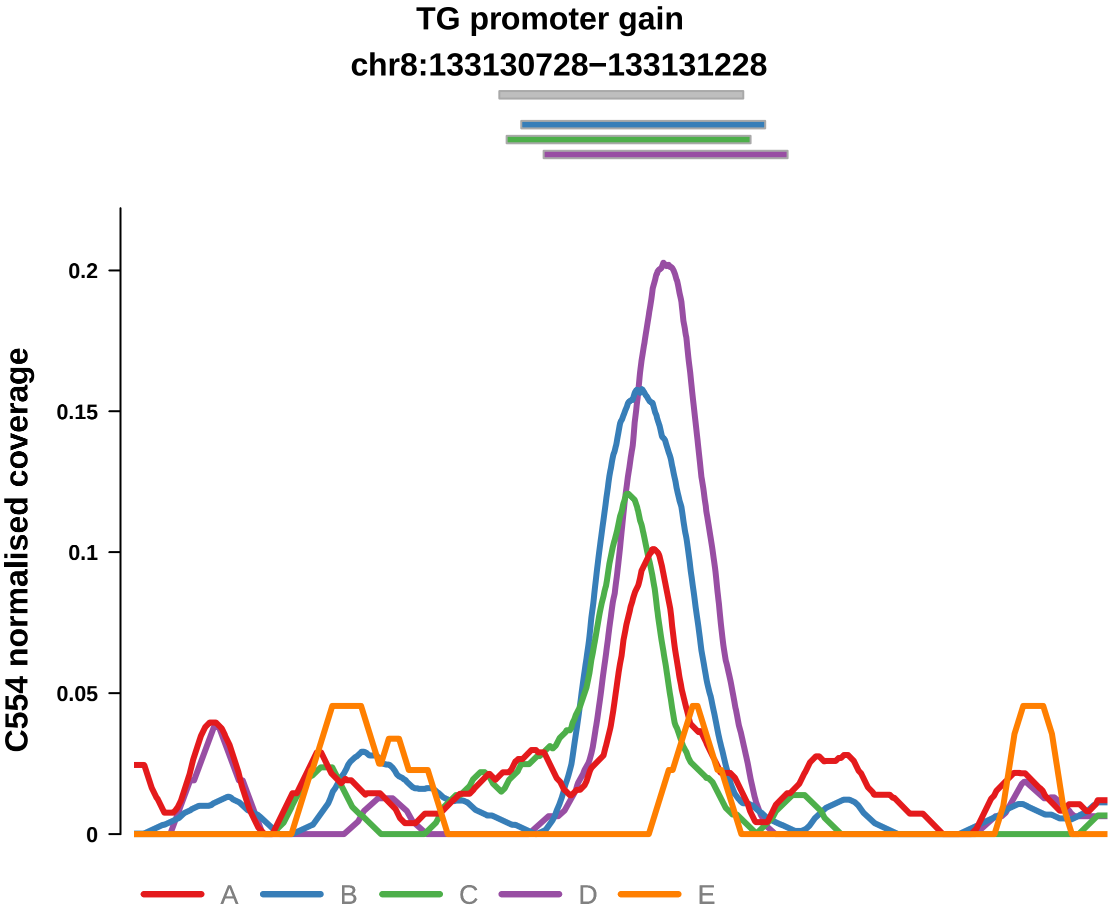


Putatively Clonal (logFC = 1.95)

| | P. | Adjusted P. |
|---|---|---|
| *~region vs. ~* | 2.94e−03 ** | 1.05e−02 * |
| *~purity vs ~* | 1.33e−02 * | 7.76e−02 |
| *~purity+region vs ~purity* | 6.72e−02 | 1.71e−01 |
| *Carcinoma vs. normal* | 4.69e−04 *** | 4.69e−04 *** |

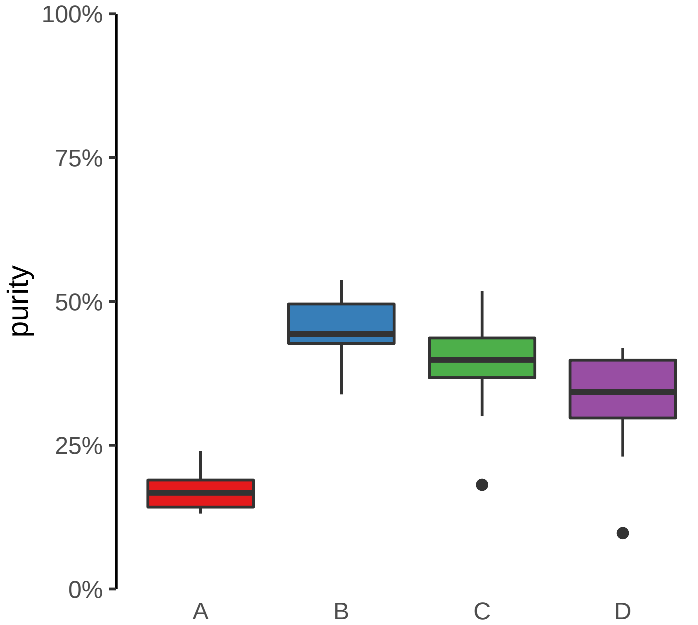

**ARHGEF17 promoter gain**
**chr11:73362044−73362544**

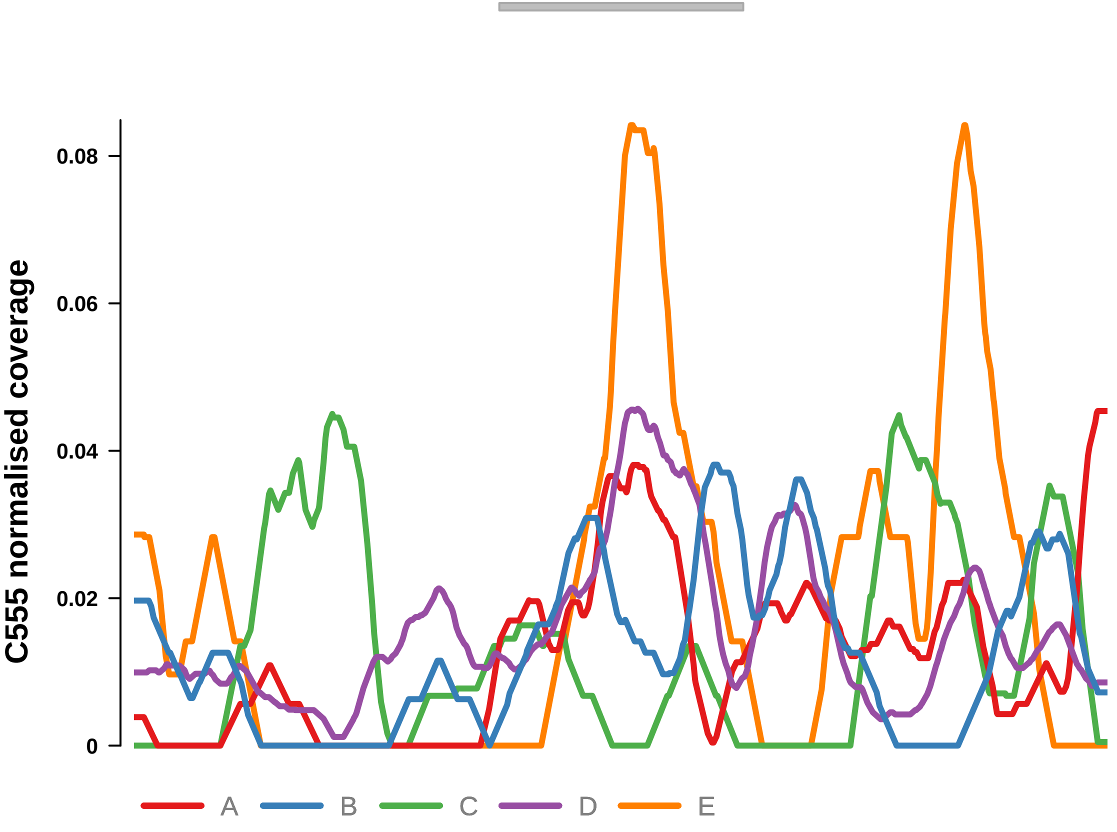


Putatively Clonal (logFC = 2.18)

| | **P.** | **Adjusted P.** |
|---|---|---|
| *~region vs. ~* | 8.63e−05 **** | 2.53e−03 ** |
| *~purity vs ~* | 2.27e−01 | 4.65e−01 |
| *~purity+region vs ~purity* | 2.16e−04 *** | 9.48e−03 ** |
| *Carcinoma vs. normal* | 4.72e−04 *** | 4.72e−04 *** |

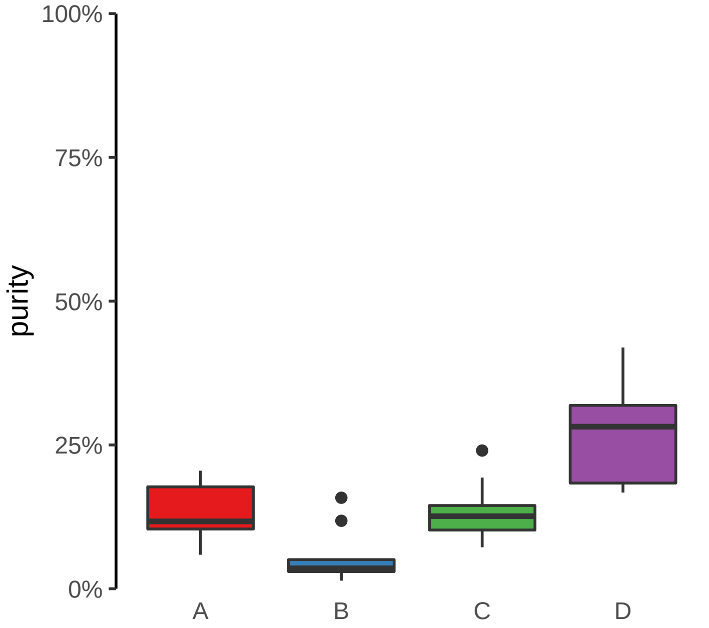

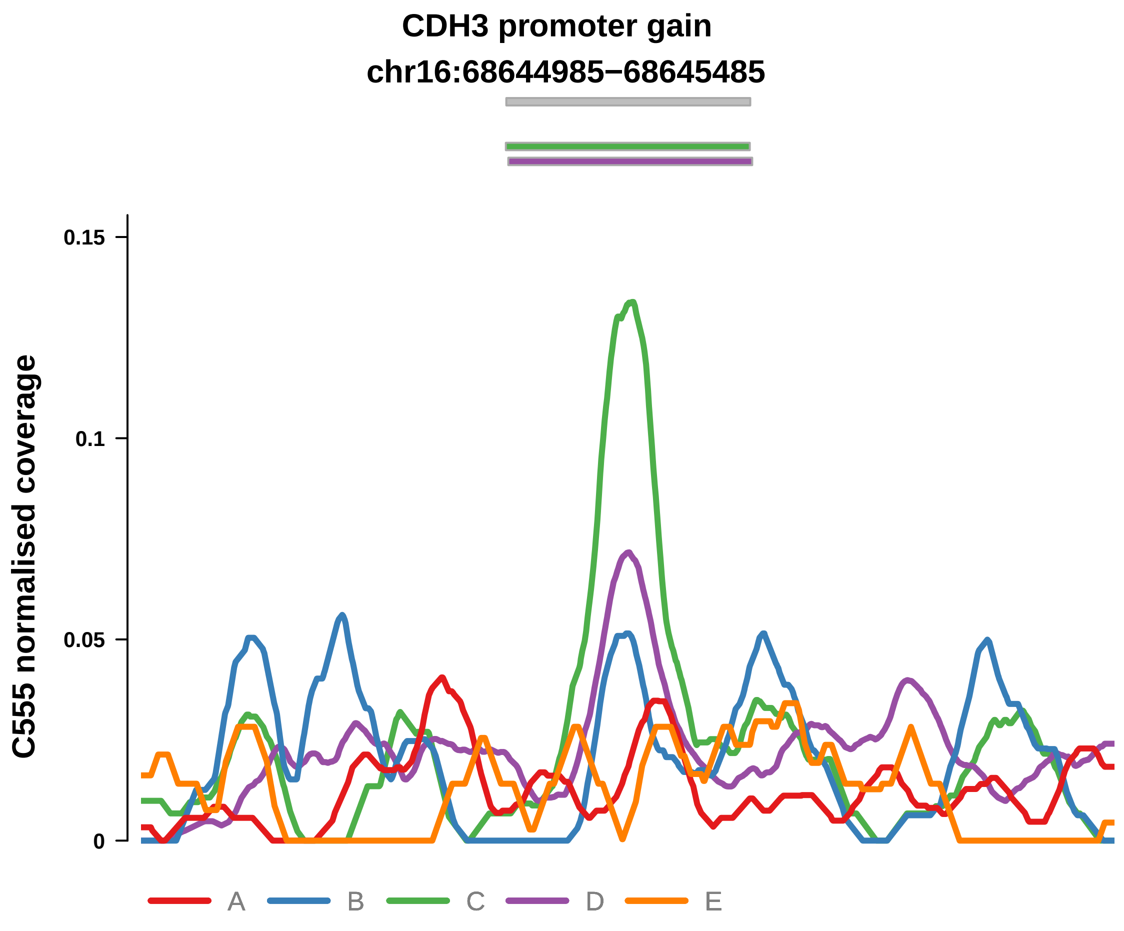


Putatively Clonal (logFC = 1.55)

| | P. | Adjusted P. |
|---|---|---|
| *~region vs. ~* | 1.58e−01 | 3.32e−01 |
| *~purity vs ~* | 5.14e−02 | 2.26e−01 |
| *~purity+region vs ~purity* | 6.33e−01 | 8.44e−01 |
| *Carcinoma vs. normal* | 9.18e−03 ** | 9.18e−03 ** |

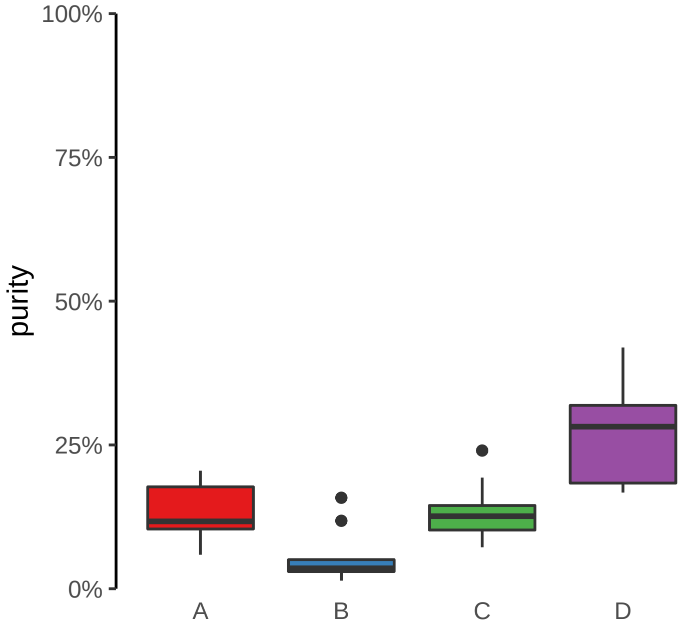

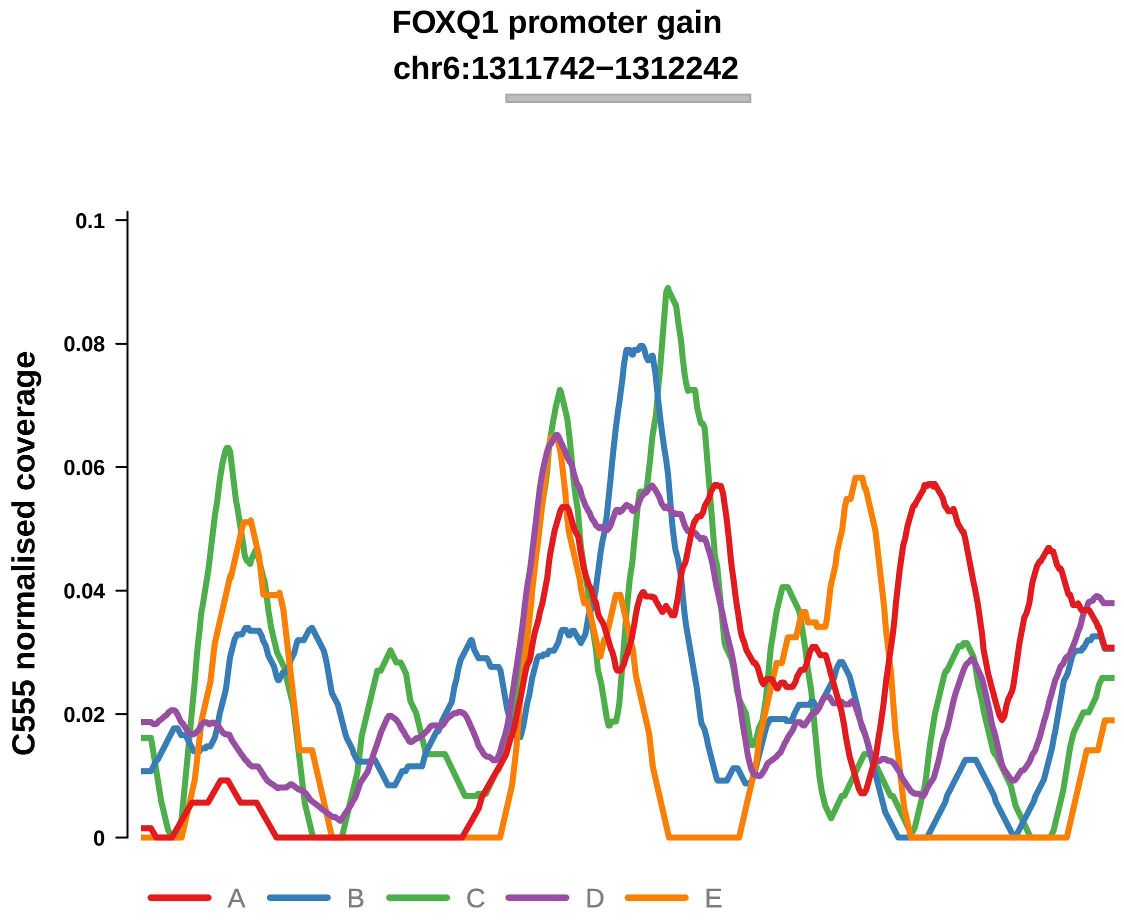


Putatively Clonal (logFC = 2.28)

| | P. | Adjusted P. |
|---|---|---|
| *~region vs. ~* | 7e−02 | 2.05e−01 |
| *~purity vs ~* | 4.78e−02 * | 2.26e−01 |
| *~purity+region vs ~purity* | 2.03e−01 | 4.81e−01 |
| *Carcinoma vs. normal* | 7.76e−05 **** | 7.76e−05 **** |

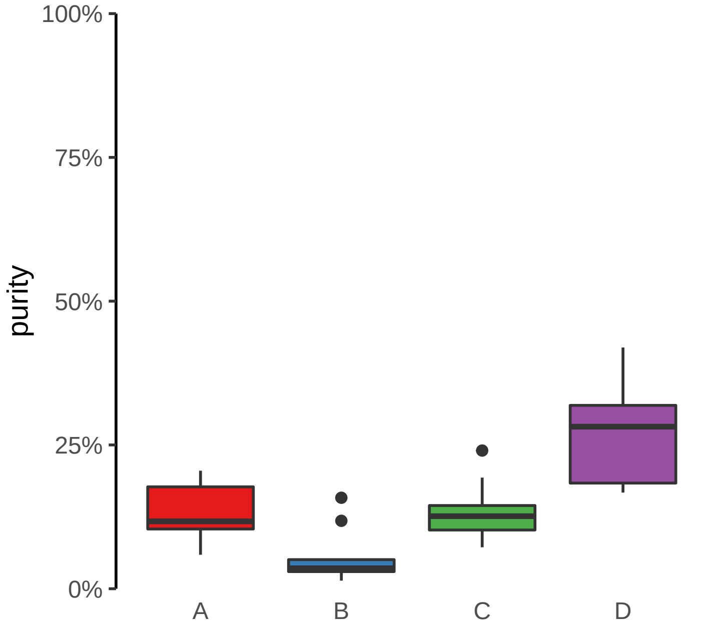

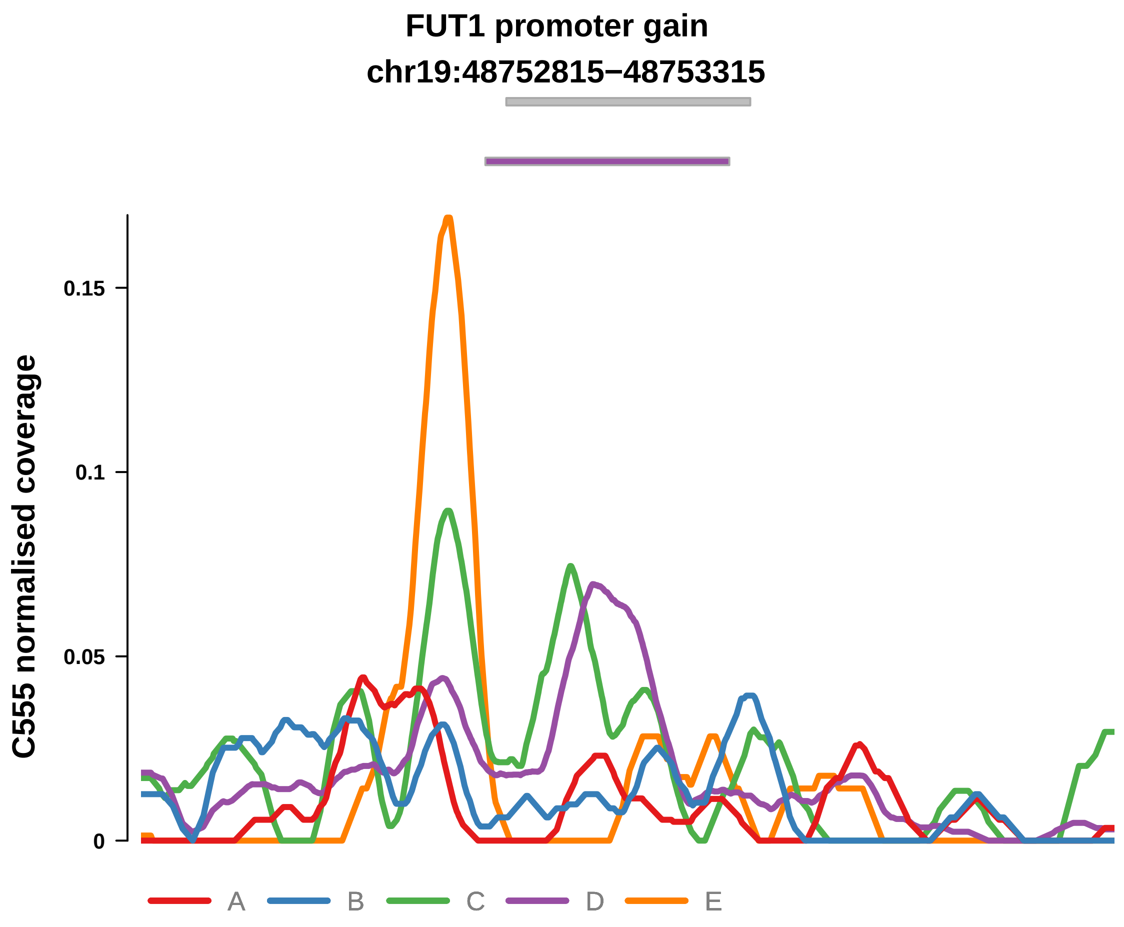


Putatively Clonal (logFC = 1.81)

| | P. | Adjusted P. |
|---|---|---|
| *~region vs. ~* | 2.14e−01 | 4.19e−01 |
| *~purity vs ~* | 6.13e−02 | 2.28e−01 |
| *~purity+region vs ~purity* | 7.02e−01 | 8.87e−01 |
| *Carcinoma vs. normal* | 2.16e−03 ** | 2.16e−03 ** |

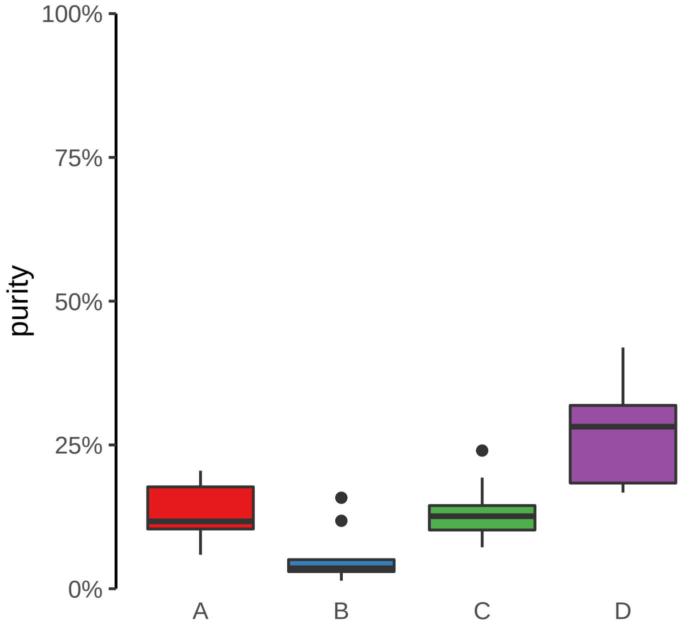

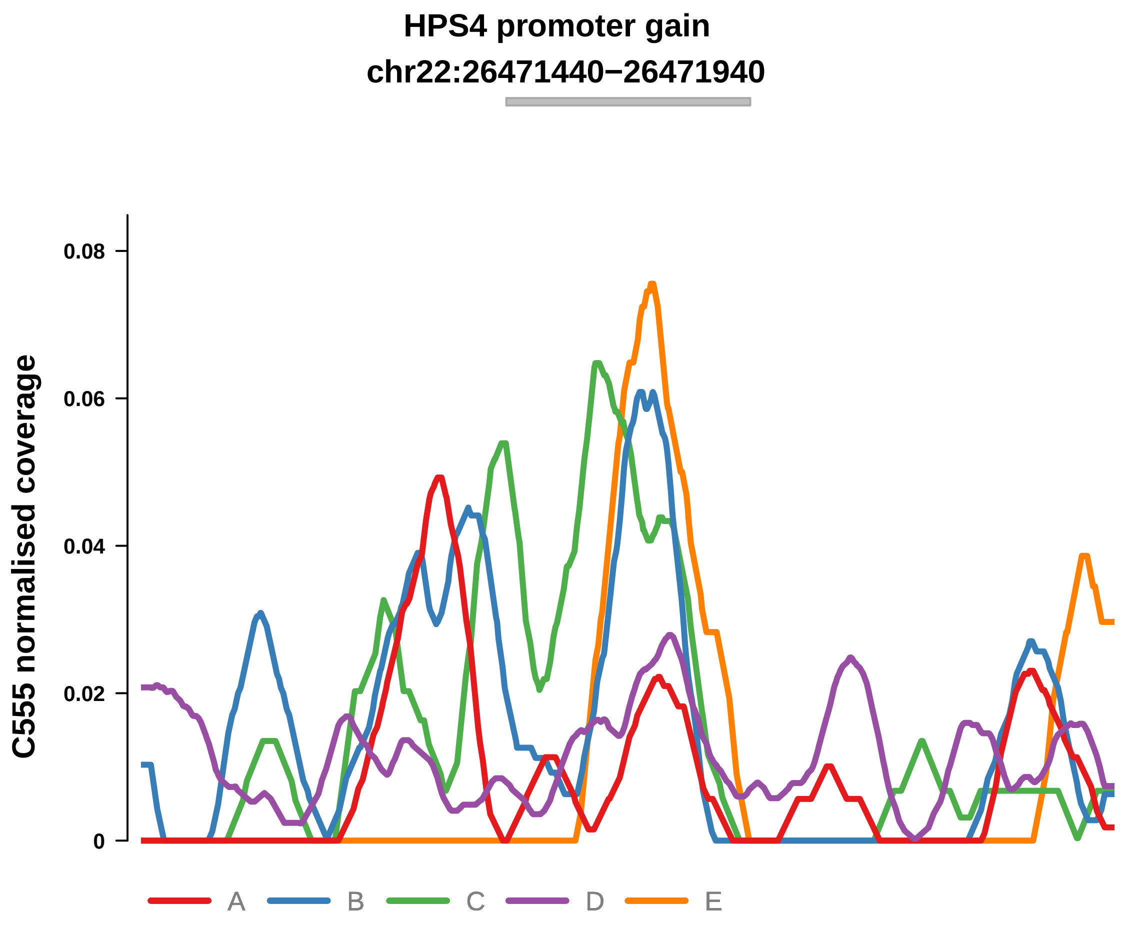


Putatively Clonal (logFC = 1.94)

| | P. | Adjusted P. |
|---|---|---|
| *~region vs. ~* | 6.73e−01 | 8.23e−01 |
| *~purity vs ~* | 5.88e−01 | 7.43e−01 |
| *~purity+region vs ~purity* | 4.51e−01 | 7.43e−01 |
| *Carcinoma vs. normal* | 3.69e−03 ** | 3.69e−03 ** |

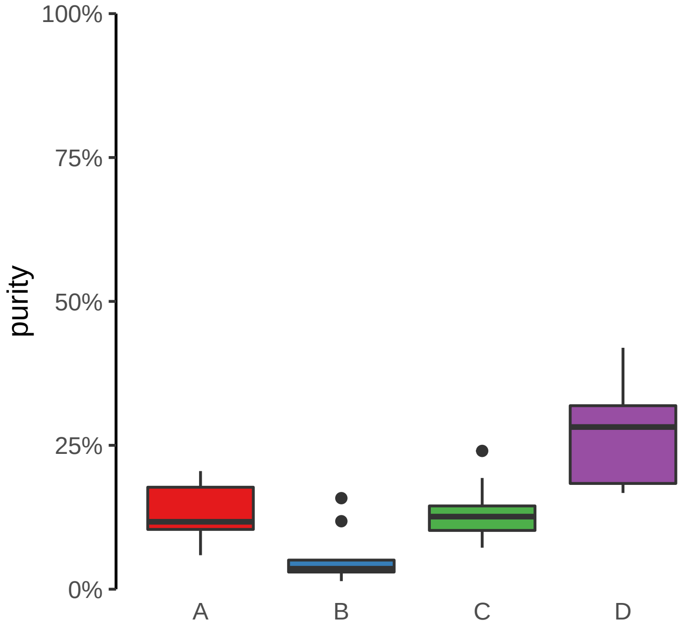

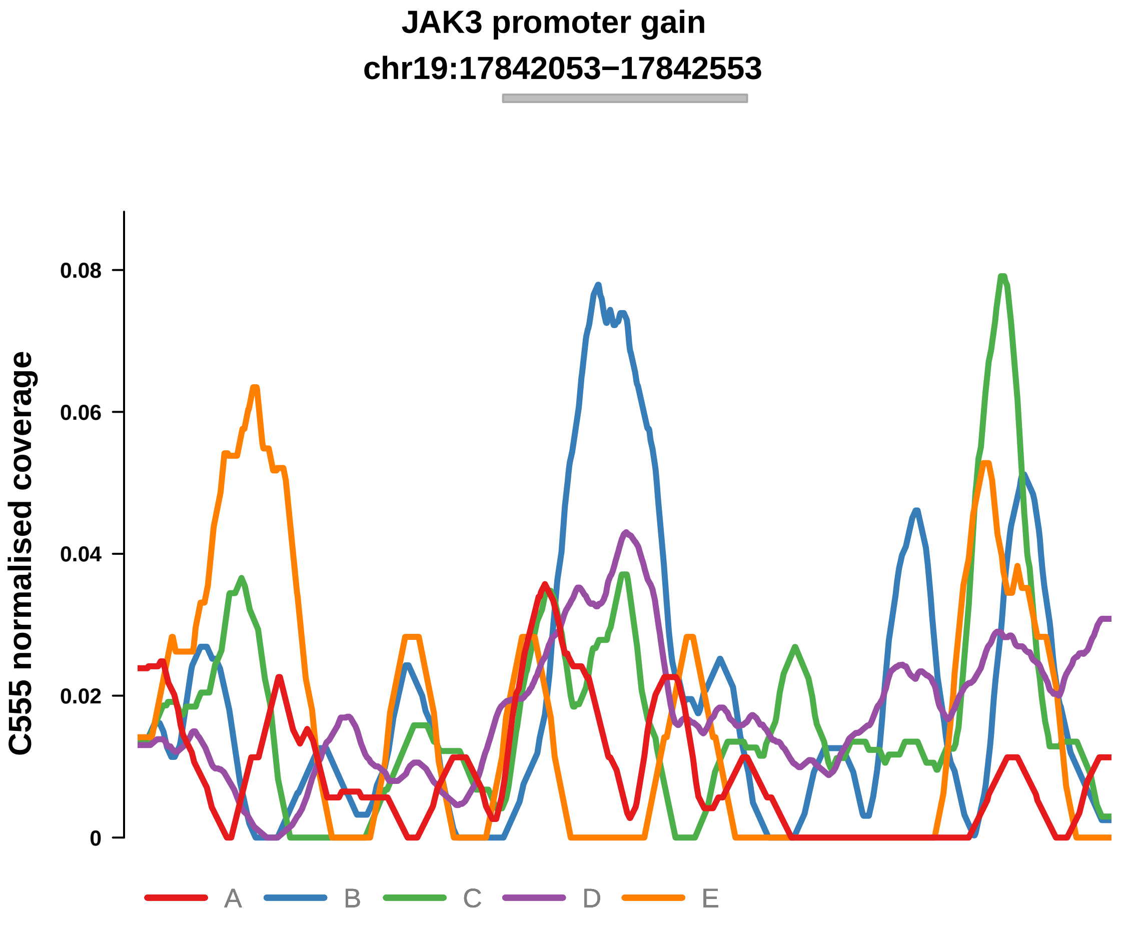


Putatively Clonal (logFC = 1.56)

| | P. | Adjusted P. |
|---|---|---|
| *~region vs. ~* | 1.45e−02 * | 6.97e−02 |
| *~purity vs ~* | 3.81e−01 | 6.32e−01 |
| *~purity+region vs ~purity* | 2.72e−02 * | 1.41e−01 |
| *Carcinoma vs. normal* | 8.85e−03 ** | 8.85e−03 ** |

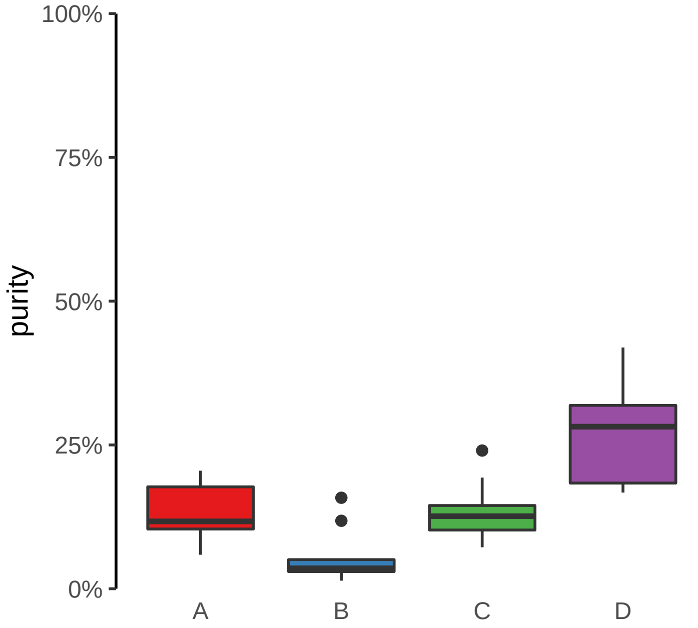

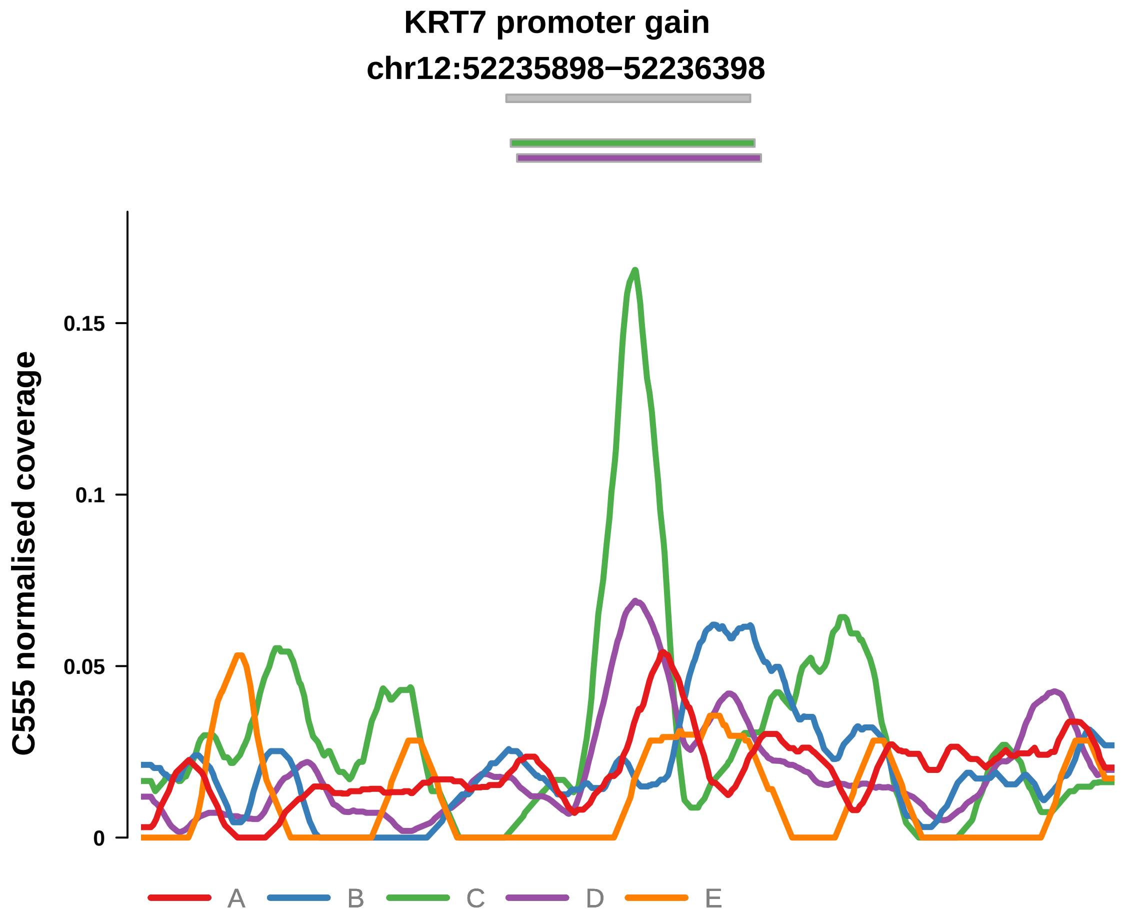


Putatively Clonal (logFC = 2)

| | P. | Adjusted P. |
|---|---|---|
| *~region vs. ~* | 6.66e−01 | 8.23e−01 |
| *~purity vs ~* | 2.26e−01 | 4.65e−01 |
| *~purity+region vs ~purity* | 9.15e−01 | 9.59e−01 |
| *Carcinoma vs. normal* | 6.96e−04 *** | 6.96e−04 *** |

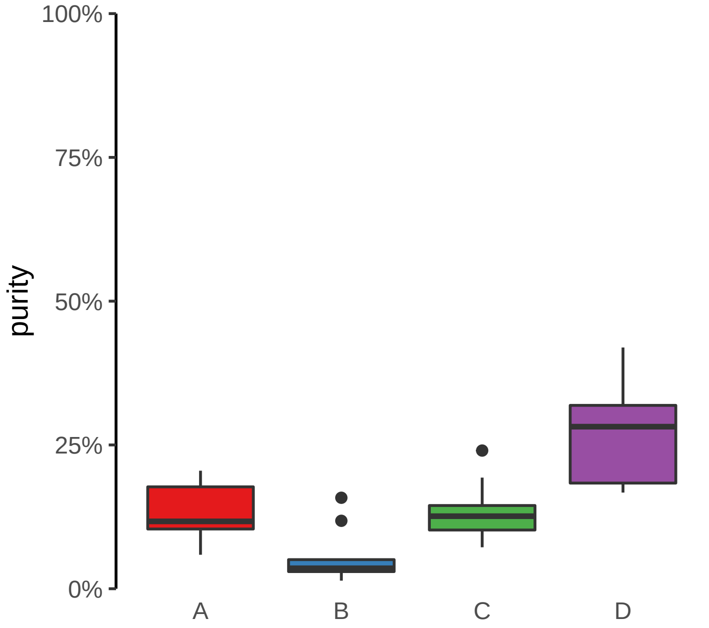

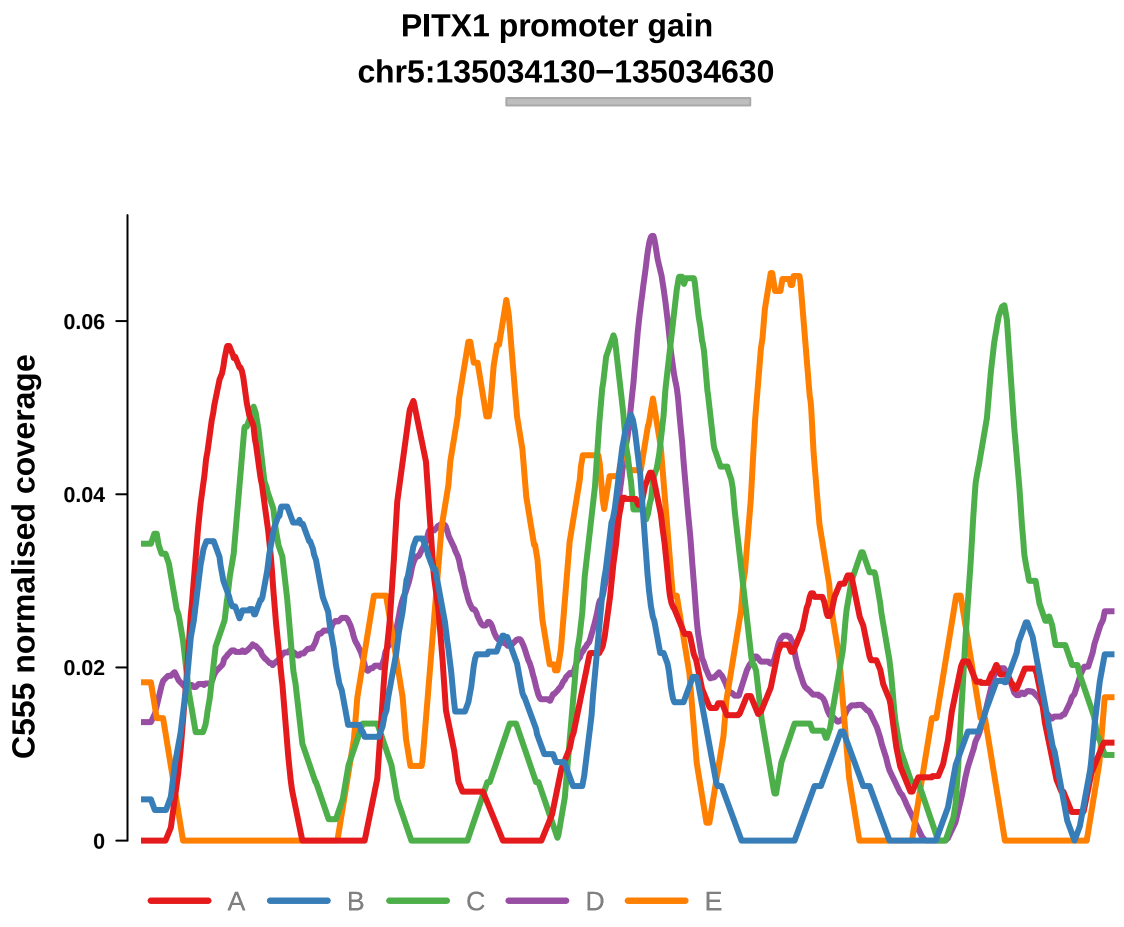


Putatively Clonal (logFC = 1.66)

| | P. | Adjusted P. |
|---|---|---|
| *~region vs. ~* | 9.63e−02 | 2.49e−01 |
| *~purity vs ~* | 7.88e−02 | 2.67e−01 |
| *~purity+region vs ~purity* | 4.11e−01 | 6.97e−01 |
| *Carcinoma vs. normal* | 5.46e−03 ** | 5.46e−03 ** |

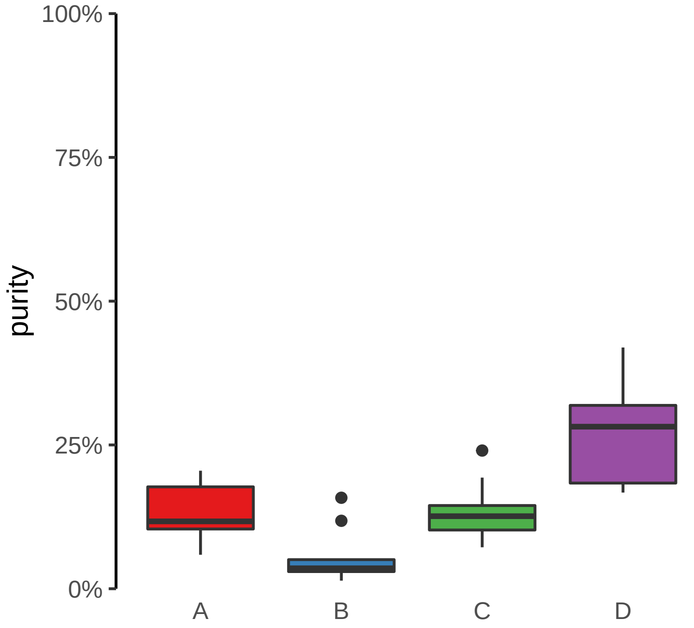

**PREX1 promoter gain**
**chr20:48656821−48657321**

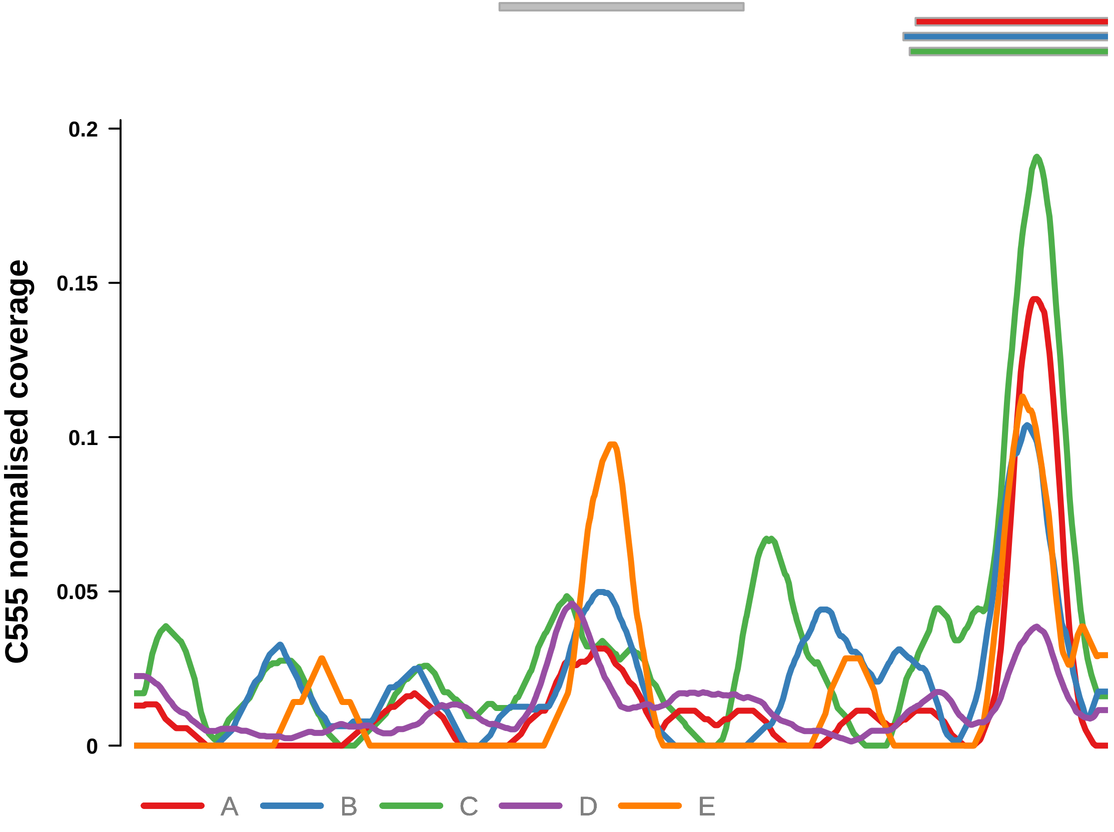


Putatively Clonal (logFC = 1.91)

| | P. | Adjusted P. |
| --- | --- | --- |
| *~region vs. ~* | 5.39e−01 | 7.13e−01 |
| *~purity vs ~* | 1.65e−01 | 4.14e−01 |
| *~purity+region vs ~purity* | 9.67e−01 | 9.78e−01 |
| *Carcinoma vs. normal* | 3.56e−03 ** | 3.56e−03 ** |

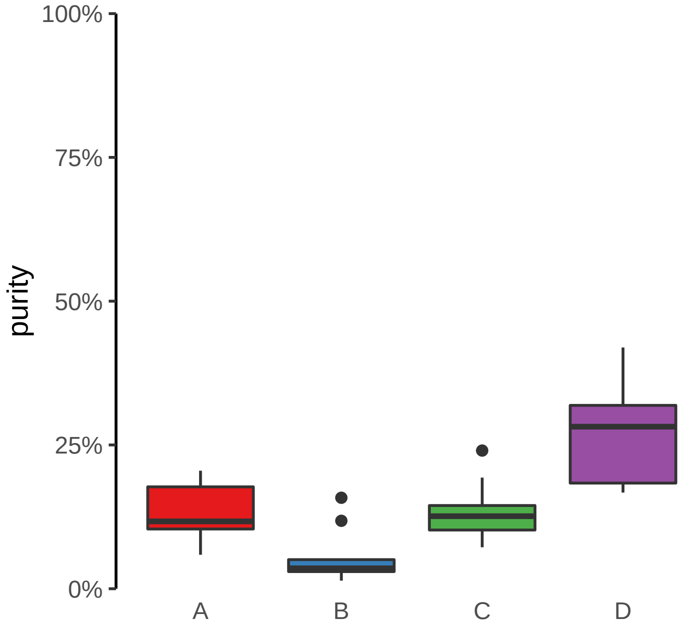

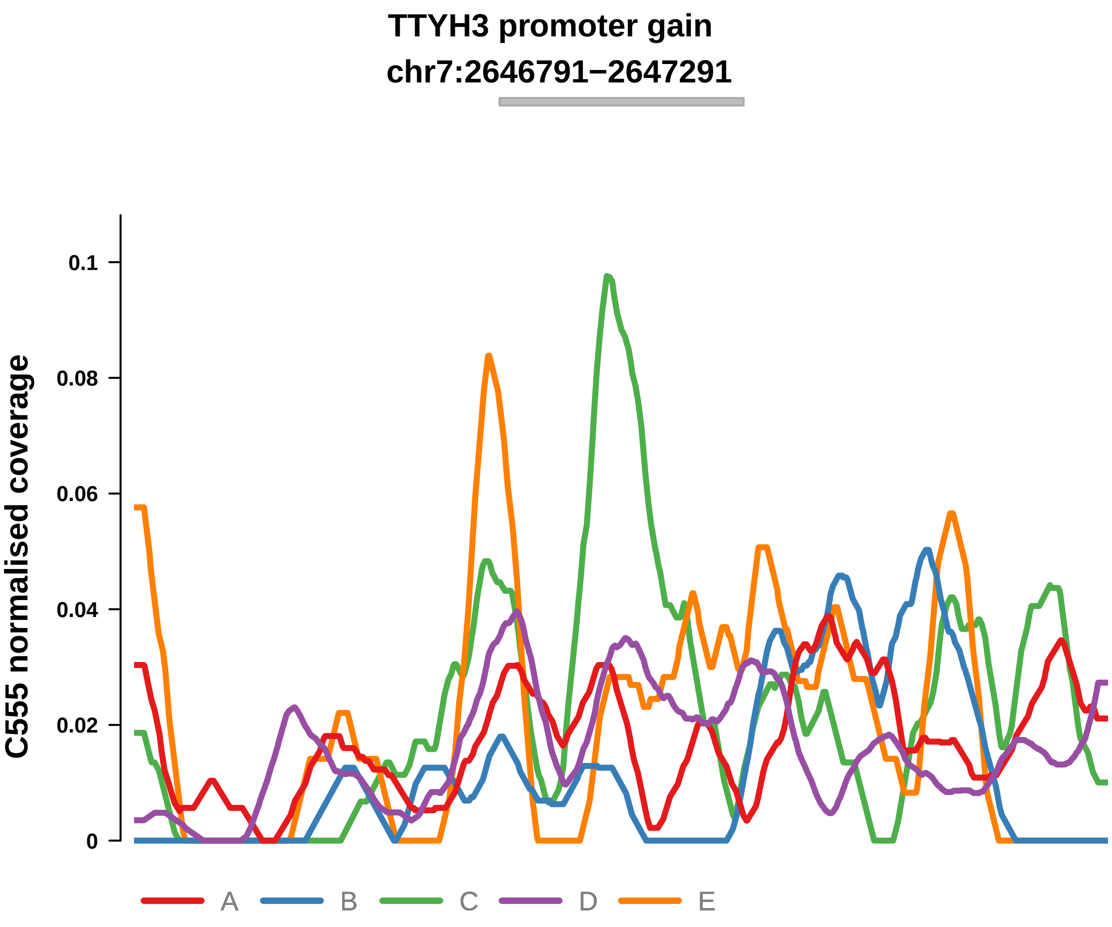


Putatively Clonal (logFC = 1.76)

| | P. | Adjusted P. |
|---|---|---|
| *~region vs. ~* | 2.68e−02 * | 1.07e−01 |
| *~purity vs ~* | 1.33e−02 * | 1.67e−01 |
| *~purity+region vs ~purity* | 3.96e−01 | 6.97e−01 |
| *Carcinoma vs. normal* | 5.18e−03 ** | 5.18e−03 ** |

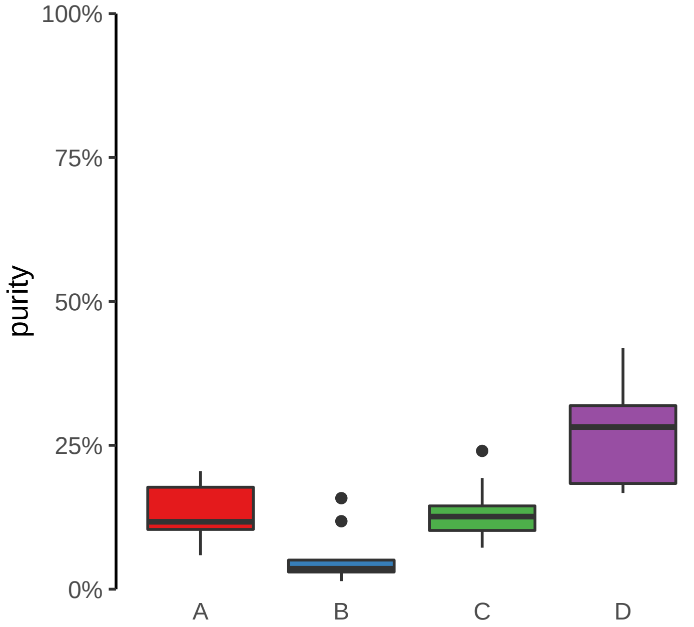

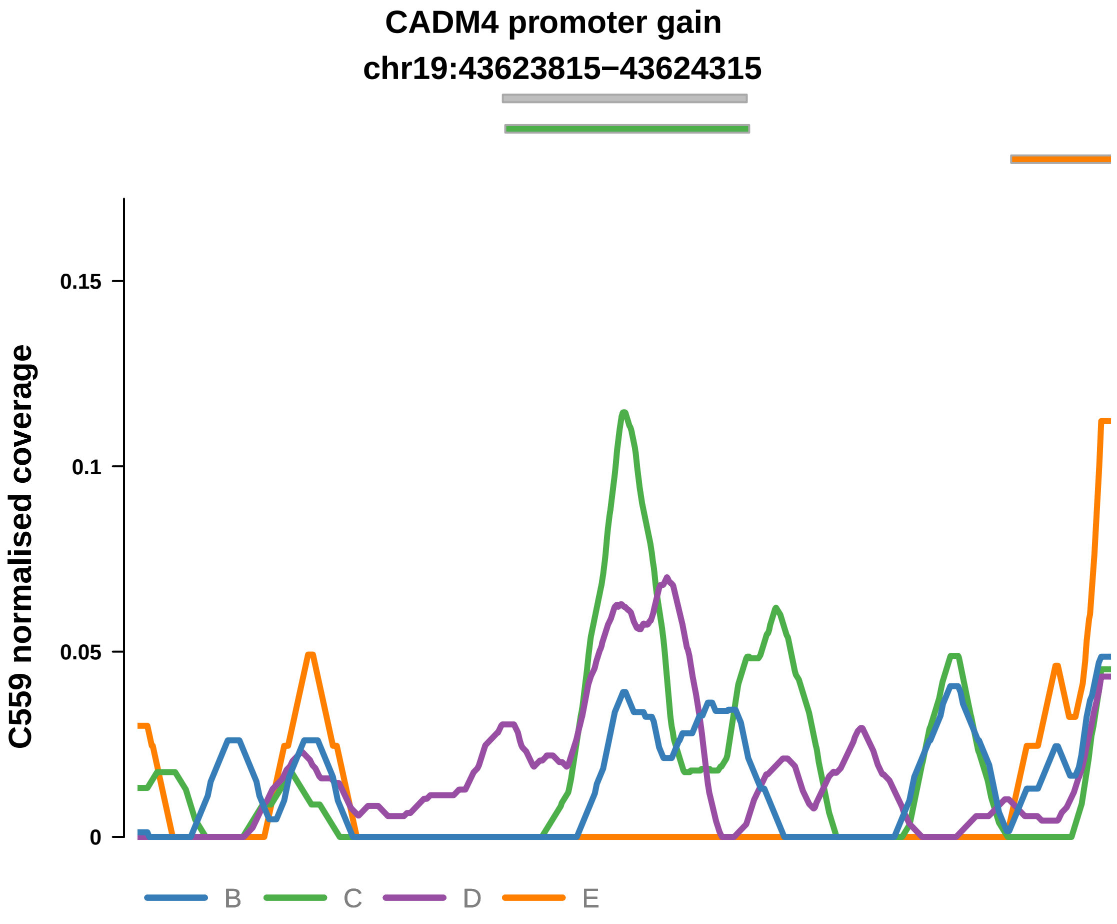


Putatively Clonal (logFC = 2.38)

| | P. | Adjusted P. |
|---|---|---|
| *~region vs. ~* | 1.14e−01 | 4.36e−01 |
| *~purity vs ~* | 1.79e−01 | 6.32e−01 |
| *~purity+region vs ~purity* | 2.63e−01 | 7.28e−01 |
| *Carcinoma vs. normal* | 2.62e−04 *** | 2.62e−04 *** |

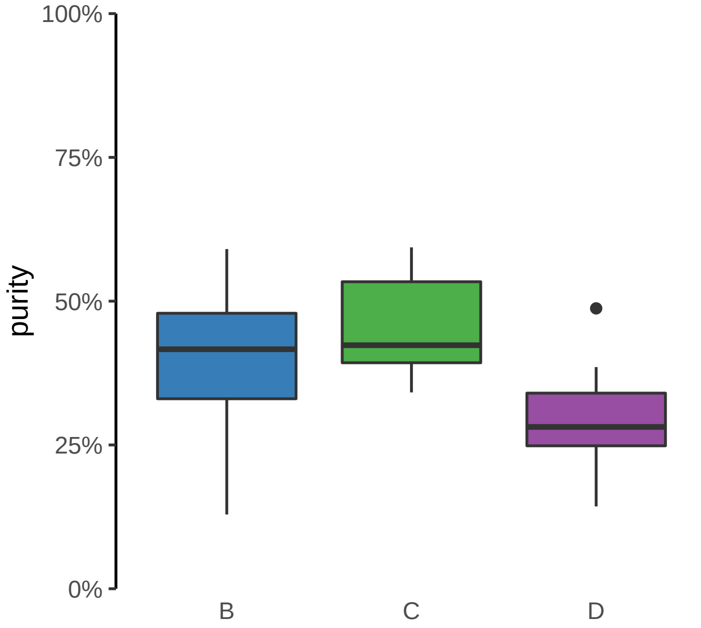

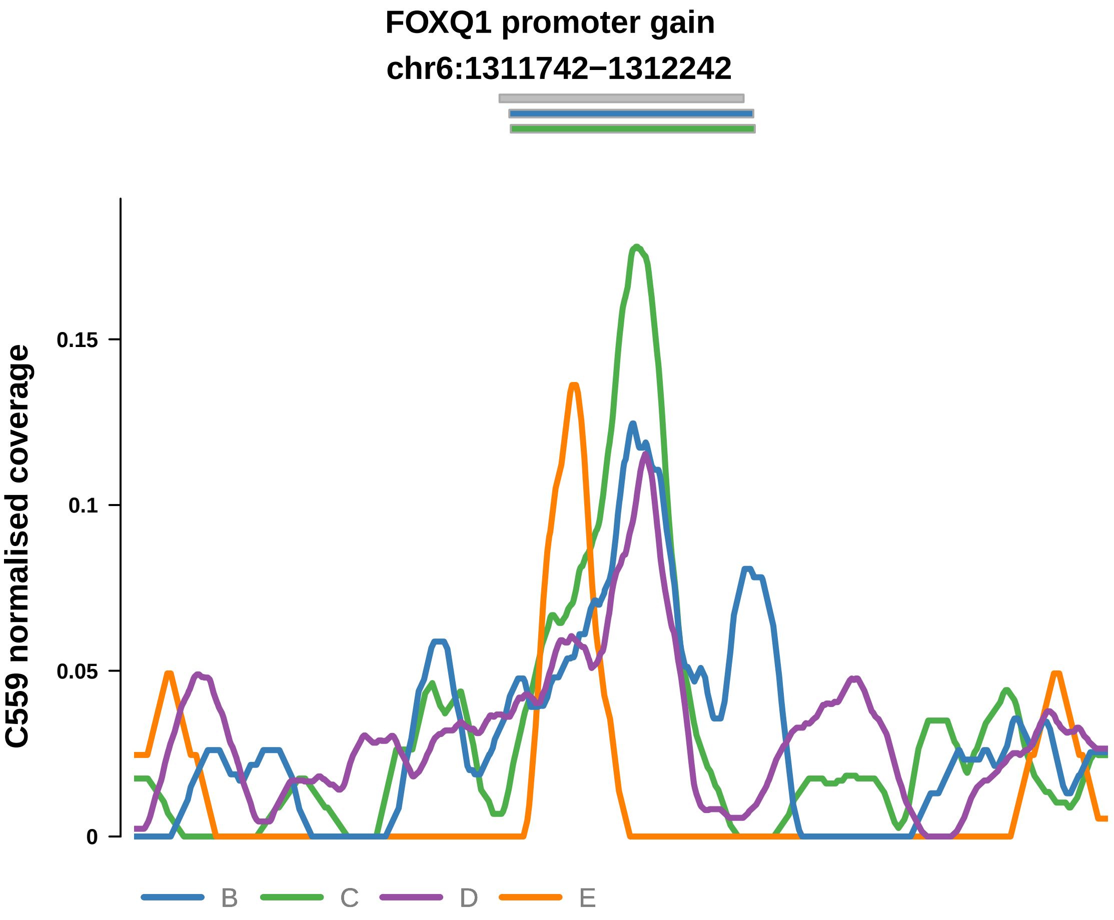


Putatively Clonal (logFC = 1.68)

| | P. | Adjusted P. |
|---|---|---|
| *~region vs. ~* | 5.58e−02 | 3.63e−01 |
| *~purity vs ~* | 4.24e−01 | 8.29e−01 |
| *~purity+region vs ~purity* | 5.7e−02 | 5.21e−01 |
| *Carcinoma vs. normal* | 5.18e−03 ** | 5.18e−03 ** |

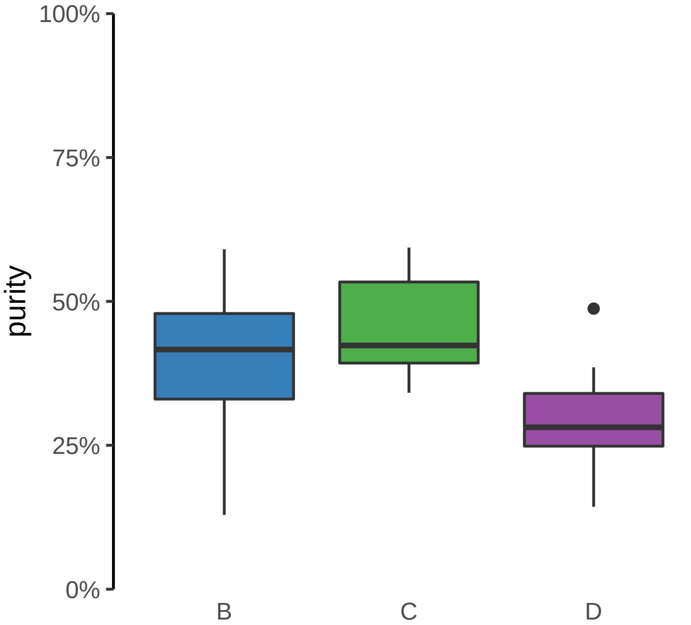

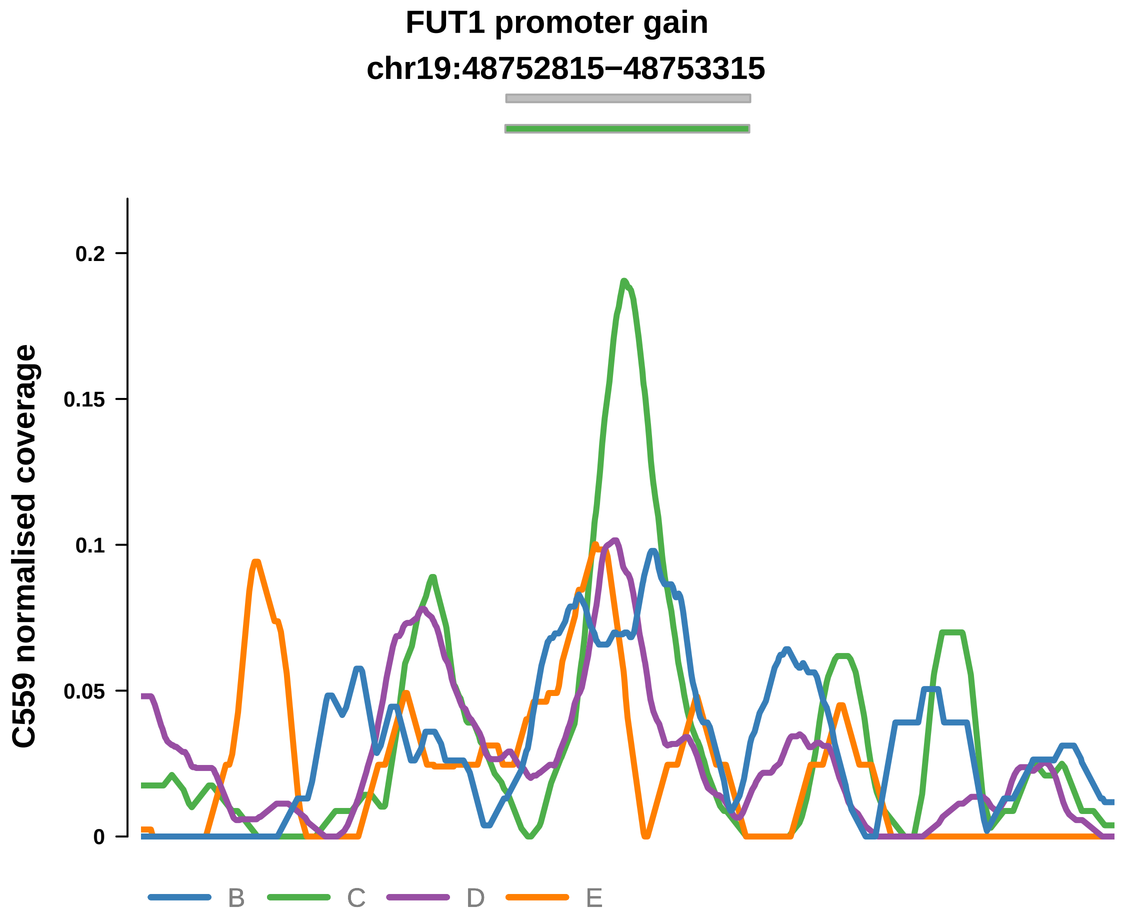


Putatively Clonal (logFC = 1.71)

| | P. | Adjusted P. |
|---|---|---|
| *~region vs. ~* | 5.25e−02 | 3.63e−01 |
| *~purity vs ~* | 2.11e−01 | 6.55e−01 |
| *~purity+region vs ~purity* | 1.11e−01 | 5.21e−01 |
| *Carcinoma vs. normal* | 5.51e−03 ** | 5.51e−03 ** |

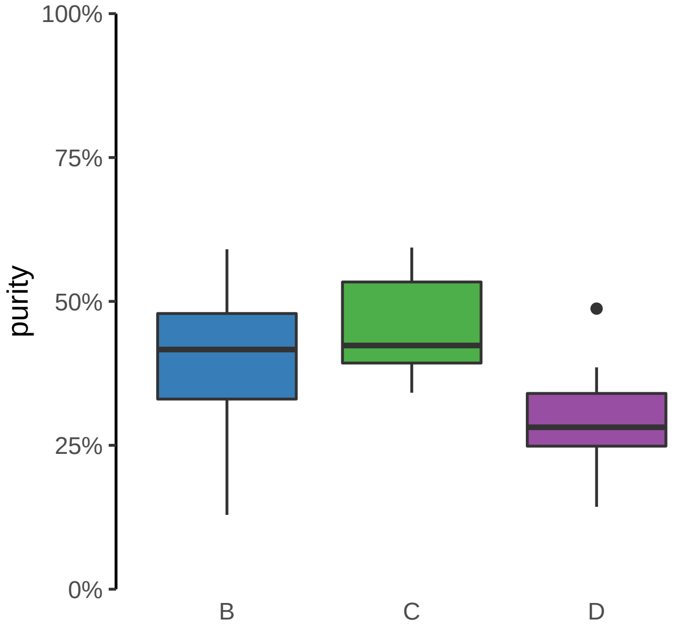

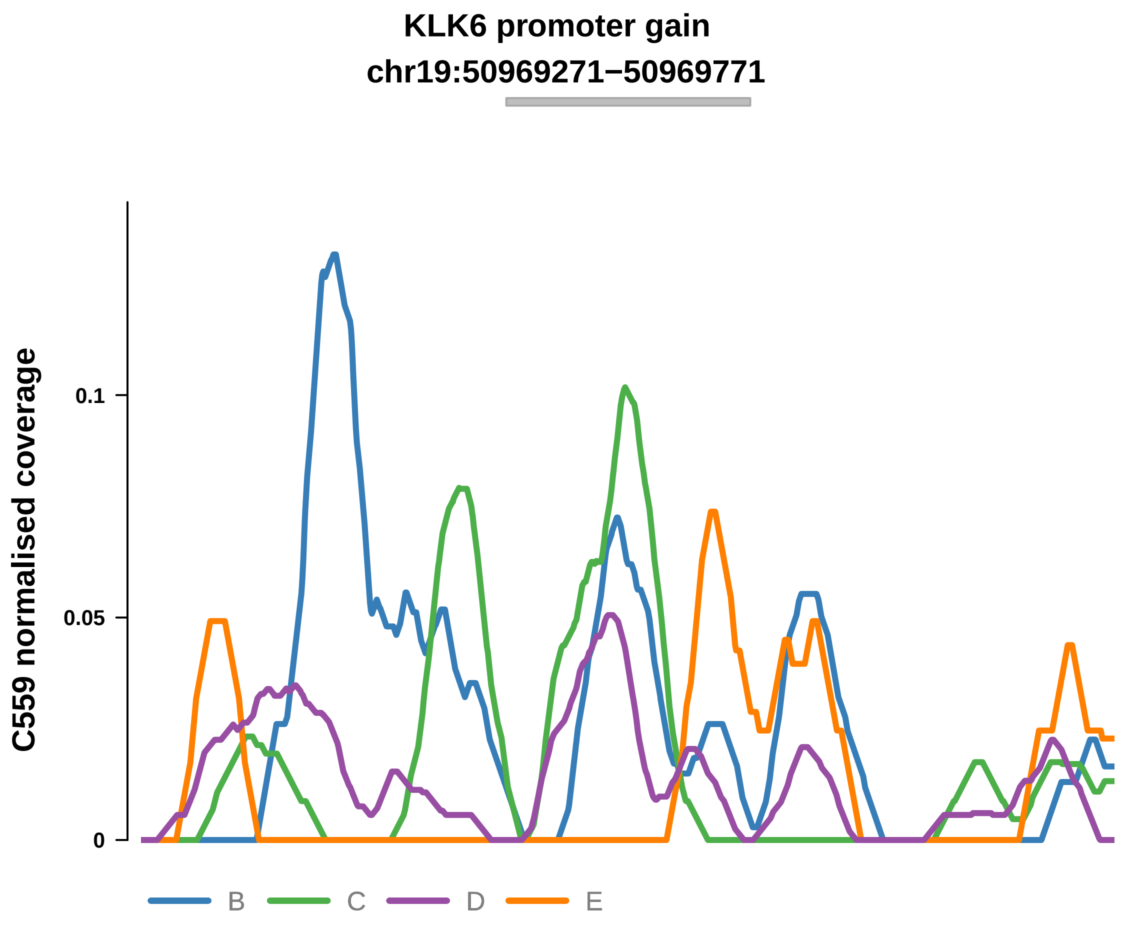


Putatively Clonal (logFC = 2)

| | P. | Adjusted P. |
|---|---|---|
| *~region vs. ~* | 1.89e−02 * | 3.63e−01 |
| *~purity vs ~* | 8.66e−02 | 5.51e−01 |
| *~purity+region vs ~purity* | 1e−01 | 5.21e−01 |
| *Carcinoma vs. normal* | 2.82e−03 ** | 2.82e−03 ** |

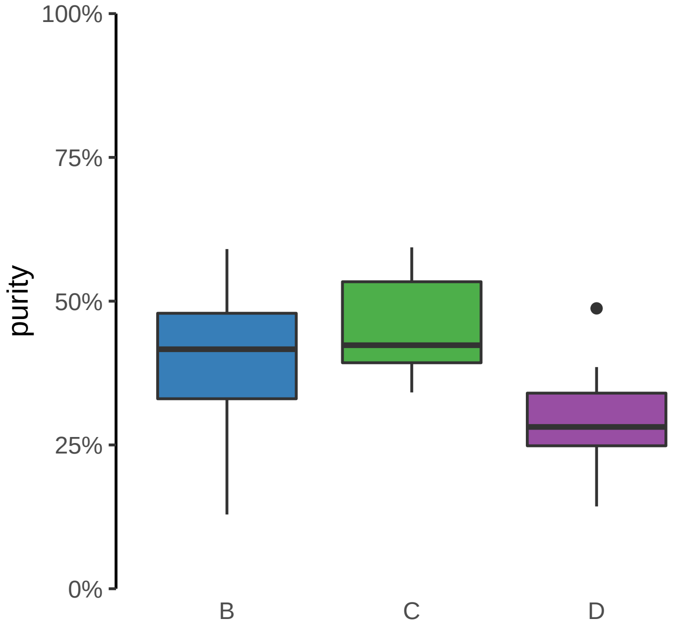

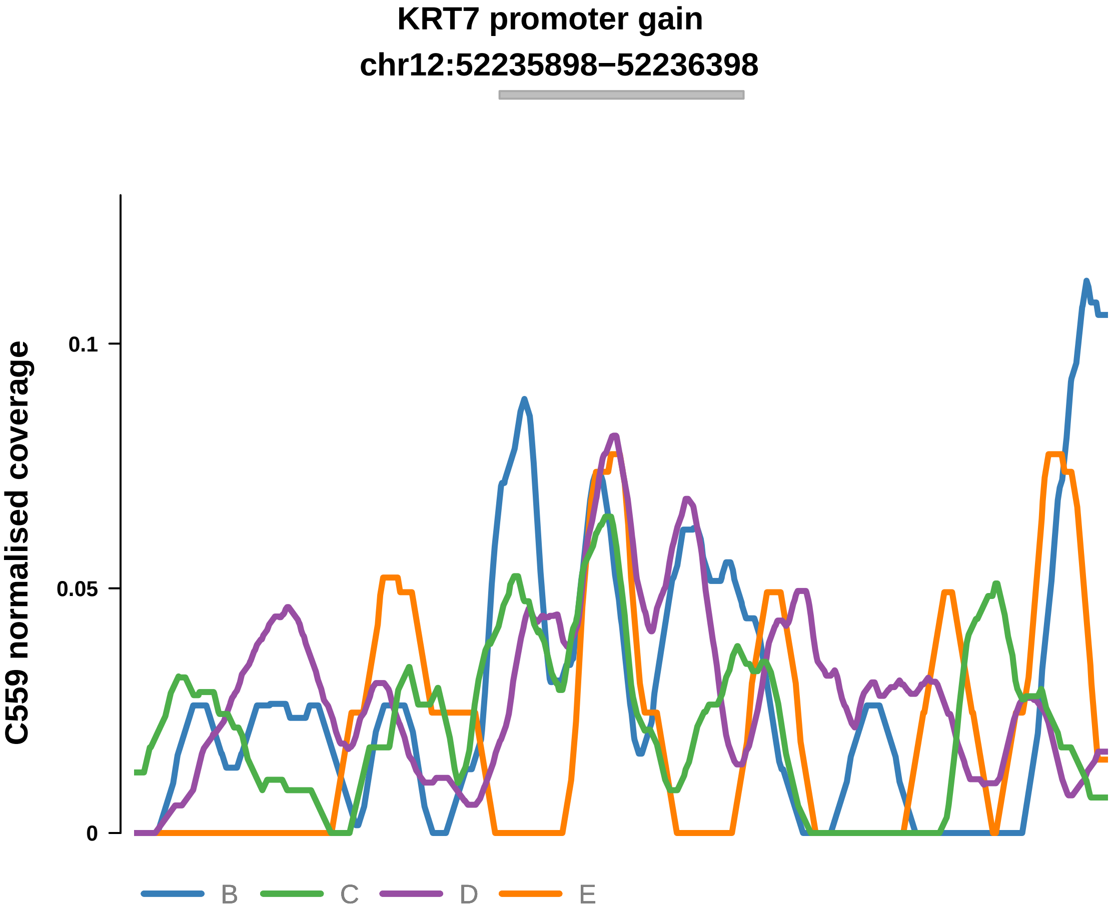


Putatively Clonal (logFC = 1.65)

| | P. | Adjusted P. |
|---|---|---|
| *~region vs. ~* | 2.26e−01 | 5.55e−01 |
| *~purity vs ~* | 8.35e−01 | 9.26e−01 |
| *~purity+region vs ~purity* | 6.85e−02 | 5.21e−01 |
| *Carcinoma vs. normal* | 7.56e−03 ** | 7.56e−03 ** |

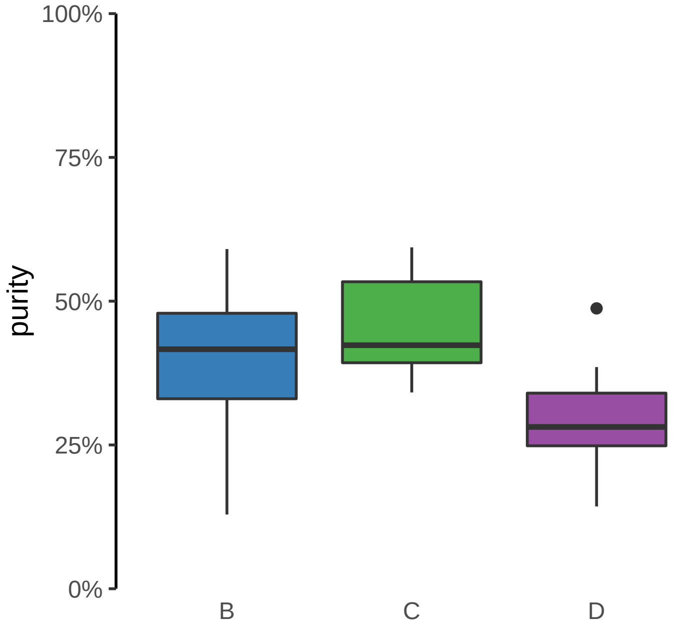

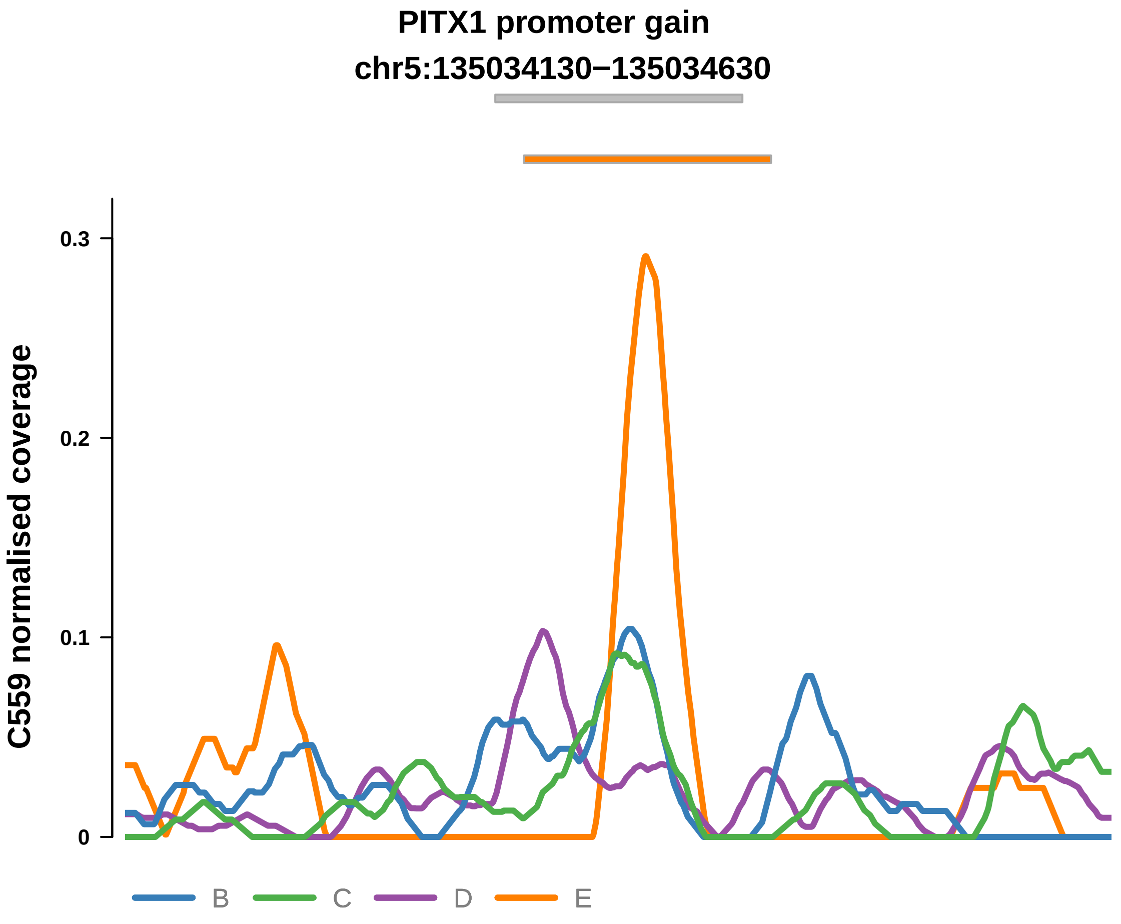


Putatively Clonal (logFC = 1.6)

| | P. | Adjusted P. |
|---|---|---|
| *~region vs. ~* | 2.31e−01 | 5.55e−01 |
| *~purity vs ~* | 7.1e−02 | 5.51e−01 |
| *~purity+region vs ~purity* | 7.71e−01 | 9.61e−01 |
| *Carcinoma vs. normal* | 9.51e−03 ** | 9.51e−03 ** |

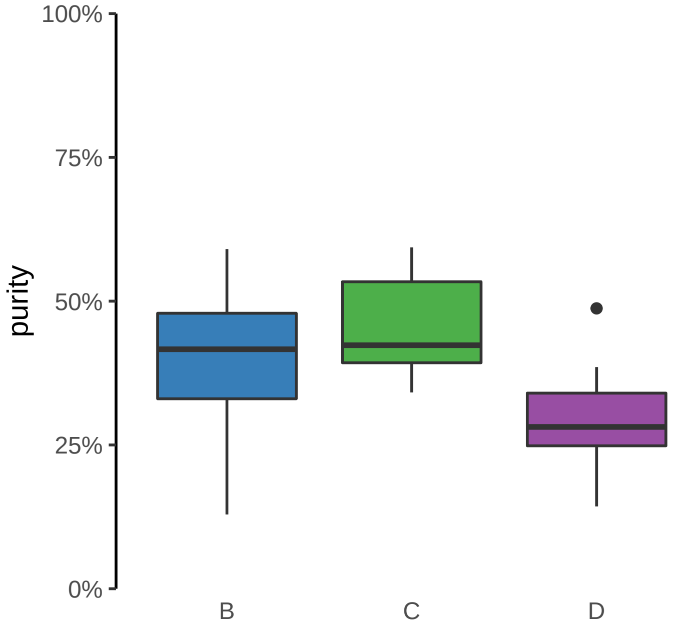

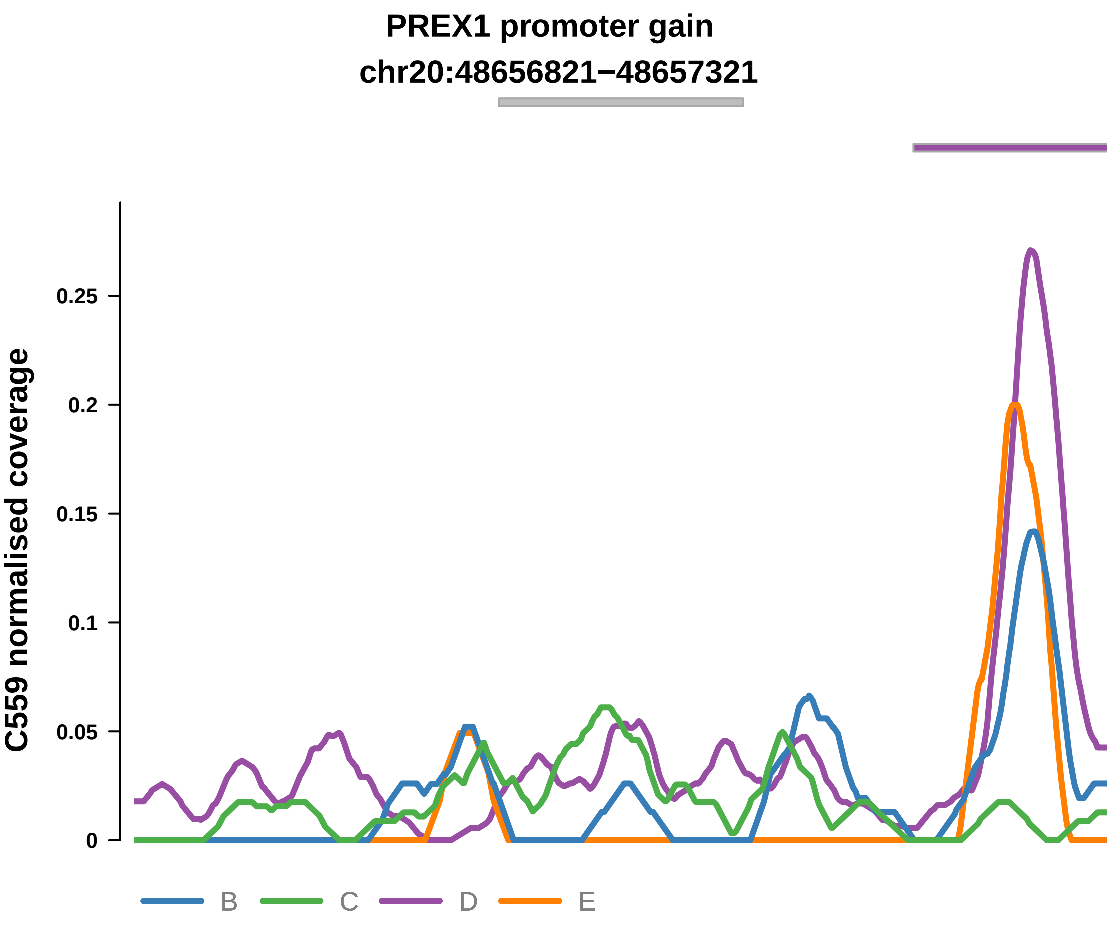


Putatively Clonal (logFC = 1.98)

| | P. | Adjusted P. |
|---|---|---|
| *~region vs. ~* | 5.06e−02 | 3.63e−01 |
| *~purity vs ~* | 8.89e−01 | 9.42e−01 |
| *~purity+region vs ~purity* | 4.23e−02 * | 5.21e−01 |
| *Carcinoma vs. normal* | 4.37e−03 ** | 4.37e−03 ** |

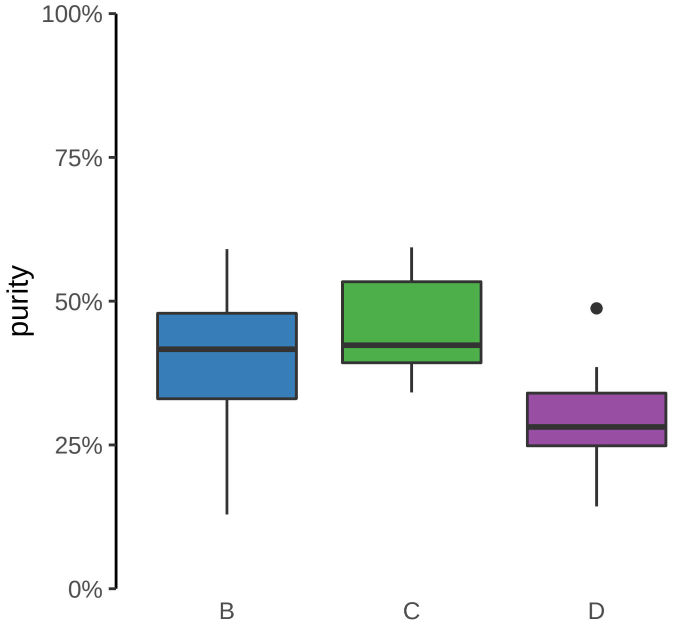

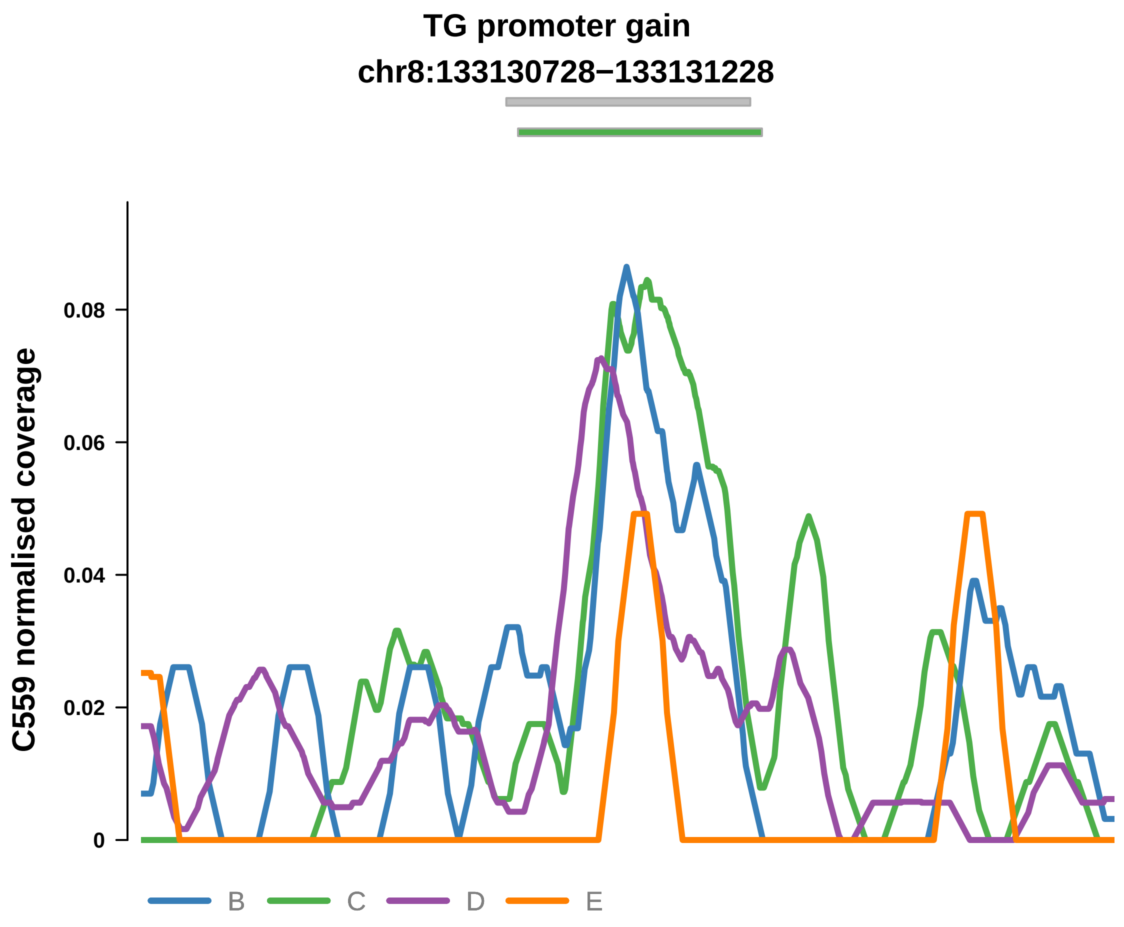


Putatively Clonal (logFC = 2.18)

| | P. | Adjusted P. |
|---|---|---|
| *~region vs. ~* | 4.57e−02 * | 3.63e−01 |
| *~purity vs ~* | 2.16e−01 | 6.55e−01 |
| *~purity+region vs ~purity* | 9.19e−02 | 5.21e−01 |
| *Carcinoma vs. normal* | 4.6e−04 *** | 4.6e−04 *** |

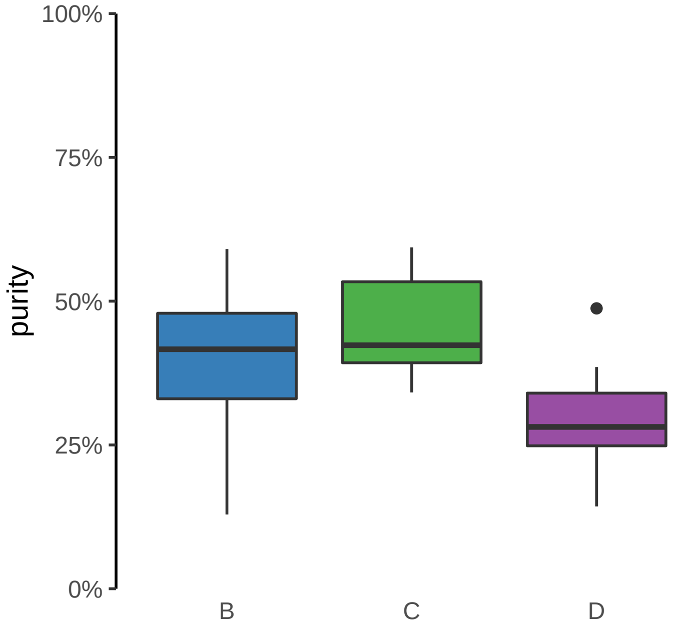

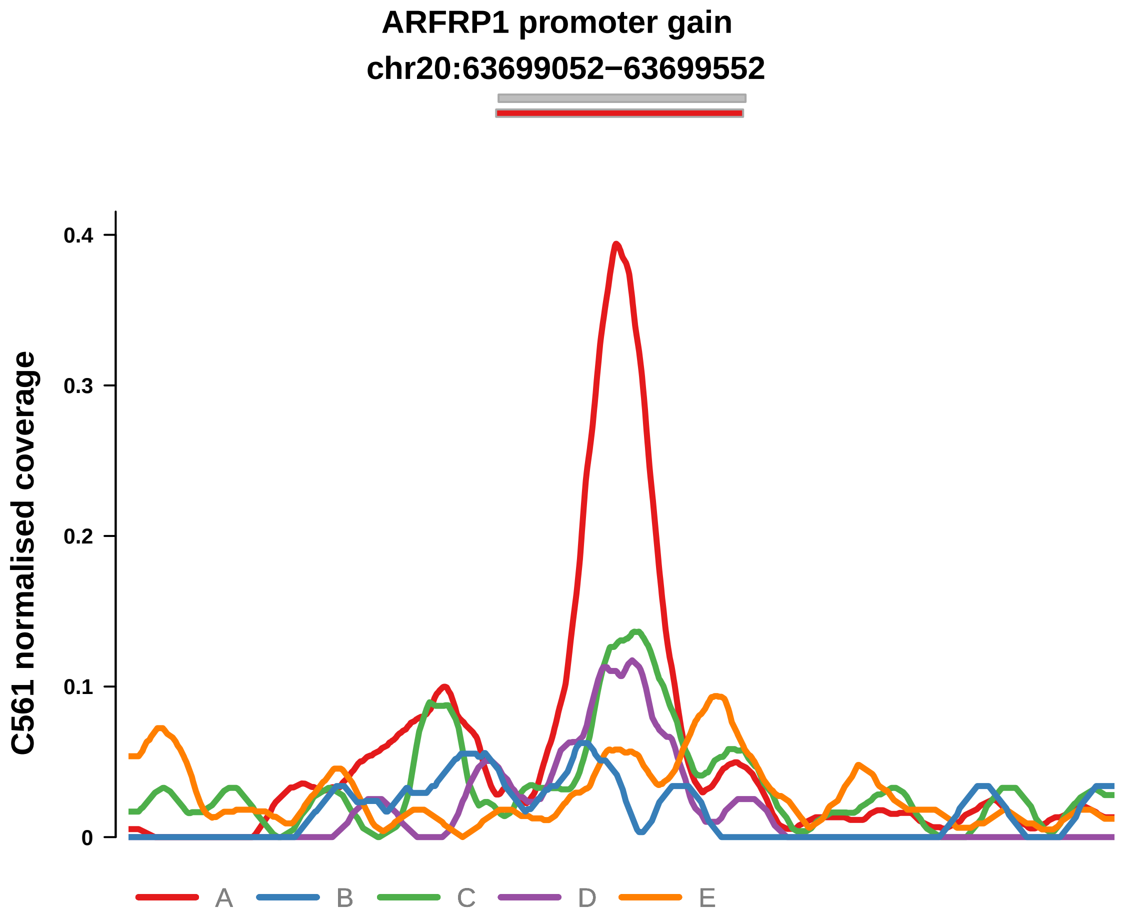


Putatively Clonal (logFC = 2.24)

| | P. | Adjusted P. |
|---|---|---|
| *~region vs. ~* | 9.63e−02 | 2.81e−01 |
| *~purity vs ~* | 4.71e−01 | 7.83e−01 |
| *~purity+region vs ~purity* | 5.5e−02 | 4.2e−01 |
| *Carcinoma vs. normal* | 5.54e−05 **** | 5.54e−05 **** |

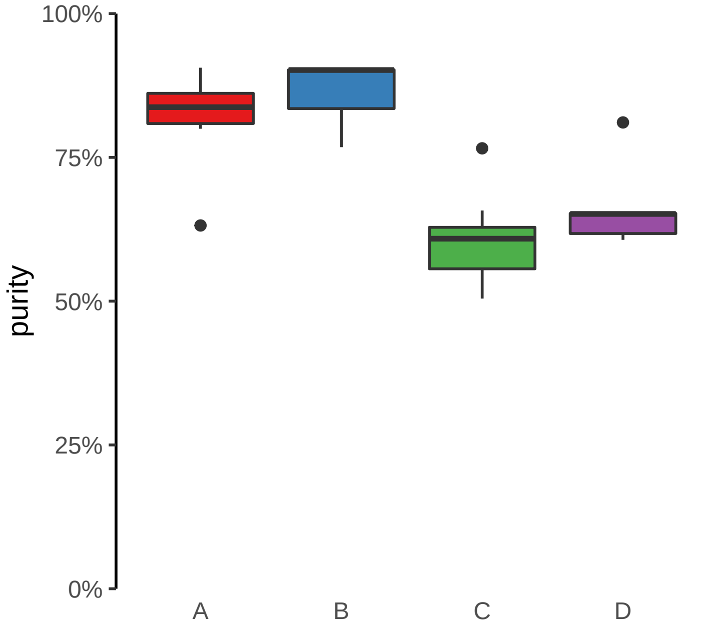

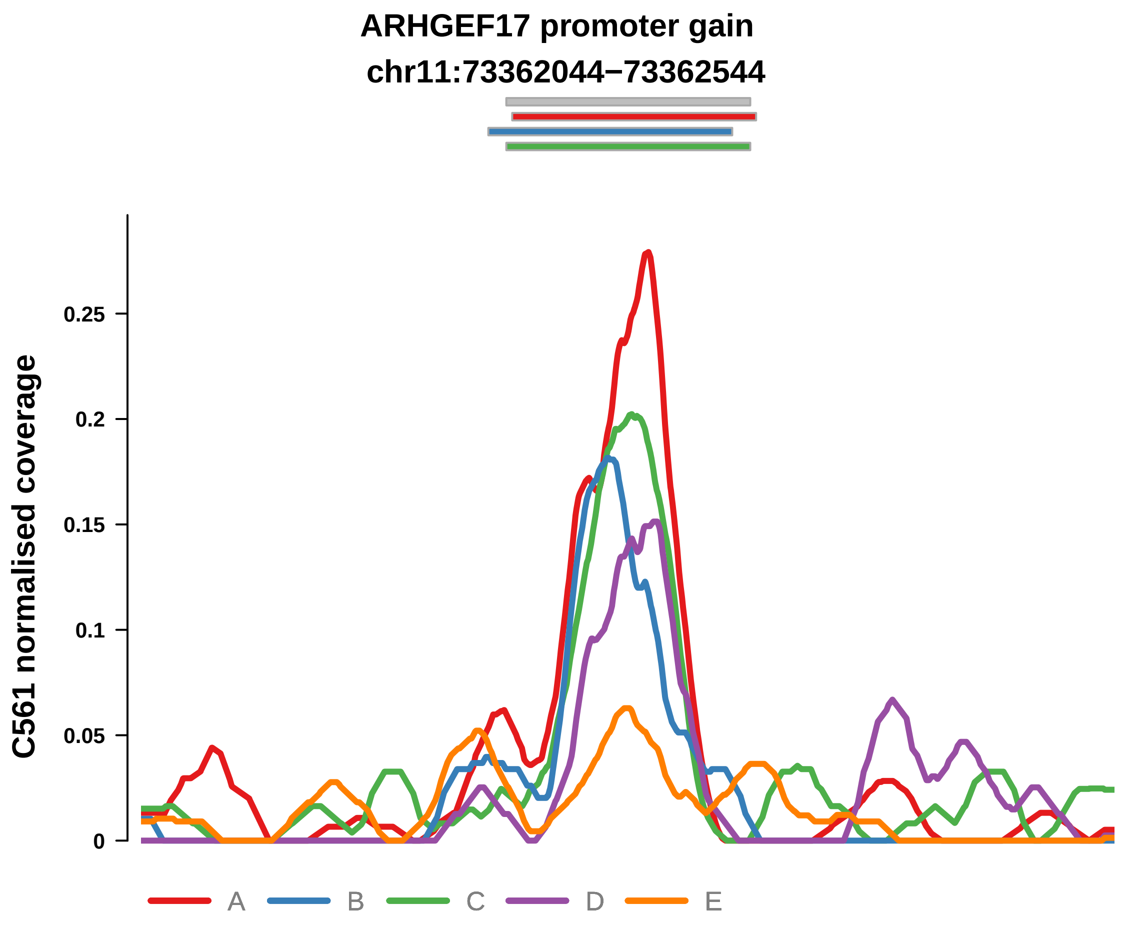


Putatively Clonal (logFC = 2.65)

| | P. | Adjusted P. |
|---|---|---|
| *~region vs. ~* | 9.56e−01 | 9.76e−01 |
| *~purity vs ~* | 5.46e−01 | 8.51e−01 |
| *~purity+region vs ~purity* | 2.25e−01 | 7.31e−01 |
| *Carcinoma vs. normal* | 3.07e−06 **** | 3.07e−06 **** |

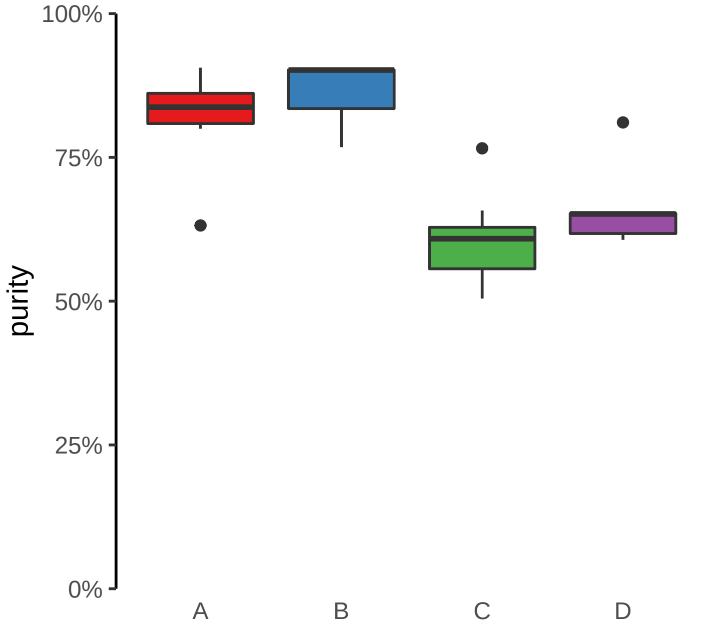

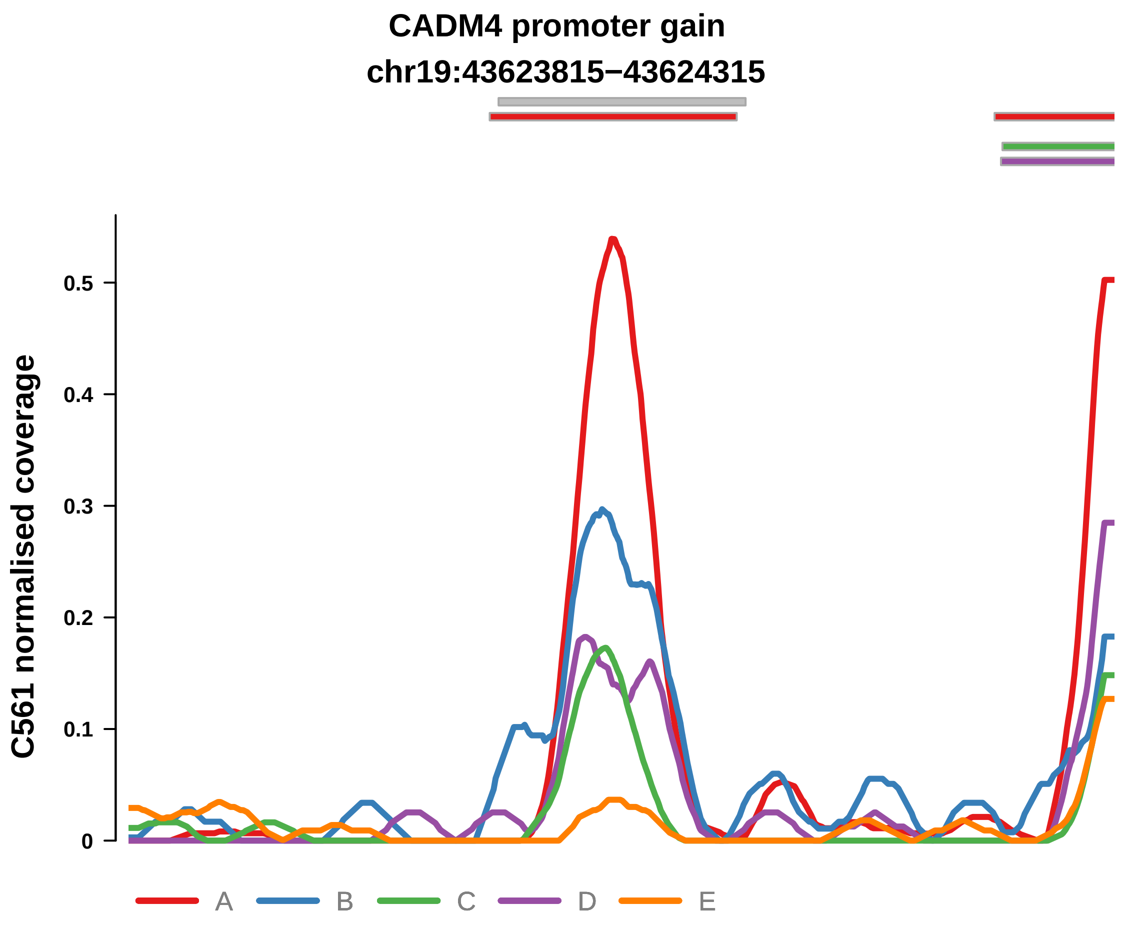


Putatively Clonal (logFC = 3.09)

| | P. | Adjusted P. |
|---|---|---|
| *~region vs. ~* | 3.09e−02 * | 1.43e−01 |
| *~purity vs ~* | 3.62e−02 * | 3.18e−01 |
| *~purity+region vs ~purity* | 2.93e−01 | 7.68e−01 |
| *Carcinoma vs. normal* | 7.38e−08 **** | 7.38e−08 **** |

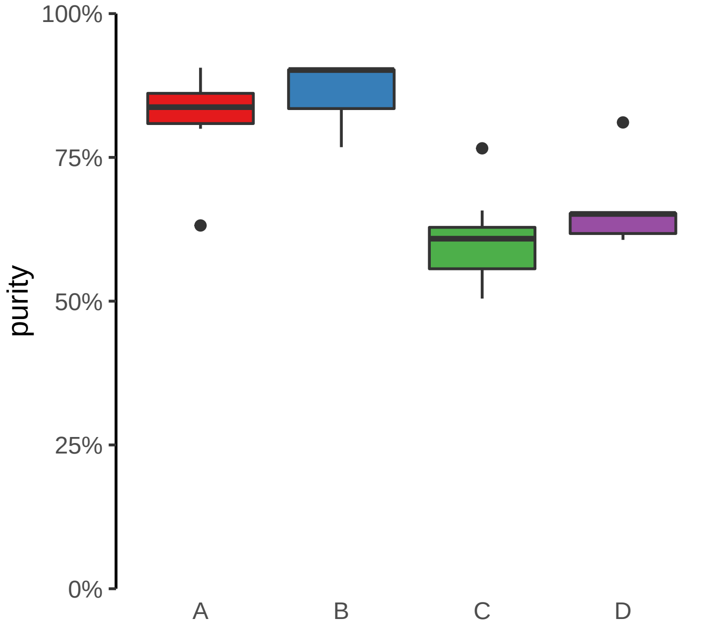

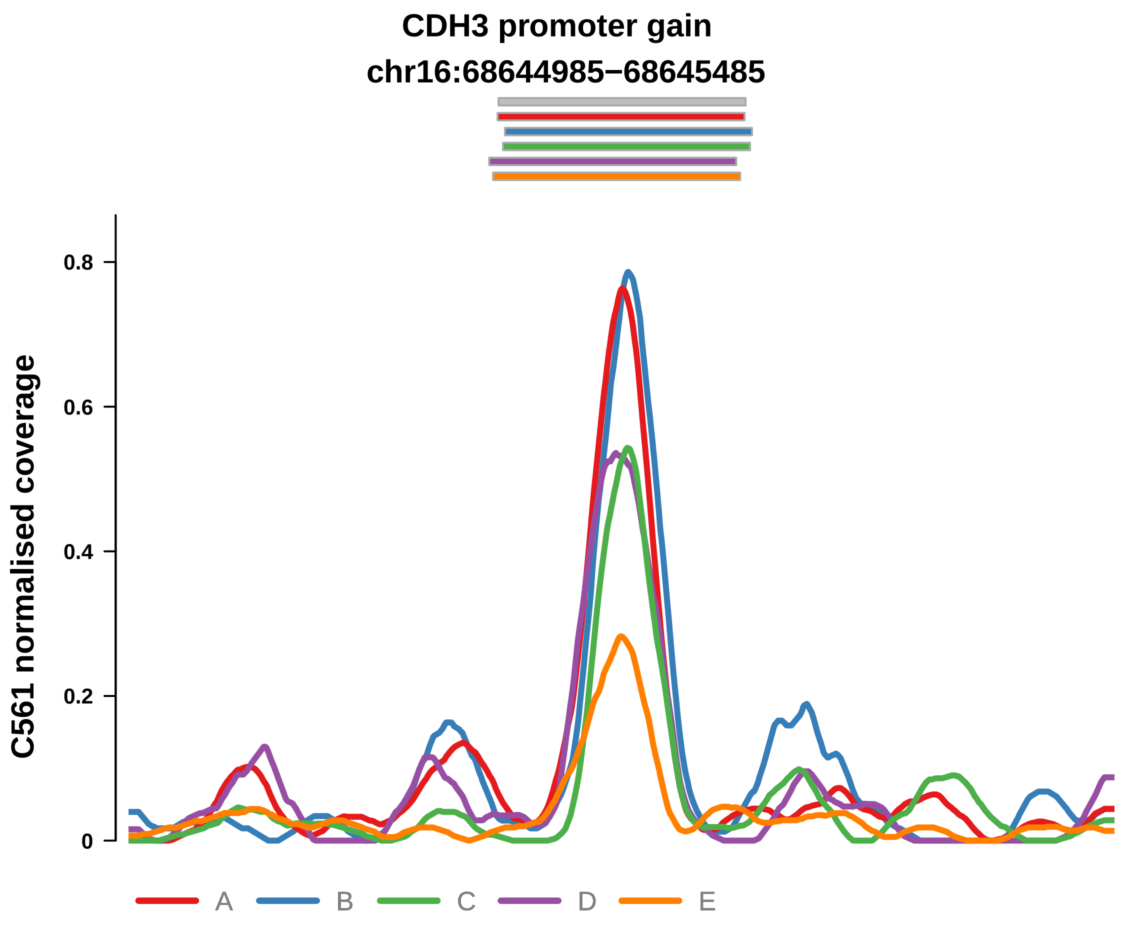


Putatively Clonal (logFC = 2.74)

| | P. | Adjusted P. |
|---|---|---|
| *~region vs. ~* | 1.14e−01 | 3e−01 |
| *~purity vs ~* | 3.53e−01 | 7.66e−01 |
| *~purity+region vs ~purity* | 1.83e−01 | 6.21e−01 |
| *Carcinoma vs. normal* | 9.05e−07 **** | 9.05e−07 **** |

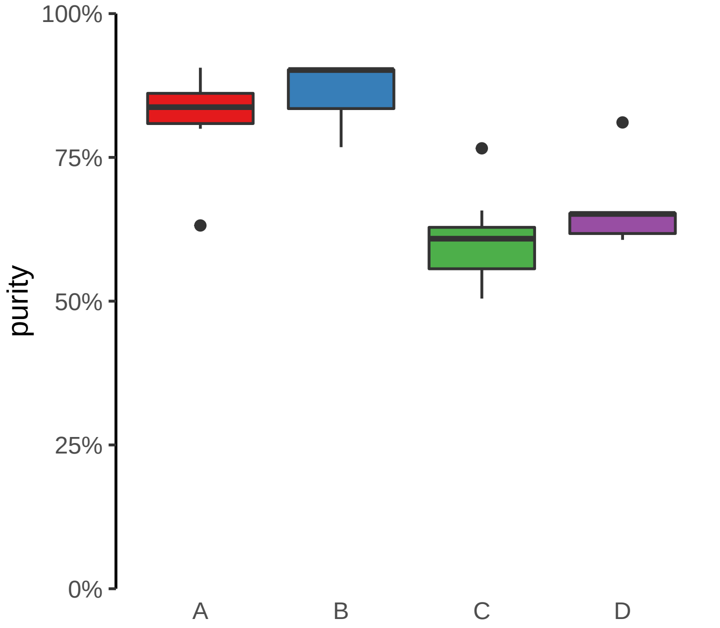

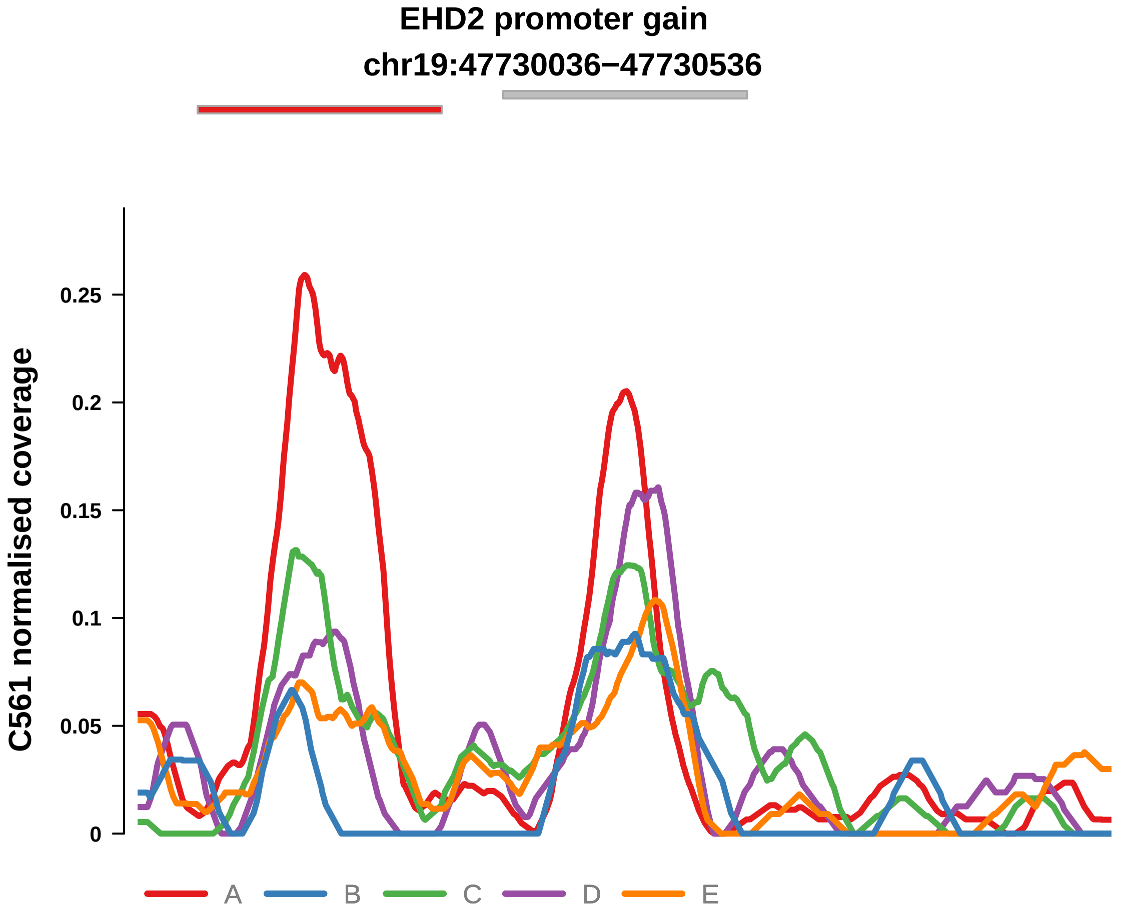


Putatively Clonal (logFC = 1.45)

| | P. | Adjusted P. |
|---|---|---|
| *~region vs. ~* | 4.01e−01 | 5.91e−01 |
| *~purity vs ~* | 1.27e−01 | 4.47e−01 |
| *~purity+region vs ~purity* | 8.45e−01 | 9.76e−01 |
| *Carcinoma vs. normal* | 8.13e−03 ** | 8.13e−03 ** |

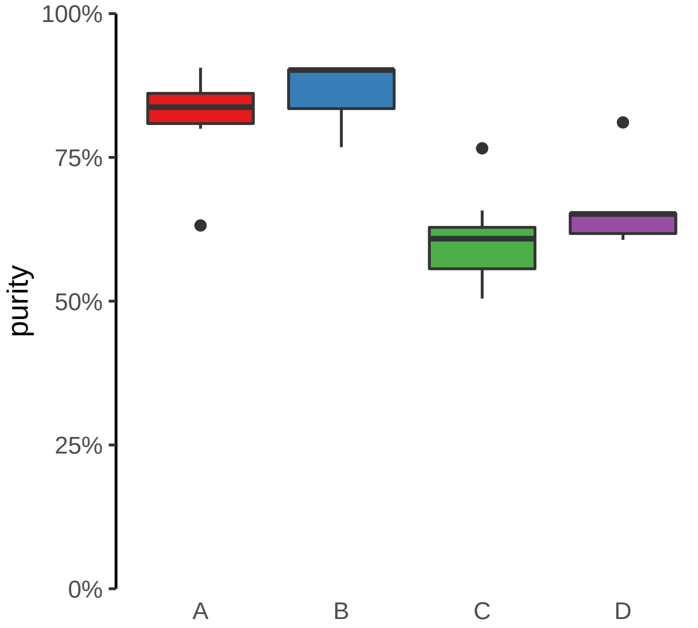

**FOXQ1 promoter gain**

**chr6:1311742−1312242**

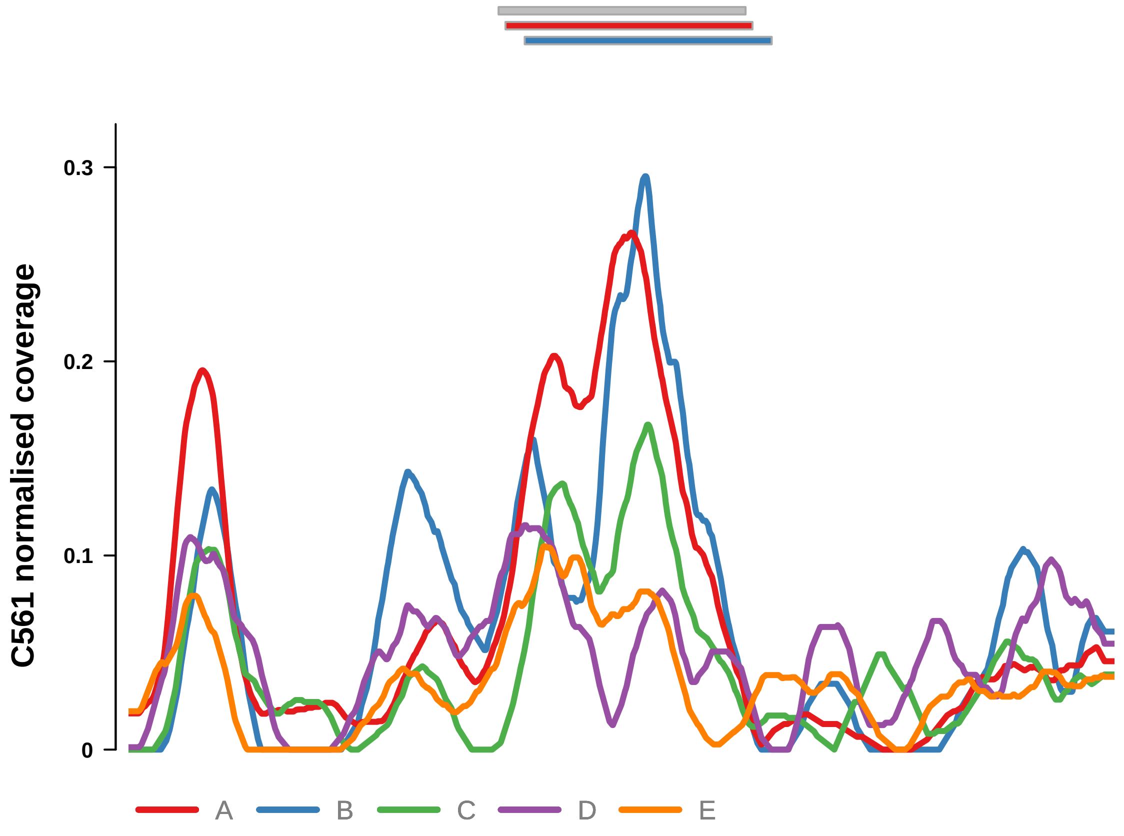


Putatively Clonal (logFC = 2.04)

| | **P.** | **Adjusted P.** |
|---|---|---|
| *~region vs. ~* | 1.43e−01 | 3.43e−01 |
| *~purity vs ~* | 8.68e−02 | 3.98e−01 |
| *~purity+region vs ~purity* | 4.95e−01 | 8.71e−01 |
| *Carcinoma vs. normal* | 1.89e−04 *** | 1.89e−04 *** |

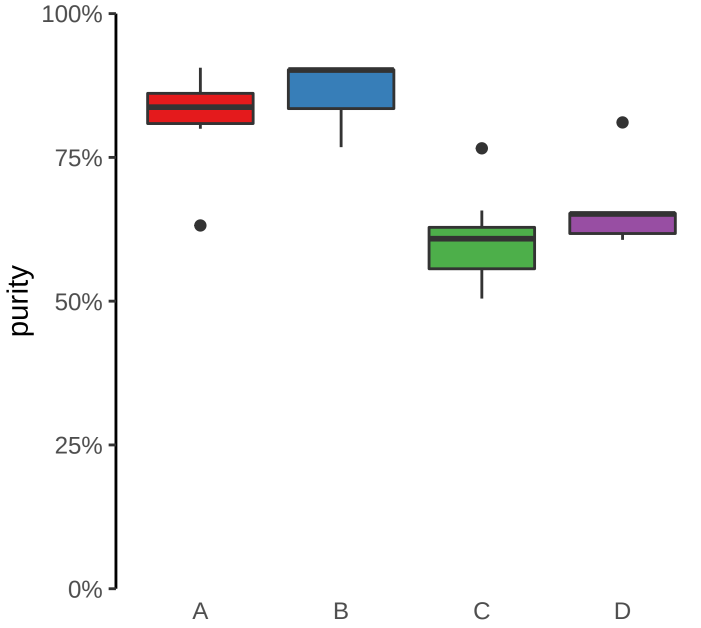

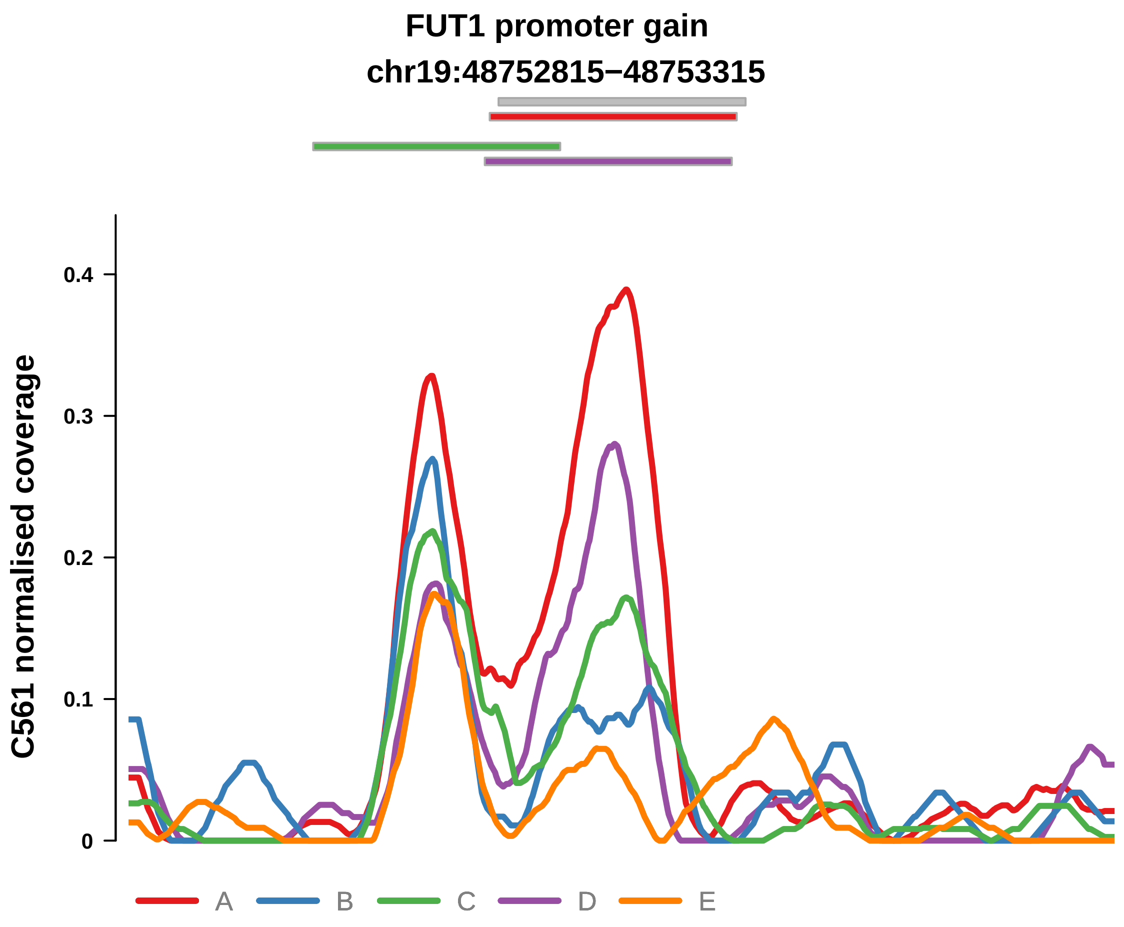


Putatively Clonal (logFC = 2.17)

| | P. | Adjusted P. |
|---|---|---|
| *~region vs. ~* | 4.87e−02 * | 1.81e−01 |
| *~purity vs ~* | 5.68e−02 | 3.73e−01 |
| *~purity+region vs ~purity* | 7.34e−03 ** | 2.35e−01 |
| *Carcinoma vs. normal* | 7.89e−05 **** | 7.89e−05 **** |

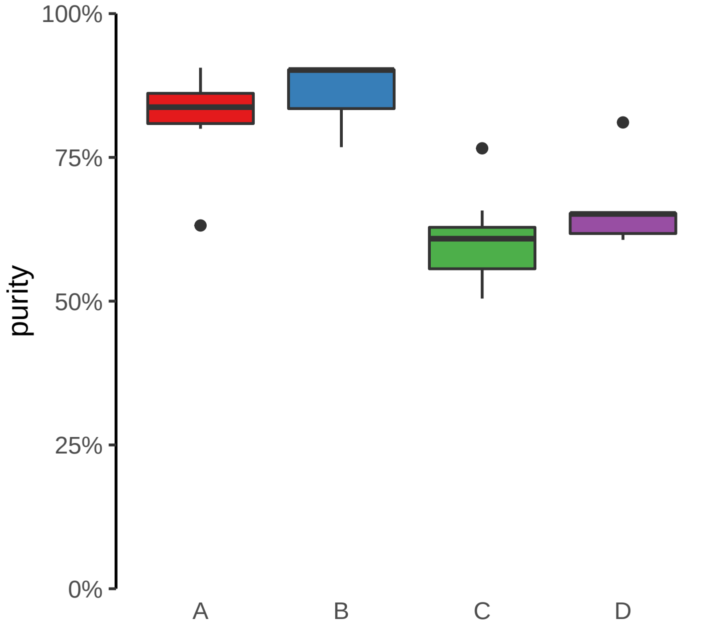

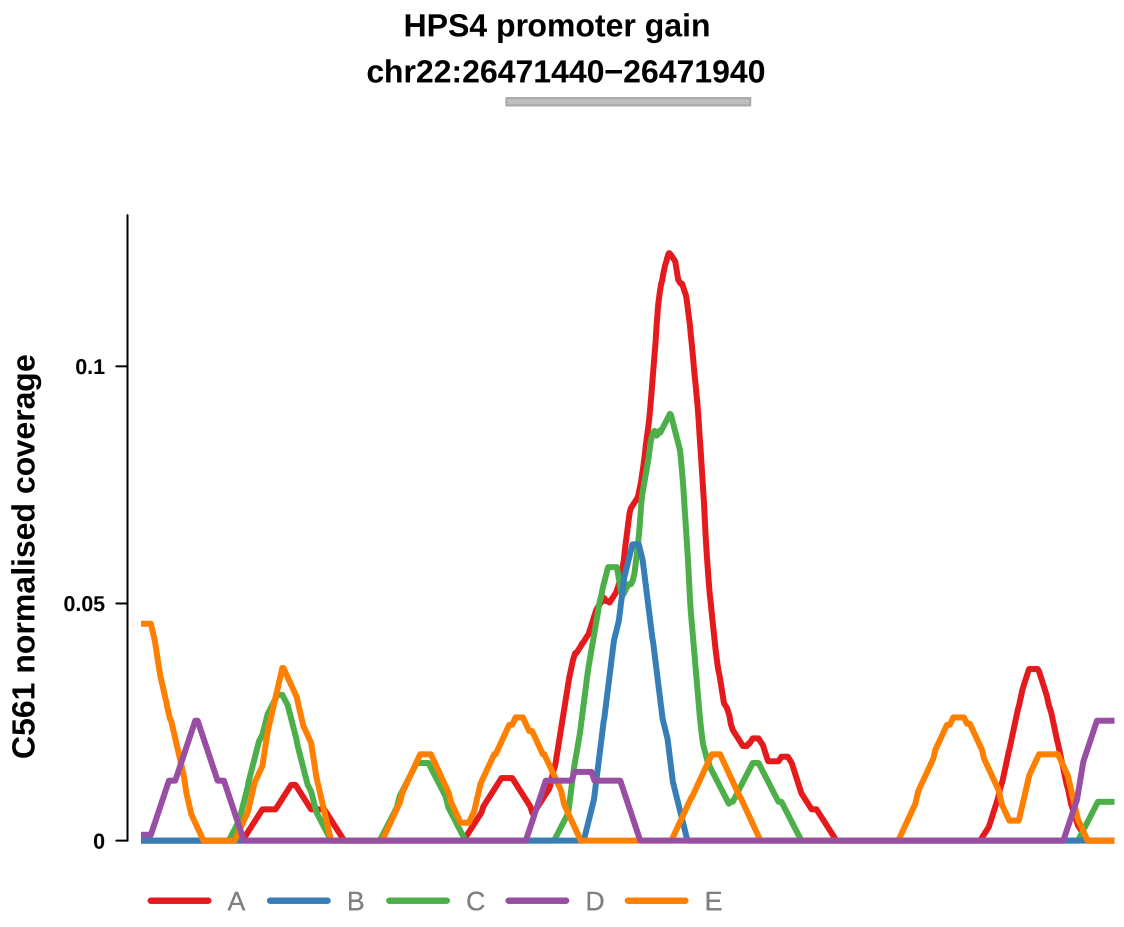


Putatively Clonal (logFC = 1.63)

| | P. | Adjusted P. |
|---|---|---|
| *~region vs. ~* | 1.35e−02 * | 8.23e−02 |
| *~purity vs ~* | 3.08e−01 | 7.53e−01 |
| *~purity+region vs ~purity* | 1.07e−02 * | 2.35e−01 |
| *Carcinoma vs. normal* | 4.16e−03 ** | 4.16e−03 ** |

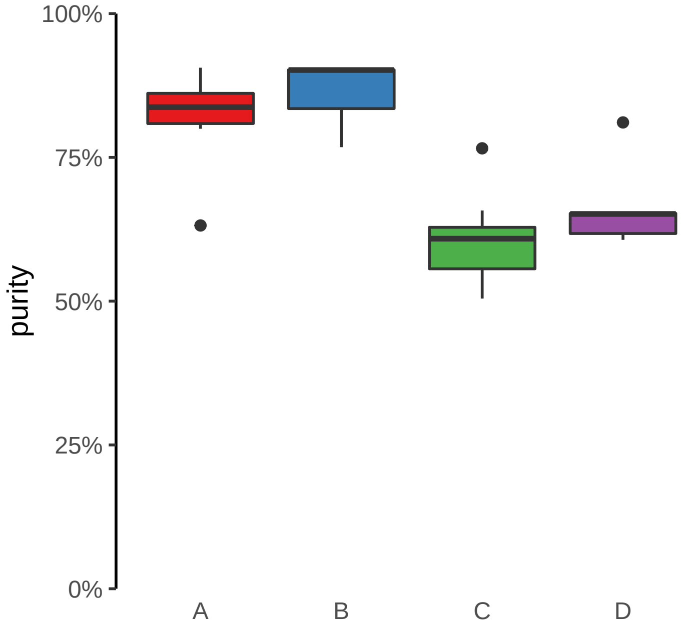

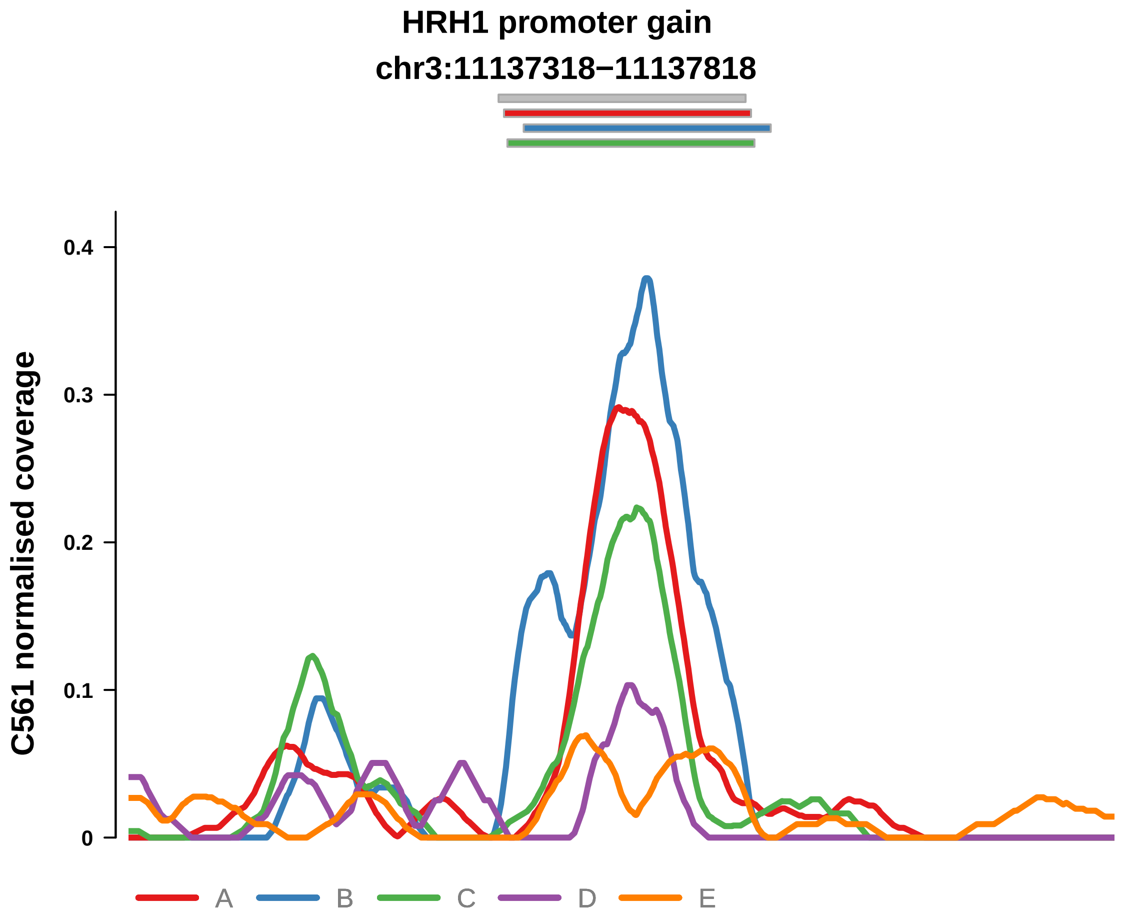


Putatively Clonal (logFC = 2.69)

| | P. | Adjusted P. |
|---|---|---|
| *~region vs. ~* | 9.51e−03 ** | 6.83e−02 |
| *~purity vs ~* | 1.55e−02 * | 1.52e−01 |
| *~purity+region vs ~purity* | 1.05e−01 | 5.15e−01 |
| *Carcinoma vs. normal* | 1.83e−06 **** | 1.83e−06 **** |

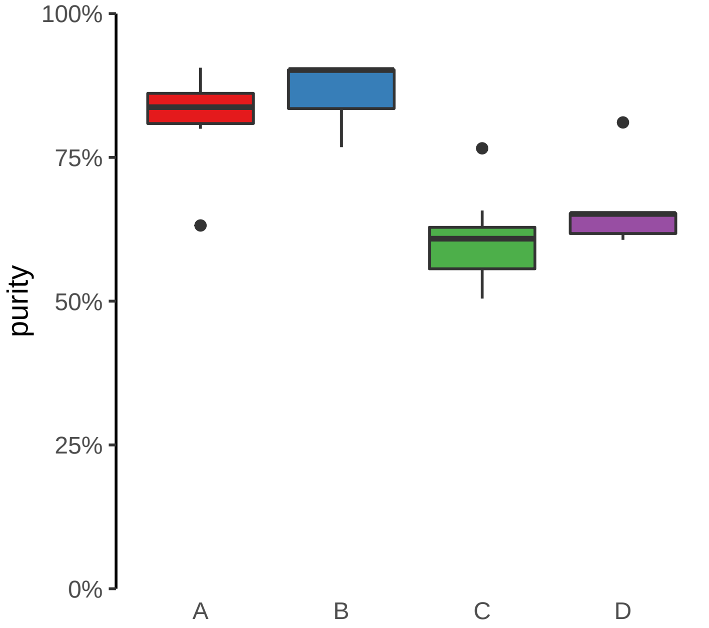

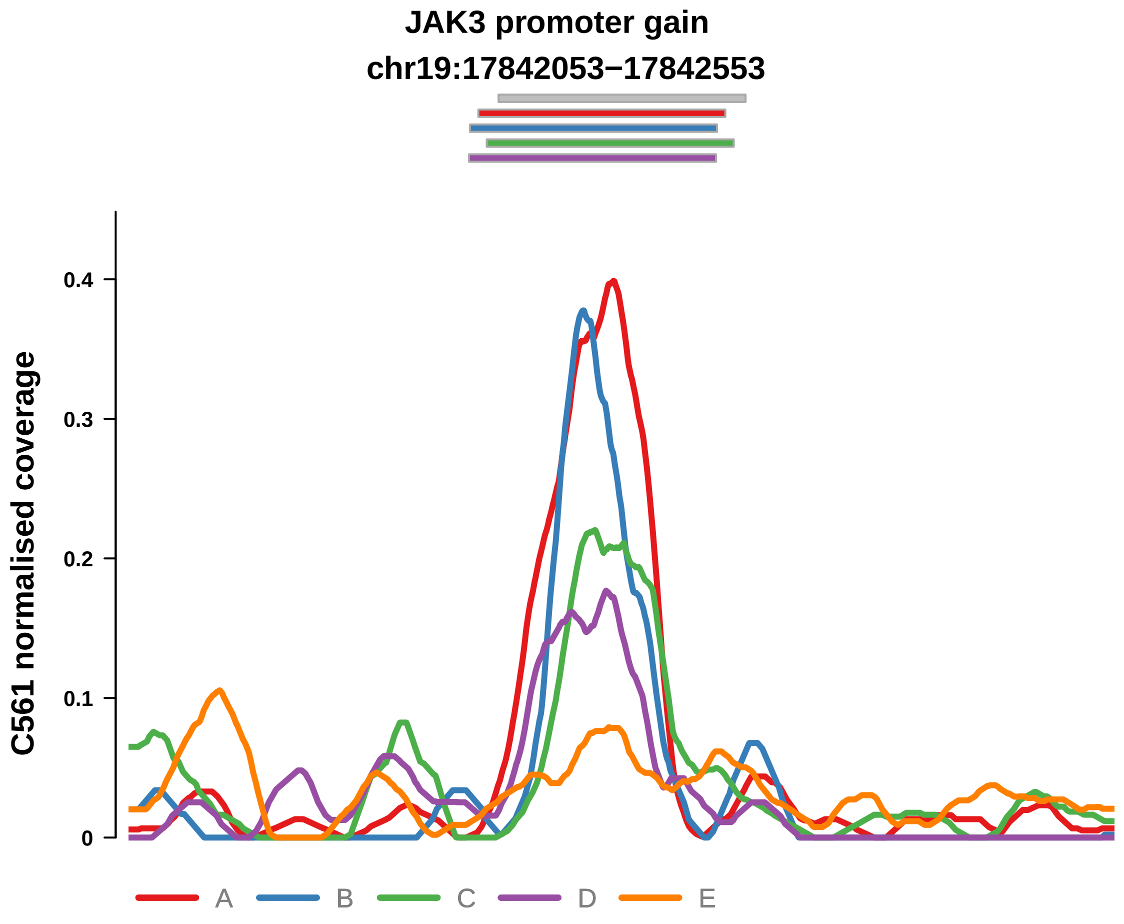


Putatively Clonal (logFC = 2.4)

| | P. | Adjusted P. |
|---|---|---|
| *~region vs. ~* | 9.14e−01 | 9.5e−01 |
| *~purity vs ~* | 4.12e−01 | 7.66e−01 |
| *~purity+region vs ~purity* | 8.55e−01 | 9.76e−01 |
| *Carcinoma vs. normal* | 1.48e−05 **** | 1.48e−05 **** |

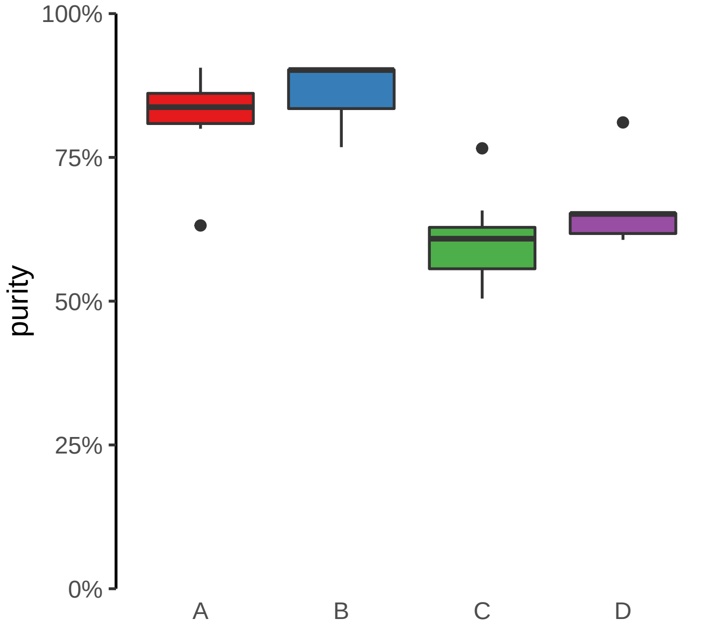

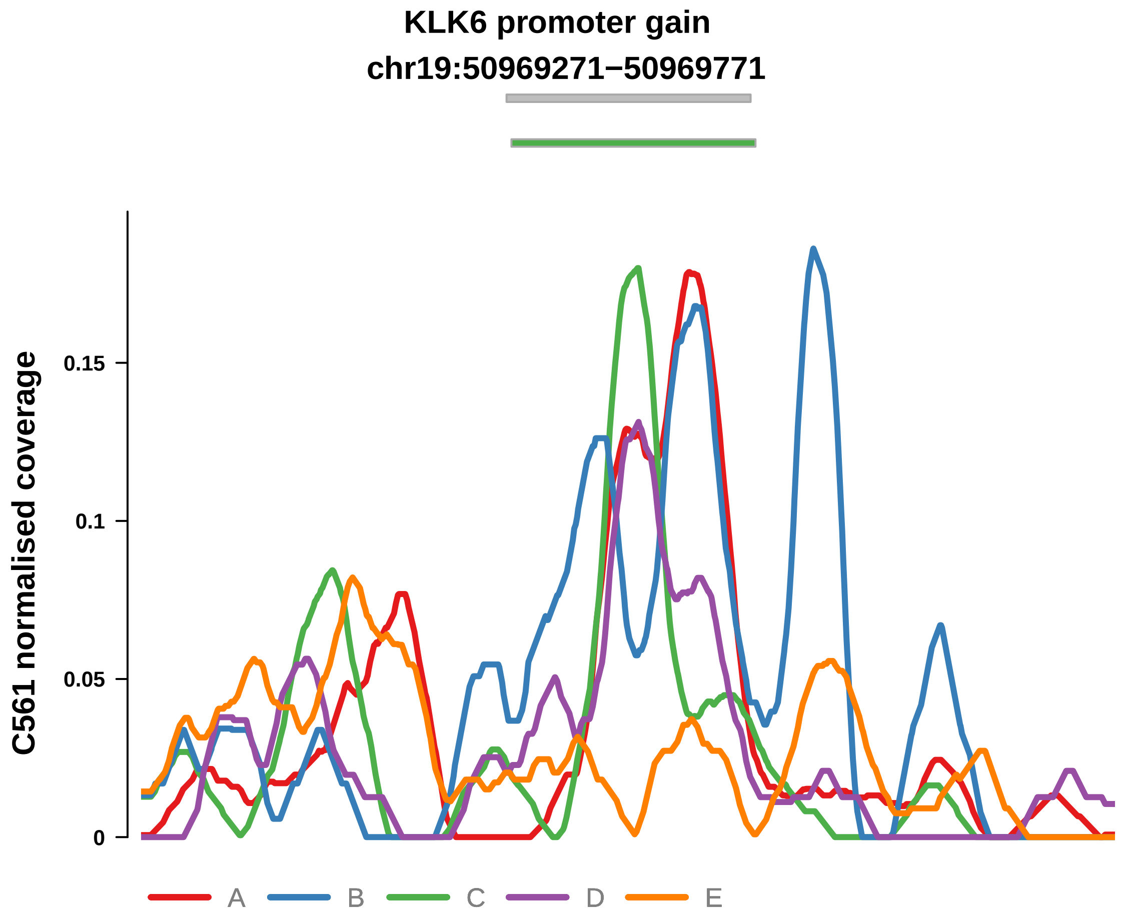


Putatively Clonal (logFC = 2.41)

| | P. | Adjusted P. |
|---|---|---|
| *~region vs. ~* | 2.25e−01 | 4.44e−01 |
| *~purity vs ~* | 7.36e−01 | 8.81e−01 |
| *~purity+region vs ~purity* | 2.36e−01 | 7.31e−01 |
| *Carcinoma vs. normal* | 1.99e−05 **** | 1.99e−05 **** |

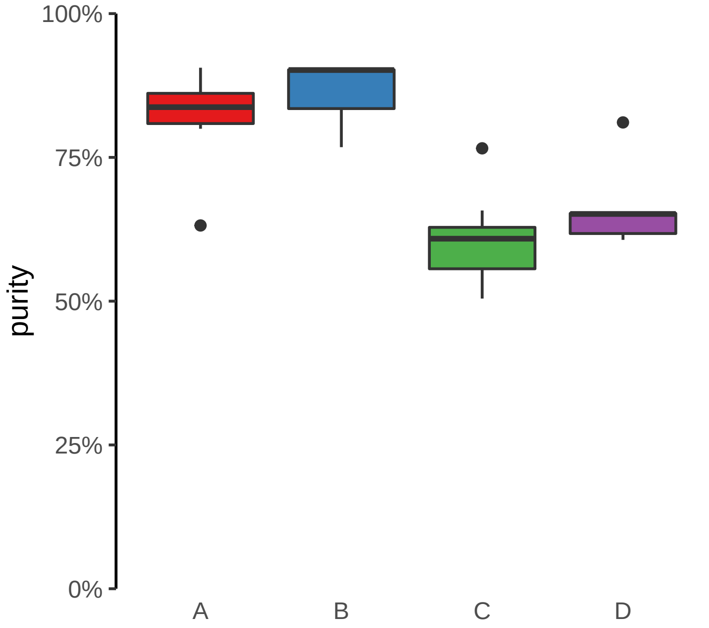

**KRT7 promoter gain**
**chr12:52235898−52236398**

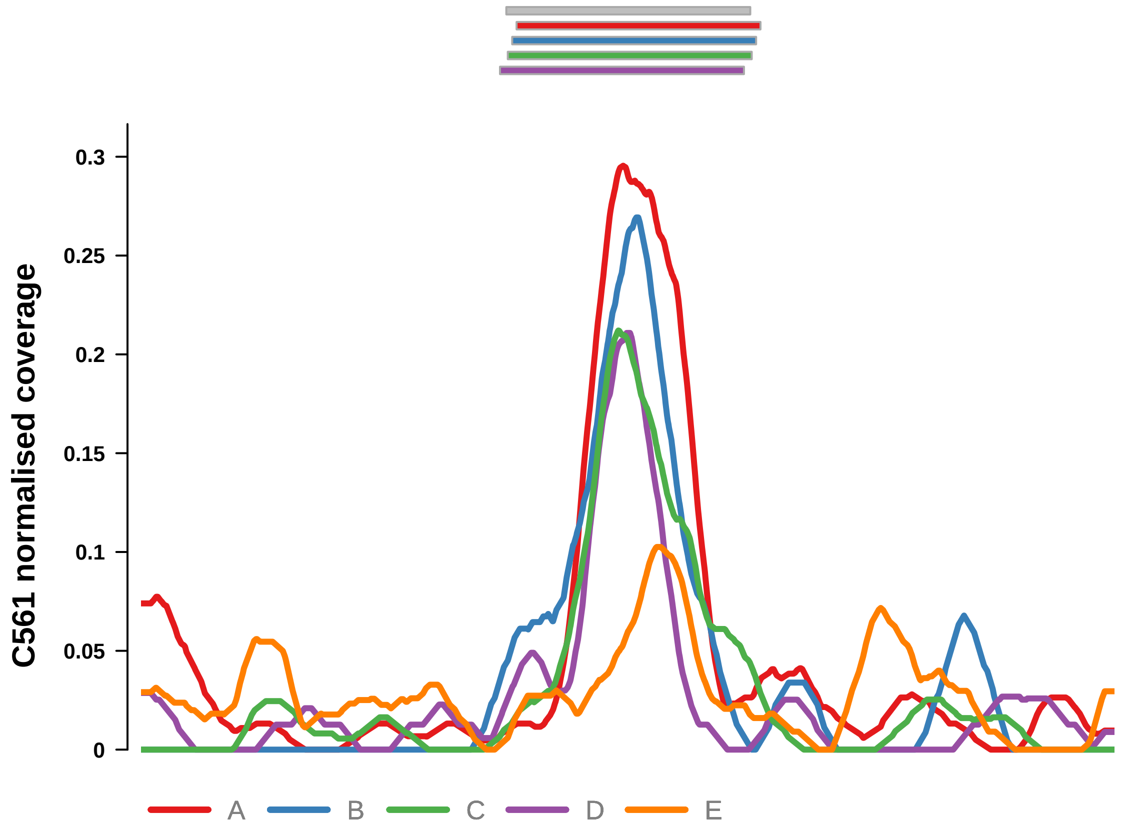


Putatively Clonal (logFC = 2.22)

| | P. | Adjusted P. |
|---|---|---|
| *~region vs. ~* | 8.56e−01 | 9.14e−01 |
| *~purity vs ~* | 1.63e−01 | 5.16e−01 |
| *~purity+region vs ~purity* | 7.64e−01 | 9.16e−01 |
| *Carcinoma vs. normal* | 5.74e−05 **** | 5.74e−05 **** |

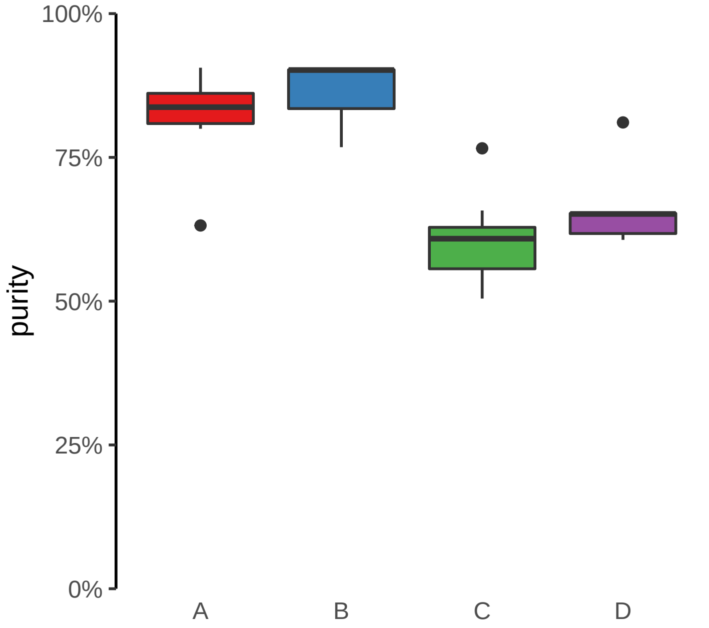

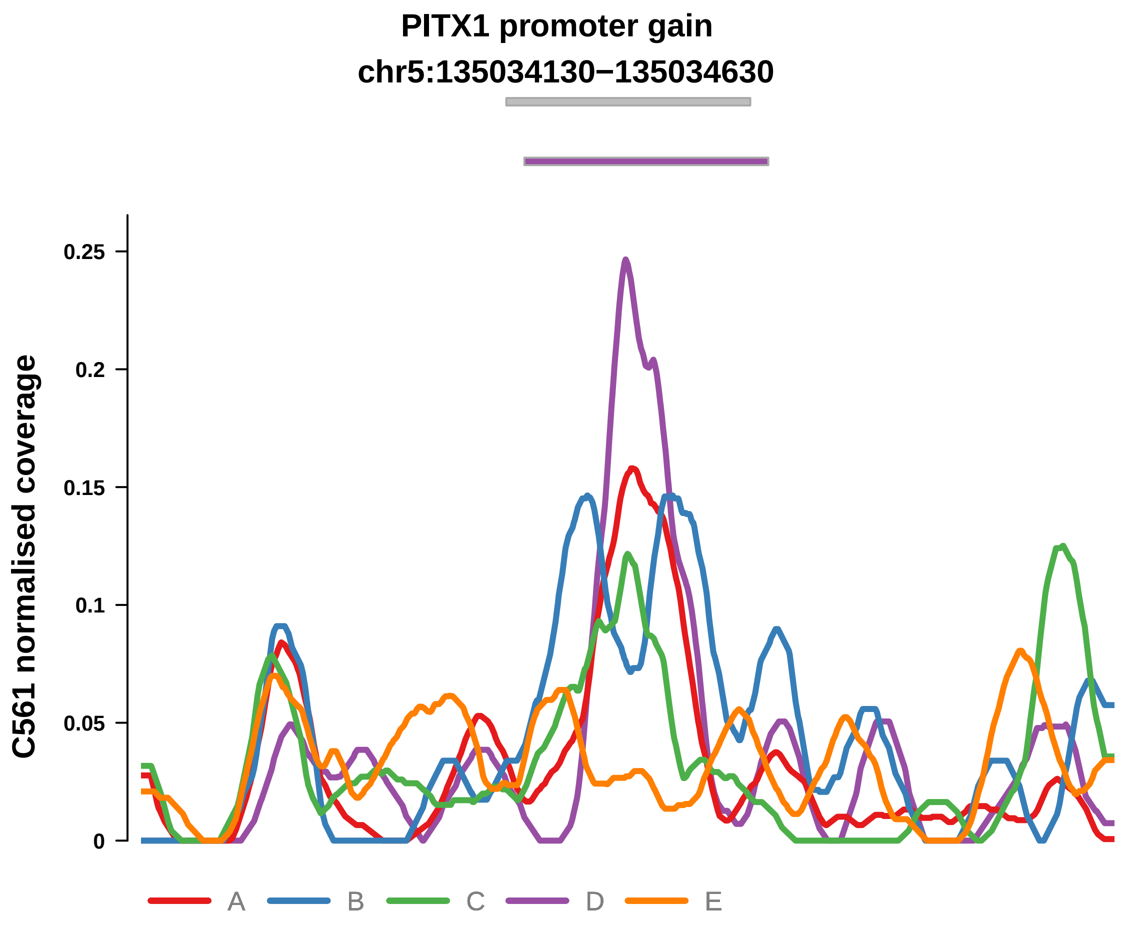


Putatively Clonal (logFC = 1.43)

| | P. | Adjusted P. |
|---|---|---|
| *~region vs. ~* | 4.12e−01 | 5.91e−01 |
| *~purity vs ~* | 6.3e−01 | 8.53e−01 |
| *~purity+region vs ~purity* | 3.75e−01 | 7.68e−01 |
| *Carcinoma vs. normal* | 8.78e−03 ** | 8.78e−03 ** |

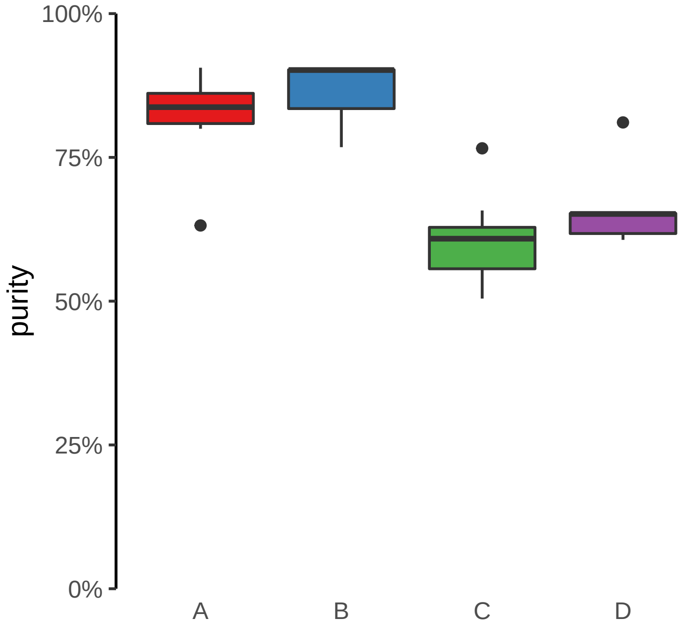

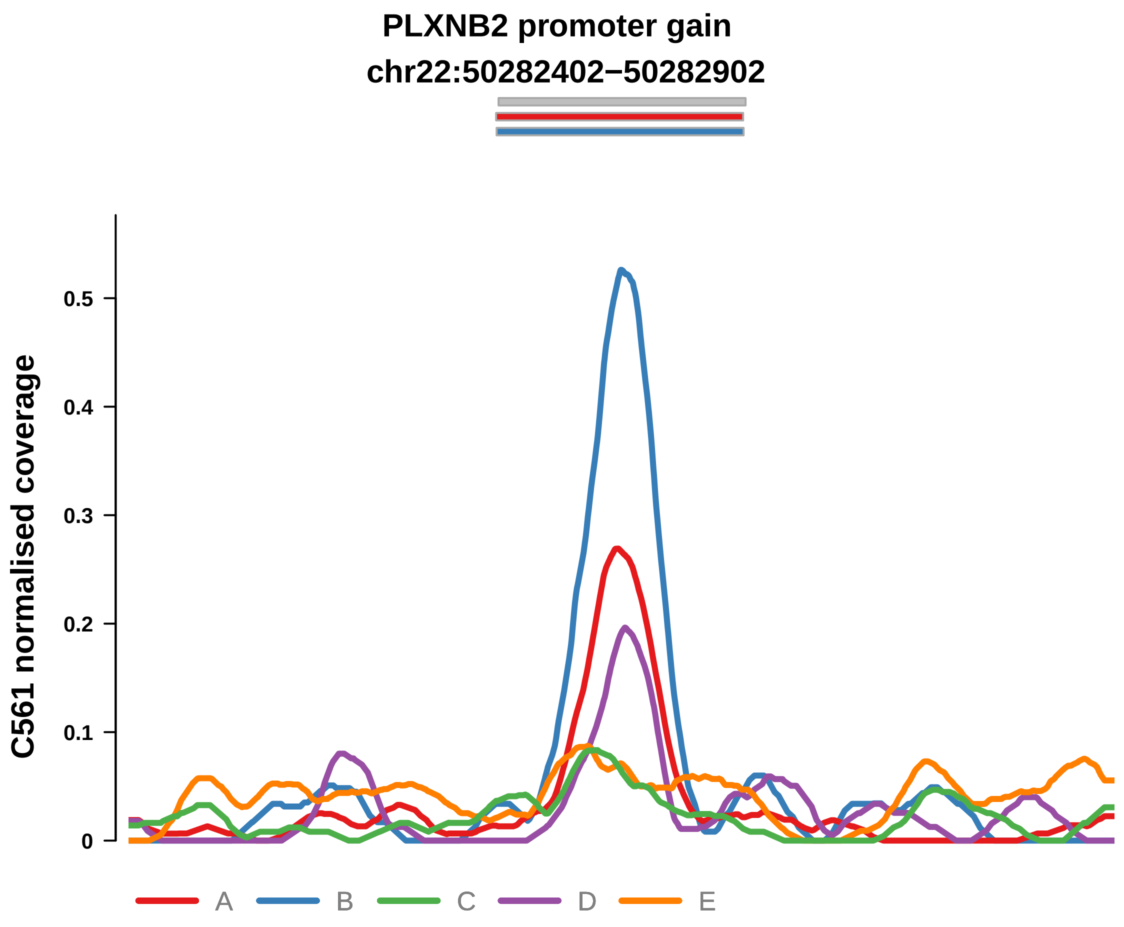


Putatively Clonal (logFC = 2.1)

| | P. | Adjusted P. |
|---|---|---|
| *~region vs. ~* | 4.29e−03 ** | 4.85e−02 * |
| *~purity vs ~* | 2.07e−03 ** | 4.55e−02 * |
| *~purity+region vs ~purity* | 3.15e−01 | 7.68e−01 |
| *Carcinoma vs. normal* | 1.55e−04 *** | 1.55e−04 *** |

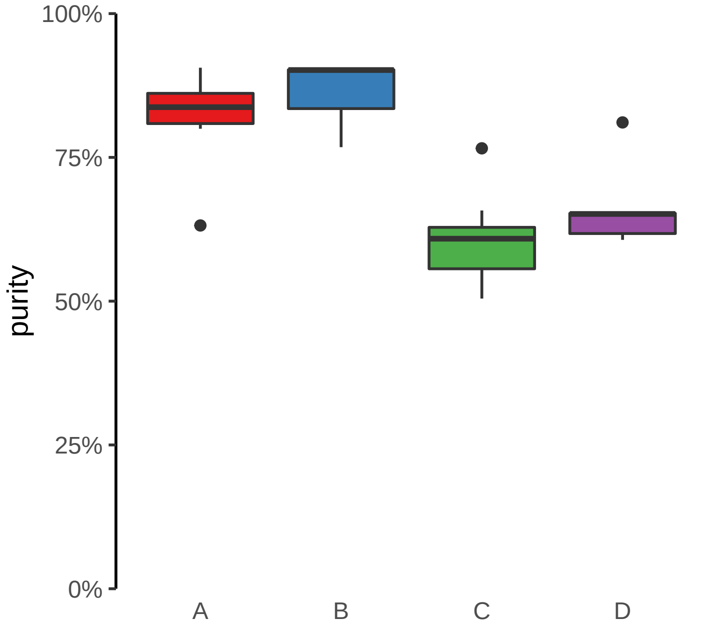

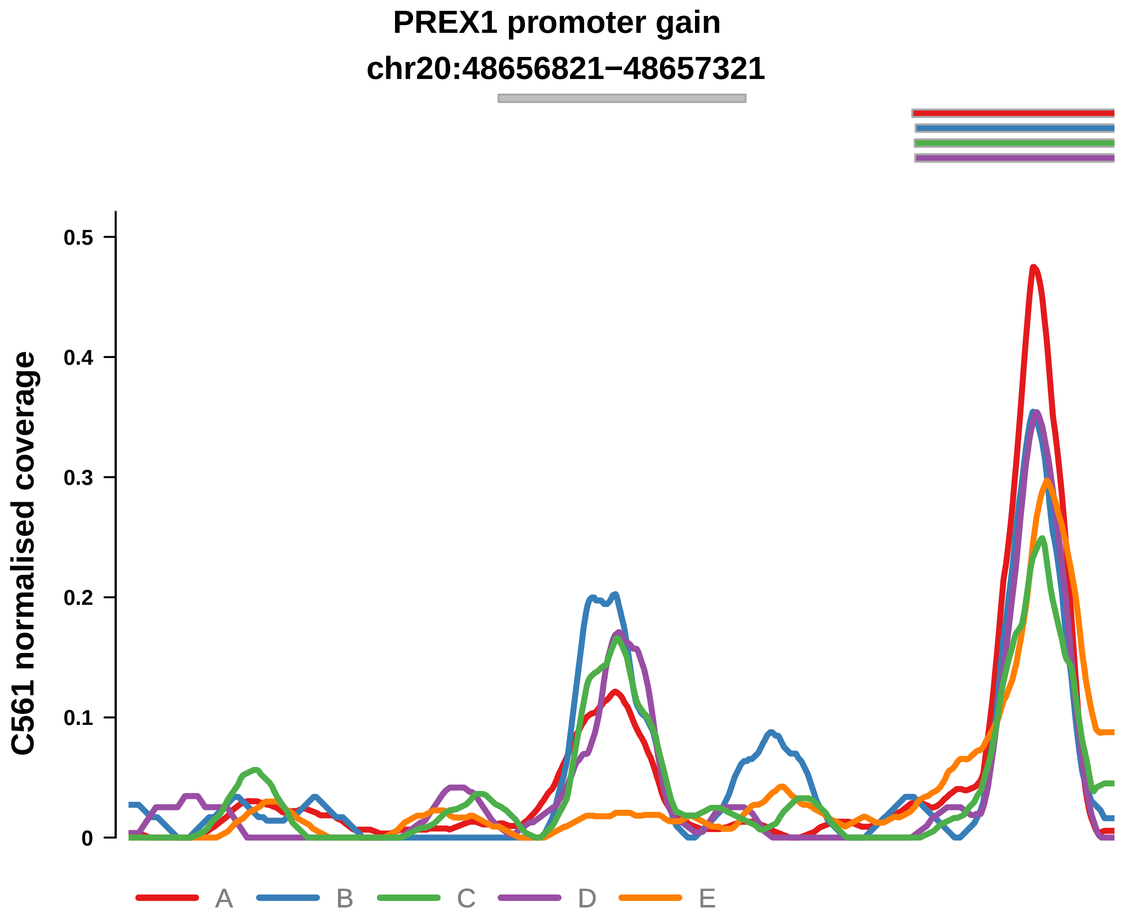


Putatively Clonal (logFC = 2.17)

| | P. | Adjusted P. |
|---|---|---|
| *~region vs. ~* | 4.32e−02 * | 1.77e−01 |
| *~purity vs ~* | 3.25e−01 | 7.66e−01 |
| *~purity+region vs ~purity* | 7.71e−02 | 4.24e−01 |
| *Carcinoma vs. normal* | 1.27e−04 *** | 1.27e−04 *** |

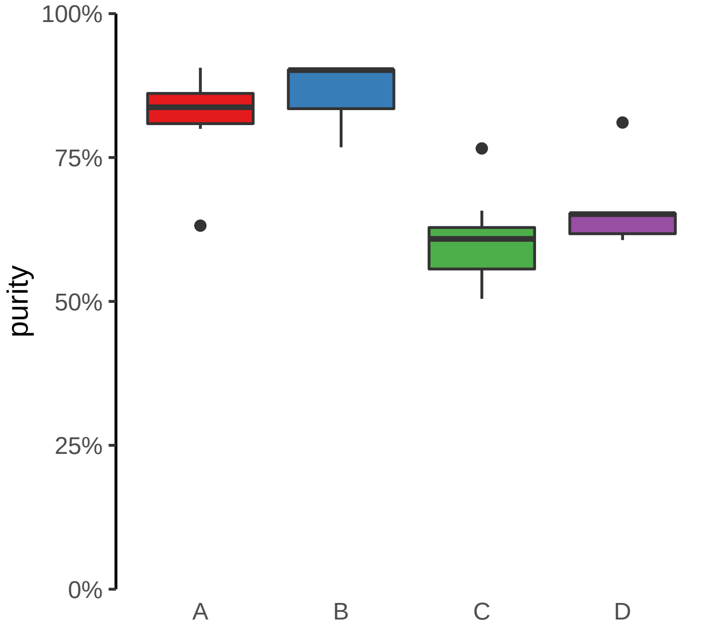

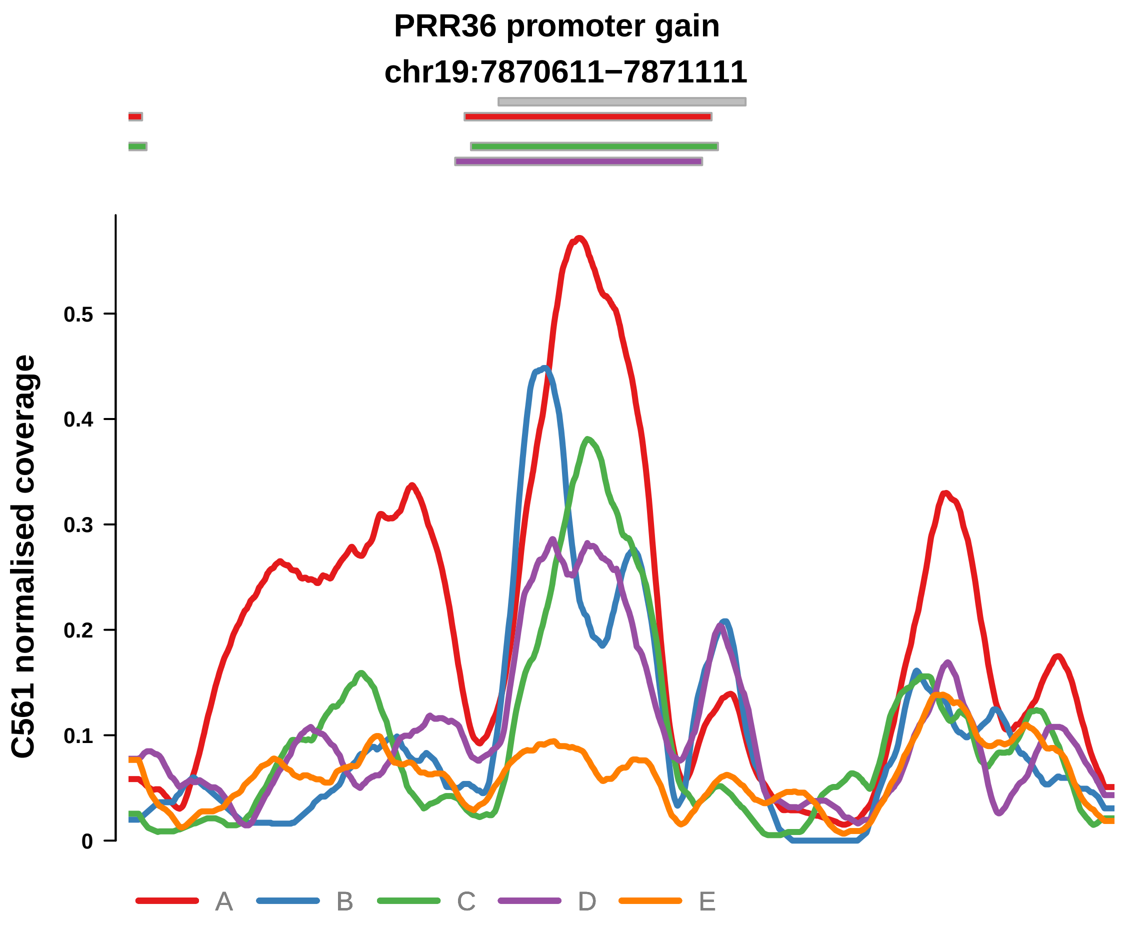


Putatively Clonal (logFC = 3.09)

| | P. | Adjusted P. |
|---|---|---|
| *~region vs. ~* | 1.77e−01 | 3.88e−01 |
| *~purity vs ~* | 8.7e−01 | 9.45e−01 |
| *~purity+region vs ~purity* | 1.47e−01 | 5.88e−01 |
| *Carcinoma vs. normal* | 4.65e−08 **** | 4.65e−08 **** |

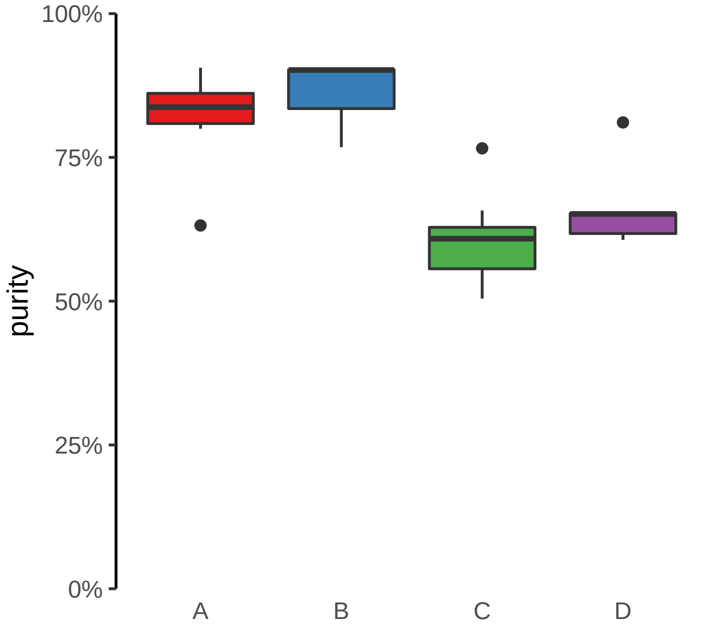

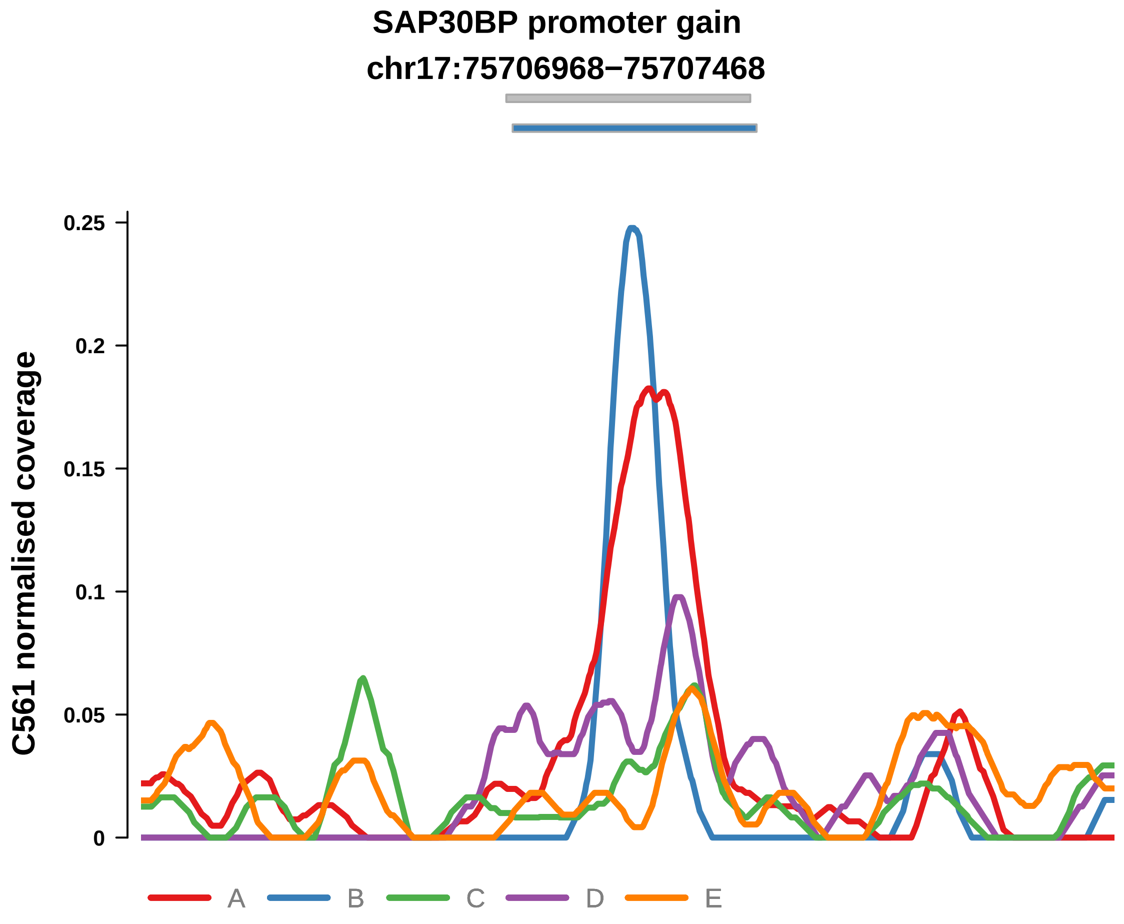


Putatively Clonal (logFC = 2.1)

| | P. | Adjusted P. |
|---|---|---|
| *~region vs. ~* | 5.26e−02 | 1.81e−01 |
| *~purity vs ~* | 6.1e−01 | 8.53e−01 |
| *~purity+region vs ~purity* | 2.48e−02 * | 2.73e−01 |
| *Carcinoma vs. normal* | 1.95e−04 *** | 1.95e−04 *** |

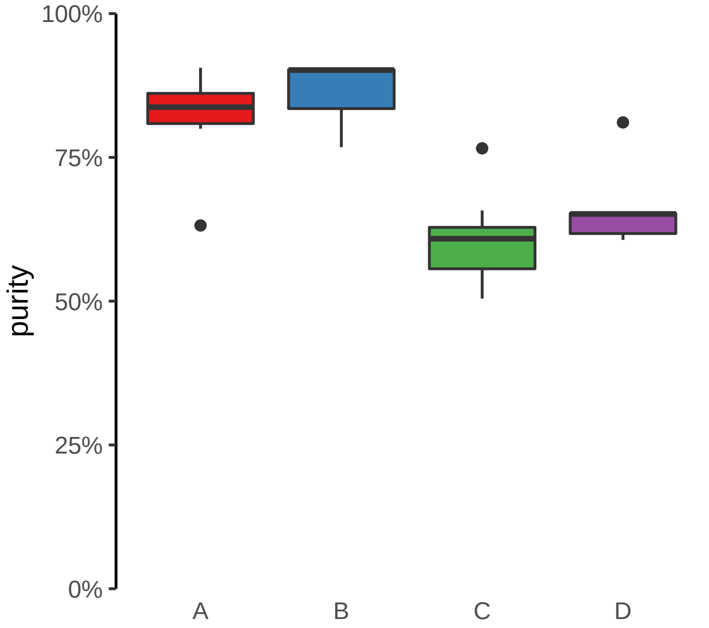

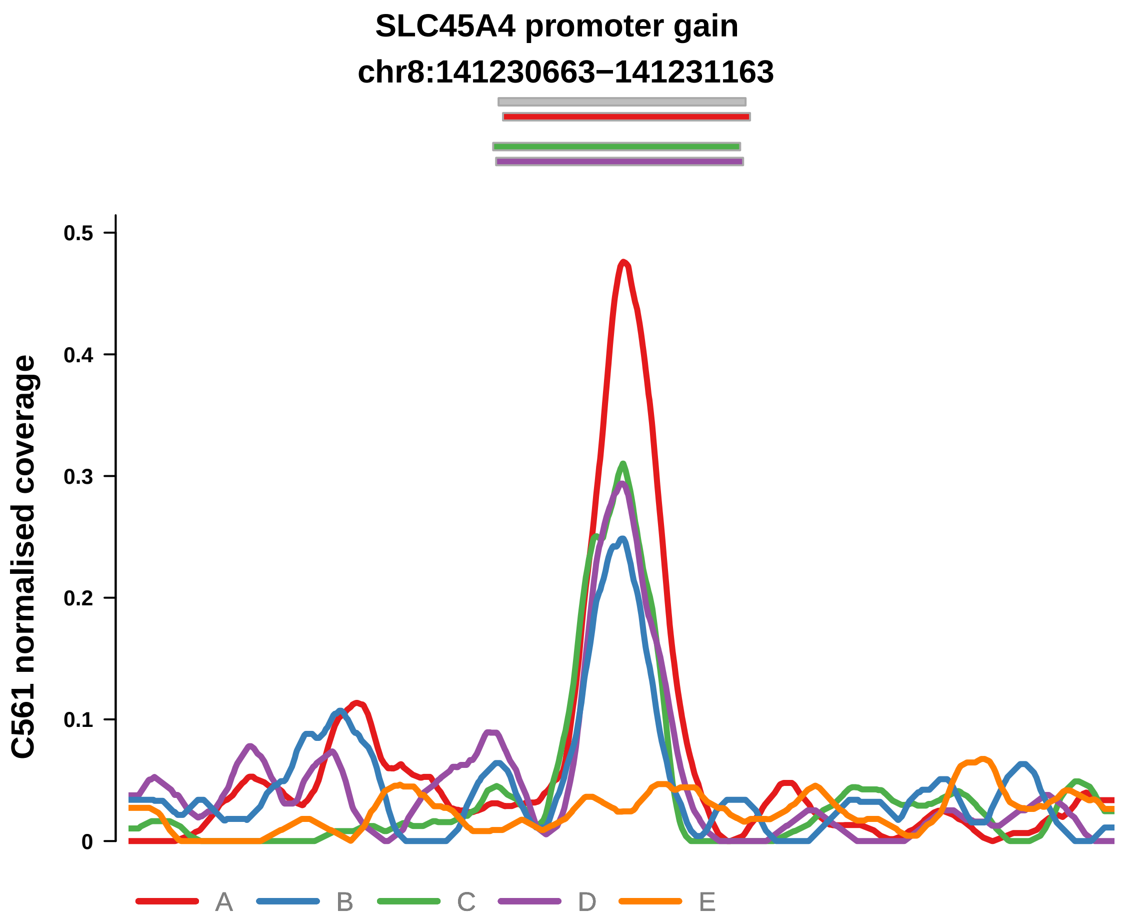


Putatively Clonal (logFC = 3.14)

| | P. | Adjusted P. |
|---|---|---|
| *~region vs. ~* | 5.8e−01 | 7.05e−01 |
| *~purity vs ~* | 2.22e−01 | 6.1e−01 |
| *~purity+region vs ~purity* | 3.33e−01 | 7.68e−01 |
| *Carcinoma vs. normal* | 4.49e−08 **** | 4.49e−08 **** |

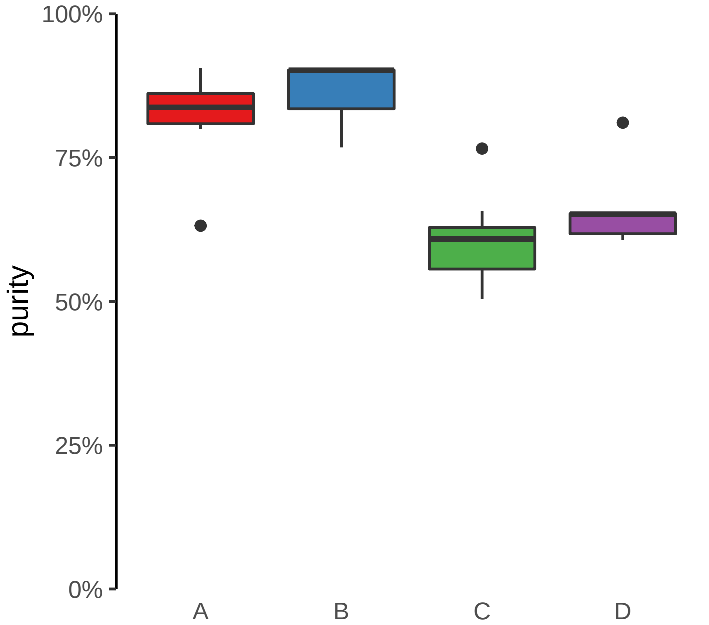

**TG promoter gain**
**chr8:133130728−133131228**

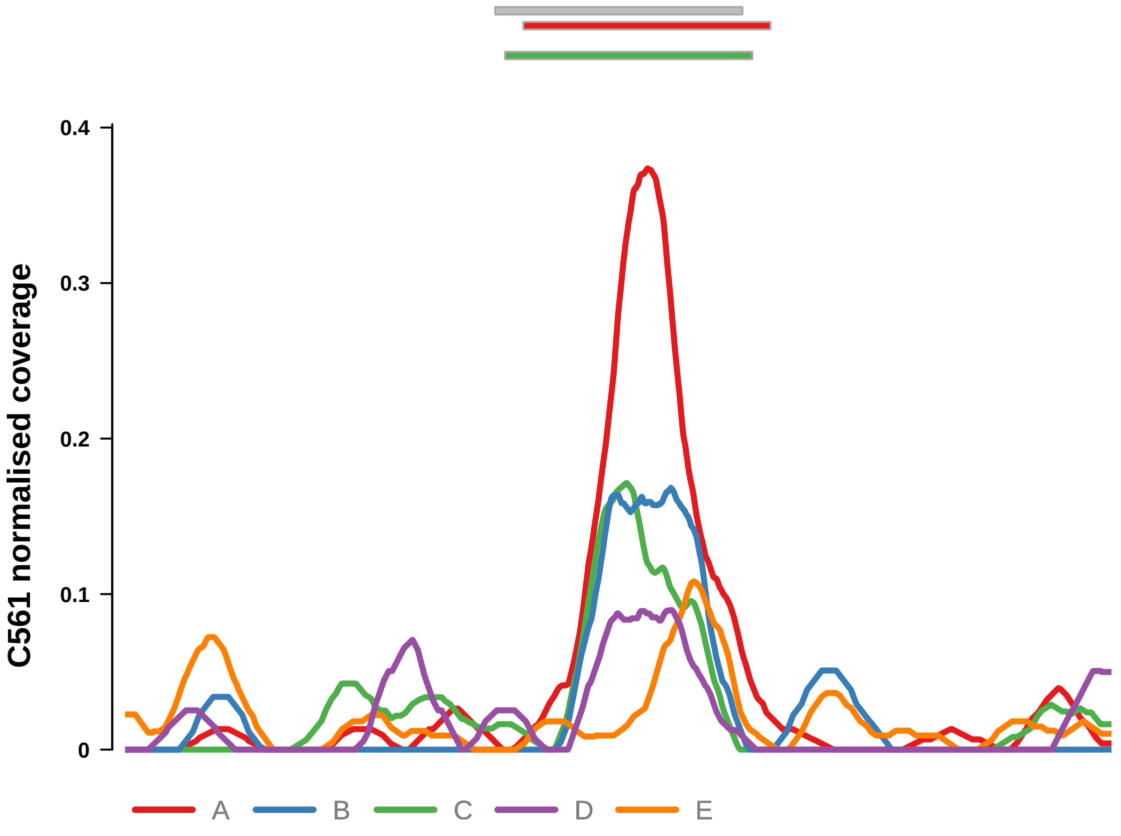


Putatively Clonal (logFC = 2.21)

| | P. | Adjusted P. |
|---|---|---|
| *~region vs. ~* | 4.19e−01 | 5.91e−01 |
| *~purity vs ~* | 2.93e−01 | 7.53e−01 |
| *~purity+region vs ~purity* | 6.88e−01 | 9.16e−01 |
| *Carcinoma vs. normal* | 6.69e−05 **** | 6.69e−05 **** |

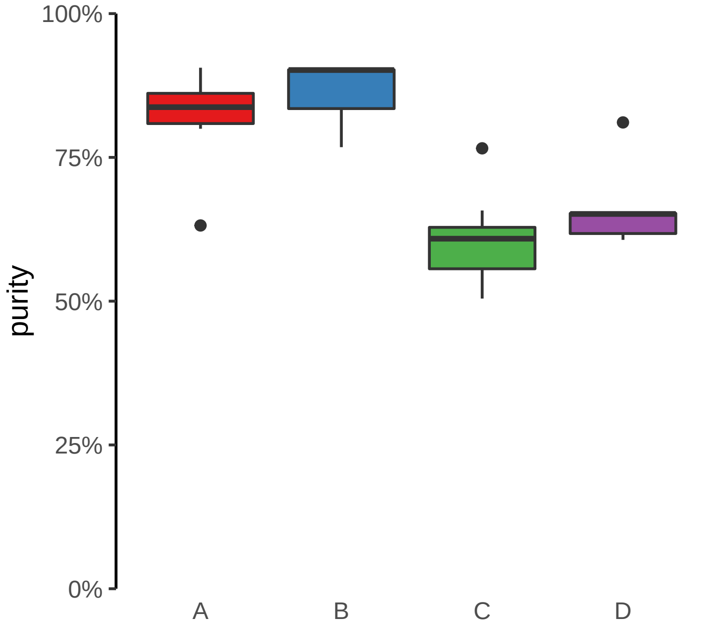

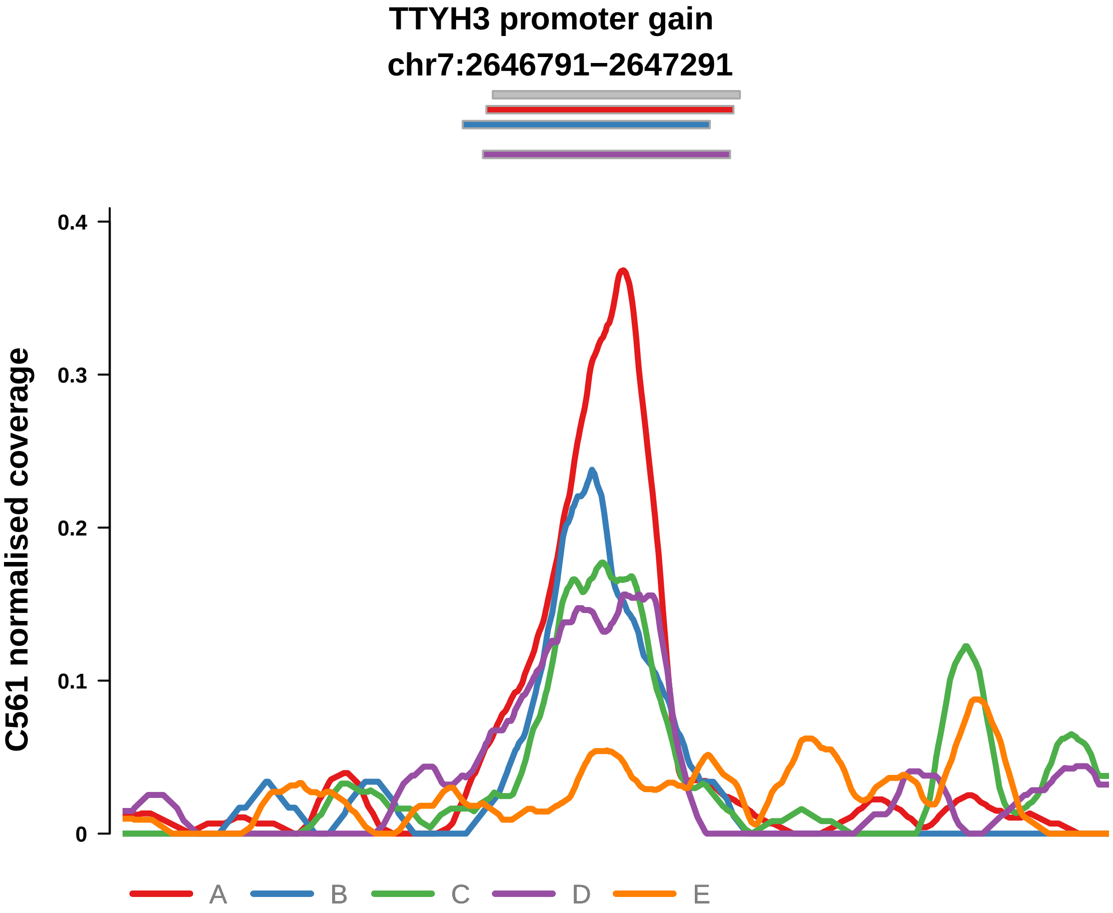


Putatively Clonal (logFC = 3.09)

| | P. | Adjusted P. |
|---|---|---|
| *~region vs. ~* | 5.77e−01 | 7.05e−01 |
| *~purity vs ~* | 8.33e−01 | 9.37e−01 |
| *~purity+region vs ~purity* | 4.13e−01 | 7.74e−01 |
| *Carcinoma vs. normal* | 6.39e−08 **** | 6.39e−08 **** |

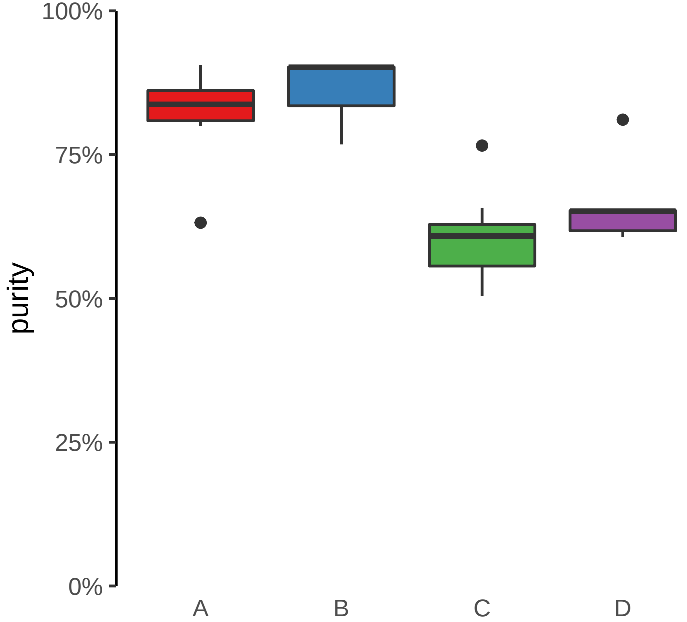

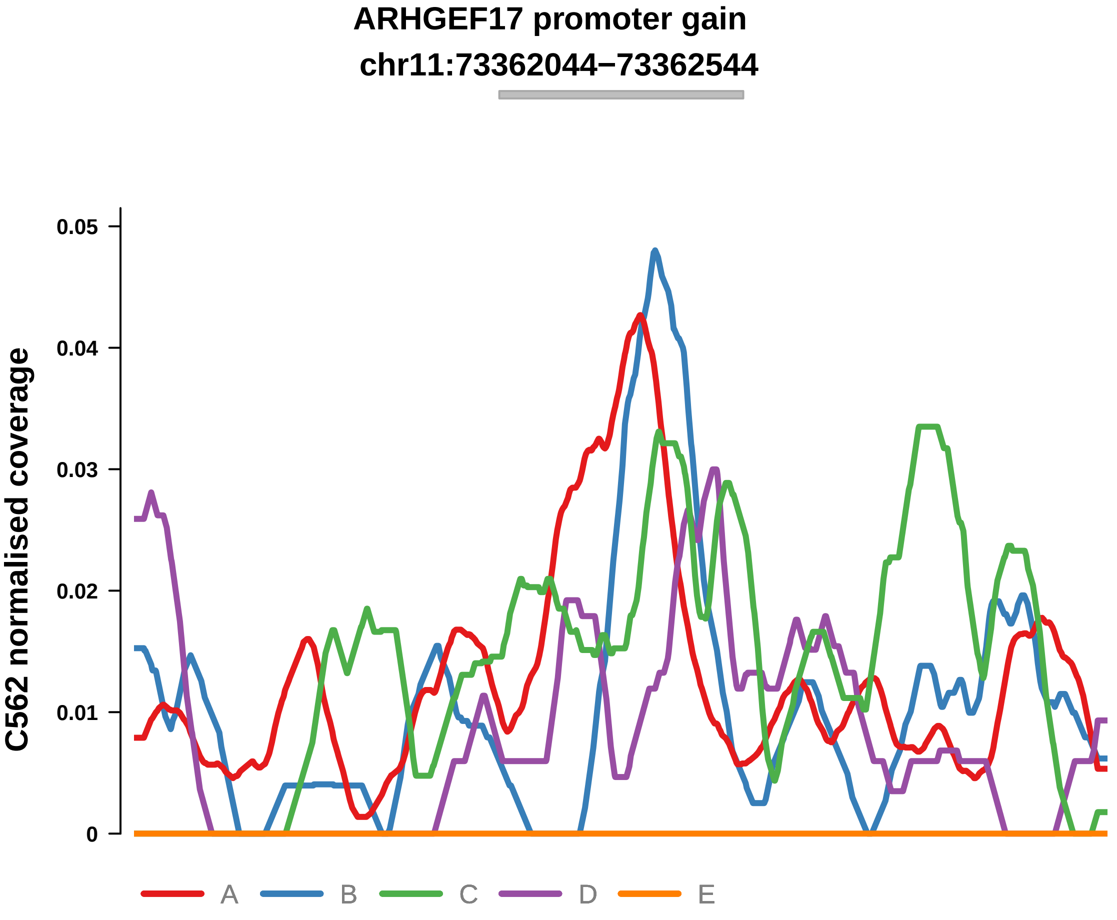


Putatively Clonal (logFC = 1.91)

| | P. | Adjusted P. |
|---|---|---|
| *~region vs. ~* | 2.79e−01 | 6.13e−01 |
| *~purity vs ~* | 1.63e−02 * | 7.56e−02 |
| *~purity+region vs ~purity* | 8.56e−01 | 9.41e−01 |
| *Carcinoma vs. normal* | 1.3e−03 ** | 1.3e−03 ** |

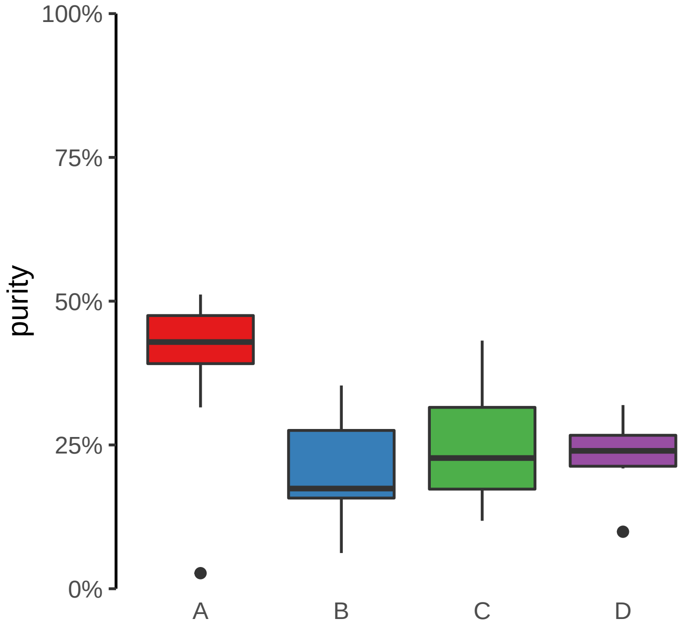

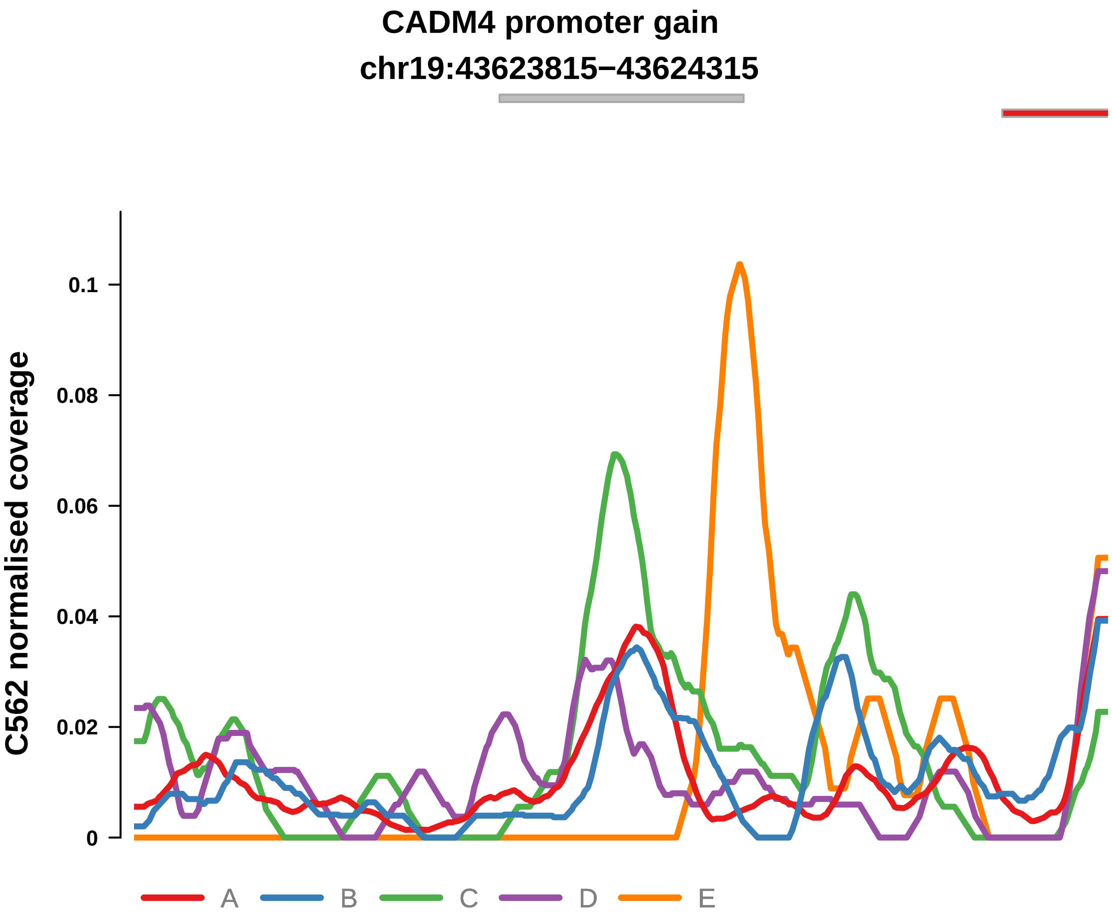


Putatively Clonal (logFC = 2.12)

| | P. | Adjusted P. |
|---|---|---|
| *~region vs. ~* | 5.89e−01 | 7.87e−01 |
| *~purity vs ~* | 7.45e−03 ** | 4.37e−02 * |
| *~purity+region vs ~purity* | 3.68e−01 | 7.81e−01 |
| *Carcinoma vs. normal* | 4.35e−04 *** | 4.35e−04 *** |

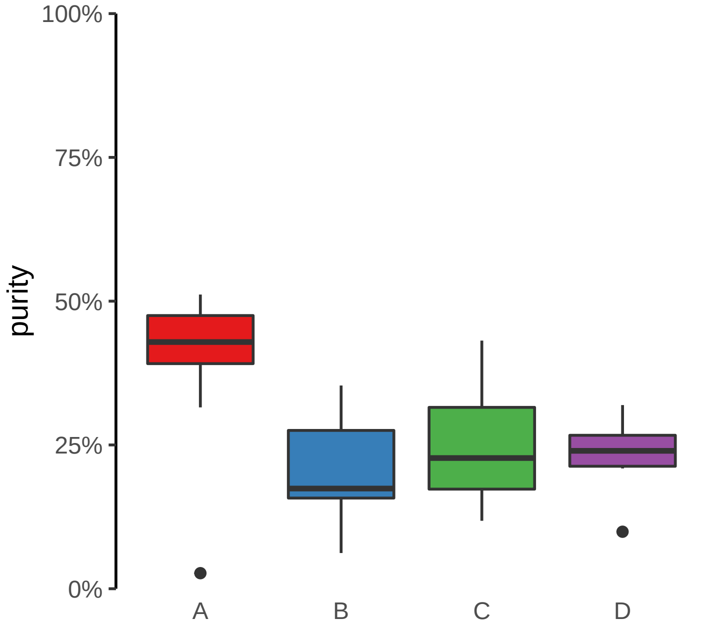

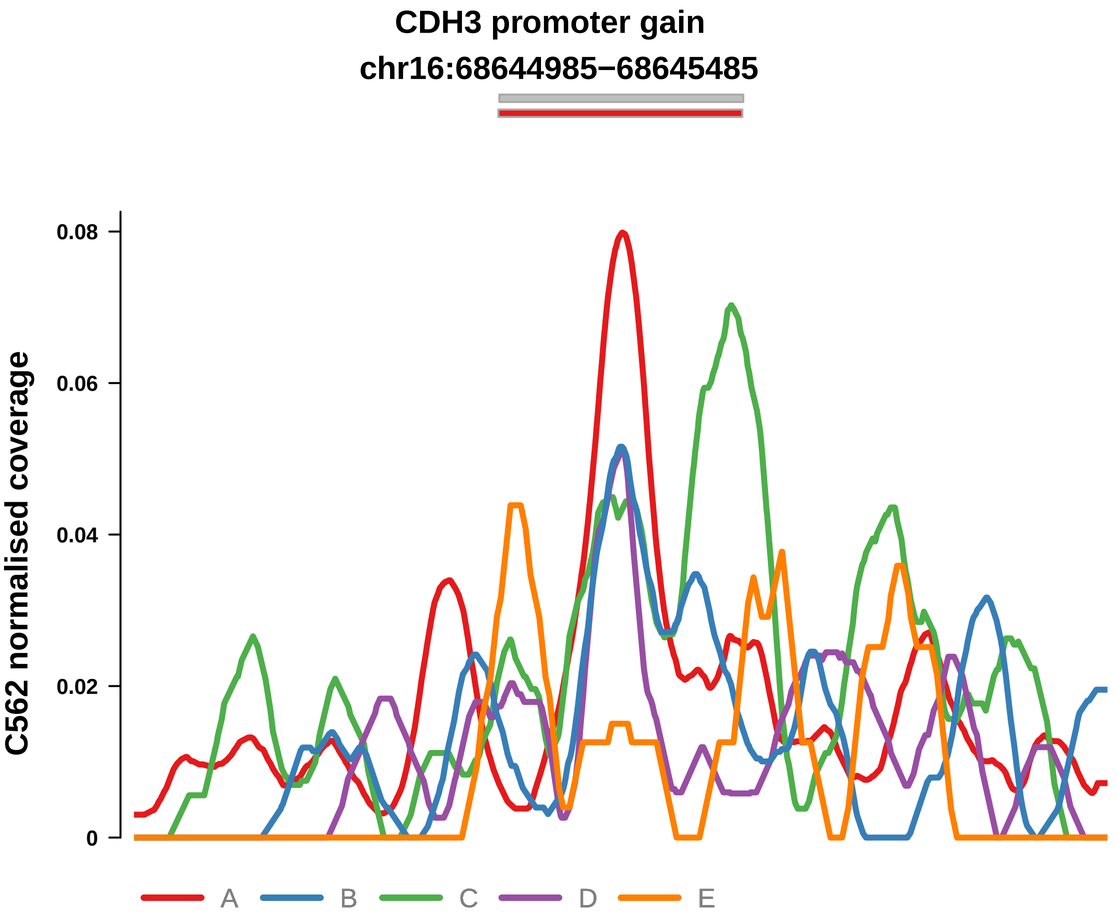


Putatively Clonal (logFC = 1.56)

| | P. | Adjusted P. |
|---|---|---|
| *~region vs. ~* | 1.43e−01 | 5.03e−01 |
| *~purity vs ~* | 1.13e−03 ** | 1.42e−02 * |
| *~purity+region vs ~purity* | 9.73e−01 | 9.85e−01 |
| *Carcinoma vs. normal* | 5.64e−03 ** | 5.64e−03 ** |

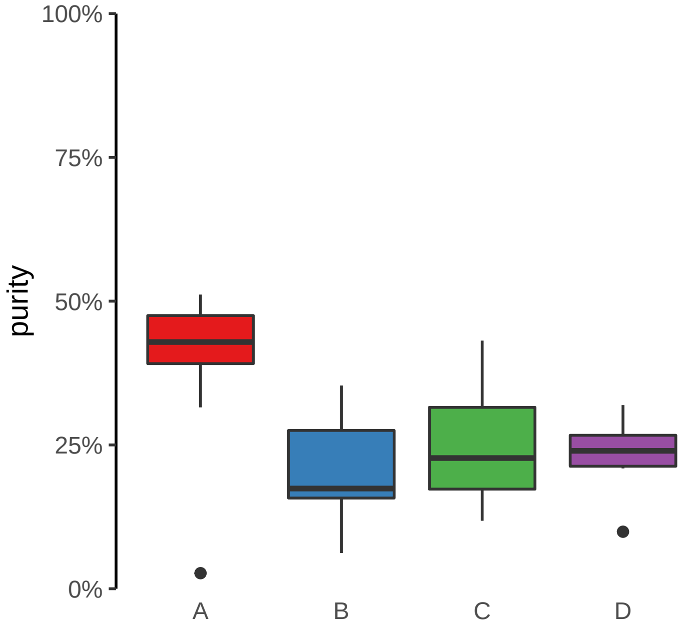

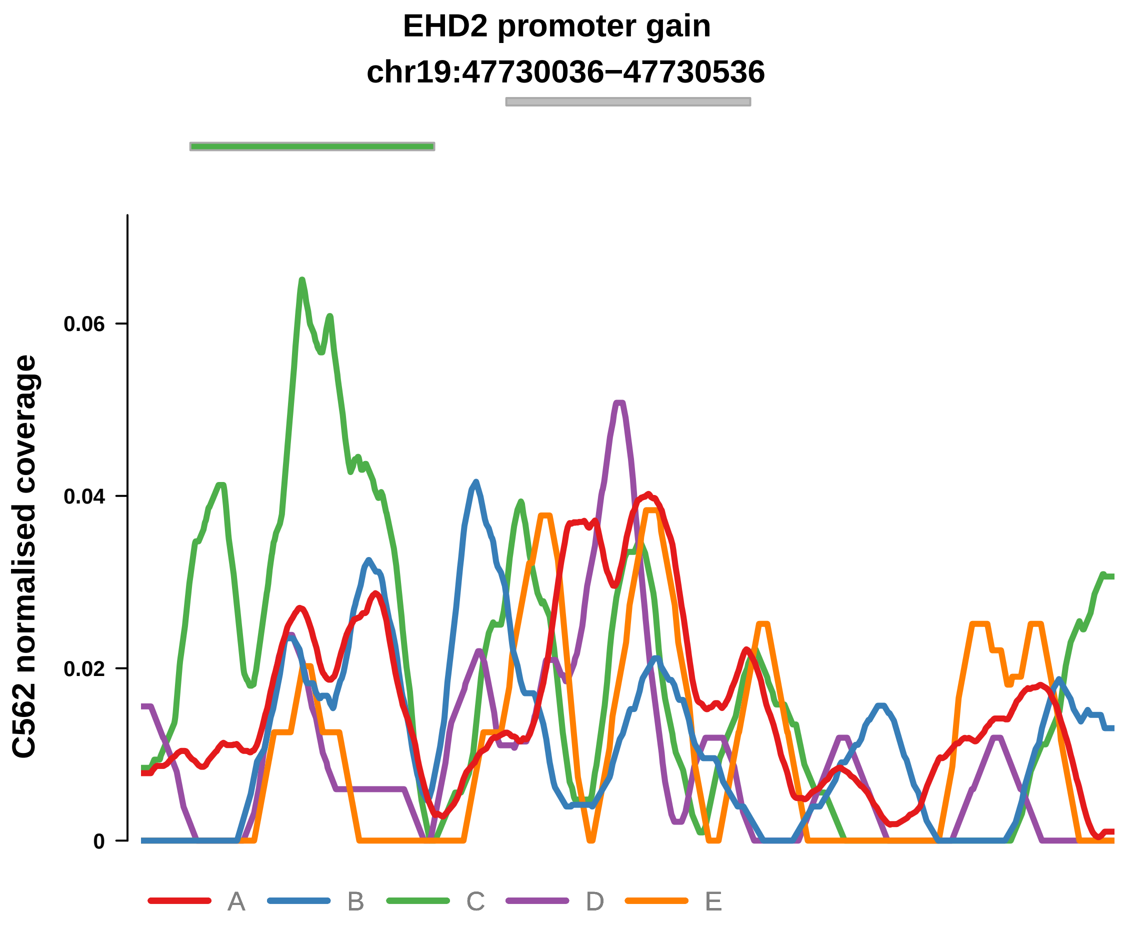


Putatively Clonal (logFC = 1.59)

| | P. | Adjusted P. |
|---|---|---|
| *~region vs. ~* | 9.57e−02 | 4.01e−01 |
| *~purity vs ~* | 5.37e−03 ** | 3.63e−02 * |
| *~purity+region vs ~purity* | 8.48e−01 | 9.41e−01 |
| *Carcinoma vs. normal* | 5.68e−03 ** | 5.68e−03 ** |

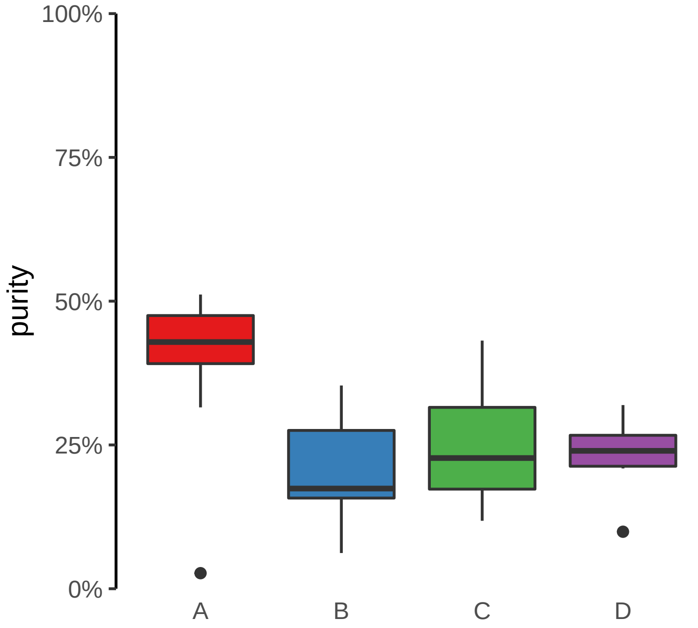

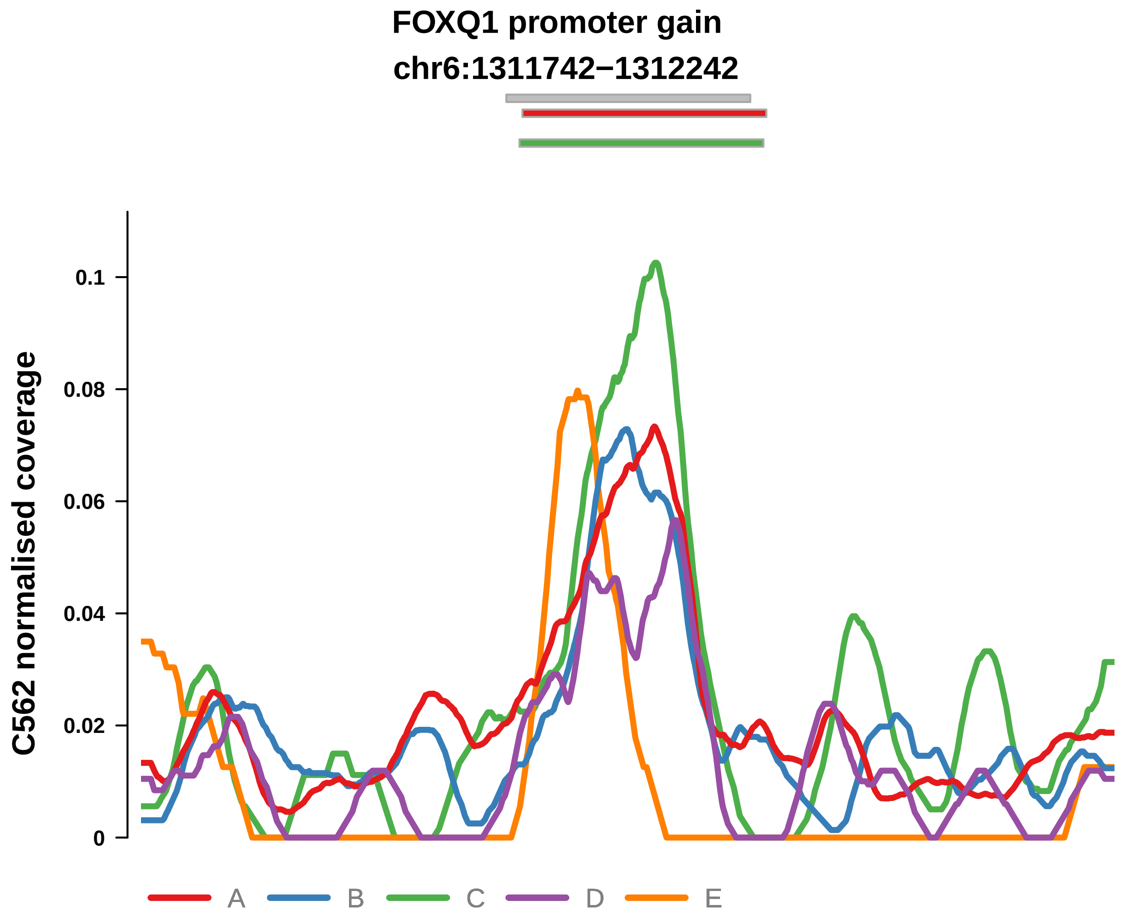


Putatively Clonal (logFC = 2.16)

| | P. | Adjusted P. |
|---|---|---|
| *~region vs. ~* | 7.38e−02 | 3.54e−01 |
| *~purity vs ~* | 1.13e−03 ** | 1.42e−02 * |
| *~purity+region vs ~purity* | 7.89e−01 | 9.41e−01 |
| *Carcinoma vs. normal* | 1.18e−04 *** | 1.18e−04 *** |

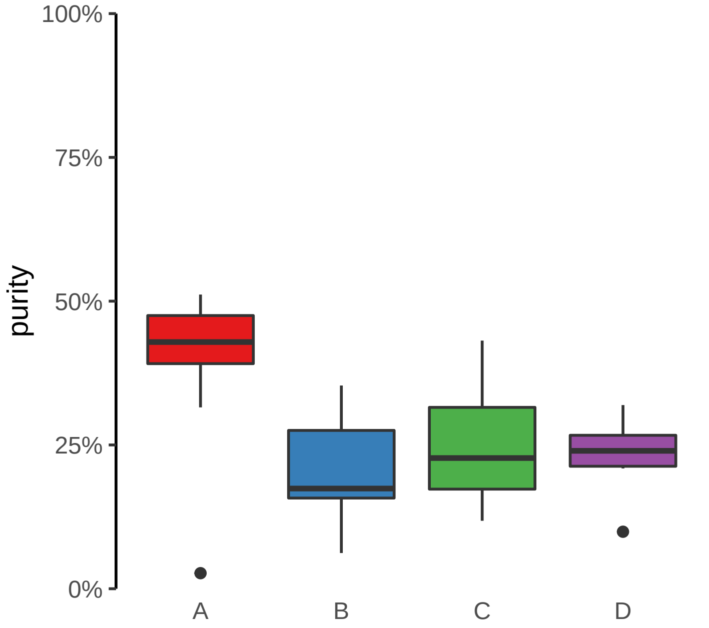

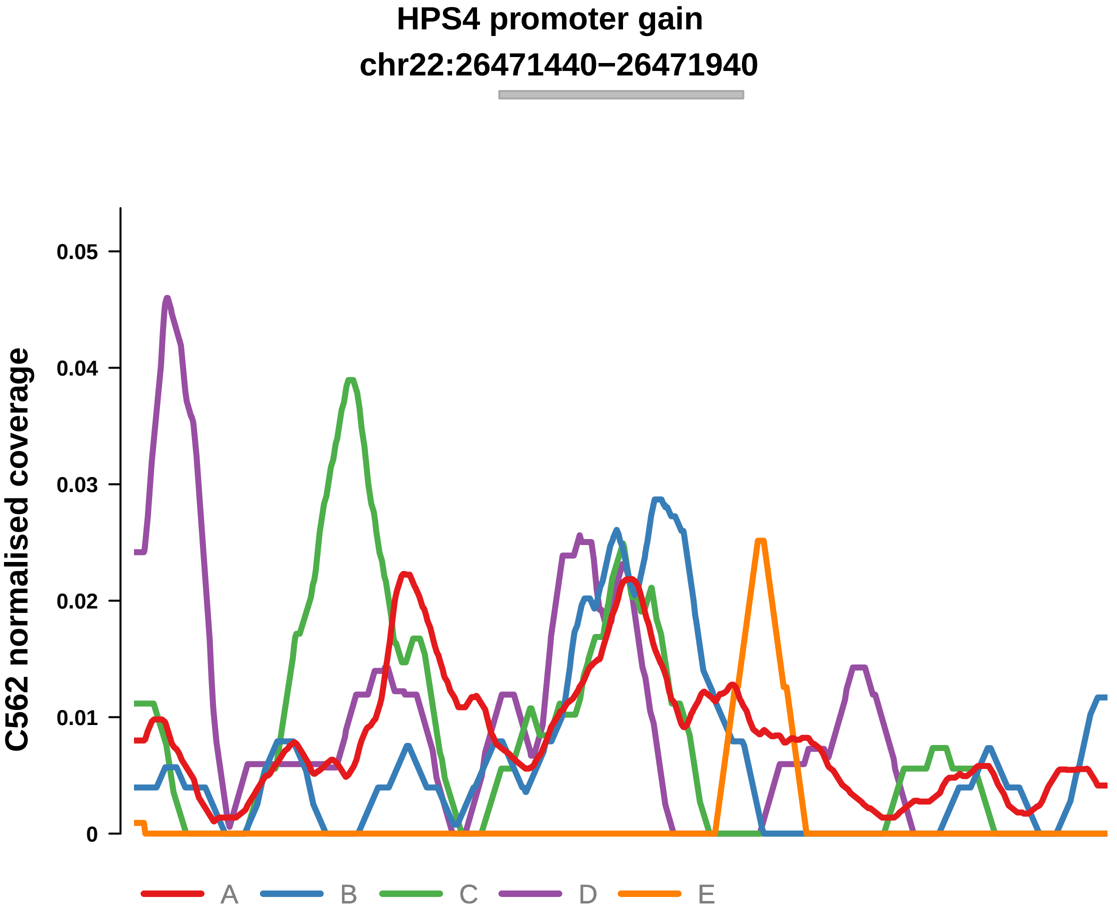


Putatively Clonal (logFC = 2.19)

| | P. | Adjusted P. |
|---|---|---|
| *~region vs. ~* | 2.93e−01 | 6.13e−01 |
| *~purity vs ~* | 2.62e−02 * | 1.05e−01 |
| *~purity+region vs ~purity* | 5.74e−01 | 9.36e−01 |
| *Carcinoma vs. normal* | 2.81e−04 *** | 2.81e−04 *** |

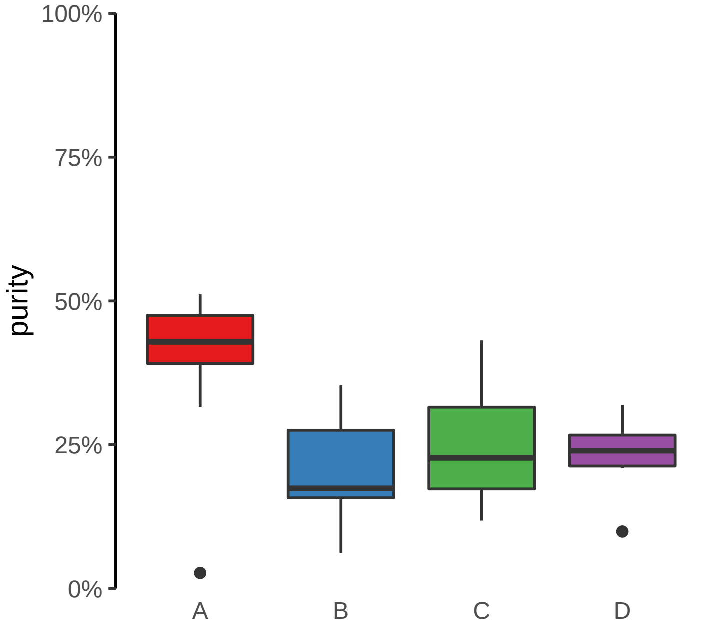

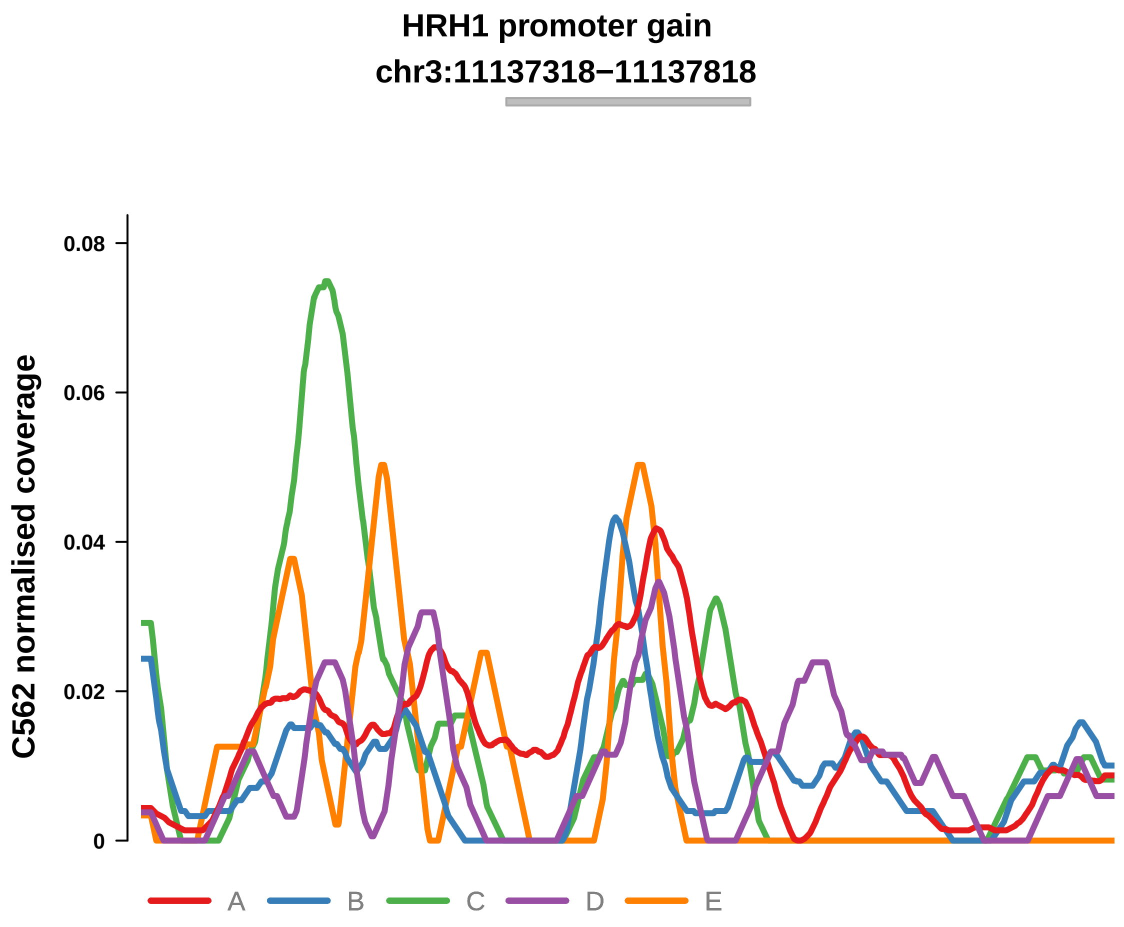


Putatively Clonal (logFC = 1.9)

| | P. | Adjusted P. |
|---|---|---|
| *~region vs. ~* | 9.93e−04 *** | 2.91e−02 * |
| *~purity vs ~* | 4.58e−04 *** | 8.05e−03 ** |
| *~purity+region vs ~purity* | 1.15e−01 | 5.08e−01 |
| *Carcinoma vs. normal* | 1.24e−03 ** | 1.24e−03 ** |

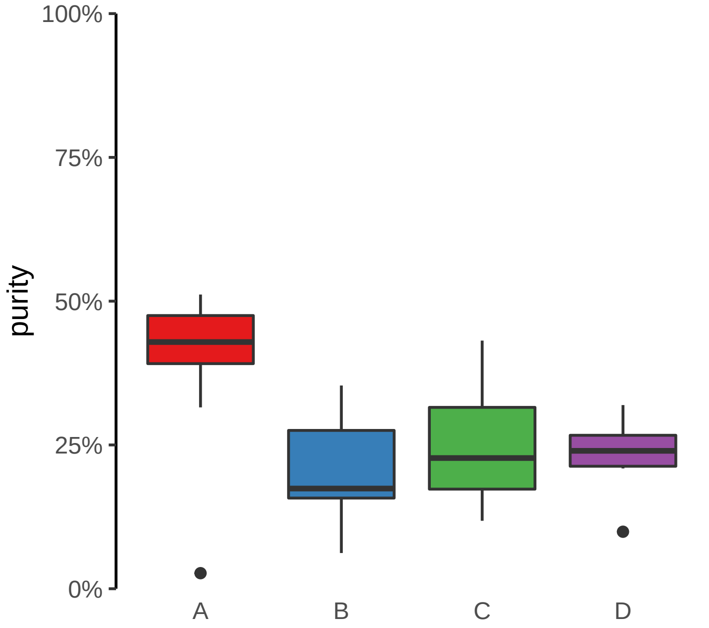

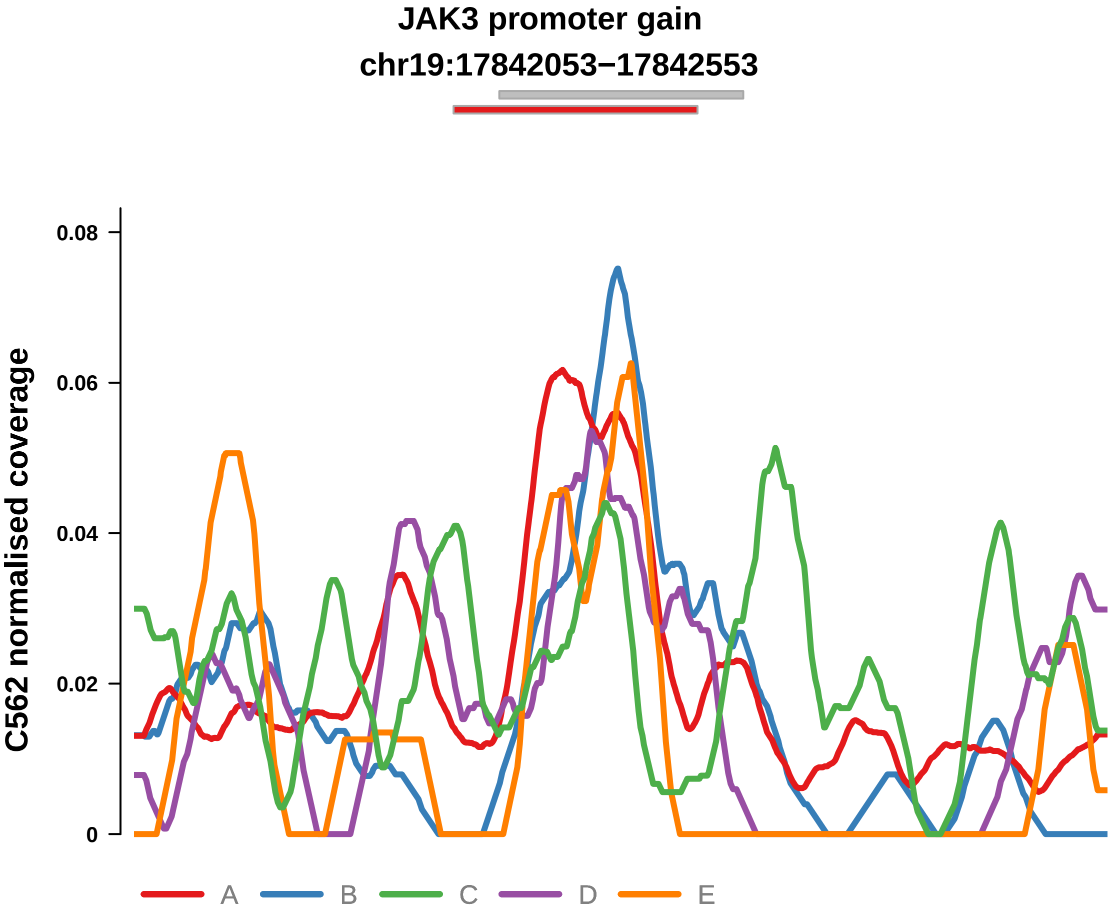


Putatively Clonal (logFC = 1.96)

| | P. | Adjusted P. |
|---|---|---|
| *~region vs. ~* | 2.24e−03 ** | 3.29e−02 * |
| *~purity vs ~* | 9.1e−03 ** | 4.71e−02 * |
| *~purity+region vs ~purity* | 3.07e−02 * | 3.16e−01 |
| *Carcinoma vs. normal* | 5.8e−04 *** | 5.8e−04 *** |

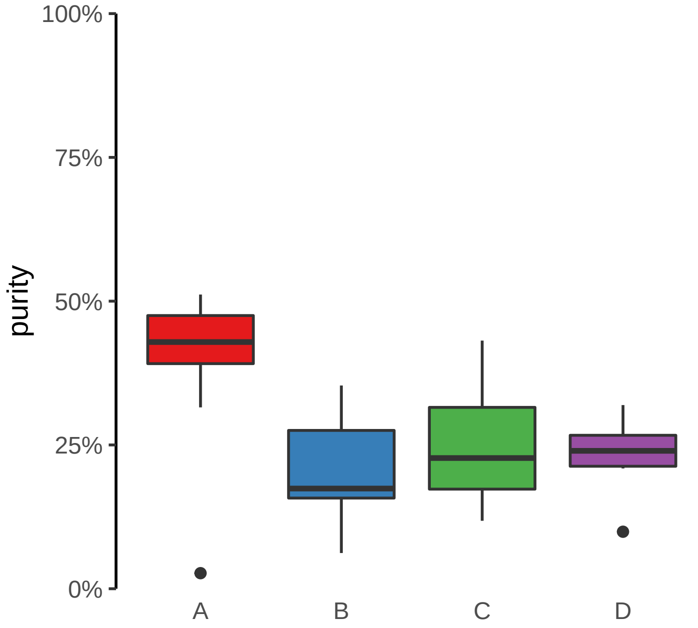

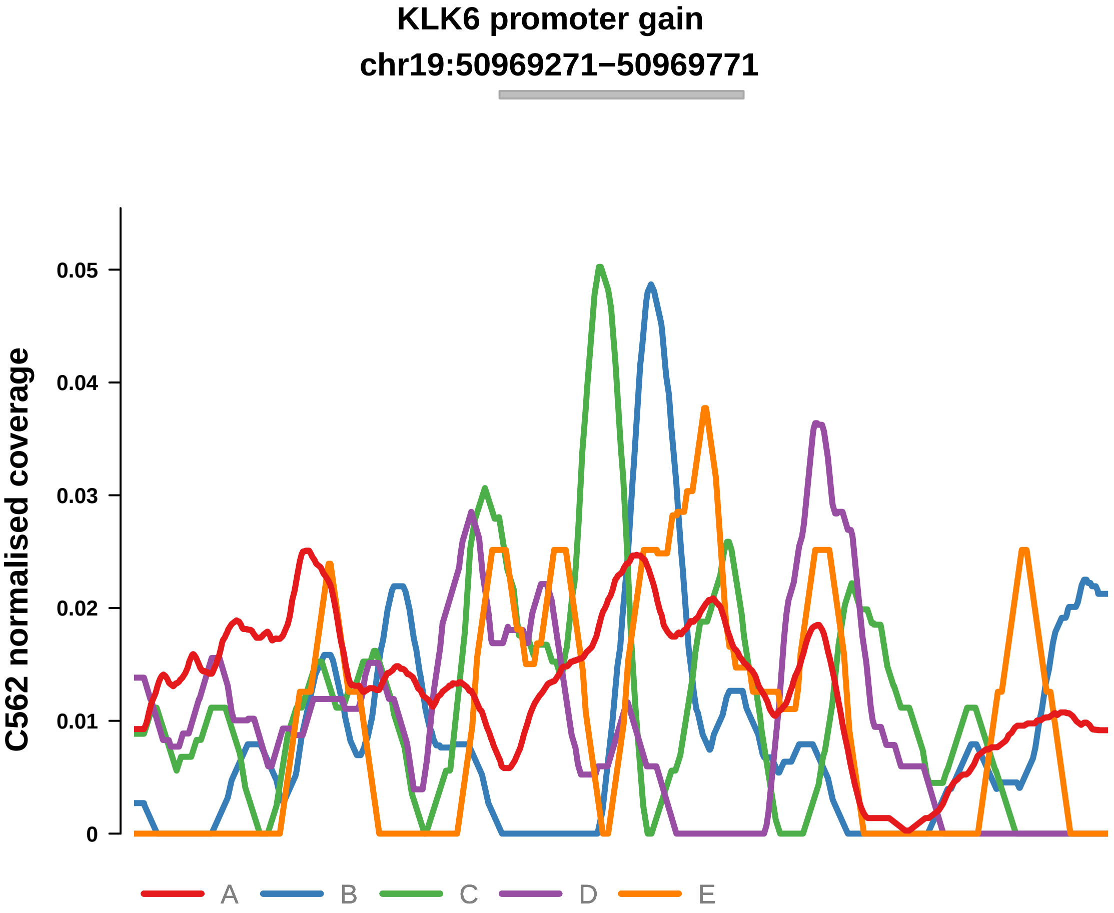


Putatively Clonal (logFC = 2)

| | P. | Adjusted P. |
|---|---|---|
| *~region vs. ~* | 3.21e−01 | 6.56e−01 |
| *~purity vs ~* | 5.43e−02 | 1.71e−01 |
| *~purity+region vs ~purity* | 3.01e−01 | 7.81e−01 |
| *Carcinoma vs. normal* | 7.01e−04 *** | 7.01e−04 *** |

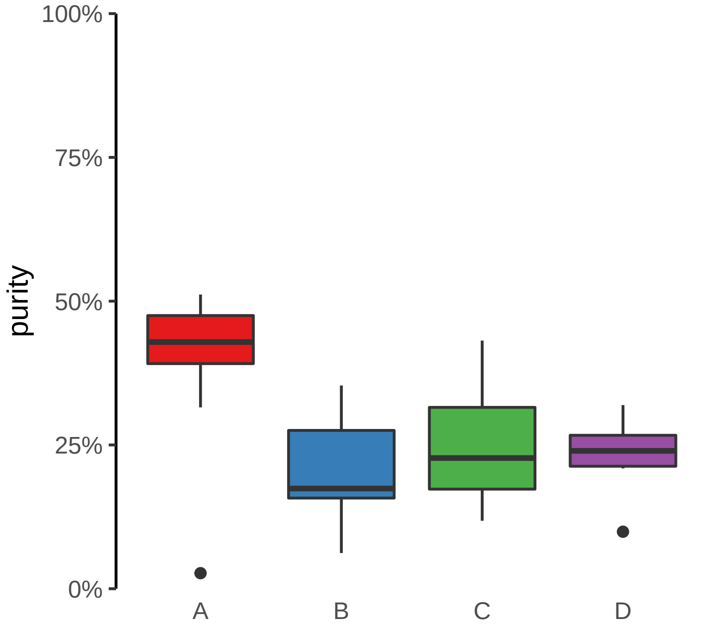

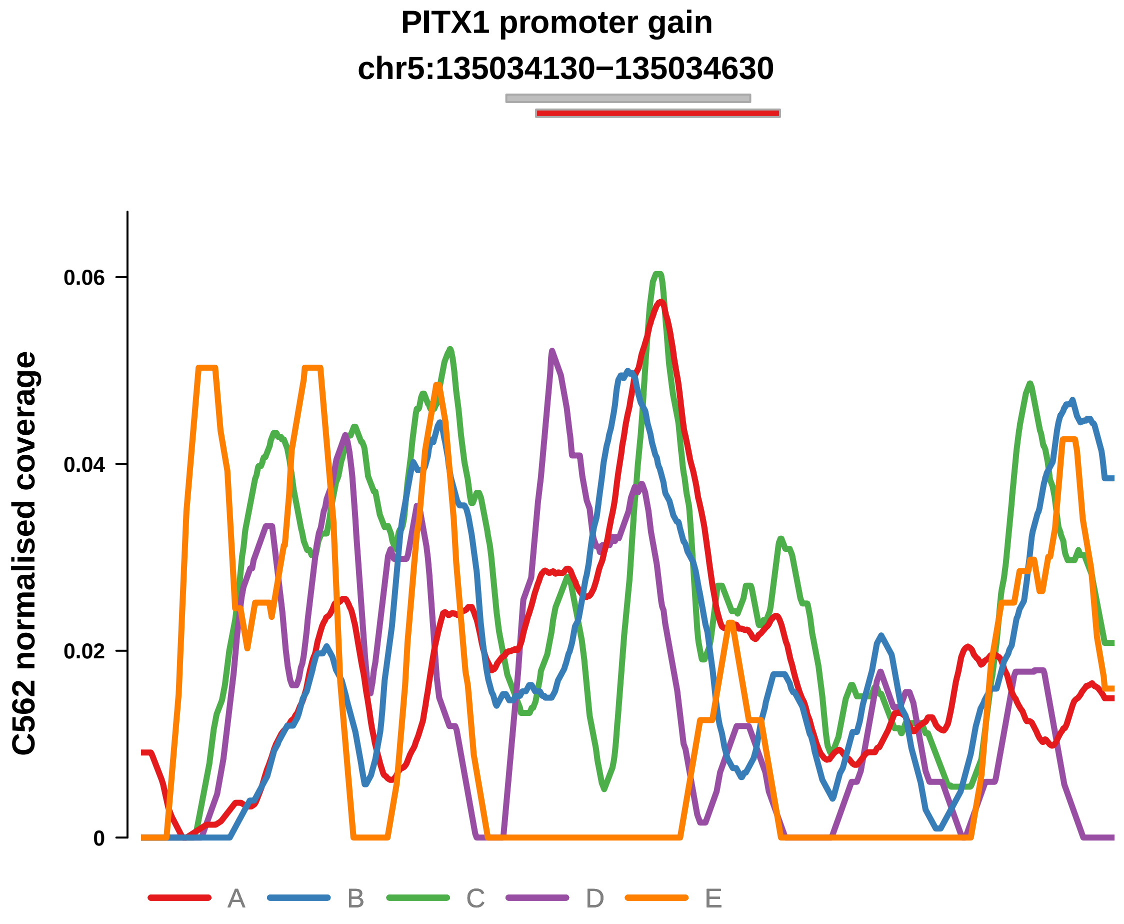


Putatively Clonal (logFC = 1.79)

| | P. | Adjusted P. |
|---|---|---|
| *~region vs. ~* | 1.29e−02 * | 1.26e−01 |
| *~purity vs ~* | 4.16e−03 ** | 3.41e−02 * |
| *~purity+region vs ~purity* | 3.34e−01 | 7.81e−01 |
| *Carcinoma vs. normal* | 1.6e−03 ** | 1.6e−03 ** |

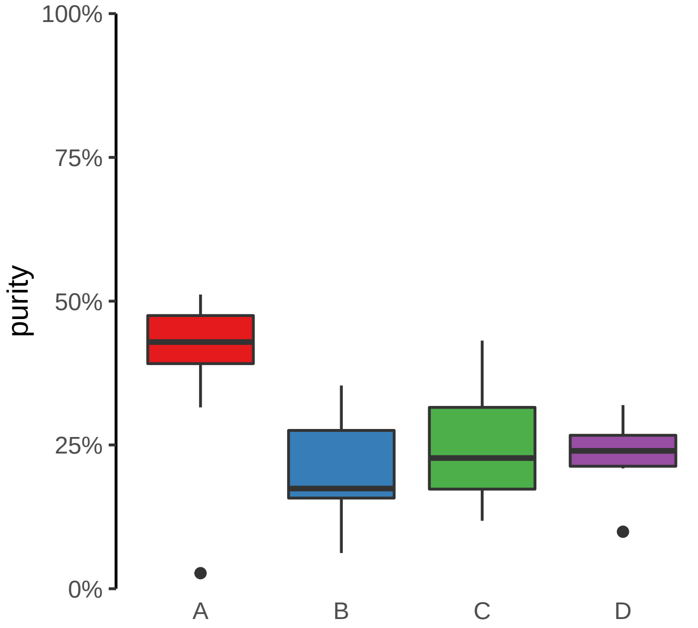

**PLXNB2 promoter gain**
**chr22:50282402−50282902**

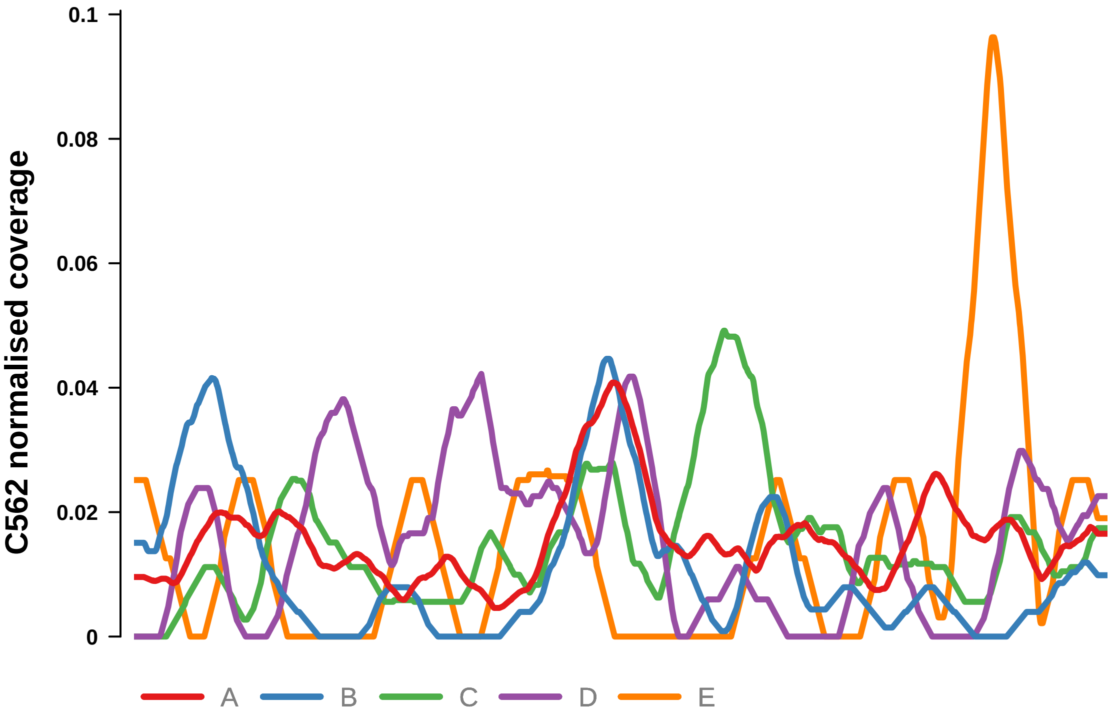


Putatively Clonal (logFC = 1.66)

| | P. | Adjusted P. |
|---|---|---|
| *~region vs. ~* | 4.78e−01 | 7.13e−01 |
| *~purity vs ~* | 4.59e−03 ** | 3.41e−02 * |
| *~purity+region vs ~purity* | 8.93e−01 | 9.7e−01 |
| *Carcinoma vs. normal* | 4.32e−03 ** | 4.32e−03 ** |

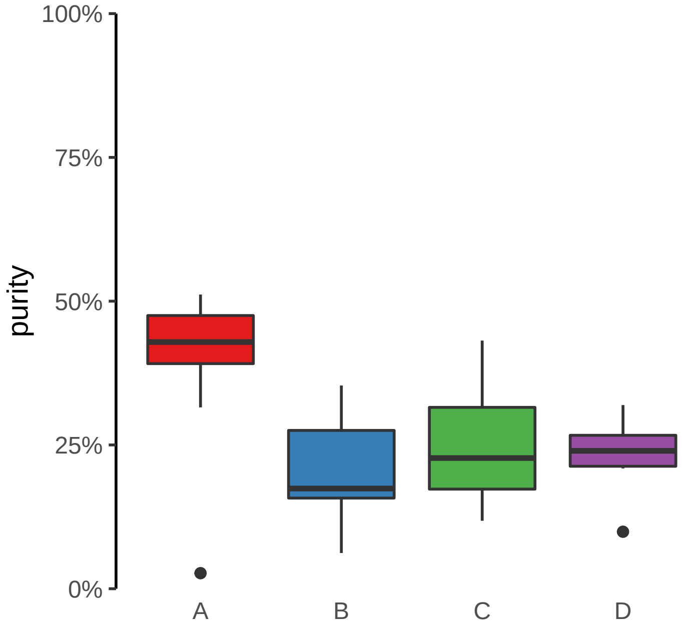

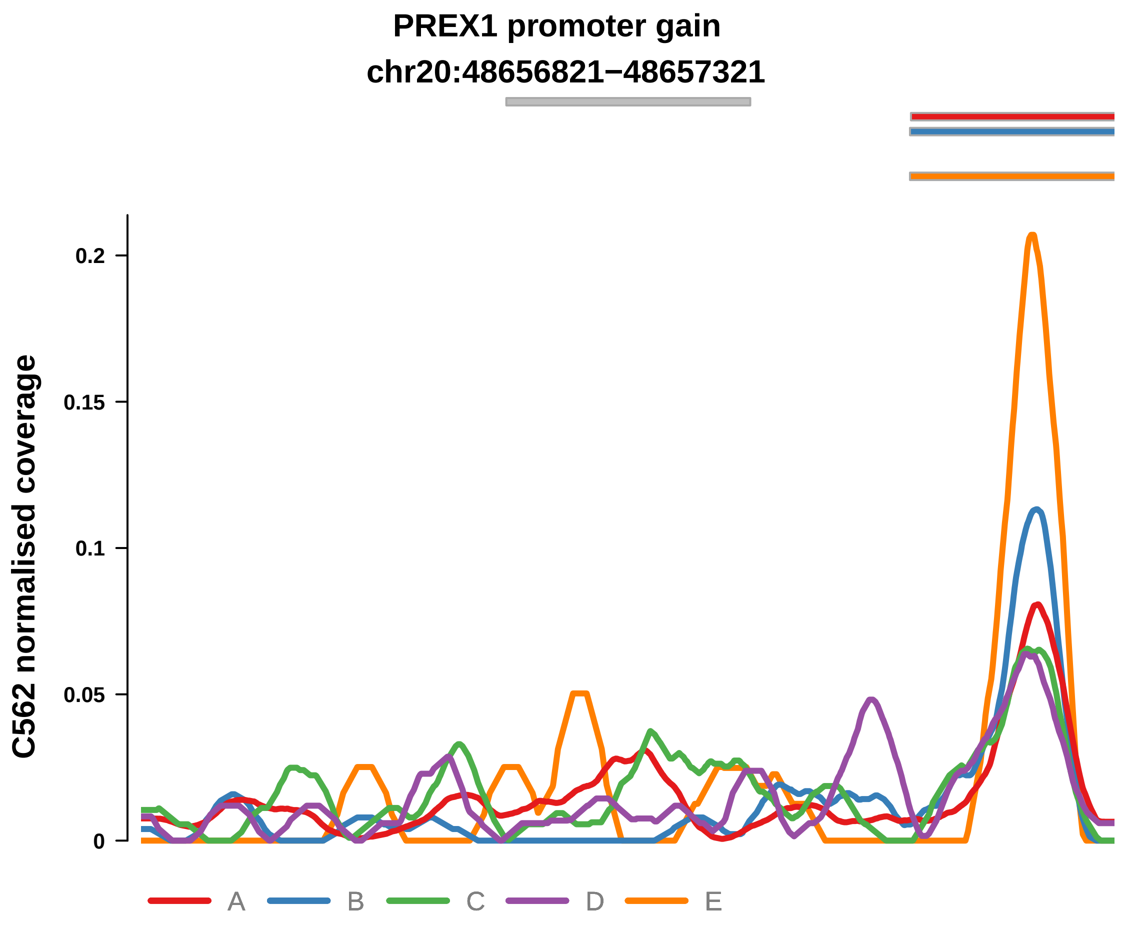


Putatively Clonal (logFC = 1.86)

| | P. | Adjusted P. |
|---|---|---|
| *~region vs. ~* | 1.84e−03 ** | 3.23e−02 * |
| *~purity vs ~* | 2.79e−03 ** | 2.87e−02 * |
| *~purity+region vs ~purity* | 4.34e−02 * | 3.82e−01 |
| *Carcinoma vs. normal* | 1.94e−03 ** | 1.94e−03 ** |

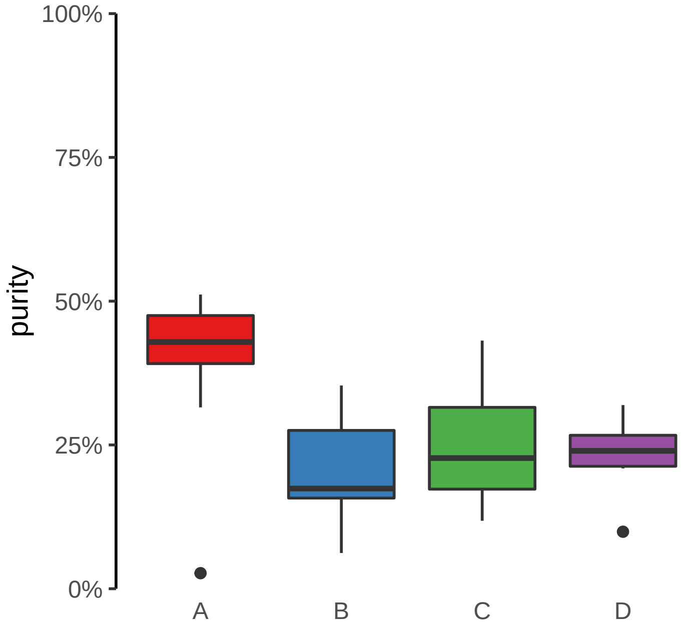

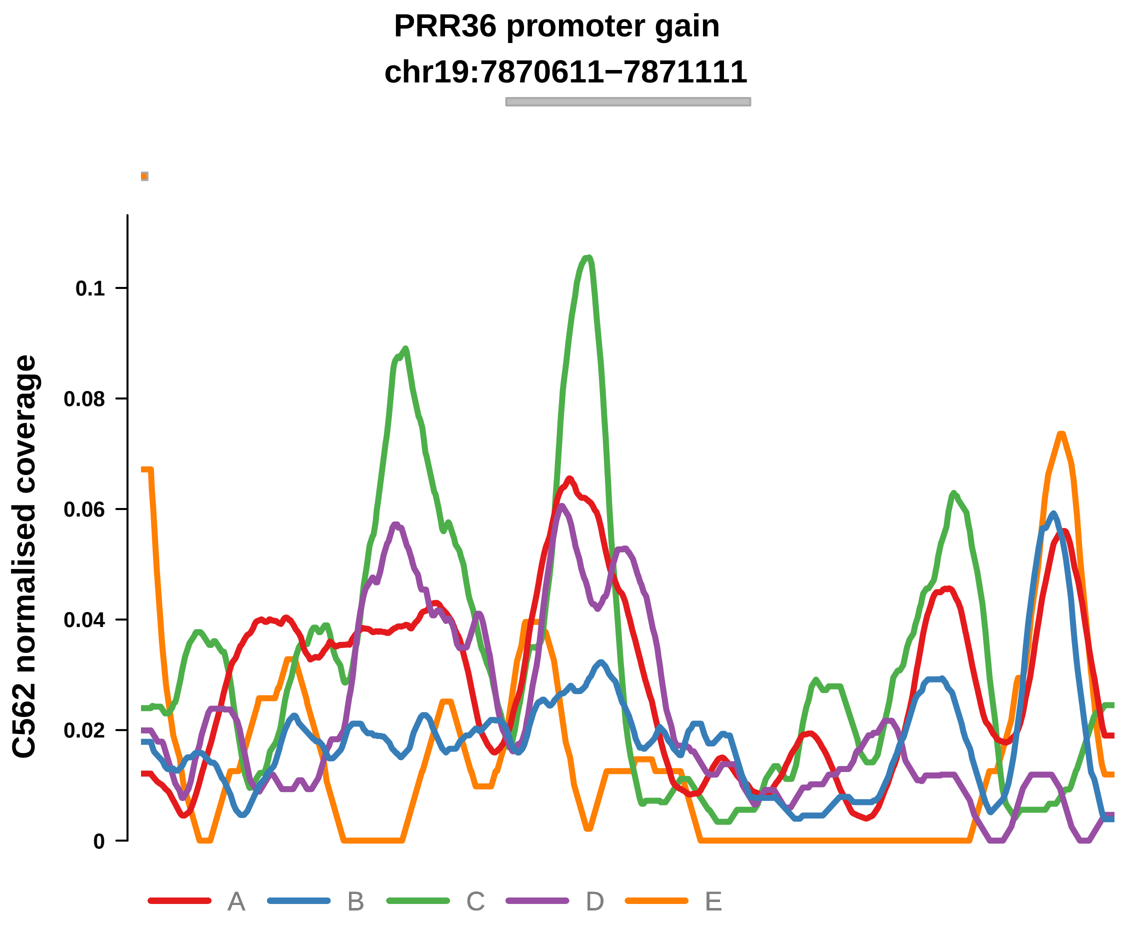


Putatively Clonal (logFC = 1.65)

| | P. | Adjusted P. |
|---|---|---|
| *~region vs. ~* | 1.21e−01 | 4.79e−01 |
| *~purity vs ~* | 1.01e−05 **** | 4.44e−04 *** |
| *~purity+region vs ~purity* | 6.34e−01 | 9.41e−01 |
| *Carcinoma vs. normal* | 3.4e−03 ** | 3.4e−03 ** |

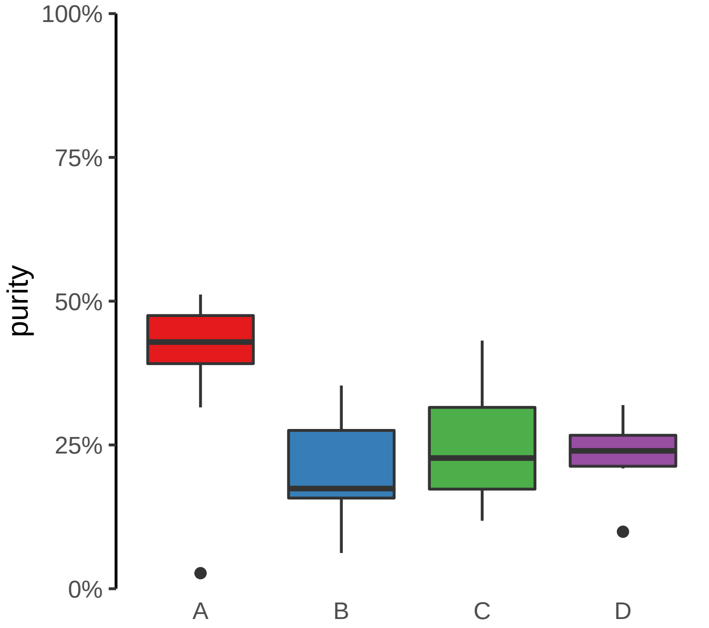

**SAP30BP promoter gain**

**chr17:75706968−75707468**

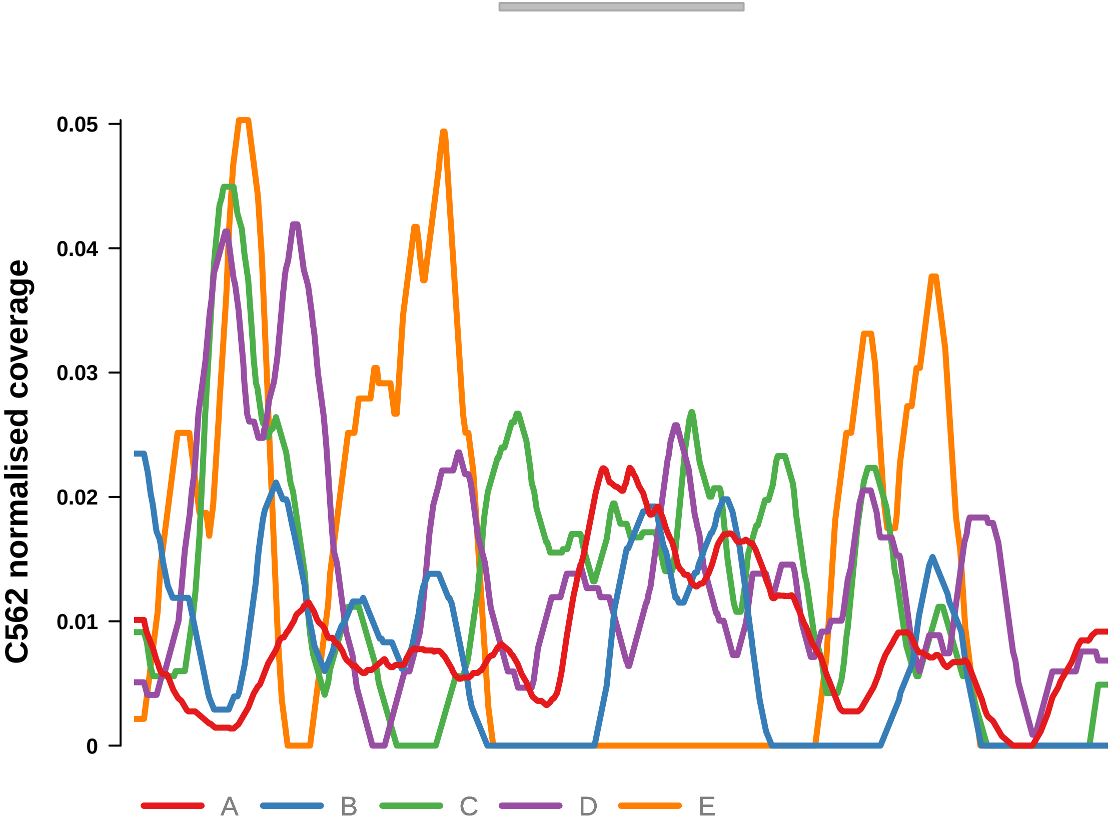


Putatively Clonal (logFC = 1.71)

| | P. | Adjusted P. |
|---|---|---|
| *~region vs. ~* | 2.32e−01 | 5.99e−01 |
| *~purity vs ~* | 1.06e−01 | 2.51e−01 |
| *~purity+region vs ~purity* | 5.68e−01 | 9.36e−01 |
| *Carcinoma vs. normal* | 4.61e−03 ** | 4.61e−03 ** |

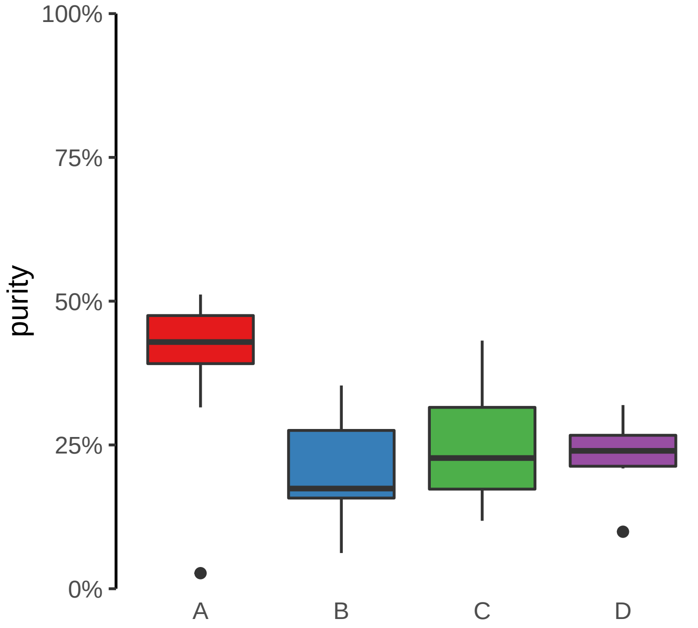

**TTYH3 promoter gain**
**chr7:2646791−2647291**

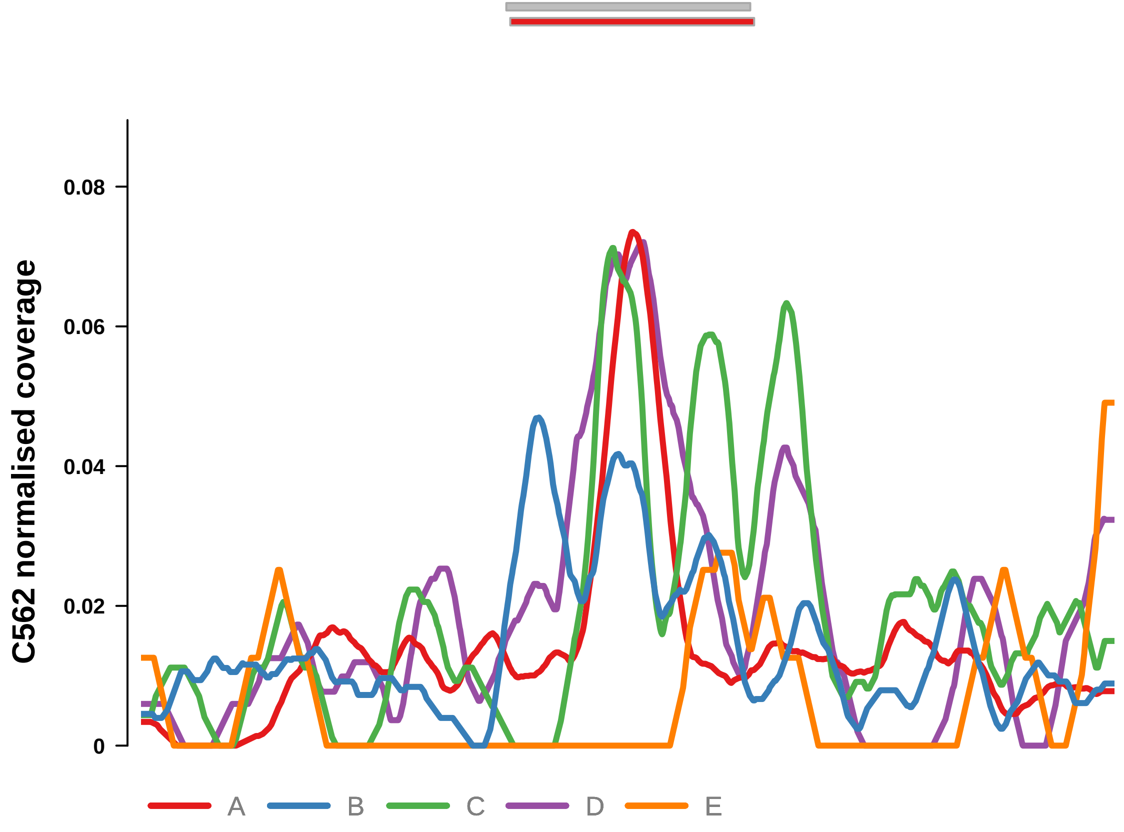


Putatively Clonal (logFC = 2.47)

| | P. | Adjusted P. |
|---|---|---|
| *~region vs. ~* | 5.07e−01 | 7.44e−01 |
| *~purity vs ~* | 1.86e−04 *** | 4.1e−03 ** |
| *~purity+region vs ~purity* | 2.51e−01 | 7.21e−01 |
| *Carcinoma vs. normal* | 2.23e−05 **** | 2.23e−05 **** |

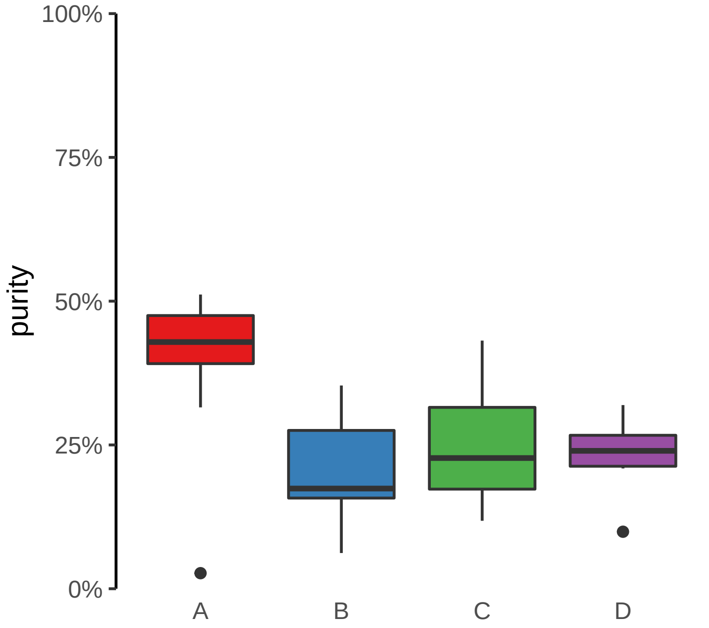
