## Supplementary Figures for "The co-evolution of the genome and epigenome in colorectal cancer": figS11_promo_loss.pdf

**CCDC6 promoter loss**
**chr10:59805006−59805506**

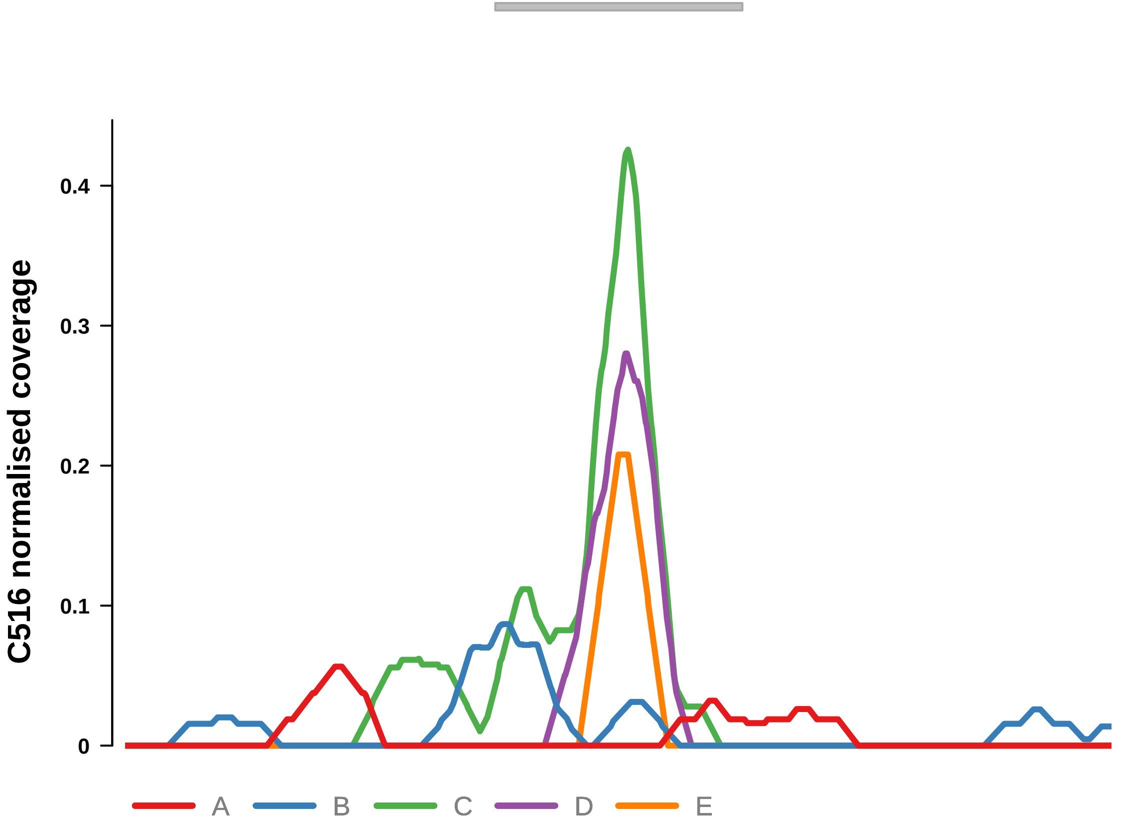


Putatively Clonal (logFC = −11.49)

| | P. | Adjusted P. |
|---|---|---|
| *~region vs. ~* | 3.69e−01 | 8.67e−01 |
| *~purity vs ~* | 8.59e−01 | 9.53e−01 |
| *~purity+region vs ~purity* | 2.09e−01 | 1e+00 |
| *Carcinoma vs. normal* | 9.77e−04 *** | 9.77e−04 *** |

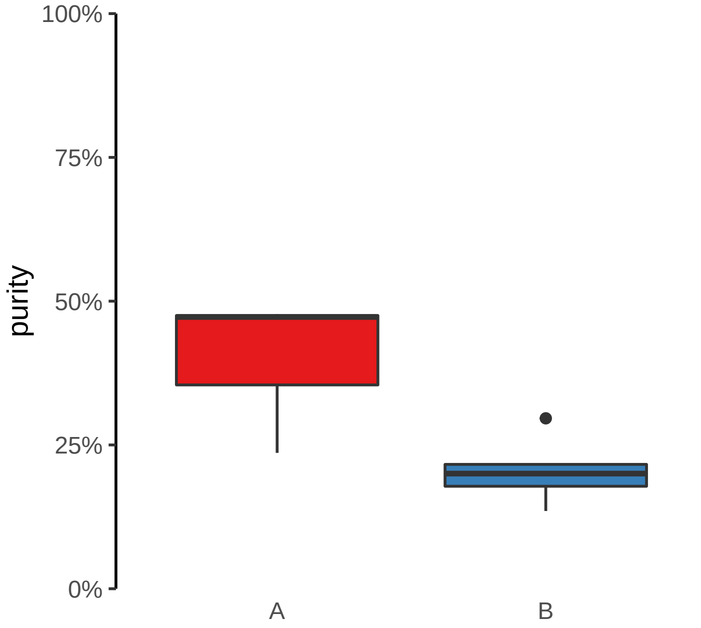

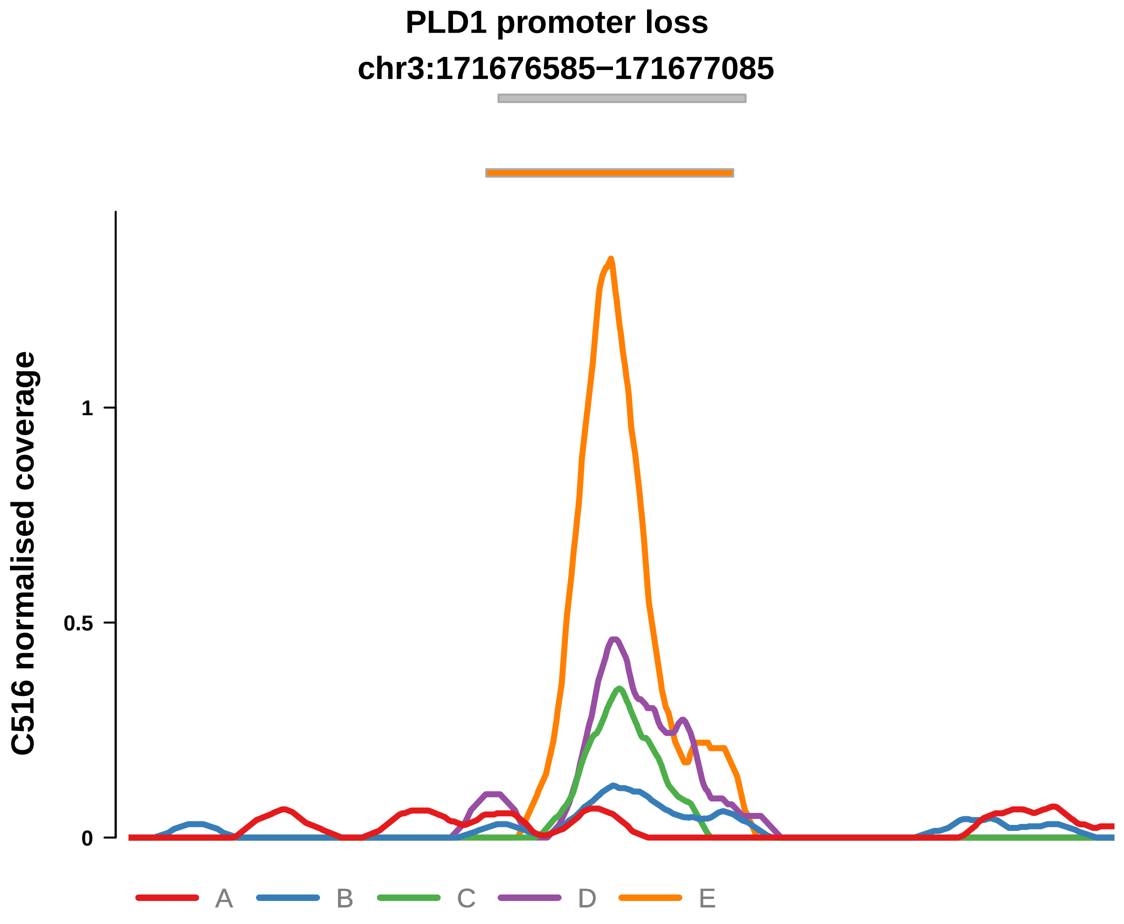


Putatively Clonal (logFC = −11.6)

| | P. | Adjusted P. |
|---|---|---|
| *~region vs. ~* | 5.55e−02 | 7.04e−01 |
| *~purity vs ~* | 1.92e−01 | 7.73e−01 |
| *~purity+region vs ~purity* | 1.51e−01 | 1e+00 |
| *Carcinoma vs. normal* | 4.95e−04 *** | 4.95e−04 *** |

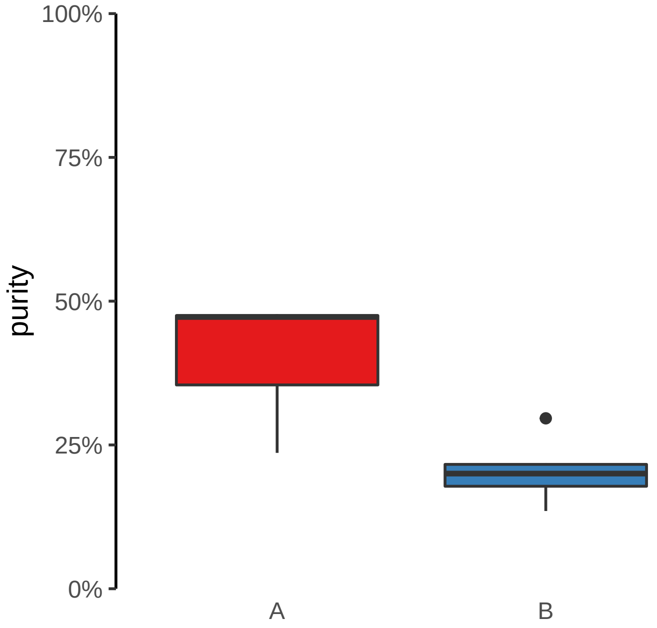

**SMARCA2 promoter loss**
**chr9:2053751−2054251**

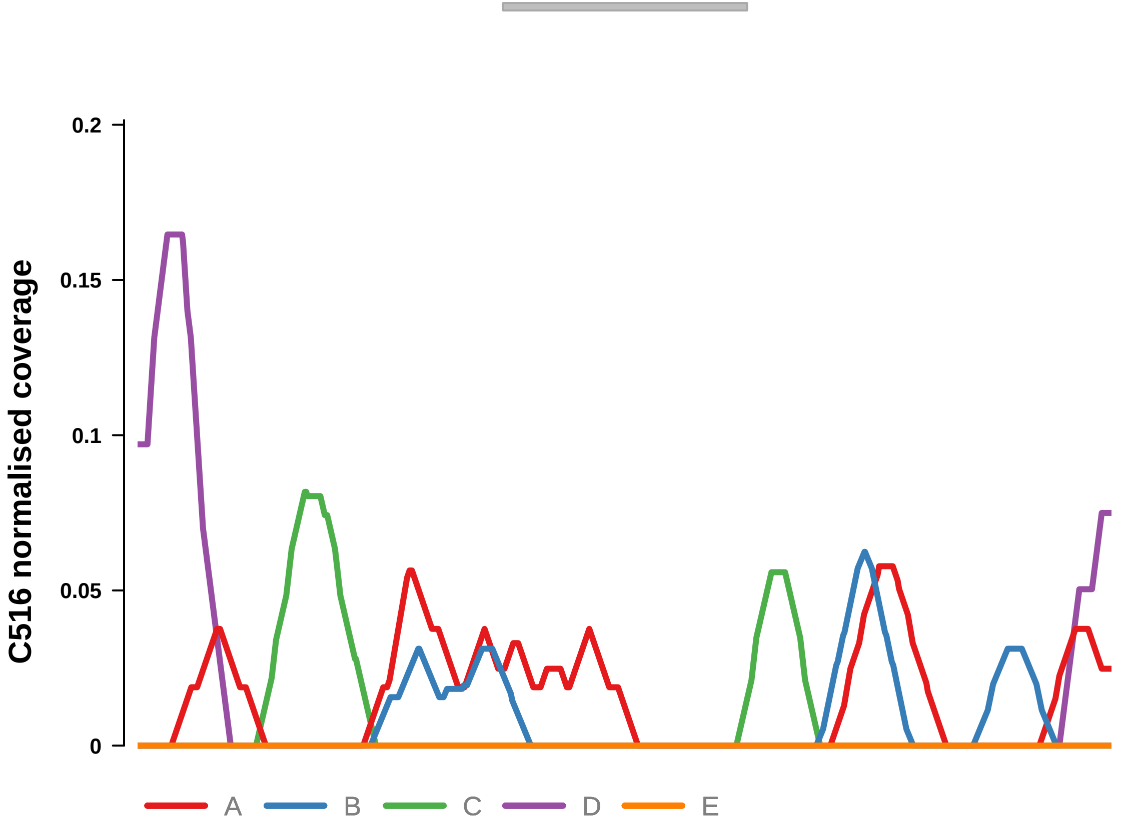


Putatively Clonal (logFC = −10.65)

| | P. | Adjusted P. |
|---|---|---|
| *~region vs. ~* | 1e+00 | 1e+00 |
| *~purity vs ~* | 9.2e−01 | 9.53e−01 |
| *~purity+region vs ~purity* | 7.01e−01 | 1e+00 |
| *Carcinoma vs. normal* | 9.01e−03 ** | 9.01e−03 ** |

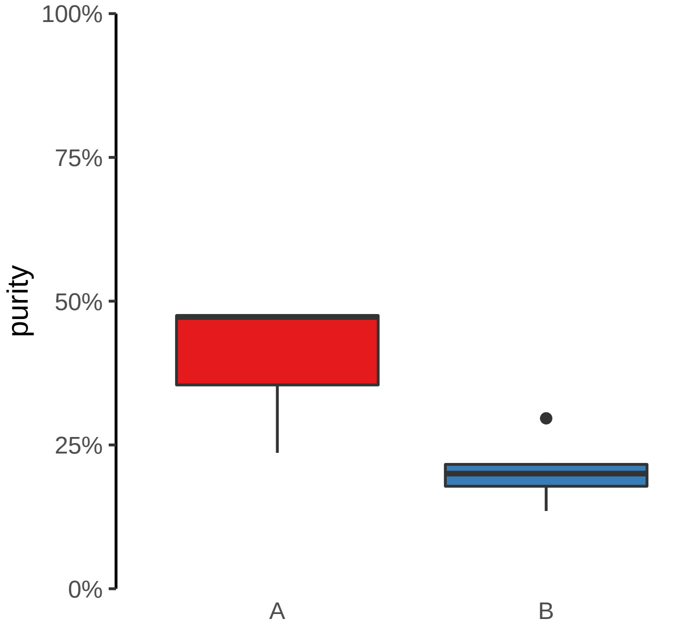

**AMBRA1 promoter loss**

**chr11:46492426−46492926**

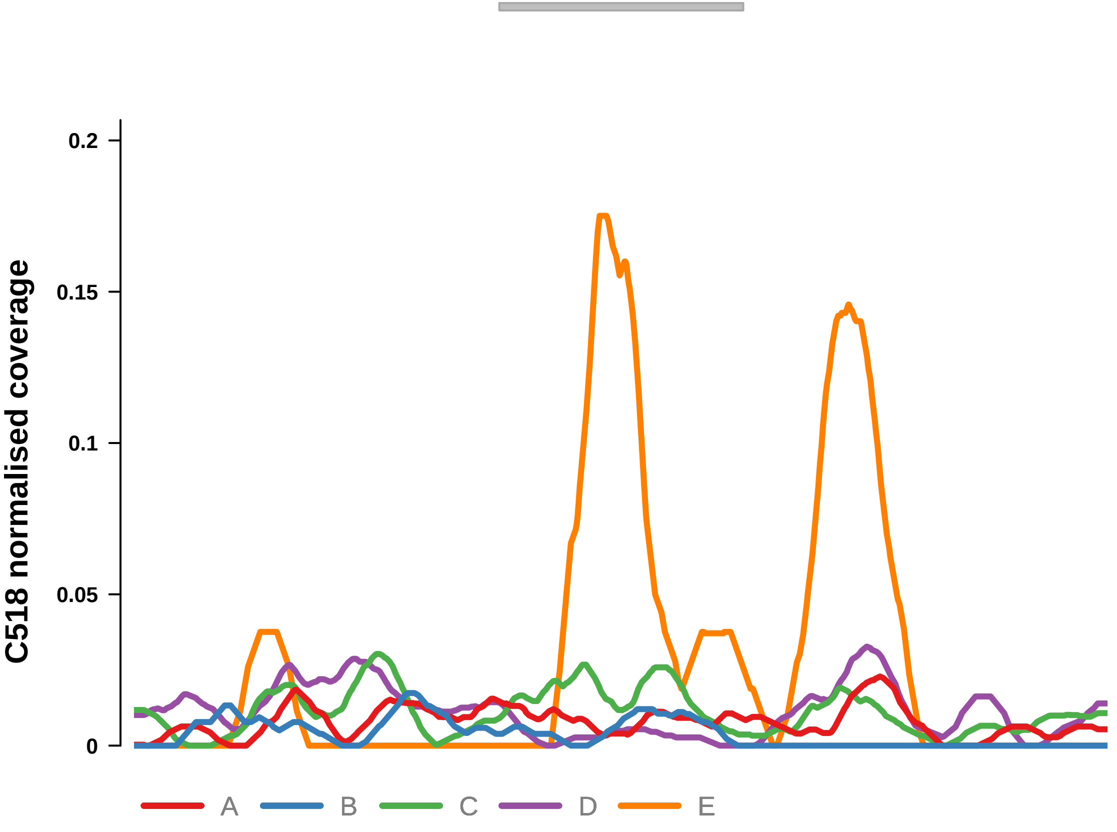


Putatively Clonal (logFC = −2.83)

| | **P.** | **Adjusted P.** |
|---|---|---|
| *~region vs. ~* | 2.55e−03 ** | 5.6e−02 |
| *~purity vs ~* | 1.5e−02 * | 3.49e−01 |
| *~purity+region vs ~purity* | 9.61e−02 | 4.7e−01 |
| *Carcinoma vs. normal* | 9.16e−05 **** | 9.16e−05 **** |

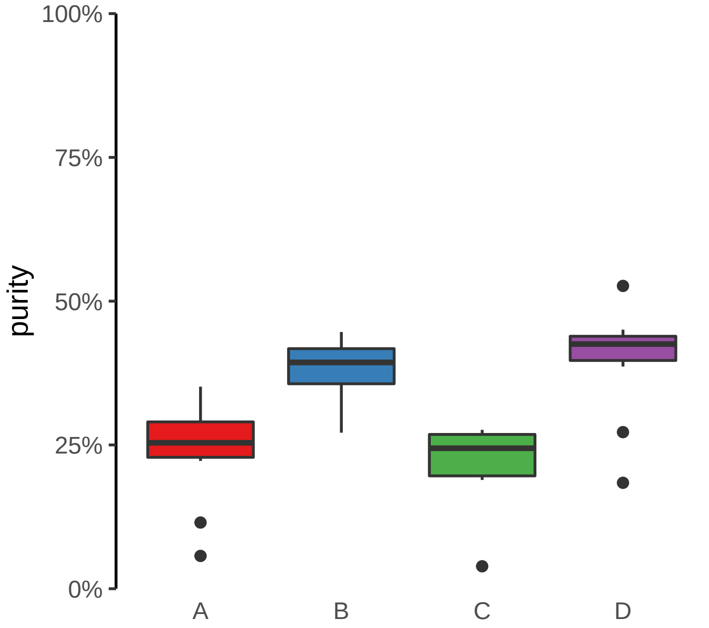

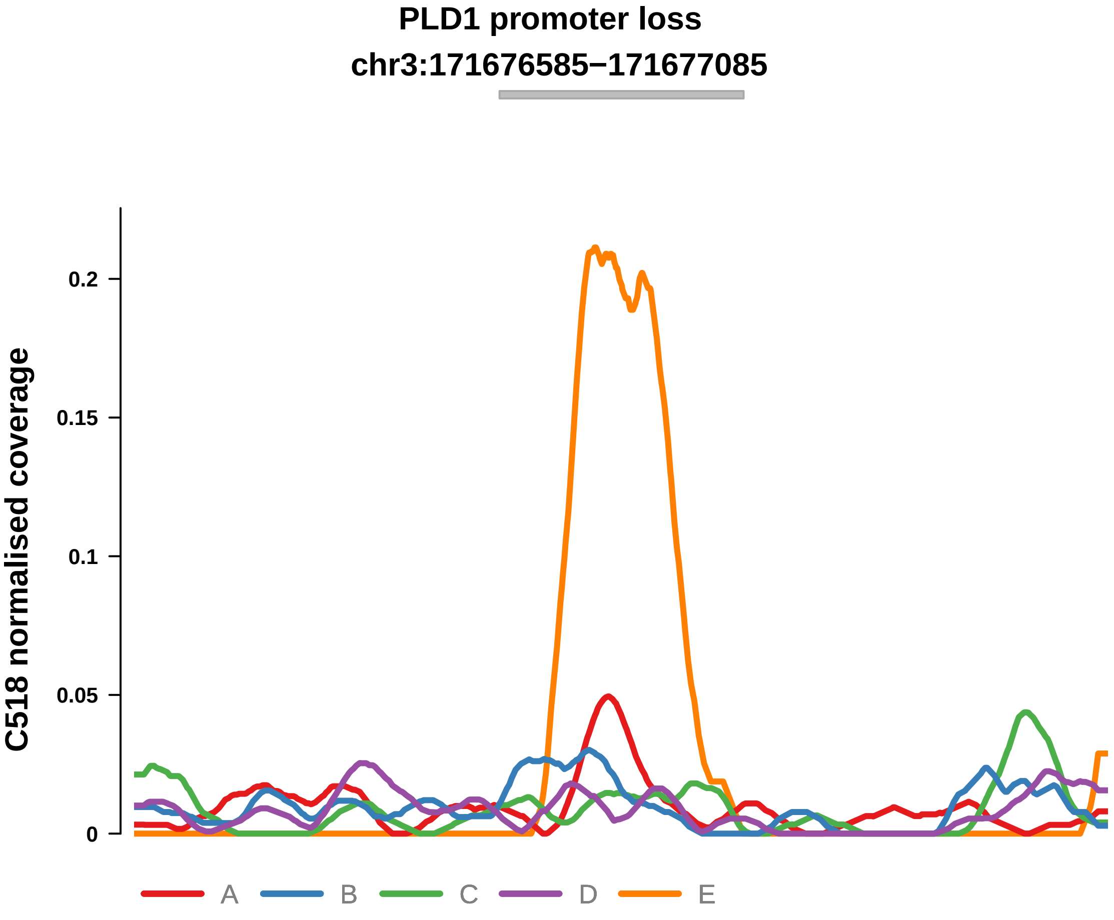


Putatively Clonal (logFC = −2.06)

| | P. | Adjusted P. |
|---|---|---|
| *~region vs. ~* | 7.14e−02 | 3.49e−01 |
| *~purity vs ~* | 1.35e−02 * | 3.49e−01 |
| *~purity+region vs ~purity* | 3.74e−01 | 7.49e−01 |
| *Carcinoma vs. normal* | 1.35e−03 ** | 1.35e−03 ** |

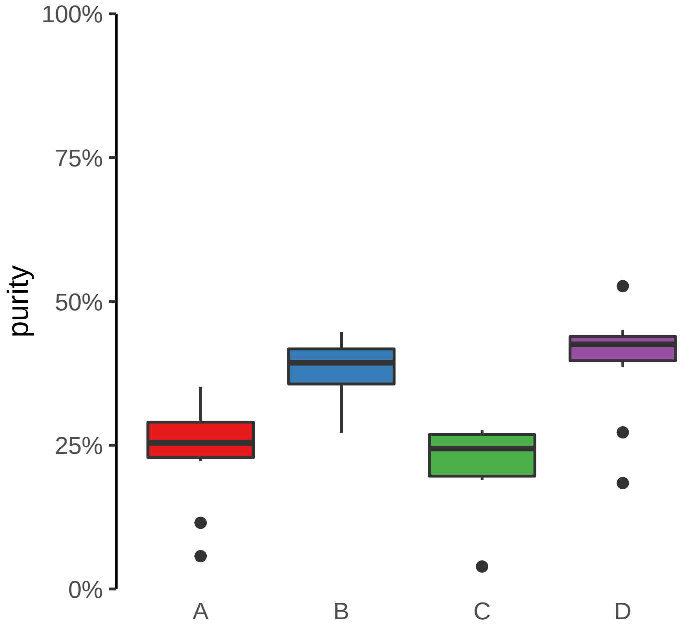

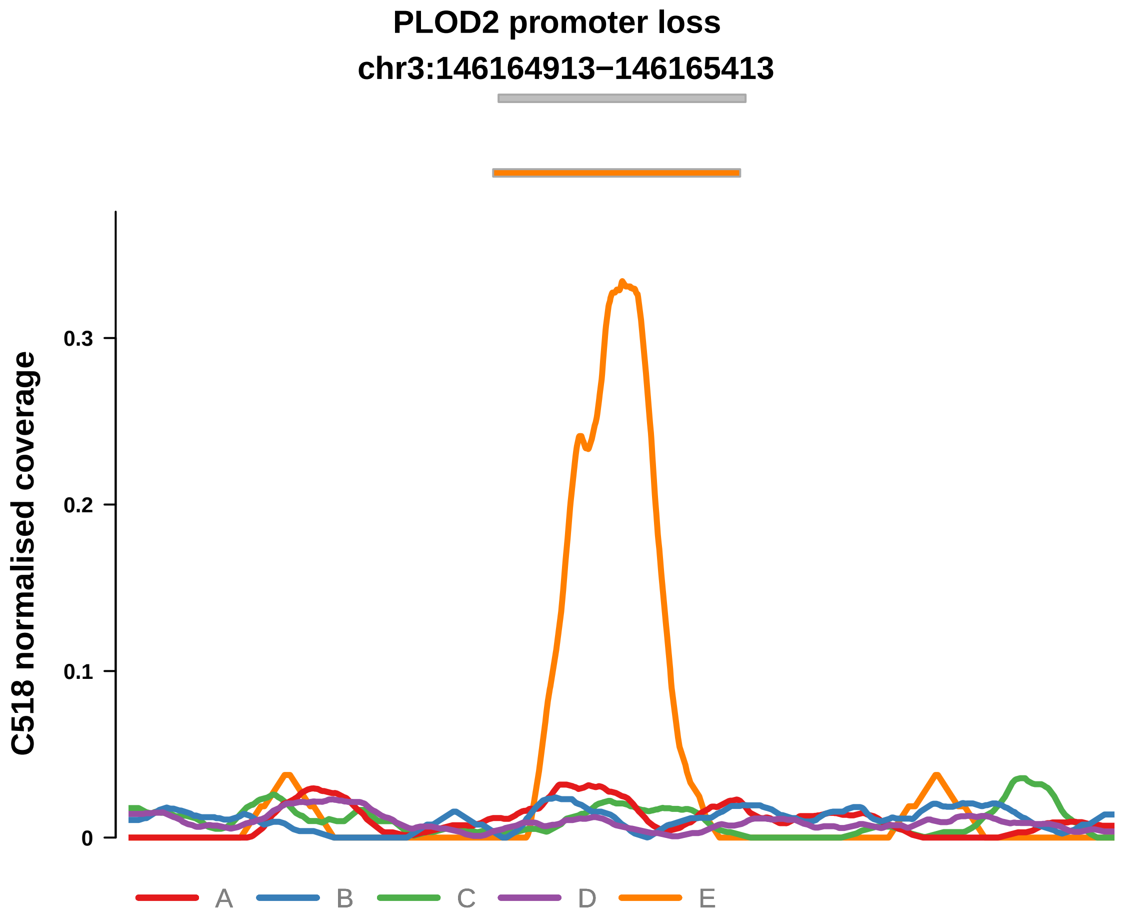


Putatively Clonal (logFC = −2.12)

| | P. | Adjusted P. |
|---|---|---|
| *~region vs. ~* | 1.74e−02 * | 1.55e−01 |
| *~purity vs ~* | 2.85e−02 * | 3.59e−01 |
| *~purity+region vs ~purity* | 1.57e−01 | 5.63e−01 |
| *Carcinoma vs. normal* | 1.5e−03 ** | 1.5e−03 ** |

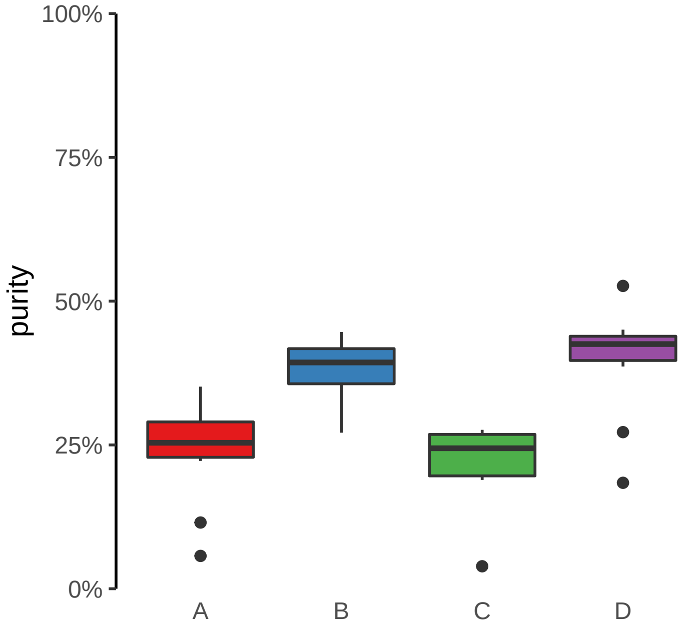

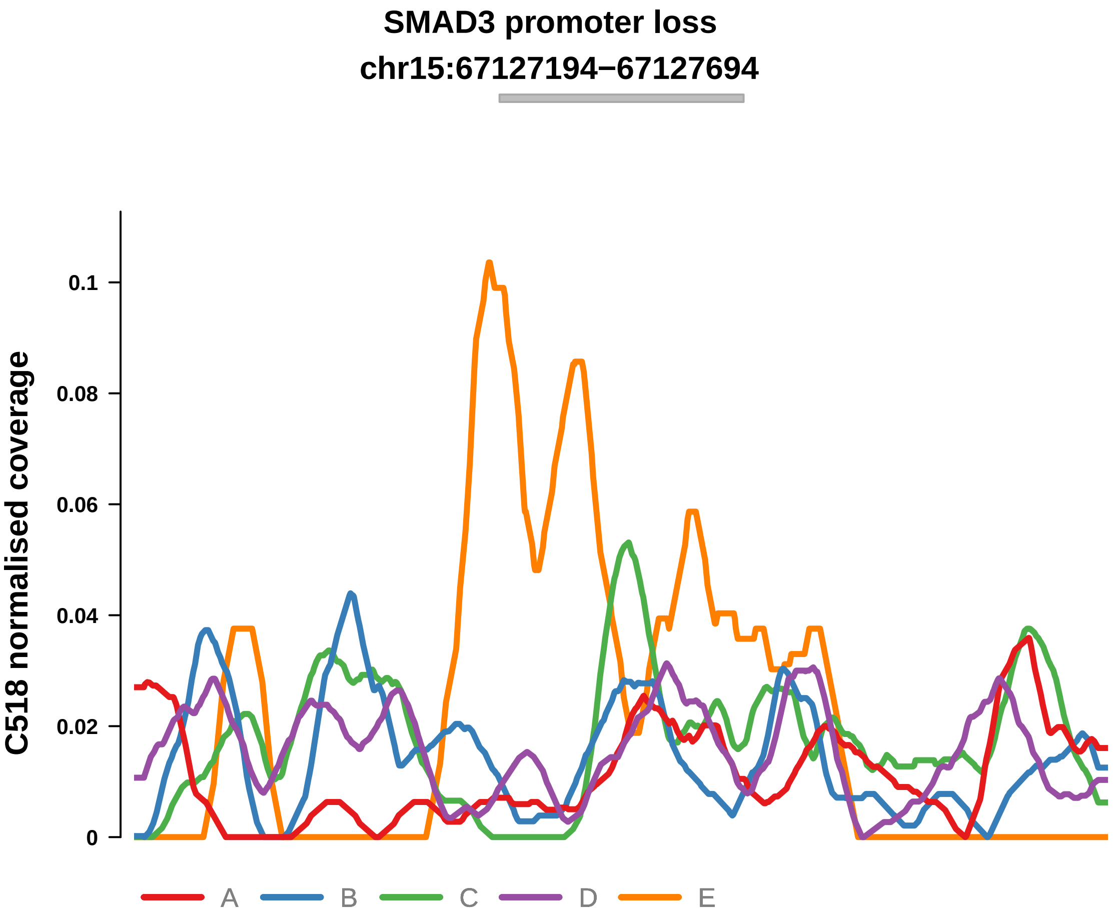


Putatively Clonal (logFC = −1.89)

| | P. | Adjusted P. |
|---|---|---|
| *~region vs. ~* | 6.23e−01 | 7.83e−01 |
| *~purity vs ~* | 2.38e−02 * | 3.49e−01 |
| *~purity+region vs ~purity* | 9.25e−01 | 9.67e−01 |
| *Carcinoma vs. normal* | 2.48e−03 ** | 2.48e−03 ** |

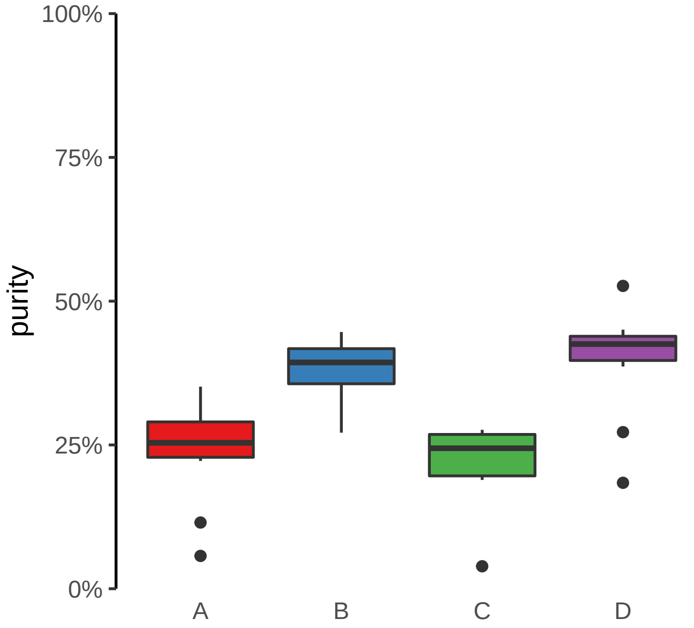

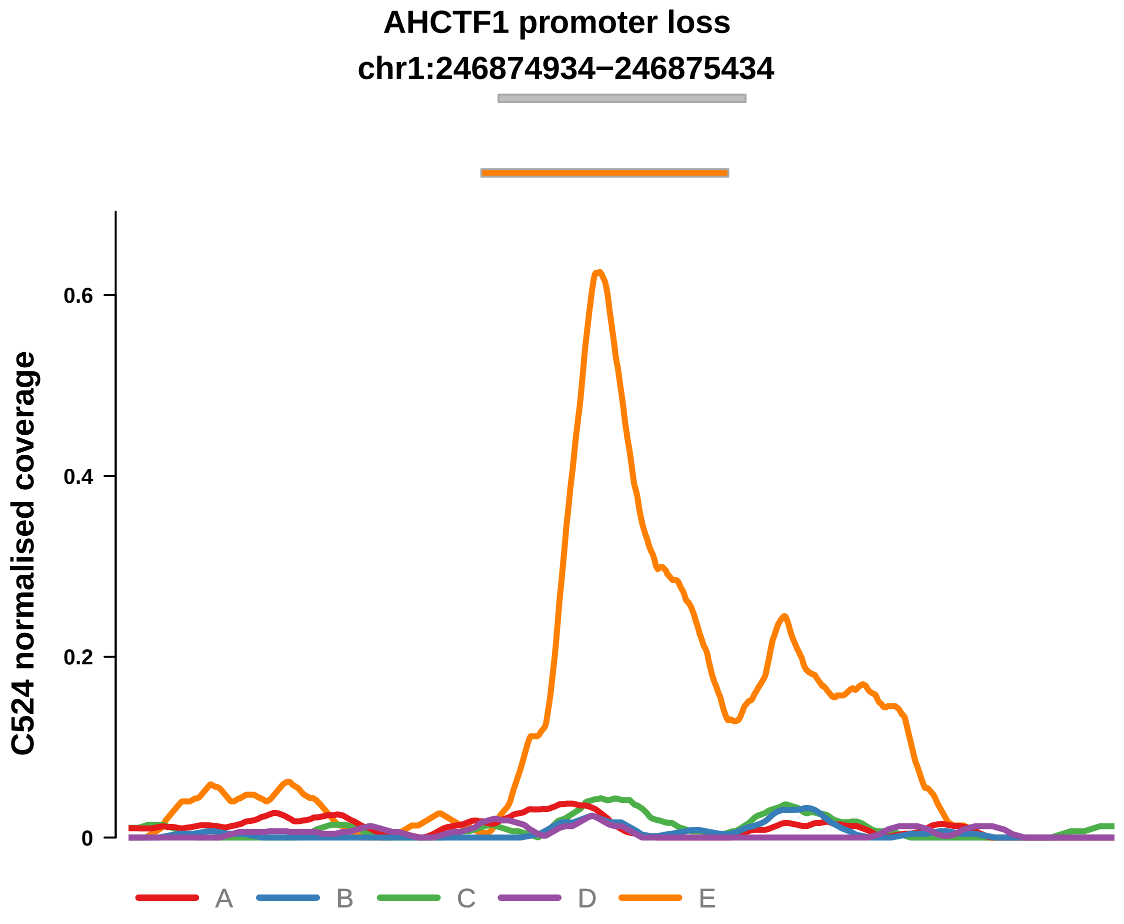


Putatively Clonal (logFC = −3.55)

| | P. | Adjusted P. |
|---|---|---|
| *~region vs. ~* | 6.33e−01 | 8.5e−01 |
| *~purity vs ~* | 2.69e−01 | 5.92e−01 |
| *~purity+region vs ~purity* | 8.76e−01 | 9.83e−01 |
| *Carcinoma vs. normal* | 8.41e−05 **** | 8.41e−05 **** |

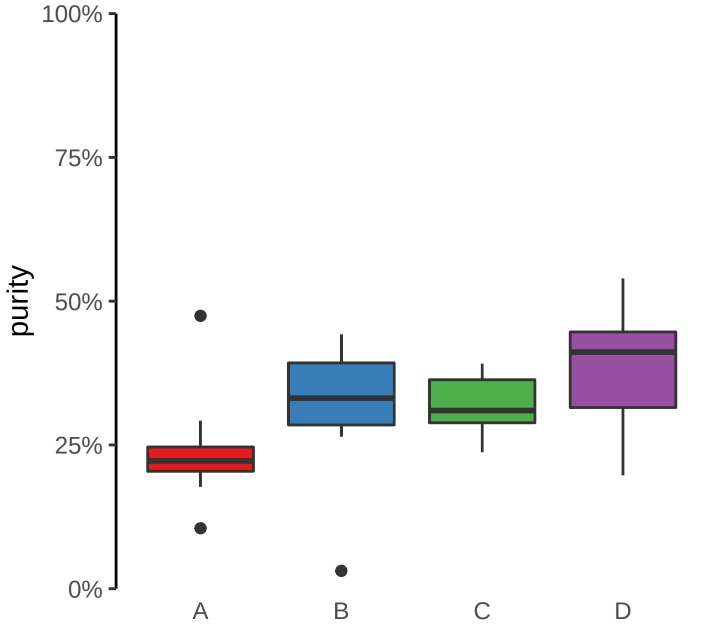

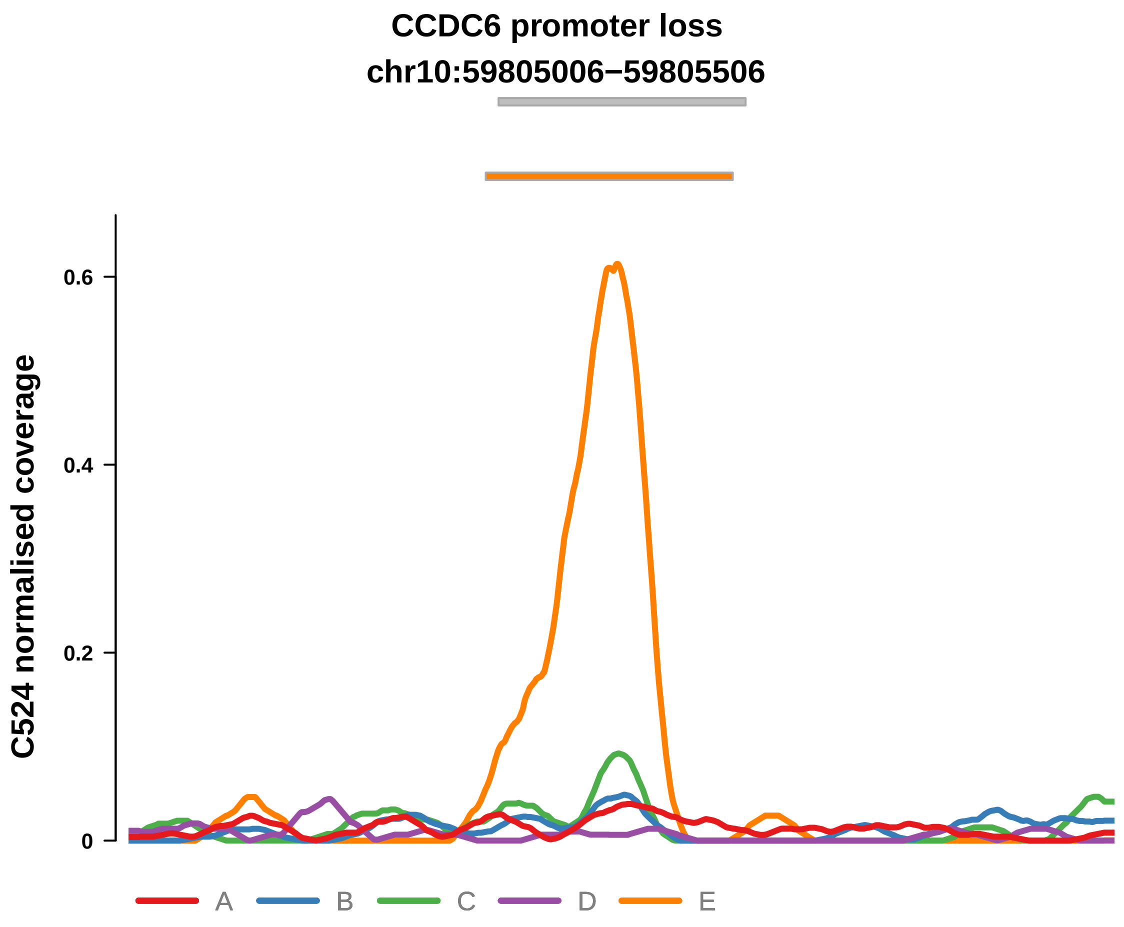


Putatively Clonal (logFC = −2.5)

| | P. | Adjusted P. |
|---|---|---|
| *~region vs. ~* | 4.71e−01 | 7.84e−01 |
| *~purity vs ~* | 8.78e−02 | 4.3e−01 |
| *~purity+region vs ~purity* | 7.9e−01 | 9.27e−01 |
| *Carcinoma vs. normal* | 1.91e−03 ** | 1.91e−03 ** |

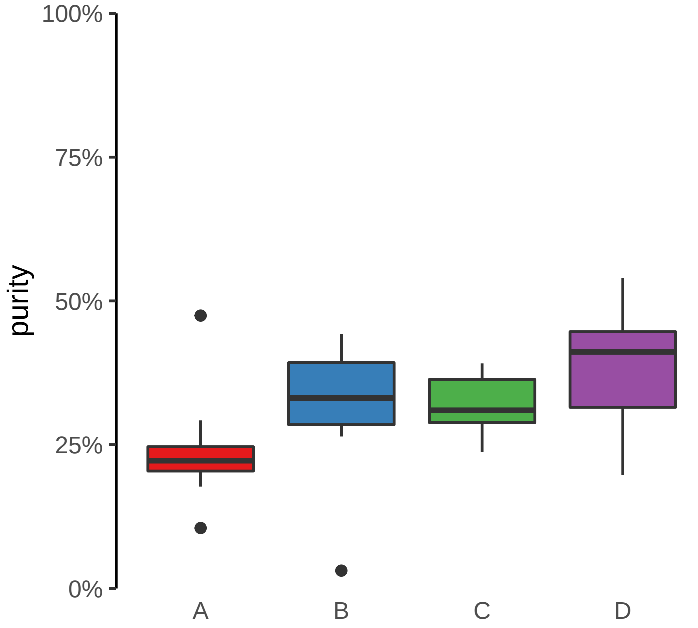

**EPB41L3 promoter loss**
**chr18:5630460−5630960**

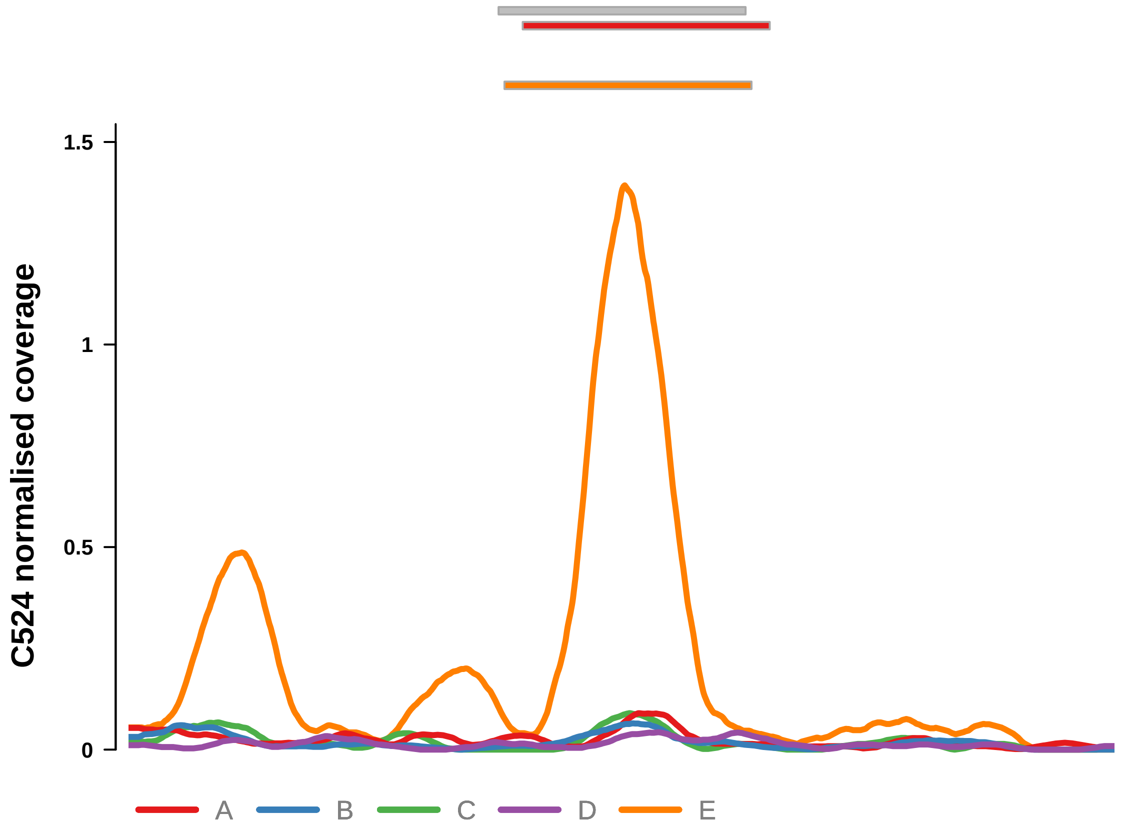


Putatively Clonal (logFC = −1.63)

| | P. | Adjusted P. |
|---|---|---|
| *~region vs. ~* | 1.22e−02 * | 2.69e−01 |
| *~purity vs ~* | 5.52e−01 | 7.12e−01 |
| *~purity+region vs ~purity* | 1.5e−02 * | 1.87e−01 |
| *Carcinoma vs. normal* | 9.82e−03 ** | 9.82e−03 ** |

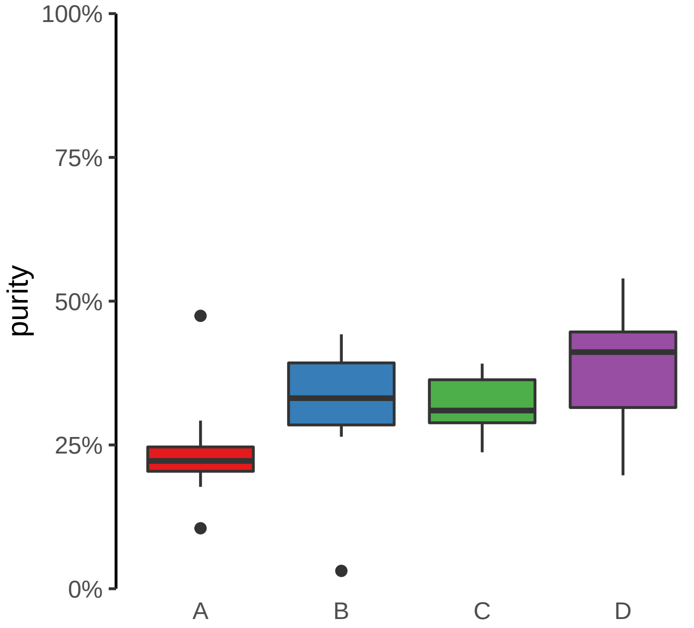

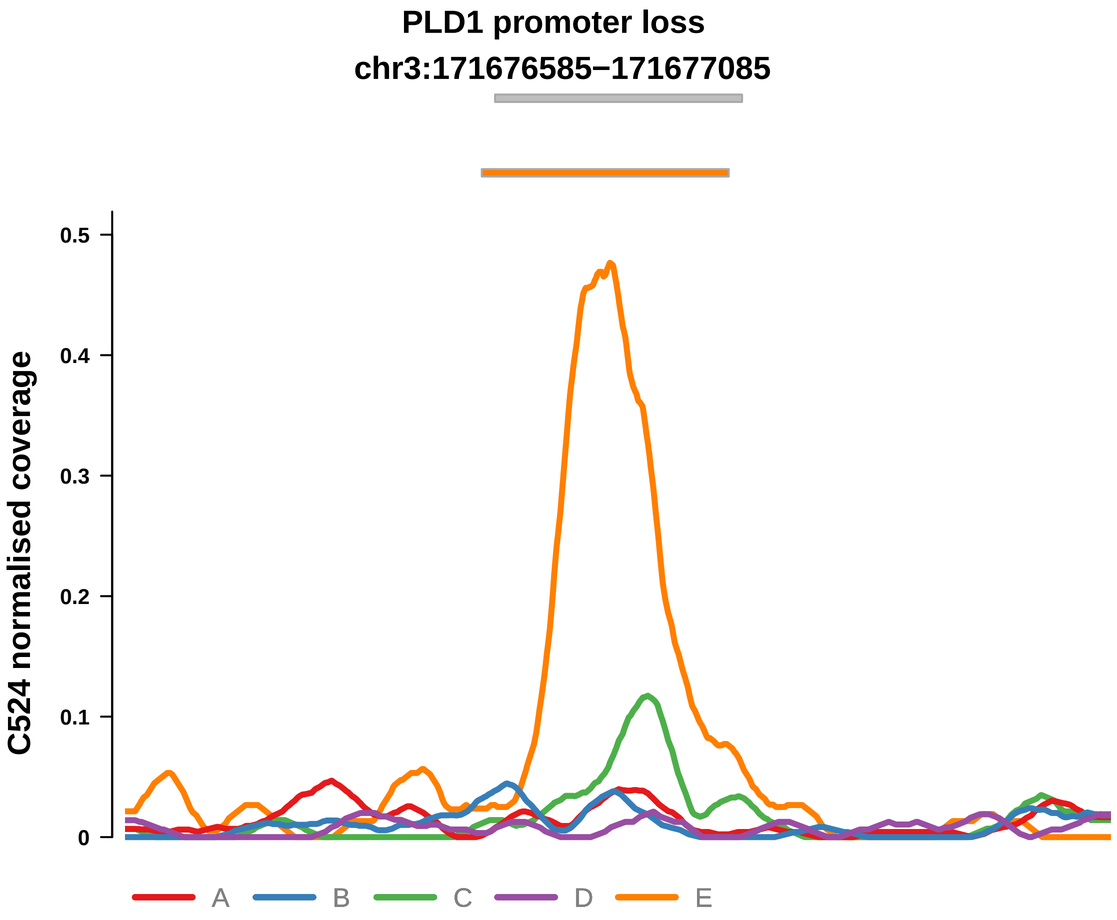


Putatively Clonal (logFC = −2.02)

| | P. | Adjusted P. |
|---|---|---|
| *~region vs. ~* | 5.26e−01 | 8.11e−01 |
| *~purity vs ~* | 3.33e−01 | 6.23e−01 |
| *~purity+region vs ~purity* | 4.74e−01 | 7.82e−01 |
| *Carcinoma vs. normal* | 6.3e−03 ** | 6.3e−03 ** |

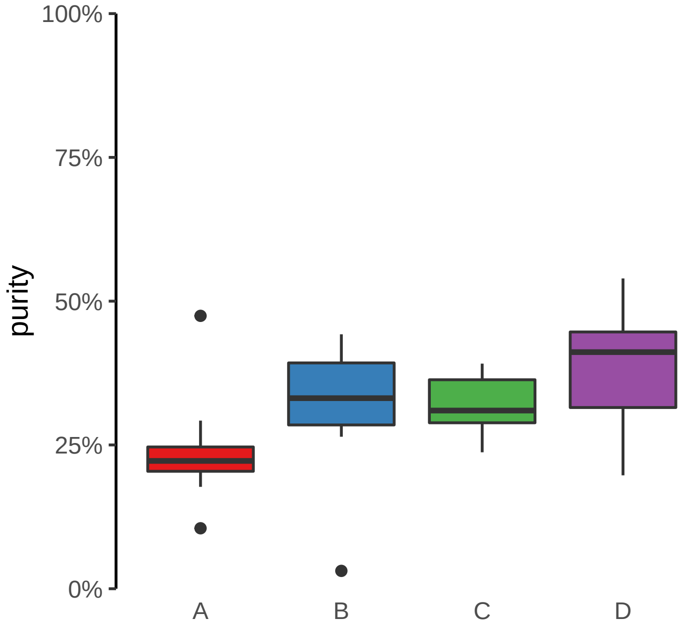

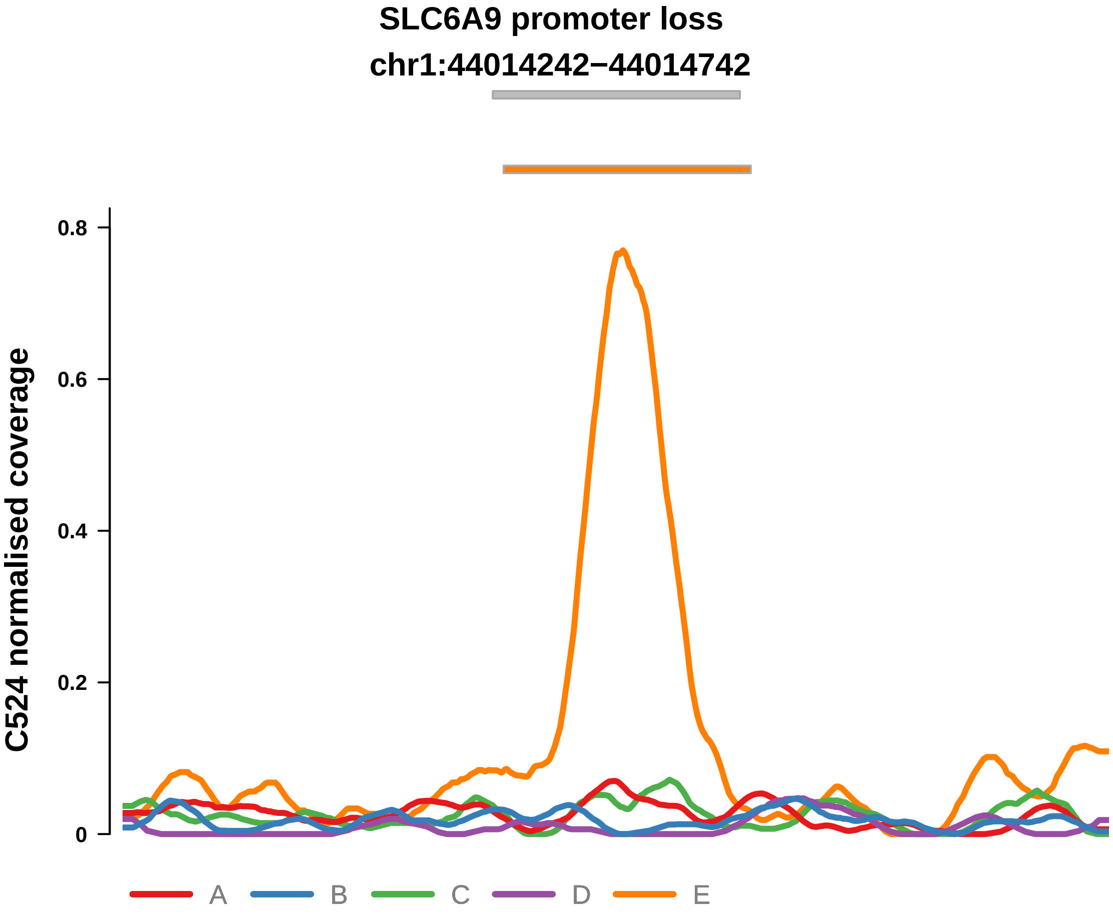


Putatively Clonal (logFC = −1.94)

| | P. | Adjusted P. |
|---|---|---|
| *~region vs. ~* | 9.03e−02 | 4.97e−01 |
| *~purity vs ~* | 1.76e−01 | 5e−01 |
| *~purity+region vs ~purity* | 2.01e−01 | 6e−01 |
| *Carcinoma vs. normal* | 7.05e−03 ** | 7.05e−03 ** |

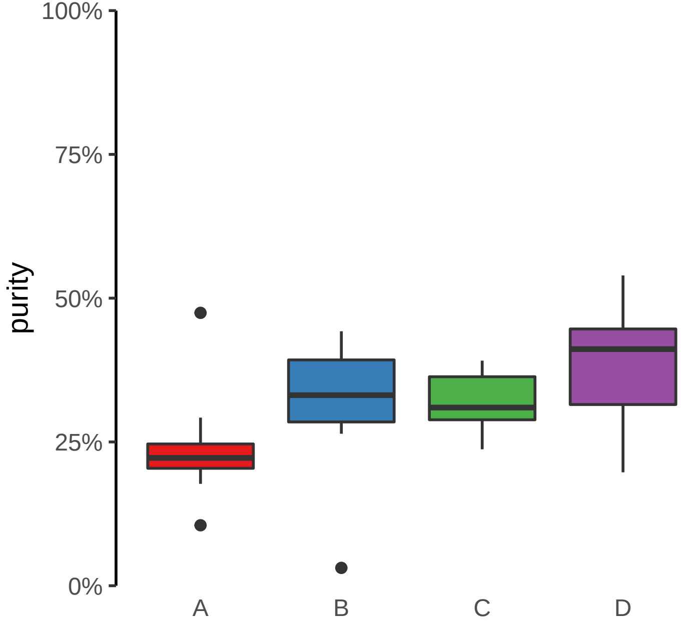

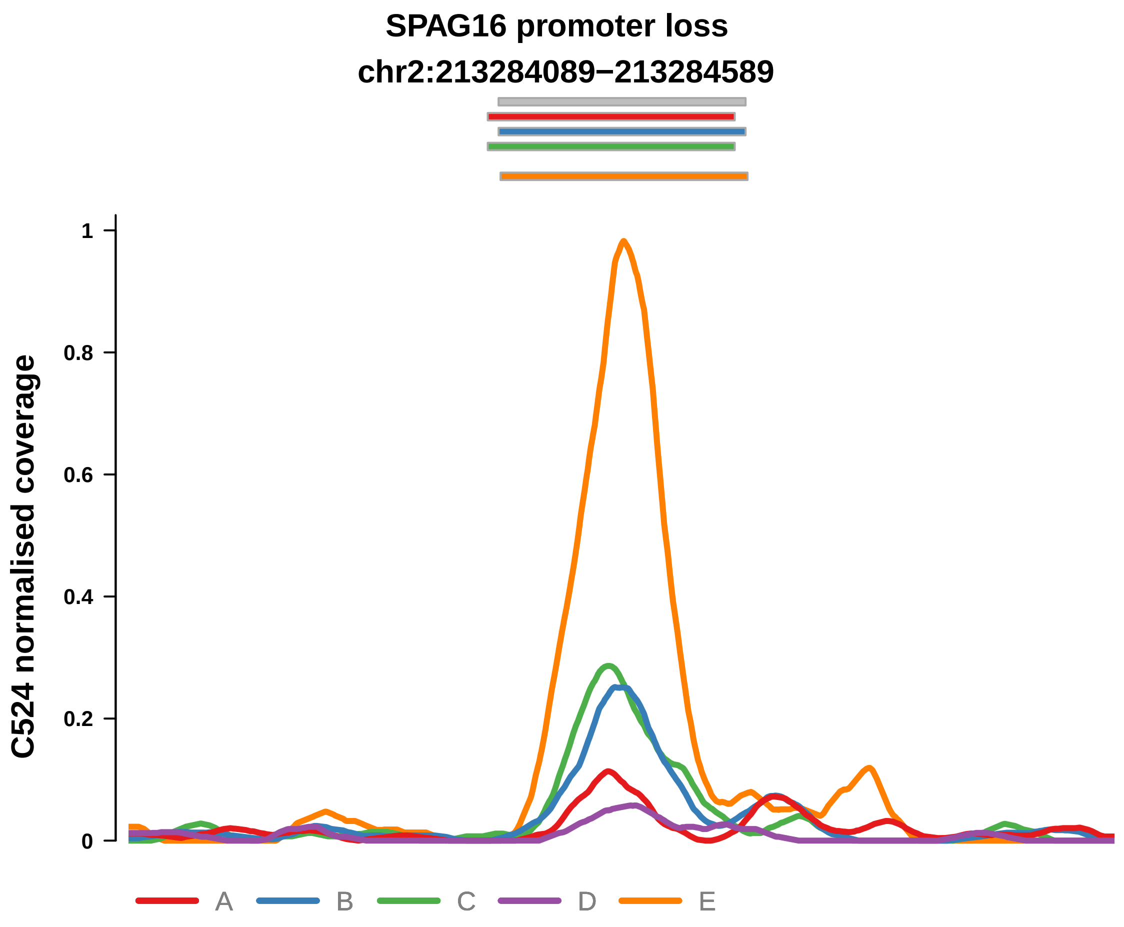


Putatively Clonal (logFC = −1.84)

| | P. | Adjusted P. |
|---|---|---|
| *~region vs. ~* | 9.18e−01 | 9.62e−01 |
| *~purity vs ~* | 2.48e−01 | 5.77e−01 |
| *~purity+region vs ~purity* | 6.26e−01 | 8.8e−01 |
| *Carcinoma vs. normal* | 2.66e−03 ** | 2.66e−03 ** |

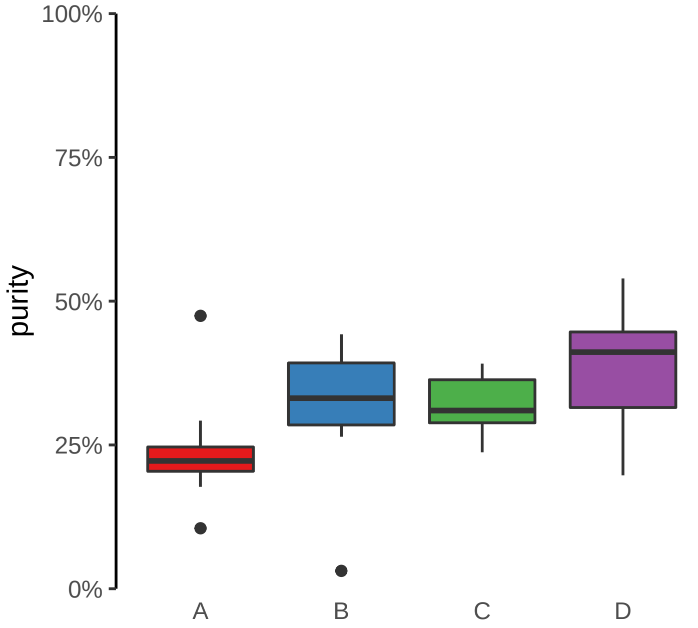

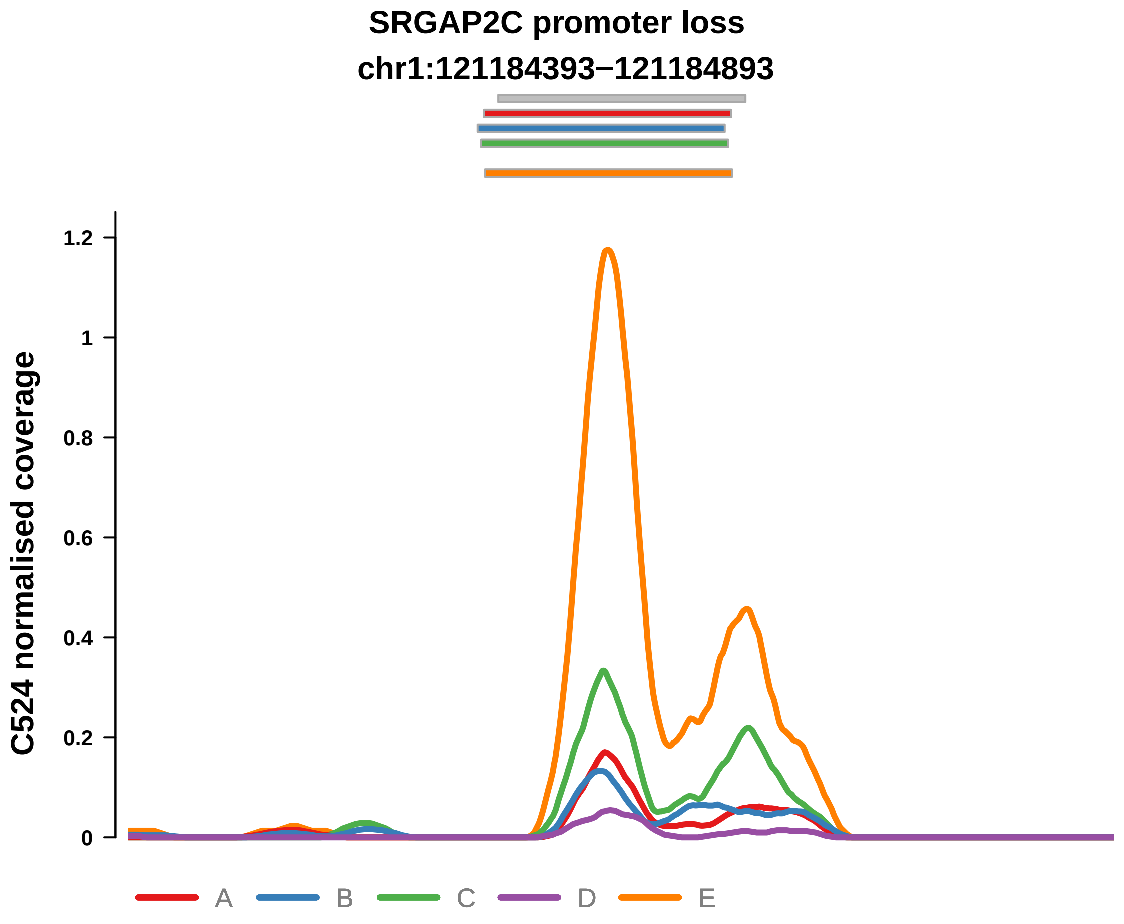


Putatively Clonal (logFC = −1.87)

| | P. | Adjusted P. |
|---|---|---|
| *~region vs. ~* | 2.05e−01 | 5.75e−01 |
| *~purity vs ~* | 3.44e−01 | 6.27e−01 |
| *~purity+region vs ~purity* | 2.49e−01 | 6e−01 |
| *Carcinoma vs. normal* | 2.18e−03 ** | 2.18e−03 ** |

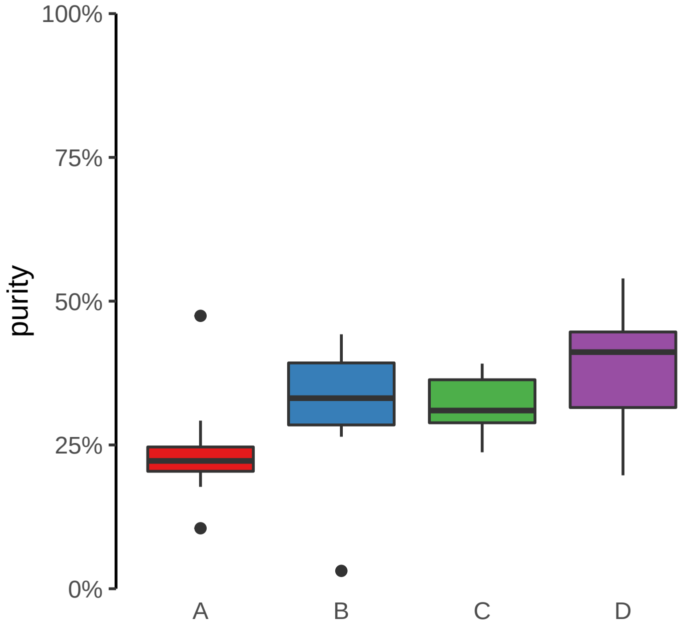

**AHCTF1 promoter loss**
**chr1:246874934−246875434**

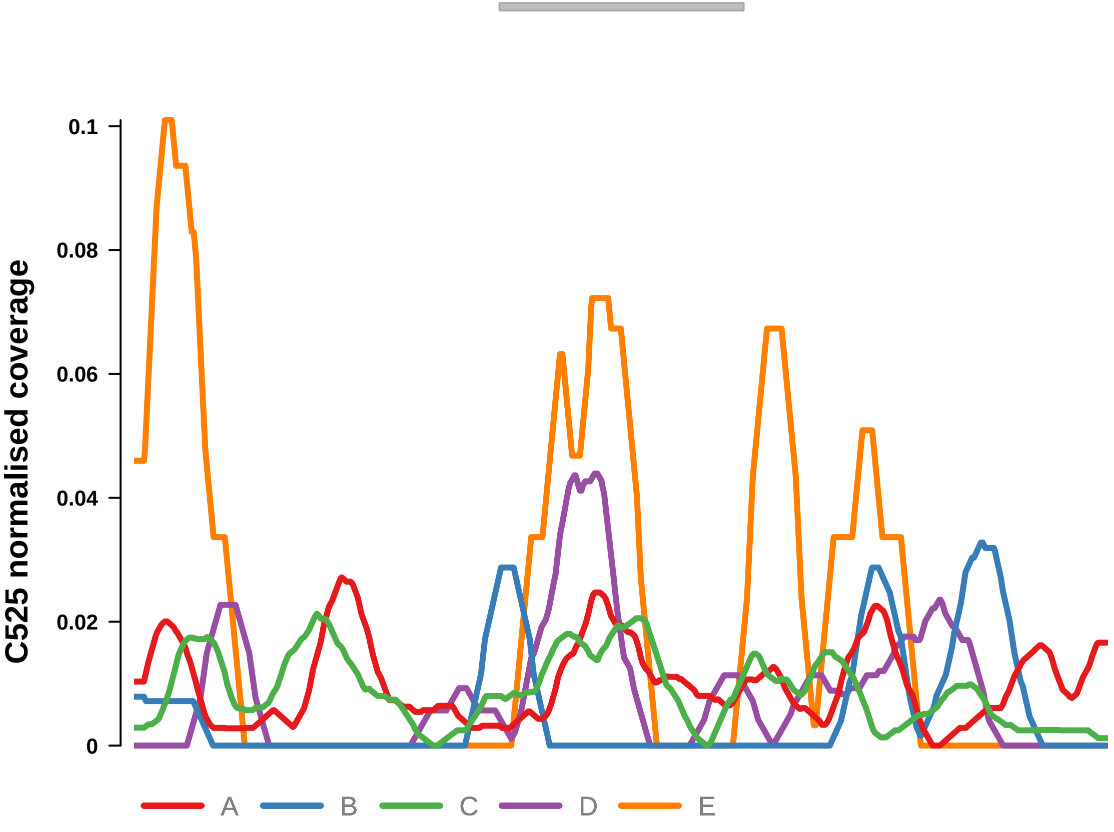


Putatively Clonal (logFC = −2.32)

| | P. | Adjusted P. |
|---|---|---|
| *~region vs. ~* | 4.77e−02 * | 9.77e−02 |
| *~purity vs ~* | 2.71e−01 | 7.48e−01 |
| *~purity+region vs ~purity* | 7.93e−02 | 1.48e−01 |
| *Carcinoma vs. normal* | 1.6e−03 ** | 1.6e−03 ** |

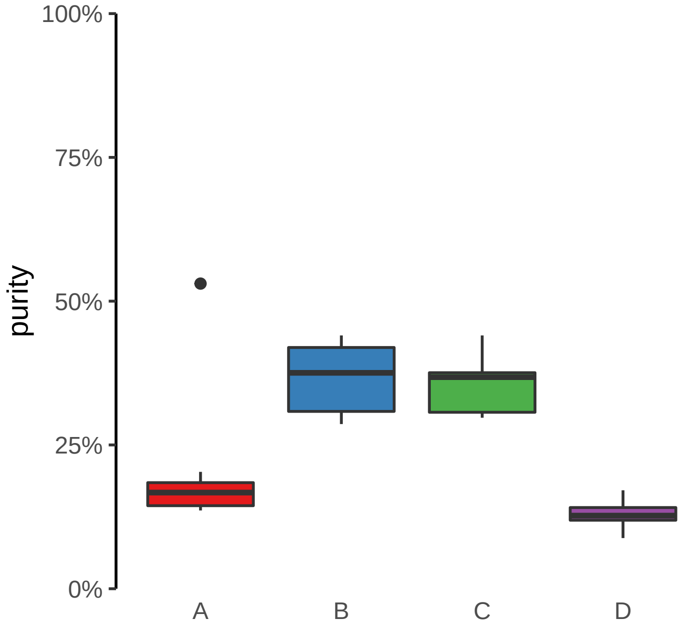

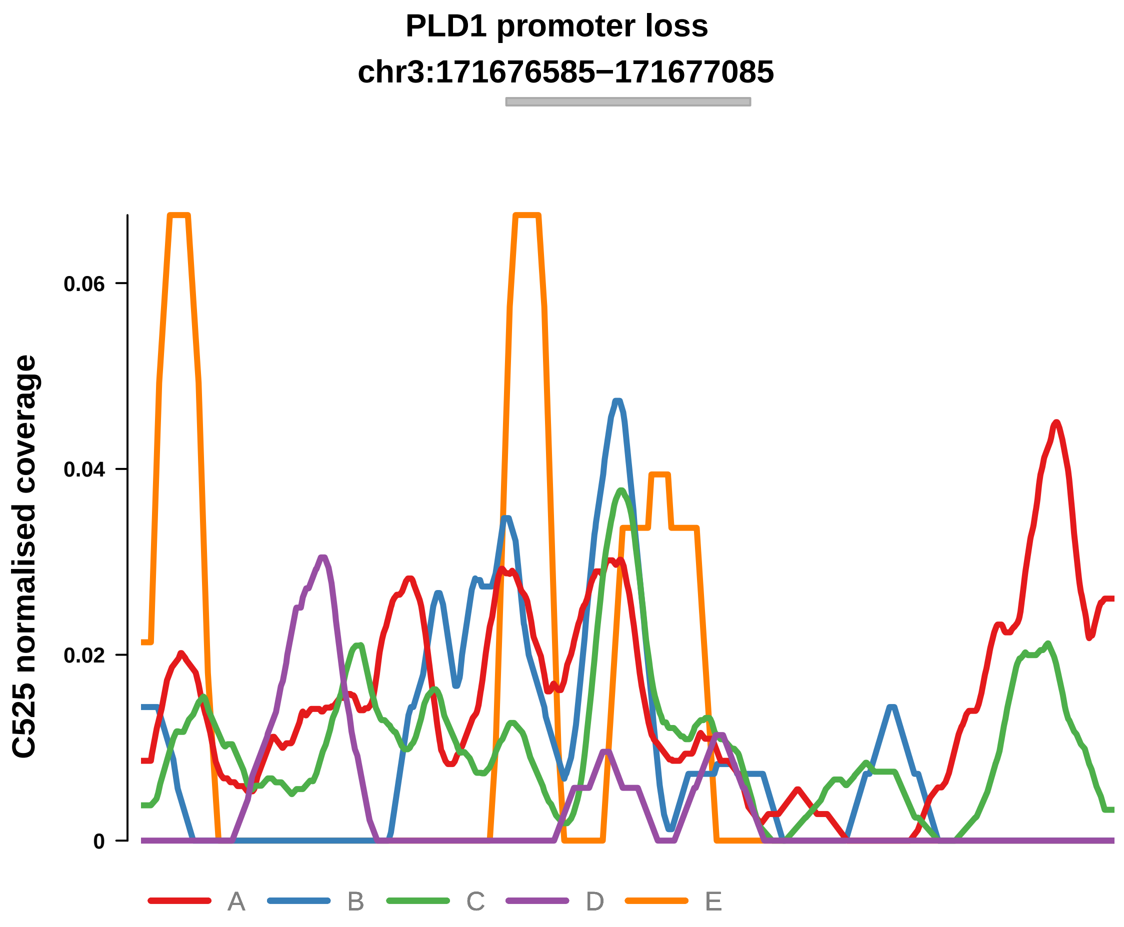


Putatively Clonal (logFC = −11.65)

| | P. | Adjusted P. |
|---|---|---|
| *~region vs. ~* | 1.22e−01 | 1.88e−01 |
| *~purity vs ~* | 6.05e−01 | 8.63e−01 |
| *~purity+region vs ~purity* | 6e−02 | 1.17e−01 |
| *Carcinoma vs. normal* | 8.49e−09 **** | 8.49e−09 **** |

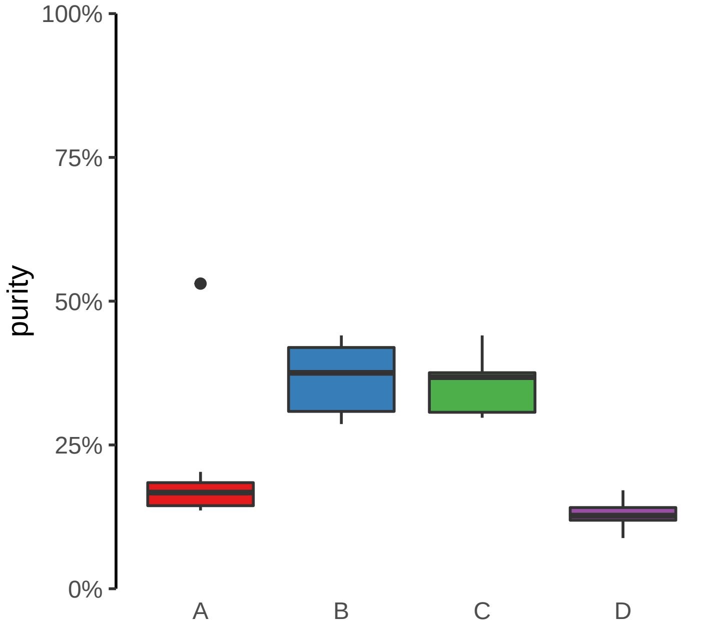

**R3HDM1 promoter loss**

**chr2:135633520−135634020**

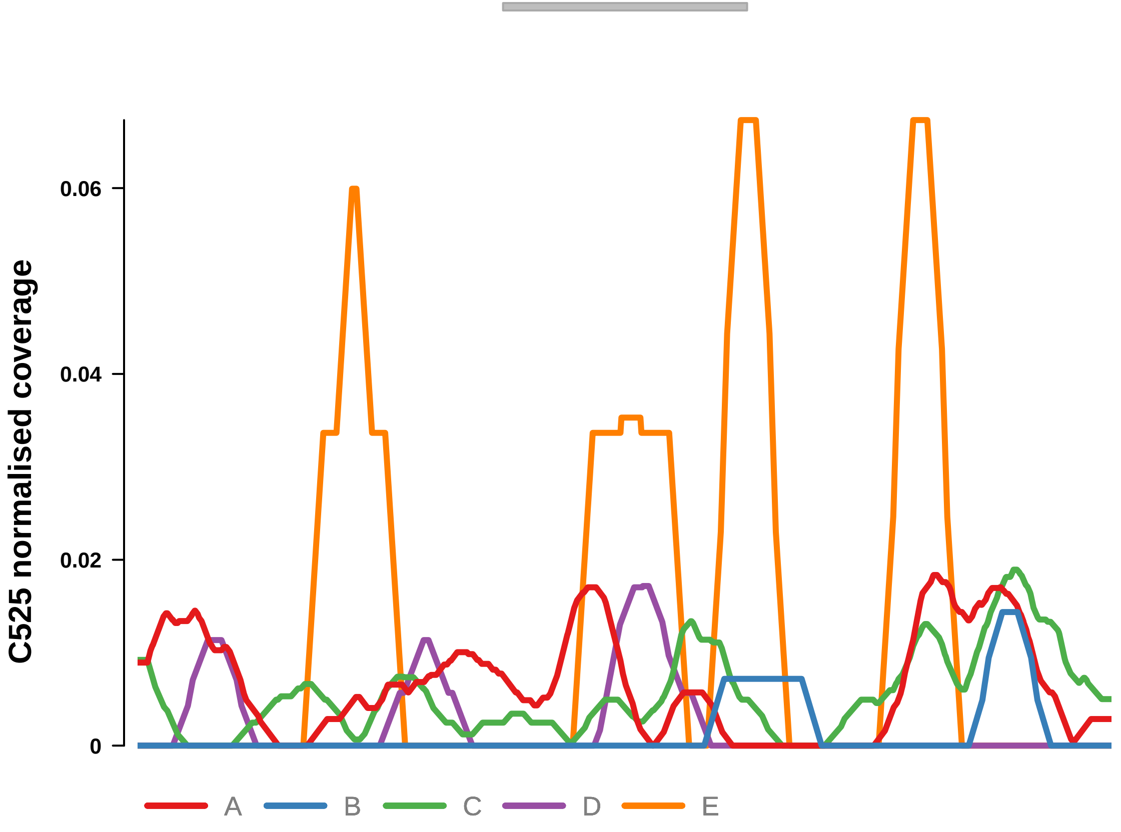


Putatively Clonal (logFC = −3.61)

| | **P.** | **Adjusted P.** |
|---|---|---|
| *~region vs. ~* | 3.2e−01 | NA |
| *~purity vs ~* | 8.5e−01 | 9.23e−01 |
| *~purity+region vs ~purity* | 3.3e−01 | 4.76e−01 |
| *Carcinoma vs. normal* | 7.12e−05 **** | 7.12e−05 **** |

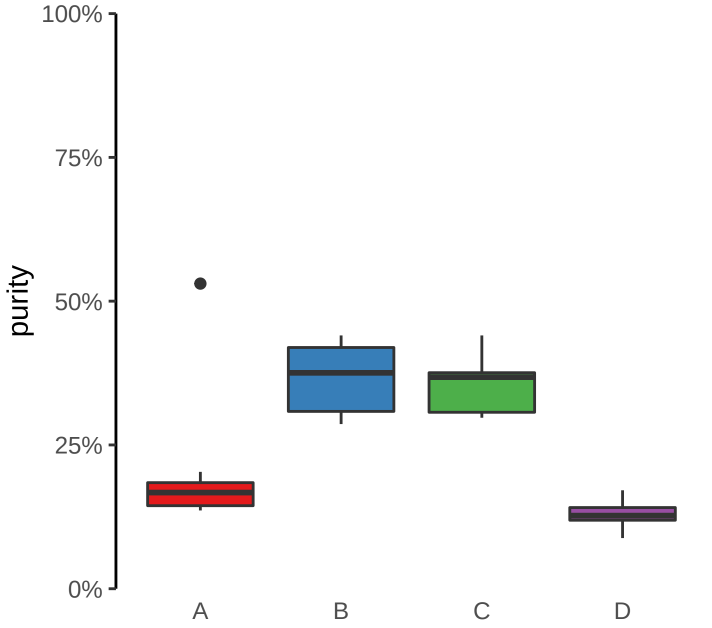

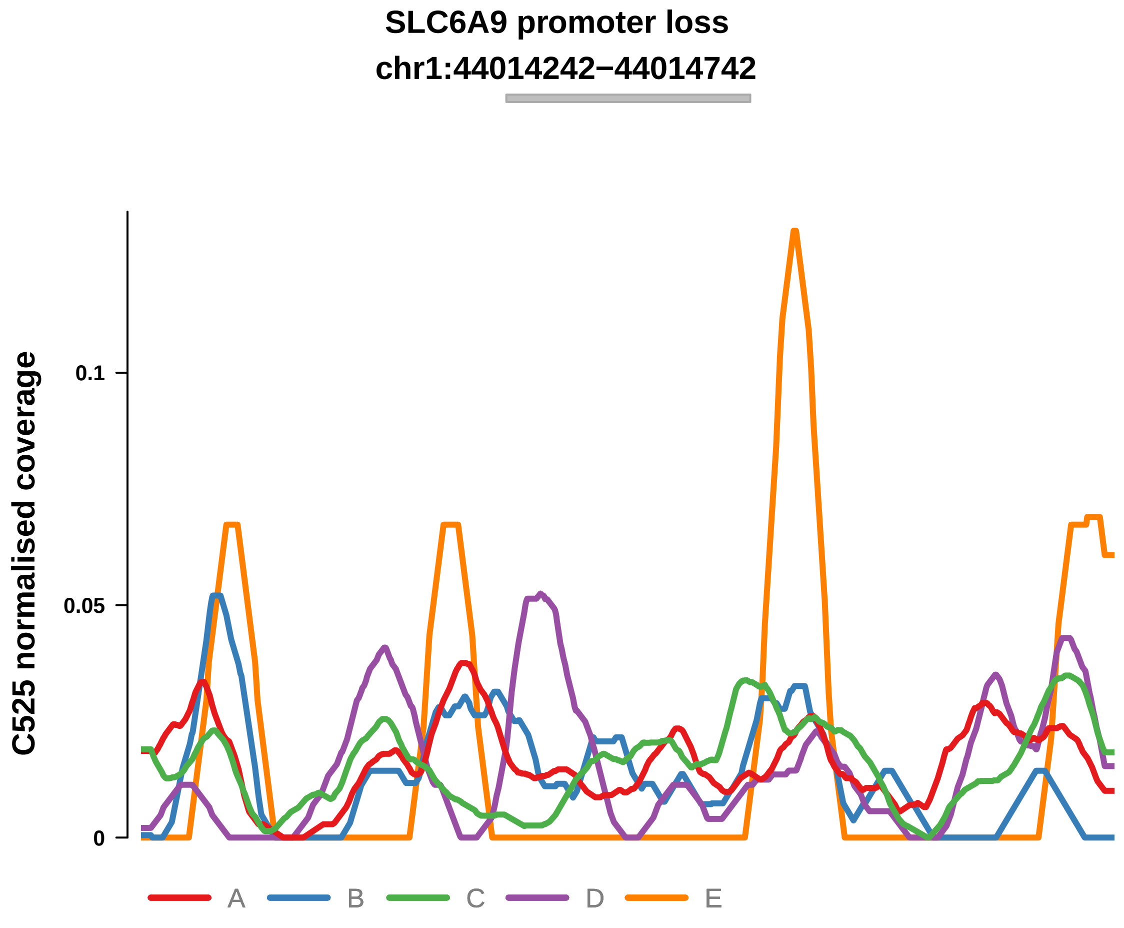


Putatively Clonal (logFC = −2.78)

| | P. | Adjusted P. |
|---|---|---|
| *~region vs. ~* | 1.83e−01 | 2.65e−01 |
| *~purity vs ~* | 5.45e−01 | 8.63e−01 |
| *~purity+region vs ~purity* | 2.17e−01 | 3.35e−01 |
| *Carcinoma vs. normal* | 2.14e−04 *** | 2.14e−04 *** |

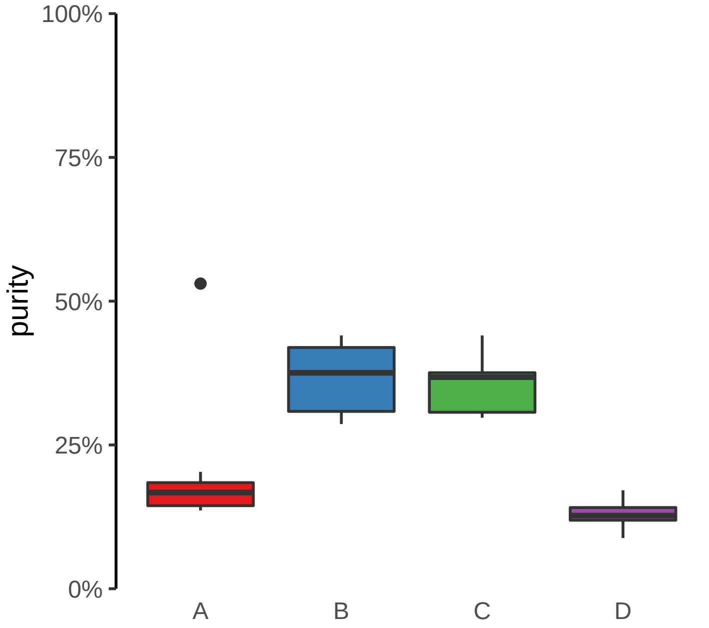

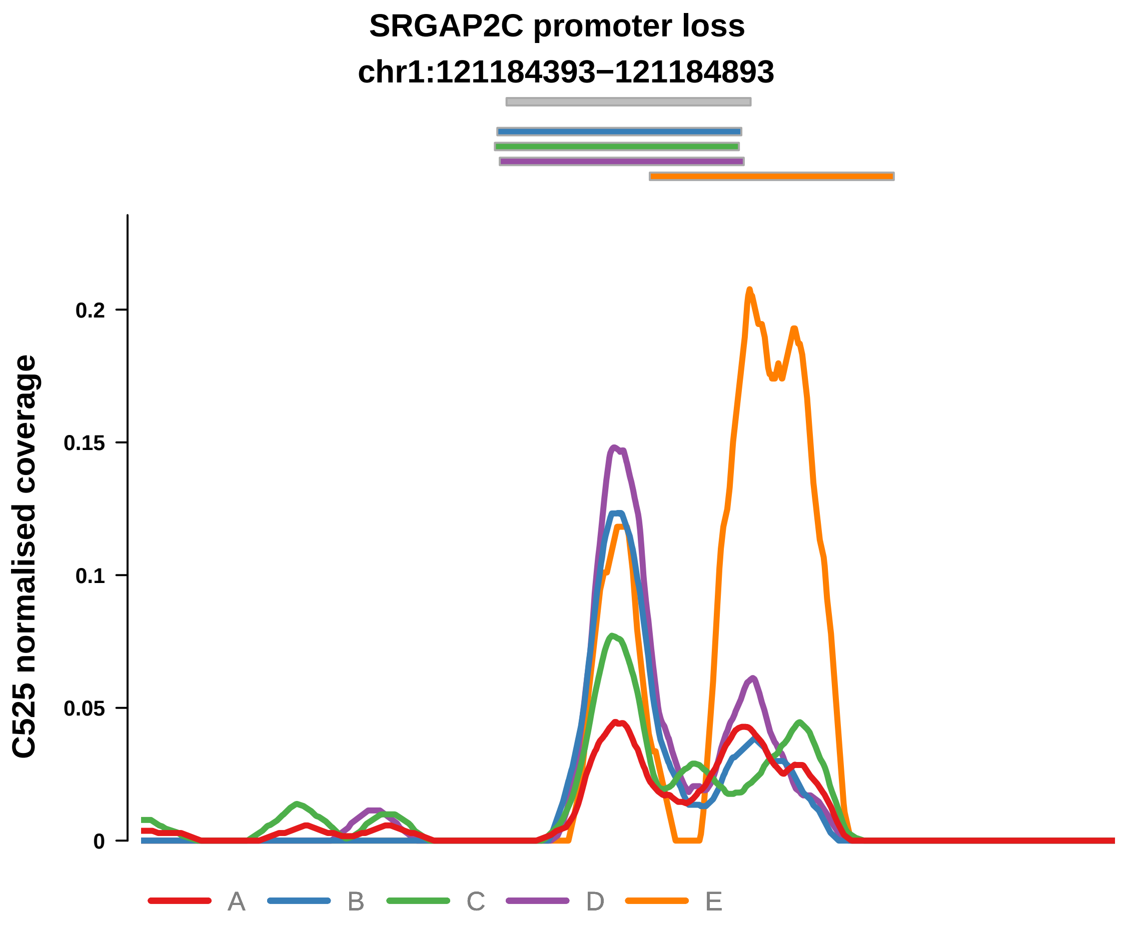


Putatively Clonal (logFC = −2.01)

| | P. | Adjusted P. |
|---|---|---|
| *~region vs. ~* | 2.12e−02 * | 5.35e−02 |
| *~purity vs ~* | 8.98e−02 | 7.18e−01 |
| *~purity+region vs ~purity* | 4.52e−02 * | 9.58e−02 |
| *Carcinoma vs. normal* | 7.76e−04 *** | 7.76e−04 *** |

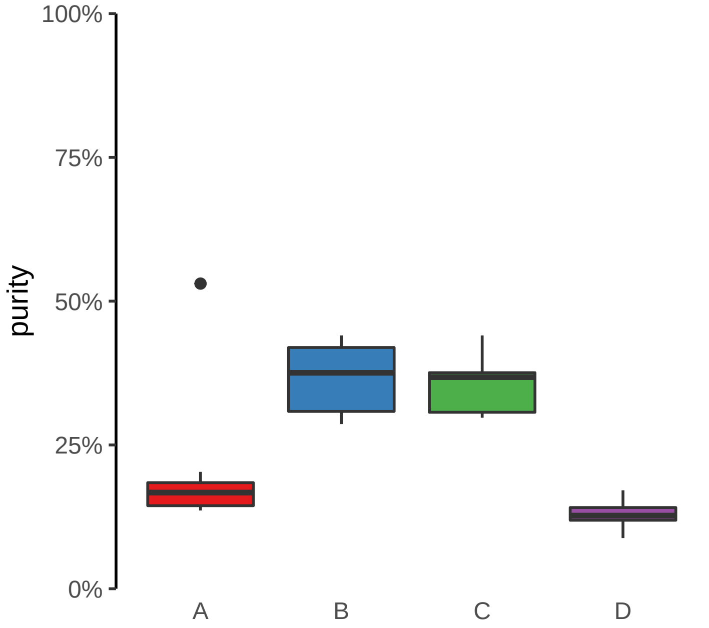

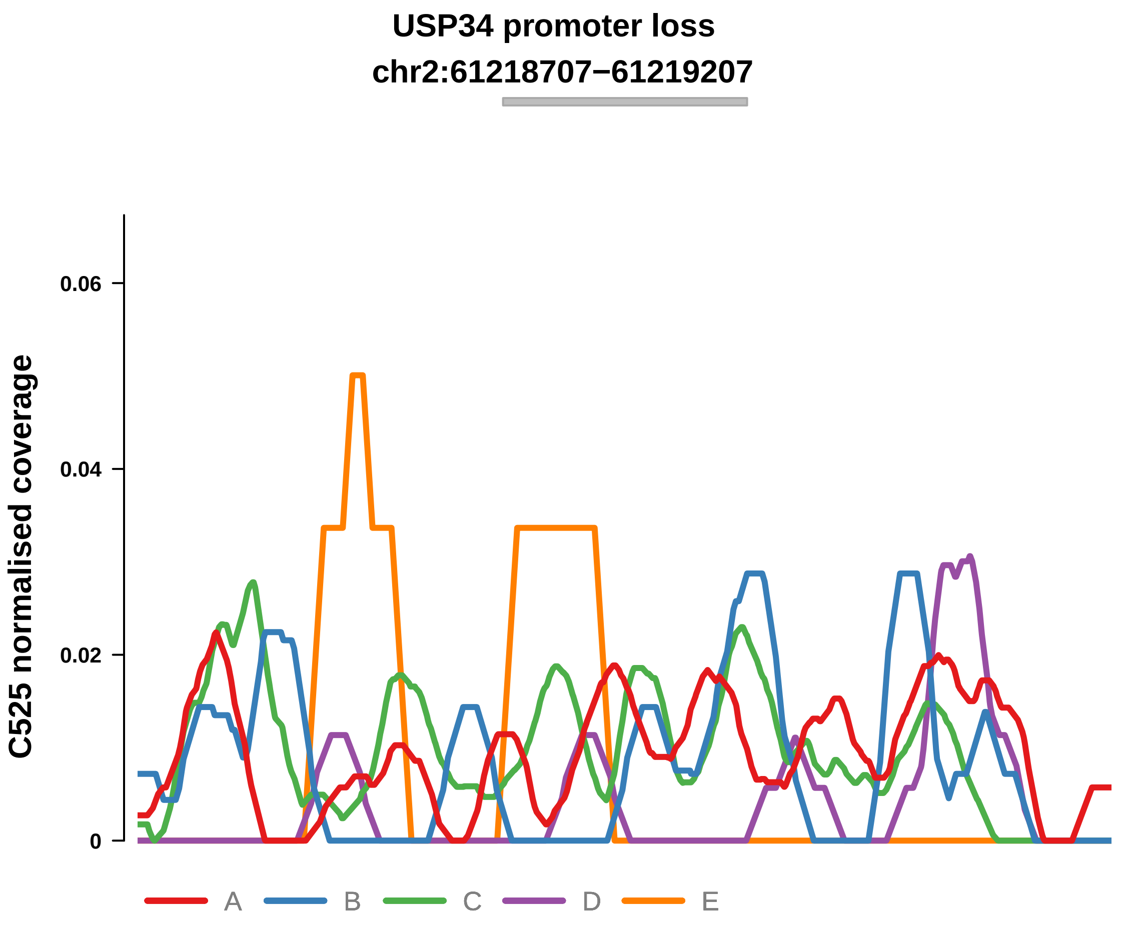


Putatively Clonal (logFC = −2.54)

| | P. | Adjusted P. |
|---|---|---|
| *~region vs. ~* | 1.55e−02 * | 4.44e−02 * |
| *~purity vs ~* | 1.74e−01 | 7.37e−01 |
| *~purity+region vs ~purity* | 4.13e−02 * | 9.32e−02 |
| *Carcinoma vs. normal* | 3.35e−03 ** | 3.35e−03 ** |

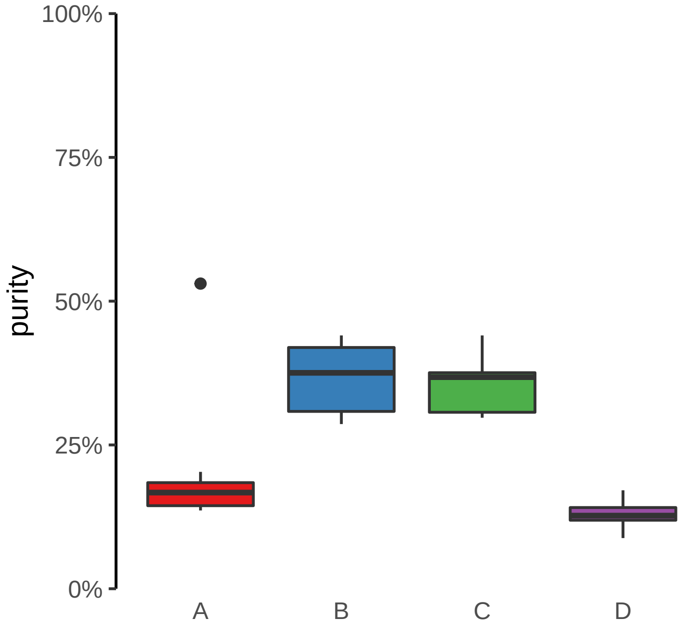

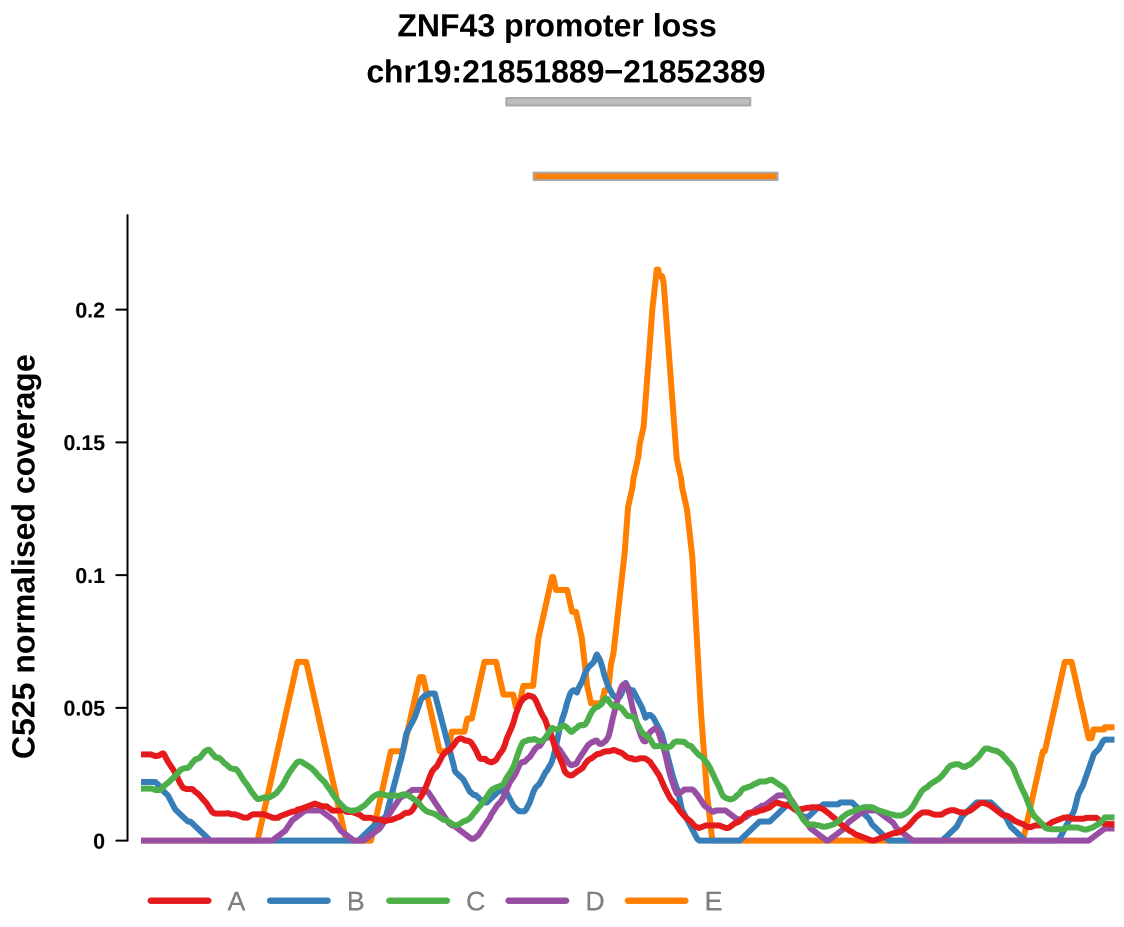


Putatively Clonal (logFC = −1.91)

| | P. | Adjusted P. |
|---|---|---|
| *~region vs. ~* | 9.58e−03 ** | 2.94e−02 * |
| *~purity vs ~* | 3.52e−01 | 7.75e−01 |
| *~purity+region vs ~purity* | 1.76e−02 * | 5.15e−02 |
| *Carcinoma vs. normal* | 1.59e−03 ** | 1.59e−03 ** |

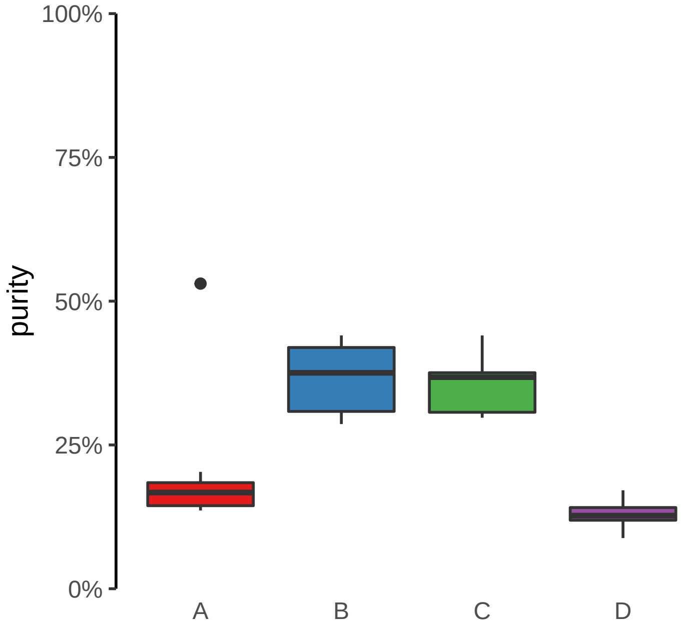

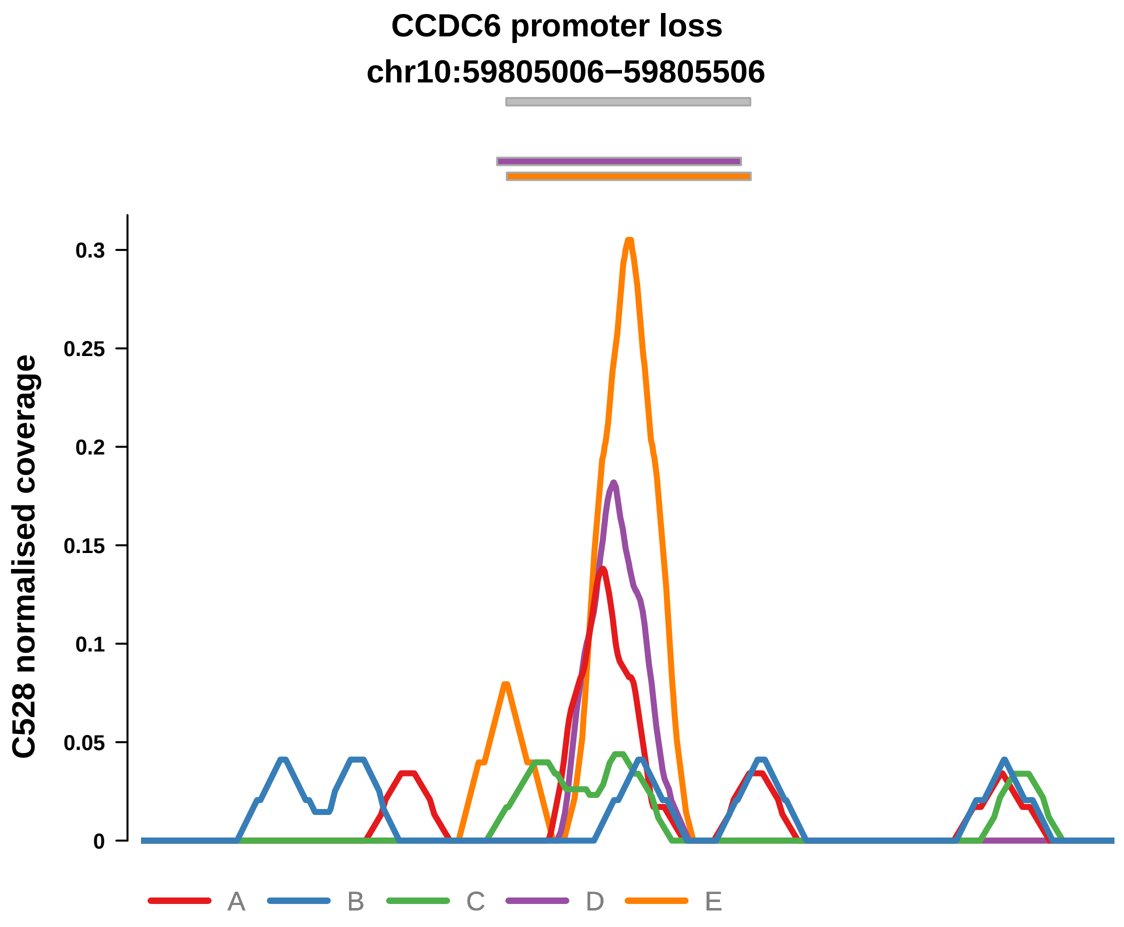


Putatively Clonal (logFC = −2.36)

| | P. | Adjusted P. |
|---|---|---|
| *~region vs. ~* | 2.67e−01 | 5.35e−01 |
| *~purity vs ~* | 5.95e−02 | 1.75e−01 |
| *~purity+region vs ~purity* | 7.94e−01 | 9.31e−01 |
| *Carcinoma vs. normal* | 3.41e−03 ** | 3.41e−03 ** |

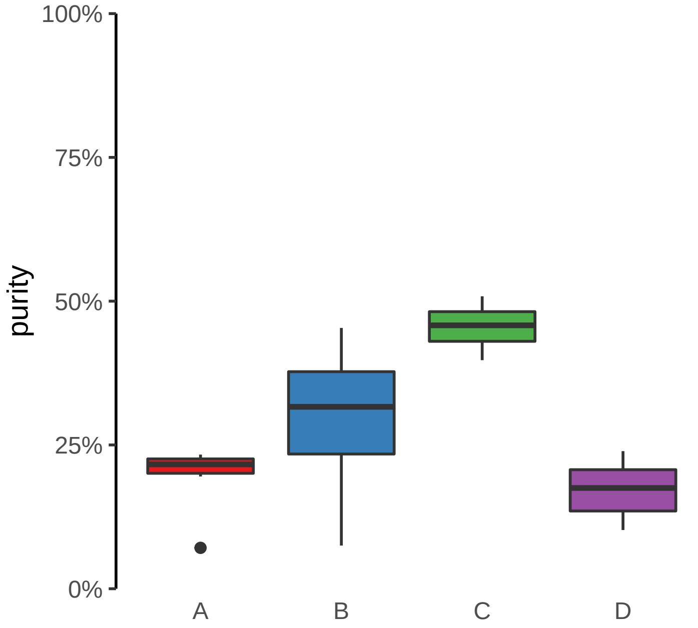

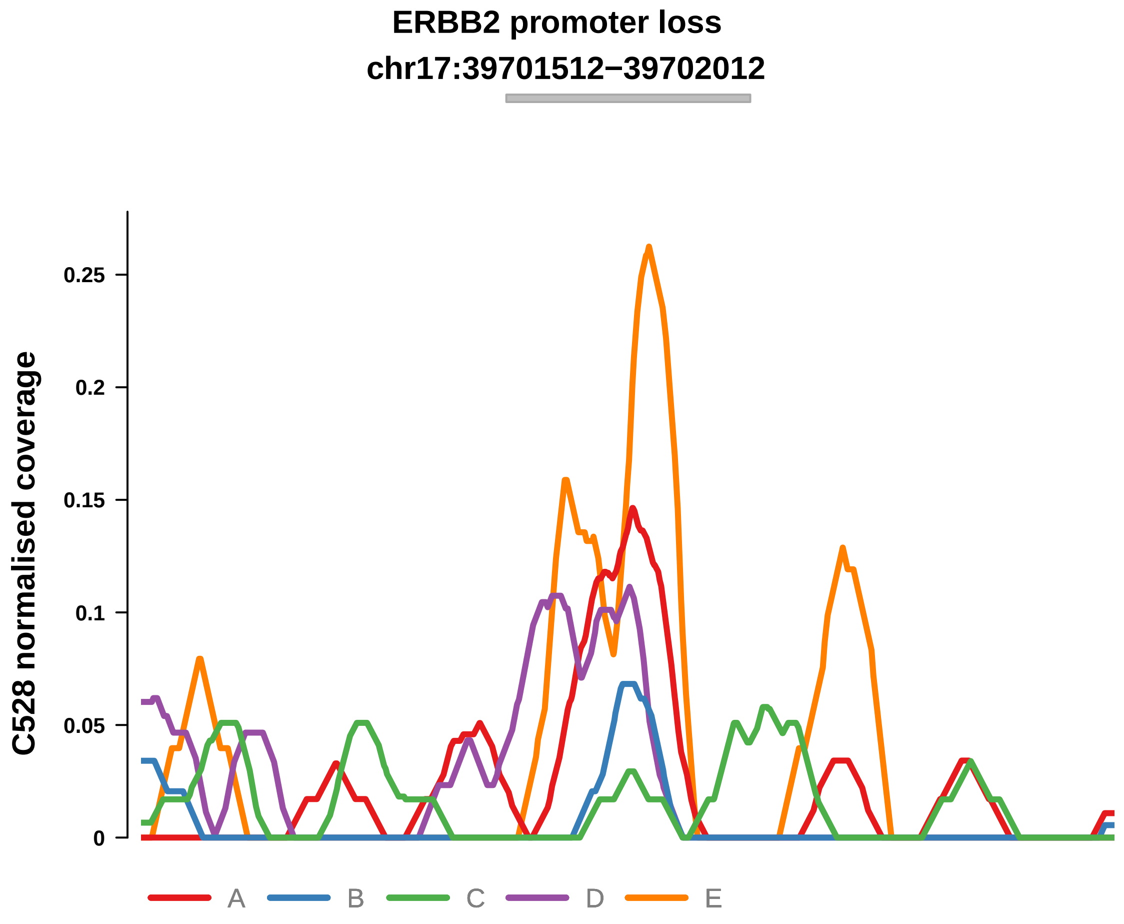


Putatively Clonal (logFC = −2.22)

| | P. | Adjusted P. |
|---|---|---|
| *~region vs. ~* | 9.82e−02 | 3.6e−01 |
| *~purity vs ~* | 4.92e−02 * | 1.62e−01 |
| *~purity+region vs ~purity* | 4.36e−01 | 7.86e−01 |
| *Carcinoma vs. normal* | 3.73e−03 ** | 3.73e−03 ** |

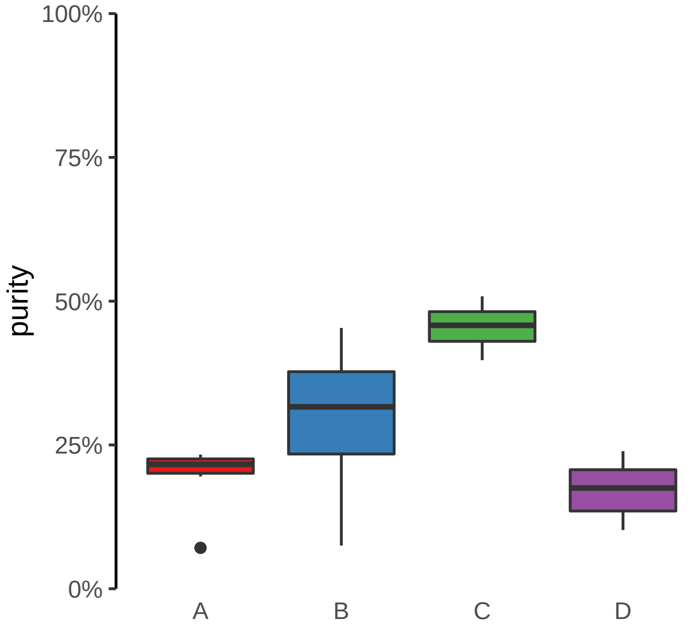

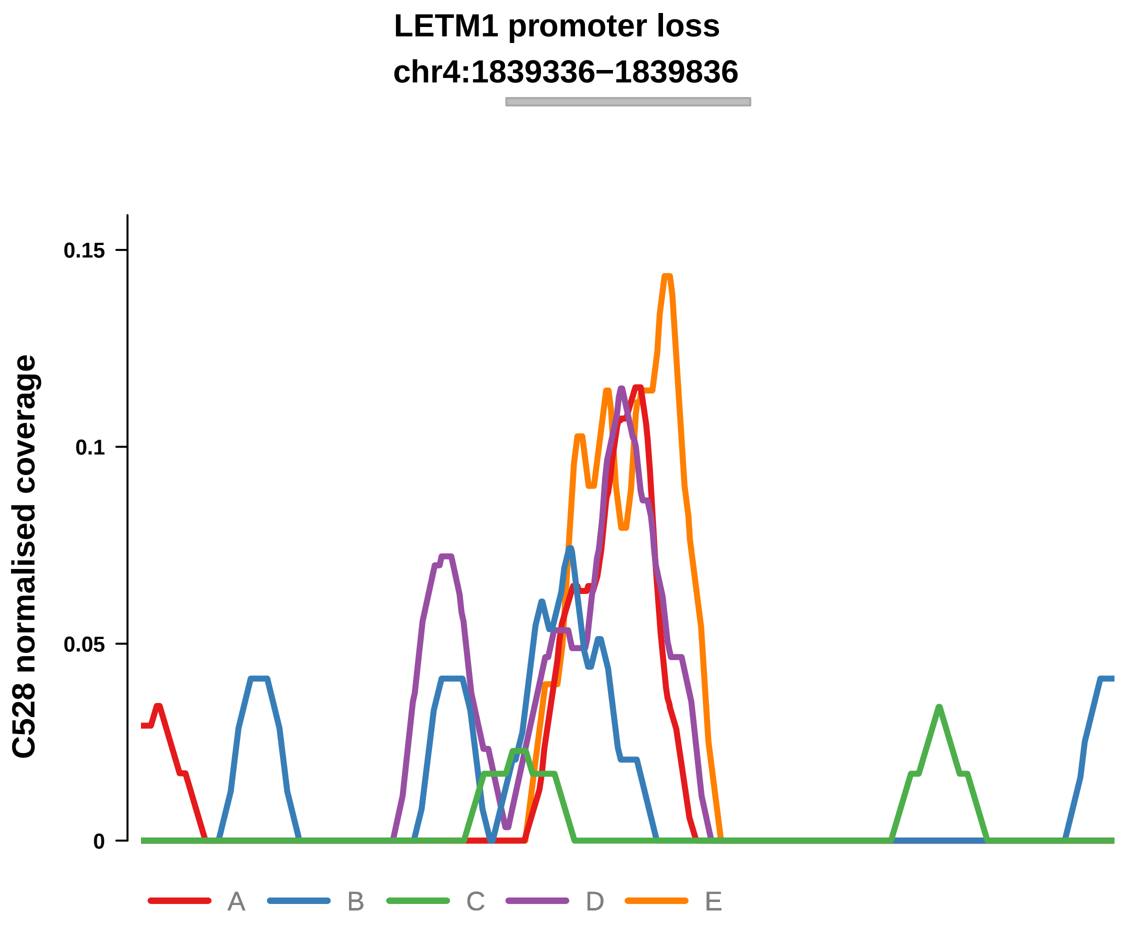


Putatively Clonal (logFC = −2.7)

| | P. | Adjusted P. |
|---|---|---|
| *~region vs. ~* | 1.9e−01 | 4.39e−01 |
| *~purity vs ~* | 1.22e−01 | 2.55e−01 |
| *~purity+region vs ~purity* | 4.1e−01 | 7.84e−01 |
| *Carcinoma vs. normal* | 1.58e−03 ** | 1.58e−03 ** |

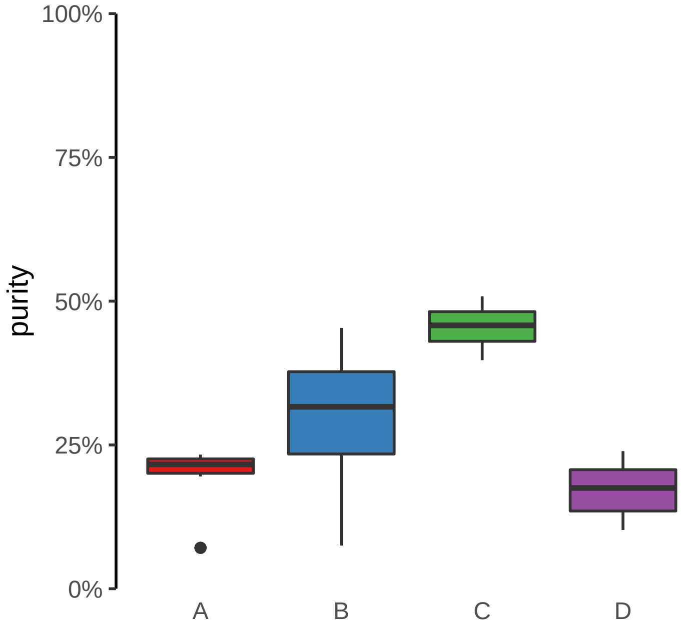

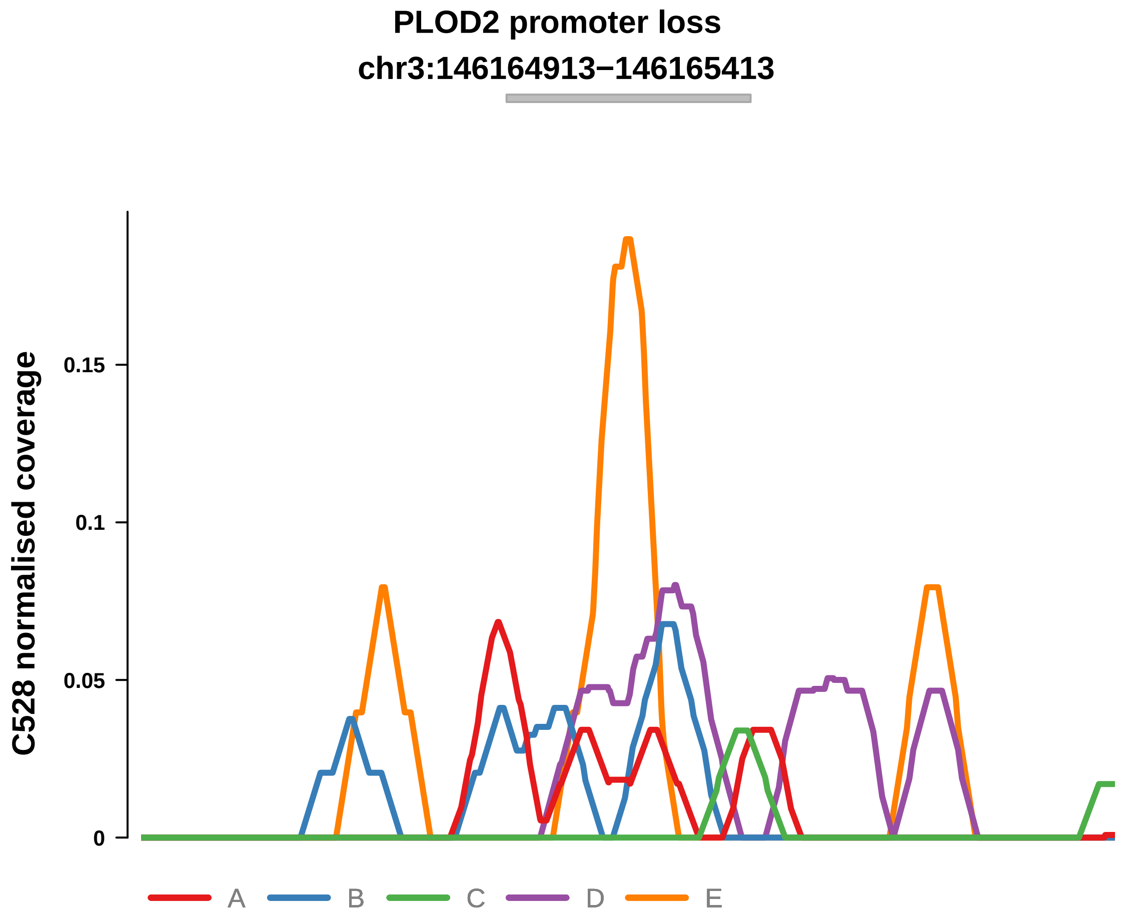


Putatively Clonal (logFC = −2.56)

| | P. | Adjusted P. |
|---|---|---|
| *~region vs. ~* | 4.31e−01 | 6.34e−01 |
| *~purity vs ~* | 3.53e−01 | NA |
| *~purity+region vs ~purity* | 5.73e−01 | 8.94e−01 |
| *Carcinoma vs. normal* | 3.52e−03 ** | 3.52e−03 ** |

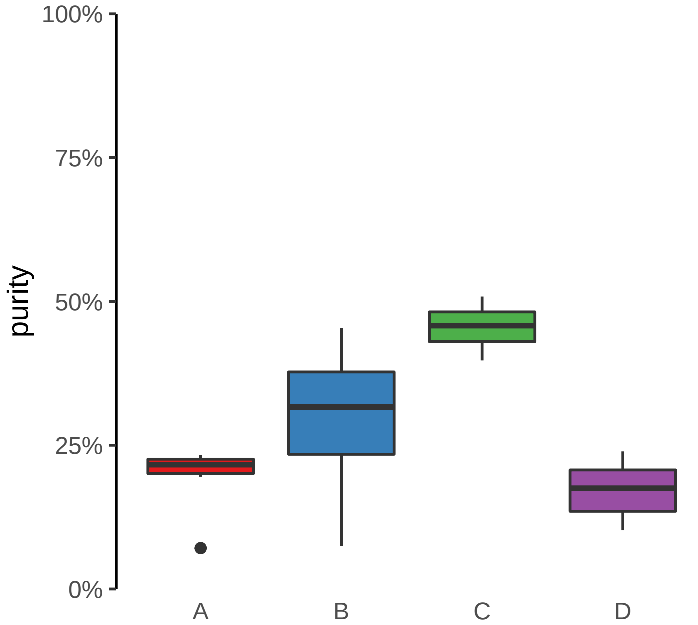

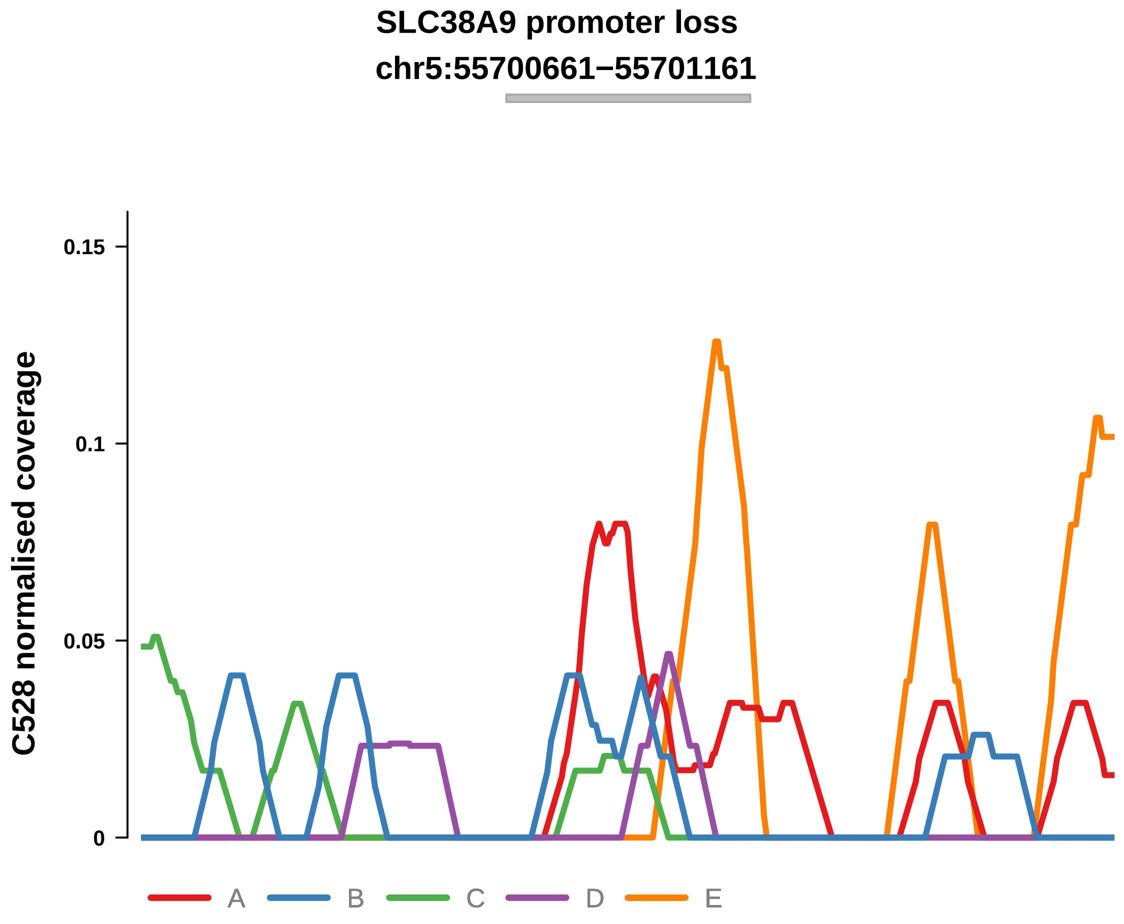


Putatively Clonal (logFC = −2.6)

| | P. | Adjusted P. |
|---|---|---|
| *~region vs. ~* | 7.48e−01 | 8.89e−01 |
| *~purity vs ~* | 3.02e−01 | NA |
| *~purity+region vs ~purity* | 7.31e−01 | 9.17e−01 |
| *Carcinoma vs. normal* | 7.97e−03 ** | 7.97e−03 ** |

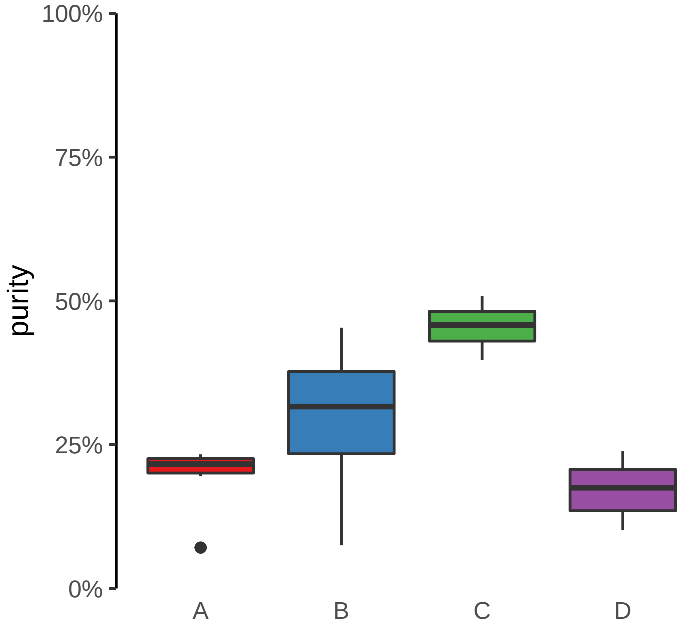

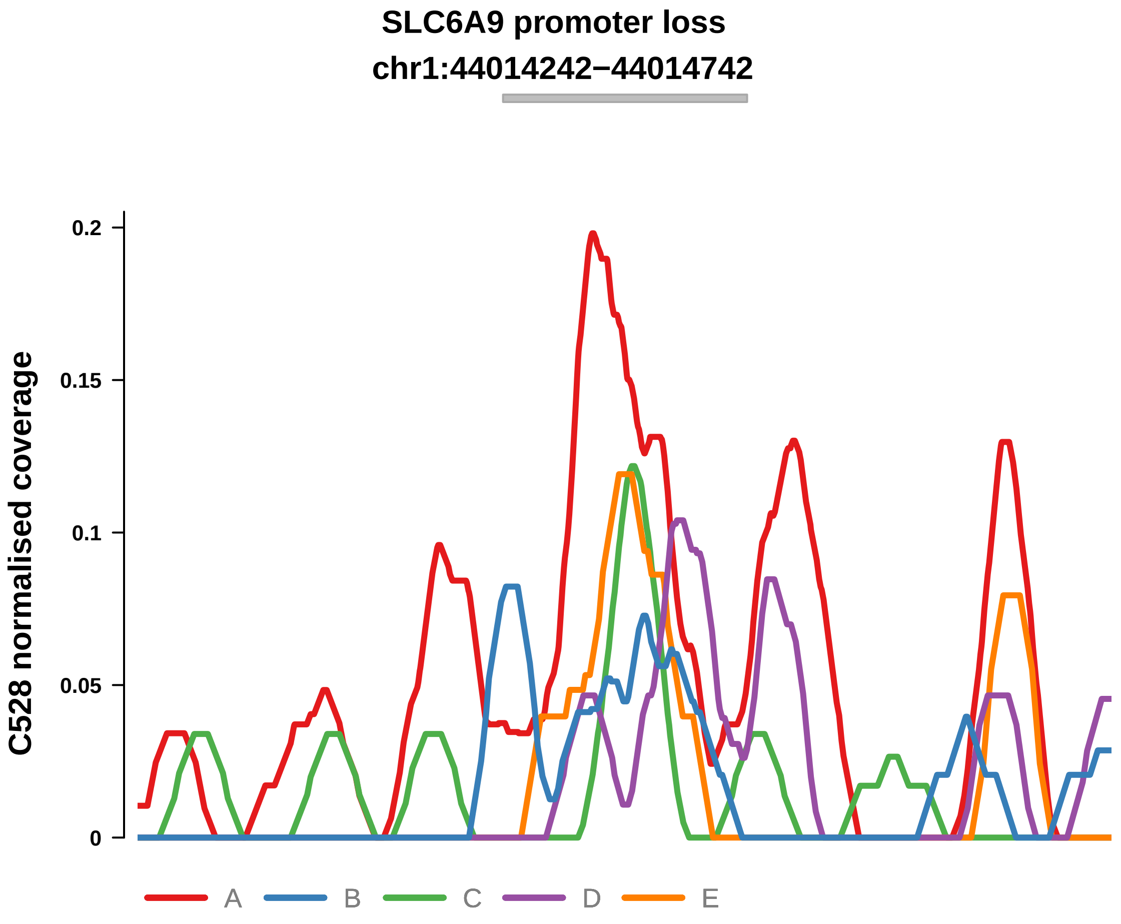


Putatively Clonal (logFC = −2.02)

| | P. | Adjusted P. |
|---|---|---|
| *~region vs. ~* | 2.21e−01 | 4.85e−01 |
| *~purity vs ~* | 1.2e−01 | 2.55e−01 |
| *~purity+region vs ~purity* | 5.48e−01 | 8.94e−01 |
| *Carcinoma vs. normal* | 6.69e−03 ** | 6.69e−03 ** |

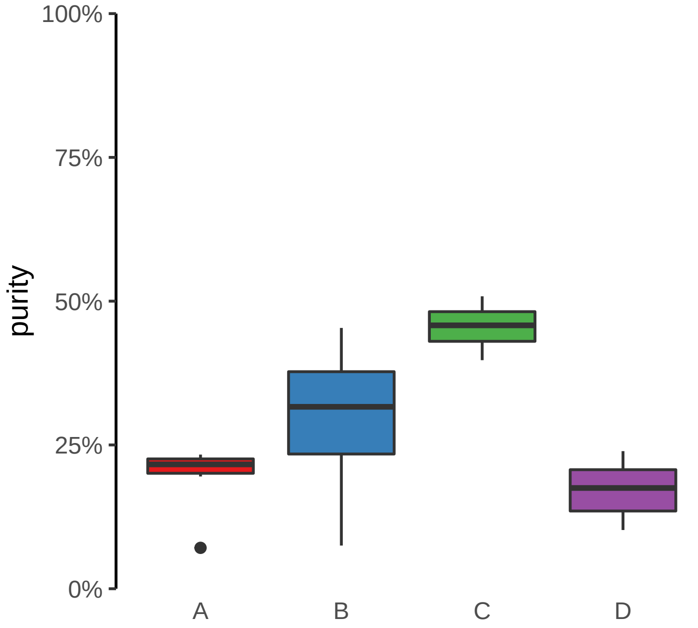

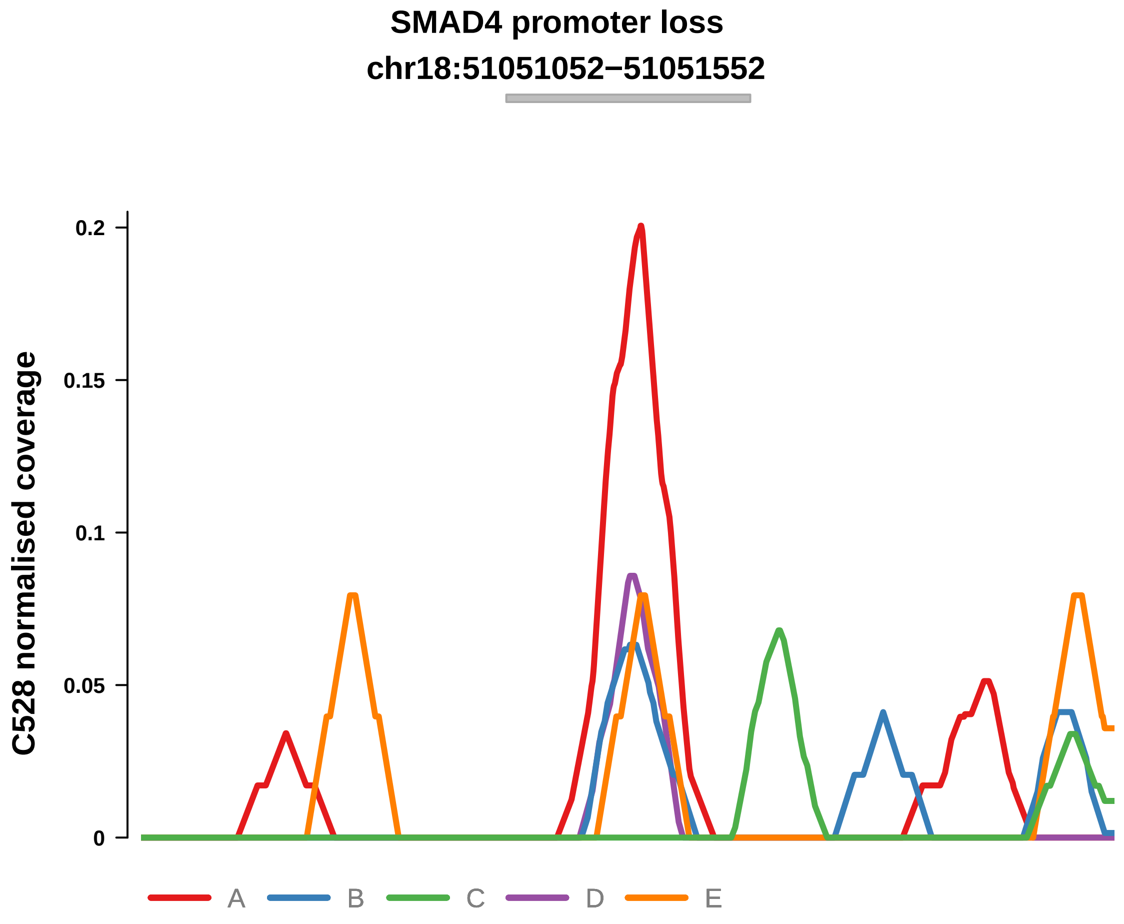


Putatively Clonal (logFC = −10.9)

| | P. | Adjusted P. |
|---|---|---|
| *~region vs. ~* | 7.2e−01 | 8.89e−01 |
| *~purity vs ~* | 2.46e−01 | NA |
| *~purity+region vs ~purity* | 9.57e−01 | 1e+00 |
| *Carcinoma vs. normal* | 9e−06 **** | 9e−06 **** |

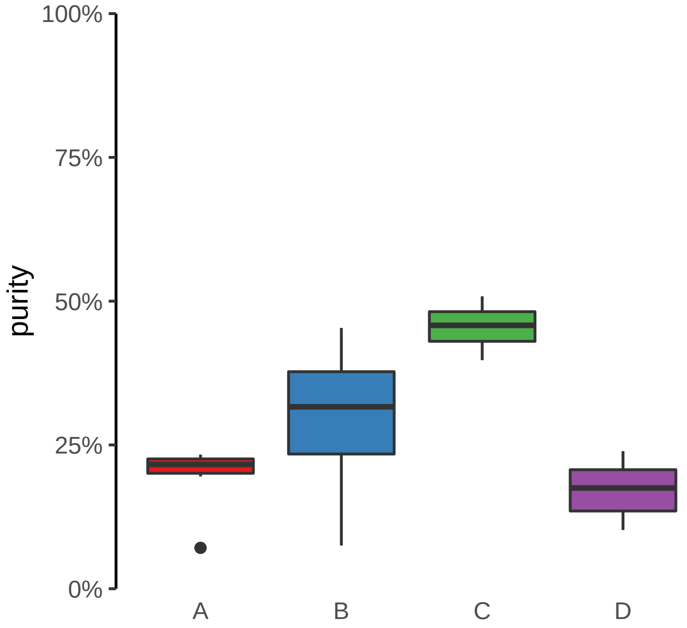

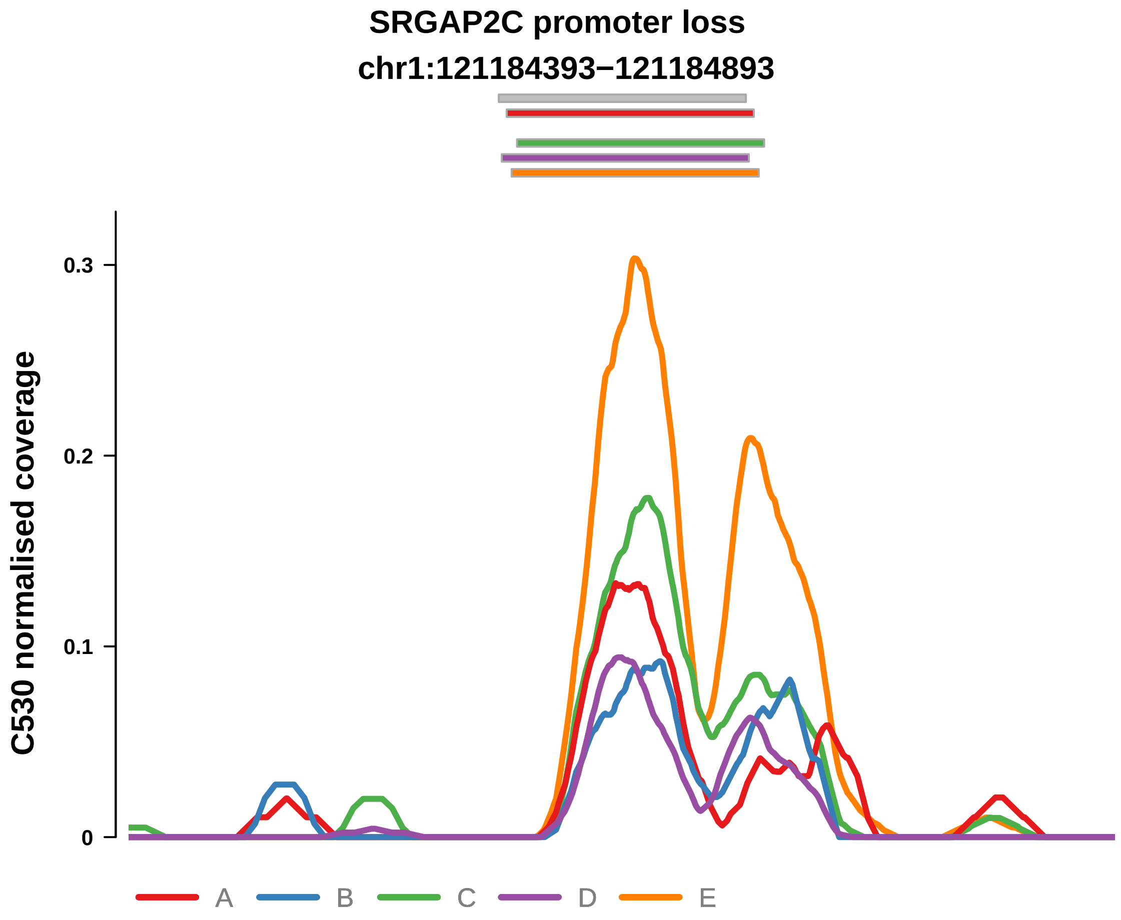


Putatively Clonal (logFC = −1.51)

| | P. | Adjusted P. |
|---|---|---|
| *~region vs. ~* | 3.9e−01 | 7.45e−01 |
| *~purity vs ~* | 3.79e−01 | 8.26e−01 |
| *~purity+region vs ~purity* | 4.89e−01 | 7.59e−01 |
| *Carcinoma vs. normal* | 6.8e−03 ** | 6.8e−03 ** |

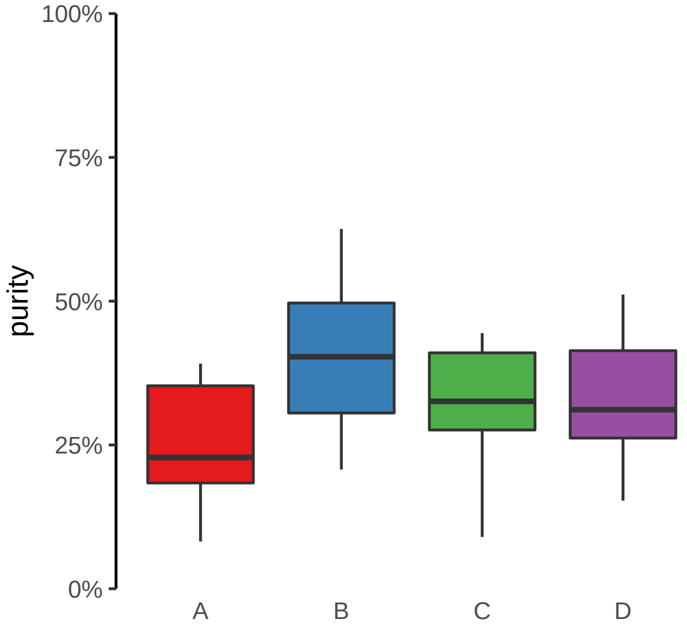

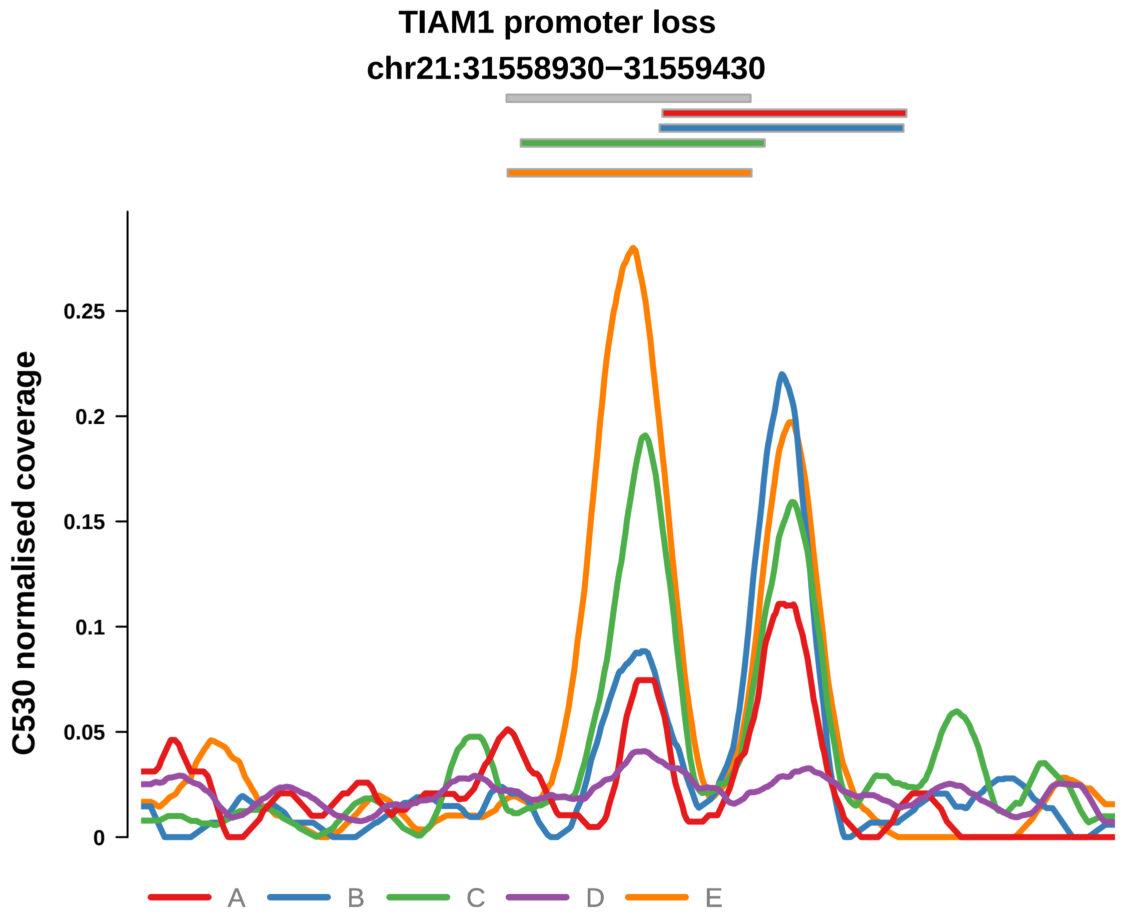


Putatively Clonal (logFC = −1.52)

| | P. | Adjusted P. |
|---|---|---|
| *~region vs. ~* | 5.09e−01 | 8.19e−01 |
| *~purity vs ~* | 4.47e−01 | 8.56e−01 |
| *~purity+region vs ~purity* | 5.2e−01 | 7.8e−01 |
| *Carcinoma vs. normal* | 6.88e−03 ** | 6.88e−03 ** |

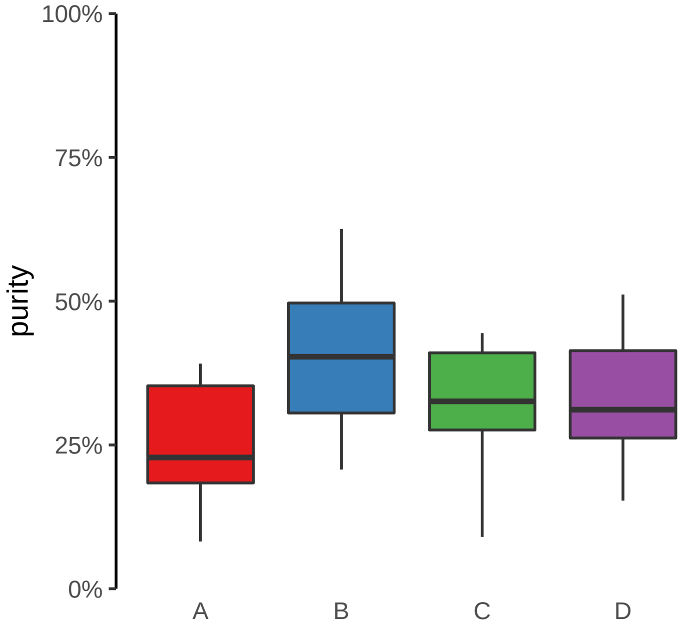

**ZNF43 promoter loss**

**chr19:21851889−21852389**

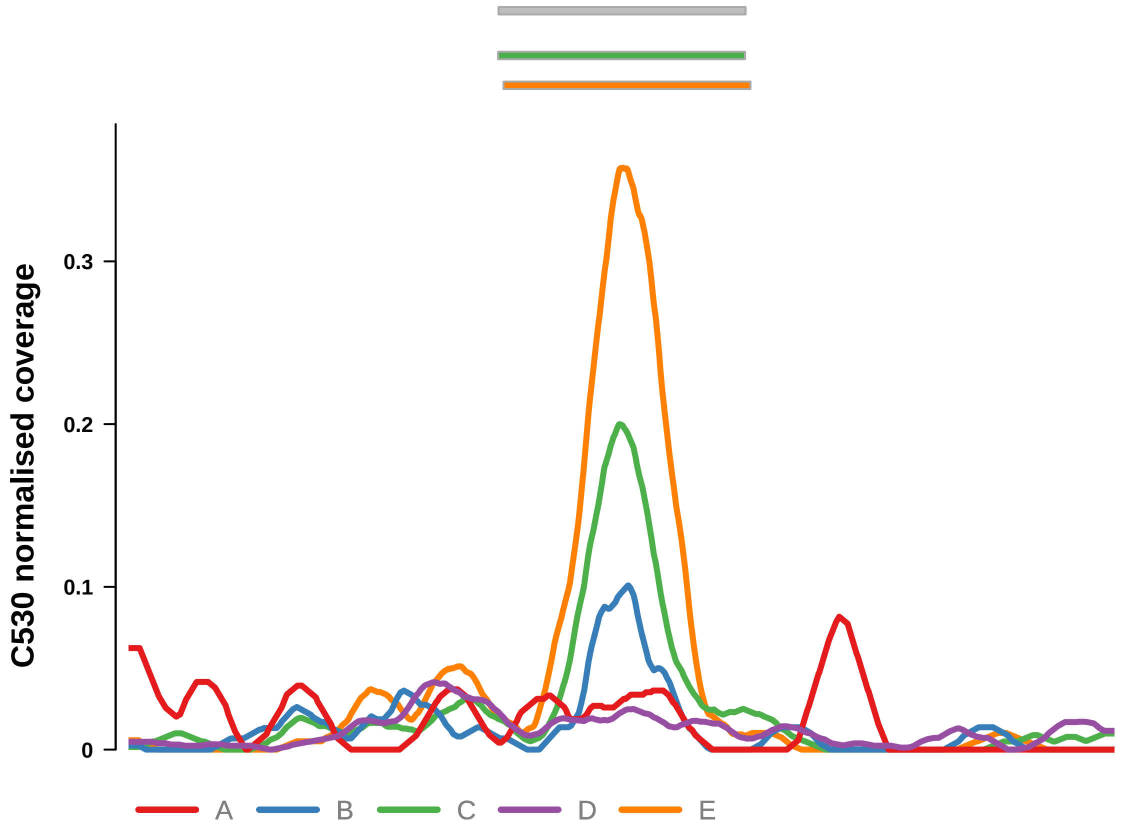


Putatively Clonal (logFC = −2.09)

| | P. | Adjusted P. |
|---|---|---|
| *~region vs. ~* | 4.87e−03 ** | 7.14e−02 |
| *~purity vs ~* | 2.84e−01 | 7.57e−01 |
| *~purity+region vs ~purity* | 5.4e−03 ** | 7.92e−02 |
| *Carcinoma vs. normal* | 3.21e−04 *** | 3.21e−04 *** |

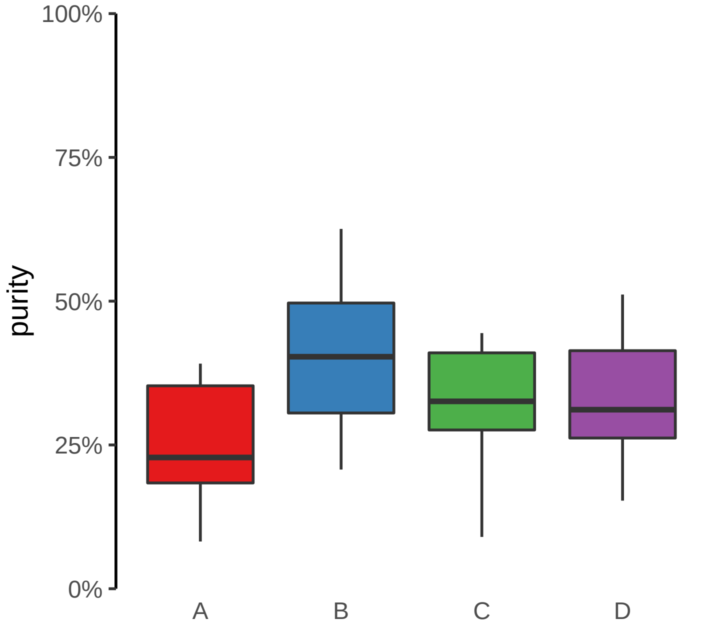

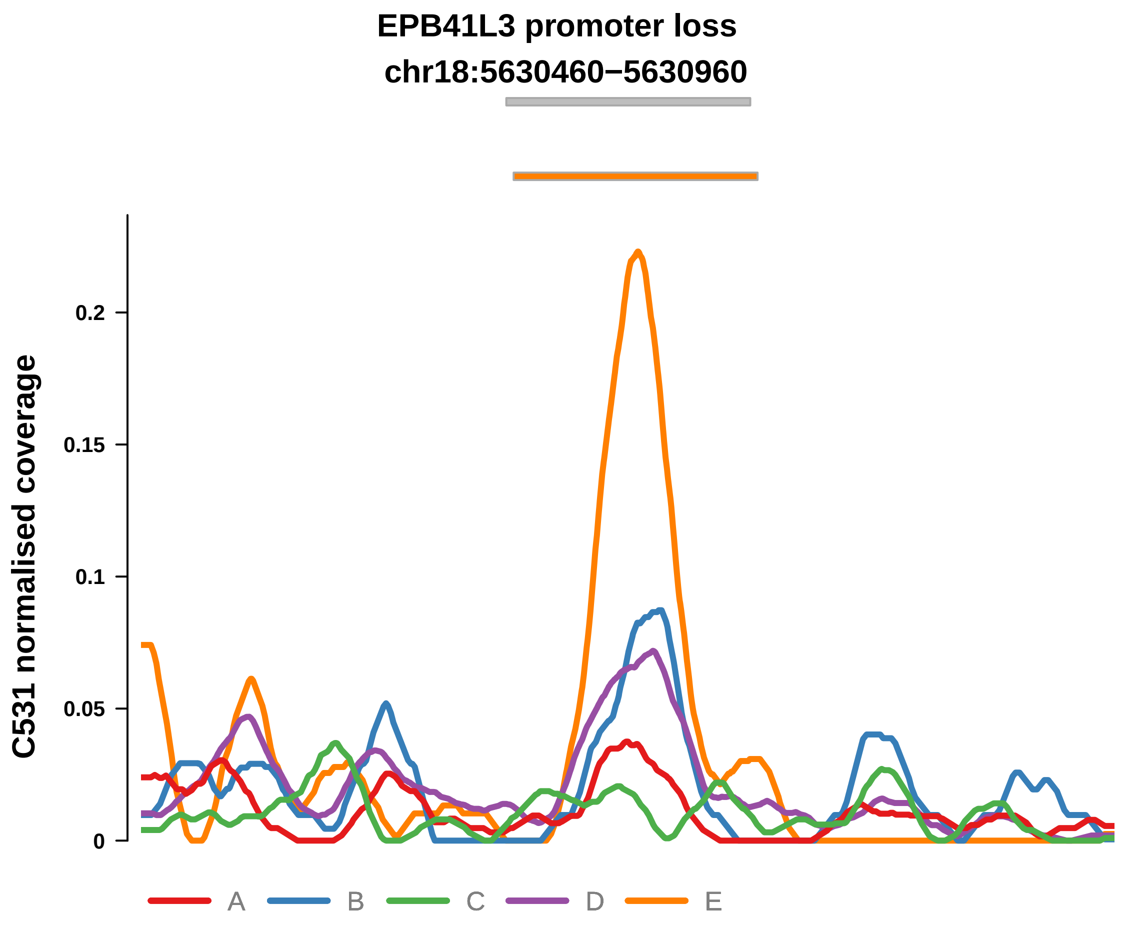


Putatively Clonal (logFC = −2.06)

| | P. | Adjusted P. |
|---|---|---|
| *~region vs. ~* | 6.61e−01 | 7.47e−01 |
| *~purity vs ~* | 2.22e−01 | 3.83e−01 |
| *~purity+region vs ~purity* | 9.5e−01 | 9.61e−01 |
| *Carcinoma vs. normal* | 2.98e−03 ** | 2.98e−03 ** |

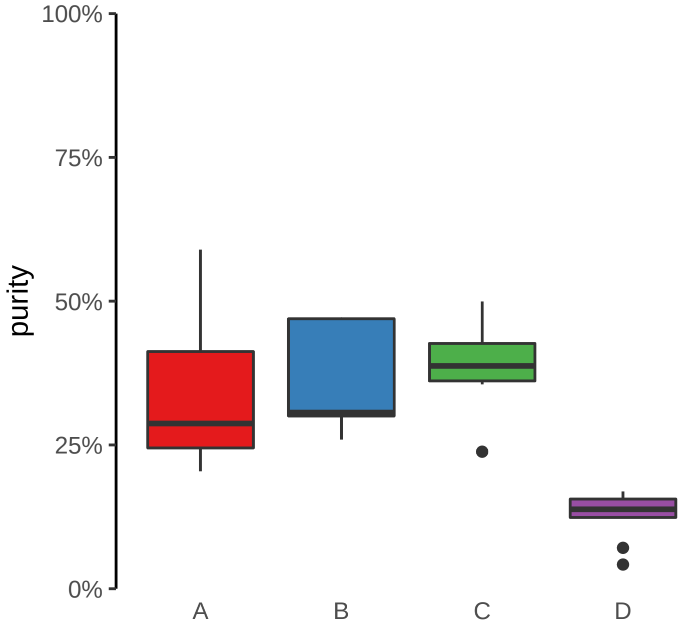

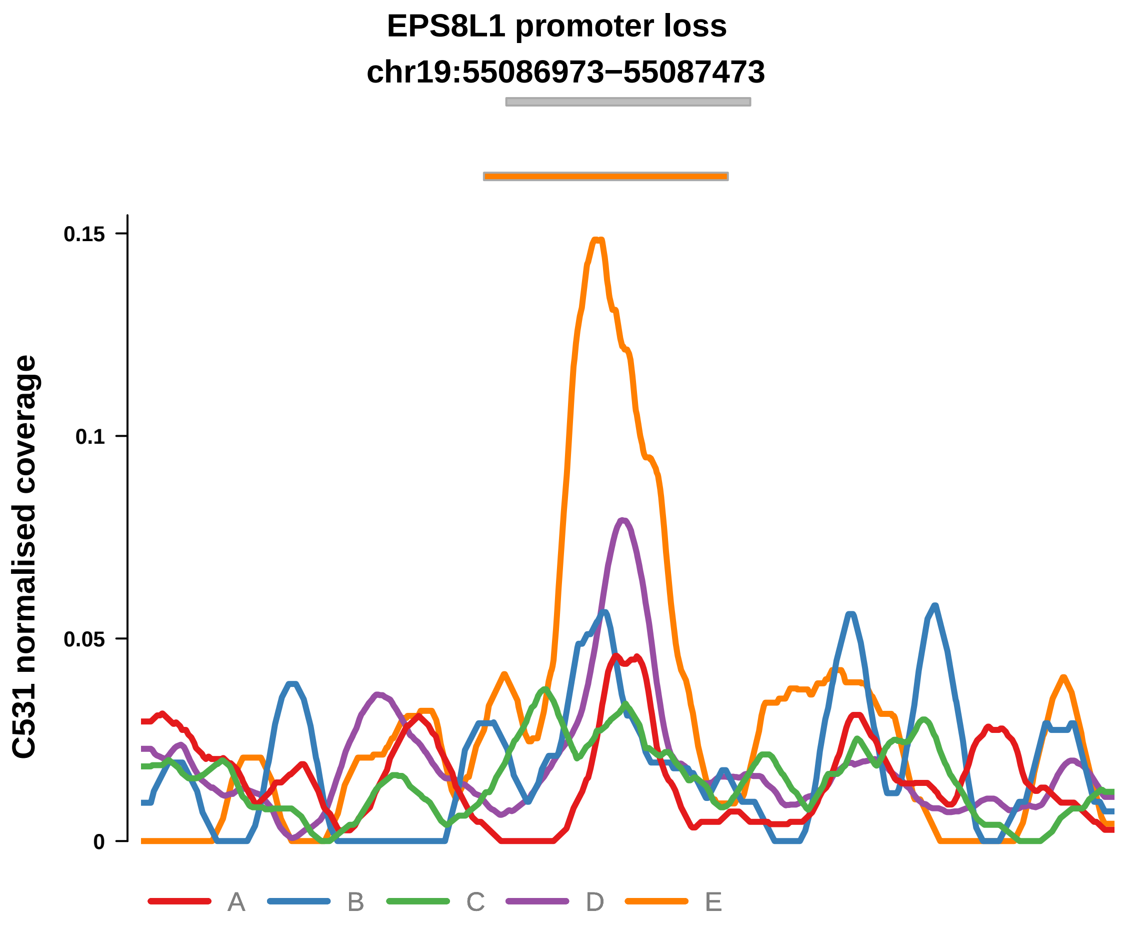


Putatively Clonal (logFC = −2)

| | P. | Adjusted P. |
|---|---|---|
| *~region vs. ~* | 4.03e−01 | 5.52e−01 |
| *~purity vs ~* | 6.18e−01 | 7.52e−01 |
| *~purity+region vs ~purity* | 4.28e−01 | 7.11e−01 |
| *Carcinoma vs. normal* | 7.05e−03 ** | 7.05e−03 ** |

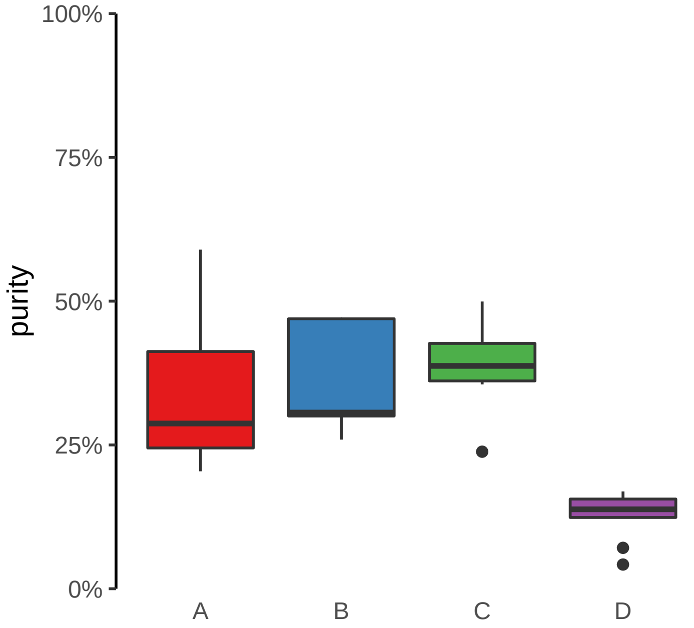

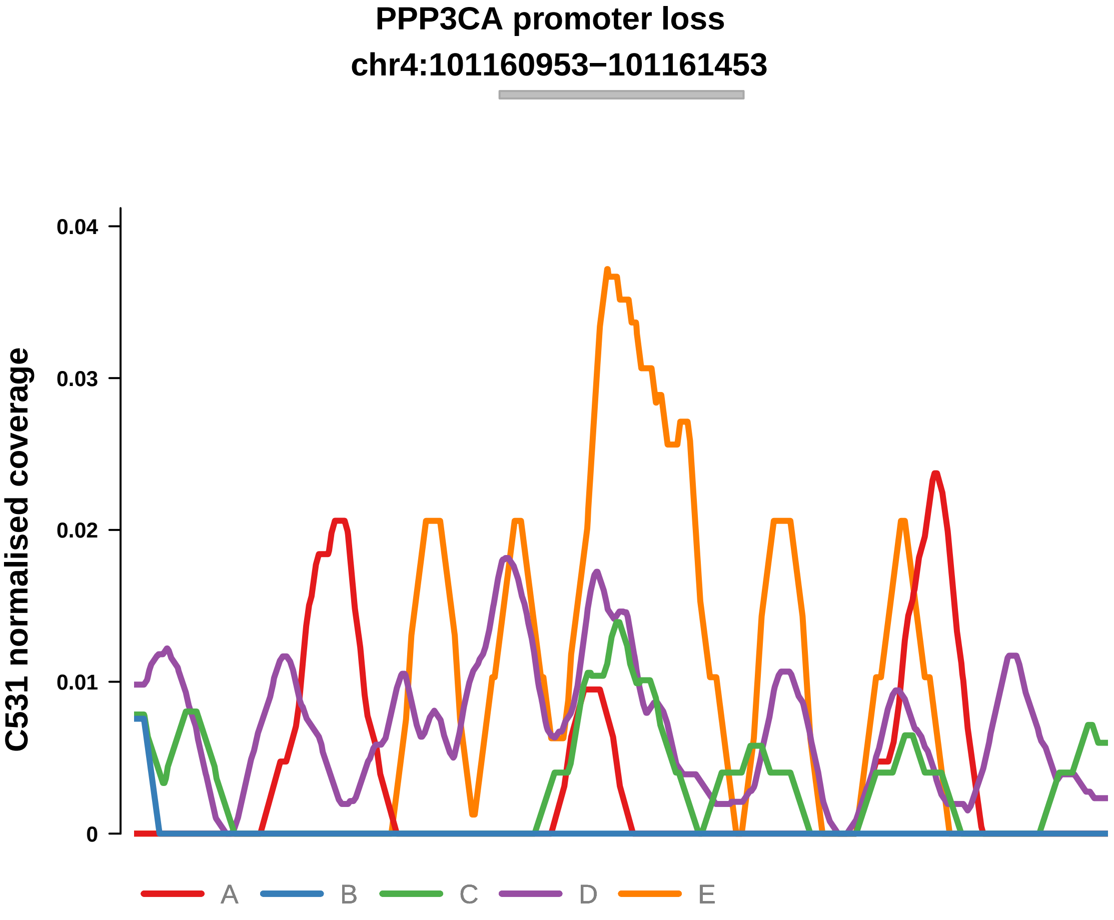


Putatively Clonal (logFC = −3.34)

| | P. | Adjusted P. |
|---|---|---|
| *~region vs. ~* | 3.24e−01 | NA |
| *~purity vs ~* | 1.67e−01 | 3.27e−01 |
| *~purity+region vs ~purity* | 6.16e−01 | 8.52e−01 |
| *Carcinoma vs. normal* | 4.28e−03 ** | 4.28e−03 ** |

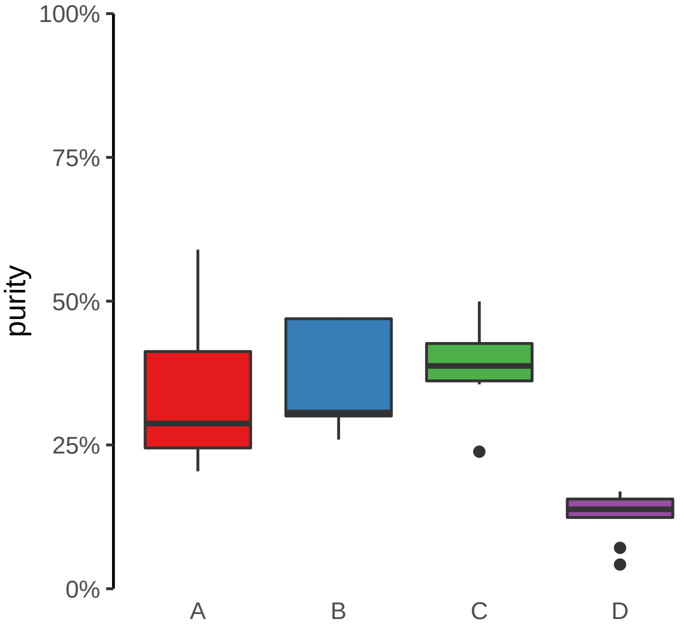

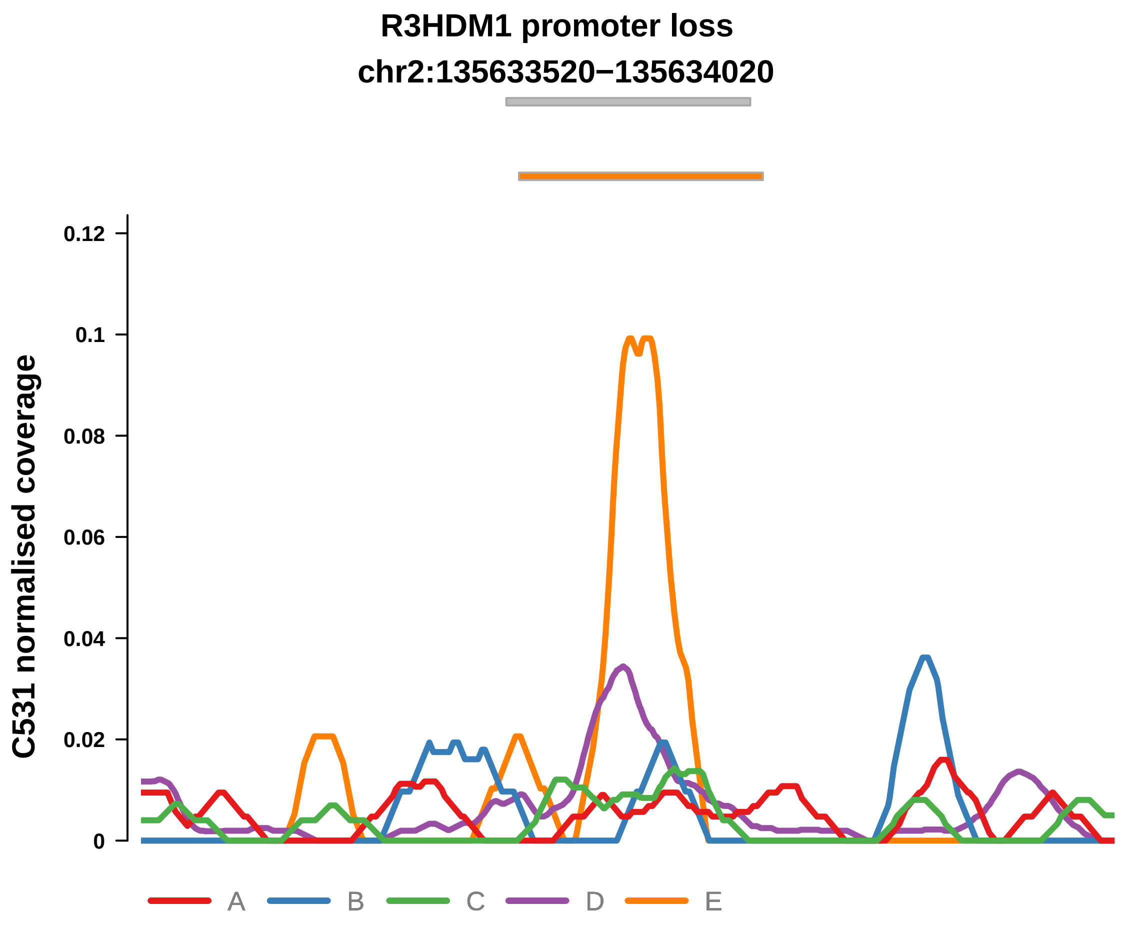


Putatively Clonal (logFC = −3.77)

| | P. | Adjusted P. |
|---|---|---|
| *~region vs. ~* | 4.01e−01 | NA |
| *~purity vs ~* | 2.69e−01 | 4.07e−01 |
| *~purity+region vs ~purity* | 6.04e−01 | 8.52e−01 |
| *Carcinoma vs. normal* | 7.28e−04 *** | 7.28e−04 *** |

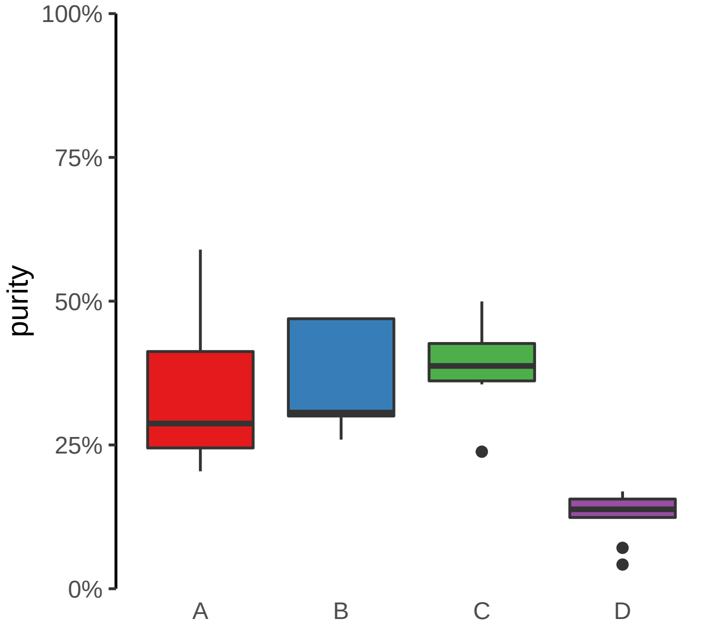

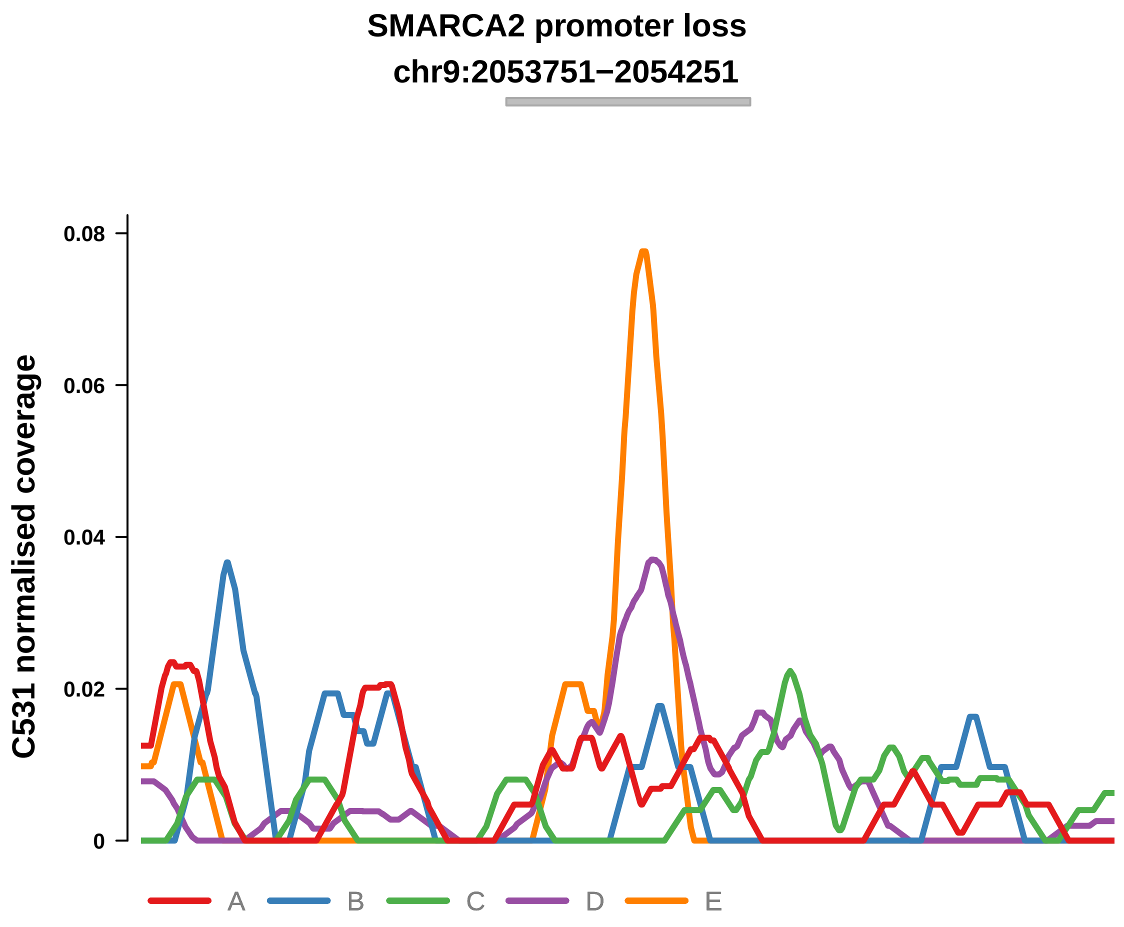


Putatively Clonal (logFC = −10.67)

| | P. | Adjusted P. |
|---|---|---|
| *~region vs. ~* | 1.46e−01 | NA |
| *~purity vs ~* | 1.18e−02 * | 5.87e−02 |
| *~purity+region vs ~purity* | 4.45e−01 | 7.26e−01 |
| *Carcinoma vs. normal* | 6.13e−05 **** | 6.13e−05 **** |

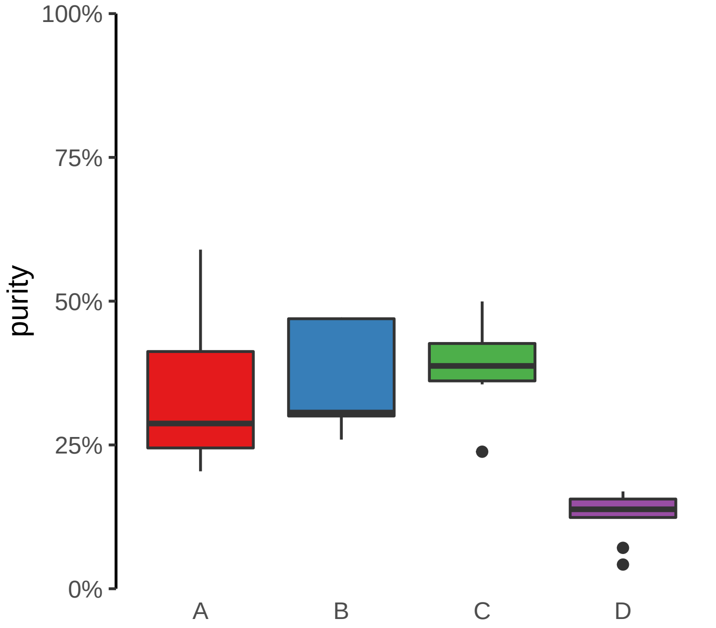

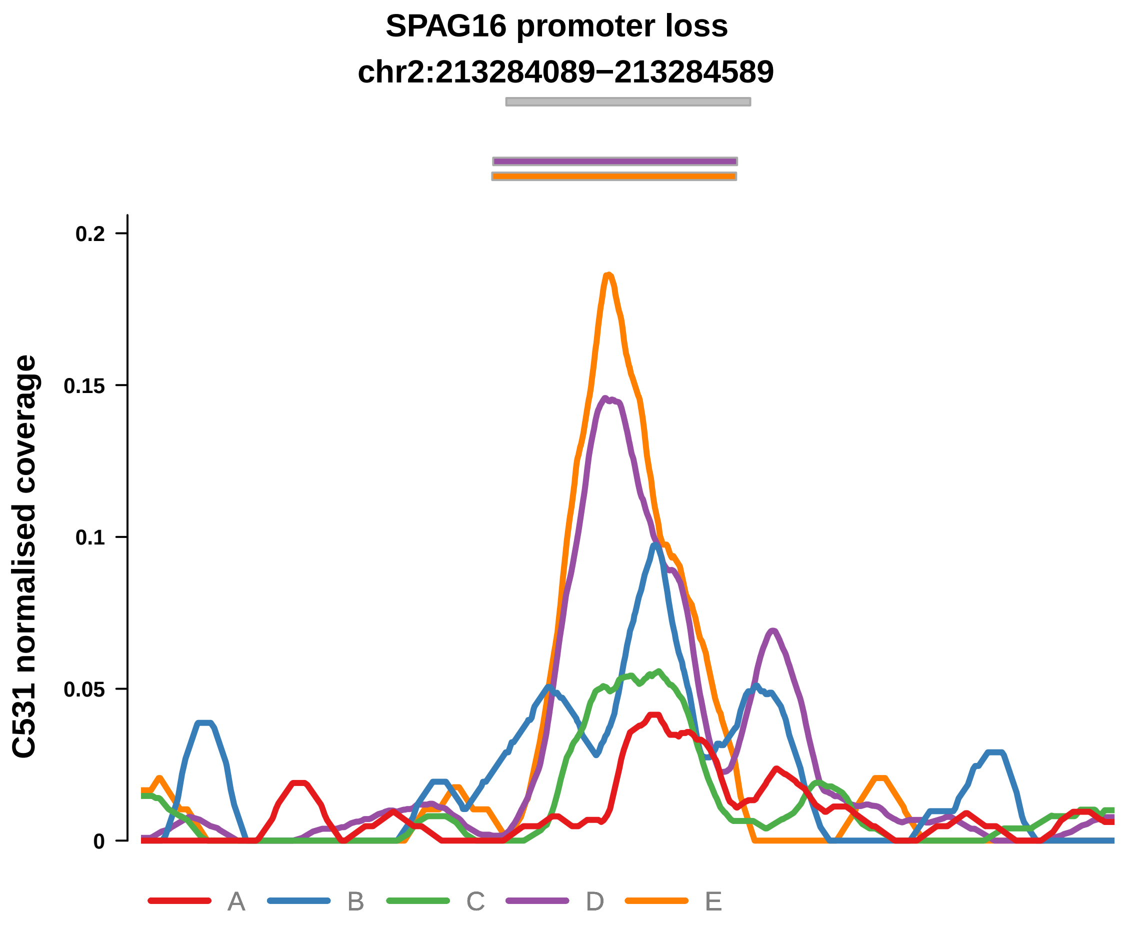


Putatively Clonal (logFC = −1.94)

| | P. | Adjusted P. |
|---|---|---|
| *~region vs. ~* | 3.74e−02 * | 9.11e−02 |
| *~purity vs ~* | 1.41e−04 *** | 4.12e−03 ** |
| *~purity+region vs ~purity* | 2.08e−01 | 6.13e−01 |
| *Carcinoma vs. normal* | 2.81e−03 ** | 2.81e−03 ** |

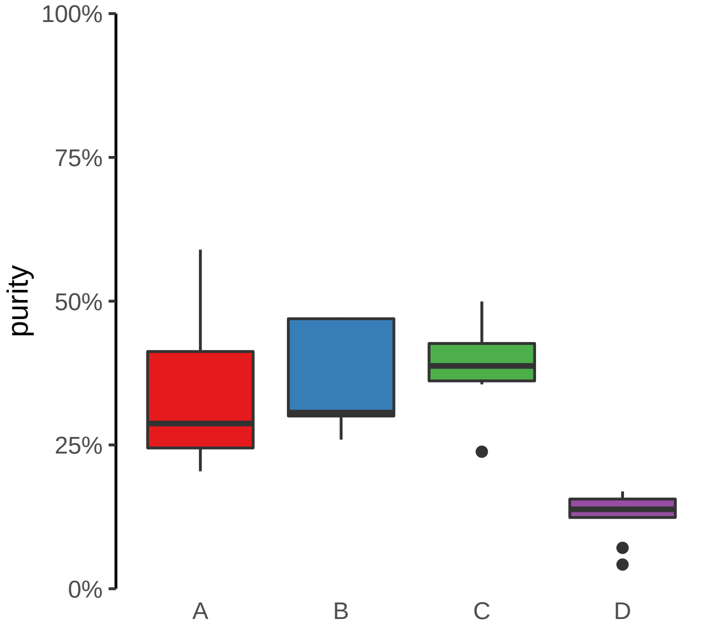

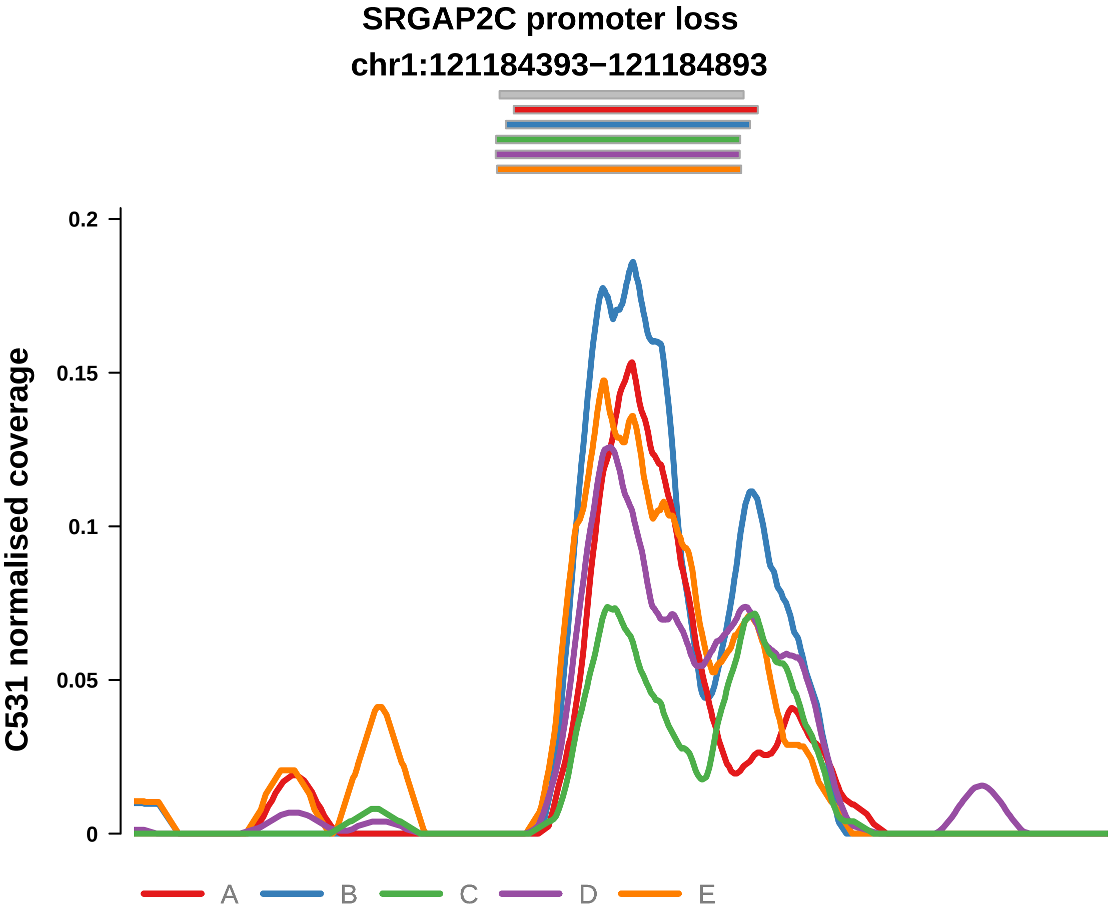


Putatively Clonal (logFC = −1.82)

| | P. | Adjusted P. |
|---|---|---|
| *~region vs. ~* | 3.85e−02 * | 9.11e−02 |
| *~purity vs ~* | 8.24e−03 ** | 4.83e−02 * |
| *~purity+region vs ~purity* | 1.09e−01 | 5.23e−01 |
| *Carcinoma vs. normal* | 4.12e−03 ** | 4.12e−03 ** |

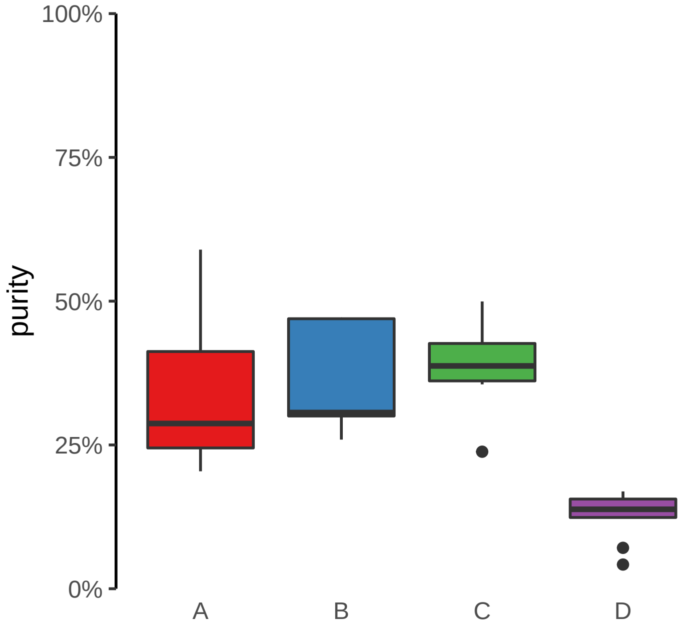

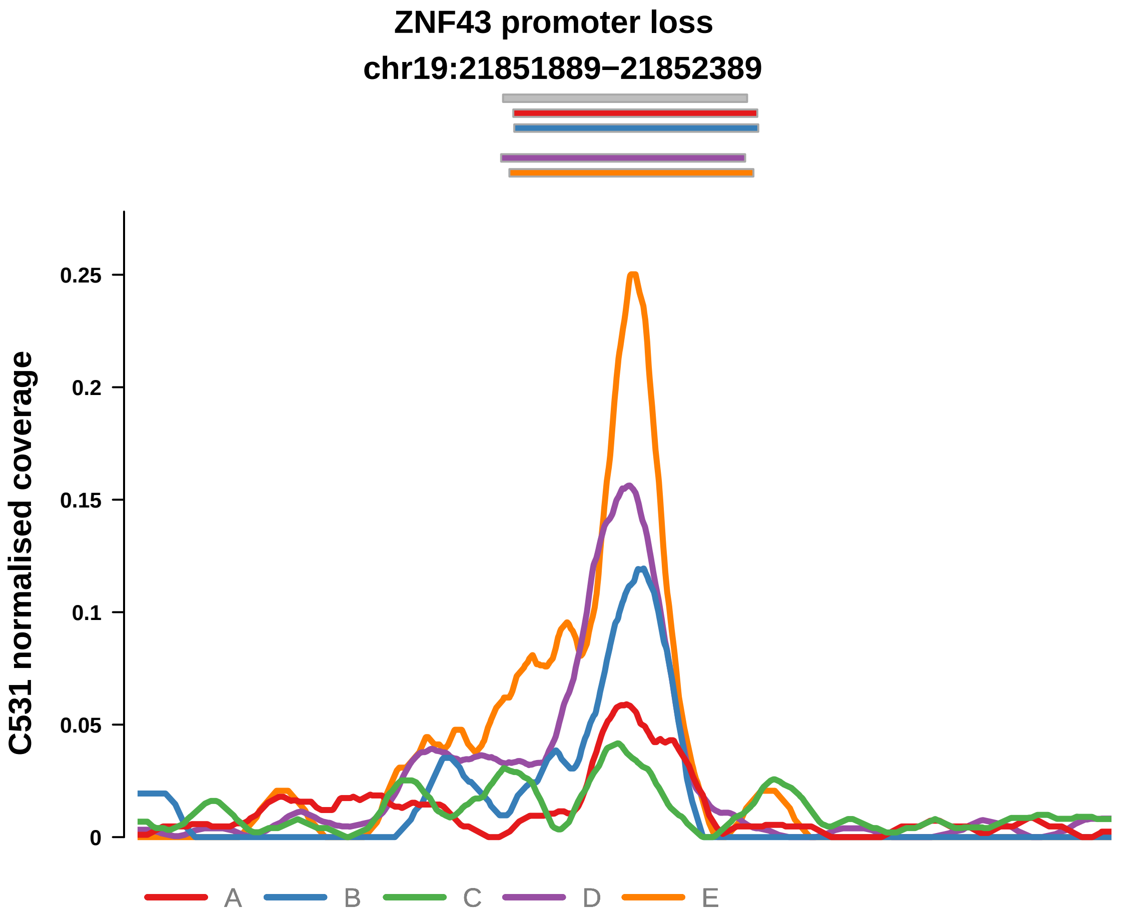


Putatively Clonal (logFC = −3.14)

| | P. | Adjusted P. |
|---|---|---|
| *~region vs. ~* | 6.14e−03 ** | 2.13e−02 * |
| *~purity vs ~* | 5.1e−04 *** | 8.97e−03 ** |
| *~purity+region vs ~purity* | 8.04e−01 | 9.16e−01 |
| *Carcinoma vs. normal* | 1.88e−05 **** | 1.88e−05 **** |

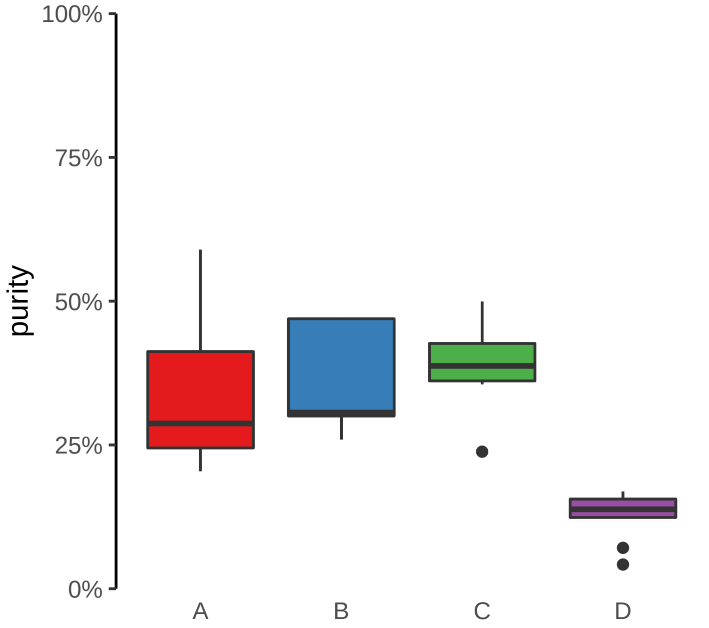

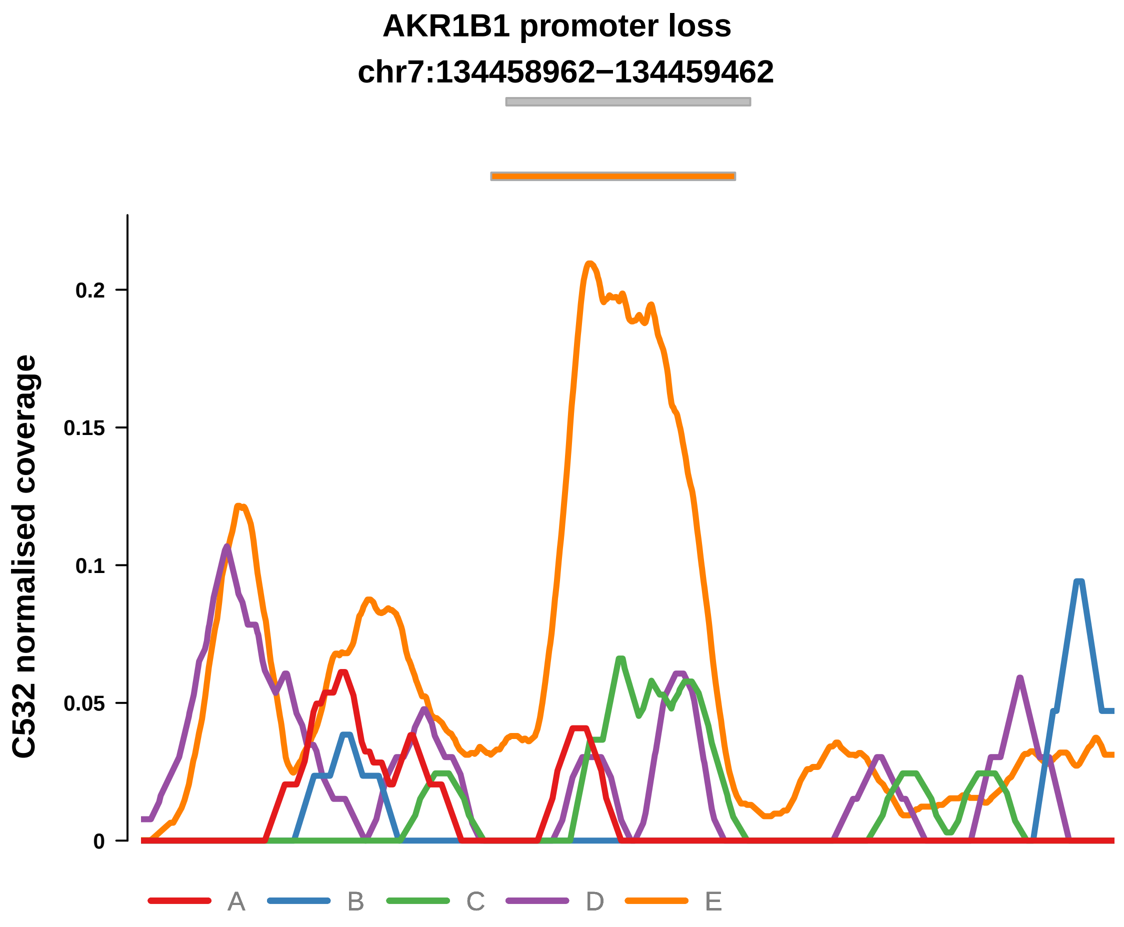


Putatively Clonal (logFC = −2.59)

| | P. | Adjusted P. |
|---|---|---|
| *~region vs. ~* | 1.06e−01 | 4.53e−01 |
| *~purity vs ~* | 4.57e−01 | 9.05e−01 |
| *~purity+region vs ~purity* | 1.64e−01 | 6.56e−01 |
| *Carcinoma vs. normal* | 5.74e−05 **** | 5.74e−05 **** |

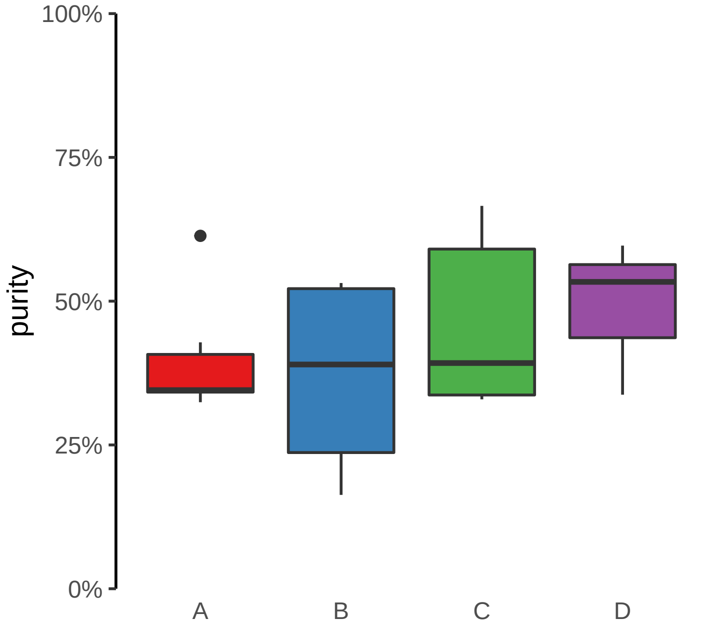

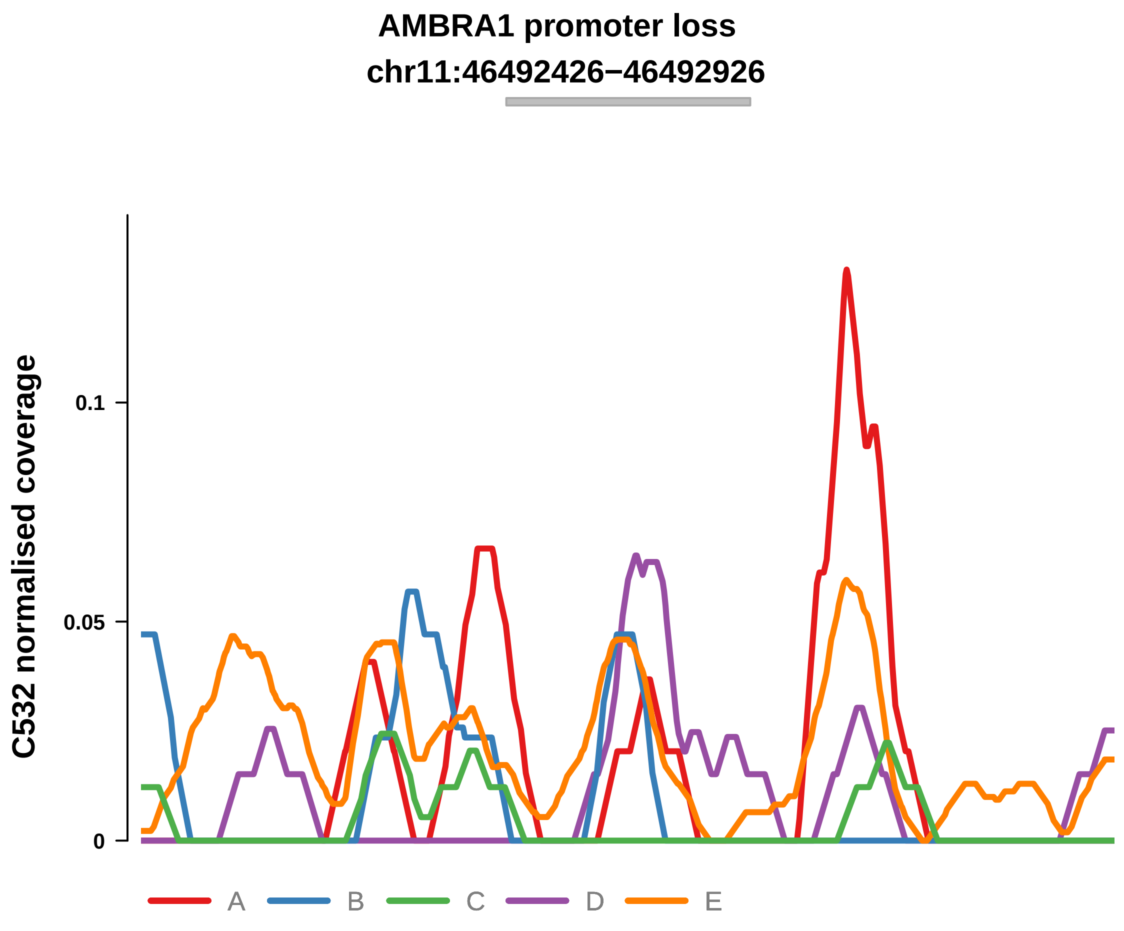


Putatively Clonal (logFC = −1.78)

| | P. | Adjusted P. |
|---|---|---|
| *~region vs. ~* | 3.29e−01 | 7.41e−01 |
| *~purity vs ~* | 8.88e−01 | 9.86e−01 |
| *~purity+region vs ~purity* | 4.15e−01 | 9.12e−01 |
| *Carcinoma vs. normal* | 7.65e−03 ** | 7.65e−03 ** |

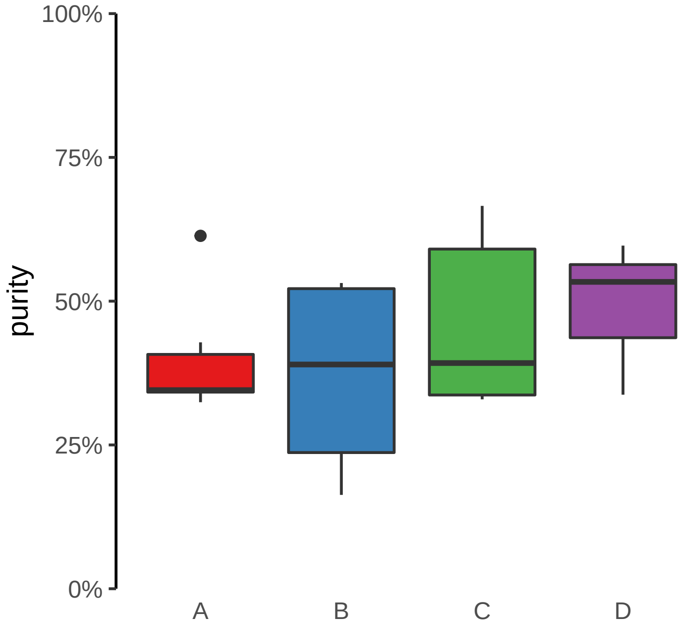

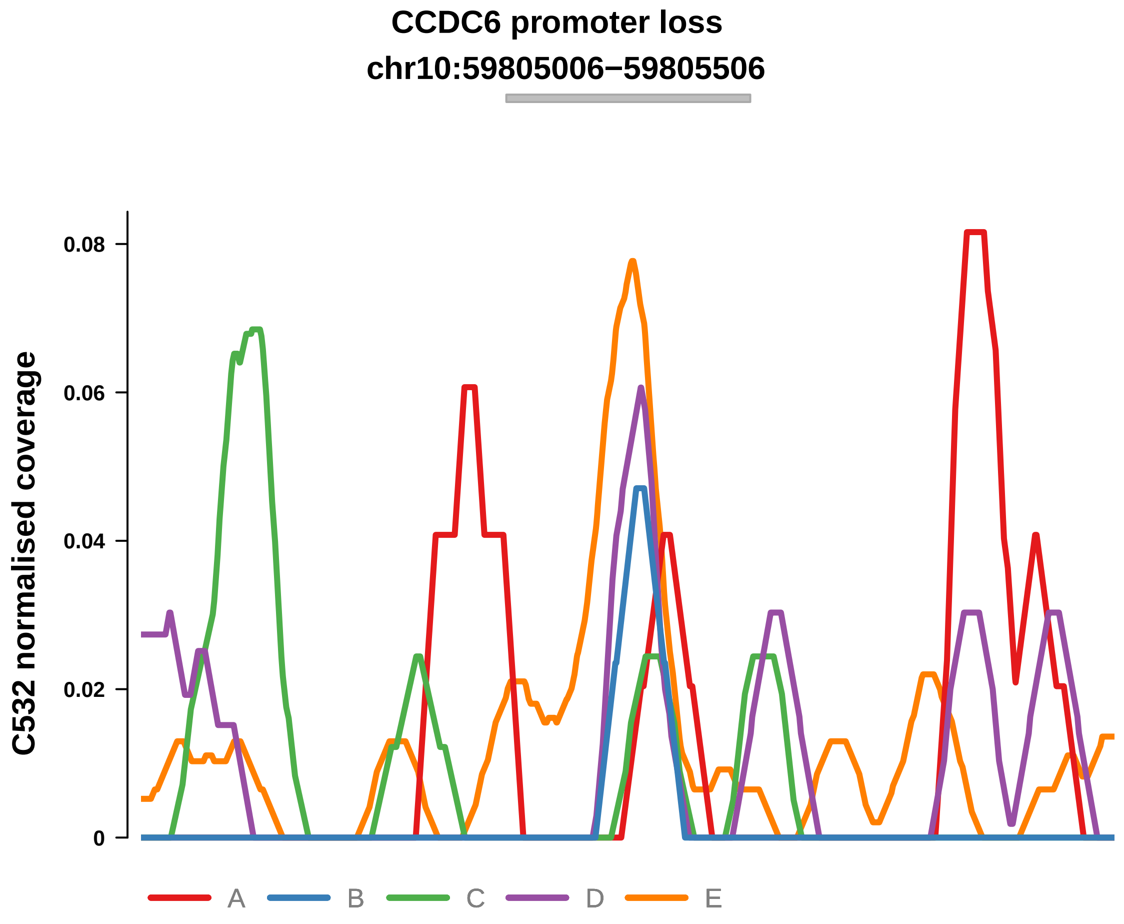


Putatively Clonal (logFC = −2.16)

| | P. | Adjusted P. |
|---|---|---|
| *~region vs. ~* | 9.59e−01 | 9.9e−01 |
| *~purity vs ~* | 9.08e−01 | 9.86e−01 |
| *~purity+region vs ~purity* | 9.79e−01 | 1e+00 |
| *Carcinoma vs. normal* | 1.29e−03 ** | 1.29e−03 ** |

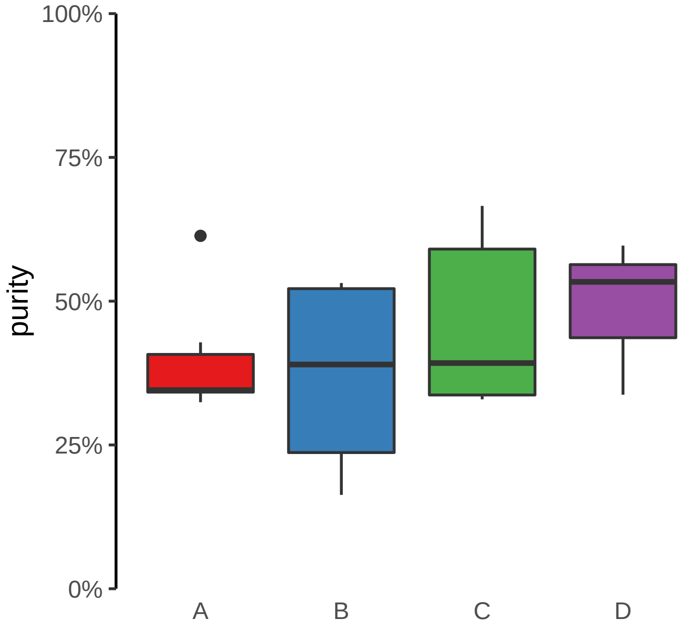

**EPS8L1 promoter loss**
**chr19:55086973−55087473**

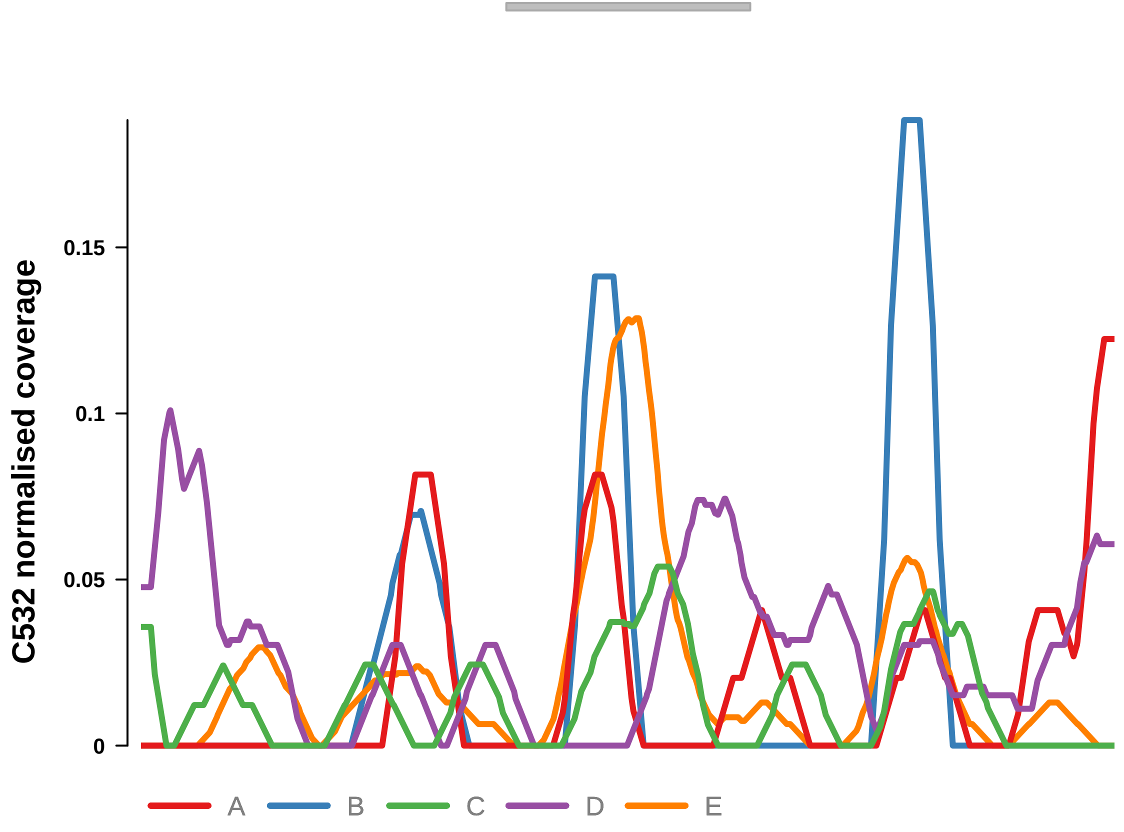


Putatively Clonal (logFC = −2.34)

| | **P.** | **Adjusted P.** |
|---|---|---|
| *~region vs. ~* | 8.37e−01 | 9.57e−01 |
| *~purity vs ~* | 2.73e−01 | 8.56e−01 |
| *~purity+region vs ~purity* | 7.4e−01 | 9.42e−01 |
| *Carcinoma vs. normal* | 2.61e−04 *** | 2.61e−04 *** |

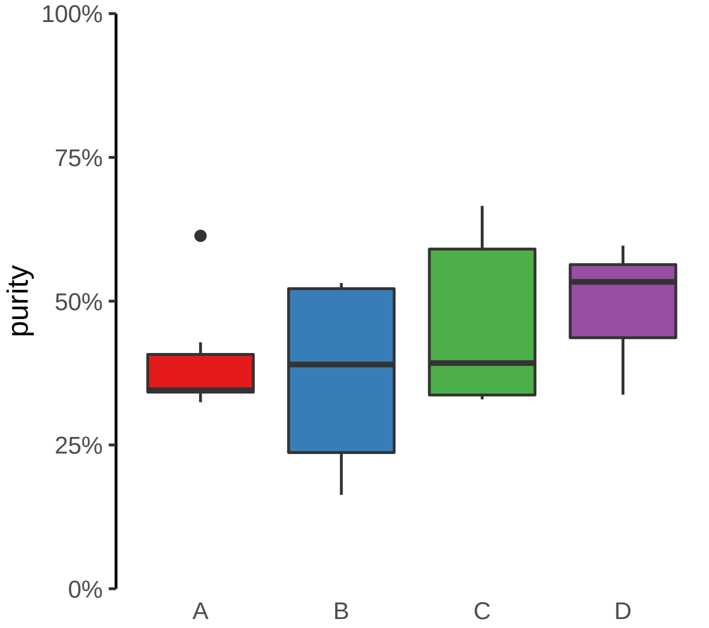

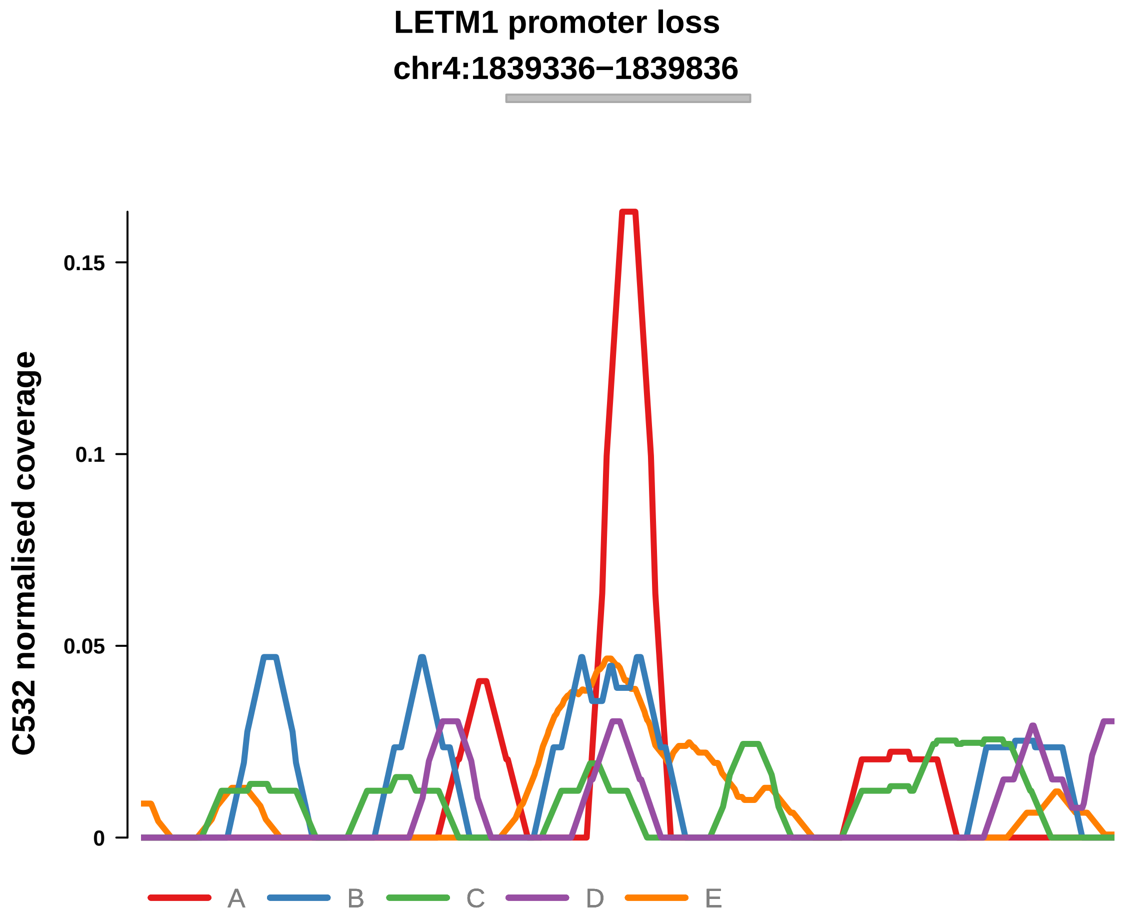


Putatively Clonal (logFC = −1.95)

| | P. | Adjusted P. |
|---|---|---|
| *~region vs. ~* | 5.69e−01 | 8.54e−01 |
| *~purity vs ~* | 7.02e−01 | 9.86e−01 |
| *~purity+region vs ~purity* | 5.81e−01 | 9.42e−01 |
| *Carcinoma vs. normal* | 3.22e−03 ** | 3.22e−03 ** |

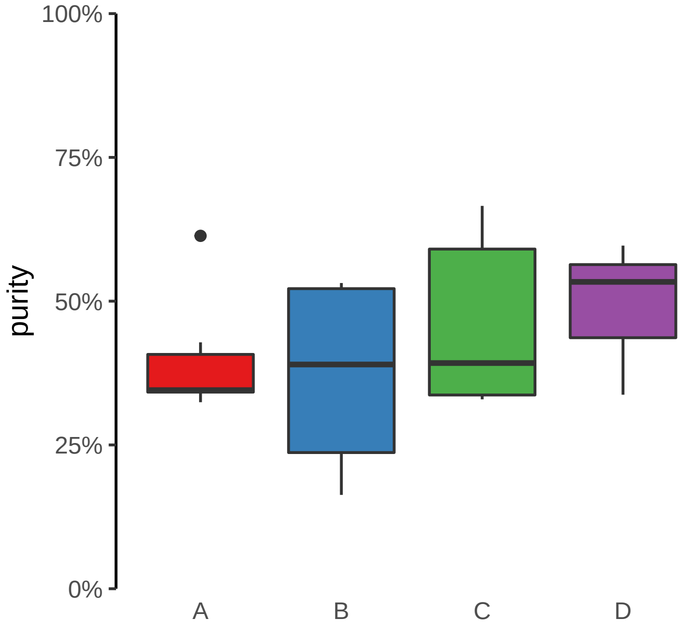

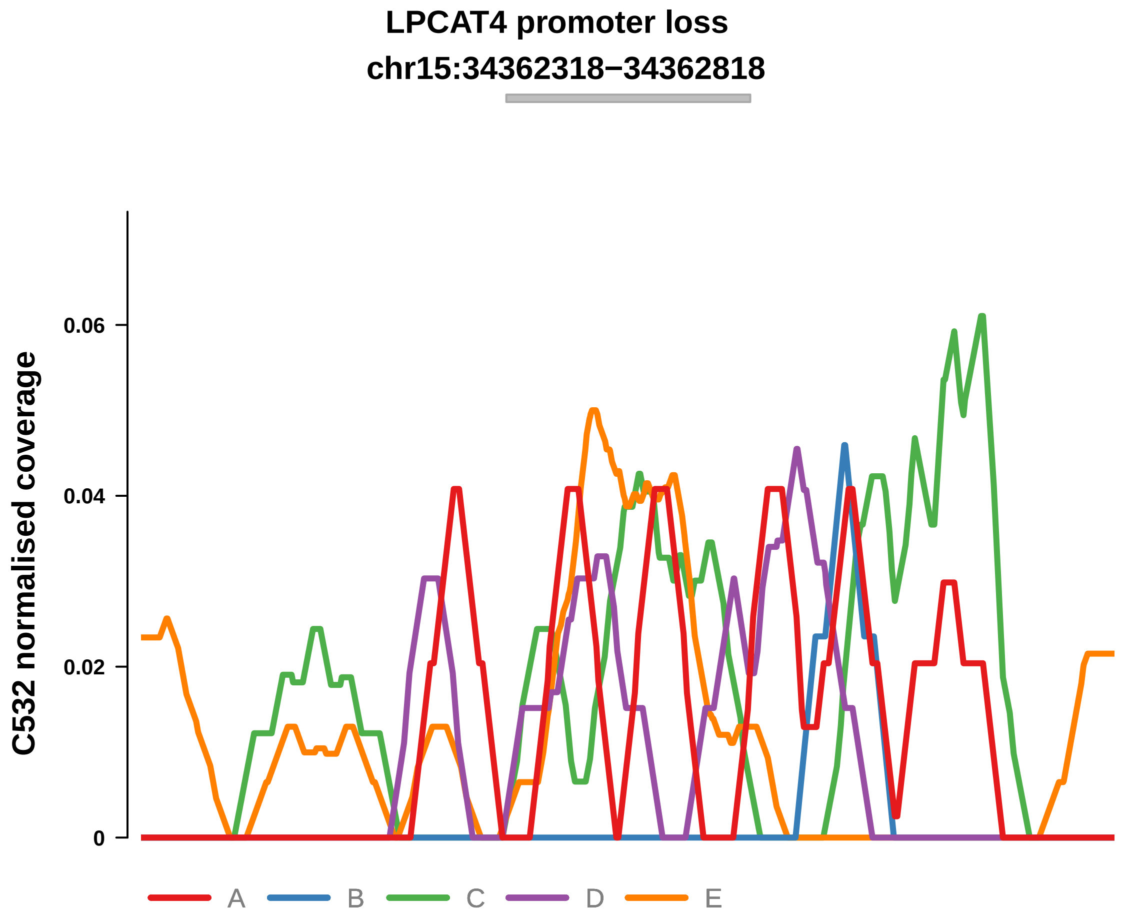


Putatively Clonal (logFC = −1.78)

| | **P.** | **Adjusted P.** |
|---|---|---|
| *~region vs. ~* | 8.26e−01 | 9.56e−01 |
| *~purity vs ~* | 6.3e−01 | 9.86e−01 |
| *~purity+region vs ~purity* | 1e+00 | 1e+00 |
| *Carcinoma vs. normal* | 4.4e−03 ** | 4.4e−03 ** |

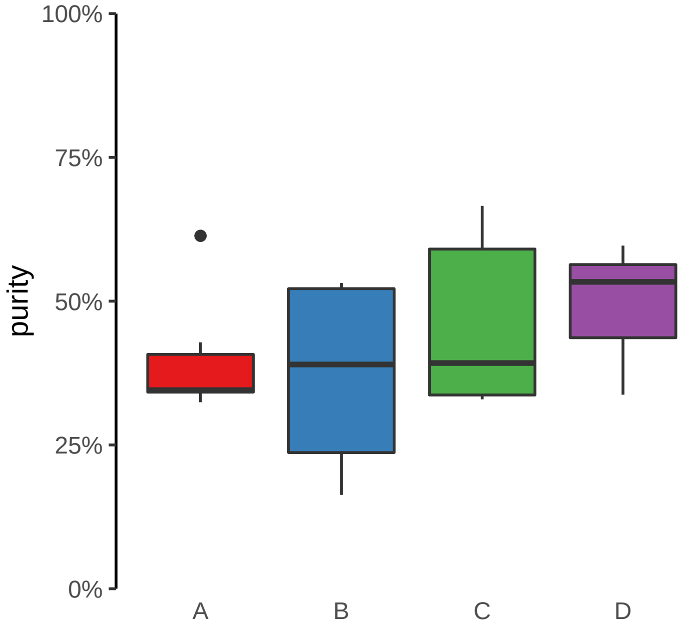

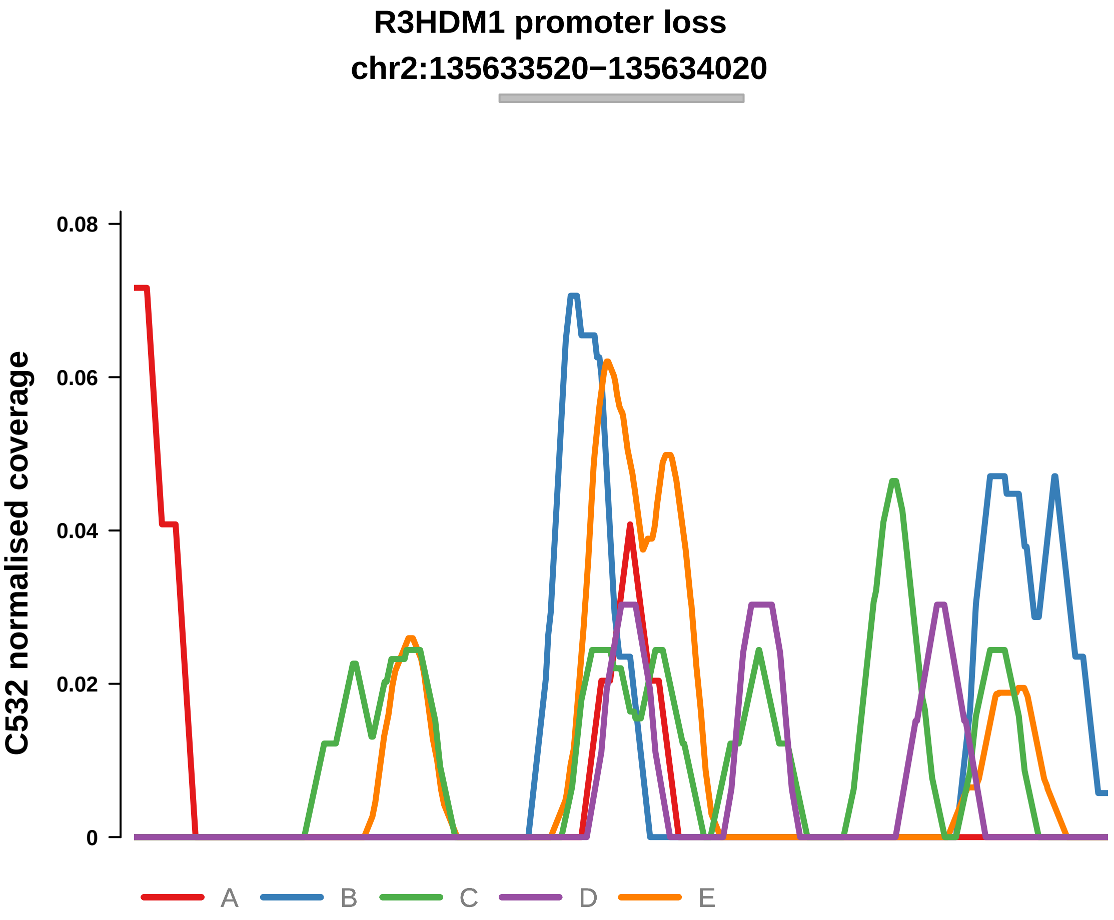


Putatively Clonal (logFC = −2.65)

| | P. | Adjusted P. |
|---|---|---|
| *~region vs. ~* | 9.49e−01 | 9.9e−01 |
| *~purity vs ~* | 7.22e−01 | 9.86e−01 |
| *~purity+region vs ~purity* | 9.52e−01 | 1e+00 |
| *Carcinoma vs. normal* | 2.76e−04 *** | 2.76e−04 *** |

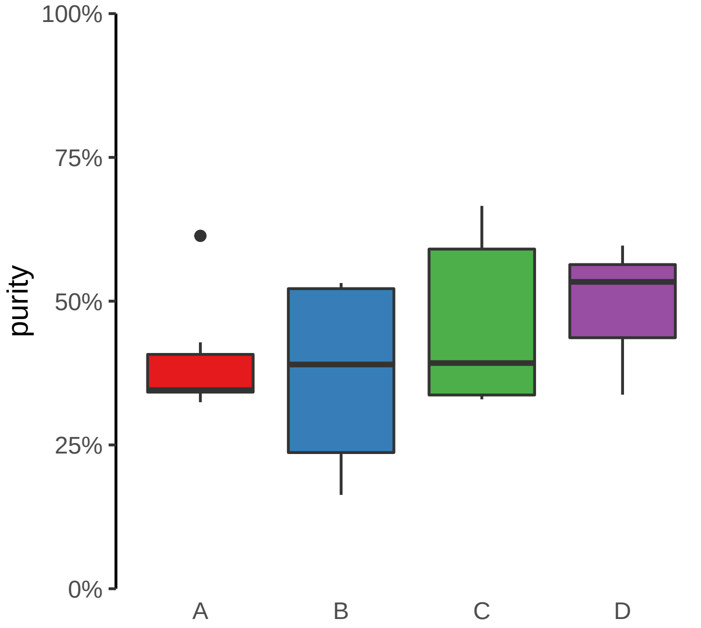

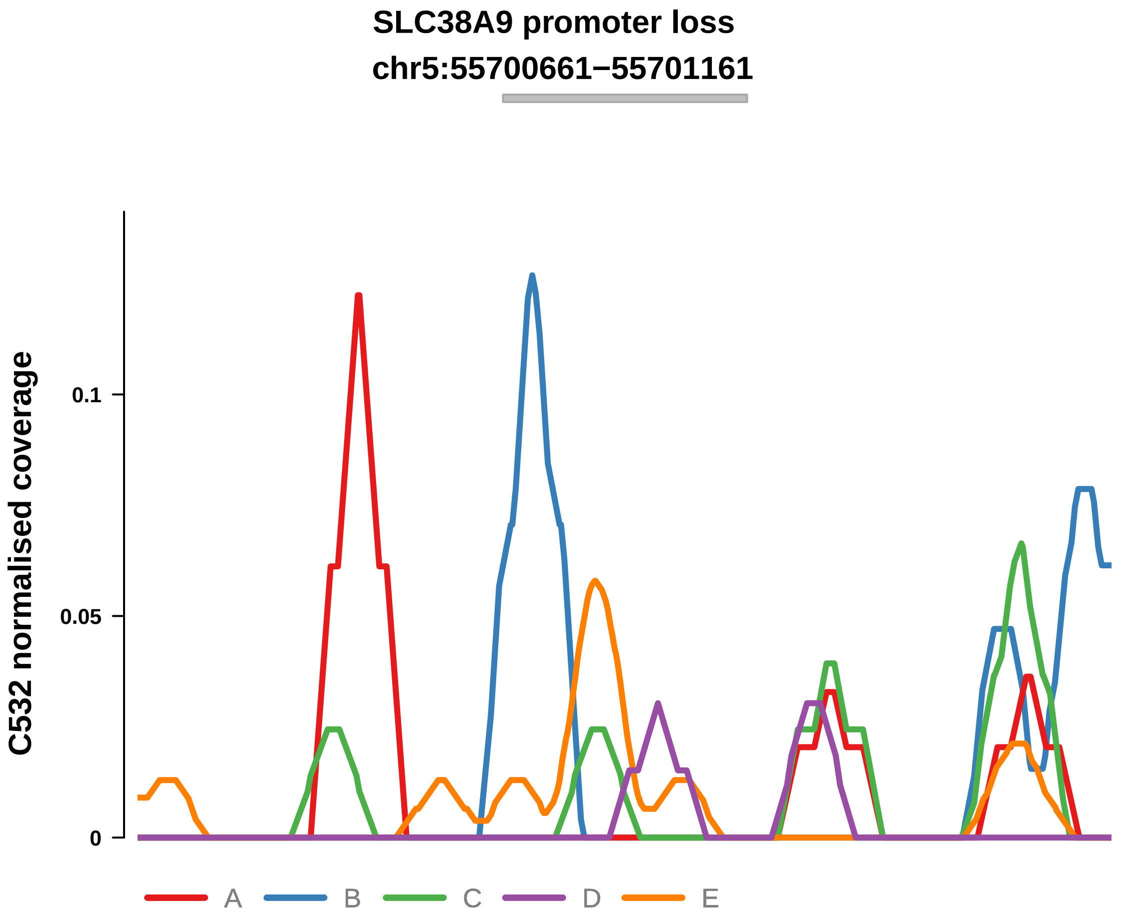


Putatively Clonal (logFC = −2.14)

| | P. | Adjusted P. |
|---|---|---|
| *~region vs. ~* | 8.04e−01 | 9.43e−01 |
| *~purity vs ~* | 7.04e−01 | 9.86e−01 |
| *~purity+region vs ~purity* | 1e+00 | 1e+00 |
| *Carcinoma vs. normal* | 3.77e−03 ** | 3.77e−03 ** |

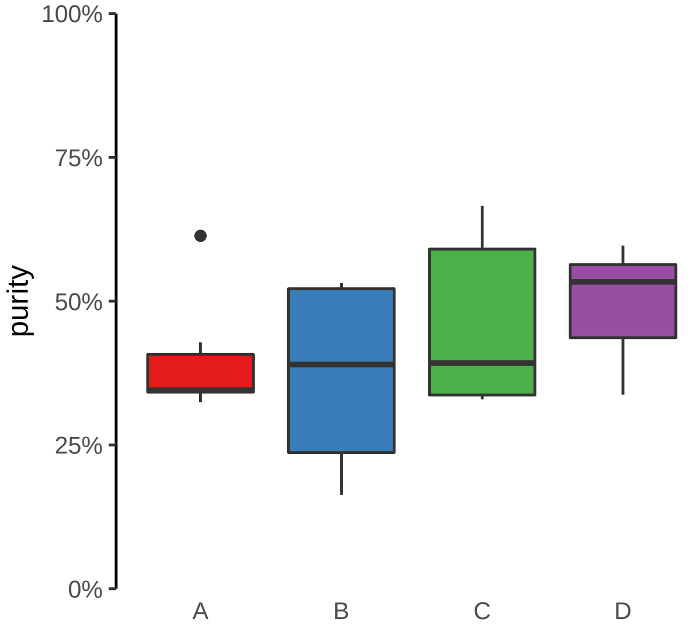

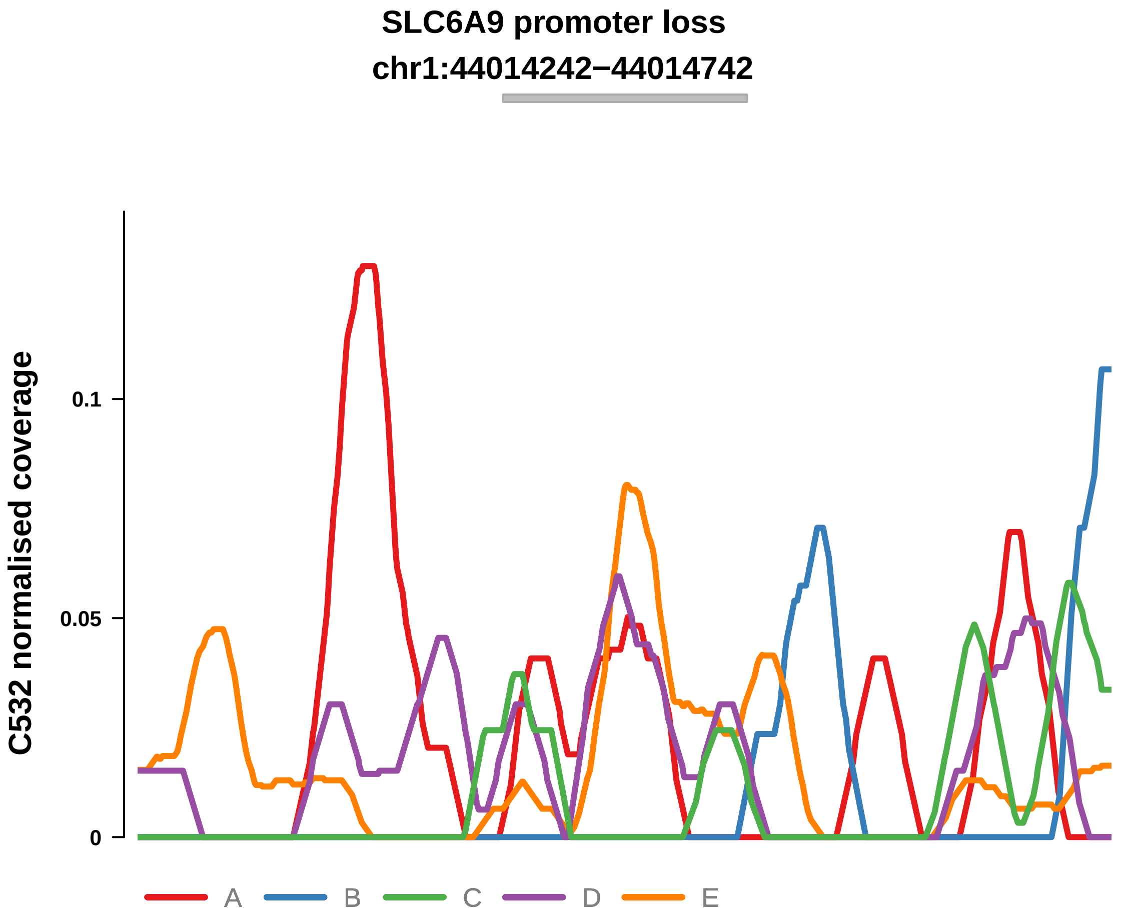


Putatively Clonal (logFC = −1.73)

| | P. | Adjusted P. |
|---|---|---|
| *~region vs. ~* | 7.32e−01 | 9.07e−01 |
| *~purity vs ~* | 4.61e−01 | 9.05e−01 |
| *~purity+region vs ~purity* | 7.71e−01 | 9.42e−01 |
| *Carcinoma vs. normal* | 5.88e−03 ** | 5.88e−03 ** |

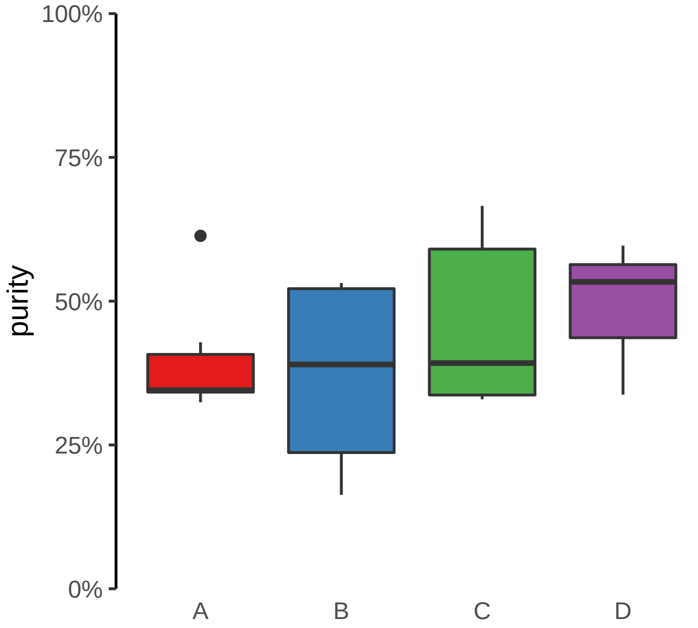

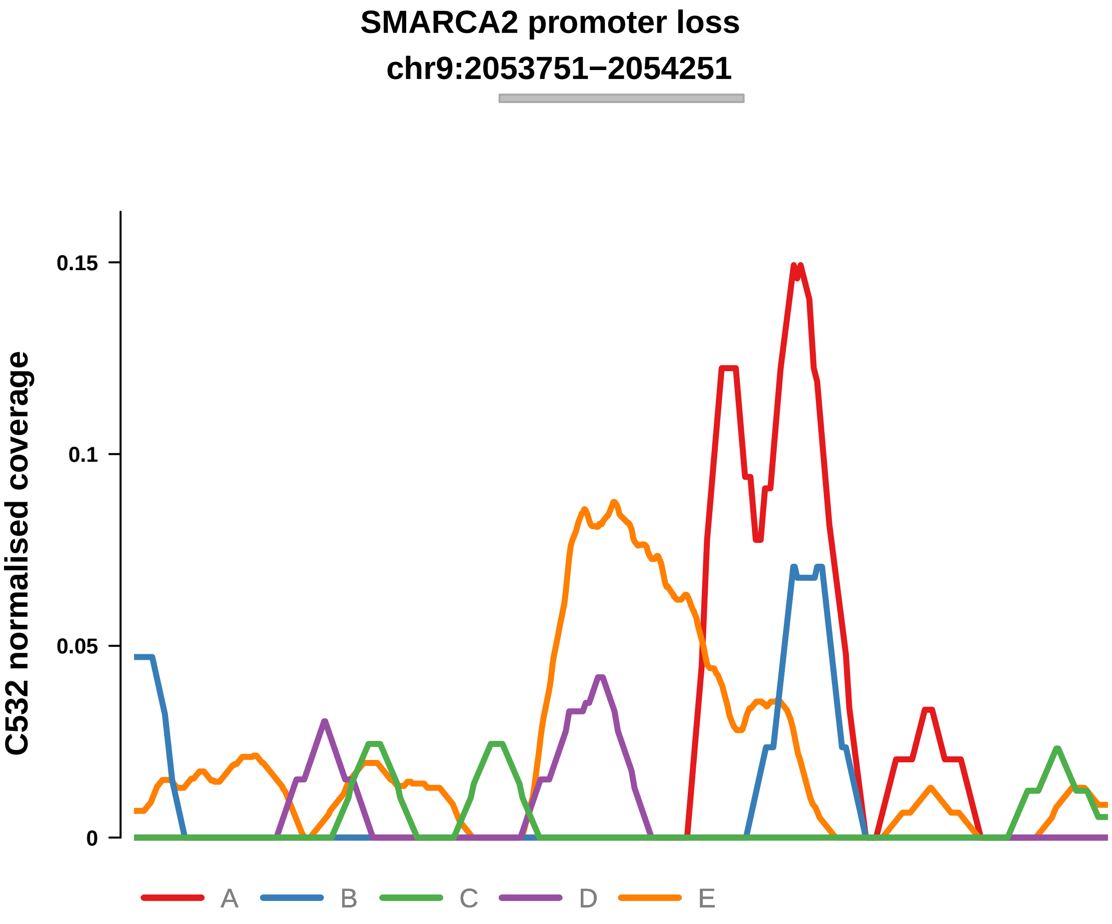


Putatively Clonal (logFC = −3.05)

| | P. | Adjusted P. |
|---|---|---|
| *~region vs. ~* | 6.63e−01 | 8.56e−01 |
| *~purity vs ~* | 7.57e−01 | 9.86e−01 |
| *~purity+region vs ~purity* | 6.94e−01 | 9.42e−01 |
| *Carcinoma vs. normal* | 3.2e−04 *** | 3.2e−04 *** |

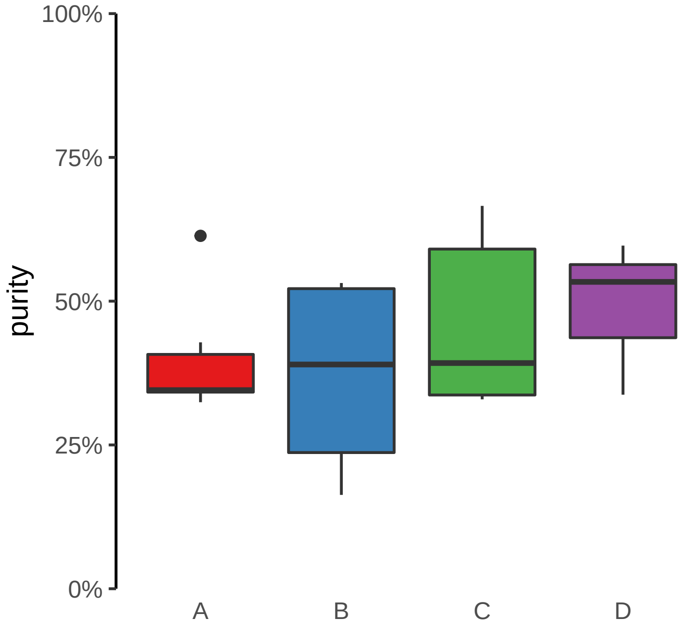

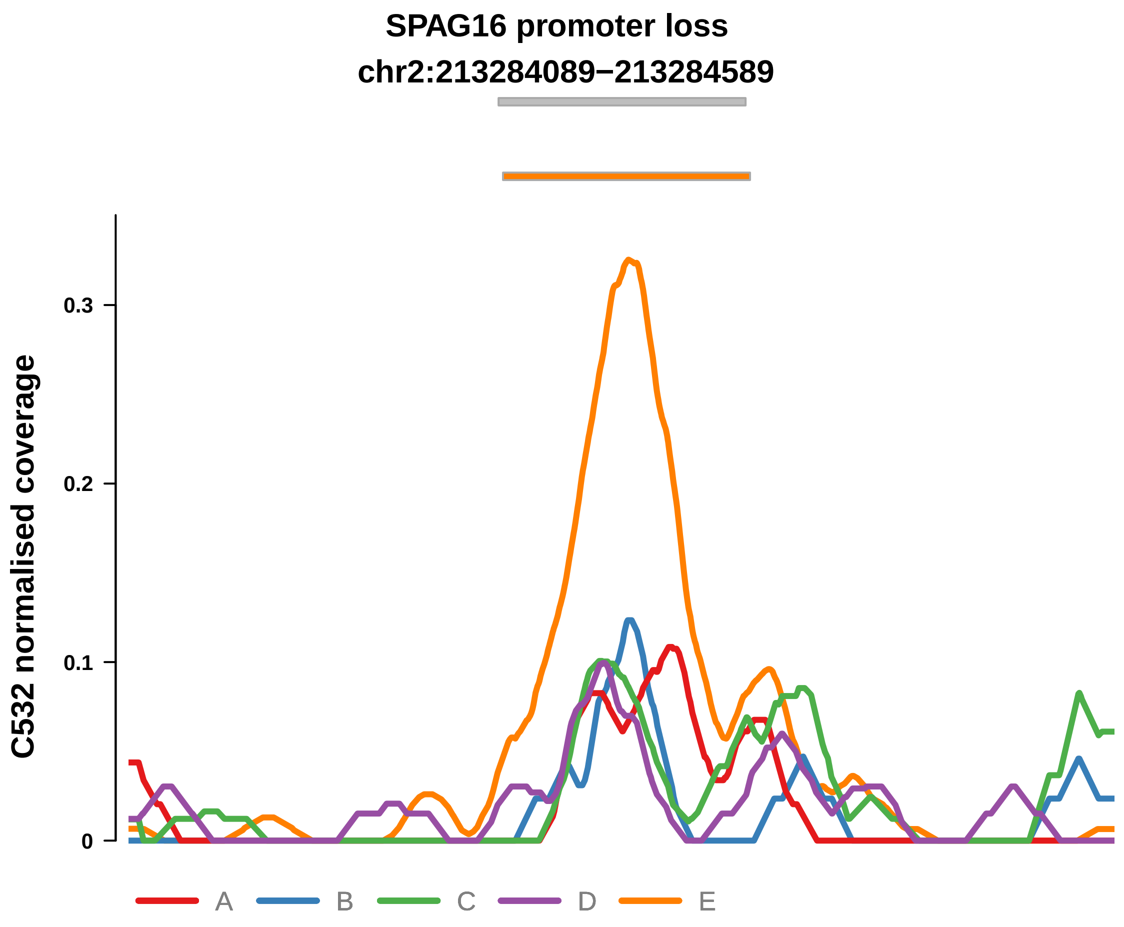


Putatively Clonal (logFC = −2.34)

| | P. | Adjusted P. |
|---|---|---|
| *~region vs. ~* | 9.49e−01 | 9.9e−01 |
| *~purity vs ~* | 8.16e−01 | 9.86e−01 |
| *~purity+region vs ~purity* | 8.71e−01 | 1e+00 |
| *Carcinoma vs. normal* | 8.37e−05 **** | 8.37e−05 **** |

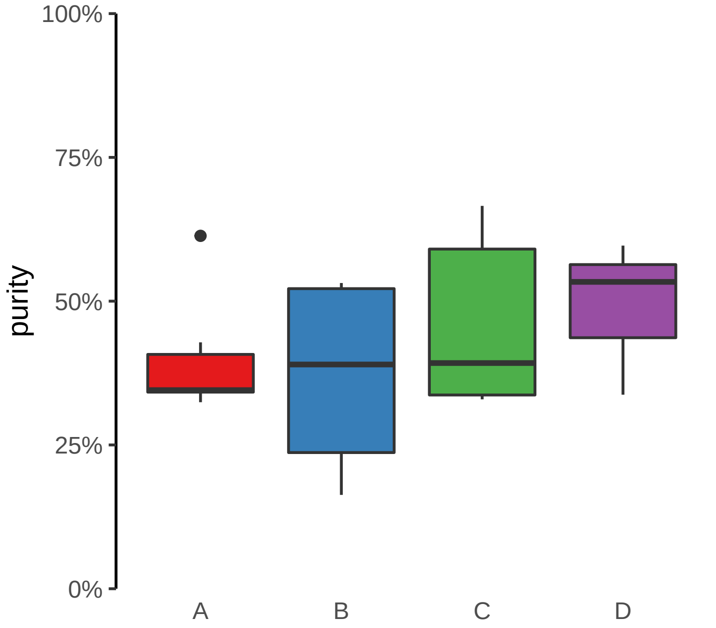

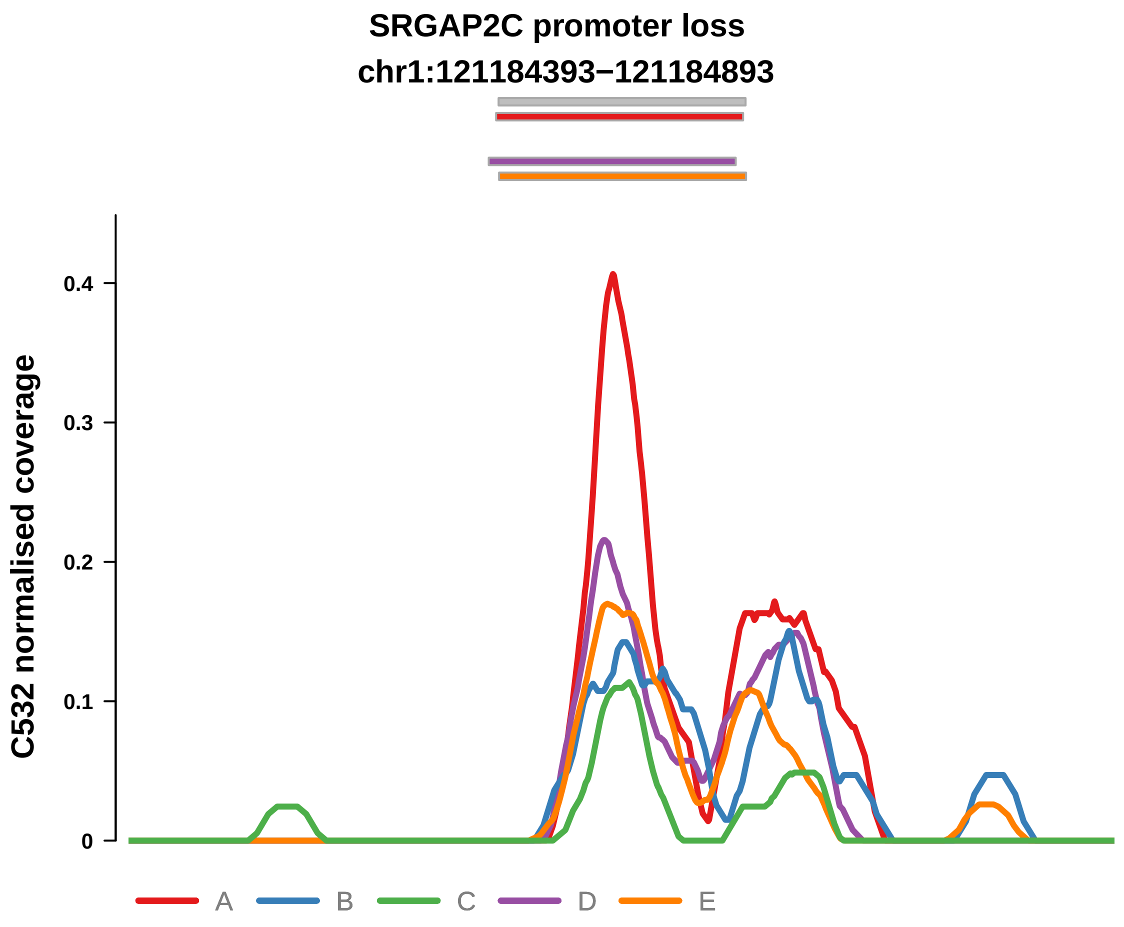


Putatively Clonal (logFC = −1.57)

| | P. | Adjusted P. |
|---|---|---|
| *~region vs. ~* | 8.08e−02 | 4.44e−01 |
| *~purity vs ~* | 3.52e−01 | 8.63e−01 |
| *~purity+region vs ~purity* | 8.33e−02 | 5.15e−01 |
| *Carcinoma vs. normal* | 5.45e−03 ** | 5.45e−03 ** |

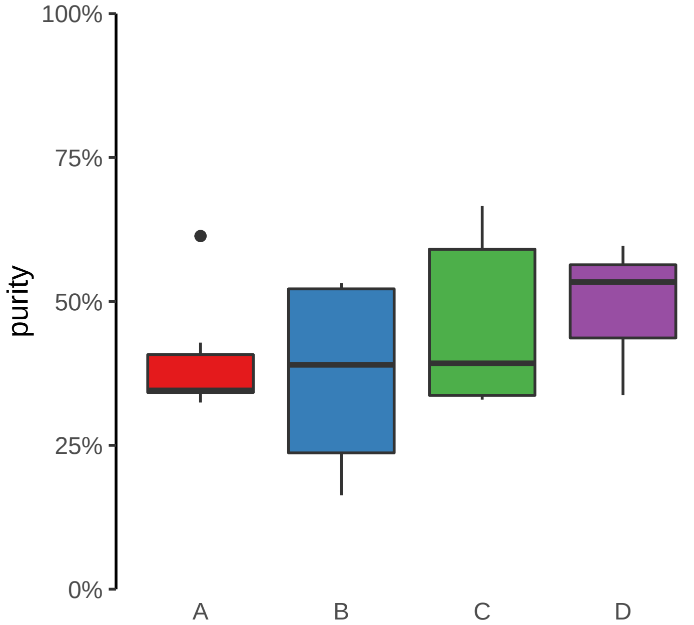

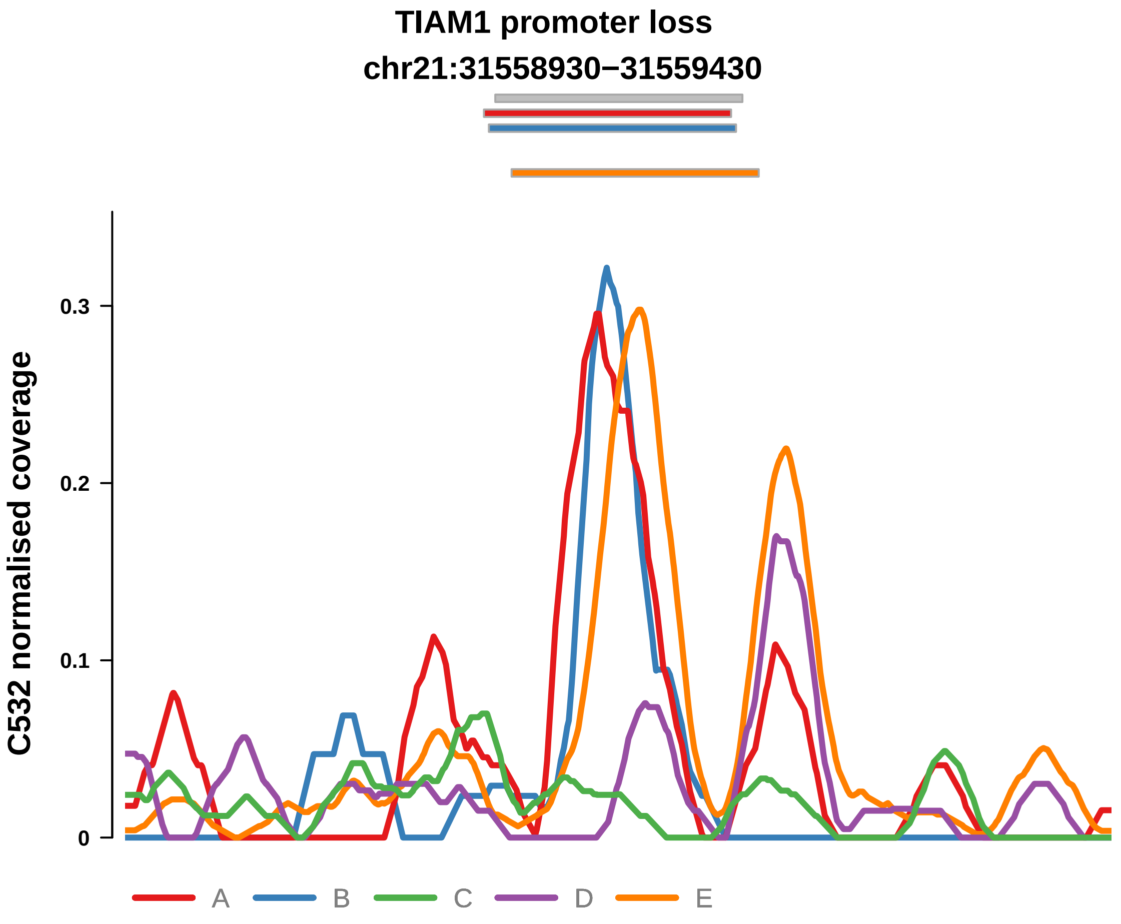


Putatively Clonal (logFC = −2.02)

| | P. | Adjusted P. |
|---|---|---|
| *~region vs. ~* | 5.65e−03 ** | 1.24e−01 |
| *~purity vs ~* | 2.42e−02 * | 3.05e−01 |
| *~purity+region vs ~purity* | 5.59e−02 | 5.15e−01 |
| *Carcinoma vs. normal* | 5.73e−04 *** | 5.73e−04 *** |

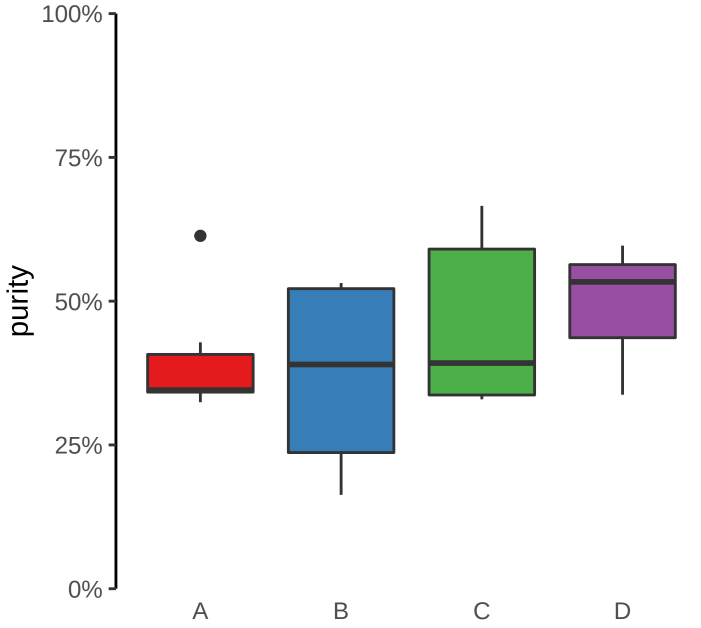

**ZNF43 promoter loss**
**chr19:21851889−21852389**

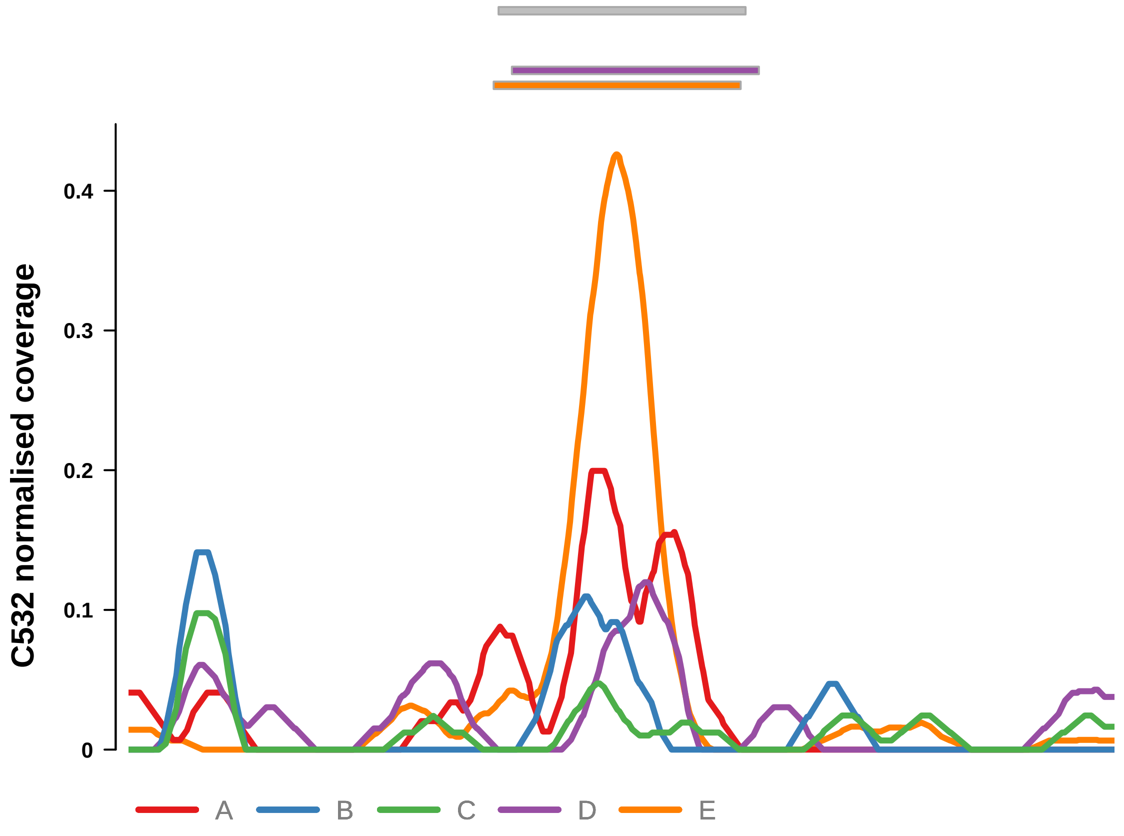


Putatively Clonal (logFC = −2.28)

| | P. | Adjusted P. |
|---|---|---|
| *~region vs. ~* | 1.09e−01 | 4.53e−01 |
| *~purity vs ~* | 3.59e−01 | 8.63e−01 |
| *~purity+region vs ~purity* | 1.73e−01 | 6.56e−01 |
| *Carcinoma vs. normal* | 1.31e−04 *** | 1.31e−04 *** |

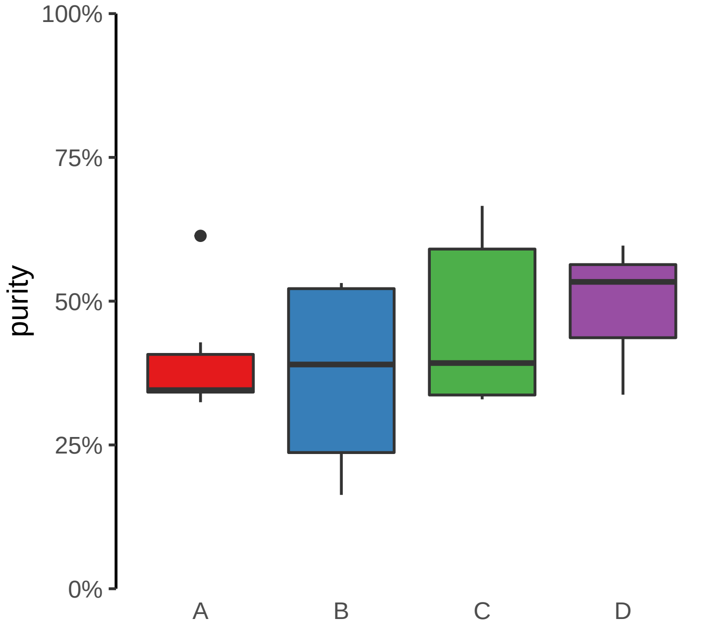

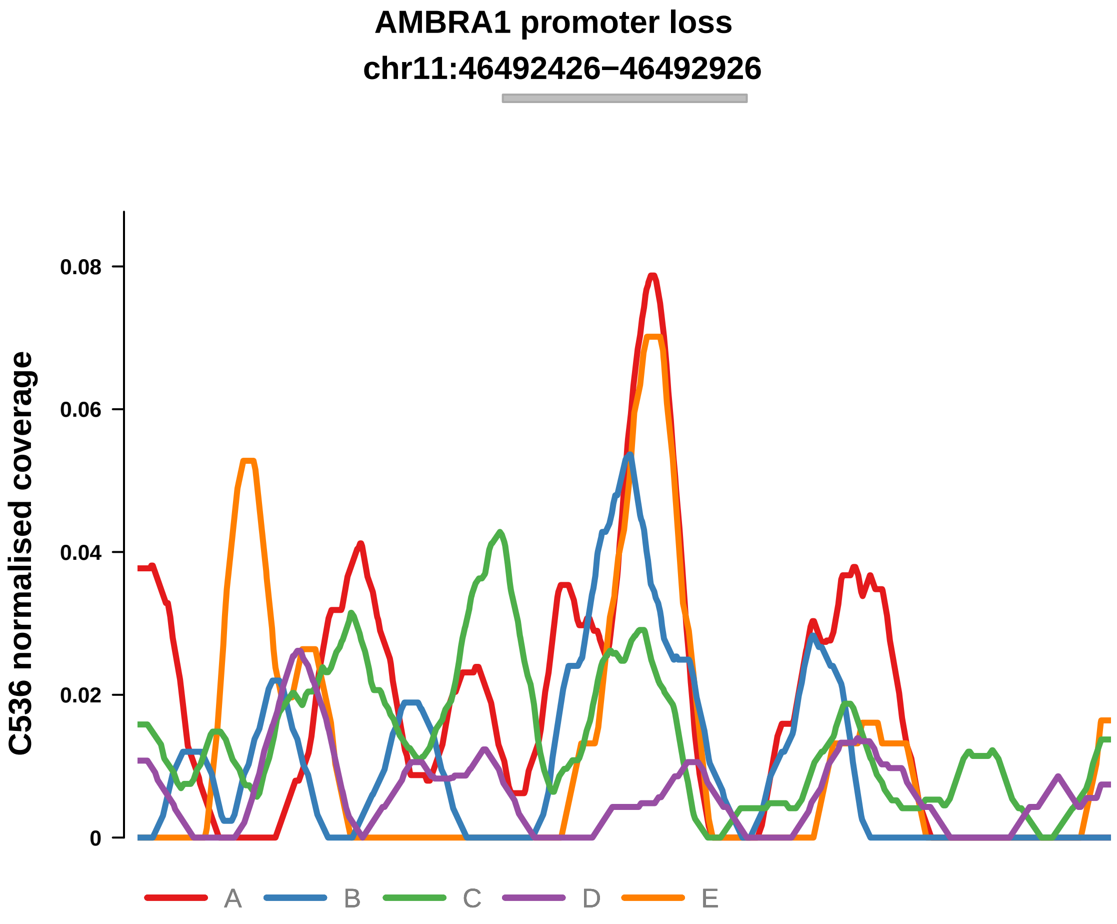


Putatively Clonal (logFC = −2.89)

| | P. | Adjusted P. |
|---|---|---|
| *~region vs. ~* | 6.45e−01 | NA |
| *~purity vs ~* | 5.8e−02 | 2.13e−01 |
| *~purity+region vs ~purity* | 6.92e−01 | NA |
| *Carcinoma vs. normal* | 2.35e−04 *** | 2.35e−04 *** |

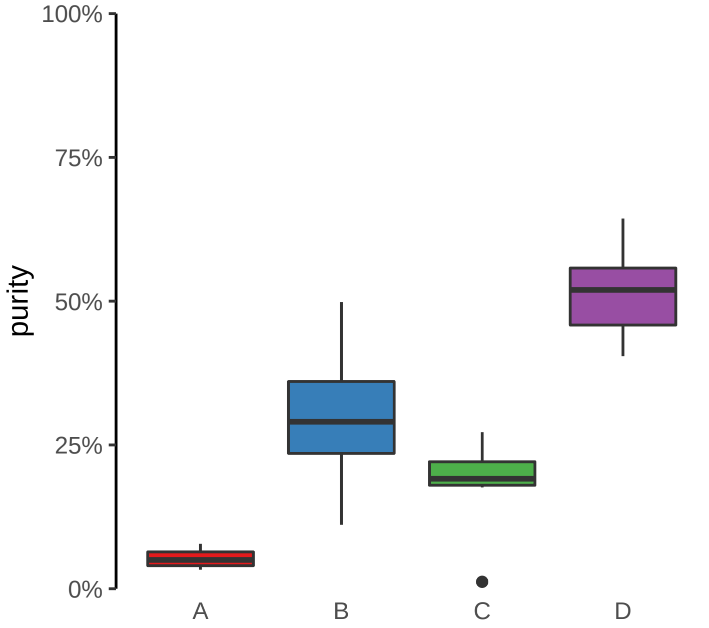

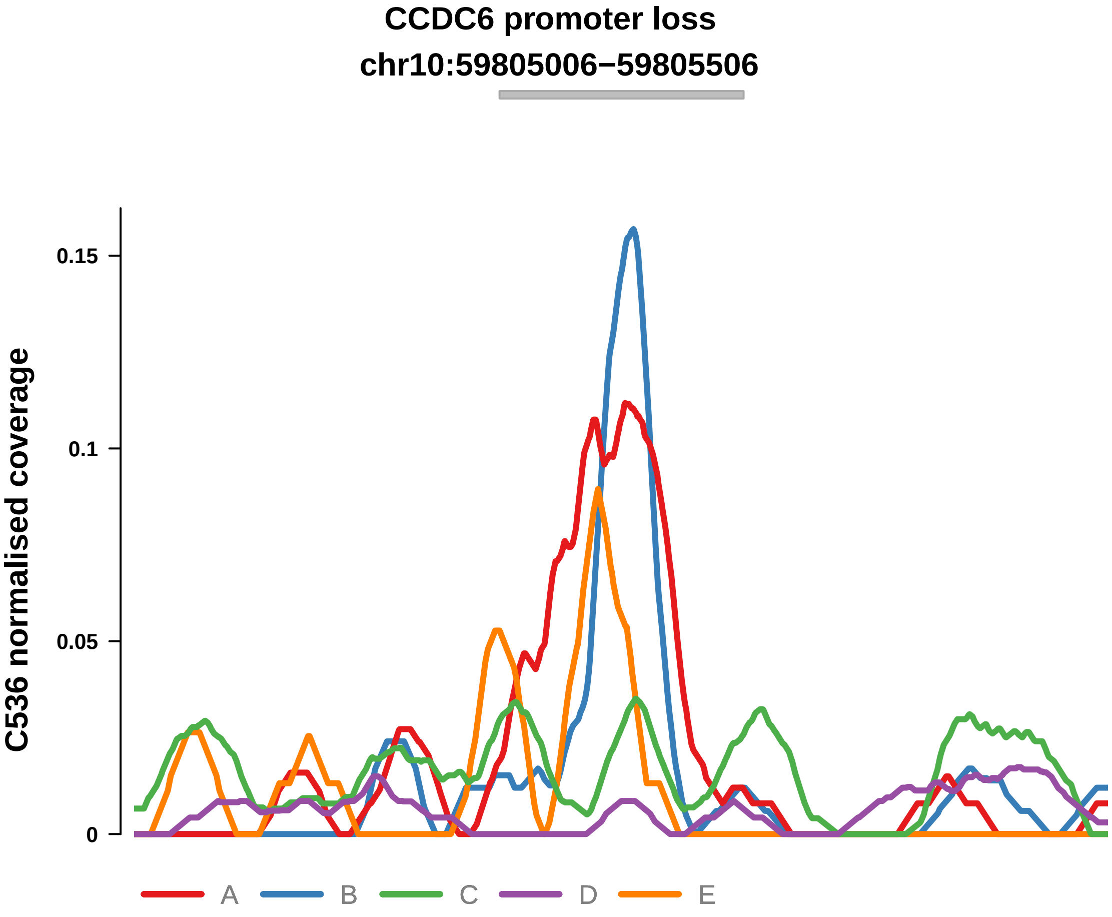


Putatively Clonal (logFC = −3.54)

| | P. | Adjusted P. |
|---|---|---|
| *~region vs. ~* | 1.23e−01 | 2.42e−01 |
| *~purity vs ~* | 1.08e−03 ** | 2.37e−02 * |
| *~purity+region vs ~purity* | 9.26e−01 | 9.39e−01 |
| *Carcinoma vs. normal* | 1.3e−05 **** | 1.3e−05 **** |

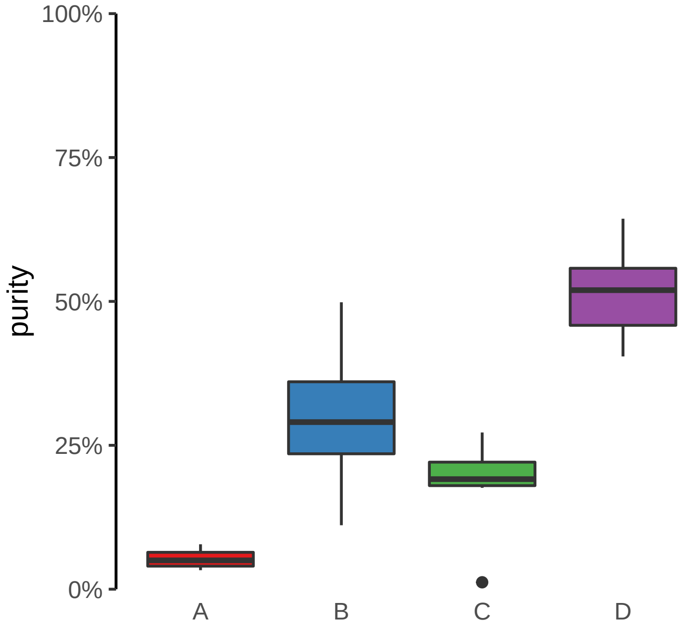

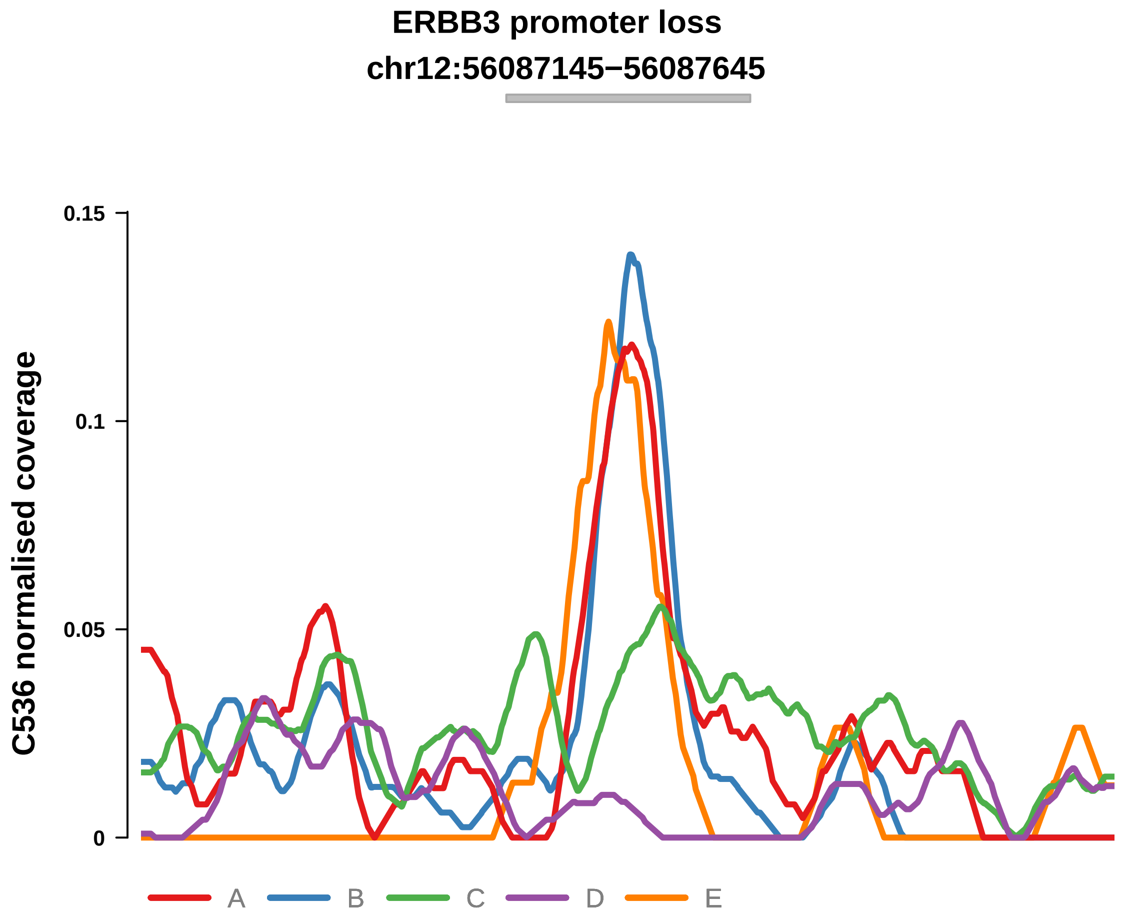


Putatively Clonal (logFC = −2.42)

| | P. | Adjusted P. |
|---|---|---|
| *~region vs. ~* | 1.09e−01 | 2.39e−01 |
| *~purity vs ~* | 4.8e−02 * | 2.04e−01 |
| *~purity+region vs ~purity* | 4.72e−01 | 6.04e−01 |
| *Carcinoma vs. normal* | 4.56e−04 *** | 4.56e−04 *** |

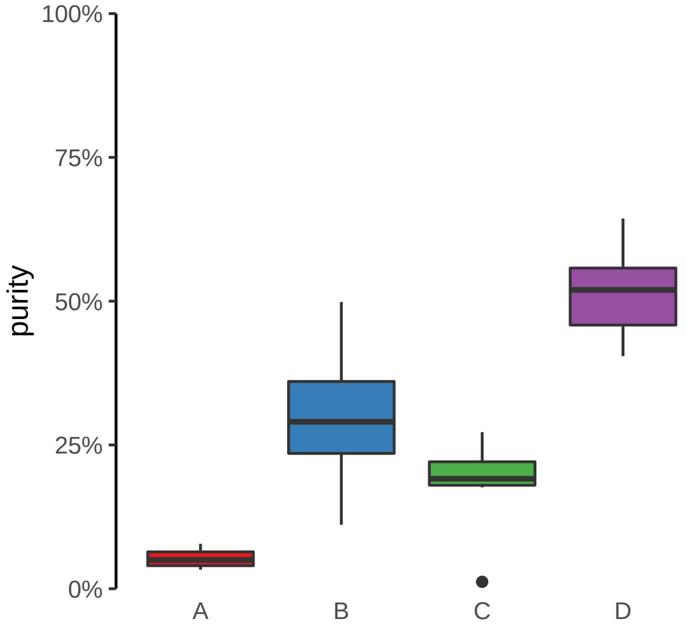

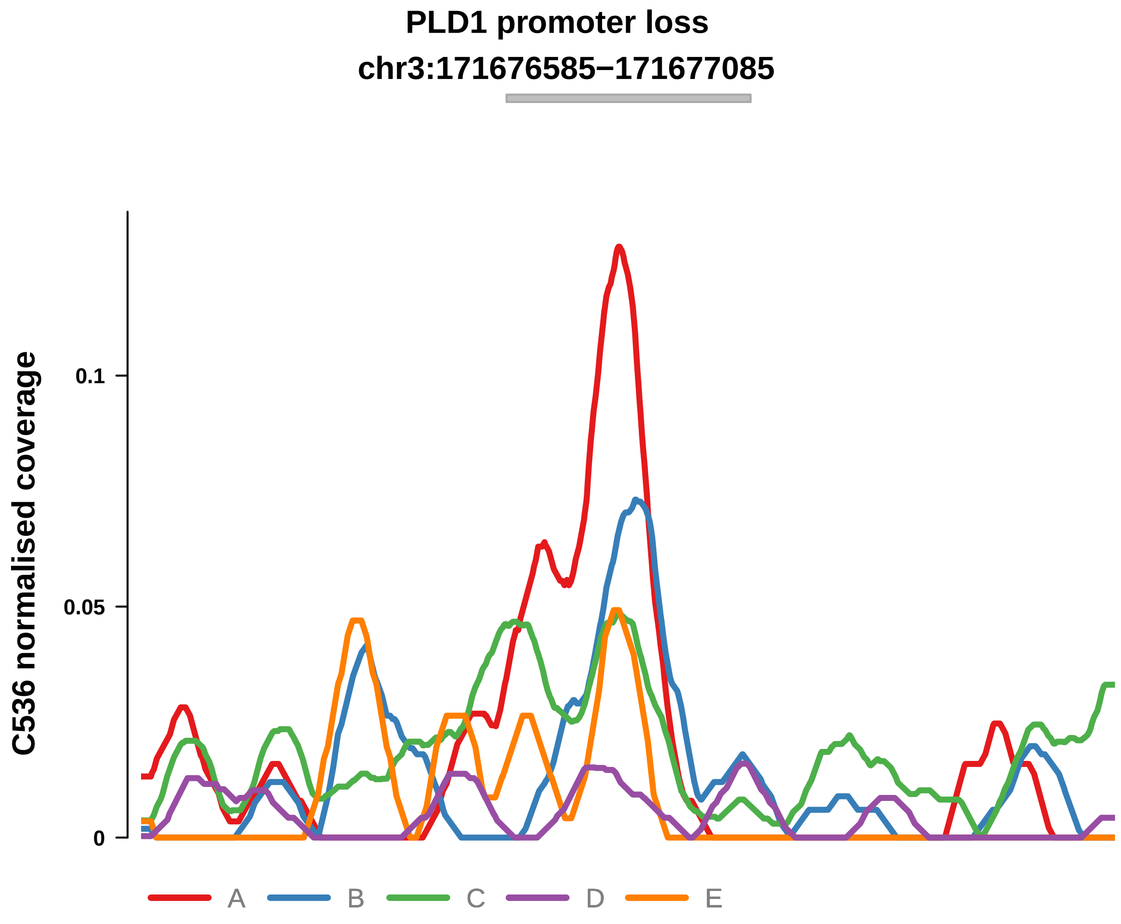


Putatively Clonal (logFC = −2.24)

| | P. | Adjusted P. |
|---|---|---|
| *~region vs. ~* | 3.53e−01 | 4.53e−01 |
| *~purity vs ~* | 2.05e−02 * | 1.39e−01 |
| *~purity+region vs ~purity* | 1.28e−01 | 2.87e−01 |
| *Carcinoma vs. normal* | 1.24e−03 ** | 1.24e−03 ** |

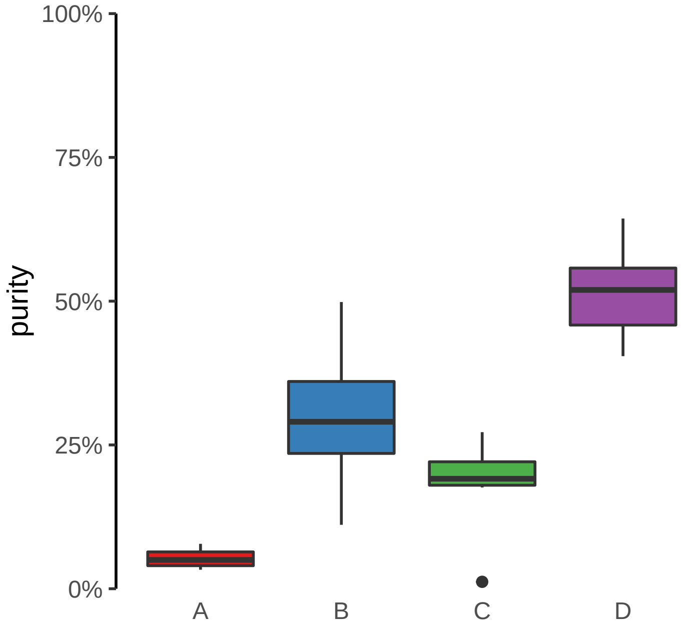

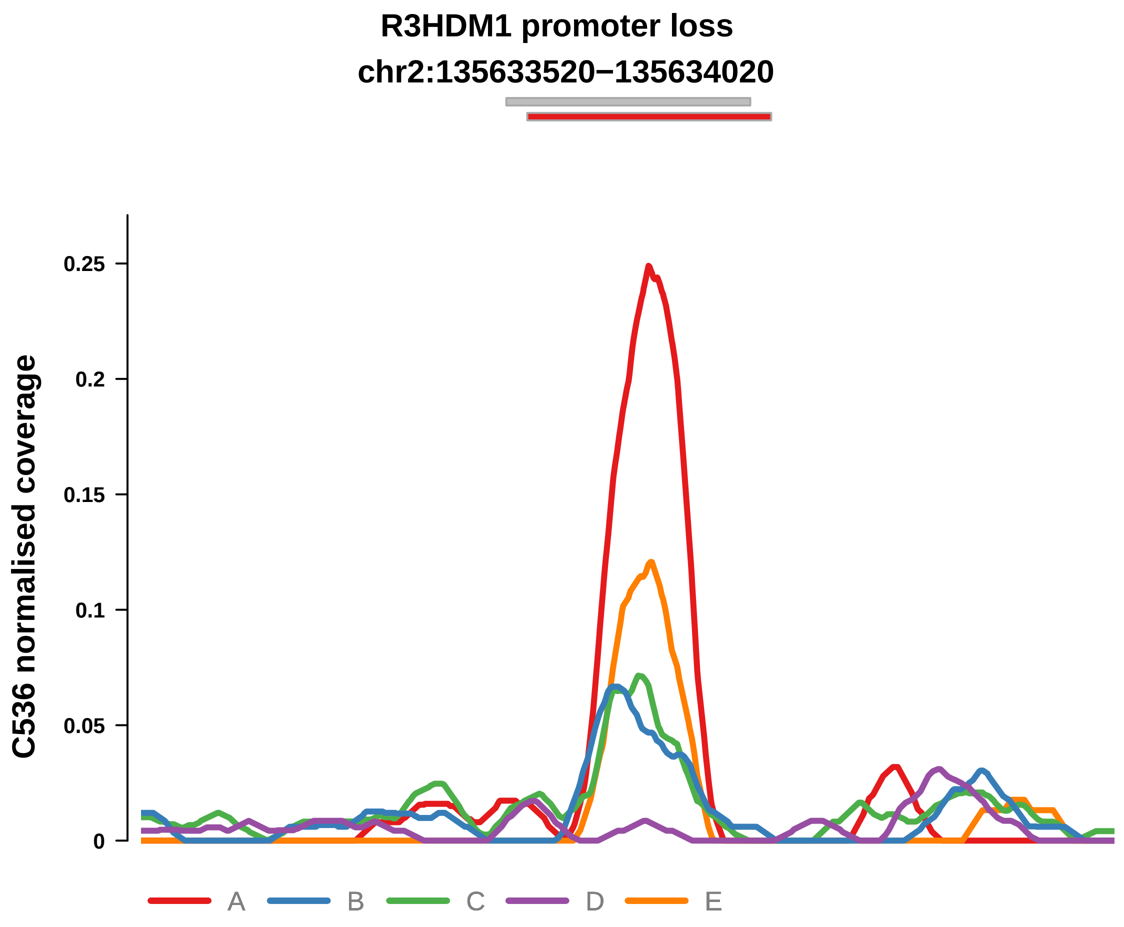


Putatively Clonal (logFC = −2.3)

| | P. | Adjusted P. |
|---|---|---|
| *~region vs. ~* | 7.33e−02 | 1.88e−01 |
| *~purity vs ~* | 3.33e−02 * | 1.83e−01 |
| *~purity+region vs ~purity* | 4.31e−01 | 5.61e−01 |
| *Carcinoma vs. normal* | 1.94e−03 ** | 1.94e−03 ** |

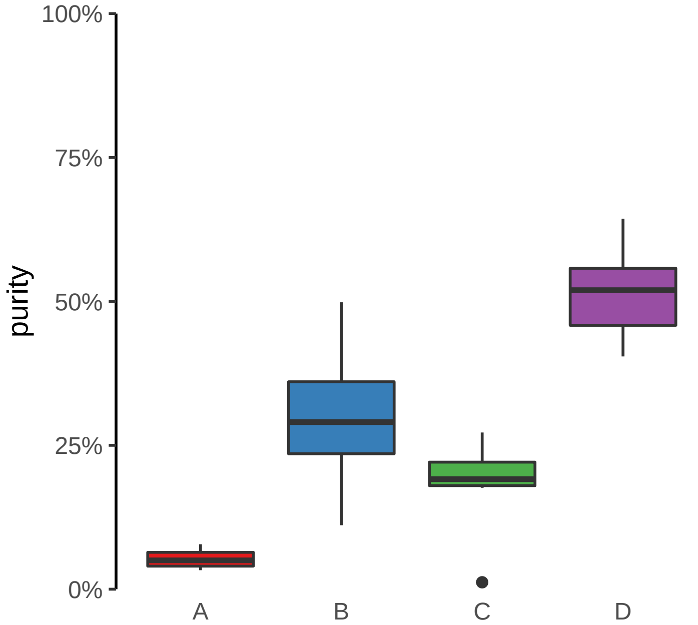

**SLC38A9 promoter loss**
**chr5:55700661−55701161**

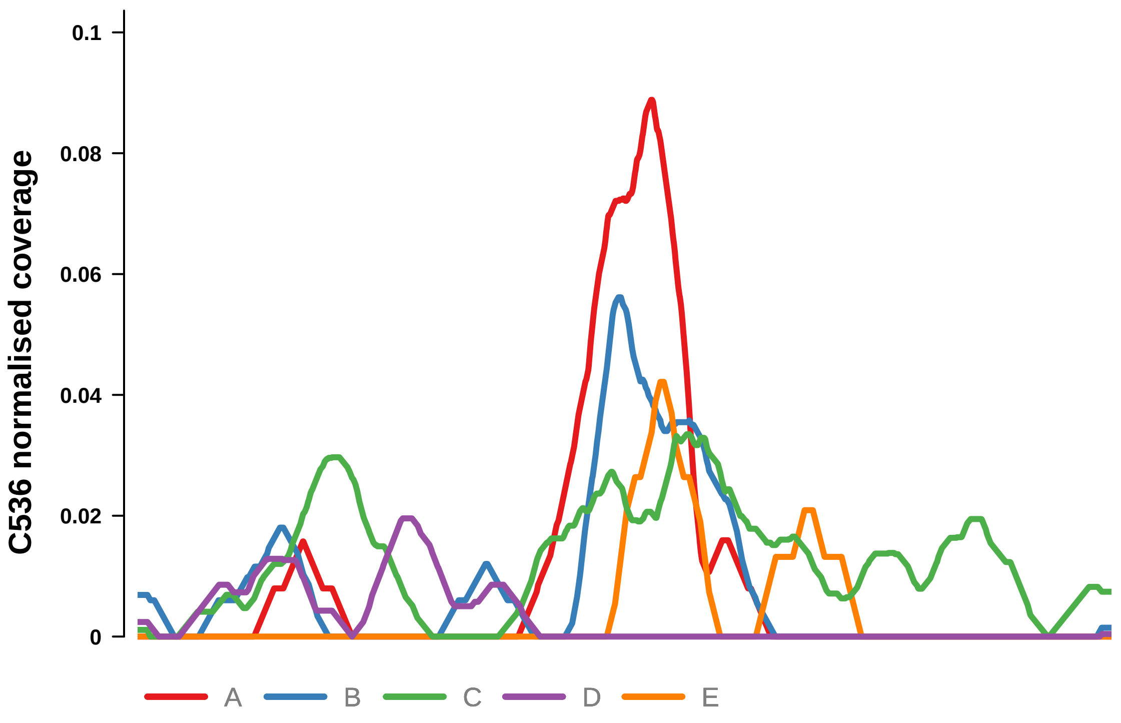


Putatively Clonal (logFC = −4.36)

| | P. | Adjusted P. |
|---|---|---|
| *~region vs. ~* | 1.72e−01 | NA |
| *~purity vs ~* | 1.89e−03 ** | 3.32e−02 * |
| *~purity+region vs ~purity* | 3.44e−01 | NA |
| *Carcinoma vs. normal* | 2.83e−05 **** | 2.83e−05 **** |

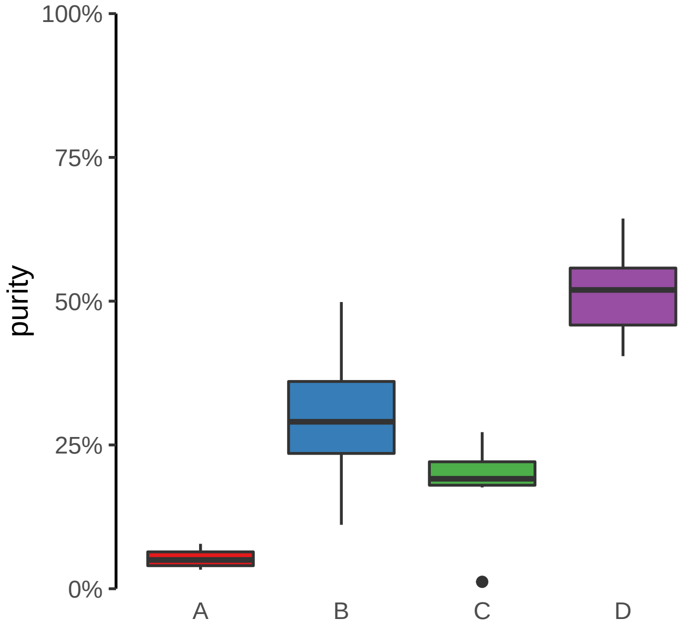

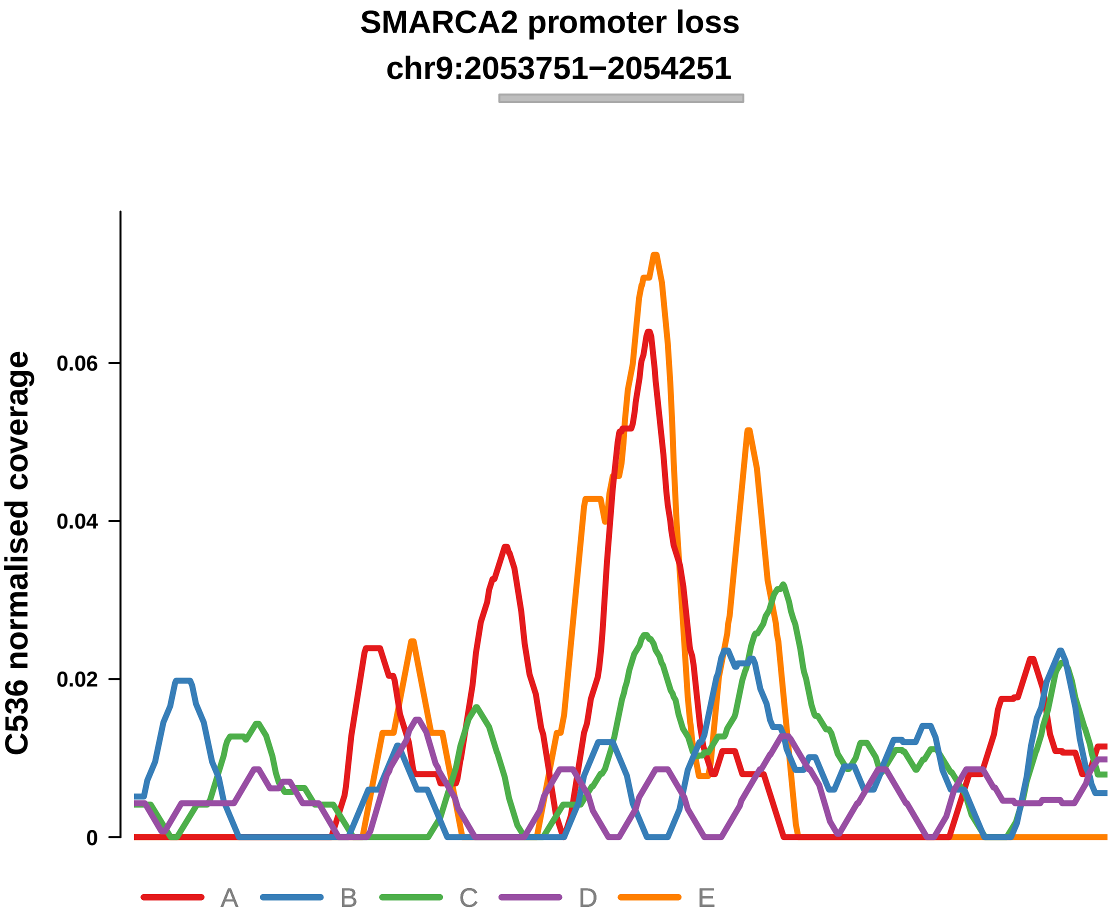


Putatively Clonal (logFC = −2.29)

| | P. | Adjusted P. |
|---|---|---|
| *~region vs. ~* | 9.77e−02 | NA |
| *~purity vs ~* | 7.44e−03 ** | 8.18e−02 |
| *~purity+region vs ~purity* | 7.87e−01 | NA |
| *Carcinoma vs. normal* | 5.04e−03 ** | 5.04e−03 ** |

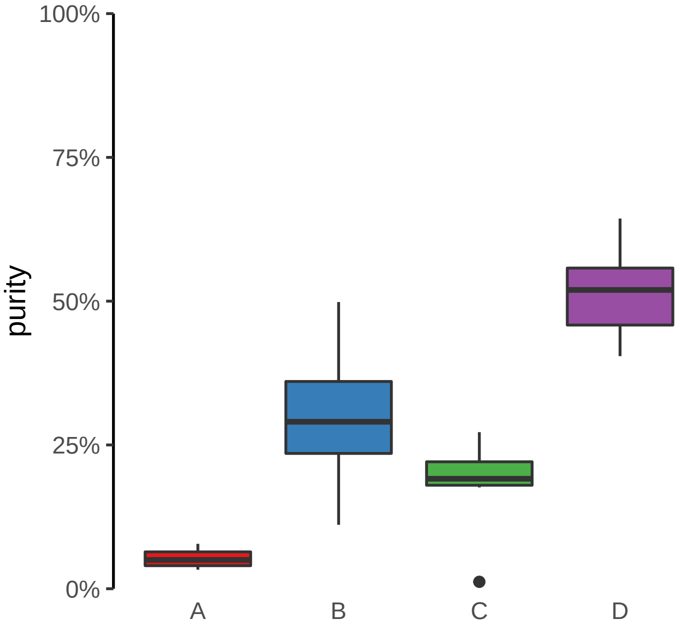

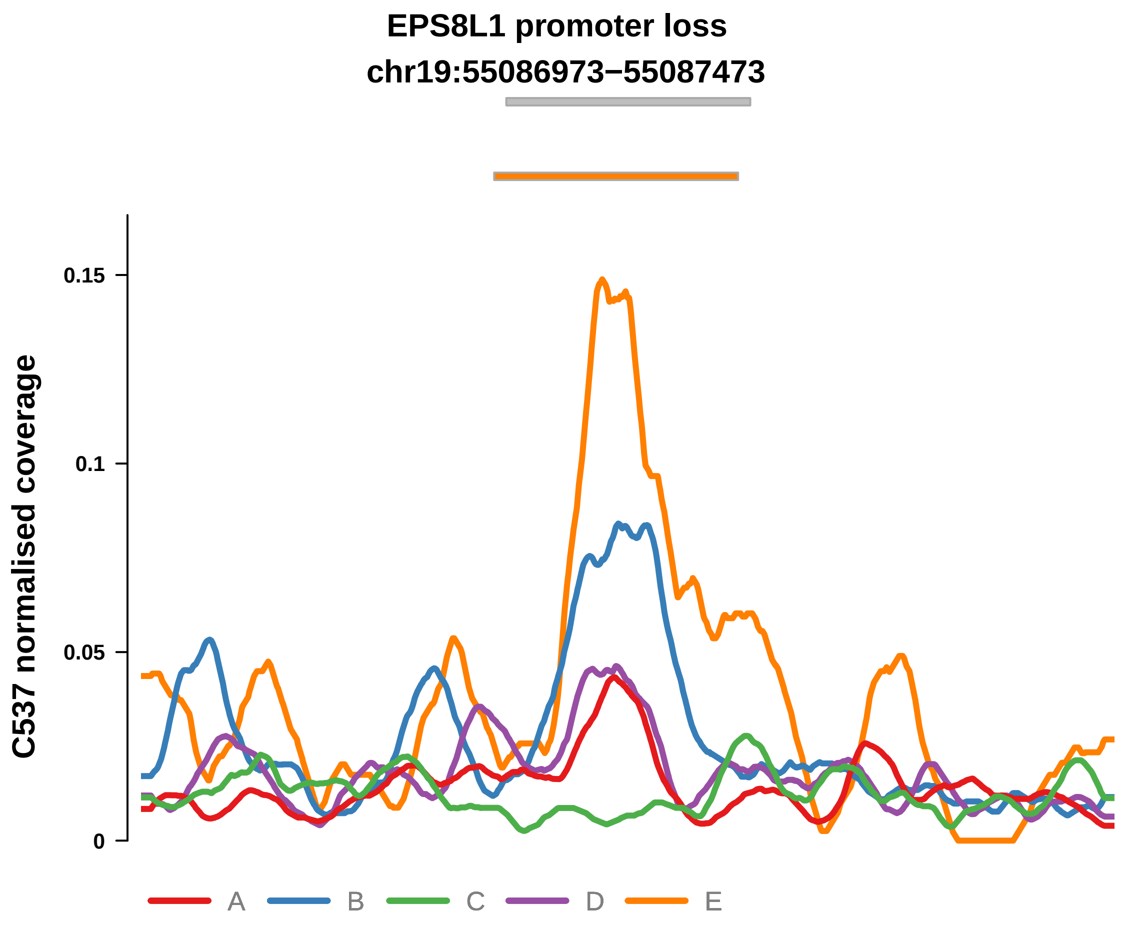


Putatively Clonal (logFC = −2.19)

| | P. | Adjusted P. |
|---|---|---|
| *~region vs. ~* | 2.49e−02 * | 7.46e−02 |
| *~purity vs ~* | 3.14e−02 * | 9.84e−02 |
| *~purity+region vs ~purity* | 8.6e−02 | 2.12e−01 |
| *Carcinoma vs. normal* | 3.94e−04 *** | 3.94e−04 *** |

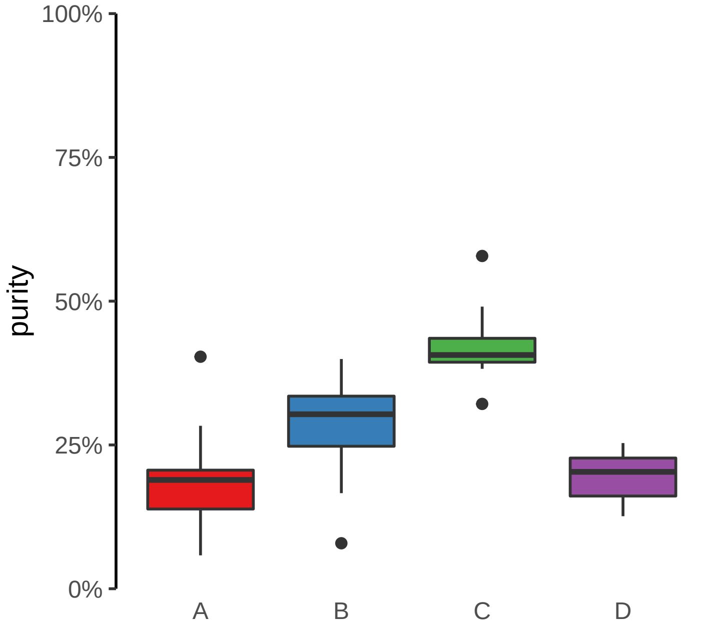

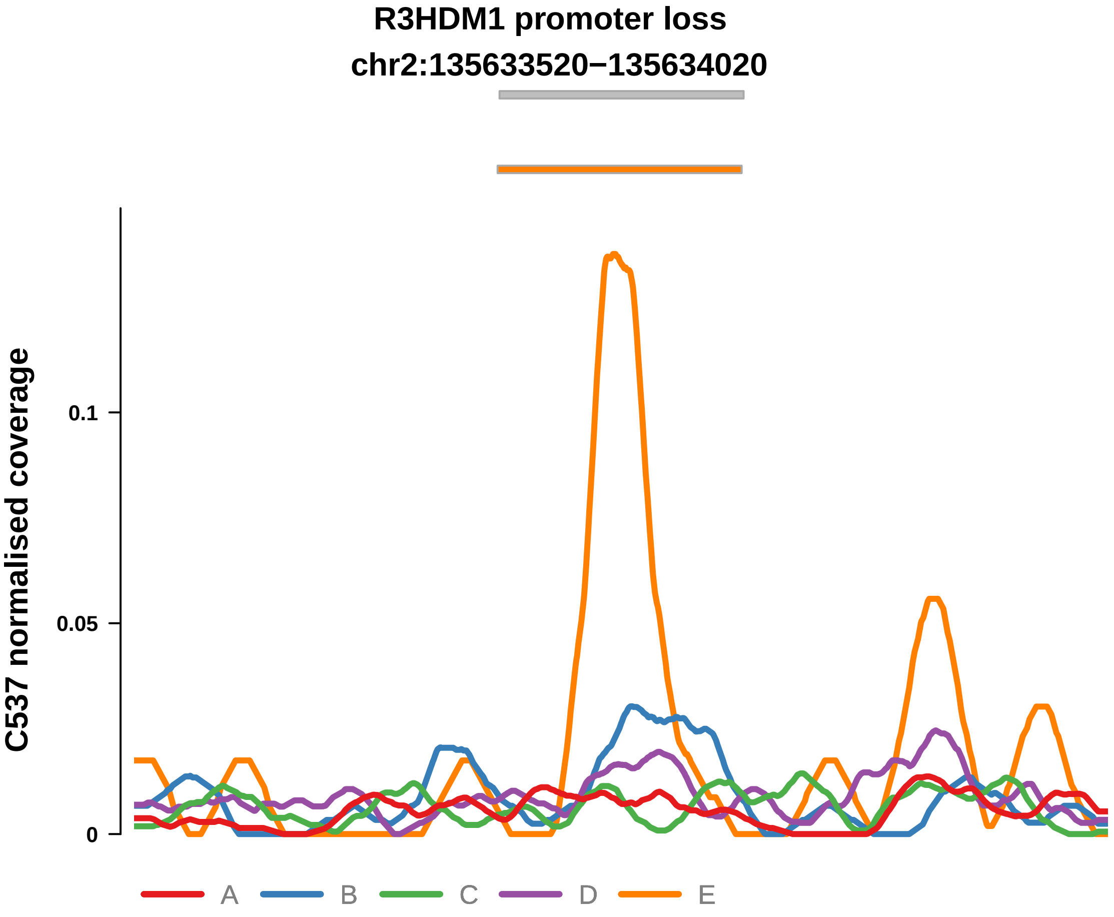


Putatively Clonal (logFC = −1.66)

| | P. | Adjusted P. |
|---|---|---|
| *~region vs. ~* | 5.71e−01 | 6.58e−01 |
| *~purity vs ~* | 6.27e−01 | NA |
| *~purity+region vs ~purity* | 6.15e−01 | NA |
| *Carcinoma vs. normal* | 9.09e−03 ** | 9.09e−03 ** |

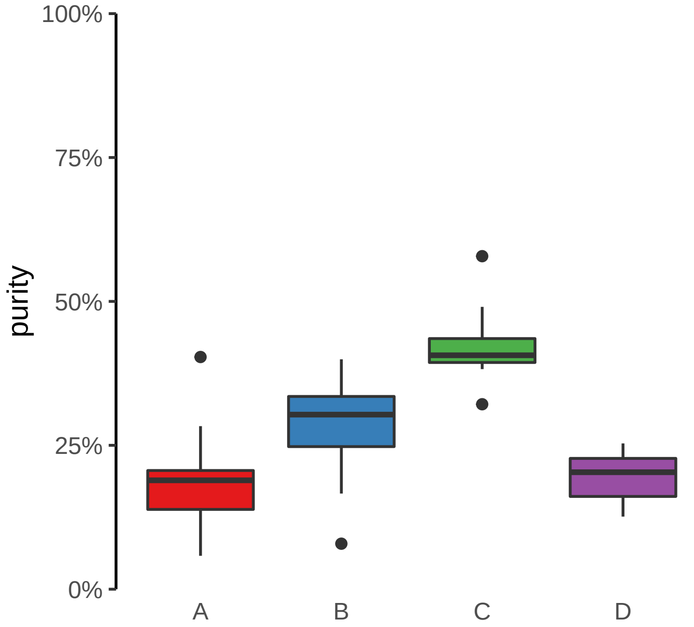

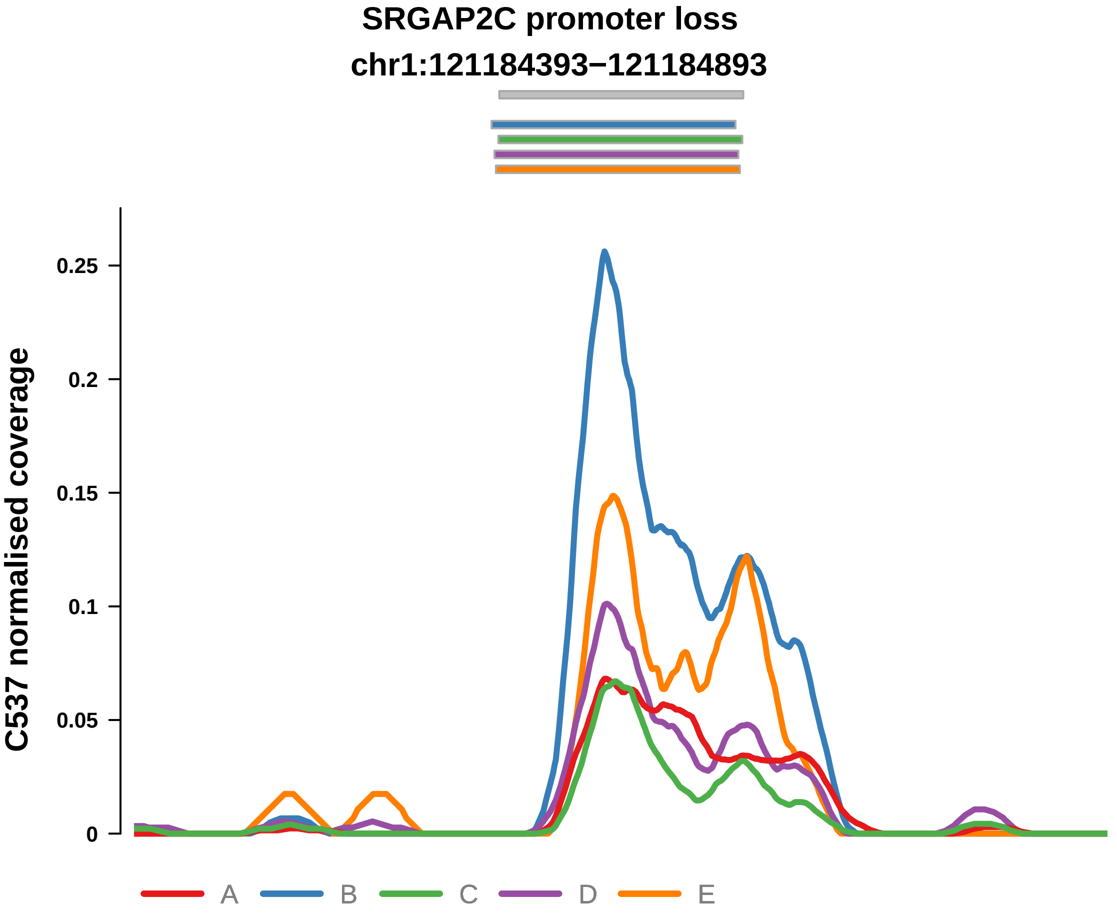


Putatively Clonal (logFC = −1.44)

| | P. | Adjusted P. |
|---|---|---|
| *~region vs. ~* | 2.83e−02 * | 7.46e−02 |
| *~purity vs ~* | 1.47e−01 | 2.66e−01 |
| *~purity+region vs ~purity* | 6.05e−02 | 1.77e−01 |
| *Carcinoma vs. normal* | 9.3e−03 ** | 9.3e−03 ** |

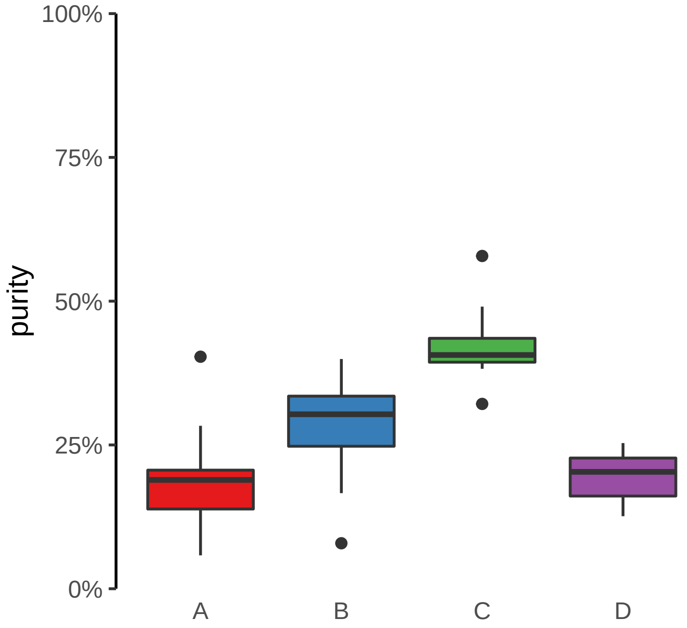

**AHCTF1 promoter loss**
**chr1:246874934−246875434**

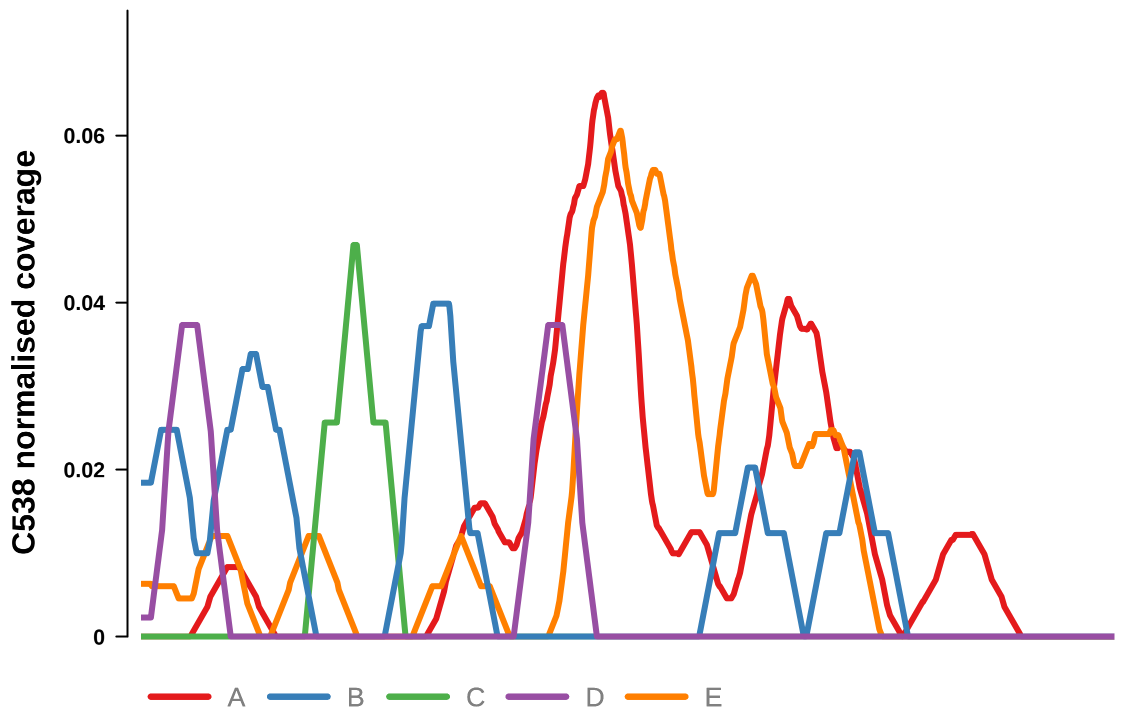


Putatively Clonal (logFC = −3.97)

| | P. | Adjusted P. |
|---|---|---|
| *~region vs. ~* | 2.66e−03 ** | 1.67e−02 * |
| *~purity vs ~* | 5.4e−04 *** | 4.32e−03 ** |
| *~purity+region vs ~purity* | 3.26e−01 | 8.7e−01 |
| *Carcinoma vs. normal* | 6.84e−07 **** | 6.84e−07 **** |

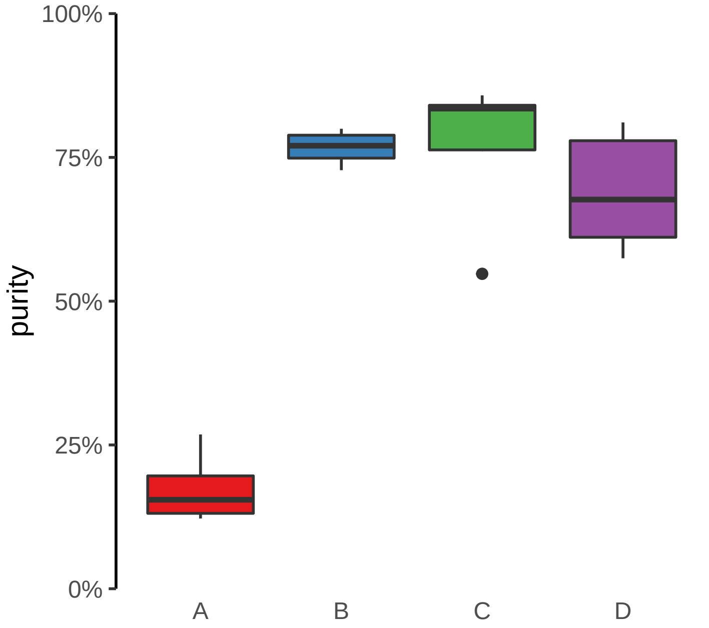

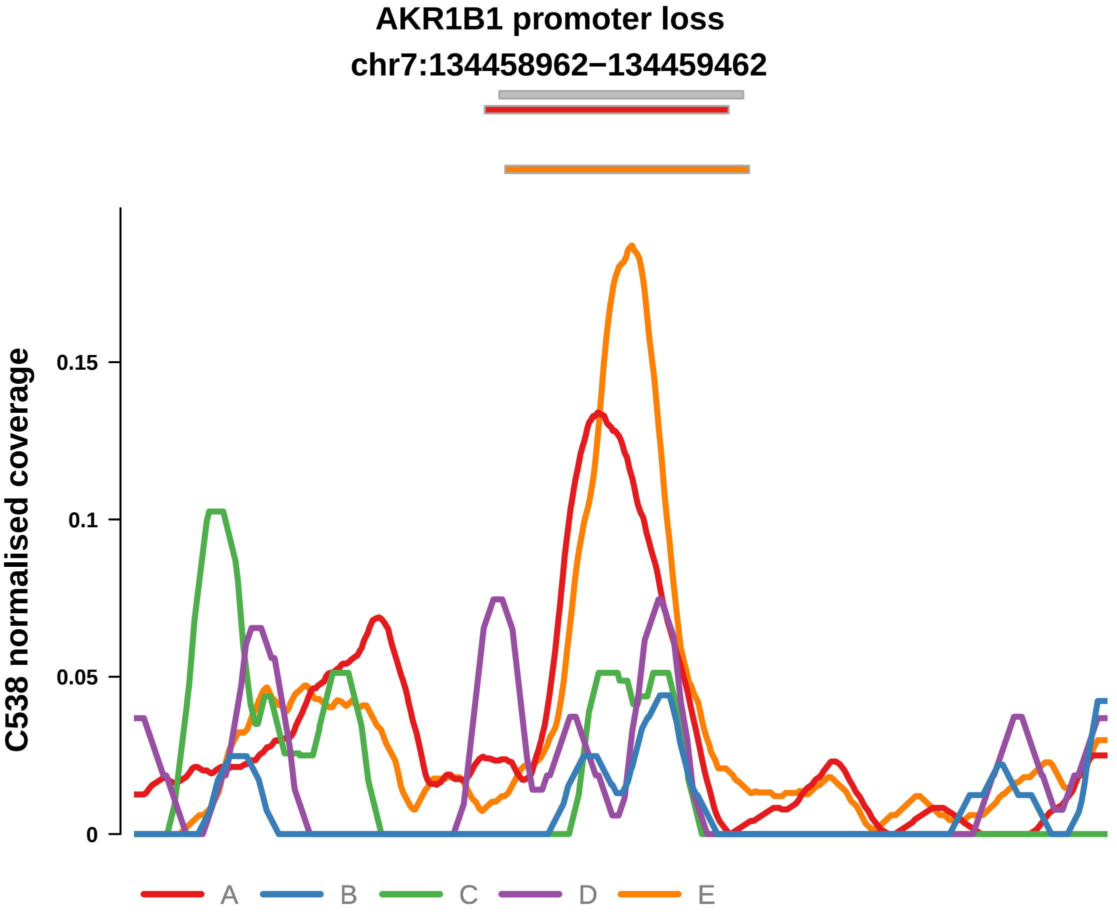


Putatively Clonal (logFC = −1.96)

| | P. | Adjusted P. |
|---|---|---|
| *~region vs. ~* | 7.41e−06 **** | 1.63e−04 *** |
| *~purity vs ~* | 1e−05 **** | 1.76e−04 *** |
| *~purity+region vs ~purity* | 2.28e−01 | 8.7e−01 |
| *Carcinoma vs. normal* | 1.29e−03 ** | 1.29e−03 ** |

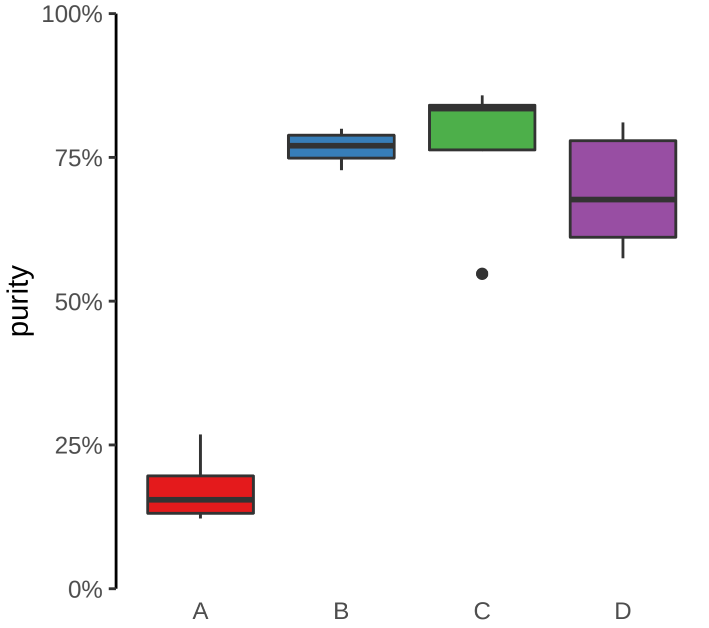

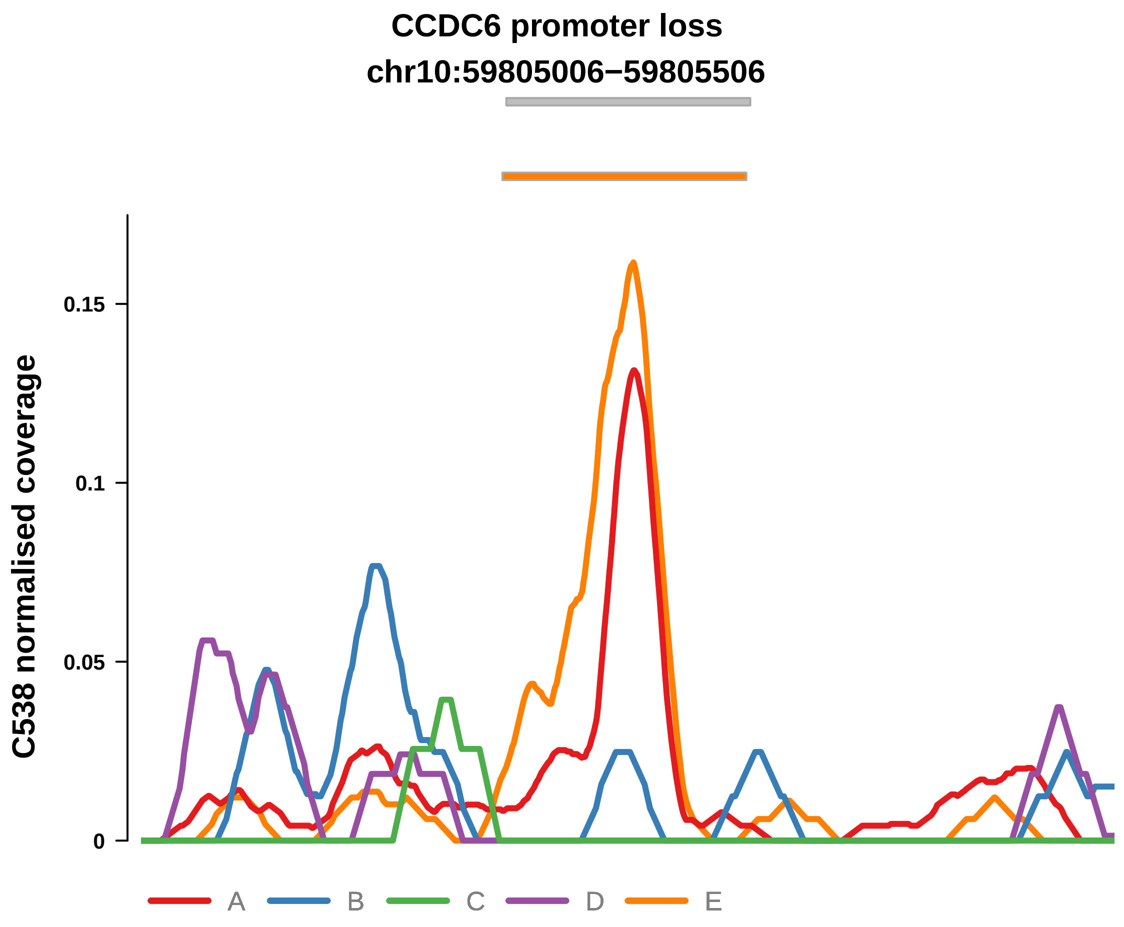


Putatively Clonal (logFC = −3.92)

| | P. | Adjusted P. |
|---|---|---|
| *~region vs. ~* | 2.97e−02 * | 9.07e−02 |
| *~purity vs ~* | 2.81e−03 ** | 1.19e−02 * |
| *~purity+region vs ~purity* | 5.67e−01 | 8.7e−01 |
| *Carcinoma vs. normal* | 9.88e−07 **** | 9.88e−07 **** |

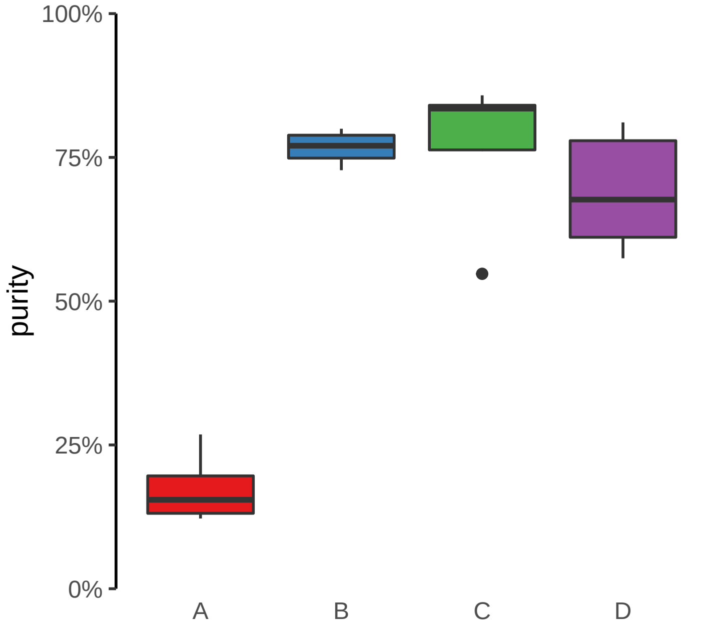

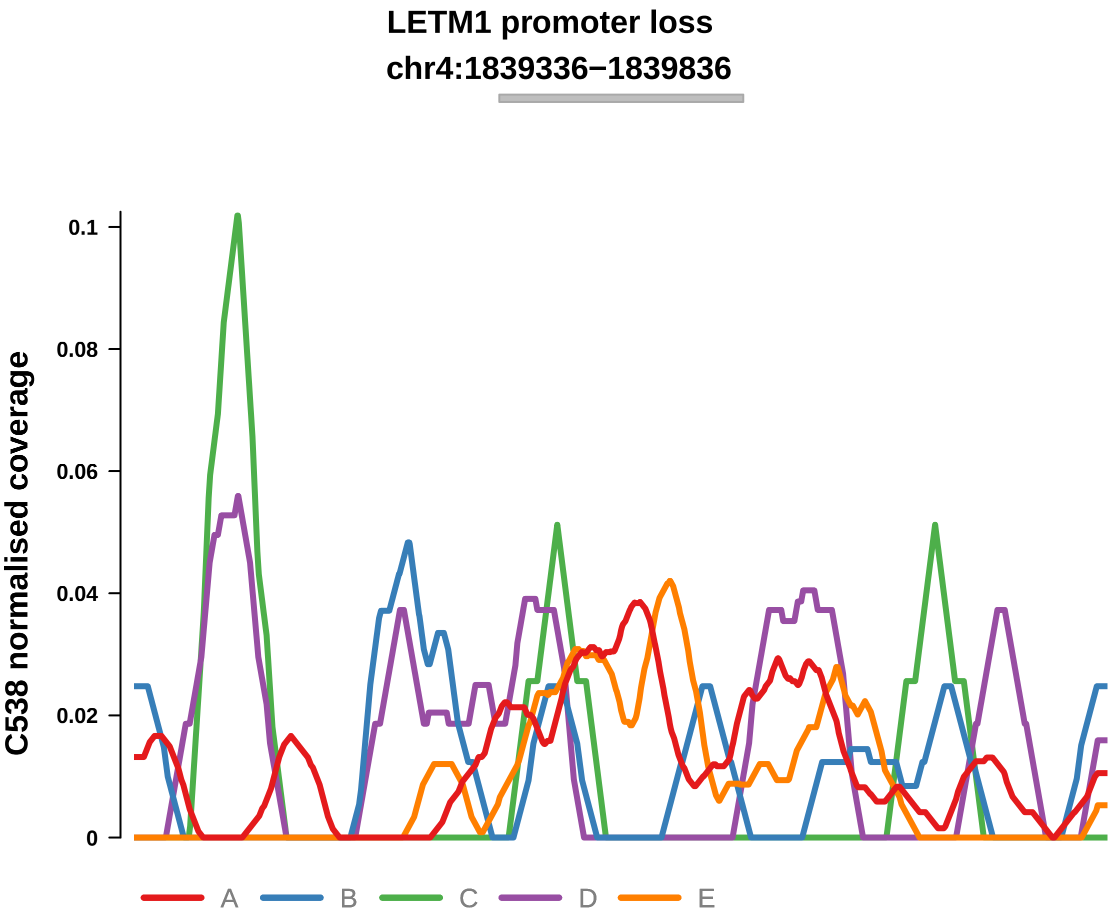


Putatively Clonal (logFC = −2.27)

| | P. | Adjusted P. |
|---|---|---|
| *~region vs. ~* | 2.54e−02 * | 8.95e−02 |
| *~purity vs ~* | 1.74e−03 ** | 8.89e−03 ** |
| *~purity+region vs ~purity* | 1e+00 | 1e+00 |
| *Carcinoma vs. normal* | 8.37e−04 *** | 8.37e−04 *** |

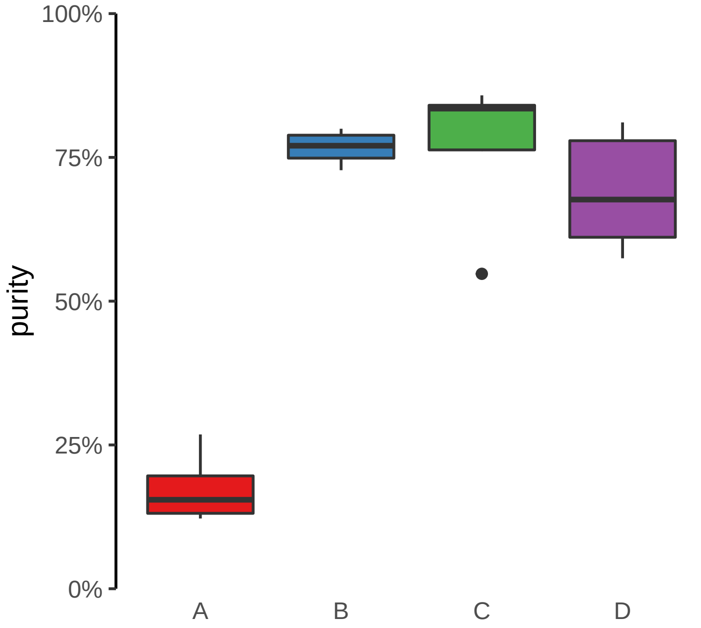

**LPCAT4 promoter loss**

**chr15:34362318−34362818**

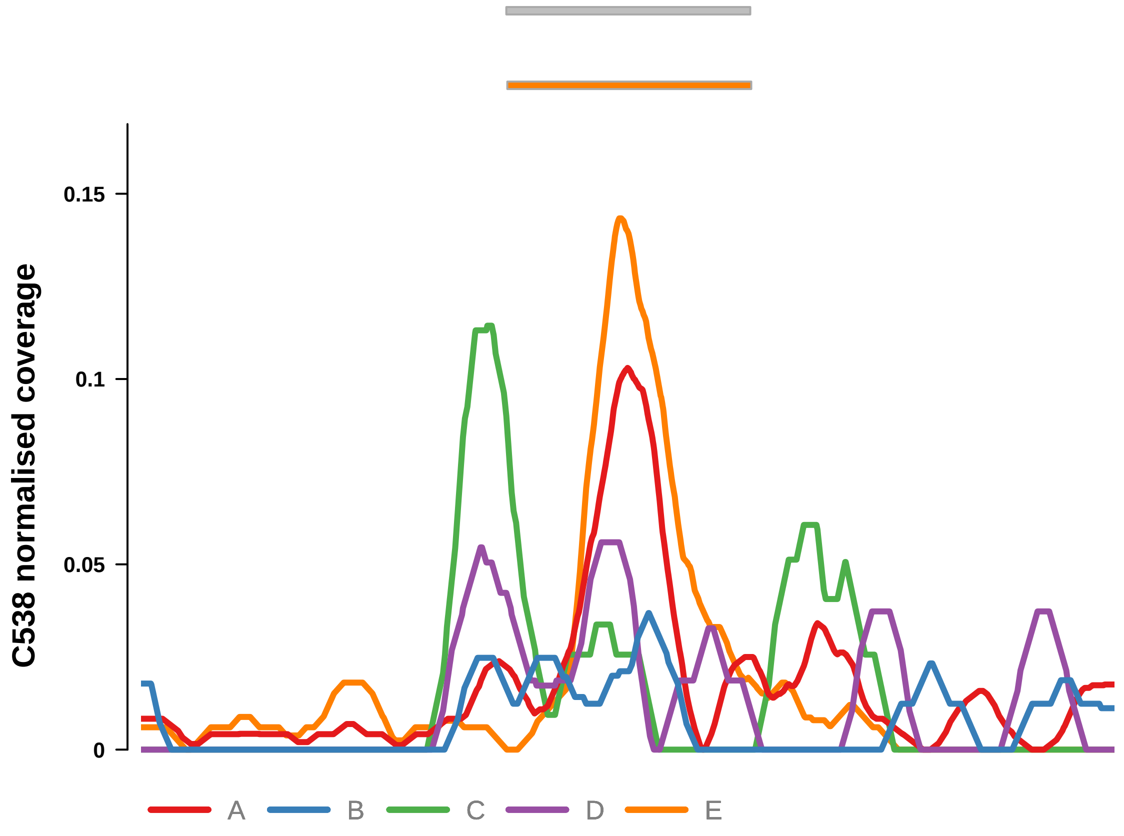


Putatively Clonal (logFC = −1.81)

| | **P.** | **Adjusted P.** |
|---|---|---|
| *~region vs. ~* | 1.42e−02 * | 6.57e−02 |
| *~purity vs ~* | 4.64e−03 ** | 1.77e−02 * |
| *~purity+region vs ~purity* | 5.56e−01 | 8.7e−01 |
| *Carcinoma vs. normal* | 3.95e−03 ** | 3.95e−03 ** |

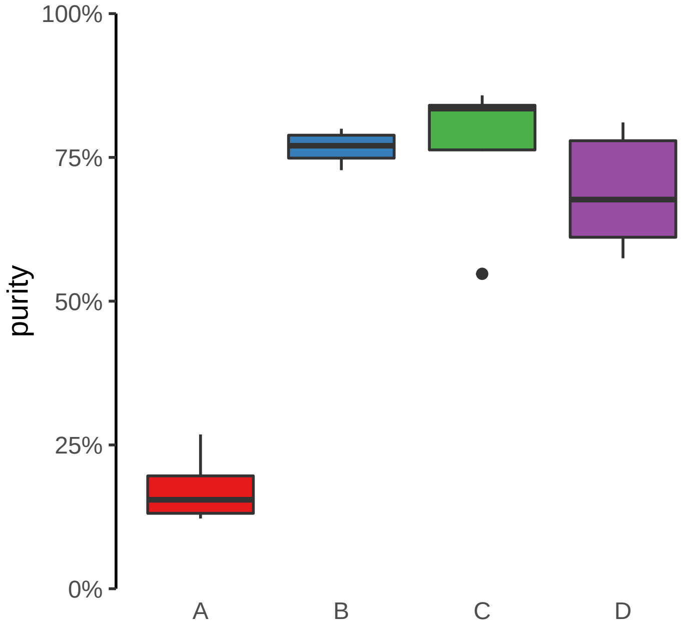

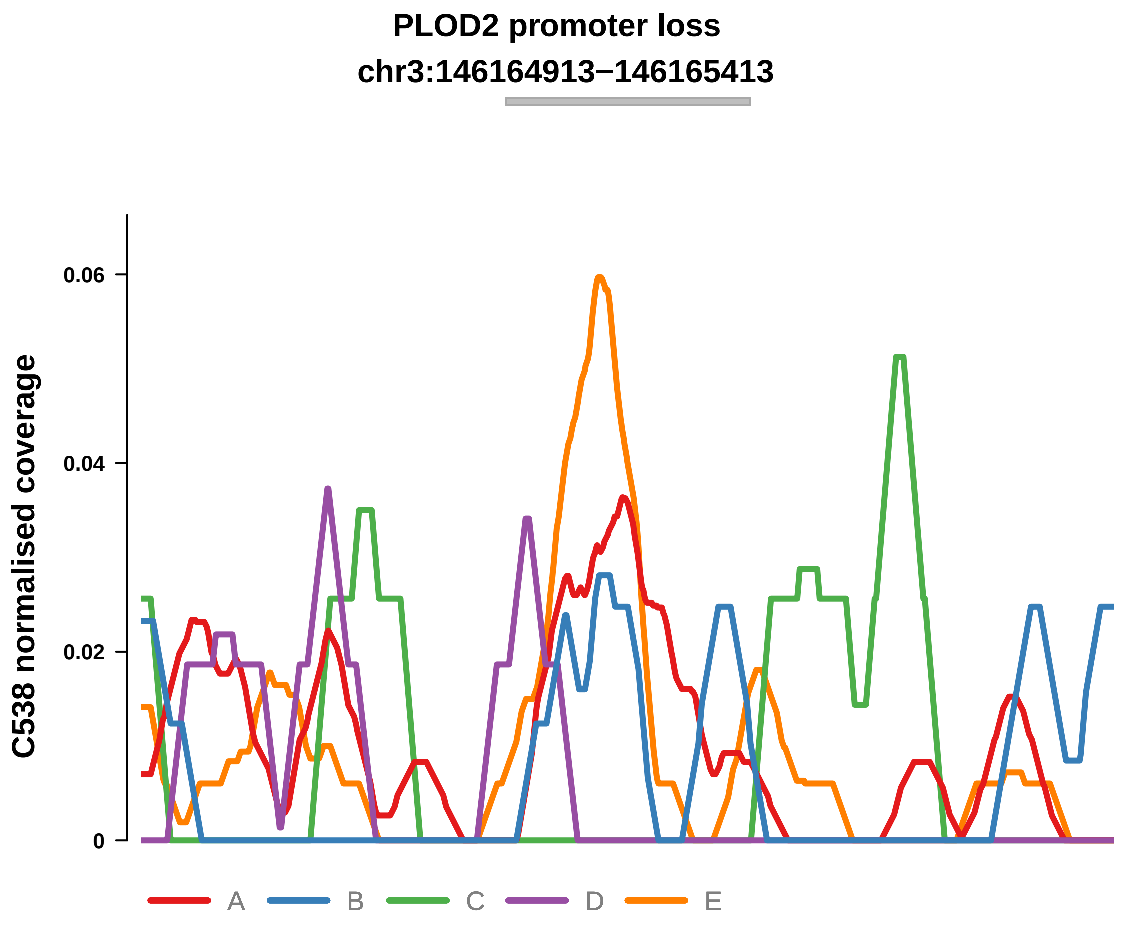


Putatively Clonal (logFC = −2.29)

| | P. | Adjusted P. |
|---|---|---|
| *~region vs. ~* | 4.15e−01 | 5.99e−01 |
| *~purity vs ~* | 1.45e−01 | 2.74e−01 |
| *~purity+region vs ~purity* | 5.67e−01 | 8.7e−01 |
| *Carcinoma vs. normal* | 9.51e−04 *** | 9.51e−04 *** |

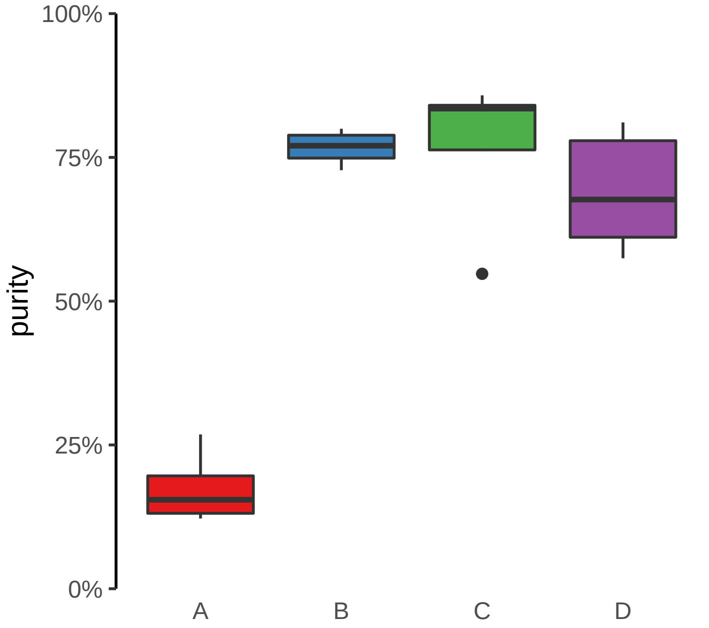

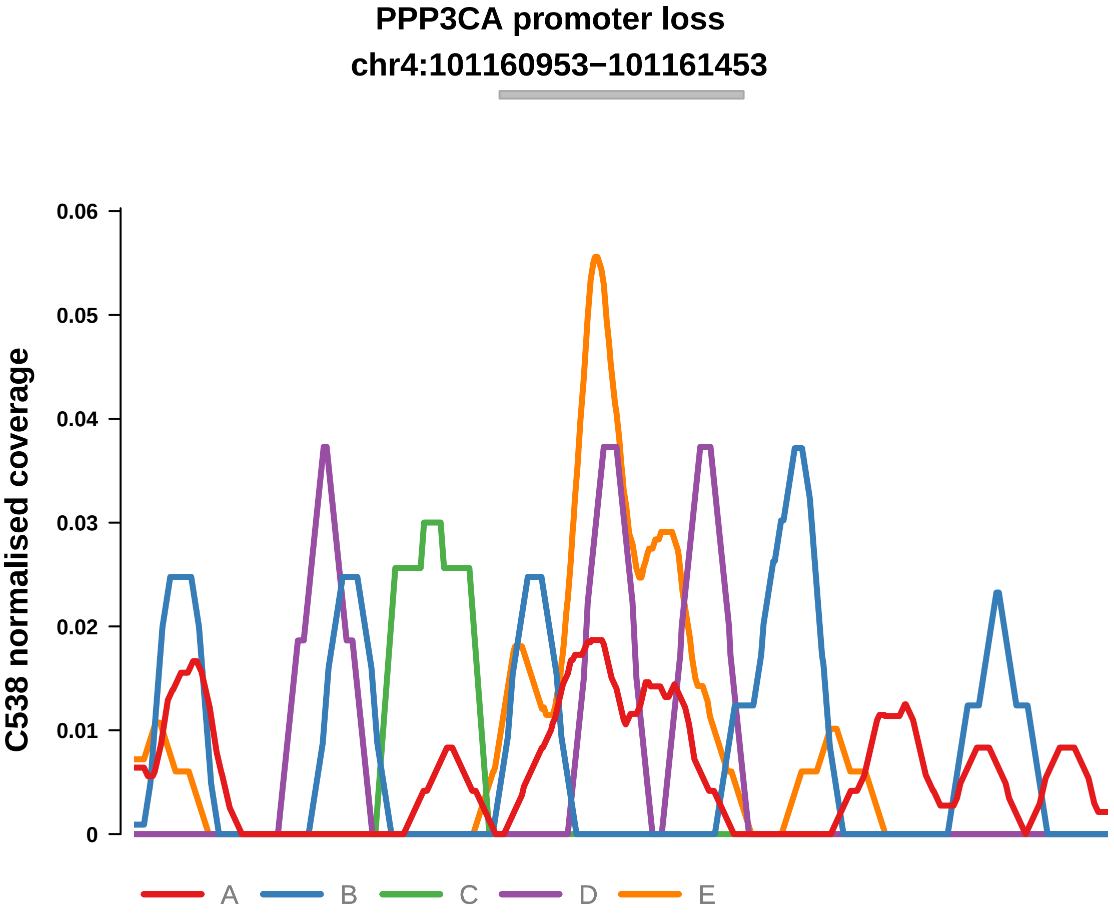


Putatively Clonal (logFC = −2.25)

| | P. | Adjusted P. |
|---|---|---|
| *~region vs. ~* | 6.26e−01 | 7.76e−01 |
| *~purity vs ~* | 3.39e−01 | 5.05e−01 |
| *~purity+region vs ~purity* | 7.44e−01 | 9.36e−01 |
| *Carcinoma vs. normal* | 2.12e−03 ** | 2.12e−03 ** |

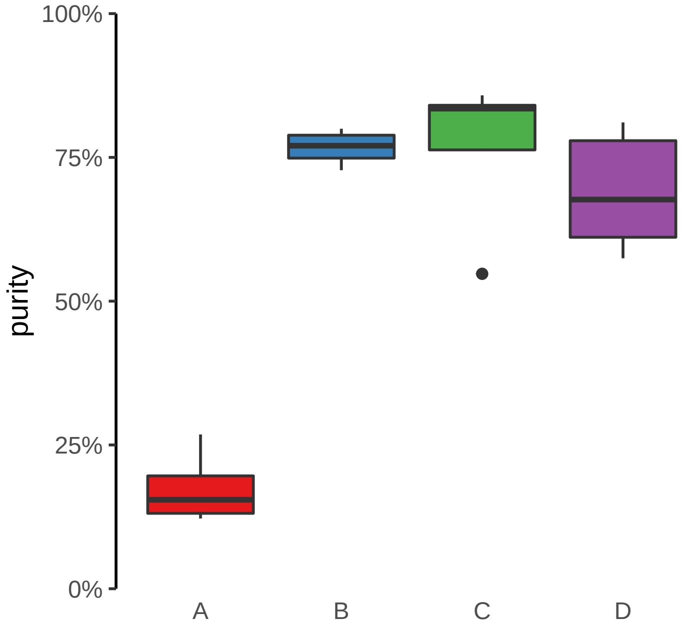

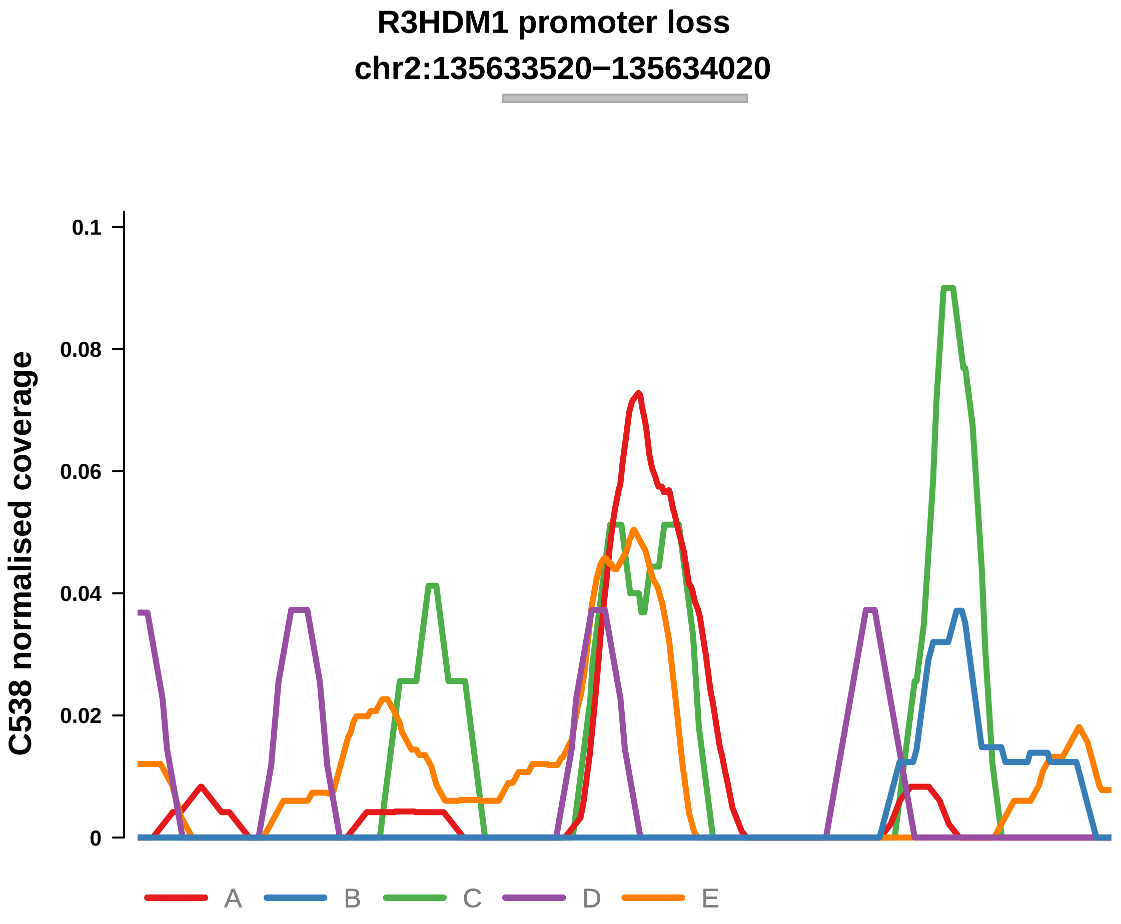


Putatively Clonal (logFC = −2.68)

| | P. | Adjusted P. |
|---|---|---|
| *~region vs. ~* | 5.18e−02 | 1.46e−01 |
| *~purity vs ~* | 9.21e−03 ** | 3e−02 * |
| *~purity+region vs ~purity* | 4.84e−01 | 8.7e−01 |
| *Carcinoma vs. normal* | 2.59e−04 *** | 2.59e−04 *** |

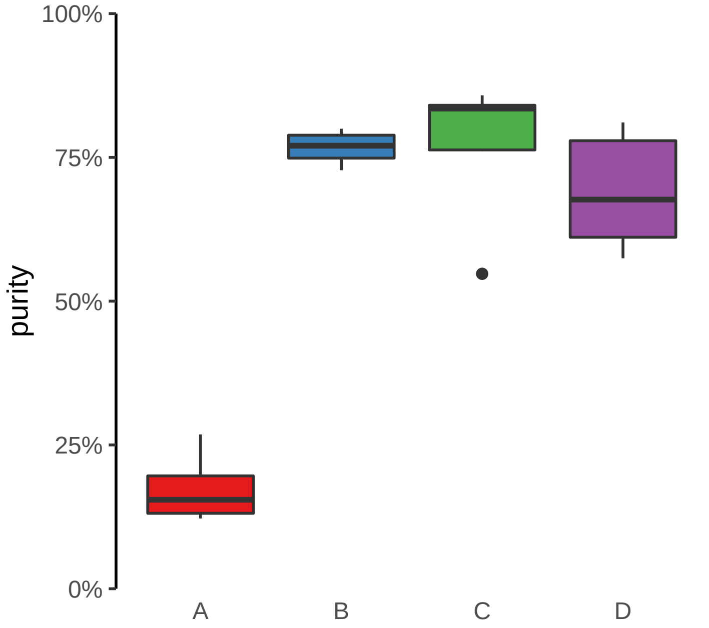

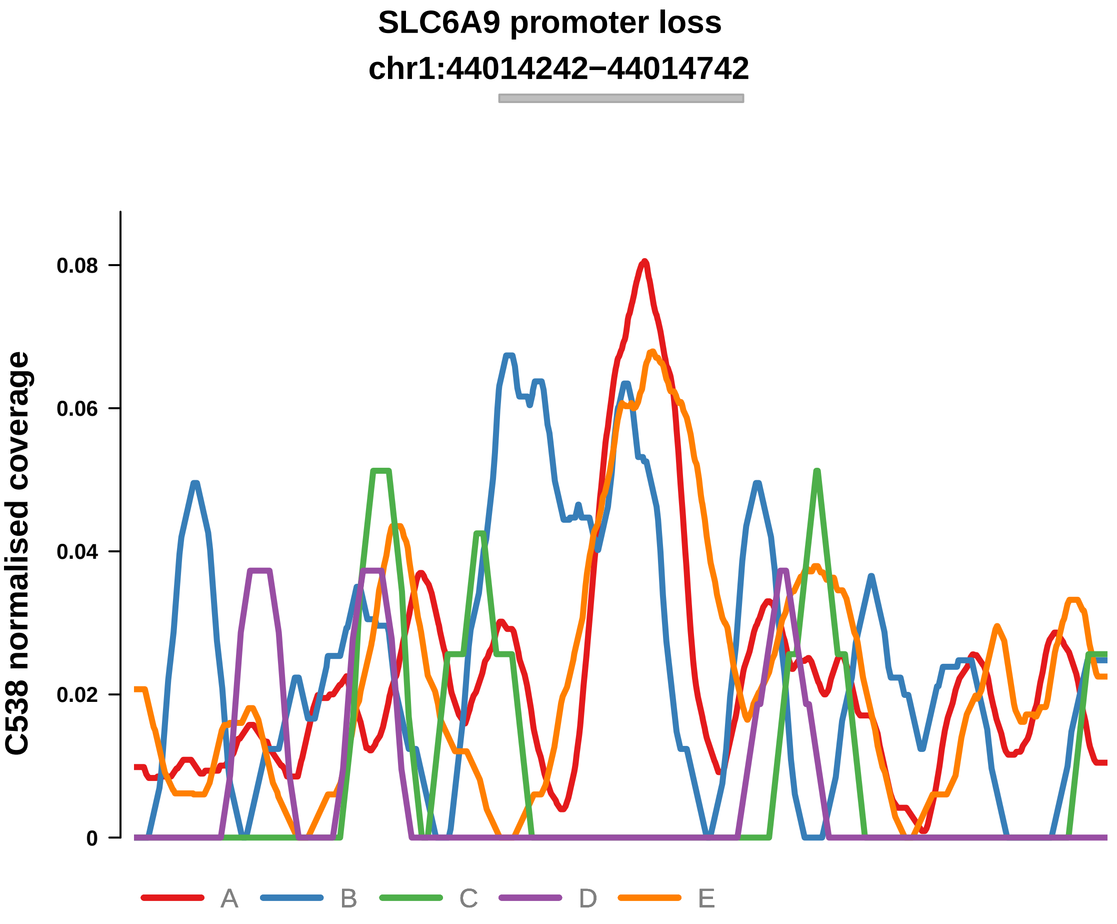


Putatively Clonal (logFC = −1.85)

| | P. | Adjusted P. |
|---|---|---|
| *~region vs. ~* | 3.89e−04 *** | 4.28e−03 ** |
| *~purity vs ~* | 1.76e−03 ** | 8.89e−03 ** |
| *~purity+region vs ~purity* | 6.63e−02 | 6.8e−01 |
| *Carcinoma vs. normal* | 3.47e−03 ** | 3.47e−03 ** |

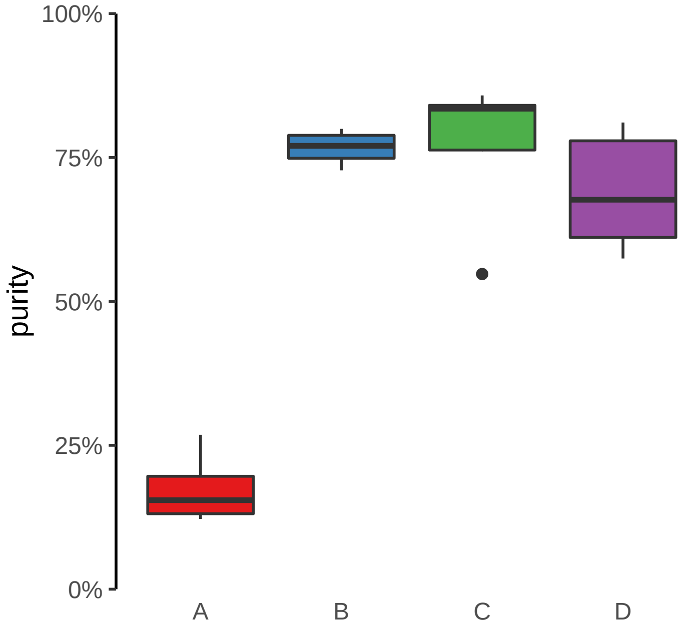

**SMAD3 promoter loss**
**chr15:67127194−67127694**

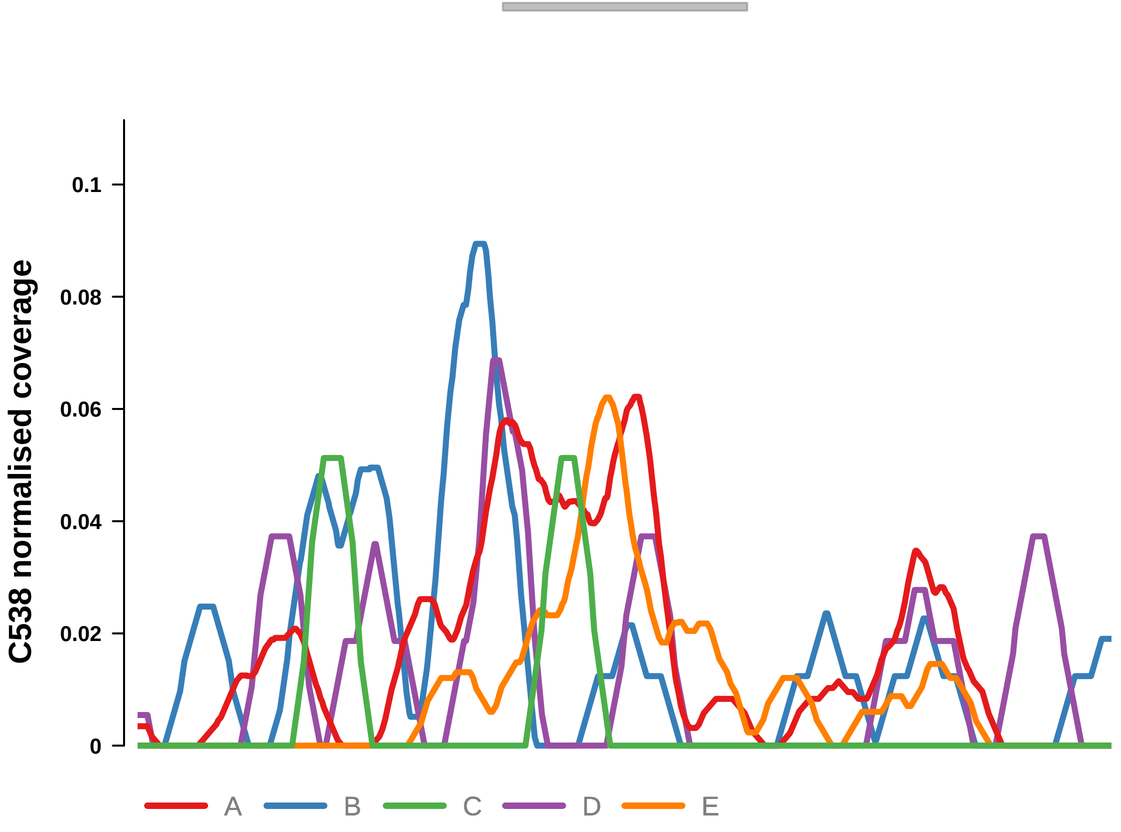


Putatively Clonal (logFC = −2.75)

| | P. | Adjusted P. |
|---|---|---|
| *~region vs. ~* | 1.73e−03 ** | 1.25e−02 * |
| *~purity vs ~* | 1.77e−04 *** | 1.77e−03 ** |
| *~purity+region vs ~purity* | 6.59e−01 | 9.21e−01 |
| *Carcinoma vs. normal* | 5.96e−05 **** | 5.96e−05 **** |

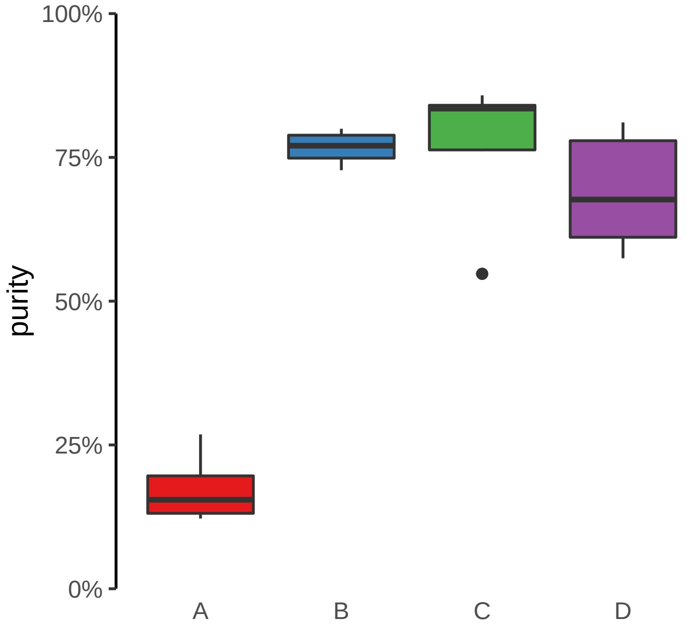

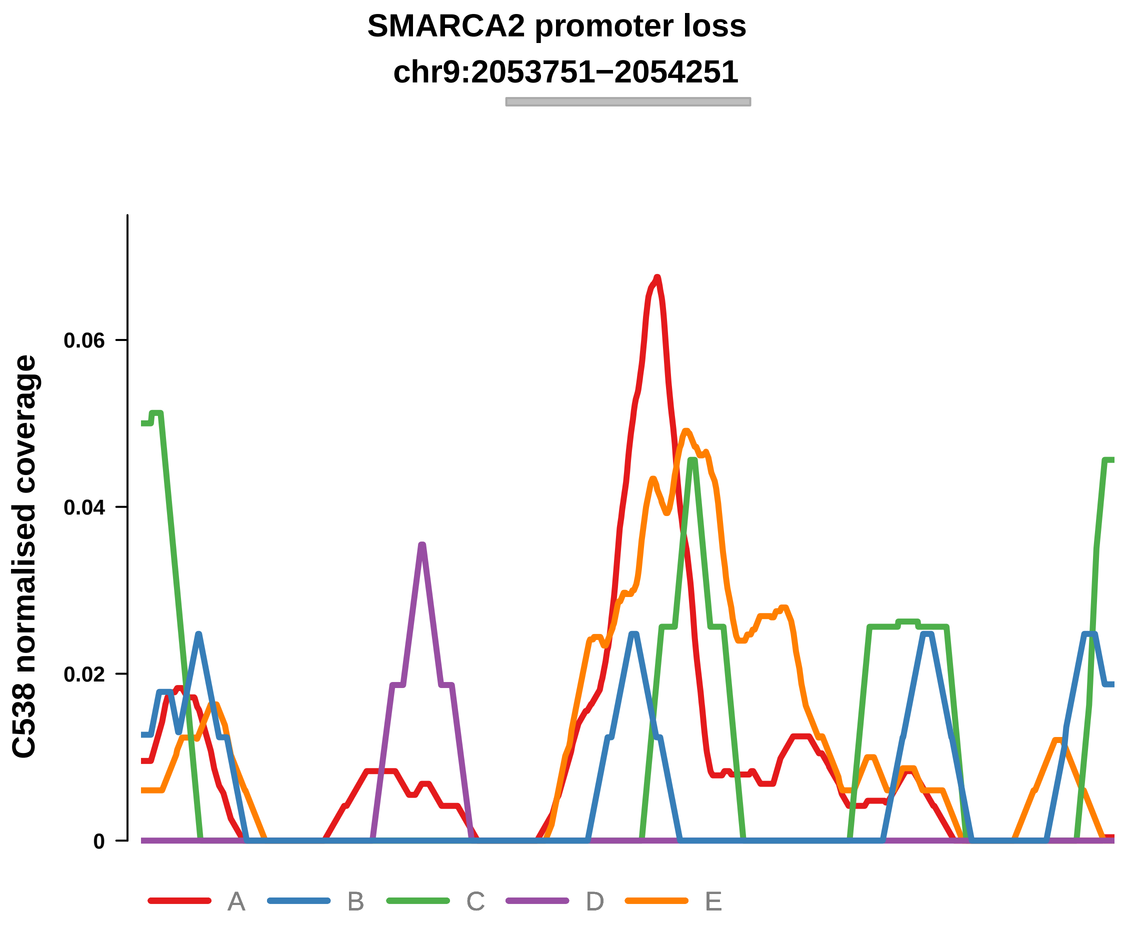


Putatively Clonal (logFC = −2.67)

| | P. | Adjusted P. |
|---|---|---|
| *~region vs. ~* | 7e−02 | 1.76e−01 |
| *~purity vs ~* | 4.44e−03 ** | 1.77e−02 * |
| *~purity+region vs ~purity* | 9.48e−01 | 9.79e−01 |
| *Carcinoma vs. normal* | 1.01e−03 ** | 1.01e−03 ** |

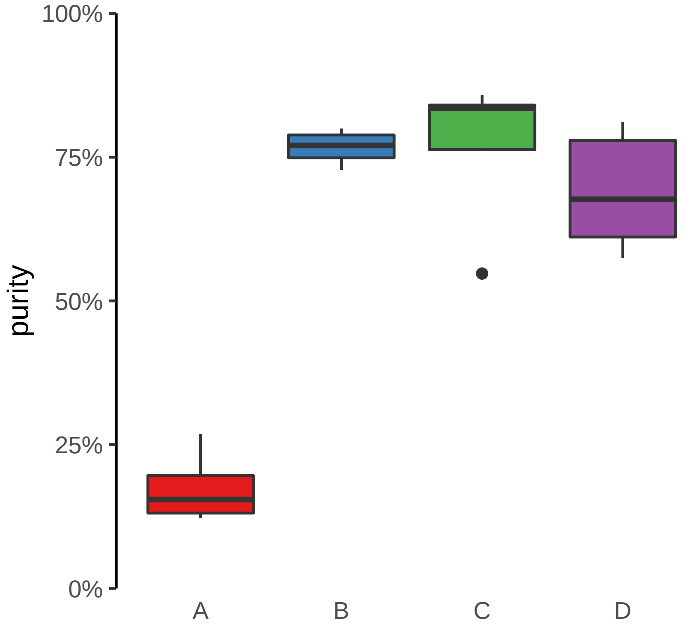

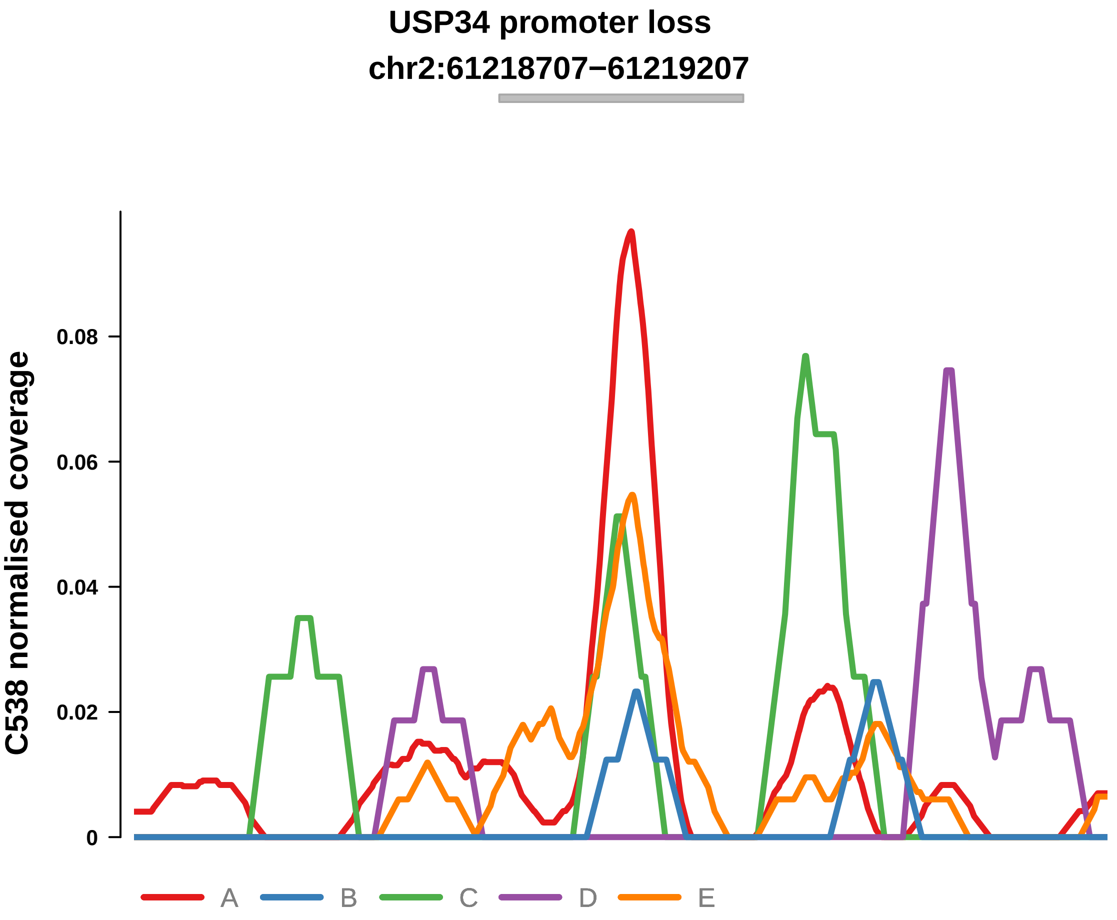


Putatively Clonal (logFC = −2.78)

| | P. | Adjusted P. |
|---|---|---|
| *~region vs. ~* | 5.32e−02 | 1.46e−01 |
| *~purity vs ~* | 8.08e−03 ** | 2.74e−02 * |
| *~purity+region vs ~purity* | 6.4e−01 | 9.12e−01 |
| *Carcinoma vs. normal* | 5.28e−04 *** | 5.28e−04 *** |

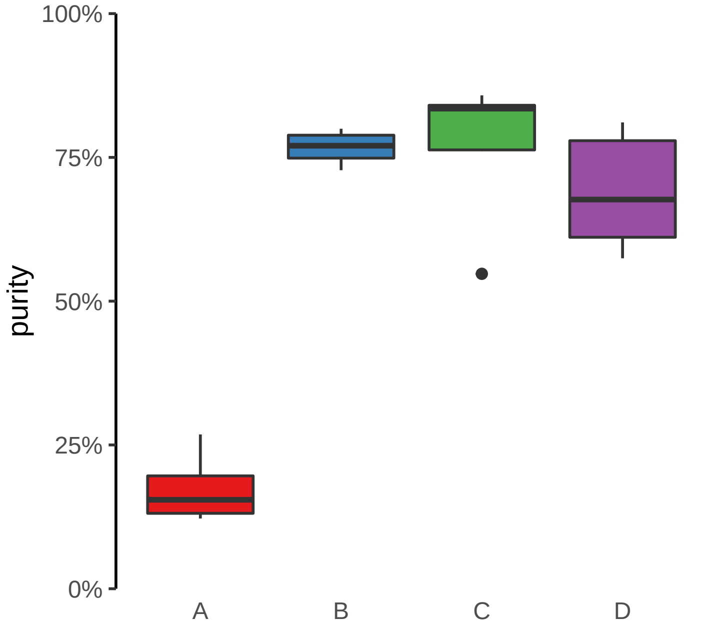

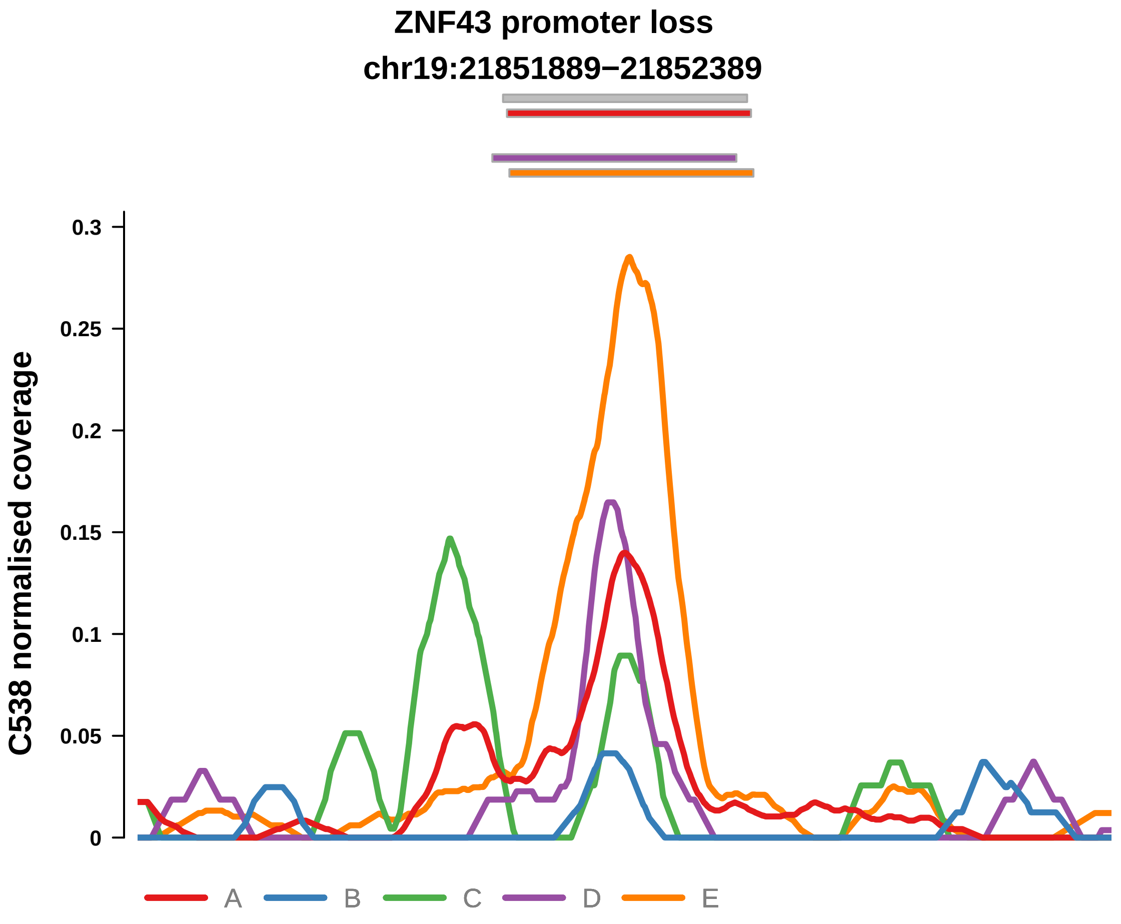


Putatively Clonal (logFC = −2.56)

| | P. | Adjusted P. |
|---|---|---|
| *~region vs. ~* | 9.24e−08 **** | 8.13e−06 **** |
| *~purity vs ~* | 1.49e−06 **** | 4.38e−05 **** |
| *~purity+region vs ~purity* | 3.93e−02 * | 5.76e−01 |
| *Carcinoma vs. normal* | 2.36e−05 **** | 2.36e−05 **** |

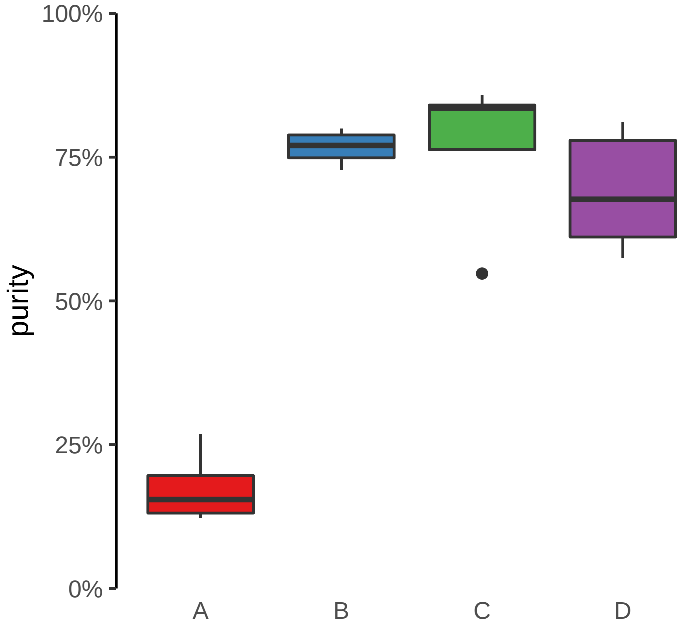

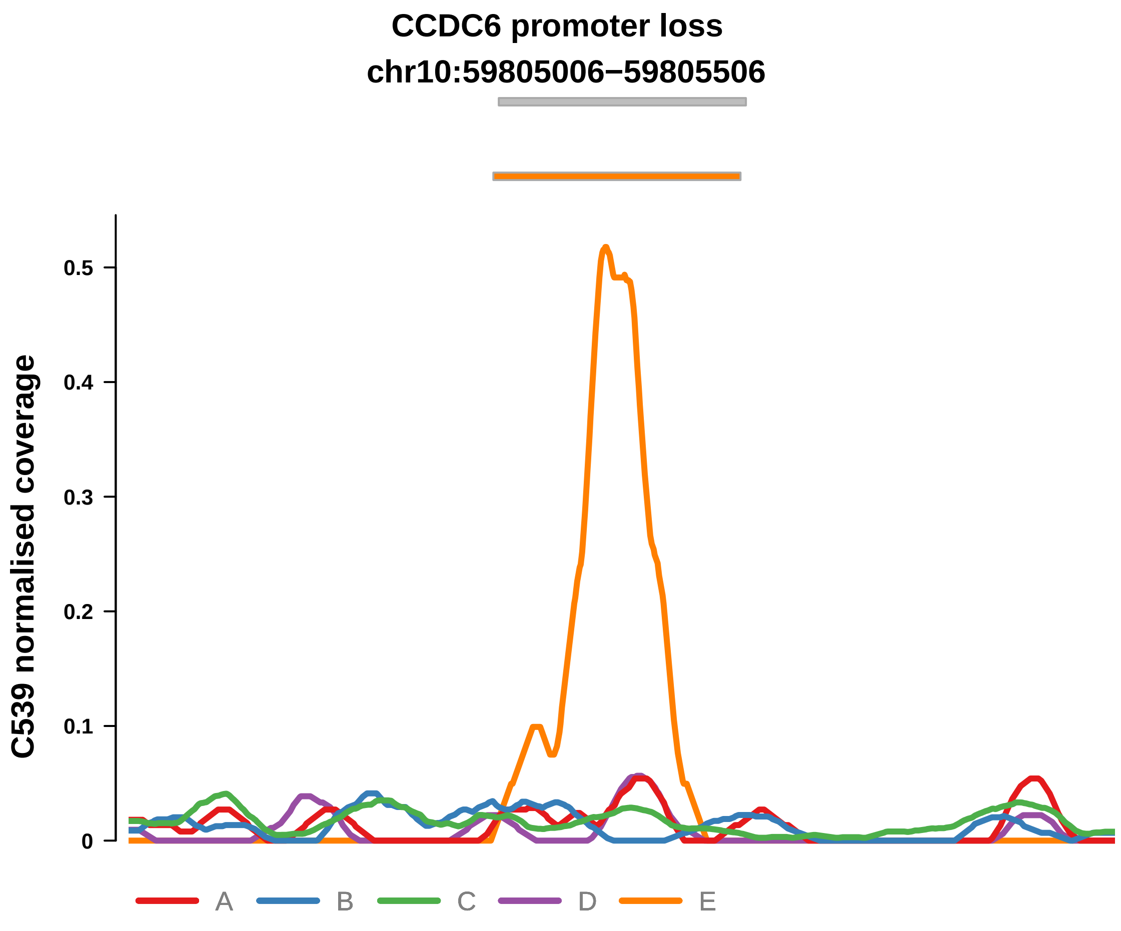


Putatively Clonal (logFC = −3.4)

| | P. | Adjusted P. |
|---|---|---|
| *~region vs. ~* | 1.1e−03 ** | 2.76e−03 ** |
| *~purity vs ~* | 5.09e−04 *** | 2.36e−03 ** |
| *~purity+region vs ~purity* | 3.82e−01 | 4.44e−01 |
| *Carcinoma vs. normal* | 3.35e−03 ** | 3.35e−03 ** |

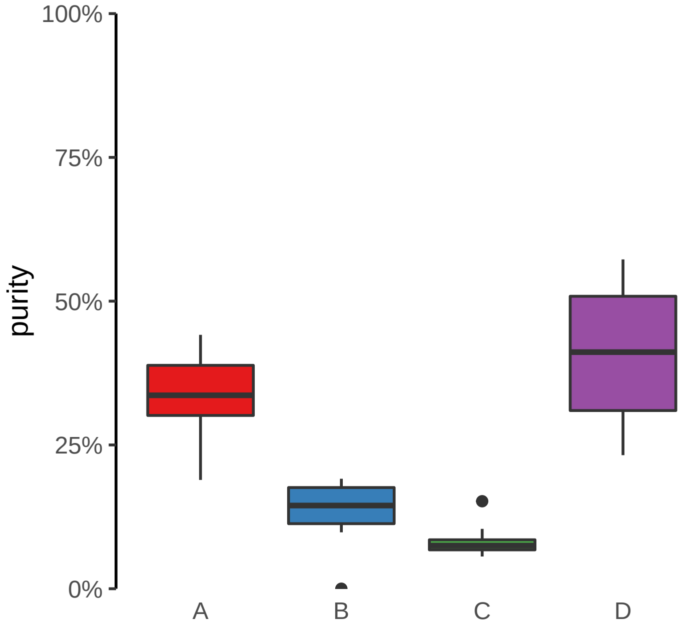

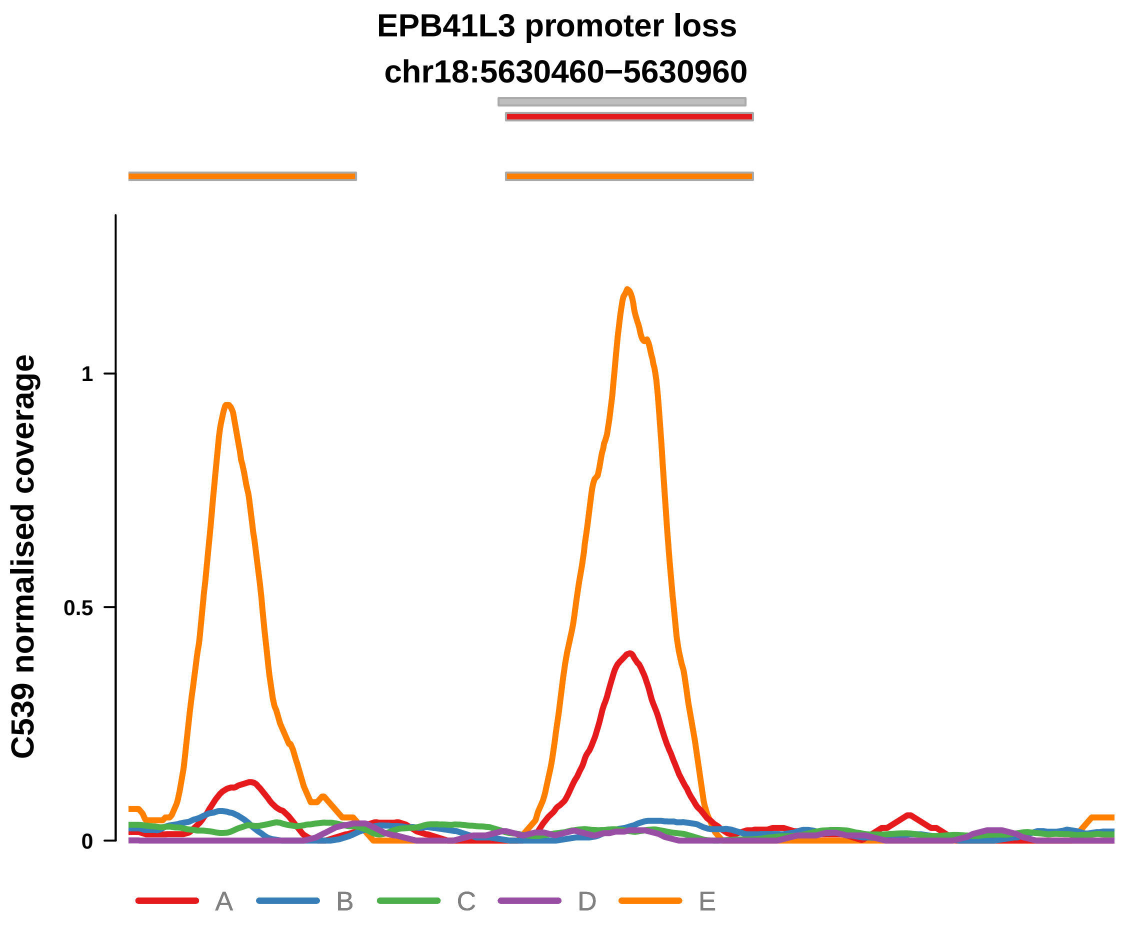


Putatively Clonal (logFC = −2.44)

| | P. | Adjusted P. |
|---|---|---|
| *~region vs. ~* | 2.91e−03 ** | 6.24e−03 ** |
| *~purity vs ~* | 2.52e−03 ** | 5.98e−03 ** |
| *~purity+region vs ~purity* | 2.37e−01 | 3.39e−01 |
| *Carcinoma vs. normal* | 2.43e−03 ** | 2.43e−03 ** |

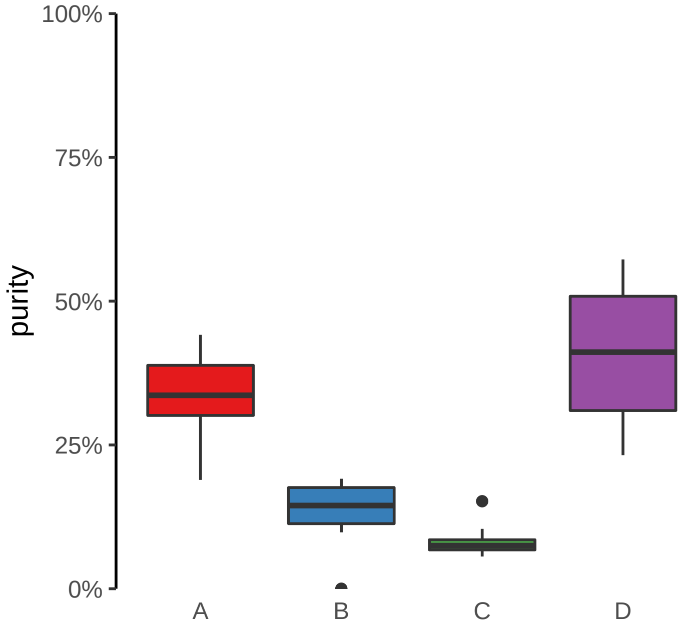

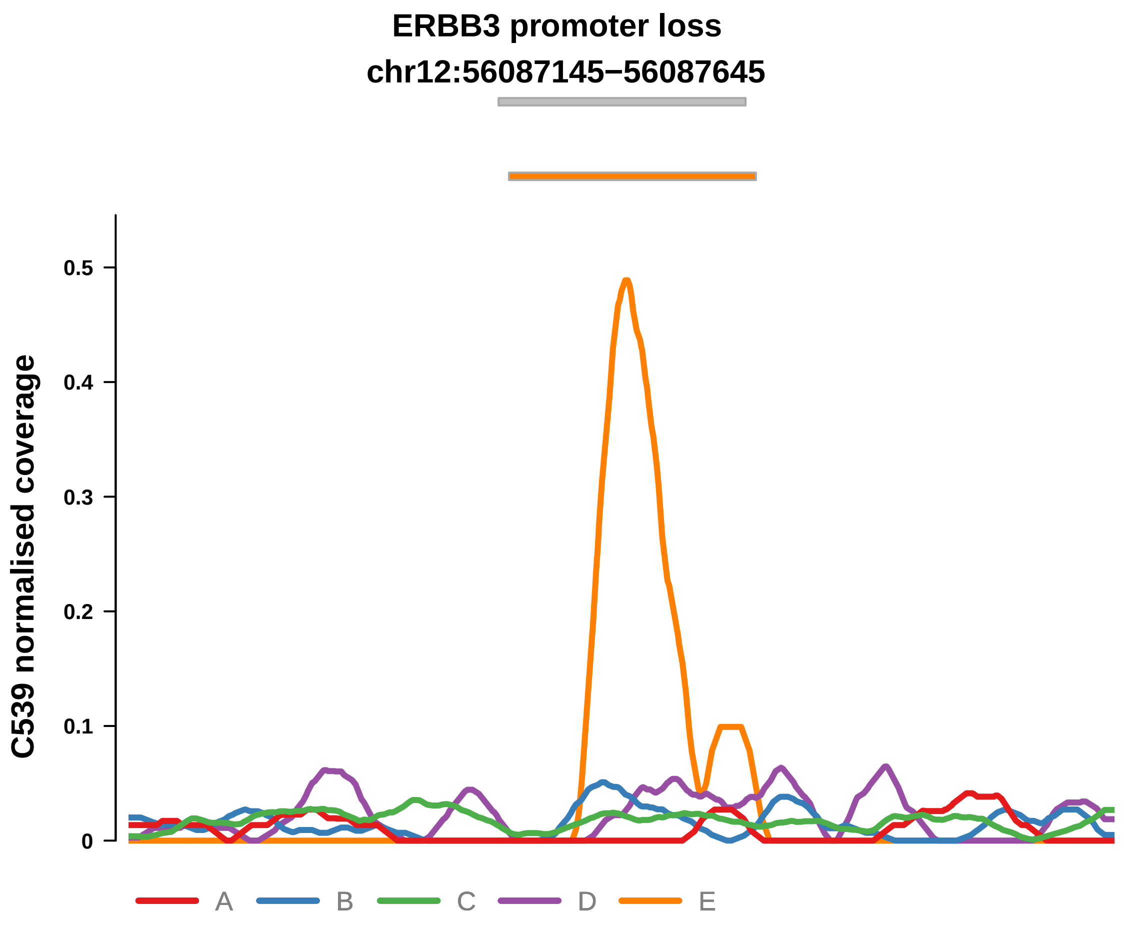


Putatively Clonal (logFC = −2.69)

| | P. | Adjusted P. |
|---|---|---|
| *~region vs. ~* | 1.17e−04 *** | 6.05e−04 *** |
| *~purity vs ~* | 1.13e−03 ** | 3.72e−03 ** |
| *~purity+region vs ~purity* | 3.6e−02 * | 1.02e−01 |
| *Carcinoma vs. normal* | 5.26e−03 ** | 5.26e−03 ** |

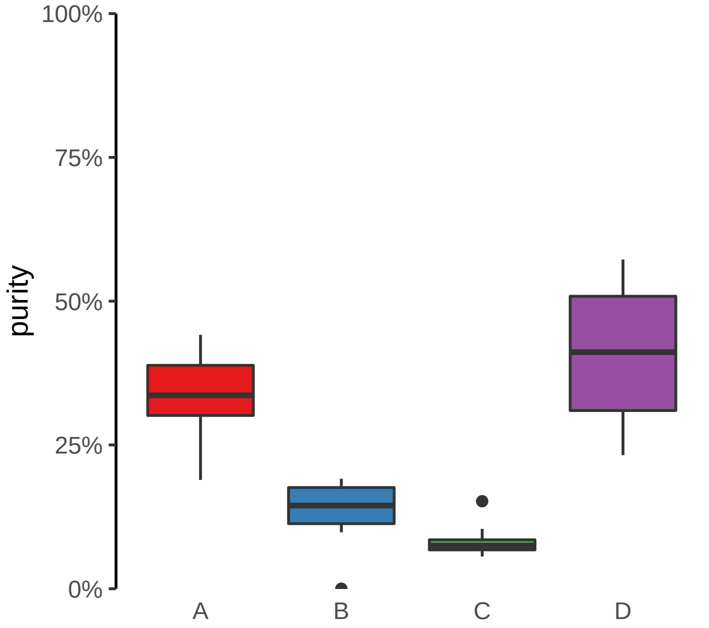

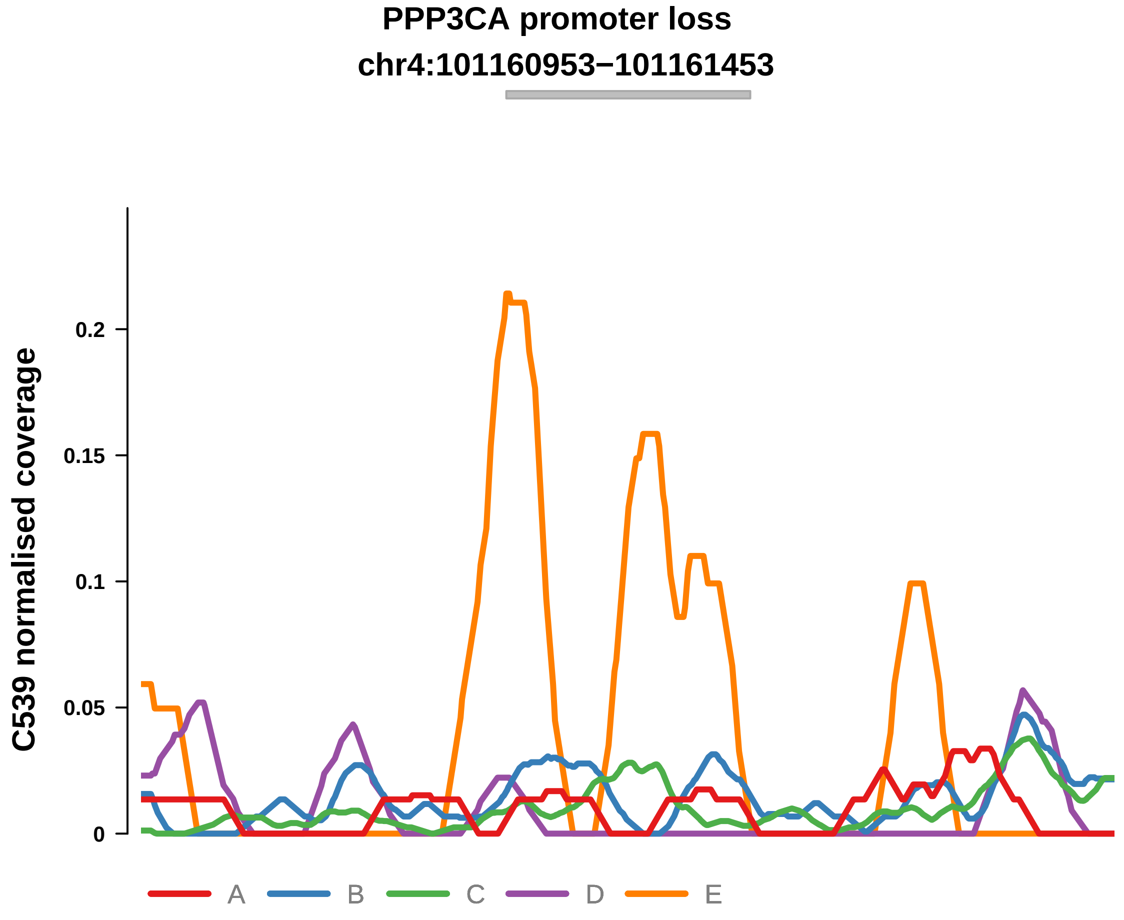


Putatively Clonal (logFC = −10.83)

| | P. | Adjusted P. |
|---|---|---|
| *~region vs. ~* | 5.94e−04 *** | 1.69e−03 ** |
| *~purity vs ~* | 1.19e−04 *** | 6.99e−04 *** |
| *~purity+region vs ~purity* | 6.28e−01 | 6.71e−01 |
| *Carcinoma vs. normal* | 2.97e−04 *** | 2.97e−04 *** |

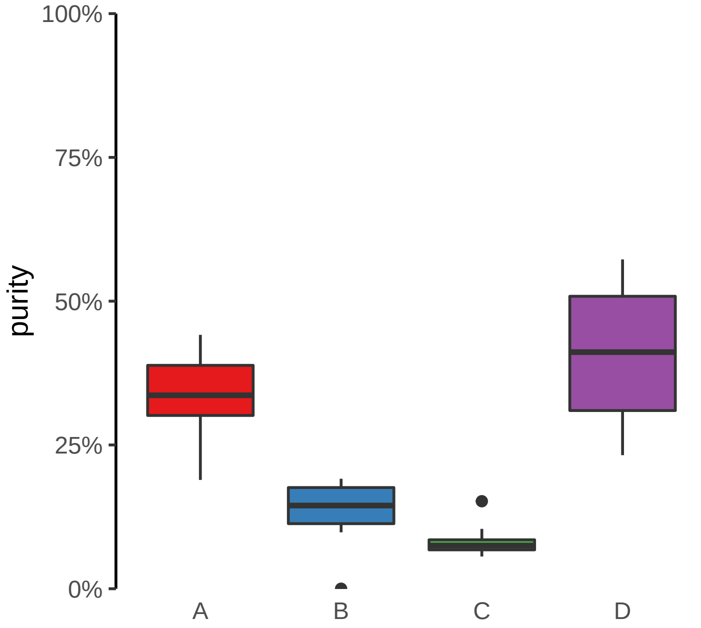

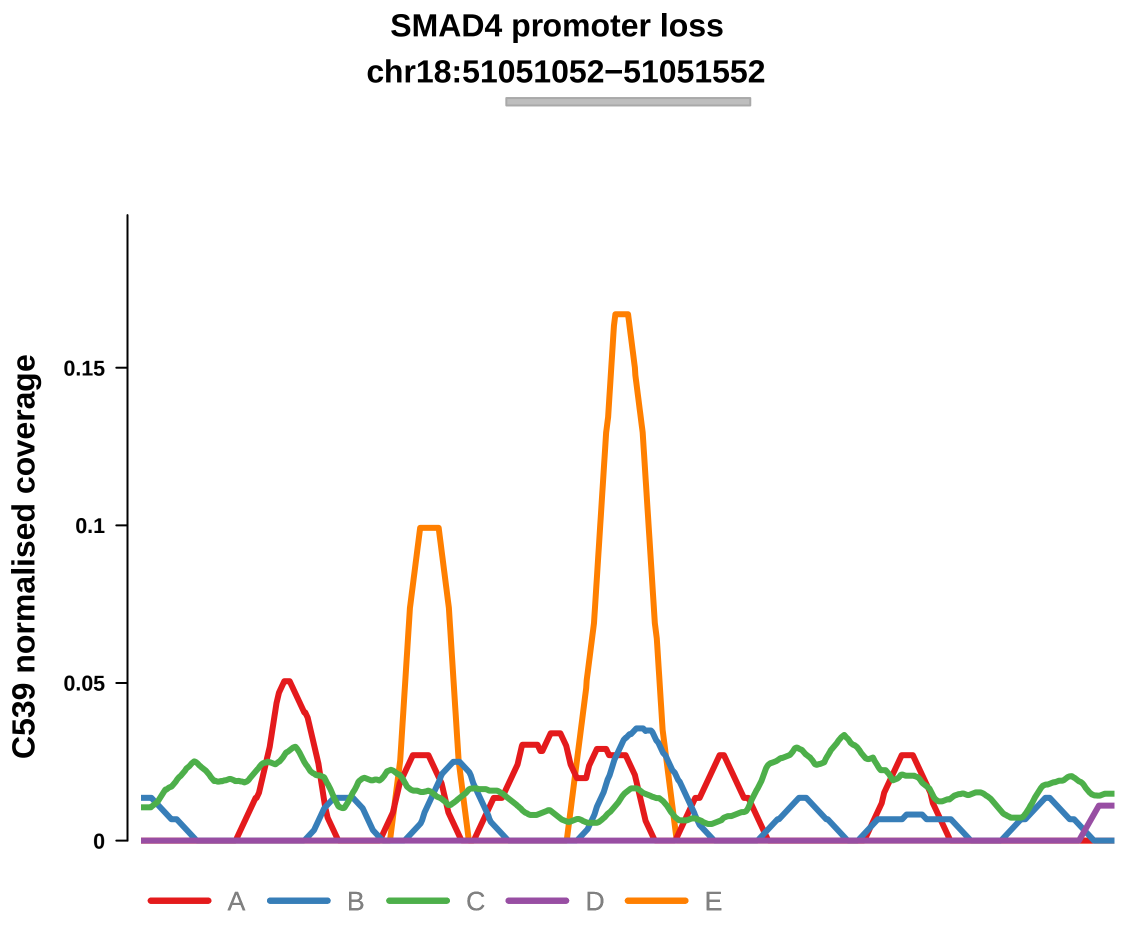


Putatively Clonal (logFC = −10.88)

| | P. | Adjusted P. |
|---|---|---|
| *~region vs. ~* | 1.37e−03 ** | 3.35e−03 ** |
| *~purity vs ~* | 8.84e−04 *** | 3.53e−03 ** |
| *~purity+region vs ~purity* | 2.88e−01 | NA |
| *Carcinoma vs. normal* | 2.19e−04 *** | 2.19e−04 *** |

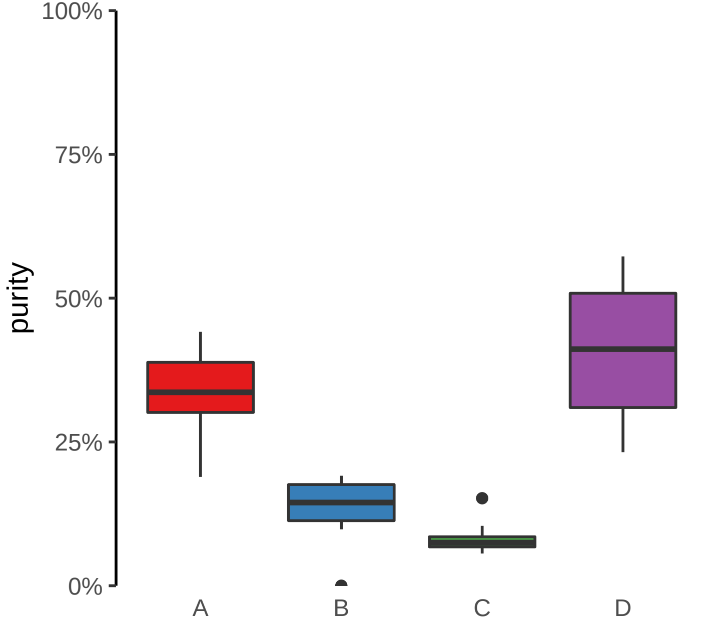

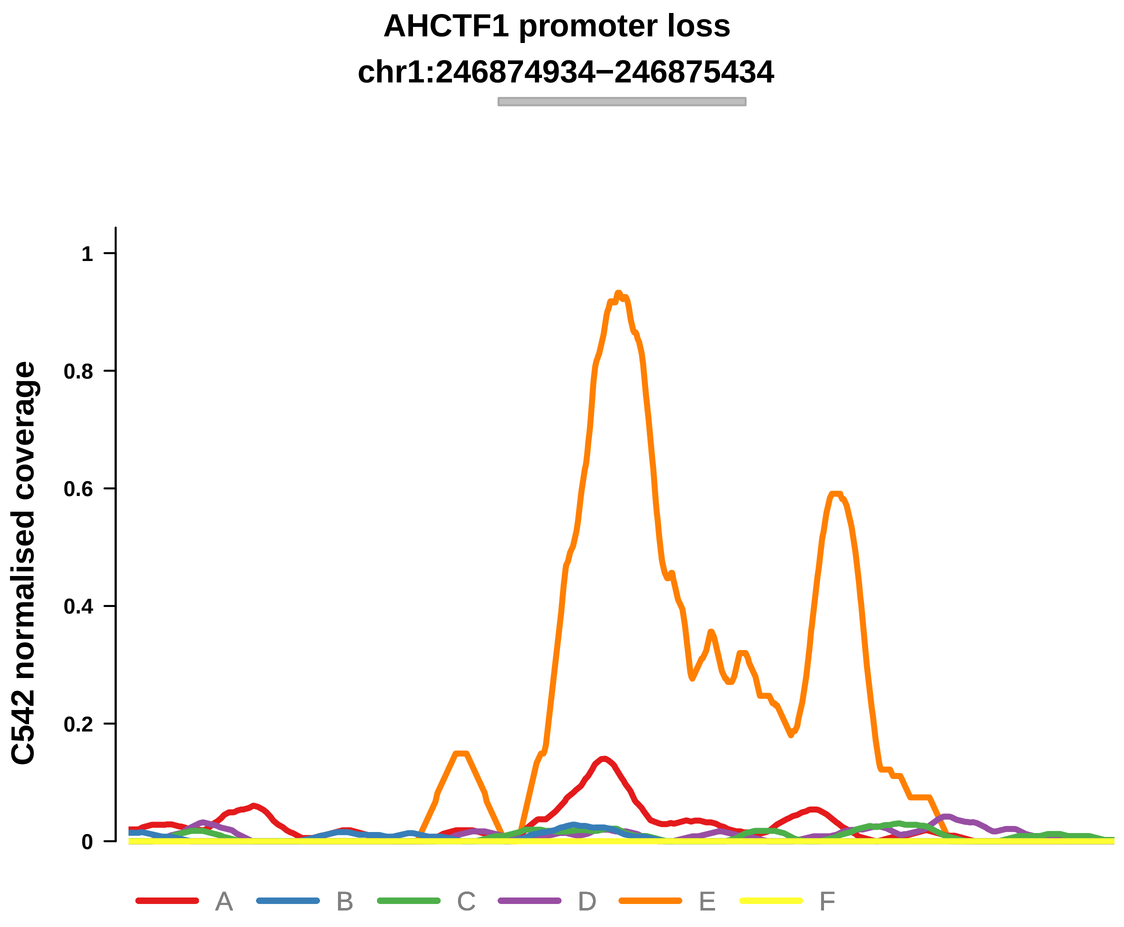


Putatively Clonal (logFC = −11.54)

| | P. | Adjusted P. |
|---|---|---|
| *~region vs. ~* | 2.72e−02 * | 4e−01 |
| *~purity vs ~* | 1.78e−03 ** | 5.22e−02 |
| *~purity+region vs ~purity* | 4.4e−01 | 8.56e−01 |
| *Carcinoma vs. normal* | 1.12e−03 ** | 1.12e−03 ** |

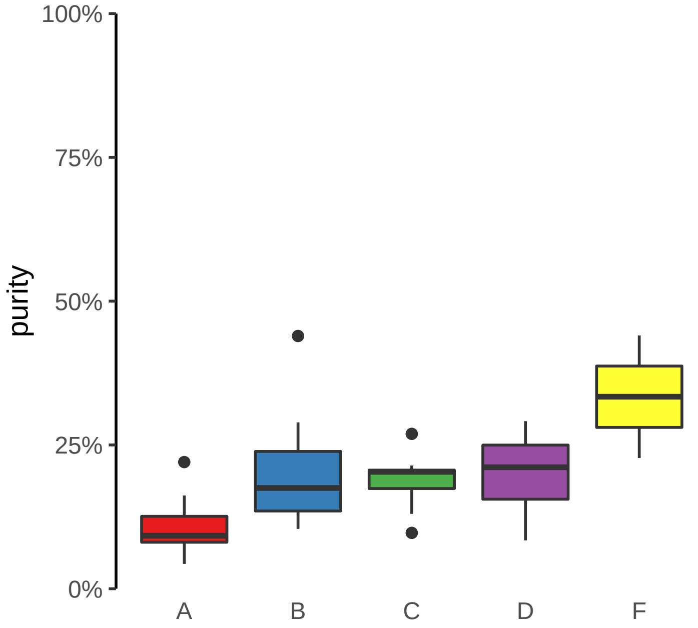

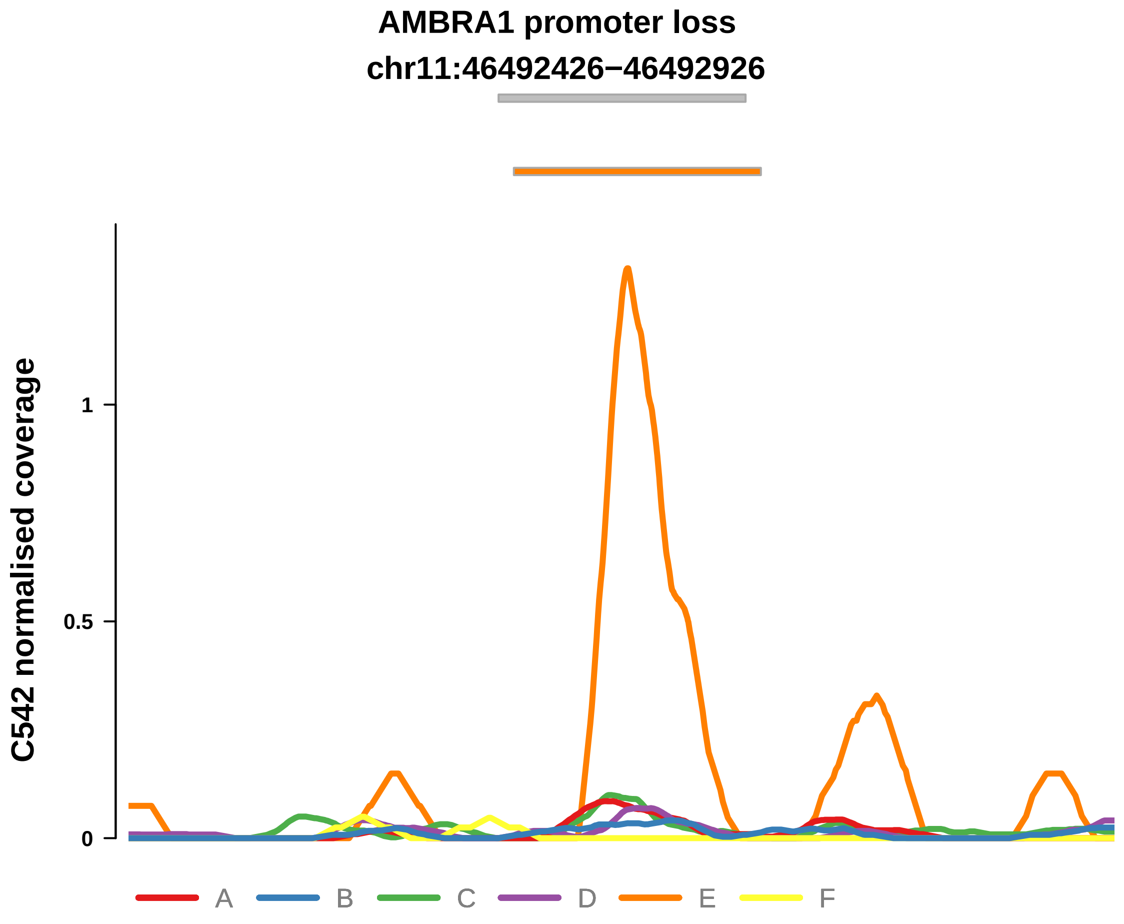


Putatively Clonal (logFC = −11.25)

| | P. | Adjusted P. |
|---|---|---|
| *~region vs. ~* | 6.27e−01 | 8.3e−01 |
| *~purity vs ~* | 4.55e−01 | 9.67e−01 |
| *~purity+region vs ~purity* | 6.95e−01 | 8.56e−01 |
| *Carcinoma vs. normal* | 2.16e−03 ** | 2.16e−03 ** |

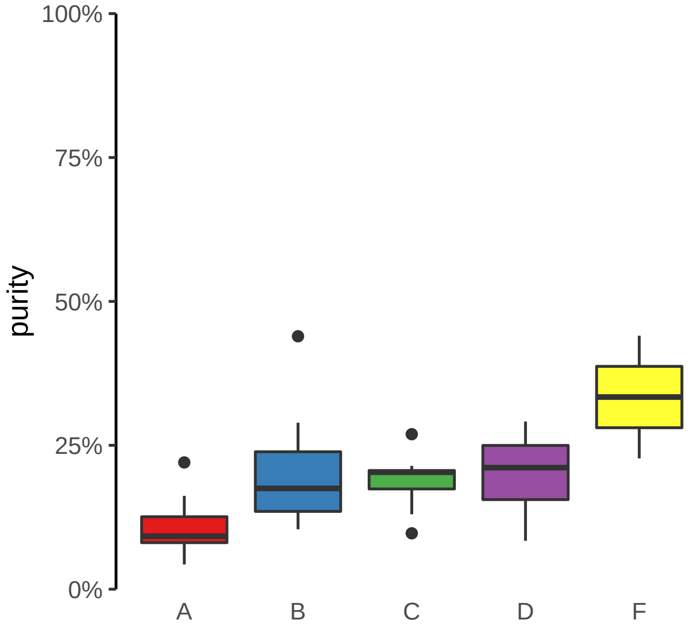

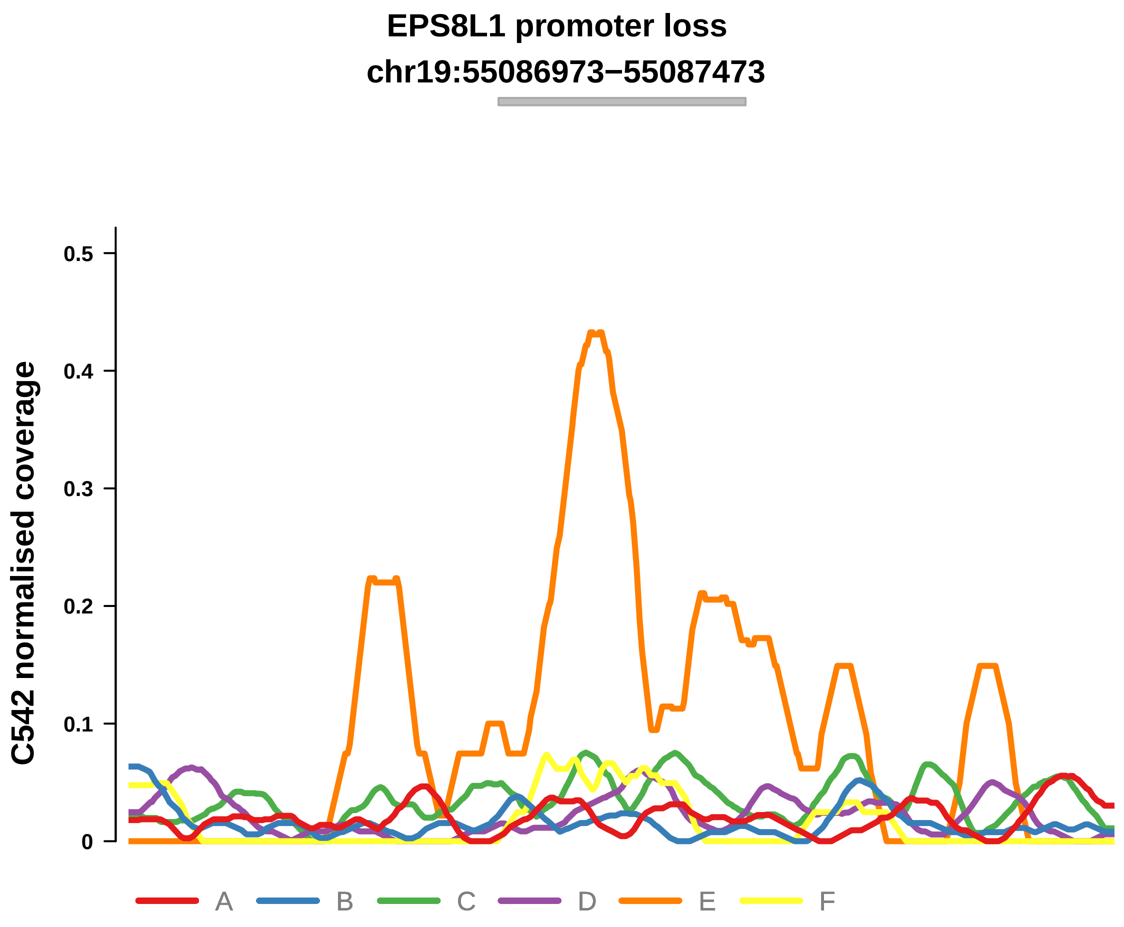


Putatively Clonal (logFC = −12.05)

| | P. | Adjusted P. |
|---|---|---|
| *~region vs. ~* | 5.38e−01 | 8.01e−01 |
| *~purity vs ~* | 1e+00 | 1e+00 |
| *~purity+region vs ~purity* | 5.47e−01 | 8.56e−01 |
| *Carcinoma vs. normal* | 2.27e−04 *** | 2.27e−04 *** |

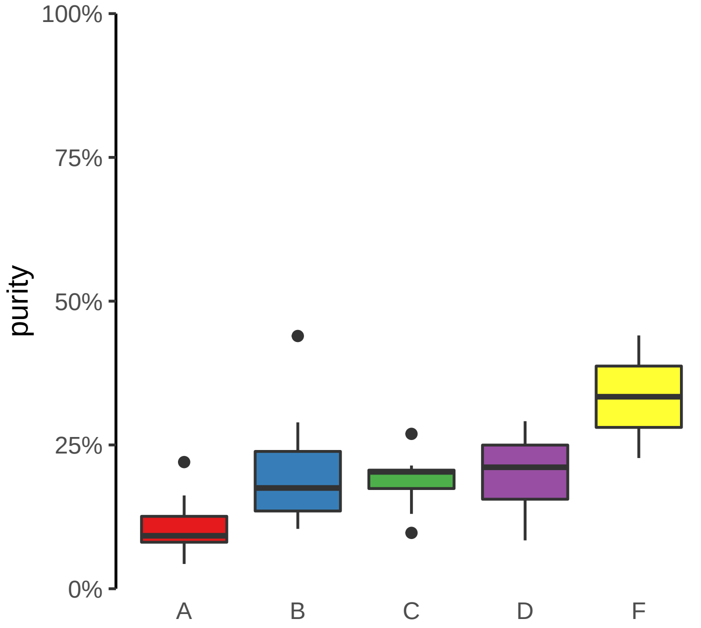

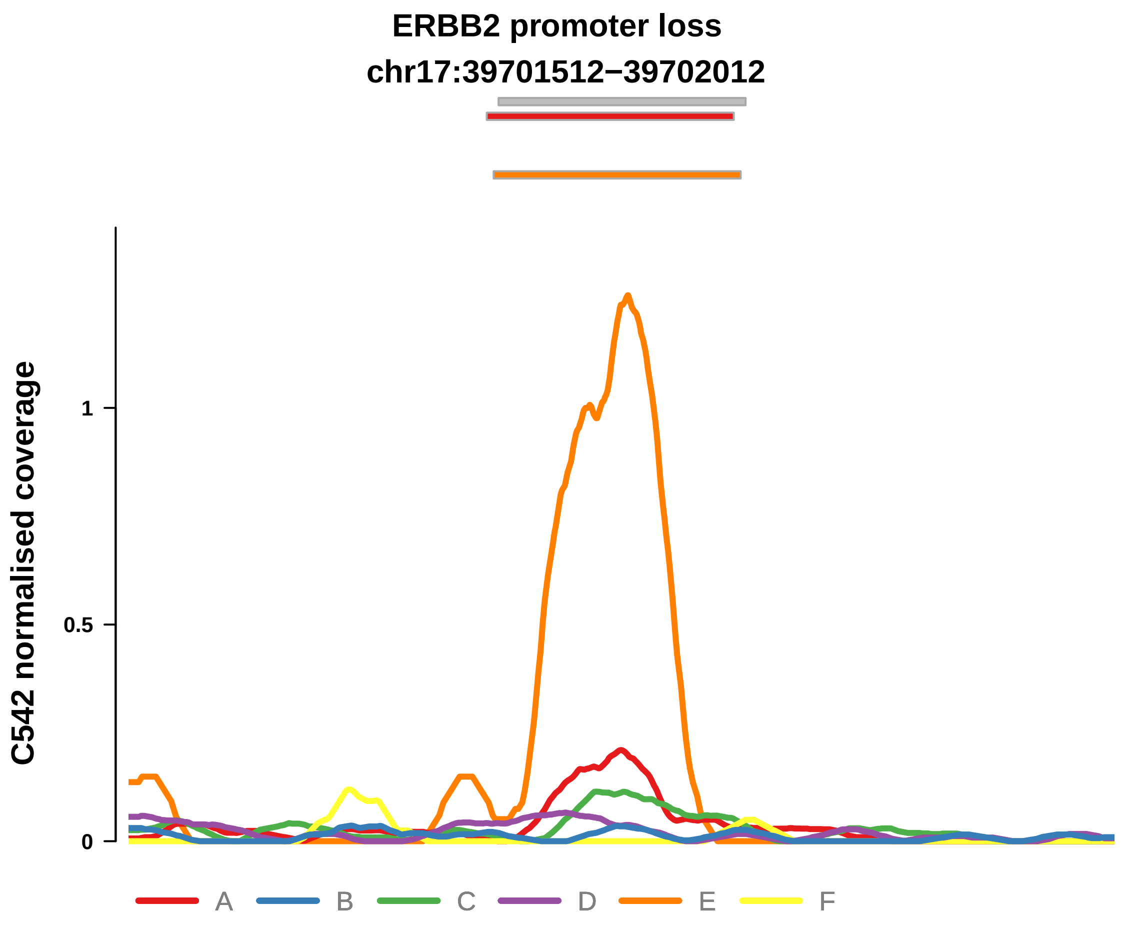


Putatively Clonal (logFC = −11.67)

| | P. | Adjusted P. |
|---|---|---|
| *~region vs. ~* | 1.62e−02 * | 4e−01 |
| *~purity vs ~* | 5.8e−02 | 5.28e−01 |
| *~purity+region vs ~purity* | 8.14e−02 | 5.5e−01 |
| *Carcinoma vs. normal* | 9.1e−04 *** | 9.1e−04 *** |

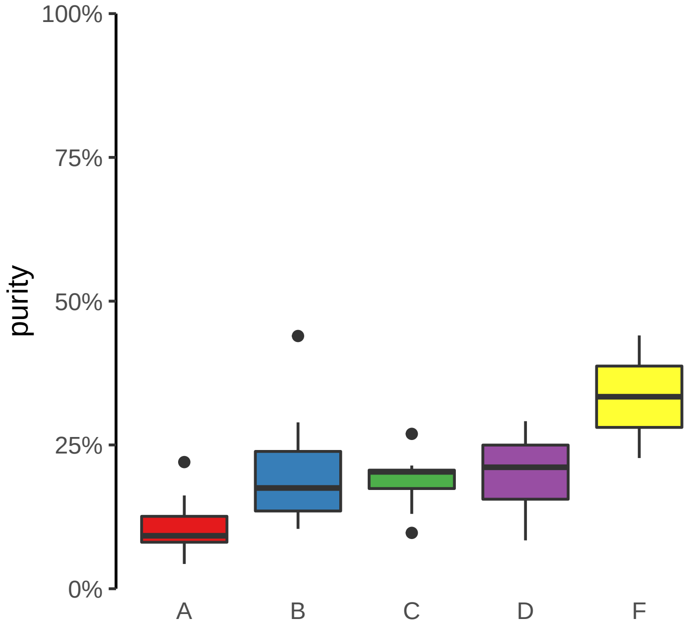

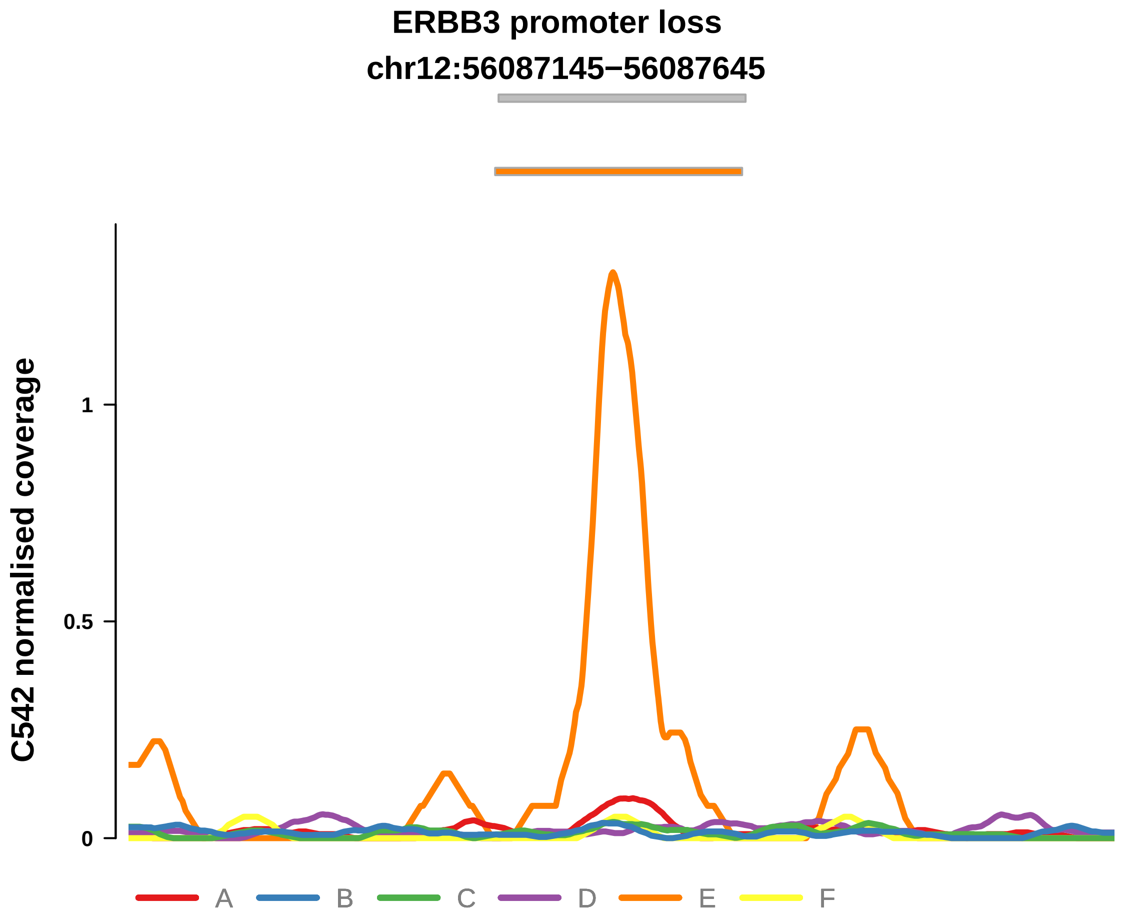


Putatively Clonal (logFC = −11.78)

| | P. | Adjusted P. |
|---|---|---|
| *~region vs. ~* | 4.39e−01 | 7.33e−01 |
| *~purity vs ~* | 1.6e−01 | 6.38e−01 |
| *~purity+region vs ~purity* | 7.21e−01 | 8.56e−01 |
| *Carcinoma vs. normal* | 4.78e−04 *** | 4.78e−04 *** |

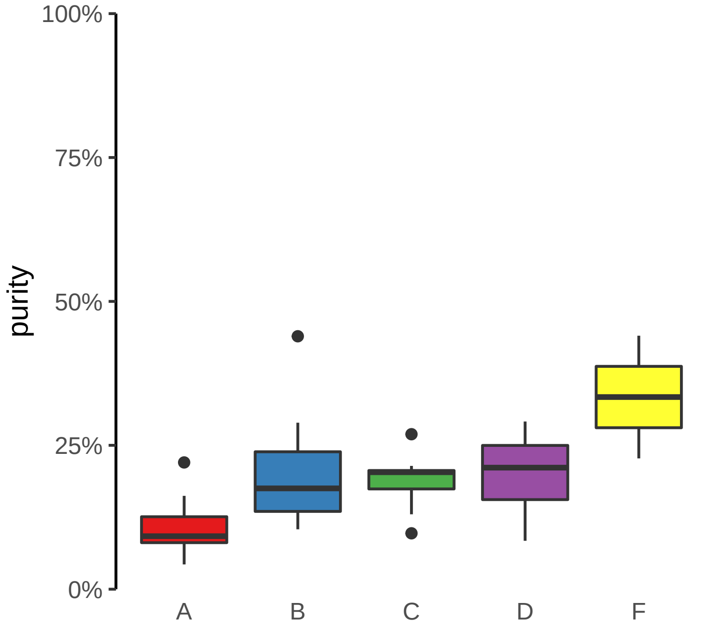

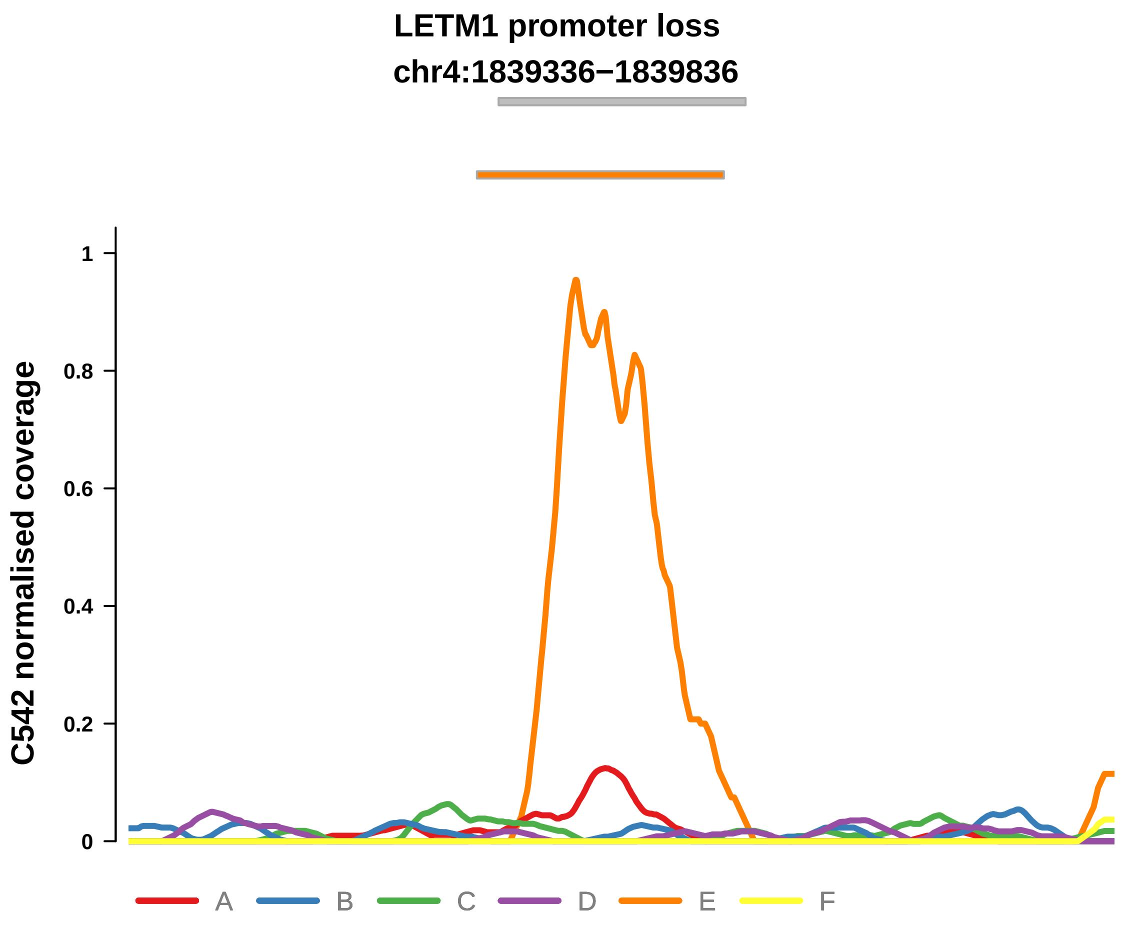


Putatively Clonal (logFC = −11.42)

| | P. | Adjusted P. |
|---|---|---|
| *~region vs. ~* | 1.45e−01 | 5.9e−01 |
| *~purity vs ~* | 2.77e−01 | 8.11e−01 |
| *~purity+region vs ~purity* | 2e−01 | 7.3e−01 |
| *Carcinoma vs. normal* | 1.42e−03 ** | 1.42e−03 ** |

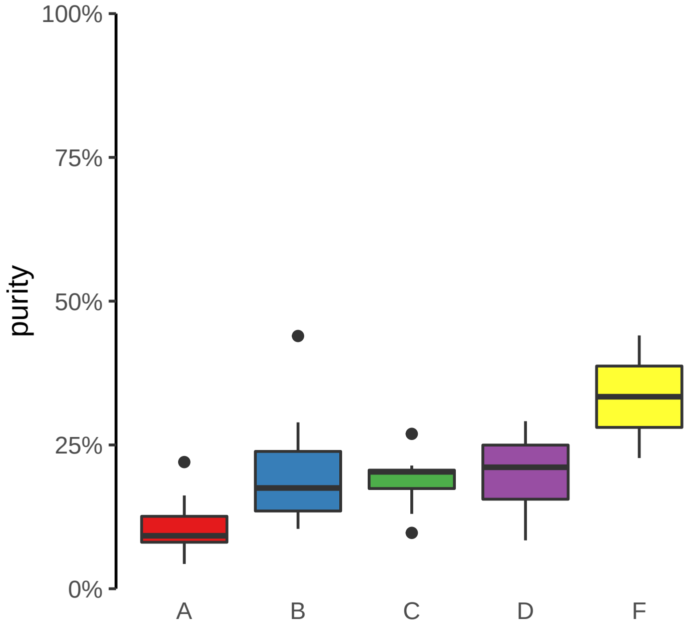

**LPCAT4 promoter loss**
**chr15:34362318−34362818**

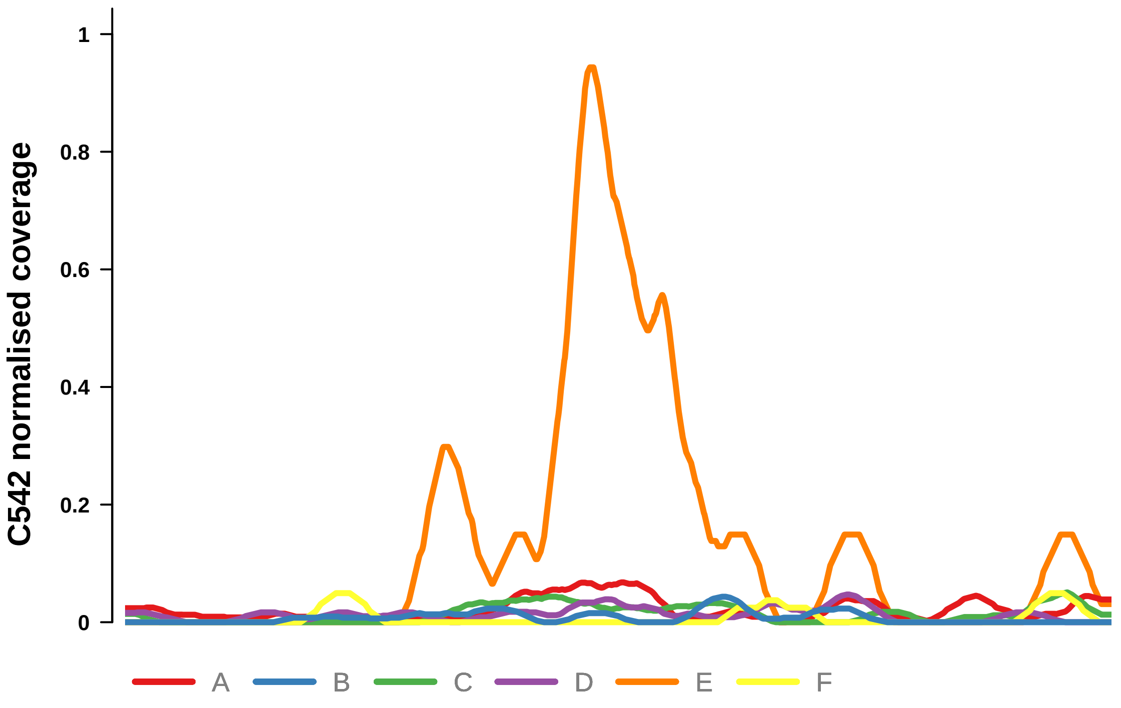


Putatively Clonal (logFC = −11.79)

| | P. | Adjusted P. |
|---|---|---|
| *~region vs. ~* | 1.38e−01 | 5.9e−01 |
| *~purity vs ~* | 2.07e−01 | 7.01e−01 |
| *~purity+region vs ~purity* | 2.6e−01 | 7.3e−01 |
| *Carcinoma vs. normal* | 4.21e−04 *** | 4.21e−04 *** |

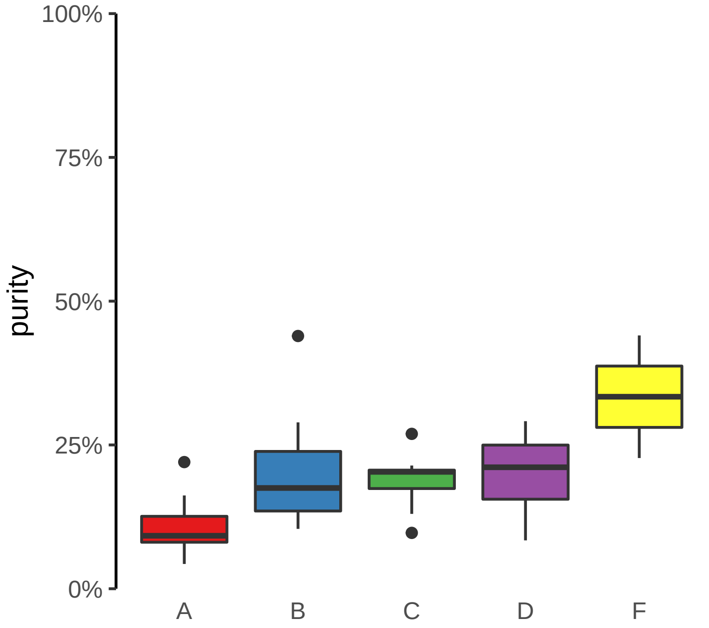

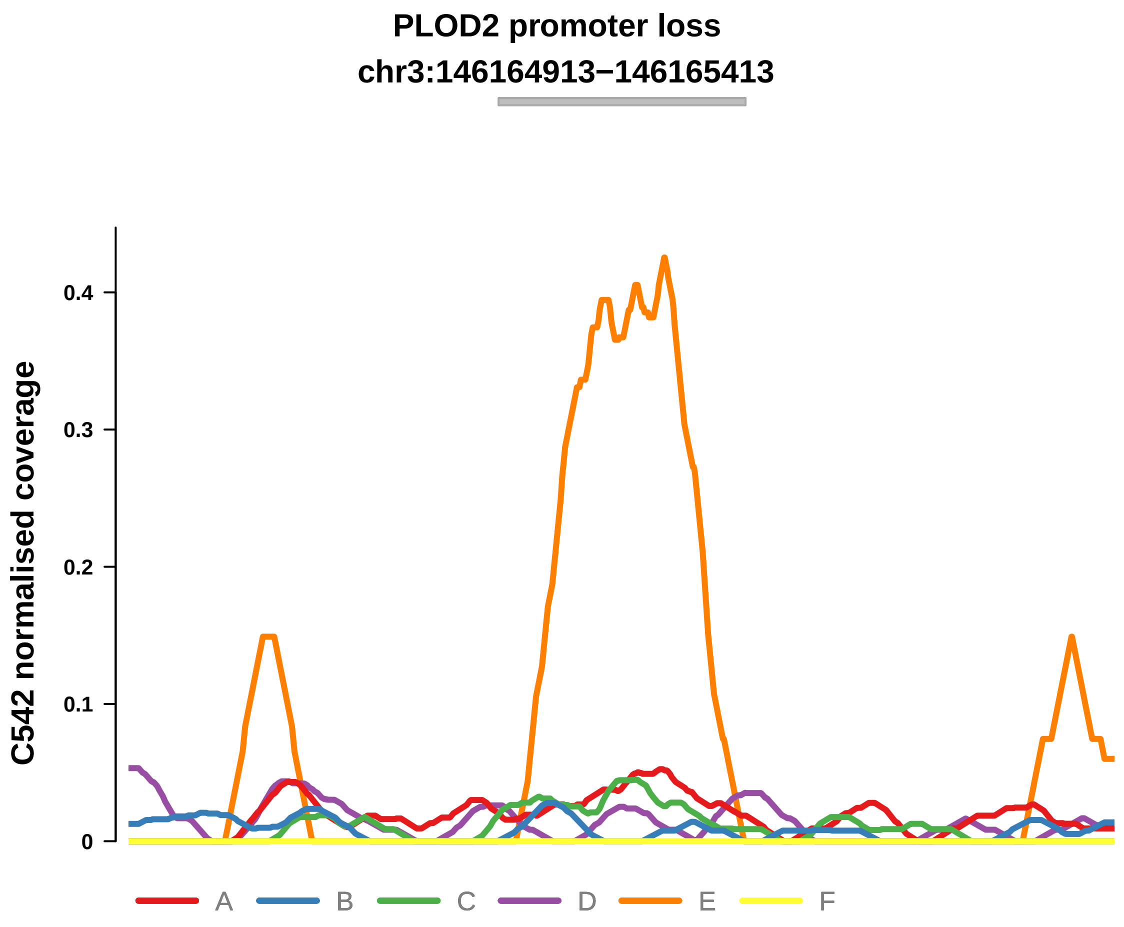


Putatively Clonal (logFC = −11.27)

| | P. | Adjusted P. |
|---|---|---|
| *~region vs. ~* | 8.98e−02 | 5.9e−01 |
| *~purity vs ~* | 3.77e−01 | 9.67e−01 |
| *~purity+region vs ~purity* | 1.29e−01 | 6.68e−01 |
| *Carcinoma vs. normal* | 3.03e−03 ** | 3.03e−03 ** |

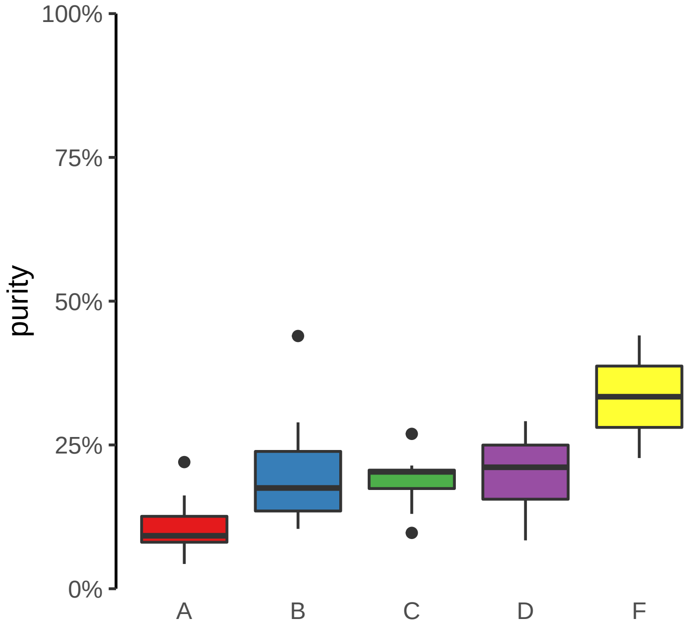

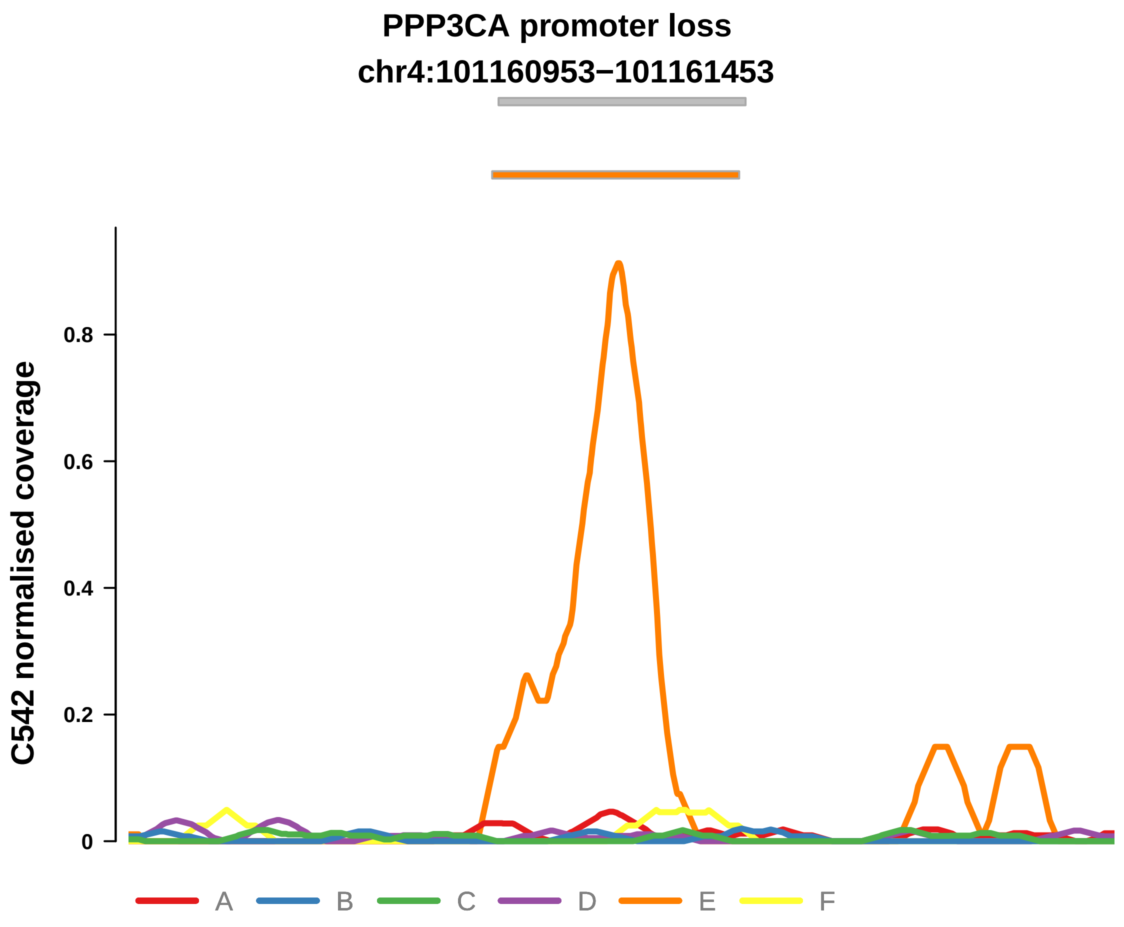


Putatively Clonal (logFC = −10.82)

| | P. | Adjusted P. |
|---|---|---|
| *~region vs. ~* | 4.58e−01 | 7.33e−01 |
| *~purity vs ~* | 2.02e−01 | 7.01e−01 |
| *~purity+region vs ~purity* | 6.87e−01 | 8.56e−01 |
| *Carcinoma vs. normal* | 6.13e−03 ** | 6.13e−03 ** |

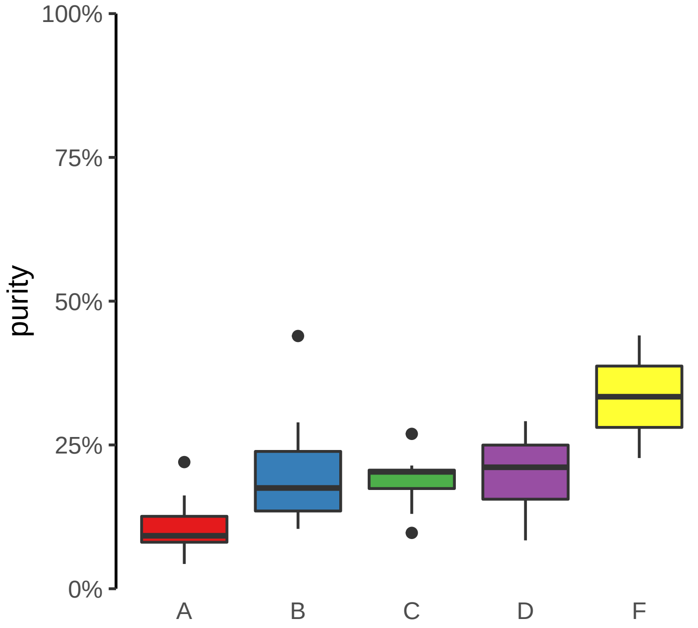

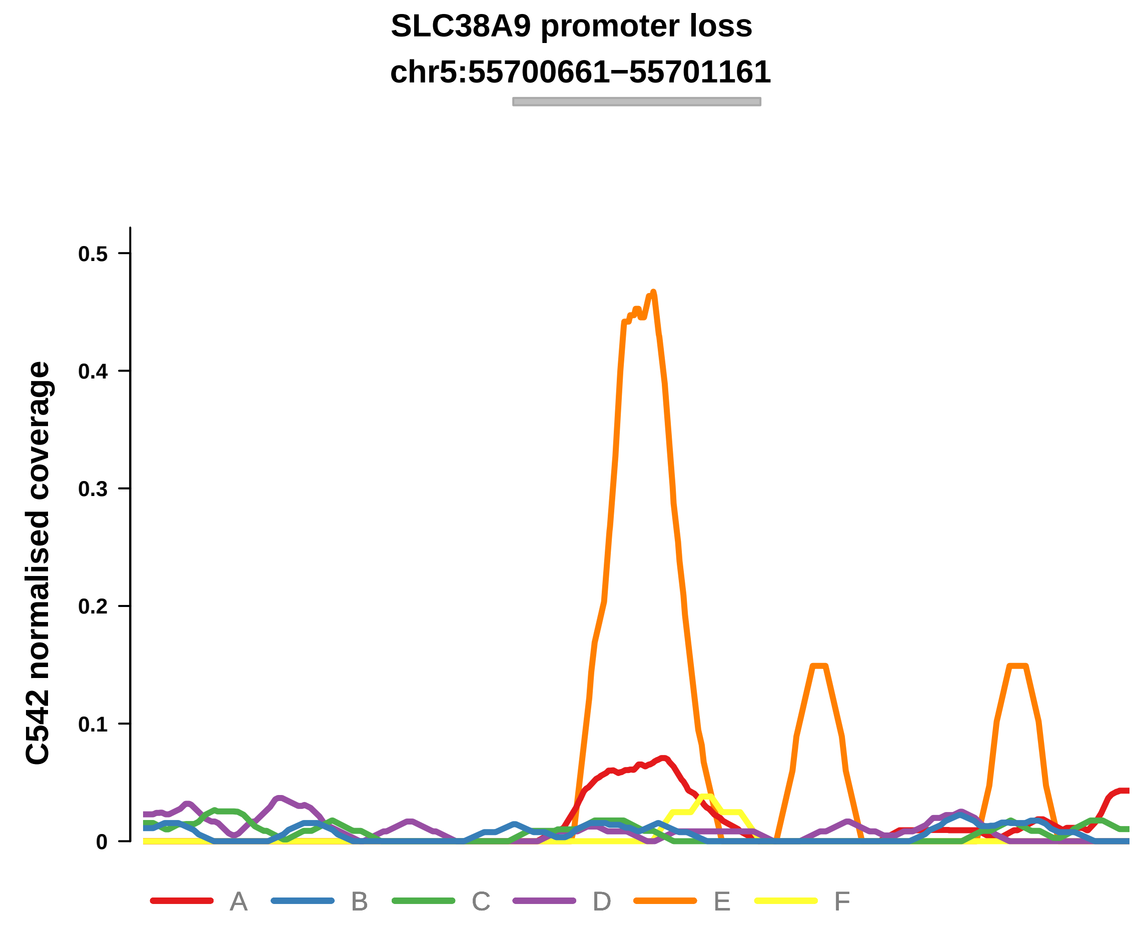


Putatively Clonal (logFC = −10.74)

| | P. | Adjusted P. |
|---|---|---|
| *~region vs. ~* | 5.48e−01 | 8.01e−01 |
| *~purity vs ~* | 2.46e−01 | 7.45e−01 |
| *~purity+region vs ~purity* | 7.53e−01 | 8.56e−01 |
| *Carcinoma vs. normal* | 9.01e−03 ** | 9.01e−03 ** |

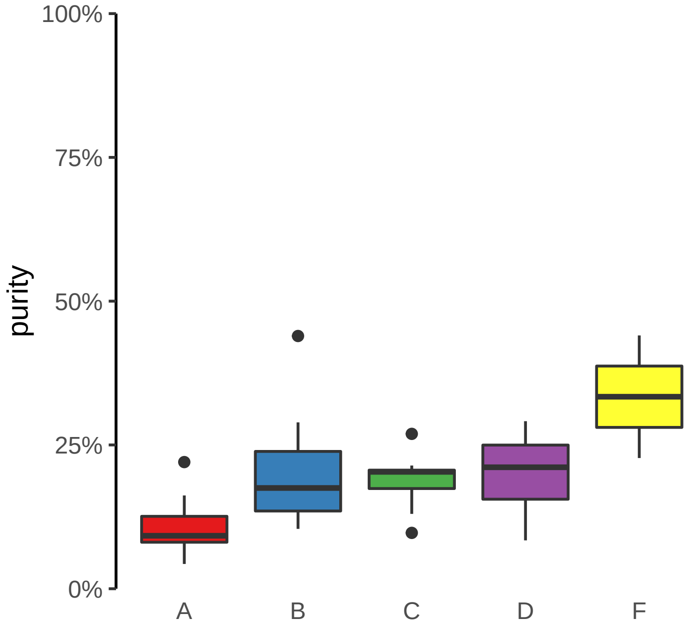

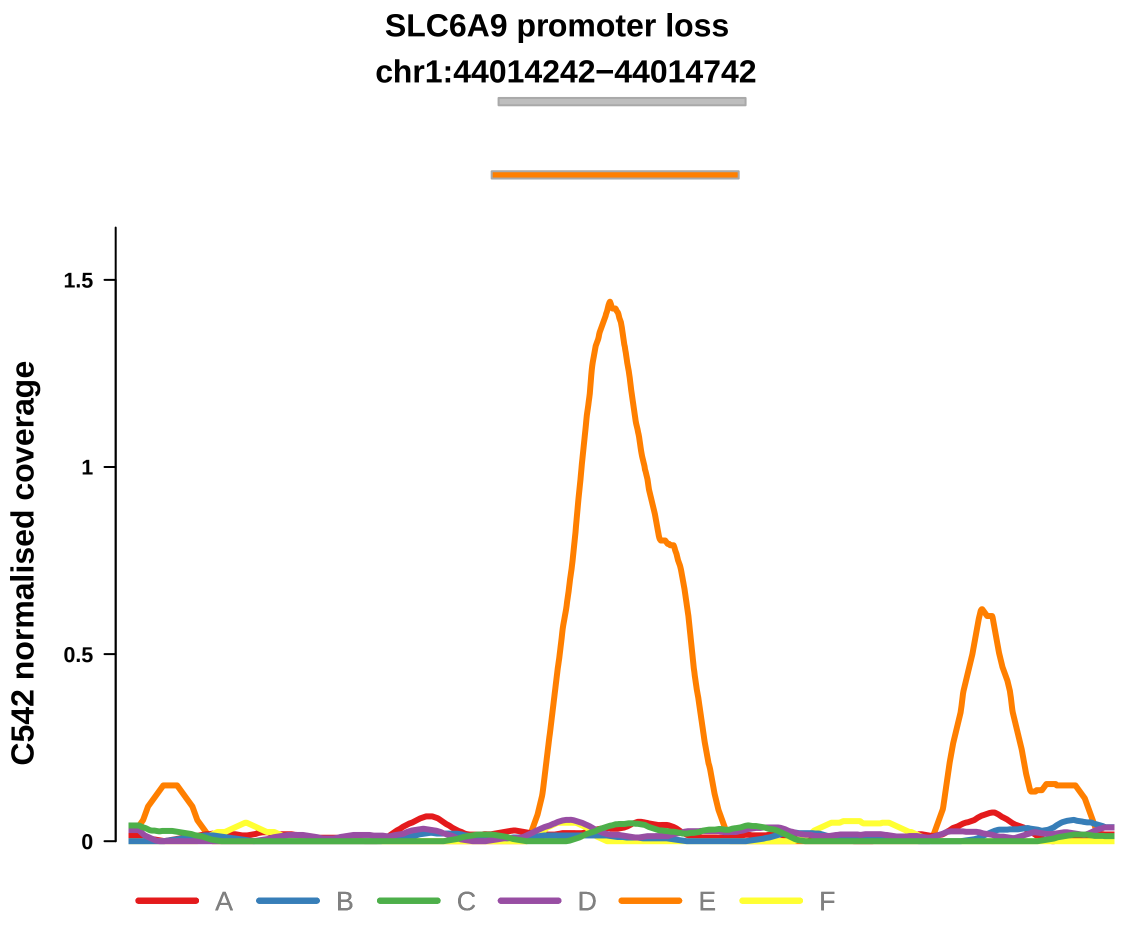


Putatively Clonal (logFC = −11.74)

| | P. | Adjusted P. |
|---|---|---|
| *~region vs. ~* | 1.29e−01 | 5.9e−01 |
| *~purity vs ~* | 4.82e−01 | 9.67e−01 |
| *~purity+region vs ~purity* | 1.6e−01 | 6.99e−01 |
| *Carcinoma vs. normal* | 6.17e−04 *** | 6.17e−04 *** |

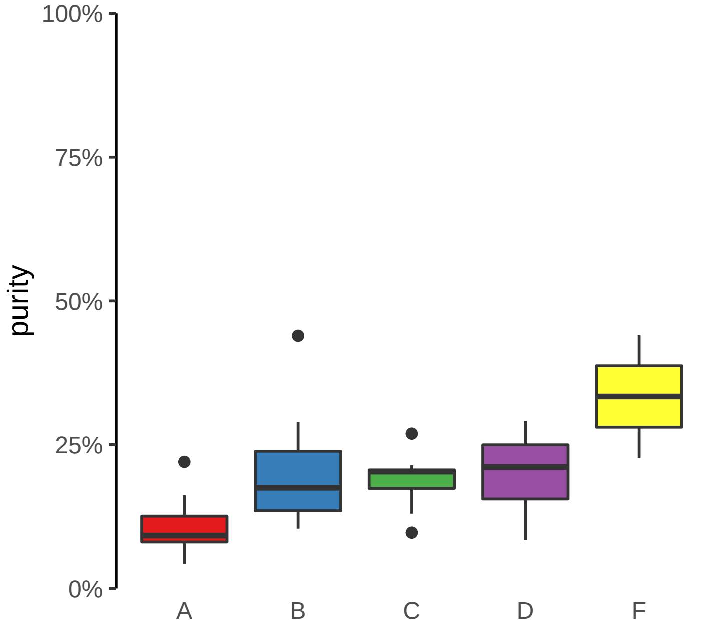

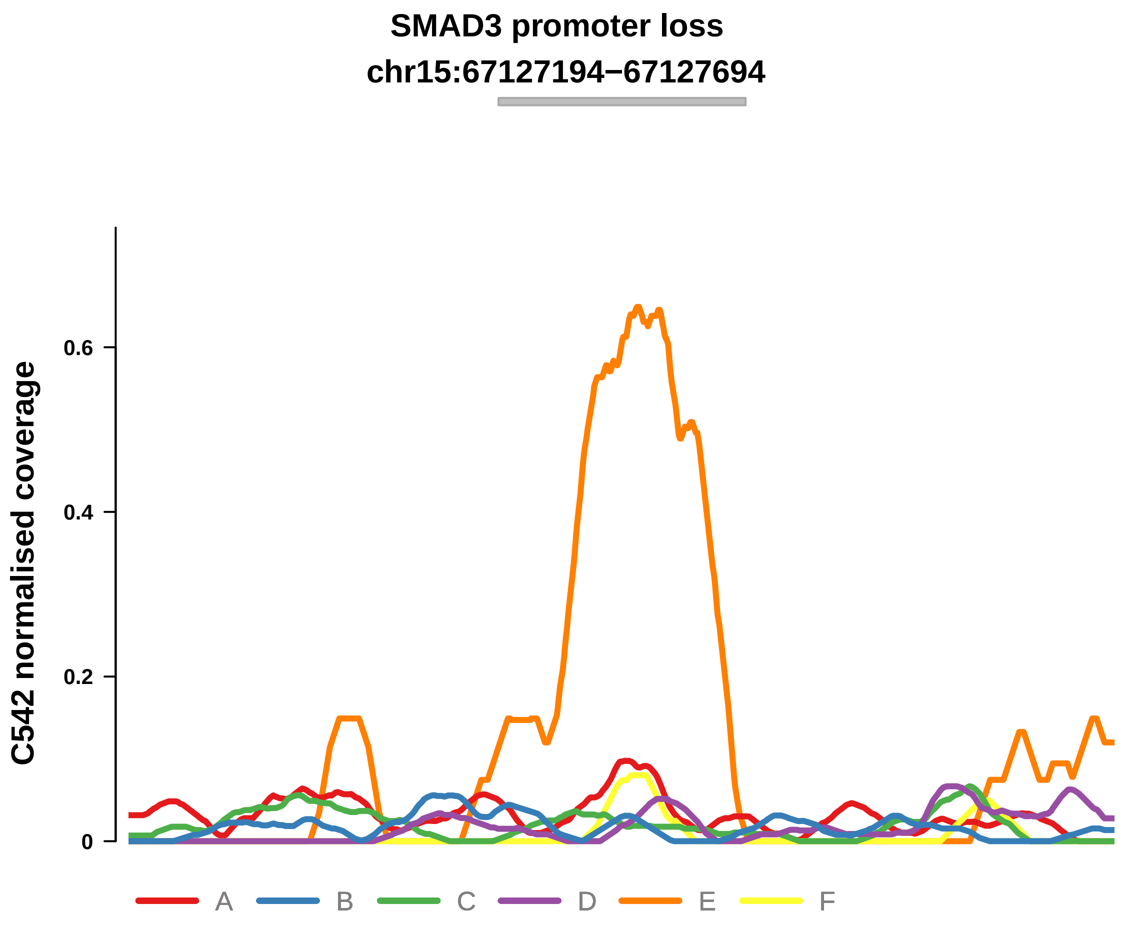


Putatively Clonal (logFC = −11.73)

| | P. | Adjusted P. |
|---|---|---|
| *~region vs. ~* | 2.27e−01 | 6.21e−01 |
| *~purity vs ~* | 1.26e−02 * | 2.13e−01 |
| *~purity+region vs ~purity* | 6.43e−01 | 8.56e−01 |
| *Carcinoma vs. normal* | 6.17e−04 *** | 6.17e−04 *** |

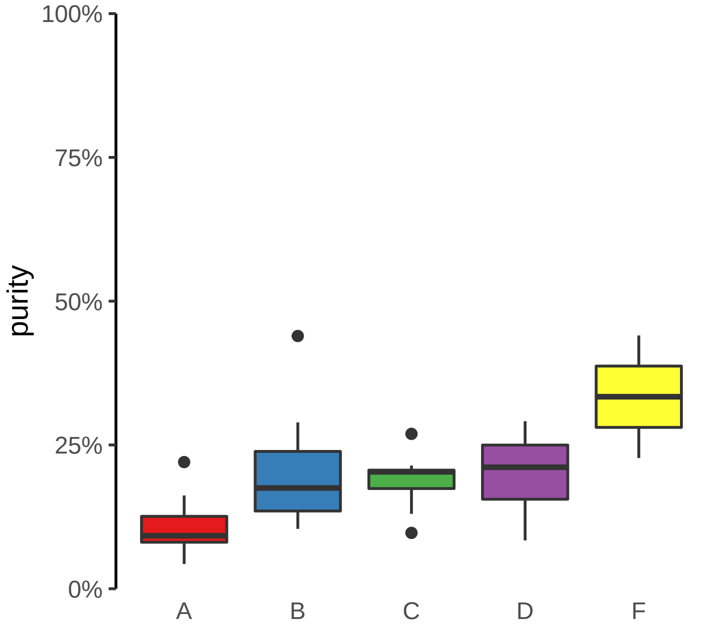

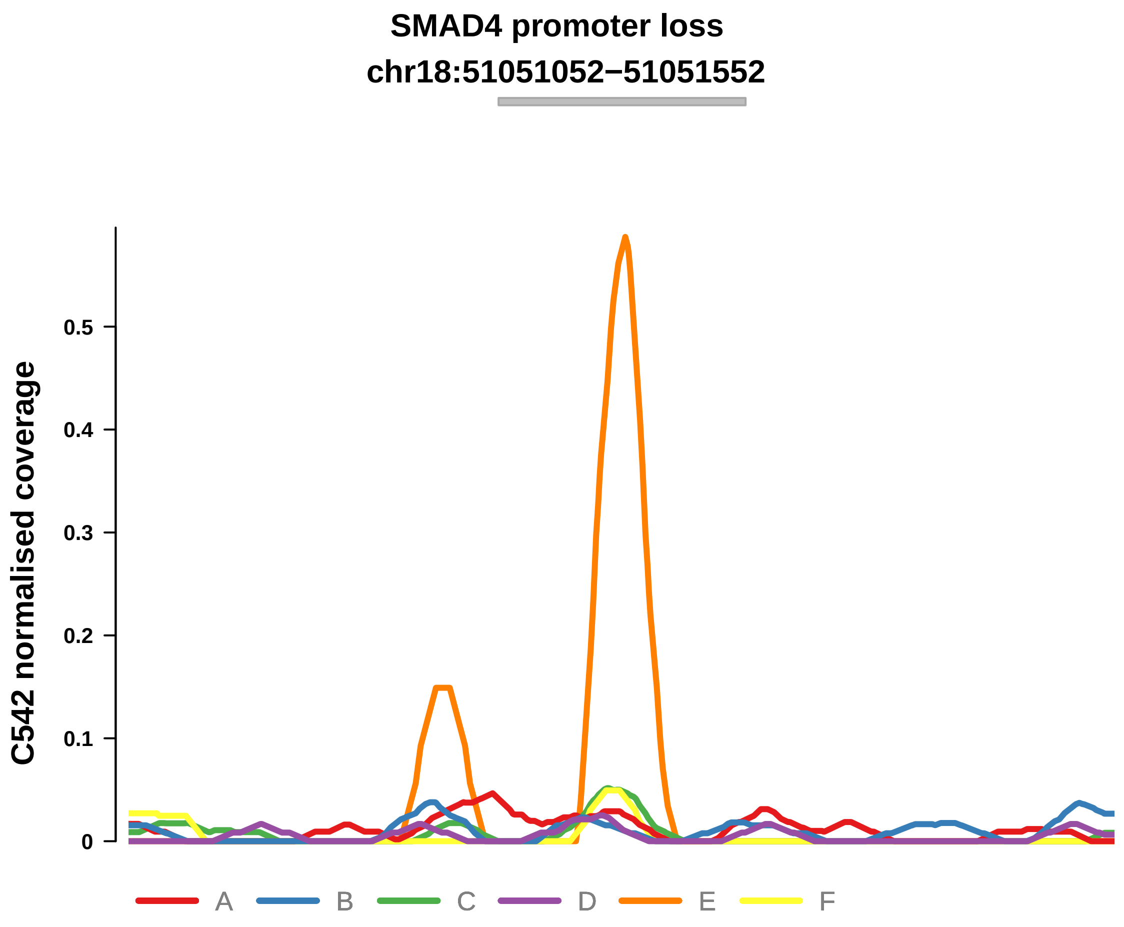


Putatively Clonal (logFC = −10.87)

| | P. | Adjusted P. |
|---|---|---|
| *~region vs. ~* | 8.96e−01 | 9.42e−01 |
| *~purity vs ~* | 7.57e−02 | 5.28e−01 |
| *~purity+region vs ~purity* | 9.96e−01 | 9.96e−01 |
| *Carcinoma vs. normal* | 8.55e−03 ** | 8.55e−03 ** |

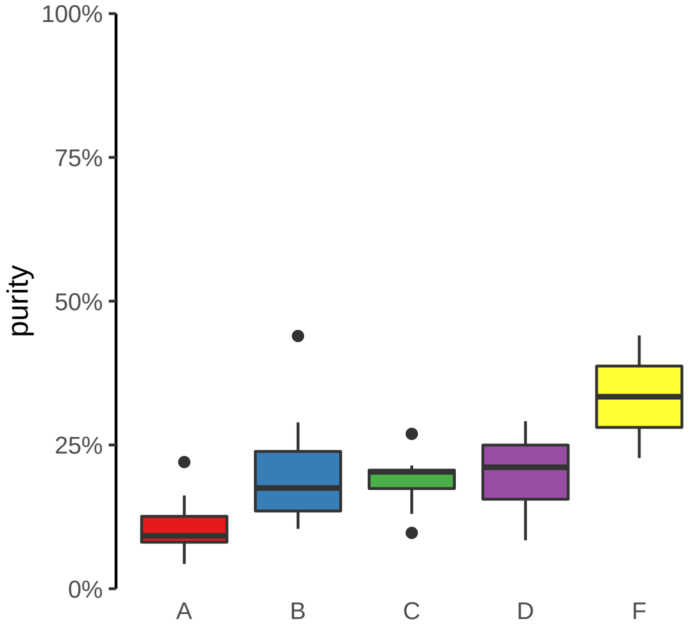

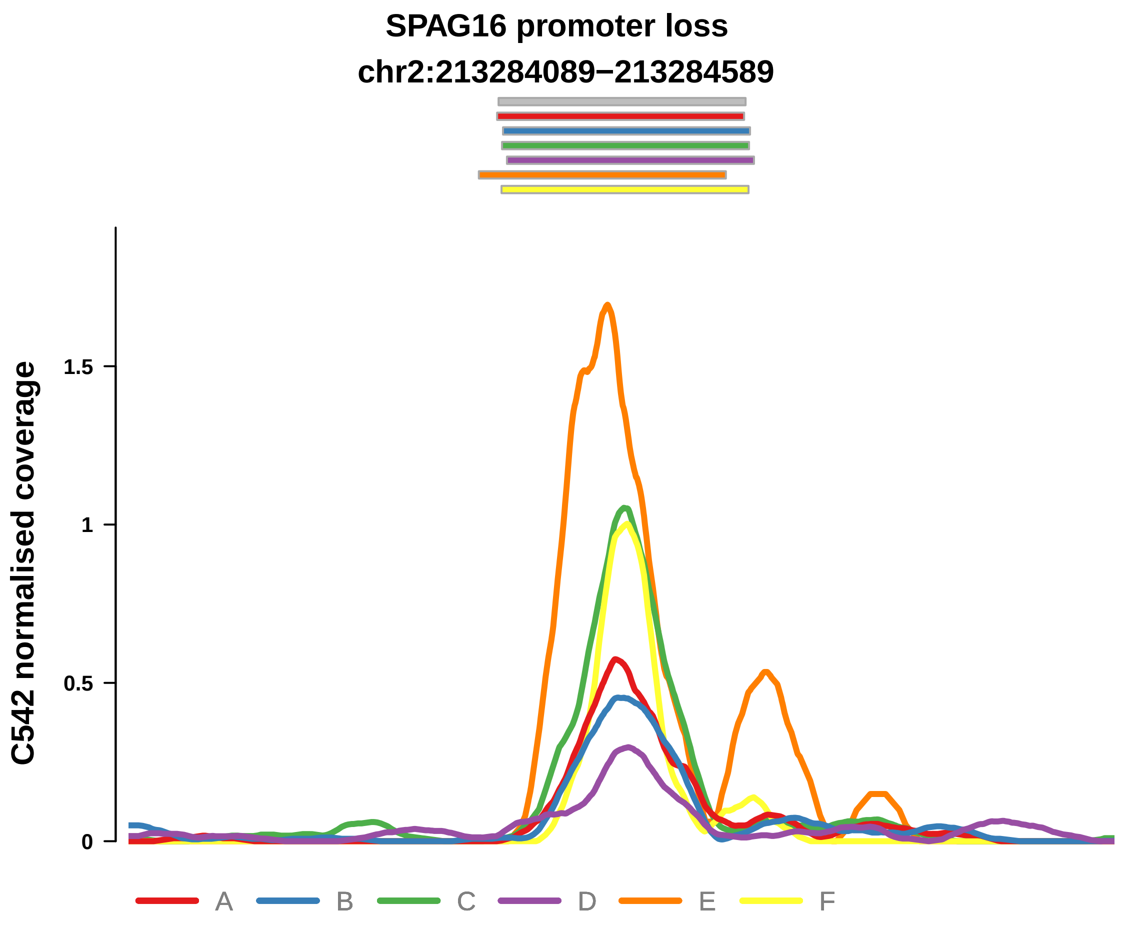


Putatively Clonal (logFC = 1.64)

| | P. | Adjusted P. |
|---|---|---|
| *~region vs. ~* | 7.81e−01 | 8.83e−01 |
| *~purity vs ~* | 2.8e−04 *** | 2.25e−02 * |
| *~purity+region vs ~purity* | 1.13e−02 * | 3.94e−01 |
| *Carcinoma vs. normal* | 5.3e−03 ** | 5.3e−03 ** |

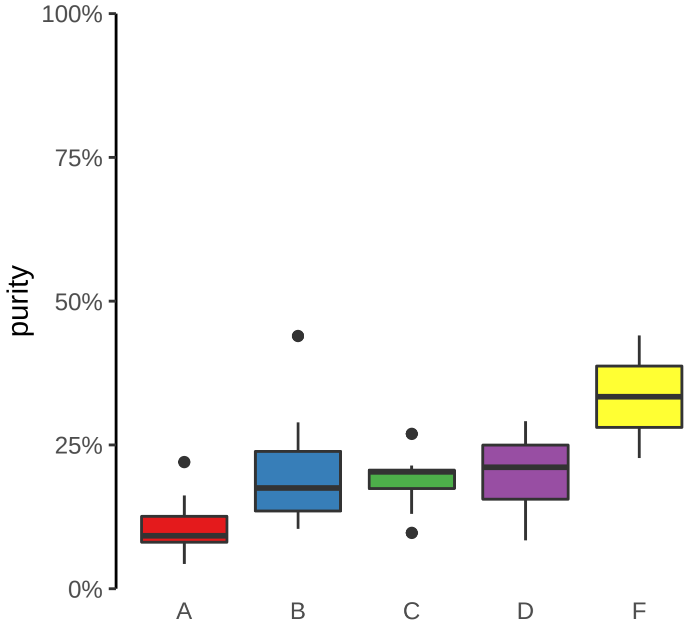

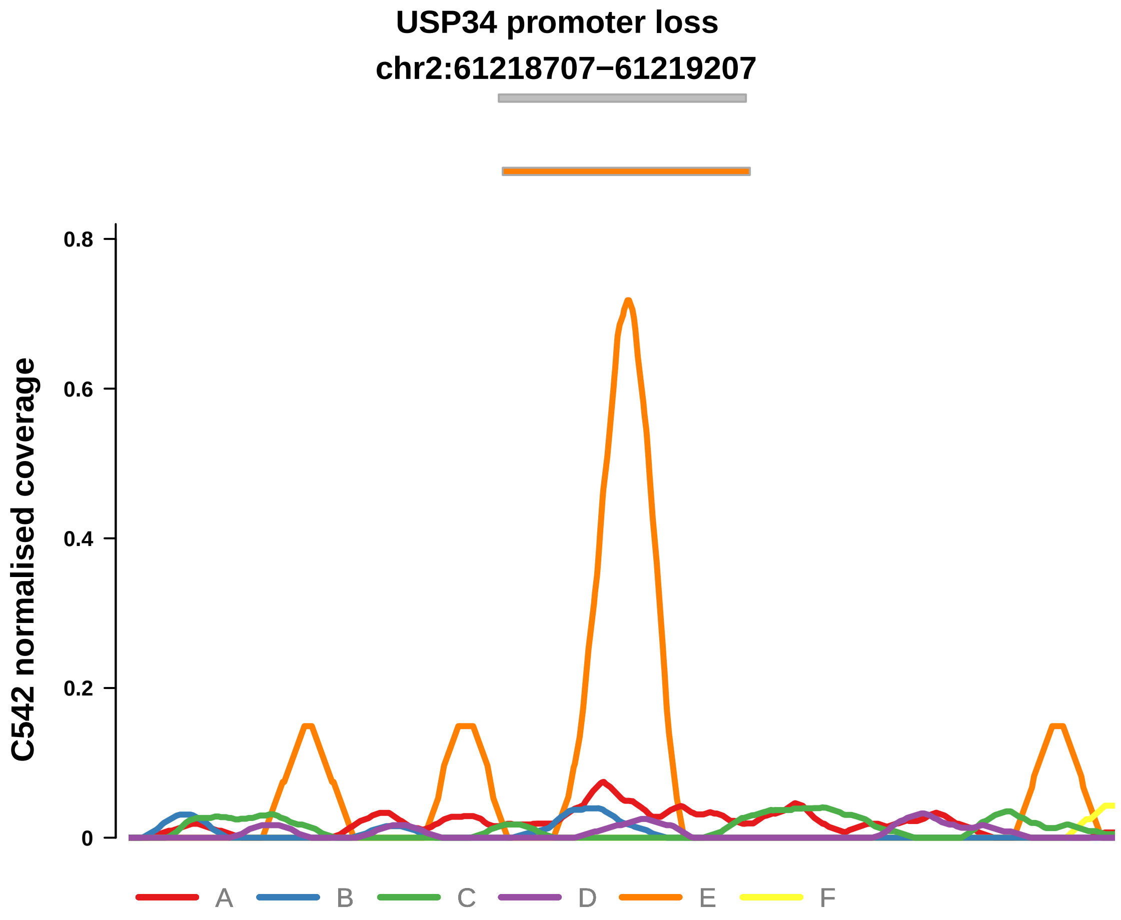


Putatively Clonal (logFC = −10.76)

| | P. | Adjusted P. |
|---|---|---|
| *~region vs. ~* | 1.54e−01 | 5.9e−01 |
| *~purity vs ~* | 1.12e−01 | 5.28e−01 |
| *~purity+region vs ~purity* | 4.13e−01 | 8.46e−01 |
| *Carcinoma vs. normal* | 7.42e−03 ** | 7.42e−03 ** |

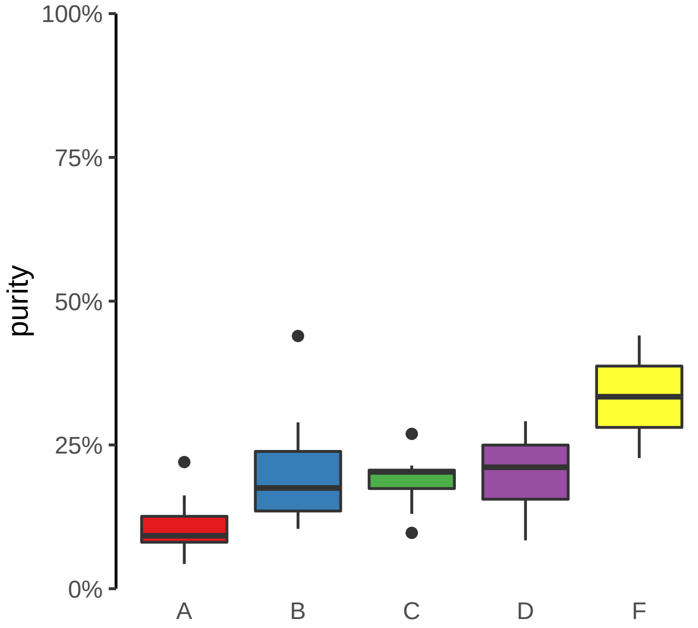

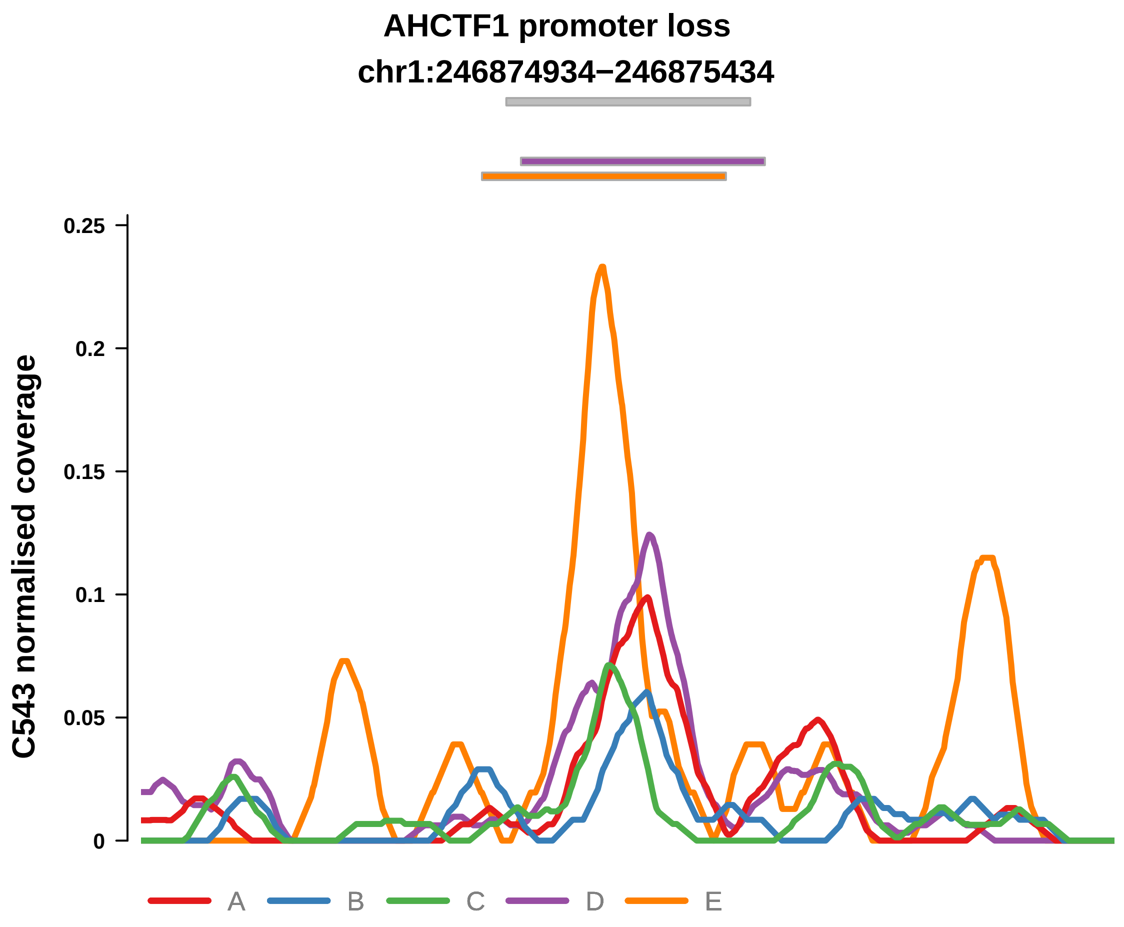


Putatively Clonal (logFC = −11.54)

| | P. | Adjusted P. |
|---|---|---|
| *~region vs. ~* | 6.44e−03 ** | 1.46e−01 |
| *~purity vs ~* | 1.6e−01 | NA |
| *~purity+region vs ~purity* | 1.31e−02 * | 2.31e−01 |
| *Carcinoma vs. normal* | 9.14e−03 ** | 9.14e−03 ** |

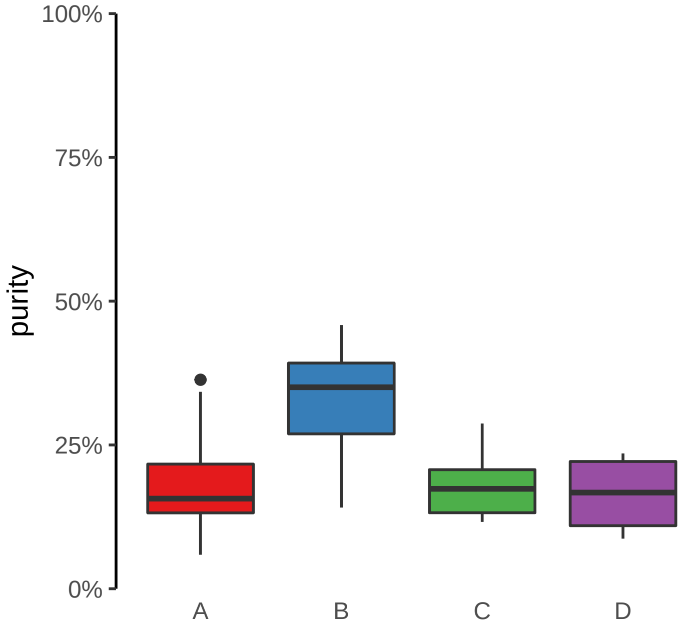

**AKR1B1 promoter loss**
**chr7:134458962−134459462**

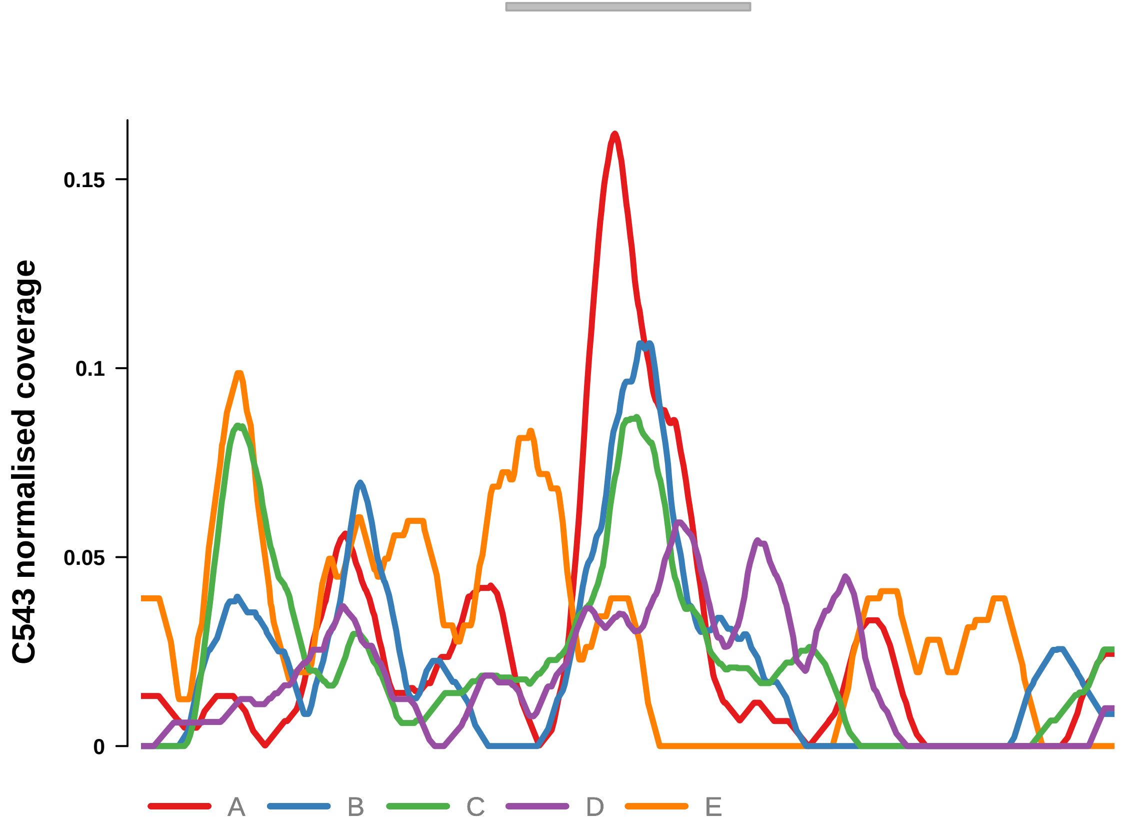


Putatively Clonal (logFC = −12.18)

| | **P.** | **Adjusted P.** |
|---|---|---|
| *~region vs. ~* | 6.14e−01 | 8.12e−01 |
| *~purity vs ~* | 1.05e−01 | 1.95e−01 |
| *~purity+region vs ~purity* | 9.32e−01 | 9.72e−01 |
| *Carcinoma vs. normal* | 1.66e−03 ** | 1.66e−03 ** |

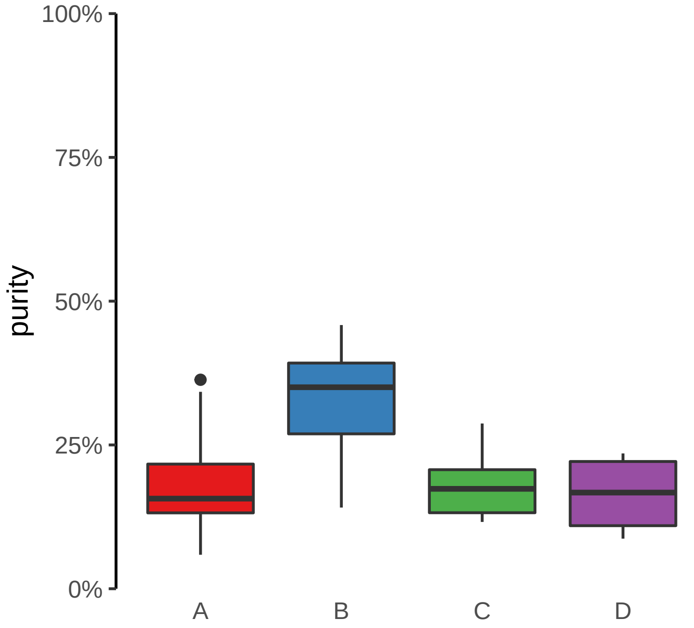

**ERBB3 promoter loss**
**chr12:56087145−56087645**

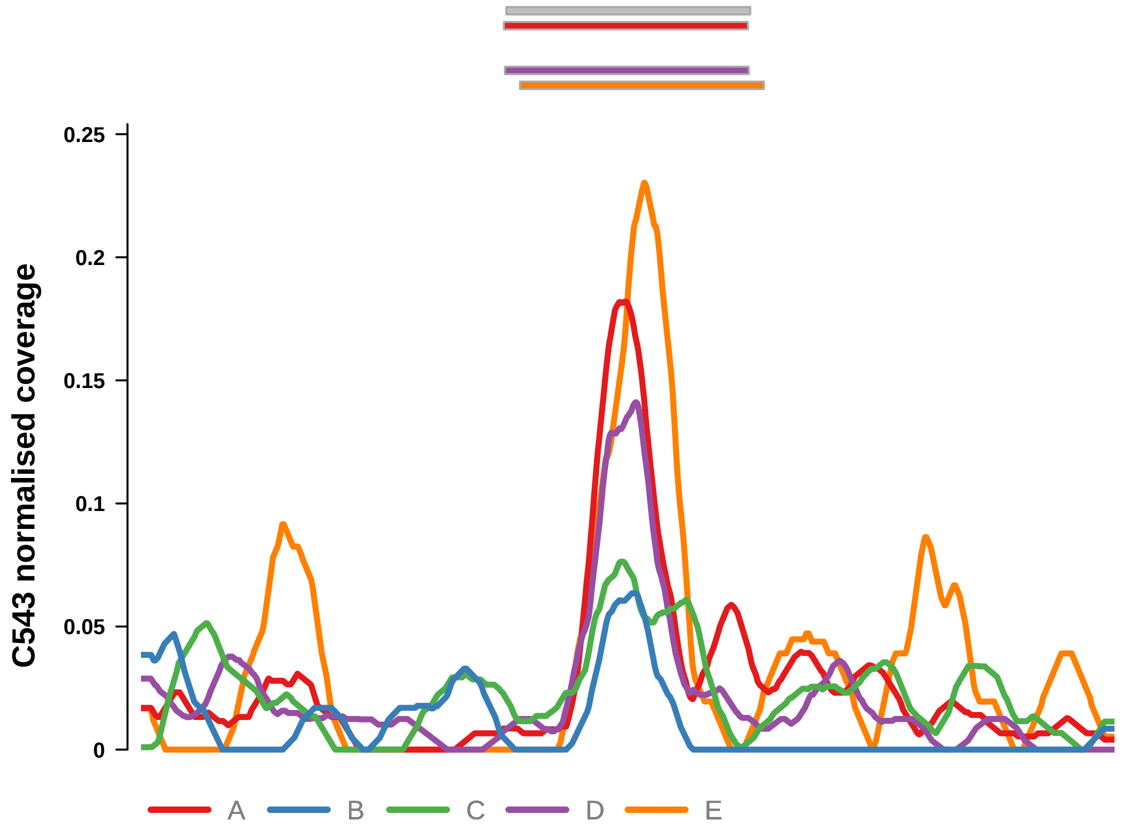


Putatively Clonal (logFC = −11.78)

| | **P.** | **Adjusted P.** |
|---|---|---|
| *~region vs. ~* | 1.47e−01 | 5.16e−01 |
| *~purity vs ~* | 5.66e−03 ** | 7.93e−02 |
| *~purity+region vs ~purity* | 6.22e−01 | 8.43e−01 |
| *Carcinoma vs. normal* | 4.1e−03 ** | 4.1e−03 ** |

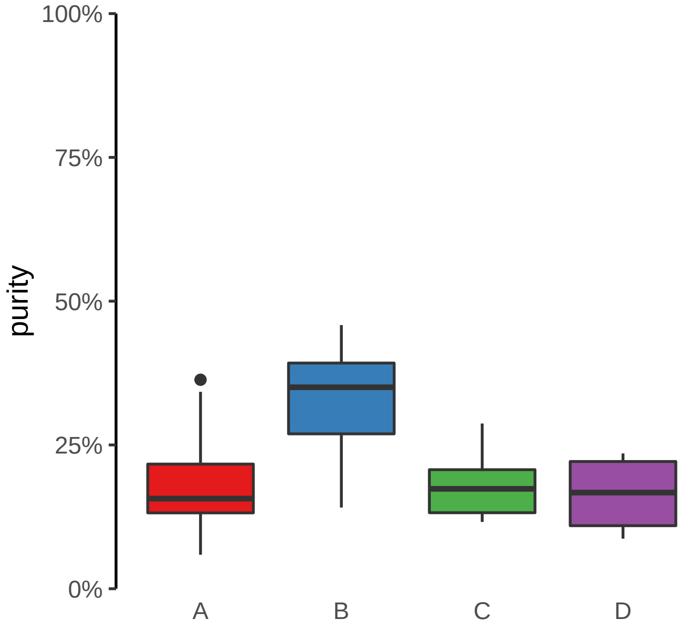

**LETM1 promoter loss**

**chr4:1839336−1839836**

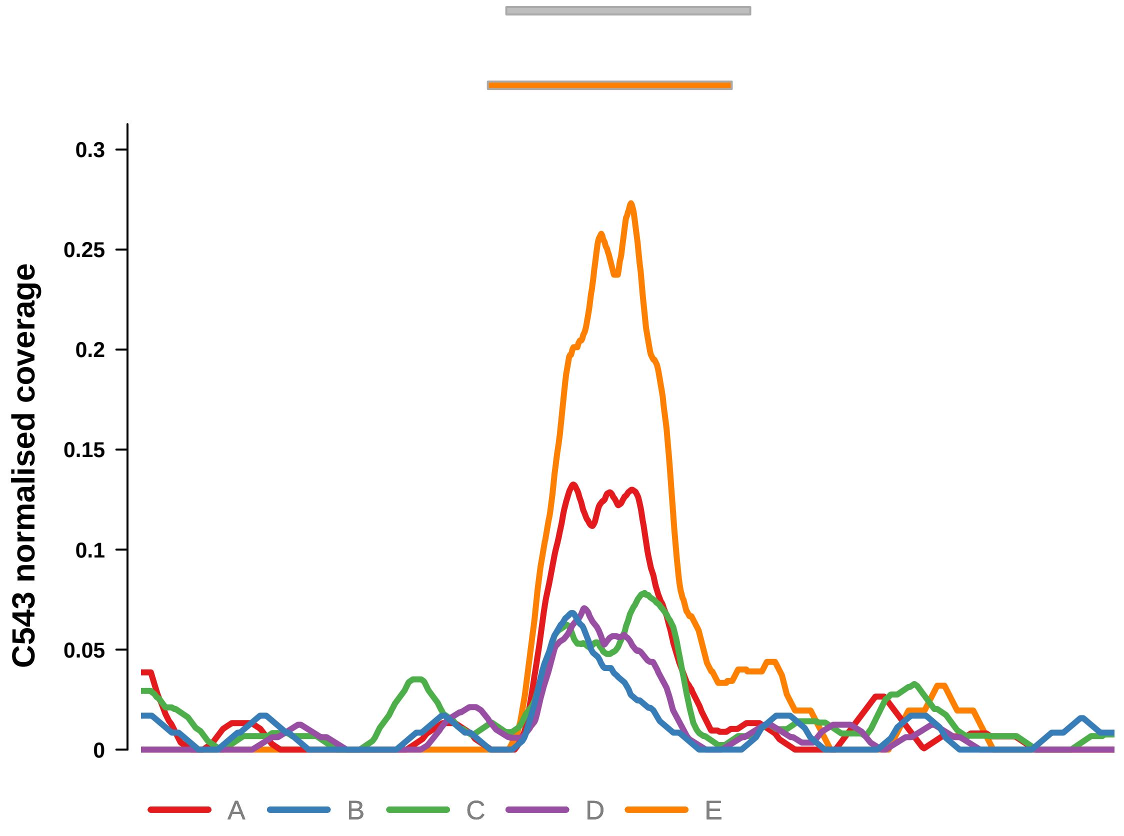


Putatively Clonal (logFC = −11.41)

| | P. | Adjusted P. |
|---|---|---|
| *~region vs. ~* | 3.24e−01 | 7.13e−01 |
| *~purity vs ~* | 1.26e−01 | 2.2e−01 |
| *~purity+region vs ~purity* | 6.32e−01 | 8.43e−01 |
| *Carcinoma vs. normal* | 9.66e−03 ** | 9.66e−03 ** |

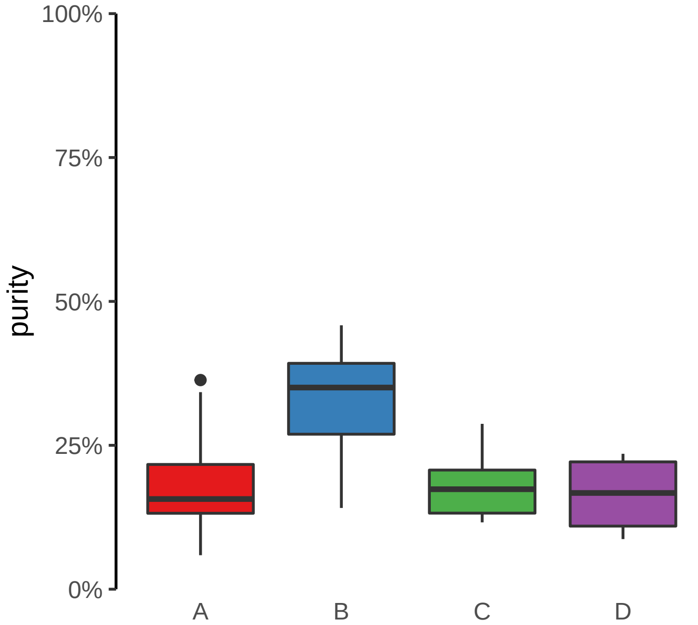

**LPCAT4 promoter loss**
**chr15:34362318−34362818**

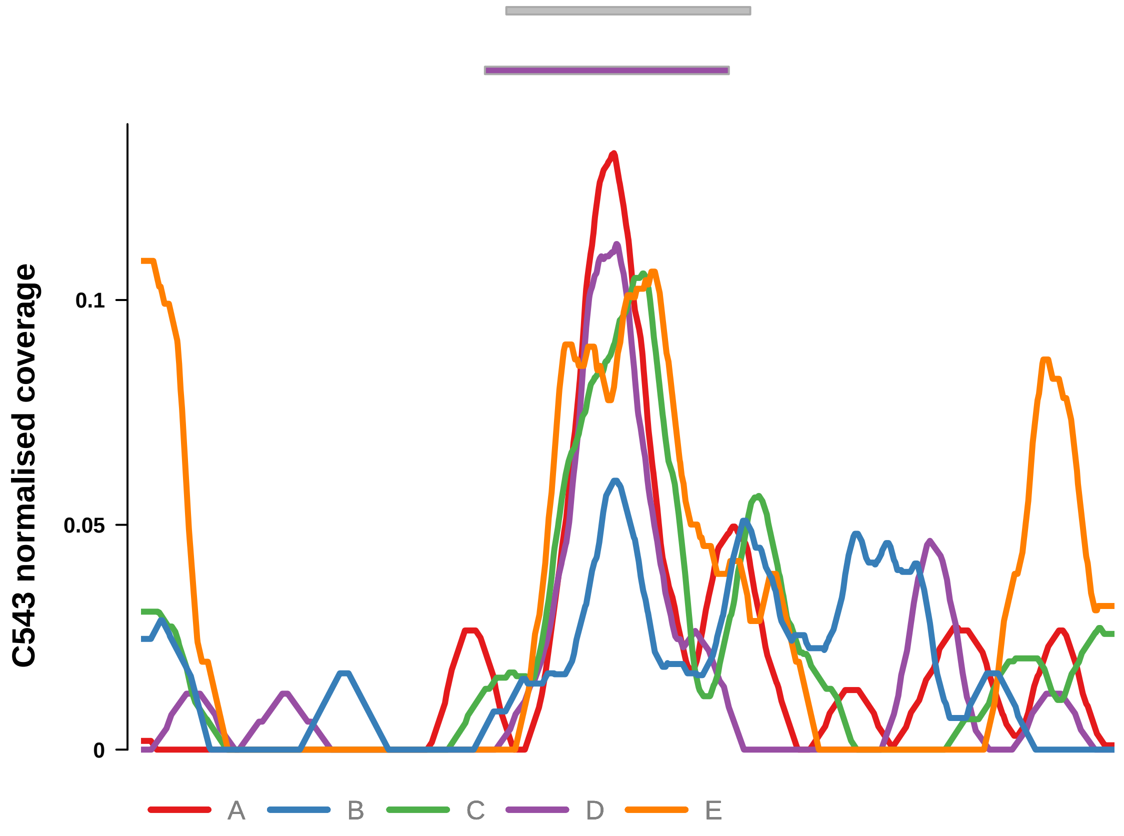


Putatively Clonal (logFC = −11.78)

| | P. | Adjusted P. |
|---|---|---|
| *~region vs. ~* | 3.79e−01 | 7.5e−01 |
| *~purity vs ~* | 2.45e−02 * | 9.57e−02 |
| *~purity+region vs ~purity* | 5.64e−01 | 8.43e−01 |
| *Carcinoma vs. normal* | 4.1e−03 ** | 4.1e−03 ** |

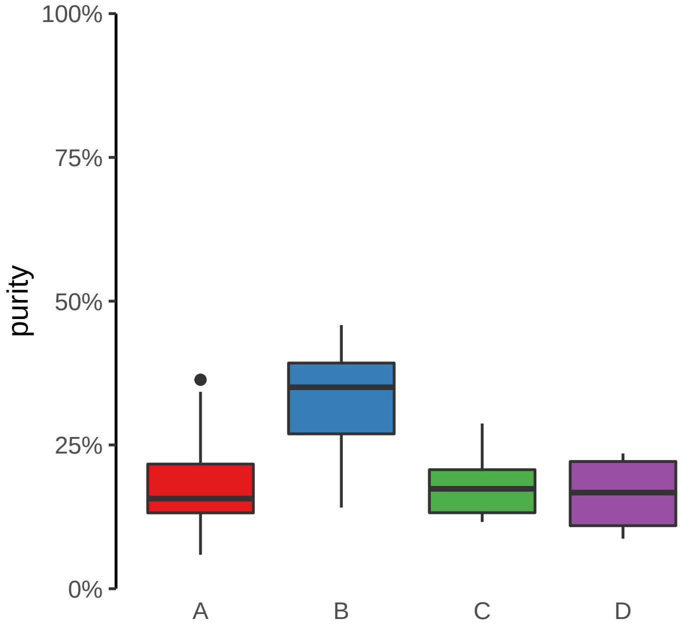

**SMAD3 promoter loss**
**chr15:67127194−67127694**

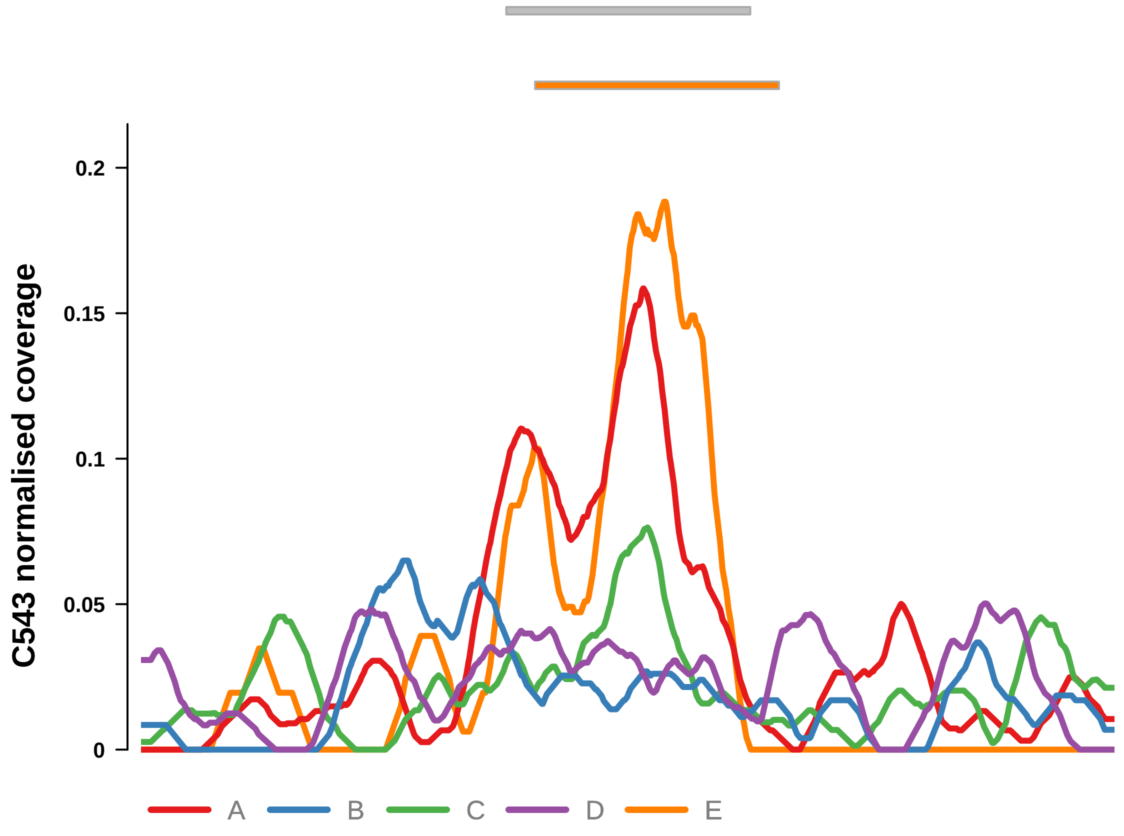


Putatively Clonal (logFC = −11.72)

| | P. | Adjusted P. |
|---|---|---|
| *~region vs. ~* | 4.23e−02 * | 3.47e−01 |
| *~purity vs ~* | 8.95e−01 | 9.51e−01 |
| *~purity+region vs ~purity* | 2.31e−02 * | 3.27e−01 |
| *Carcinoma vs. normal* | 5.52e−03 ** | 5.52e−03 ** |

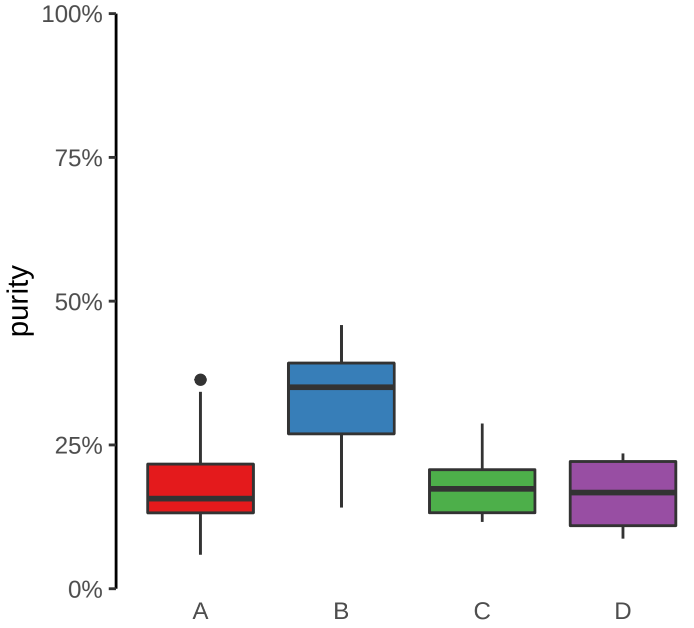

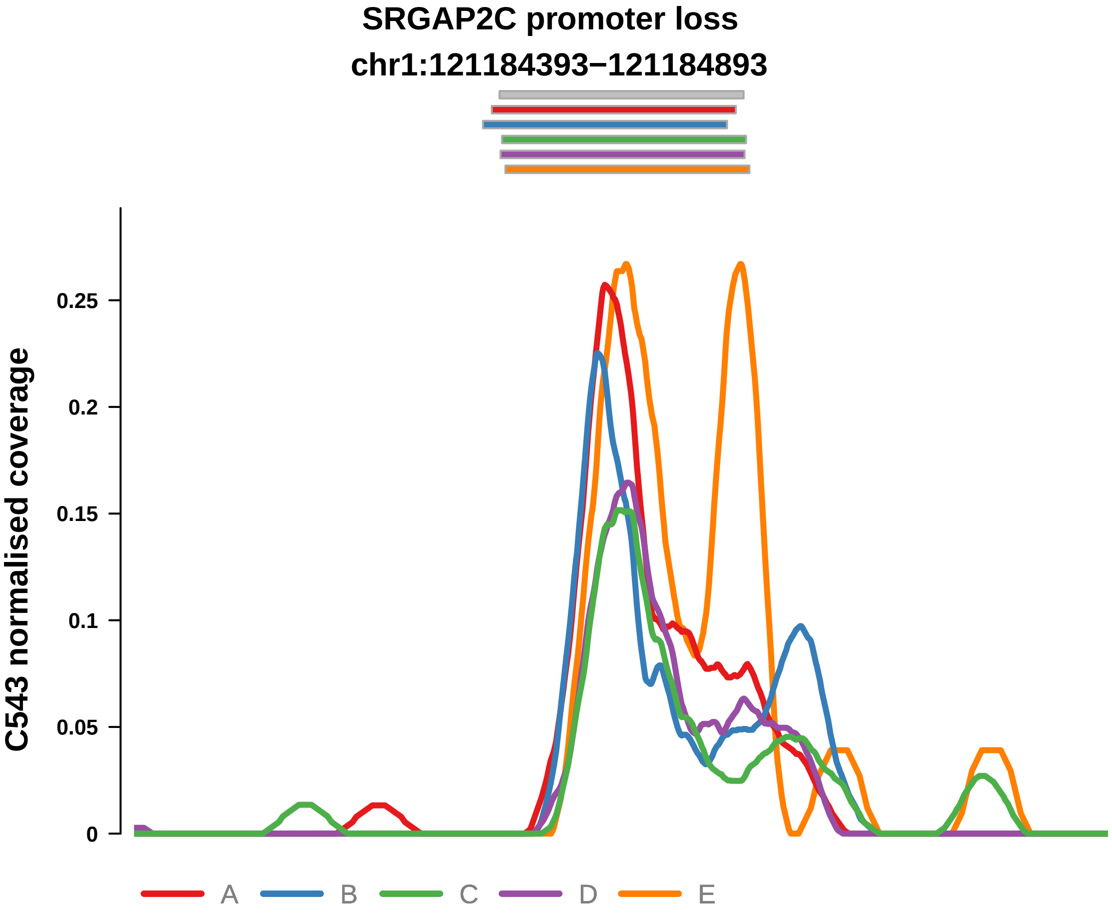


Putatively Clonal (logFC = −13.1)

| | P. | Adjusted P. |
|---|---|---|
| *~region vs. ~* | 4.51e−02 * | 3.47e−01 |
| *~purity vs ~* | 3.27e−02 * | 9.57e−02 |
| *~purity+region vs ~purity* | 1.72e−01 | 6.14e−01 |
| *Carcinoma vs. normal* | 8.35e−05 **** | 8.35e−05 **** |

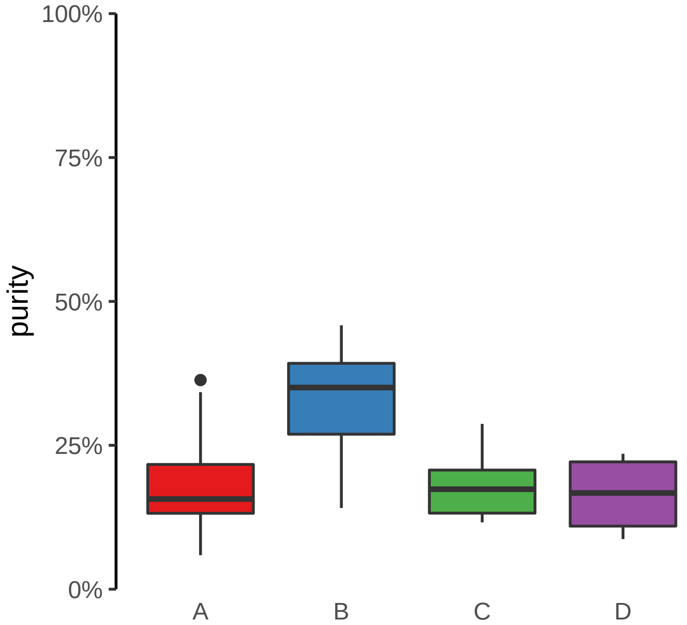

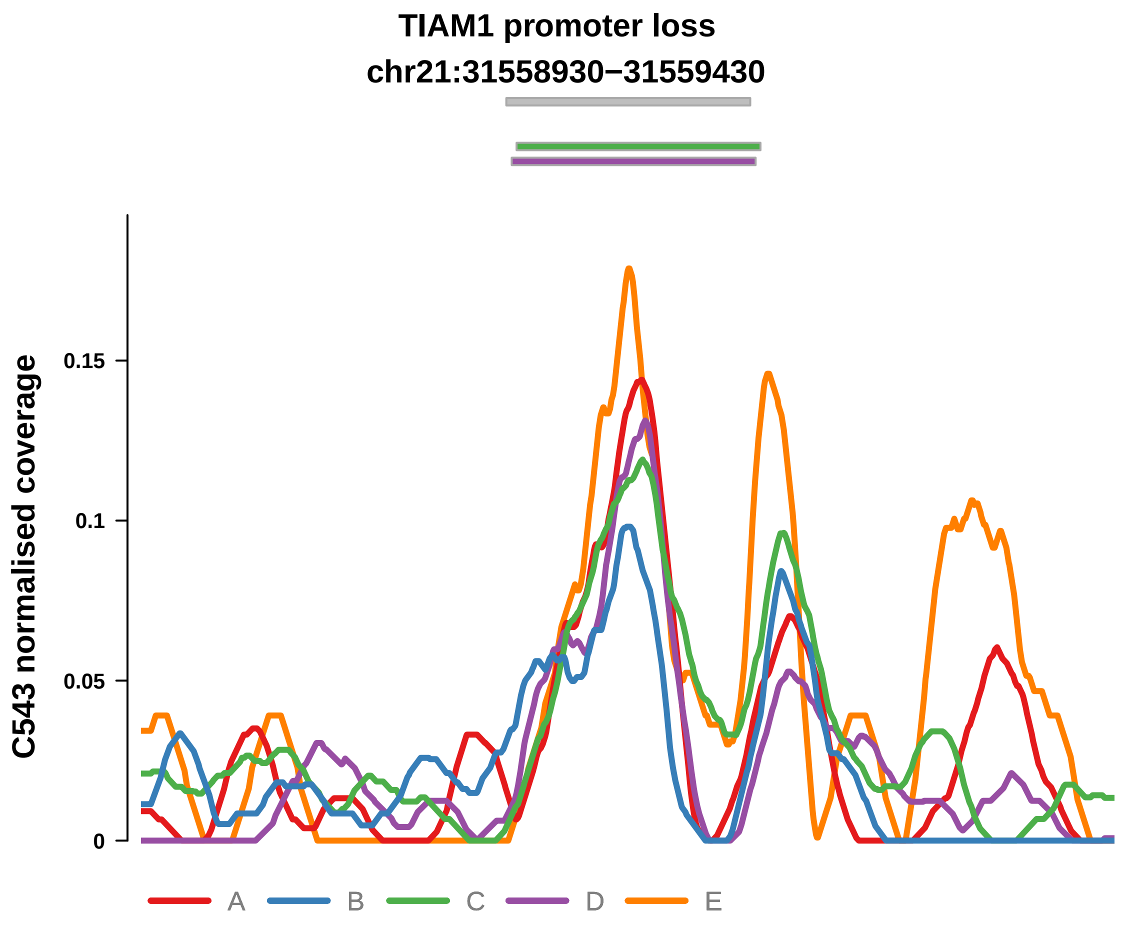


Putatively Clonal (logFC = −12.88)

| | P. | Adjusted P. |
|---|---|---|
| *~region vs. ~* | 8.05e−02 | 4.17e−01 |
| *~purity vs ~* | 1.78e−02 * | 8.29e−02 |
| *~purity+region vs ~purity* | 2.6e−01 | 7.73e−01 |
| *Carcinoma vs. normal* | 1.72e−04 *** | 1.72e−04 *** |

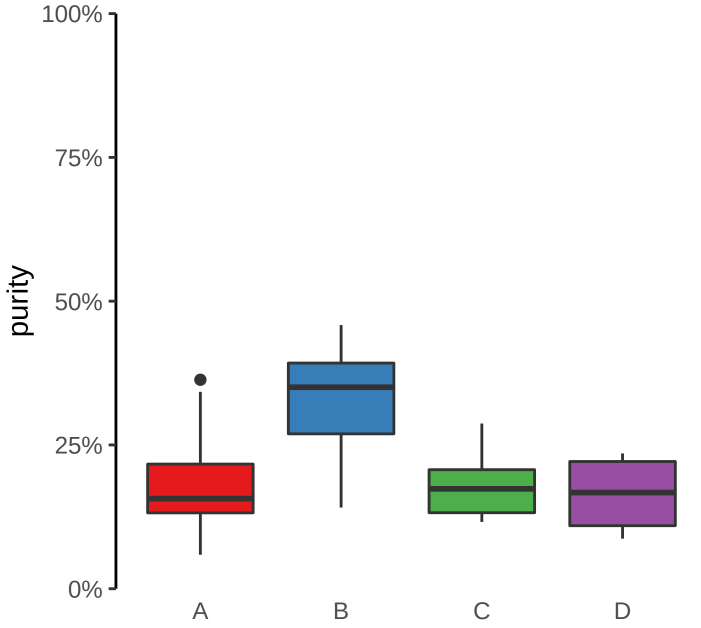

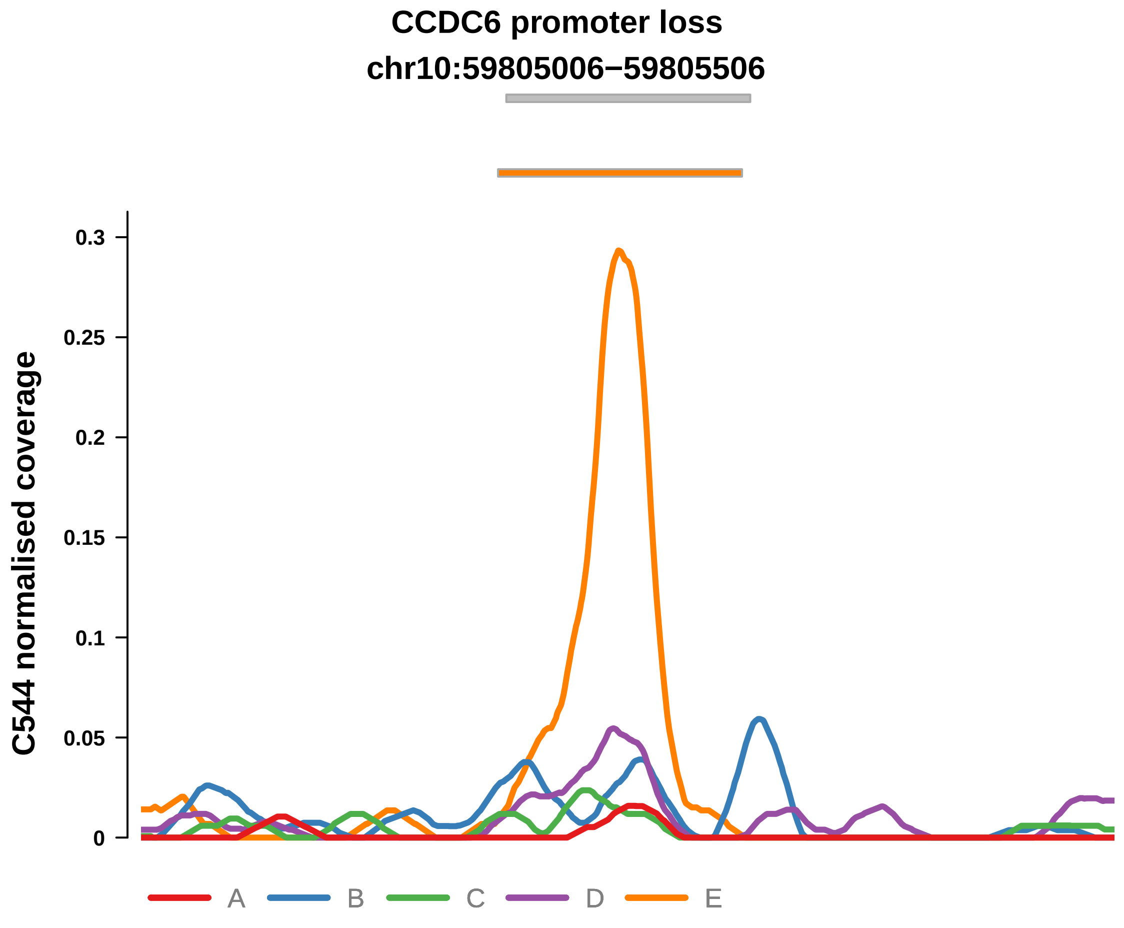


Putatively Clonal (logFC = −1.68)

| | P. | Adjusted P. |
|---|---|---|
| *~region vs. ~* | 4.01e−03 ** | 1.76e−02 * |
| *~purity vs ~* | 1.34e−01 | 2.79e−01 |
| *~purity+region vs ~purity* | 6.65e−03 ** | NA |
| *Carcinoma vs. normal* | 9.28e−03 ** | 9.28e−03 ** |

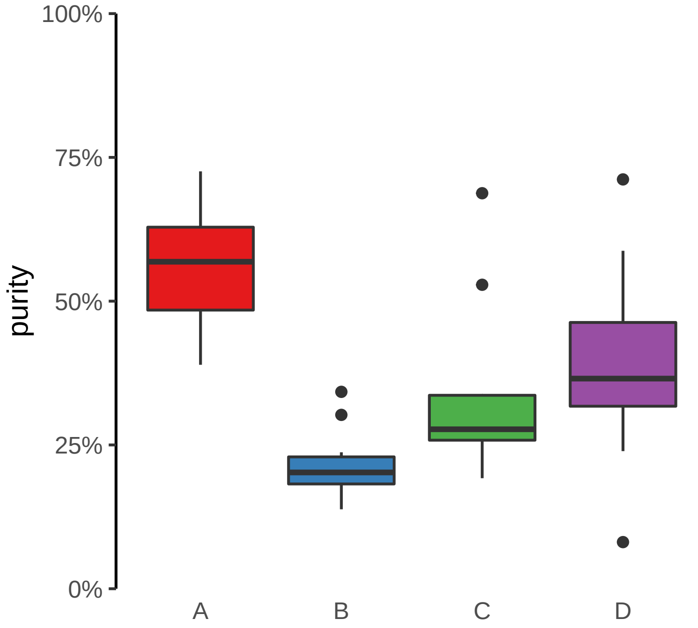

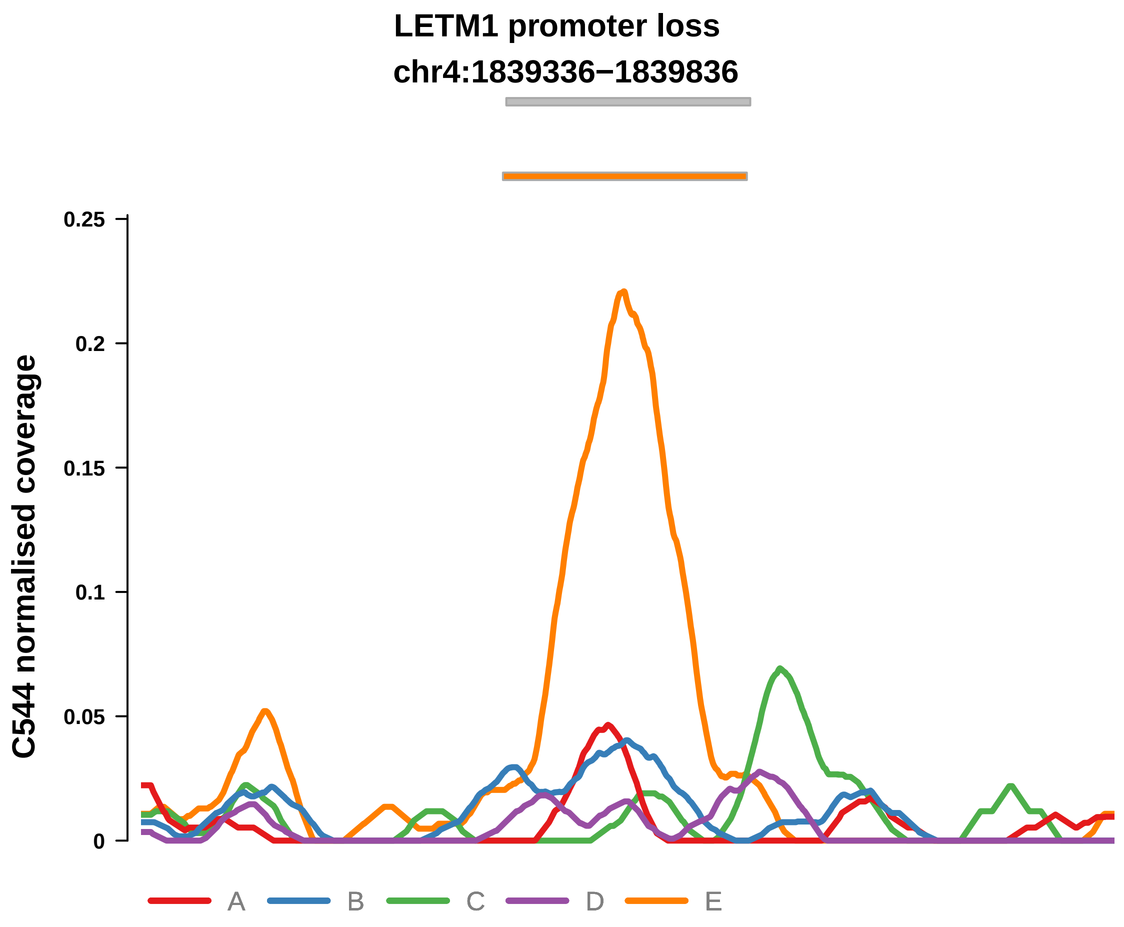


Putatively Clonal (logFC = −1.71)

| | P. | Adjusted P. |
|---|---|---|
| *~region vs. ~* | 2.84e−01 | 3.95e−01 |
| *~purity vs ~* | 4.11e−01 | 5.69e−01 |
| *~purity+region vs ~purity* | 3.71e−01 | NA |
| *Carcinoma vs. normal* | 8.77e−03 ** | 8.77e−03 ** |

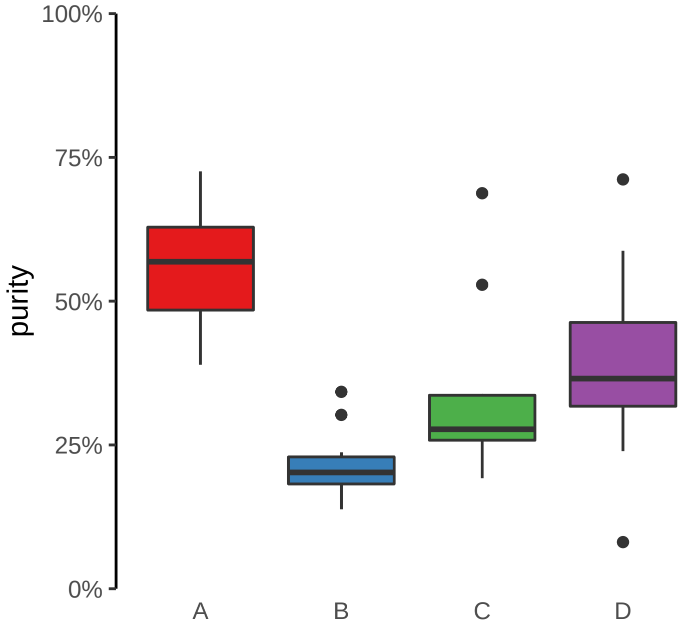

**PLOD2 promoter loss**
**chr3:146164913−146165413**

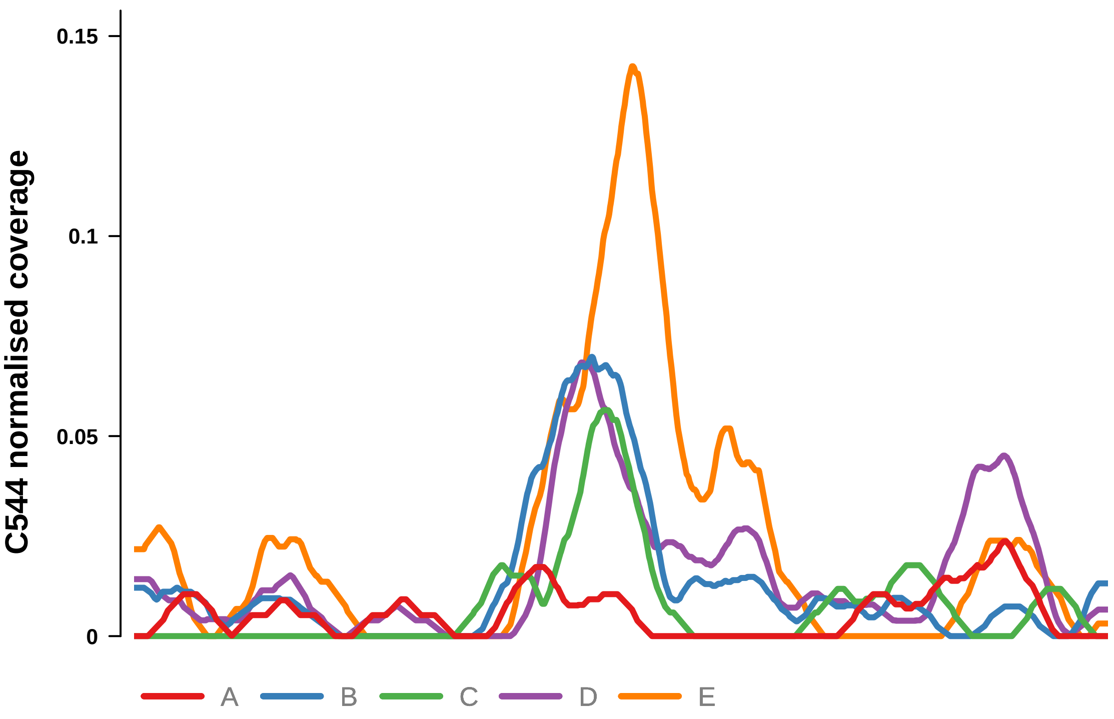


Putatively Clonal (logFC = −1.94)

| | P. | Adjusted P. |
|---|---|---|
| *~region vs. ~* | 8.94e−02 | 1.64e−01 |
| *~purity vs ~* | 1.75e−02 * | 7.18e−02 |
| *~purity+region vs ~purity* | 6.11e−01 | 7e−01 |
| *Carcinoma vs. normal* | 3.99e−03 ** | 3.99e−03 ** |

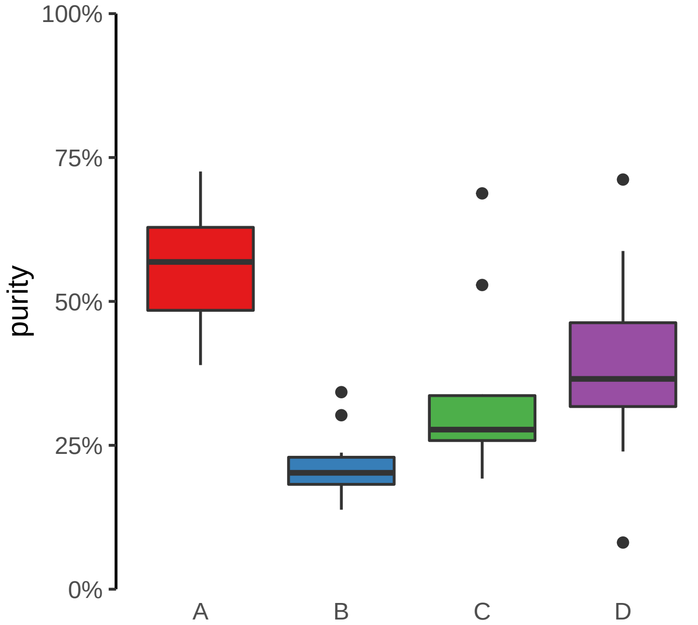

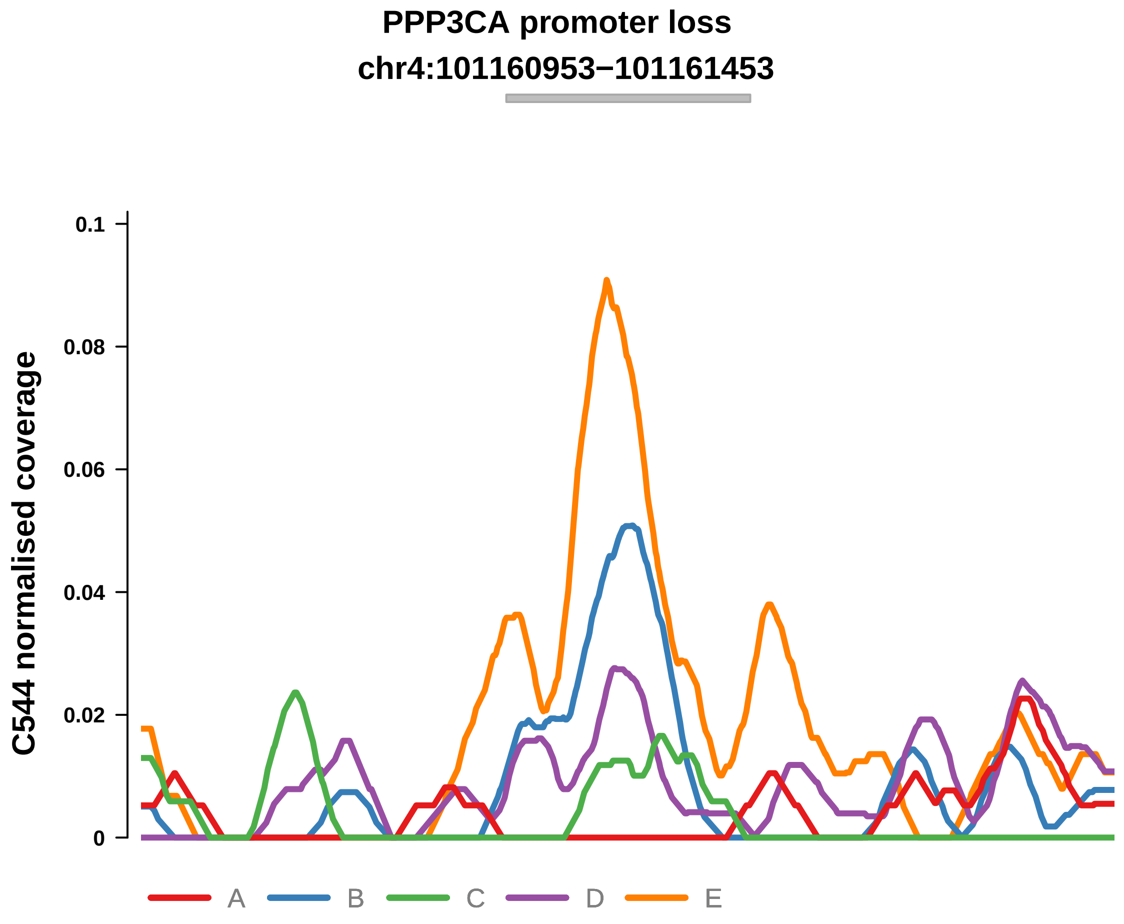


Putatively Clonal (logFC = −2.22)

| | P. | Adjusted P. |
|---|---|---|
| *~region vs. ~* | 1.3e−01 | 2.08e−01 |
| *~purity vs ~* | 1.2e−01 | NA |
| *~purity+region vs ~purity* | 3.62e−01 | NA |
| *Carcinoma vs. normal* | 3.03e−03 ** | 3.03e−03 ** |

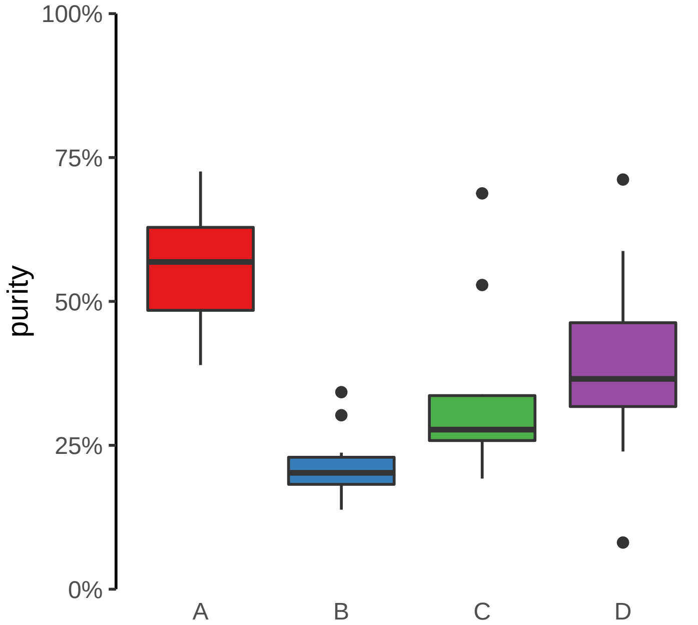

**SRGAP2C promoter loss**
**chr1:121184393−121184893**

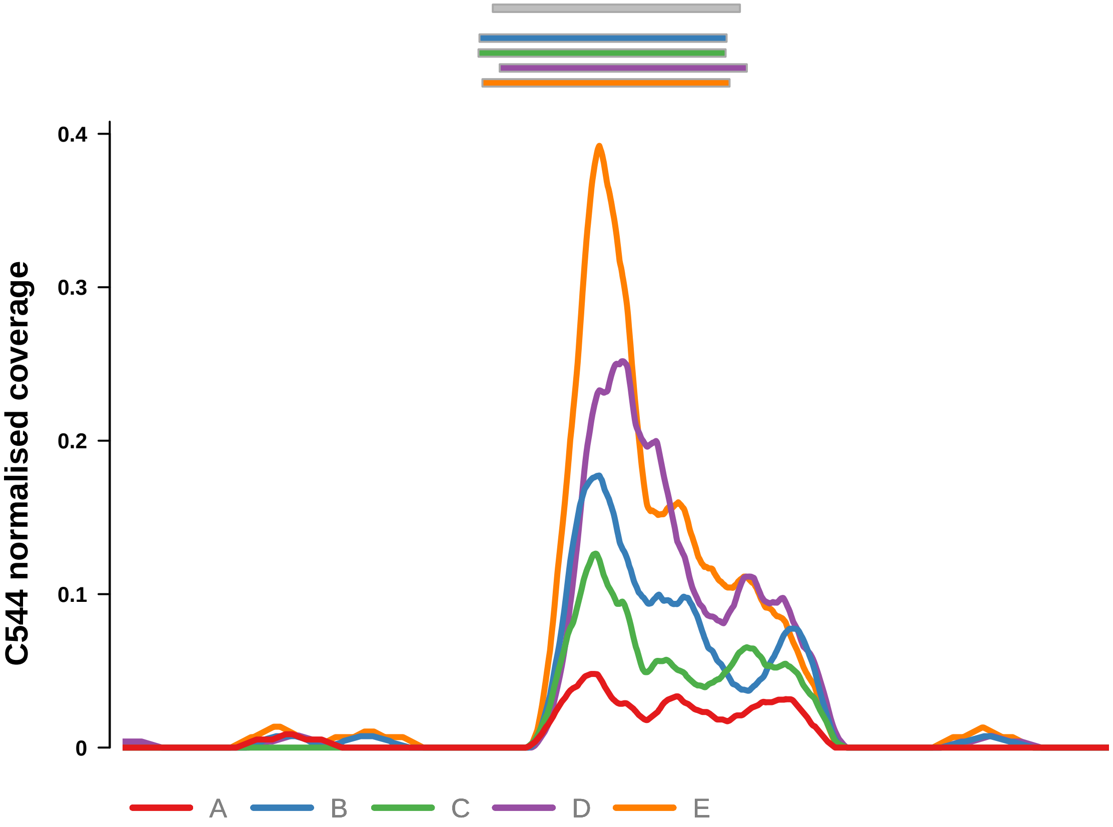


Putatively Clonal (logFC = −1.74)

| | P. | Adjusted P. |
|---|---|---|
| *~region vs. ~* | 1.01e−03 ** | 5.9e−03 ** |
| *~purity vs ~* | 1.55e−03 ** | 1.57e−02 * |
| *~purity+region vs ~purity* | 1.18e−01 | 2.23e−01 |
| *Carcinoma vs. normal* | 2.27e−03 ** | 2.27e−03 ** |

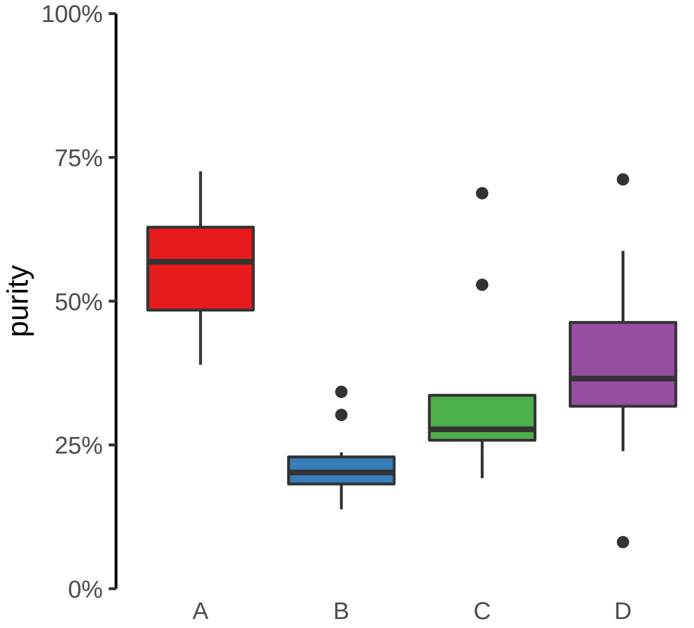

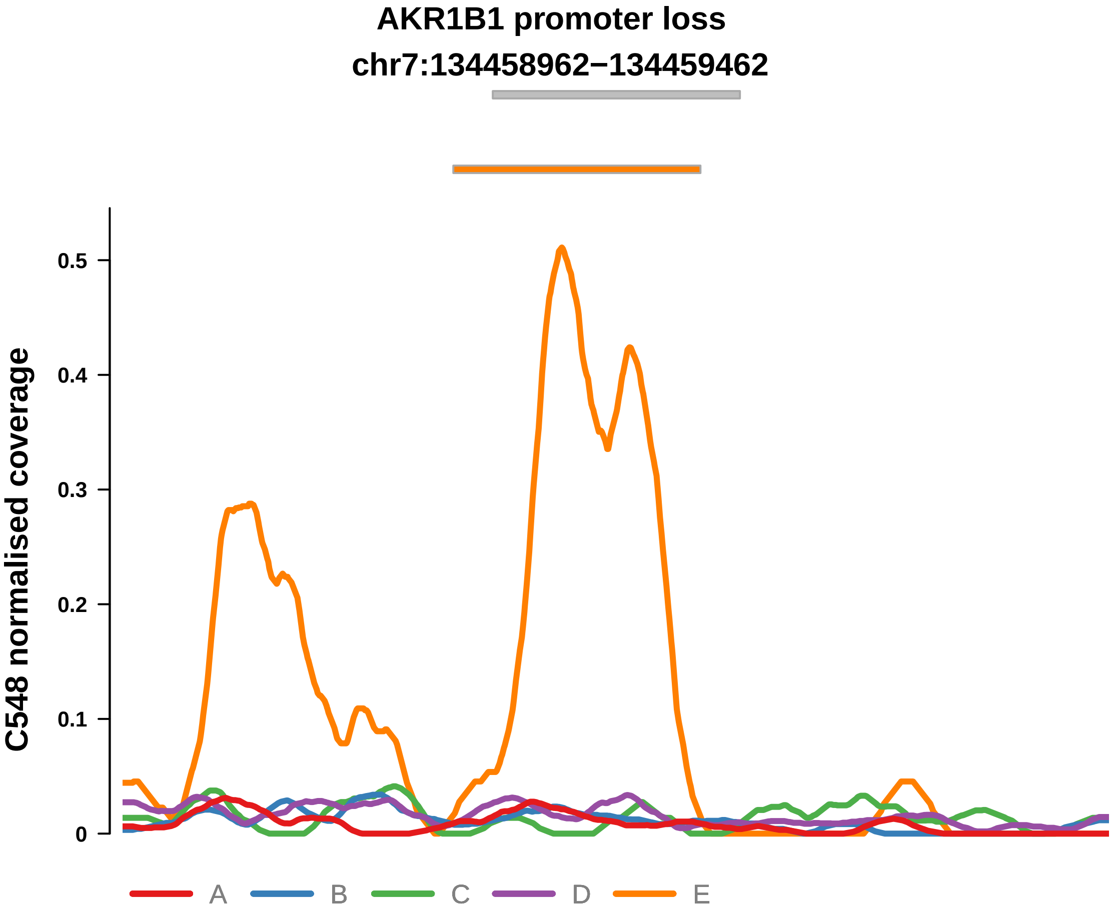


Putatively Clonal (logFC = −2.16)

| | P. | Adjusted P. |
|---|---|---|
| *~region vs. ~* | 8.09e−02 | 1.98e−01 |
| *~purity vs ~* | 1.62e−01 | 3.57e−01 |
| *~purity+region vs ~purity* | 2.44e−01 | 5.31e−01 |
| *Carcinoma vs. normal* | 1.83e−03 ** | 1.83e−03 ** |

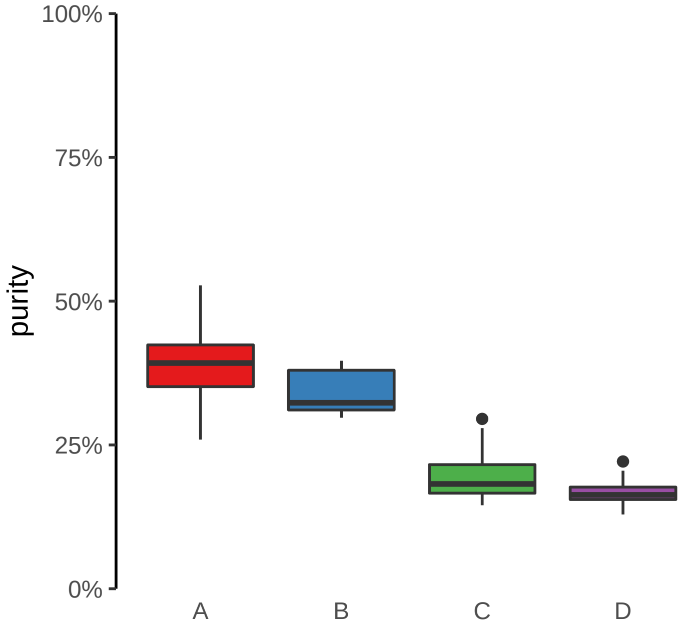

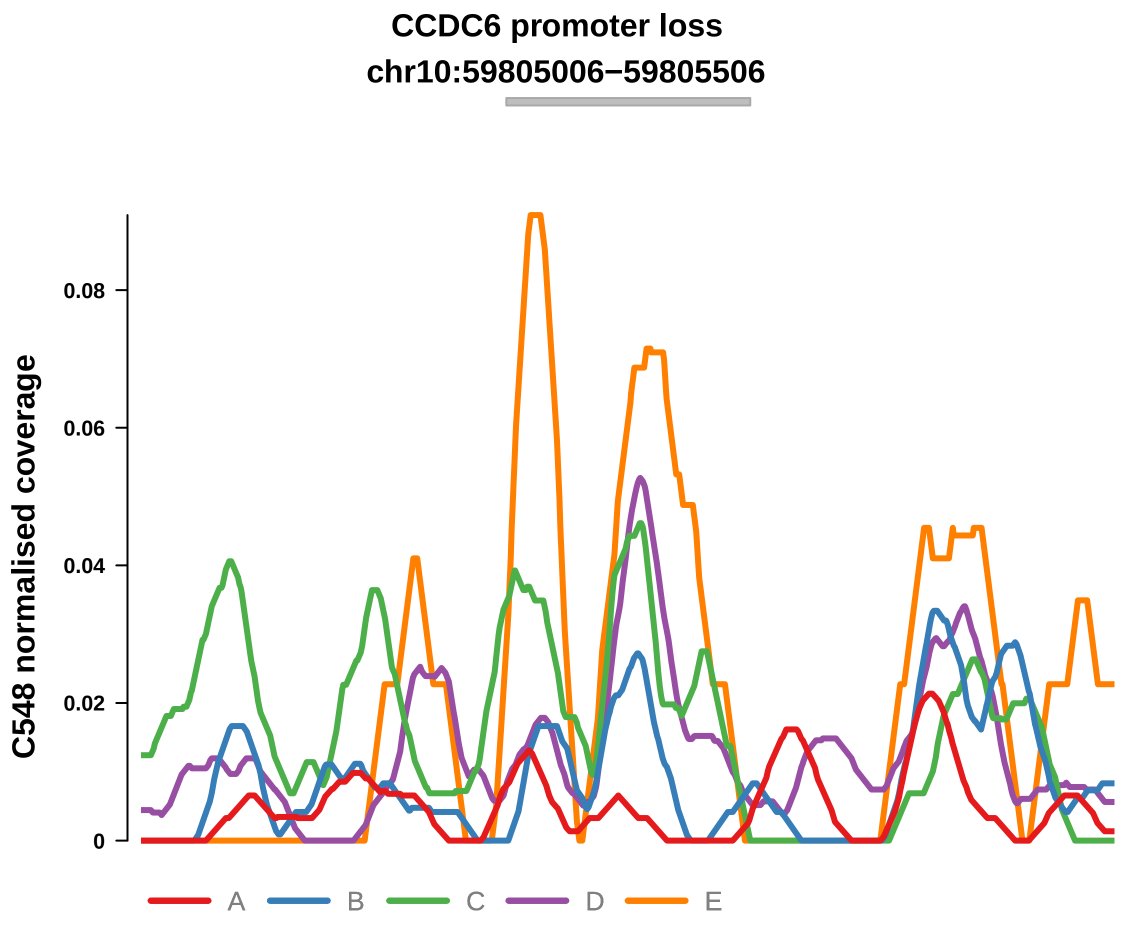


Putatively Clonal (logFC = −3.46)

| | P. | Adjusted P. |
|---|---|---|
| *~region vs. ~* | 1.43e−03 ** | 1.8e−02 * |
| *~purity vs ~* | 1.44e−05 **** | 1.27e−03 ** |
| *~purity+region vs ~purity* | 9.64e−01 | 9.75e−01 |
| *Carcinoma vs. normal* | 1.31e−04 *** | 1.31e−04 *** |

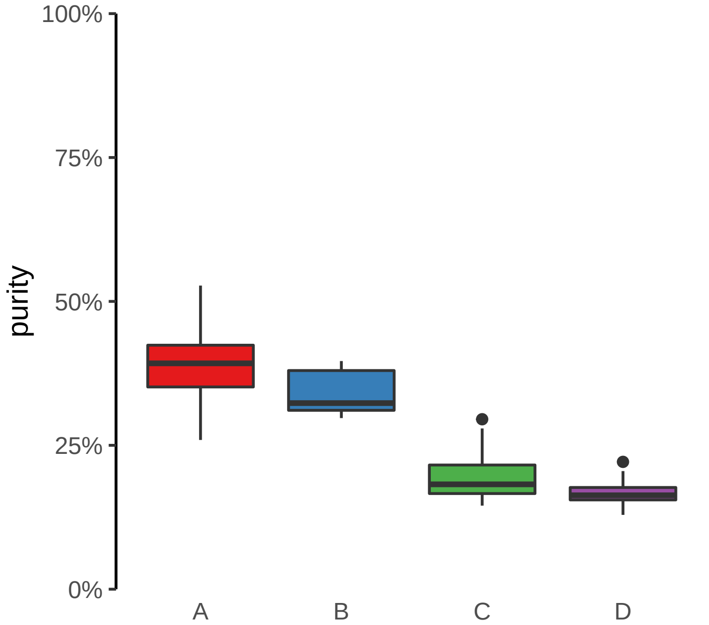

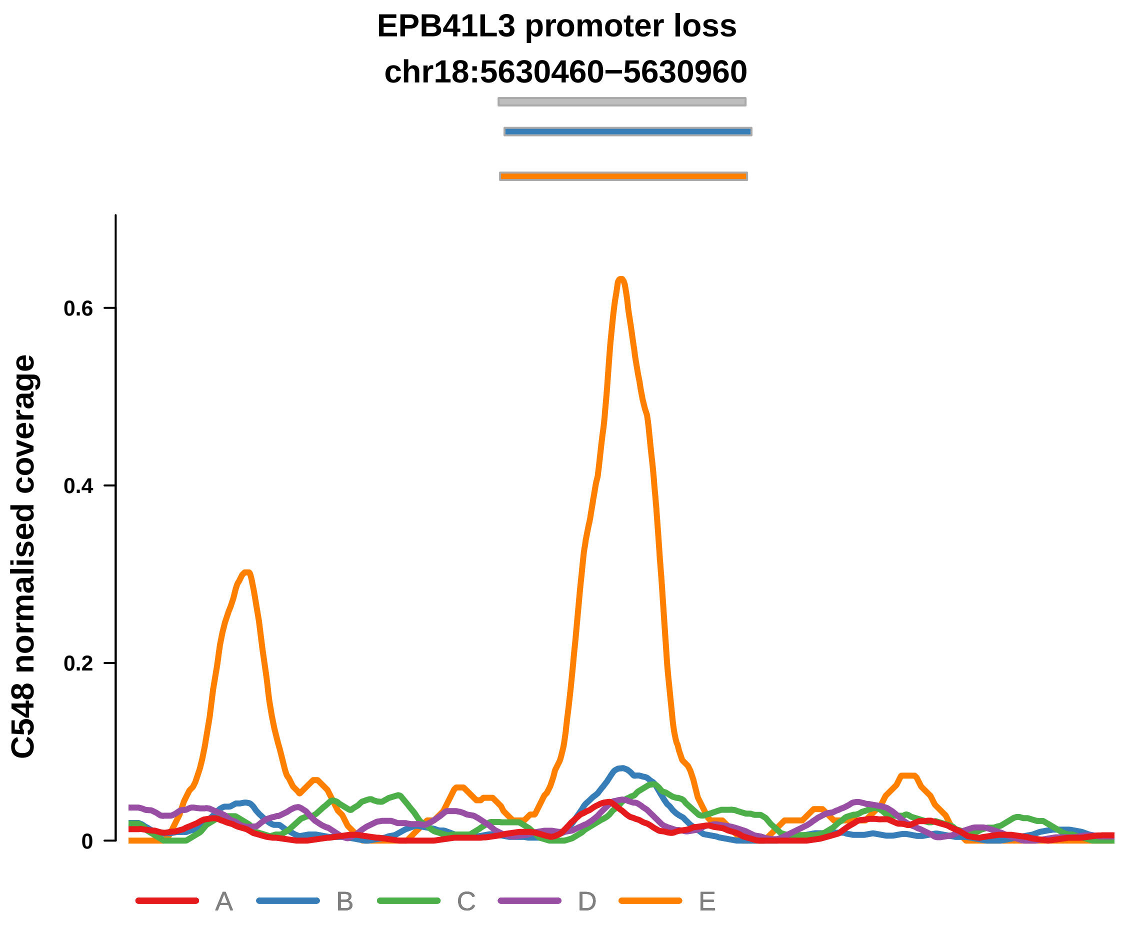


Putatively Clonal (logFC = −1.69)

| | P. | Adjusted P. |
|---|---|---|
| *~region vs. ~* | 7.67e−01 | 8.76e−01 |
| *~purity vs ~* | 2.25e−01 | 3.97e−01 |
| *~purity+region vs ~purity* | 9.95e−01 | 9.95e−01 |
| *Carcinoma vs. normal* | 7.94e−03 ** | 7.94e−03 ** |

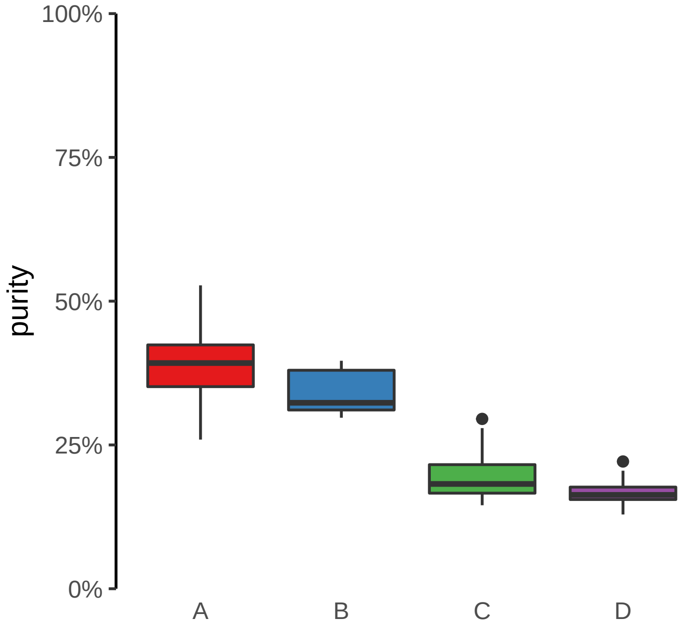

**LPCAT4 promoter loss**
**chr15:34362318−34362818**

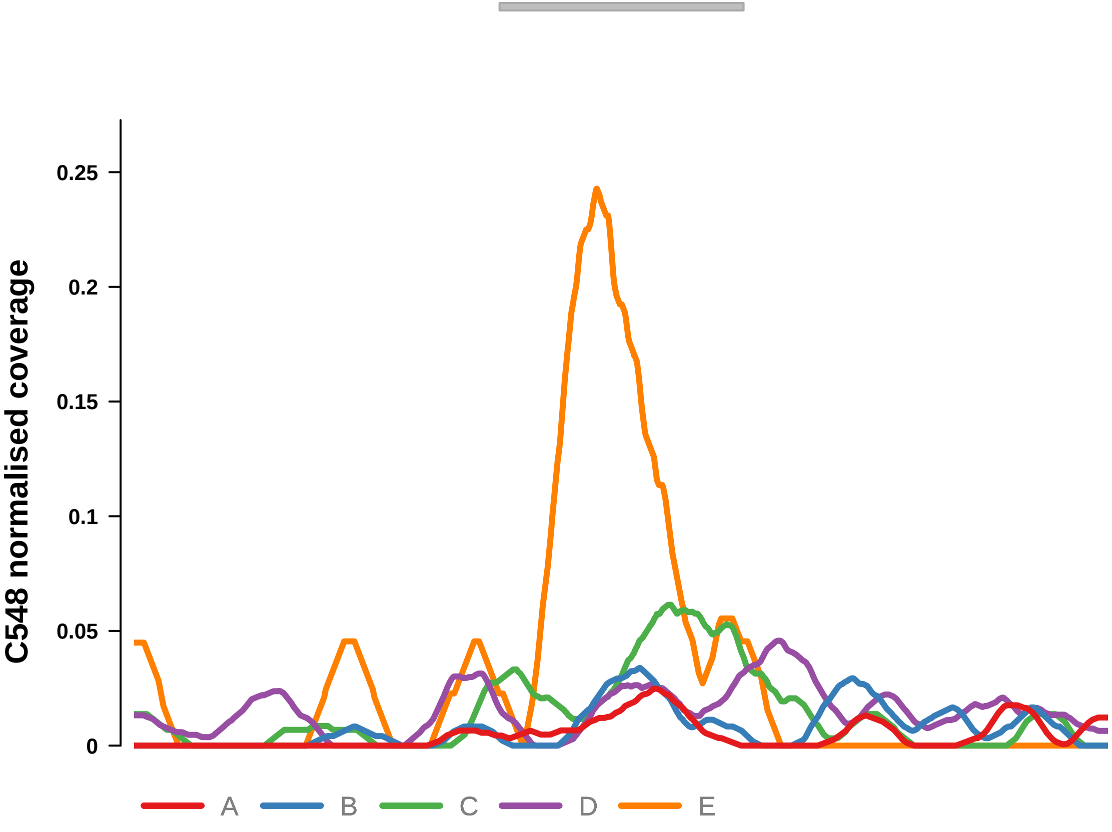


Putatively Clonal (logFC = −2.18)

| | P. | Adjusted P. |
|---|---|---|
| *~region vs. ~* | 3.24e−03 ** | 2.38e−02 * |
| *~purity vs ~* | 4.96e−04 *** | 8.73e−03 ** |
| *~purity+region vs ~purity* | 3.94e−01 | 6.09e−01 |
| *Carcinoma vs. normal* | 3.27e−03 ** | 3.27e−03 ** |

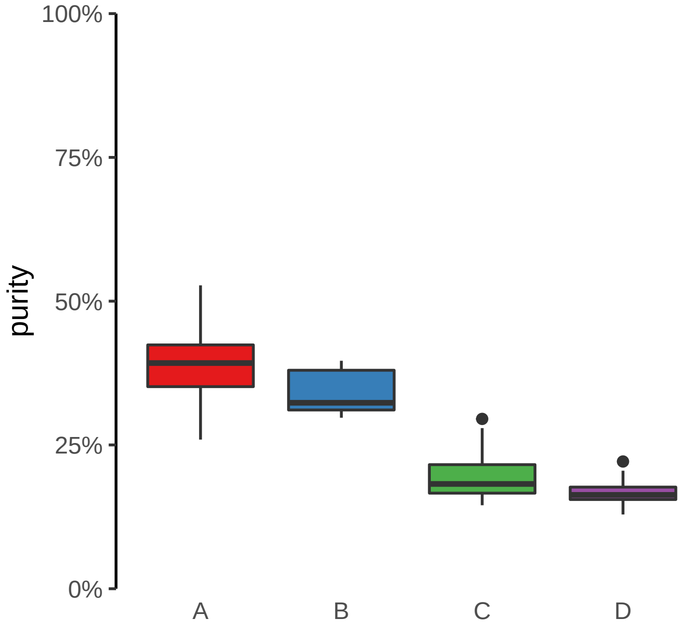

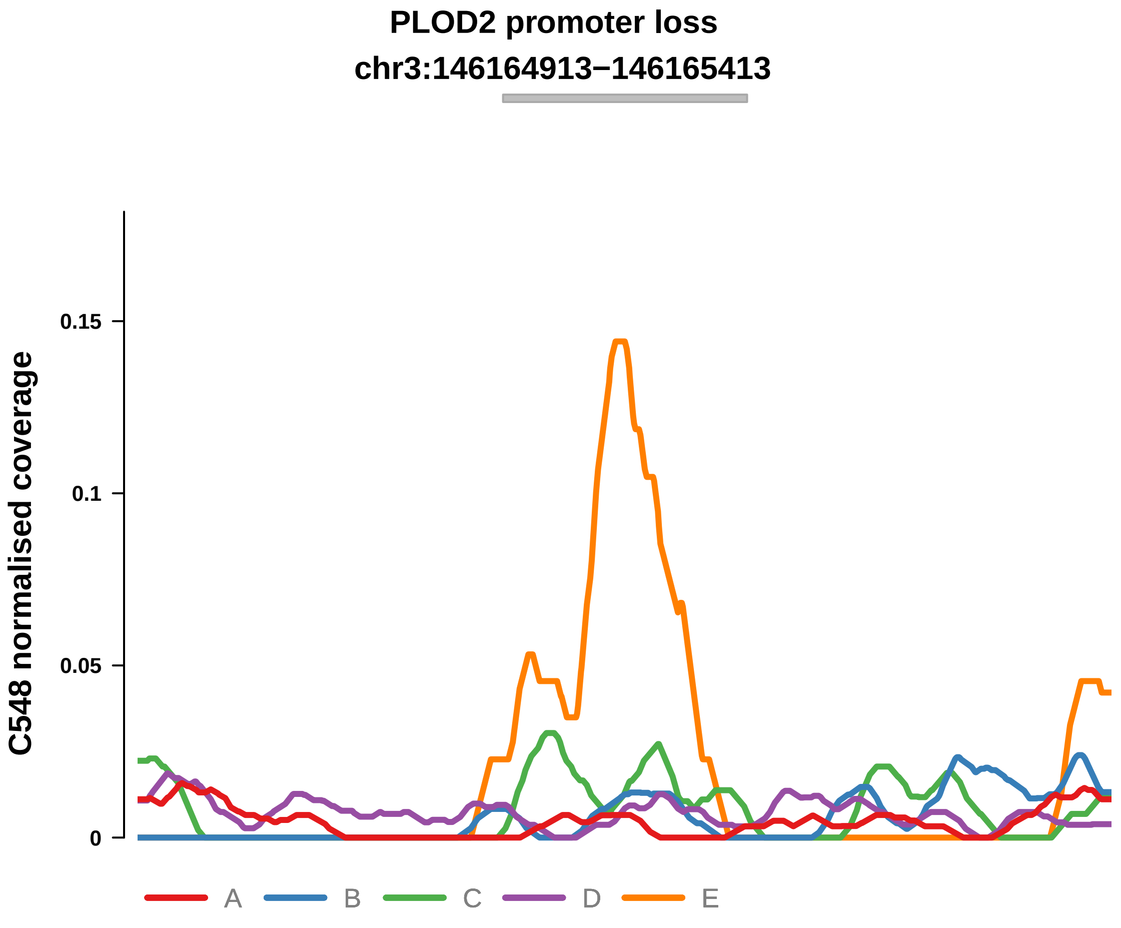


Putatively Clonal (logFC = −11.31)

| | P. | Adjusted P. |
|---|---|---|
| *~region vs. ~* | 2.14e−01 | 3.19e−01 |
| *~purity vs ~* | 7.8e−02 | 2.37e−01 |
| *~purity+region vs ~purity* | 6.91e−01 | 8.44e−01 |
| *Carcinoma vs. normal* | 6.09e−07 **** | 6.09e−07 **** |

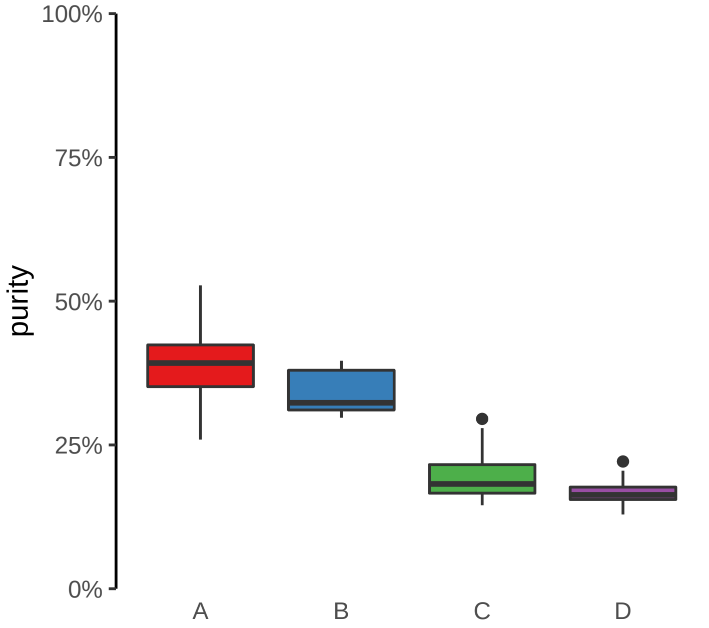

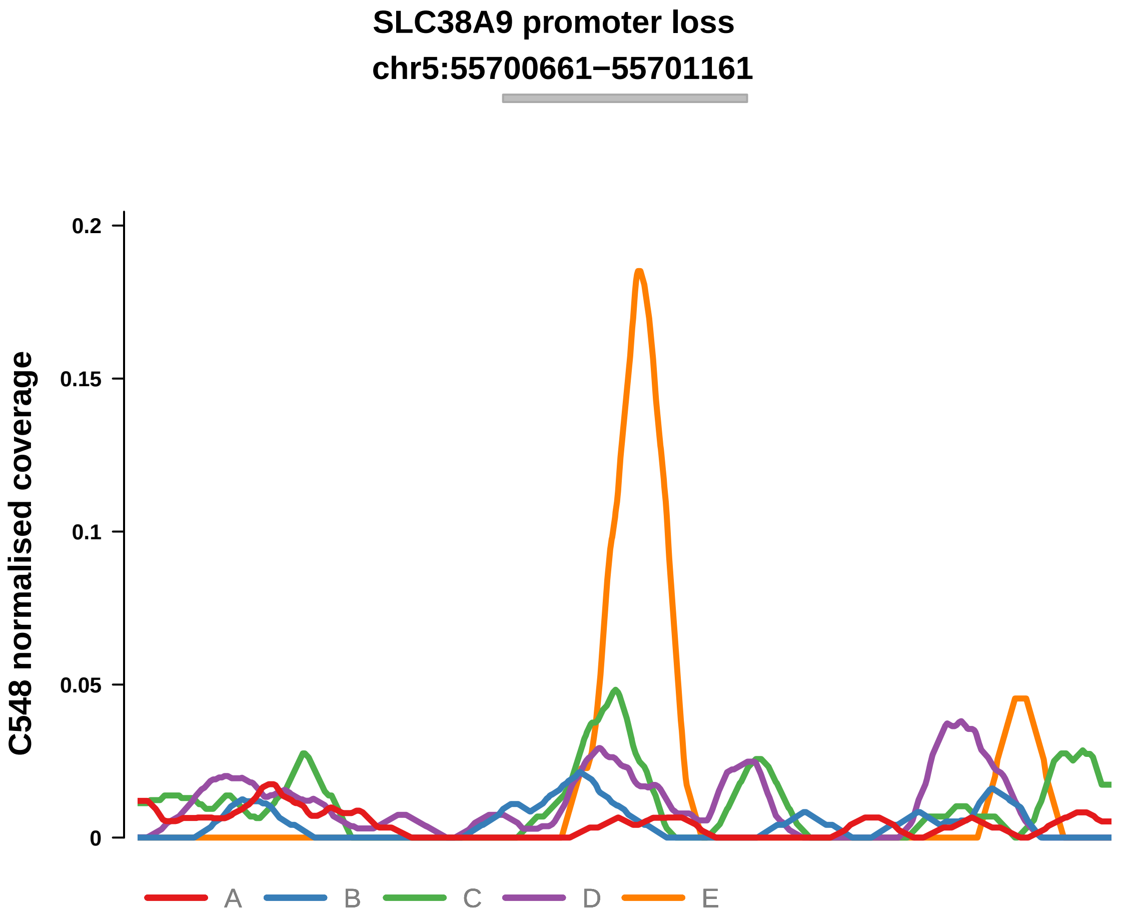


Putatively Clonal (logFC = −2.71)

| | P. | Adjusted P. |
|---|---|---|
| *~region vs. ~* | 7.4e−04 *** | 1.63e−02 * |
| *~purity vs ~* | 3.57e−04 *** | 8.73e−03 ** |
| *~purity+region vs ~purity* | 4.96e−01 | 6.93e−01 |
| *Carcinoma vs. normal* | 6.57e−03 ** | 6.57e−03 ** |

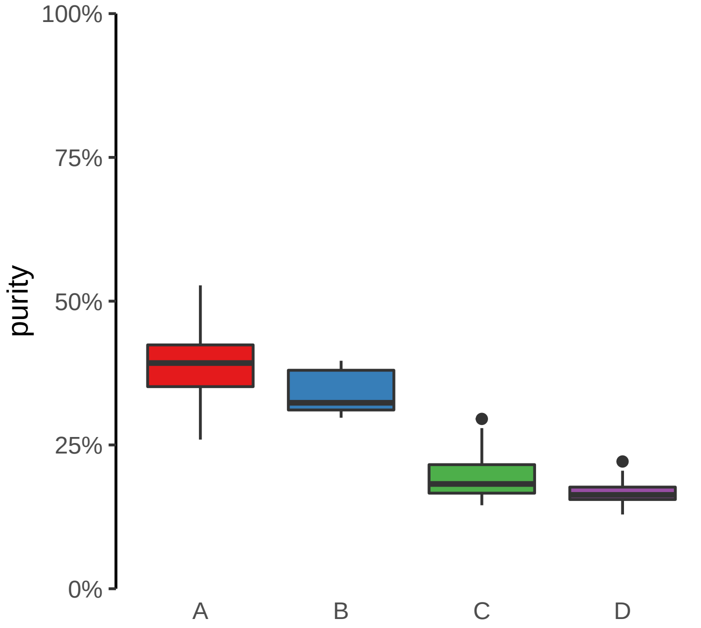

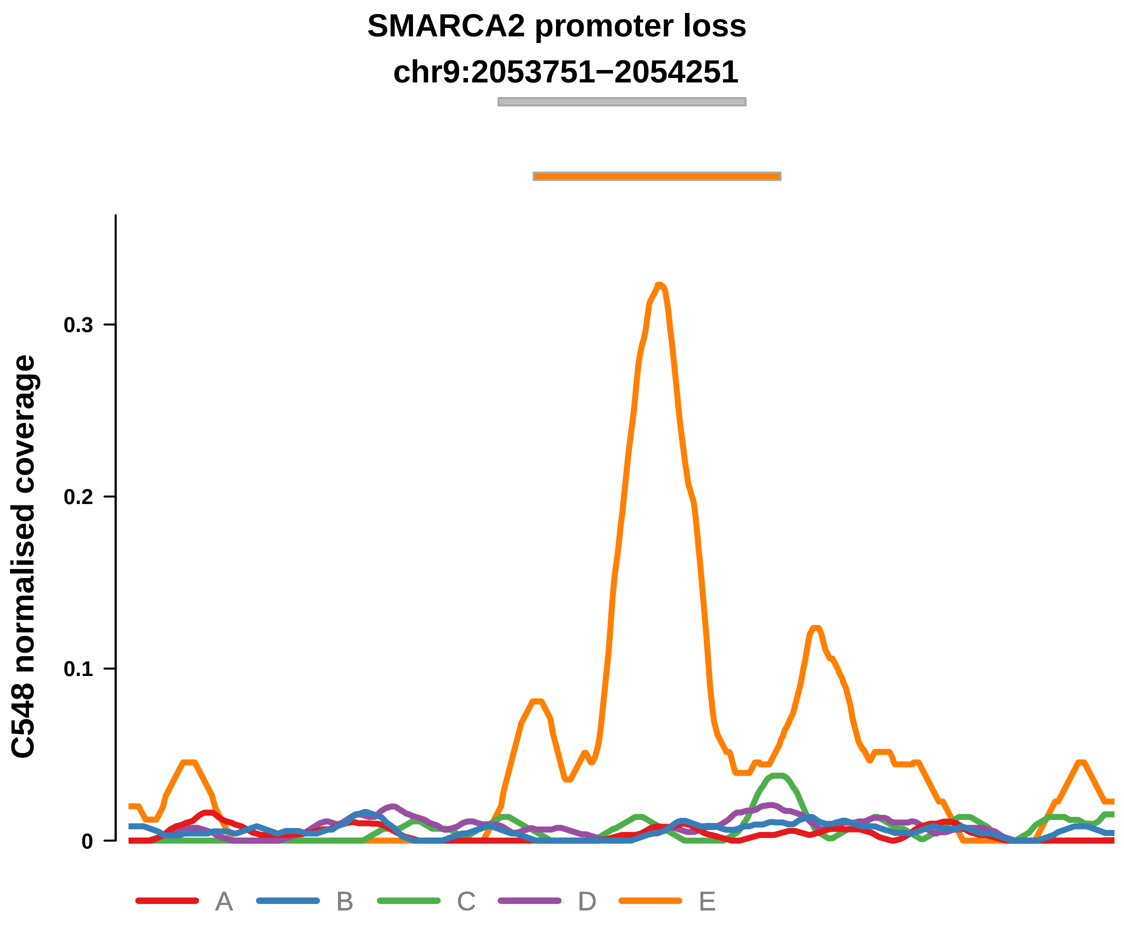


Putatively Clonal (logFC = −2.62)

| | P. | Adjusted P. |
|---|---|---|
| *~region vs. ~* | 2.55e−01 | 3.62e−01 |
| *~purity vs ~* | 5.92e−02 | 2.27e−01 |
| *~purity+region vs ~purity* | 8.44e−01 | 9.31e−01 |
| *Carcinoma vs. normal* | 7.46e−03 ** | 7.46e−03 ** |

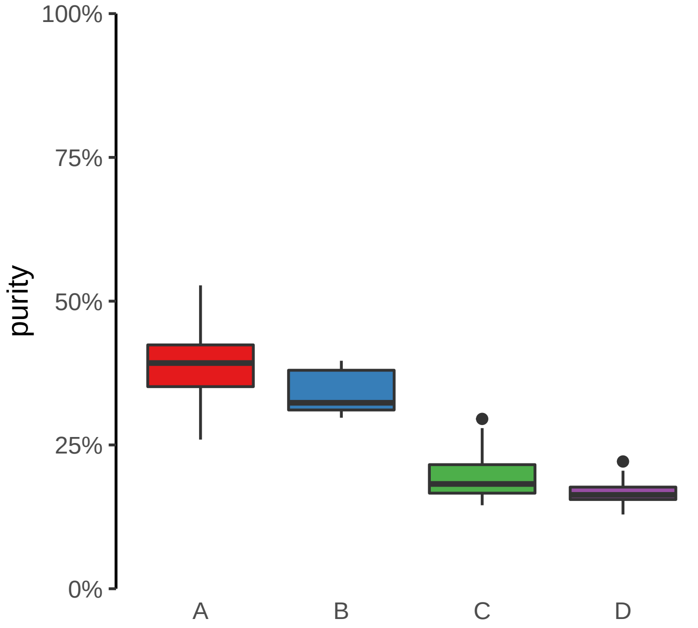

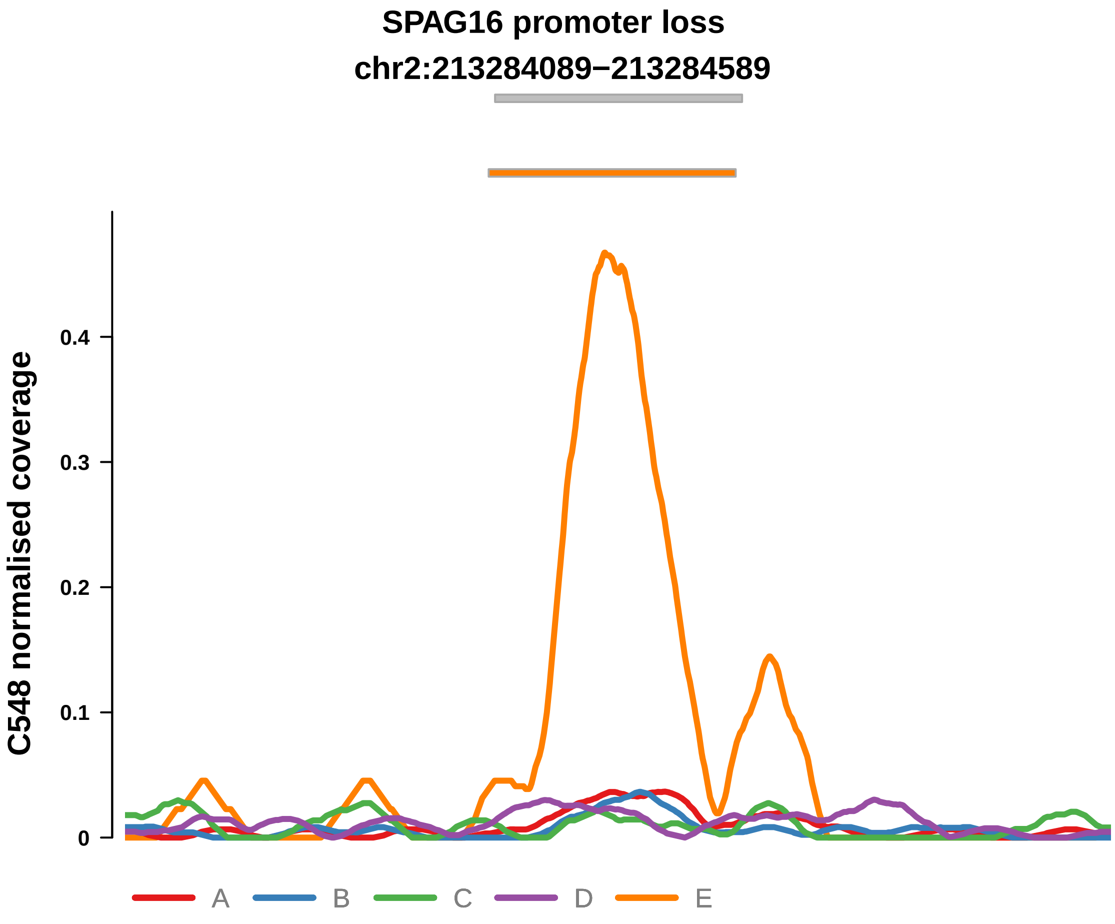


Putatively Clonal (logFC = −2.38)

| | P. | Adjusted P. |
|---|---|---|
| *~region vs. ~* | 1.07e−01 | 2.24e−01 |
| *~purity vs ~* | 7.83e−01 | 8.72e−01 |
| *~purity+region vs ~purity* | 1.08e−01 | 3.66e−01 |
| *Carcinoma vs. normal* | 2.09e−04 *** | 2.09e−04 *** |

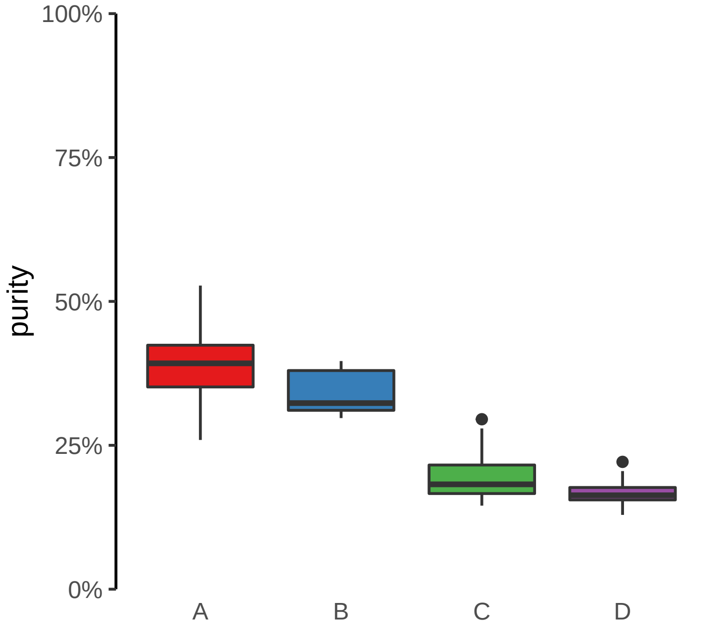

**SRGAP2C promoter loss**
**chr1:121184393−121184893**

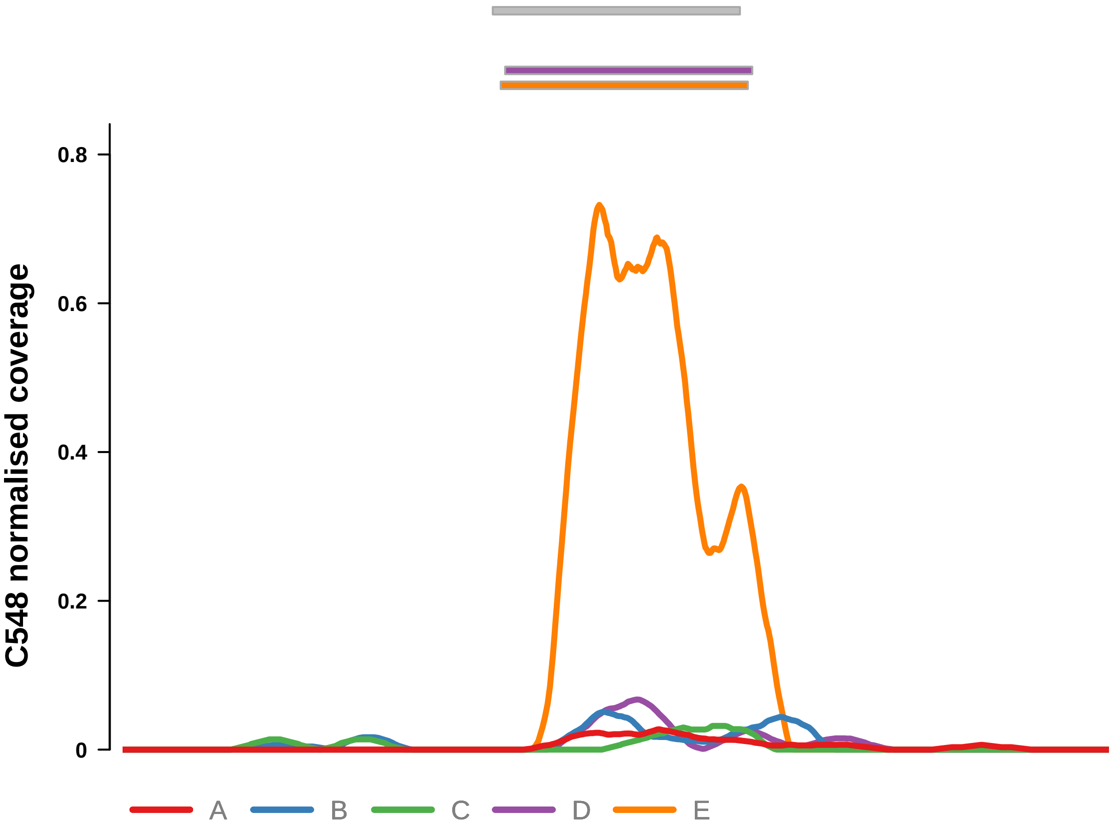


Putatively Clonal (logFC = −2.75)

| | P. | Adjusted P. |
|---|---|---|
| *~region vs. ~* | 1.54e−01 | 2.63e−01 |
| *~purity vs ~* | 1.17e−01 | 3.07e−01 |
| *~purity+region vs ~purity* | 3.15e−01 | 5.77e−01 |
| *Carcinoma vs. normal* | 2.99e−05 **** | 2.99e−05 **** |

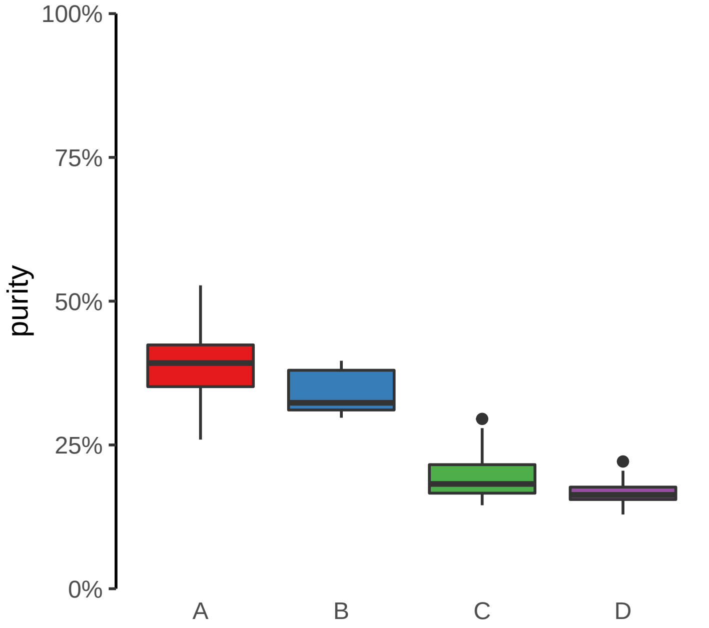

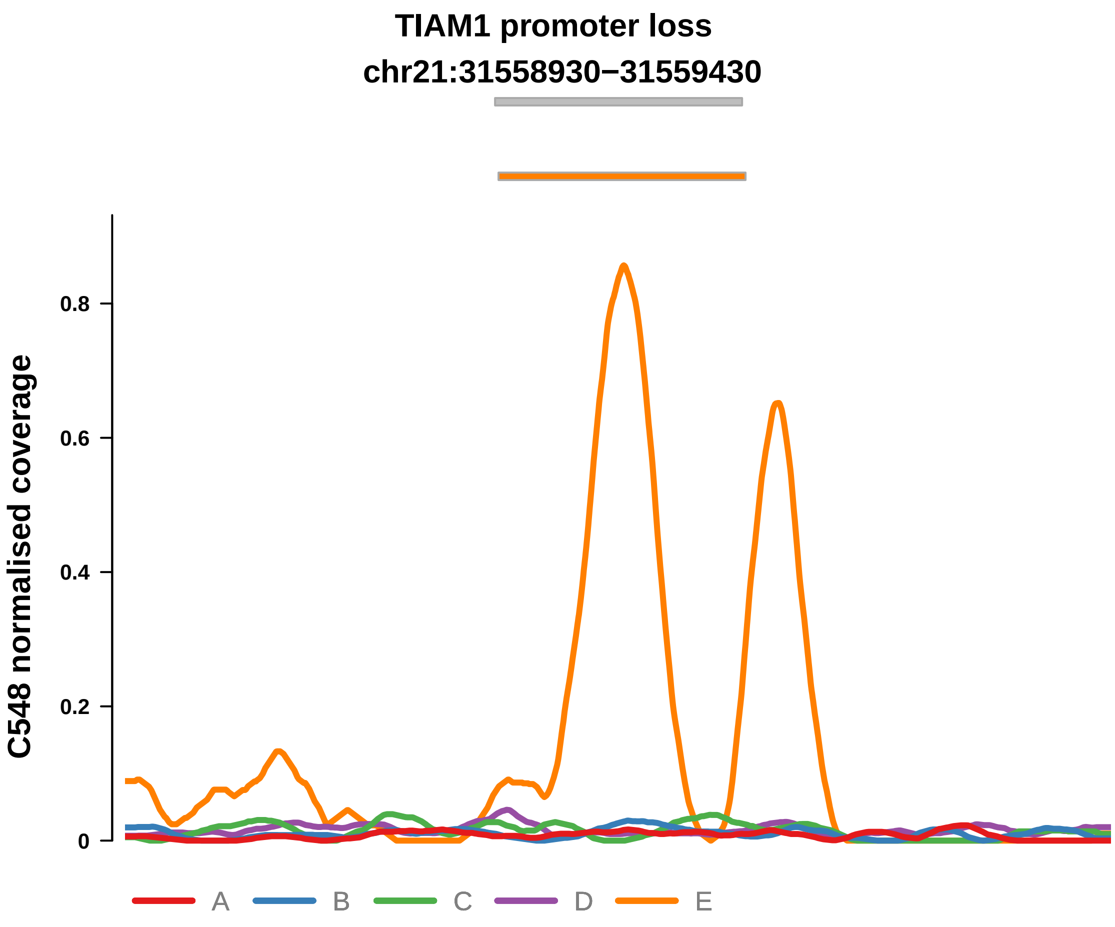


Putatively Clonal (logFC = −2.4)

| | P. | Adjusted P. |
|---|---|---|
| *~region vs. ~* | 1.13e−01 | 2.26e−01 |
| *~purity vs ~* | 1.9e−01 | 3.97e−01 |
| *~purity+region vs ~purity* | 7.21e−02 | 3.52e−01 |
| *Carcinoma vs. normal* | 2.24e−04 *** | 2.24e−04 *** |

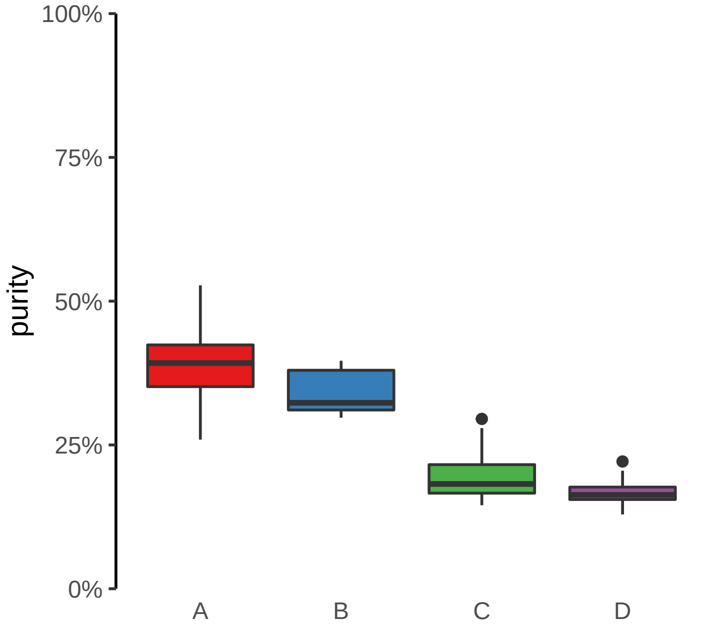

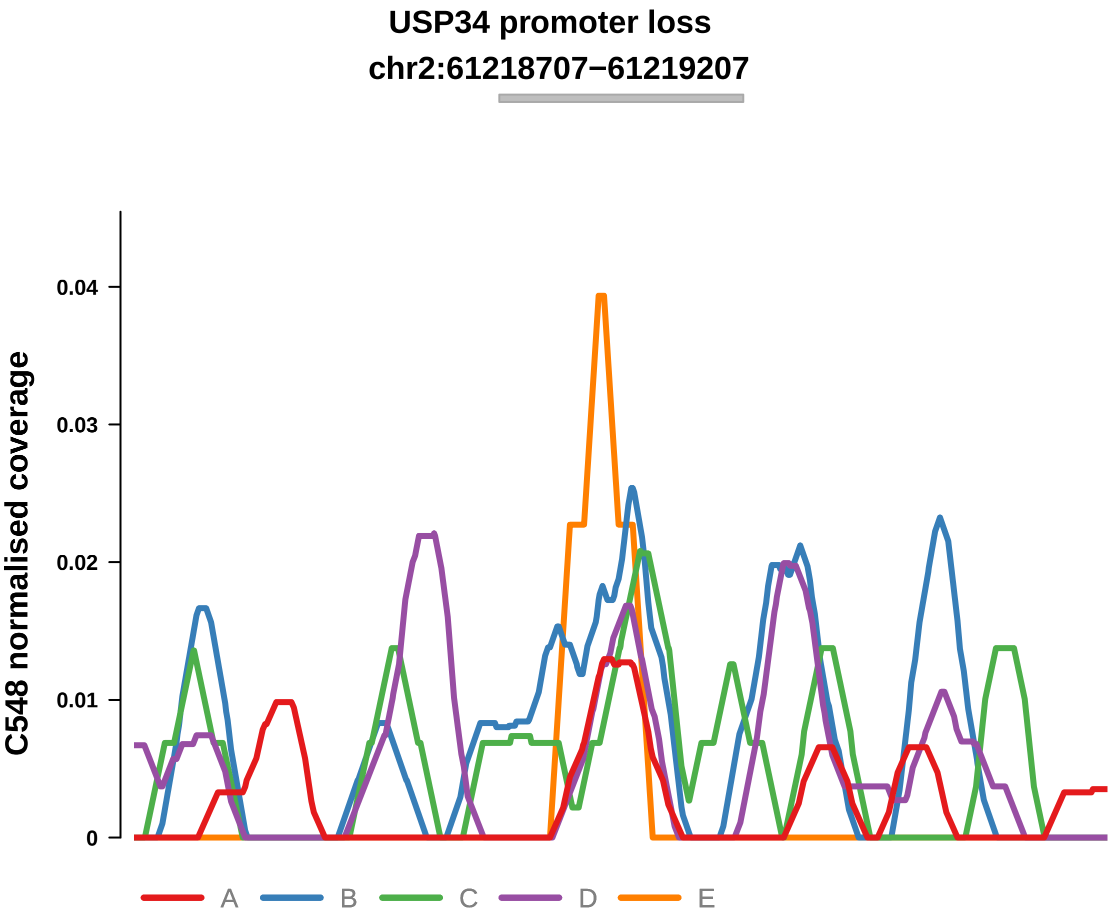


Putatively Clonal (logFC = −2.73)

| | P. | Adjusted P. |
|---|---|---|
| *~region vs. ~* | 8.14e−01 | 9.18e−01 |
| *~purity vs ~* | 3.78e−01 | 5.64e−01 |
| *~purity+region vs ~purity* | 2.85e−01 | 5.33e−01 |
| *Carcinoma vs. normal* | 5.43e−03 ** | 5.43e−03 ** |

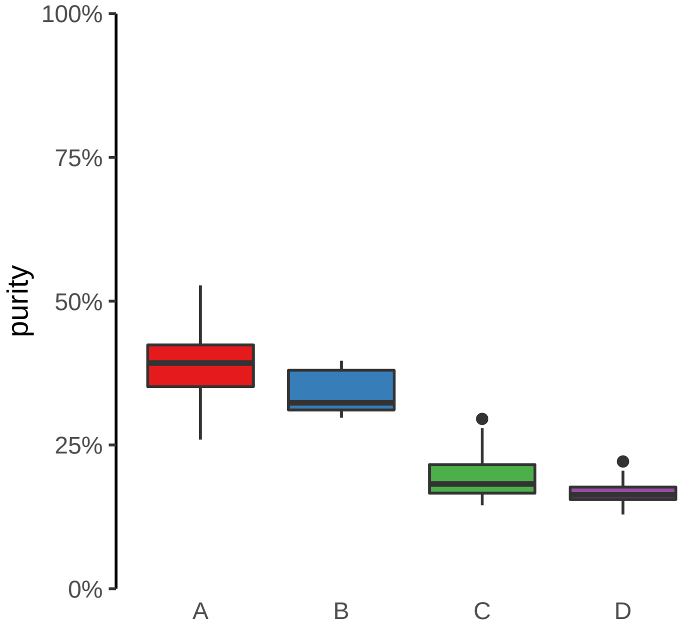

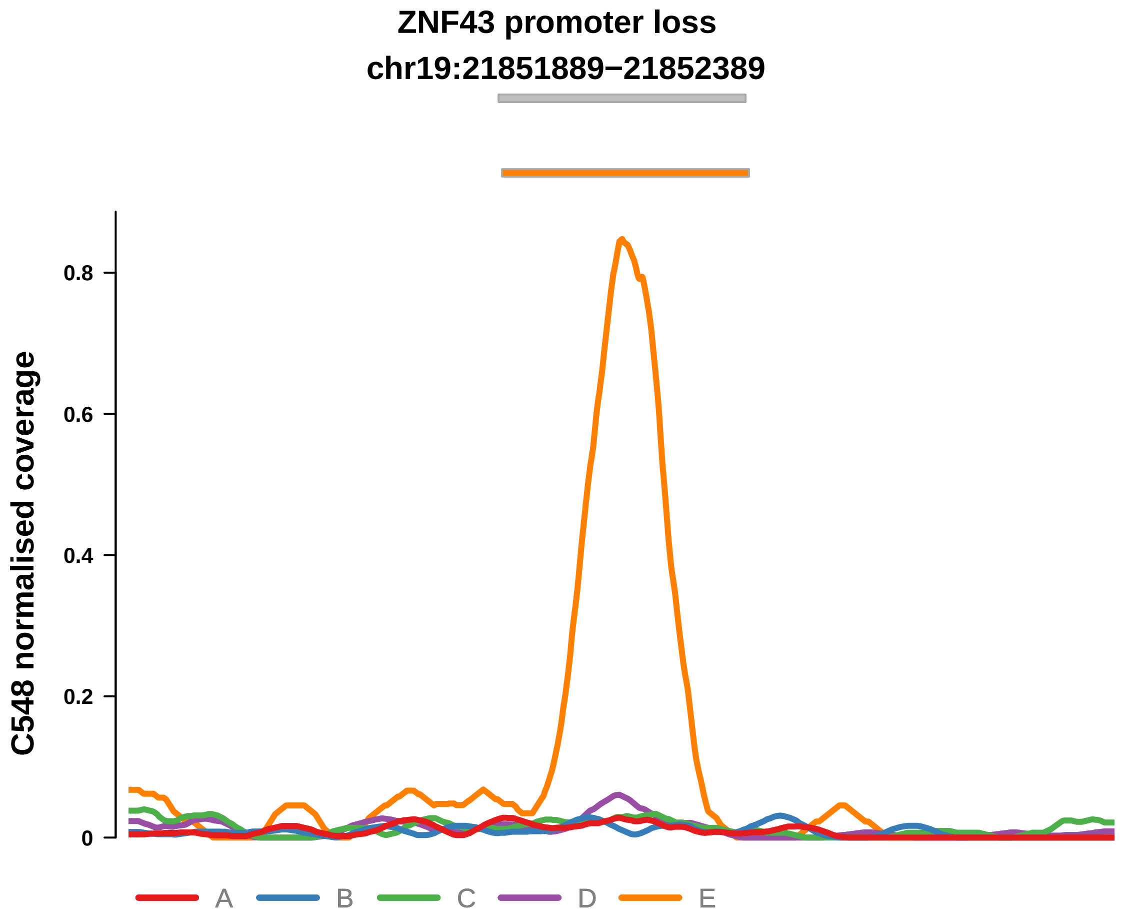


Putatively Clonal (logFC = −1.66)

| | P. | Adjusted P. |
|---|---|---|
| *~region vs. ~* | 7.19e−02 | 1.82e−01 |
| *~purity vs ~* | 7.67e−02 | 2.37e−01 |
| *~purity+region vs ~purity* | 2.46e−01 | 5.31e−01 |
| *Carcinoma vs. normal* | 6.79e−03 ** | 6.79e−03 ** |

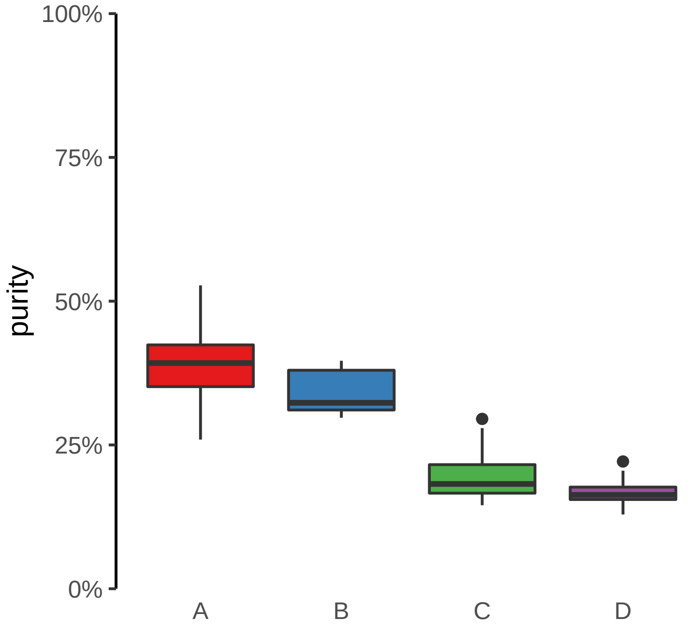

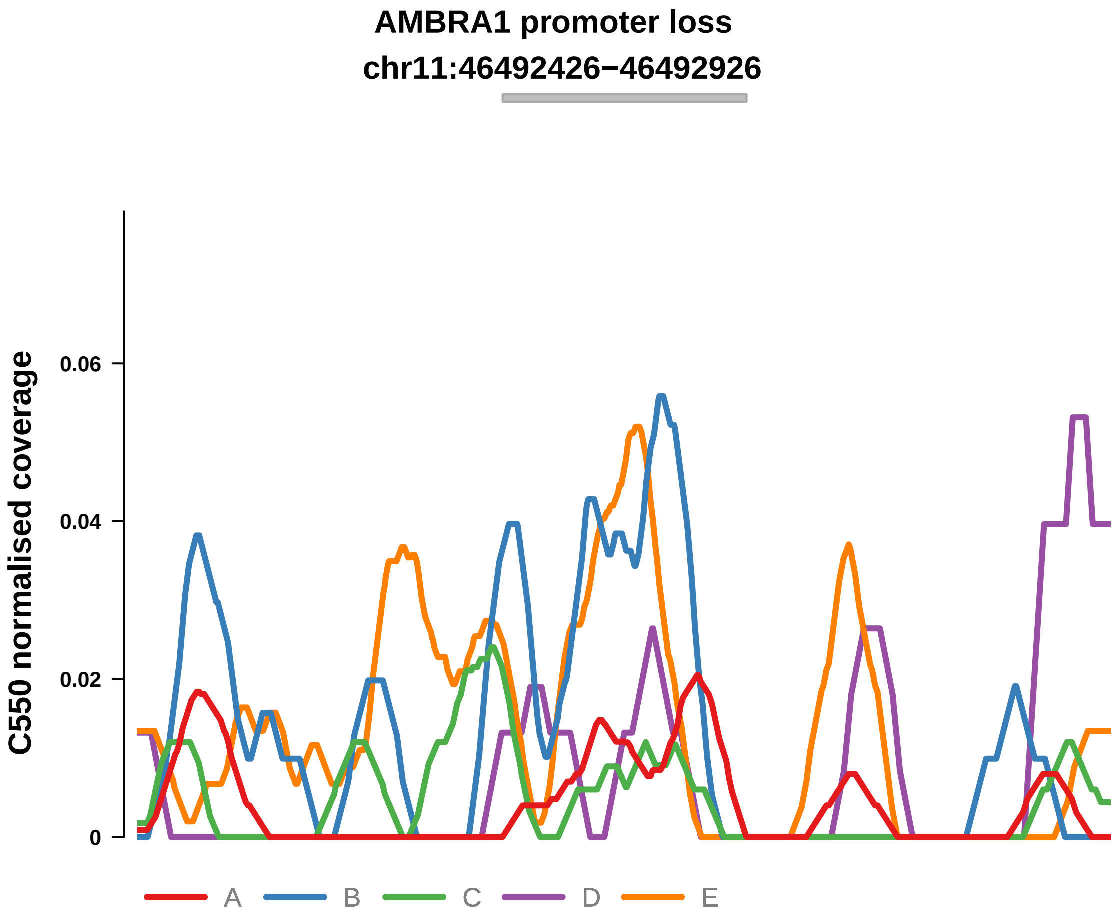


Putatively Clonal (logFC = −2.7)

| | P. | Adjusted P. |
|---|---|---|
| *~region vs. ~* | 4.37e−01 | 9.11e−01 |
| *~purity vs ~* | 7.84e−02 | 4.59e−01 |
| *~purity+region vs ~purity* | 7.91e−01 | 1e+00 |
| *Carcinoma vs. normal* | 5.85e−03 ** | 5.85e−03 ** |

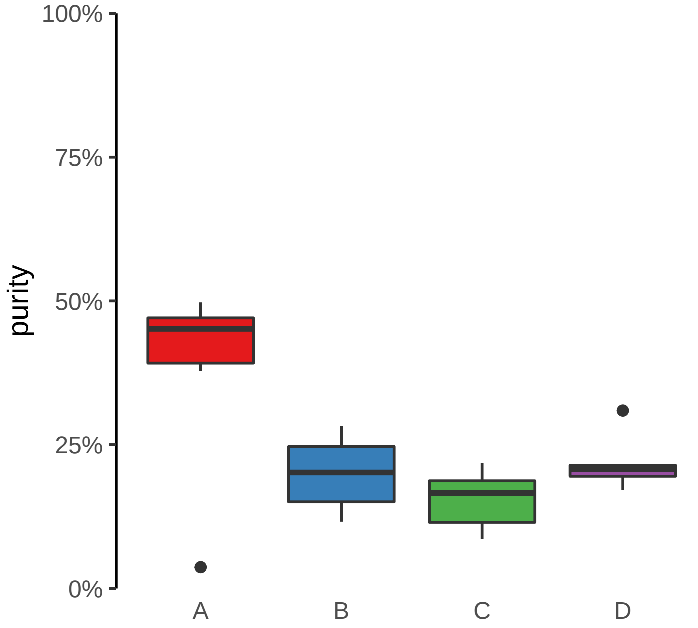

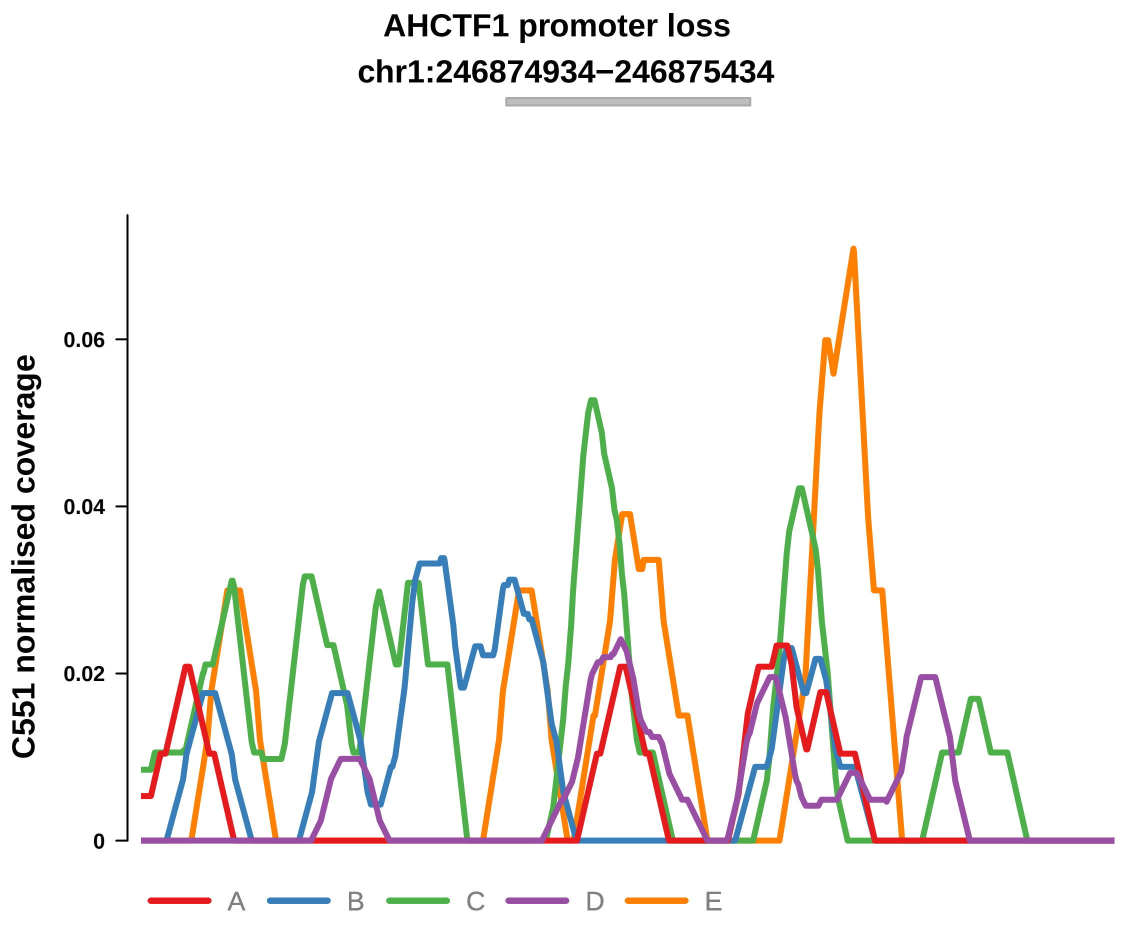


Putatively Clonal (logFC = −2.91)

| | P. | Adjusted P. |
|---|---|---|
| *~region vs. ~* | 9.52e−01 | 9.74e−01 |
| *~purity vs ~* | 6.01e−01 | 9.06e−01 |
| *~purity+region vs ~purity* | 9.69e−01 | 9.99e−01 |
| *Carcinoma vs. normal* | 3.14e−06 **** | 3.14e−06 **** |

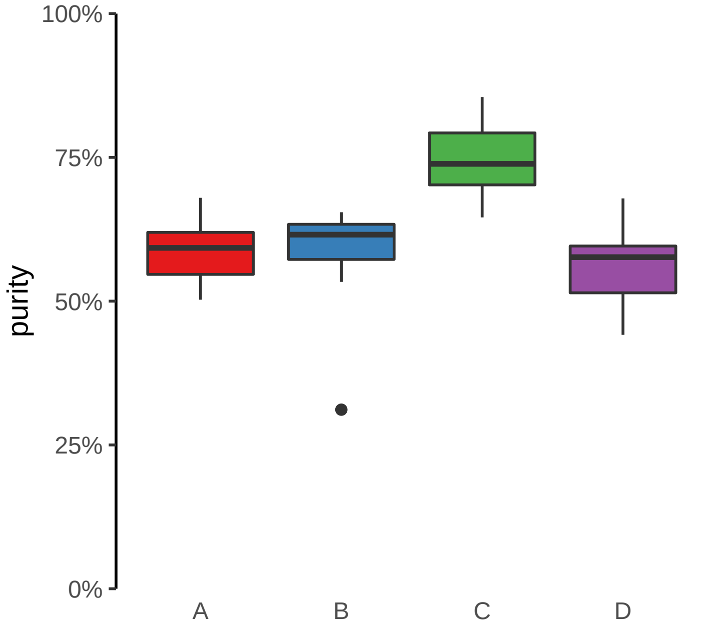

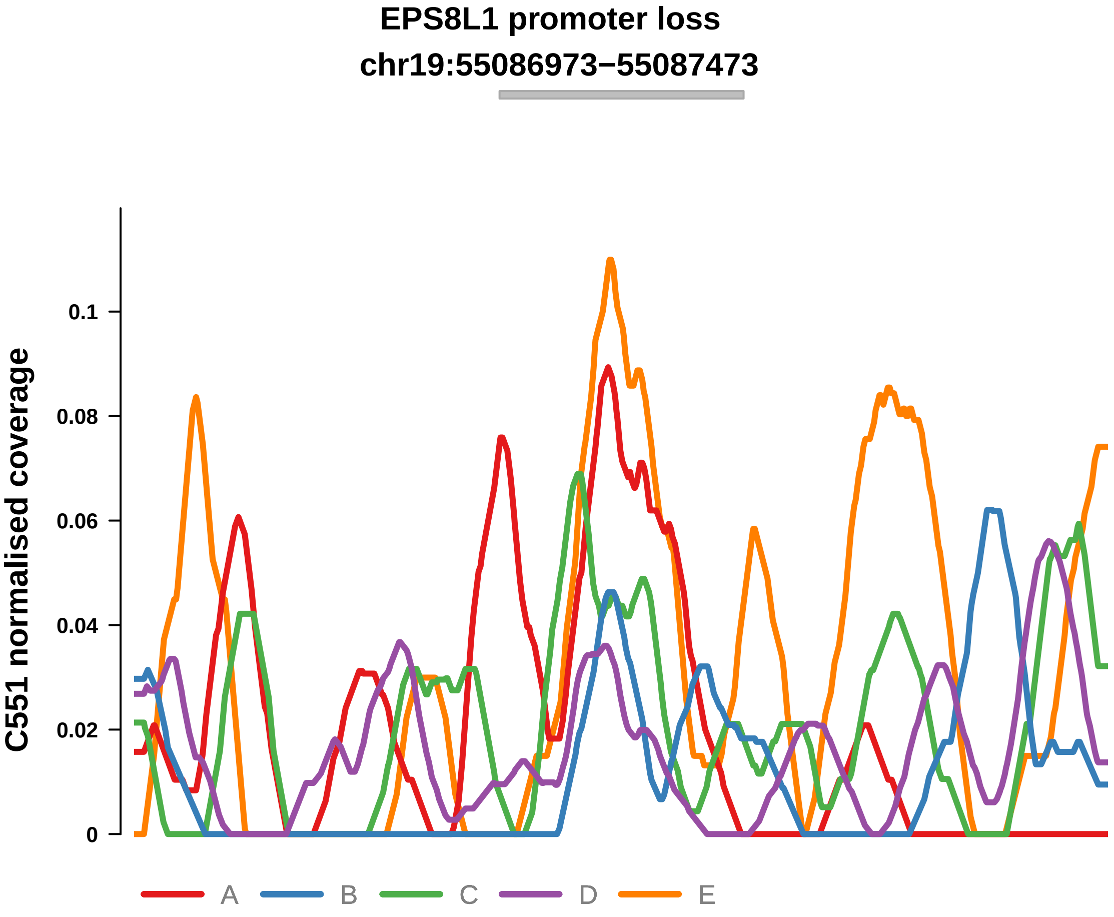


Putatively Clonal (logFC = −1.68)

| | P. | Adjusted P. |
|---|---|---|
| *~region vs. ~* | 1.47e−01 | 4.79e−01 |
| *~purity vs ~* | 2.45e−01 | 9.06e−01 |
| *~purity+region vs ~purity* | 2.56e−01 | 7.26e−01 |
| *Carcinoma vs. normal* | 2.61e−03 ** | 2.61e−03 ** |

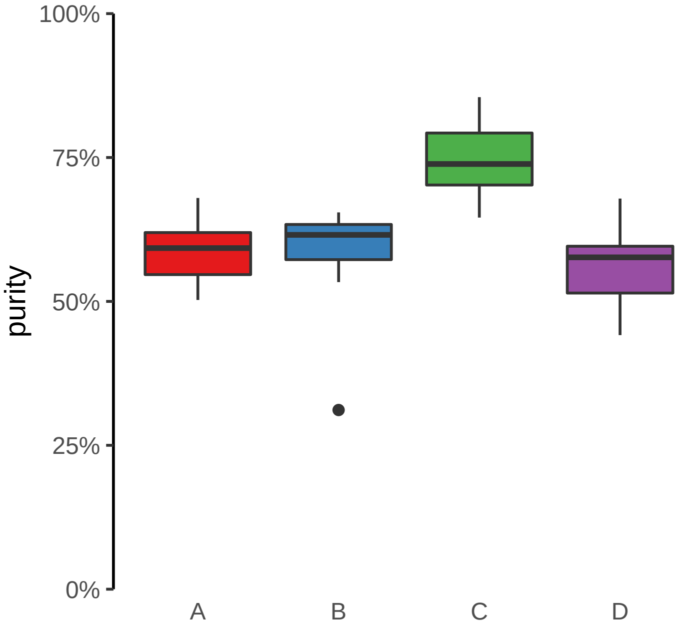

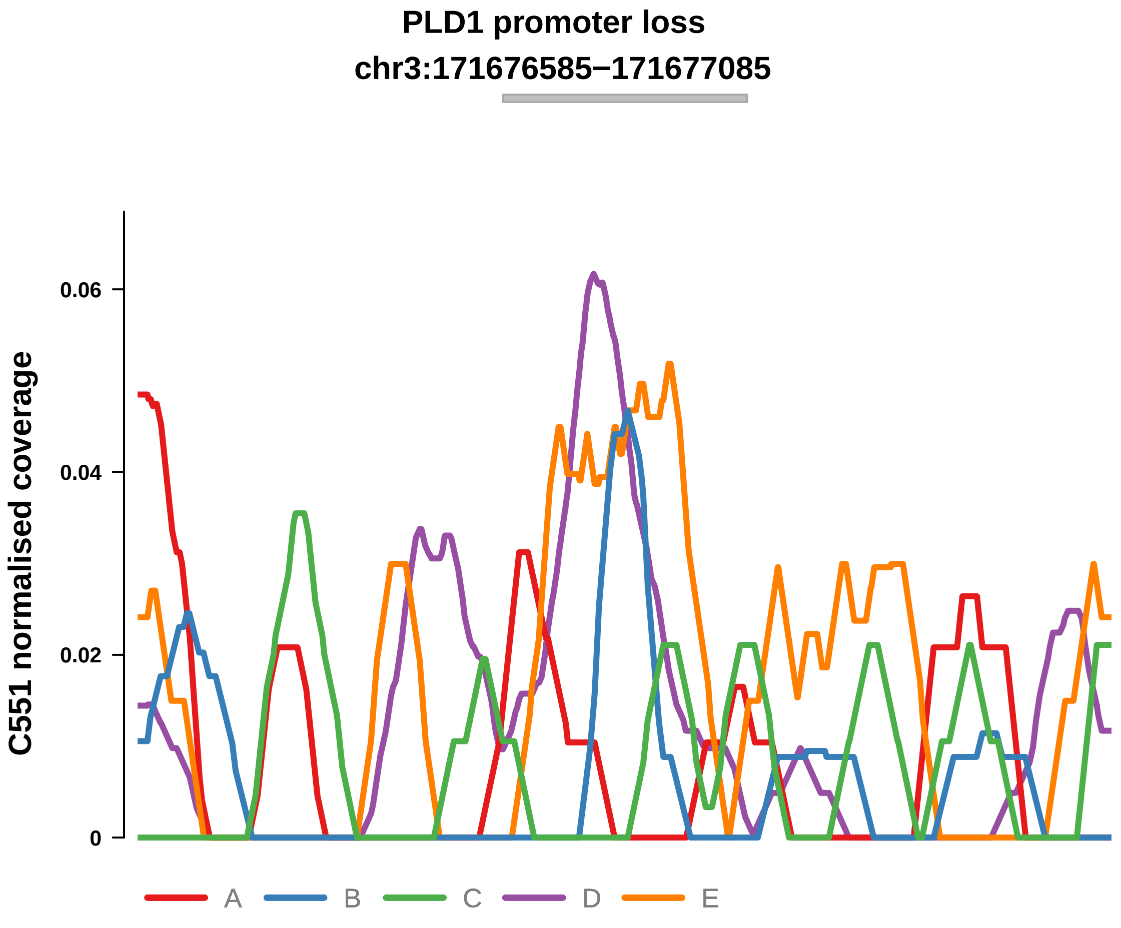


Putatively Clonal (logFC = −1.74)

| | P. | Adjusted P. |
|---|---|---|
| *~region vs. ~* | 1.13e−01 | 4.16e−01 |
| *~purity vs ~* | 8.11e−02 | 7.93e−01 |
| *~purity+region vs ~purity* | 4.59e−01 | 9.48e−01 |
| *Carcinoma vs. normal* | 2.34e−03 ** | 2.34e−03 ** |

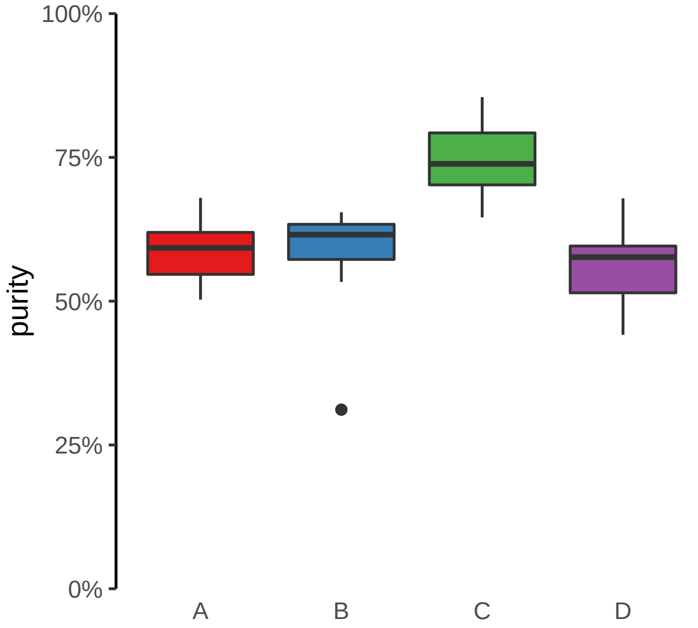

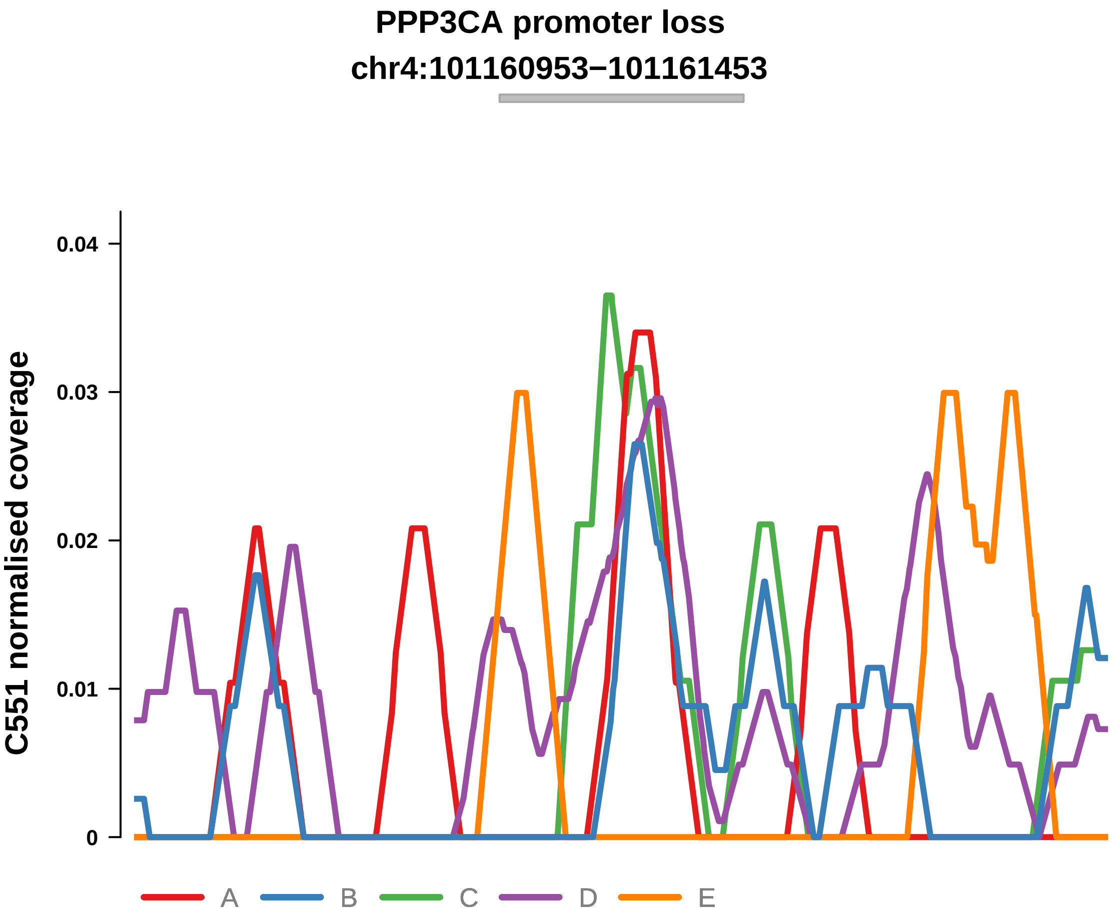


Putatively Clonal (logFC = −1.65)

| | P. | Adjusted P. |
|---|---|---|
| *~region vs. ~* | 8.26e−01 | 9.65e−01 |
| *~purity vs ~* | 6.11e−01 | 9.06e−01 |
| *~purity+region vs ~purity* | 7.2e−01 | 9.78e−01 |
| *Carcinoma vs. normal* | 5.69e−03 ** | 5.69e−03 ** |

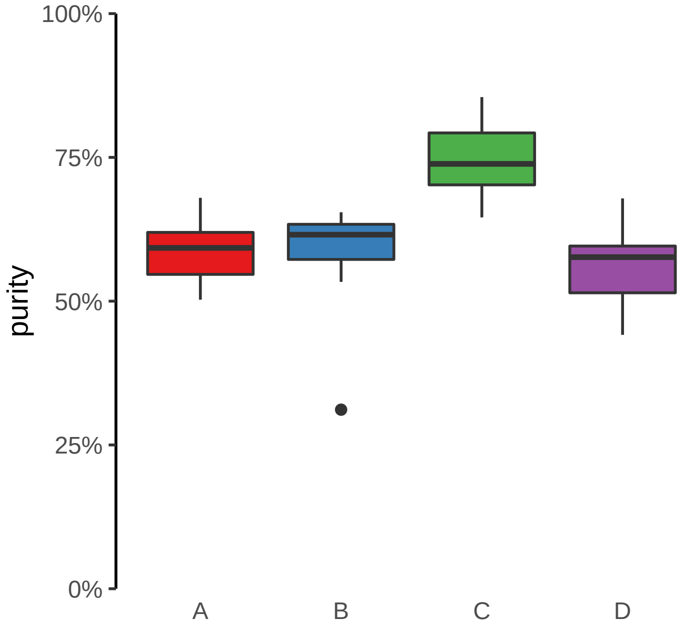

**R3HDM1 promoter loss**
**chr2:135633520−135634020**

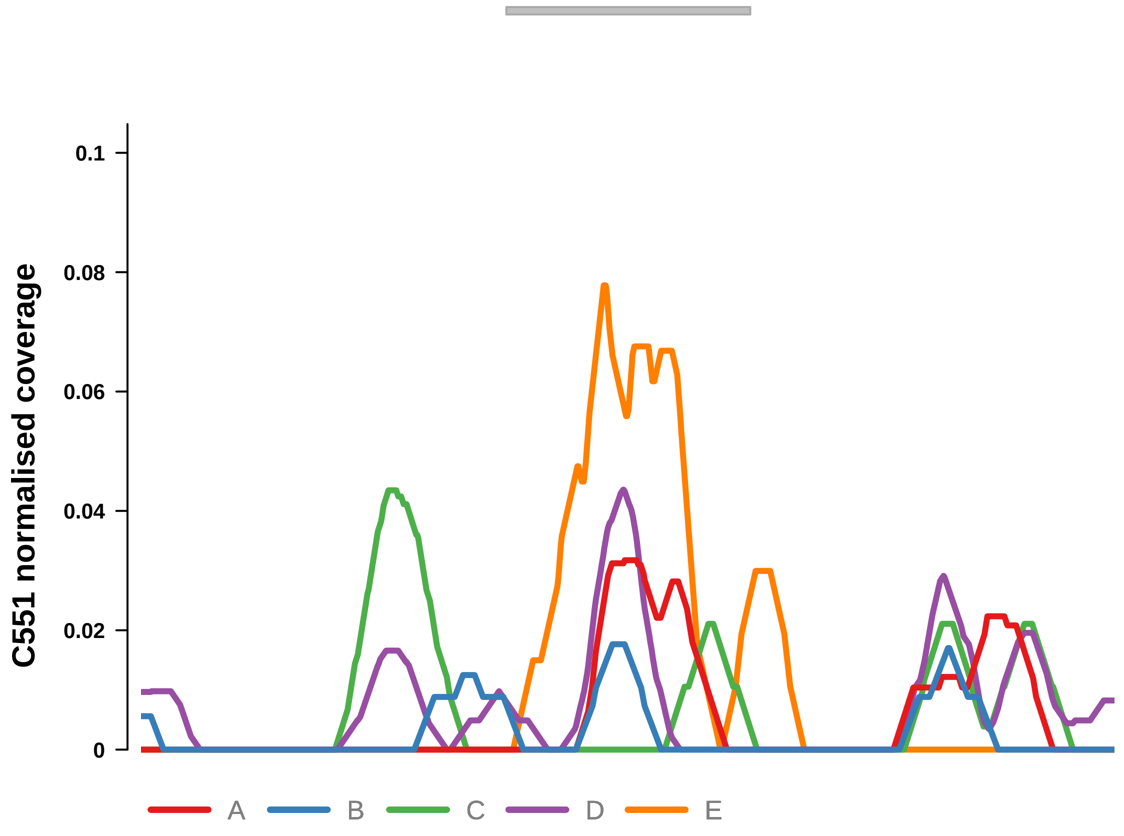


Putatively Clonal (logFC = −2.53)

| | P. | Adjusted P. |
|---|---|---|
| *~region vs. ~* | 9e−01 | 9.74e−01 |
| *~purity vs ~* | 2.8e−01 | 9.06e−01 |
| *~purity+region vs ~purity* | 1e+00 | 1e+00 |
| *Carcinoma vs. normal* | 4.11e−05 **** | 4.11e−05 **** |

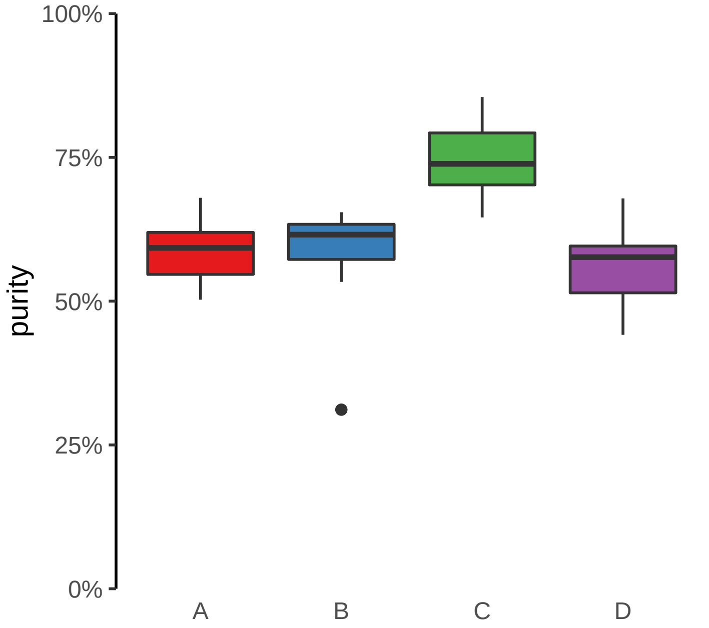

**SLC38A9 promoter loss**
**chr5:55700661−55701161**

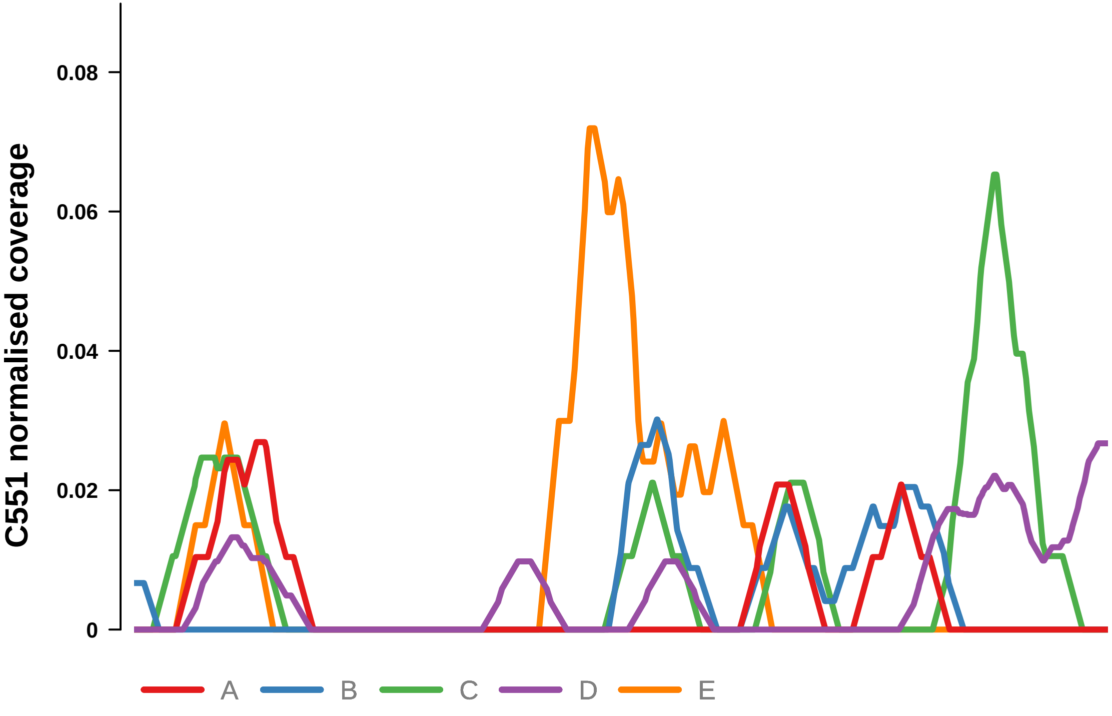


Putatively Clonal (logFC = −2.95)

| | P. | Adjusted P. |
|---|---|---|
| *~region vs. ~* | 9.52e−01 | 9.74e−01 |
| *~purity vs ~* | 9.15e−01 | 9.69e−01 |
| *~purity+region vs ~purity* | 9.51e−01 | 9.99e−01 |
| *Carcinoma vs. normal* | 1.21e−05 **** | 1.21e−05 **** |

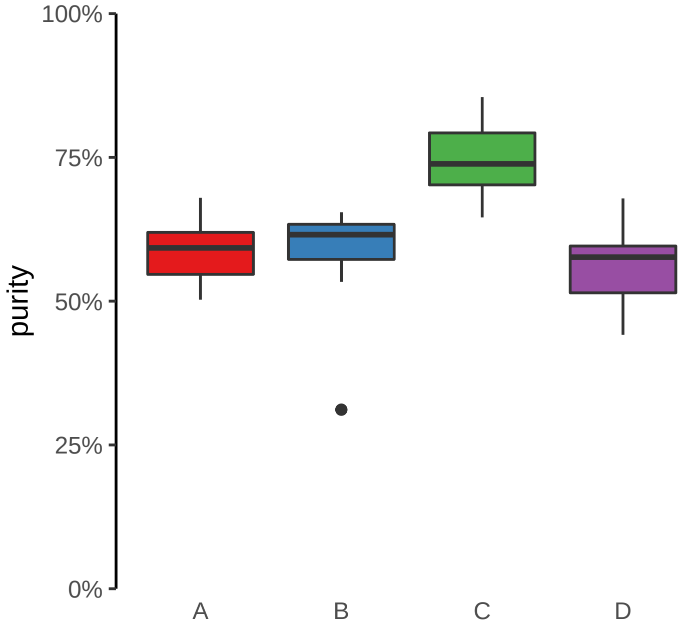

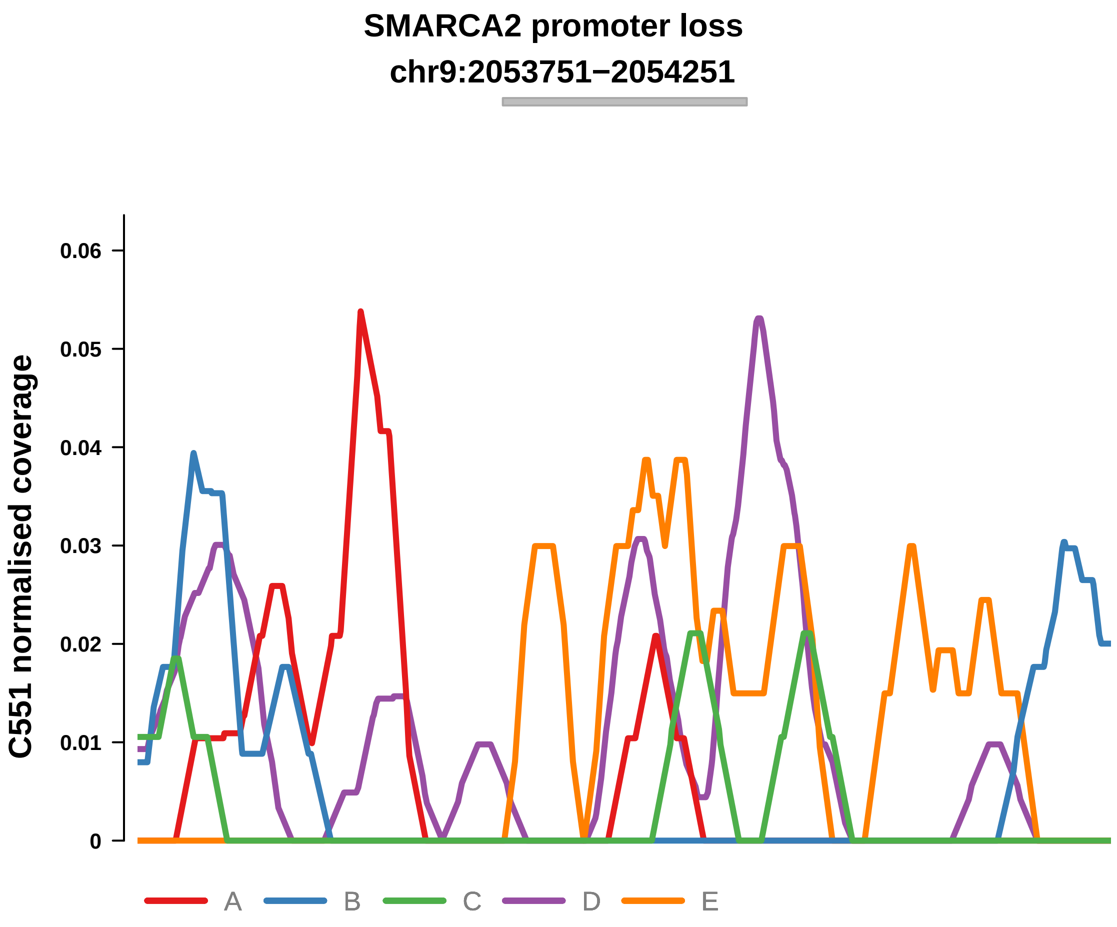


Putatively Clonal (logFC = −2.1)

| | P. | Adjusted P. |
|---|---|---|
| *~region vs. ~* | 4.63e−01 | 8.39e−01 |
| *~purity vs ~* | 5.61e−01 | 9.06e−01 |
| *~purity+region vs ~purity* | 5.56e−01 | 9.48e−01 |
| *Carcinoma vs. normal* | 7.38e−04 *** | 7.38e−04 *** |

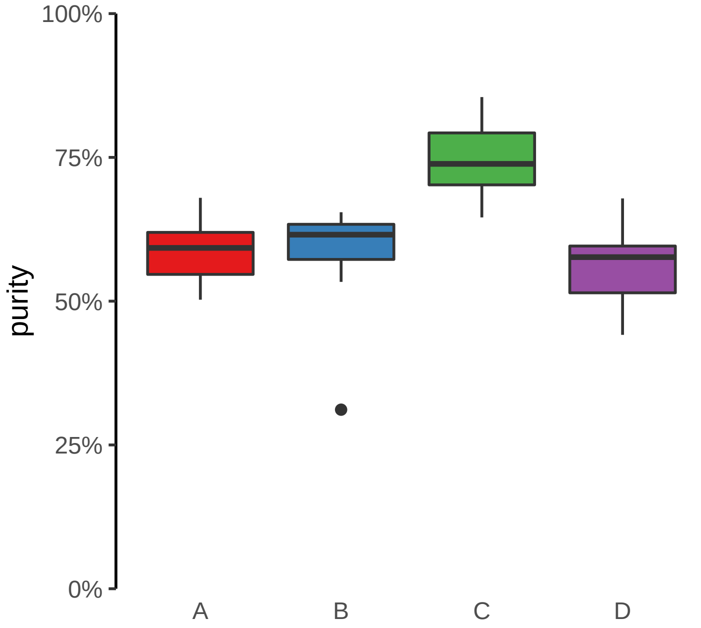

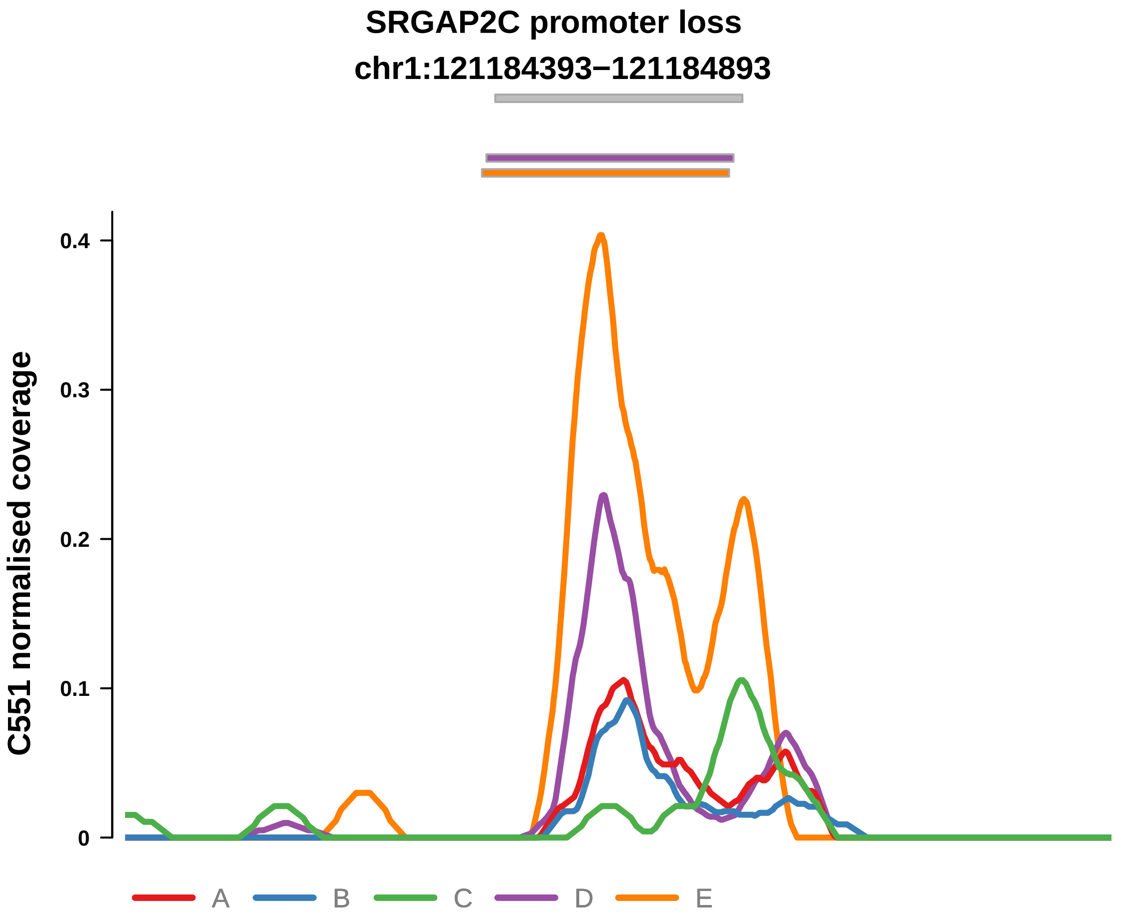


Putatively Clonal (logFC = −1.76)

| | P. | Adjusted P. |
|---|---|---|
| *~region vs. ~* | 5.55e−02 | 3.05e−01 |
| *~purity vs ~* | 1.16e−01 | 7.97e−01 |
| *~purity+region vs ~purity* | 1.9e−01 | 5.76e−01 |
| *Carcinoma vs. normal* | 1.26e−03 ** | 1.26e−03 ** |

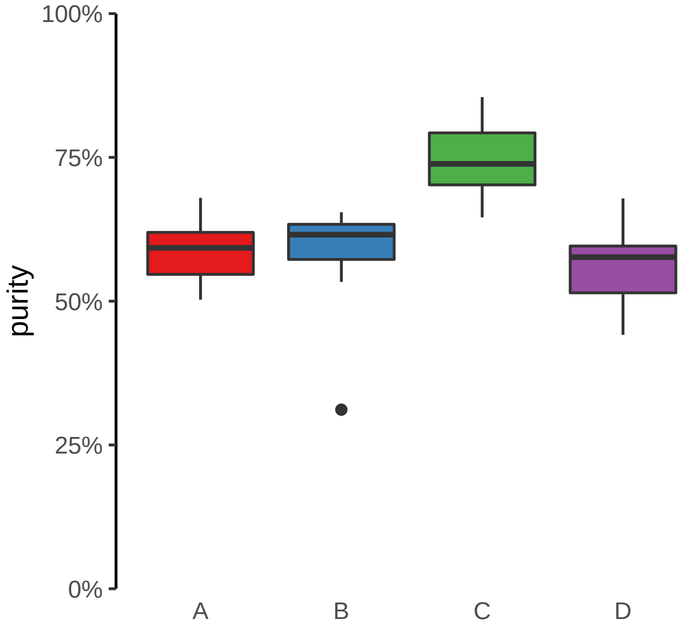

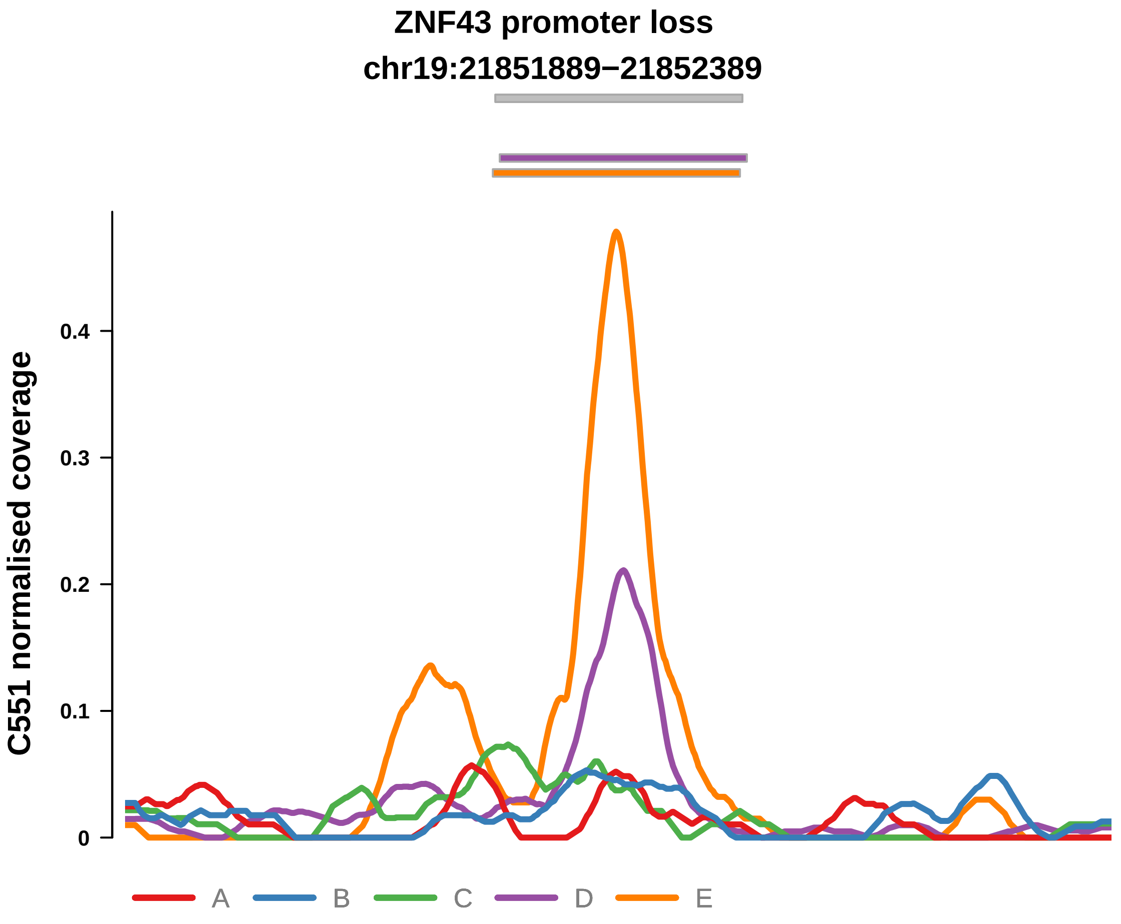


Putatively Clonal (logFC = −1.55)

| | P. | Adjusted P. |
|---|---|---|
| *~region vs. ~* | 4.58e−02 * | 2.69e−01 |
| *~purity vs ~* | 4.99e−01 | 9.06e−01 |
| *~purity+region vs ~purity* | 6.55e−02 | 5.24e−01 |
| *Carcinoma vs. normal* | 4.27e−03 ** | 4.27e−03 ** |

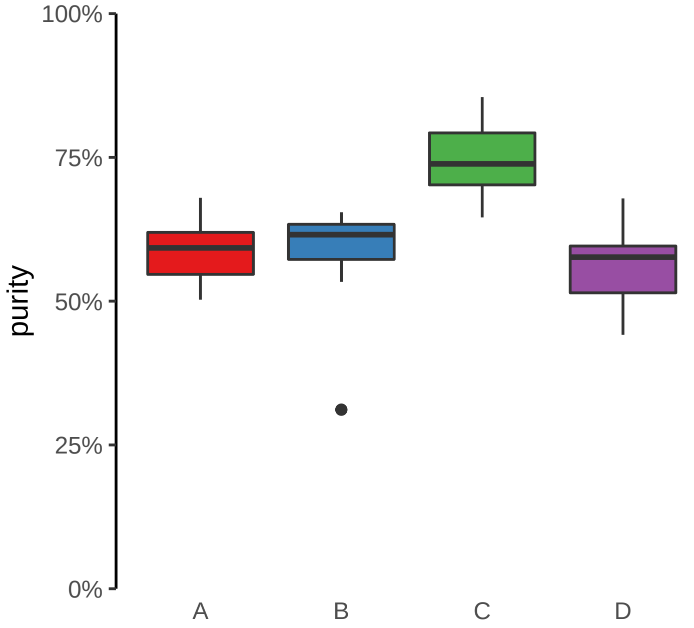

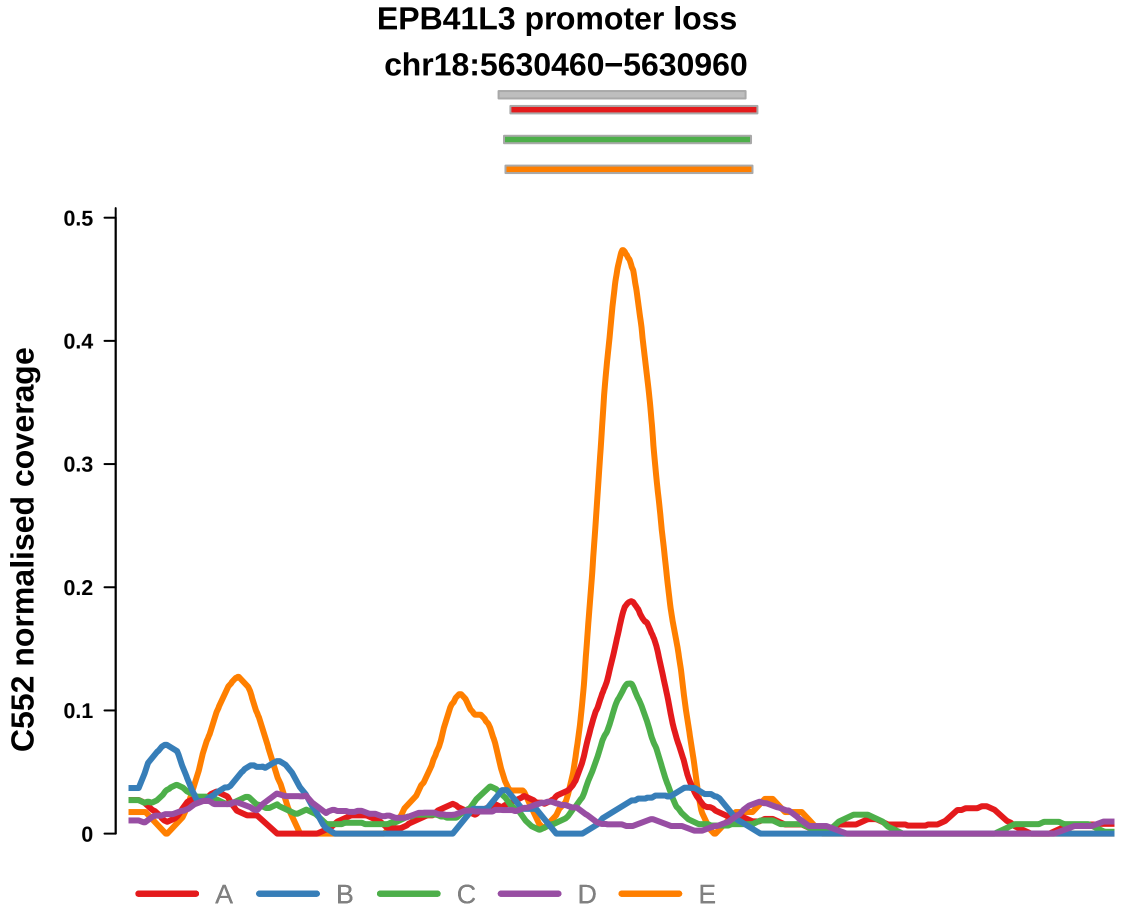


Putatively Clonal (logFC = −1.78)

| | P. | Adjusted P. |
|---|---|---|
| *~region vs. ~* | 8.74e−02 | 1.96e−01 |
| *~purity vs ~* | 1.01e−01 | 1.62e−01 |
| *~purity+region vs ~purity* | 1.29e−01 | 9.1e−01 |
| *Carcinoma vs. normal* | 7.9e−03 ** | 7.9e−03 ** |

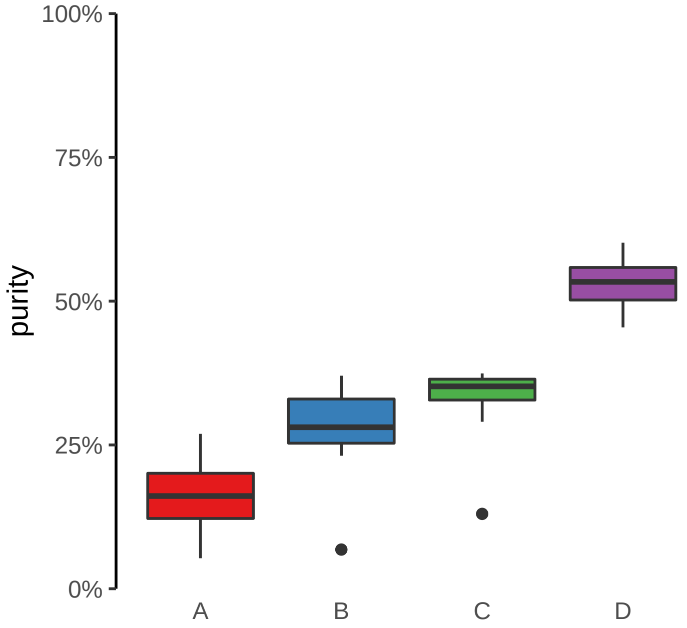

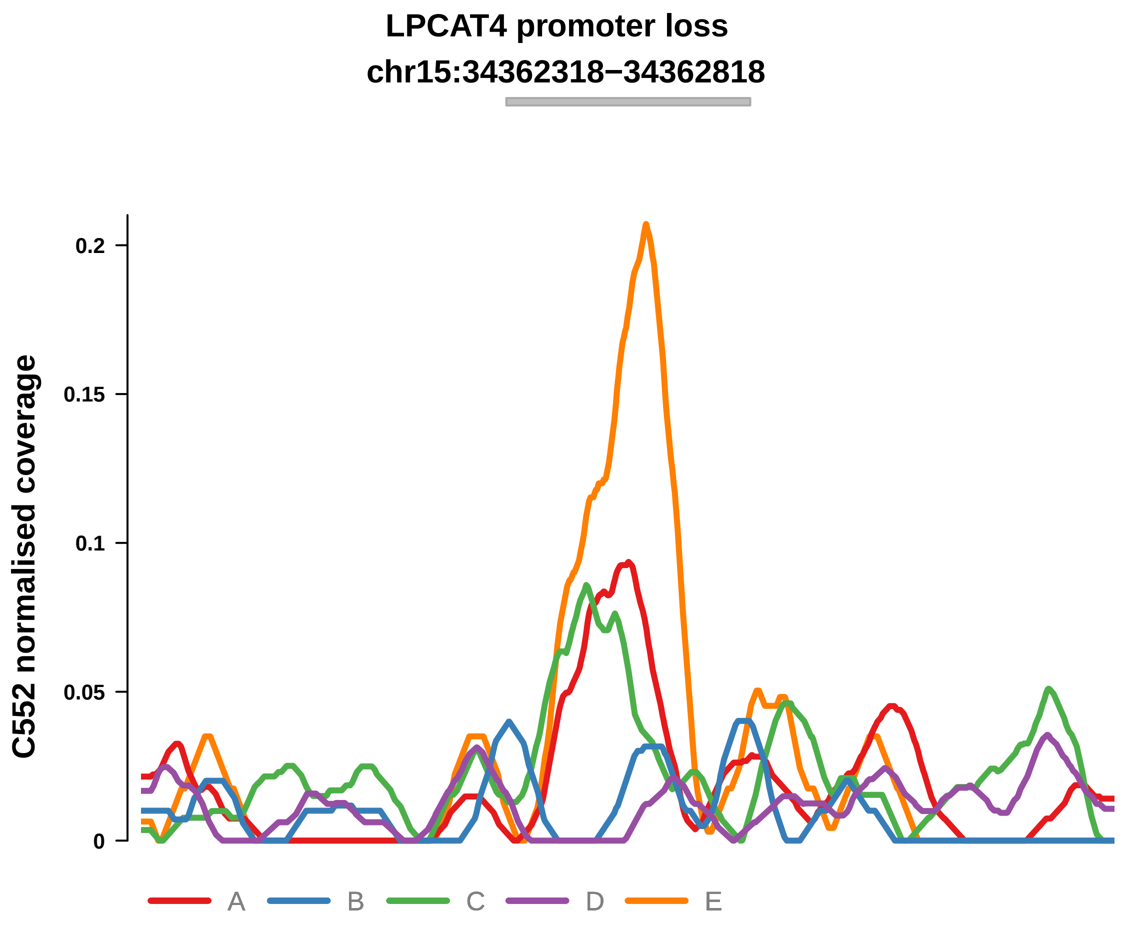


Putatively Clonal (logFC = −2.36)

| | P. | Adjusted P. |
|---|---|---|
| *~region vs. ~* | 7.62e−01 | 8.14e−01 |
| *~purity vs ~* | 9.78e−01 | 9.78e−01 |
| *~purity+region vs ~purity* | 4.47e−01 | 9.85e−01 |
| *Carcinoma vs. normal* | 3.16e−03 ** | 3.16e−03 ** |

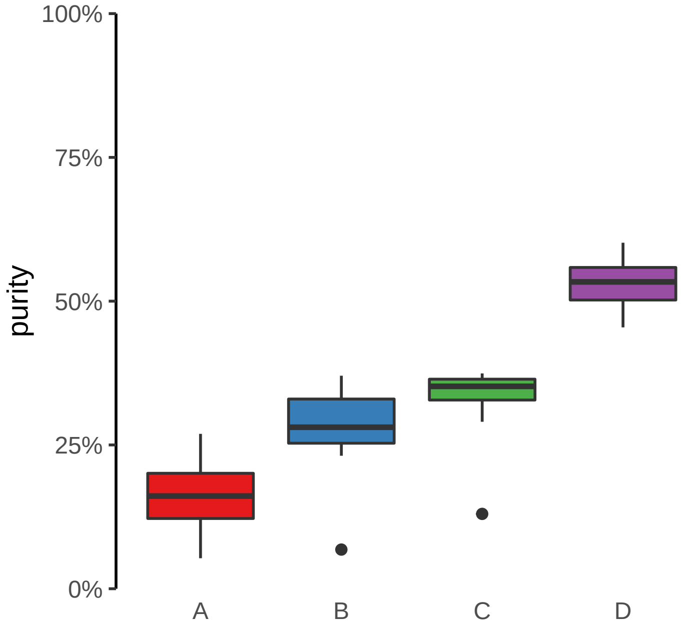

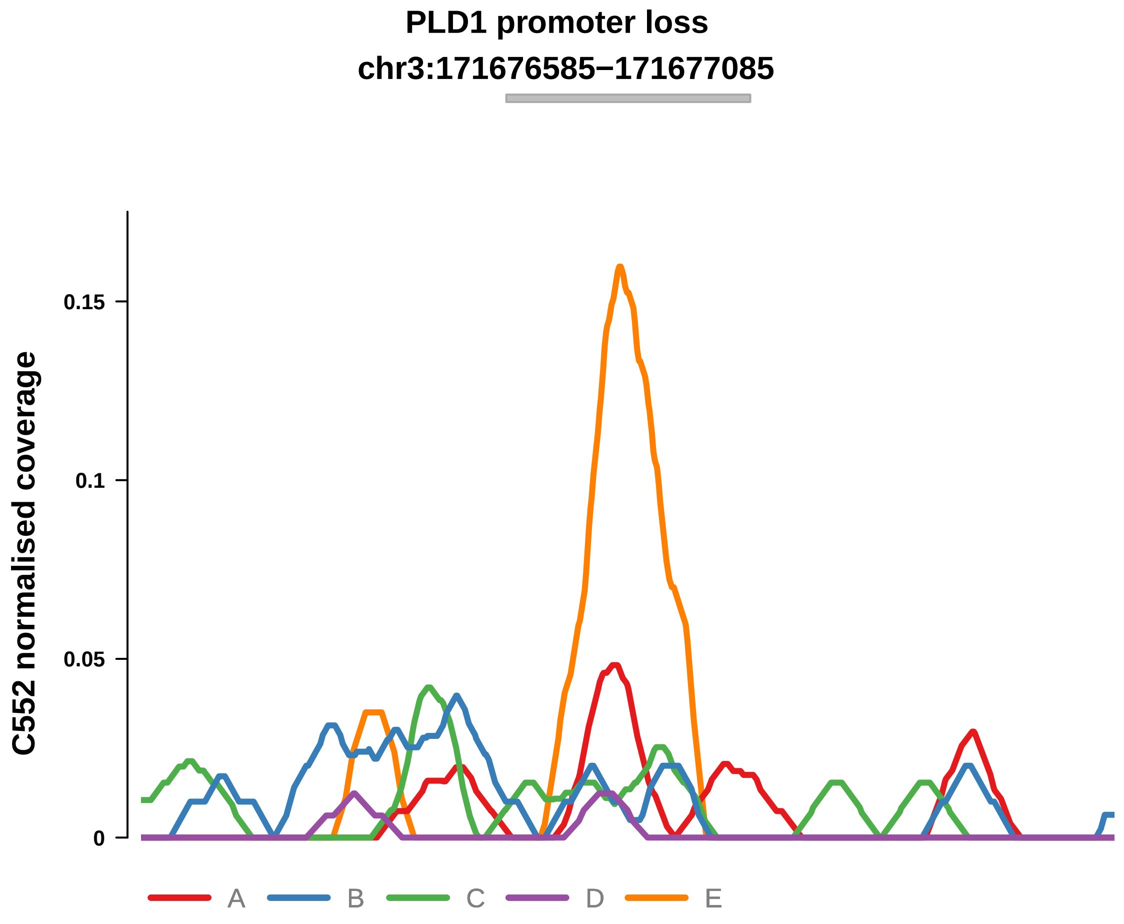


Putatively Clonal (logFC = −3.16)

| | P. | Adjusted P. |
|---|---|---|
| *~region vs. ~* | 9.28e−01 | NA |
| *~purity vs ~* | 3.03e−01 | NA |
| *~purity+region vs ~purity* | 9.45e−01 | 9.85e−01 |
| *Carcinoma vs. normal* | 6.05e−04 *** | 6.05e−04 *** |

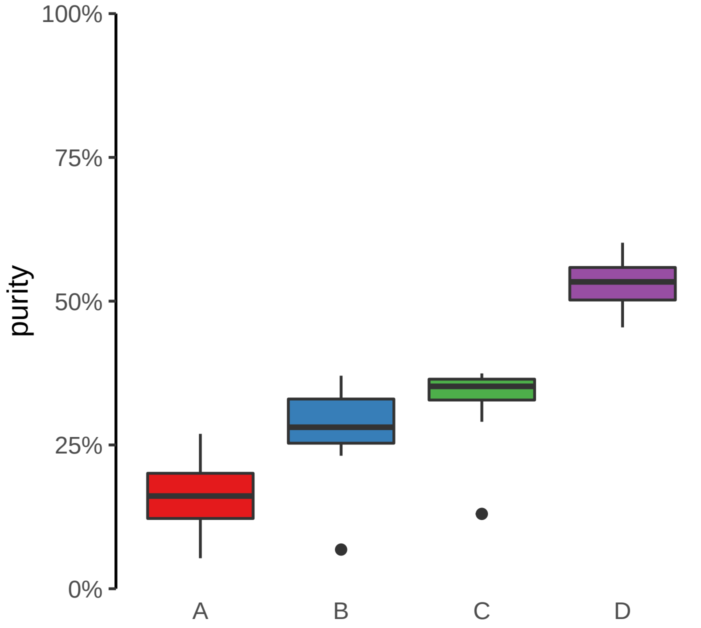

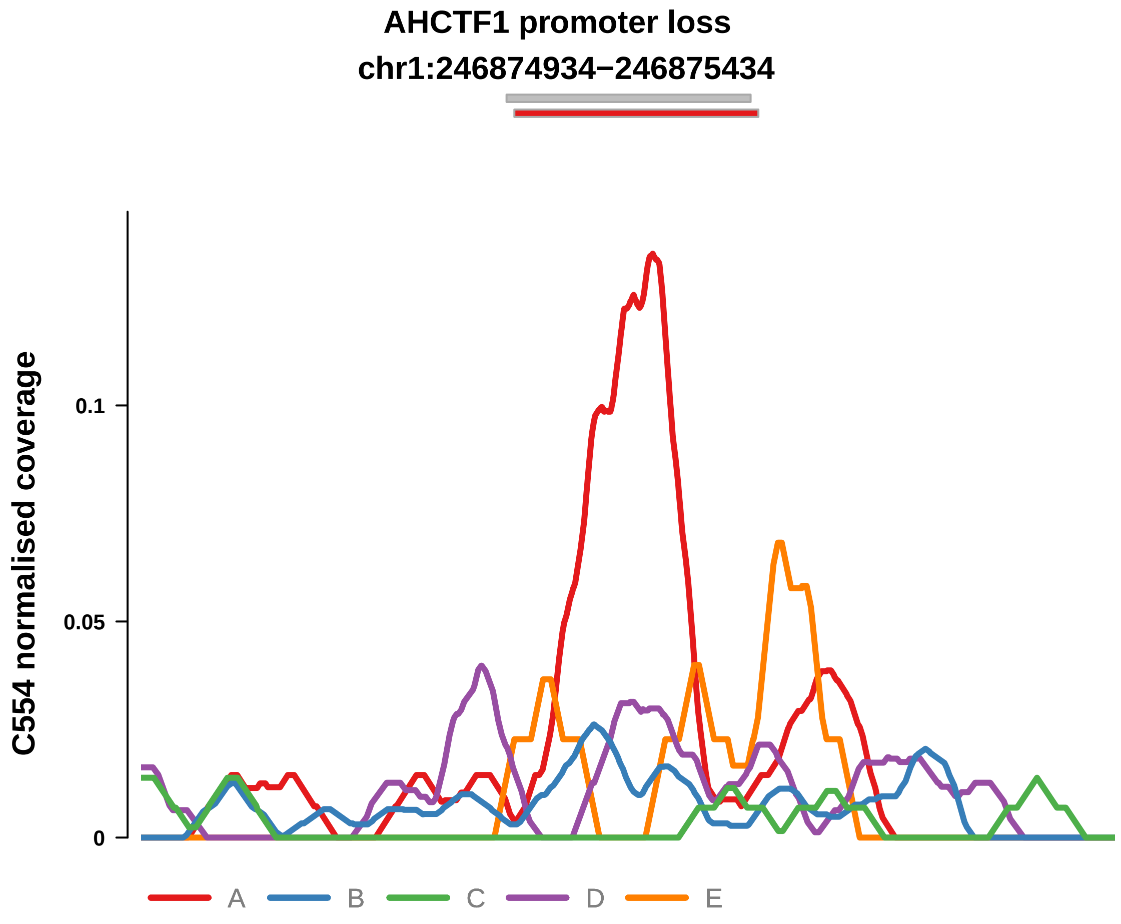


Putatively Clonal (logFC = −2.68)

| | P. | Adjusted P. |
|---|---|---|
| *~region vs. ~* | 5.66e−02 | 1.29e−01 |
| *~purity vs ~* | 5.68e−02 | 1.72e−01 |
| *~purity+region vs ~purity* | 3.47e−01 | 4.58e−01 |
| *Carcinoma vs. normal* | 2.04e−05 **** | 2.04e−05 **** |

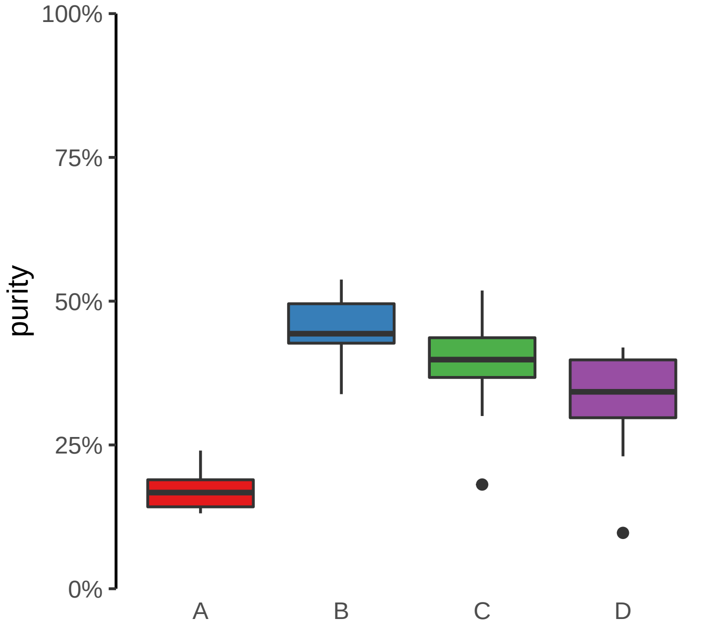

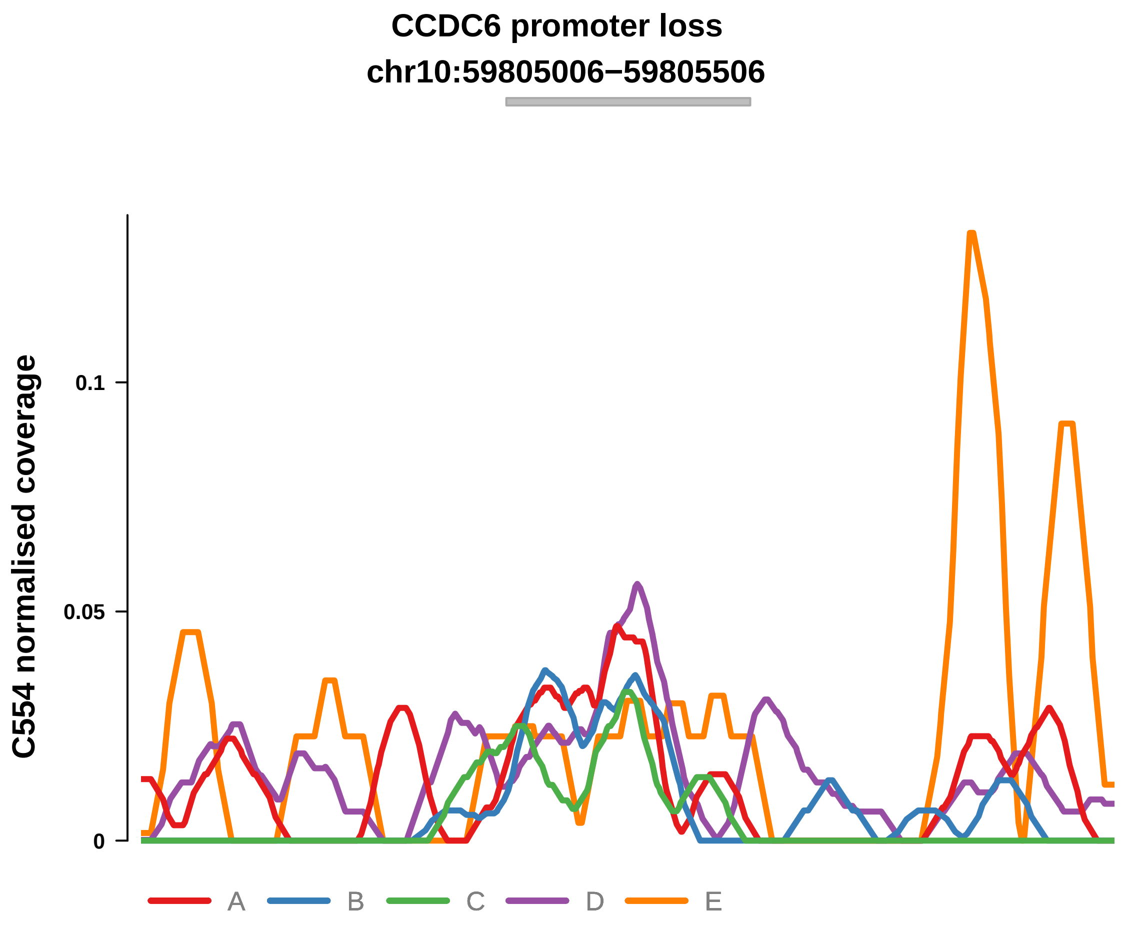


Putatively Clonal (logFC = −1.68)

| | P. | Adjusted P. |
|---|---|---|
| *~region vs. ~* | 5.22e−01 | 5.86e−01 |
| *~purity vs ~* | 8.11e−01 | 9.11e−01 |
| *~purity+region vs ~purity* | 1.82e−01 | 3.11e−01 |
| *Carcinoma vs. normal* | 4.17e−03 ** | 4.17e−03 ** |

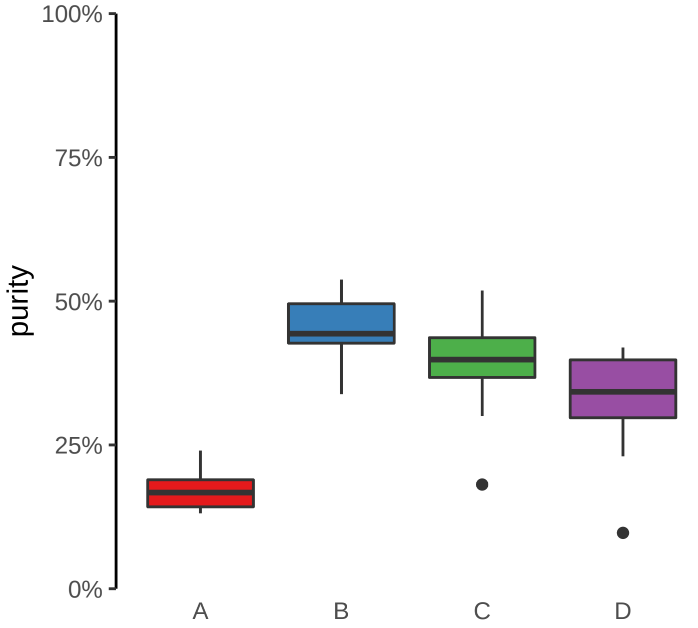

**ERBB3 promoter loss**
**chr12:56087145−56087645**

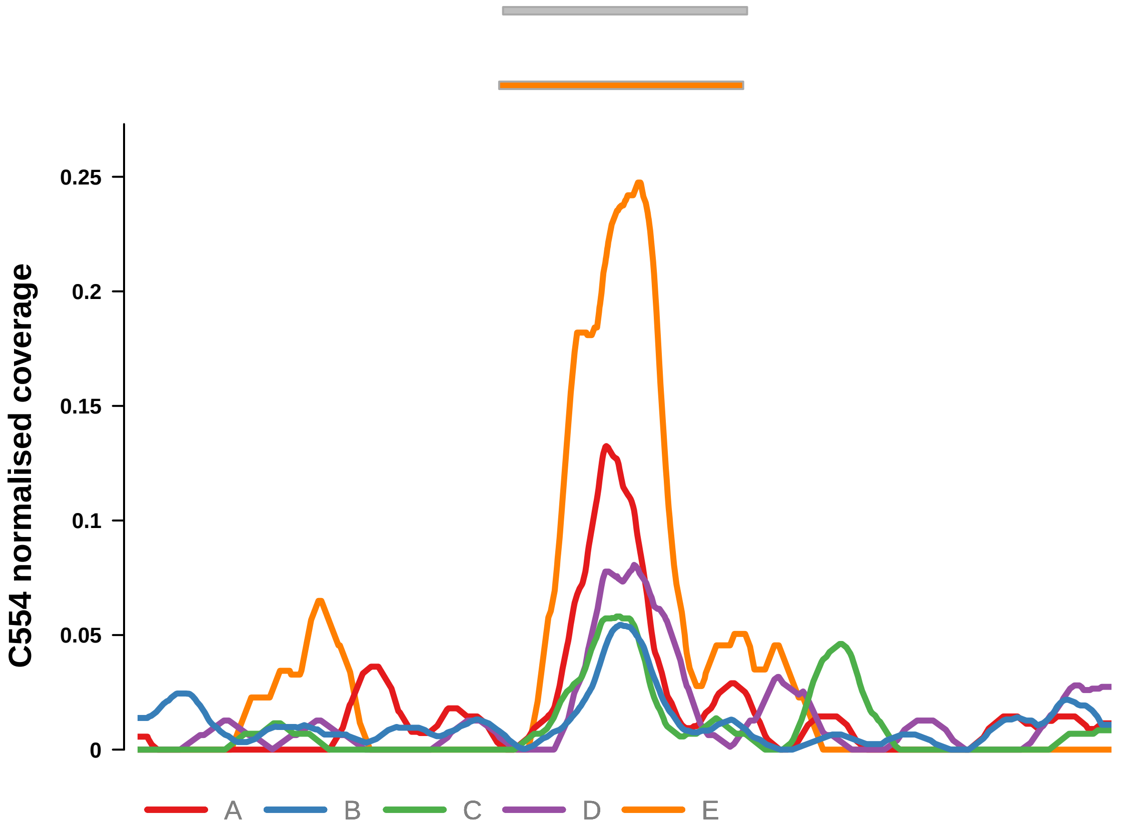


Putatively Clonal (logFC = −1.93)

| | P. | Adjusted P. |
|---|---|---|
| *~region vs. ~* | 1.73e−01 | 2.73e−01 |
| *~purity vs ~* | 1.07e−01 | 2.57e−01 |
| *~purity+region vs ~purity* | 1.94e−01 | 3.12e−01 |
| *Carcinoma vs. normal* | 9.74e−04 *** | 9.74e−04 *** |

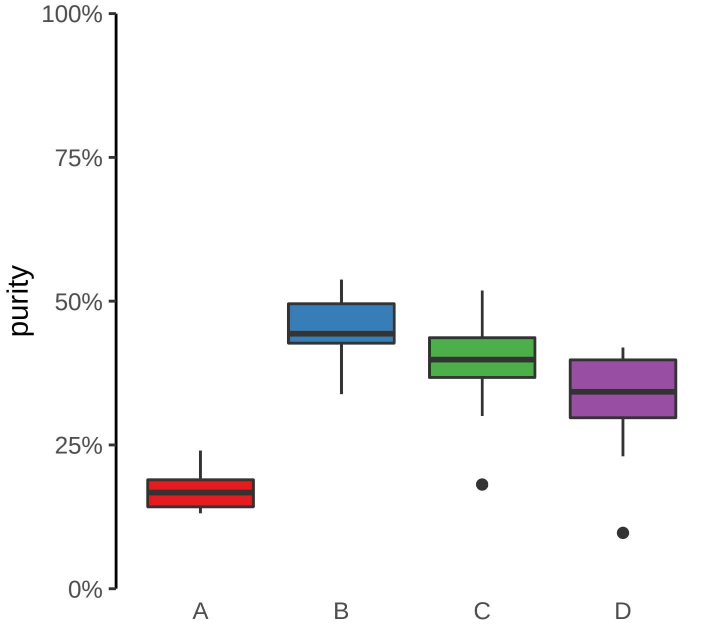

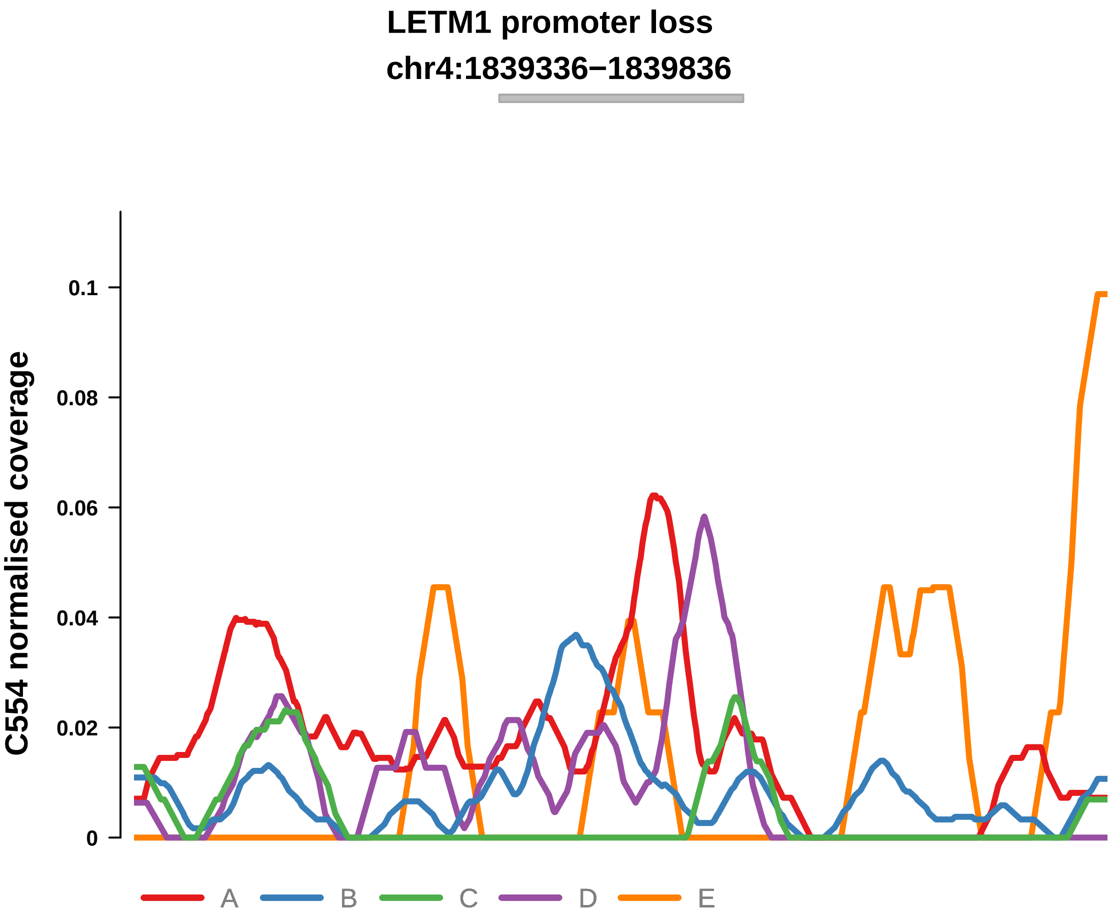


Putatively Clonal (logFC = −1.75)

| | P. | Adjusted P. |
|---|---|---|
| *~region vs. ~* | 6.41e−01 | 6.92e−01 |
| *~purity vs ~* | 3.98e−01 | 5.93e−01 |
| *~purity+region vs ~purity* | 7.99e−01 | NA |
| *Carcinoma vs. normal* | 3.04e−03 ** | 3.04e−03 ** |

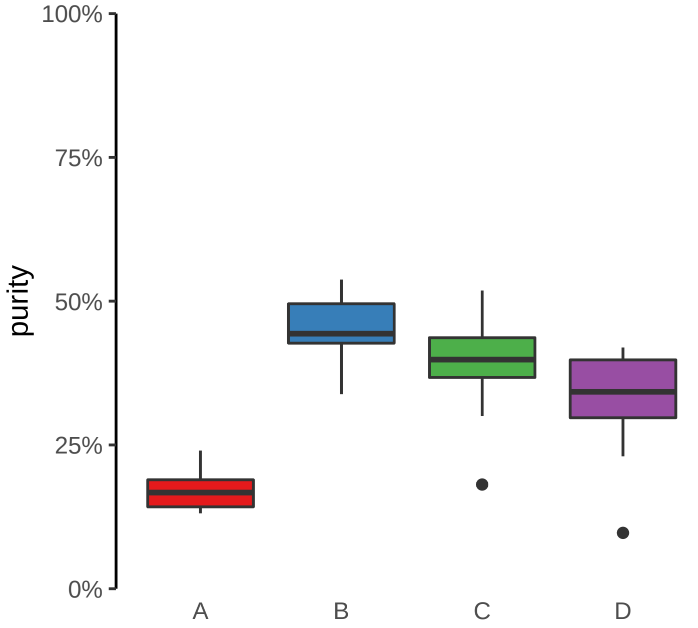

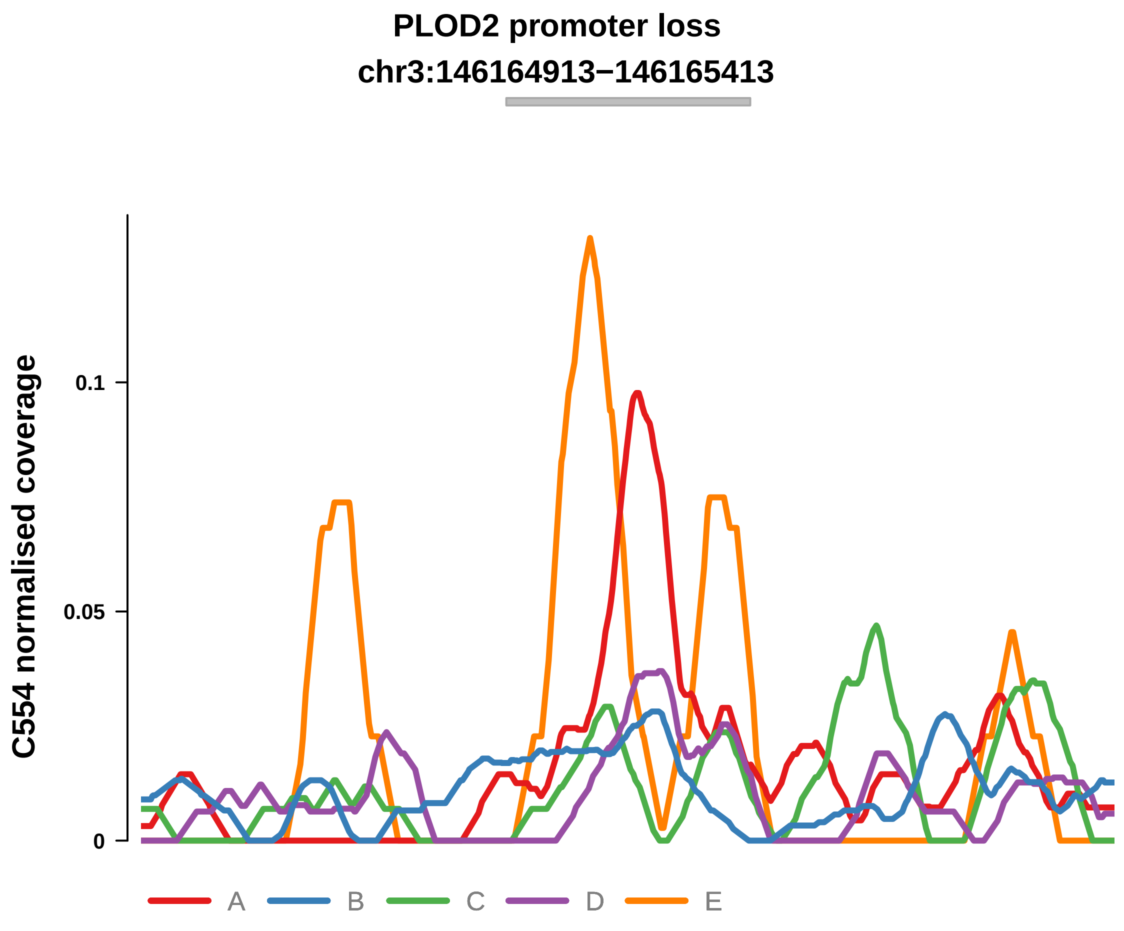


Putatively Clonal (logFC = −2.09)

| | P. | Adjusted P. |
|---|---|---|
| *~region vs. ~* | 1.23e−01 | 2.28e−01 |
| *~purity vs ~* | 5.05e−02 | 1.66e−01 |
| *~purity+region vs ~purity* | 1.93e−02 * | 6.01e−02 |
| *Carcinoma vs. normal* | 6.42e−04 *** | 6.42e−04 *** |

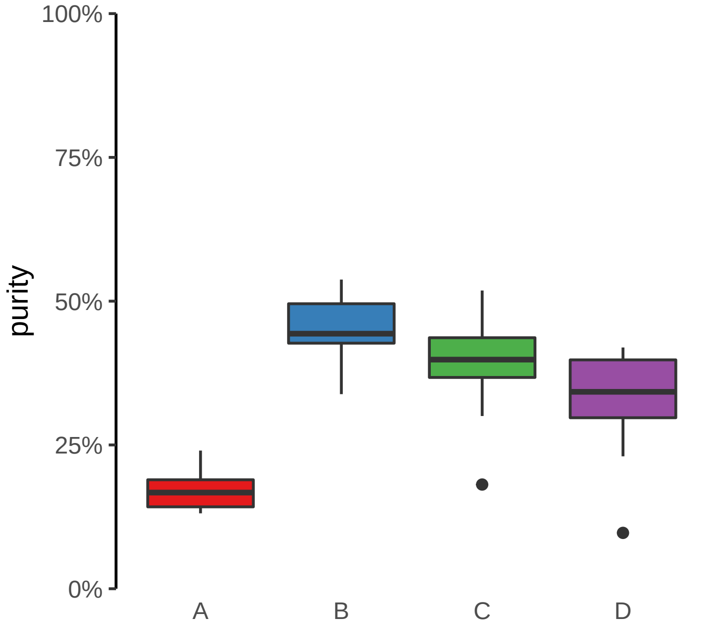

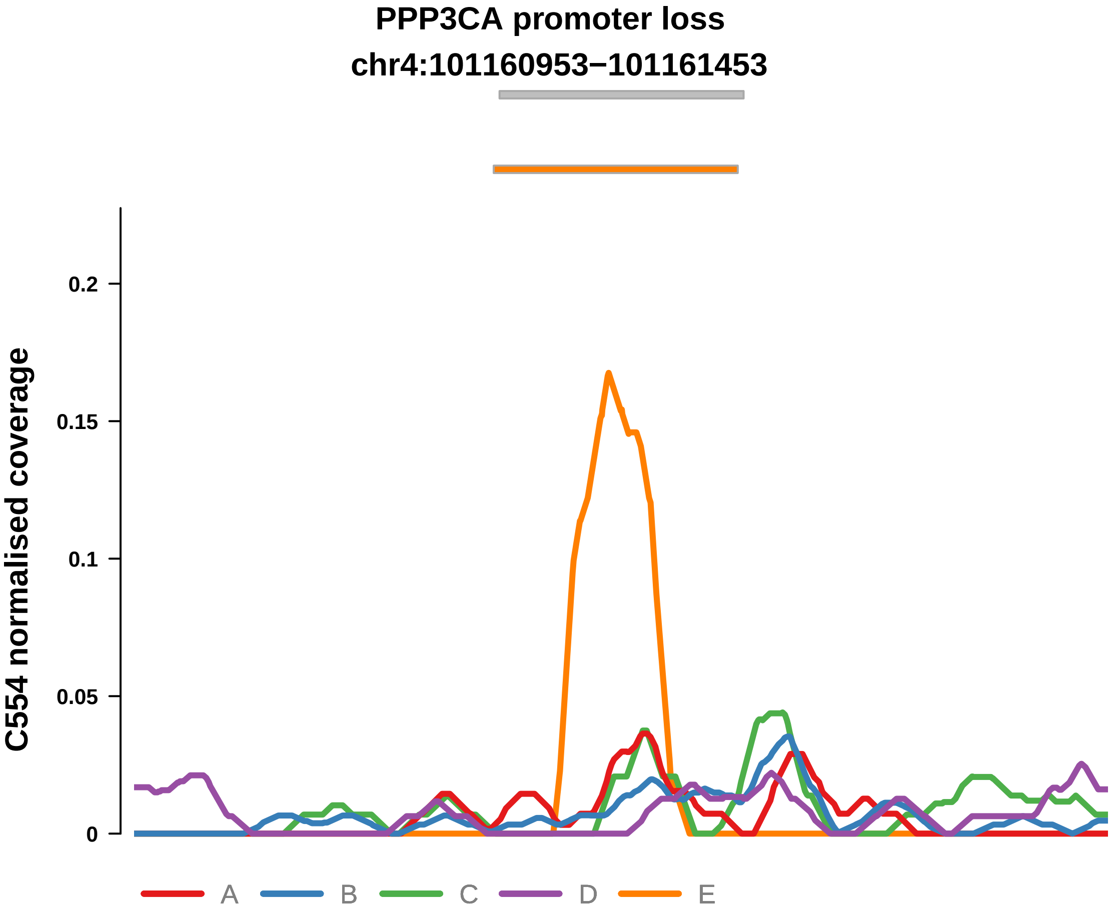


Putatively Clonal (logFC = −2.26)

| | P. | Adjusted P. |
|---|---|---|
| *~region vs. ~* | 3.05e−01 | NA |
| *~purity vs ~* | 7.18e−01 | NA |
| *~purity+region vs ~purity* | 2.77e−01 | NA |
| *Carcinoma vs. normal* | 4.93e−04 *** | 4.93e−04 *** |

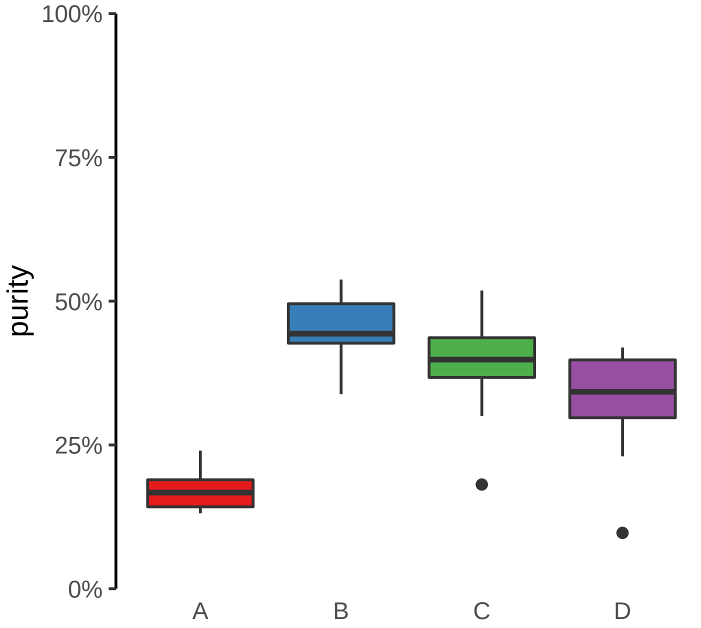

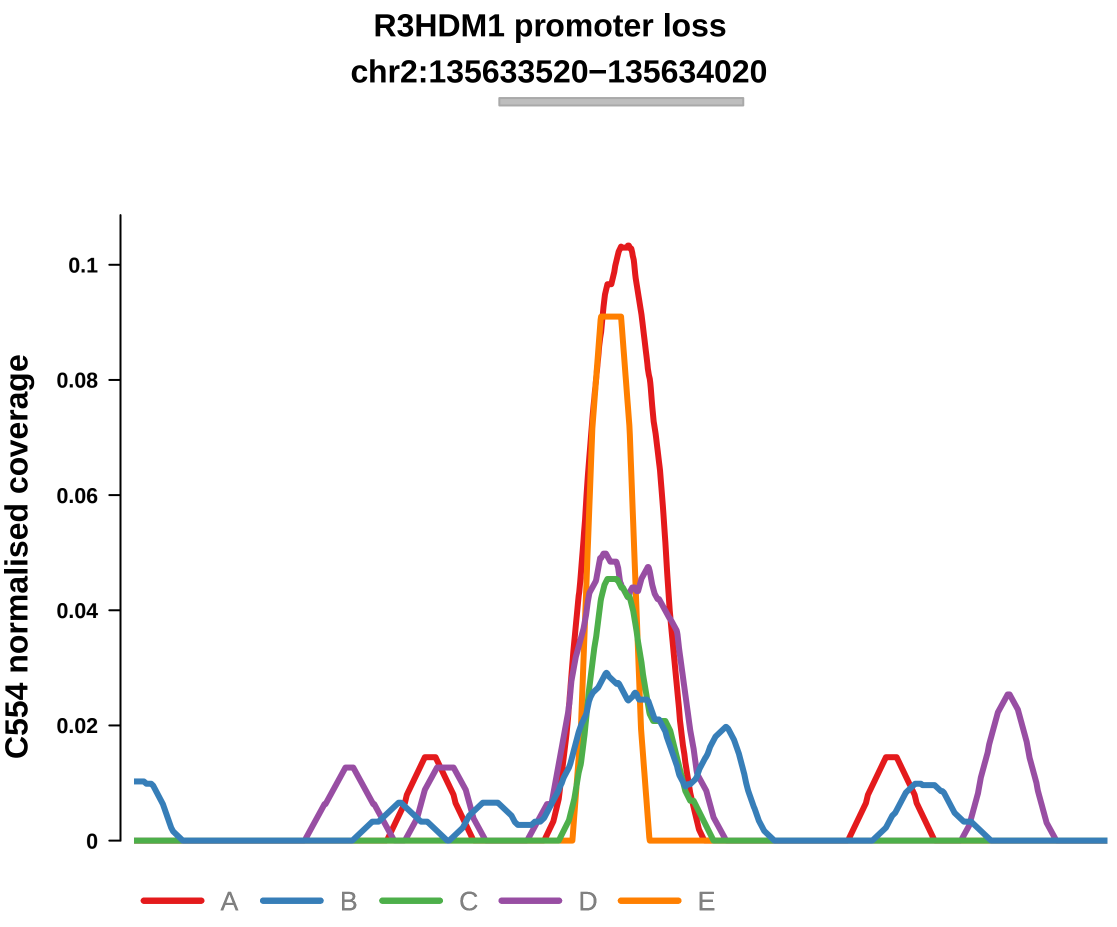


Putatively Clonal (logFC = −1.69)

| | P. | Adjusted P. |
|---|---|---|
| *~region vs. ~* | 8.14e−01 | 8.34e−01 |
| *~purity vs ~* | 8.37e−01 | 9.15e−01 |
| *~purity+region vs ~purity* | 7.68e−01 | 8.12e−01 |
| *Carcinoma vs. normal* | 4.5e−03 ** | 4.5e−03 ** |

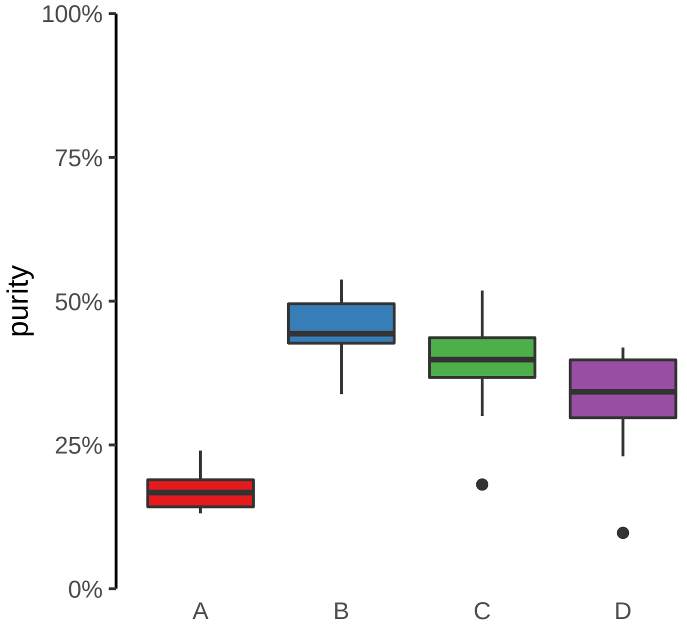

**SLC38A9 promoter loss**
**chr5:55700661−55701161**

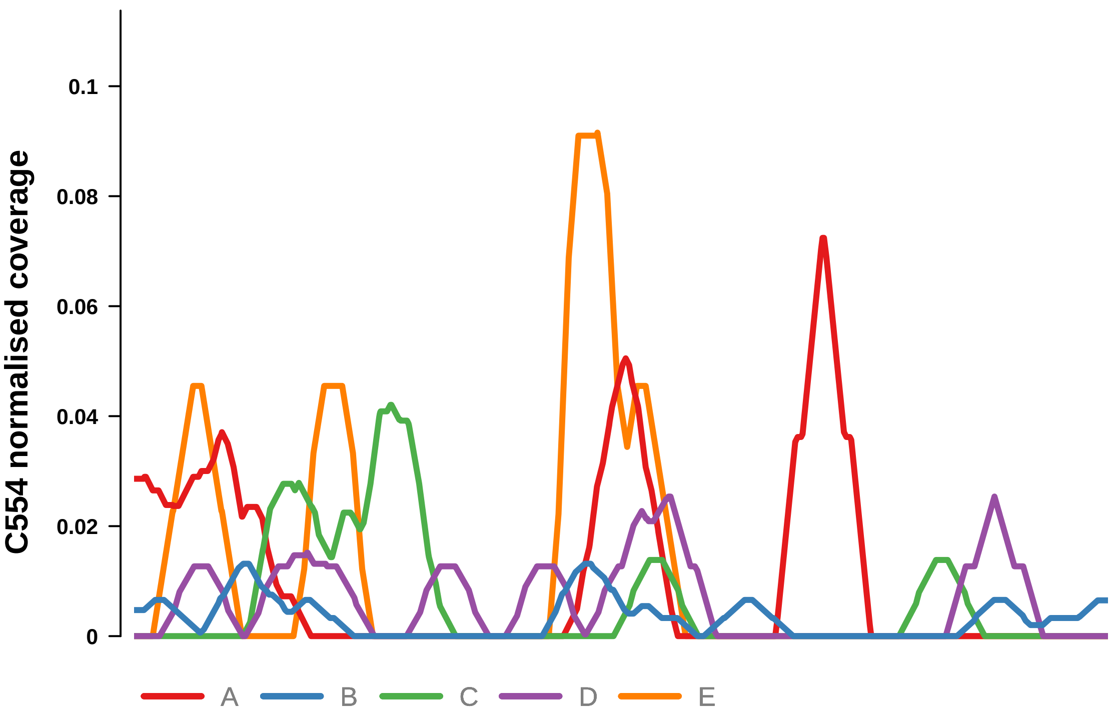


Putatively Clonal (logFC = −2.71)

| | **P.** | **Adjusted P.** |
|---|---|---|
| *~region vs. ~* | 1e+00 | NA |
| *~purity vs ~* | 7.74e−01 | NA |
| *~purity+region vs ~purity* | 1e+00 | NA |
| *Carcinoma vs. normal* | 6.44e−05 **** | 6.44e−05 **** |

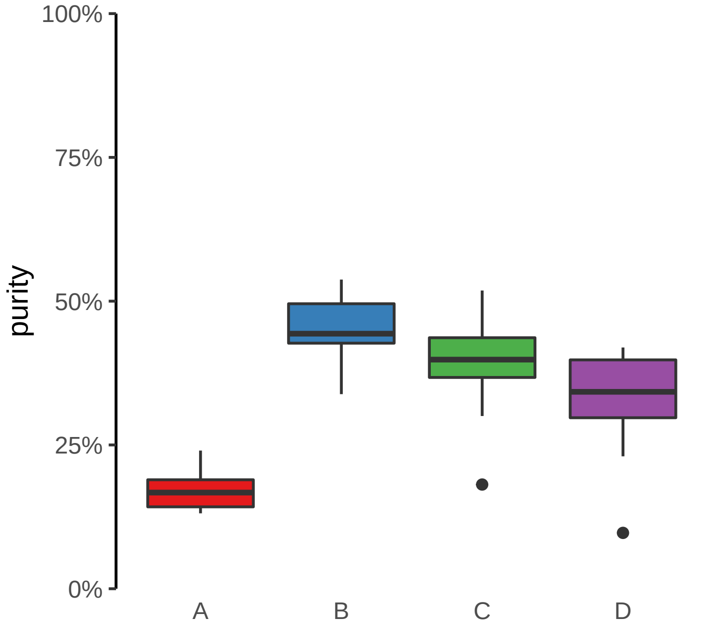

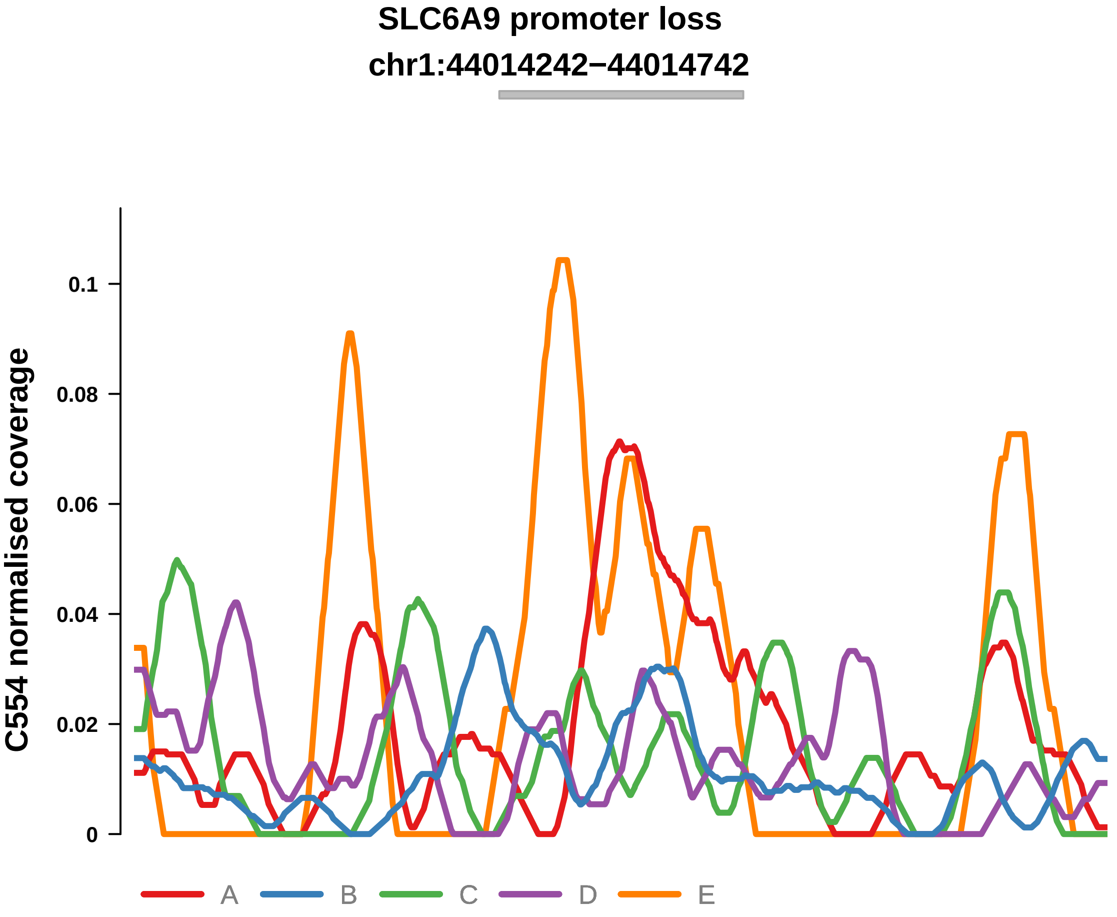


Putatively Clonal (logFC = −2.13)

| | P. | Adjusted P. |
|---|---|---|
| *~region vs. ~* | 1.46e−01 | 2.45e−01 |
| *~purity vs ~* | 3.64e−01 | 5.75e−01 |
| *~purity+region vs ~purity* | 1.05e−01 | 2.21e−01 |
| *Carcinoma vs. normal* | 3.48e−04 *** | 3.48e−04 *** |

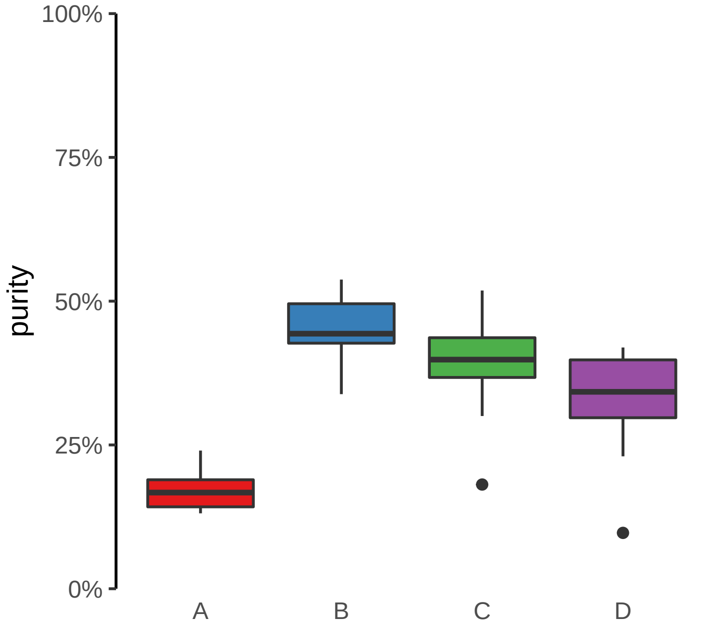

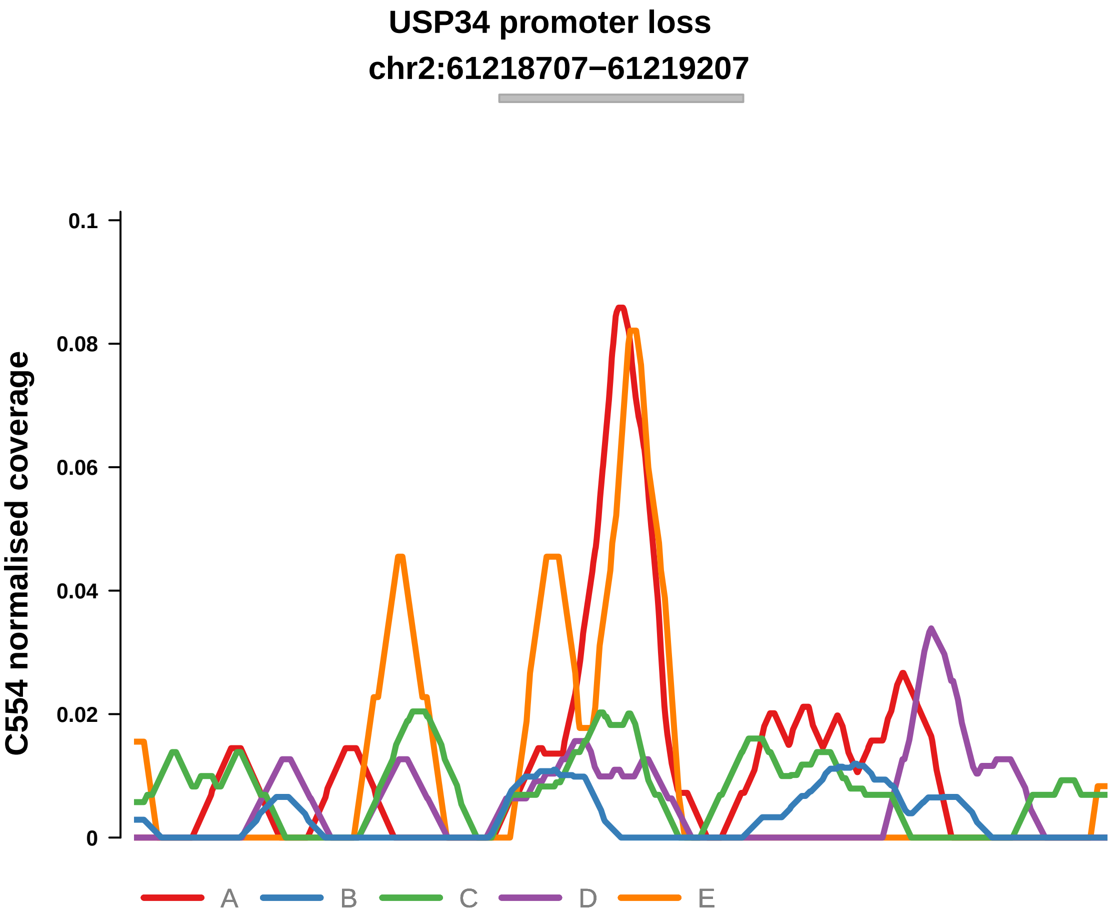


Putatively Clonal (logFC = −2.9)

| | P. | Adjusted P. |
|---|---|---|
| *~region vs. ~* | 4.44e−02 * | NA |
| *~purity vs ~* | 7.14e−02 | NA |
| *~purity+region vs ~purity* | 5.22e−02 | NA |
| *Carcinoma vs. normal* | 3.19e−05 **** | 3.19e−05 **** |

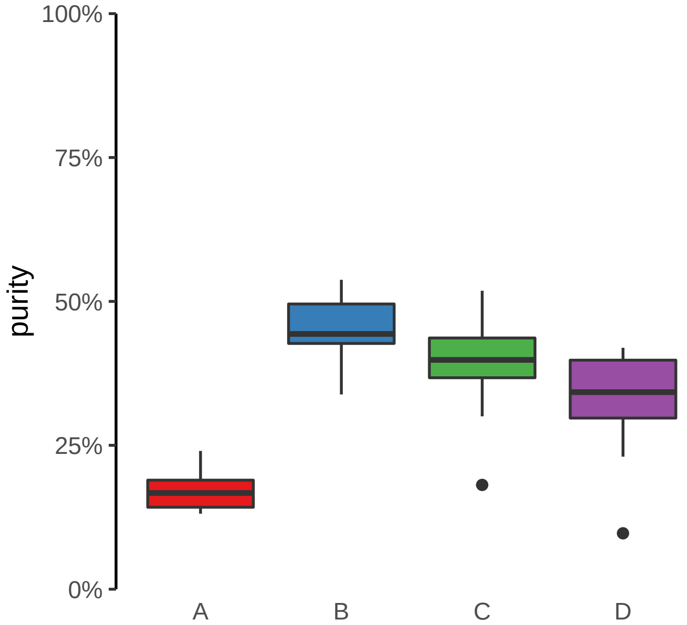

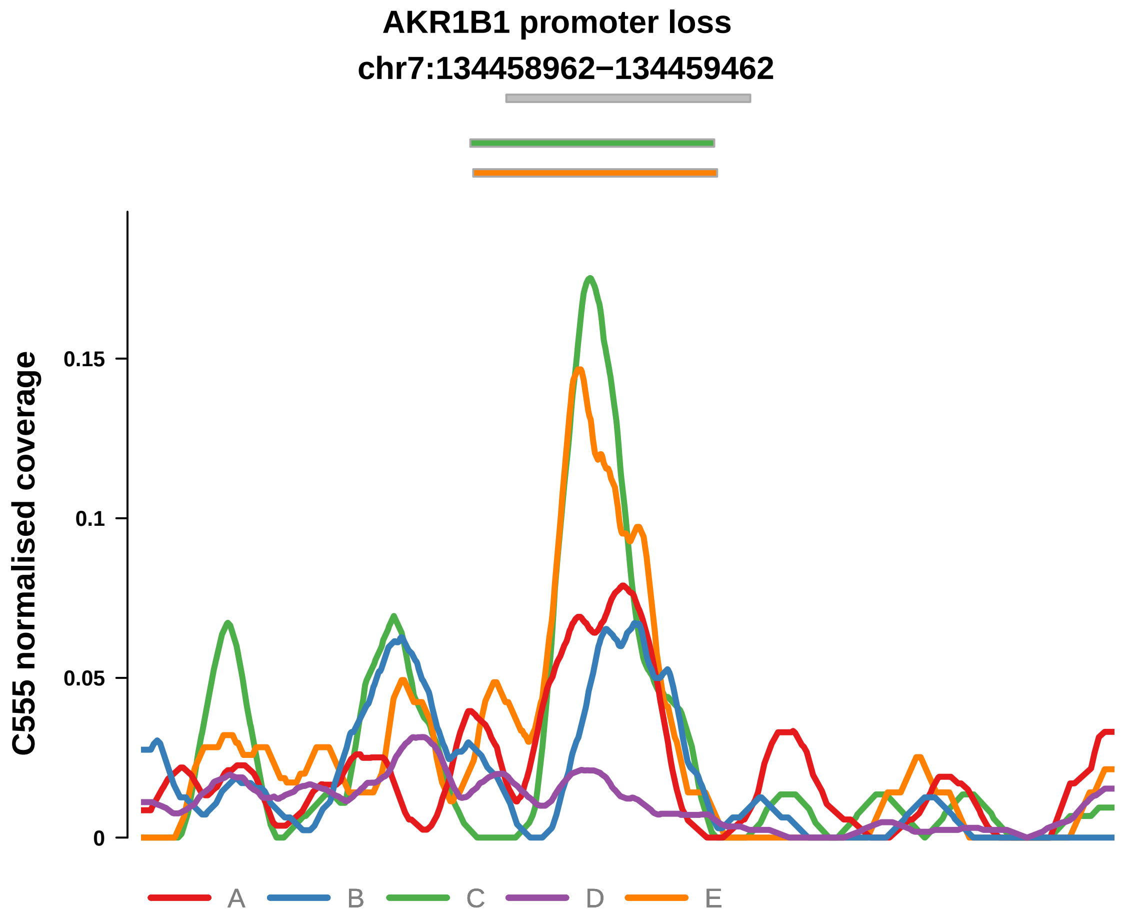


Putatively Clonal (logFC = −3.14)

| | P. | Adjusted P. |
|---|---|---|
| *~region vs. ~* | 7.3e−01 | 8.45e−01 |
| *~purity vs ~* | 4.58e−01 | 6.95e−01 |
| *~purity+region vs ~purity* | 4.12e−01 | 6.97e−01 |
| *Carcinoma vs. normal* | 2e−05 **** | 2e−05 **** |

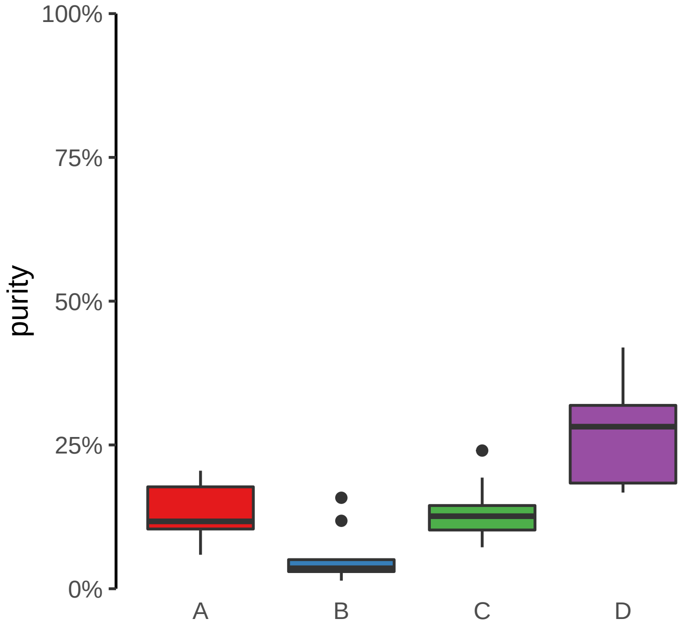

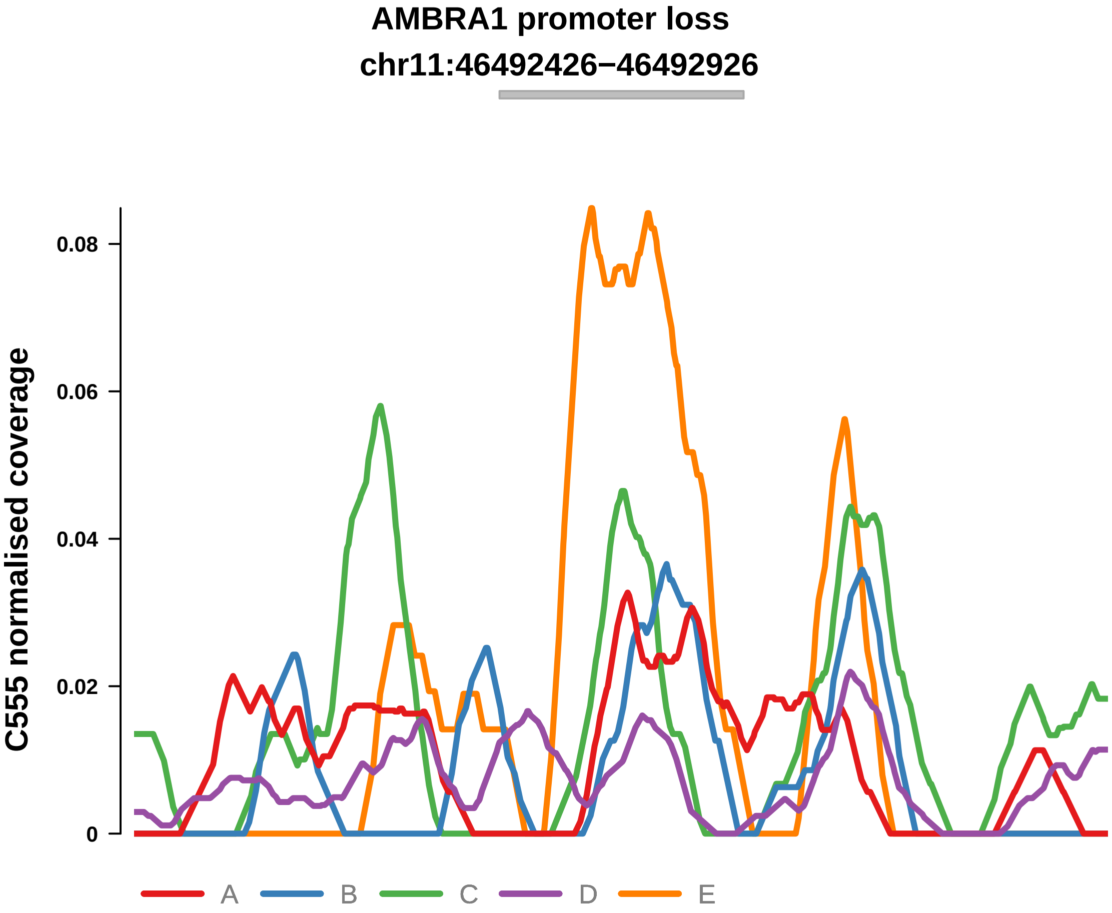


Putatively Clonal (logFC = −2.2)

| | P. | Adjusted P. |
|---|---|---|
| *~region vs. ~* | 3.33e−01 | 5.32e−01 |
| *~purity vs ~* | 7.43e−01 | 8.38e−01 |
| *~purity+region vs ~purity* | 3.73e−01 | 6.97e−01 |
| *Carcinoma vs. normal* | 4.42e−03 ** | 4.42e−03 ** |

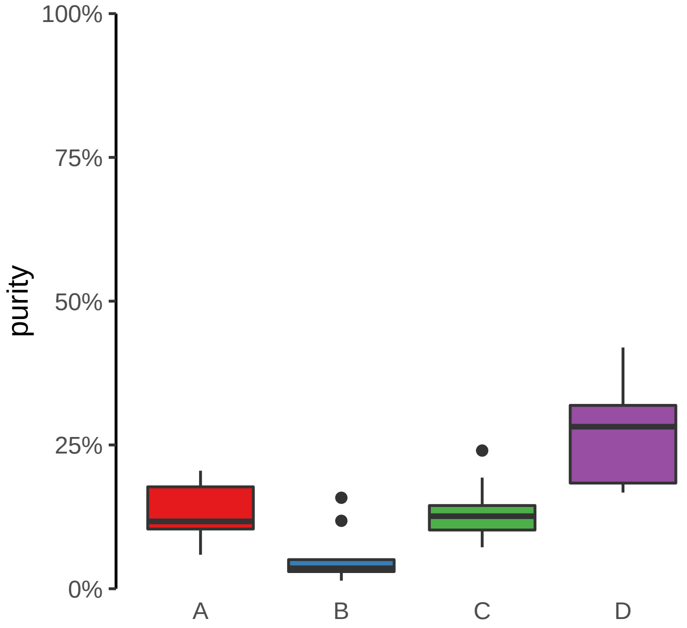

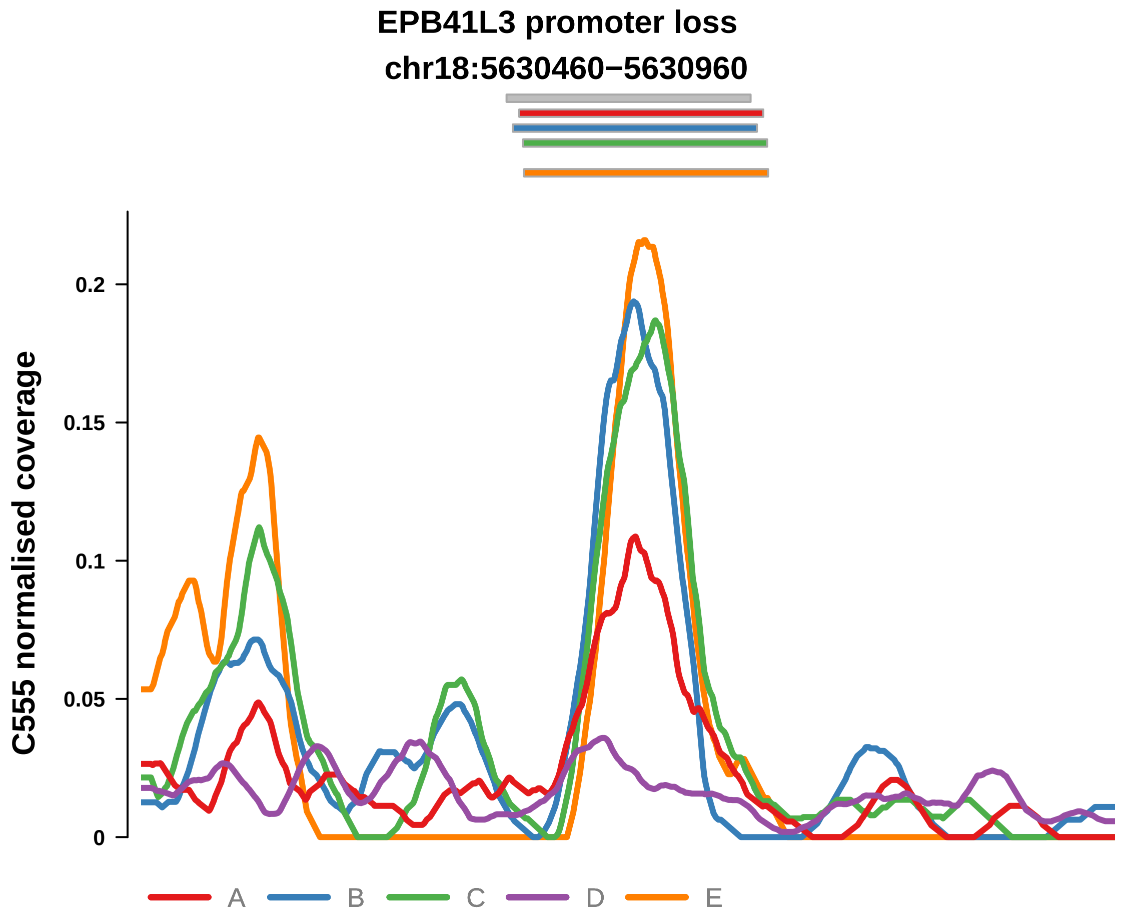


Putatively Clonal (logFC = −1.72)

| | P. | Adjusted P. |
|---|---|---|
| *~region vs. ~* | 3.98e−01 | 6.22e−01 |
| *~purity vs ~* | 4.1e−01 | 6.34e−01 |
| *~purity+region vs ~purity* | 4.71e−01 | 7.43e−01 |
| *Carcinoma vs. normal* | 5.66e−03 ** | 5.66e−03 ** |

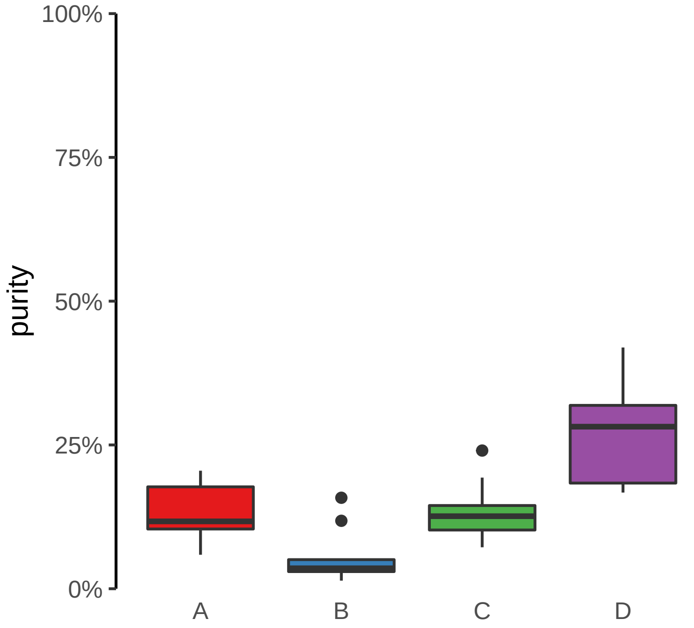

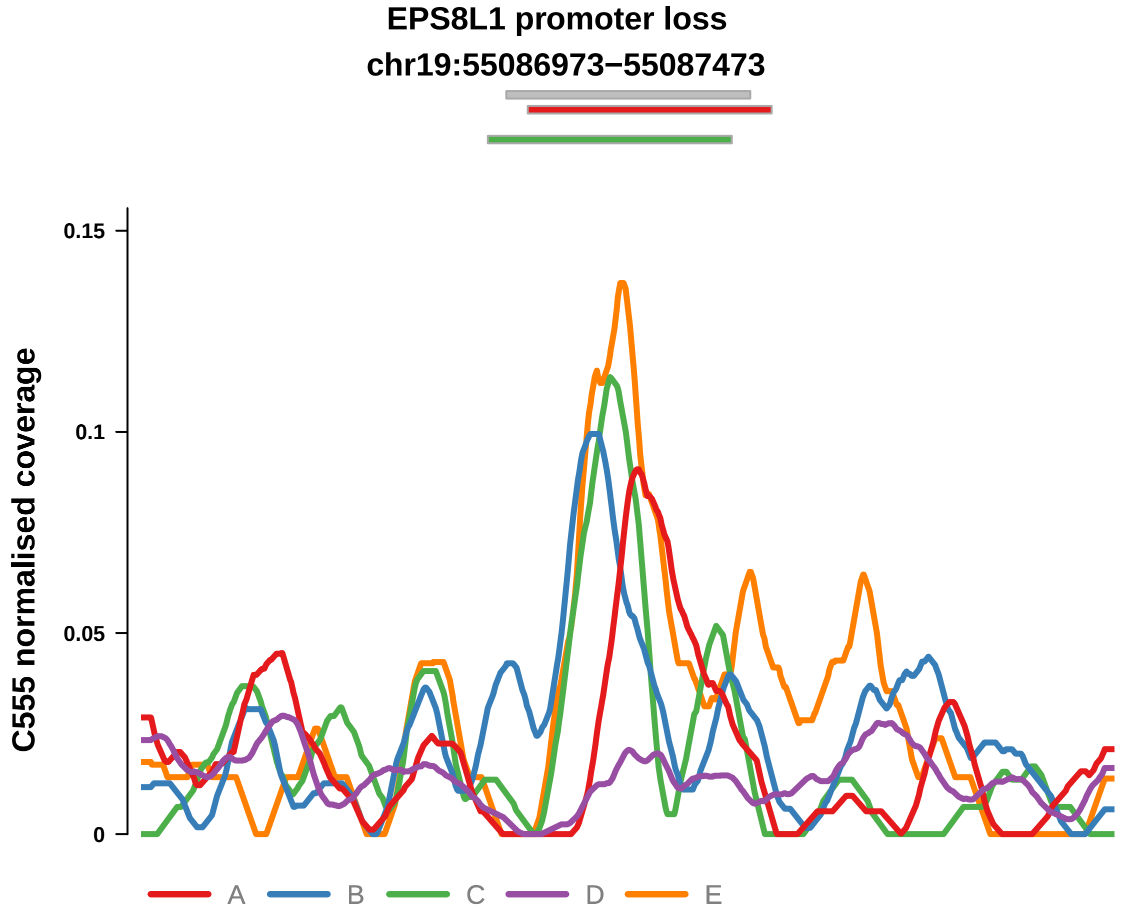


Putatively Clonal (logFC = −3.01)

| | P. | Adjusted P. |
|---|---|---|
| *~region vs. ~* | 3.22e−01 | 5.32e−01 |
| *~purity vs ~* | 6.17e−01 | 7.43e−01 |
| *~purity+region vs ~purity* | 3.35e−01 | 6.86e−01 |
| *Carcinoma vs. normal* | 5.42e−05 **** | 5.42e−05 **** |

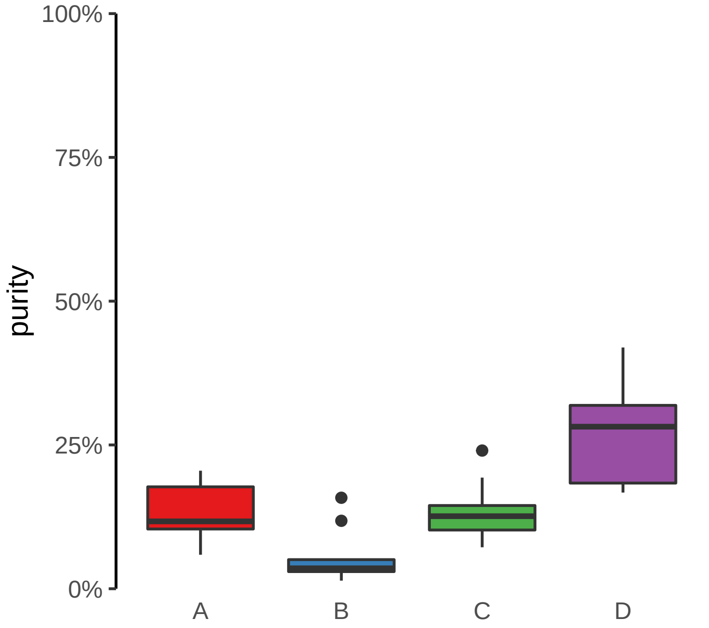

**SPAG16 promoter loss**
**chr2:213284089−213284589**

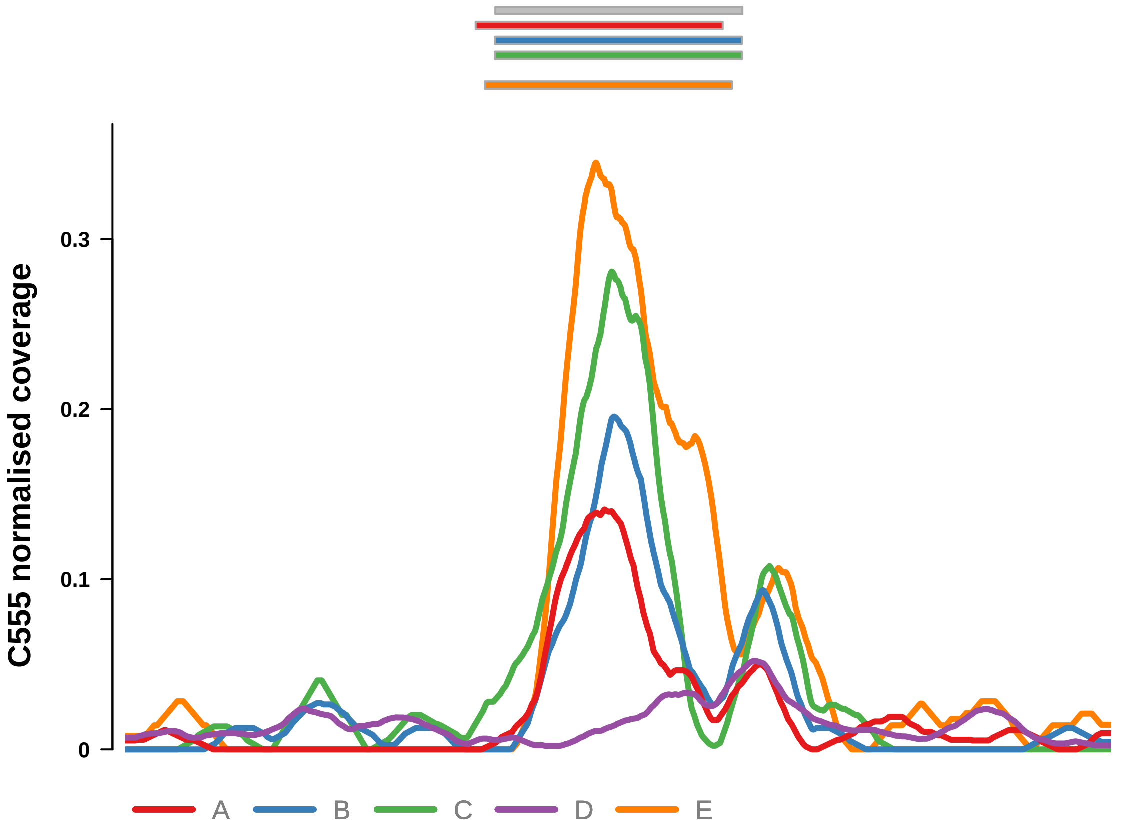


Putatively Clonal (logFC = −1.75)

| | P. | Adjusted P. |
|---|---|---|
| *~region vs. ~* | 6.76e−03 ** | 3.96e−02 * |
| *~purity vs ~* | 8.67e−03 ** | 1.53e−01 |
| *~purity+region vs ~purity* | 2.06e−01 | 4.81e−01 |
| *Carcinoma vs. normal* | 3.34e−03 ** | 3.34e−03 ** |

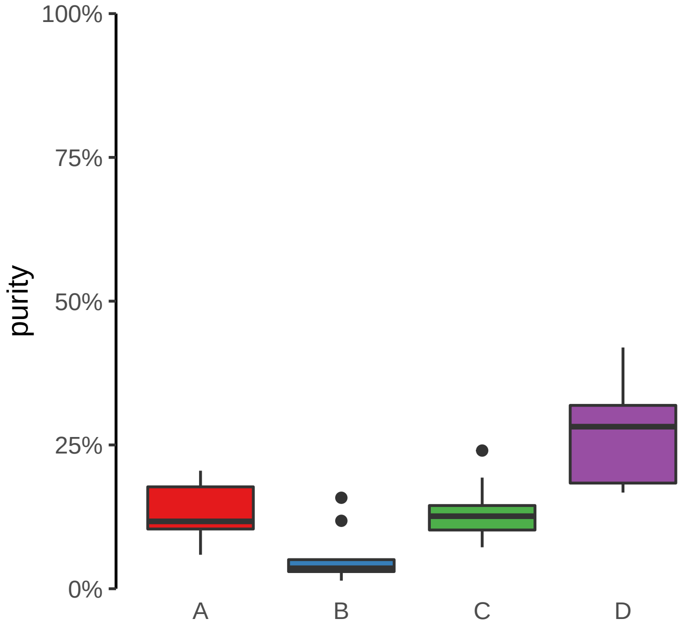

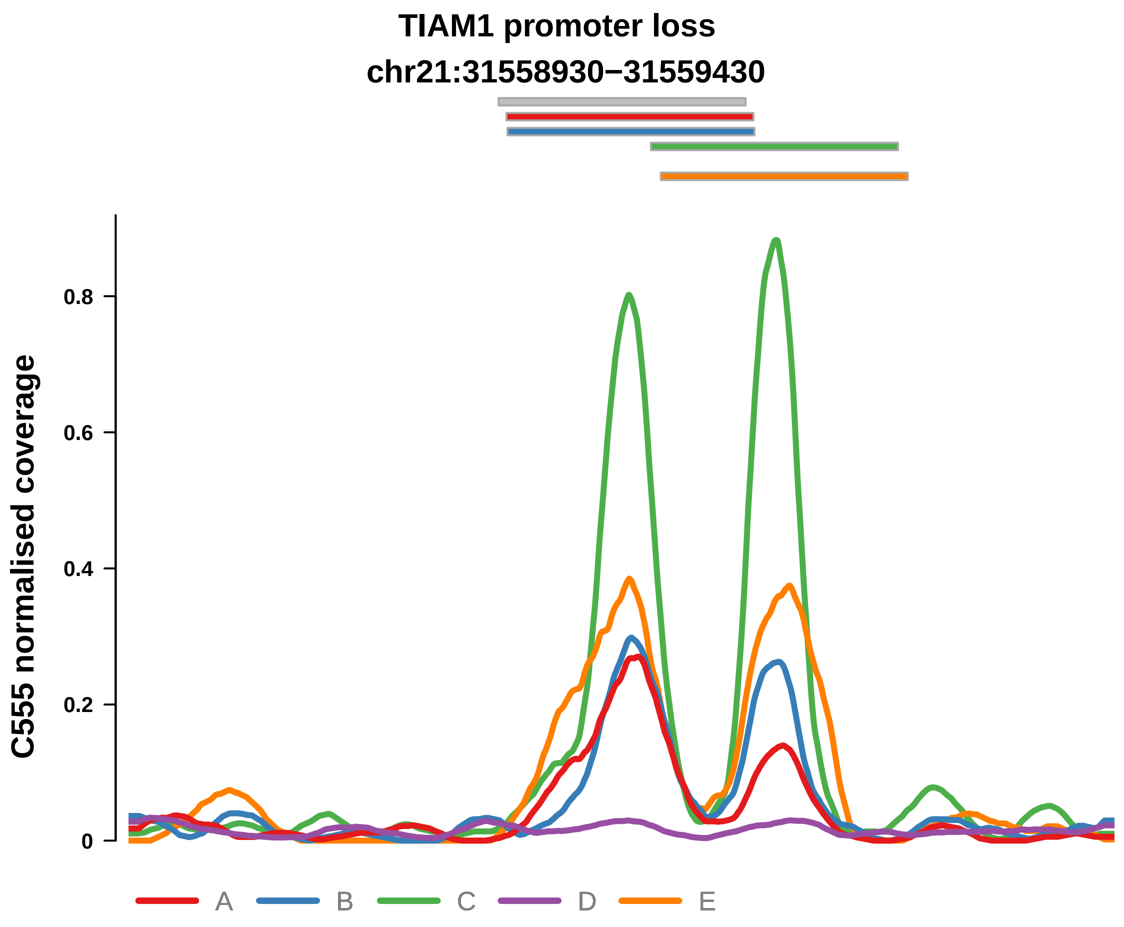


Putatively Clonal (logFC = −2.07)

| | P. | Adjusted P. |
|---|---|---|
| *~region vs. ~* | 1.9e−05 **** | 8.37e−04 *** |
| *~purity vs ~* | 2e−03 ** | 1.1e−01 |
| *~purity+region vs ~purity* | 7.23e−03 ** | 5.9e−02 |
| *Carcinoma vs. normal* | 8.11e−04 *** | 8.11e−04 *** |

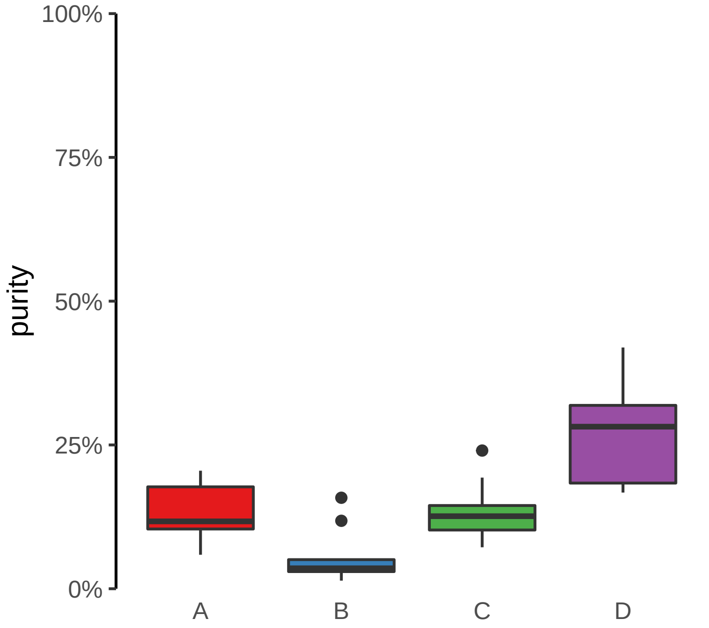

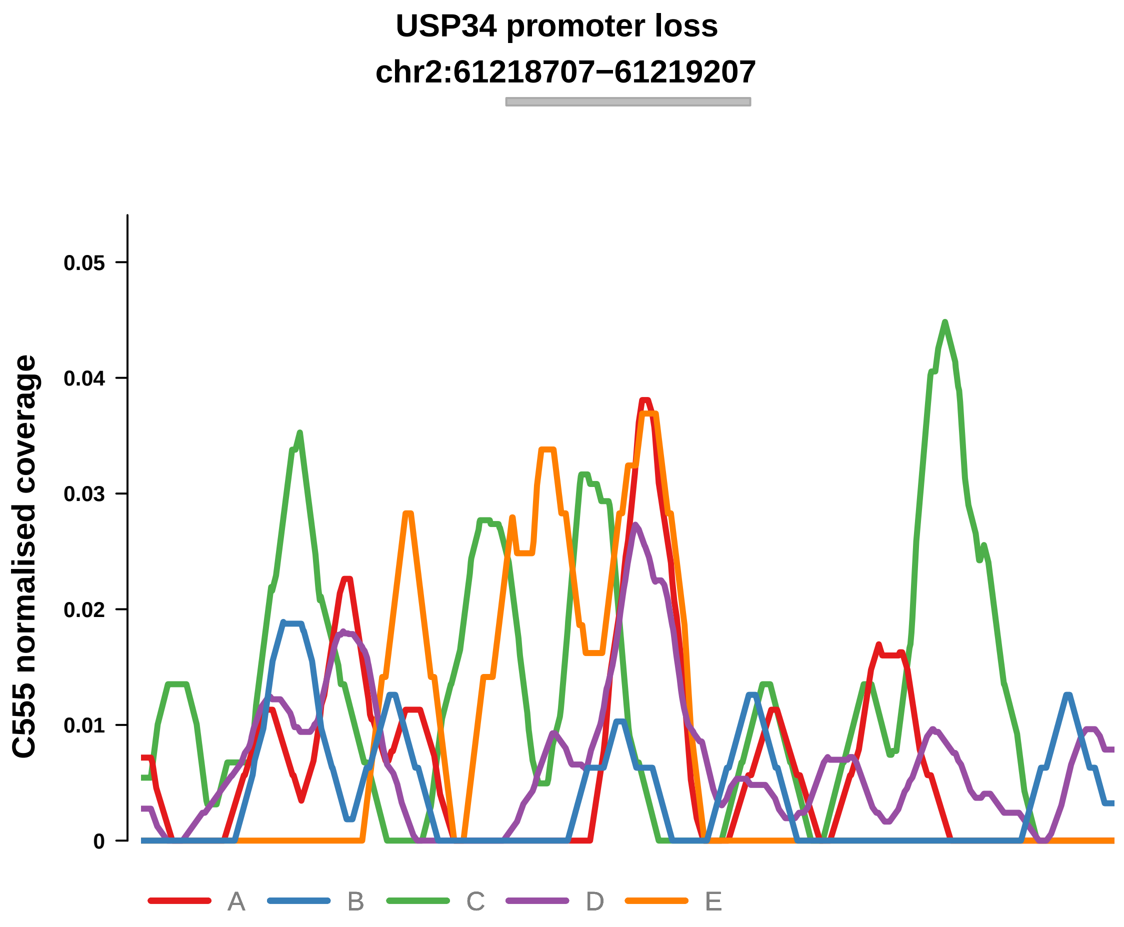


Putatively Clonal (logFC = −3.03)

| | P. | Adjusted P. |
|---|---|---|
| *~region vs. ~* | 2.95e−01 | 4.99e−01 |
| *~purity vs ~* | 4.66e−02 * | 2.26e−01 |
| *~purity+region vs ~purity* | 9.12e−01 | 9.59e−01 |
| *Carcinoma vs. normal* | 1.41e−03 ** | 1.41e−03 ** |

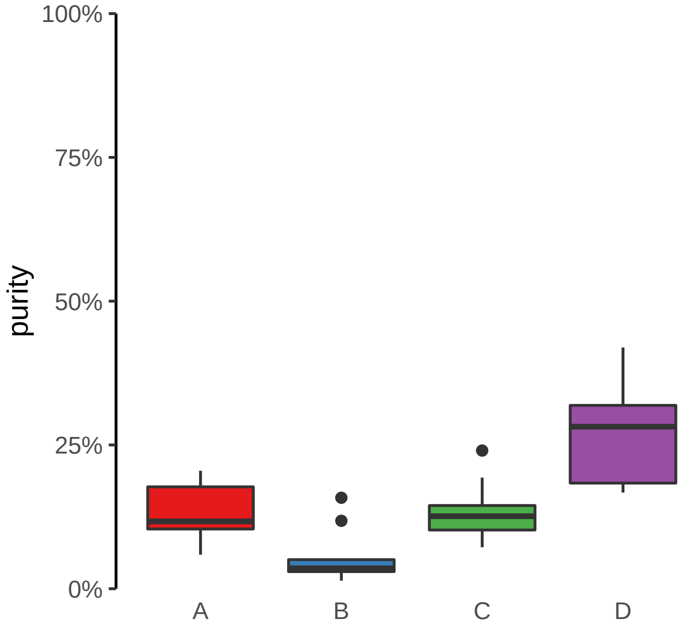

**ZNF43 promoter loss**
**chr19:21851889−21852389**

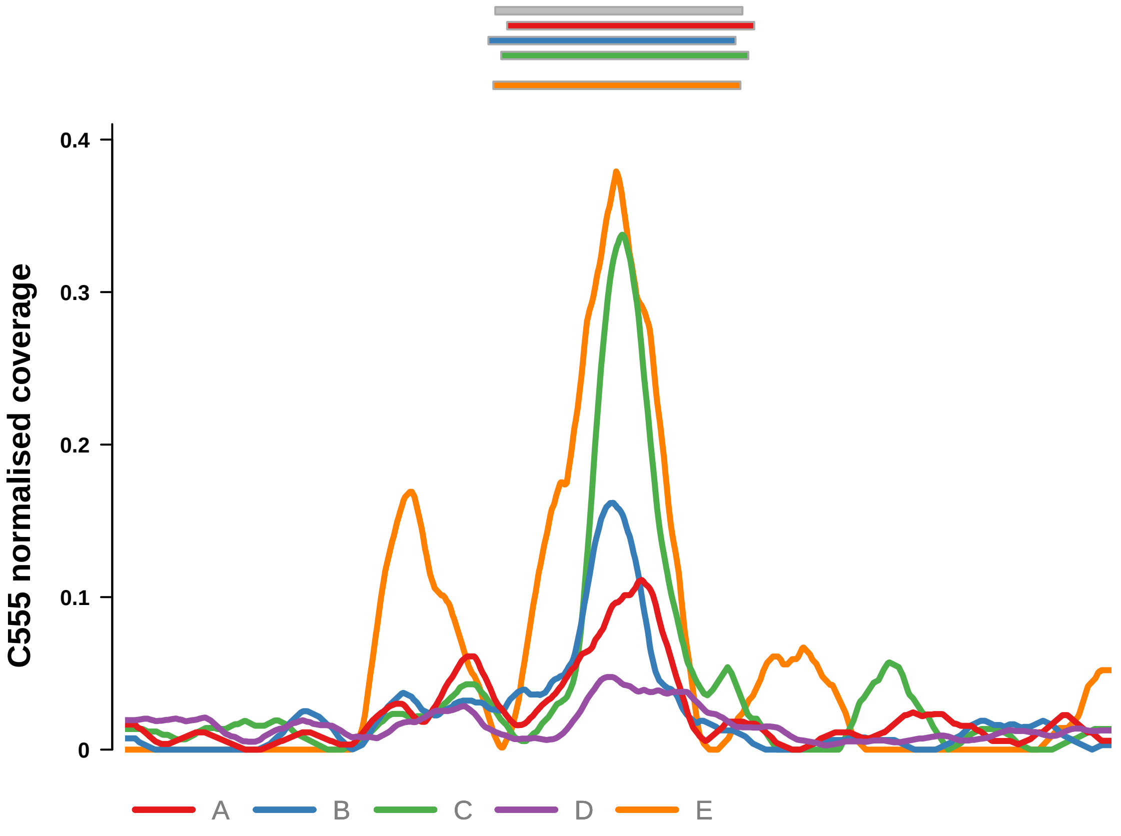


Putatively Clonal (logFC = −1.68)

| | P. | Adjusted P. |
|---|---|---|
| *~region vs. ~* | 9.17e−01 | 9.41e−01 |
| *~purity vs ~* | 3.92e−01 | 6.34e−01 |
| *~purity+region vs ~purity* | 9.87e−01 | 9.87e−01 |
| *Carcinoma vs. normal* | 4.77e−03 ** | 4.77e−03 ** |

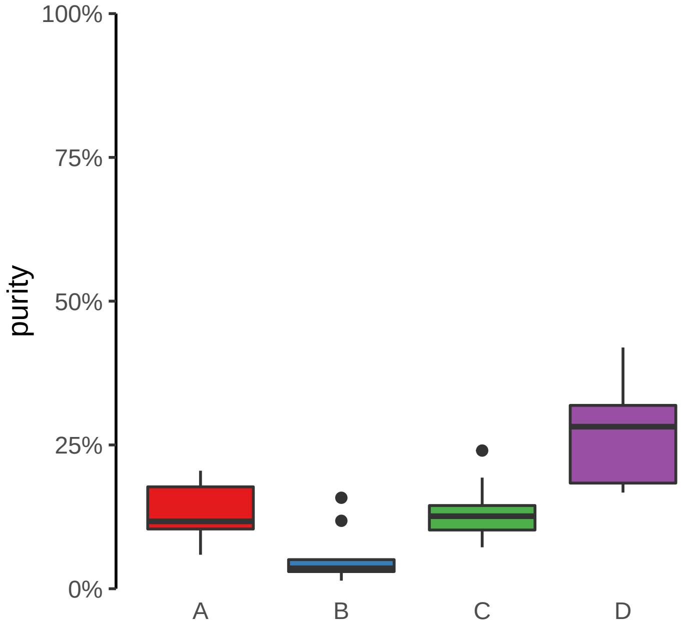

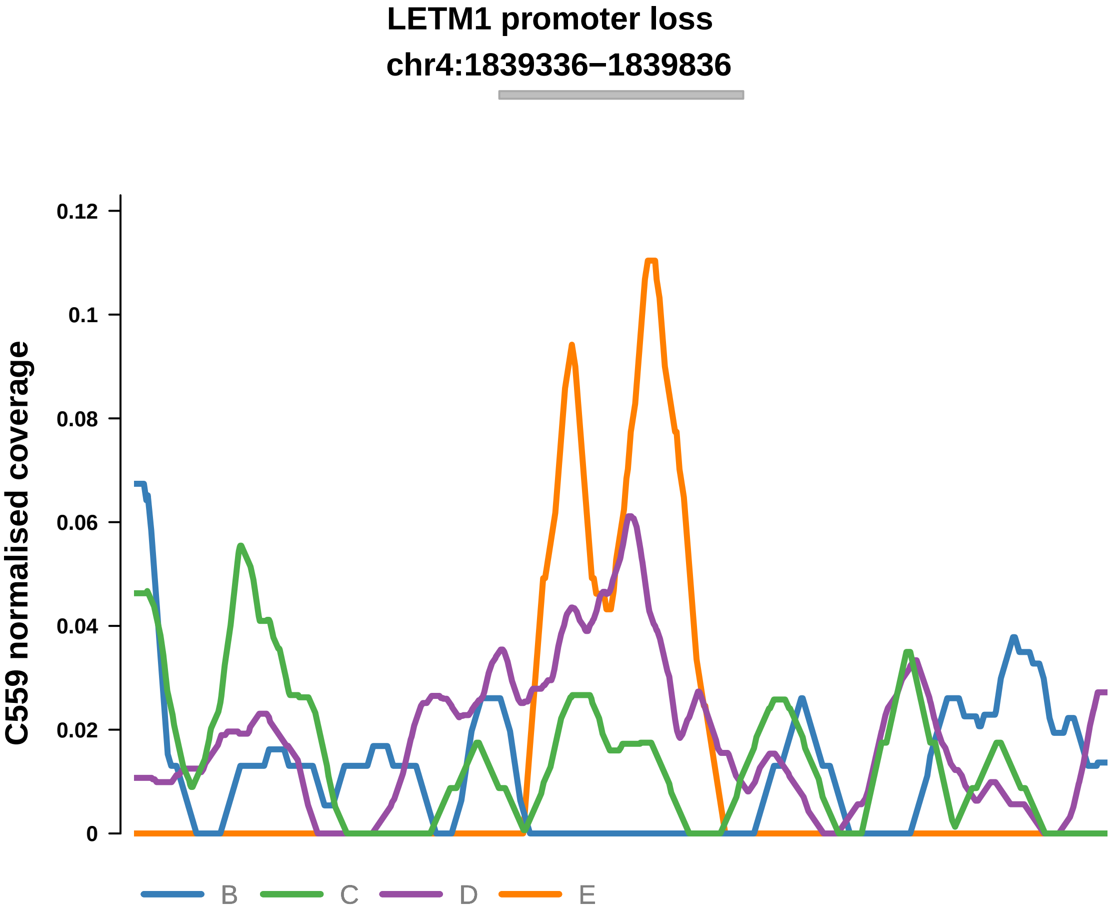


Putatively Clonal (logFC = −2.27)

| | P. | Adjusted P. |
|---|---|---|
| *~region vs. ~* | 9.88e−02 | 4.14e−01 |
| *~purity vs ~* | 1.31e−01 | 5.51e−01 |
| *~purity+region vs ~purity* | 1.37e−01 | 5.29e−01 |
| *Carcinoma vs. normal* | 5.38e−03 ** | 5.38e−03 ** |

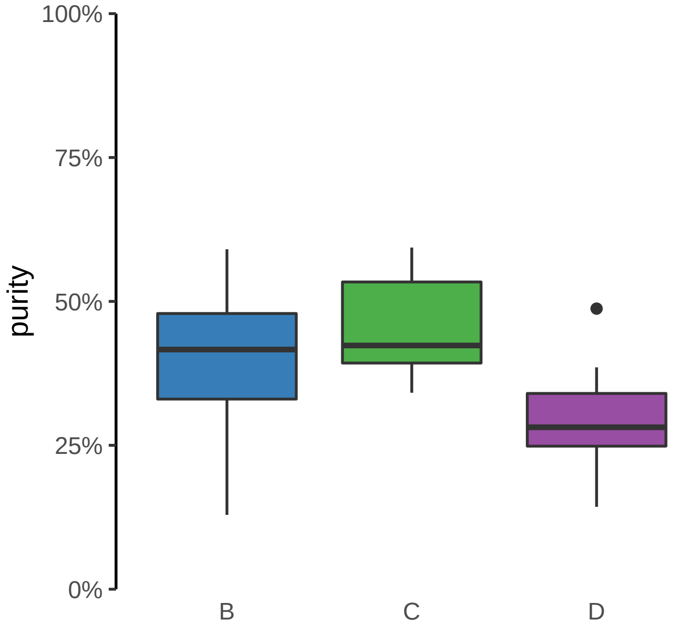

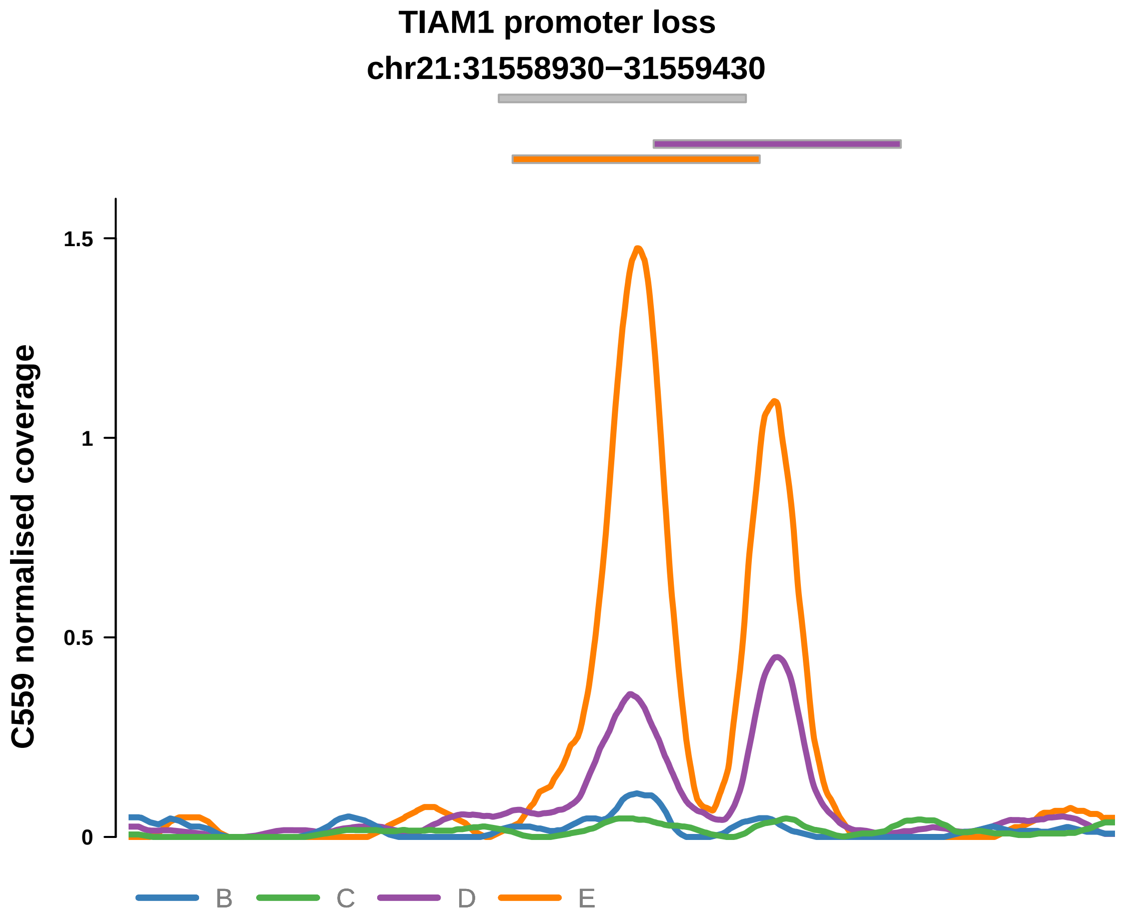


Putatively Clonal (logFC = −2.42)

| | P. | Adjusted P. |
|---|---|---|
| *~region vs. ~* | 5.49e−04 *** | 4.43e−02 * |
| *~purity vs ~* | 6.56e−04 *** | 2.89e−02 * |
| *~purity+region vs ~purity* | 1.17e−01 | 5.21e−01 |
| *Carcinoma vs. normal* | 2.61e−04 *** | 2.61e−04 *** |

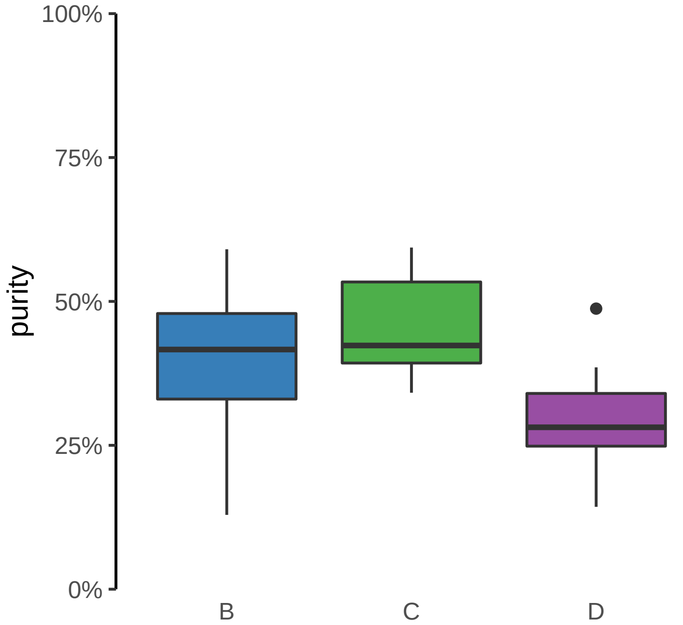

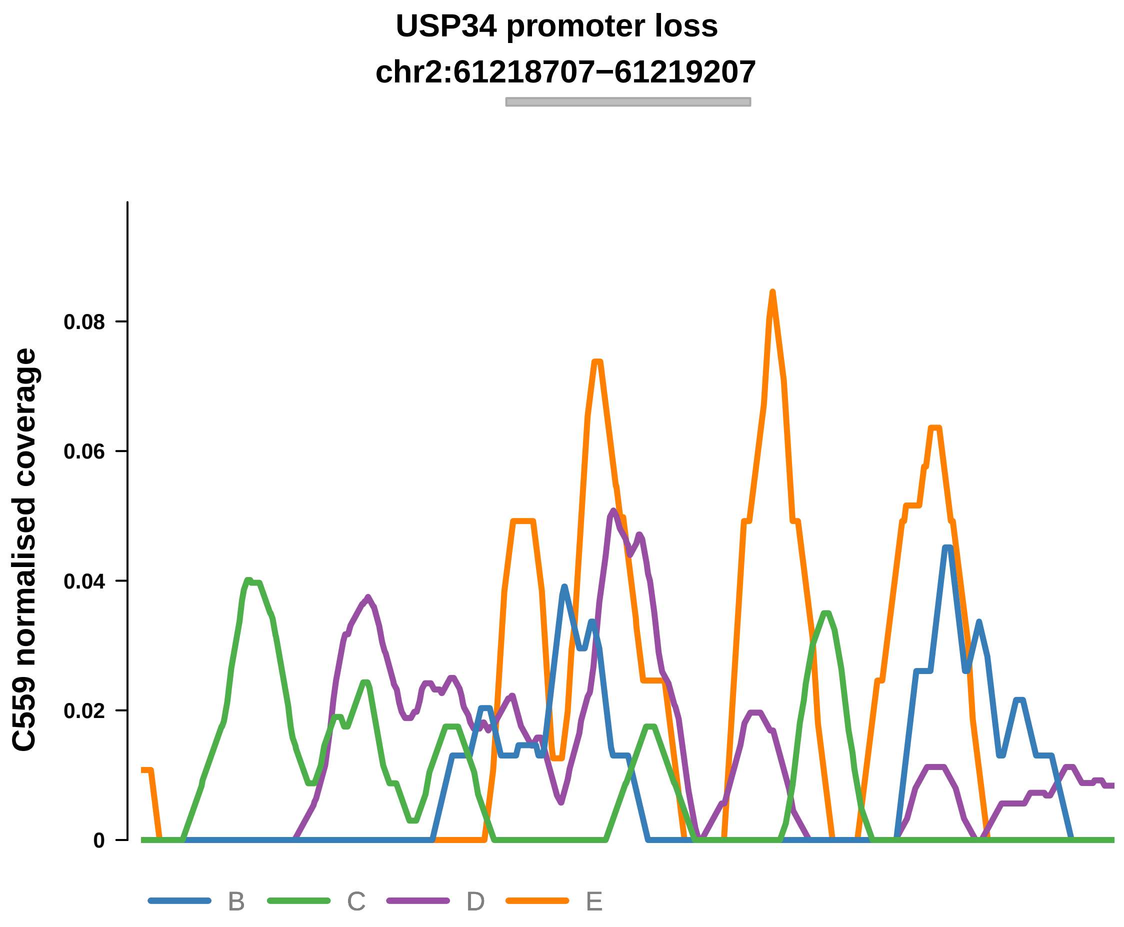


Putatively Clonal (logFC = −10.79)

| | P. | Adjusted P. |
|---|---|---|
| *~region vs. ~* | 2.83e−01 | 5.75e−01 |
| *~purity vs ~* | 1.07e−01 | 5.51e−01 |
| *~purity+region vs ~purity* | 7.08e−01 | 9.61e−01 |
| *Carcinoma vs. normal* | 9.54e−06 **** | 9.54e−06 **** |

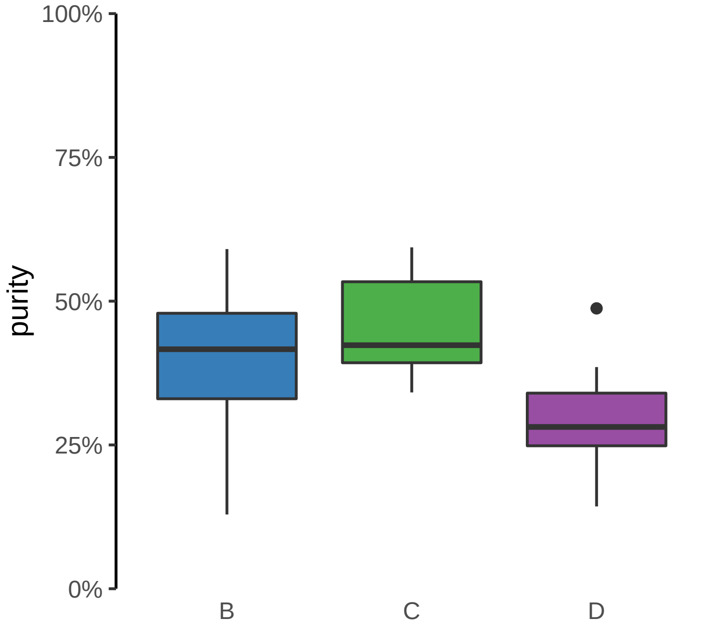

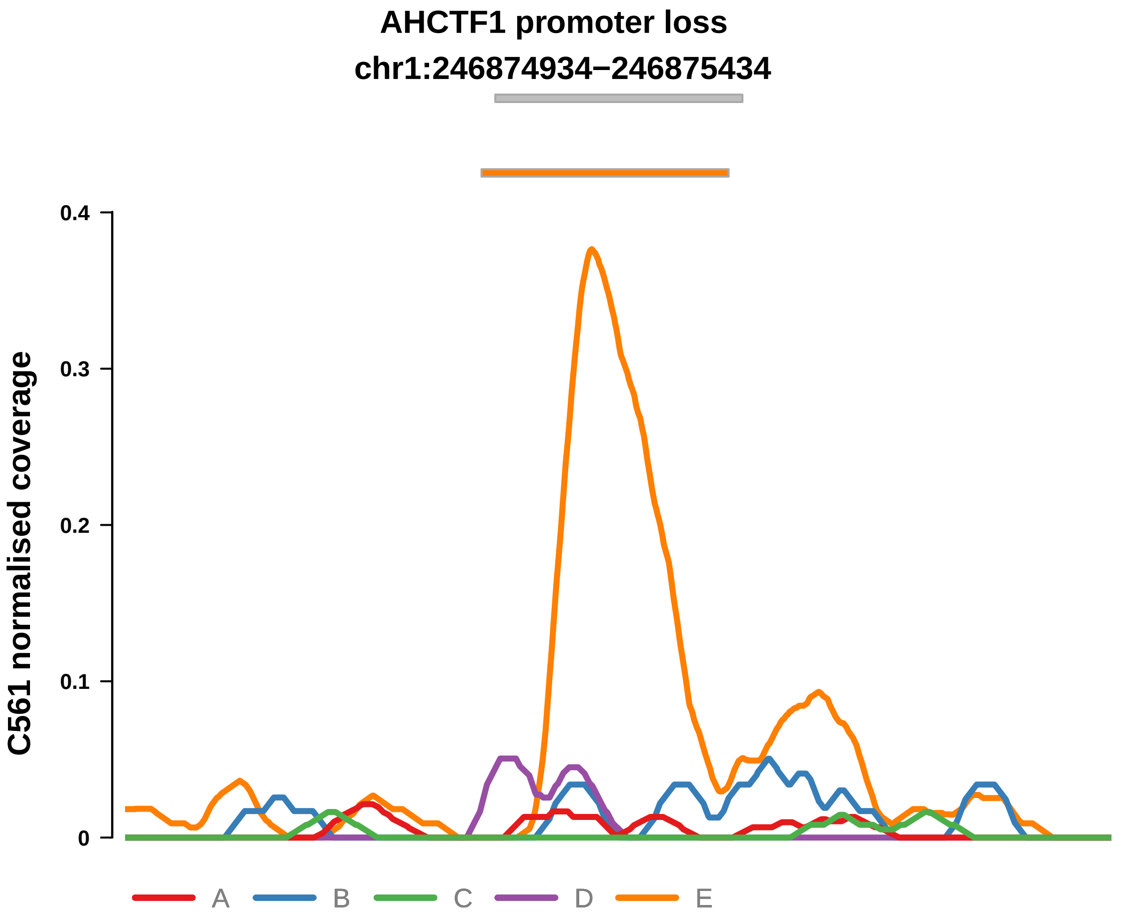


Putatively Clonal (logFC = −2.64)

| | P. | Adjusted P. |
|---|---|---|
| *~region vs. ~* | 1.5e−01 | NA |
| *~purity vs ~* | 4.47e−01 | 7.66e−01 |
| *~purity+region vs ~purity* | 2.8e−01 | 7.68e−01 |
| *Carcinoma vs. normal* | 1.29e−05 **** | 1.29e−05 **** |

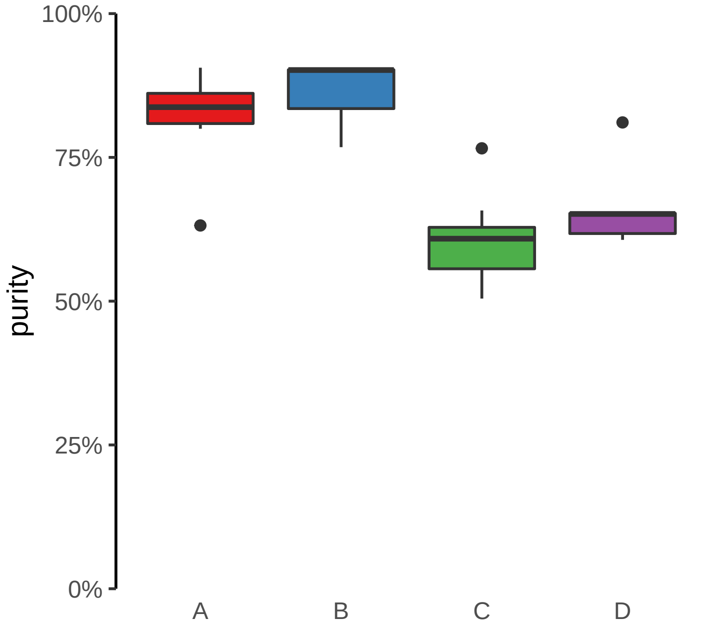

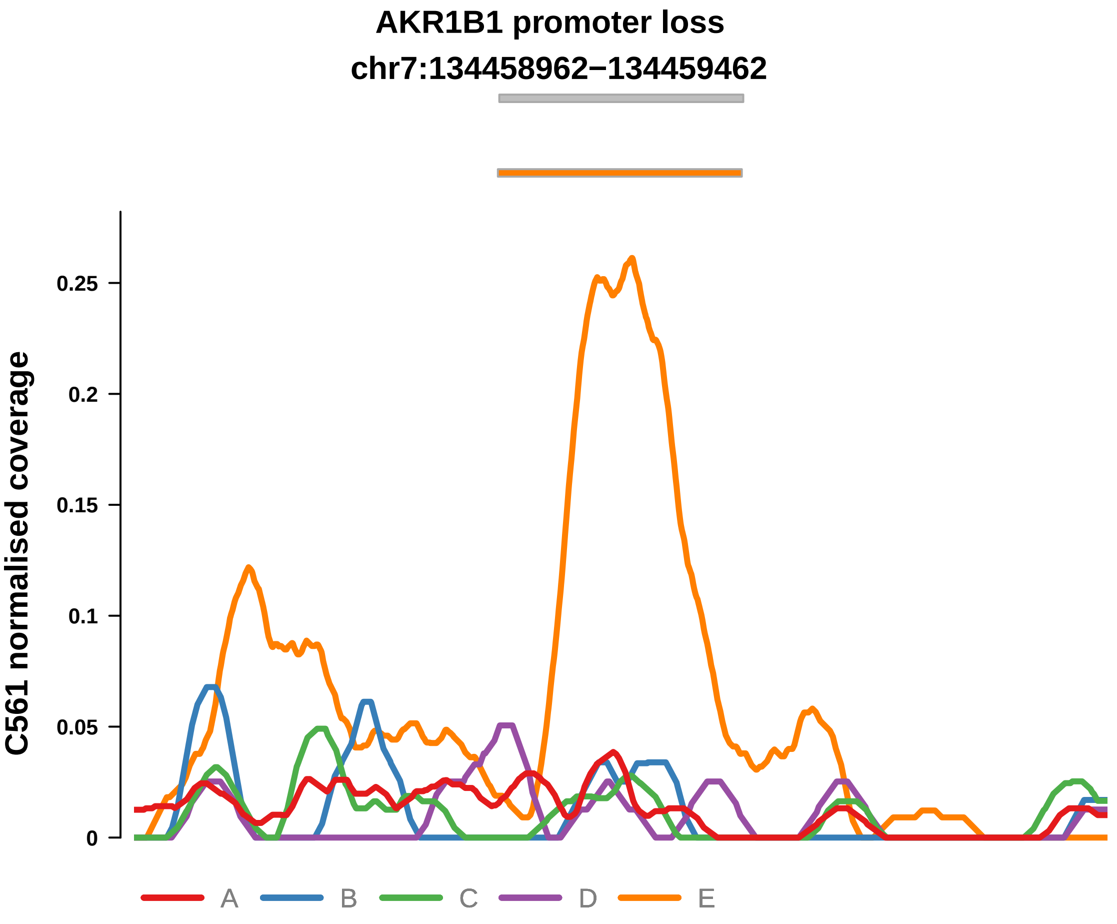


Putatively Clonal (logFC = −2.79)

| | P. | Adjusted P. |
|---|---|---|
| *~region vs. ~* | 5.14e−01 | 6.77e−01 |
| *~purity vs ~* | 7.92e−01 | 9.05e−01 |
| *~purity+region vs ~purity* | 5.49e−01 | 9.16e−01 |
| *Carcinoma vs. normal* | 2.58e−06 **** | 2.58e−06 **** |

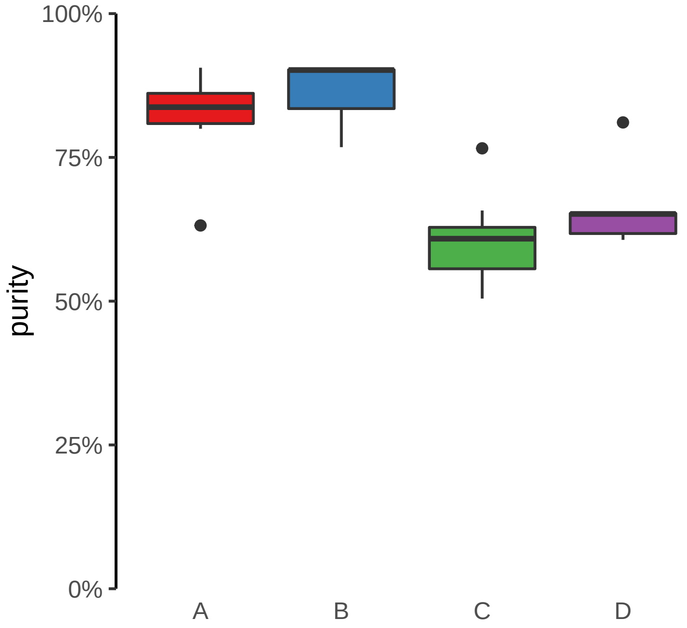

**CCDC6 promoter loss**
**chr10:59805006−59805506**

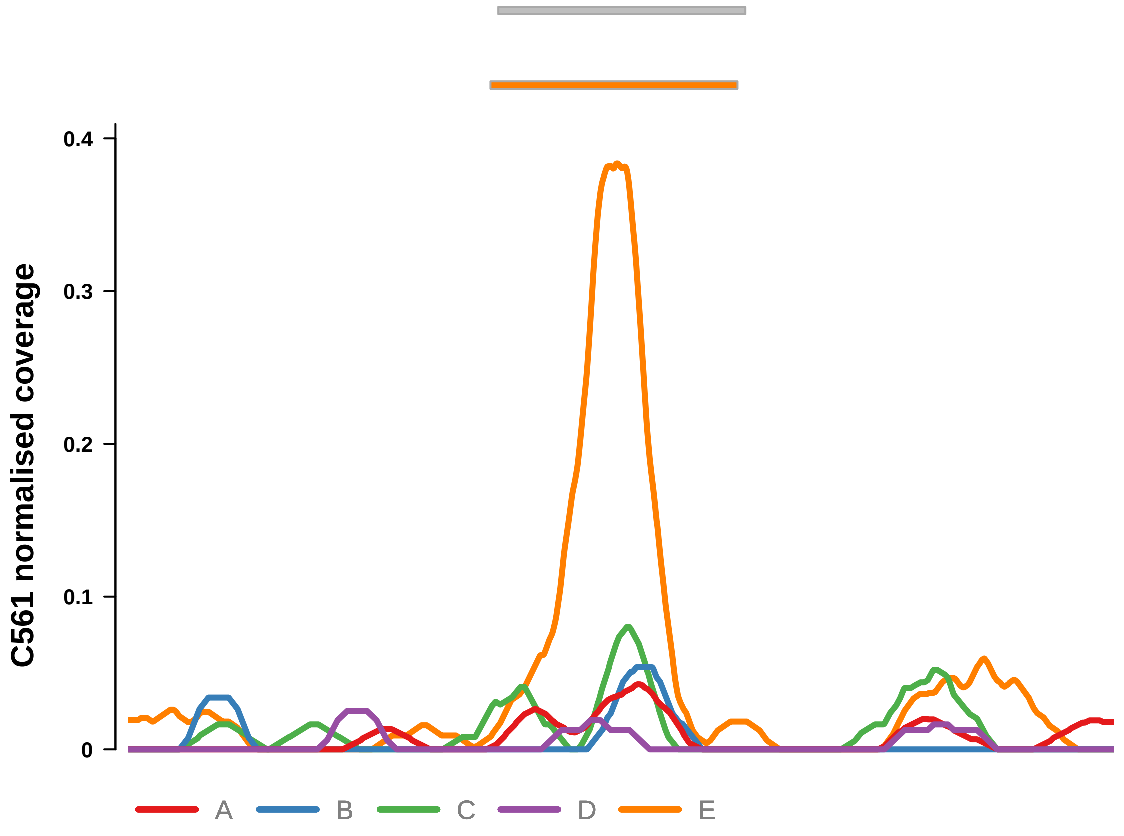


Putatively Clonal (logFC = −1.5)

| | P. | Adjusted P. |
|---|---|---|
| *~region vs. ~* | 3.94e−01 | 5.91e−01 |
| *~purity vs ~* | 4.42e−02 * | 3.24e−01 |
| *~purity+region vs ~purity* | 7.03e−01 | 9.16e−01 |
| *Carcinoma vs. normal* | 8.11e−03 ** | 8.11e−03 ** |

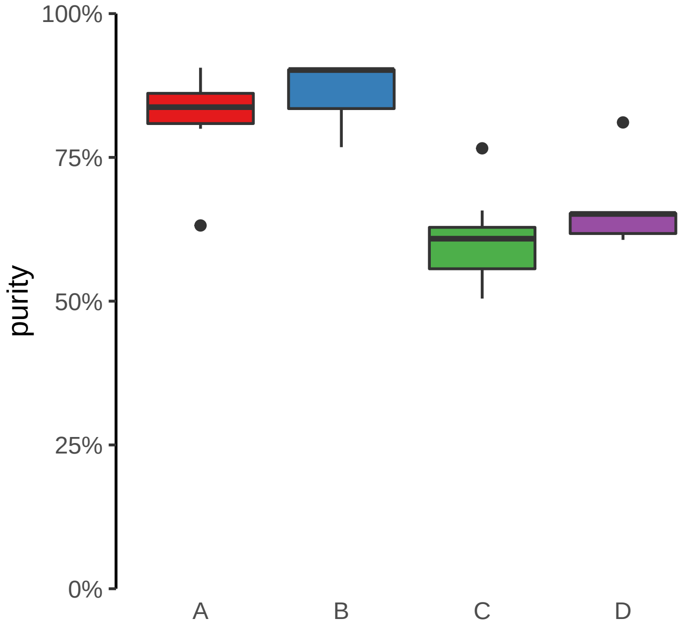

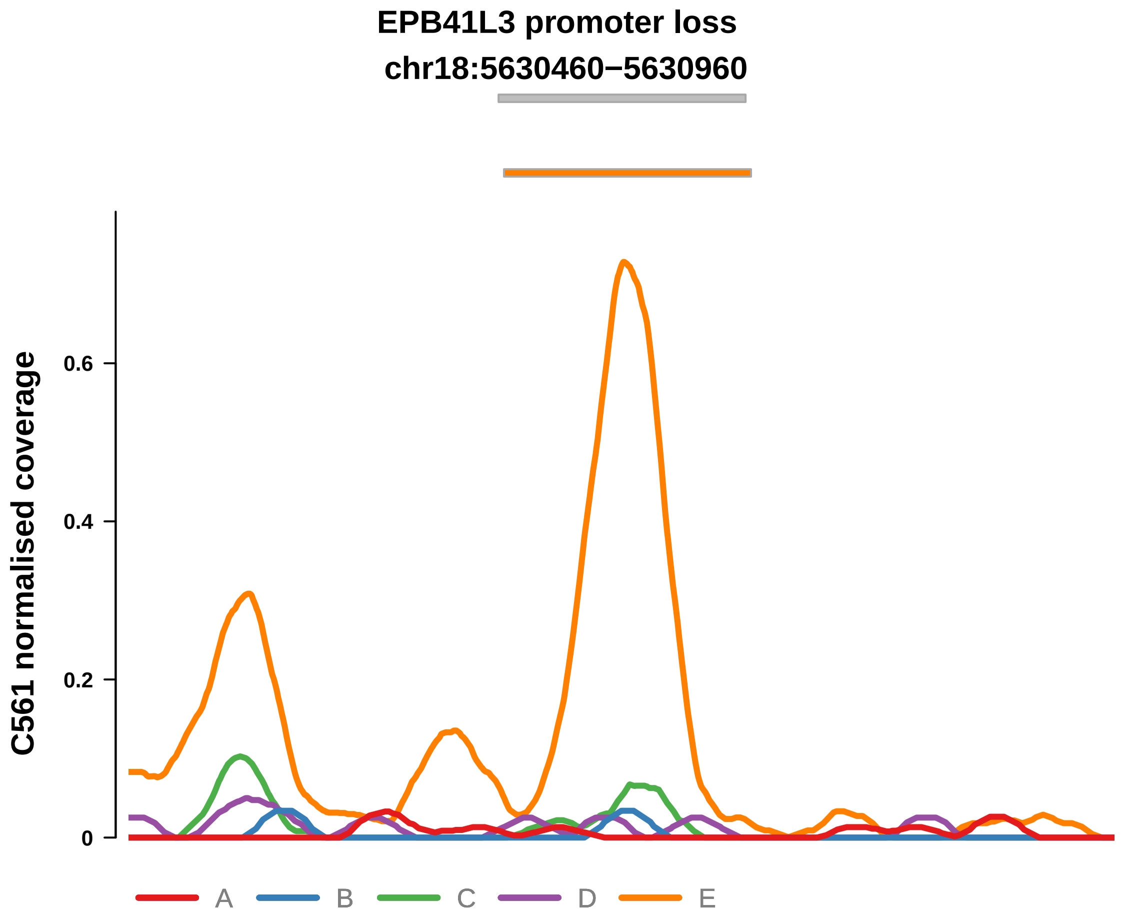


Putatively Clonal (logFC = −3.5)

| | P. | Adjusted P. |
|---|---|---|
| *~region vs. ~* | 4.85e−01 | 6.49e−01 |
| *~purity vs ~* | 1.74e−01 | 5.28e−01 |
| *~purity+region vs ~purity* | 8.78e−01 | 9.76e−01 |
| *Carcinoma vs. normal* | 1.13e−08 **** | 1.13e−08 **** |

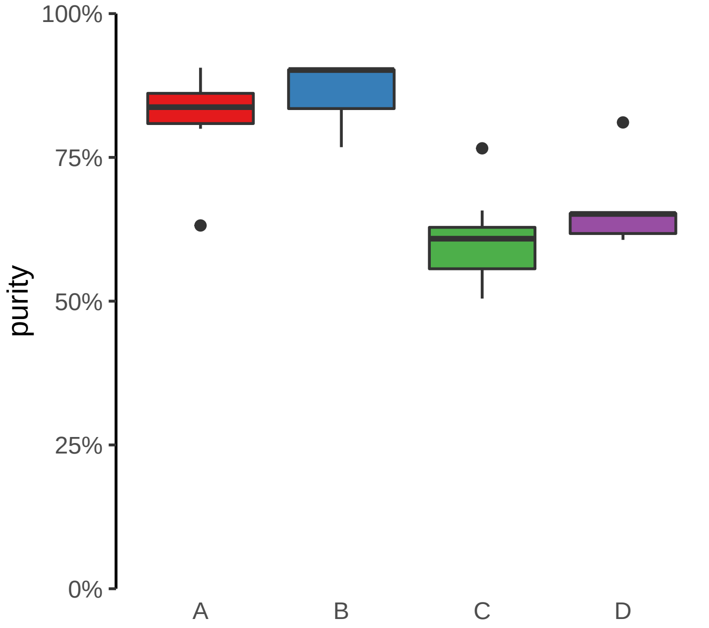

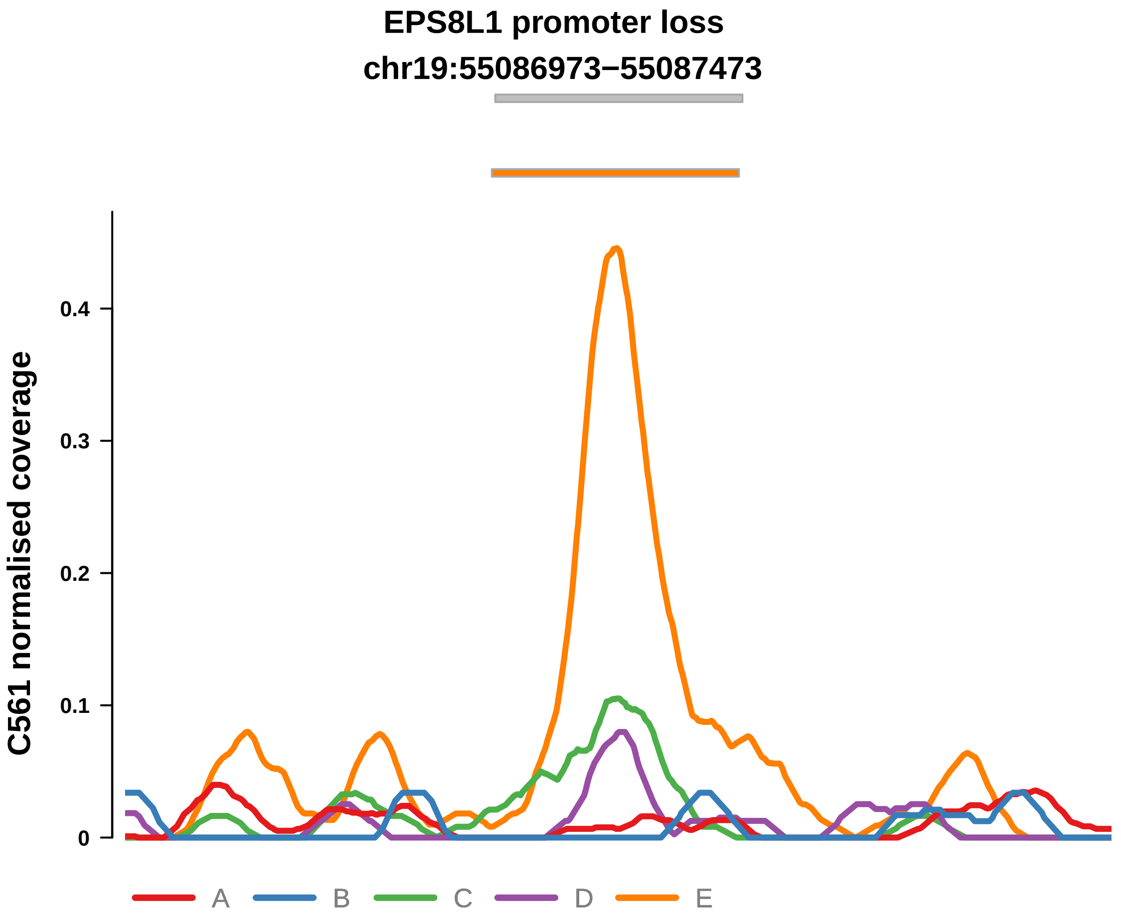


Putatively Clonal (logFC = −2.03)

| | P. | Adjusted P. |
|---|---|---|
| *~region vs. ~* | 1.56e−03 ** | 3.09e−02 * |
| *~purity vs ~* | 1.79e−04 *** | 5.26e−03 ** |
| *~purity+region vs ~purity* | 8.77e−01 | 9.76e−01 |
| *Carcinoma vs. normal* | 3.8e−04 *** | 3.8e−04 *** |

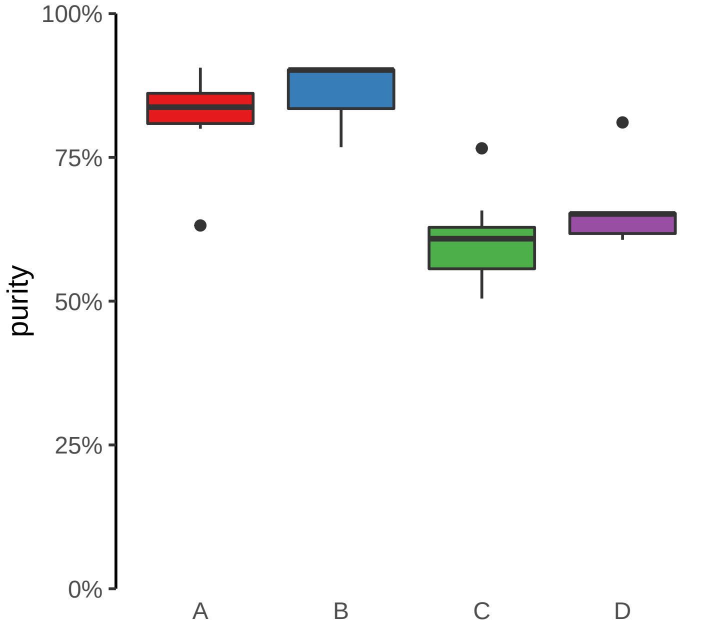

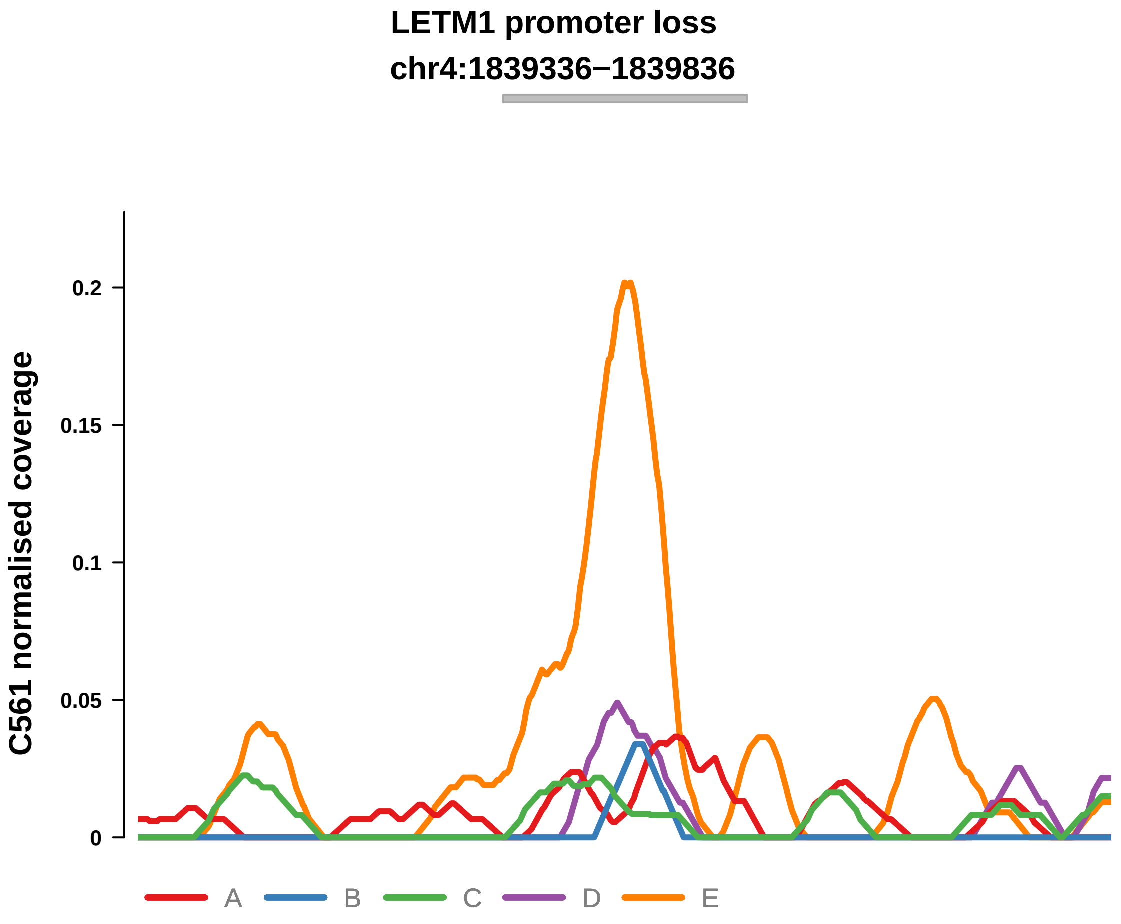


Putatively Clonal (logFC = −1.89)

| | P. | Adjusted P. |
|---|---|---|
| *~region vs. ~* | 8.45e−01 | 9.14e−01 |
| *~purity vs ~* | 6.13e−01 | 8.53e−01 |
| *~purity+region vs ~purity* | 9.1e−01 | 9.76e−01 |
| *Carcinoma vs. normal* | 1.21e−03 ** | 1.21e−03 ** |

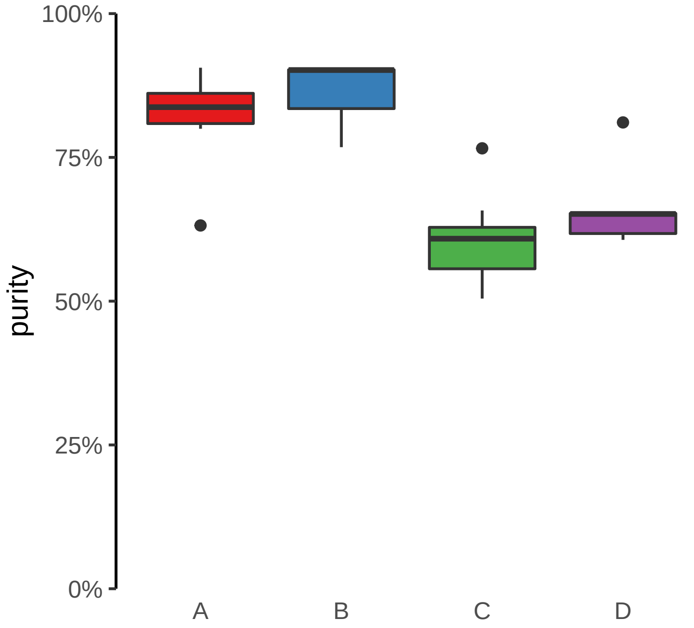

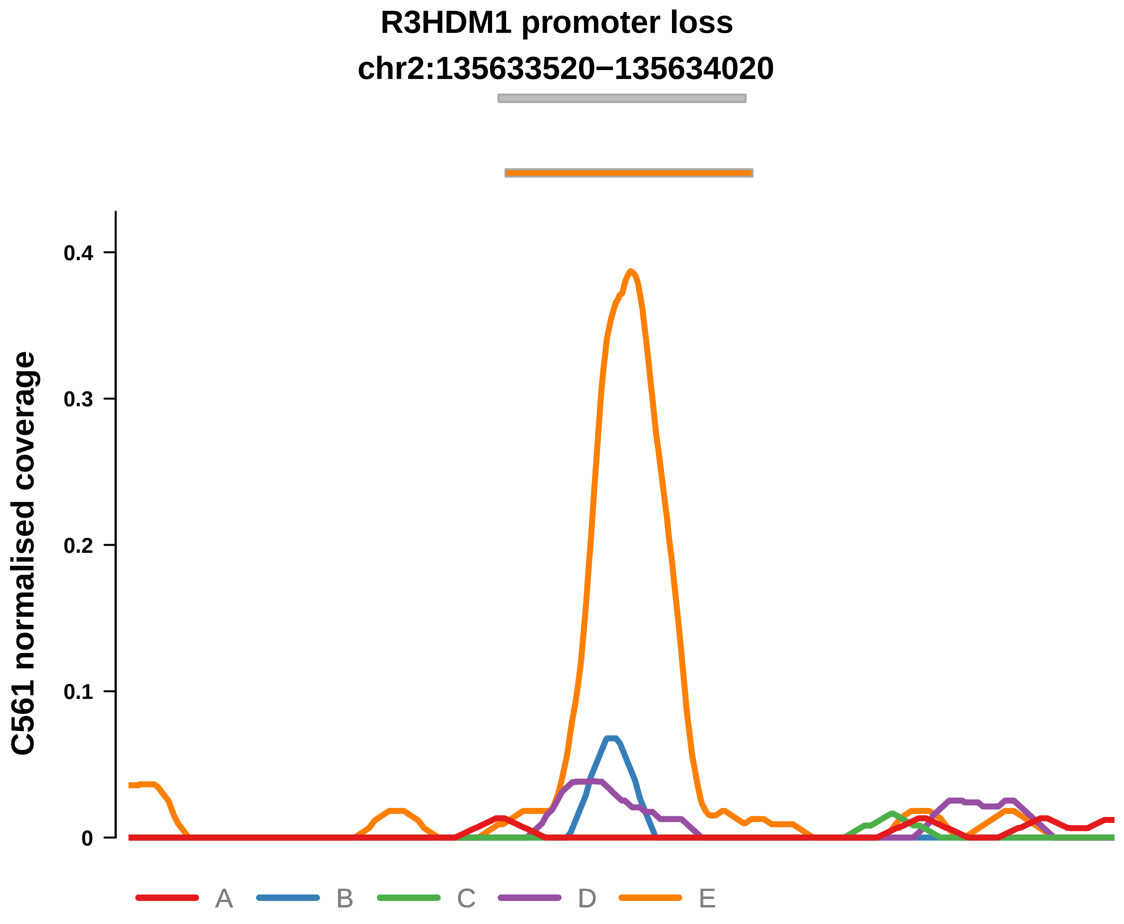


Putatively Clonal (logFC = −2.72)

| | P. | Adjusted P. |
| --- | --- | --- |
| *~region vs. ~* | 6.88e−01 | NA |
| *~purity vs ~* | 7.28e−01 | 8.81e−01 |
| *~purity+region vs ~purity* | 7.6e−01 | 9.16e−01 |
| *Carcinoma vs. normal* | 1.42e−05 **** | 1.42e−05 **** |

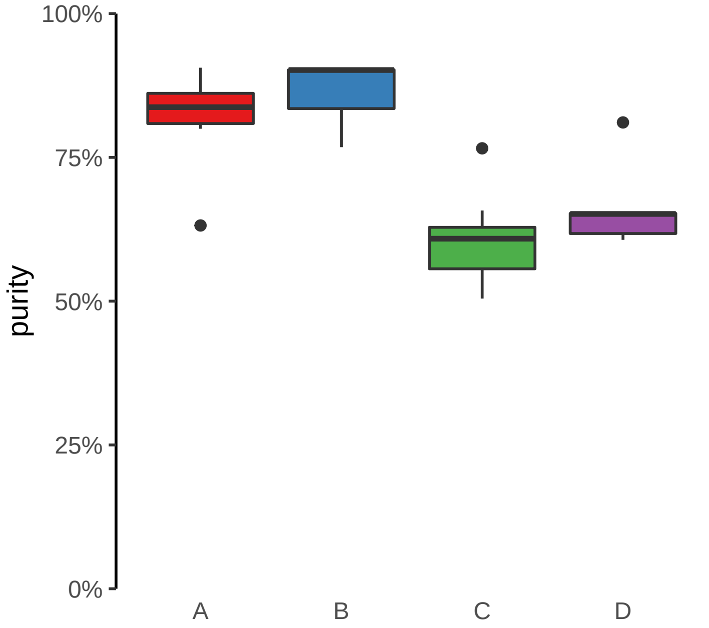

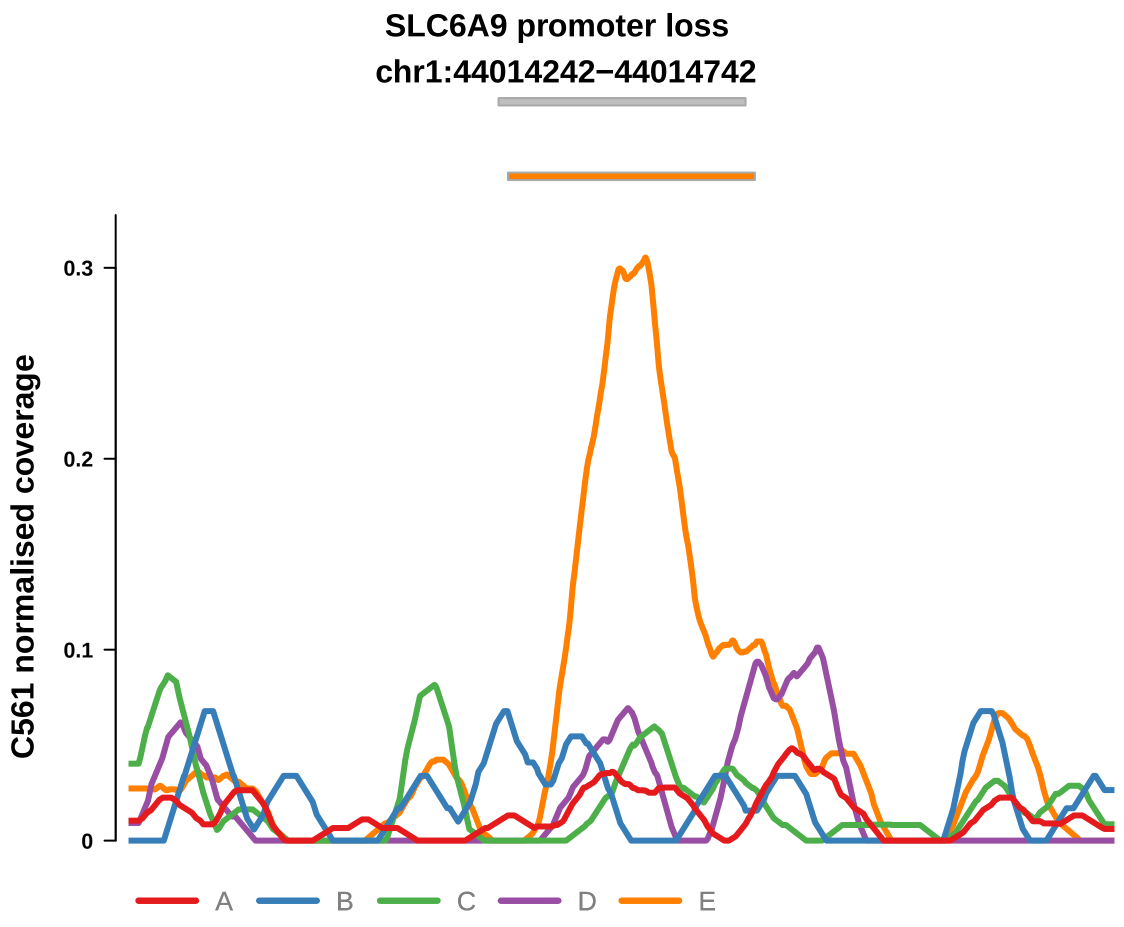


Putatively Clonal (logFC = −1.52)

| | P. | Adjusted P. |
|---|---|---|
| *~region vs. ~* | 5.42e−01 | 6.91e−01 |
| *~purity vs ~* | 7.56e−02 | 3.98e−01 |
| *~purity+region vs ~purity* | 3.17e−01 | 7.68e−01 |
| *Carcinoma vs. normal* | 6.95e−03 ** | 6.95e−03 ** |

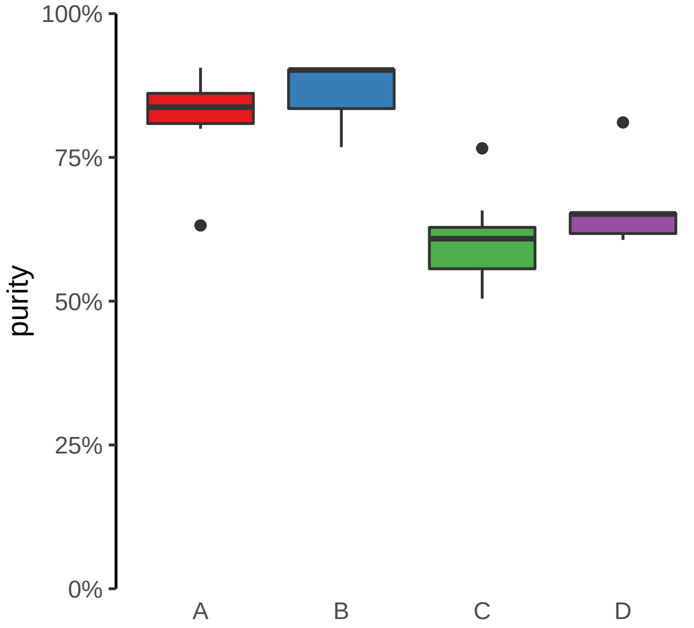

**SMAD4 promoter loss**
**chr18:51051052−51051552**

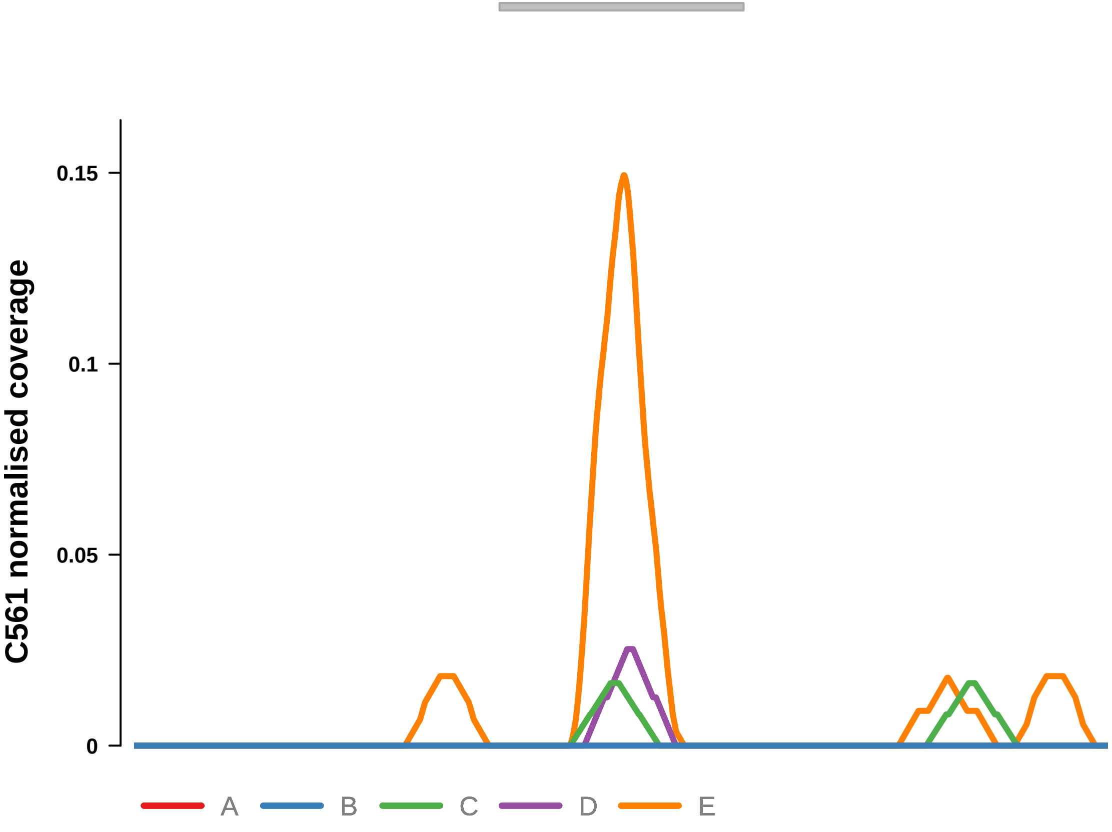


Putatively Clonal (logFC = −4.42)

| | P. | Adjusted P. |
|---|---|---|
| *~region vs. ~* | 8.86e−01 | NA |
| *~purity vs ~* | 4.15e−01 | 7.66e−01 |
| *~purity+region vs ~purity* | 9.67e−01 | 9.79e−01 |
| *Carcinoma vs. normal* | 5.69e−09 **** | 5.69e−09 **** |

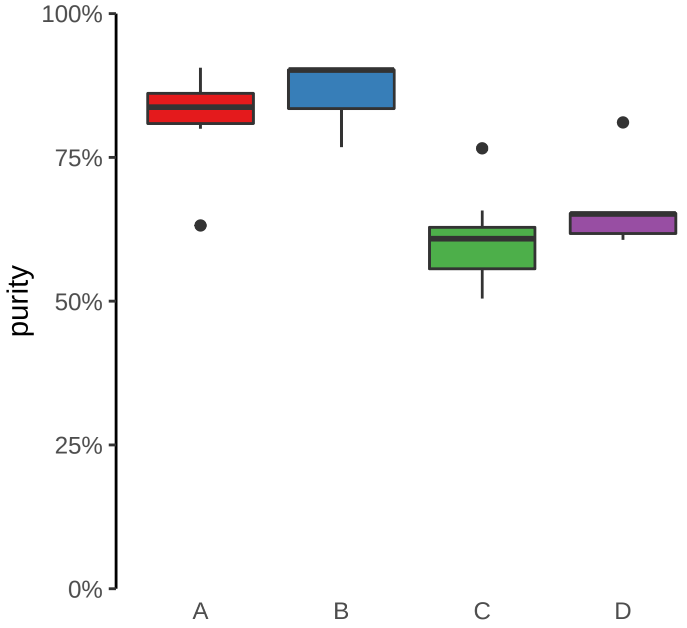

**SMARCA2 promoter loss**
**chr9:2053751−2054251**

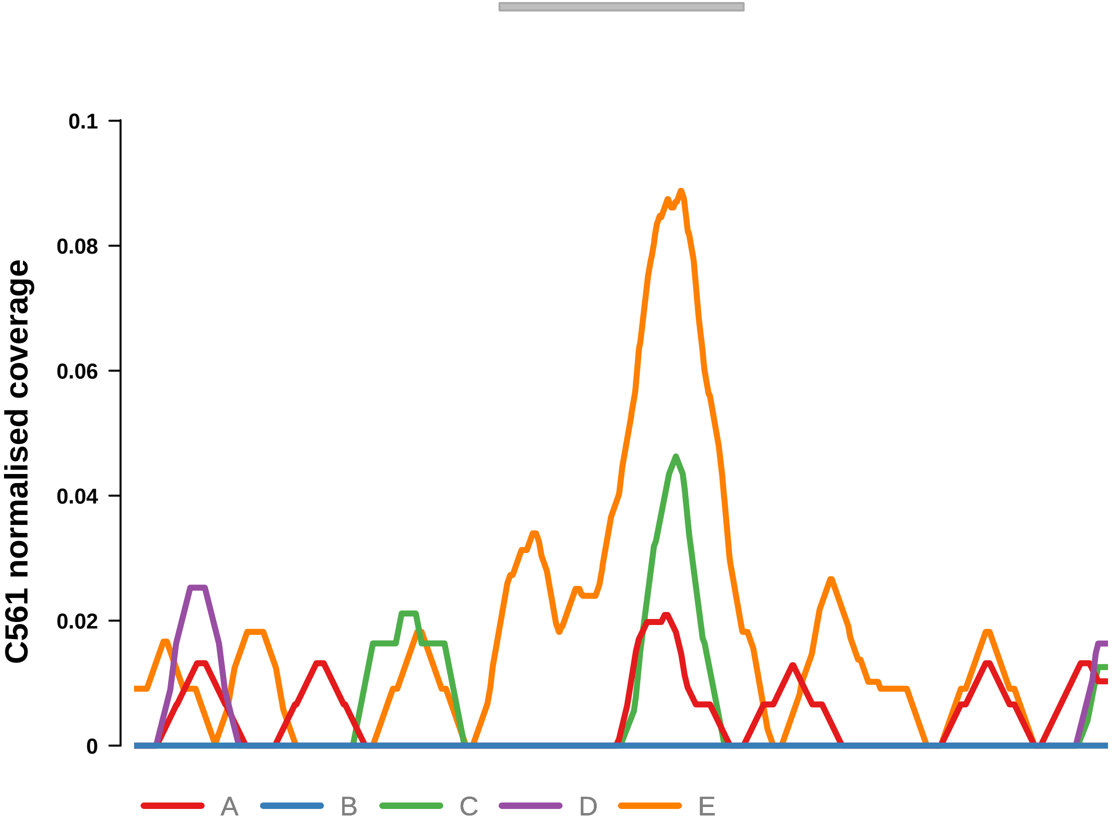


Putatively Clonal (logFC = −2.88)

| | P. | Adjusted P. |
|---|---|---|
| *~region vs. ~* | 8.64e−01 | NA |
| *~purity vs ~* | 3.89e−01 | 7.66e−01 |
| *~purity+region vs ~purity* | 9.47e−01 | 9.79e−01 |
| *Carcinoma vs. normal* | 1.52e−05 **** | 1.52e−05 **** |

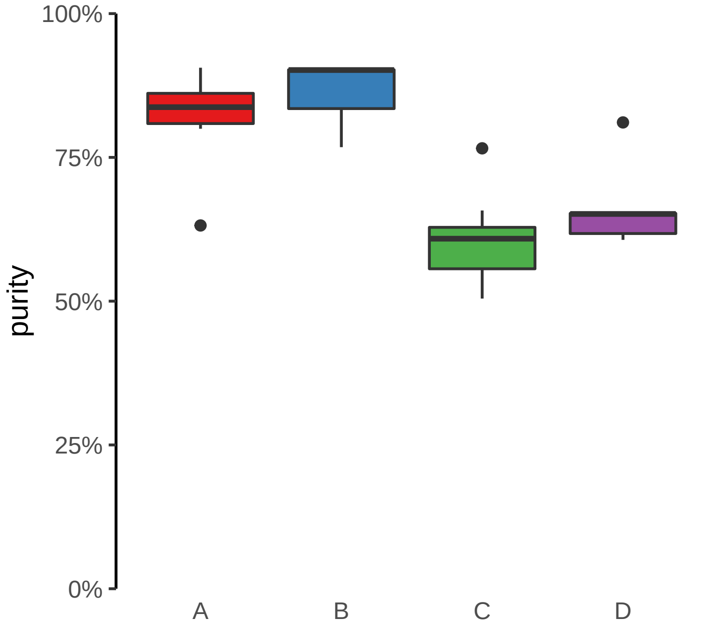

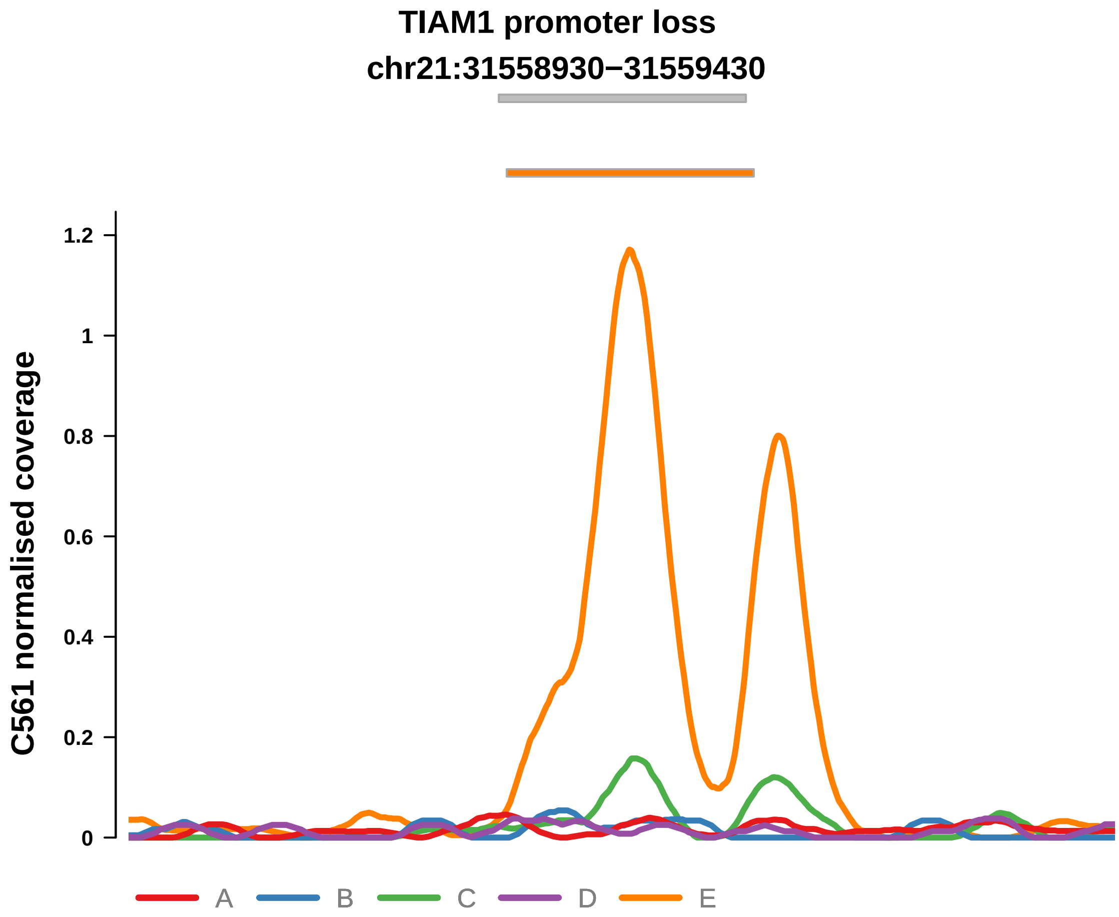


Putatively Clonal (logFC = −2.18)

| | P. | Adjusted P. |
|---|---|---|
| *~region vs. ~* | 1.26e−02 * | 8.23e−02 |
| *~purity vs ~* | 1.33e−02 * | 1.46e−01 |
| *~purity+region vs ~purity* | 3.84e−01 | 7.68e−01 |
| *Carcinoma vs. normal* | 9.4e−05 **** | 9.4e−05 **** |

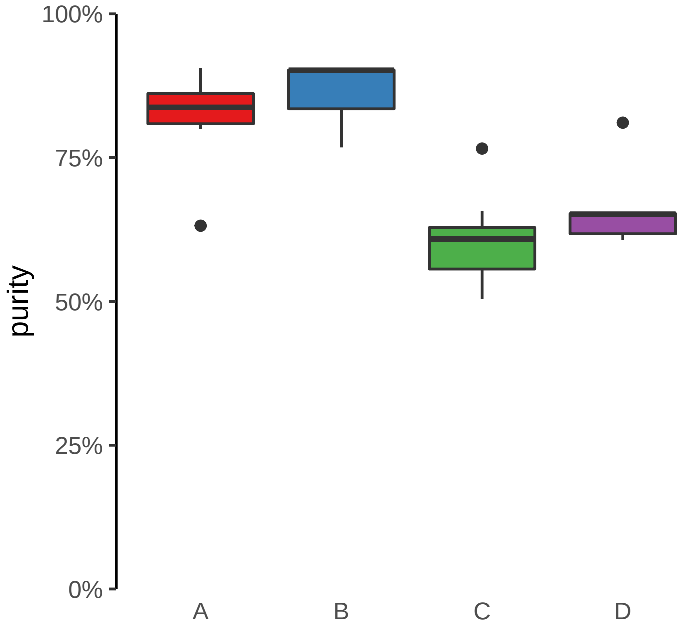

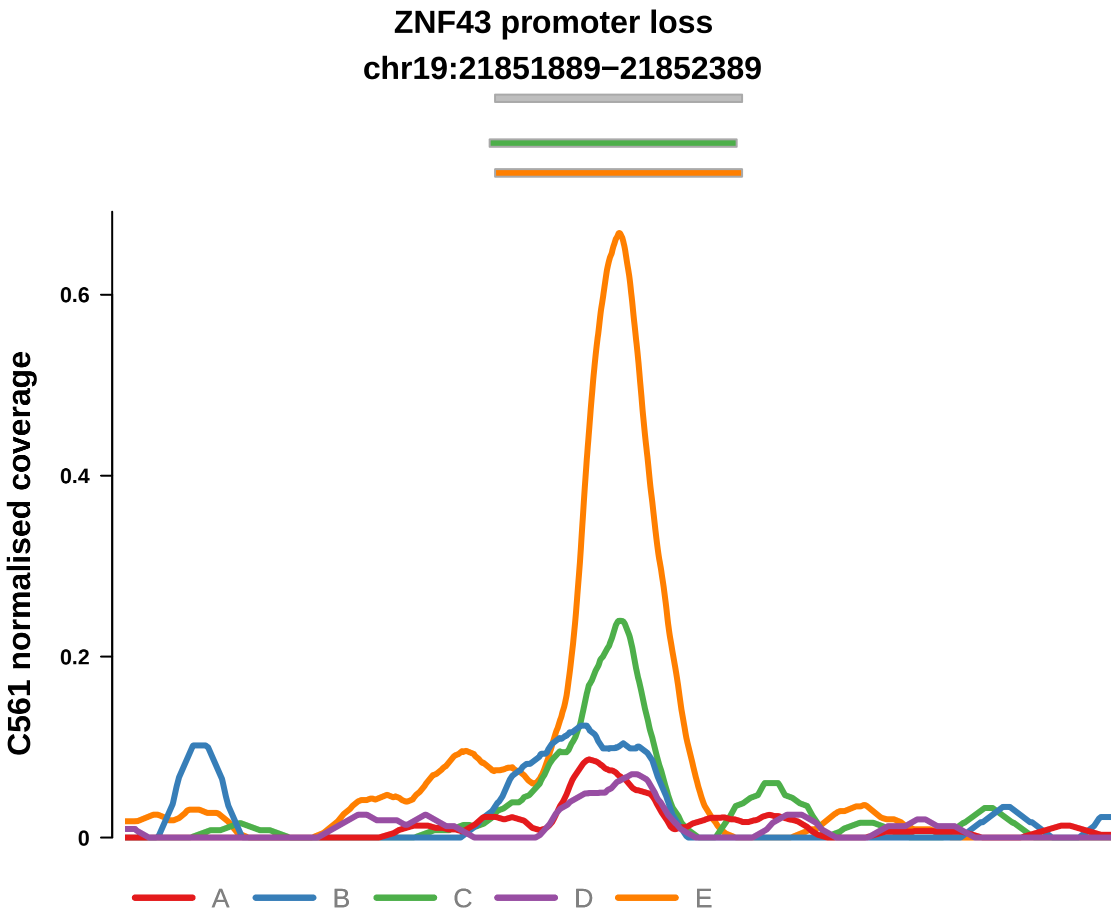


Putatively Clonal (logFC = −1.58)

| | P. | Adjusted P. |
|---|---|---|
| *~region vs. ~* | 6.69e−03 ** | 5.52e−02 |
| *~purity vs ~* | 1.11e−01 | 4.2e−01 |
| *~purity+region vs ~purity* | 3.51e−02 * | 3.43e−01 |
| *Carcinoma vs. normal* | 3.49e−03 ** | 3.49e−03 ** |

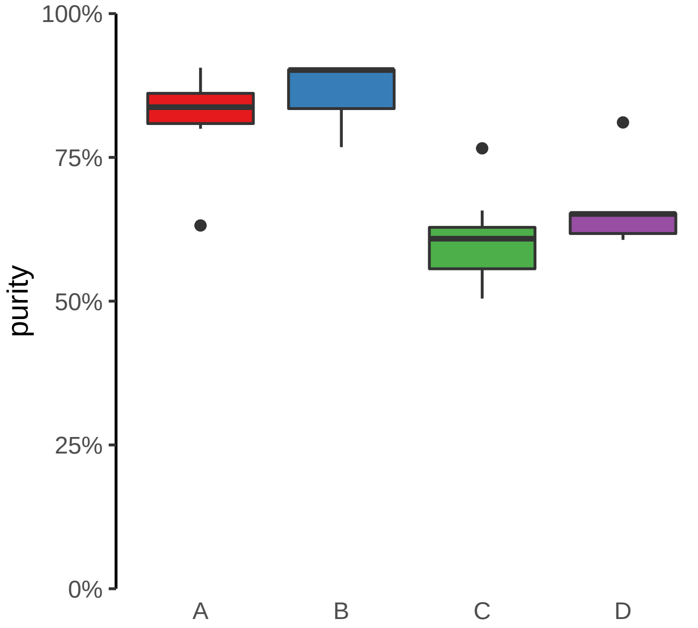

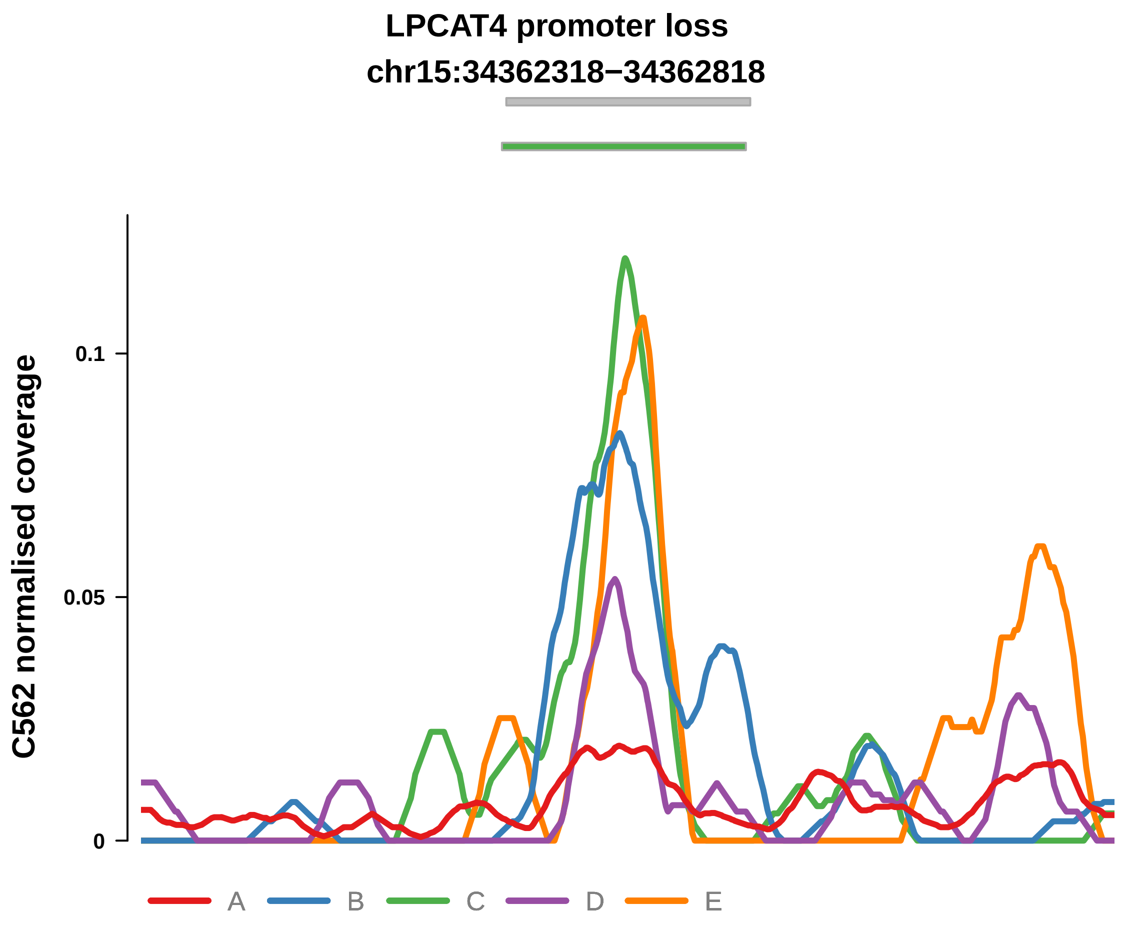


Putatively Clonal (logFC = −1.6)

| | P. | Adjusted P. |
|---|---|---|
| *~region vs. ~* | 3.98e−04 *** | 1.75e−02 * |
| *~purity vs ~* | 6.27e−02 | 1.78e−01 |
| *~purity+region vs ~purity* | 7.57e−04 *** | 5.3e−02 |
| *Carcinoma vs. normal* | 9.39e−03 ** | 9.39e−03 ** |

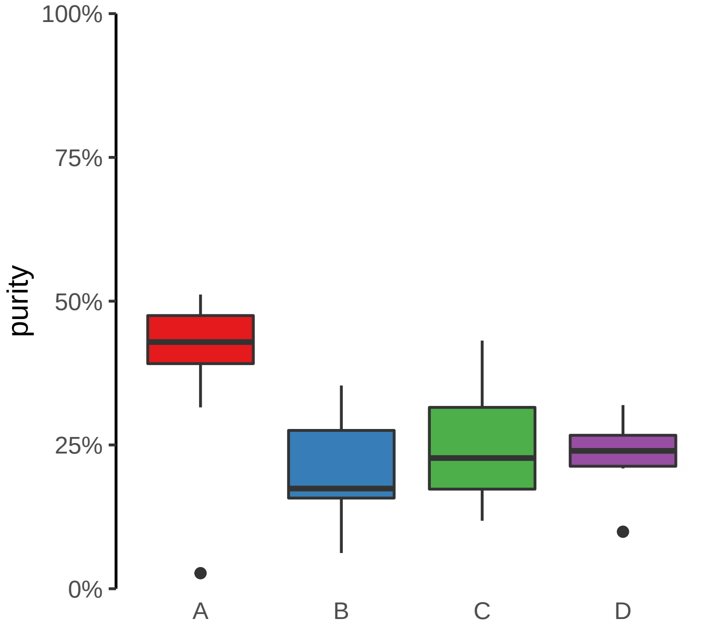

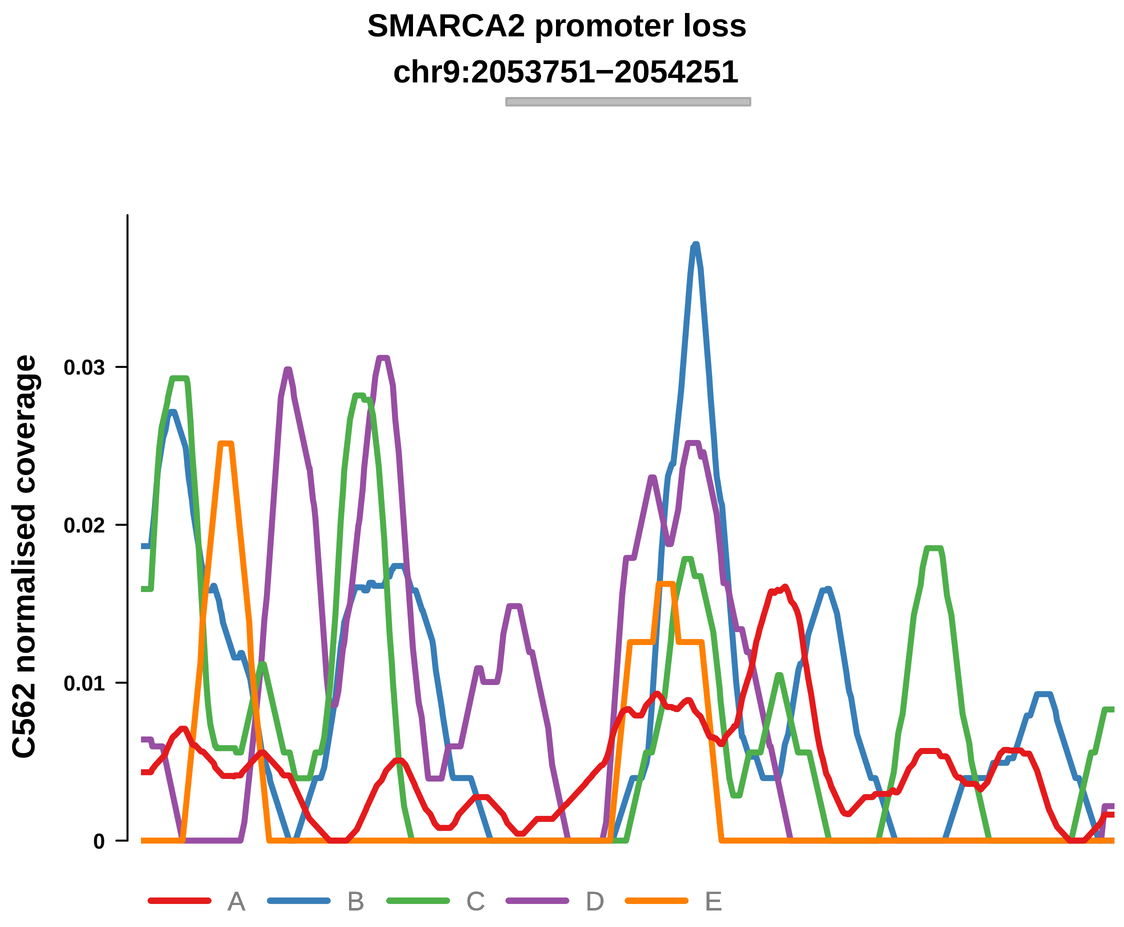


Putatively Clonal (logFC = −2.12)

| | P. | Adjusted P. |
|---|---|---|
| *~region vs. ~* | 1.81e−01 | 5.14e−01 |
| *~purity vs ~* | 7.9e−01 | 9.58e−01 |
| *~purity+region vs ~purity* | 1.83e−01 | 5.73e−01 |
| *Carcinoma vs. normal* | 5.04e−03 ** | 5.04e−03 ** |

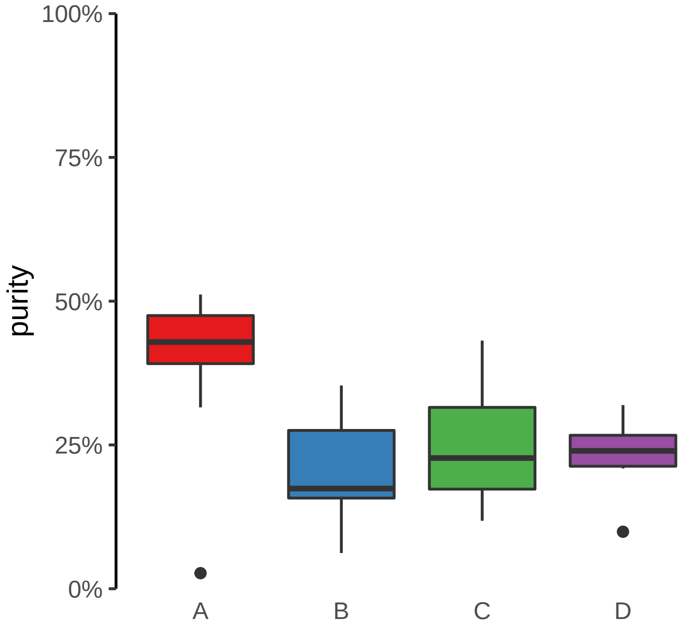

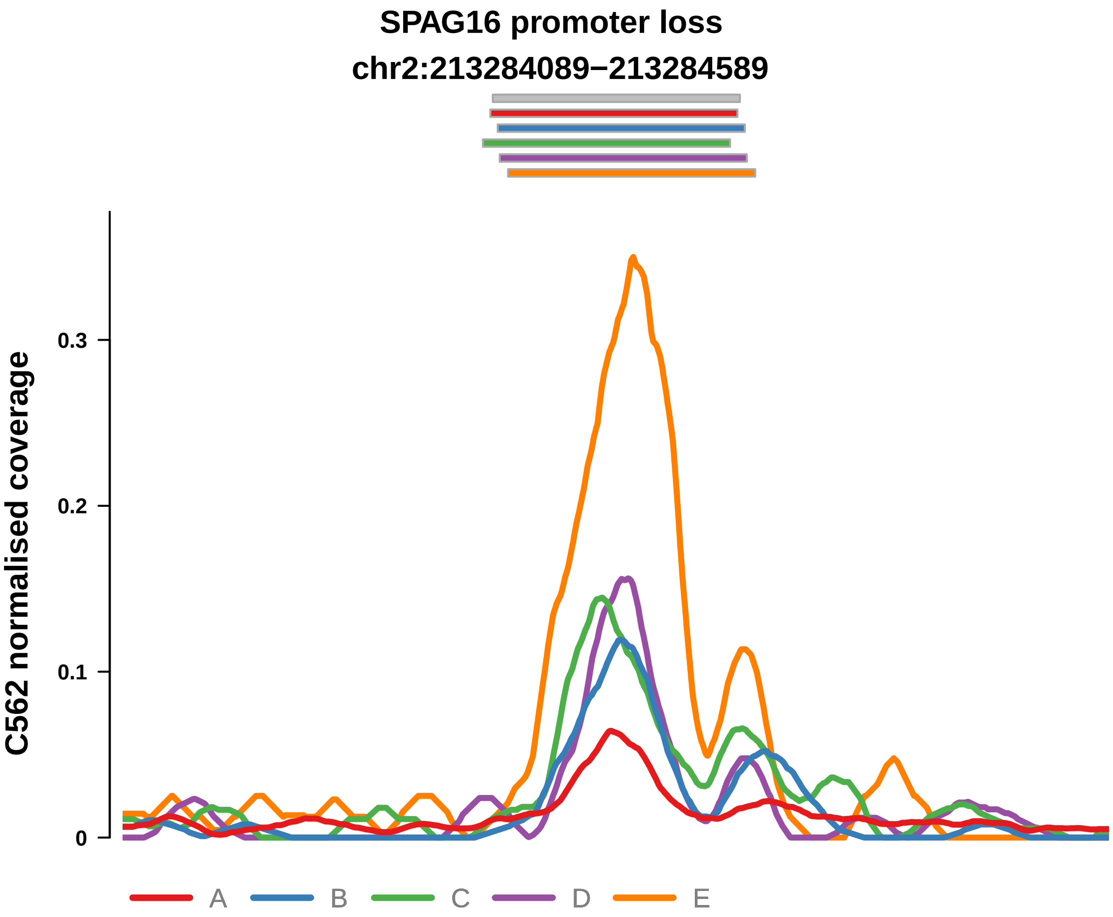


Putatively Clonal (logFC = −1.66)

| | P. | Adjusted P. |
|---|---|---|
| *~region vs. ~* | 4.25e−02 * | 2.87e−01 |
| *~purity vs ~* | 5.85e−02 | 1.71e−01 |
| *~purity+region vs ~purity* | 7.8e−02 | 4.58e−01 |
| *Carcinoma vs. normal* | 3.47e−03 ** | 3.47e−03 ** |

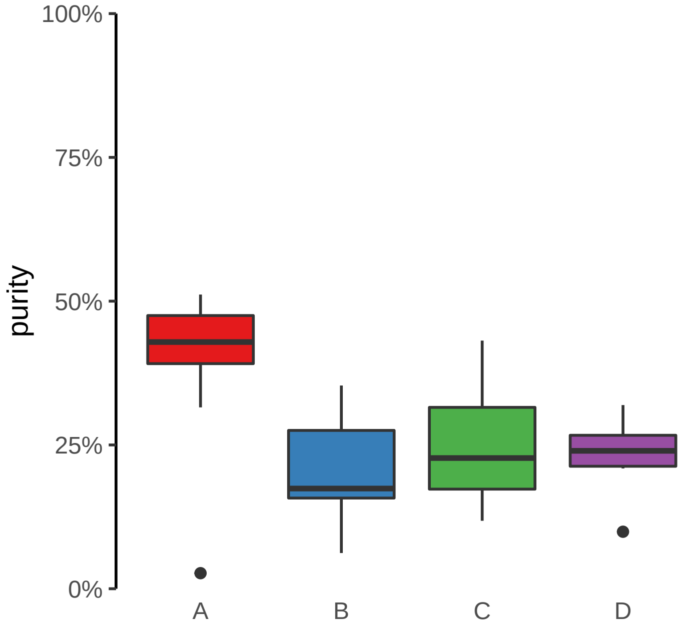
