## Supplementary Figures for "The co-evolution of the genome and epigenome in colorectal cancer": figS12_enhancer_gain.pdf

**TPRA1, ABTB1, MGLL (GH03J127766) enhancer gain**

**chr3:127767157−127767657**

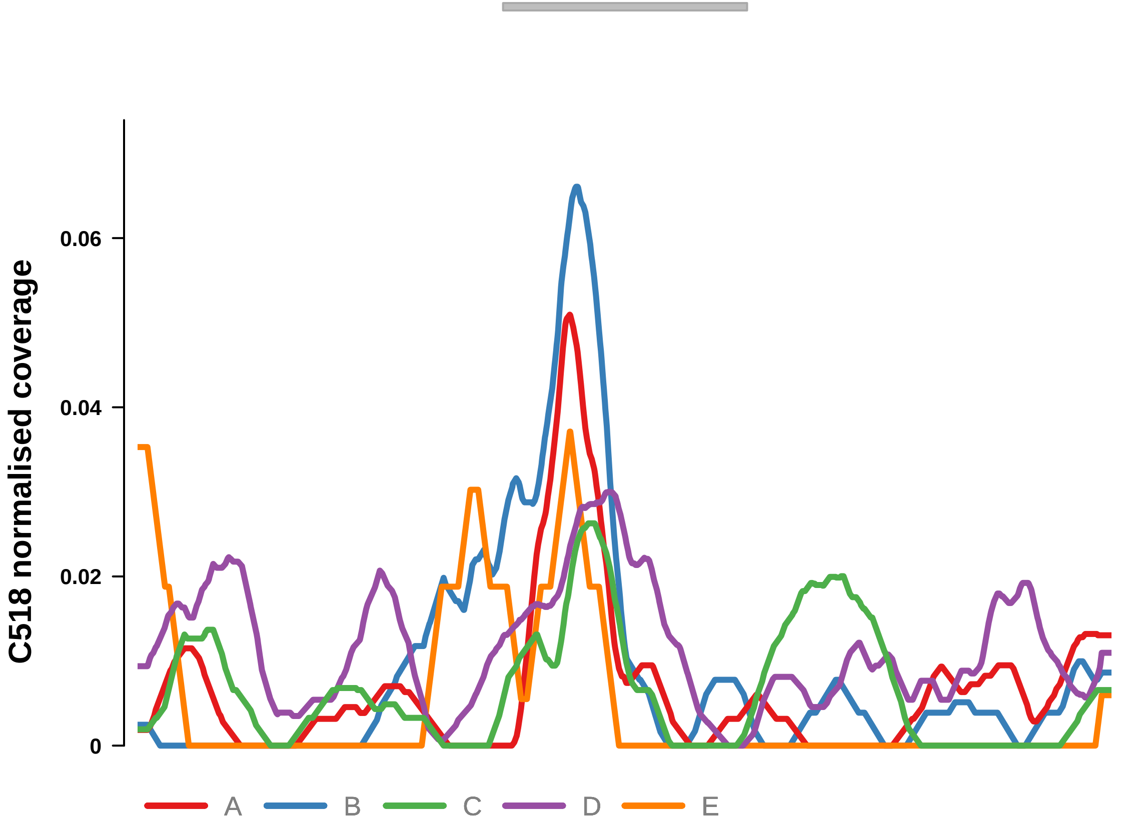


Putatively Clonal (logFC = 1.99)

| | P. | Adjusted P. |
|---|---|---|
| *~region vs. ~* | 3.93e−01 | 6.53e−01 |
| *~purity vs ~* | 9.99e−01 | 9.99e−01 |
| *~purity+region vs ~purity* | 4e−01 | 7.49e−01 |
| *Carcinoma vs. normal* | 8.54e−03 ** | 8.54e−03 ** |

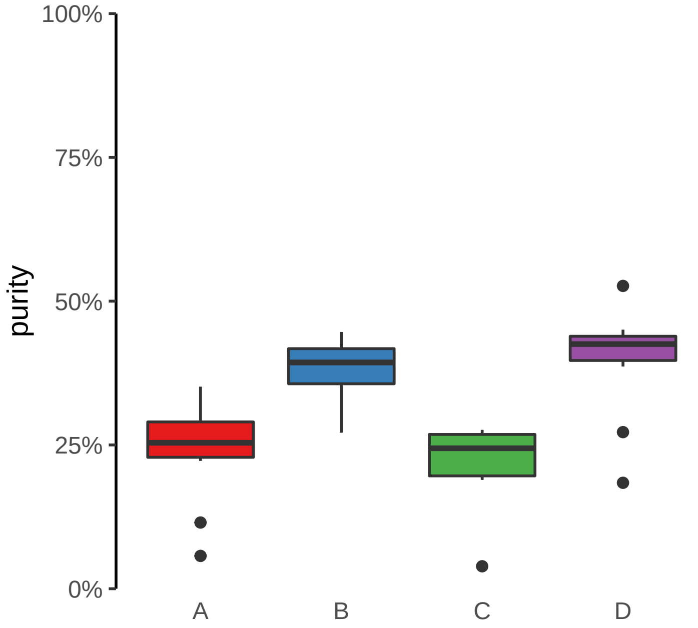

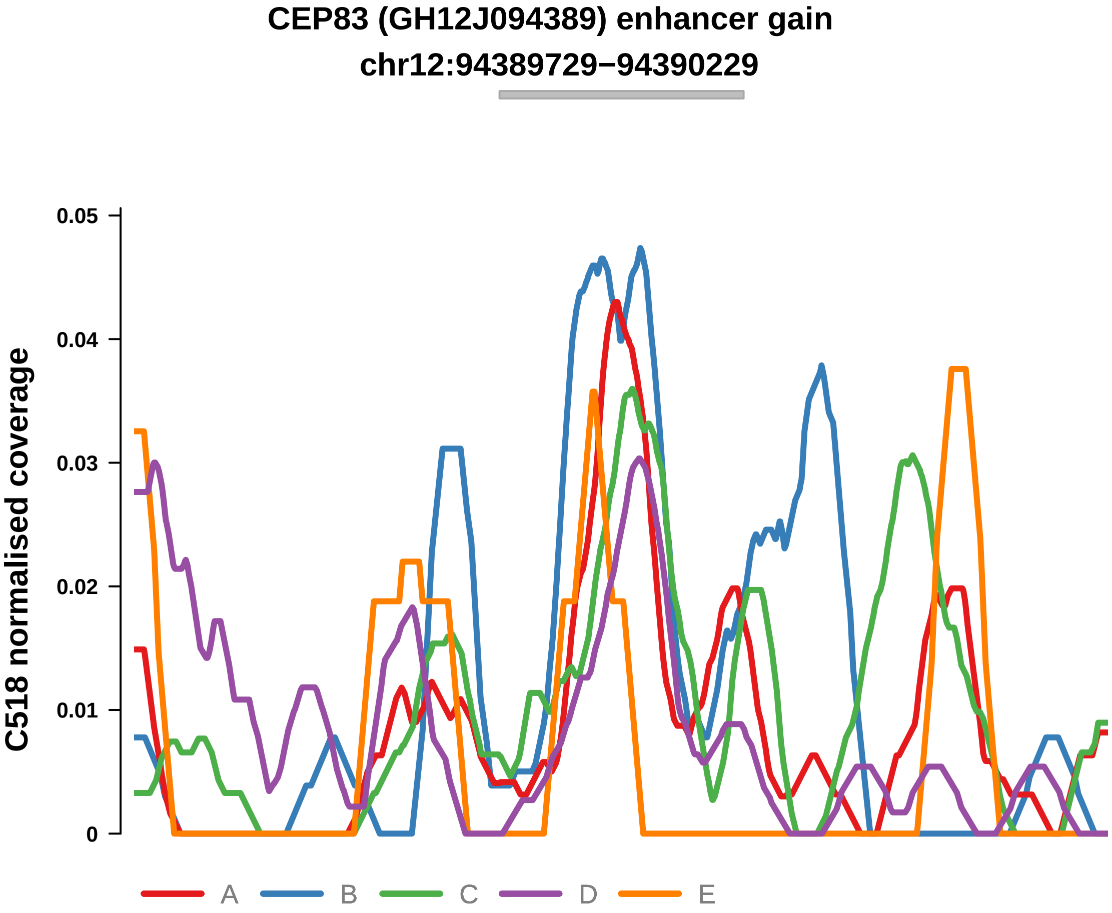


Putatively Clonal (logFC = 2.3)

| | P. | Adjusted P. |
|---|---|---|
| *~region vs. ~* | 2.35e−01 | 6.27e−01 |
| *~purity vs ~* | 2.88e−01 | 7.38e−01 |
| *~purity+region vs ~purity* | 3.85e−01 | 7.49e−01 |
| *Carcinoma vs. normal* | 2.58e−03 ** | 2.58e−03 ** |

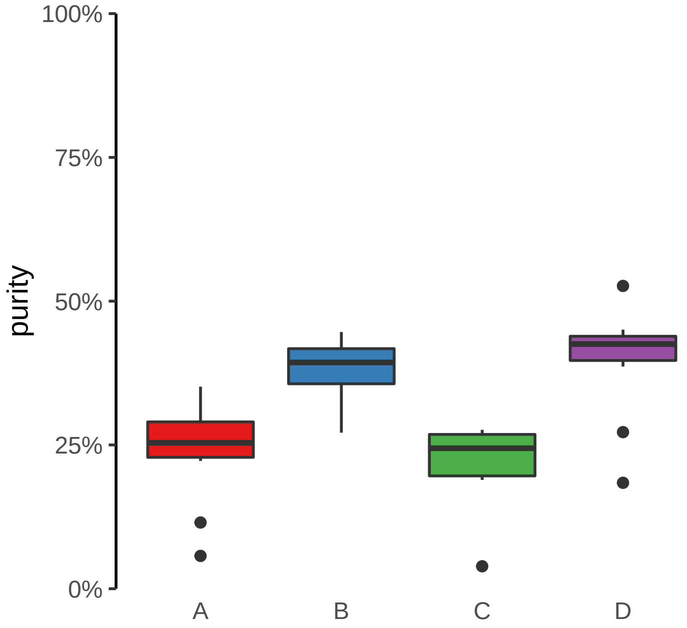

**CLASRP, MARK4 (GH19J045307) enhancer gain**
**chr19:45308035−45308535**

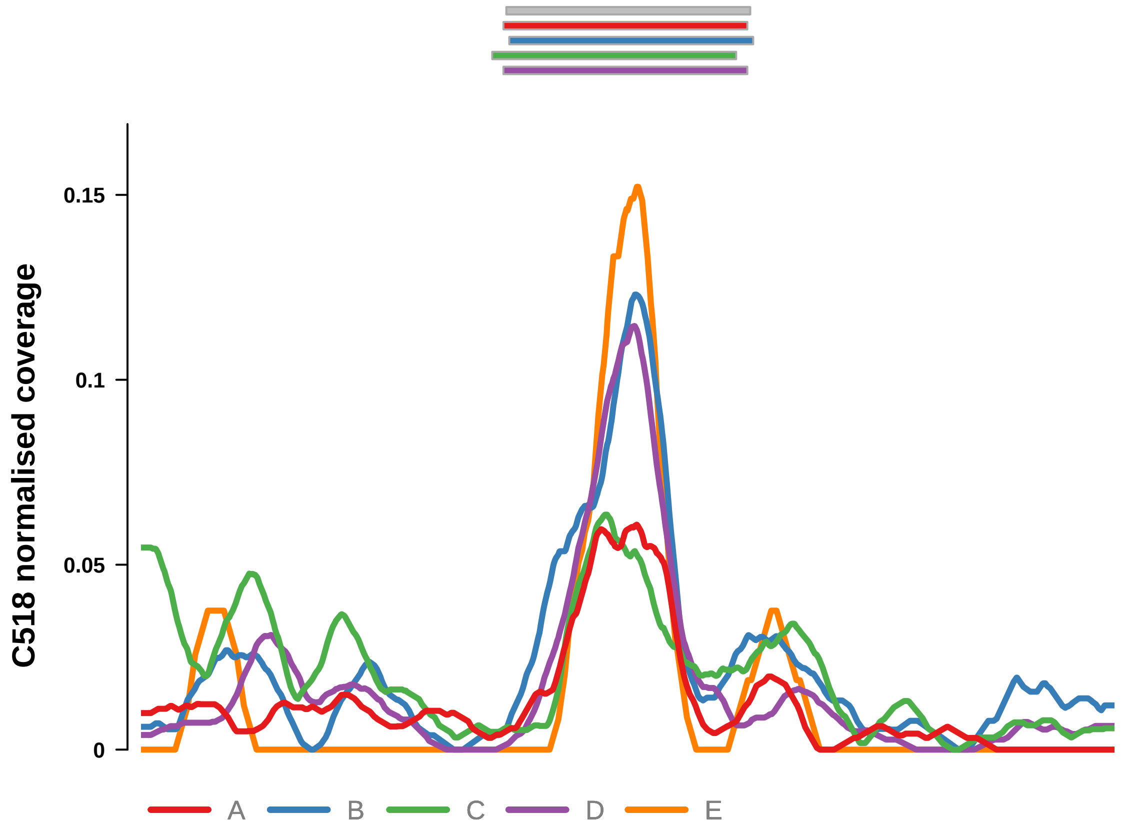


Putatively Clonal (logFC = 2.25)

| | P. | Adjusted P. |
|---|---|---|
| *~region vs. ~* | 6.89e−01 | 8.17e−01 |
| *~purity vs ~* | 8.13e−03 ** | 3.49e−01 |
| *~purity+region vs ~purity* | 4.59e−01 | 7.51e−01 |
| *Carcinoma vs. normal* | 1.4e−03 ** | 1.4e−03 ** |

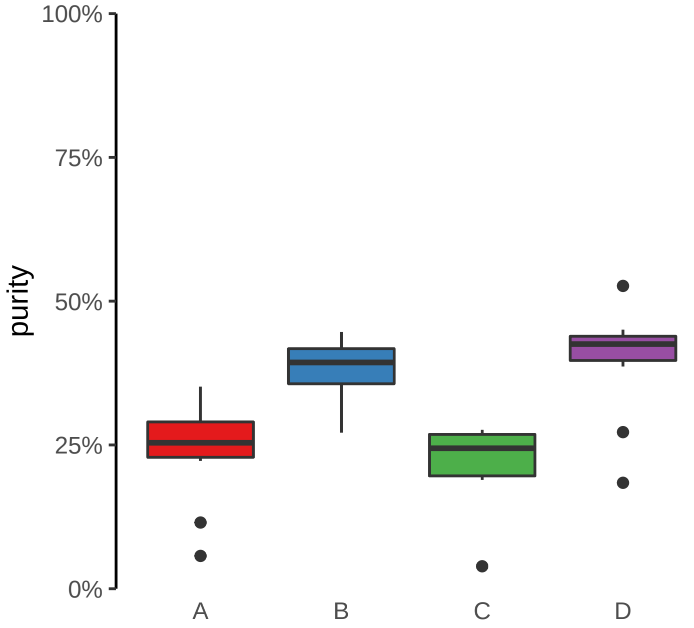

**FGD2 (GH06J037022) enhancer gain**
**chr6:37024608−37025108**

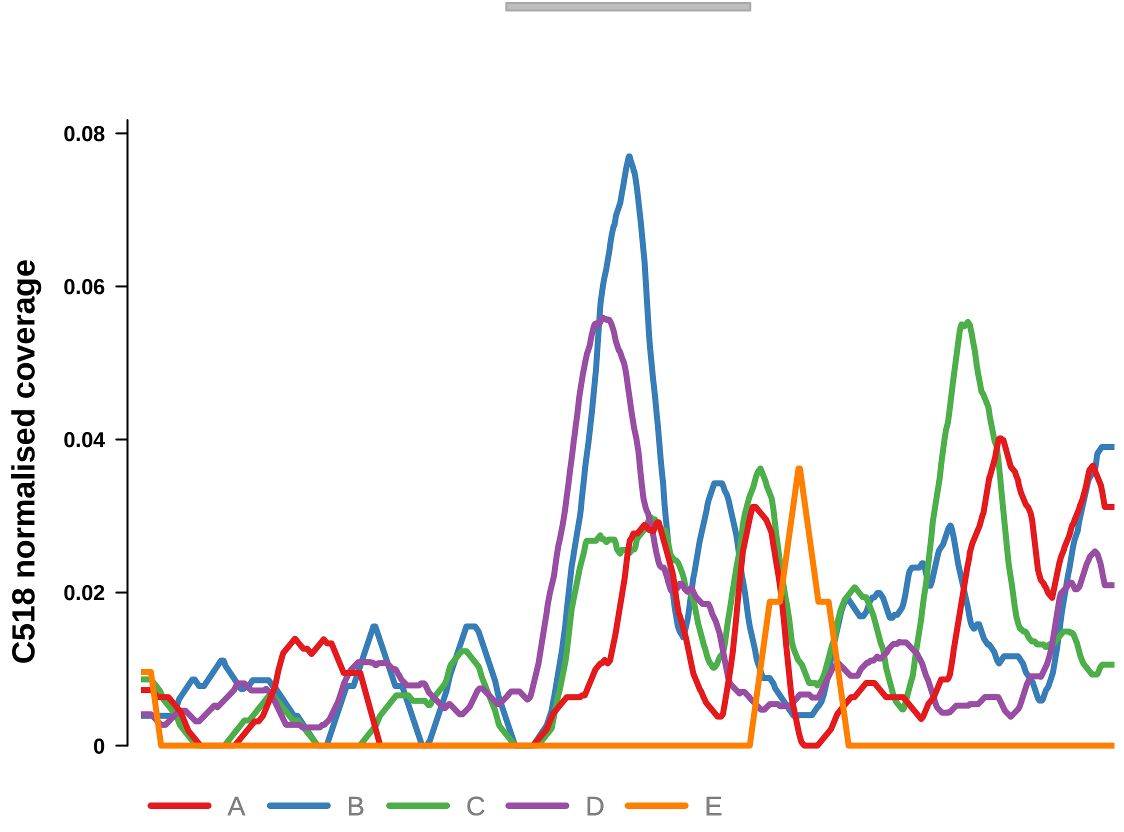


Putatively Clonal (logFC = 2.04)

| | P. | Adjusted P. |
|---|---|---|
| *~region vs. ~* | 8.64e−01 | 9.27e−01 |
| *~purity vs ~* | 6.87e−01 | 9.13e−01 |
| *~purity+region vs ~purity* | 9.02e−01 | 9.57e−01 |
| *Carcinoma vs. normal* | 4.54e−03 ** | 4.54e−03 ** |

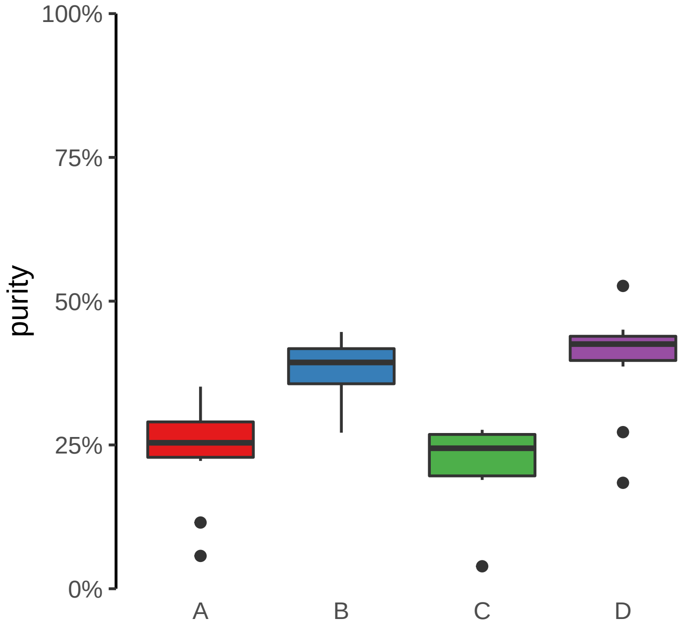

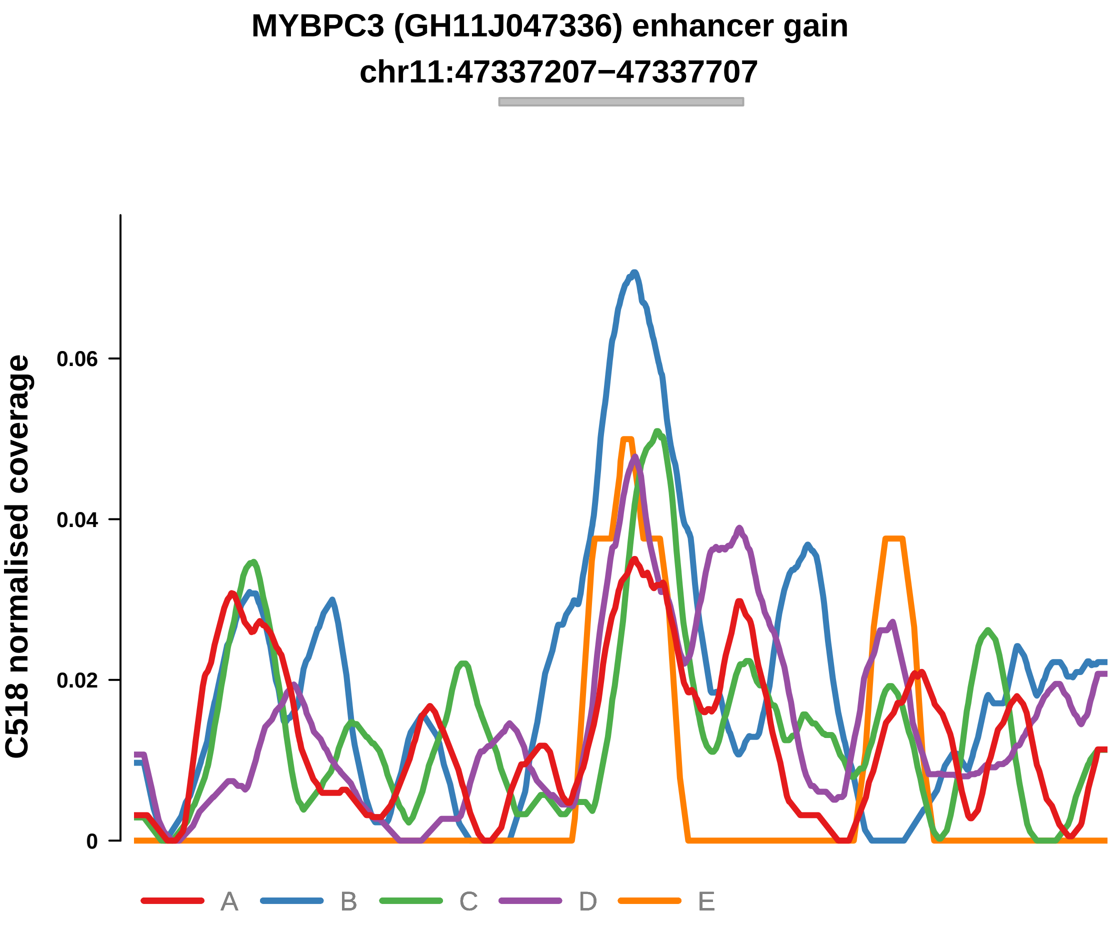


Putatively Clonal (logFC = 2.39)

| | P. | Adjusted P. |
|---|---|---|
| *~region vs. ~* | 5.26e−01 | 7.35e−01 |
| *~purity vs ~* | 6.87e−01 | 9.13e−01 |
| *~purity+region vs ~purity* | 3.48e−01 | 7.49e−01 |
| *Carcinoma vs. normal* | 1.08e−03 ** | 1.08e−03 ** |

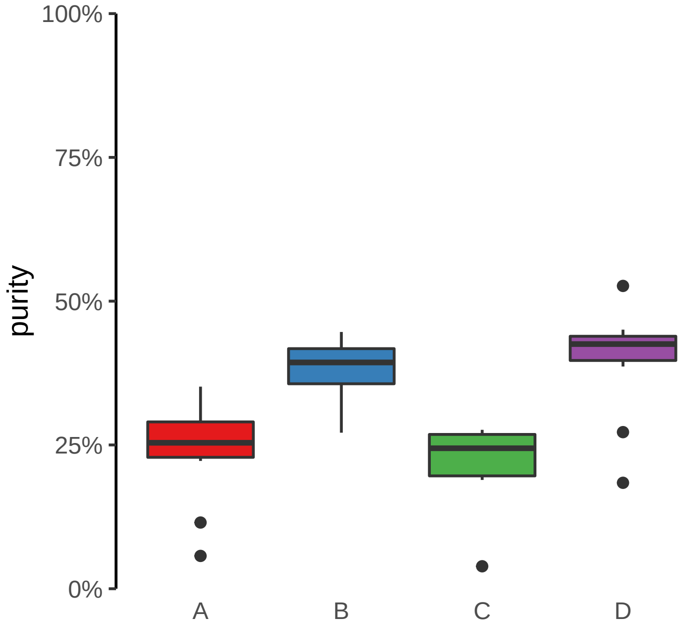

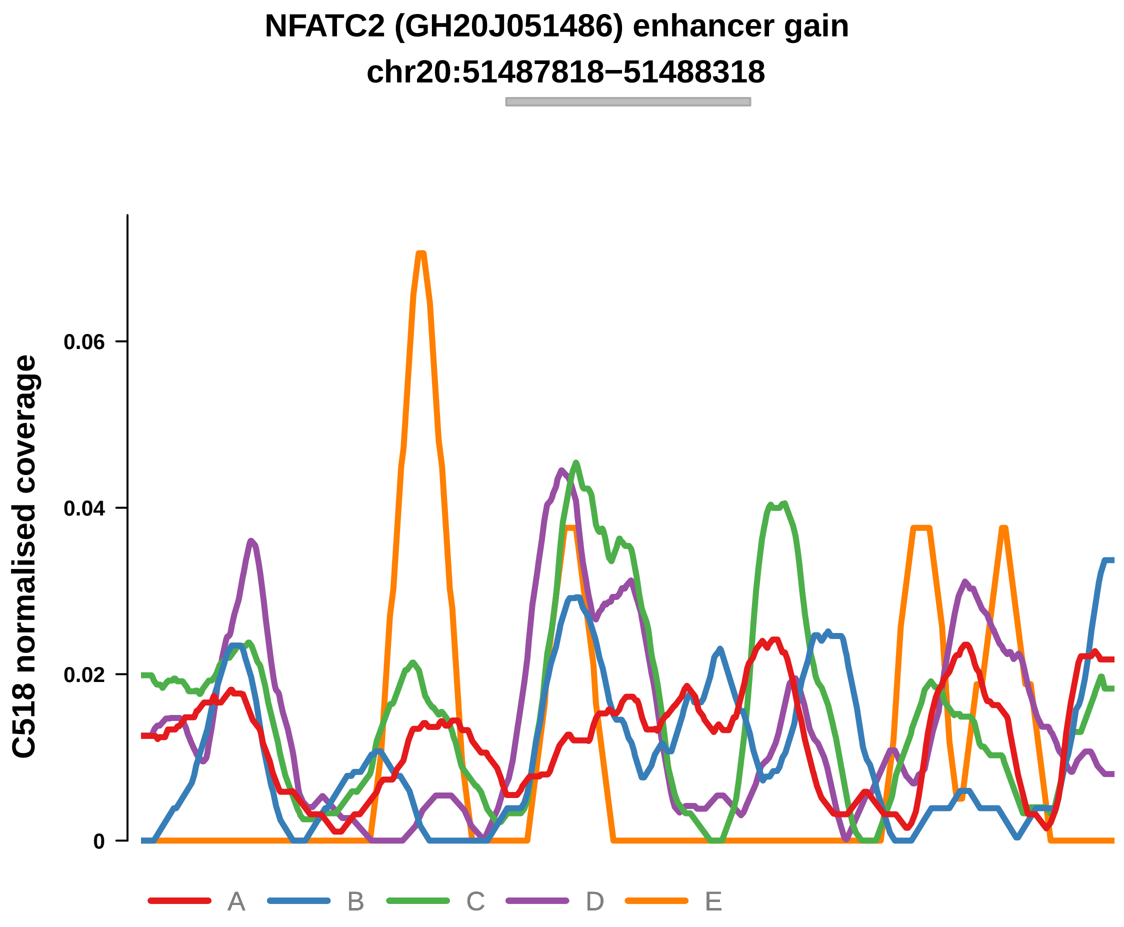


Putatively Clonal (logFC = 2.04)

| | P. | Adjusted P. |
|---|---|---|
| *~region vs. ~* | 2.67e−01 | 6.53e−01 |
| *~purity vs ~* | 8e−02 | 5.03e−01 |
| *~purity+region vs ~purity* | 5.86e−01 | 8.52e−01 |
| *Carcinoma vs. normal* | 6.02e−03 ** | 6.02e−03 ** |

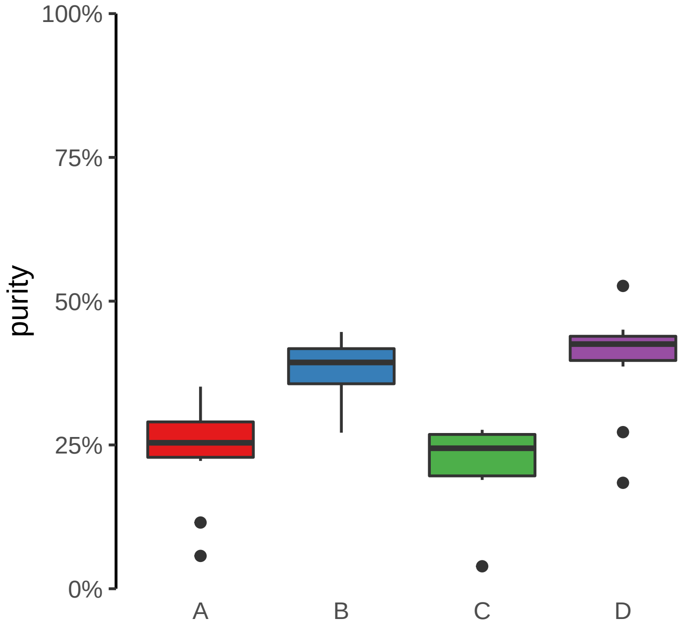

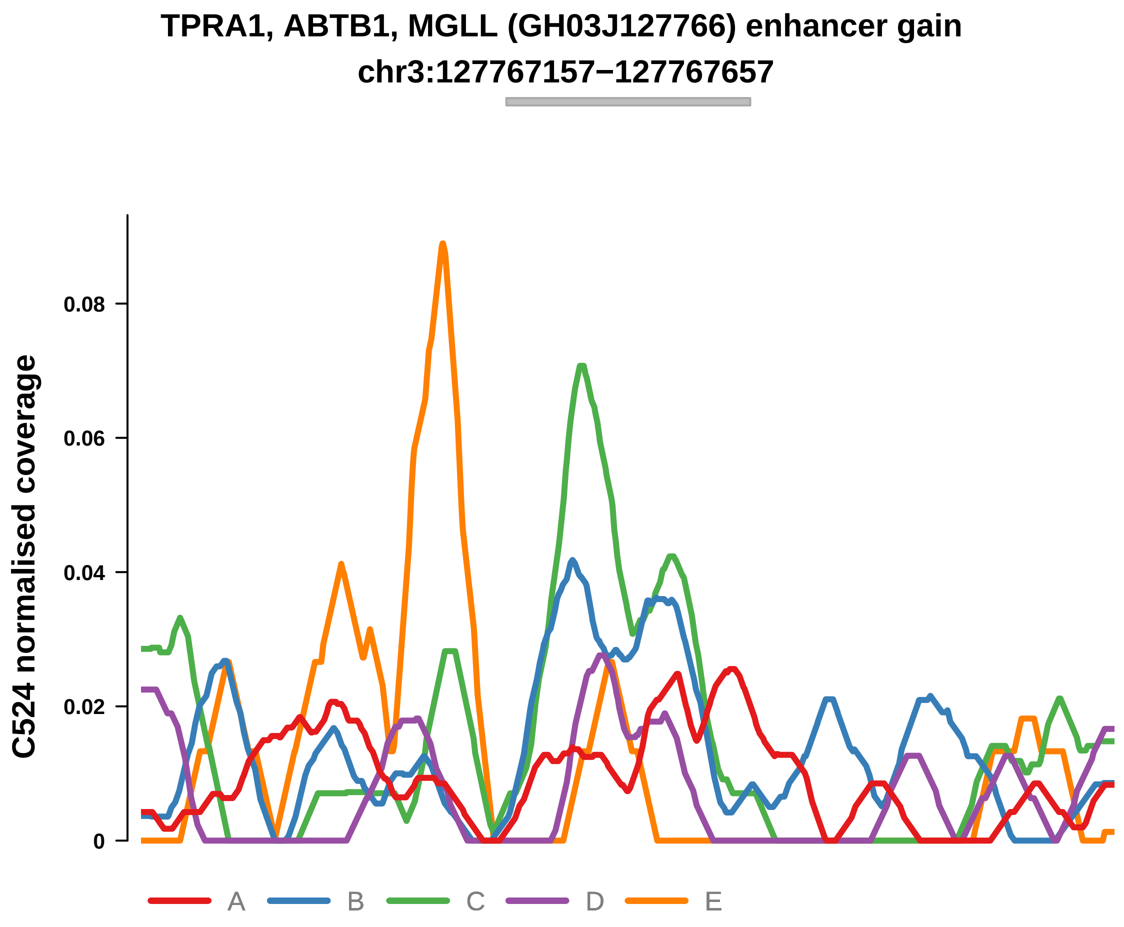


Putatively Clonal (logFC = 2.31)

| | P. | Adjusted P. |
|---|---|---|
| *~region vs. ~* | 2.83e−01 | 6.57e−01 |
| *~purity vs ~* | 3.74e−02 * | 2.8e−01 |
| *~purity+region vs ~purity* | 3.94e−01 | 7.48e−01 |
| *Carcinoma vs. normal* | 5.05e−03 ** | 5.05e−03 ** |

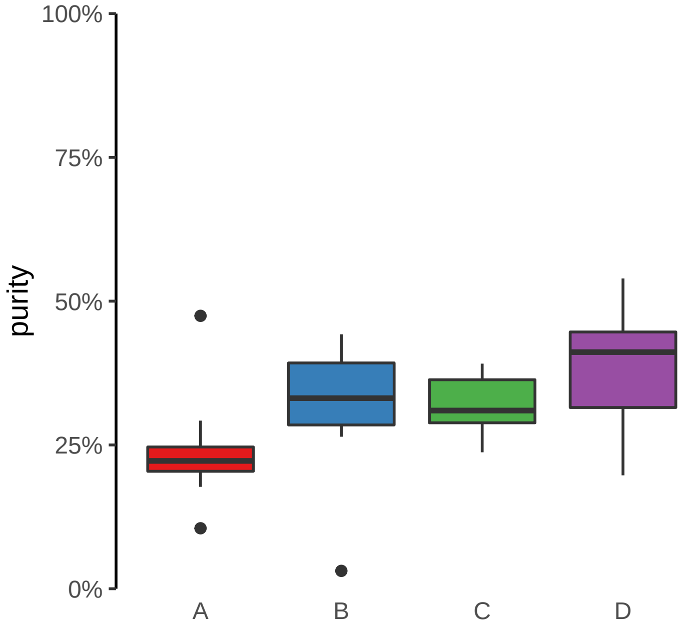

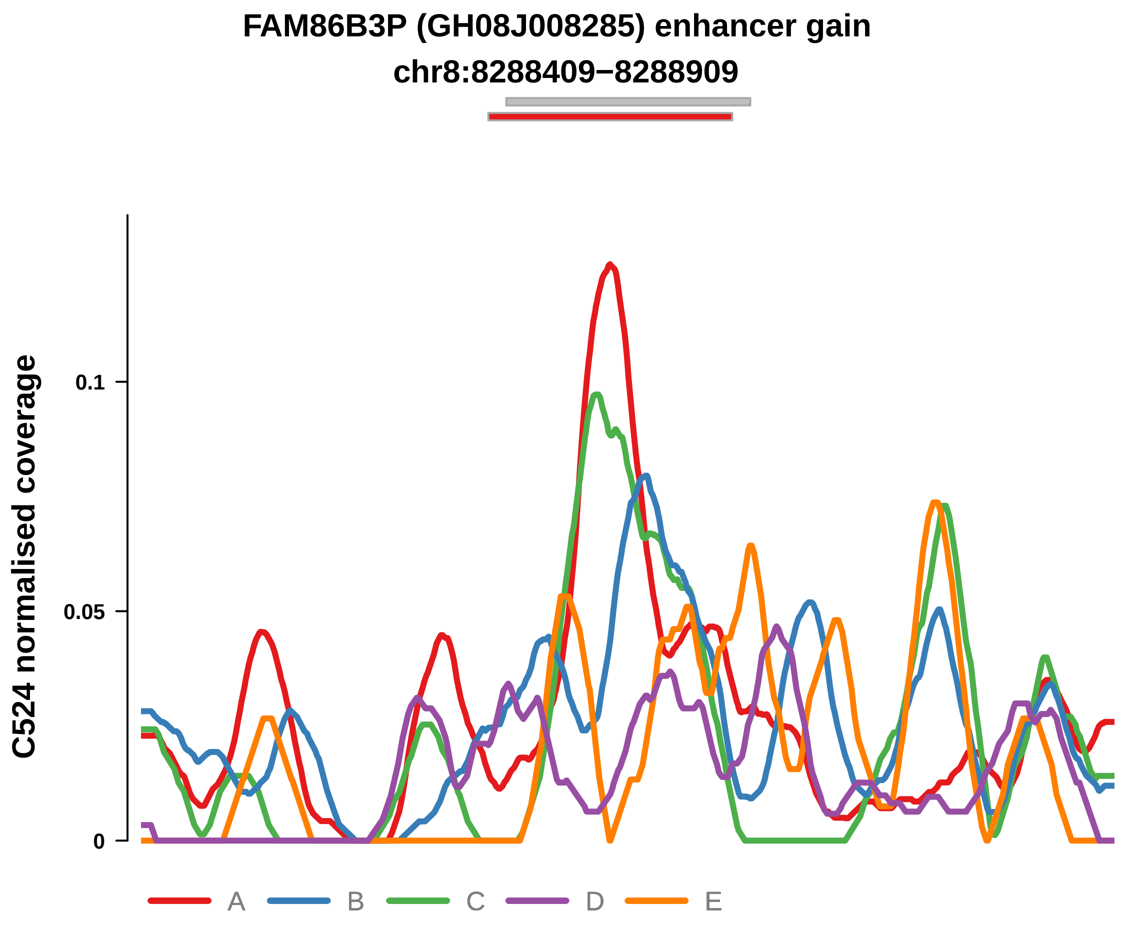


Putatively Clonal (logFC = 3.2)

| | P. | Adjusted P. |
|---|---|---|
| *~region vs. ~* | 1.32e−01 | 5.53e−01 |
| *~purity vs ~* | 1.68e−01 | 5e−01 |
| *~purity+region vs ~purity* | 9.33e−02 | 3.98e−01 |
| *Carcinoma vs. normal* | 2.44e−05 **** | 2.44e−05 **** |

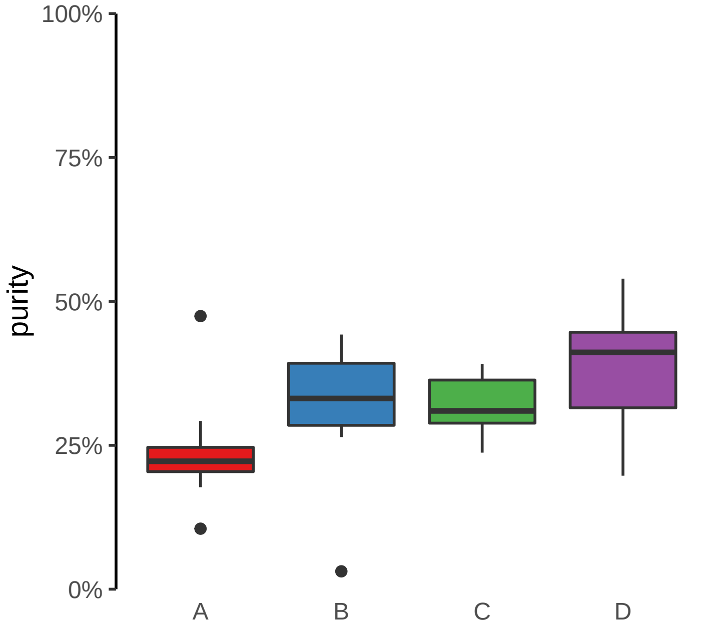

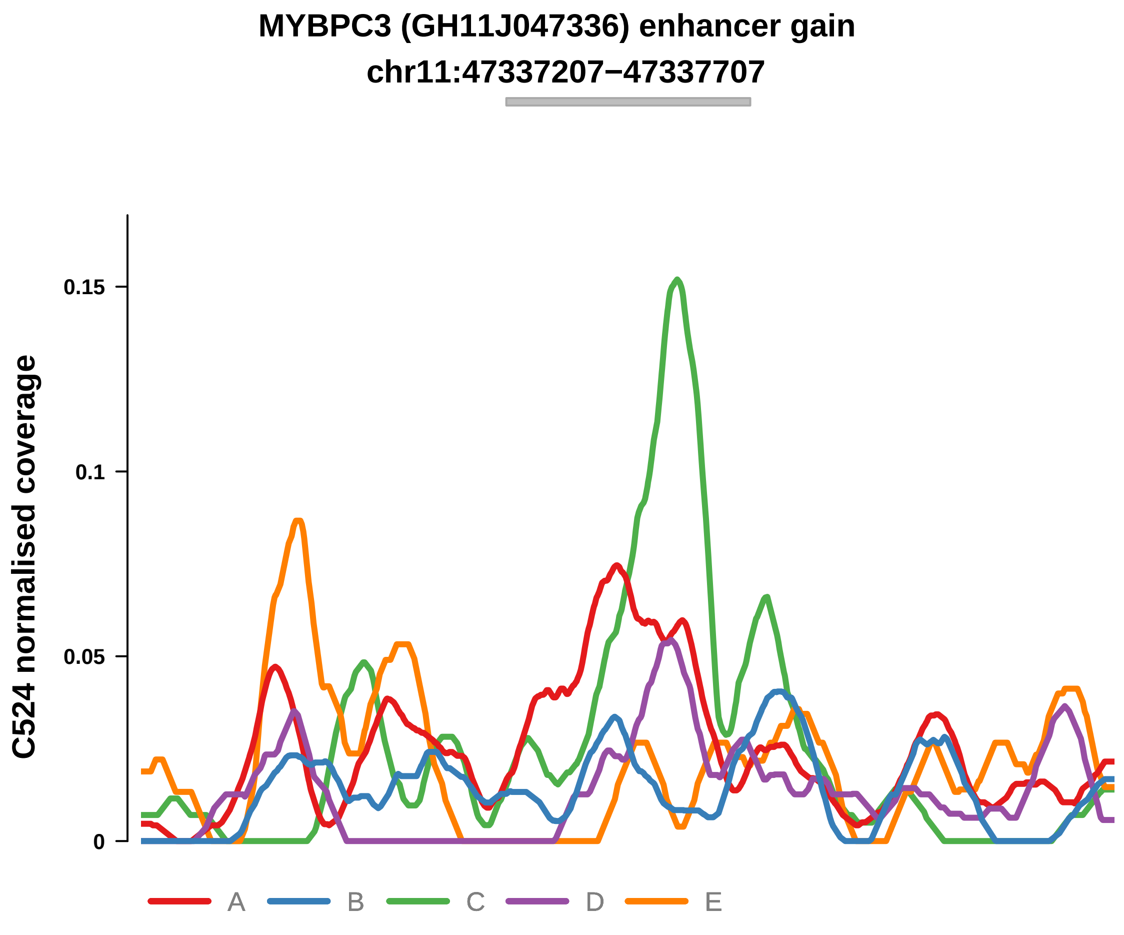


Putatively Clonal (logFC = 2.66)

| | P. | Adjusted P. |
|---|---|---|
| *~region vs. ~* | 2.98e−01 | 6.57e−01 |
| *~purity vs ~* | 6.68e−01 | 7.74e−01 |
| *~purity+region vs ~purity* | 2.39e−01 | 6e−01 |
| *Carcinoma vs. normal* | 5.34e−04 *** | 5.34e−04 *** |

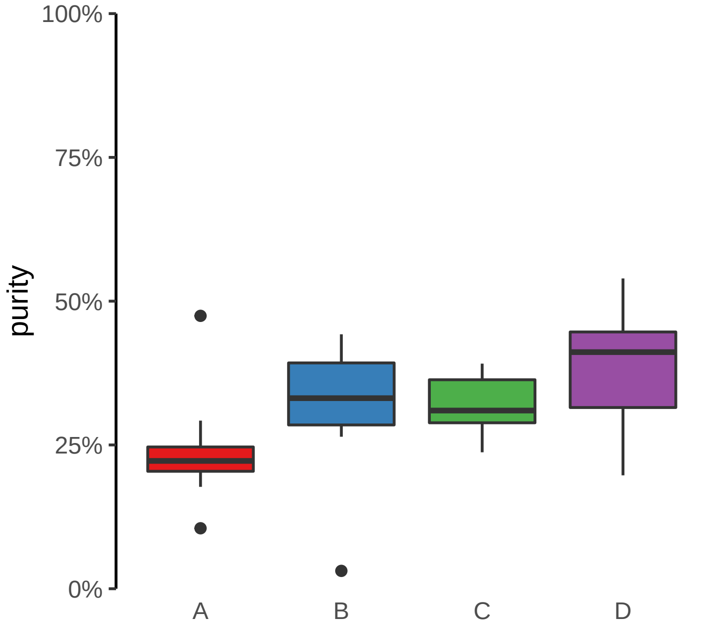

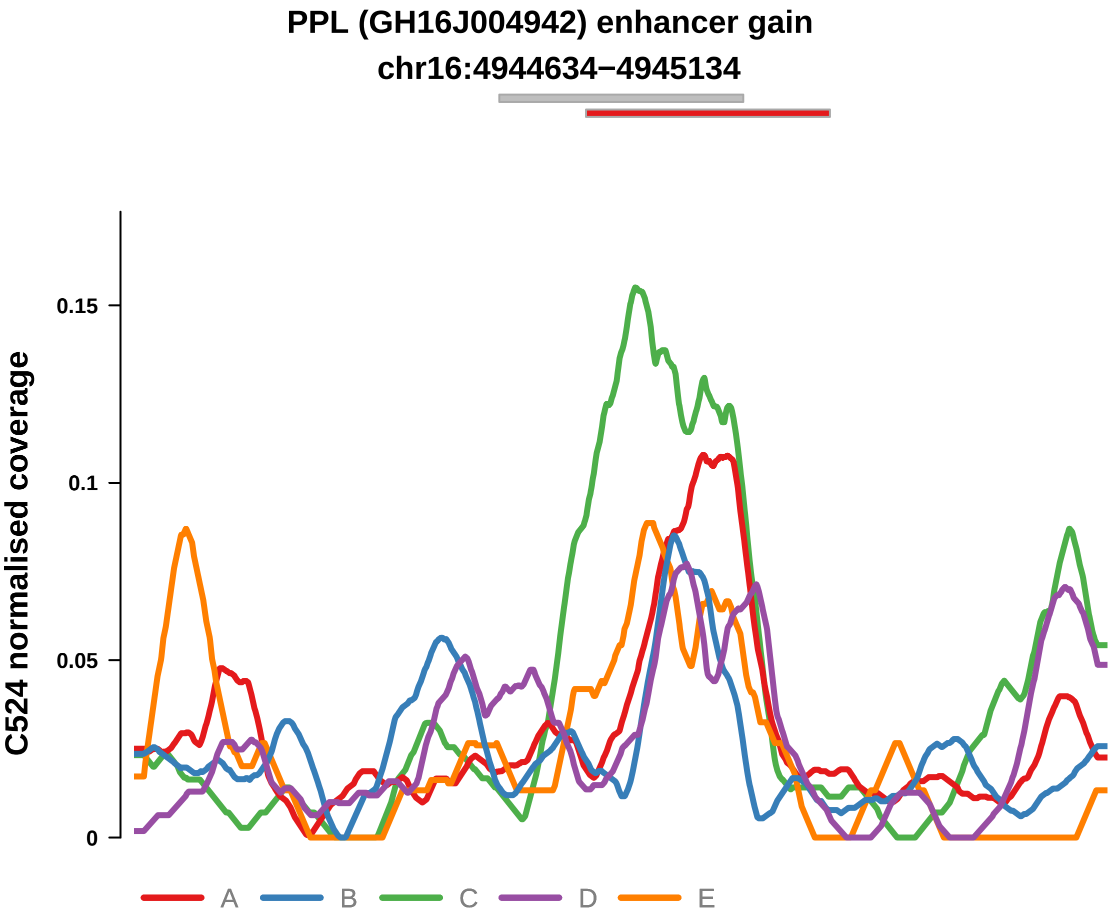


Putatively Clonal (logFC = 2.22)

| | P. | Adjusted P. |
|---|---|---|
| *~region vs. ~* | 5.66e−02 | 4.53e−01 |
| *~purity vs ~* | 1.75e−02 * | 2.2e−01 |
| *~purity+region vs ~purity* | 3.24e−02 * | 2.85e−01 |
| *Carcinoma vs. normal* | 2.24e−03 ** | 2.24e−03 ** |

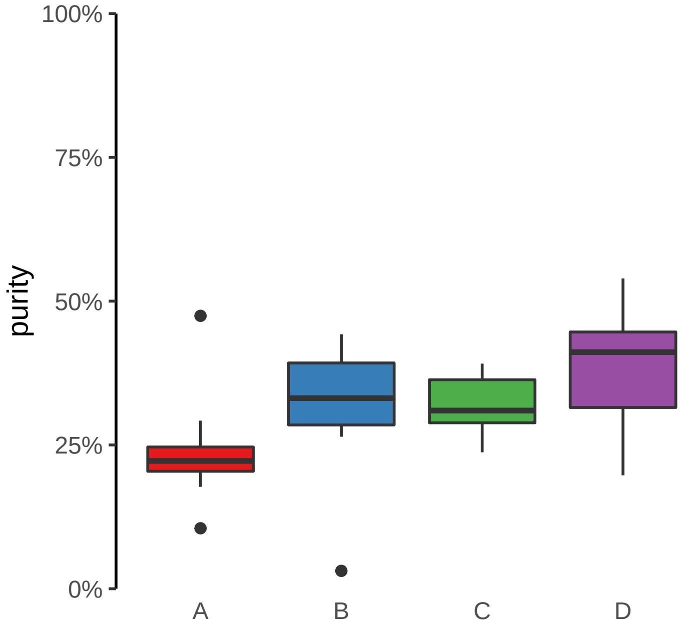

**RNU7−124P, SHB, ENSG00000255872 (GH09J037962) enhancer gain**
**chr9:37966483−37966983**

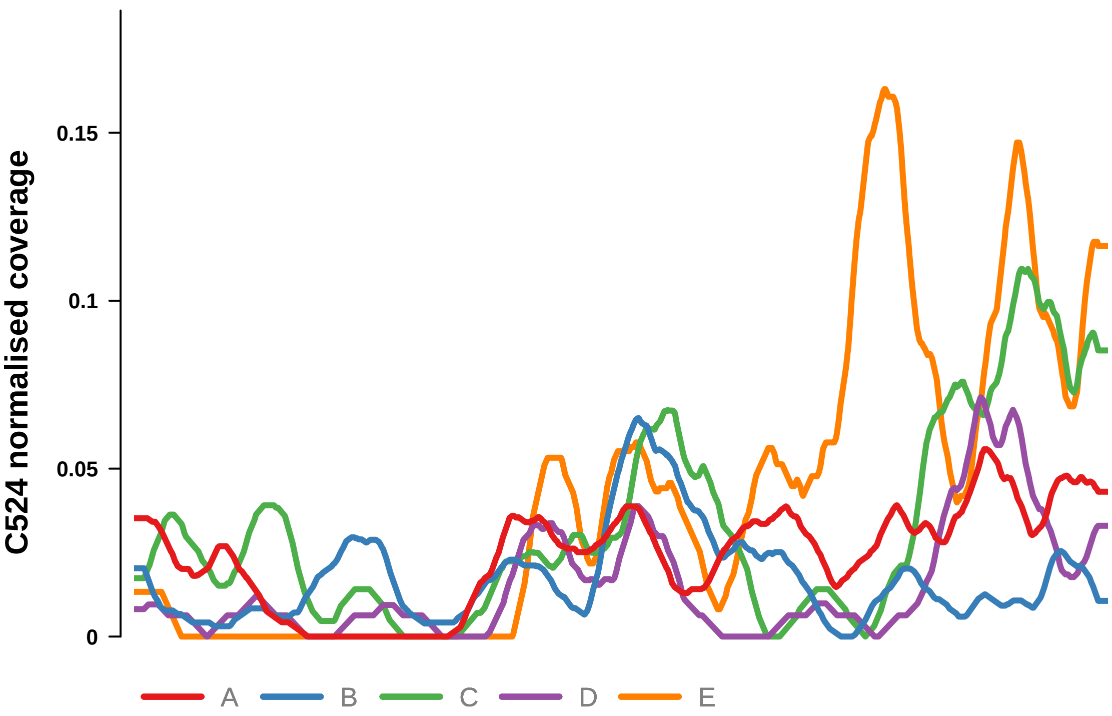


Putatively Clonal (logFC = 2.23)

| | P. | Adjusted P. |
|---|---|---|
| *~region vs. ~* | 2.09e−01 | 5.75e−01 |
| *~purity vs ~* | 3.64e−02 * | 2.8e−01 |
| *~purity+region vs ~purity* | 9.51e−02 | 3.98e−01 |
| *Carcinoma vs. normal* | 3.08e−03 ** | 3.08e−03 ** |

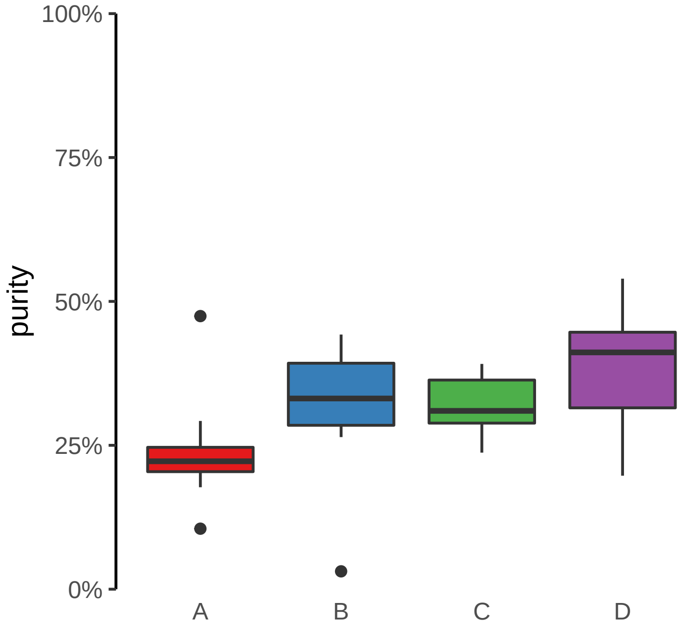

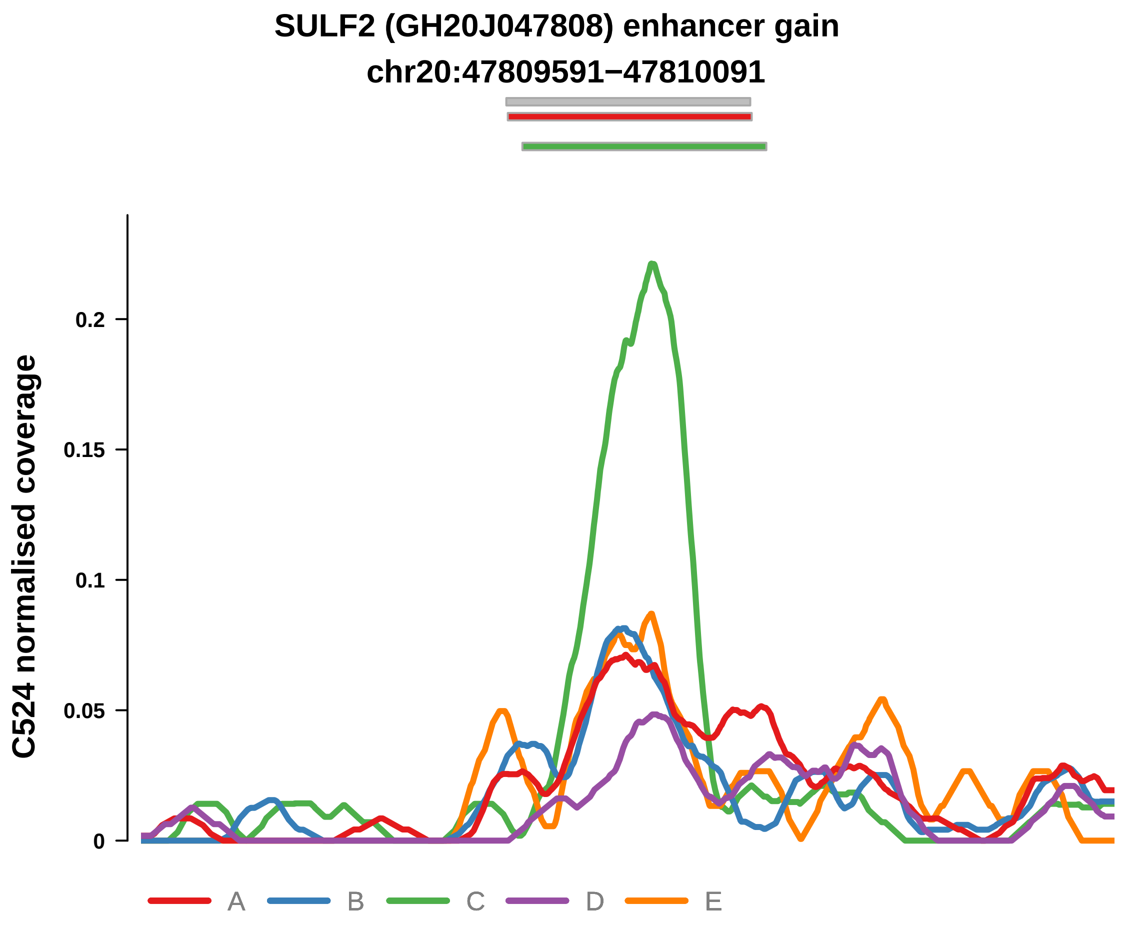


Putatively Clonal (logFC = 1.95)

| | P. | Adjusted P. |
|---|---|---|
| *~region vs. ~* | 8.38e−01 | 9.34e−01 |
| *~purity vs ~* | 5.31e−01 | 7.12e−01 |
| *~purity+region vs ~purity* | 7.68e−01 | 9.26e−01 |
| *Carcinoma vs. normal* | 7.02e−03 ** | 7.02e−03 ** |

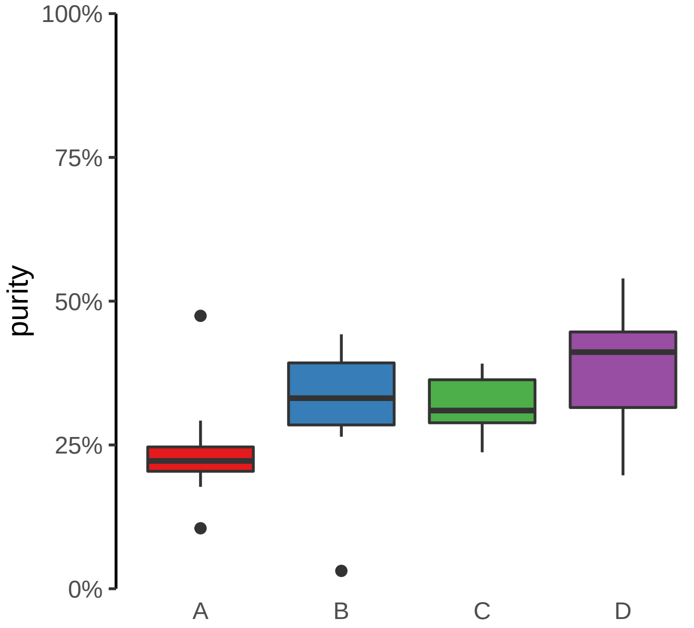

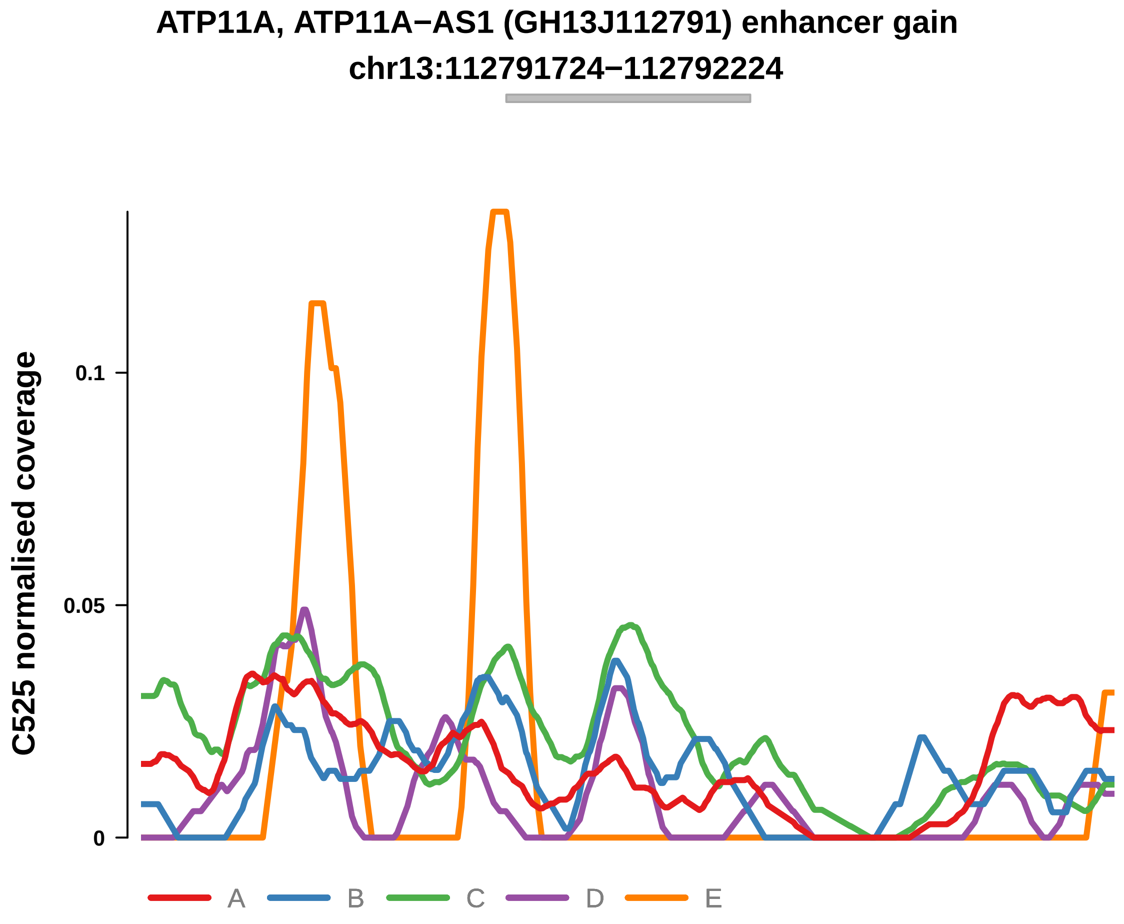


Putatively Clonal (logFC = 2.28)

| | P. | Adjusted P. |
|---|---|---|
| *~region vs. ~* | 8.1e−05 **** | 7.83e−04 *** |
| *~purity vs ~* | 1.48e−02 * | 3.25e−01 |
| *~purity+region vs ~purity* | 5.39e−03 ** | 2.21e−02 * |
| *Carcinoma vs. normal* | 3.69e−03 ** | 3.69e−03 ** |

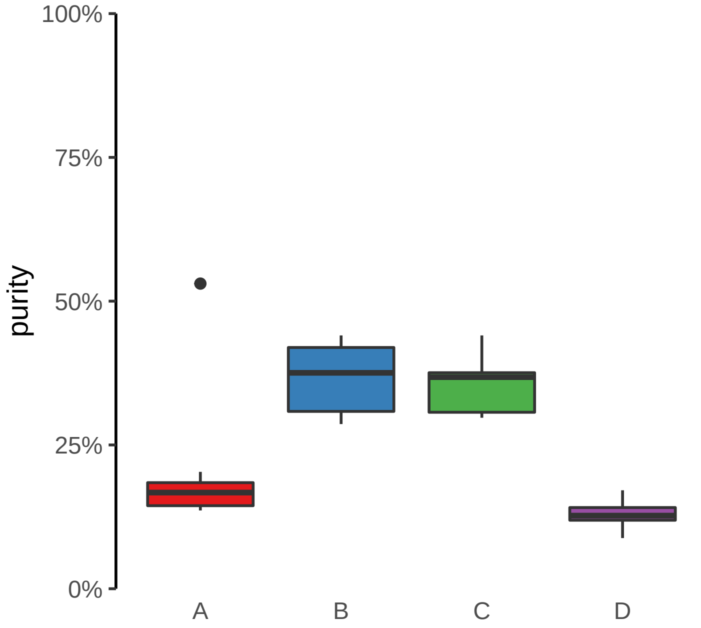

**FAM86B3P (GH08J008285) enhancer gain**
**chr8:8288409−8288909**

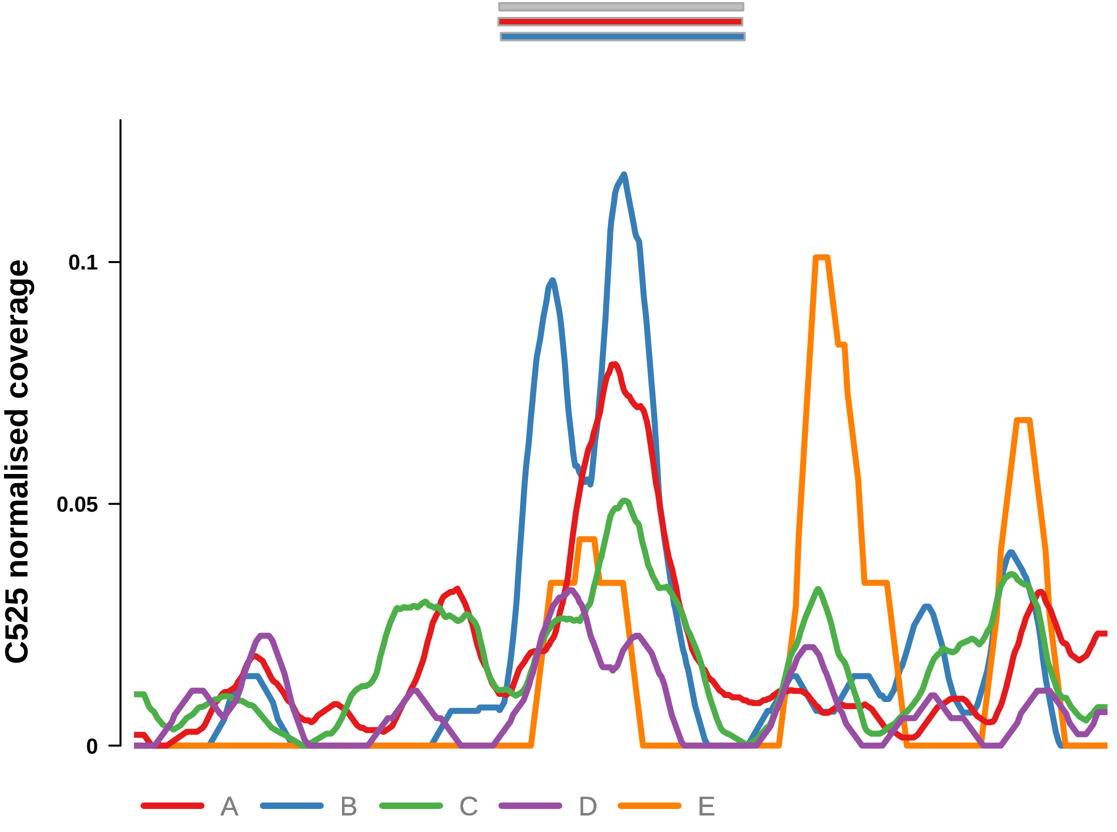


Putatively Clonal (logFC = 2.38)

| | P. | Adjusted P. |
|---|---|---|
| *~region vs. ~* | 2.7e−01 | 3.48e−01 |
| *~purity vs ~* | 1.05e−01 | 7.37e−01 |
| *~purity+region vs ~purity* | 4.9e−01 | 5.68e−01 |
| *Carcinoma vs. normal* | 1.83e−03 ** | 1.83e−03 ** |

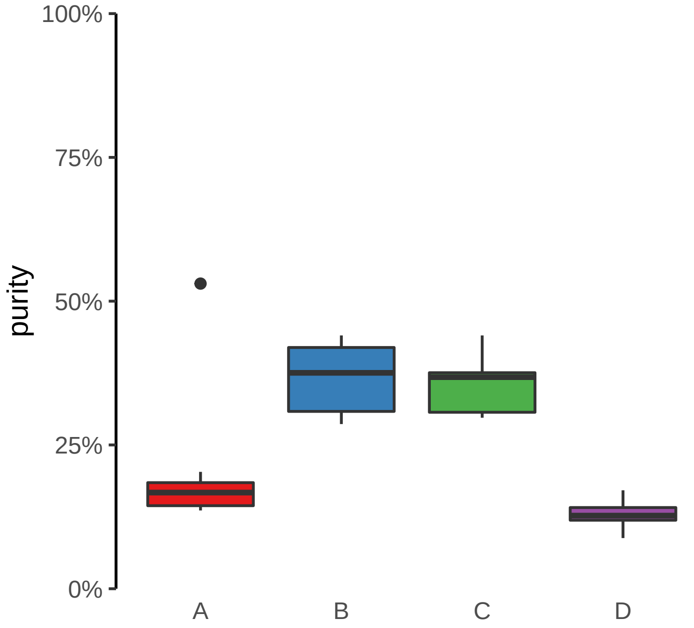

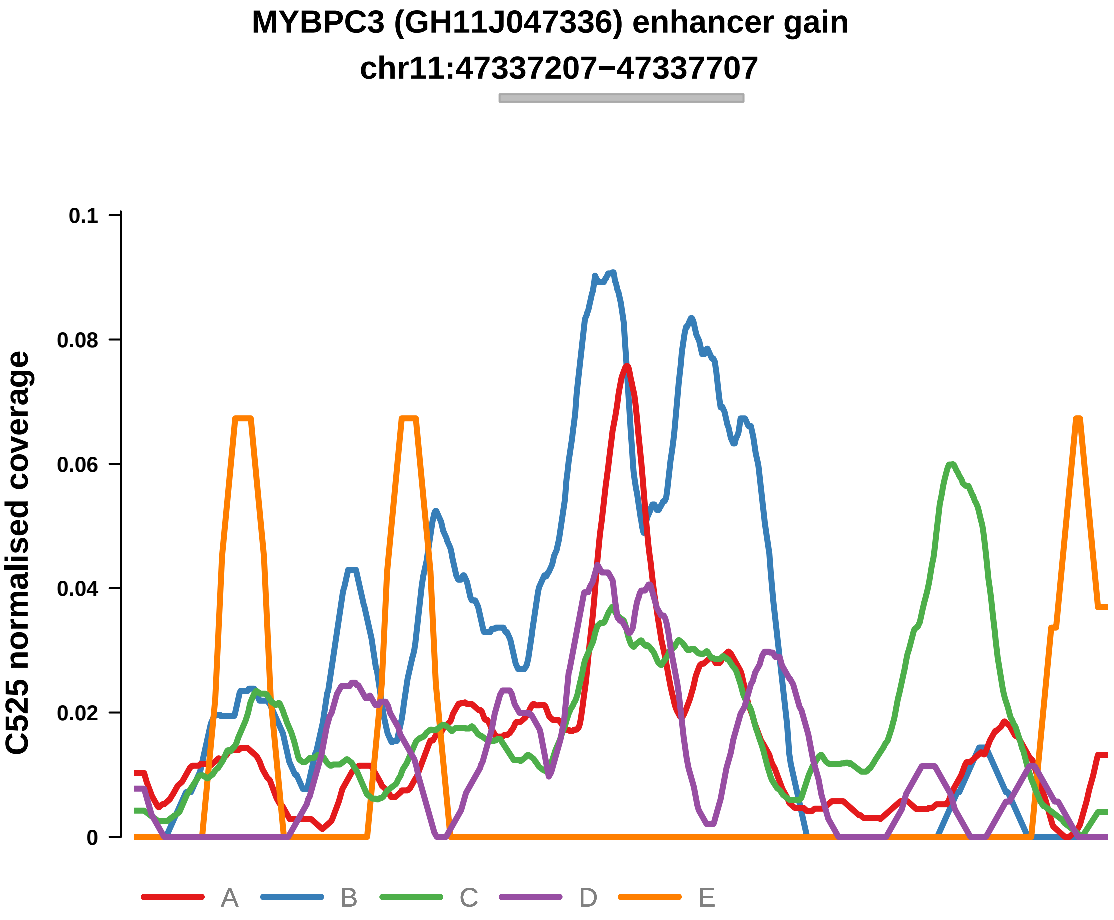


Putatively Clonal (logFC = 2.25)

| | P. | Adjusted P. |
|---|---|---|
| *~region vs. ~* | 9.14e−01 | 9.14e−01 |
| *~purity vs ~* | 4.47e−01 | 8.63e−01 |
| *~purity+region vs ~purity* | 5.45e−02 | 1.09e−01 |
| *Carcinoma vs. normal* | 4.05e−03 ** | 4.05e−03 ** |

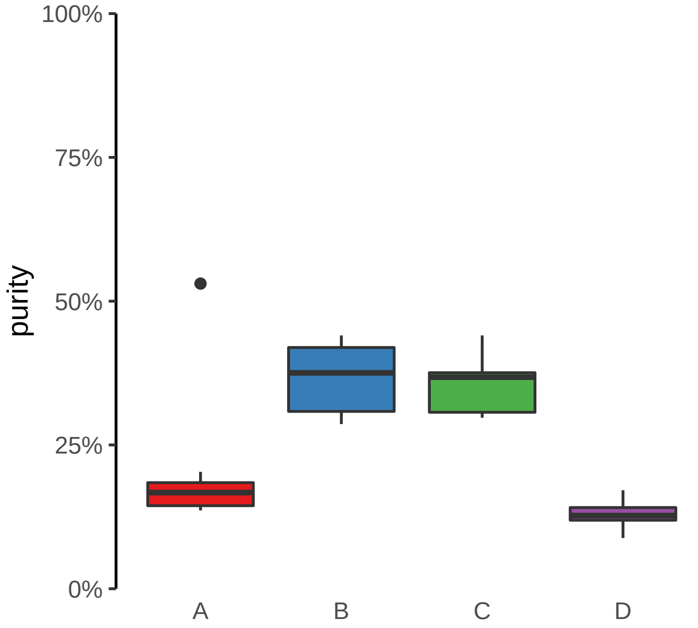

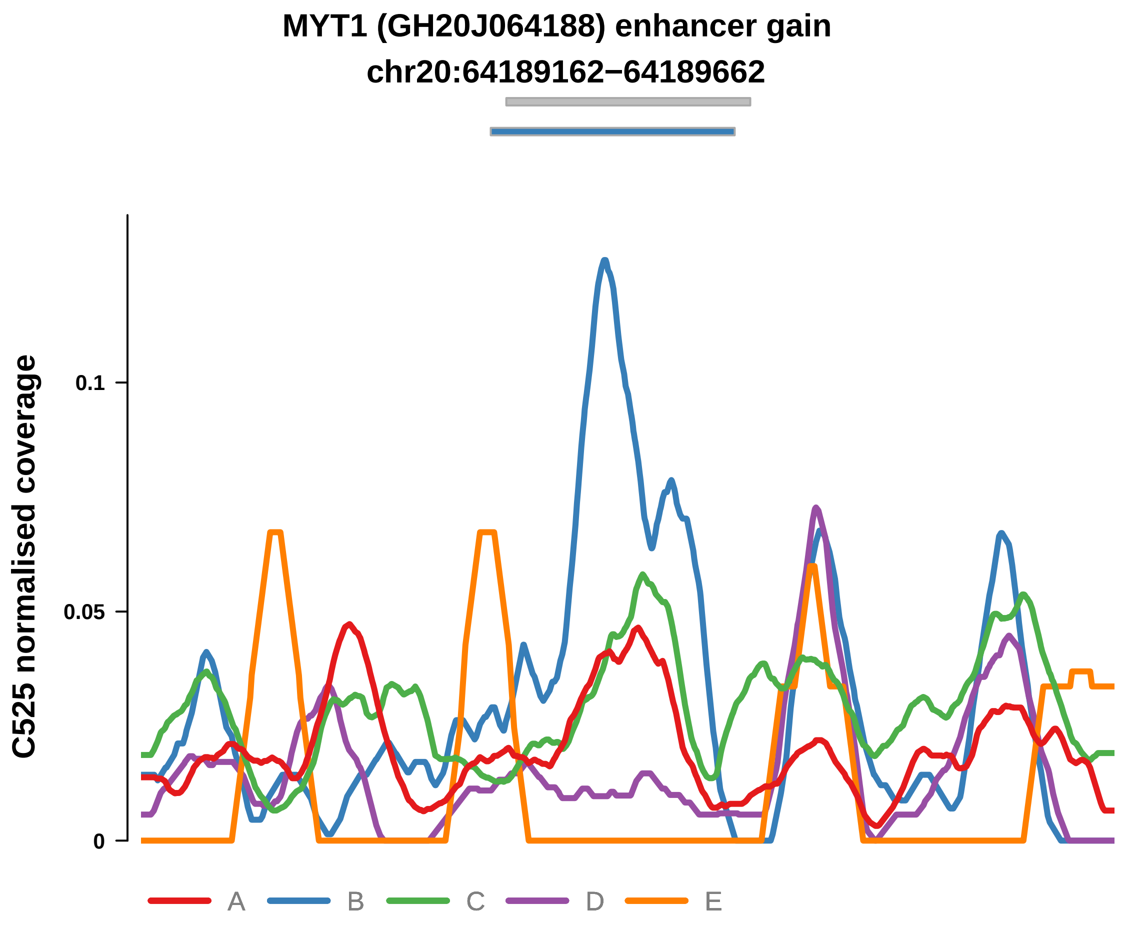


Putatively Clonal (logFC = 3.3)

| | P. | Adjusted P. |
|---|---|---|
| *~region vs. ~* | 1.77e−02 * | 4.62e−02 * |
| *~purity vs ~* | 2.44e−03 ** | 1.07e−01 |
| *~purity+region vs ~purity* | 5.8e−01 | 6.3e−01 |
| *Carcinoma vs. normal* | 1.49e−05 **** | 1.49e−05 **** |

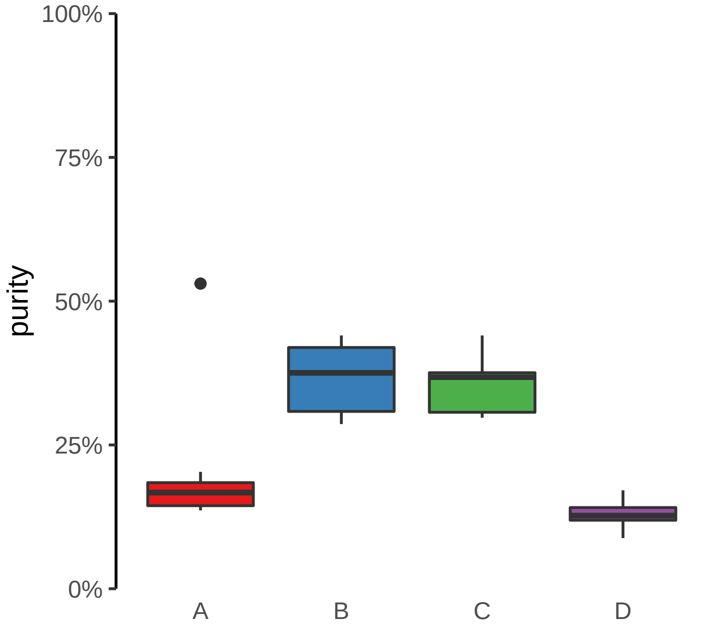

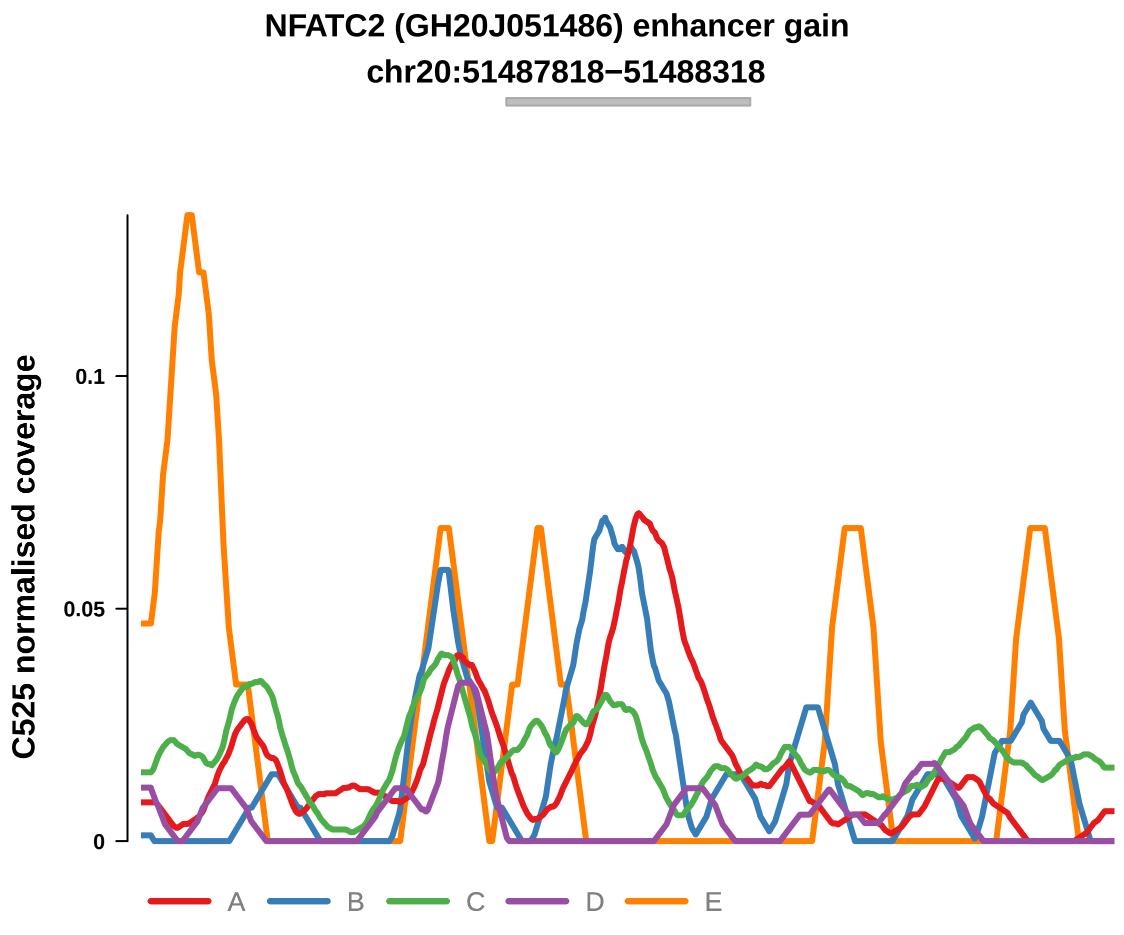


Putatively Clonal (logFC = 2.3)

| | P. | Adjusted P. |
|---|---|---|
| *~region vs. ~* | 8.19e−05 **** | 7.83e−04 *** |
| *~purity vs ~* | 6.43e−01 | 8.63e−01 |
| *~purity+region vs ~purity* | 7.85e−05 **** | 2.3e−03 ** |
| *Carcinoma vs. normal* | 3.44e−03 ** | 3.44e−03 ** |

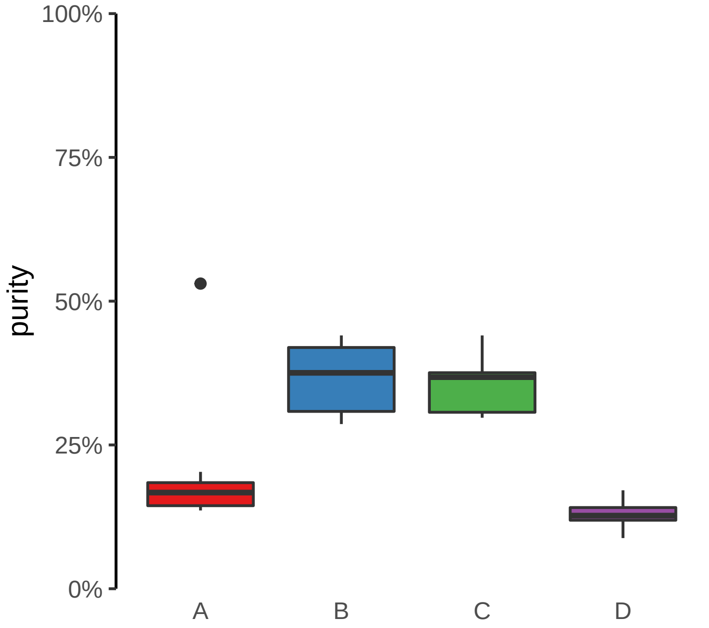

**RNU7−124P, SHB, ENSG00000255872 (GH09J037962) enhancer gain**
**chr9:37966483−37966983**

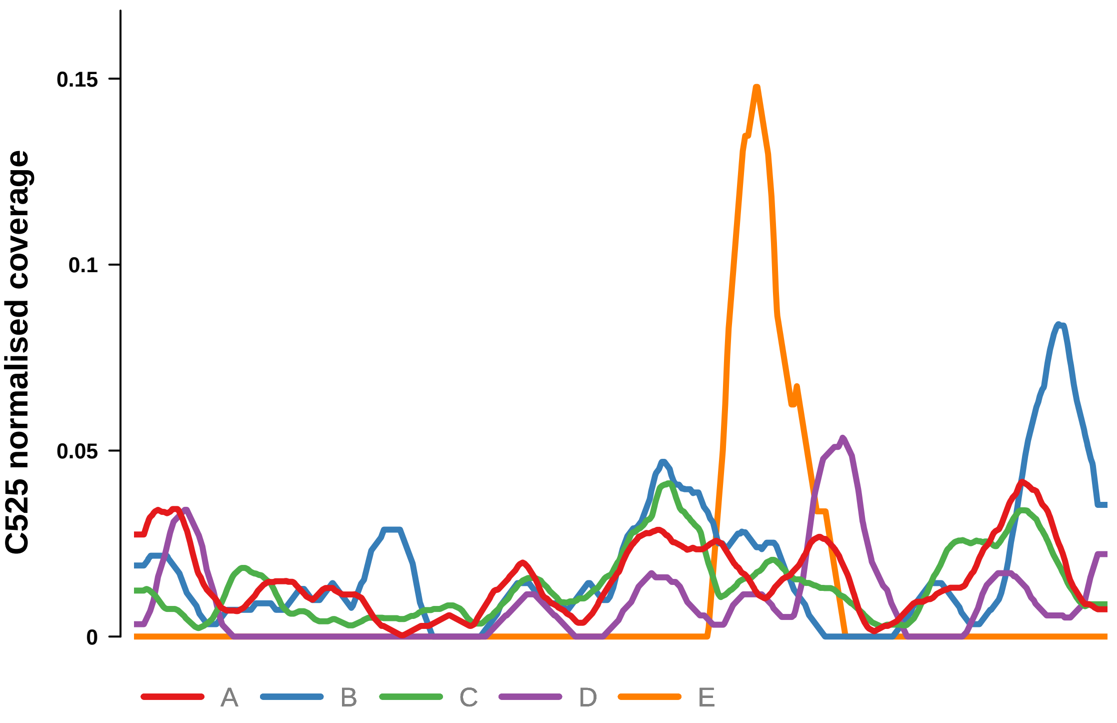


Putatively Clonal (logFC = 2.18)

| | P. | Adjusted P. |
|---|---|---|
| *~region vs. ~* | 1.18e−01 | 1.88e−01 |
| *~purity vs ~* | 1.99e−01 | 7.37e−01 |
| *~purity+region vs ~purity* | 2.85e−01 | 4.19e−01 |
| *Carcinoma vs. normal* | 3.71e−03 ** | 3.71e−03 ** |

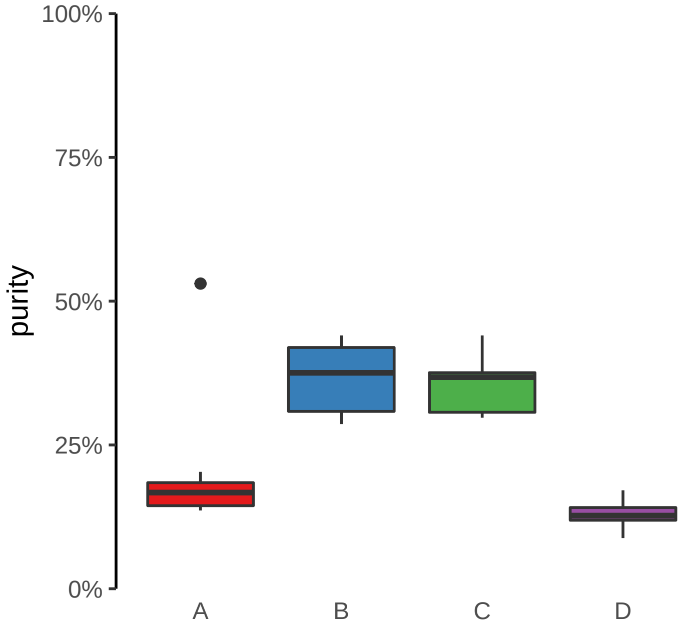

**HOXC11, HOXC10, HOTAIR (GH12J053971) enhancer gain**
**chr12:53978873−53979373**

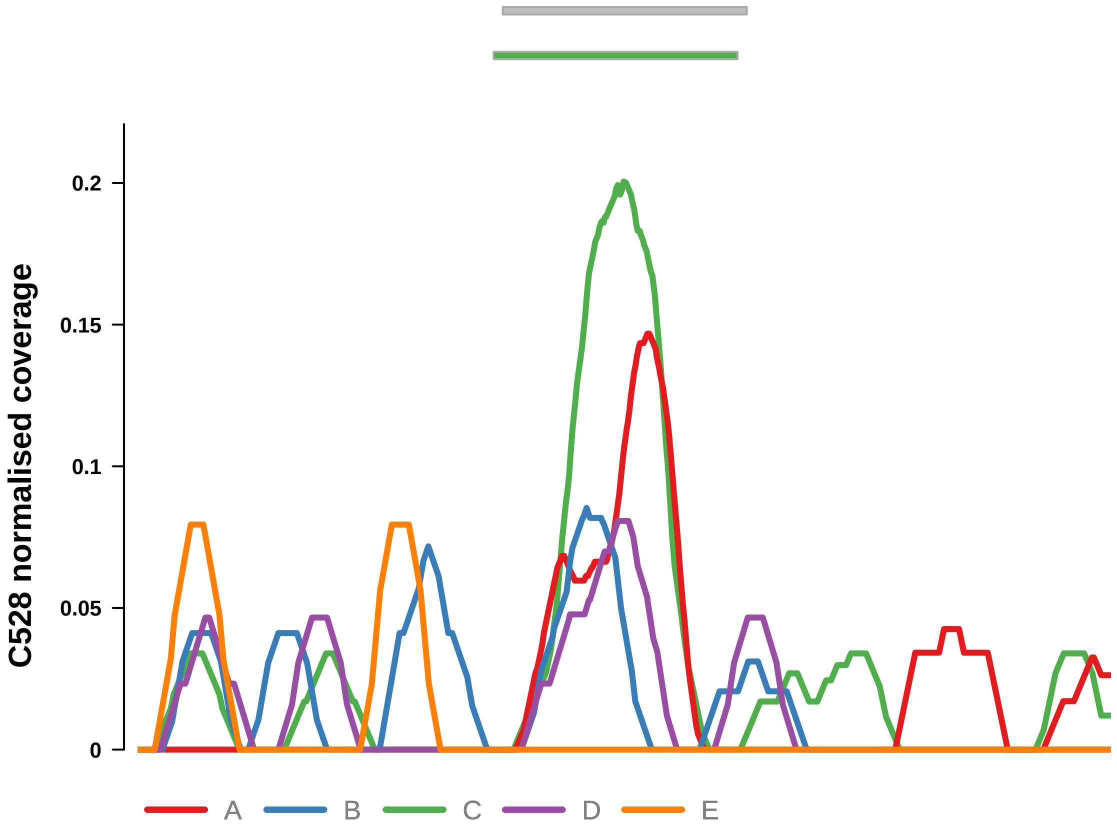


Putatively Clonal (logFC = 2.5)

| | **P.** | **Adjusted P.** |
|---|---|---|
| *~region vs. ~* | 7.41e−01 | 8.89e−01 |
| *~purity vs ~* | 2.38e−01 | 4e−01 |
| *~purity+region vs ~purity* | 5.51e−01 | 8.94e−01 |
| *Carcinoma vs. normal* | 1.24e−03 ** | 1.24e−03 ** |

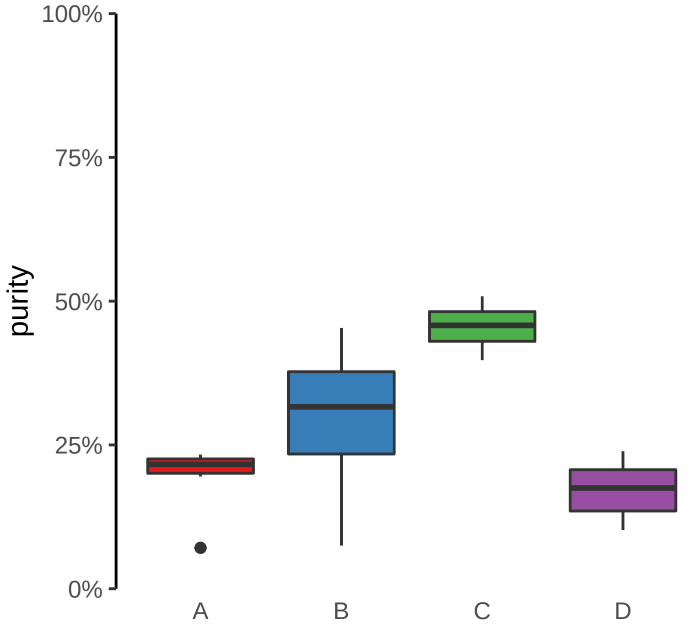

**ATP11A, ATP11A−AS1 (GH13J112791) enhancer gain**
**chr13:112791724−112792224**

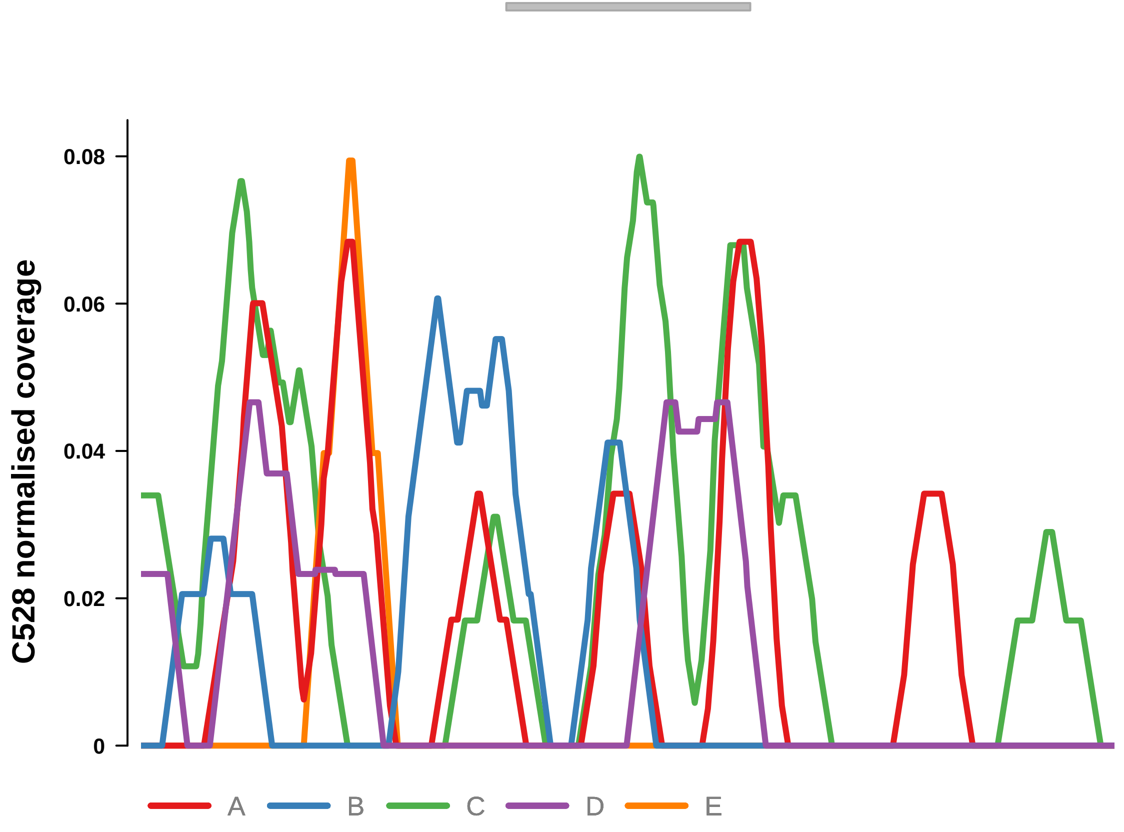


Putatively Clonal (logFC = 2.28)

| | P. | Adjusted P. |
|---|---|---|
| *~region vs. ~* | 9.72e−01 | 9.95e−01 |
| *~purity vs ~* | 5.32e−01 | NA |
| *~purity+region vs ~purity* | 7.28e−01 | 9.17e−01 |
| *Carcinoma vs. normal* | 8.07e−03 ** | 8.07e−03 ** |

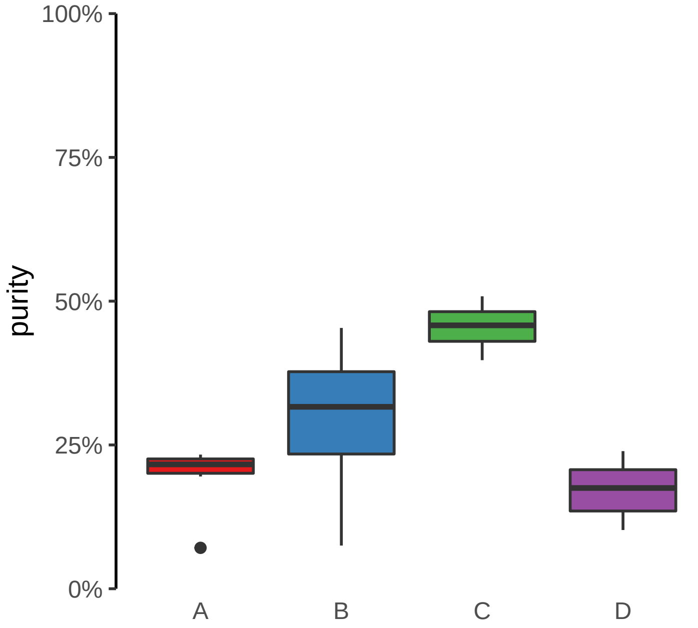

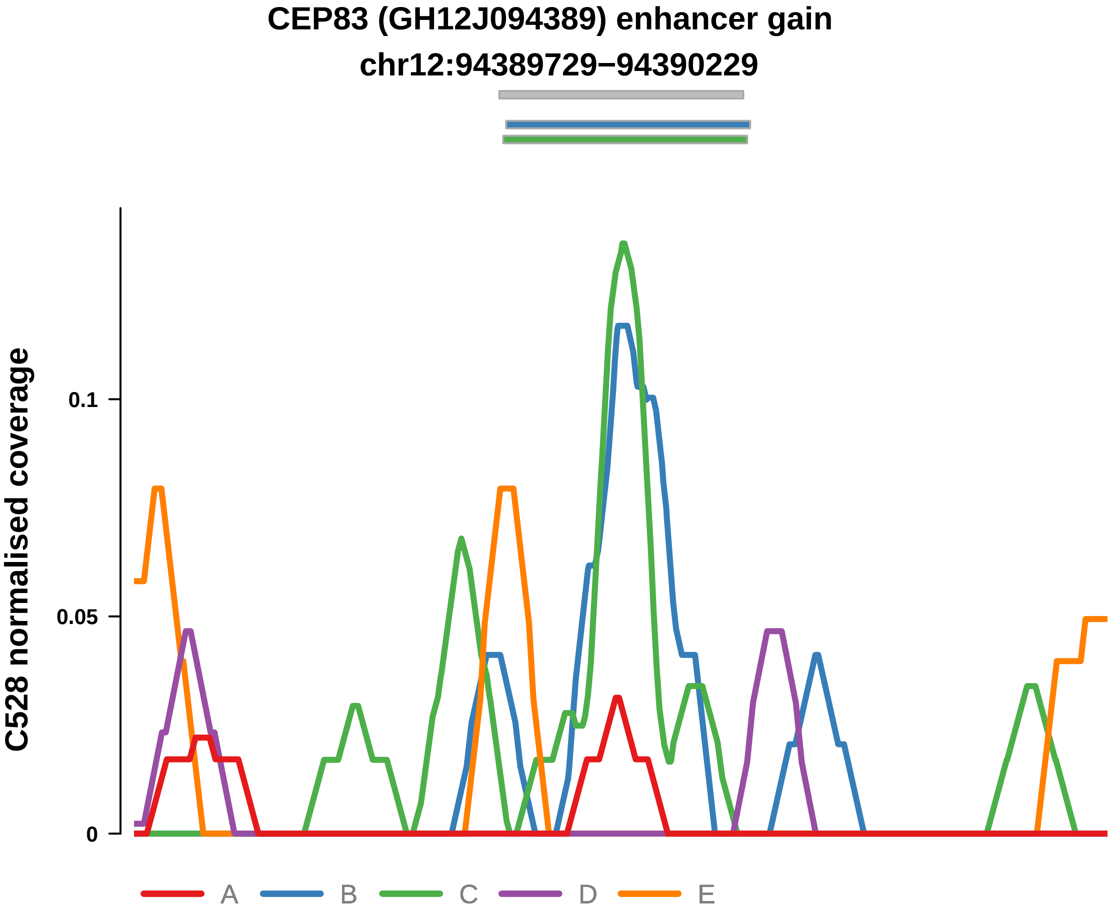


Putatively Clonal (logFC = 2.93)

| | P. | Adjusted P. |
|---|---|---|
| *~region vs. ~* | 3.52e−02 * | 2.21e−01 |
| *~purity vs ~* | 1.74e−02 * | 9.87e−02 |
| *~purity+region vs ~purity* | 2.52e−01 | 6.8e−01 |
| *Carcinoma vs. normal* | 5.53e−04 *** | 5.53e−04 *** |

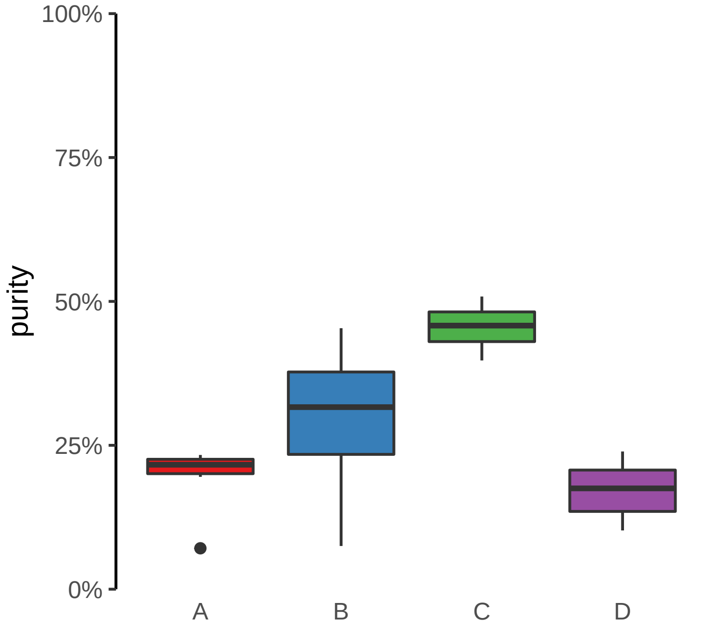

**CSGALNACT1 (GH08J019660) enhancer gain**
**chr8:19661066−19661566**

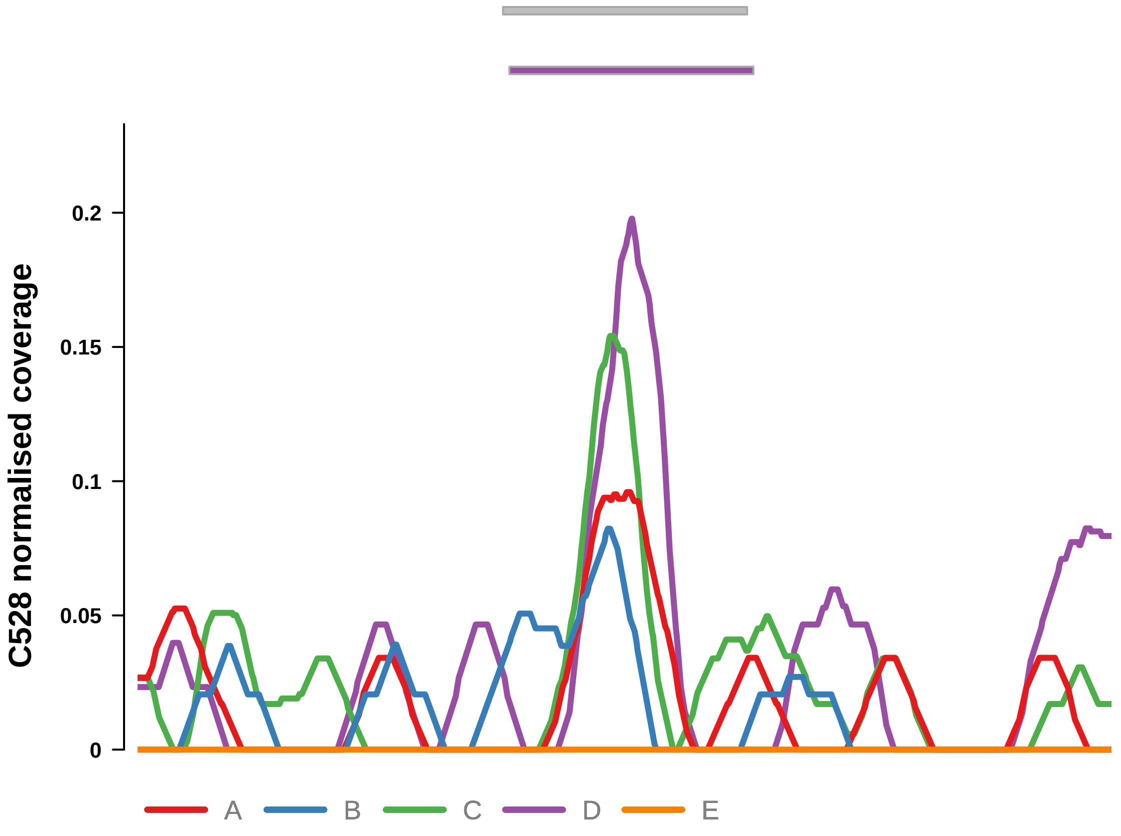


Putatively Clonal (logFC = 2.34)

| | P. | Adjusted P. |
|---|---|---|
| *~region vs. ~* | 6.91e−01 | 8.68e−01 |
| *~purity vs ~* | 6.62e−01 | 7.45e−01 |
| *~purity+region vs ~purity* | 1.28e−01 | 6.24e−01 |
| *Carcinoma vs. normal* | 3.24e−03 ** | 3.24e−03 ** |

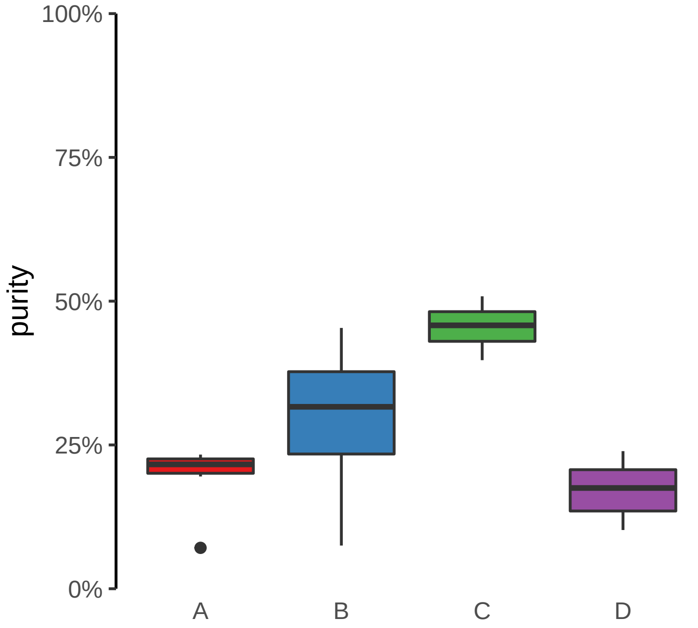

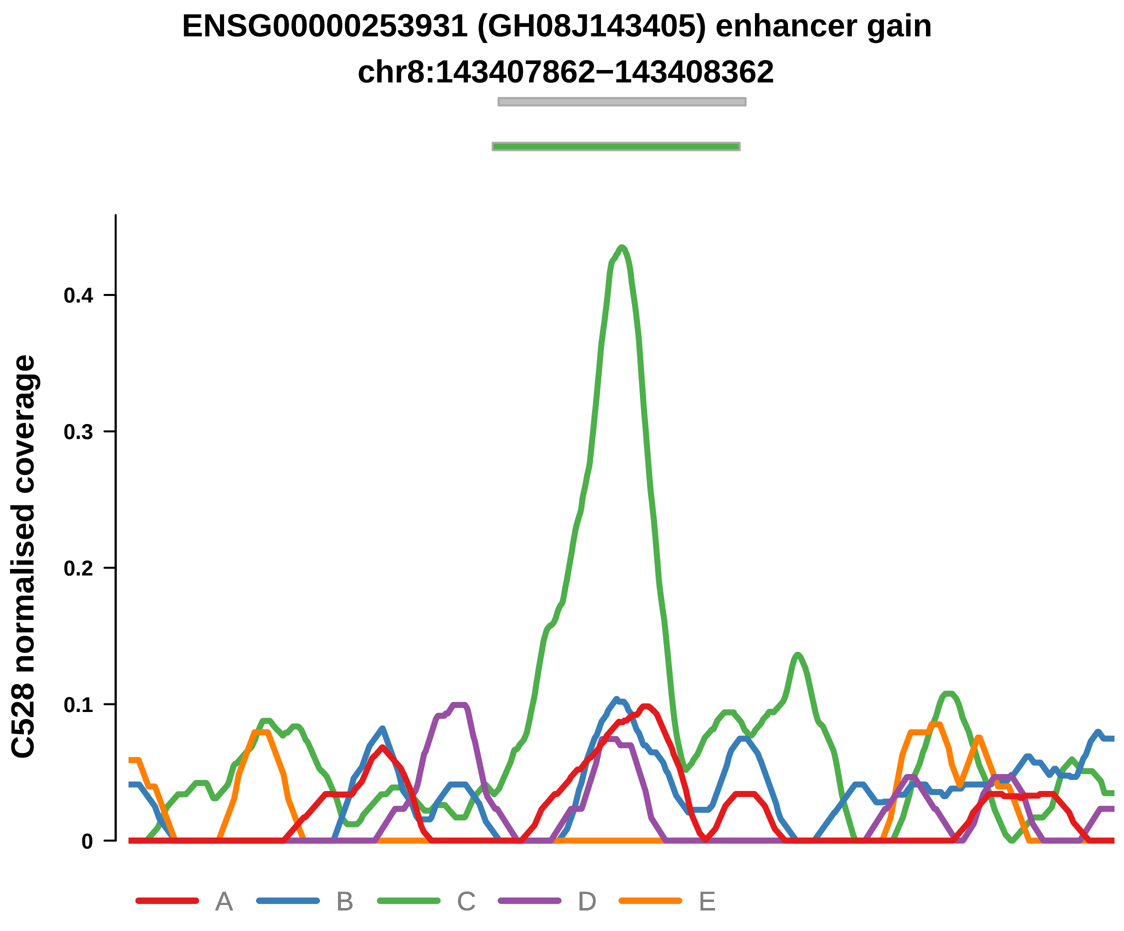


Putatively Clonal (logFC = 2.21)

| | P. | Adjusted P. |
|---|---|---|
| *~region vs. ~* | 2.86e−04 *** | 2.52e−02 * |
| *~purity vs ~* | 2.58e−05 **** | 8.92e−04 *** |
| *~purity+region vs ~purity* | 6.64e−01 | 9.17e−01 |
| *Carcinoma vs. normal* | 2.19e−03 ** | 2.19e−03 ** |

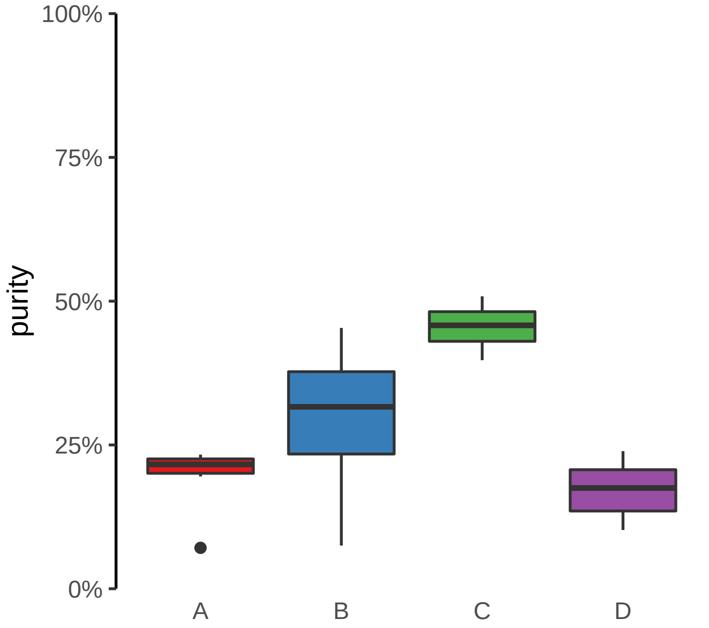

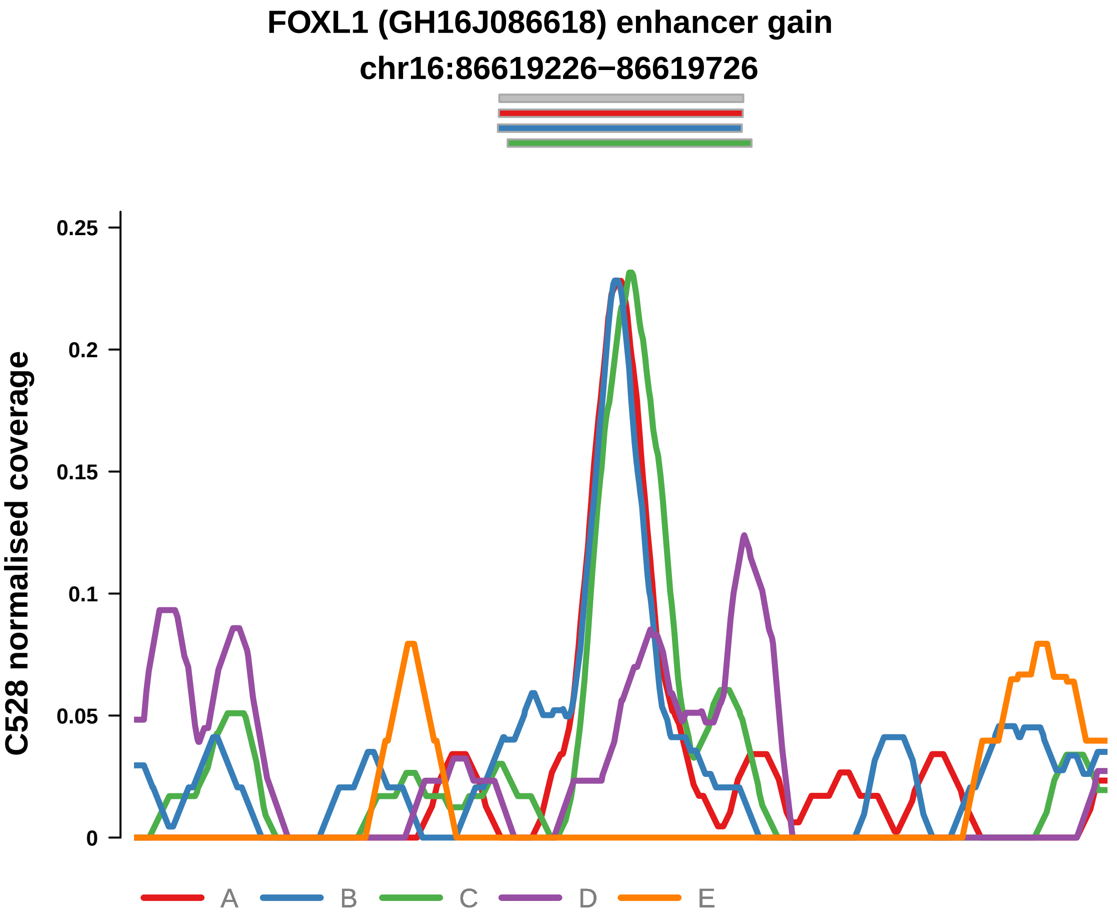


Putatively Clonal (logFC = 2.73)

| | P. | Adjusted P. |
|---|---|---|
| *~region vs. ~* | 8.87e−01 | 9.65e−01 |
| *~purity vs ~* | 3.69e−01 | 5.31e−01 |
| *~purity+region vs ~purity* | 6.06e−02 | 6.11e−01 |
| *Carcinoma vs. normal* | 3.14e−04 *** | 3.14e−04 *** |

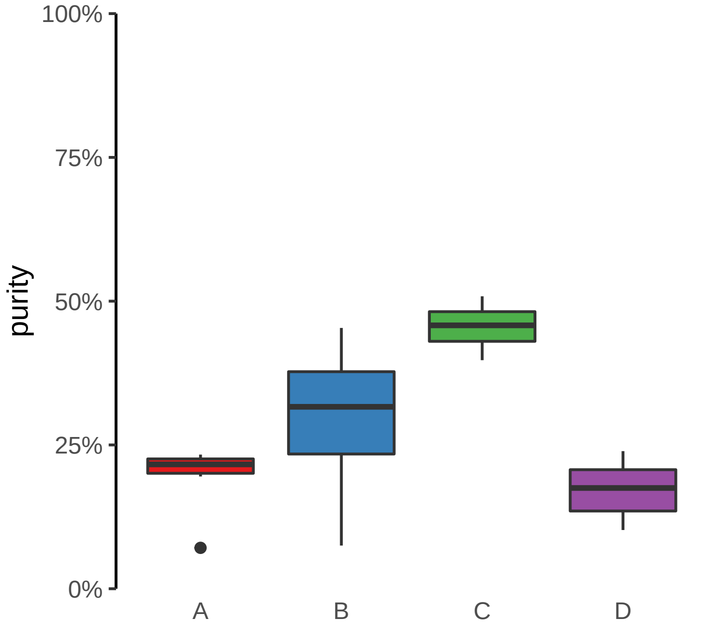

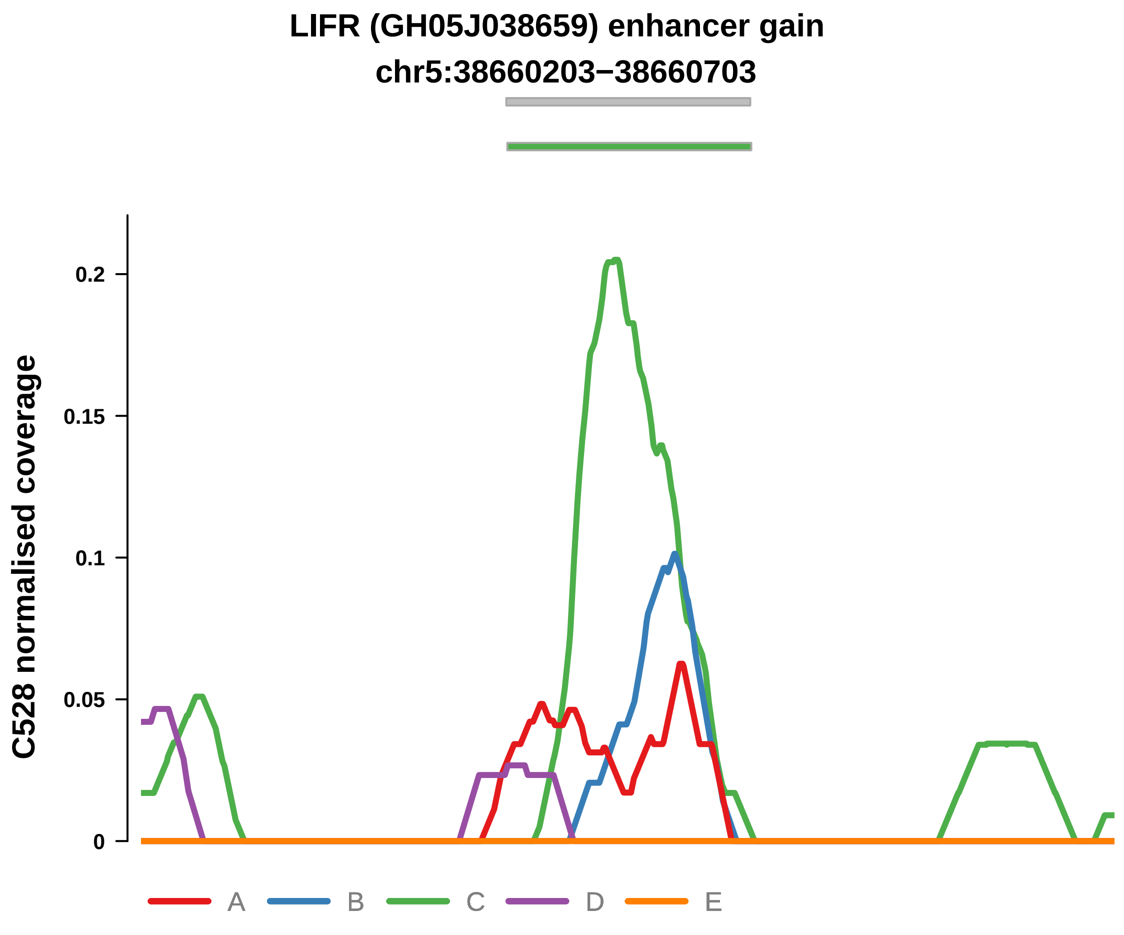


Putatively Clonal (logFC = 2.63)

| | P. | Adjusted P. |
|---|---|---|
| *~region vs. ~* | 6.1e−01 | 7.9e−01 |
| *~purity vs ~* | 7.8e−02 | 1.92e−01 |
| *~purity+region vs ~purity* | 3.43e−01 | 7.26e−01 |
| *Carcinoma vs. normal* | 4.87e−04 *** | 4.87e−04 *** |

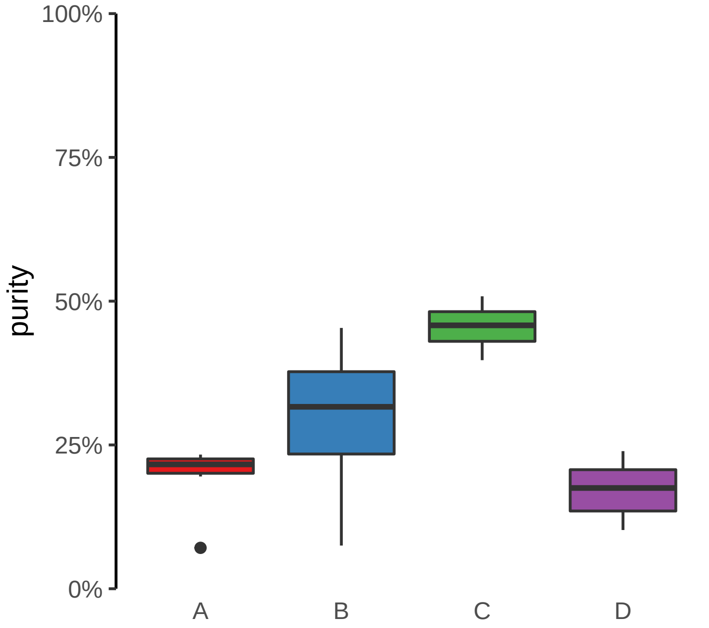

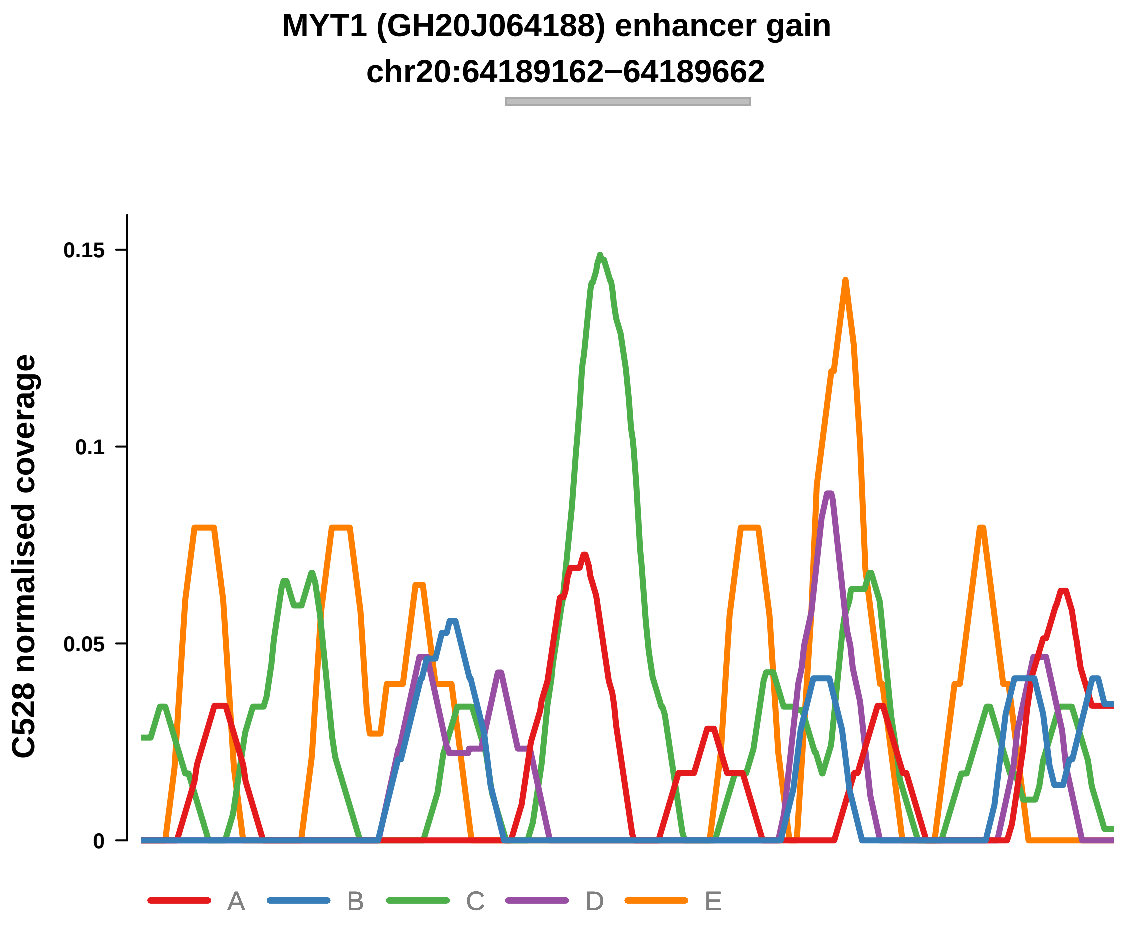


Putatively Clonal (logFC = 2.63)

| | P. | Adjusted P. |
|---|---|---|
| *~region vs. ~* | 4.2e−01 | 6.34e−01 |
| *~purity vs ~* | 3.75e−01 | NA |
| *~purity+region vs ~purity* | 3.1e−01 | 7.26e−01 |
| *Carcinoma vs. normal* | 1.39e−03 ** | 1.39e−03 ** |

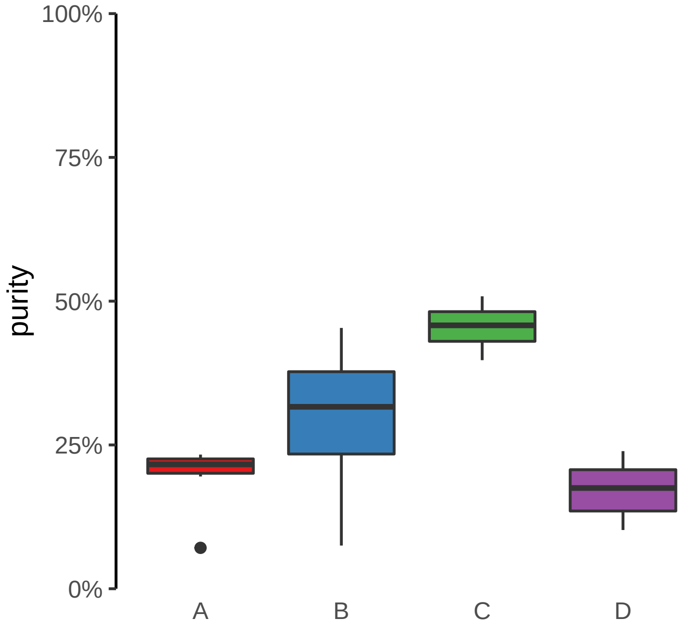

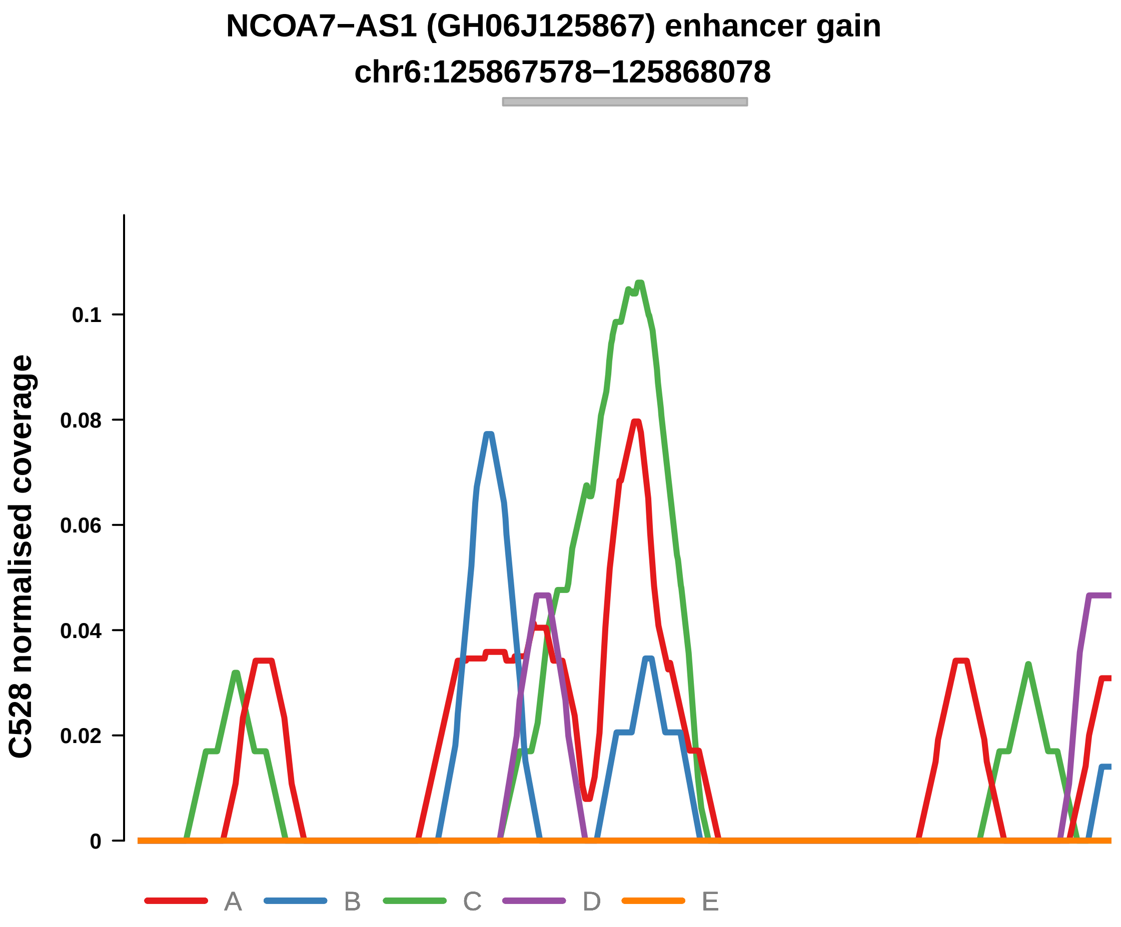


Putatively Clonal (logFC = 2.25)

| | P. | Adjusted P. |
|---|---|---|
| *~region vs. ~* | 9.41e−01 | 9.91e−01 |
| *~purity vs ~* | 7.5e−01 | NA |
| *~purity+region vs ~purity* | 9.1e−01 | 1e+00 |
| *Carcinoma vs. normal* | 5.6e−03 ** | 5.6e−03 ** |

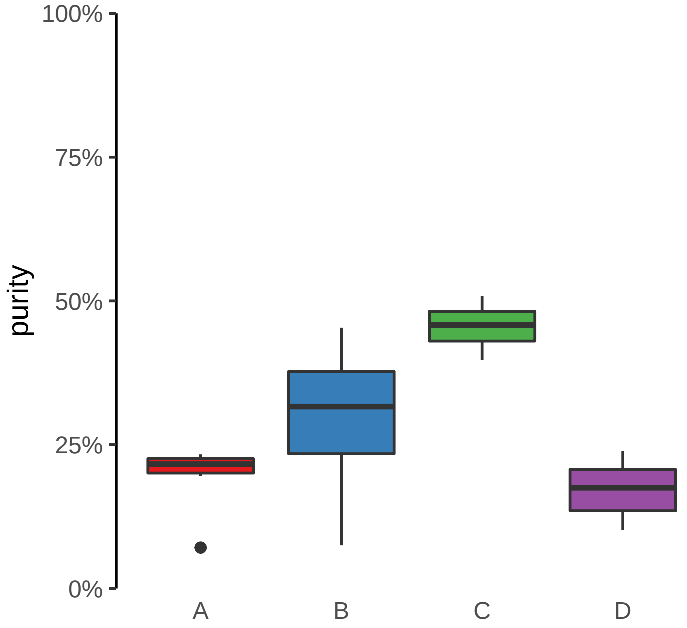

**NFATC2 (GH20J051486) enhancer gain**

**chr20:51487818−51488318**

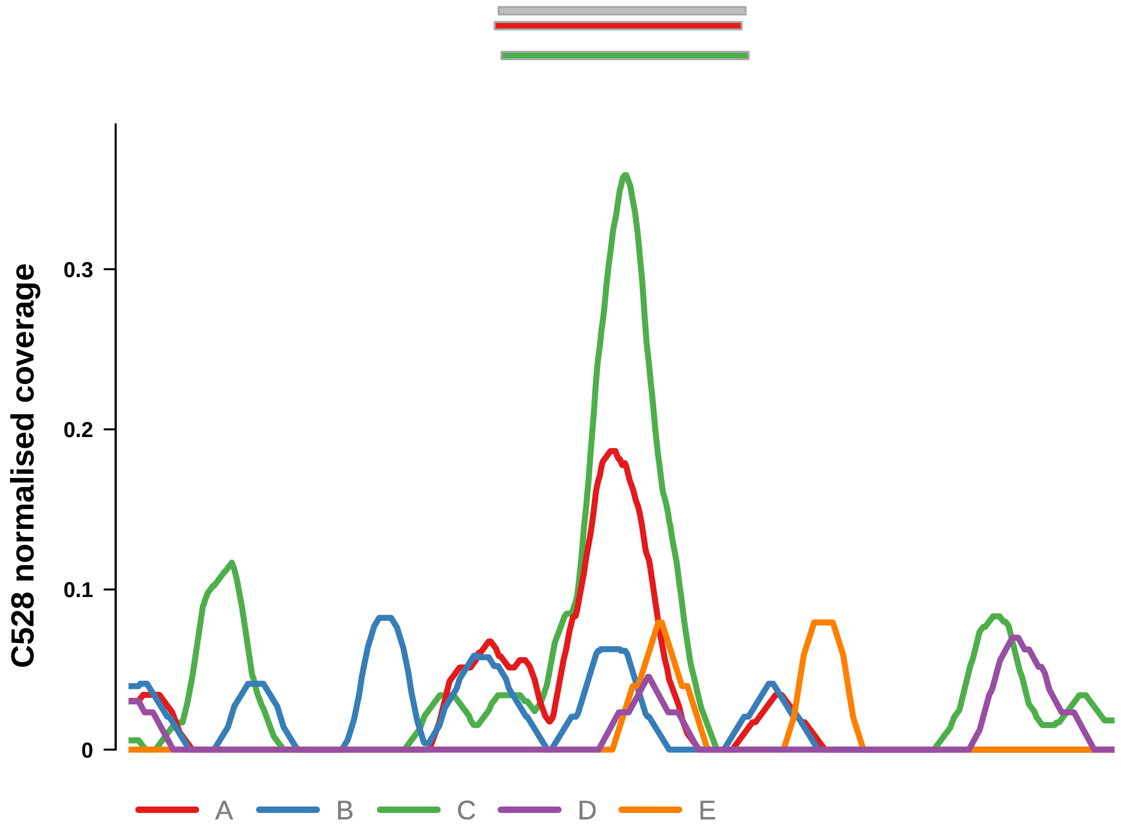


Putatively Clonal (logFC = 3.47)

| | P. | Adjusted P. |
|---|---|---|
| *~region vs. ~* | 7.84e−03 ** | 7.67e−02 |
| *~purity vs ~* | 4.94e−02 * | 1.62e−01 |
| *~purity+region vs ~purity* | 5.45e−02 | 6.11e−01 |
| *Carcinoma vs. normal* | 1.17e−05 **** | 1.17e−05 **** |

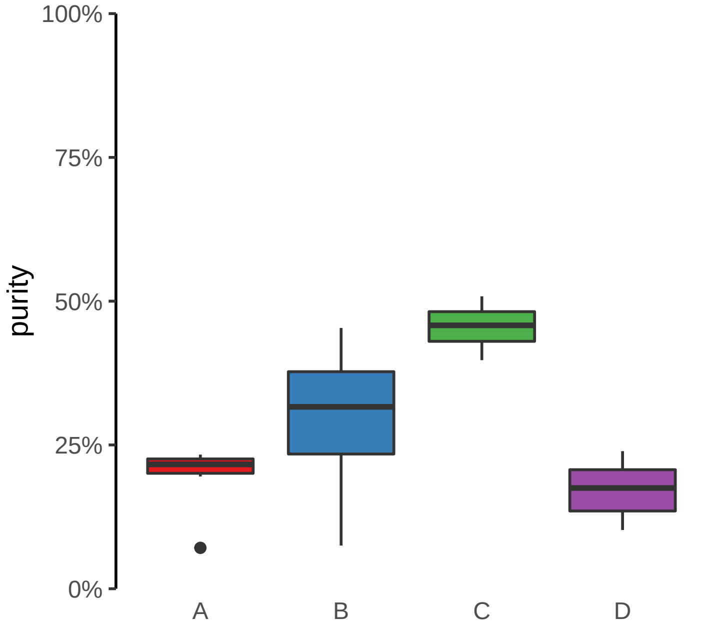

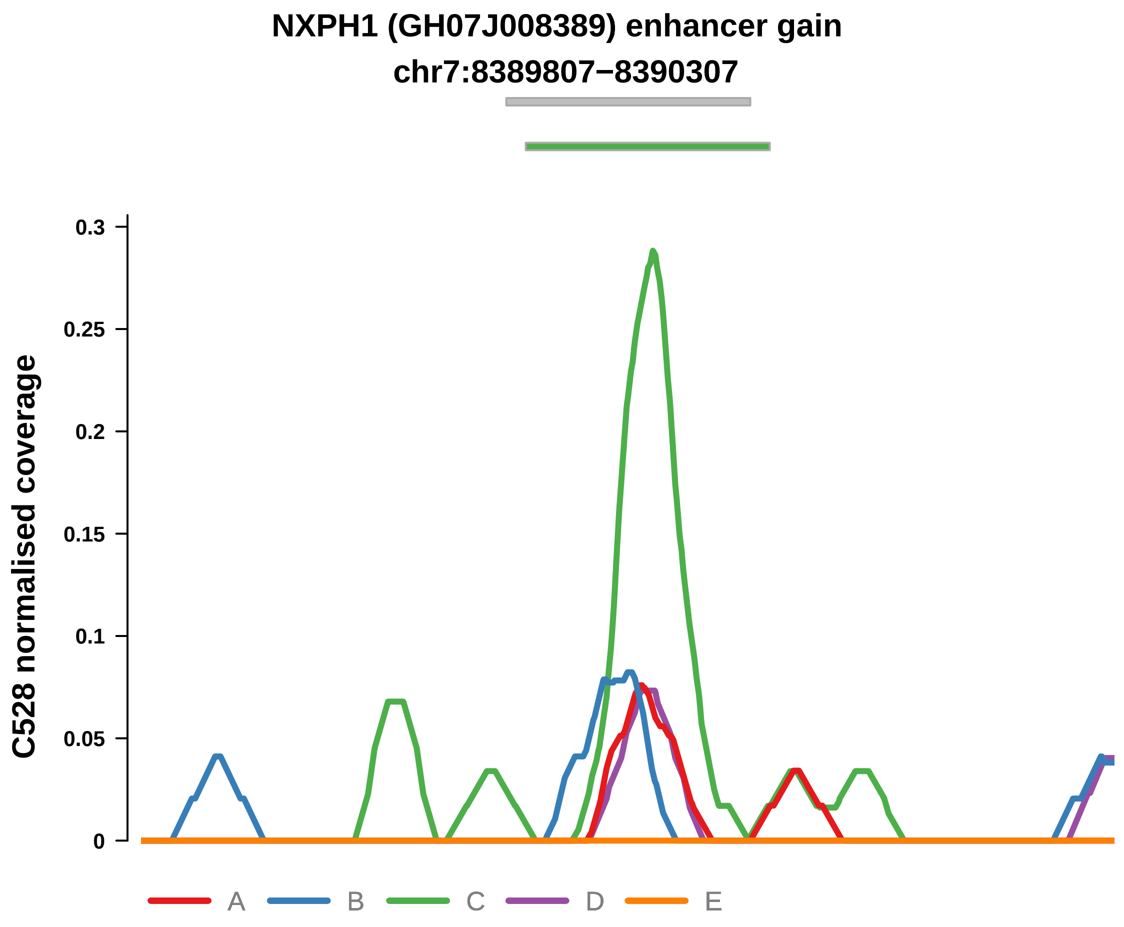


Putatively Clonal (logFC = 2.07)

| | P. | Adjusted P. |
|---|---|---|
| *~region vs. ~* | 3.47e−02 * | 2.21e−01 |
| *~purity vs ~* | 3.68e−02 * | 1.59e−01 |
| *~purity+region vs ~purity* | 1.43e−01 | 6.24e−01 |
| *Carcinoma vs. normal* | 7.42e−03 ** | 7.42e−03 ** |

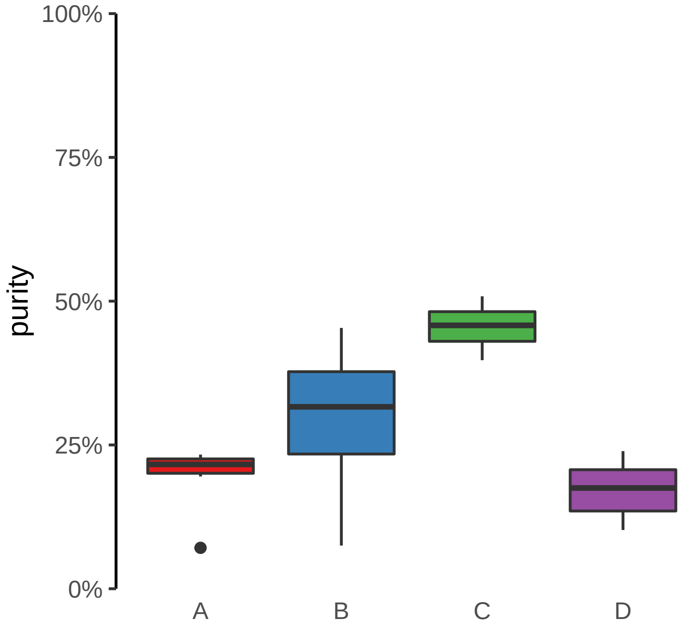

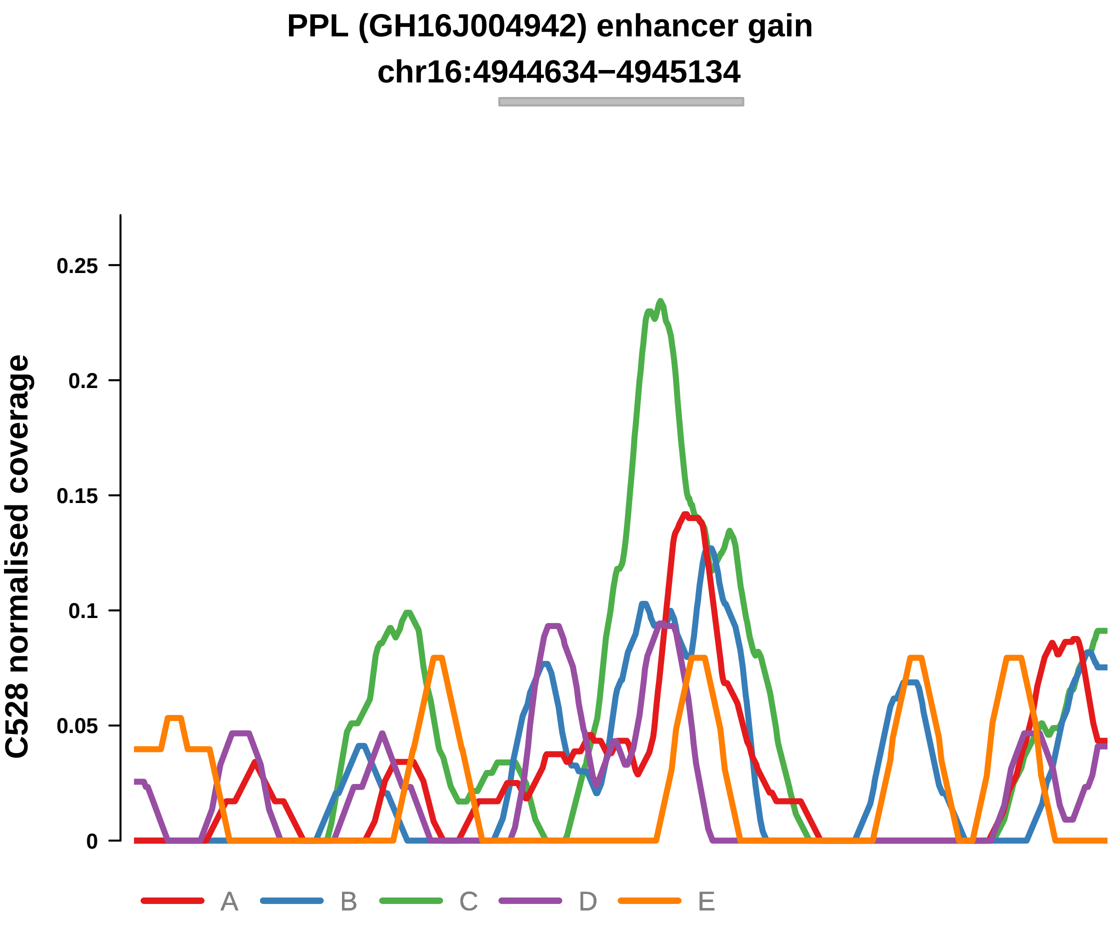


Putatively Clonal (logFC = 2.18)

| | P. | Adjusted P. |
|---|---|---|
| *~region vs. ~* | 4.09e−01 | 6.31e−01 |
| *~purity vs ~* | 4.83e−02 * | 1.62e−01 |
| *~purity+region vs ~purity* | 9.64e−01 | 1e+00 |
| *Carcinoma vs. normal* | 3.21e−03 ** | 3.21e−03 ** |

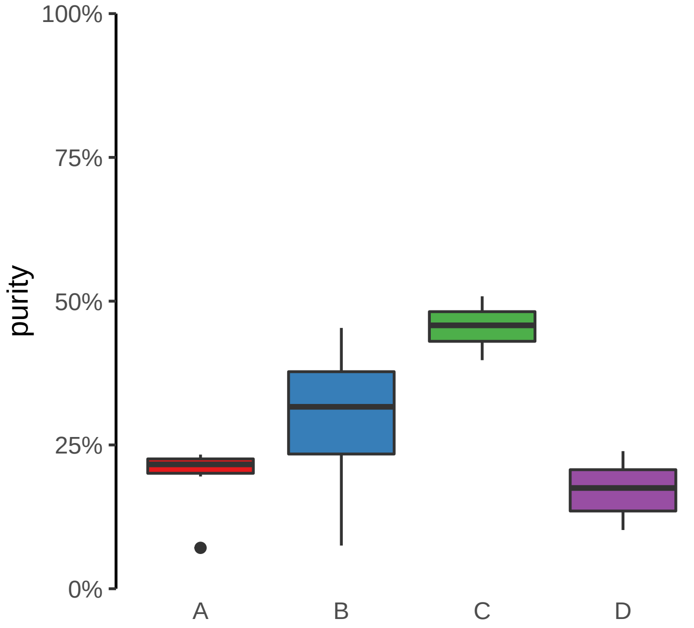

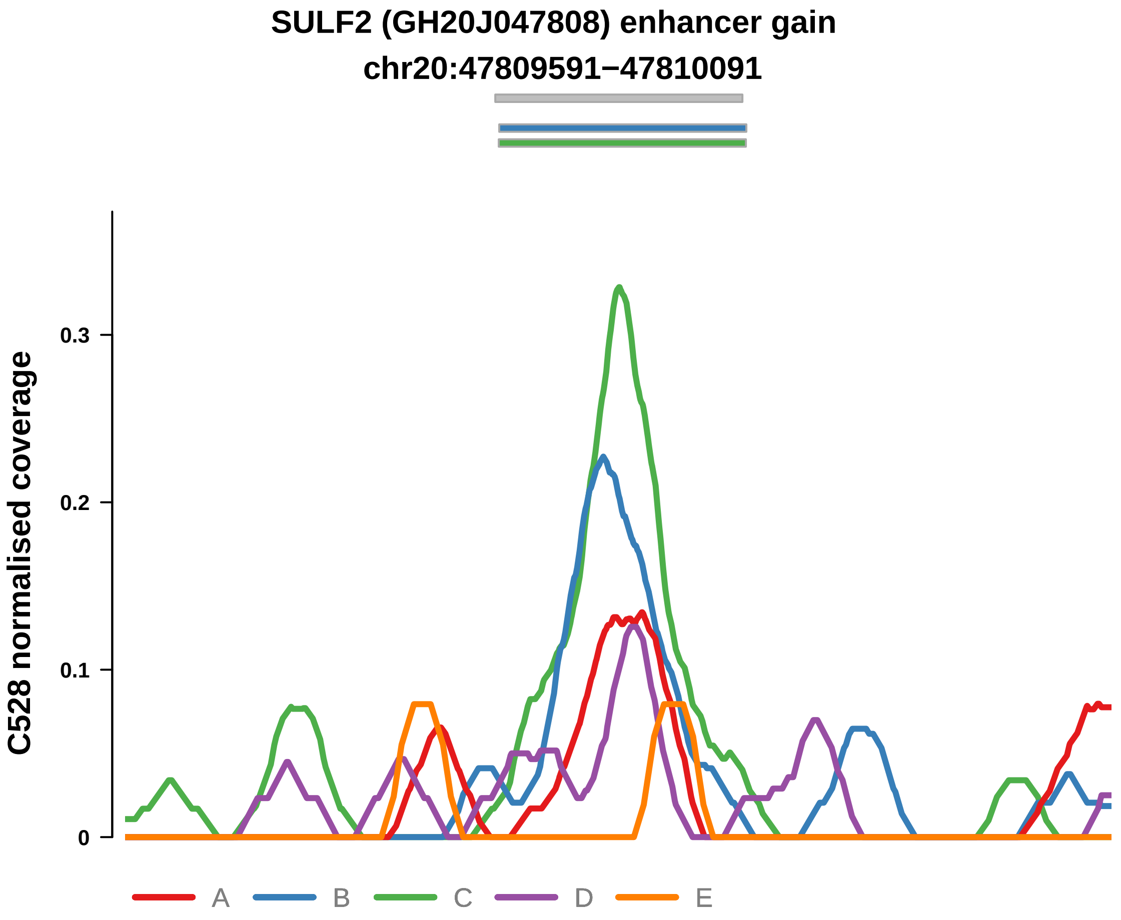


Putatively Clonal (logFC = 2.76)

| | P. | Adjusted P. |
|---|---|---|
| *~region vs. ~* | 5.04e−02 | 2.46e−01 |
| *~purity vs ~* | 1.41e−02 * | 9.7e−02 |
| *~purity+region vs ~purity* | 3.33e−01 | 7.26e−01 |
| *Carcinoma vs. normal* | 1.77e−04 *** | 1.77e−04 *** |

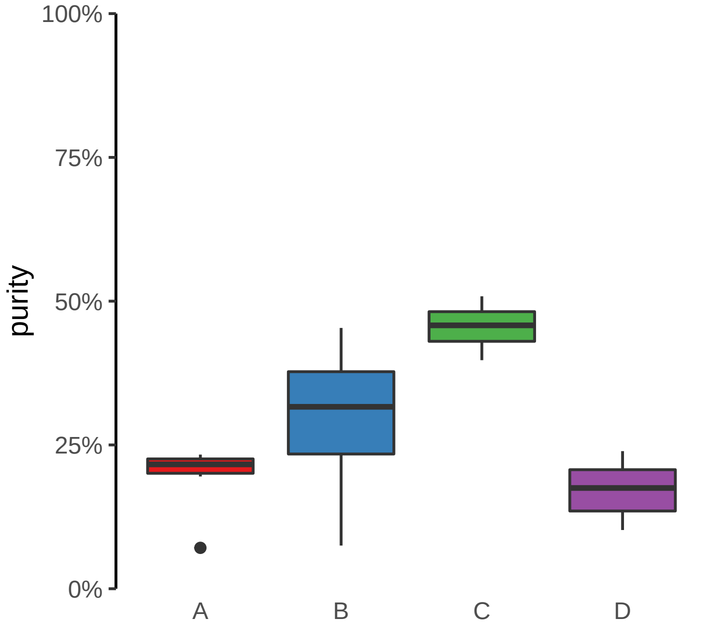

**TEX26 (GH13J030867) enhancer gain**
**chr13:30870352−30870852**

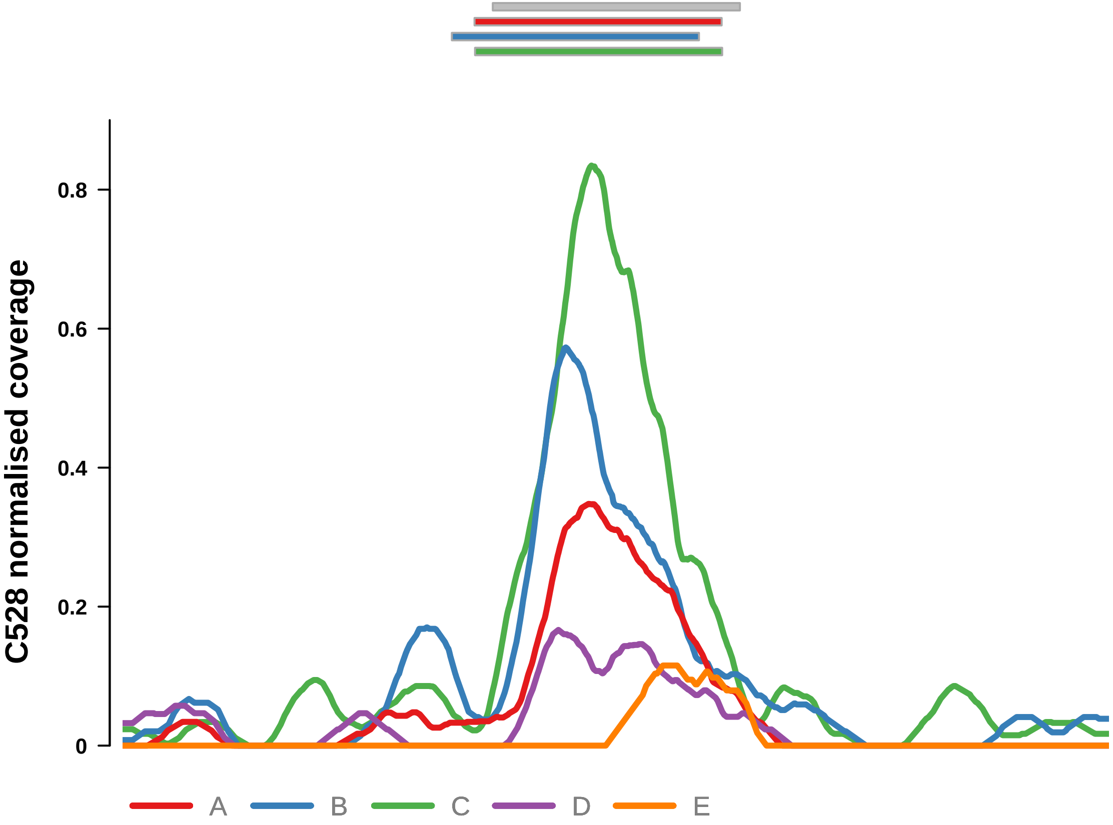


Putatively Clonal (logFC = 3.22)

| | P. | Adjusted P. |
|---|---|---|
| *~region vs. ~* | 2.6e−03 ** | 5.14e−02 |
| *~purity vs ~* | 9.89e−03 ** | 8.53e−02 |
| *~purity+region vs ~purity* | 9.73e−02 | 6.11e−01 |
| *Carcinoma vs. normal* | 8.46e−06 **** | 8.46e−06 **** |

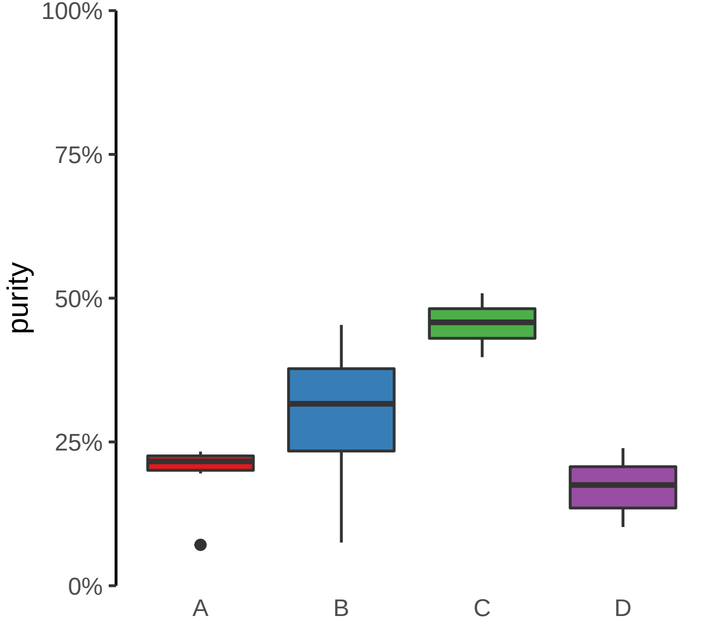

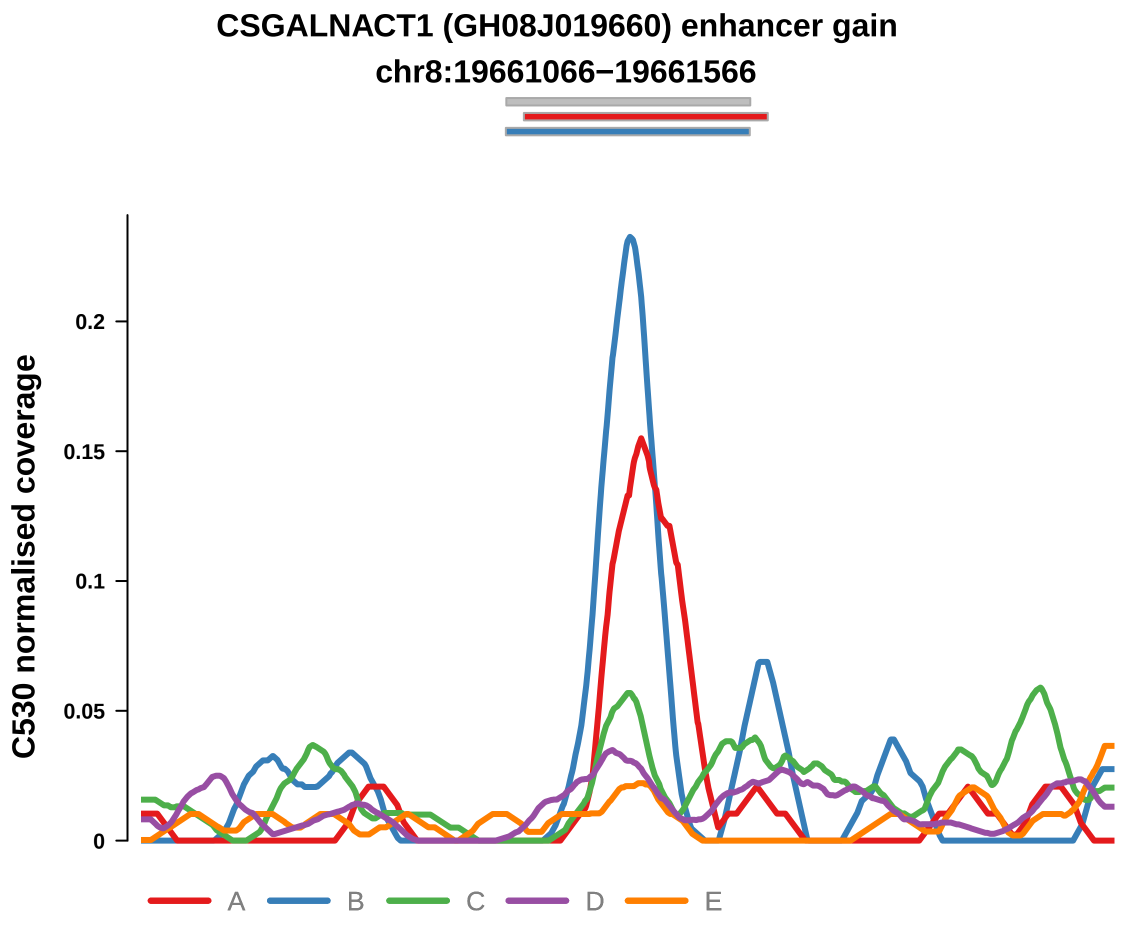


Putatively Clonal (logFC = 2.1)

| | P. | Adjusted P. |
|---|---|---|
| *~region vs. ~* | 6.88e−02 | 3.55e−01 |
| *~purity vs ~* | 1.76e−01 | 5.54e−01 |
| *~purity+region vs ~purity* | 6.01e−02 | 3.5e−01 |
| *Carcinoma vs. normal* | 3.44e−03 ** | 3.44e−03 ** |

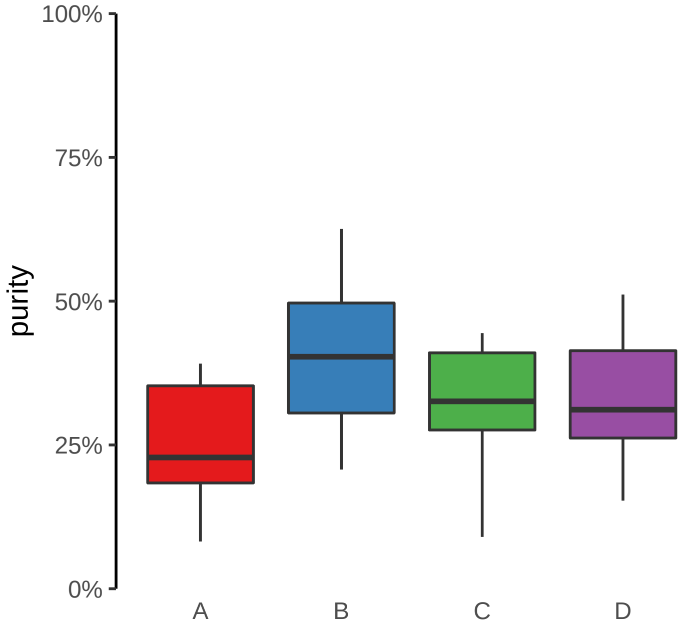

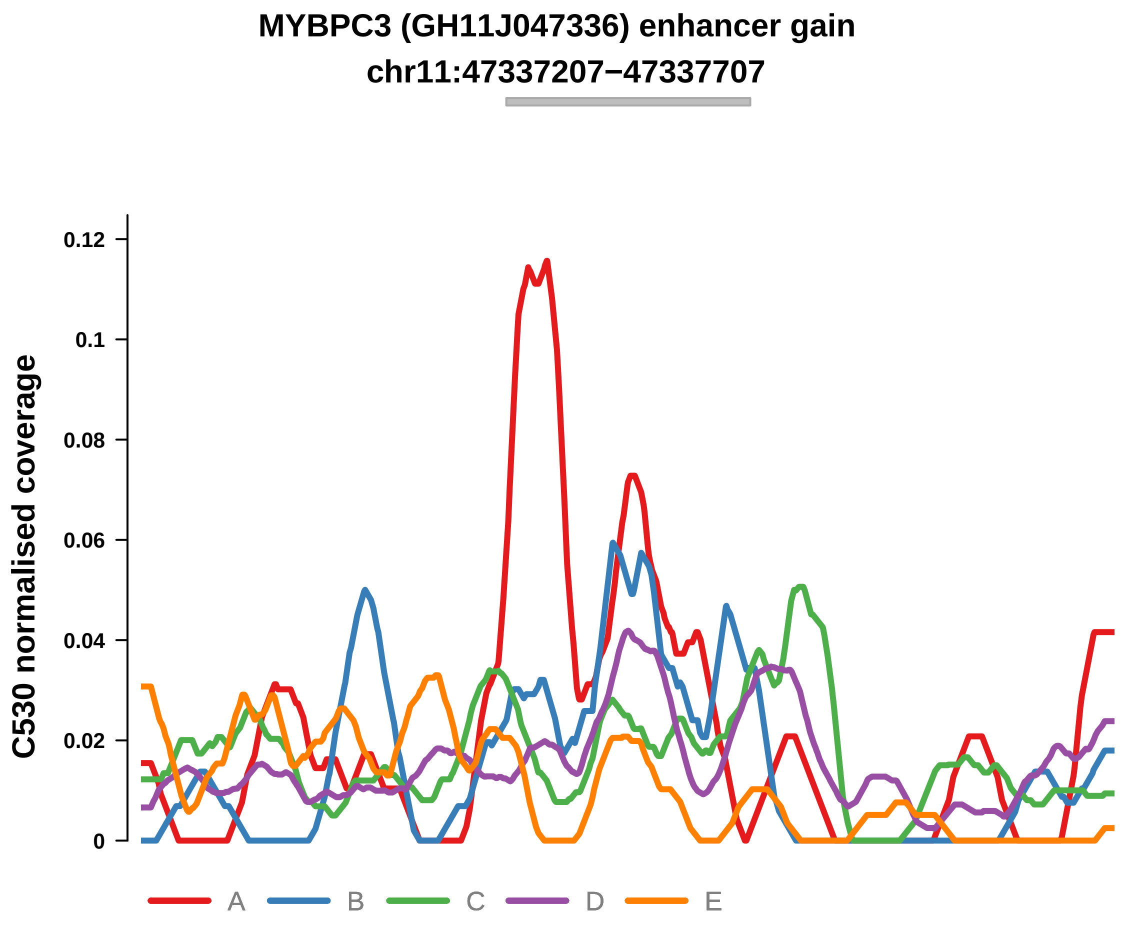


Putatively Clonal (logFC = 1.97)

| | P. | Adjusted P. |
|---|---|---|
| *~region vs. ~* | 7.57e−01 | 8.86e−01 |
| *~purity vs ~* | 1.18e−01 | 4.33e−01 |
| *~purity+region vs ~purity* | 4.85e−01 | 7.59e−01 |
| *Carcinoma vs. normal* | 7.1e−03 ** | 7.1e−03 ** |

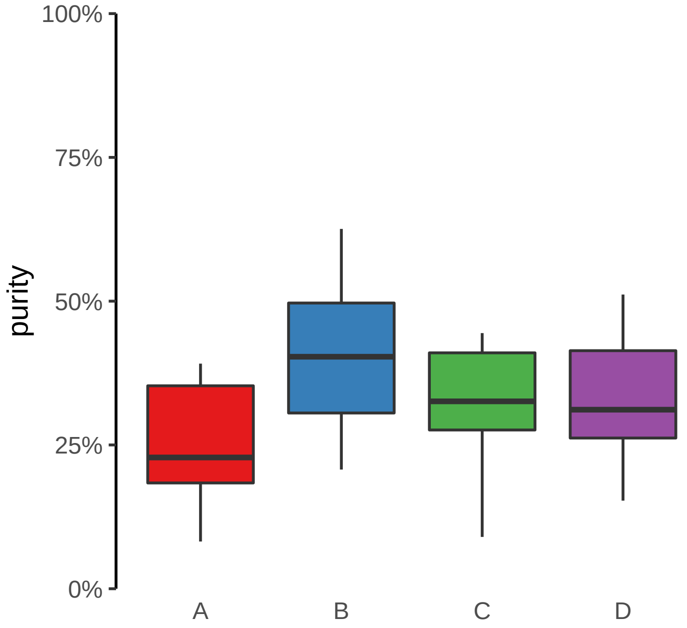

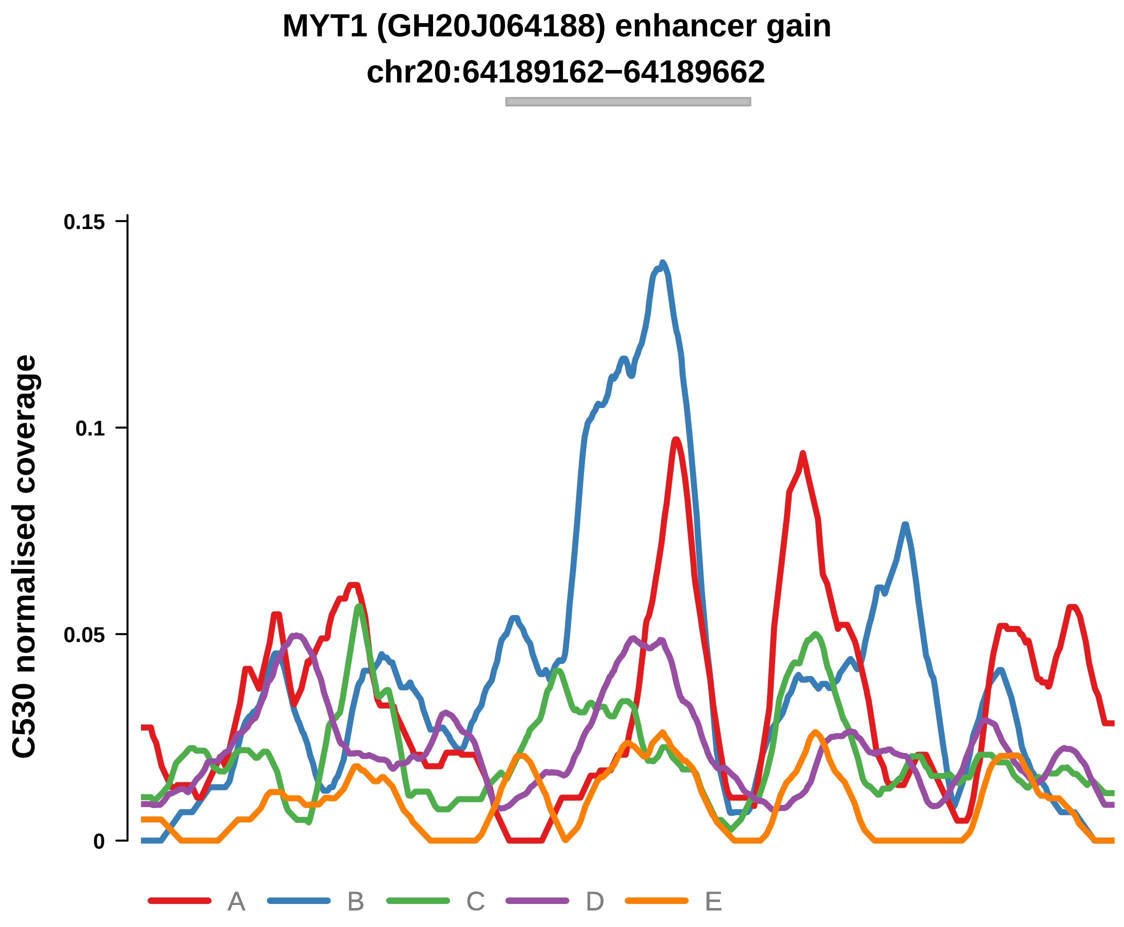


Putatively Clonal (logFC = 2.55)

| | P. | Adjusted P. |
|---|---|---|
| *~region vs. ~* | 3.89e−02 * | 2.52e−01 |
| *~purity vs ~* | 1.18e−01 | 4.33e−01 |
| *~purity+region vs ~purity* | 9.89e−02 | 3.5e−01 |
| *Carcinoma vs. normal* | 4.97e−04 *** | 4.97e−04 *** |

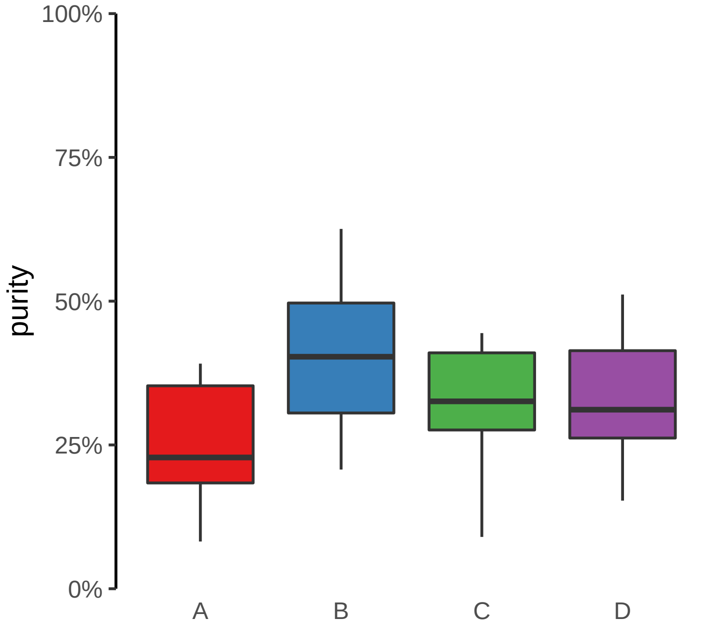

**NFATC2 (GH20J051486) enhancer gain**
**chr20:51487818−51488318**

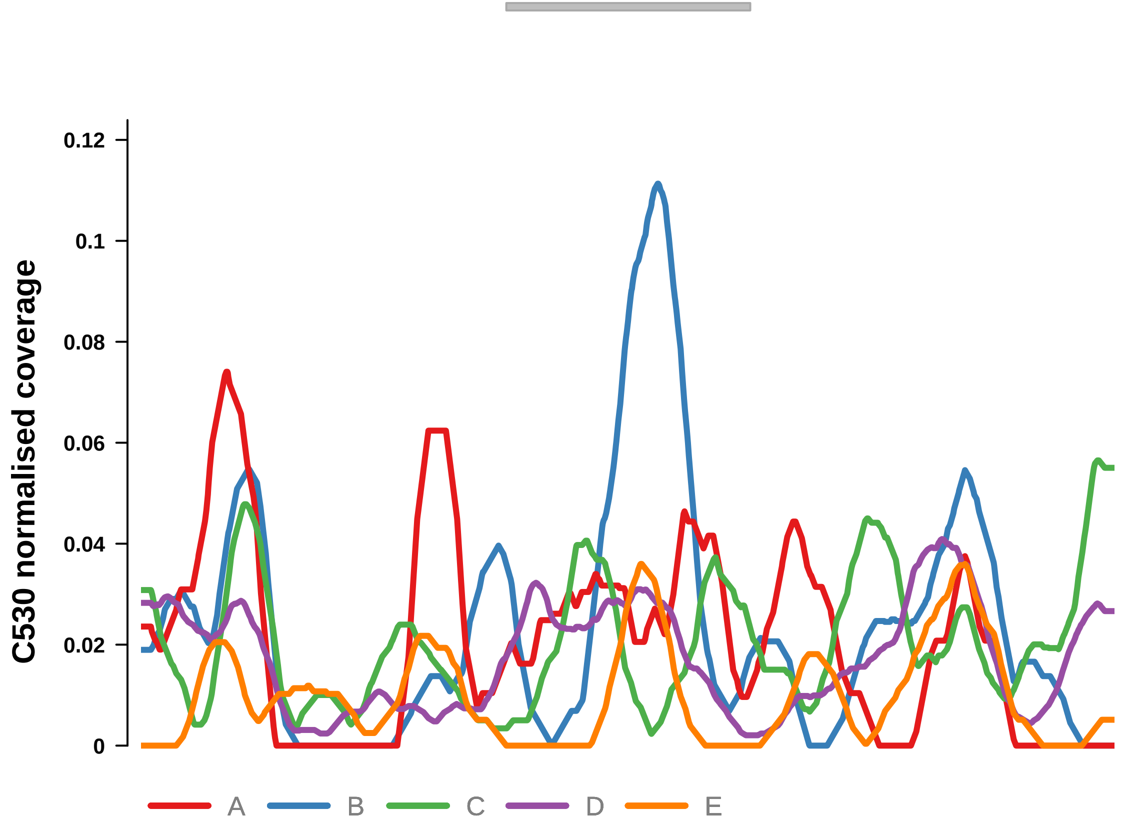


Putatively Clonal (logFC = 2.08)

| | P. | Adjusted P. |
|---|---|---|
| *~region vs. ~* | 6.77e−01 | 8.86e−01 |
| *~purity vs ~* | 5.85e−03 ** | 1.72e−01 |
| *~purity+region vs ~purity* | 9.81e−01 | 9.84e−01 |
| *Carcinoma vs. normal* | 4.5e−03 ** | 4.5e−03 ** |

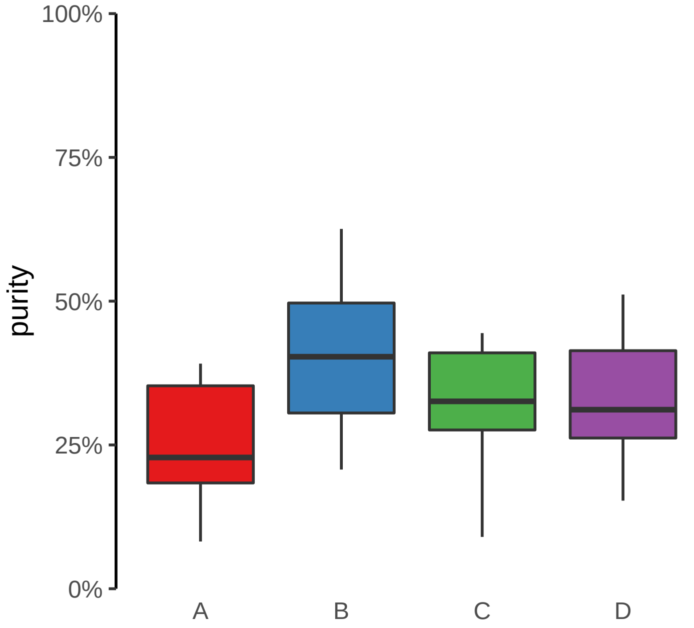

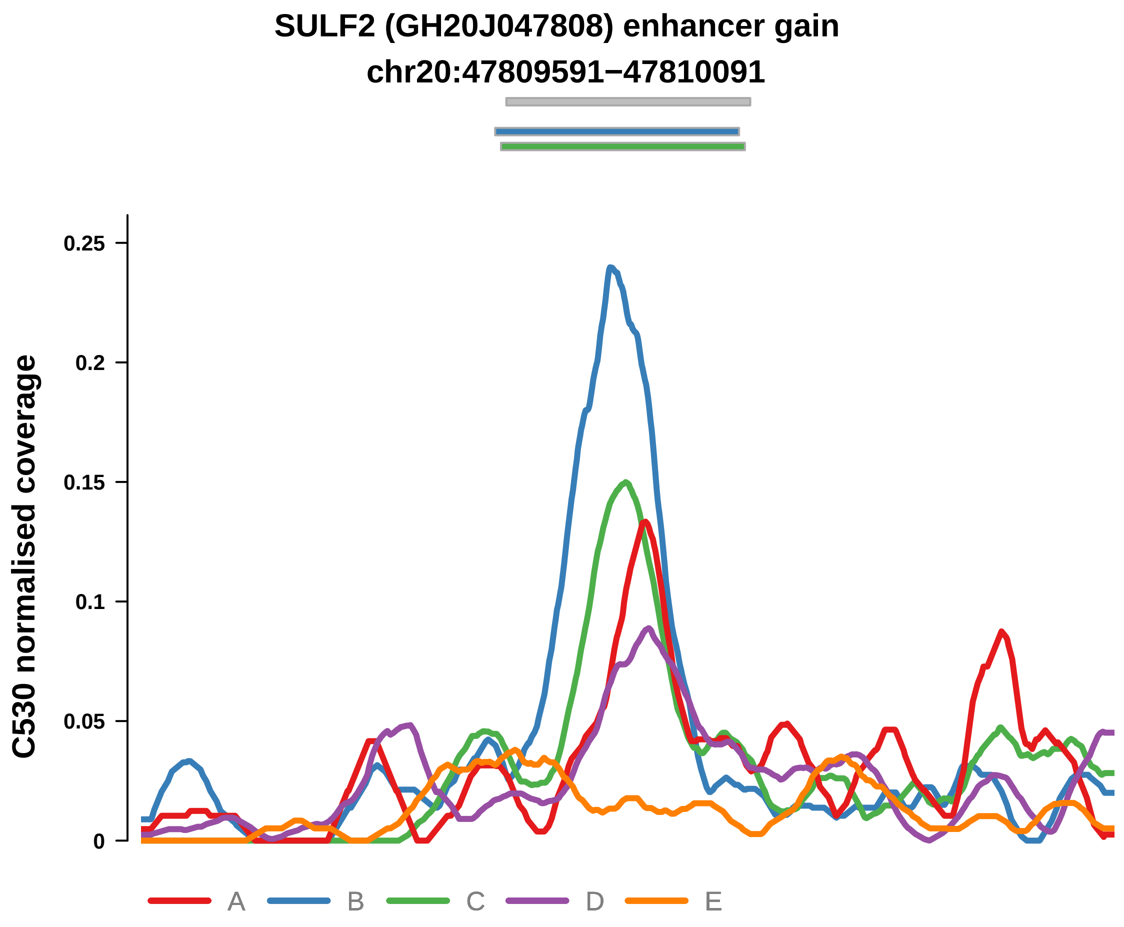


Putatively Clonal (logFC = 2.2)

| | P. | Adjusted P. |
|---|---|---|
| *~region vs. ~* | 3.55e−01 | 7.02e−01 |
| *~purity vs ~* | 4.29e−03 ** | 1.72e−01 |
| *~purity+region vs ~purity* | 8.16e−01 | 8.58e−01 |
| *Carcinoma vs. normal* | 1.71e−03 ** | 1.71e−03 ** |

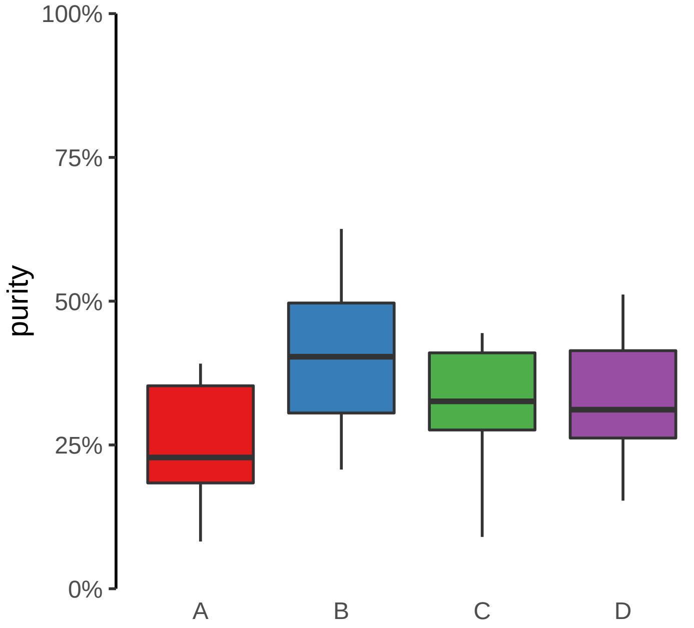

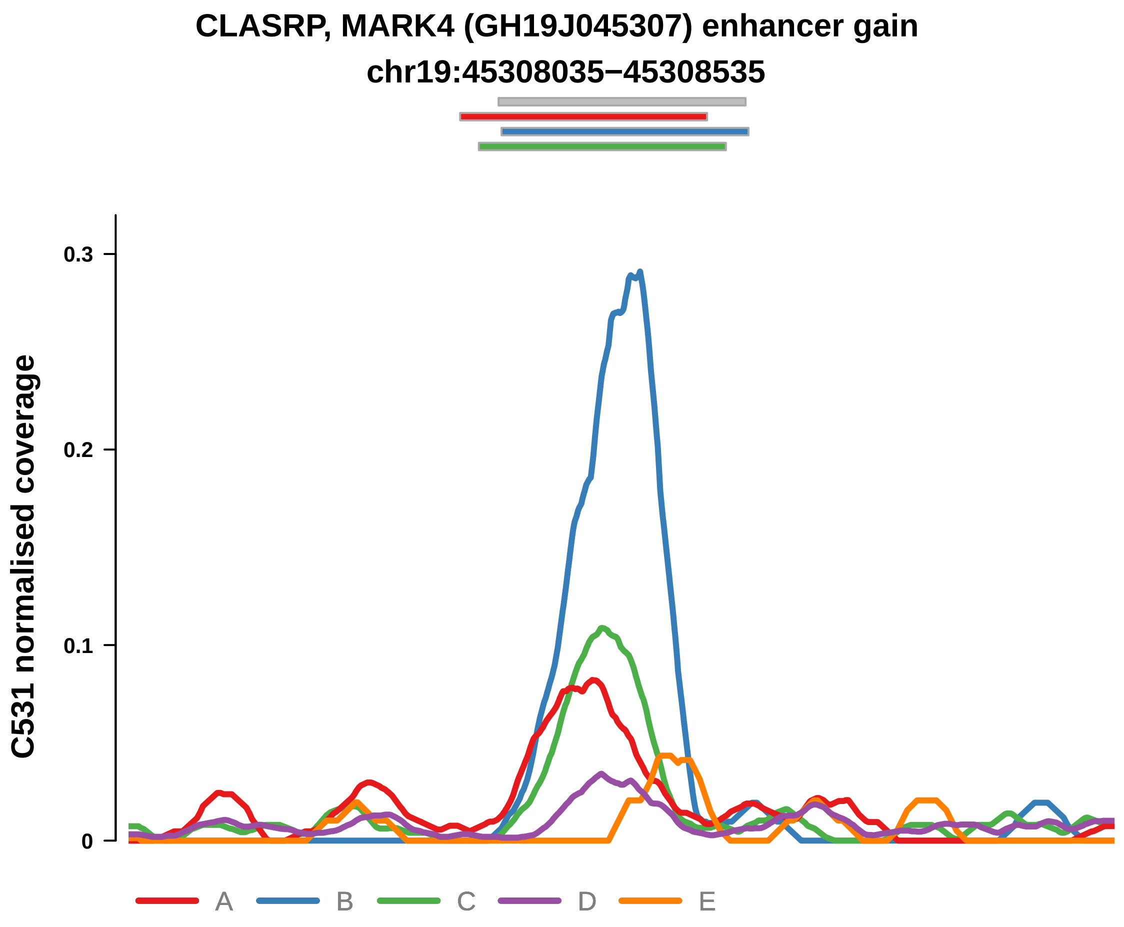


Putatively Clonal (logFC = 2.1)

| | P. | Adjusted P. |
|---|---|---|
| *~region vs. ~* | 3.6e−06 **** | 8e−05 **** |
| *~purity vs ~* | 5.13e−03 ** | 3.76e−02 * |
| *~purity+region vs ~purity* | 1.73e−03 ** | 6.55e−02 |
| *Carcinoma vs. normal* | 5.28e−03 ** | 5.28e−03 ** |

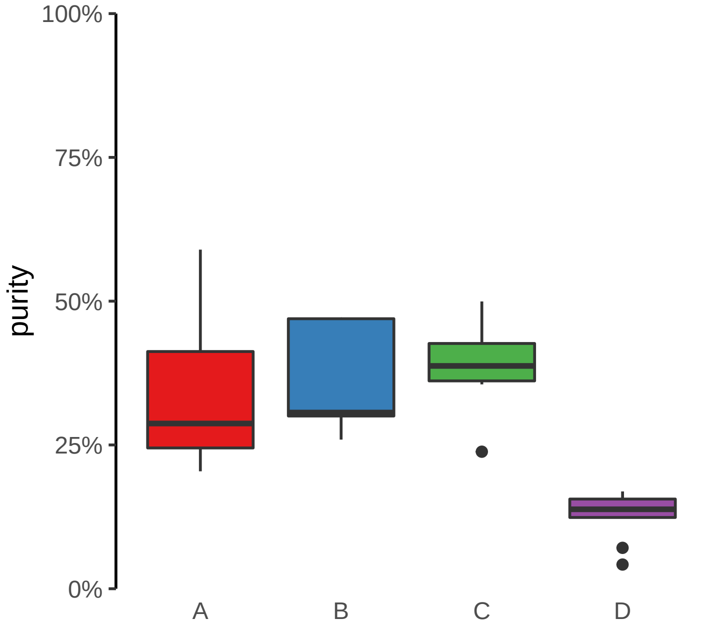

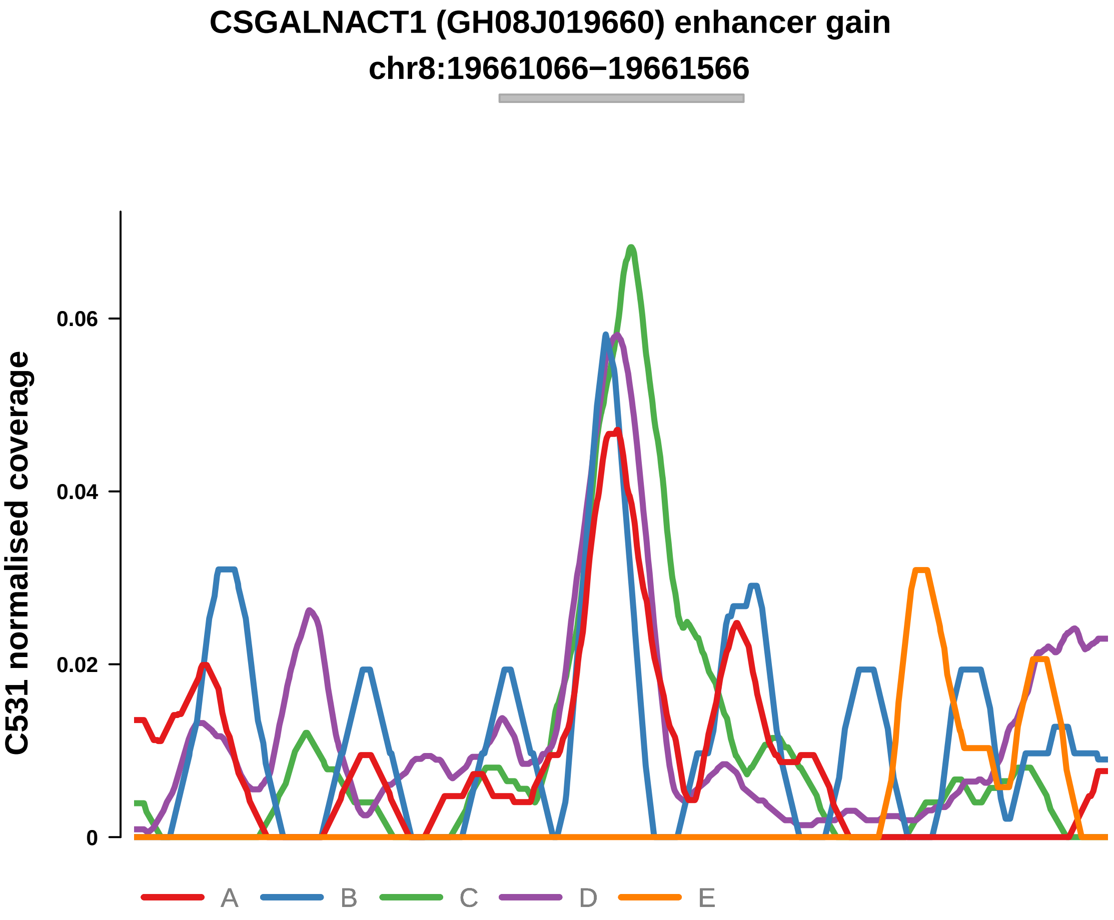


Putatively Clonal (logFC = 2.09)

| | P. | Adjusted P. |
|---|---|---|
| *~region vs. ~* | 2.41e−01 | 3.91e−01 |
| *~purity vs ~* | 4.9e−01 | 6.35e−01 |
| *~purity+region vs ~purity* | 2.56e−01 | 6.13e−01 |
| *Carcinoma vs. normal* | 9.72e−03 ** | 9.72e−03 ** |

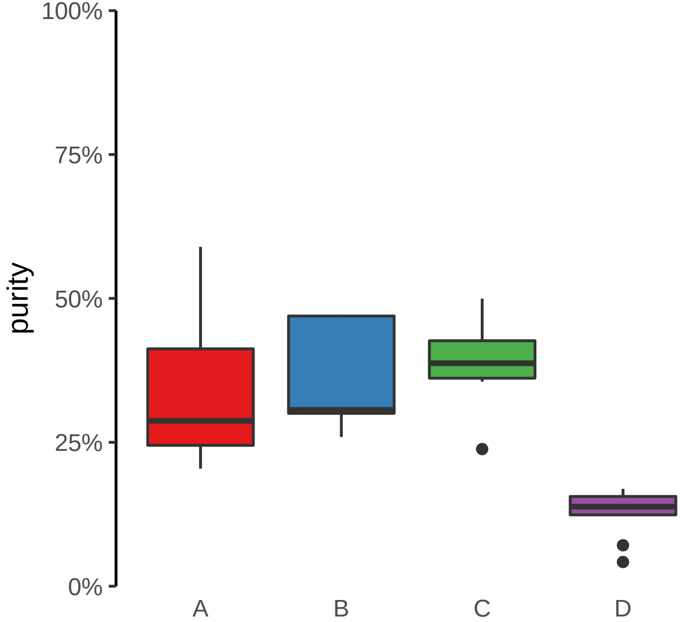

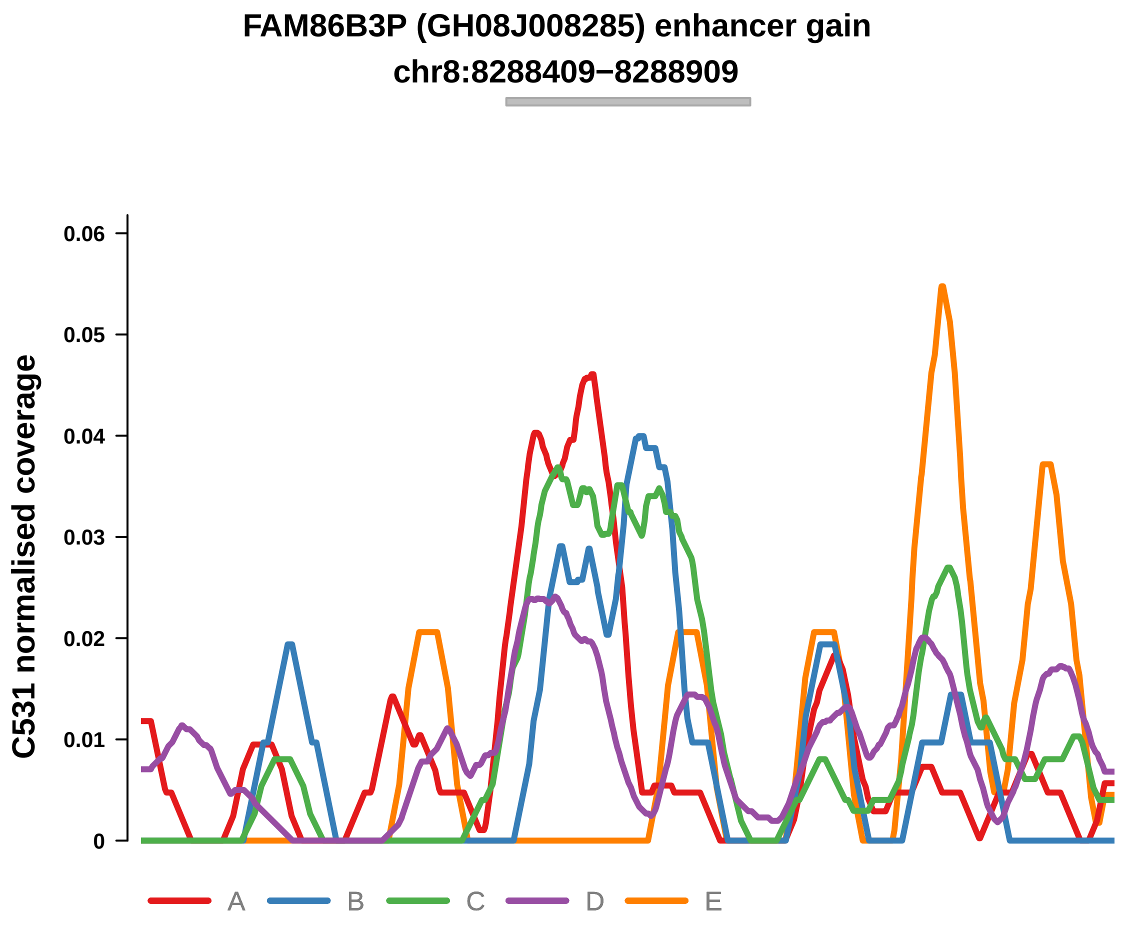


Putatively Clonal (logFC = 2.2)

| | P. | Adjusted P. |
|---|---|---|
| *~region vs. ~* | 1.5e−01 | NA |
| *~purity vs ~* | 1.72e−01 | 3.27e−01 |
| *~purity+region vs ~purity* | 3.11e−01 | 6.13e−01 |
| *Carcinoma vs. normal* | 6.45e−03 ** | 6.45e−03 ** |

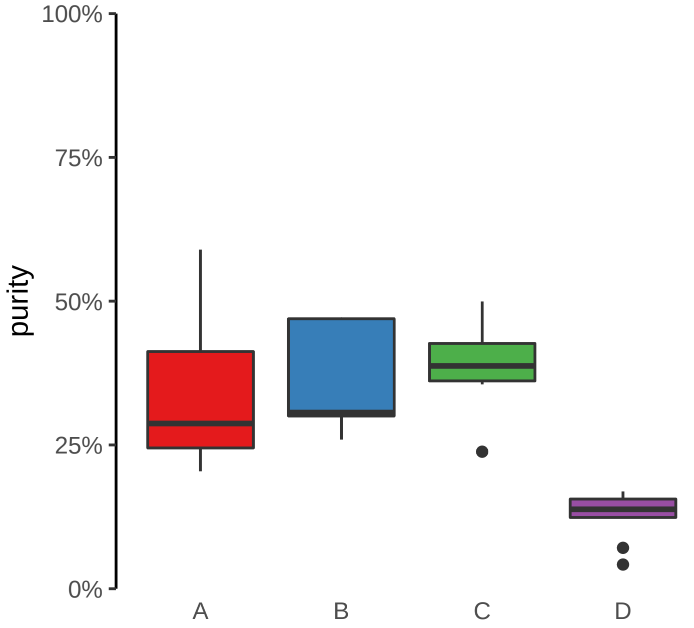

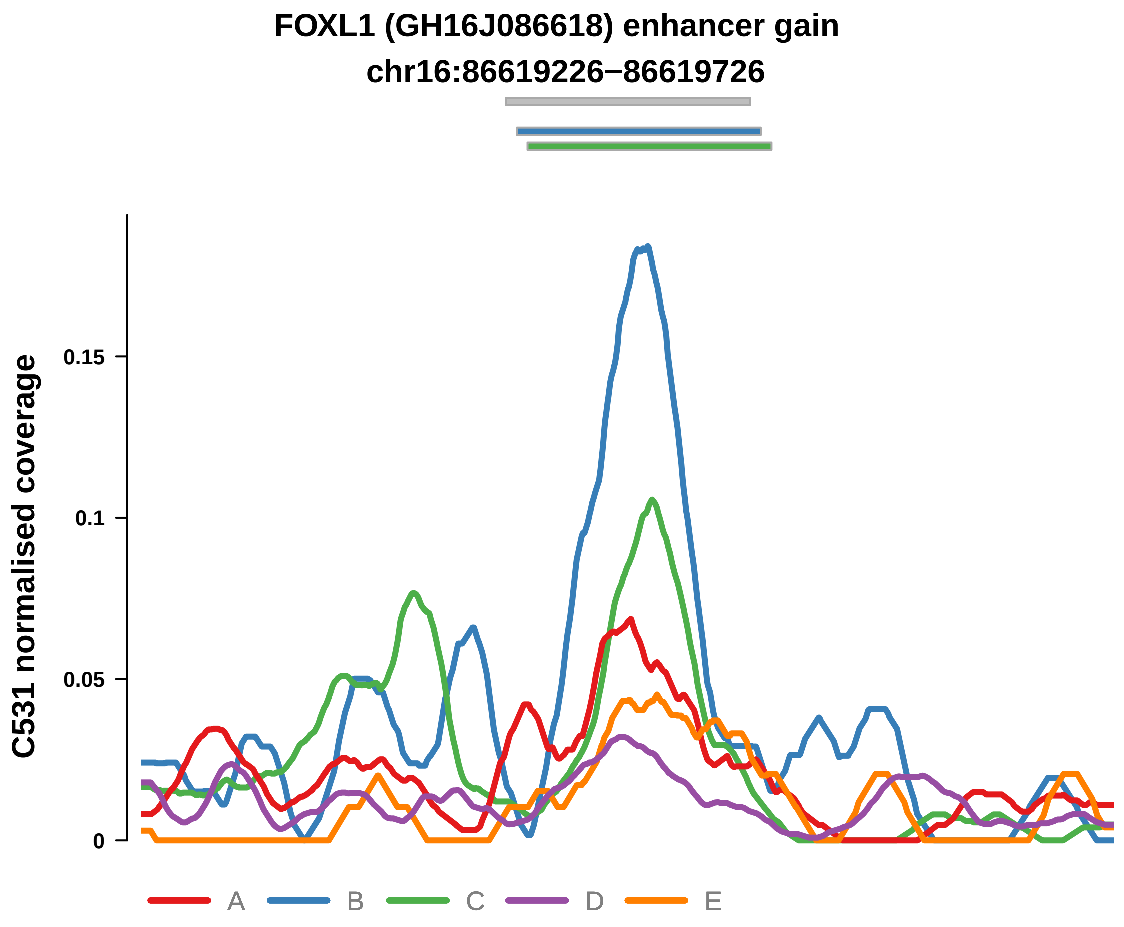


Putatively Clonal (logFC = 2.78)

| | P. | Adjusted P. |
|---|---|---|
| *~region vs. ~* | 7.11e−06 **** | 9.24e−05 **** |
| *~purity vs ~* | 1.53e−05 **** | 6.75e−04 *** |
| *~purity+region vs ~purity* | 1.19e−01 | 5.23e−01 |
| *Carcinoma vs. normal* | 2.4e−04 *** | 2.4e−04 *** |

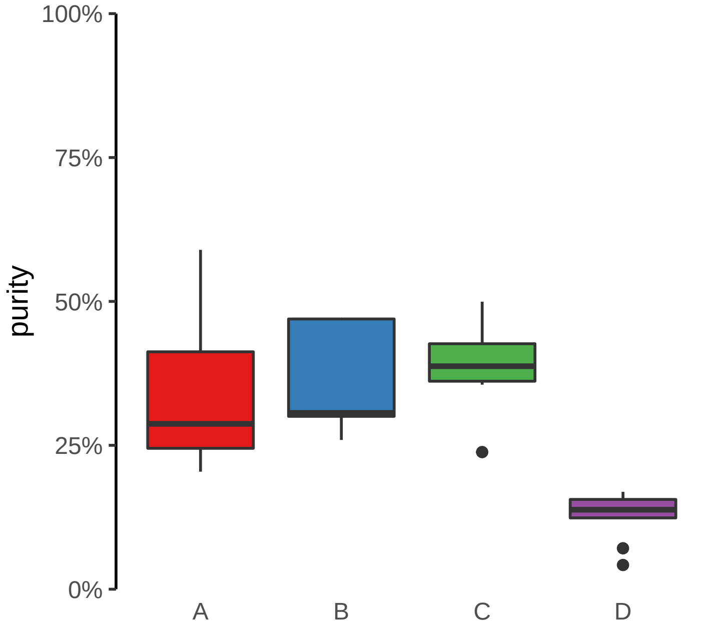

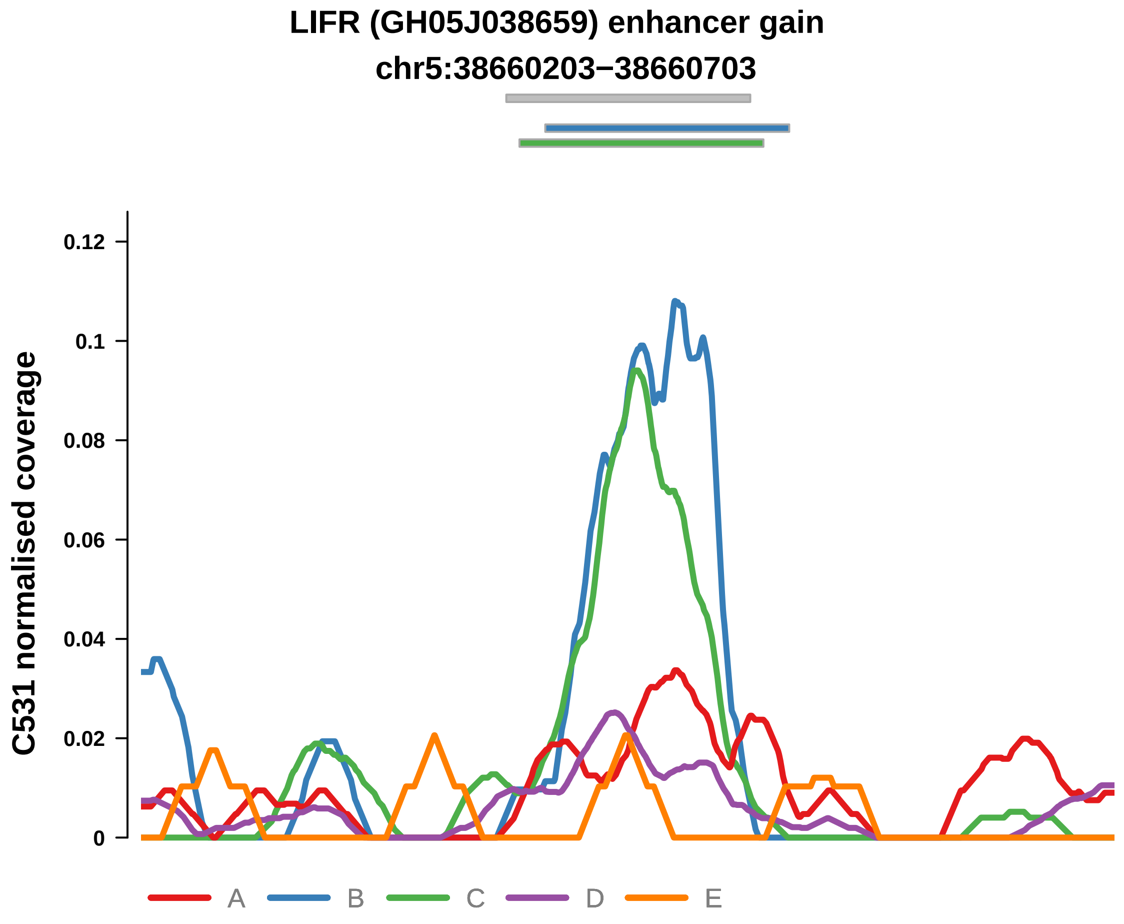


Putatively Clonal (logFC = 2.07)

| | P. | Adjusted P. |
|---|---|---|
| *~region vs. ~* | 2.58e−02 * | 7.06e−02 |
| *~purity vs ~* | 1.72e−03 ** | 2.52e−02 * |
| *~purity+region vs ~purity* | 6.94e−01 | 8.85e−01 |
| *Carcinoma vs. normal* | 7.59e−03 ** | 7.59e−03 ** |

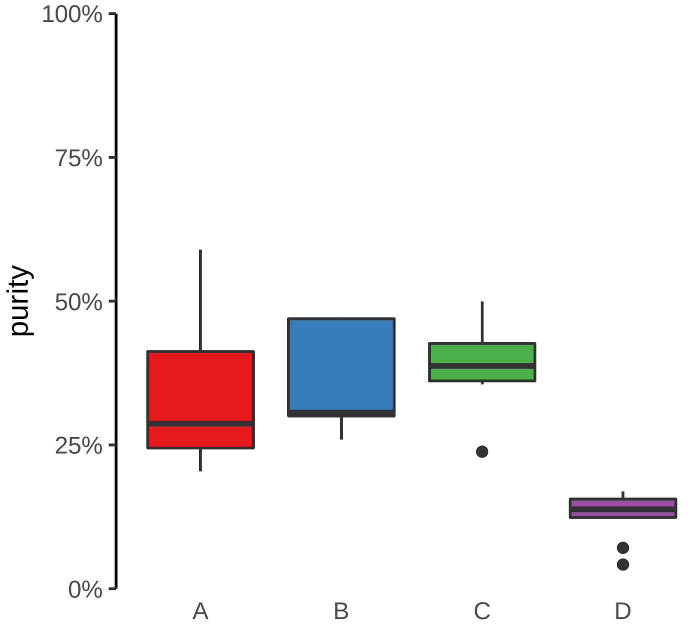

**MYBPC3 (GH11J047336) enhancer gain**
**chr11:47337207−47337707**

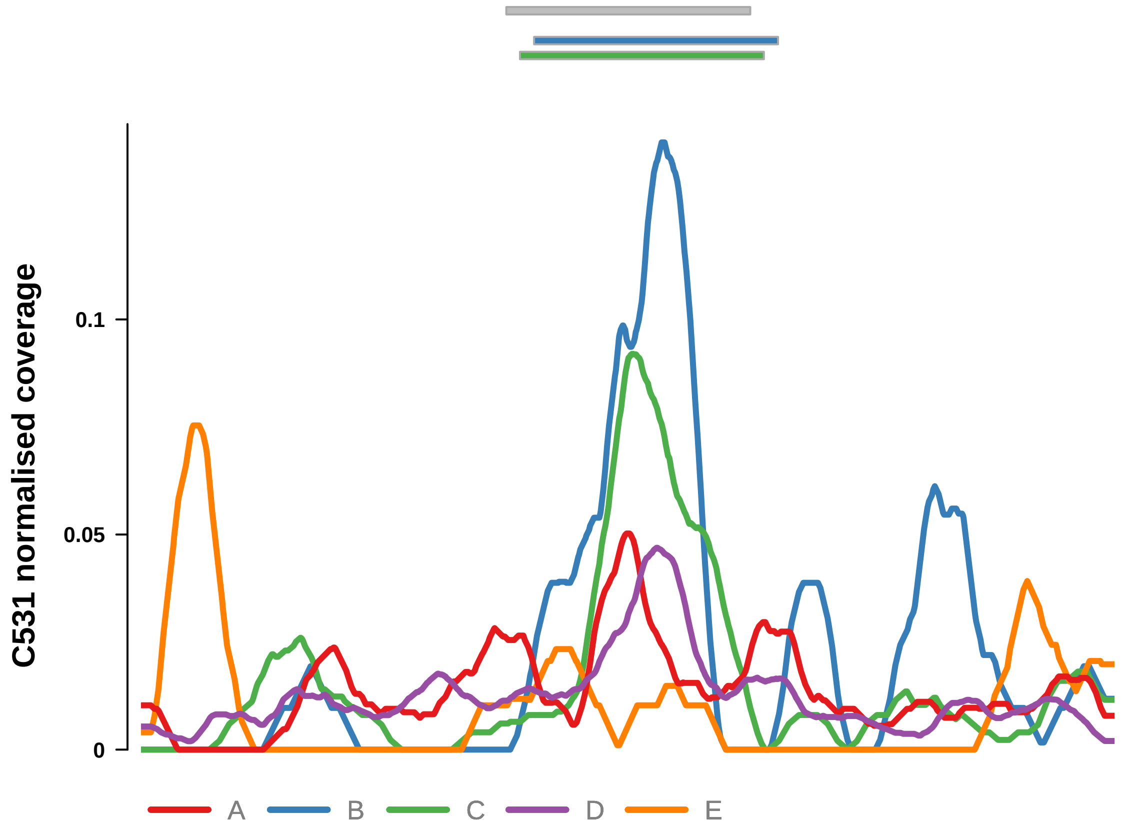


Putatively Clonal (logFC = 2.87)

| | P. | Adjusted P. |
|---|---|---|
| *~region vs. ~* | 7.63e−01 | 8.1e−01 |
| *~purity vs ~* | 6.58e−01 | 7.72e−01 |
| *~purity+region vs ~purity* | 5.41e−01 | 7.95e−01 |
| *Carcinoma vs. normal* | 3.03e−04 *** | 3.03e−04 *** |

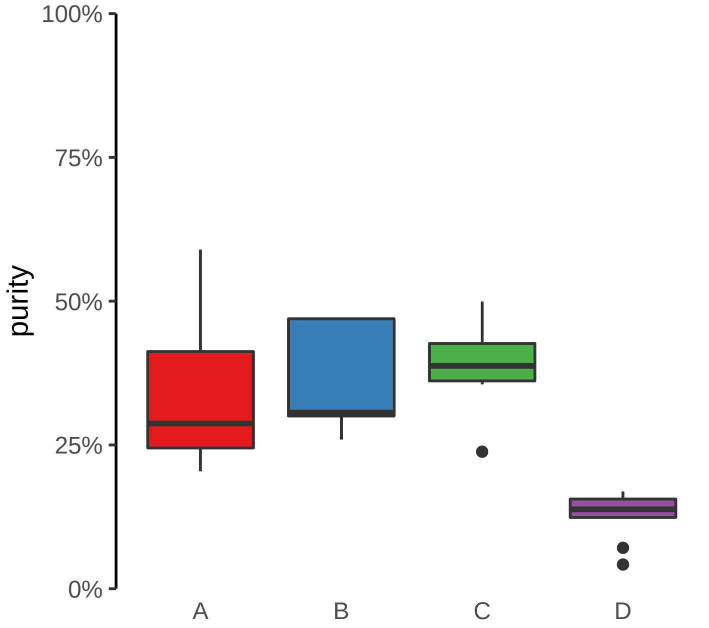

**MYT1 (GH20J064188) enhancer gain**
**chr20:64189162−64189662**

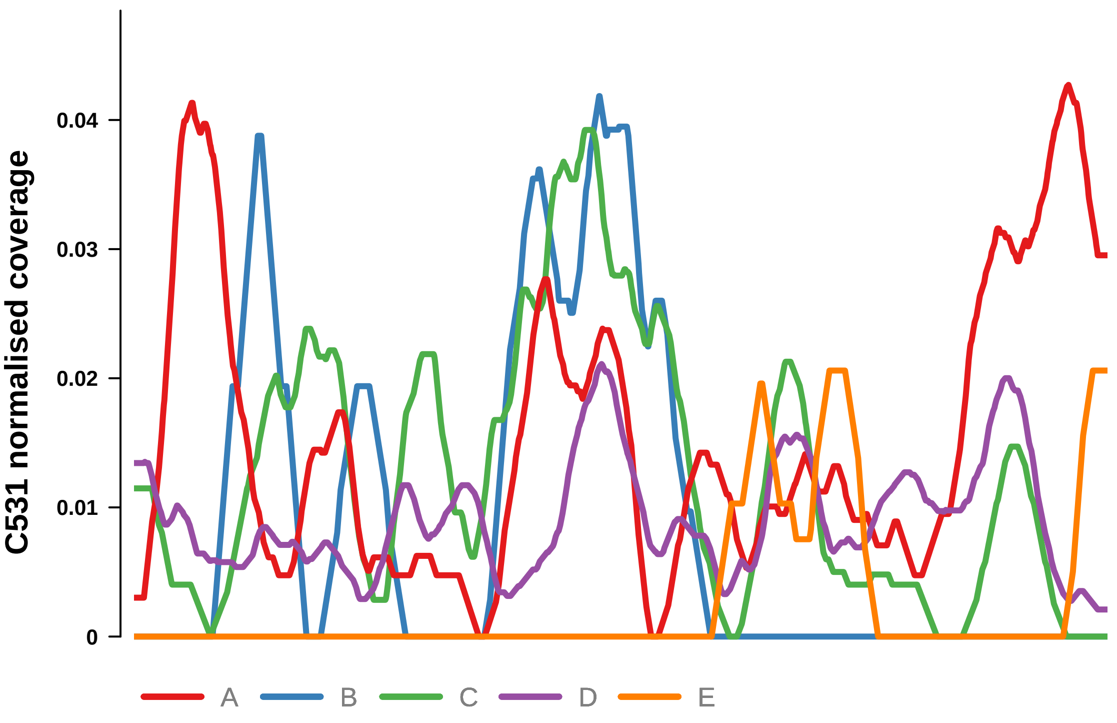


Putatively Clonal (logFC = 2.55)

| | P. | Adjusted P. |
|---|---|---|
| *~region vs. ~* | 2.63e−01 | NA |
| *~purity vs ~* | 2.55e−01 | 3.97e−01 |
| *~purity+region vs ~purity* | 3.13e−01 | 6.13e−01 |
| *Carcinoma vs. normal* | 2.07e−03 ** | 2.07e−03 ** |

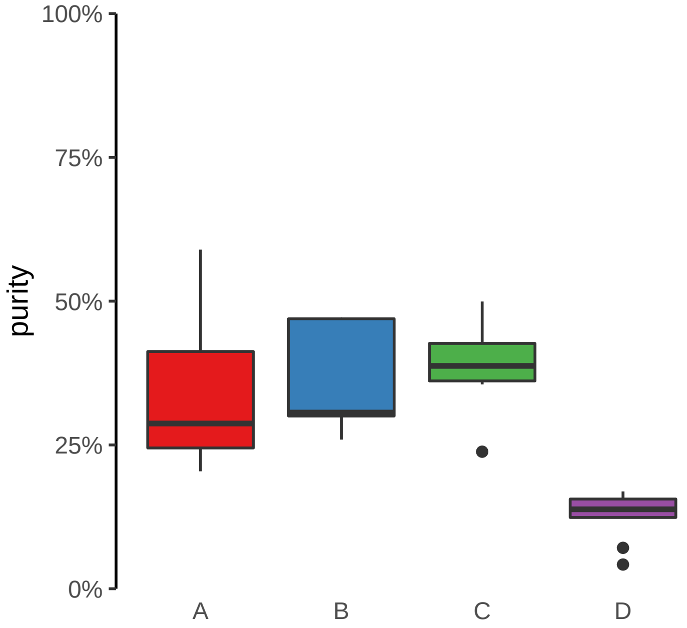

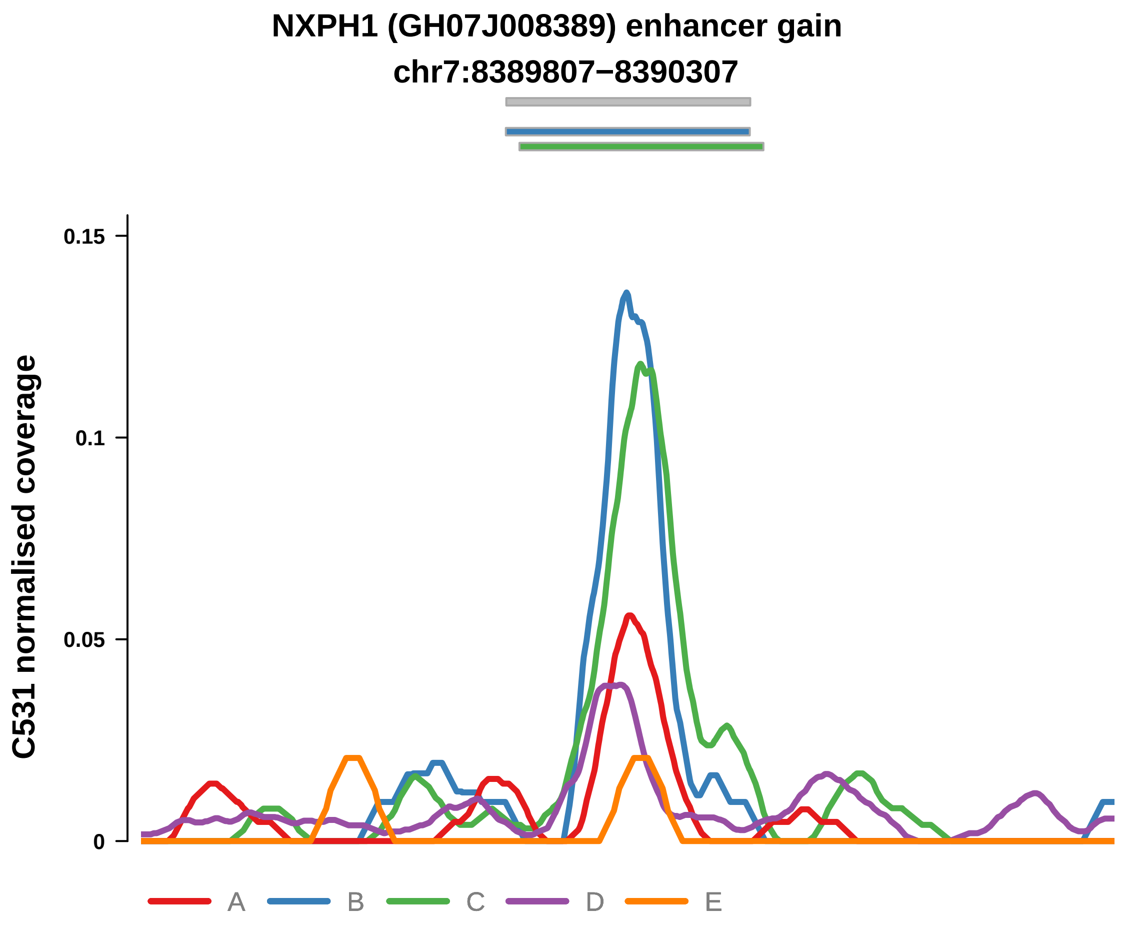


Putatively Clonal (logFC = 2.11)

| | P. | Adjusted P. |
|---|---|---|
| *~region vs. ~* | 5.43e−03 ** | 2.02e−02 * |
| *~purity vs ~* | 3.46e−03 ** | 2.95e−02 * |
| *~purity+region vs ~purity* | 2.83e−01 | 6.13e−01 |
| *Carcinoma vs. normal* | 5.75e−03 ** | 5.75e−03 ** |

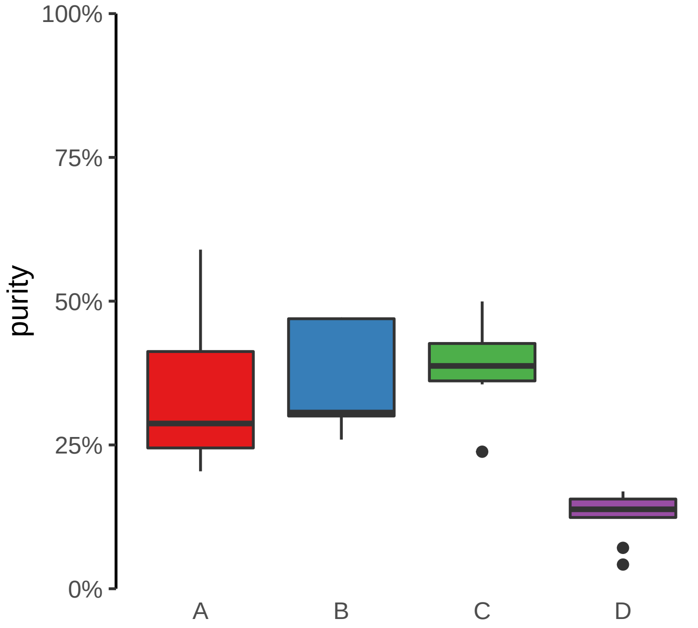

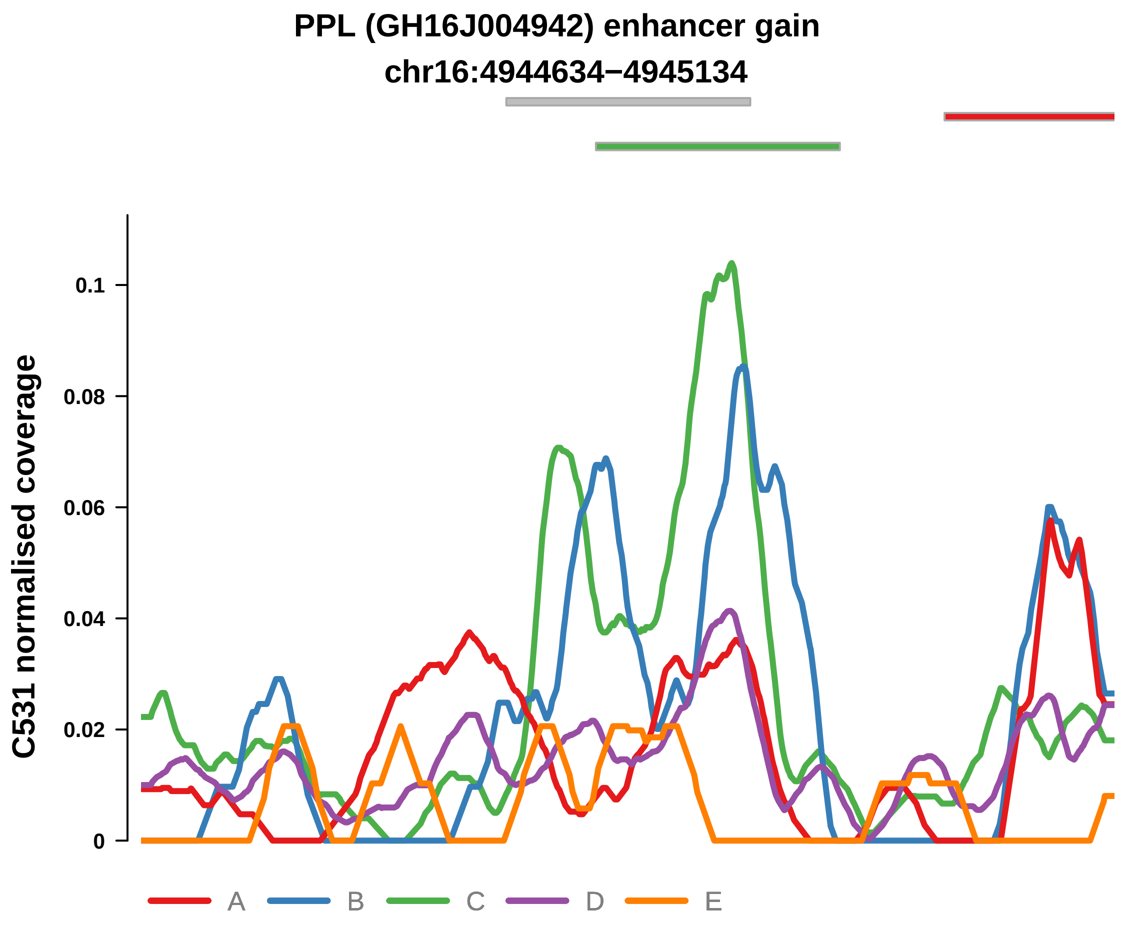


Putatively Clonal (logFC = 2.56)

| | P. | Adjusted P. |
|---|---|---|
| *~region vs. ~* | 3.32e−03 ** | 1.44e−02 * |
| *~purity vs ~* | 3.08e−03 ** | 2.95e−02 * |
| *~purity+region vs ~purity* | 2.16e−01 | 6.13e−01 |
| *Carcinoma vs. normal* | 4.79e−04 *** | 4.79e−04 *** |

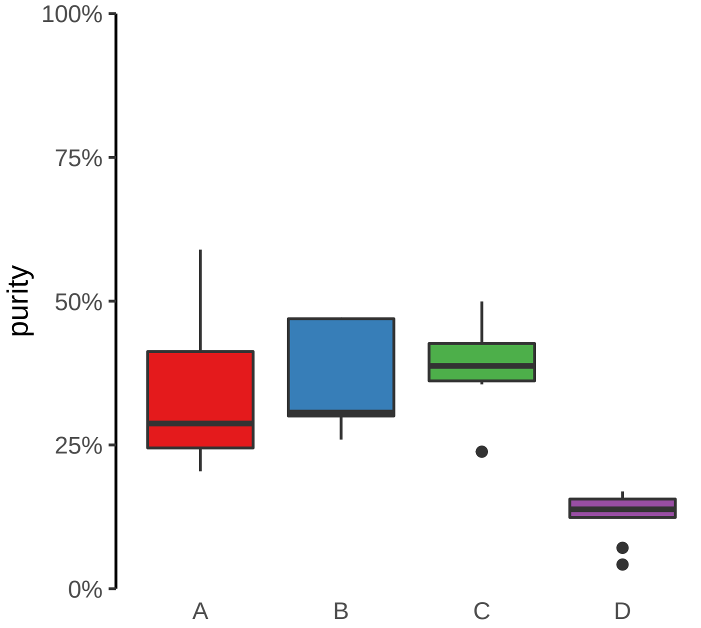

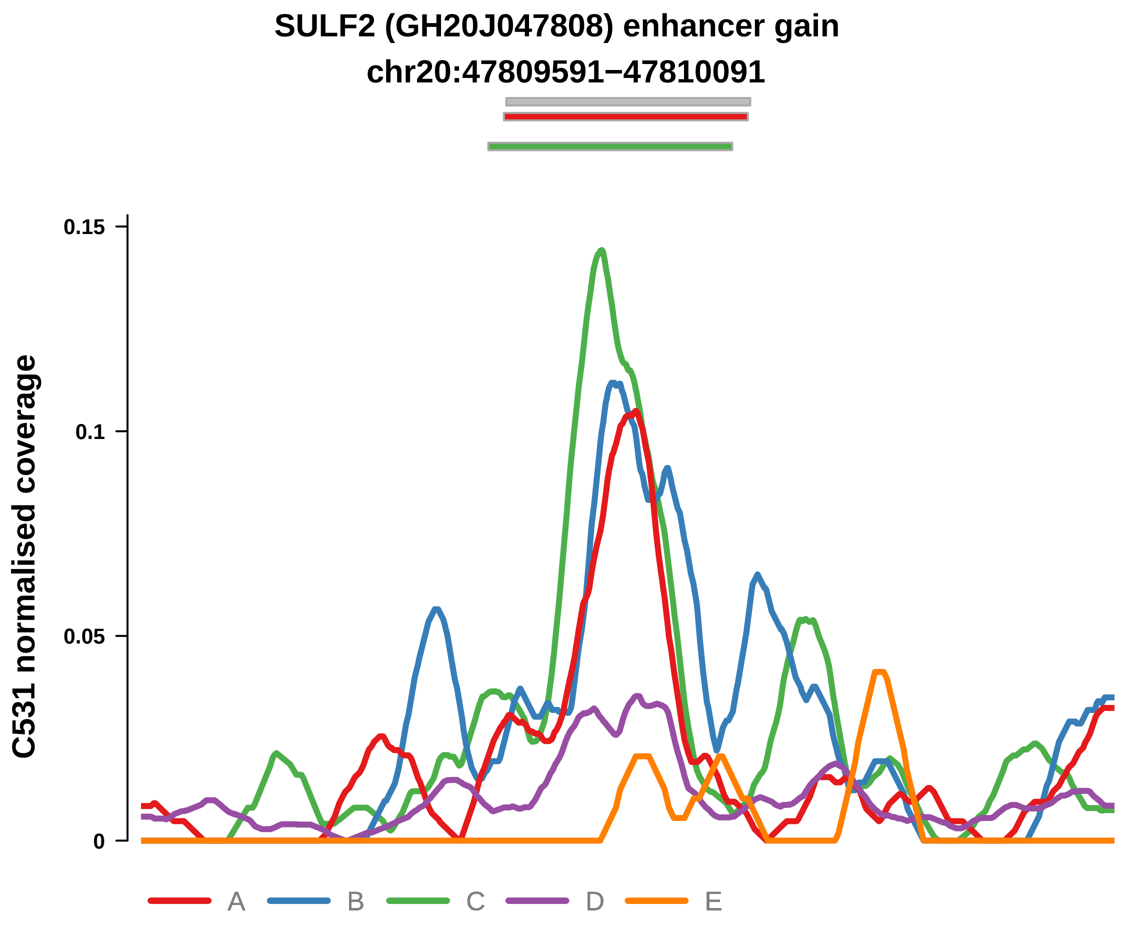


Putatively Clonal (logFC = 2.93)

| | P. | Adjusted P. |
|---|---|---|
| *~region vs. ~* | 4.62e−06 **** | 8e−05 **** |
| *~purity vs ~* | 1.06e−07 **** | 9.36e−06 **** |
| *~purity+region vs ~purity* | 2.52e−01 | 6.13e−01 |
| *Carcinoma vs. normal* | 6.52e−05 **** | 6.52e−05 **** |

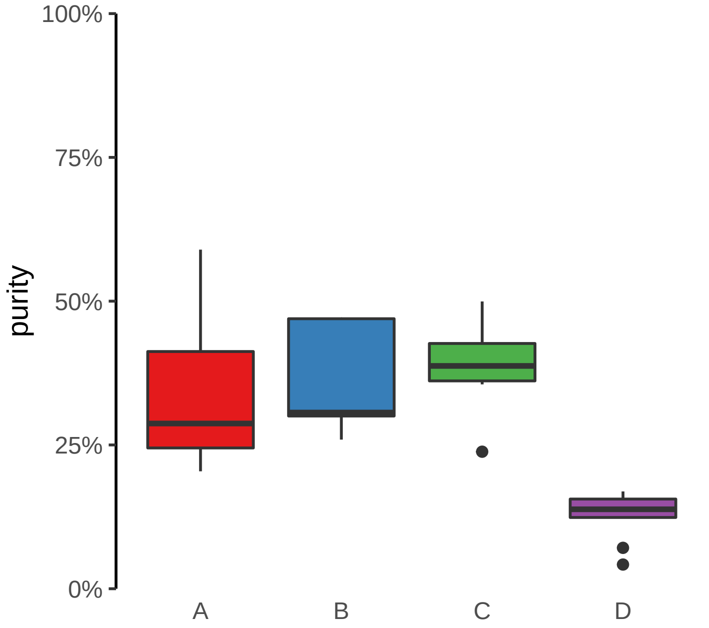

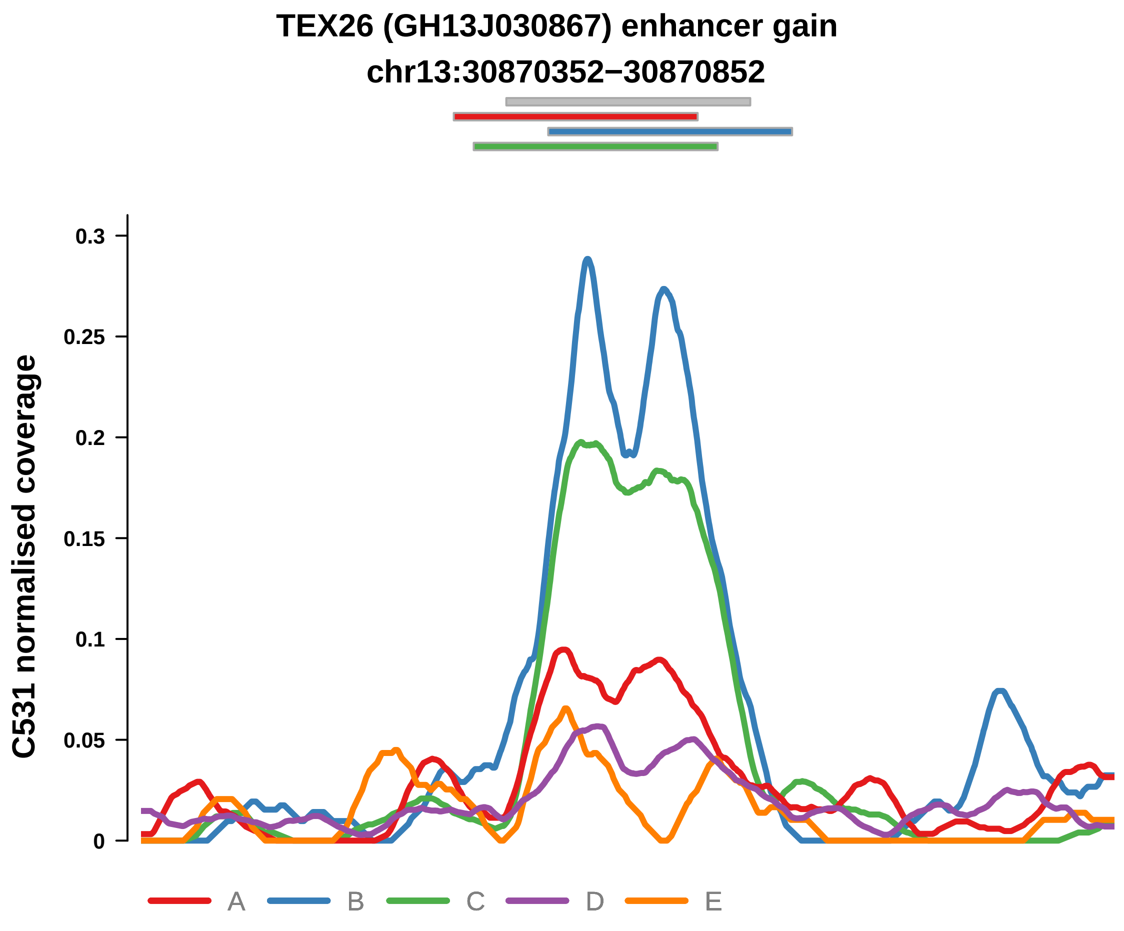


Putatively Clonal (logFC = 2.41)

| | P. | Adjusted P. |
|---|---|---|
| *~region vs. ~* | 8.45e−08 **** | 4.4e−06 **** |
| *~purity vs ~* | 2.94e−04 *** | 6.46e−03 ** |
| *~purity+region vs ~purity* | 3.72e−03 ** | 6.55e−02 |
| *Carcinoma vs. normal* | 6.81e−04 *** | 6.81e−04 *** |

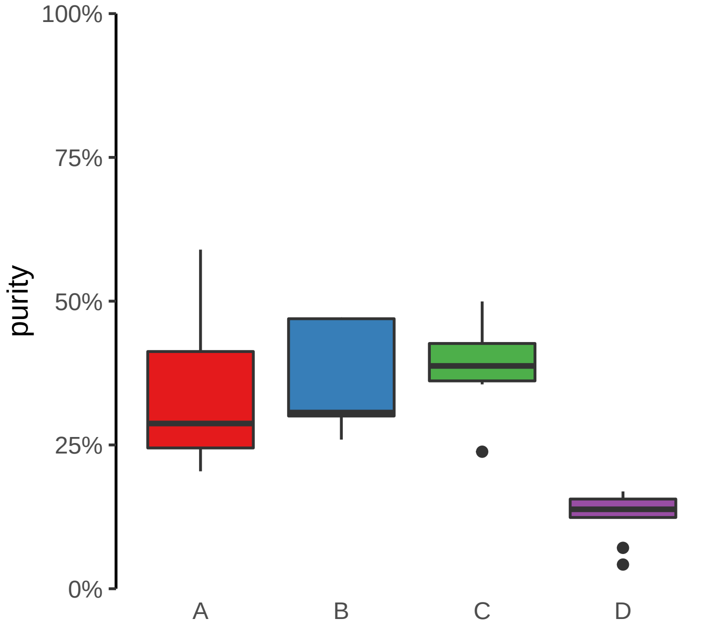

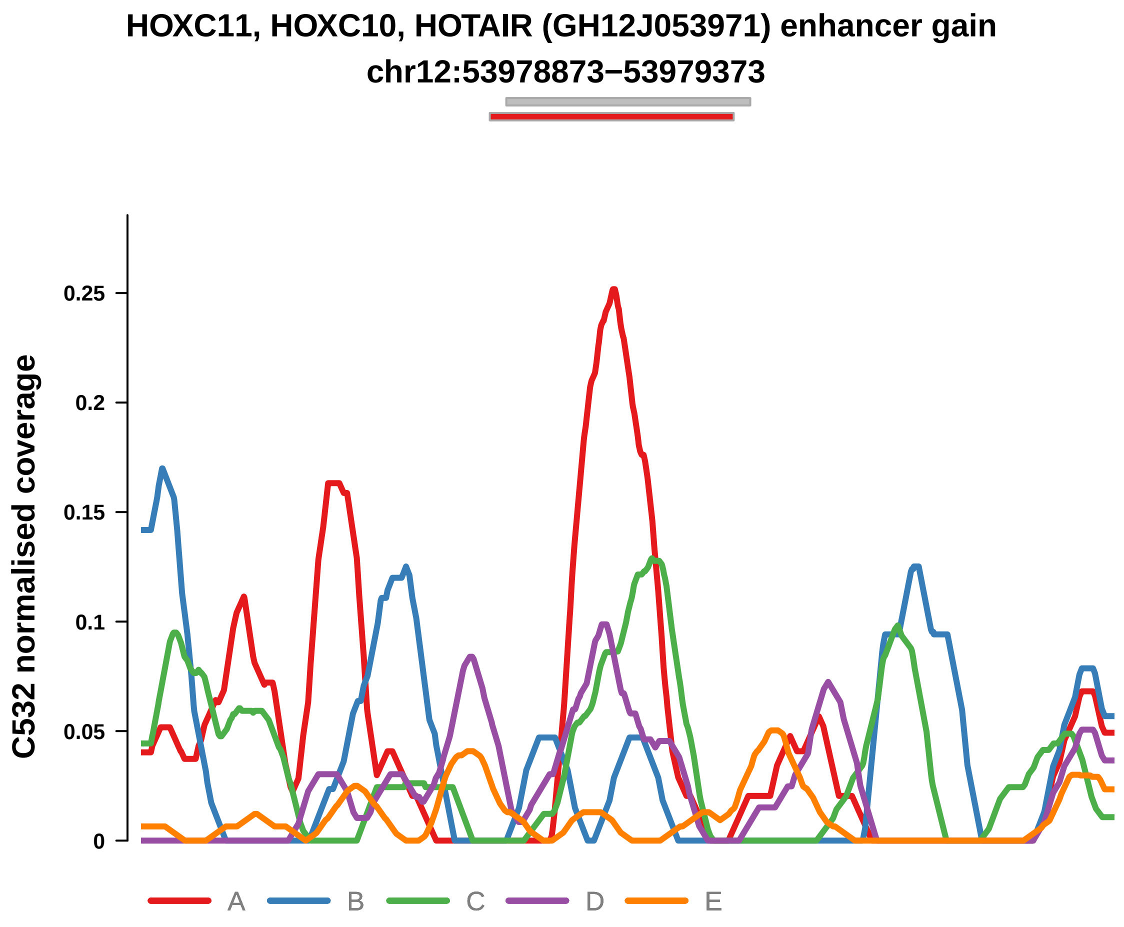


Putatively Clonal (logFC = 2.01)

| | P. | Adjusted P. |
|---|---|---|
| *~region vs. ~* | 2.47e−01 | 6.78e−01 |
| *~purity vs ~* | 3.72e−01 | 8.63e−01 |
| *~purity+region vs ~purity* | 1.42e−01 | 6.56e−01 |
| *Carcinoma vs. normal* | 5.96e−03 ** | 5.96e−03 ** |

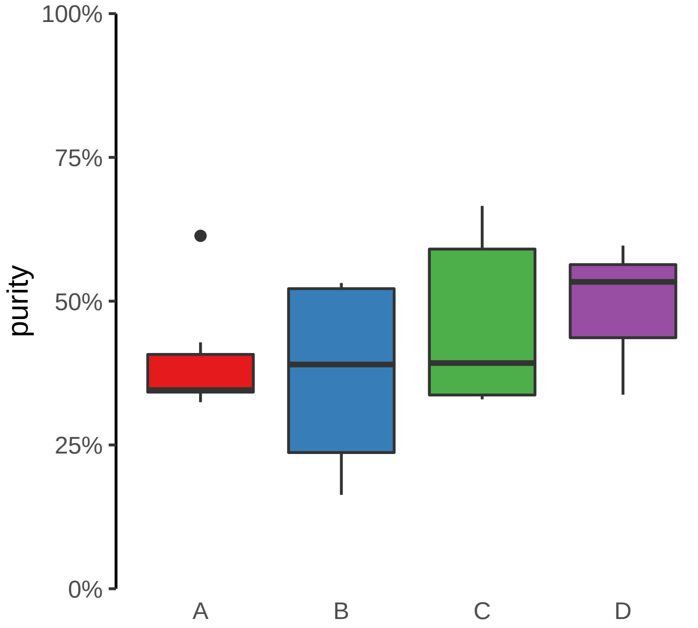

**CEP83 (GH12J094389) enhancer gain**
**chr12:94389729−94390229**

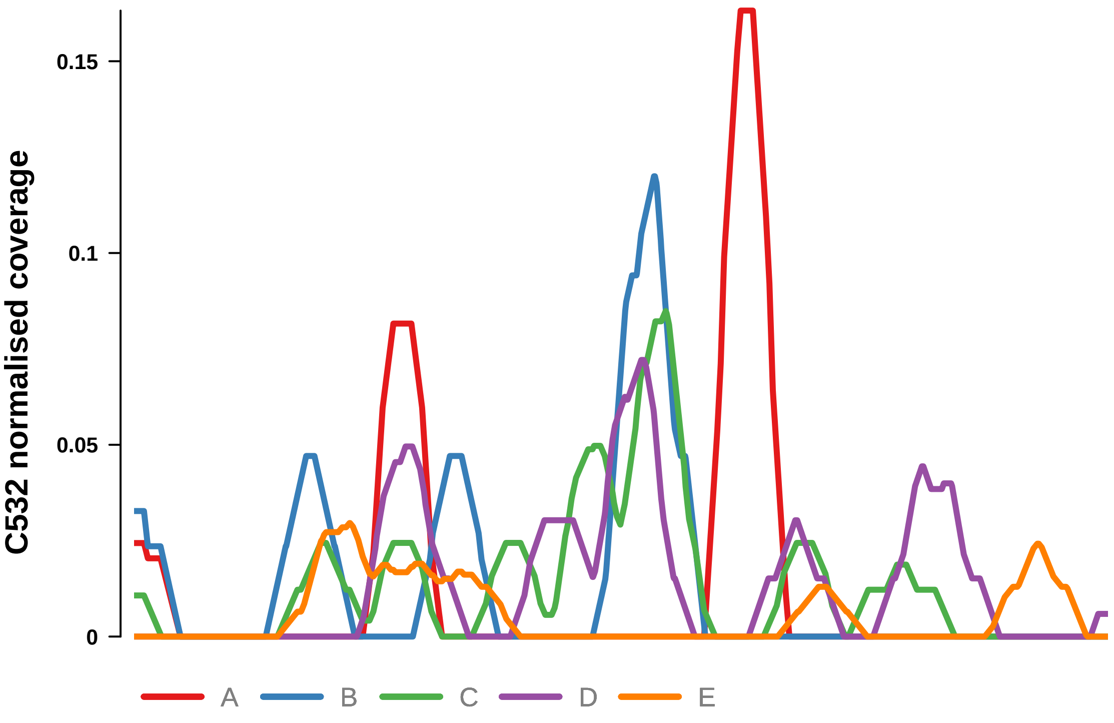


Putatively Clonal (logFC = 2.38)

| | P. | Adjusted P. |
|---|---|---|
| *~region vs. ~* | 9.46e−01 | 9.9e−01 |
| *~purity vs ~* | 1.5e−01 | 8.27e−01 |
| *~purity+region vs ~purity* | 8.69e−01 | 1e+00 |
| *Carcinoma vs. normal* | 1.88e−03 ** | 1.88e−03 ** |

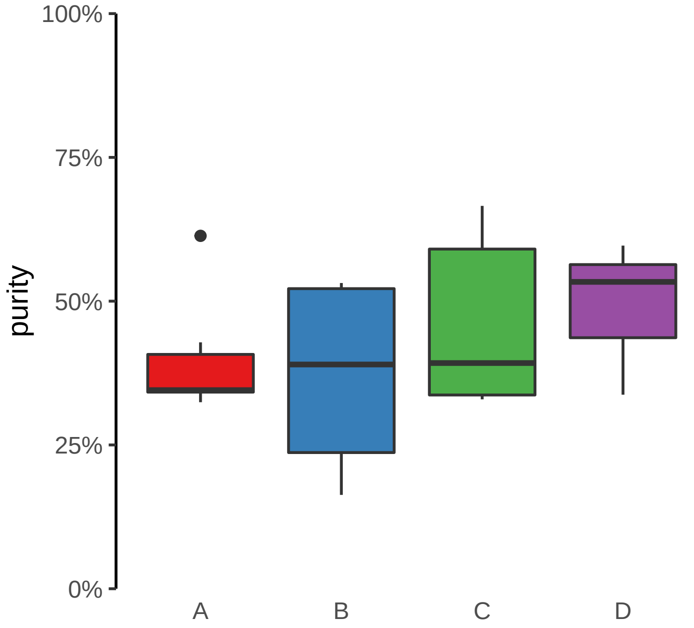

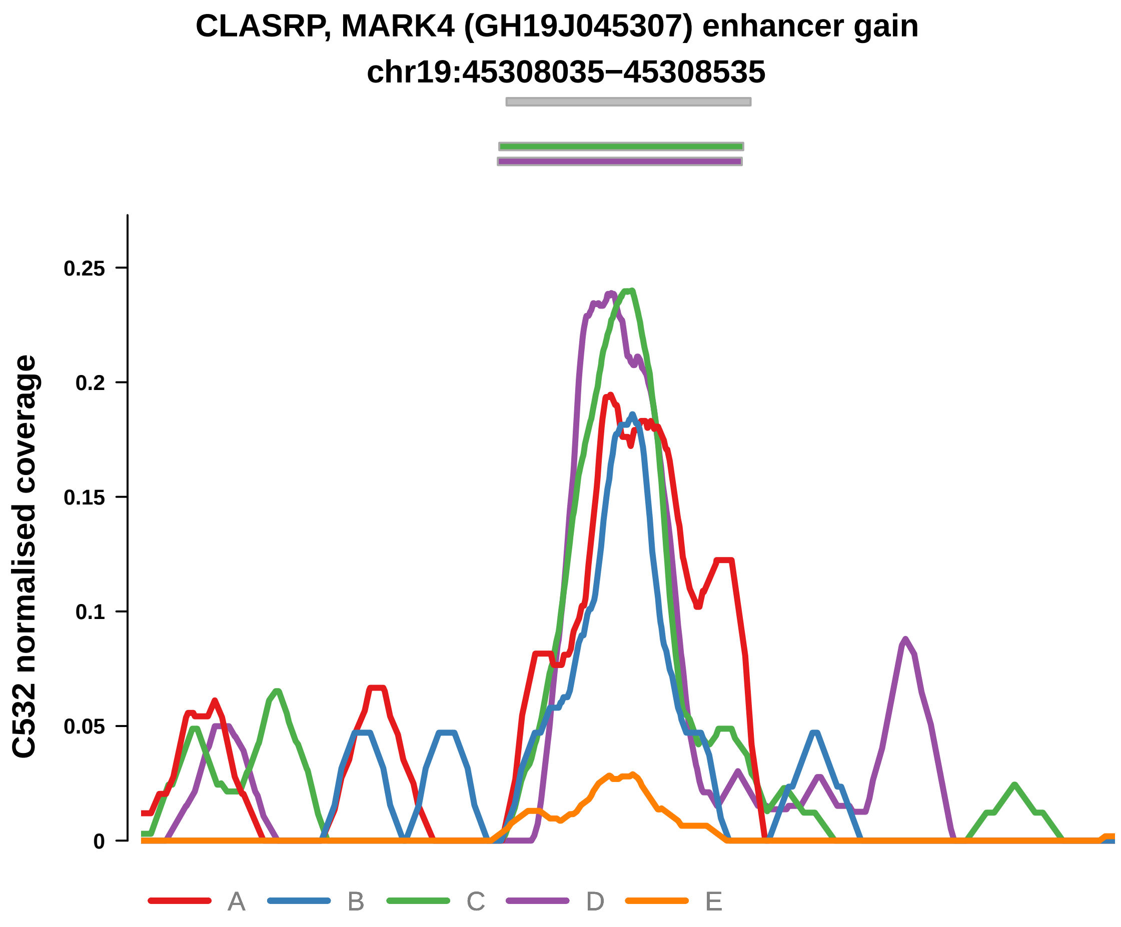


Putatively Clonal (logFC = 2.84)

| | P. | Adjusted P. |
|---|---|---|
| *~region vs. ~* | 6.93e−01 | 8.72e−01 |
| *~purity vs ~* | 7.53e−03 ** | 2.21e−01 |
| *~purity+region vs ~purity* | 8.55e−01 | 1e+00 |
| *Carcinoma vs. normal* | 8.39e−05 **** | 8.39e−05 **** |

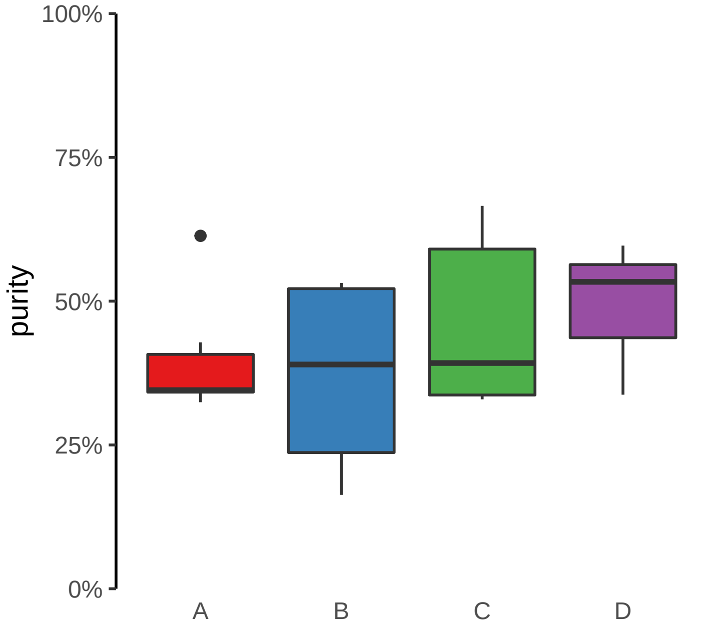

**CSGALNACT1 (GH08J019660) enhancer gain**
**chr8:19661066−19661566**

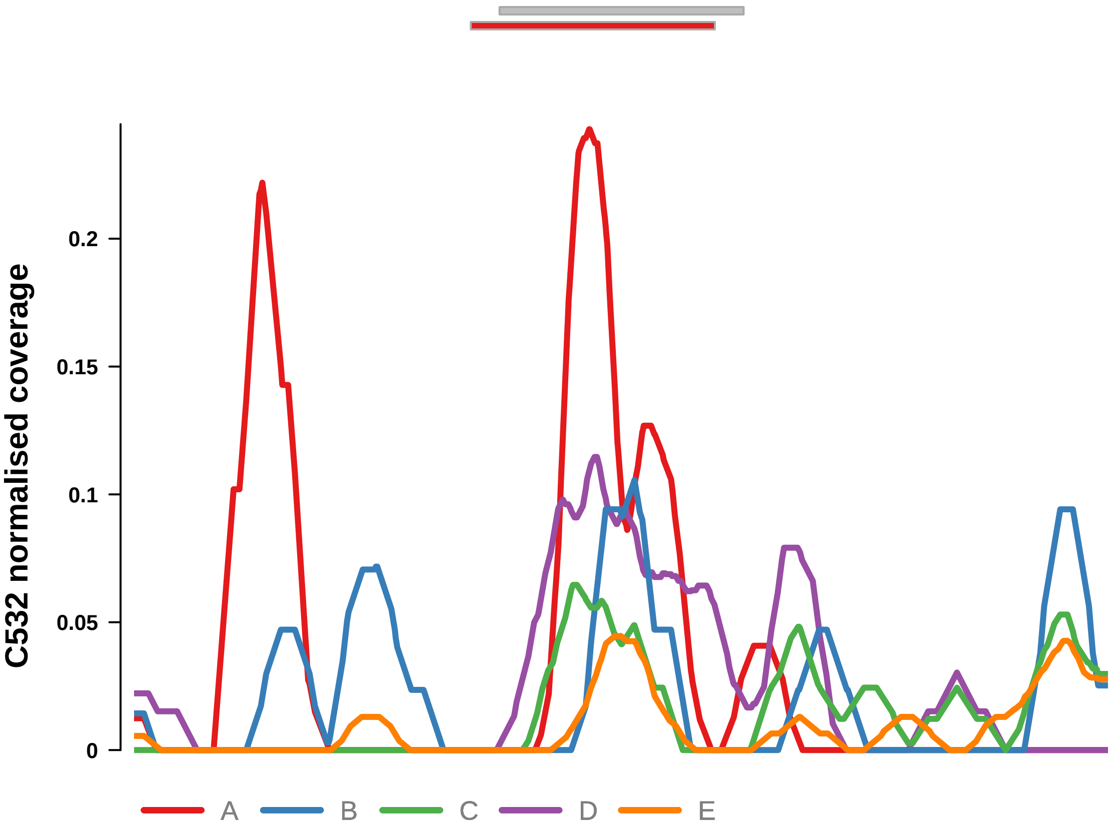


Putatively Clonal (logFC = 2.01)

| | P. | Adjusted P. |
|---|---|---|
| *~region vs. ~* | 1.89e−01 | 5.85e−01 |
| *~purity vs ~* | 6.54e−01 | 9.86e−01 |
| *~purity+region vs ~purity* | 1.91e−01 | 6.56e−01 |
| *Carcinoma vs. normal* | 7.08e−03 ** | 7.08e−03 ** |

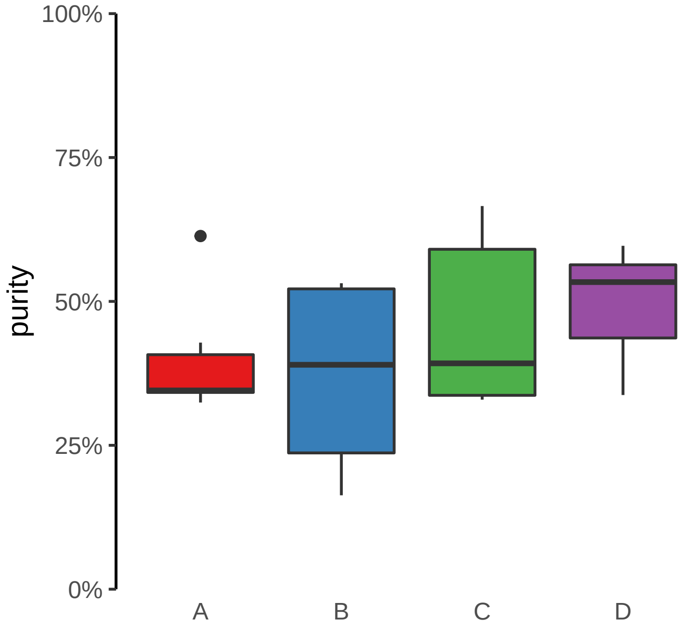

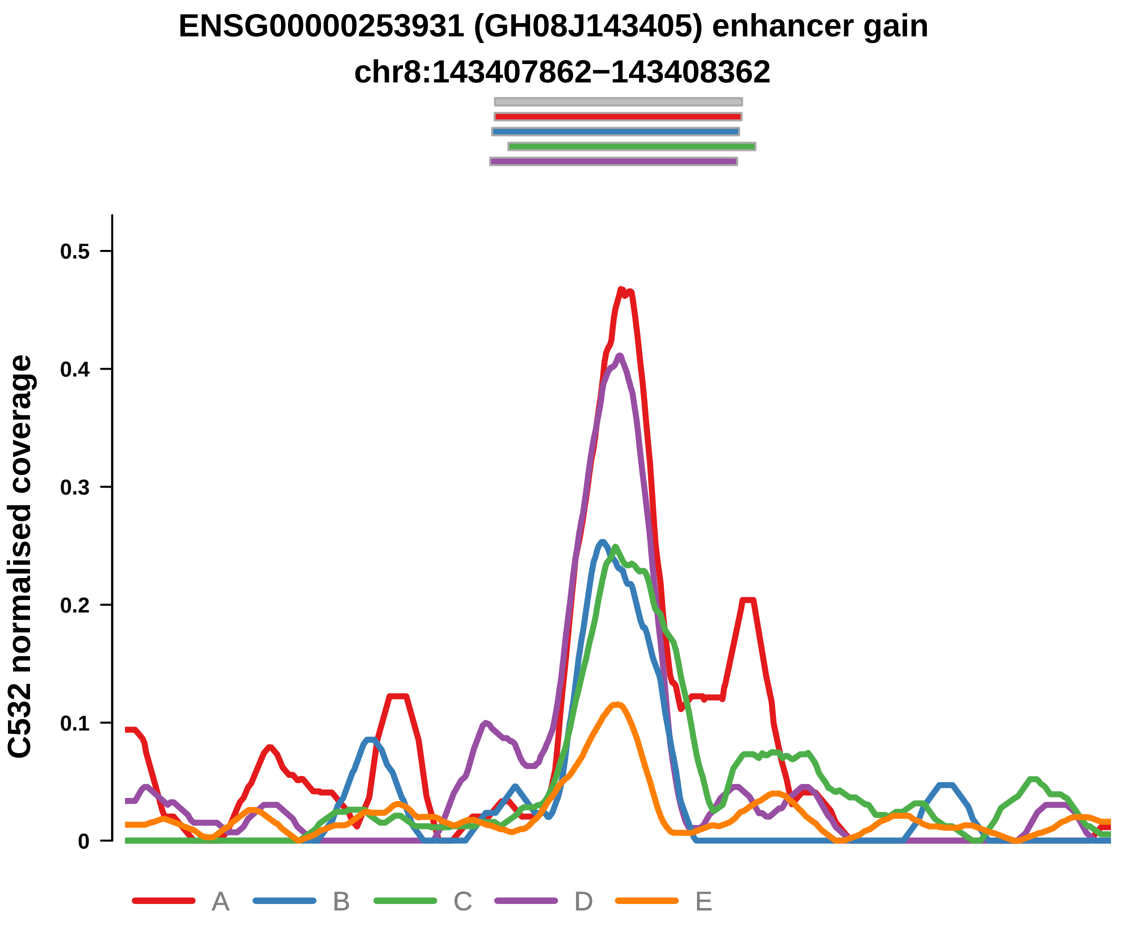


Putatively Clonal (logFC = 2.12)

| | P. | Adjusted P. |
|---|---|---|
| *~region vs. ~* | 2.05e−01 | 5.98e−01 |
| *~purity vs ~* | 2.27e−01 | 8.56e−01 |
| *~purity+region vs ~purity* | 2.45e−01 | 6.74e−01 |
| *Carcinoma vs. normal* | 2.4e−03 ** | 2.4e−03 ** |

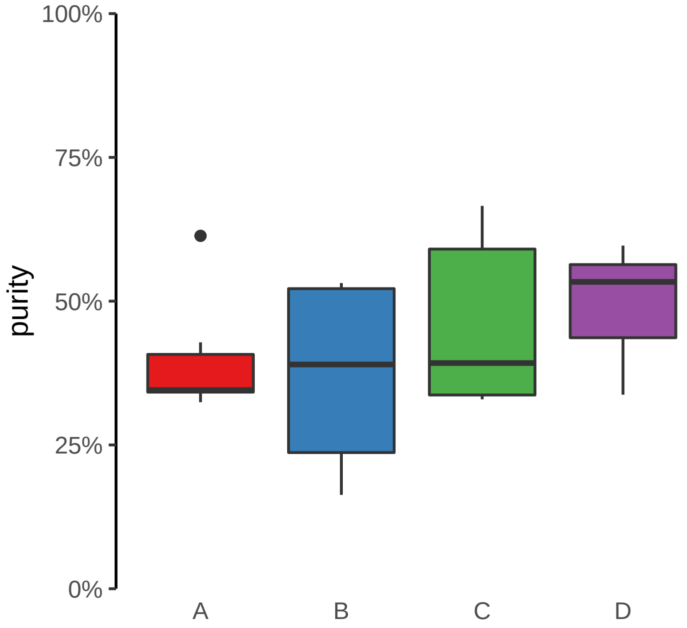

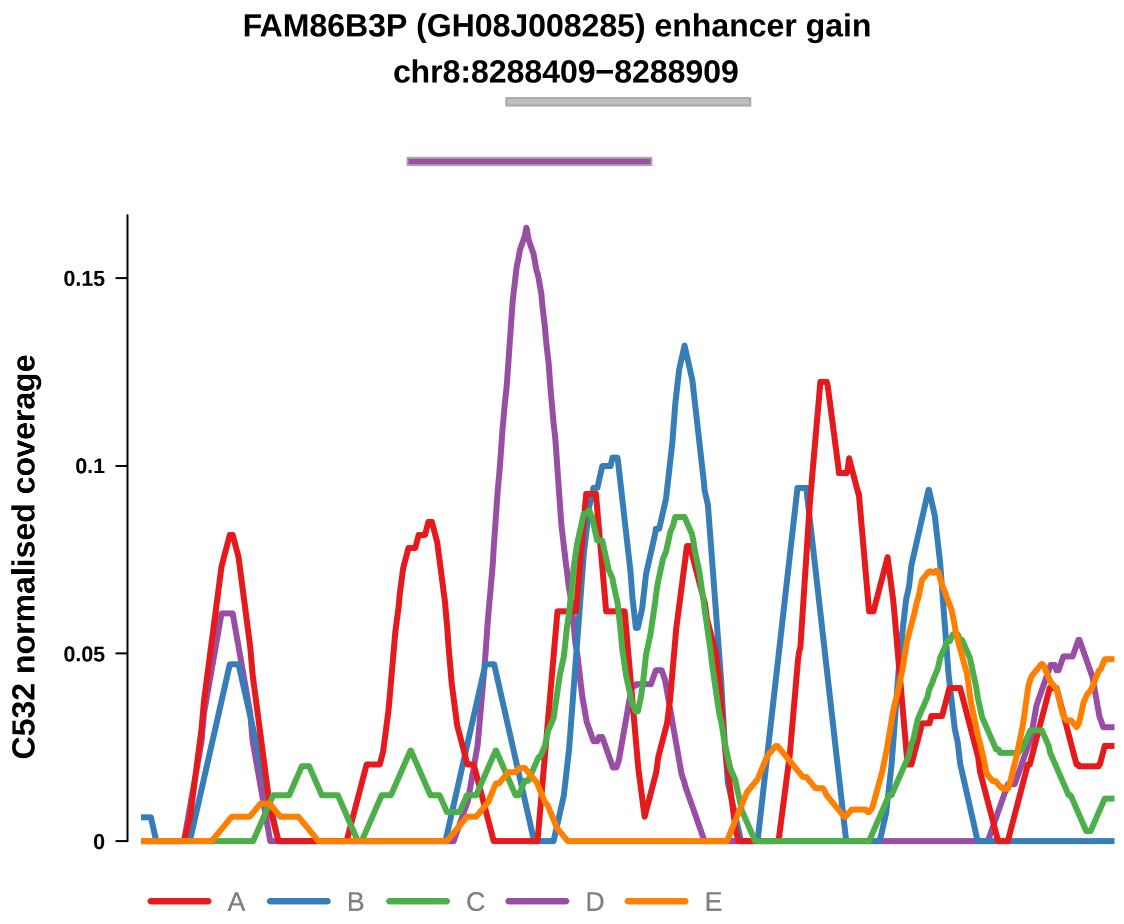


Putatively Clonal (logFC = 2.53)

| | P. | Adjusted P. |
|---|---|---|
| *~region vs. ~* | 5.39e−01 | 8.54e−01 |
| *~purity vs ~* | 3.02e−01 | 8.56e−01 |
| *~purity+region vs ~purity* | 6.57e−01 | 9.42e−01 |
| *Carcinoma vs. normal* | 6.66e−04 *** | 6.66e−04 *** |

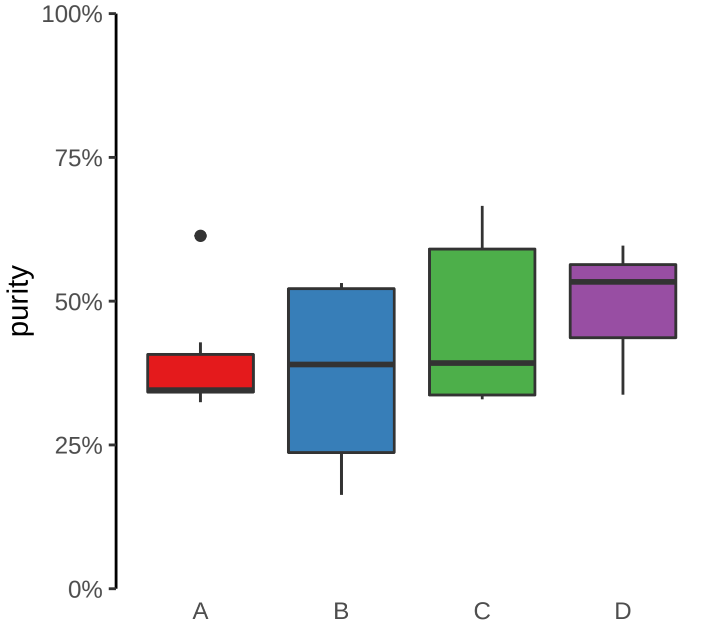

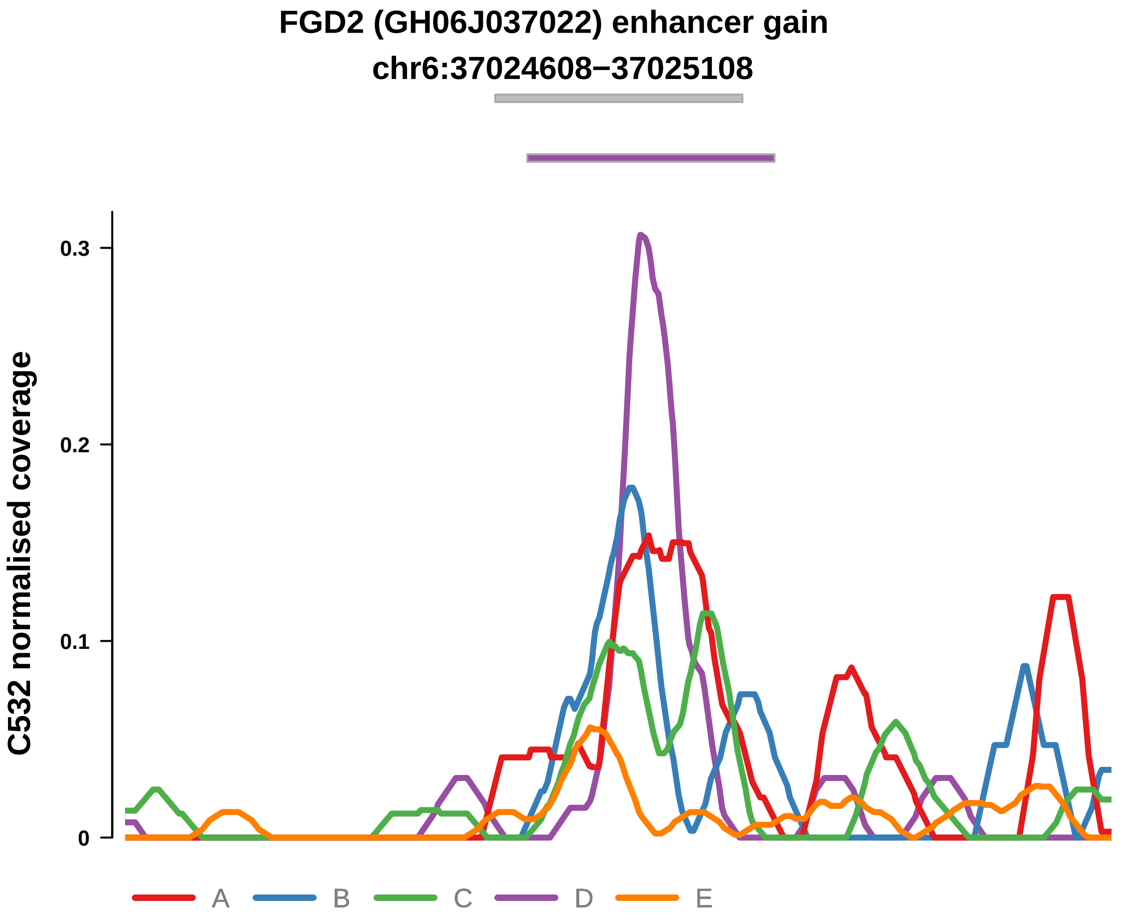


Putatively Clonal (logFC = 2.7)

| | P. | Adjusted P. |
|---|---|---|
| *~region vs. ~* | 6.5e−01 | 8.54e−01 |
| *~purity vs ~* | 5.04e−01 | 9.05e−01 |
| *~purity+region vs ~purity* | 7.24e−01 | 9.42e−01 |
| *Carcinoma vs. normal* | 2.4e−04 *** | 2.4e−04 *** |

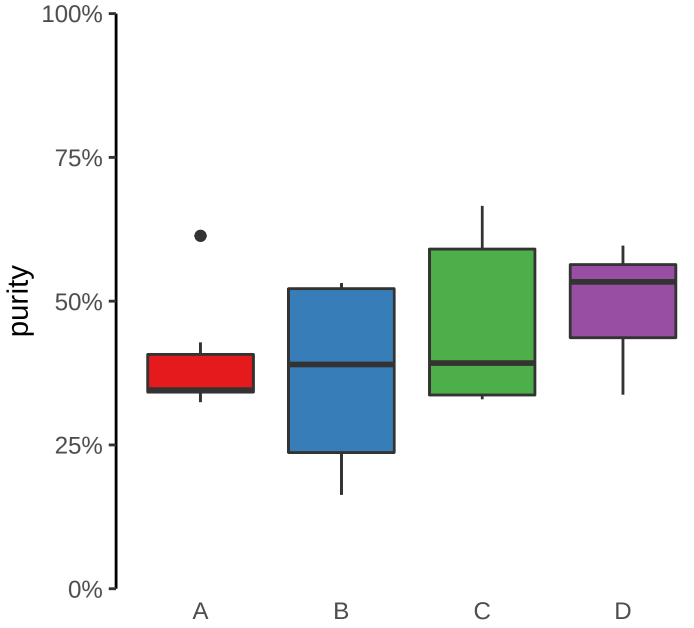

**FOXL1 (GH16J086618) enhancer gain**
**chr16:86619226−86619726**

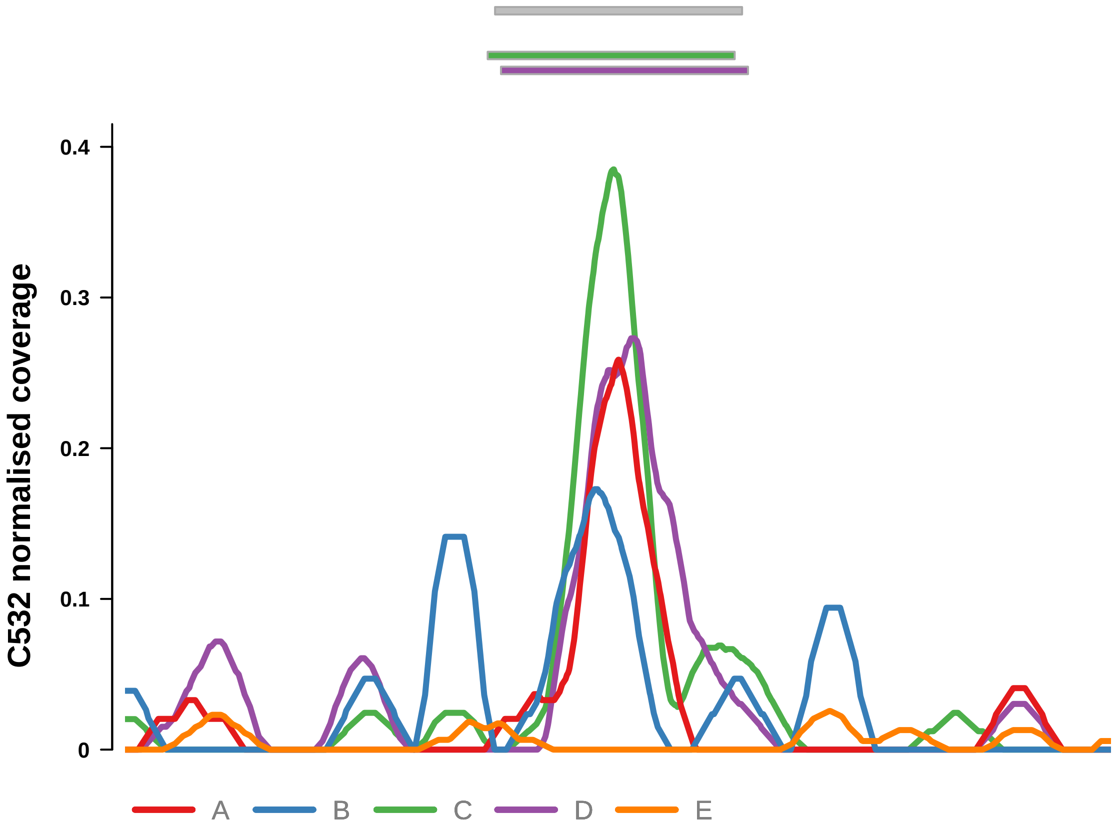


Putatively Clonal (logFC = 3.13)

| | P. | Adjusted P. |
|---|---|---|
| *~region vs. ~* | 5.75e−02 | 3.62e−01 |
| *~purity vs ~* | 2.11e−02 * | 3.05e−01 |
| *~purity+region vs ~purity* | 1.92e−01 | 6.56e−01 |
| *Carcinoma vs. normal* | 1.91e−05 **** | 1.91e−05 **** |

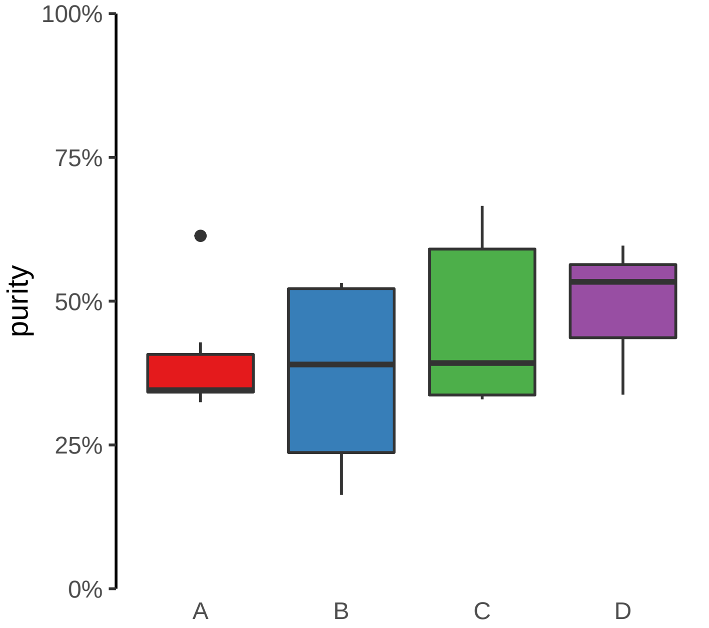

**LIFR (GH05J038659) enhancer gain**
**chr5:38660203−38660703**

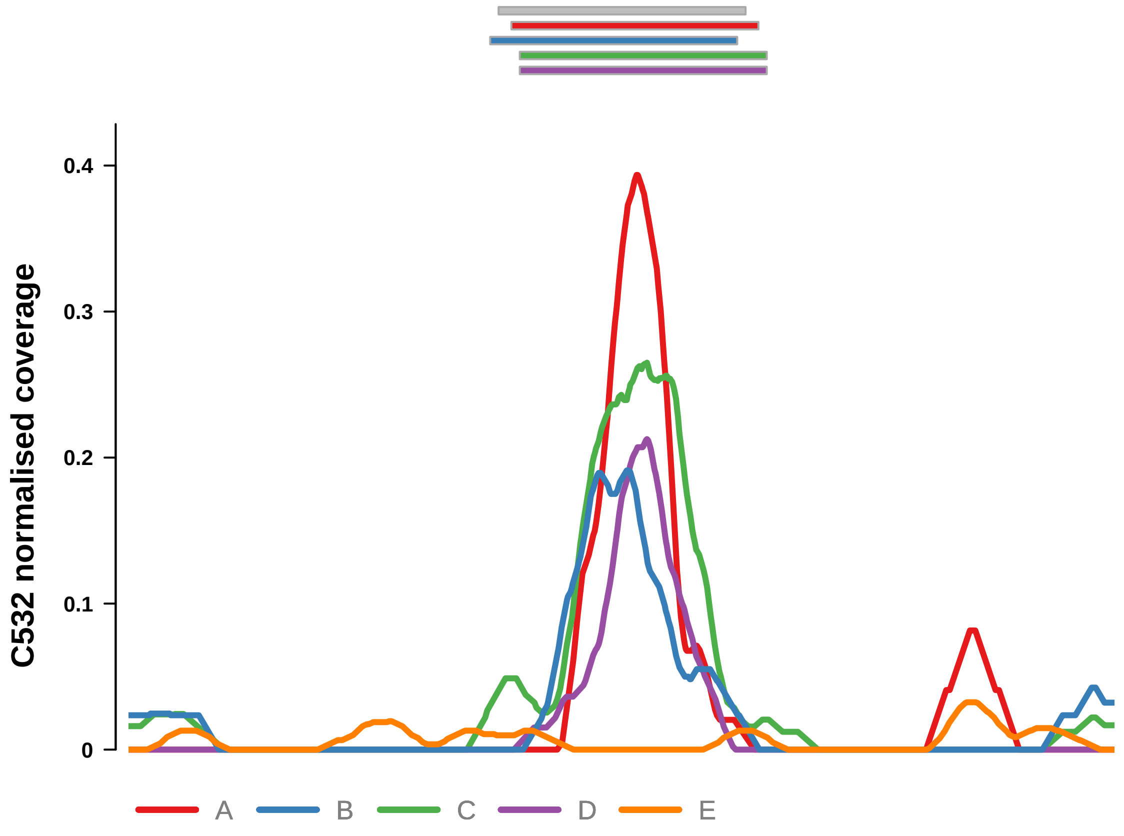


Putatively Clonal (logFC = 2.84)

| | P. | Adjusted P. |
|---|---|---|
| *~region vs. ~* | 2.34e−02 * | 2.06e−01 |
| *~purity vs ~* | 6.05e−01 | 9.86e−01 |
| *~purity+region vs ~purity* | 1.9e−02 * | 2.6e−01 |
| *Carcinoma vs. normal* | 9.05e−05 **** | 9.05e−05 **** |

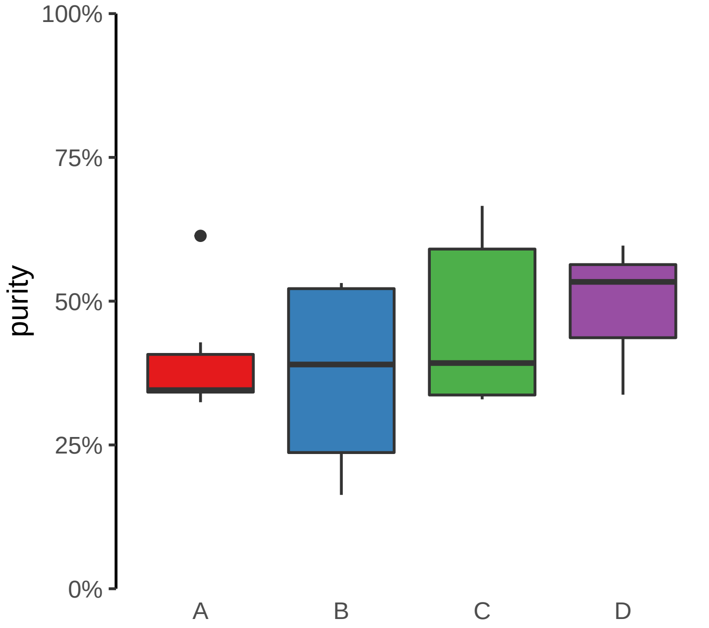

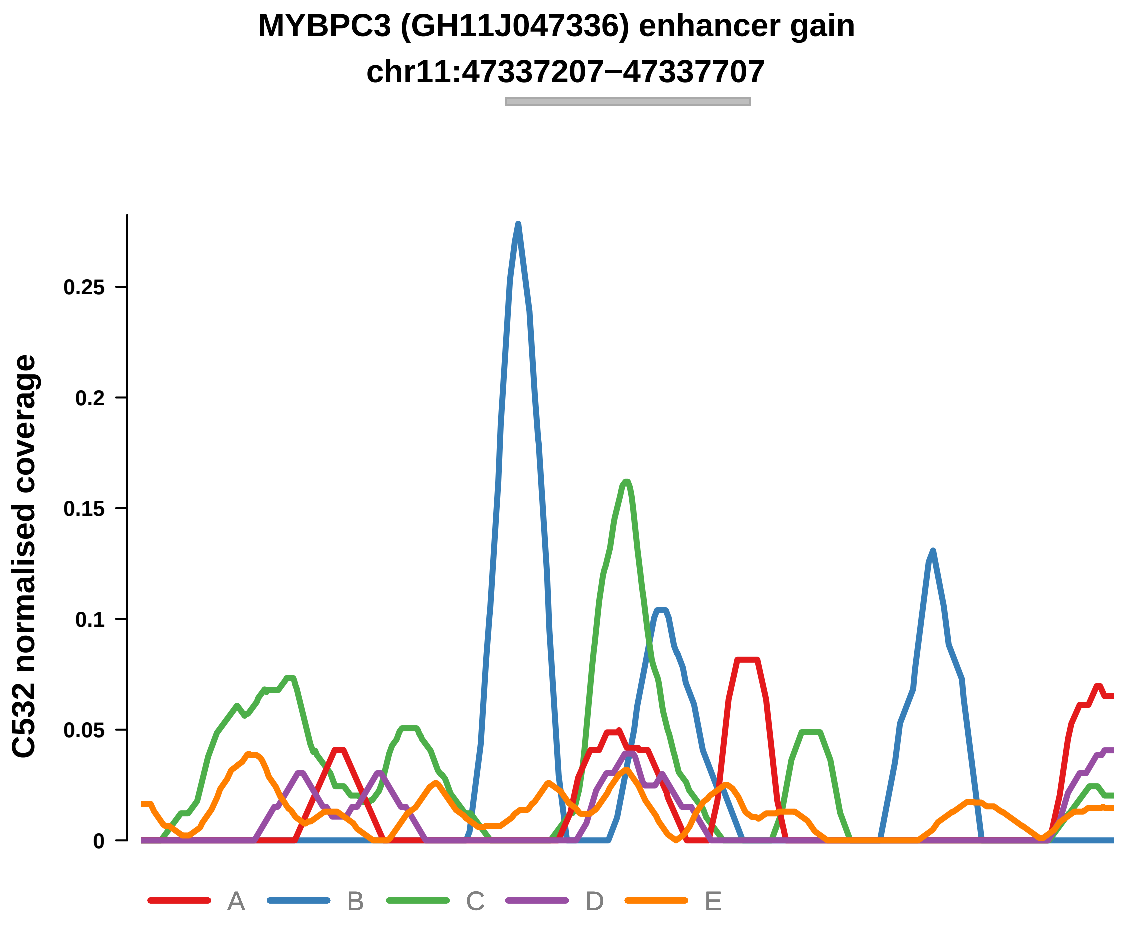


Putatively Clonal (logFC = 2.25)

| | P. | Adjusted P. |
|---|---|---|
| *~region vs. ~* | 4.97e−02 * | 3.62e−01 |
| *~purity vs ~* | 8.91e−01 | 9.86e−01 |
| *~purity+region vs ~purity* | 6.6e−02 | 5.15e−01 |
| *Carcinoma vs. normal* | 2.54e−03 ** | 2.54e−03 ** |

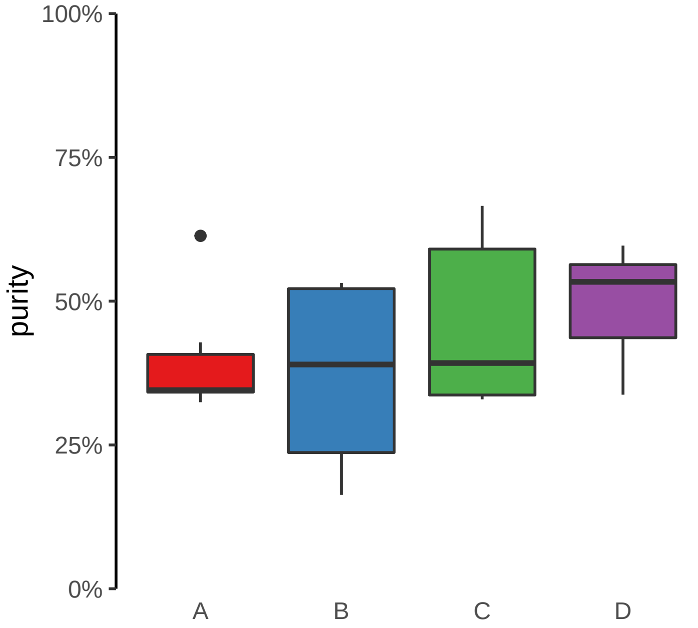

**MYT1 (GH20J064188) enhancer gain**
**chr20:64189162−64189662**

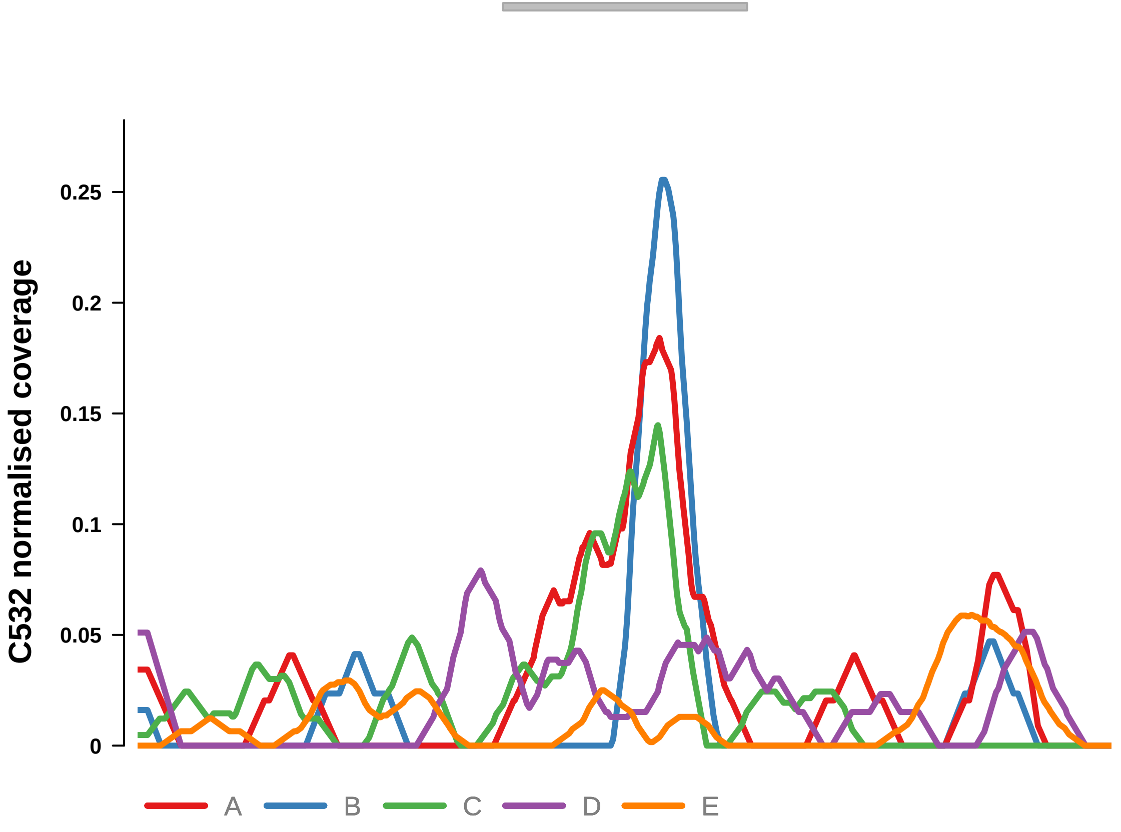


Putatively Clonal (logFC = 2.88)

| | **P.** | **Adjusted P.** |
|---|---|---|
| *~region vs. ~* | 8.93e−01 | 9.8e−01 |
| *~purity vs ~* | 2.01e−01 | 8.56e−01 |
| *~purity+region vs ~purity* | 8.99e−01 | 1e+00 |
| *Carcinoma vs. normal* | 1.4e−04 *** | 1.4e−04 *** |

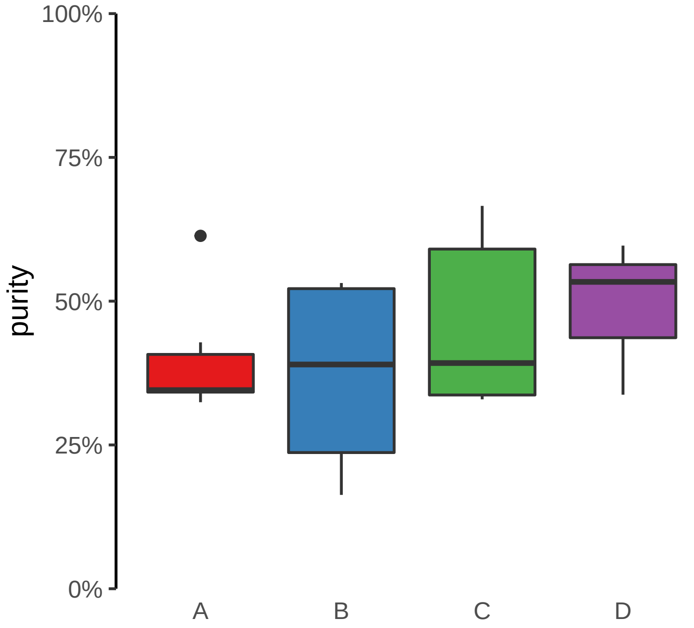

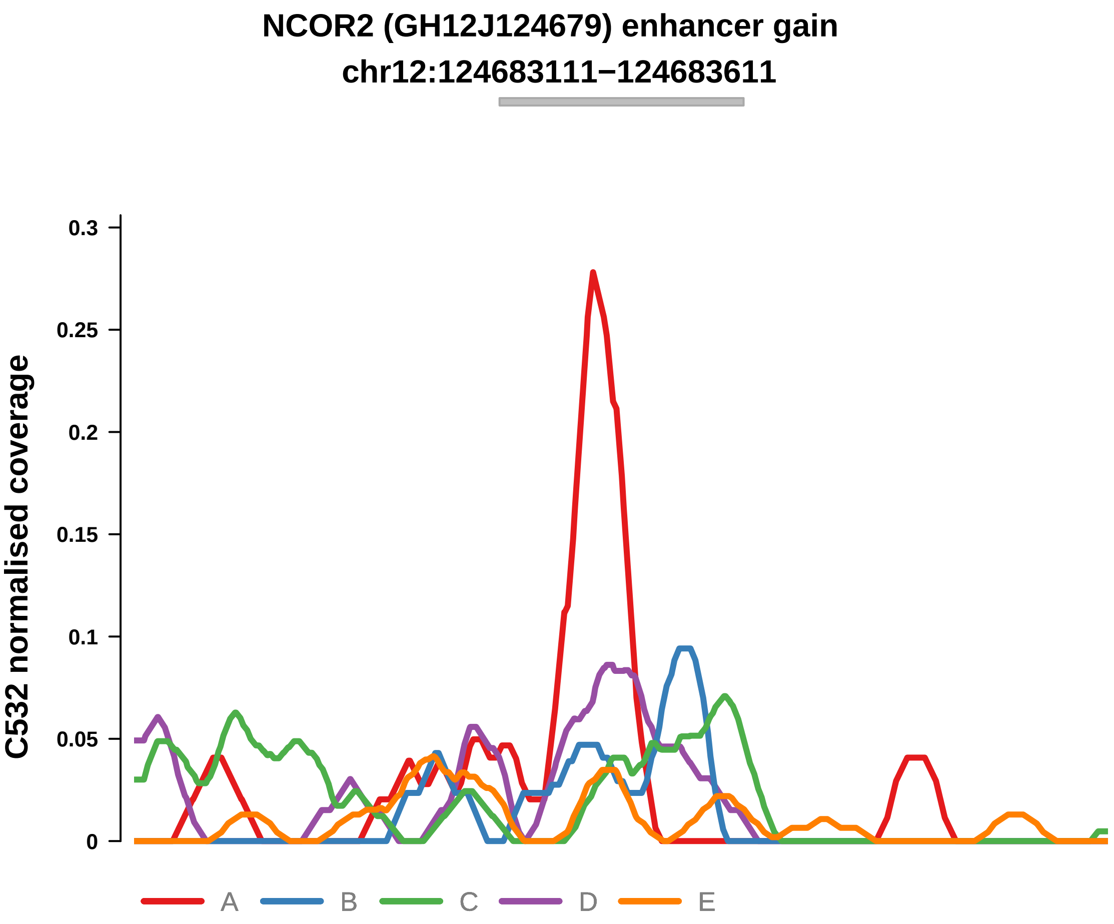


Putatively Clonal (logFC = 2.08)

| | P. | Adjusted P. |
|---|---|---|
| *~region vs. ~* | 5.36e−01 | 8.54e−01 |
| *~purity vs ~* | 8.53e−01 | 9.86e−01 |
| *~purity+region vs ~purity* | 4.6e−01 | 9.41e−01 |
| *Carcinoma vs. normal* | 4.7e−03 ** | 4.7e−03 ** |

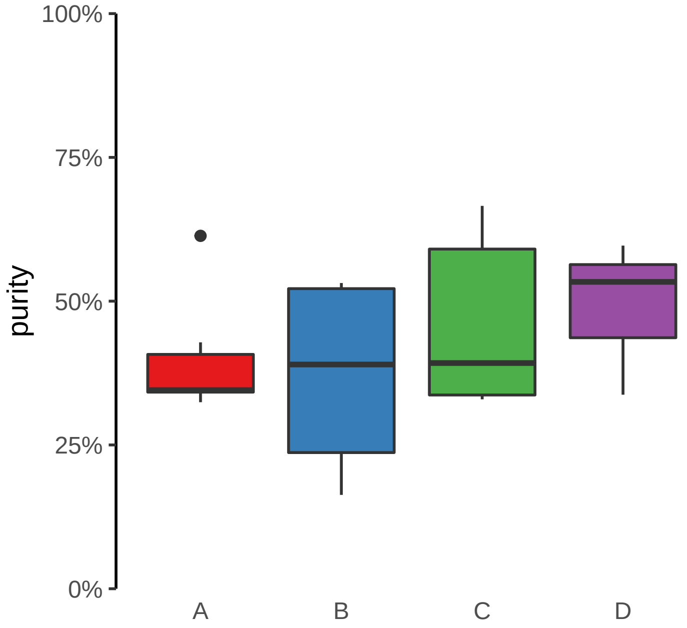

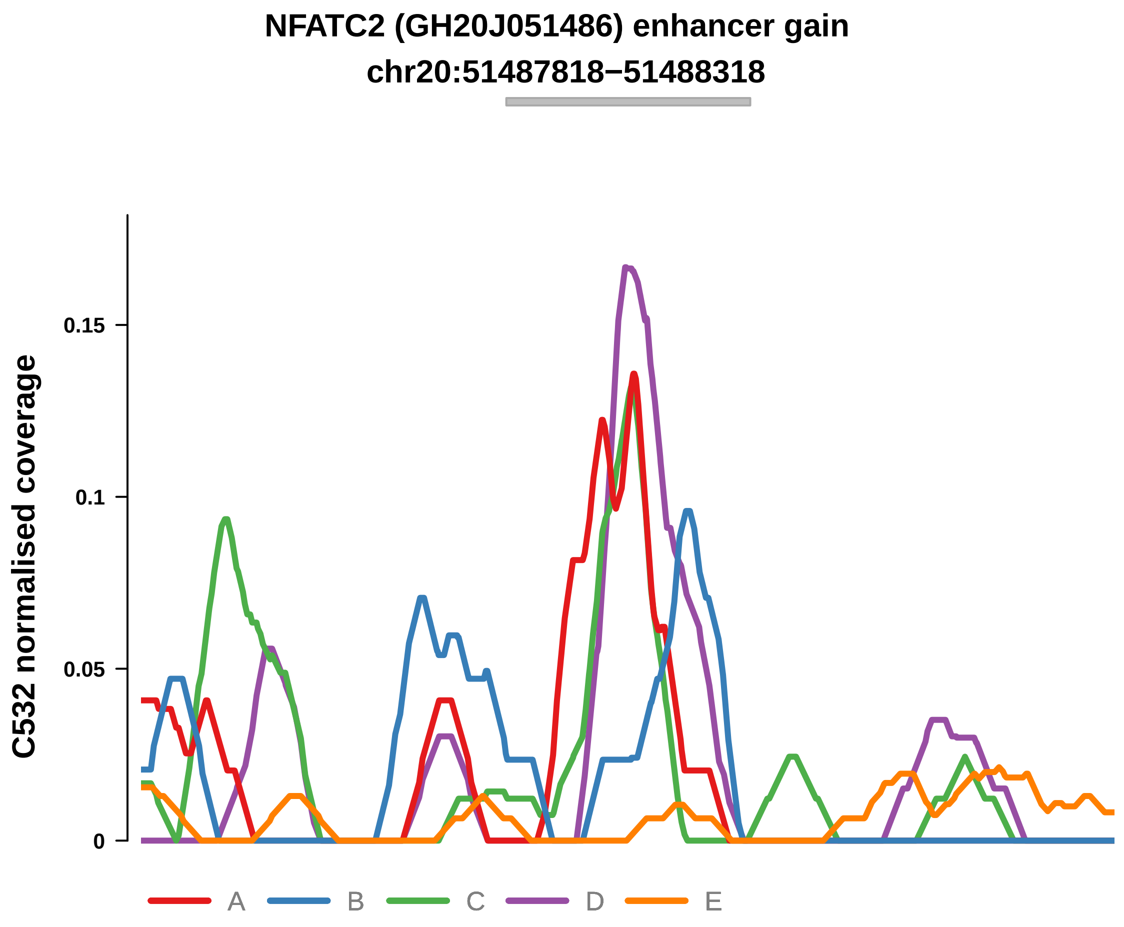


Putatively Clonal (logFC = 2.65)

| | P. | Adjusted P. |
|---|---|---|
| *~region vs. ~* | 6.43e−01 | 8.54e−01 |
| *~purity vs ~* | 1.16e−01 | 7.63e−01 |
| *~purity+region vs ~purity* | 9.04e−01 | 1e+00 |
| *Carcinoma vs. normal* | 4.09e−04 *** | 4.09e−04 *** |

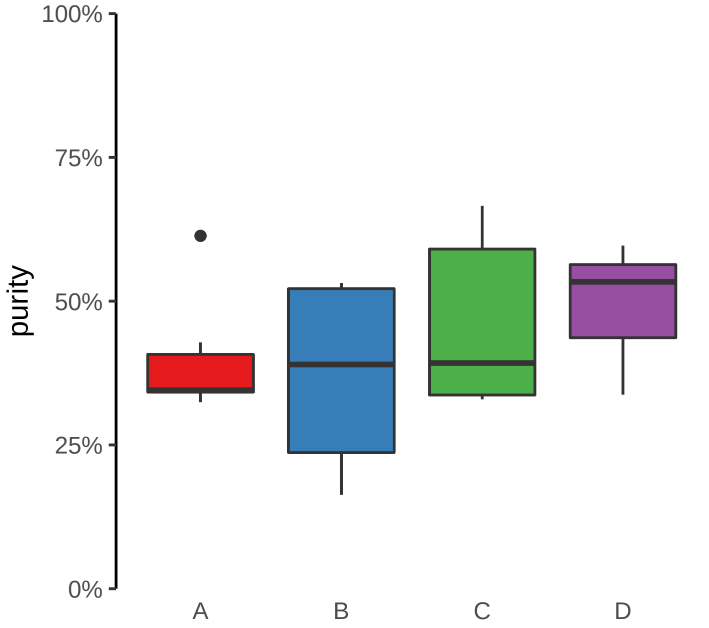

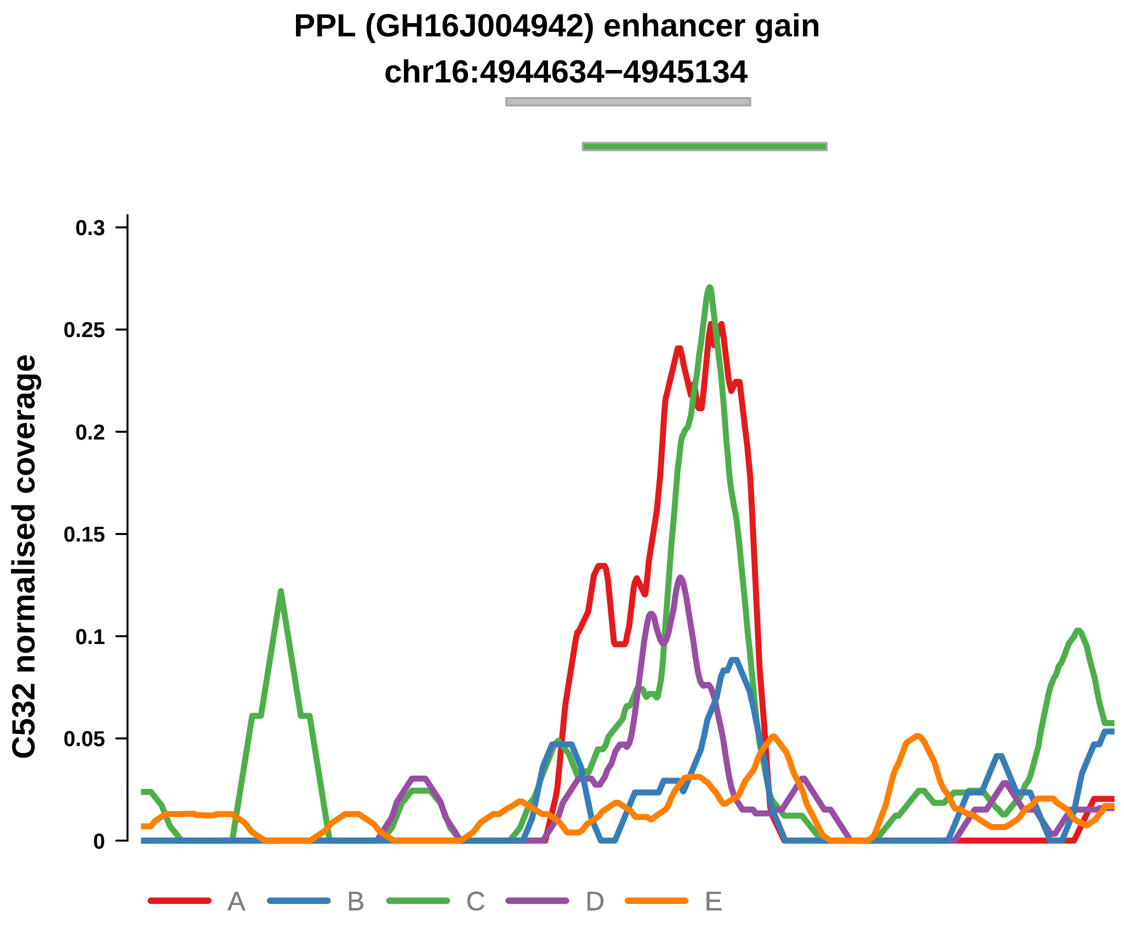


Putatively Clonal (logFC = 1.96)

| | P. | Adjusted P. |
|---|---|---|
| *~region vs. ~* | 2e−02 * | 2.06e−01 |
| *~purity vs ~* | 6.14e−01 | 9.86e−01 |
| *~purity+region vs ~purity* | 2.06e−02 * | 2.6e−01 |
| *Carcinoma vs. normal* | 5.55e−03 ** | 5.55e−03 ** |

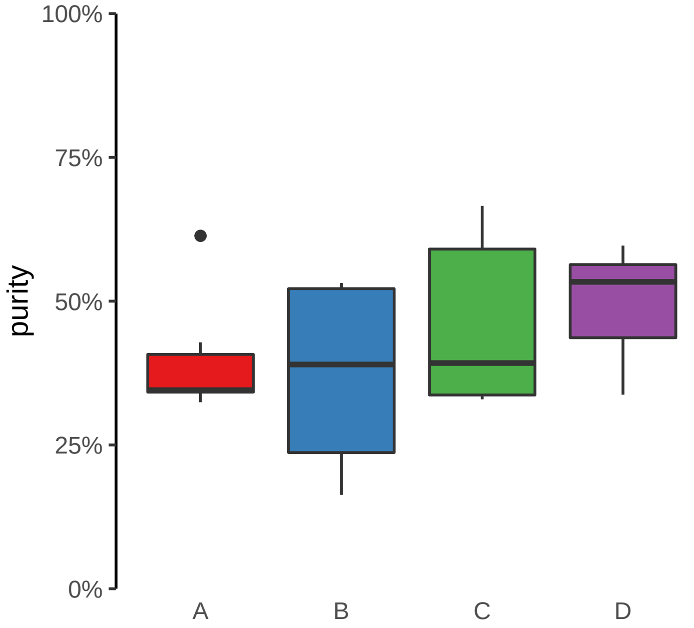

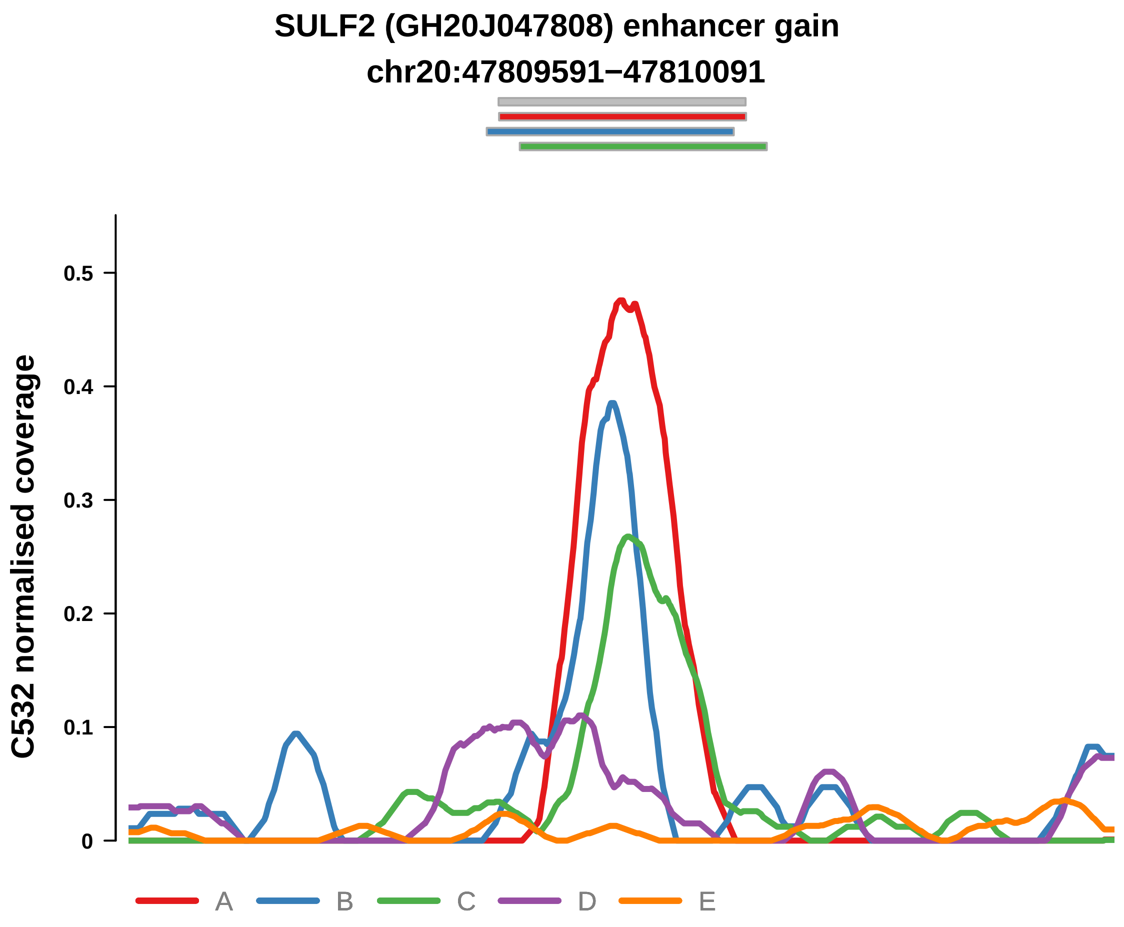


Putatively Clonal (logFC = 2.56)

| | P. | Adjusted P. |
|---|---|---|
| *~region vs. ~* | 1.42e−01 | 4.99e−01 |
| *~purity vs ~* | 8.18e−01 | 9.86e−01 |
| *~purity+region vs ~purity* | 8.78e−02 | 5.15e−01 |
| *Carcinoma vs. normal* | 3.32e−04 *** | 3.32e−04 *** |

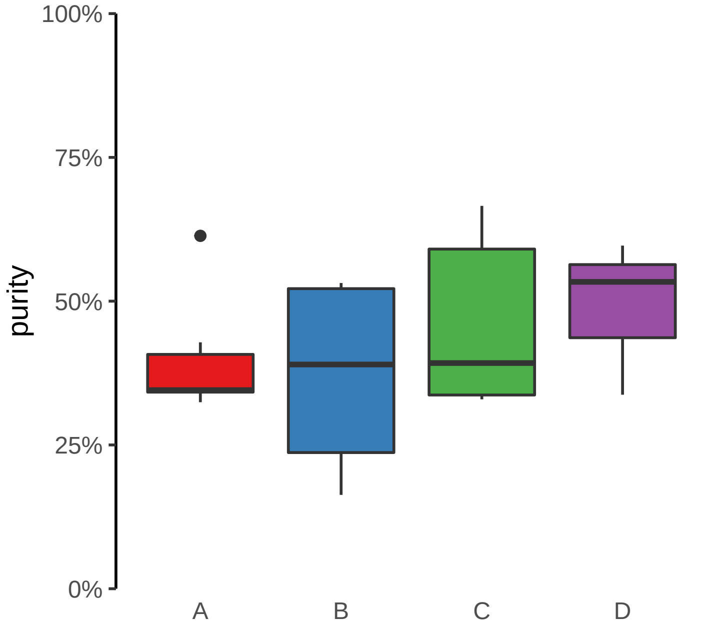

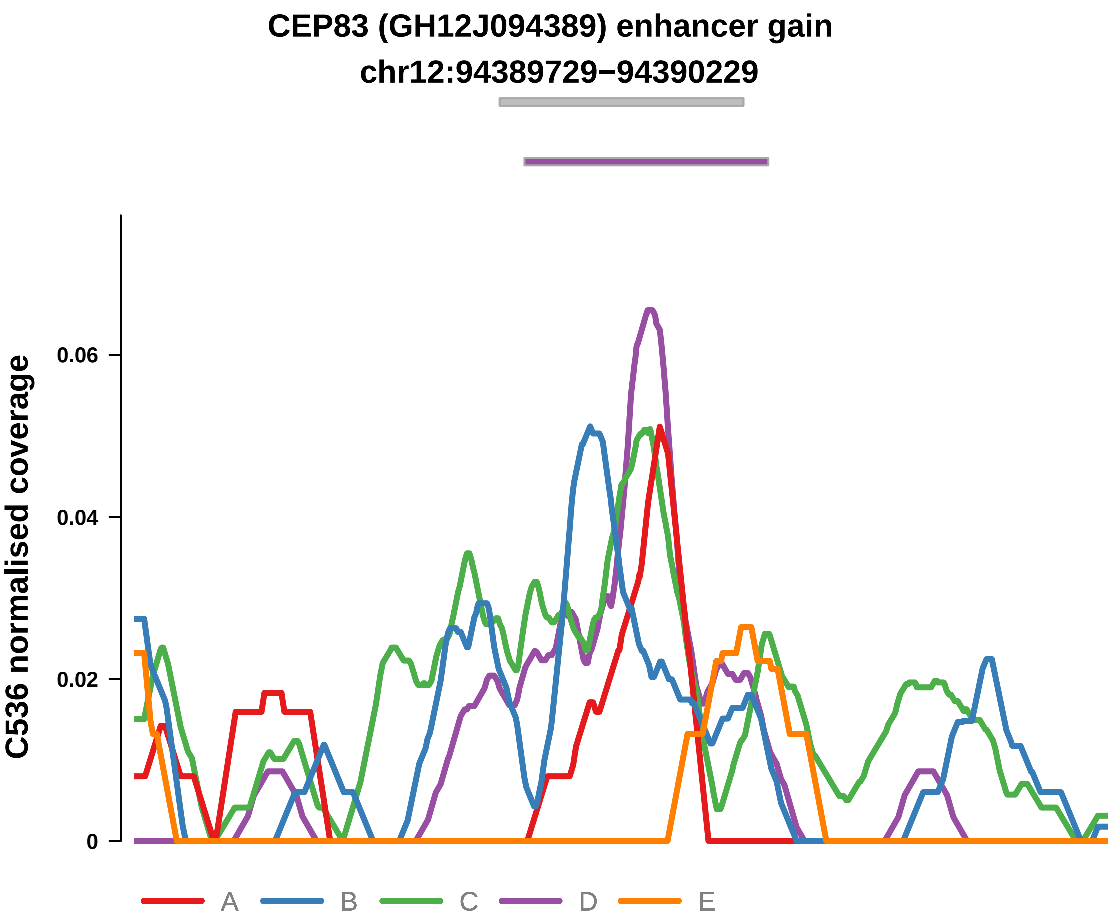


Putatively Clonal (logFC = 3.48)

| | P. | Adjusted P. |
|---|---|---|
| *~region vs. ~* | 1.99e−01 | 3.45e−01 |
| *~purity vs ~* | 3.08e−01 | 5.93e−01 |
| *~purity+region vs ~purity* | 3.39e−01 | 4.87e−01 |
| *Carcinoma vs. normal* | 7.21e−06 **** | 7.21e−06 **** |

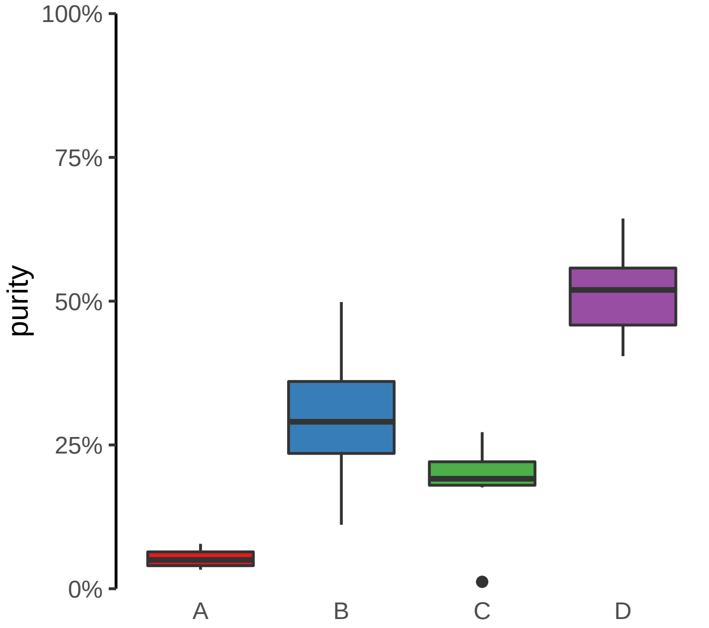

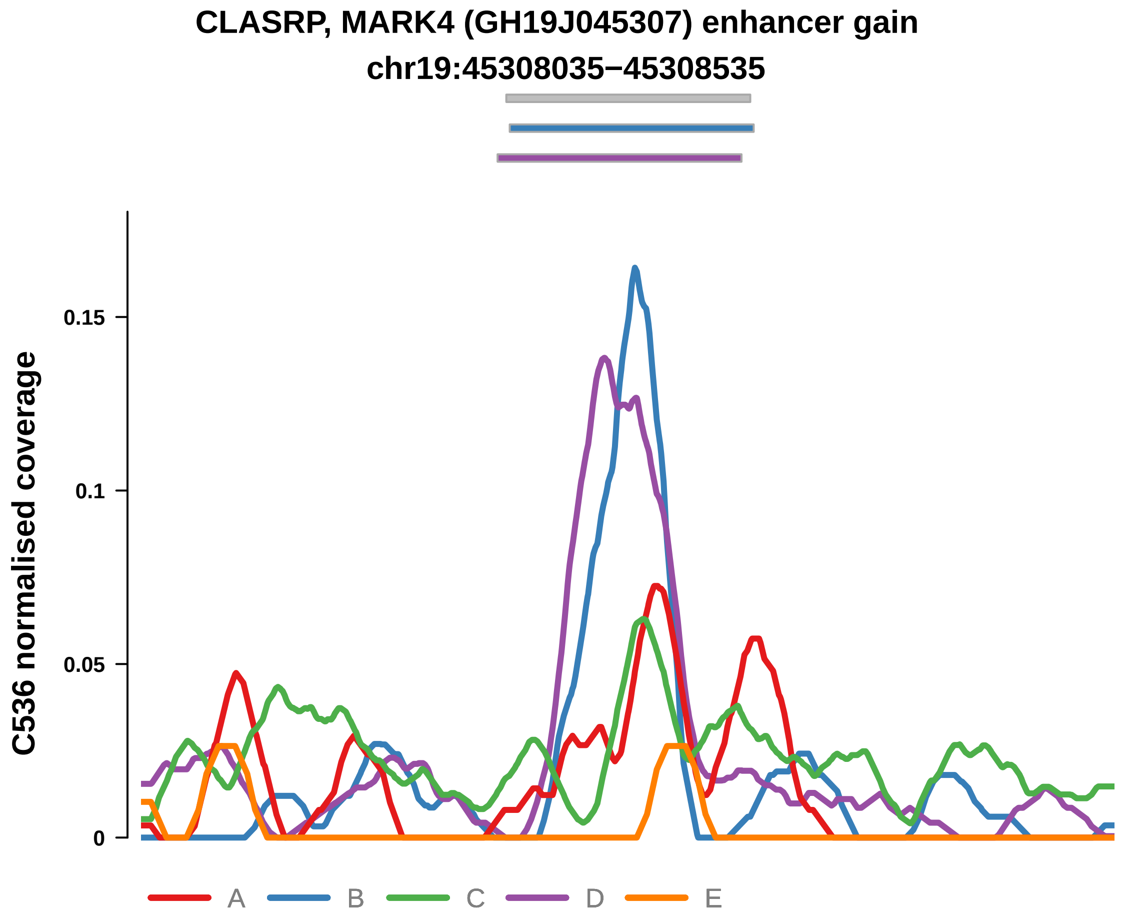


Putatively Clonal (logFC = 2.83)

| | P. | Adjusted P. |
|---|---|---|
| *~region vs. ~* | 6.7e−02 | 1.8e−01 |
| *~purity vs ~* | 5.46e−02 | 2.09e−01 |
| *~purity+region vs ~purity* | 3.5e−01 | 4.9e−01 |
| *Carcinoma vs. normal* | 8.82e−05 **** | 8.82e−05 **** |

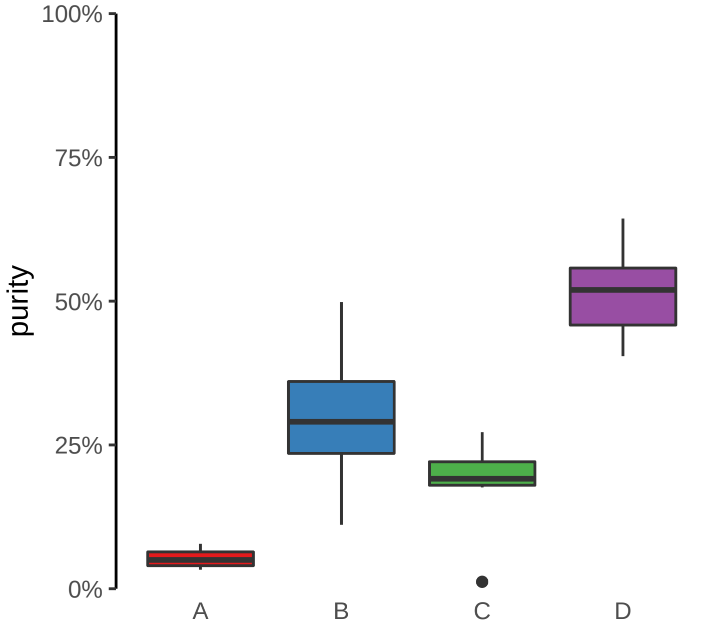

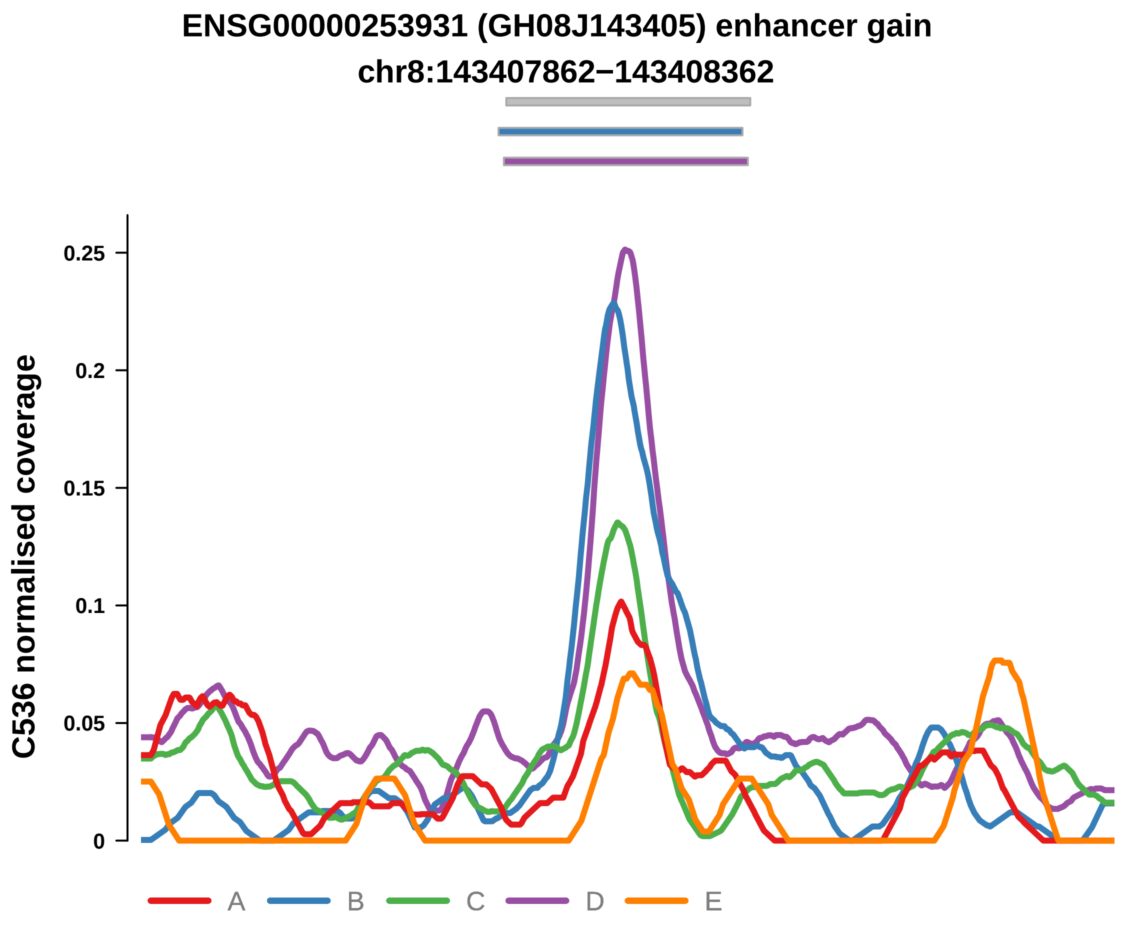


Putatively Clonal (logFC = 2.45)

| | P. | Adjusted P. |
|---|---|---|
| *~region vs. ~* | 1.86e−03 ** | 2.75e−02 * |
| *~purity vs ~* | 4.88e−04 *** | 2.37e−02 * |
| *~purity+region vs ~purity* | 4.17e−01 | 5.55e−01 |
| *Carcinoma vs. normal* | 5.03e−04 *** | 5.03e−04 *** |

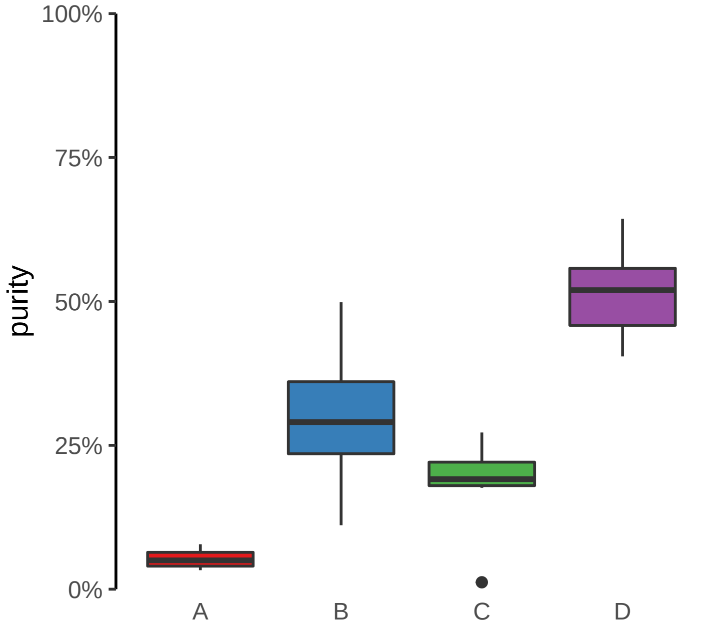

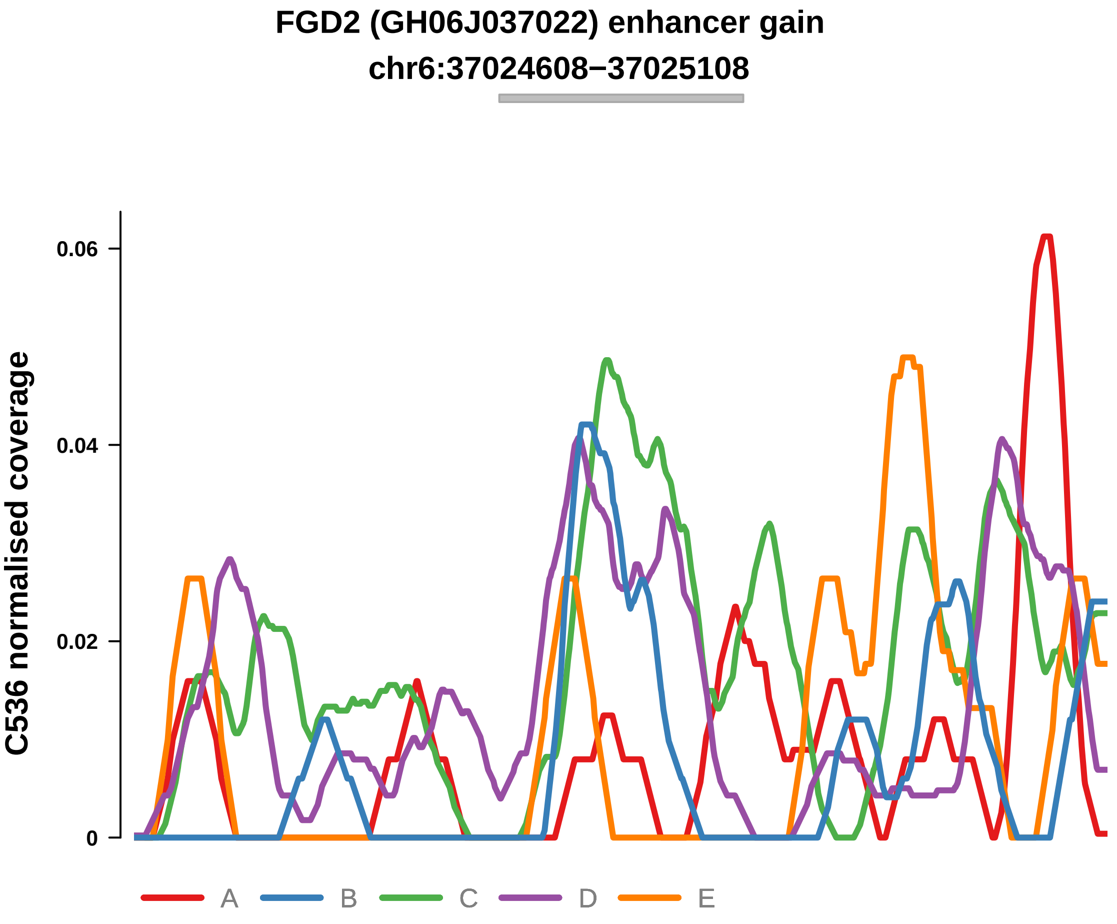


Putatively Clonal (logFC = 1.94)

| | P. | Adjusted P. |
|---|---|---|
| *~region vs. ~* | 2.86e−02 * | NA |
| *~purity vs ~* | 5.75e−01 | 7.44e−01 |
| *~purity+region vs ~purity* | 6.81e−03 ** | 5.22e−02 |
| *Carcinoma vs. normal* | 9.44e−03 ** | 9.44e−03 ** |

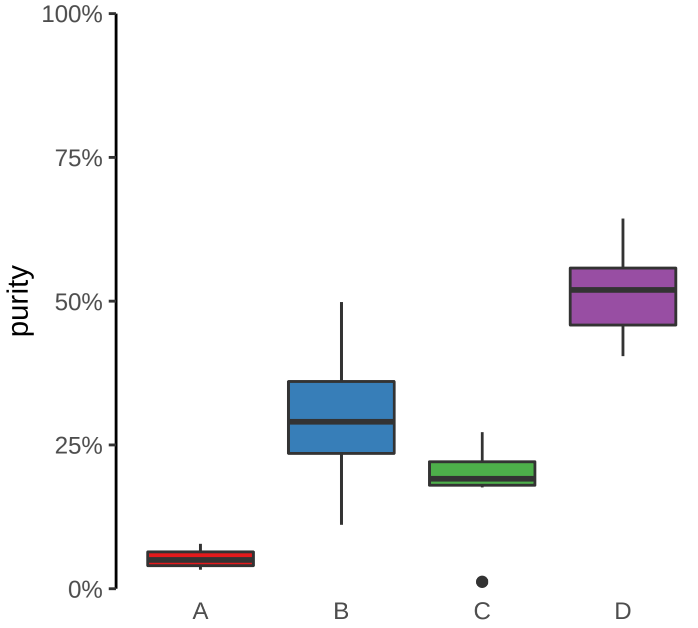

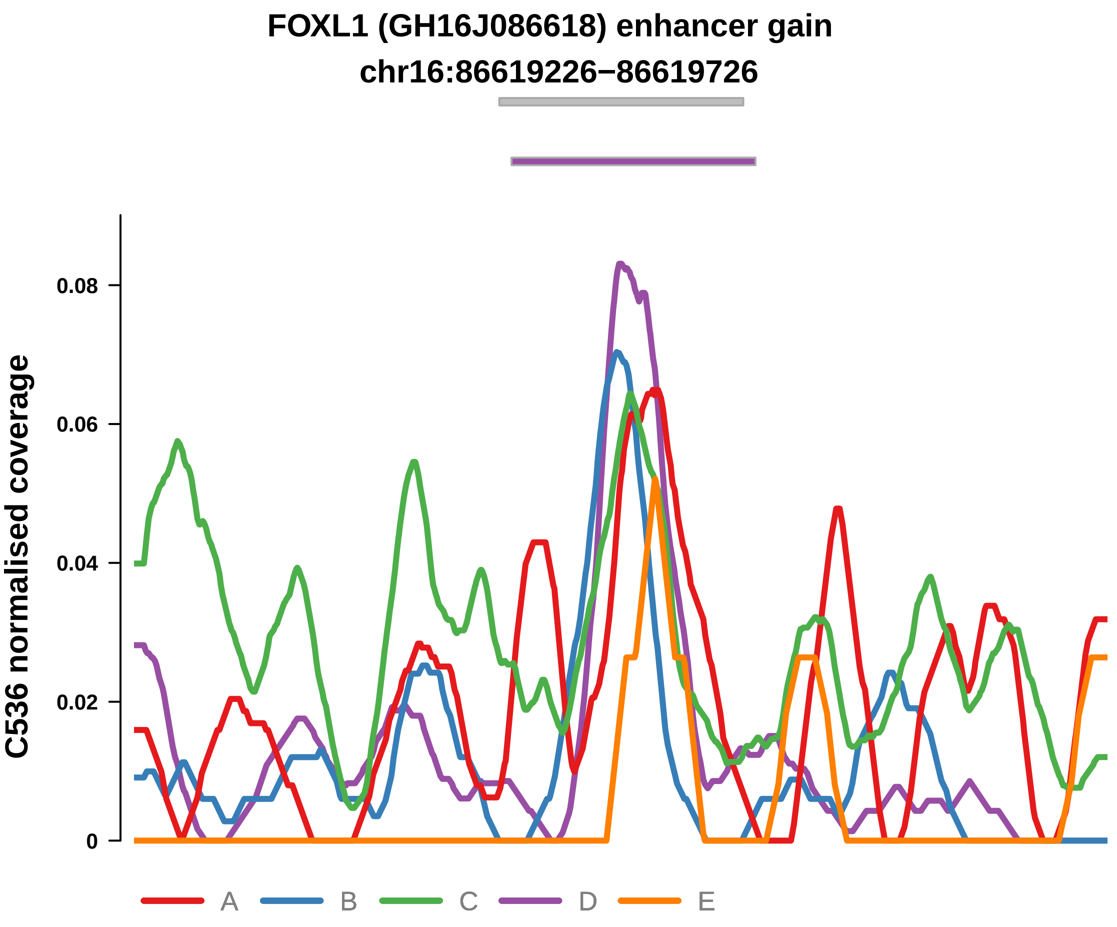


Putatively Clonal (logFC = 2)

| | P. | Adjusted P. |
|---|---|---|
| *~region vs. ~* | 2.89e−01 | 4.06e−01 |
| *~purity vs ~* | 9.57e−01 | 9.68e−01 |
| *~purity+region vs ~purity* | 2.29e−01 | 3.96e−01 |
| *Carcinoma vs. normal* | 6.13e−03 ** | 6.13e−03 ** |

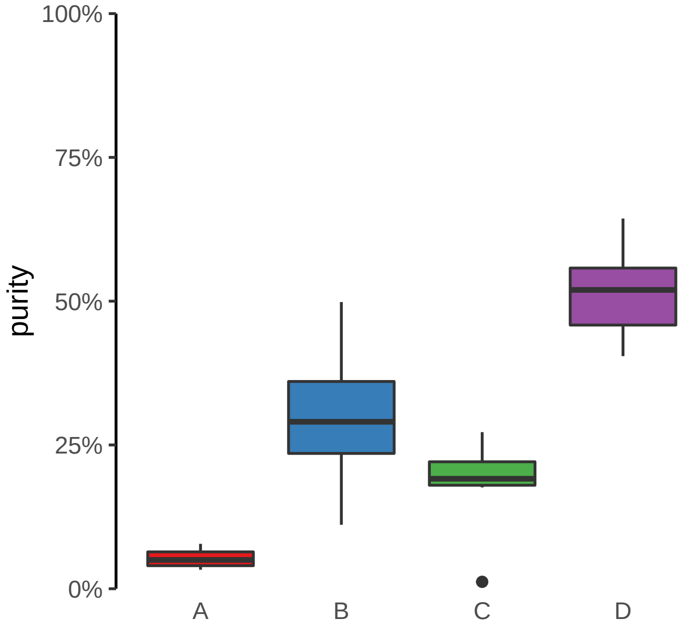

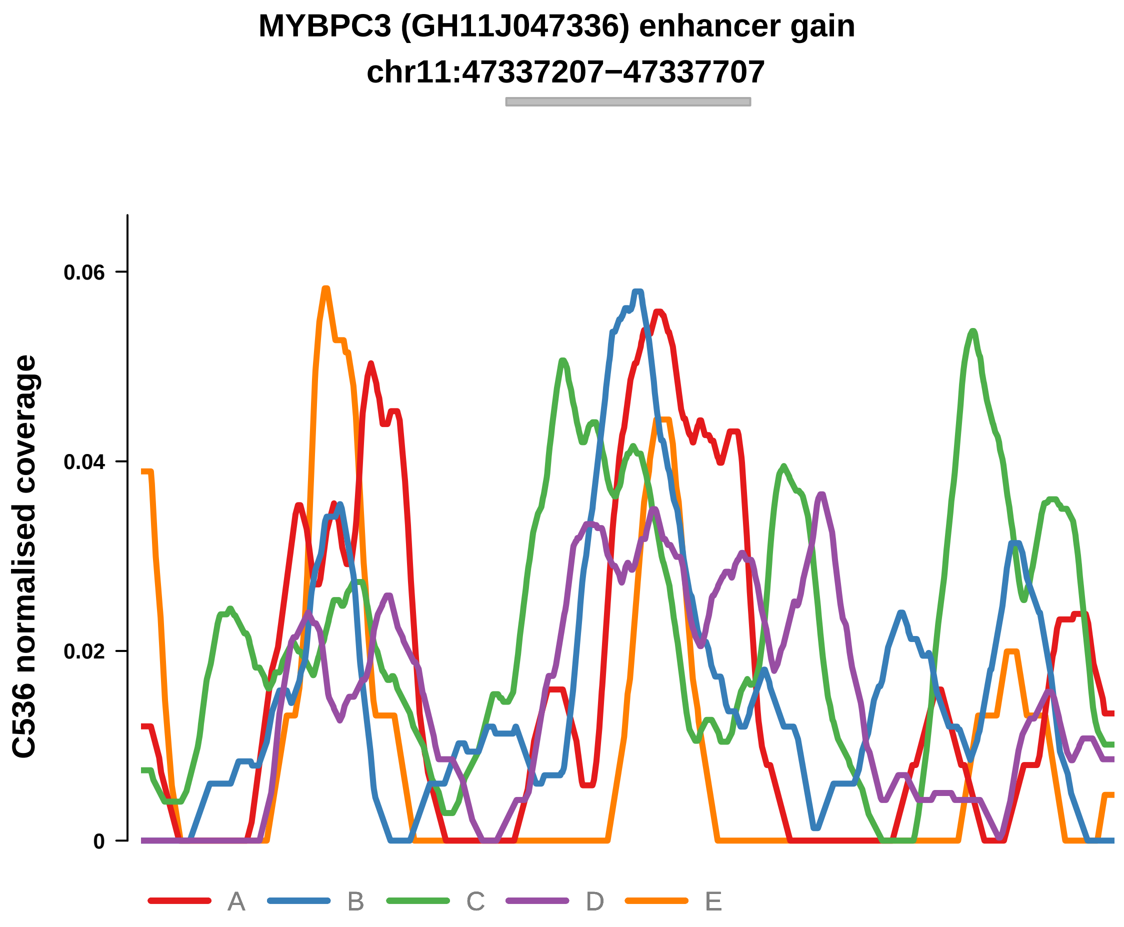


Putatively Clonal (logFC = 2.38)

| | P. | Adjusted P. |
|---|---|---|
| *~region vs. ~* | 4.34e−01 | 5.08e−01 |
| *~purity vs ~* | 6.3e−01 | 7.7e−01 |
| *~purity+region vs ~purity* | 1.33e−02 * | 7.08e−02 |
| *Carcinoma vs. normal* | 1.65e−03 ** | 1.65e−03 ** |

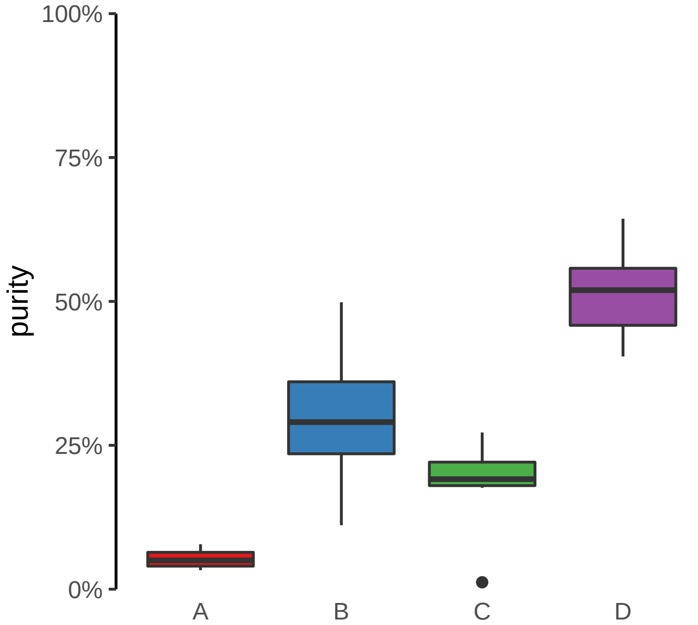

**TPRA1, ABTB1, MGLL (GH03J127766) enhancer gain**
**chr3:127767157−127767657**

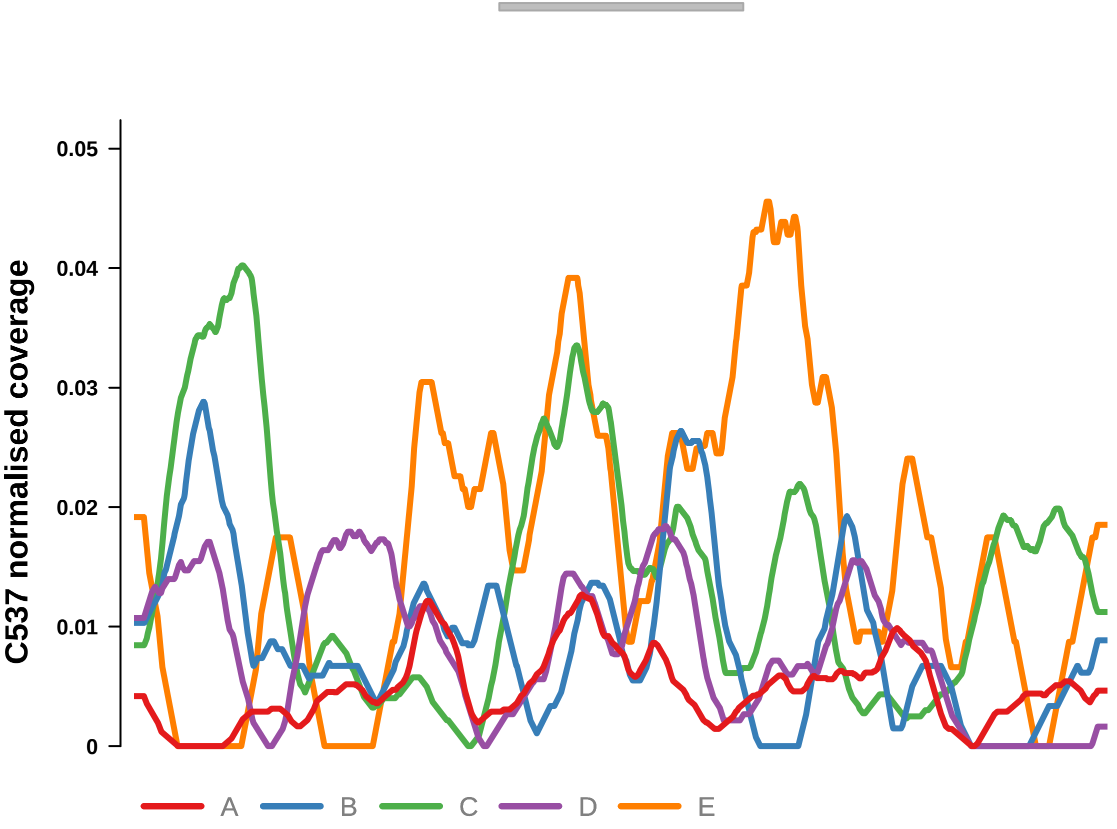


Putatively Clonal (logFC = 1.98)

| | P. | Adjusted P. |
|---|---|---|
| *~region vs. ~* | 7.1e−02 | 1.36e−01 |
| *~purity vs ~* | 5.94e−02 | NA |
| *~purity+region vs ~purity* | 3.72e−01 | NA |
| *Carcinoma vs. normal* | 6.87e−03 ** | 6.87e−03 ** |

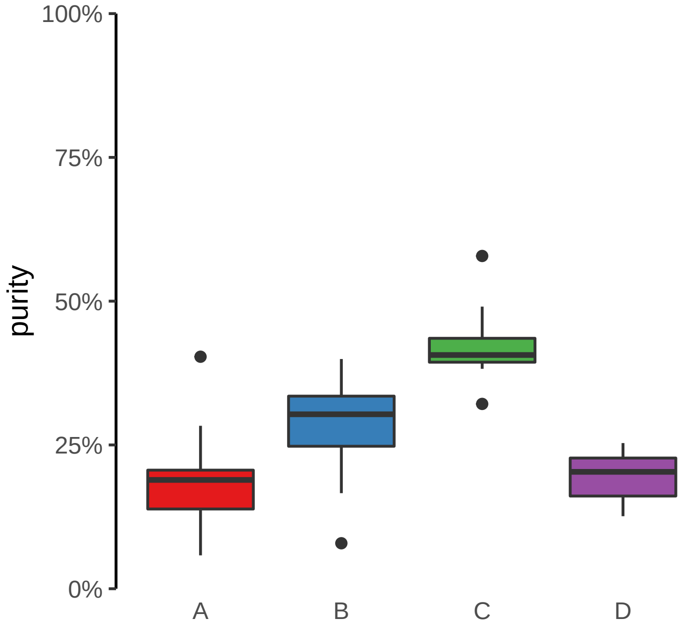

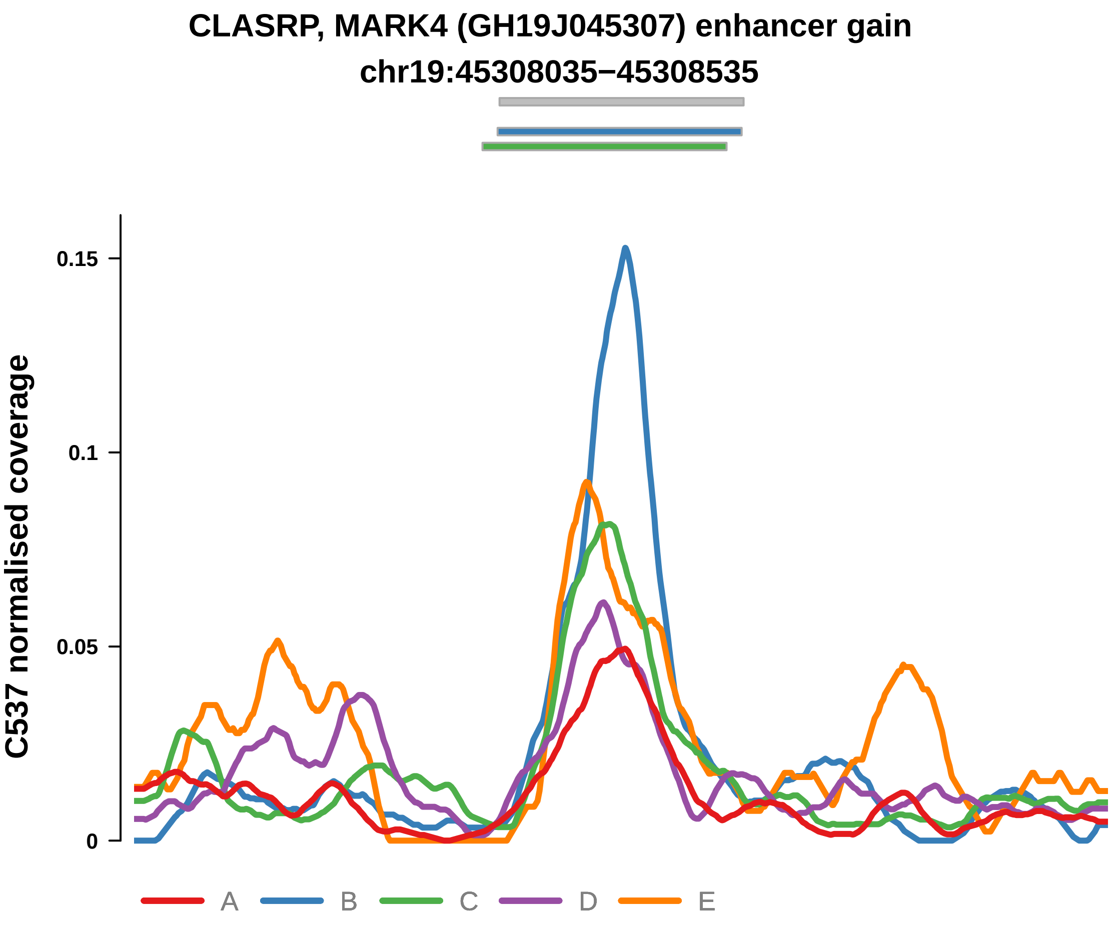


Putatively Clonal (logFC = 1.84)

| | P. | Adjusted P. |
|---|---|---|
| *~region vs. ~* | 1.62e−01 | 2.44e−01 |
| *~purity vs ~* | 1.88e−04 *** | 4.32e−03 ** |
| *~purity+region vs ~purity* | 2.11e−01 | 3.48e−01 |
| *Carcinoma vs. normal* | 8.57e−03 ** | 8.57e−03 ** |

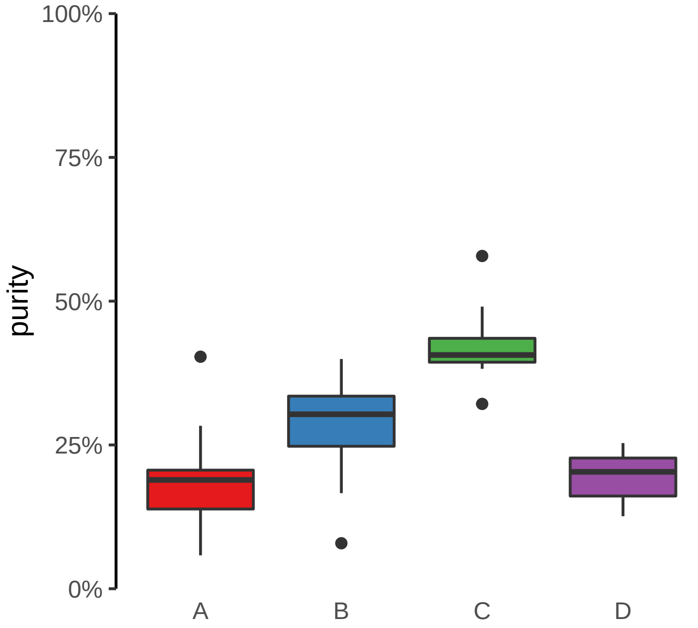

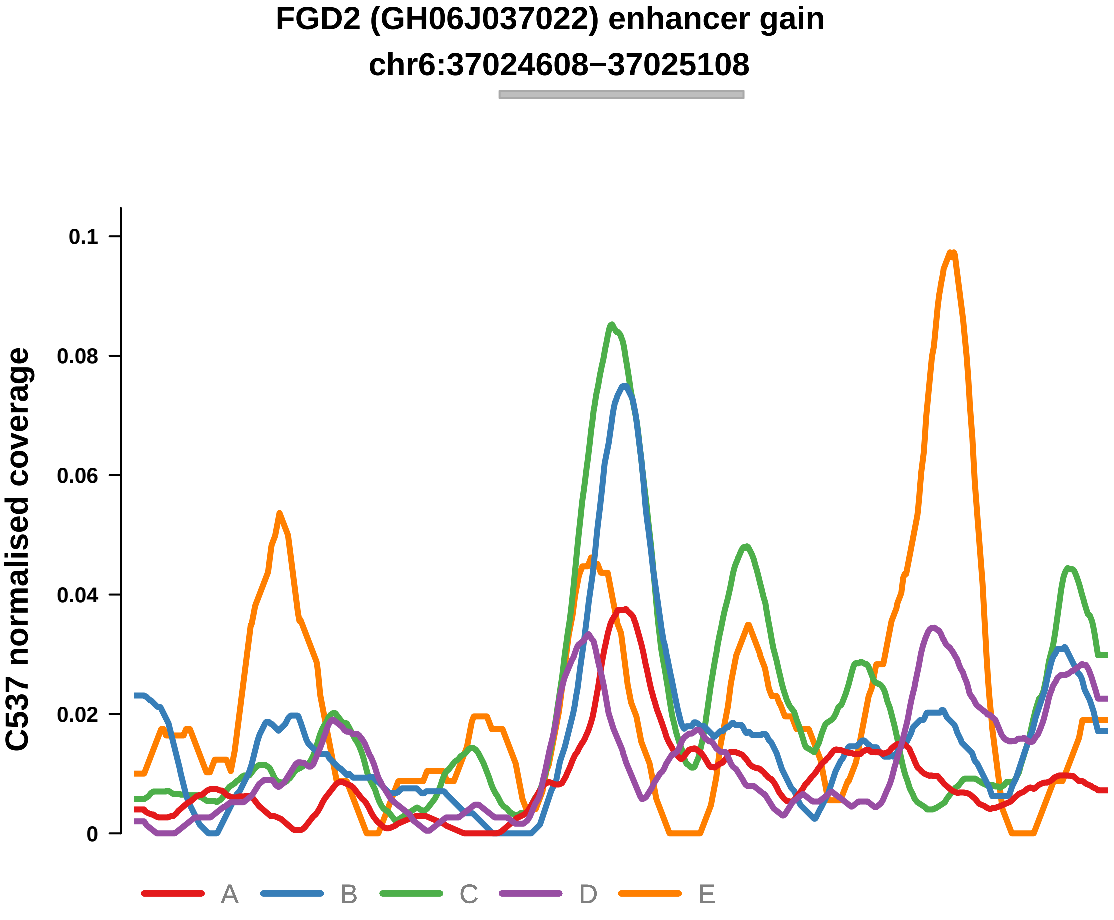


Putatively Clonal (logFC = 2.37)

| | P. | Adjusted P. |
|---|---|---|
| *~region vs. ~* | 1.42e−03 ** | 2.08e−02 * |
| *~purity vs ~* | 1.06e−05 **** | 7.3e−04 *** |
| *~purity+region vs ~purity* | 4.34e−01 | 6.01e−01 |
| *Carcinoma vs. normal* | 8.91e−04 *** | 8.91e−04 *** |

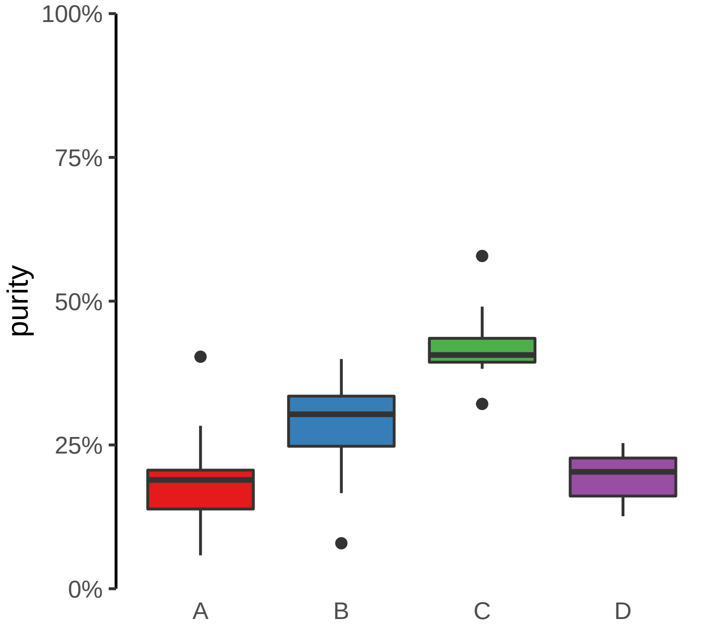

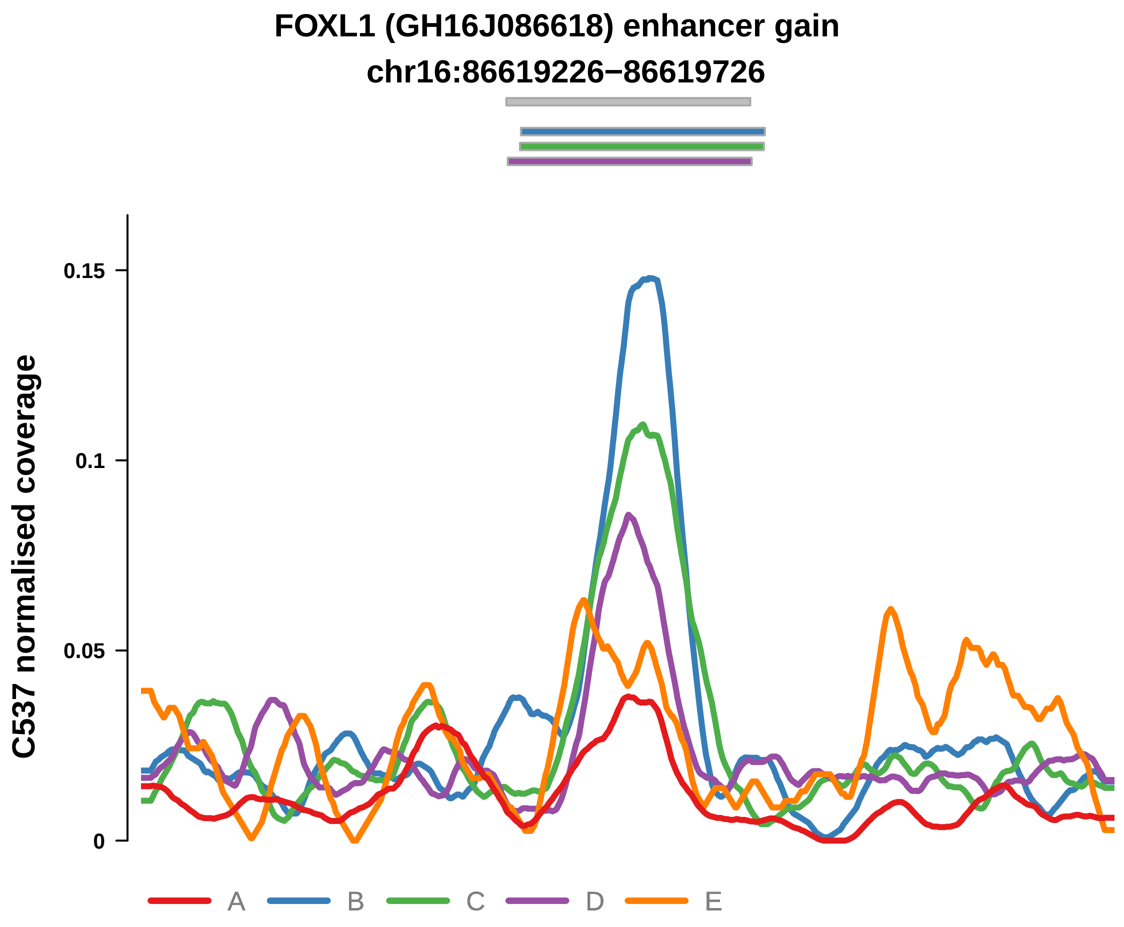


Putatively Clonal (logFC = 2.48)

| | P. | Adjusted P. |
|---|---|---|
| *~region vs. ~* | 5.91e−03 ** | 4.57e−02 * |
| *~purity vs ~* | 2.33e−03 ** | 1.73e−02 * |
| *~purity+region vs ~purity* | 1.59e−01 | 3.04e−01 |
| *Carcinoma vs. normal* | 4.96e−04 *** | 4.96e−04 *** |

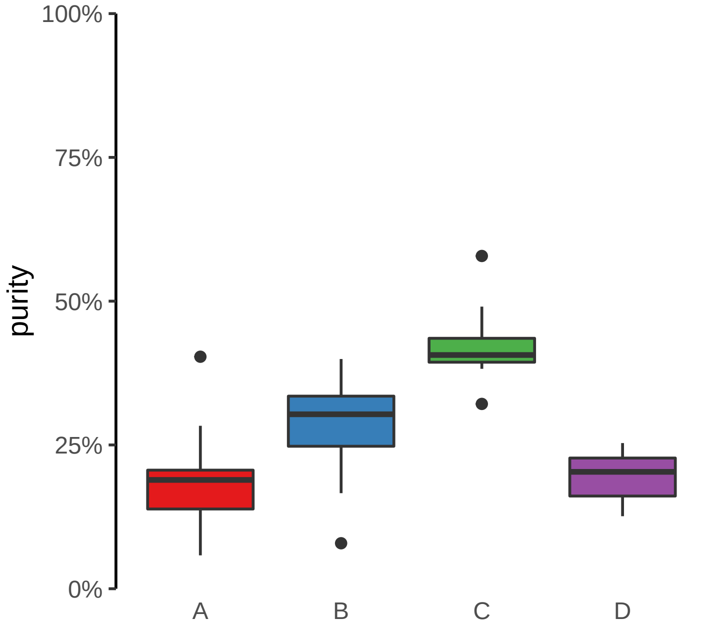

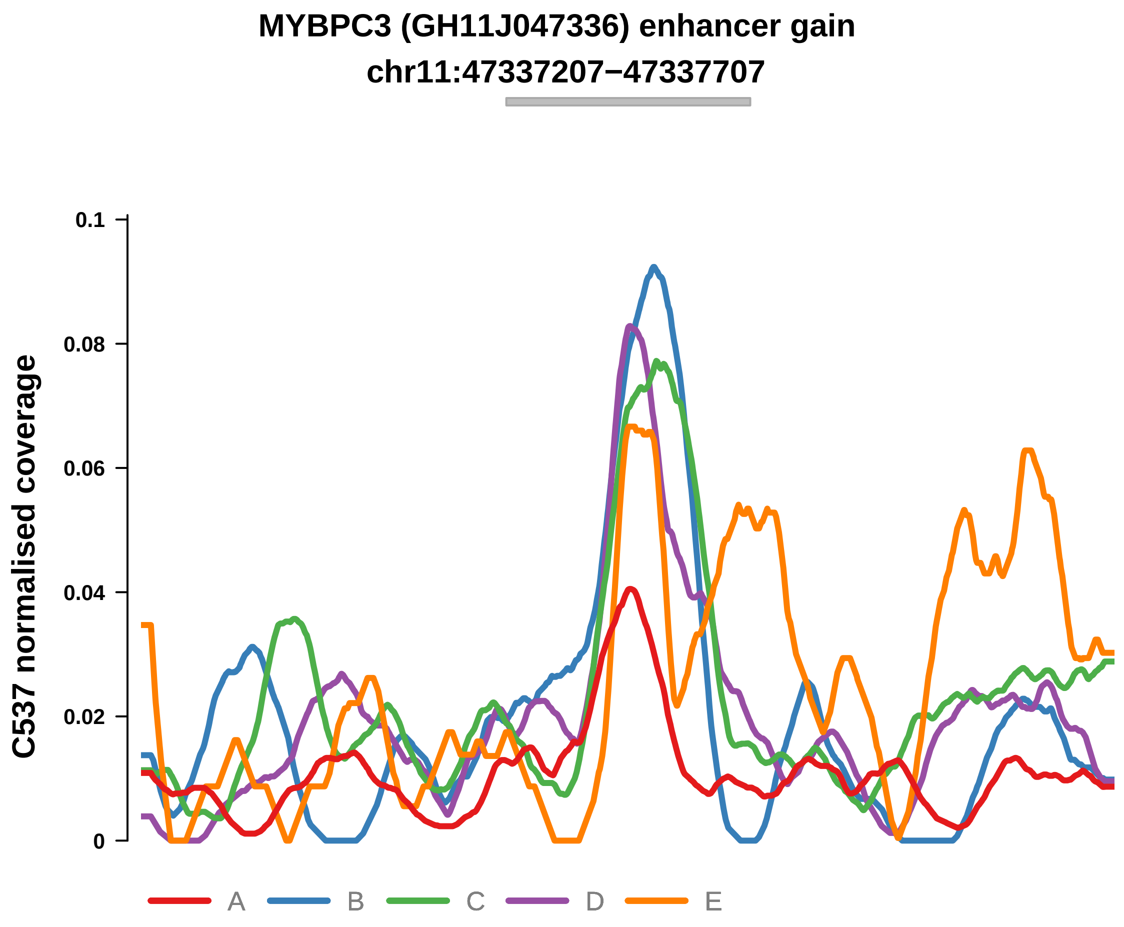


Putatively Clonal (logFC = 2.65)

| | P. | Adjusted P. |
|---|---|---|
| *~region vs. ~* | 6.87e−04 *** | 1.27e−02 * |
| *~purity vs ~* | 1.33e−02 * | 5.39e−02 |
| *~purity+region vs ~purity* | 2.12e−03 ** | 3.88e−02 * |
| *Carcinoma vs. normal* | 2.52e−04 *** | 2.52e−04 *** |

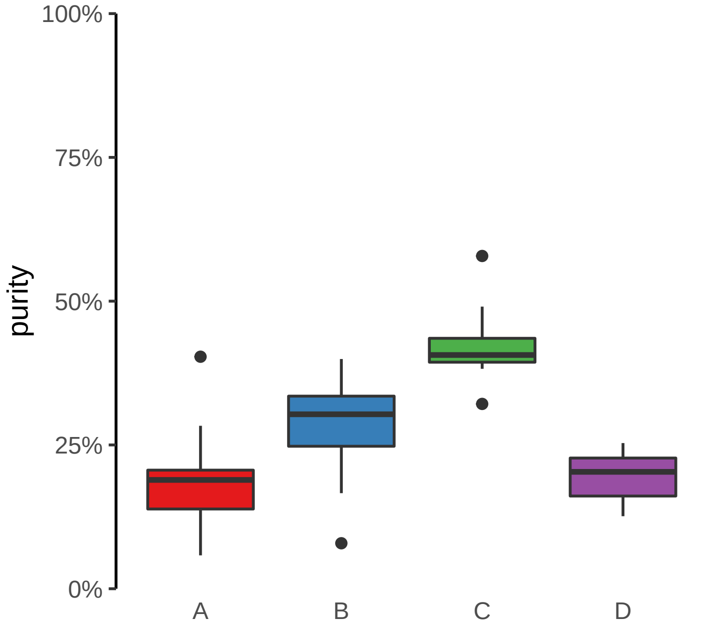

**MYT1 (GH20J064188) enhancer gain**
**chr20:64189162−64189662**

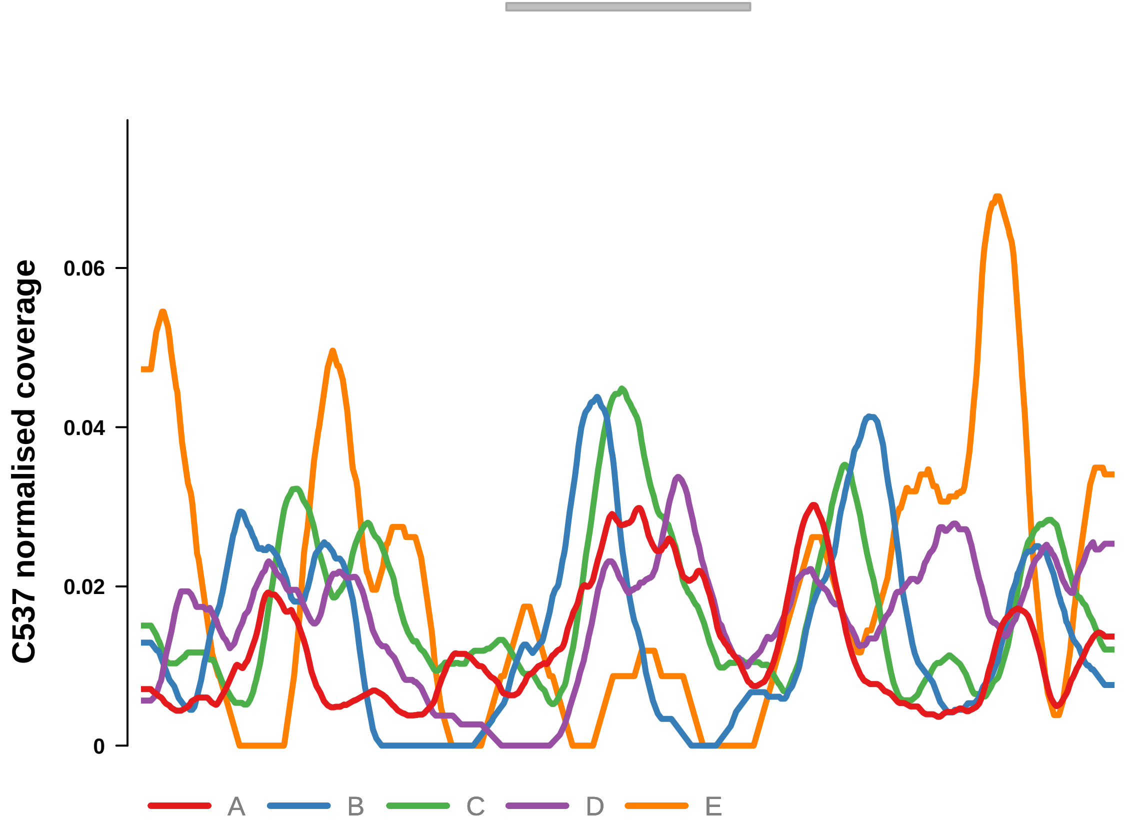


Putatively Clonal (logFC = 2.11)

| | P. | Adjusted P. |
|---|---|---|
| *~region vs. ~* | 6.87e−02 | 1.36e−01 |
| *~purity vs ~* | 5.06e−01 | 6.47e−01 |
| *~purity+region vs ~purity* | 5.69e−02 | 1.77e−01 |
| *Carcinoma vs. normal* | 3.82e−03 ** | 3.82e−03 ** |

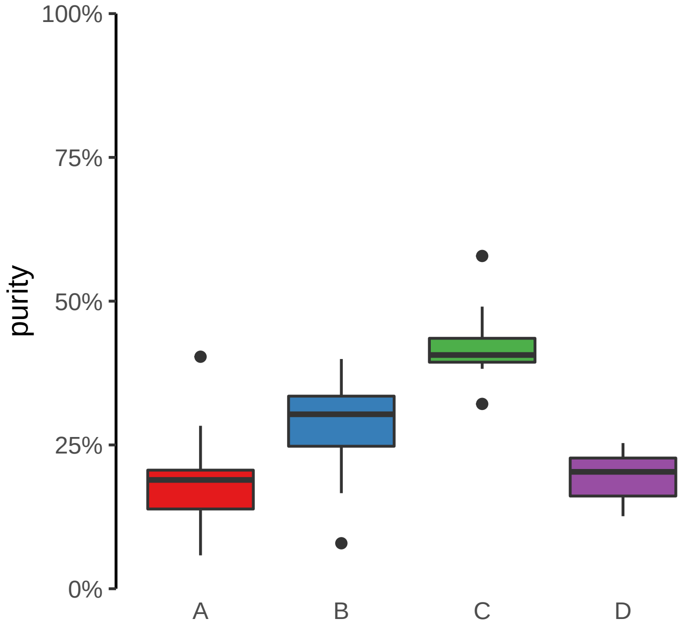

**NFATC2 (GH20J051486) enhancer gain**
**chr20:51487818−51488318**

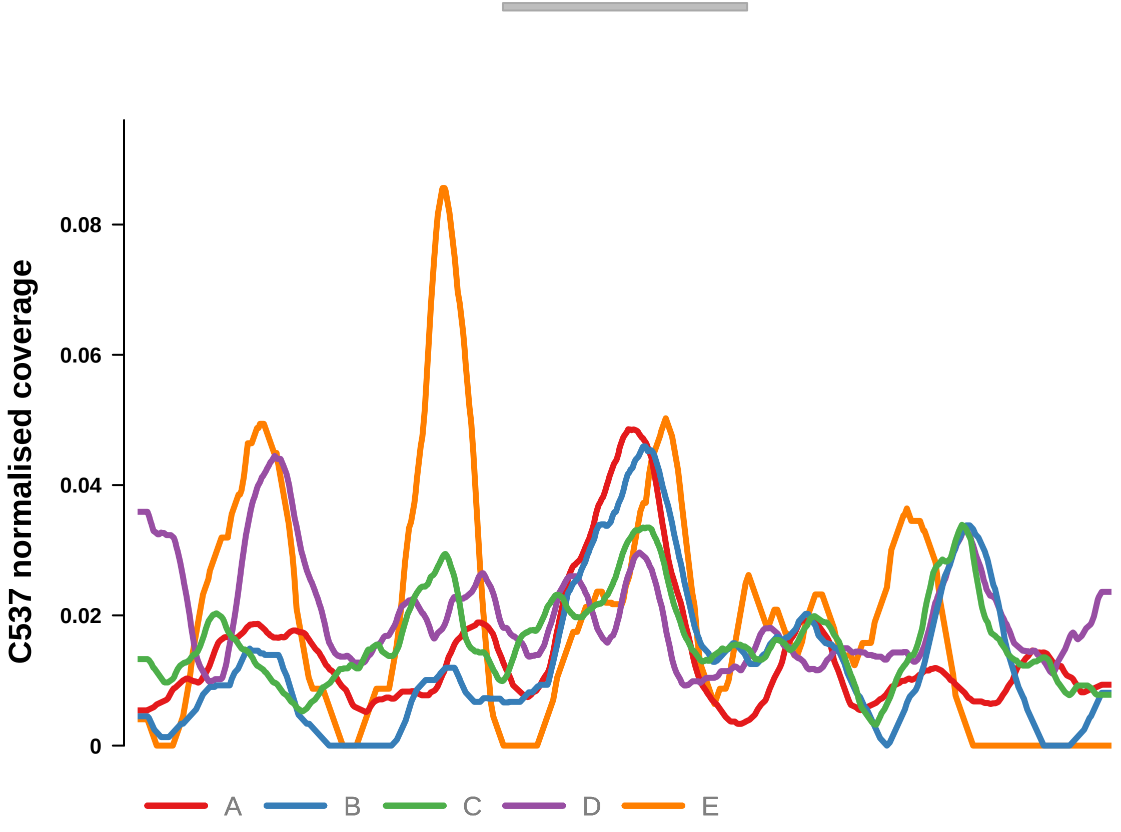


Putatively Clonal (logFC = 1.98)

| | P. | Adjusted P. |
|---|---|---|
| *~region vs. ~* | 2.04e−01 | 2.9e−01 |
| *~purity vs ~* | 7.62e−01 | 8.34e−01 |
| *~purity+region vs ~purity* | 2.12e−01 | 3.48e−01 |
| *Carcinoma vs. normal* | 6.79e−03 ** | 6.79e−03 ** |

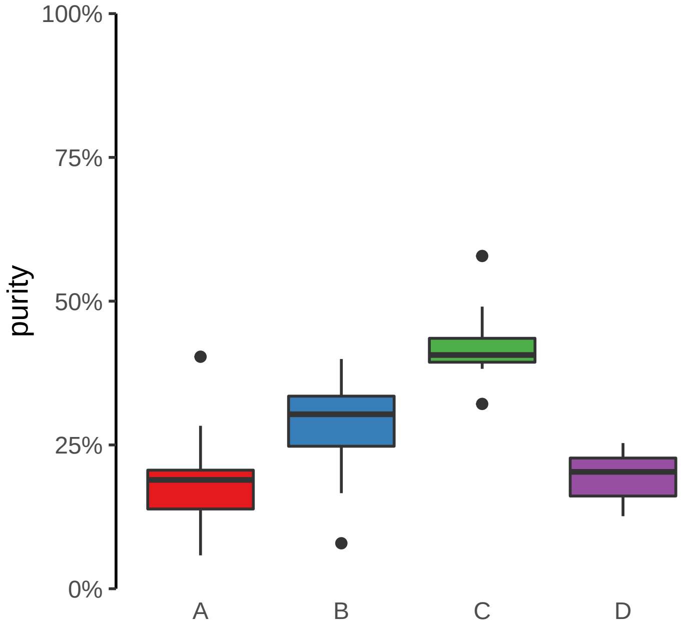

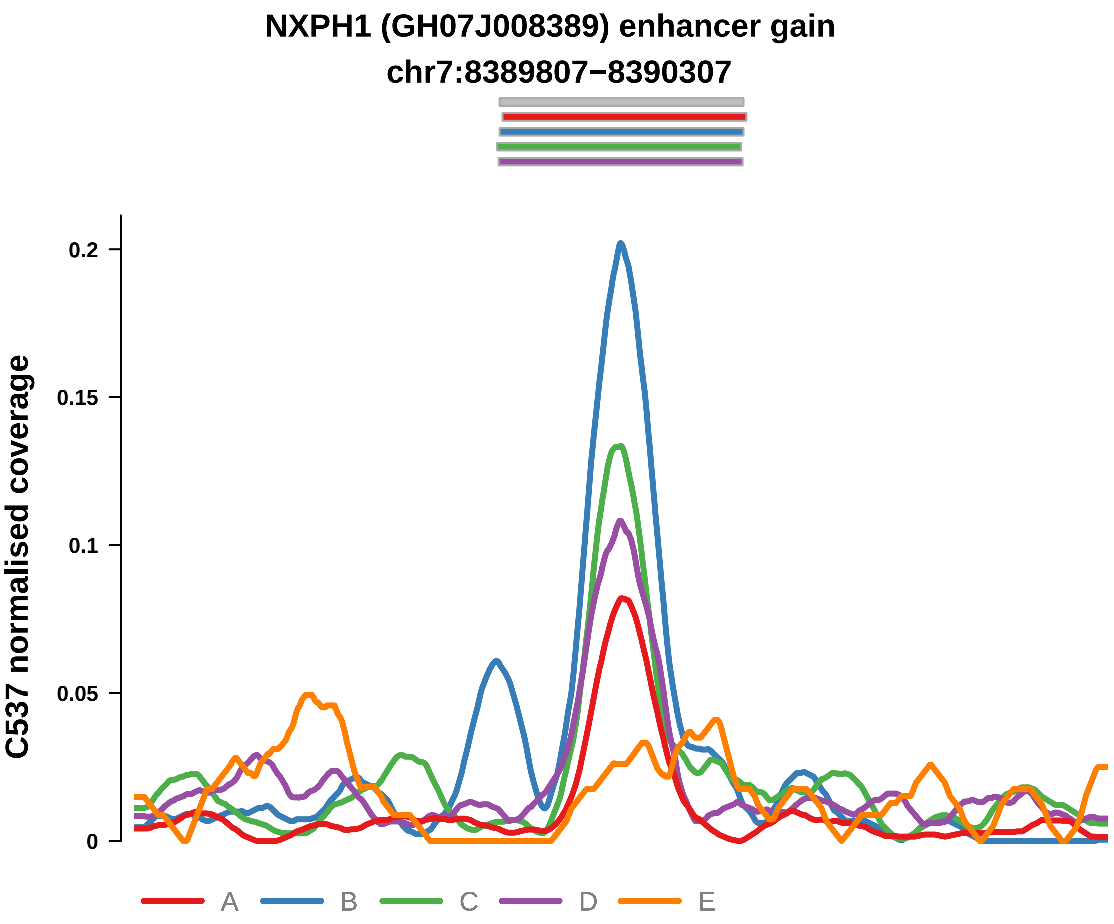


Putatively Clonal (logFC = 2.29)

| | P. | Adjusted P. |
|---|---|---|
| *~region vs. ~* | 1.26e−01 | 2.05e−01 |
| *~purity vs ~* | 1.29e−02 * | 5.39e−02 |
| *~purity+region vs ~purity* | 2.37e−01 | 3.78e−01 |
| *Carcinoma vs. normal* | 1.26e−03 ** | 1.26e−03 ** |

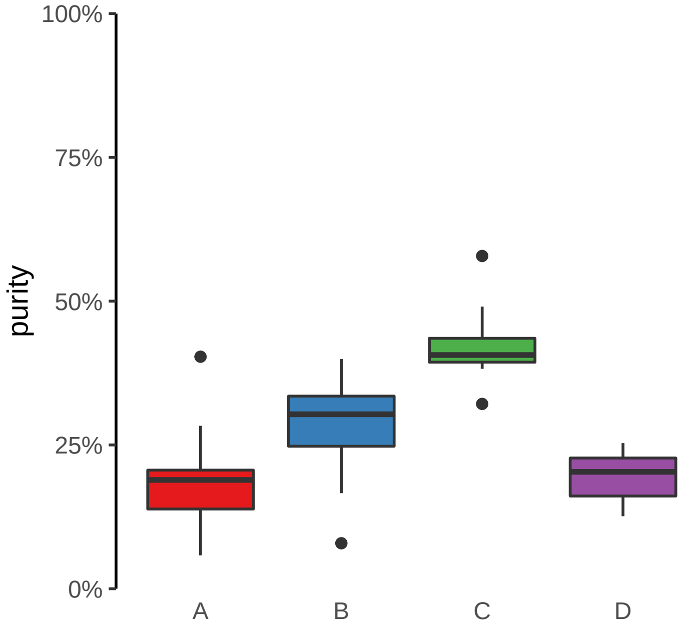

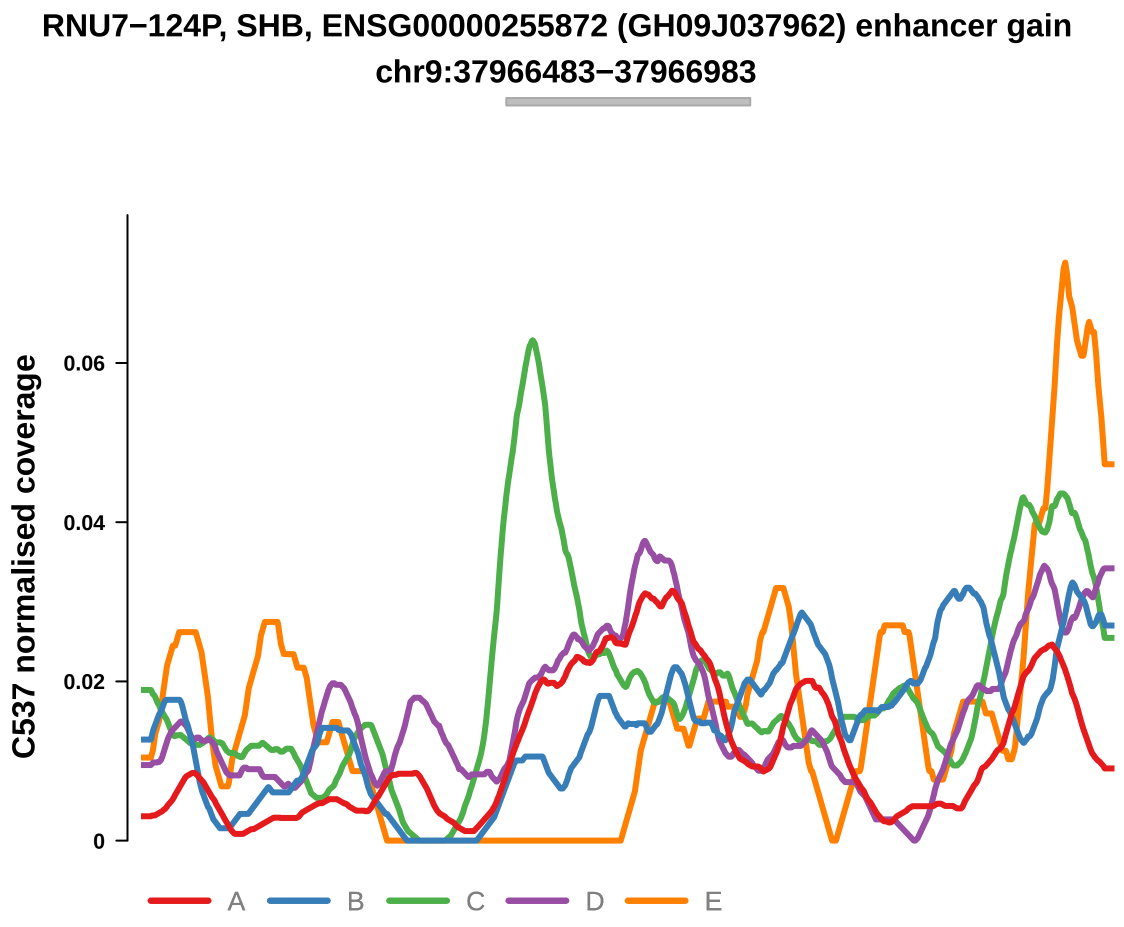


Putatively Clonal (logFC = 2.06)

| | P. | Adjusted P. |
|---|---|---|
| *~region vs. ~* | 2.77e−03 ** | 3.48e−02 * |
| *~purity vs ~* | 2.93e−01 | 4.39e−01 |
| *~purity+region vs ~purity* | 2.24e−03 ** | 3.88e−02 * |
| *Carcinoma vs. normal* | 4.09e−03 ** | 4.09e−03 ** |

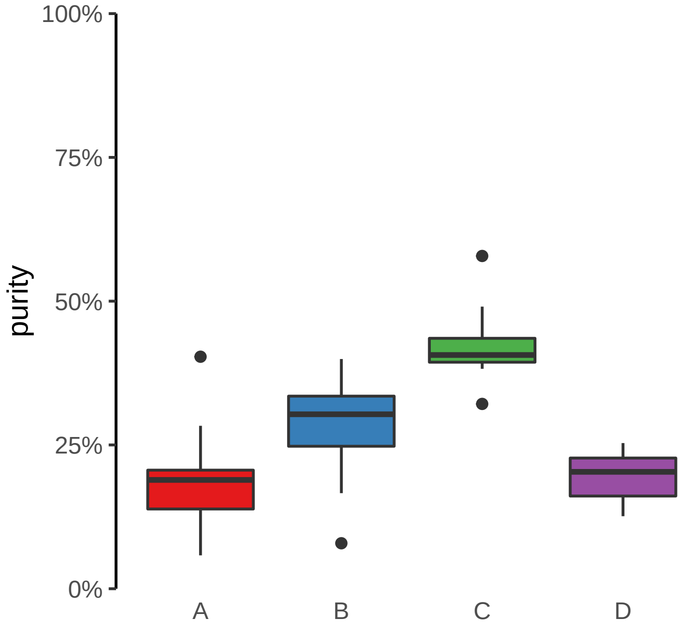

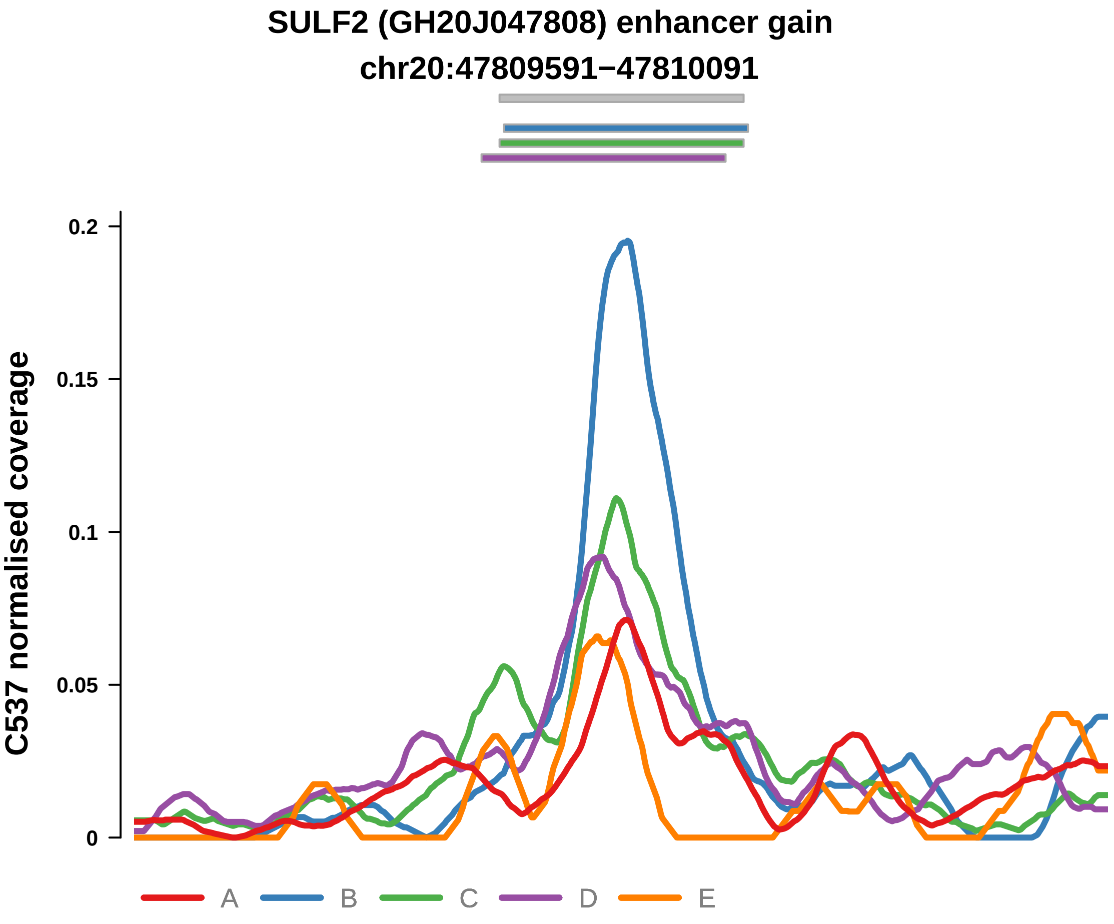


Putatively Clonal (logFC = 2)

| | P. | Adjusted P. |
|---|---|---|
| *~region vs. ~* | 7.85e−03 ** | 4.57e−02 * |
| *~purity vs ~* | 6.32e−02 | 1.62e−01 |
| *~purity+region vs ~purity* | 2.97e−02 * | 1.13e−01 |
| *Carcinoma vs. normal* | 4.21e−03 ** | 4.21e−03 ** |

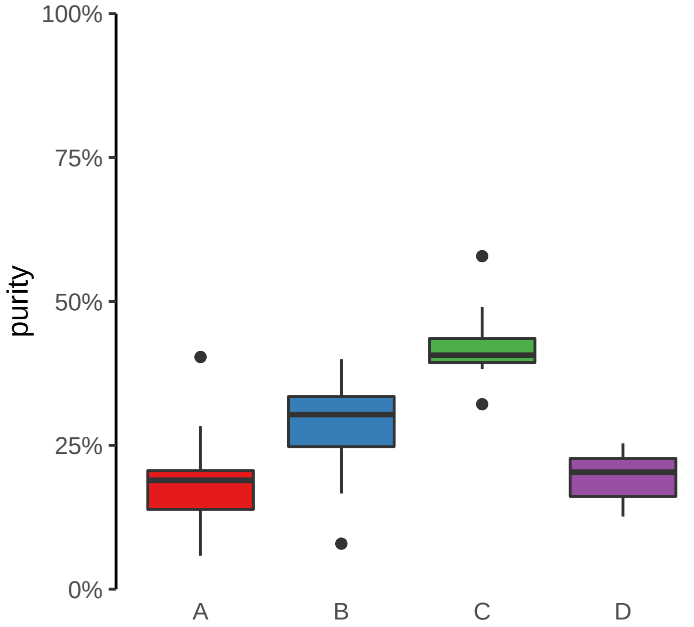

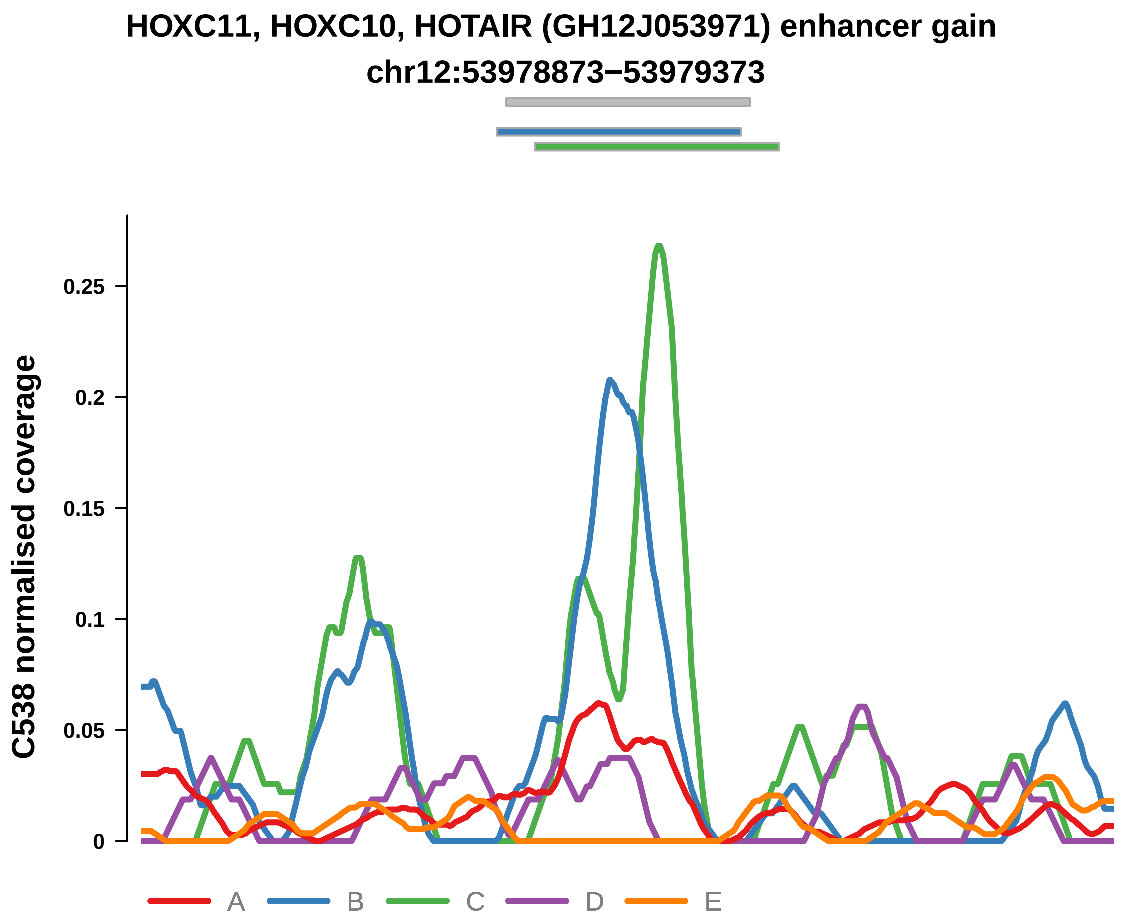


Putatively Clonal (logFC = 2.29)

| | P. | Adjusted P. |
|---|---|---|
| *~region vs. ~* | 2.48e−02 * | 8.95e−02 |
| *~purity vs ~* | 5.95e−01 | 7.28e−01 |
| *~purity+region vs ~purity* | 7.72e−03 ** | 3.4e−01 |
| *Carcinoma vs. normal* | 1.65e−03 ** | 1.65e−03 ** |

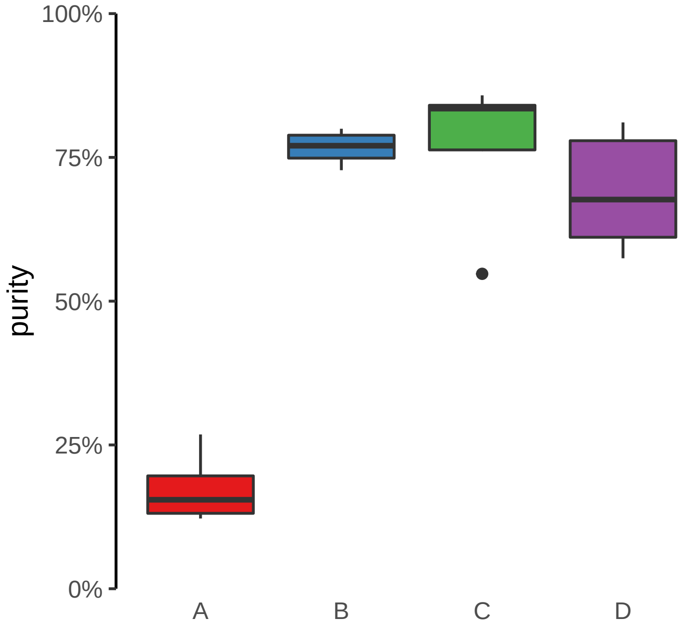

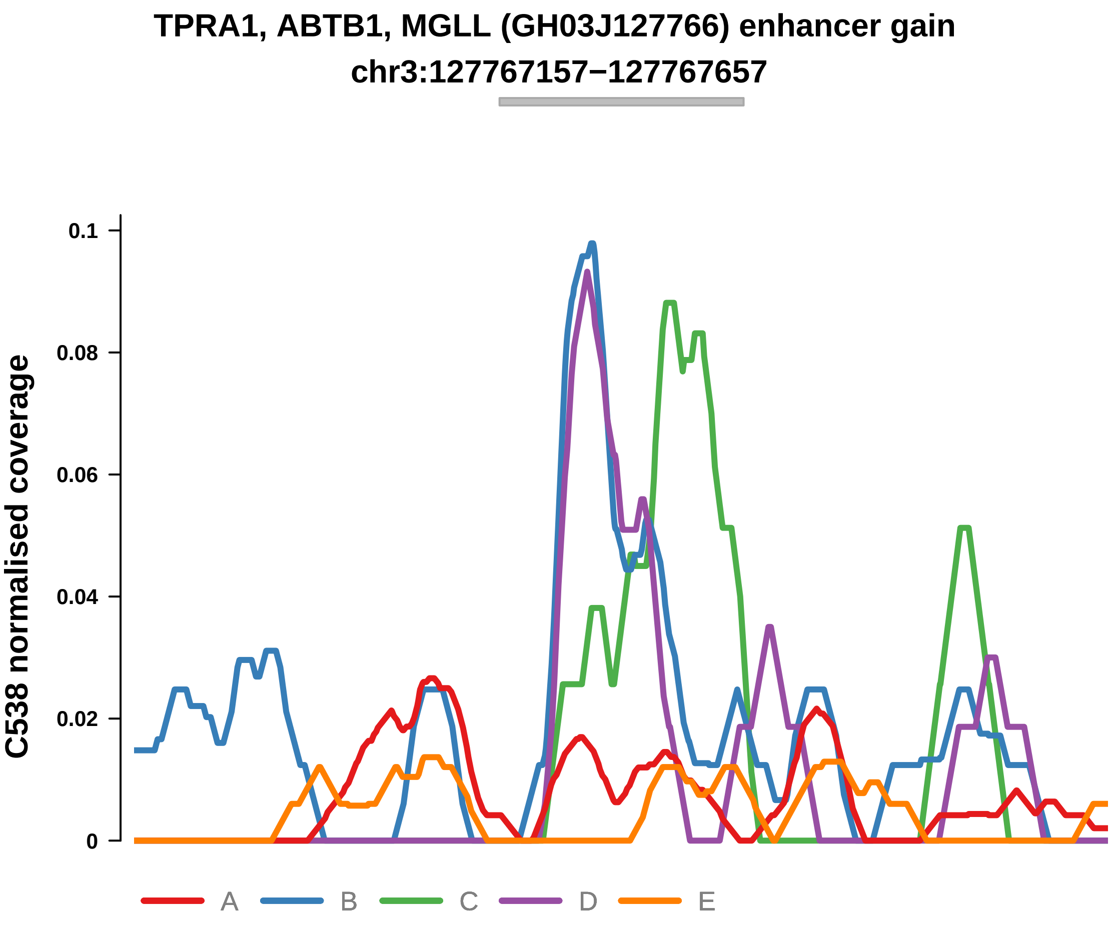


Putatively Clonal (logFC = 2.49)

| | P. | Adjusted P. |
|---|---|---|
| *~region vs. ~* | 4.03e−01 | 5.91e−01 |
| *~purity vs ~* | 1.55e−01 | 2.79e−01 |
| *~purity+region vs ~purity* | 7.02e−01 | 9.36e−01 |
| *Carcinoma vs. normal* | 1.19e−03 ** | 1.19e−03 ** |

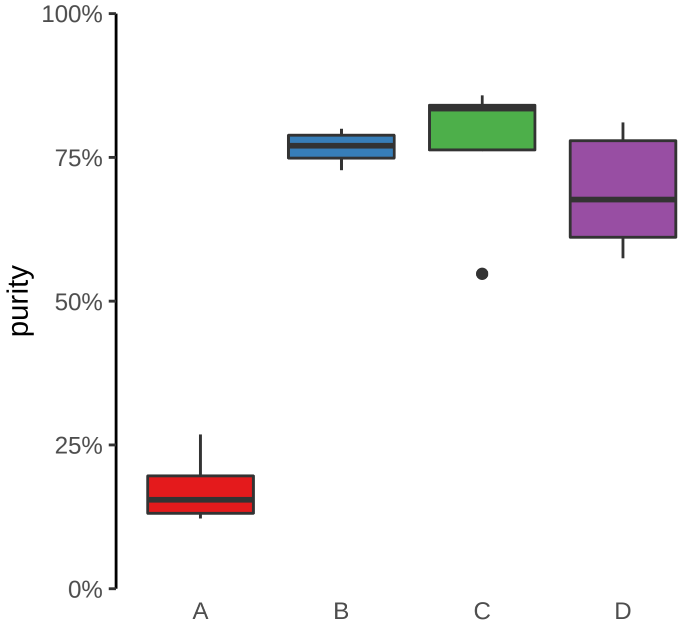

**CLASRP, MARK4 (GH19J045307) enhancer gain**
**chr19:45308035−45308535**

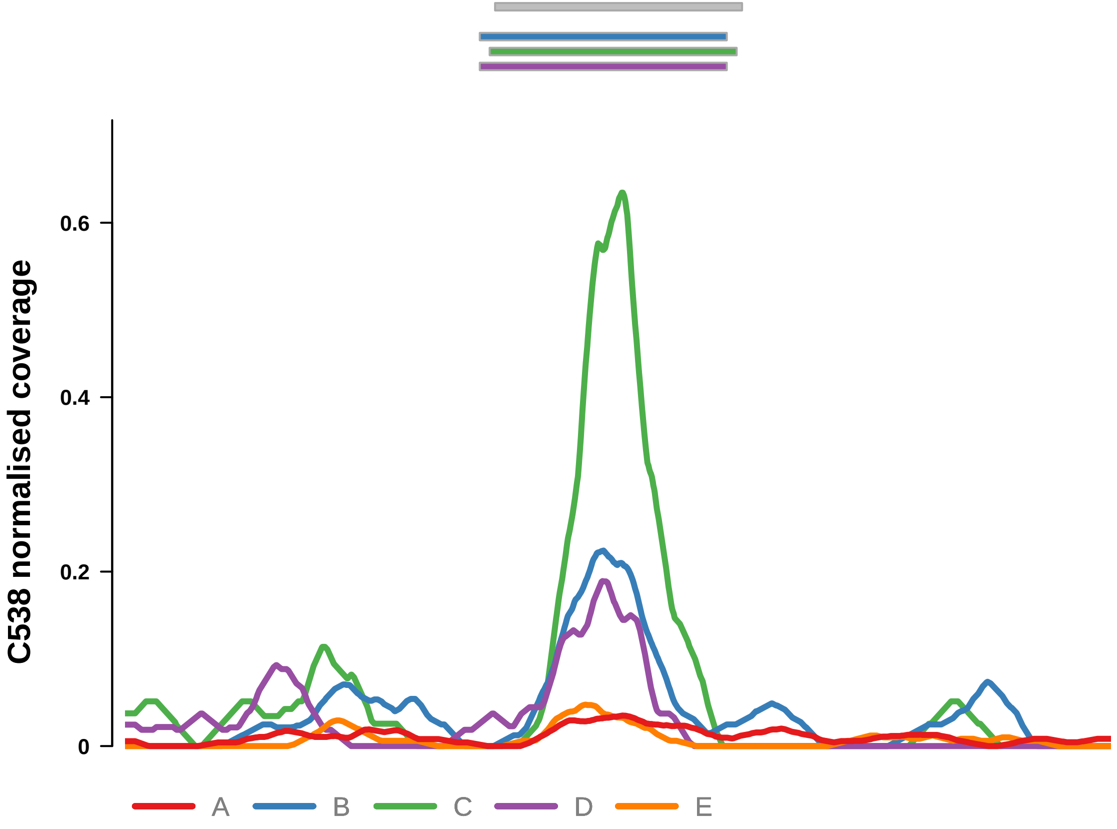


Putatively Clonal (logFC = 2.85)

| | P. | Adjusted P. |
|---|---|---|
| *~region vs. ~* | 2.88e−02 * | 9.07e−02 |
| *~purity vs ~* | 2.84e−03 ** | 1.19e−02 * |
| *~purity+region vs ~purity* | 9.24e−01 | 9.79e−01 |
| *Carcinoma vs. normal* | 7.85e−05 **** | 7.85e−05 **** |

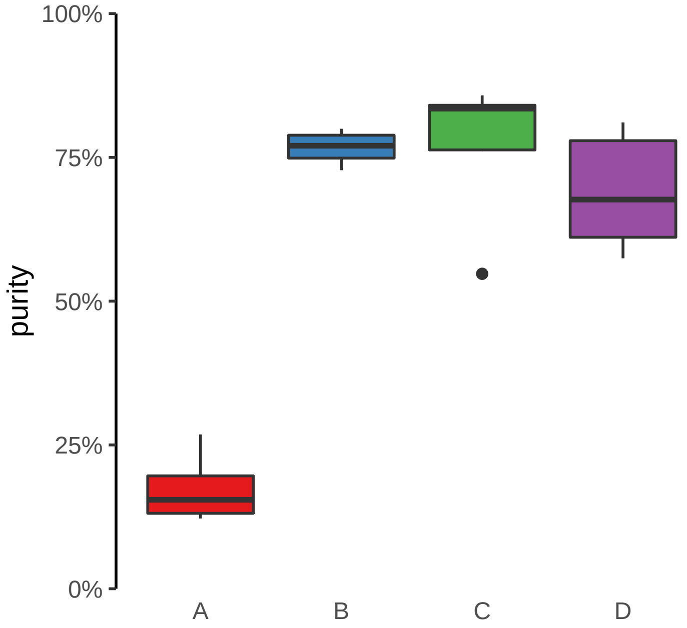

**ENSG00000253931 (GH08J143405) enhancer gain**
**chr8:143407862−143408362**

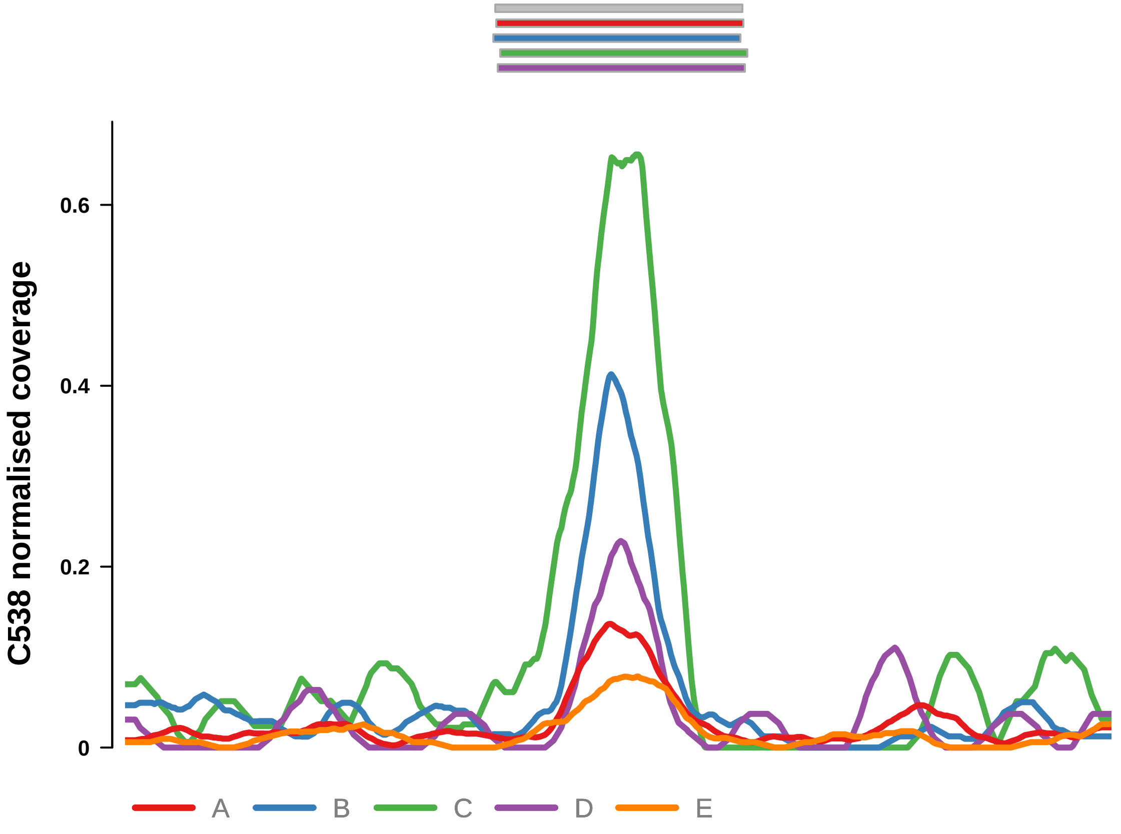


Putatively Clonal (logFC = 2.22)

| | P. | Adjusted P. |
|---|---|---|
| *~region vs. ~* | 1.38e−01 | 2.82e−01 |
| *~purity vs ~* | 4.59e−02 * | 1.44e−01 |
| *~purity+region vs ~purity* | 4.35e−01 | 8.7e−01 |
| *Carcinoma vs. normal* | 1.49e−03 ** | 1.49e−03 ** |

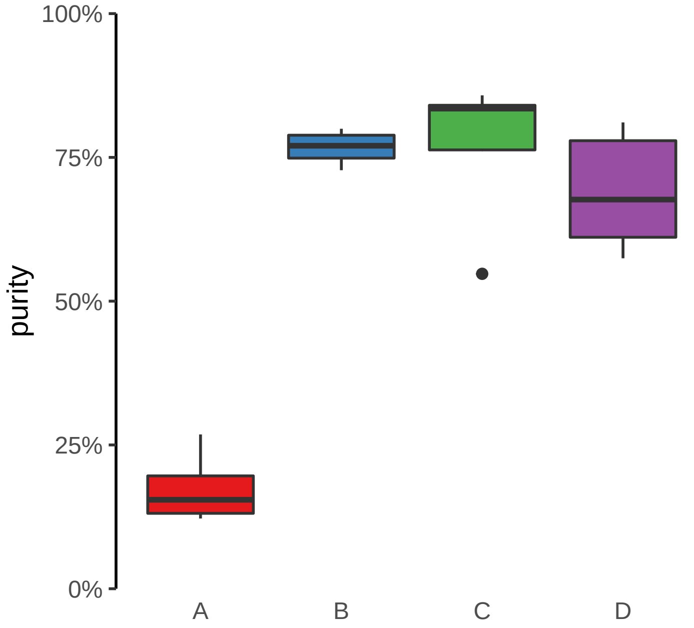

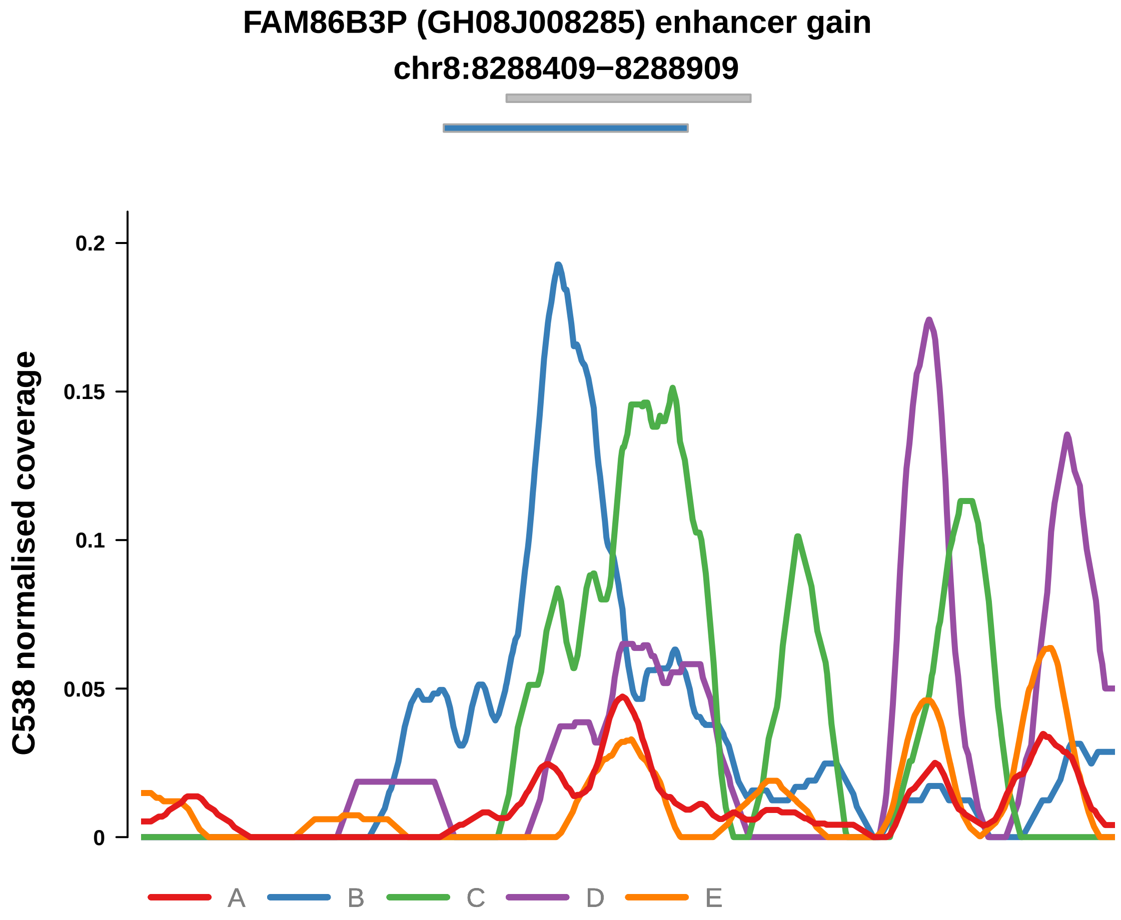


Putatively Clonal (logFC = 3)

| | P. | Adjusted P. |
|---|---|---|
| *~region vs. ~* | 6.24e−02 | 1.66e−01 |
| *~purity vs ~* | 1.47e−01 | 2.74e−01 |
| *~purity+region vs ~purity* | 1.6e−01 | 8.28e−01 |
| *Carcinoma vs. normal* | 5.11e−05 **** | 5.11e−05 **** |

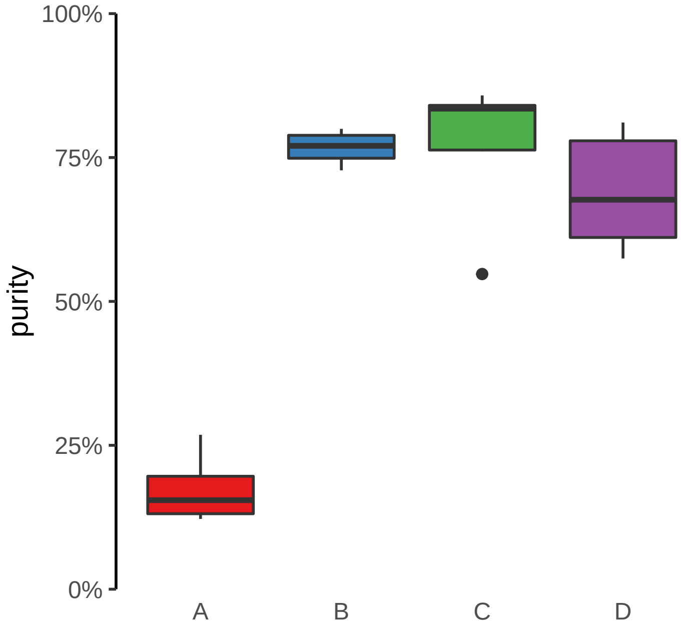

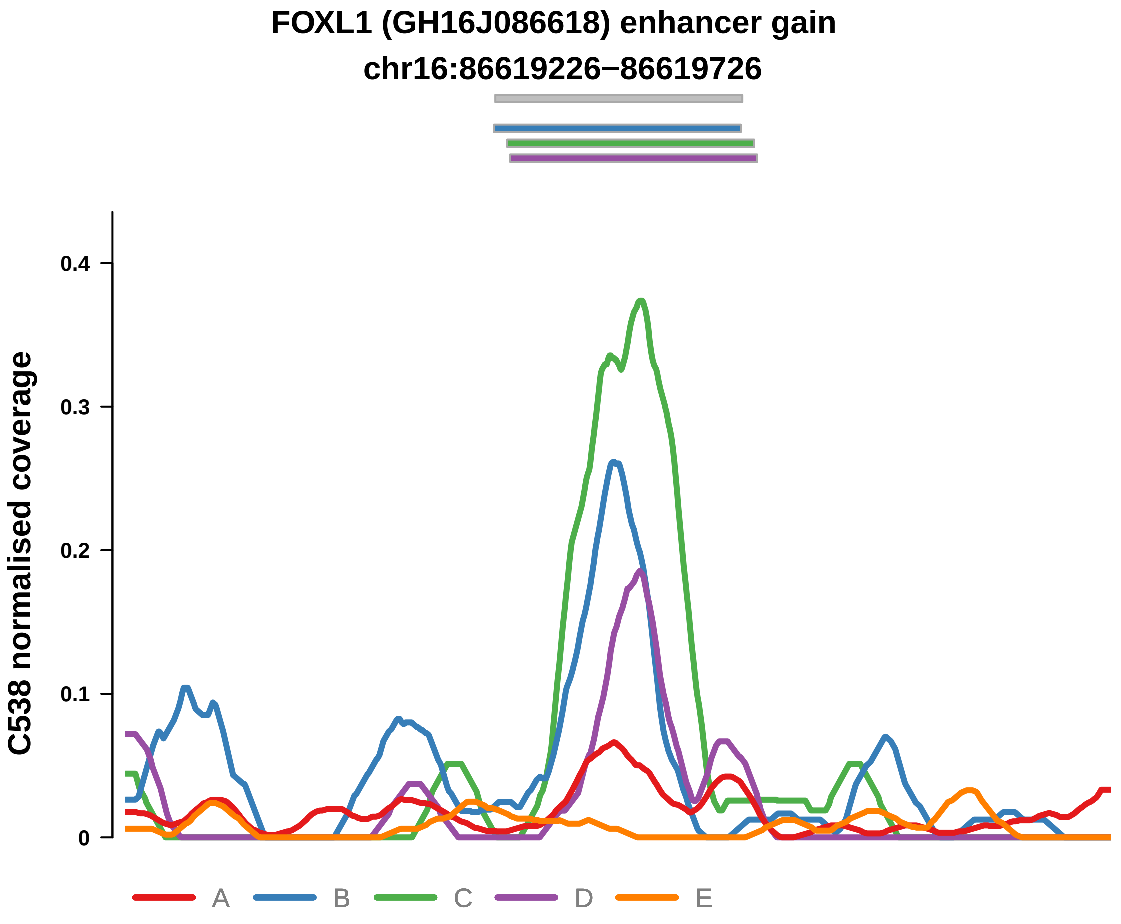


Putatively Clonal (logFC = 2.93)

| | P. | Adjusted P. |
|---|---|---|
| *~region vs. ~* | 9.87e−01 | 9.87e−01 |
| *~purity vs ~* | 9.14e−01 | 9.35e−01 |
| *~purity+region vs ~purity* | 7.43e−01 | 9.36e−01 |
| *Carcinoma vs. normal* | 5.3e−05 **** | 5.3e−05 **** |

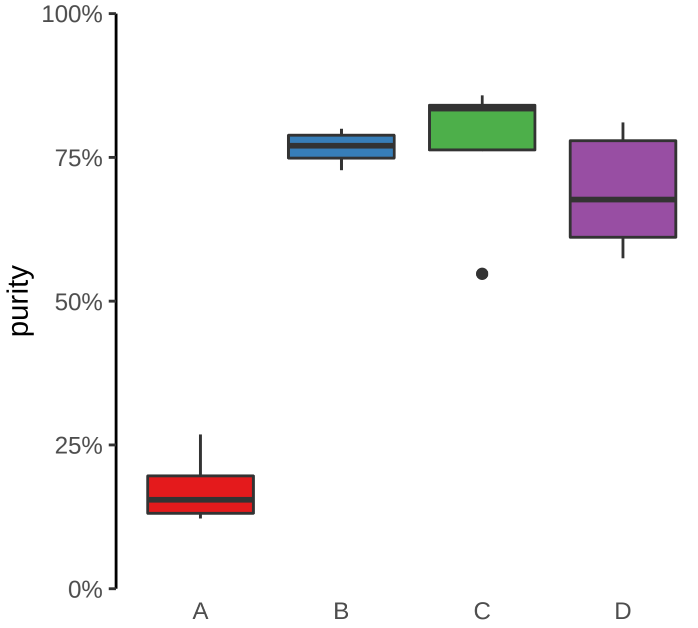

**LIFR (GH05J038659) enhancer gain**
**chr5:38660203−38660703**

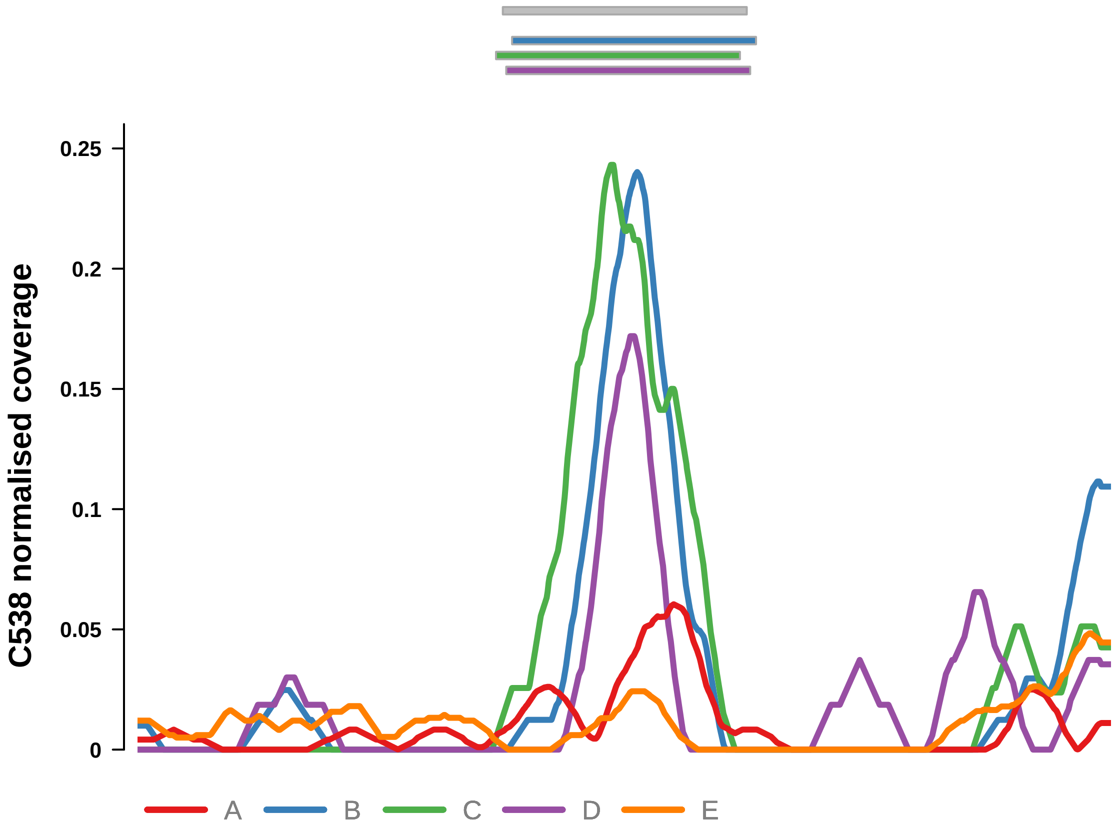


Putatively Clonal (logFC = 2.5)

| | P. | Adjusted P. |
|---|---|---|
| *~region vs. ~* | 6.89e−01 | 7.98e−01 |
| *~purity vs ~* | 6.99e−01 | 8.09e−01 |
| *~purity+region vs ~purity* | 4.94e−01 | 8.7e−01 |
| *Carcinoma vs. normal* | 4.92e−04 *** | 4.92e−04 *** |

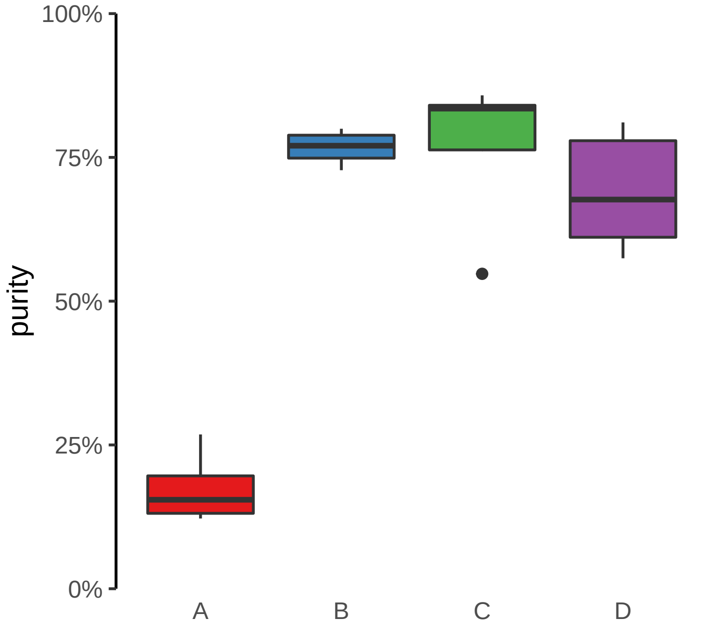

**MYBPC3 (GH11J047336) enhancer gain**
**chr11:47337207−47337707**

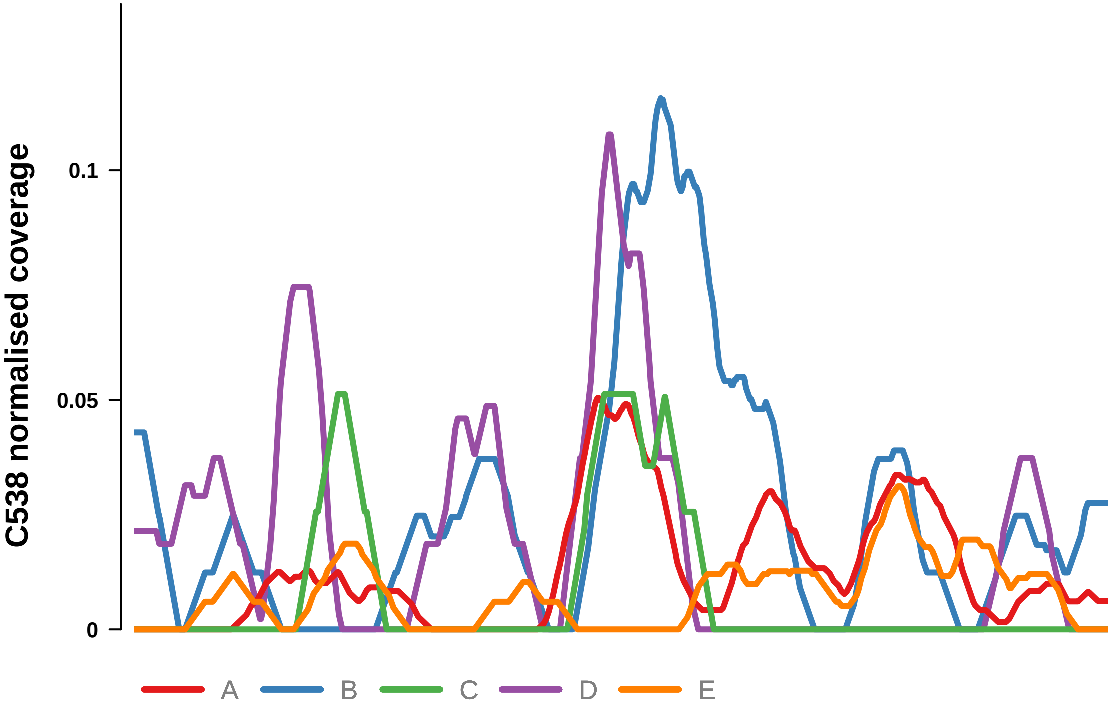


Putatively Clonal (logFC = 2.24)

| | P. | Adjusted P. |
|---|---|---|
| *~region vs. ~* | 2.51e−02 * | 8.95e−02 |
| *~purity vs ~* | 8.99e−01 | 9.31e−01 |
| *~purity+region vs ~purity* | 2.74e−02 * | 4.83e−01 |
| *Carcinoma vs. normal* | 2.94e−03 ** | 2.94e−03 ** |

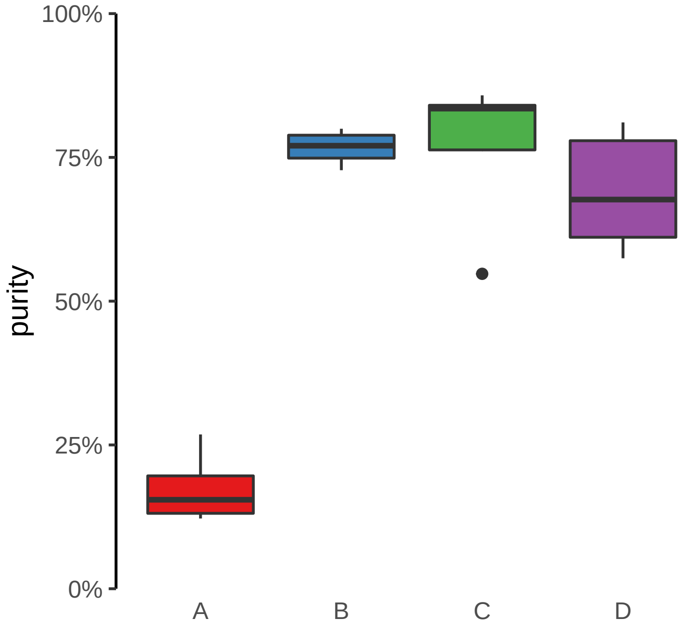

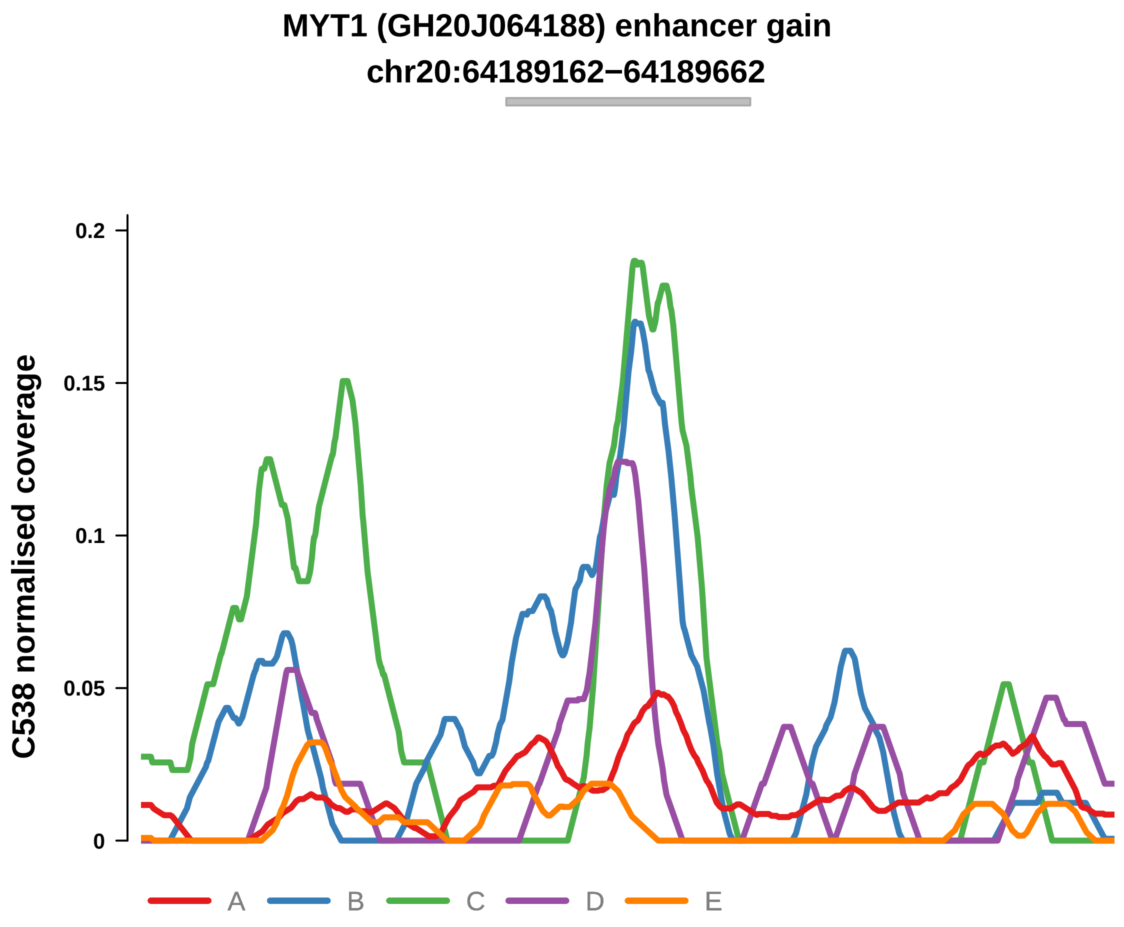


Putatively Clonal (logFC = 3.19)

| | P. | Adjusted P. |
|---|---|---|
| *~region vs. ~* | 7.07e−01 | 8e−01 |
| *~purity vs ~* | 8.51e−01 | 9.13e−01 |
| *~purity+region vs ~purity* | 3e−01 | 8.7e−01 |
| *Carcinoma vs. normal* | 2.26e−05 **** | 2.26e−05 **** |

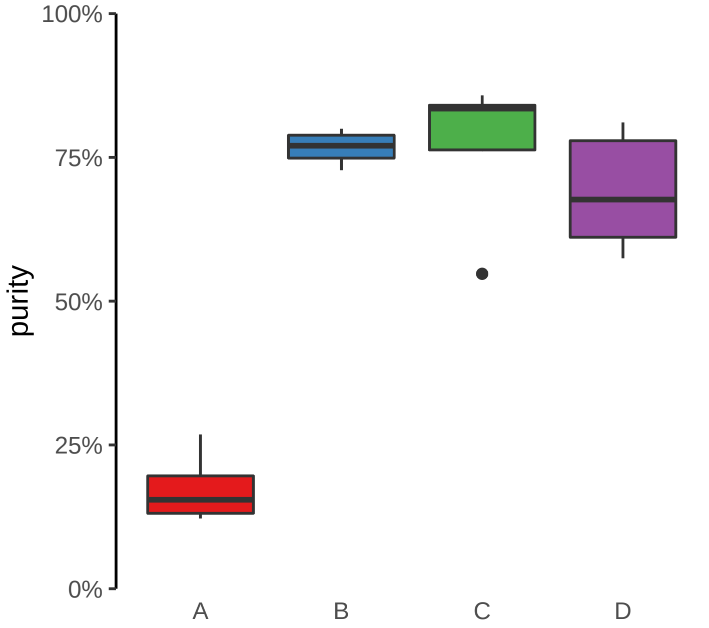

**NCOA7−AS1 (GH06J125867) enhancer gain**
**chr6:125867578−125868078**

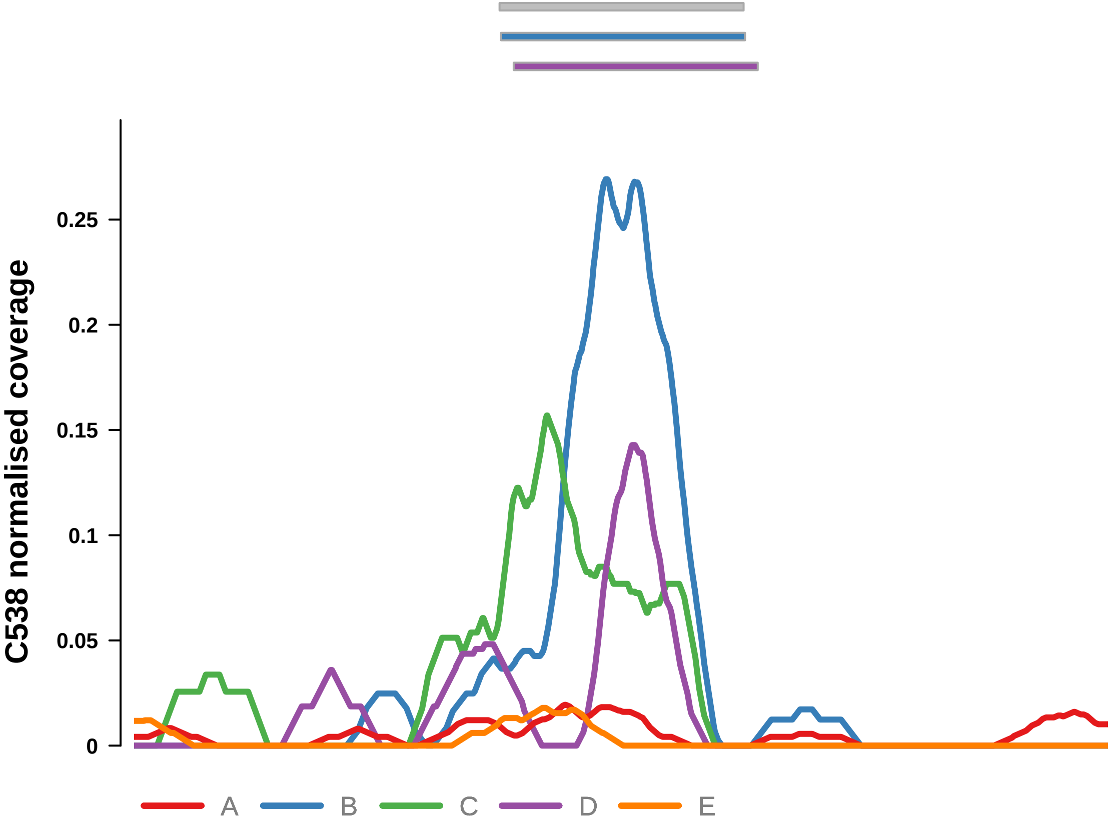


Putatively Clonal (logFC = 3.31)

| | P. | Adjusted P. |
|---|---|---|
| *~region vs. ~* | 2.46e−01 | 4.71e−01 |
| *~purity vs ~* | 2.24e−01 | 3.78e−01 |
| *~purity+region vs ~purity* | 5.07e−01 | 8.7e−01 |
| *Carcinoma vs. normal* | 9.45e−06 **** | 9.45e−06 **** |

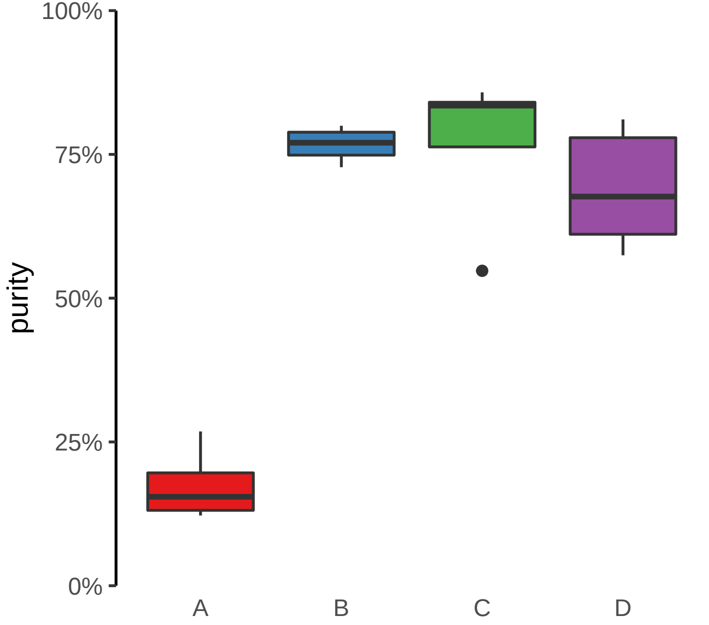

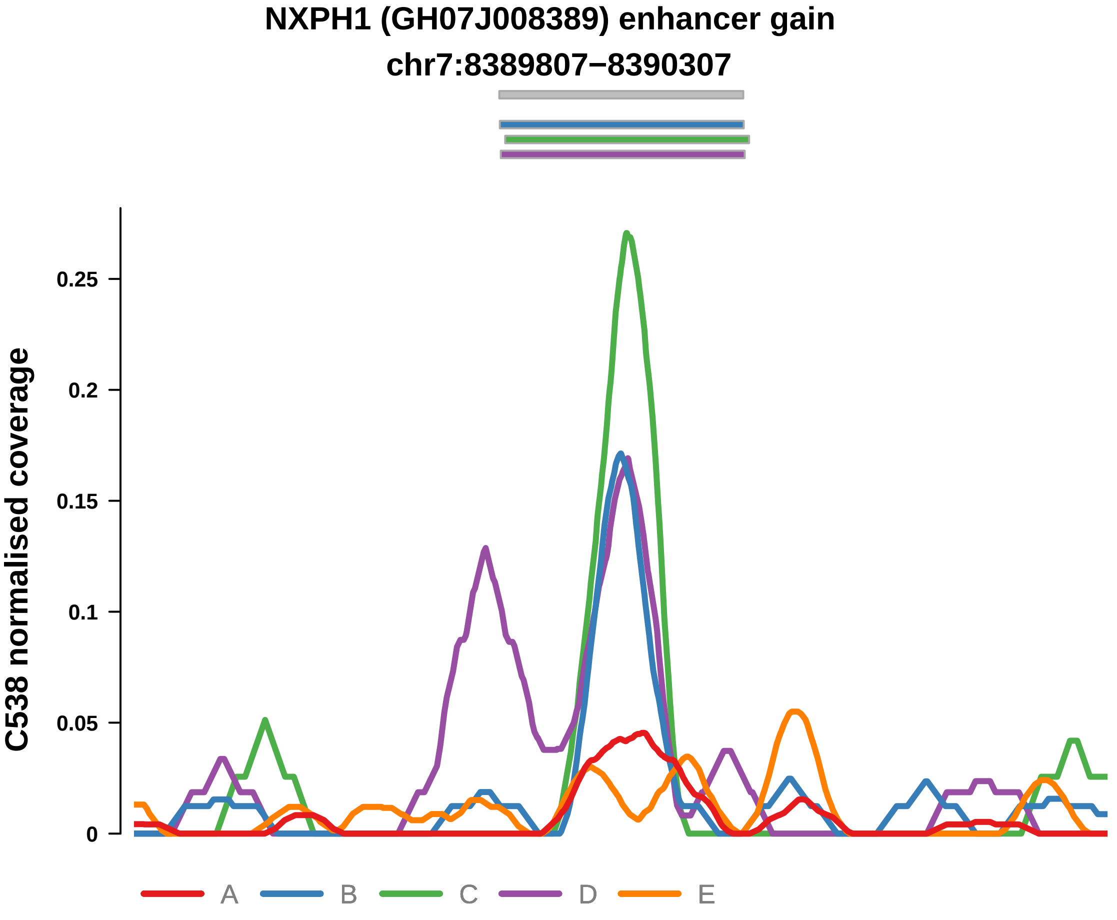


Putatively Clonal (logFC = 2.1)

| | P. | Adjusted P. |
|---|---|---|
| *~region vs. ~* | 3.53e−01 | 5.55e−01 |
| *~purity vs ~* | 3.54e−01 | 5.1e−01 |
| *~purity+region vs ~purity* | 4.93e−01 | 8.7e−01 |
| *Carcinoma vs. normal* | 3.51e−03 ** | 3.51e−03 ** |

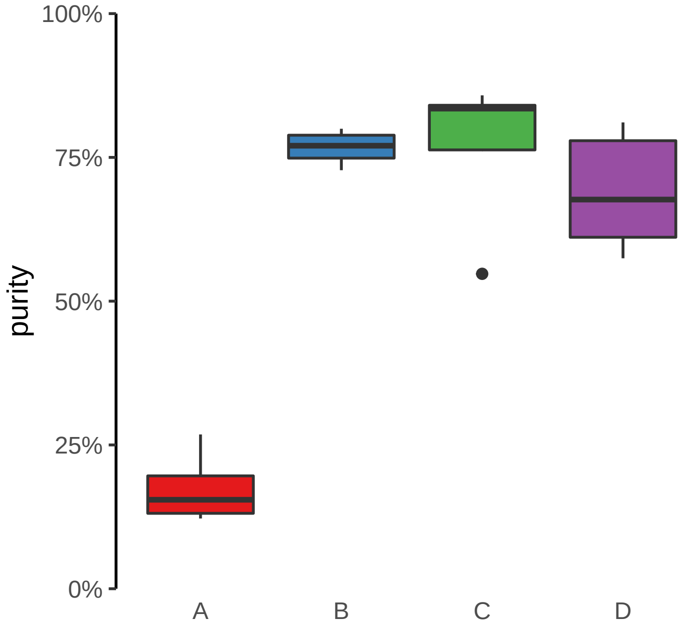

**PPL (GH16J004942) enhancer gain**
**chr16:4944634−4945134**

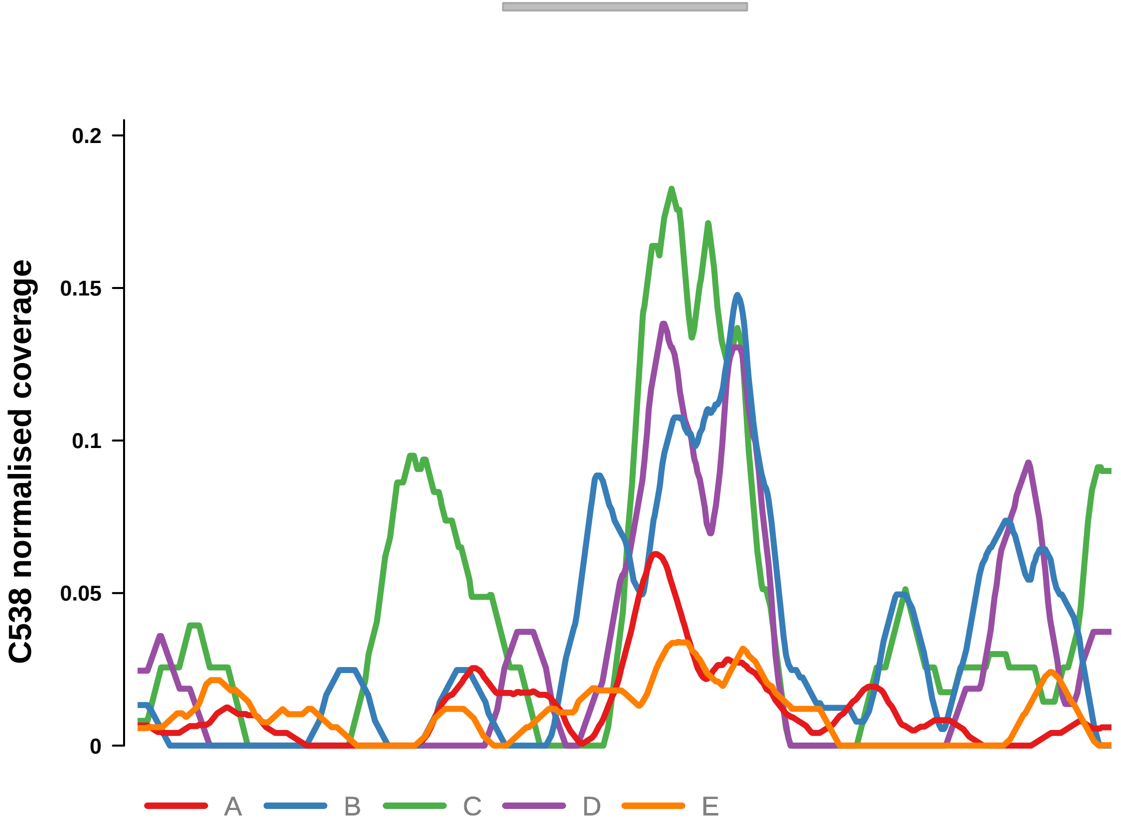


Putatively Clonal (logFC = 1.86)

| | P. | Adjusted P. |
|---|---|---|
| *~region vs. ~* | 5.45e−01 | 7.15e−01 |
| *~purity vs ~* | 5.01e−01 | 6.35e−01 |
| *~purity+region vs ~purity* | 6.42e−01 | 9.12e−01 |
| *Carcinoma vs. normal* | 9.24e−03 ** | 9.24e−03 ** |

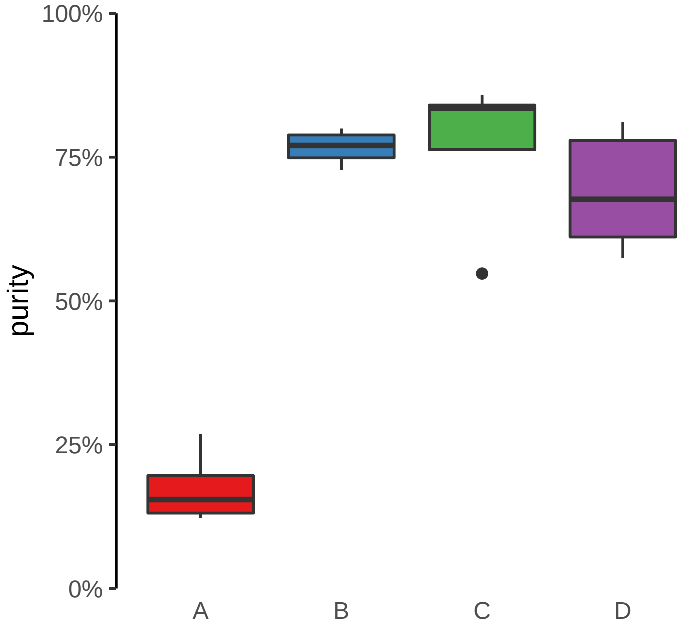

**RNU7−124P, SHB, ENSG00000255872 (GH09J037962) enhancer gain**
**chr9:37966483−37966983**

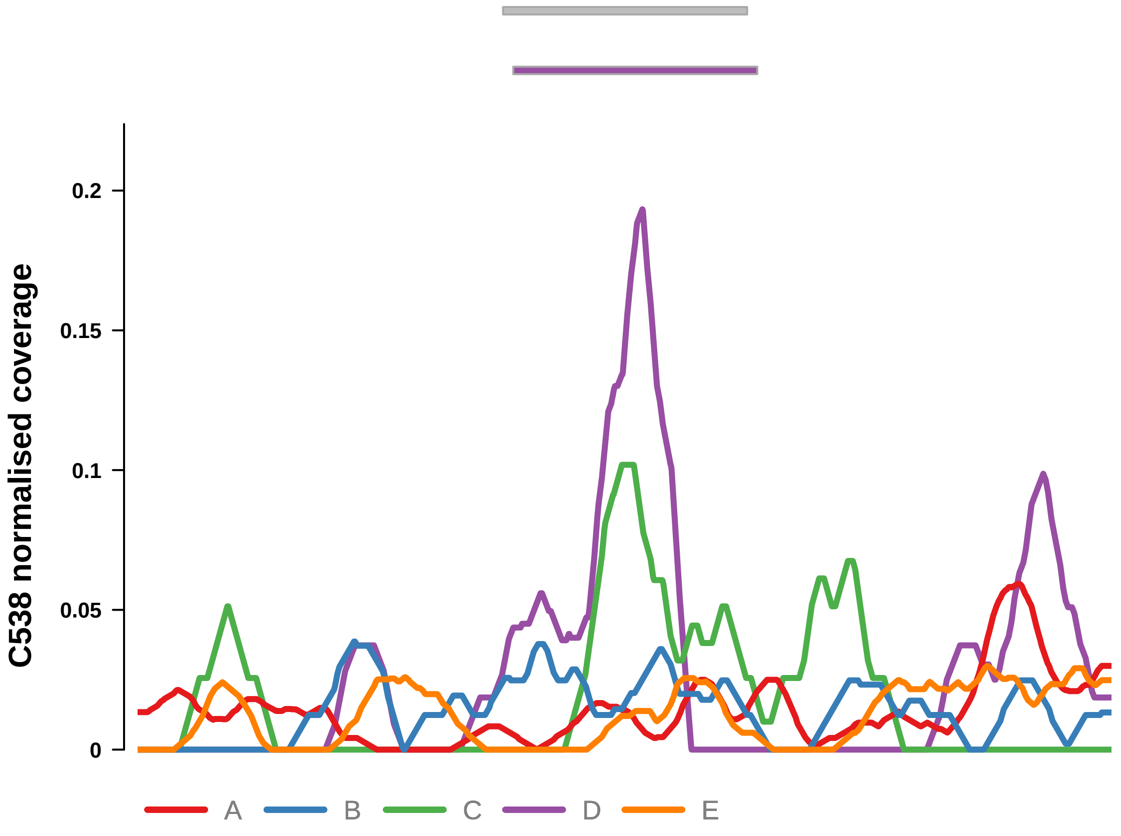


Putatively Clonal (logFC = 1.95)

| | P. | Adjusted P. |
|---|---|---|
| *~region vs. ~* | 7.02e−02 | 1.76e−01 |
| *~purity vs ~* | 4.56e−01 | 6.07e−01 |
| *~purity+region vs ~purity* | 9.79e−02 | 7.83e−01 |
| *Carcinoma vs. normal* | 8.58e−03 ** | 8.58e−03 ** |

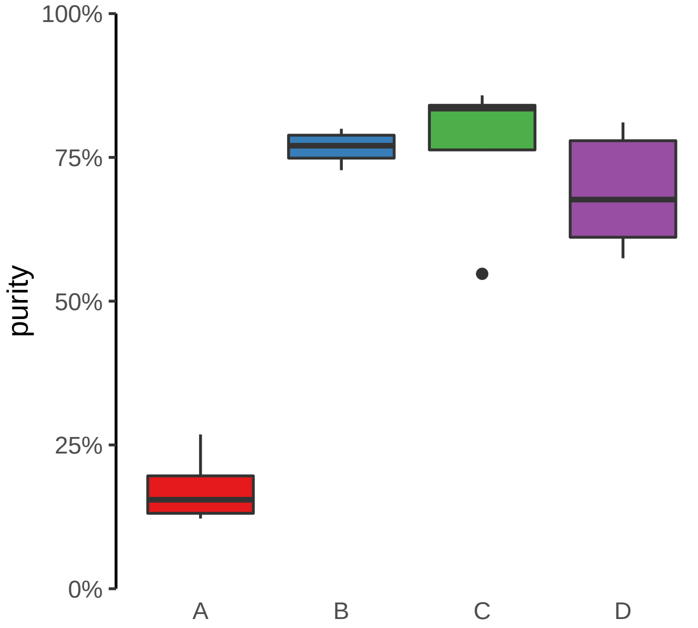

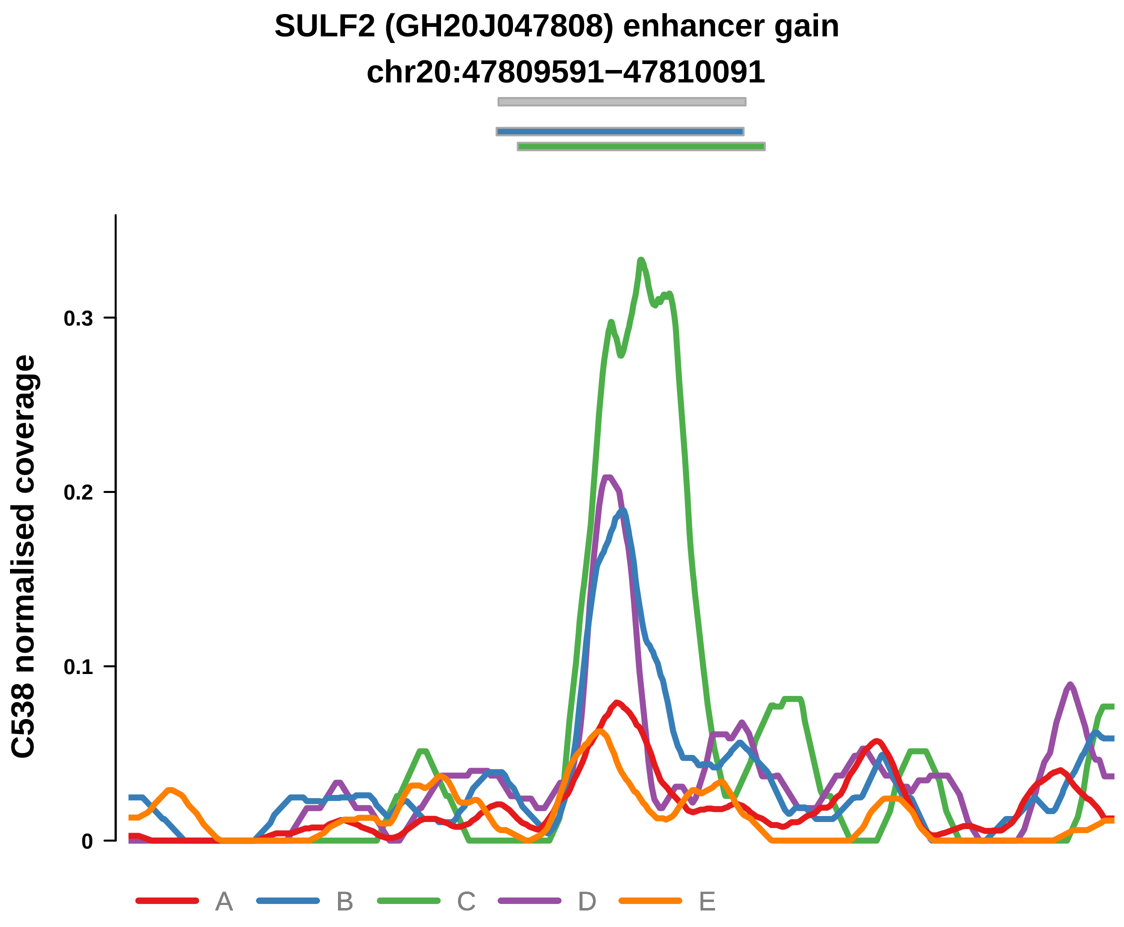


Putatively Clonal (logFC = 2.26)

| | P. | Adjusted P. |
|---|---|---|
| *~region vs. ~* | 9.21e−01 | 9.43e−01 |
| *~purity vs ~* | 2.97e−01 | 4.58e−01 |
| *~purity+region vs ~purity* | 5.16e−01 | 8.7e−01 |
| *Carcinoma vs. normal* | 1.41e−03 ** | 1.41e−03 ** |

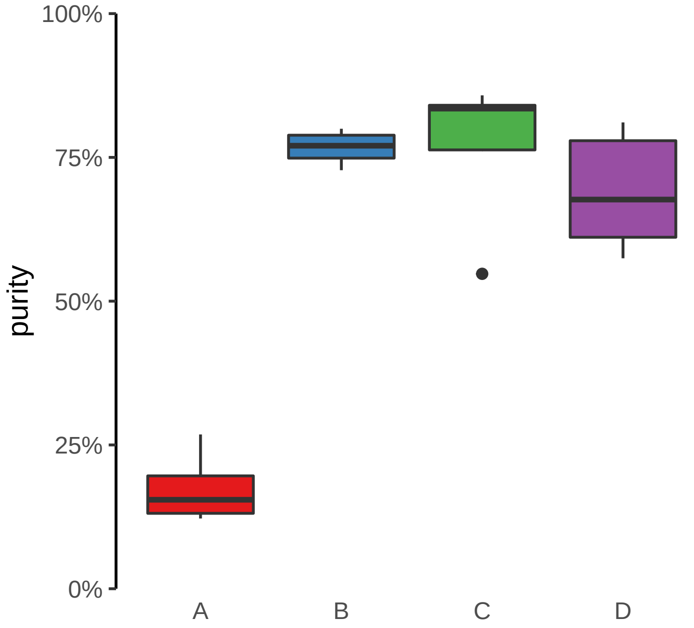

**TEX26 (GH13J030867) enhancer gain**
**chr13:30870352−30870852**

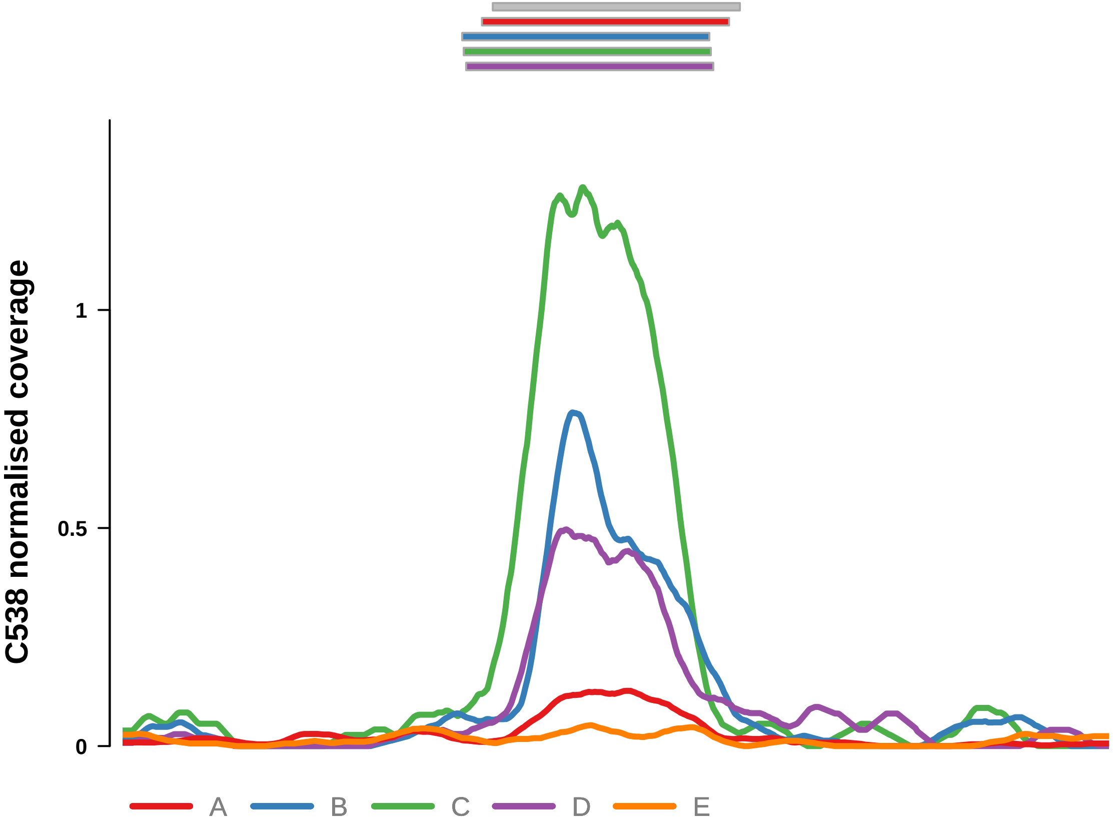


Putatively Clonal (logFC = 3.53)

| | P. | Adjusted P. |
|---|---|---|
| *~region vs. ~* | 3.45e−07 **** | 1.52e−05 **** |
| *~purity vs ~* | 8.71e−08 **** | 7.66e−06 **** |
| *~purity+region vs ~purity* | 4.46e−01 | 8.7e−01 |
| *Carcinoma vs. normal* | 1.19e−06 **** | 1.19e−06 **** |

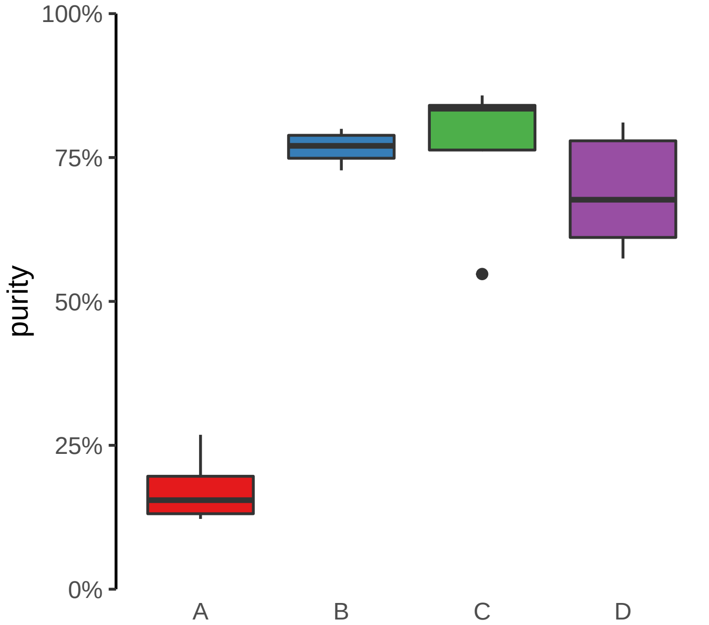

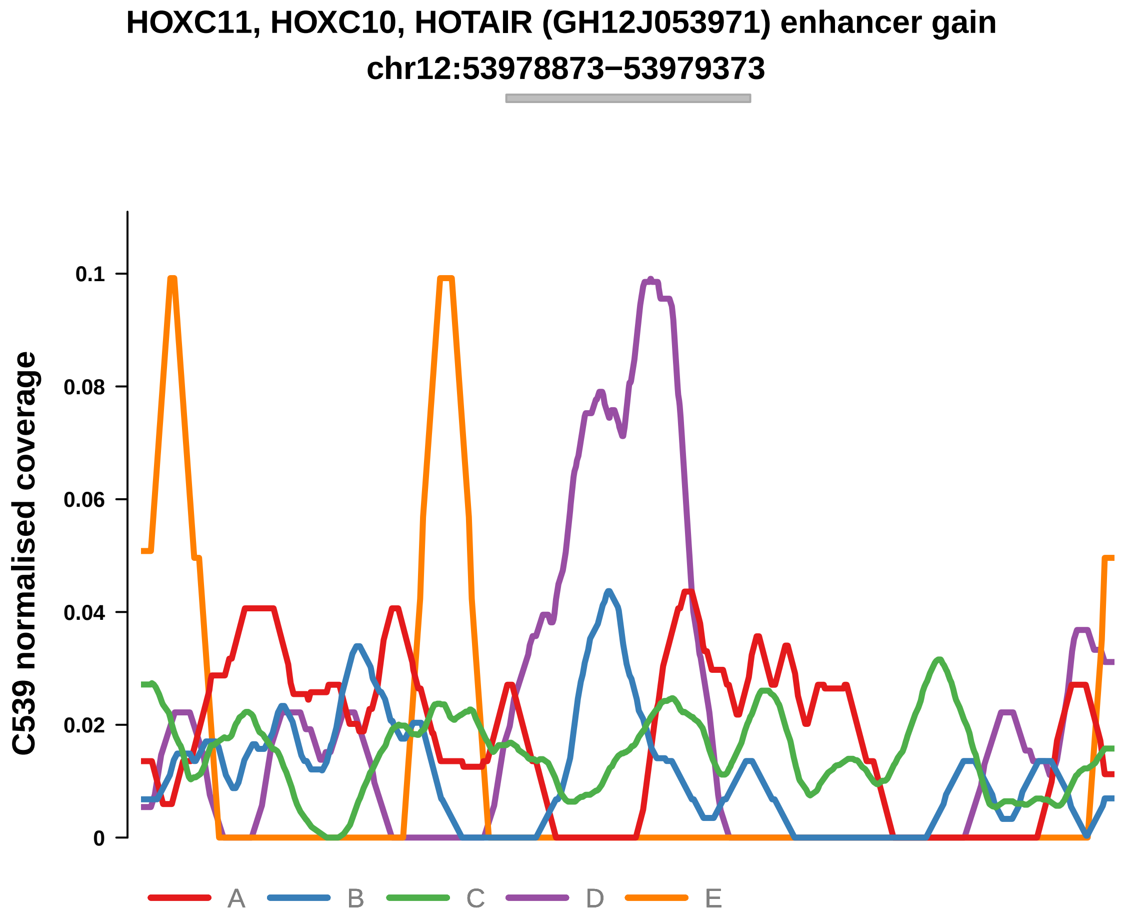


Putatively Clonal (logFC = 2.42)

| | P. | Adjusted P. |
|---|---|---|
| *~region vs. ~* | 1.34e−01 | 1.4e−01 |
| *~purity vs ~* | 1.98e−01 | 2.07e−01 |
| *~purity+region vs ~purity* | 1.68e−01 | 2.78e−01 |
| *Carcinoma vs. normal* | 3.91e−03 ** | 3.91e−03 ** |

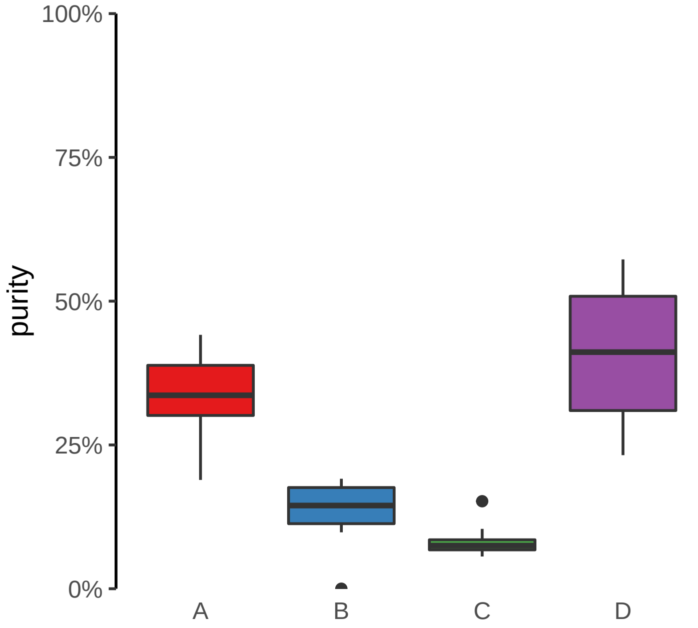

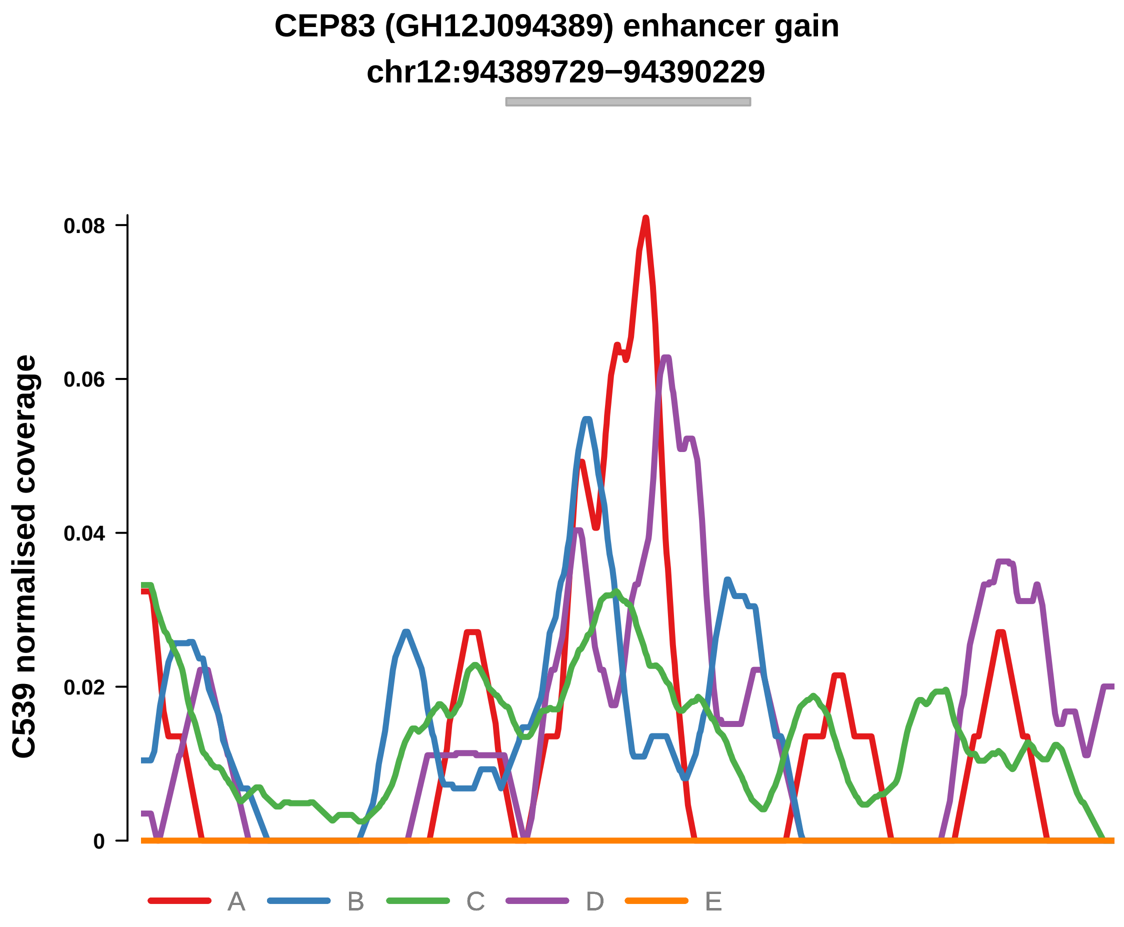


Putatively Clonal (logFC = 2.96)

| | P. | Adjusted P. |
|---|---|---|
| *~region vs. ~* | 2.43e−03 ** | 5.47e−03 ** |
| *~purity vs ~* | 4.56e−03 ** | 9.55e−03 ** |
| *~purity+region vs ~purity* | 1.39e−01 | 2.61e−01 |
| *Carcinoma vs. normal* | 1.52e−03 ** | 1.52e−03 ** |

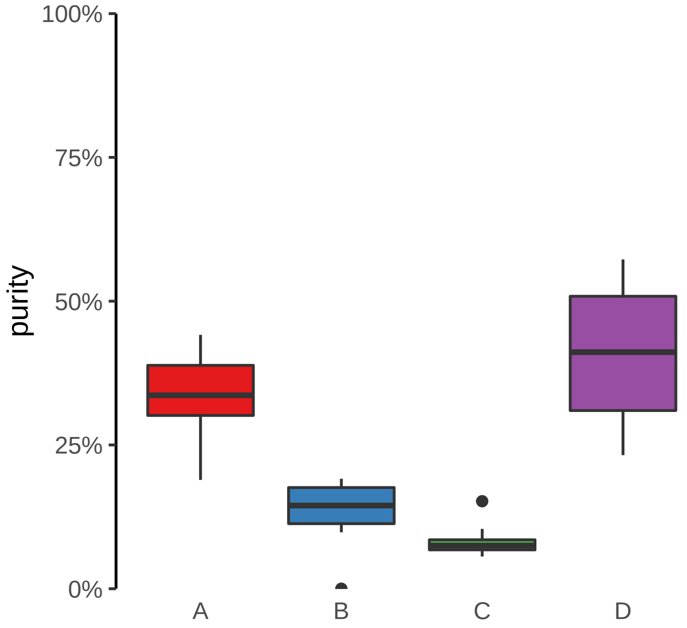

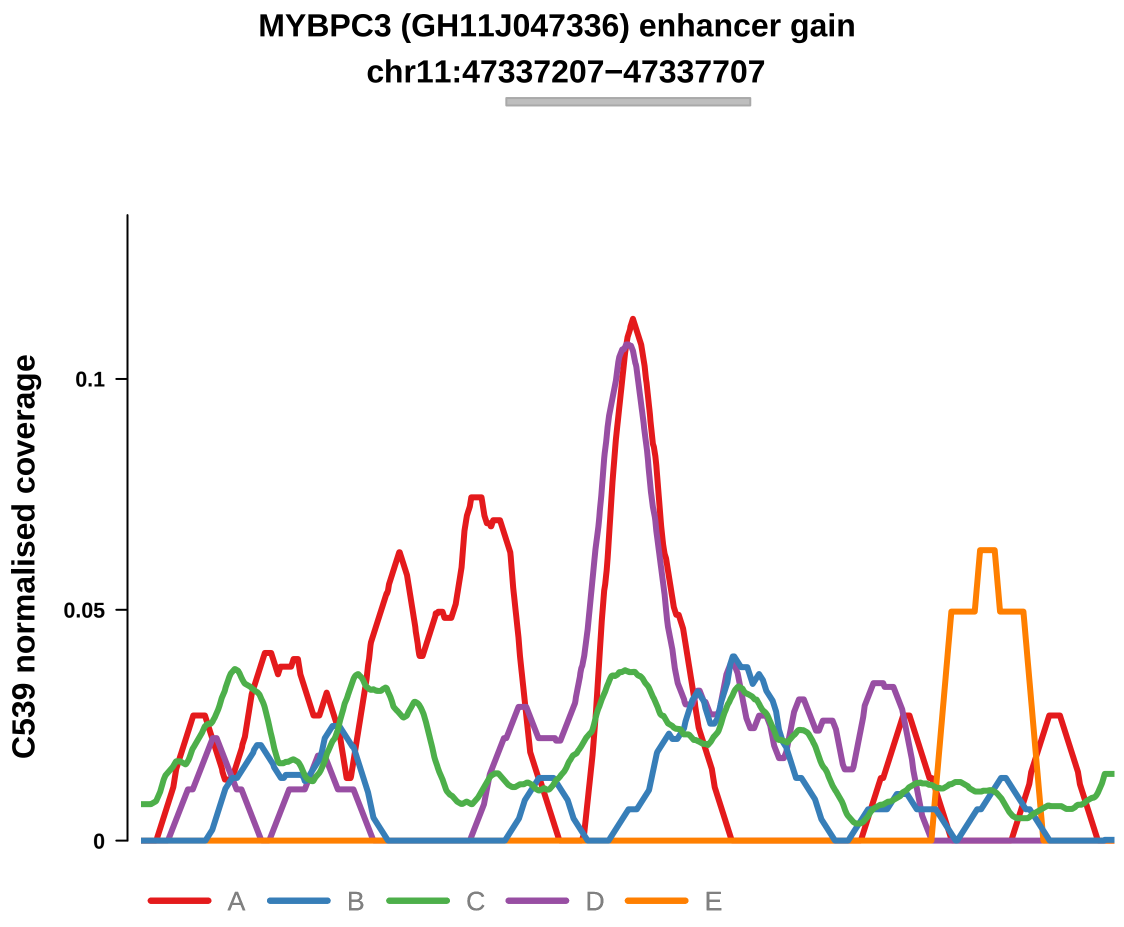


Putatively Clonal (logFC = 3.07)

| | P. | Adjusted P. |
|---|---|---|
| *~region vs. ~* | 3.9e−02 * | 4.57e−02 * |
| *~purity vs ~* | 2.29e−02 * | 3.56e−02 * |
| *~purity+region vs ~purity* | 4.06e−01 | 4.52e−01 |
| *Carcinoma vs. normal* | 2.75e−04 *** | 2.75e−04 *** |

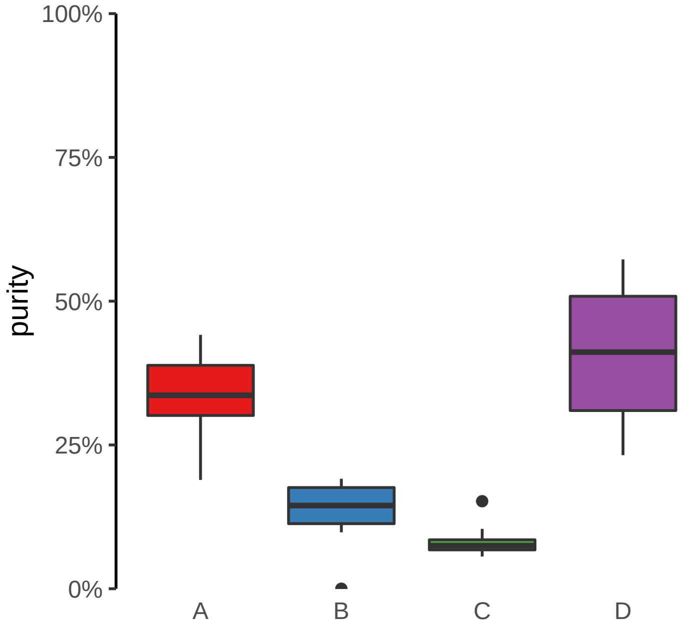

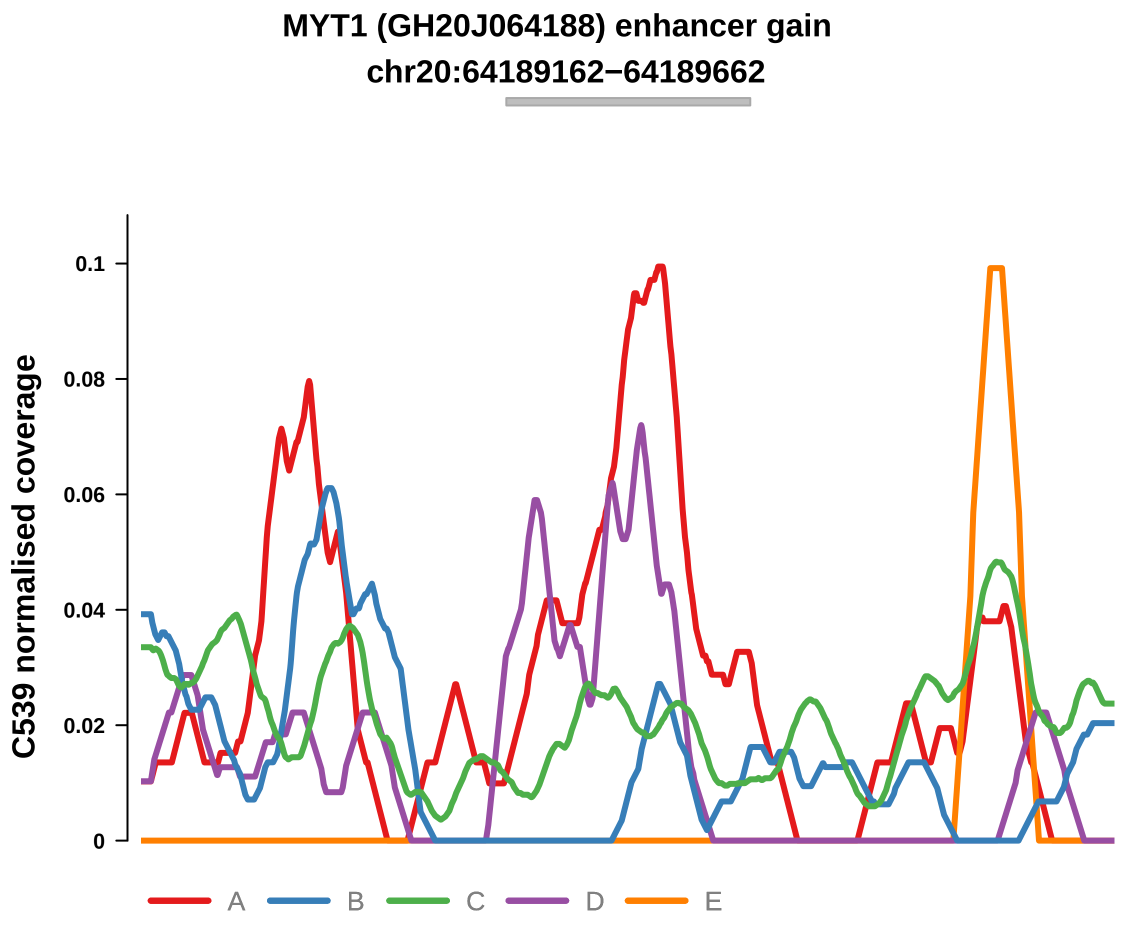


Putatively Clonal (logFC = 2.83)

| | P. | Adjusted P. |
|---|---|---|
| *~region vs. ~* | 6.74e−03 ** | 1.04e−02 * |
| *~purity vs ~* | 7.03e−02 | 8.36e−02 |
| *~purity+region vs ~purity* | 3.29e−02 * | 9.99e−02 |
| *Carcinoma vs. normal* | 1.85e−03 ** | 1.85e−03 ** |

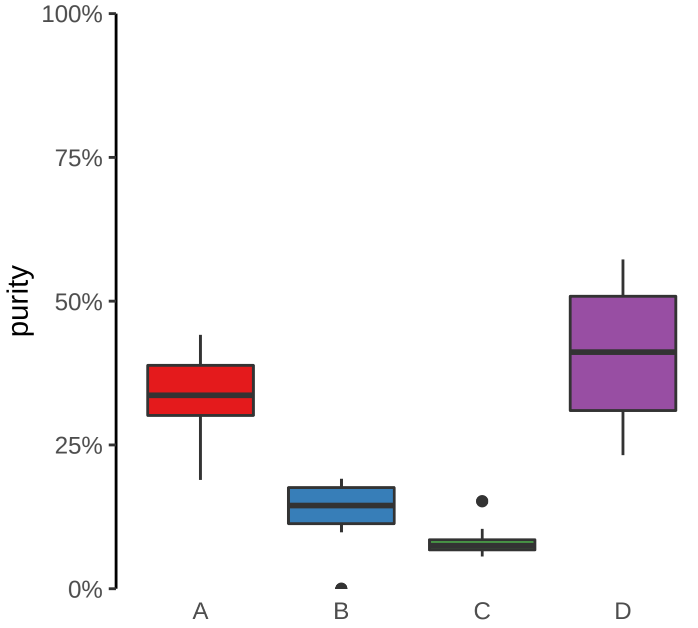

**NCOA7−AS1 (GH06J125867) enhancer gain**
**chr6:125867578−125868078**

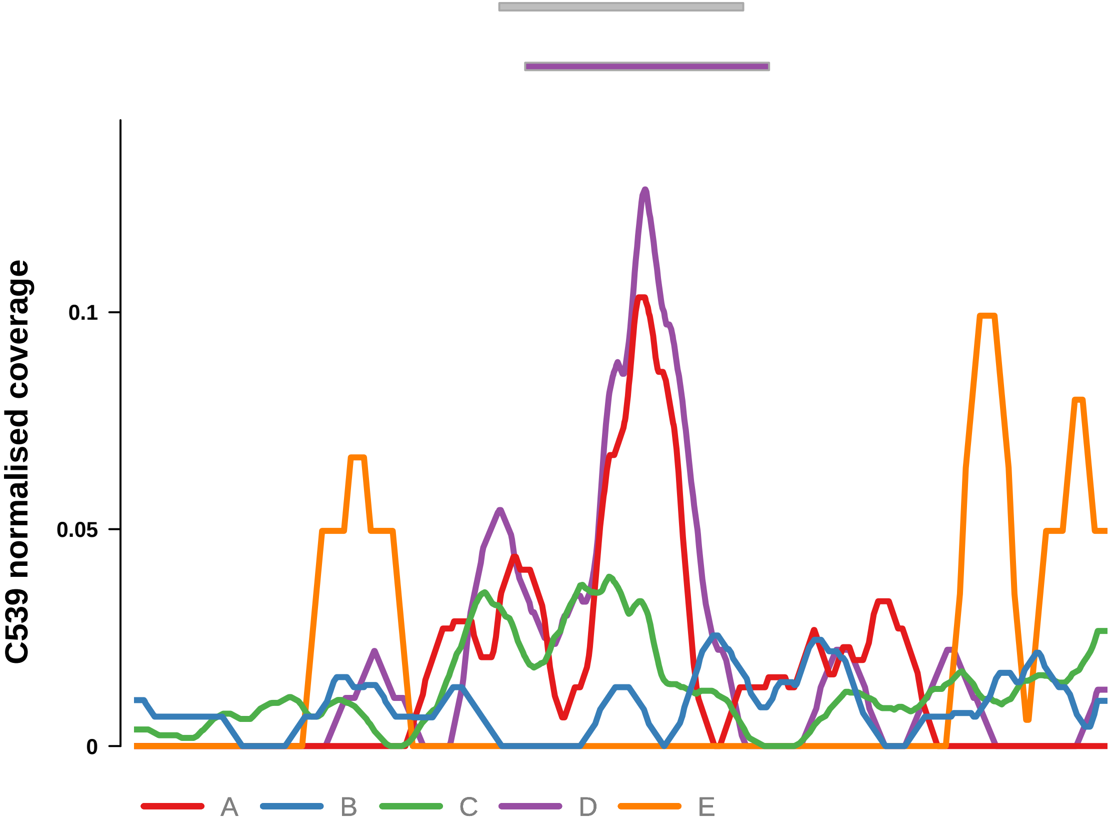


Putatively Clonal (logFC = 2.55)

| | P. | Adjusted P. |
|---|---|---|
| *~region vs. ~* | 2.57e−03 ** | 5.66e−03 ** |
| *~purity vs ~* | 6.96e−02 | 8.36e−02 |
| *~purity+region vs ~purity* | 1.56e−02 * | 5.6e−02 |
| *Carcinoma vs. normal* | 4.47e−03 ** | 4.47e−03 ** |

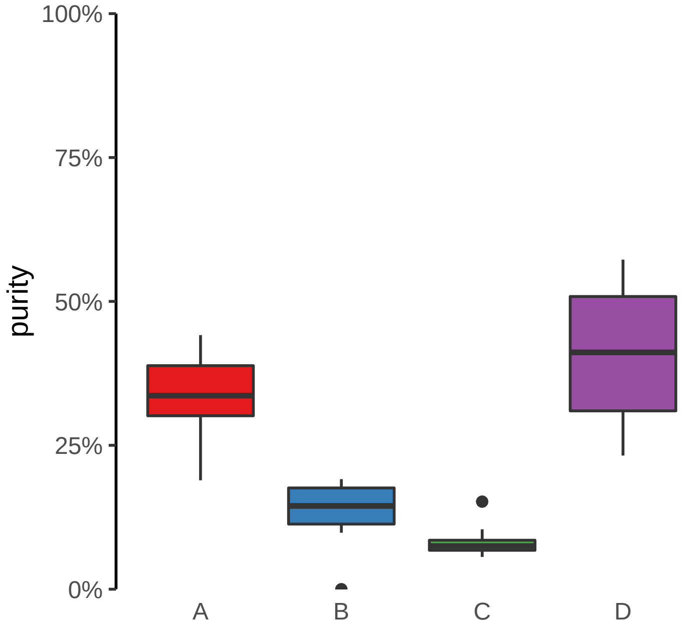

**NFATC2 (GH20J051486) enhancer gain**
**chr20:51487818−51488318**

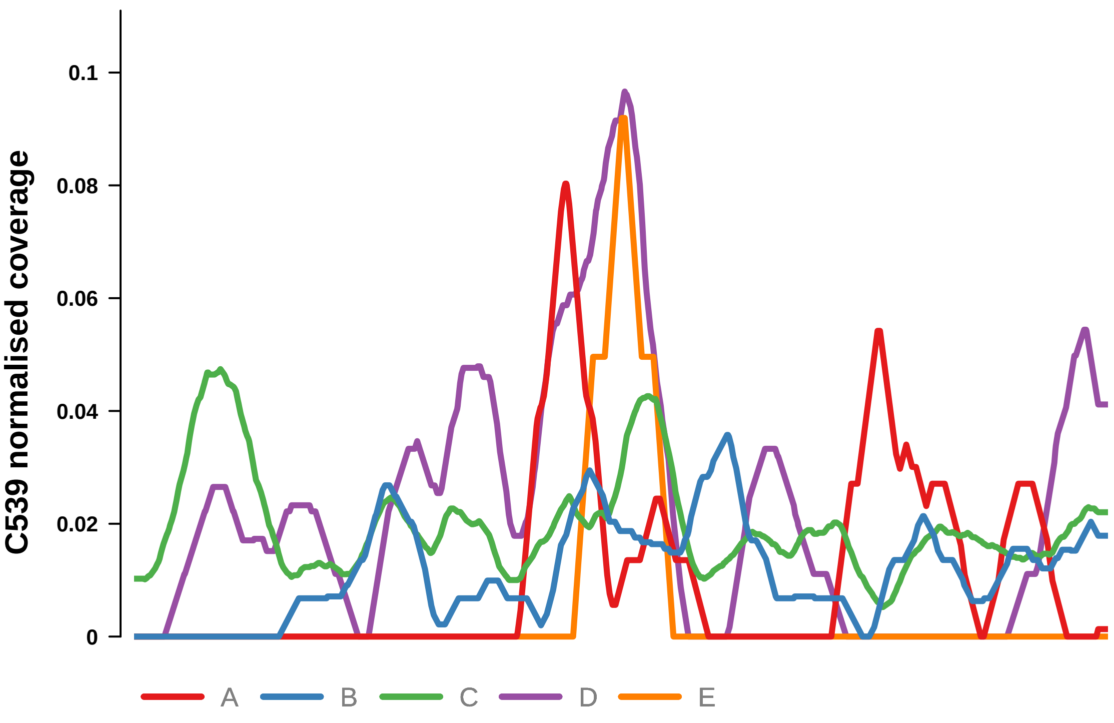


Putatively Clonal (logFC = 2.77)

| | P. | Adjusted P. |
|---|---|---|
| *~region vs. ~* | 3.39e−03 ** | 6.71e−03 ** |
| *~purity vs ~* | 1.18e−02 * | 2e−02 * |
| *~purity+region vs ~purity* | 9.43e−02 | 2.01e−01 |
| *Carcinoma vs. normal* | 1.85e−03 ** | 1.85e−03 ** |

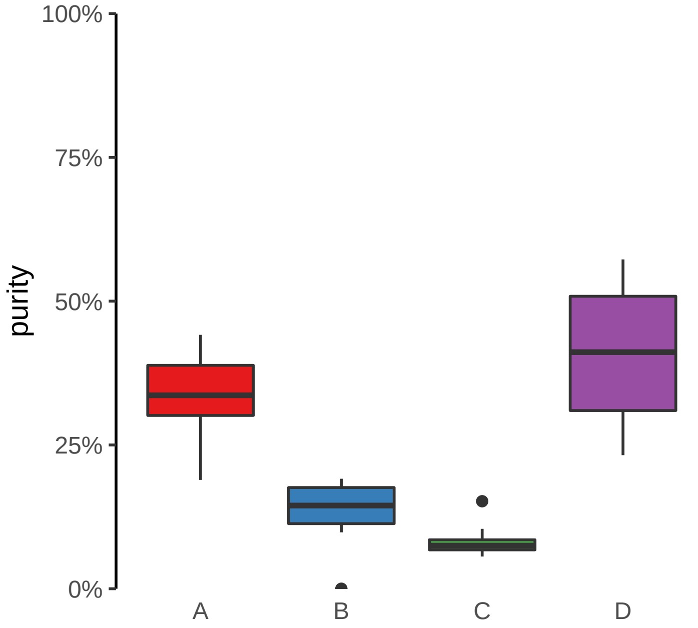

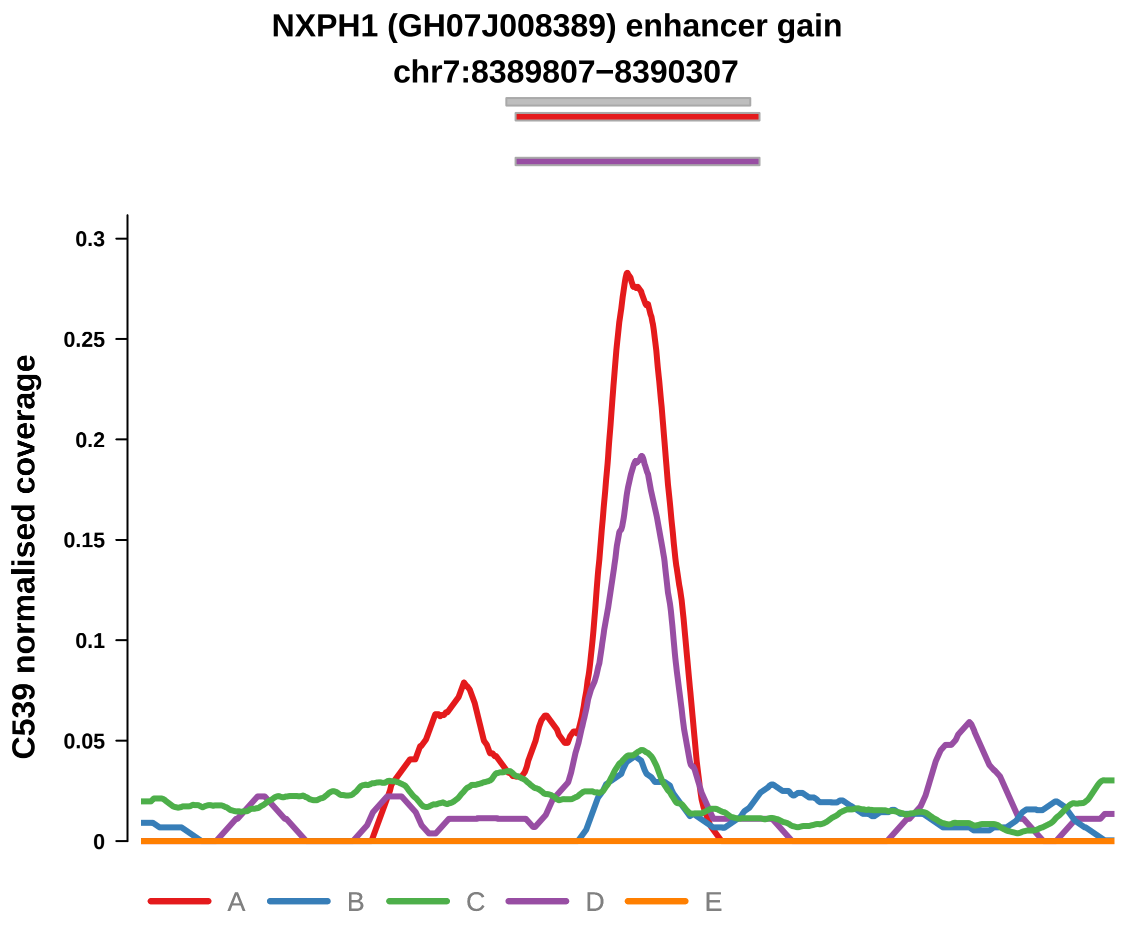


Putatively Clonal (logFC = 2.78)

| | P. | Adjusted P. |
|---|---|---|
| *~region vs. ~* | 1.84e−02 * | 2.48e−02 * |
| *~purity vs ~* | 3.53e−02 * | 4.92e−02 * |
| *~purity+region vs ~purity* | 1.66e−01 | 2.78e−01 |
| *Carcinoma vs. normal* | 4.96e−04 *** | 4.96e−04 *** |

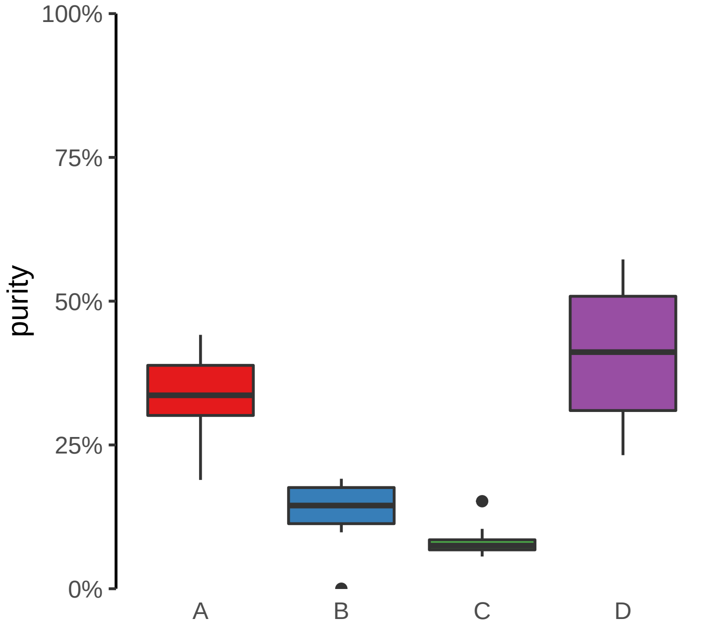

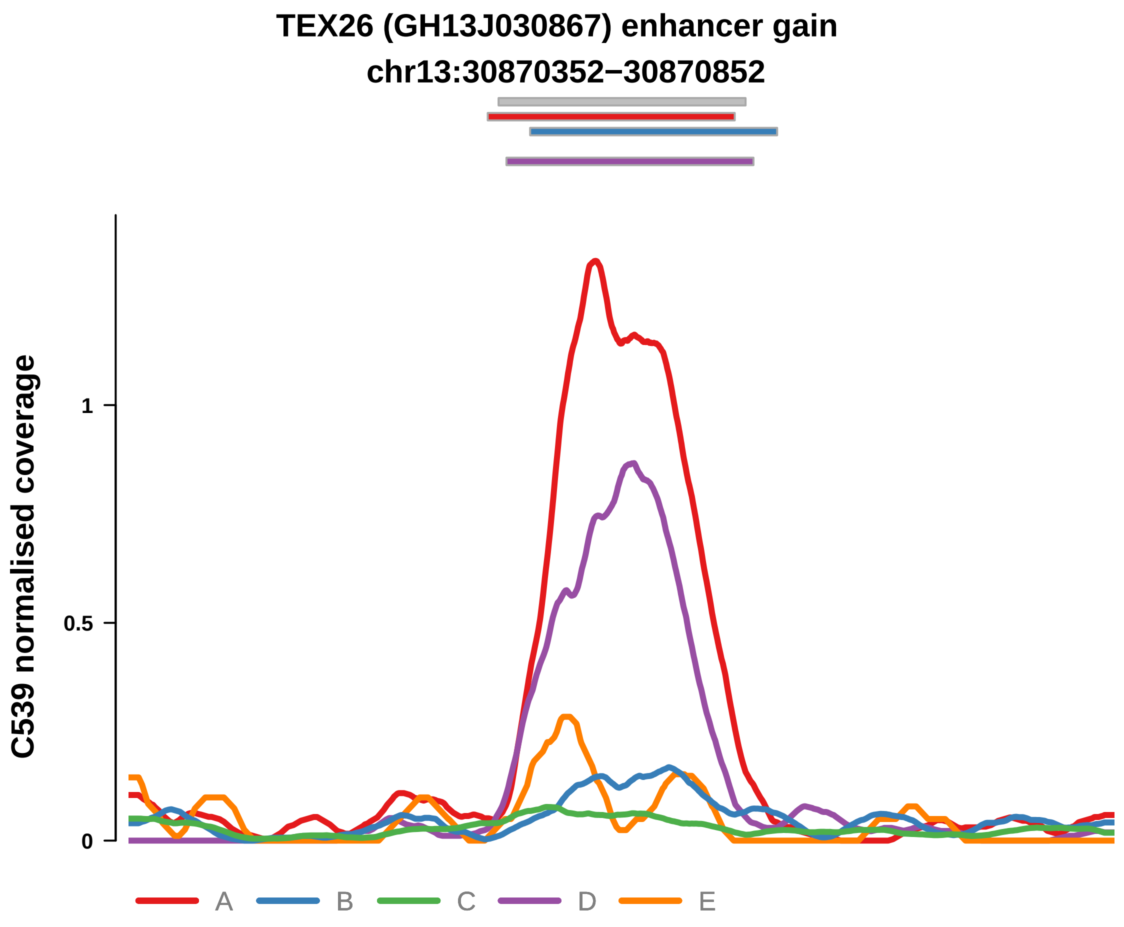


Putatively Clonal (logFC = 4.3)

| | P. | Adjusted P. |
|---|---|---|
| *~region vs. ~* | 1.5e−01 | 1.54e−01 |
| *~purity vs ~* | 5.47e−02 | 6.97e−02 |
| *~purity+region vs ~purity* | 5.38e−02 | 1.25e−01 |
| *Carcinoma vs. normal* | 1.39e−08 **** | 1.39e−08 **** |

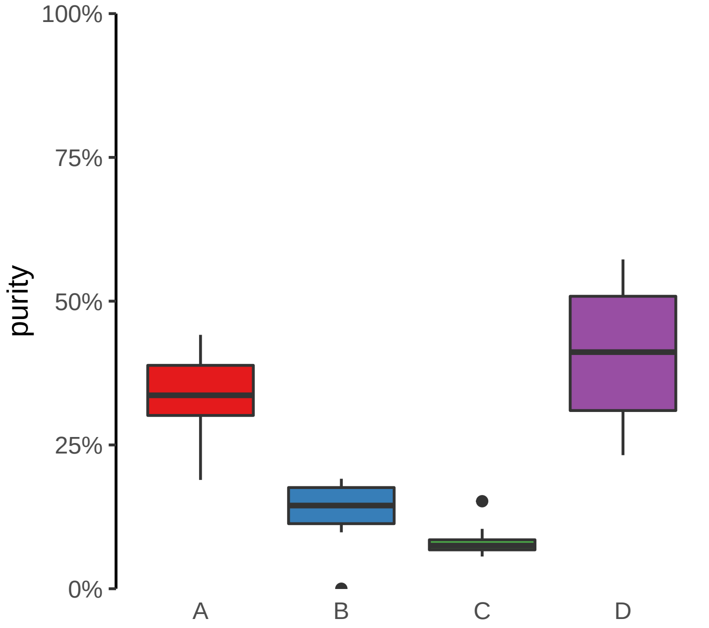

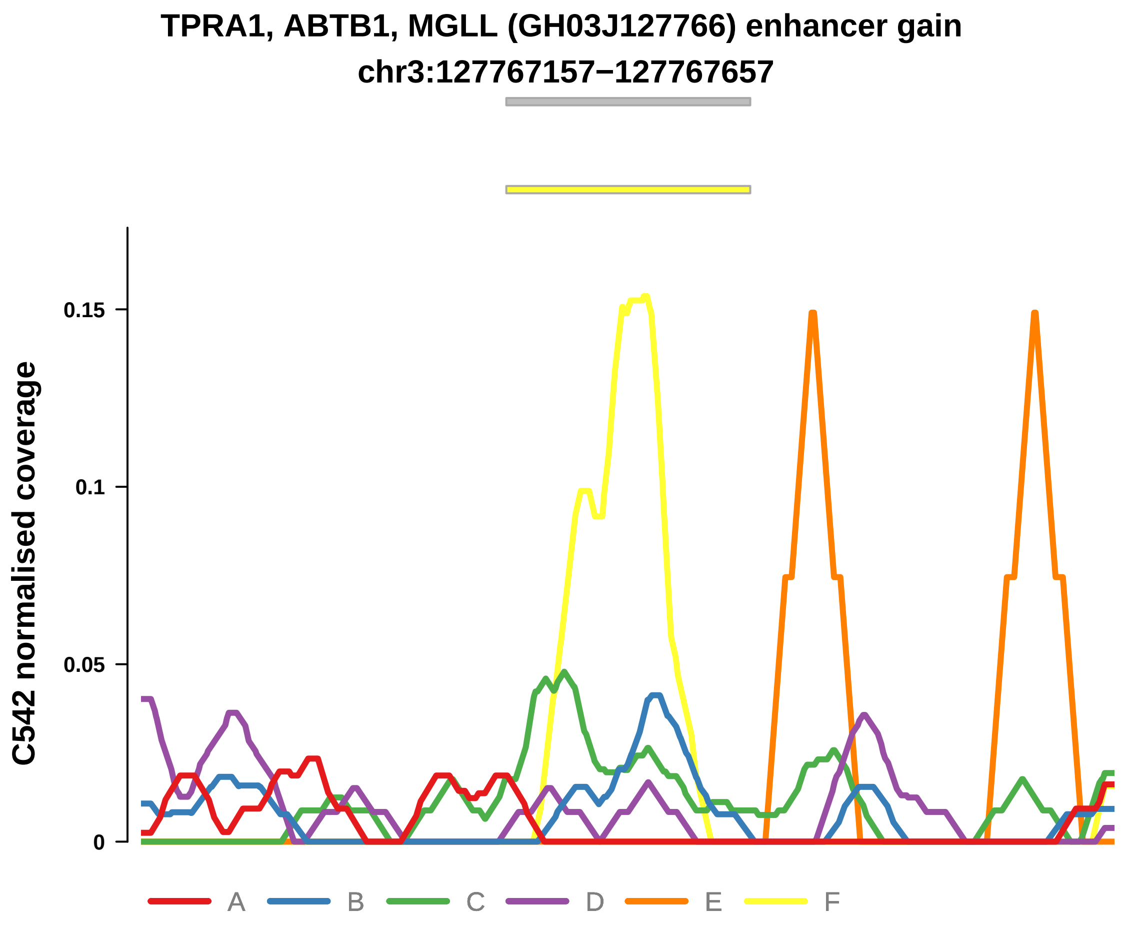


Putatively Clonal (logFC = 2.97)

| | P. | Adjusted P. |
|---|---|---|
| *~region vs. ~* | 9.1e−01 | 9.42e−01 |
| *~purity vs ~* | 2.37e−01 | 7.45e−01 |
| *~purity+region vs ~purity* | 9.95e−01 | 9.96e−01 |
| *Carcinoma vs. normal* | 9.33e−03 ** | 9.33e−03 ** |

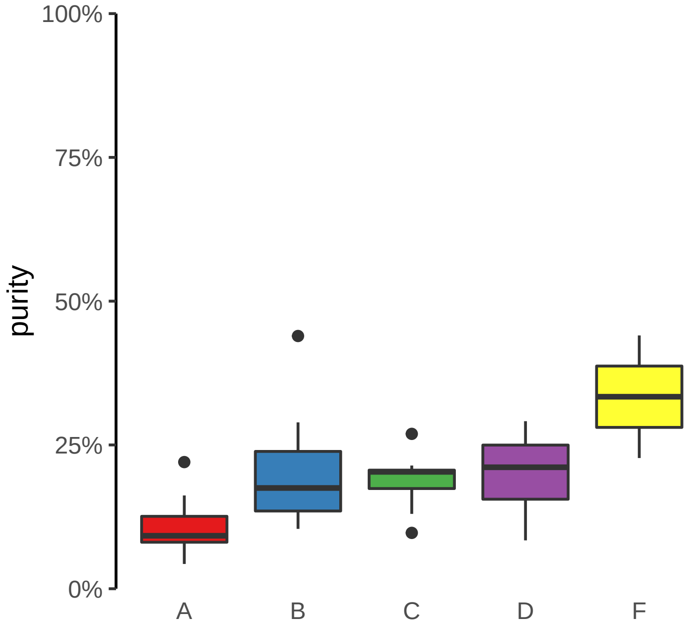

**ATP11A, ATP11A−AS1 (GH13J112791) enhancer gain**
**chr13:112791724−112792224**

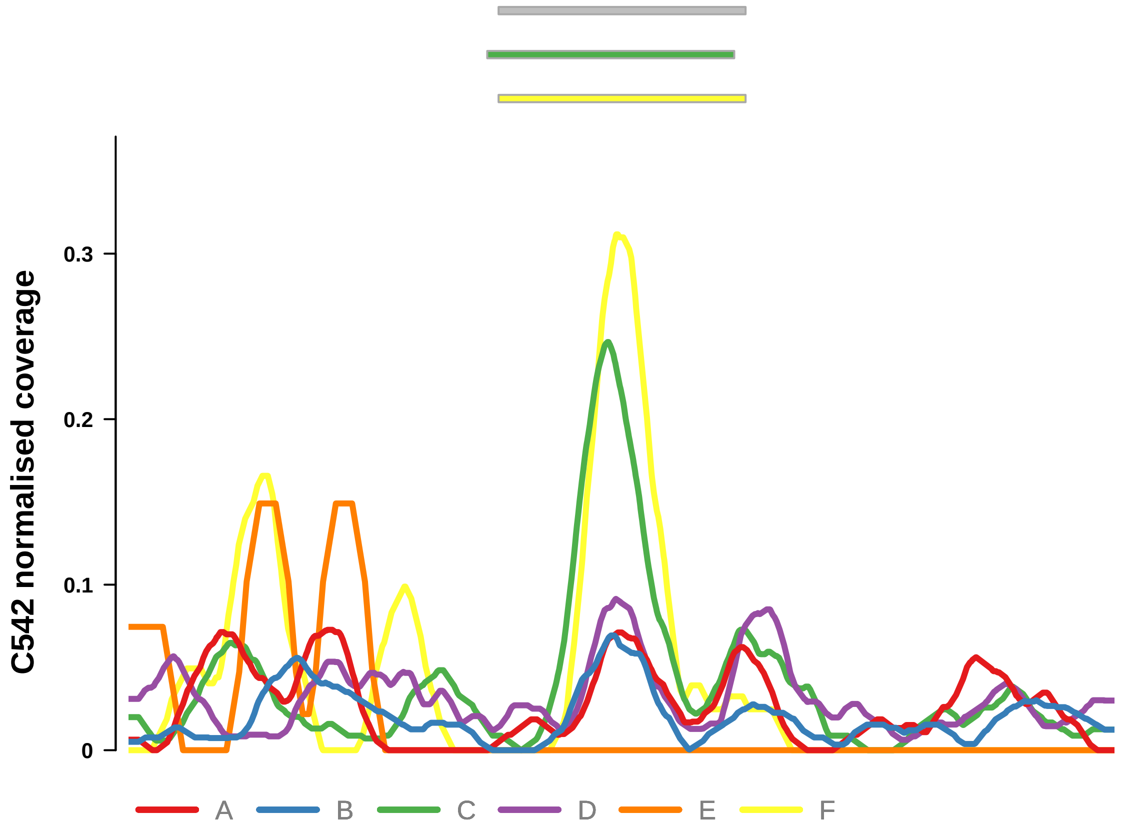


Putatively Clonal (logFC = 4.12)

| | P. | Adjusted P. |
|---|---|---|
| *~region vs. ~* | 2.49e−01 | 6.21e−01 |
| *~purity vs ~* | 1.04e−01 | 5.28e−01 |
| *~purity+region vs ~purity* | 3e−01 | 7.3e−01 |
| *Carcinoma vs. normal* | 4.35e−06 **** | 4.35e−06 **** |

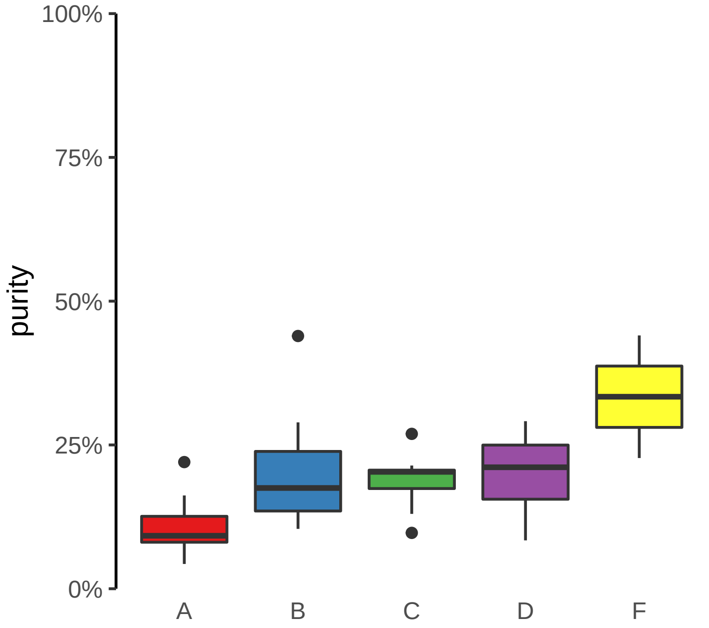

**FAM86B3P (GH08J008285) enhancer gain**
**chr8:8288409−8288909**

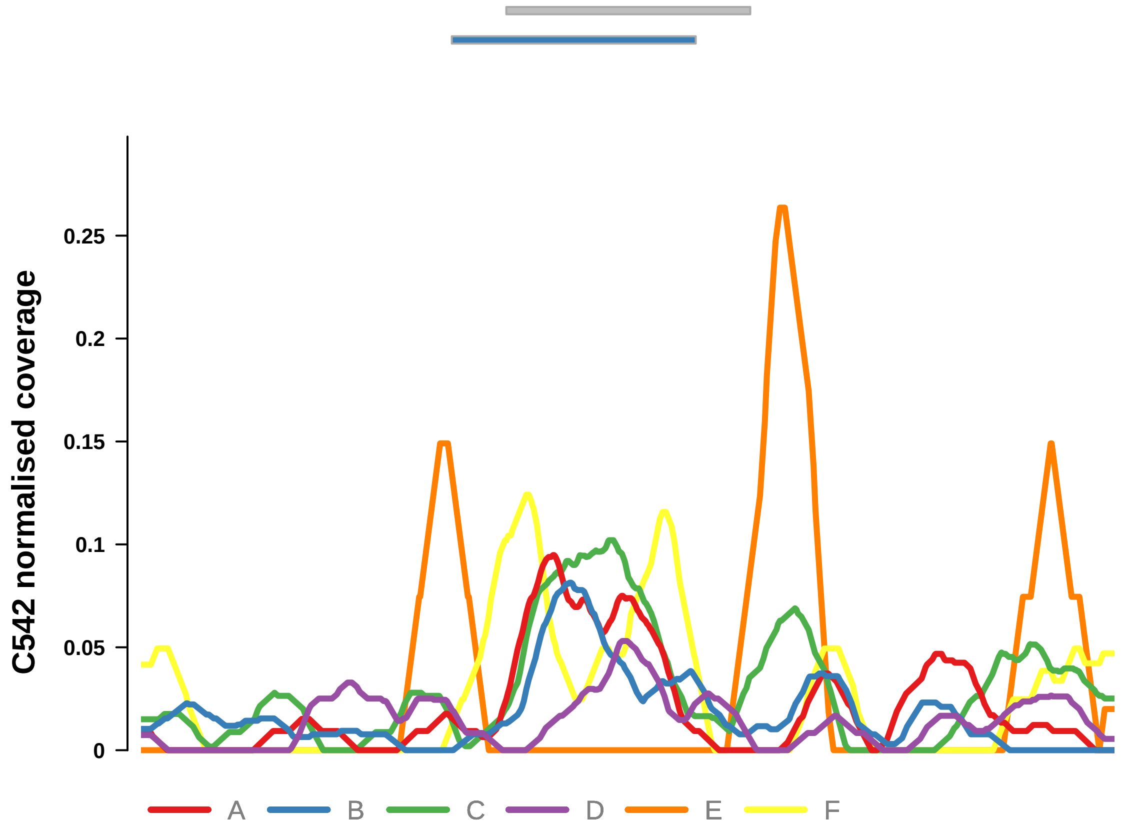


Putatively Clonal (logFC = 2.73)

| | P. | Adjusted P. |
|---|---|---|
| *~region vs. ~* | 8.62e−01 | 9.36e−01 |
| *~purity vs ~* | 8.22e−01 | 9.67e−01 |
| *~purity+region vs ~purity* | 8.07e−01 | 8.56e−01 |
| *Carcinoma vs. normal* | 9.69e−03 ** | 9.69e−03 ** |

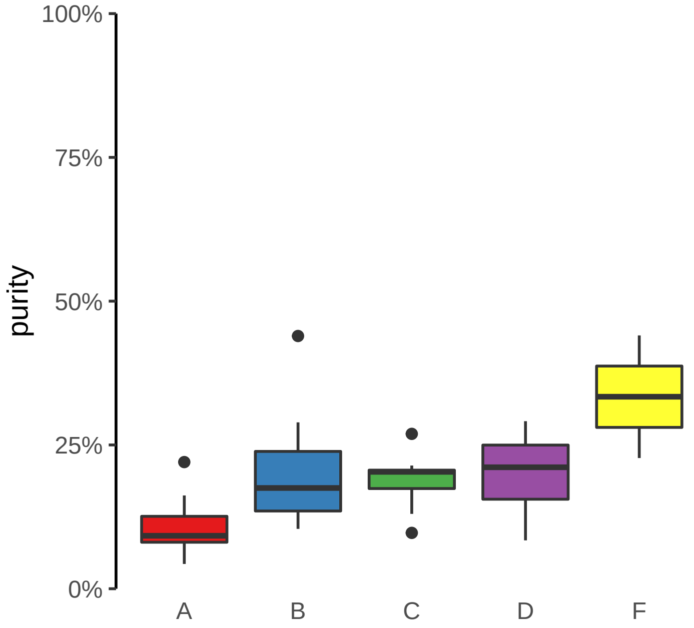

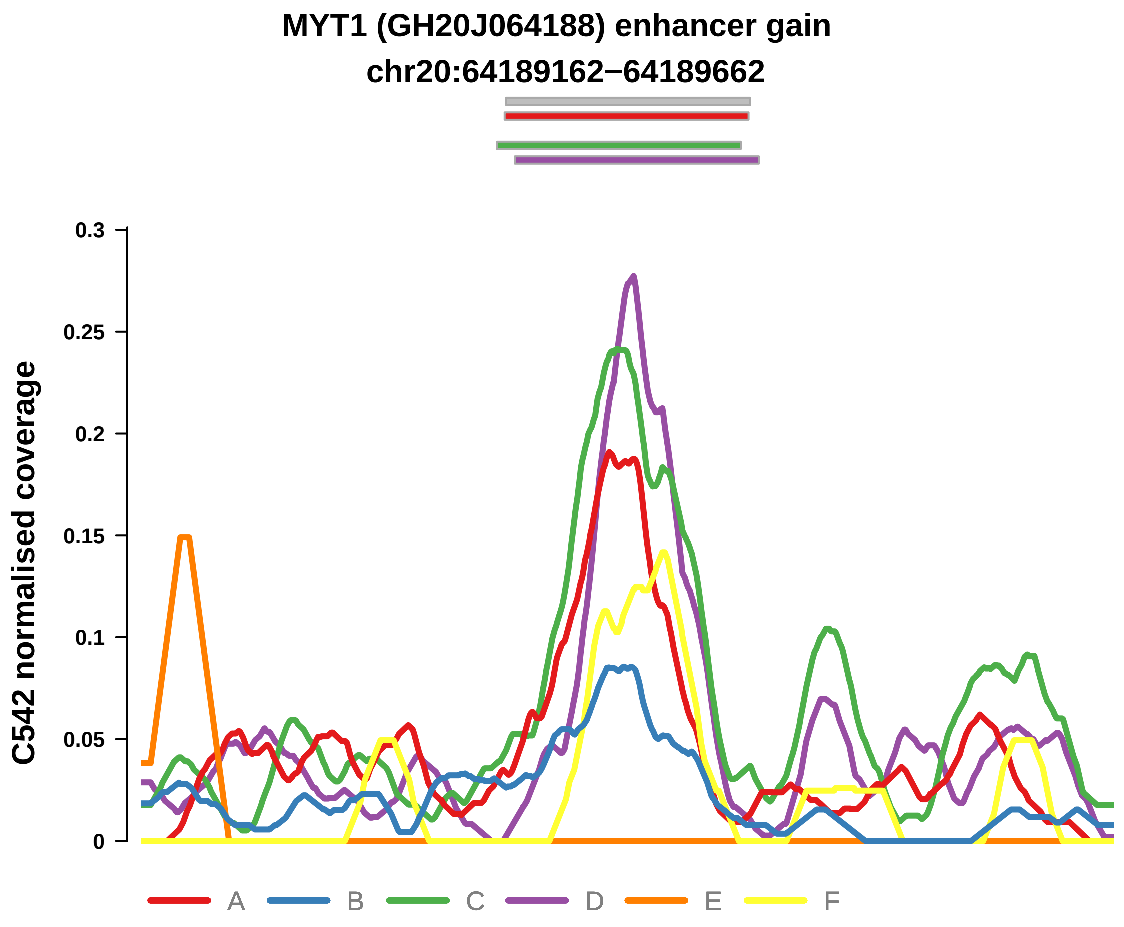


Putatively Clonal (logFC = 3.17)

| | P. | Adjusted P. |
|---|---|---|
| *~region vs. ~* | 3.99e−02 * | 5.01e−01 |
| *~purity vs ~* | 4.07e−01 | 9.67e−01 |
| *~purity+region vs ~purity* | 1.24e−02 * | 3.94e−01 |
| *Carcinoma vs. normal* | 2.16e−03 ** | 2.16e−03 ** |

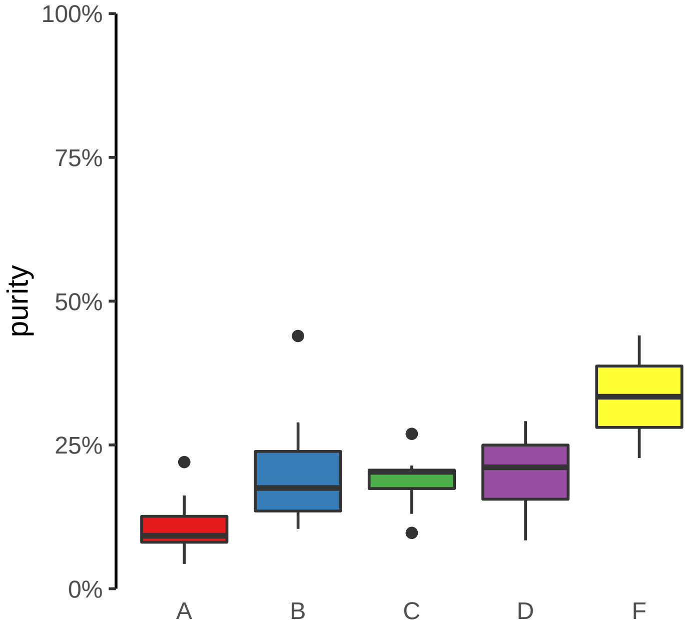

**NCOA7−AS1 (GH06J125867) enhancer gain**
**chr6:125867578−125868078**

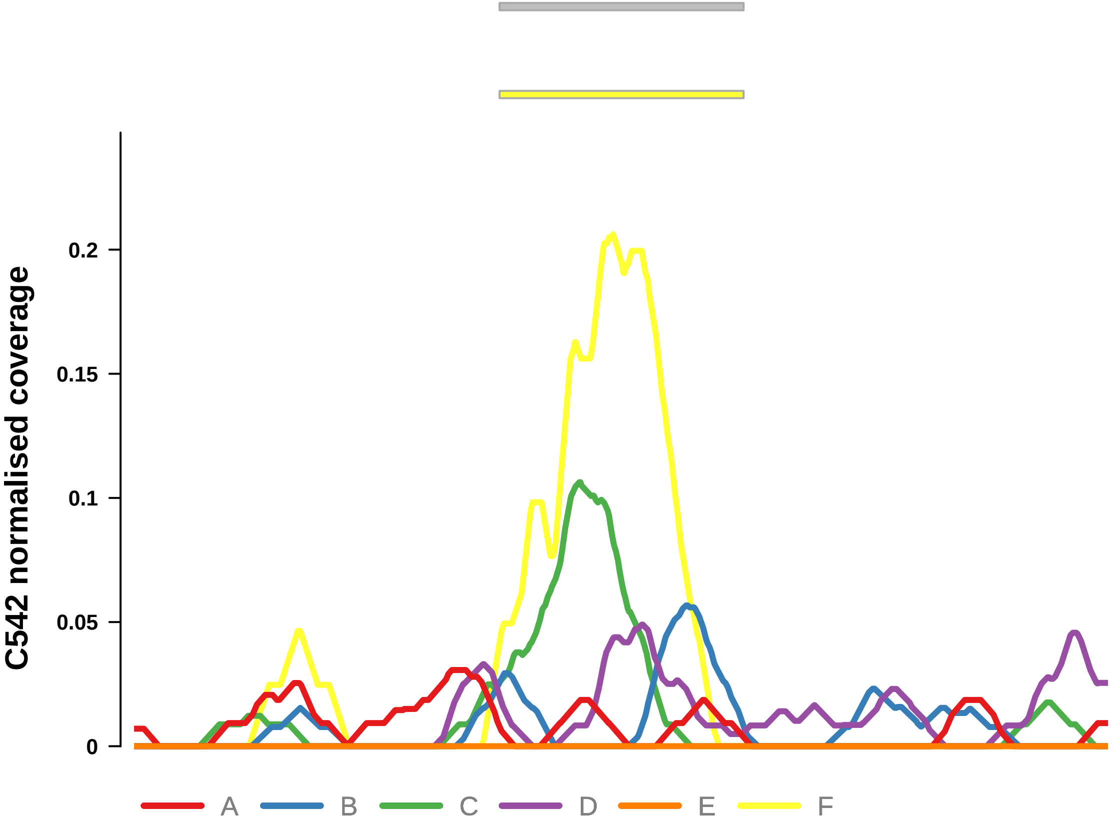


Putatively Clonal (logFC = 2.89)

| | P. | Adjusted P. |
|---|---|---|
| *~region vs. ~* | 2.98e−01 | 6.34e−01 |
| *~purity vs ~* | 6.21e−01 | 9.67e−01 |
| *~purity+region vs ~purity* | 2.65e−01 | 7.3e−01 |
| *Carcinoma vs. normal* | 3.93e−03 ** | 3.93e−03 ** |

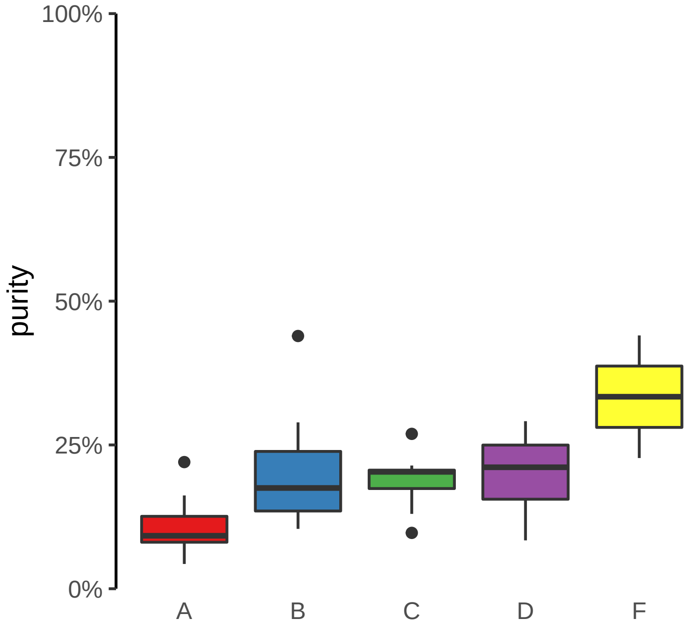

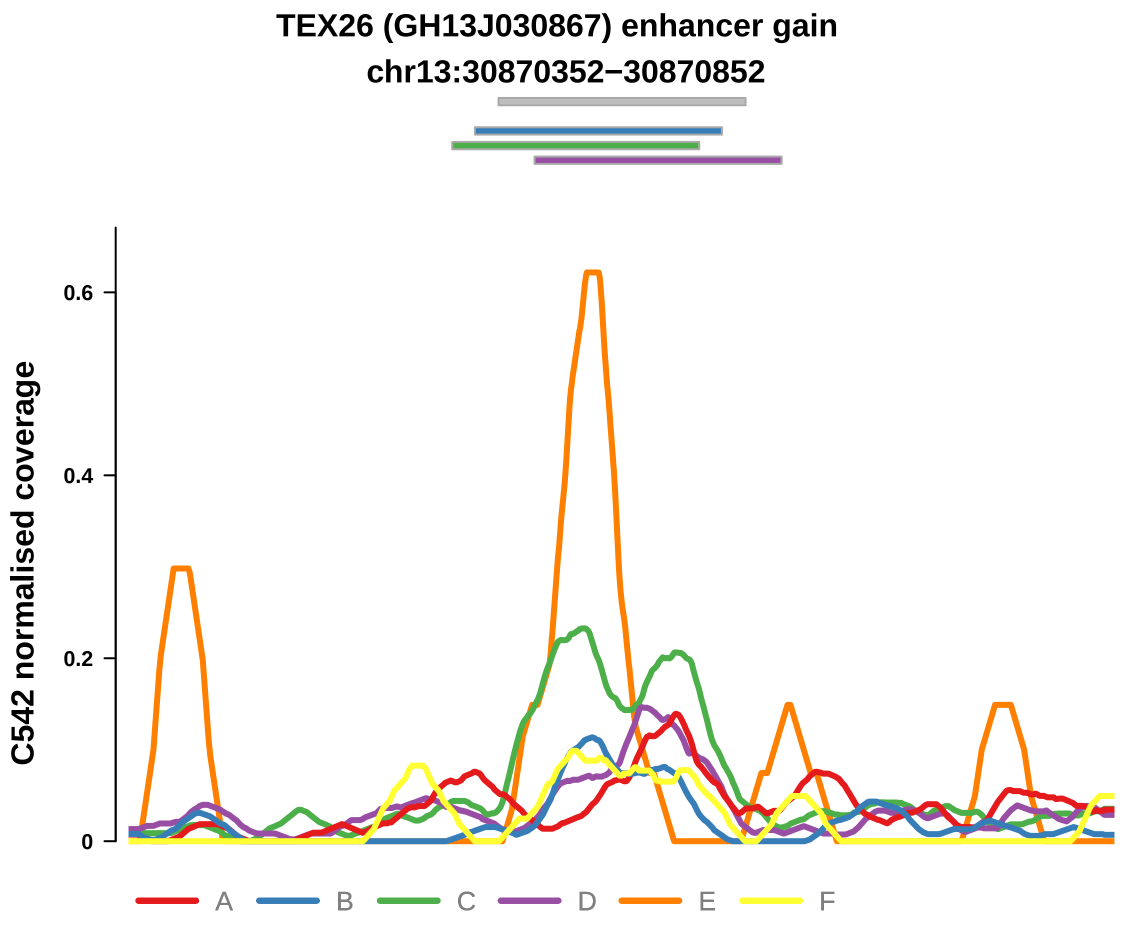


Putatively Clonal (logFC = −11.18)

| | P. | Adjusted P. |
|---|---|---|
| *~region vs. ~* | 2.74e−01 | 6.34e−01 |
| *~purity vs ~* | 5.64e−01 | 9.67e−01 |
| *~purity+region vs ~purity* | 1.72e−01 | 6.99e−01 |
| *Carcinoma vs. normal* | 2.8e−03 ** | 2.8e−03 ** |

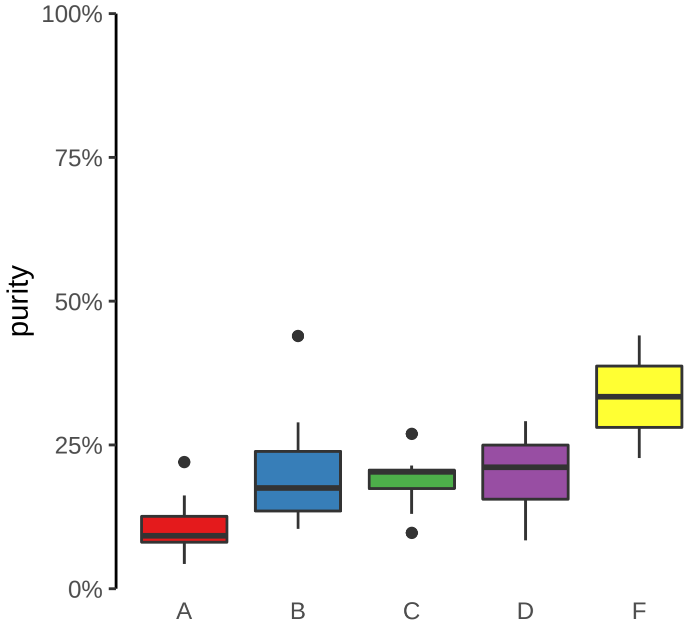

**MYT1 (GH20J064188) enhancer gain**

**chr20:64189162−64189662**

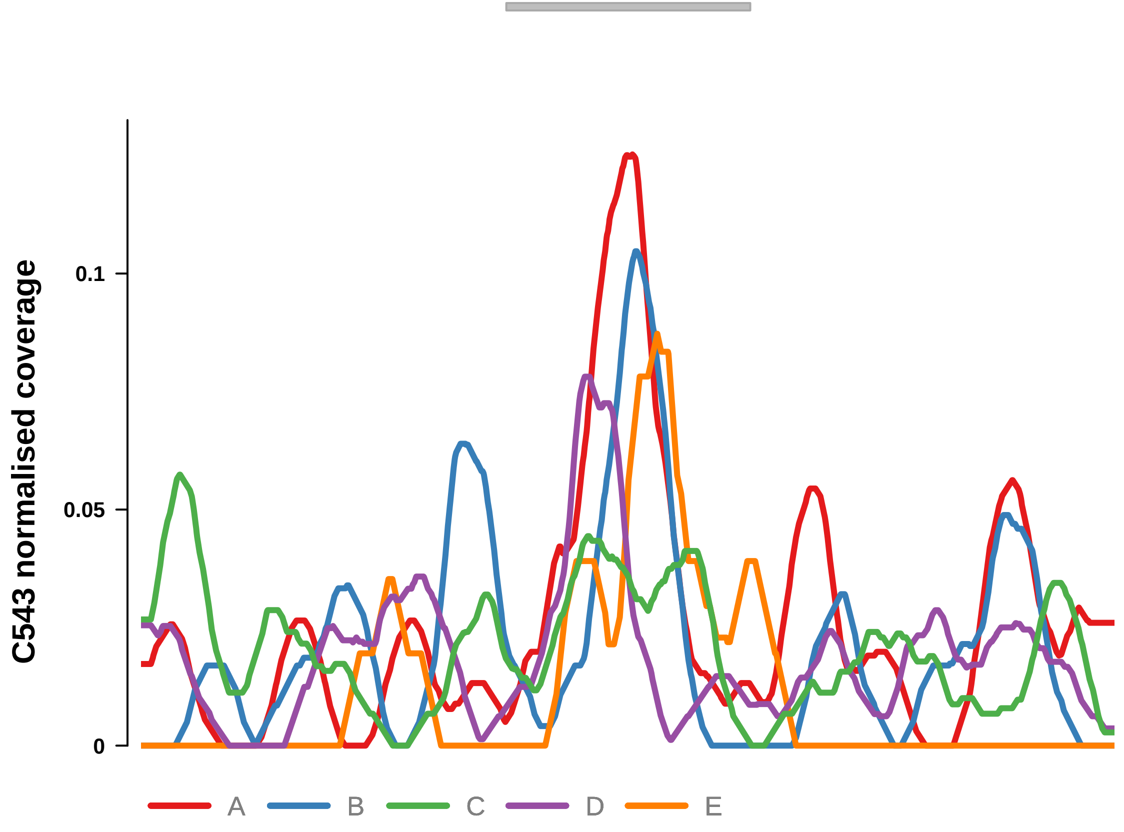


Putatively Clonal (logFC = 3.3)

| | P. | Adjusted P. |
|---|---|---|
| *~region vs. ~* | 7.82e−01 | 8.54e−01 |
| *~purity vs ~* | 2.92e−01 | NA |
| *~purity+region vs ~purity* | 2.97e−01 | 7.91e−01 |
| *Carcinoma vs. normal* | 6.05e−03 ** | 6.05e−03 ** |

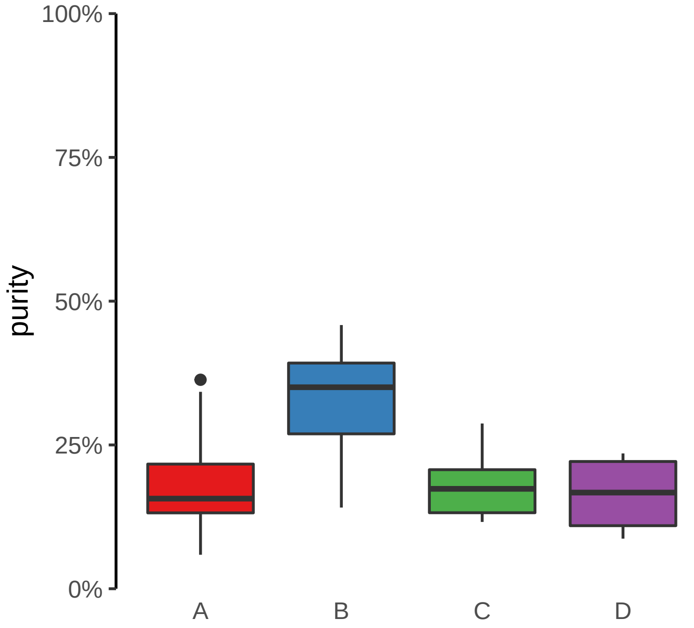

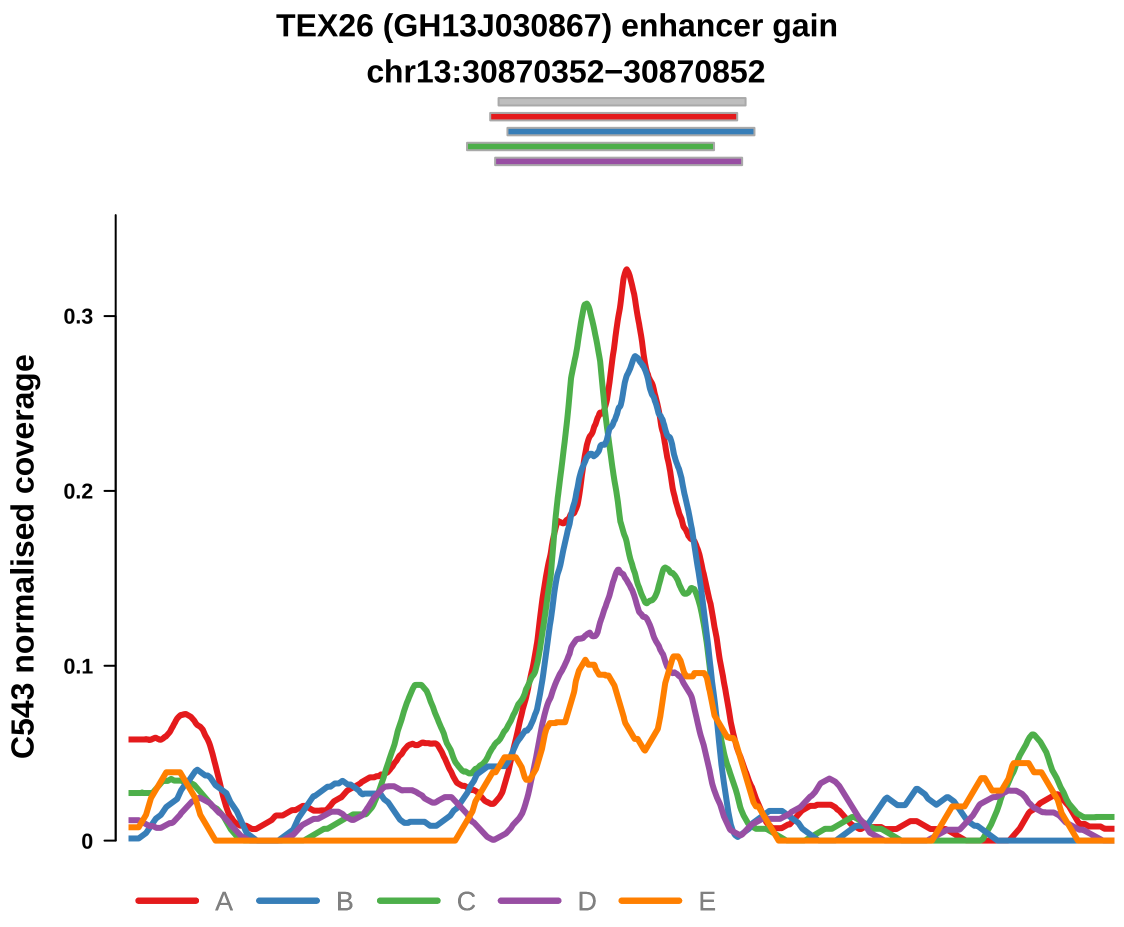


Putatively Clonal (logFC = 2.32)

| | P. | Adjusted P. |
|---|---|---|
| *~region vs. ~* | 1.38e−01 | 5.16e−01 |
| *~purity vs ~* | 1.79e−03 ** | 5.01e−02 |
| *~purity+region vs ~purity* | 5.34e−01 | 8.43e−01 |
| *Carcinoma vs. normal* | 4.26e−03 ** | 4.26e−03 ** |

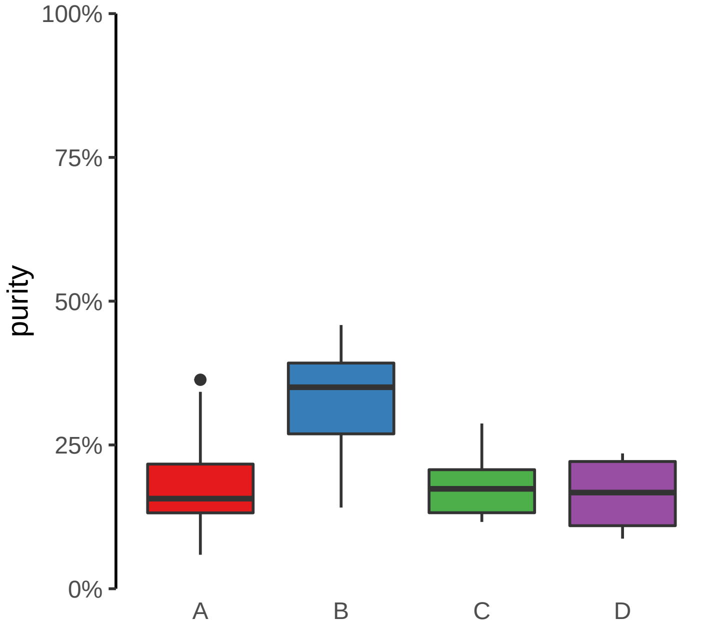

**FGD2 (GH06J037022) enhancer gain**
**chr6:37024608−37025108**

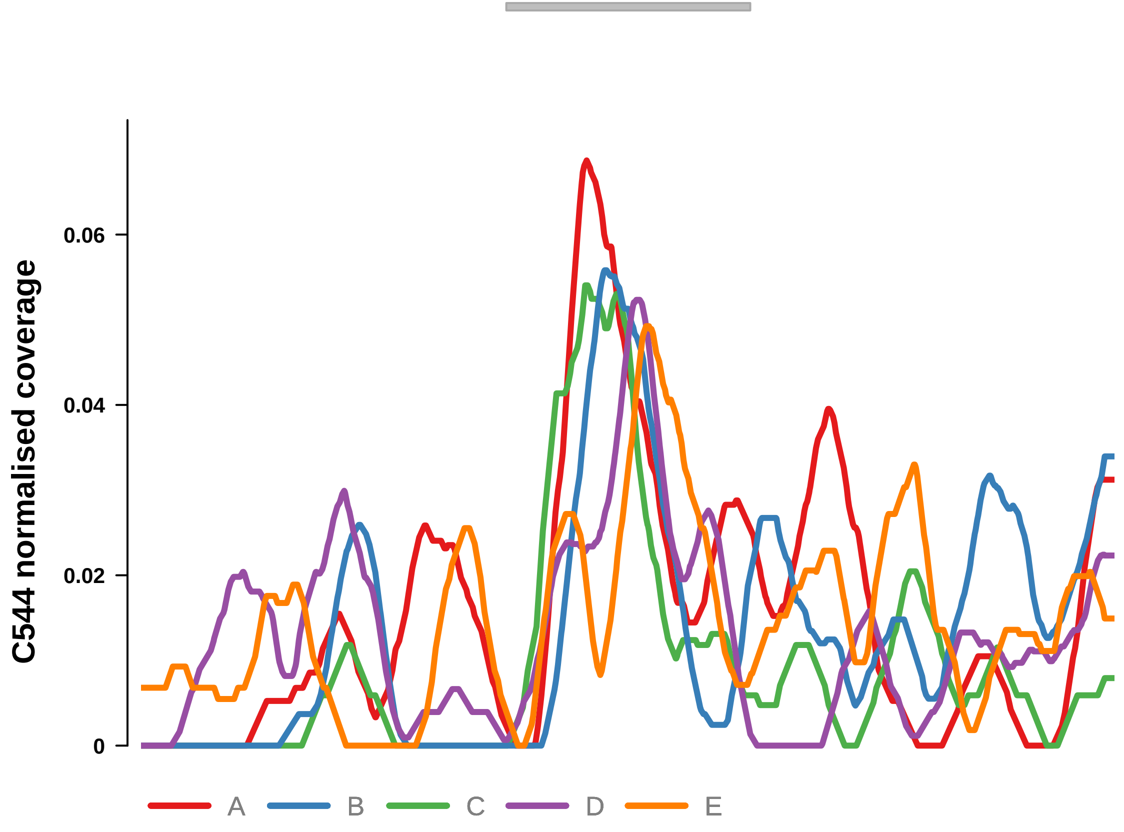


Putatively Clonal (logFC = 2.11)

| | P. | Adjusted P. |
|---|---|---|
| *~region vs. ~* | 5.99e−01 | 6.5e−01 |
| *~purity vs ~* | 3.28e−03 ** | 2.16e−02 * |
| *~purity+region vs ~purity* | 5.25e−01 | 6.82e−01 |
| *Carcinoma vs. normal* | 3.45e−03 ** | 3.45e−03 ** |

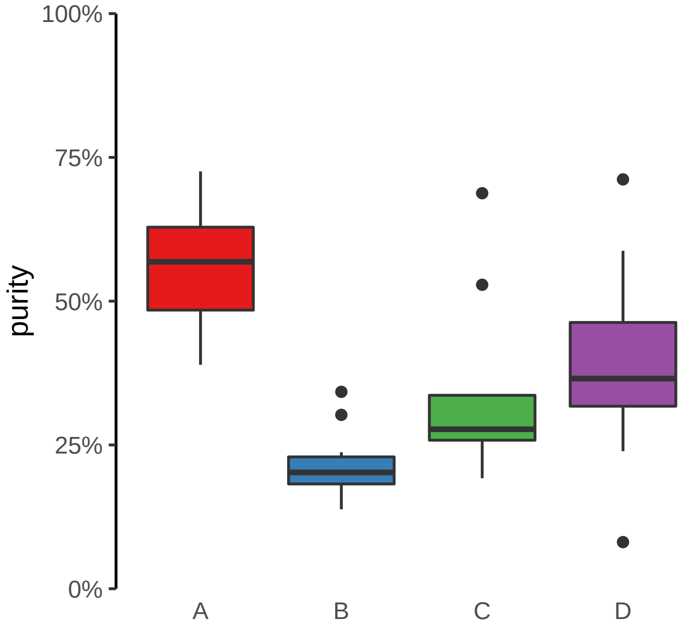

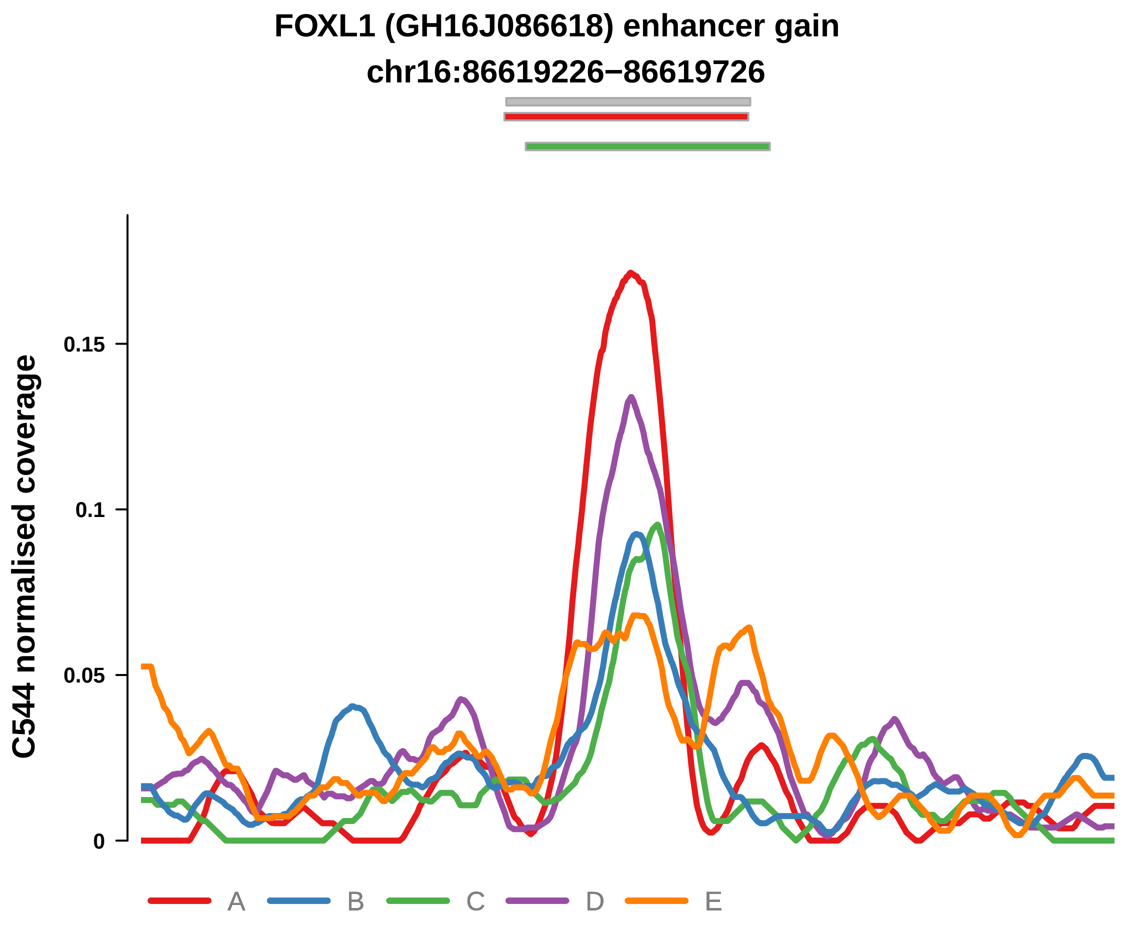


Putatively Clonal (logFC = 2.45)

| | P. | Adjusted P. |
|---|---|---|
| *~region vs. ~* | 2.56e−02 * | 6.42e−02 |
| *~purity vs ~* | 3.47e−03 ** | 2.16e−02 * |
| *~purity+region vs ~purity* | 6.04e−01 | 7e−01 |
| *Carcinoma vs. normal* | 6.3e−04 *** | 6.3e−04 *** |

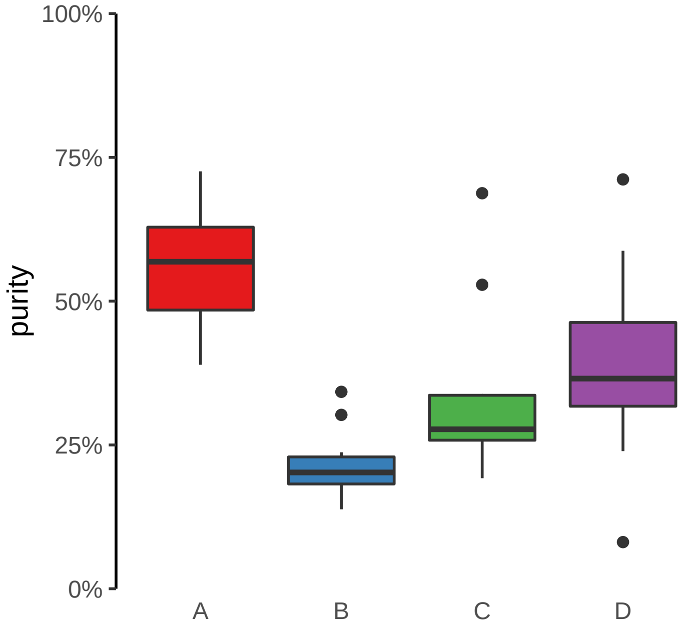

**LIFR (GH05J038659) enhancer gain**
**chr5:38660203−38660703**

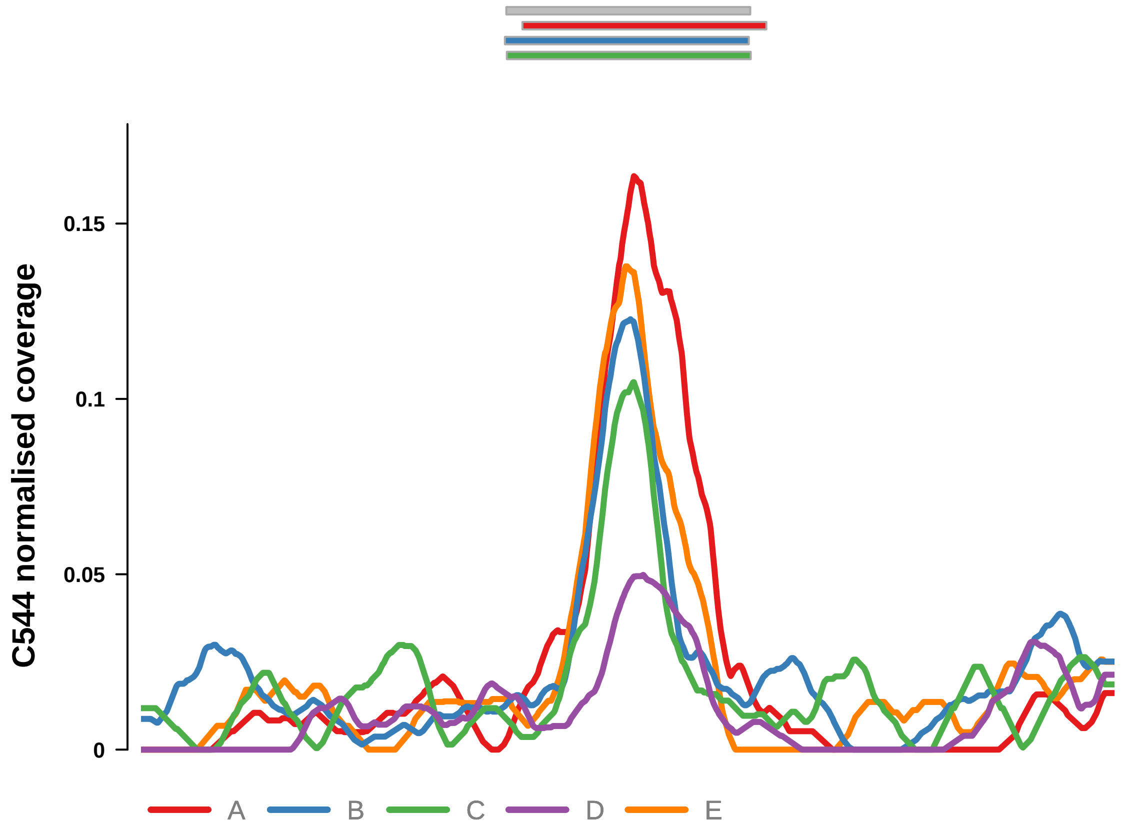


Putatively Clonal (logFC = 2.36)

| | P. | Adjusted P. |
|---|---|---|
| *~region vs. ~* | 6.09e−03 ** | 2.06e−02 * |
| *~purity vs ~* | 1.91e−02 * | 7.18e−02 |
| *~purity+region vs ~purity* | 4.9e−02 * | 1.18e−01 |
| *Carcinoma vs. normal* | 9.44e−04 *** | 9.44e−04 *** |

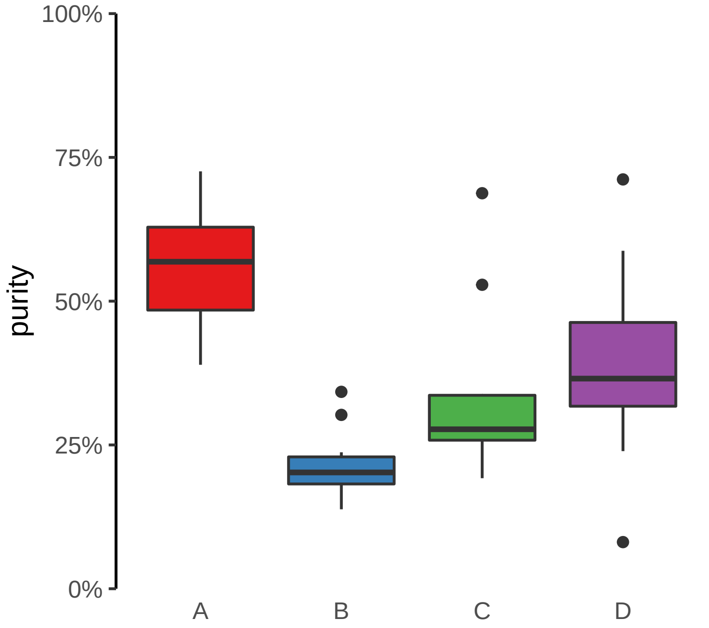

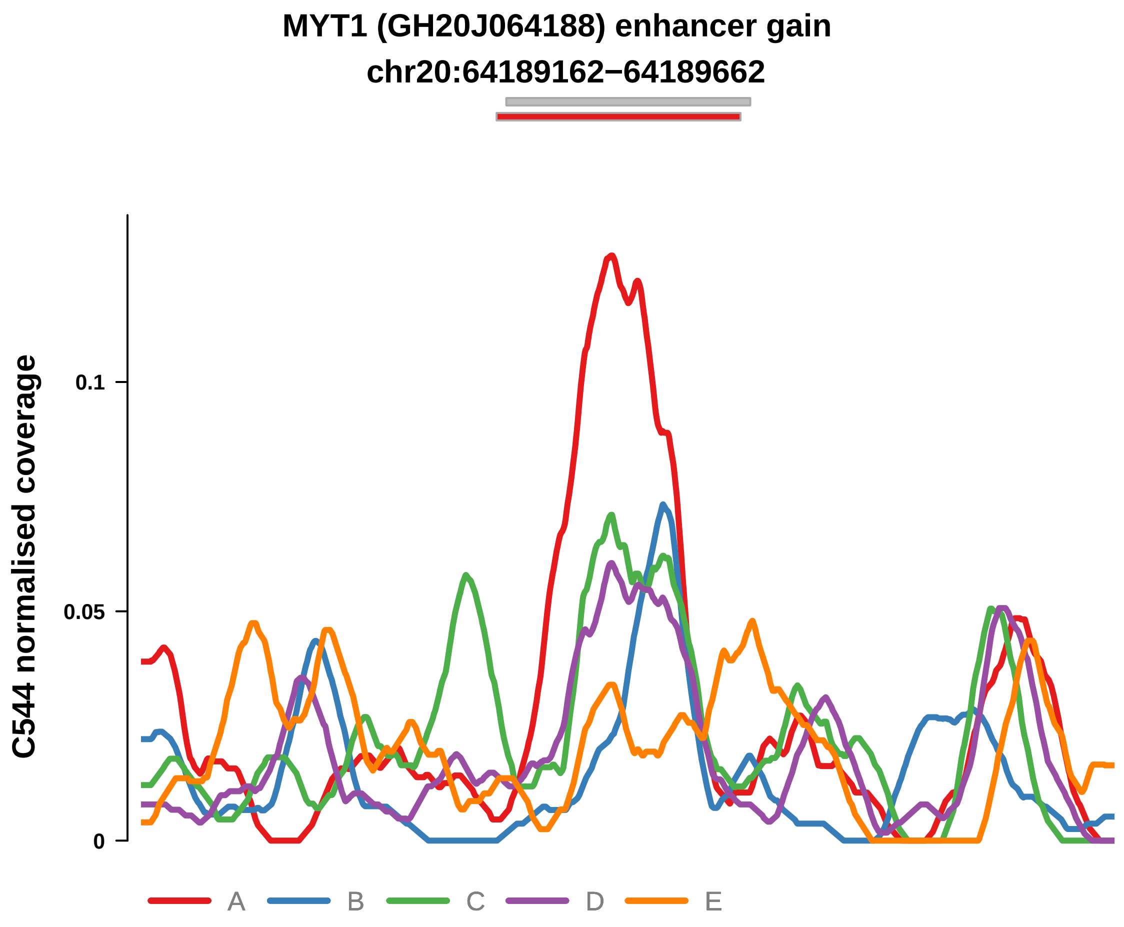


Putatively Clonal (logFC = 3.21)

| | P. | Adjusted P. |
|---|---|---|
| *~region vs. ~* | 2.12e−02 * | 5.84e−02 |
| *~purity vs ~* | 4.03e−04 *** | 6.53e−03 ** |
| *~purity+region vs ~purity* | 6.59e−01 | 7.4e−01 |
| *Carcinoma vs. normal* | 1.51e−05 **** | 1.51e−05 **** |

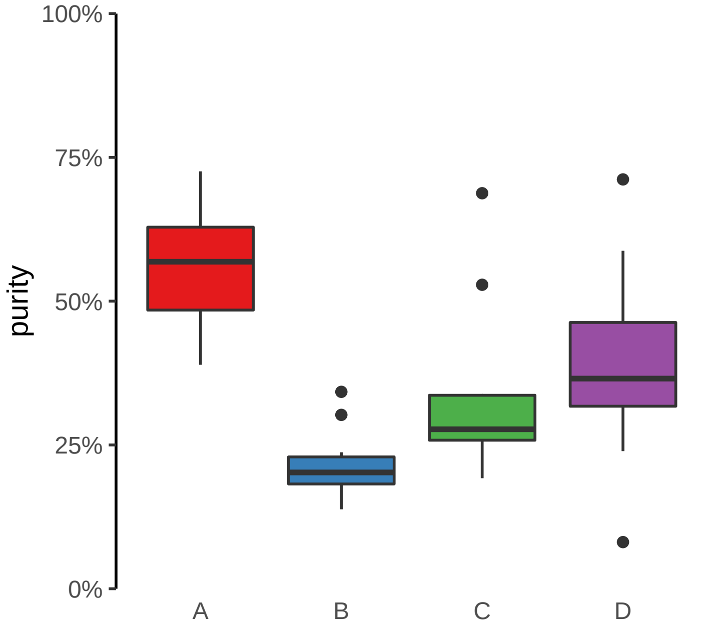

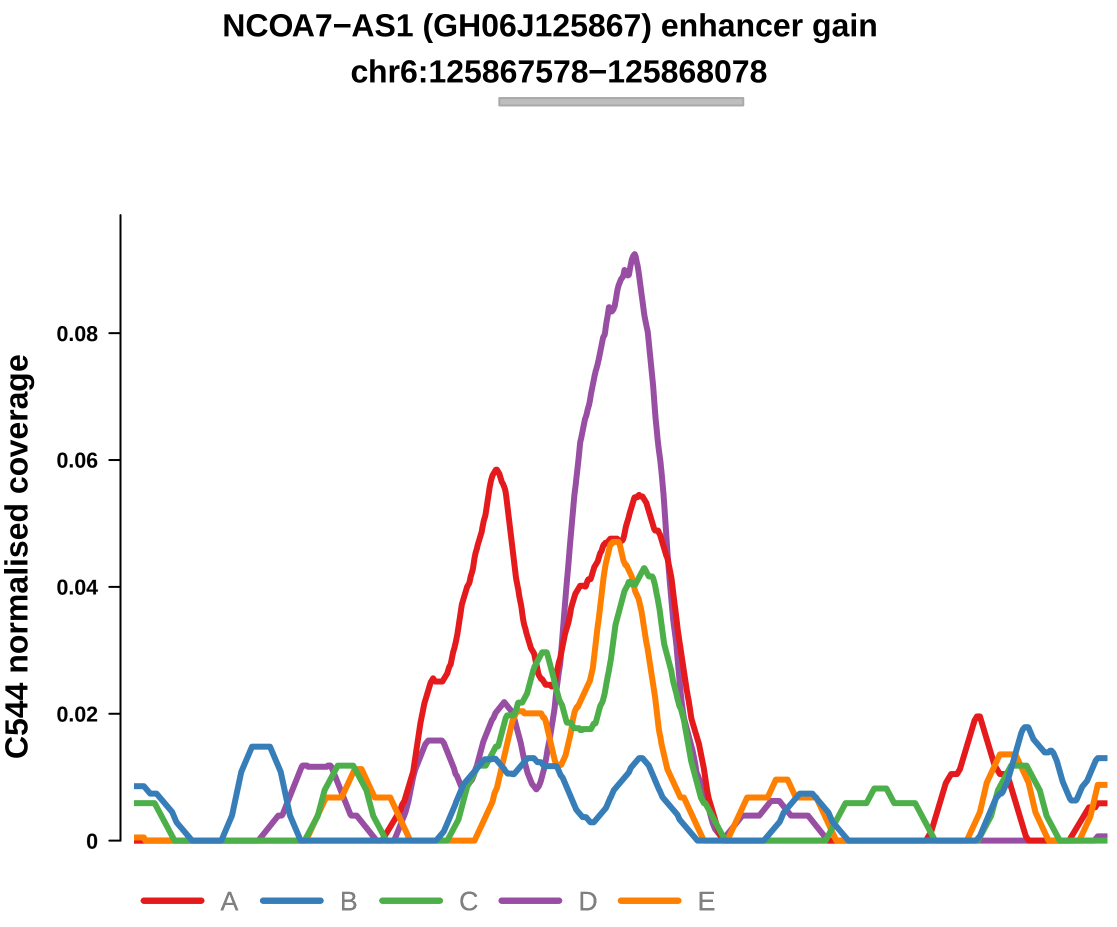


Putatively Clonal (logFC = 2.14)

| | P. | Adjusted P. |
|---|---|---|
| *~region vs. ~* | 1.58e−03 ** | 7.73e−03 ** |
| *~purity vs ~* | 7.5e−03 ** | 3.79e−02 * |
| *~purity+region vs ~purity* | 3.98e−02 * | 1.07e−01 |
| *Carcinoma vs. normal* | 3.22e−03 ** | 3.22e−03 ** |

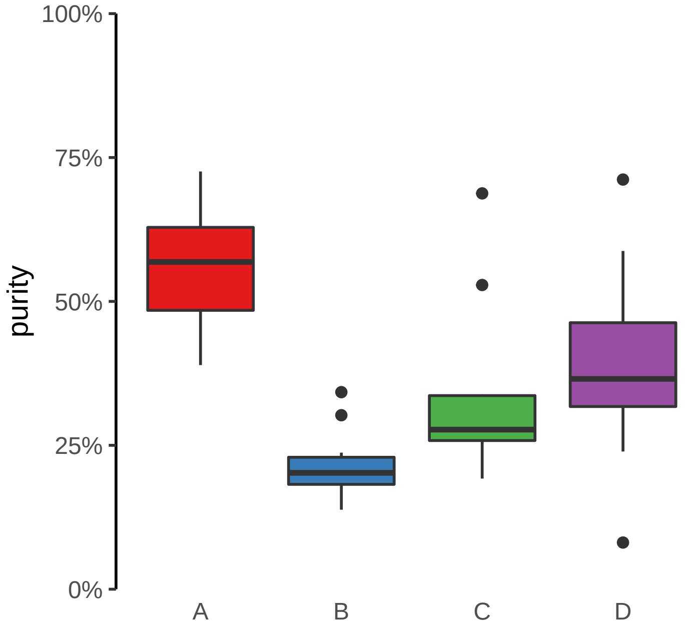

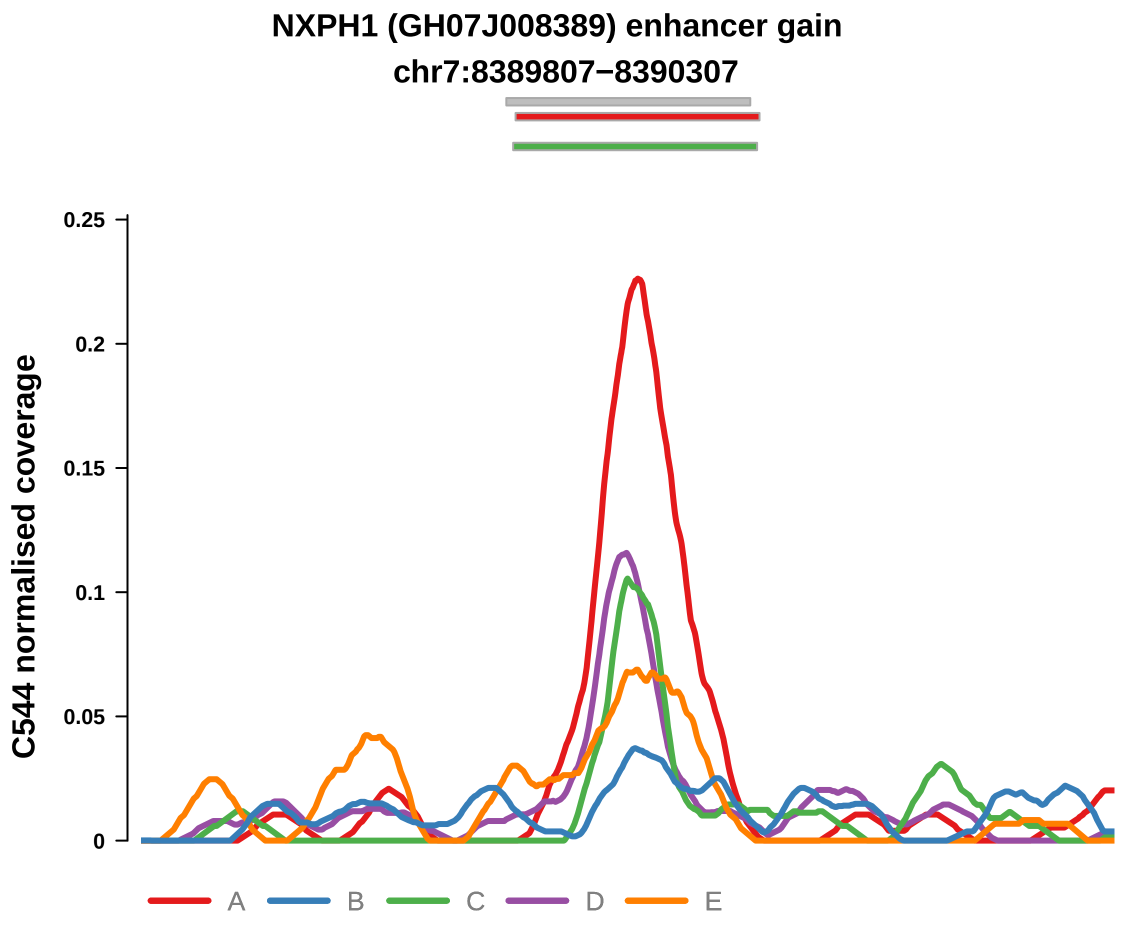


Putatively Clonal (logFC = 2.59)

| | P. | Adjusted P. |
|---|---|---|
| *~region vs. ~* | 1.07e−06 **** | 9.46e−05 **** |
| *~purity vs ~* | 7.91e−06 **** | 3.2e−04 *** |
| *~purity+region vs ~purity* | 4.03e−02 * | 1.07e−01 |
| *Carcinoma vs. normal* | 2.93e−04 *** | 2.93e−04 *** |

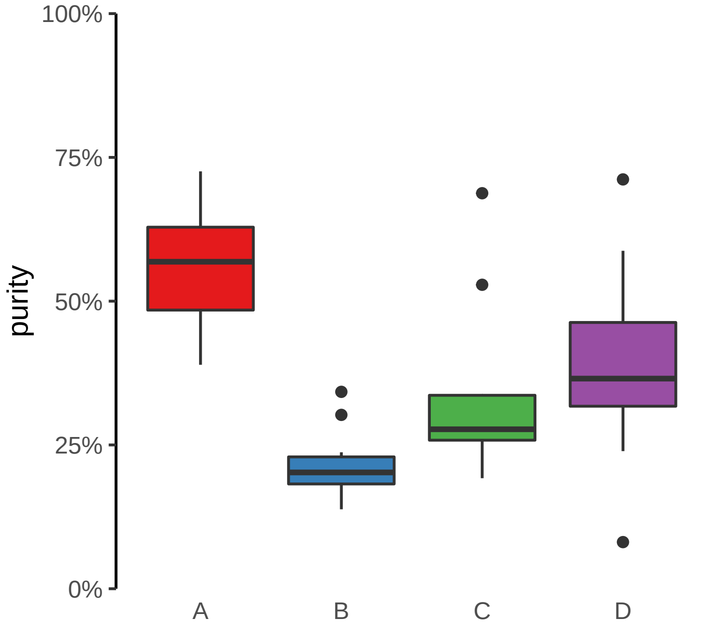

**HOXC11, HOXC10, HOTAIR (GH12J053971) enhancer gain**
**chr12:53978873−53979373**

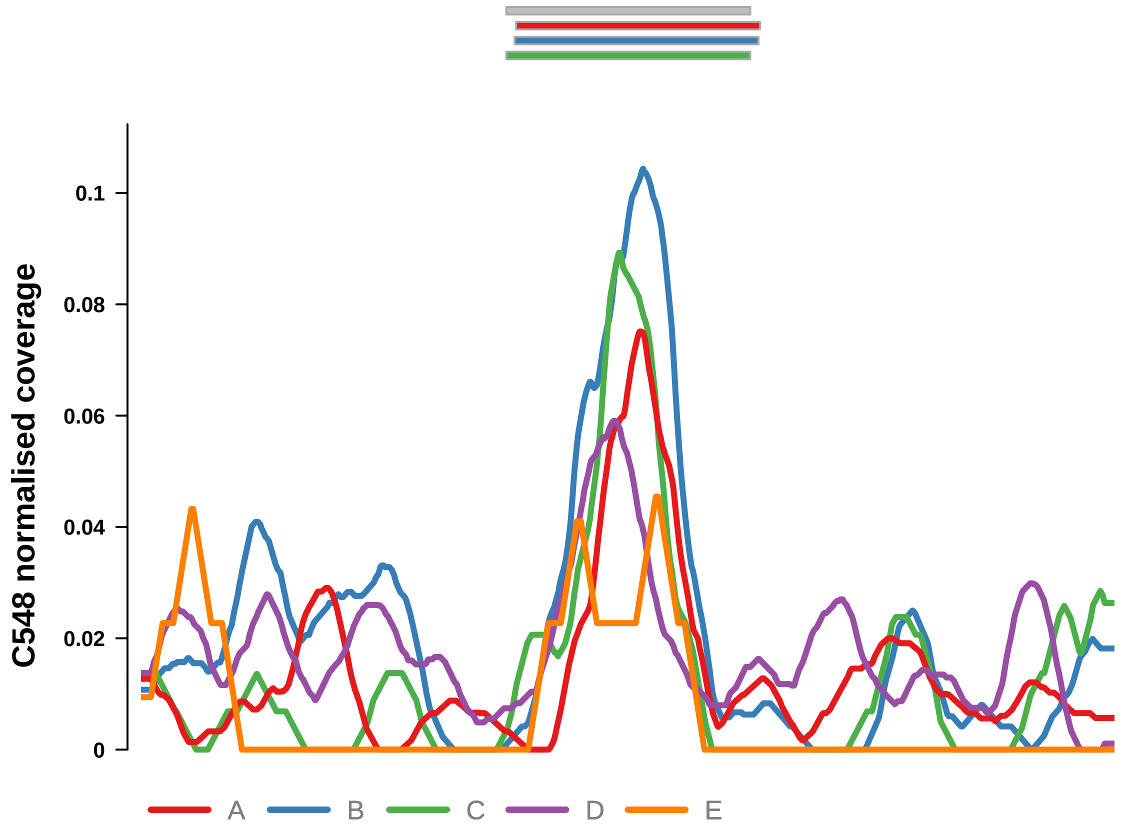


Putatively Clonal (logFC = 2.3)

| | P. | Adjusted P. |
|---|---|---|
| *~region vs. ~* | 8.77e−01 | 9.21e−01 |
| *~purity vs ~* | 8.18e−01 | 8.81e−01 |
| *~purity+region vs ~purity* | 4.5e−01 | 6.41e−01 |
| *Carcinoma vs. normal* | 2.46e−03 ** | 2.46e−03 ** |

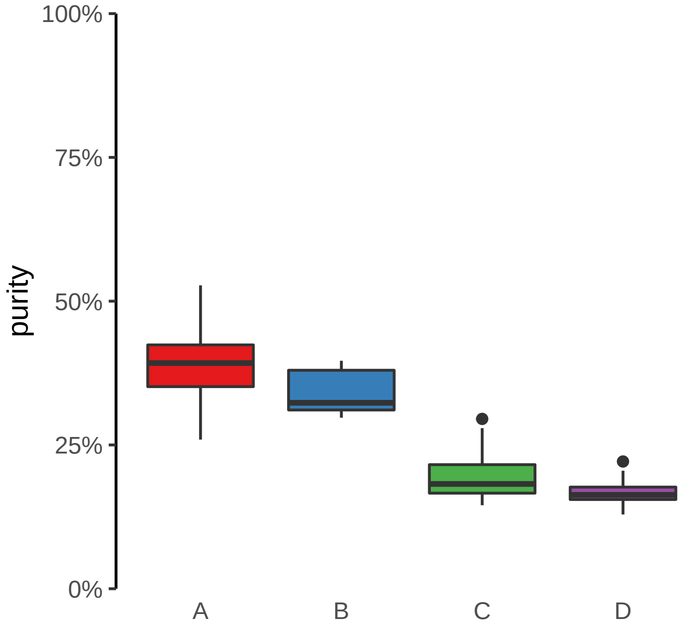

**TPRA1, ABTB1, MGLL (GH03J127766) enhancer gain**
**chr3:127767157−127767657**

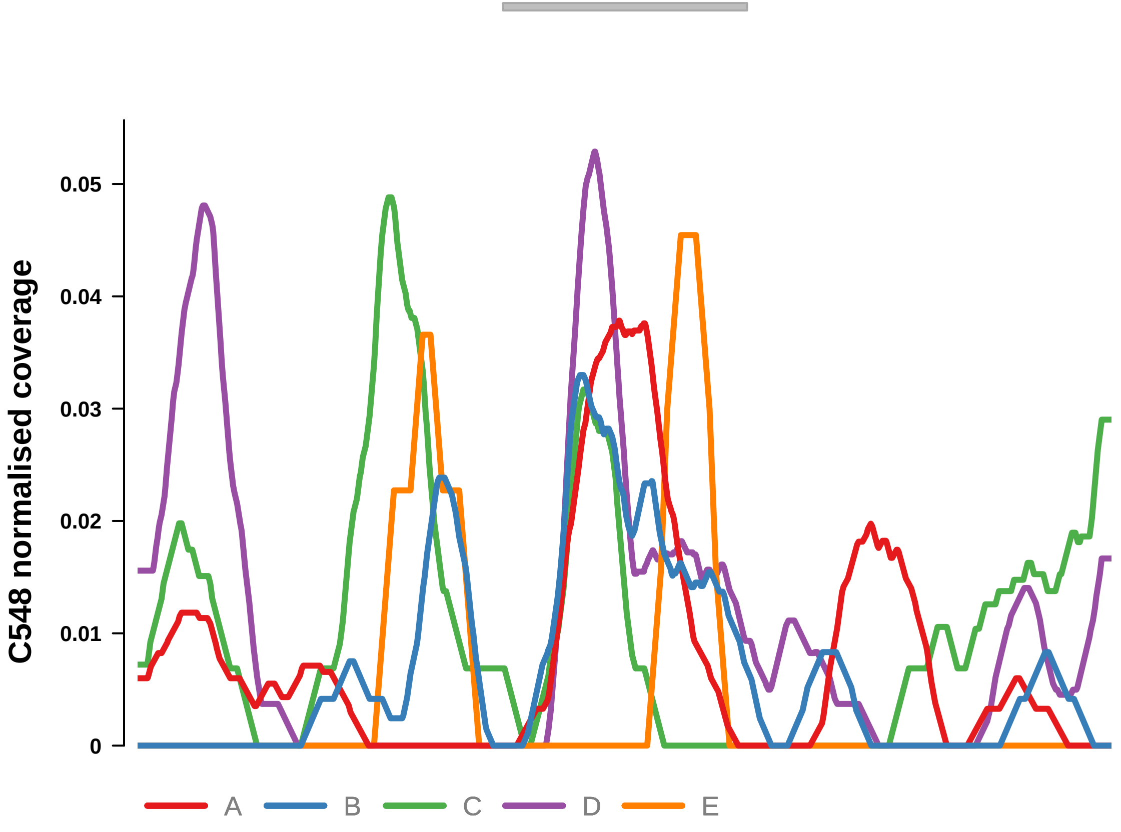


Putatively Clonal (logFC = 2.44)

| | **P.** | **Adjusted P.** |
|---|---|---|
| *~region vs. ~* | 3.93e−01 | 5.24e−01 |
| *~purity vs ~* | 9.48e−01 | 9.7e−01 |
| *~purity+region vs ~purity* | 2.84e−01 | 5.33e−01 |
| *Carcinoma vs. normal* | 3.5e−03 ** | 3.5e−03 ** |

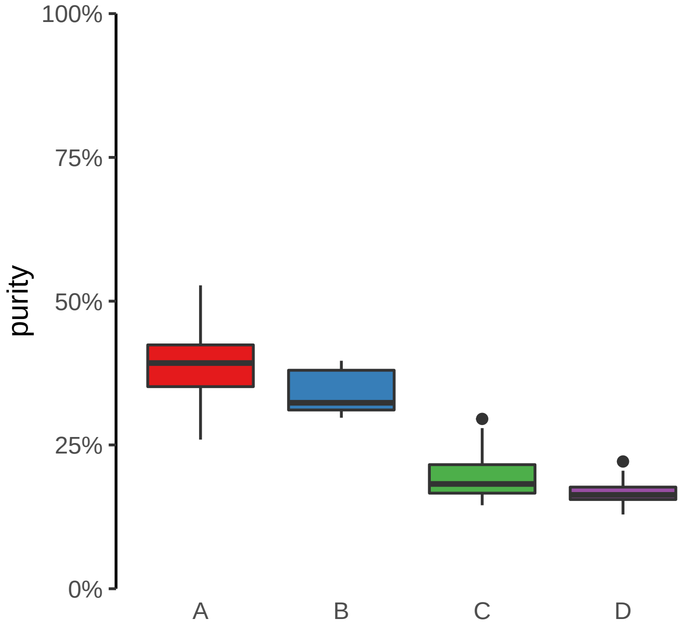

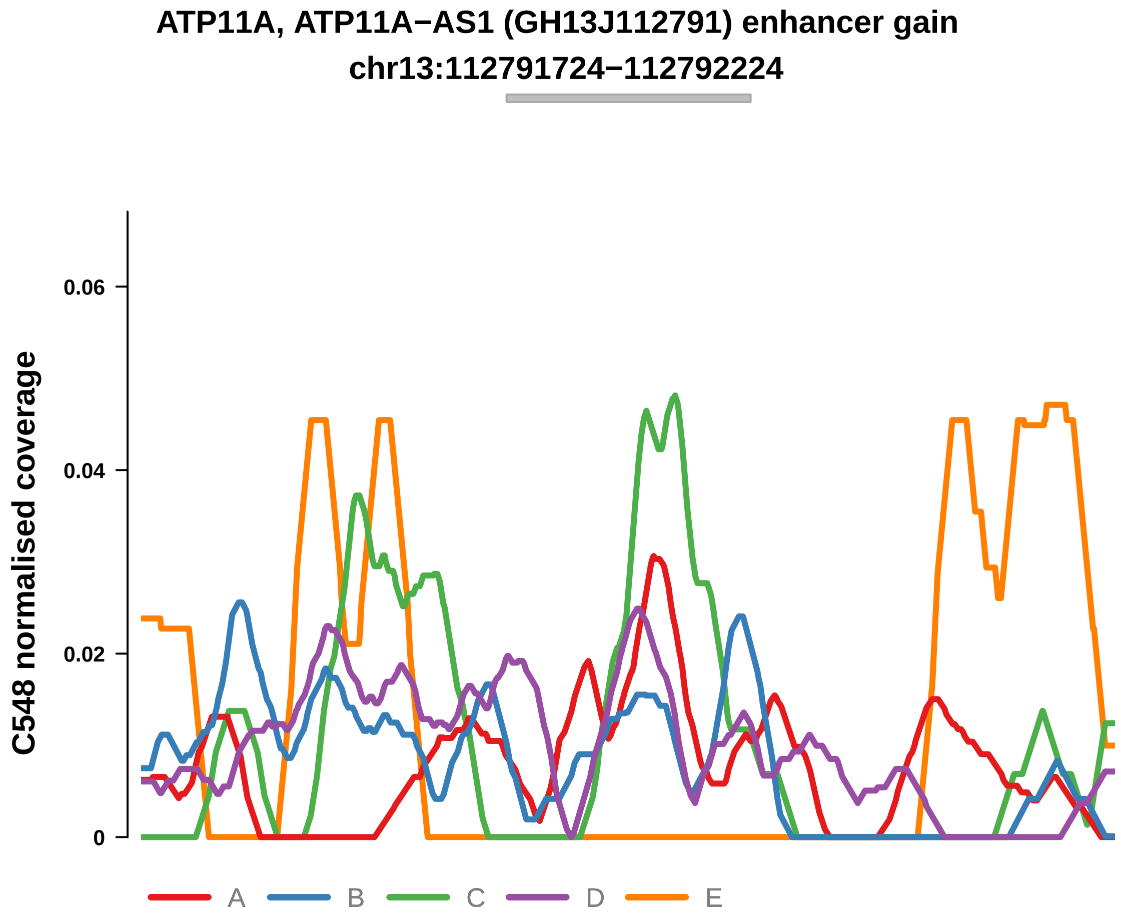


Putatively Clonal (logFC = 2.44)

| | P. | Adjusted P. |
|---|---|---|
| *~region vs. ~* | 4.68e−01 | 5.88e−01 |
| *~purity vs ~* | 5.29e−01 | 6.94e−01 |
| *~purity+region vs ~purity* | 2.65e−01 | 5.31e−01 |
| *Carcinoma vs. normal* | 2.8e−03 ** | 2.8e−03 ** |

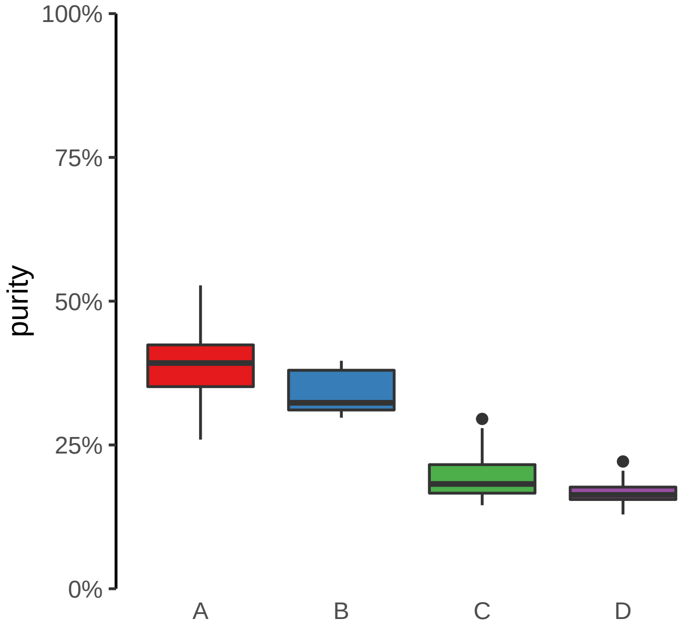

**CLASRP, MARK4 (GH19J045307) enhancer gain**

**chr19:45308035−45308535**

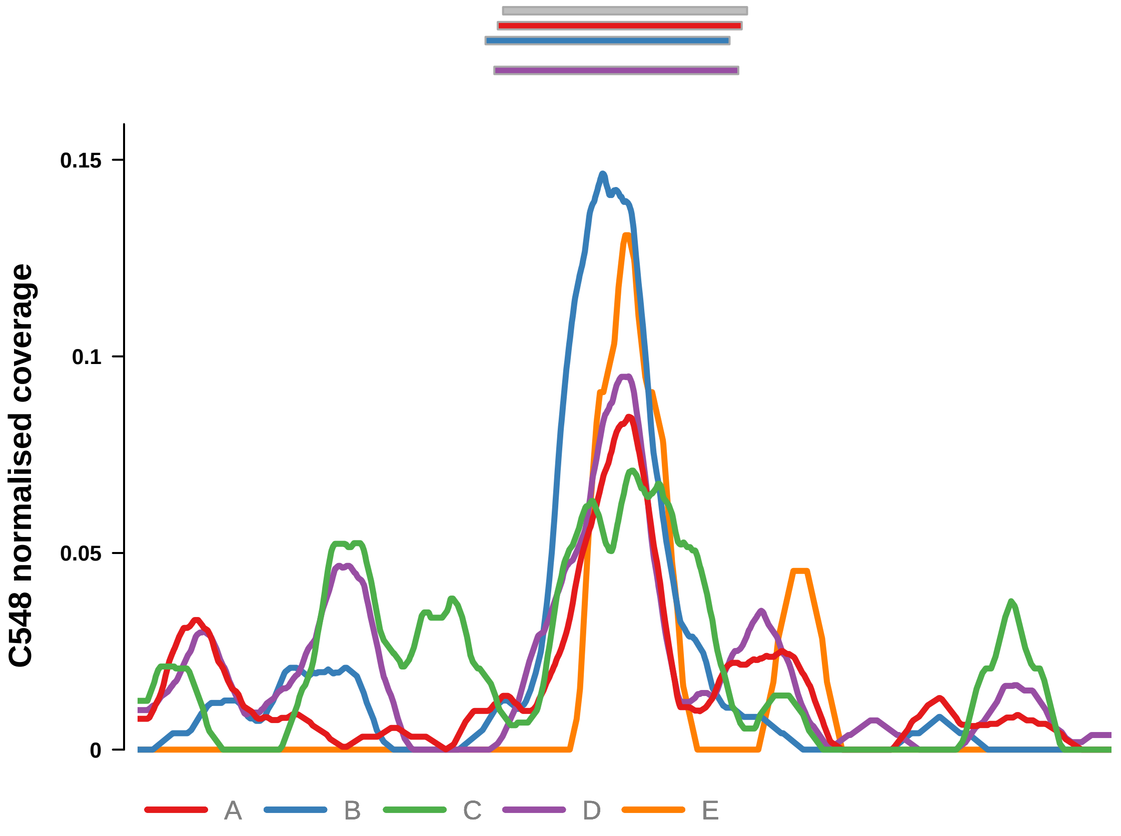


Putatively Clonal (logFC = 2.35)

| | P. | Adjusted P. |
| --- | --- | --- |
| *~region vs. ~* | 4.87e−01 | 6.04e−01 |
| *~purity vs ~* | 6.12e−01 | 7.58e−01 |
| *~purity+region vs ~purity* | 1.49e−01 | 4.18e−01 |
| *Carcinoma vs. normal* | 1.43e−03 ** | 1.43e−03 ** |

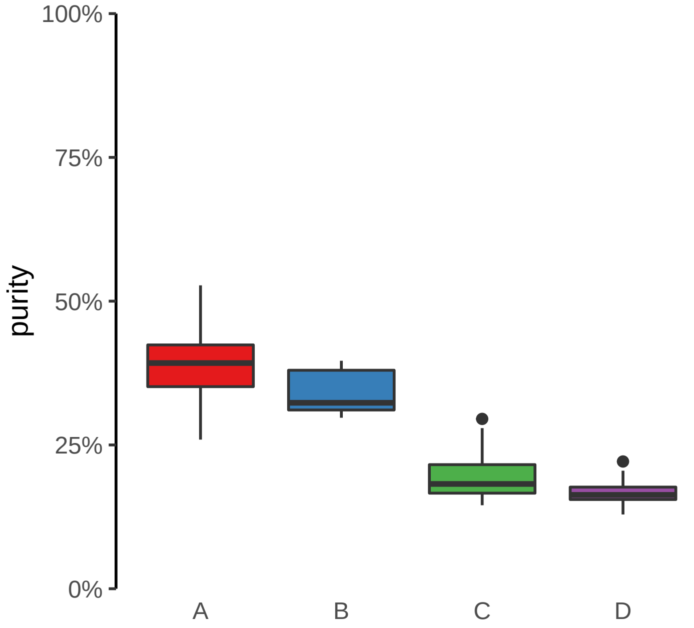

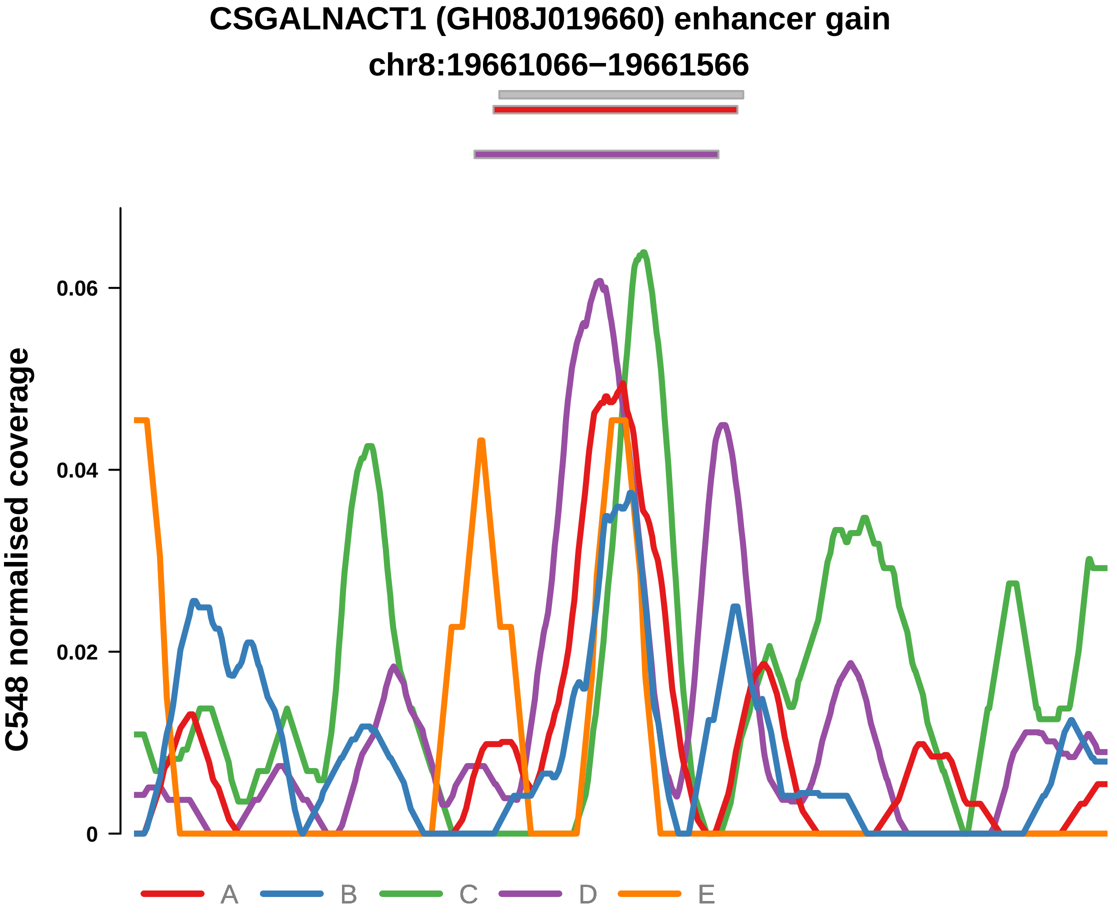


Putatively Clonal (logFC = 2.11)

| | P. | Adjusted P. |
|---|---|---|
| *~region vs. ~* | 6.93e−02 | 1.82e−01 |
| *~purity vs ~* | 2.39e−01 | 3.97e−01 |
| *~purity+region vs ~purity* | 4.5e−02 * | 2.77e−01 |
| *Carcinoma vs. normal* | 7.91e−03 ** | 7.91e−03 ** |

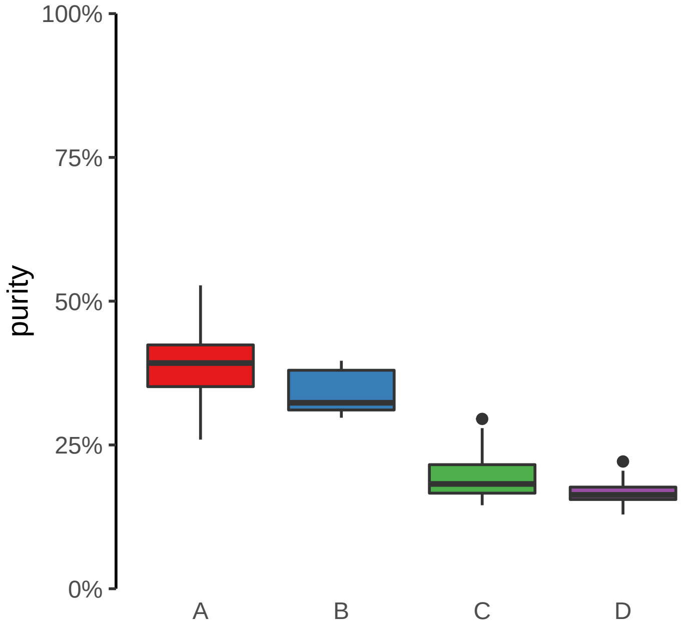

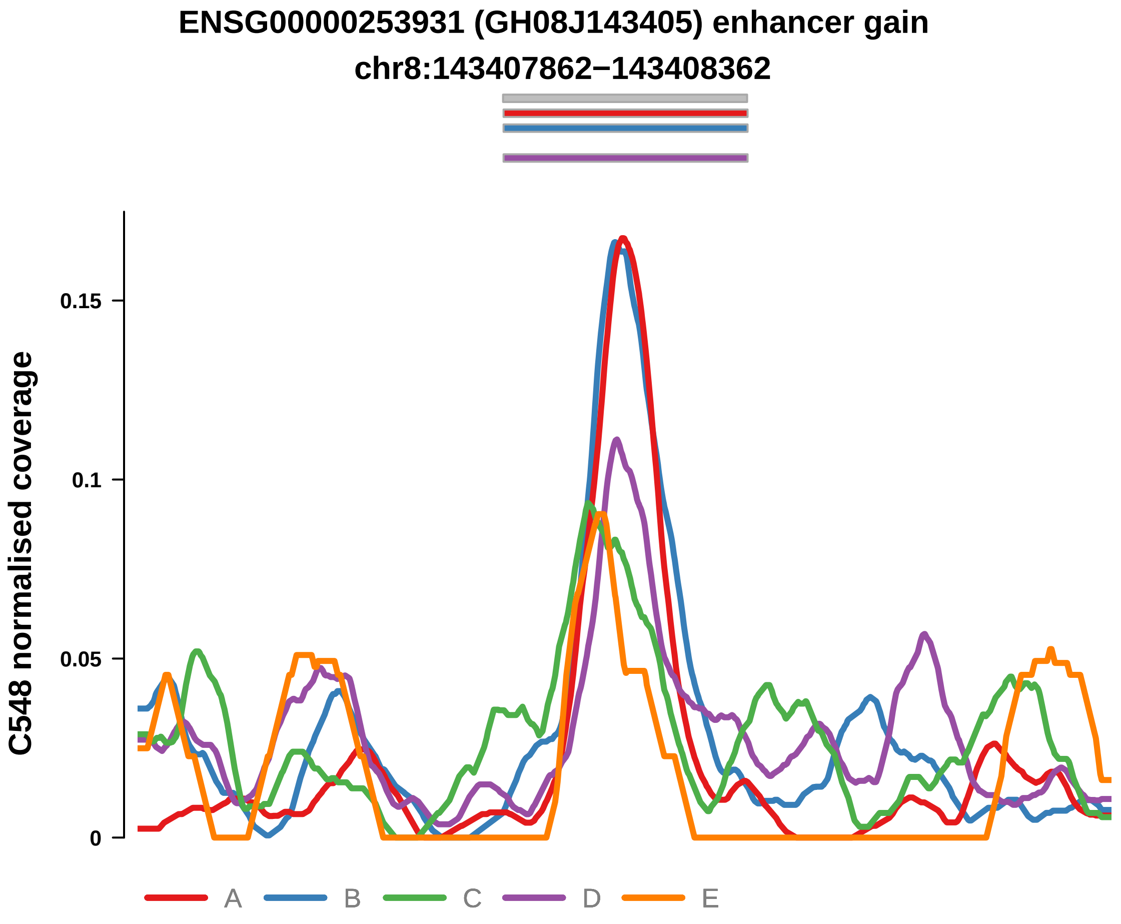


Putatively Clonal (logFC = 1.87)

| | P. | Adjusted P. |
|---|---|---|
| *~region vs. ~* | 4.68e−01 | 5.88e−01 |
| *~purity vs ~* | 7.45e−01 | 8.4e−01 |
| *~purity+region vs ~purity* | 4.51e−01 | 6.41e−01 |
| *Carcinoma vs. normal* | 8.5e−03 ** | 8.5e−03 ** |

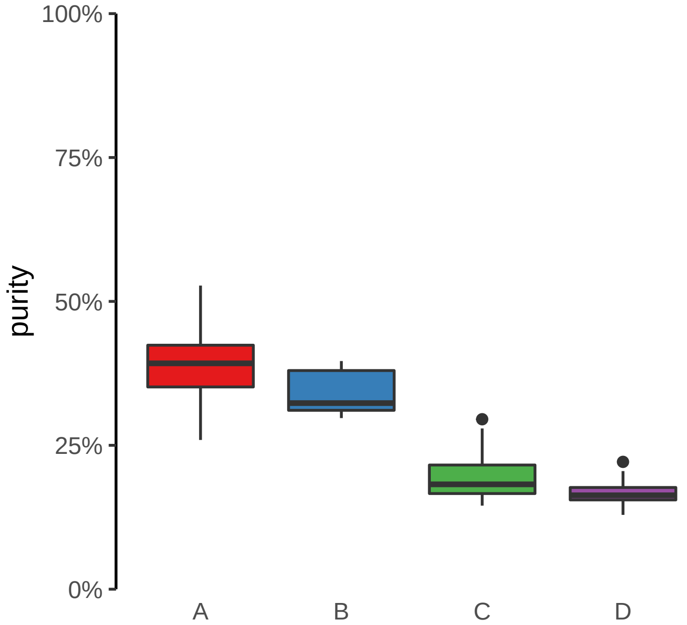

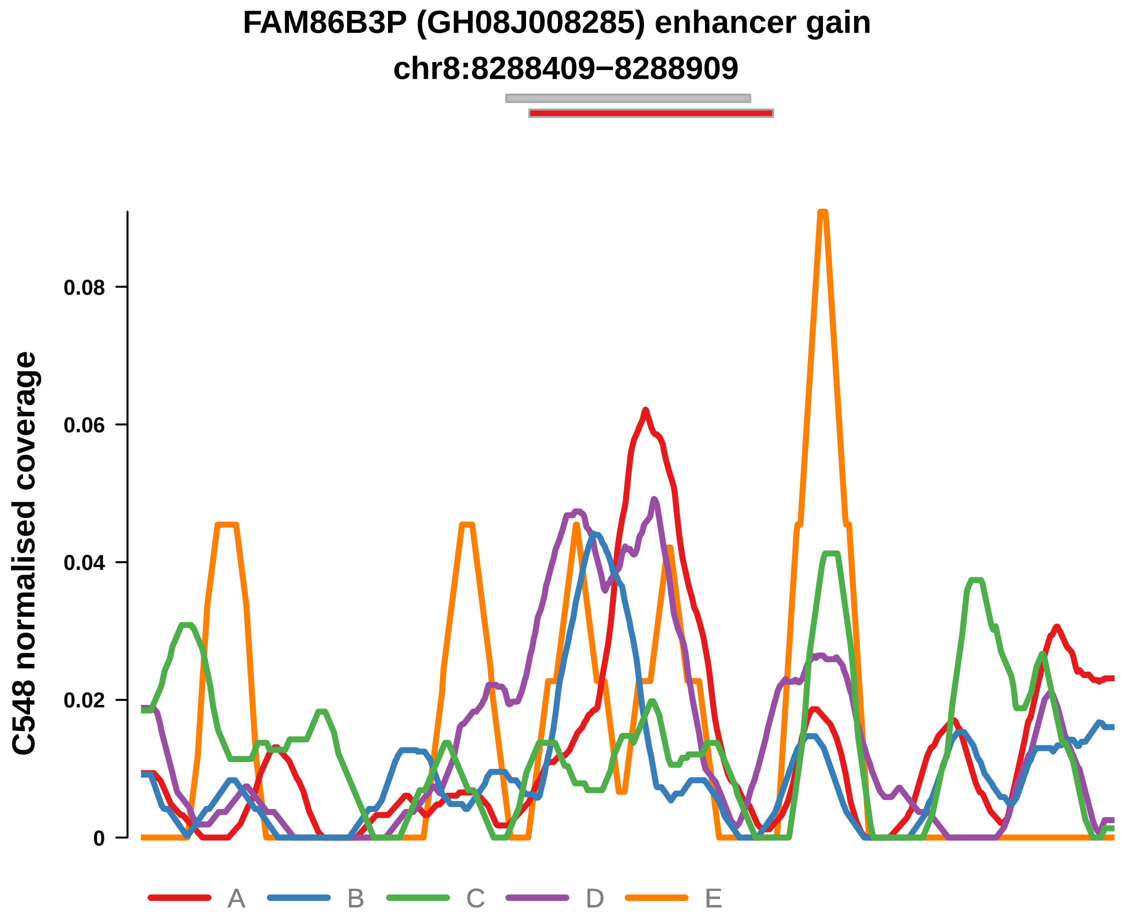


Putatively Clonal (logFC = 2.78)

| | P. | Adjusted P. |
|---|---|---|
| *~region vs. ~* | 9.78e−03 ** | 5.74e−02 |
| *~purity vs ~* | 4.85e−01 | 6.67e−01 |
| *~purity+region vs ~purity* | 4.52e−03 ** | 6.63e−02 |
| *Carcinoma vs. normal* | 3.51e−04 *** | 3.51e−04 *** |

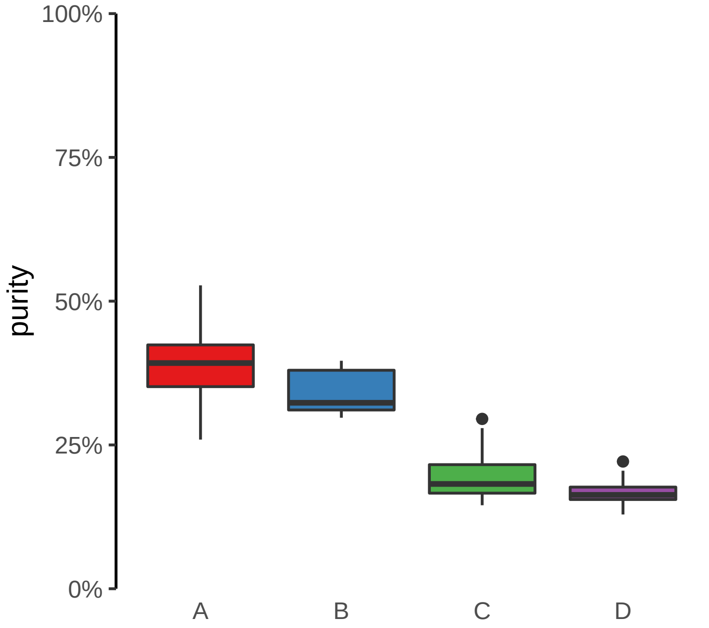

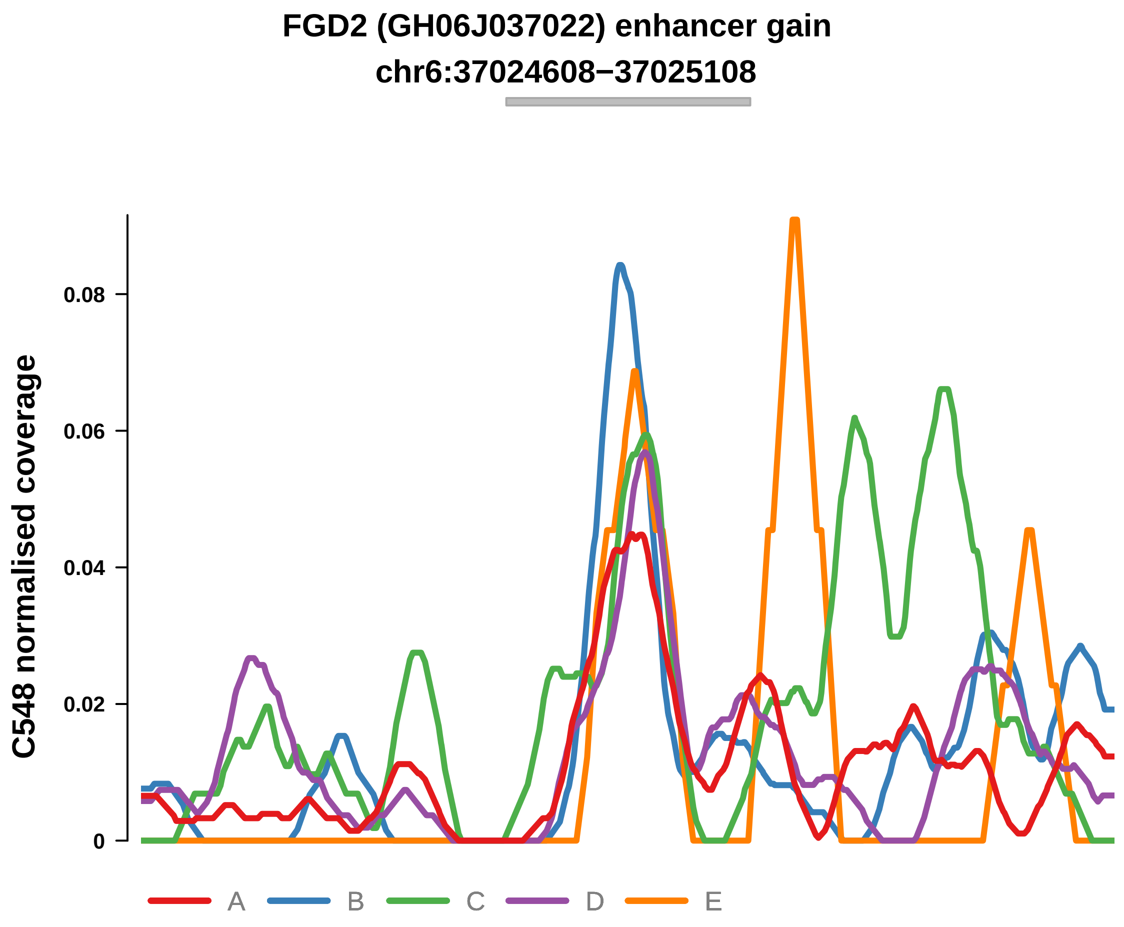


Putatively Clonal (logFC = 2.22)

| | P. | Adjusted P. |
|---|---|---|
| *~region vs. ~* | 2.73e−01 | 3.7e−01 |
| *~purity vs ~* | 3.68e−01 | 5.59e−01 |
| *~purity+region vs ~purity* | 1.39e−01 | 4.07e−01 |
| *Carcinoma vs. normal* | 4.74e−03 ** | 4.74e−03 ** |

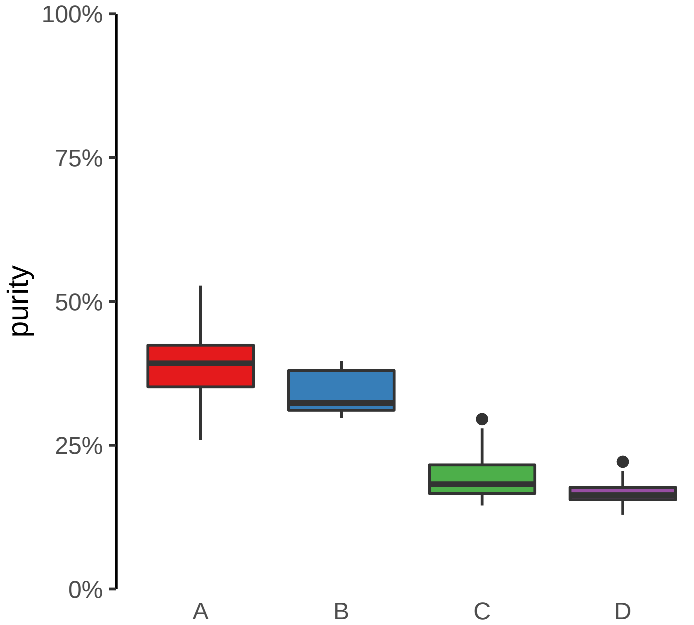

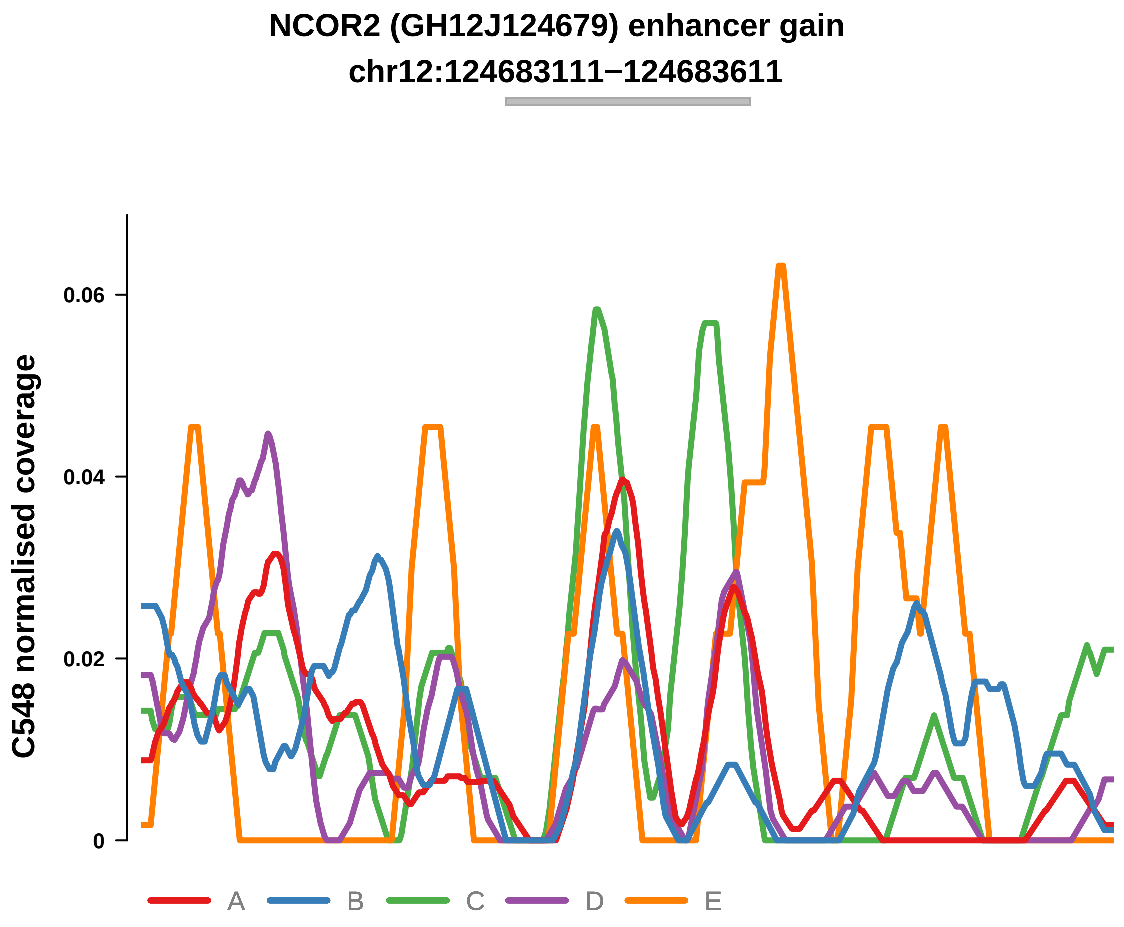


Putatively Clonal (logFC = 2.05)

| | P. | Adjusted P. |
|---|---|---|
| *~region vs. ~* | 1.66e−01 | 2.63e−01 |
| *~purity vs ~* | 5.27e−01 | 6.94e−01 |
| *~purity+region vs ~purity* | 2.38e−01 | 5.31e−01 |
| *Carcinoma vs. normal* | 8.74e−03 ** | 8.74e−03 ** |

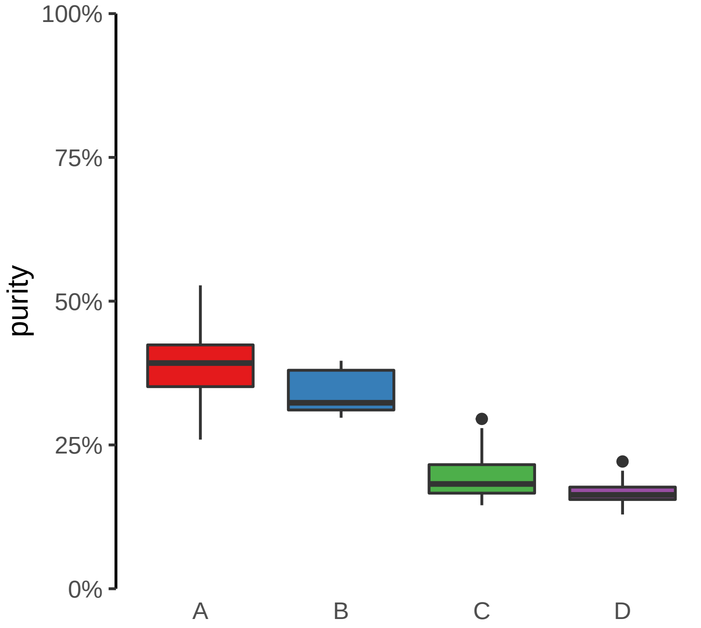

**PPL (GH16J004942) enhancer gain**
**chr16:4944634−4945134**

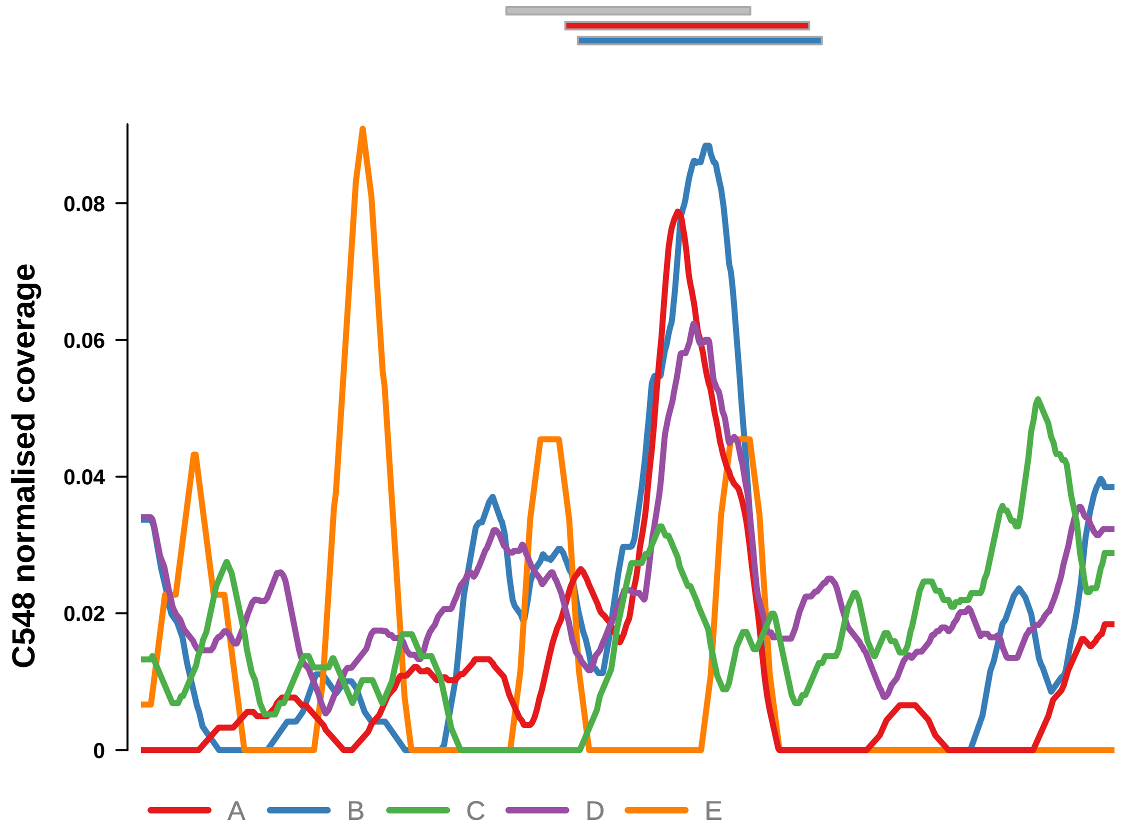


Putatively Clonal (logFC = 1.92)

| | P. | Adjusted P. |
|---|---|---|
| *~region vs. ~* | 1.89e−02 * | 8.48e−02 |
| *~purity vs ~* | 4.96e−01 | 6.71e−01 |
| *~purity+region vs ~purity* | 1.64e−02 * | 1.32e−01 |
| *Carcinoma vs. normal* | 8.7e−03 ** | 8.7e−03 ** |

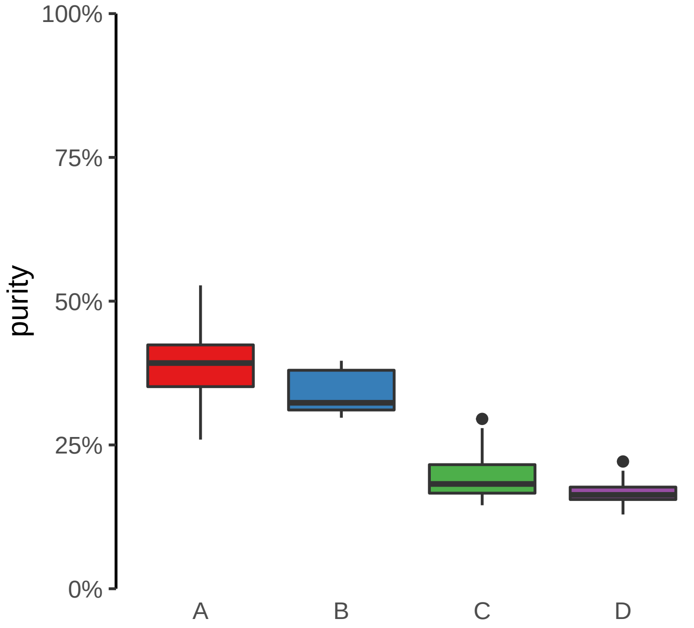

**ATP11A, ATP11A−AS1 (GH13J112791) enhancer gain**
**chr13:112791724−112792224**

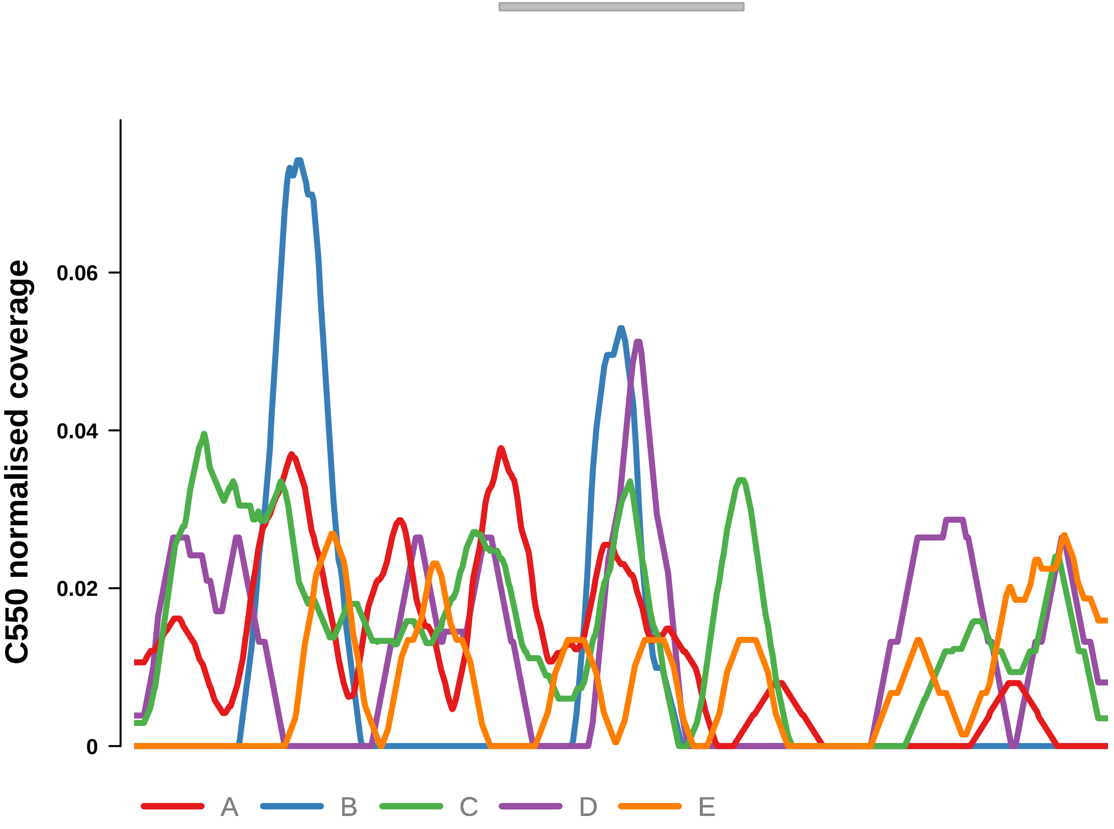


Putatively Clonal (logFC = 2.26)

| | P. | Adjusted P. |
|---|---|---|
| *~region vs. ~* | 7.81e−01 | 9.39e−01 |
| *~purity vs ~* | 1.39e−01 | 5.11e−01 |
| *~purity+region vs ~purity* | 9.92e−01 | 1e+00 |
| *Carcinoma vs. normal* | 6.05e−03 ** | 6.05e−03 ** |

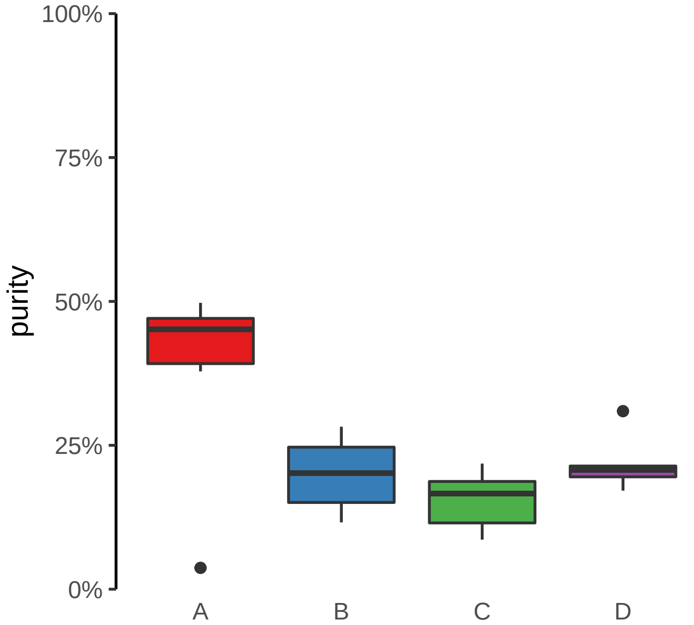

**CSGALNACT1 (GH08J019660) enhancer gain**
**chr8:19661066−19661566**

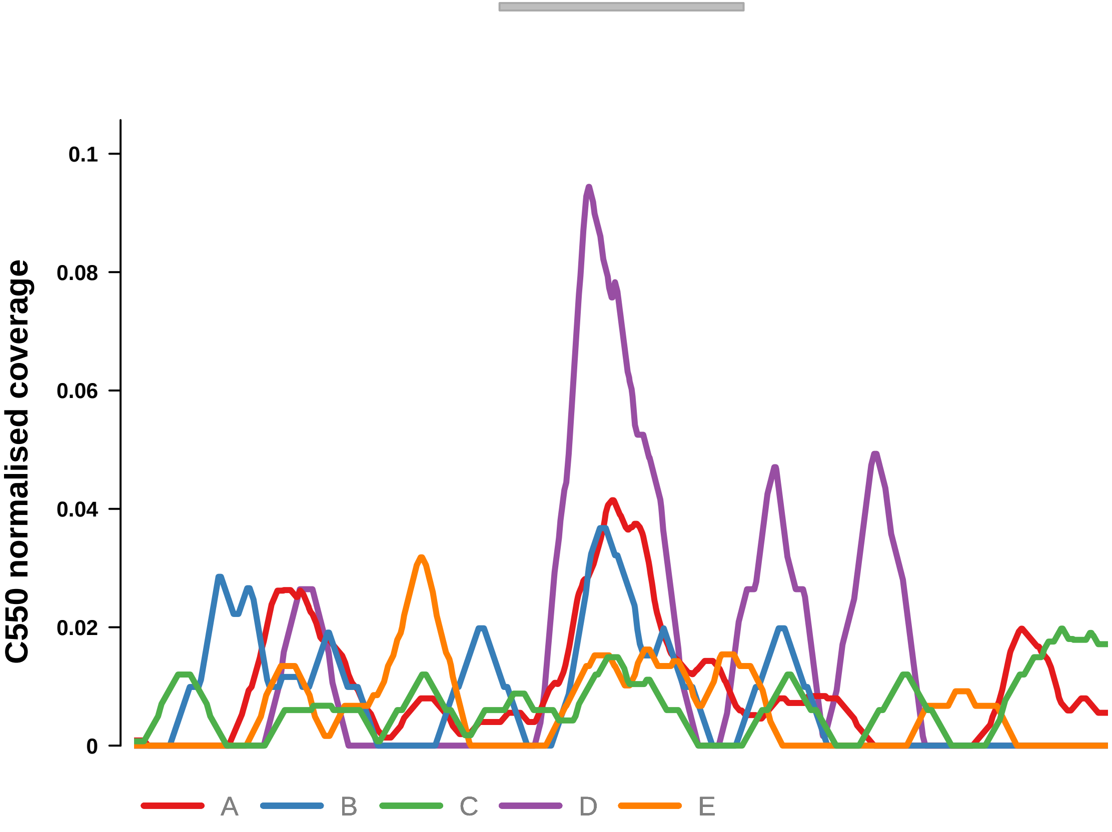


Putatively Clonal (logFC = 2.11)

| | P. | Adjusted P. |
|---|---|---|
| *~region vs. ~* | 5.38e−02 | 5.8e−01 |
| *~purity vs ~* | 1.55e−01 | 5.46e−01 |
| *~purity+region vs ~purity* | 1.36e−01 | 7.48e−01 |
| *Carcinoma vs. normal* | 7.68e−03 ** | 7.68e−03 ** |

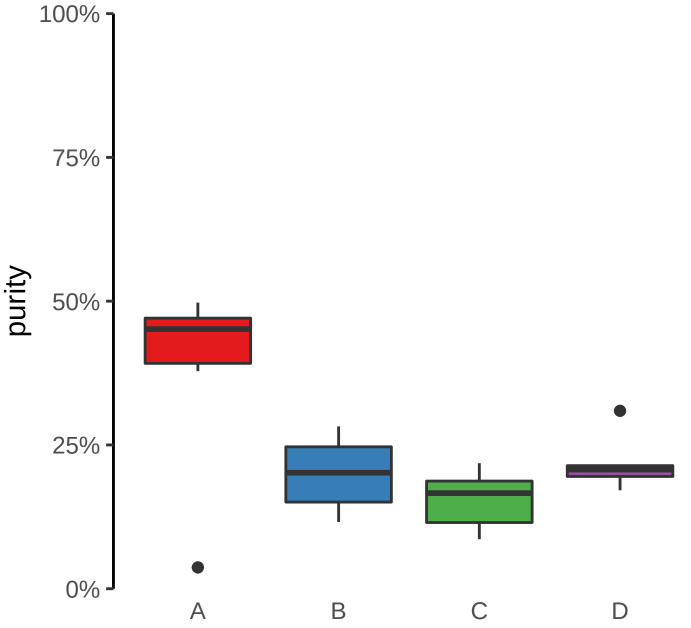

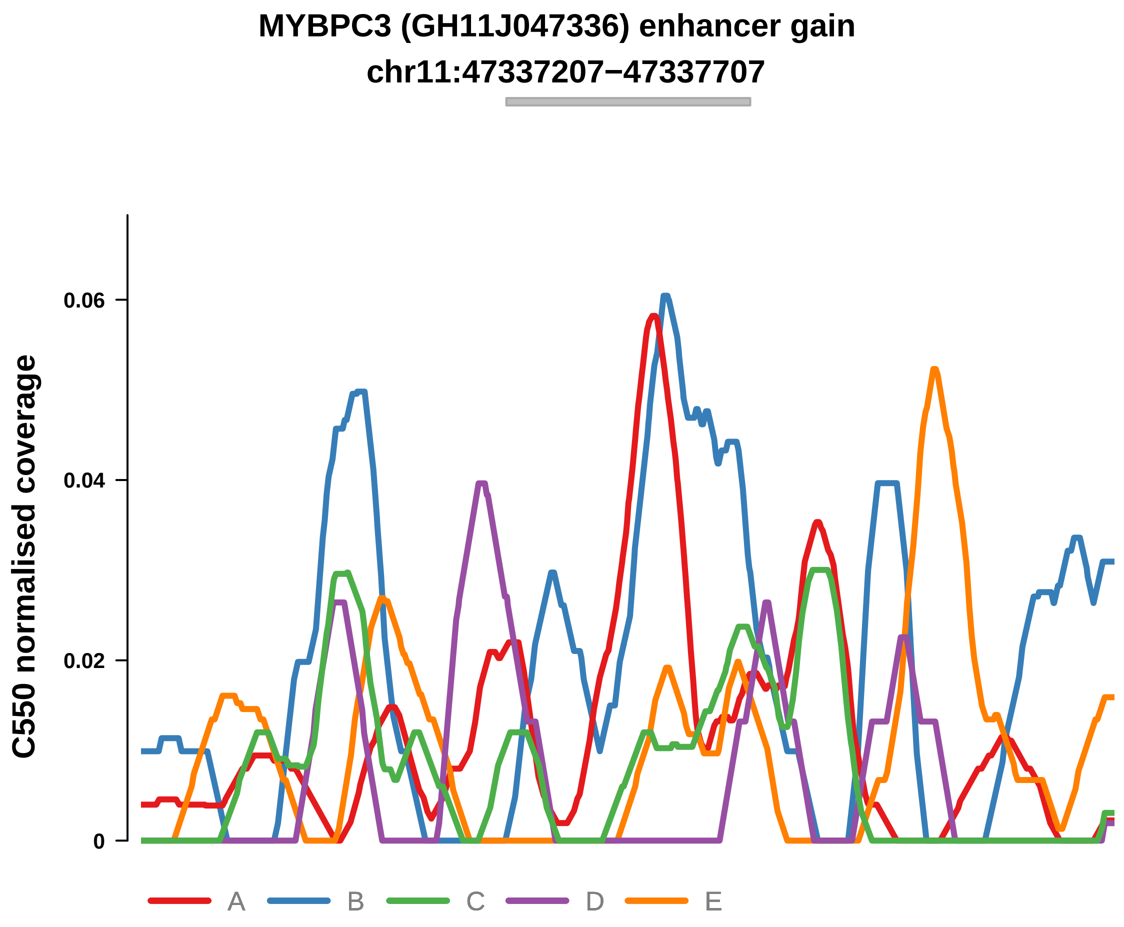


Putatively Clonal (logFC = 2.14)

| | P. | Adjusted P. |
|---|---|---|
| *~region vs. ~* | 1.32e−01 | 7.77e−01 |
| *~purity vs ~* | 2.72e−02 * | 3.42e−01 |
| *~purity+region vs ~purity* | 3.78e−01 | 9.78e−01 |
| *Carcinoma vs. normal* | 7.99e−03 ** | 7.99e−03 ** |

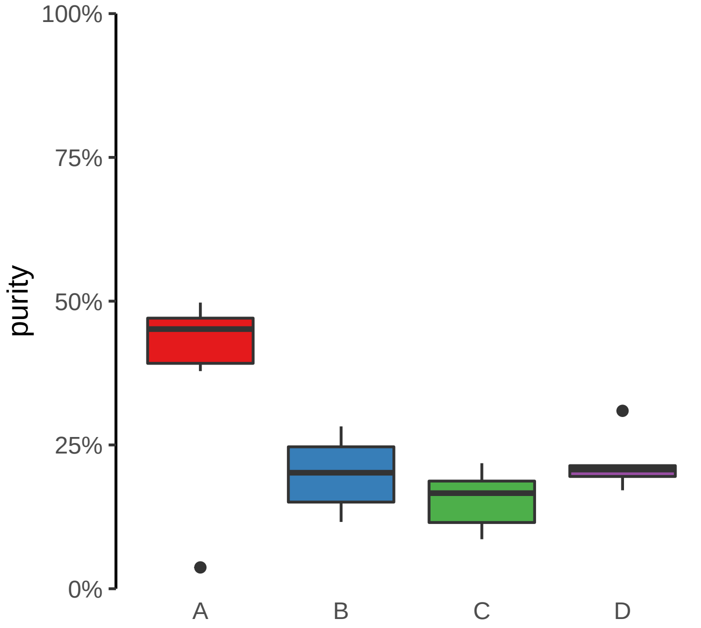

**MYT1 (GH20J064188) enhancer gain**
**chr20:64189162−64189662**

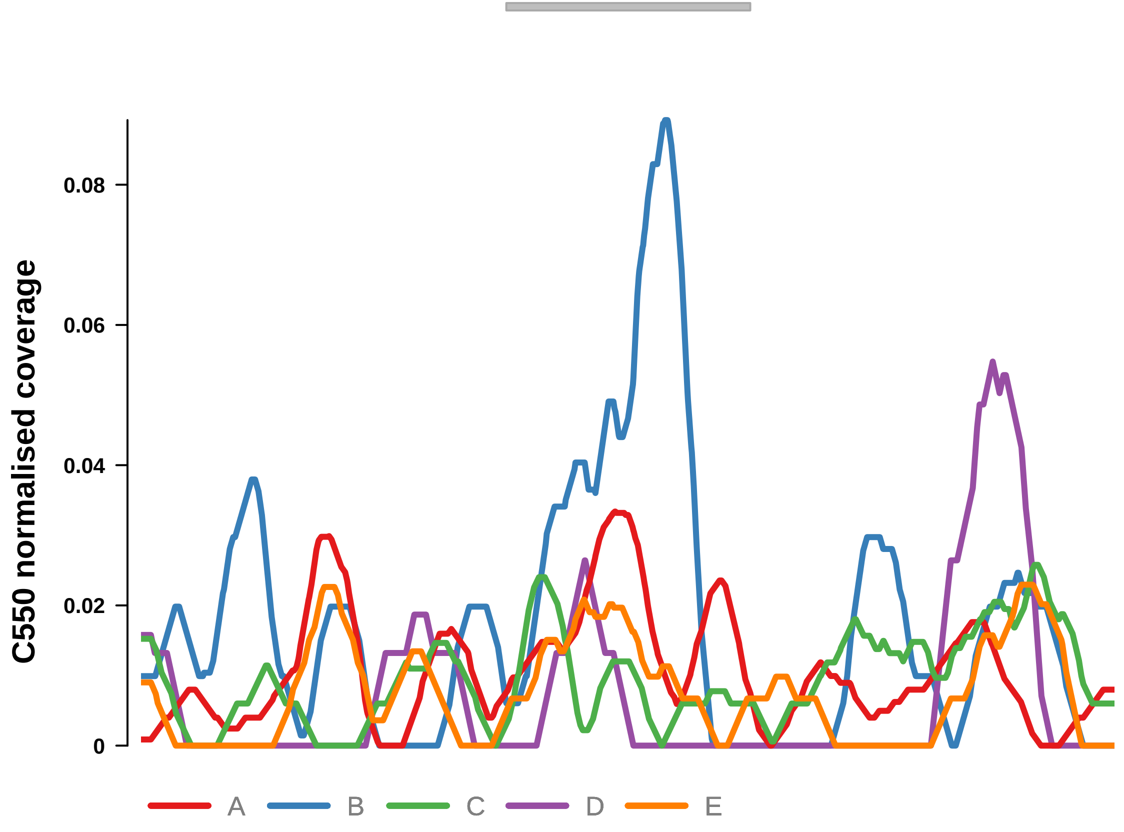


Putatively Clonal (logFC = 2.4)

| | P. | Adjusted P. |
|---|---|---|
| *~region vs. ~* | 5.26e−01 | 9.11e−01 |
| *~purity vs ~* | 5.76e−01 | 8.71e−01 |
| *~purity+region vs ~purity* | 6.29e−01 | 9.87e−01 |
| *Carcinoma vs. normal* | 3.48e−03 ** | 3.48e−03 ** |

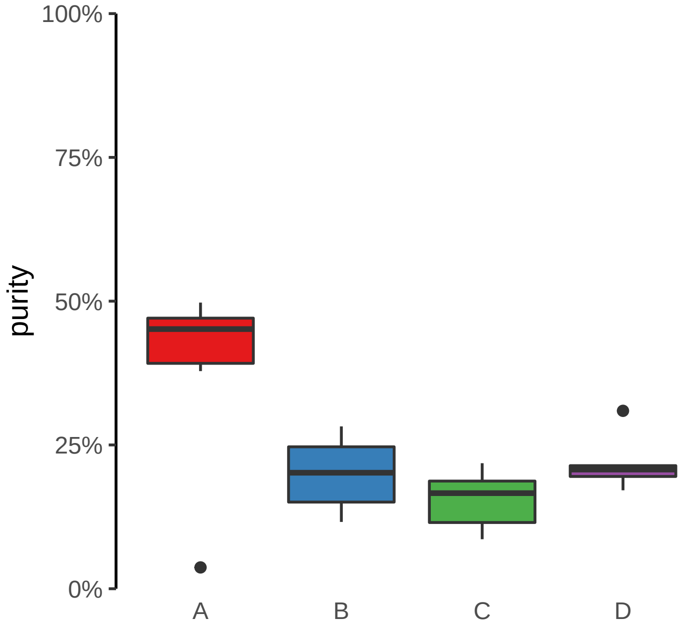

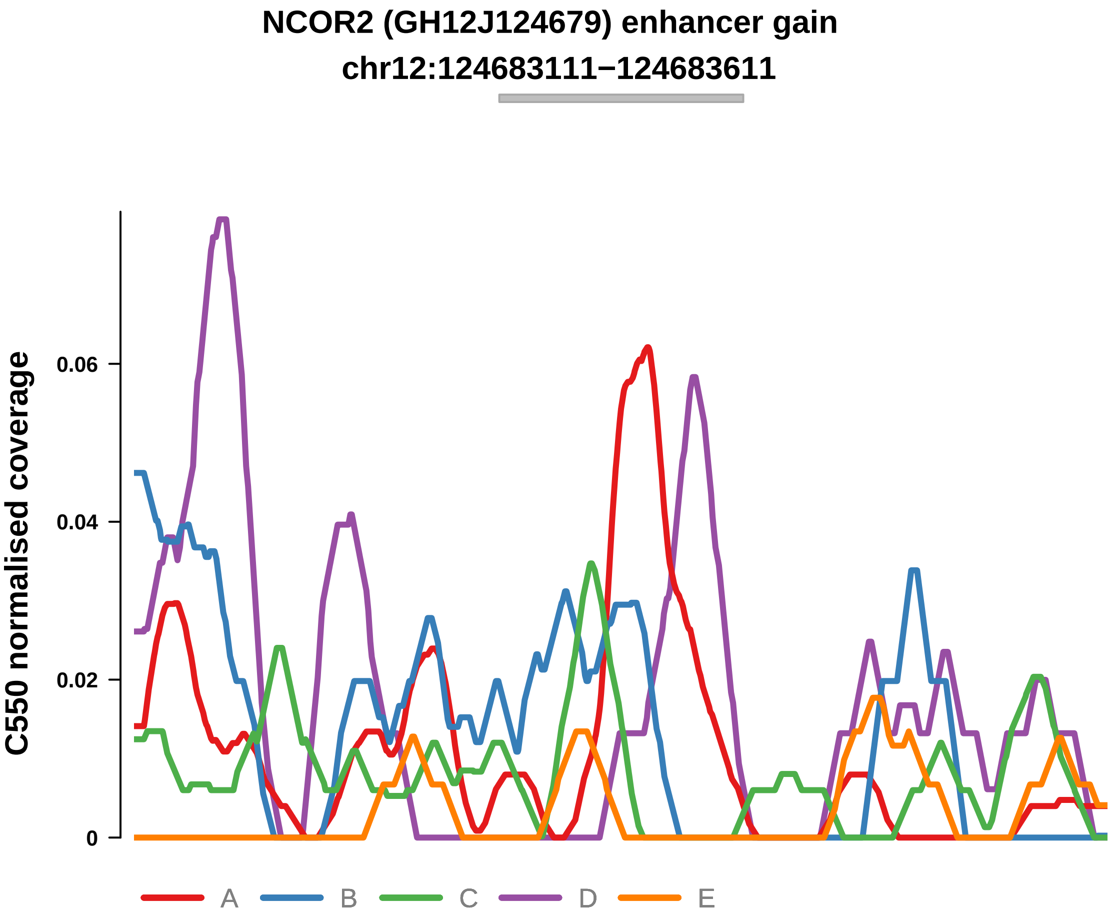


Putatively Clonal (logFC = 2.2)

| | P. | Adjusted P. |
|---|---|---|
| *~region vs. ~* | 3.02e−01 | 8.27e−01 |
| *~purity vs ~* | 1.27e−02 * | 2.79e−01 |
| *~purity+region vs ~purity* | 4.95e−01 | 9.87e−01 |
| *Carcinoma vs. normal* | 4.93e−03 ** | 4.93e−03 ** |

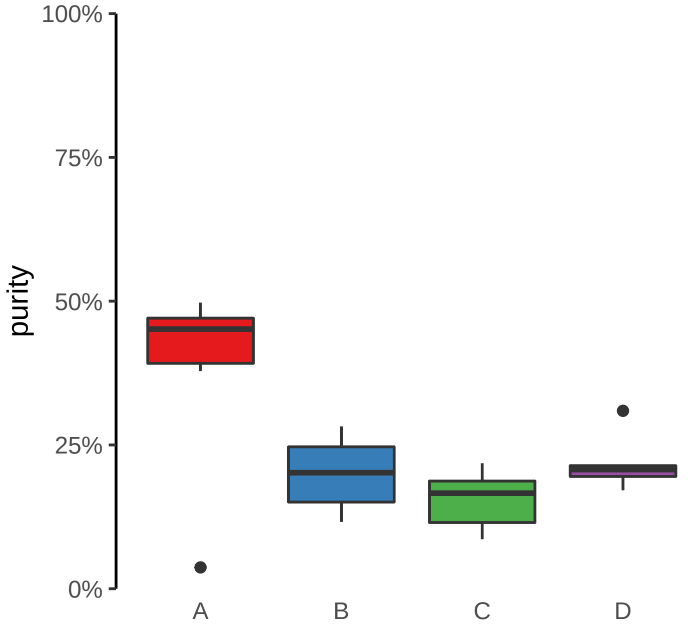

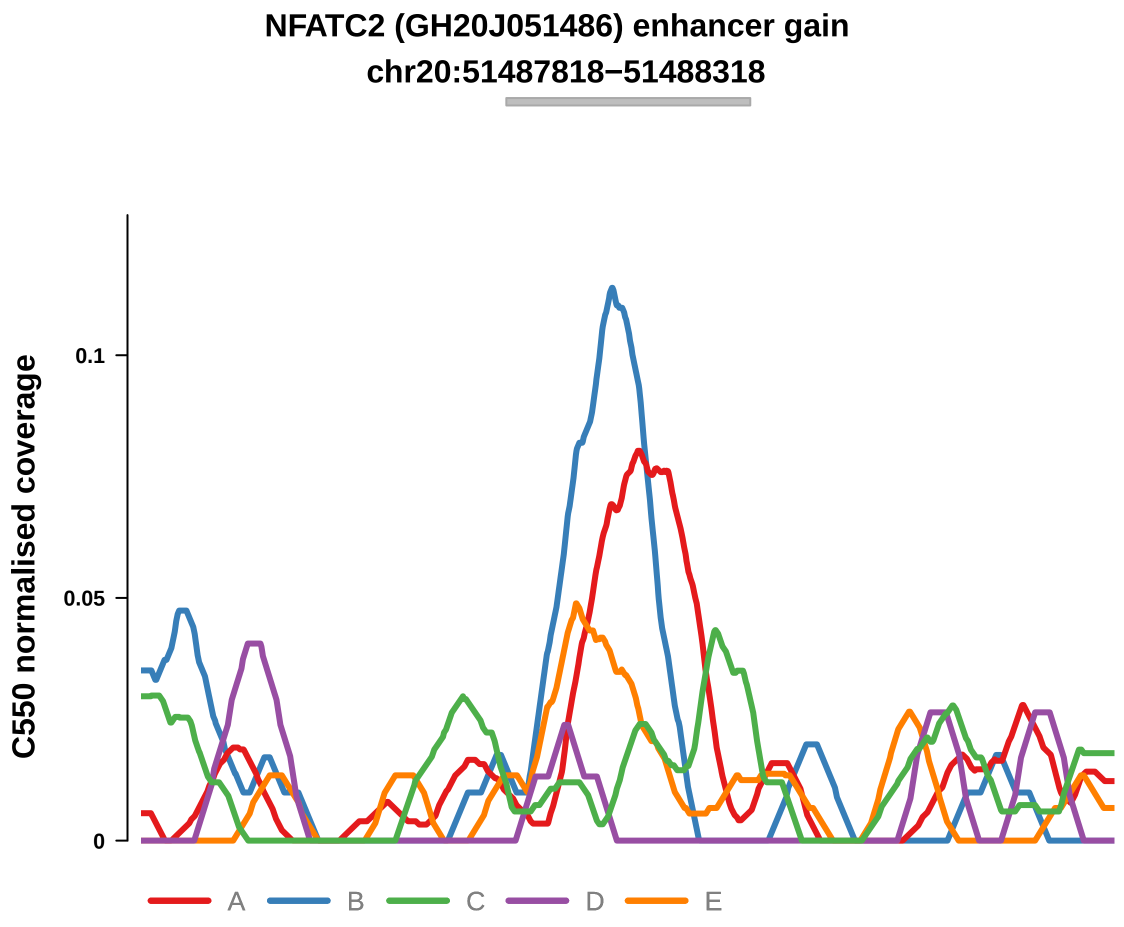


Putatively Clonal (logFC = 3.16)

| | P. | Adjusted P. |
|---|---|---|
| *~region vs. ~* | 2.89e−03 ** | 1.31e−01 |
| *~purity vs ~* | 1.26e−02 * | 2.79e−01 |
| *~purity+region vs ~purity* | 5.93e−02 | 7.48e−01 |
| *Carcinoma vs. normal* | 4.98e−05 **** | 4.98e−05 **** |

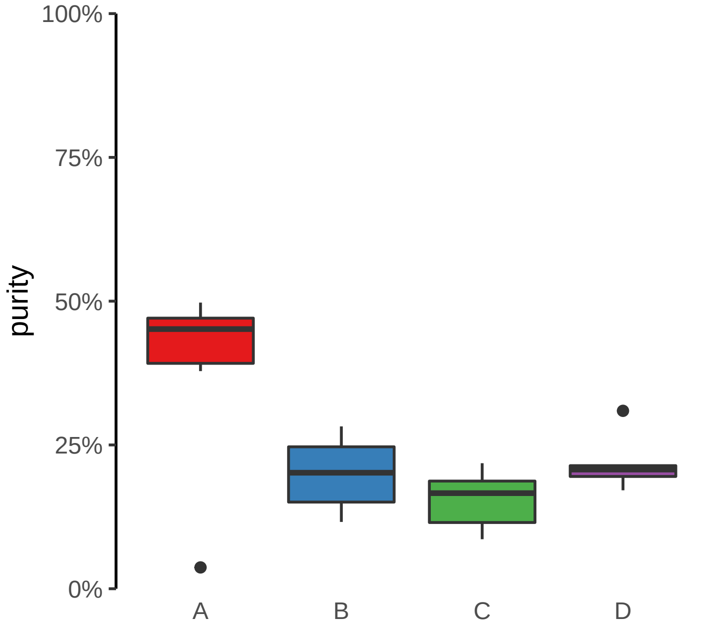

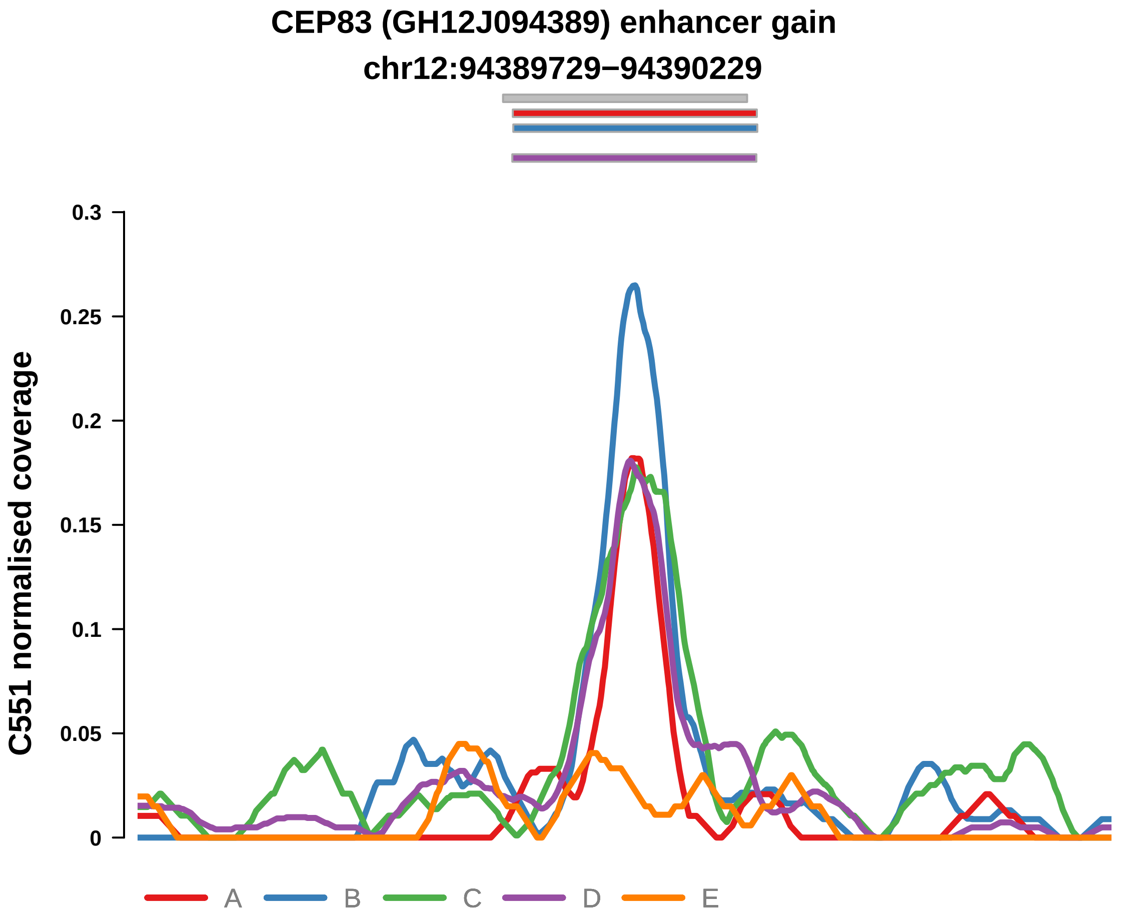


Putatively Clonal (logFC = 3.28)

| | P. | Adjusted P. |
|---|---|---|
| *~region vs. ~* | 9.73e−02 | 4.16e−01 |
| *~purity vs ~* | 3.13e−01 | 9.06e−01 |
| *~purity+region vs ~purity* | 1.62e−01 | 5.7e−01 |
| *Carcinoma vs. normal* | 8.17e−06 **** | 8.17e−06 **** |

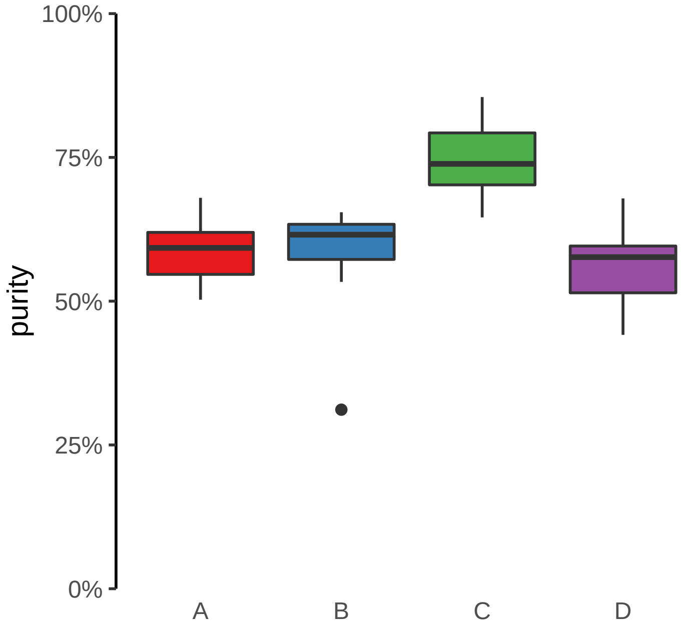

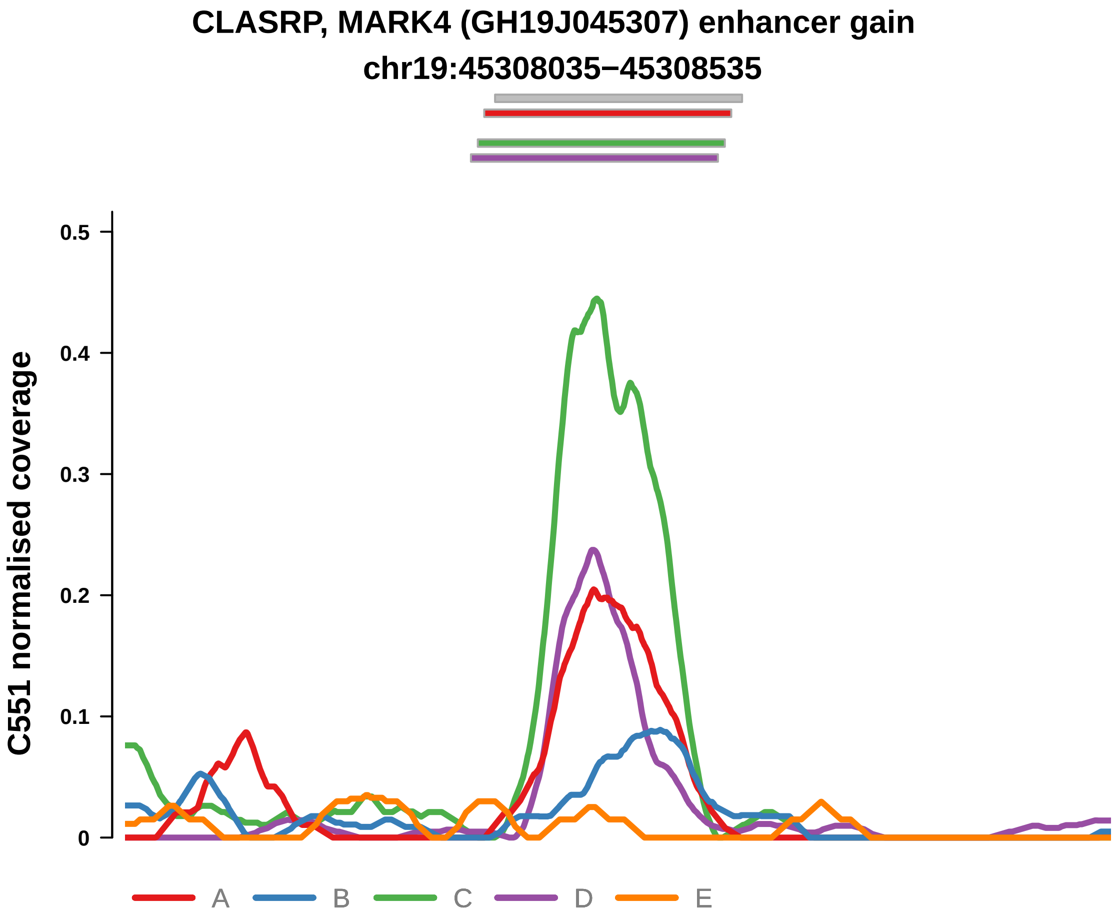


Putatively Clonal (logFC = 2.2)

| | P. | Adjusted P. |
|---|---|---|
| *~region vs. ~* | 4.25e−02 * | 2.67e−01 |
| *~purity vs ~* | 1.16e−01 | 7.97e−01 |
| *~purity+region vs ~purity* | 1.48e−01 | 5.7e−01 |
| *Carcinoma vs. normal* | 1.5e−03 ** | 1.5e−03 ** |

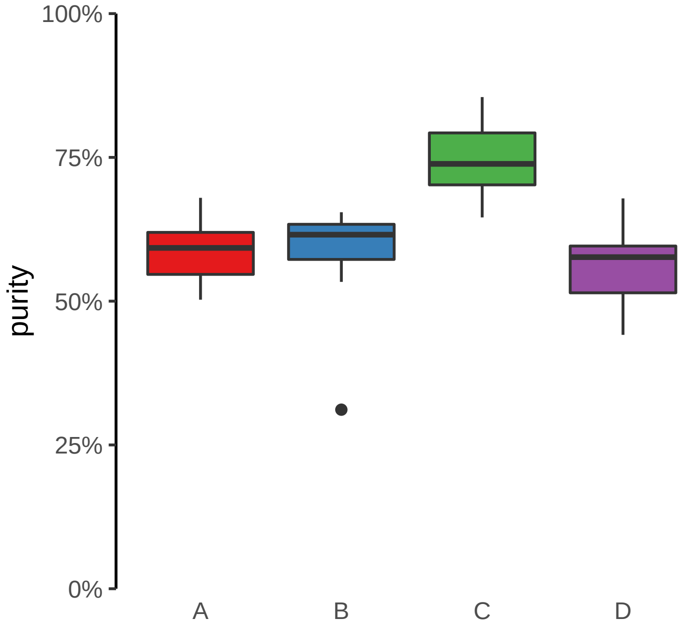

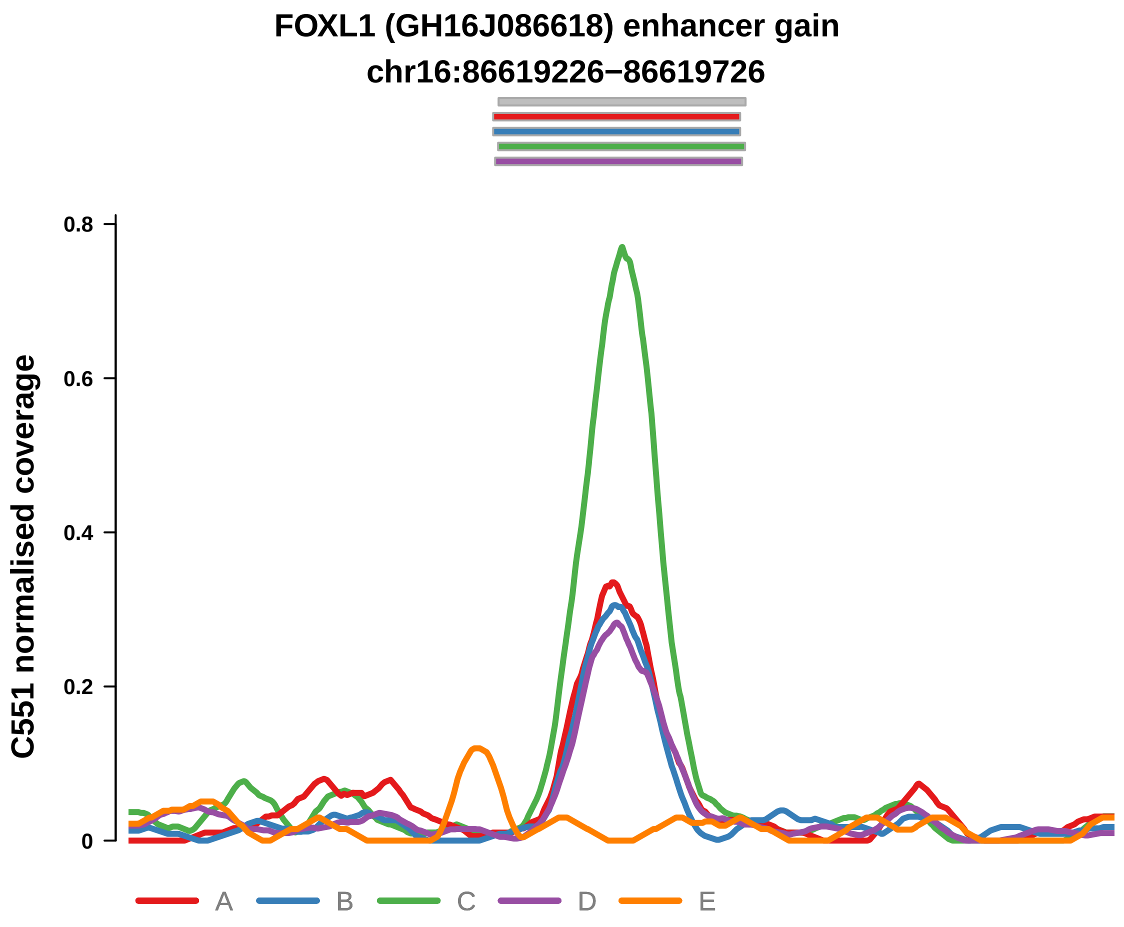


Putatively Clonal (logFC = 3.05)

| | P. | Adjusted P. |
|---|---|---|
| *~region vs. ~* | 3.53e−01 | 7.4e−01 |
| *~purity vs ~* | 3.22e−02 * | 5.38e−01 |
| *~purity+region vs ~purity* | 9.46e−01 | 9.99e−01 |
| *Carcinoma vs. normal* | 2.04e−05 **** | 2.04e−05 **** |

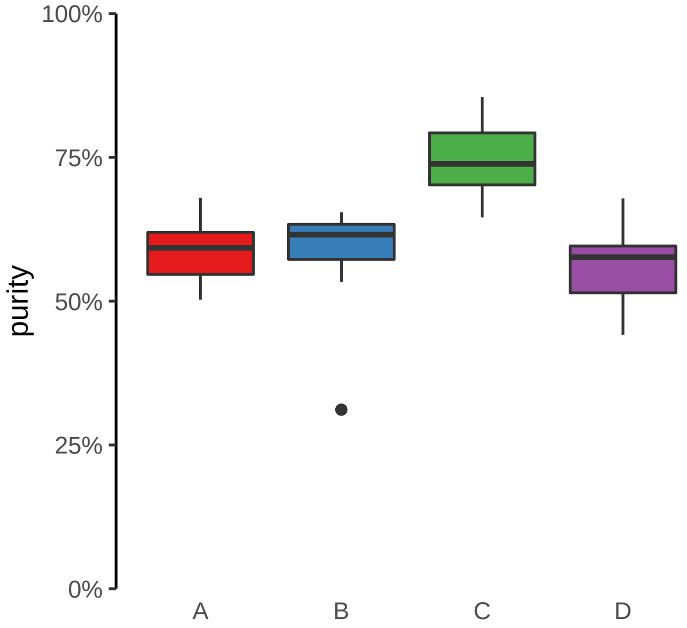

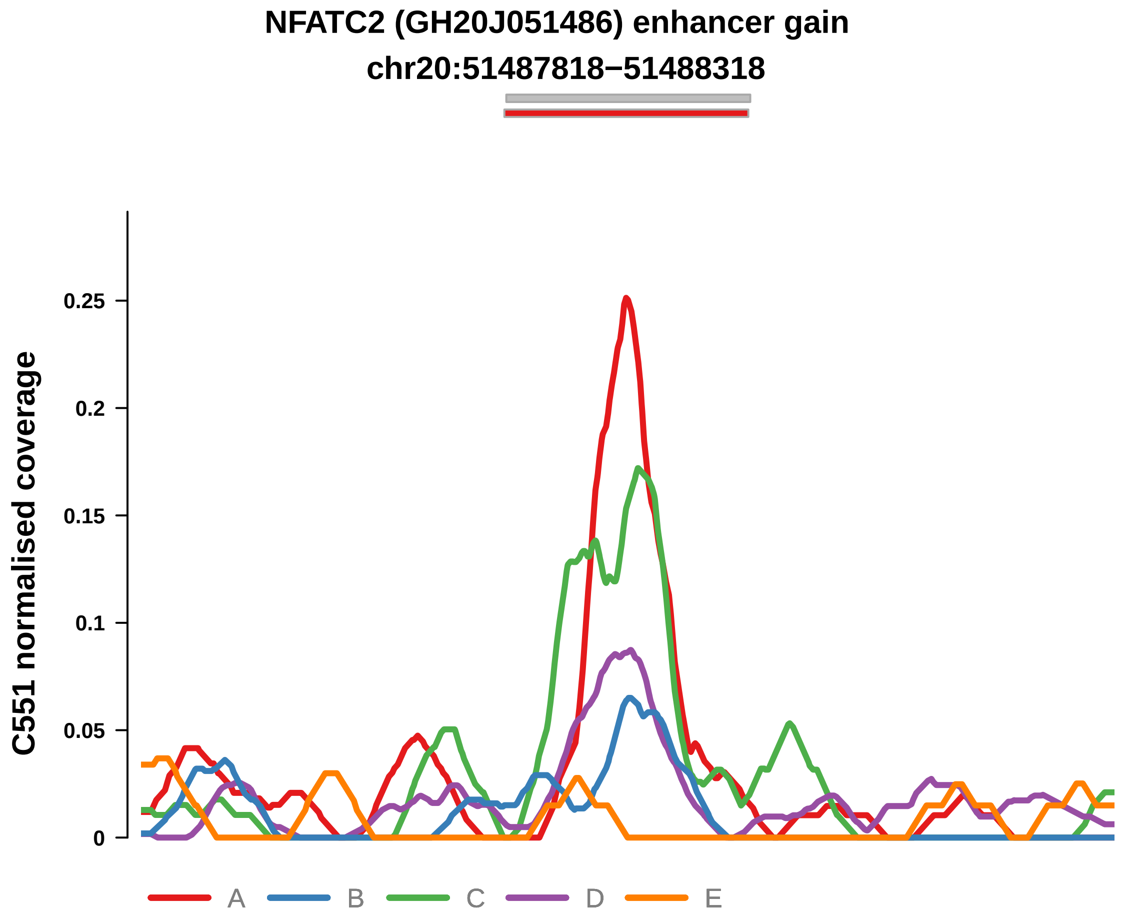


Putatively Clonal (logFC = 2.3)

| | P. | Adjusted P. |
|---|---|---|
| *~region vs. ~* | 1.23e−01 | 4.32e−01 |
| *~purity vs ~* | 2.82e−01 | 9.06e−01 |
| *~purity+region vs ~purity* | 1.82e−01 | 5.7e−01 |
| *Carcinoma vs. normal* | 1.17e−03 ** | 1.17e−03 ** |

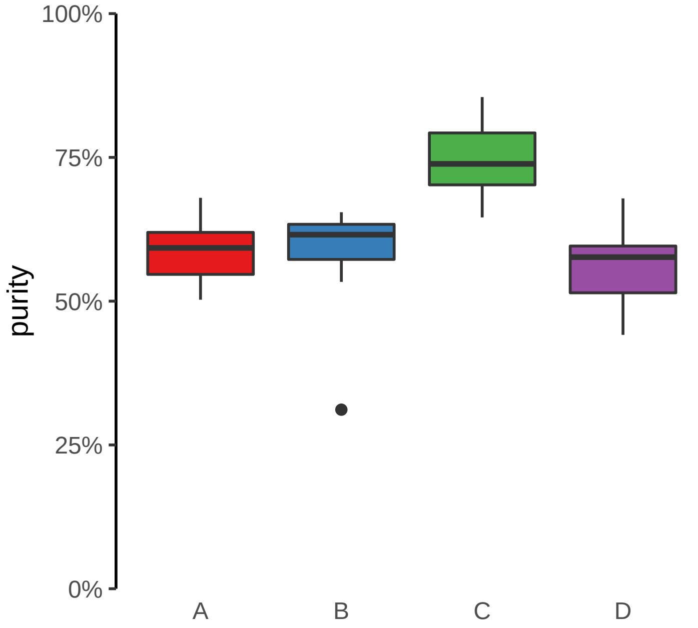

**TEX26 (GH13J030867) enhancer gain**
**chr13:30870352−30870852**

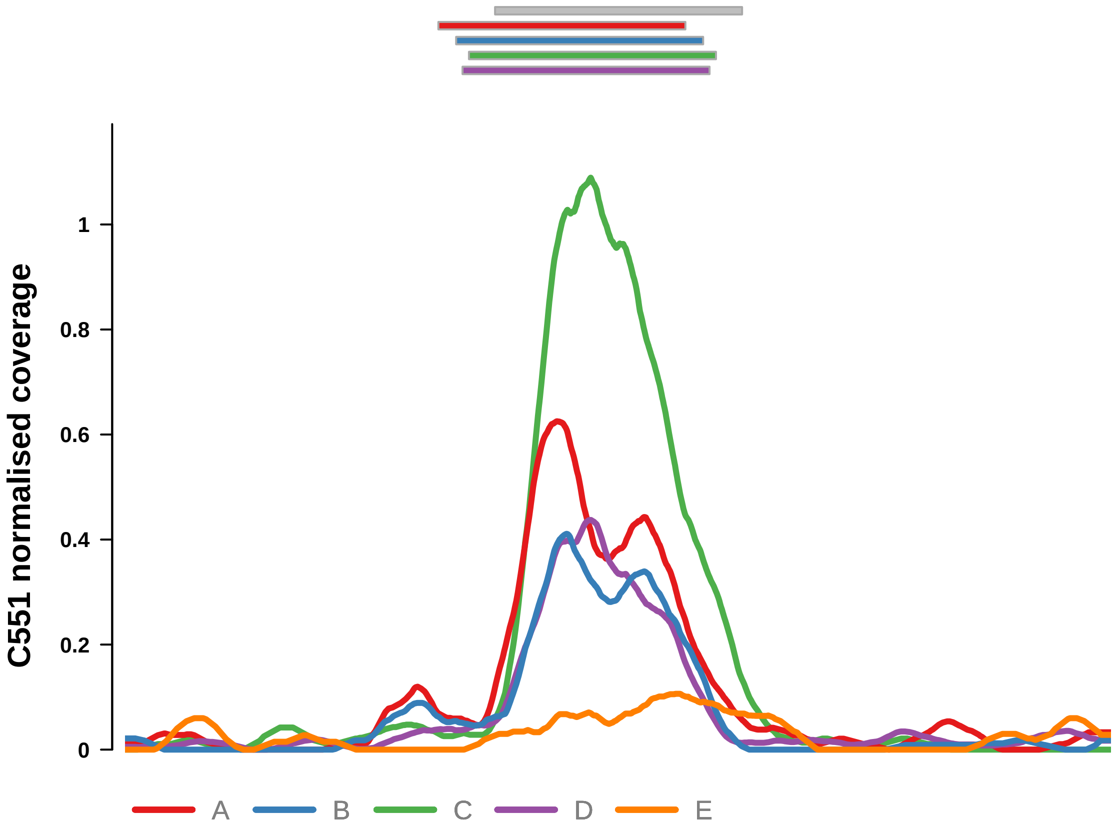


Putatively Clonal (logFC = 2.69)

| | **P.** | **Adjusted P.** |
|---|---|---|
| *~region vs. ~* | 2.37e−02 * | 2.61e−01 |
| *~purity vs ~* | 9.75e−03 ** | 4.89e−01 |
| *~purity+region vs ~purity* | 1.8e−01 | 5.7e−01 |
| *Carcinoma vs. normal* | 1.19e−04 *** | 1.19e−04 *** |

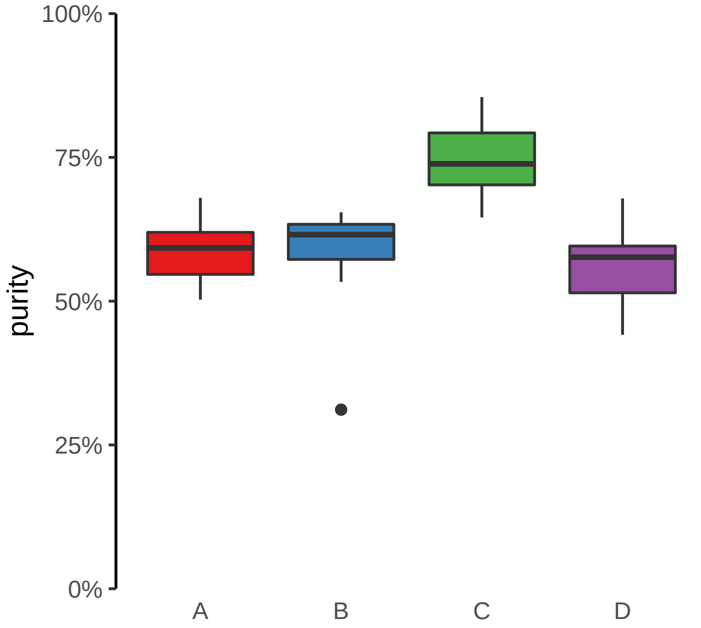

**ATP11A, ATP11A−AS1 (GH13J112791) enhancer gain**
**chr13:112791724−112792224**

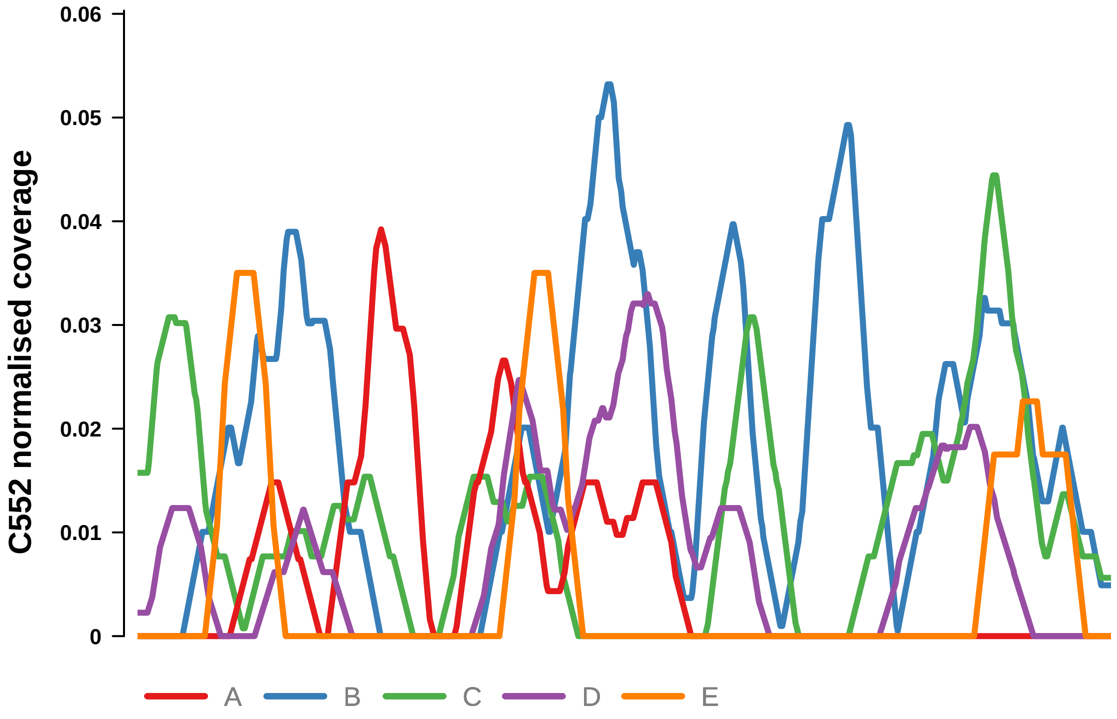


Putatively Clonal (logFC = 2.39)

| | P. | Adjusted P. |
|---|---|---|
| *~region vs. ~* | 1.04e−02 * | NA |
| *~purity vs ~* | 1.68e−01 | NA |
| *~purity+region vs ~purity* | 2.41e−02 * | 3.73e−01 |
| *Carcinoma vs. normal* | 3.77e−03 ** | 3.77e−03 ** |

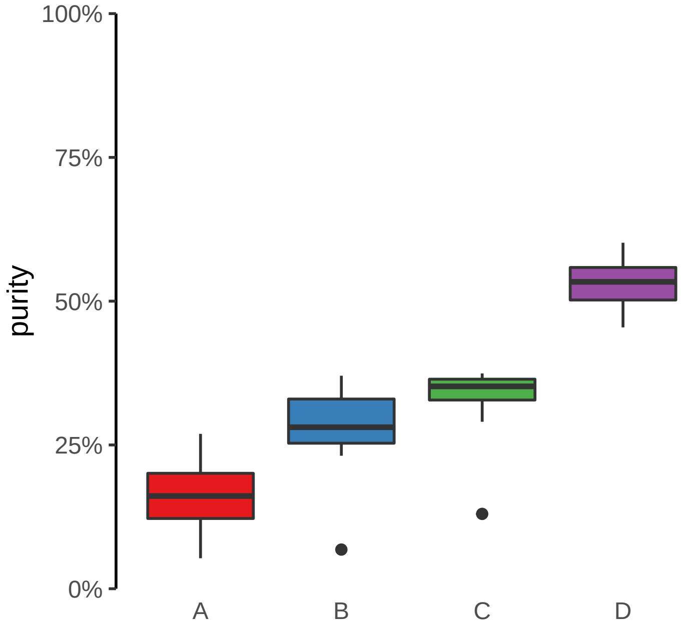

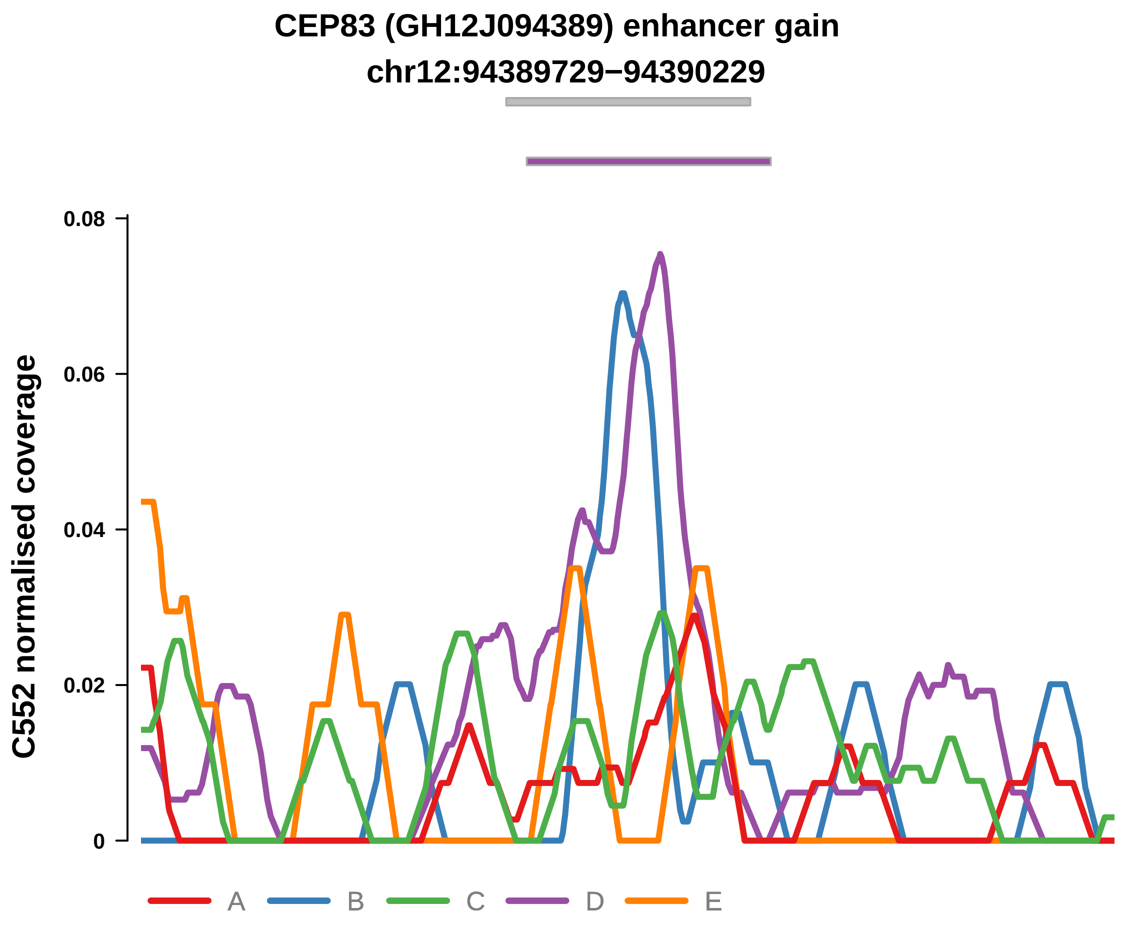


Putatively Clonal (logFC = 3.6)

| | P. | Adjusted P. |
|---|---|---|
| *~region vs. ~* | 1.98e−03 ** | 1.28e−02 * |
| *~purity vs ~* | 9.33e−04 *** | 7.77e−03 ** |
| *~purity+region vs ~purity* | 2.42e−01 | 9.1e−01 |
| *Carcinoma vs. normal* | 8.62e−06 **** | 8.62e−06 **** |

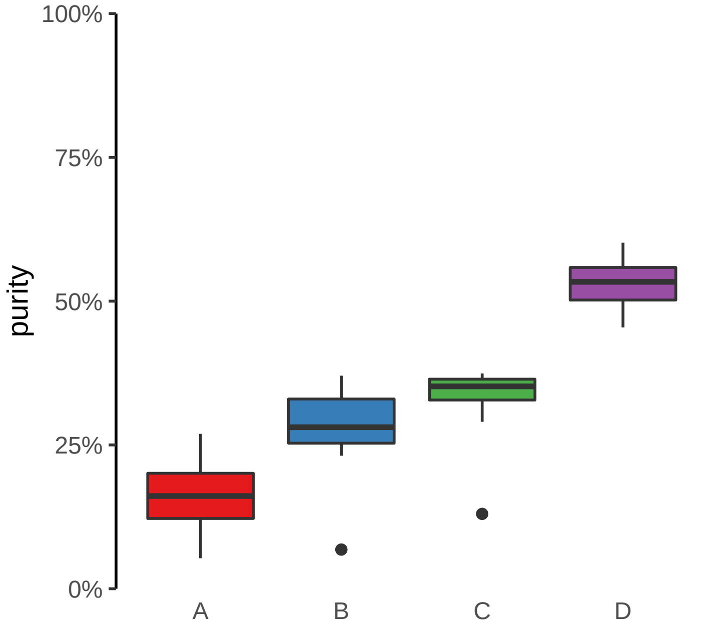

**CLASRP, MARK4 (GH19J045307) enhancer gain**
**chr19:45308035−45308535**

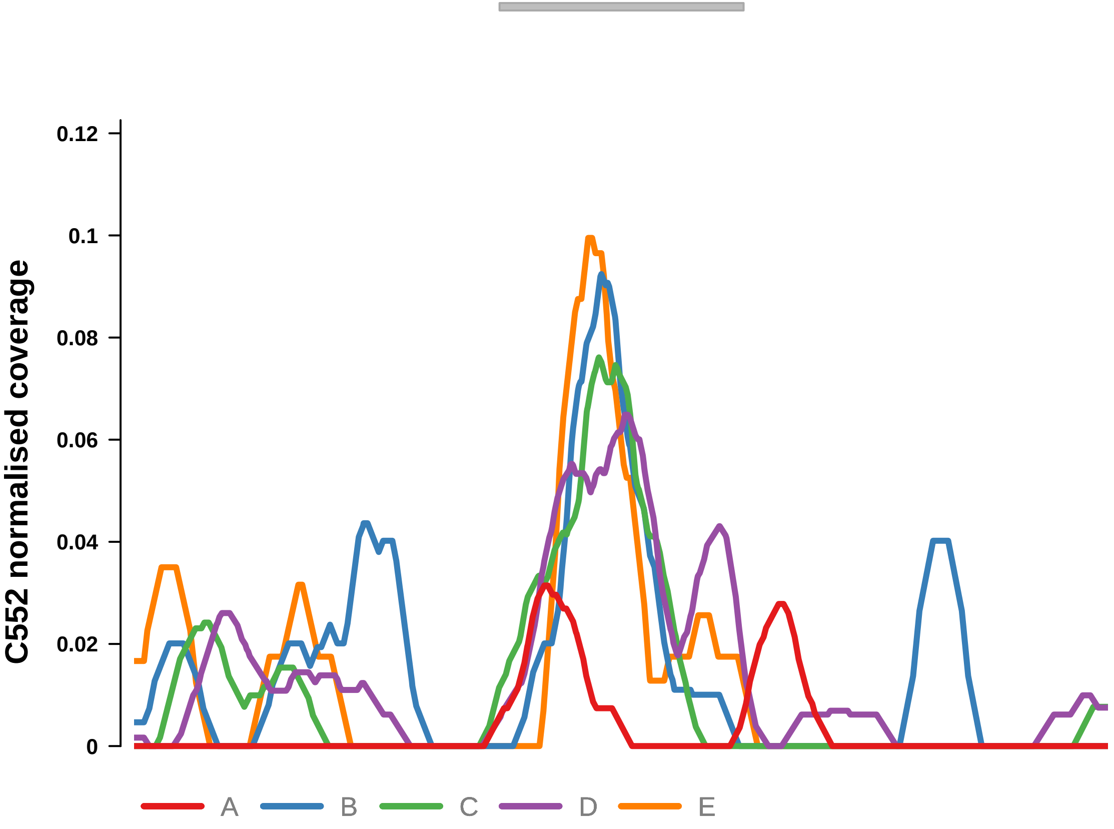


Putatively Clonal (logFC = 2.19)

| | P. | Adjusted P. |
|---|---|---|
| *~region vs. ~* | 1.89e−01 | 2.86e−01 |
| *~purity vs ~* | 2.2e−02 * | 6.46e−02 |
| *~purity+region vs ~purity* | 9.29e−01 | 9.85e−01 |
| *Carcinoma vs. normal* | 3.21e−03 ** | 3.21e−03 ** |

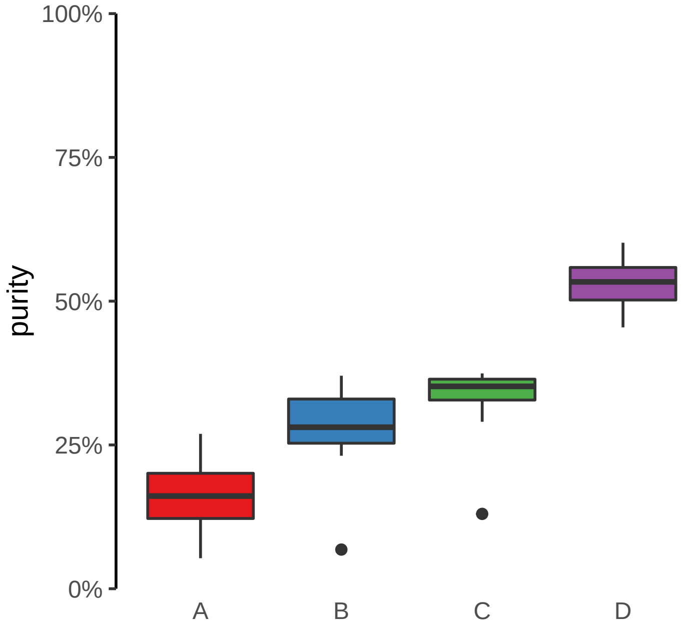

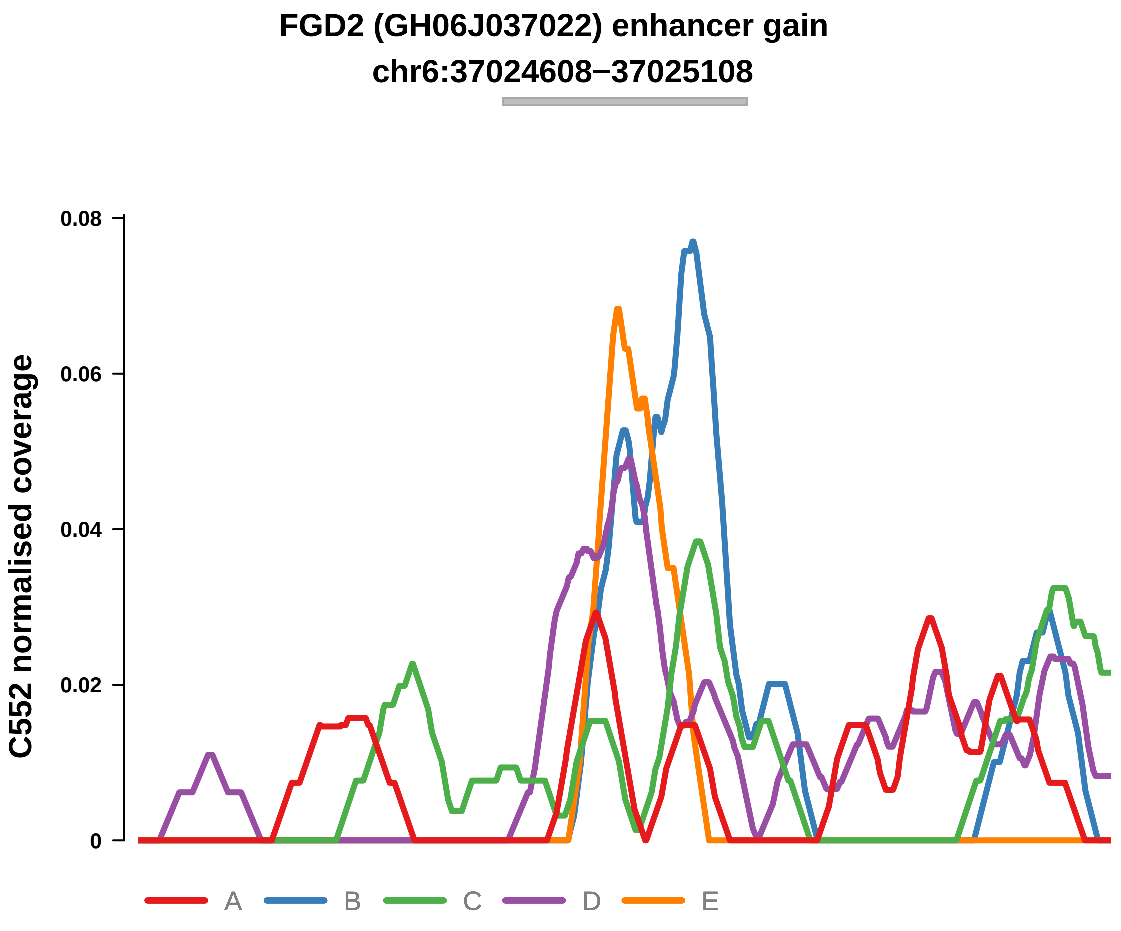


Putatively Clonal (logFC = 2.2)

| | P. | Adjusted P. |
|---|---|---|
| *~region vs. ~* | 2.52e−02 * | 9.88e−02 |
| *~purity vs ~* | 5.24e−03 ** | 2.39e−02 * |
| *~purity+region vs ~purity* | 2.2e−01 | 9.1e−01 |
| *Carcinoma vs. normal* | 5.71e−03 ** | 5.71e−03 ** |

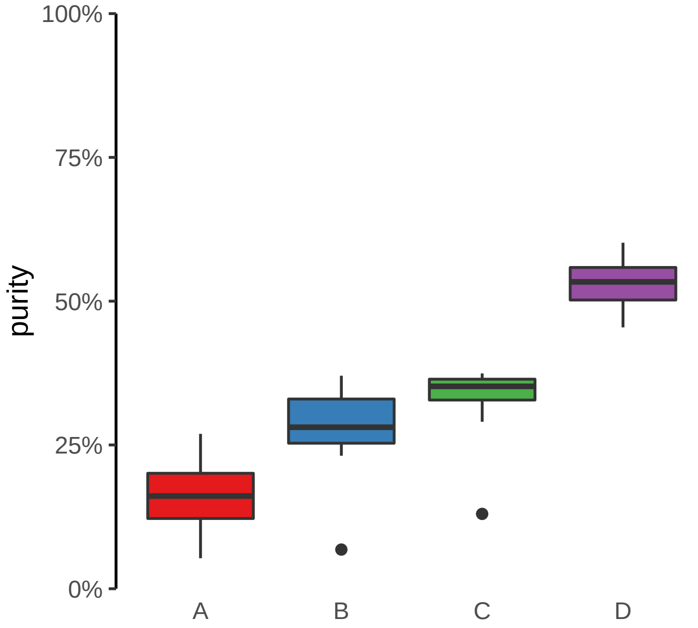

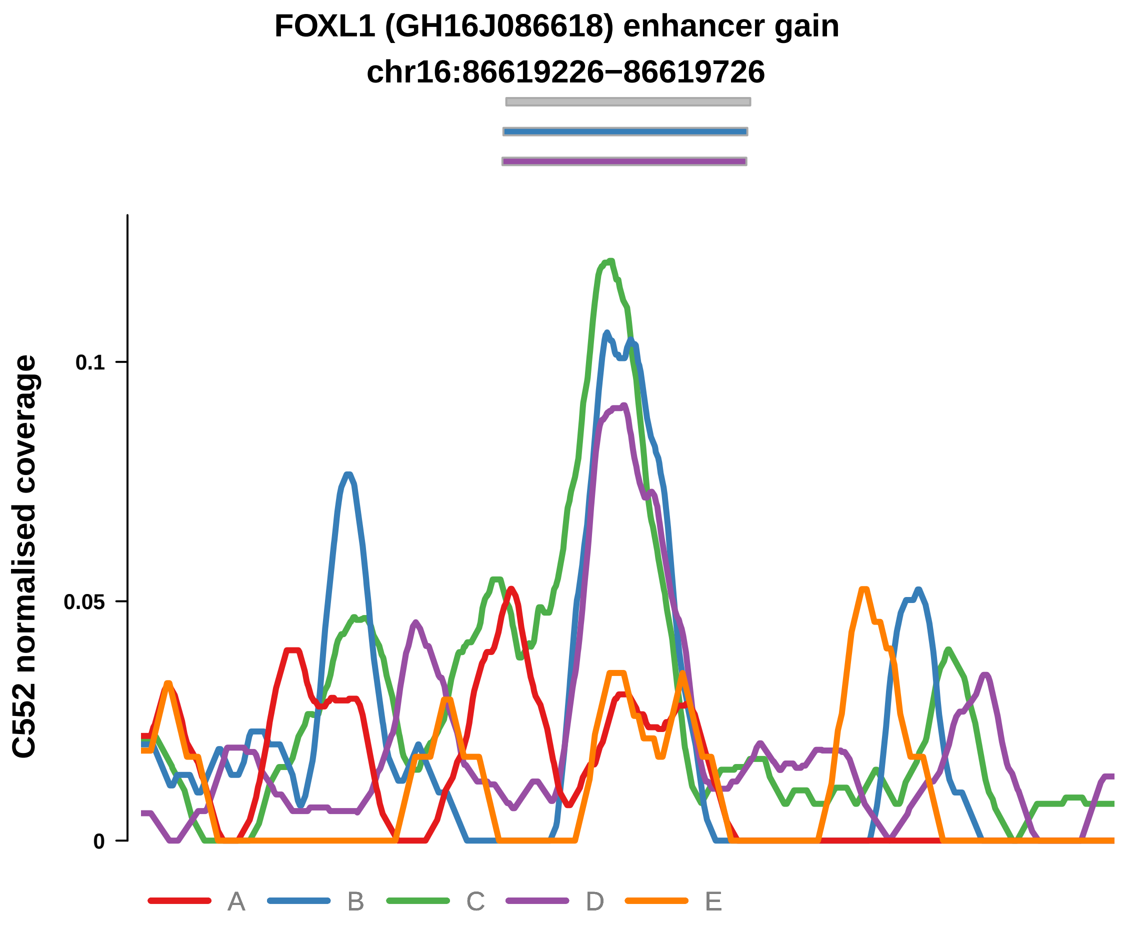


Putatively Clonal (logFC = 2.54)

| | P. | Adjusted P. |
|---|---|---|
| *~region vs. ~* | 4.76e−02 * | 1.24e−01 |
| *~purity vs ~* | 5.29e−03 ** | 2.39e−02 * |
| *~purity+region vs ~purity* | 2.89e−01 | 9.1e−01 |
| *Carcinoma vs. normal* | 7.21e−04 *** | 7.21e−04 *** |

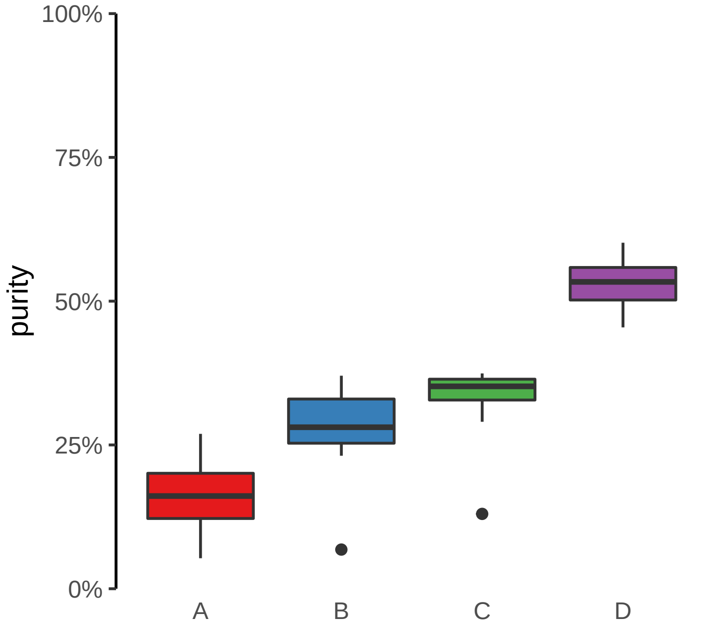

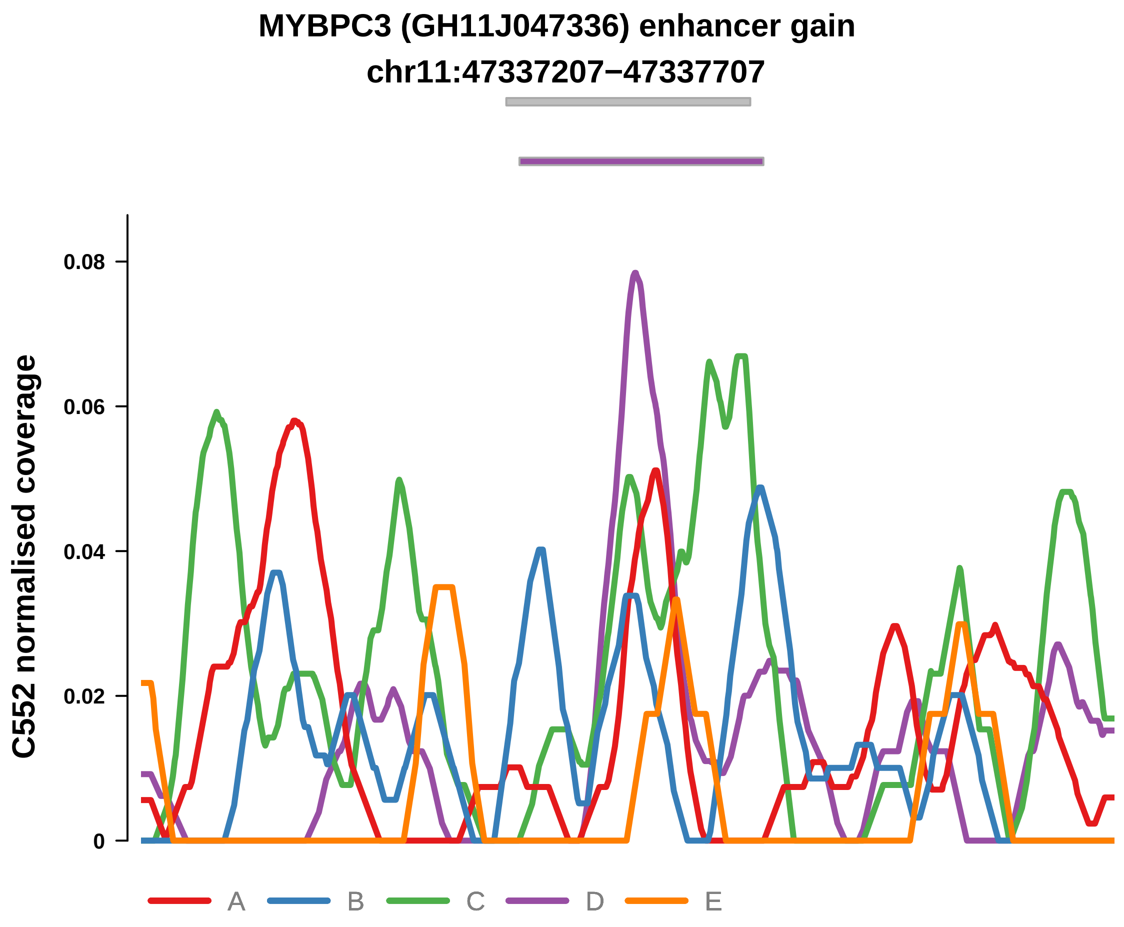


Putatively Clonal (logFC = 2.34)

| | P. | Adjusted P. |
|---|---|---|
| *~region vs. ~* | 1.67e−01 | 2.82e−01 |
| *~purity vs ~* | 1.51e−02 * | 5.79e−02 |
| *~purity+region vs ~purity* | 5.95e−01 | 9.85e−01 |
| *Carcinoma vs. normal* | 3.71e−03 ** | 3.71e−03 ** |

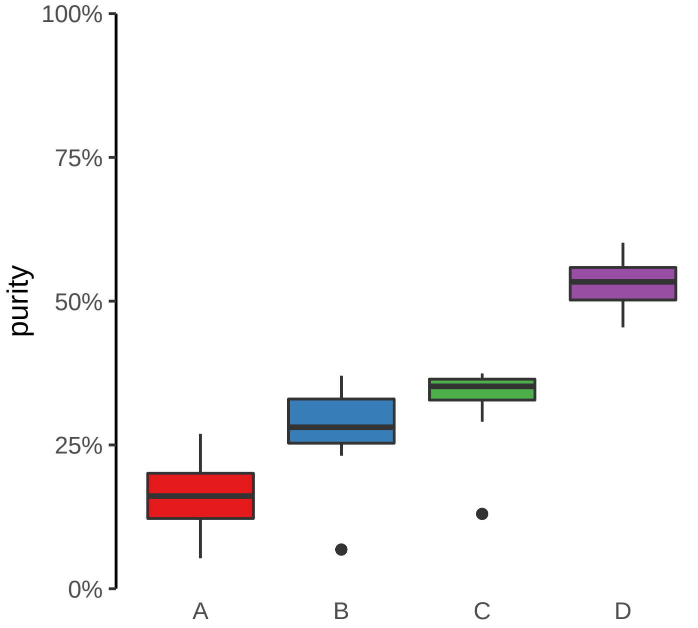

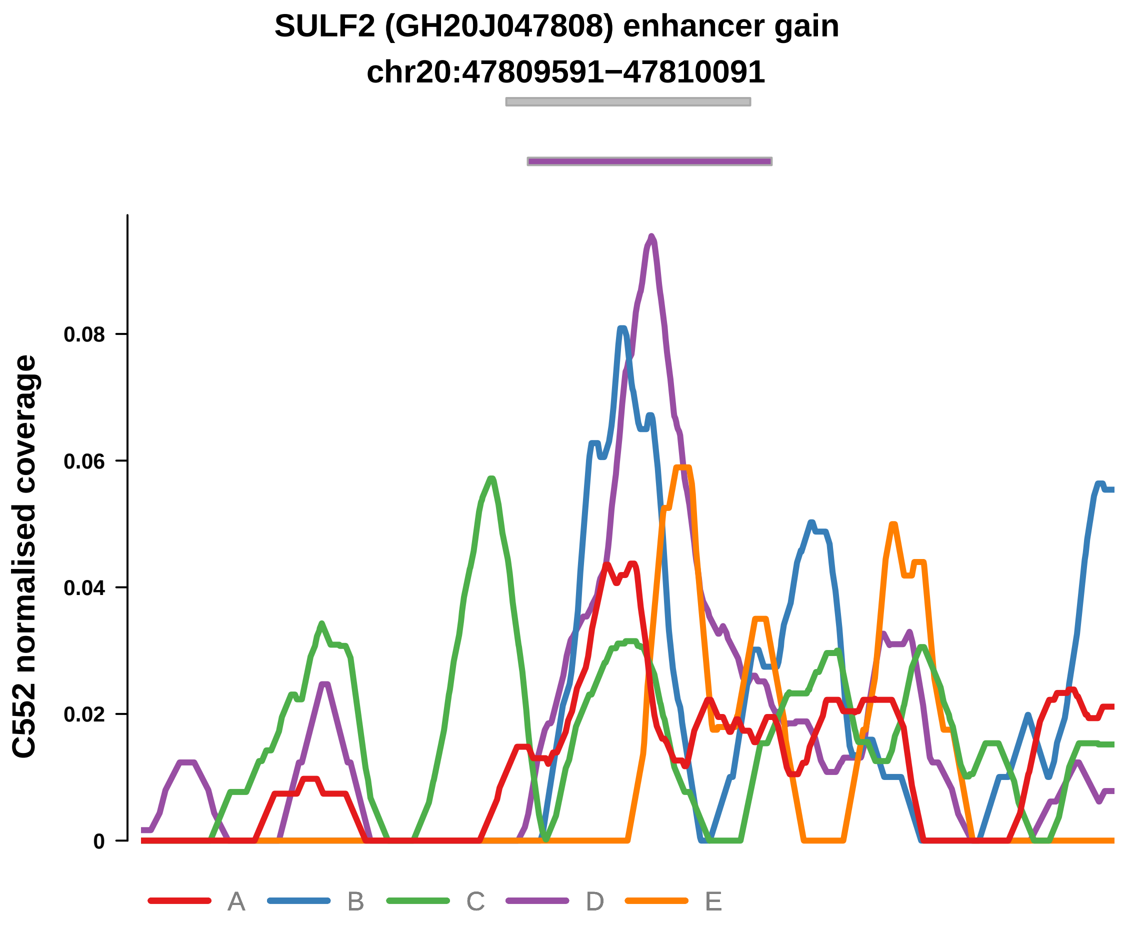


Putatively Clonal (logFC = 2.1)

| | P. | Adjusted P. |
|---|---|---|
| *~region vs. ~* | 5.87e−02 | 1.43e−01 |
| *~purity vs ~* | 3.36e−02 * | 8.57e−02 |
| *~purity+region vs ~purity* | 4.61e−01 | 9.85e−01 |
| *Carcinoma vs. normal* | 4.59e−03 ** | 4.59e−03 ** |

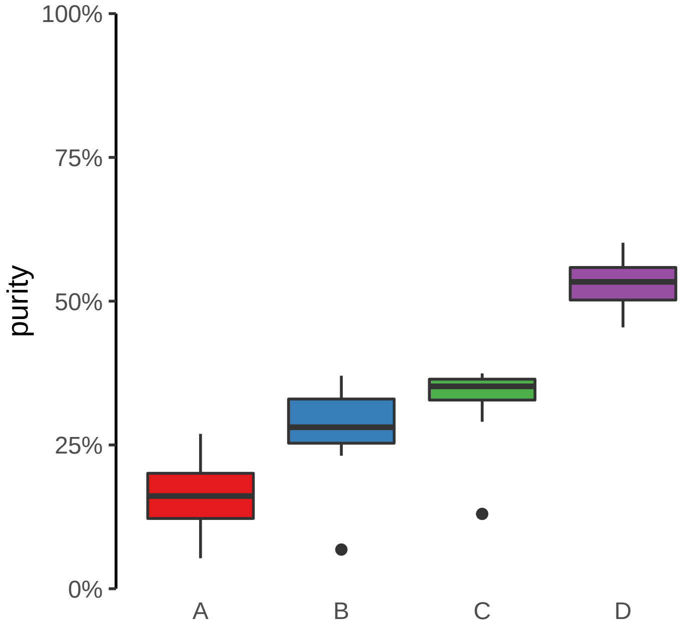

**CSGALNACT1 (GH08J019660) enhancer gain**

**chr8:19661066−19661566**

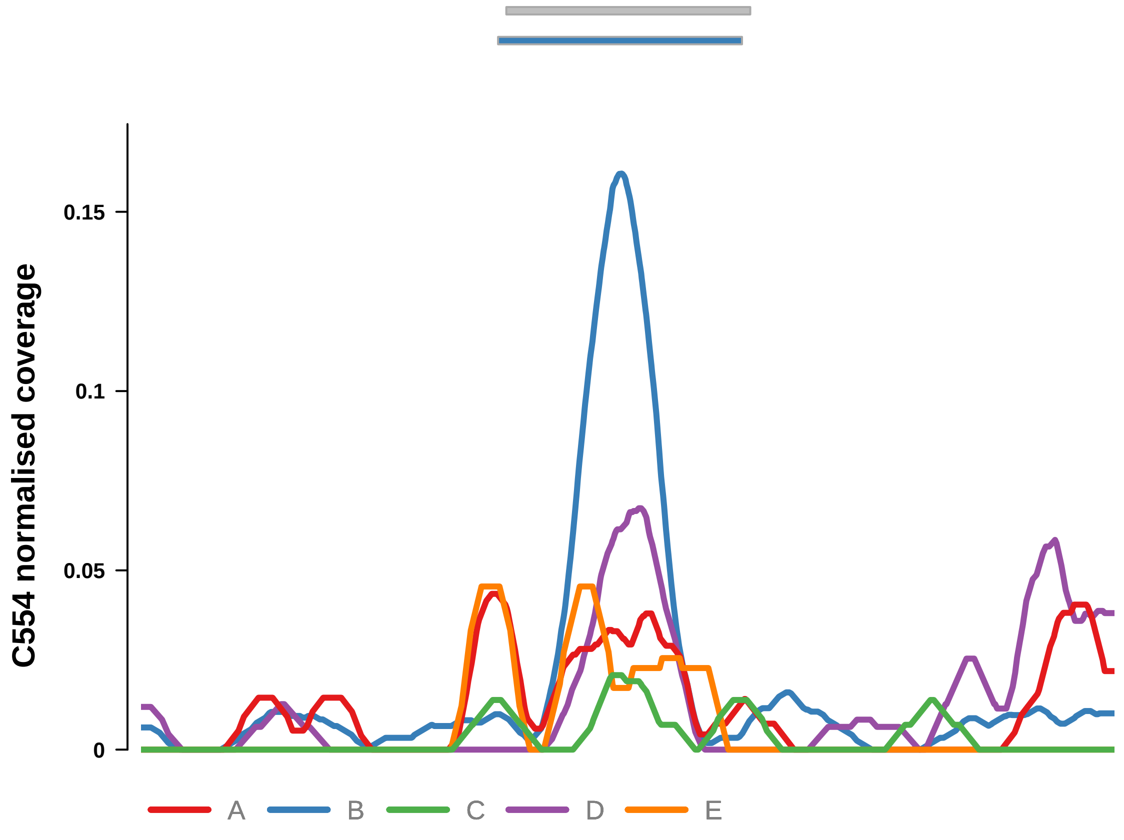


Putatively Clonal (logFC = 1.96)

| | P. | Adjusted P. |
|---|---|---|
| *~region vs. ~* | 1.74e−03 ** | 7.35e−03 ** |
| *~purity vs ~* | 1.62e−02 * | 7.81e−02 |
| *~purity+region vs ~purity* | 3.63e−02 * | 9.94e−02 |
| *Carcinoma vs. normal* | 6.04e−03 ** | 6.04e−03 ** |

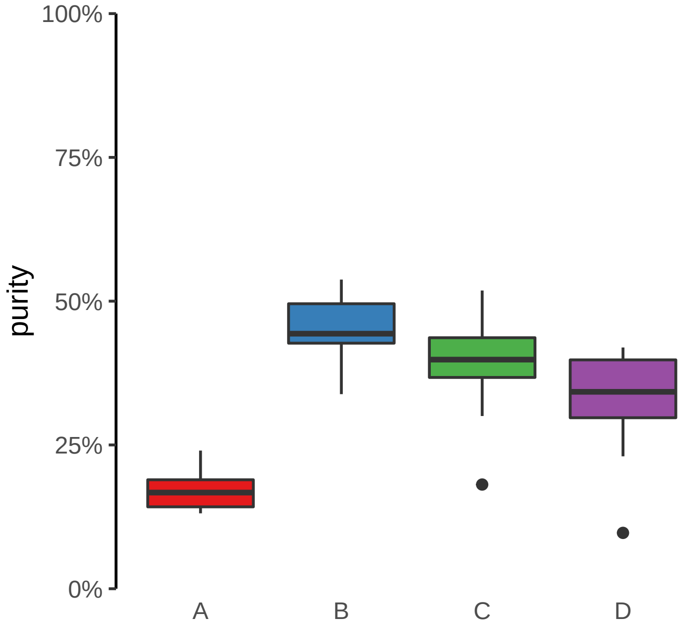

**MYBPC3 (GH11J047336) enhancer gain**
**chr11:47337207−47337707**

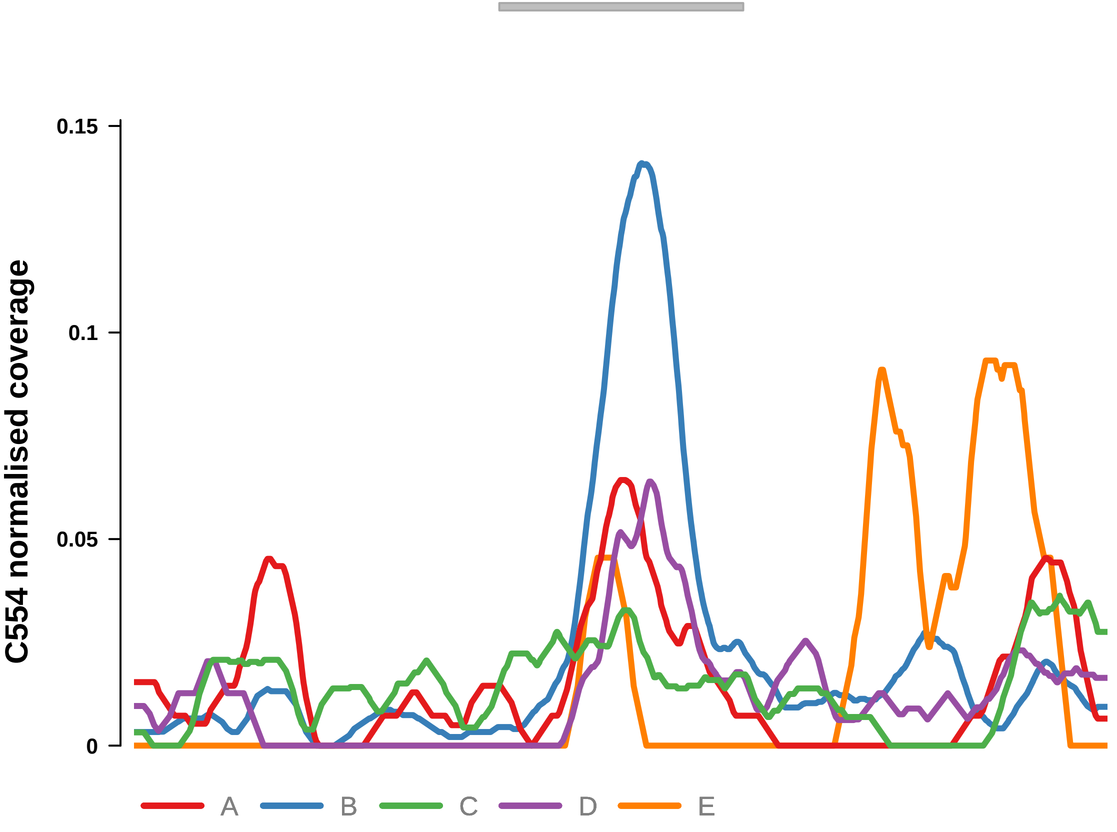


Putatively Clonal (logFC = 2.48)

| | P. | Adjusted P. |
|---|---|---|
| *~region vs. ~* | 1.71e−03 ** | 7.35e−03 ** |
| *~purity vs ~* | 8.01e−05 **** | 1.31e−03 ** |
| *~purity+region vs ~purity* | 2.67e−01 | 3.96e−01 |
| *Carcinoma vs. normal* | 6.04e−04 *** | 6.04e−04 *** |

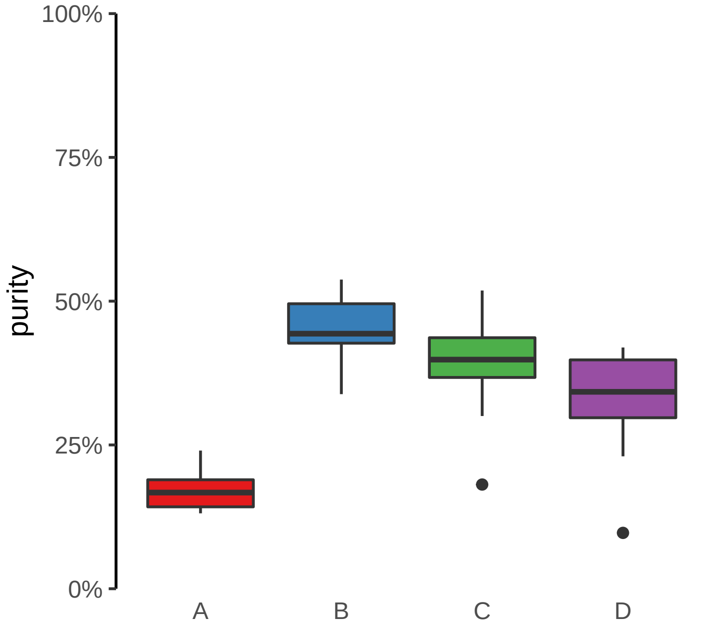

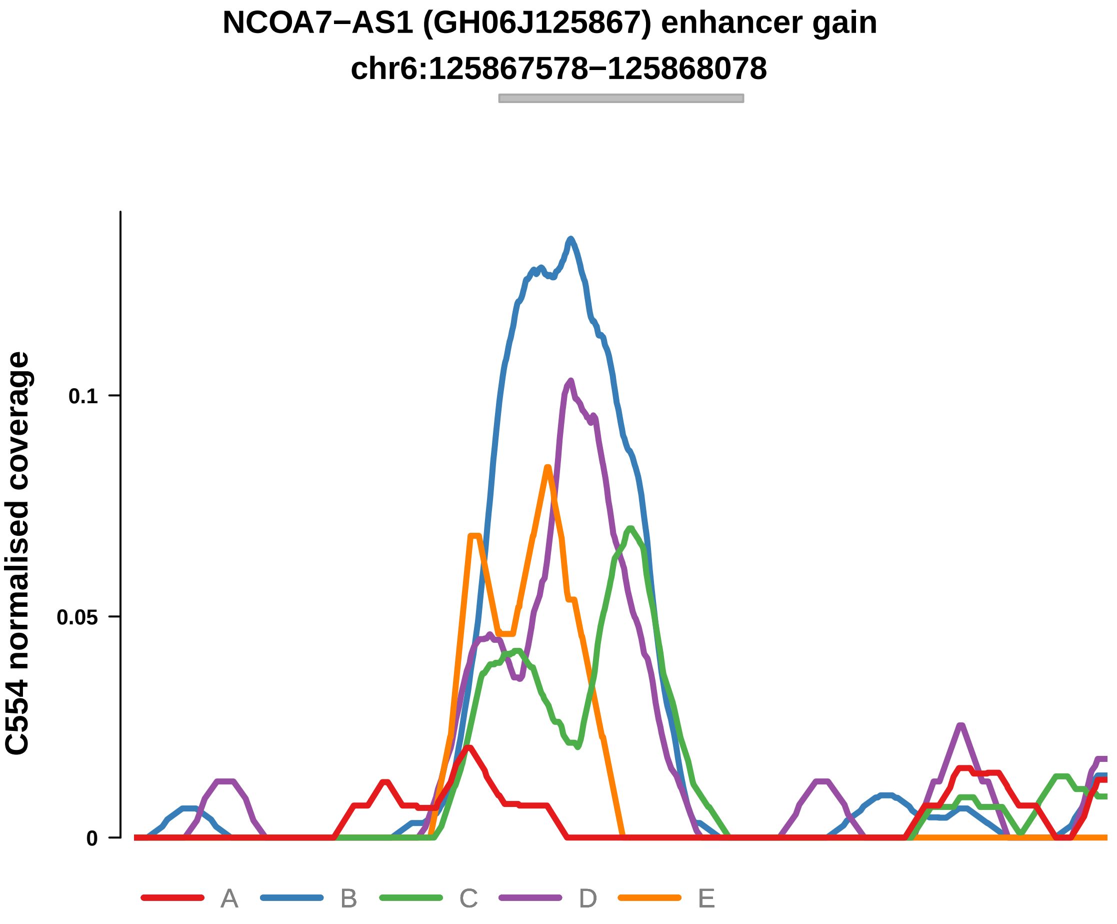


Putatively Clonal (logFC = 2.74)

| | P. | Adjusted P. |
|---|---|---|
| *~region vs. ~* | 7.49e−11 **** | 2.05e−09 **** |
| *~purity vs ~* | 2.25e−08 **** | 1.04e−06 **** |
| *~purity+region vs ~purity* | 2.31e−03 ** | 1.9e−02 * |
| *Carcinoma vs. normal* | 1.38e−04 *** | 1.38e−04 *** |

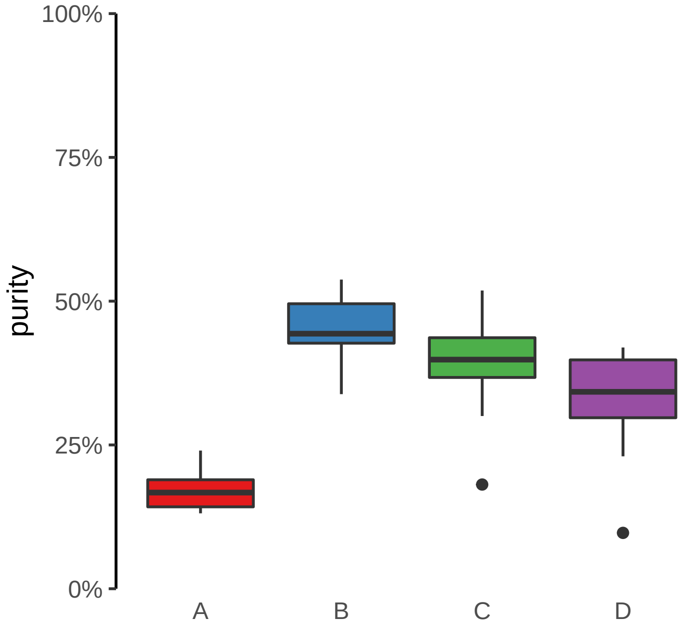

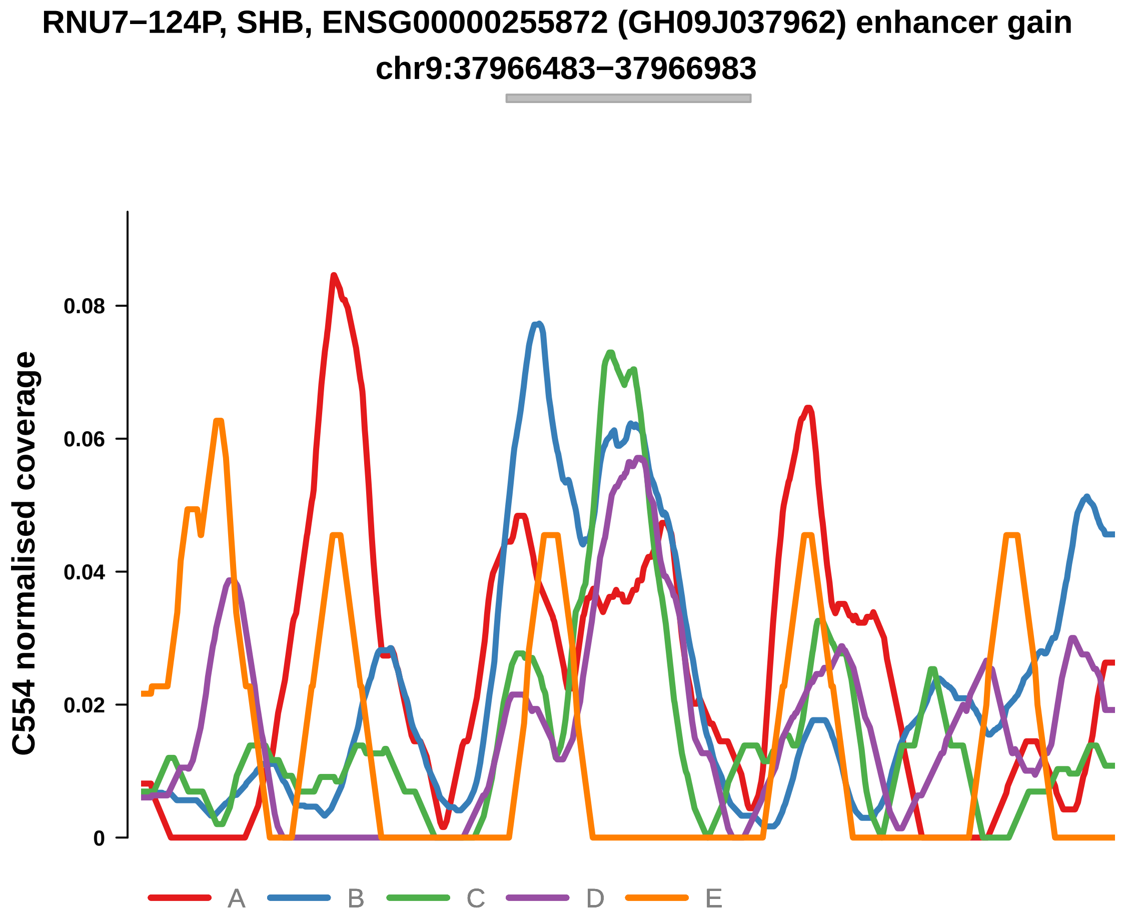


Putatively Clonal (logFC = 1.86)

| | P. | Adjusted P. |
|---|---|---|
| *~region vs. ~* | 6.3e−02 | 1.36e−01 |
| *~purity vs ~* | 2.06e−01 | 4.11e−01 |
| *~purity+region vs ~purity* | 1.51e−01 | 2.8e−01 |
| *Carcinoma vs. normal* | 8.44e−03 ** | 8.44e−03 ** |

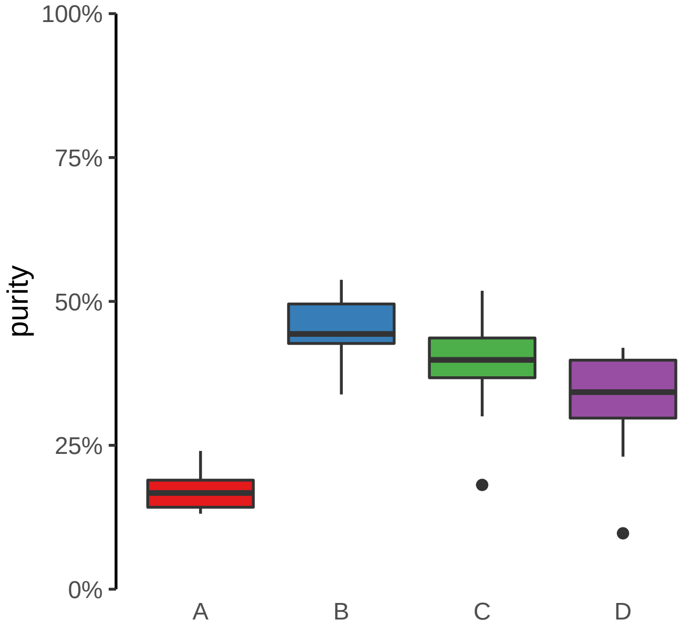

**SULF2 (GH20J047808) enhancer gain**
**chr20:47809591−47810091**

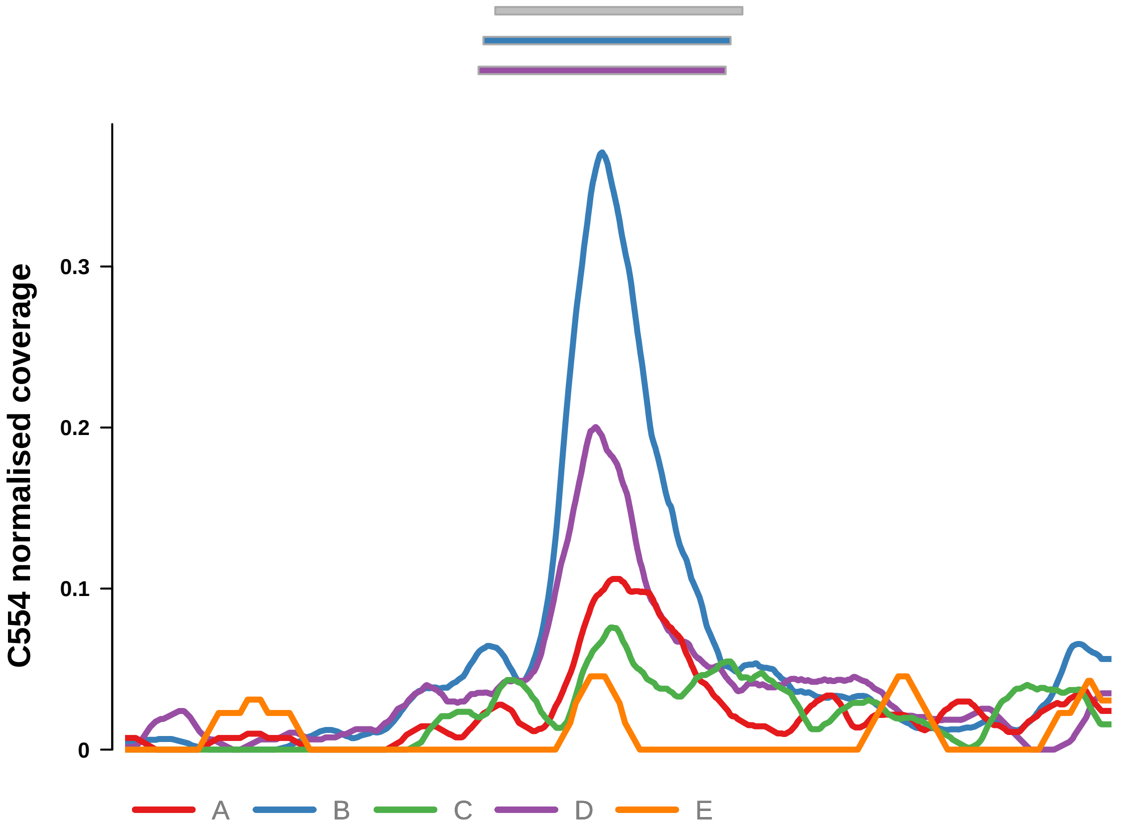


Putatively Clonal (logFC = 2.69)

| | P. | Adjusted P. |
|---|---|---|
| *~region vs. ~* | 3.81e−08 **** | 5.2e−07 **** |
| *~purity vs ~* | 1.69e−06 **** | 3.46e−05 **** |
| *~purity+region vs ~purity* | 2.2e−02 * | 6.28e−02 |
| *Carcinoma vs. normal* | 1.43e−04 *** | 1.43e−04 *** |

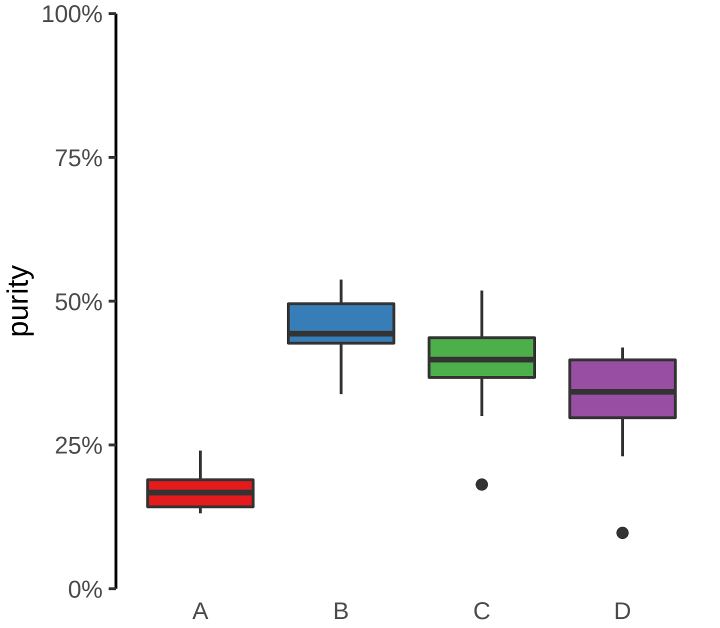

**HOXC11, HOXC10, HOTAIR (GH12J053971) enhancer gain**
**chr12:53978873−53979373**

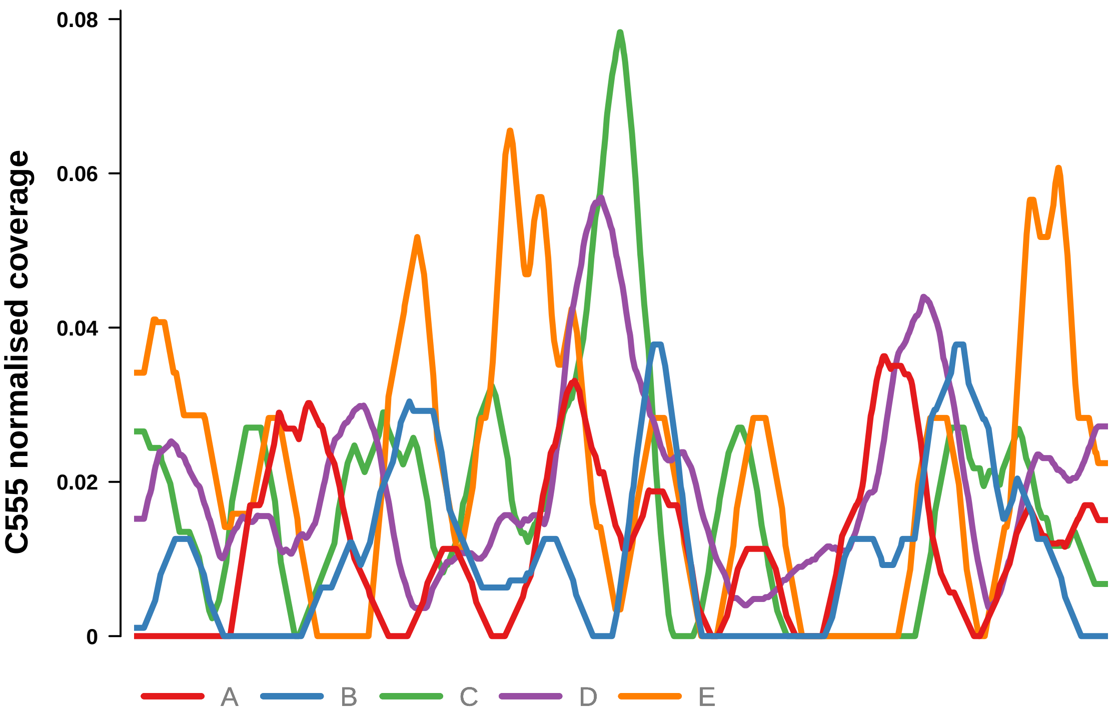


Putatively Clonal (logFC = 2.18)

| | P. | Adjusted P. |
|---|---|---|
| *~region vs. ~* | 8.56e−02 | 2.43e−01 |
| *~purity vs ~* | 4.65e−03 ** | 1.36e−01 |
| *~purity+region vs ~purity* | 9.59e−01 | 9.78e−01 |
| *Carcinoma vs. normal* | 3.35e−03 ** | 3.35e−03 ** |

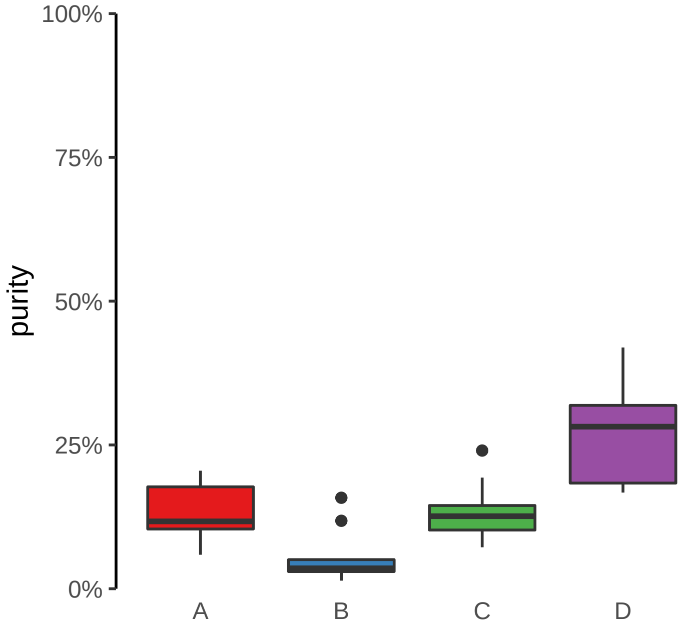

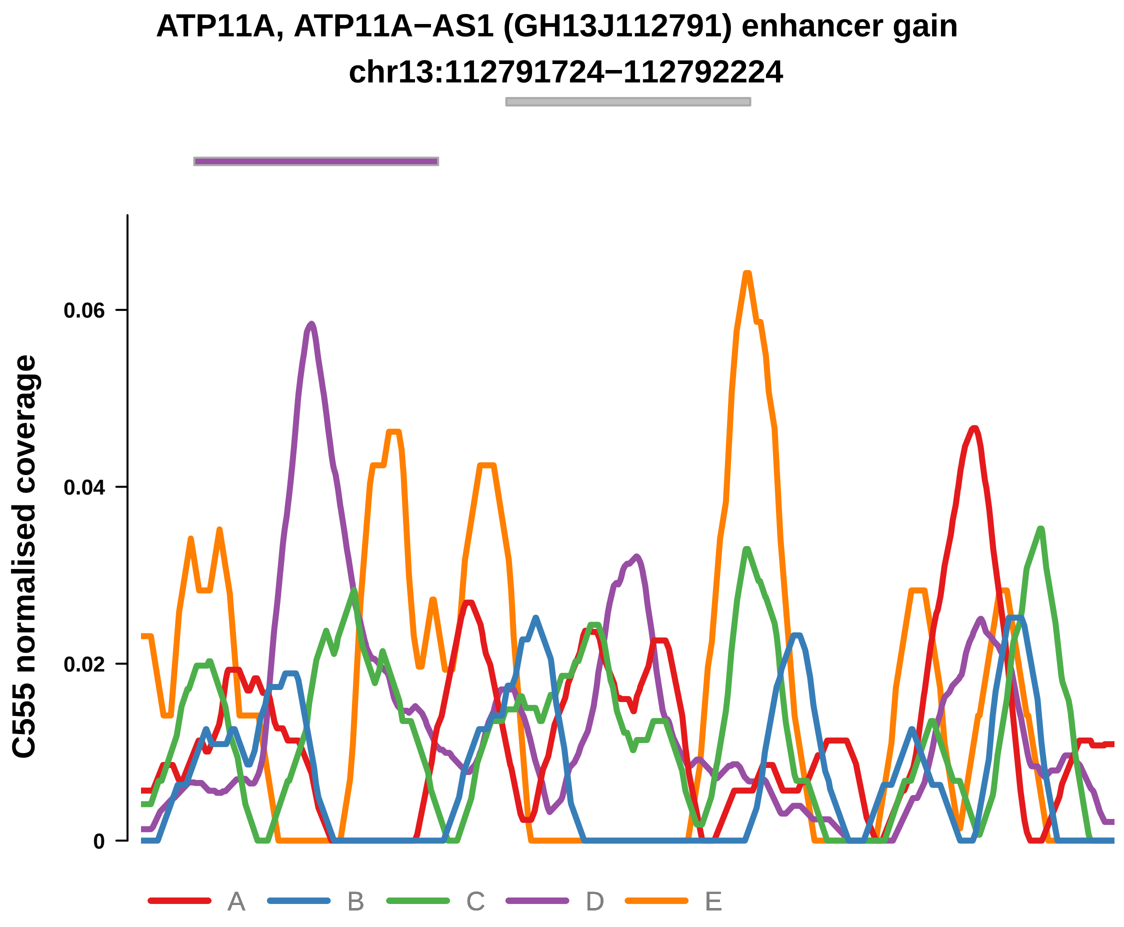


Putatively Clonal (logFC = 2.33)

| | P. | Adjusted P. |
|---|---|---|
| *~region vs. ~* | 2.69e−01 | 4.73e−01 |
| *~purity vs ~* | 2.67e−02 * | 2.09e−01 |
| *~purity+region vs ~purity* | 7.18e−01 | 8.87e−01 |
| *Carcinoma vs. normal* | 3.46e−03 ** | 3.46e−03 ** |

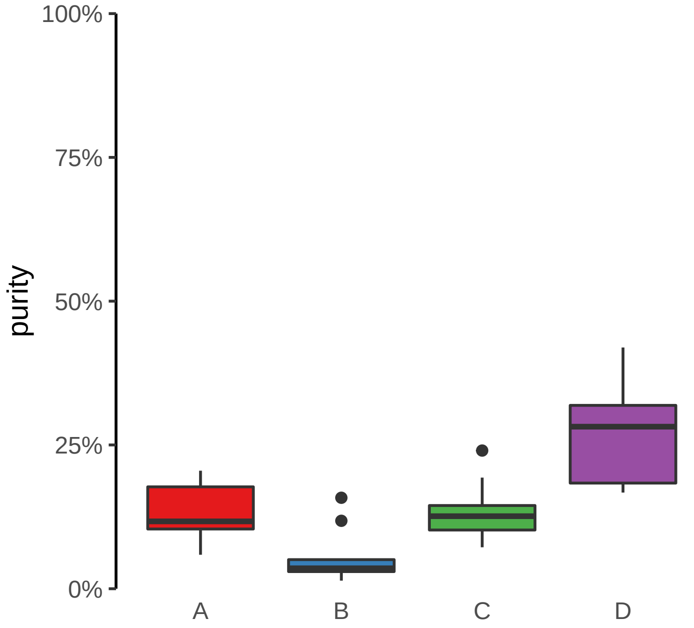

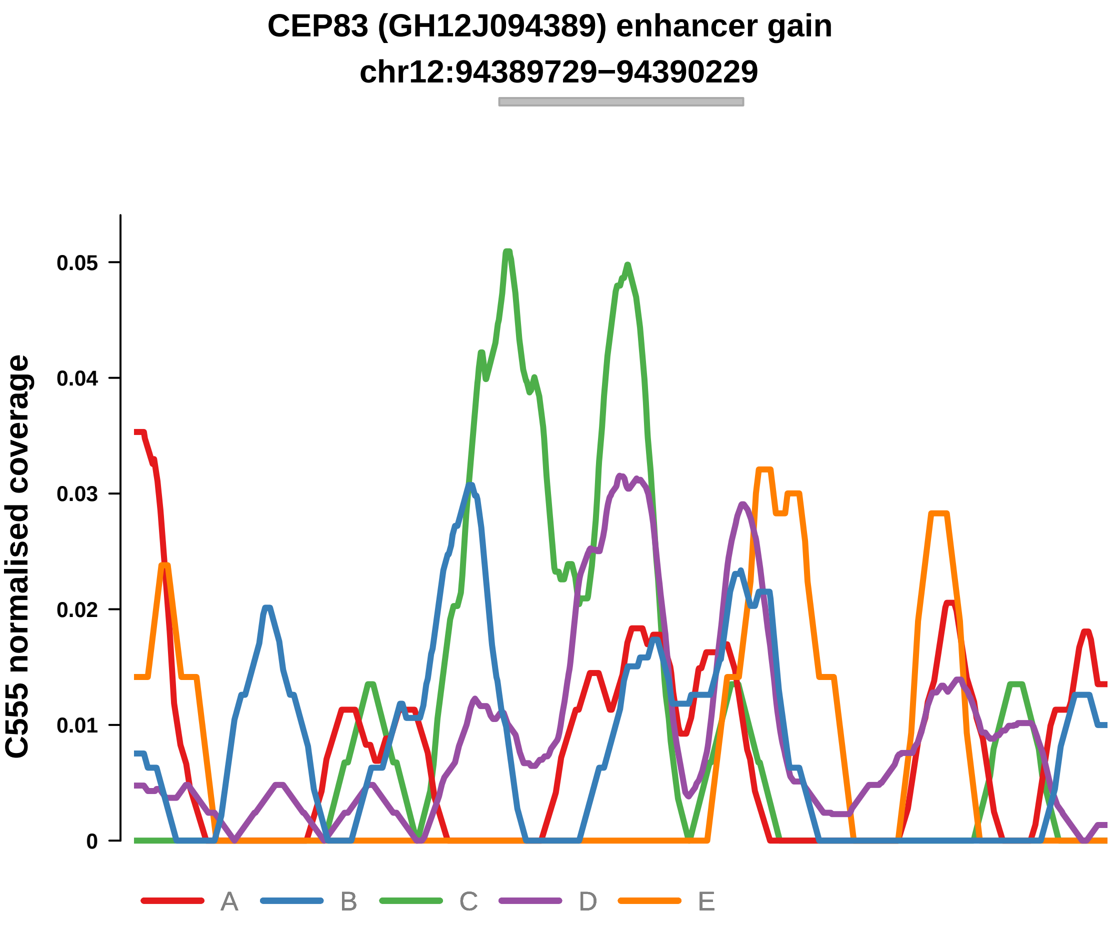


Putatively Clonal (logFC = 2.31)

| | P. | Adjusted P. |
|---|---|---|
| *~region vs. ~* | 9.33e−02 | 2.49e−01 |
| *~purity vs ~* | 1.31e−02 * | 1.67e−01 |
| *~purity+region vs ~purity* | 8.1e−01 | 9.43e−01 |
| *Carcinoma vs. normal* | 4.85e−03 ** | 4.85e−03 ** |

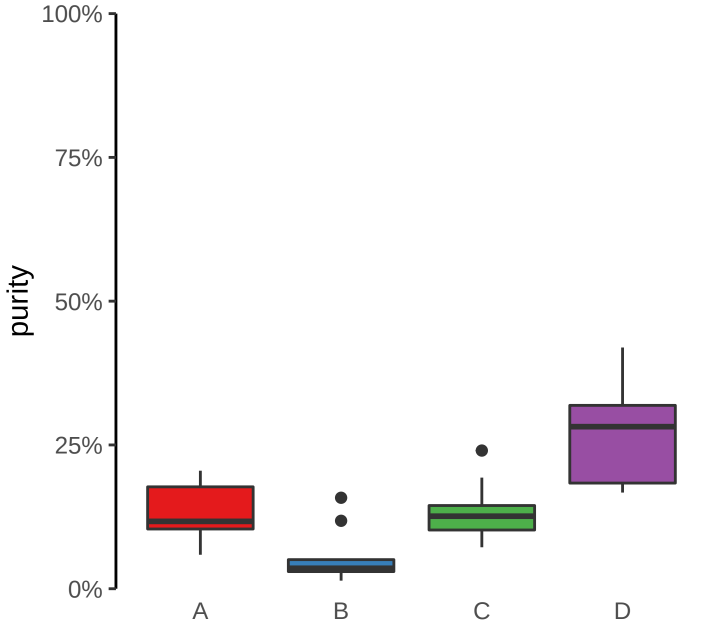

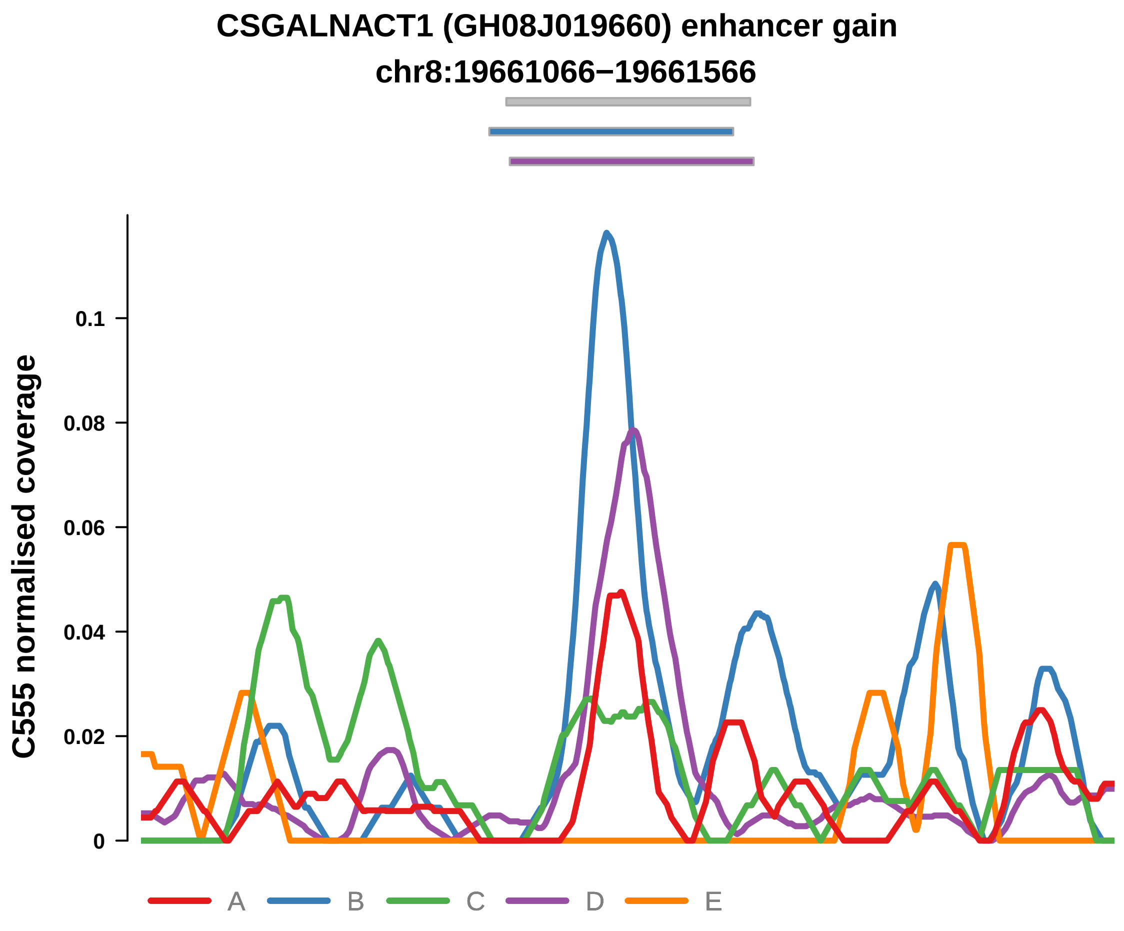


Putatively Clonal (logFC = 2.71)

| | P. | Adjusted P. |
|---|---|---|
| *~region vs. ~* | 5.89e−03 ** | 3.7e−02 * |
| *~purity vs ~* | 7.83e−01 | 8.4e−01 |
| *~purity+region vs ~purity* | 6.65e−03 ** | 5.9e−02 |
| *Carcinoma vs. normal* | 2.92e−04 *** | 2.92e−04 *** |

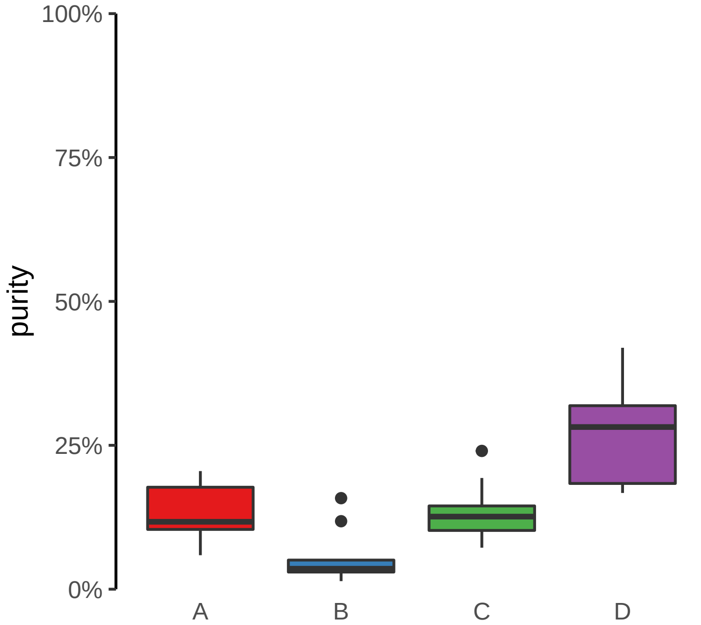

**FAM86B3P (GH08J008285) enhancer gain**
**chr8:8288409−8288909**

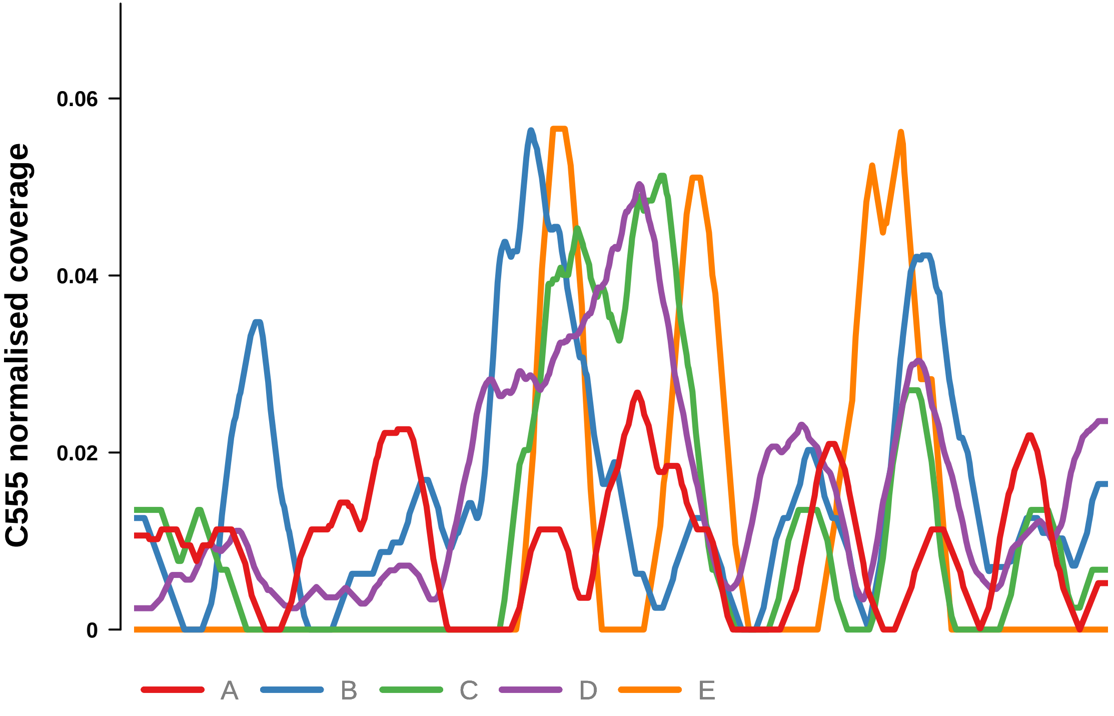


Putatively Clonal (logFC = 2.87)

| | P. | Adjusted P. |
|---|---|---|
| *~region vs. ~* | 1.84e−01 | 3.68e−01 |
| *~purity vs ~* | 2.51e−03 ** | 1.1e−01 |
| *~purity+region vs ~purity* | 2.2e−01 | 4.97e−01 |
| *Carcinoma vs. normal* | 1.51e−04 *** | 1.51e−04 *** |

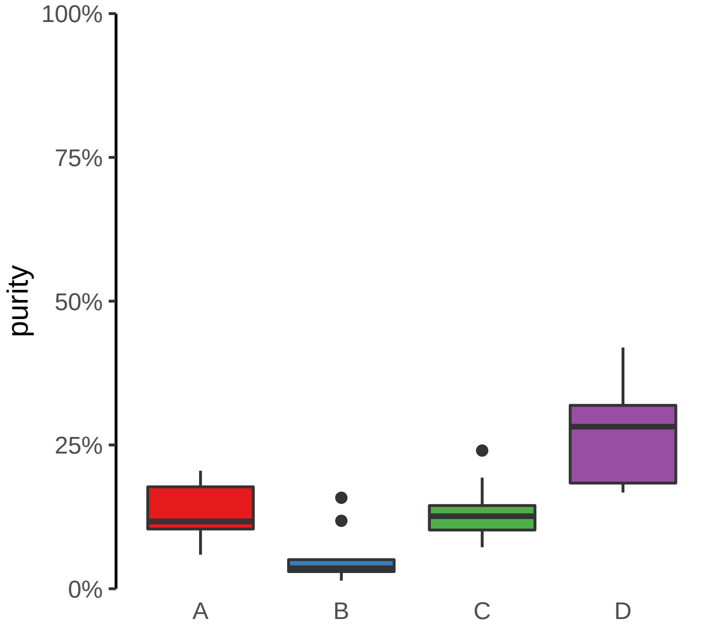

**MYBPC3 (GH11J047336) enhancer gain**
**chr11:47337207−47337707**

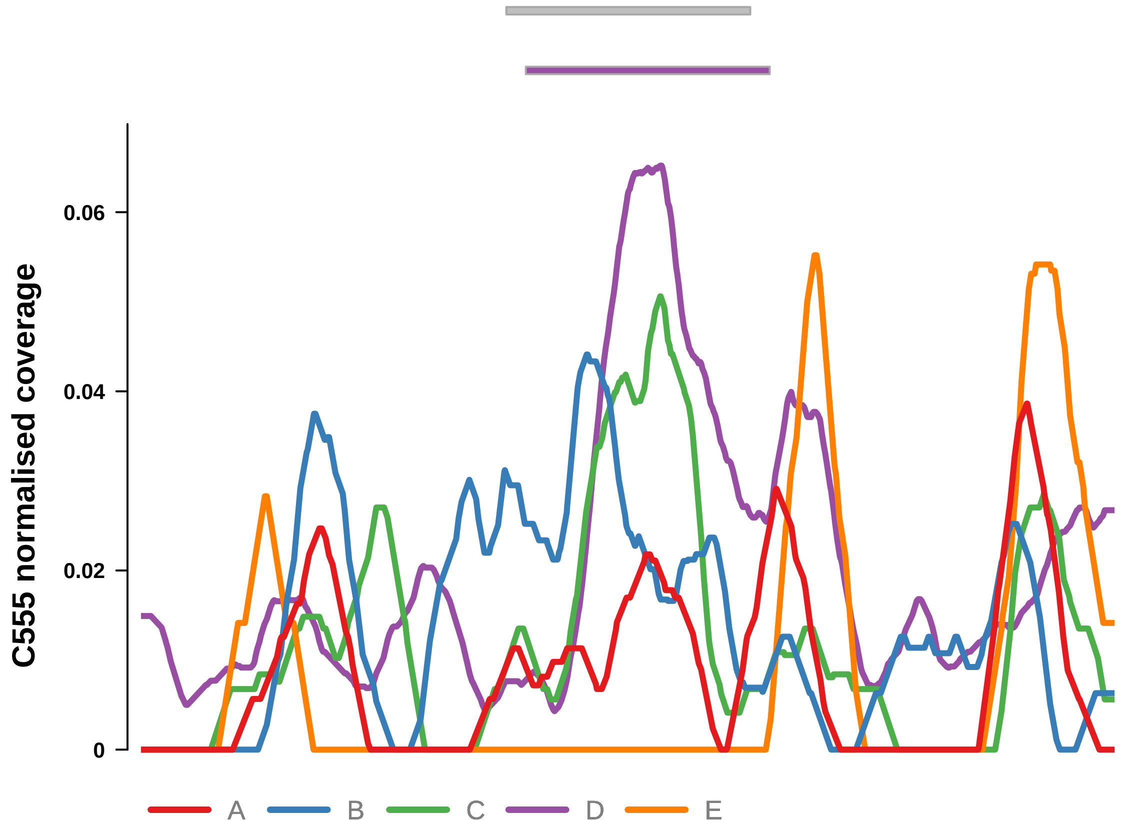


Putatively Clonal (logFC = 3.27)

| | P. | Adjusted P. |
|---|---|---|
| *~region vs. ~* | 1.8e−01 | 3.68e−01 |
| *~purity vs ~* | 2.14e−01 | 4.65e−01 |
| *~purity+region vs ~purity* | 3.96e−01 | 6.97e−01 |
| *Carcinoma vs. normal* | 1.83e−05 **** | 1.83e−05 **** |

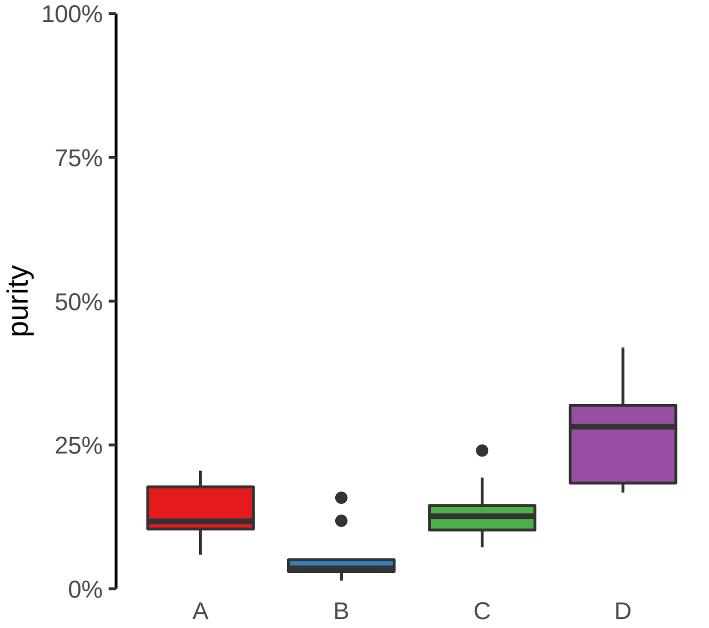

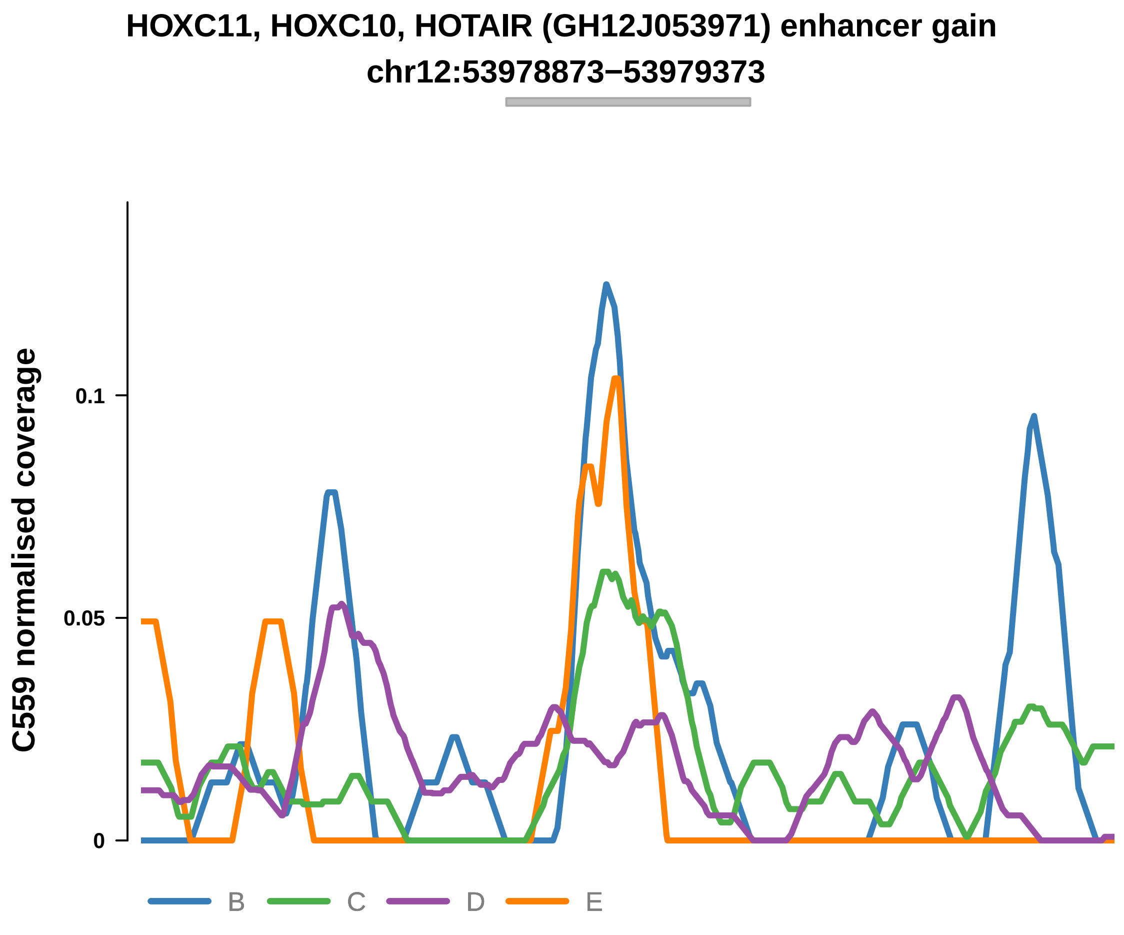


Putatively Clonal (logFC = 2.11)

| | P. | Adjusted P. |
|---|---|---|
| *~region vs. ~* | 8.3e−02 | 3.85e−01 |
| *~purity vs ~* | 1.99e−04 *** | 1.75e−02 * |
| *~purity+region vs ~purity* | 8.27e−01 | 9.61e−01 |
| *Carcinoma vs. normal* | 5.33e−03 ** | 5.33e−03 ** |

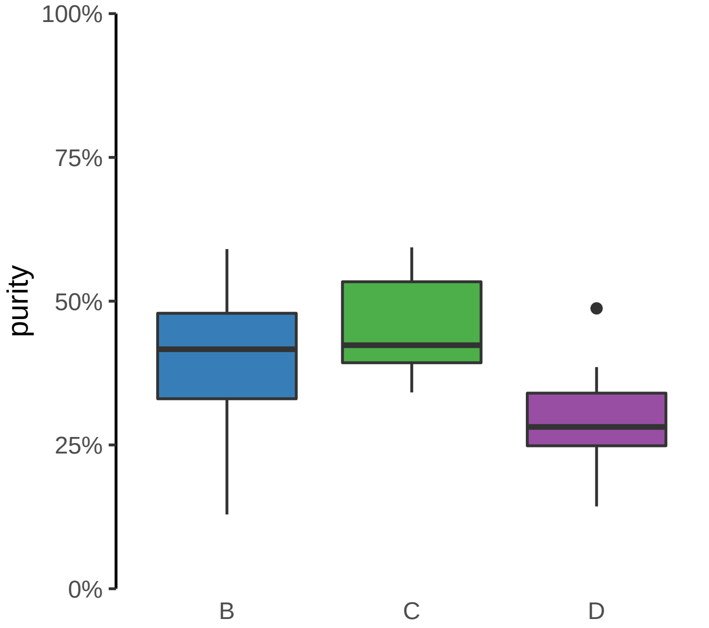

**ATP11A, ATP11A−AS1 (GH13J112791) enhancer gain**
**chr13:112791724−112792224**

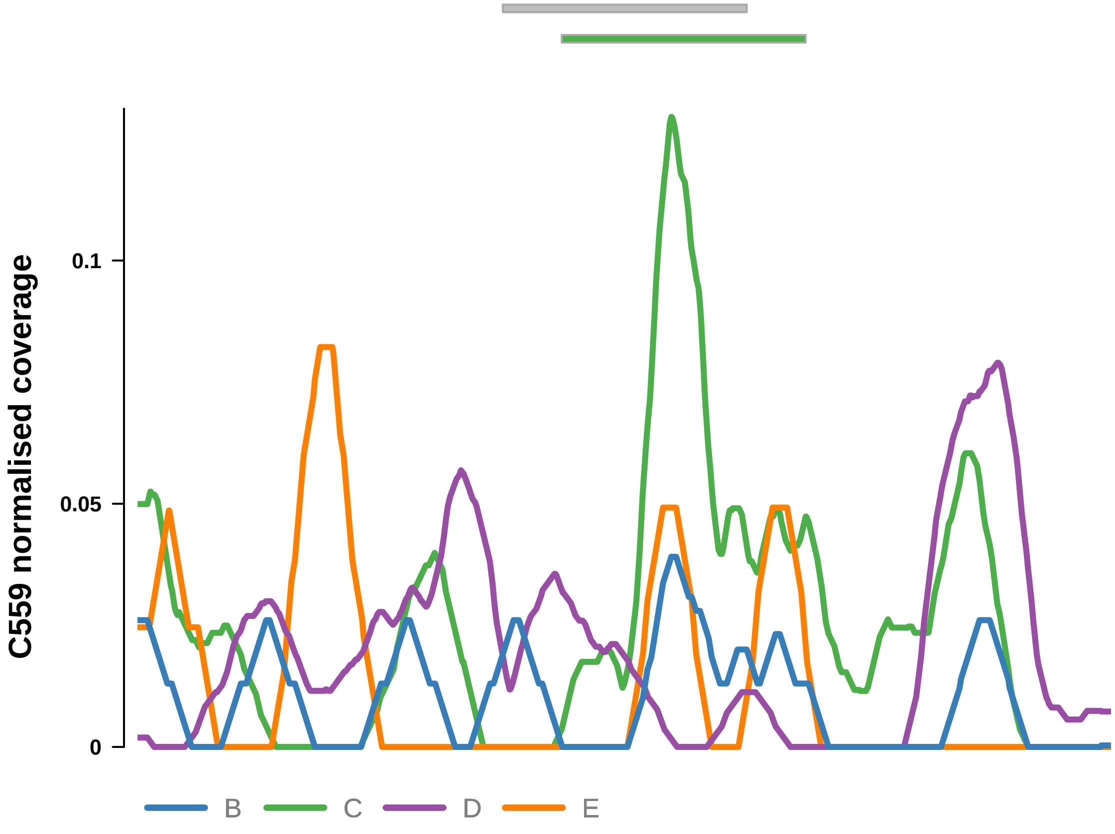


Putatively Clonal (logFC = 2.58)

| | P. | Adjusted P. |
|---|---|---|
| *~region vs. ~* | 1.01e−03 ** | 4.43e−02 * |
| *~purity vs ~* | 2.58e−03 ** | 7.56e−02 |
| *~purity+region vs ~purity* | 4.64e−02 * | 5.21e−01 |
| *Carcinoma vs. normal* | 1.14e−03 ** | 1.14e−03 ** |

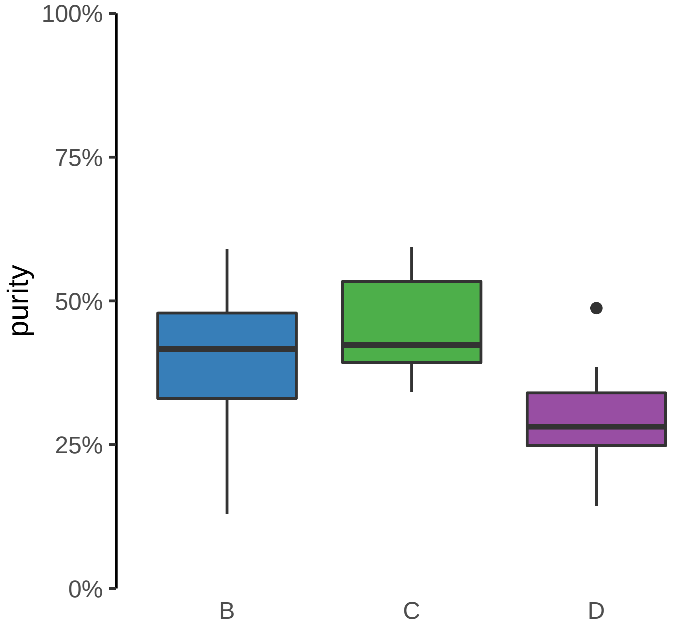

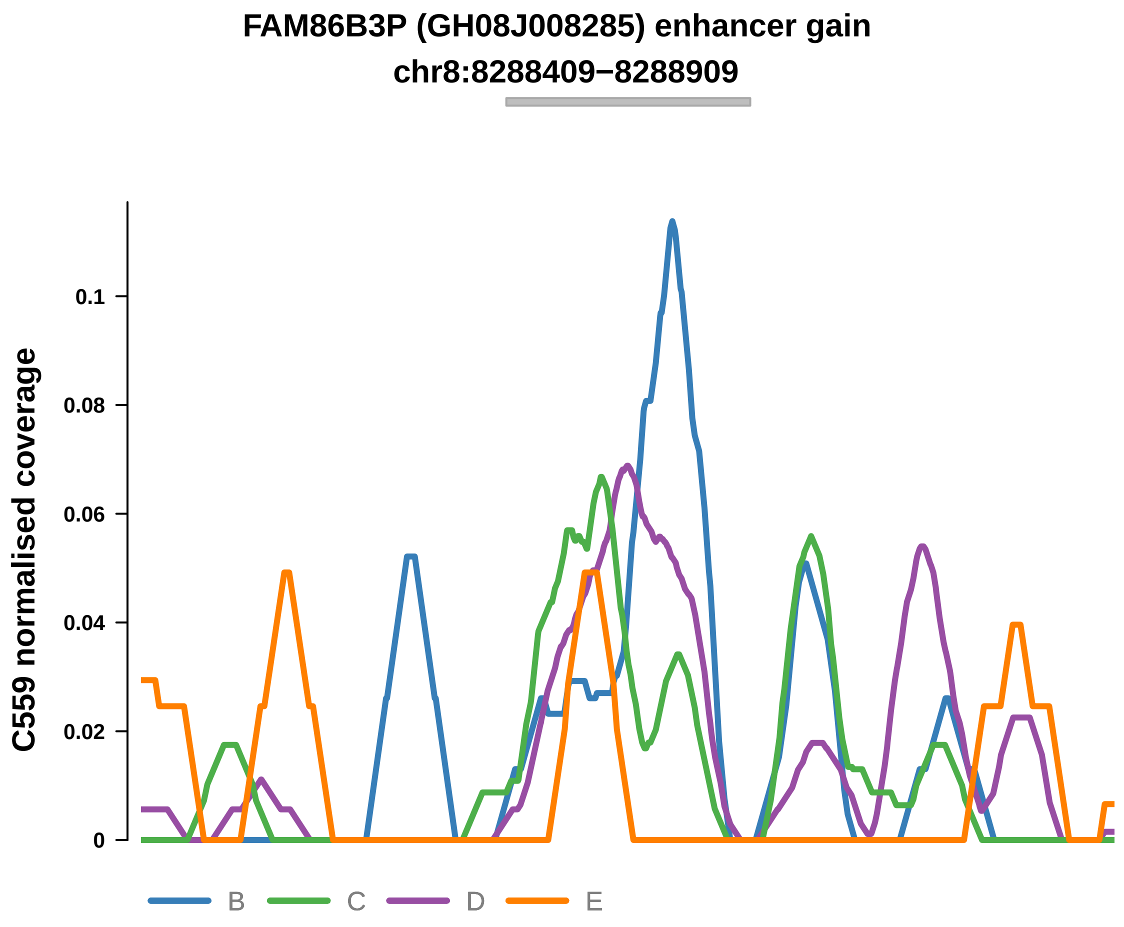


Putatively Clonal (logFC = 2.29)

| | P. | Adjusted P. |
|---|---|---|
| *~region vs. ~* | 5.36e−01 | 8.09e−01 |
| *~purity vs ~* | 6.48e−01 | 9.03e−01 |
| *~purity+region vs ~purity* | 5.44e−01 | 9.31e−01 |
| *Carcinoma vs. normal* | 3.88e−03 ** | 3.88e−03 ** |

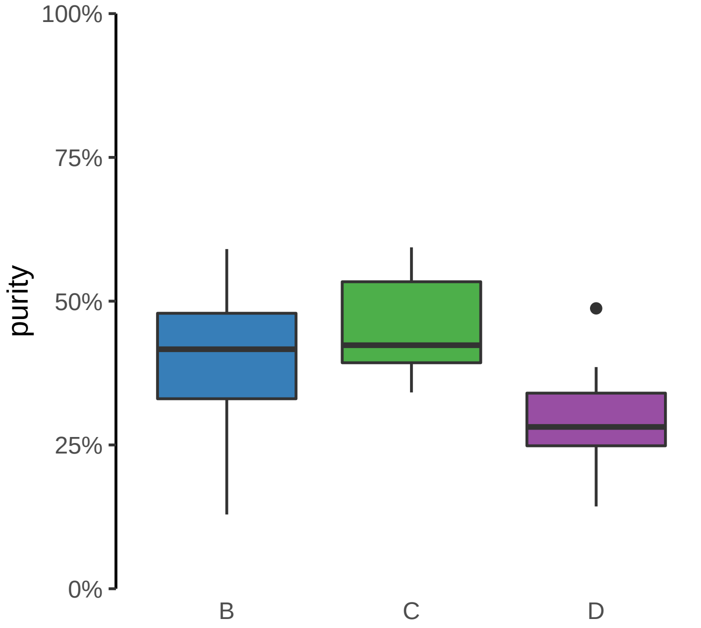

**FOXL1 (GH16J086618) enhancer gain**
**chr16:86619226−86619726**

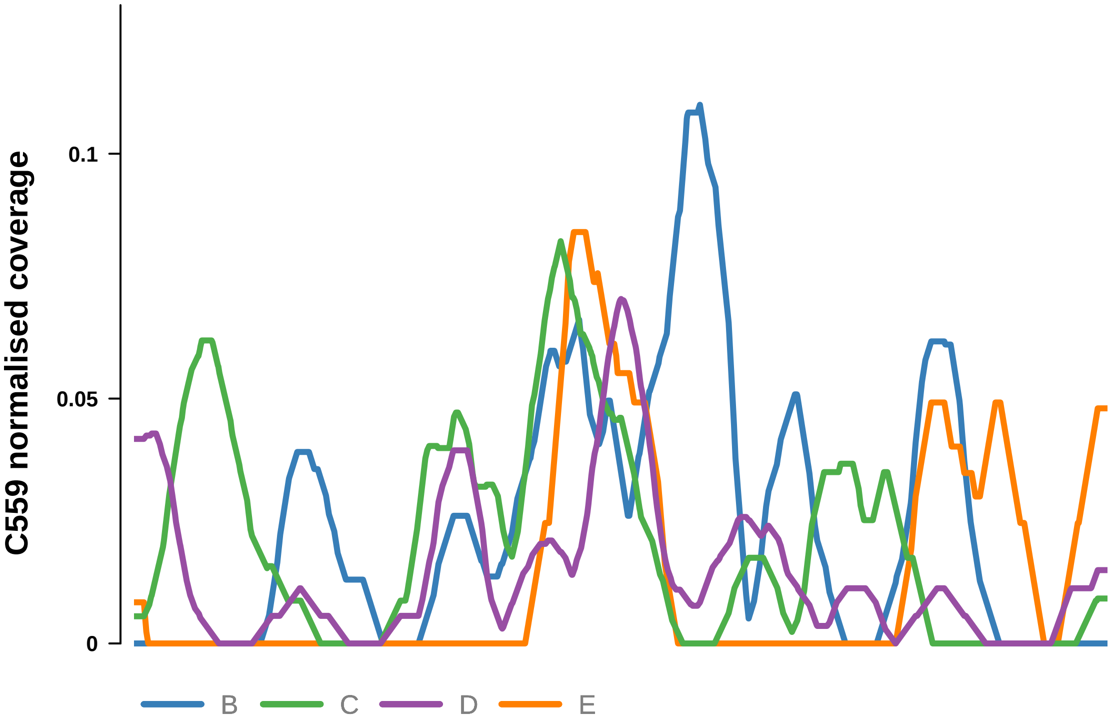


Putatively Clonal (logFC = 2.25)

| | P. | Adjusted P. |
|---|---|---|
| *~region vs. ~* | 8.99e−02 | 3.95e−01 |
| *~purity vs ~* | 7.04e−02 | 5.51e−01 |
| *~purity+region vs ~purity* | 3.15e−01 | 7.87e−01 |
| *Carcinoma vs. normal* | 2.85e−03 ** | 2.85e−03 ** |

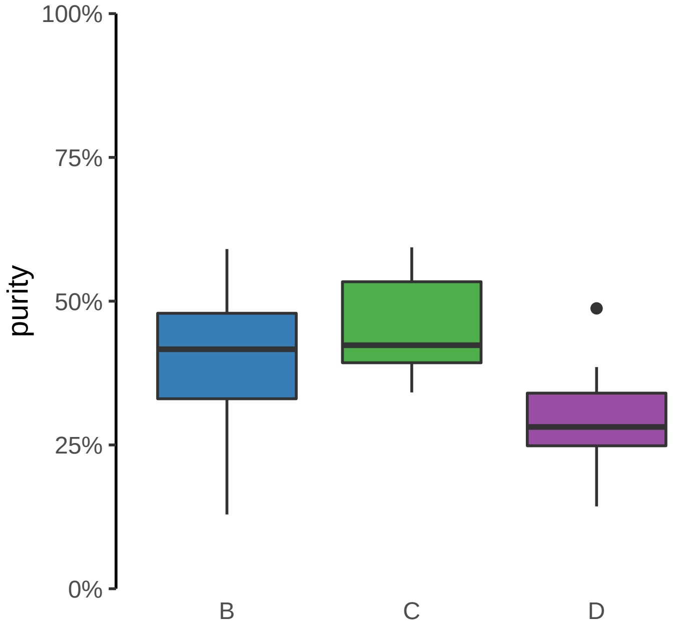

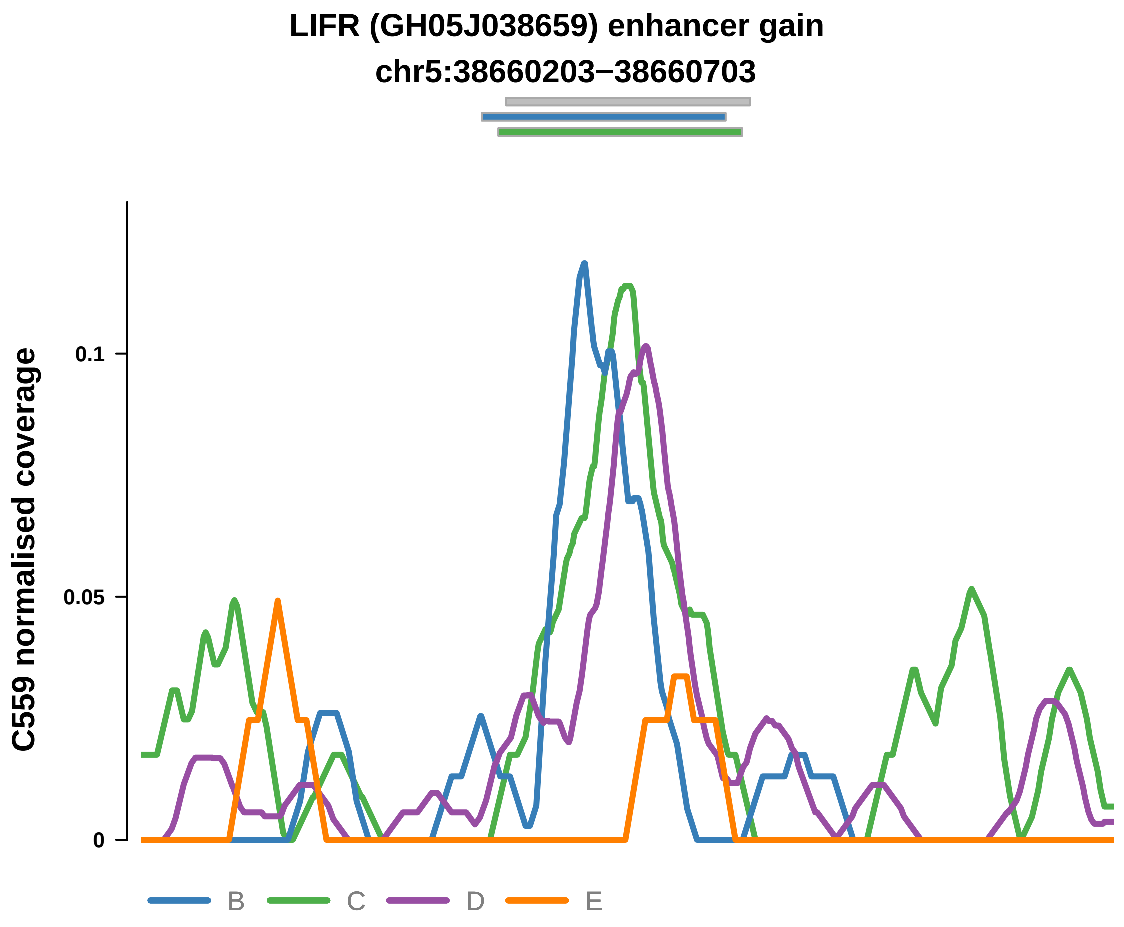


Putatively Clonal (logFC = 2.15)

| | P. | Adjusted P. |
|---|---|---|
| *~region vs. ~* | 7.16e−02 | 3.63e−01 |
| *~purity vs ~* | 7.64e−01 | 9.03e−01 |
| *~purity+region vs ~purity* | 1.78e−02 * | 5.21e−01 |
| *Carcinoma vs. normal* | 3.94e−03 ** | 3.94e−03 ** |

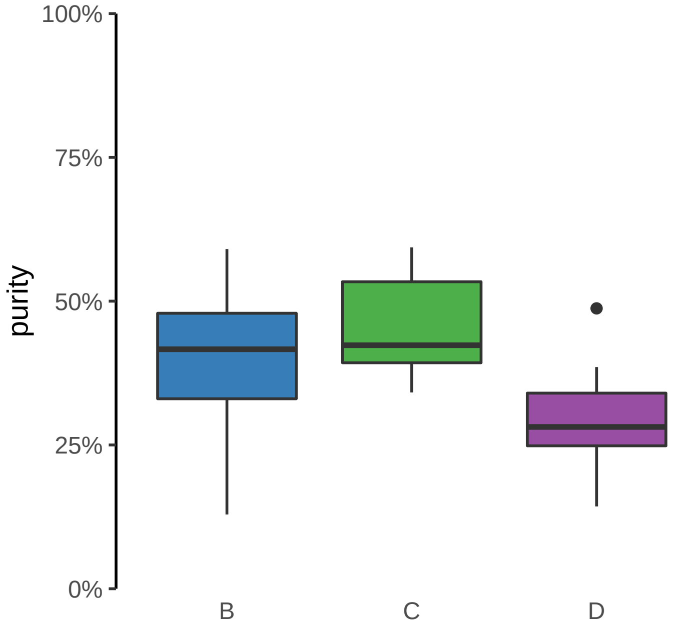

**MYT1 (GH20J064188) enhancer gain**
**chr20:64189162−64189662**

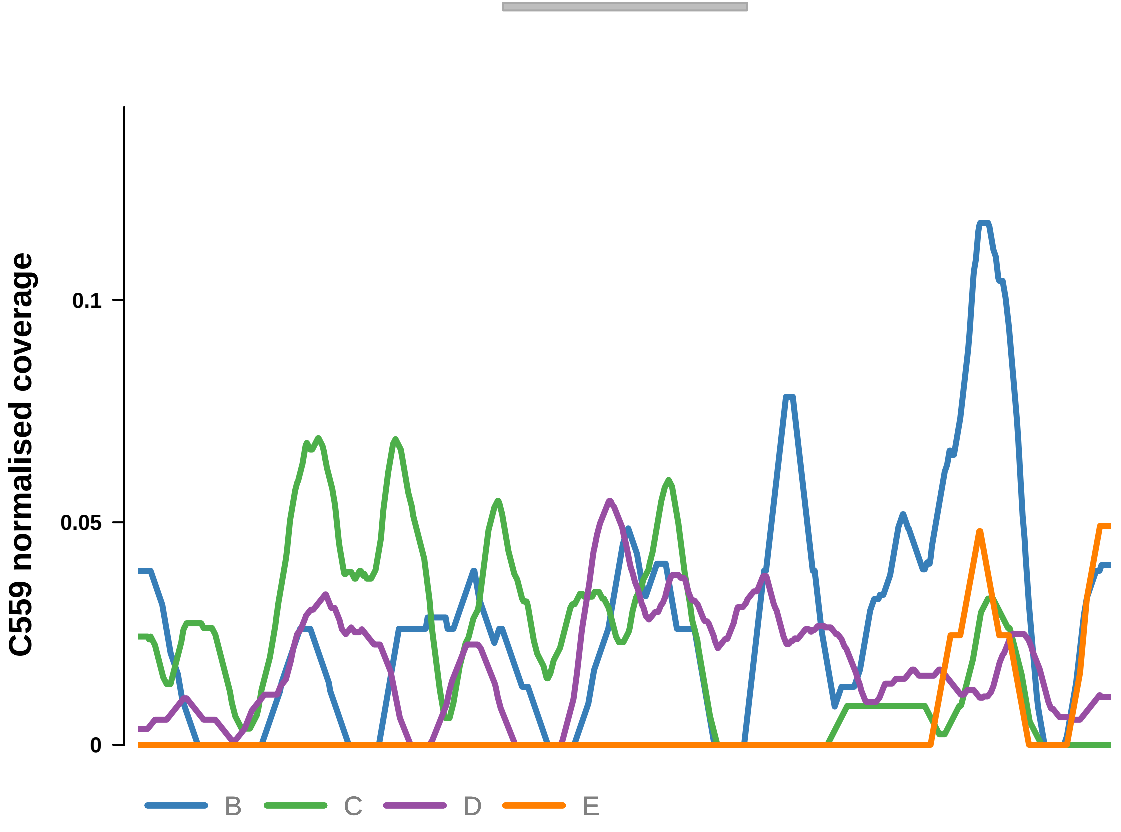


Putatively Clonal (logFC = 2.54)

| | P. | Adjusted P. |
|---|---|---|
| *~region vs. ~* | 4.57e−01 | 7.45e−01 |
| *~purity vs ~* | 2.02e−01 | 6.55e−01 |
| *~purity+region vs ~purity* | 5.82e−01 | 9.31e−01 |
| *Carcinoma vs. normal* | 1.55e−03 ** | 1.55e−03 ** |

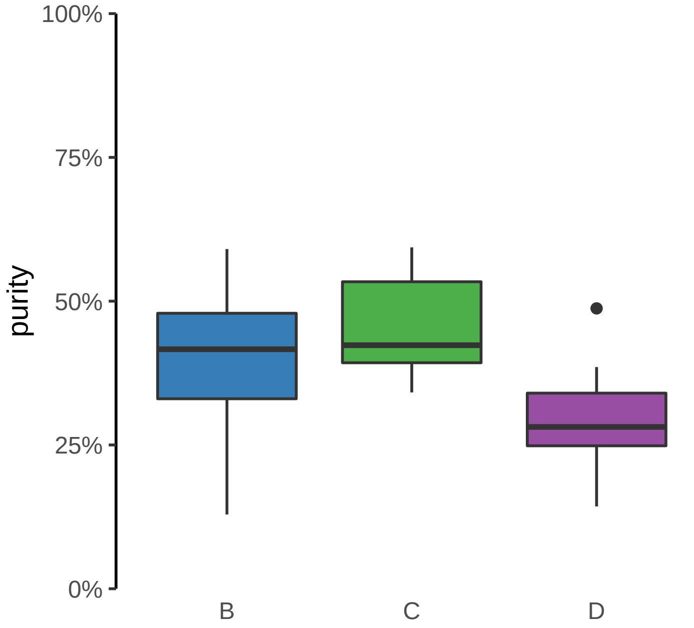

**RNU7−124P, SHB, ENSG00000255872 (GH09J037962) enhancer gain**
**chr9:37966483−37966983**

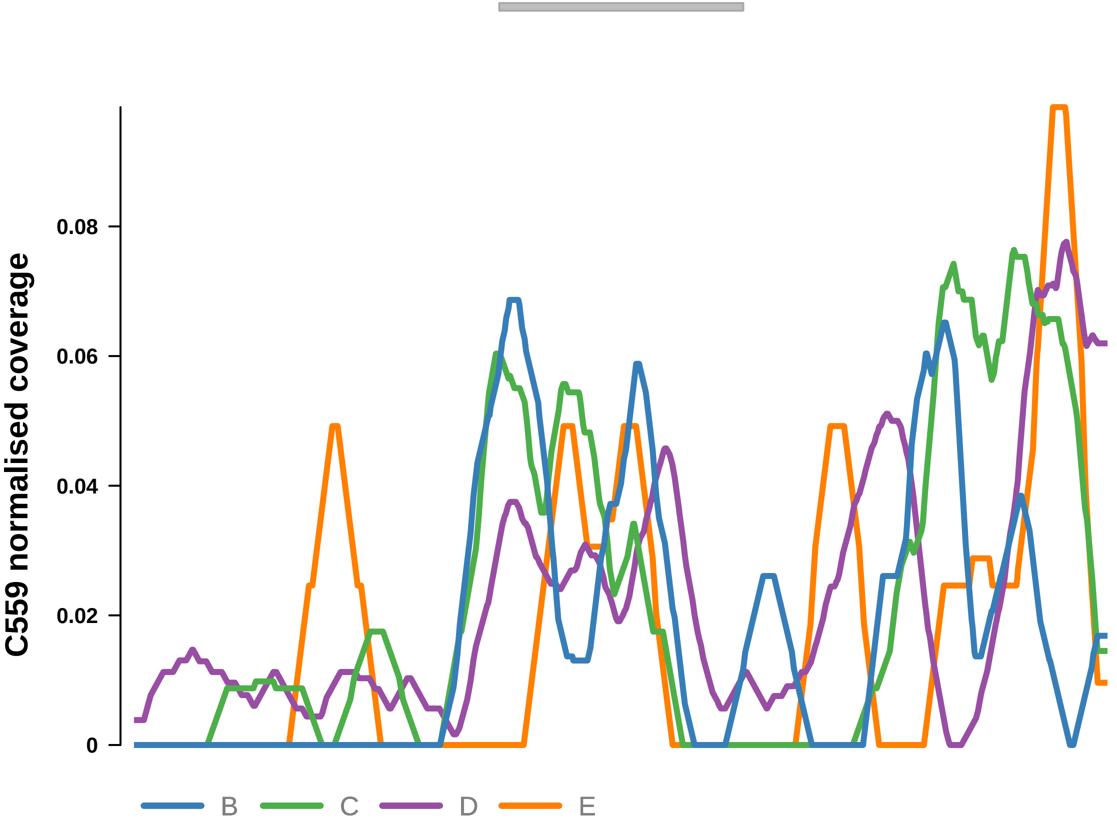


Putatively Clonal (logFC = 2.04)

| | P. | Adjusted P. |
|---|---|---|
| *~region vs. ~* | 2.87e−01 | 5.75e−01 |
| *~purity vs ~* | 1.22e−01 | 5.51e−01 |
| *~purity+region vs ~purity* | 7.49e−01 | 9.61e−01 |
| *Carcinoma vs. normal* | 8.54e−03 ** | 8.54e−03 ** |

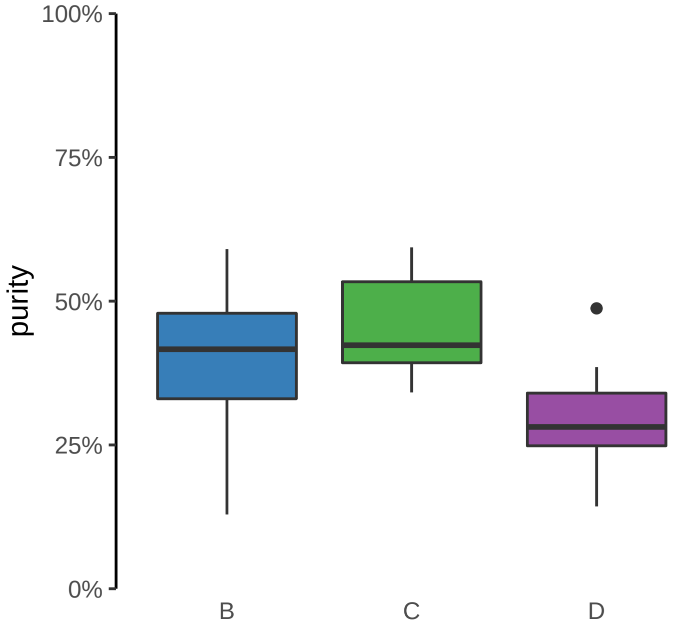

**TPRA1, ABTB1, MGLL (GH03J127766) enhancer gain**
**chr3:127767157−127767657**

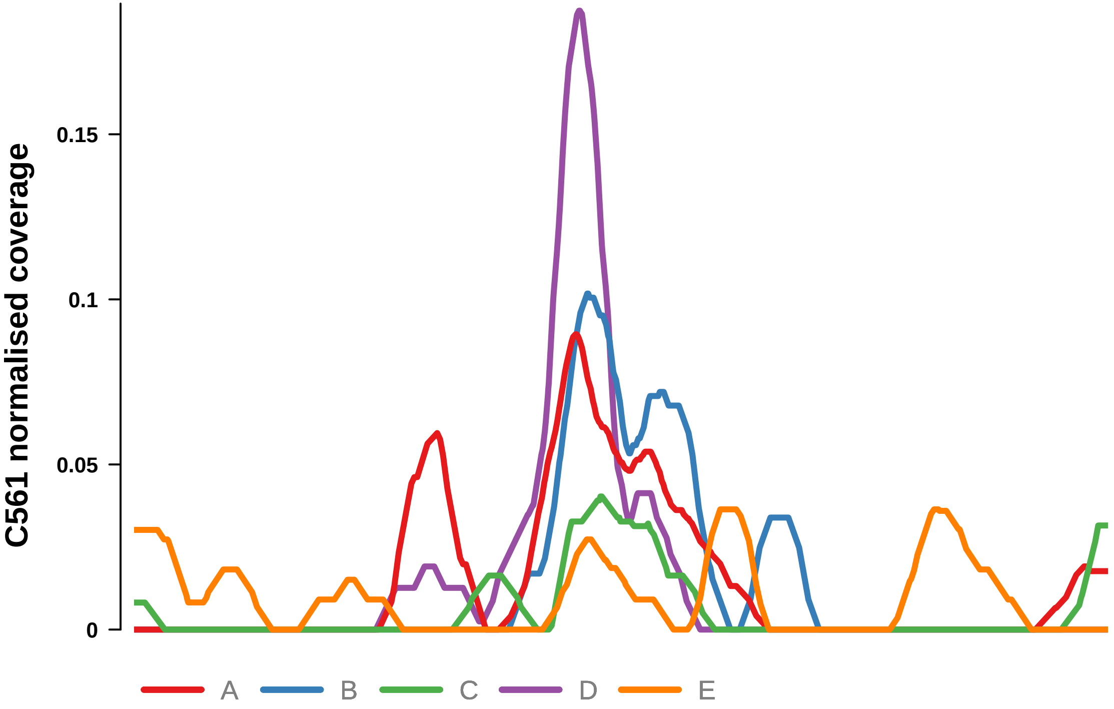


Putatively Clonal (logFC = 2.42)

| | P. | Adjusted P. |
|---|---|---|
| *~region vs. ~* | 1.83e−02 * | 9.94e−02 |
| *~purity vs ~* | 3.01e−01 | 7.53e−01 |
| *~purity+region vs ~purity* | 8.72e−03 ** | 2.35e−01 |
| *Carcinoma vs. normal* | 8.38e−04 *** | 8.38e−04 *** |

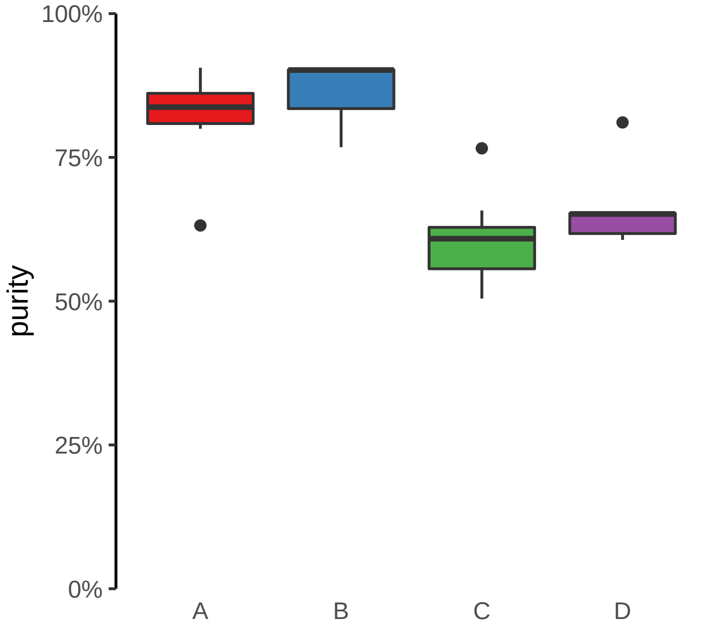

**CLASRP, MARK4 (GH19J045307) enhancer gain**
**chr19:45308035−45308535**

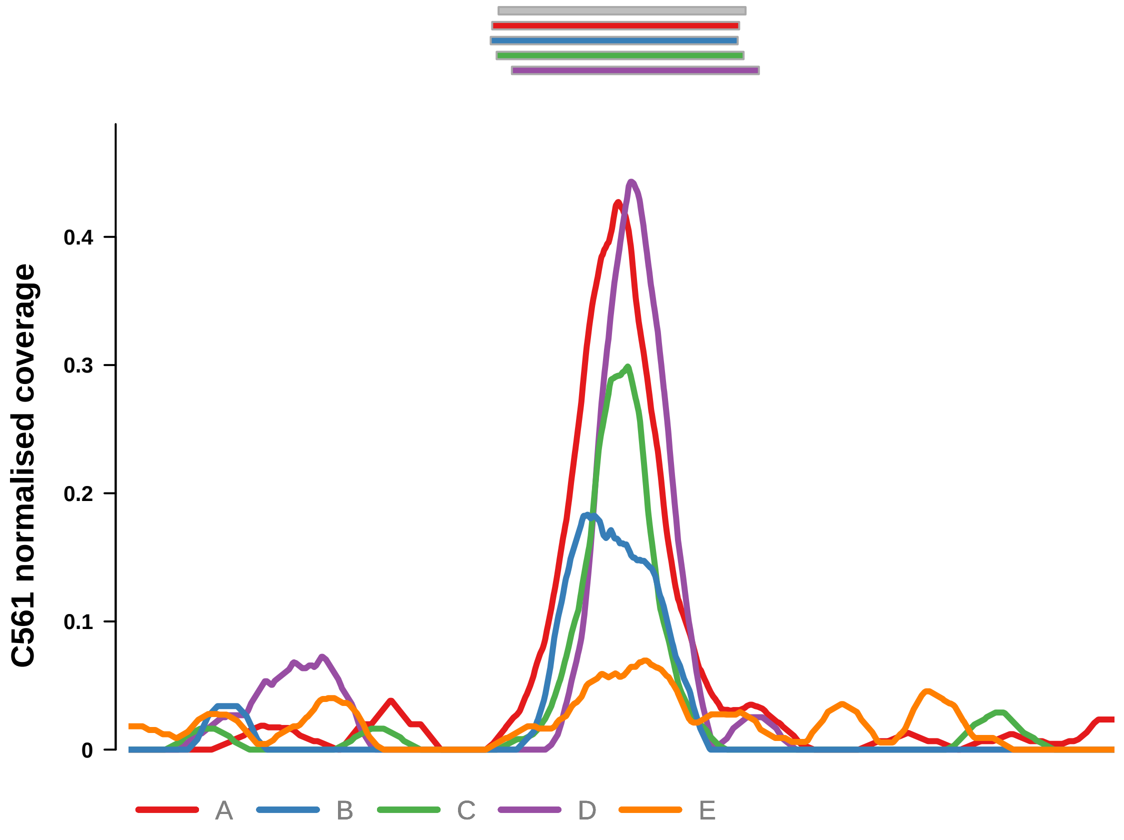


Putatively Clonal (logFC = 2.6)

| | P. | Adjusted P. |
|---|---|---|
| *~region vs. ~* | 1.31e−01 | 3.23e−01 |
| *~purity vs ~* | 4.07e−01 | 7.66e−01 |
| *~purity+region vs ~purity* | 1.62e−01 | 6.21e−01 |
| *Carcinoma vs. normal* | 2.23e−04 *** | 2.23e−04 *** |

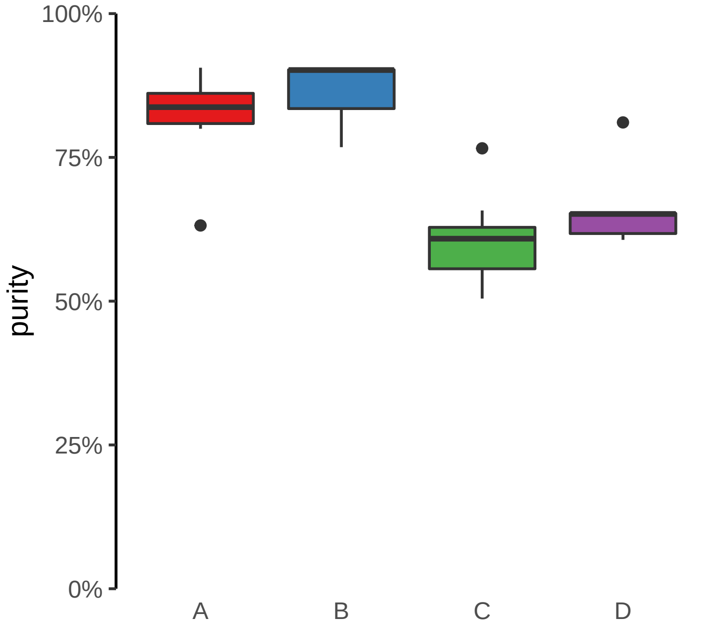

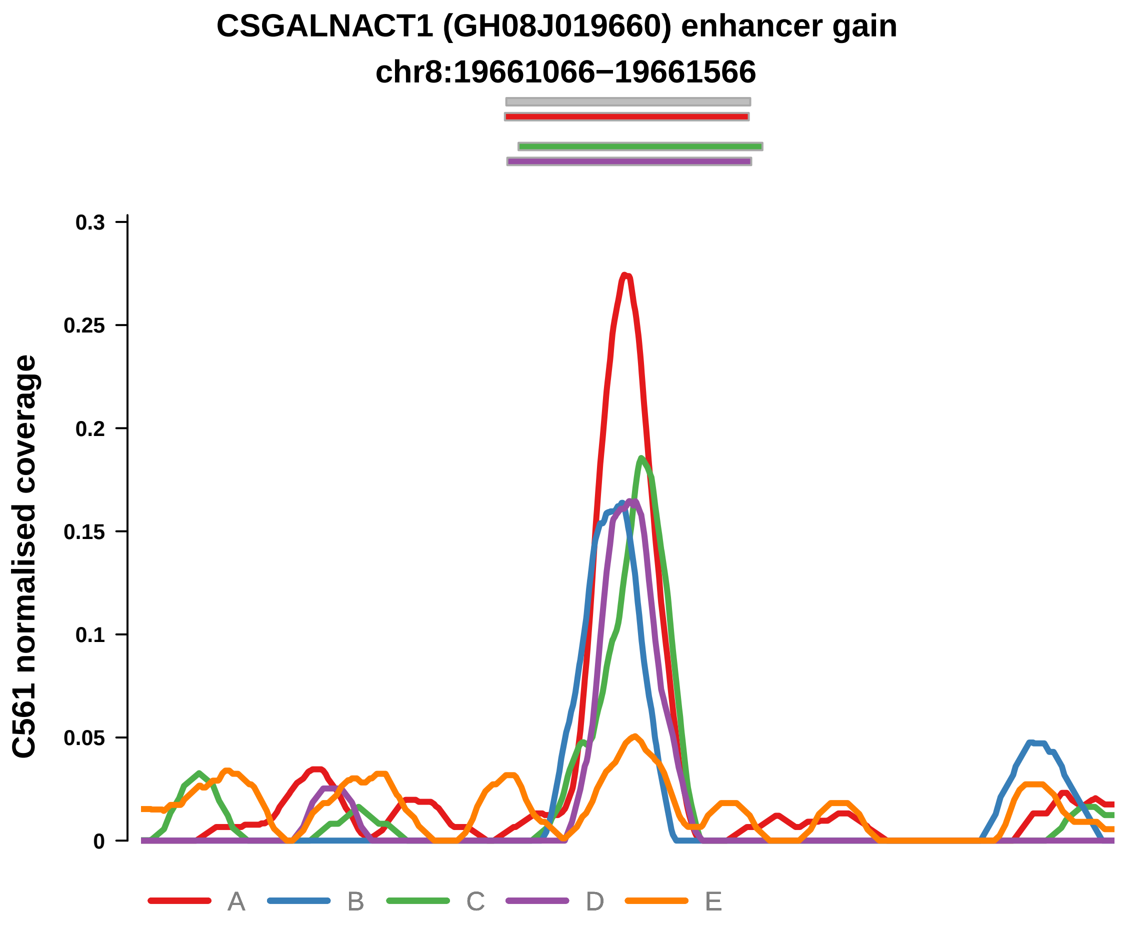


Putatively Clonal (logFC = 2.28)

| | P. | Adjusted P. |
|---|---|---|
| *~region vs. ~* | 9.76e−01 | 9.76e−01 |
| *~purity vs ~* | 9.65e−01 | 1e+00 |
| *~purity+region vs ~purity* | 9.63e−01 | 9.79e−01 |
| *Carcinoma vs. normal* | 1.23e−03 ** | 1.23e−03 ** |

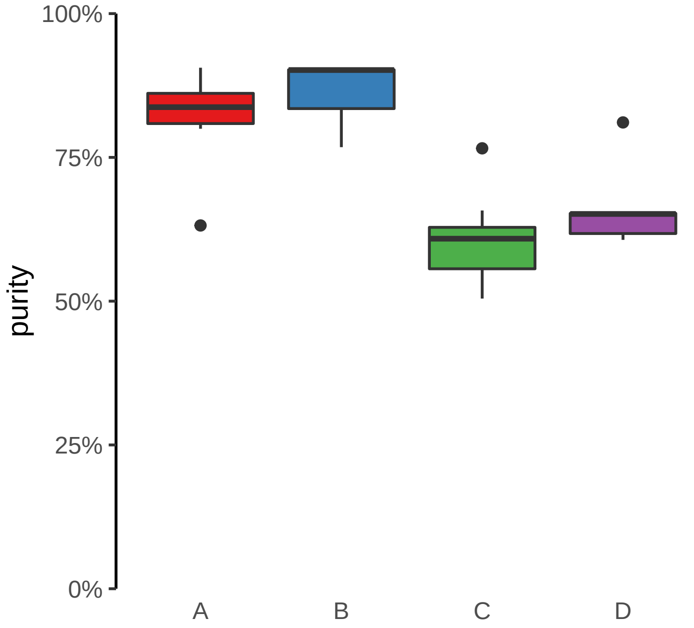

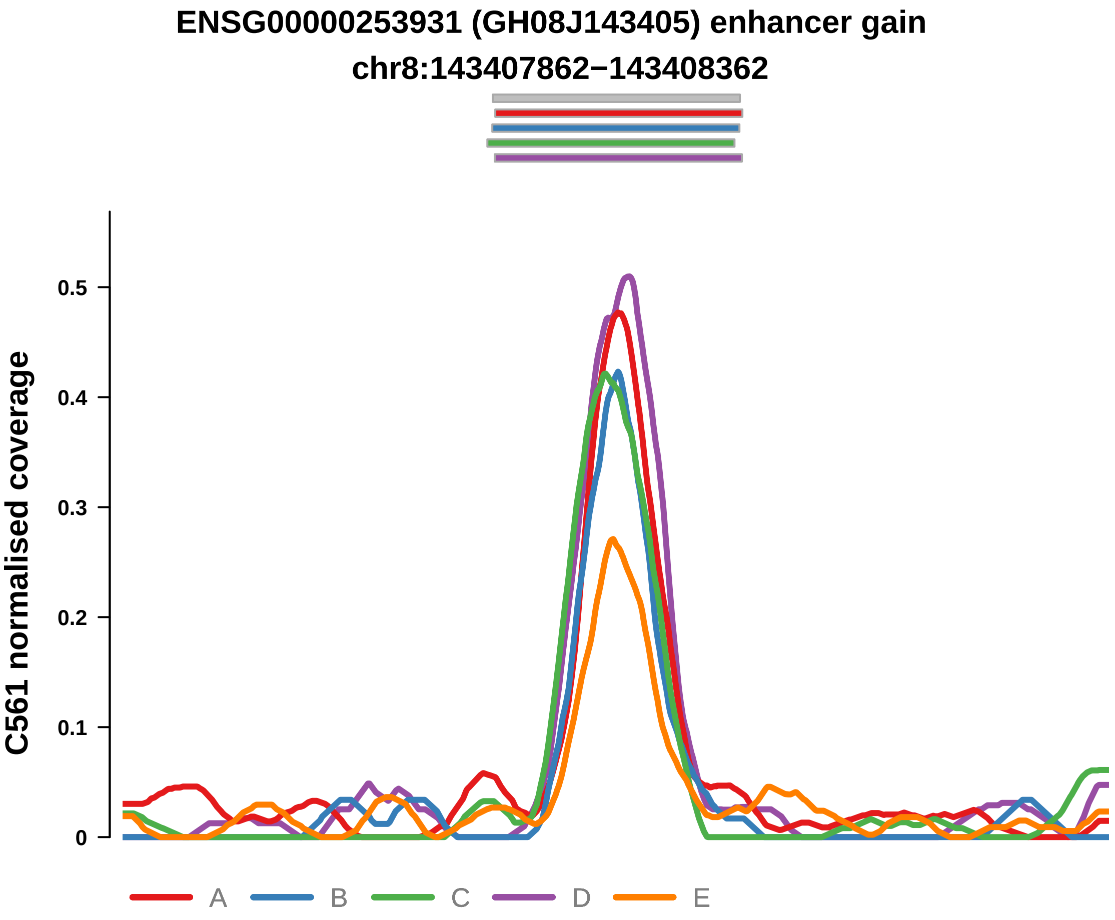


Putatively Clonal (logFC = 2.1)

| | P. | Adjusted P. |
|---|---|---|
| *~region vs. ~* | 8.28e−02 | 2.62e−01 |
| *~purity vs ~* | 3.98e−02 * | 3.18e−01 |
| *~purity+region vs ~purity* | 1.44e−01 | 5.88e−01 |
| *Carcinoma vs. normal* | 2.19e−03 ** | 2.19e−03 ** |

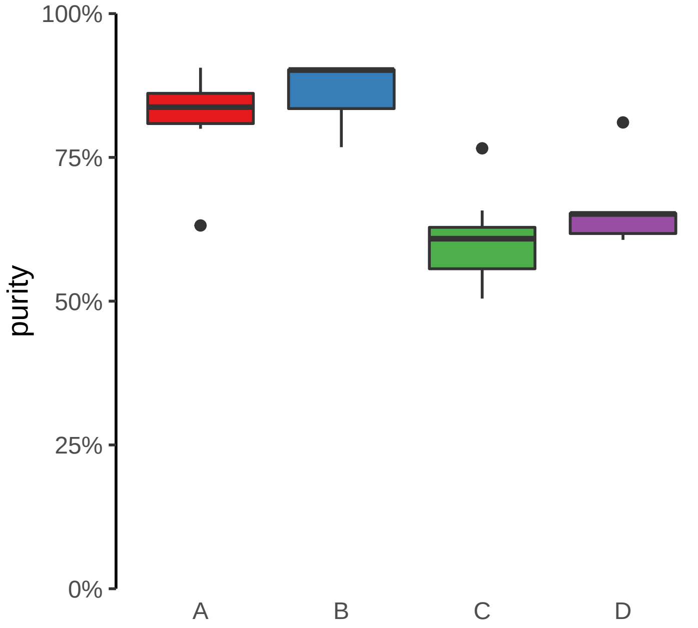

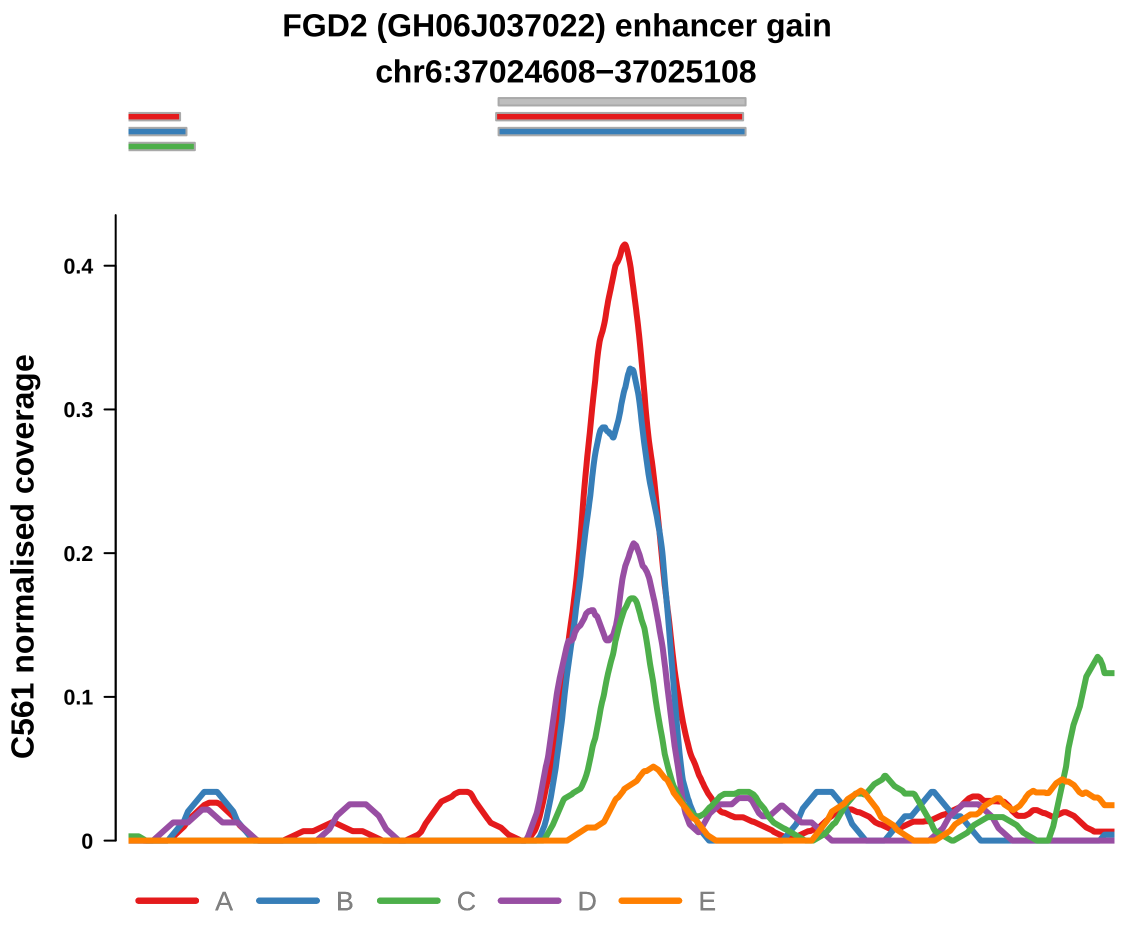


Putatively Clonal (logFC = 2.97)

| | P. | Adjusted P. |
|---|---|---|
| *~region vs. ~* | 2.51e−01 | 4.58e−01 |
| *~purity vs ~* | 9.04e−02 | 3.98e−01 |
| *~purity+region vs ~purity* | 6.67e−01 | 9.16e−01 |
| *Carcinoma vs. normal* | 3.67e−05 **** | 3.67e−05 **** |

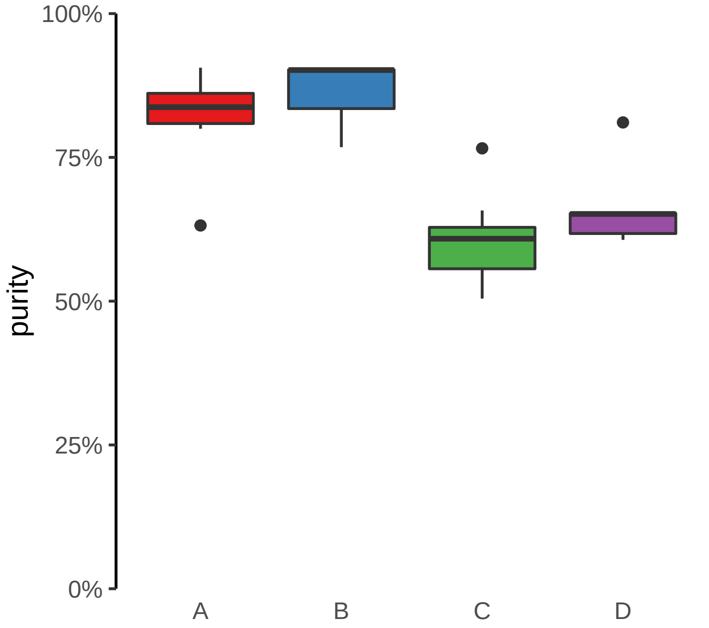

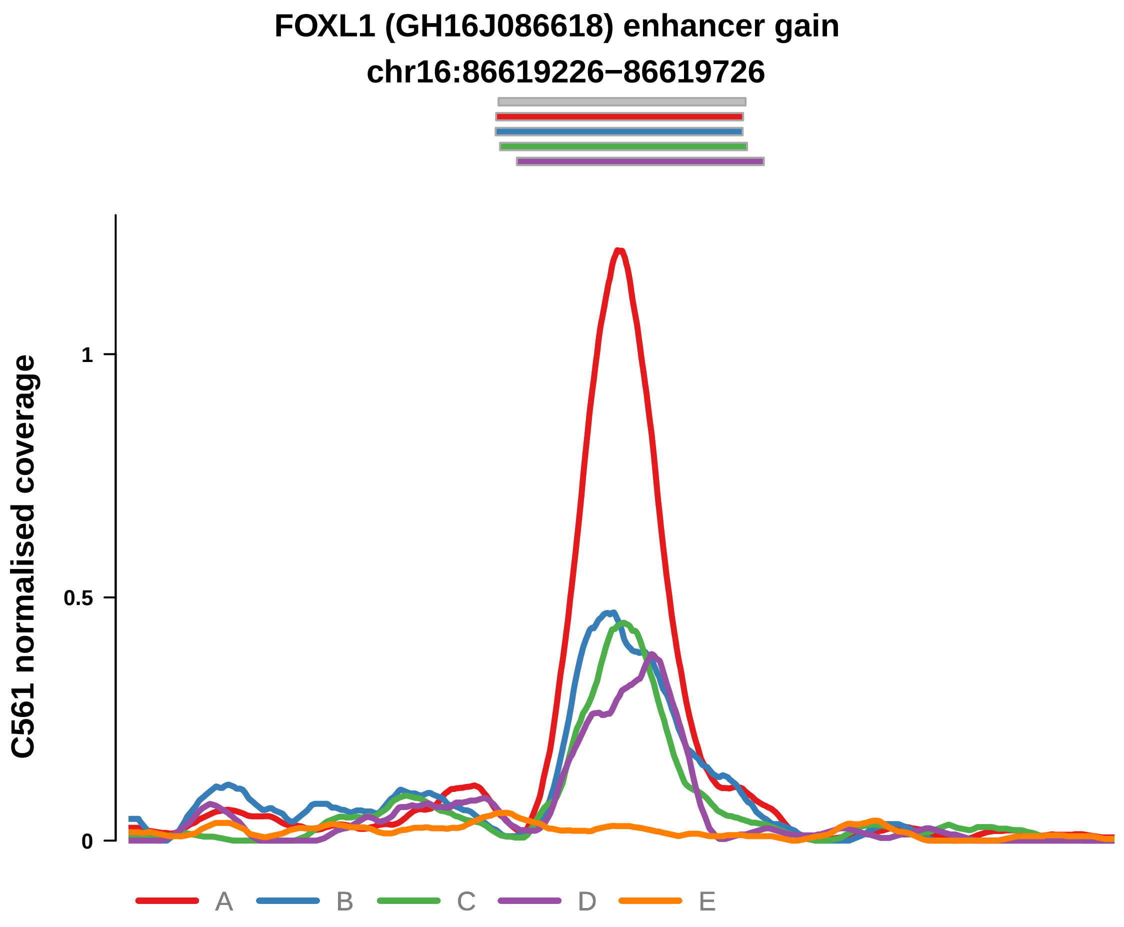


Putatively Clonal (logFC = 4.09)

| | P. | Adjusted P. |
|---|---|---|
| *~region vs. ~* | 9.95e−02 | 2.81e−01 |
| *~purity vs ~* | 6.91e−02 | 3.98e−01 |
| *~purity+region vs ~purity* | 6.23e−01 | 9.16e−01 |
| *Carcinoma vs. normal* | 4.38e−08 **** | 4.38e−08 **** |

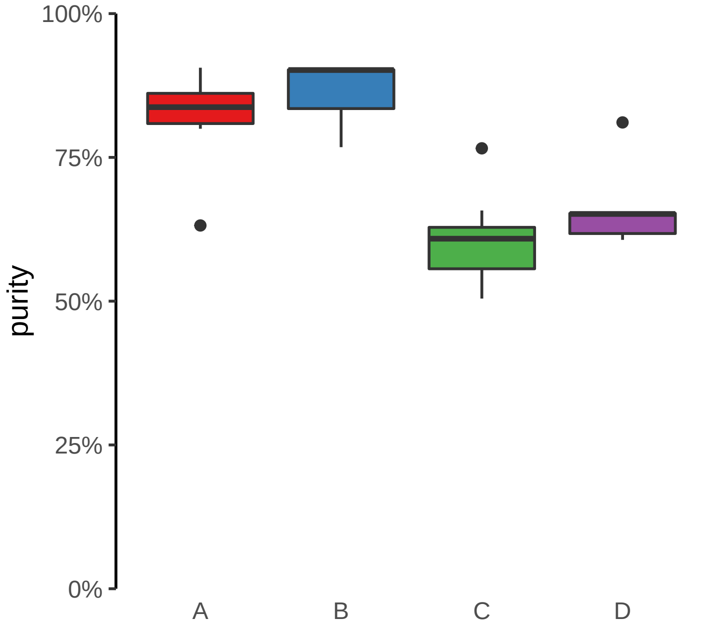

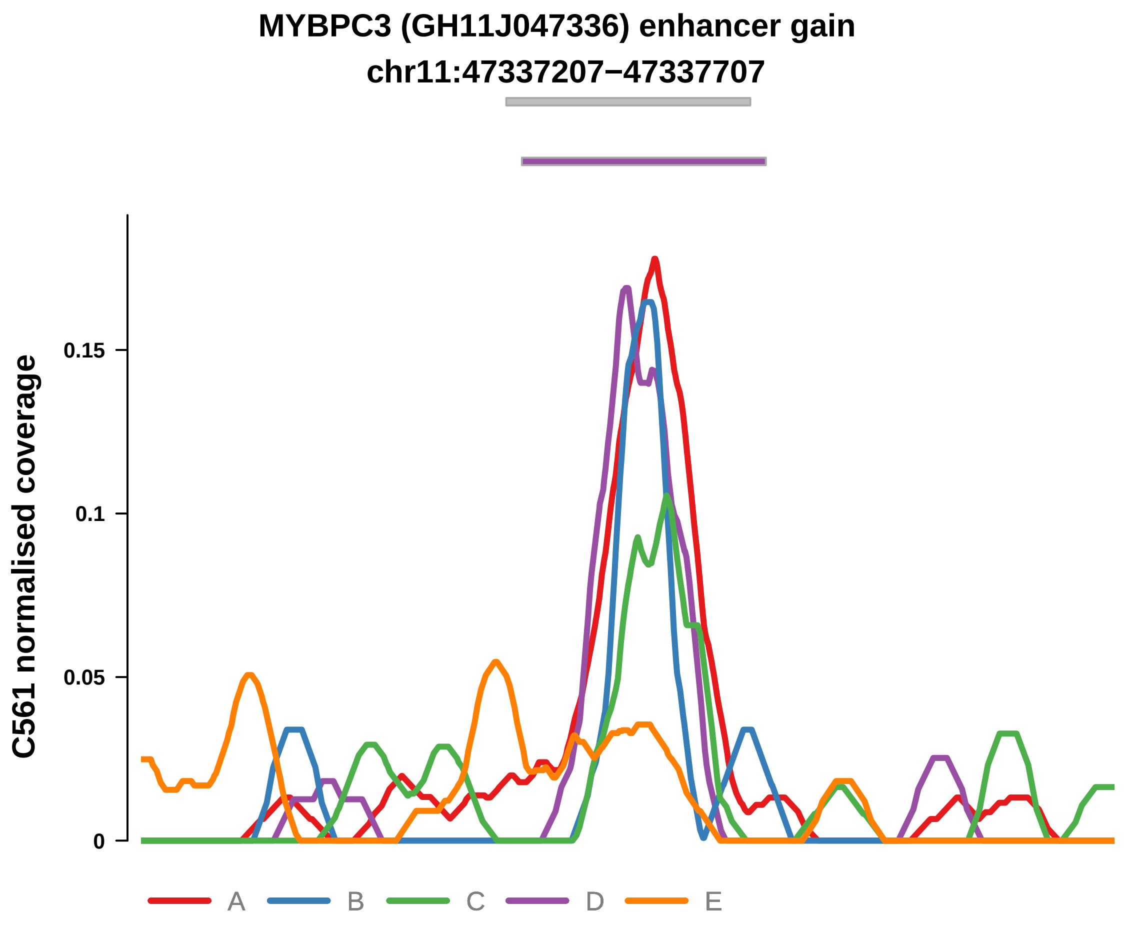


Putatively Clonal (logFC = 2.41)

| | P. | Adjusted P. |
|---|---|---|
| *~region vs. ~* | 3.13e−01 | 5.29e−01 |
| *~purity vs ~* | 8.82e−01 | 9.47e−01 |
| *~purity+region vs ~purity* | 2.8e−01 | 7.68e−01 |
| *Carcinoma vs. normal* | 7.83e−04 *** | 7.83e−04 *** |

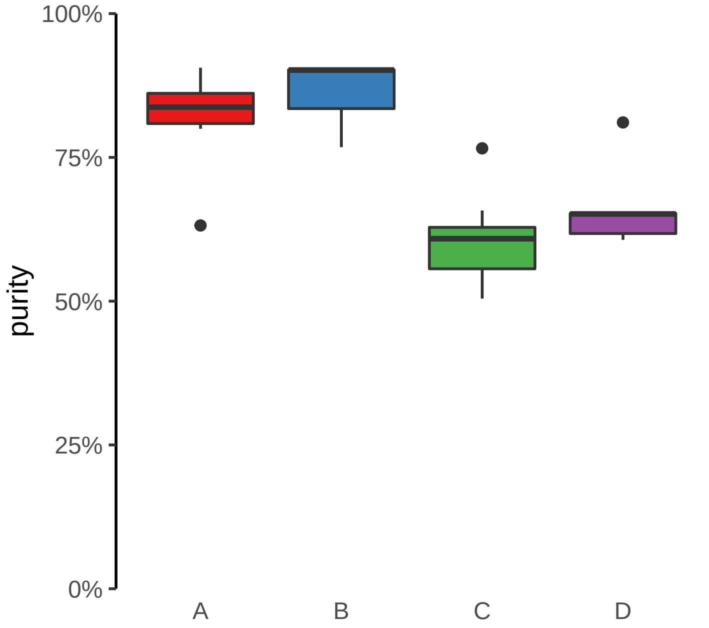

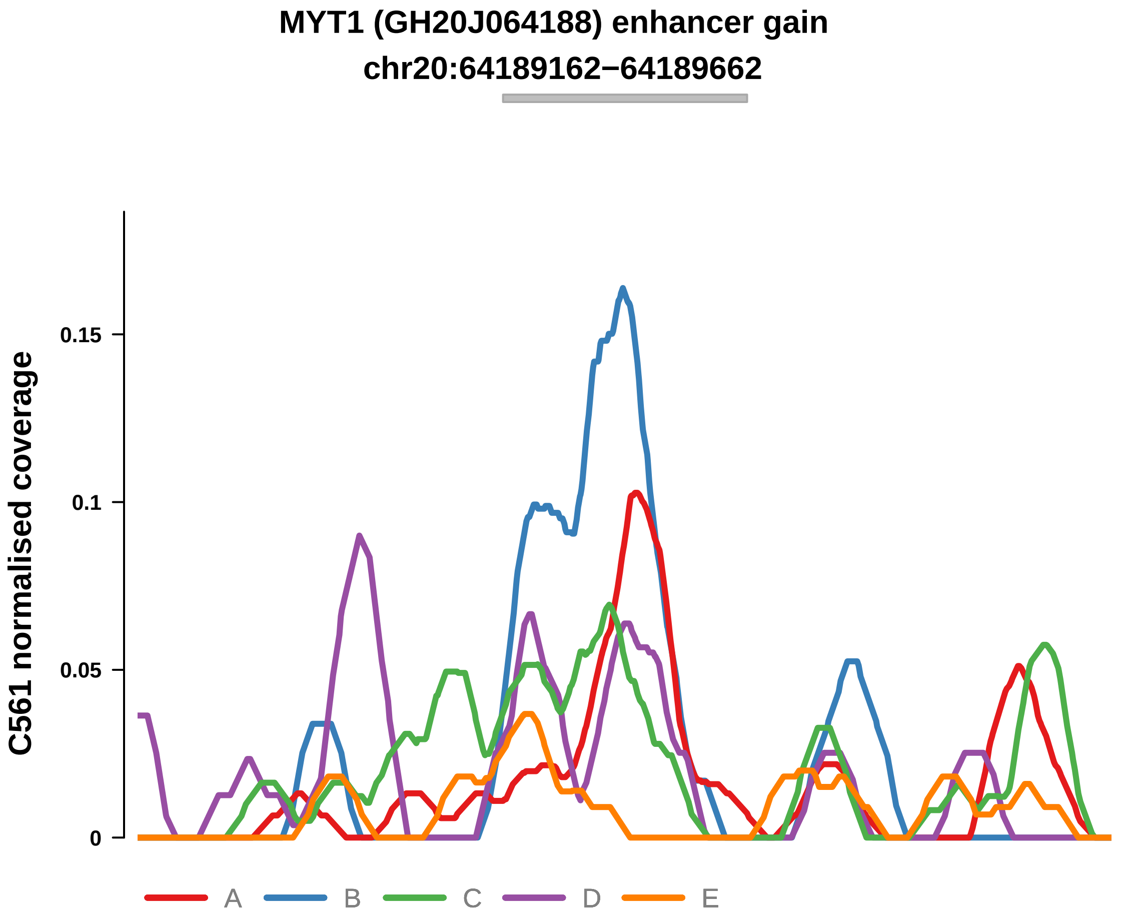


Putatively Clonal (logFC = 2.3)

| | P. | Adjusted P. |
|---|---|---|
| *~region vs. ~* | 3.77e−01 | 5.91e−01 |
| *~purity vs ~* | 9.63e−02 | 4.04e−01 |
| *~purity+region vs ~purity* | 7.45e−02 | 4.24e−01 |
| *Carcinoma vs. normal* | 1.22e−03 ** | 1.22e−03 ** |

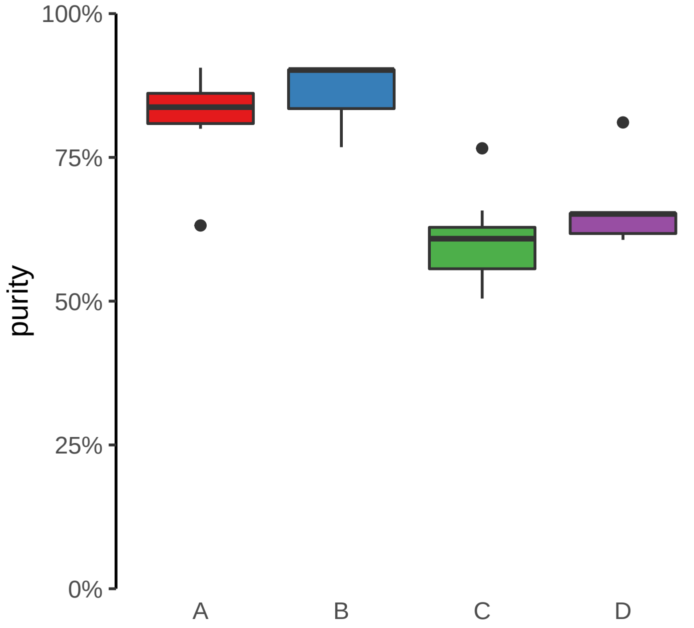

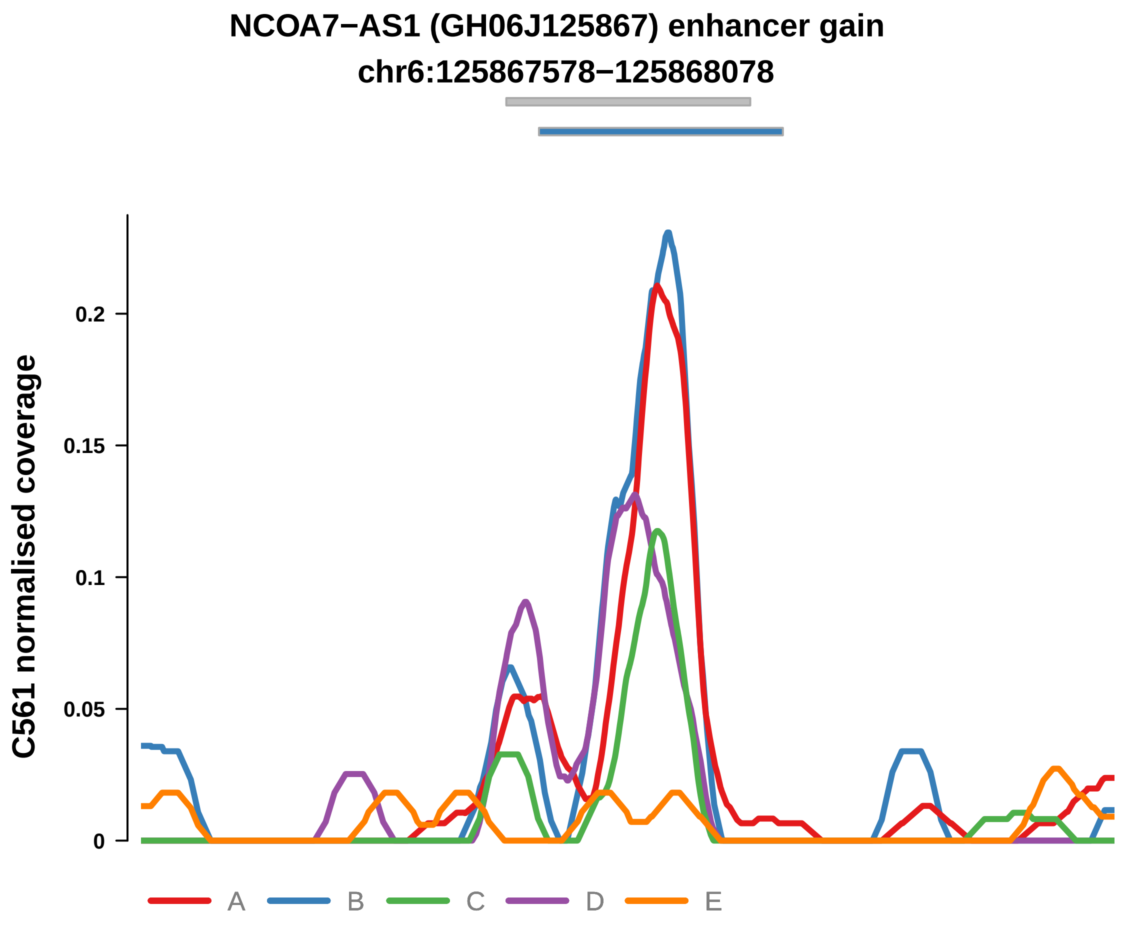


Putatively Clonal (logFC = 2.49)

| | P. | Adjusted P. |
|---|---|---|
| *~region vs. ~* | 7.91e−02 | 2.6e−01 |
| *~purity vs ~* | 5.92e−01 | 8.53e−01 |
| *~purity+region vs ~purity* | 4.46e−02 * | 3.92e−01 |
| *Carcinoma vs. normal* | 4.86e−04 *** | 4.86e−04 *** |

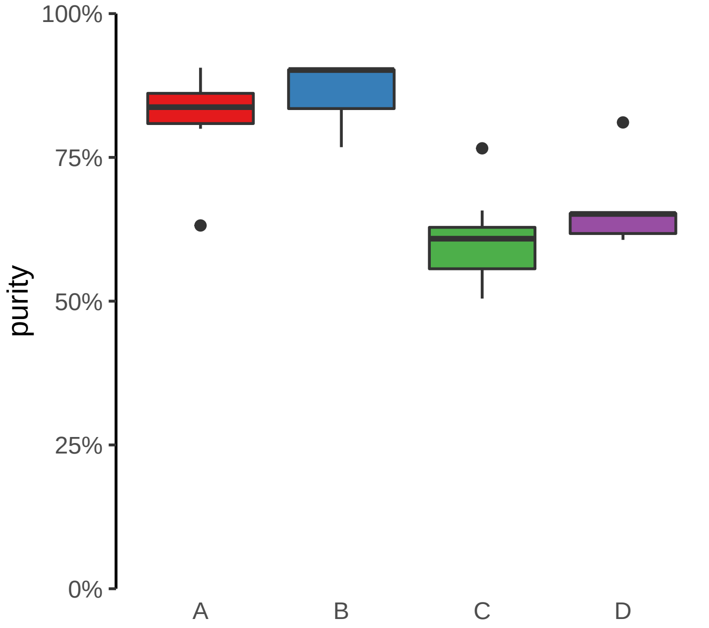

**NFATC2 (GH20J051486) enhancer gain**
**chr20:51487818−51488318**

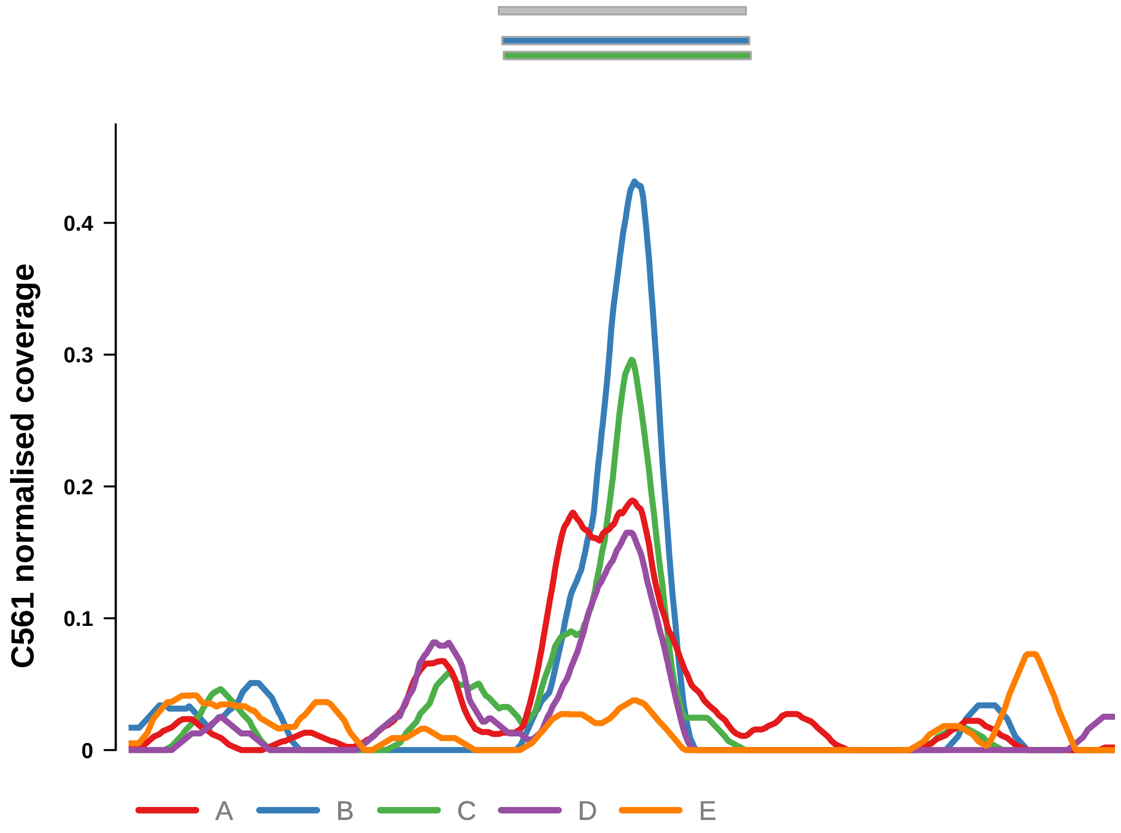


Putatively Clonal (logFC = 3.24)

| | P. | Adjusted P. |
|---|---|---|
| *~region vs. ~* | 2.43e−01 | 4.58e−01 |
| *~purity vs ~* | 6.66e−01 | 8.74e−01 |
| *~purity+region vs ~purity* | 1.44e−01 | 5.88e−01 |
| *Carcinoma vs. normal* | 8.64e−06 **** | 8.64e−06 **** |

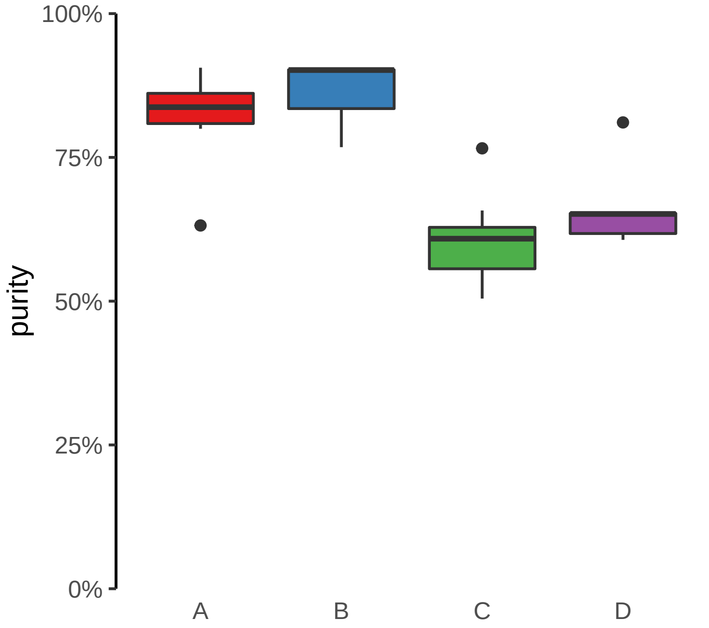

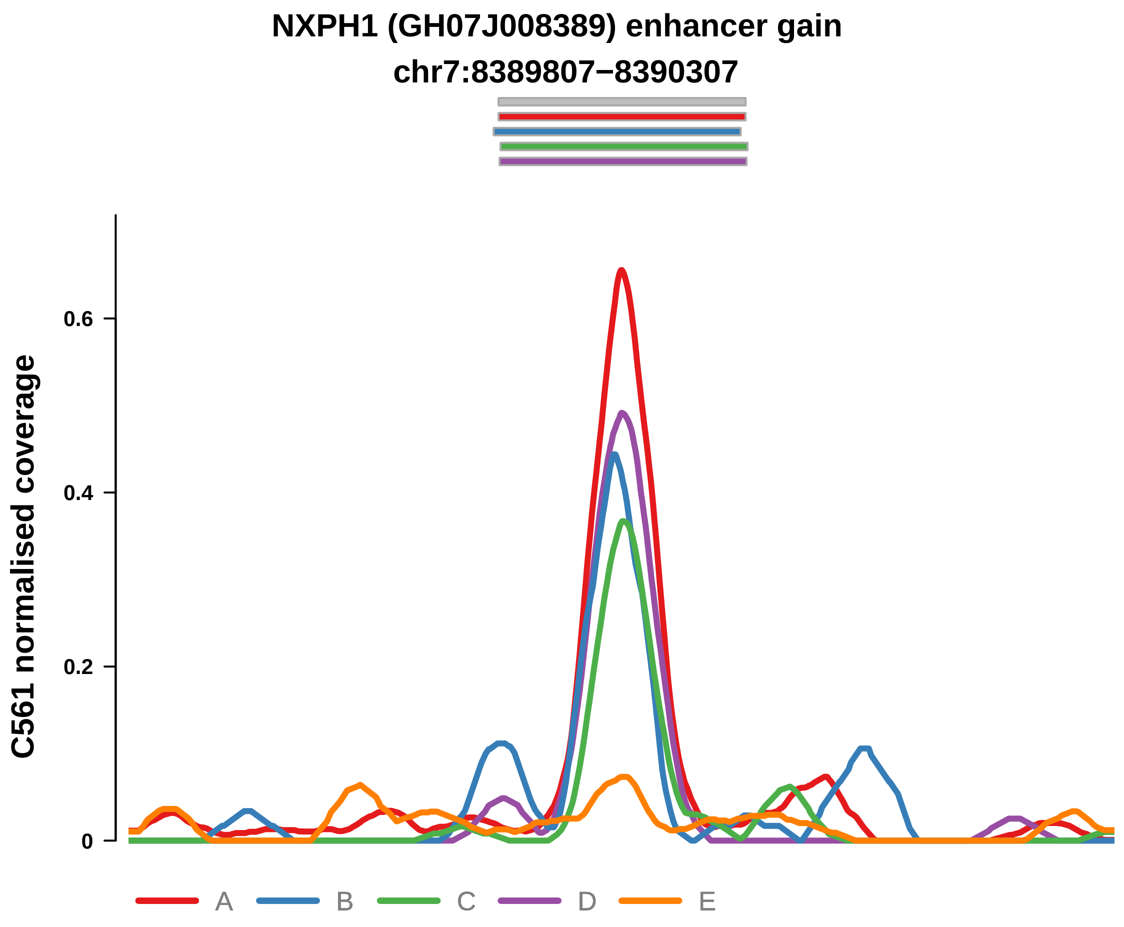


Putatively Clonal (logFC = 3.16)

| | P. | Adjusted P. |
|---|---|---|
| *~region vs. ~* | 1.18e−01 | 3.02e−01 |
| *~purity vs ~* | 9.6e−01 | 1e+00 |
| *~purity+region vs ~purity* | 5.73e−02 | 4.2e−01 |
| *Carcinoma vs. normal* | 1.16e−05 **** | 1.16e−05 **** |

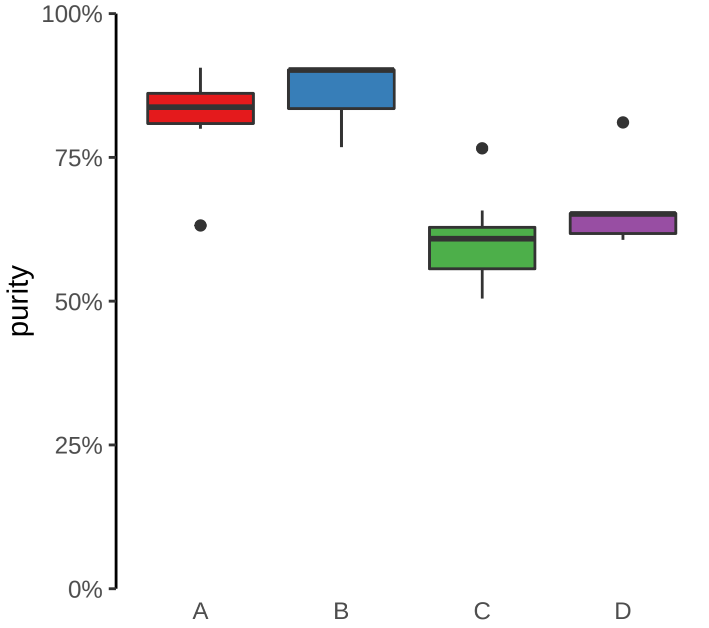

**PPL (GH16J004942) enhancer gain**
**chr16:4944634−4945134**

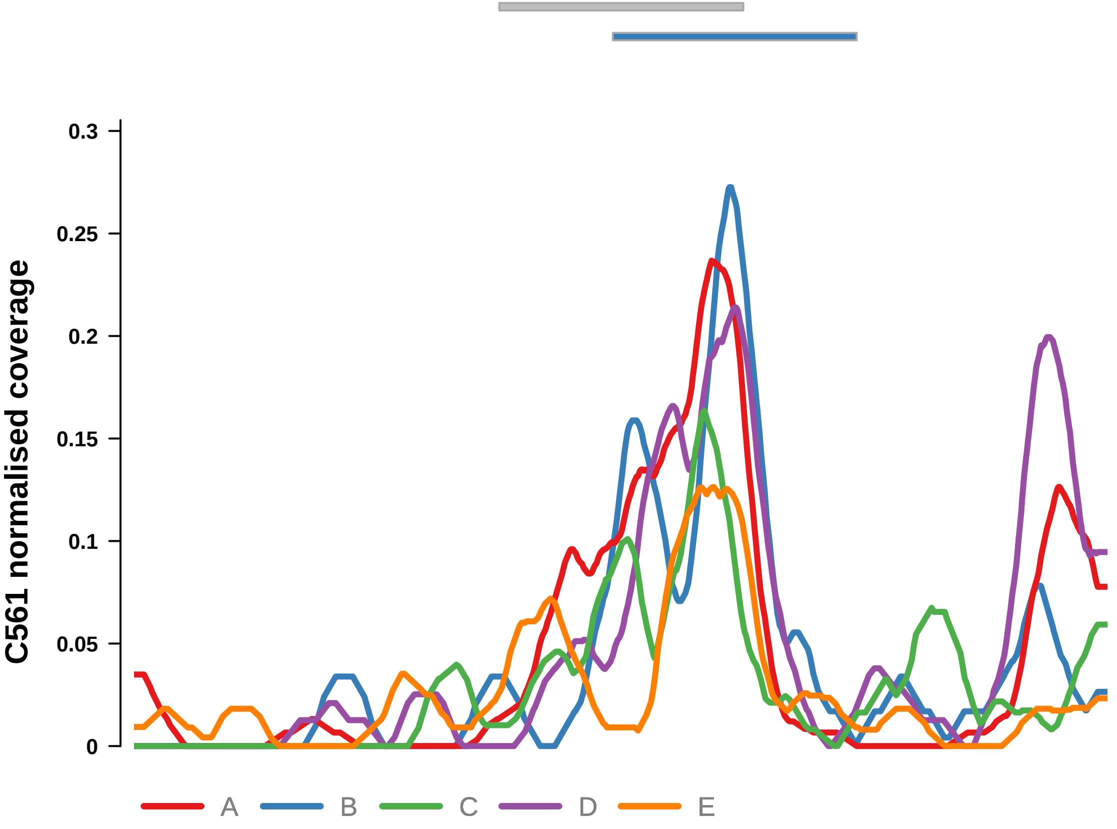


Putatively Clonal (logFC = 2.1)

| | P. | Adjusted P. |
|---|---|---|
| *~region vs. ~* | 6.41e−01 | 7.67e−01 |
| *~purity vs ~* | 4.47e−01 | 7.66e−01 |
| *~purity+region vs ~purity* | 7.48e−01 | 9.16e−01 |
| *Carcinoma vs. normal* | 2.38e−03 ** | 2.38e−03 ** |

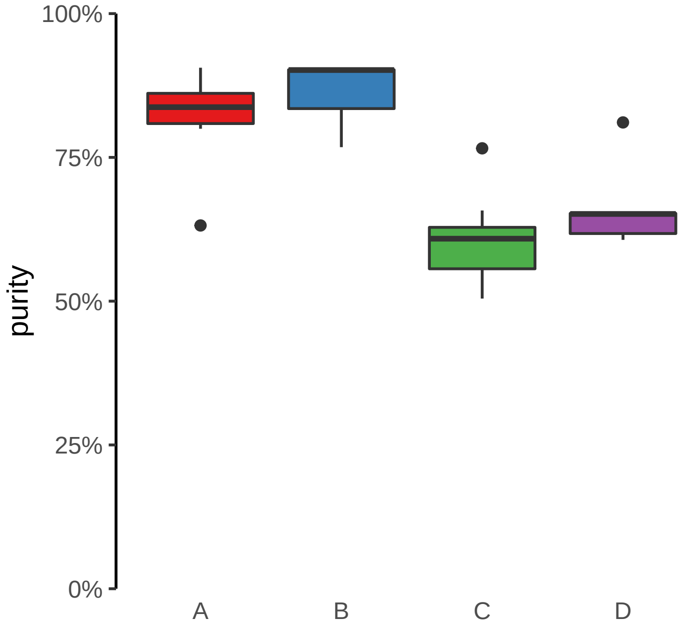

**RNU7−124P, SHB, ENSG00000255872 (GH09J037962) enhancer gain**
**chr9:37966483−37966983**

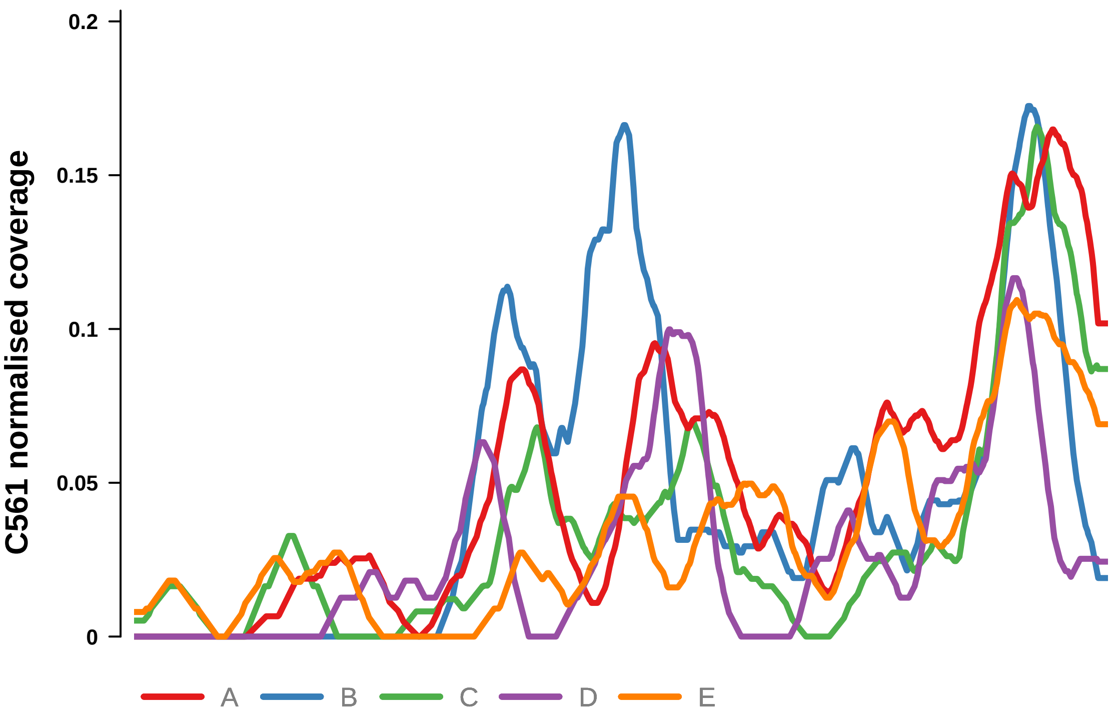


Putatively Clonal (logFC = 2.08)

| | P. | Adjusted P. |
|---|---|---|
| *~region vs. ~* | 2.16e−01 | 4.37e−01 |
| *~purity vs ~* | 3.81e−01 | 7.66e−01 |
| *~purity+region vs ~purity* | 3.61e−01 | 7.68e−01 |
| *Carcinoma vs. normal* | 3.11e−03 ** | 3.11e−03 ** |

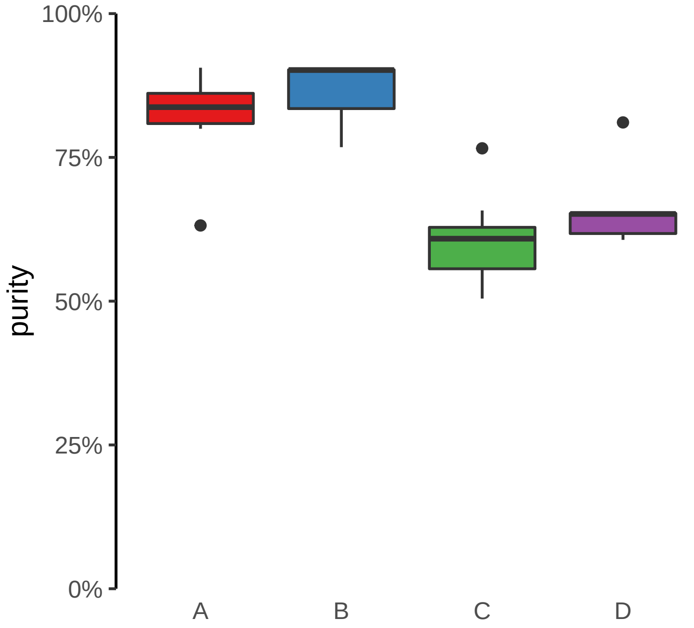

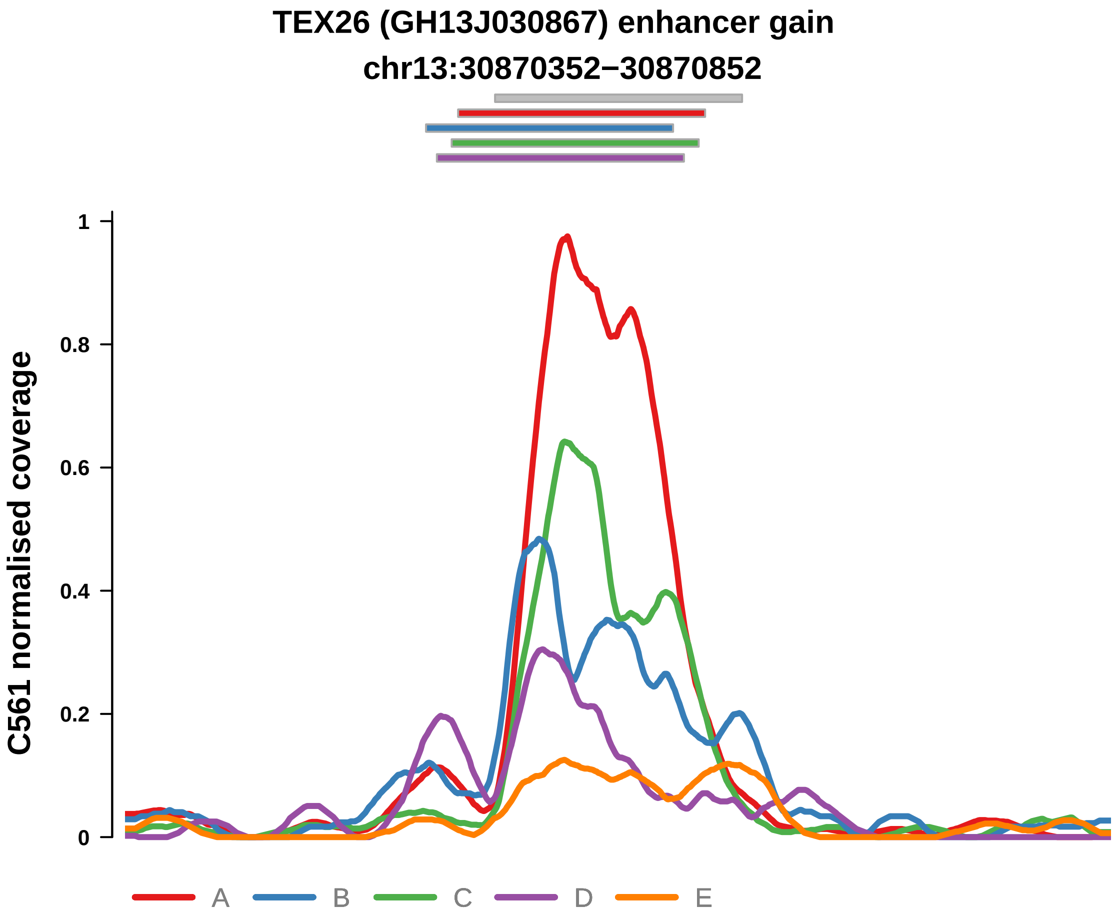


Putatively Clonal (logFC = 2.94)

| | P. | Adjusted P. |
|---|---|---|
| *~region vs. ~* | 4.41e−02 * | 1.77e−01 |
| *~purity vs ~* | 4.34e−01 | 7.66e−01 |
| *~purity+region vs ~purity* | 6.63e−02 | 4.24e−01 |
| *Carcinoma vs. normal* | 3.1e−05 **** | 3.1e−05 **** |

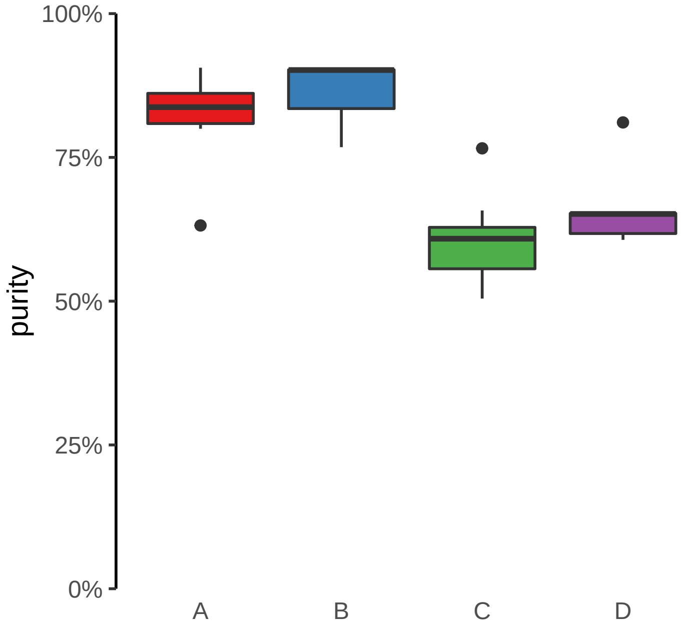

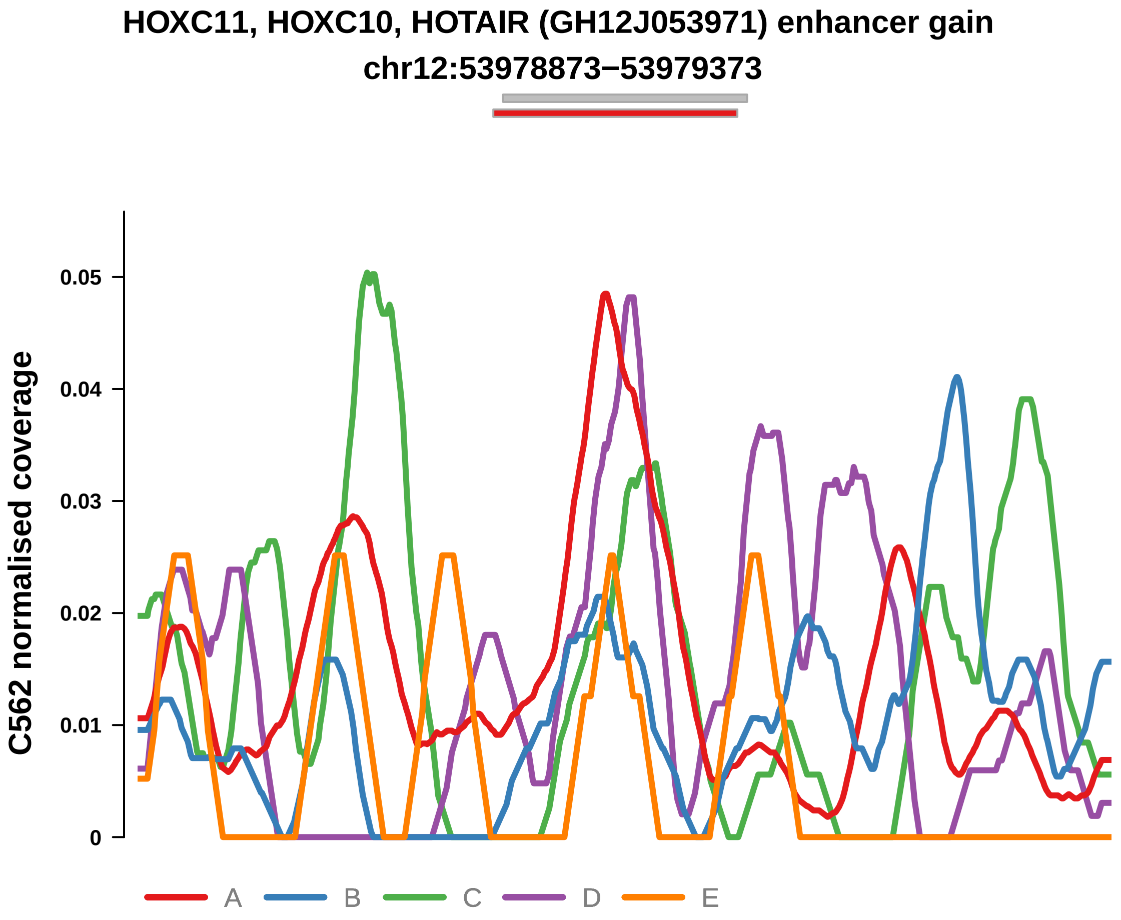


Putatively Clonal (logFC = 1.86)

| | P. | Adjusted P. |
|---|---|---|
| *~region vs. ~* | 2.02e−02 * | 1.48e−01 |
| *~purity vs ~* | 1.62e−04 *** | 4.1e−03 ** |
| *~purity+region vs ~purity* | 1.09e−01 | 5.06e−01 |
| *Carcinoma vs. normal* | 9.03e−03 ** | 9.03e−03 ** |

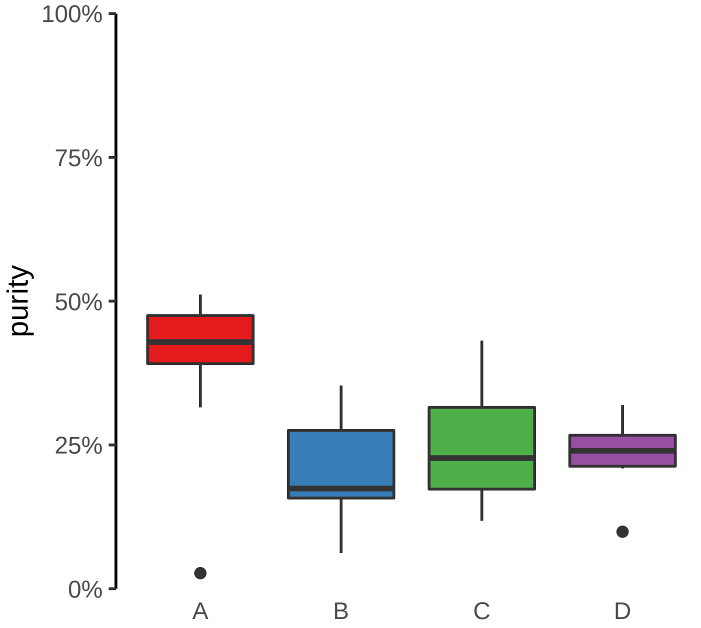

**CLASRP, MARK4 (GH19J045307) enhancer gain**
**chr19:45308035−45308535**

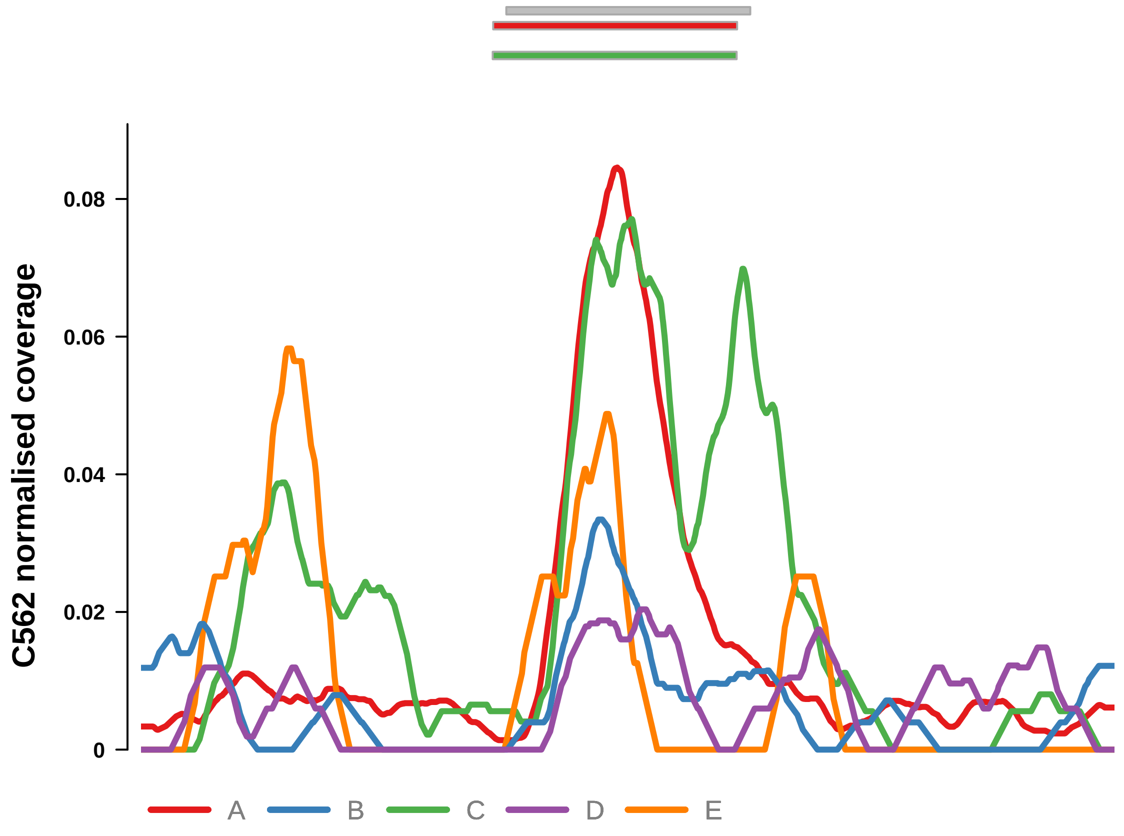


Putatively Clonal (logFC = 2.31)

| | P. | Adjusted P. |
|---|---|---|
| *~region vs. ~* | 2.05e−05 **** | 1.8e−03 ** |
| *~purity vs ~* | 1.84e−08 **** | 1.62e−06 **** |
| *~purity+region vs ~purity* | 6.37e−02 | 4.01e−01 |
| *Carcinoma vs. normal* | 1.11e−03 ** | 1.11e−03 ** |

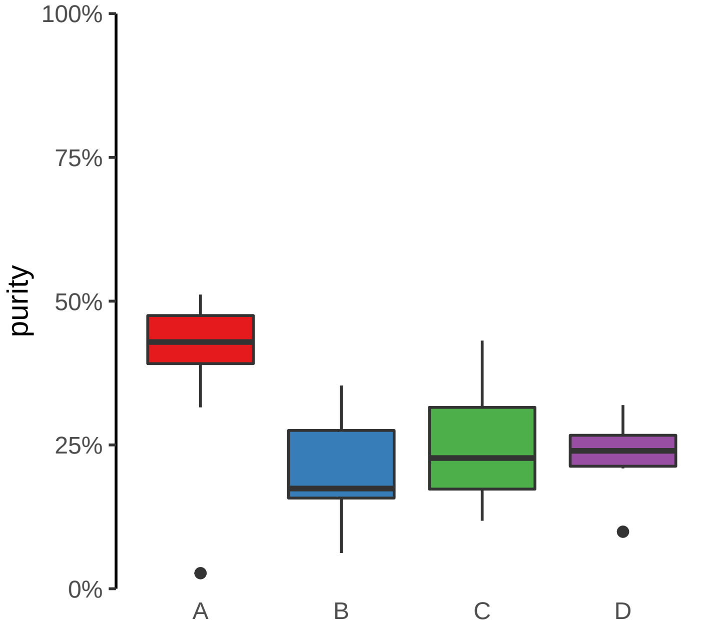

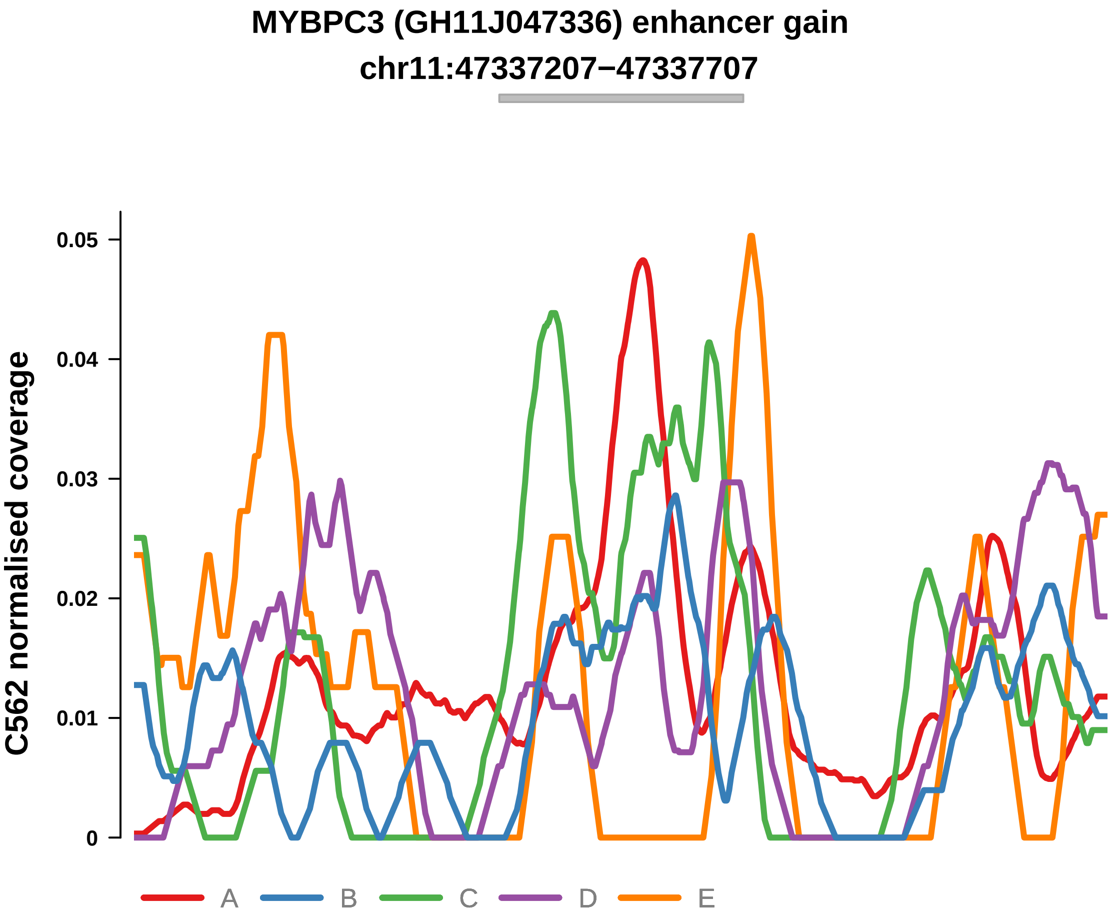


Putatively Clonal (logFC = 2.55)

| | P. | Adjusted P. |
|---|---|---|
| *~region vs. ~* | 1.55e−01 | 5.14e−01 |
| *~purity vs ~* | 4.65e−03 ** | 3.41e−02 * |
| *~purity+region vs ~purity* | 7.77e−01 | 9.41e−01 |
| *Carcinoma vs. normal* | 4.84e−04 *** | 4.84e−04 *** |

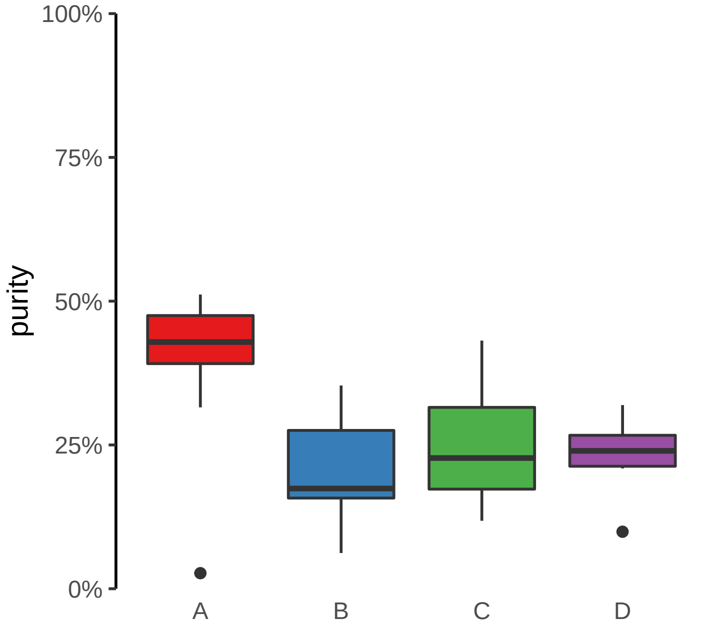
