## Supplementary Figures for "The co-evolution of the genome and epigenome in colorectal cancer": figS13_enhancer_loss.pdf

**ANTXR2 (GH04J080057) enhancer loss**
**chr4:80058687−80059187**

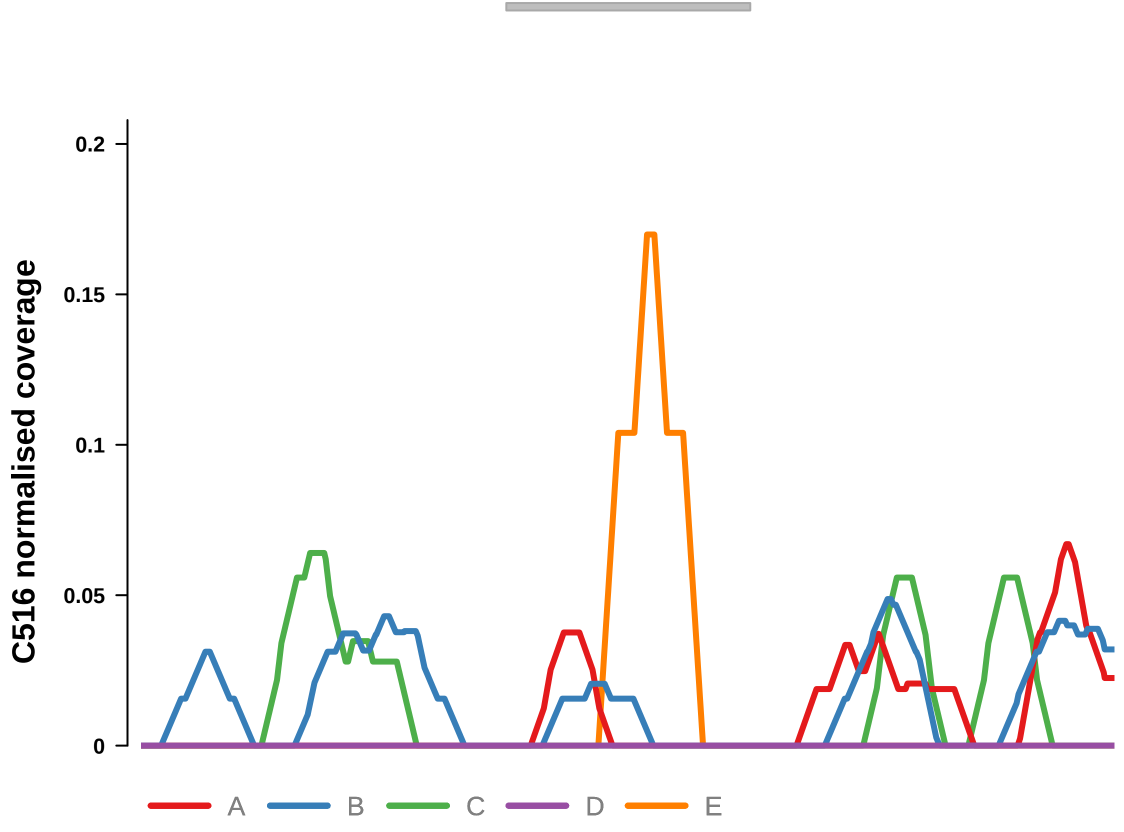


Putatively Clonal (logFC = −10.72)

| | P. | Adjusted P. |
|---|---|---|
| *~region vs. ~* | 8.49e−01 | 9.59e−01 |
| *~purity vs ~* | 8.77e−01 | 9.53e−01 |
| *~purity+region vs ~purity* | 4.87e−01 | 1e+00 |
| *Carcinoma vs. normal* | 2.67e−03 ** | 2.67e−03 ** |

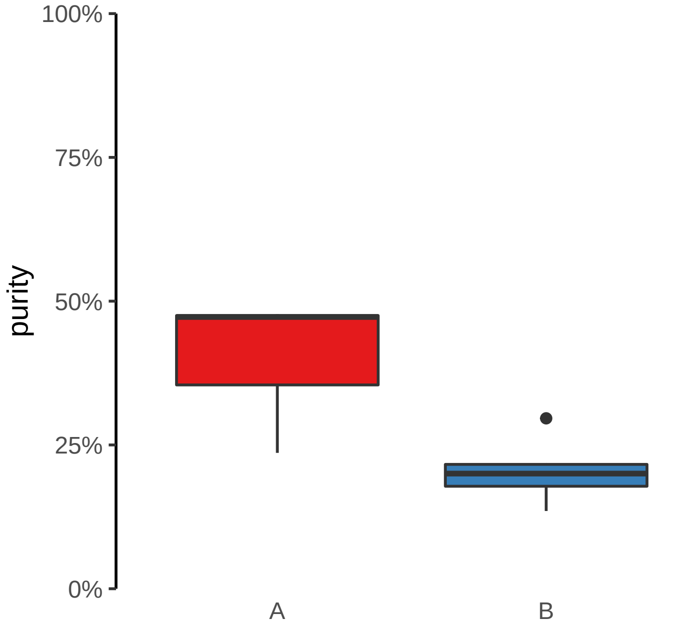

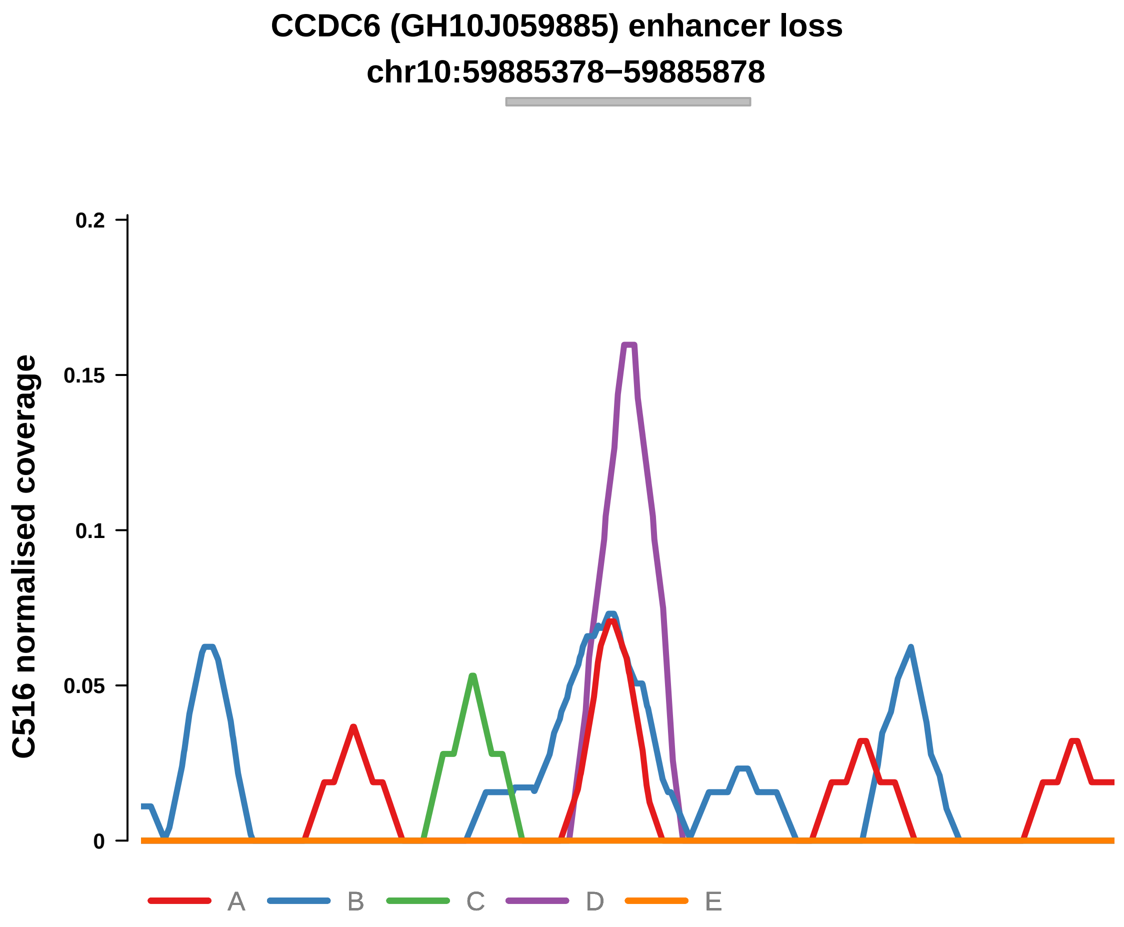


Putatively Clonal (logFC = −10.69)

| | P. | Adjusted P. |
|---|---|---|
| *~region vs. ~* | 1.37e−01 | 7.59e−01 |
| *~purity vs ~* | 9.27e−02 | 6.69e−01 |
| *~purity+region vs ~purity* | 8.86e−01 | 1e+00 |
| *Carcinoma vs. normal* | 3.05e−03 ** | 3.05e−03 ** |

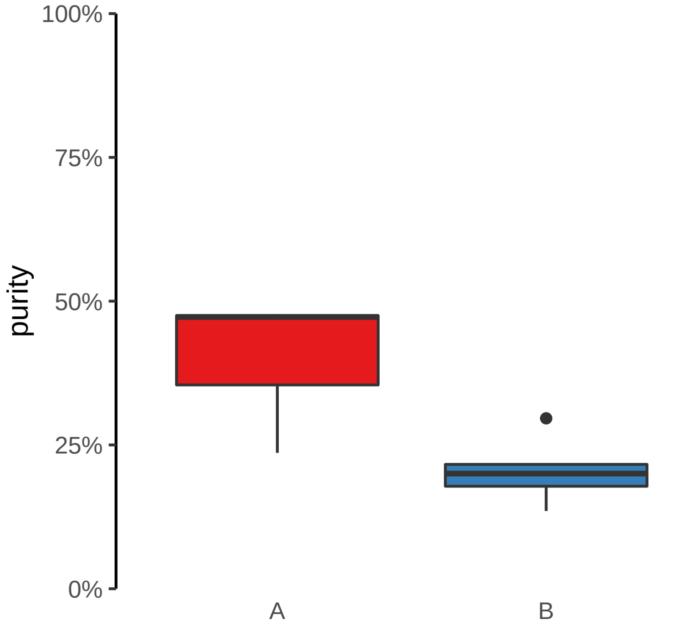

**CLASP2, ENSG00000271324 (GH03J033790) enhancer loss**
**chr3:33791629−33792129**

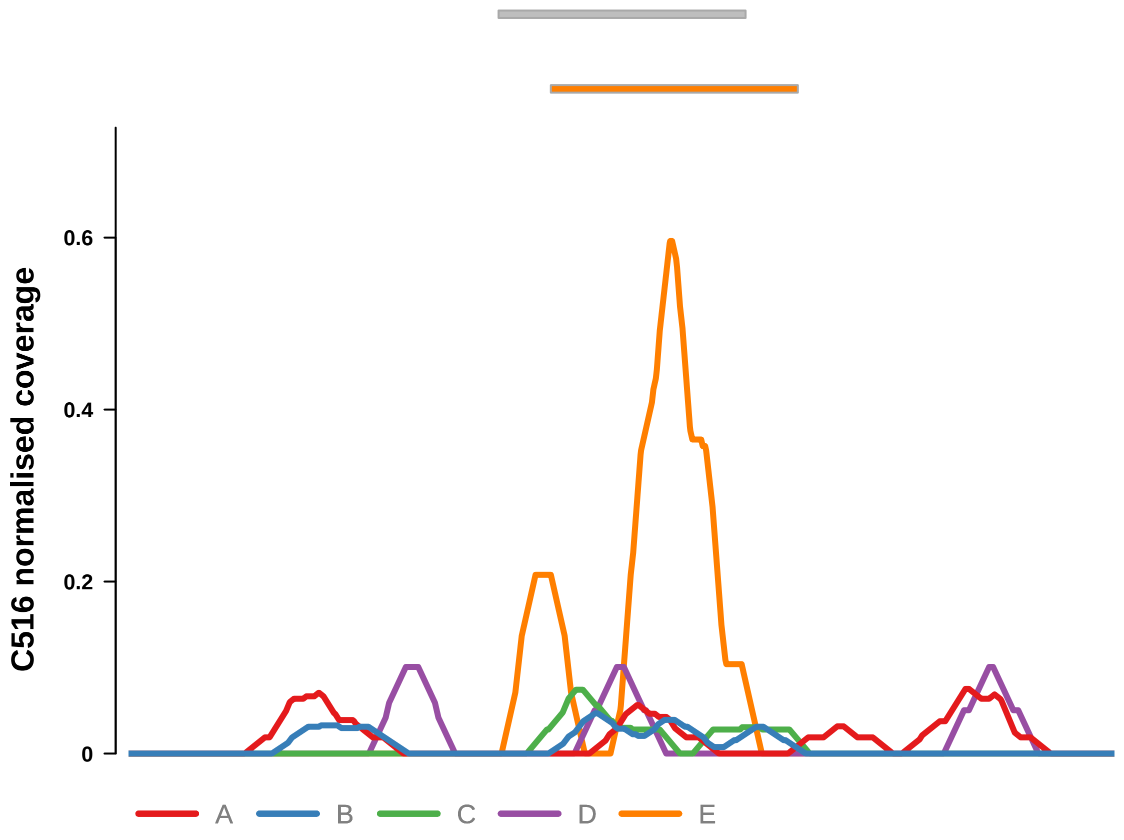


Putatively Clonal (logFC = −12.11)

| | P. | Adjusted P. |
|---|---|---|
| *~region vs. ~* | 4.42e−01 | 8.67e−01 |
| *~purity vs ~* | 3.73e−01 | 8.86e−01 |
| *~purity+region vs ~purity* | 6.96e−01 | 1e+00 |
| *Carcinoma vs. normal* | 3.29e−05 **** | 3.29e−05 **** |

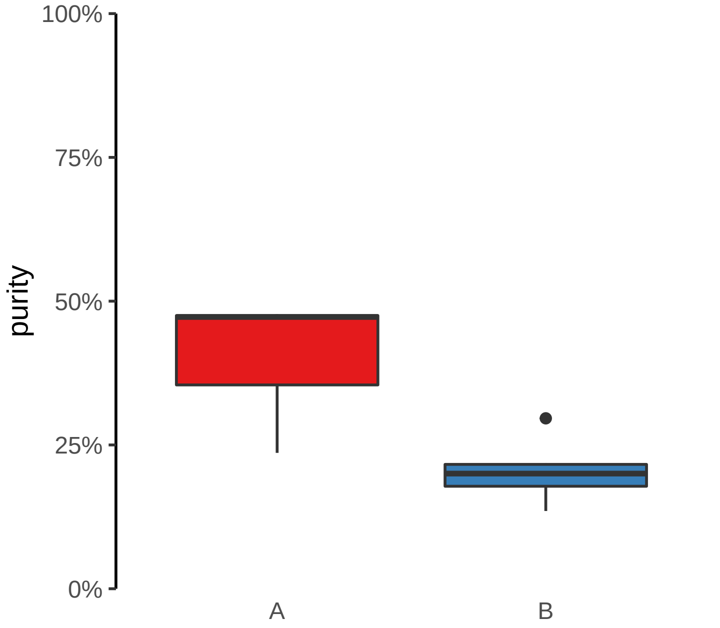

**DNAH7, ENSG00000272211 (GH02J196100) enhancer loss**
**chr2:196101142−196101642**

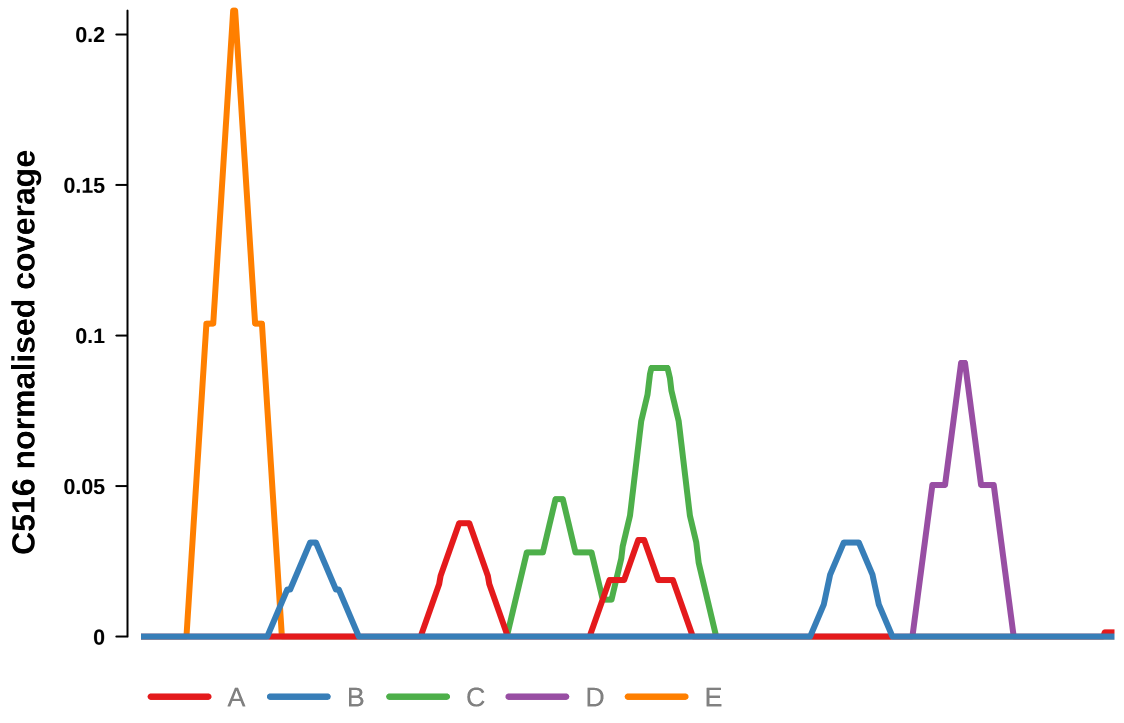


Putatively Clonal (logFC = −10.88)

| | P. | Adjusted P. |
|---|---|---|
| *~region vs. ~* | 9.23e−01 | 9.83e−01 |
| *~purity vs ~* | 9.56e−01 | 9.67e−01 |
| *~purity+region vs ~purity* | 8.96e−01 | 1e+00 |
| *Carcinoma vs. normal* | 2.43e−03 ** | 2.43e−03 ** |

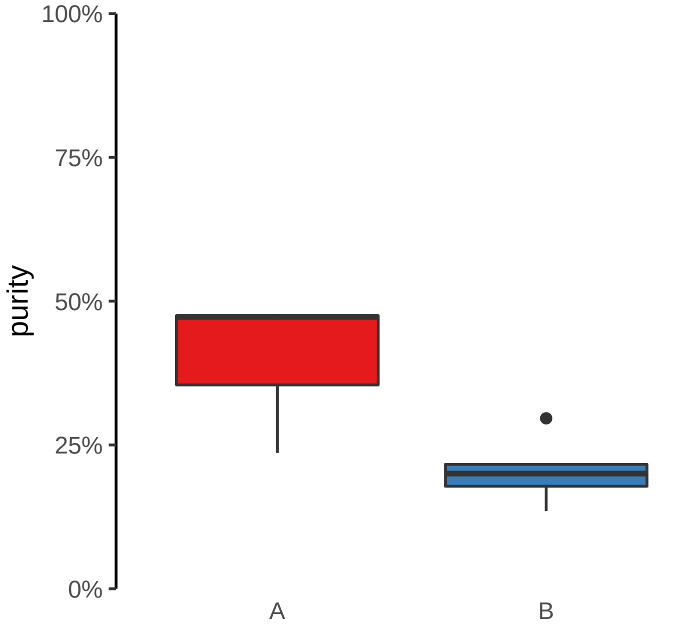

**MAP3K1, LINC01948 (GH05J056411) enhancer loss**
**chr5:56412924−56413424**

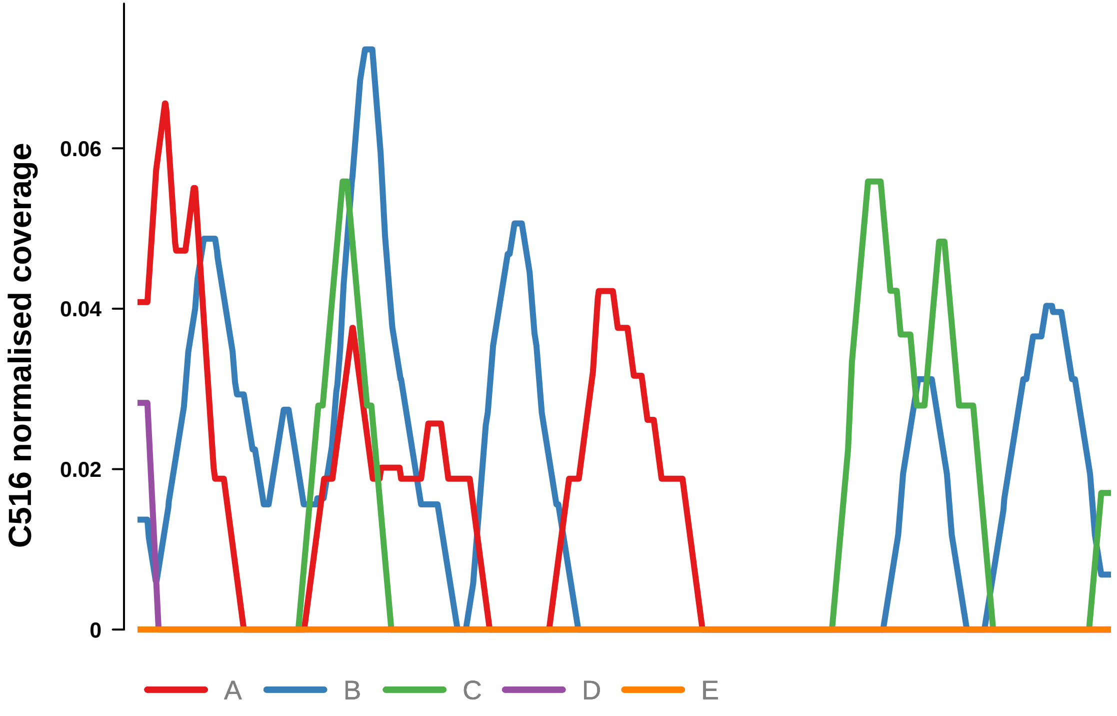


Putatively Clonal (logFC = −10.61)

| | P. | Adjusted P. |
|---|---|---|
| *~region vs. ~* | 5.77e−01 | 8.94e−01 |
| *~purity vs ~* | 7.62e−01 | 9.12e−01 |
| *~purity+region vs ~purity* | 5.99e−01 | 1e+00 |
| *Carcinoma vs. normal* | 3.99e−03 ** | 3.99e−03 ** |

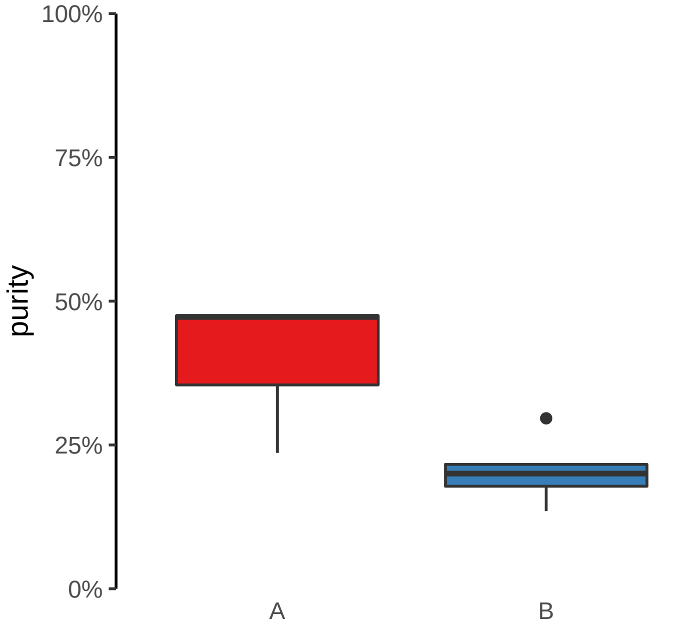

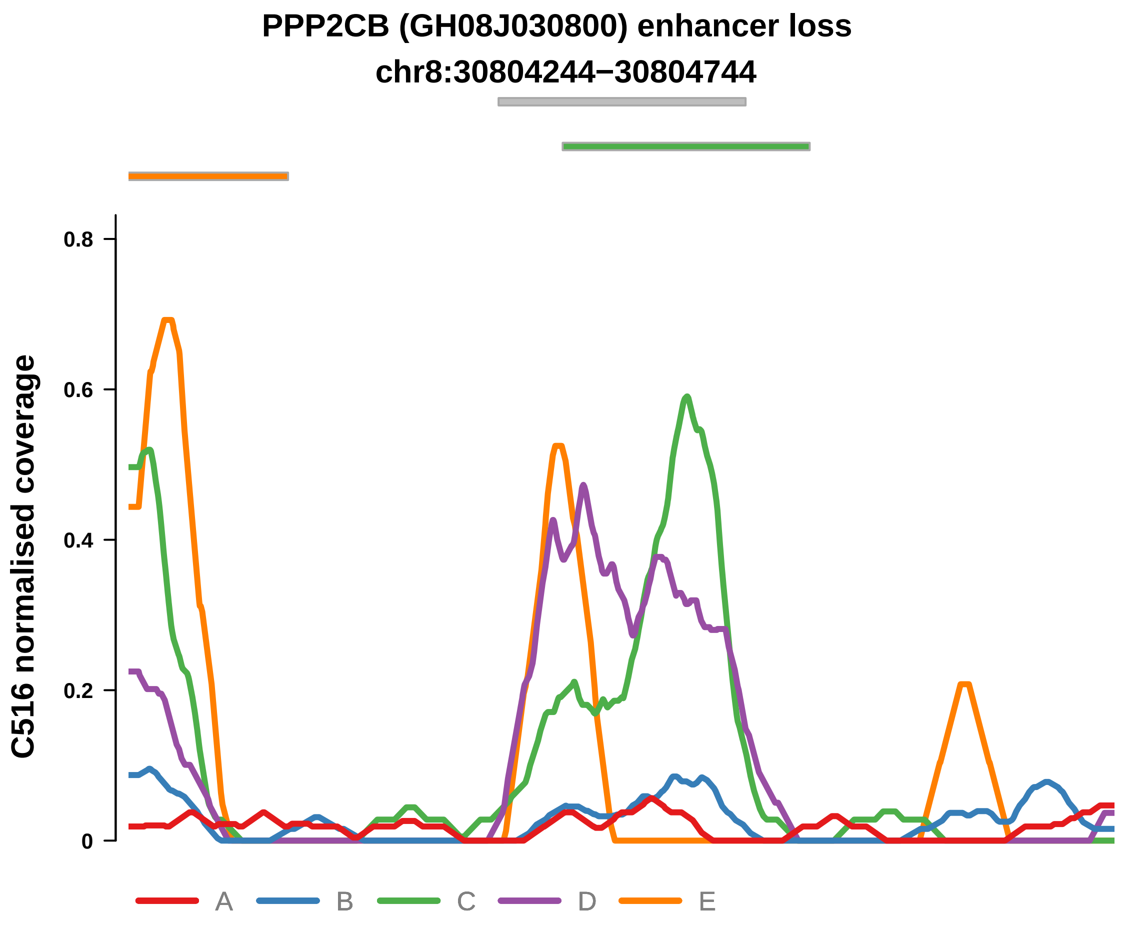


Putatively Clonal (logFC = −11.75)

| | P. | Adjusted P. |
|---|---|---|
| *~region vs. ~* | 1.64e−01 | 7.59e−01 |
| *~purity vs ~* | 4.59e−01 | 8.86e−01 |
| *~purity+region vs ~purity* | 1.49e−01 | 1e+00 |
| *Carcinoma vs. normal* | 1.18e−04 *** | 1.18e−04 *** |

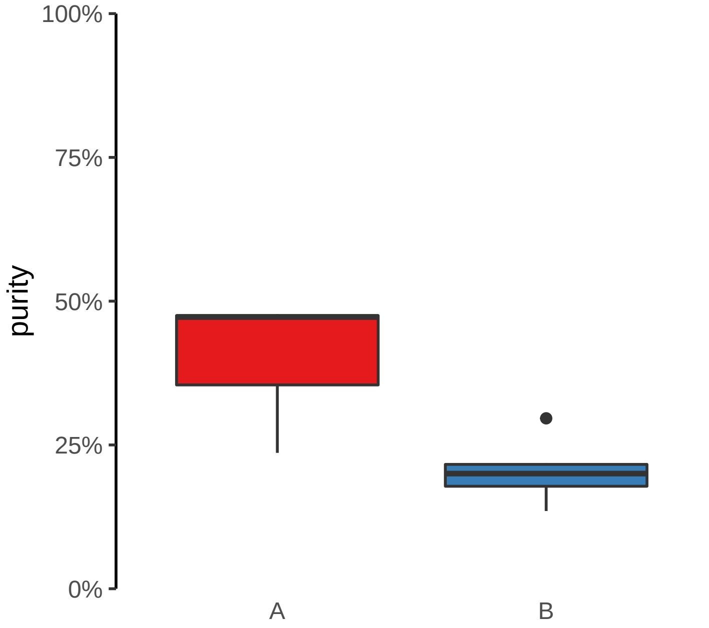

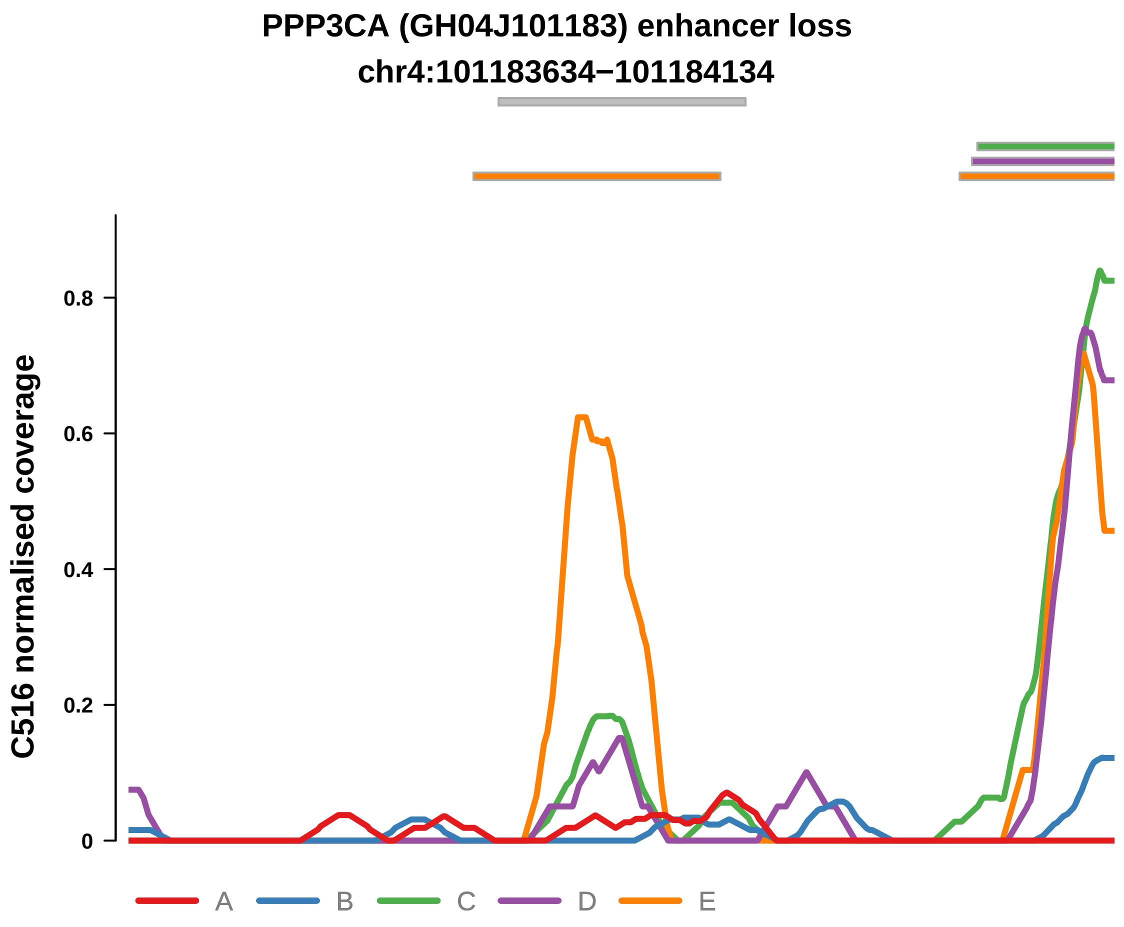


Putatively Clonal (logFC = −10.73)

| | P. | Adjusted P. |
|---|---|---|
| *~region vs. ~* | 5.26e−01 | 8.94e−01 |
| *~purity vs ~* | 5.24e−01 | 8.86e−01 |
| *~purity+region vs ~purity* | 1e+00 | 1e+00 |
| *Carcinoma vs. normal* | 2.34e−03 ** | 2.34e−03 ** |

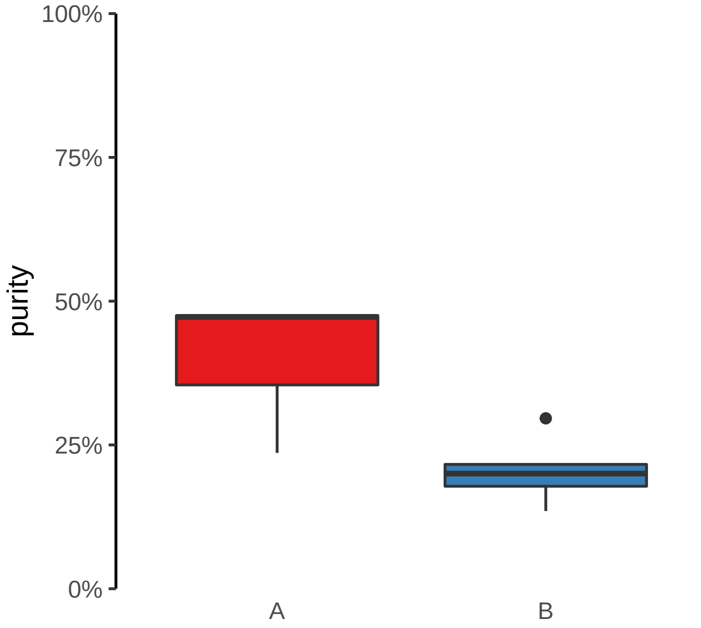

**RNLS, LIPJ (GH10J088553) enhancer loss**
**chr10:88553102−88553602**

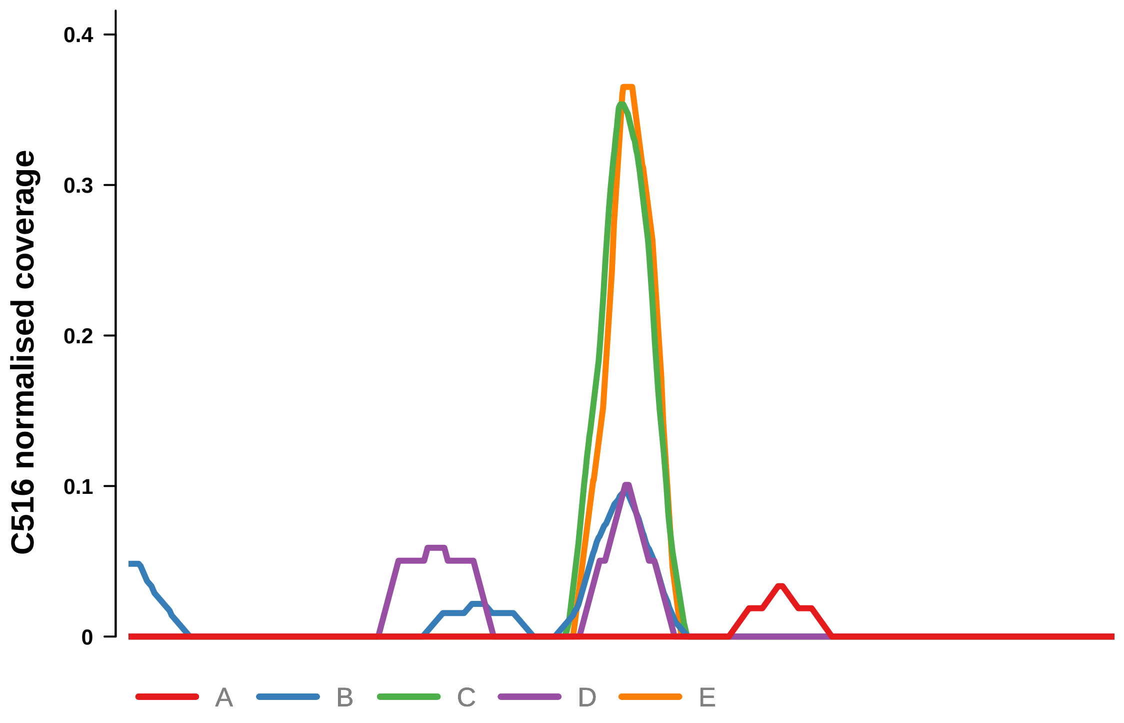


Putatively Clonal (logFC = −11.43)

| | P. | Adjusted P. |
|---|---|---|
| *~region vs. ~* | 3.19e−01 | 8.67e−01 |
| *~purity vs ~* | 3.15e−01 | 8.53e−01 |
| *~purity+region vs ~purity* | 8.28e−01 | 1e+00 |
| *Carcinoma vs. normal* | 3.27e−04 *** | 3.27e−04 *** |

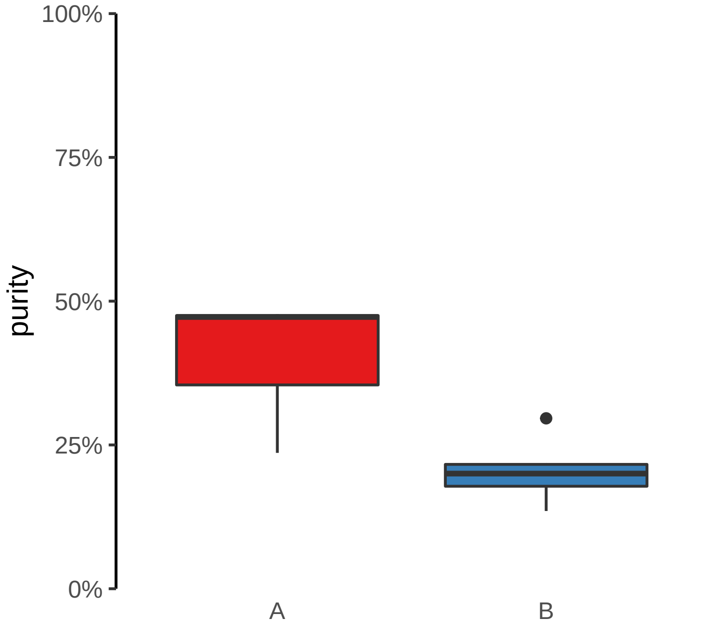

**SMAD3 (GH15J067079) enhancer loss**
**chr15:67081050−67081550**

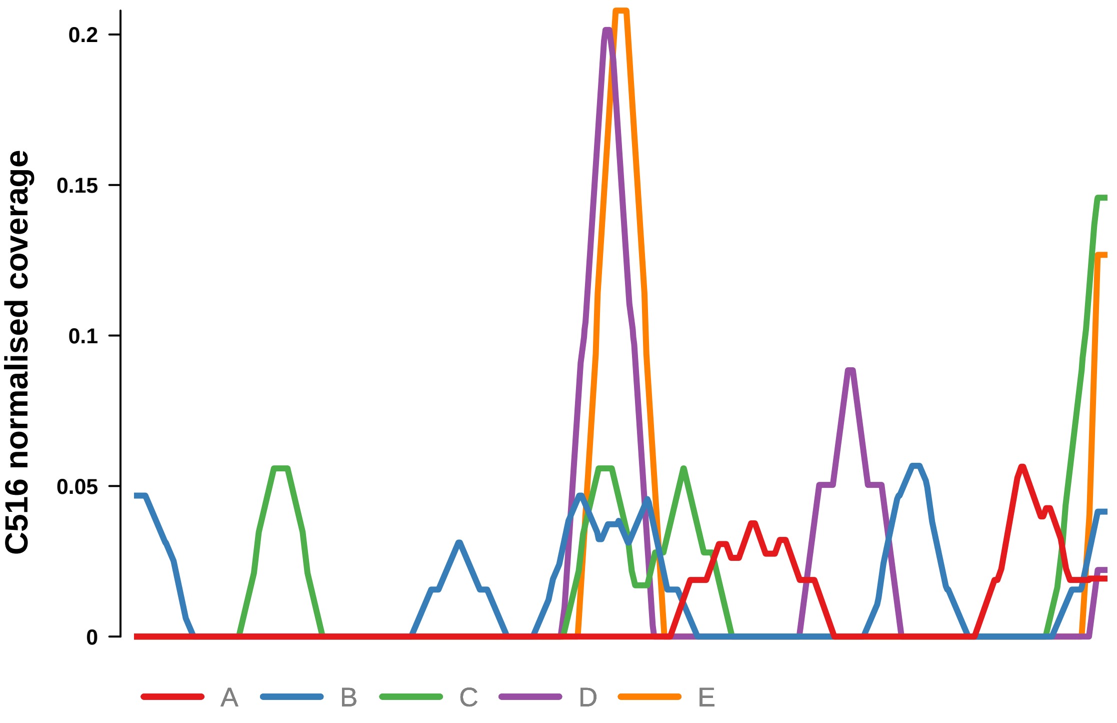


Putatively Clonal (logFC = −10.65)

| | P. | Adjusted P. |
|---|---|---|
| *~region vs. ~* | 4.87e−01 | 8.67e−01 |
| *~purity vs ~* | 7.97e−01 | 9.12e−01 |
| *~purity+region vs ~purity* | 4.45e−01 | 1e+00 |
| *Carcinoma vs. normal* | 3.48e−03 ** | 3.48e−03 ** |

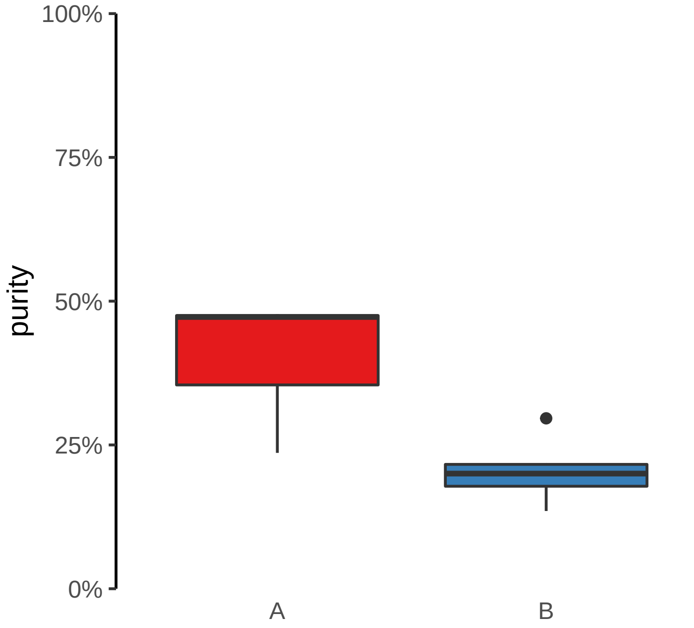

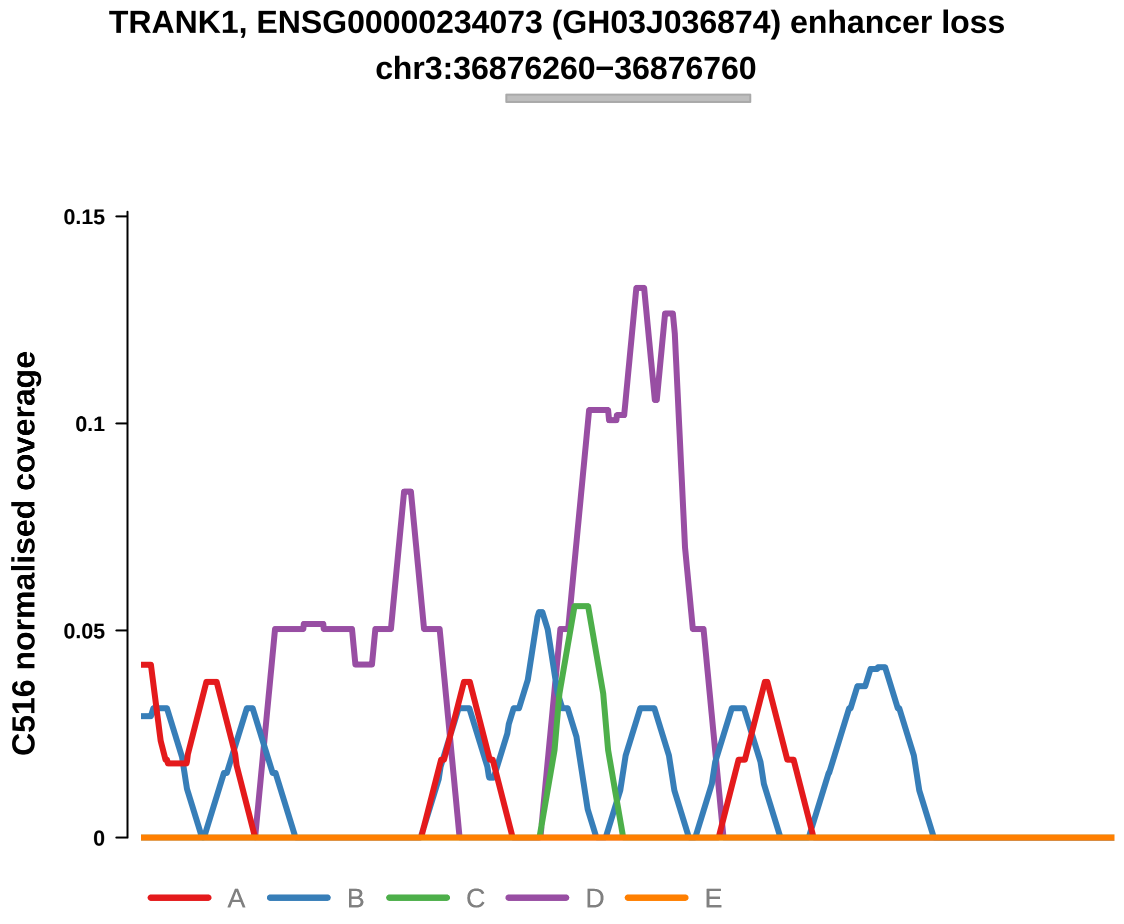


Putatively Clonal (logFC = −10.94)

| | P. | Adjusted P. |
|---|---|---|
| *~region vs. ~* | 3.47e−01 | 8.67e−01 |
| *~purity vs ~* | 4.63e−01 | 8.86e−01 |
| *~purity+region vs ~purity* | 5.7e−01 | 1e+00 |
| *Carcinoma vs. normal* | 1.91e−03 ** | 1.91e−03 ** |

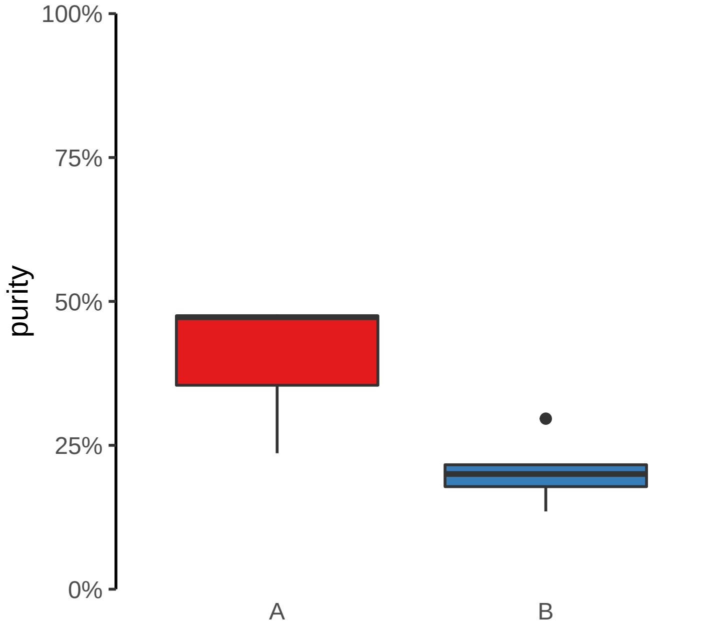

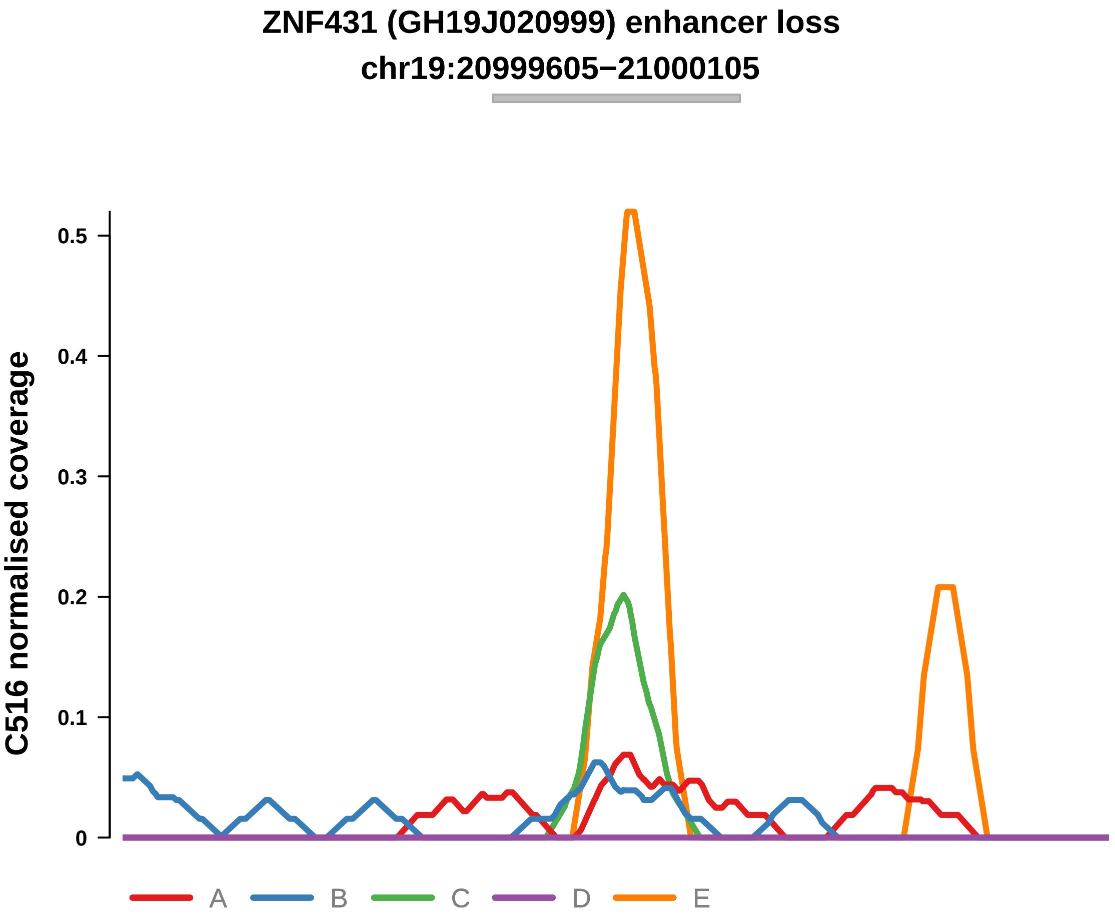


Putatively Clonal (logFC = −2.78)

| | P. | Adjusted P. |
|---|---|---|
| *~region vs. ~* | 9.98e−01 | 1e+00 |
| *~purity vs ~* | 7.61e−01 | 9.12e−01 |
| *~purity+region vs ~purity* | 5.31e−01 | 1e+00 |
| *Carcinoma vs. normal* | 8.81e−03 ** | 8.81e−03 ** |

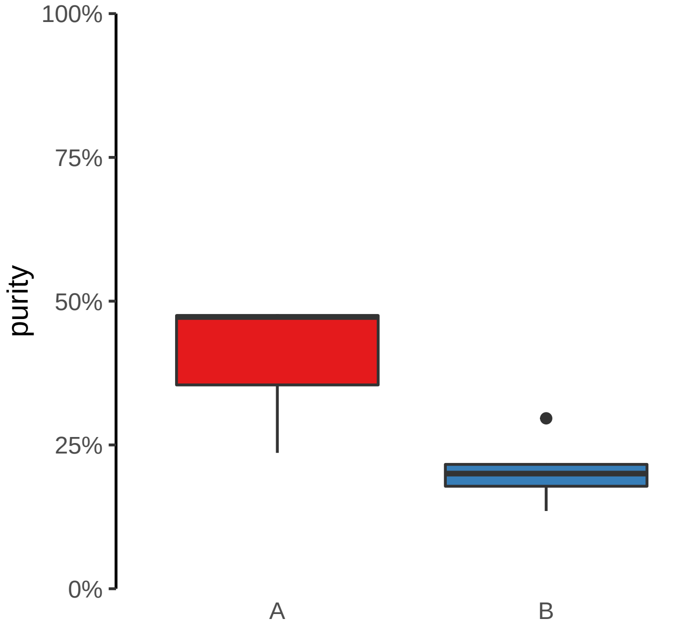

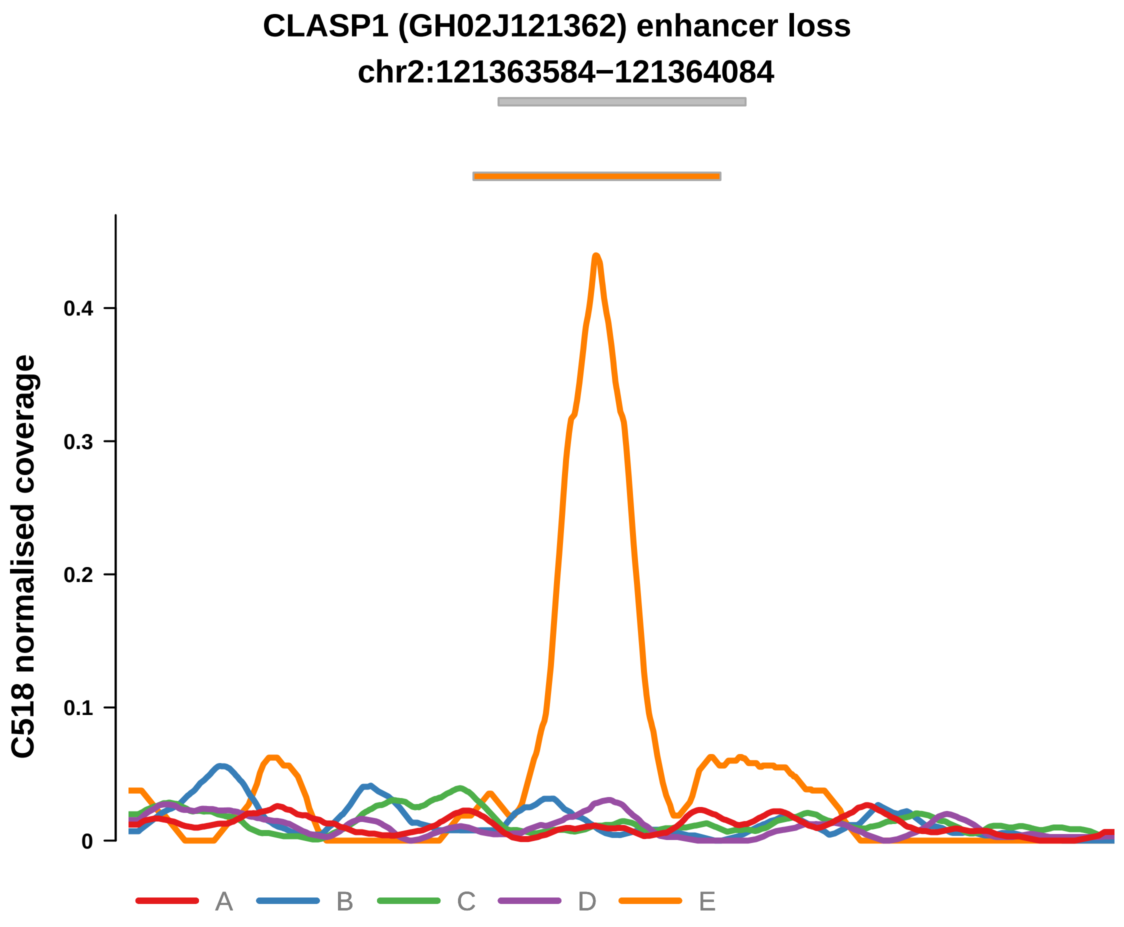


Putatively Clonal (logFC = −2.04)

| | P. | Adjusted P. |
|---|---|---|
| *~region vs. ~* | 9.67e−01 | 9.78e−01 |
| *~purity vs ~* | 9.34e−01 | 9.78e−01 |
| *~purity+region vs ~purity* | 9.68e−01 | 9.8e−01 |
| *Carcinoma vs. normal* | 7.13e−03 ** | 7.13e−03 ** |

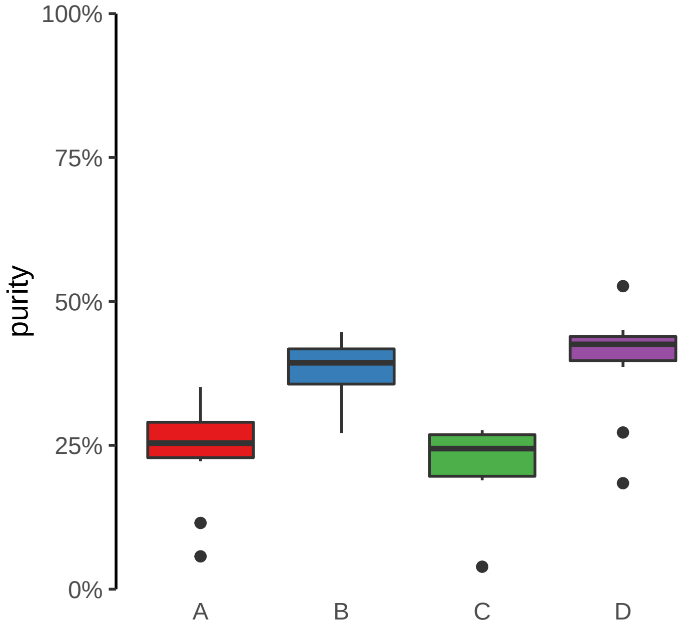

**ENSG00000228035, NGF (GH01J115322) enhancer loss**
**chr1:115323237−115323737**

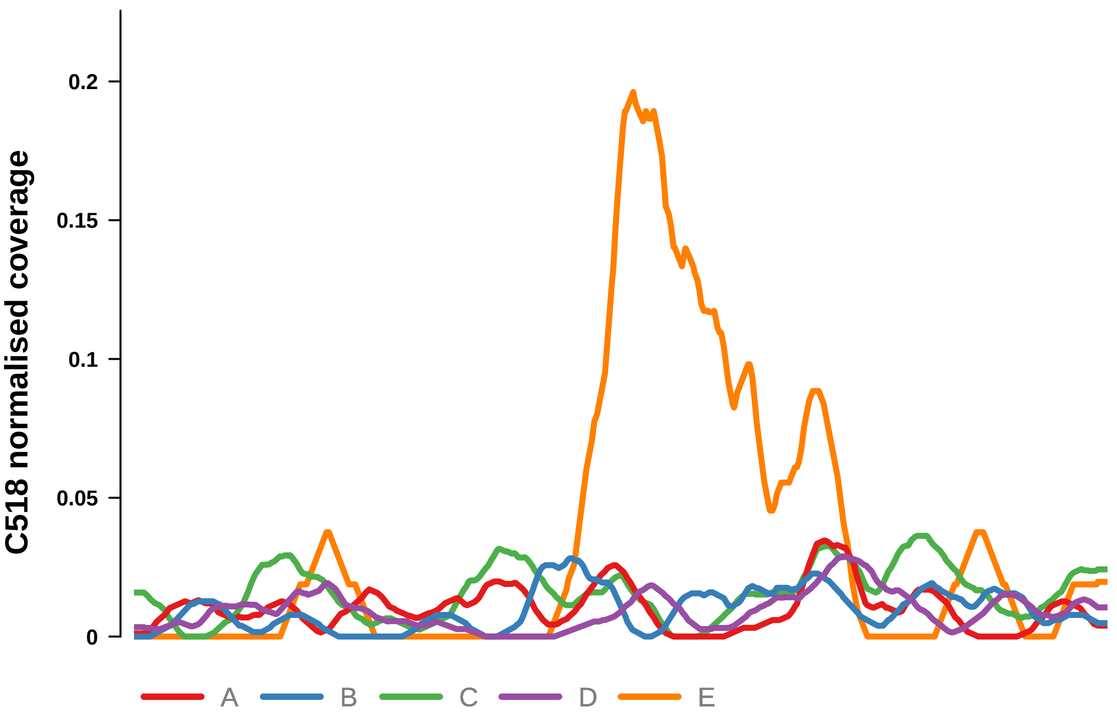


Putatively Clonal (logFC = −2.73)

| | P. | Adjusted P. |
|---|---|---|
| *~region vs. ~* | 6.4e−01 | 7.83e−01 |
| *~purity vs ~* | 1.2e−01 | 6.62e−01 |
| *~purity+region vs ~purity* | 9.54e−01 | 9.76e−01 |
| *Carcinoma vs. normal* | 7.47e−04 *** | 7.47e−04 *** |

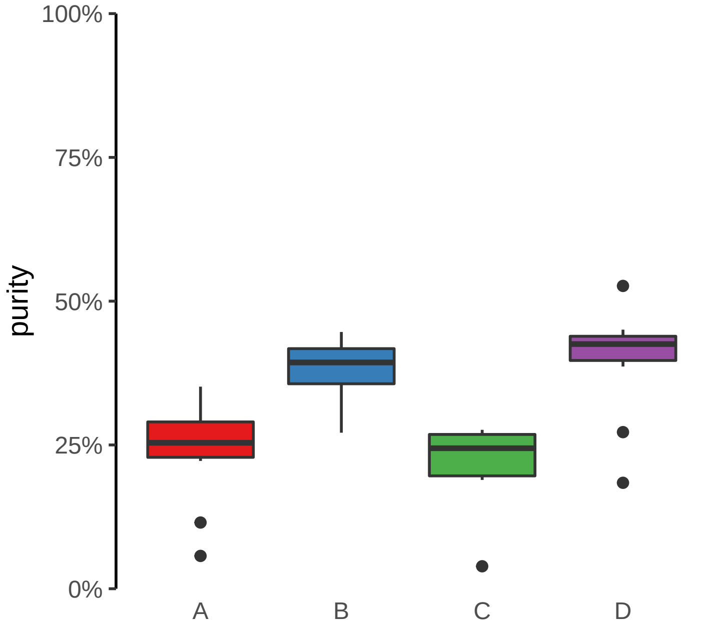

**FABP1, THNSL2 (GH02J088144) enhancer loss**
**chr2:88144817−88145317**

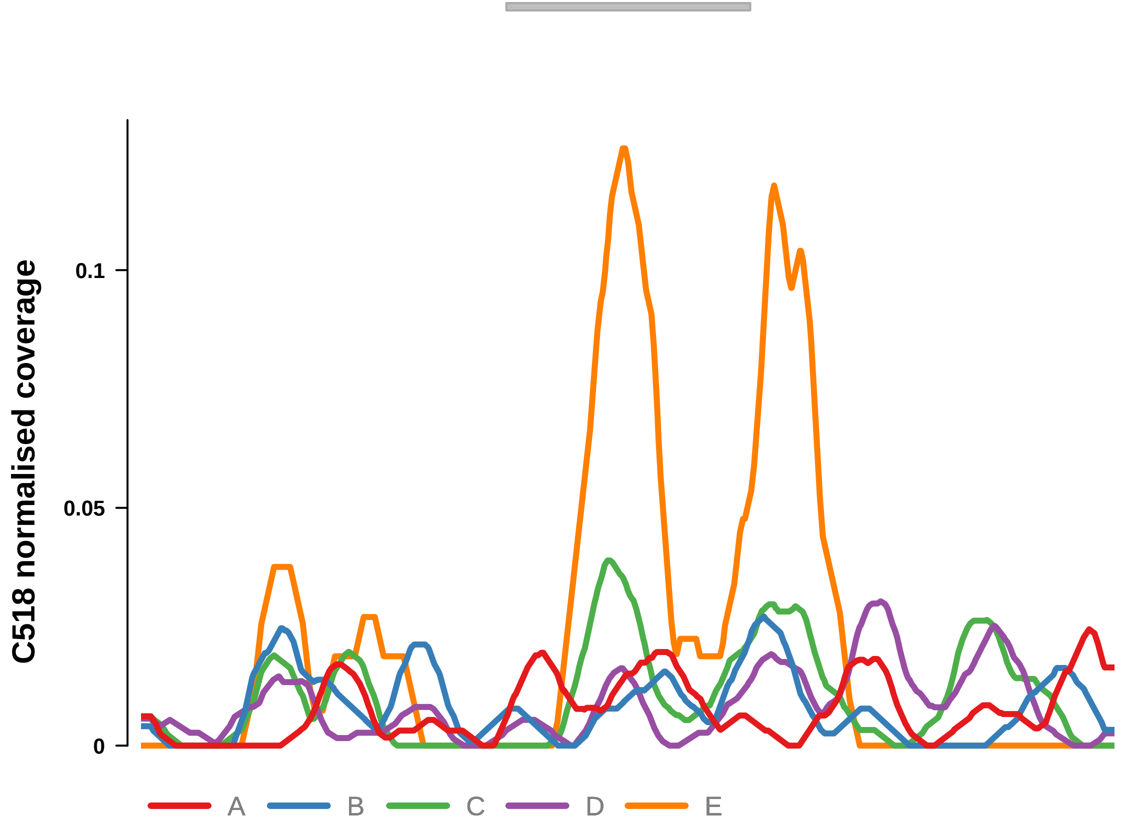


Putatively Clonal (logFC = −2.32)

| | P. | Adjusted P. |
|---|---|---|
| *~region vs. ~* | 2.97e−01 | 6.53e−01 |
| *~purity vs ~* | 5.29e−02 | 4.16e−01 |
| *~purity+region vs ~purity* | 8.69e−01 | 9.5e−01 |
| *Carcinoma vs. normal* | 2.12e−03 ** | 2.12e−03 ** |

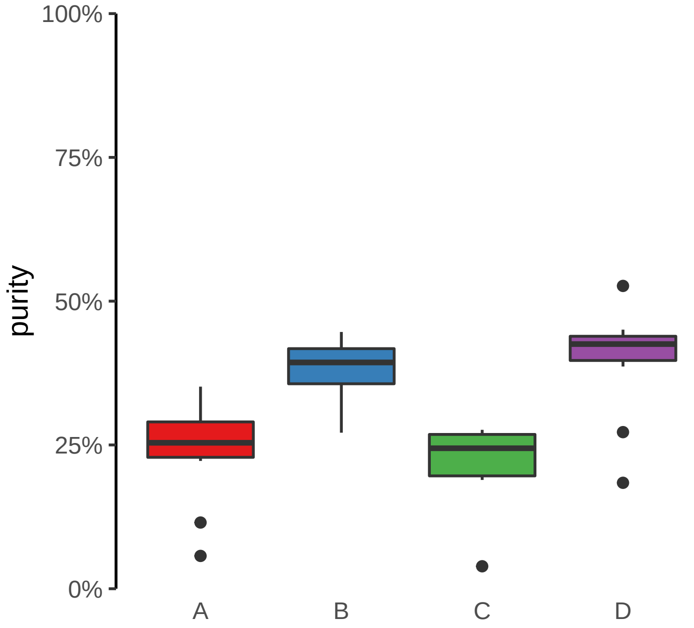

**ACAP2, PPP1R2 (GH03J195537) enhancer loss**
**chr3:195537917−195538417**

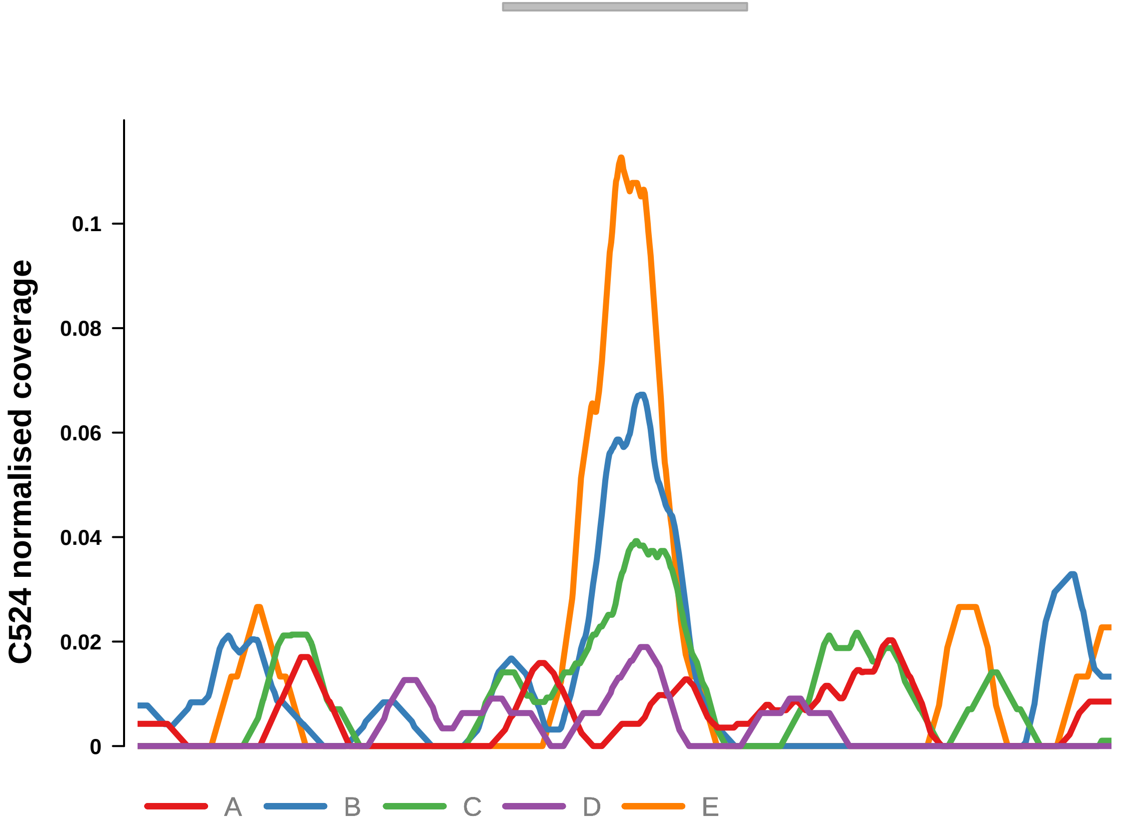


Putatively Clonal (logFC = −10.4)

| | P. | Adjusted P. |
|---|---|---|
| *~region vs. ~* | 9.65e−01 | 9.75e−01 |
| *~purity vs ~* | 1.49e−01 | 5e−01 |
| *~purity+region vs ~purity* | 7e−01 | 8.8e−01 |
| *Carcinoma vs. normal* | 6.06e−05 **** | 6.06e−05 **** |

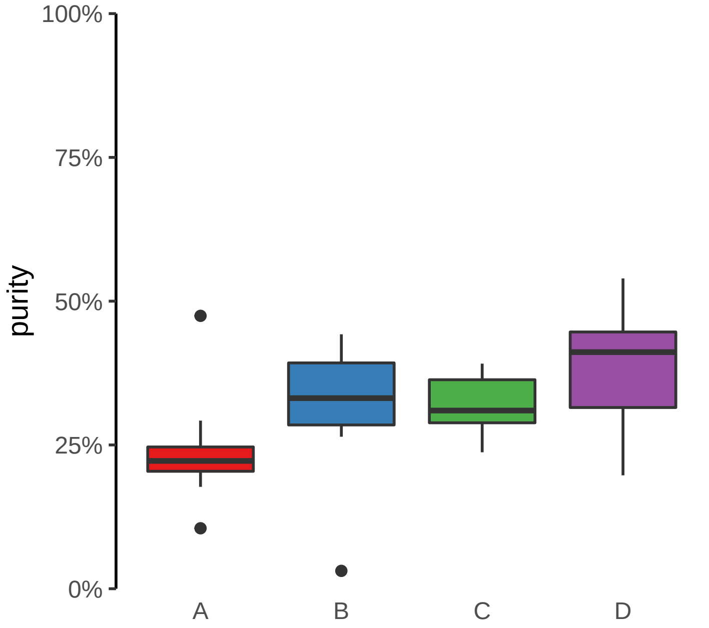

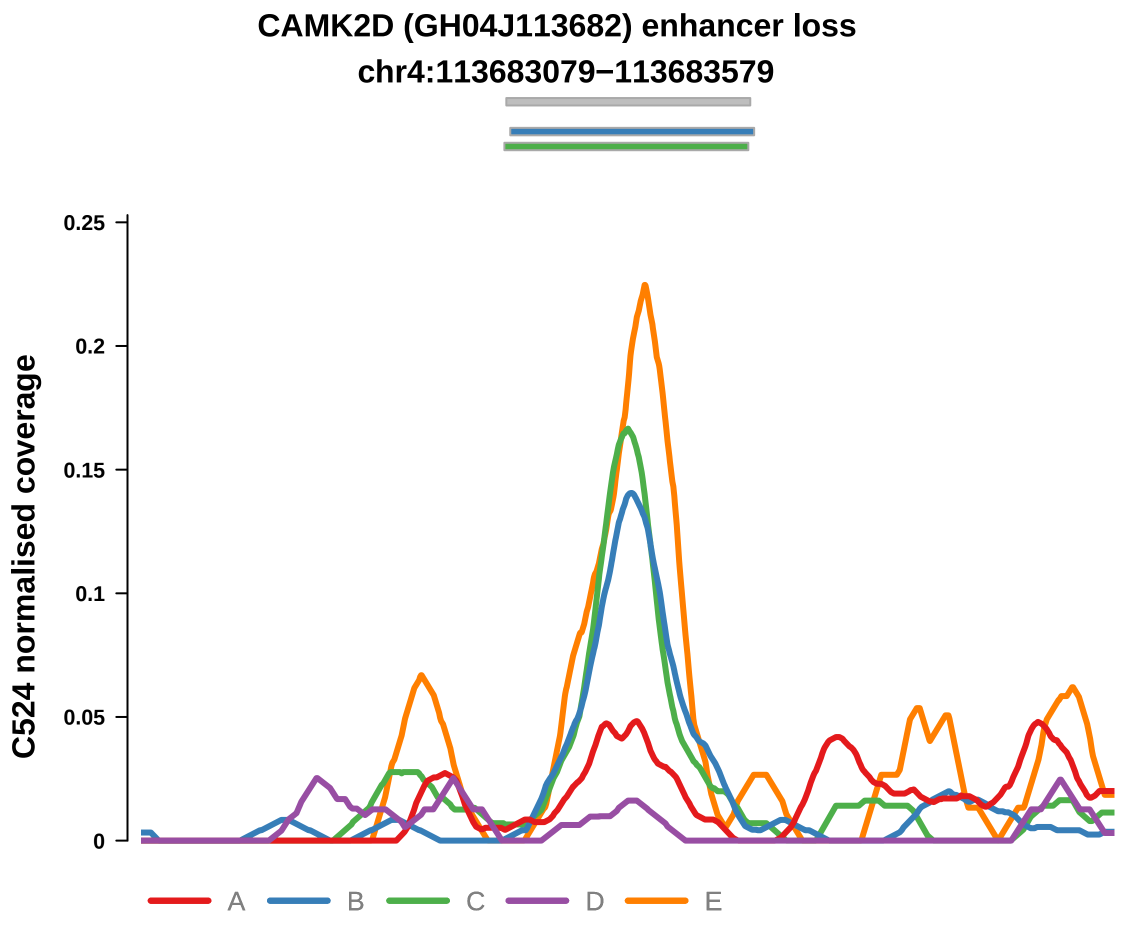


Putatively Clonal (logFC = −3.98)

| | P. | Adjusted P. |
|---|---|---|
| *~region vs. ~* | 6.5e−01 | 8.5e−01 |
| *~purity vs ~* | 9.28e−02 | 4.3e−01 |
| *~purity+region vs ~purity* | 6.26e−01 | 8.8e−01 |
| *Carcinoma vs. normal* | 9.13e−04 *** | 9.13e−04 *** |

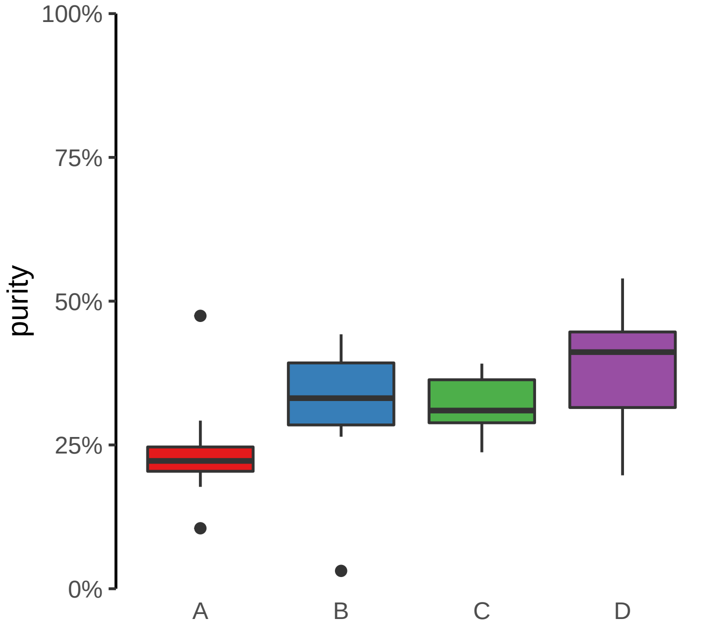

**CCDC6 (GH10J059885) enhancer loss**

**chr10:59885378−59885878**

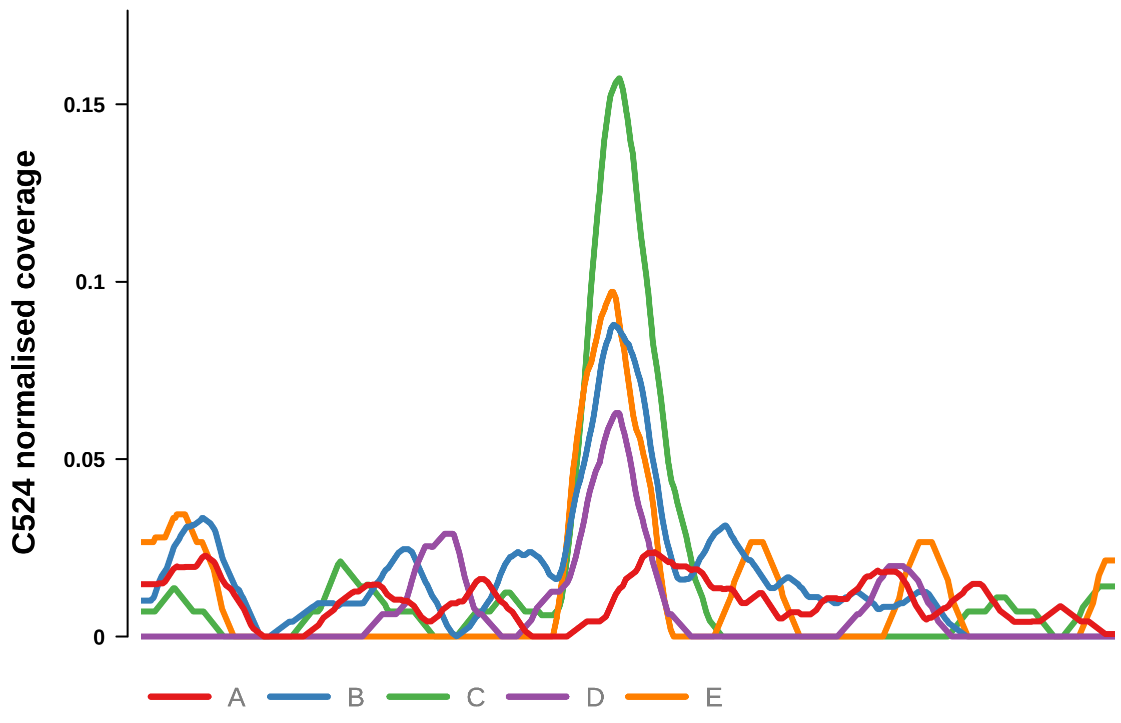


Putatively Clonal (logFC = −2.97)

| | **P.** | **Adjusted P.** |
|---|---|---|
| *~region vs. ~* | 4.72e−01 | 7.84e−01 |
| *~purity vs ~* | 8.57e−02 | 4.3e−01 |
| *~purity+region vs ~purity* | 5.56e−02 | 3.98e−01 |
| *Carcinoma vs. normal* | 4.96e−03 ** | 4.96e−03 ** |

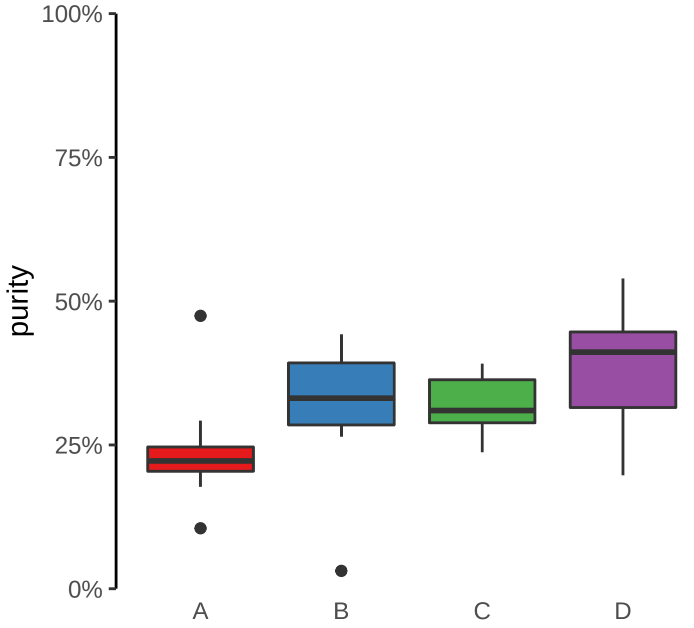

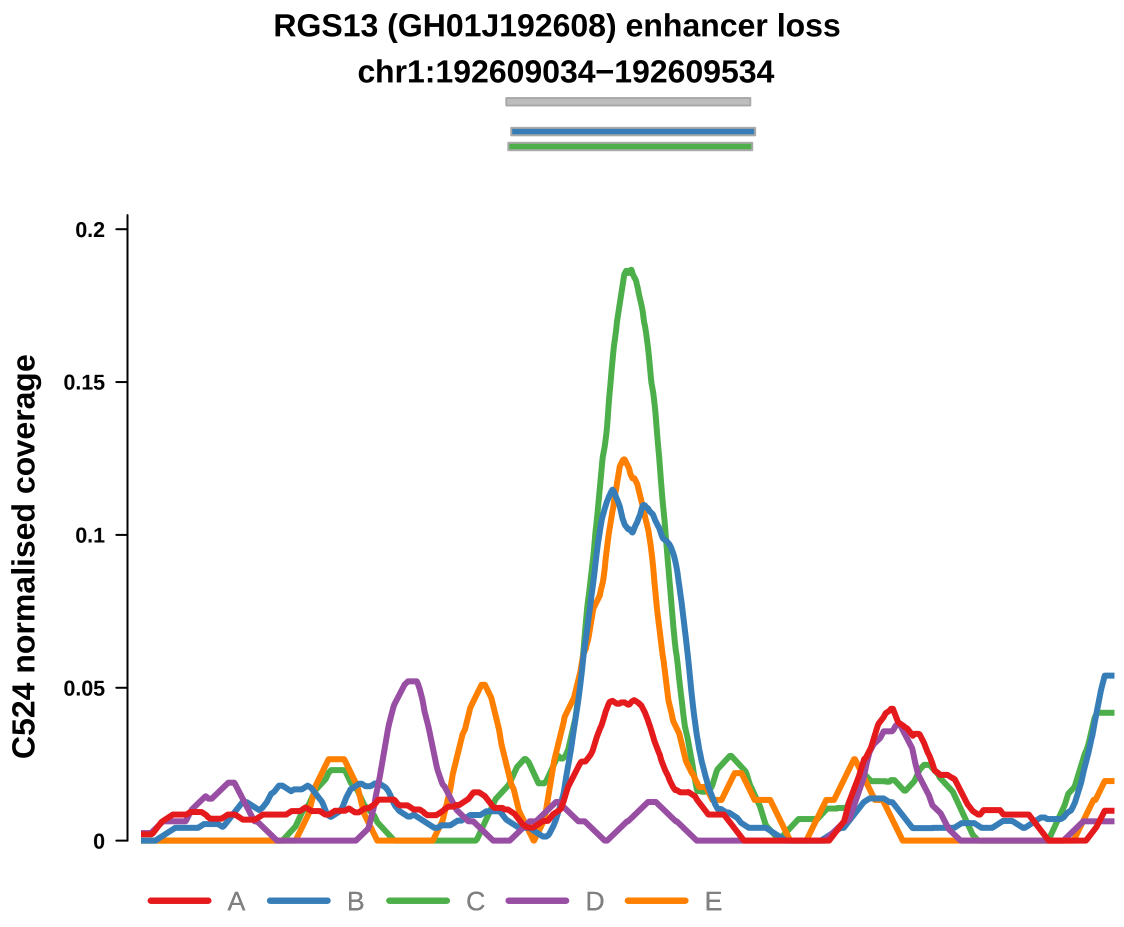


Putatively Clonal (logFC = −2.42)

| | P. | Adjusted P. |
|---|---|---|
| *~region vs. ~* | 1.46e−01 | 5.65e−01 |
| *~purity vs ~* | 4.66e−03 ** | 1.69e−01 |
| *~purity+region vs ~purity* | 9.15e−02 | 3.98e−01 |
| *Carcinoma vs. normal* | 9.1e−03 ** | 9.1e−03 ** |

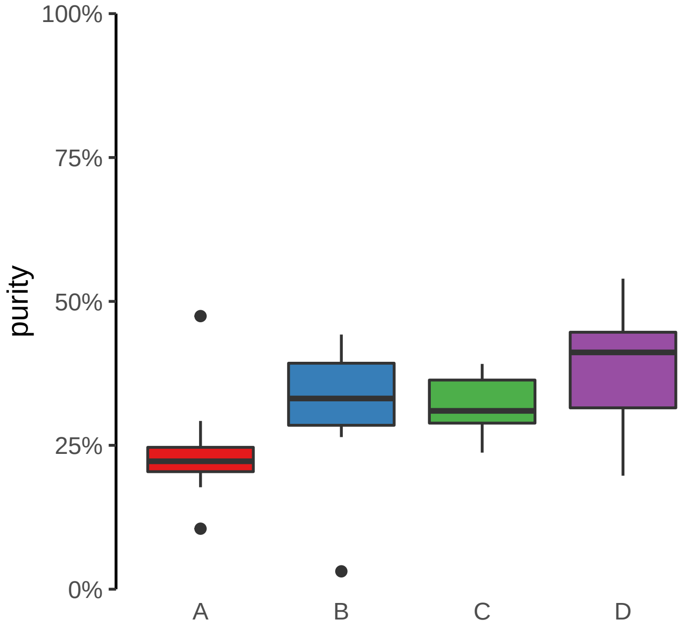

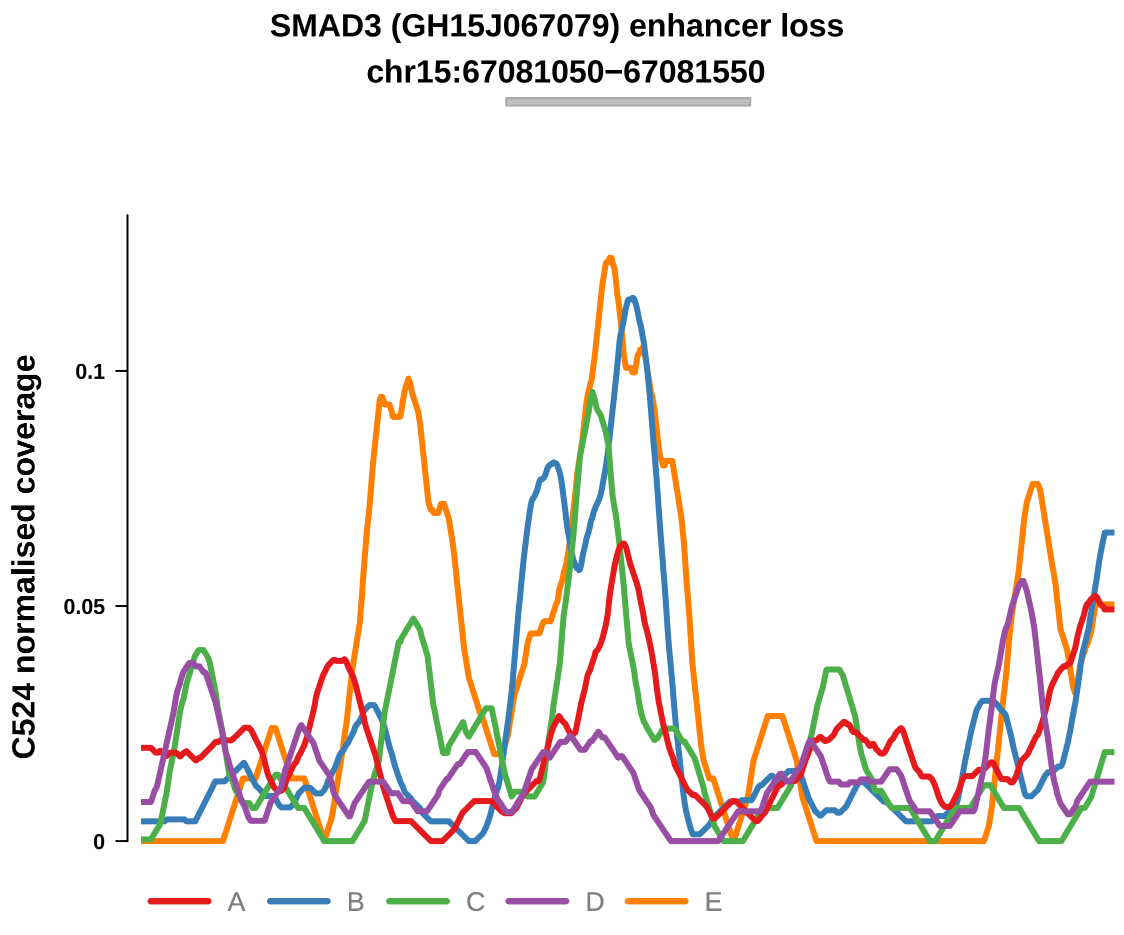


Putatively Clonal (logFC = −2.93)

| | P. | Adjusted P. |
|---|---|---|
| *~region vs. ~* | 8.82e−01 | 9.58e−01 |
| *~purity vs ~* | 3.06e−03 ** | 1.69e−01 |
| *~purity+region vs ~purity* | 9.66e−01 | 9.87e−01 |
| *Carcinoma vs. normal* | 5.74e−03 ** | 5.74e−03 ** |

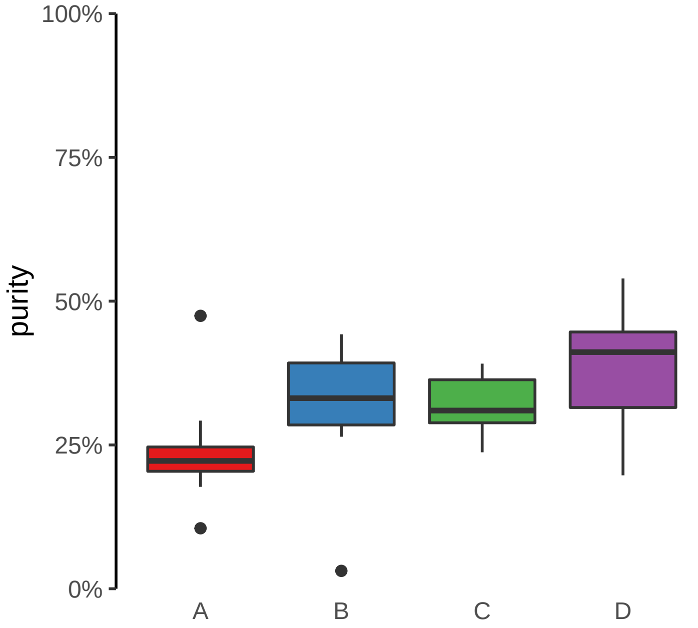

**ANTXR2 (GH04J080057) enhancer loss**
**chr4:80058687−80059187**

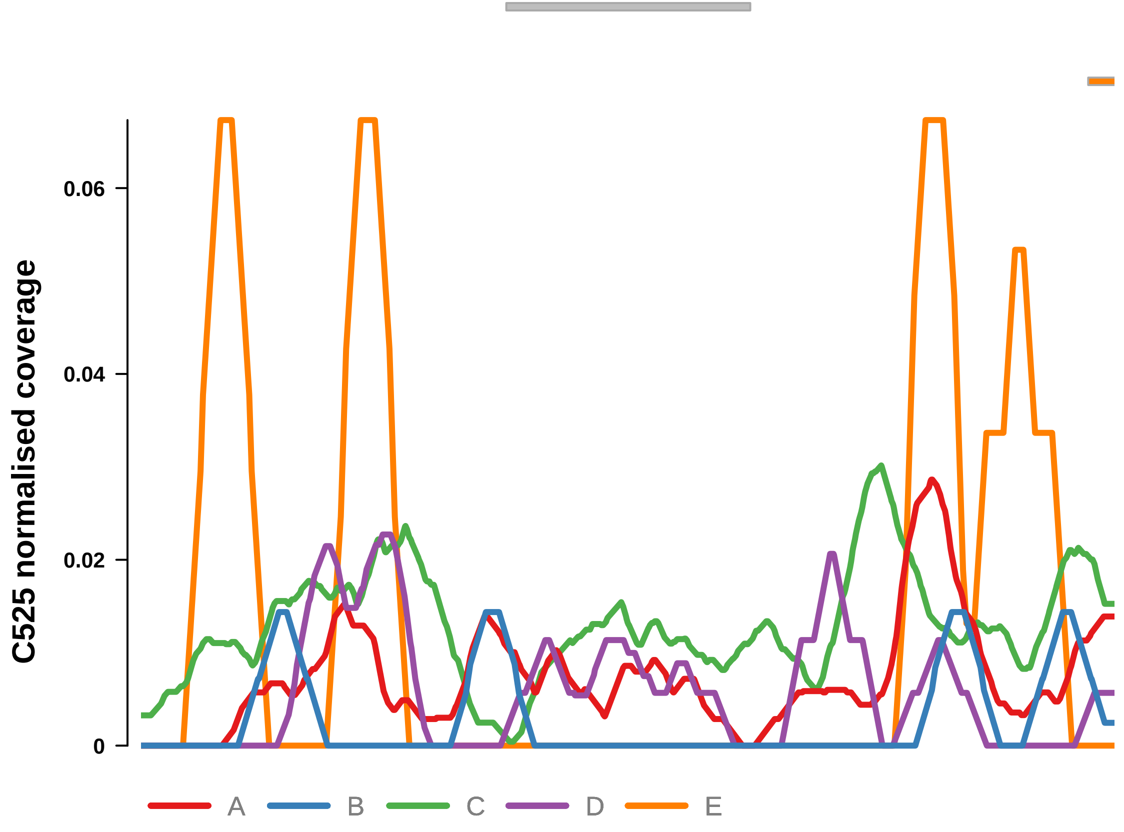


Putatively Clonal (logFC = −3.15)

| | P. | Adjusted P. |
|---|---|---|
| *~region vs. ~* | 2.65e−01 | NA |
| *~purity vs ~* | 5.38e−01 | 8.63e−01 |
| *~purity+region vs ~purity* | 3.58e−01 | 4.93e−01 |
| *Carcinoma vs. normal* | 2.43e−03 ** | 2.43e−03 ** |

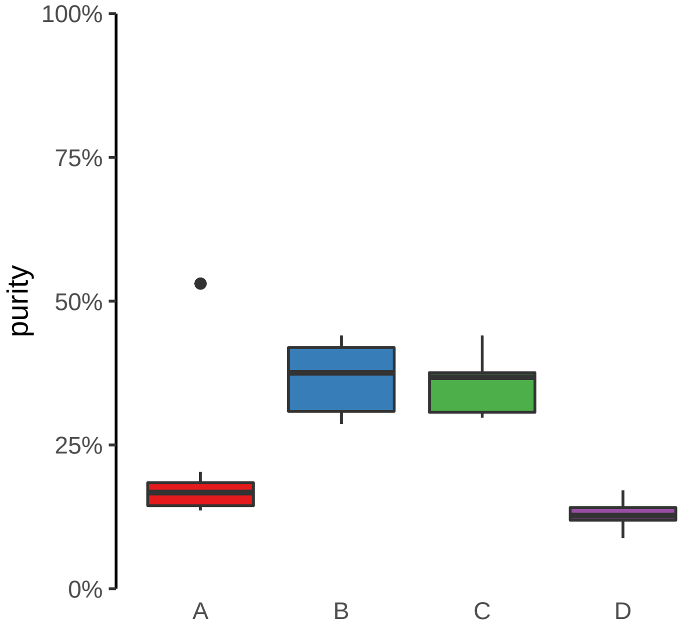

**CLASP2, ENSG00000271324 (GH03J033790) enhancer loss**
**chr3:33791629−33792129**

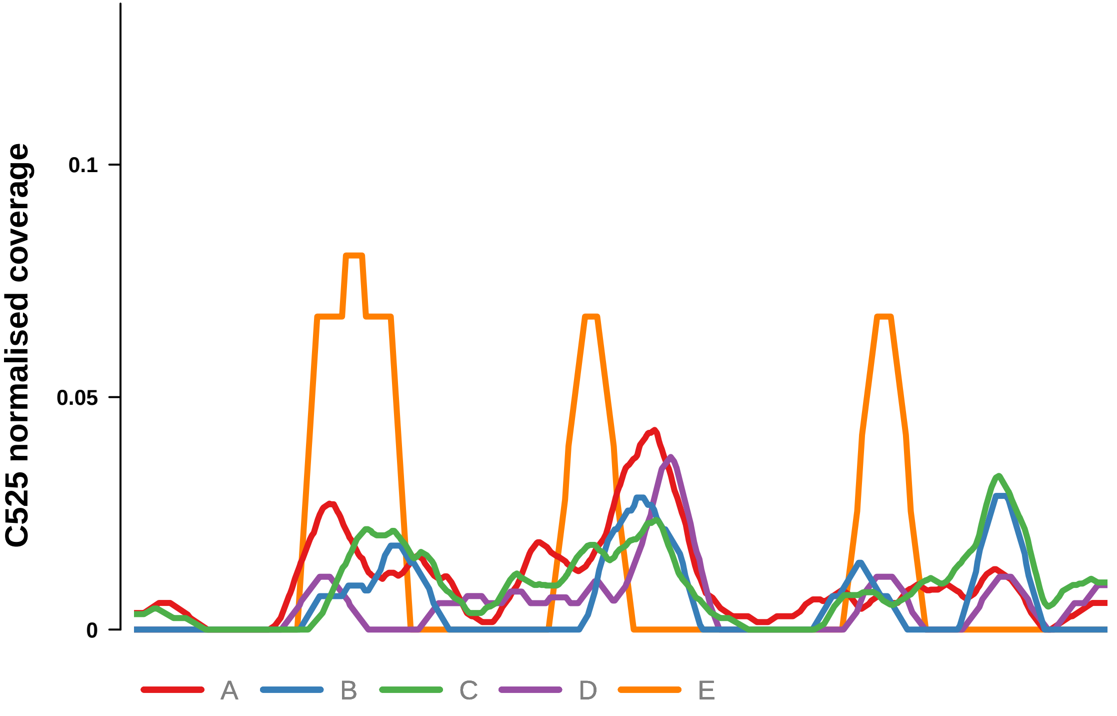


Putatively Clonal (logFC = −2.54)

| | P. | Adjusted P. |
|---|---|---|
| *~region vs. ~* | 7.16e−02 | 1.26e−01 |
| *~purity vs ~* | 2.33e−01 | 7.37e−01 |
| *~purity+region vs ~purity* | 4.68e−02 * | 9.58e−02 |
| *Carcinoma vs. normal* | 1.77e−03 ** | 1.77e−03 ** |

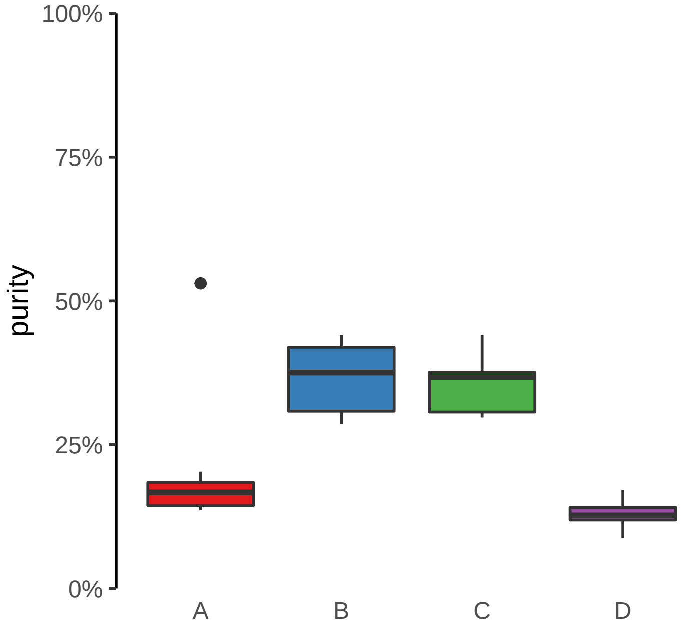

**FAM83G (GH17J019010) enhancer loss**
**chr17:19011073−19011573**

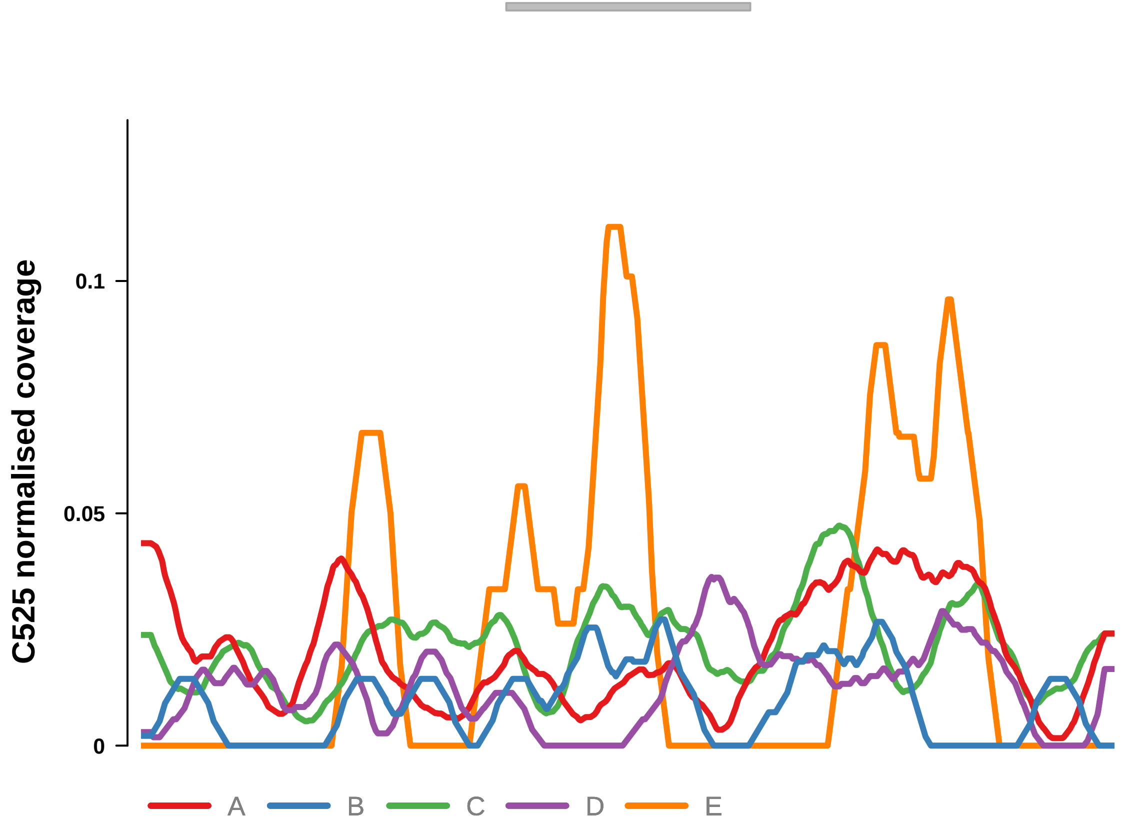


Putatively Clonal (logFC = −2.18)

| | P. | Adjusted P. |
|---|---|---|
| *~region vs. ~* | 1.77e−02 * | 4.62e−02 * |
| *~purity vs ~* | 6.53e−01 | 8.63e−01 |
| *~purity+region vs ~purity* | 2.57e−02 * | 6.85e−02 |
| *Carcinoma vs. normal* | 4.9e−03 ** | 4.9e−03 ** |

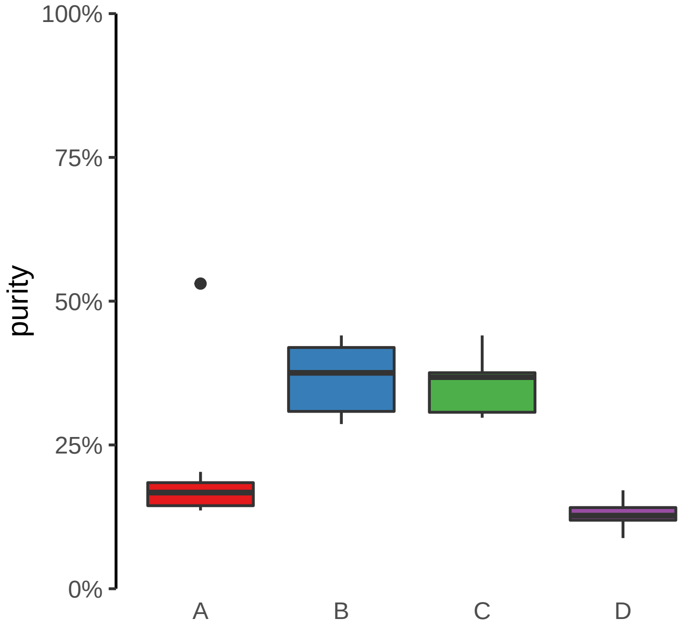

**PLAGL1 (GH06J144054) enhancer loss**
**chr6:144055046−144055546**

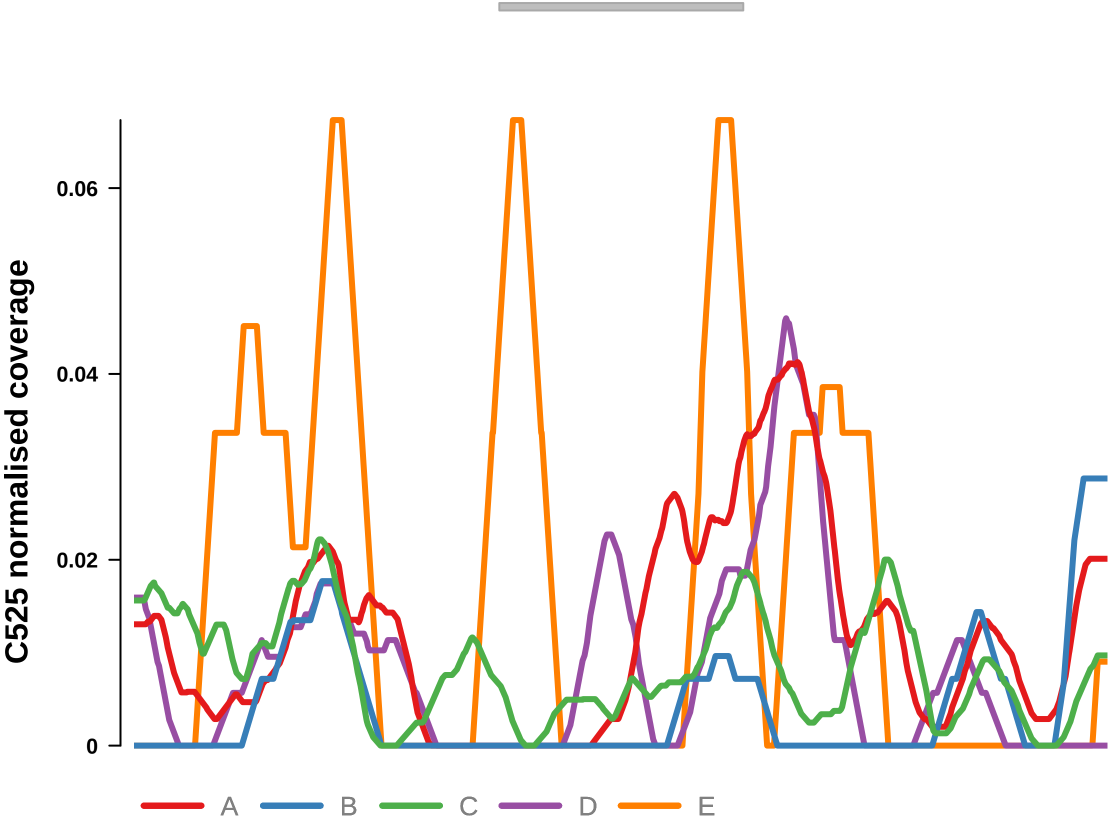


Putatively Clonal (logFC = −2.81)

| | P. | Adjusted P. |
|---|---|---|
| *~region vs. ~* | 1.79e−01 | 2.65e−01 |
| *~purity vs ~* | 5.53e−01 | 8.63e−01 |
| *~purity+region vs ~purity* | 2.07e−01 | 3.25e−01 |
| *Carcinoma vs. normal* | 3.24e−03 ** | 3.24e−03 ** |

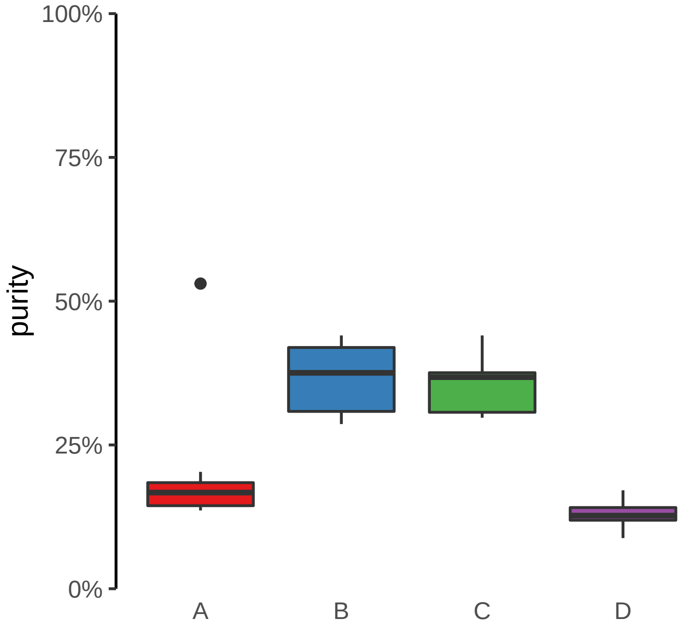

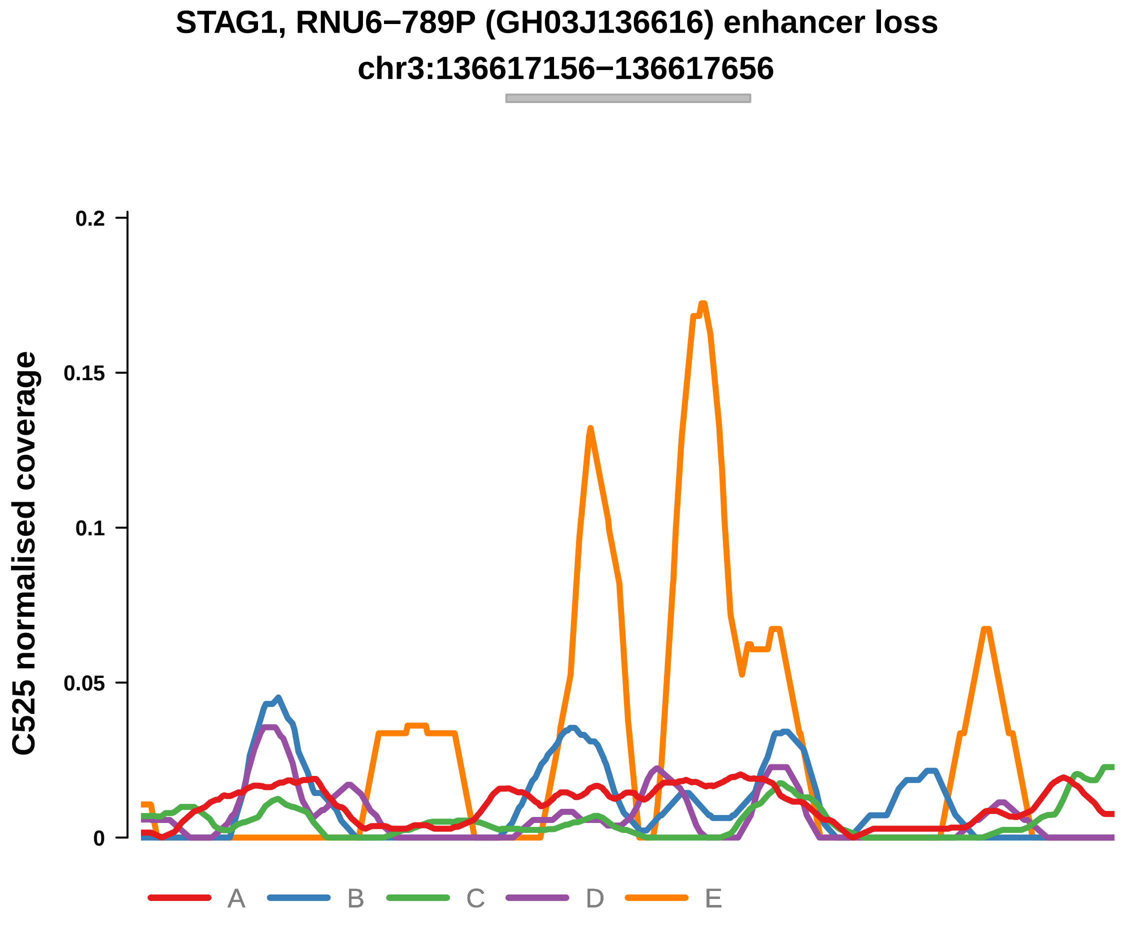


Putatively Clonal (logFC = −2.76)

| | P. | Adjusted P. |
|---|---|---|
| *~region vs. ~* | 1.76e−01 | 2.65e−01 |
| *~purity vs ~* | 1.72e−01 | 7.37e−01 |
| *~purity+region vs ~purity* | 4.45e−01 | 5.5e−01 |
| *Carcinoma vs. normal* | 1.55e−03 ** | 1.55e−03 ** |

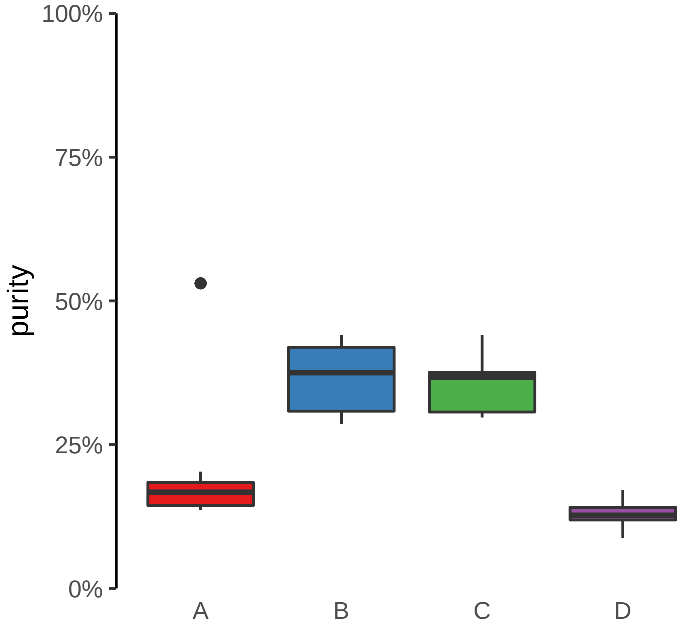

**CAMK2D (GH04J113682) enhancer loss**

**chr4:113683079−113683579**

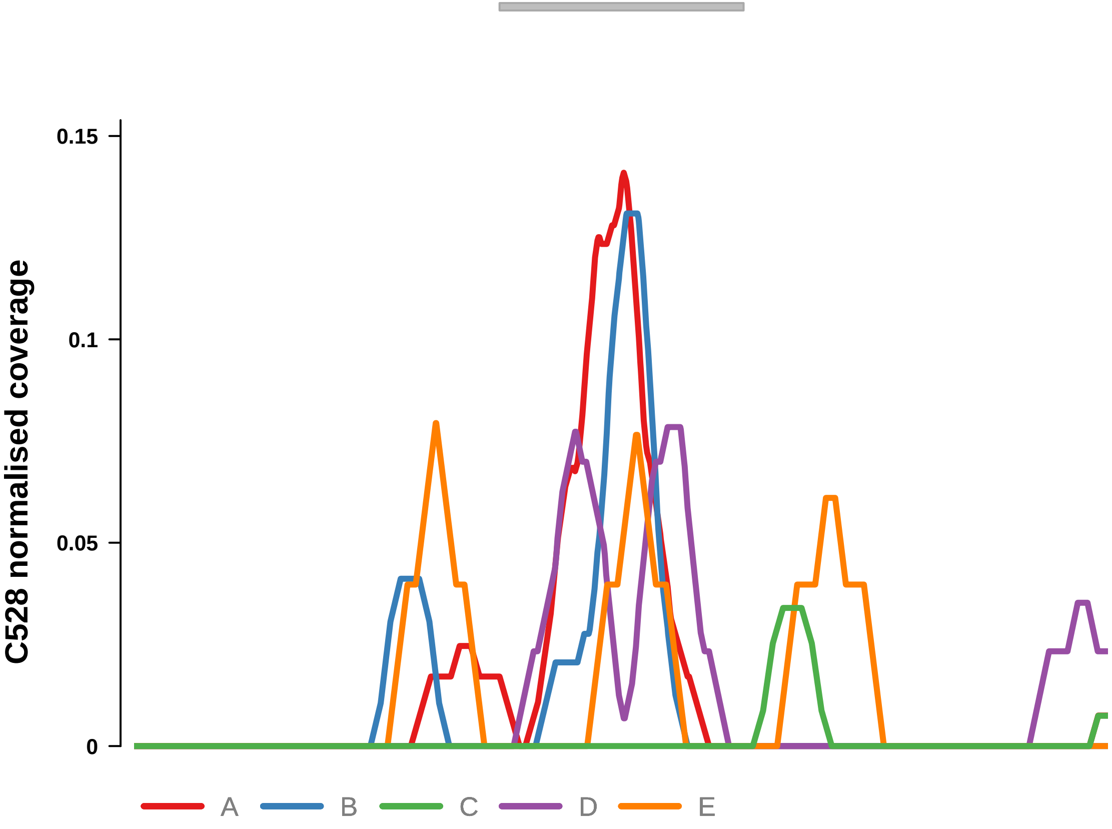


Putatively Clonal (logFC = −10.72)

| | P. | Adjusted P. |
|---|---|---|
| *~region vs. ~* | 2.96e−01 | 5.55e−01 |
| *~purity vs ~* | 7.81e−02 | NA |
| *~purity+region vs ~purity* | 7.48e−01 | 9.17e−01 |
| *Carcinoma vs. normal* | 2.05e−04 *** | 2.05e−04 *** |

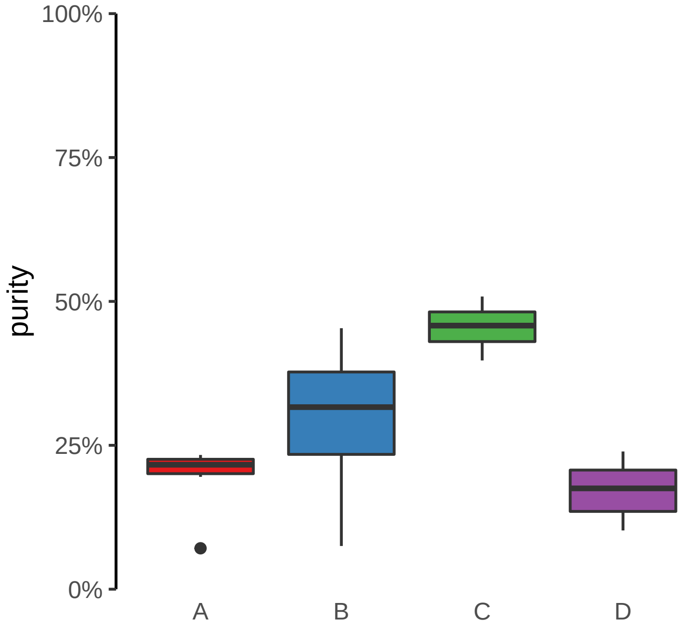

**CNST (GH01J246591) enhancer loss**
**chr1:246591917−246592417**

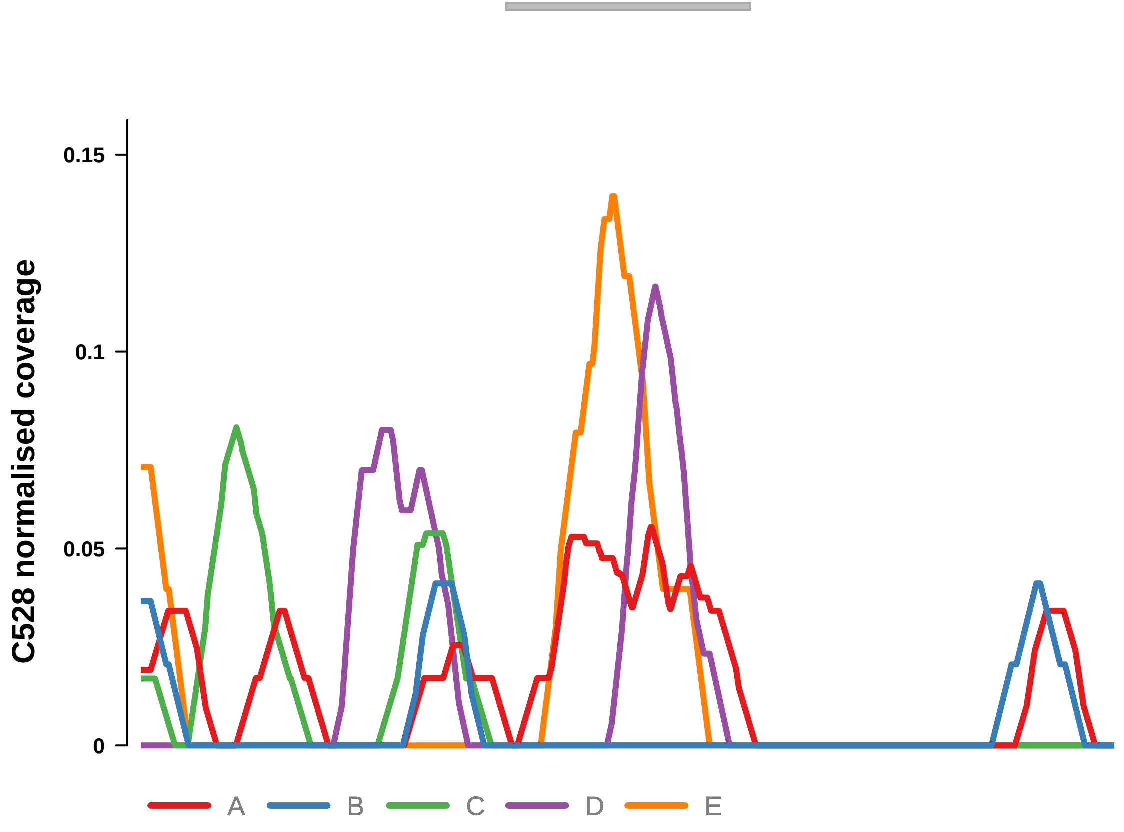


Putatively Clonal (logFC = −11.12)

| | P. | Adjusted P. |
|---|---|---|
| *~region vs. ~* | 5.31e−01 | 7.3e−01 |
| *~purity vs ~* | 1.9e−01 | NA |
| *~purity+region vs ~purity* | 8.48e−01 | 9.69e−01 |
| *Carcinoma vs. normal* | 2.84e−05 **** | 2.84e−05 **** |

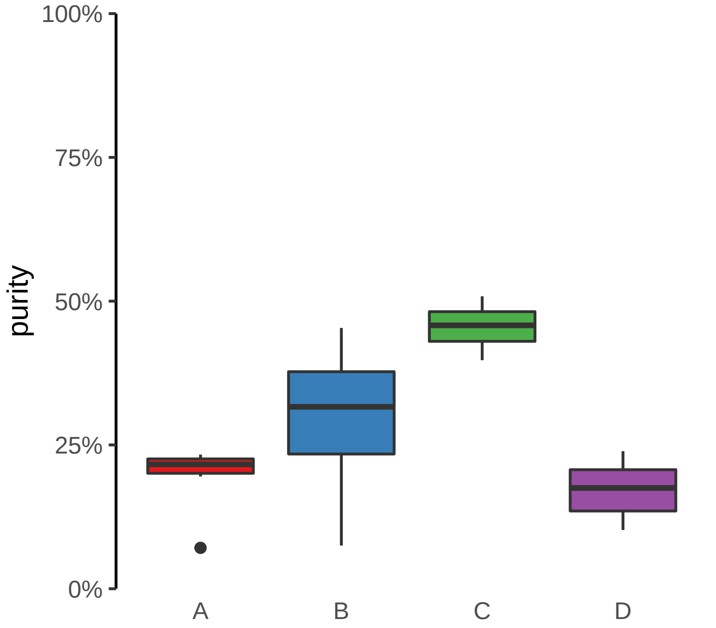

**FABP1, THNSL2 (GH02J088144) enhancer loss**
**chr2:88144817−88145317**

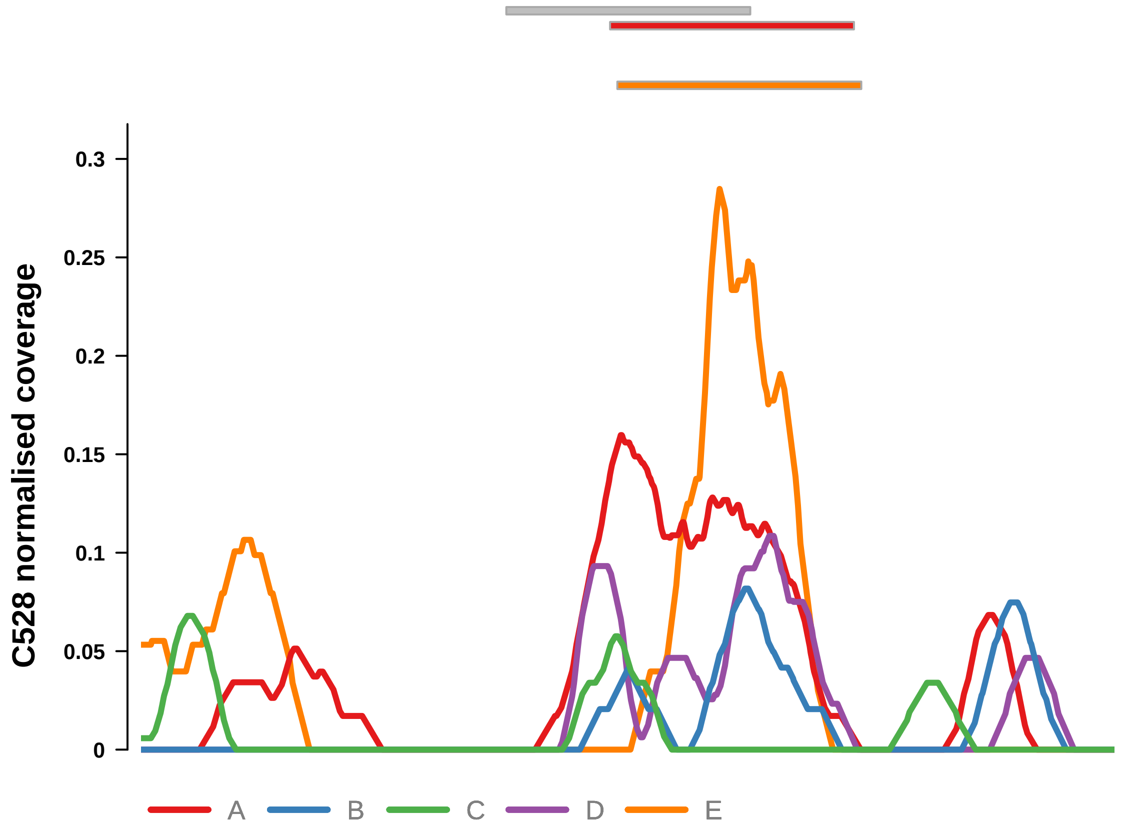


Putatively Clonal (logFC = −3.47)

| | P. | Adjusted P. |
|---|---|---|
| *~region vs. ~* | 2.3e−02 * | 2.02e−01 |
| *~purity vs ~* | 3.26e−03 ** | 3.75e−02 * |
| *~purity+region vs ~purity* | 6.39e−01 | 9.07e−01 |
| *Carcinoma vs. normal* | 6.91e−04 *** | 6.91e−04 *** |

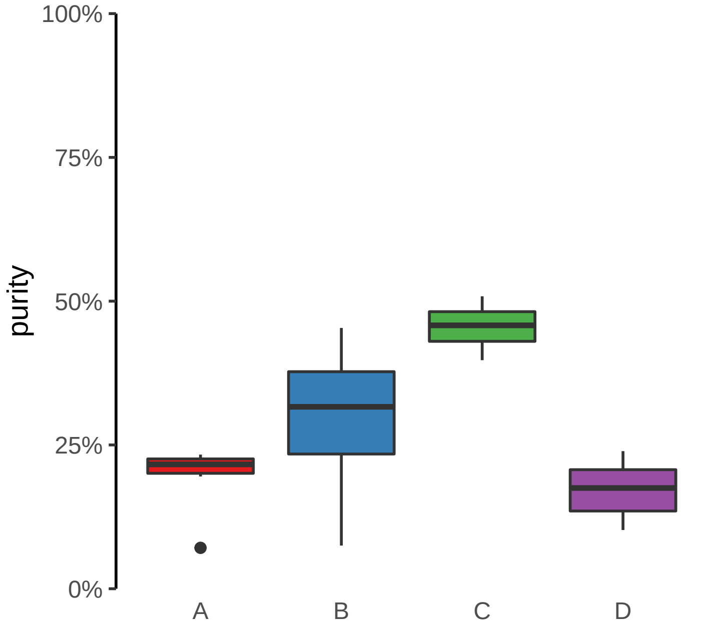

**PPP2CB (GH08J030800) enhancer loss**
**chr8:30804244−30804744**

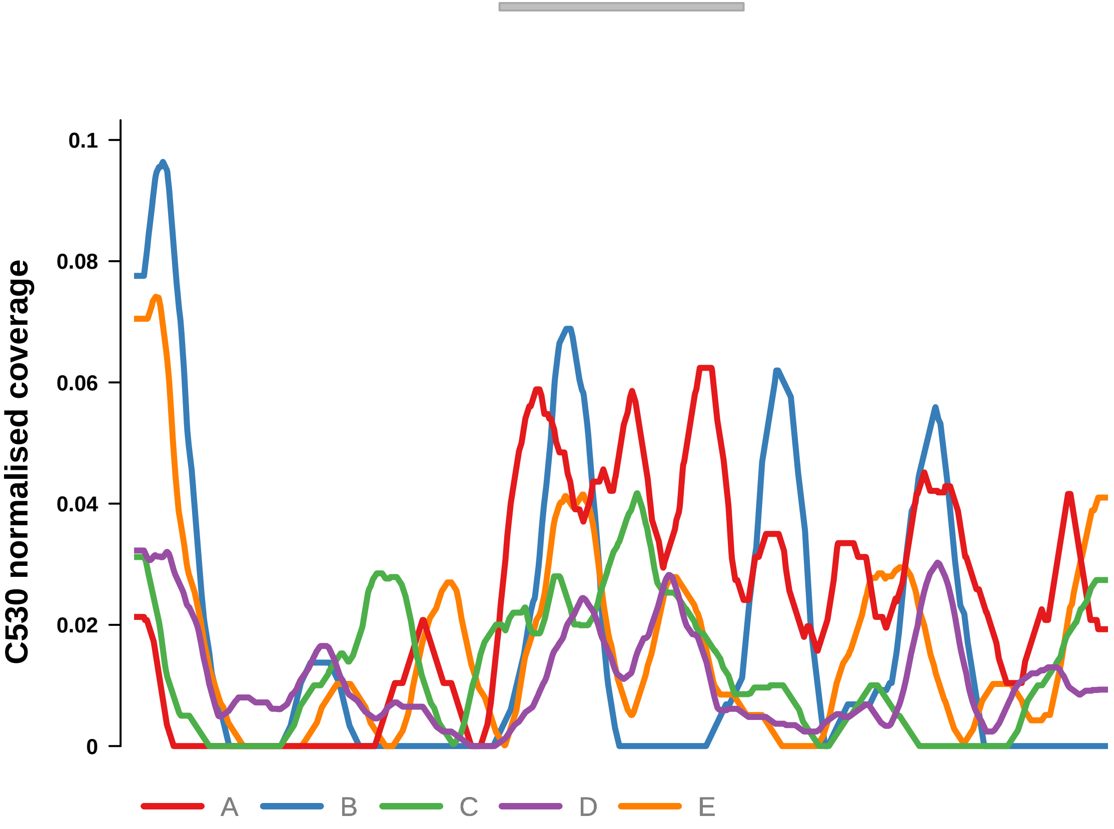


Putatively Clonal (logFC = −2.05)

| | P. | Adjusted P. |
|---|---|---|
| *~region vs. ~* | 6.42e−01 | 8.86e−01 |
| *~purity vs ~* | 3.56e−01 | 8.26e−01 |
| *~purity+region vs ~purity* | 6.82e−01 | 8.58e−01 |
| *Carcinoma vs. normal* | 5.42e−03 ** | 5.42e−03 ** |

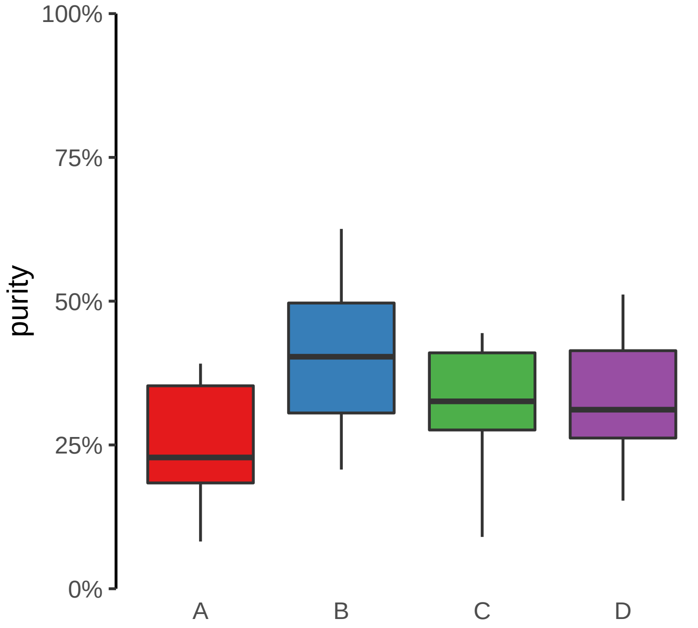

**ZNF431 (GH19J020999) enhancer loss**
**chr19:20999605−21000105**

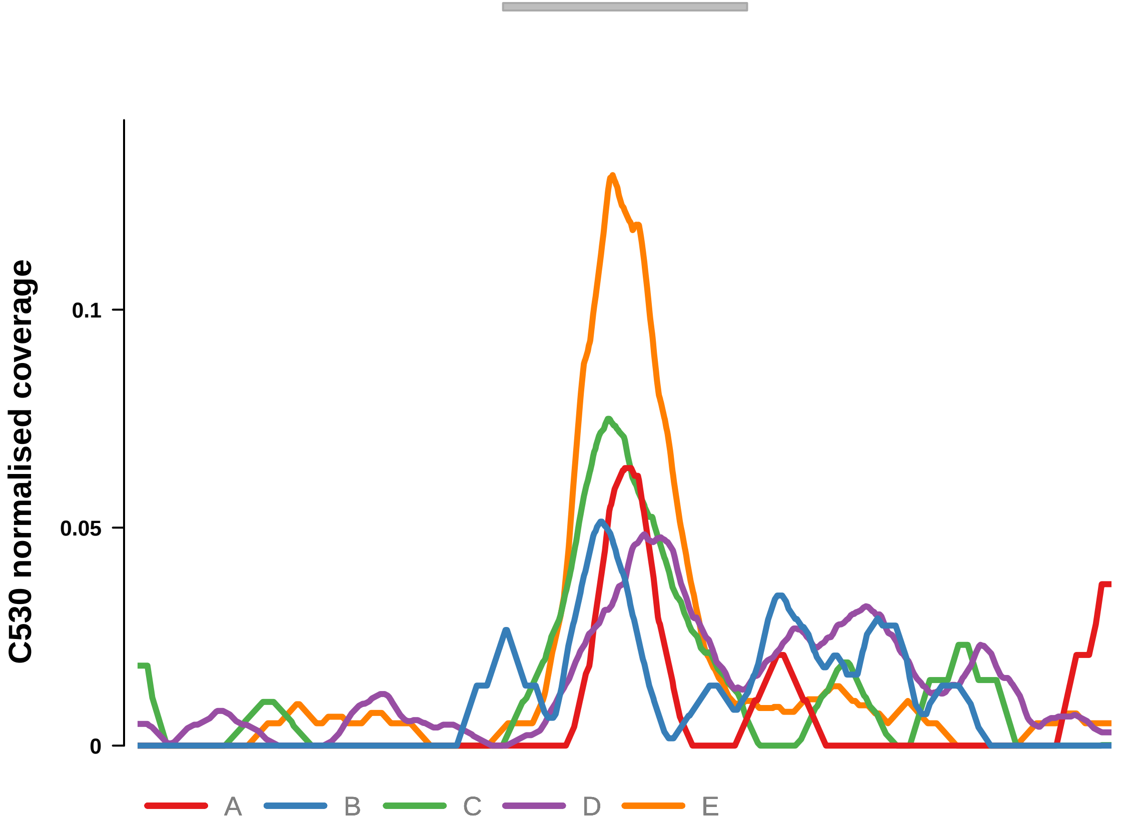


Putatively Clonal (logFC = −1.91)

| | P. | Adjusted P. |
|---|---|---|
| *~region vs. ~* | 5.24e−01 | 8.24e−01 |
| *~purity vs ~* | 9.25e−01 | 9.83e−01 |
| *~purity+region vs ~purity* | 5.39e−01 | 7.8e−01 |
| *Carcinoma vs. normal* | 7.77e−03 ** | 7.77e−03 ** |

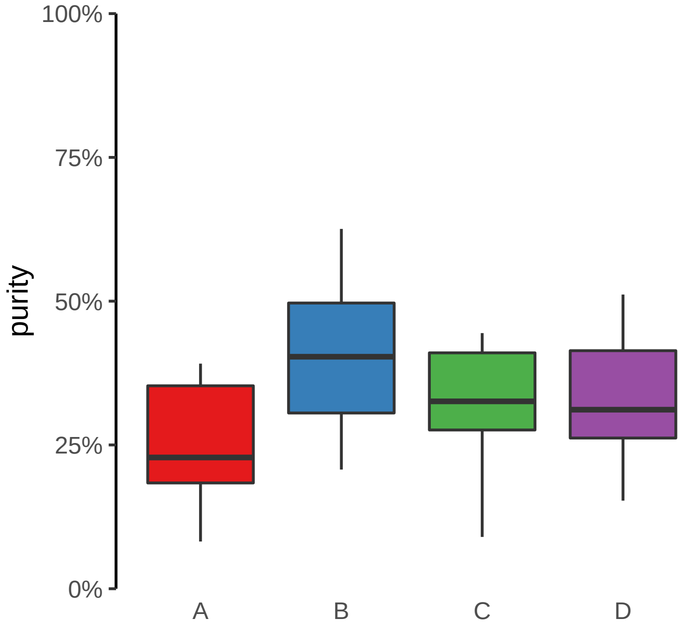

**ACAP2, PPP1R2 (GH03J195537) enhancer loss**
**chr3:195537917−195538417**

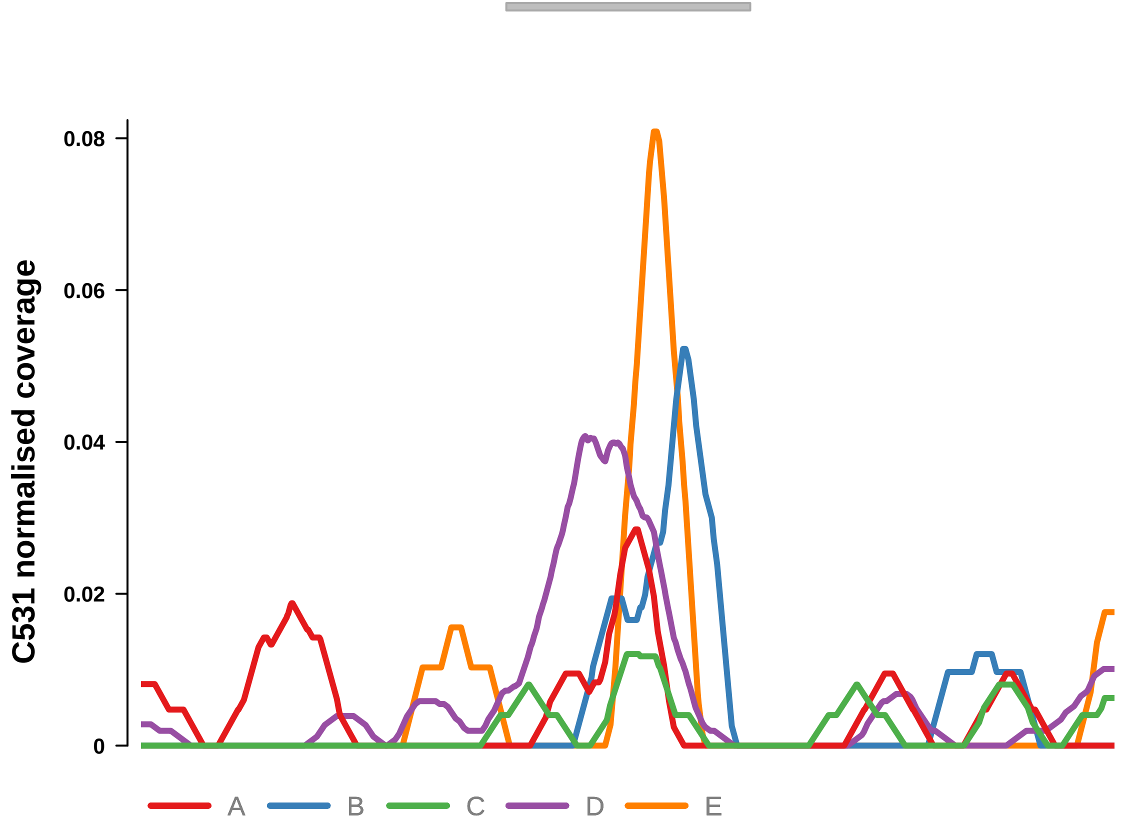


Putatively Clonal (logFC = −10.38)

| | P. | Adjusted P. |
|---|---|---|
| *~region vs. ~* | 5.39e−01 | NA |
| *~purity vs ~* | 5.83e−02 | 1.6e−01 |
| *~purity+region vs ~purity* | 9.3e−01 | 9.52e−01 |
| *Carcinoma vs. normal* | 4.6e−04 *** | 4.6e−04 *** |

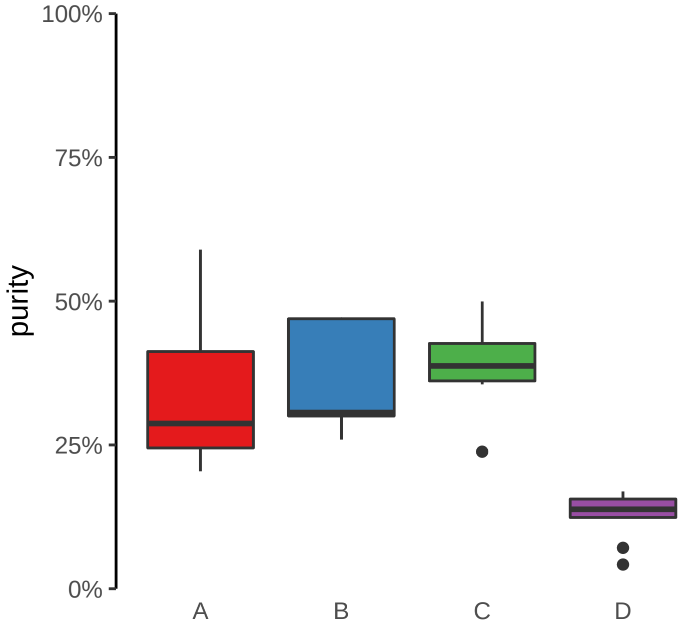

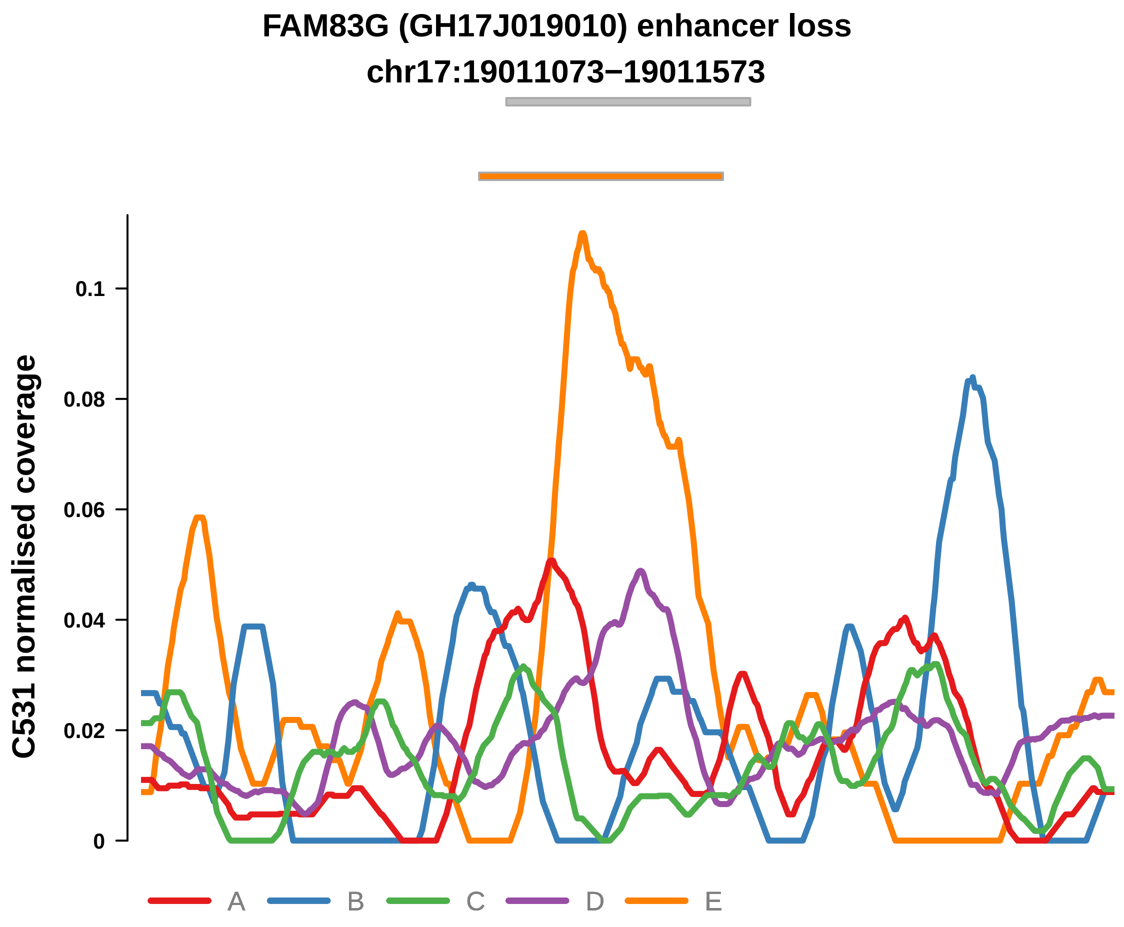


Putatively Clonal (logFC = −2.28)

| | P. | Adjusted P. |
|---|---|---|
| *~region vs. ~* | 1.63e−01 | 2.92e−01 |
| *~purity vs ~* | 2.8e−02 * | 9.85e−02 |
| *~purity+region vs ~purity* | 5.29e−02 | 4.16e−01 |
| *Carcinoma vs. normal* | 5.91e−03 ** | 5.91e−03 ** |

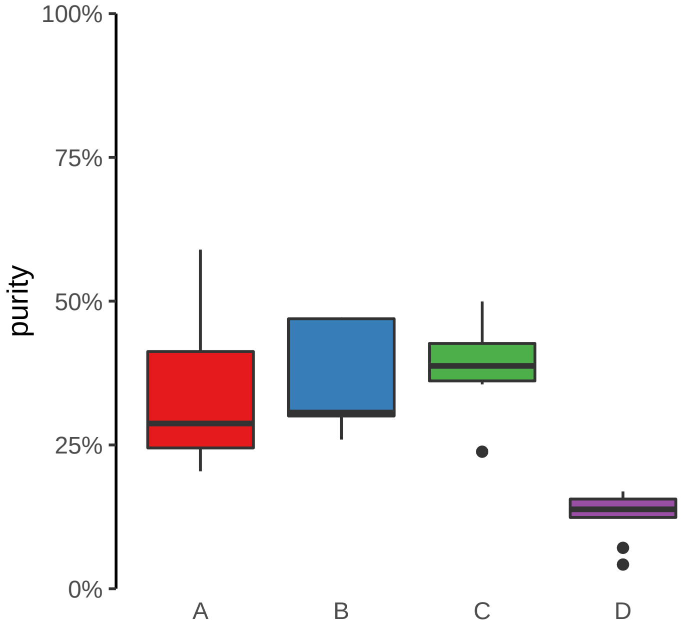

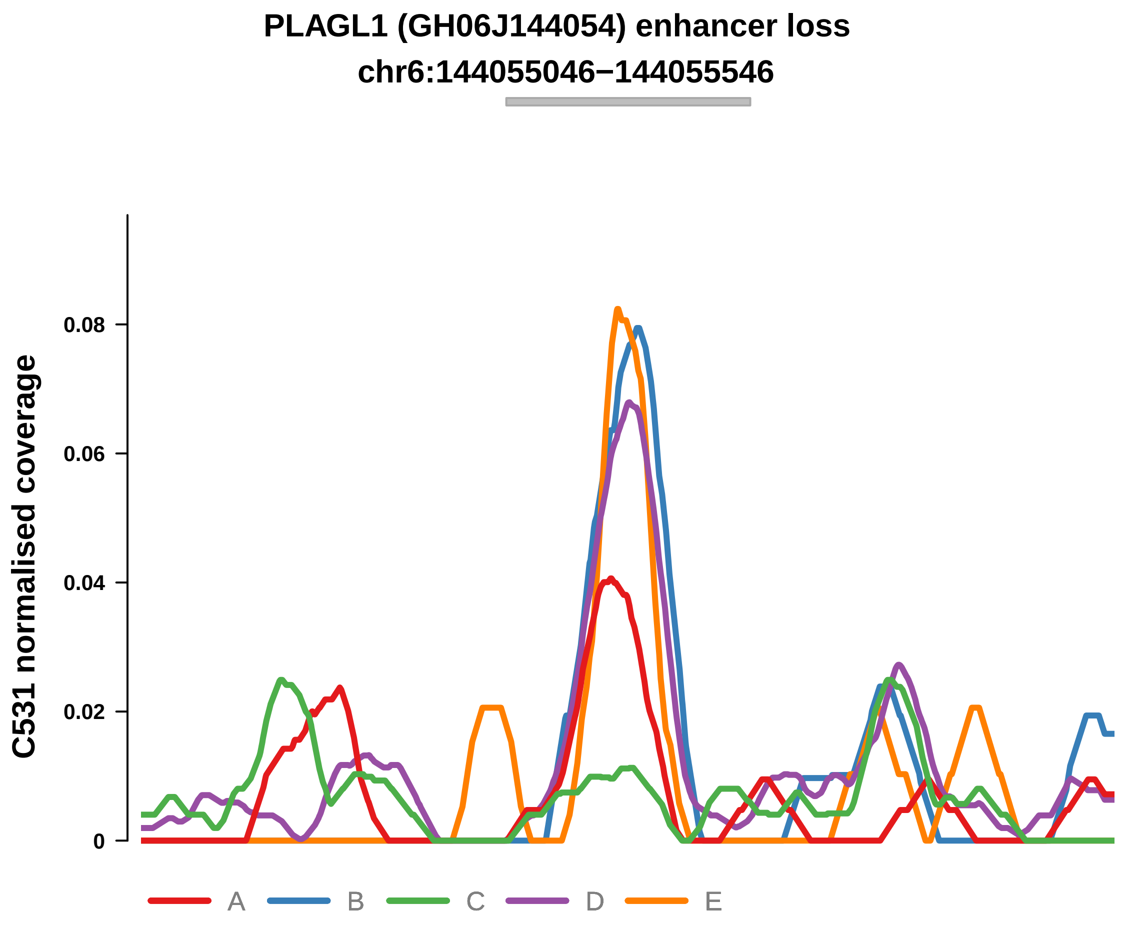


Putatively Clonal (logFC = −10.99)

| | P. | Adjusted P. |
|---|---|---|
| *~region vs. ~* | 2.3e−01 | NA |
| *~purity vs ~* | 5.15e−02 | 1.51e−01 |
| *~purity+region vs ~purity* | 5.05e−01 | 7.7e−01 |
| *Carcinoma vs. normal* | 4.39e−05 **** | 4.39e−05 **** |

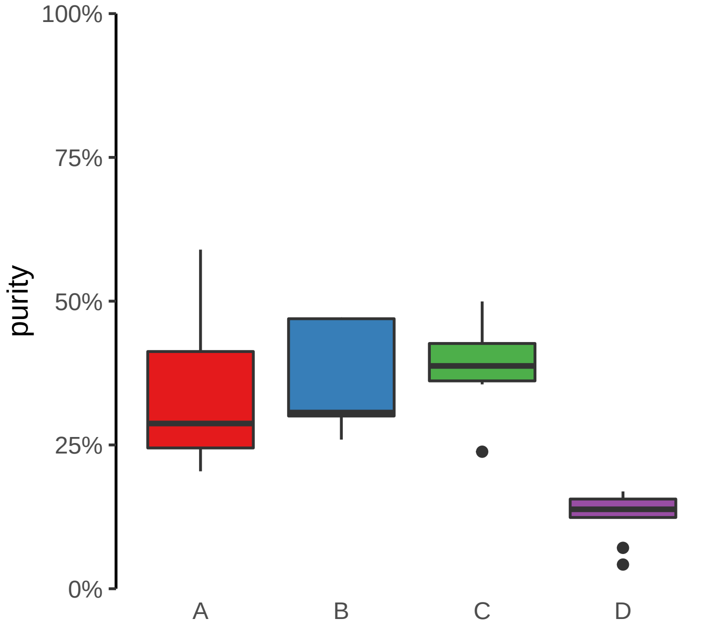

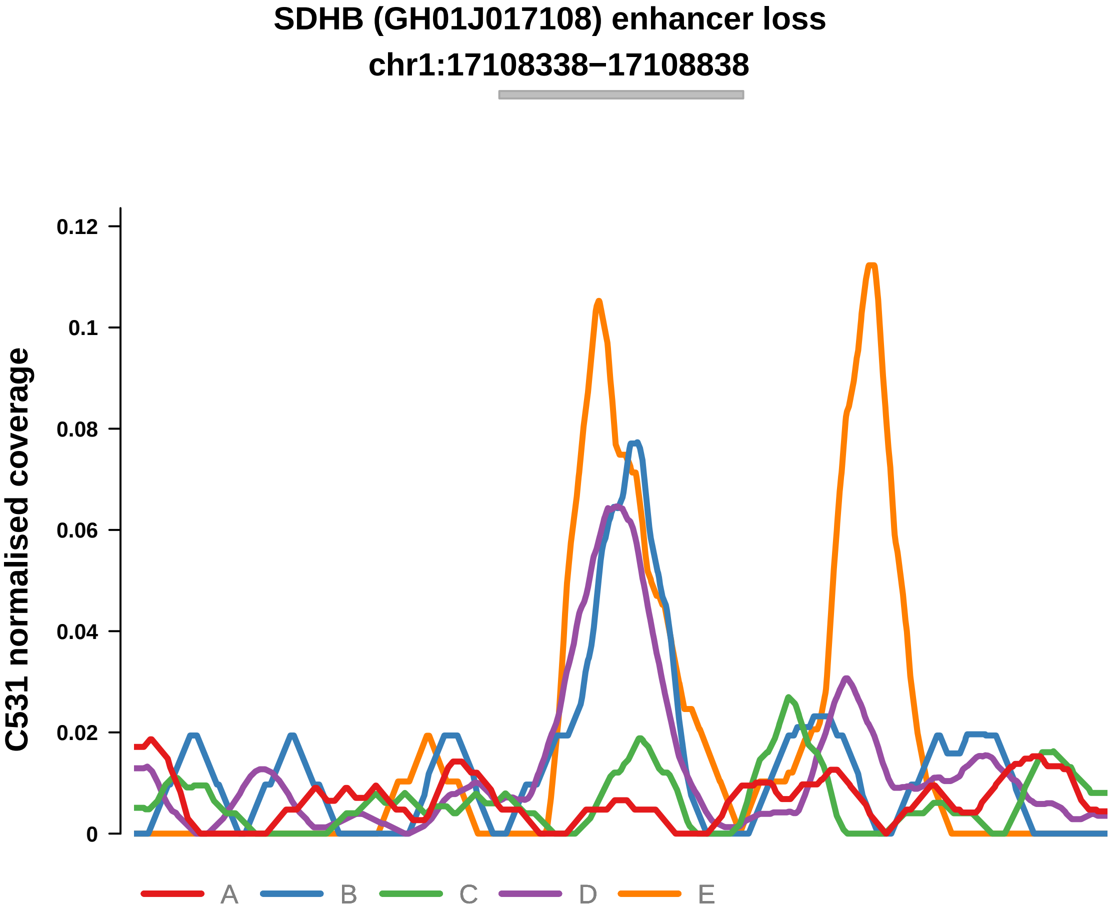


Putatively Clonal (logFC = −3.32)

| | P. | Adjusted P. |
|---|---|---|
| *~region vs. ~* | 2.1e−01 | NA |
| *~purity vs ~* | 2.04e−02 * | 8.14e−02 |
| *~purity+region vs ~purity* | 1e+00 | 1e+00 |
| *Carcinoma vs. normal* | 1.34e−03 ** | 1.34e−03 ** |

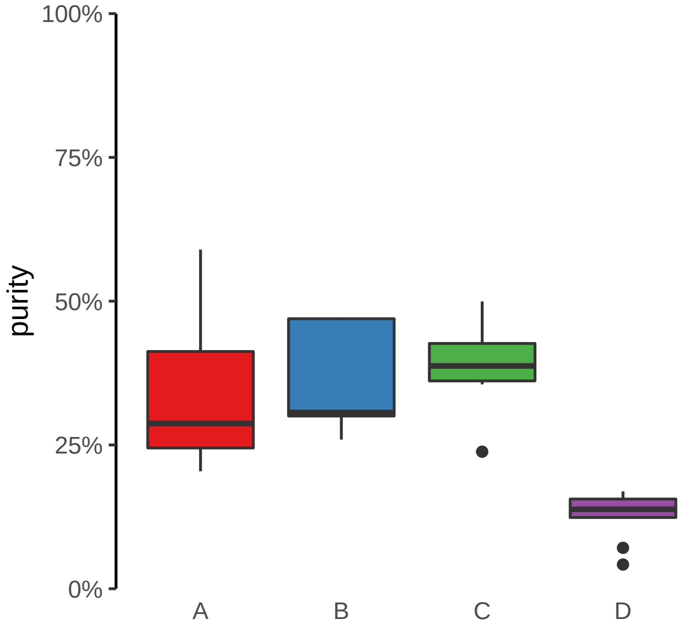

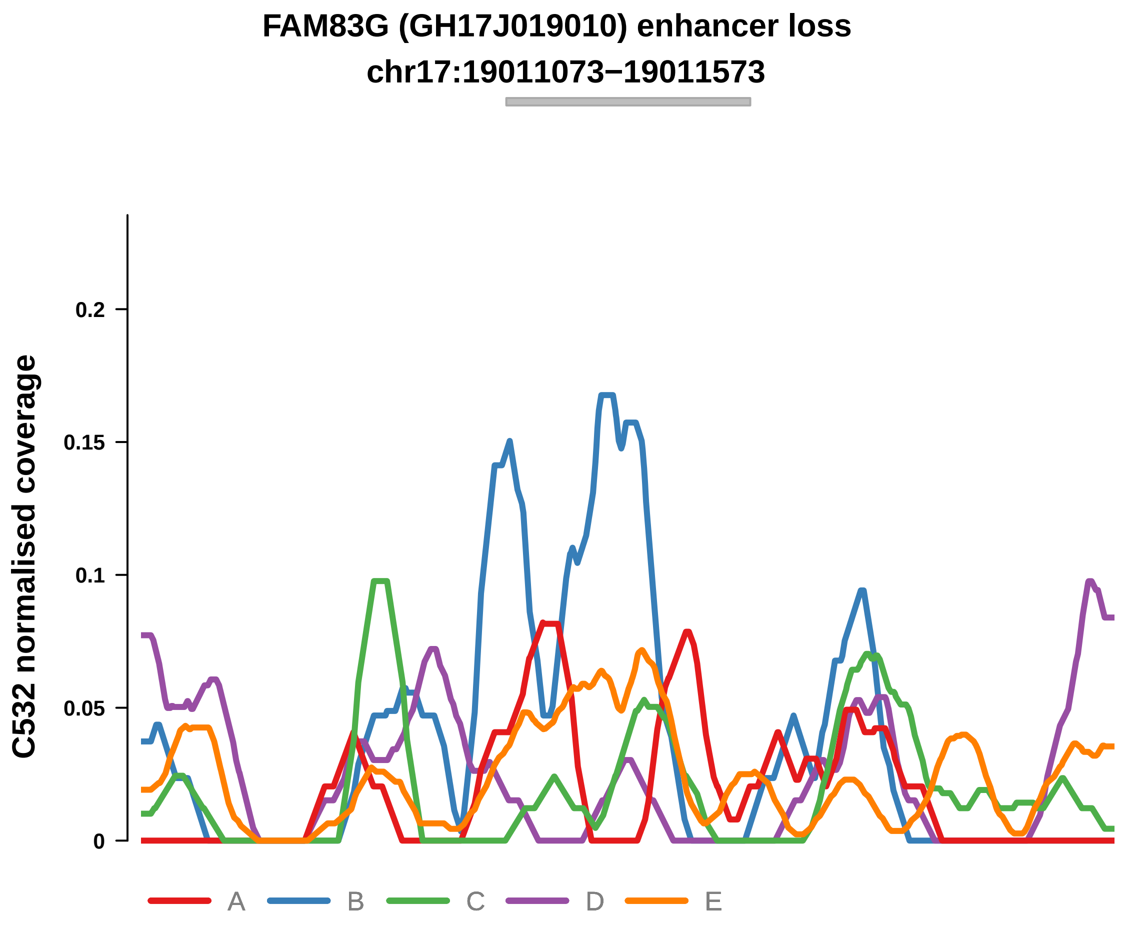


Putatively Clonal (logFC = −2.79)

| | P. | Adjusted P. |
|---|---|---|
| *~region vs. ~* | 1.91e−02 * | 2.06e−01 |
| *~purity vs ~* | 2.11e−02 * | 3.05e−01 |
| *~purity+region vs ~purity* | 2.06e−01 | 6.56e−01 |
| *Carcinoma vs. normal* | 3.68e−04 *** | 3.68e−04 *** |

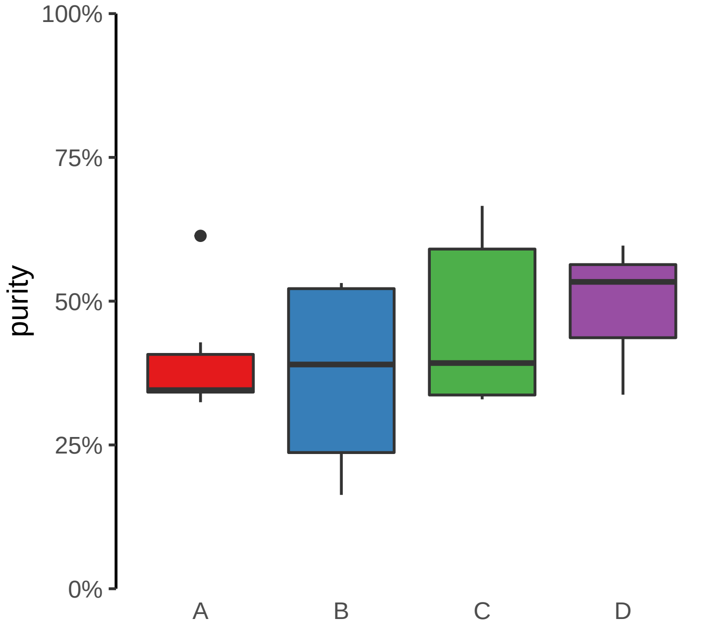

**HMGN2, ARID1A, PIGV (GH01J026739) enhancer loss**
**chr1:26743976−26744476**

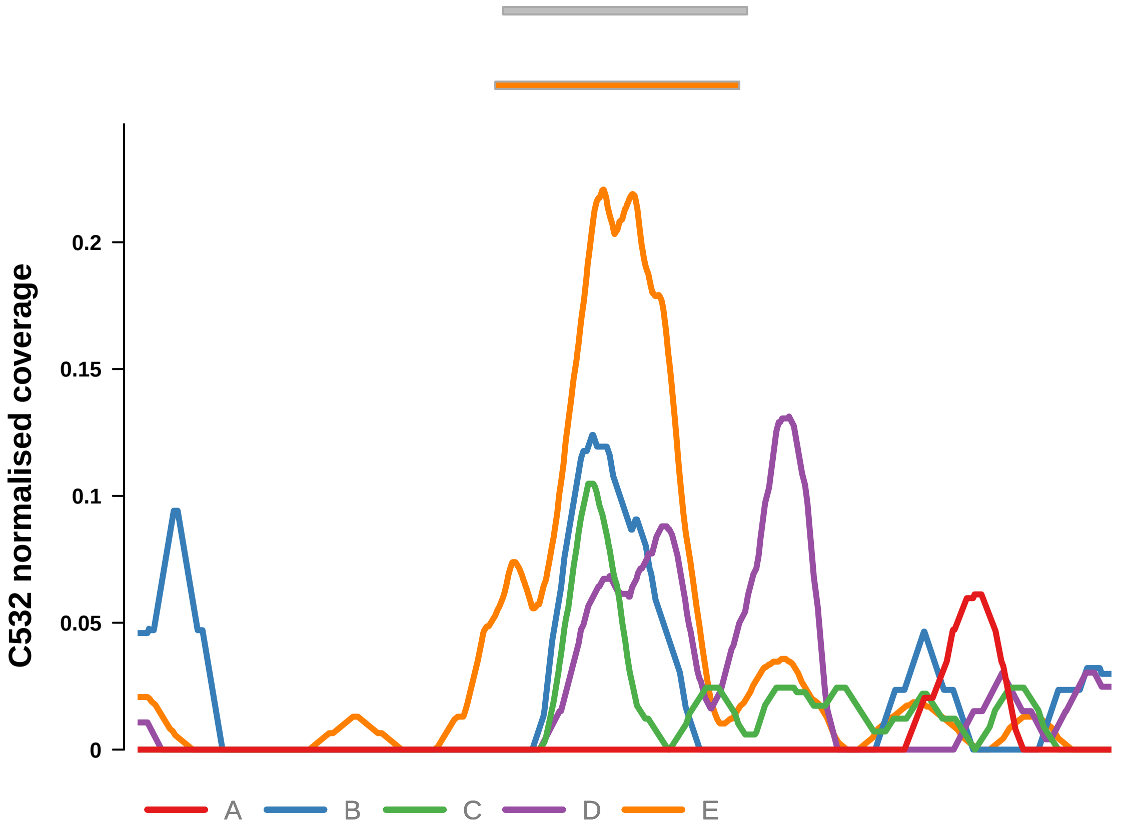


Putatively Clonal (logFC = −2.29)

| | P. | Adjusted P. |
|---|---|---|
| *~region vs. ~* | 9.03e−03 ** | 1.59e−01 |
| *~purity vs ~* | 2e−01 | 8.56e−01 |
| *~purity+region vs ~purity* | 5.13e−03 ** | 1.13e−01 |
| *Carcinoma vs. normal* | 2.38e−03 ** | 2.38e−03 ** |

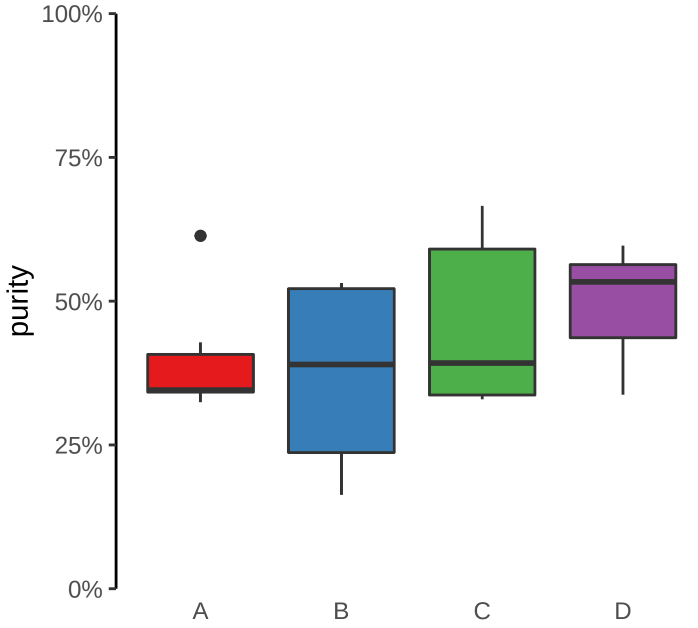

**CLASP1 (GH02J121362) enhancer loss**
**chr2:121363584−121364084**

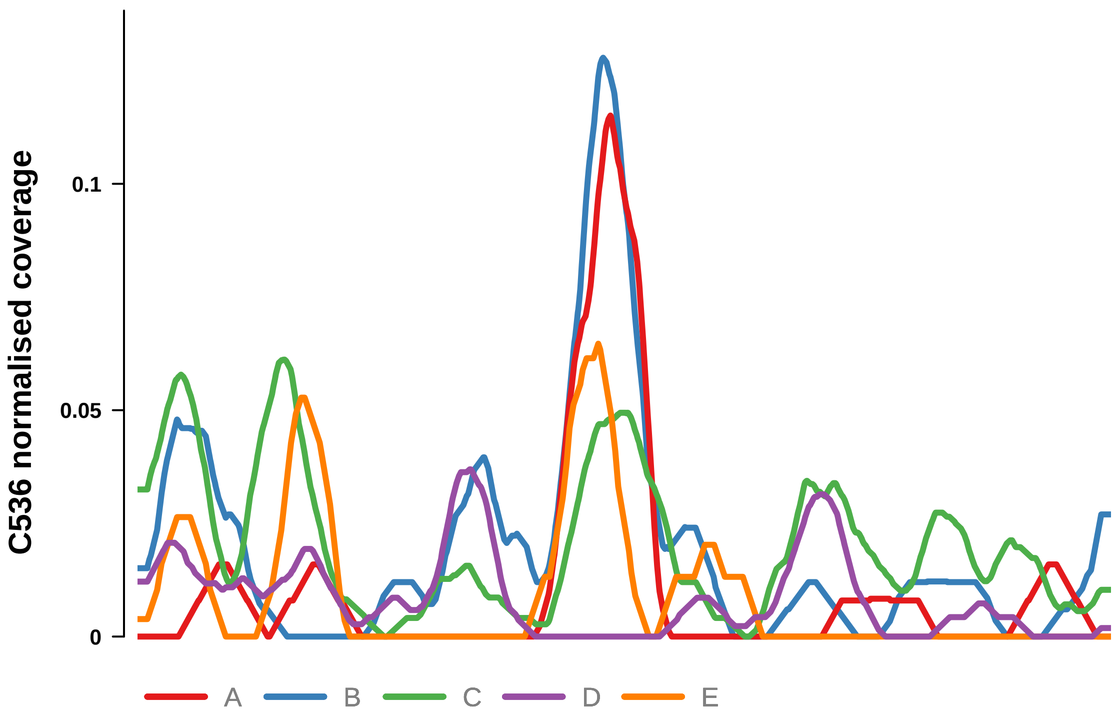


Putatively Clonal (logFC = −4.02)

| | P. | Adjusted P. |
|---|---|---|
| *~region vs. ~* | 5.13e−01 | NA |
| *~purity vs ~* | 4.68e−02 * | 2.04e−01 |
| *~purity+region vs ~purity* | 3.37e−01 | 4.87e−01 |
| *Carcinoma vs. normal* | 5.78e−05 **** | 5.78e−05 **** |

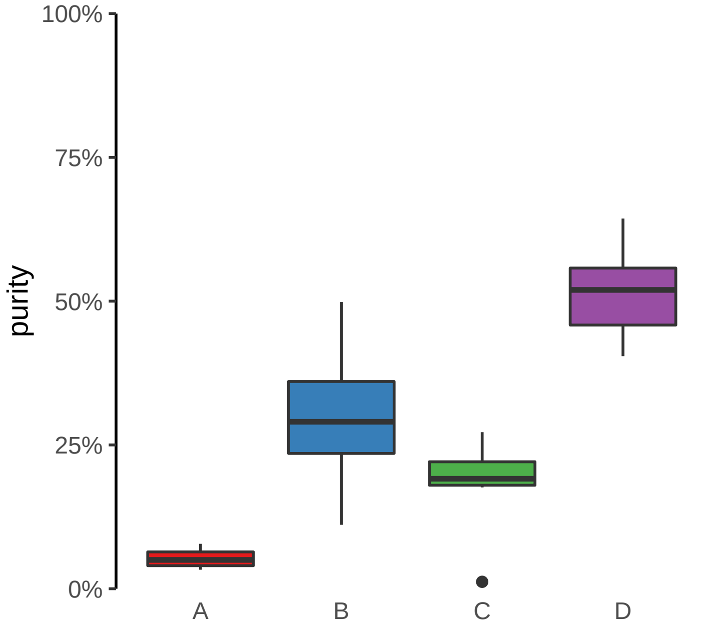

**CLASP2, ENSG00000271324 (GH03J033790) enhancer loss**
**chr3:33791629−33792129**

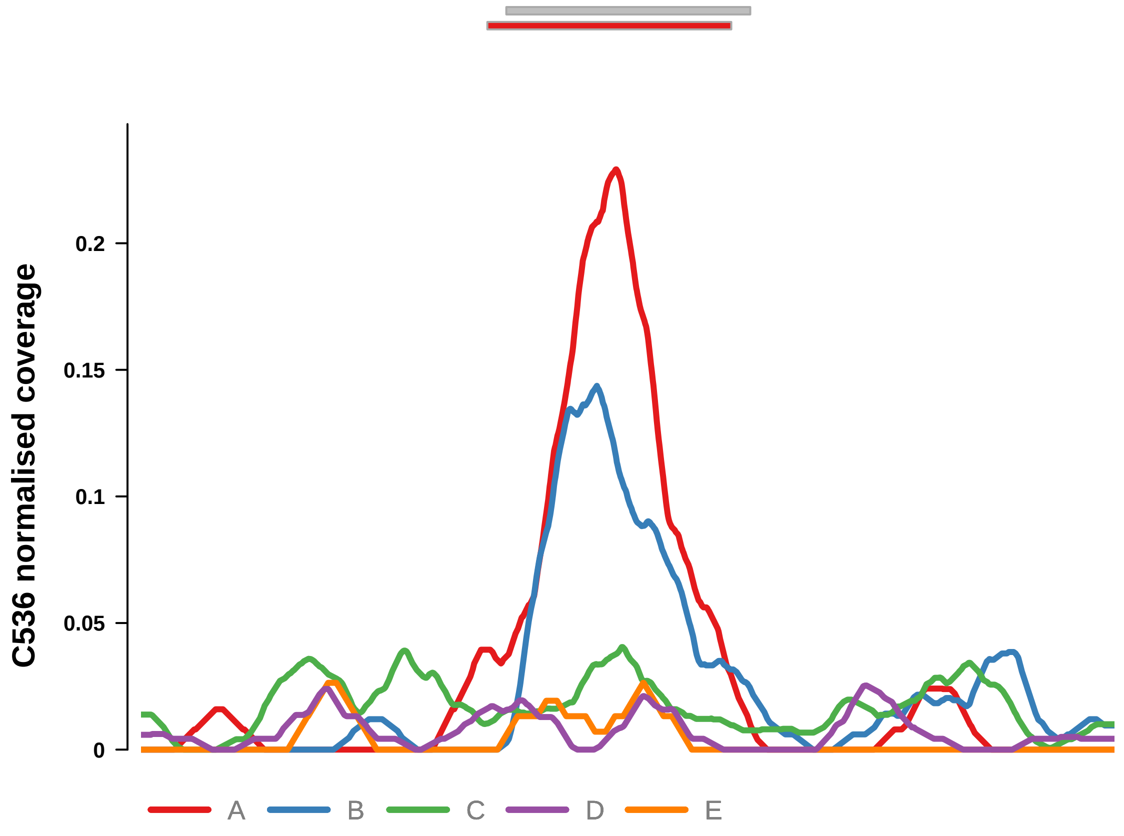


Putatively Clonal (logFC = −2.02)

| | P. | Adjusted P. |
|---|---|---|
| *~region vs. ~* | 3.66e−01 | 4.53e−01 |
| *~purity vs ~* | 1.15e−01 | 2.81e−01 |
| *~purity+region vs ~purity* | 8.73e−01 | 9.13e−01 |
| *Carcinoma vs. normal* | 8.45e−03 ** | 8.45e−03 ** |

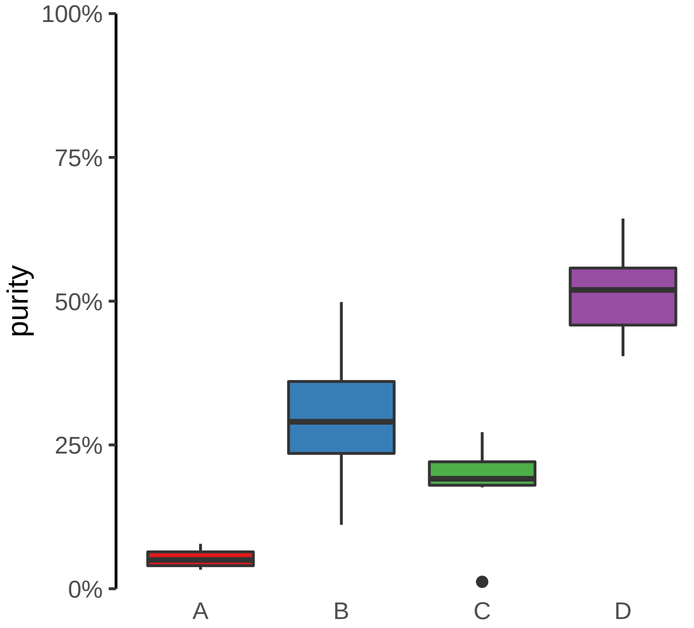

**FABP1, THNSL2 (GH02J088144) enhancer loss**
**chr2:88144817−88145317**

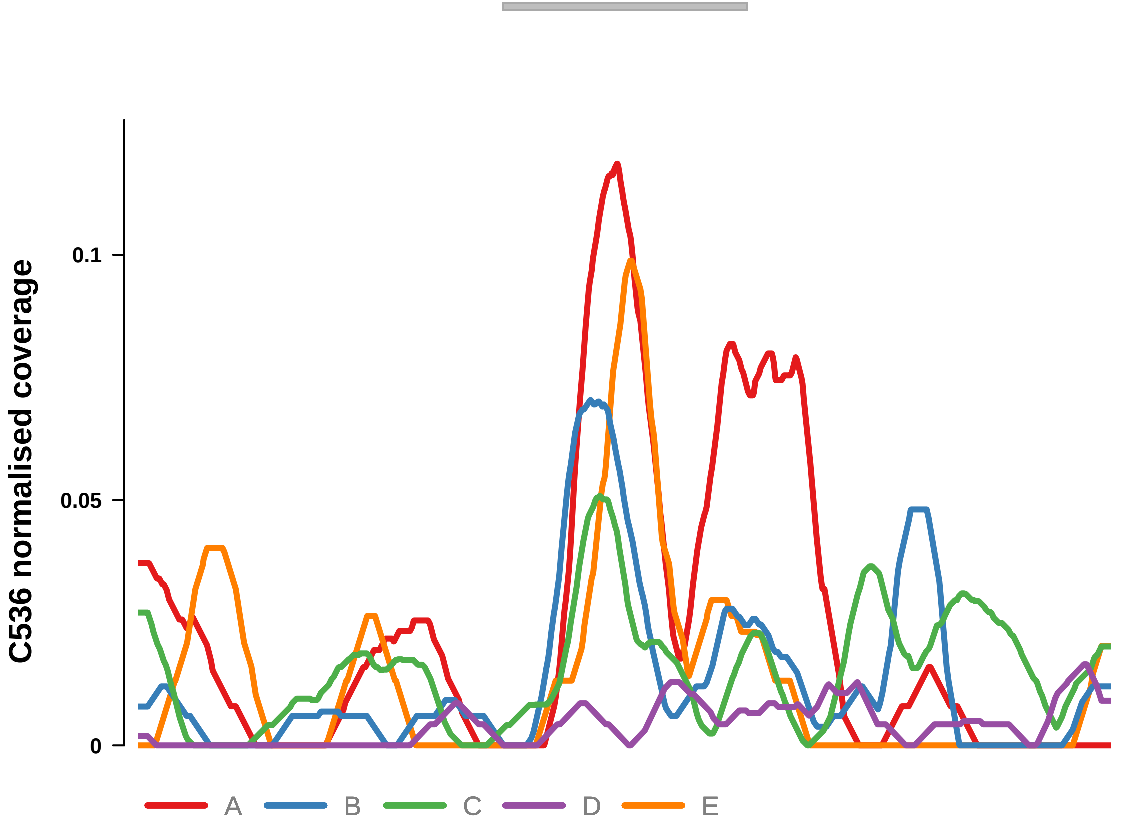


Putatively Clonal (logFC = −2.83)

| | P. | Adjusted P. |
|---|---|---|
| *~region vs. ~* | 6.05e−01 | NA |
| *~purity vs ~* | 3.01e−02 * | 1.77e−01 |
| *~purity+region vs ~purity* | 2.04e−01 | NA |
| *Carcinoma vs. normal* | 8.54e−04 *** | 8.54e−04 *** |

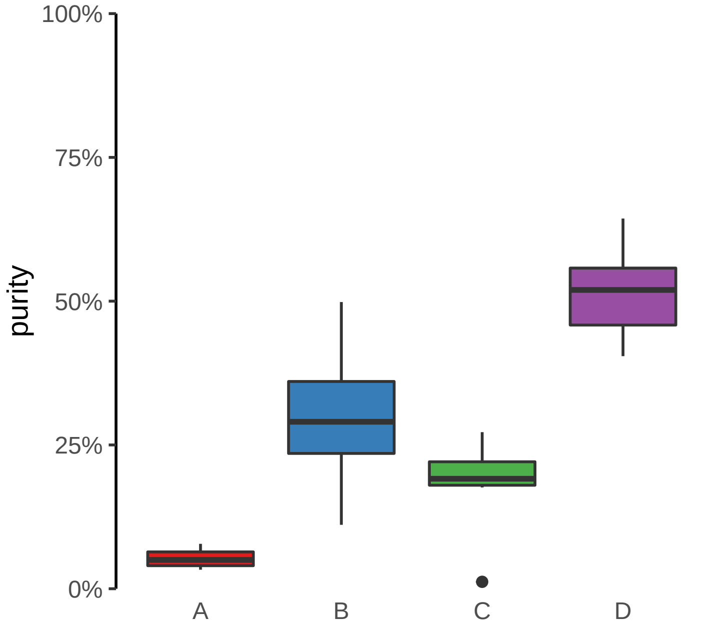

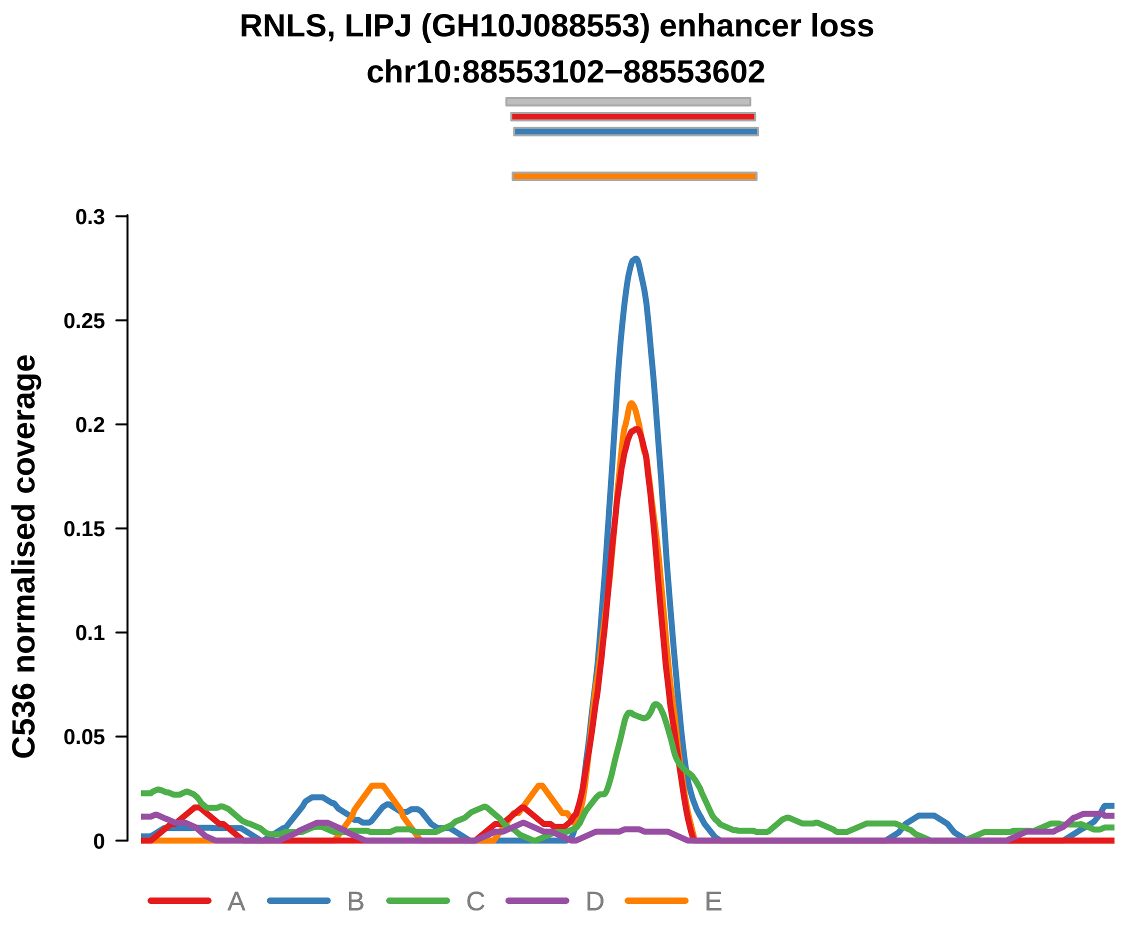


Putatively Clonal (logFC = −3.04)

| | P. | Adjusted P. |
|---|---|---|
| *~region vs. ~* | 2.62e−01 | 3.87e−01 |
| *~purity vs ~* | 1.53e−02 * | 1.22e−01 |
| *~purity+region vs ~purity* | 5.23e−01 | 6.38e−01 |
| *Carcinoma vs. normal* | 6.92e−04 *** | 6.92e−04 *** |

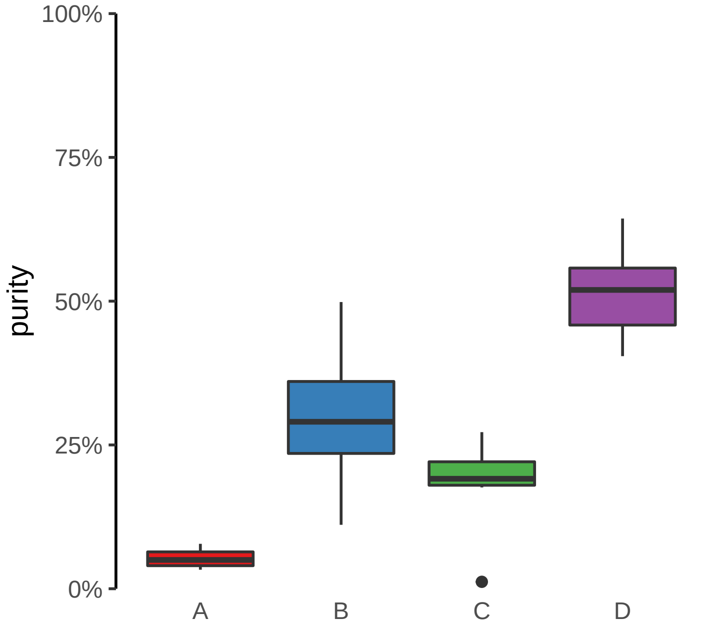

**CLASP1 (GH02J121362) enhancer loss**
**chr2:121363584−121364084**

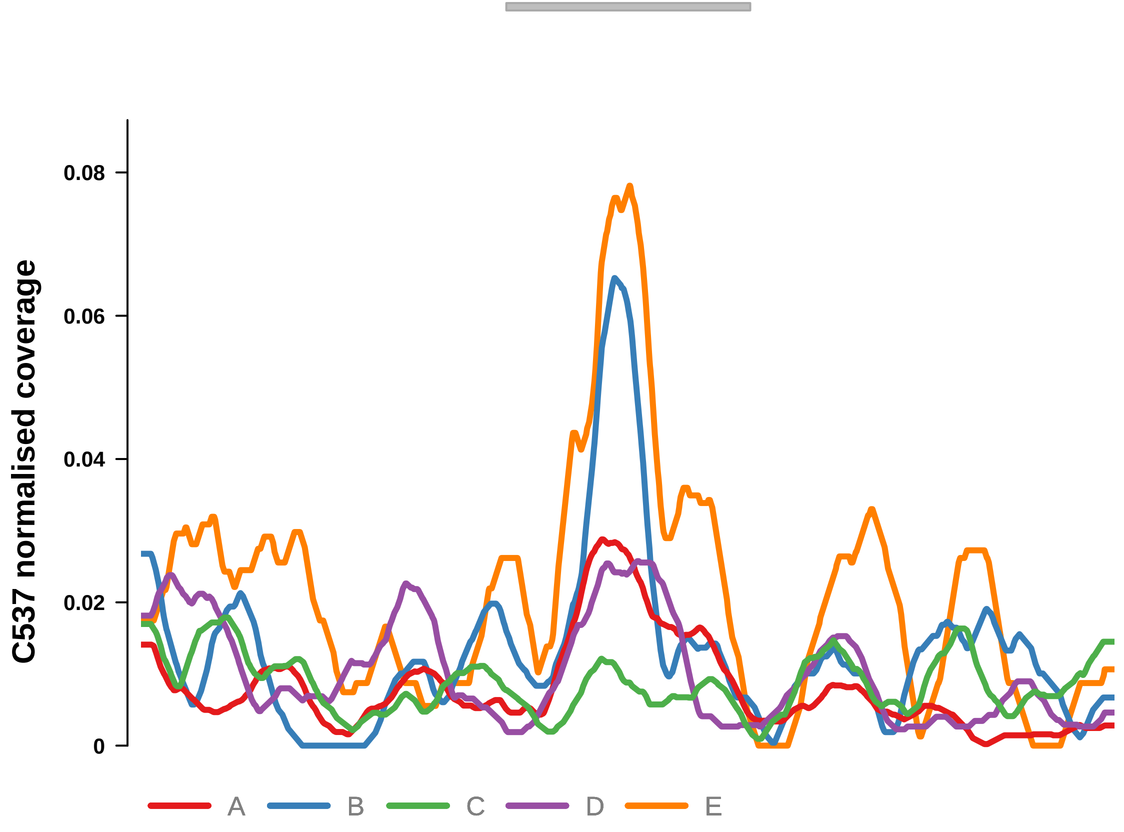


Putatively Clonal (logFC = −2.65)

| | **P.** | **Adjusted P.** |
|---|---|---|
| *~region vs. ~* | 5.83e−01 | 6.58e−01 |
| *~purity vs ~* | 2.08e−01 | 3.42e−01 |
| *~purity+region vs ~purity* | 9.38e−01 | 9.5e−01 |
| *Carcinoma vs. normal* | 7.87e−04 *** | 7.87e−04 *** |

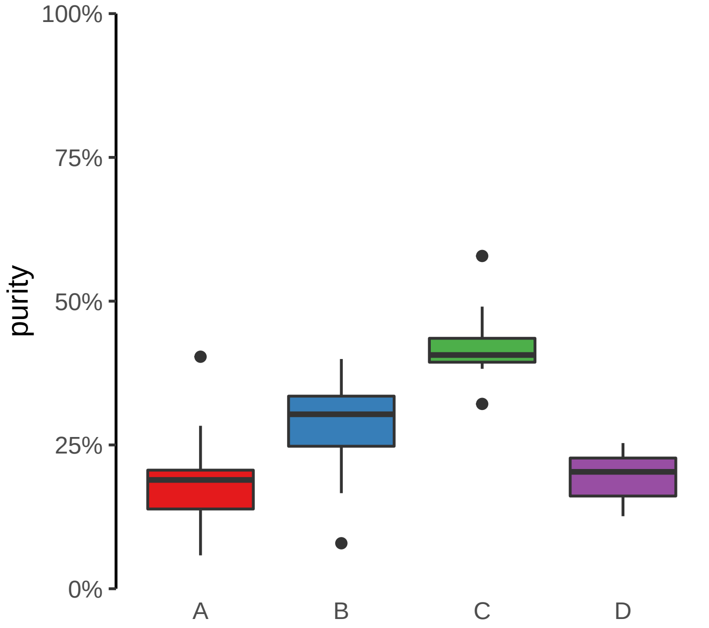

**CLASP2, ENSG00000271324 (GH03J033790) enhancer loss**
**chr3:33791629−33792129**

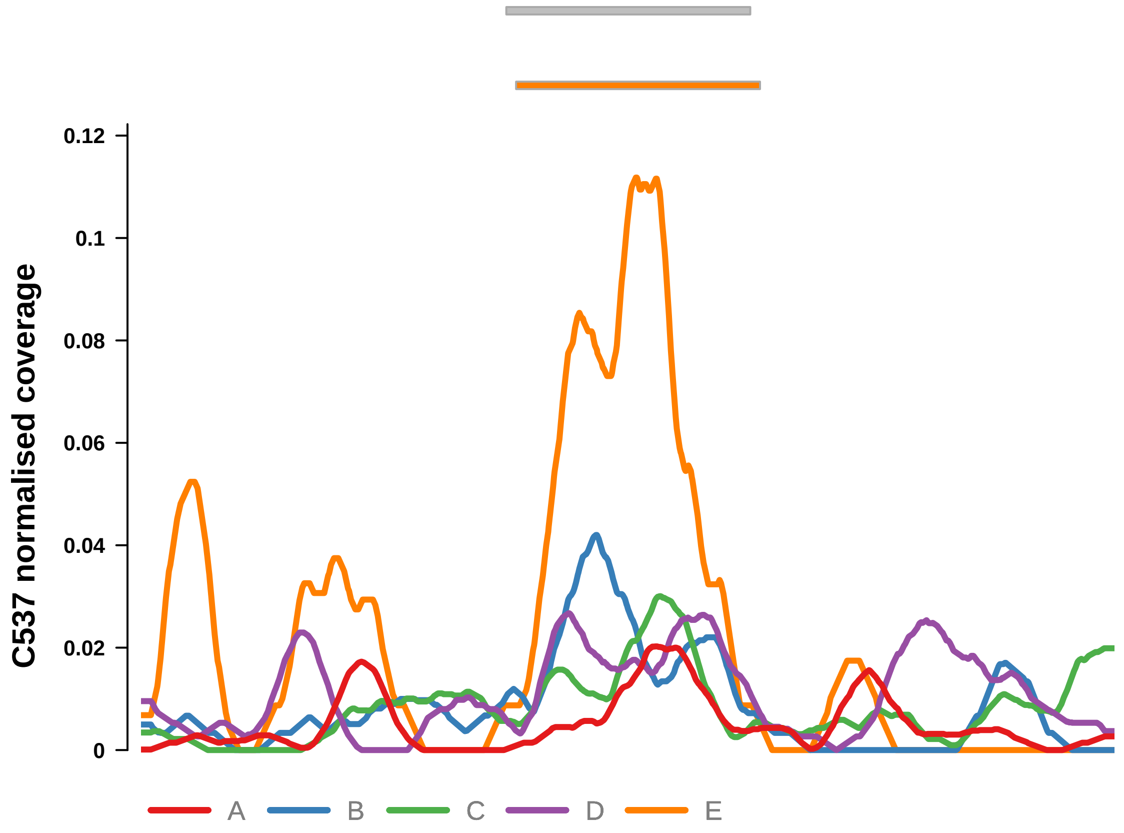


Putatively Clonal (logFC = −2.04)

| | P. | Adjusted P. |
|---|---|---|
| *~region vs. ~* | 1.76e−01 | 2.54e−01 |
| *~purity vs ~* | 9.2e−01 | NA |
| *~purity+region vs ~purity* | 1.48e−01 | 2.99e−01 |
| *Carcinoma vs. normal* | 4.86e−03 ** | 4.86e−03 ** |

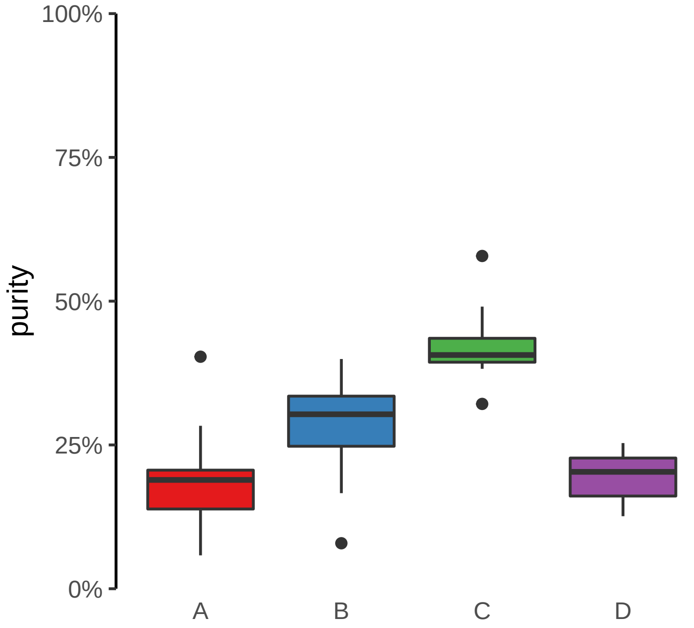

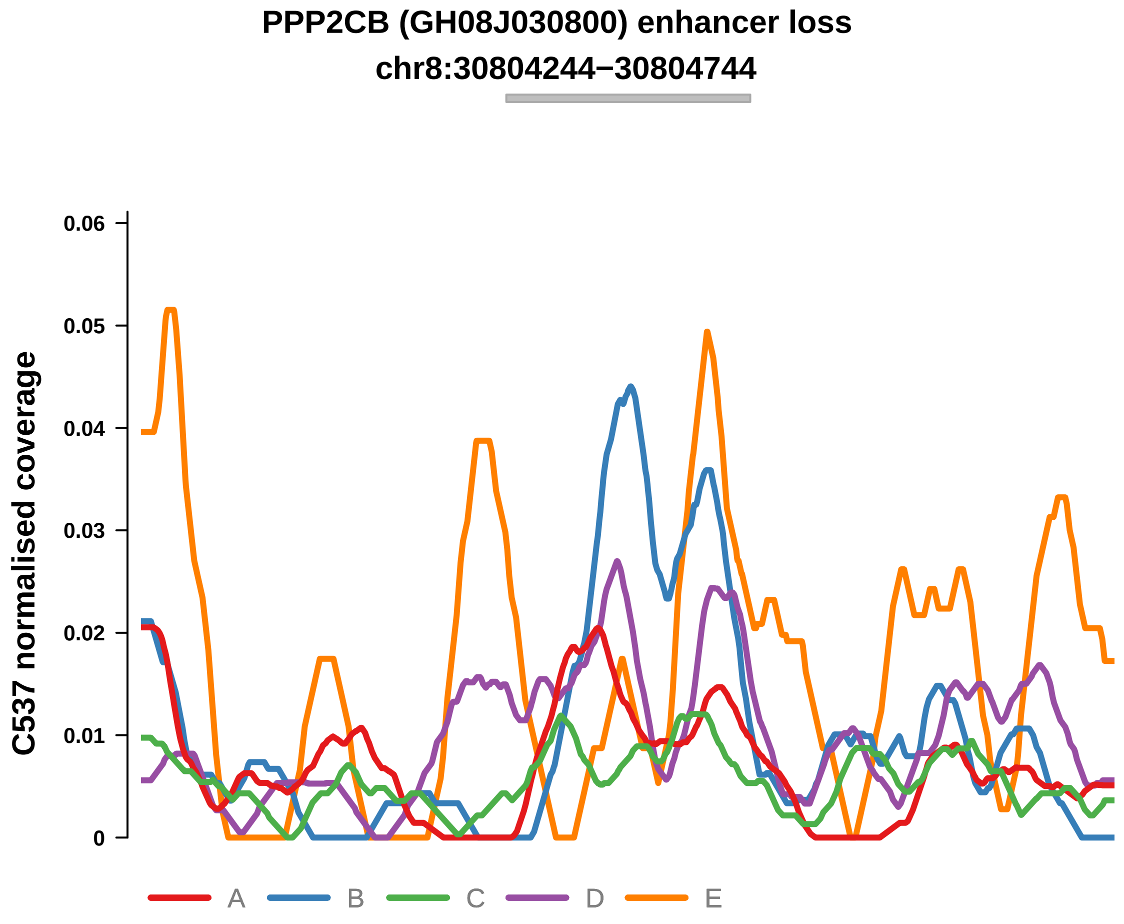


Putatively Clonal (logFC = −2.12)

| | P. | Adjusted P. |
|---|---|---|
| *~region vs. ~* | 2.31e−01 | 3.23e−01 |
| *~purity vs ~* | 2.18e−01 | NA |
| *~purity+region vs ~purity* | 3.58e−01 | 5.23e−01 |
| *Carcinoma vs. normal* | 4.24e−03 ** | 4.24e−03 ** |

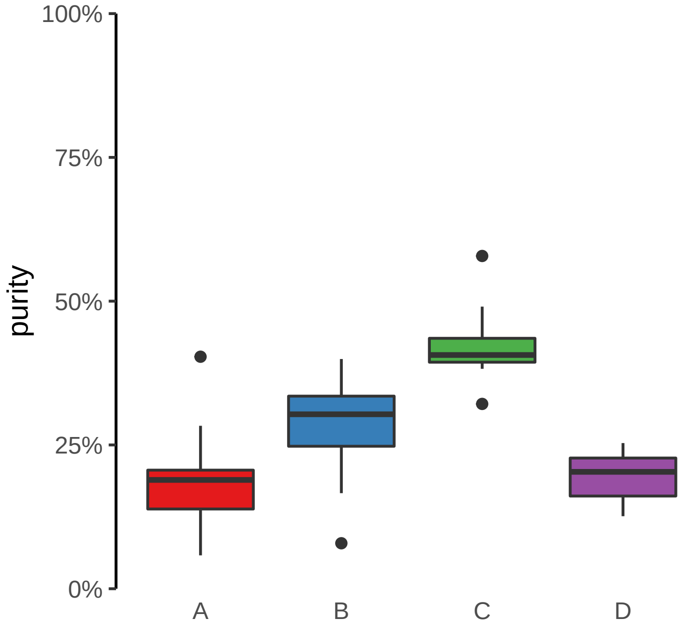

**STAG1, RNU6−789P (GH03J136616) enhancer loss**
**chr3:136617156−136617656**

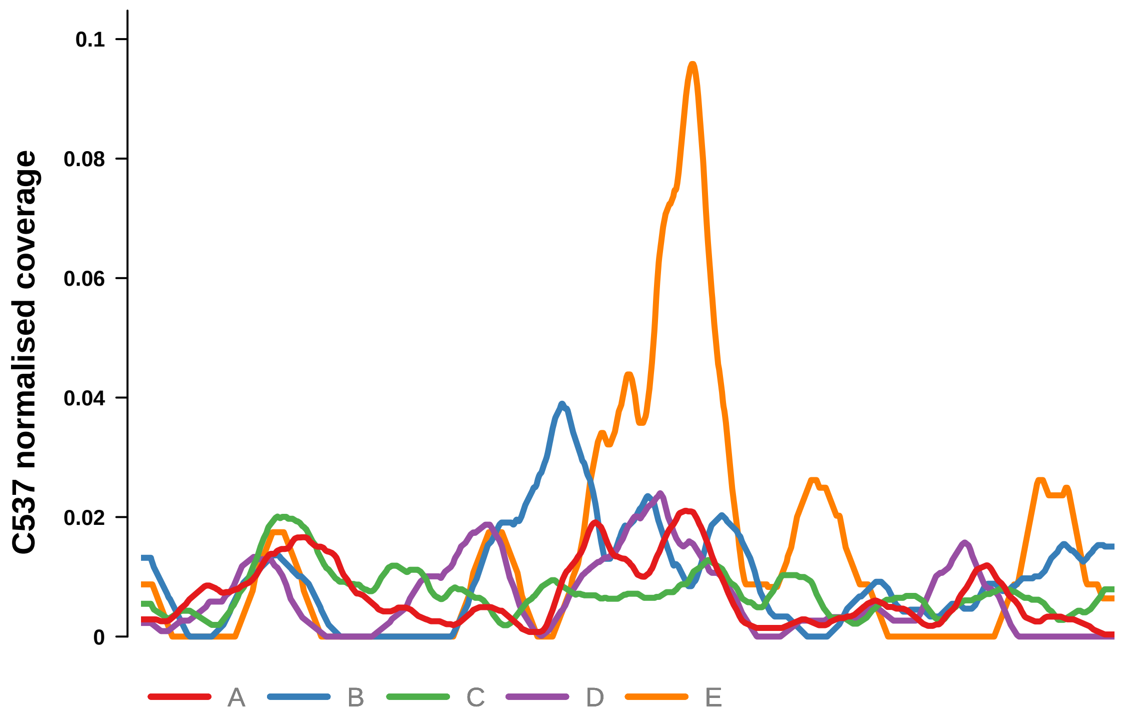


Putatively Clonal (logFC = −2.17)

| | P. | Adjusted P. |
|---|---|---|
| *~region vs. ~* | 4.85e−01 | 5.84e−01 |
| *~purity vs ~* | 5.11e−01 | NA |
| *~purity+region vs ~purity* | 4.67e−01 | NA |
| *Carcinoma vs. normal* | 3.6e−03 ** | 3.6e−03 ** |

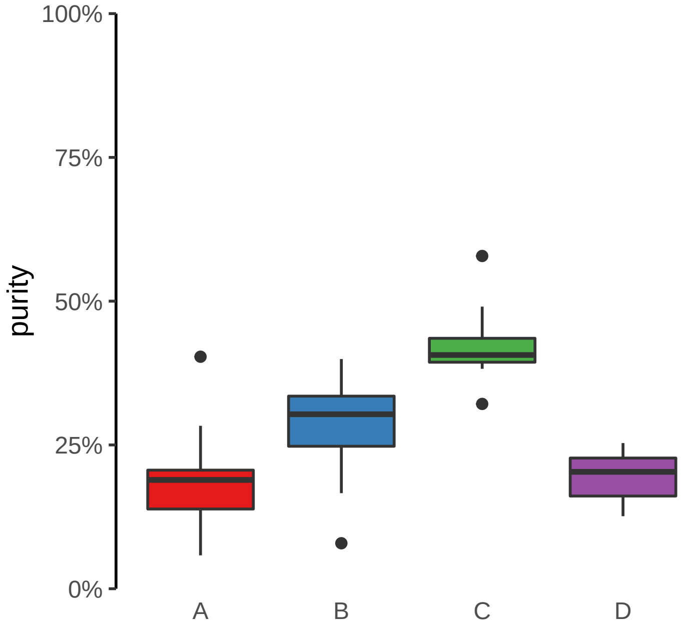

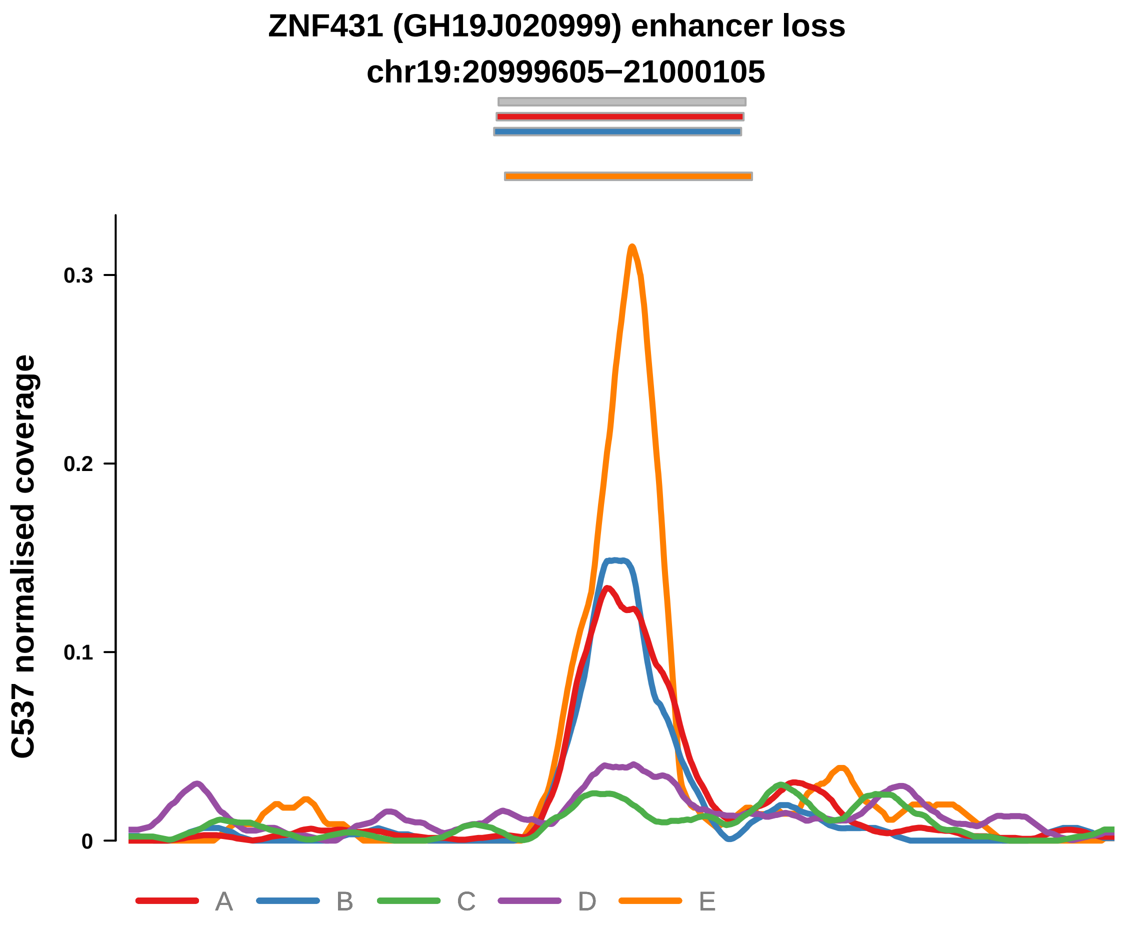


Putatively Clonal (logFC = −2.17)

| | P. | Adjusted P. |
|---|---|---|
| *~region vs. ~* | 7.23e−04 *** | 1.27e−02 * |
| *~purity vs ~* | 6.19e−04 *** | 7.12e−03 ** |
| *~purity+region vs ~purity* | 6.77e−02 | 1.91e−01 |
| *Carcinoma vs. normal* | 2.77e−03 ** | 2.77e−03 ** |

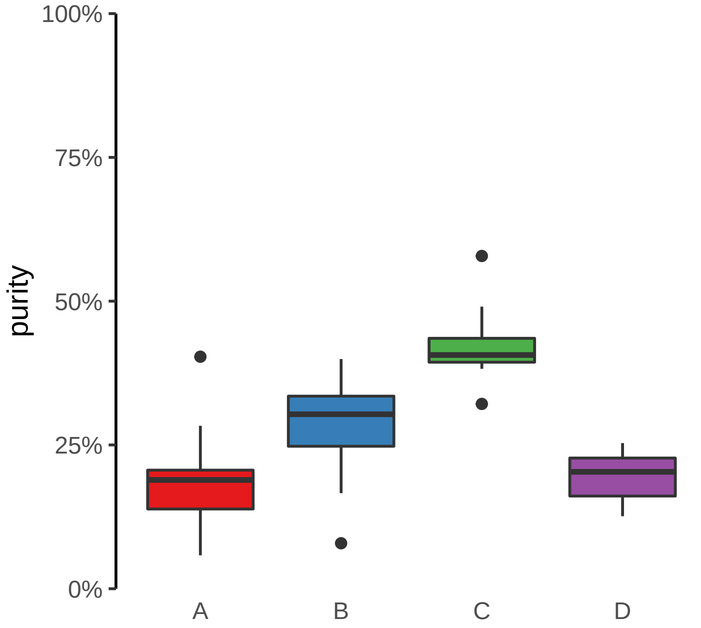

**ANTXR2 (GH04J080057) enhancer loss**
**chr4:80058687−80059187**

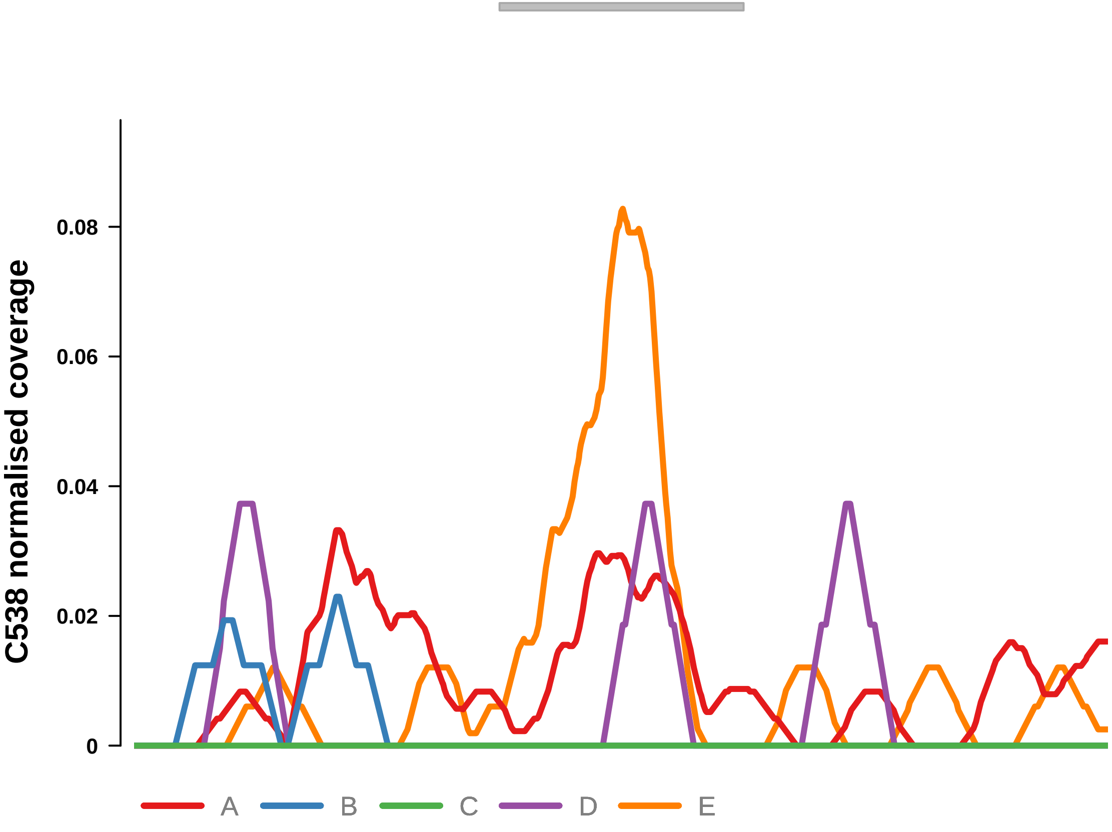


Putatively Clonal (logFC = −3.59)

| | P. | Adjusted P. |
|---|---|---|
| *~region vs. ~* | 4.5e−01 | 6.29e−01 |
| *~purity vs ~* | 8.1e−02 | 1.93e−01 |
| *~purity+region vs ~purity* | 9.34e−01 | 9.79e−01 |
| *Carcinoma vs. normal* | 4.49e−04 *** | 4.49e−04 *** |

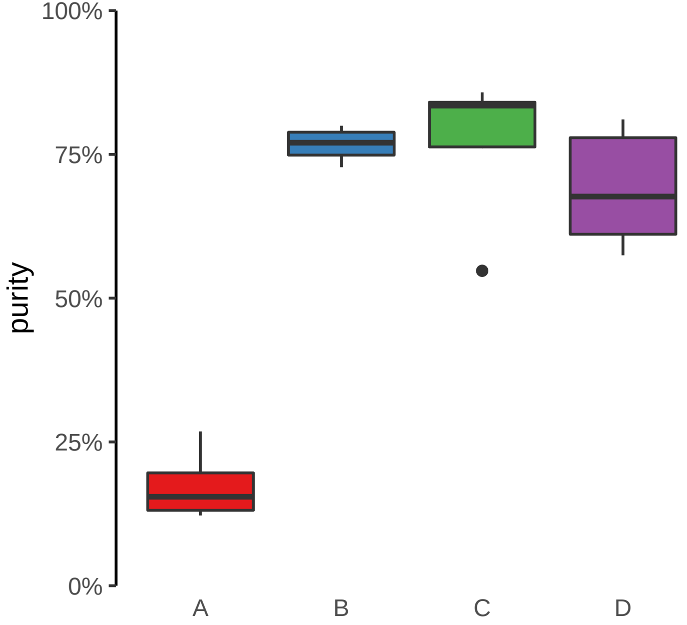

**CAMK2D (GH04J113682) enhancer loss**
**chr4:113683079−113683579**

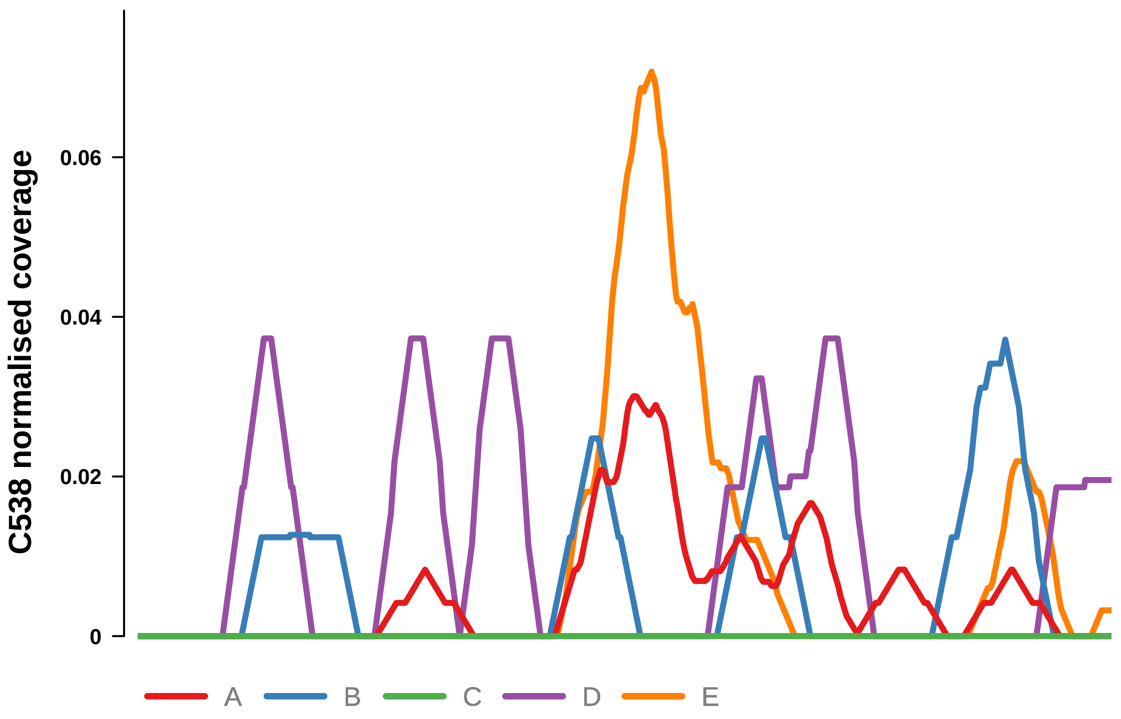


Putatively Clonal (logFC = −2.58)

| | P. | Adjusted P. |
|---|---|---|
| *~region vs. ~* | 2.31e−01 | 4.51e−01 |
| *~purity vs ~* | 6.25e−02 | 1.62e−01 |
| *~purity+region vs ~purity* | 7.23e−01 | 9.36e−01 |
| *Carcinoma vs. normal* | 4.87e−03 ** | 4.87e−03 ** |

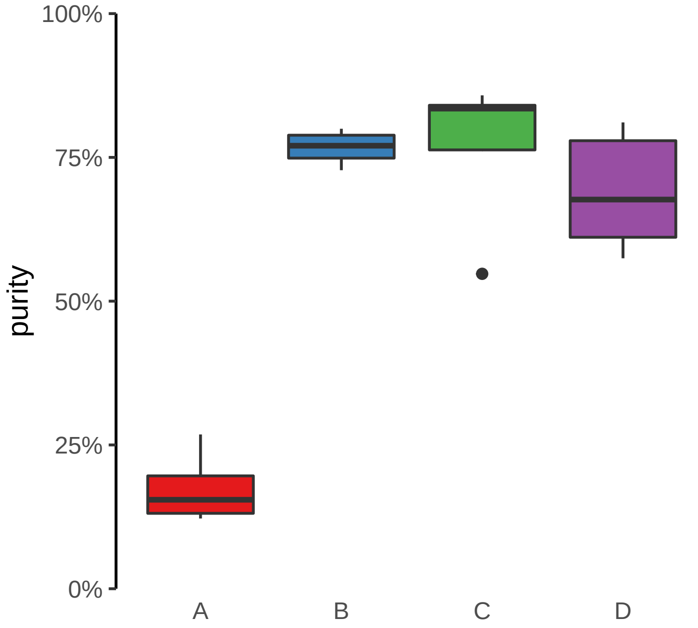

**CLASP2, ENSG00000271324 (GH03J033790) enhancer loss**
**chr3:33791629−33792129**

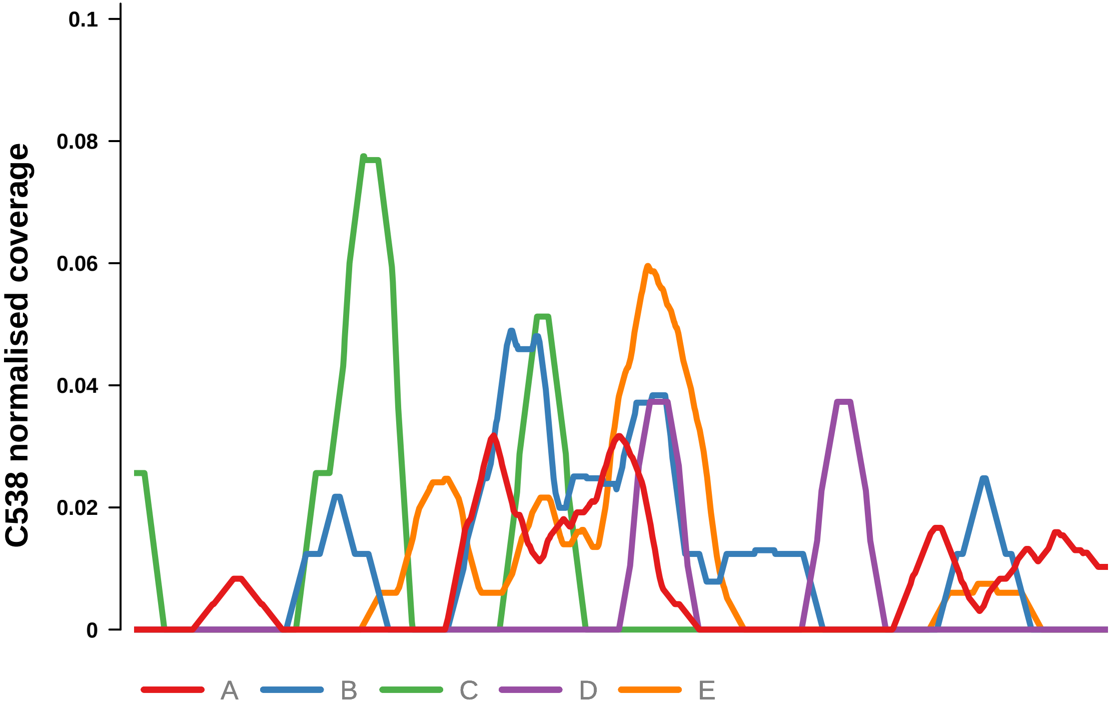


Putatively Clonal (logFC = −2.09)

| | P. | Adjusted P. |
|---|---|---|
| *~region vs. ~* | 3.9e−01 | 5.81e−01 |
| *~purity vs ~* | 2.66e−01 | 4.18e−01 |
| *~purity+region vs ~purity* | 2.99e−01 | 8.7e−01 |
| *Carcinoma vs. normal* | 5.67e−03 ** | 5.67e−03 ** |

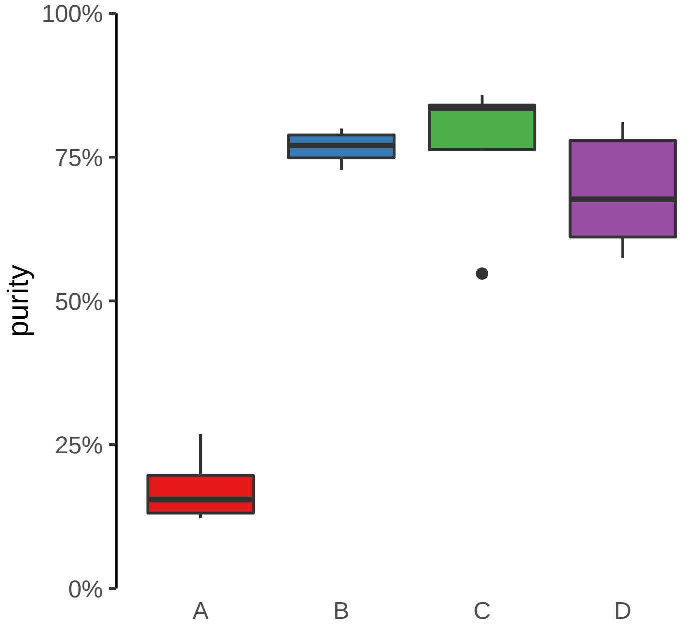

**CNST (GH01J246591) enhancer loss**
**chr1:246591917−246592417**

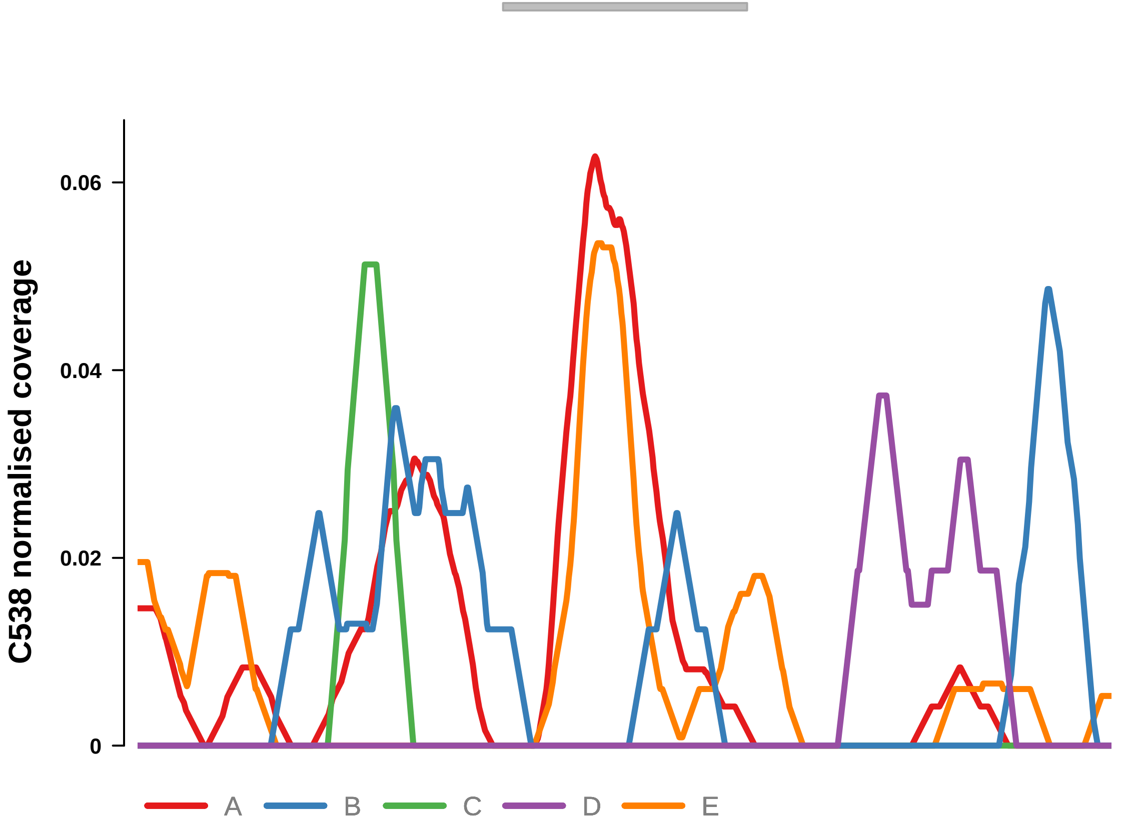


Putatively Clonal (logFC = −3.97)

| | **P.** | **Adjusted P.** |
|---|---|---|
| *~region vs. ~* | 8.26e−03 ** | 4.04e−02 * |
| *~purity vs ~* | 1.82e−03 ** | 8.89e−03 ** |
| *~purity+region vs ~purity* | 4.16e−01 | 8.7e−01 |
| *Carcinoma vs. normal* | 7.17e−05 **** | 7.17e−05 **** |

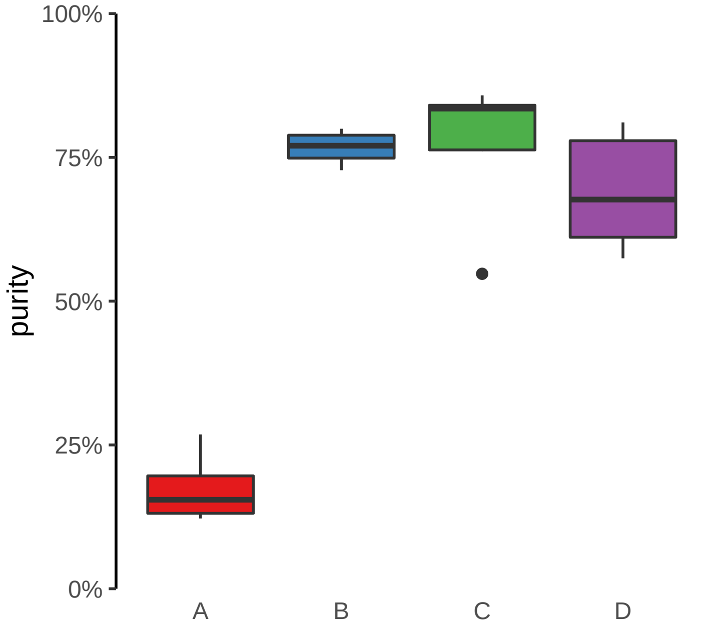

**DNAH7, ENSG00000272211 (GH02J196100) enhancer loss**
**chr2:196101142−196101642**

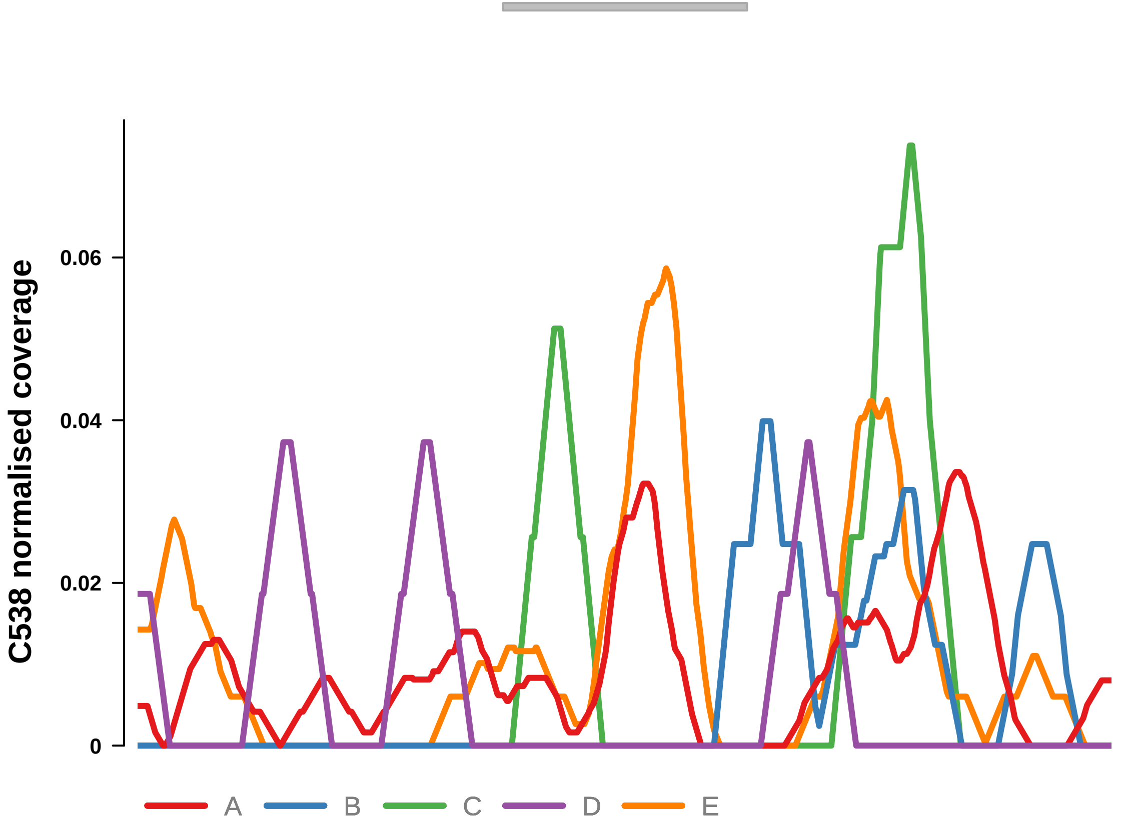


Putatively Clonal (logFC = −3.76)

| | P. | Adjusted P. |
|---|---|---|
| *~region vs. ~* | 5.49e−01 | 7.15e−01 |
| *~purity vs ~* | 1.1e−01 | 2.31e−01 |
| *~purity+region vs ~purity* | 9.77e−01 | 9.88e−01 |
| *Carcinoma vs. normal* | 2.07e−04 *** | 2.07e−04 *** |

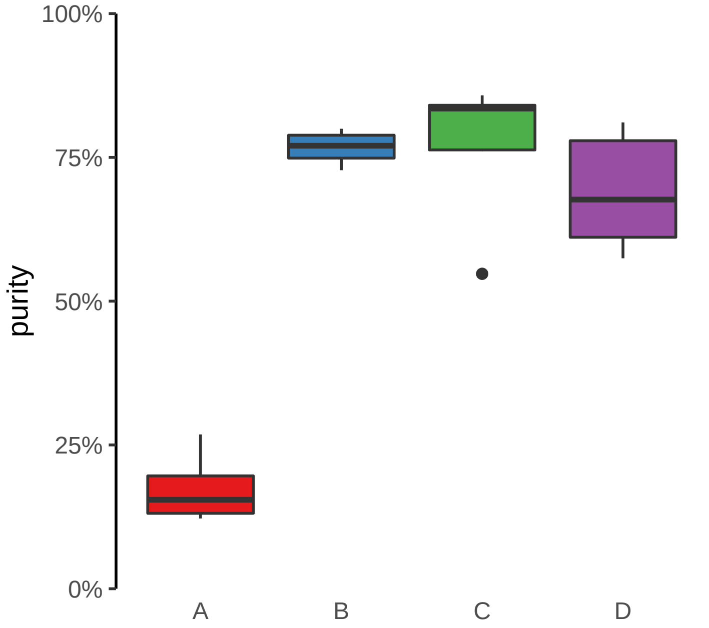

**ENSG00000228035, NGF (GH01J115322) enhancer loss**
**chr1:115323237−115323737**

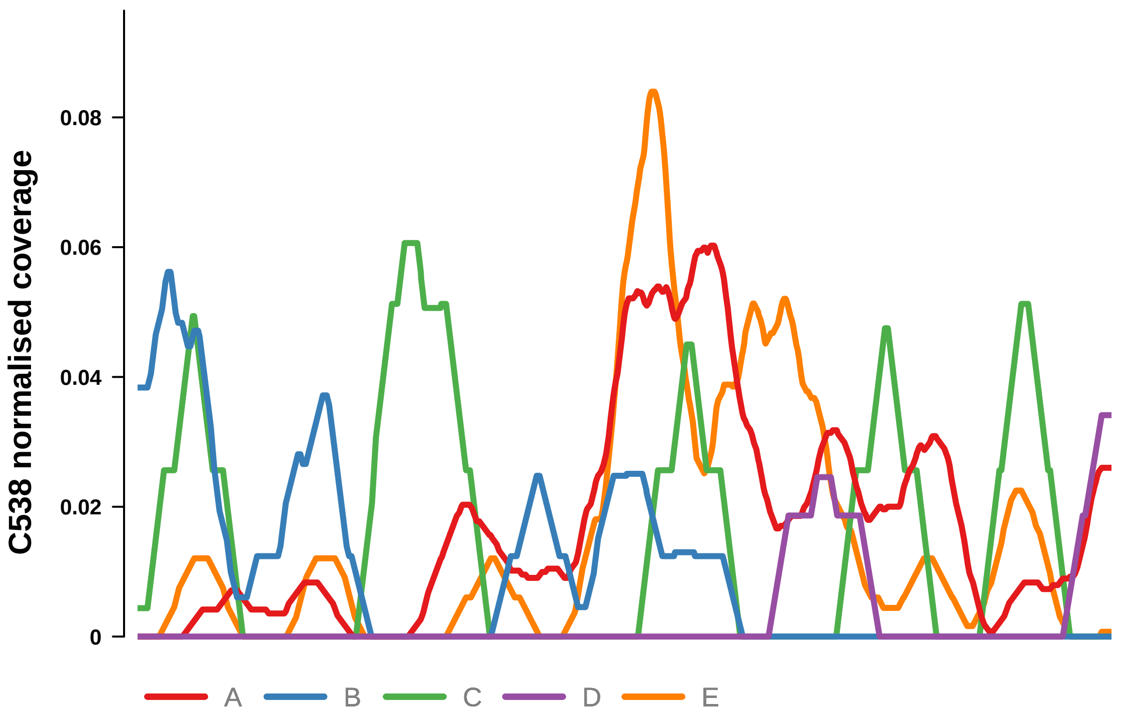


Putatively Clonal (logFC = −2.19)

| | P. | Adjusted P. |
|---|---|---|
| *~region vs. ~* | 1.35e−03 ** | 1.19e−02 * |
| *~purity vs ~* | 1.81e−04 *** | 1.77e−03 ** |
| *~purity+region vs ~purity* | 4.1e−01 | 8.7e−01 |
| *Carcinoma vs. normal* | 7.53e−03 ** | 7.53e−03 ** |

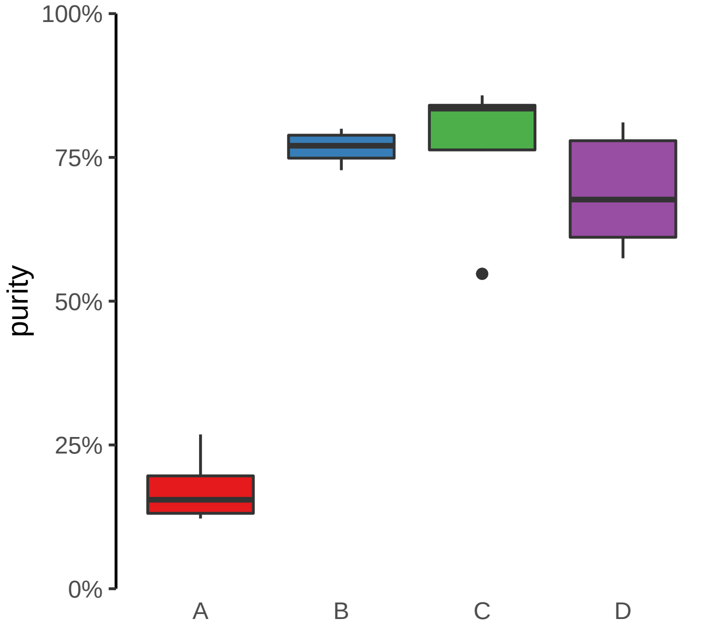

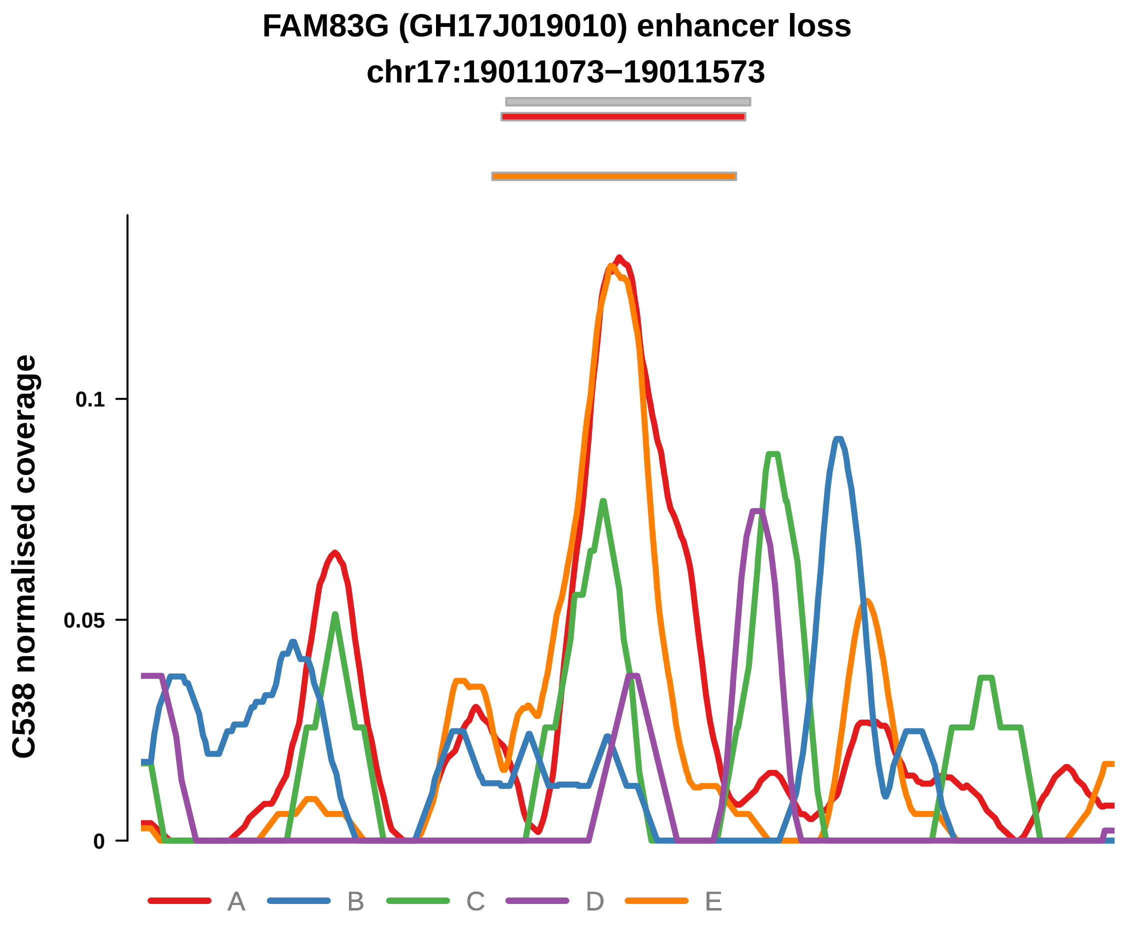


Putatively Clonal (logFC = −2.31)

| | P. | Adjusted P. |
|---|---|---|
| *~region vs. ~* | 1.85e−03 ** | 1.25e−02 * |
| *~purity vs ~* | 1.75e−03 ** | 8.89e−03 ** |
| *~purity+region vs ~purity* | 1.25e−01 | 8.28e−01 |
| *Carcinoma vs. normal* | 2.36e−03 ** | 2.36e−03 ** |

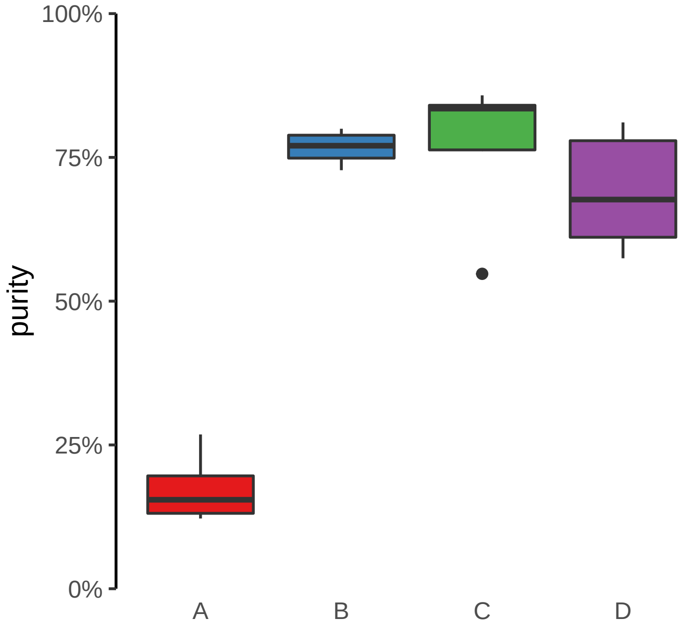

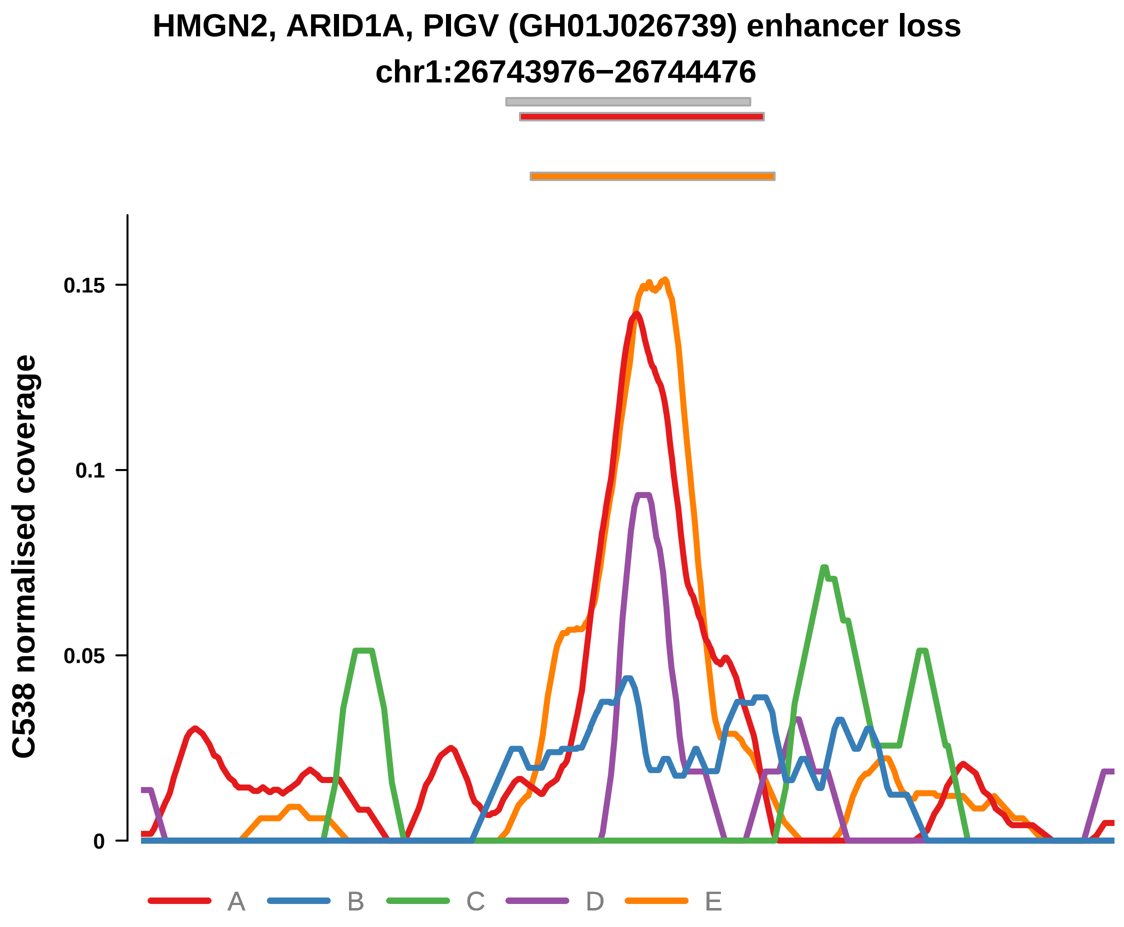


Putatively Clonal (logFC = −2.25)

| | P. | Adjusted P. |
|---|---|---|
| *~region vs. ~* | 7.27e−07 **** | 2.13e−05 **** |
| *~purity vs ~* | 2.49e−05 **** | 3.13e−04 *** |
| *~purity+region vs ~purity* | 2.11e−02 * | 4.63e−01 |
| *Carcinoma vs. normal* | 2.81e−03 ** | 2.81e−03 ** |

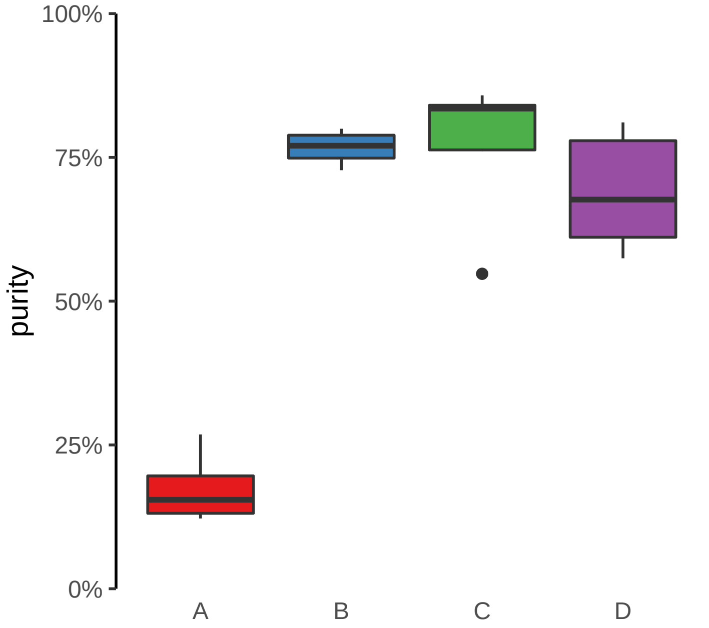

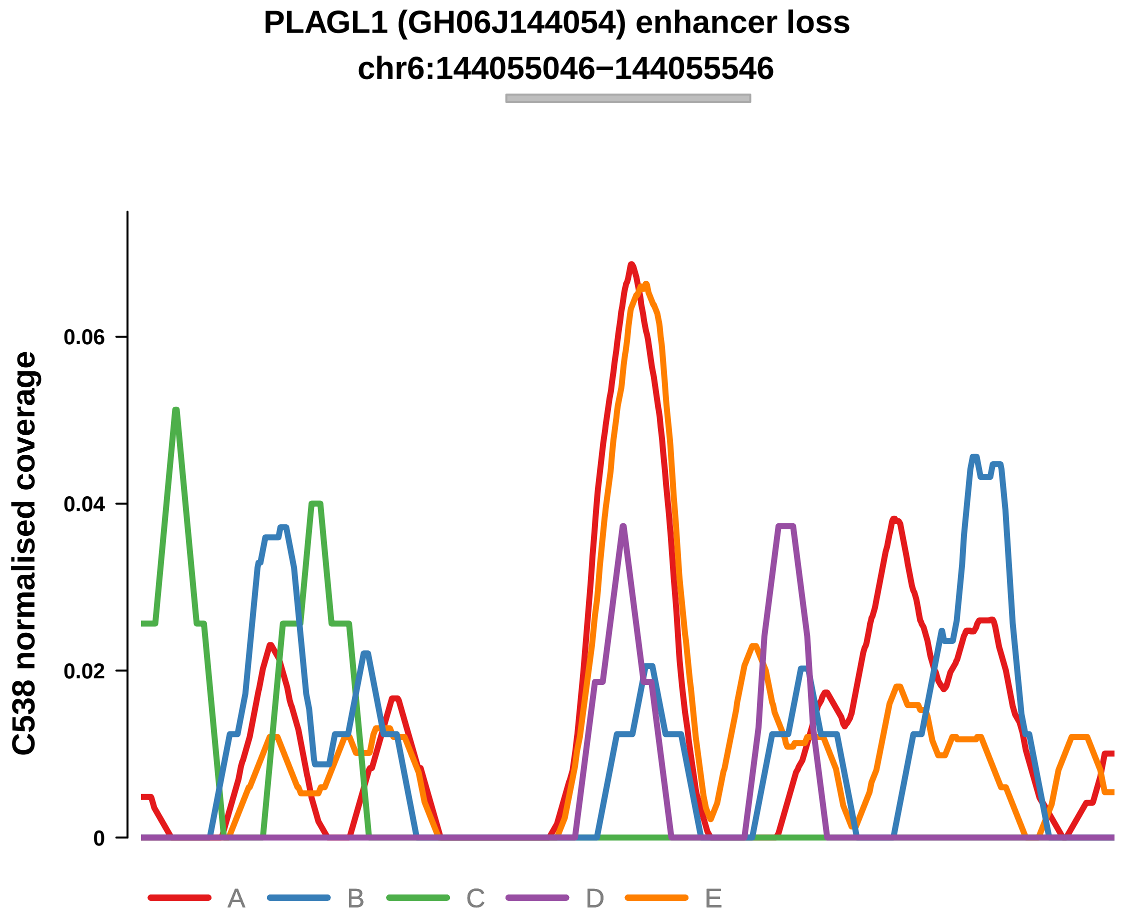


Putatively Clonal (logFC = −2.85)

| | P. | Adjusted P. |
|---|---|---|
| *~region vs. ~* | 2.99e−02 * | 9.07e−02 |
| *~purity vs ~* | 2.38e−03 ** | 1.1e−02 * |
| *~purity+region vs ~purity* | 8.96e−01 | 9.79e−01 |
| *Carcinoma vs. normal* | 1.99e−03 ** | 1.99e−03 ** |

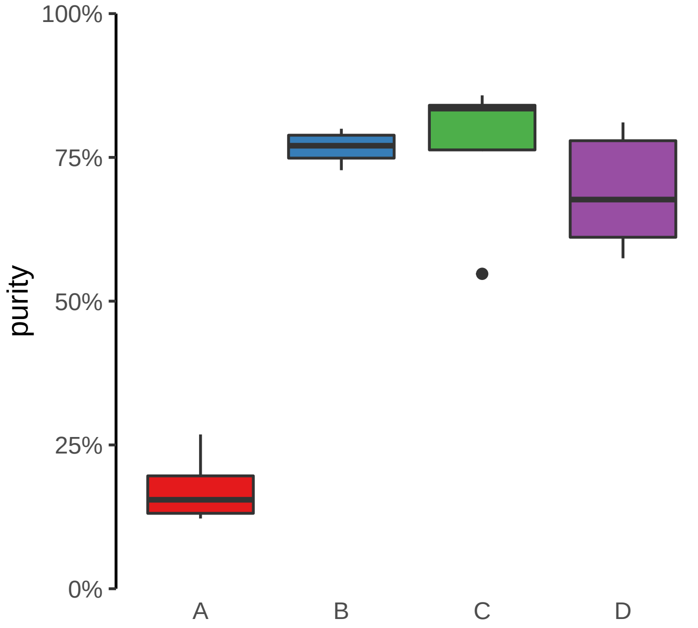

**PPP3CA (GH04J101183) enhancer loss**
**chr4:101183634−101184134**

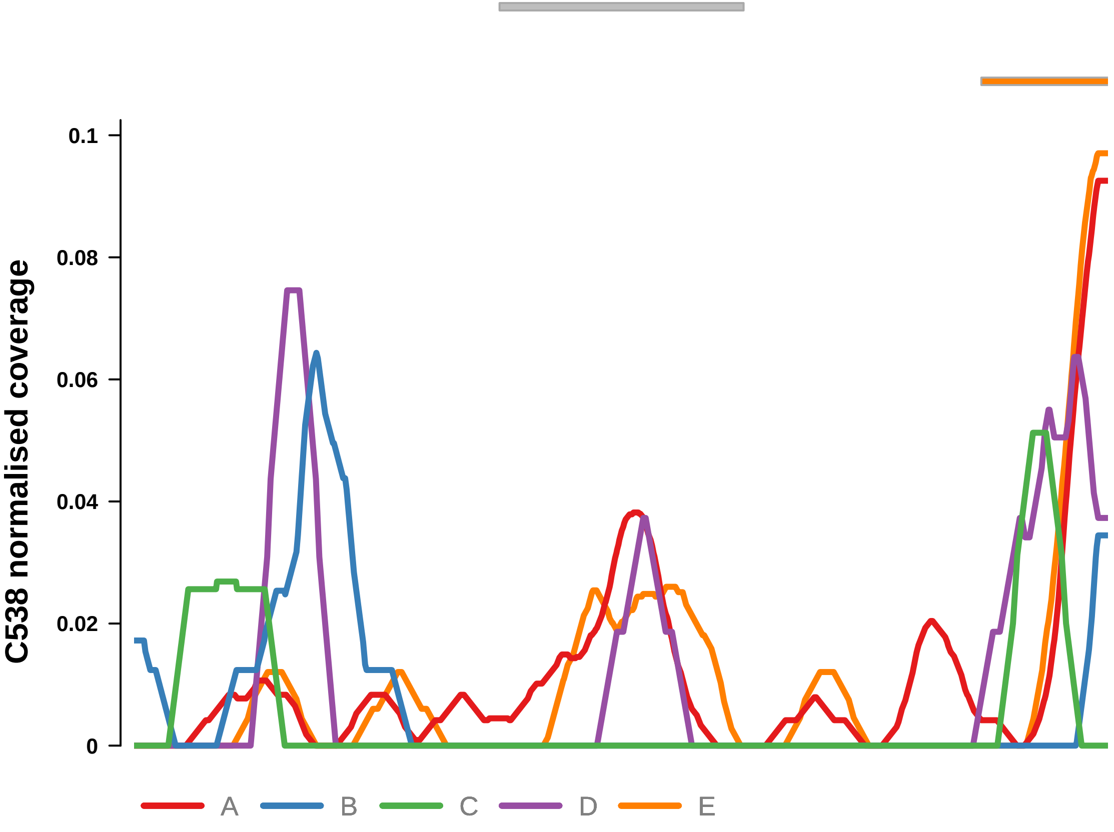


Putatively Clonal (logFC = −3.61)

| | P. | Adjusted P. |
|---|---|---|
| *~region vs. ~* | 4.5e−01 | 6.29e−01 |
| *~purity vs ~* | 6.98e−02 | 1.76e−01 |
| *~purity+region vs ~purity* | 9.56e−01 | 9.79e−01 |
| *Carcinoma vs. normal* | 4.2e−04 *** | 4.2e−04 *** |

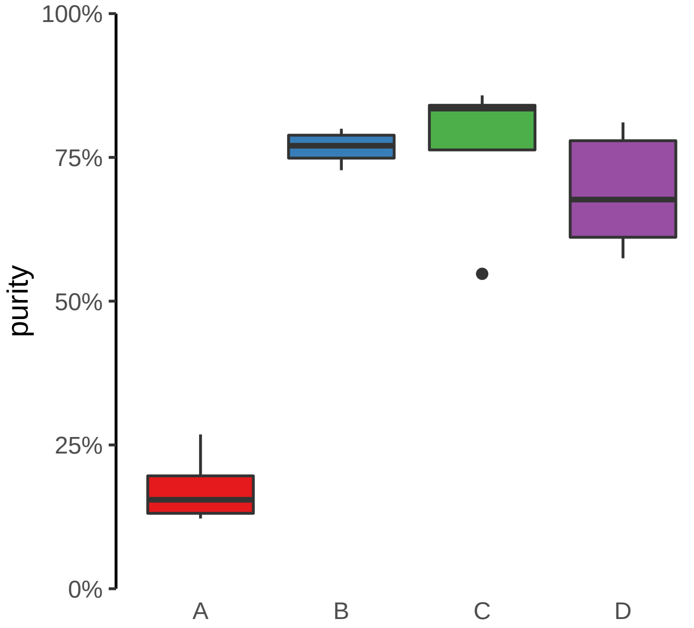

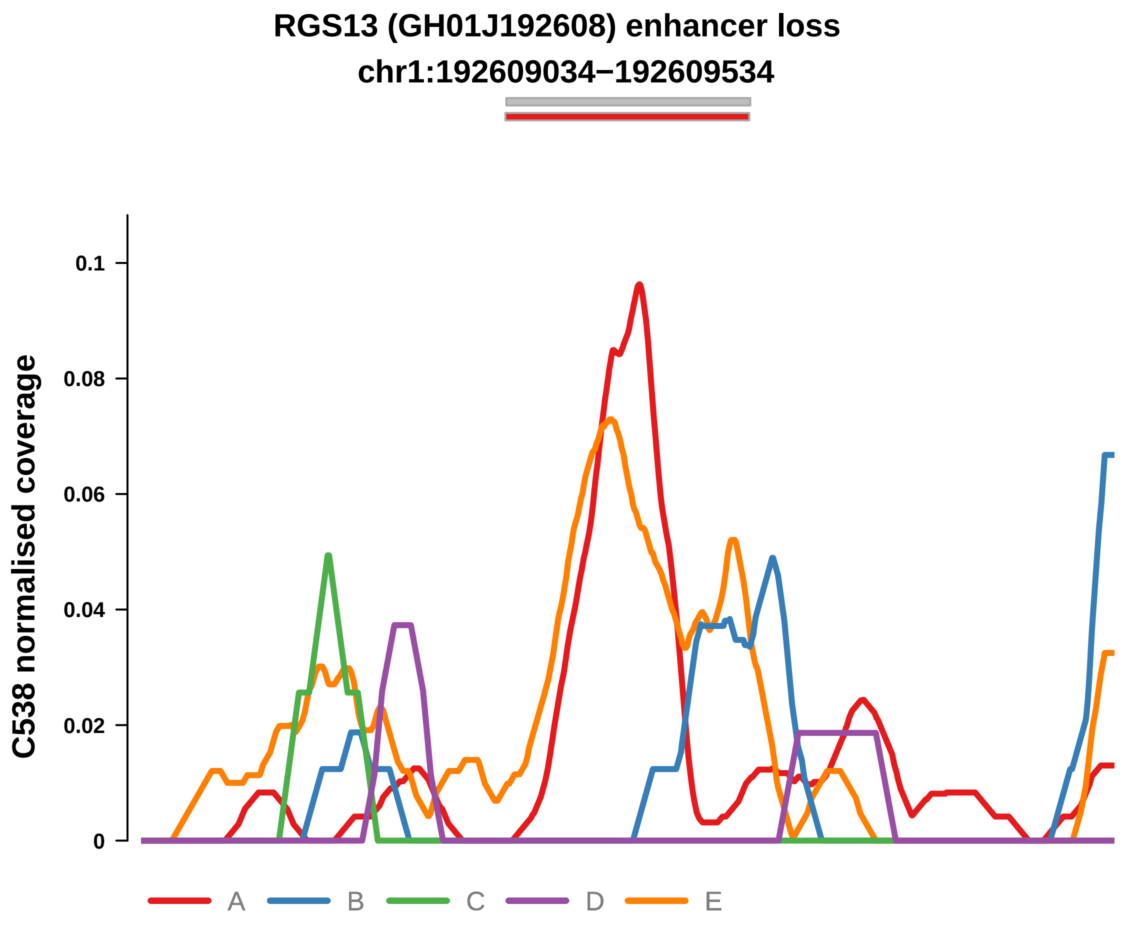


Putatively Clonal (logFC = −3.02)

| | P. | Adjusted P. |
|---|---|---|
| *~region vs. ~* | 1.74e−03 ** | 1.25e−02 * |
| *~purity vs ~* | 8.17e−04 *** | 5.99e−03 ** |
| *~purity+region vs ~purity* | 2.74e−01 | 8.7e−01 |
| *Carcinoma vs. normal* | 8.81e−04 *** | 8.81e−04 *** |

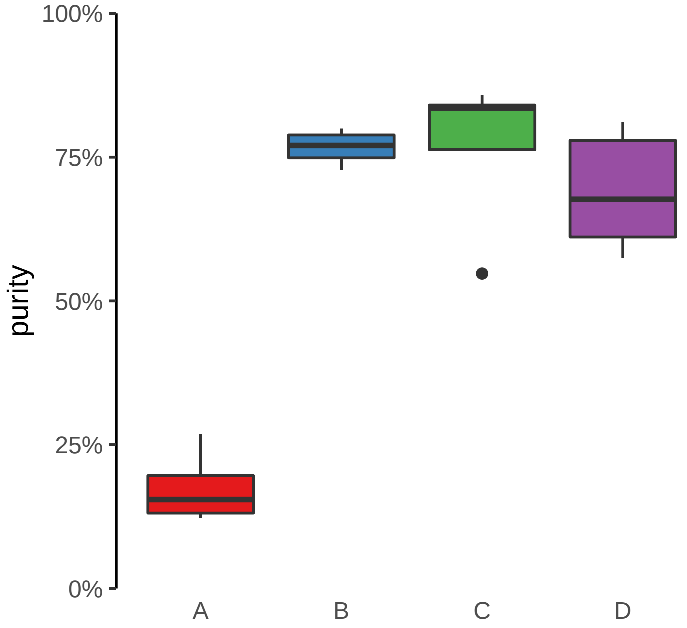

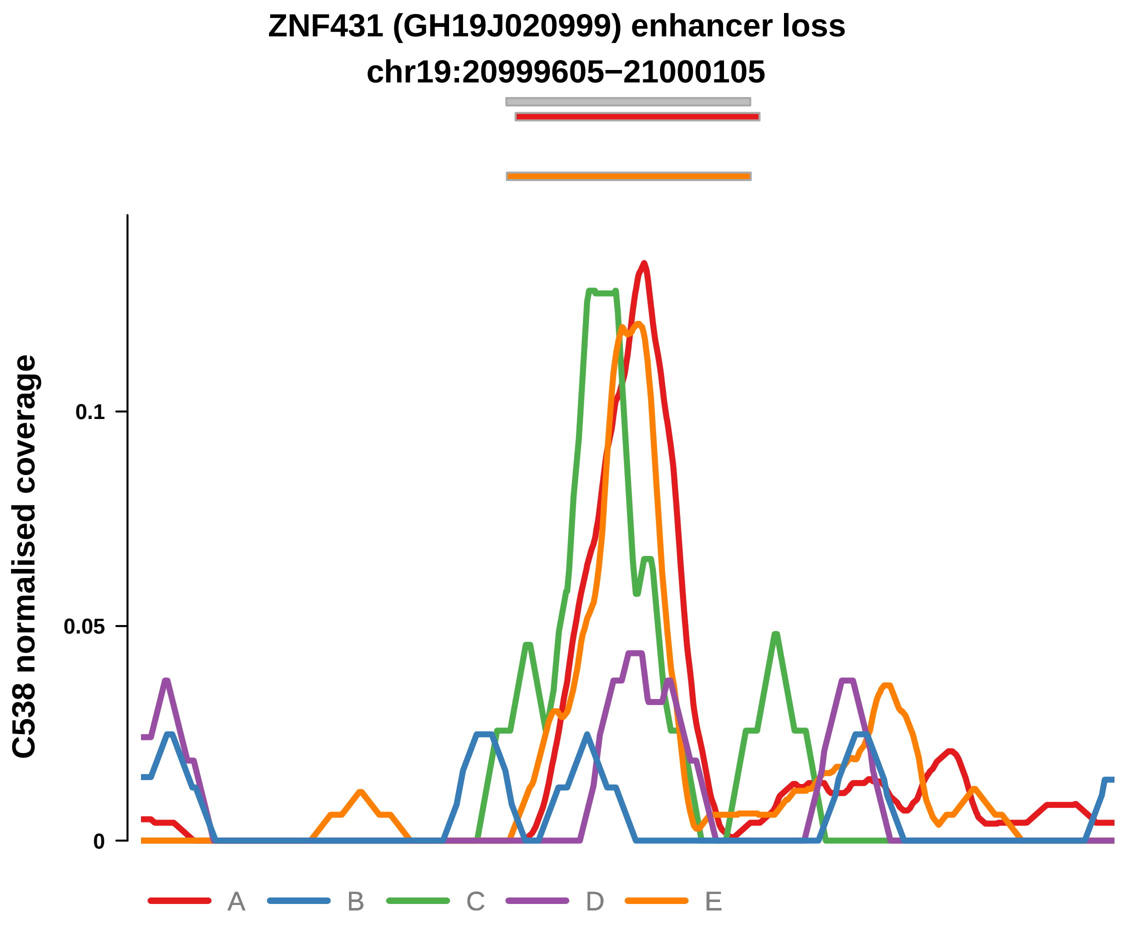


Putatively Clonal (logFC = −2.17)

| | P. | Adjusted P. |
|---|---|---|
| *~region vs. ~* | 5.77e−03 ** | 3.17e−02 * |
| *~purity vs ~* | 2.21e−04 *** | 1.94e−03 ** |
| *~purity+region vs ~purity* | 4.1e−01 | 8.7e−01 |
| *Carcinoma vs. normal* | 4.06e−03 ** | 4.06e−03 ** |

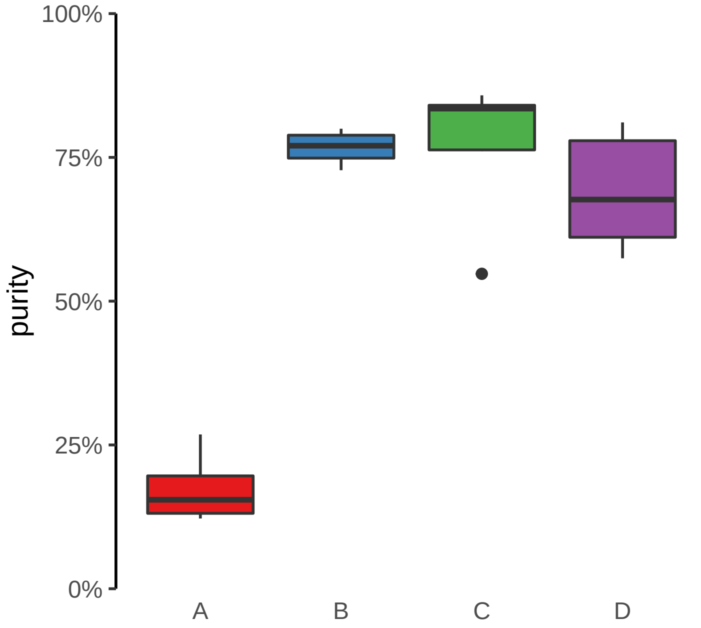

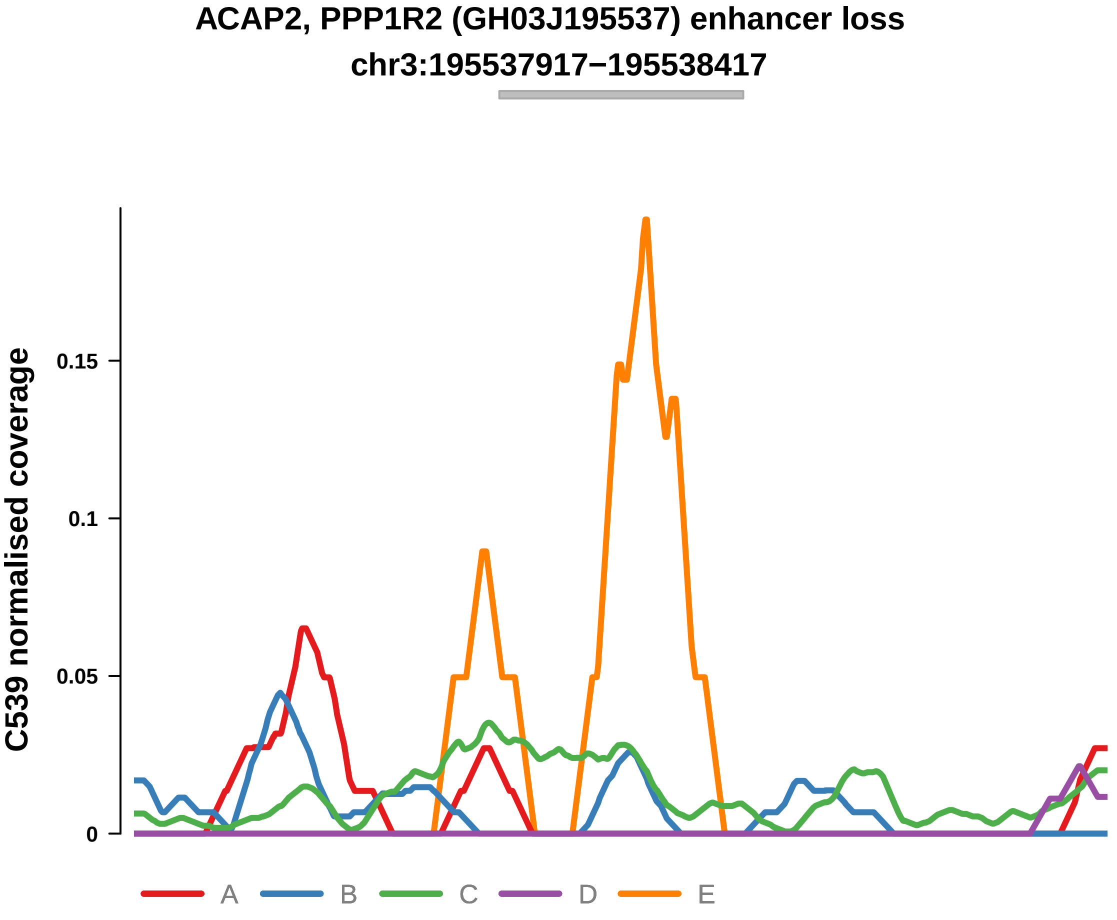


Putatively Clonal (logFC = −10.36)

| | P. | Adjusted P. |
|---|---|---|
| *~region vs. ~* | 1.43e−04 *** | 6.63e−04 *** |
| *~purity vs ~* | 6.81e−05 **** | 6.48e−04 *** |
| *~purity+region vs ~purity* | 3.75e−01 | 4.44e−01 |
| *Carcinoma vs. normal* | 2.57e−03 ** | 2.57e−03 ** |

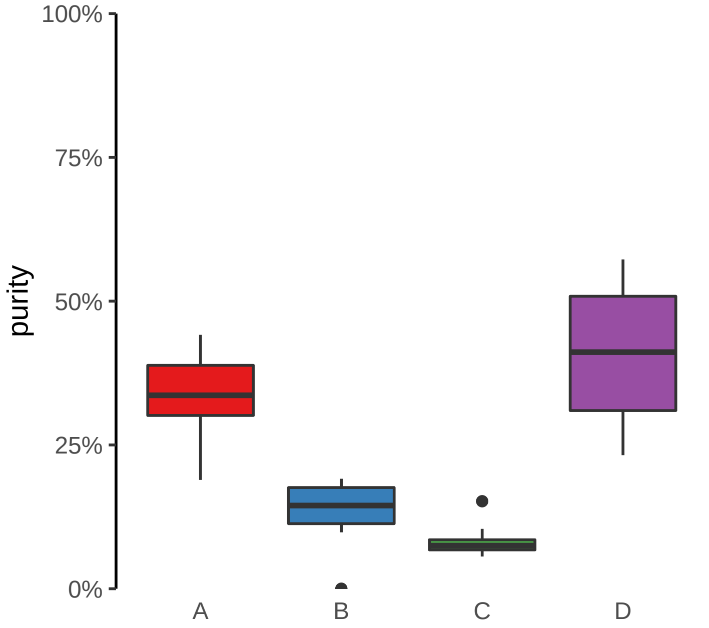

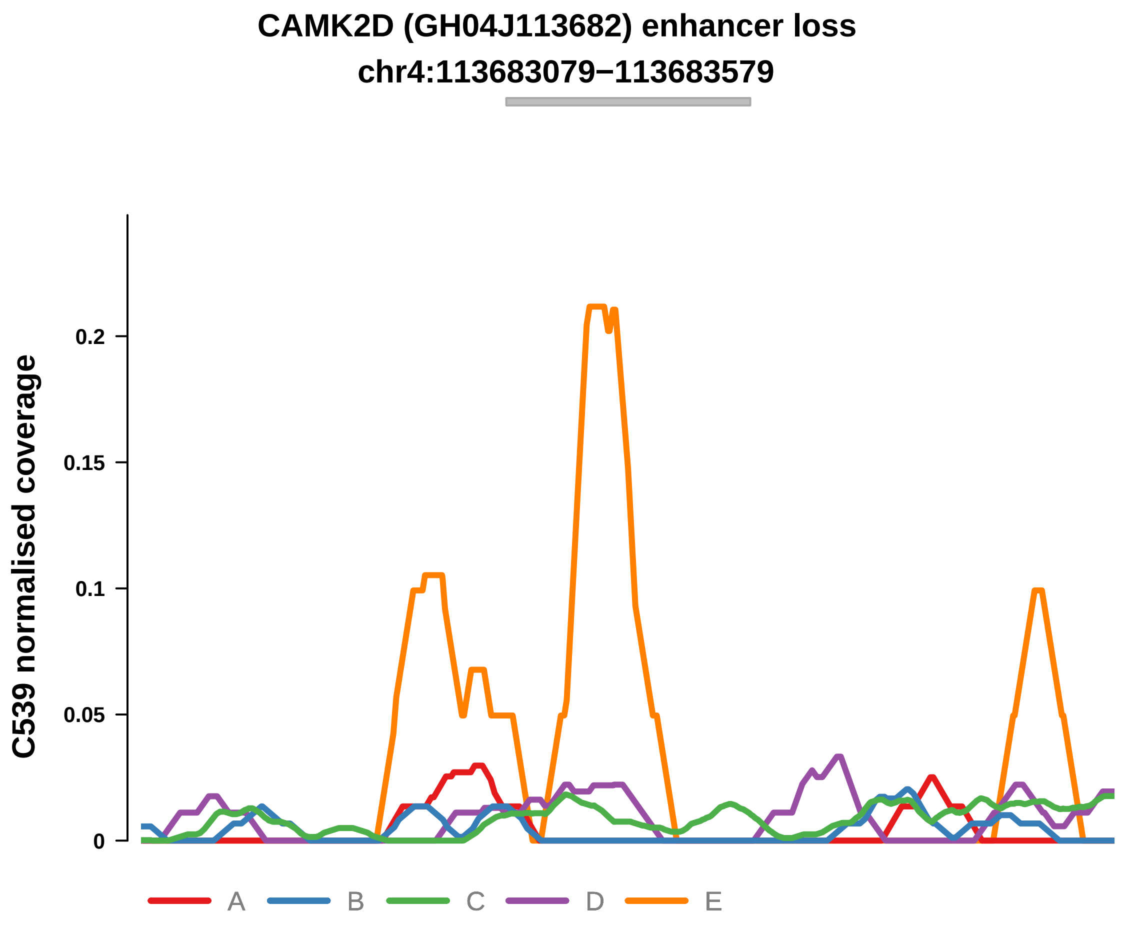


Putatively Clonal (logFC = −10.71)

| | P. | Adjusted P. |
|---|---|---|
| *~region vs. ~* | 2.17e−03 ** | 5.17e−03 ** |
| *~purity vs ~* | 1.23e−03 ** | 3.72e−03 ** |
| *~purity+region vs ~purity* | 2.32e−01 | NA |
| *Carcinoma vs. normal* | 1.1e−03 ** | 1.1e−03 ** |

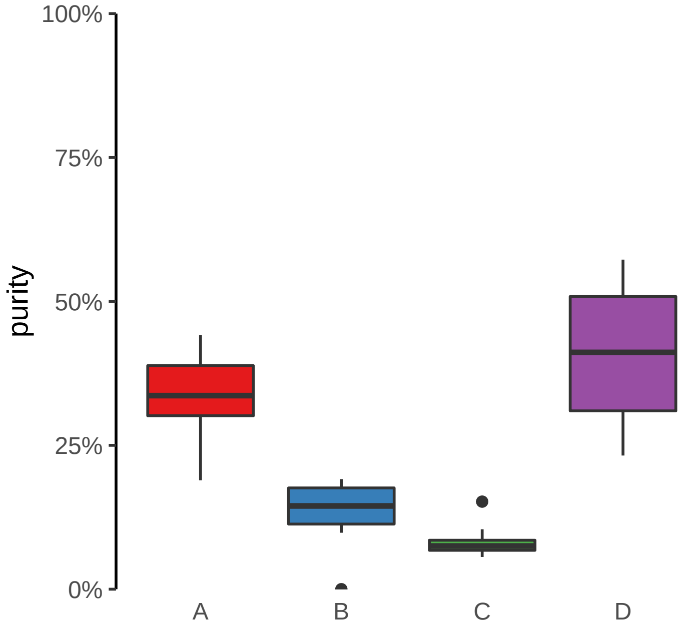

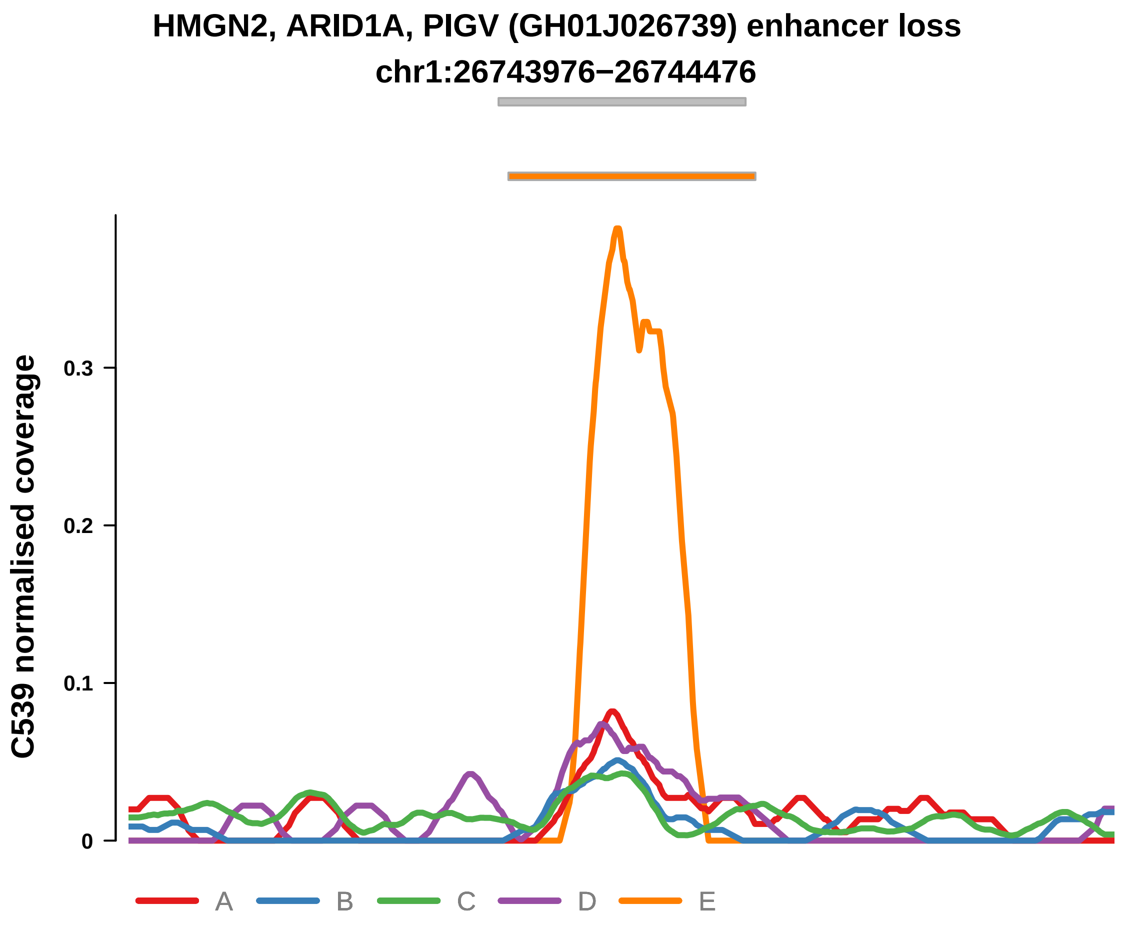


Putatively Clonal (logFC = −3.33)

| | P. | Adjusted P. |
|---|---|---|
| *~region vs. ~* | 5.95e−04 *** | 1.69e−03 ** |
| *~purity vs ~* | 2.91e−06 **** | 6.41e−05 **** |
| *~purity+region vs ~purity* | 3.2e−01 | 4.01e−01 |
| *Carcinoma vs. normal* | 1.12e−03 ** | 1.12e−03 ** |

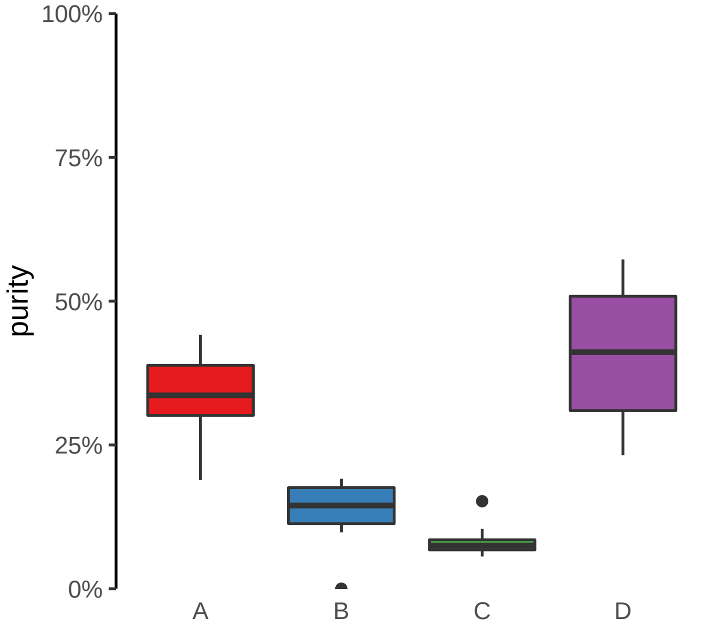

**RNLS, LIPJ (GH10J088553) enhancer loss**
**chr10:88553102−88553602**

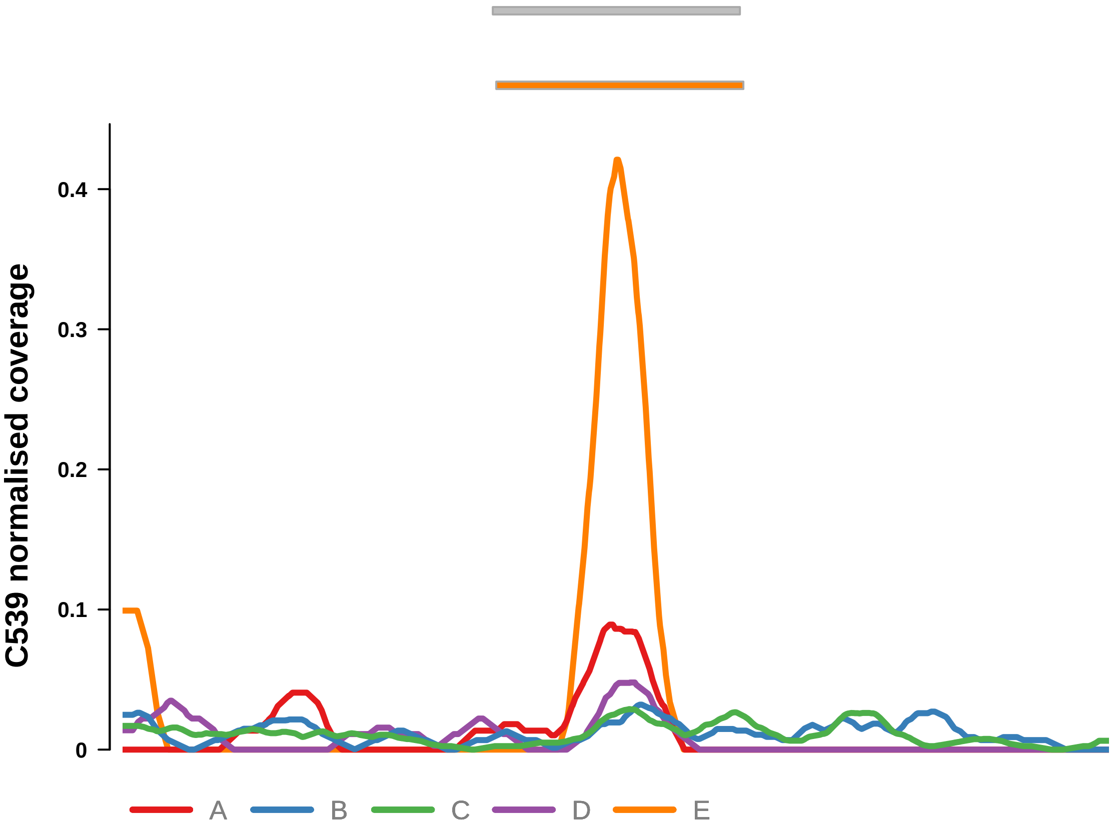


Putatively Clonal (logFC = −11.44)

| | P. | Adjusted P. |
|---|---|---|
| *~region vs. ~* | 8.52e−02 | 9.44e−02 |
| *~purity vs ~* | 3.14e−03 ** | 6.92e−03 ** |
| *~purity+region vs ~purity* | 8.95e−01 | 8.95e−01 |
| *Carcinoma vs. normal* | 1.04e−04 *** | 1.04e−04 *** |

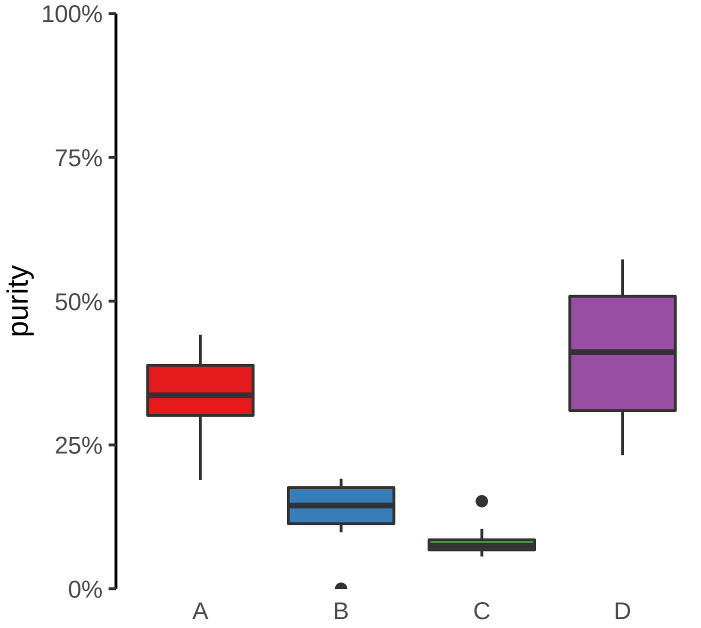

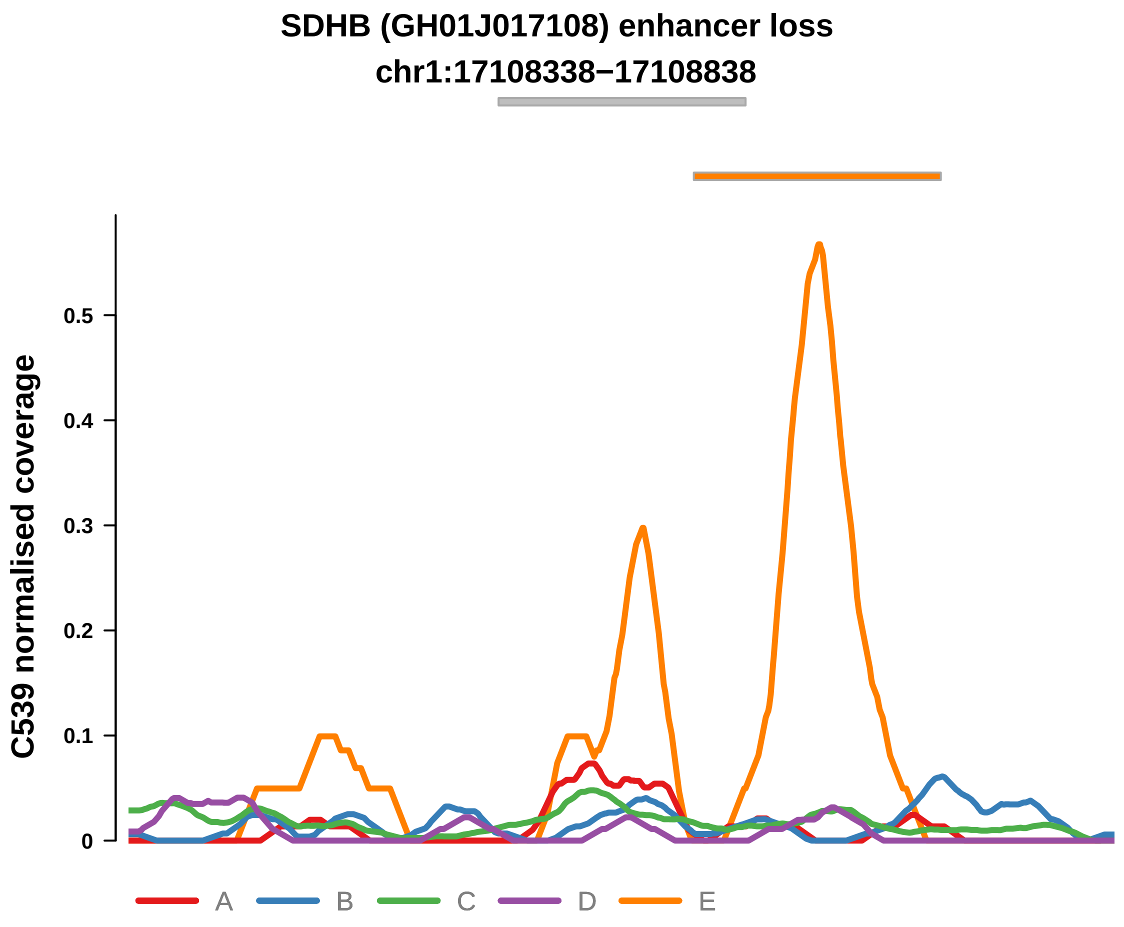


Putatively Clonal (logFC = −11.57)

| | P. | Adjusted P. |
|---|---|---|
| *~region vs. ~* | 1.72e−04 *** | 6.86e−04 *** |
| *~purity vs ~* | 7.62e−05 **** | 6.48e−04 *** |
| *~purity+region vs ~purity* | 4e−01 | 4.51e−01 |
| *Carcinoma vs. normal* | 5.48e−05 **** | 5.48e−05 **** |

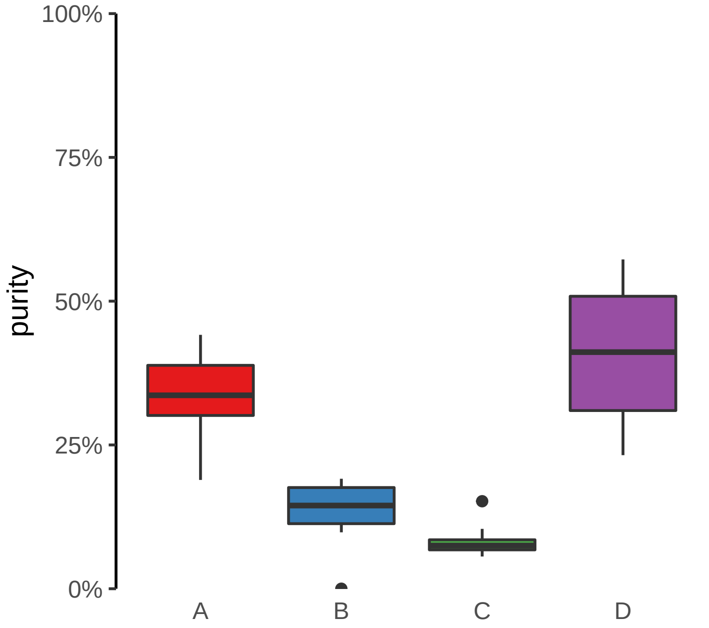

**TRANK1, ENSG00000234073 (GH03J036874) enhancer loss**
**chr3:36876260−36876760**

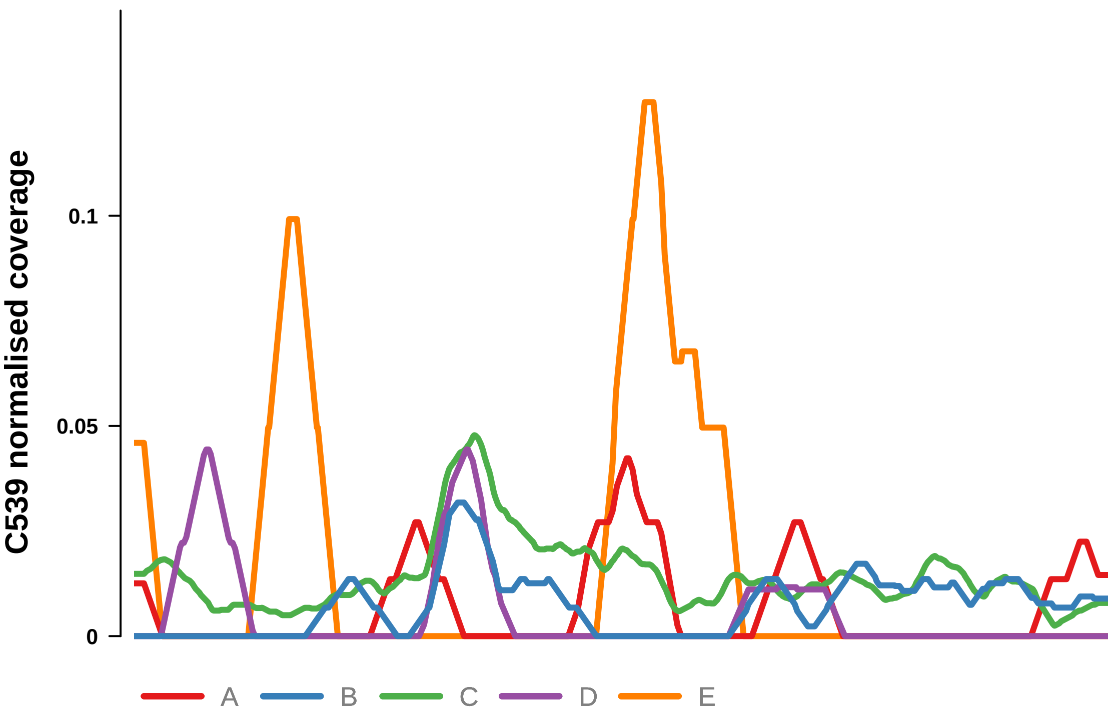


Putatively Clonal (logFC = −10.94)

| | P. | Adjusted P. |
|---|---|---|
| *~region vs. ~* | 7.85e−05 **** | 4.32e−04 *** |
| *~purity vs ~* | 7.69e−04 *** | 3.22e−03 ** |
| *~purity+region vs ~purity* | 3.42e−02 * | 9.99e−02 |
| *Carcinoma vs. normal* | 6.26e−04 *** | 6.26e−04 *** |

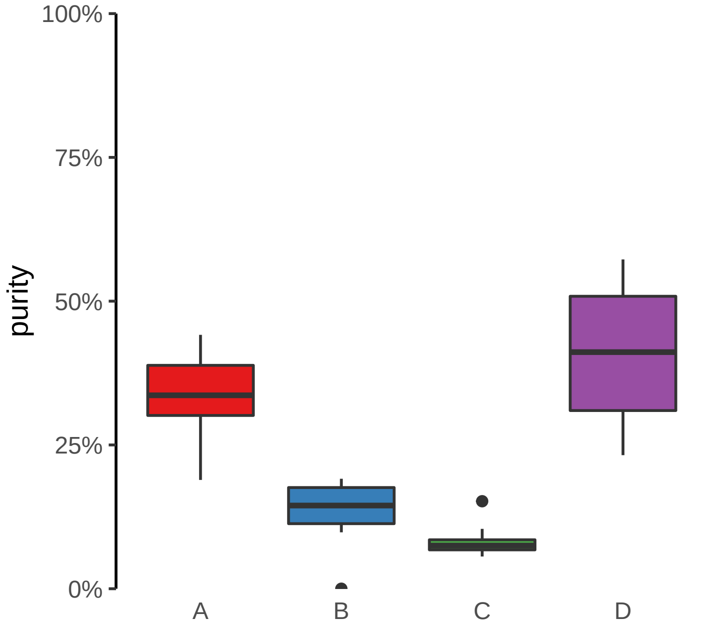

**CAMK2D (GH04J113682) enhancer loss**
**chr4:113683079−113683579**

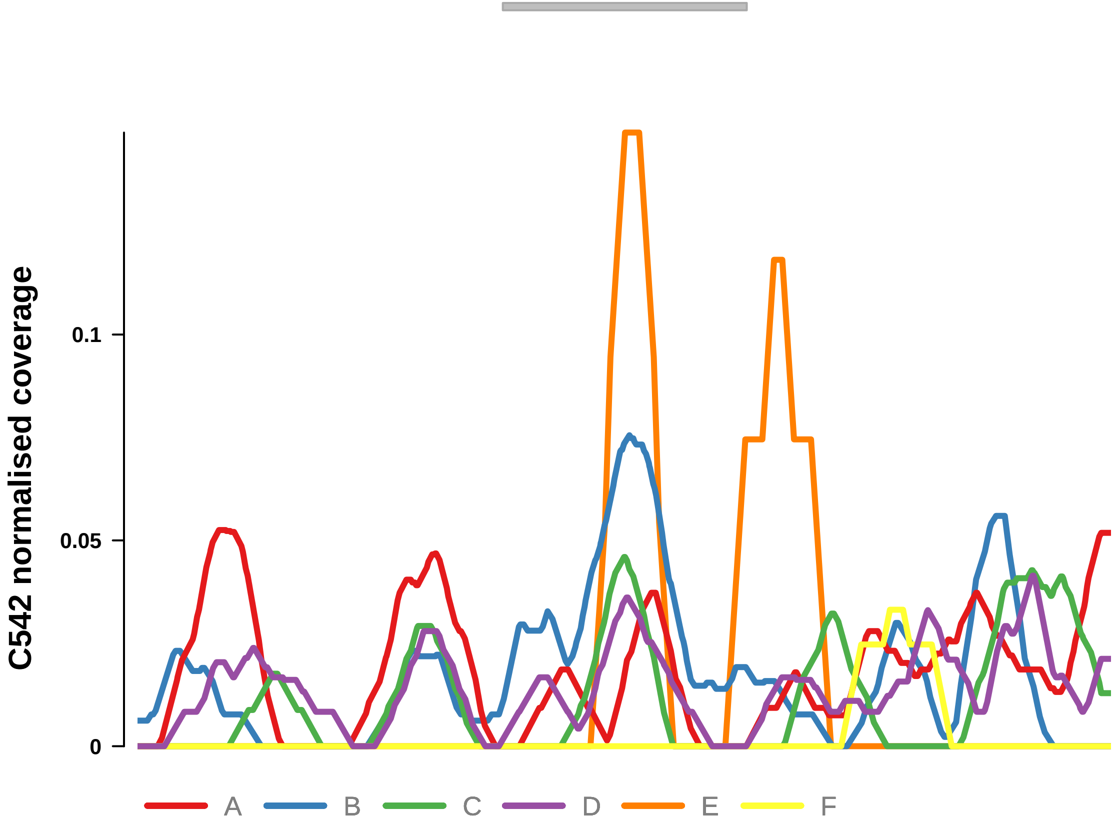


Putatively Clonal (logFC = −10.7)

| | P. | Adjusted P. |
|---|---|---|
| *~region vs. ~* | 1.86e−01 | 6.17e−01 |
| *~purity vs ~* | 8.78e−01 | 9.67e−01 |
| *~purity+region vs ~purity* | 2.12e−01 | 7.3e−01 |
| *Carcinoma vs. normal* | 6.98e−03 ** | 6.98e−03 ** |

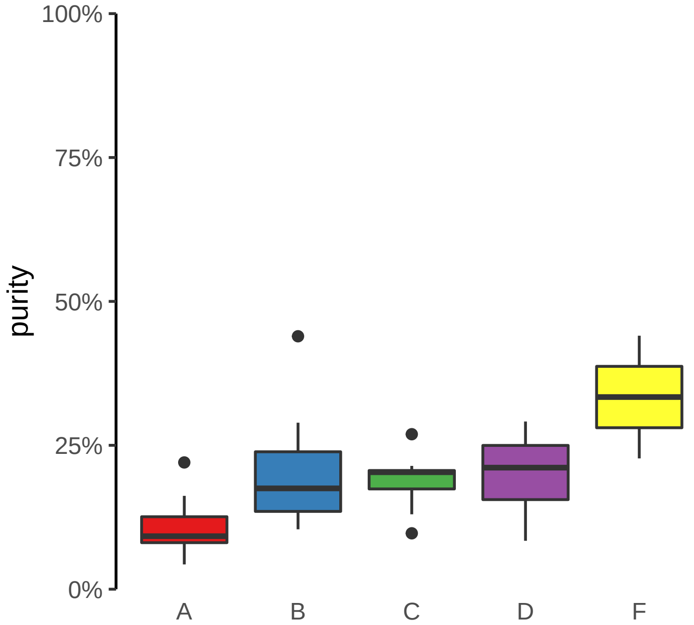

**CLASP1 (GH02J121362) enhancer loss**
**chr2:121363584−121364084**

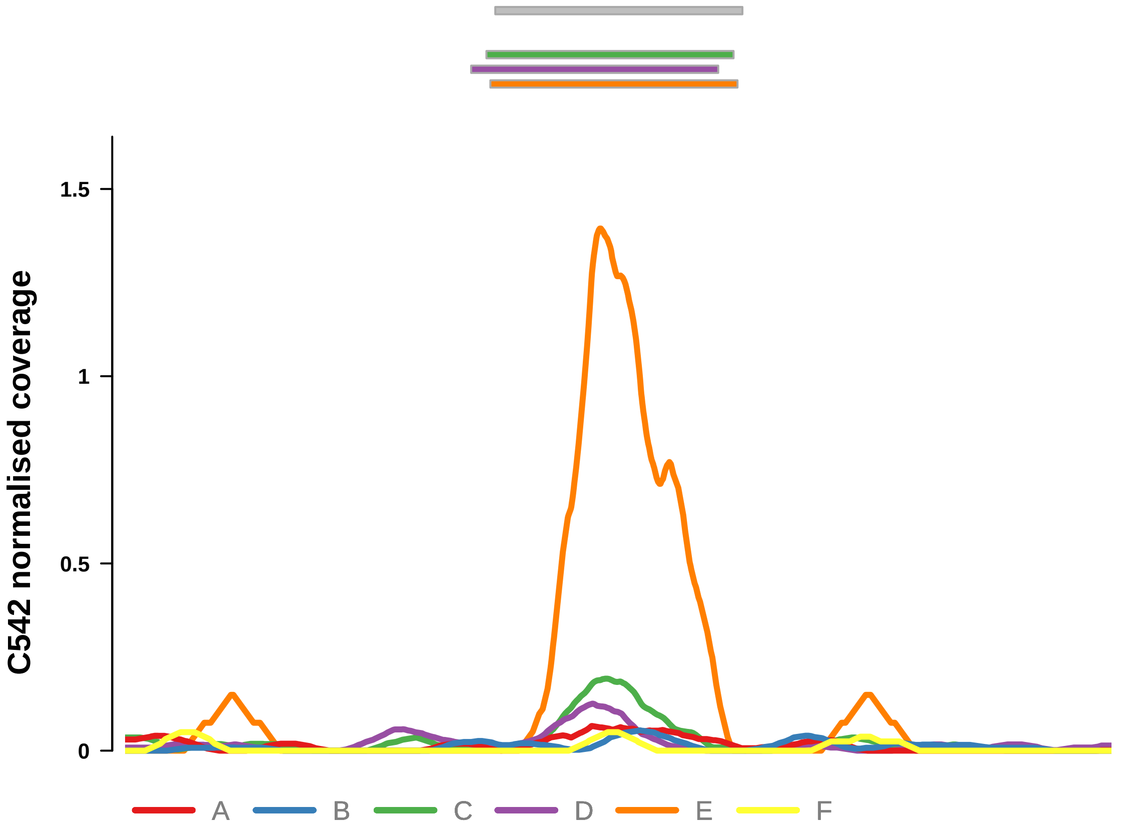


Putatively Clonal (logFC = −11.42)

| | P. | Adjusted P. |
|---|---|---|
| *~region vs. ~* | 1.11e−01 | 5.9e−01 |
| *~purity vs ~* | 6.61e−01 | 9.67e−01 |
| *~purity+region vs ~purity* | 1.25e−01 | 6.68e−01 |
| *Carcinoma vs. normal* | 1.06e−03 ** | 1.06e−03 ** |

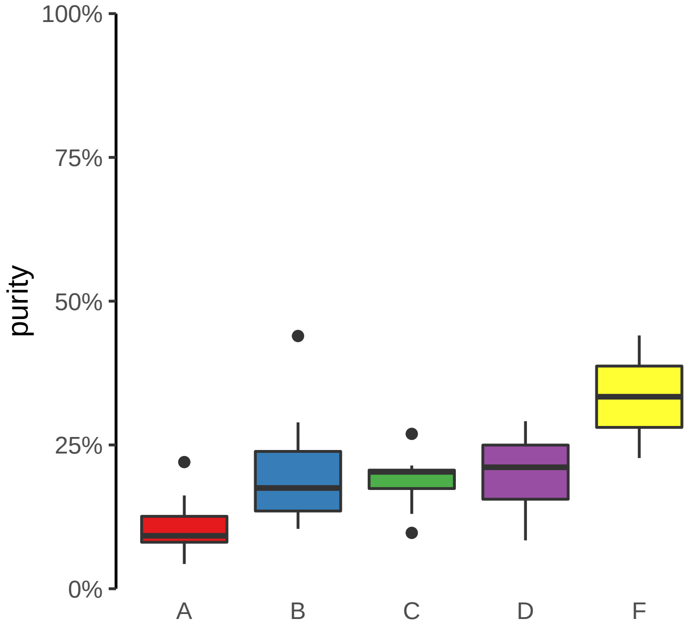

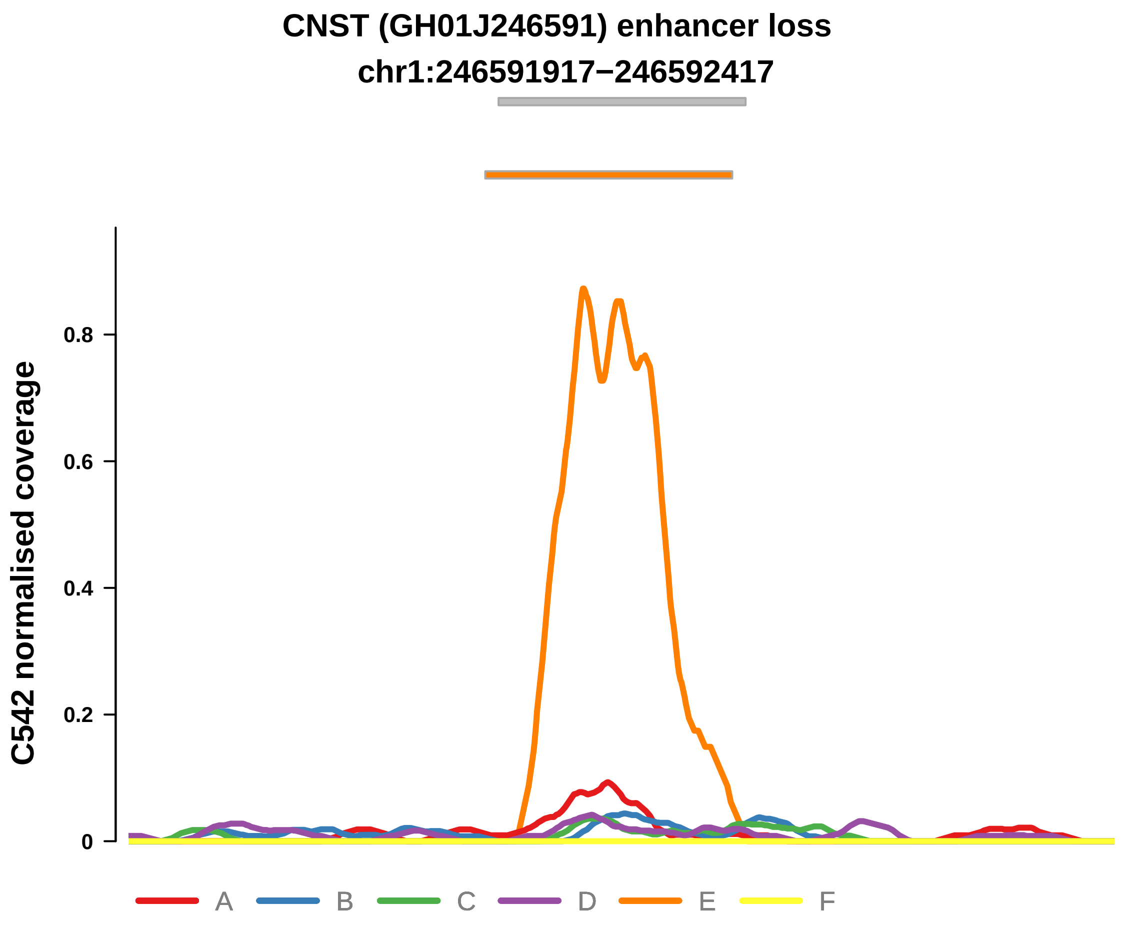


Putatively Clonal (logFC = −11.09)

| | P. | Adjusted P. |
|---|---|---|
| *~region vs. ~* | 4.96e−01 | 7.66e−01 |
| *~purity vs ~* | 7.08e−02 | 5.28e−01 |
| *~purity+region vs ~purity* | 6.92e−01 | 8.56e−01 |
| *Carcinoma vs. normal* | 2.43e−03 ** | 2.43e−03 ** |

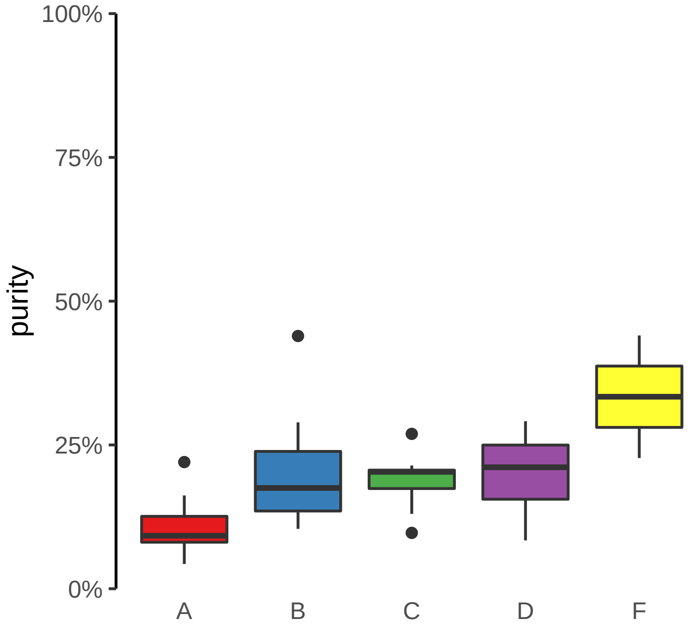

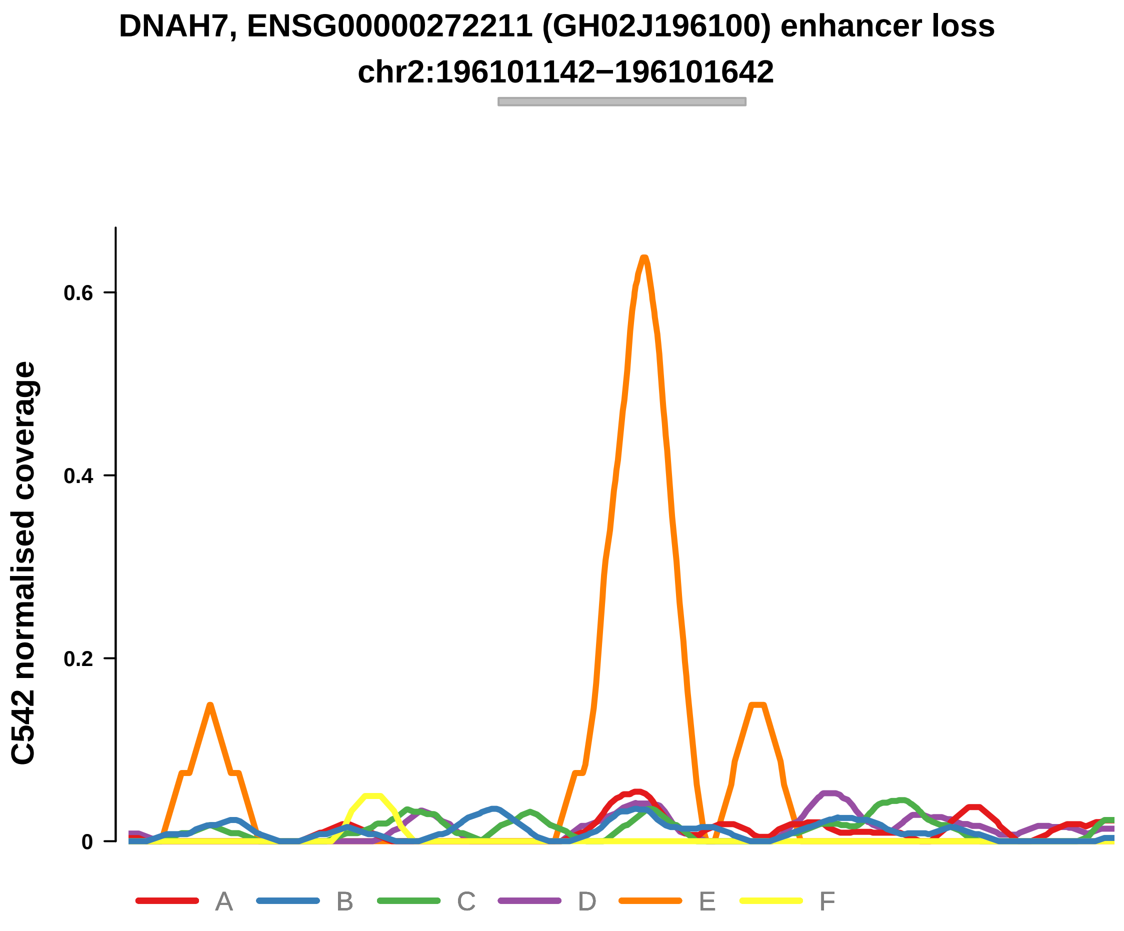


Putatively Clonal (logFC = −10.88)

| | P. | Adjusted P. |
|---|---|---|
| *~region vs. ~* | 7.22e−01 | 8.78e−01 |
| *~purity vs ~* | 5.5e−01 | 9.67e−01 |
| *~purity+region vs ~purity* | 7.89e−01 | 8.56e−01 |
| *Carcinoma vs. normal* | 5.83e−03 ** | 5.83e−03 ** |

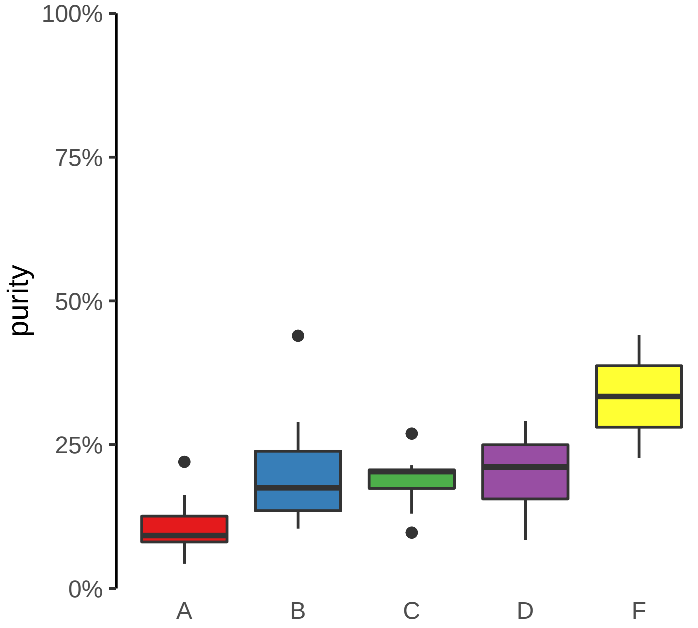

**ENSG00000228035, NGF (GH01J115322) enhancer loss**
**chr1:115323237−115323737**

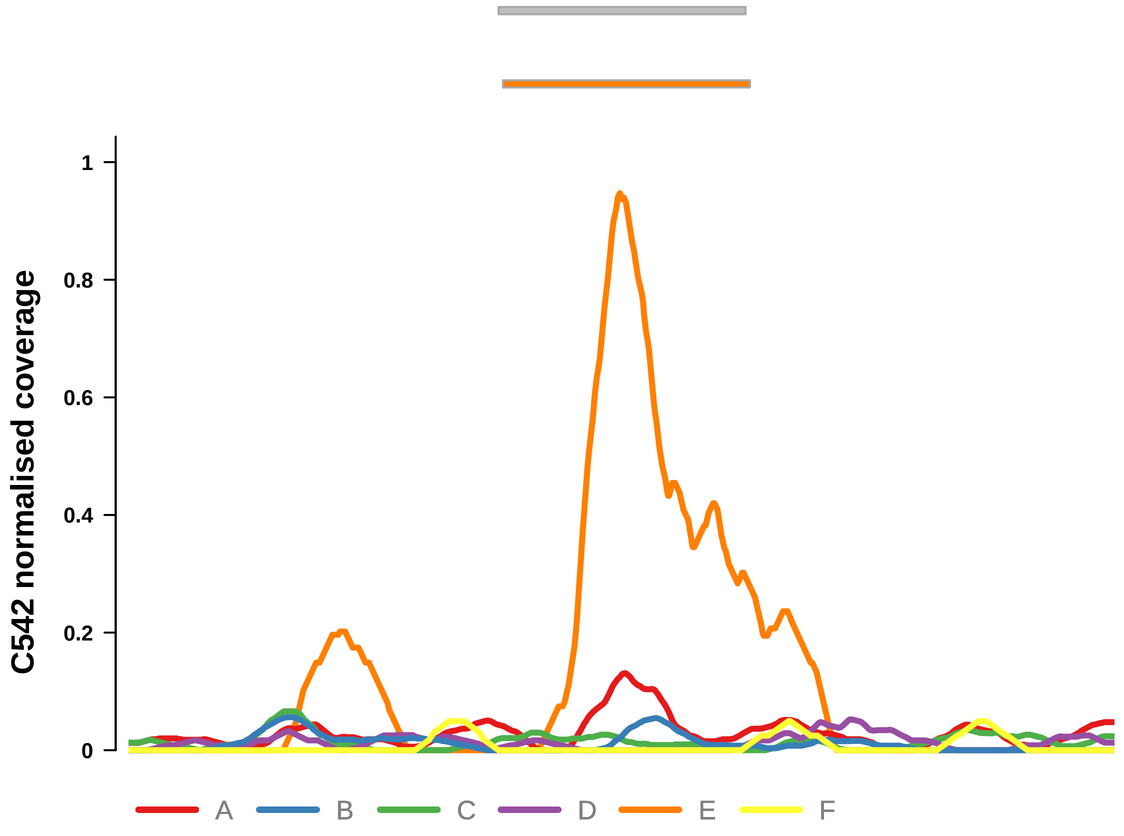


Putatively Clonal (logFC = −11.3)

| | P. | Adjusted P. |
|---|---|---|
| *~region vs. ~* | 4.56e−02 * | 5.01e−01 |
| *~purity vs ~* | 5.12e−04 *** | 2.25e−02 * |
| *~purity+region vs ~purity* | 6.51e−01 | 8.56e−01 |
| *Carcinoma vs. normal* | 1.61e−03 ** | 1.61e−03 ** |

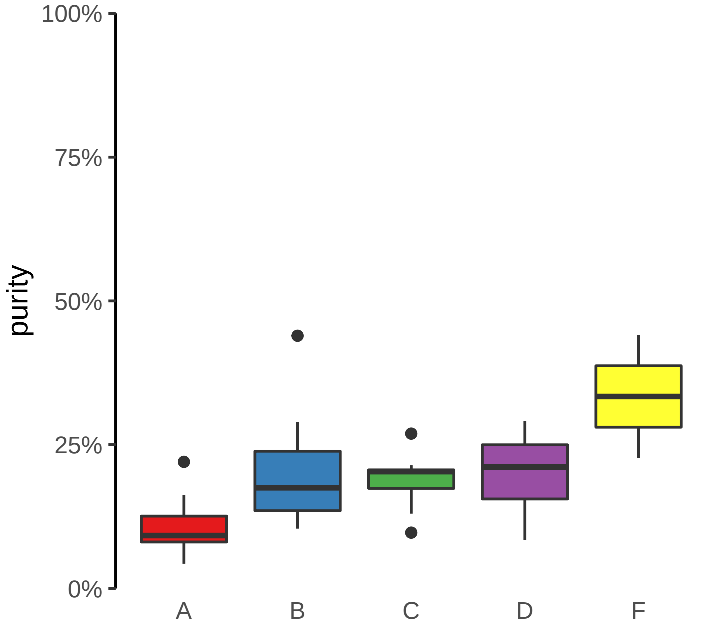

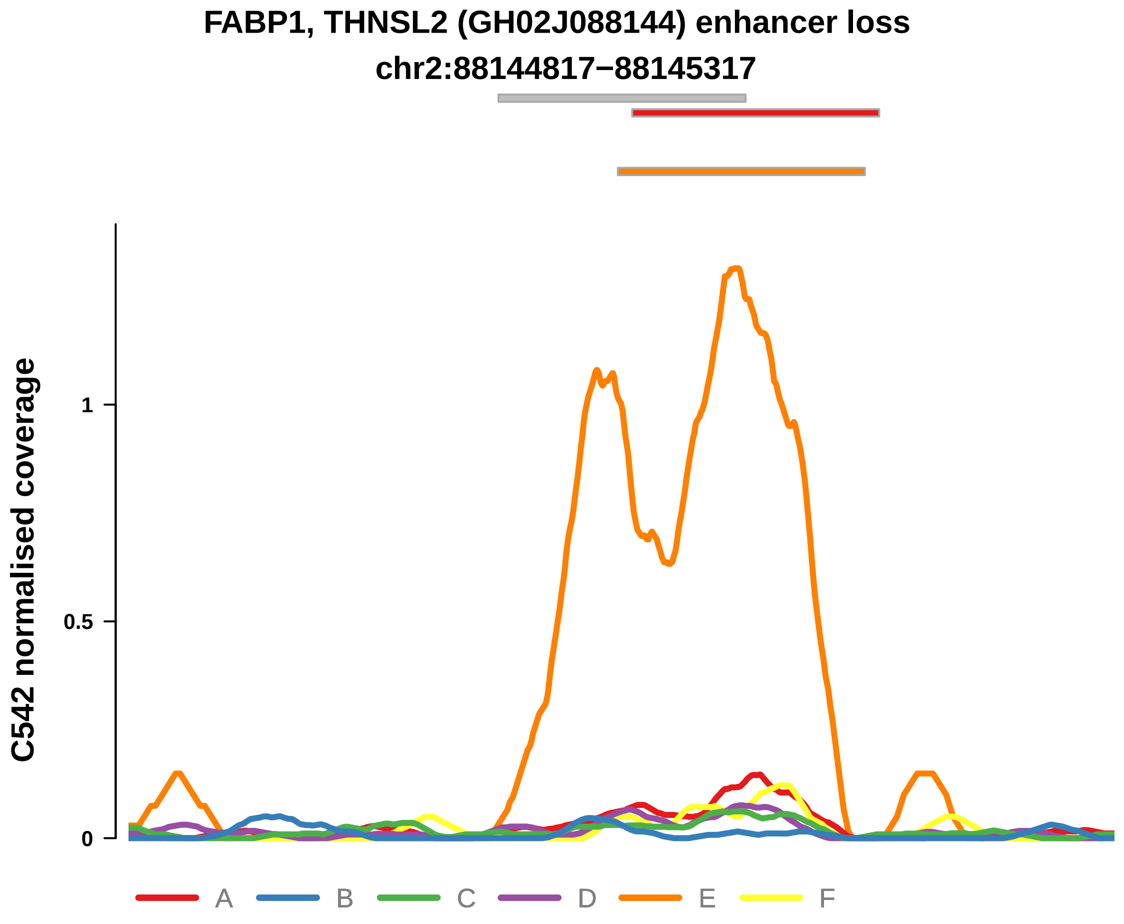


Putatively Clonal (logFC = −11.8)

| | P. | Adjusted P. |
|---|---|---|
| *~region vs. ~* | 8.39e−03 ** | 4e−01 |
| *~purity vs ~* | 4.21e−01 | 9.67e−01 |
| *~purity+region vs ~purity* | 1.34e−02 * | 3.94e−01 |
| *Carcinoma vs. normal* | 4.67e−04 *** | 4.67e−04 *** |

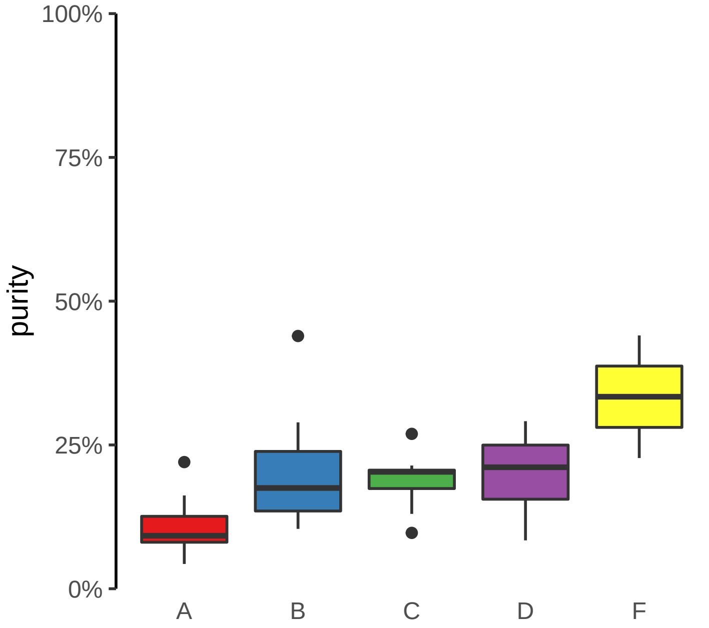

**MAP3K1, LINC01948 (GH05J056411) enhancer loss**
**chr5:56412924−56413424**

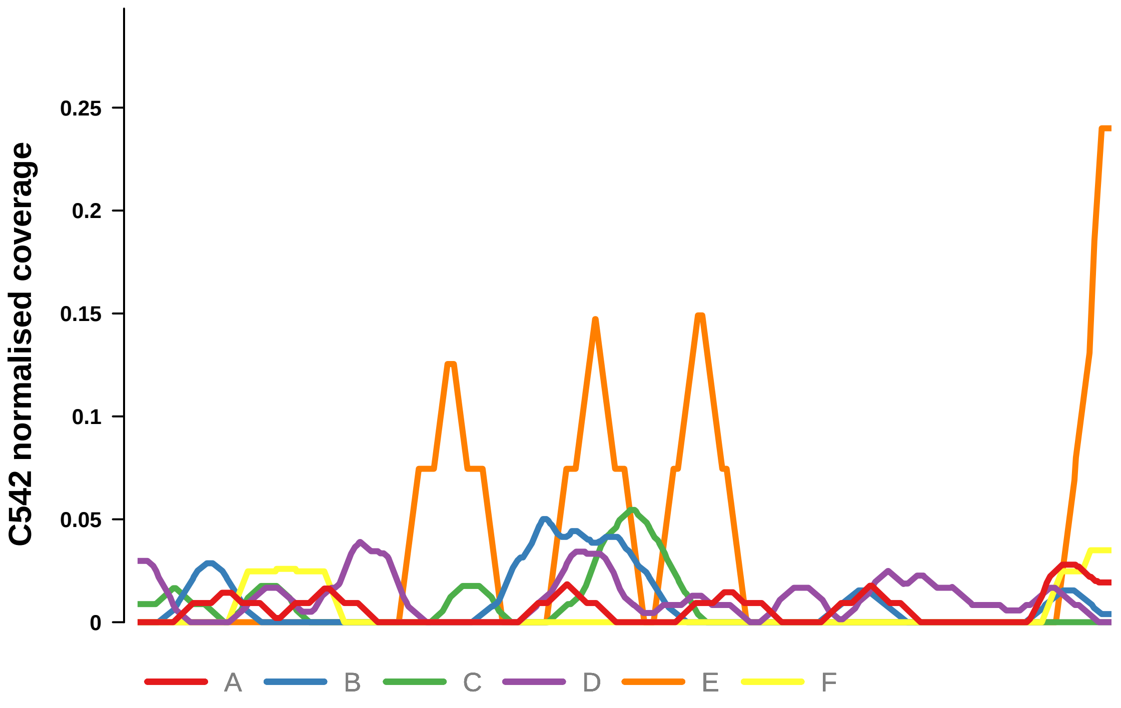


Putatively Clonal (logFC = −10.61)

| | P. | Adjusted P. |
|---|---|---|
| *~region vs. ~* | 3.51e−01 | 6.71e−01 |
| *~purity vs ~* | 3.52e−01 | 9.67e−01 |
| *~purity+region vs ~purity* | 1.75e−01 | 6.99e−01 |
| *Carcinoma vs. normal* | 9.33e−03 ** | 9.33e−03 ** |

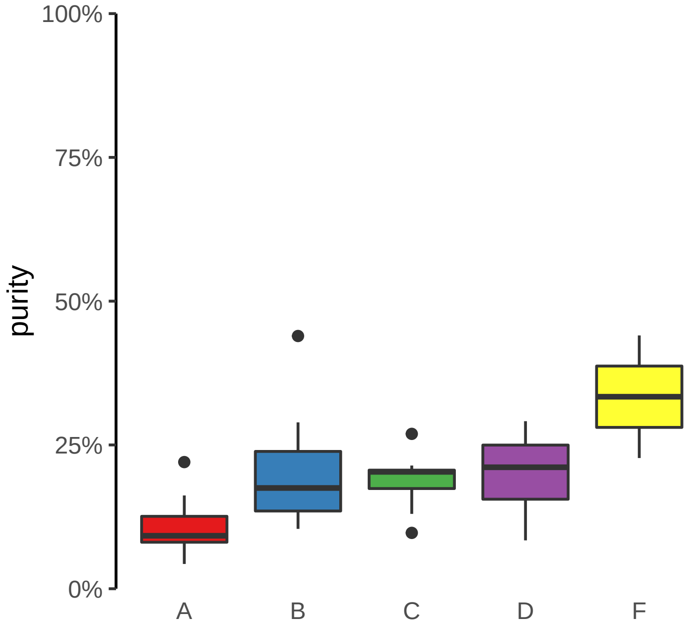

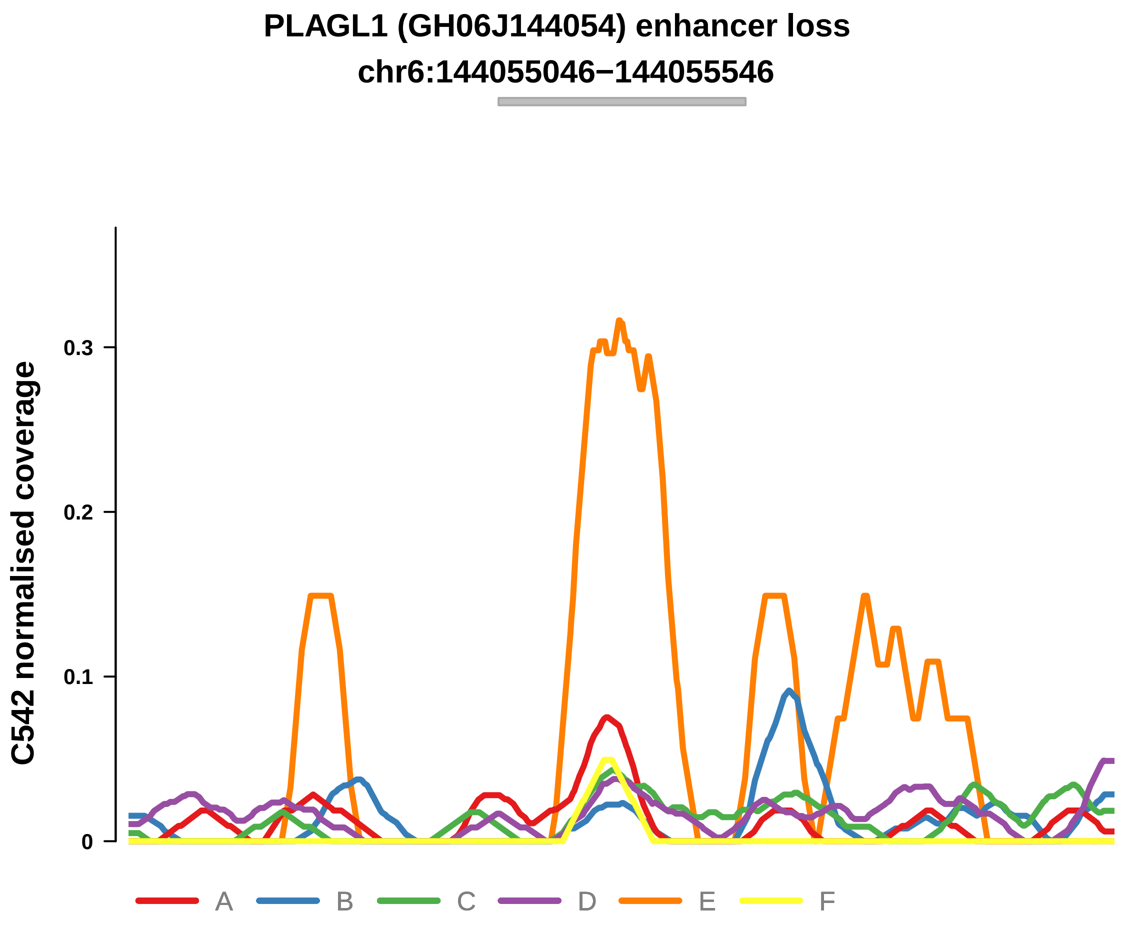


Putatively Clonal (logFC = −10.96)

| | P. | Adjusted P. |
|---|---|---|
| *~region vs. ~* | 7.04e−01 | 8.78e−01 |
| *~purity vs ~* | 7.76e−01 | 9.67e−01 |
| *~purity+region vs ~purity* | 7.23e−01 | 8.56e−01 |
| *Carcinoma vs. normal* | 3.97e−03 ** | 3.97e−03 ** |

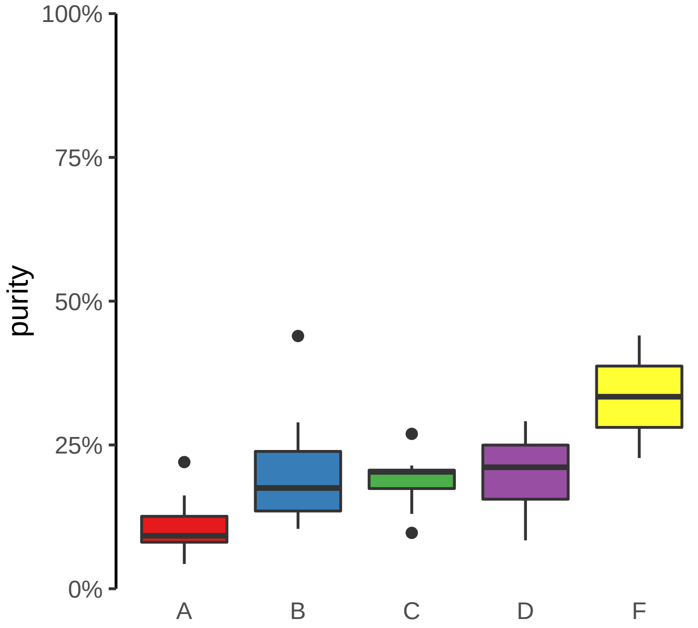

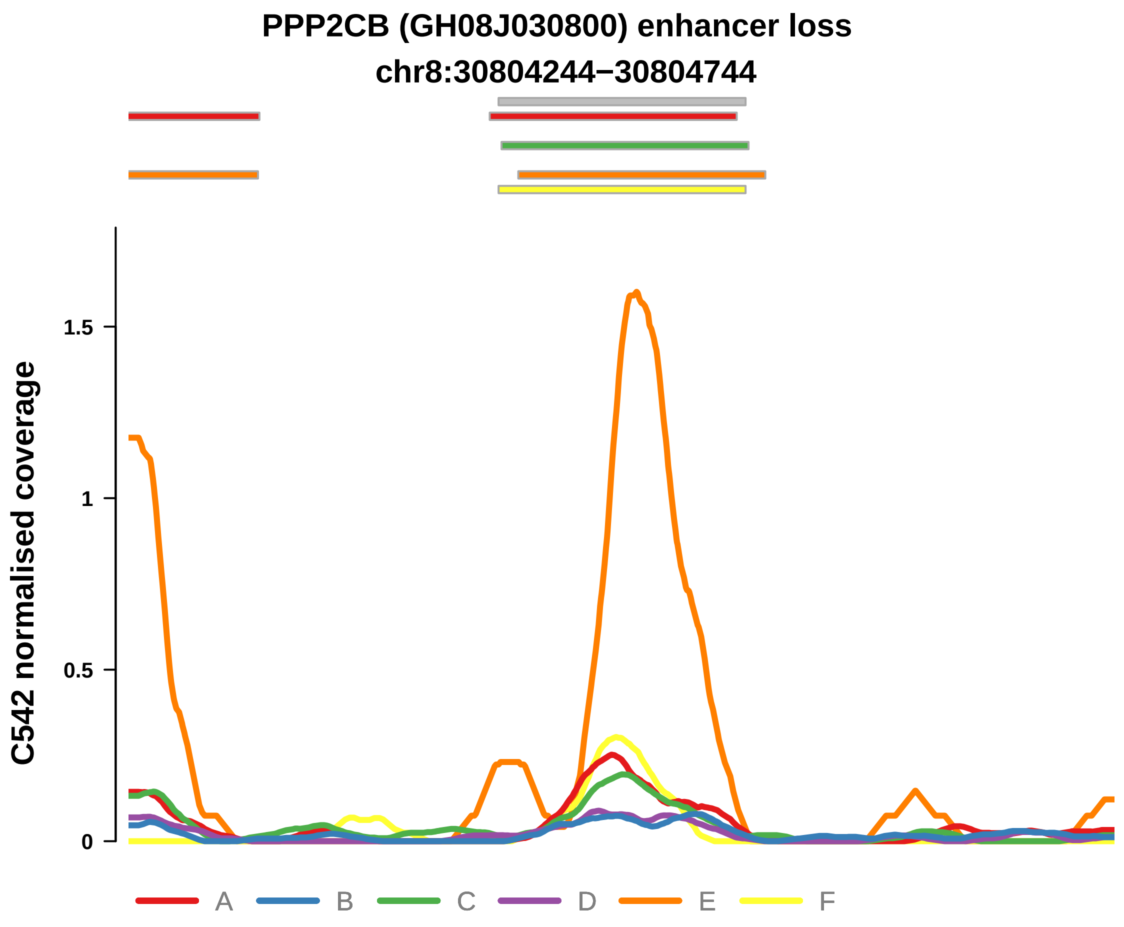


Putatively Clonal (logFC = −11.75)

| | P. | Adjusted P. |
|---|---|---|
| *~region vs. ~* | 4.29e−01 | 7.33e−01 |
| *~purity vs ~* | 4.85e−01 | 9.67e−01 |
| *~purity+region vs ~purity* | 5e−01 | 8.56e−01 |
| *Carcinoma vs. normal* | 6.02e−04 *** | 6.02e−04 *** |

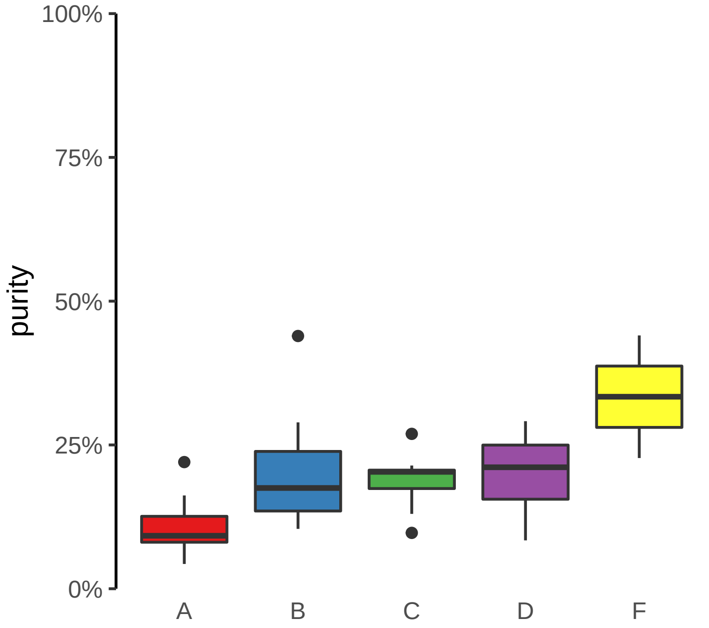

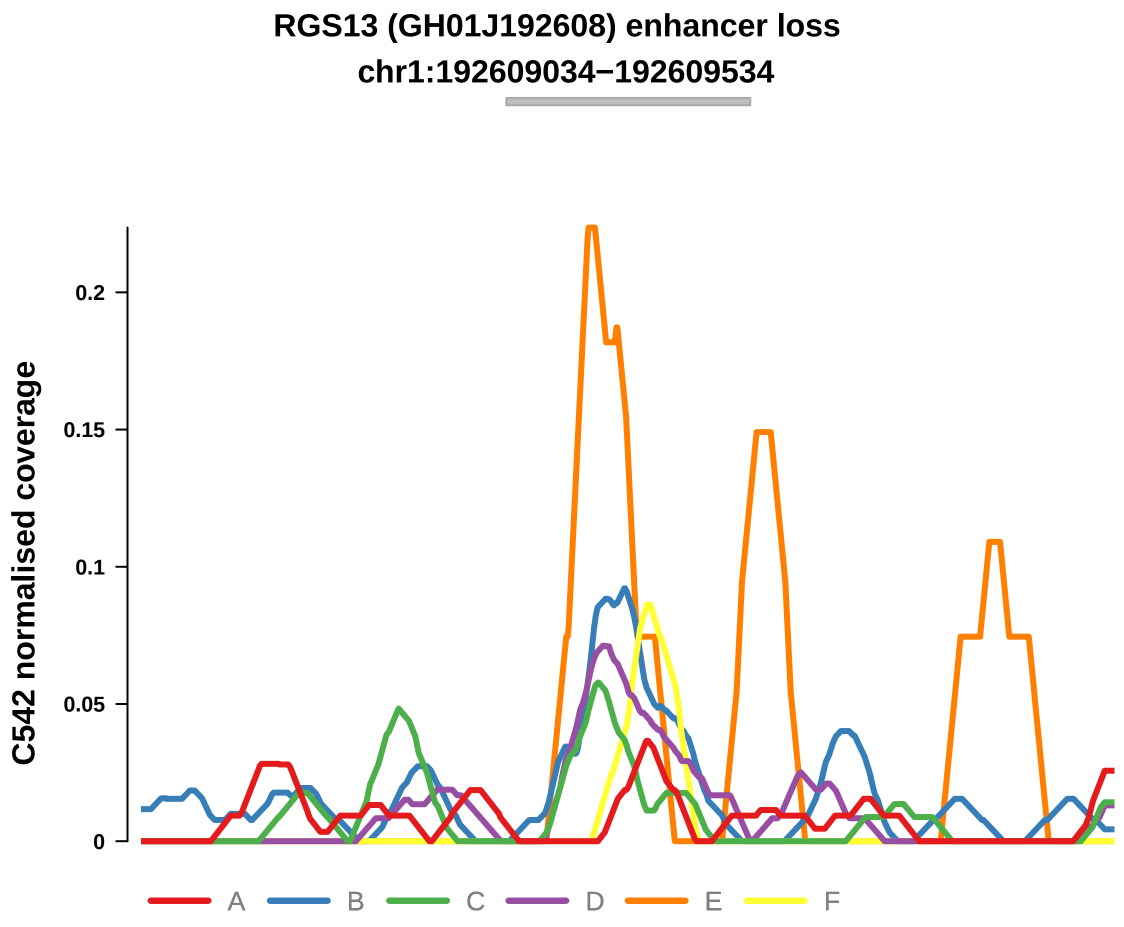


Putatively Clonal (logFC = −11.14)

| | P. | Adjusted P. |
|---|---|---|
| *~region vs. ~* | 8.54e−02 | 5.9e−01 |
| *~purity vs ~* | 8.22e−01 | 9.67e−01 |
| *~purity+region vs ~purity* | 6.33e−02 | 5.5e−01 |
| *Carcinoma vs. normal* | 2.16e−03 ** | 2.16e−03 ** |

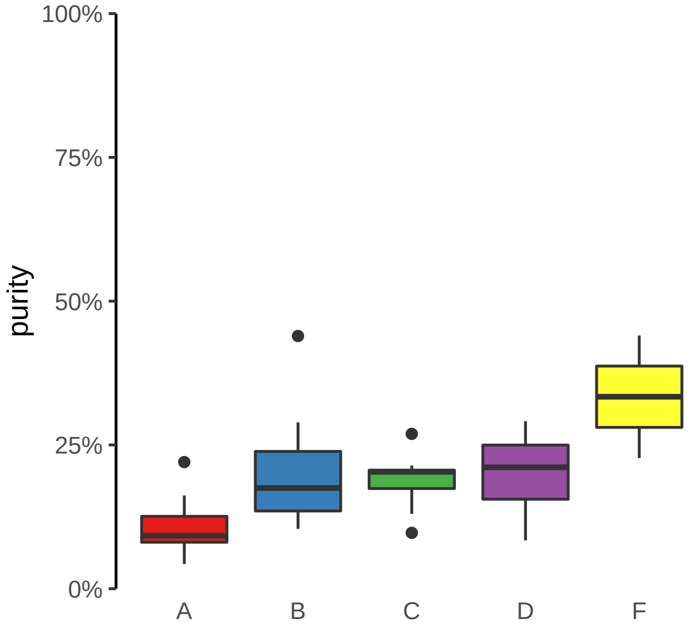

**RNLS, LIPJ (GH10J088553) enhancer loss**
**chr10:88553102−88553602**

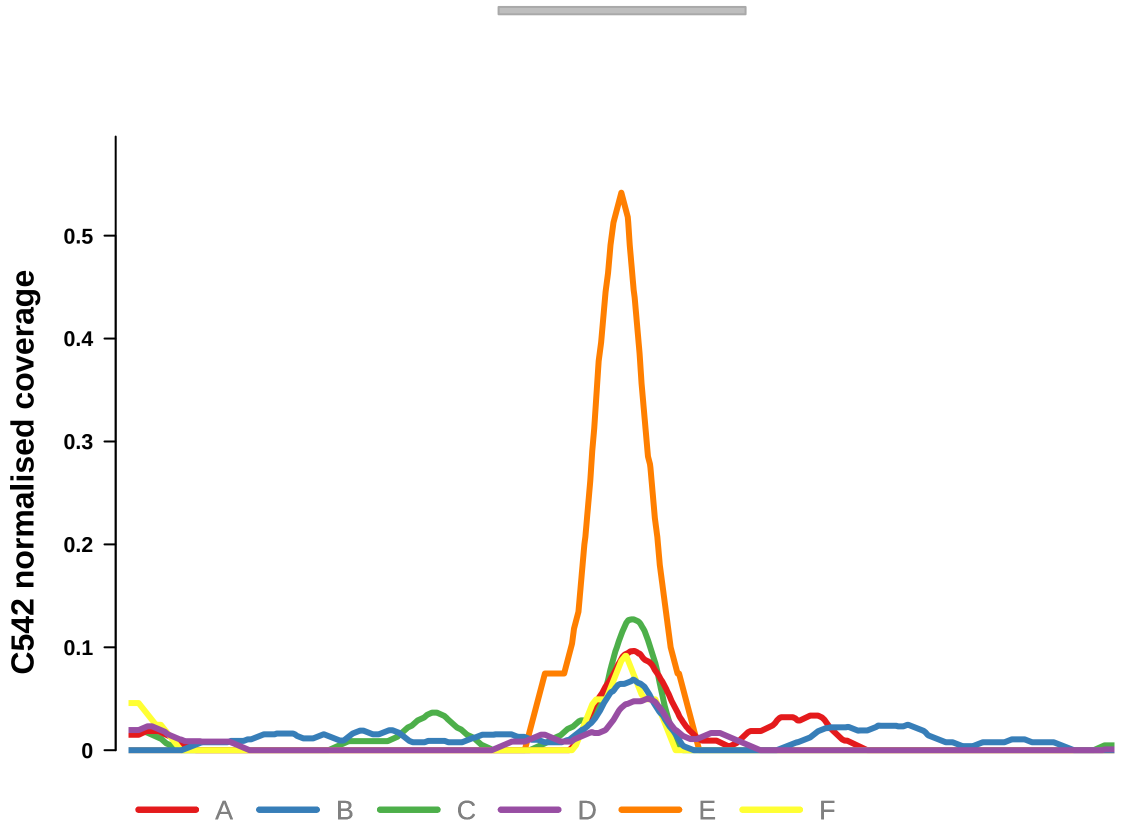


Putatively Clonal (logFC = −11.43)

| | P. | Adjusted P. |
|---|---|---|
| *~region vs. ~* | 6.3e−01 | 8.3e−01 |
| *~purity vs ~* | 9.06e−01 | 9.84e−01 |
| *~purity+region vs ~purity* | 6.35e−01 | 8.56e−01 |
| *Carcinoma vs. normal* | 9.55e−04 *** | 9.55e−04 *** |

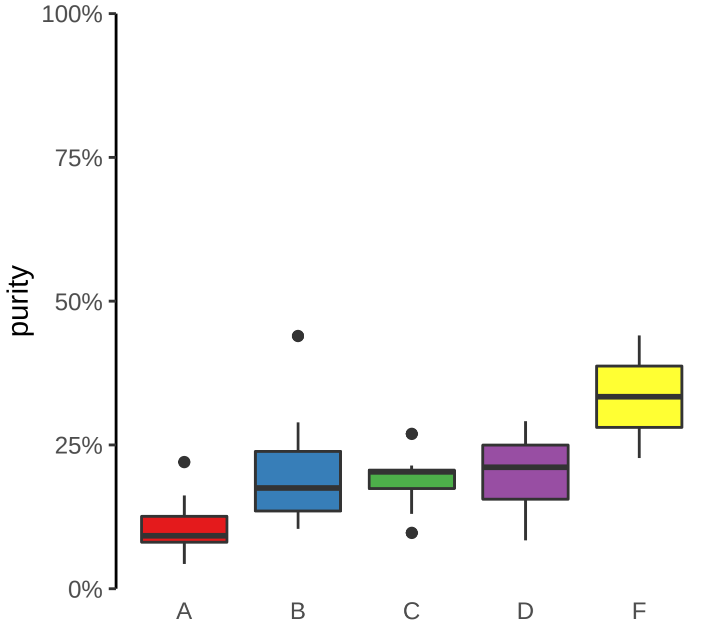

**SMAD3 (GH15J067079) enhancer loss**
**chr15:67081050−67081550**

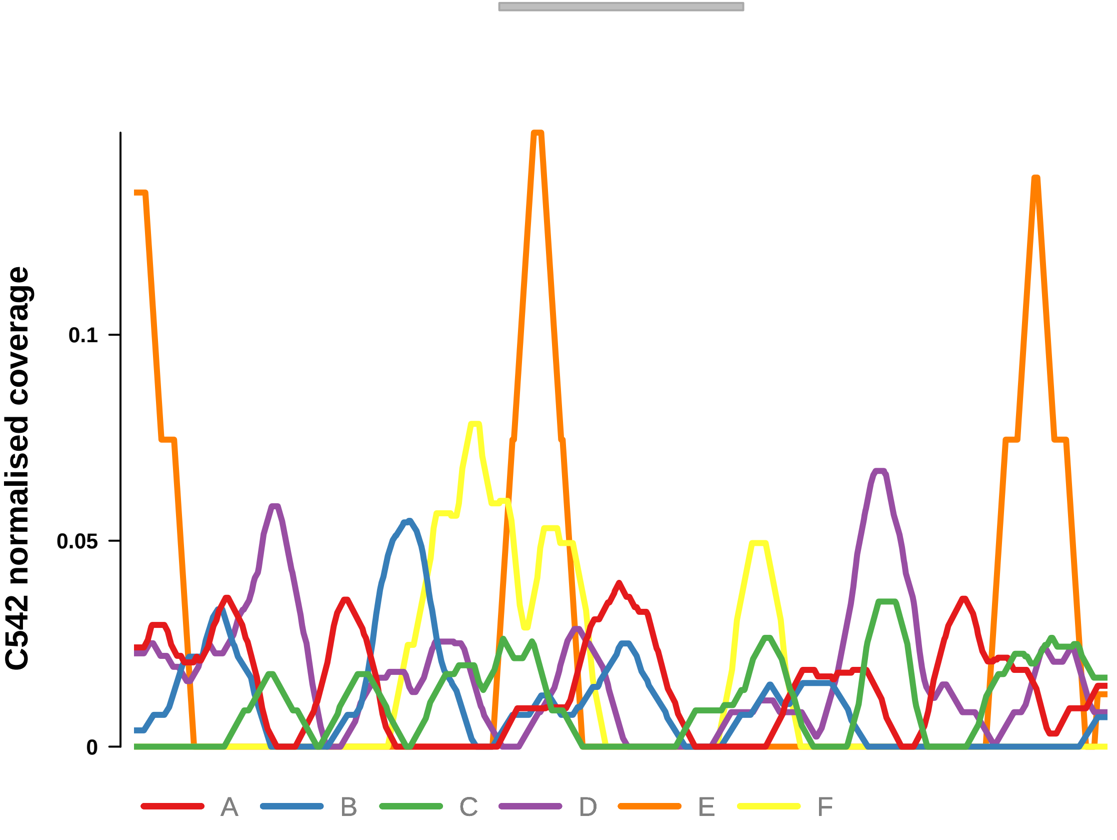


Putatively Clonal (logFC = −10.65)

| | **P.** | **Adjusted P.** |
|---|---|---|
| *~region vs. ~* | 7.58e−01 | 8.78e−01 |
| *~purity vs ~* | 6.85e−01 | 9.67e−01 |
| *~purity+region vs ~purity* | 8.08e−01 | 8.56e−01 |
| *Carcinoma vs. normal* | 8.06e−03 ** | 8.06e−03 ** |

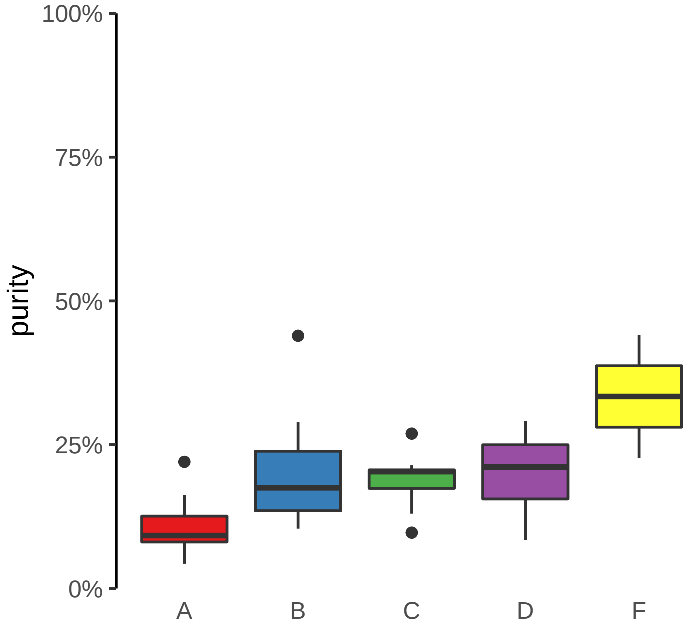

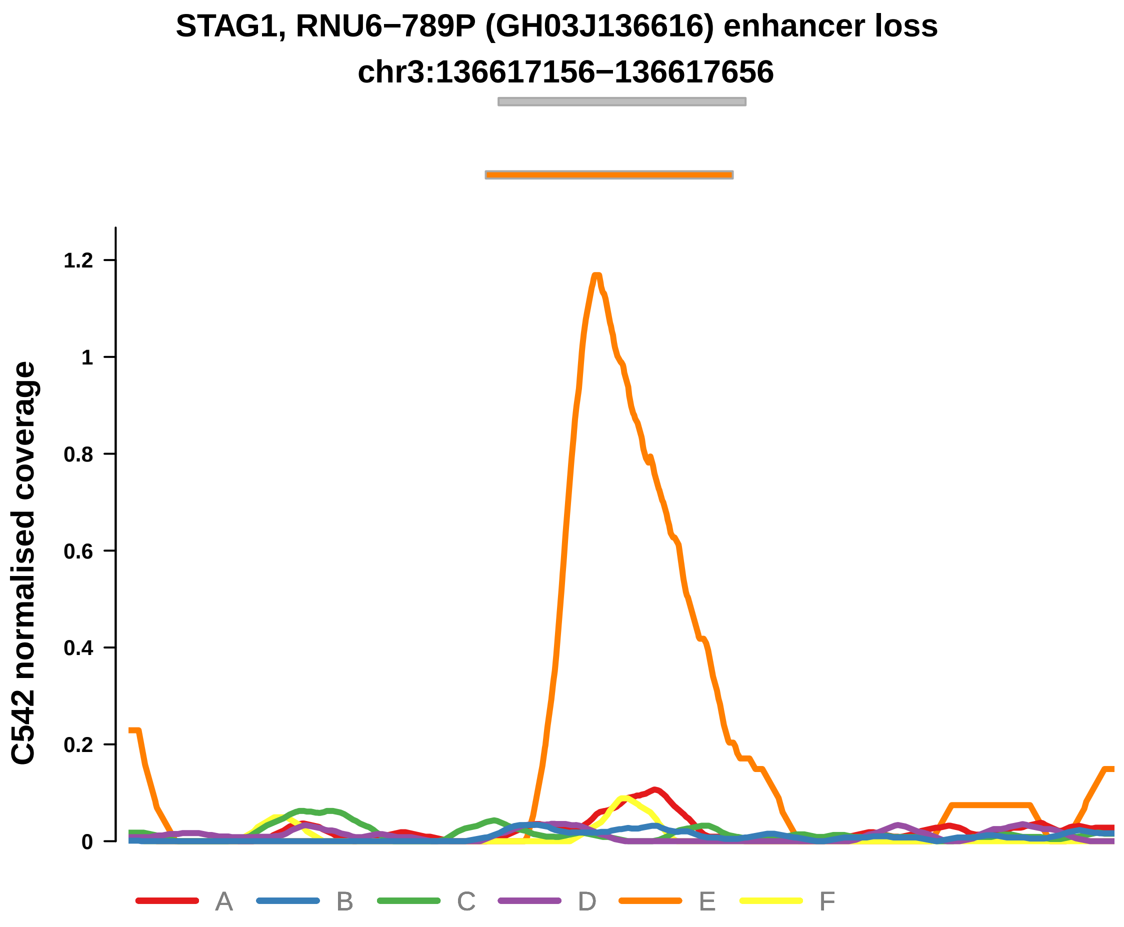


Putatively Clonal (logFC = −11.64)

| | P. | Adjusted P. |
|---|---|---|
| *~region vs. ~* | 3.62e−01 | 6.78e−01 |
| *~purity vs ~* | 3.44e−02 * | 4.32e−01 |
| *~purity+region vs ~purity* | 7.76e−01 | 8.56e−01 |
| *Carcinoma vs. normal* | 5.92e−04 *** | 5.92e−04 *** |

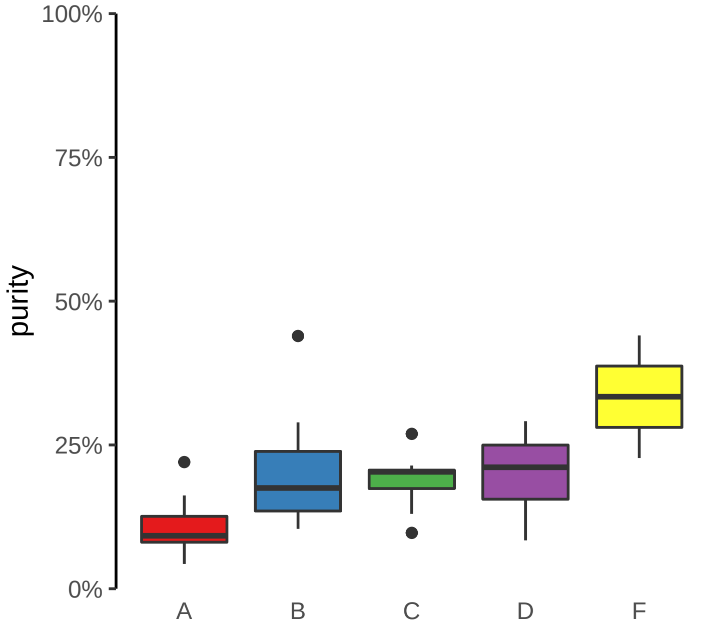

**TRANK1, ENSG00000234073 (GH03J036874) enhancer loss**
**chr3:36876260−36876760**

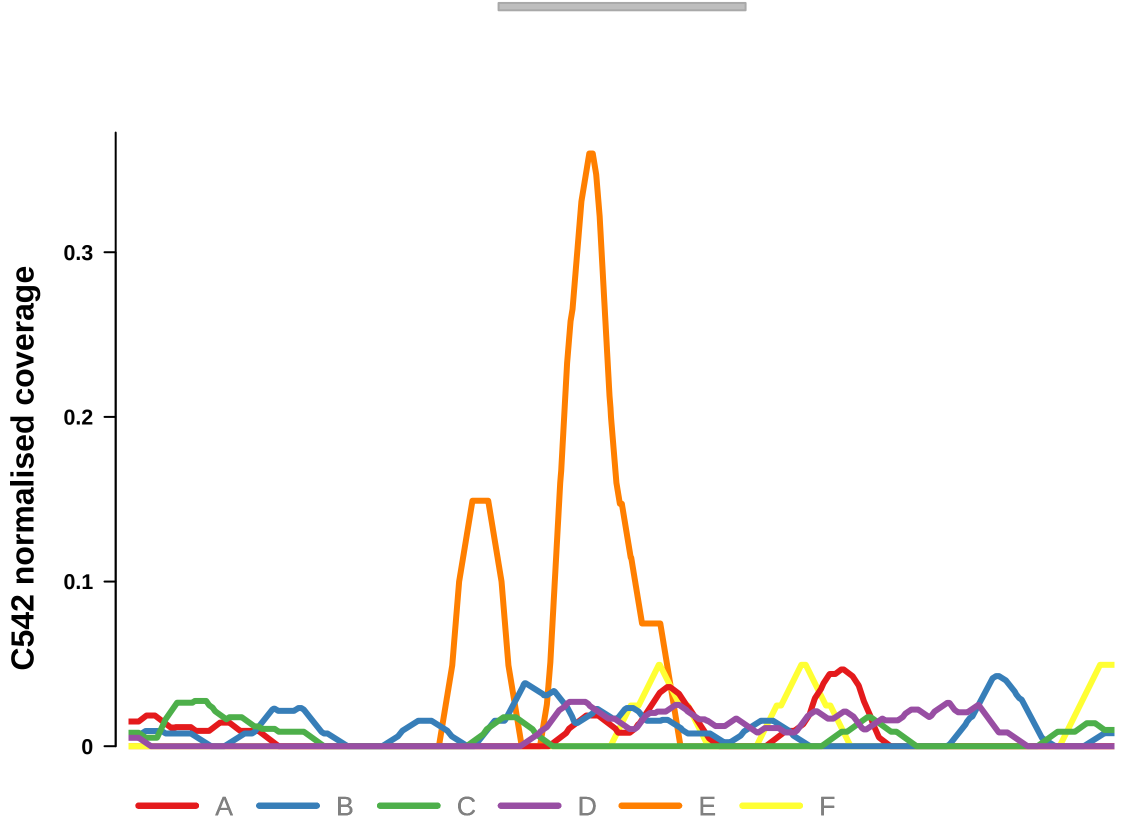


Putatively Clonal (logFC = −10.93)

| | P. | Adjusted P. |
|---|---|---|
| *~region vs. ~* | 2.53e−01 | 6.21e−01 |
| *~purity vs ~* | 1e+00 | 1e+00 |
| *~purity+region vs ~purity* | 2.52e−01 | 7.3e−01 |
| *Carcinoma vs. normal* | 4.51e−03 ** | 4.51e−03 ** |

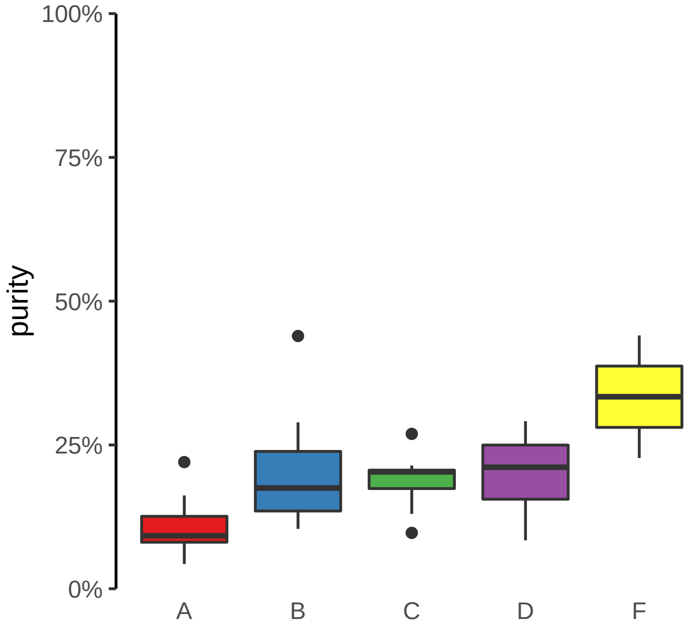

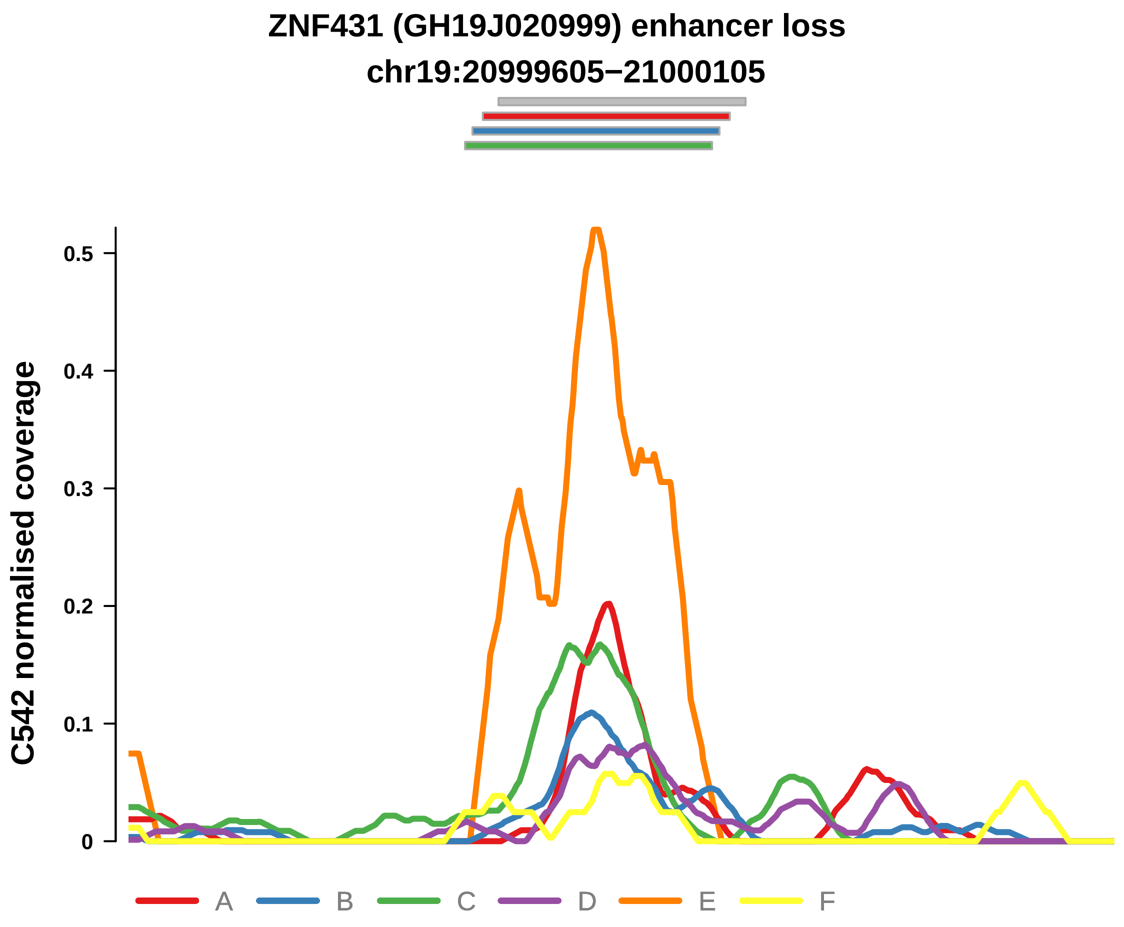


Putatively Clonal (logFC = −12.28)

| | P. | Adjusted P. |
|---|---|---|
| *~region vs. ~* | 6.32e−01 | 8.3e−01 |
| *~purity vs ~* | 1.45e−02 * | 2.13e−01 |
| *~purity+region vs ~purity* | 3.8e−01 | 7.96e−01 |
| *Carcinoma vs. normal* | 6.88e−05 **** | 6.88e−05 **** |

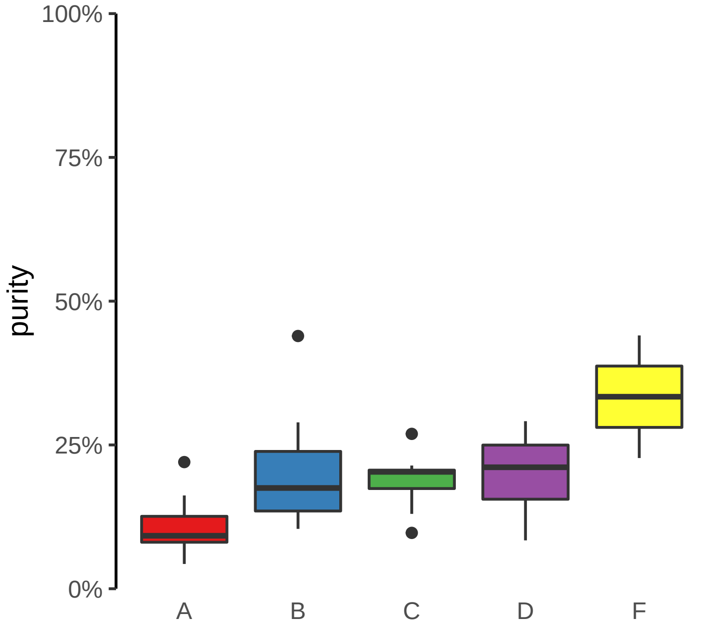

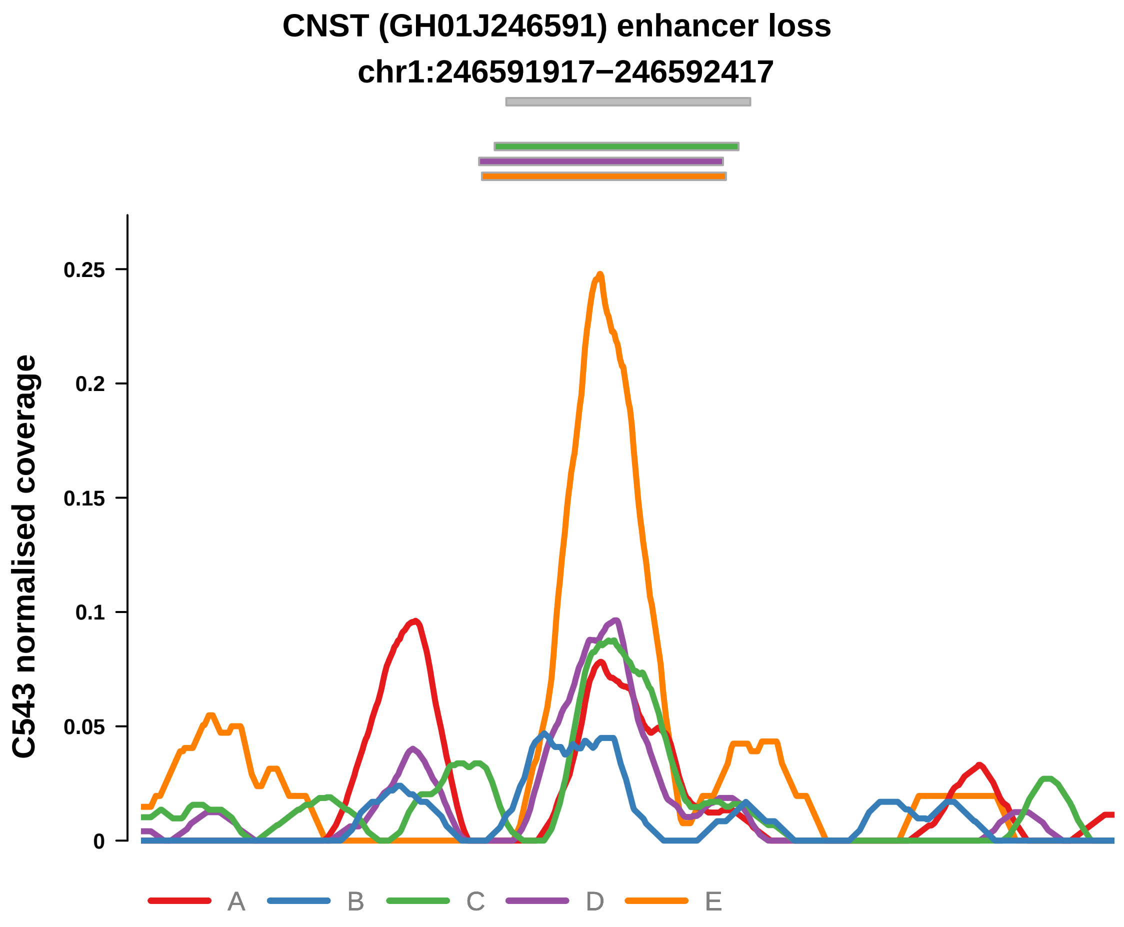


Putatively Clonal (logFC = −11.09)

| | P. | Adjusted P. |
|---|---|---|
| *~region vs. ~* | 2.98e−01 | 6.76e−01 |
| *~purity vs ~* | 9.74e−01 | NA |
| *~purity+region vs ~purity* | 2.66e−01 | 7.73e−01 |
| *Carcinoma vs. normal* | 7.58e−03 ** | 7.58e−03 ** |

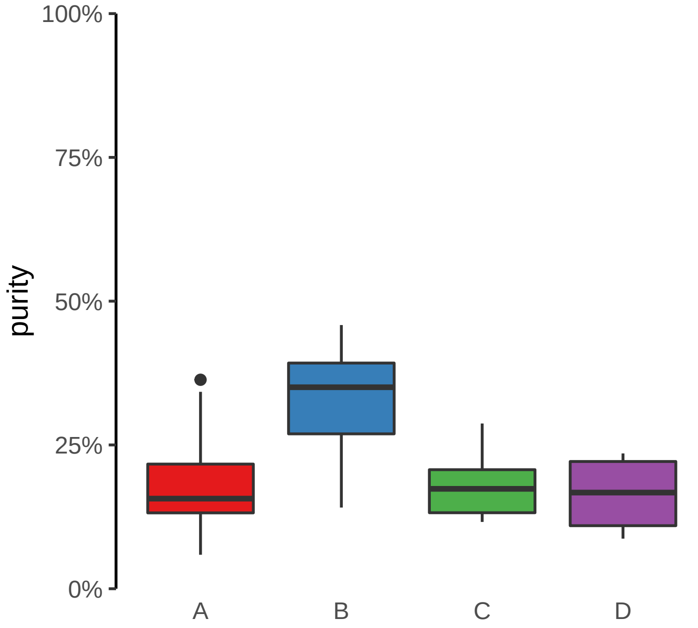

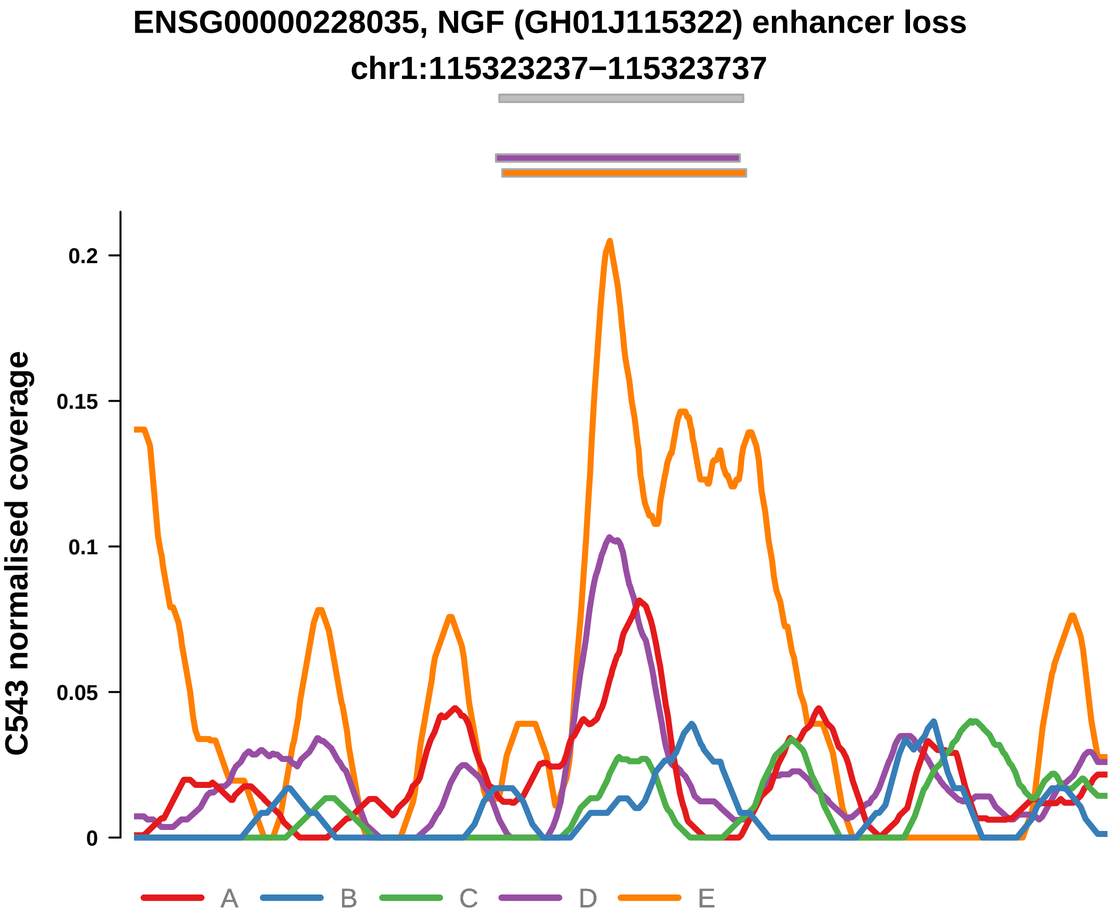


Putatively Clonal (logFC = −11.3)

| | P. | Adjusted P. |
|---|---|---|
| *~region vs. ~* | 5.34e−01 | 8.11e−01 |
| *~purity vs ~* | 6.82e−01 | NA |
| *~purity+region vs ~purity* | 4.31e−01 | 8.43e−01 |
| *Carcinoma vs. normal* | 5e−03 ** | 5e−03 ** |

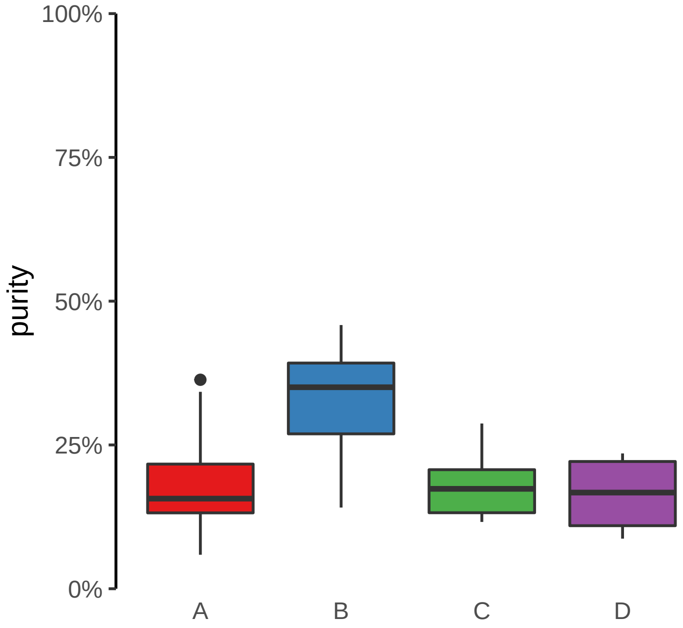

HMGN2, ARID1A, PIGV (GH01J026739) enhancer loss
chr1:26743976−26744476

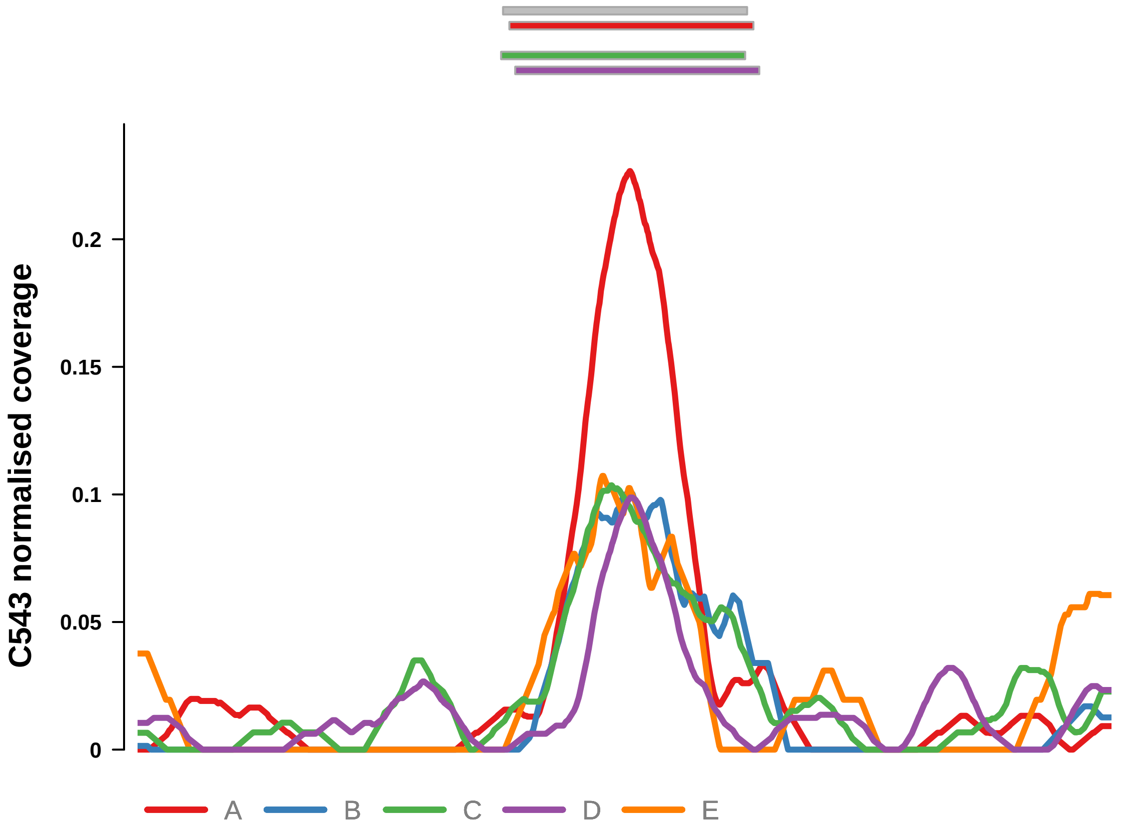


Putatively Clonal (logFC = −12.36)

| | P. | Adjusted P. |
|---|---|---|
| *~region vs. ~* | 1.88e−01 | 5.46e−01 |
| *~purity vs ~* | 3.76e−02 * | 9.57e−02 |
| *~purity+region vs ~purity* | 4.41e−01 | 8.43e−01 |
| *Carcinoma vs. normal* | 2.75e−04 *** | 2.75e−04 *** |

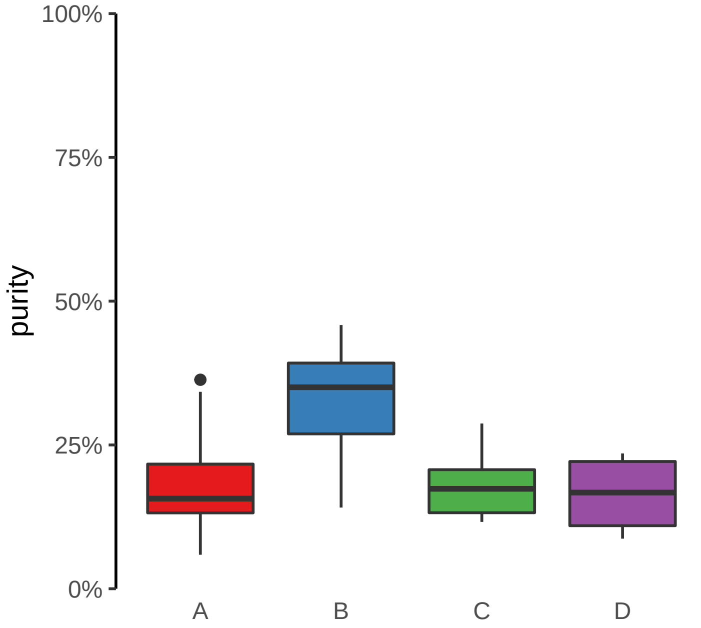

**PPP2CB (GH08J030800) enhancer loss**
**chr8:30804244−30804744**

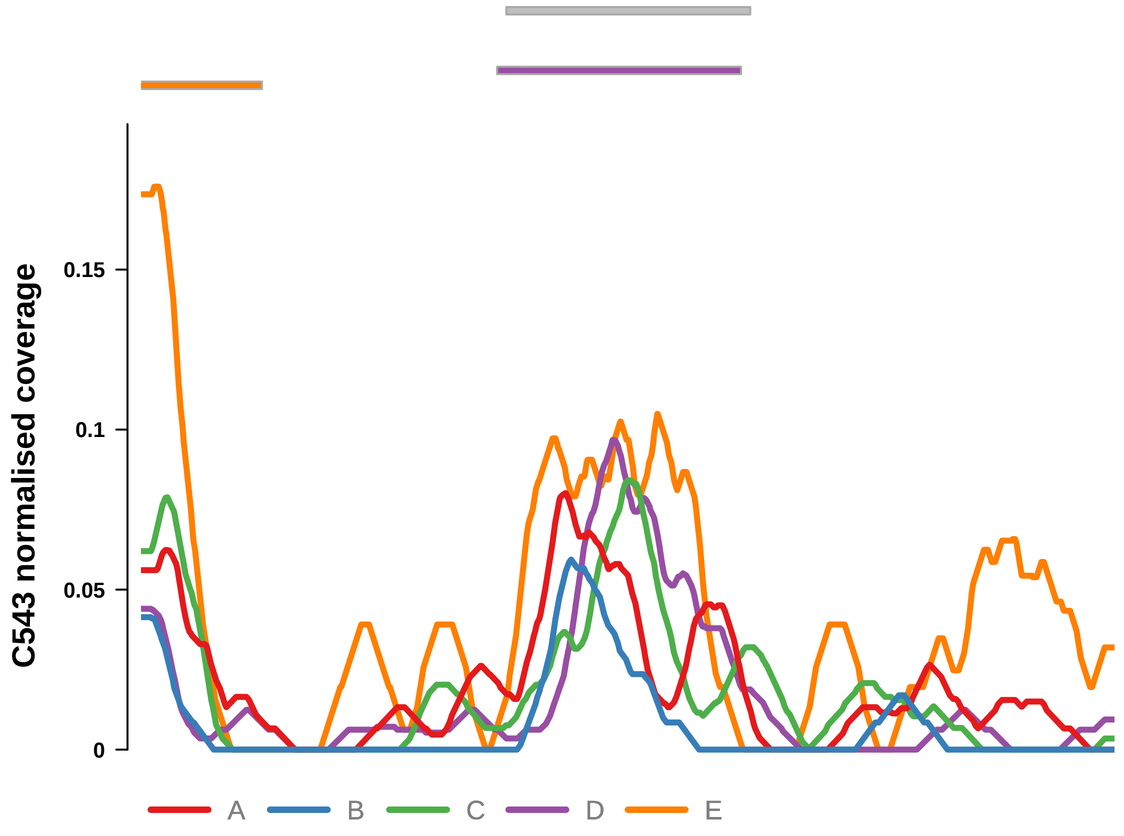


Putatively Clonal (logFC = −11.75)

| | P. | Adjusted P. |
|---|---|---|
| *~region vs. ~* | 3.87e−01 | 7.5e−01 |
| *~purity vs ~* | 9.18e−02 | NA |
| *~purity+region vs ~purity* | 6.53e−01 | 8.58e−01 |
| *Carcinoma vs. normal* | 1.96e−03 ** | 1.96e−03 ** |

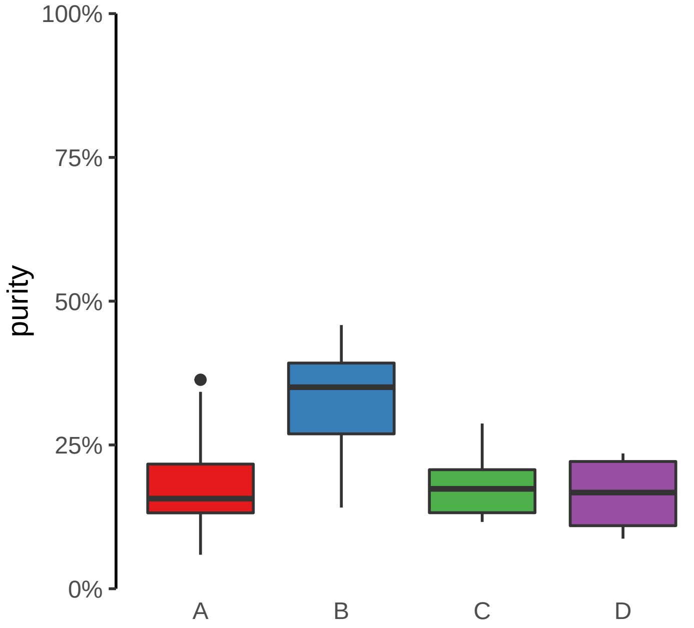

**RGS13 (GH01J192608) enhancer loss**
**chr1:192609034−192609534**

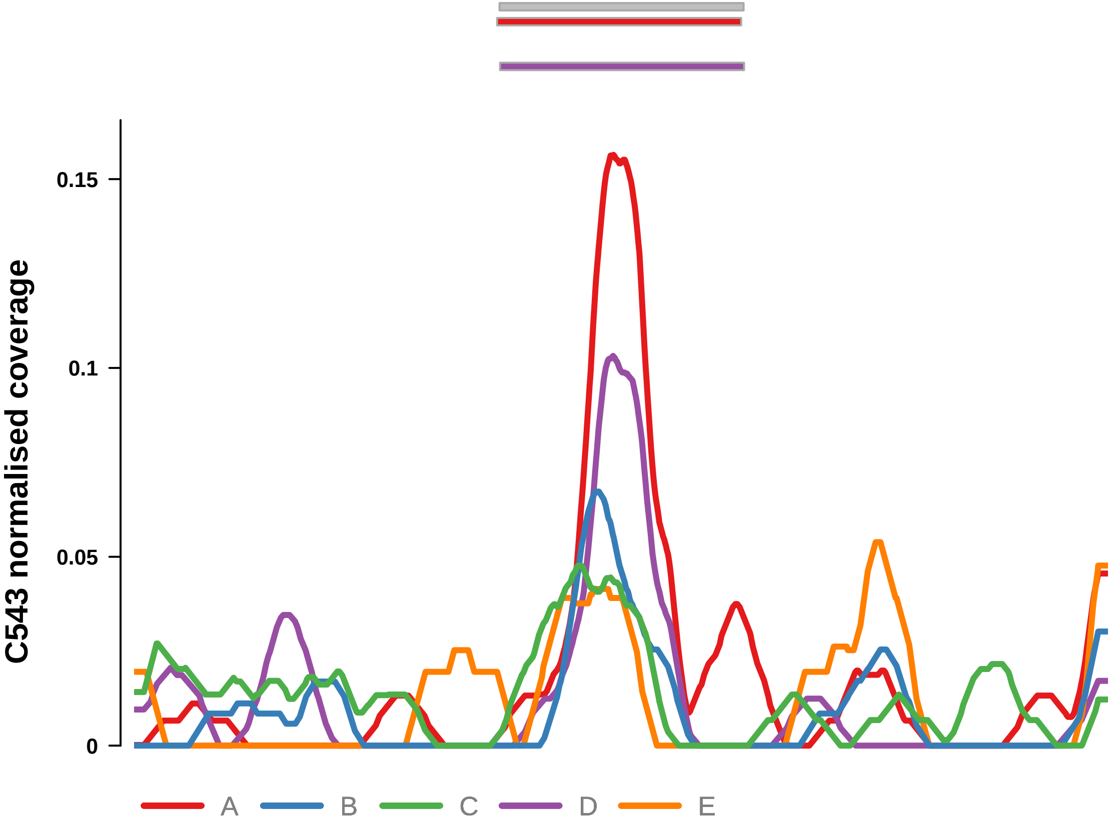


Putatively Clonal (logFC = −11.13)

| | P. | Adjusted P. |
|---|---|---|
| *~region vs. ~* | 5.01e−03 ** | 1.46e−01 |
| *~purity vs ~* | 8.44e−02 | 1.82e−01 |
| *~purity+region vs ~purity* | 4.38e−02 * | 4.28e−01 |
| *Carcinoma vs. normal* | 6.64e−03 ** | 6.64e−03 ** |

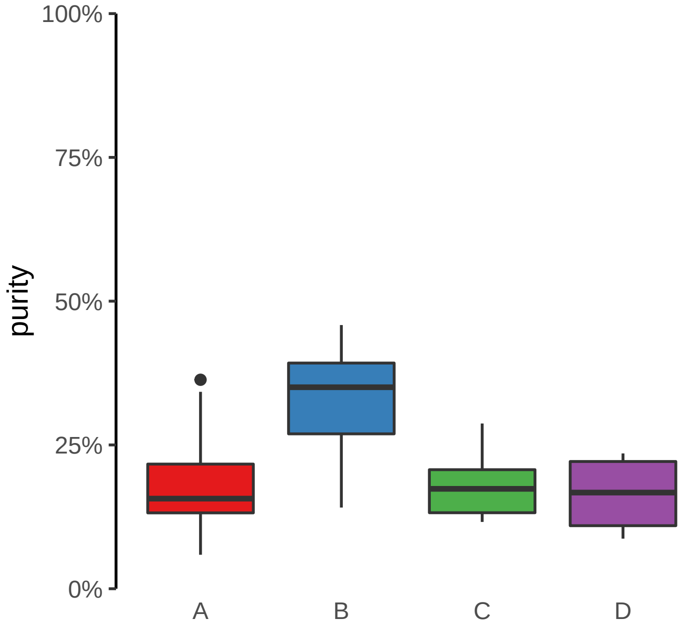

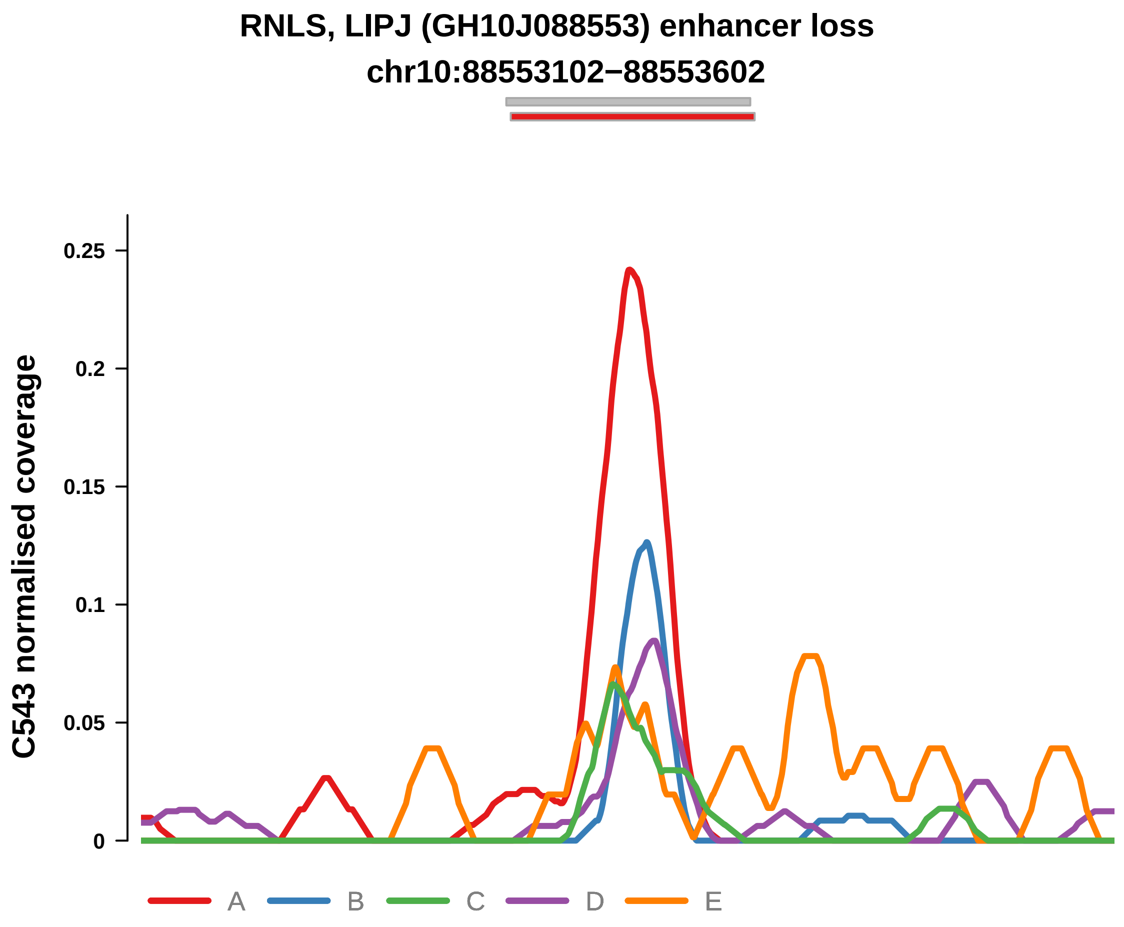


Putatively Clonal (logFC = −11.42)

| | P. | Adjusted P. |
|---|---|---|
| *~region vs. ~* | 8.55e−03 ** | 1.5e−01 |
| *~purity vs ~* | 2.54e−02 * | NA |
| *~purity+region vs ~purity* | 3.66e−02 * | 4.02e−01 |
| *Carcinoma vs. normal* | 3.14e−03 ** | 3.14e−03 ** |

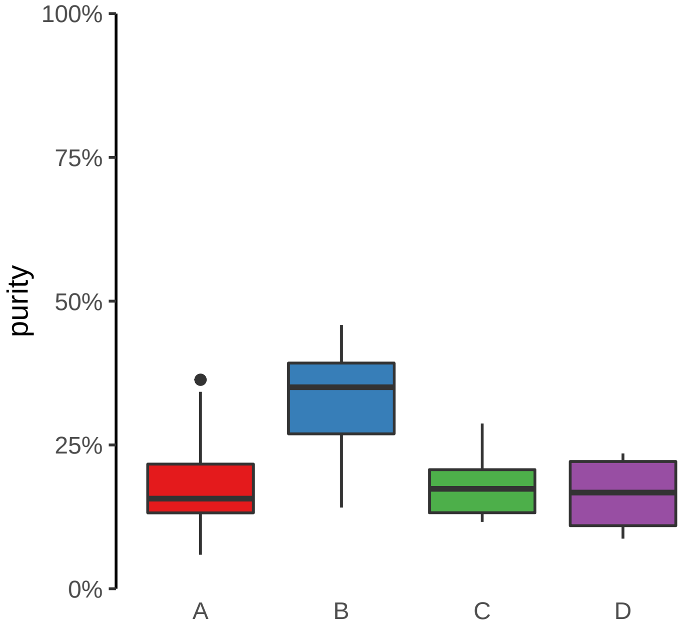

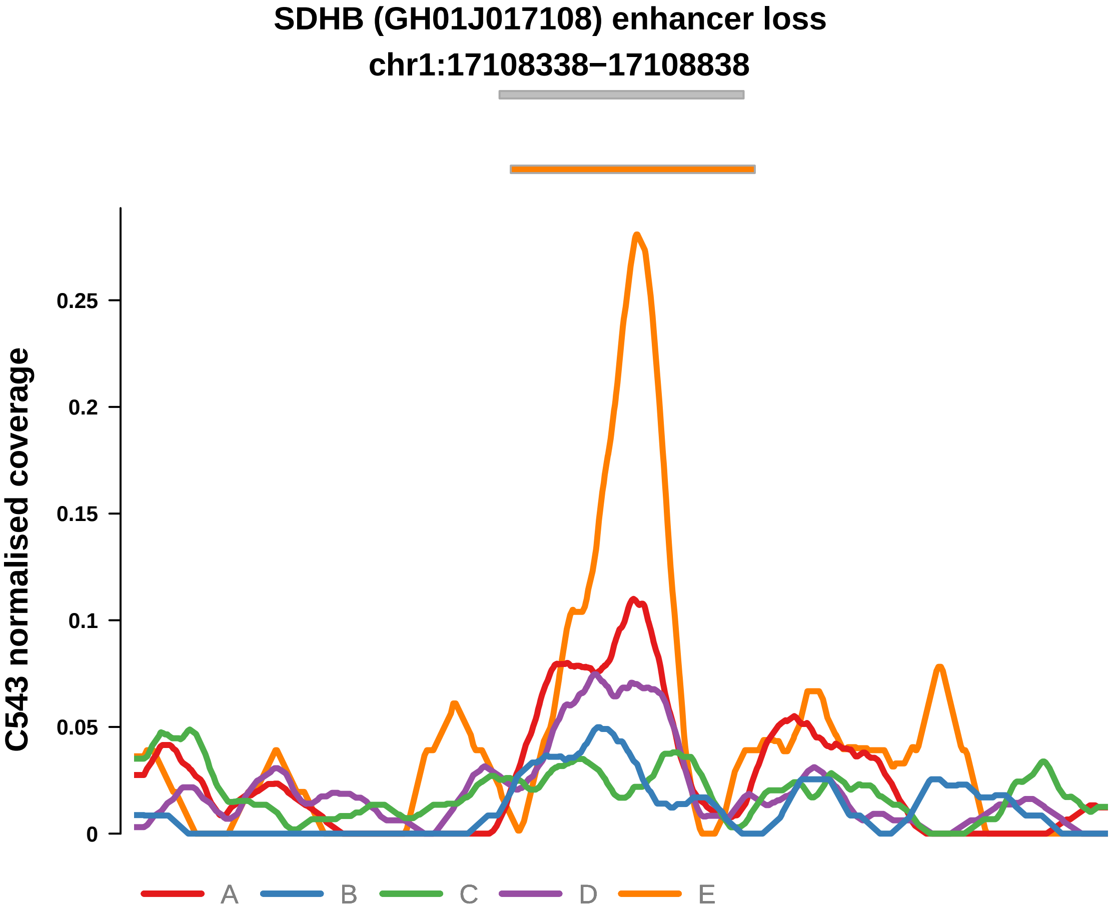


Putatively Clonal (logFC = −11.56)

| | P. | Adjusted P. |
|---|---|---|
| *~region vs. ~* | 2.28e−01 | 6.07e−01 |
| *~purity vs ~* | 1.1e−01 | NA |
| *~purity+region vs ~purity* | 4.34e−01 | 8.43e−01 |
| *Carcinoma vs. normal* | 2.85e−03 ** | 2.85e−03 ** |

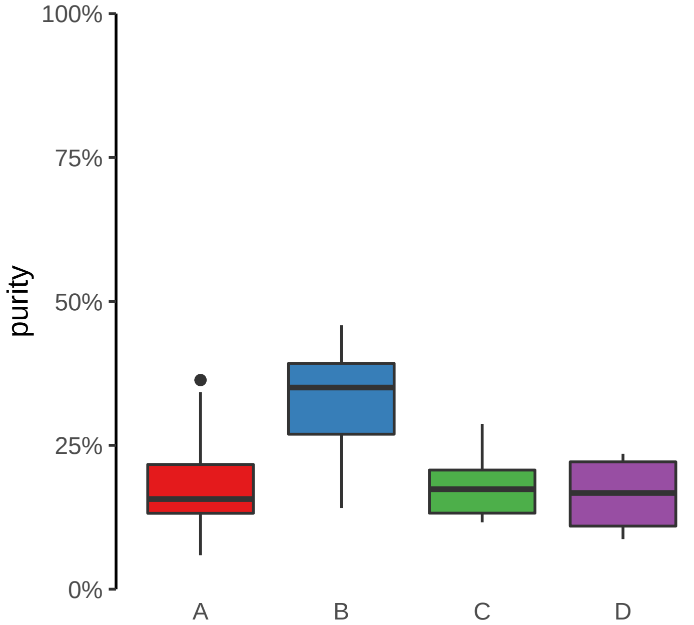

**ZNF431 (GH19J020999) enhancer loss**
**chr19:20999605−21000105**

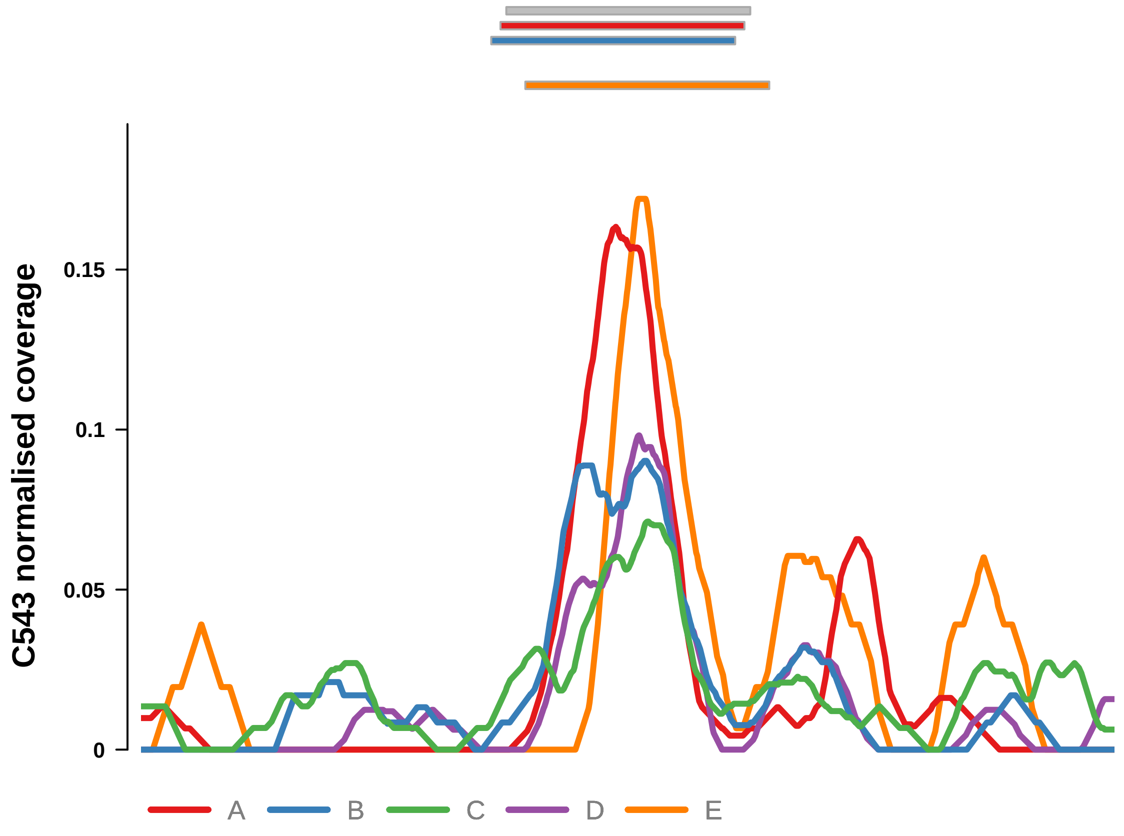


Putatively Clonal (logFC = −12.28)

| | P. | Adjusted P. |
|---|---|---|
| *~region vs. ~* | 1.22e−01 | 5.16e−01 |
| *~purity vs ~* | 1.13e−02 * | 8.29e−02 |
| *~purity+region vs ~purity* | 4.82e−01 | 8.43e−01 |
| *Carcinoma vs. normal* | 3.95e−04 *** | 3.95e−04 *** |

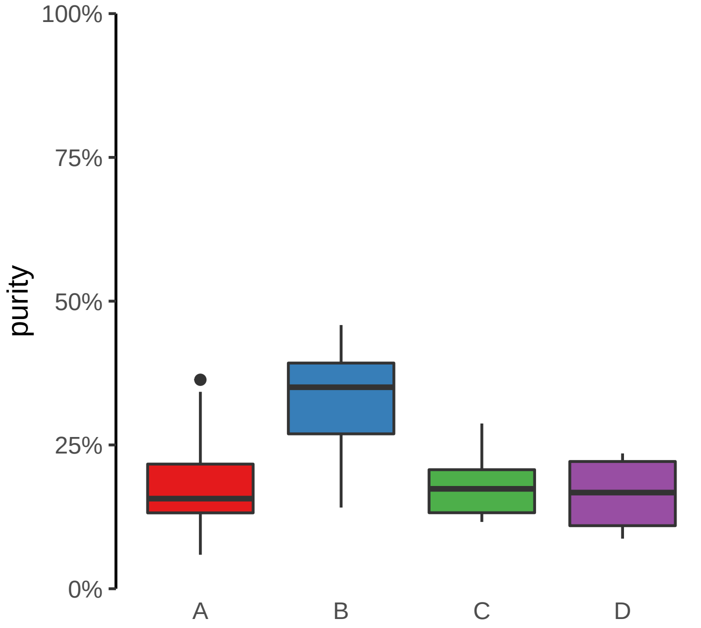

**ANTXR2 (GH04J080057) enhancer loss**
**chr4:80058687−80059187**

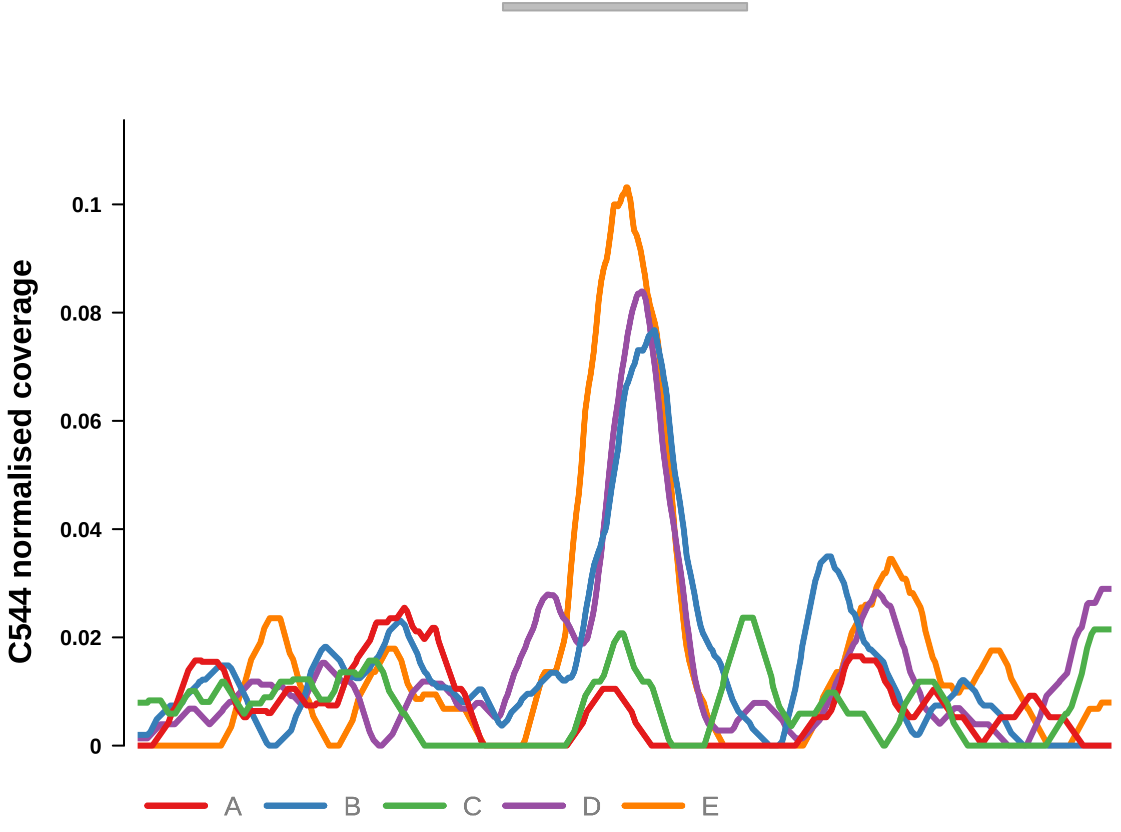


Putatively Clonal (logFC = −3.02)

| | P. | Adjusted P. |
|---|---|---|
| *~region vs. ~* | 2.35e−04 *** | 1.72e−03 ** |
| *~purity vs ~* | 6.42e−05 **** | 1.73e−03 ** |
| *~purity+region vs ~purity* | 1.59e−01 | NA |
| *Carcinoma vs. normal* | 8.6e−04 *** | 8.6e−04 *** |

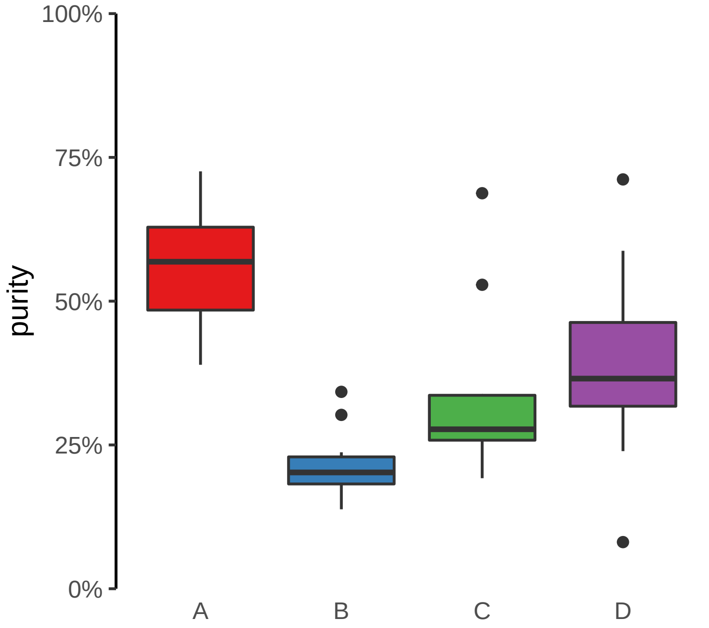

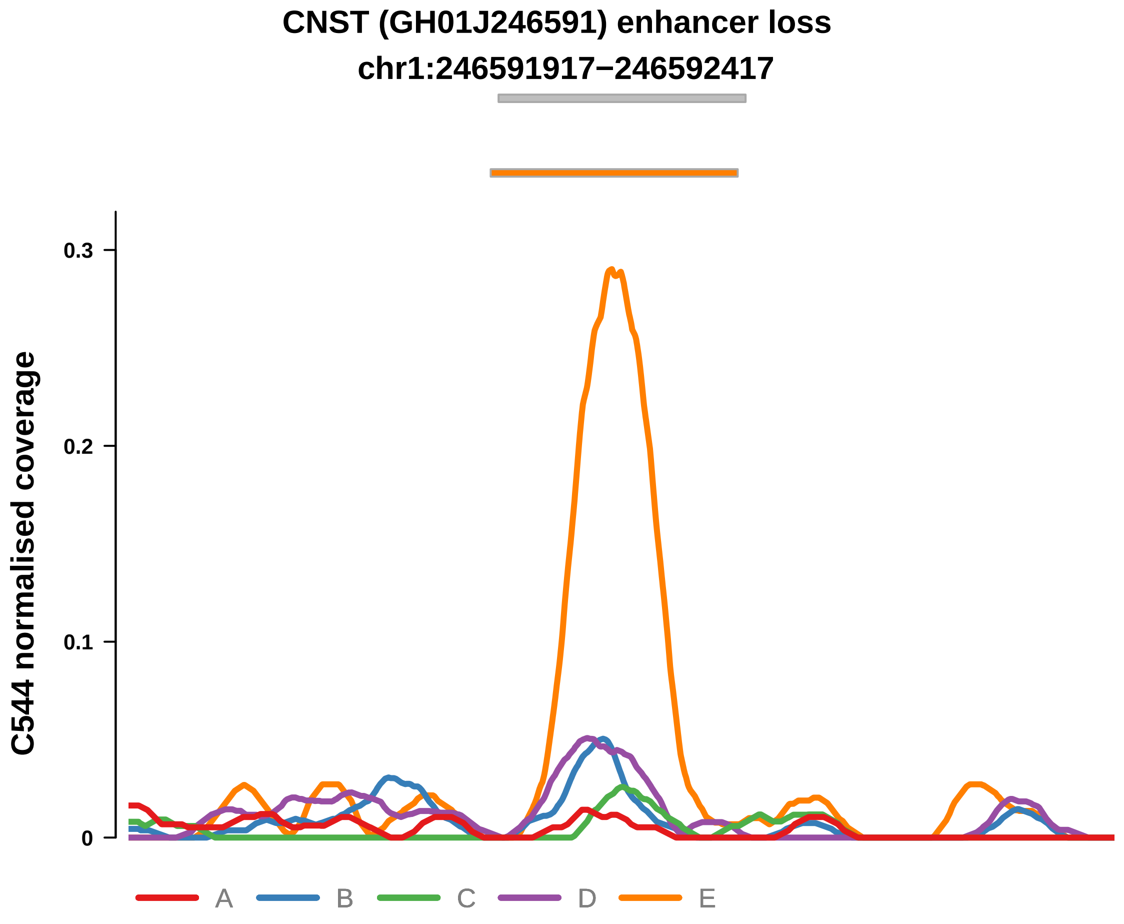


Putatively Clonal (logFC = −2.08)

| | P. | Adjusted P. |
|---|---|---|
| *~region vs. ~* | 3.37e−01 | 4.36e−01 |
| *~purity vs ~* | 5.48e−02 | NA |
| *~purity+region vs ~purity* | 6.18e−01 | NA |
| *Carcinoma vs. normal* | 8.58e−03 ** | 8.58e−03 ** |

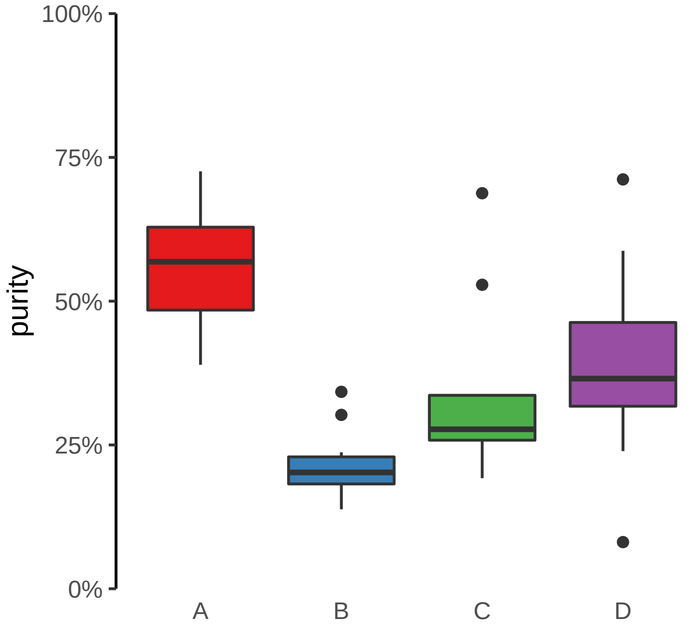

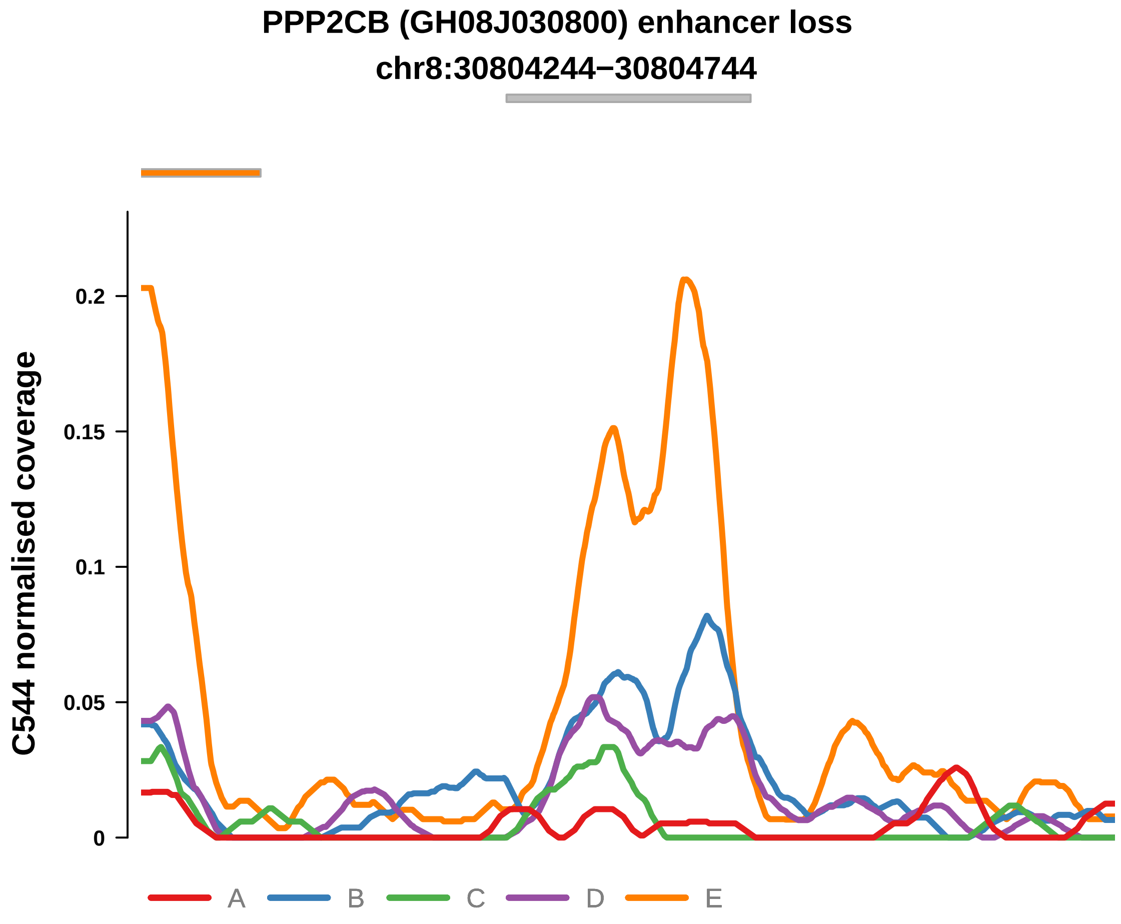


Putatively Clonal (logFC = −2.25)

| | P. | Adjusted P. |
|---|---|---|
| *~region vs. ~* | 6.67e−04 *** | 4.19e−03 ** |
| *~purity vs ~* | 6.19e−03 ** | 3.4e−02 * |
| *~purity+region vs ~purity* | 3.36e−02 * | 9.79e−02 |
| *Carcinoma vs. normal* | 3.11e−03 ** | 3.11e−03 ** |

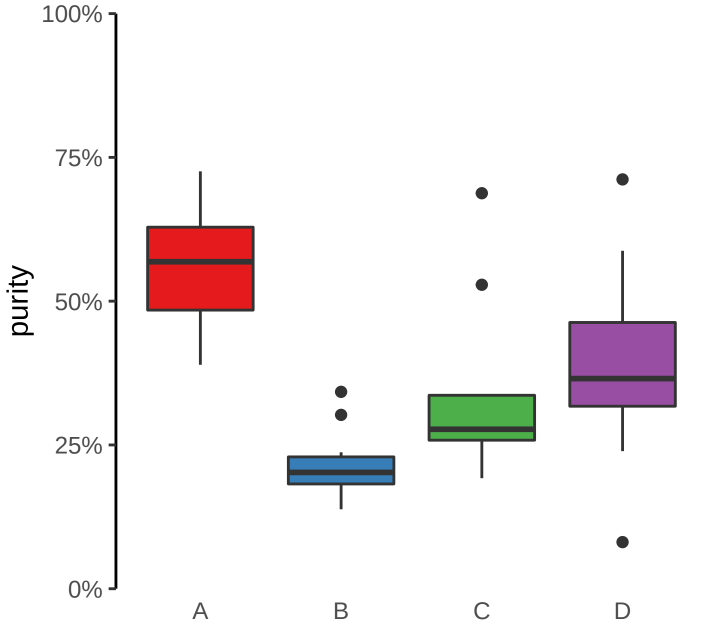

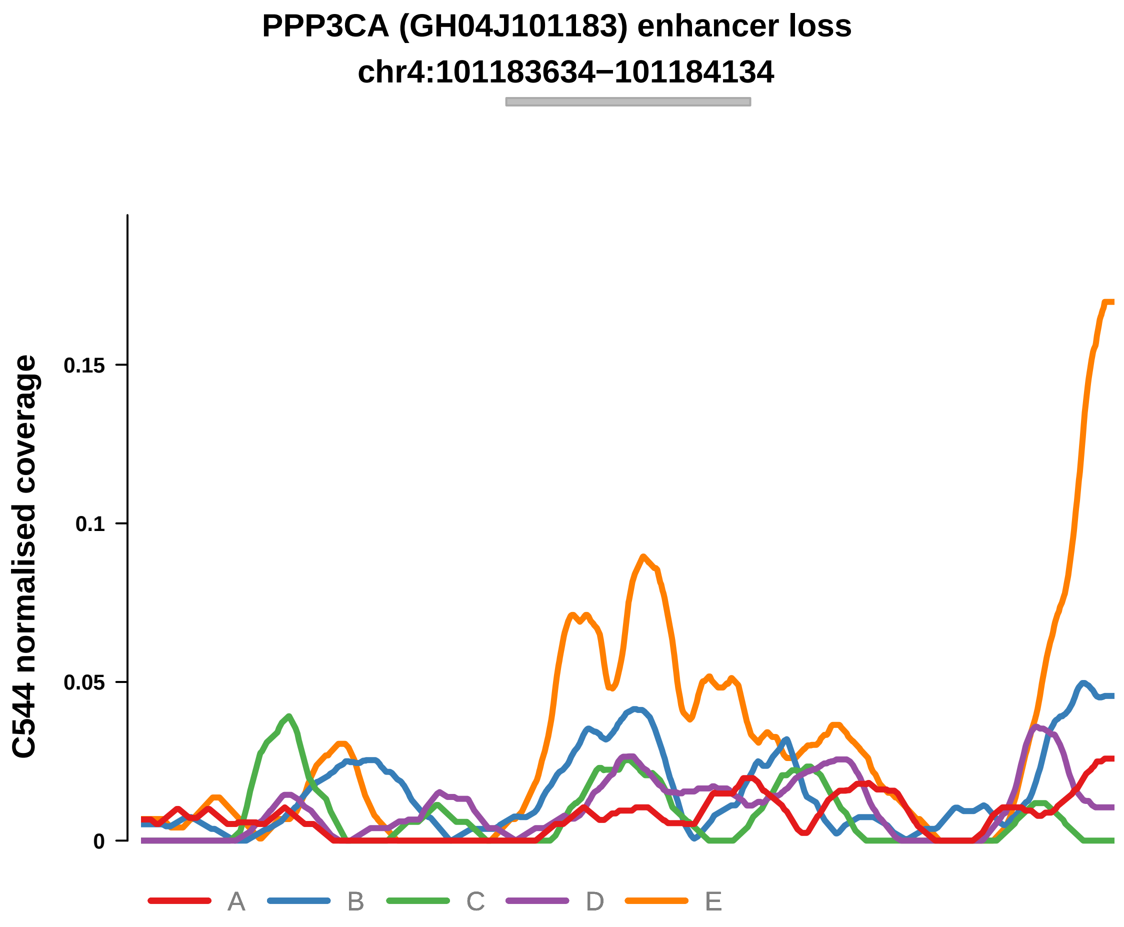


Putatively Clonal (logFC = −2.23)

| | P. | Adjusted P. |
|---|---|---|
| *~region vs. ~* | 7.11e−01 | 7.45e−01 |
| *~purity vs ~* | 2.12e−01 | 3.73e−01 |
| *~purity+region vs ~purity* | 7.26e−01 | NA |
| *Carcinoma vs. normal* | 8.57e−03 ** | 8.57e−03 ** |

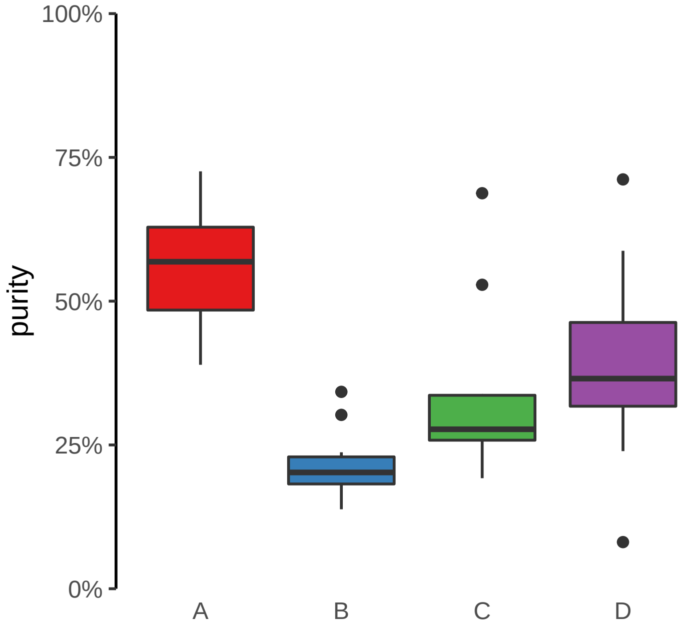

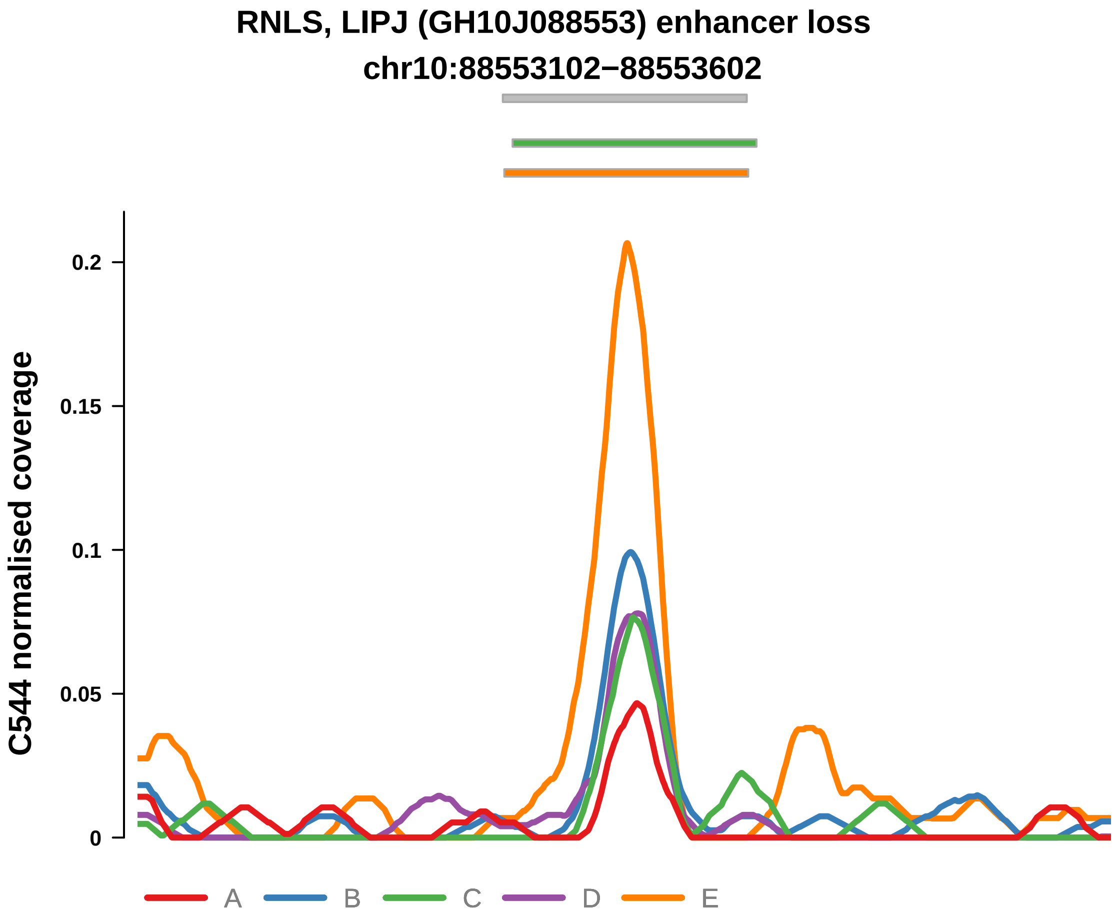


Putatively Clonal (logFC = −2.15)

| | P. | Adjusted P. |
|---|---|---|
| *~region vs. ~* | 4.83e−01 | 5.52e−01 |
| *~purity vs ~* | 1.82e−01 | 3.42e−01 |
| *~purity+region vs ~purity* | 4.4e−01 | NA |
| *Carcinoma vs. normal* | 5.71e−03 ** | 5.71e−03 ** |

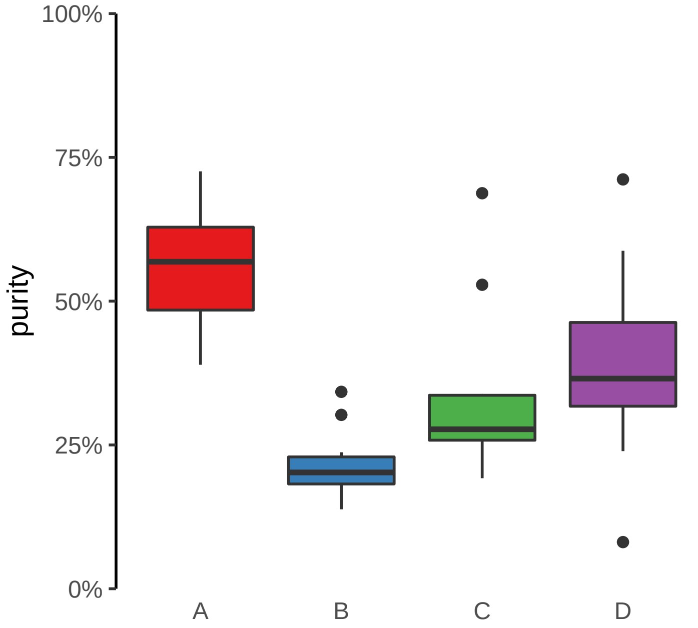

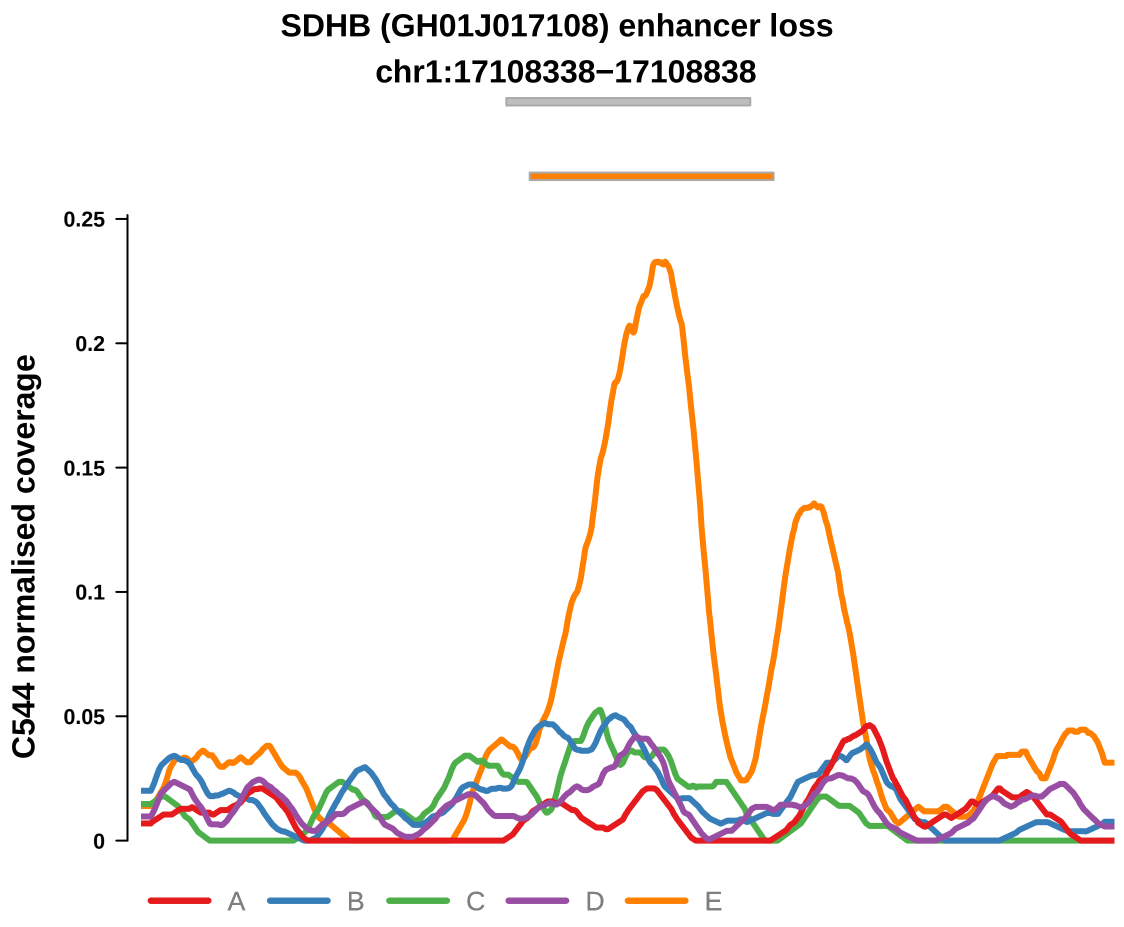


Putatively Clonal (logFC = −2.28)

| | P. | Adjusted P. |
|---|---|---|
| *~region vs. ~* | 1.51e−02 * | 4.42e−02 * |
| *~purity vs ~* | 1.95e−02 * | 7.18e−02 |
| *~purity+region vs ~purity* | 1.39e−01 | NA |
| *Carcinoma vs. normal* | 3.29e−03 ** | 3.29e−03 ** |

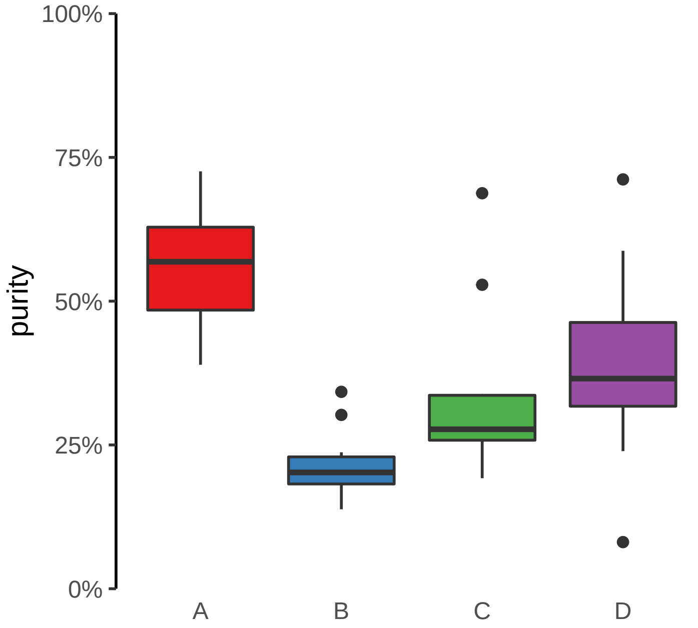

**STAG1, RNU6−789P (GH03J136616) enhancer loss**
**chr3:136617156−136617656**

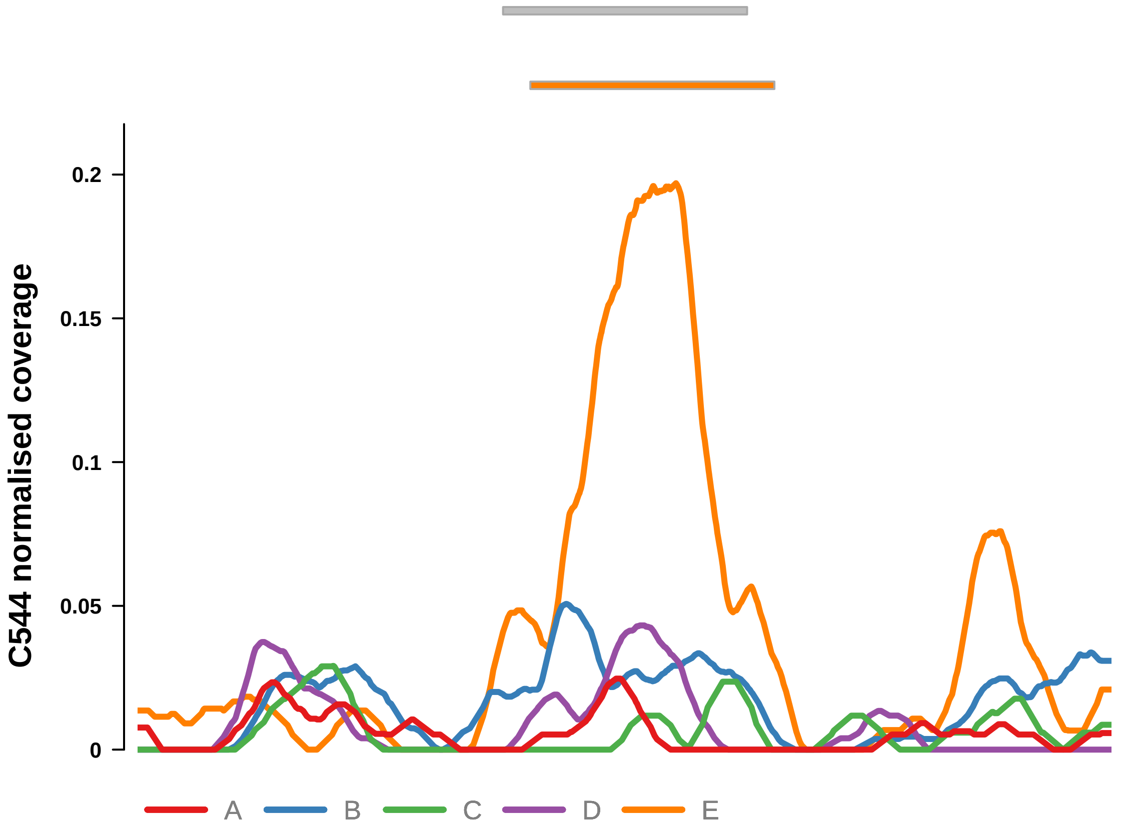


Putatively Clonal (logFC = −3.36)

| | P. | Adjusted P. |
|---|---|---|
| *~region vs. ~* | 5.38e−03 ** | 2.04e−02 * |
| *~purity vs ~* | 2.81e−03 ** | 2.16e−02 * |
| *~purity+region vs ~purity* | 2.45e−01 | NA |
| *Carcinoma vs. normal* | 6.66e−05 **** | 6.66e−05 **** |

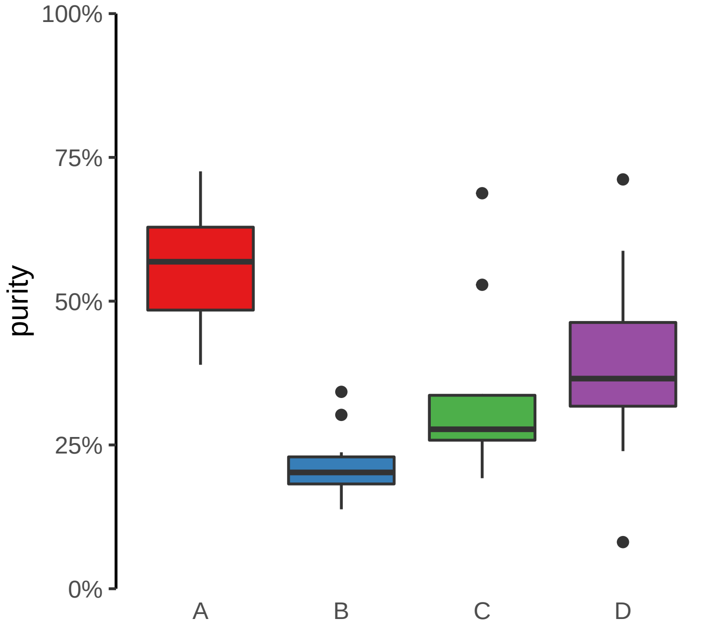

**ACAP2, PPP1R2 (GH03J195537) enhancer loss**
**chr3:195537917−195538417**

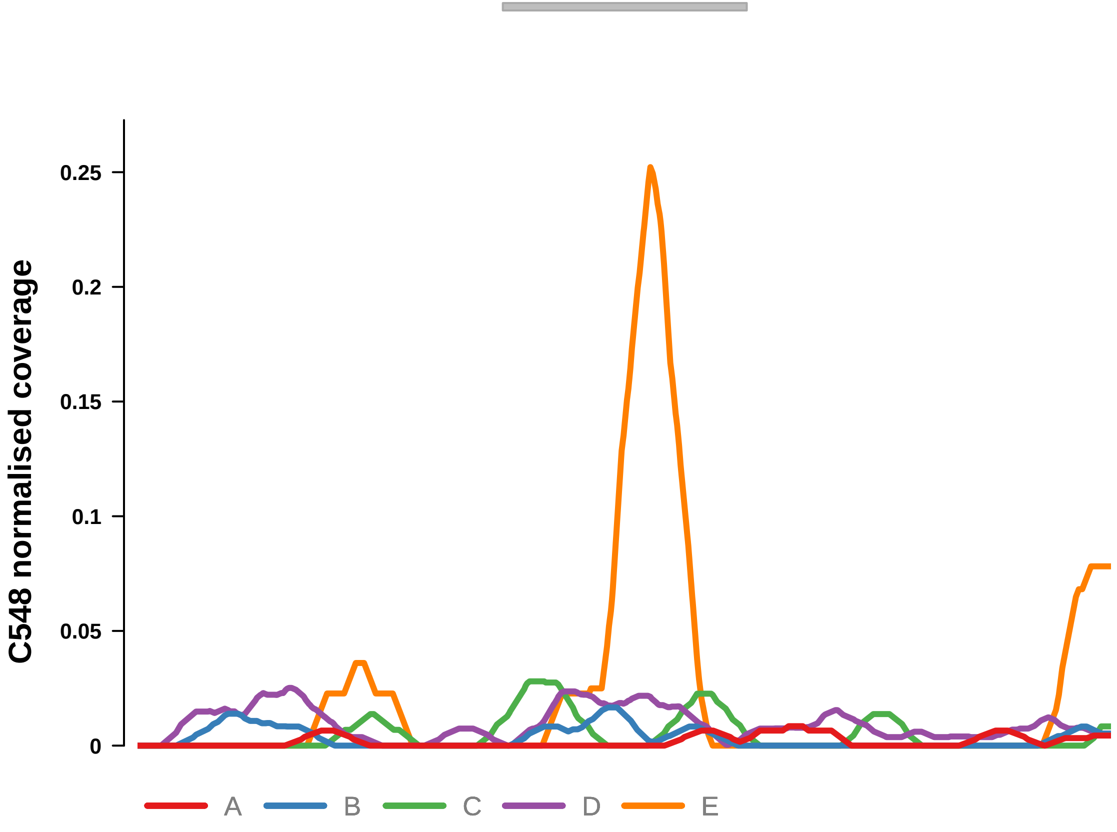


Putatively Clonal (logFC = −10.39)

| | P. | Adjusted P. |
|---|---|---|
| *~region vs. ~* | 1.93e−01 | 2.97e−01 |
| *~purity vs ~* | 2.78e−02 * | 1.44e−01 |
| *~purity+region vs ~purity* | 6.54e−01 | 8.22e−01 |
| *Carcinoma vs. normal* | 2.53e−04 *** | 2.53e−04 *** |

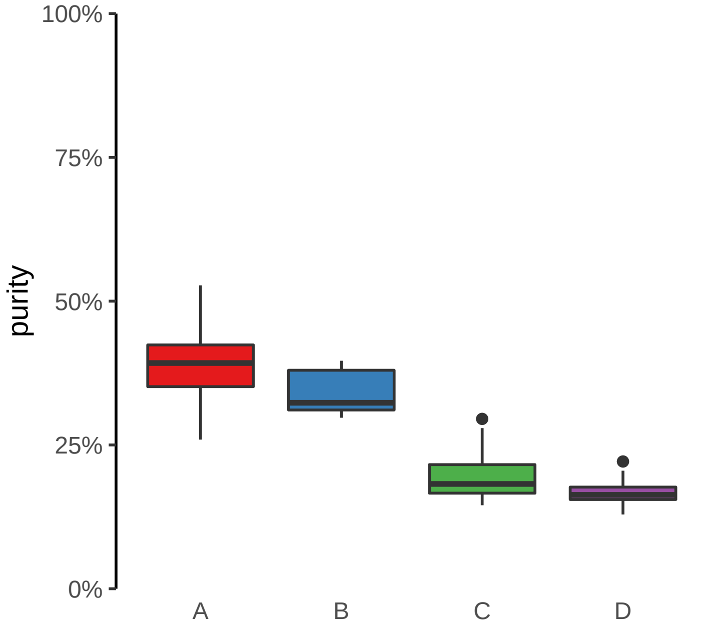

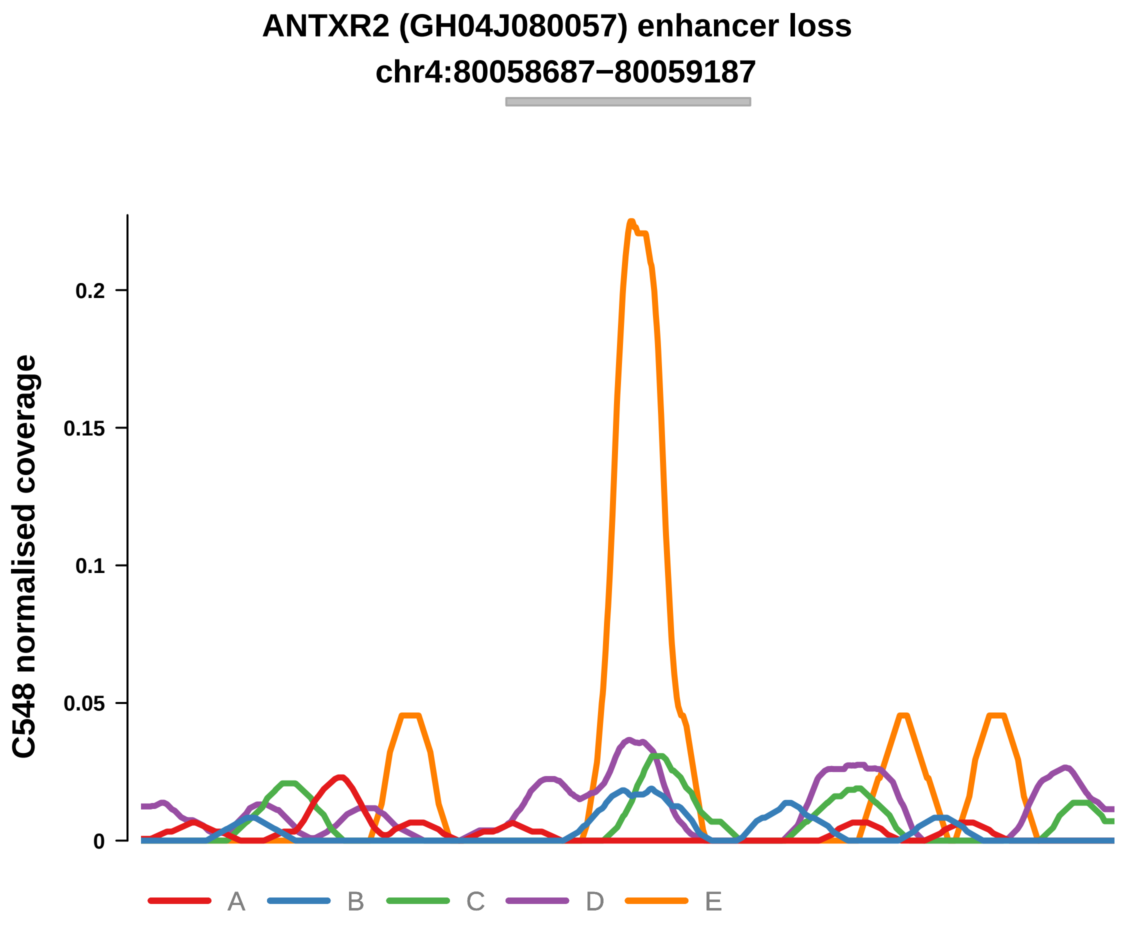


Putatively Clonal (logFC = −3.69)

| | P. | Adjusted P. |
|---|---|---|
| *~region vs. ~* | 4.63e−02 * | 1.46e−01 |
| *~purity vs ~* | 8.51e−03 ** | 7.49e−02 |
| *~purity+region vs ~purity* | 5.33e−01 | 7.33e−01 |
| *Carcinoma vs. normal* | 2.51e−03 ** | 2.51e−03 ** |

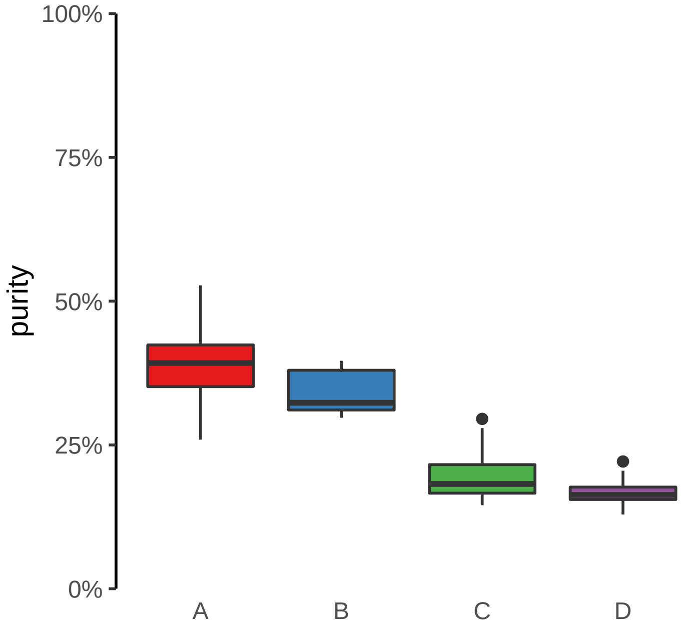

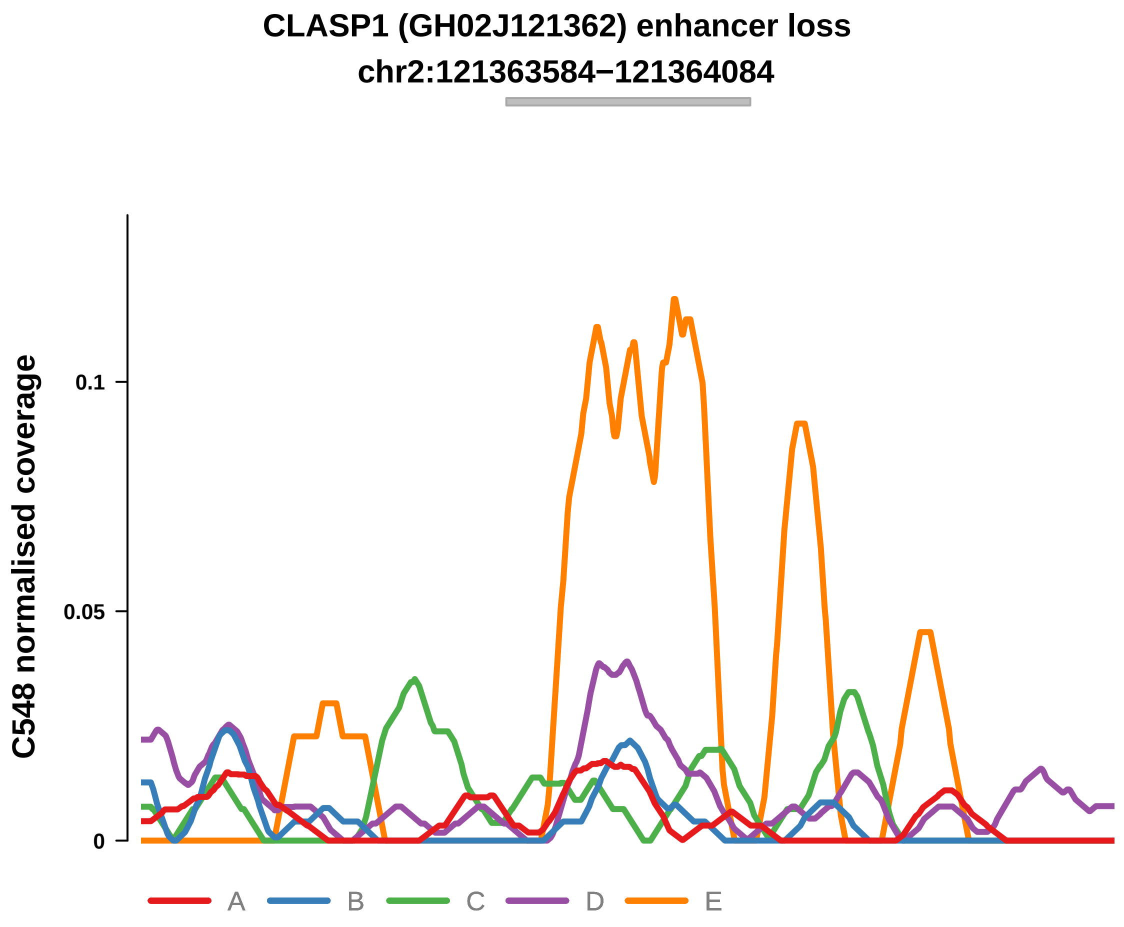


Putatively Clonal (logFC = −2.4)

| | P. | Adjusted P. |
| --- | --- | --- |
| *~region vs. ~* | 1.1e−02 * | 6.03e−02 |
| *~purity vs ~* | 1.52e−02 * | 1.11e−01 |
| *~purity+region vs ~purity* | 2.03e−01 | 4.97e−01 |
| *Carcinoma vs. normal* | 9.16e−03 ** | 9.16e−03 ** |

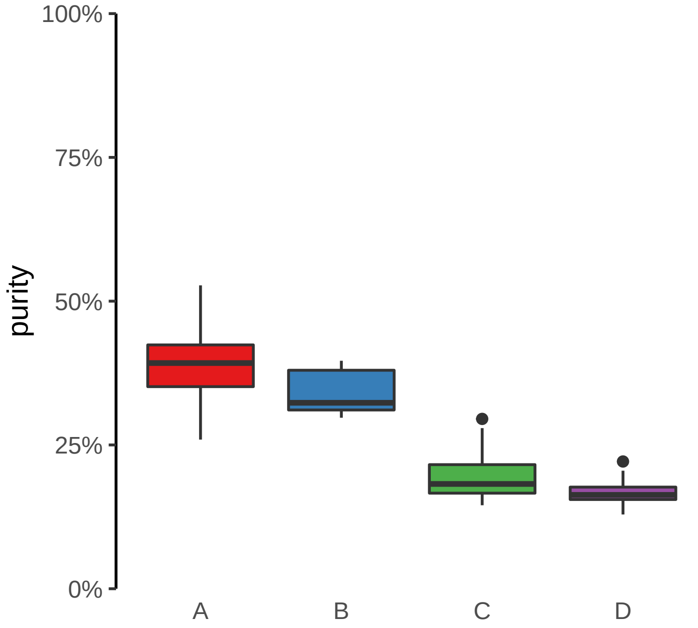

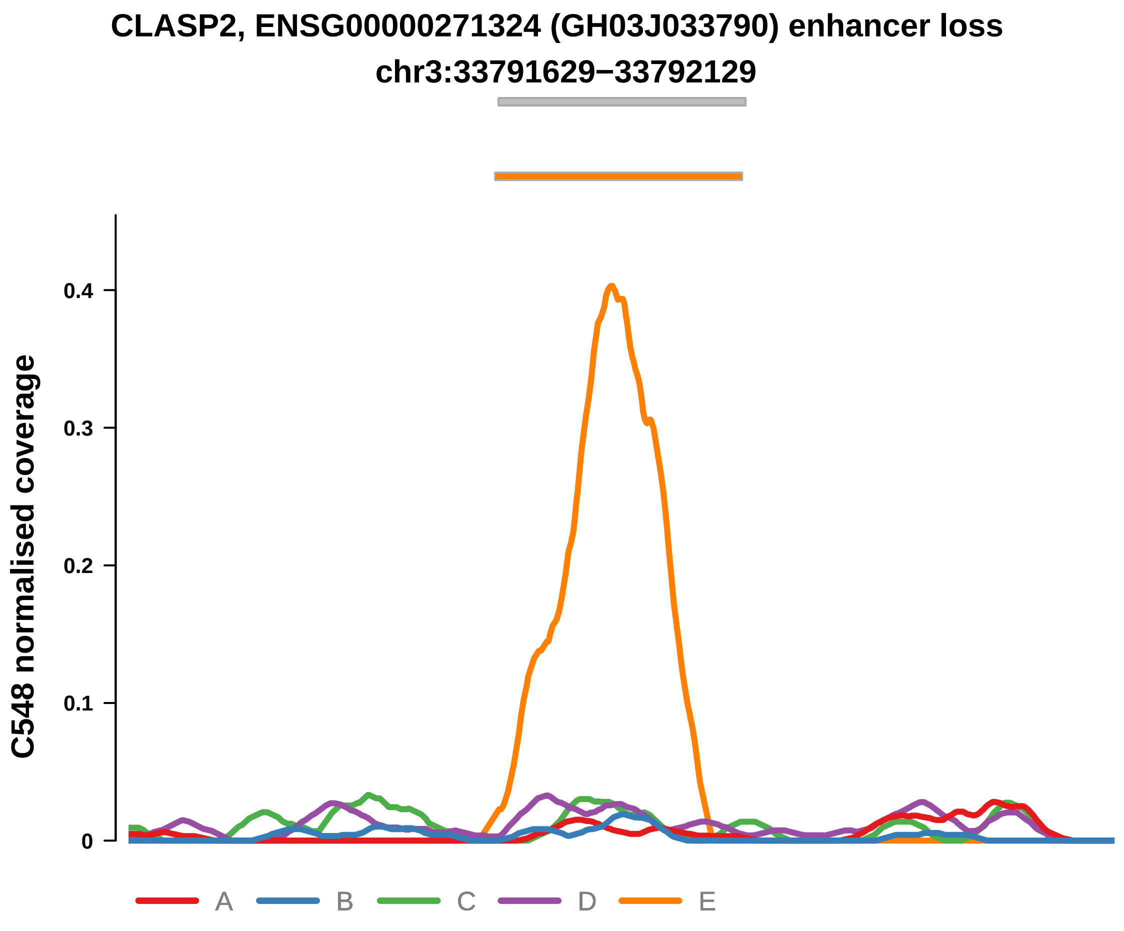


Putatively Clonal (logFC = −2.51)

| | P. | Adjusted P. |
|---|---|---|
| *~region vs. ~* | 8.27e−03 ** | 5.6e−02 |
| *~purity vs ~* | 1.5e−03 ** | 1.88e−02 * |
| *~purity+region vs ~purity* | 8.19e−01 | 9.31e−01 |
| *Carcinoma vs. normal* | 3.31e−03 ** | 3.31e−03 ** |

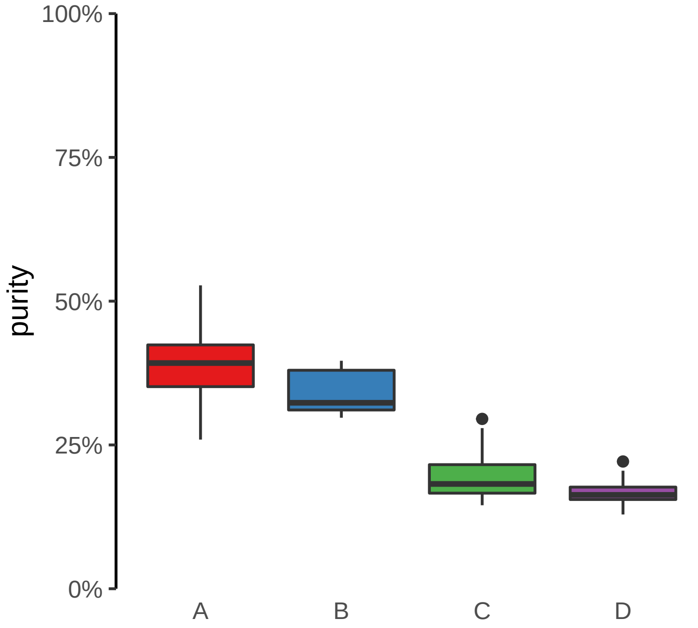

**CNST (GH01J246591) enhancer loss**
**chr1:246591917−246592417**

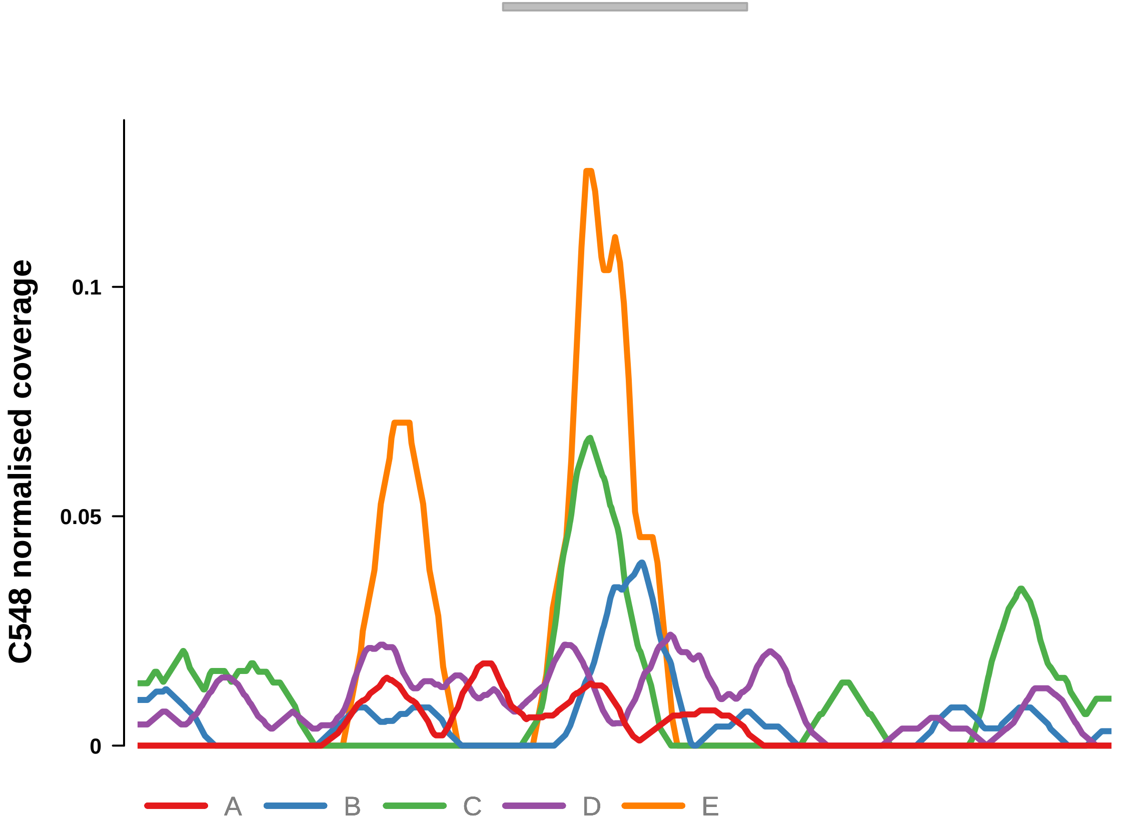


Putatively Clonal (logFC = −3.07)

| | P. | Adjusted P. |
|---|---|---|
| *~region vs. ~* | 1.23e−01 | 2.3e−01 |
| *~purity vs ~* | 1.85e−02 * | 1.16e−01 |
| *~purity+region vs ~purity* | 9.42e−01 | 9.75e−01 |
| *Carcinoma vs. normal* | 3.91e−03 ** | 3.91e−03 ** |

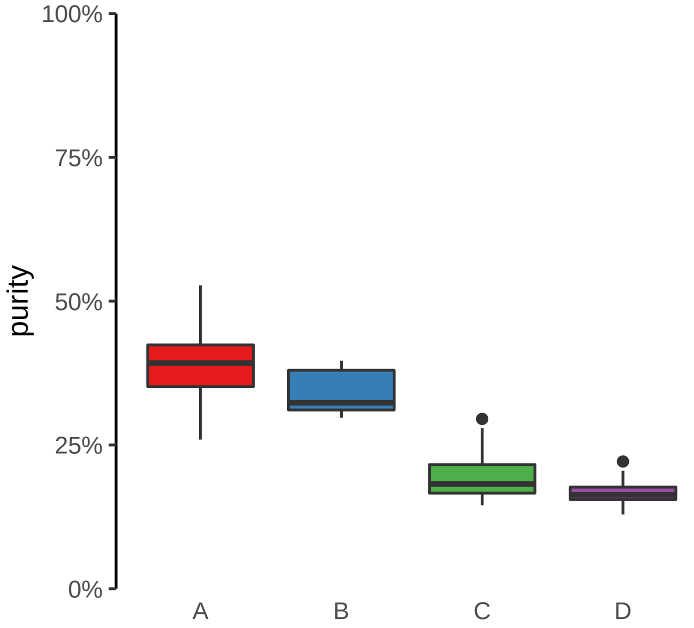

**DNAH7, ENSG00000272211 (GH02J196100) enhancer loss**
**chr2:196101142−196101642**

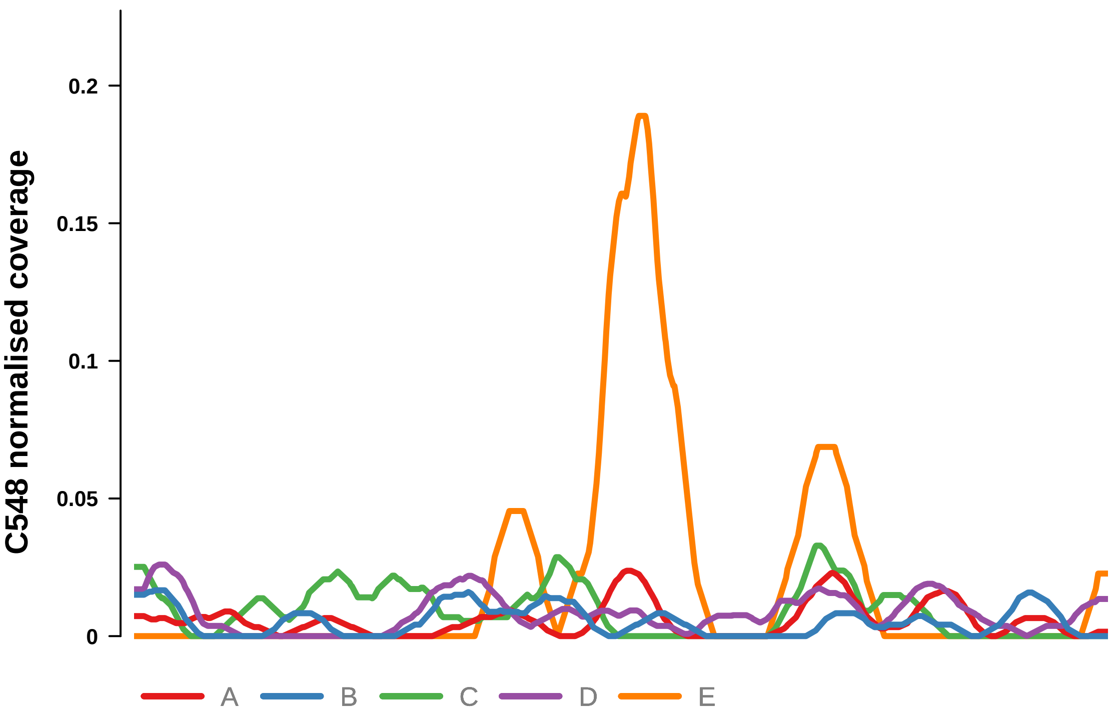


Putatively Clonal (logFC = −2.86)

| | P. | Adjusted P. |
|---|---|---|
| *~region vs. ~* | 8.37e−01 | 9.21e−01 |
| *~purity vs ~* | 2.11e−01 | 3.97e−01 |
| *~purity+region vs ~purity* | 8.46e−01 | 9.31e−01 |
| *Carcinoma vs. normal* | 7.18e−03 ** | 7.18e−03 ** |

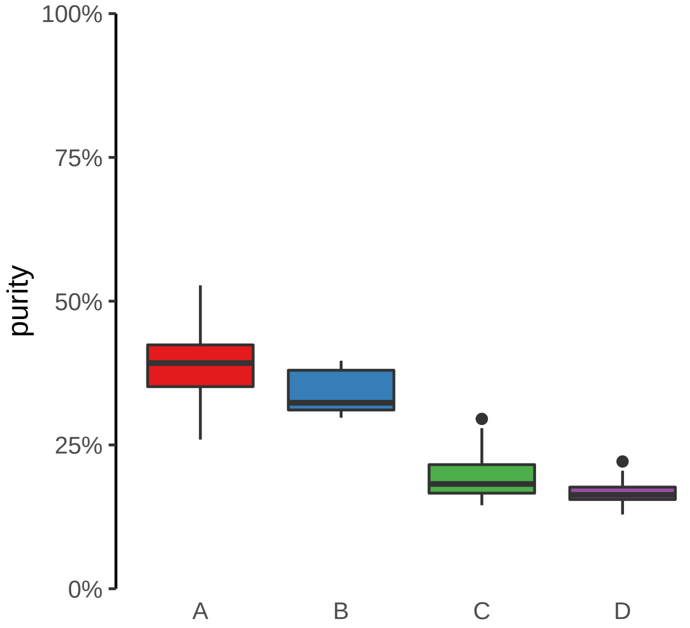

**MAP3K1, LINC01948 (GH05J056411) enhancer loss**
**chr5:56412924−56413424**

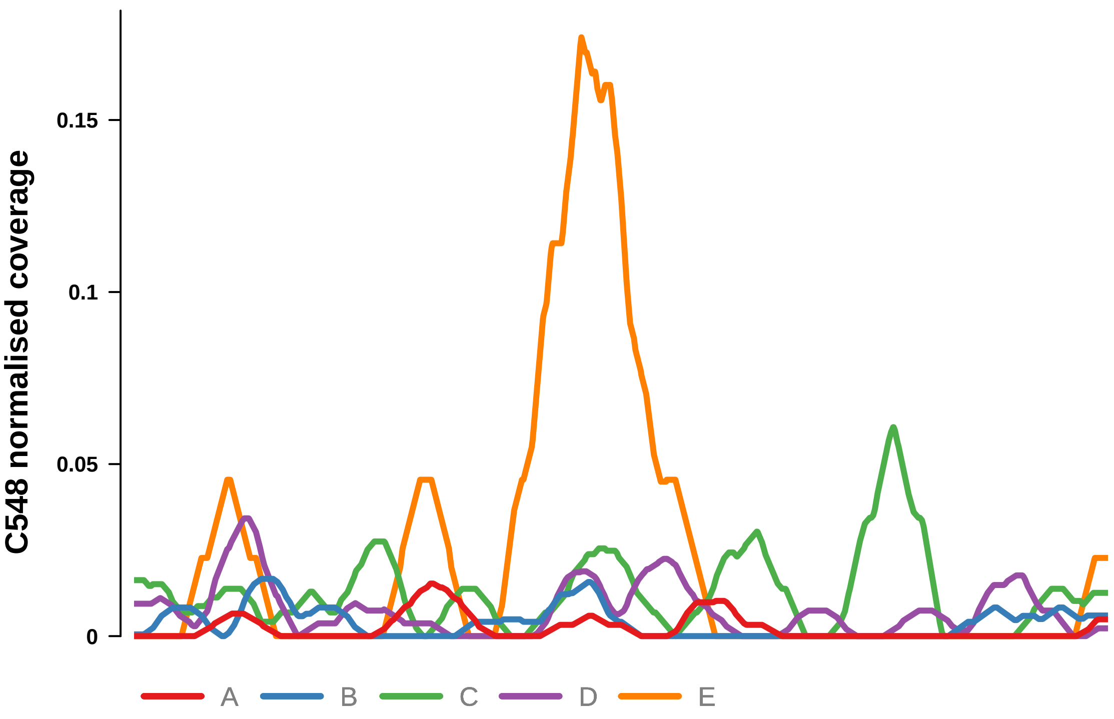


Putatively Clonal (logFC = −10.64)

| | P. | Adjusted P. |
|---|---|---|
| *~region vs. ~* | 1.67e−01 | 2.63e−01 |
| *~purity vs ~* | 2.07e−02 * | 1.21e−01 |
| *~purity+region vs ~purity* | 7.9e−01 | 9.15e−01 |
| *Carcinoma vs. normal* | 7.63e−05 **** | 7.63e−05 **** |

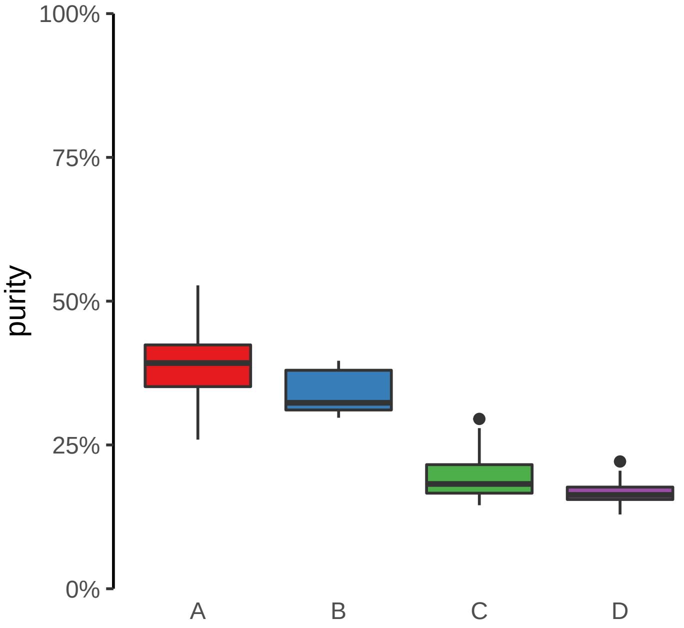

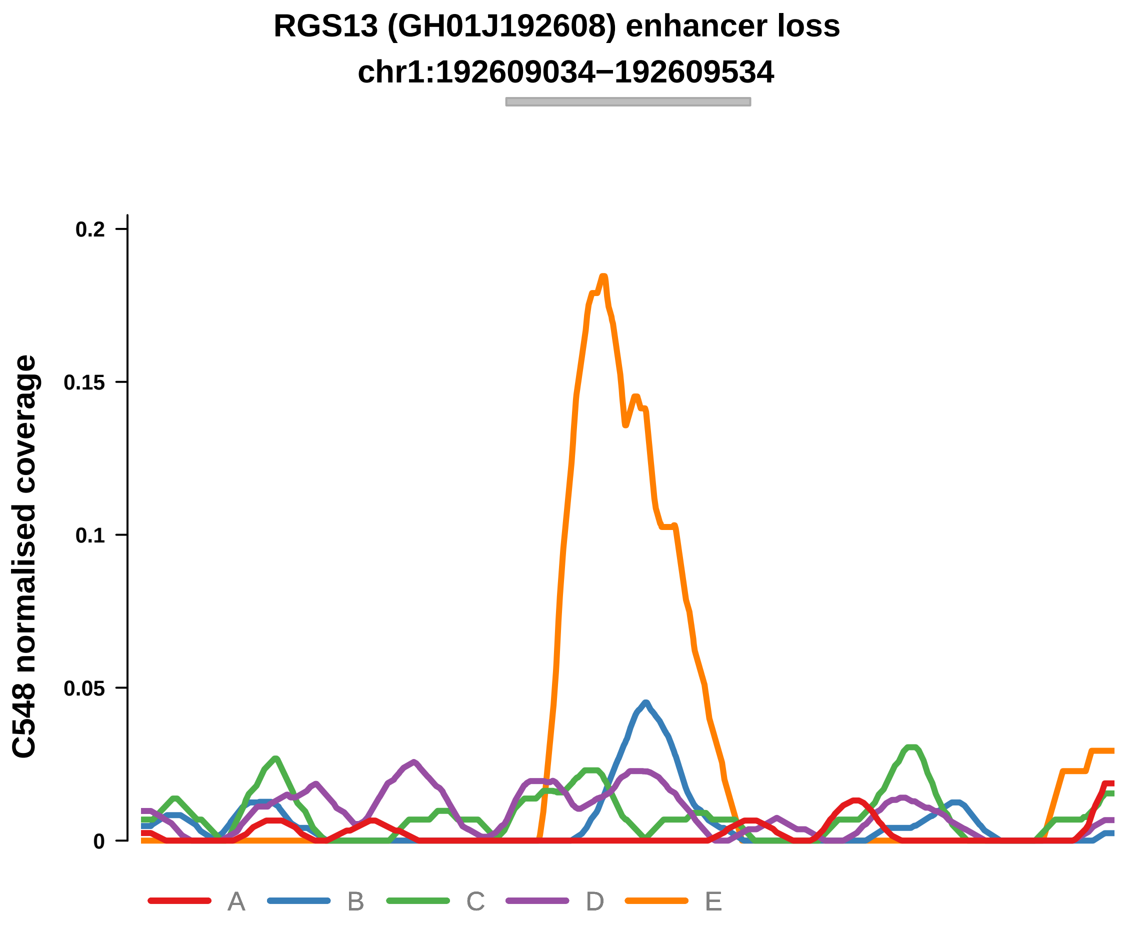


Putatively Clonal (logFC = −4.11)

| | P. | Adjusted P. |
|---|---|---|
| *~region vs. ~* | 2.76e−02 * | 1.16e−01 |
| *~purity vs ~* | 1.19e−01 | 3.07e−01 |
| *~purity+region vs ~purity* | 8.91e−02 | 3.58e−01 |
| *Carcinoma vs. normal* | 5.14e−04 *** | 5.14e−04 *** |

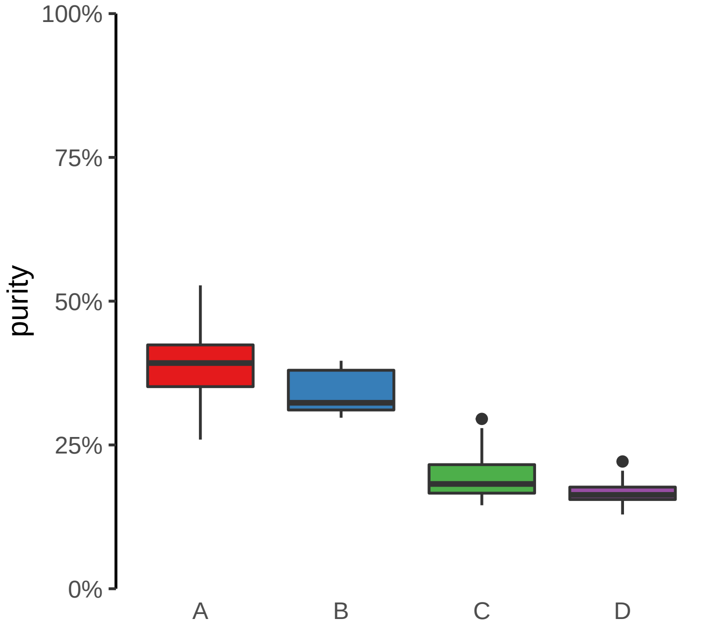

**STAG1, RNU6−789P (GH03J136616) enhancer loss**
**chr3:136617156−136617656**

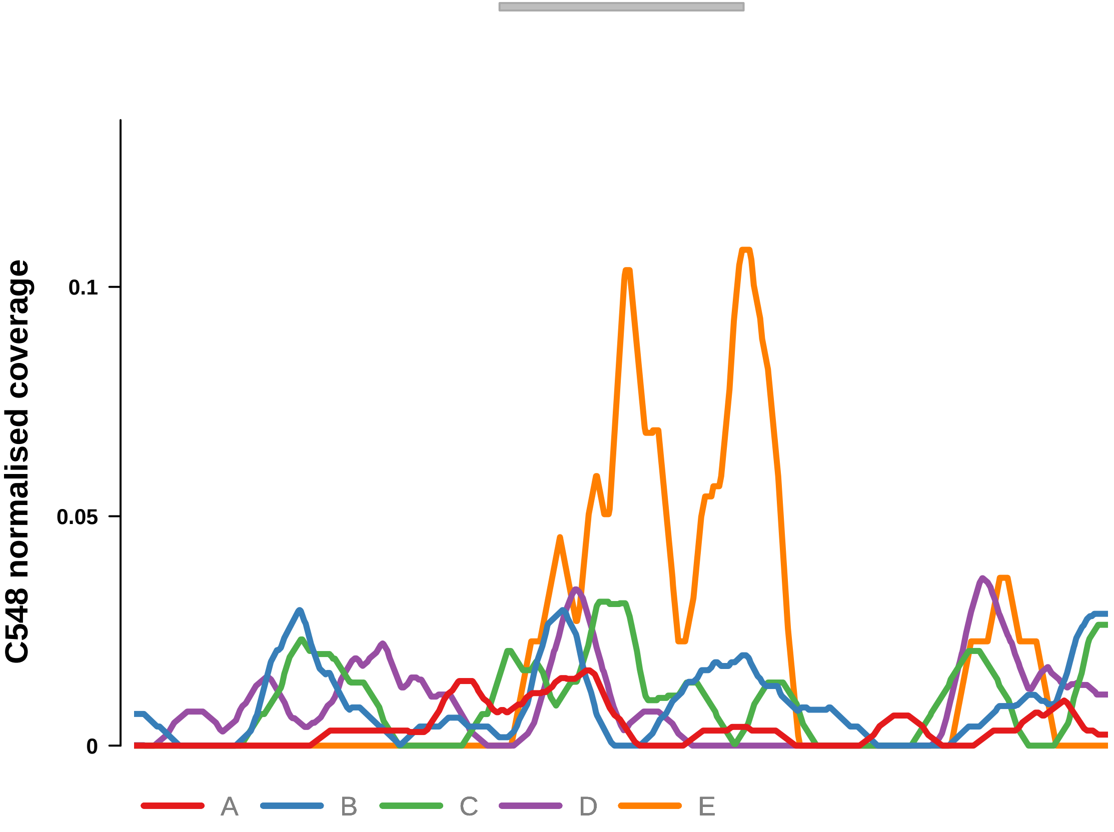


Putatively Clonal (logFC = −2.31)

| | P. | Adjusted P. |
|---|---|---|
| *~region vs. ~* | 6.45e−01 | 7.67e−01 |
| *~purity vs ~* | 7.1e−01 | 8.11e−01 |
| *~purity+region vs ~purity* | 3.4e−01 | 5.86e−01 |
| *Carcinoma vs. normal* | 8.87e−03 ** | 8.87e−03 ** |

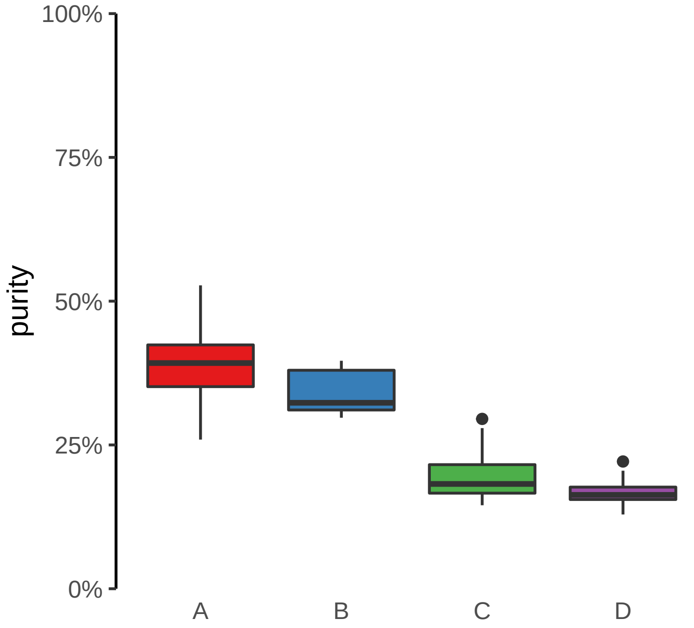

**ENSG00000228035, NGF (GH01J115322) enhancer loss**
**chr1:115323237−115323737**

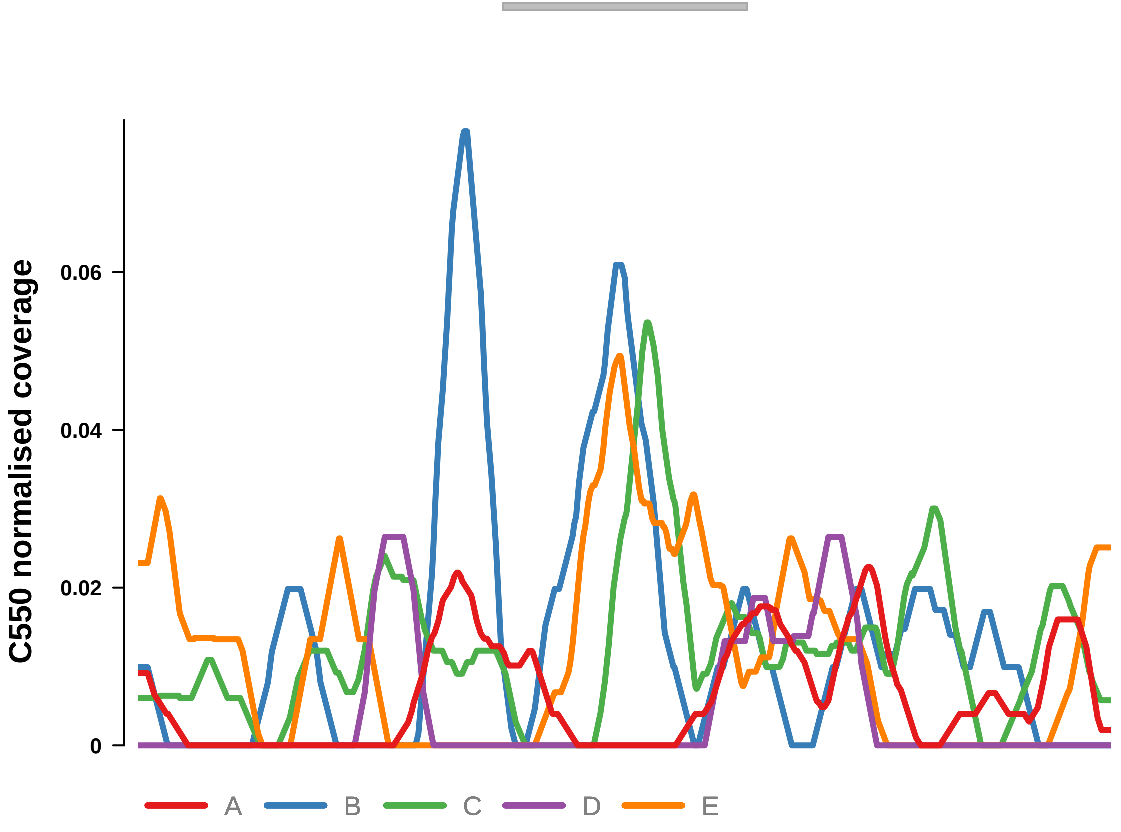


Putatively Clonal (logFC = −2.62)

| | P. | Adjusted P. |
|---|---|---|
| *~region vs. ~* | 3.95e−01 | 8.74e−01 |
| *~purity vs ~* | 4.77e−01 | 8.23e−01 |
| *~purity+region vs ~purity* | 5.03e−01 | 9.87e−01 |
| *Carcinoma vs. normal* | 7.04e−03 ** | 7.04e−03 ** |

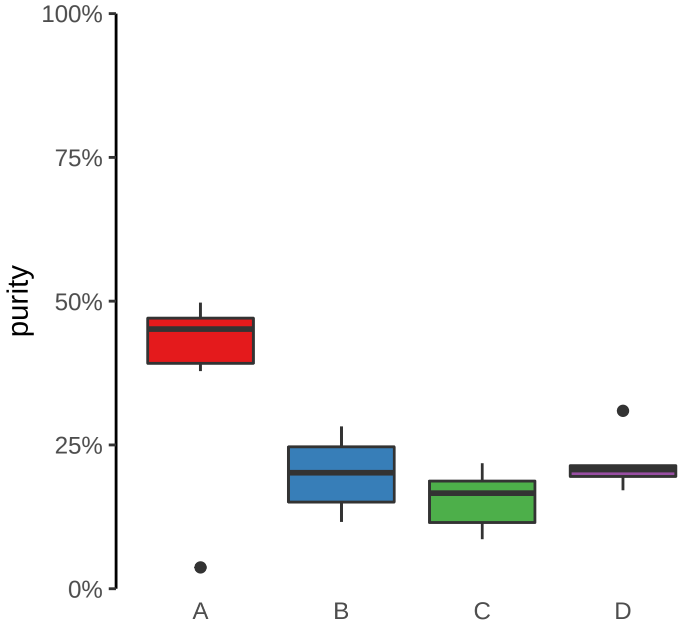

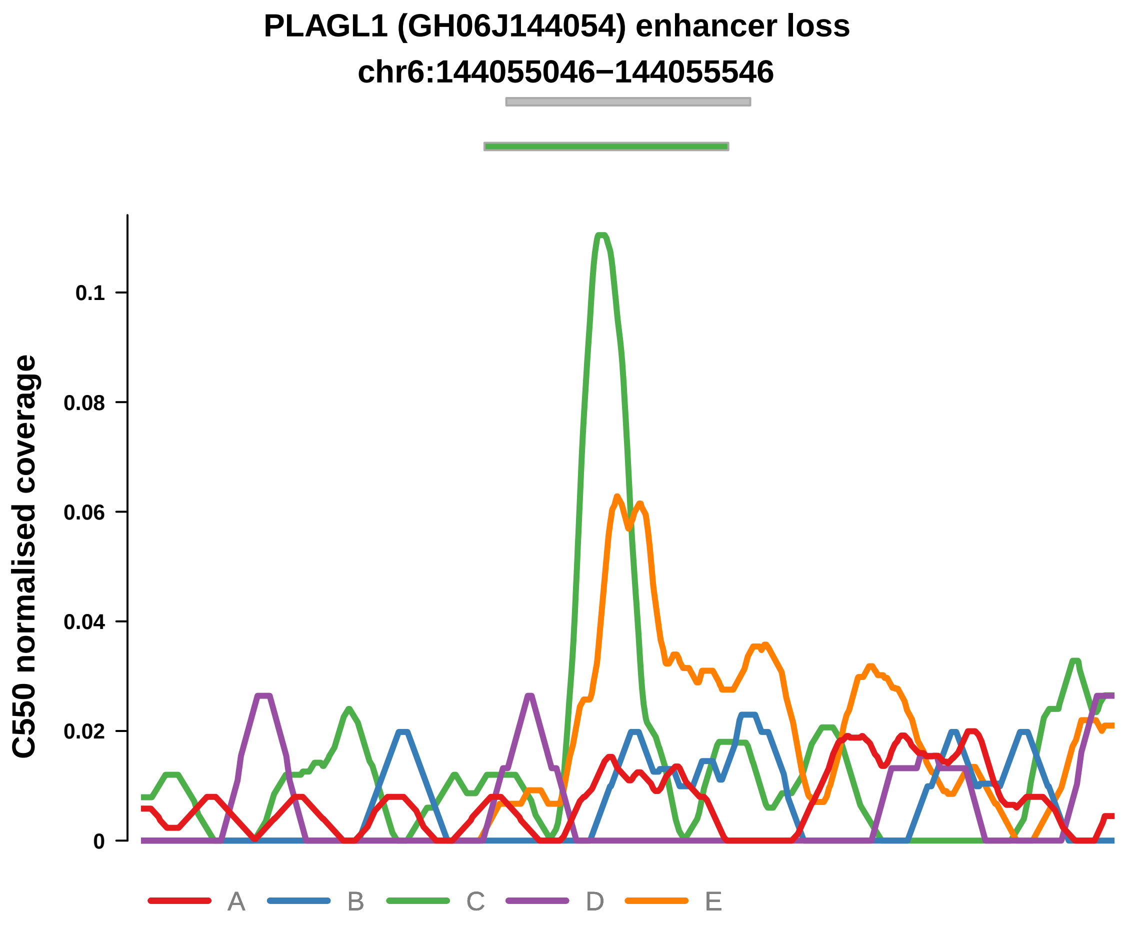


Putatively Clonal (logFC = −10.99)

| | P. | Adjusted P. |
|---|---|---|
| *~region vs. ~* | 1.72e−01 | 7.94e−01 |
| *~purity vs ~* | 4.4e−02 * | 3.57e−01 |
| *~purity+region vs ~purity* | 5.85e−01 | 9.87e−01 |
| *Carcinoma vs. normal* | 3.09e−05 **** | 3.09e−05 **** |

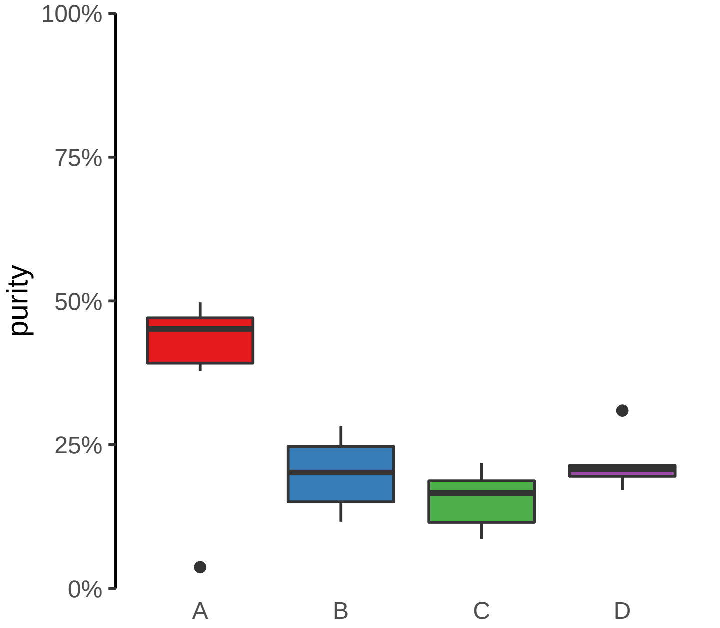

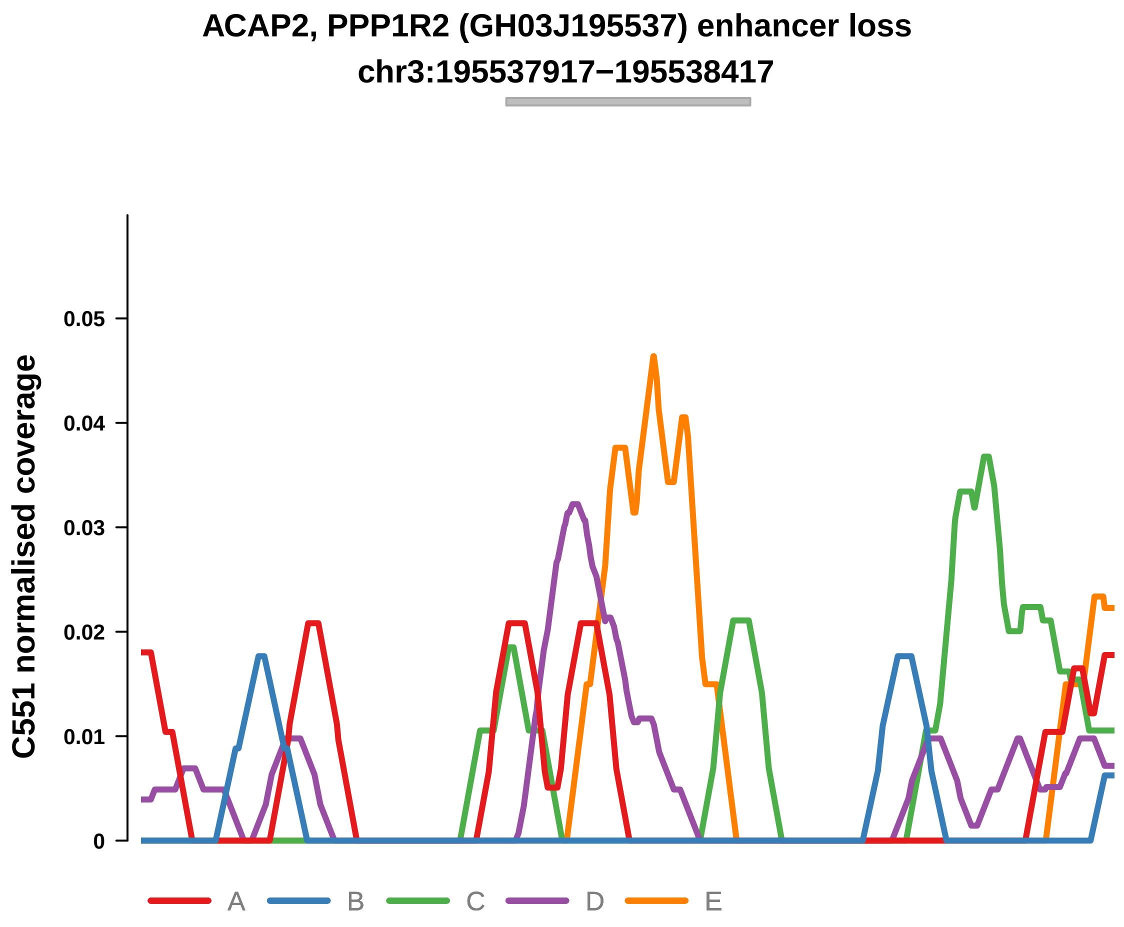


Putatively Clonal (logFC = −1.96)

| | P. | Adjusted P. |
|---|---|---|
| *~region vs. ~* | 5.28e−01 | 8.69e−01 |
| *~purity vs ~* | 6.59e−01 | 9.06e−01 |
| *~purity+region vs ~purity* | 5.92e−01 | 9.48e−01 |
| *Carcinoma vs. normal* | 8.4e−03 ** | 8.4e−03 ** |

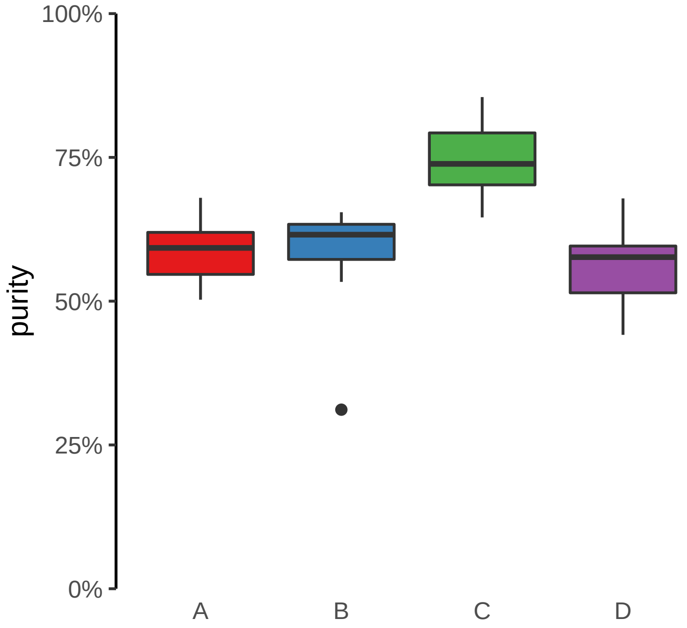

**ANTXR2 (GH04J080057) enhancer loss**
**chr4:80058687−80059187**

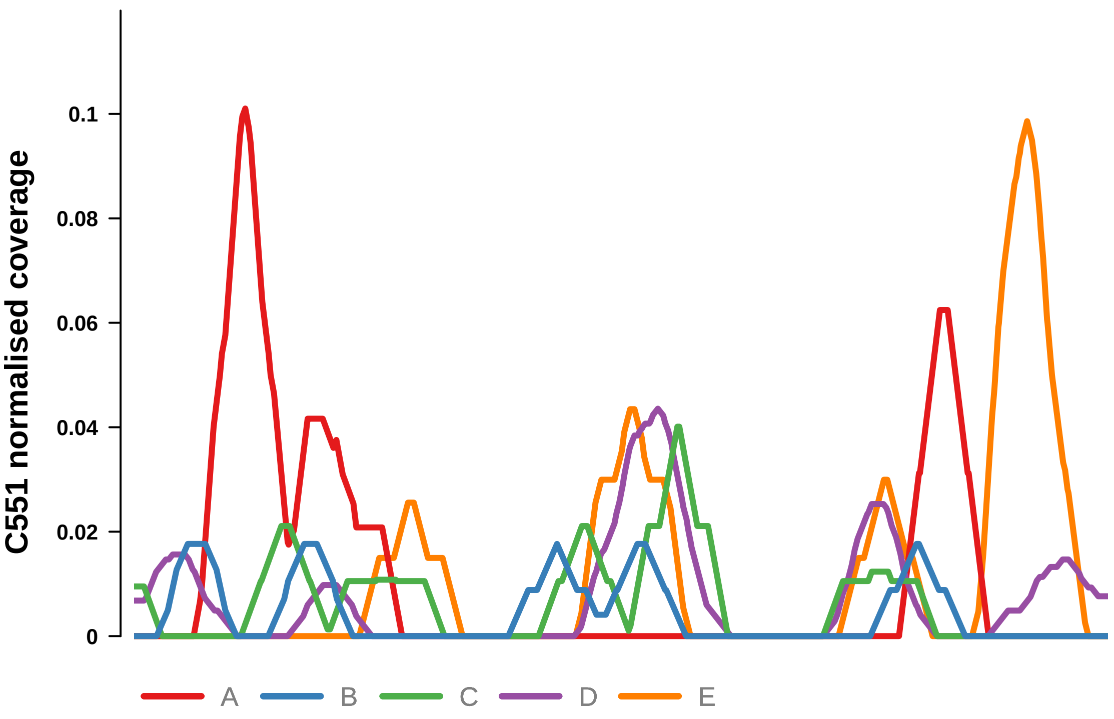


Putatively Clonal (logFC = −1.99)

| | P. | Adjusted P. |
|---|---|---|
| *~region vs. ~* | 5.01e−01 | 8.69e−01 |
| *~purity vs ~* | 8.26e−01 | 9.44e−01 |
| *~purity+region vs ~purity* | 5.55e−01 | 9.48e−01 |
| *Carcinoma vs. normal* | 6.92e−03 ** | 6.92e−03 ** |

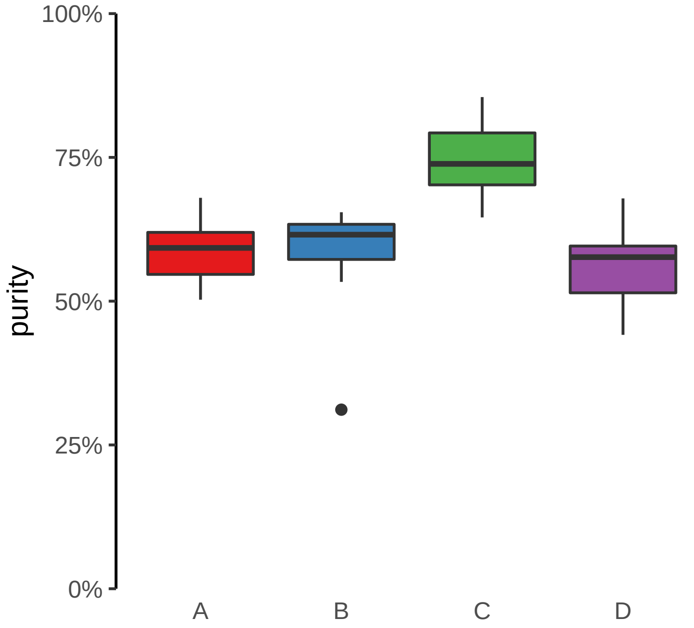

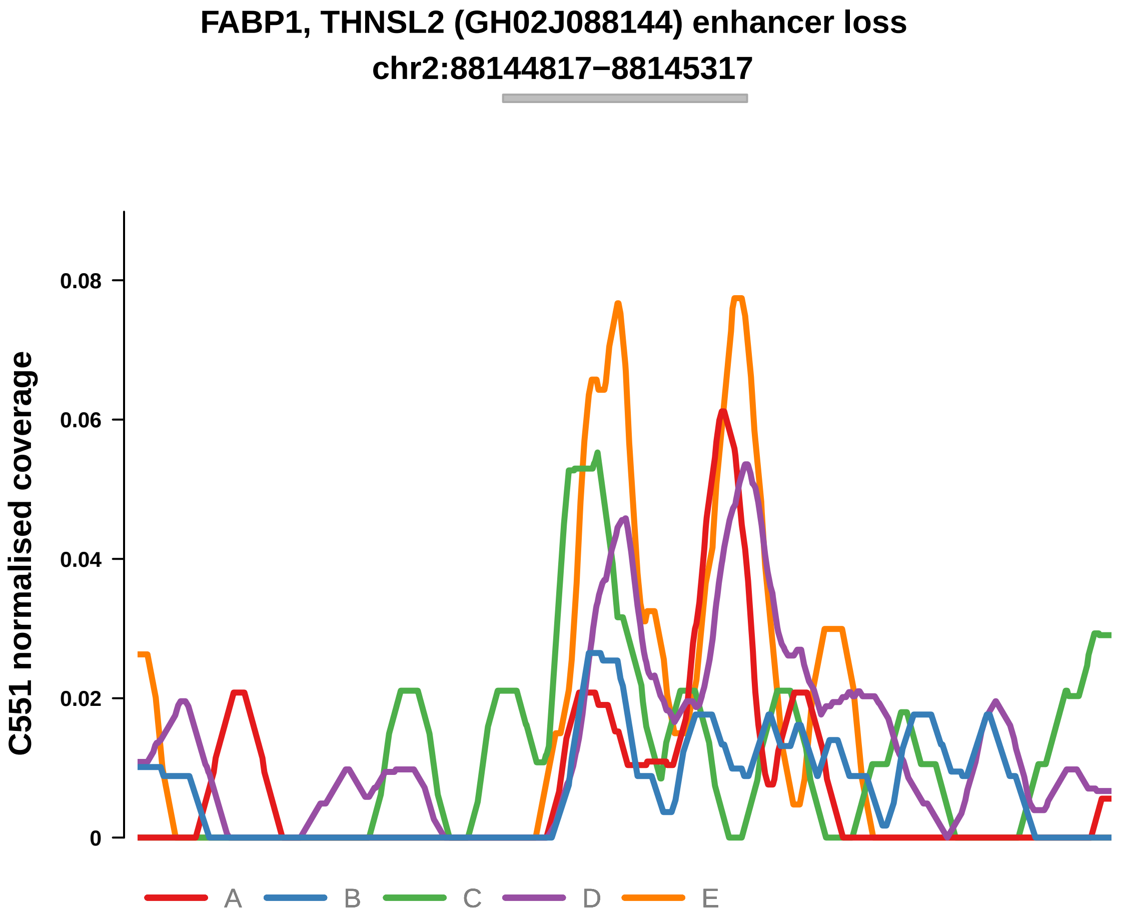


Putatively Clonal (logFC = −2.02)

| | P. | Adjusted P. |
|---|---|---|
| *~region vs. ~* | 9.13e−01 | 9.74e−01 |
| *~purity vs ~* | 8.95e−01 | 9.69e−01 |
| *~purity+region vs ~purity* | 8.85e−01 | 9.99e−01 |
| *Carcinoma vs. normal* | 4.21e−03 ** | 4.21e−03 ** |

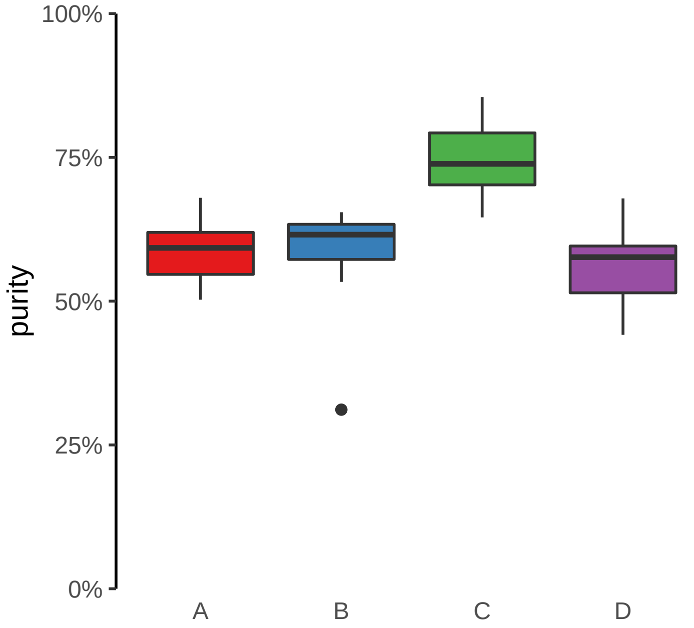

**FAM83G (GH17J019010) enhancer loss**
**chr17:19011073−19011573**

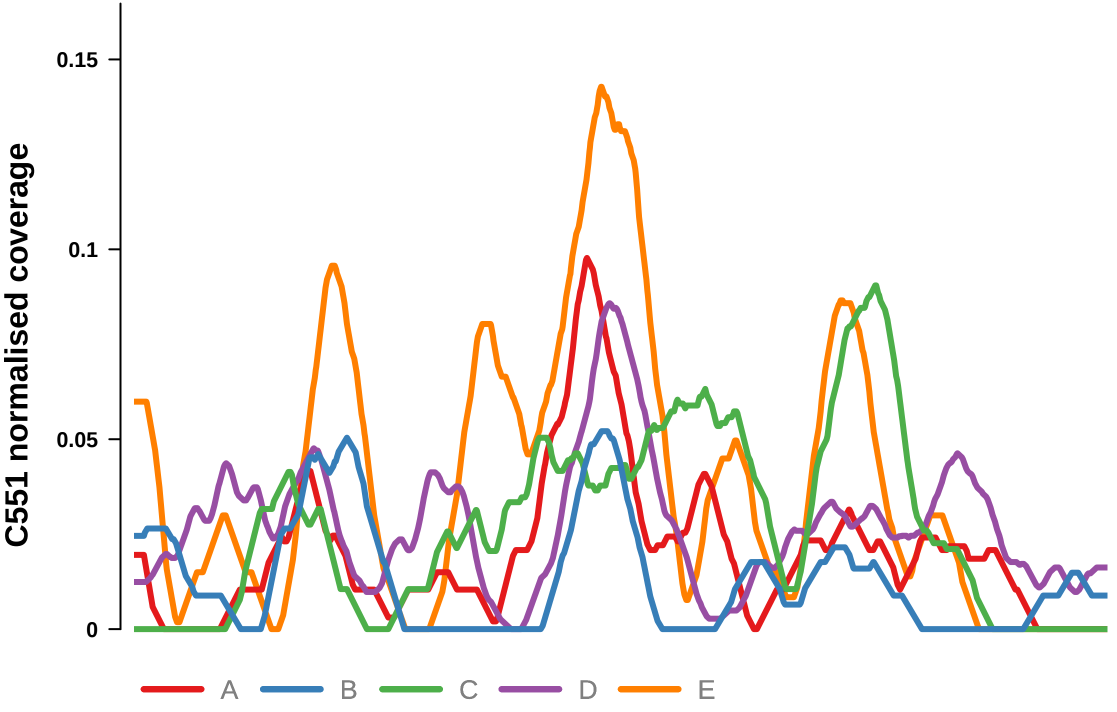


Putatively Clonal (logFC = −1.8)

| | P. | Adjusted P. |
|---|---|---|
| *~region vs. ~* | 6.63e−01 | 9.65e−01 |
| *~purity vs ~* | 2.14e−01 | 8.57e−01 |
| *~purity+region vs ~purity* | 7.08e−01 | 9.78e−01 |
| *Carcinoma vs. normal* | 9.2e−03 ** | 9.2e−03 ** |

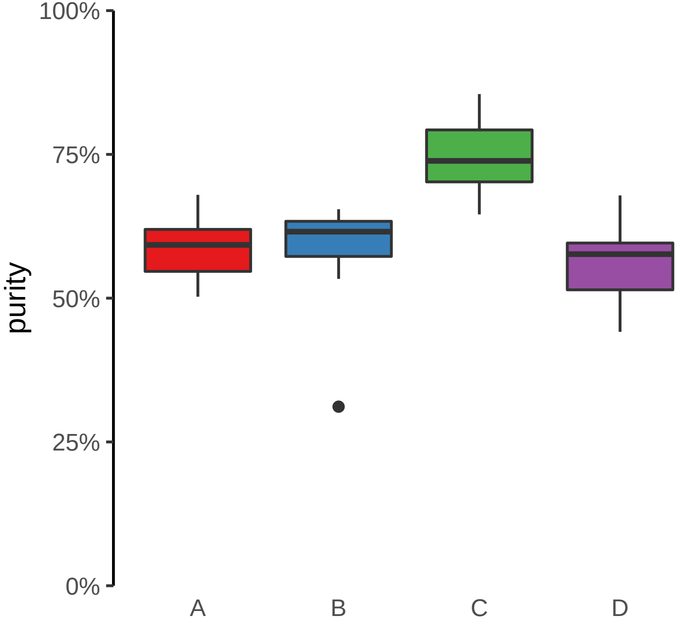

**PLAGL1 (GH06J144054) enhancer loss**
**chr6:144055046−144055546**

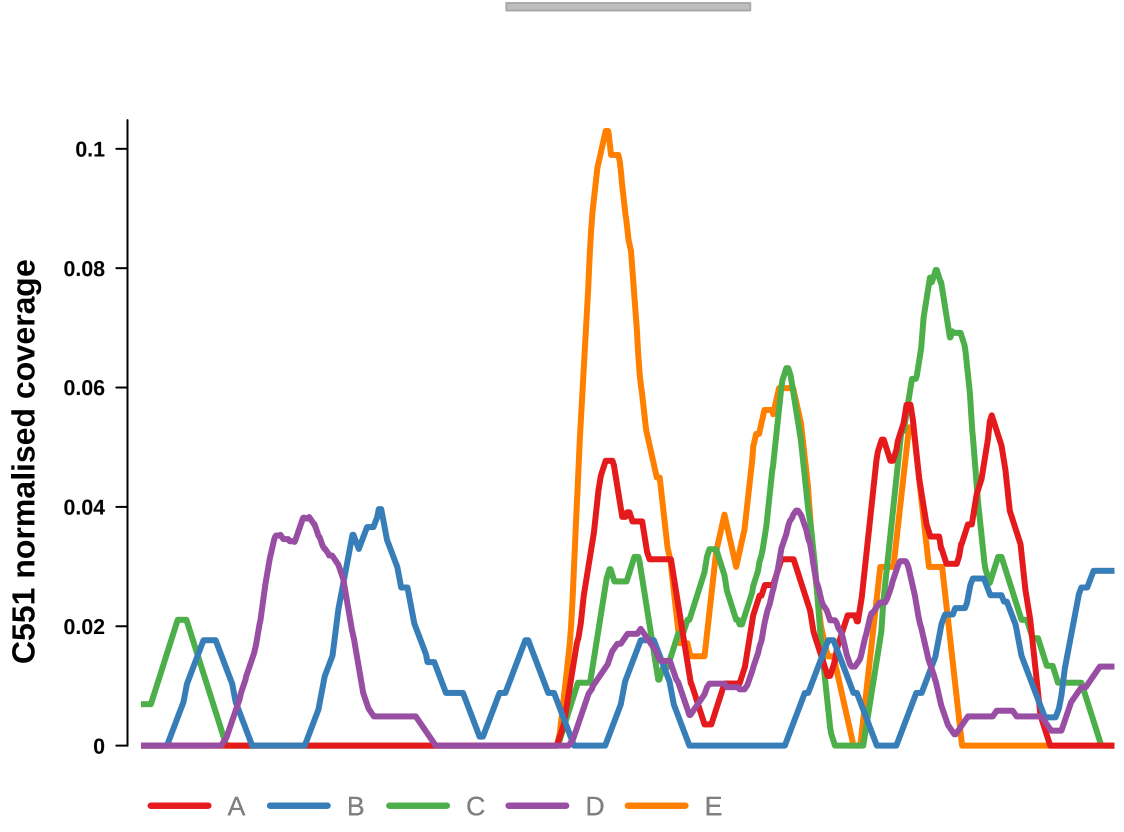


Putatively Clonal (logFC = −1.92)

| | P. | Adjusted P. |
|---|---|---|
| *~region vs. ~* | 8.35e−01 | 9.65e−01 |
| *~purity vs ~* | 8.3e−01 | 9.44e−01 |
| *~purity+region vs ~purity* | 7.22e−01 | 9.78e−01 |
| *Carcinoma vs. normal* | 7.88e−03 ** | 7.88e−03 ** |

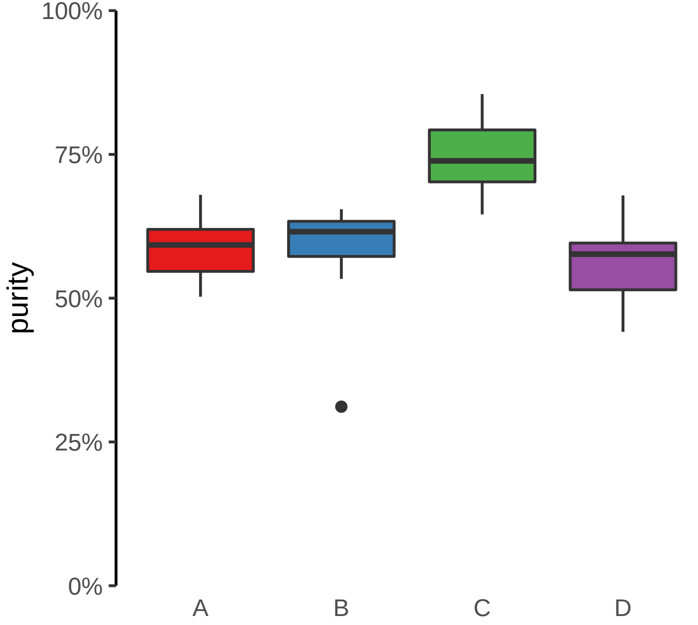

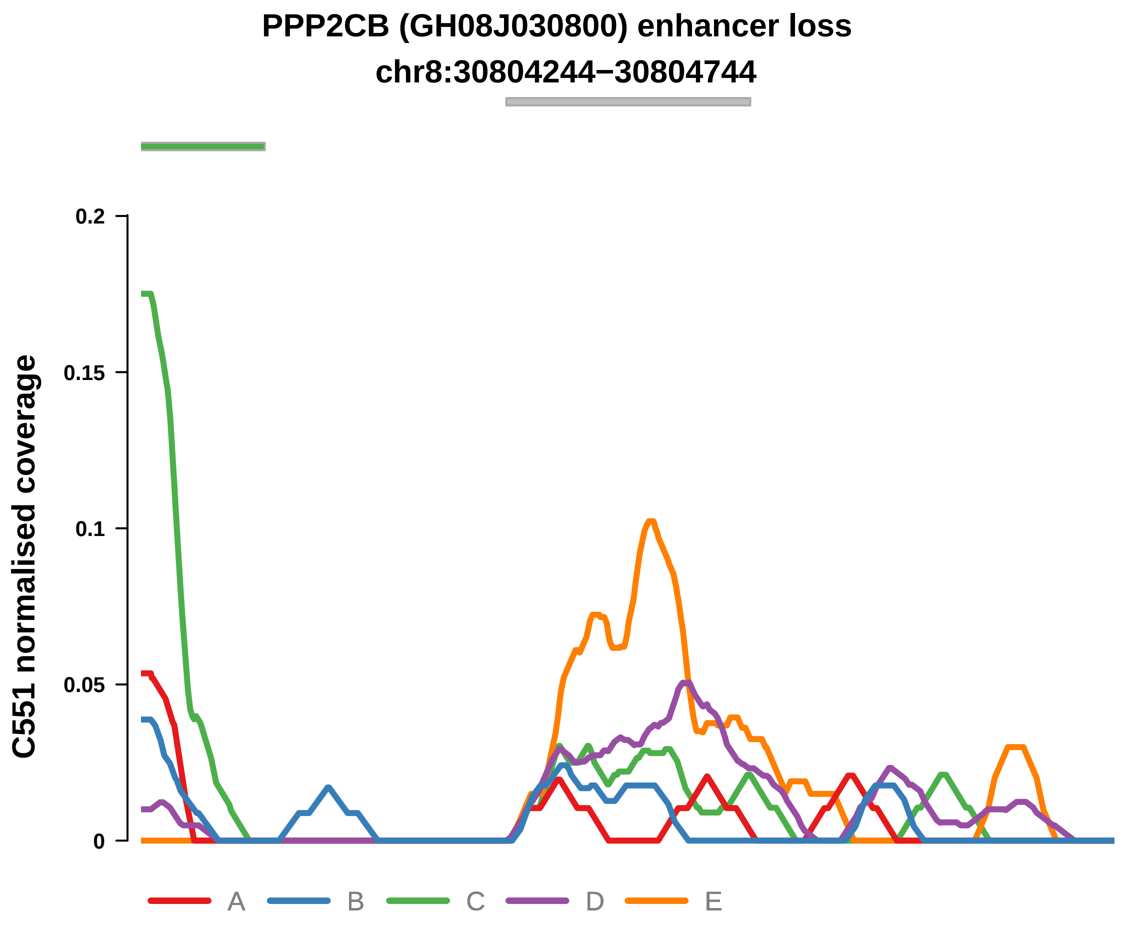


Putatively Clonal (logFC = −2.06)

| | P. | Adjusted P. |
|---|---|---|
| *~region vs. ~* | 2.49e−01 | 6.08e−01 |
| *~purity vs ~* | 8.23e−01 | 9.44e−01 |
| *~purity+region vs ~purity* | 3.22e−01 | 7.87e−01 |
| *Carcinoma vs. normal* | 3.62e−03 ** | 3.62e−03 ** |

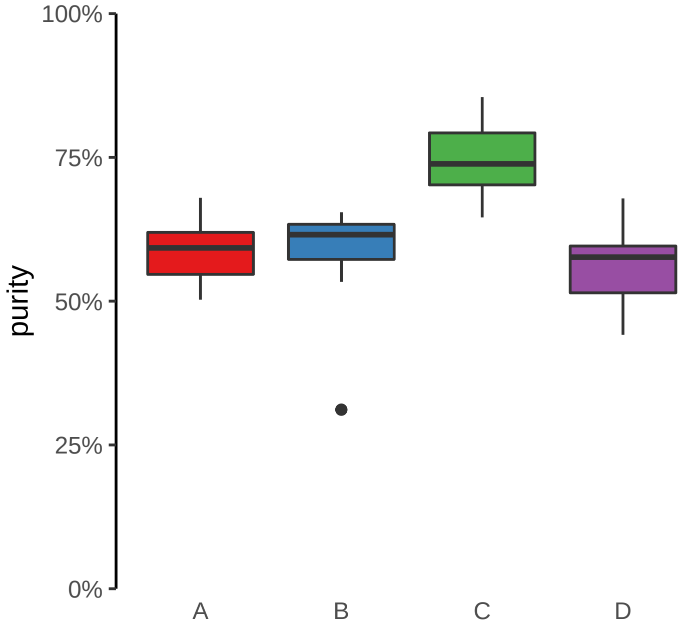

**PPP3CA (GH04J101183) enhancer loss**
**chr4:101183634−101184134**

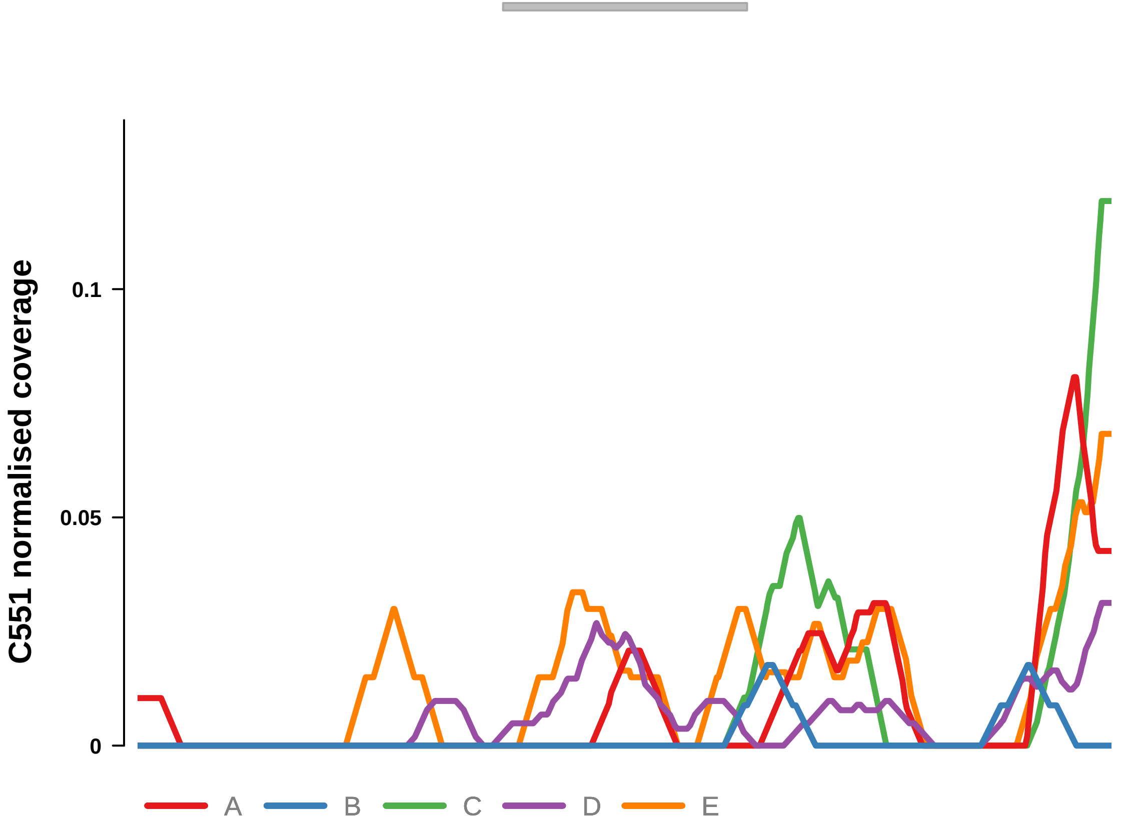


Putatively Clonal (logFC = −2.78)

| | **P.** | **Adjusted P.** |
|---|---|---|
| *~region vs. ~* | 8.35e−01 | 9.65e−01 |
| *~purity vs ~* | 6.21e−01 | 9.06e−01 |
| *~purity+region vs ~purity* | 8.95e−01 | 9.99e−01 |
| *Carcinoma vs. normal* | 3.23e−04 *** | 3.23e−04 *** |

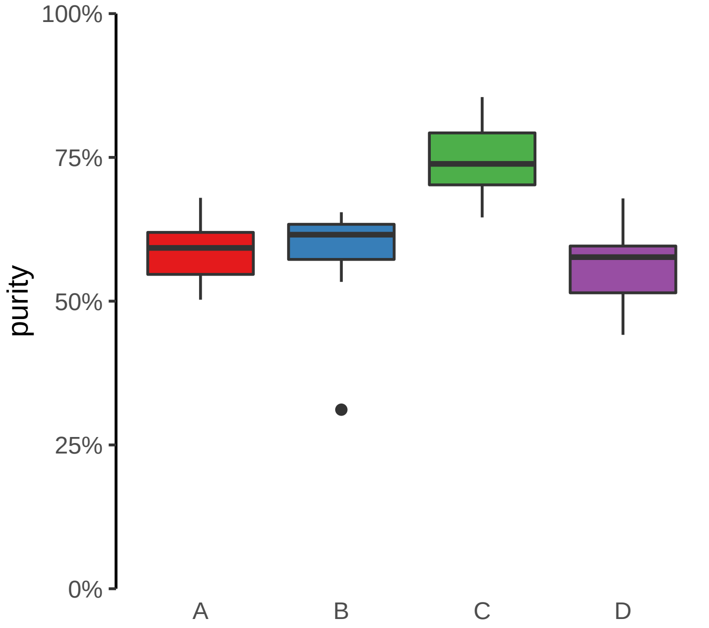

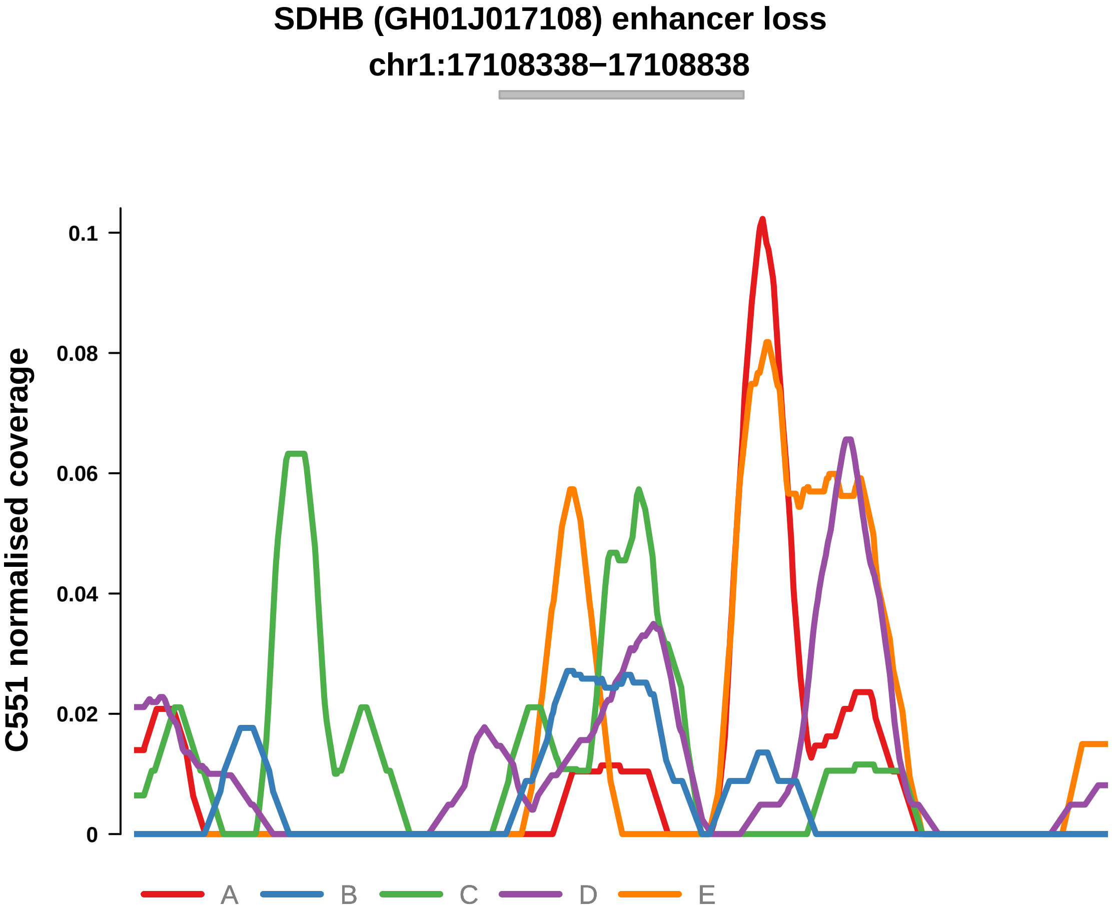


Putatively Clonal (logFC = −2.34)

| | P. | Adjusted P. |
|---|---|---|
| *~region vs. ~* | 9.41e−01 | 9.74e−01 |
| *~purity vs ~* | 5.78e−01 | 9.06e−01 |
| *~purity+region vs ~purity* | 9.86e−01 | 9.99e−01 |
| *Carcinoma vs. normal* | 1.24e−03 ** | 1.24e−03 ** |

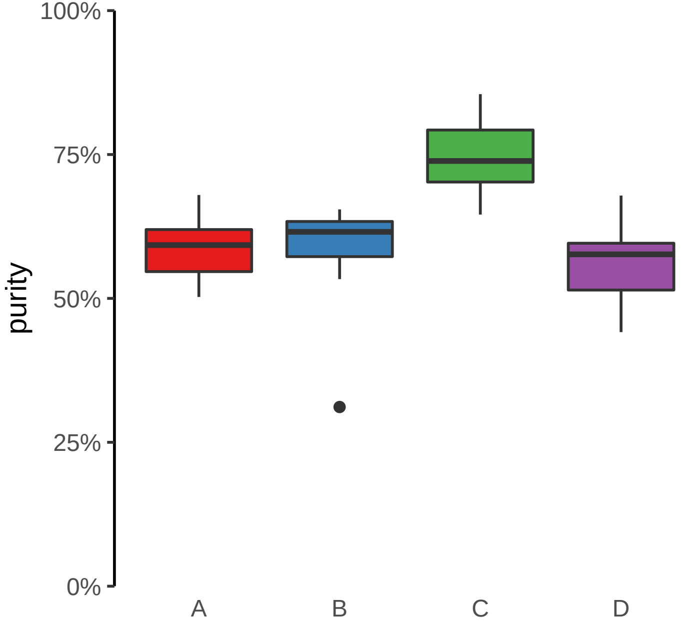

**TRANK1, ENSG00000234073 (GH03J036874) enhancer loss**
**chr3:36876260−36876760**

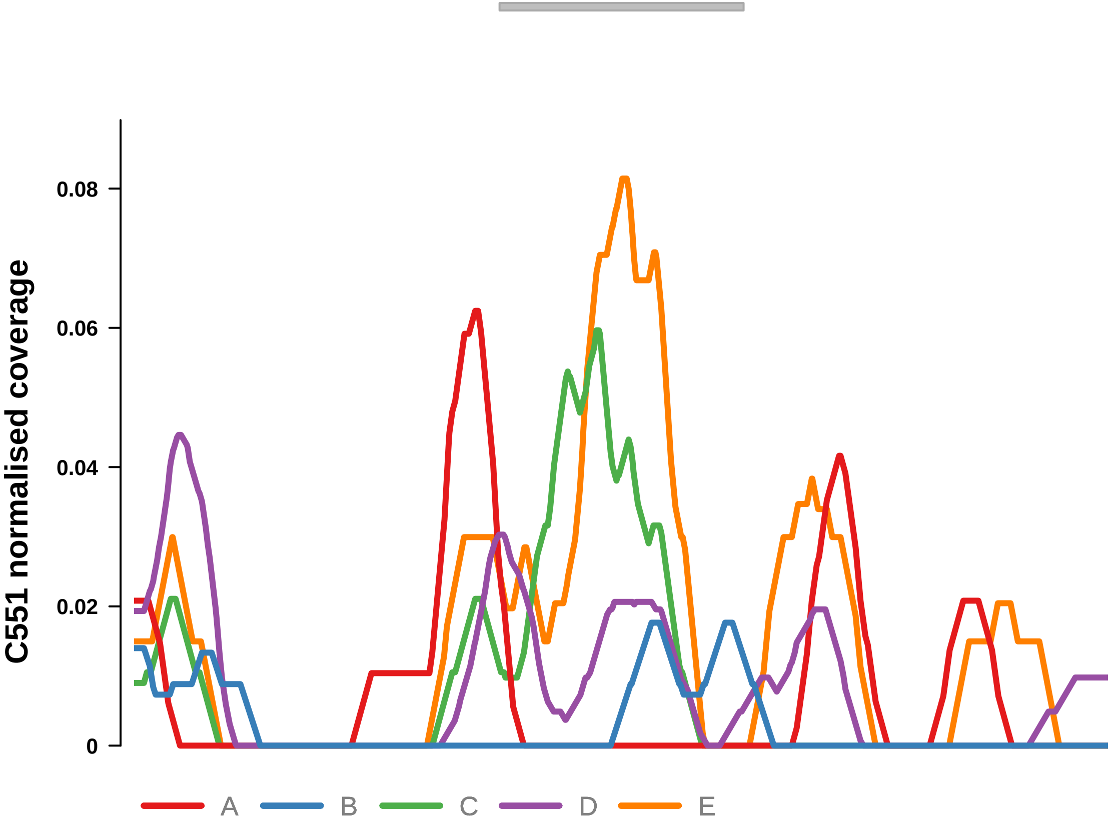


Putatively Clonal (logFC = −1.94)

| | P. | Adjusted P. |
|---|---|---|
| *~region vs. ~* | 5.28e−01 | 8.69e−01 |
| *~purity vs ~* | 9.08e−01 | 9.69e−01 |
| *~purity+region vs ~purity* | 5.03e−01 | 9.48e−01 |
| *Carcinoma vs. normal* | 7.32e−03 ** | 7.32e−03 ** |

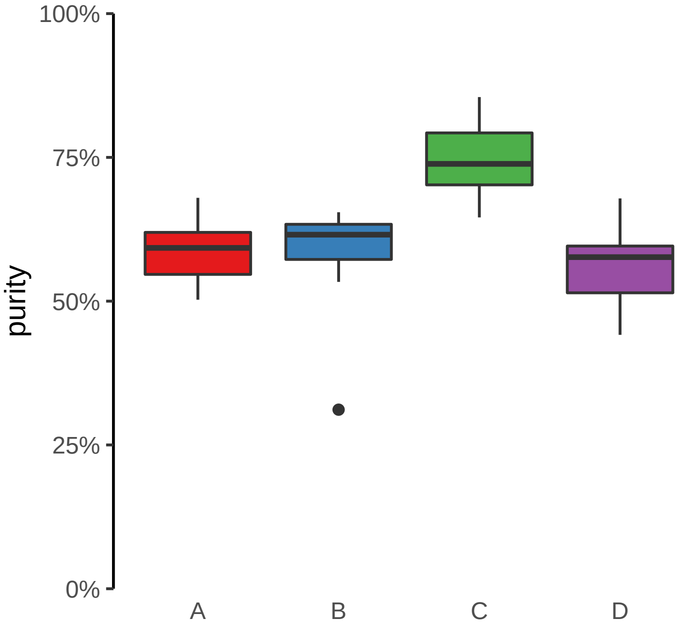

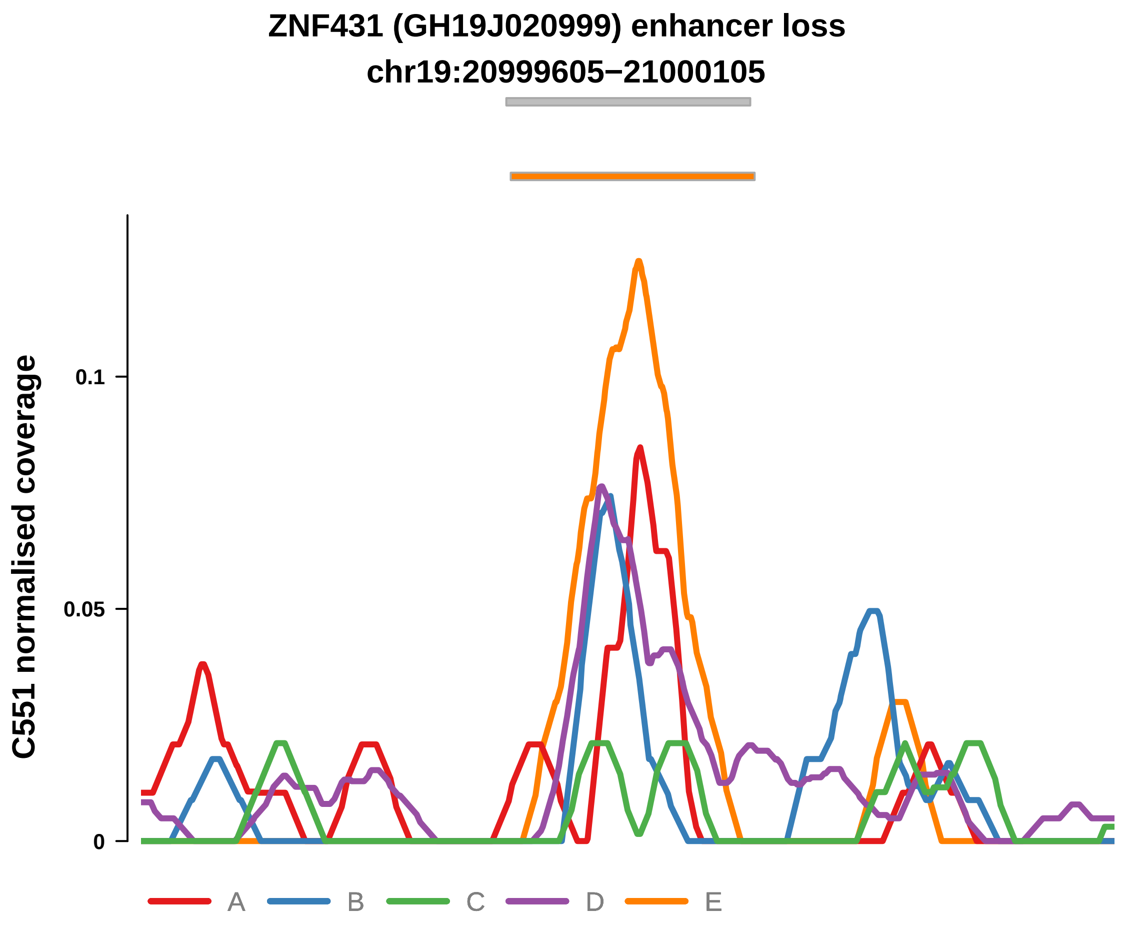


Putatively Clonal (logFC = −2.34)

| | P. | Adjusted P. |
|---|---|---|
| *~region vs. ~* | 1.7e−01 | 4.98e−01 |
| *~purity vs ~* | 1.87e−01 | 8.37e−01 |
| *~purity+region vs ~purity* | 4.03e−01 | 8.86e−01 |
| *Carcinoma vs. normal* | 9.6e−04 *** | 9.6e−04 *** |

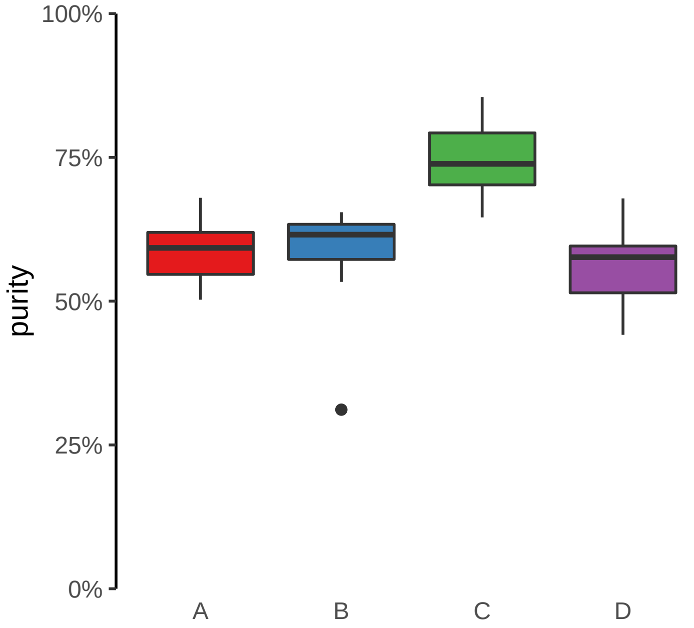

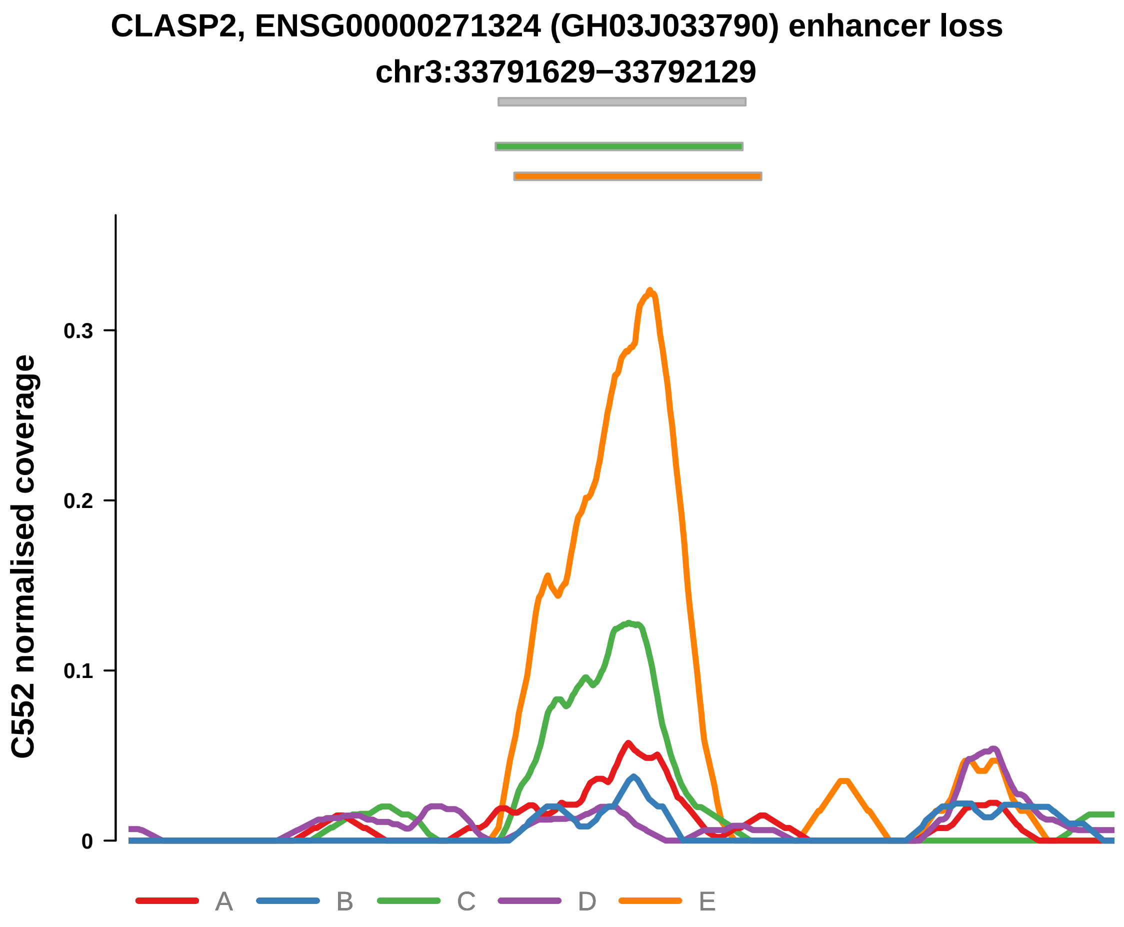


Putatively Clonal (logFC = −2.17)

| | P. | Adjusted P. |
|---|---|---|
| *~region vs. ~* | 8.04e−01 | 8.4e−01 |
| *~purity vs ~* | 2.93e−01 | 3.96e−01 |
| *~purity+region vs ~purity* | 2e−01 | 9.1e−01 |
| *Carcinoma vs. normal* | 9.53e−03 ** | 9.53e−03 ** |

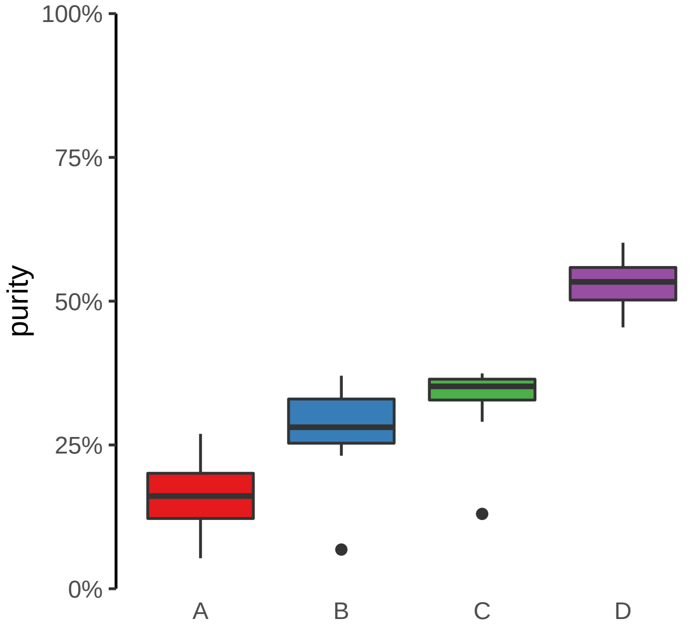

**DNAH7, ENSG00000272211 (GH02J196100) enhancer loss**
**chr2:196101142−196101642**

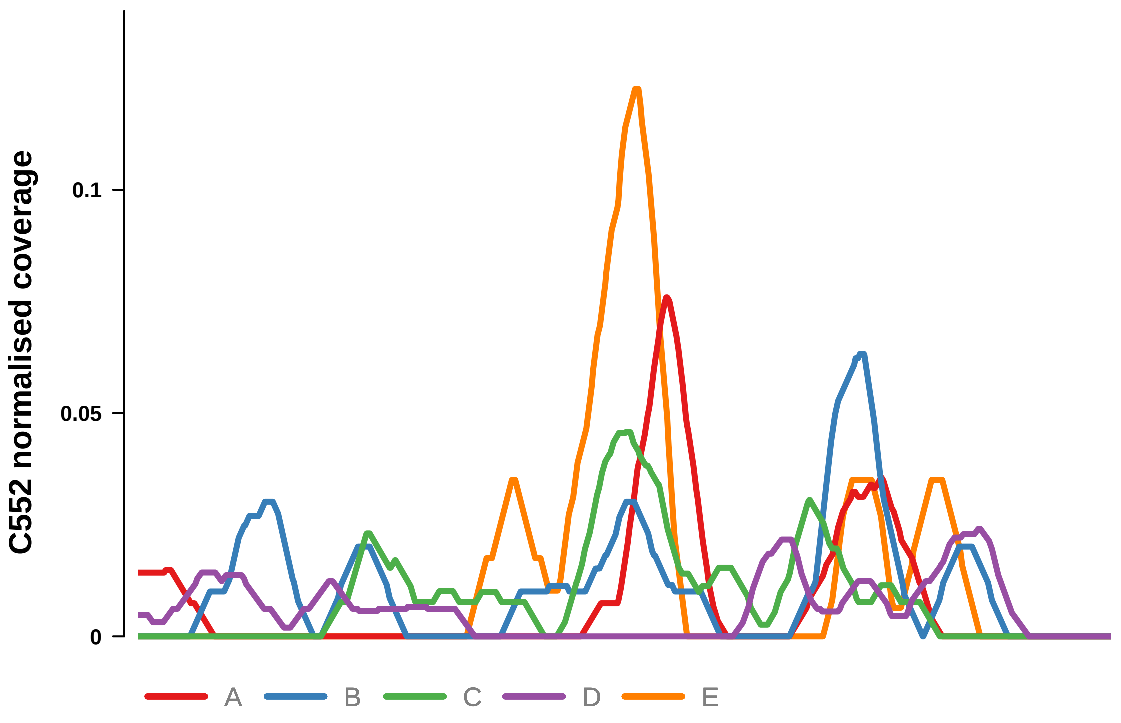


Putatively Clonal (logFC = −10.91)

| | P. | Adjusted P. |
|---|---|---|
| *~region vs. ~* | 5.99e−01 | NA |
| *~purity vs ~* | 4.26e−01 | NA |
| *~purity+region vs ~purity* | 6.71e−01 | 9.85e−01 |
| *Carcinoma vs. normal* | 5.26e−05 **** | 5.26e−05 **** |

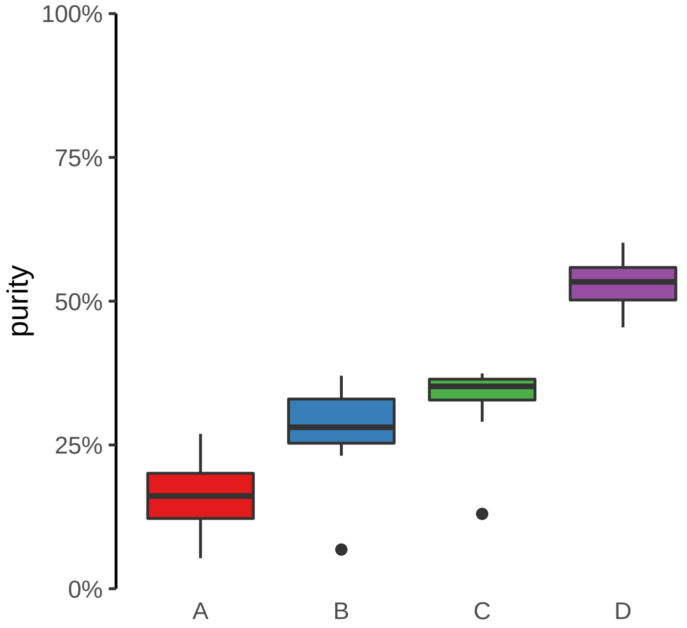

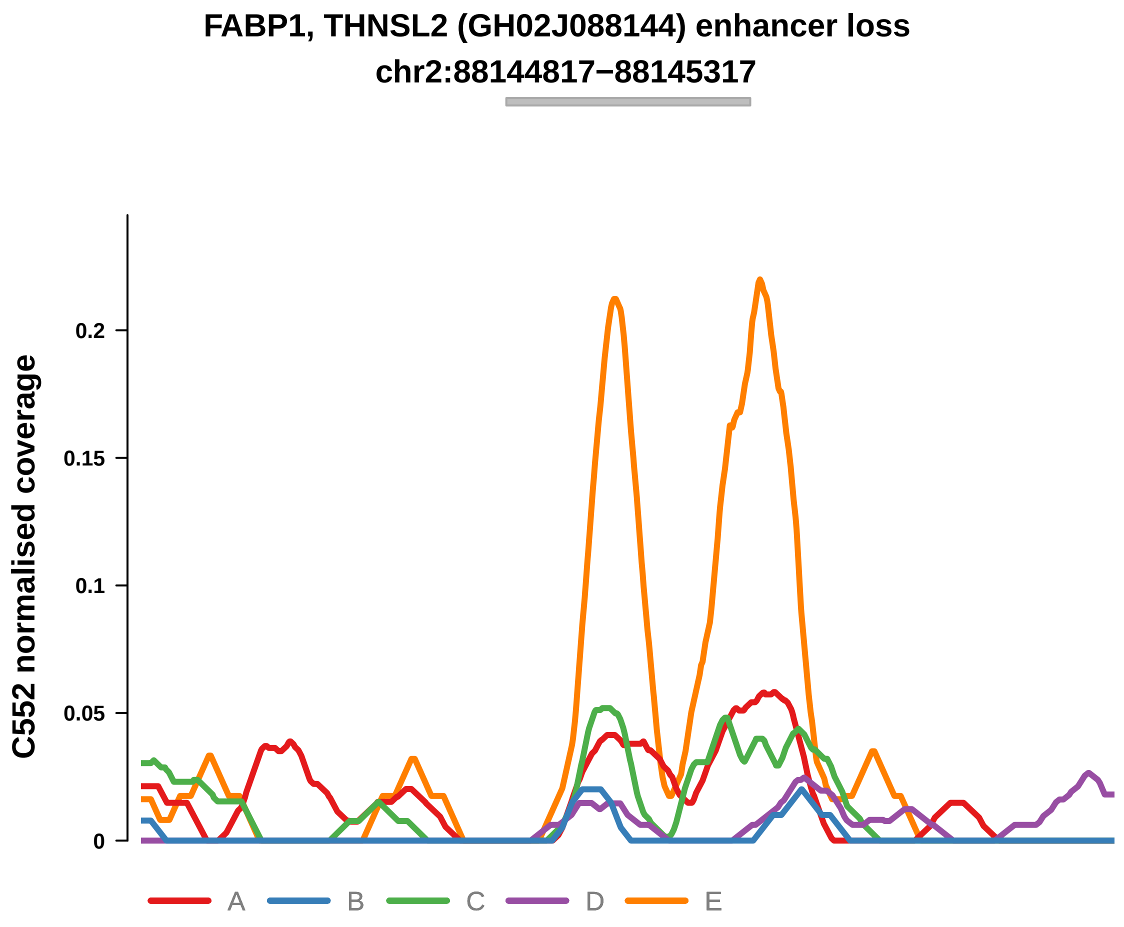


Putatively Clonal (logFC = −2.66)

| | P. | Adjusted P. |
|---|---|---|
| *~region vs. ~* | 9.36e−01 | NA |
| *~purity vs ~* | 5.59e−01 | NA |
| *~purity+region vs ~purity* | 7.85e−01 | 9.85e−01 |
| *Carcinoma vs. normal* | 3.22e−03 ** | 3.22e−03 ** |

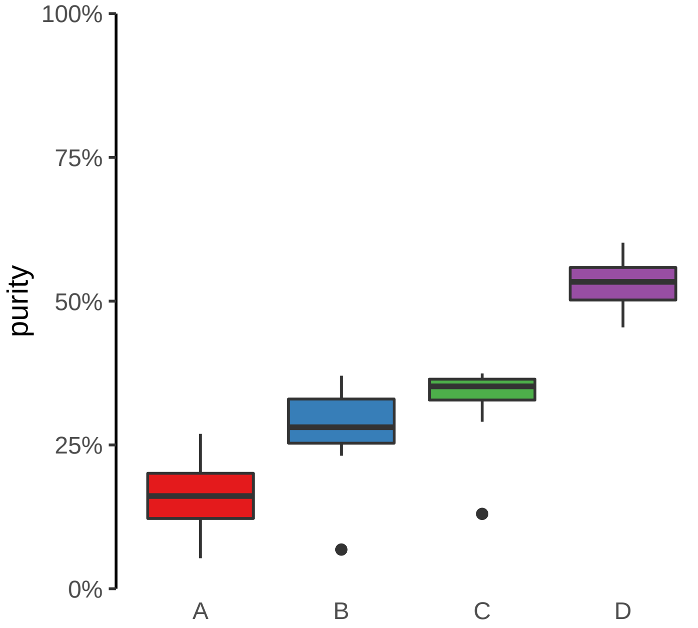

**HMGN2, ARID1A, PIGV (GH01J026739) enhancer loss**
**chr1:26743976−26744476**

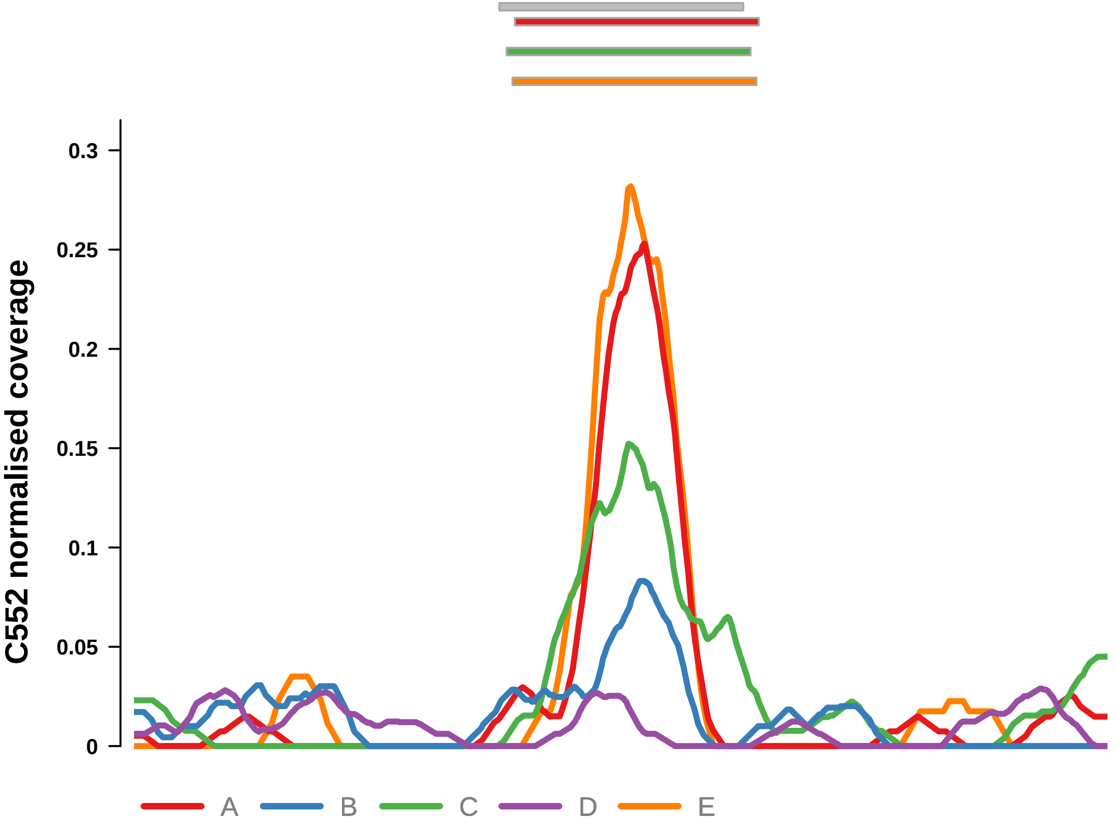


Putatively Clonal (logFC = −2.65)

| | P. | Adjusted P. |
|---|---|---|
| *~region vs. ~* | 2.2e−03 ** | 1.28e−02 * |
| *~purity vs ~* | 2.74e−03 ** | 1.52e−02 * |
| *~purity+region vs ~purity* | 1.42e−01 | 9.1e−01 |
| *Carcinoma vs. normal* | 1.75e−03 ** | 1.75e−03 ** |

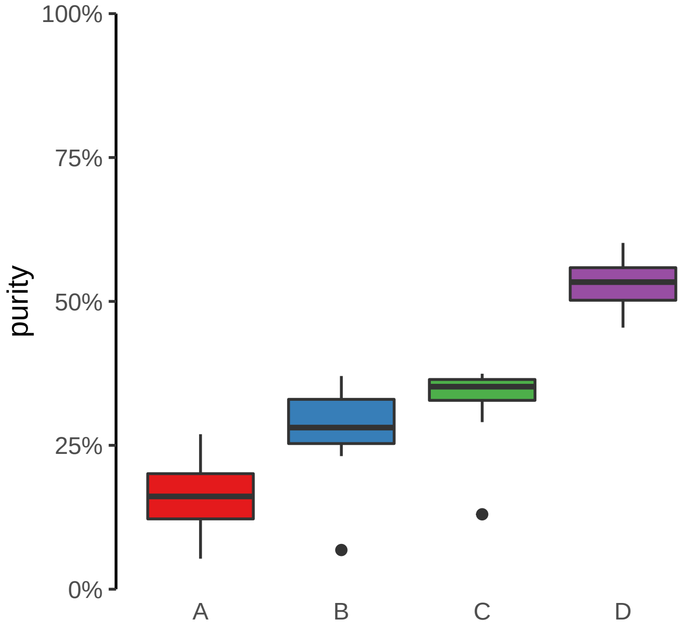

**PPP3CA (GH04J101183) enhancer loss**
**chr4:101183634−101184134**

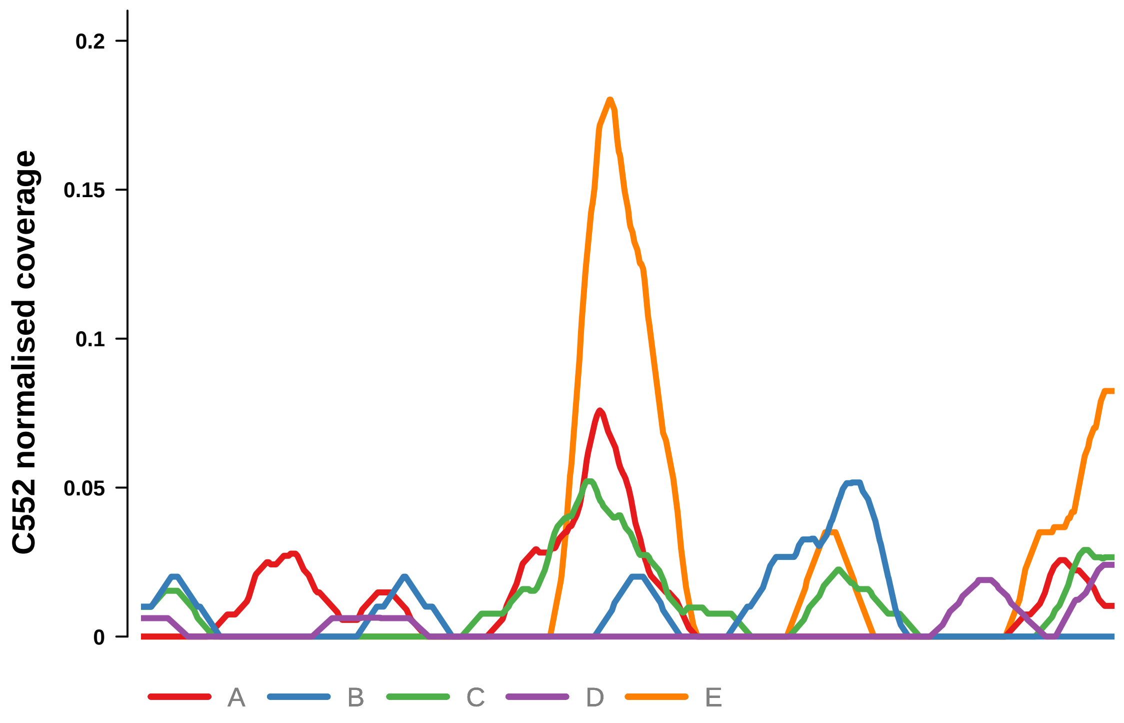


Putatively Clonal (logFC = −10.76)

| | P. | Adjusted P. |
|---|---|---|
| *~region vs. ~* | 5.75e−01 | NA |
| *~purity vs ~* | 1.59e−01 | NA |
| *~purity+region vs ~purity* | 9.5e−01 | 9.85e−01 |
| *Carcinoma vs. normal* | 5.63e−05 **** | 5.63e−05 **** |

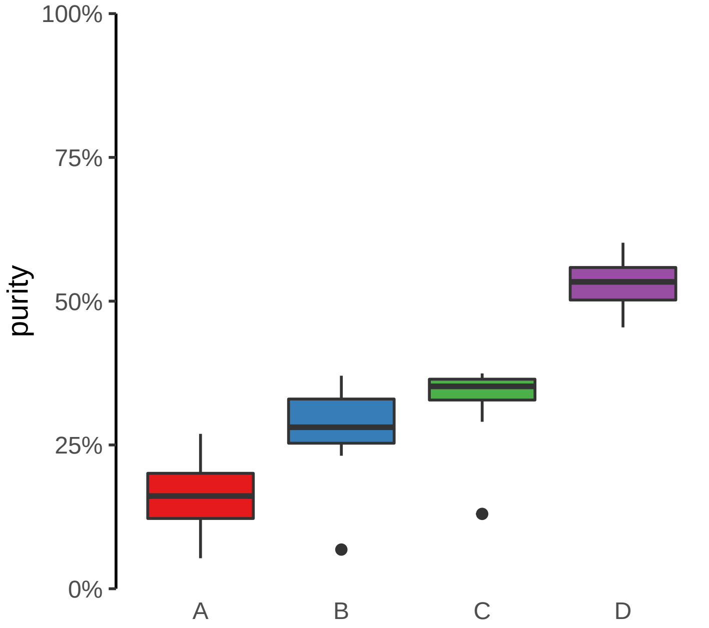

**TRANRK1, ENSG00000234073 (GH03J036874) enhancer loss**
**chr3:36876260−36876760**

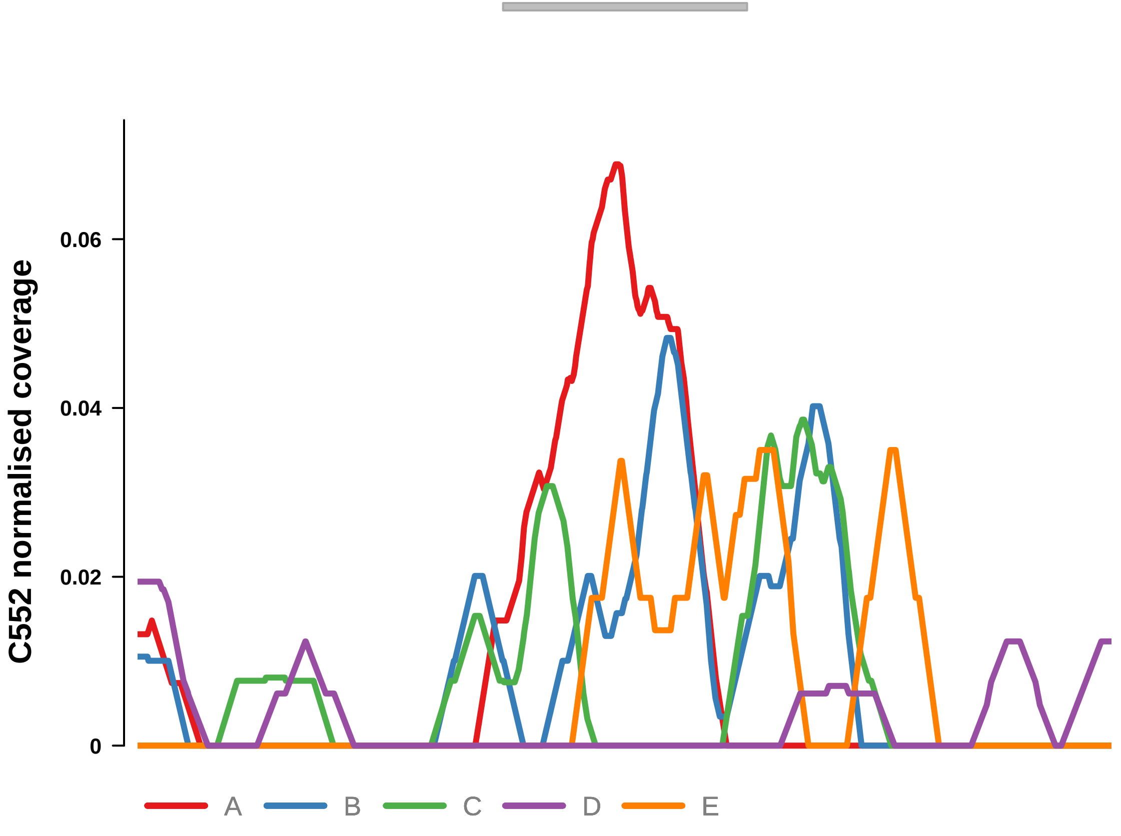


Putatively Clonal (logFC = −10.96)

| | P. | Adjusted P. |
|---|---|---|
| *~region vs. ~* | 1.41e−01 | NA |
| *~purity vs ~* | 1.36e−02 * | NA |
| *~purity+region vs ~purity* | 8.91e−01 | 9.85e−01 |
| *Carcinoma vs. normal* | 4.02e−05 **** | 4.02e−05 **** |

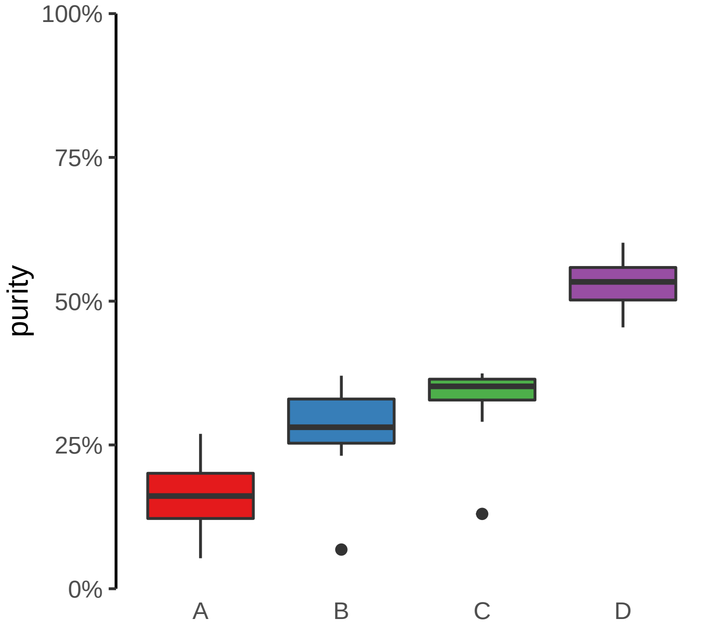

**ACAP2, PPP1R2 (GH03J195537) enhancer loss**
**chr3:195537917−195538417**

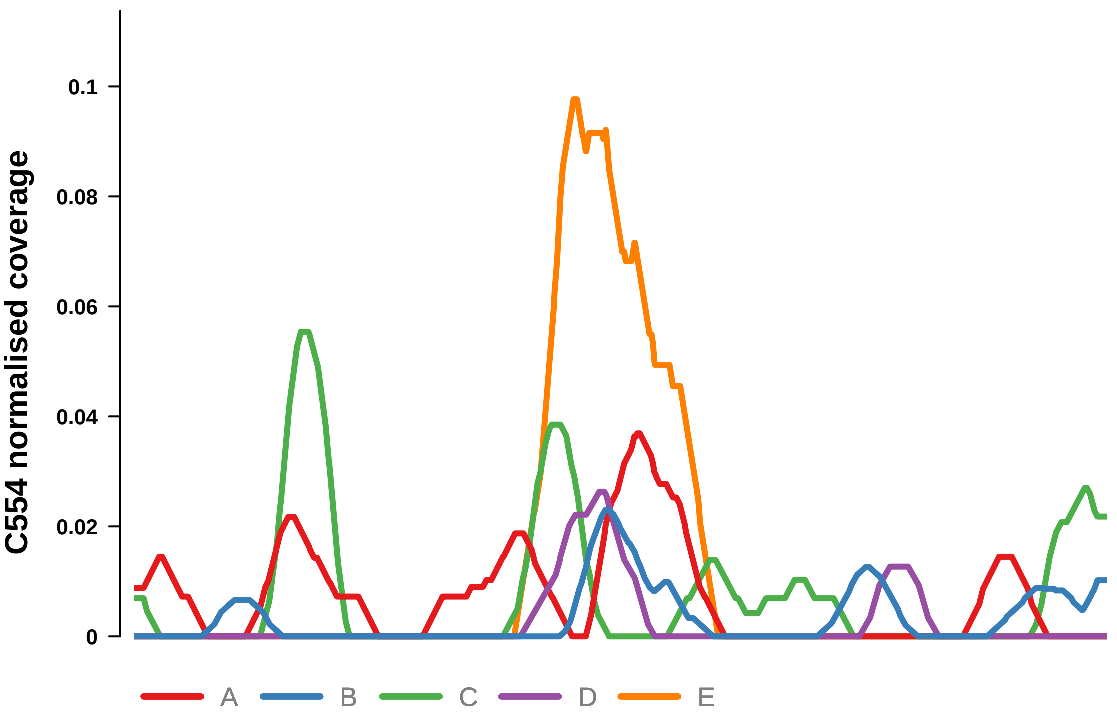


Putatively Clonal (logFC = −2.19)

| | P. | Adjusted P. |
|---|---|---|
| *~region vs. ~* | 3.19e−01 | NA |
| *~purity vs ~* | 8.29e−01 | NA |
| *~purity+region vs ~purity* | 3.89e−01 | NA |
| *Carcinoma vs. normal* | 5.68e−03 ** | 5.68e−03 ** |

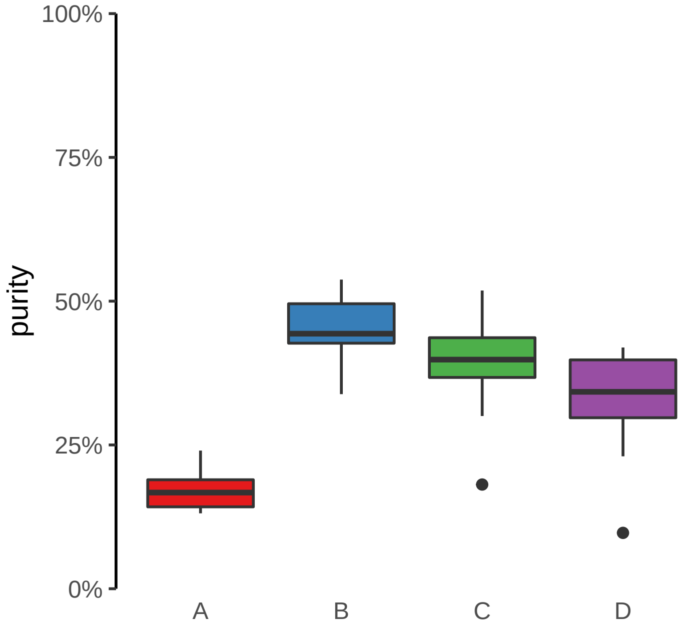

**ANTXR2 (GH04J080057) enhancer loss**
**chr4:80058687−80059187**

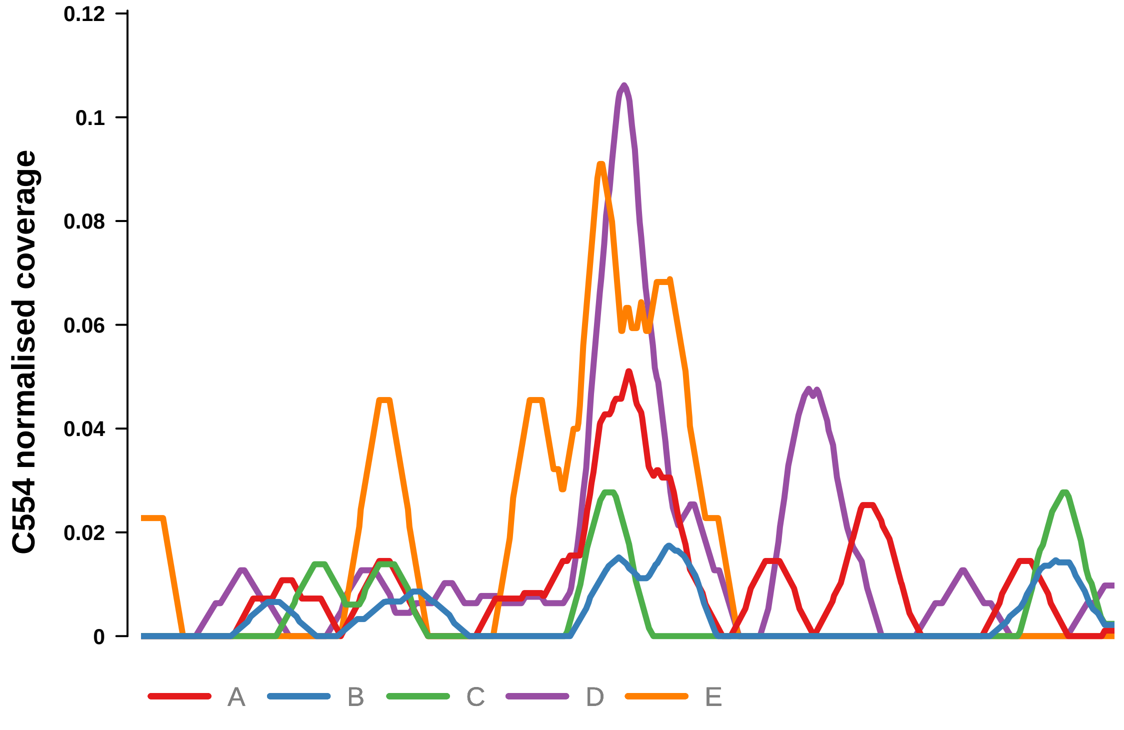


Putatively Clonal (logFC = −2.07)

| | P. | Adjusted P. |
|---|---|---|
| *~region vs. ~* | 2.42e−01 | 3.36e−01 |
| *~purity vs ~* | 4.26e−01 | 6.24e−01 |
| *~purity+region vs ~purity* | 3.41e−01 | NA |
| *Carcinoma vs. normal* | 7.11e−03 ** | 7.11e−03 ** |

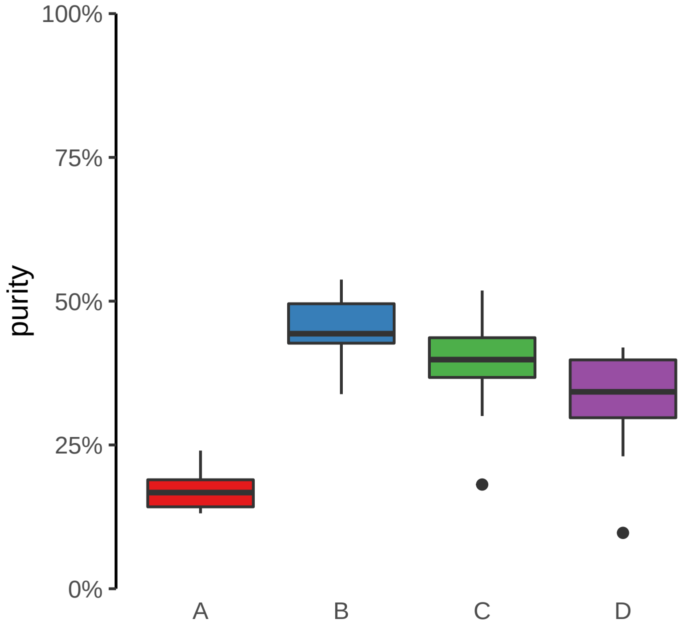

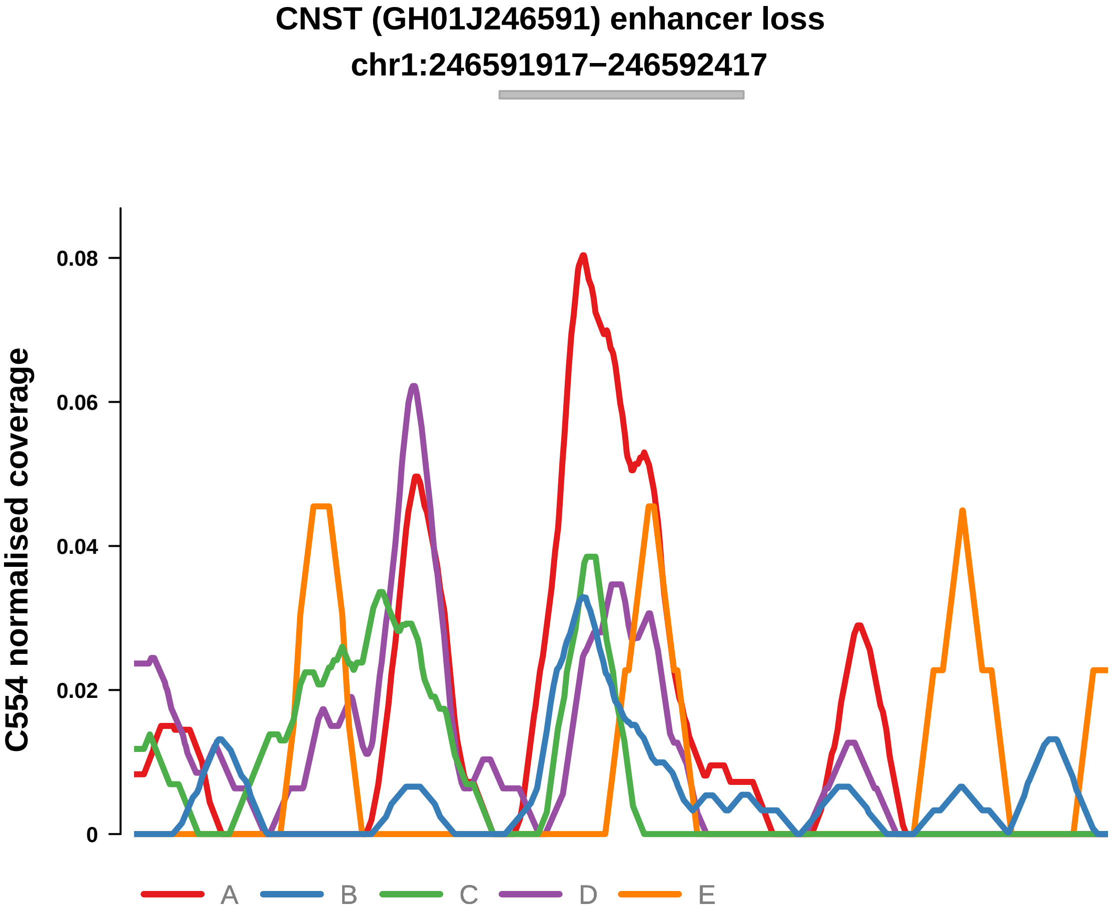


Putatively Clonal (logFC = −2.01)

| | P. | Adjusted P. |
|---|---|---|
| *~region vs. ~* | 3.96e−01 | 4.71e−01 |
| *~purity vs ~* | 2.88e−01 | 5.02e−01 |
| *~purity+region vs ~purity* | 6.46e−01 | 7.4e−01 |
| *Carcinoma vs. normal* | 6.83e−03 ** | 6.83e−03 ** |

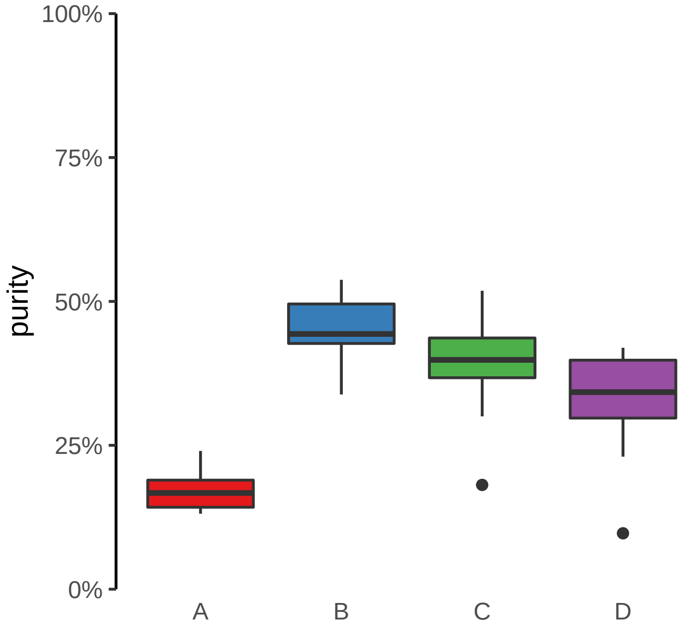

**FAM83G (GH17J019010) enhancer loss**

**chr17:19011073−19011573**

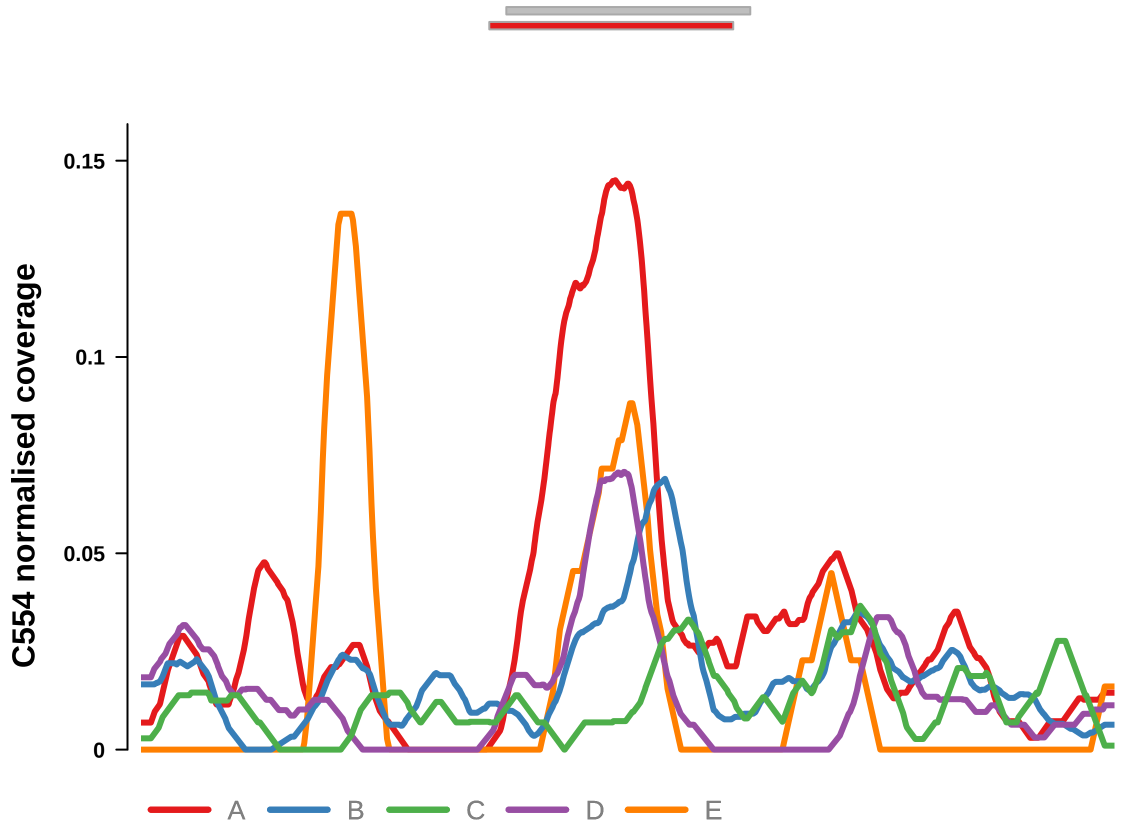


Putatively Clonal (logFC = −2)

| | **P.** | **Adjusted P.** |
|---|---|---|
| *~region vs. ~* | 1.44e−01 | 2.45e−01 |
| *~purity vs ~* | 2.39e−01 | 4.35e−01 |
| *~purity+region vs ~purity* | 2.76e−01 | 4.01e−01 |
| *Carcinoma vs. normal* | 4.66e−03 ** | 4.66e−03 ** |

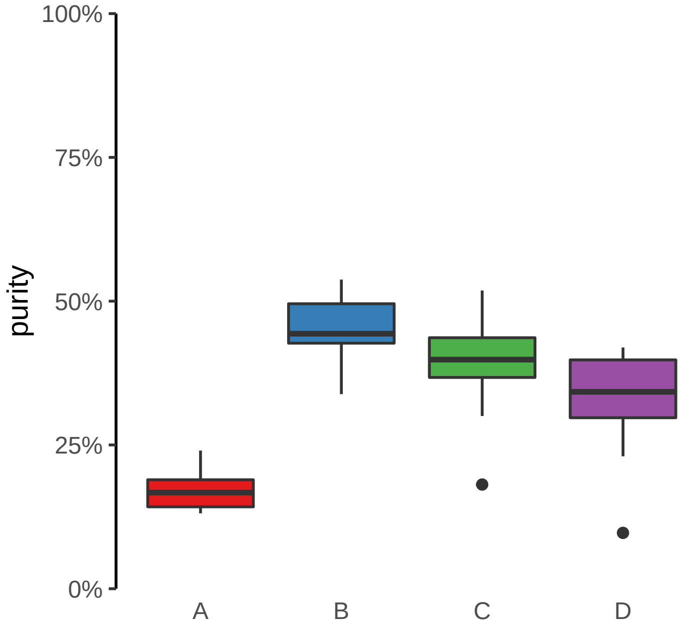

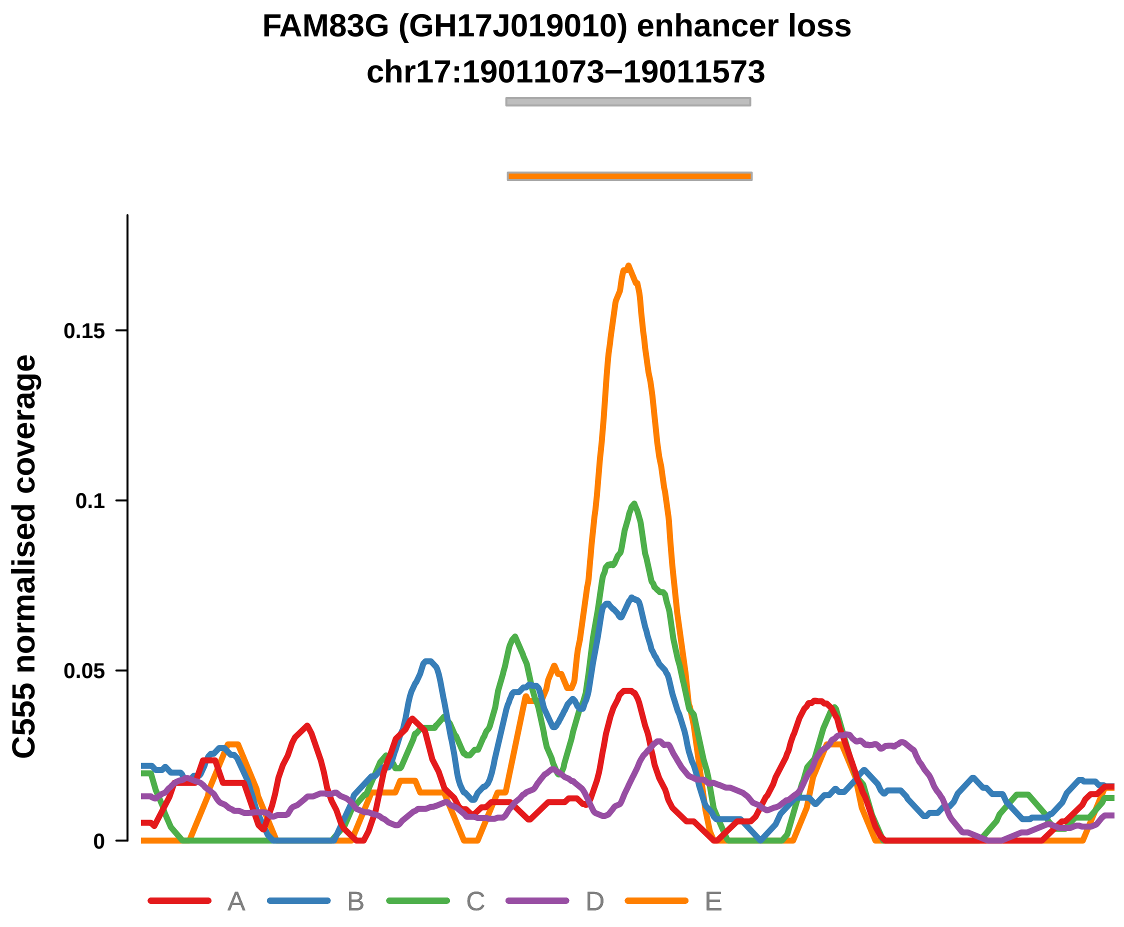


Putatively Clonal (logFC = −2.9)

| | P. | Adjusted P. |
|---|---|---|
| *~region vs. ~* | 4.24e−01 | 6.34e−01 |
| *~purity vs ~* | 5.87e−01 | 7.43e−01 |
| *~purity+region vs ~purity* | 1.66e−01 | 4.3e−01 |
| *Carcinoma vs. normal* | 4.09e−04 *** | 4.09e−04 *** |

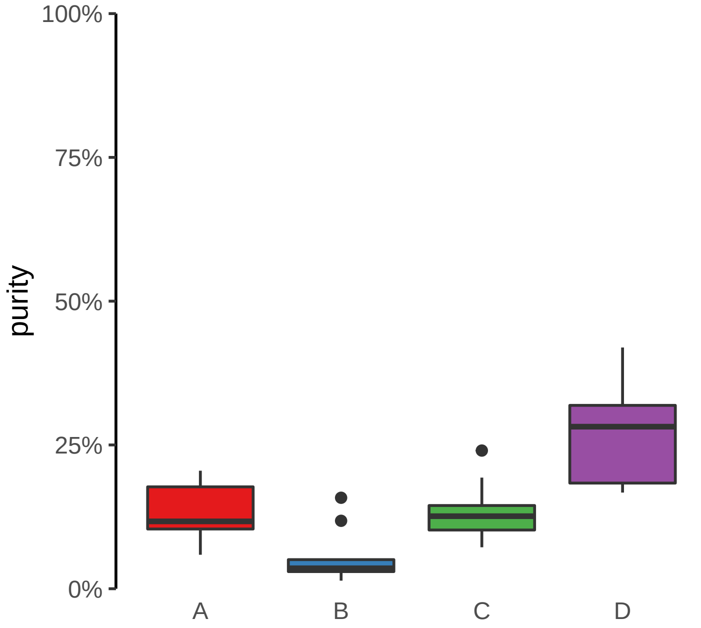

**RNLS, LIPJ (GH10J088553) enhancer loss**
**chr10:88553102−88553602**

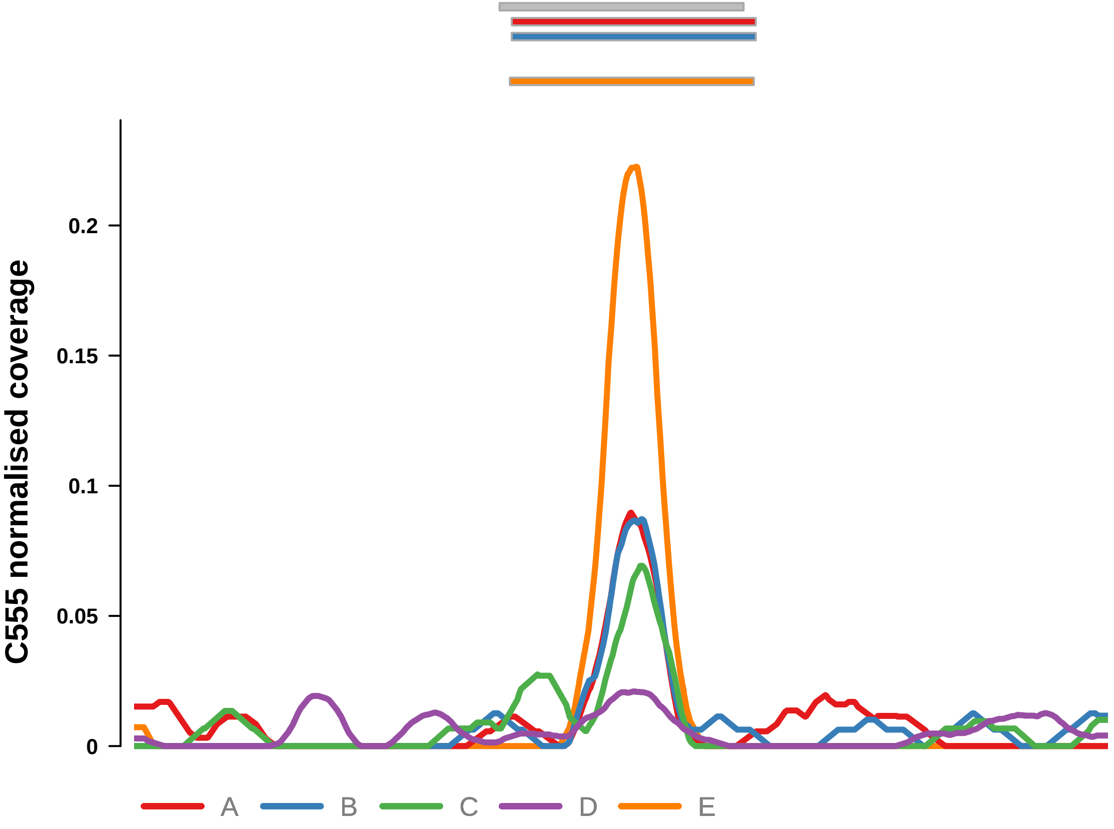


Putatively Clonal (logFC = −2.22)

| | P. | Adjusted P. |
|---|---|---|
| *~region vs. ~* | 5.51e−01 | 7.13e−01 |
| *~purity vs ~* | 6.11e−01 | 7.43e−01 |
| *~purity+region vs ~purity* | 5.64e−01 | 7.88e−01 |
| *Carcinoma vs. normal* | 9.62e−03 ** | 9.62e−03 ** |

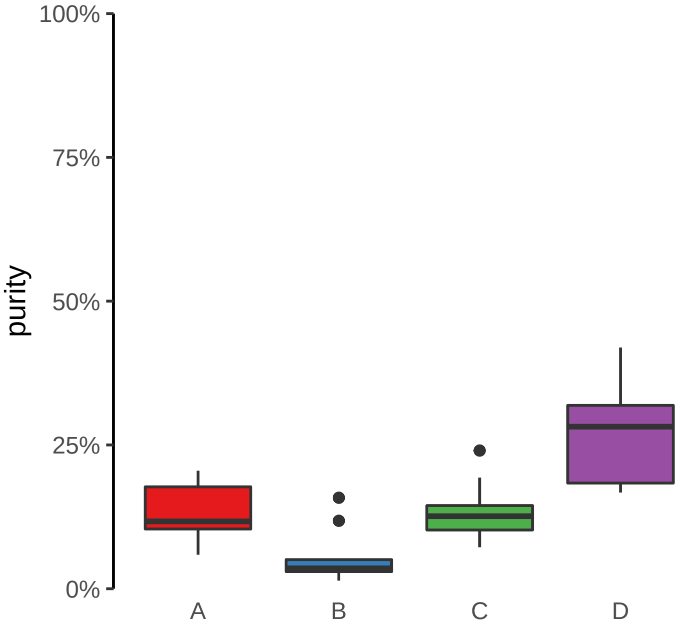

**CLASP1 (GH02J121362) enhancer loss**
**chr2:121363584−121364084**

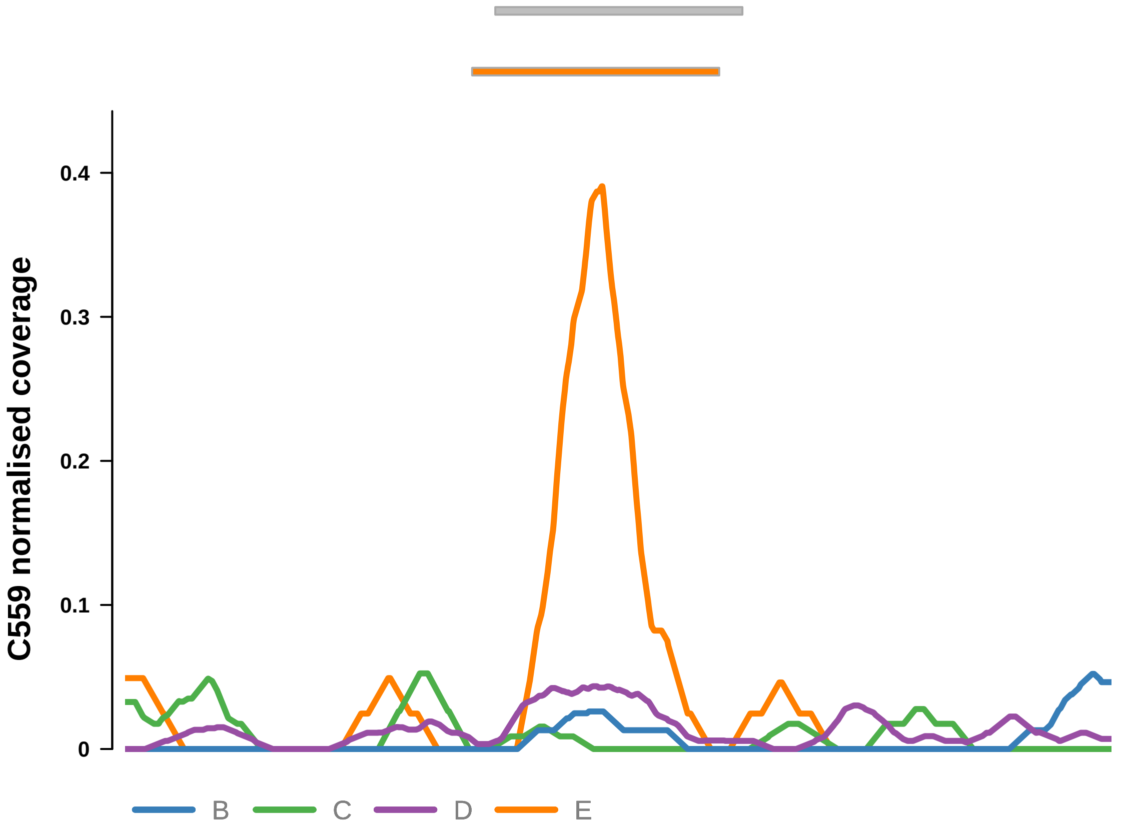


Putatively Clonal (logFC = −2.48)

| | P. | Adjusted P. |
|---|---|---|
| *~region vs. ~* | 2.3e−01 | 5.55e−01 |
| *~purity vs ~* | 9.36e−02 | 5.51e−01 |
| *~purity+region vs ~purity* | 7.95e−01 | 9.61e−01 |
| *Carcinoma vs. normal* | 6.43e−03 ** | 6.43e−03 ** |

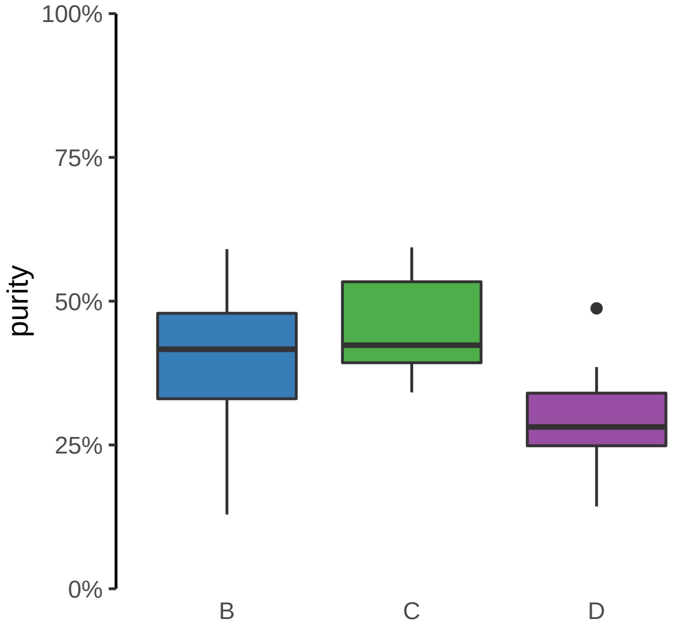

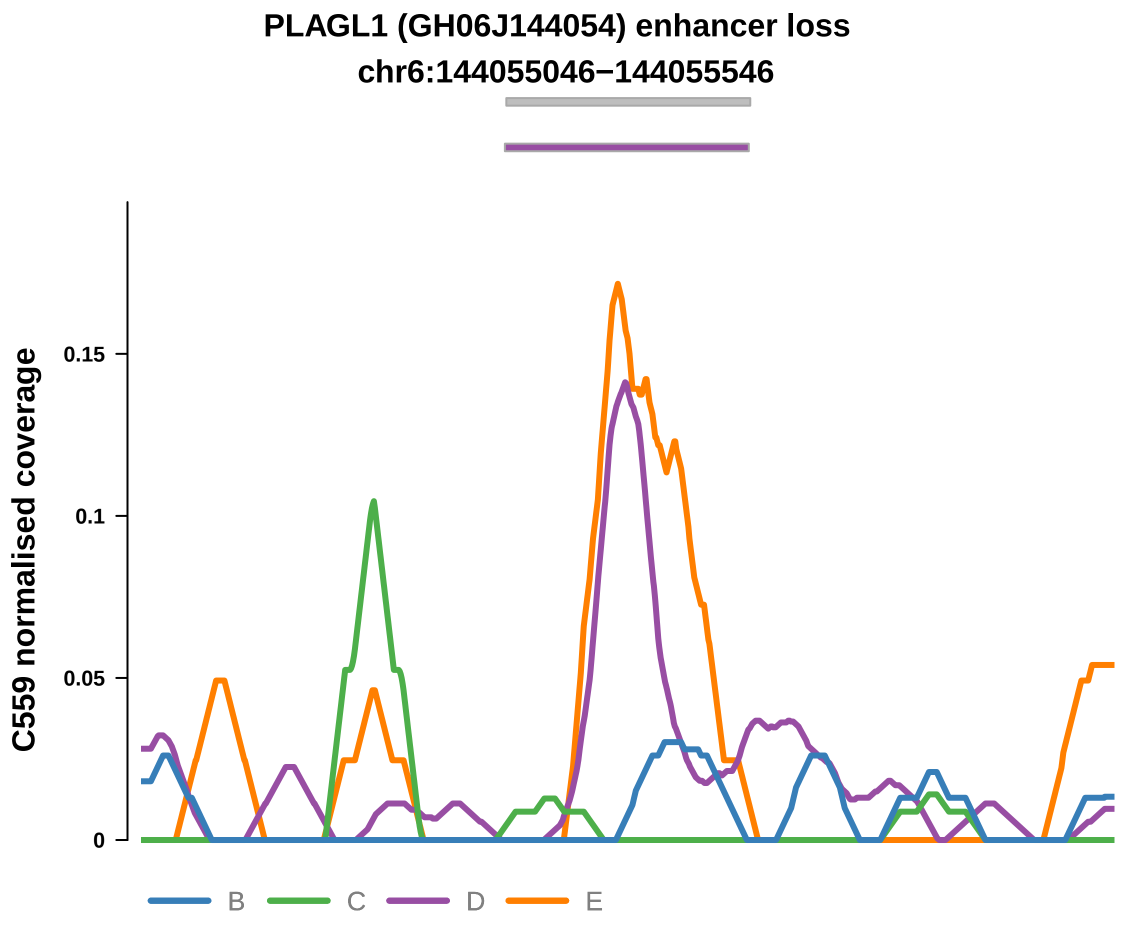


Putatively Clonal (logFC = −3.02)

| | P. | Adjusted P. |
|---|---|---|
| *~region vs. ~* | 2.41e−02 * | 3.63e−01 |
| *~purity vs ~* | 5.29e−03 ** | 1.16e−01 |
| *~purity+region vs ~purity* | 4.59e−01 | 9.11e−01 |
| *Carcinoma vs. normal* | 4.49e−03 ** | 4.49e−03 ** |

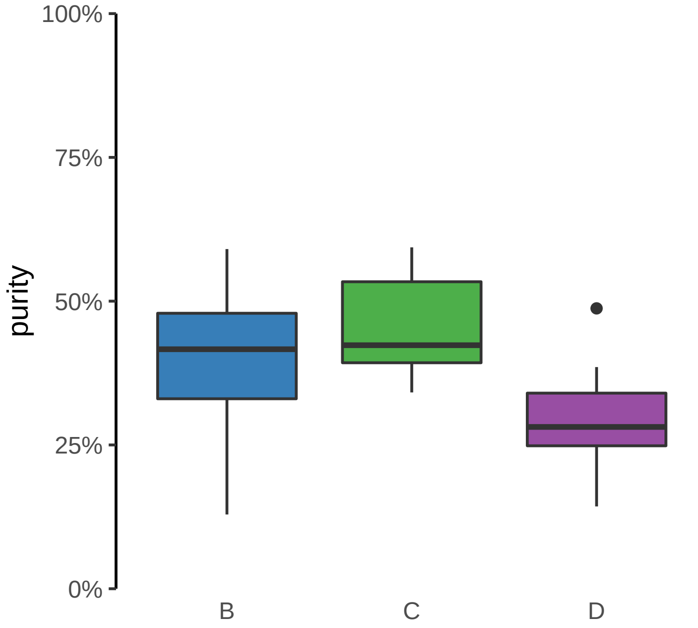

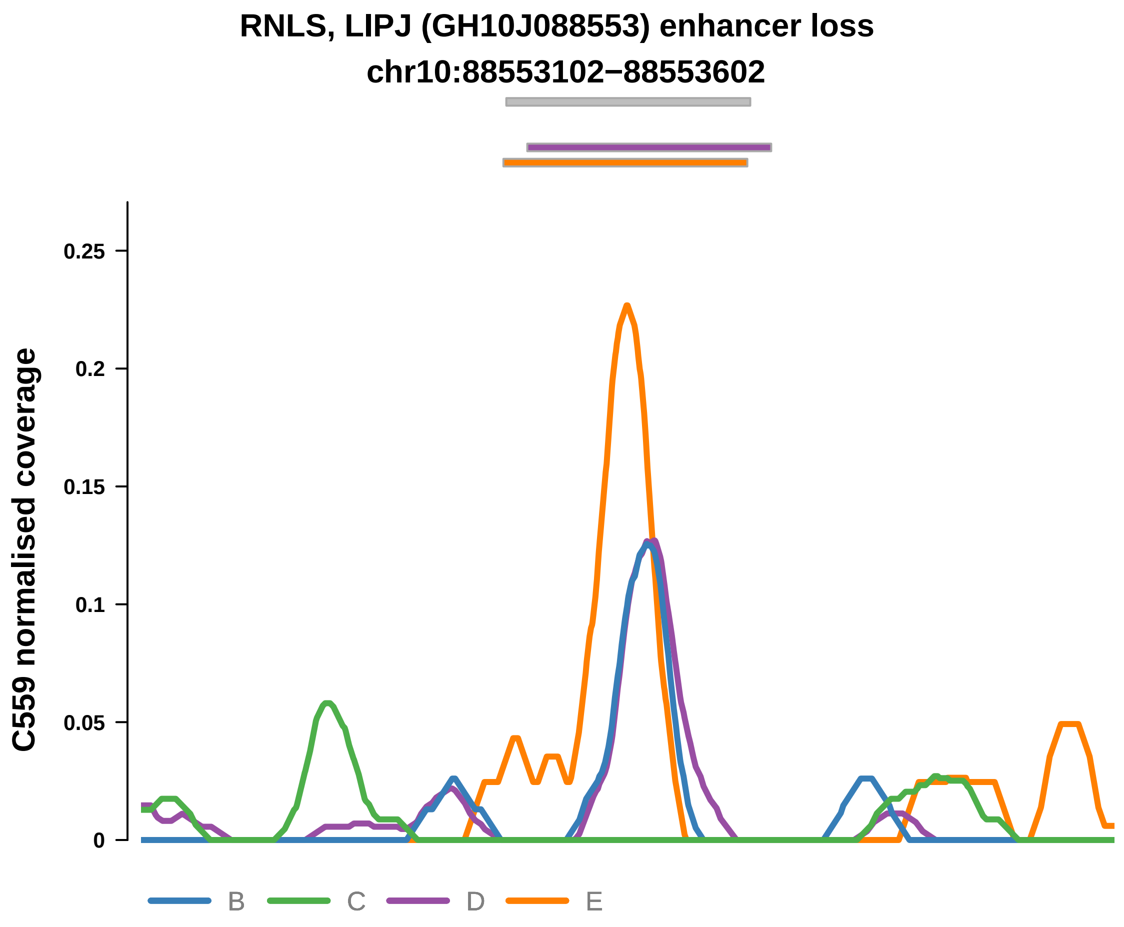


Putatively Clonal (logFC = −2.49)

| | P. | Adjusted P. |
|---|---|---|
| *~region vs. ~* | 1.14e−02 * | 3.35e−01 |
| *~purity vs ~* | 1.7e−02 * | 2.5e−01 |
| *~purity+region vs ~purity* | 1.18e−01 | 5.21e−01 |
| *Carcinoma vs. normal* | 6.13e−03 ** | 6.13e−03 ** |

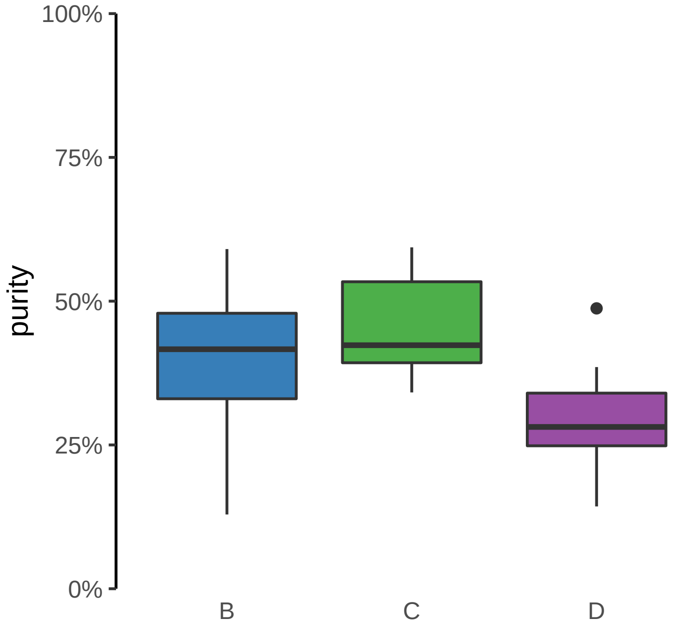

**ANTXR2 (GH04J080057) enhancer loss**
**chr4:80058687−80059187**

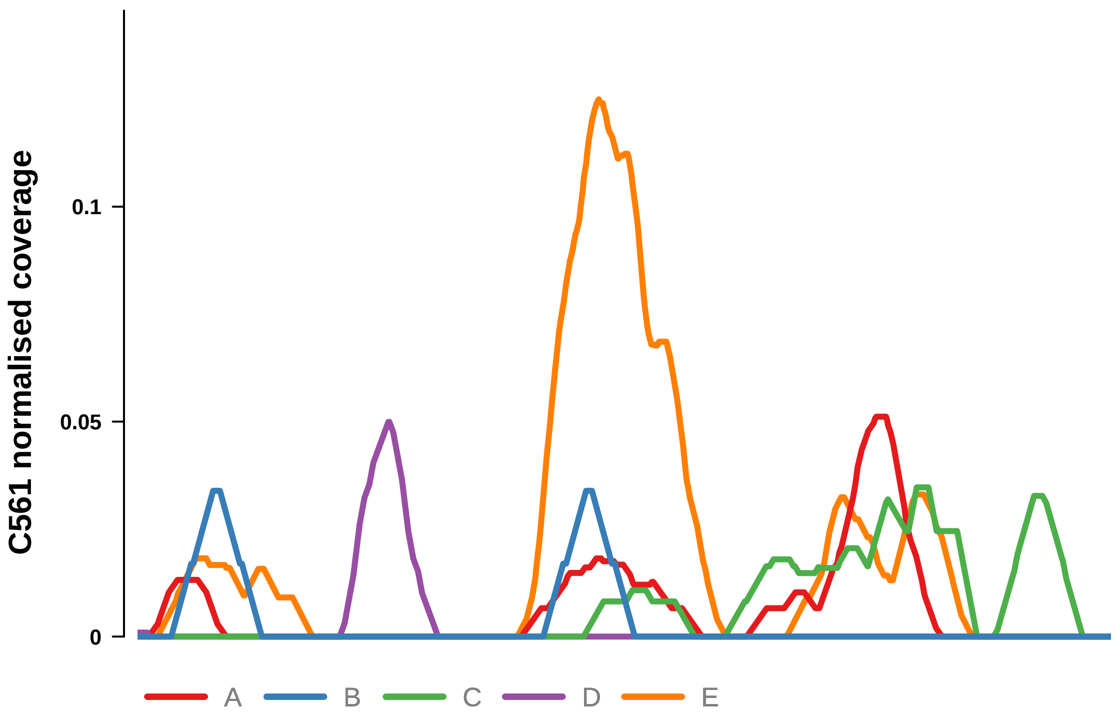


Putatively Clonal (logFC = −2.74)

| | P. | Adjusted P. |
|---|---|---|
| *~region vs. ~* | 9.48e−01 | NA |
| *~purity vs ~* | 6.93e−01 | 8.81e−01 |
| *~purity+region vs ~purity* | 9.68e−01 | 9.79e−01 |
| *Carcinoma vs. normal* | 5.11e−04 *** | 5.11e−04 *** |

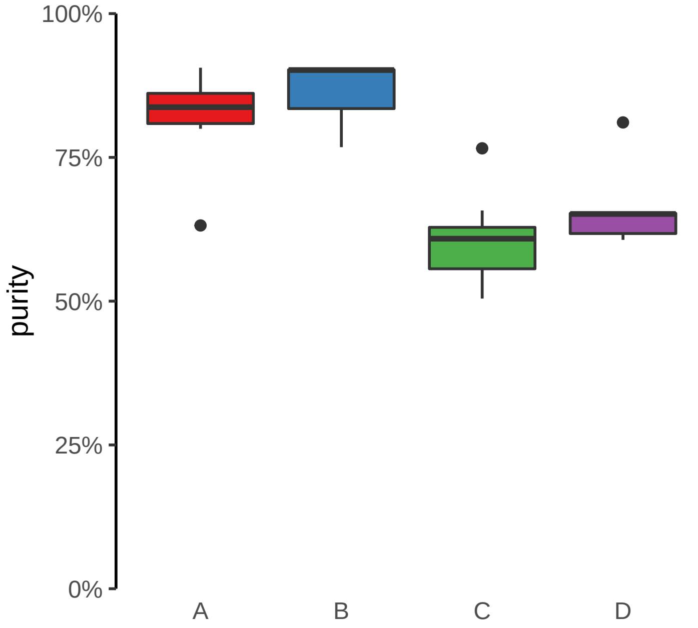

**CAMK2D (GH04J113682) enhancer loss**
**chr4:113683079−113683579**

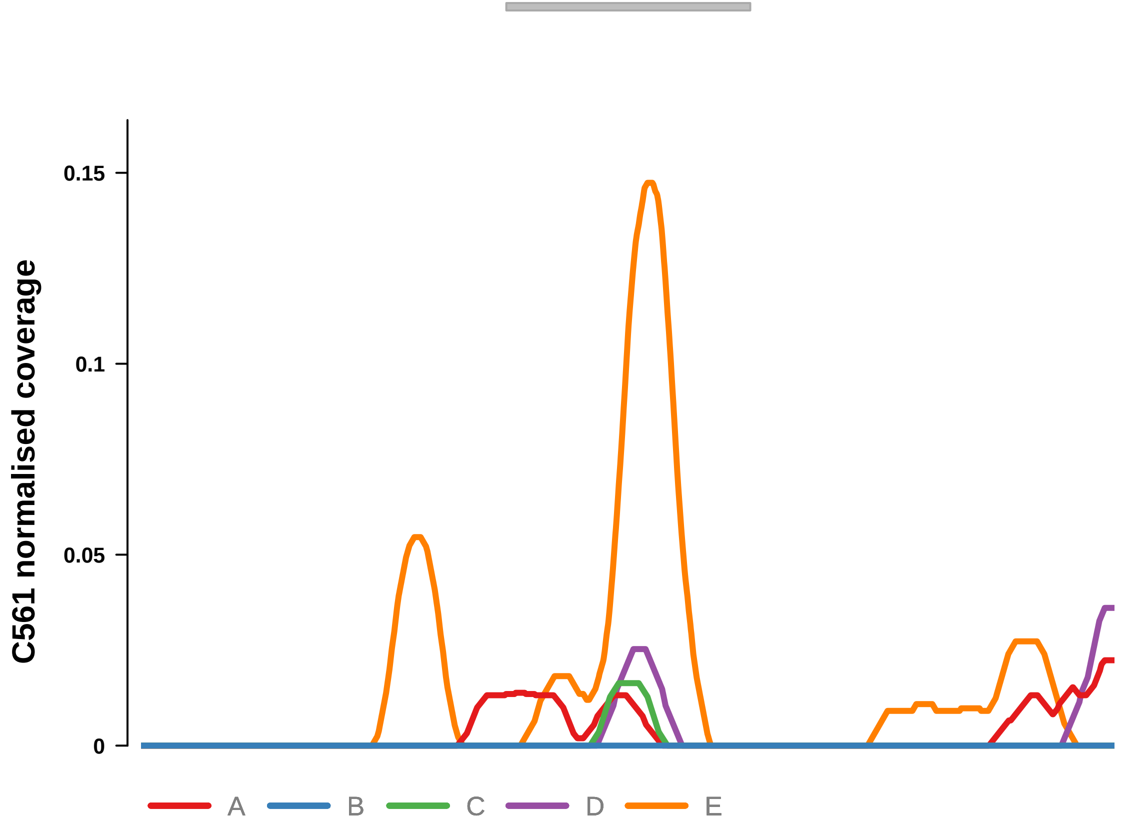


Putatively Clonal (logFC = −2.99)

| | P. | Adjusted P. |
|---|---|---|
| *~region vs. ~* | 9.86e−01 | NA |
| *~purity vs ~* | 1e+00 | 1e+00 |
| *~purity+region vs ~purity* | 9.83e−01 | 9.83e−01 |
| *Carcinoma vs. normal* | 1.87e−04 *** | 1.87e−04 *** |

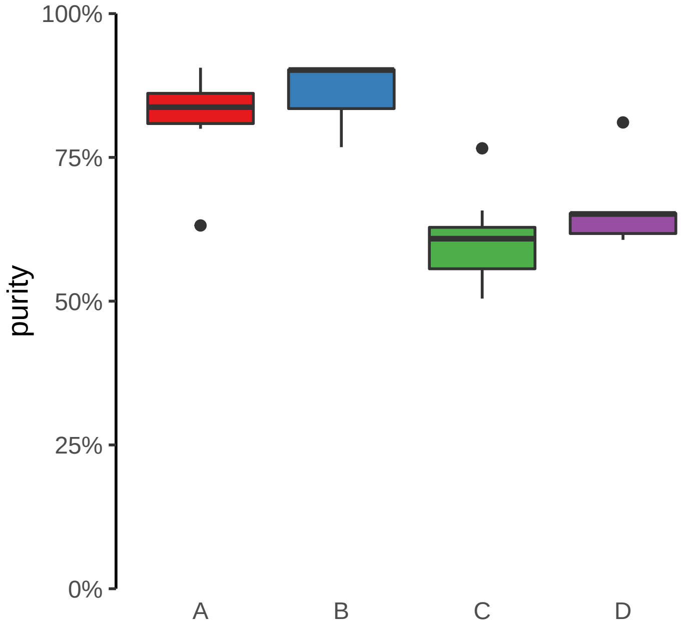

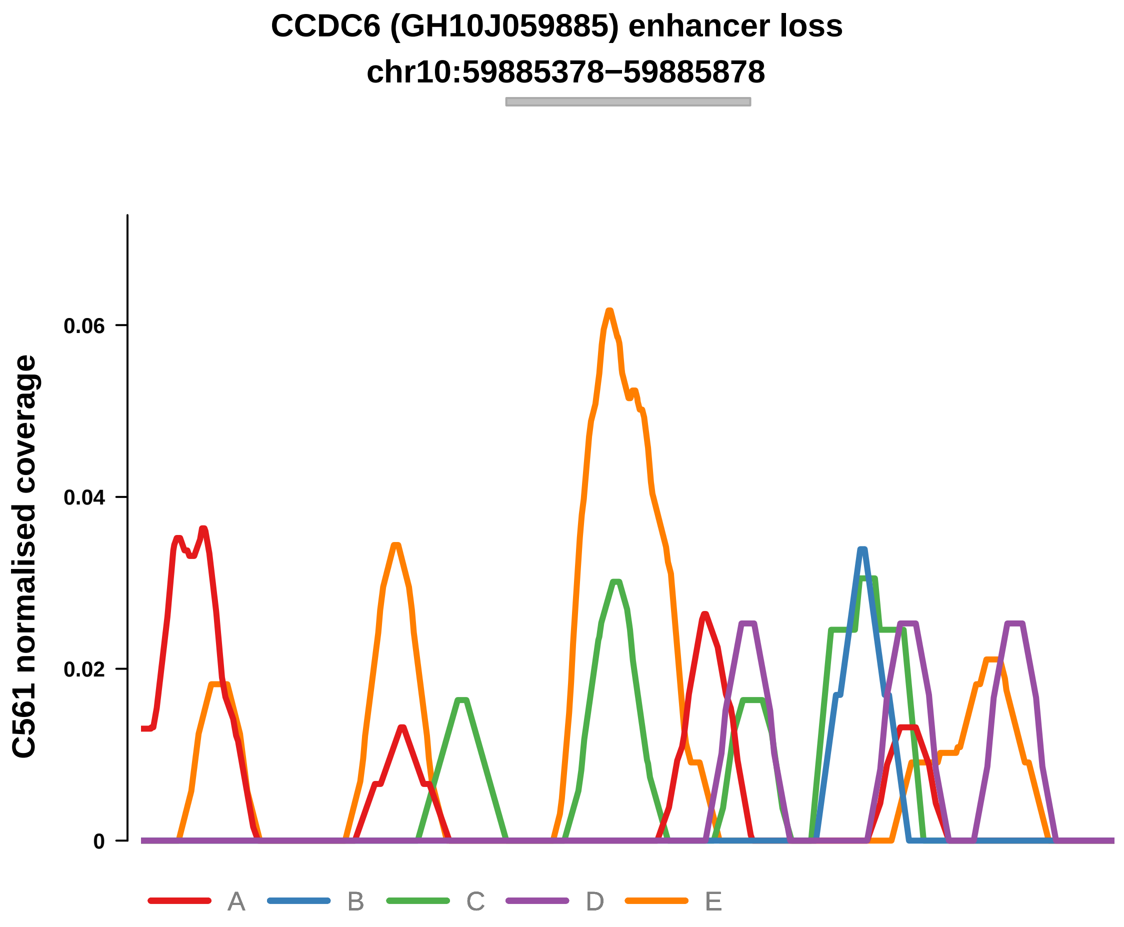


Putatively Clonal (logFC = −2.71)

| | P. | Adjusted P. |
|---|---|---|
| *~region vs. ~* | 8.68e−01 | NA |
| *~purity vs ~* | 4.07e−01 | 7.66e−01 |
| *~purity+region vs ~purity* | 9.57e−01 | 9.79e−01 |
| *Carcinoma vs. normal* | 5.95e−04 *** | 5.95e−04 *** |

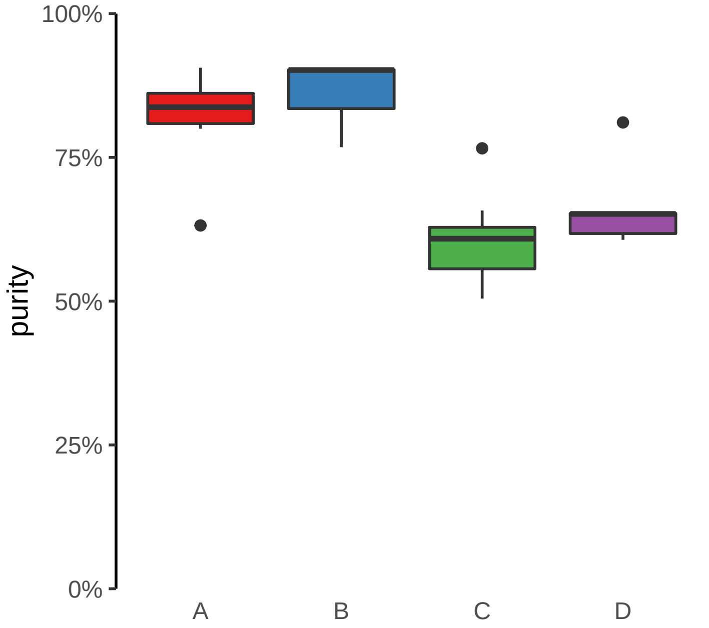

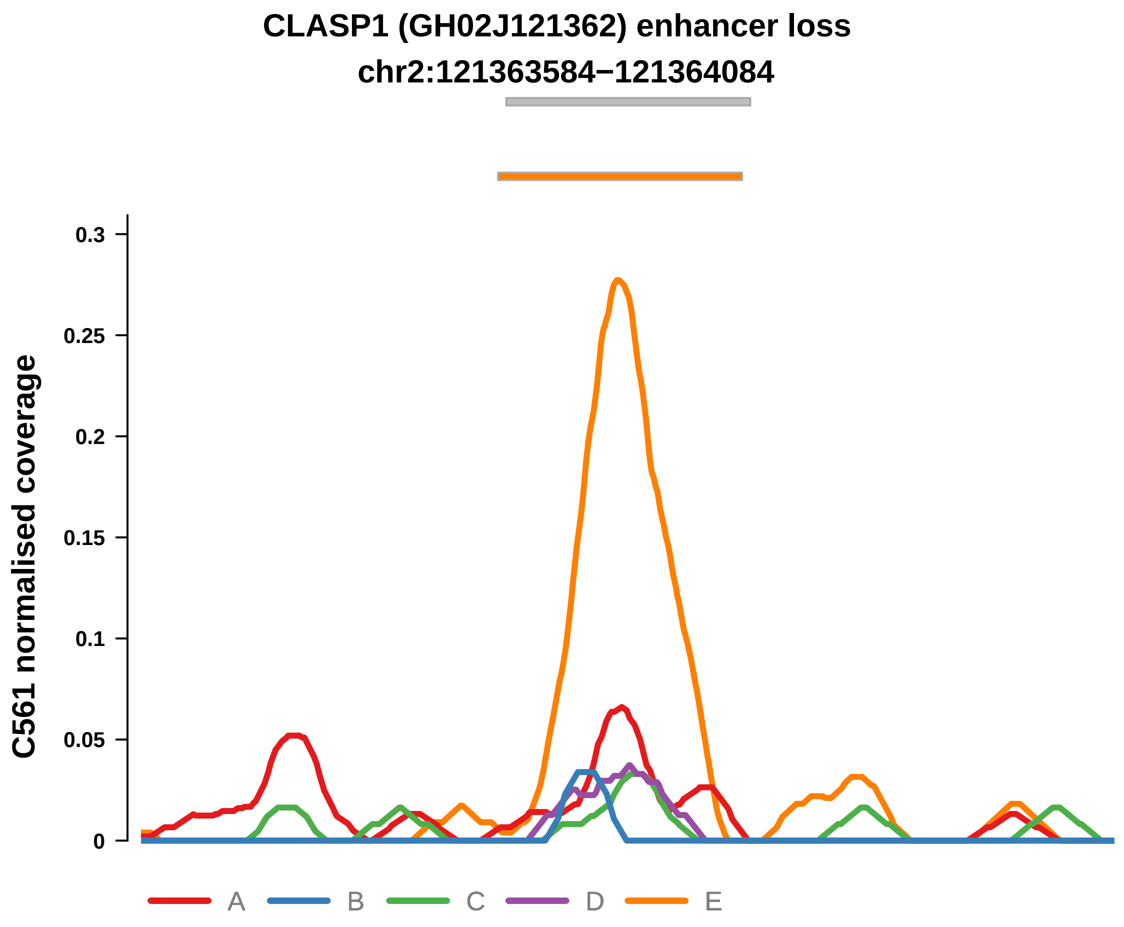


Putatively Clonal (logFC = −1.86)

| | P. | Adjusted P. |
|---|---|---|
| *~region vs. ~* | 6.72e−01 | 7.8e−01 |
| *~purity vs ~* | 1e+00 | 1e+00 |
| *~purity+region vs ~purity* | 7.14e−01 | 9.16e−01 |
| *Carcinoma vs. normal* | 9.19e−03 ** | 9.19e−03 ** |

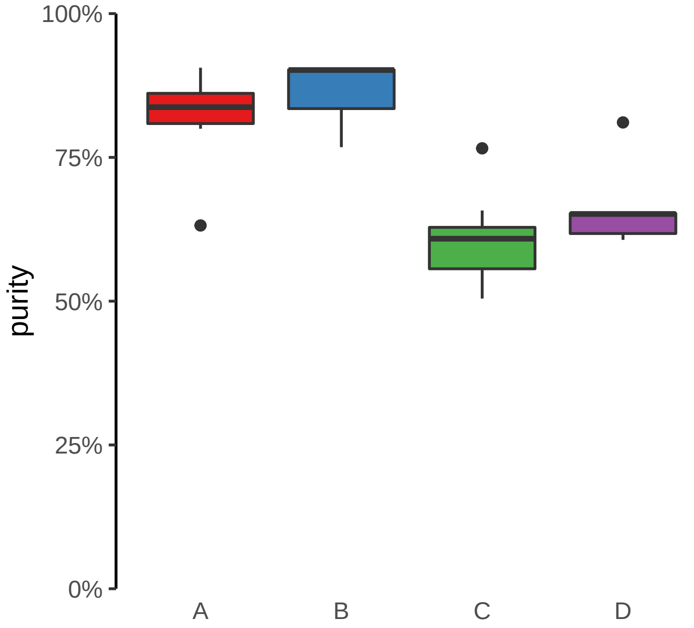

**DNAH7, ENSG00000272211 (GH02J196100) enhancer loss**
**chr2:196101142−196101642**

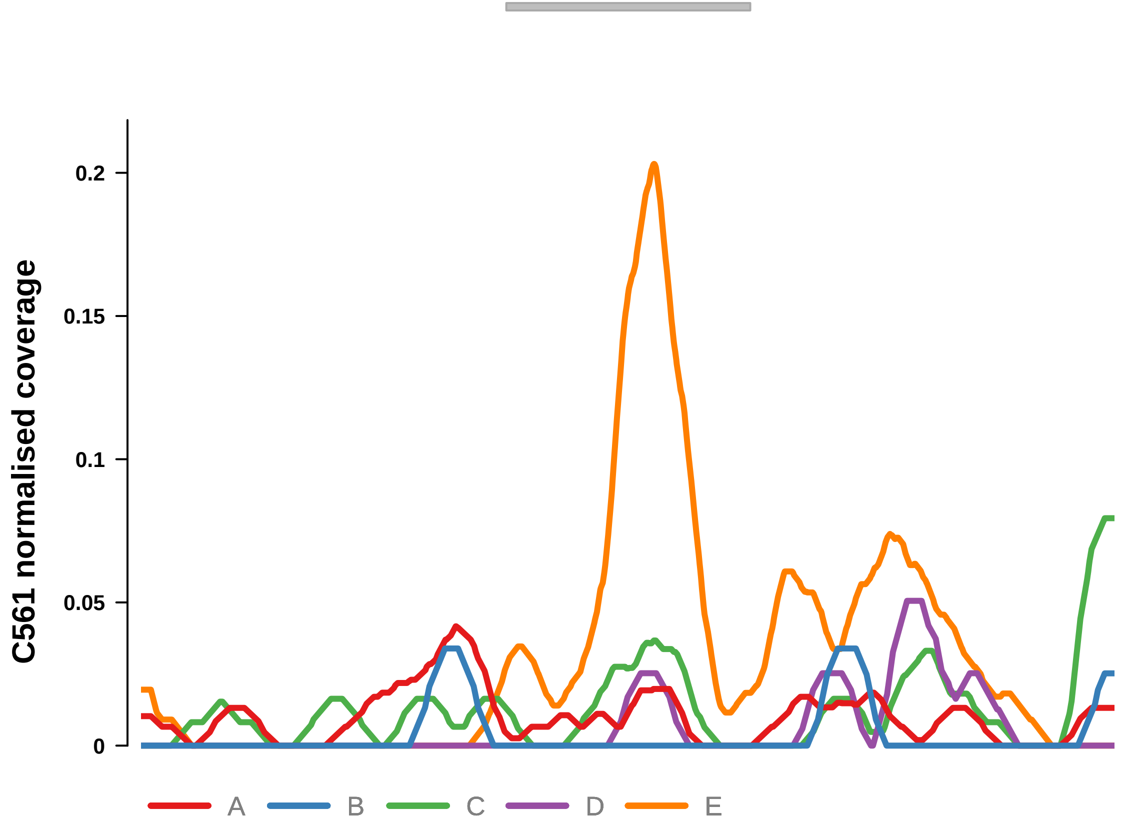


Putatively Clonal (logFC = −2.49)

| | P. | Adjusted P. |
|---|---|---|
| *~region vs. ~* | 4.28e−01 | 5.93e−01 |
| *~purity vs ~* | 1.64e−01 | 5.16e−01 |
| *~purity+region vs ~purity* | 8.57e−01 | 9.76e−01 |
| *Carcinoma vs. normal* | 1.16e−03 ** | 1.16e−03 ** |

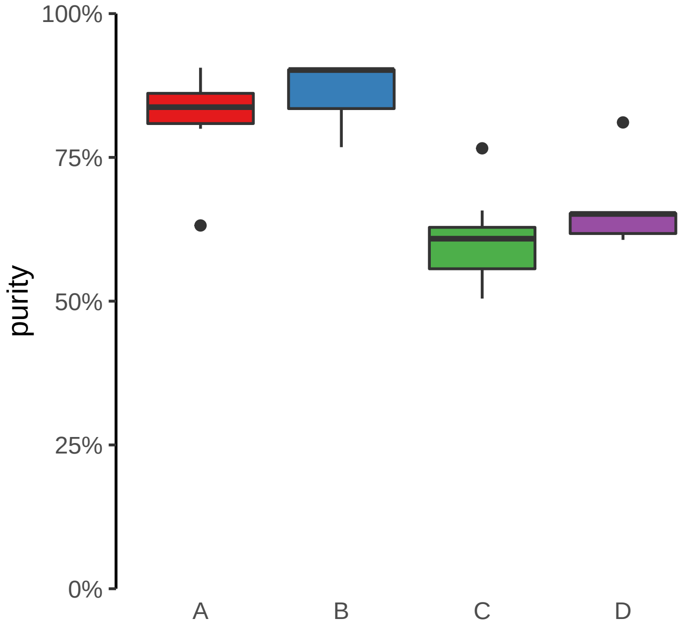

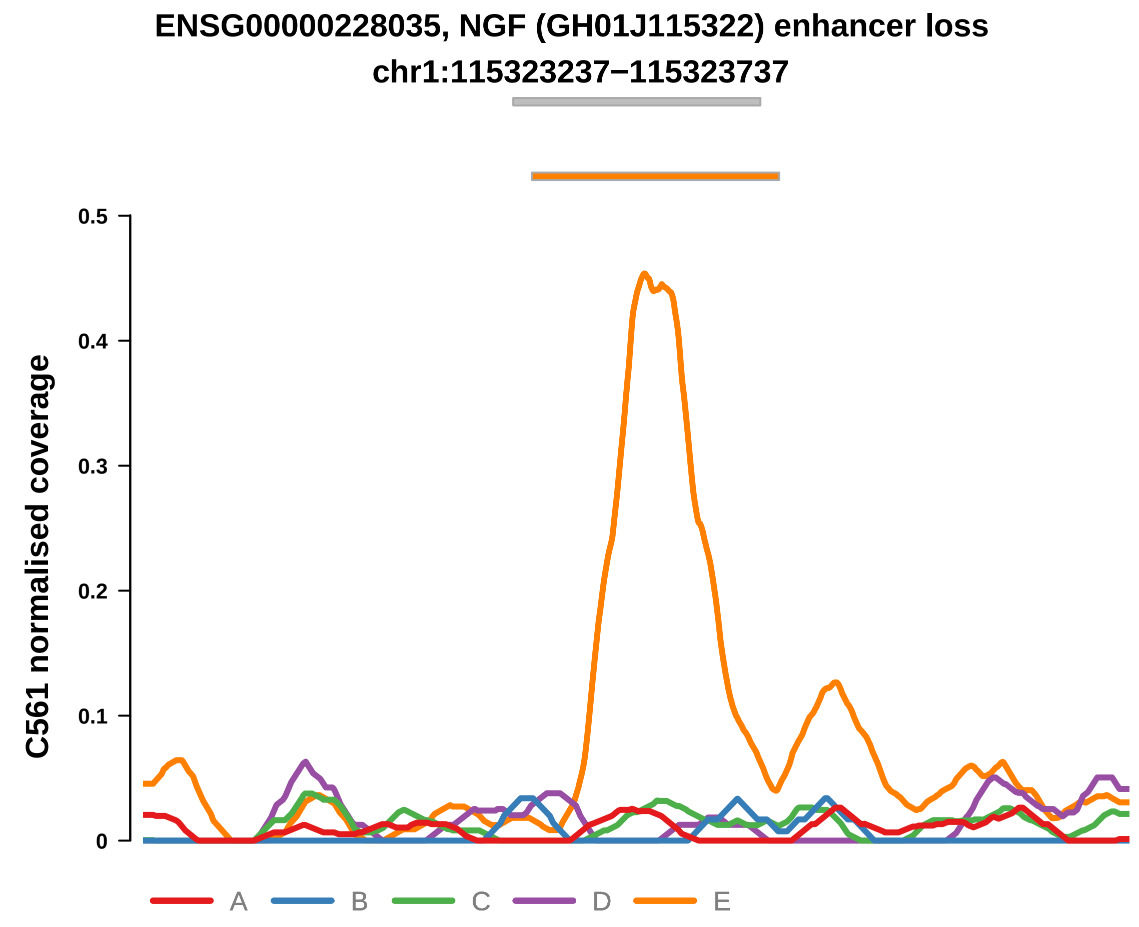


Putatively Clonal (logFC = −2.01)

| | P. | Adjusted P. |
|---|---|---|
| *~region vs. ~* | 4.51e−01 | 6.14e−01 |
| *~purity vs ~* | 5.06e−01 | 8.24e−01 |
| *~purity+region vs ~purity* | 5.29e−01 | 9.12e−01 |
| *Carcinoma vs. normal* | 5.34e−03 ** | 5.34e−03 ** |

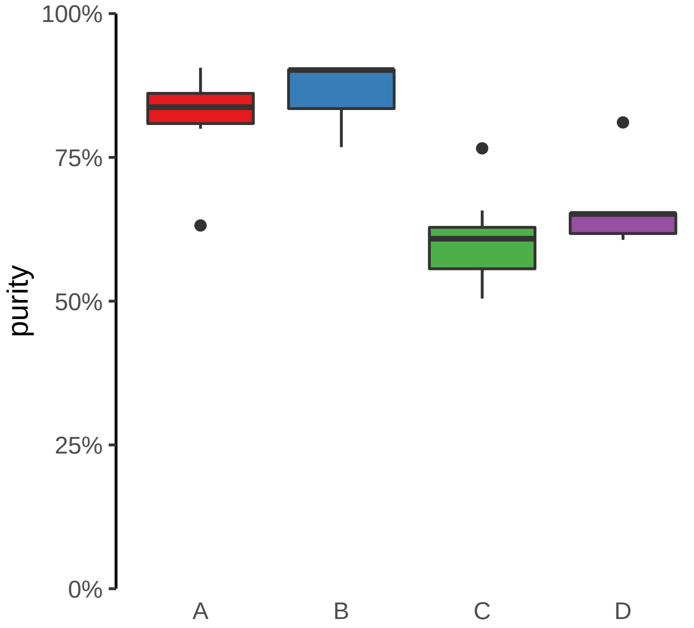

**FABP1, THNSL2 (GH02J088144) enhancer loss**
**chr2:88144817−88145317**

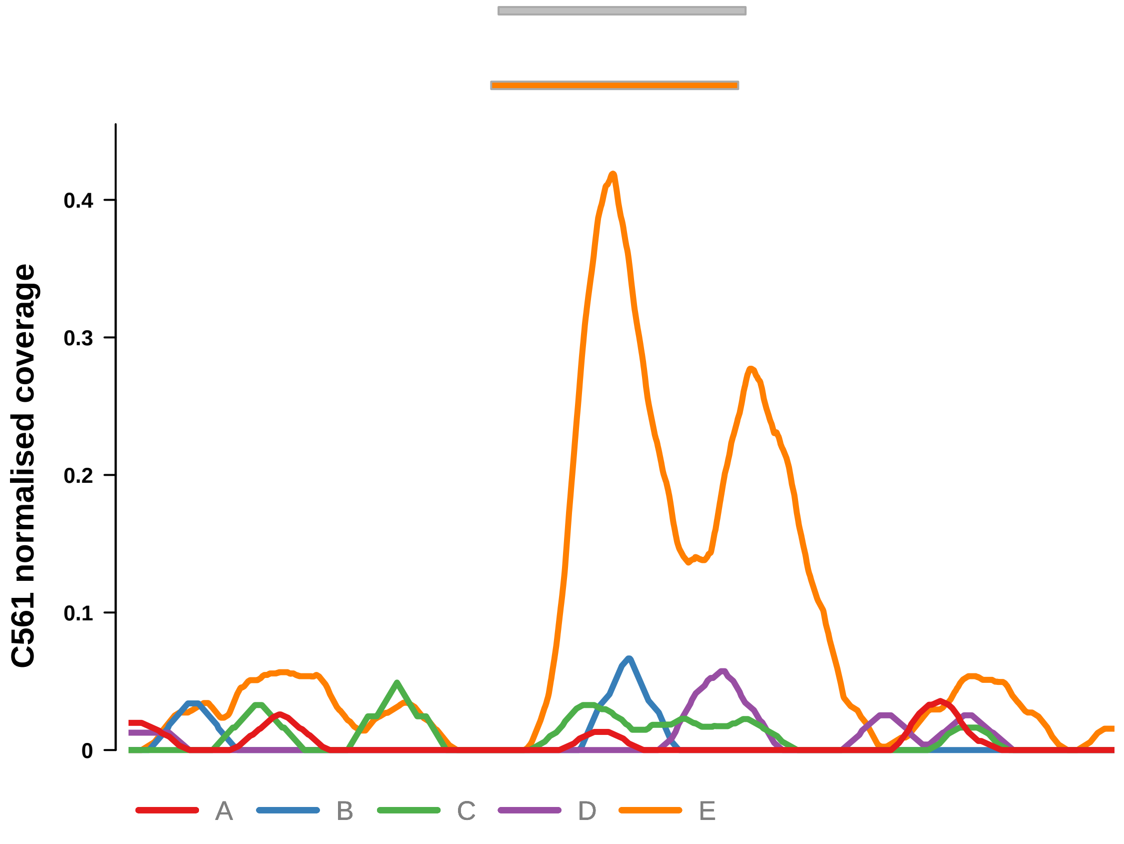


Putatively Clonal (logFC = −2.71)

| | P. | Adjusted P. |
|---|---|---|
| *~region vs. ~* | 2.55e−01 | 4.58e−01 |
| *~purity vs ~* | 8.79e−02 | 3.98e−01 |
| *~purity+region vs ~purity* | 7.01e−01 | 9.16e−01 |
| *Carcinoma vs. normal* | 2.5e−04 *** | 2.5e−04 *** |

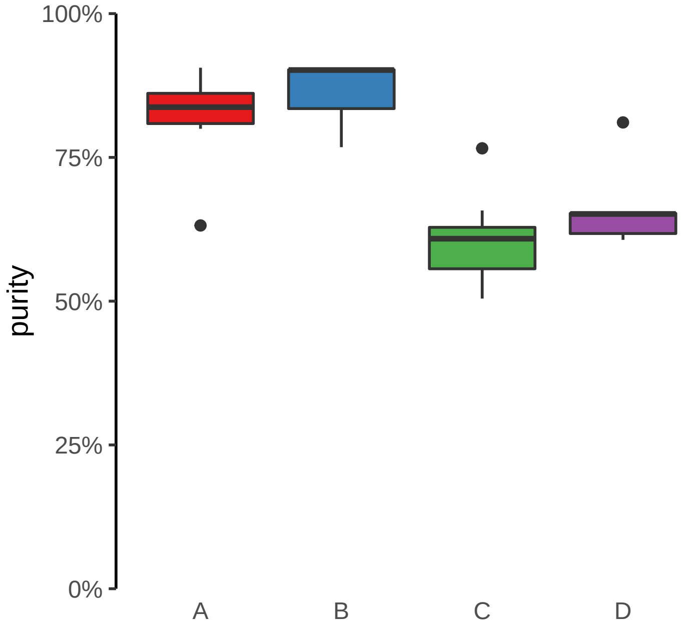

**HMGN2, ARID1A, PIGV (GH01J026739) enhancer loss**
**chr1:26743976−26744476**

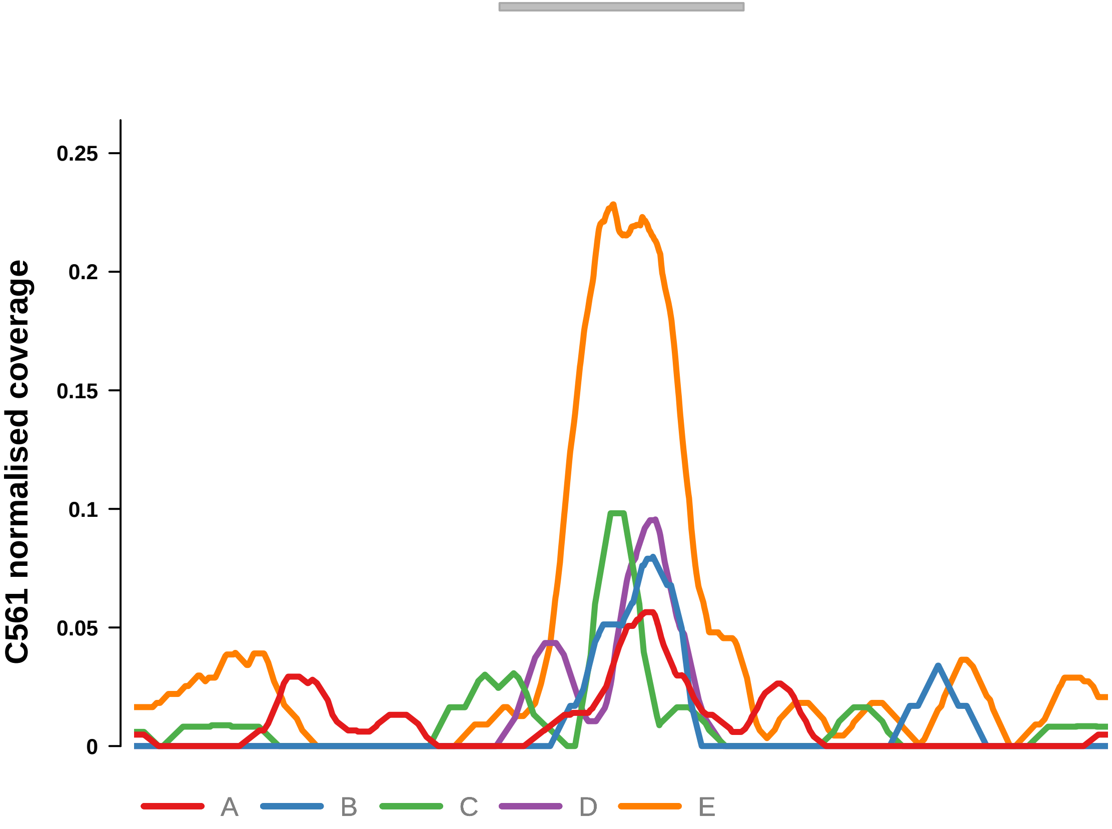


Putatively Clonal (logFC = −2.12)

| | **P.** | **Adjusted P.** |
|---|---|---|
| *~region vs. ~* | 4.05e−01 | 5.91e−01 |
| *~purity vs ~* | 2.12e−01 | 6.08e−01 |
| *~purity+region vs ~purity* | 7.19e−01 | 9.16e−01 |
| *Carcinoma vs. normal* | 2.47e−03 ** | 2.47e−03 ** |

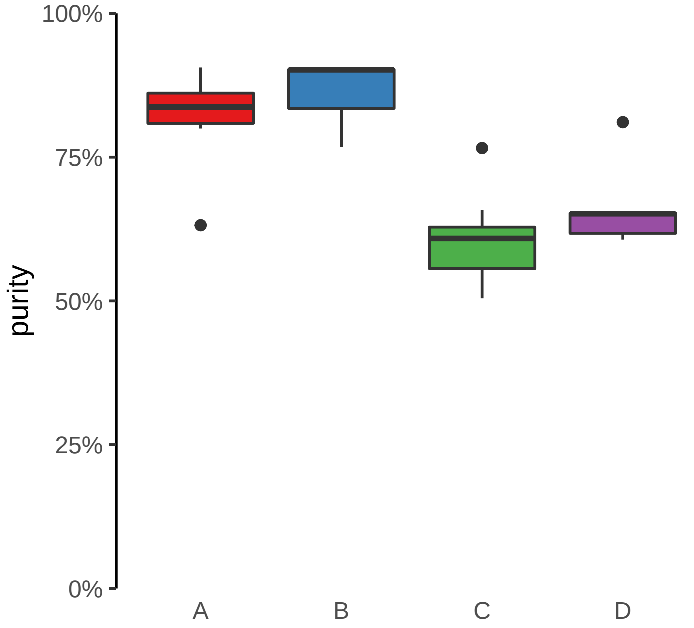

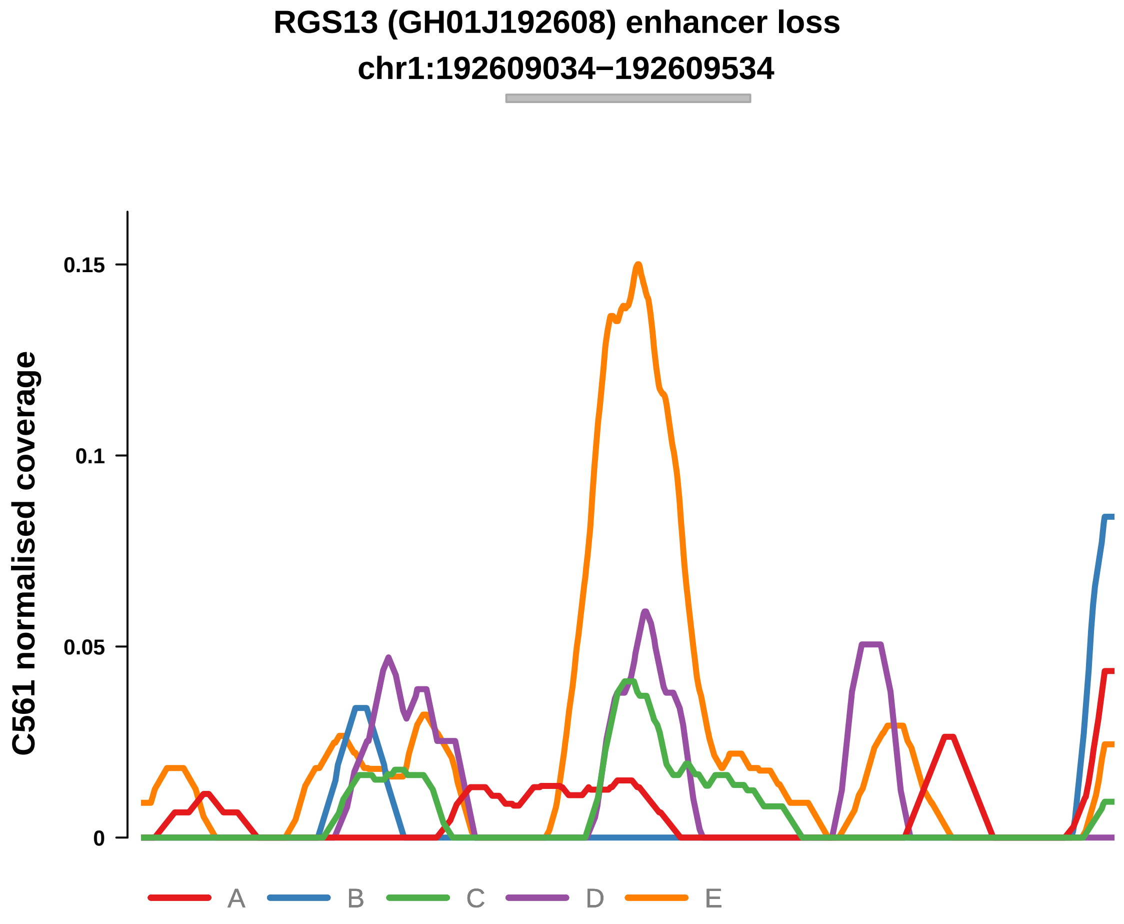


Putatively Clonal (logFC = −2.36)

| | P. | Adjusted P. |
|---|---|---|
| *~region vs. ~* | 3.14e−01 | 5.29e−01 |
| *~purity vs ~* | 1.43e−01 | 4.84e−01 |
| *~purity+region vs ~purity* | 6.97e−01 | 9.16e−01 |
| *Carcinoma vs. normal* | 1.51e−03 ** | 1.51e−03 ** |

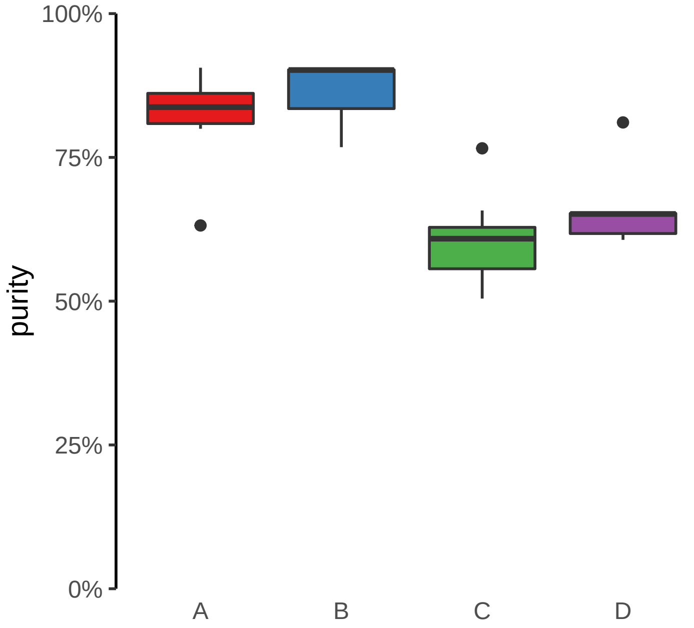

**SMAD3 (GH15J067079) enhancer loss**
**chr15:67081050−67081550**

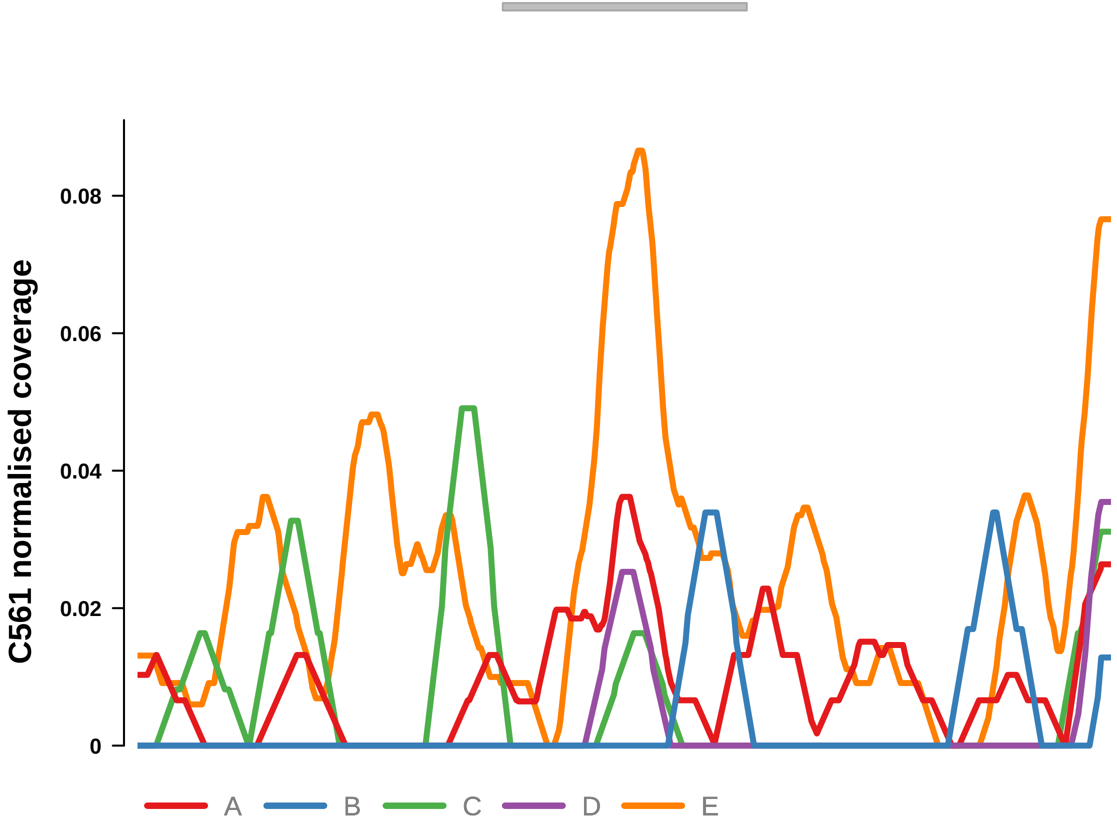


Putatively Clonal (logFC = −2.17)

| | P. | Adjusted P. |
|---|---|---|
| *~region vs. ~* | 8.12e−01 | 8.91e−01 |
| *~purity vs ~* | 6.76e−01 | 8.75e−01 |
| *~purity+region vs ~purity* | 7.7e−01 | 9.16e−01 |
| *Carcinoma vs. normal* | 4.23e−03 ** | 4.23e−03 ** |

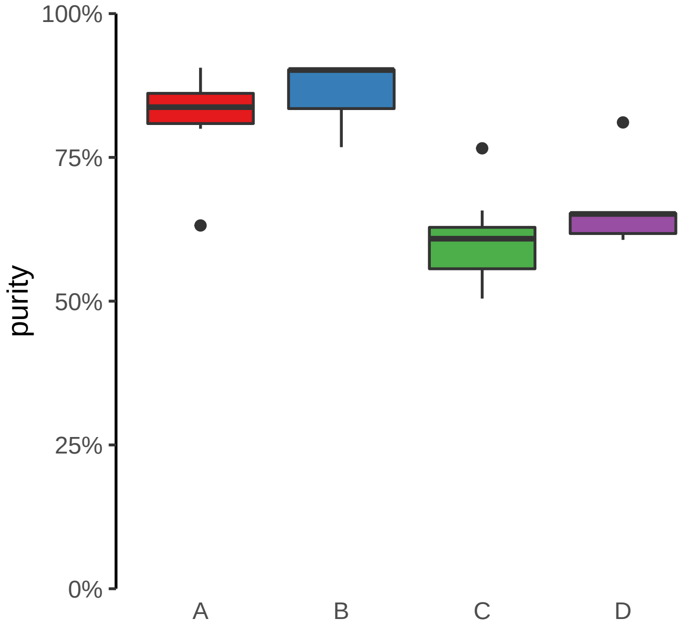

**STAG1, RNU6−789P (GH03J136616) enhancer loss**
**chr3:136617156−136617656**

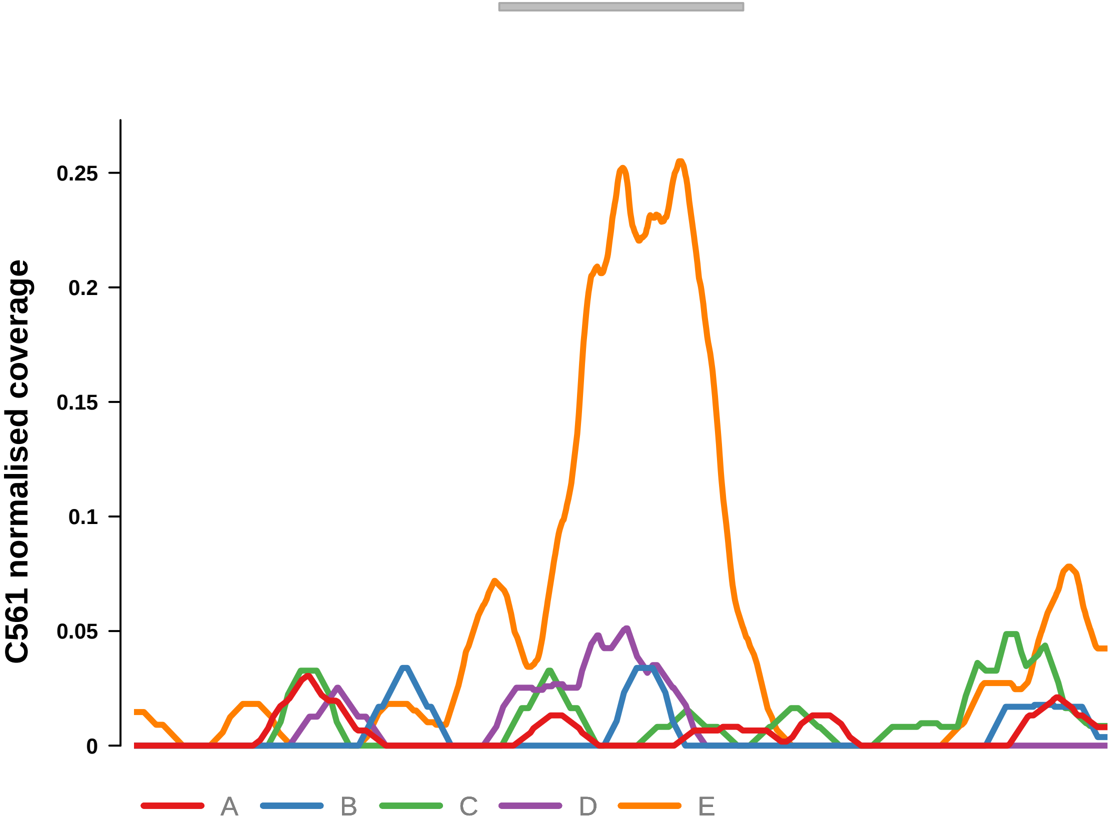


Putatively Clonal (logFC = −2.73)

| | P. | Adjusted P. |
|---|---|---|
| *~region vs. ~* | 3.59e−01 | 5.79e−01 |
| *~purity vs ~* | 7.91e−01 | 9.05e−01 |
| *~purity+region vs ~purity* | 2.41e−01 | 7.31e−01 |
| *Carcinoma vs. normal* | 2.43e−04 *** | 2.43e−04 *** |

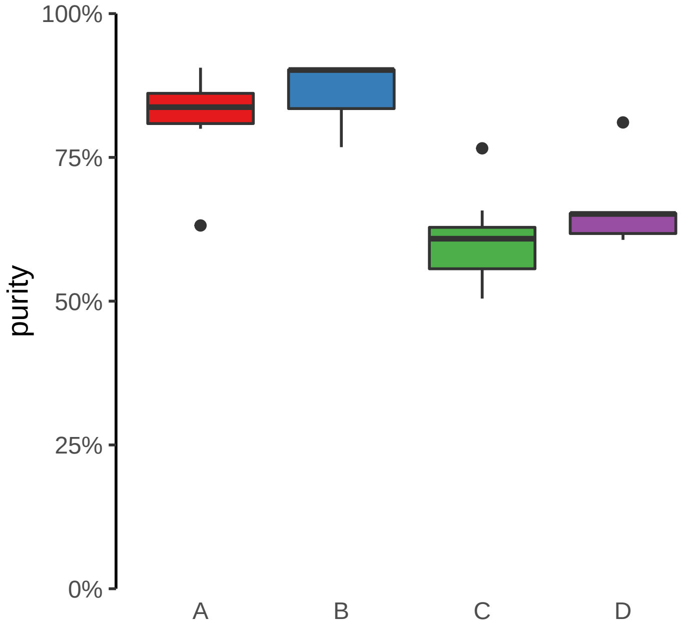

**TRANK1, ENSG00000234073 (GH03J036874) enhancer loss**
**chr3:36876260−36876760**

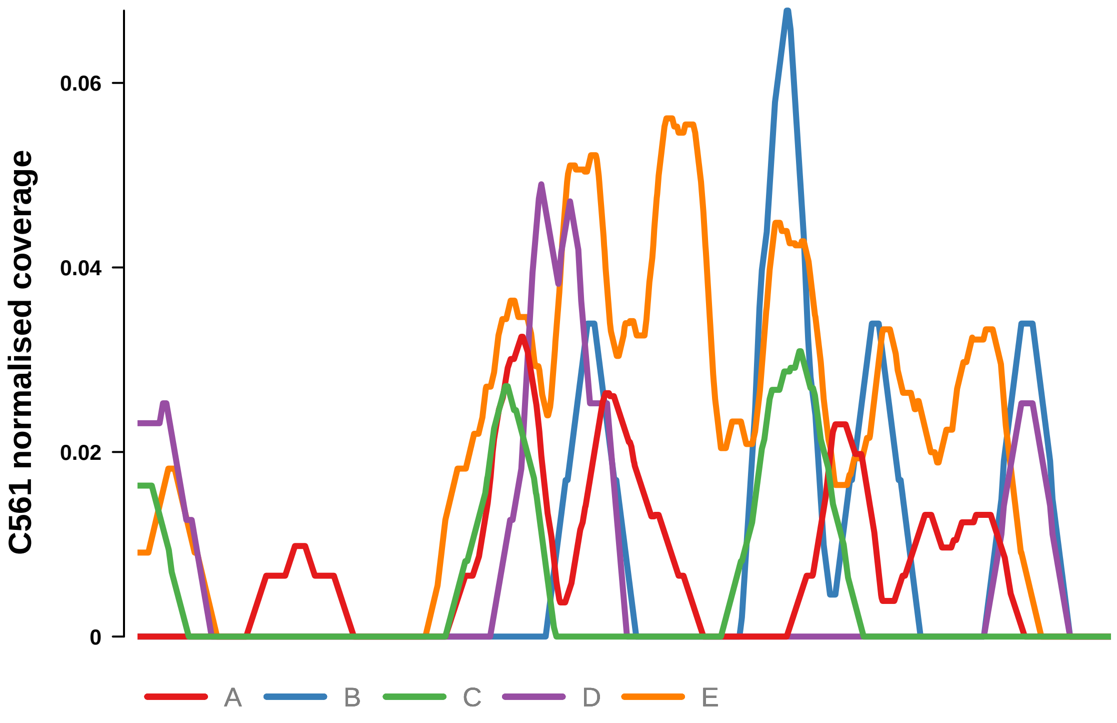


Putatively Clonal (logFC = −2.15)

| | P. | Adjusted P. |
|---|---|---|
| *~region vs. ~* | 5.31e−01 | 6.87e−01 |
| *~purity vs ~* | 4.16e−01 | 7.66e−01 |
| *~purity+region vs ~purity* | 6.11e−01 | 9.16e−01 |
| *Carcinoma vs. normal* | 3.69e−03 ** | 3.69e−03 ** |

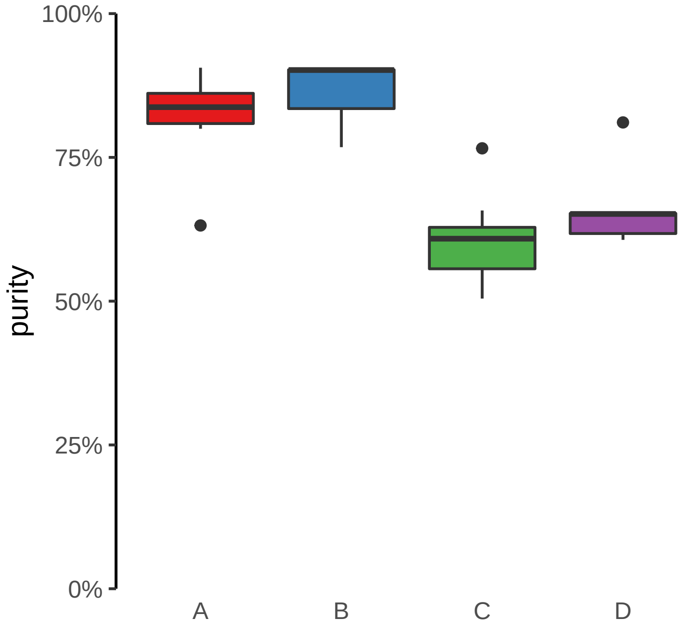
