## Supplementary Figures for "The co-evolution of the genome and epigenome in colorectal cancer": figS14_subclonal_adenoma.pdf

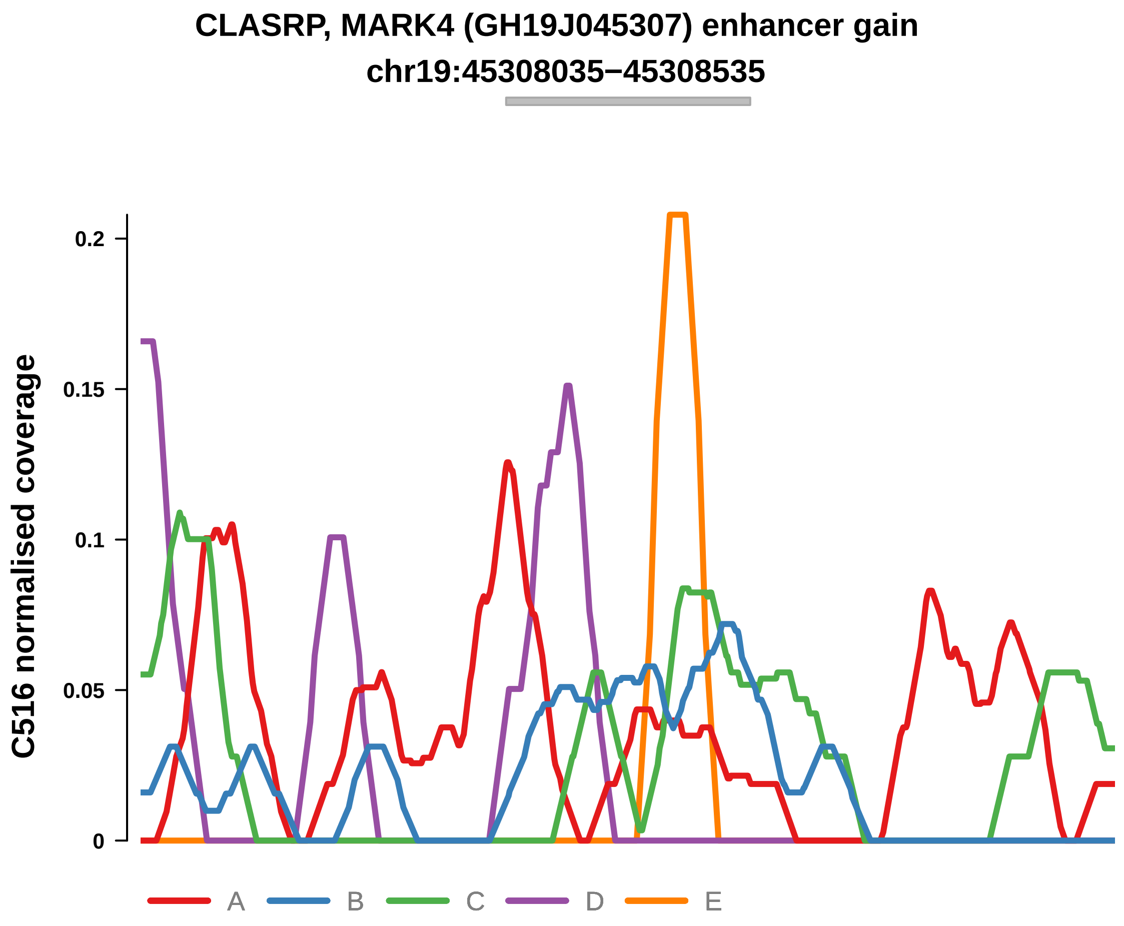


Putatively Subclonal

| | LogFC | P. | Adjusted P. |
|---|---|---|---|
| *Adenoma vs. normal* | −0.28 | NA | NA |
| *Adenoma vs. Cancer ~purity+type* | −3.57 | 3.55e−03 ** | 3.91e−02 * |

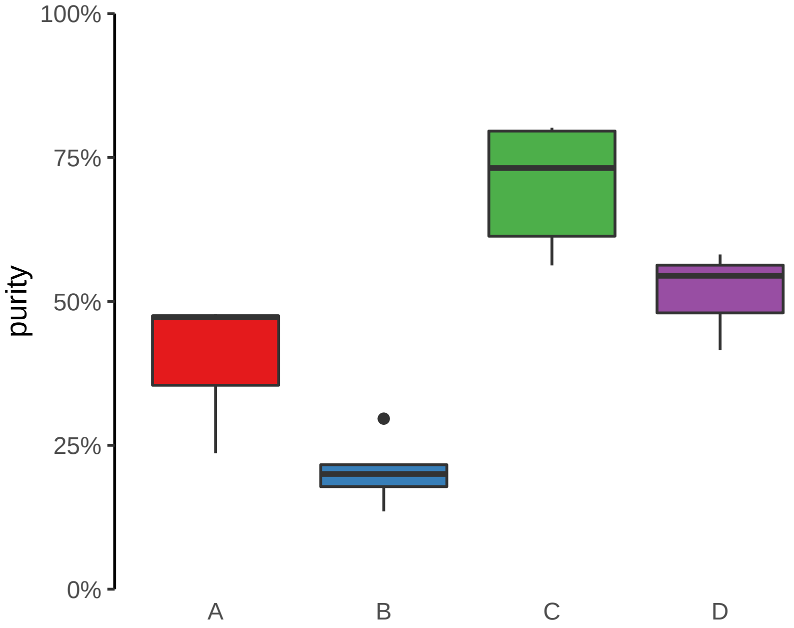

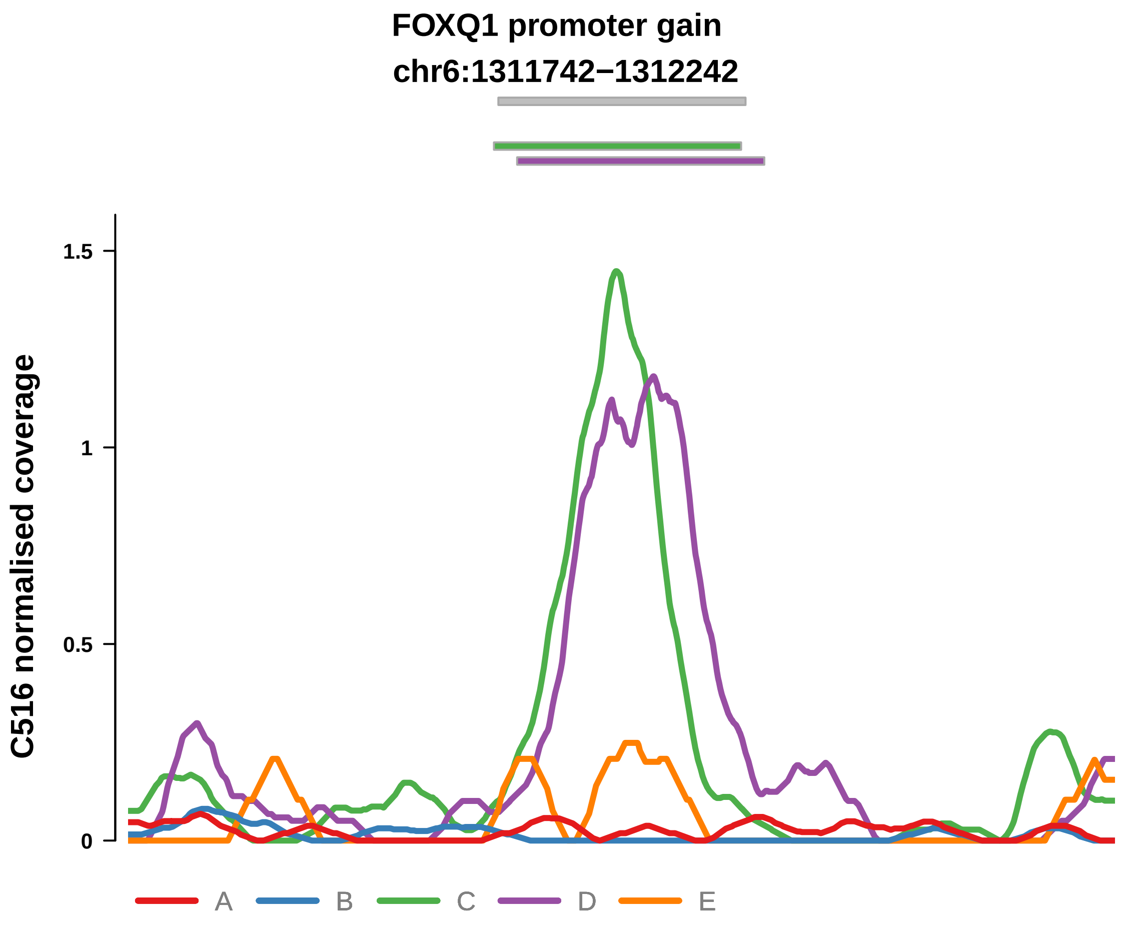


Putatively Subclonal

| | LogFC | P. | Adjusted P. |
|---|---|---|---|
| *Adenoma vs. normal* | 2.85 | 5.24e−07 **** | 5.24e−07 **** |
| *Adenoma vs. Cancer ~purity+type* | 2.57 | 1.49e−03 ** | 2.88e−02 * |

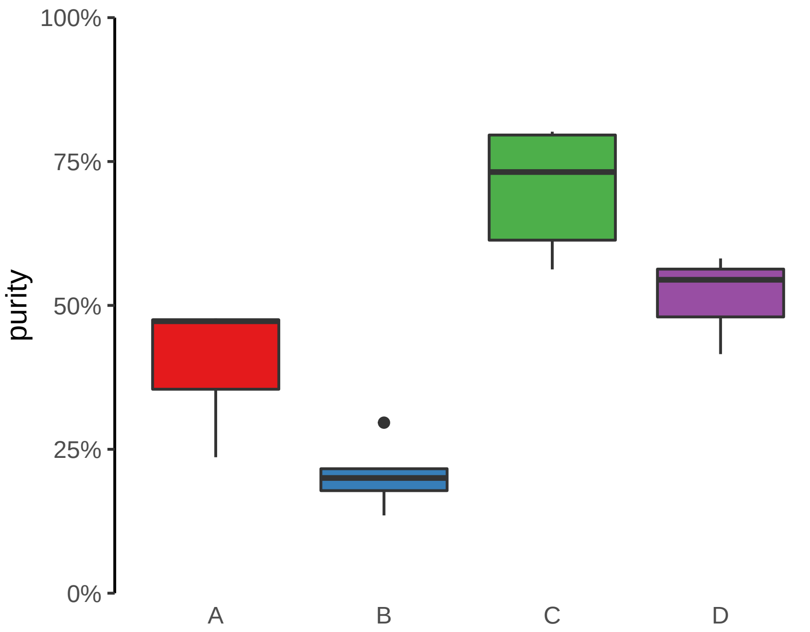

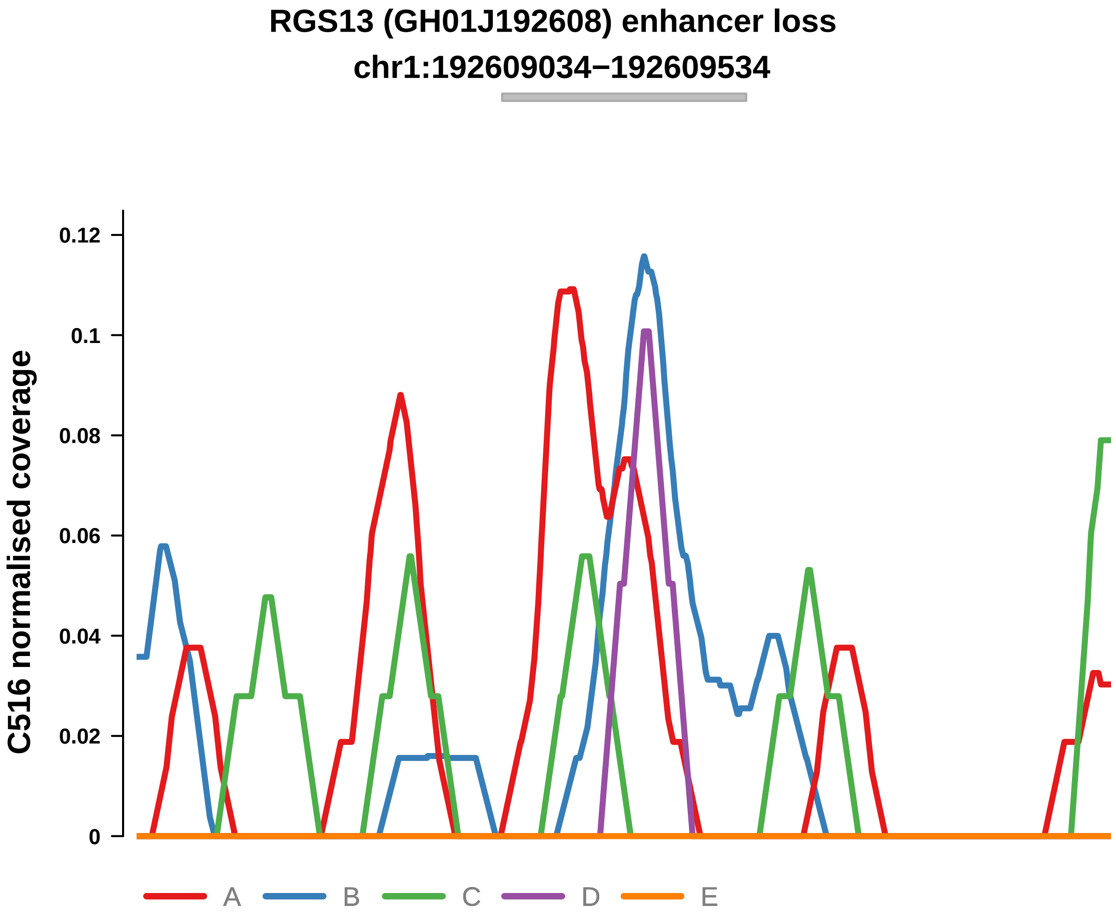


Putatively Subclonal

| | LogFC | P. | Adjusted P. |
|---|---|---|---|
| *Adenoma vs. normal* | −2.93 | 1.42e−03 ** | 1.42e−03 ** |
| *Adenoma vs. Cancer ~purity+type* | −4.53 | 3e−03 ** | 3.84e−02 * |

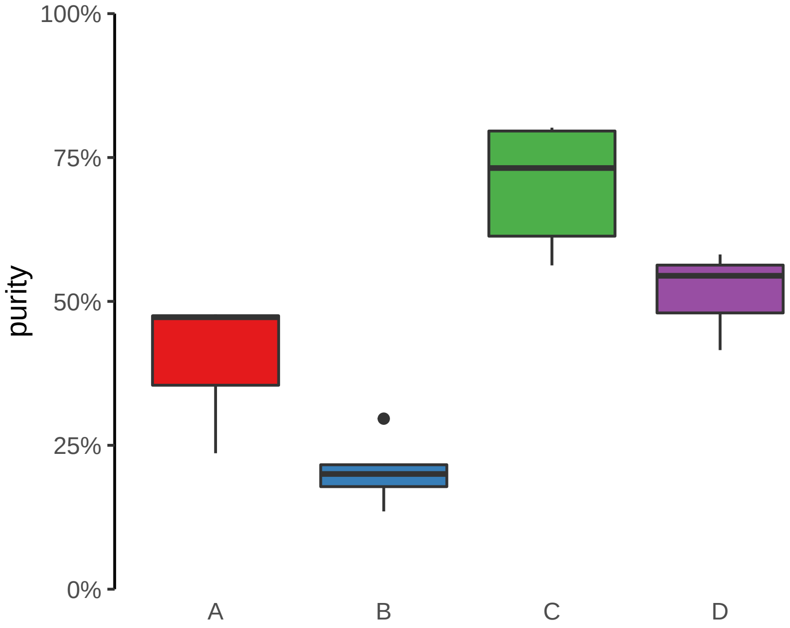

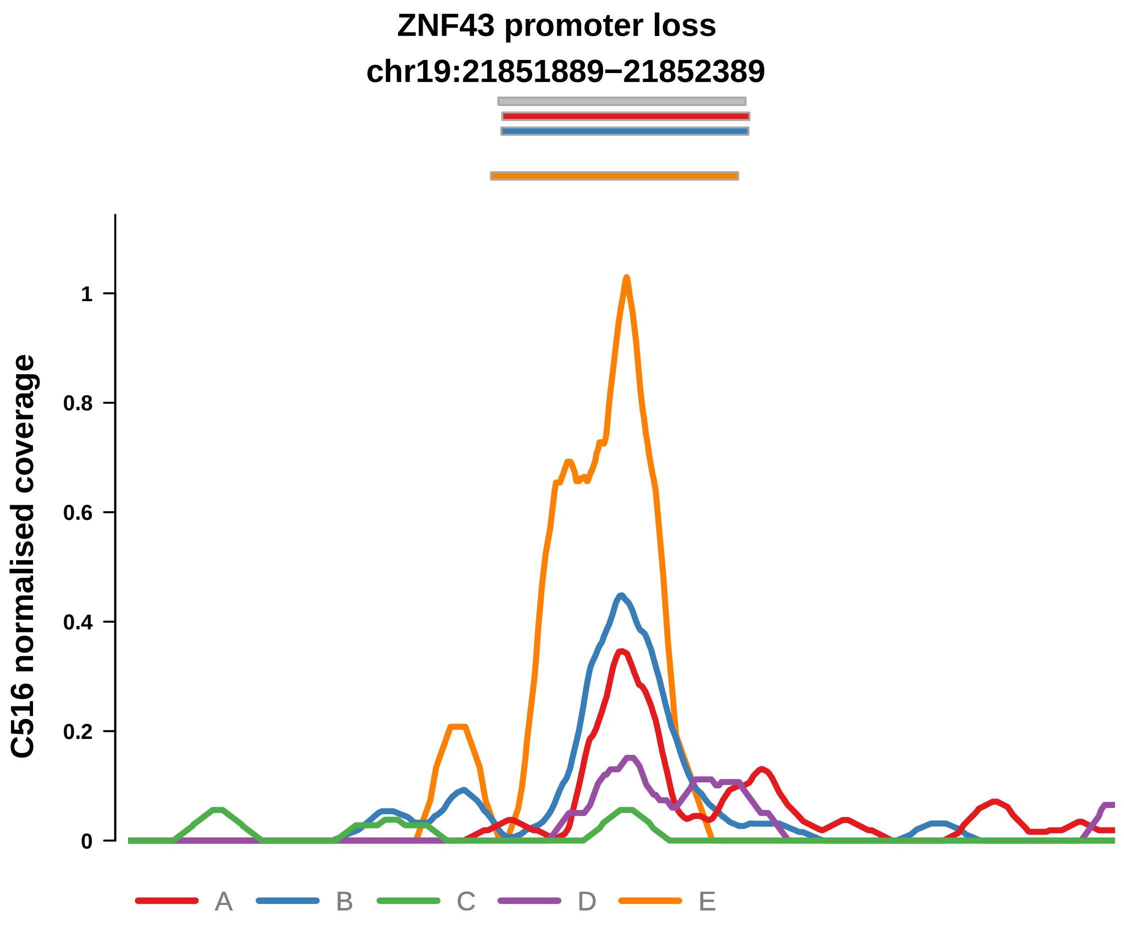


Putatively Subclonal

| | LogFC | P. | Adjusted P. |
|---|---|---|---|
| *Adenoma vs. normal* | −3.83 | 1e−08 **** | 1e−08 **** |
| *Adenoma vs. Cancer ~purity+type* | −3.63 | 3.49e−05 **** | 8.96e−04 *** |

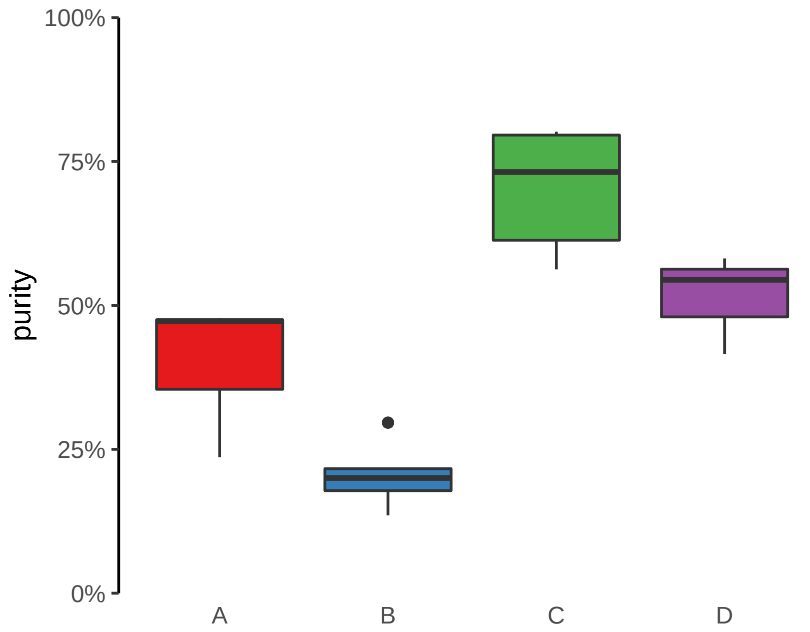

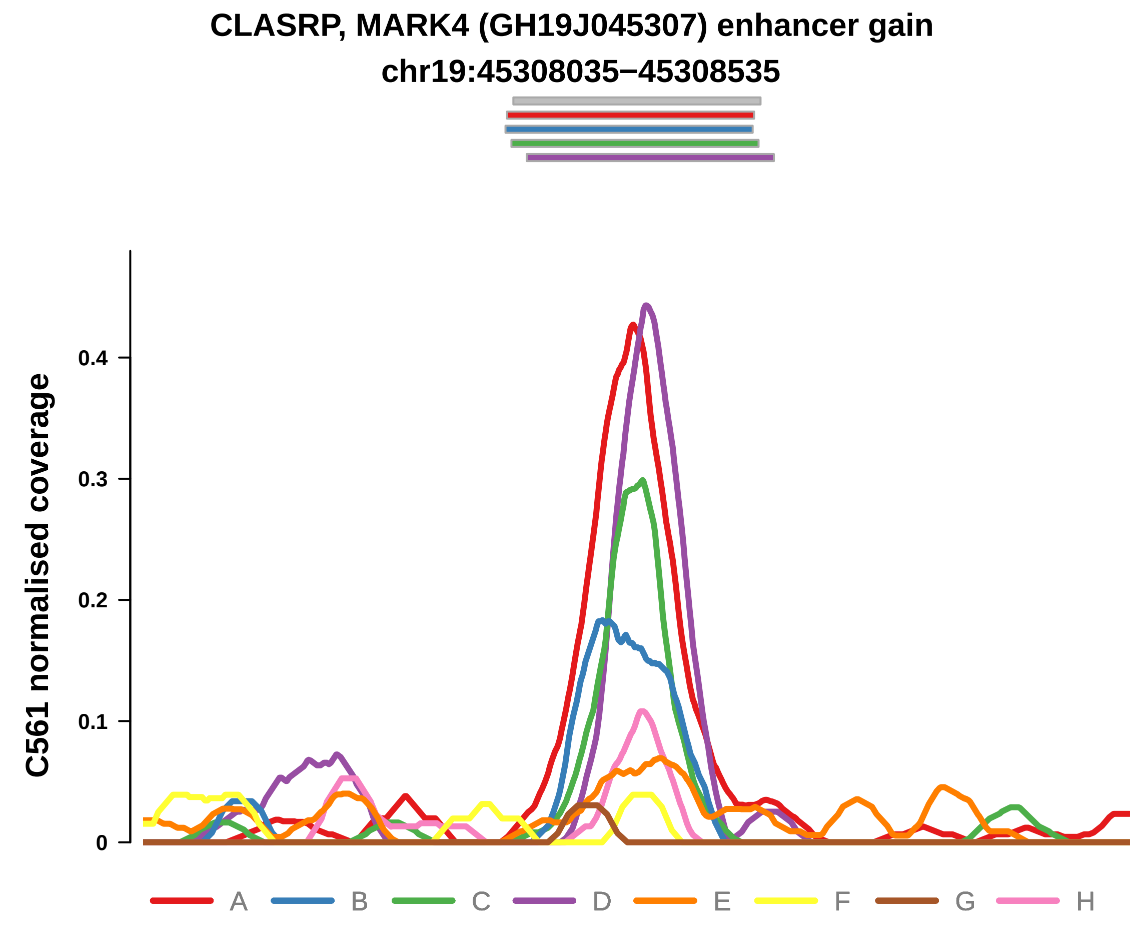


Putatively Subclonal

| | LogFC | P. | Adjusted P. |
|---|---|---|---|
| *Adenoma vs. normal* | −0.94 | 4.98e−02 * | 4.98e−02 * |
| *F vs.Cancer ~purity+type* | −4.19 | 2.87e−04 *** | 6.12e−03 ** |
| *G vs.Cancer ~purity+type* | −5.53 | 2.28e−05 **** | 1.39e−03 ** |

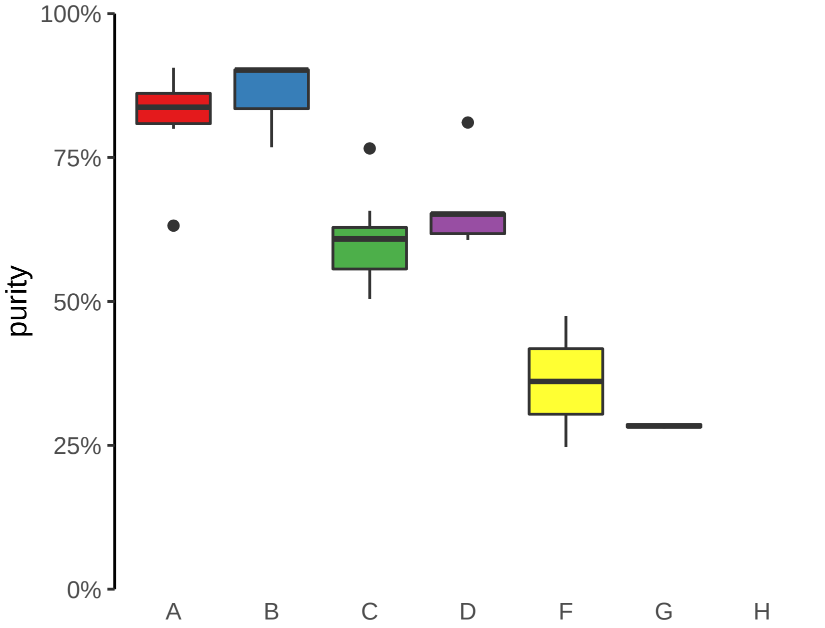

**ENSG00000253931 (GH08J143405) enhancer gain**
**chr8:143407862−143408362**

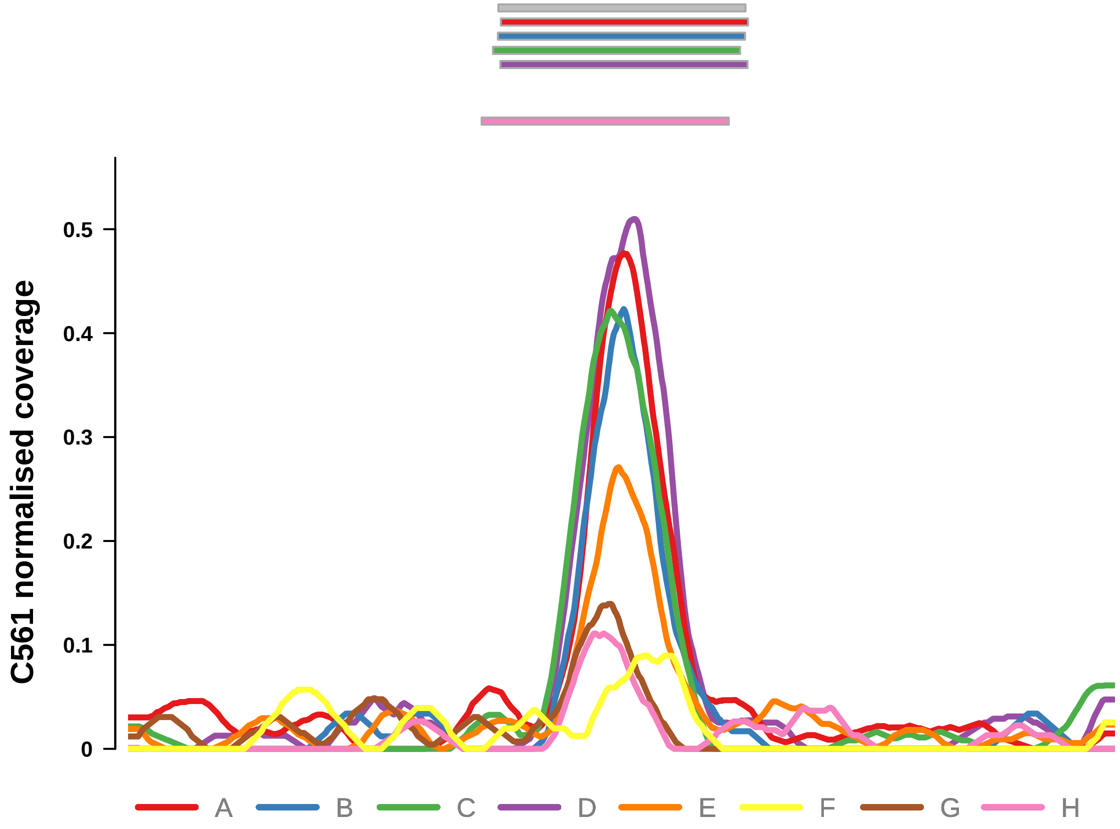


Putatively Subclonal

| | LogFC | P. | Adjusted P. |
|---|---|---|---|
| *Adenoma vs. normal* | −0.94 | 4.98e−02 * | 4.98e−02 * |
| *F vs.Cancer ~purity+type* | −3.64 | 1.05e−04 *** | 3.35e−03 ** |
| *G vs.Cancer ~purity+type* | −4.03 | 6.69e−05 **** | 2.04e−03 ** |

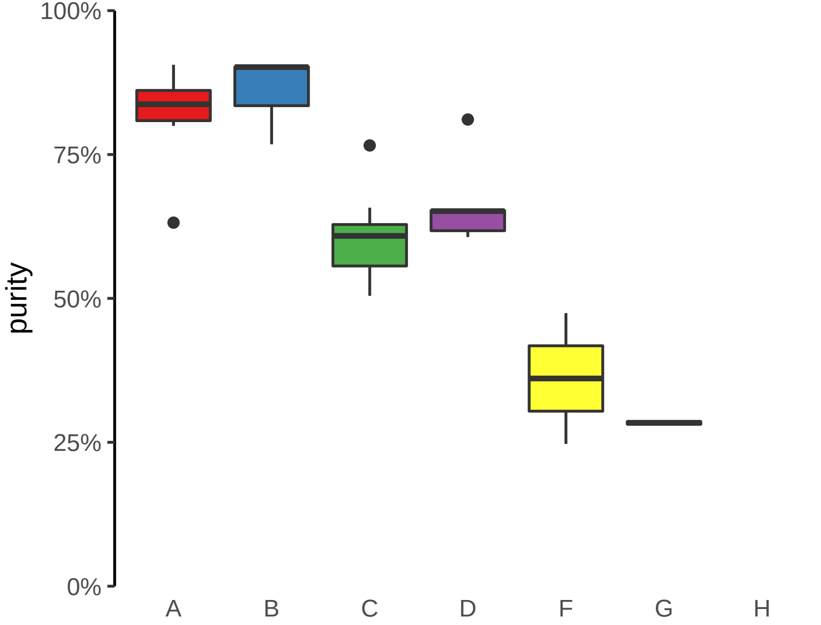

**NXPH1 (GH07J008389) enhancer gain**
**chr7:8389807−8390307**

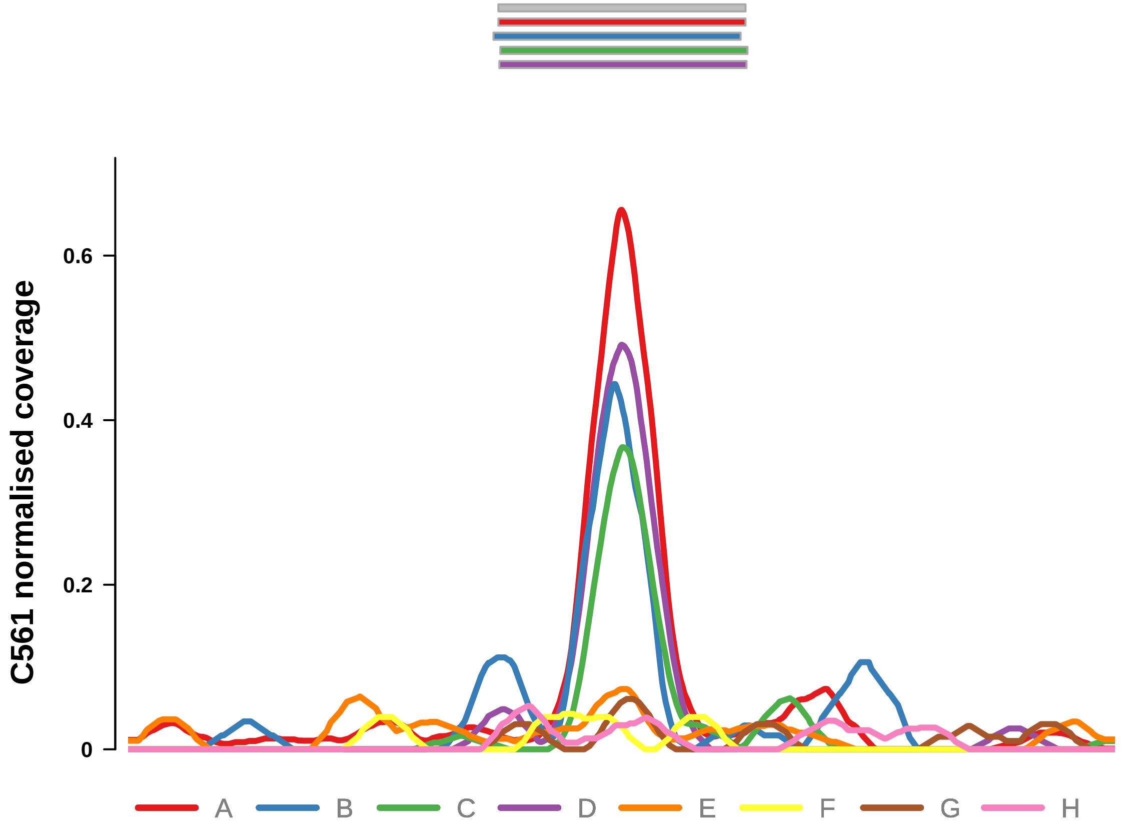


Putatively Subclonal

| | LogFC | P. | Adjusted P. |
|---|---|---|---|
| *Adenoma vs. normal* | −0.94 | 4.98e−02 * | 4.98e−02 * |
| *F vs.Cancer ~purity+type* | −2.99 | 4.27e−03 ** | 5.47e−02 |
| *G vs.Cancer ~purity+type* | −3.23 | 3.58e−03 ** | 3.64e−02 * |

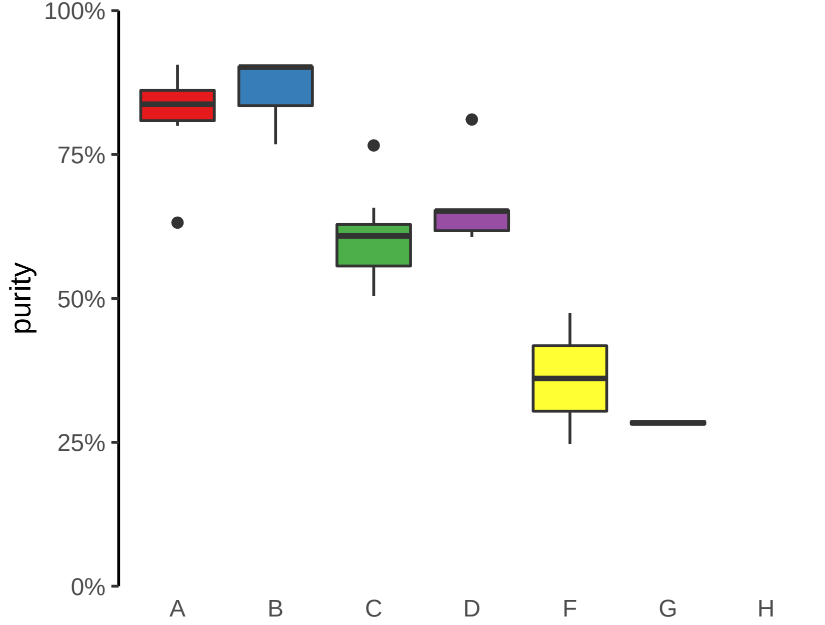

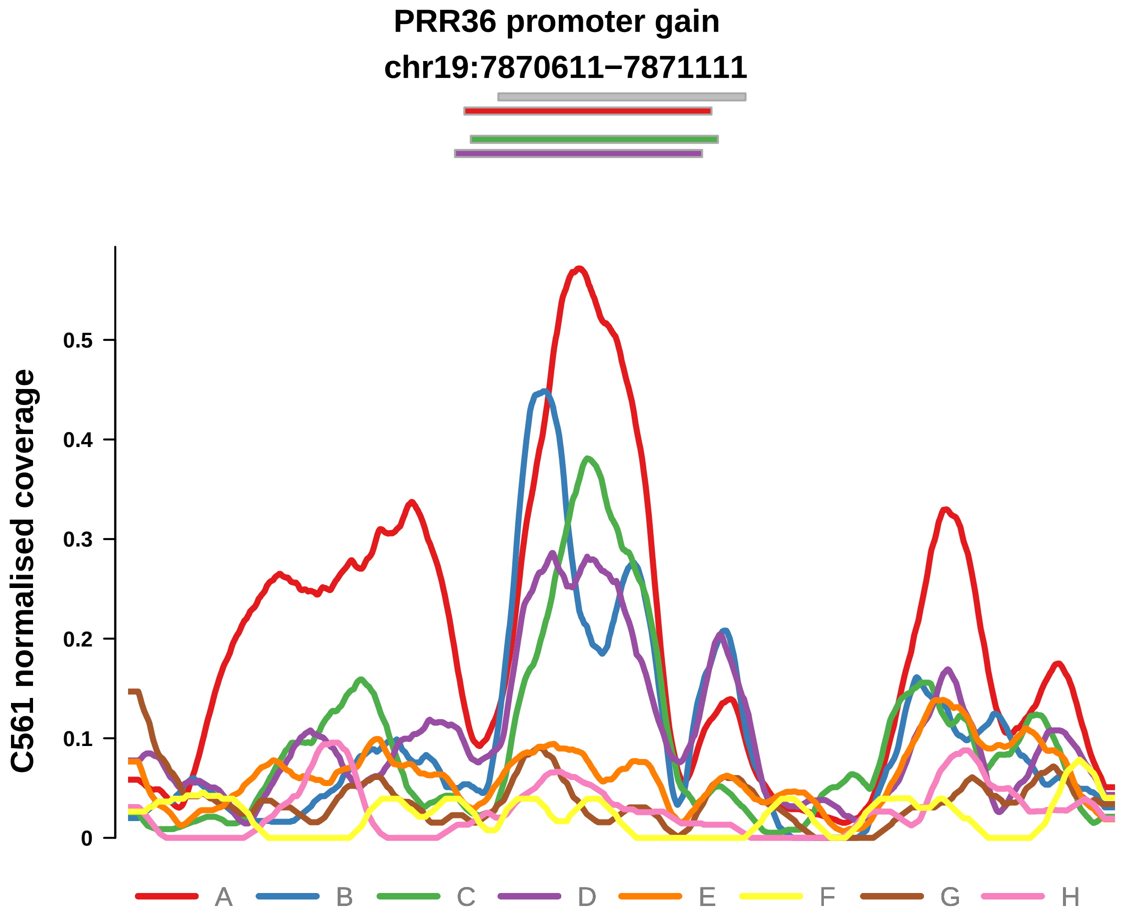


Putatively Subclonal

| | LogFC | P. | Adjusted P. |
|---|---|---|---|
| *Adenoma vs. normal* | −0.94 | 4.98e−02 * | 4.98e−02 * |
| *F vs.Cancer ~purity+type* | −3.79 | 2.59e−05 **** | 1.66e−03 ** |
| *G vs.Cancer ~purity+type* | −2.49 | 2.58e−03 ** | 3.14e−02 * |

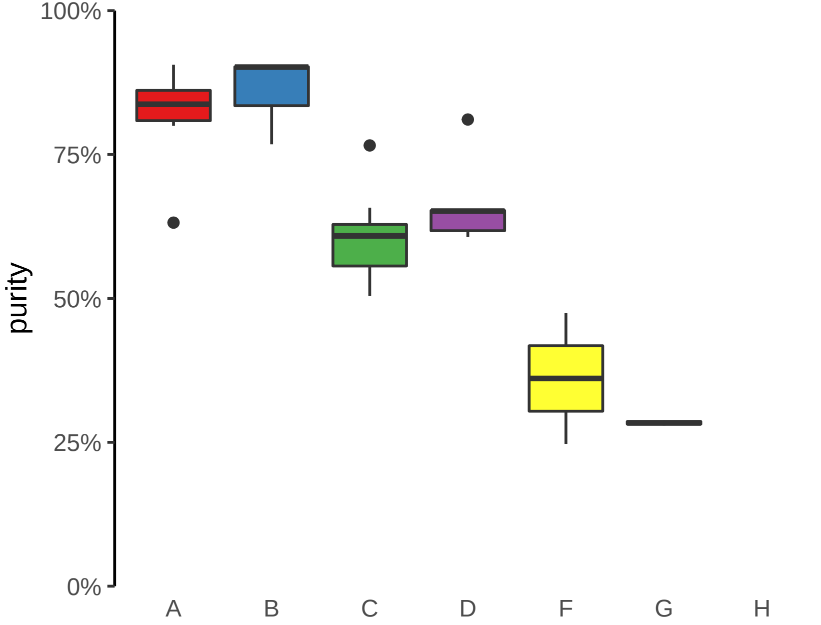

**RNLS, LIPJ (GH10J088553) enhancer loss**
**chr10:88553102−88553602**

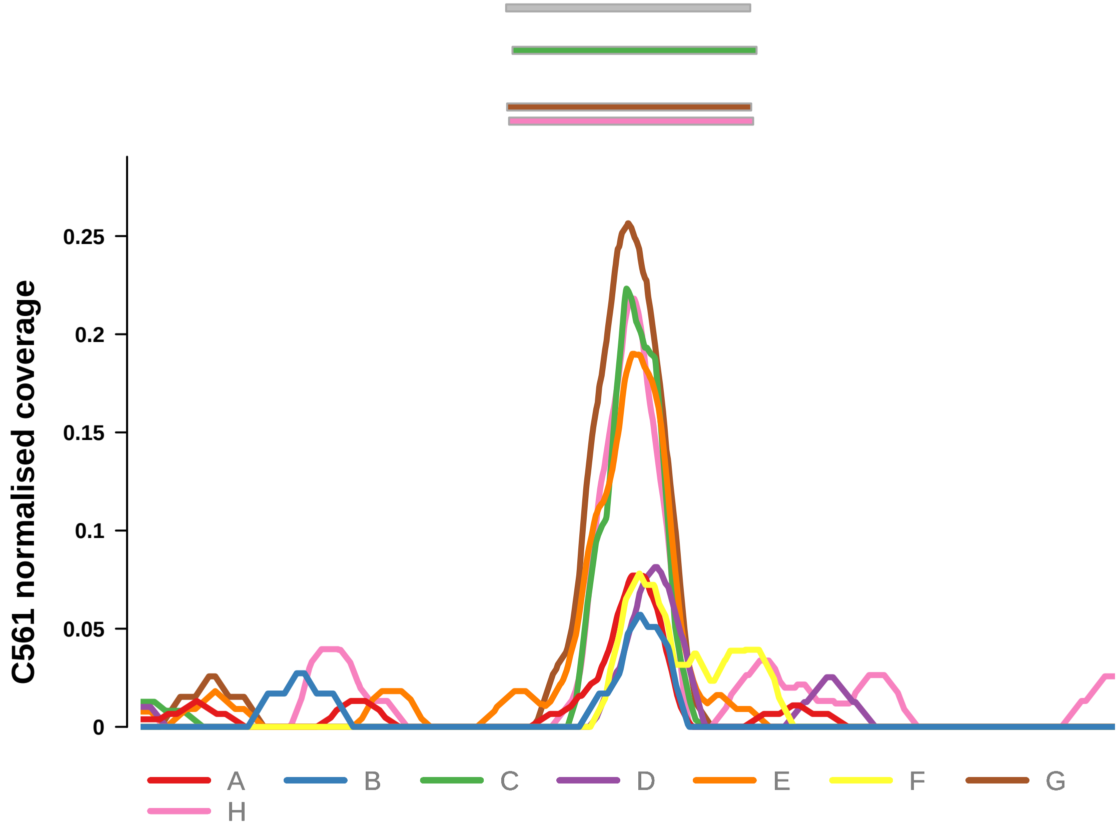


Putatively Subclonal

| | LogFC | P. | Adjusted P. |
|---|---|---|---|
| *Adenoma vs. normal* | −0.94 | 4.98e−02 * | 4.98e−02 * |
| *F vs.Cancer ~purity+type* | −4.84 | 4.02e−03 ** | 5.47e−02 |
| *G vs.Cancer ~purity+type* | −5.51 | 2.47e−03 ** | 3.14e−02 * |

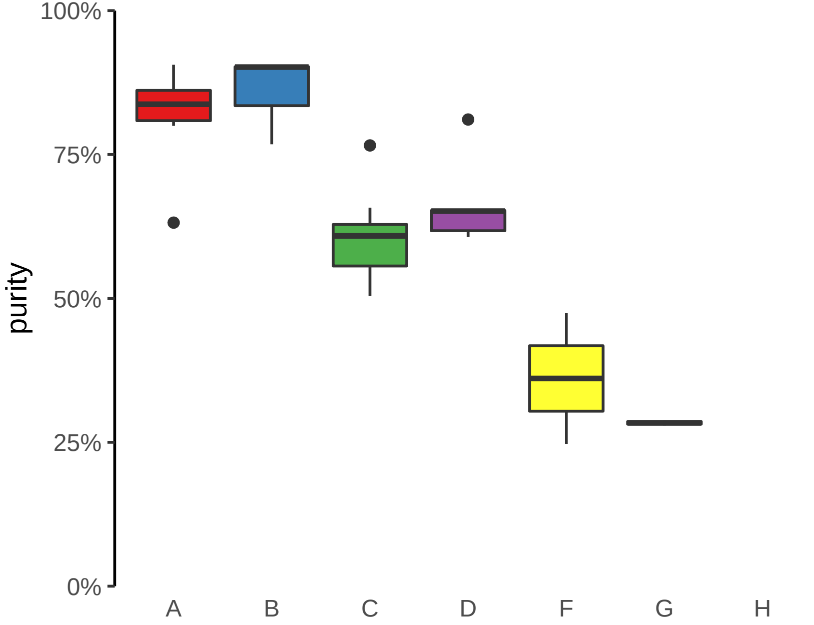

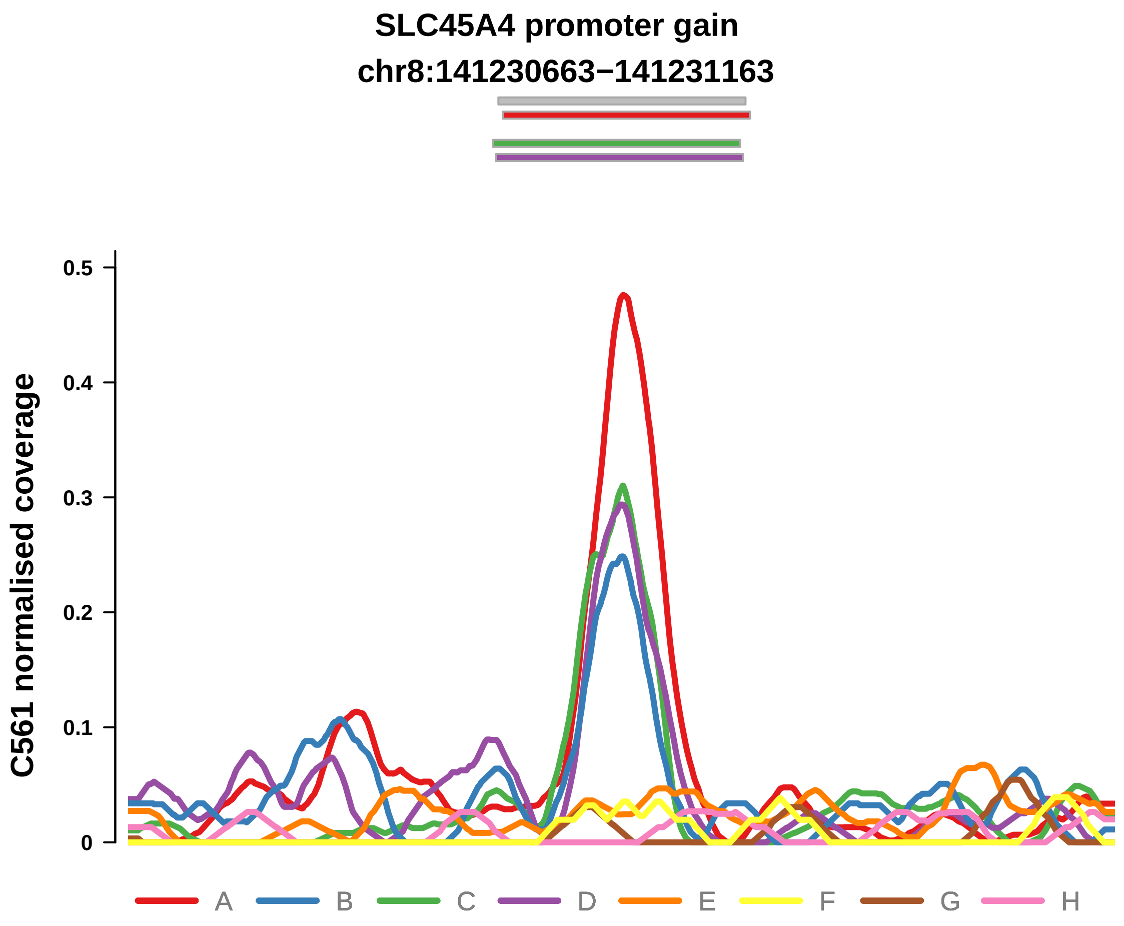


Putatively Subclonal

| | LogFC | P. | Adjusted P. |
|---|---|---|---|
| *Adenoma vs. normal* | −0.94 | 4.98e−02 * | 4.98e−02 * |
| *F vs.Cancer ~purity+type* | −2.22 | 5.79e−02 | 2.32e−01 |
| *G vs.Cancer ~purity+type* | −3.78 | 6.2e−03 ** | 5e−02 * |

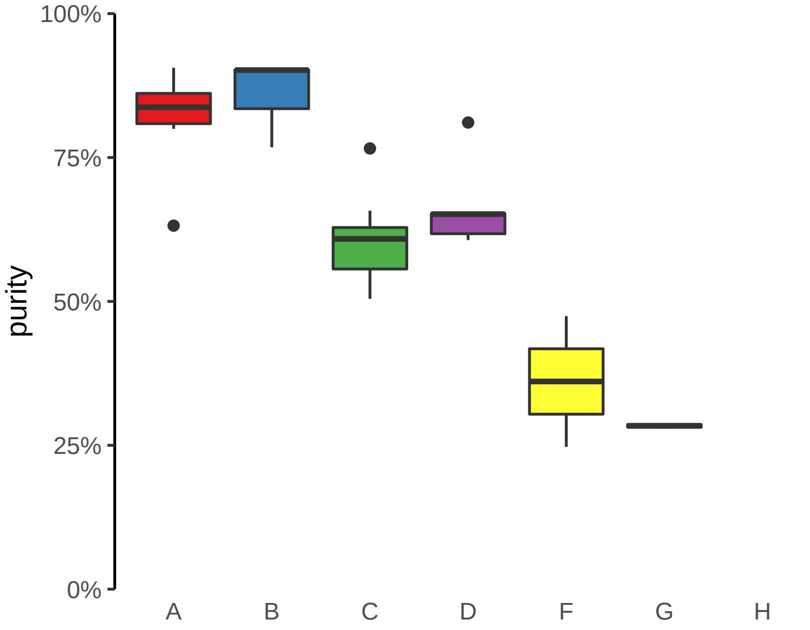

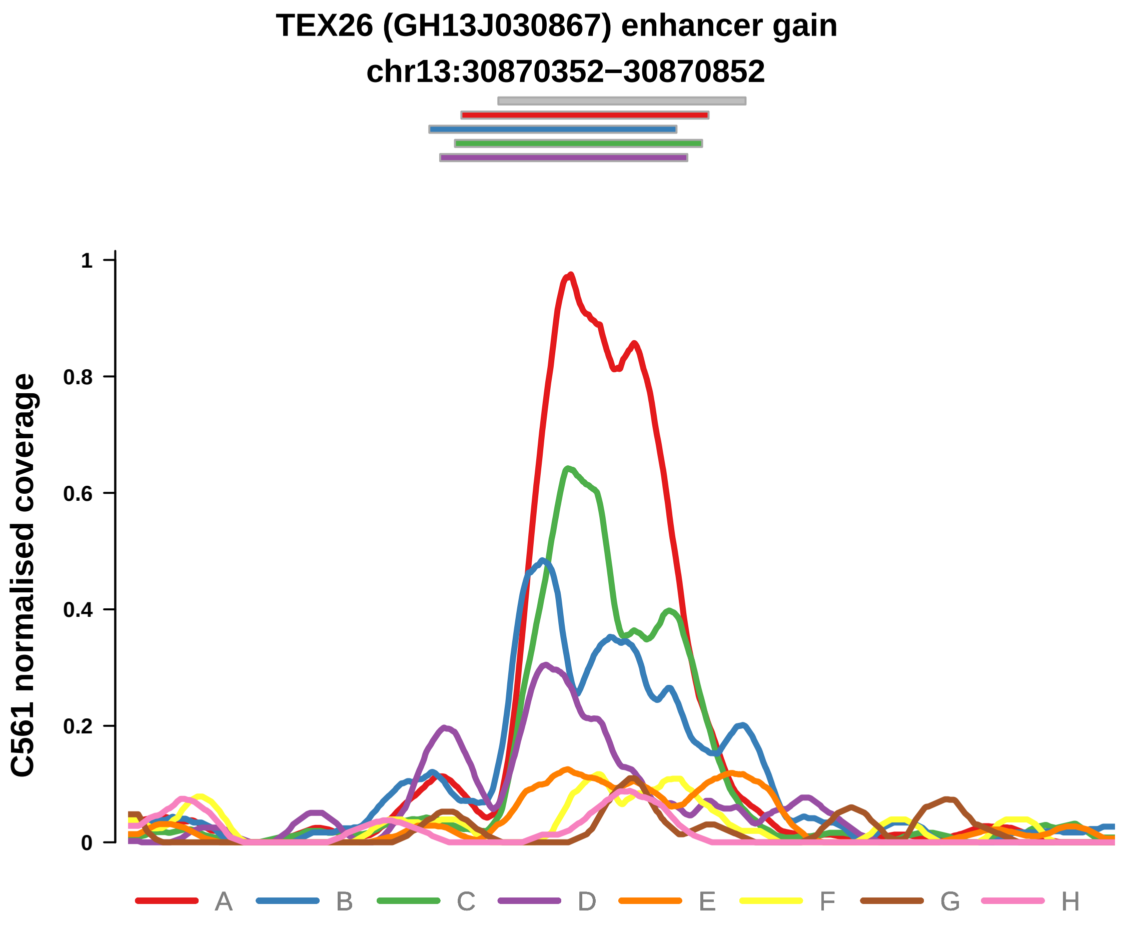


Putatively Subclonal

| | LogFC | P. | Adjusted P. |
|---|---|---|---|
| *Adenoma vs. normal* | −0.94 | 4.98e−02 * | 4.98e−02 * |
| *F vs.Cancer ~purity+type* | −1.99 | 5.43e−03 ** | 5.79e−02 |
| *G vs.Cancer ~purity+type* | −3.21 | 1.11e−04 *** | 2.26e−03 ** |

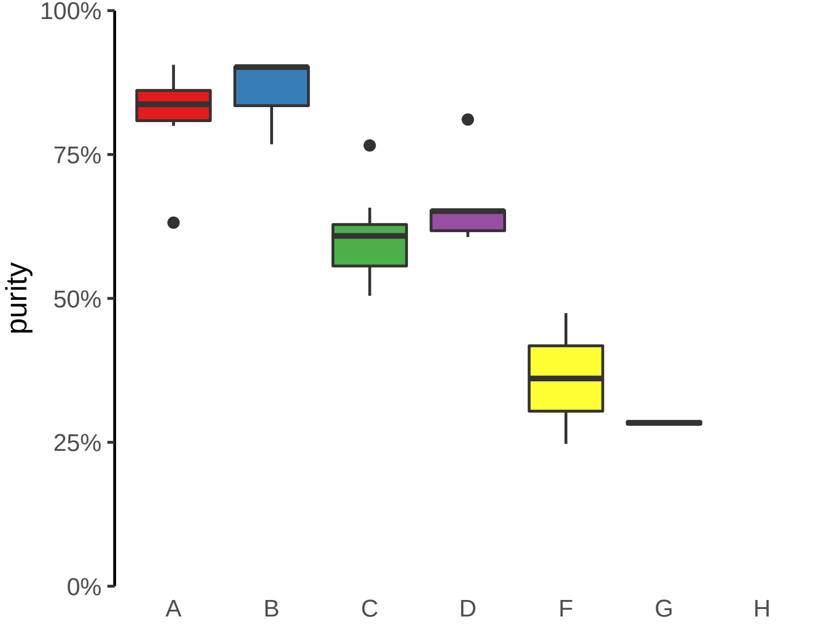

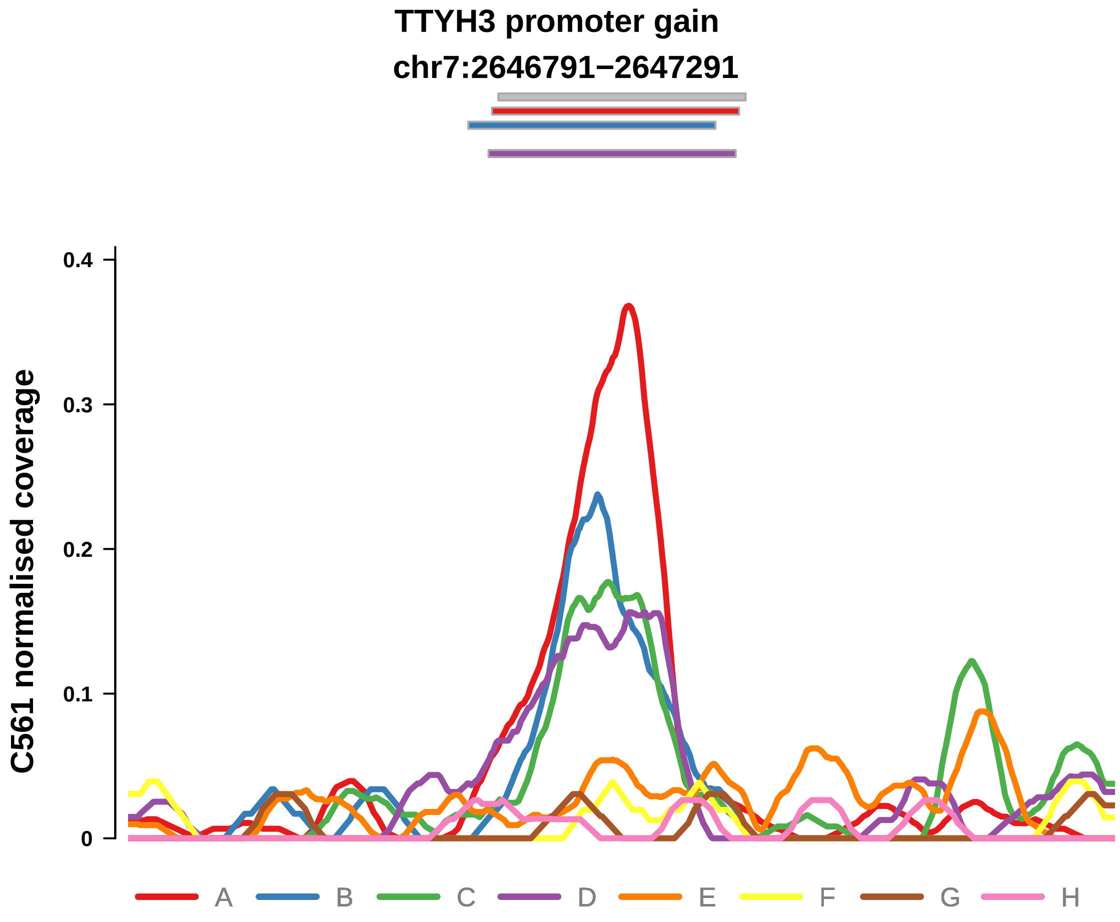


Putatively Subclonal

| | LogFC | P. | Adjusted P. |
|---|---|---|---|
| *Adenoma vs. normal* | −0.94 | 4.98e−02 * | 4.98e−02 * |
| *F vs.Cancer ~purity+type* | −2.91 | 1.16e−02 * | 8.87e−02 |
| *G vs.Cancer ~purity+type* | −3.44 | 6.55e−03 ** | 5e−02 * |

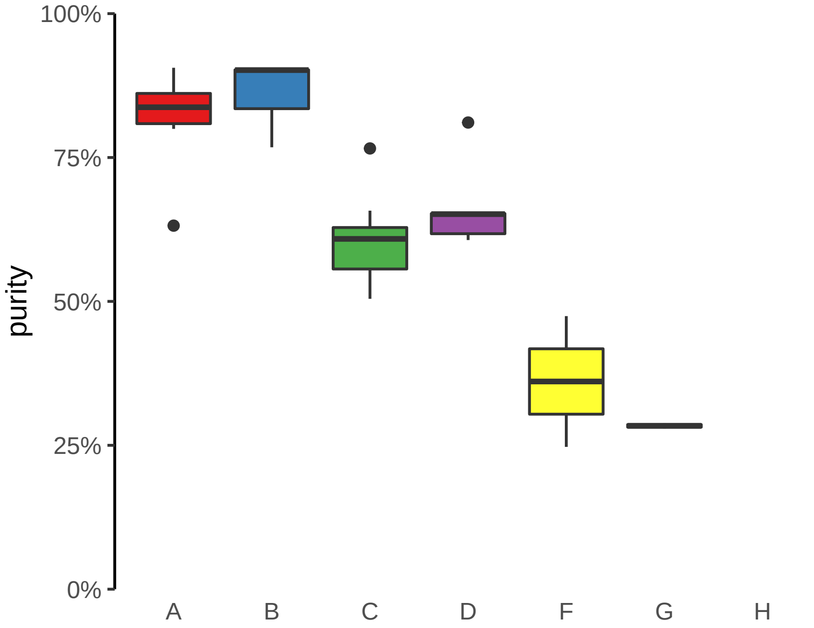
