## Supplementary Figures for "The co-evolution of the genome and epigenome in colorectal cancer": figS15_subclonal_promoter_gain.pdf

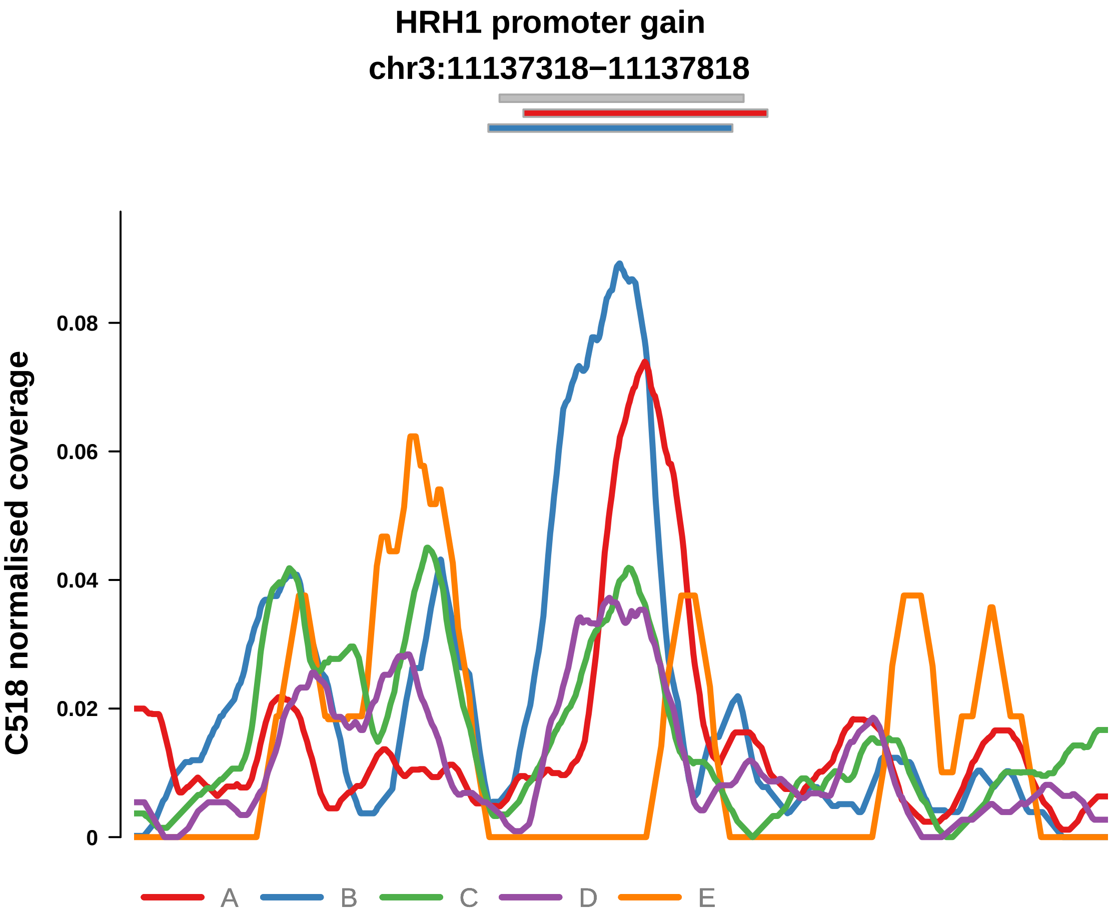


Putatively Subclonal (logFC = 1.37)

| | P. | Adjusted P. |
|---|---|---|
| *~region vs. ~* | 7.13e−04 *** | 3.14e−02 * |
| *~purity vs ~* | 2.52e−01 | 7.38e−01 |
| *~purity+region vs ~purity* | 1.14e−03 ** | 3.34e−02 * |
| *Carcinoma vs. normal* | 2.32e−02 * | 2.32e−02 * |

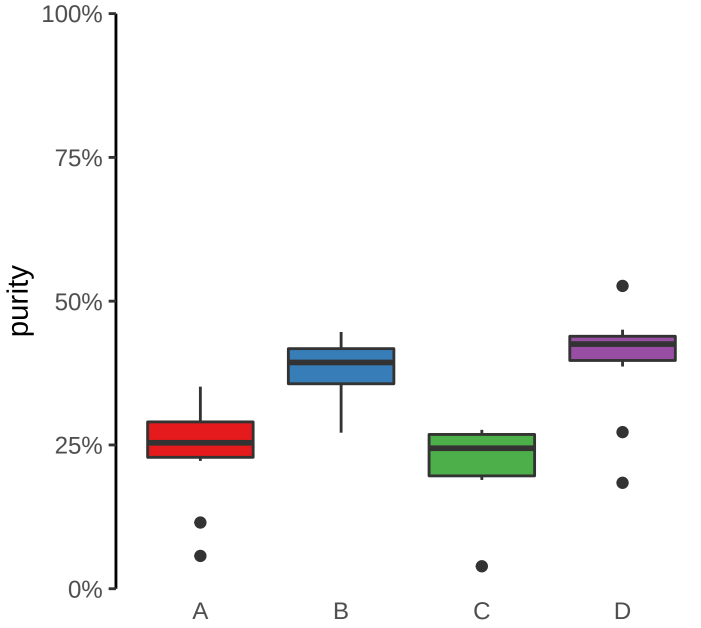

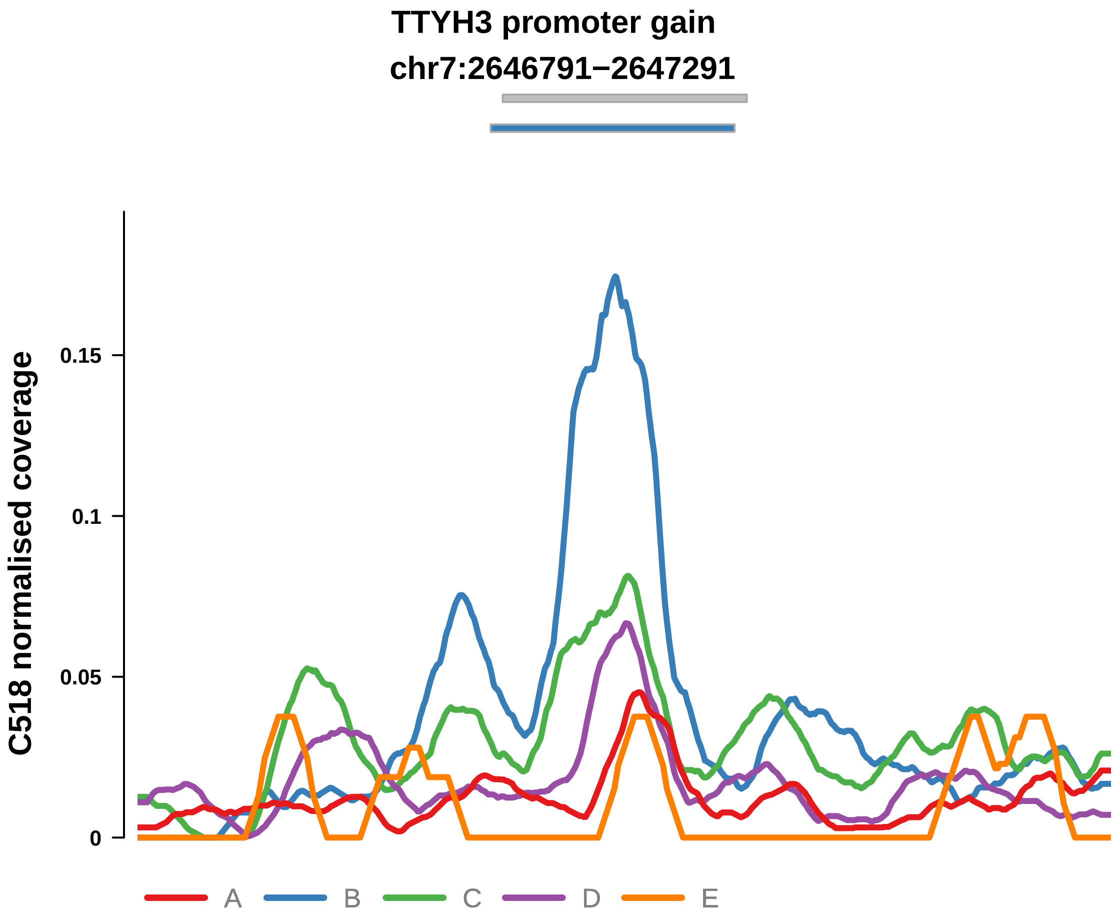


Putatively Subclonal (logFC = 2.69)

| | P. | Adjusted P. |
|---|---|---|
| *~region vs. ~* | 5.48e−05 **** | 4.82e−03 ** |
| *~purity vs ~* | 1.79e−01 | 7.38e−01 |
| *~purity+region vs ~purity* | 2.42e−05 **** | 2.13e−03 ** |
| *Carcinoma vs. normal* | 3.75e−06 **** | 3.75e−06 **** |

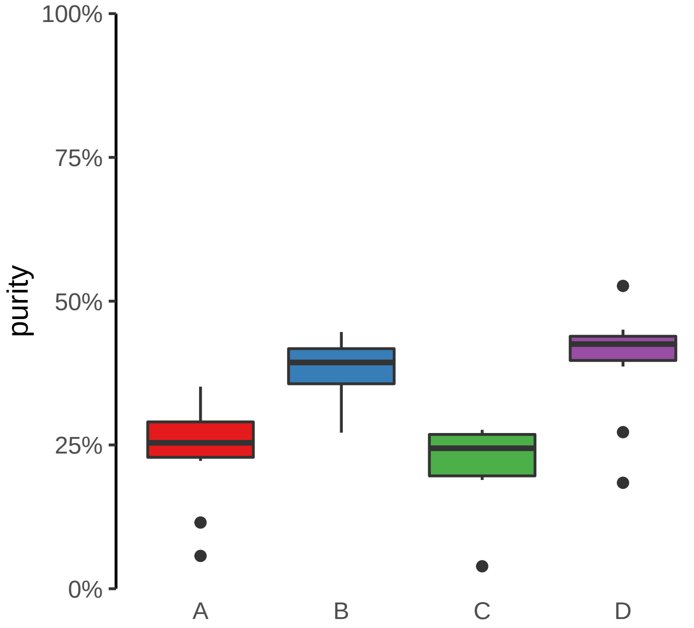

**HPS4 promoter gain**

**chr22:26471440−26471940**

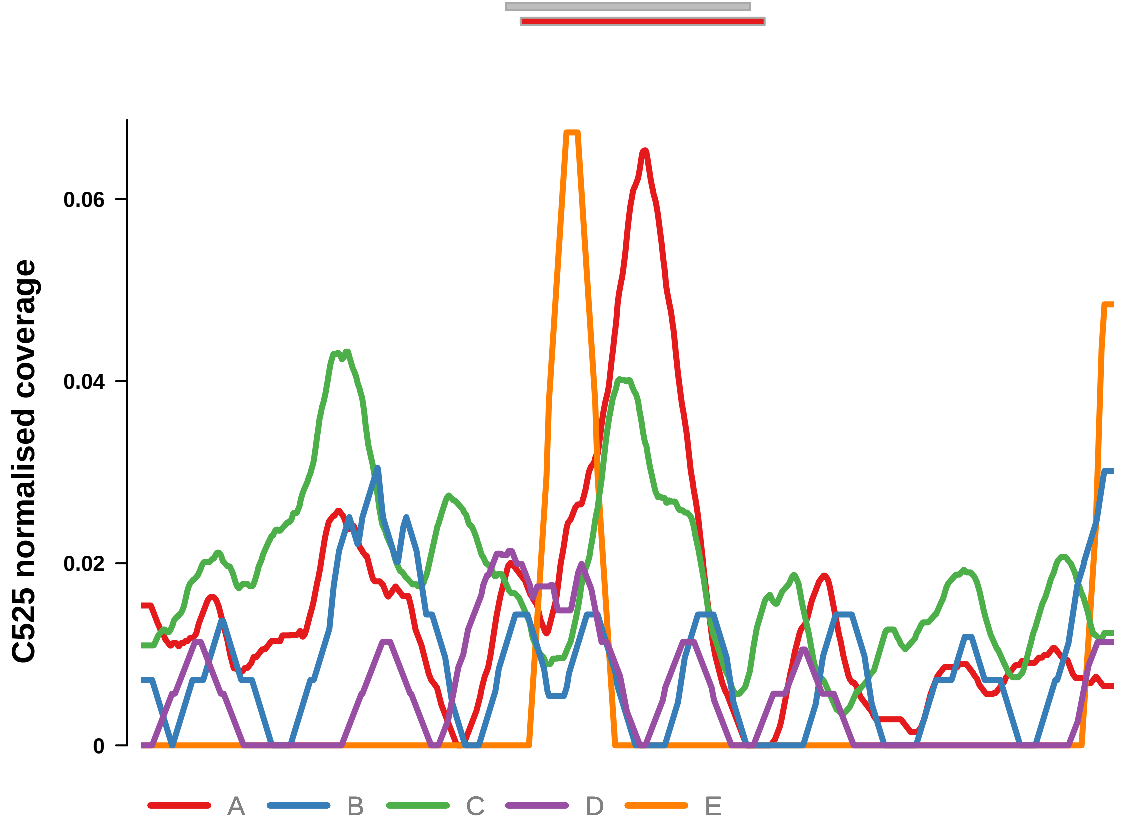


Putatively Subclonal (logFC = 0.84)

| | **P.** | **Adjusted P.** |
|---|---|---|
| *~region vs. ~* | 1.6e−04 *** | 1.14e−03 ** |
| *~purity vs ~* | 7.82e−01 | 8.93e−01 |
| *~purity+region vs ~purity* | 2.79e−04 *** | 2.95e−03 ** |
| *Carcinoma vs. normal* | NA | NA |

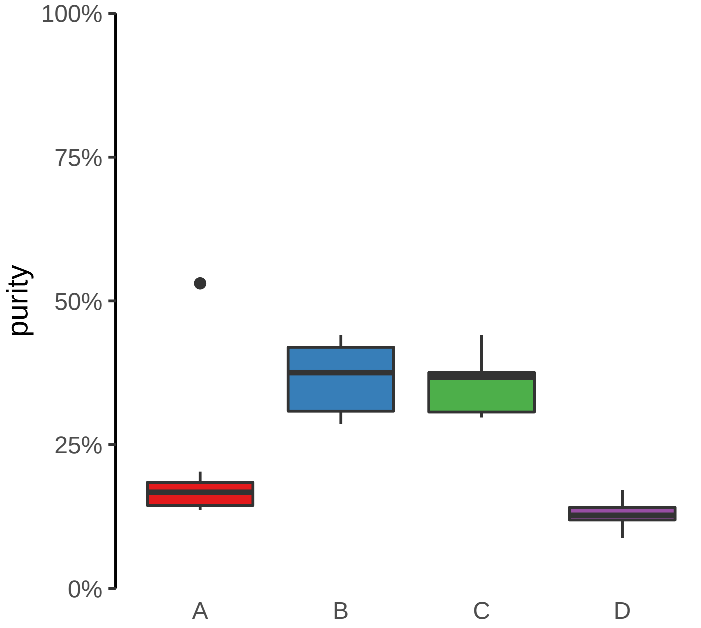

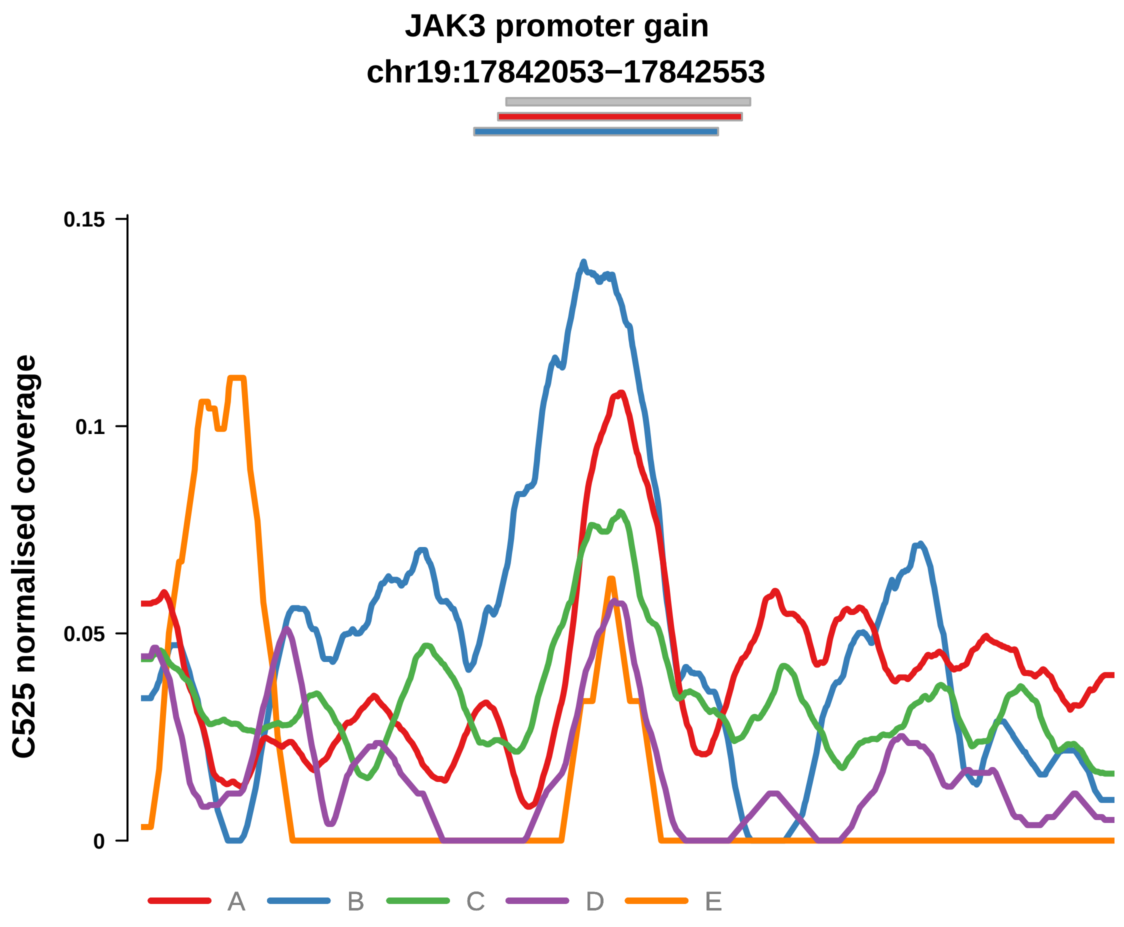


Putatively Subclonal (logFC = 1.76)

| | P. | Adjusted P. |
| --- | --- | --- |
| *~region vs. ~* | 3.66e−04 *** | 2.1e−03 ** |
| *~purity vs ~* | 4.06e−02 * | 5.96e−01 |
| *~purity+region vs ~purity* | 4.86e−03 ** | 2.21e−02 * |
| *Carcinoma vs. normal* | 2.78e−03 ** | 2.78e−03 ** |

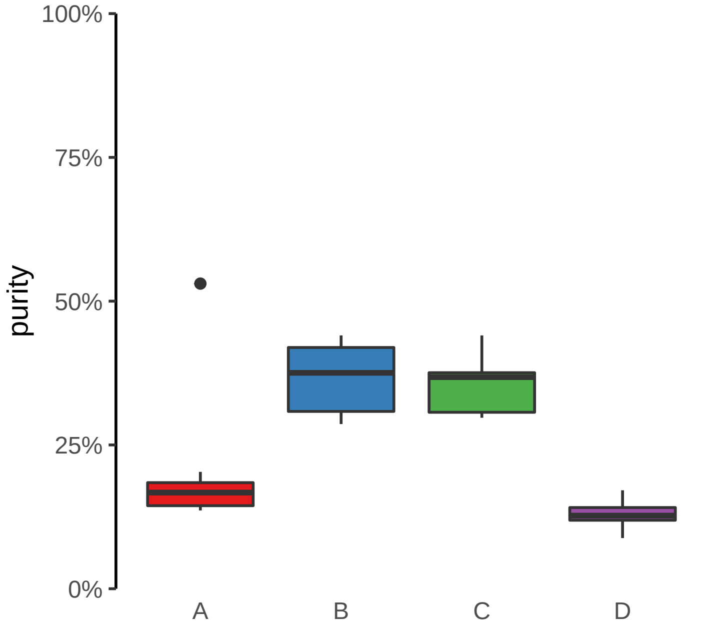

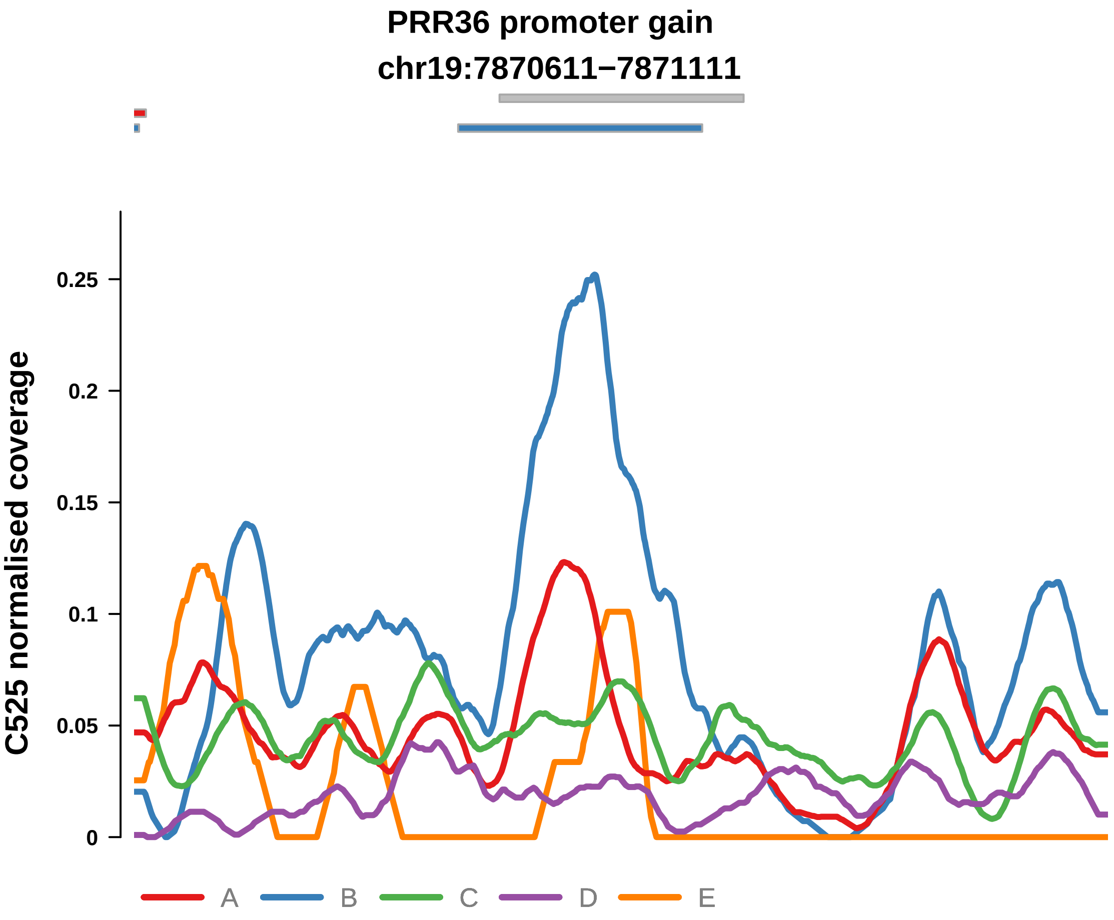


Putatively Subclonal (logFC = 2.1)

| | P. | Adjusted P. |
|---|---|---|
| *~region vs. ~* | 3.08e−06 **** | 1.32e−04 *** |
| *~purity vs ~* | 6.62e−05 **** | 5.83e−03 ** |
| *~purity+region vs ~purity* | 5.41e−03 ** | 2.21e−02 * |
| *Carcinoma vs. normal* | 3.02e−04 *** | 3.02e−04 *** |

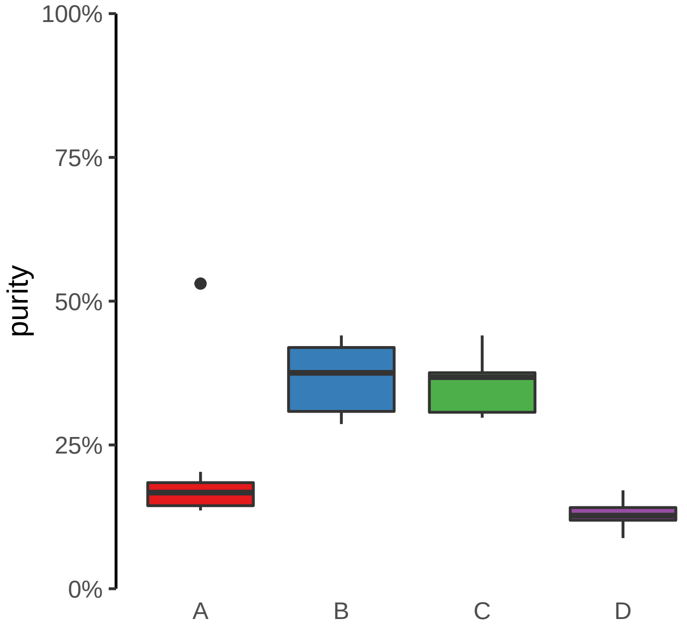

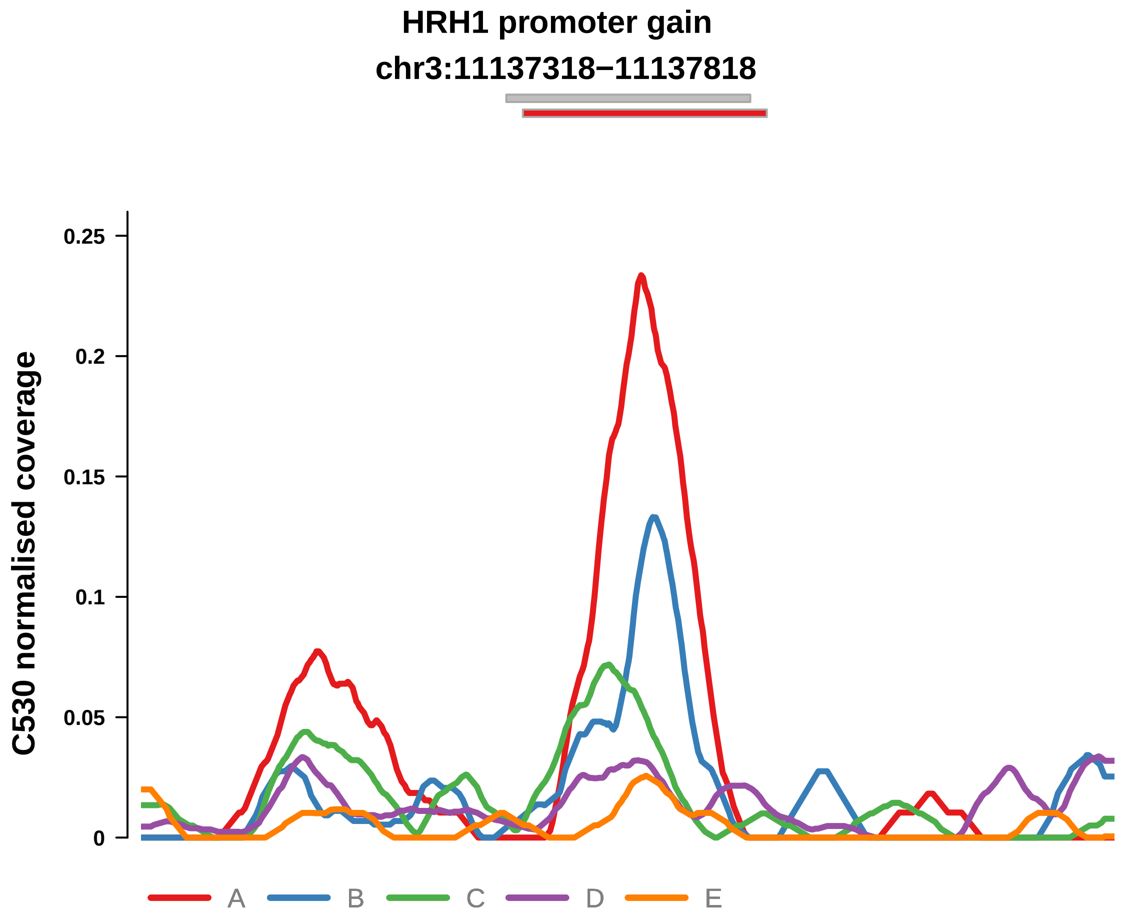


Putatively Subclonal (logFC = 1.29)

| | P. | Adjusted P. |
|---|---|---|
| *~region vs. ~* | 1.93e−03 ** | 5.12e−02 |
| *~purity vs ~* | 5.23e−01 | 9.32e−01 |
| *~purity+region vs ~purity* | 1.4e−03 ** | 4.52e−02 * |
| *Carcinoma vs. normal* | 3.3e−02 * | 3.3e−02 * |

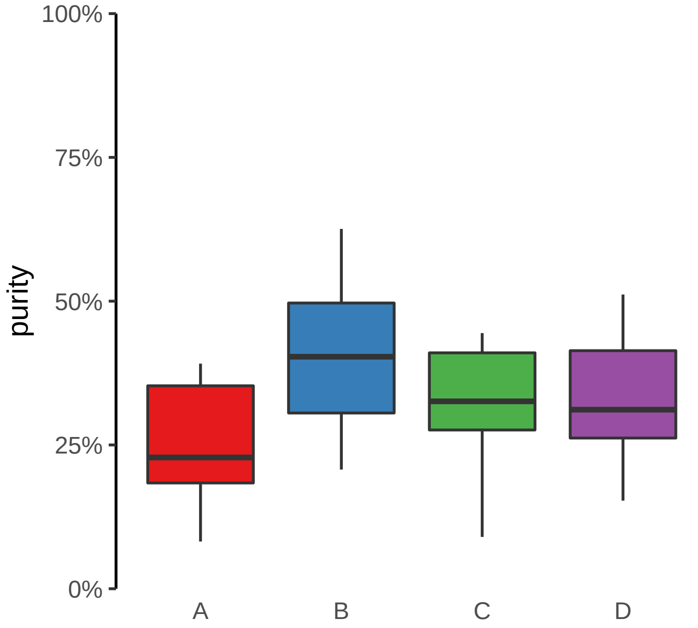

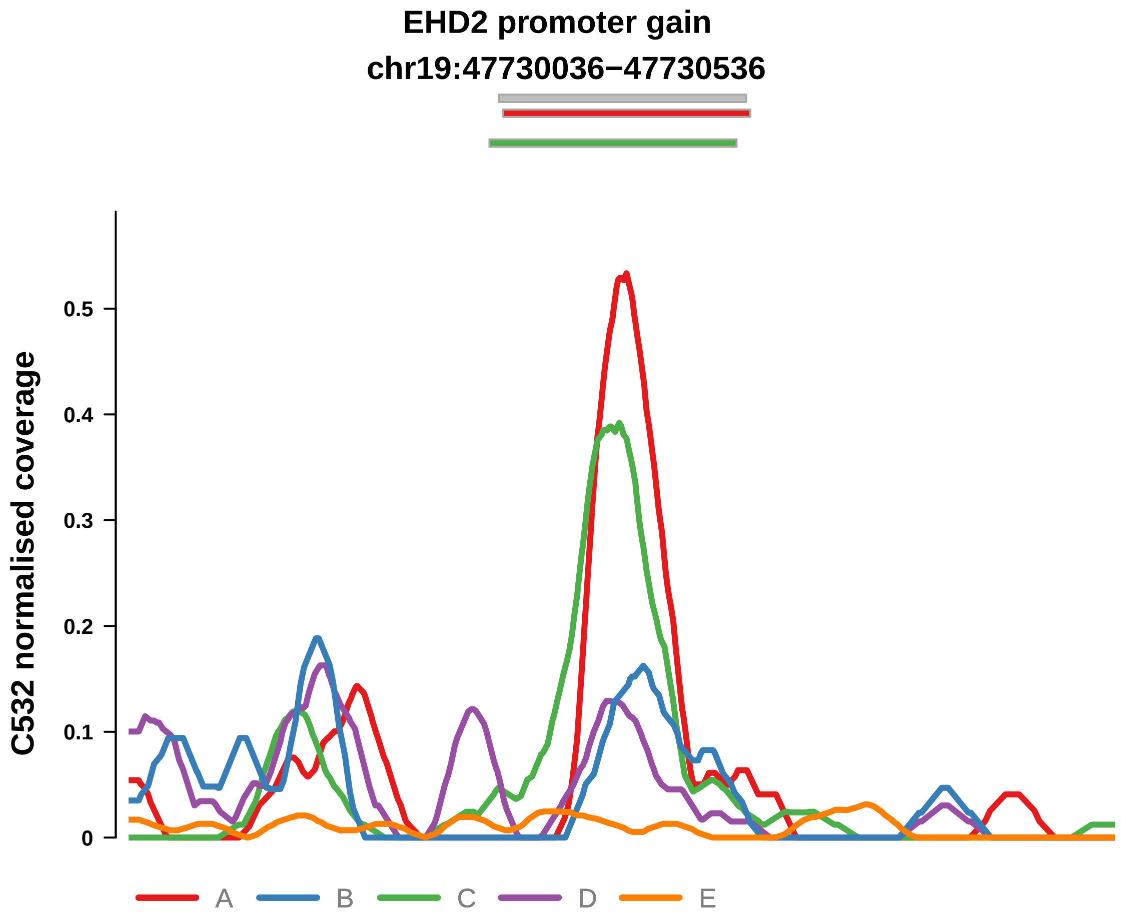


Putatively Subclonal (logFC = 2.46)

| | P. | Adjusted P. |
|---|---|---|
| *~region vs. ~* | 1.74e−03 ** | 5.11e−02 |
| *~purity vs ~* | 8.52e−01 | 9.86e−01 |
| *~purity+region vs ~purity* | 1.64e−03 ** | 4.81e−02 * |
| *Carcinoma vs. normal* | 1.85e−05 **** | 1.85e−05 **** |

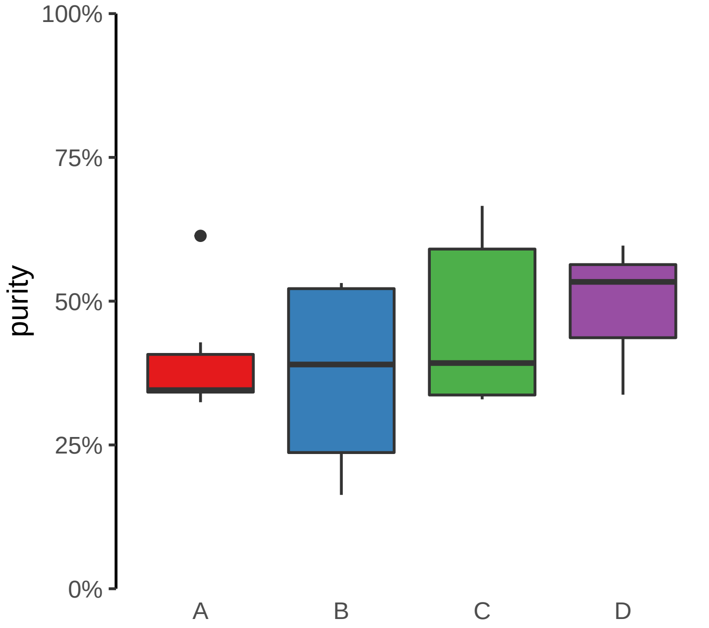

**KRT7 promoter gain**
**chr12:52235898−52236398**

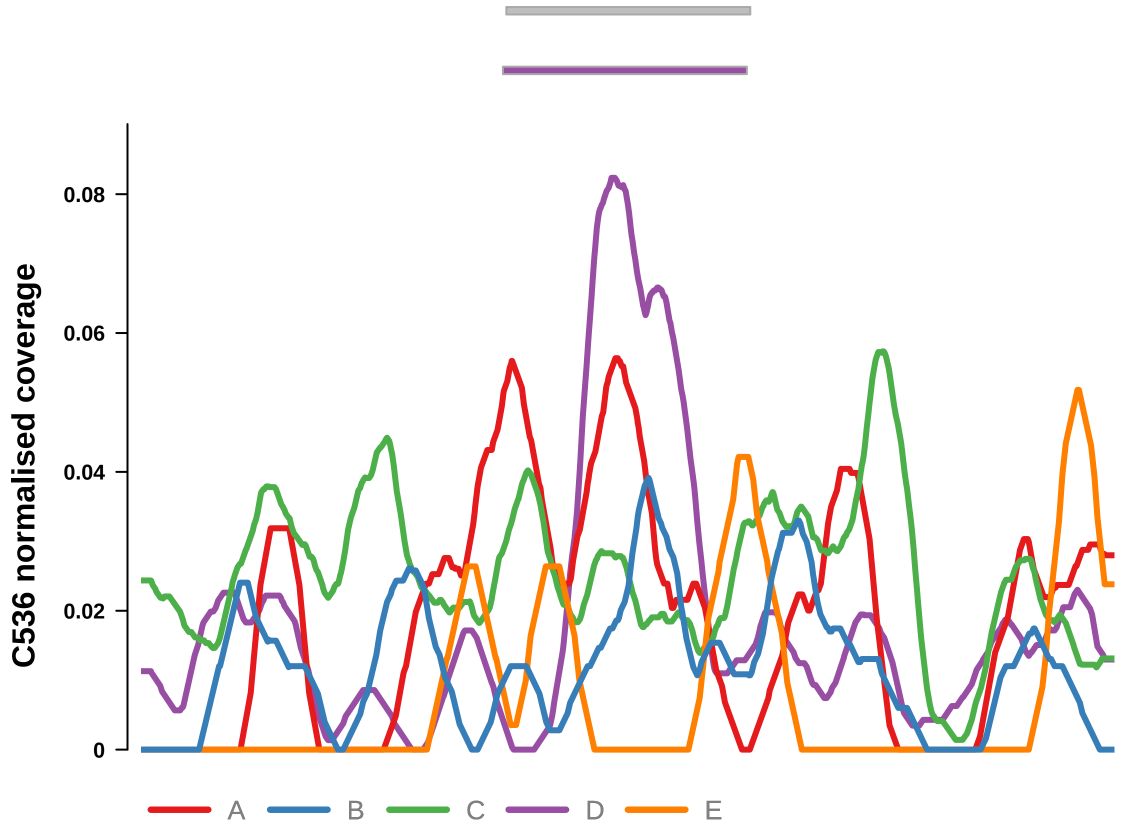


Putatively Subclonal (logFC = 1.61)

| | P. | Adjusted P. |
|---|---|---|
| *~region vs. ~* | 2.73e−02 * | 1.04e−01 |
| *~purity vs ~* | 3.93e−01 | 6.29e−01 |
| *~purity+region vs ~purity* | 3.62e−03 ** | 4.57e−02 * |
| *Carcinoma vs. normal* | 5.99e−03 ** | 5.99e−03 ** |

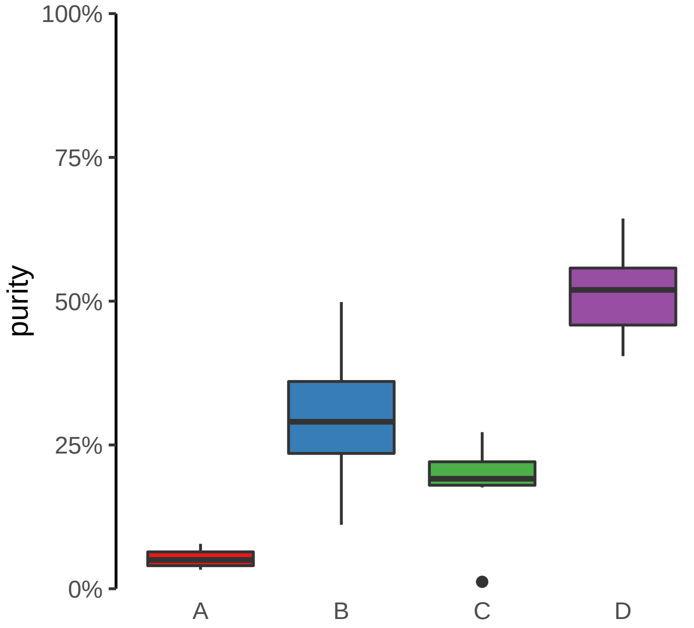

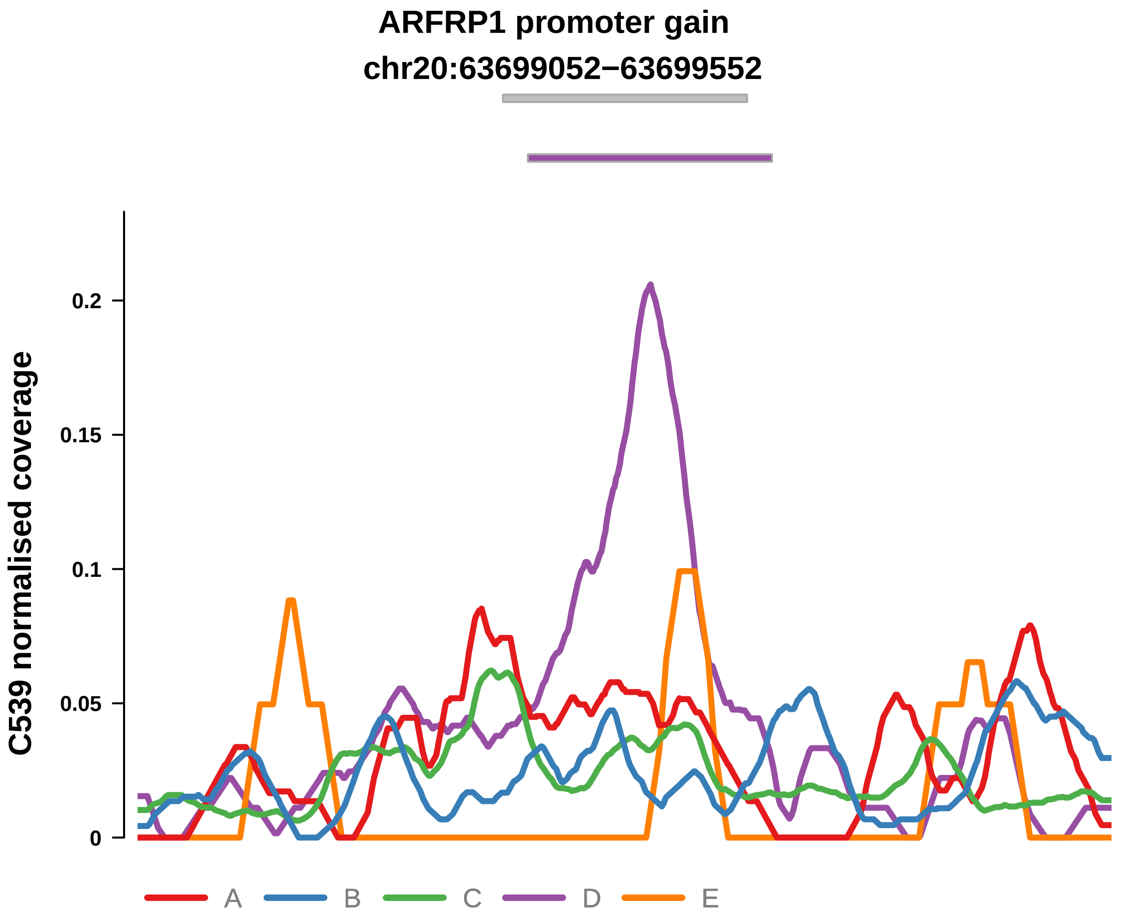


Putatively Subclonal (logFC = 3.25)

| | P. | Adjusted P. |
|---|---|---|
| *~region vs. ~* | 2.61e−04 *** | 8.5e−04 *** |
| *~purity vs ~* | 9.81e−02 | 1.09e−01 |
| *~purity+region vs ~purity* | 5.87e−05 **** | 2.32e−03 ** |
| *Carcinoma vs. normal* | 4.44e−07 **** | 4.44e−07 **** |

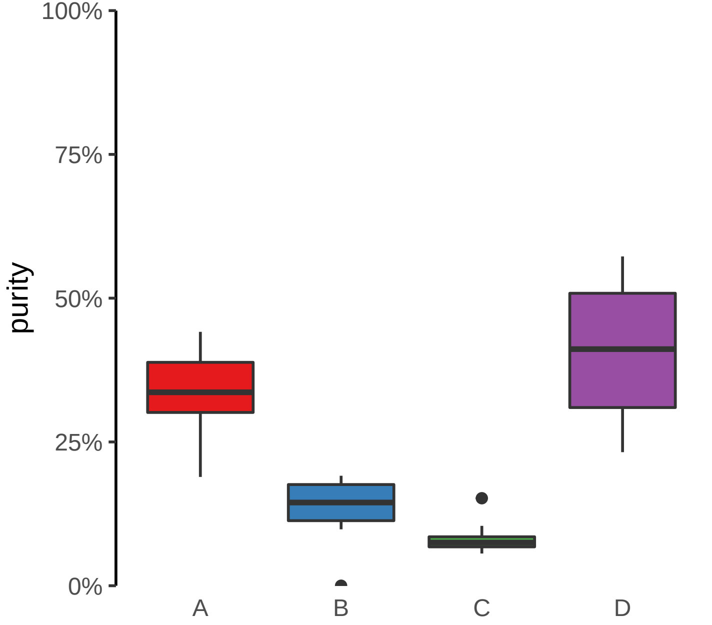

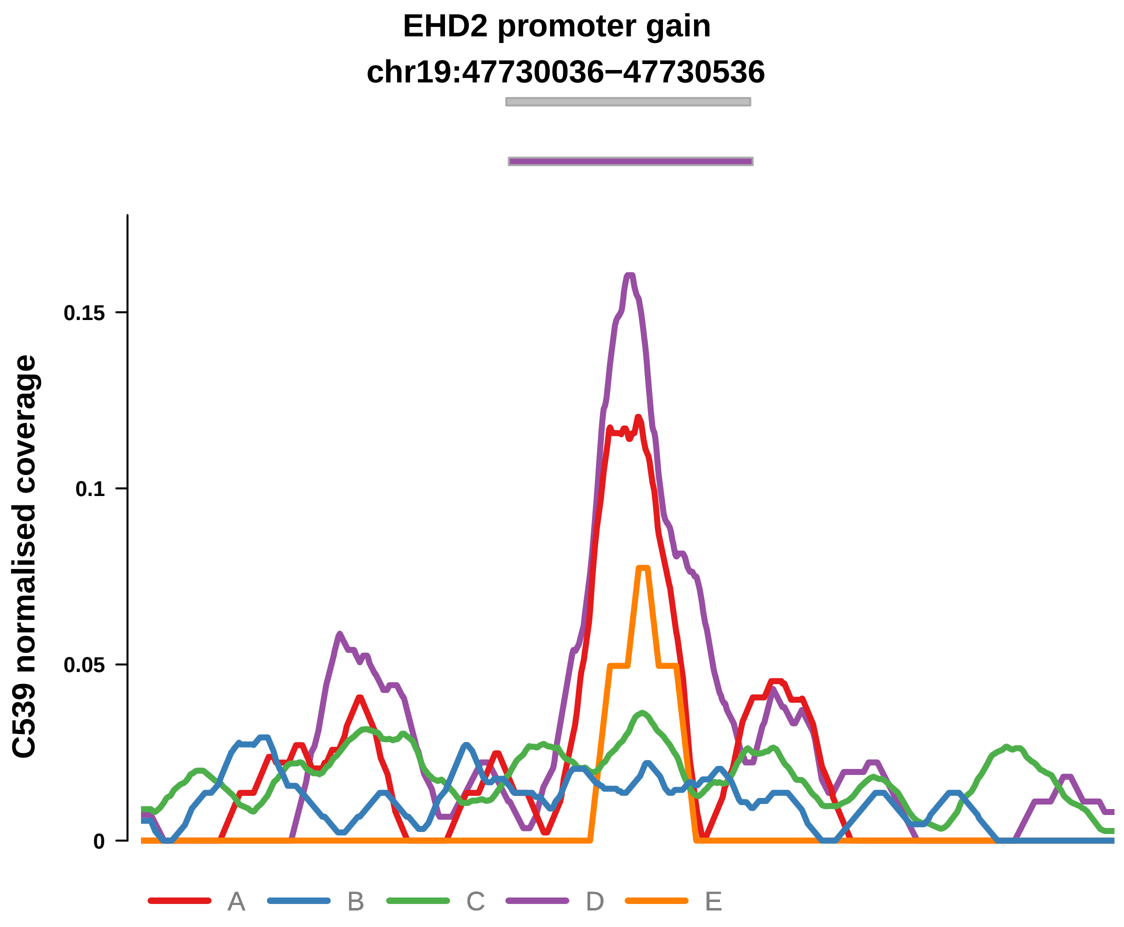


Putatively Subclonal (logFC = 2.71)

| | P. | Adjusted P. |
|---|---|---|
| *~region vs. ~* | 3e−03 ** | 6.29e−03 ** |
| *~purity vs ~* | 1.13e−01 | 1.24e−01 |
| *~purity+region vs ~purity* | 1.53e−03 ** | 1.45e−02 * |
| *Carcinoma vs. normal* | 2.44e−05 **** | 2.44e−05 **** |

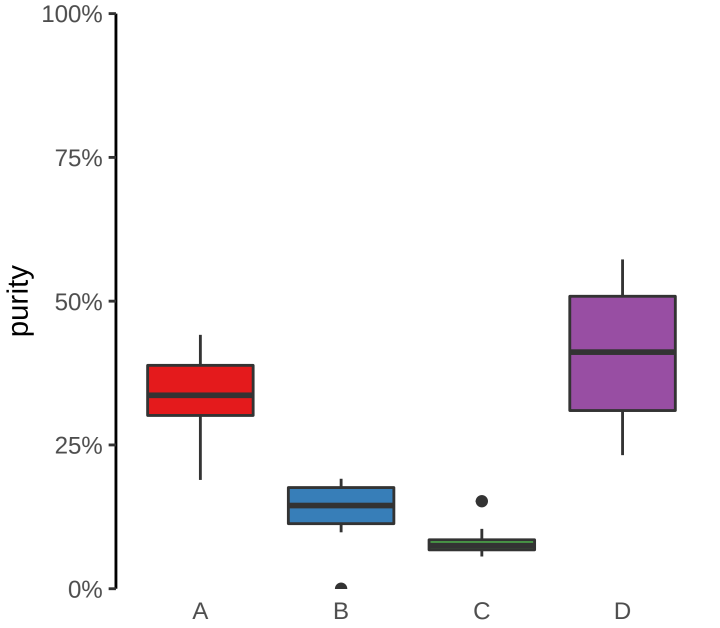

**FUT1 promoter gain**
**chr19:48752815−48753315**

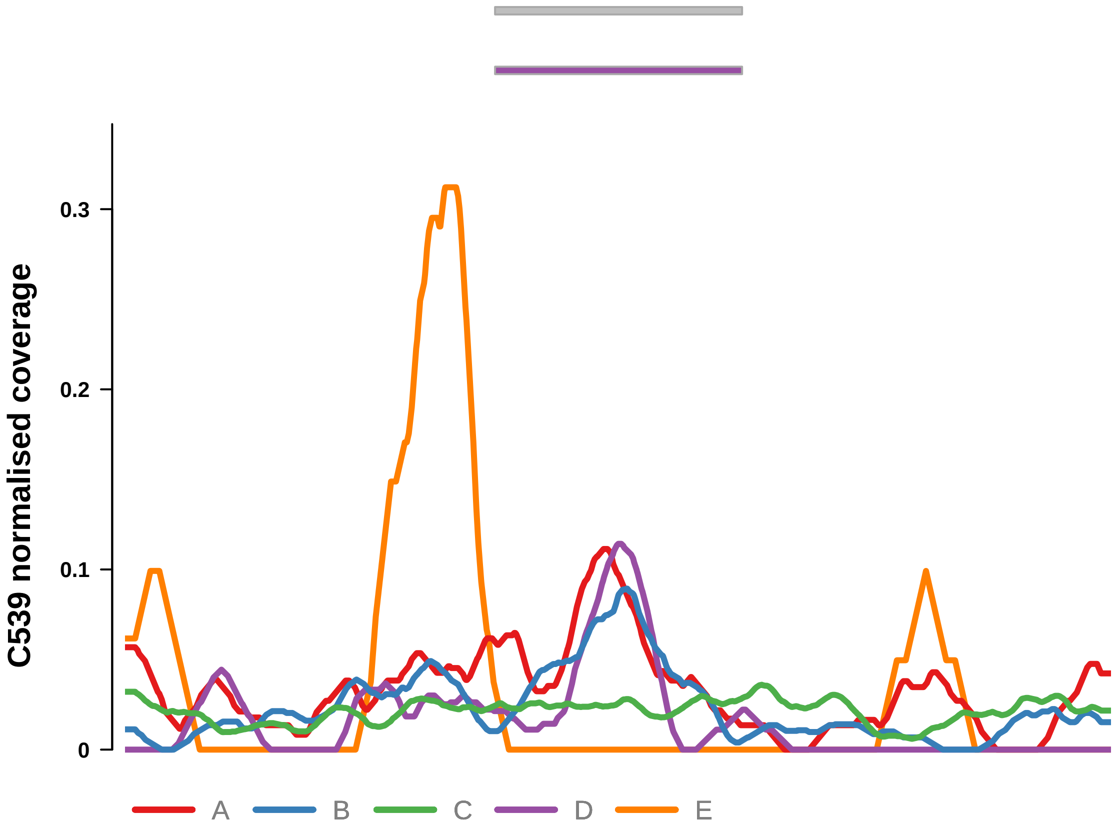


Putatively Subclonal (logFC = 1.75)

| | **P.** | **Adjusted P.** |
|---|---|---|
| *~region vs. ~* | 5.99e−03 ** | 9.59e−03 ** |
| *~purity vs ~* | 3.29e−02 * | 4.67e−02 * |
| *~purity+region vs ~purity* | 3.6e−03 ** | 2.22e−02 * |
| *Carcinoma vs. normal* | 1.19e−02 * | 1.19e−02 * |

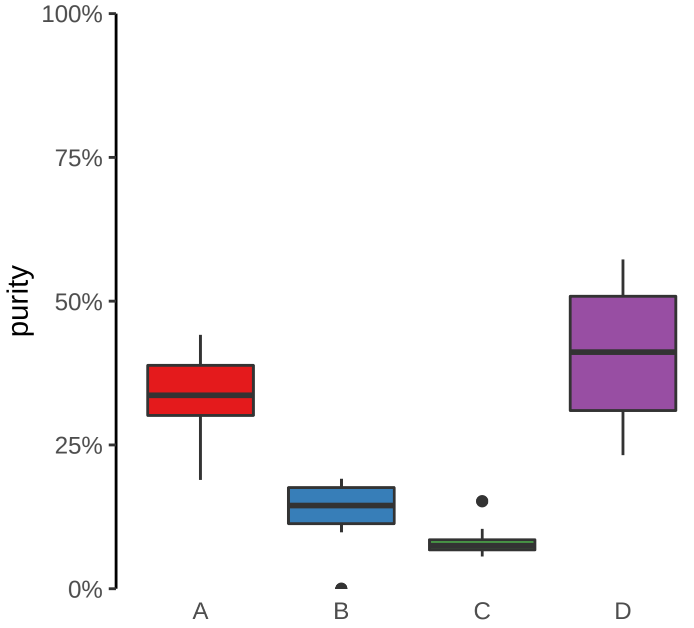

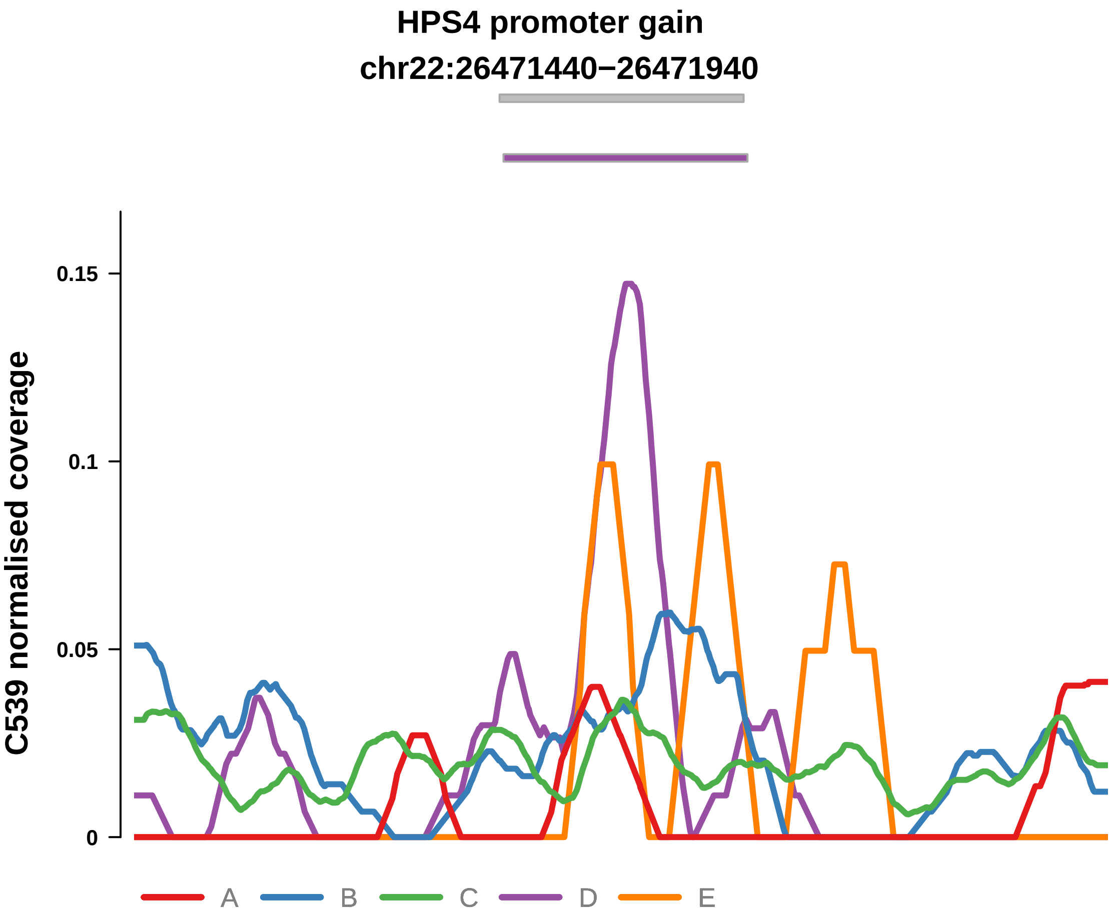


Putatively Subclonal (logFC = 2.98)

| | P. | Adjusted P. |
|---|---|---|
| *~region vs. ~* | 1.95e−04 *** | 7.42e−04 *** |
| *~purity vs ~* | 5.89e−02 | 7.41e−02 |
| *~purity+region vs ~purity* | 1.35e−04 *** | 2.66e−03 ** |
| *Carcinoma vs. normal* | 7.36e−05 **** | 7.36e−05 **** |

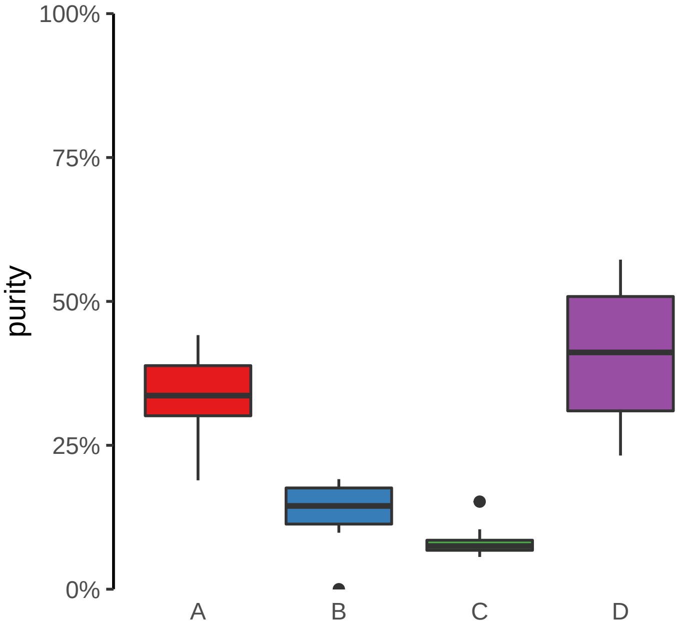

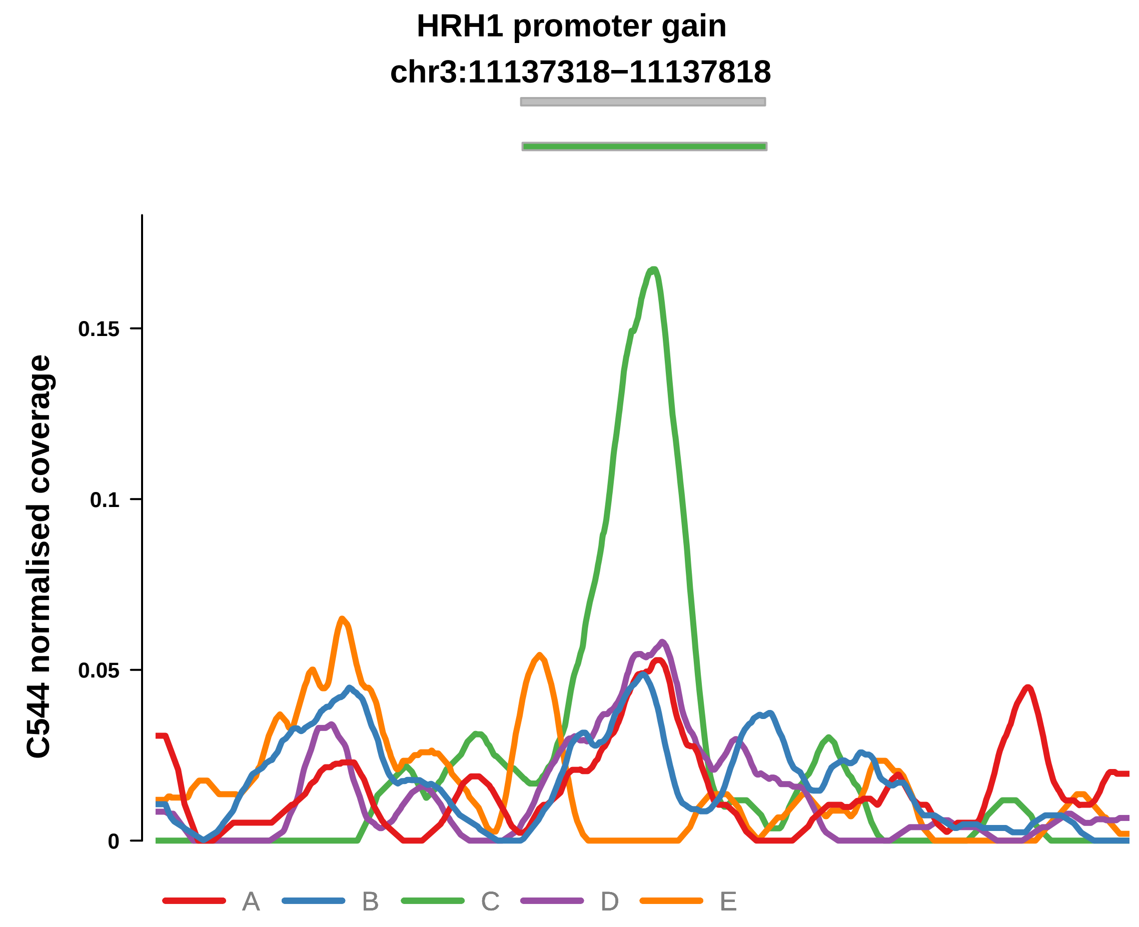


Putatively Subclonal (logFC = 1.96)

| | P. | Adjusted P. |
|---|---|---|
| *~region vs. ~* | 1.46e−02 * | 4.42e−02 * |
| *~purity vs ~* | 3.77e−01 | 5.36e−01 |
| *~purity+region vs ~purity* | 5e−03 ** | 2.83e−02 * |
| *Carcinoma vs. normal* | 8.82e−04 *** | 8.82e−04 *** |

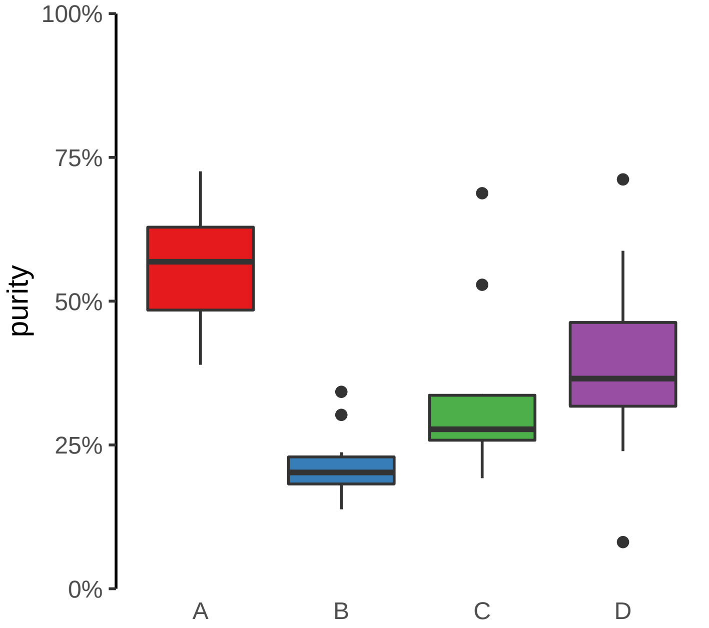

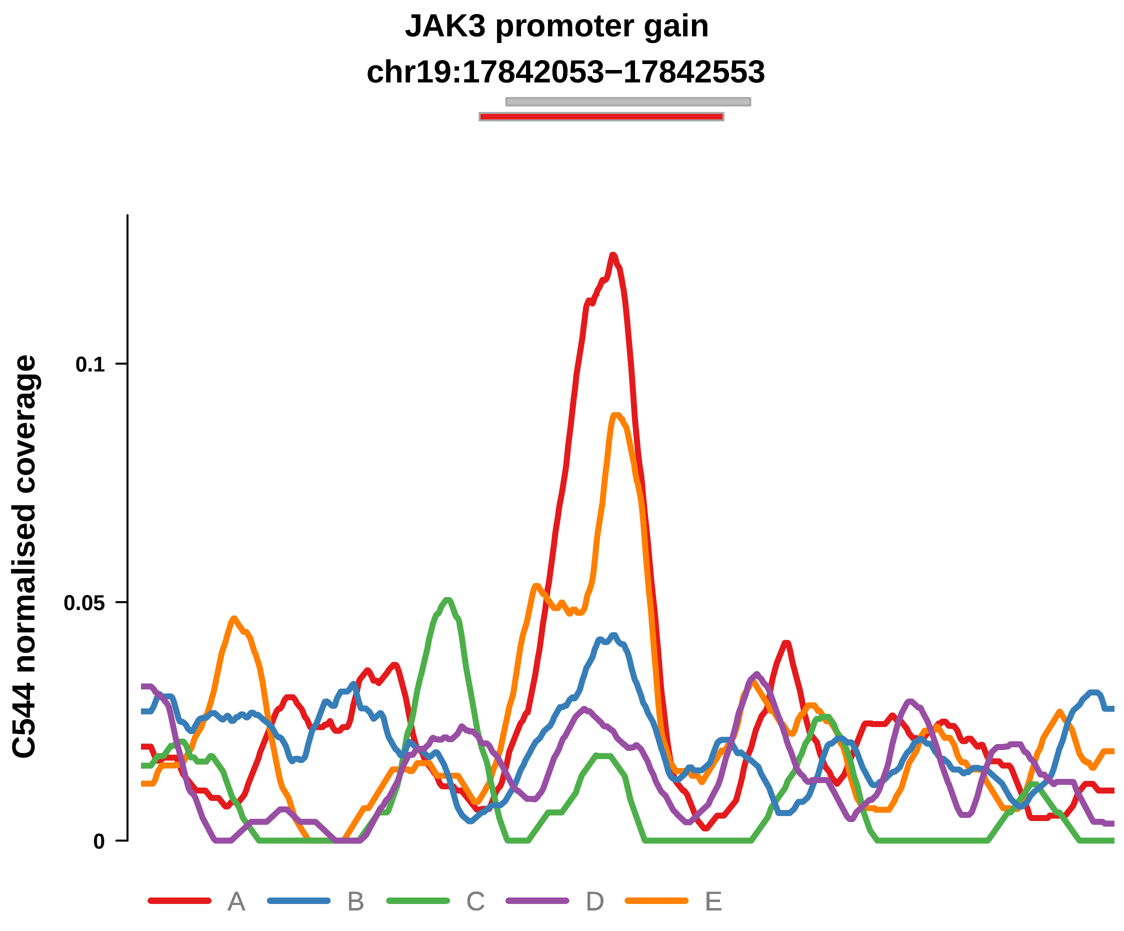


Putatively Subclonal (logFC = 1.52)

| | P. | Adjusted P. |
|---|---|---|
| *~region vs. ~* | 2.9e−05 **** | 3.48e−04 *** |
| *~purity vs ~* | 3.54e−02 * | 9.88e−02 |
| *~purity+region vs ~purity* | 6.78e−04 *** | 1.08e−02 * |
| *Carcinoma vs. normal* | 8.44e−03 ** | 8.44e−03 ** |

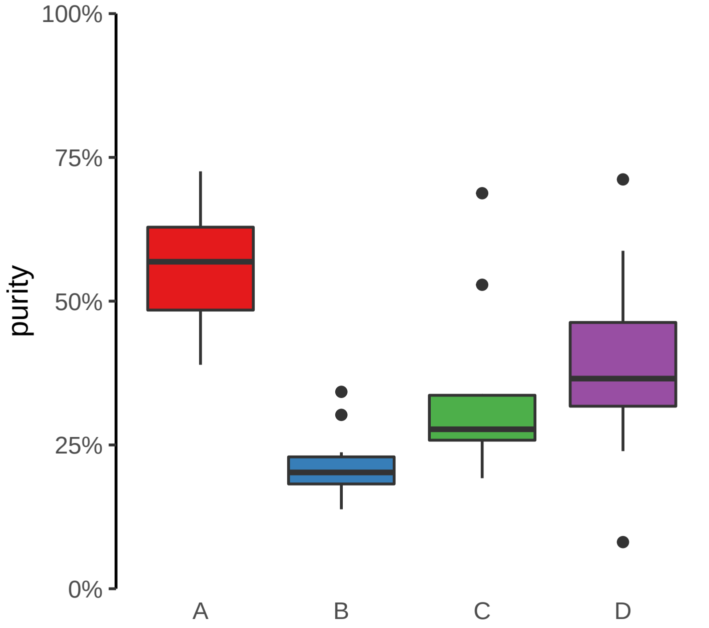

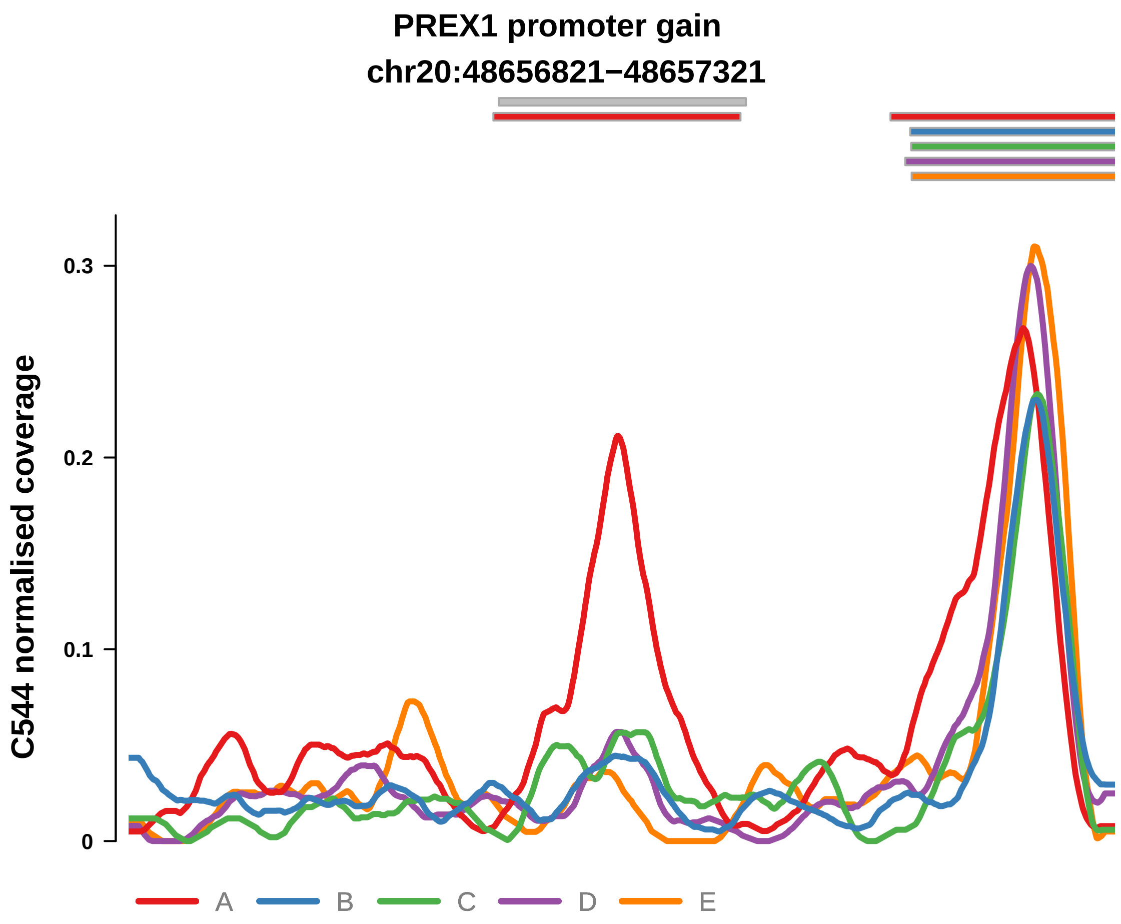


Putatively Subclonal (logFC = 3.1)

| | P. | Adjusted P. |
|---|---|---|
| *~region vs. ~* | 5.21e−05 **** | 5.1e−04 *** |
| *~purity vs ~* | 1.82e−02 * | 7.18e−02 |
| *~purity+region vs ~purity* | 3.32e−03 ** | 2.36e−02 * |
| *Carcinoma vs. normal* | 1.97e−07 **** | 1.97e−07 **** |

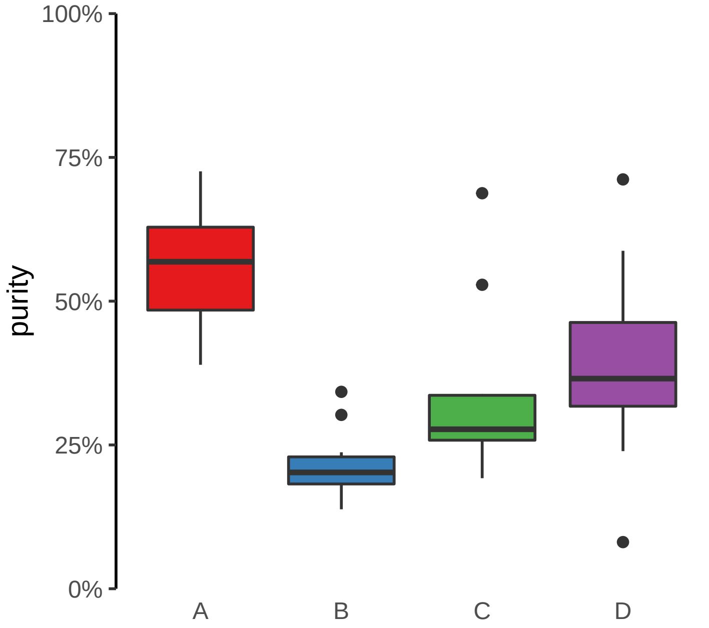

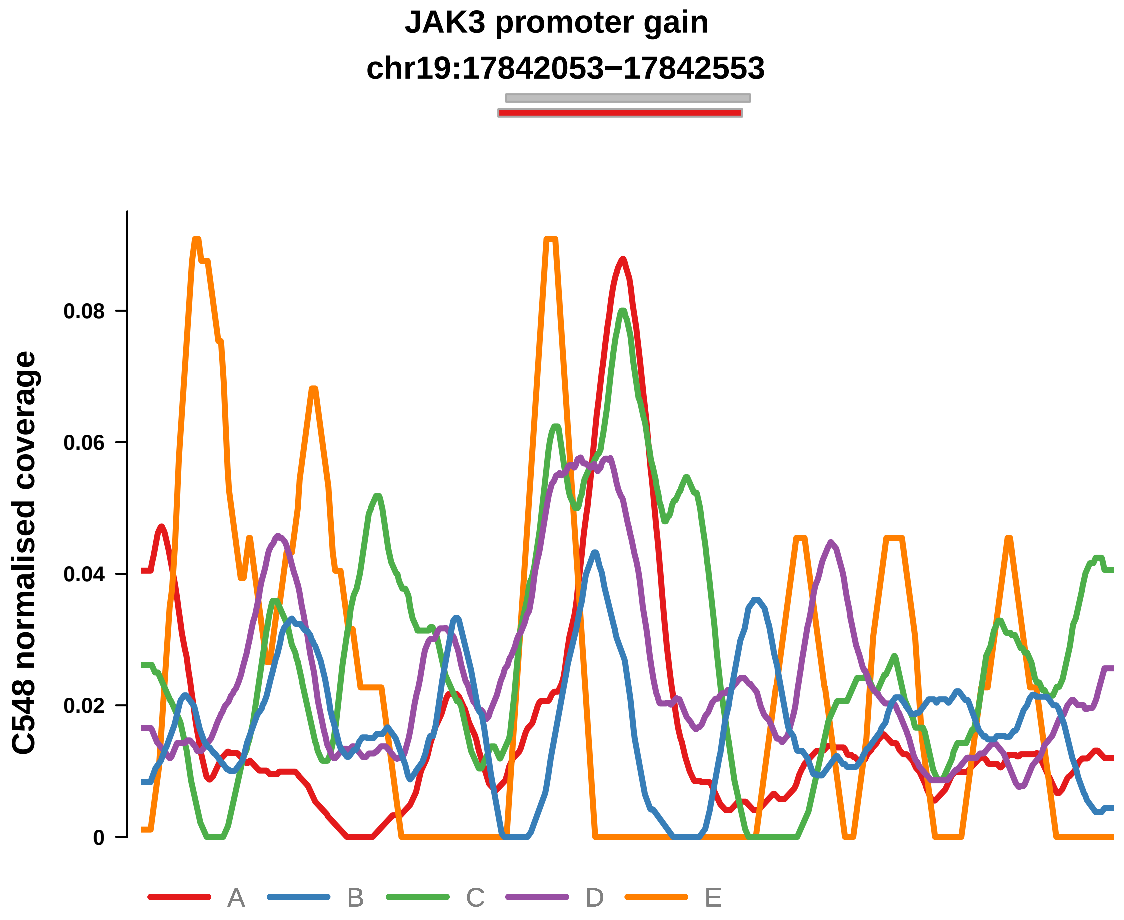


Putatively Subclonal (logFC = 1.74)

| | P. | Adjusted P. |
|---|---|---|
| *~region vs. ~* | 2.29e−04 *** | 6.72e−03 ** |
| *~purity vs ~* | 5.11e−02 | 2.05e−01 |
| *~purity+region vs ~purity* | 4.98e−04 *** | 1.46e−02 * |
| *Carcinoma vs. normal* | 4.31e−03 ** | 4.31e−03 ** |

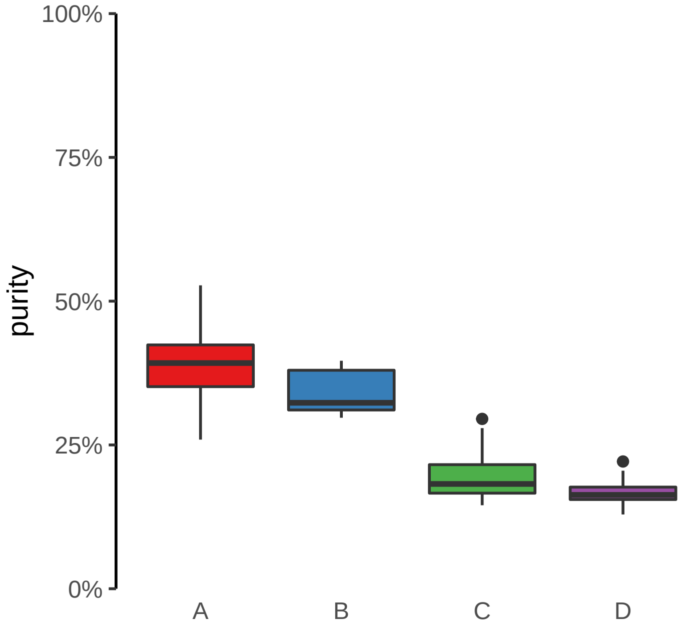

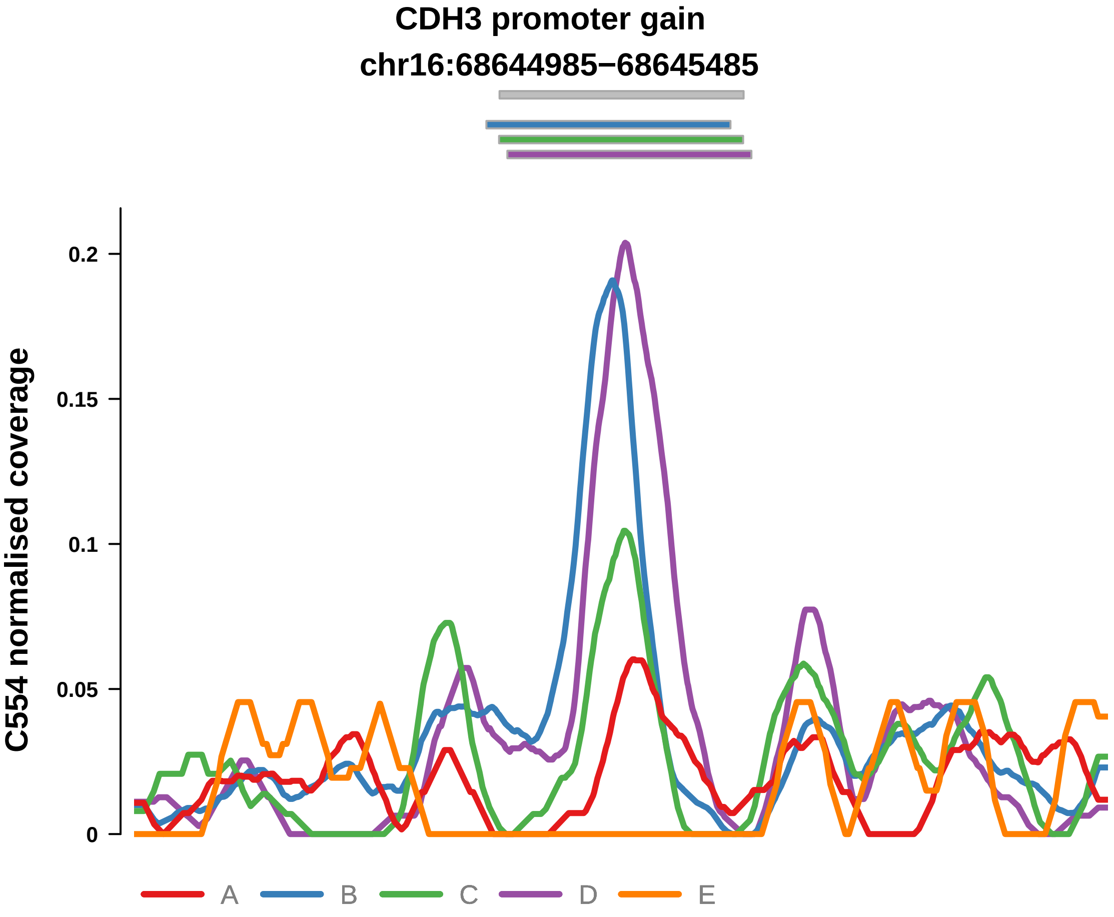


Putatively Subclonal (logFC = 1.23)

| | P. | Adjusted P. |
|---|---|---|
| *~region vs. ~* | 1.22e−04 *** | 9.98e−04 *** |
| *~purity vs ~* | 8.48e−03 ** | 5.8e−02 |
| *~purity+region vs ~purity* | 9.19e−03 ** | 3.88e−02 * |
| *Carcinoma vs. normal* | 2.5e−02 * | 2.5e−02 * |

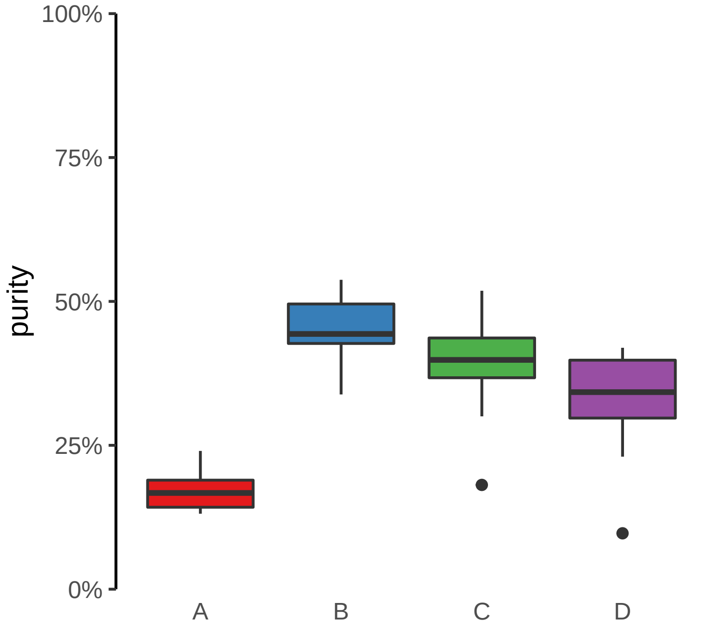

**FOXQ1 promoter gain**
**chr6:1311742−1312242**

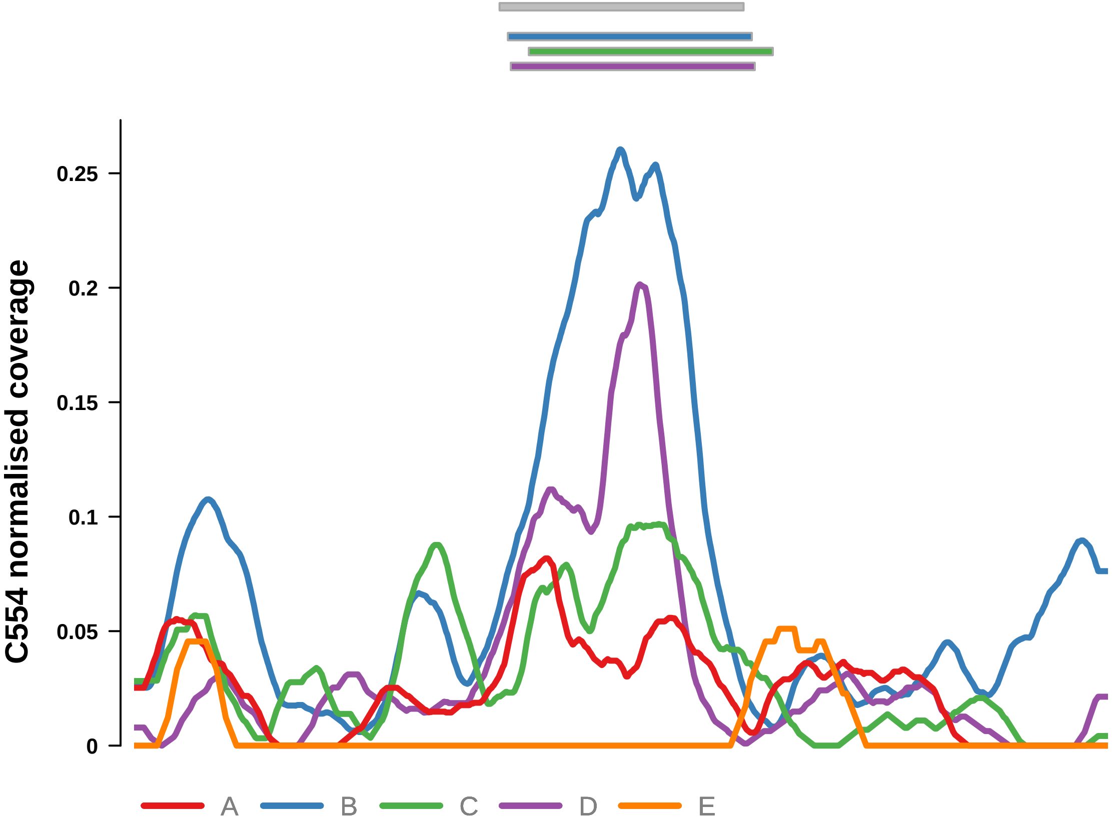


Putatively Subclonal (logFC = 2.43)

| | P. | Adjusted P. |
|---|---|---|
| *~region vs. ~* | 1.69e−10 **** | 3.46e−09 **** |
| *~purity vs ~* | 2.54e−08 **** | 1.04e−06 **** |
| *~purity+region vs ~purity* | 6.66e−03 ** | 3.52e−02 * |
| *Carcinoma vs. normal* | 1.16e−05 **** | 1.16e−05 **** |

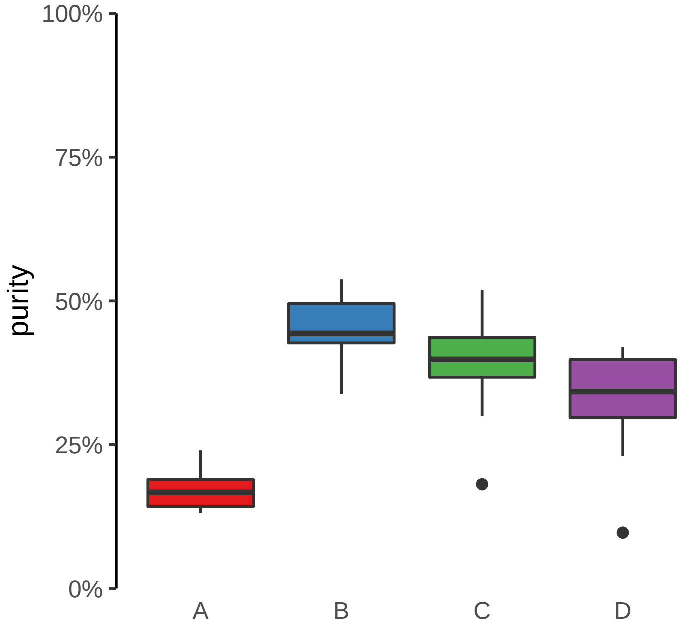

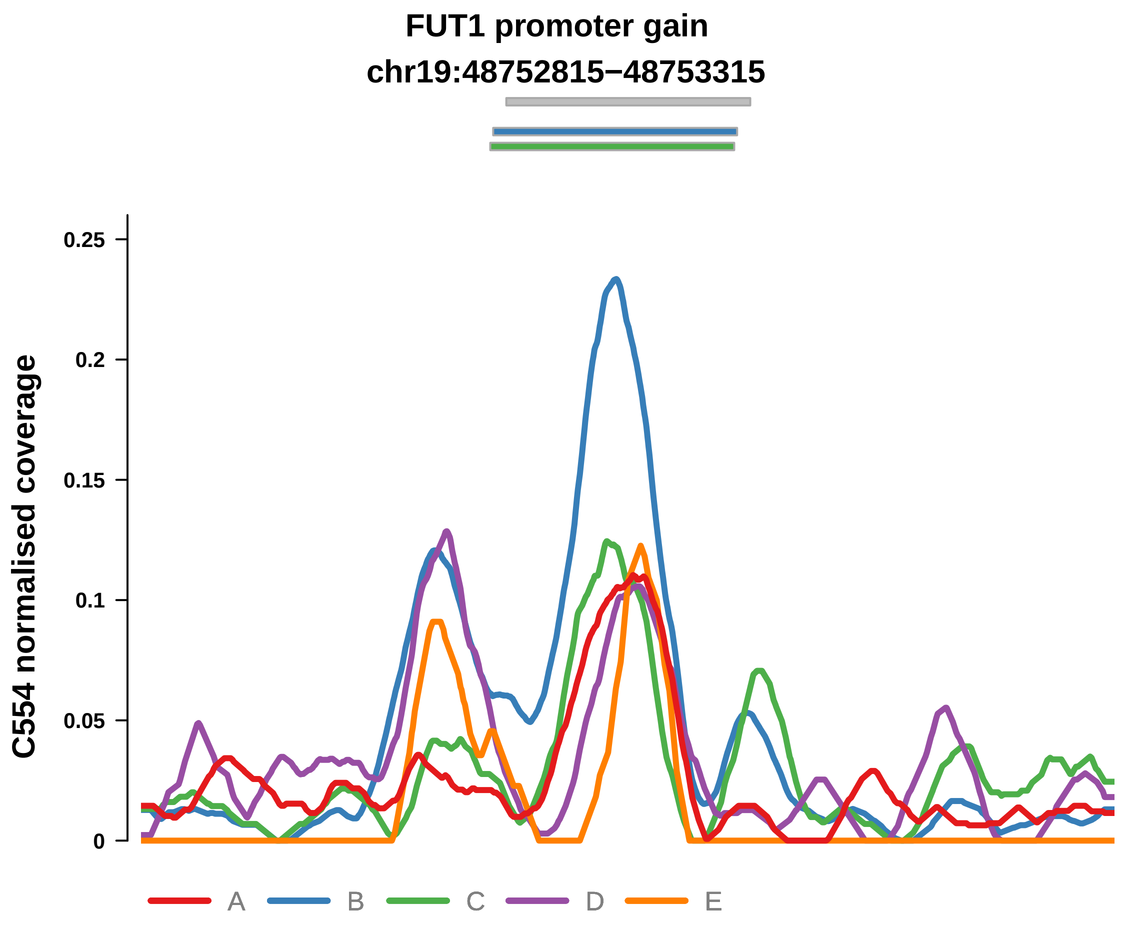


Putatively Subclonal (logFC = 1.85)

| | P. | Adjusted P. |
|---|---|---|
| *~region vs. ~* | 6.32e−06 **** | 6.48e−05 **** |
| *~purity vs ~* | 8.82e−04 *** | 8.03e−03 ** |
| *~purity+region vs ~purity* | 8.03e−03 ** | 3.71e−02 * |
| *Carcinoma vs. normal* | 7.65e−04 *** | 7.65e−04 *** |

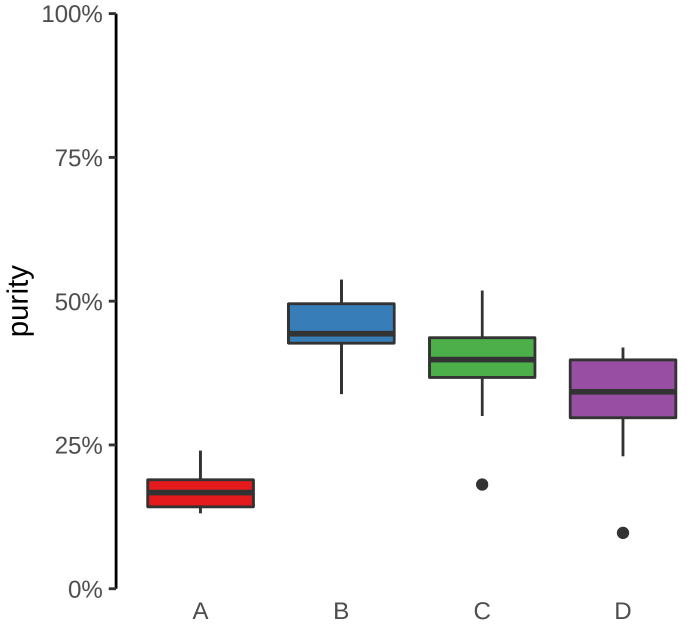

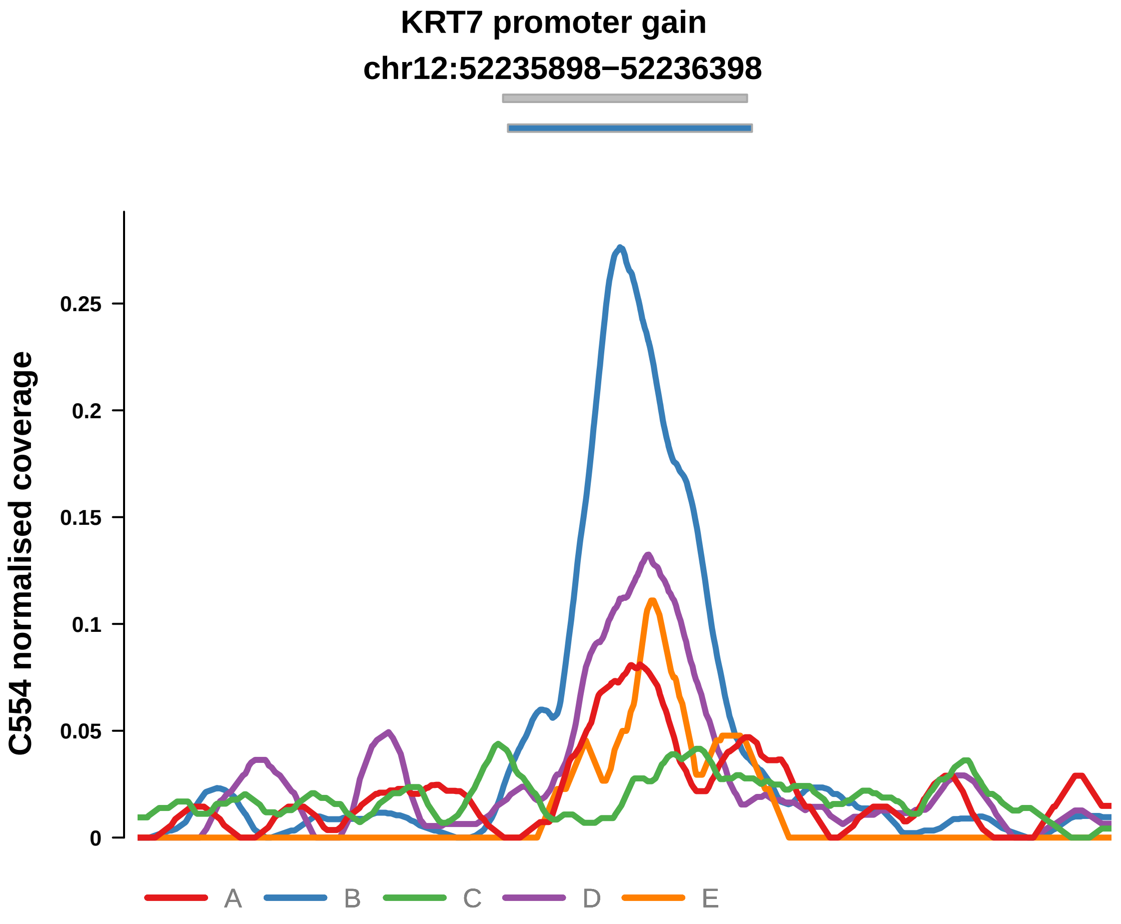


Putatively Subclonal (logFC = 2.27)

| | P. | Adjusted P. |
|---|---|---|
| *~region vs. ~* | 2.17e−08 **** | 3.56e−07 **** |
| *~purity vs ~* | 3.41e−04 *** | 4.66e−03 ** |
| *~purity+region vs ~purity* | 3.61e−04 *** | 6.67e−03 ** |
| *Carcinoma vs. normal* | 4.62e−05 **** | 4.62e−05 **** |

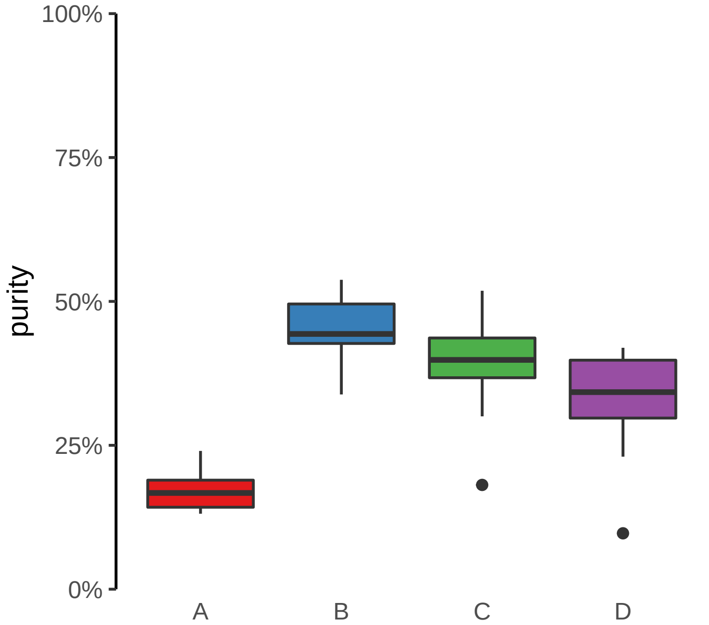
