## Supplementary Figures for "The co-evolution of the genome and epigenome in colorectal cancer": figS16_subclonal_promoter_loss.pdf

**AKR1B1 promoter loss**
**chr7:134458962−134459462**

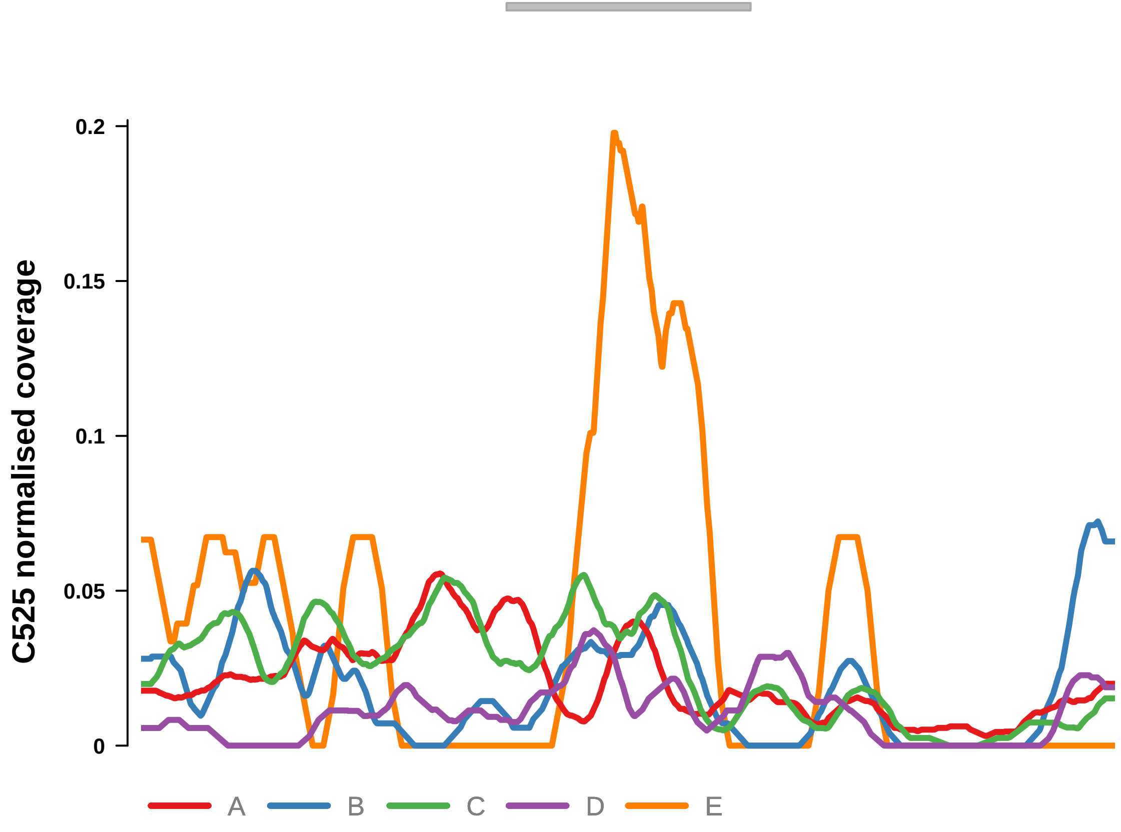


Putatively Subclonal (logFC = −0.86)

| | P. | Adjusted P. |
|---|---|---|
| *~region vs. ~* | 2.77e−03 ** | 1.04e−02 * |
| *~purity vs ~* | 4.77e−01 | 8.63e−01 |
| *~purity+region vs ~purity* | 6.74e−03 ** | 2.37e−02 * |
| *Carcinoma vs. normal* | NA | NA |

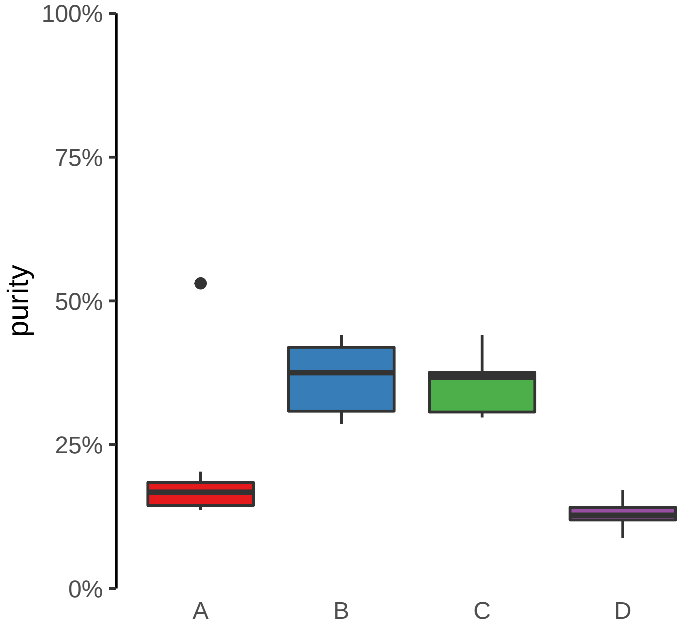

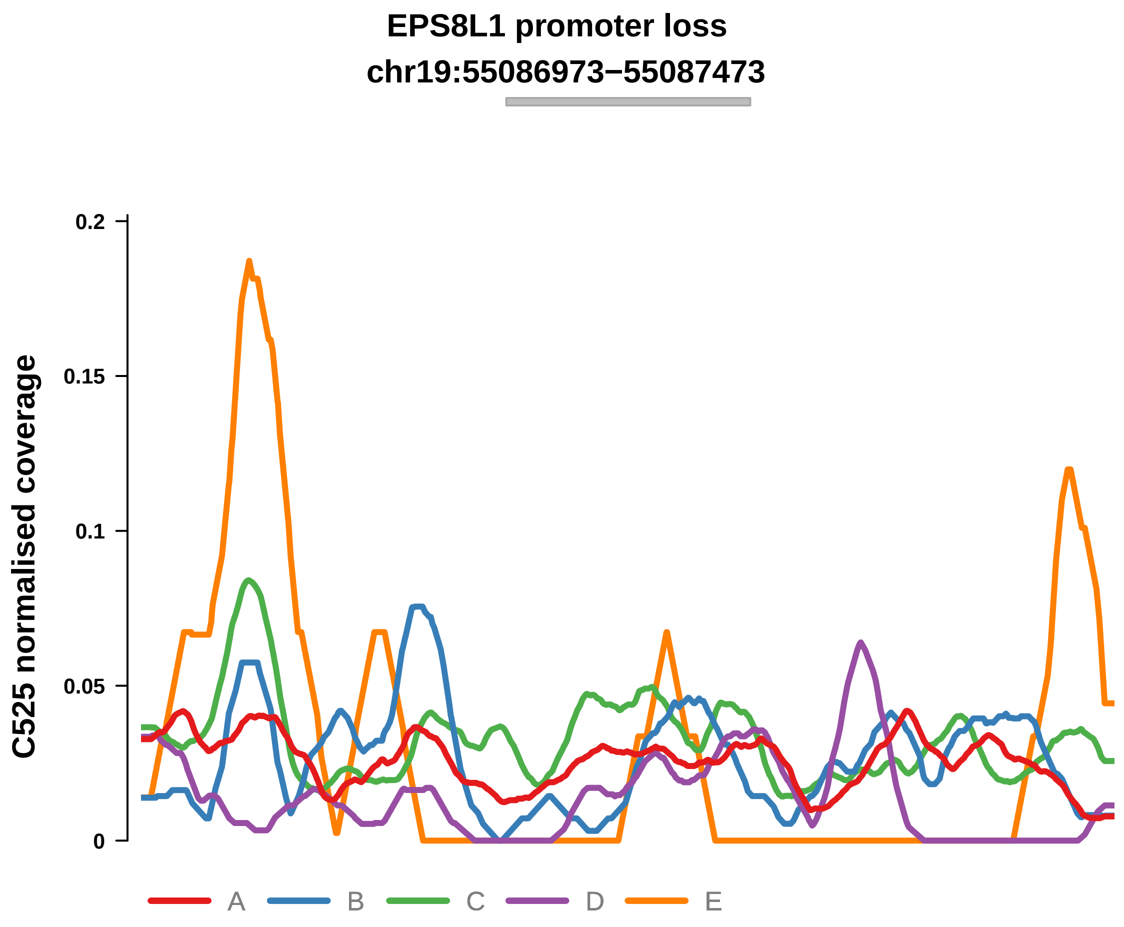


Putatively Subclonal (logFC = −1.42)

| | P. | Adjusted P. |
|---|---|---|
| *~region vs. ~* | 1.14e−04 *** | 9.8e−04 *** |
| *~purity vs ~* | 7.1e−02 | 7.18e−01 |
| *~purity+region vs ~purity* | 1.22e−03 ** | 7.66e−03 ** |
| *Carcinoma vs. normal* | 2.39e−02 * | 2.39e−02 * |

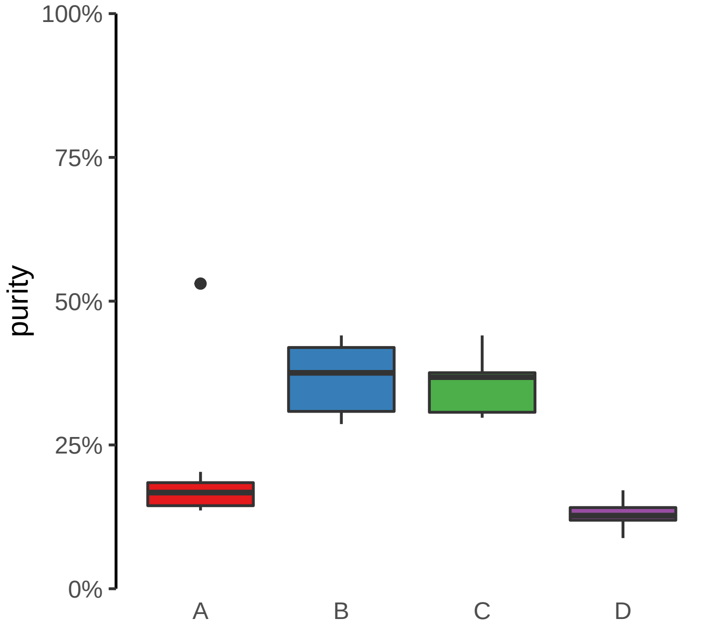

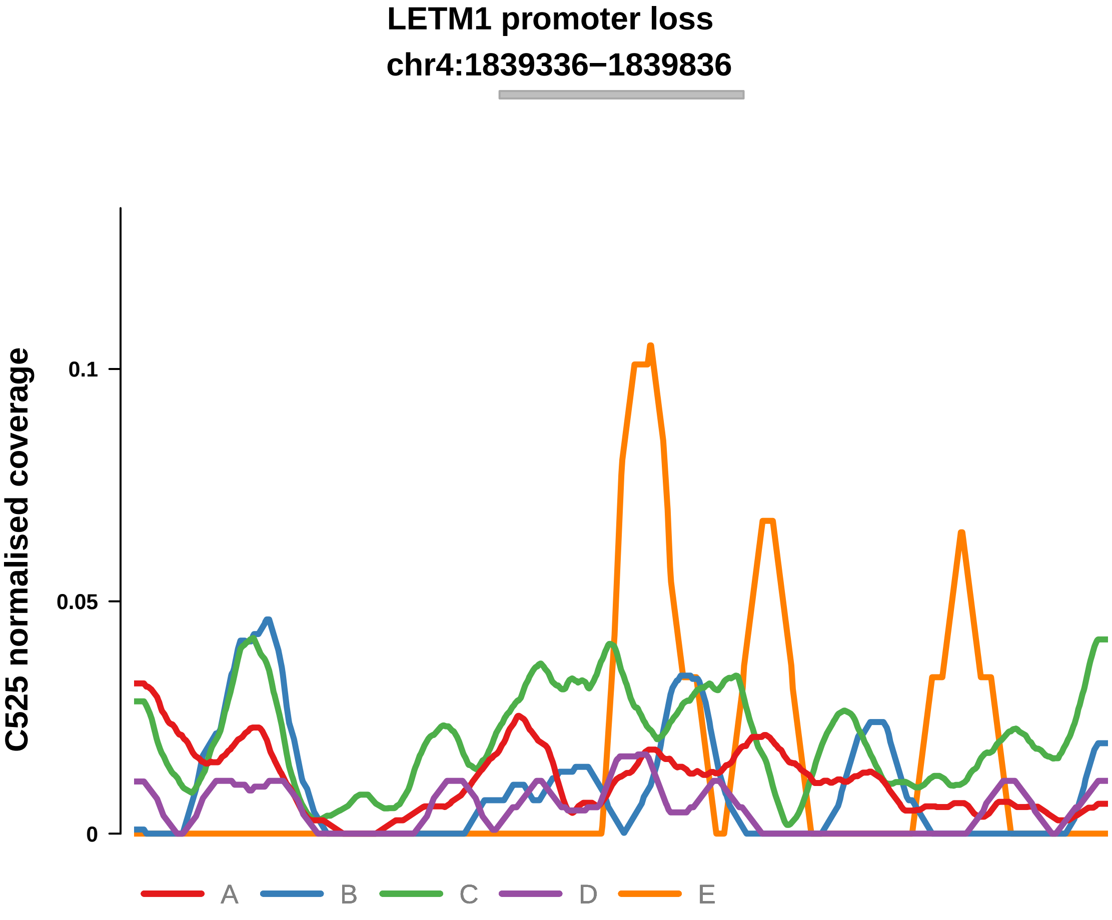


Putatively Subclonal (logFC = −0.47)

| | P. | Adjusted P. |
|---|---|---|
| *~region vs. ~* | 1.64e−05 **** | 4.7e−04 *** |
| *~purity vs ~* | 4.86e−02 * | 6.12e−01 |
| *~purity+region vs ~purity* | 4.79e−04 *** | 3.9e−03 ** |
| *Carcinoma vs. normal* | NA | NA |

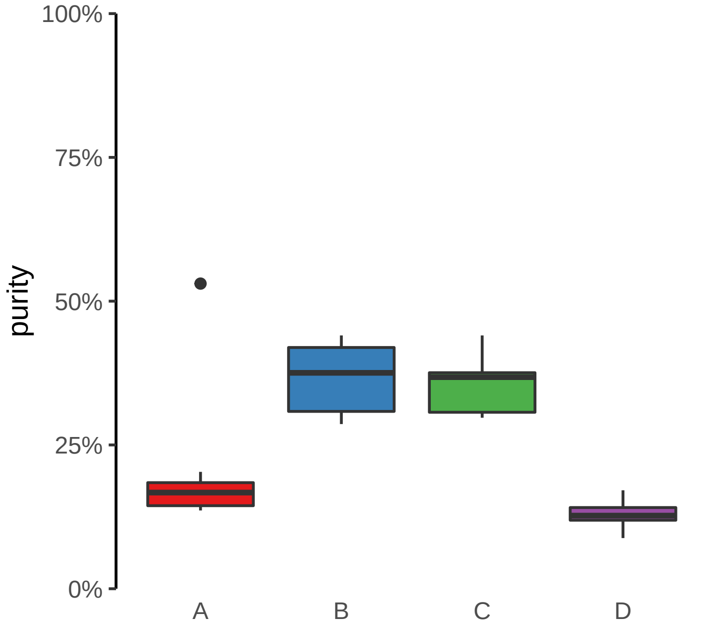

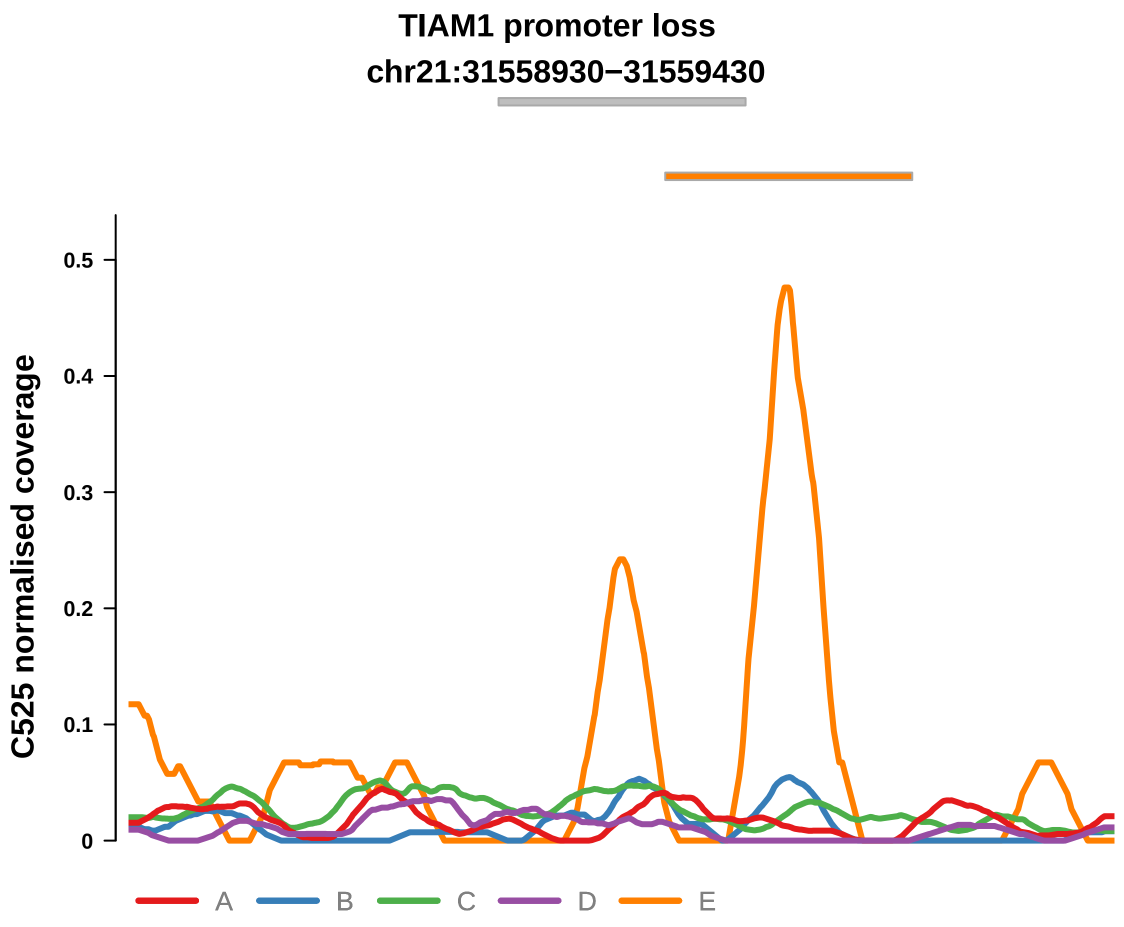


Putatively Subclonal (logFC = −1.93)

| | P. | Adjusted P. |
|---|---|---|
| *~region vs. ~* | 1.77e−03 ** | 7.63e−03 ** |
| *~purity vs ~* | 3.22e−01 | 7.48e−01 |
| *~purity+region vs ~purity* | 5.79e−03 ** | 2.21e−02 * |
| *Carcinoma vs. normal* | 1.45e−03 ** | 1.45e−03 ** |

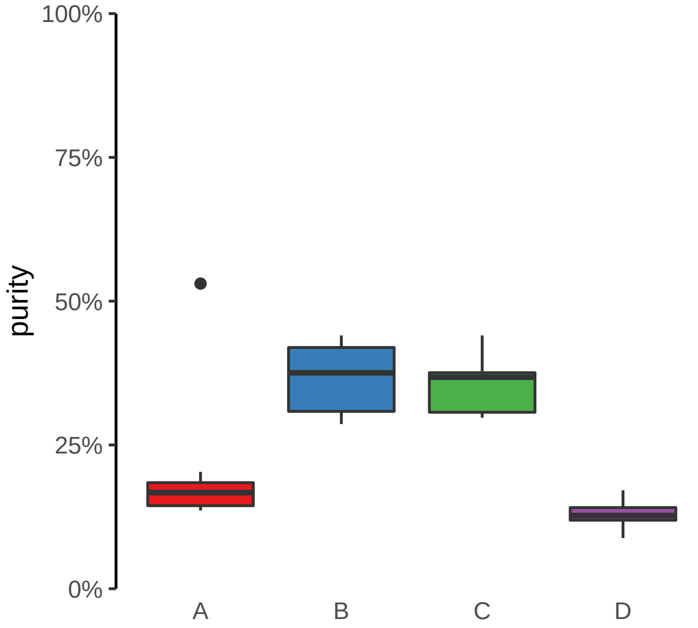

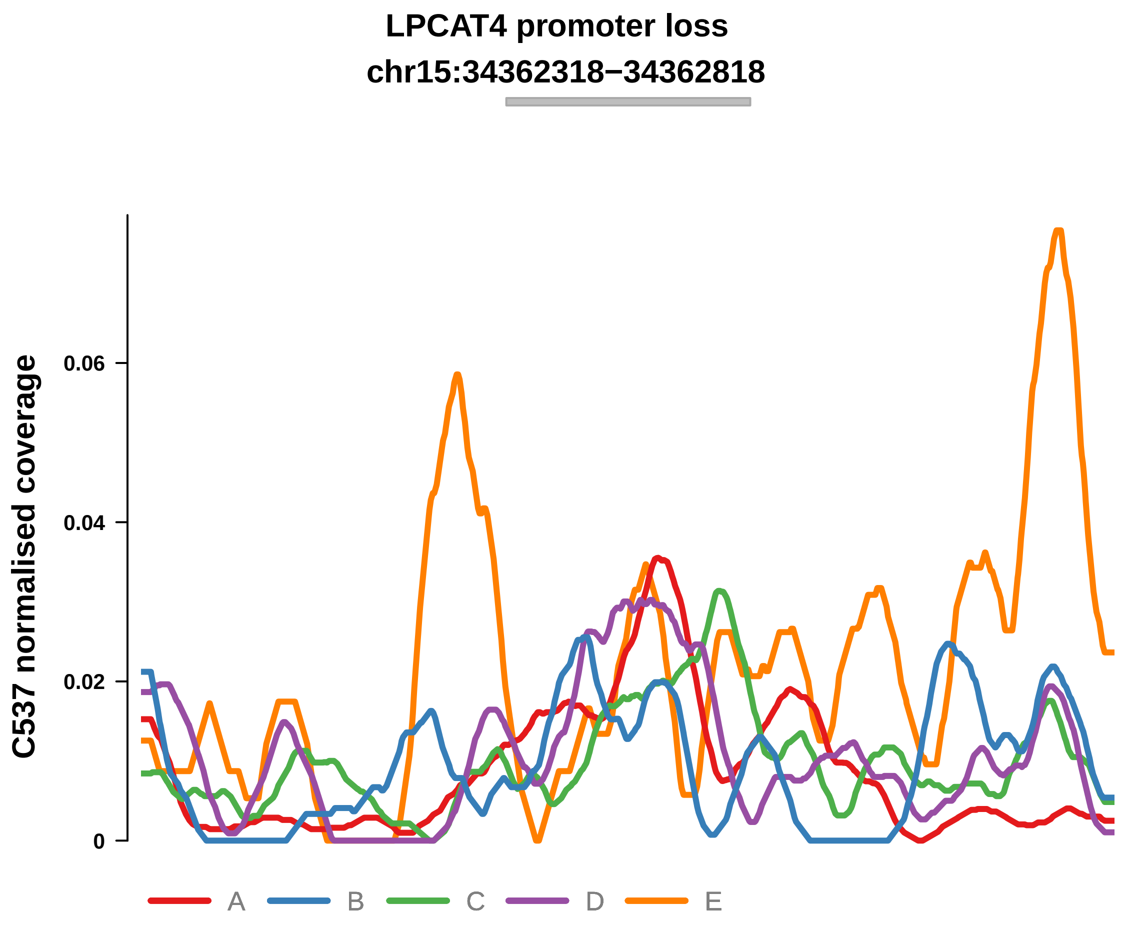


Putatively Subclonal (logFC = −0.69)

| | P. | Adjusted P. |
|---|---|---|
| *~region vs. ~* | 1.44e−02 * | 6.02e−02 |
| *~purity vs ~* | 7.33e−01 | 8.29e−01 |
| *~purity+region vs ~purity* | 3.01e−03 ** | 3.96e−02 * |
| *Carcinoma vs. normal* | NA | NA |

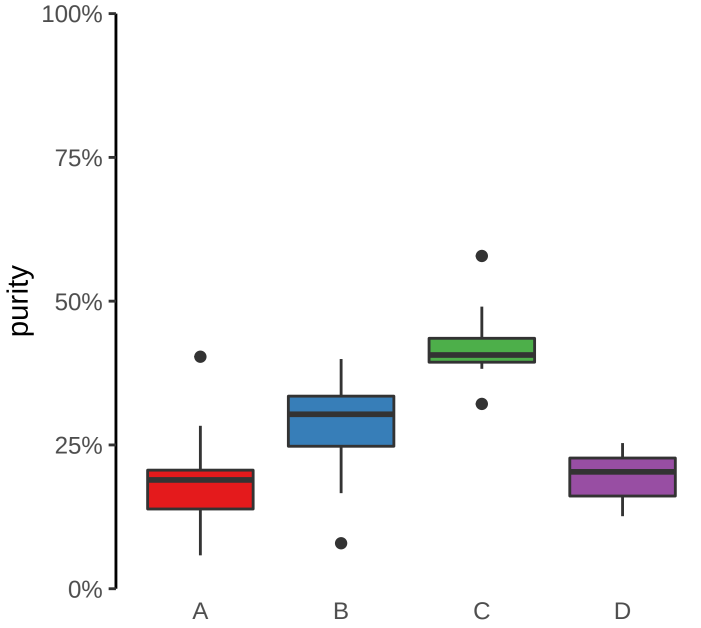

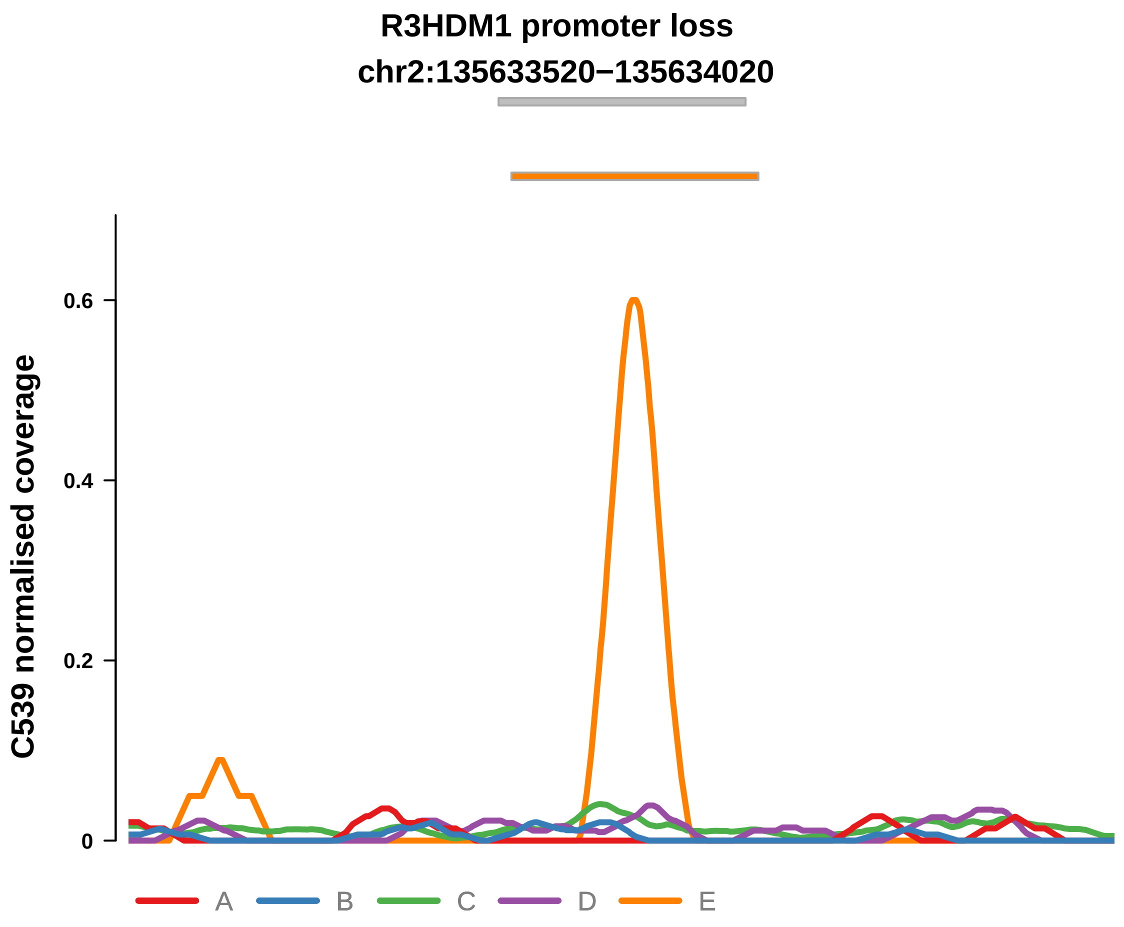


Putatively Subclonal (logFC = −0.57)

| | P. | Adjusted P. |
|---|---|---|
| *~region vs. ~* | 4.47e−05 **** | 2.62e−04 *** |
| *~purity vs ~* | 2.84e−03 ** | 6.41e−03 ** |
| *~purity+region vs ~purity* | 8.54e−03 ** | 4.22e−02 * |
| *Carcinoma vs. normal* | NA | NA |

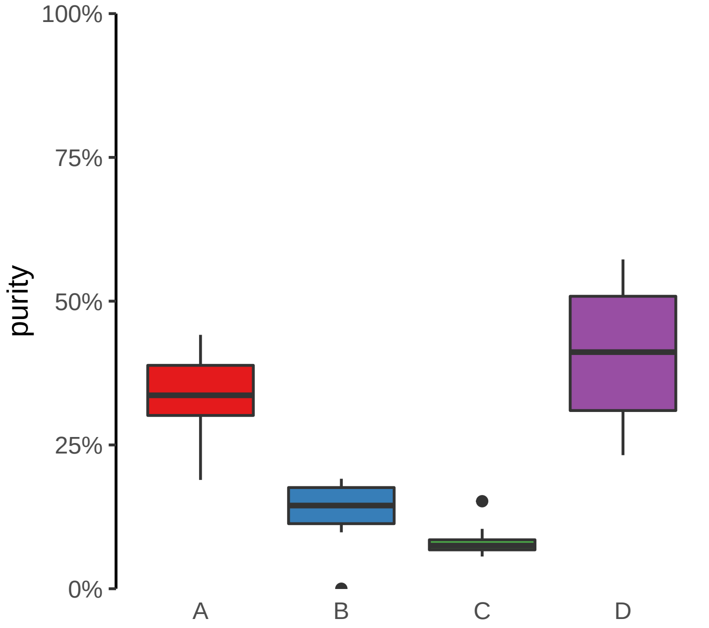

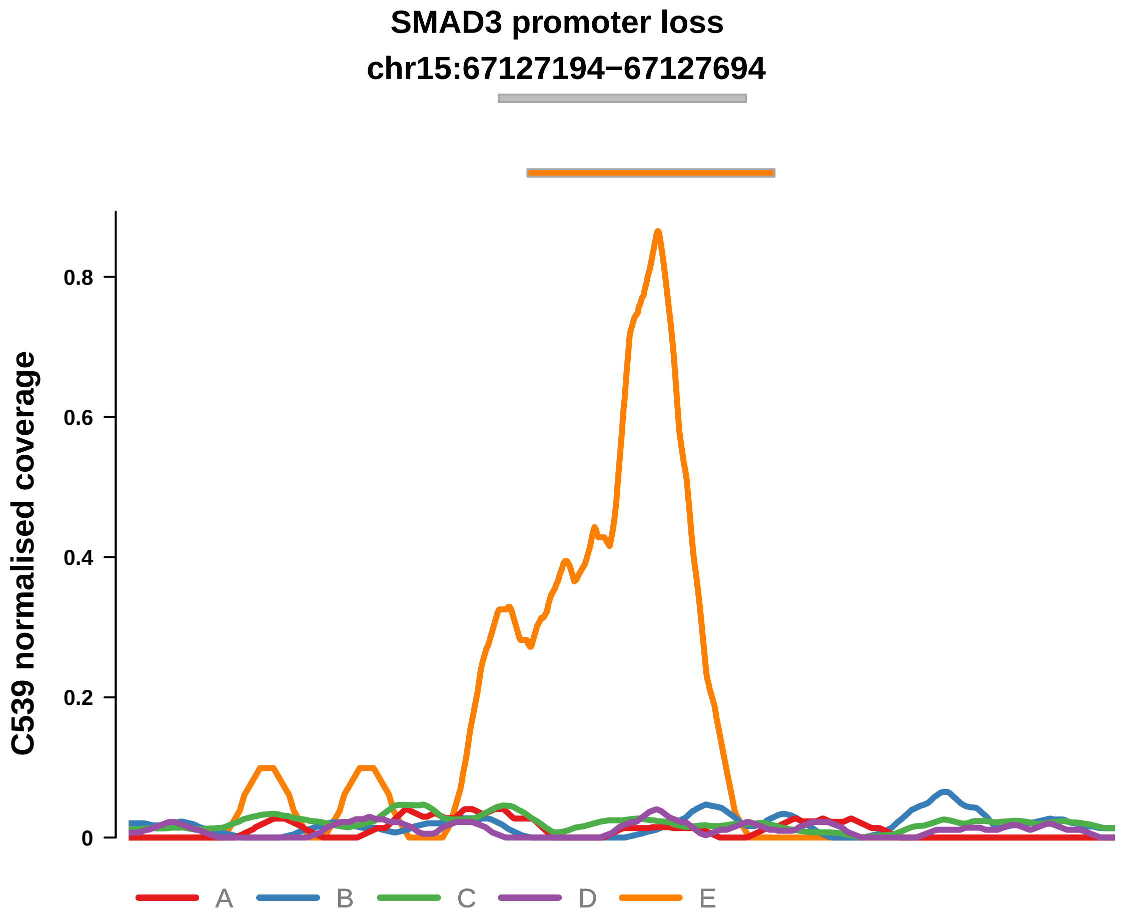


Putatively Subclonal (logFC = −1.32)

| | P. | Adjusted P. |
|---|---|---|
| *~region vs. ~* | 1.42e−07 **** | 4.16e−06 **** |
| *~purity vs ~* | 7.03e−05 **** | 6.48e−04 *** |
| *~purity+region vs ~purity* | 3.12e−04 *** | 4.93e−03 ** |
| *Carcinoma vs. normal* | 1.05e−01 | 1.05e−01 |

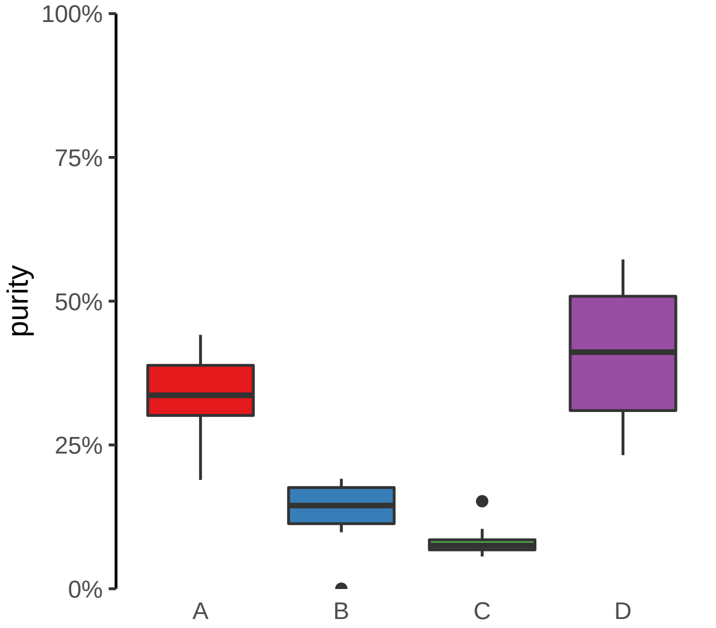

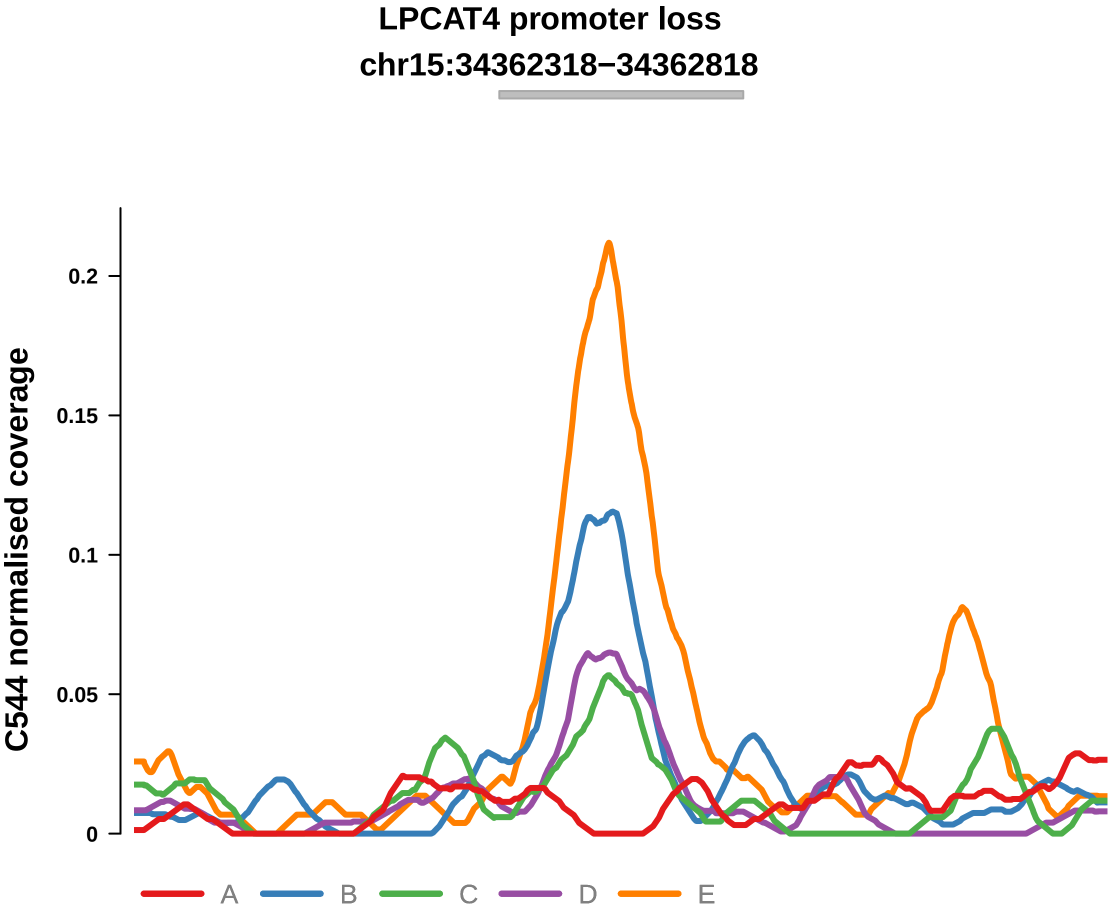


Putatively Subclonal (logFC = −1.32)

| | P. | Adjusted P. |
|---|---|---|
| *~region vs. ~* | 1.89e−05 **** | 3.48e−04 *** |
| *~purity vs ~* | 3.26e−03 ** | 2.16e−02 * |
| *~purity+region vs ~purity* | 1.8e−03 ** | 1.59e−02 * |
| *Carcinoma vs. normal* | 2.91e−02 * | 2.91e−02 * |

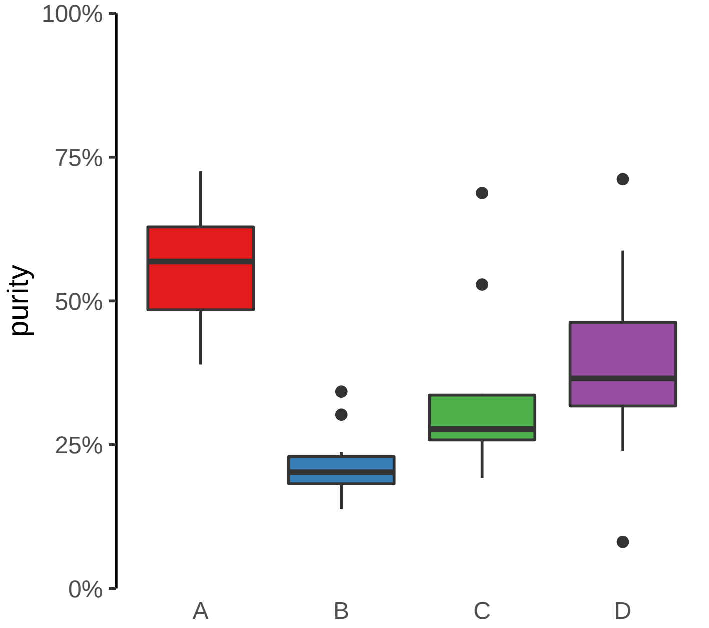

**R3HDM1 promoter loss**
**chr2:135633520−135634020**

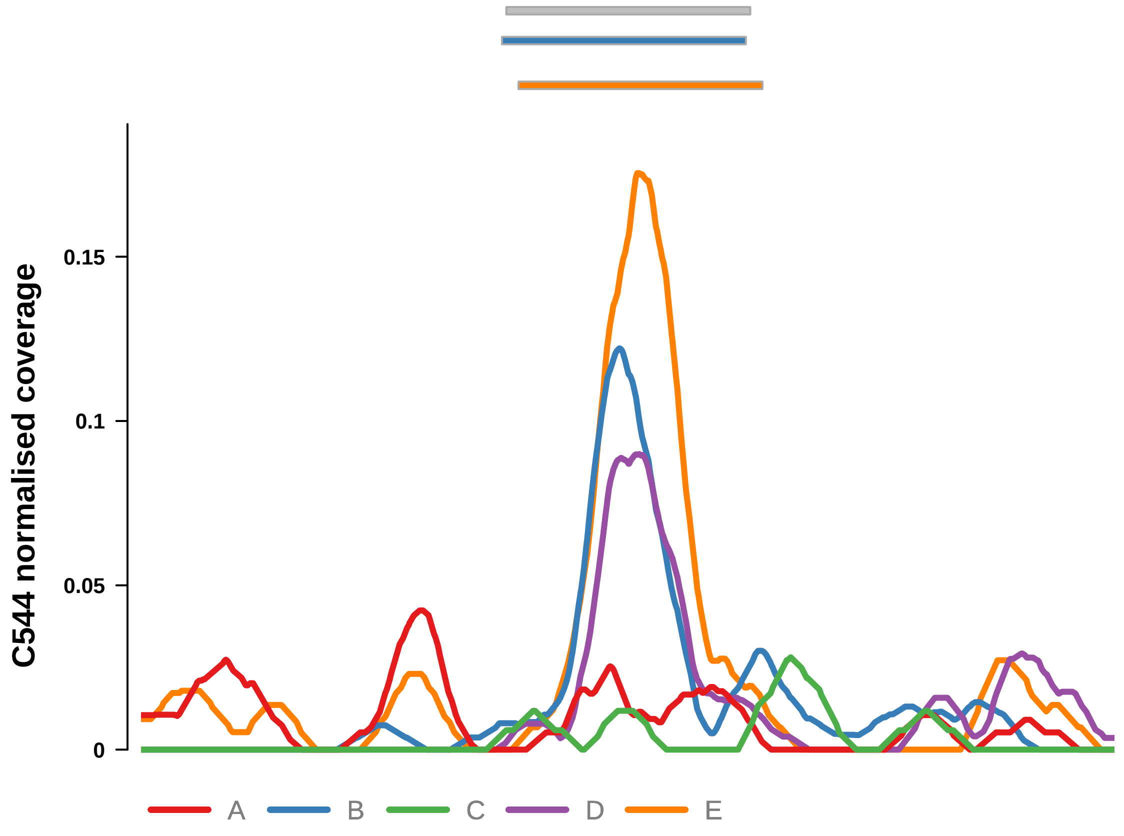


Putatively Subclonal (logFC = −1.24)

| | P. | Adjusted P. |
|---|---|---|
| *~region vs. ~* | 6.97e−05 **** | 6.14e−04 *** |
| *~purity vs ~* | 3.73e−02 * | 9.93e−02 |
| *~purity+region vs ~purity* | 9.16e−04 *** | 1.17e−02 * |
| *Carcinoma vs. normal* | 5.05e−02 | 5.05e−02 |

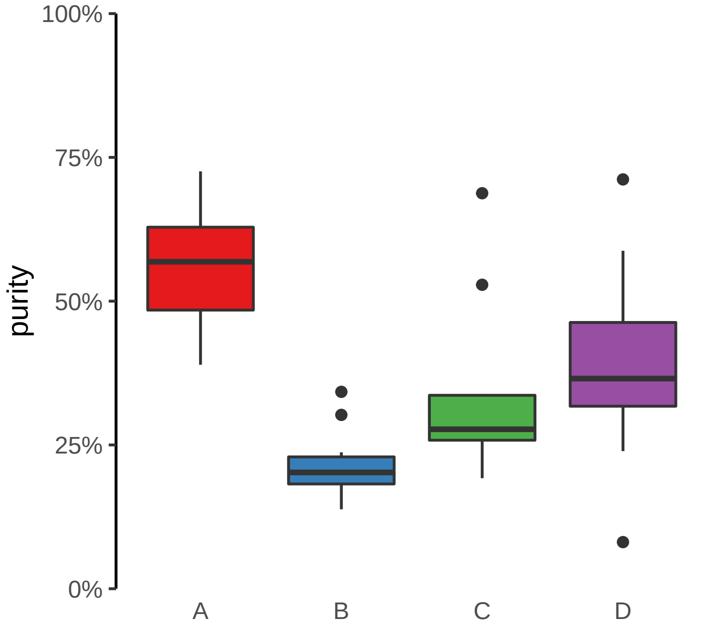

**SPAG16 promoter loss**
**chr2:213284089−213284589**

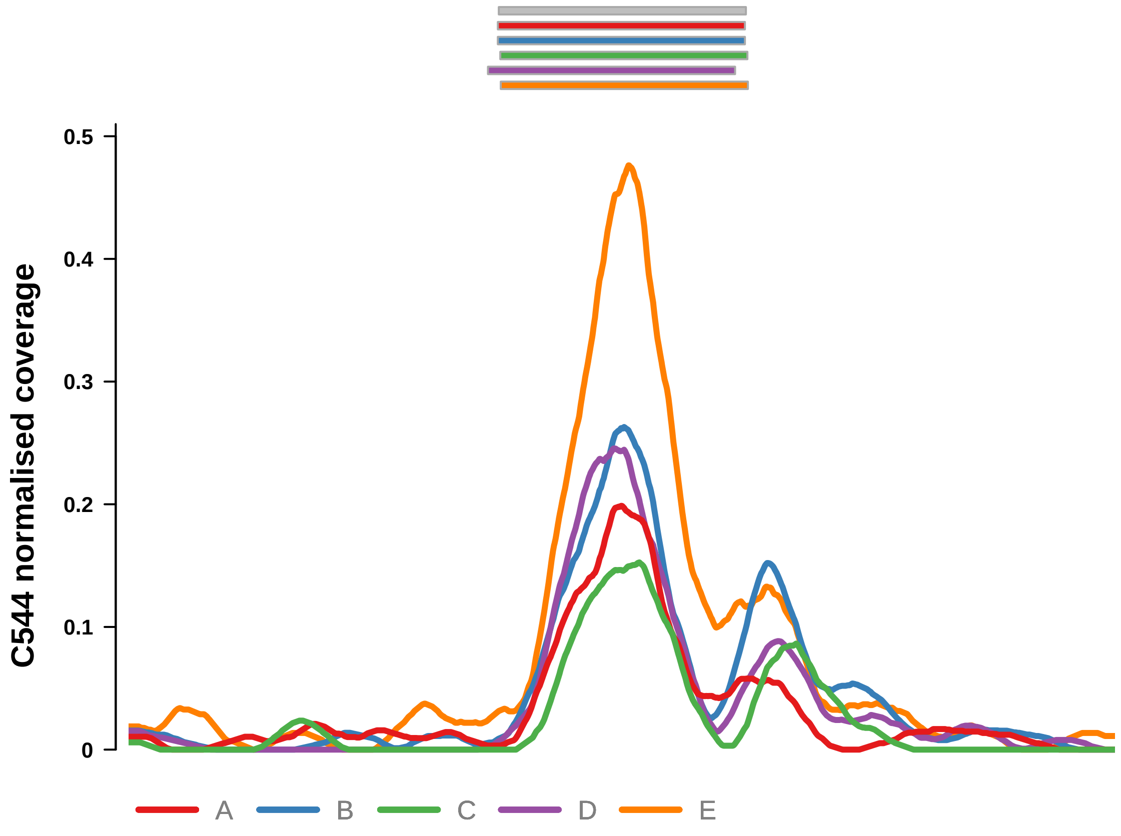


Putatively Subclonal (logFC = −0.12)

| | P. | Adjusted P. |
|---|---|---|
| *~region vs. ~* | 4.67e−03 ** | 1.96e−02 * |
| *~purity vs ~* | 7.21e−01 | 8e−01 |
| *~purity+region vs ~purity* | 5.32e−03 ** | 2.83e−02 * |
| *Carcinoma vs. normal* | NA | NA |

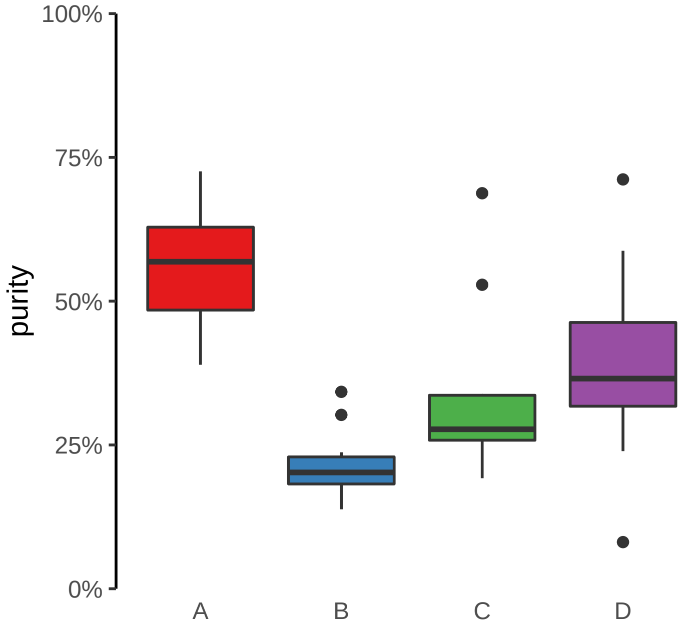

**TIAM1 promoter loss**
**chr21:31558930−31559430**

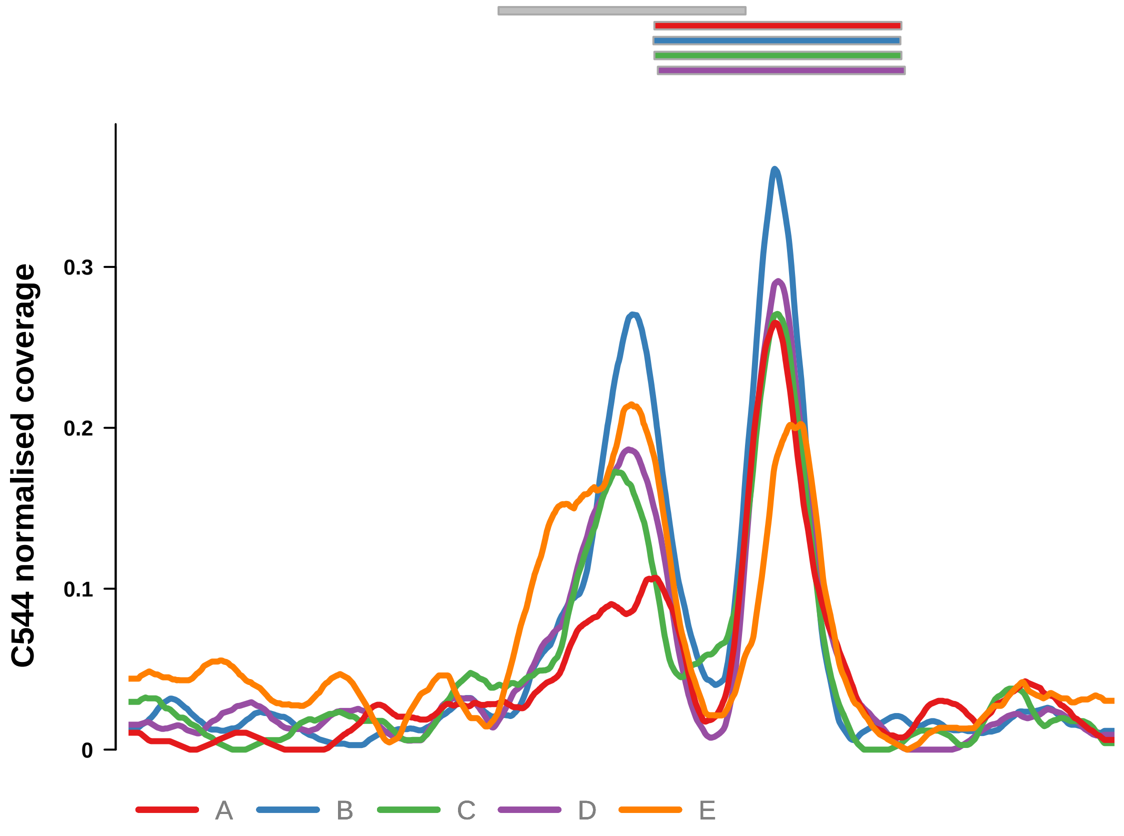


Putatively Subclonal (logFC = −0.63)

| | P. | Adjusted P. |
|---|---|---|
| *~region vs. ~* | 1.93e−04 *** | 1.55e−03 ** |
| *~purity vs ~* | 3.51e−01 | 5.18e−01 |
| *~purity+region vs ~purity* | 3.65e−04 *** | 9.1e−03 ** |
| *Carcinoma vs. normal* | NA | NA |

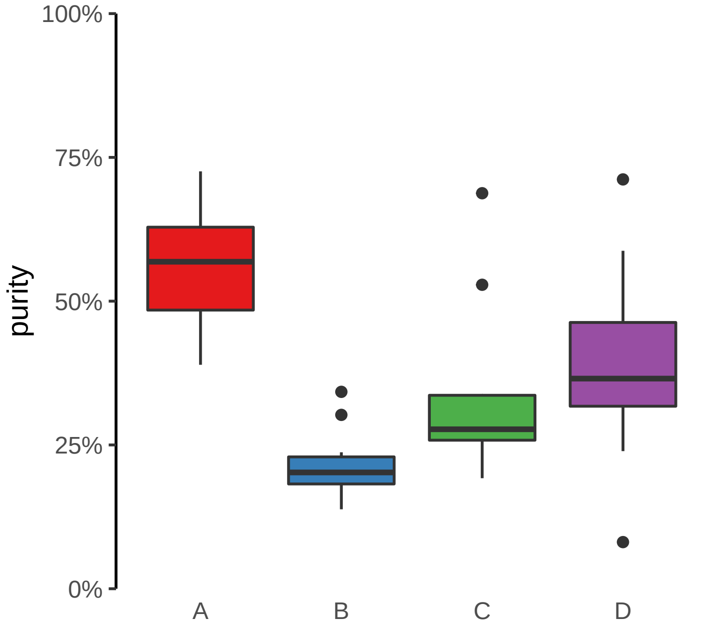

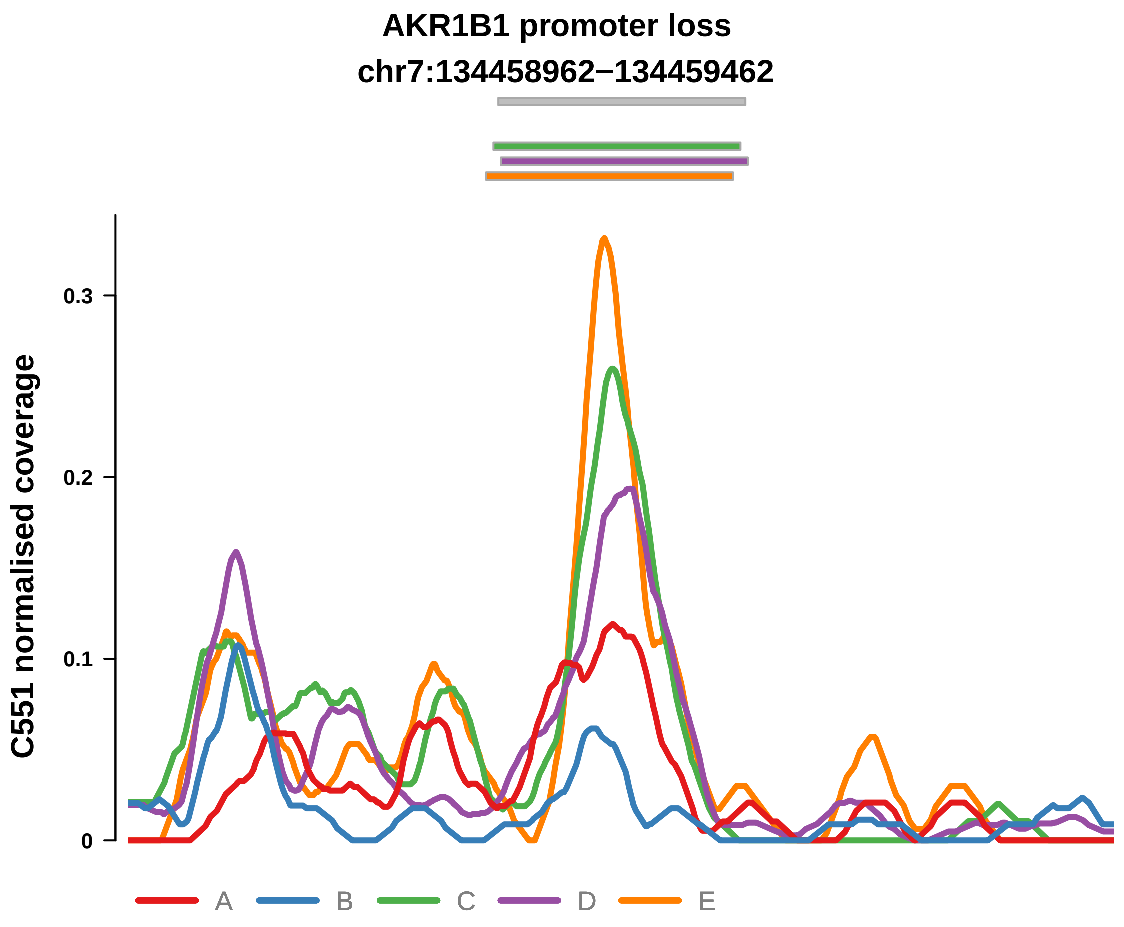


Putatively Subclonal (logFC = −0.24)

| | P. | Adjusted P. |
|---|---|---|
| *~region vs. ~* | 1.84e−04 *** | 8.09e−03 ** |
| *~purity vs ~* | 7.62e−01 | 9.44e−01 |
| *~purity+region vs ~purity* | 6.86e−05 **** | 6.03e−03 ** |
| *Carcinoma vs. normal* | NA | NA |

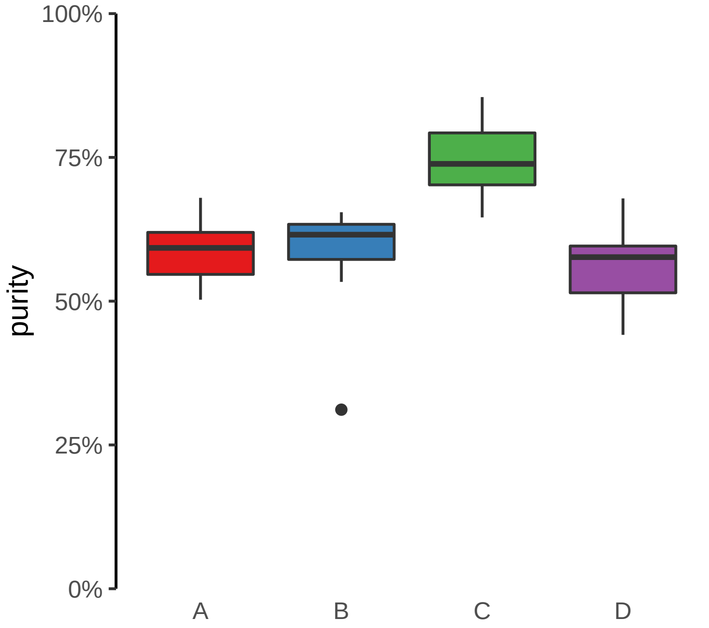

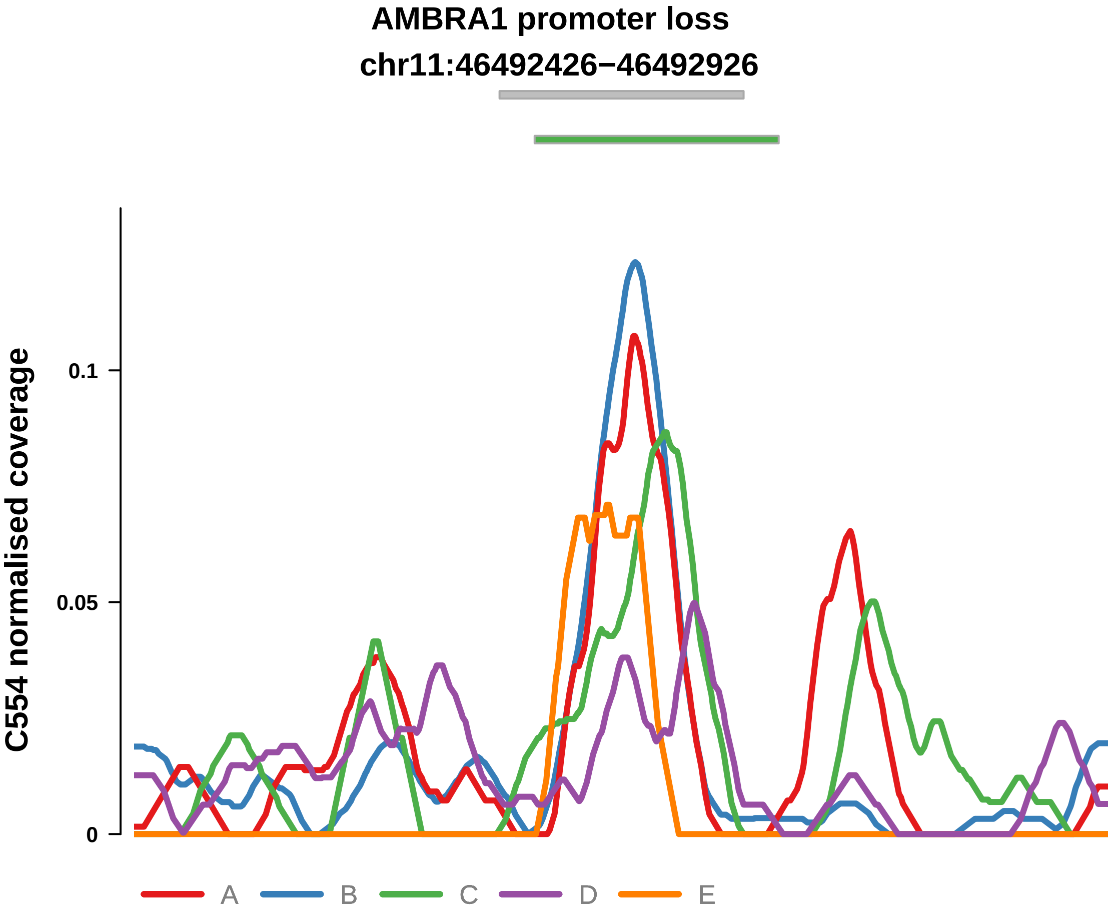


Putatively Subclonal (logFC = −0.2)

| | P. | Adjusted P. |
|---|---|---|
| *~region vs. ~* | 8.53e−04 *** | 4.37e−03 ** |
| *~purity vs ~* | 7.87e−02 | 2.22e−01 |
| *~purity+region vs ~purity* | 5.97e−03 ** | 3.52e−02 * |
| *Carcinoma vs. normal* | NA | NA |

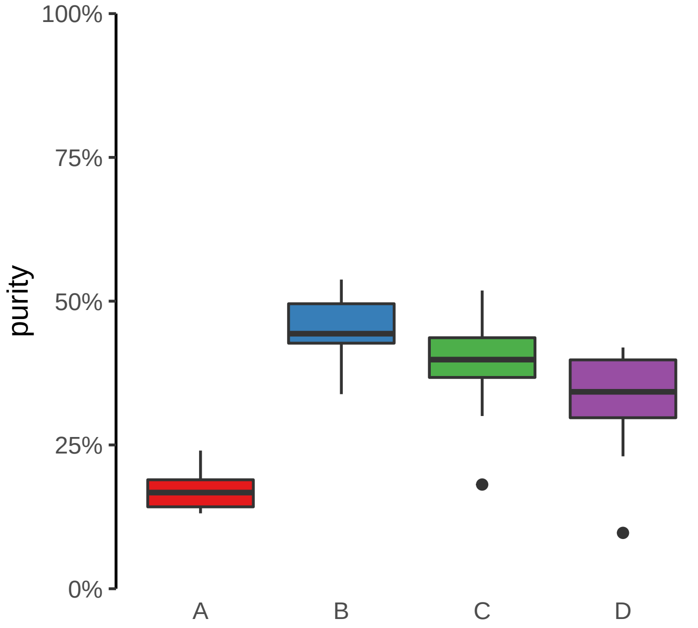

**TIAM1 promoter loss**

**chr21:31558930−31559430**

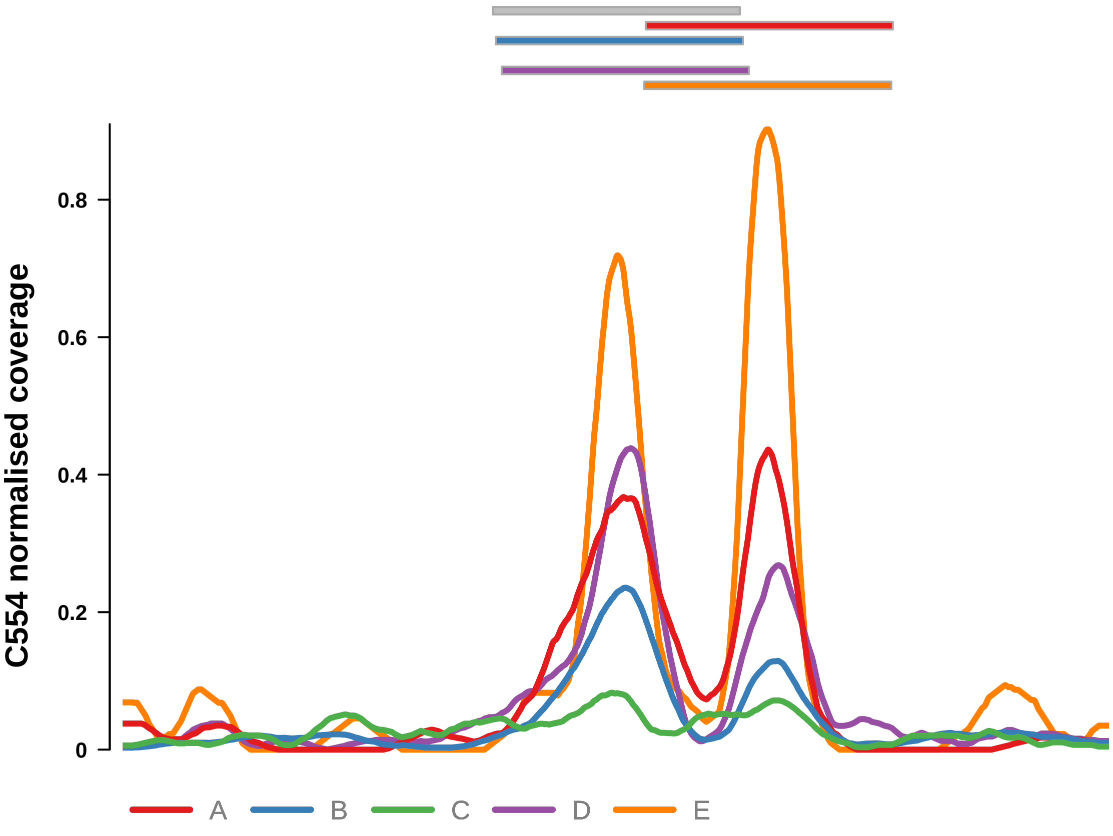


Putatively Subclonal (logFC = −0.43)

| | P. | Adjusted P. |
|---|---|---|
| *~region vs. ~* | 2.82e−01 | 3.63e−01 |
| *~purity vs ~* | 8.9e−01 | 9.43e−01 |
| *~purity+region vs ~purity* | 1.24e−02 * | 4.38e−02 * |
| *Carcinoma vs. normal* | NA | NA |

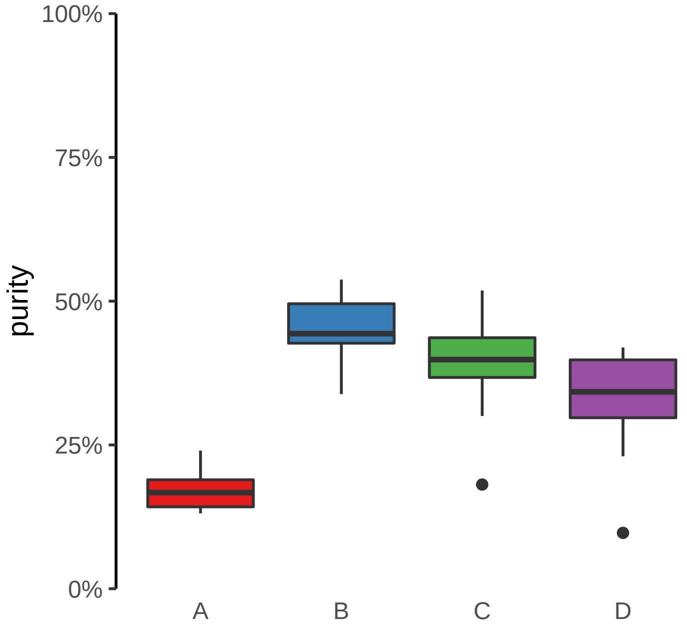
