## Supplementary Figures for "The co-evolution of the genome and epigenome in colorectal cancer": figS17_subclonal_enhancer_gain.pdf

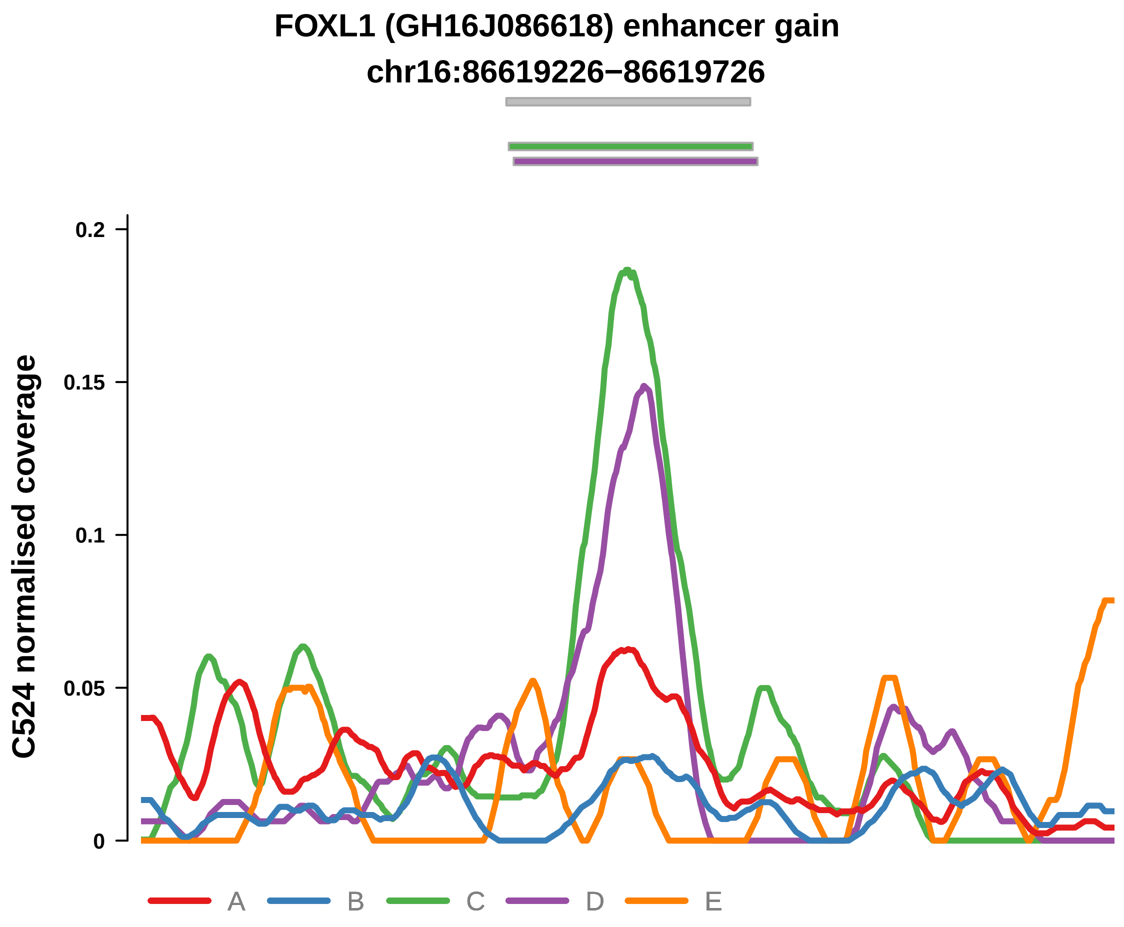


Putatively Subclonal (logFC = 1.83)

| | P. | Adjusted P. |
| --- | --- | --- |
| *~region vs. ~* | 1.48e−06 **** | 1.3e−04 *** |
| *~purity vs ~* | 1.93e−01 | 5e−01 |
| *~purity+region vs ~purity* | 4.01e−06 **** | 3.53e−04 *** |
| *Carcinoma vs. normal* | 1.52e−02 * | 1.52e−02 * |

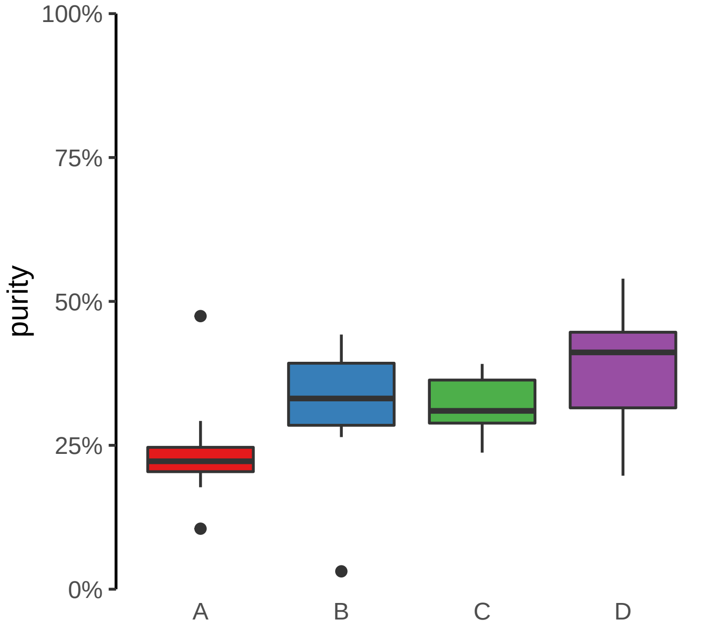

**SULF2 (GH20J047808) enhancer gain**

**chr20:47809591−47810091**

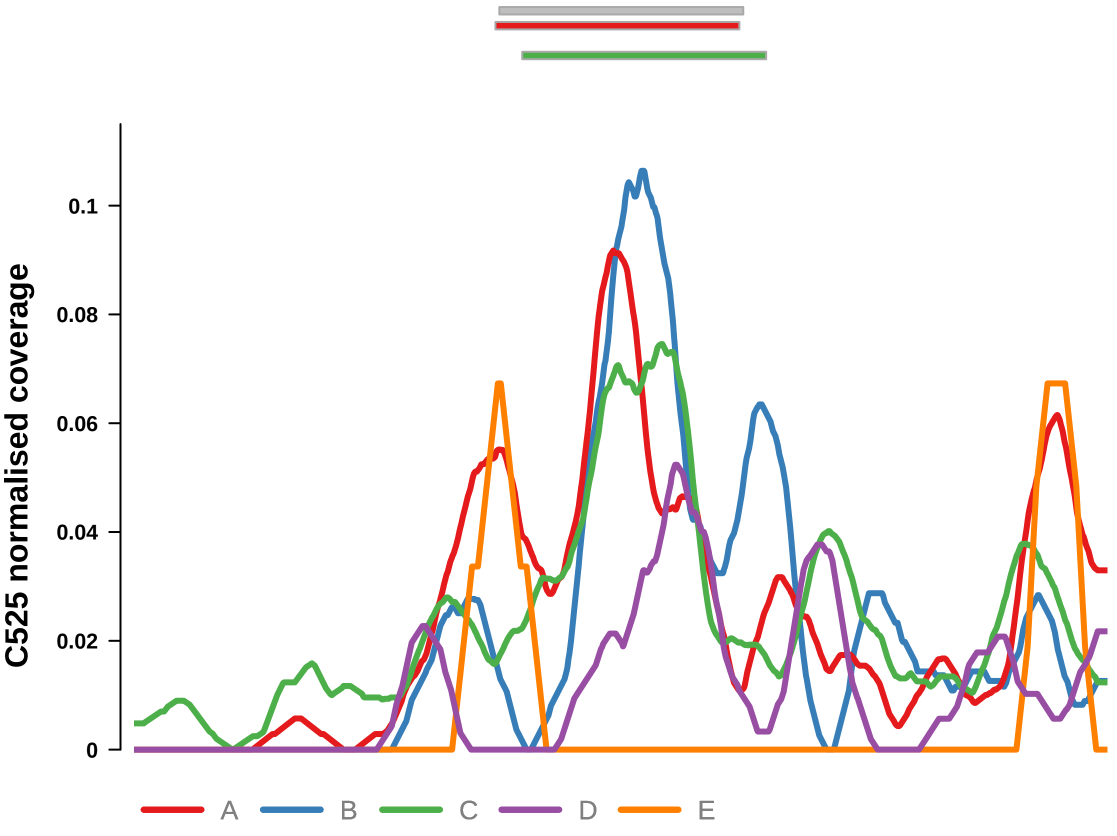


Putatively Subclonal (logFC = 1.65)

| | **P.** | **Adjusted P.** |
|---|---|---|
| *~region vs. ~* | 1.34e−03 ** | 6.78e−03 ** |
| *~purity vs ~* | 3.82e−01 | 8.2e−01 |
| *~purity+region vs ~purity* | 2.27e−03 ** | 1.25e−02 * |
| *Carcinoma vs. normal* | 2.24e−02 * | 2.24e−02 * |

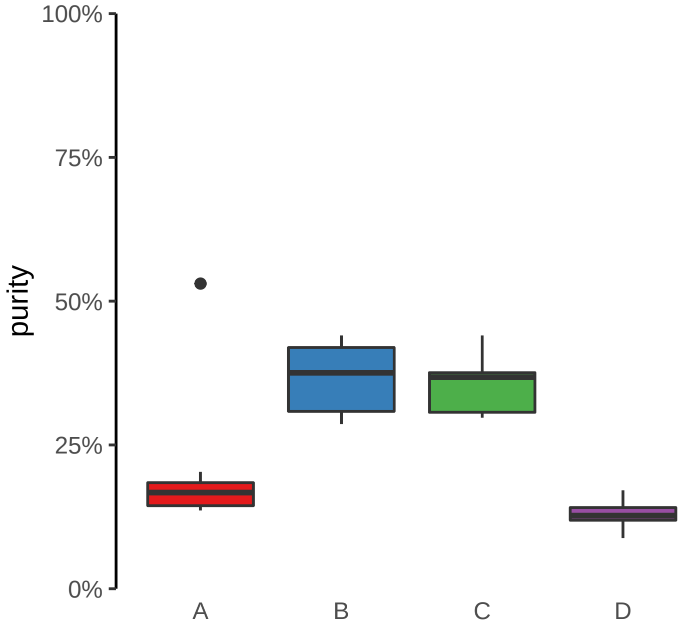

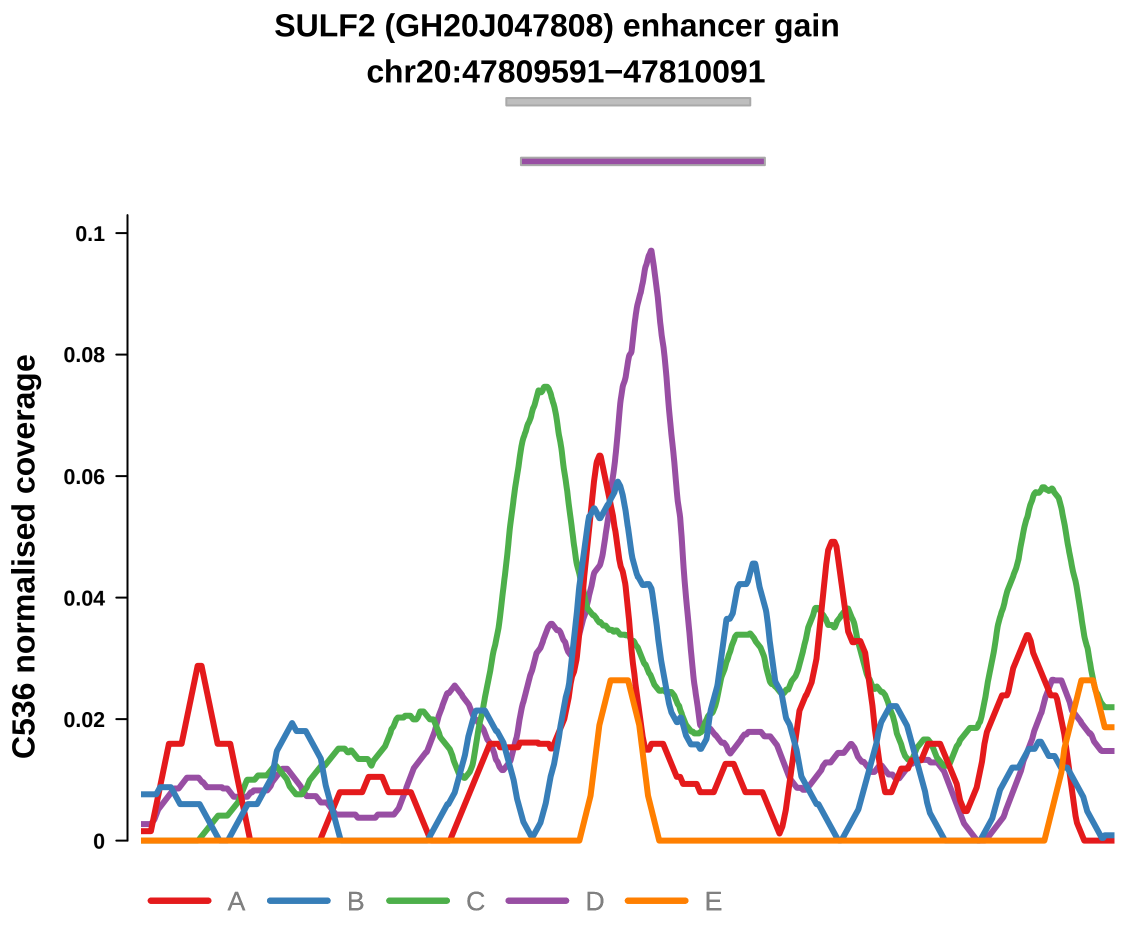


Putatively Subclonal (logFC = 1.91)

| | P. | Adjusted P. |
|---|---|---|
| *~region vs. ~* | 1.19e−01 | 2.41e−01 |
| *~purity vs ~* | 8.74e−01 | 9.5e−01 |
| *~purity+region vs ~purity* | 1.25e−03 ** | 2.4e−02 * |
| *Carcinoma vs. normal* | 7.95e−03 ** | 7.95e−03 ** |

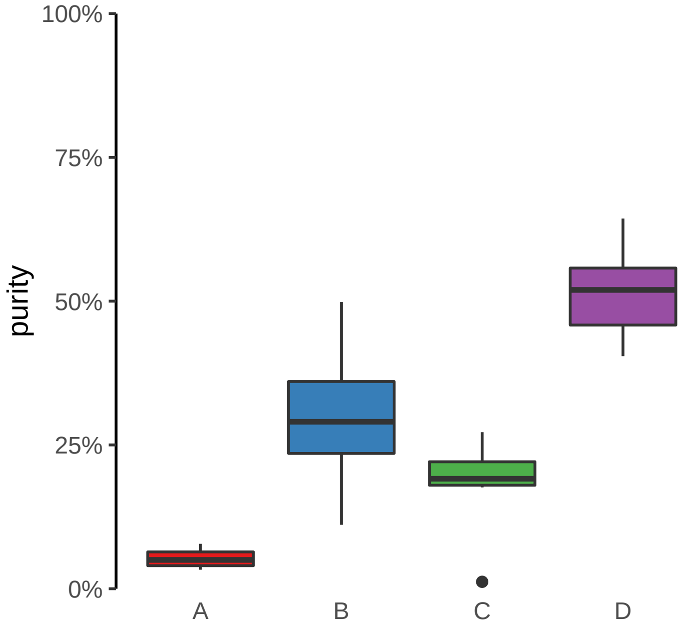

**CLASRP, MARK4 (GH19J045307) enhancer gain**
**chr19:45308035−45308535**

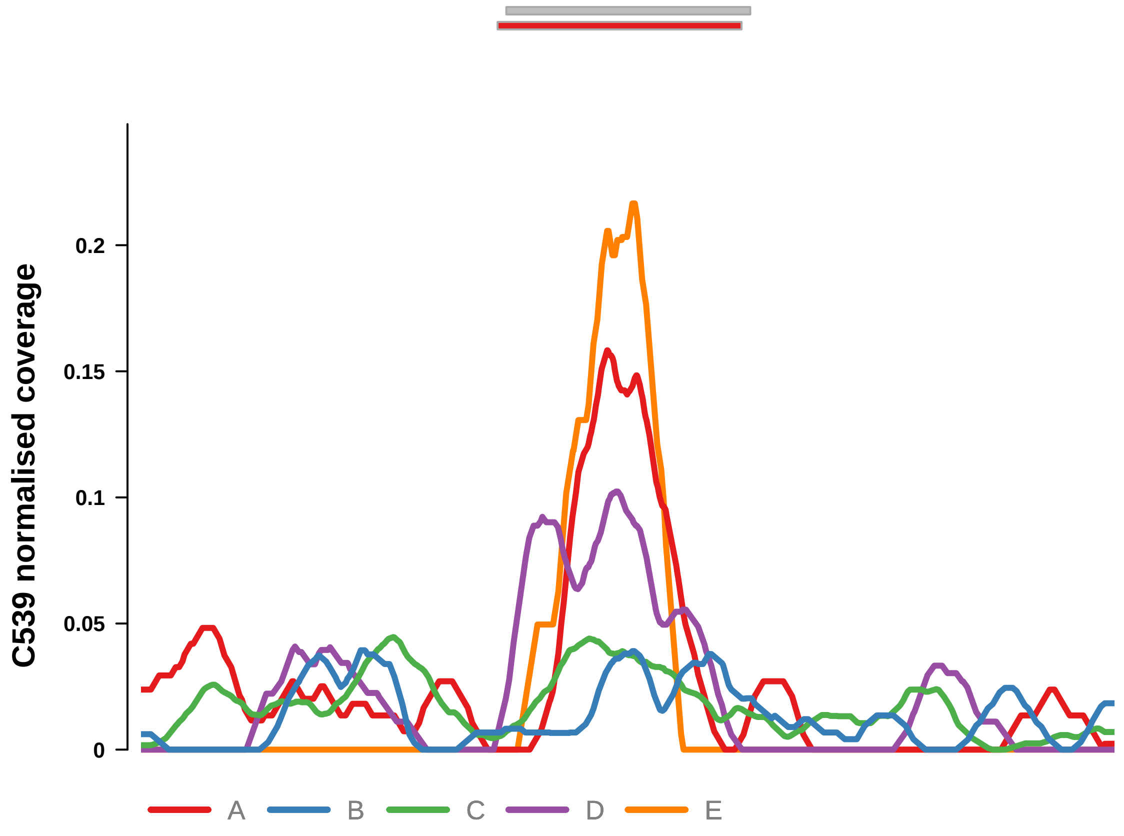


Putatively Subclonal (logFC = 2.89)

| | **P.** | **Adjusted P.** |
|---|---|---|
| *~region vs. ~* | 3.51e−03 ** | 6.71e−03 ** |
| *~purity vs ~* | 7.71e−02 | 8.92e−02 |
| *~purity+region vs ~purity* | 1.27e−04 *** | 2.66e−03 ** |
| *Carcinoma vs. normal* | 1.8e−04 *** | 1.8e−04 *** |

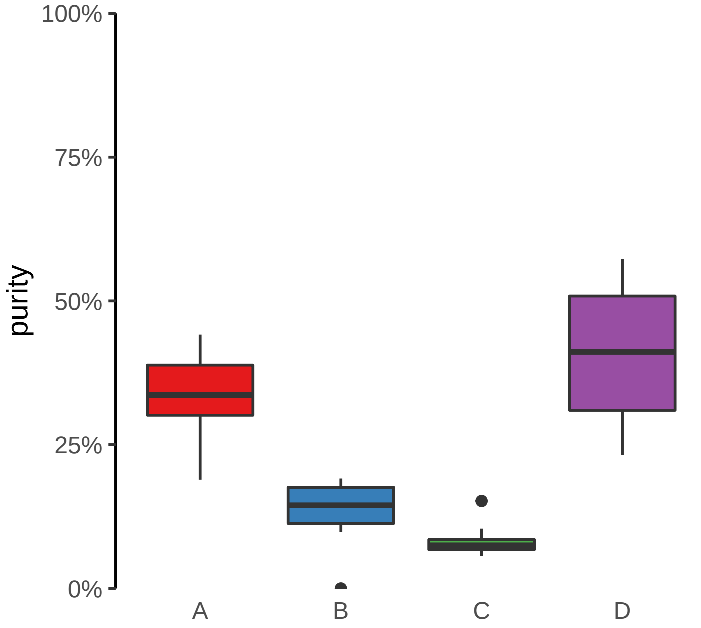

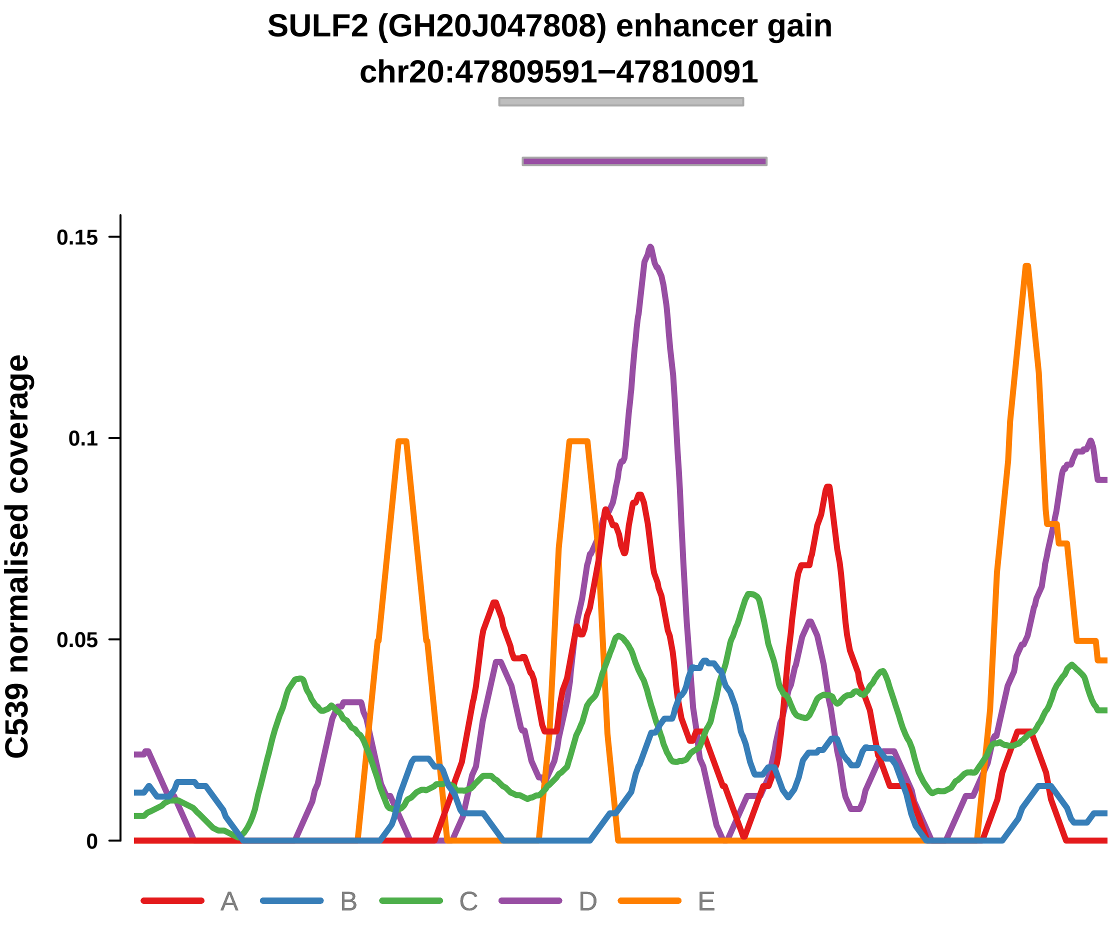


Putatively Subclonal (logFC = 2.24)

| | P. | Adjusted P. |
|---|---|---|
| *~region vs. ~* | 2.29e−03 ** | 5.3e−03 ** |
| *~purity vs ~* | 4.28e−02 * | 5.62e−02 |
| *~purity+region vs ~purity* | 2.56e−03 ** | 1.89e−02 * |
| *Carcinoma vs. normal* | 4.2e−03 ** | 4.2e−03 ** |

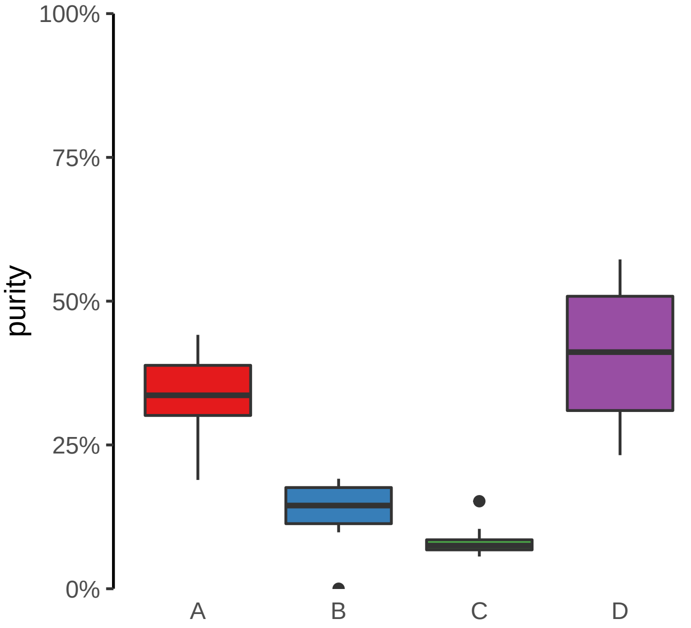

**CLASRP, MARK4 (GH19J045307) enhancer gain**

**chr19:45308035−45308535**

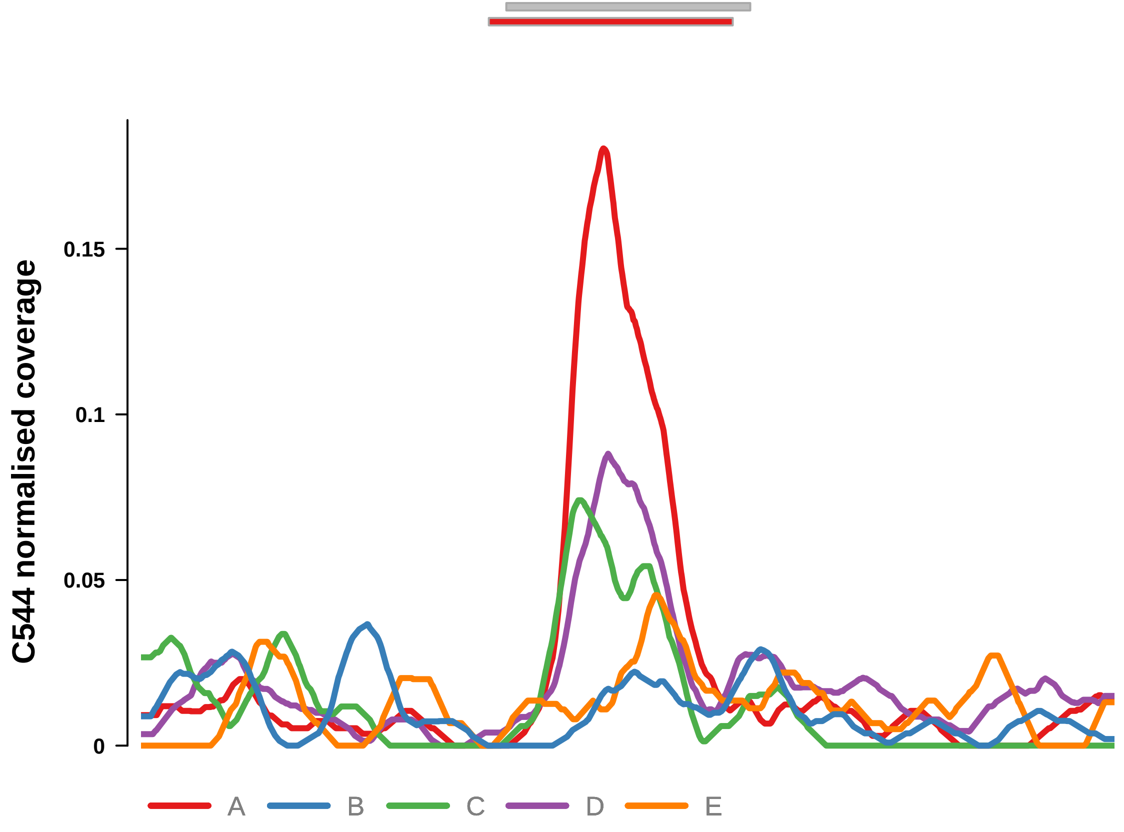


Putatively Subclonal (logFC = 2.05)

| | P. | Adjusted P. |
|---|---|---|
| *~region vs. ~* | 2.87e−06 **** | 1.07e−04 *** |
| *~purity vs ~* | 1.36e−03 ** | 1.57e−02 * |
| *~purity+region vs ~purity* | 1.98e−03 ** | 1.59e−02 * |
| *Carcinoma vs. normal* | 3.56e−03 ** | 3.56e−03 ** |

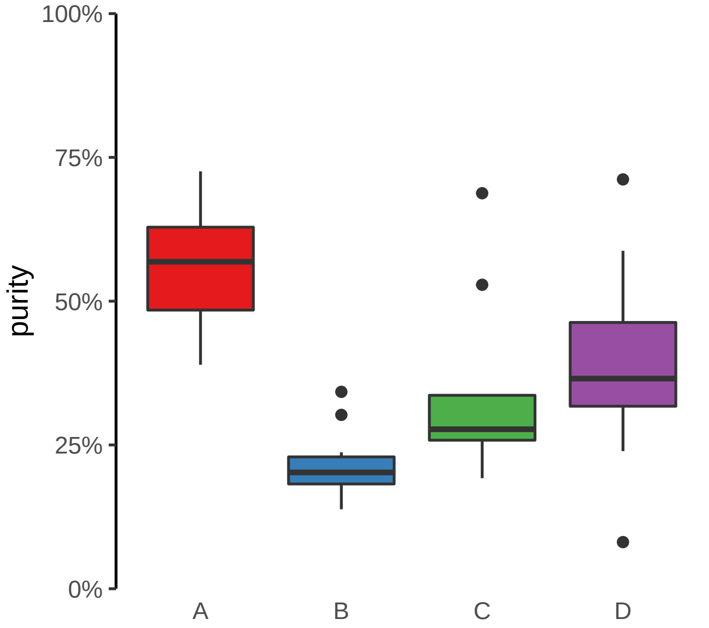

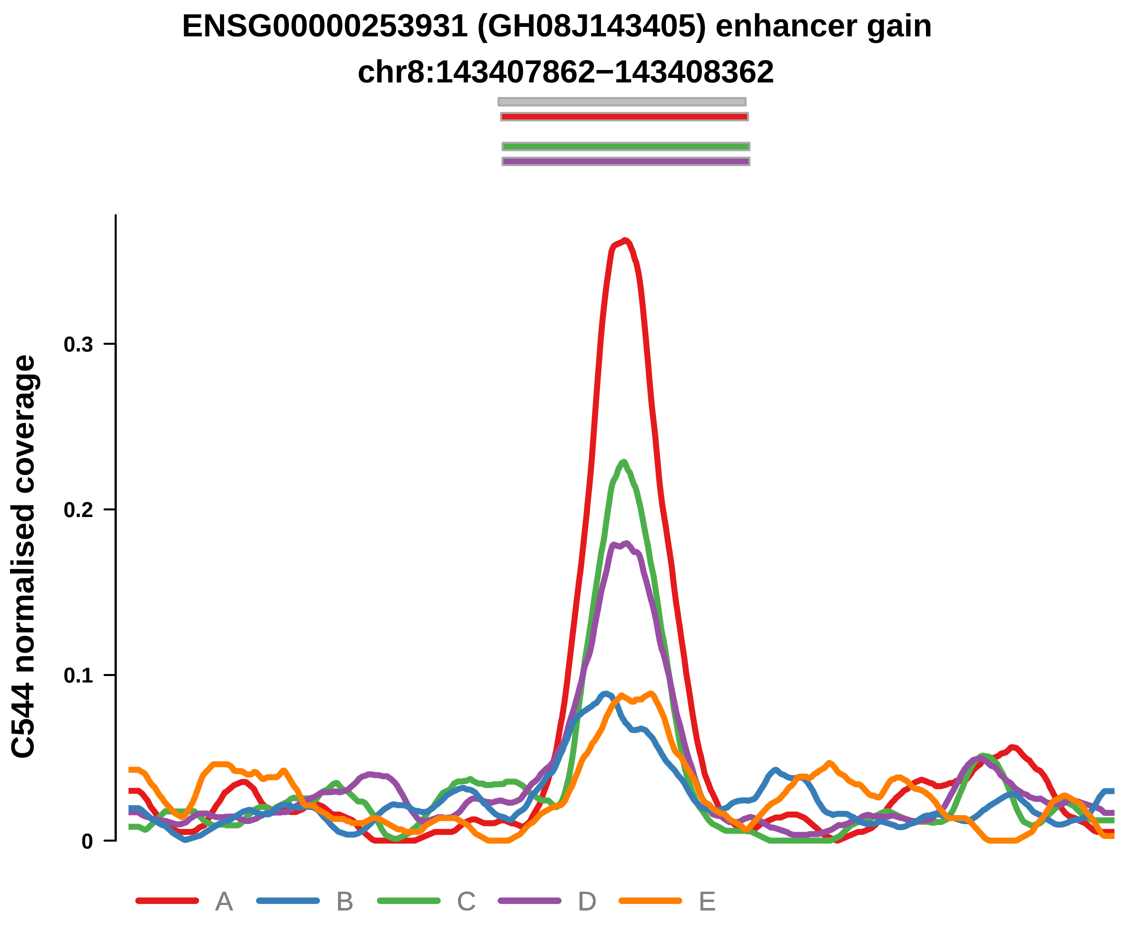


Putatively Subclonal (logFC = 2.02)

| | P. | Adjusted P. |
|---|---|---|
| *~region vs. ~* | 3.64e−06 **** | 1.07e−04 *** |
| *~purity vs ~* | 1.85e−04 *** | 3.74e−03 ** |
| *~purity+region vs ~purity* | 1.11e−02 * | 4.73e−02 * |
| *Carcinoma vs. normal* | 3.55e−03 ** | 3.55e−03 ** |

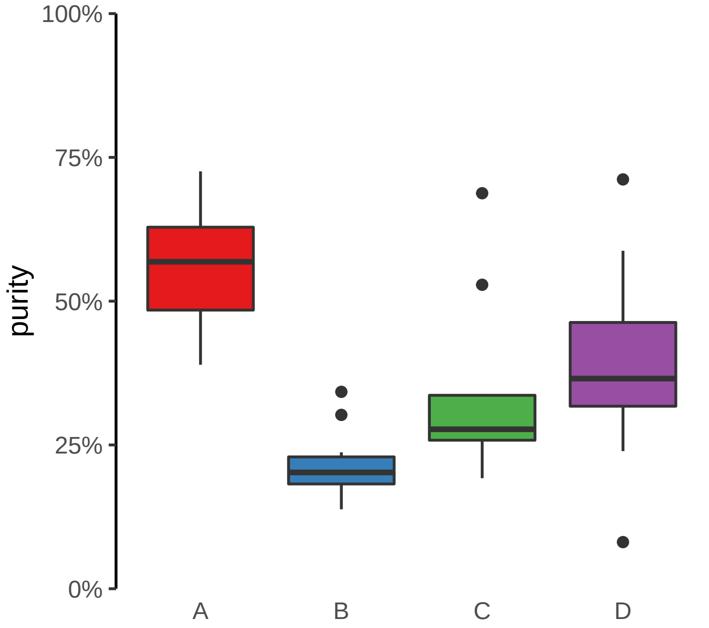

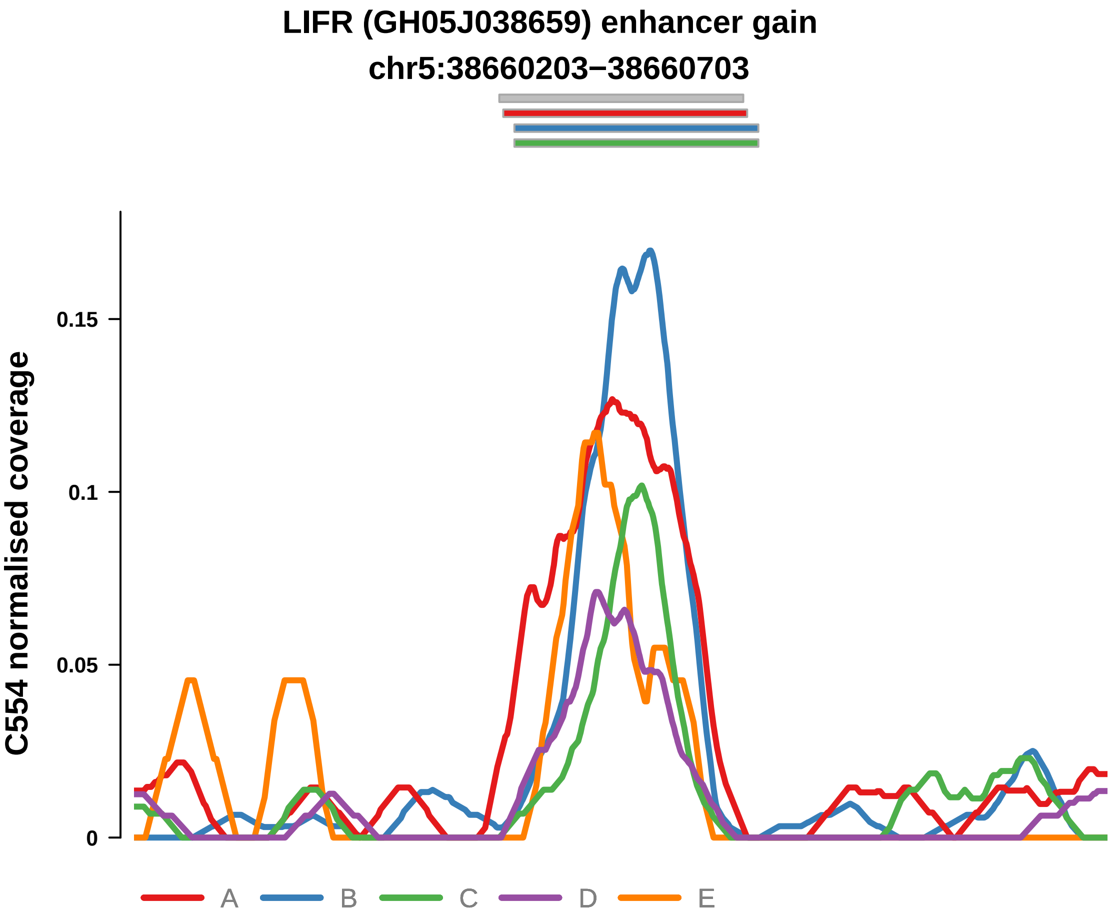


Putatively Subclonal (logFC = 2.07)

| | P. | Adjusted P. |
|---|---|---|
| *~region vs. ~* | 2.31e−03 ** | 8.62e−03 ** |
| *~purity vs ~* | 1.02e−01 | 2.57e−01 |
| *~purity+region vs ~purity* | 5.8e−03 ** | 3.52e−02 * |
| *Carcinoma vs. normal* | 3.23e−03 ** | 3.23e−03 ** |

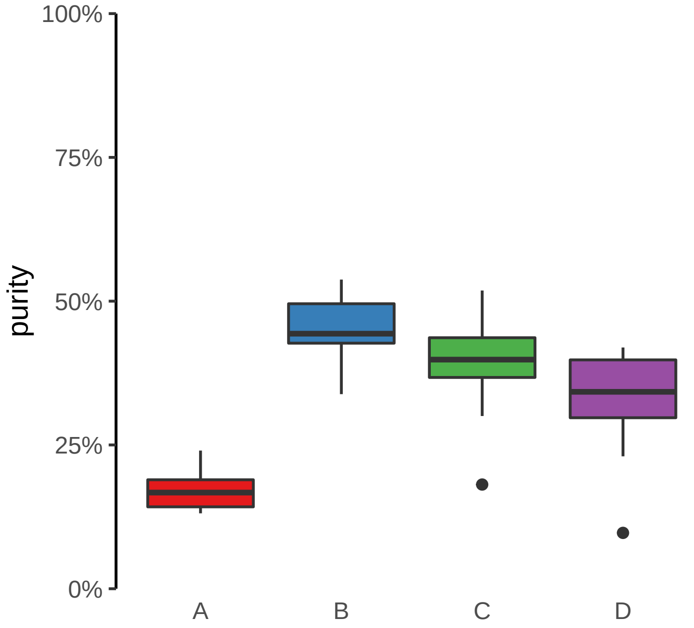

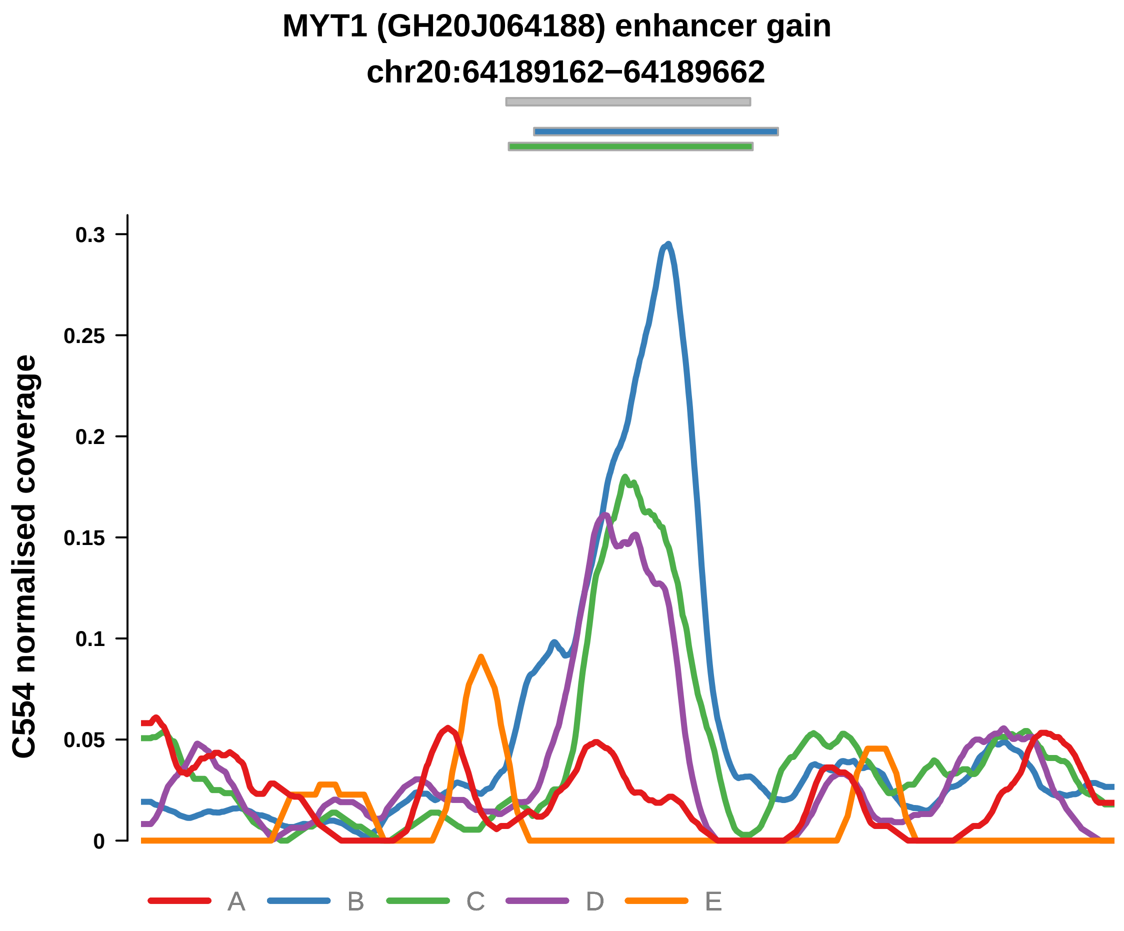


Putatively Subclonal (logFC = 3.84)

| | P. | Adjusted P. |
|---|---|---|
| *~region vs. ~* | 8.46e−16 **** | 6.94e−14 **** |
| *~purity vs ~* | 1.29e−06 **** | 3.46e−05 **** |
| *~purity+region vs ~purity* | 1.36e−06 **** | 5.03e−05 **** |
| *Carcinoma vs. normal* | 2.99e−07 **** | 2.99e−07 **** |

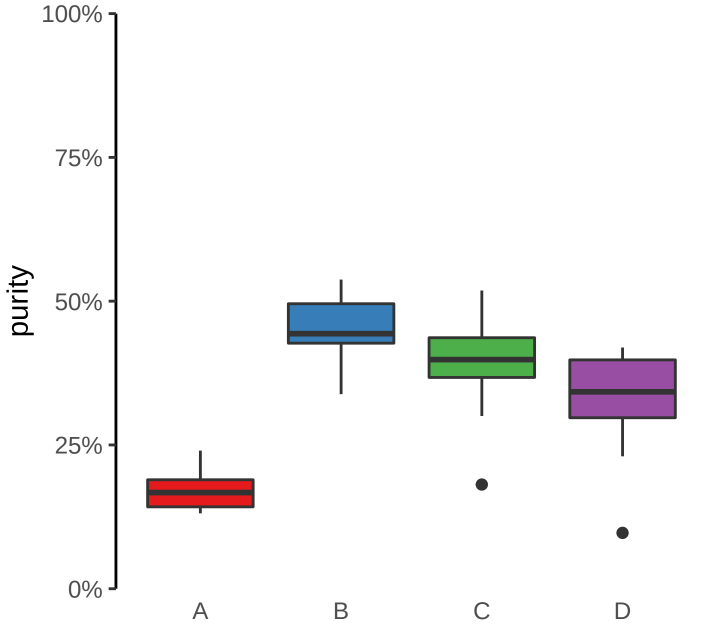

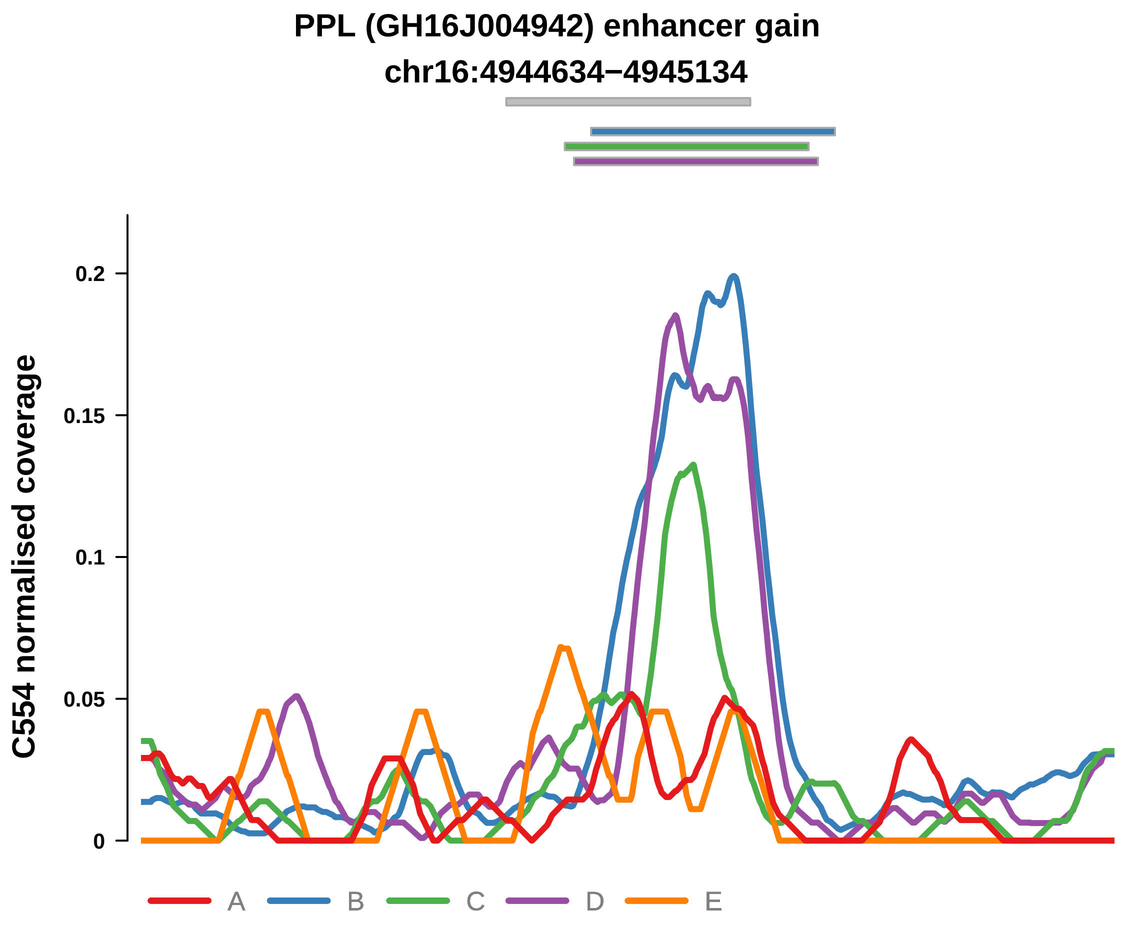


Putatively Subclonal (logFC = 2.04)

| | P. | Adjusted P. |
|---|---|---|
| *~region vs. ~* | 2.09e−06 **** | 2.45e−05 **** |
| *~purity vs ~* | 4.32e−04 *** | 5.06e−03 ** |
| *~purity+region vs ~purity* | 7.34e−03 ** | 3.62e−02 * |
| *Carcinoma vs. normal* | 3.15e−03 ** | 3.15e−03 ** |

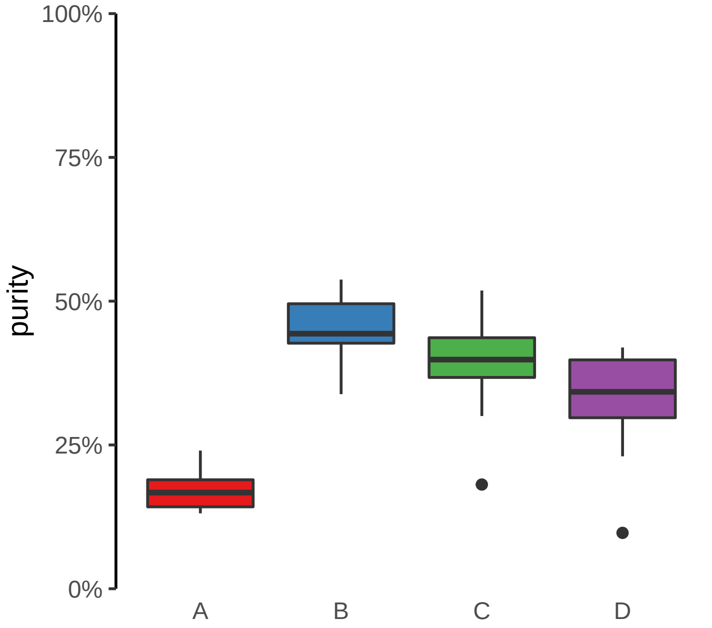

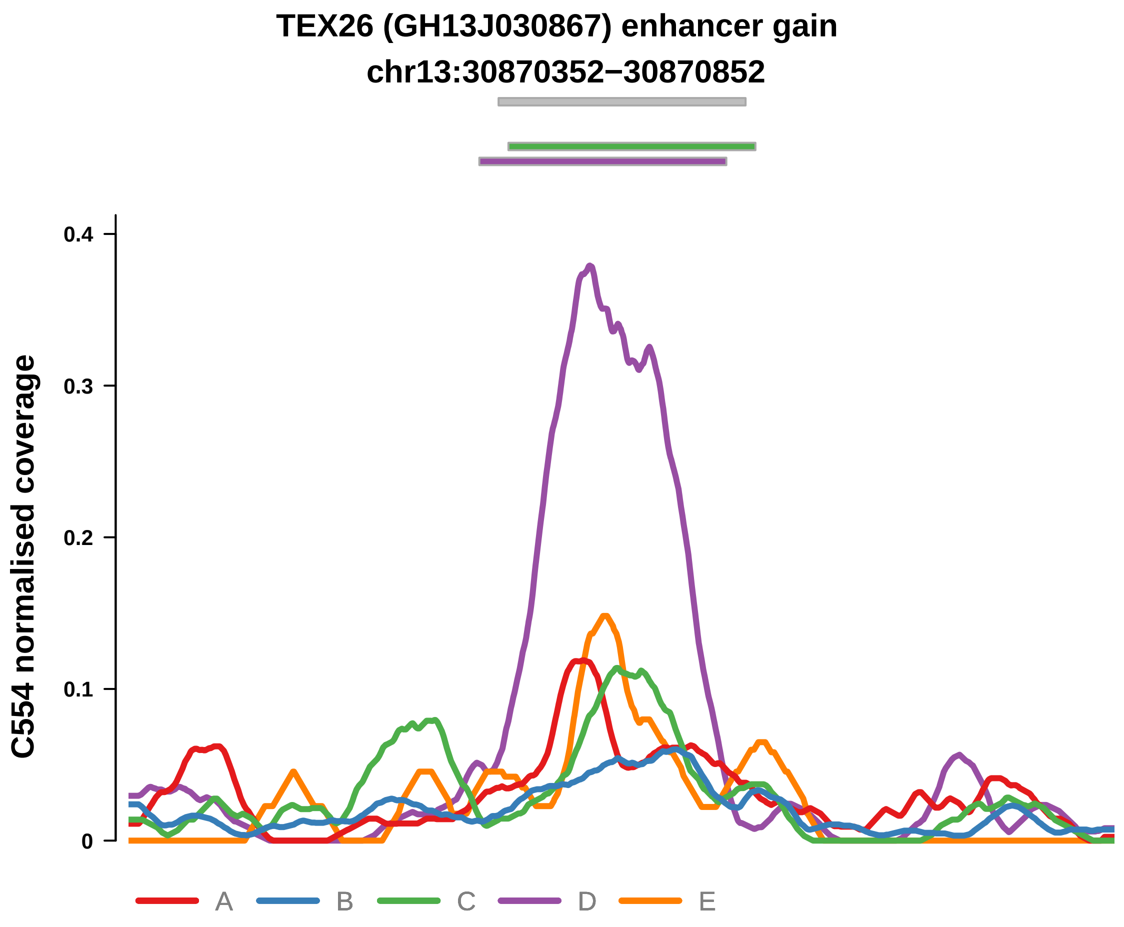


Putatively Subclonal (logFC = 0.33)

| | P. | Adjusted P. |
|---|---|---|
| *~region vs. ~* | 5.58e−14 **** | 2.29e−12 **** |
| *~purity vs ~* | 5.76e−01 | 7.62e−01 |
| *~purity+region vs ~purity* | 5.94e−14 **** | 4.4e−12 **** |
| *Carcinoma vs. normal* | NA | NA |

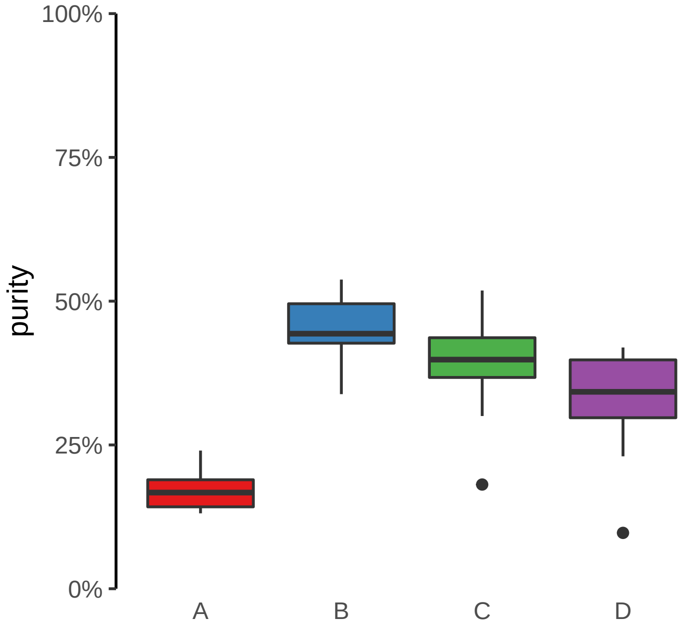

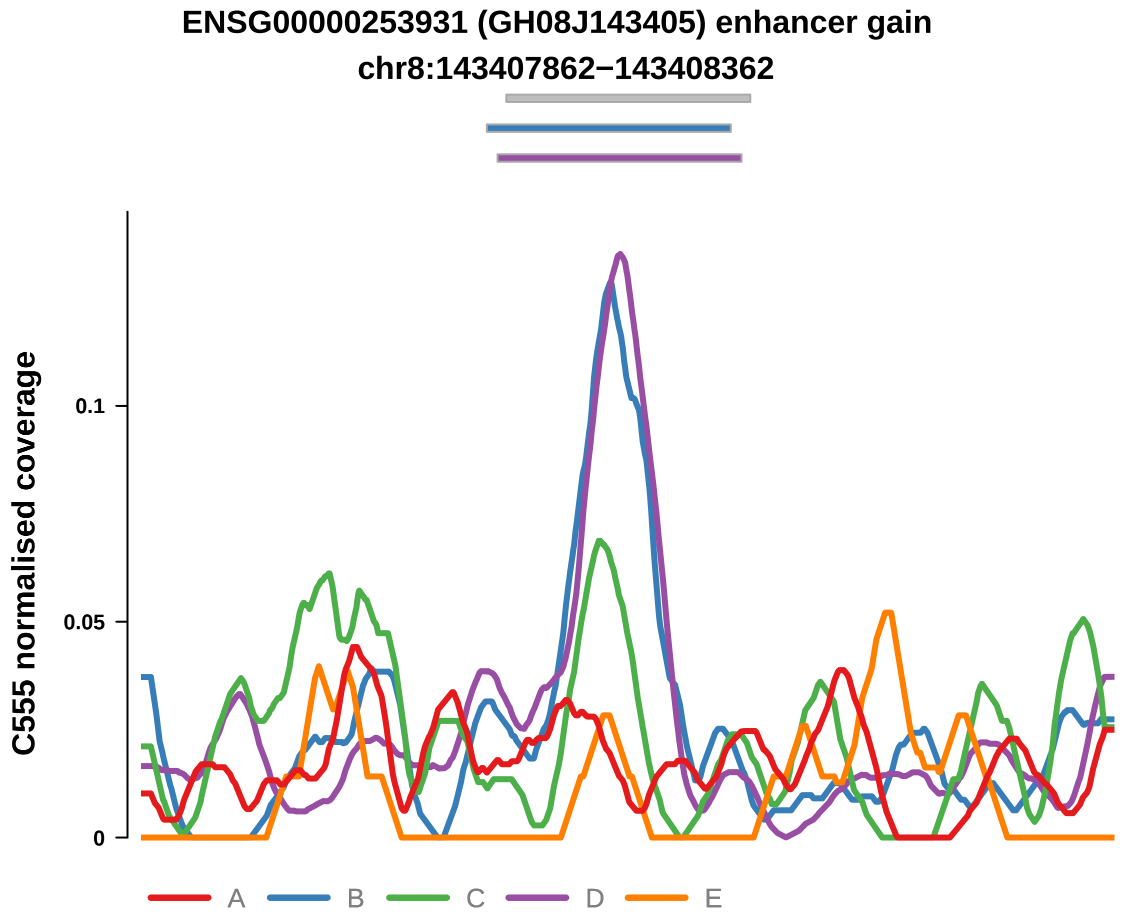


Putatively Subclonal (logFC = 1.85)

| | P. | Adjusted P. |
|---|---|---|
| *~region vs. ~* | 1.85e−05 **** | 8.37e−04 *** |
| *~purity vs ~* | 1.61e−01 | 4.14e−01 |
| *~purity+region vs ~purity* | 1.41e−04 *** | 9.48e−03 ** |
| *Carcinoma vs. normal* | 8.17e−03 ** | 8.17e−03 ** |

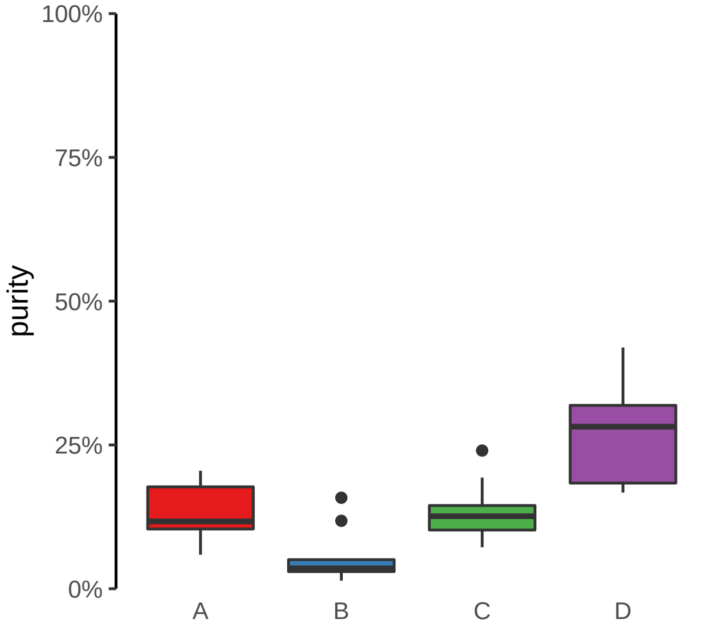
