## Supplementary Figures for "The co-evolution of the genome and epigenome in colorectal cancer": figS18_subclonal_enhancer_loss.pdf

**FAM83G (GH17J019010) enhancer loss**
**chr17:19011073−19011573**

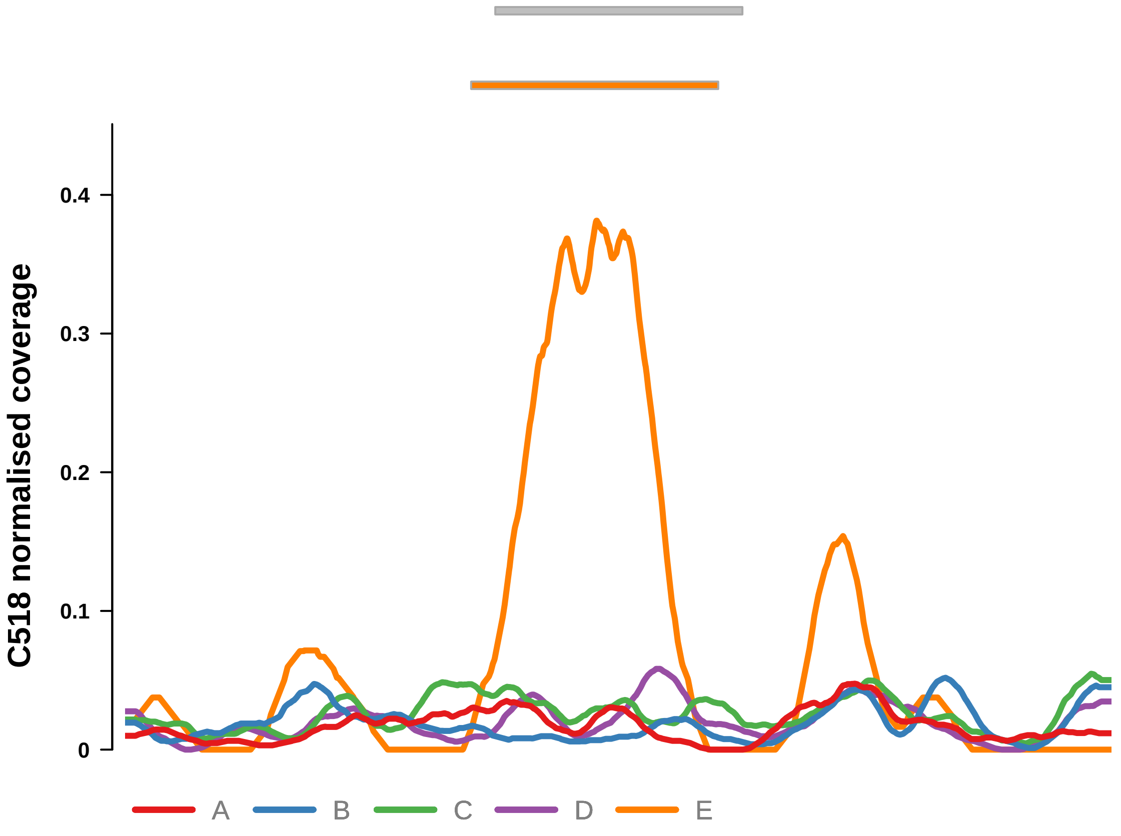


Putatively Subclonal (logFC = −1.06)

| | P. | Adjusted P. |
|---|---|---|
| *~region vs. ~* | 1.33e−03 ** | 3.91e−02 * |
| *~purity vs ~* | 4.7e−01 | 8.48e−01 |
| *~purity+region vs ~purity* | 1.06e−03 ** | 3.34e−02 * |
| *Carcinoma vs. normal* | 1.25e−01 | 1.25e−01 |

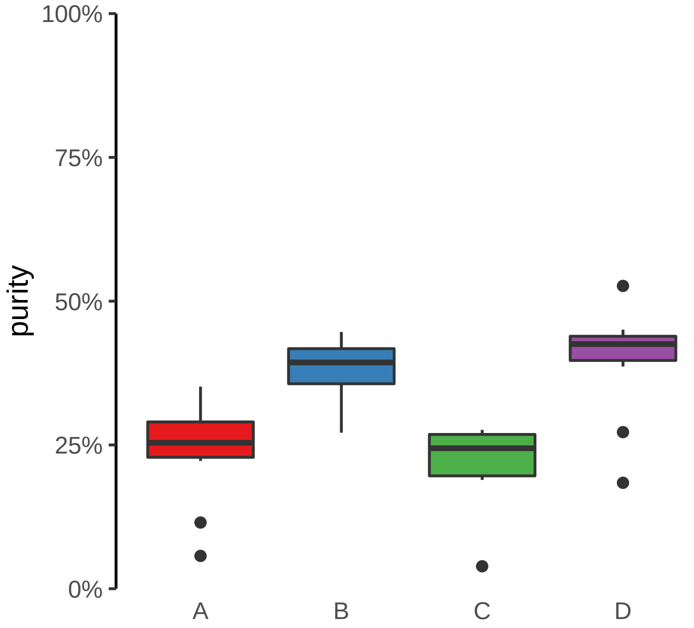

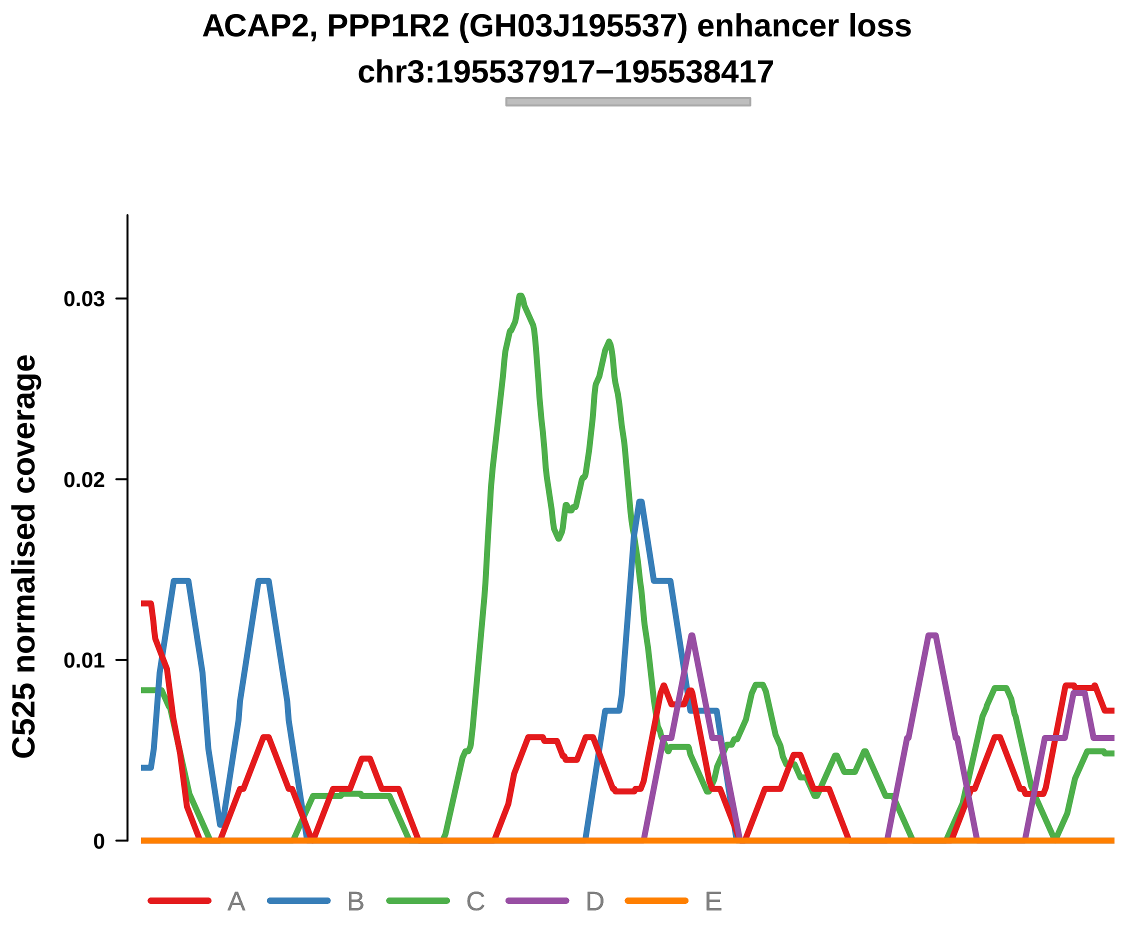


Putatively Subclonal (logFC = −1.47)

| | P. | Adjusted P. |
|---|---|---|
| *~region vs. ~* | 8.63e−04 *** | 4.64e−03 ** |
| *~purity vs ~* | 8.21e−02 | 7.18e−01 |
| *~purity+region vs ~purity* | 4.04e−03 ** | 1.98e−02 * |
| *Carcinoma vs. normal* | 1.16e−01 | 1.16e−01 |

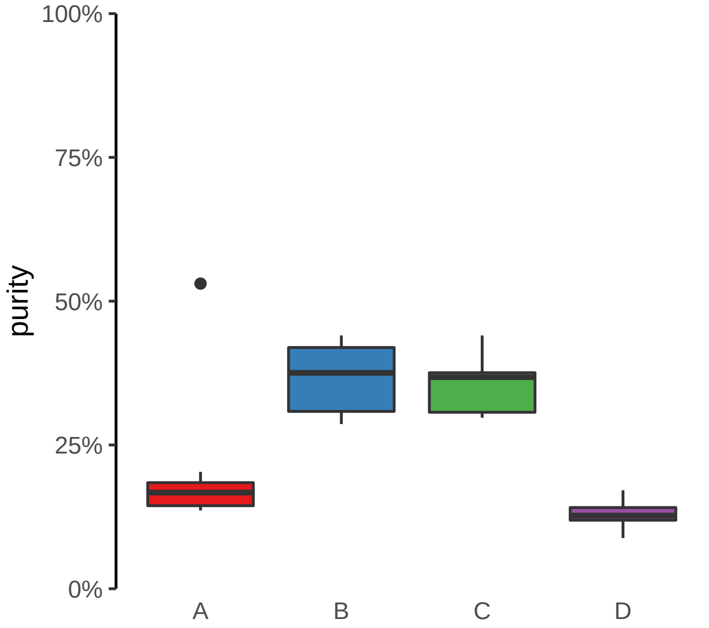

**CNST (GH01J246591) enhancer loss**
**chr1:246591917−246592417**

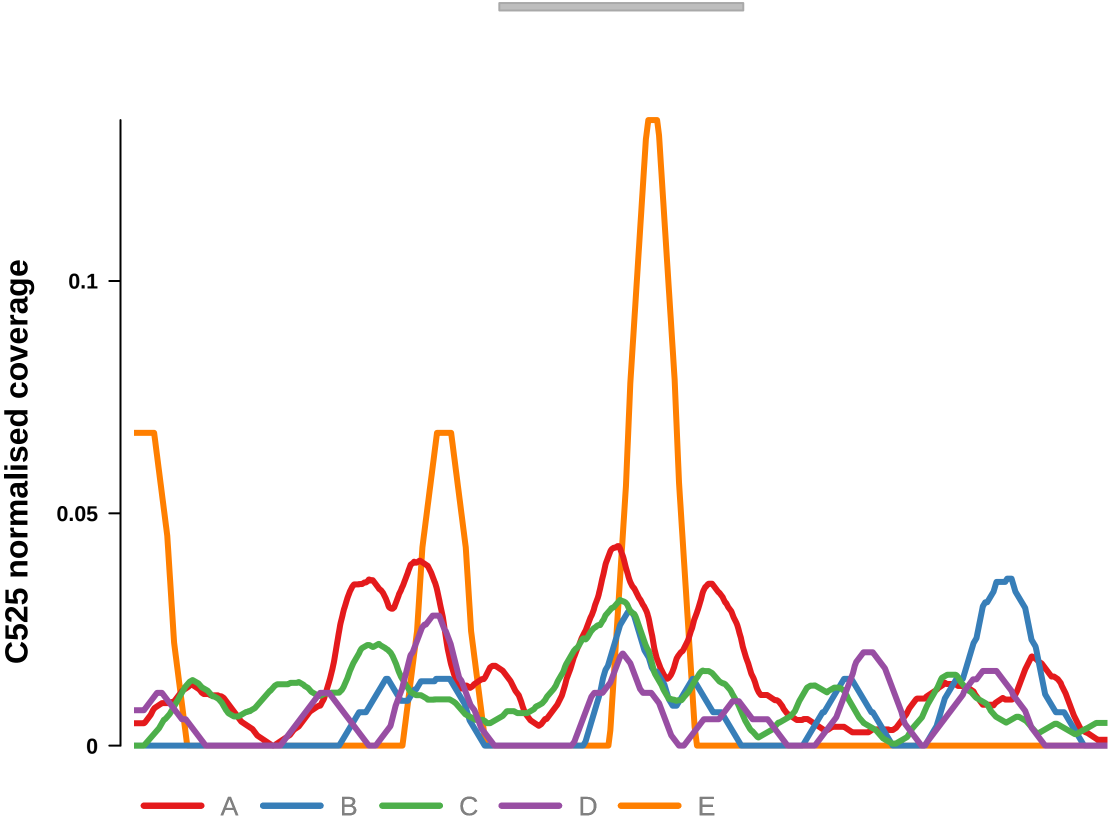


Putatively Subclonal (logFC = −0.73)

| | P. | Adjusted P. |
|---|---|---|
| *~region vs. ~* | 1.68e−02 * | 4.62e−02 * |
| *~purity vs ~* | 5.16e−01 | 8.63e−01 |
| *~purity+region vs ~purity* | 7.48e−03 ** | 2.53e−02 * |
| *Carcinoma vs. normal* | NA | NA |

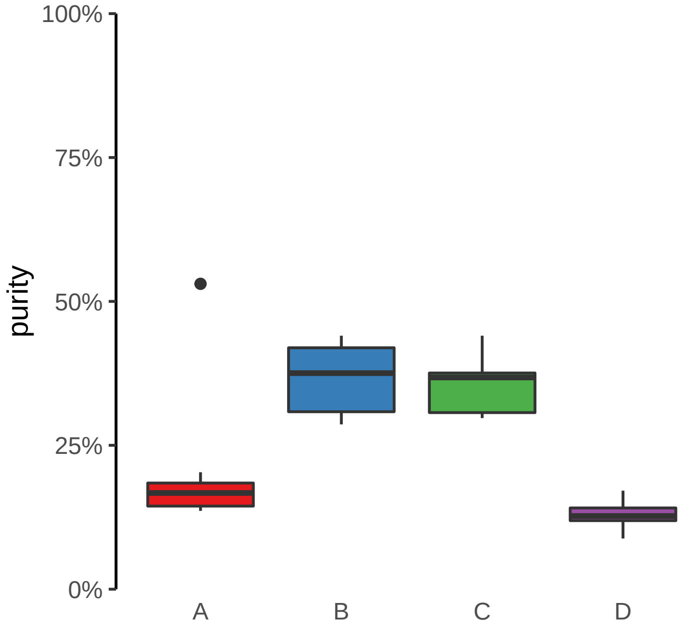

**DNAH7, ENSG00000272211 (GH02J196100) enhancer loss**
**chr2:196101142−196101642**

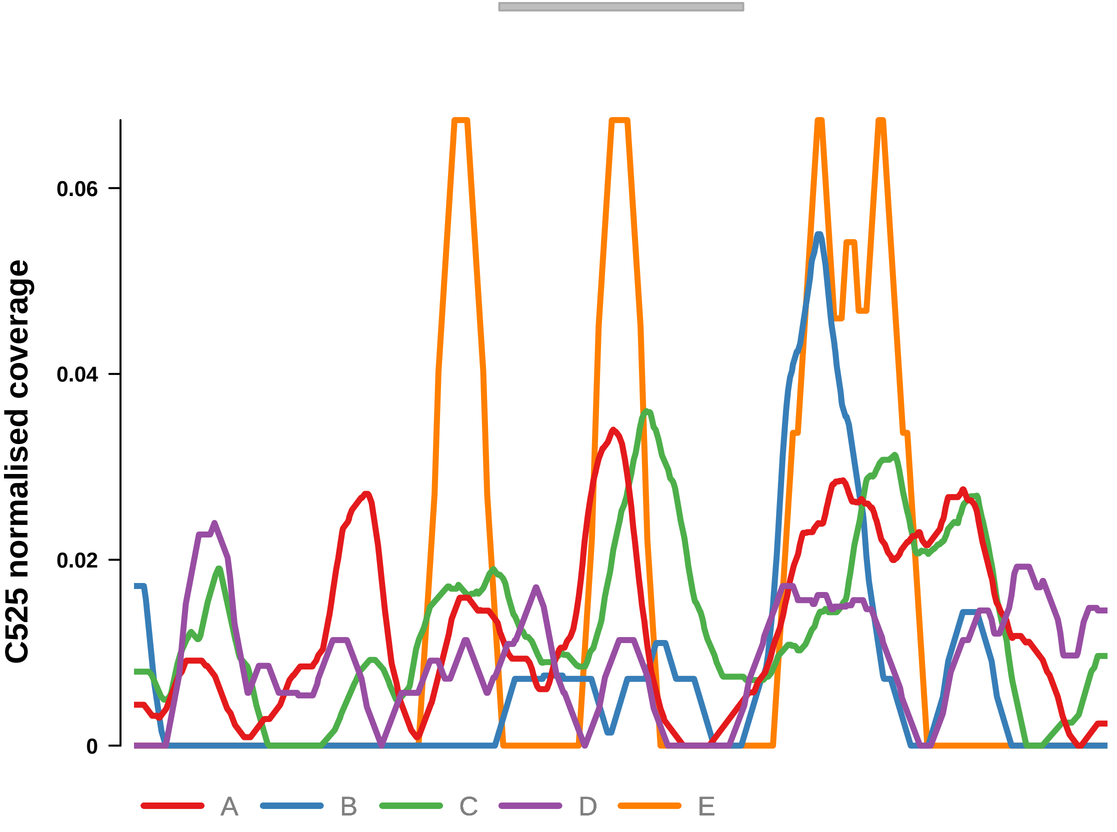


Putatively Subclonal (logFC = −0.99)

| | P. | Adjusted P. |
|---|---|---|
| *~region vs. ~* | 2.27e−03 ** | 9.28e−03 ** |
| *~purity vs ~* | 7.81e−01 | 8.93e−01 |
| *~purity+region vs ~purity* | 6.57e−04 *** | 4.82e−03 ** |
| *Carcinoma vs. normal* | NA | NA |

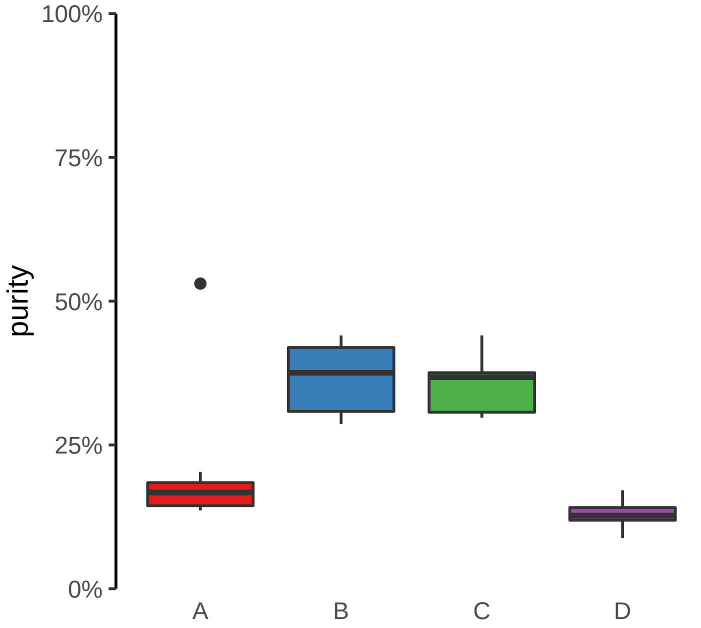

**FABP1, THNSL2 (GH02J088144) enhancer loss**
**chr2:88144817−88145317**

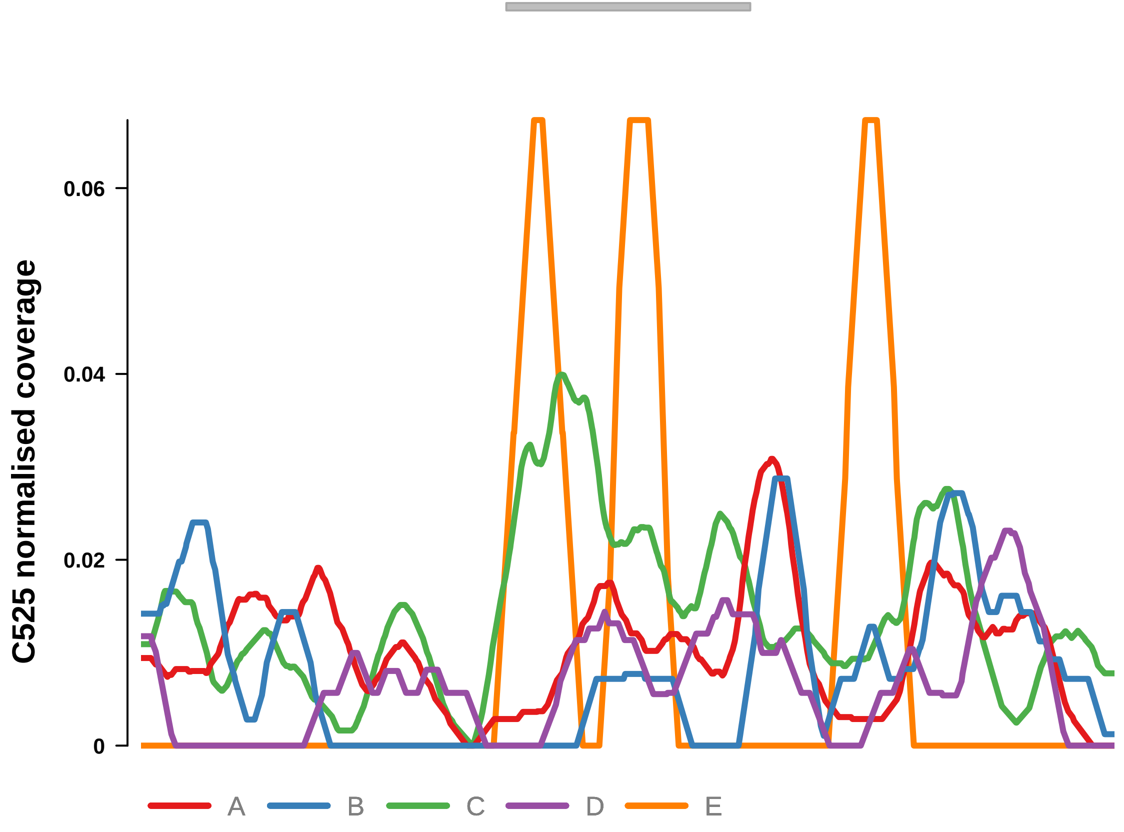


Putatively Subclonal (logFC = −1.92)

| | P. | Adjusted P. |
|---|---|---|
| *~region vs. ~* | 7.41e−05 **** | 7.83e−04 *** |
| *~purity vs ~* | 2.43e−01 | 7.37e−01 |
| *~purity+region vs ~purity* | 3.02e−04 *** | 2.95e−03 ** |
| *Carcinoma vs. normal* | 1.54e−02 * | 1.54e−02 * |

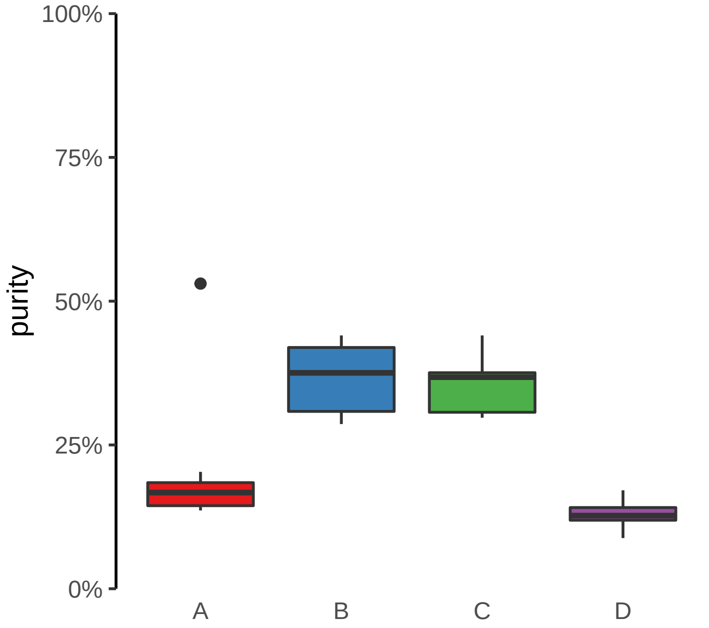

**MAP3K1, LINC01948 (GH05J056411) enhancer loss**
**chr5:56412924−56413424**

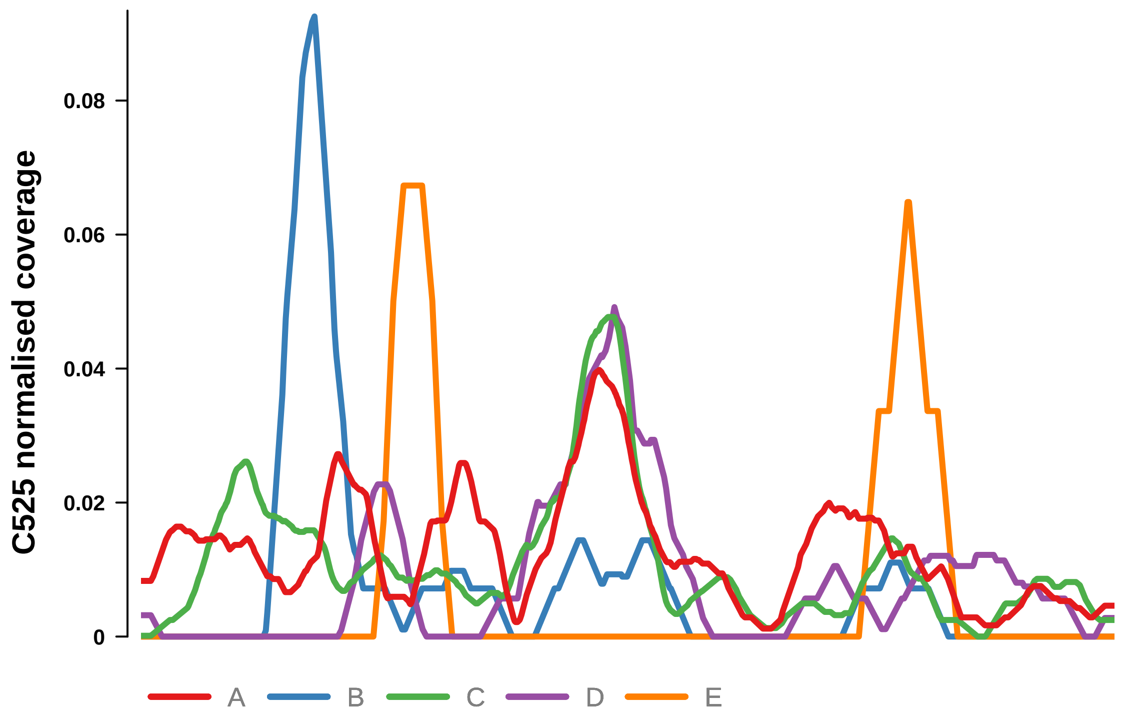


Putatively Subclonal (logFC = −1.04)

| | P. | Adjusted P. |
|---|---|---|
| *~region vs. ~* | 2.64e−03 ** | 1.03e−02 * |
| *~purity vs ~* | 4.77e−01 | 8.63e−01 |
| *~purity+region vs ~purity* | 4.02e−03 ** | 1.98e−02 * |
| *Carcinoma vs. normal* | 2.11e−01 | 2.11e−01 |

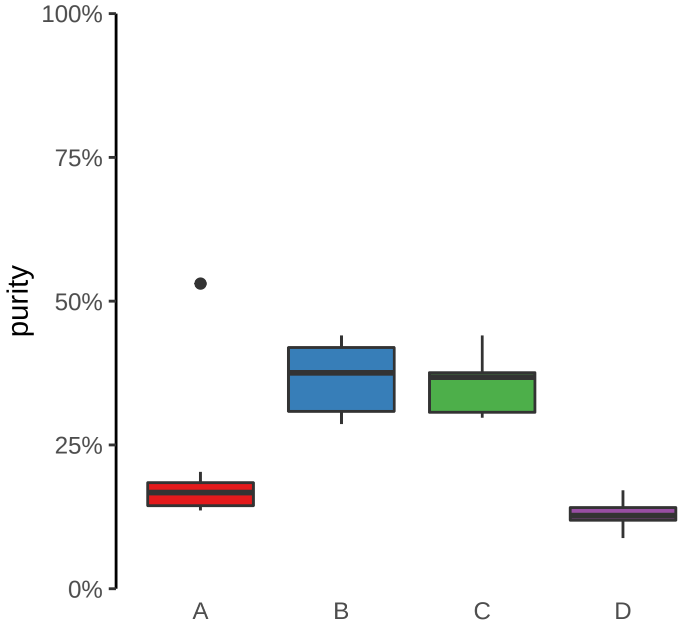

**PPP2CB (GH08J030800) enhancer loss**
**chr8:30804244−30804744**

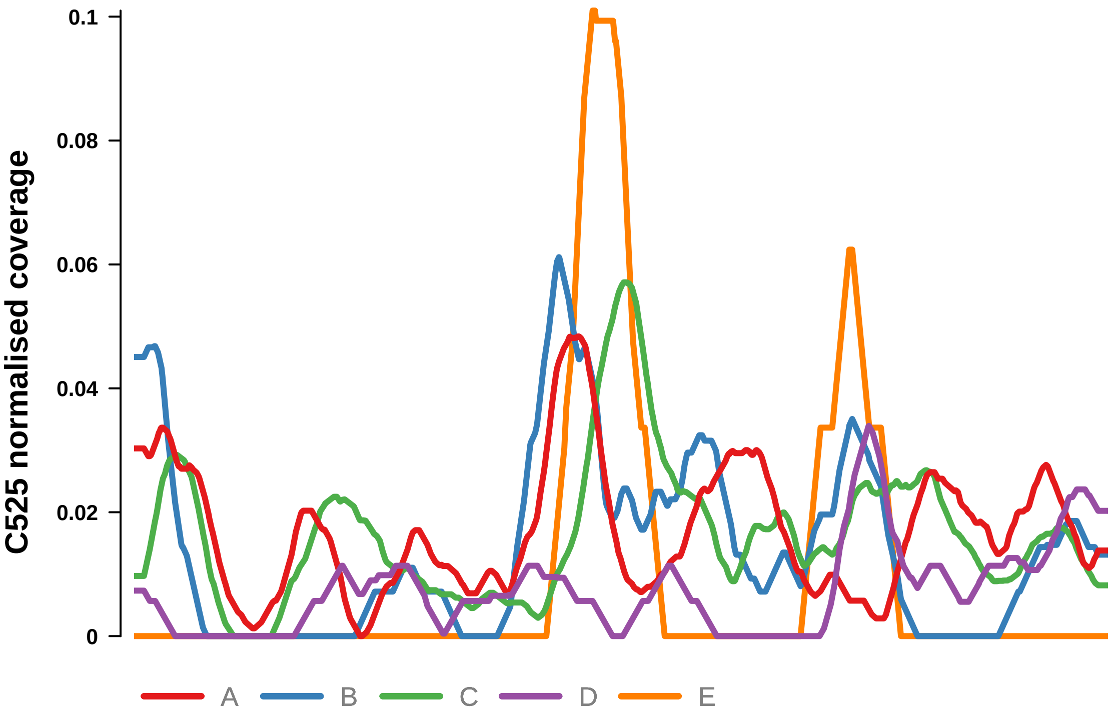


Putatively Subclonal (logFC = −0.8)

| | **P.** | **Adjusted P.** |
|---|---|---|
| *~region vs. ~* | 3.81e−03 ** | 1.37e−02 * |
| *~purity vs ~* | 3.05e−01 | 7.48e−01 |
| *~purity+region vs ~purity* | 6.71e−03 ** | 2.37e−02 * |
| *Carcinoma vs. normal* | NA | NA |

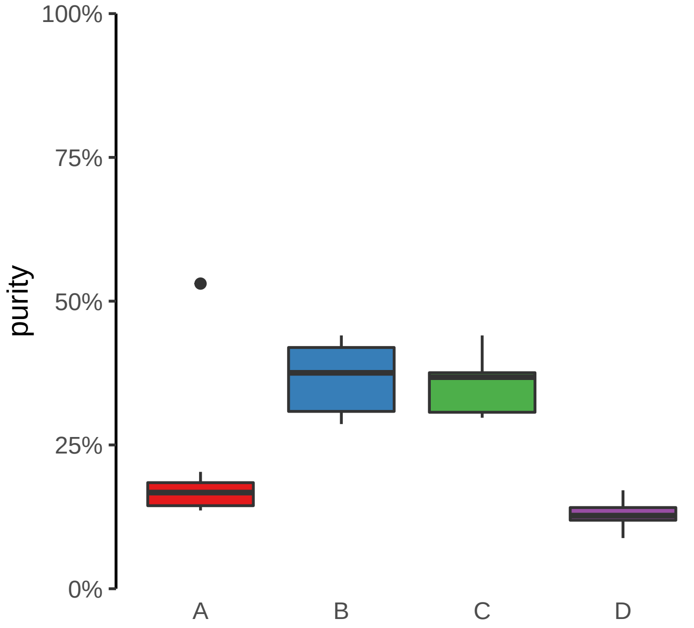

**PPP3CA (GH04J101183) enhancer loss**
**chr4:101183634−101184134**

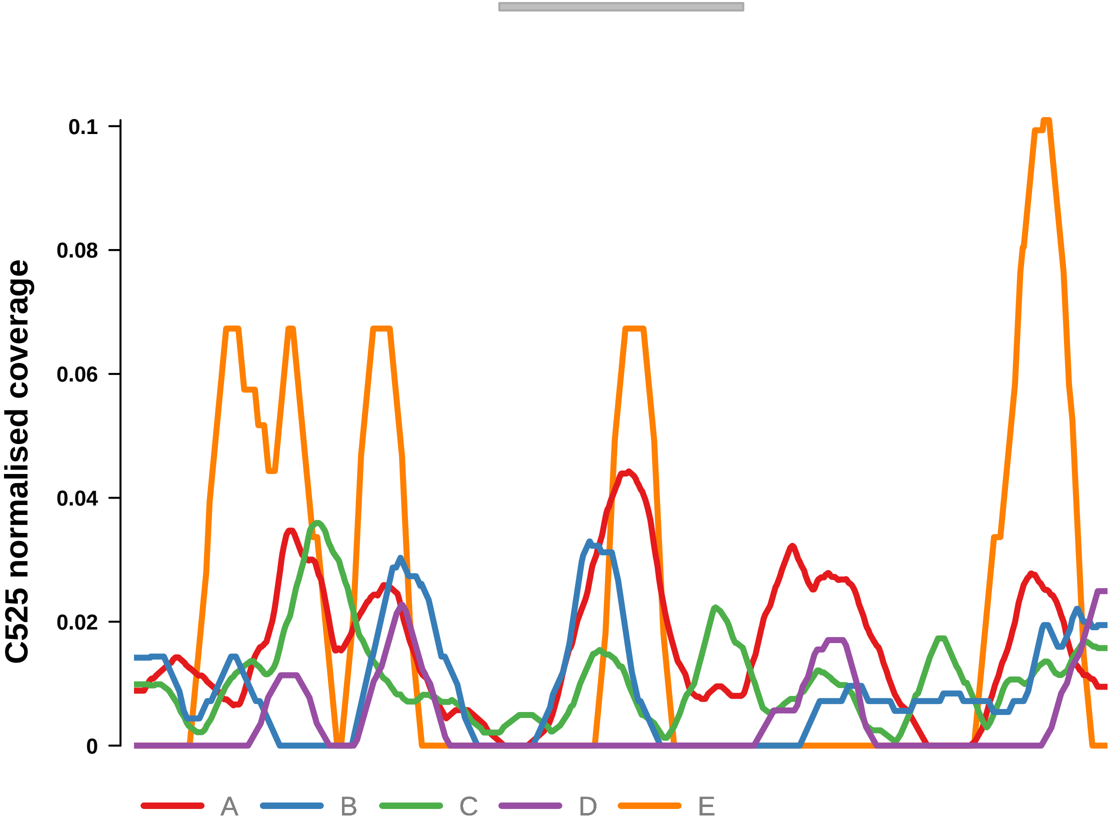


Putatively Subclonal (logFC = −0.85)

| | **P.** | **Adjusted P.** |
|---|---|---|
| *~region vs. ~* | 1.72e−03 ** | 7.63e−03 ** |
| *~purity vs ~* | 8.83e−01 | 9.48e−01 |
| *~purity+region vs ~purity* | 2.13e−03 ** | 1.25e−02 * |
| *Carcinoma vs. normal* | NA | NA |

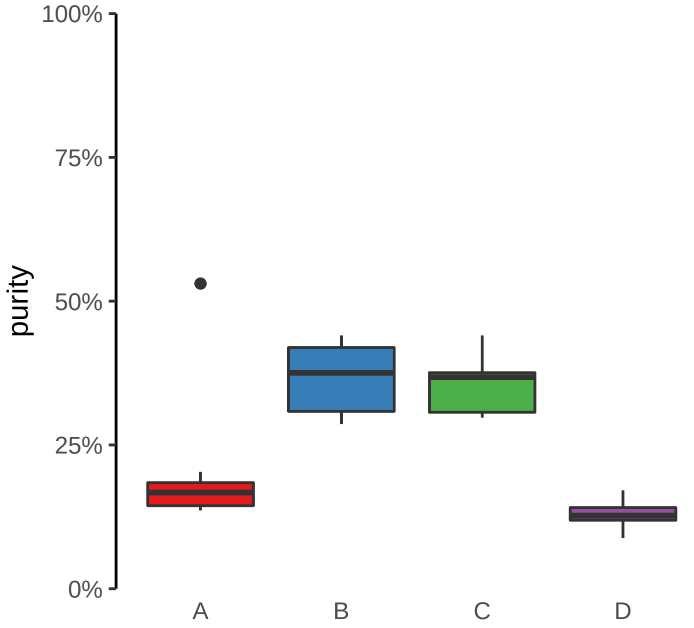

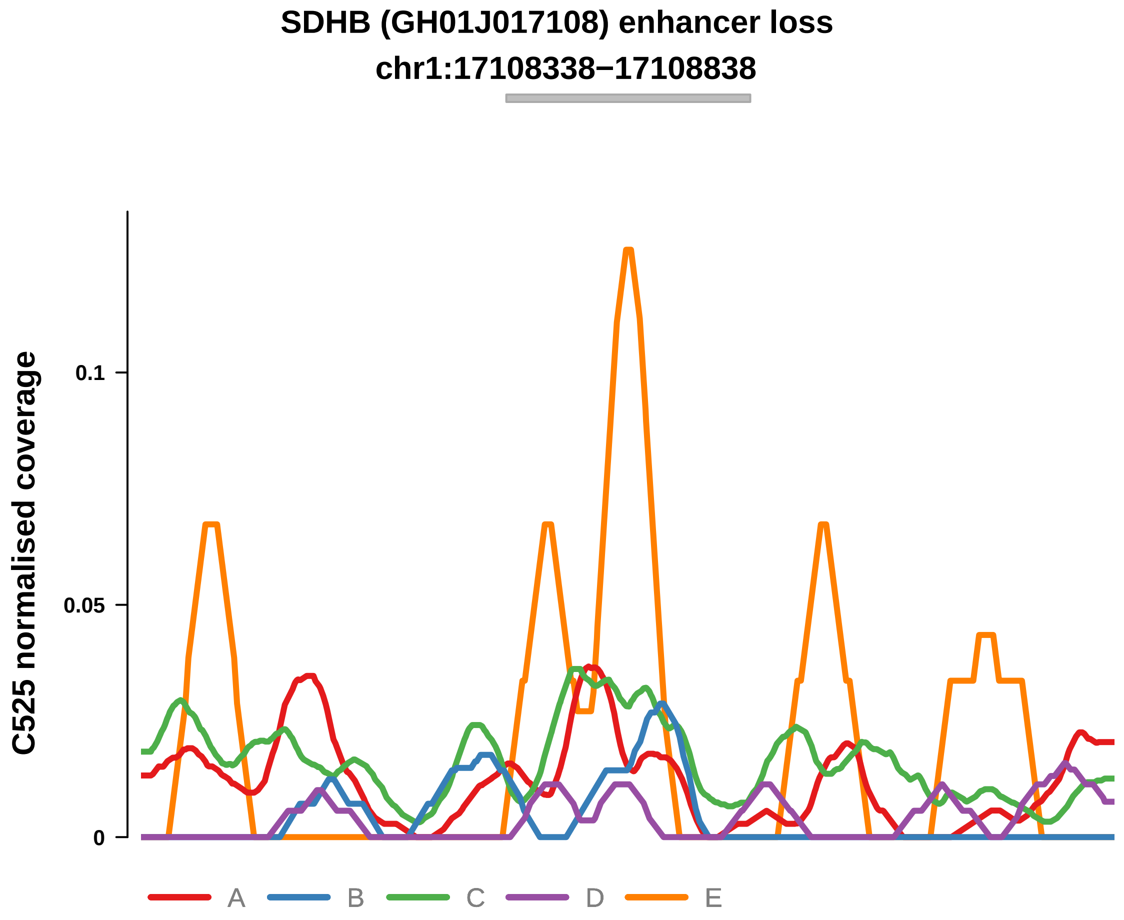


Putatively Subclonal (logFC = −1)

| | P. | Adjusted P. |
|---|---|---|
| *~region vs. ~* | 1.46e−03 ** | 6.96e−03 ** |
| *~purity vs ~* | 1.91e−01 | 7.37e−01 |
| *~purity+region vs ~purity* | 5.71e−03 ** | 2.21e−02 * |
| *Carcinoma vs. normal* | 1.9e−01 | 1.9e−01 |

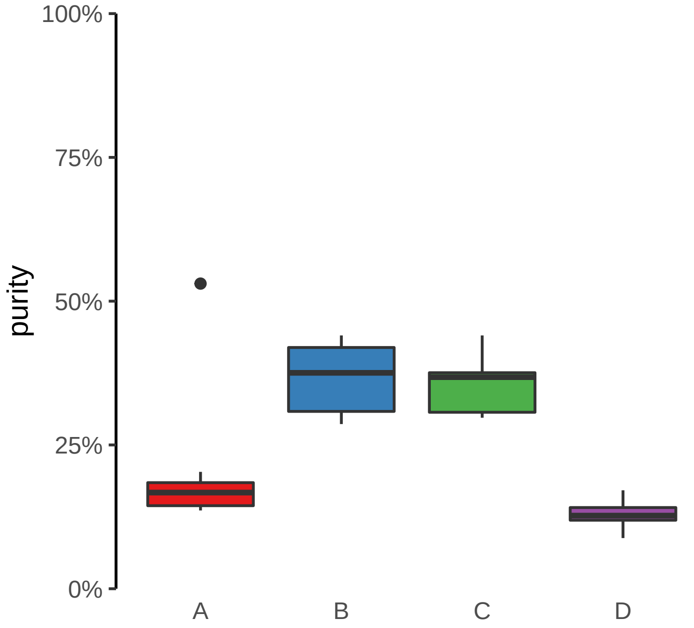

**ZNF431 (GH19J020999) enhancer loss**
**chr19:20999605−21000105**

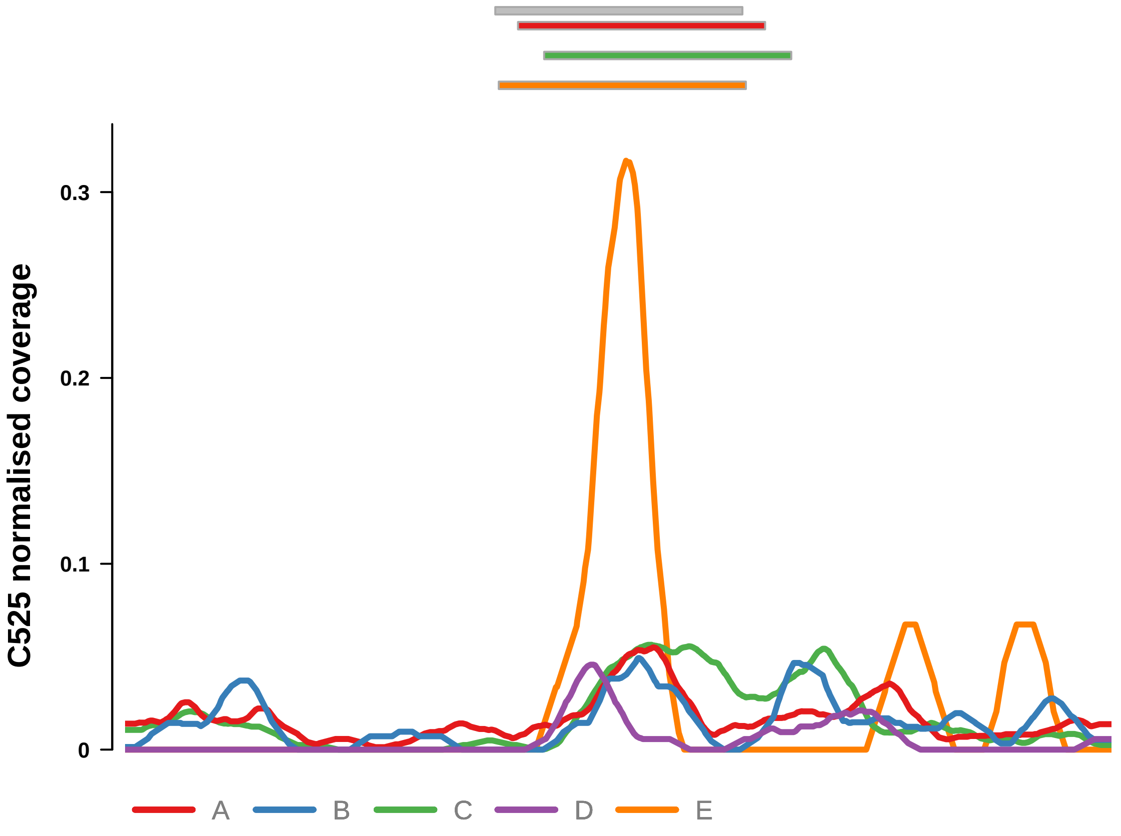


Putatively Subclonal (logFC = −0.82)

| | P. | Adjusted P. |
|---|---|---|
| *~region vs. ~* | 5.02e−05 **** | 7.46e−04 *** |
| *~purity vs ~* | 2.85e−01 | 7.48e−01 |
| *~purity+region vs ~purity* | 1.86e−04 *** | 2.95e−03 ** |
| *Carcinoma vs. normal* | NA | NA |

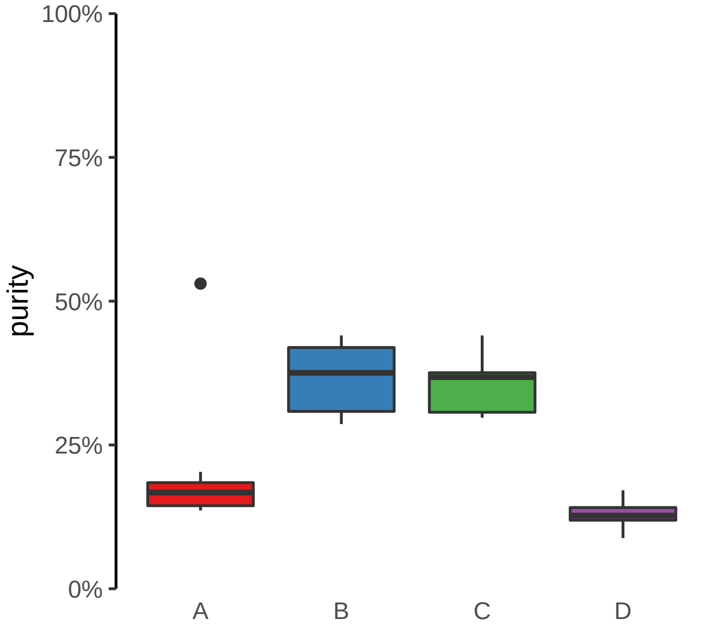

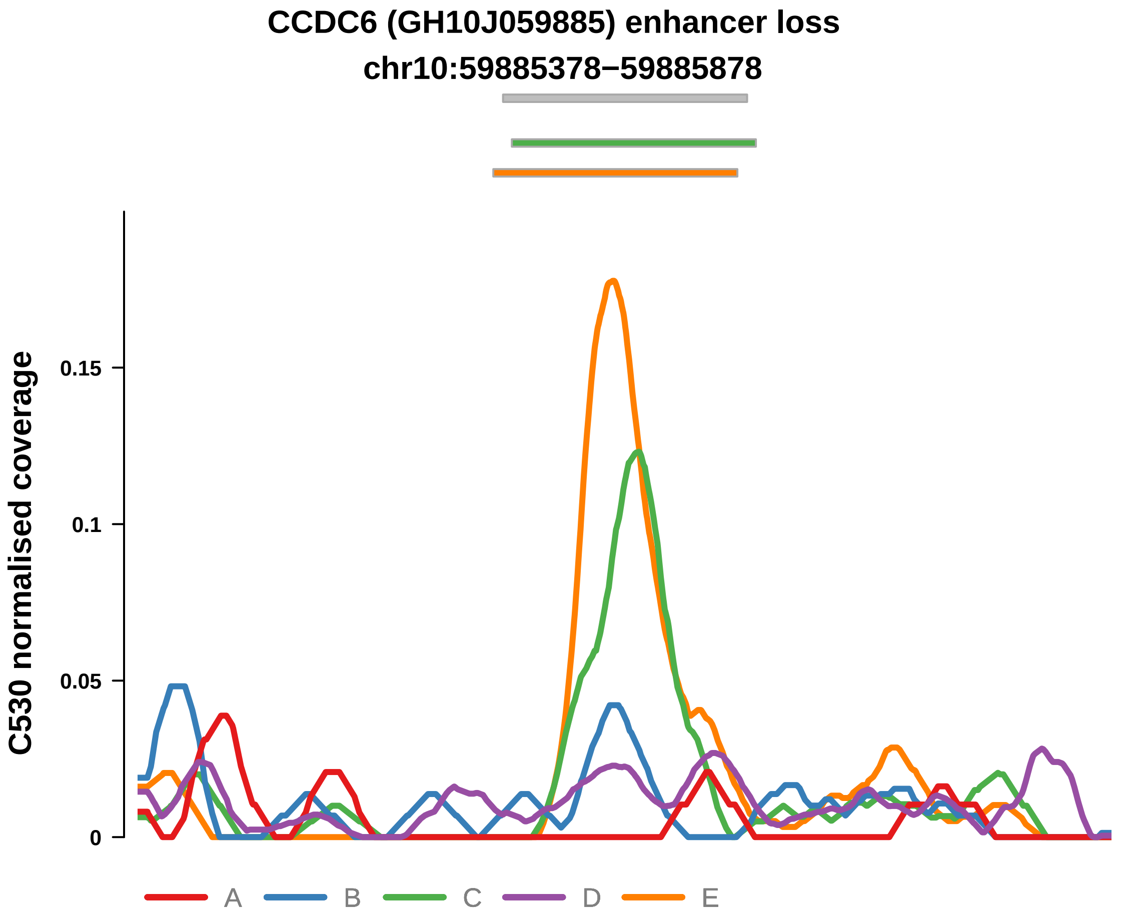


Putatively Subclonal (logFC = −0.51)

| | P. | Adjusted P. |
|---|---|---|
| *~region vs. ~* | 1.18e−03 ** | 5.12e−02 |
| *~purity vs ~* | 8.75e−01 | 9.62e−01 |
| *~purity+region vs ~purity* | 1.54e−03 ** | 4.52e−02 * |
| *Carcinoma vs. normal* | NA | NA |

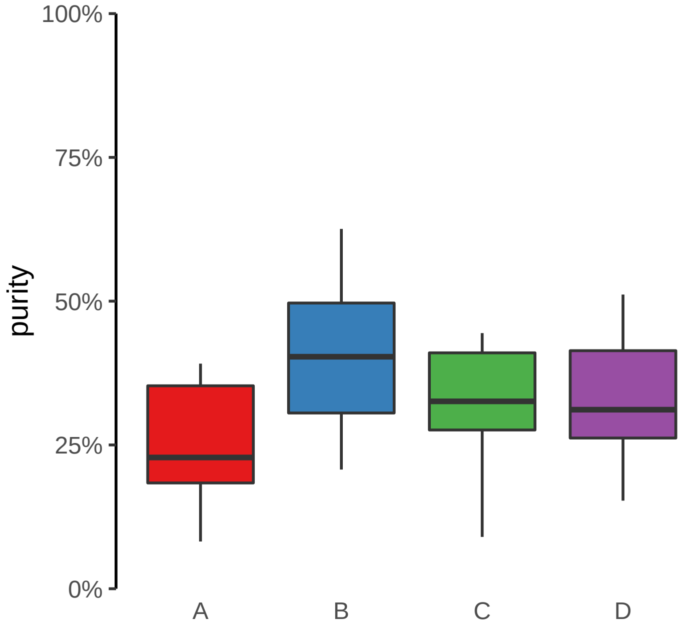

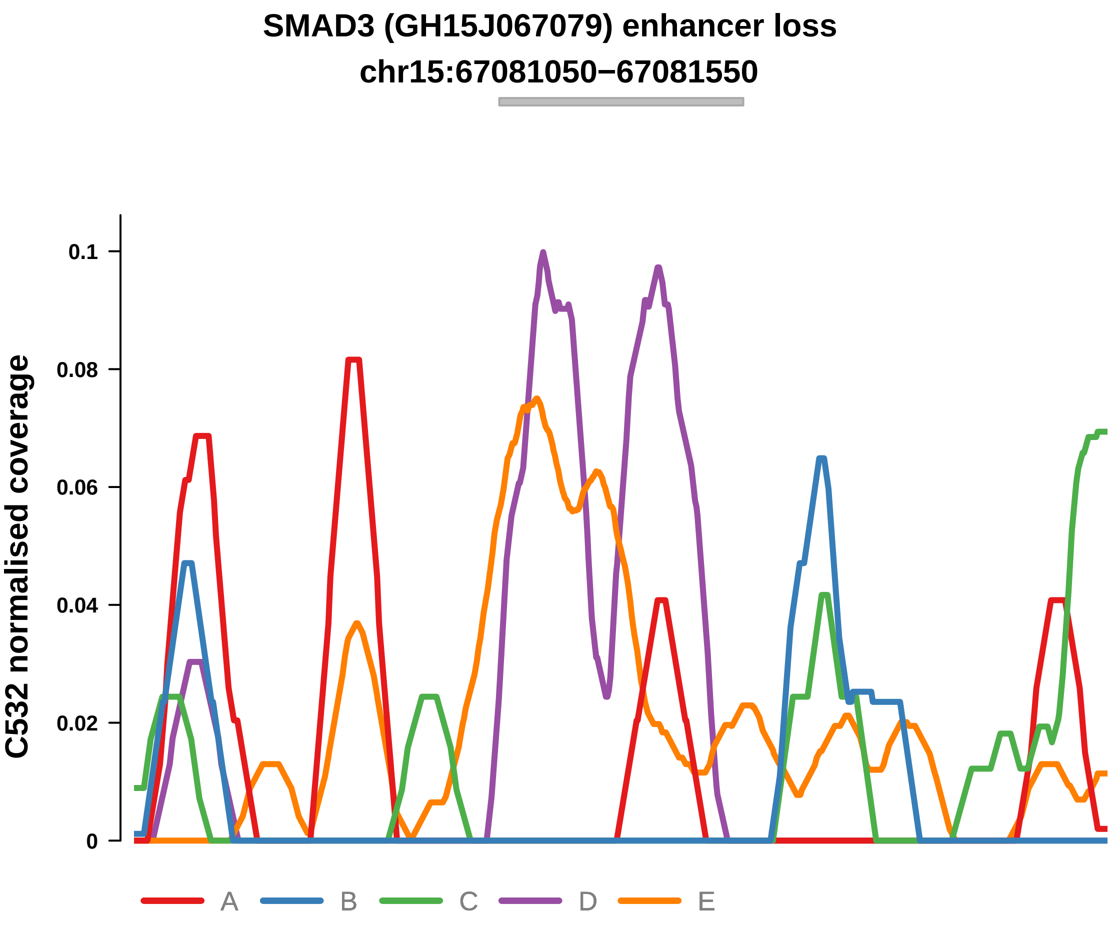


Putatively Subclonal (logFC = −0.57)

| | P. | Adjusted P. |
|---|---|---|
| *~region vs. ~* | 6.8e−05 **** | 2.99e−03 ** |
| *~purity vs ~* | 3.35e−01 | 8.63e−01 |
| *~purity+region vs ~purity* | 3.83e−04 *** | 1.69e−02 * |
| *Carcinoma vs. normal* | NA | NA |

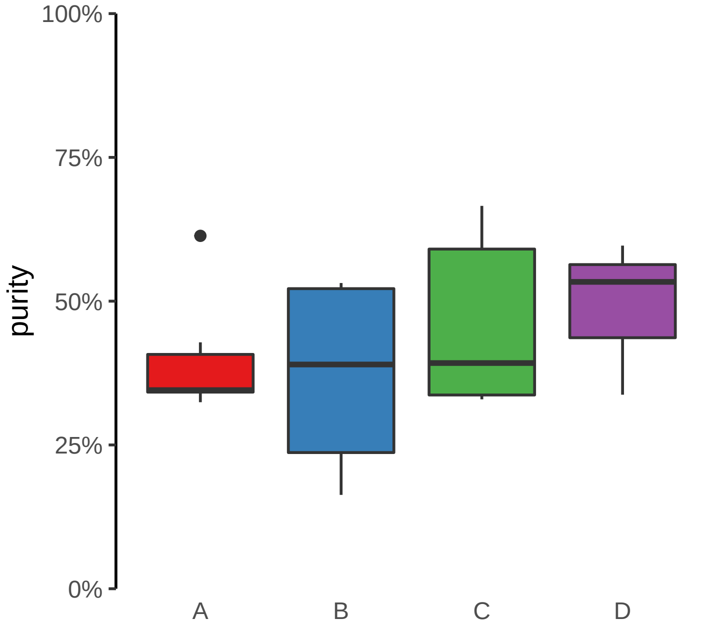

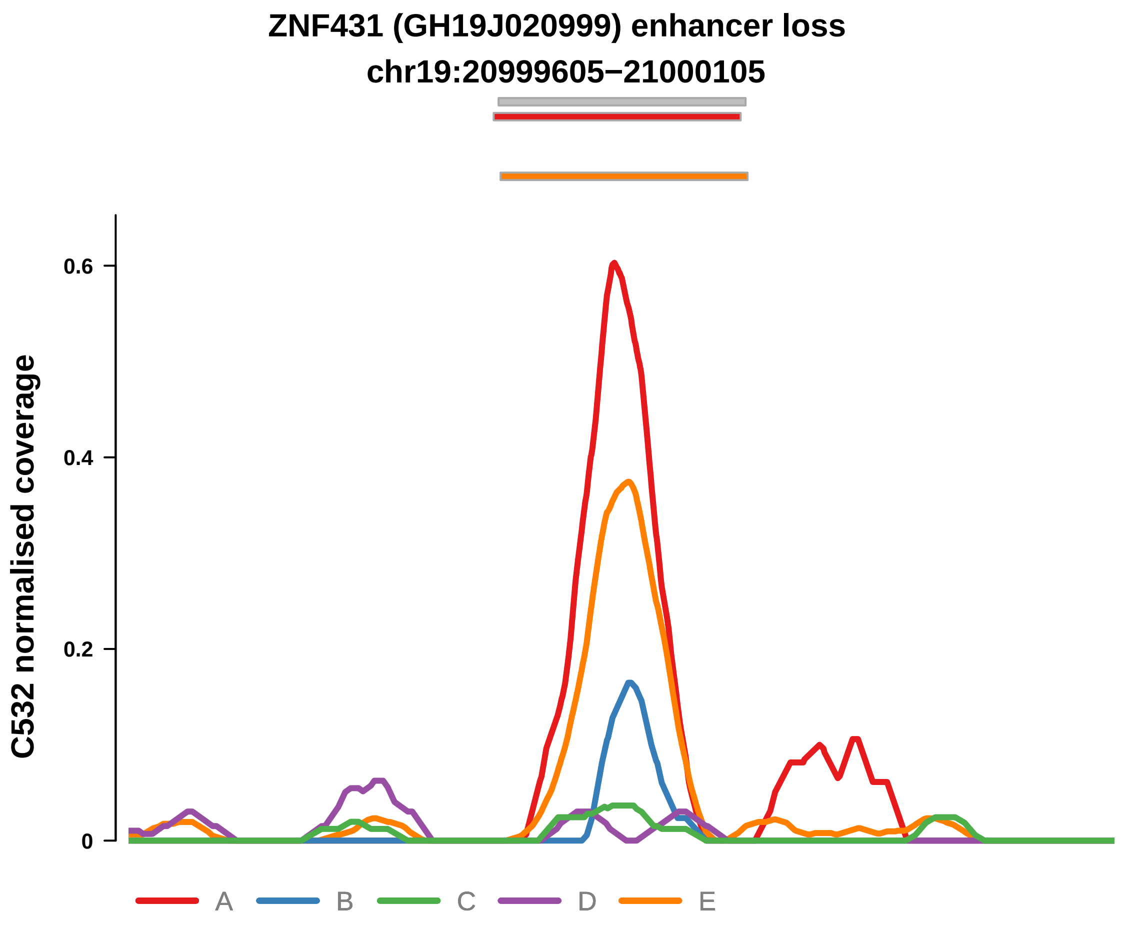


Putatively Subclonal (logFC = −1.3)

| | P. | Adjusted P. |
|---|---|---|
| *~region vs. ~* | 6.69e−07 **** | 5.88e−05 **** |
| *~purity vs ~* | 5.27e−01 | 9.09e−01 |
| *~purity+region vs ~purity* | 1.62e−06 **** | 1.42e−04 *** |
| *Carcinoma vs. normal* | 6.85e−02 | 6.85e−02 |

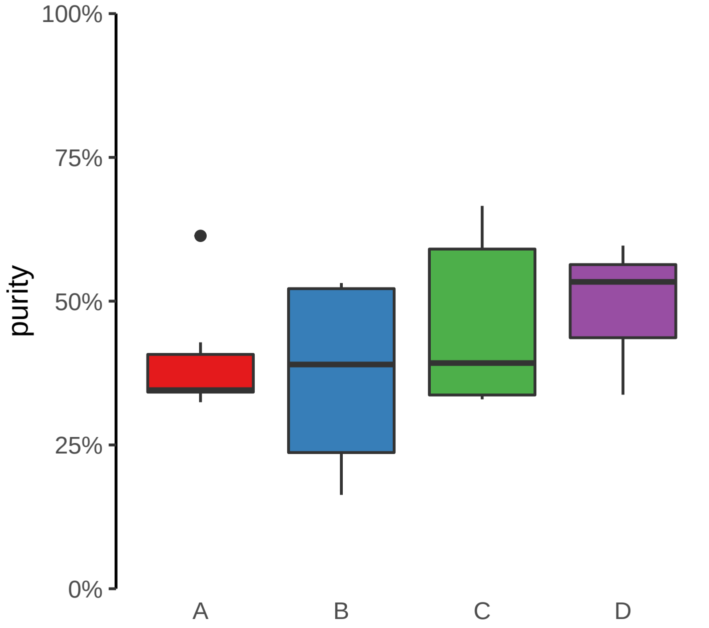

**SDHB (GH01J017108) enhancer loss**
**chr1:17108338−17108838**

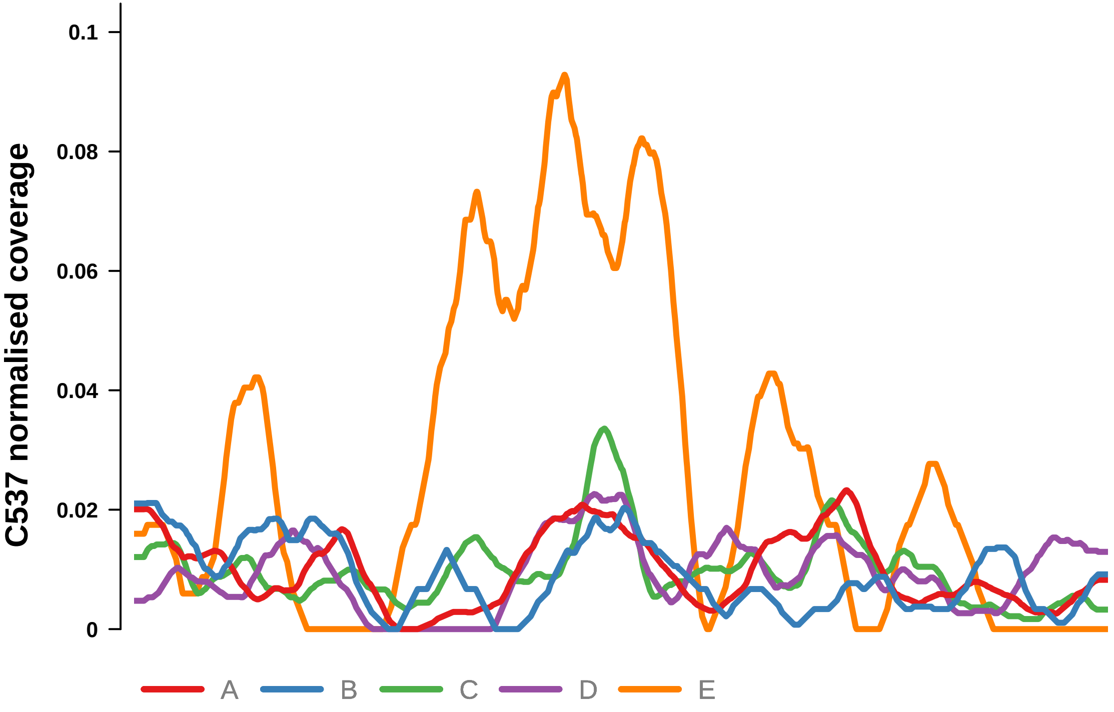


Putatively Subclonal (logFC = −1.93)

| | P. | Adjusted P. |
|---|---|---|
| *~region vs. ~* | 2.17e−02 * | 7.18e−02 |
| *~purity vs ~* | 8.42e−01 | 8.78e−01 |
| *~purity+region vs ~purity* | 2.46e−03 ** | 3.88e−02 * |
| *Carcinoma vs. normal* | 8.66e−03 ** | 8.66e−03 ** |

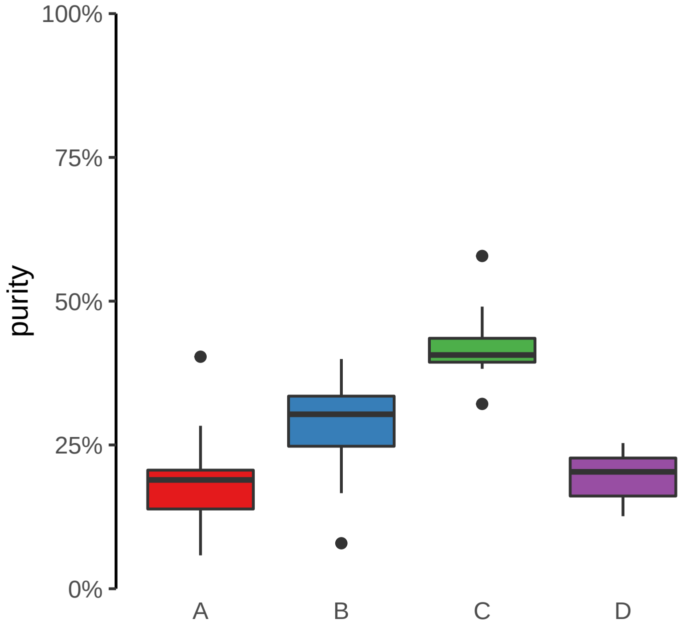

**ANTXR2 (GH04J080057) enhancer loss**
**chr4:80058687−80059187**

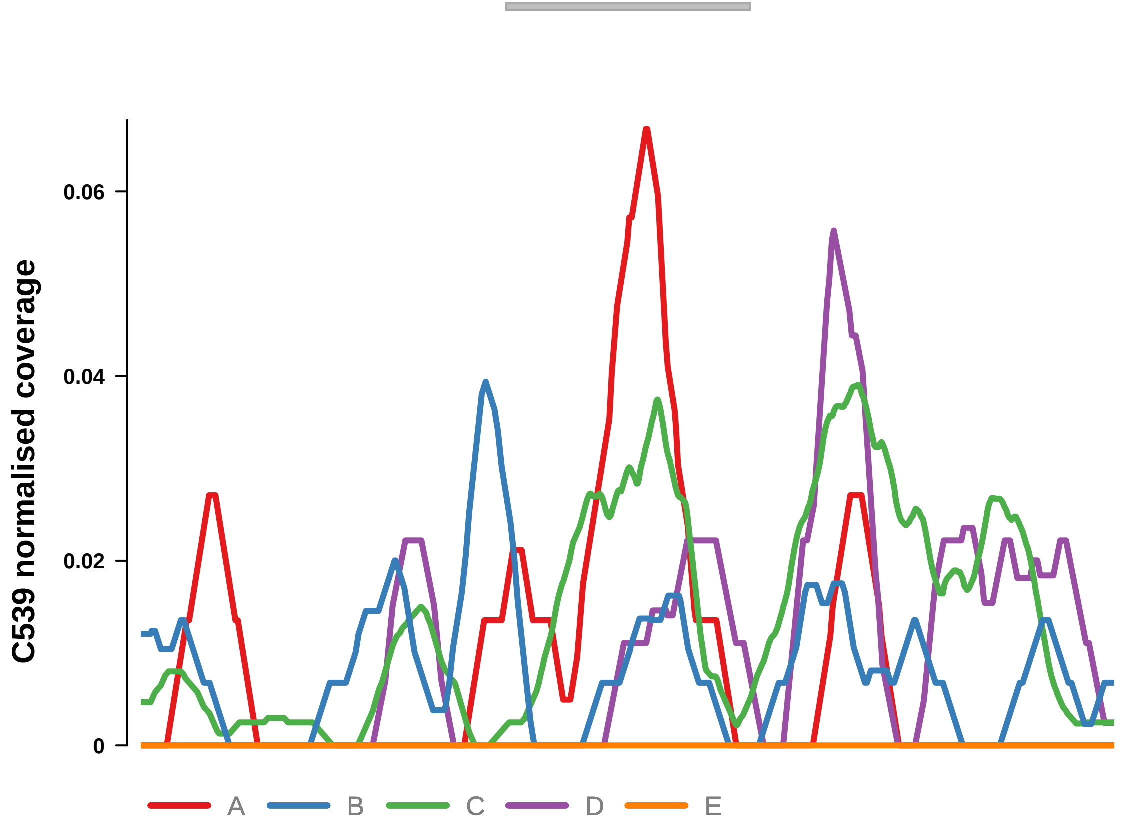


Putatively Subclonal (logFC = −0.68)

| | P. | Adjusted P. |
|---|---|---|
| *~region vs. ~* | 1.94e−05 **** | 1.71e−04 *** |
| *~purity vs ~* | 9.26e−04 *** | 3.54e−03 ** |
| *~purity+region vs ~purity* | 1.65e−03 ** | 1.45e−02 * |
| *Carcinoma vs. normal* | NA | NA |

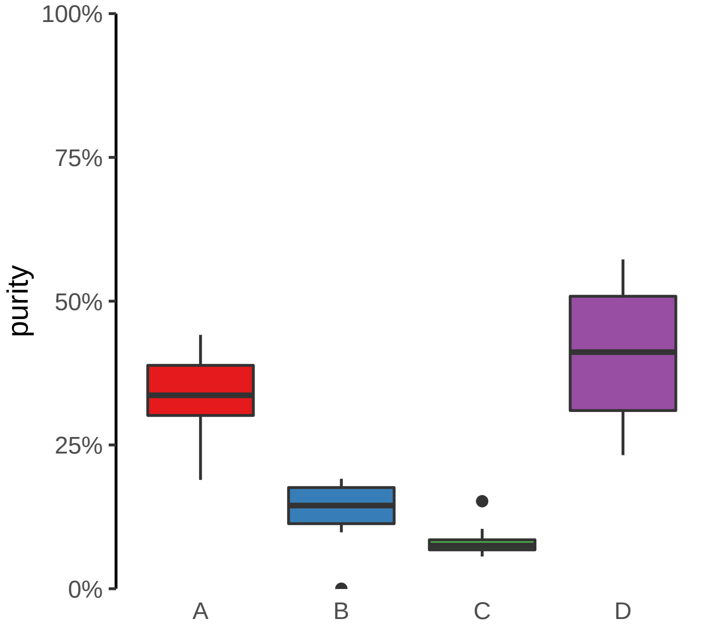

**ENSG00000228035, NGF (GH01J115322) enhancer loss**
**chr1:115323237−115323737**

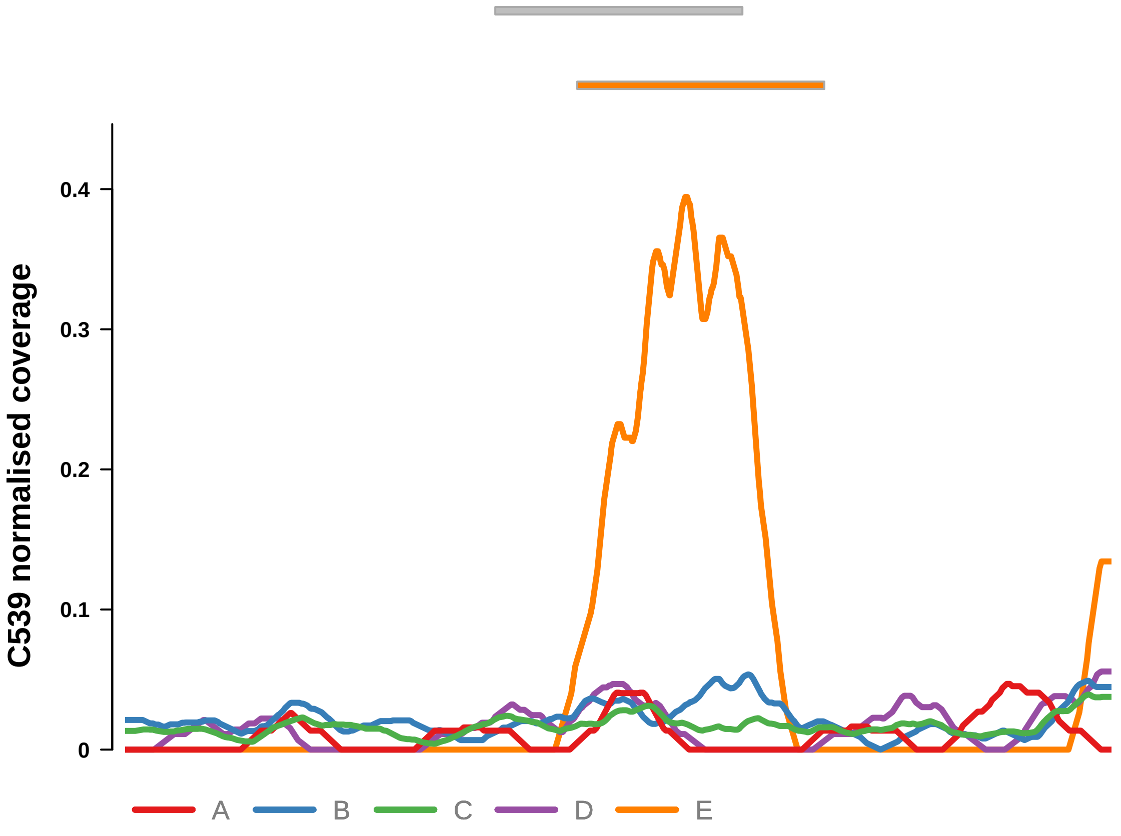


Putatively Subclonal (logFC = −0.46)

| | P. | Adjusted P. |
|---|---|---|
| *~region vs. ~* | 1.66e−06 **** | 2.92e−05 **** |
| *~purity vs ~* | 8.85e−05 **** | 6.48e−04 *** |
| *~purity+region vs ~purity* | 9.37e−03 ** | 4.35e−02 * |
| *Carcinoma vs. normal* | NA | NA |

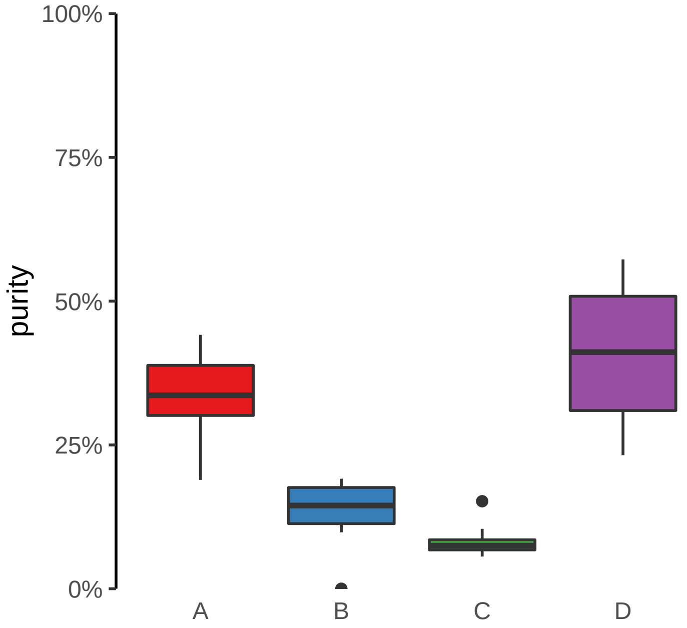

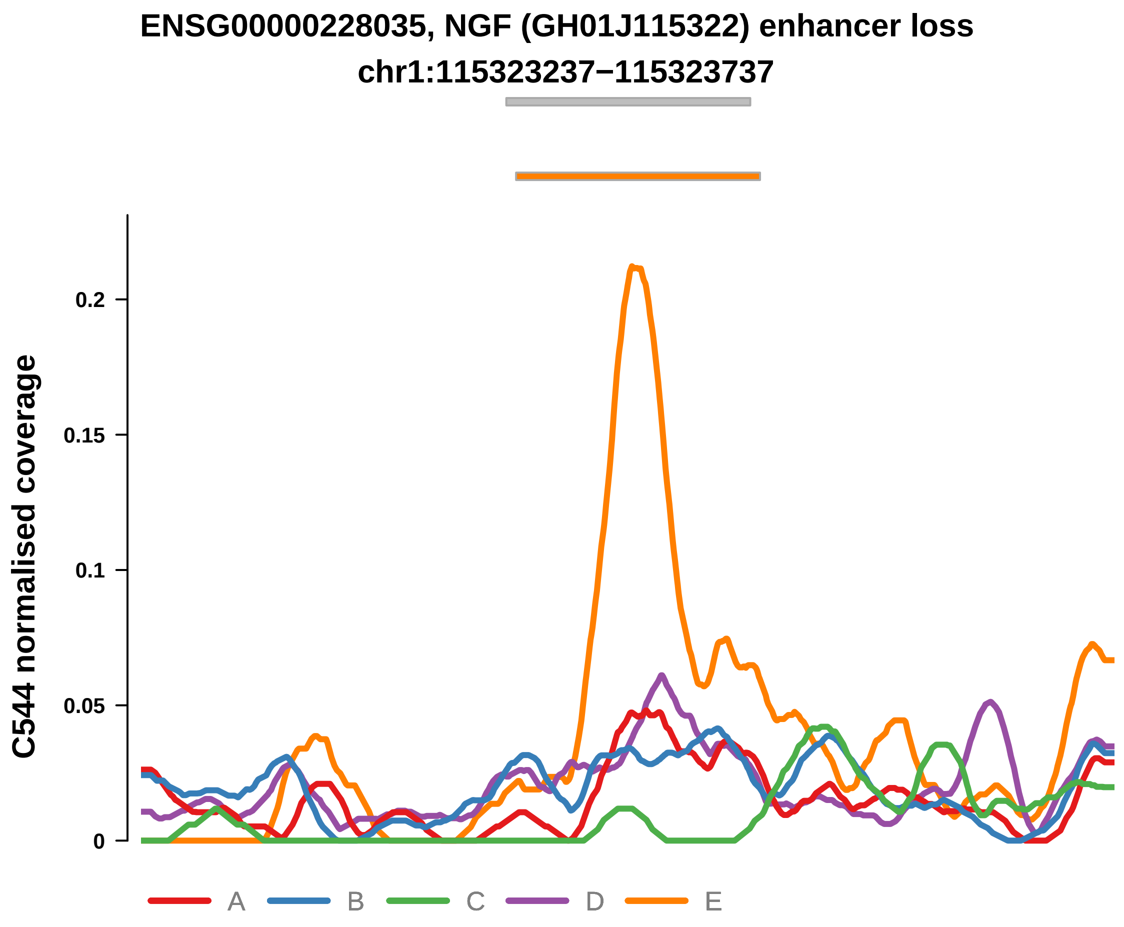


Putatively Subclonal (logFC = −0.7)

| | P. | Adjusted P. |
|---|---|---|
| *~region vs. ~* | 1.53e−03 ** | 7.73e−03 ** |
| *~purity vs ~* | 4.64e−01 | 6.09e−01 |
| *~purity+region vs ~purity* | 1.75e−04 *** | 9.1e−03 ** |
| *Carcinoma vs. normal* | NA | NA |

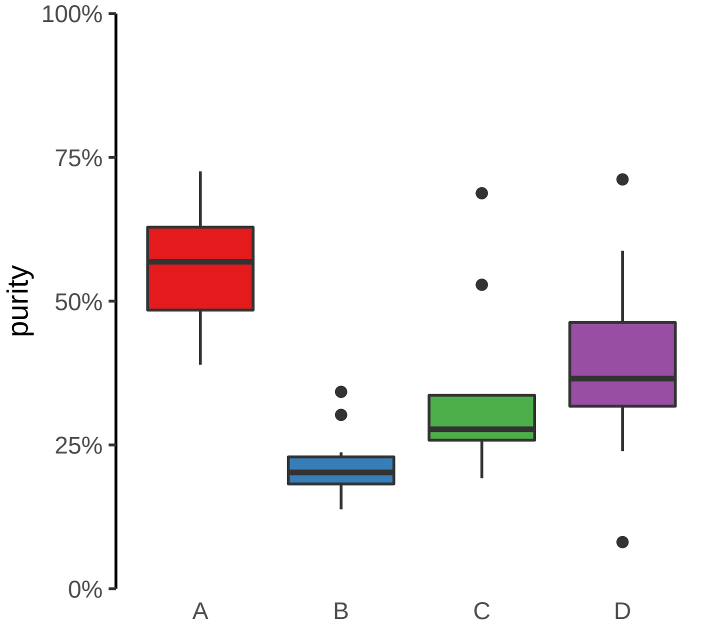

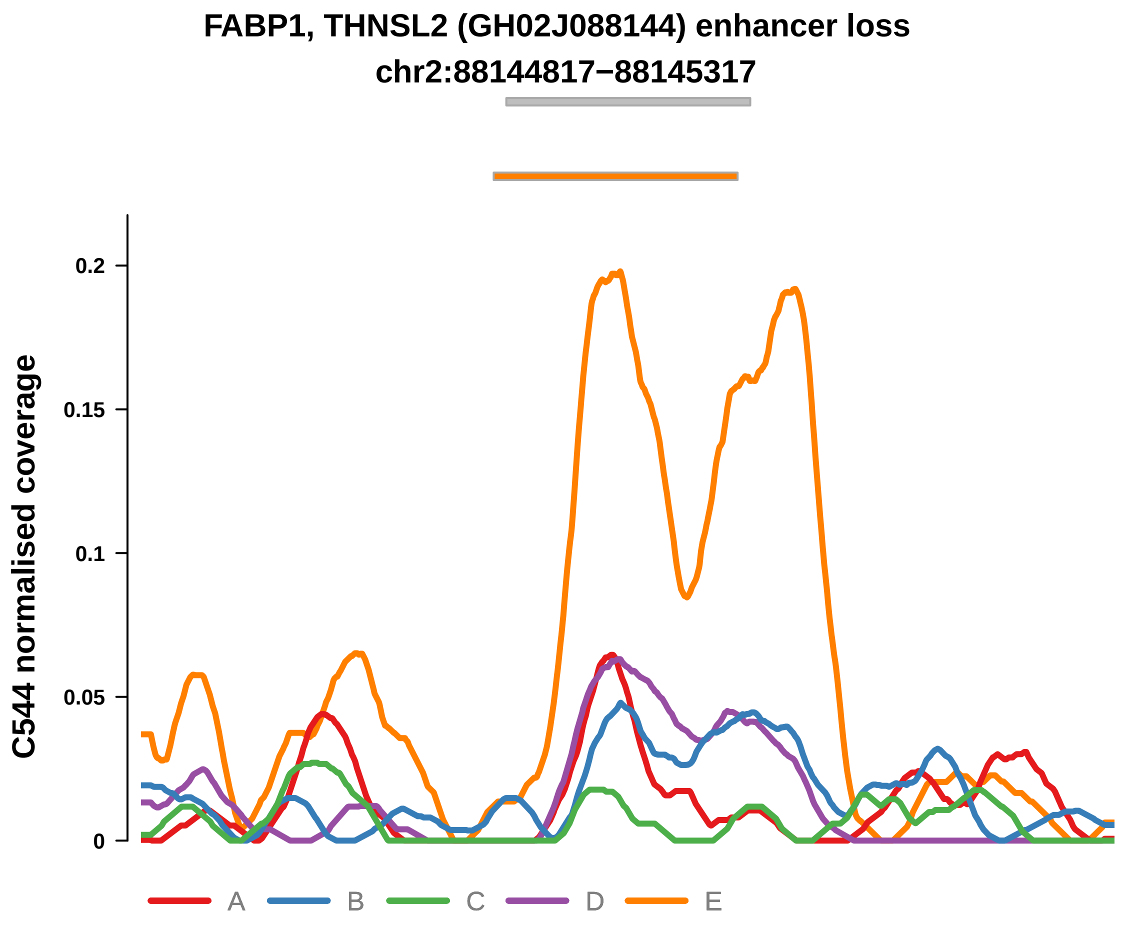


Putatively Subclonal (logFC = −1.25)

| | P. | Adjusted P. |
|---|---|---|
| *~region vs. ~* | 5.88e−03 ** | 2.06e−02 * |
| *~purity vs ~* | 8.86e−01 | 9.1e−01 |
| *~purity+region vs ~purity* | 4.97e−03 ** | 2.83e−02 * |
| *Carcinoma vs. normal* | 8.04e−02 | 8.04e−02 |

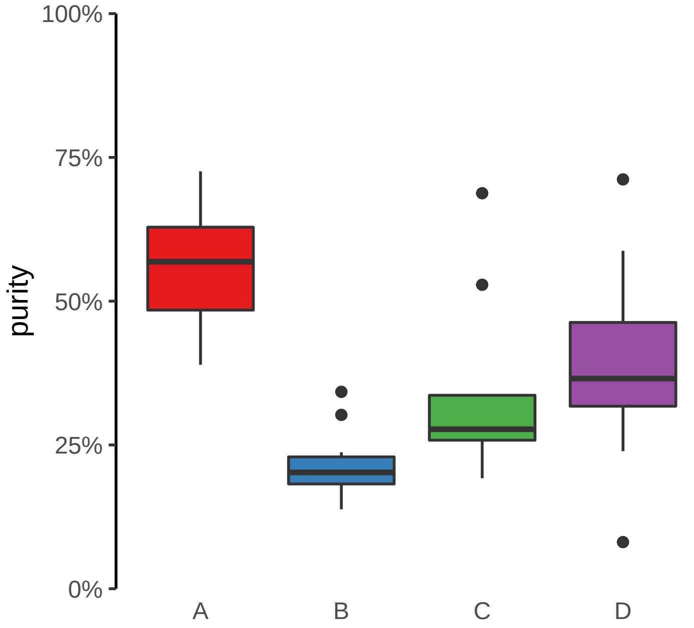

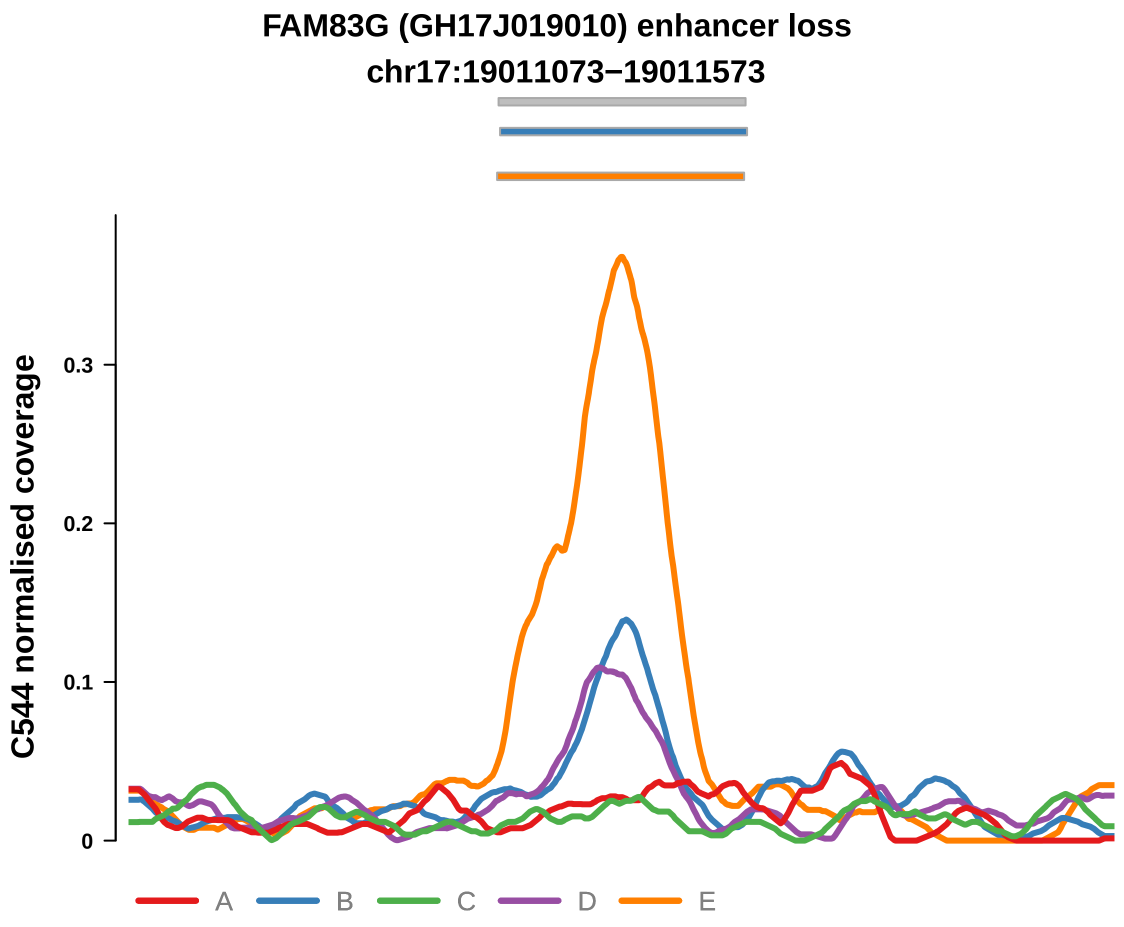


Putatively Subclonal (logFC = −1.51)

| | P. | Adjusted P. |
|---|---|---|
| *~region vs. ~* | 3.46e−03 ** | 1.6e−02 * |
| *~purity vs ~* | 1.97e−01 | 3.54e−01 |
| *~purity+region vs ~purity* | 9.87e−03 ** | 4.65e−02 * |
| *Carcinoma vs. normal* | 3.3e−02 * | 3.3e−02 * |

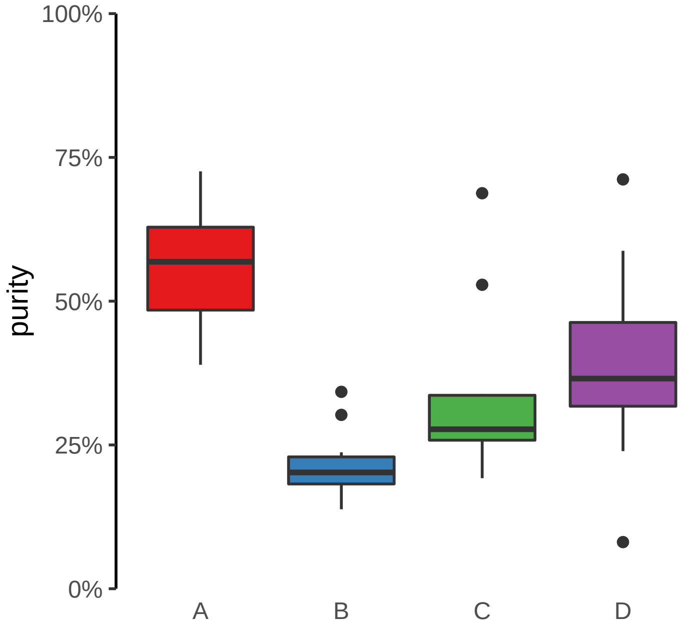

**HMGN2, ARID1A, PIGV (GH01J026739) enhancer loss**
**chr1:26743976−26744476**

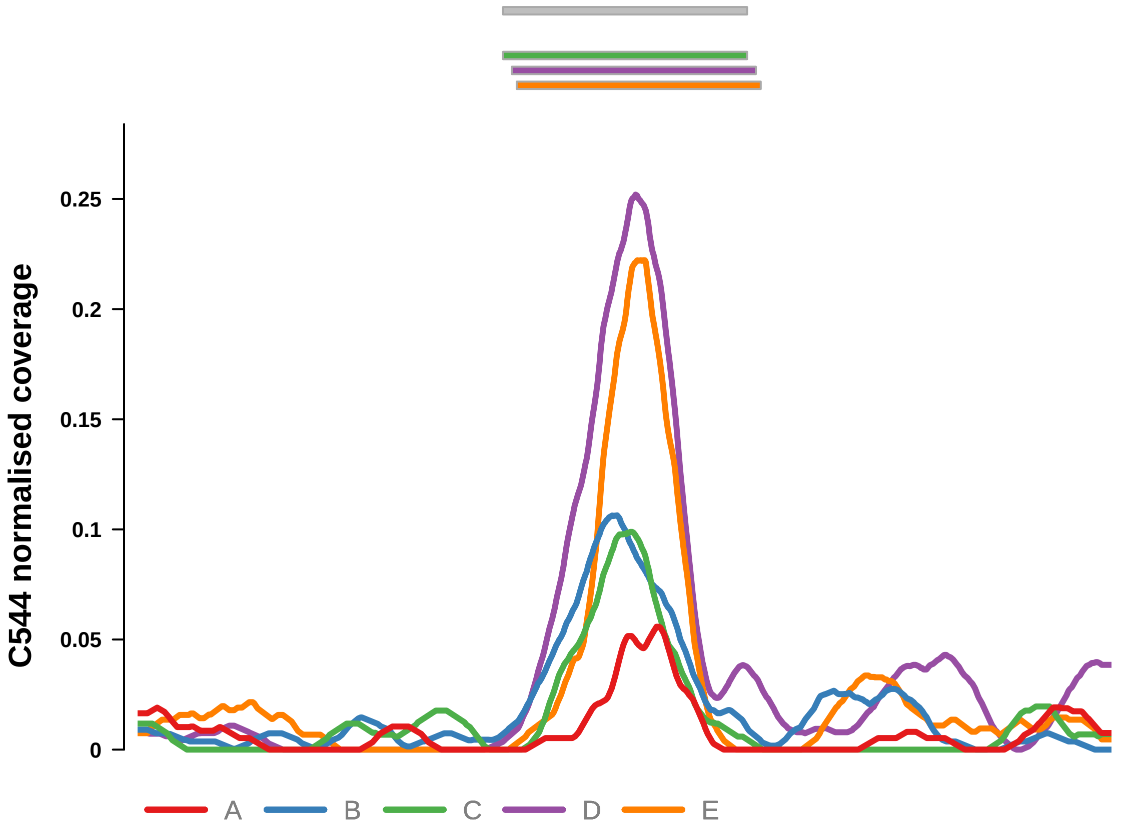


Putatively Subclonal (logFC = −1.72)

| | P. | Adjusted P. |
|---|---|---|
| *~region vs. ~* | 1.19e−03 ** | 6.55e−03 ** |
| *~purity vs ~* | 1.6e−02 * | 7.18e−02 |
| *~purity+region vs ~purity* | 1.02e−02 * | 4.65e−02 * |
| *Carcinoma vs. normal* | 1.62e−02 * | 1.62e−02 * |

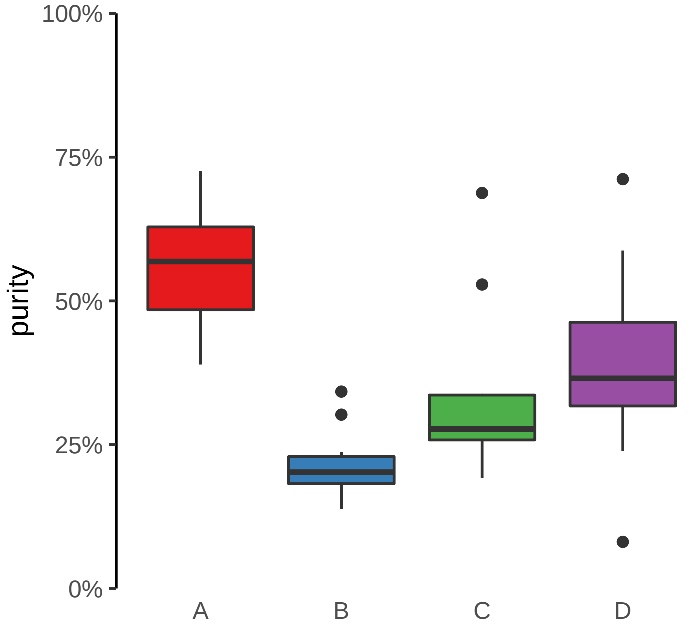

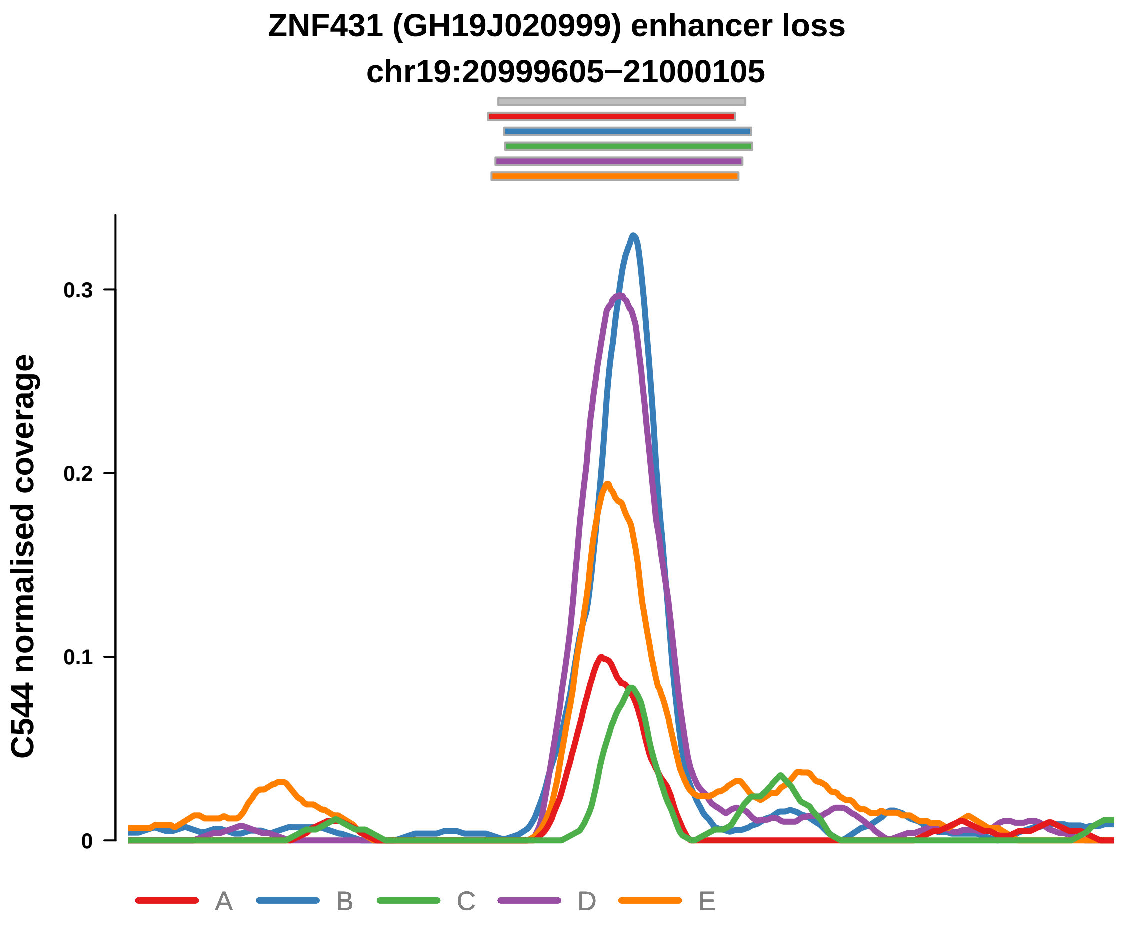


Putatively Subclonal (logFC = −1.01)

| | P. | Adjusted P. |
|---|---|---|
| *~region vs. ~* | 2.72e−05 **** | 3.48e−04 *** |
| *~purity vs ~* | 6.29e−03 ** | 3.4e−02 * |
| *~purity+region vs ~purity* | 1.1e−03 ** | 1.17e−02 * |
| *Carcinoma vs. normal* | 1.49e−01 | 1.49e−01 |

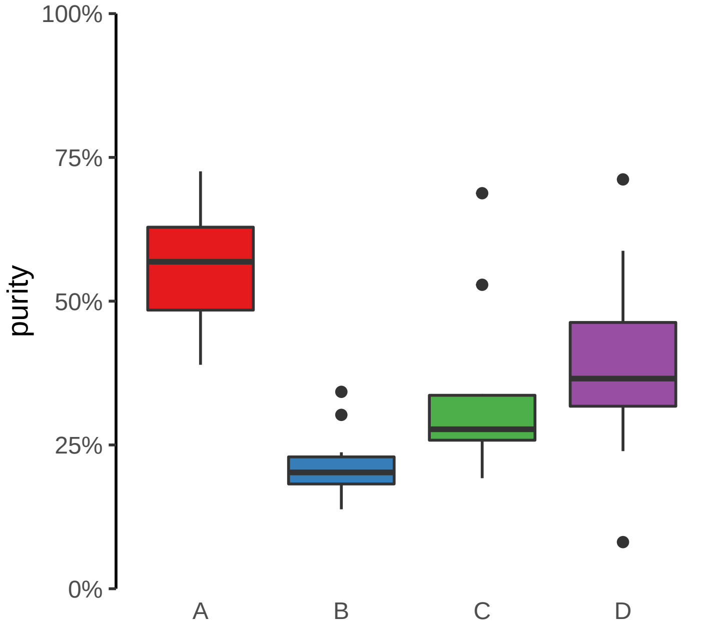

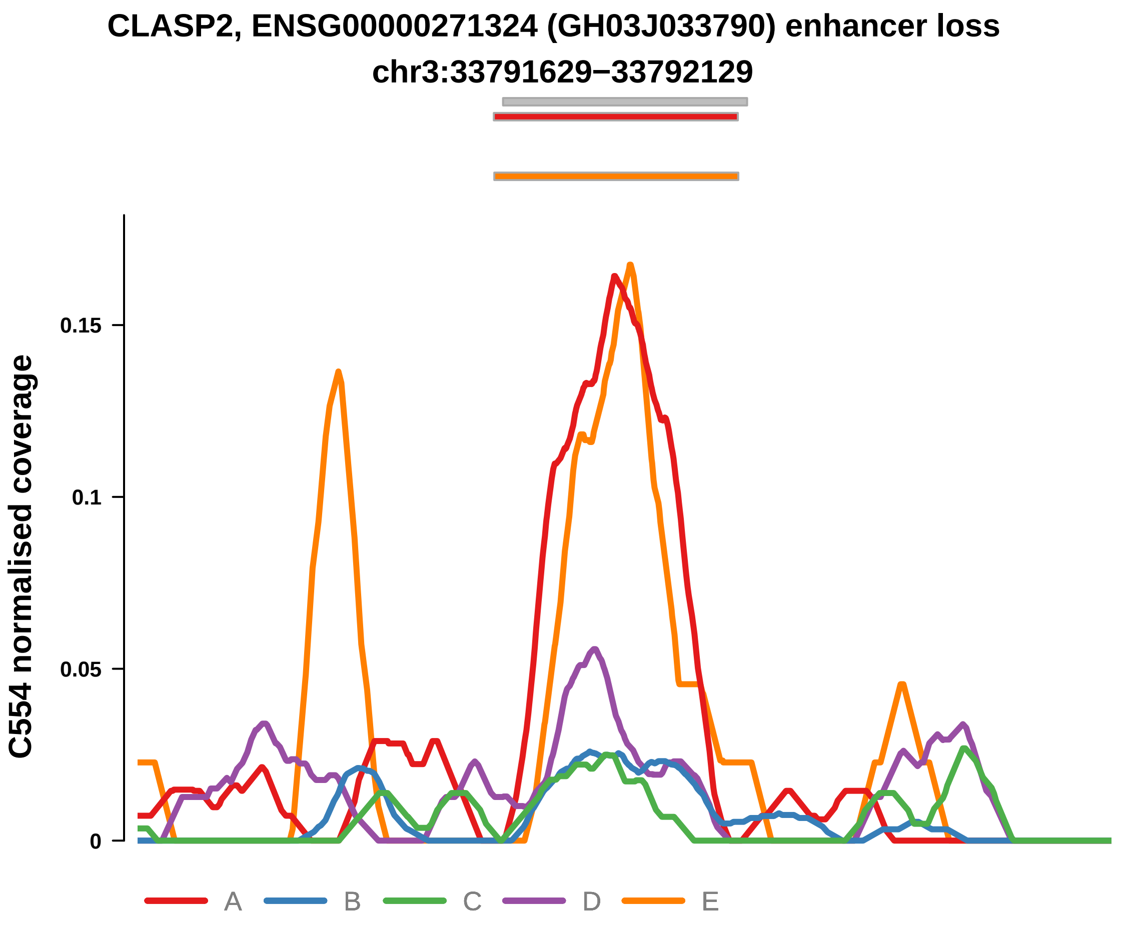


Putatively Subclonal (logFC = −2.18)

| | P. | Adjusted P. |
|---|---|---|
| *~region vs. ~* | 1.53e−03 ** | 7.35e−03 ** |
| *~purity vs ~* | 2.42e−02 * | 9.72e−02 |
| *~purity+region vs ~purity* | 1.12e−02 * | 4.35e−02 * |
| *Carcinoma vs. normal* | 2.41e−03 ** | 2.41e−03 ** |

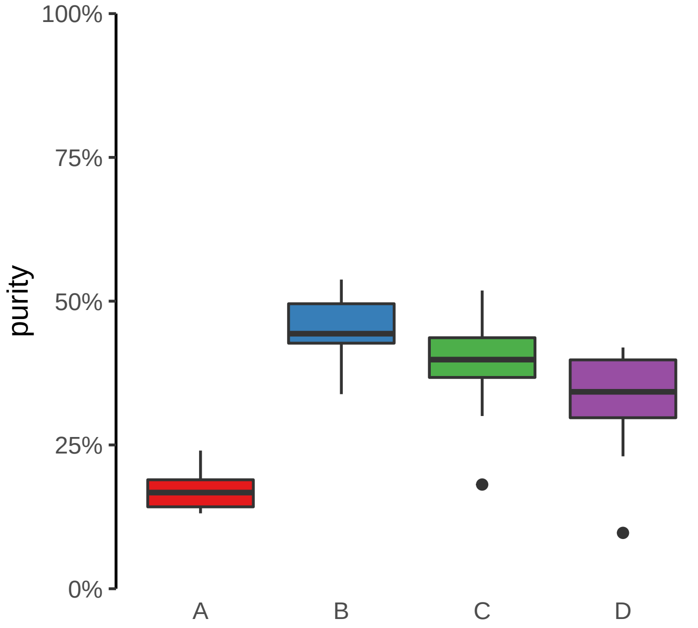

**FABP1, THNSL2 (GH02J088144) enhancer loss**
**chr2:88144817−88145317**

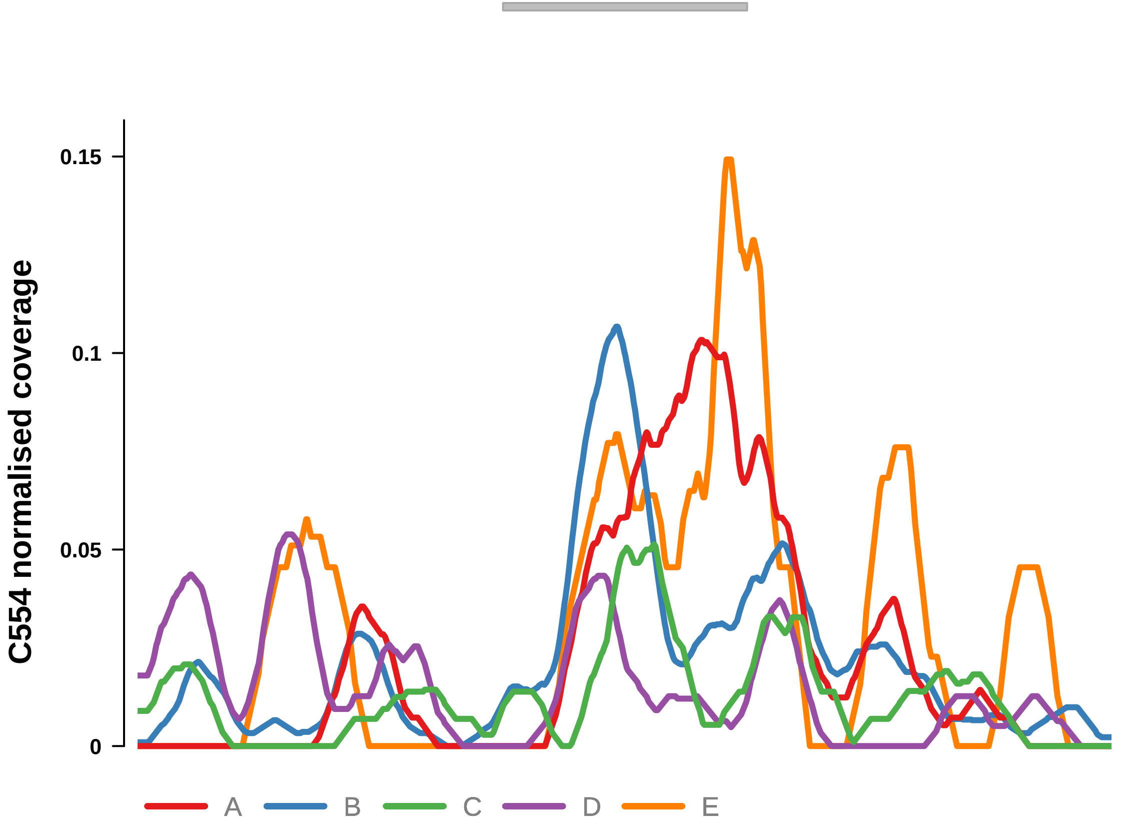


Putatively Subclonal (logFC = −0.86)

| | **P.** | **Adjusted P.** |
|---|---|---|
| *~region vs. ~* | 3.08e−03 ** | 1.05e−02 * |
| *~purity vs ~* | 6.67e−01 | 7.93e−01 |
| *~purity+region vs ~purity* | 3.78e−03 ** | 2.79e−02 * |
| *Carcinoma vs. normal* | NA | NA |

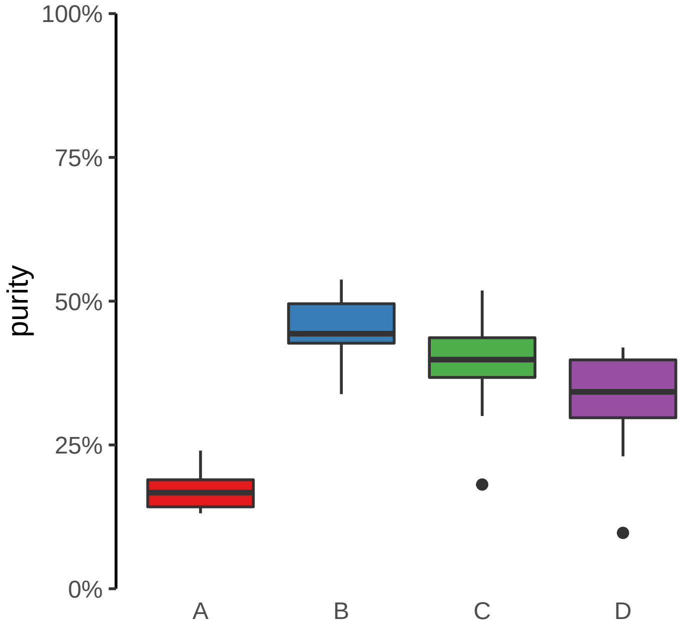

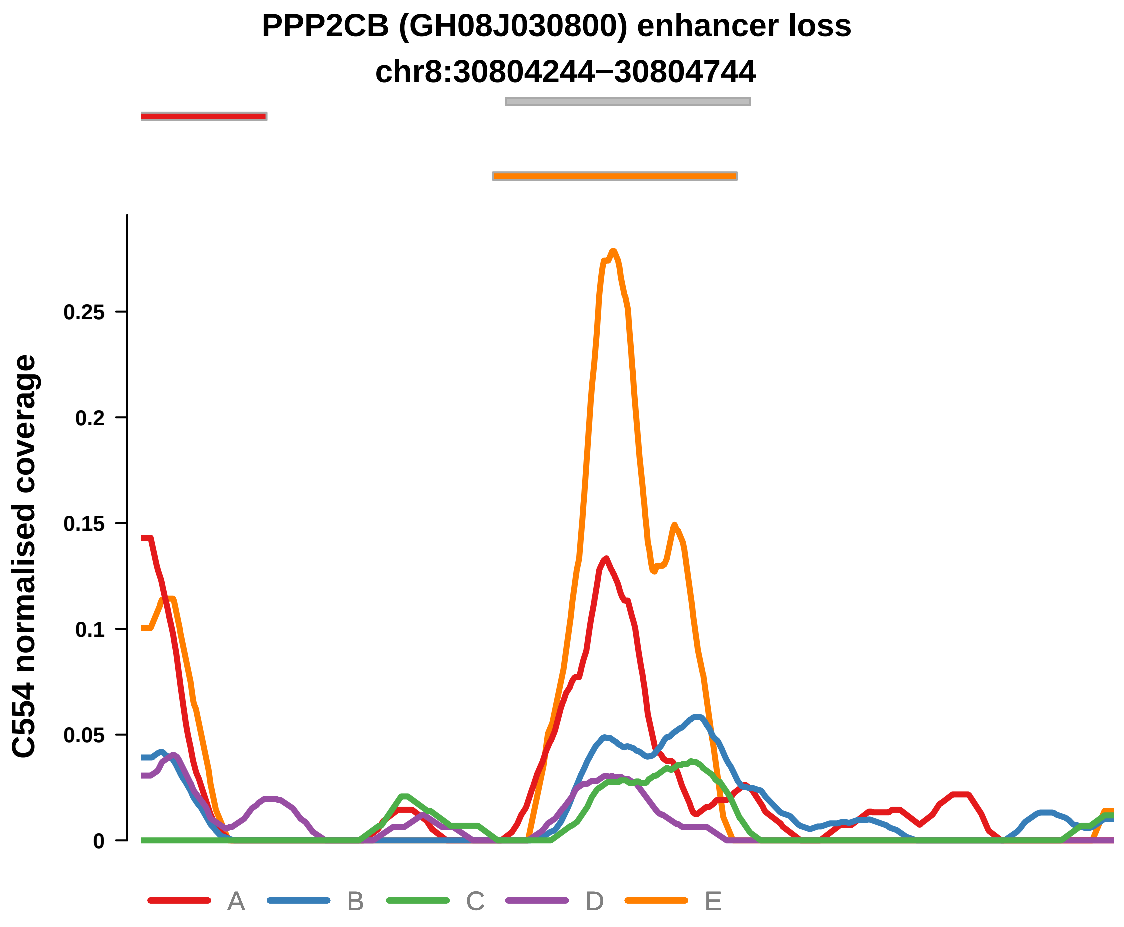


Putatively Subclonal (logFC = −1.19)

| | P. | Adjusted P. |
|---|---|---|
| *~region vs. ~* | 1.06e−02 * | 3.1e−02 * |
| *~purity vs ~* | 9.09e−01 | 9.43e−01 |
| *~purity+region vs ~purity* | 9.43e−03 ** | 3.88e−02 * |
| *Carcinoma vs. normal* | 8.76e−02 | 8.76e−02 |

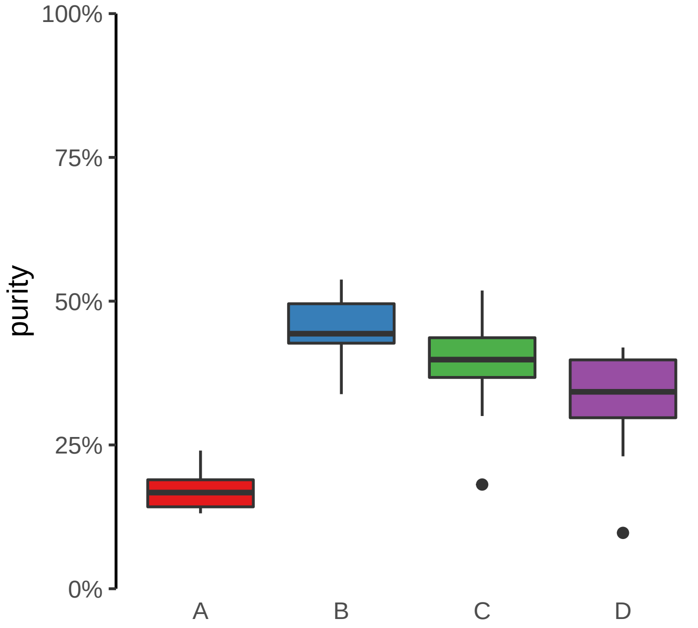

**PPP3CA (GH04J101183) enhancer loss**
**chr4:101183634−101184134**

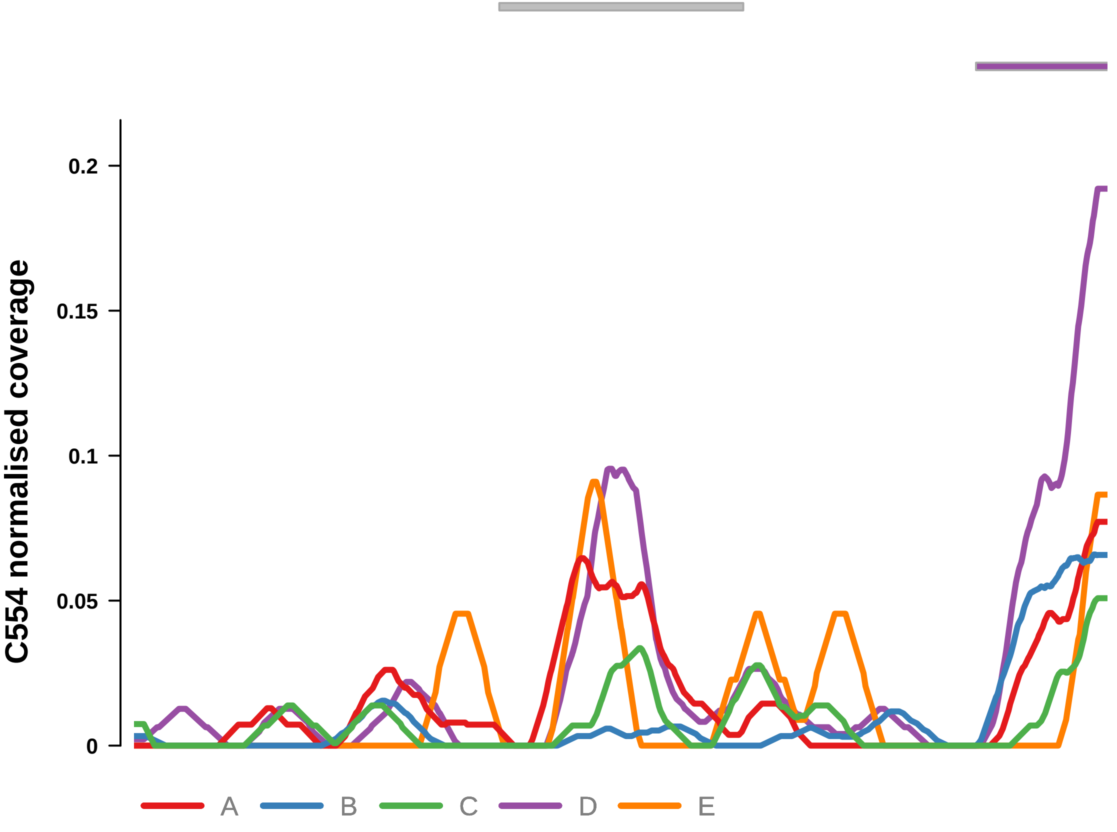


Putatively Subclonal (logFC = −2.31)

| | P. | Adjusted P. |
|---|---|---|
| *~region vs. ~* | 5.81e−04 *** | 3.17e−03 ** |
| *~purity vs ~* | 1.8e−01 | 3.72e−01 |
| *~purity+region vs ~purity* | 2.09e−03 ** | 1.9e−02 * |
| *Carcinoma vs. normal* | 3.22e−03 ** | 3.22e−03 ** |

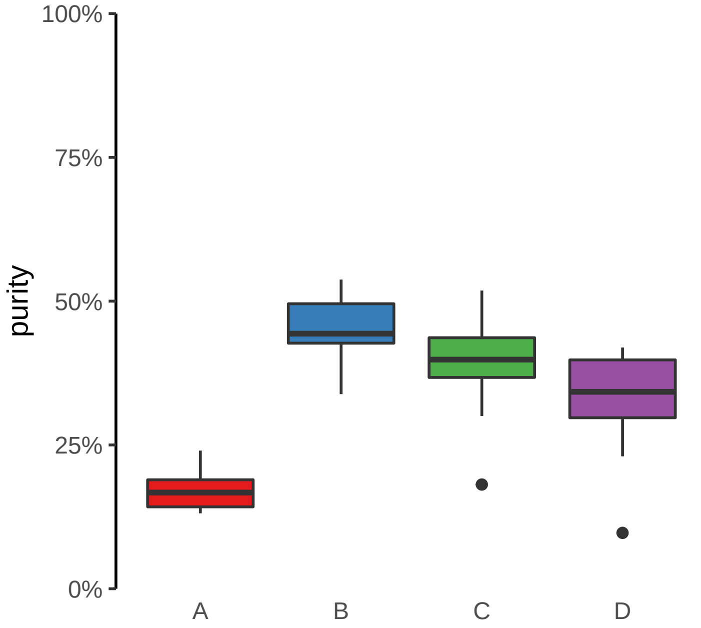

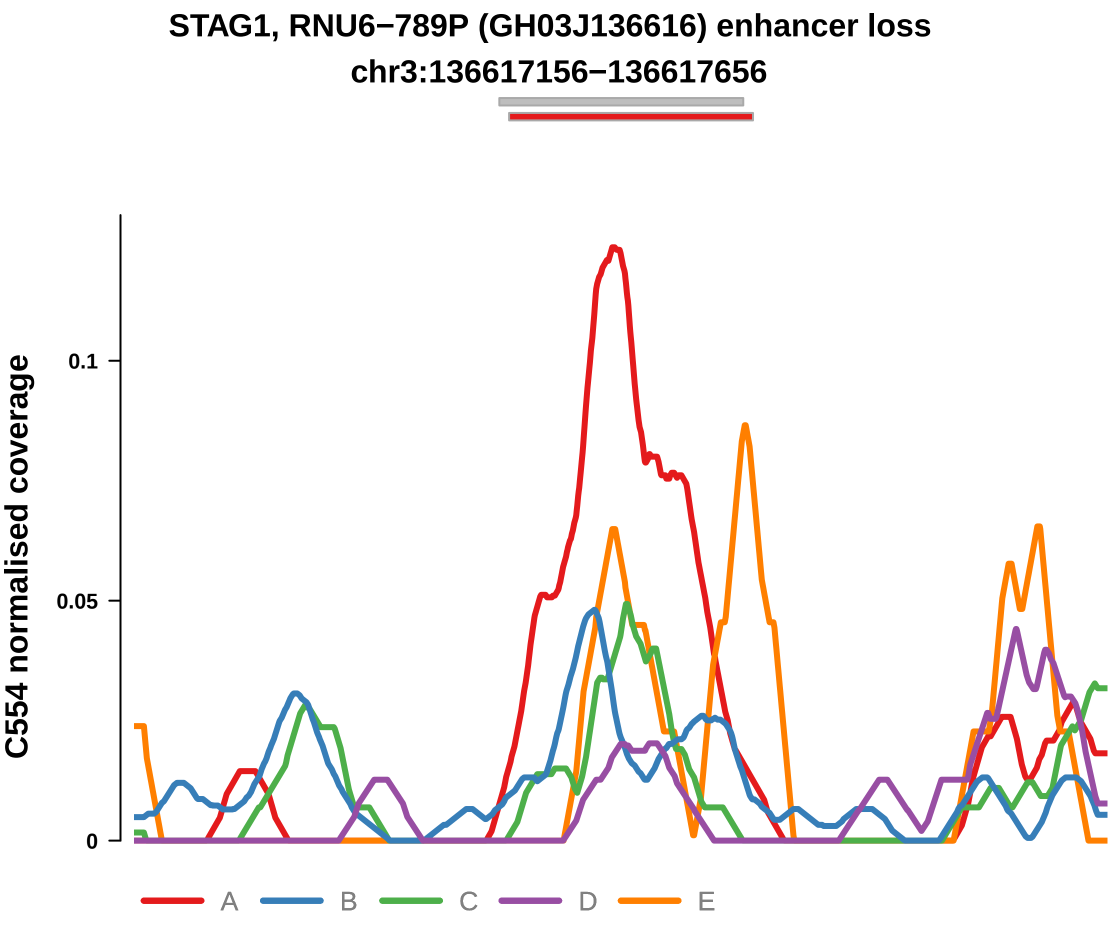


Putatively Subclonal (logFC = −1.45)

| | P. | Adjusted P. |
|---|---|---|
| *~region vs. ~* | 4.69e−04 *** | 2.75e−03 ** |
| *~purity vs ~* | 3.06e−01 | 5.23e−01 |
| *~purity+region vs ~purity* | 1.02e−03 ** | 1.08e−02 * |
| *Carcinoma vs. normal* | 3.84e−02 * | 3.84e−02 * |

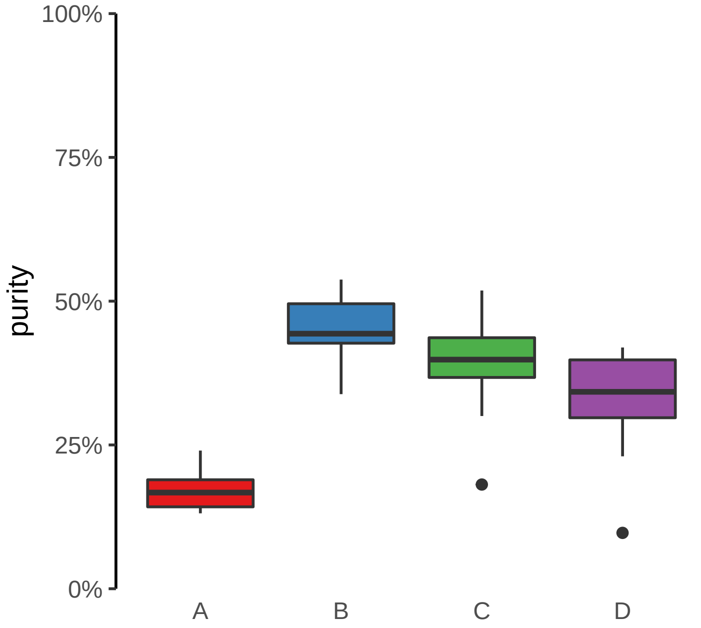
