## Supplementary Figures for "The co-evolution of the genome and epigenome in colorectal cancer": figS19.pdf

### C516 – Cancer (Pure >= 0.4) – FOXL1_uc_pTSS_oPEAK

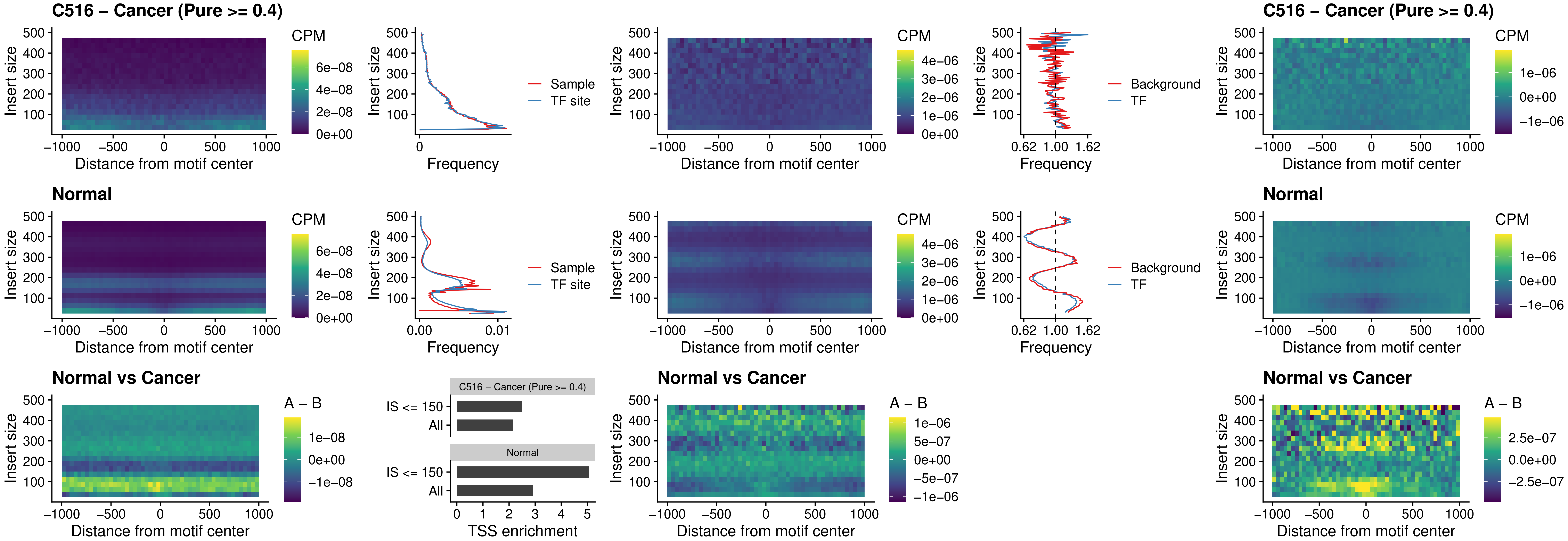

### C516 – Cancer (Pure >= 0.4) – CPEB1_uc_cTSS_oPEAK

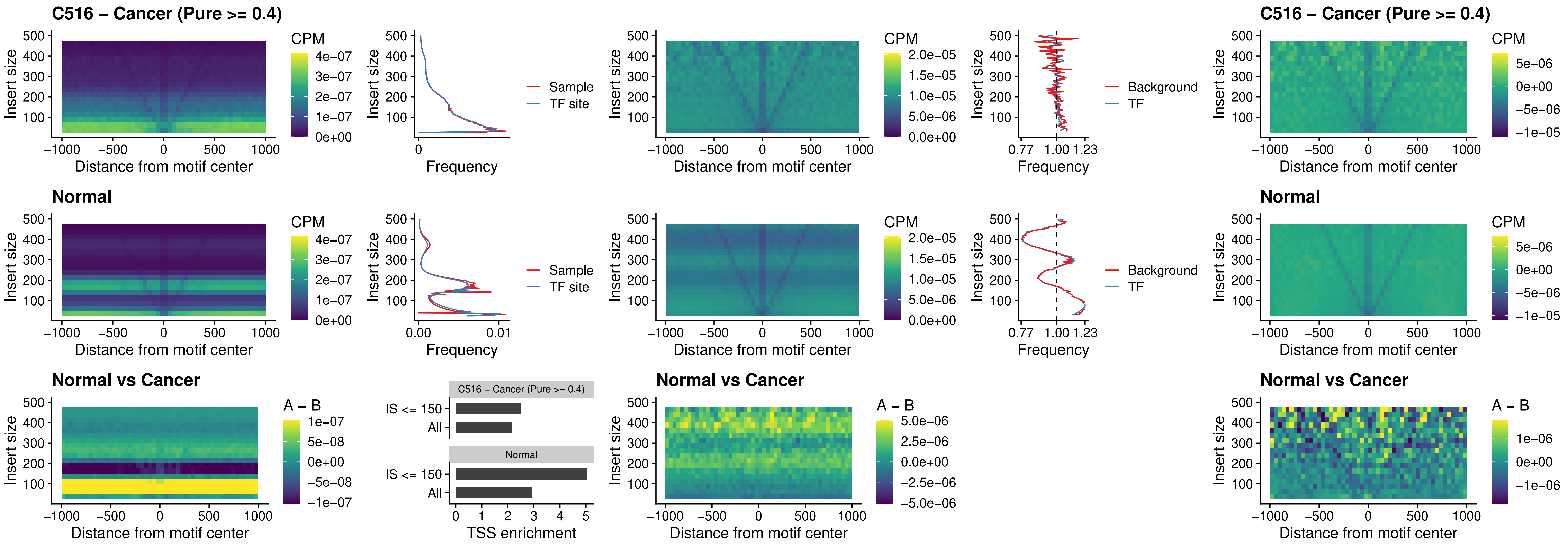

#### C516 – Cancer (Pure >= 0.4) – HMGA1_uc_pTSS_nPEAK

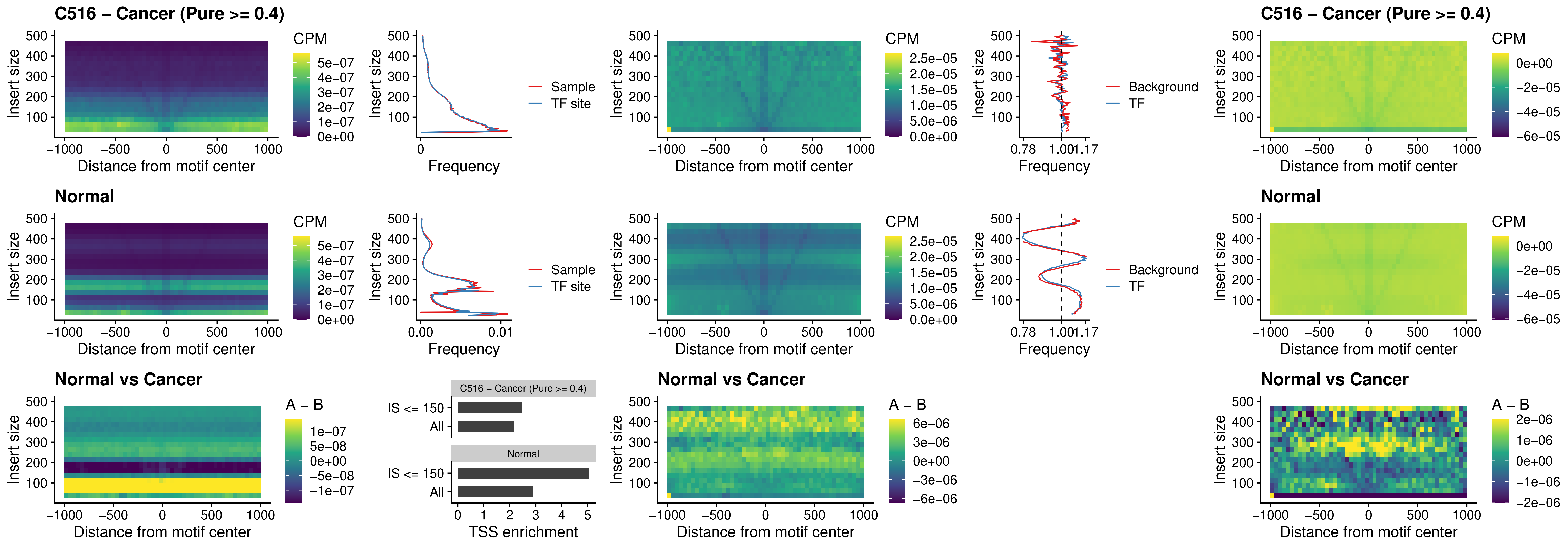

### C516 – Cancer (Pure >= 0.4) – SOX13_uc_cTSS_nPEAK

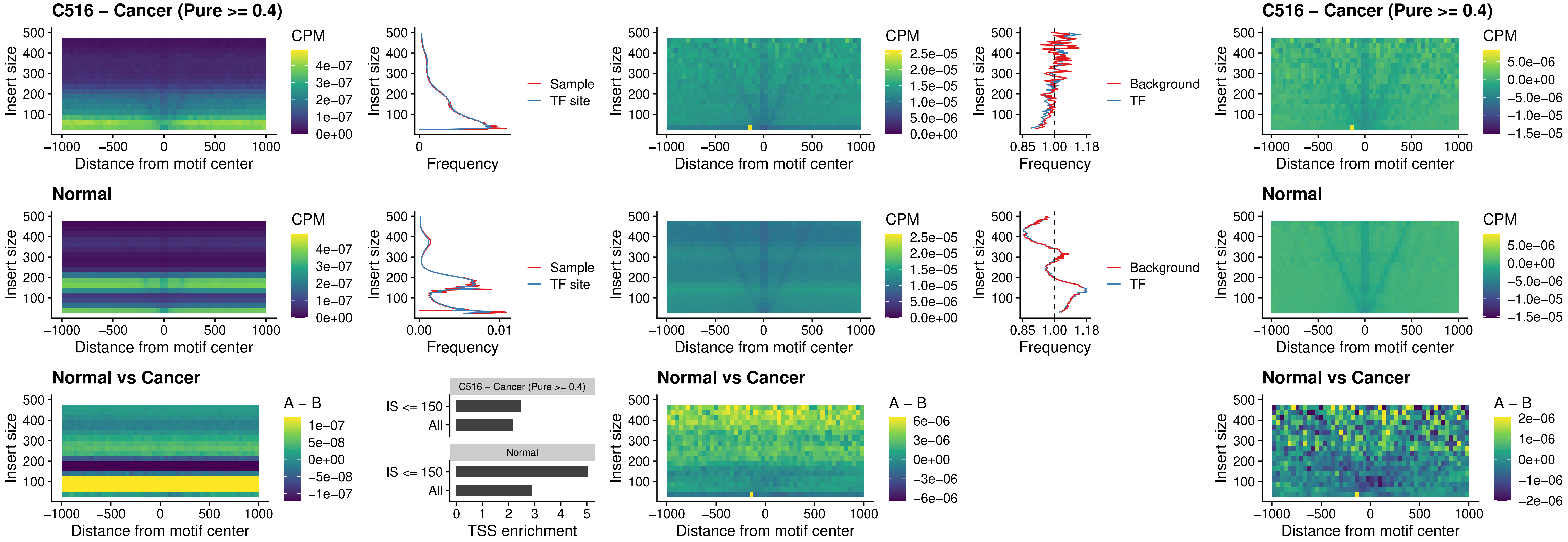

### C516 – Cancer (Pure >= 0.4) – LHX4_uc_cTSS_nPEAK

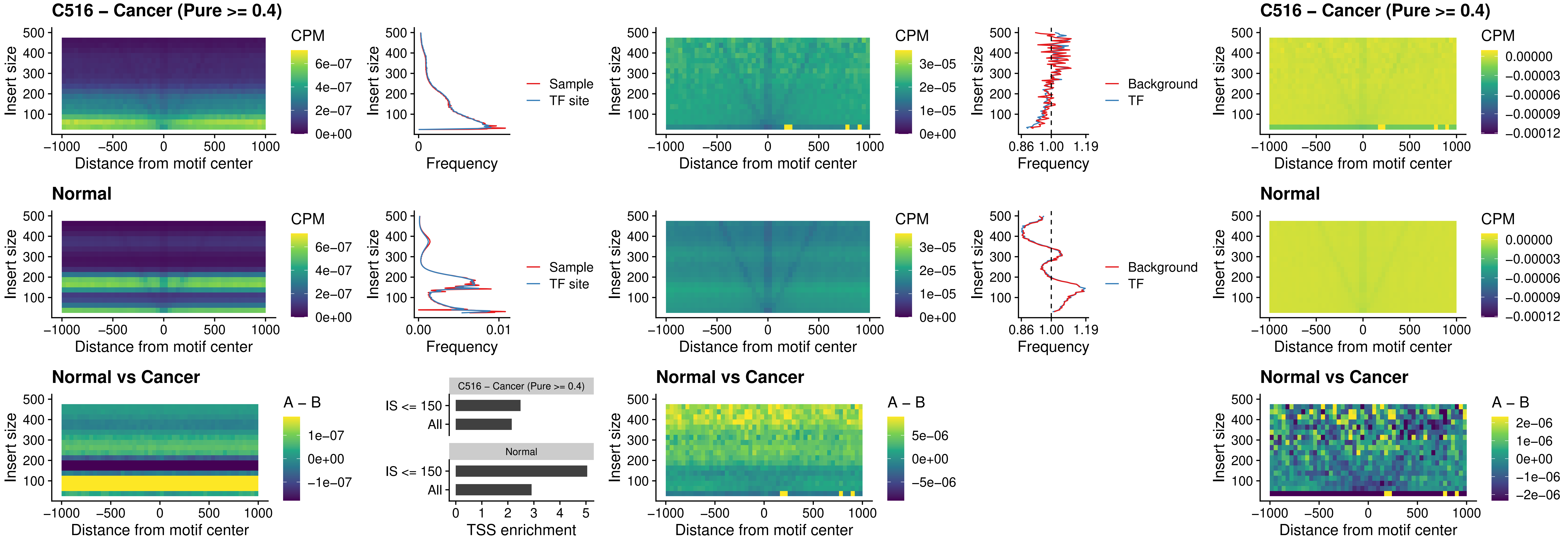

#### C516 – Cancer (Pure >= 0.4) – HOXA4_uc_cTSS_nPEAK

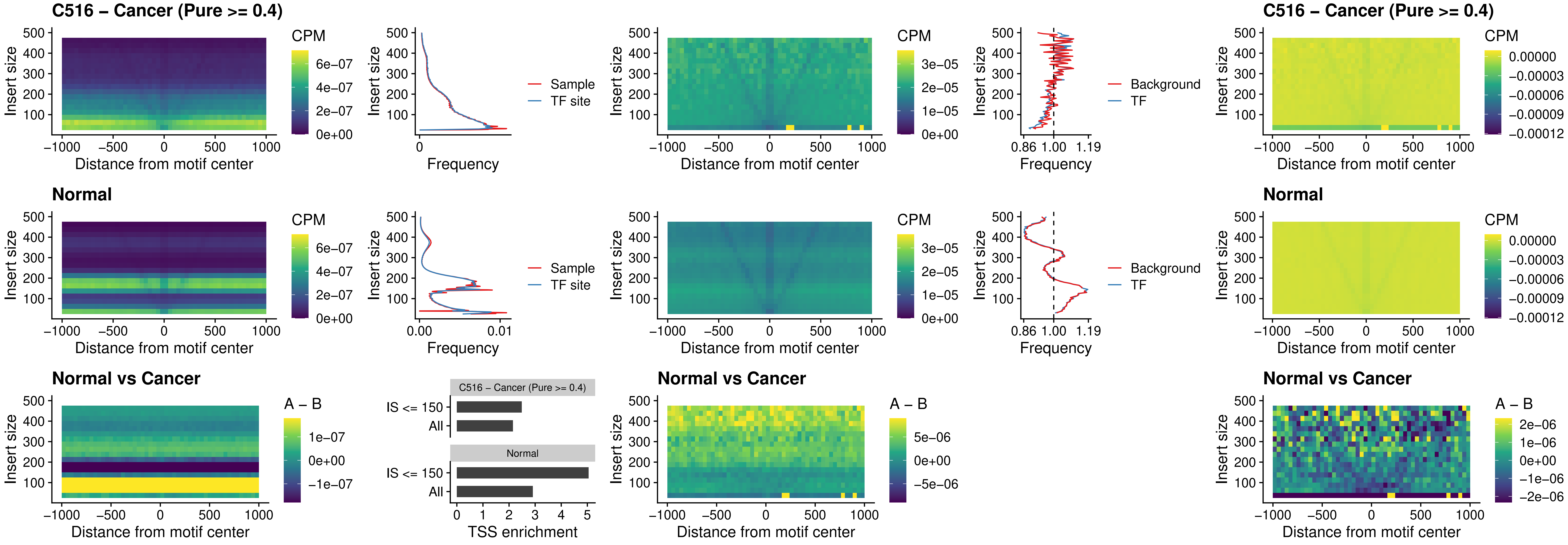

#### C516 – Cancer (Pure >= 0.4) – HLX_uc_cTSS_nPEAK

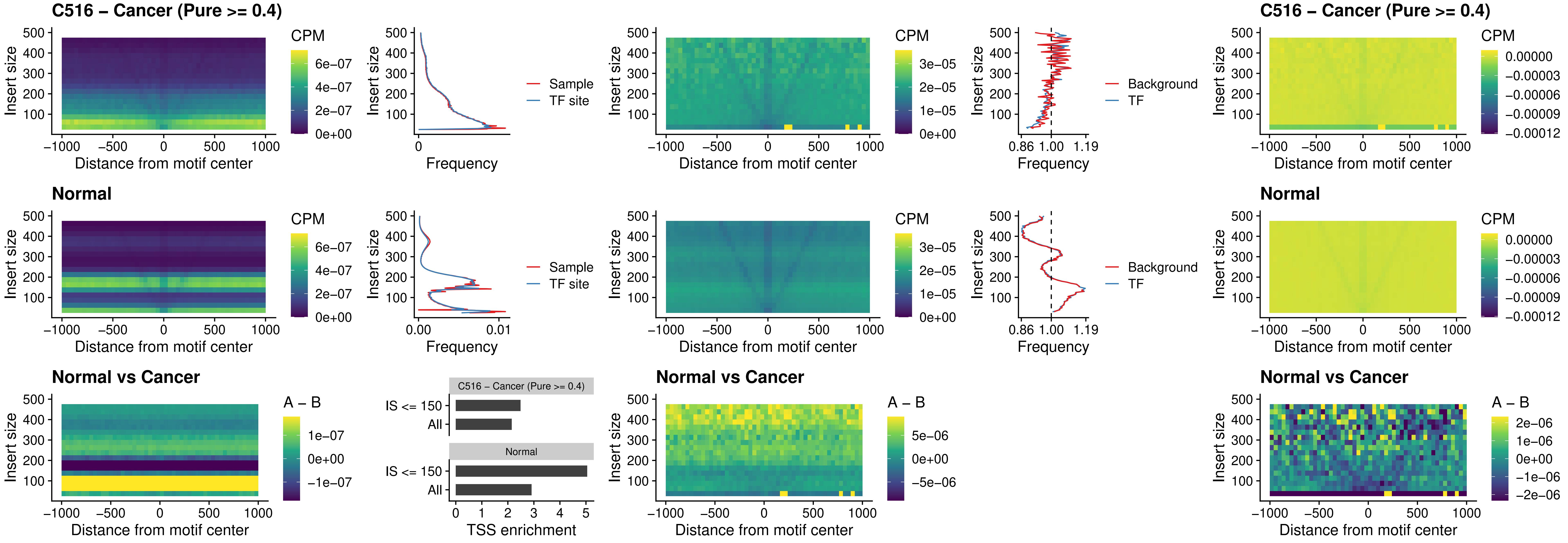

#### C516 – Cancer (Pure >= 0.4) – AC0125311_uc_cTSS_nPEAK

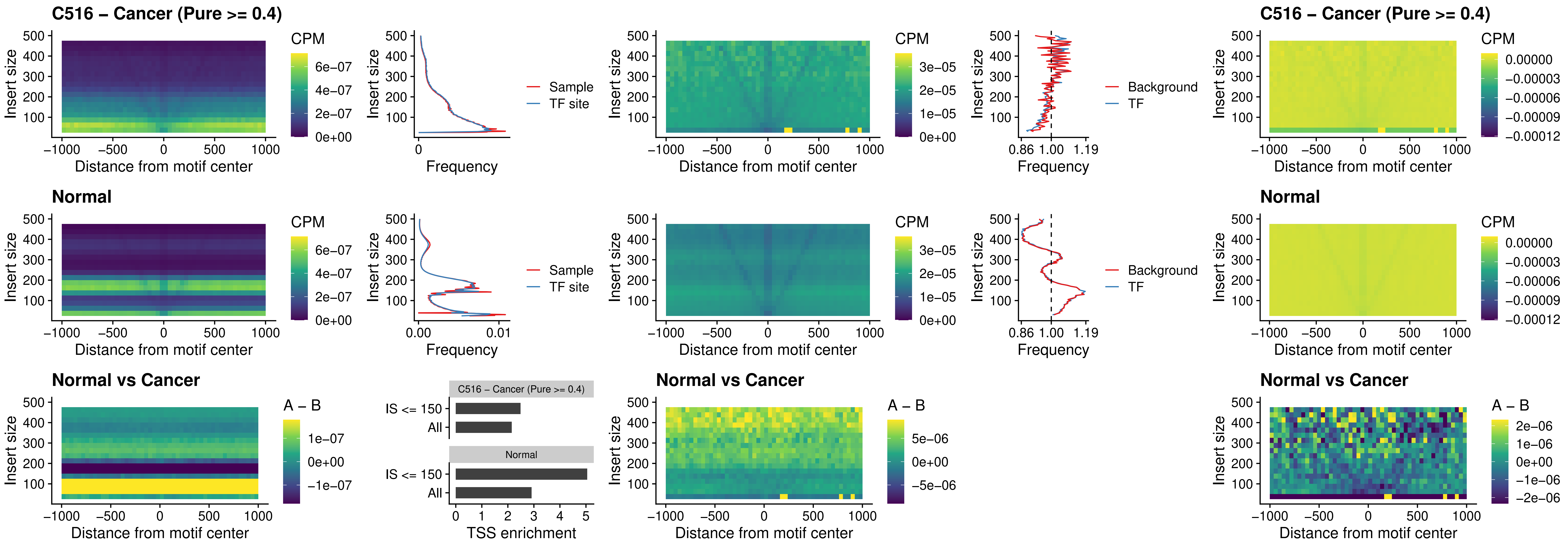

### C516 – Cancer (Pure >= 0.4) – DBX2_uc_cTSS_nPEAK

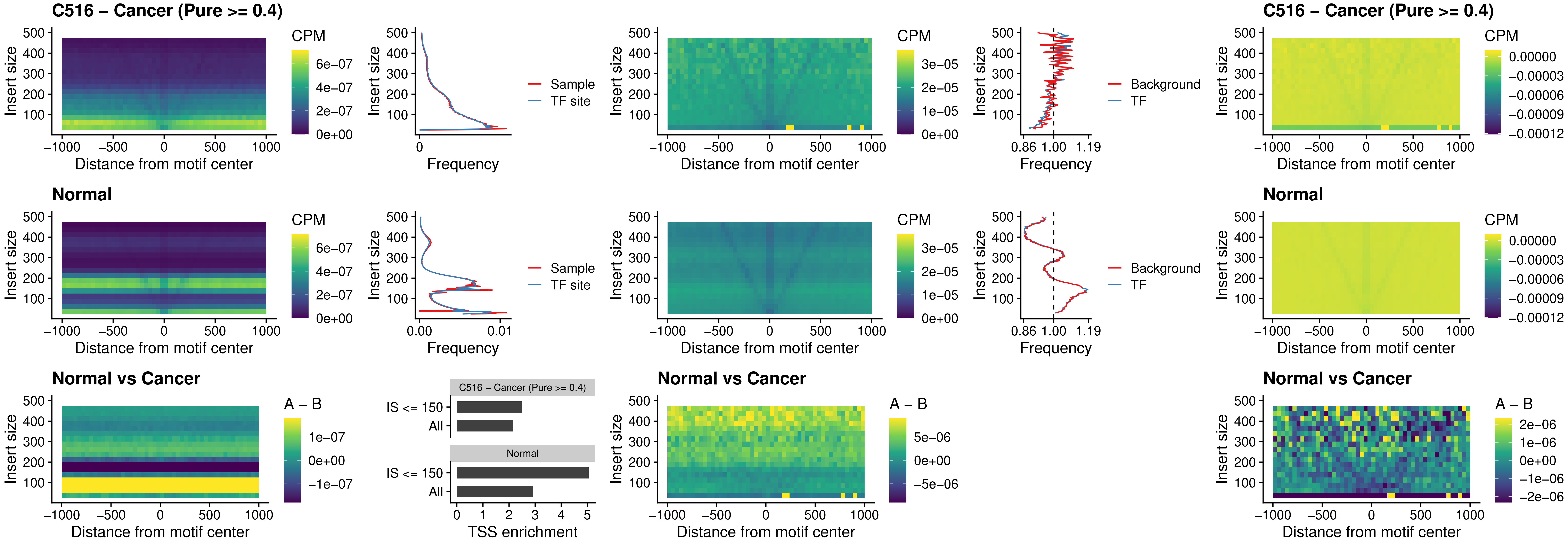

### C516 – Cancer (Pure >= 0.4) – LHX6_uc_dTSS_nPEAK

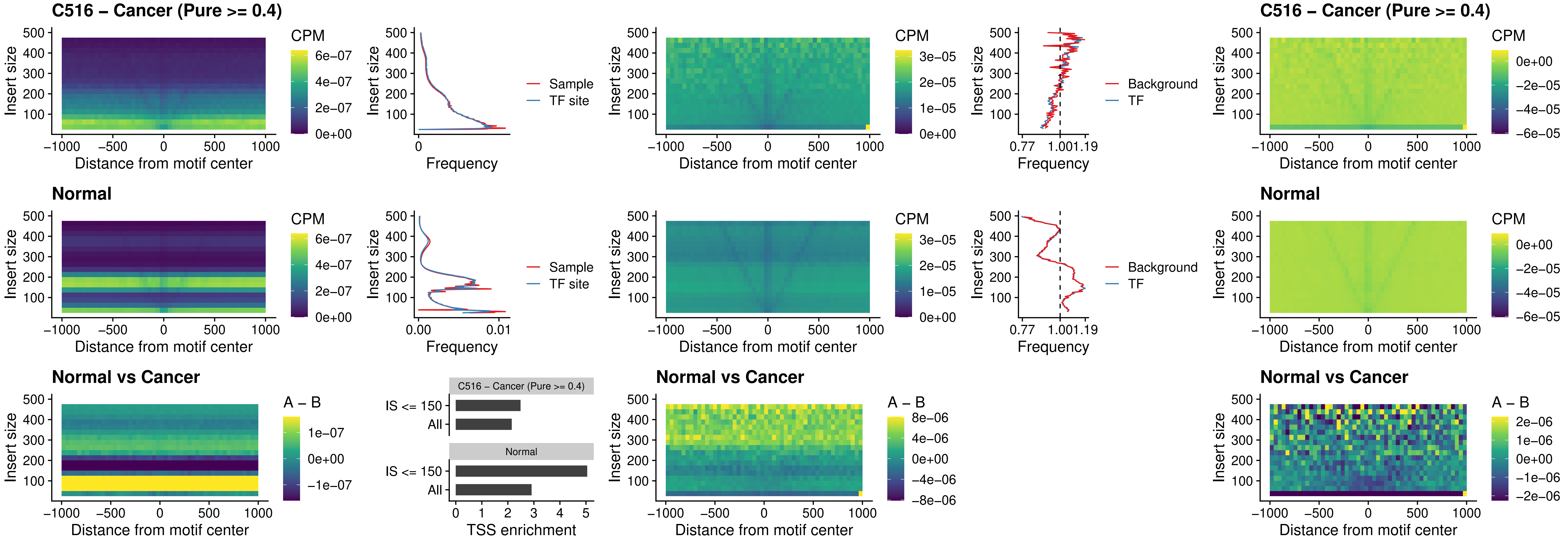

#### C516 – Cancer (Pure >= 0.4) – SOX1_uc_dTSS_nPEAK

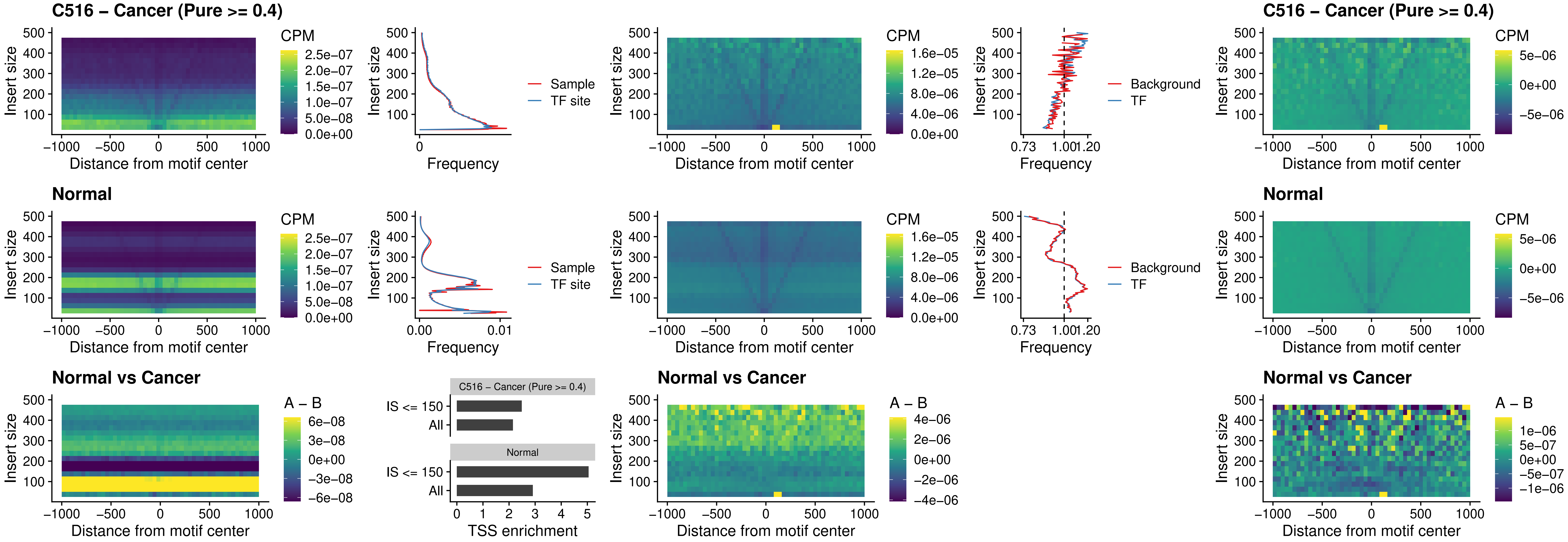

#### C516 – Cancer (Pure >= 0.4) – NKX24_uc_dTSS_nPEAK

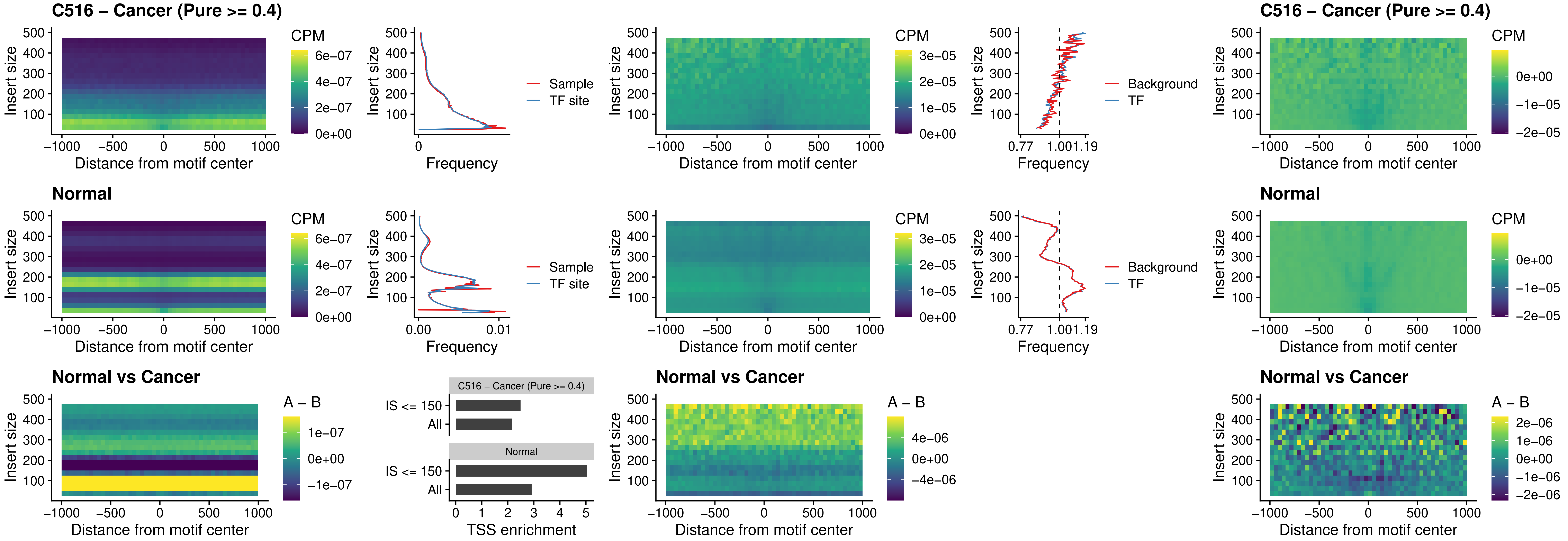

### C516 – Cancer (Pure >= 0.4) – DLX3_uc_cTSS_nPEAK

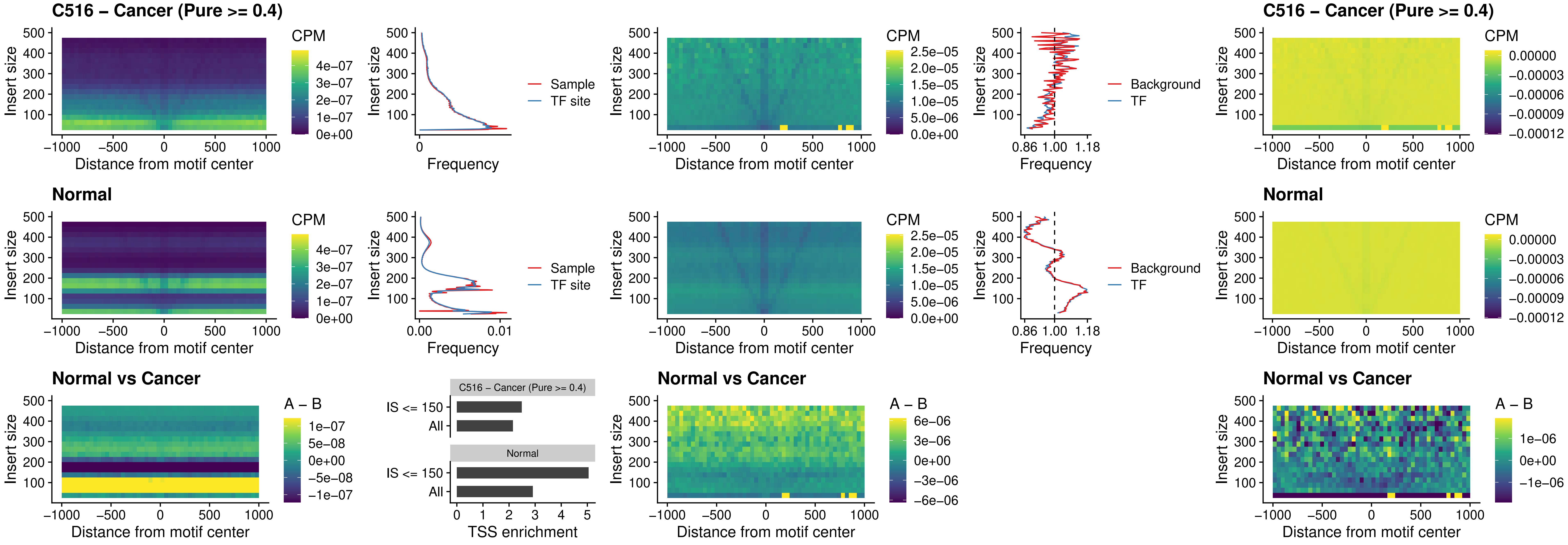

#### C516 – Cancer (Pure >= 0.4) – HOXD3_uc_cTSS_nPEAK

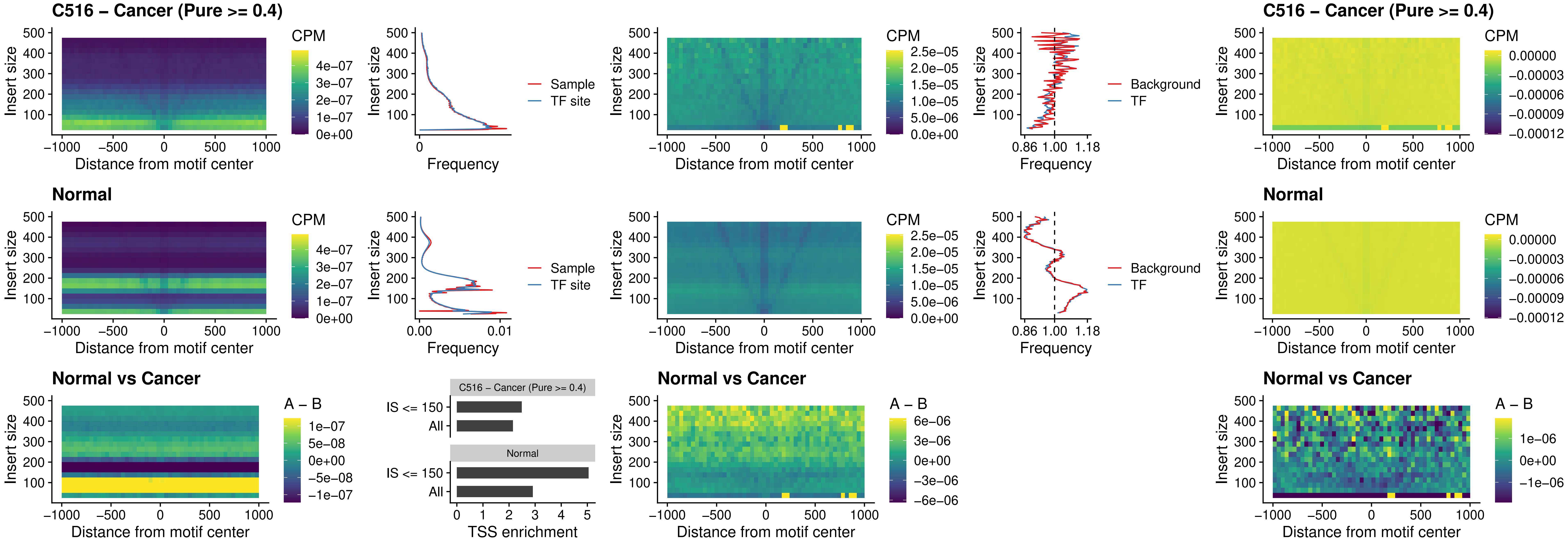

### C516 – Cancer (Pure >= 0.4) – ISL1_uc_dTSS_oPEAK

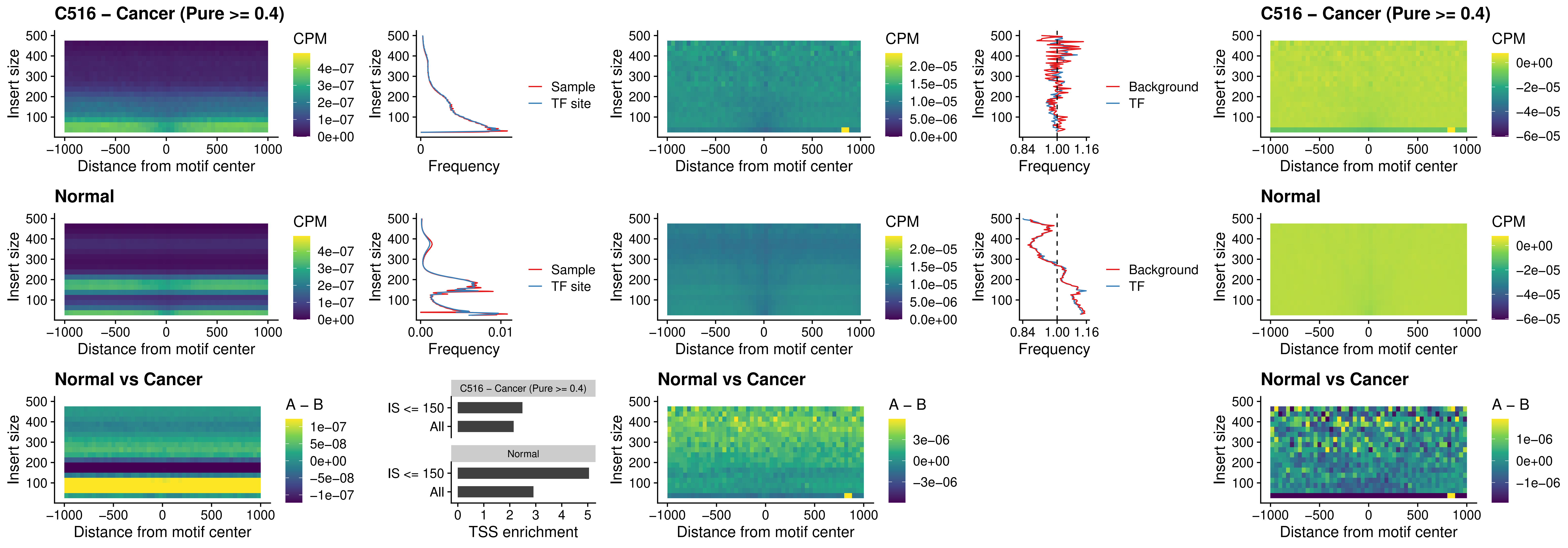

#### C516 – Cancer (Pure >= 0.4) – HOXB4_uc_dTSS_oPEAK

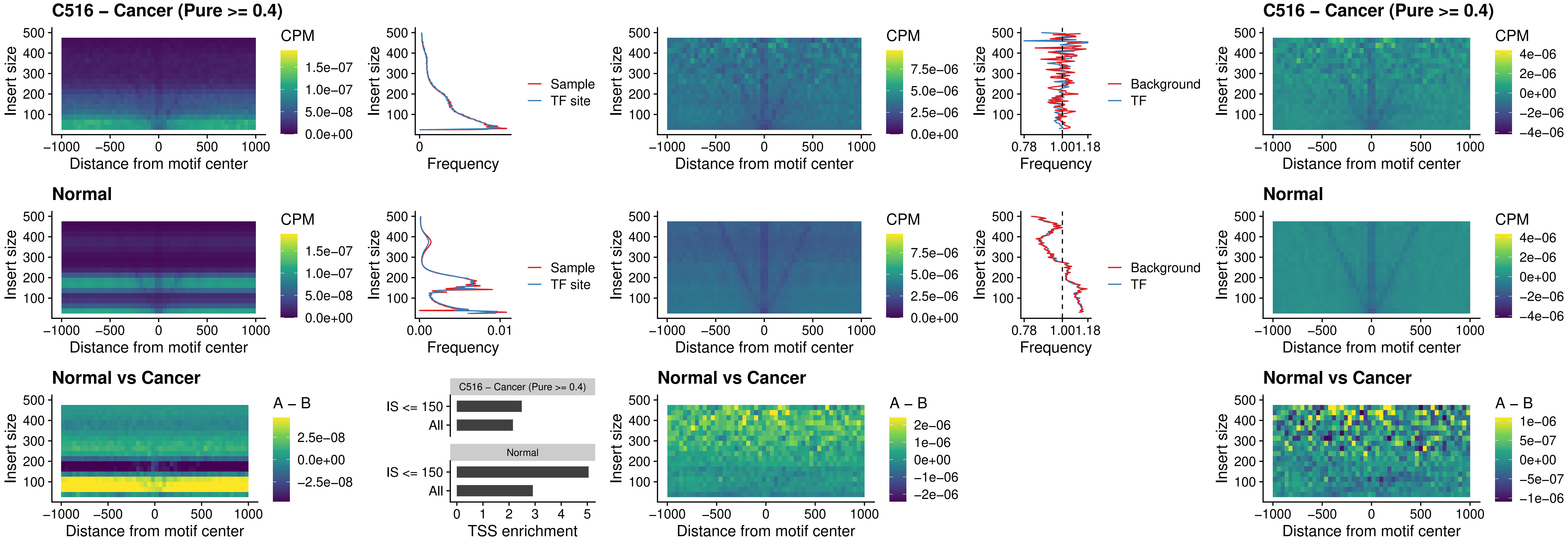

#### C516 – Cancer (Pure >= 0.4) – VAX1_uc_dTSS_oPEAK

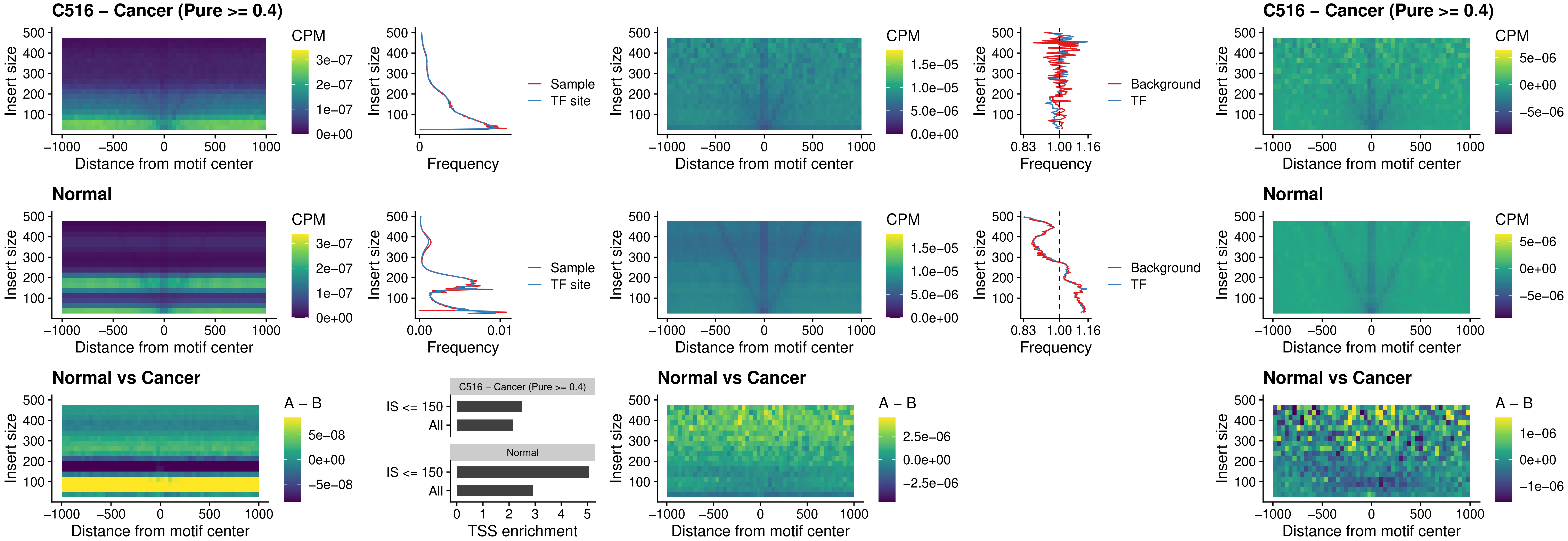

#### C516 – Cancer (Pure >= 0.4) – DLX3_uc_dTSS_oPEAK

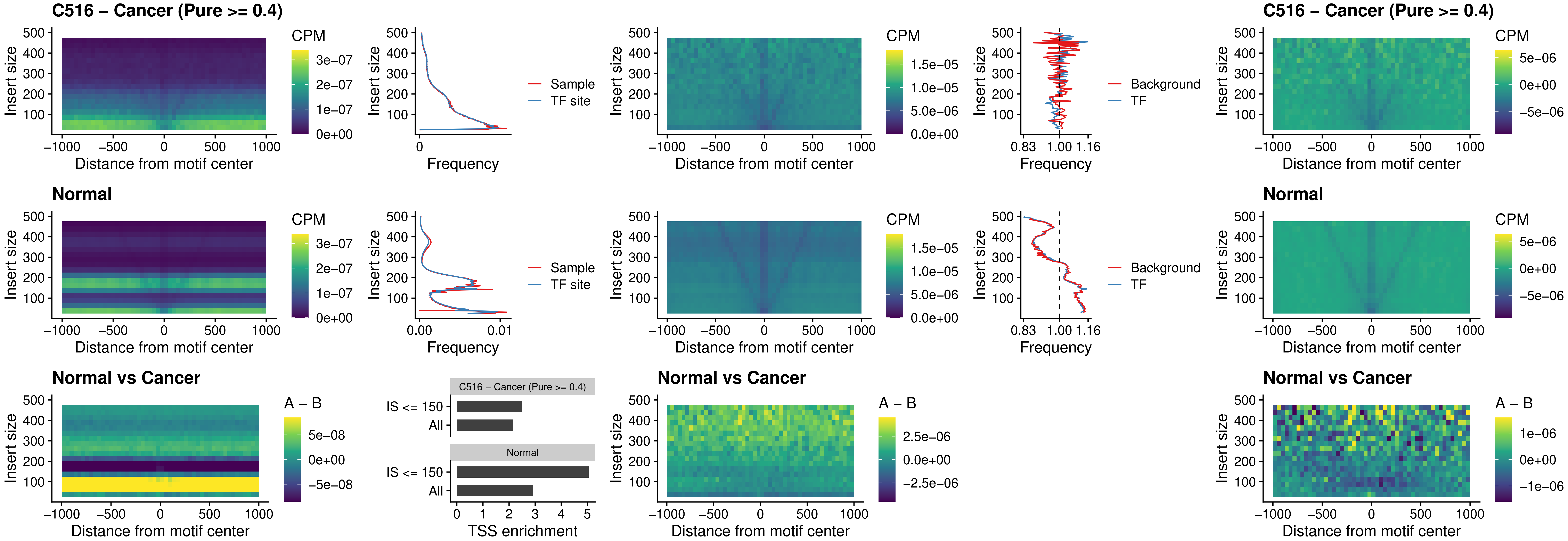

#### C516 – Cancer (Pure >= 0.4) – HOXD3_uc_dTSS_oPEAK

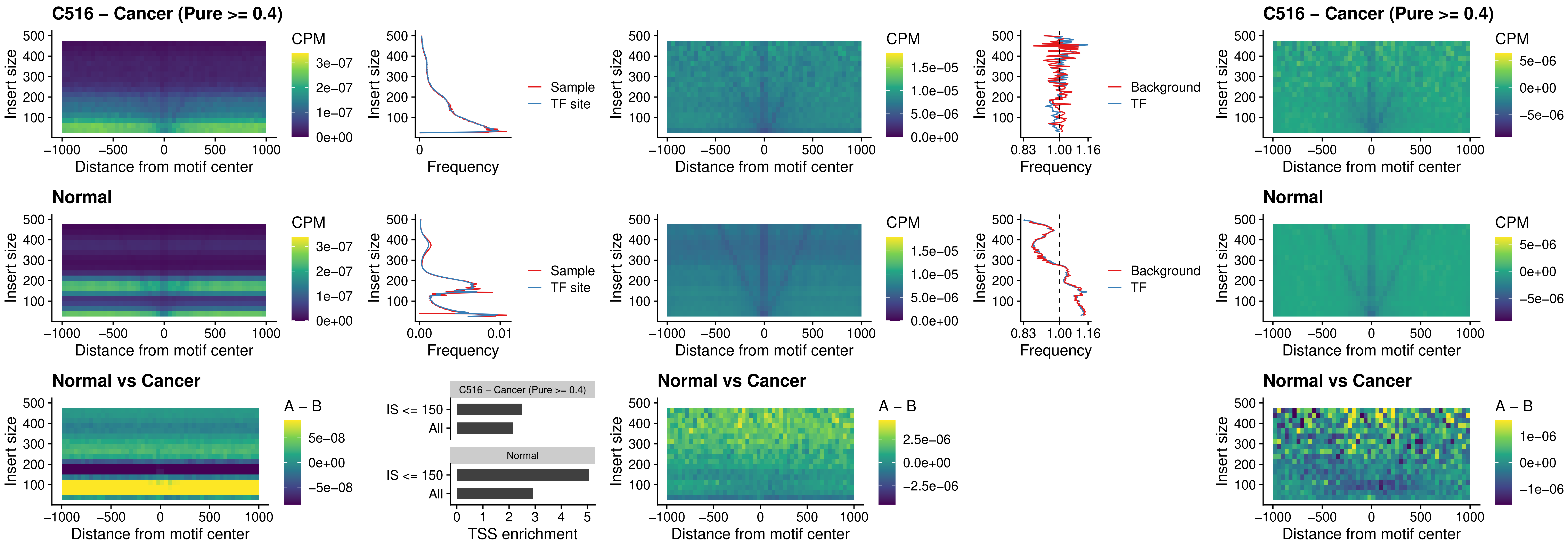

#### C516 – Cancer (Pure >= 0.4) – BBX_uc_dTSS_nPEAK

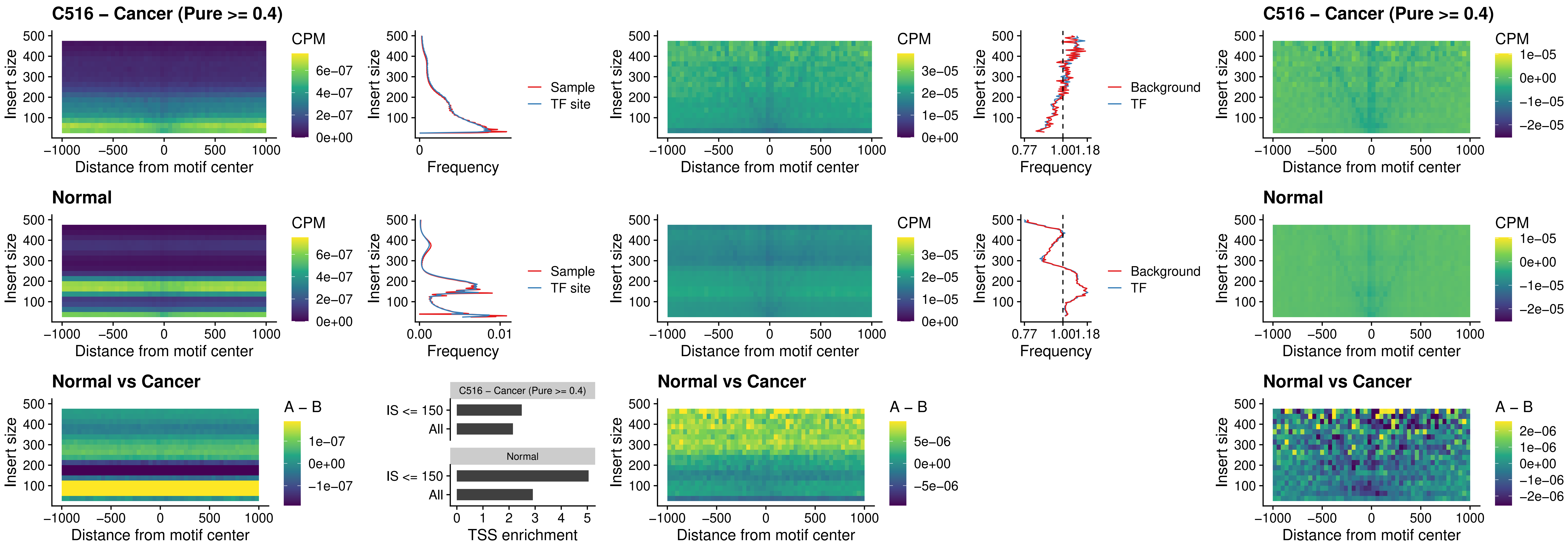

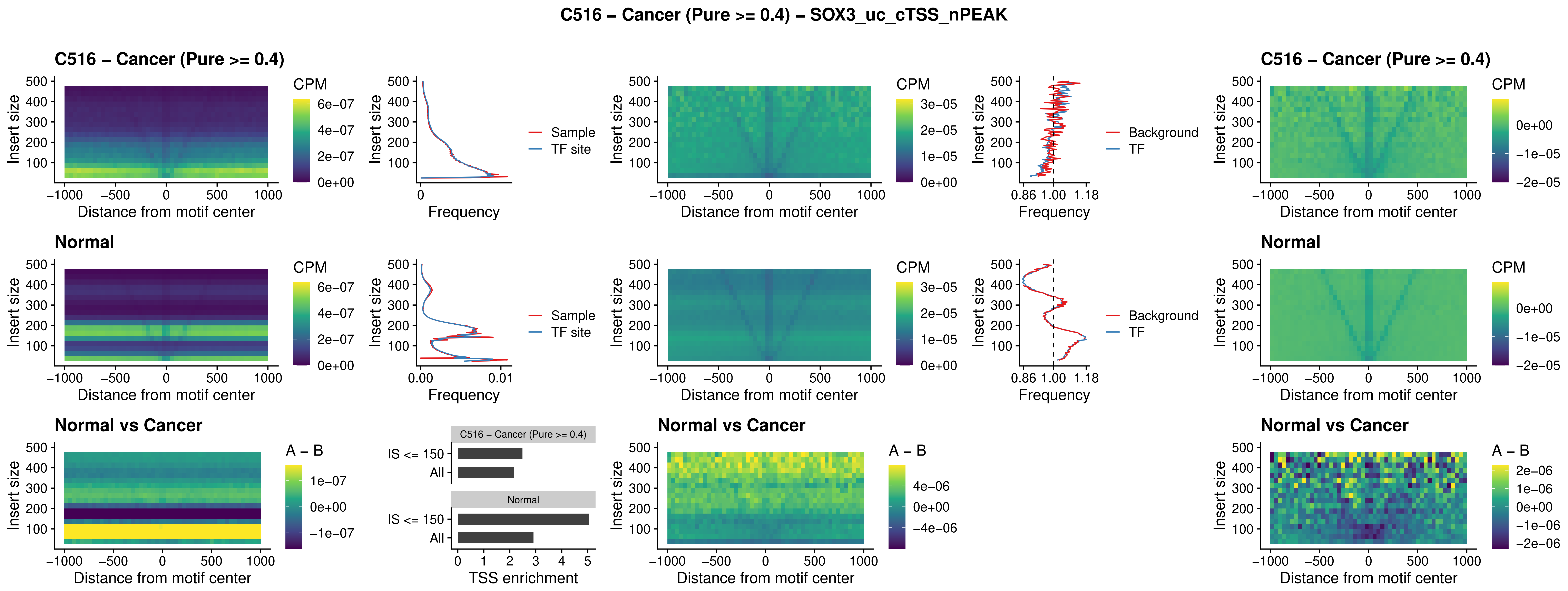
C516 – Cancer (Pure >= 0.4) – SOX3_uc_cTSS_nPEAK
C516 – Cancer (Pure >= 0.4)
Normal
Normal vs Cancer
Insert size
Distance from motif center
Frequency
CPM
A – B
Sample
TF site
Background
TF
C516 – Cancer (Pure >= 0.4)
Normal
IS <= 150
All
TSS enrichment

### C516 – Cancer (Pure >= 0.4) – TBPL2_uc_cTSS_nPEAK

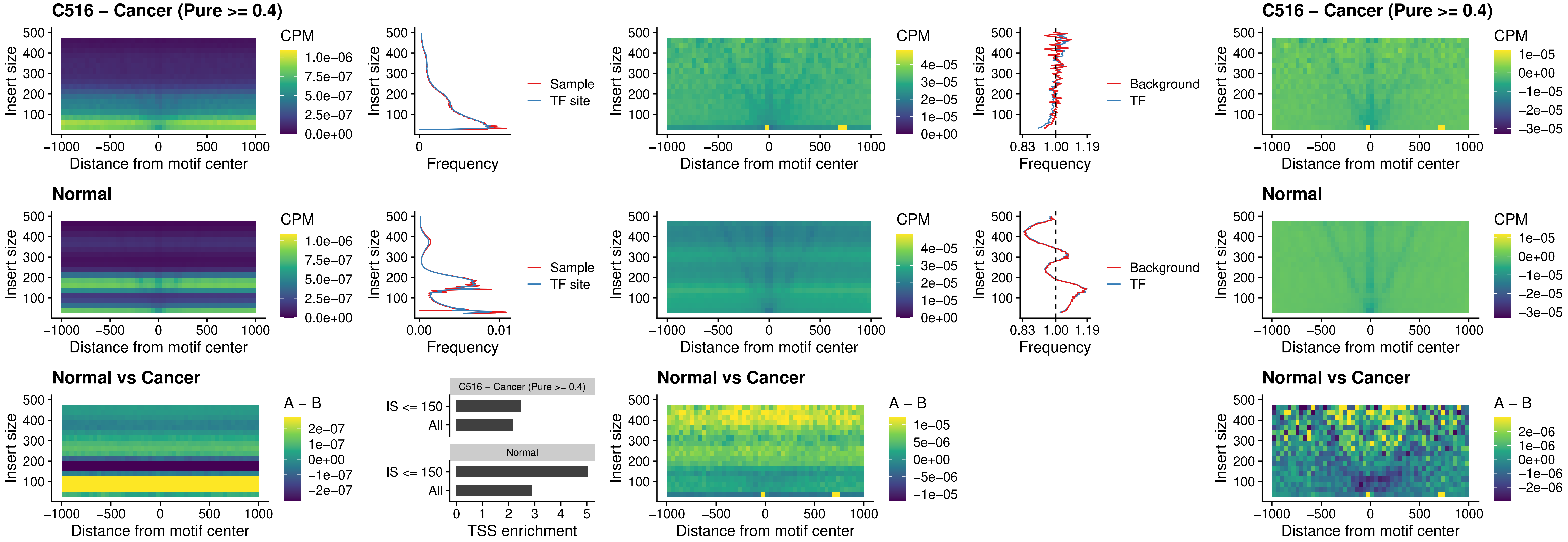

### C516 – Cancer (Pure >= 0.4) – HOXA4_uc_cTSS_oPEAK

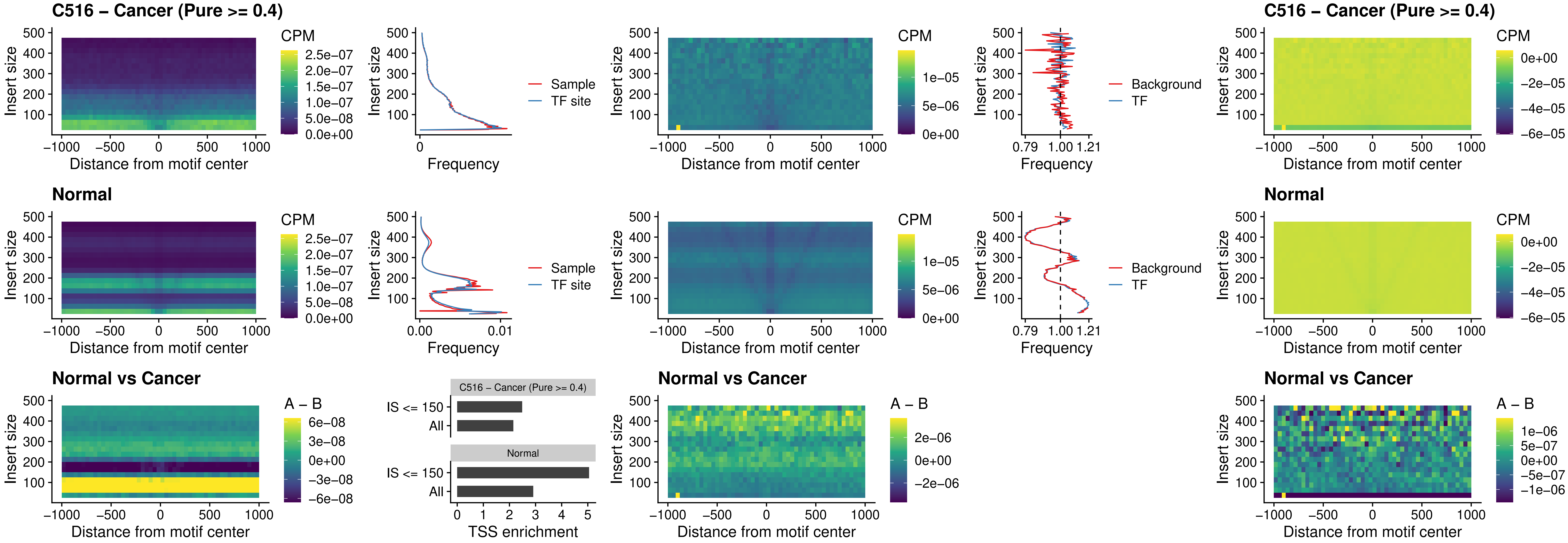

#### C516 – Cancer (Pure >= 0.4) – PDX1_uc_dTSS_oPEAK

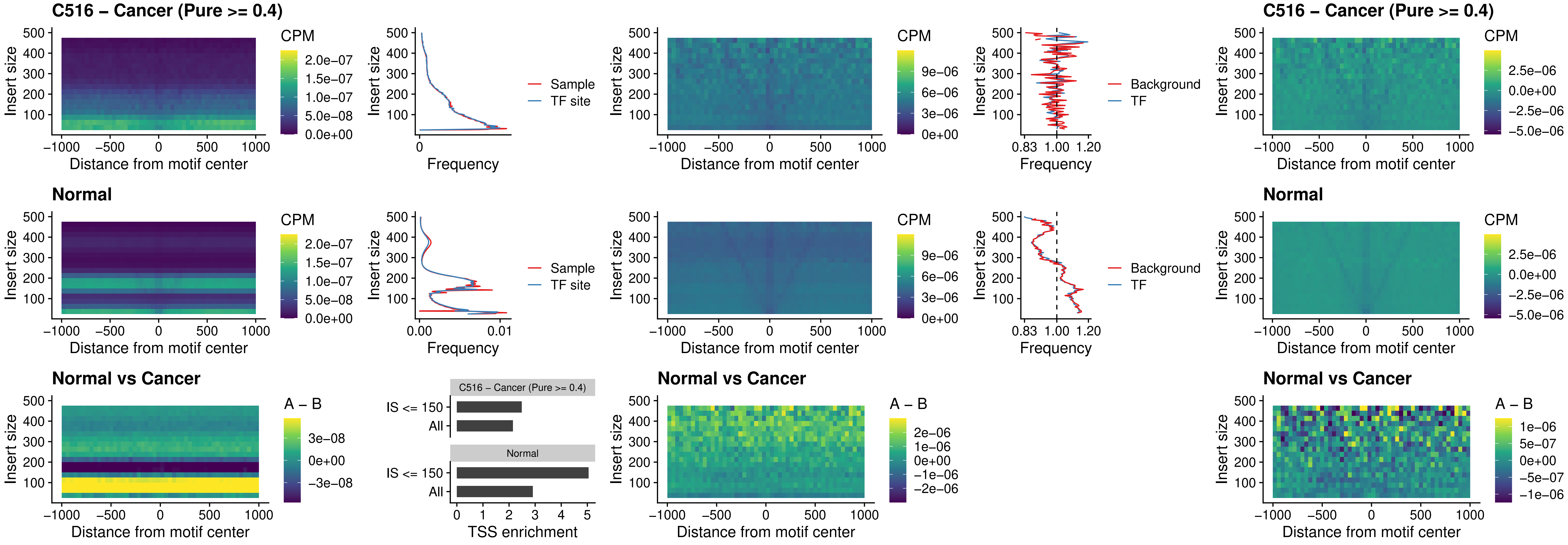

#### C516 – Cancer (Pure >= 0.4) – LHX4_uc_dTSS_oPEAK

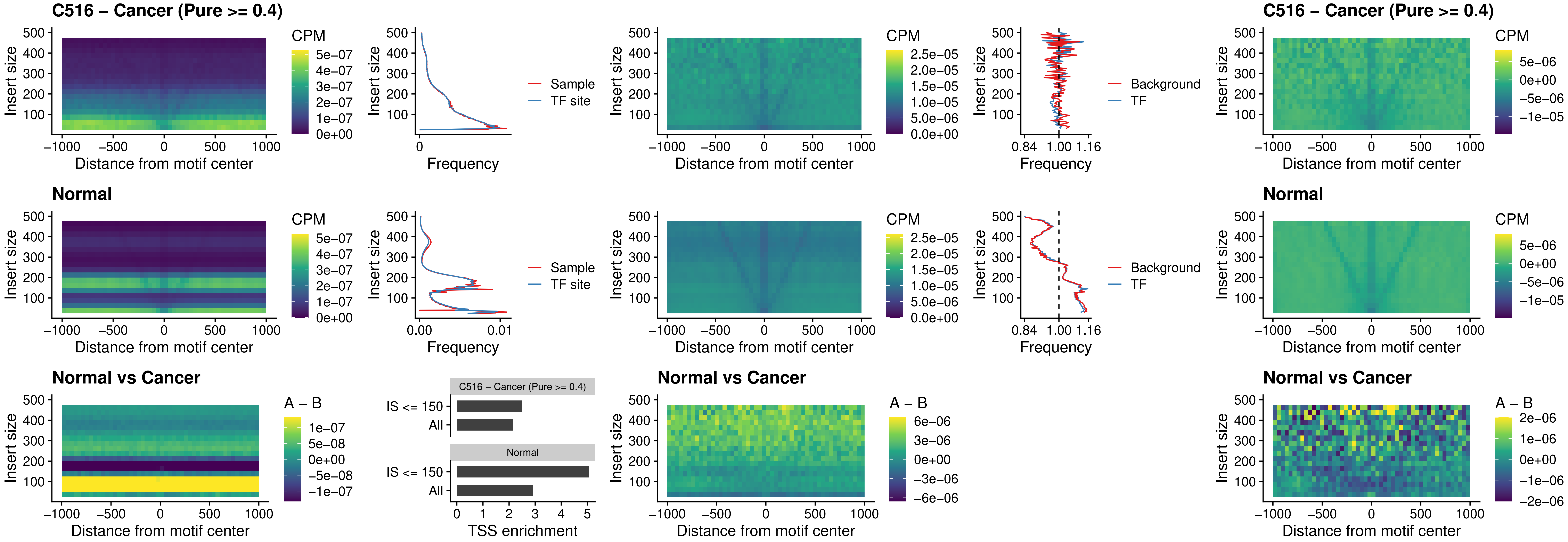

#### C516 – Cancer (Pure >= 0.4) – HOXA4_uc_dTSS_oPEAK

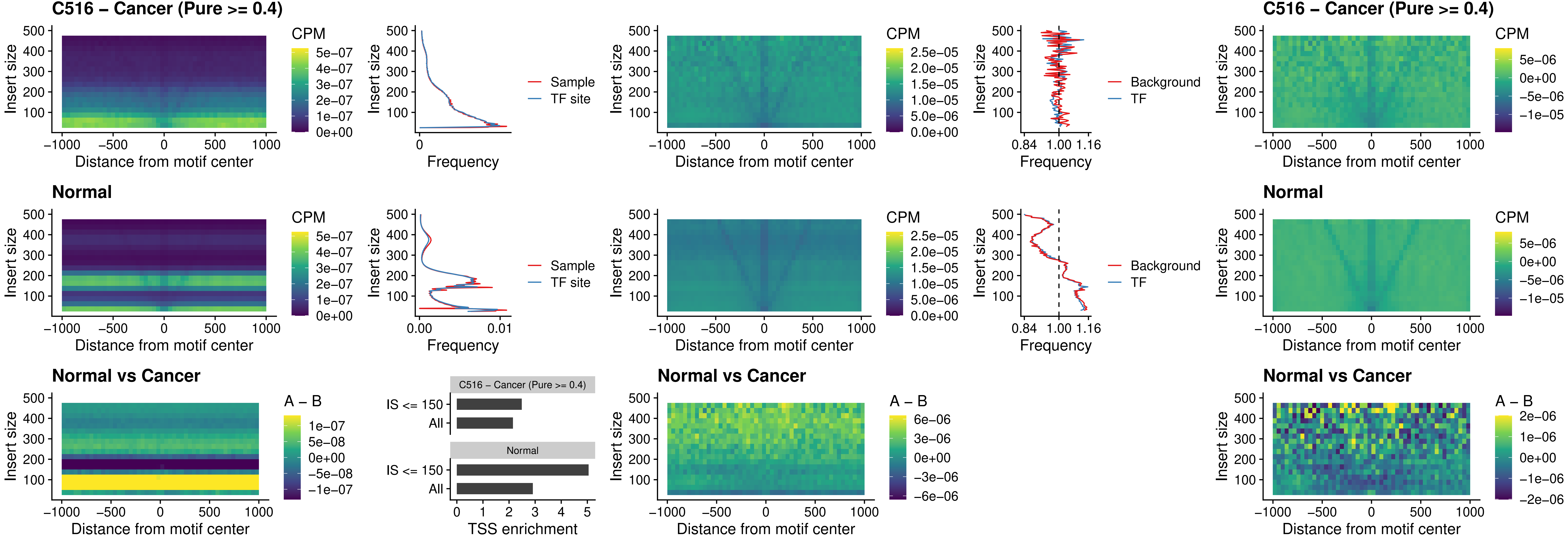

### C516 – Cancer (Pure >= 0.4) – HLX_uc_dTSS_oPEAK

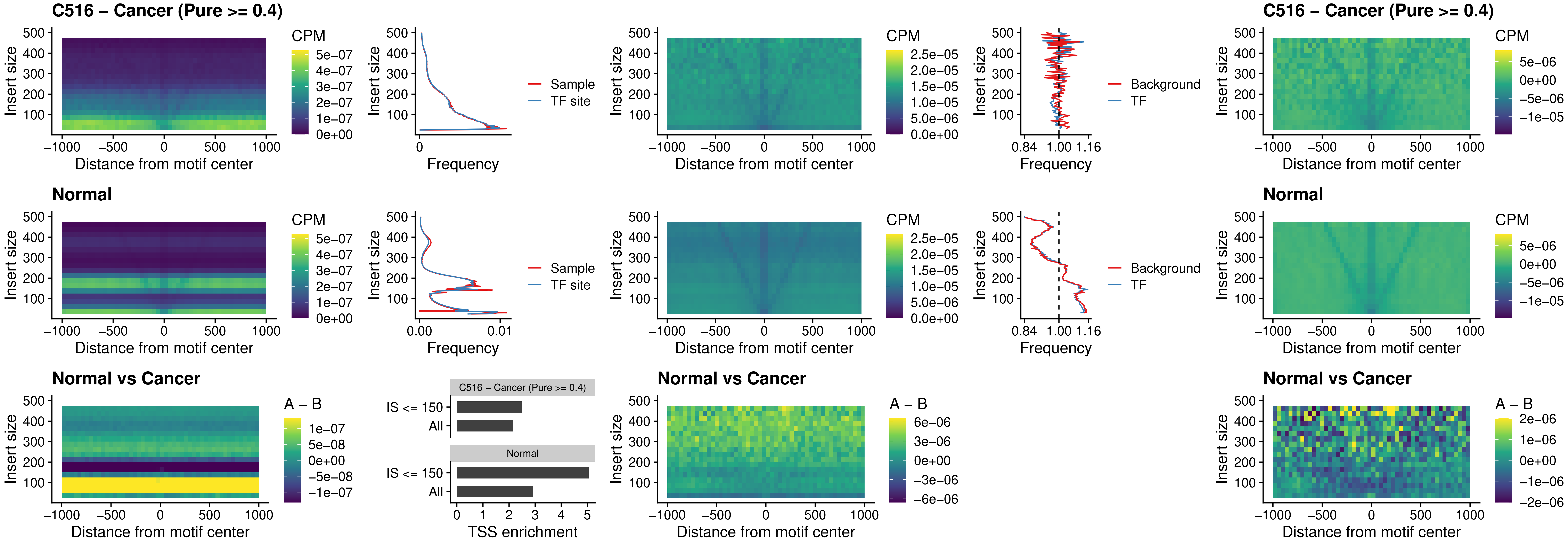

#### C516 – Cancer (Pure >= 0.4) – AC0125311_uc_dTSS_oPEAK

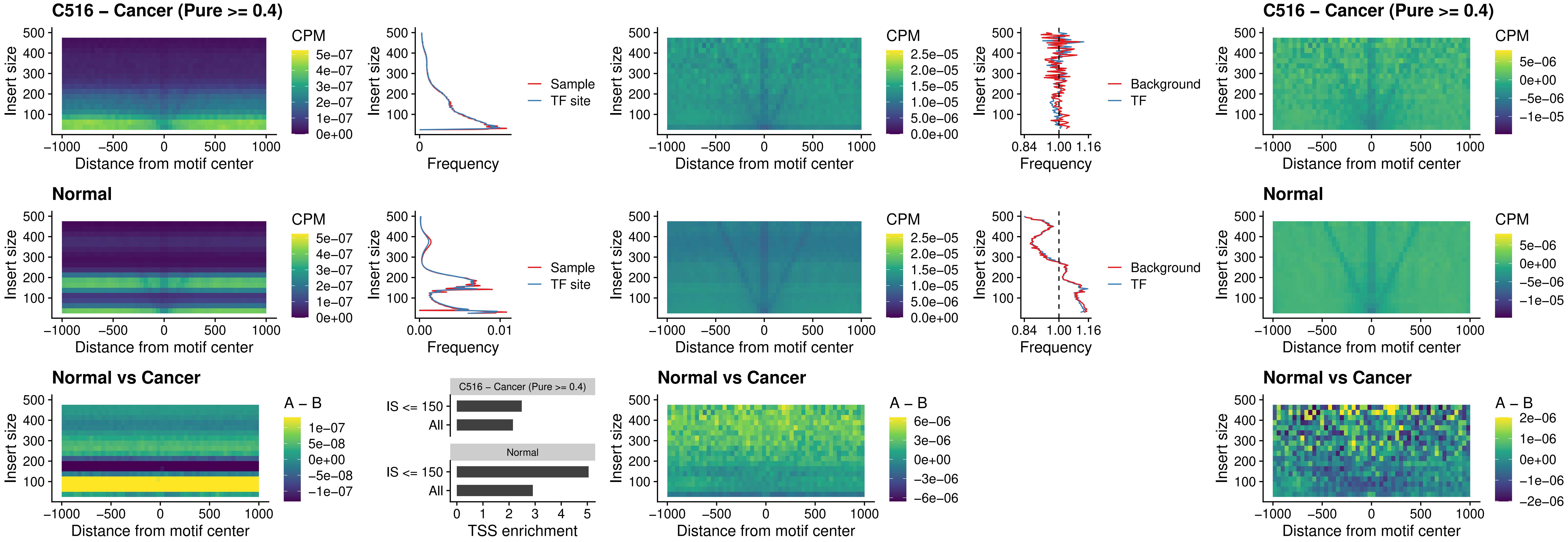

#### C516 – Cancer (Pure >= 0.4) – DBX2_uc_dTSS_oPEAK

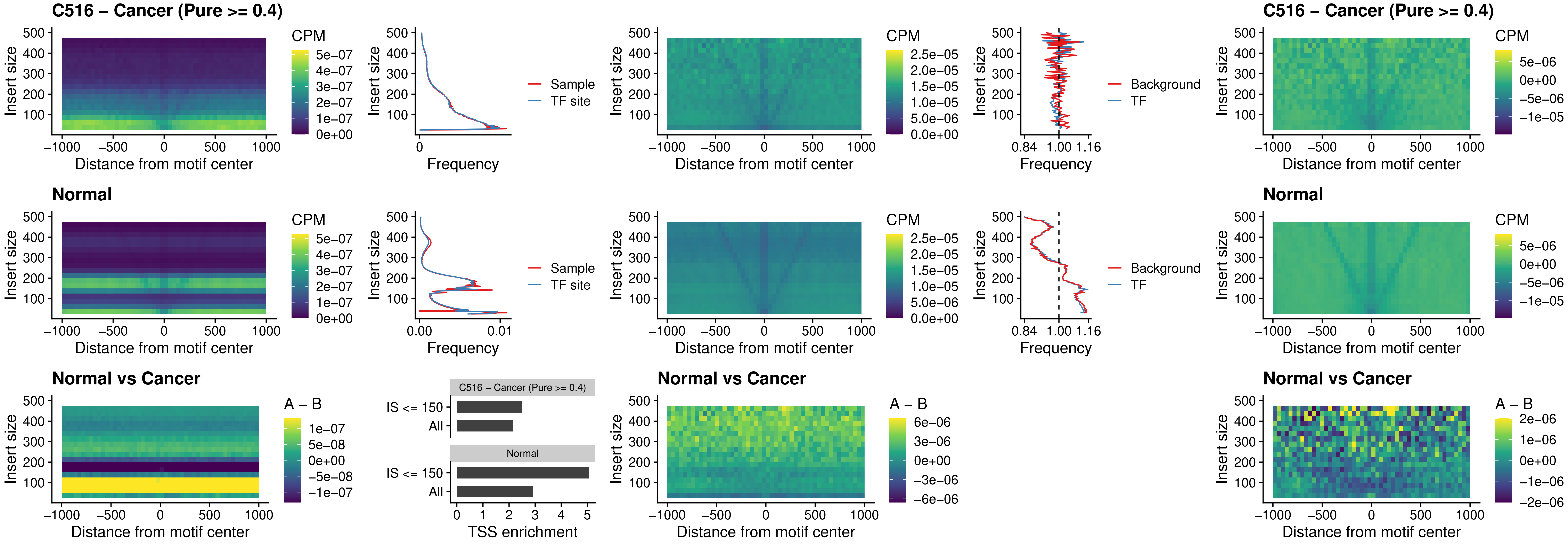

#### C516 – Cancer (Pure >= 0.4) – HOXA2_uc_dTSS_nPEAK

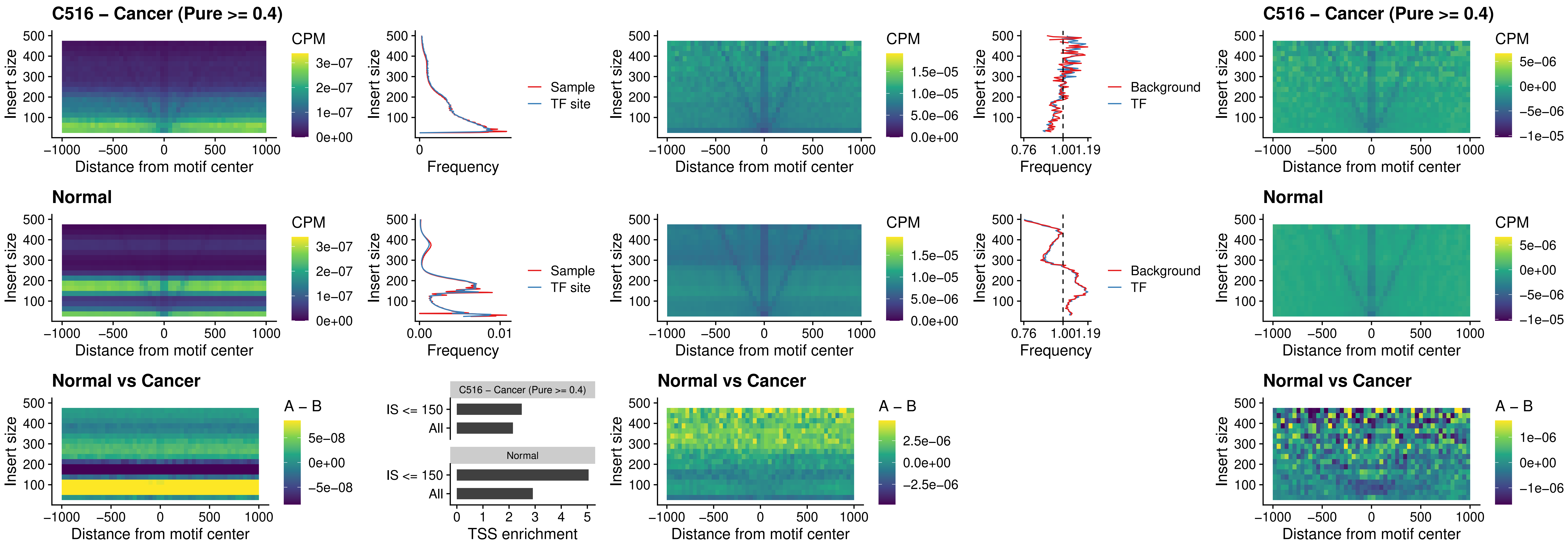

### C516 – Cancer (Pure >= 0.4) – SHOX_uc_dTSS_nPEAK

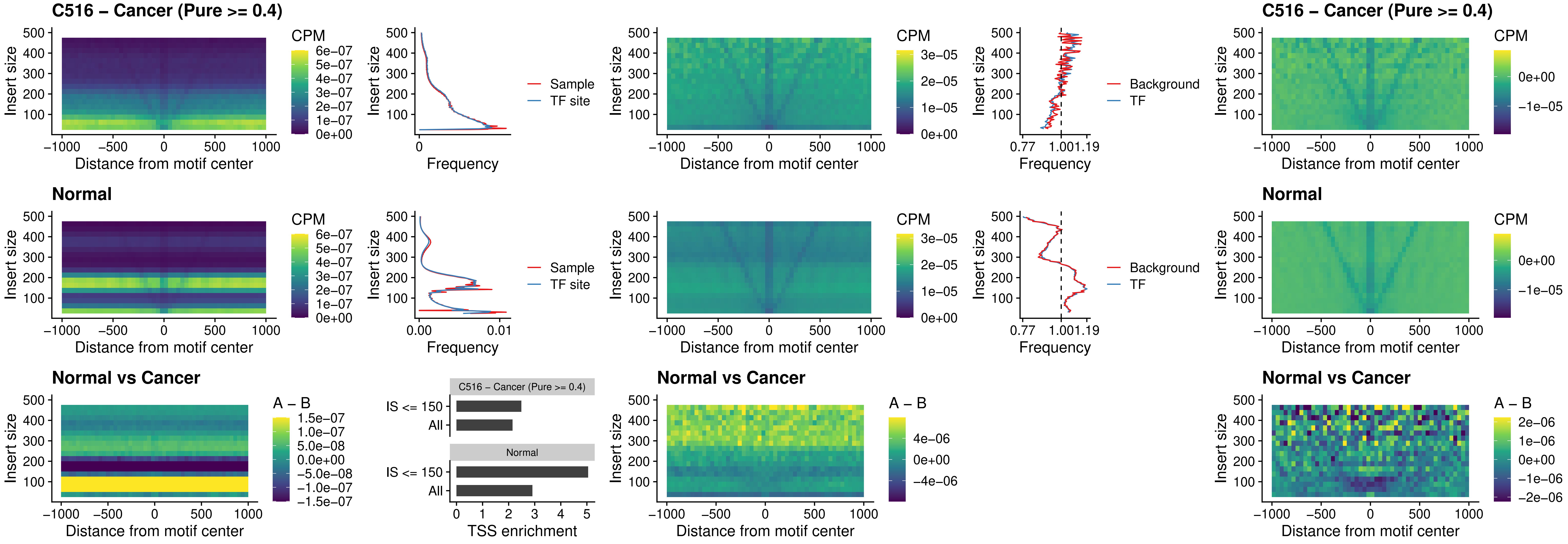

#### C516 – Cancer (Pure >= 0.4) – PRRX1_uc_dTSS_nPEAK

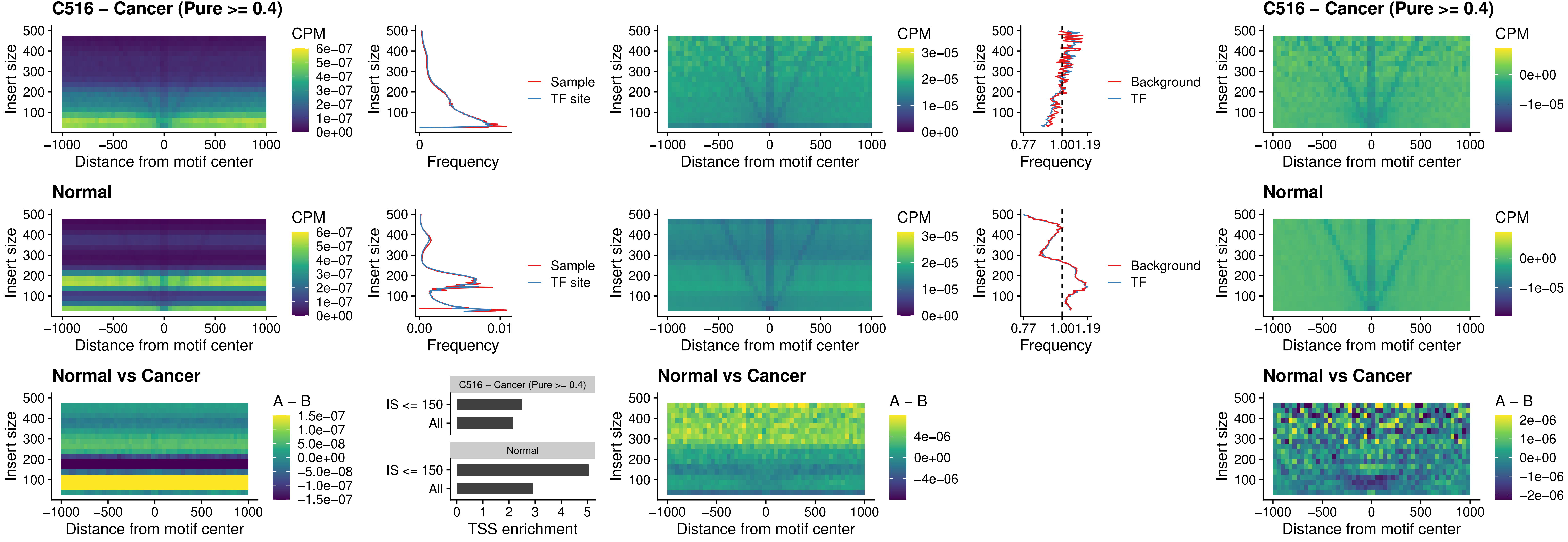

#### C516 – Cancer (Pure >= 0.4) – LMX1A_uc_dTSS_nPEAK

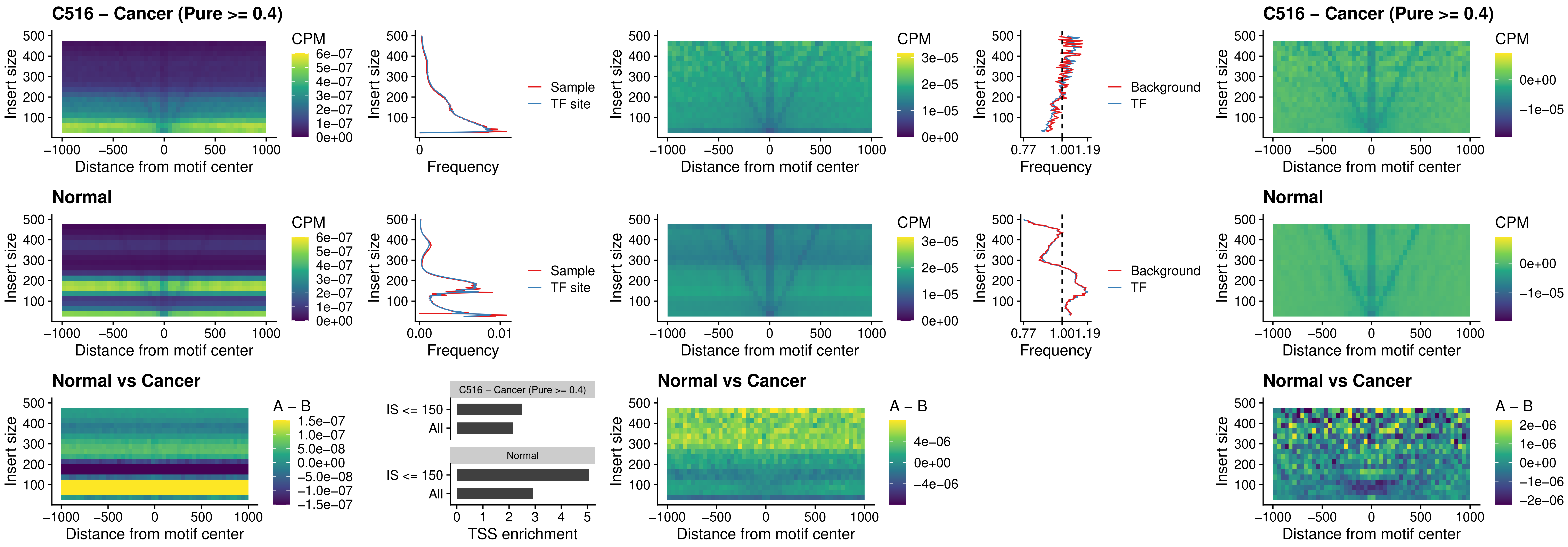

#### C516 – Cancer (Pure >= 0.4) – LMX1B_uc_dTSS_nPEAK

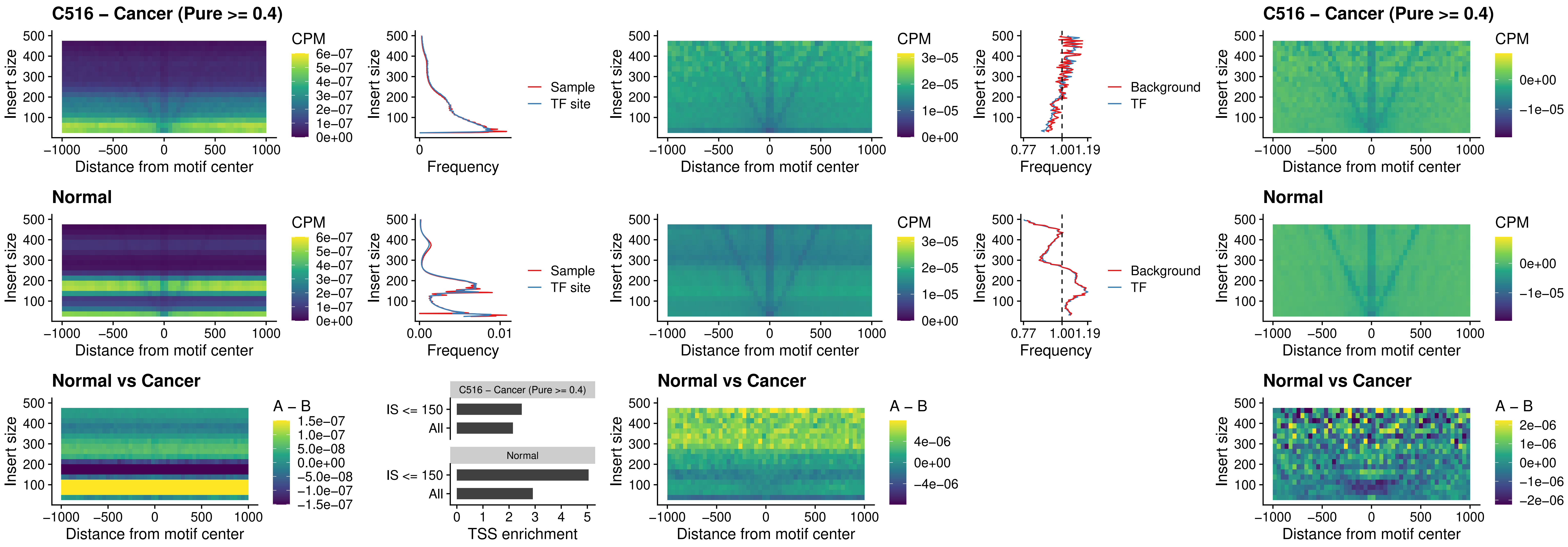

#### C516 – Cancer (Pure >= 0.4) – DLX5_uc_dTSS_oPEAK

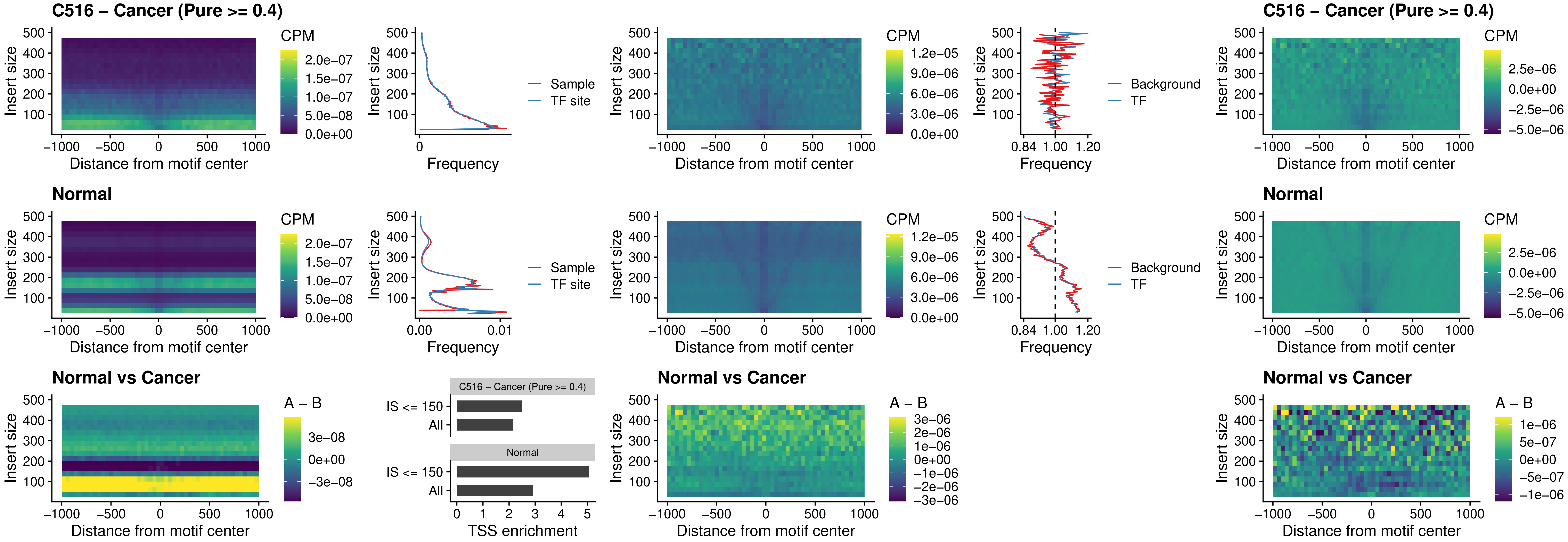

#### C516 – Cancer (Pure >= 0.4) – ARID3C_uc_dTSS_nPEAK

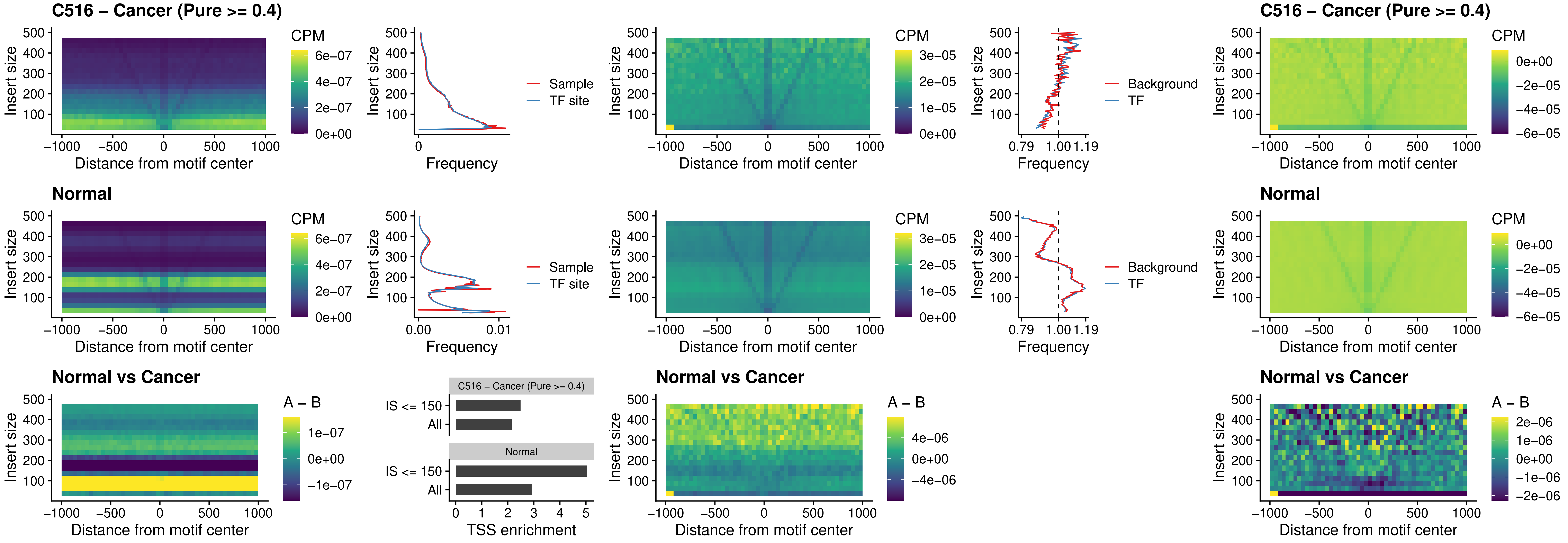

#### C516 – Cancer (Pure >= 0.4) – HOXD12_uc_dTSS_nPEAK

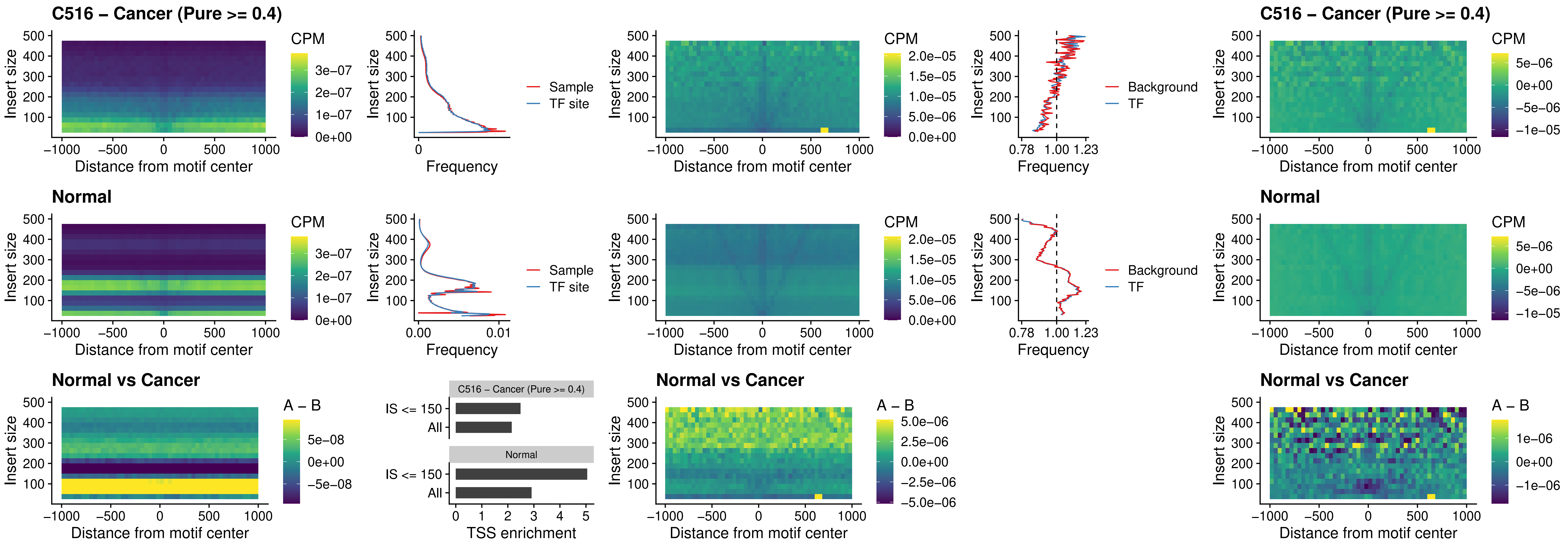

### C516 – Cancer (Pure >= 0.4) – CDX2_uc_dTSS_nPEAK

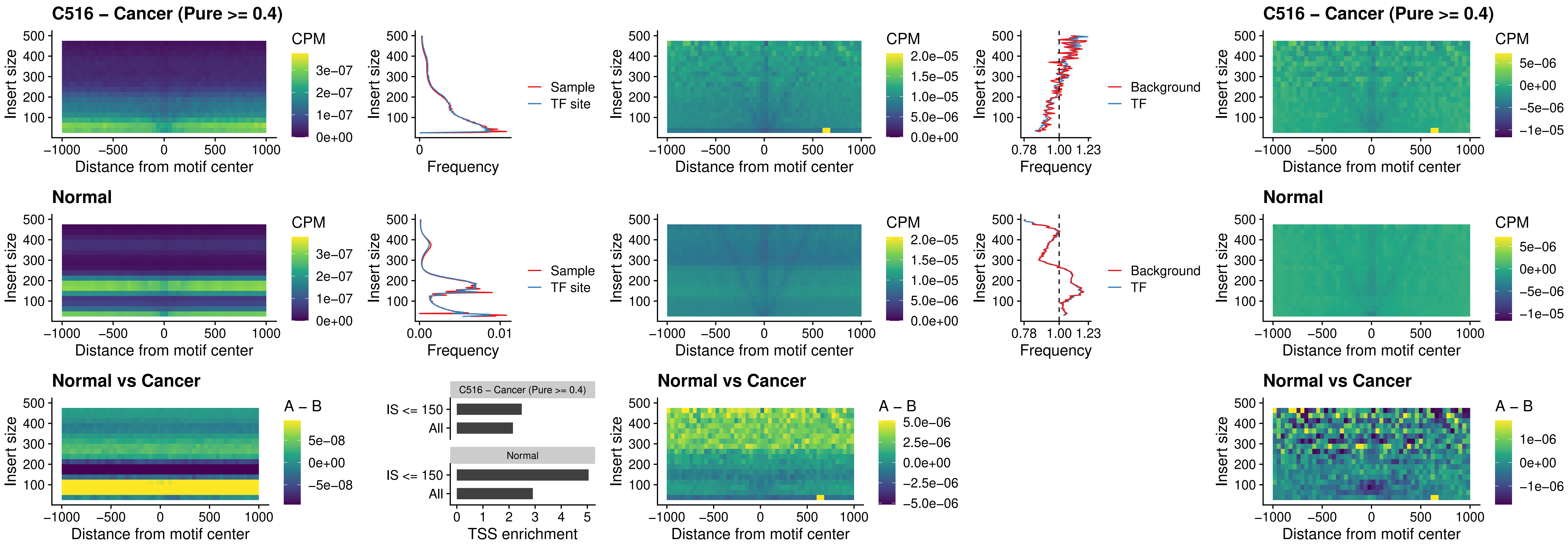

#### C516 – Cancer (Pure >= 0.4) – HOXC12_uc_dTSS_nPEAK

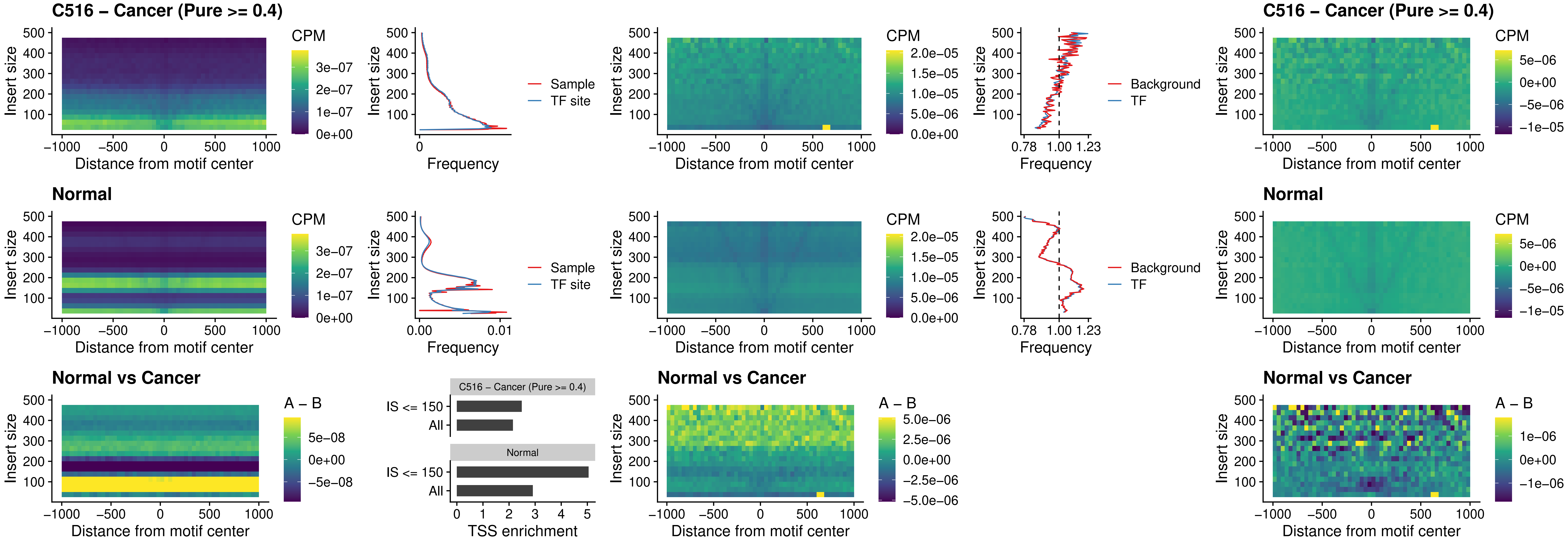

#### C516 – Cancer (Pure >= 0.4) – HHEX_uc_dTSS_oPEAK

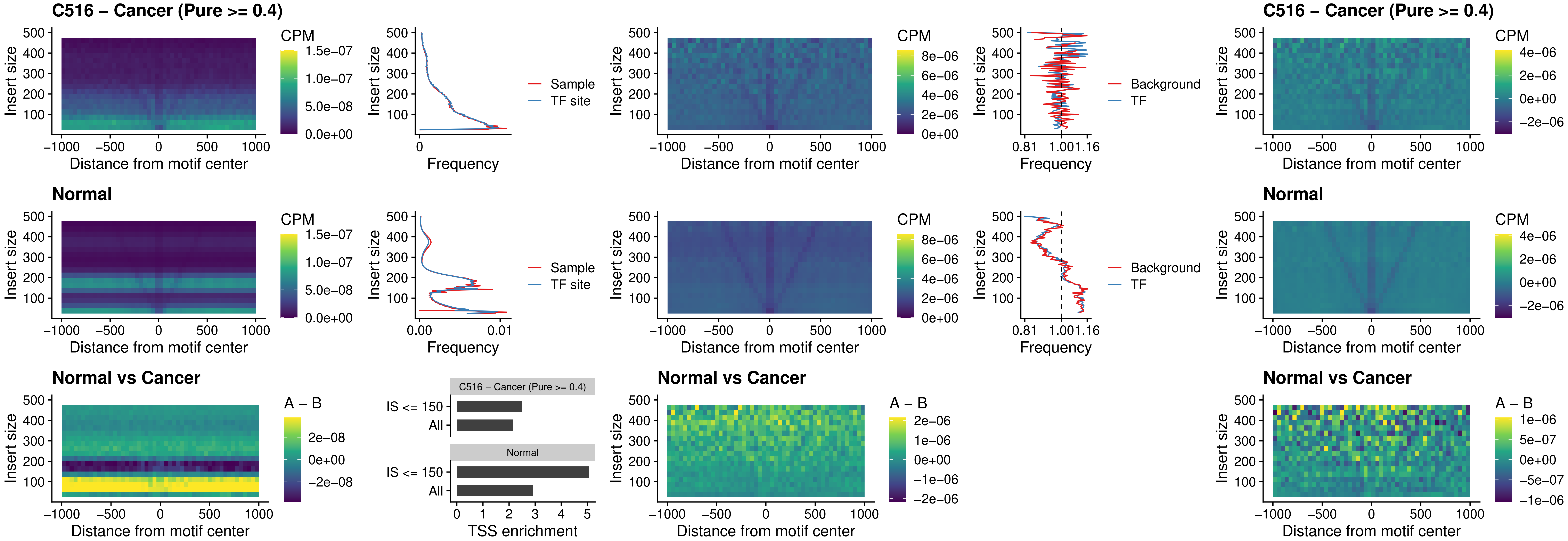

#### C516 – Cancer (Pure >= 0.4) – HOXA11_uc_dTSS_nPEAK

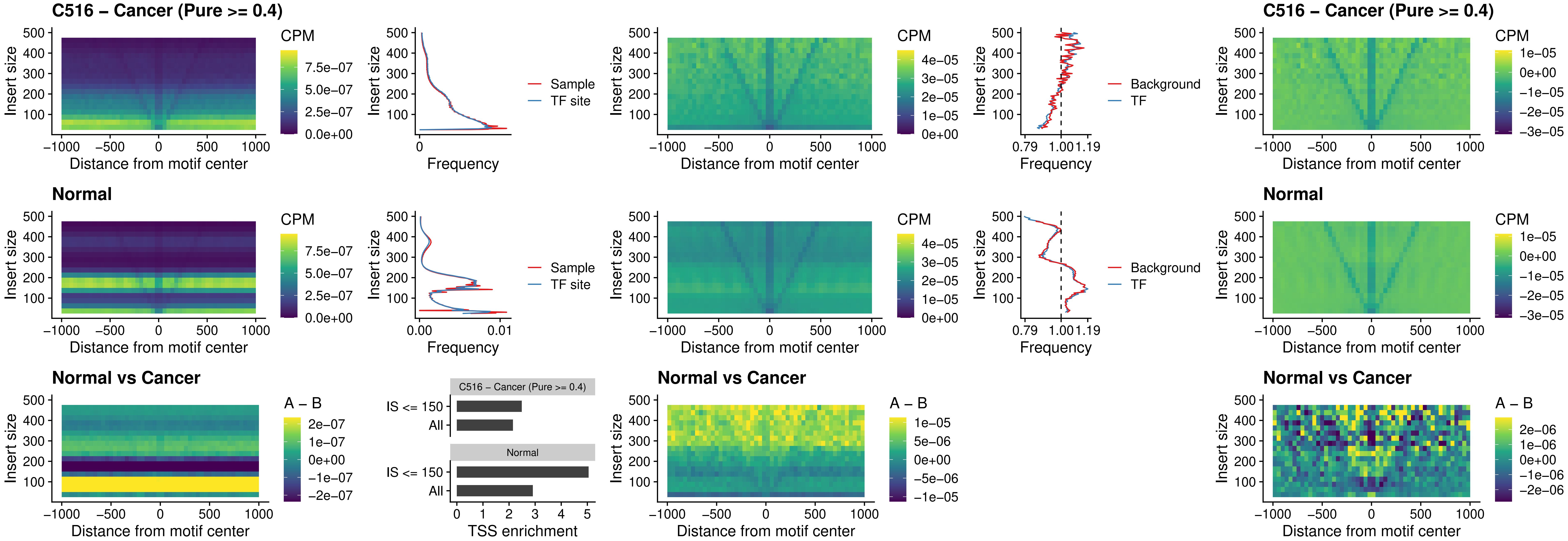

#### C516 – Cancer (Pure >= 0.4) – HOXA11_uc_cTSS_nPEAK

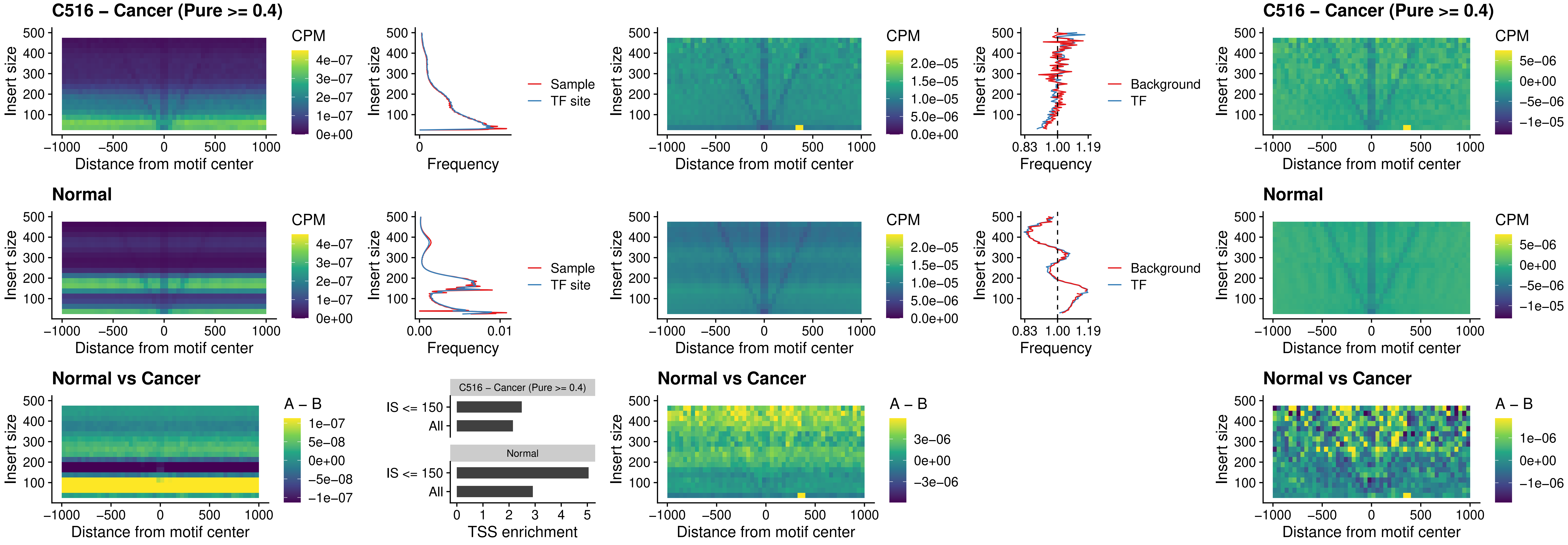

### C516 – Cancer (Pure >= 0.4) – ZFHX3_uc_dTSS_nPEAK

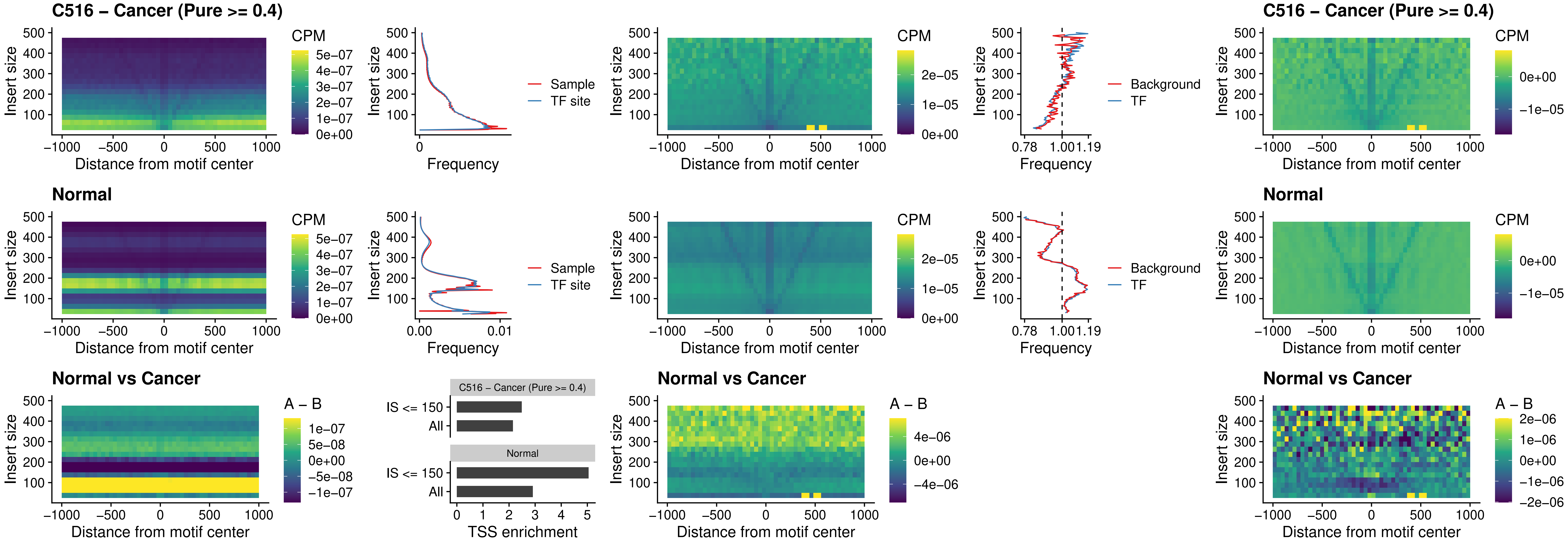

#### C516 – Cancer (Pure >= 0.4) – HOXB4_uc_dTSS_nPEAK

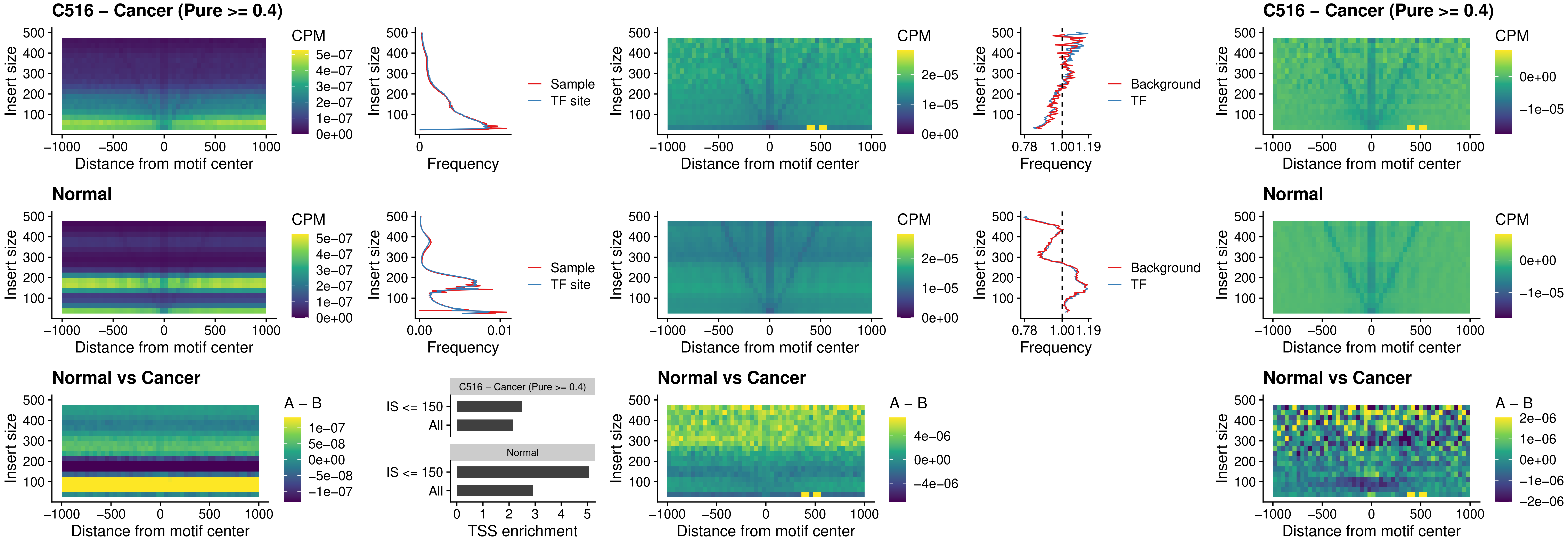

#### C516 – Cancer (Pure >= 0.4) – HOXC5_uc_dTSS_nPEAK

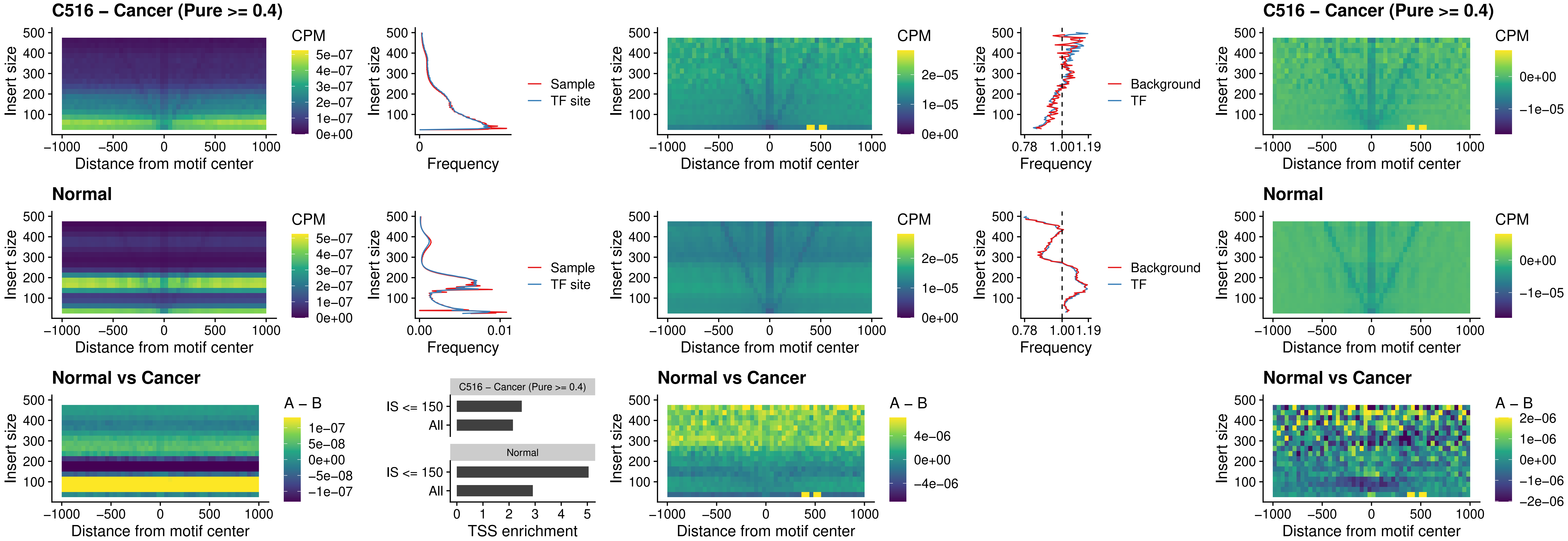

#### C516 – Cancer (Pure >= 0.4) – HOXD1_uc_dTSS_nPEAK

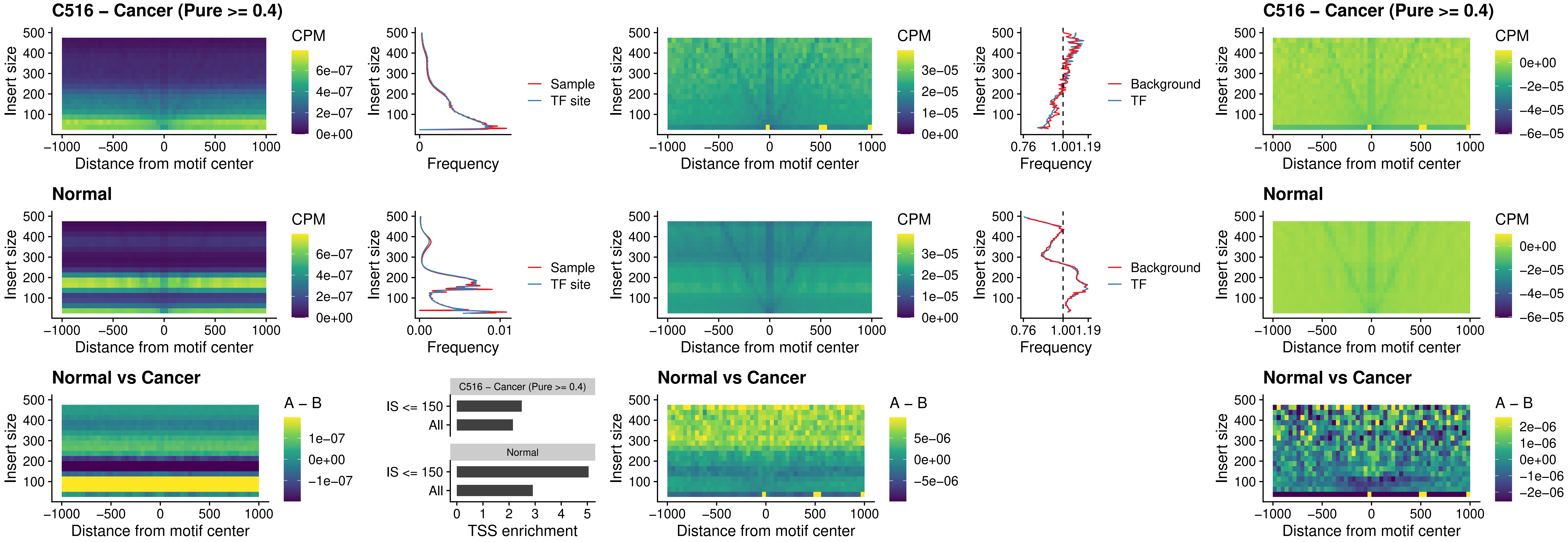

#### C516 – Cancer (Pure >= 0.4) – OTP_uc_dTSS_nPEAK

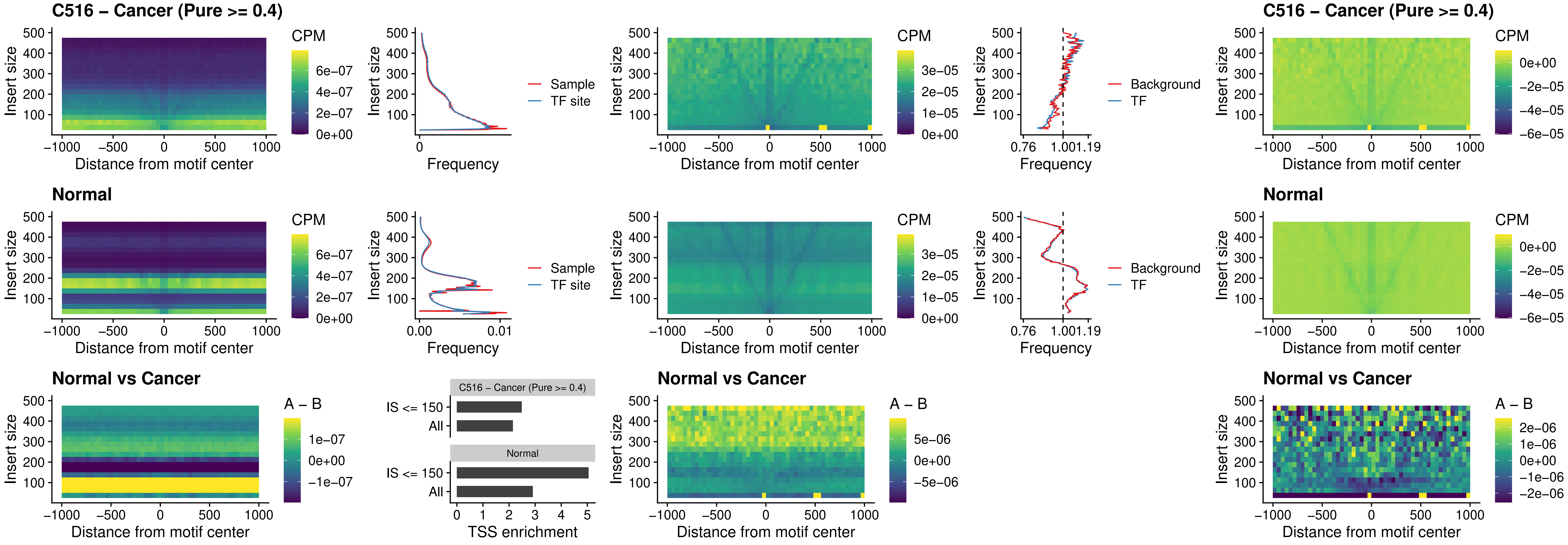

### C516 – Cancer (Pure >= 0.4) – FOXL1_uc_dTSS_oPEAK

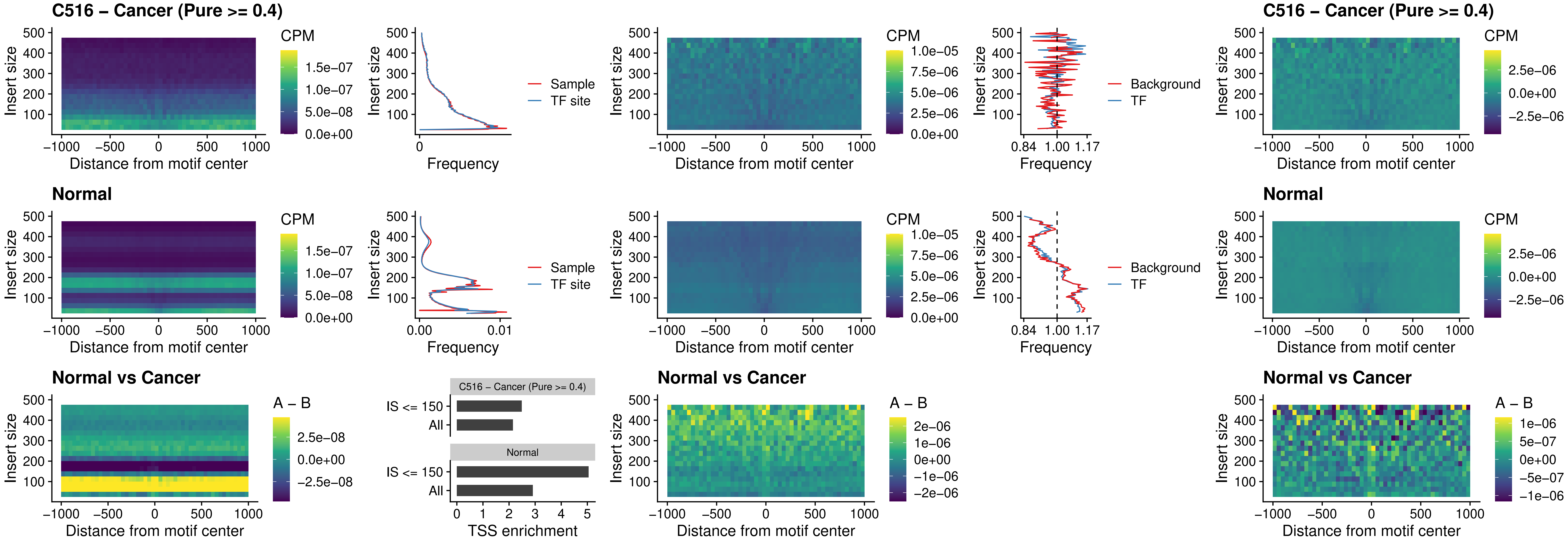

#### C516 – Cancer (Pure >= 0.4) – ZNF32_uc_dTSS_oPEAK

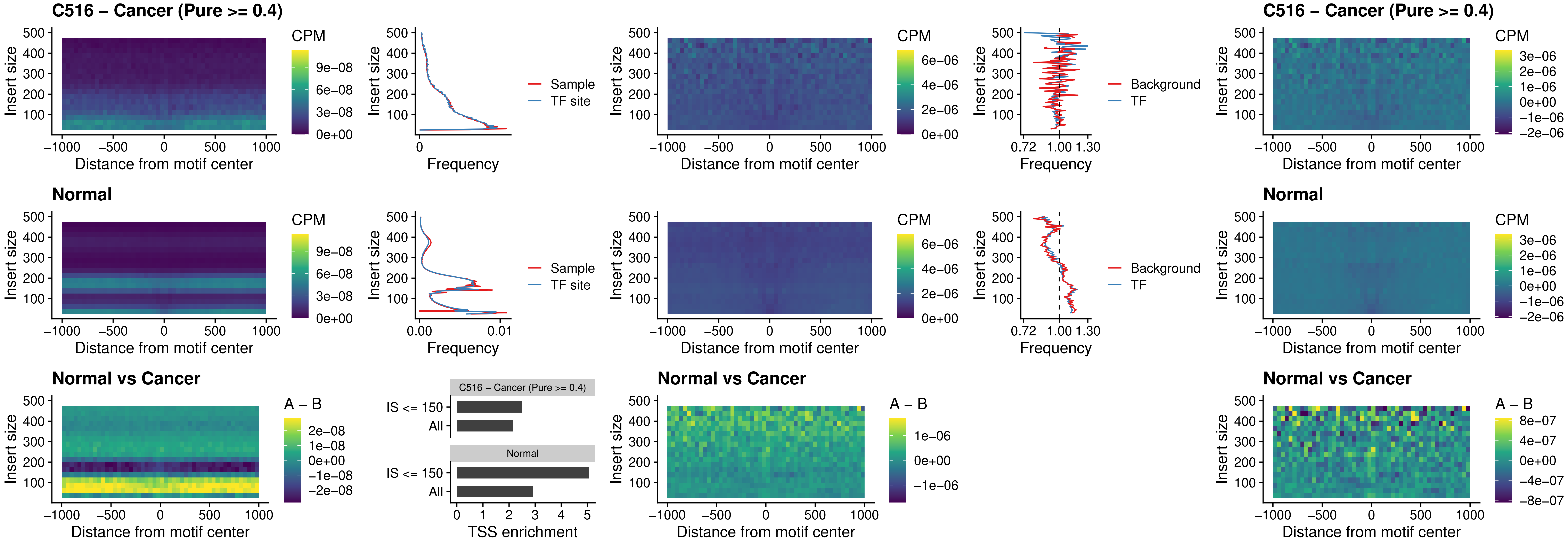

#### C516 – Cancer (Pure >= 0.4) – FOXL1_uc_dTSS_nPEAK

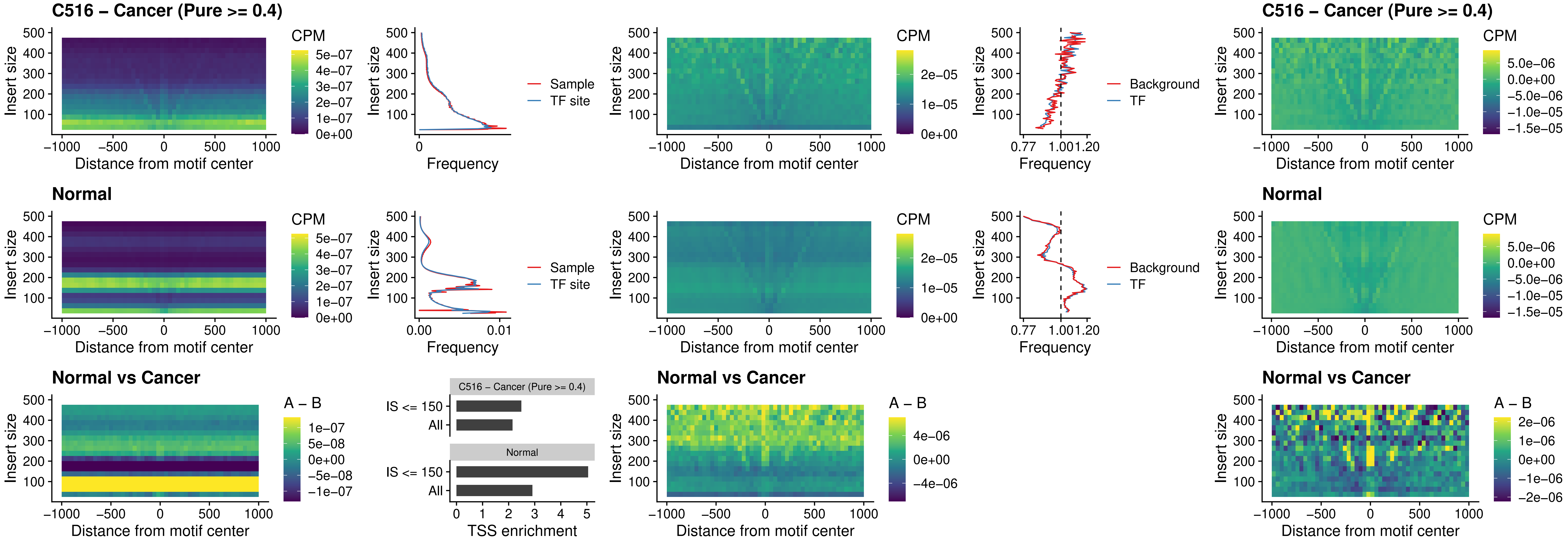

#### C516 – Cancer (Pure >= 0.4) – ZNF32_uc_dTSS_nPEAK

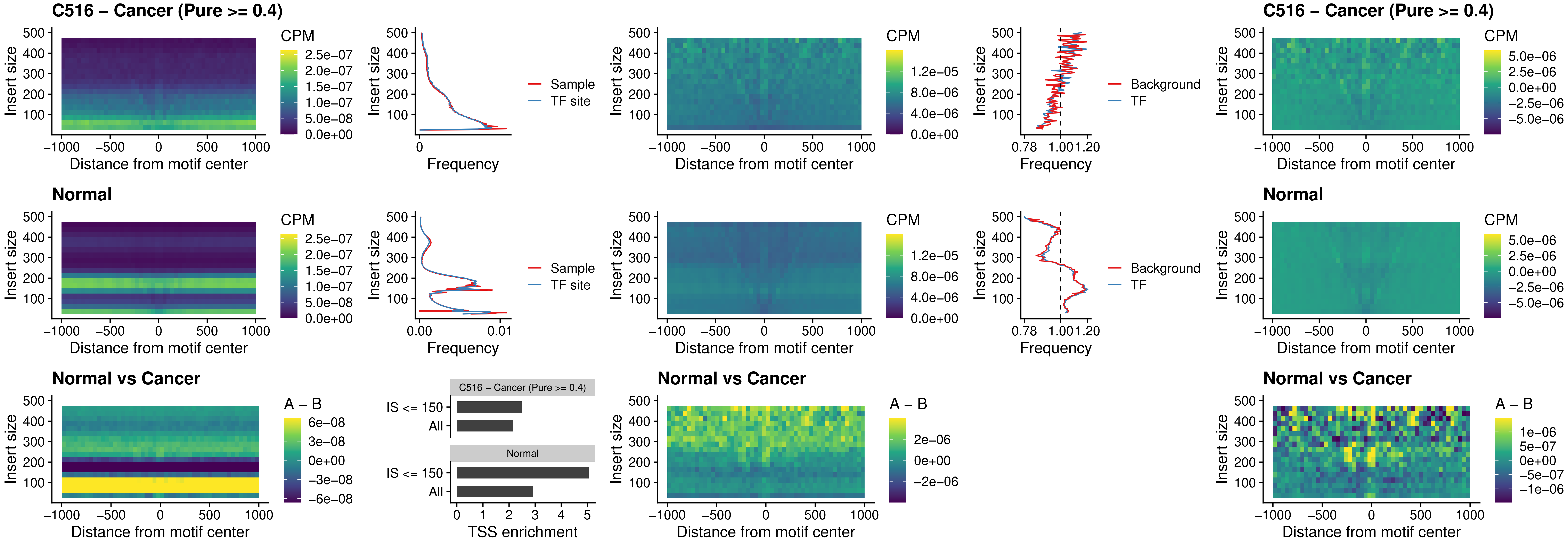

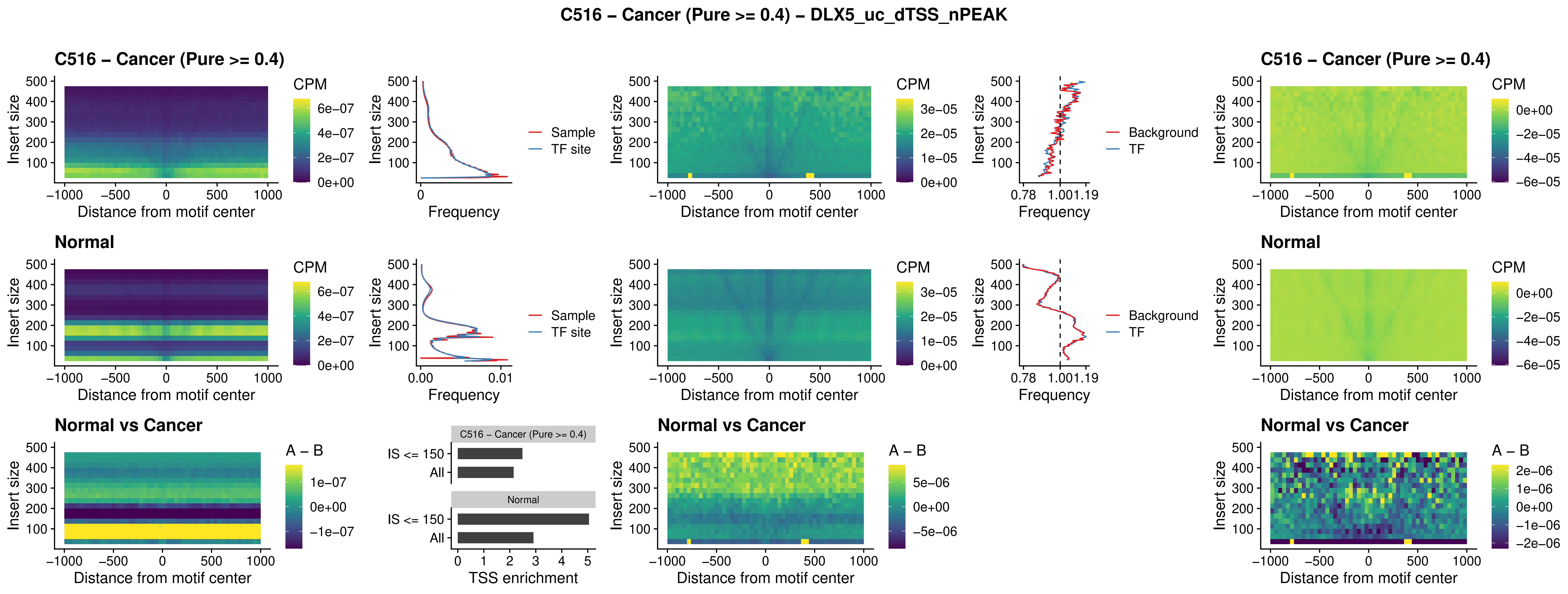

C516 – Cancer (Pure >= 0.4) – DLX5_uc_dTSS_nPEAK
C516 – Cancer (Pure >= 0.4)
Normal
Normal vs Cancer
Insert size
Distance from motif center
Frequency
CPM
A – B
Sample
TF site
Background
TF
C516 – Cancer (Pure >= 0.4)
Normal
IS <= 150
All
TSS enrichment

#### C516 – Cancer (Pure >= 0.4) – DMRT2_uc_dTSS_nPEAK

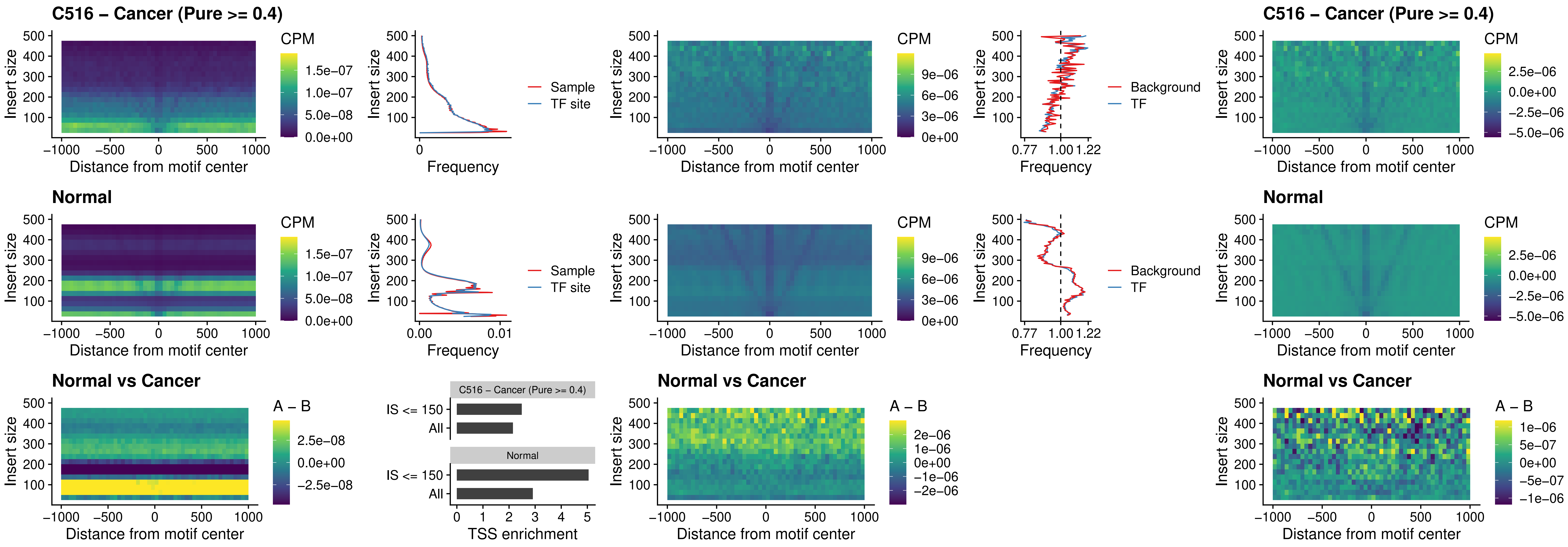

#### C516 – Cancer (Pure >= 0.4) – DMRTC2_uc_cTSS_nPEAK

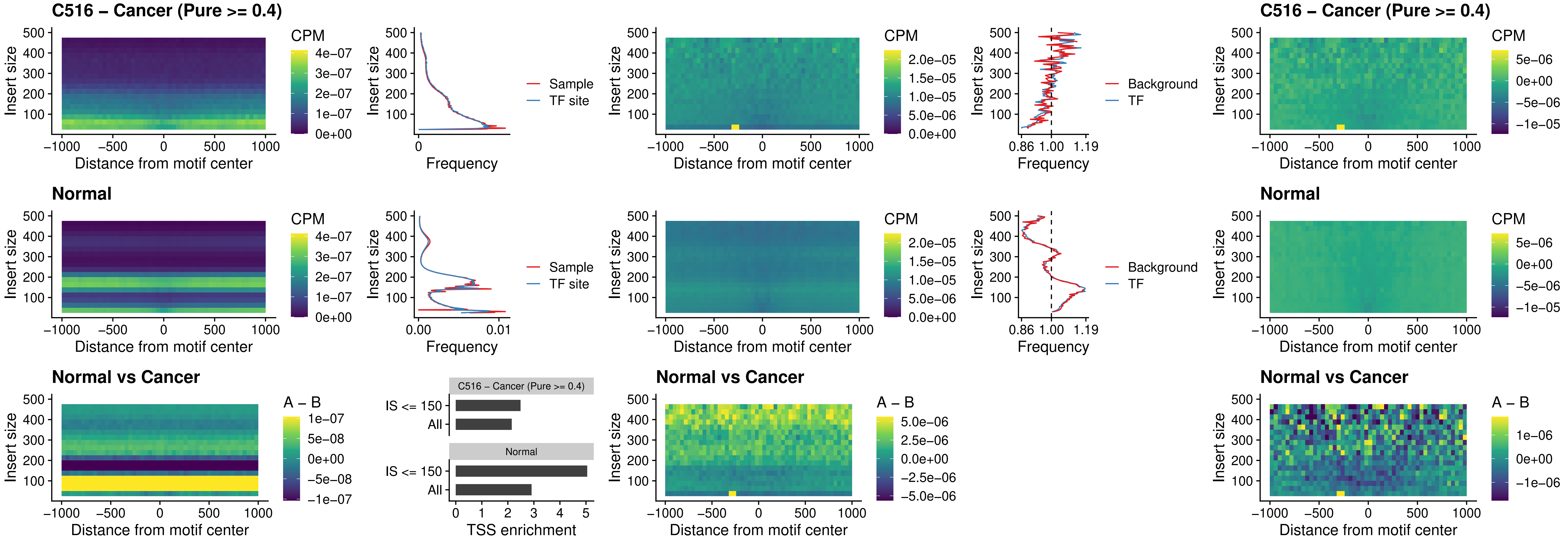

#### C516 – Cancer (Pure >= 0.4) – HOXD9_uc_dTSS_nPEAK

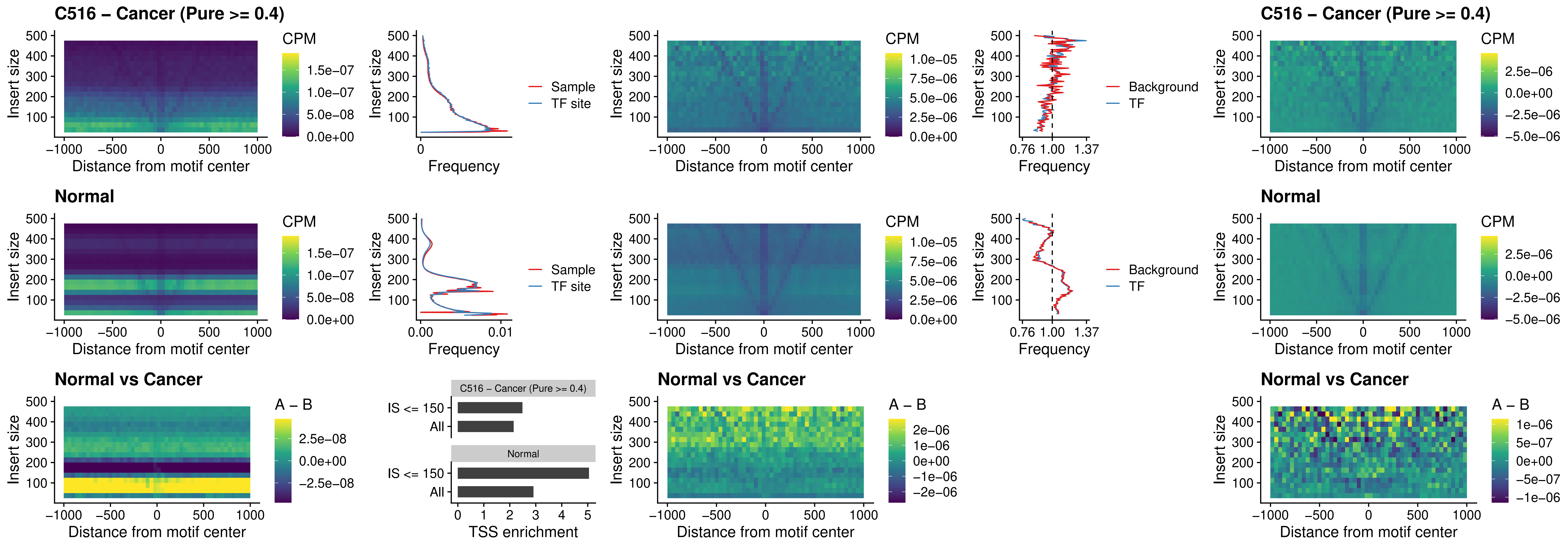

#### C516 – Cancer (Pure >= 0.4) – HOXD4_uc_dTSS_nPEAK

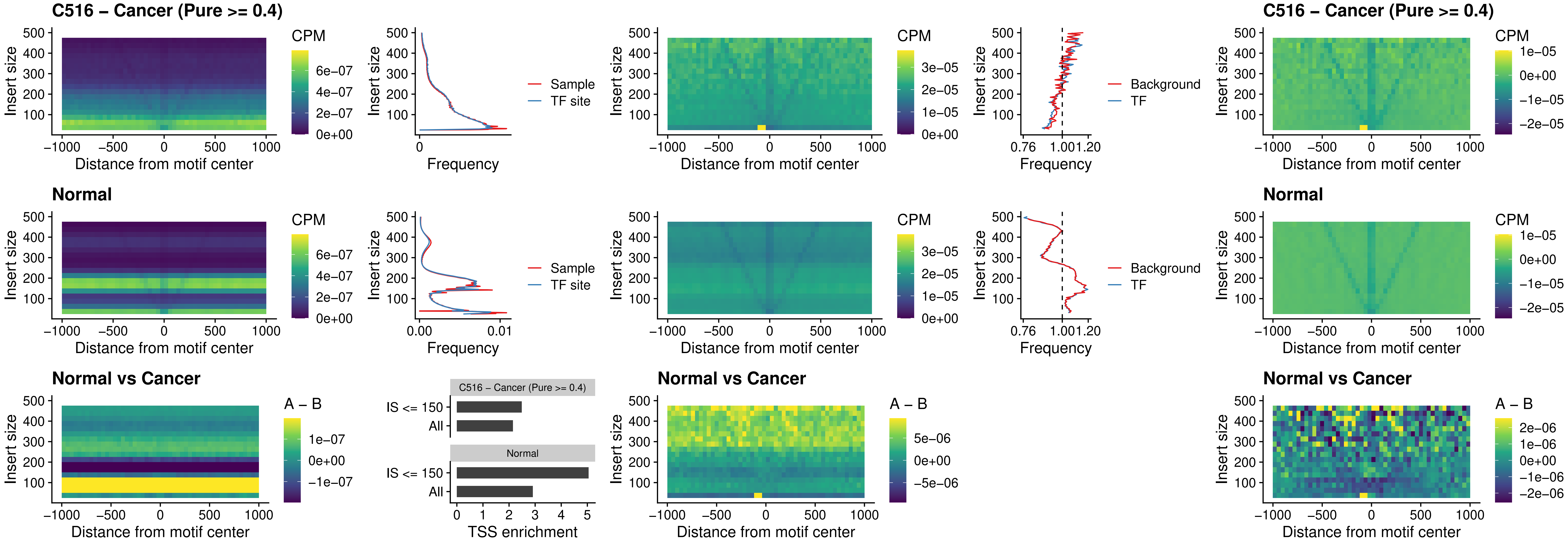

#### C516 – Cancer (Pure >= 0.4) – FOXP3_uc_cTSS_nPEAK

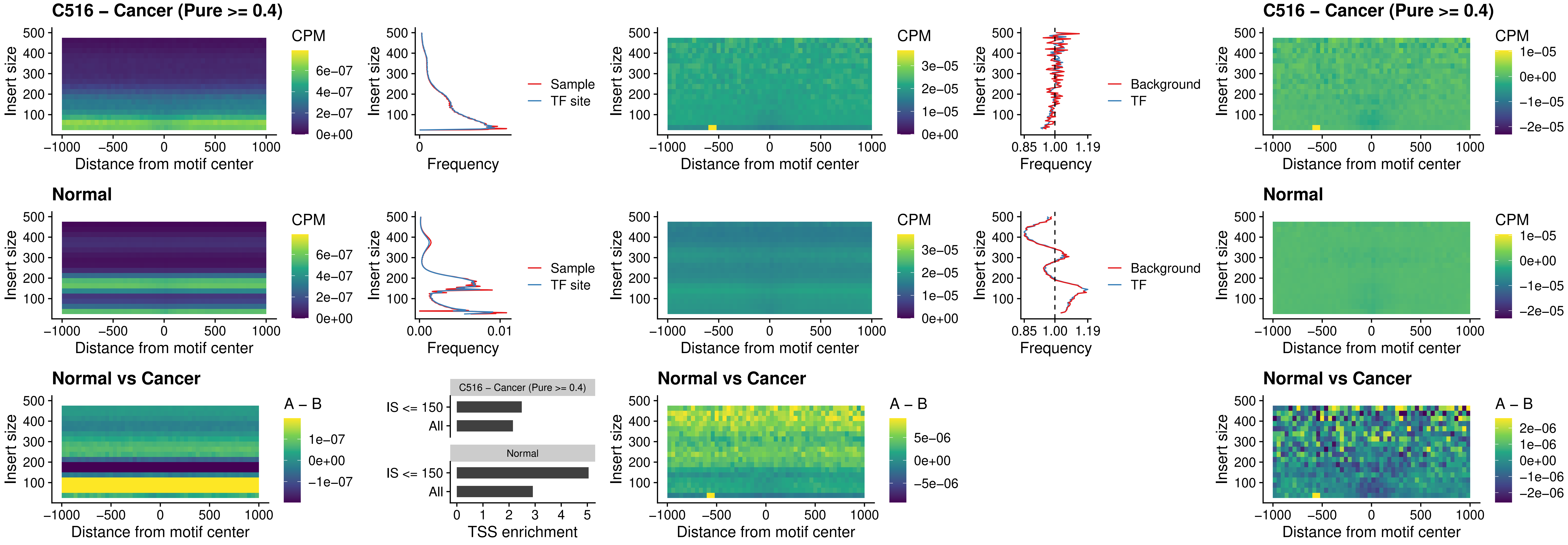

### C516 – Cancer (Pure >= 0.4) – FOXD4L6_uc_cTSS_nPEAK

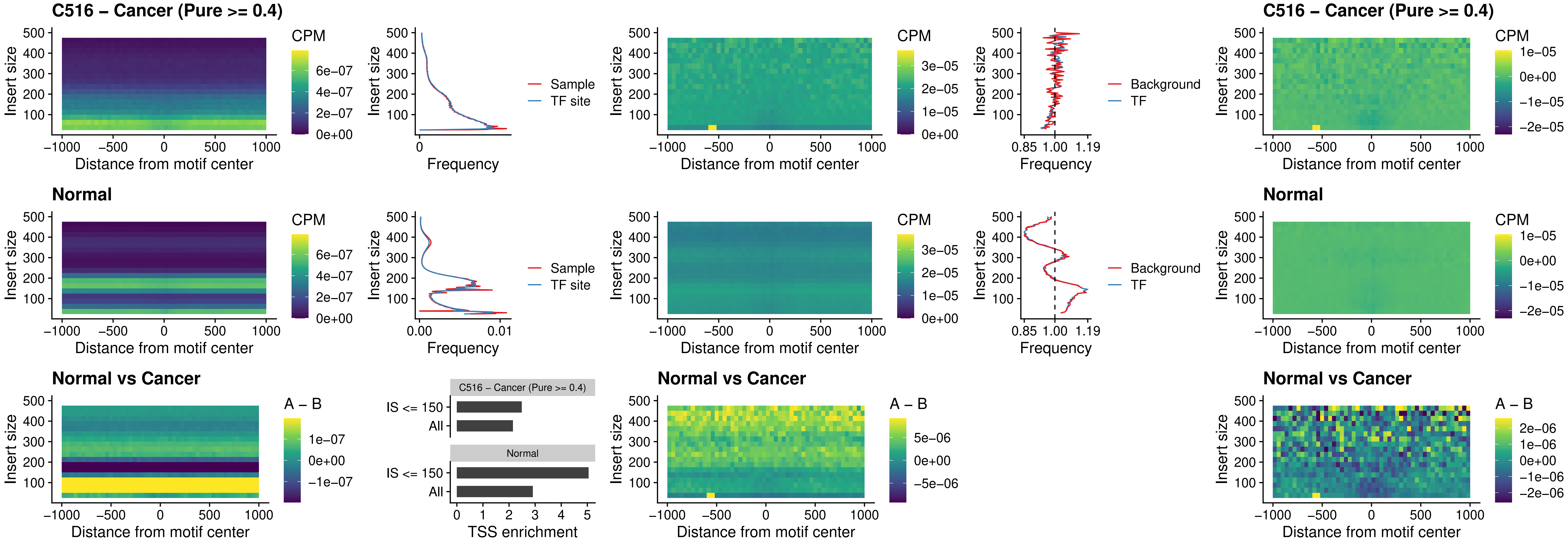

### C516 – Cancer (Pure >= 0.4) – FOXD4L5_uc_cTSS_nPEAK

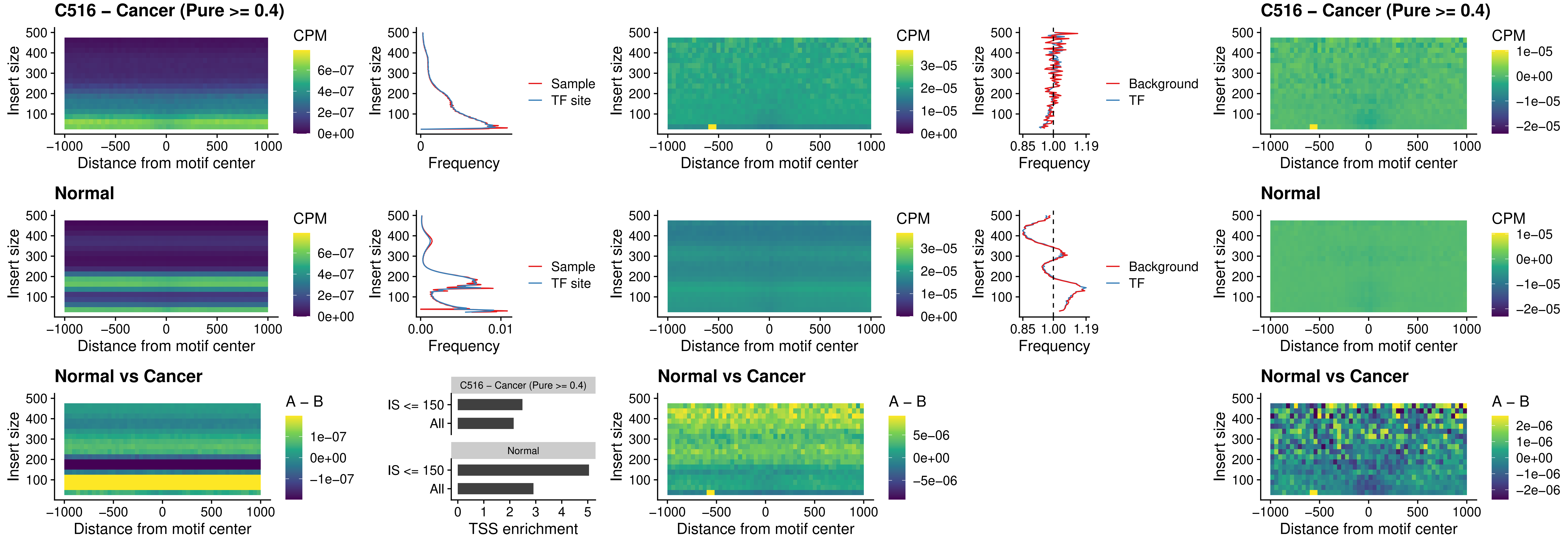

#### C516 – Cancer (Pure >= 0.4) – FOXD4L4_uc_cTSS_nPEAK

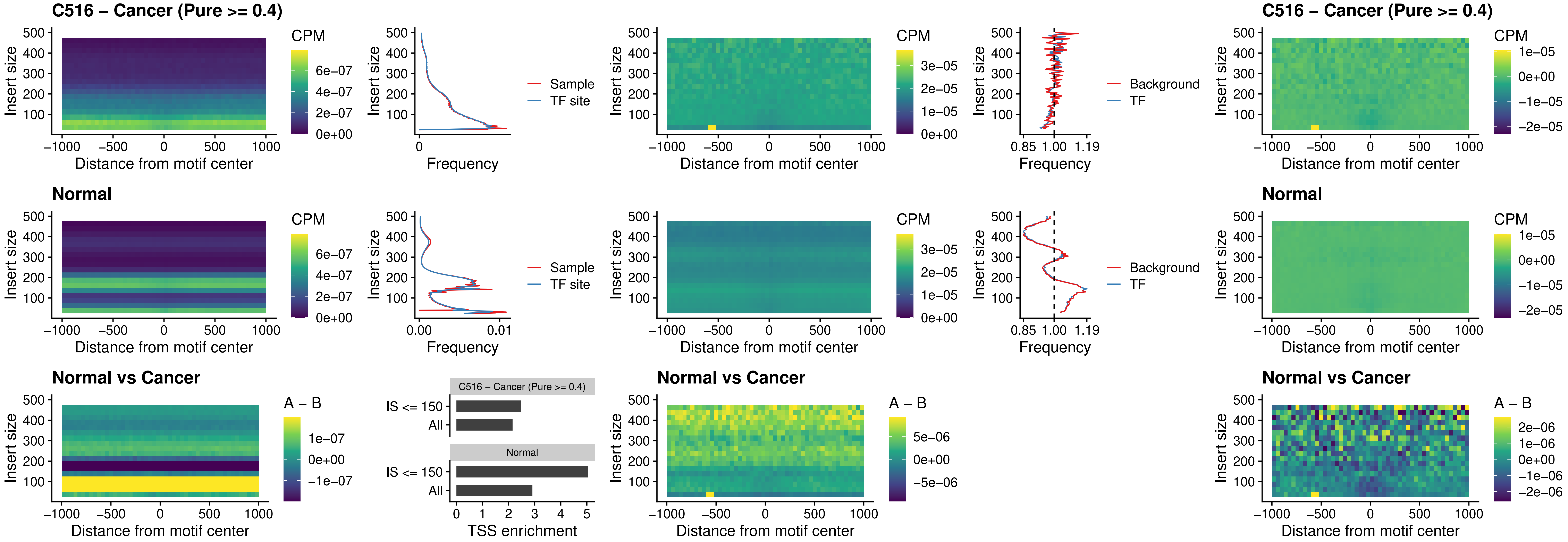

### C516 – Cancer (Pure >= 0.4) – FOXD4L2_uc_cTSS_nPEAK

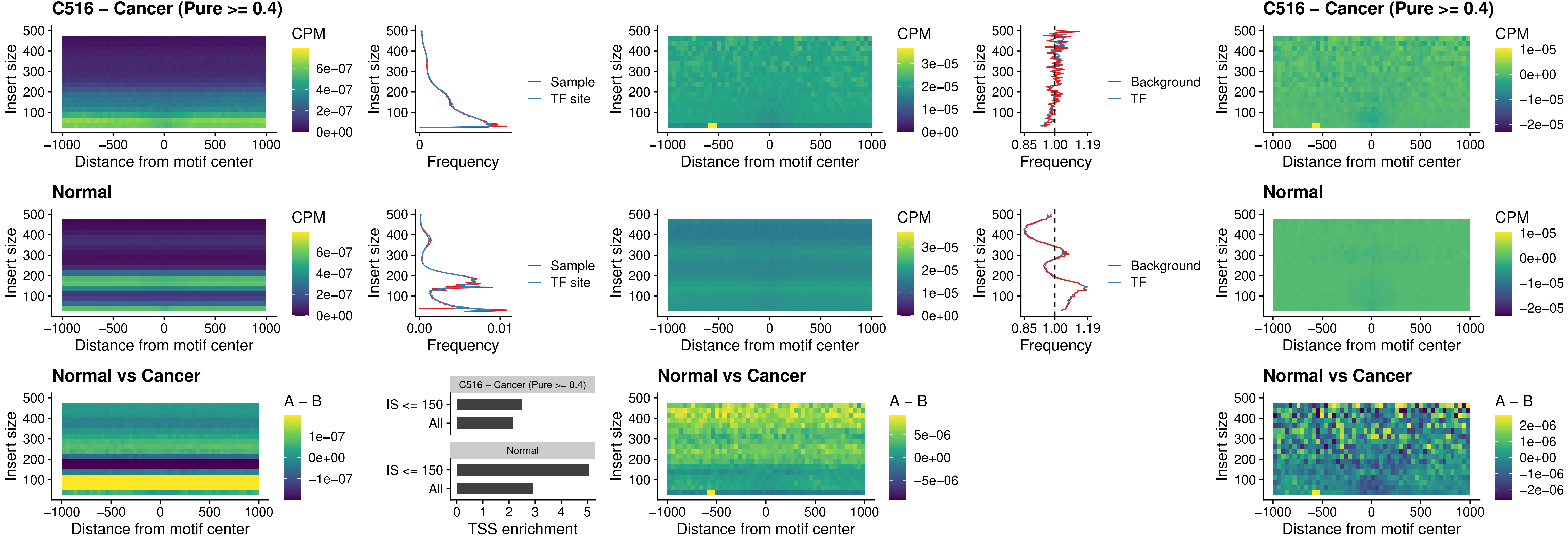

### C516 – Cancer (Pure >= 0.4) – FOXD4L3_uc_cTSS_nPEAK

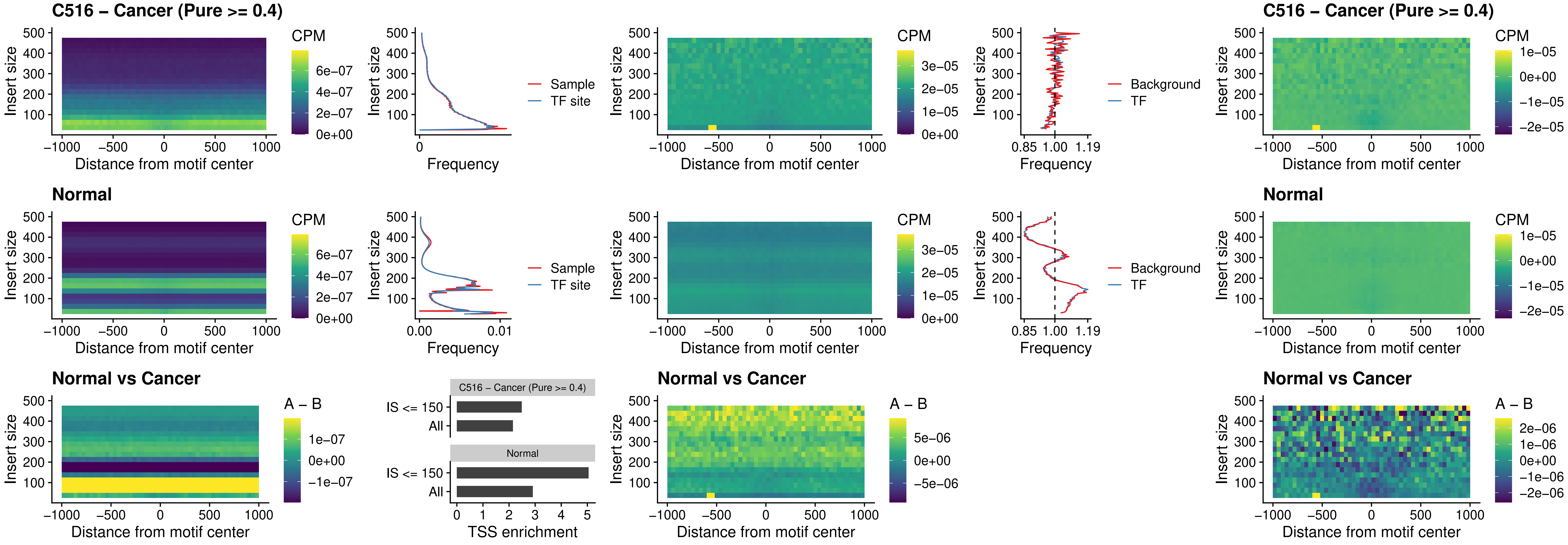

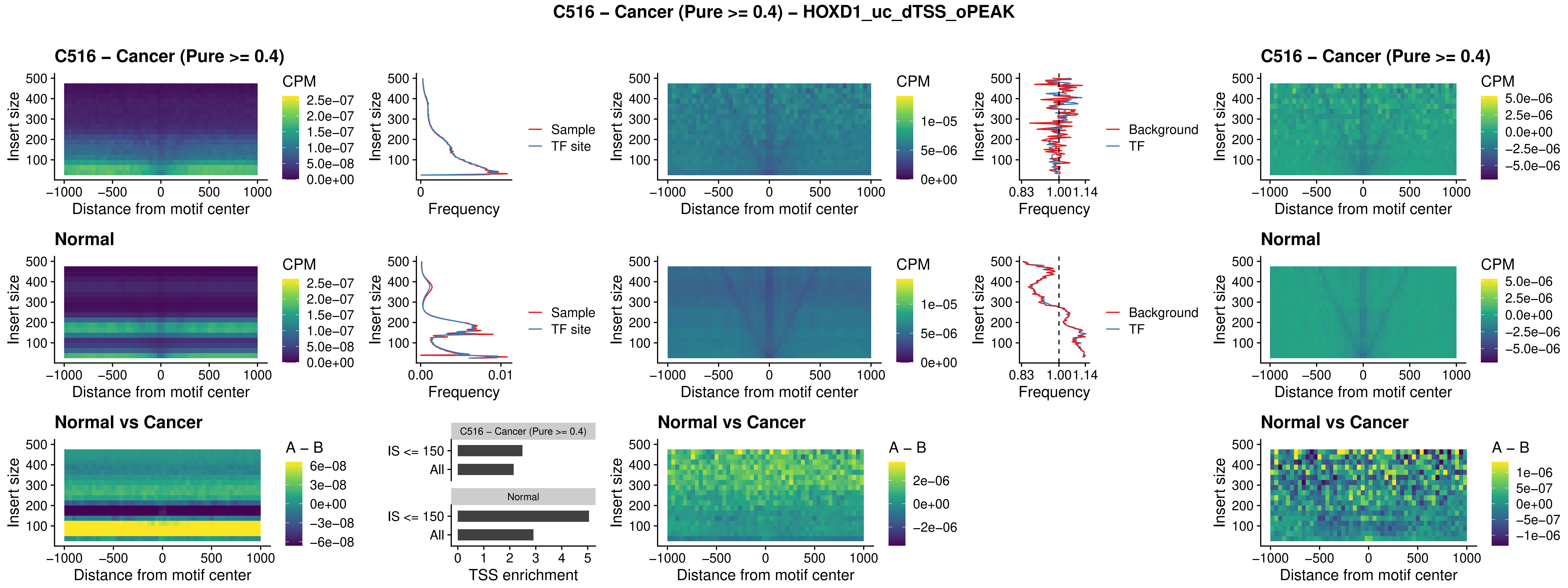
C516 – Cancer (Pure >= 0.4) – HOXD1_uc_dTSS_oPEAK
C516 – Cancer (Pure >= 0.4)
Normal
Normal vs Cancer
Insert size
Distance from motif center
CPM
A – B
Frequency
Sample
TF site
Background
TF
TSS enrichment
C516 – Cancer (Pure >= 0.4)
Normal
IS <= 150
All

#### C516 – Cancer (Pure >= 0.4) – VAX1_uc_pTSS_nPEAK

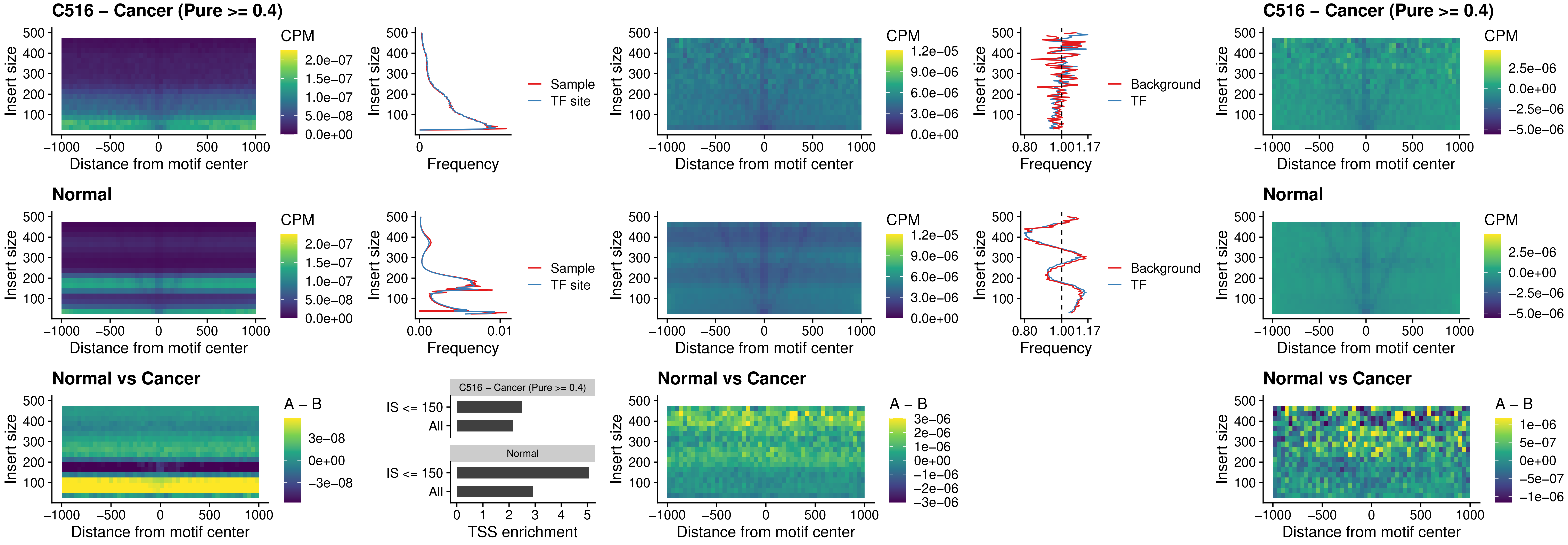

#### C516 – Cancer (Pure >= 0.4) – DLX3_uc_pTSS_nPEAK

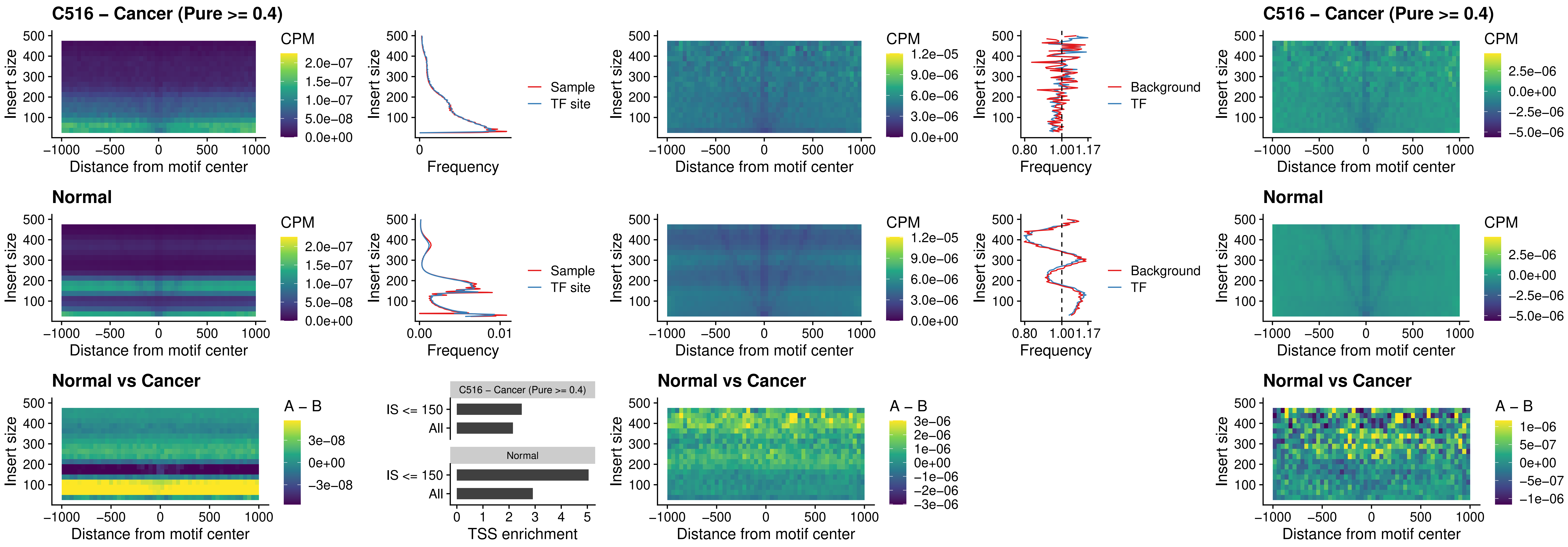

#### C516 – Cancer (Pure >= 0.4) – HOXD3_uc_pTSS_nPEAK

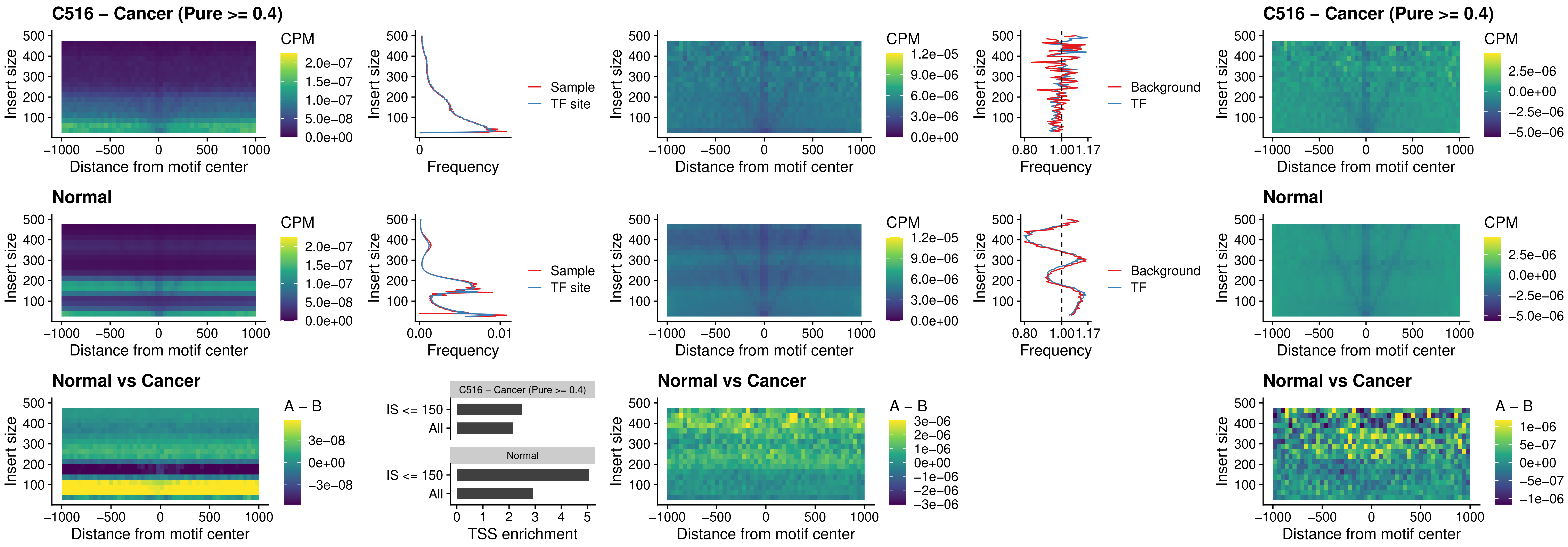

#### C516 – Cancer (Pure >= 0.4) – PDX1_uc_dTSS_nPEAK

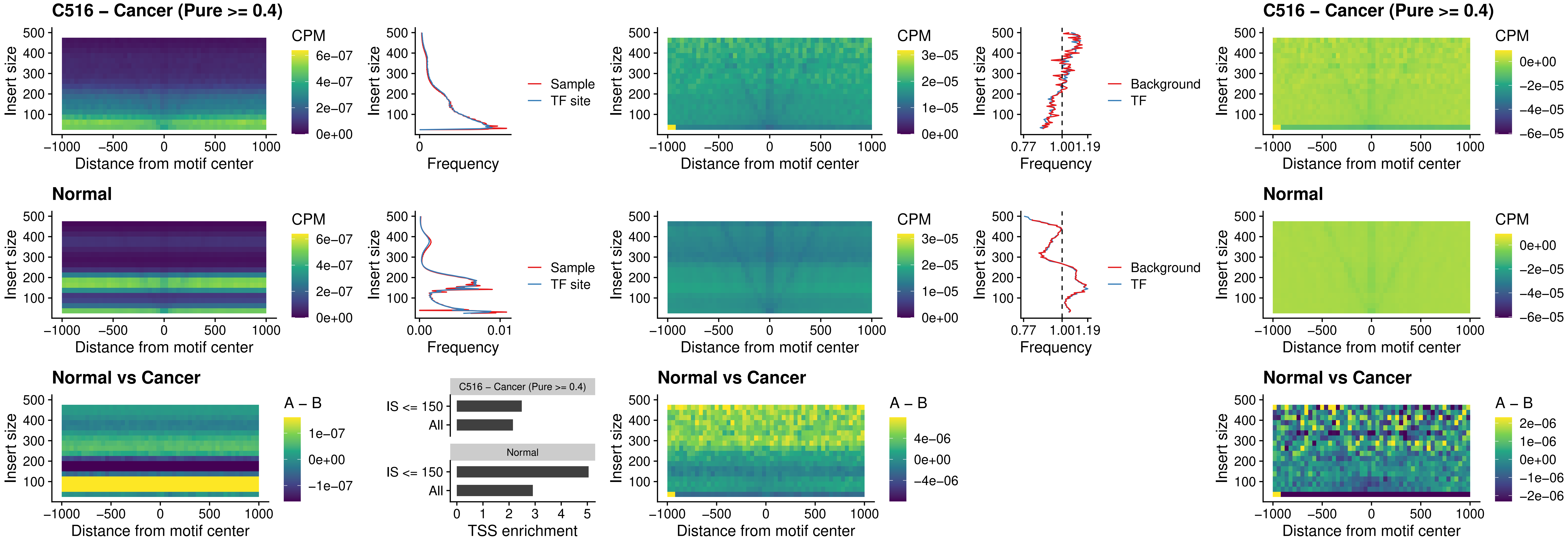

### C516 – Cancer (Pure >= 0.4) – VAX2_uc_dTSS_nPEAK

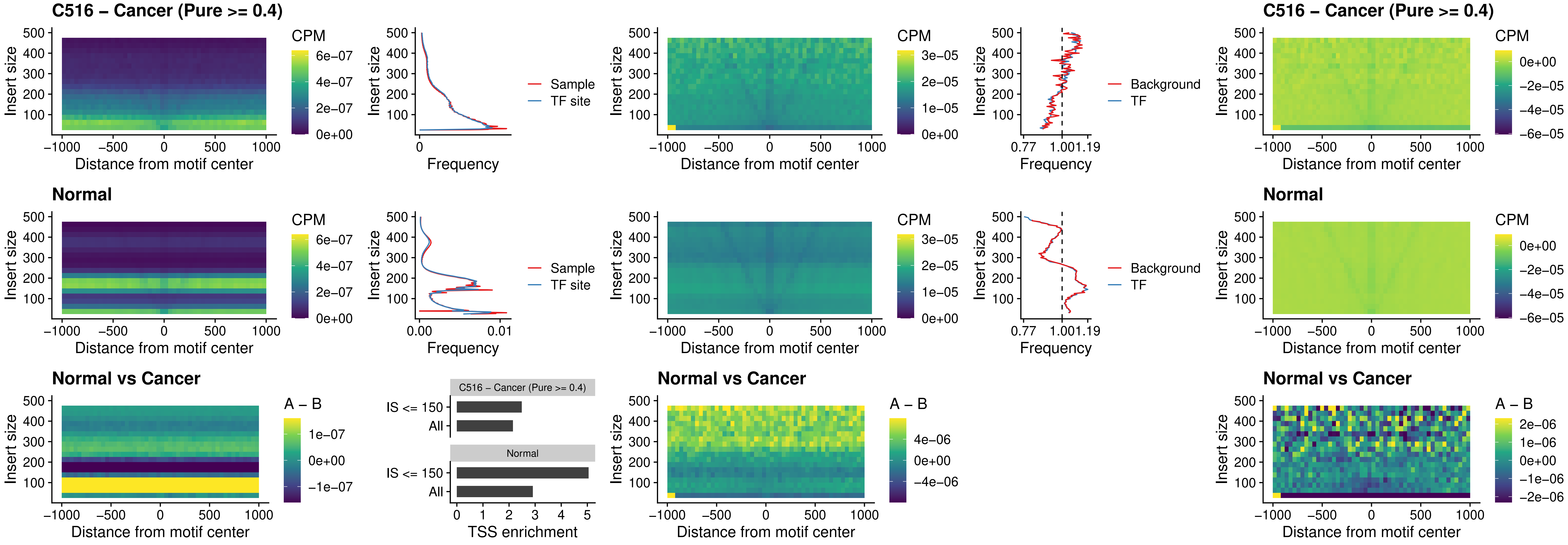

#### C516 – Cancer (Pure >= 0.4) – SOX5_uc_dTSS_nPEAK

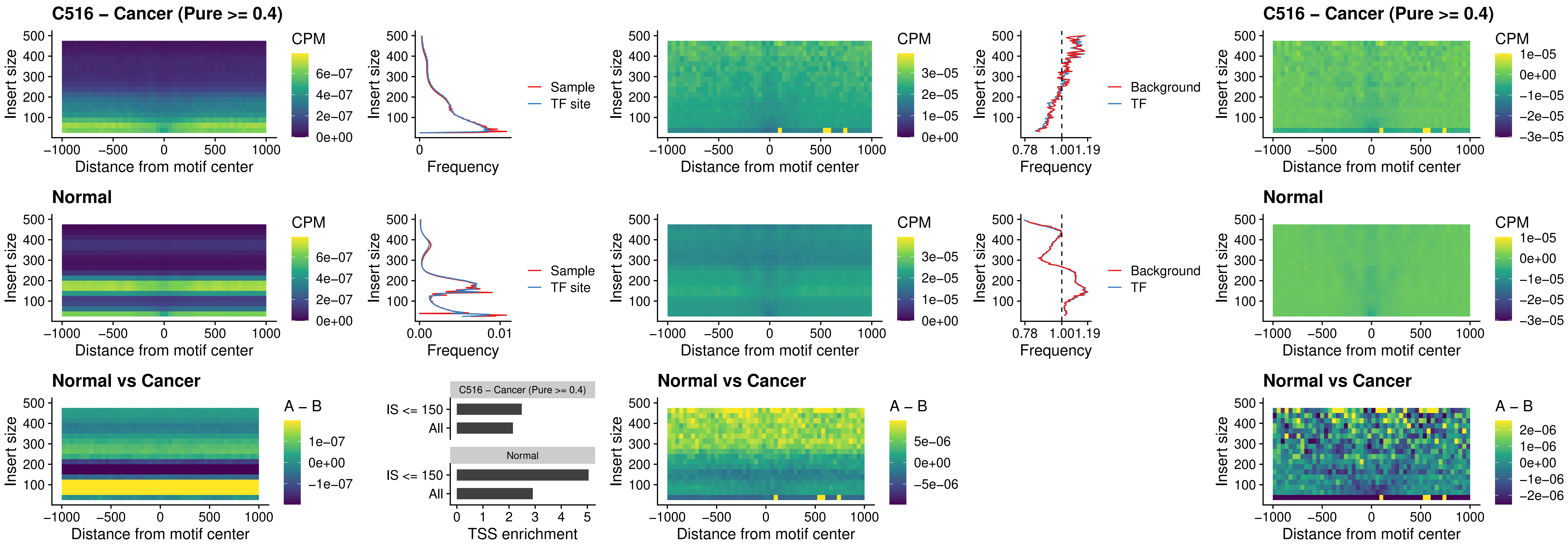

#### C516 – Cancer (Pure >= 0.4) – SOX6_uc_dTSS_nPEAK

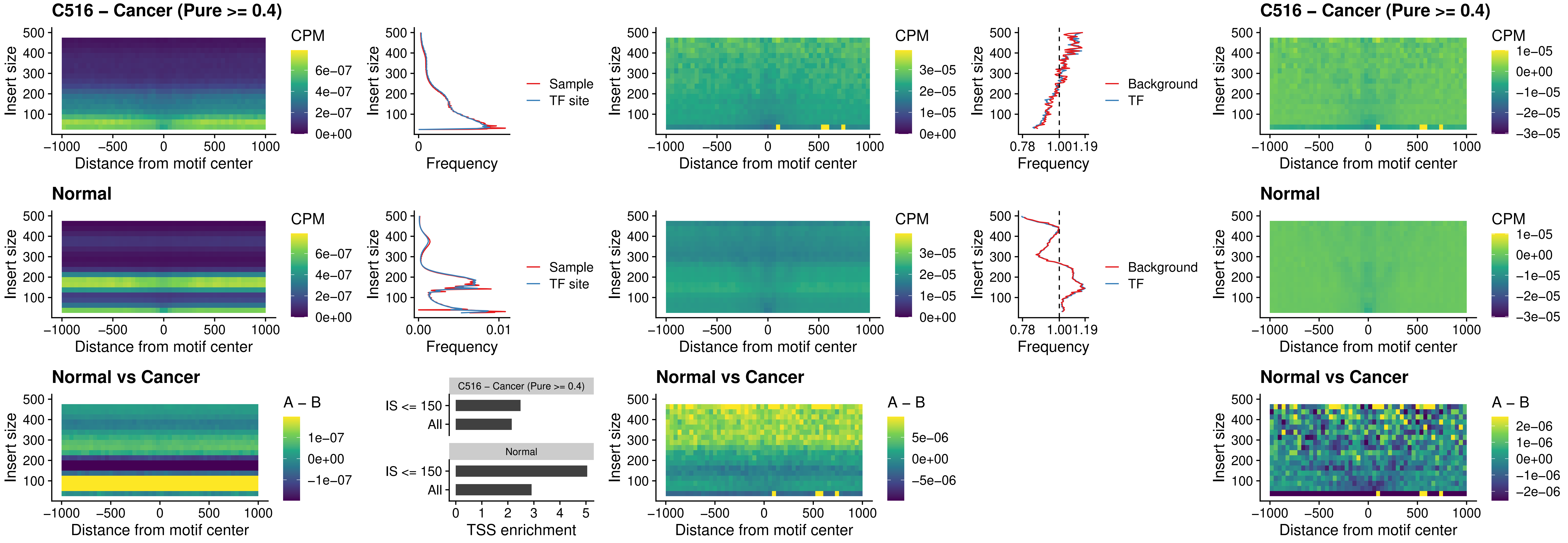

### C516 – Cancer (Pure >= 0.4) – JDP2_uc_cTSS_nPEAK

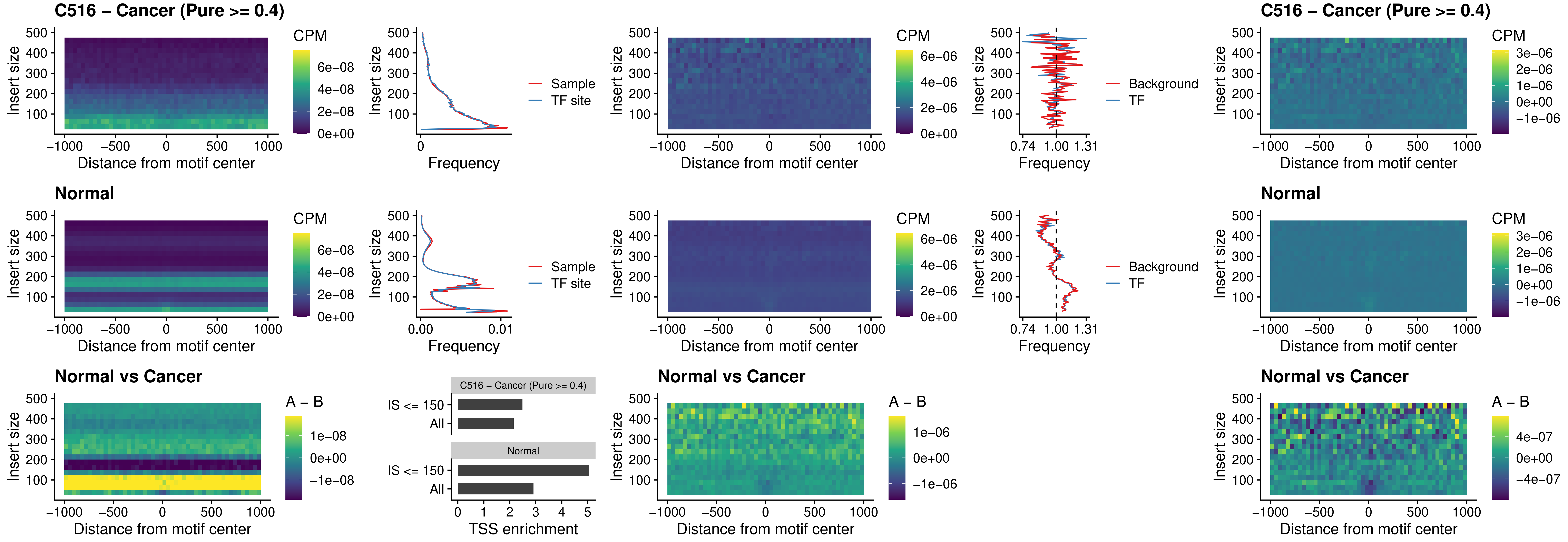

#### C516 – Cancer (Pure >= 0.4) – PURA_uc_dTSS_oPEAK

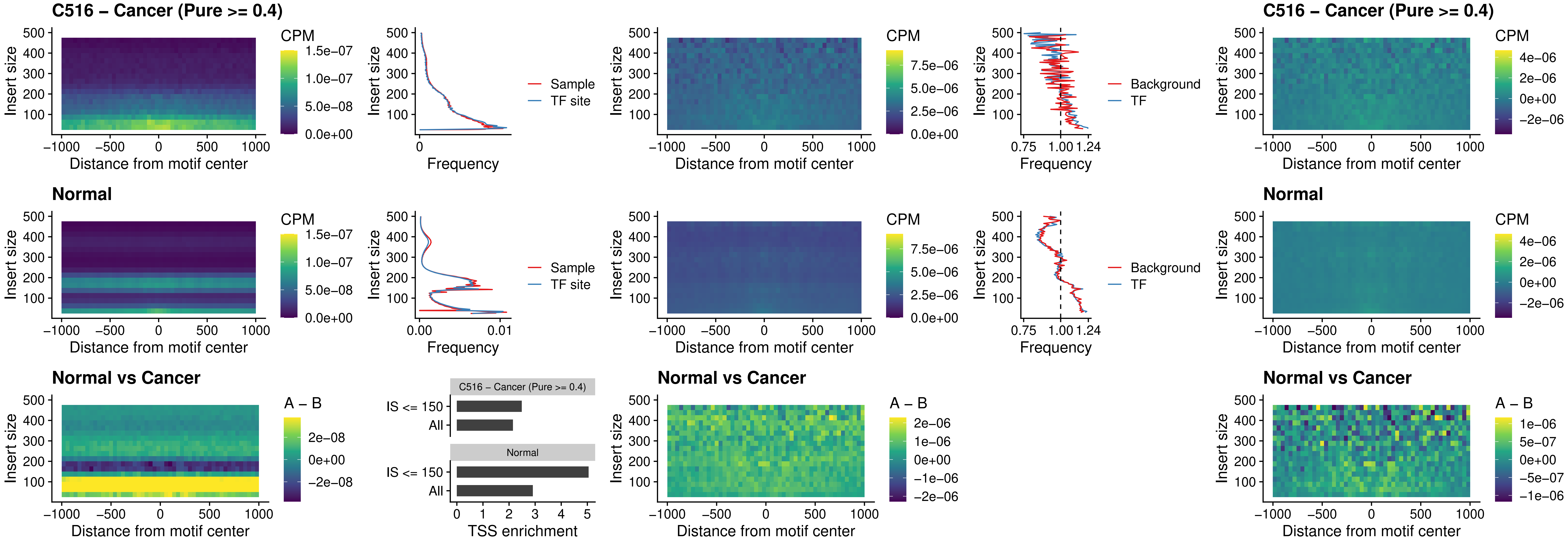

#### C516 – Cancer (Pure >= 0.4) – HHEX_uc_dTSS_nPEAK

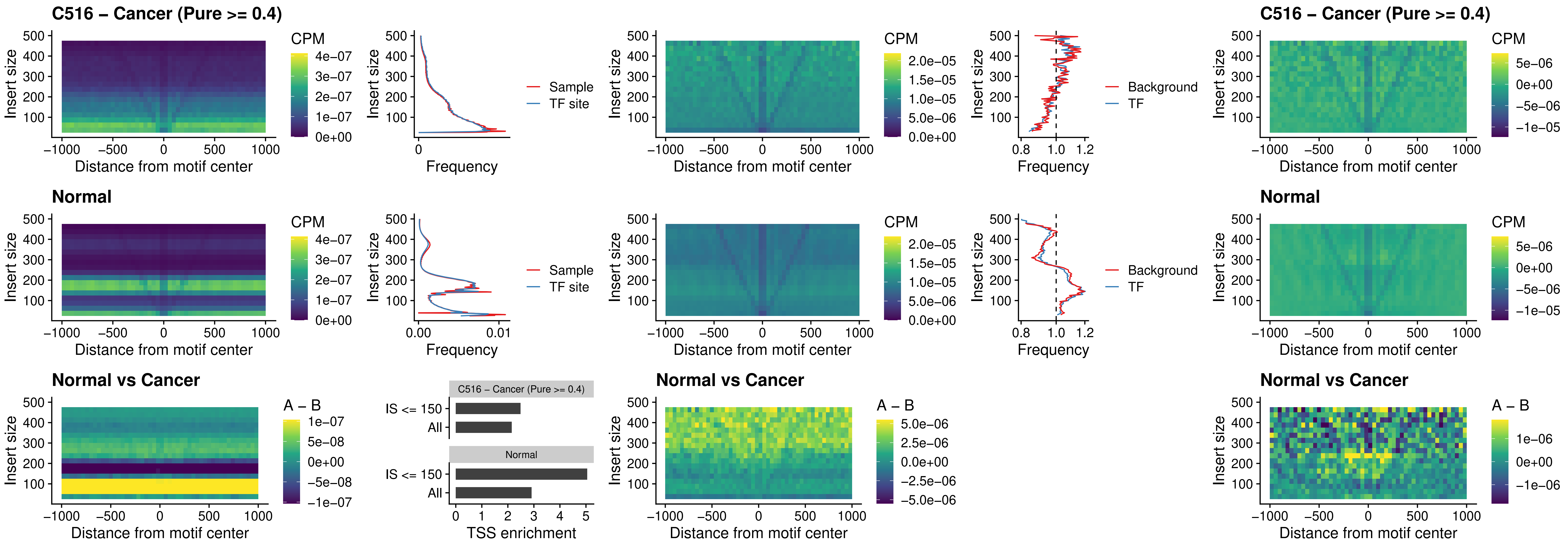

#### C516 – Cancer (Pure >= 0.4) – HMG20B_uc_dTSS_nPEAK

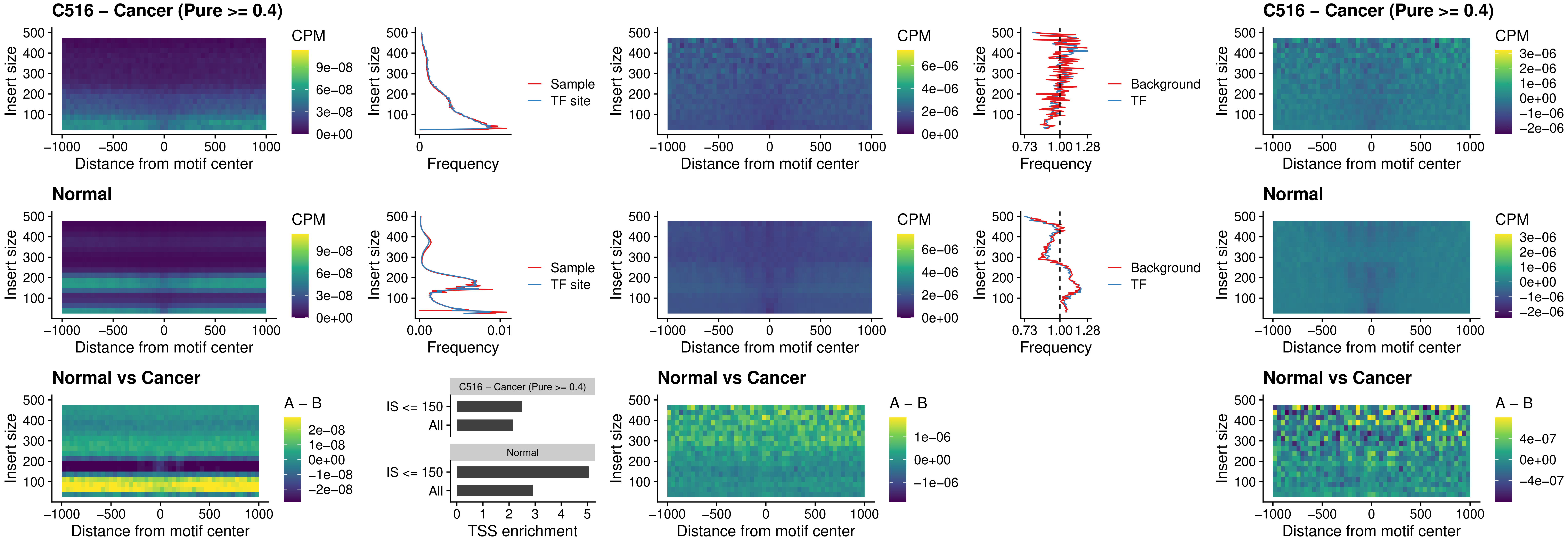

#### C516 – Cancer (Pure >= 0.4) – FOXB2_uc_dTSS_nPEAK

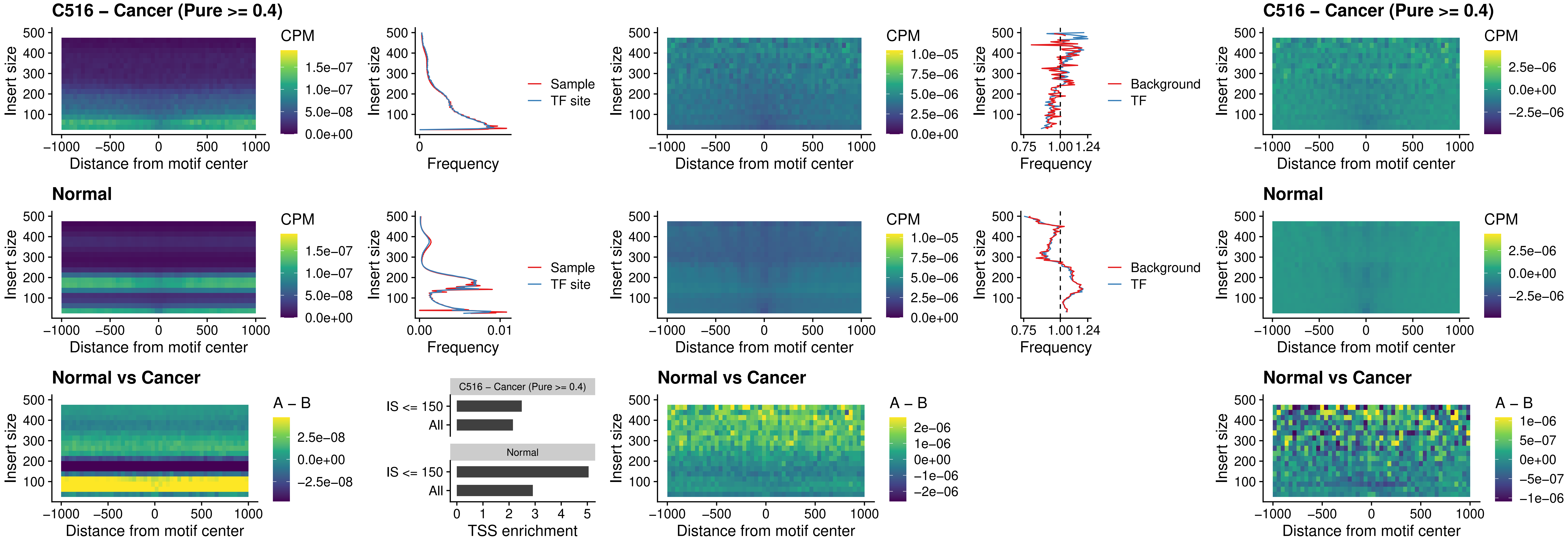

#### C516 – Cancer (Pure >= 0.4) – FOXL1_uc_cTSS_nPEAK

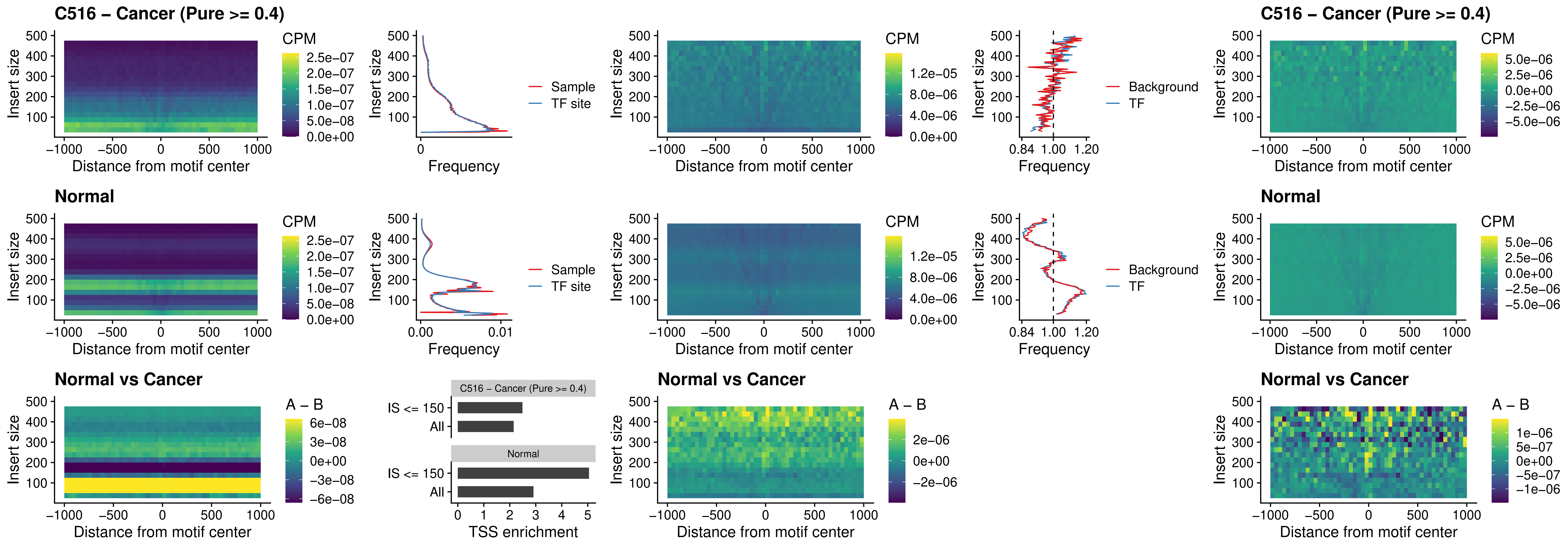

### C516 – Cancer (Pure >= 0.4) – PBX3_uc_pTSS_oPEAK

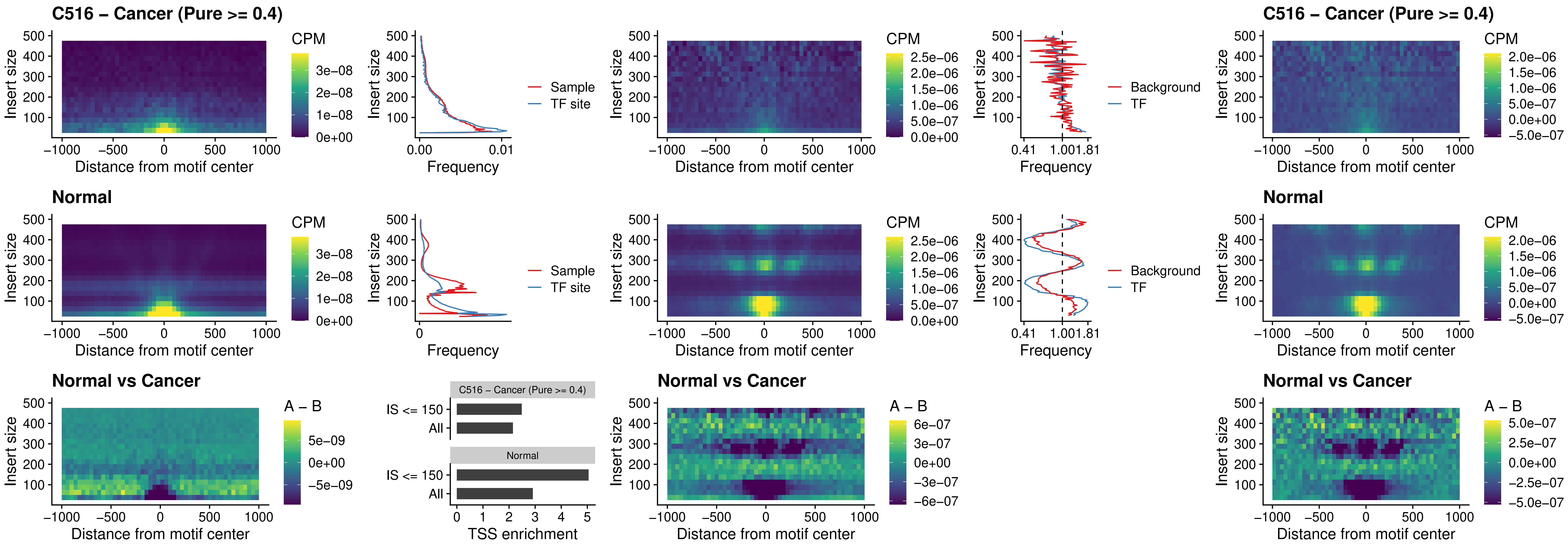

### C516 – Cancer (Pure >= 0.4) – SNAI2_uc_cTSS_nPEAK

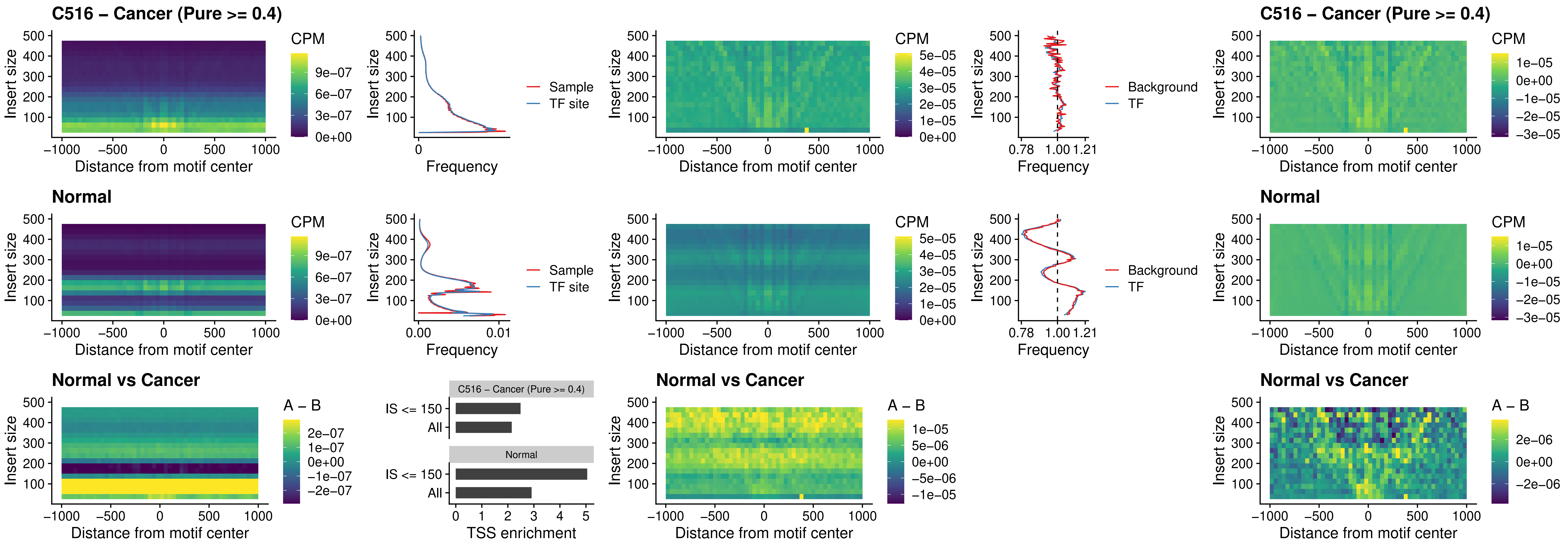

### C516 – Cancer (Pure >= 0.4) – SNAI2_uc_dTSS_oPEAK

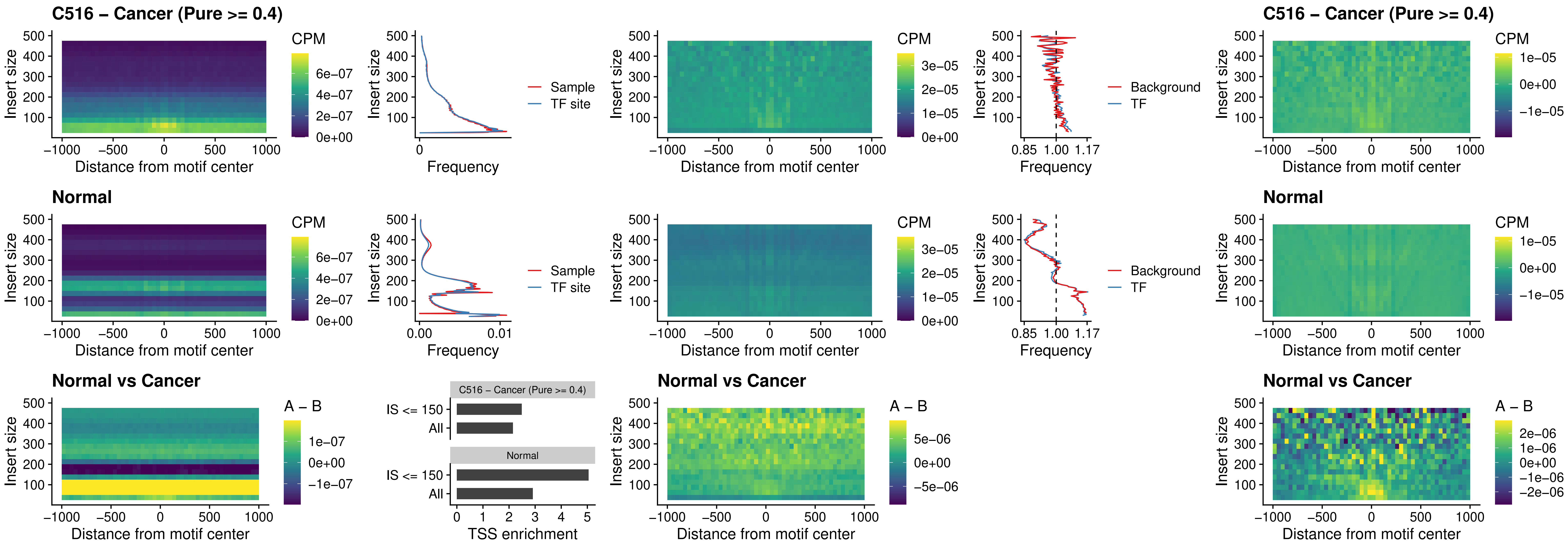

### C516 – Cancer (Pure >= 0.4) – SPI1_uc_pTSS_nPEAK

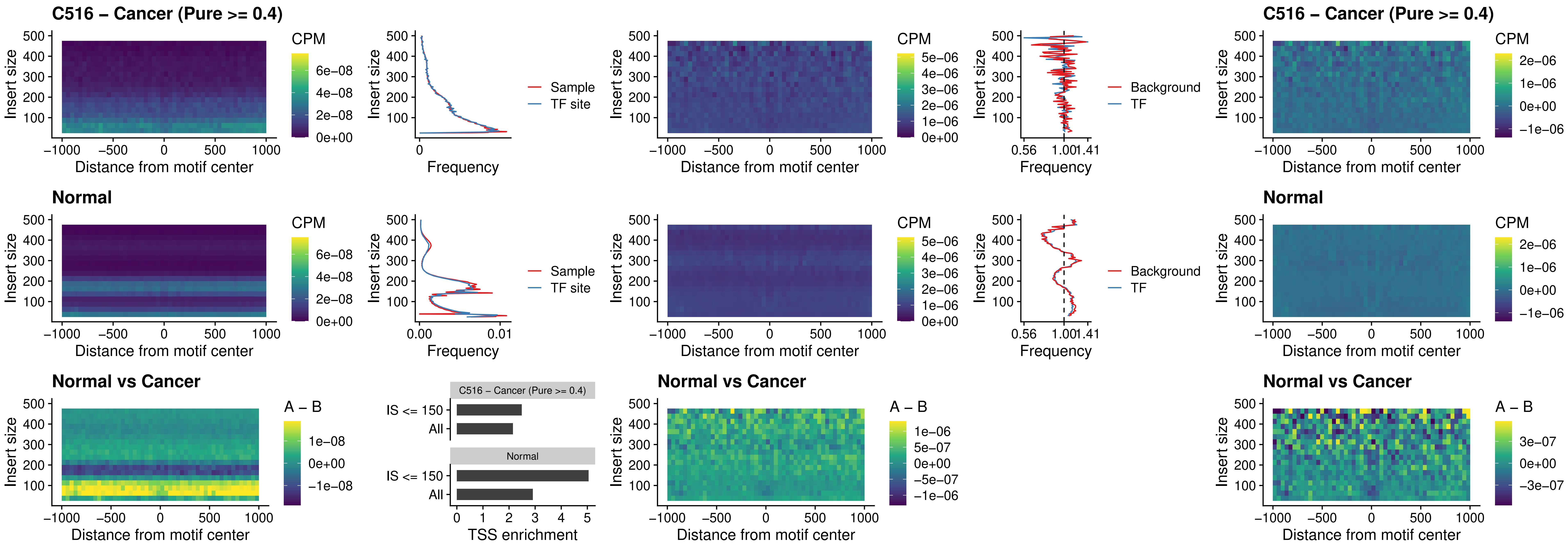

#### C516 – Cancer (Pure >= 0.4) – KLF16_uc_pTSS_nPEAK

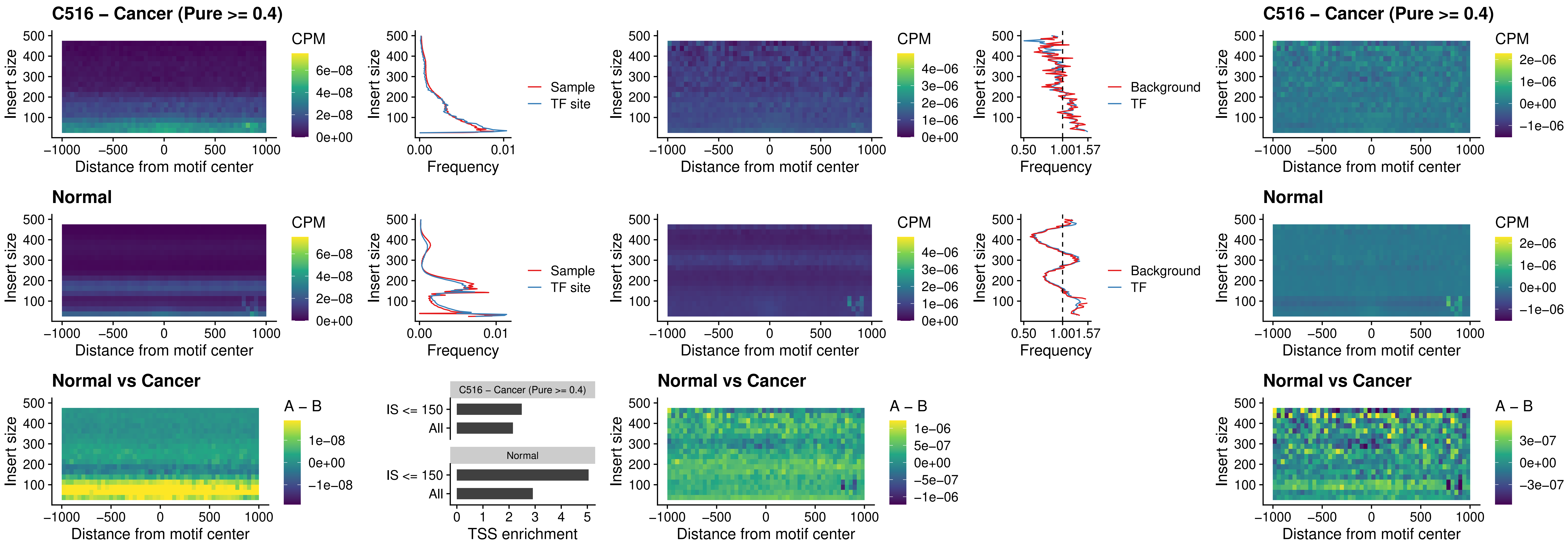

### C516 – Cancer (Pure >= 0.4) – SP3_uc_pTSS_nPEAK

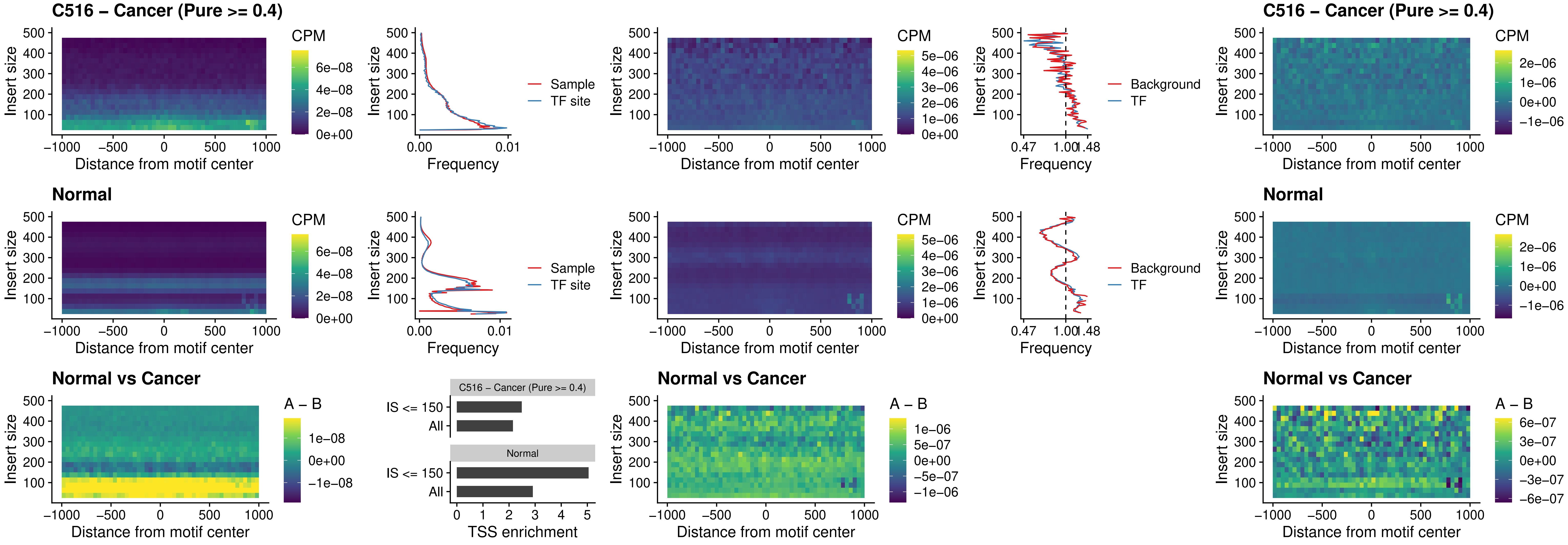

### C516 – Cancer (Pure >= 0.4) – NEUROD1_uc_dTSS_oPEAK

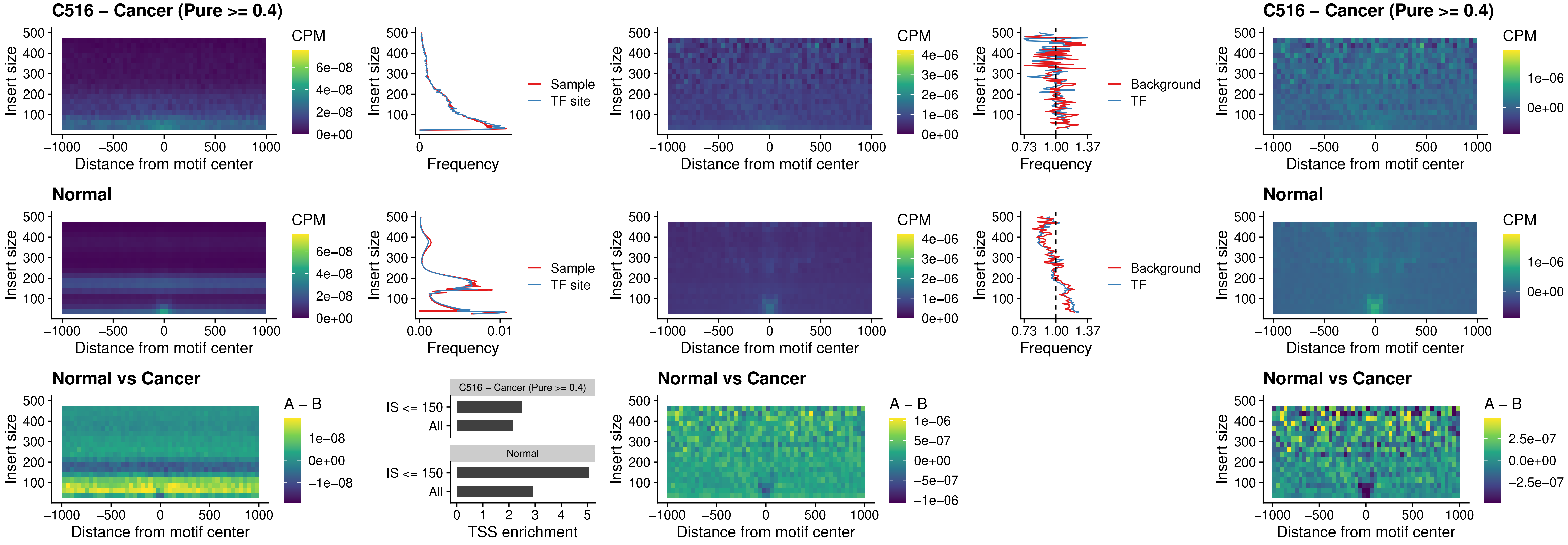

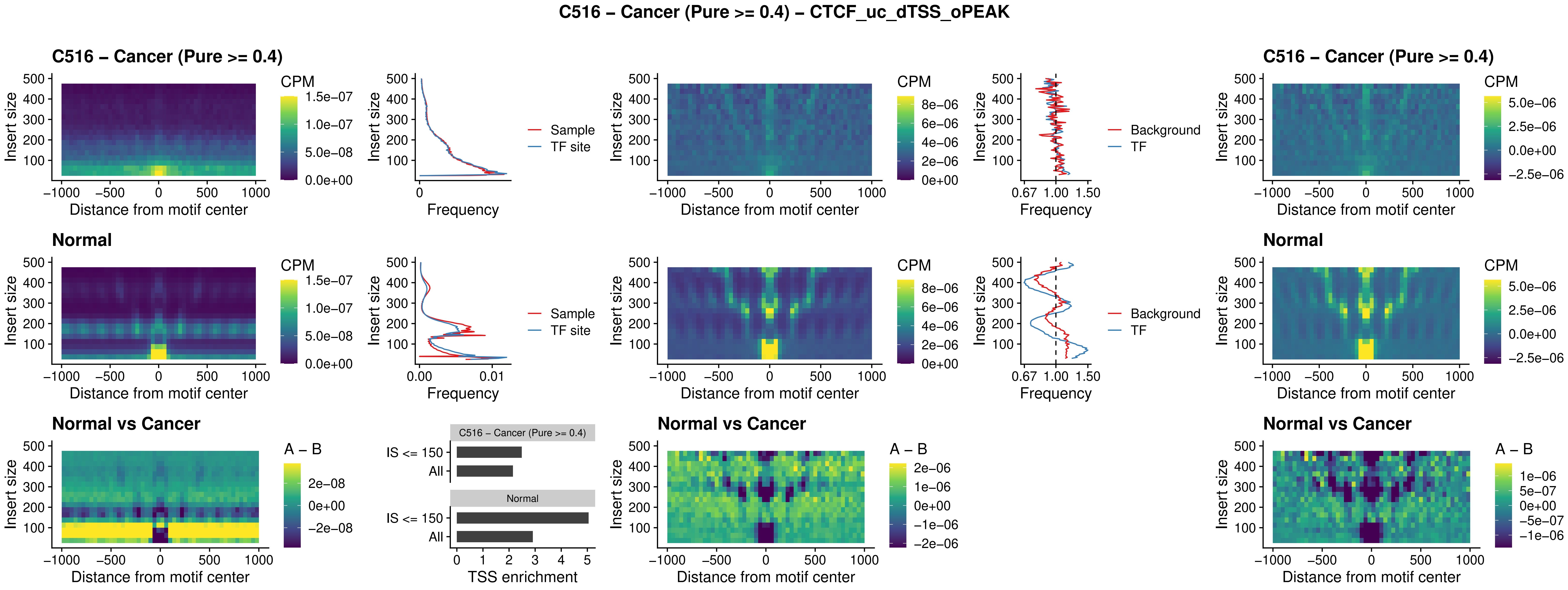
C516 – Cancer (Pure >= 0.4) – CTCF_uc_dTSS_oPEAK
C516 – Cancer (Pure >= 0.4)
Normal
Normal vs Cancer
Insert size
Distance from motif center
CPM
A – B
Frequency
Sample
TF site
Background
TF
C516 – Cancer (Pure >= 0.4)
IS <= 150
All
TSS enrichment

#### C516 – Cancer (Pure >= 0.4) – CTCF_uc_cTSS_oPEAK

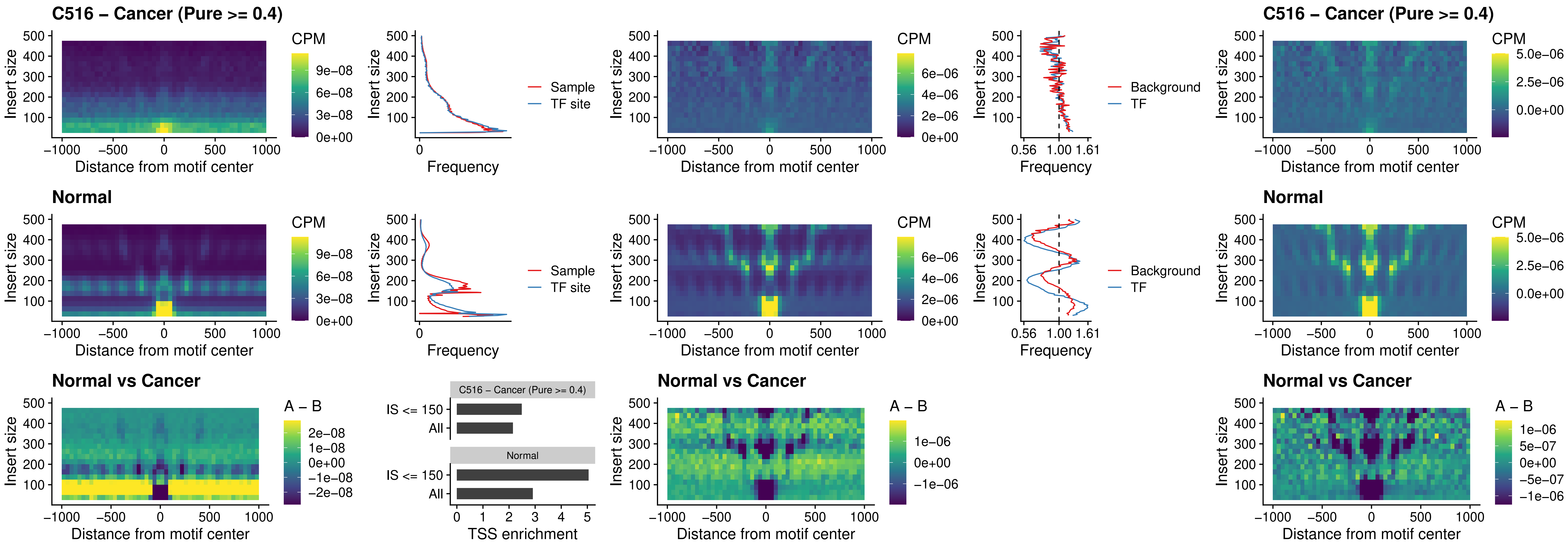

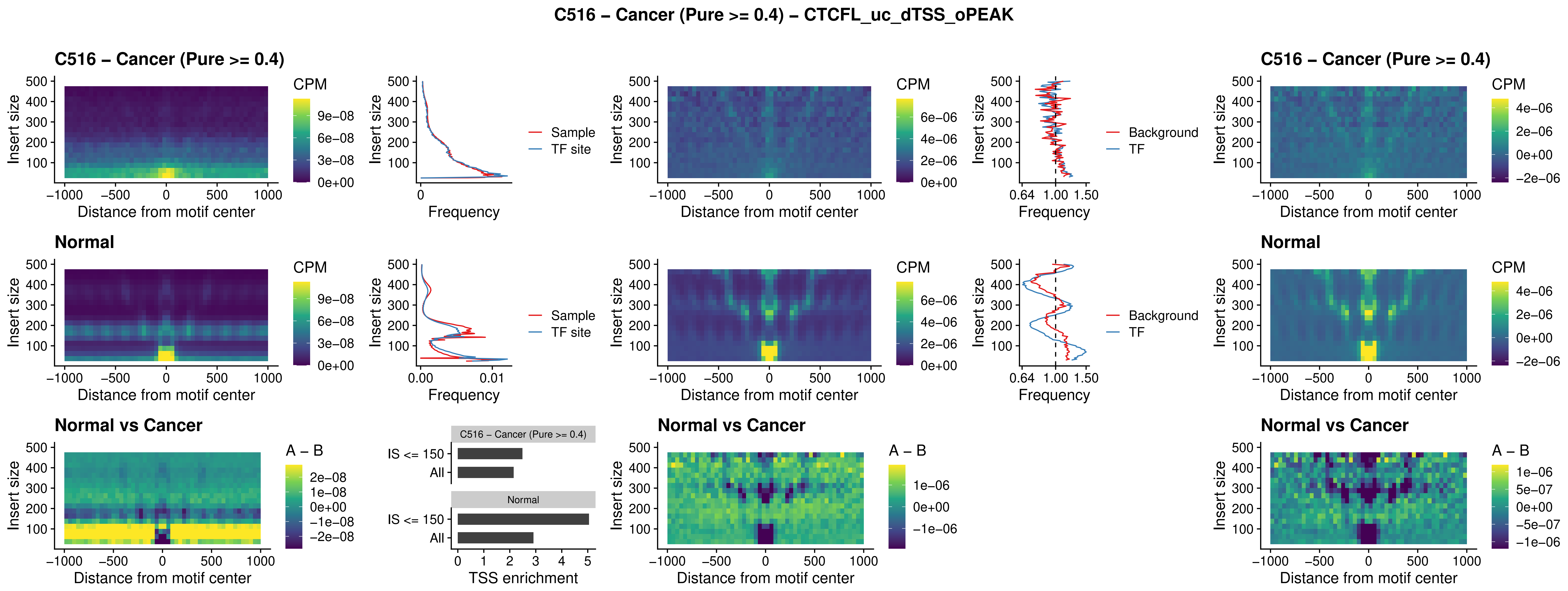
C516 – Cancer (Pure >= 0.4) – CTCFL_uc_dTSS_oPEAK
C516 – Cancer (Pure >= 0.4)
Normal
Normal vs Cancer
Insert size
Distance from motif center
Frequency
CPM
A – B
Sample
TF site
Background
TF
C516 – Cancer (Pure >= 0.4)
Normal
IS <= 150
All
TSS enrichment

#### C516 – Cancer (Pure >= 0.4) – ENCODE_CTCF_sites_COLON

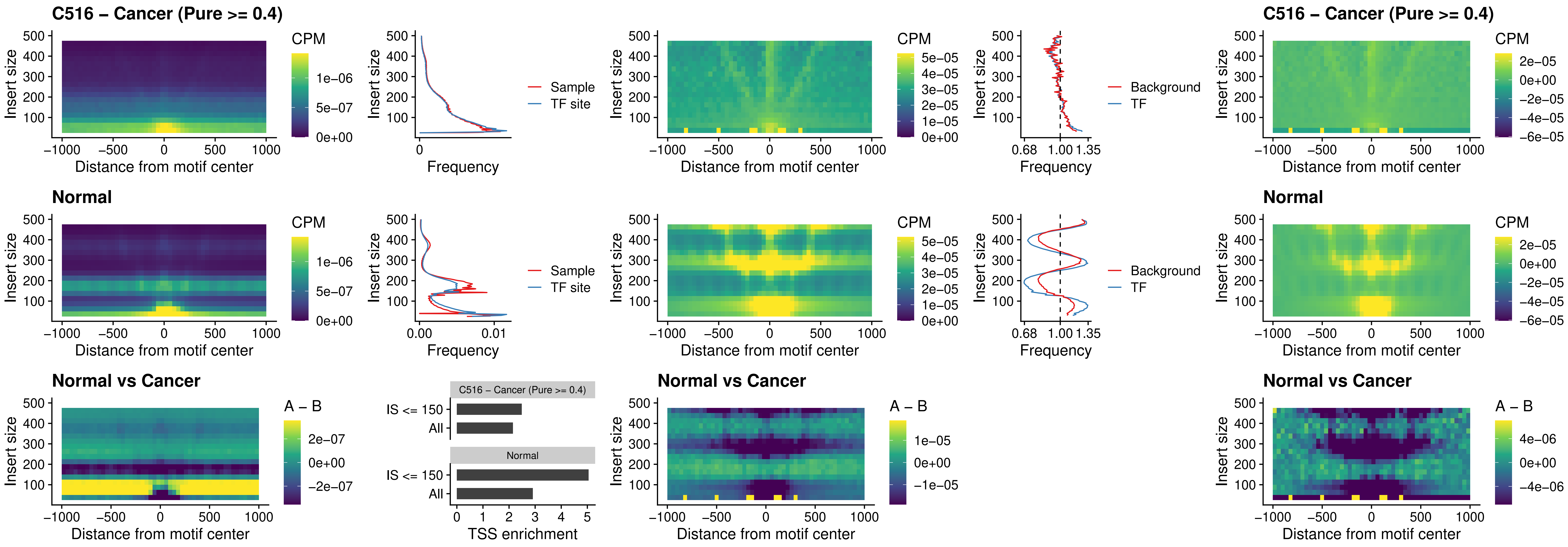

**C516 – Cancer (Pure >= 0.4) – ENCODE_CTCF_sites_COLON_loop_ctcf**

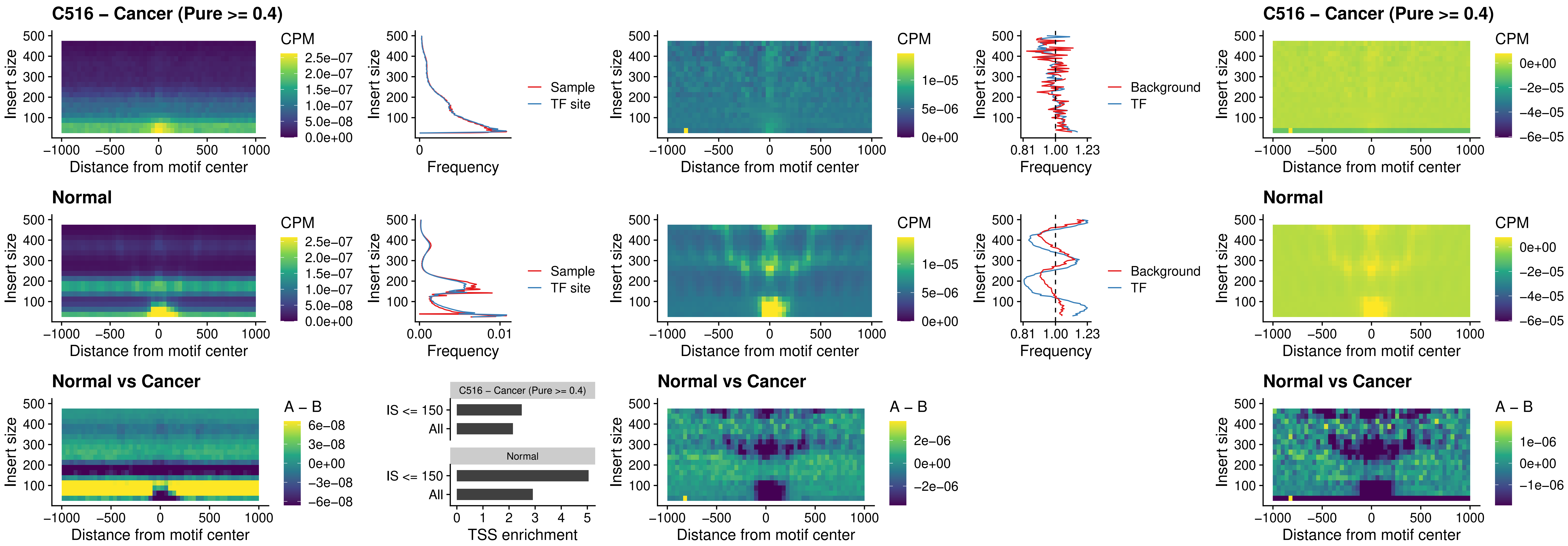

#### C516 – Cancer (Pure >= 0.4) – ENCODE_CTCF_sites_COLON_intergenic_ctcf

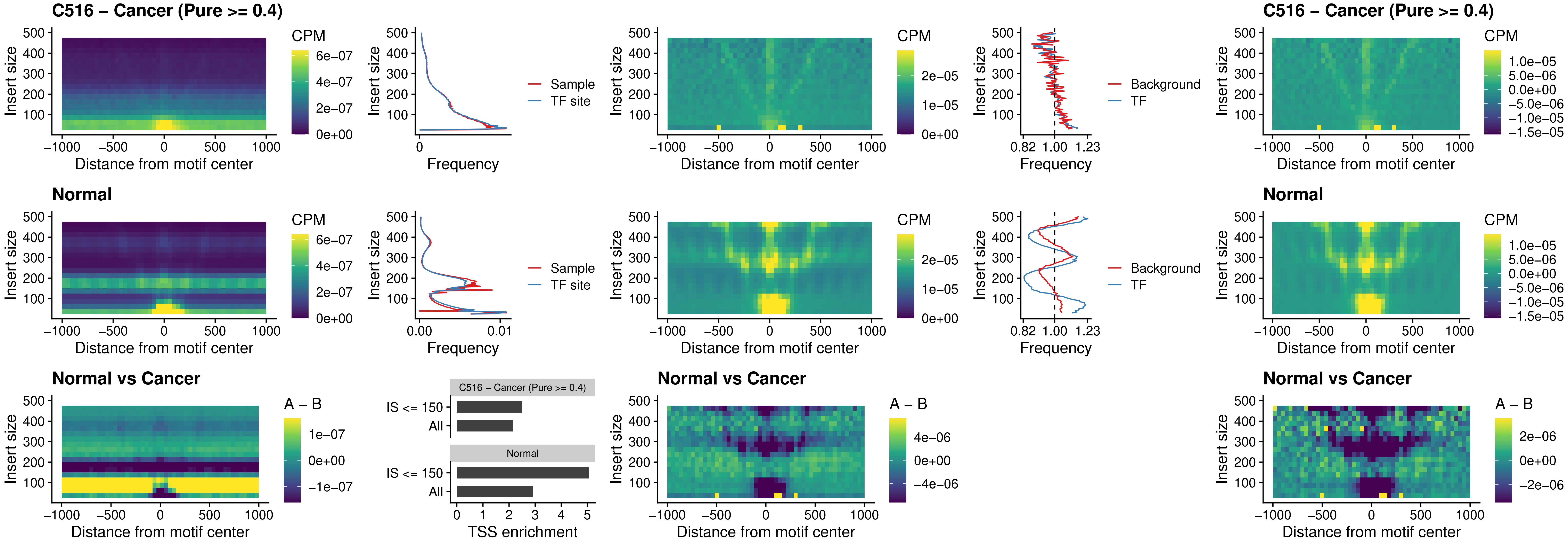

#### C516 – Cancer (Pure >= 0.4) – ENCODE_CTCF_sites_COLON_nonpromotor

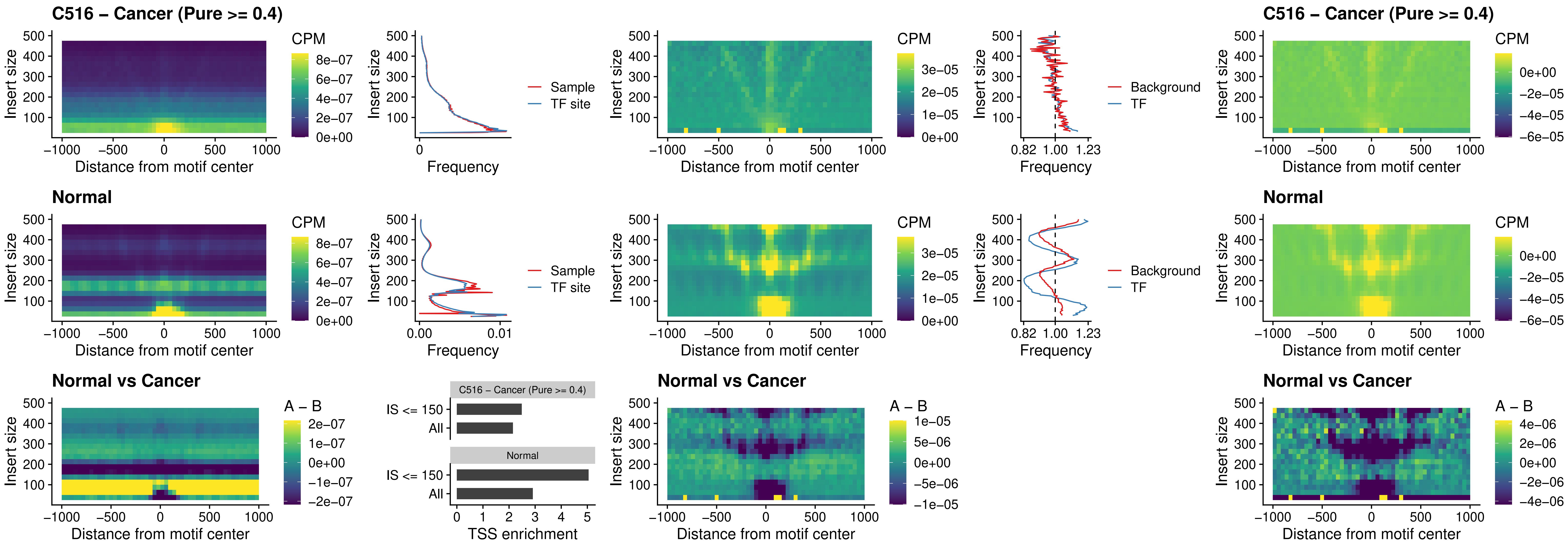

#### C516 – Cancer (Pure >= 0.4) – CTCFL_uc_cTSS_oPEAK

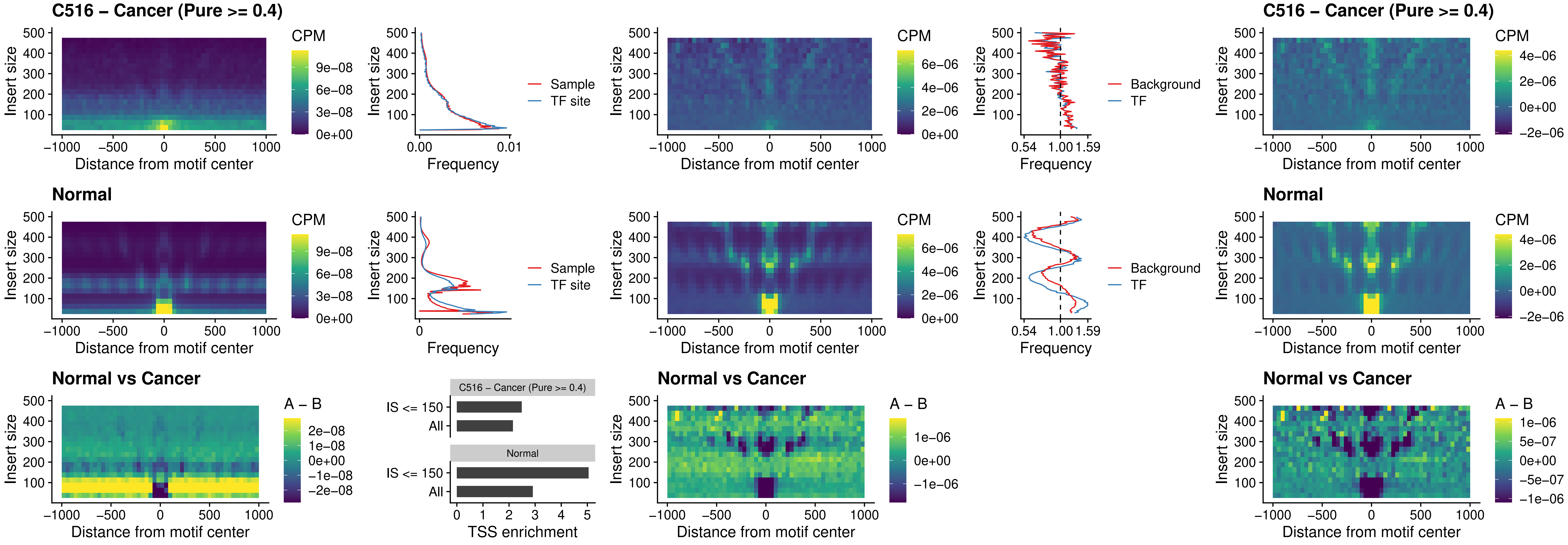

#### C516 – Cancer (Pure >= 0.4) – NEUROD1_uc_cTSS_oPEAK

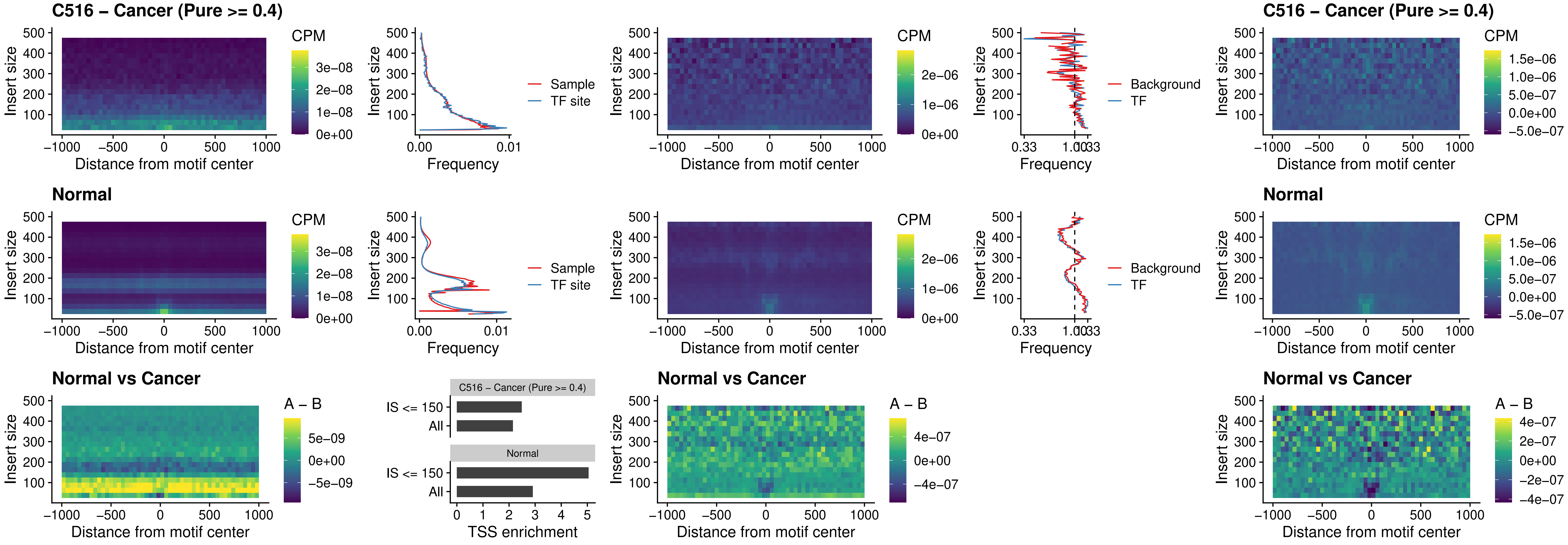

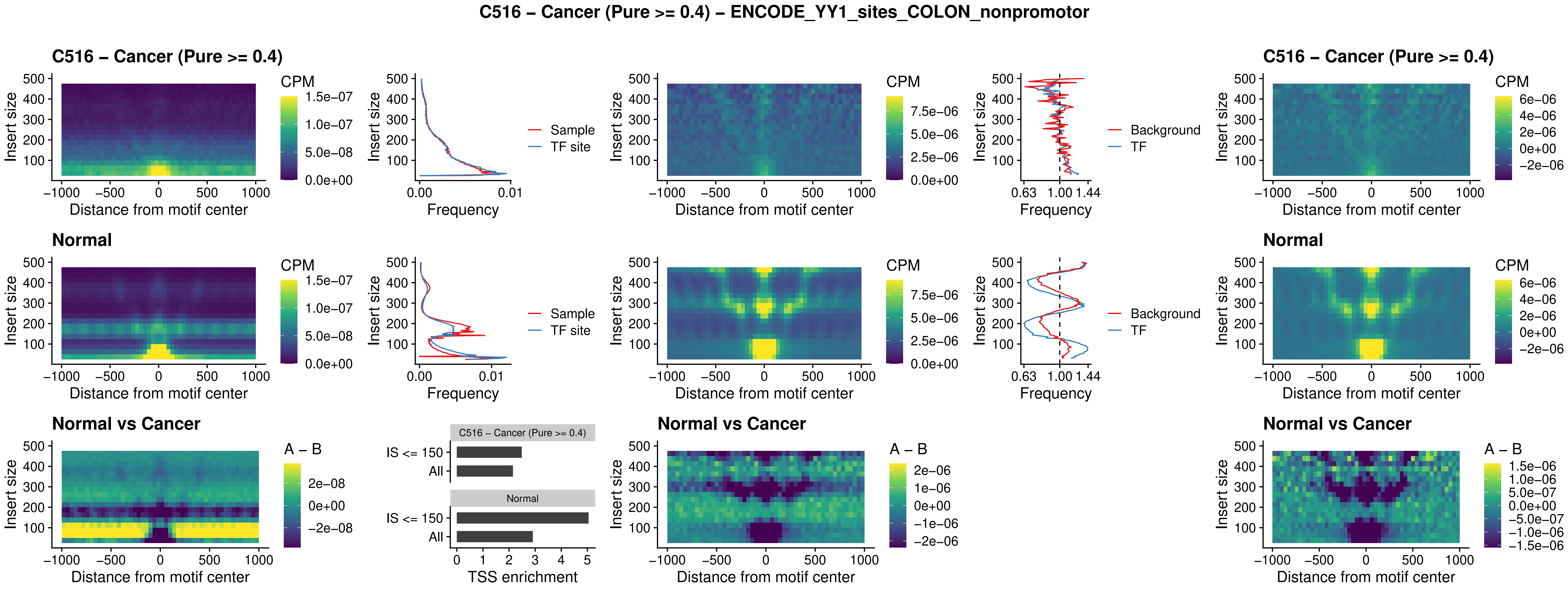
C516 – Cancer (Pure >= 0.4) – ENCODE_YY1_sites_COLON_nonpromotor
C516 – Cancer (Pure >= 0.4)
Normal
Normal vs Cancer
Insert size
Distance from motif center
CPM
A – B
Frequency
Sample
TF site
Background
TF
C516 – Cancer (Pure >= 0.4)
IS <= 150
All
TSS enrichment

#### C516 – Cancer (Pure >= 0.4) – CTCF_uc_pTSS_oPEAK

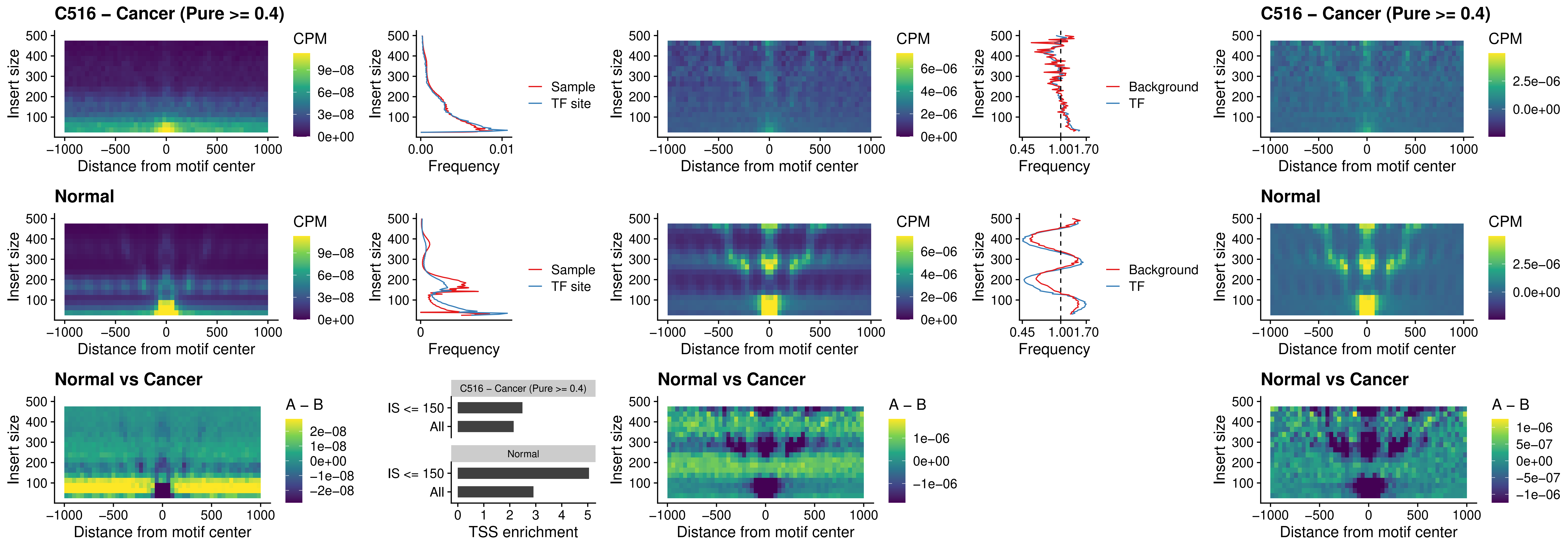

#### C516 – Cancer (Pure >= 0.4) – ENCODE_CTCF_sites_COLON_promotor

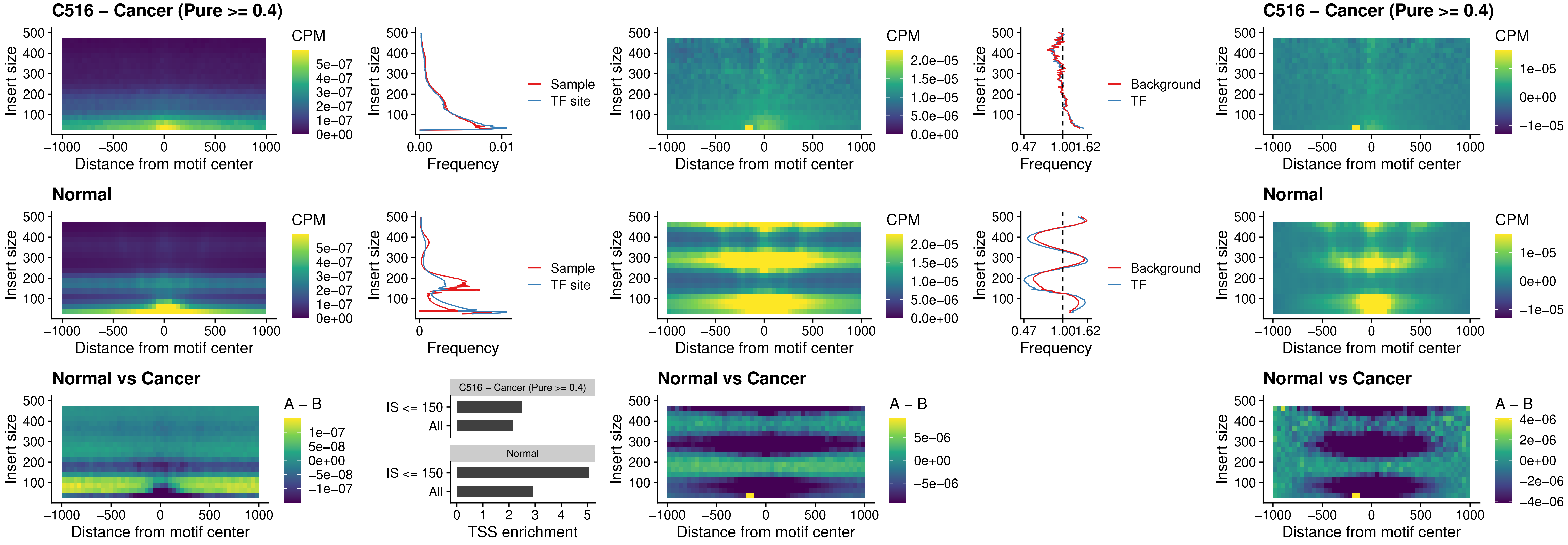

#### C516 – Cancer (Pure >= 0.4) – ENCODE_CTCF_sites_COLON_promotor_ctcf

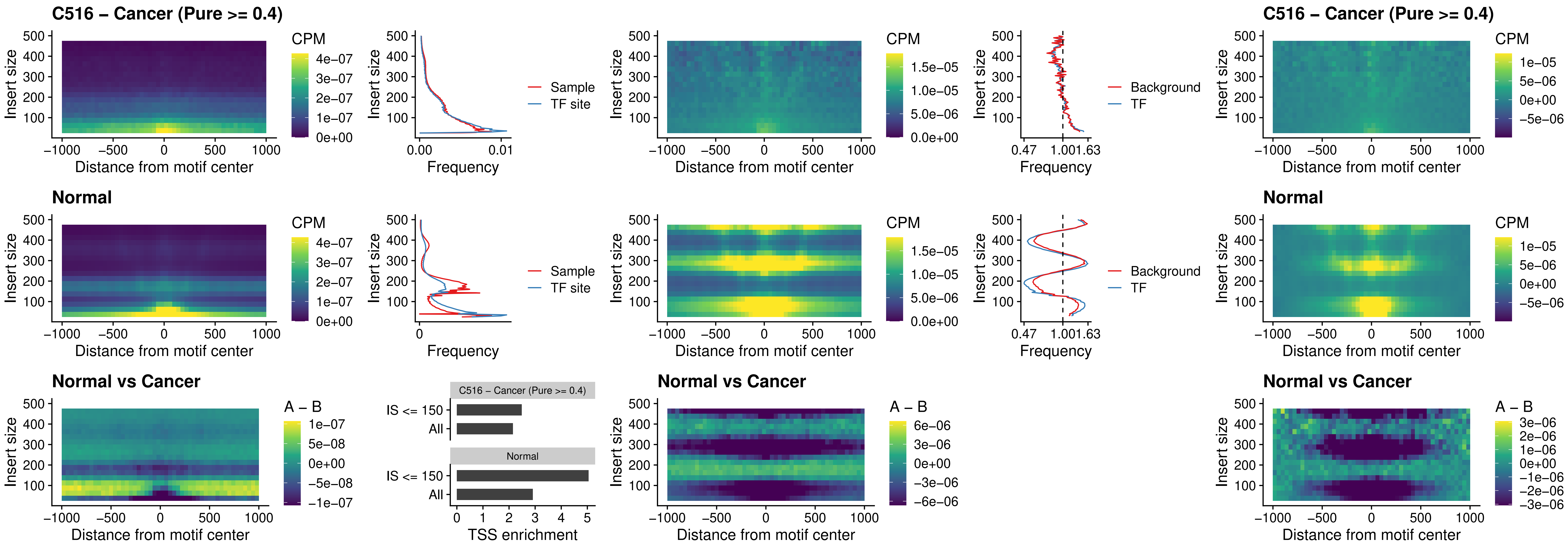

#### C516 – Cancer (Pure >= 0.4) – SPIC_uc_pTSS_oPEAK

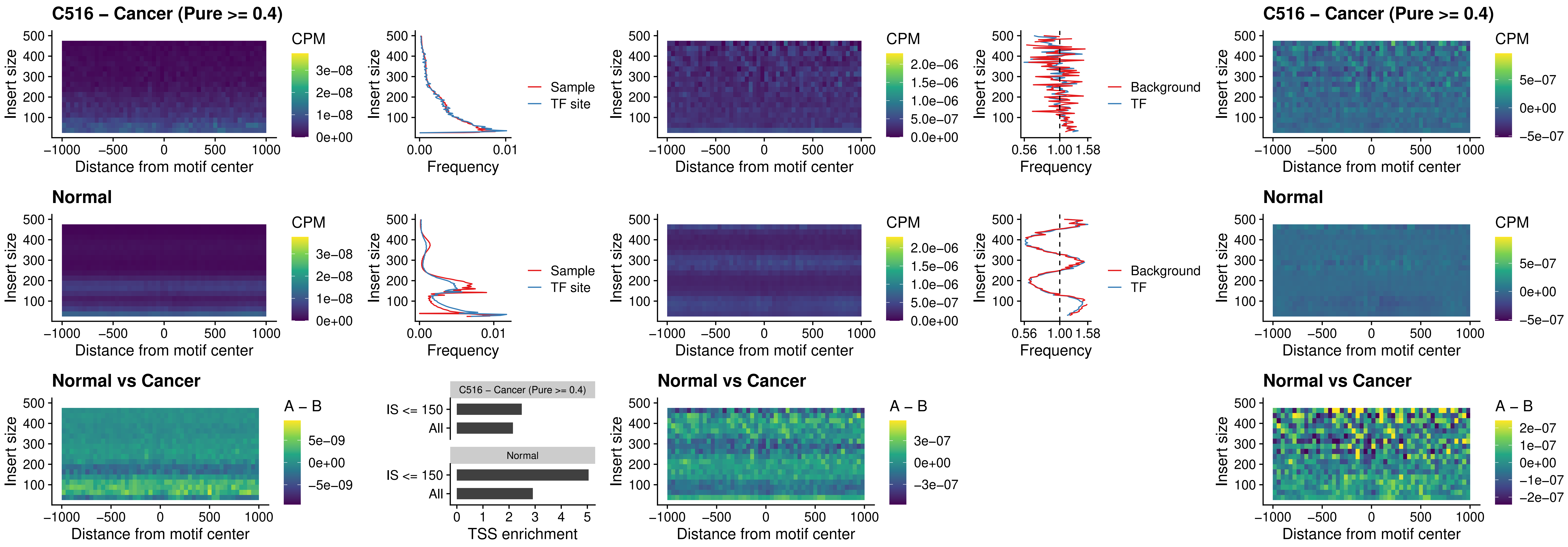

#### C516 – Cancer (Pure >= 0.4) – SPI1_uc_dTSS_nPEAK

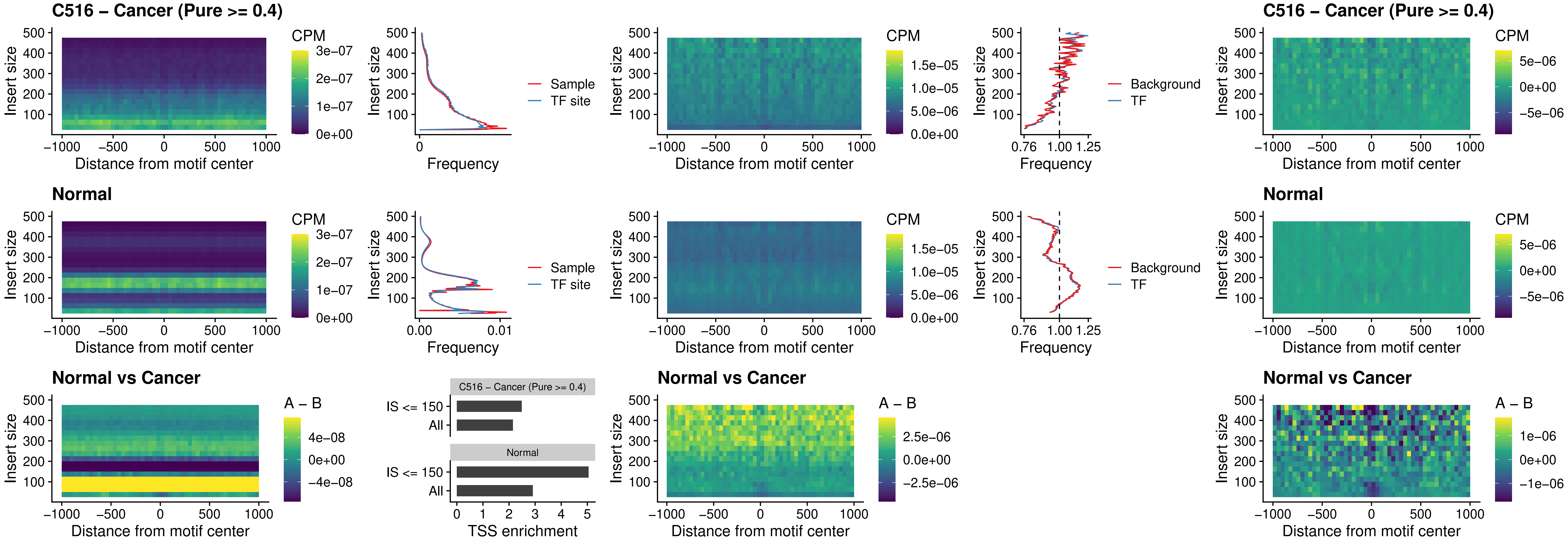

#### C516 – Cancer (Pure >= 0.4) – SPI1_uc_cTSS_nPEAK

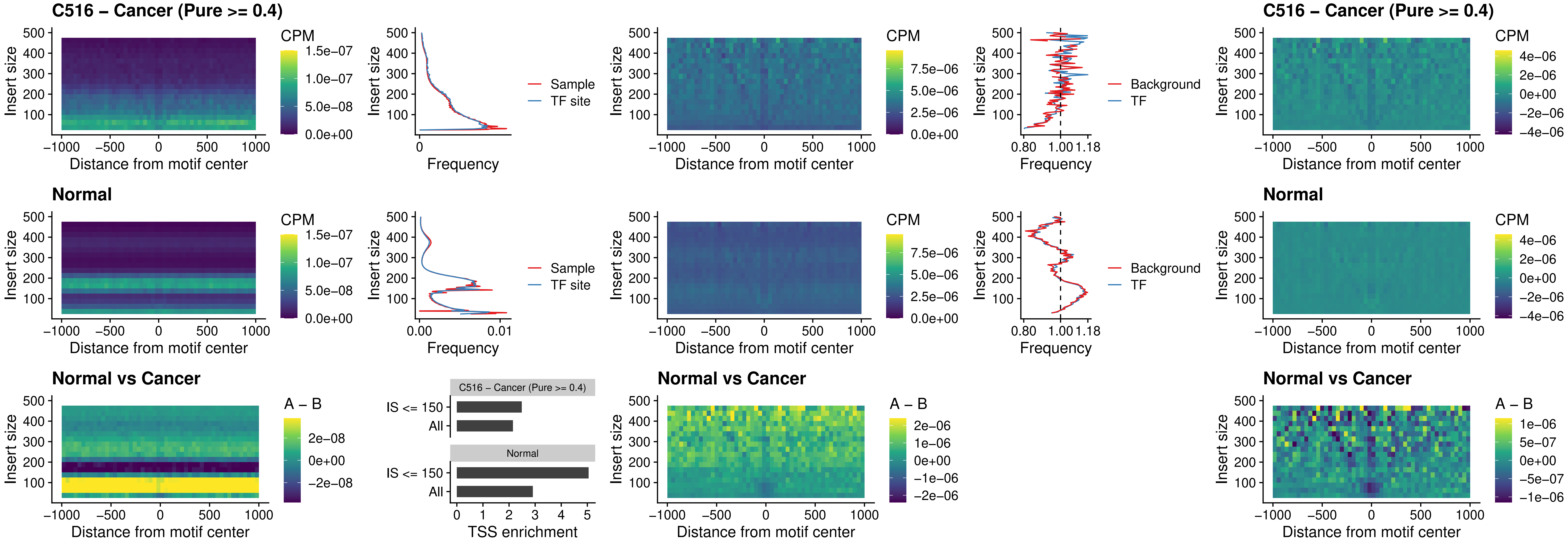

#### C516 – Cancer (Pure >= 0.4) – BCL11A_uc_pTSS_nPEAK

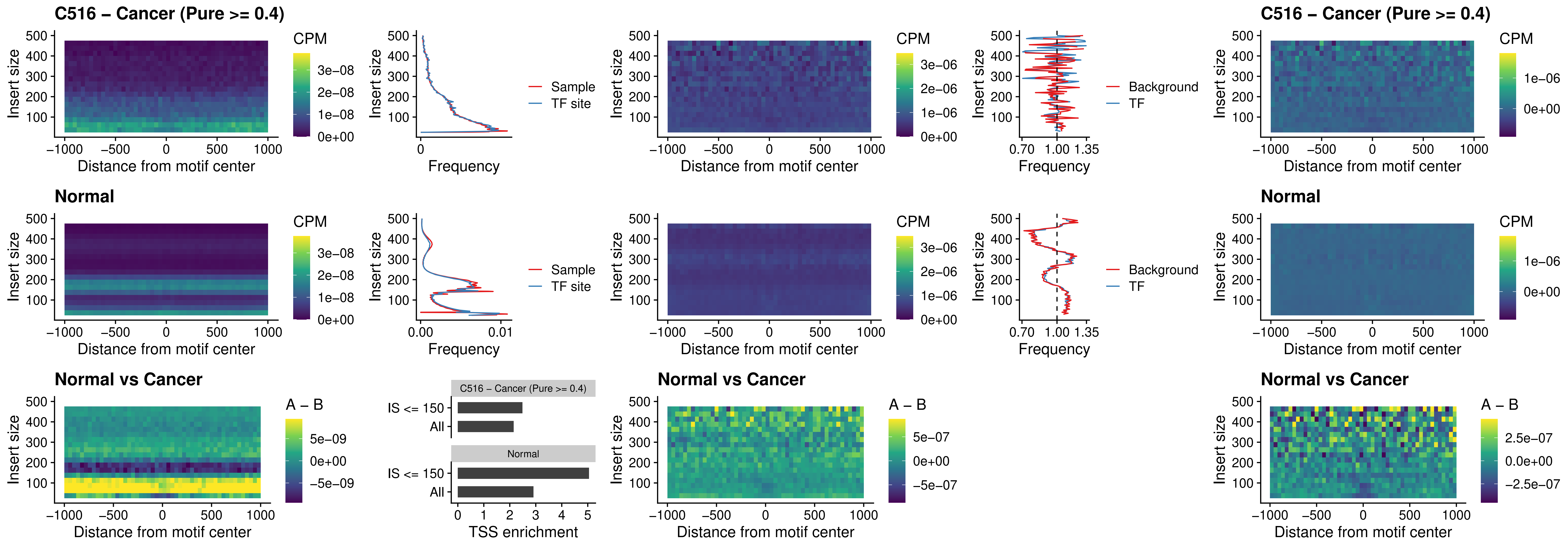

#### C516 – Cancer (Pure >= 0.4) – BCL11B_uc_pTSS_nPEAK

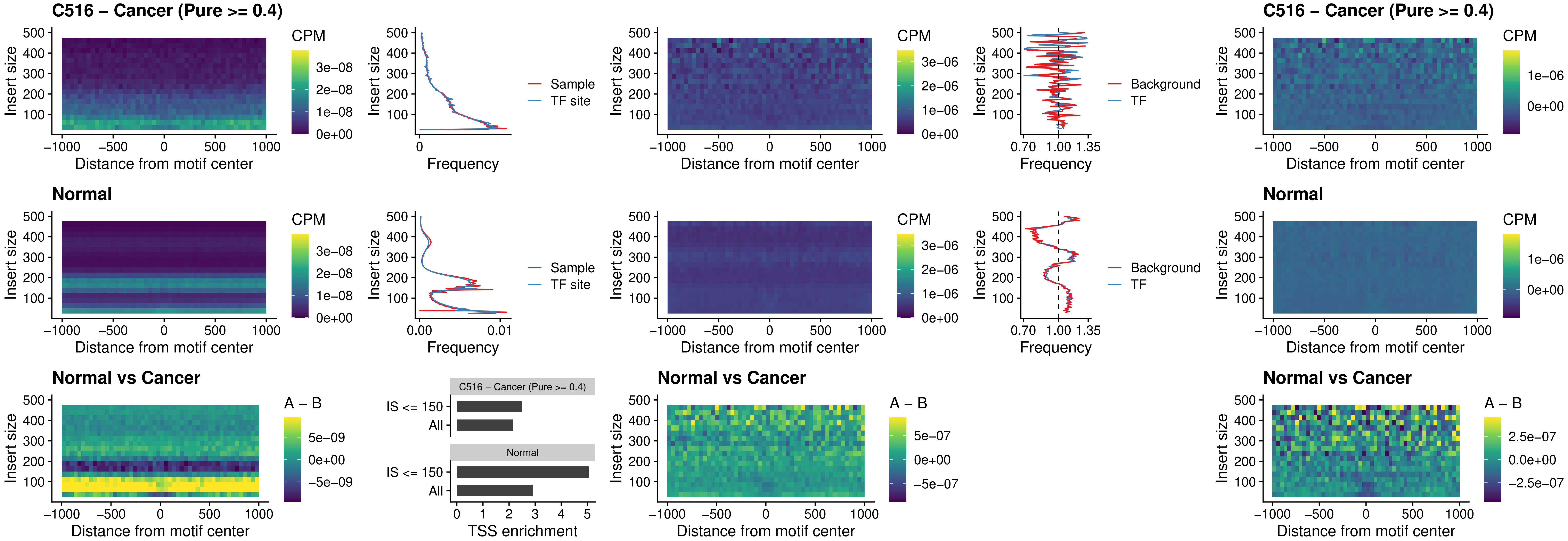

#### C516 – Cancer (Pure >= 0.4) – IRF9_uc_dTSS_oPEAK

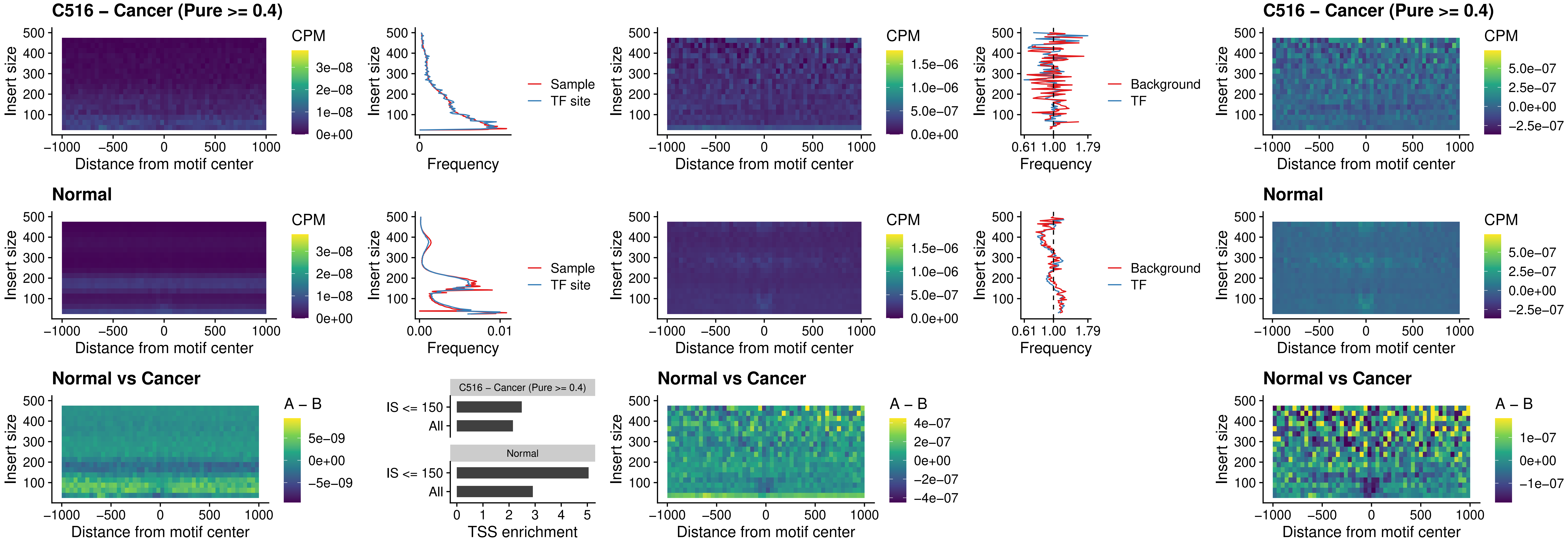

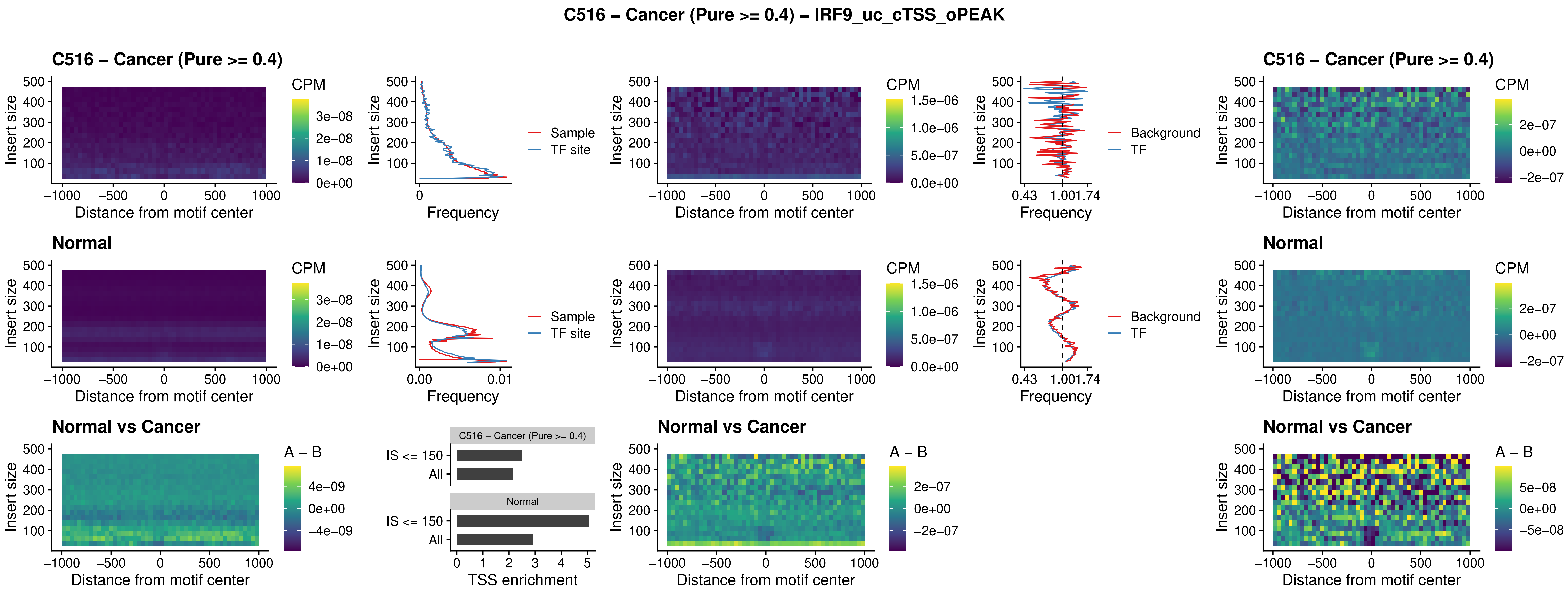
C516 – Cancer (Pure >= 0.4) – IRF9_uc_cTSS_oPEAK
C516 – Cancer (Pure >= 0.4)
Normal
Normal vs Cancer
Insert size
Distance from motif center
Frequency
CPM
A – B
Sample
TF site
Background
TF
TSS enrichment
IS <= 150
All
C516 – Cancer (Pure >= 0.4)
Normal

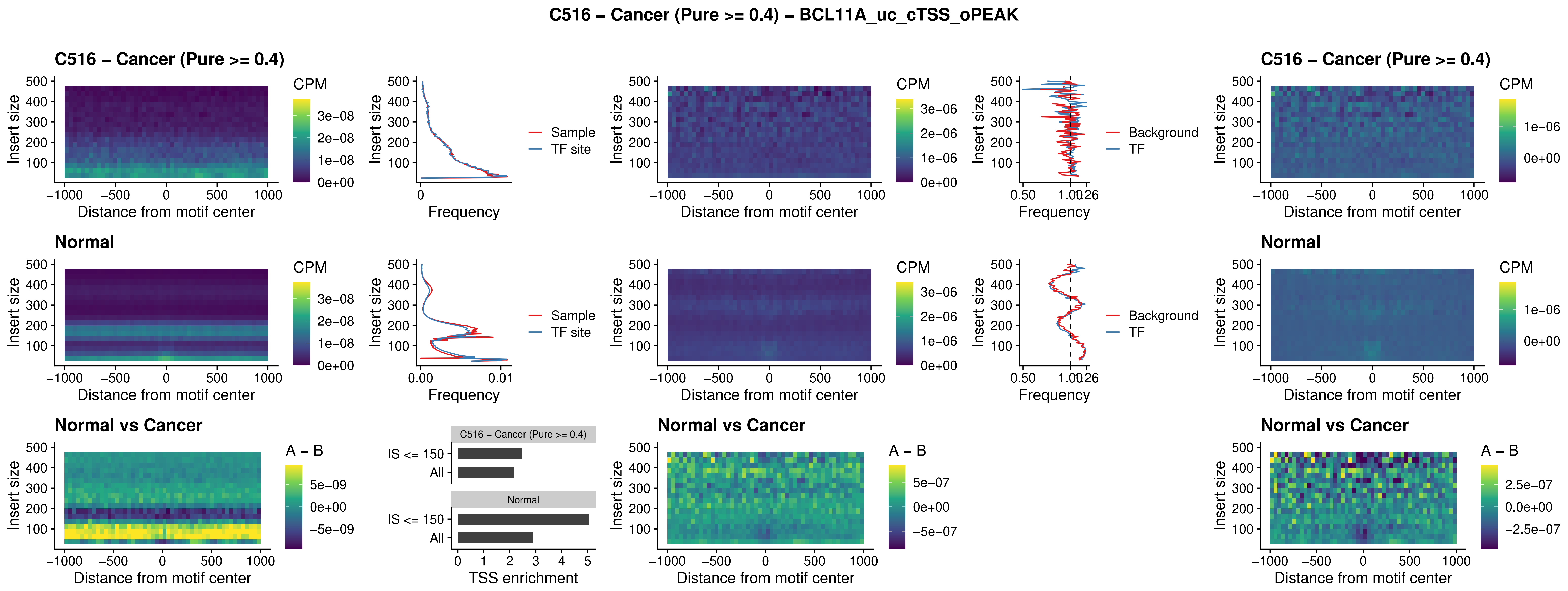
C516 – Cancer (Pure >= 0.4) – BCL11A_uc_cTSS_oPEAK
C516 – Cancer (Pure >= 0.4)
Normal
Normal vs Cancer
Insert size
Distance from motif center
Frequency
CPM
A – B
Sample
TF site
Background
TF
C516 – Cancer (Pure >= 0.4)
Normal
IS <= 150
All
TSS enrichment

#### C516 – Cancer (Pure >= 0.4) – BCL11B_uc_cTSS_oPEAK

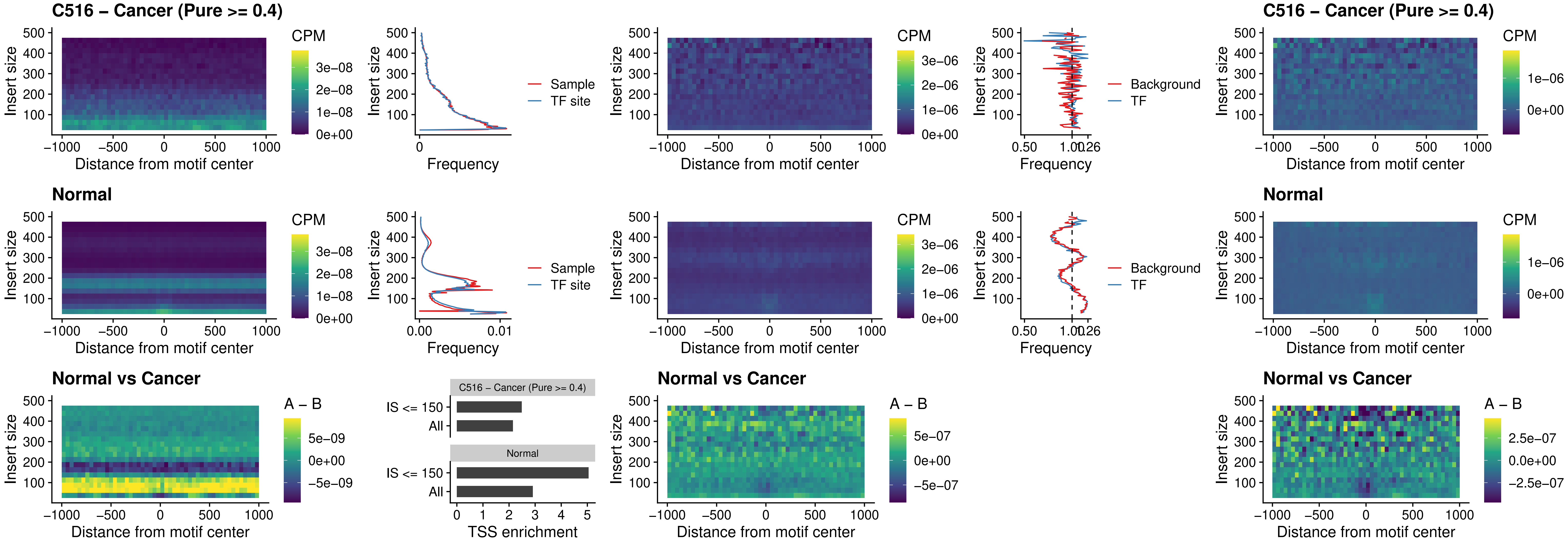

### C516 – Cancer (Pure >= 0.4) – BCL11A_uc_dTSS_oPEAK

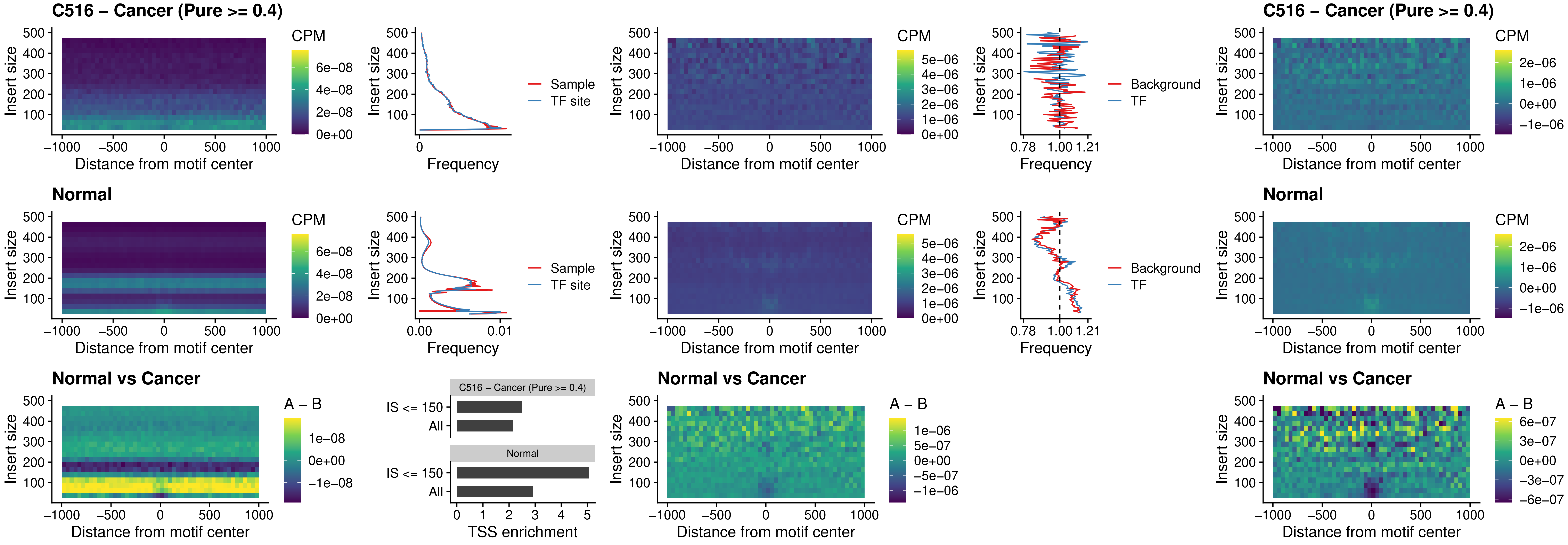

#### C516 – Cancer (Pure >= 0.4) – BCL11B_uc_dTSS_oPEAK

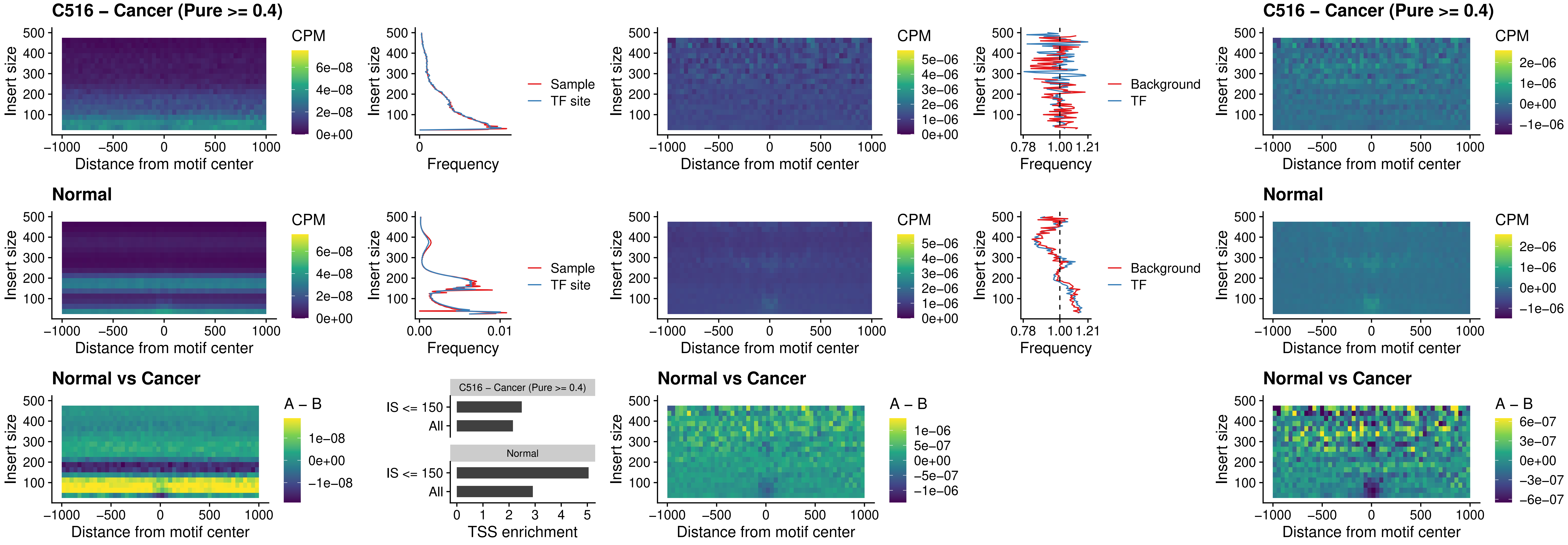

#### C516 – Cancer (Pure >= 0.4) – SPIB_uc_cTSS_nPEAK

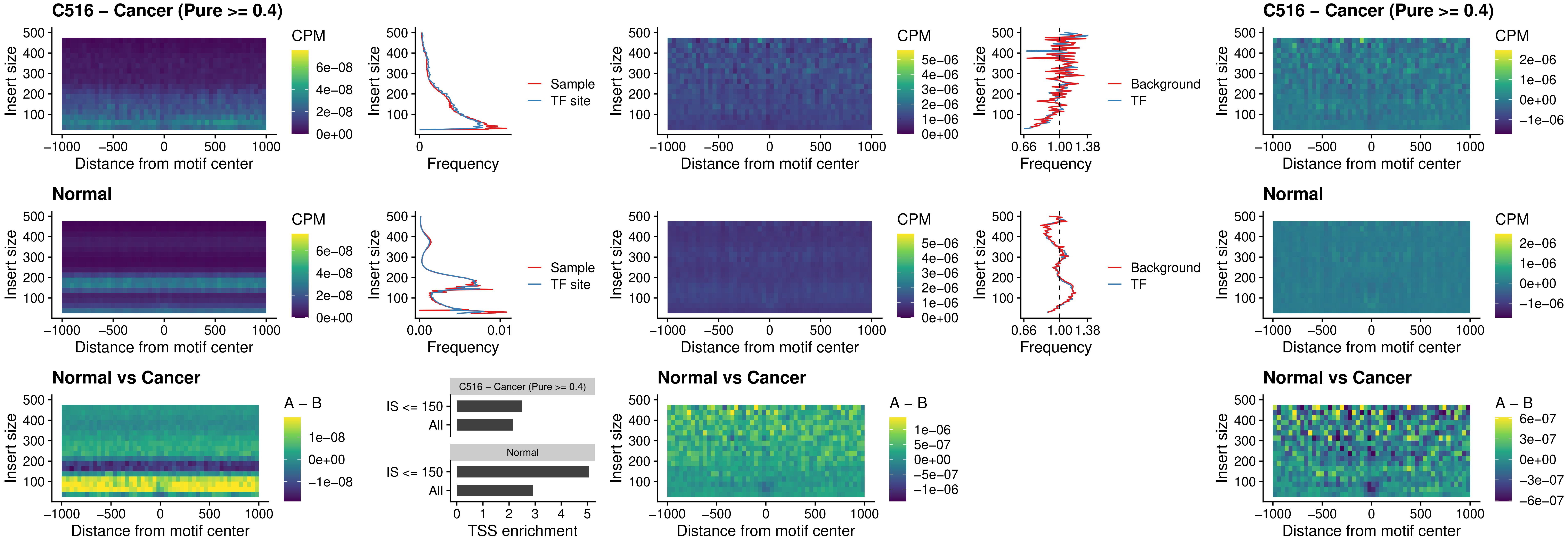

#### C516 – Cancer (Pure >= 0.4) – SPIC_uc_cTSS_nPEAK

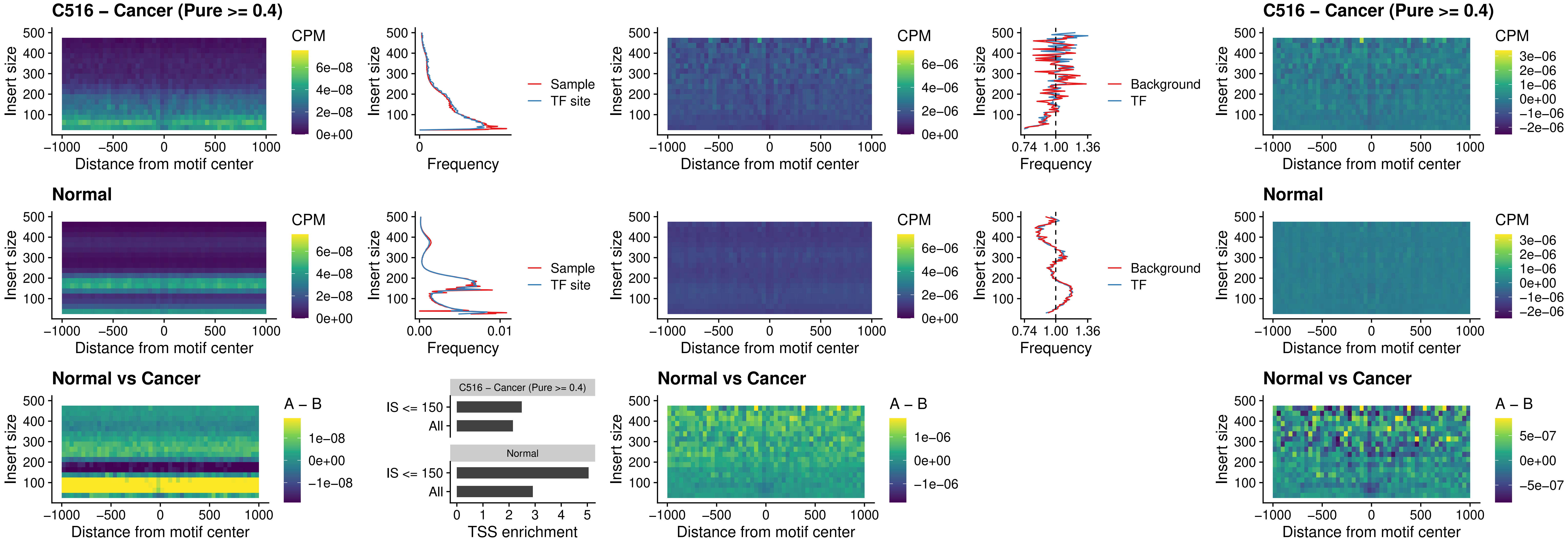

#### C516 – Cancer (Pure >= 0.4) – SPI1_uc_pTSS_oPEAK

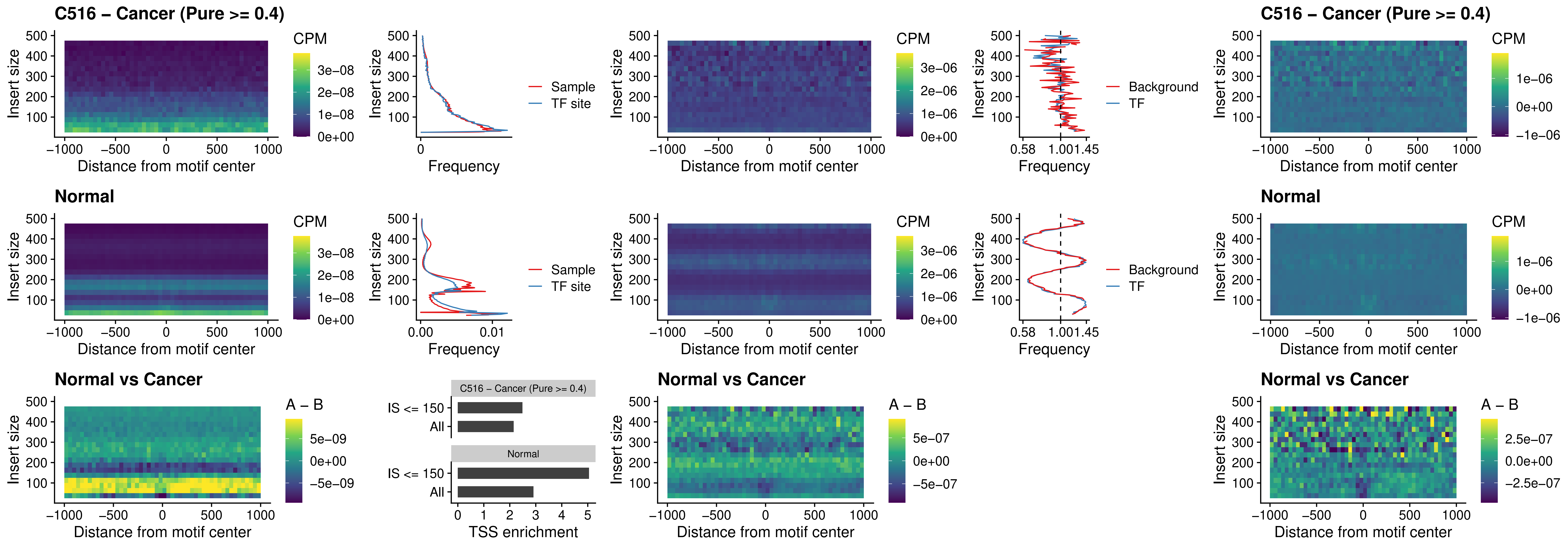

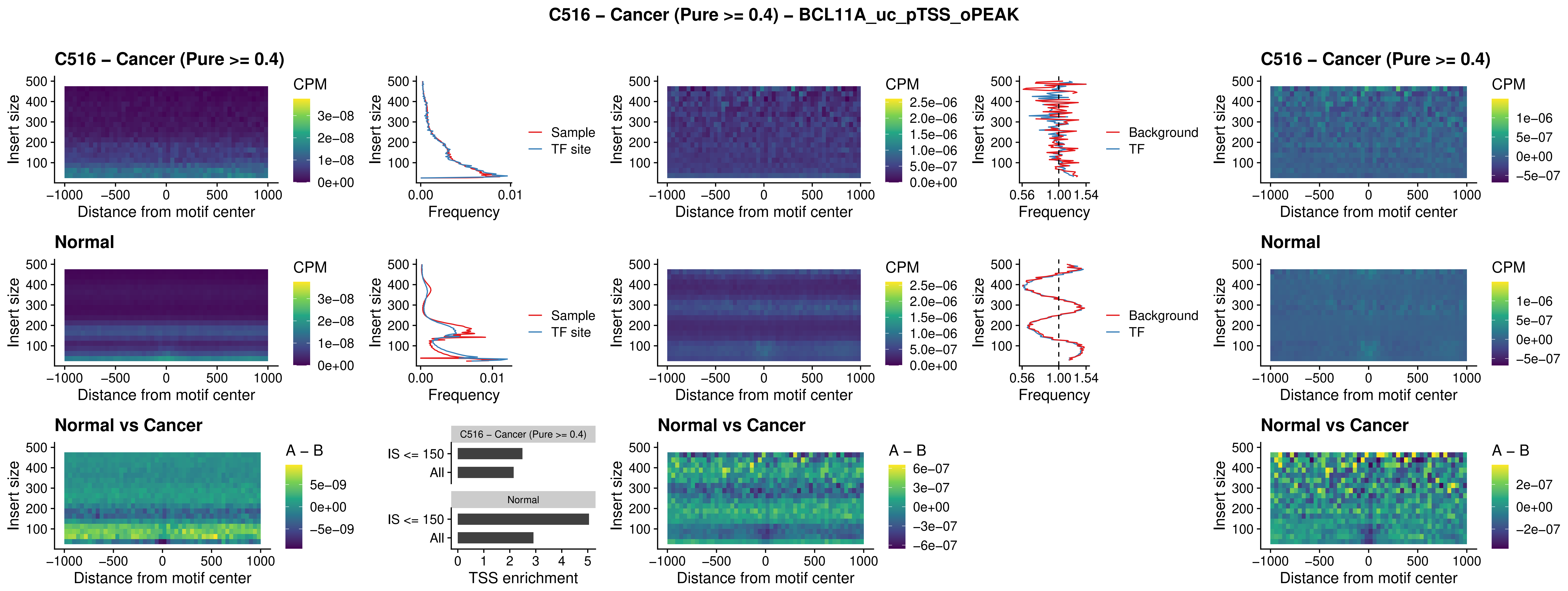

C516 – Cancer (Pure >= 0.4) – BCL11A_uc_pTSS_oPEAK
C516 – Cancer (Pure >= 0.4)
Normal
Normal vs Cancer
Insert size
Distance from motif center
Frequency
CPM
A – B
Sample
TF site
Background
TF
TSS enrichment
IS <= 150
All
C516 – Cancer (Pure >= 0.4)
Normal

#### C516 – Cancer (Pure >= 0.4) – BCL11B_uc_pTSS_oPEAK

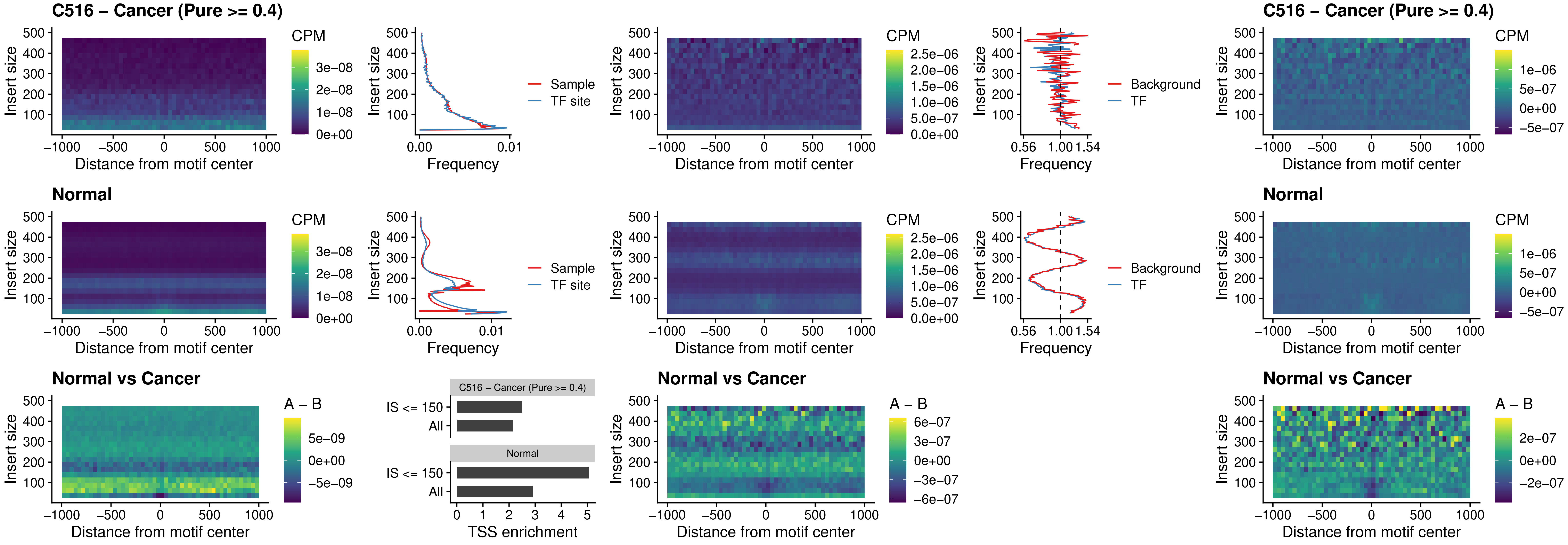

#### C516 – Cancer (Pure >= 0.4) – SPI1_uc_cTSS_oPEAK

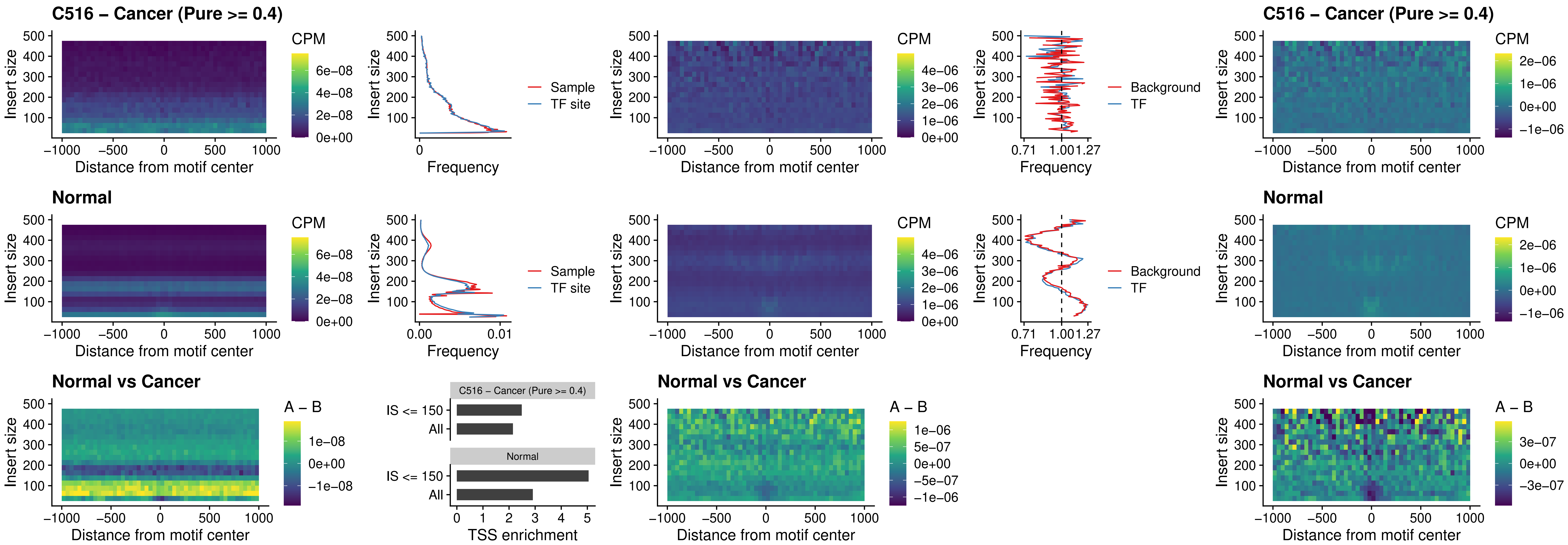

### C516 – Cancer (Pure >= 0.4) – SPIB_uc_cTSS_oPEAK

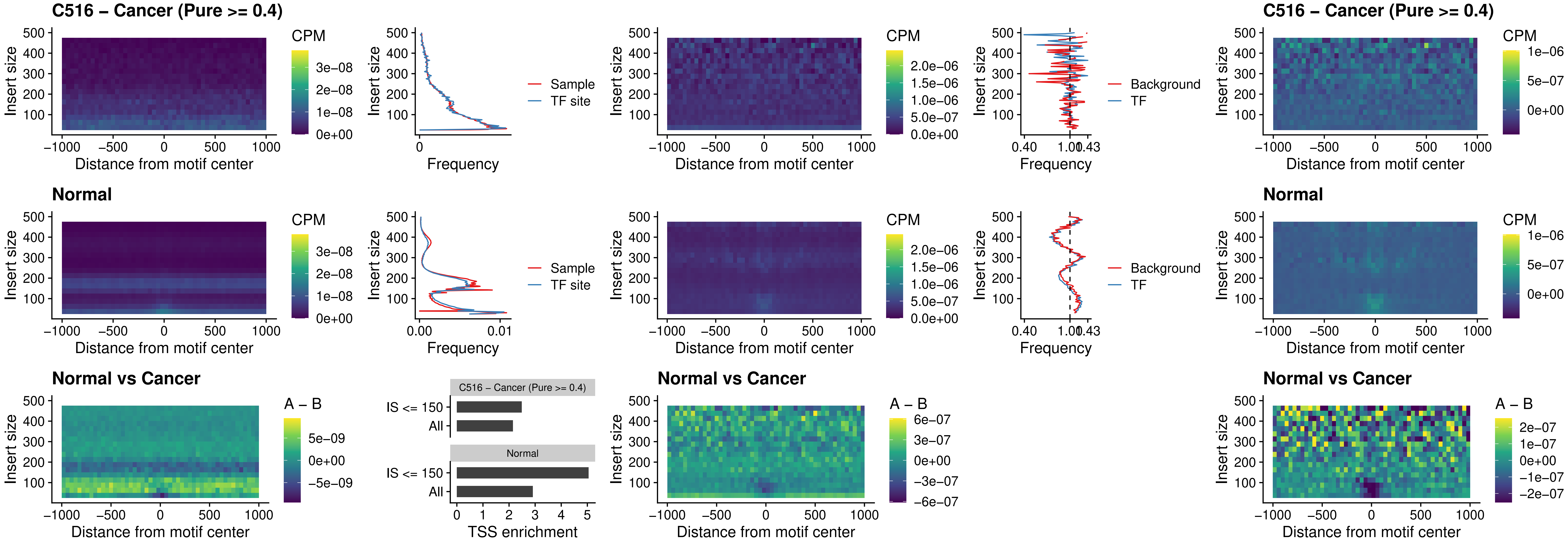

### C516 – Cancer (Pure >= 0.4) – SPIB_uc_pTSS_oPEAK

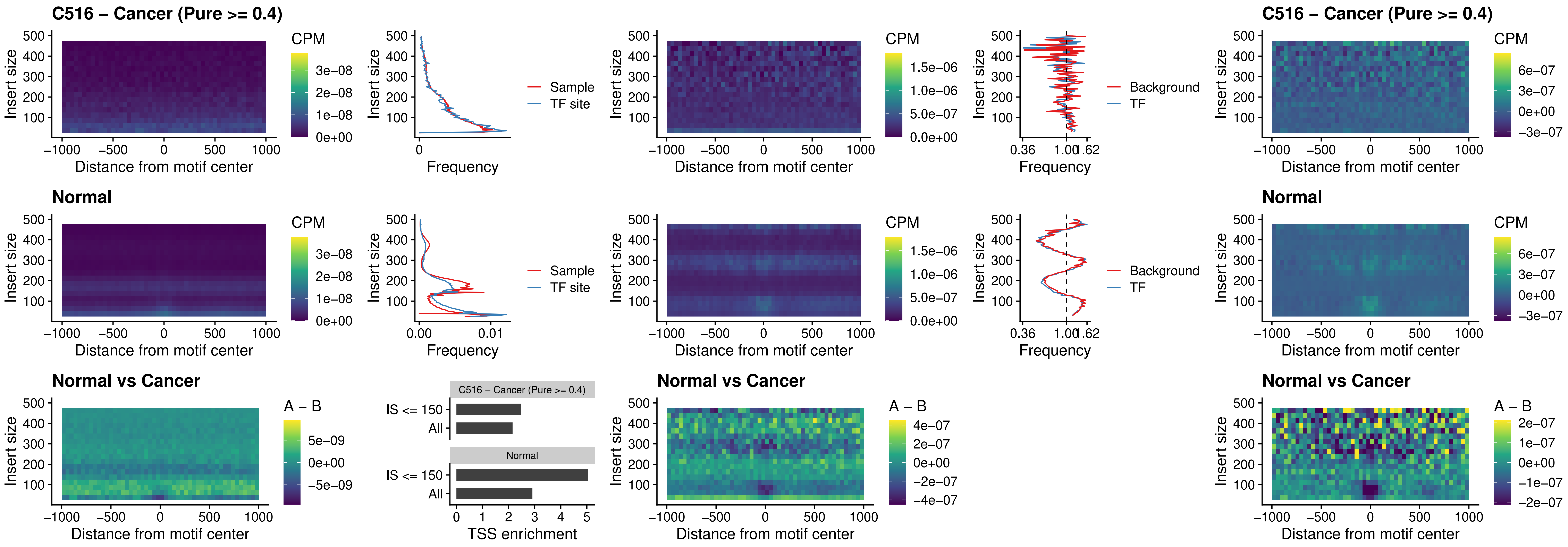

#### C516 – Cancer (Pure >= 0.4) – SPI1_uc_dTSS_oPEAK

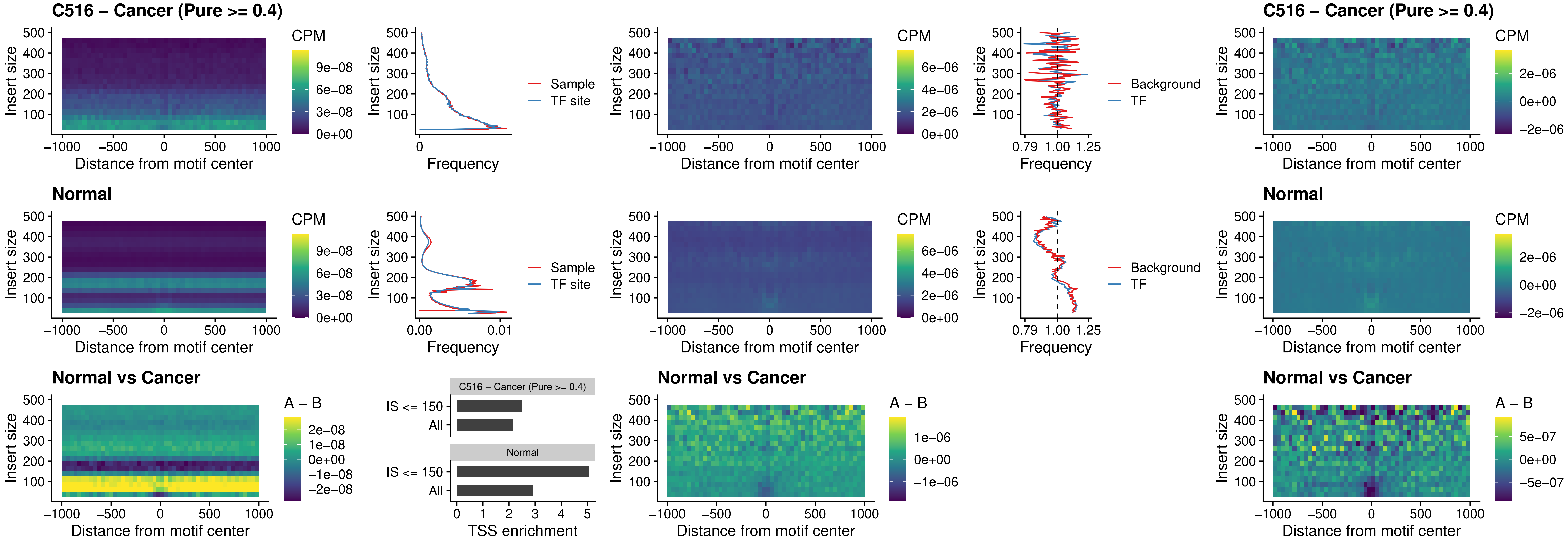

#### C516 – Cancer (Pure >= 0.4) – SPIB_uc_dTSS_oPEAK

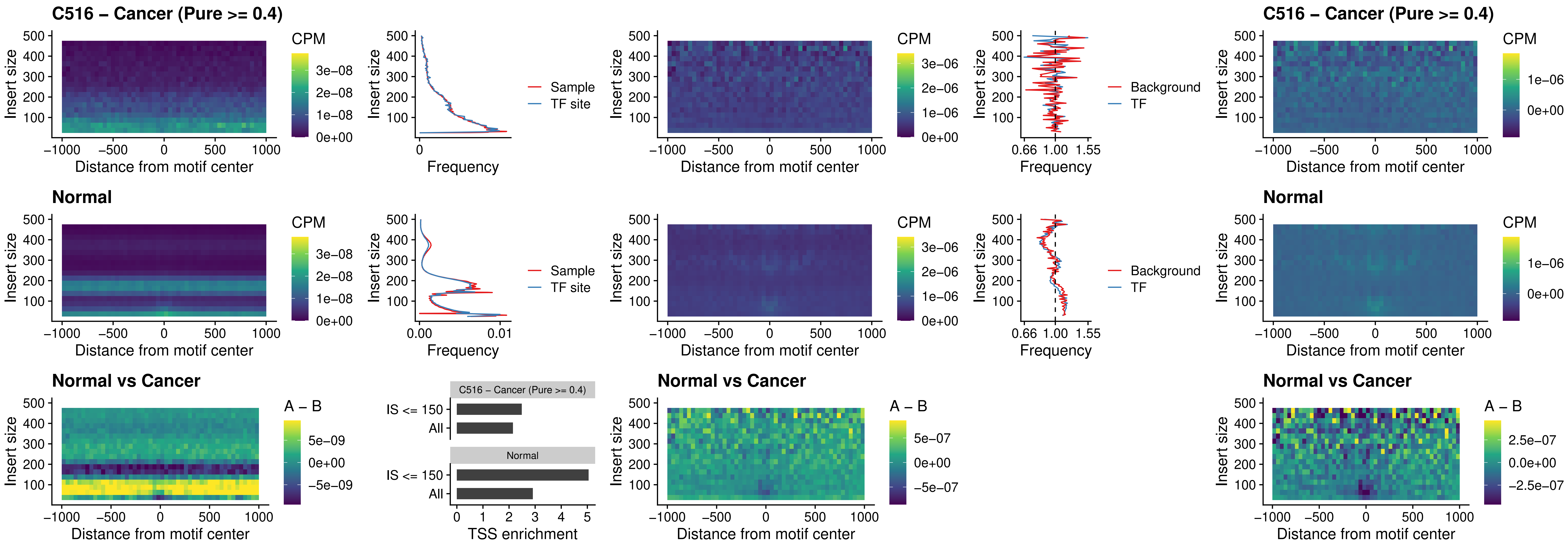

#### C516 – Cancer (Pure >= 0.4) – IRF3_uc_pTSS_oPEAK

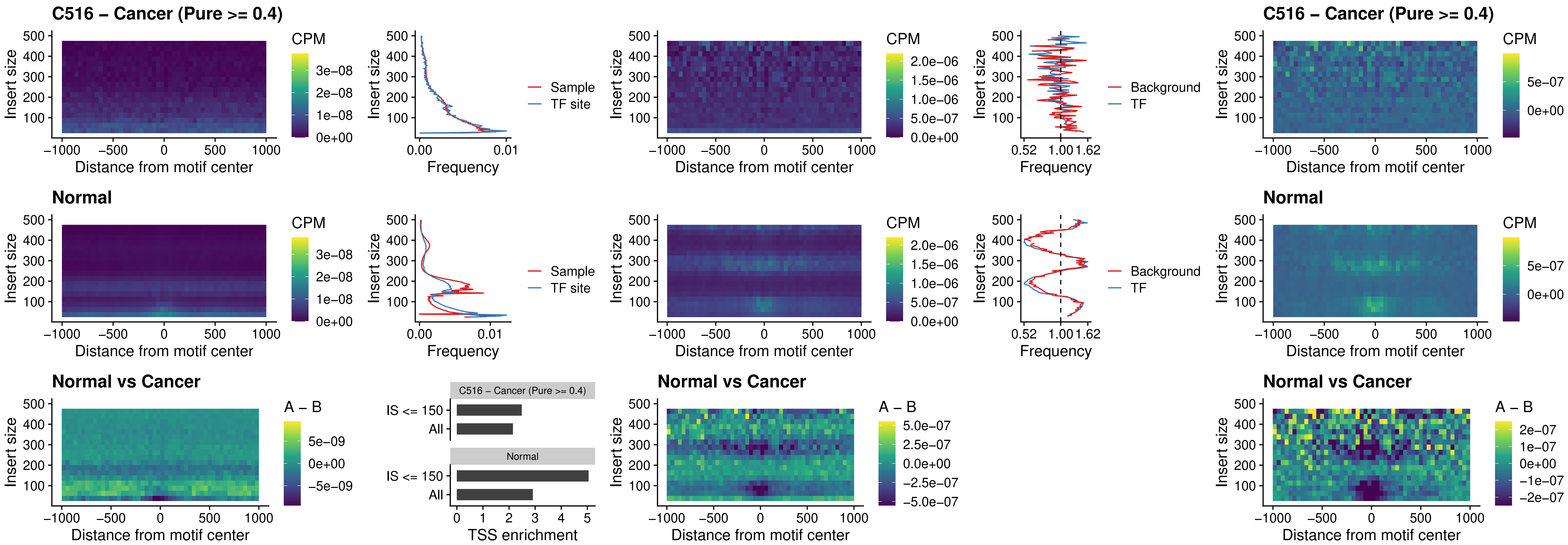

#### C516 – Cancer (Pure >= 0.4) – IRF2_uc_dTSS_oPEAK

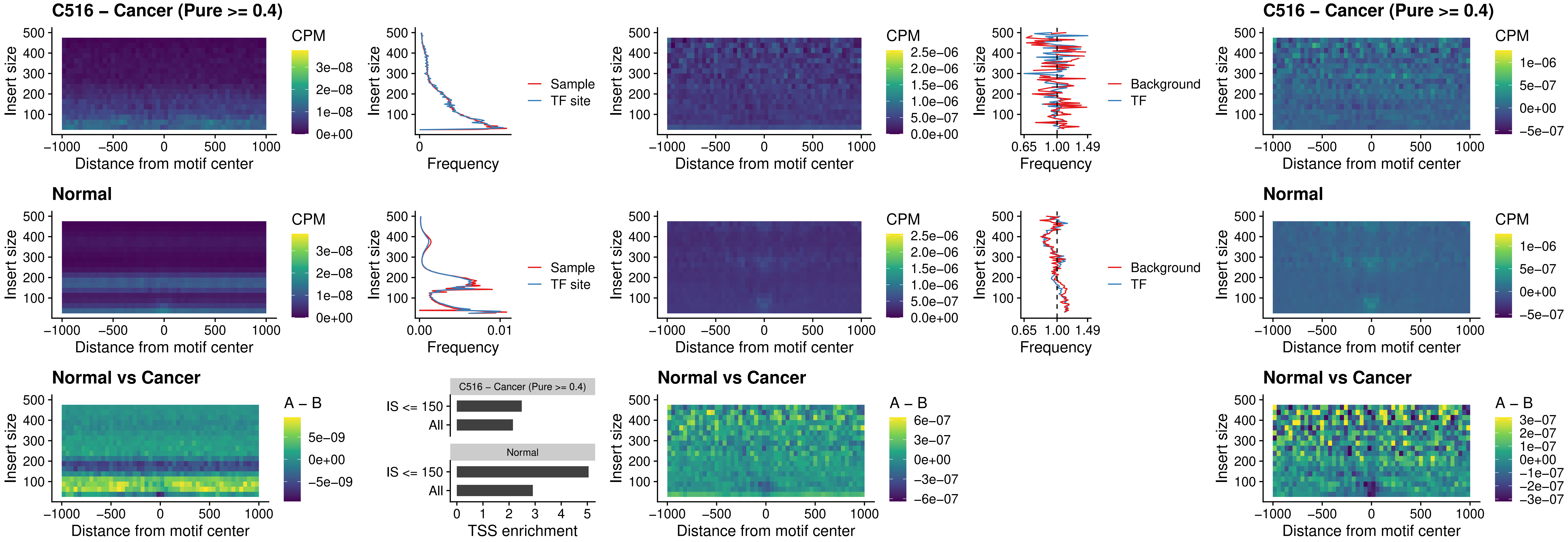

#### C516 – Cancer (Pure >= 0.4) – IRF2_uc_pTSS_oPEAK

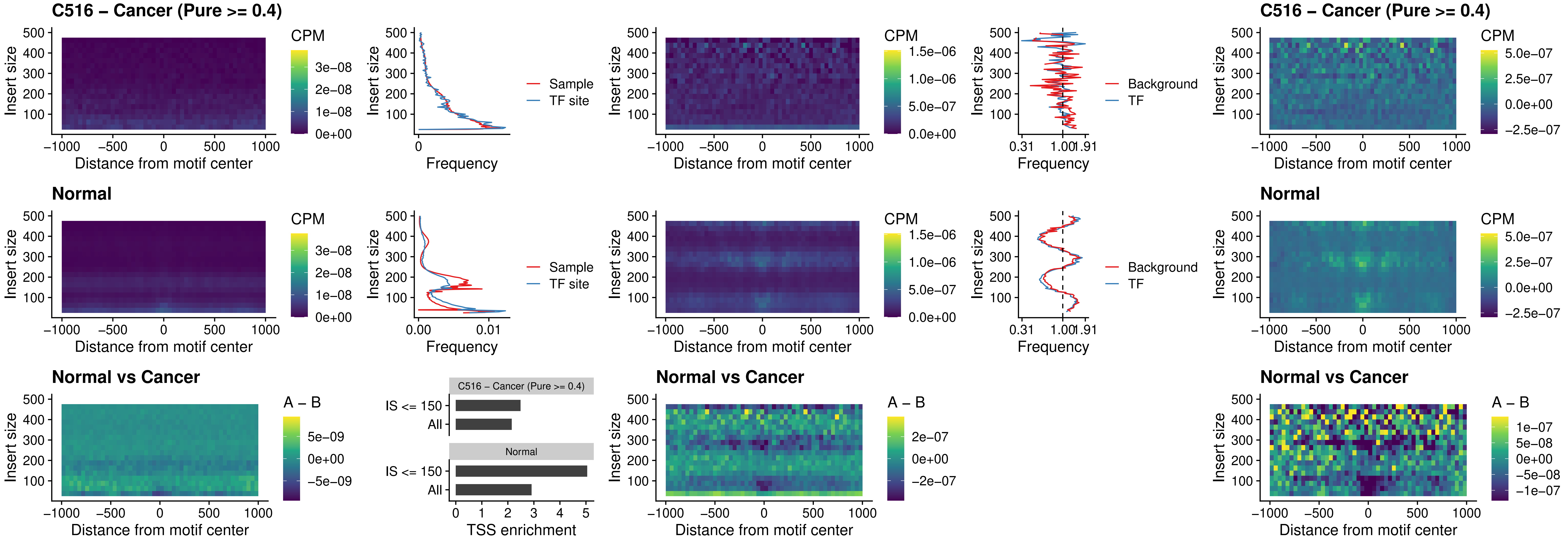

#### C516 – Cancer (Pure >= 0.4) – STAT2_uc_pTSS_oPEAK

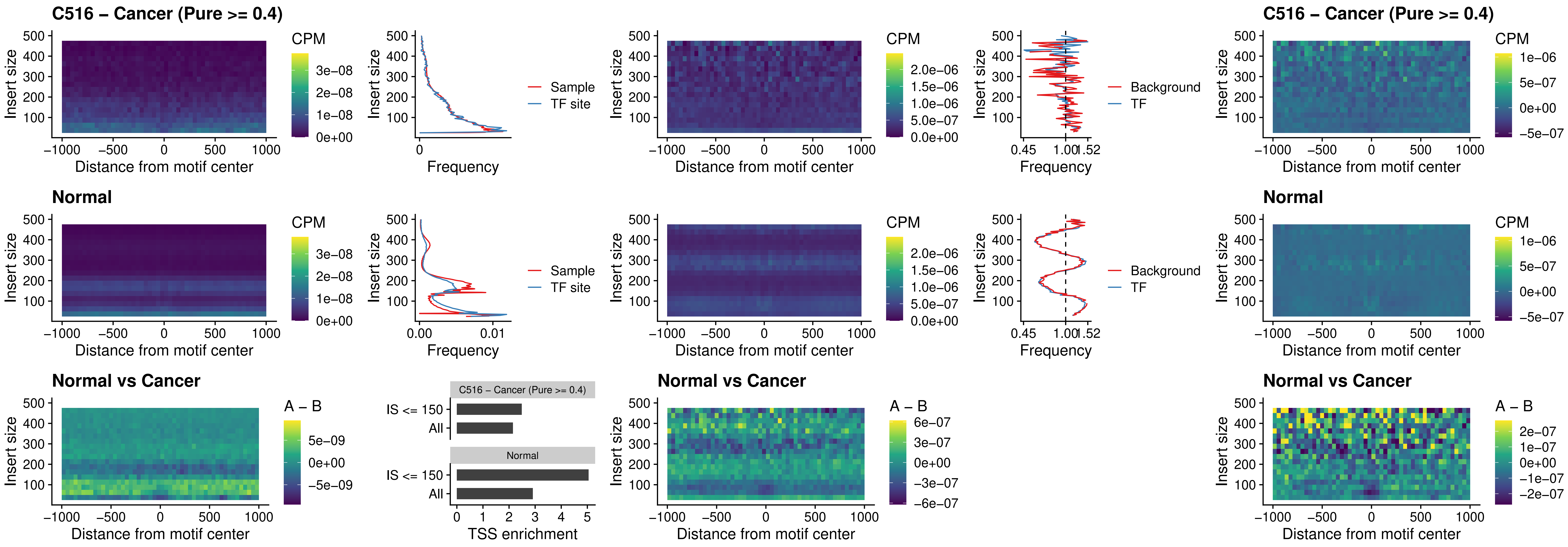

#### C516 – Cancer (Pure >= 0.4) – SPIB_uc_dTSS_nPEAK

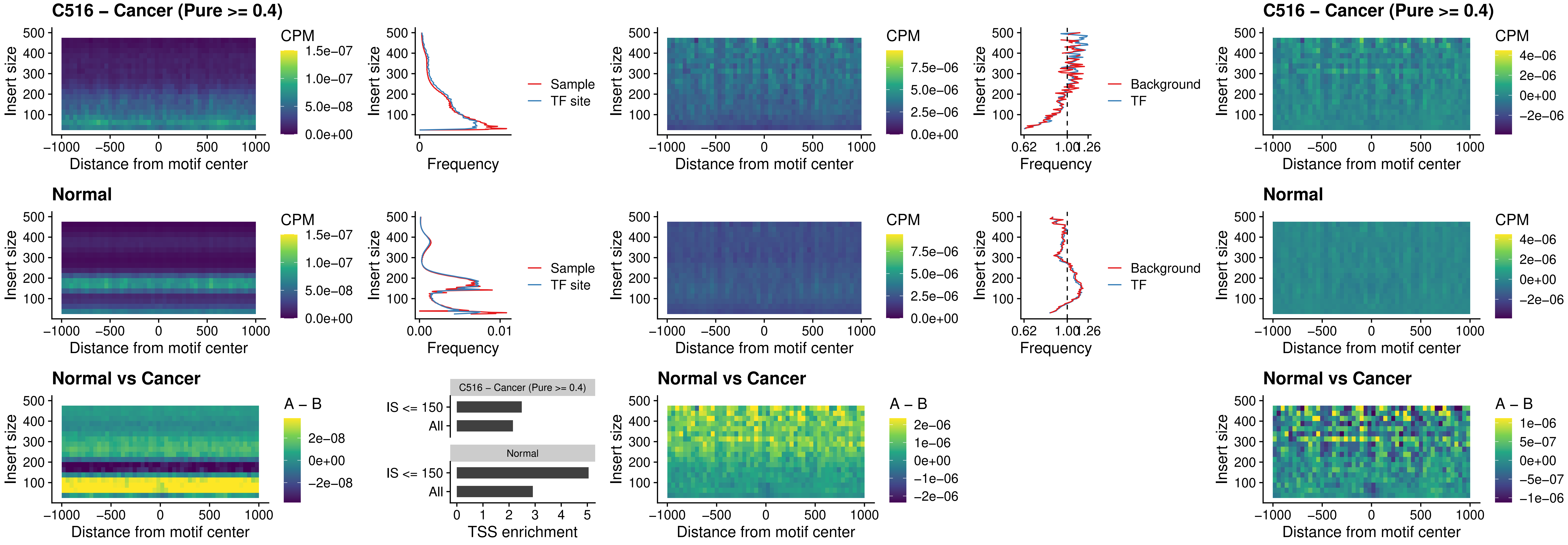

#### C516 – Cancer (Pure >= 0.4) – SPIC_uc_cTSS_oPEAK

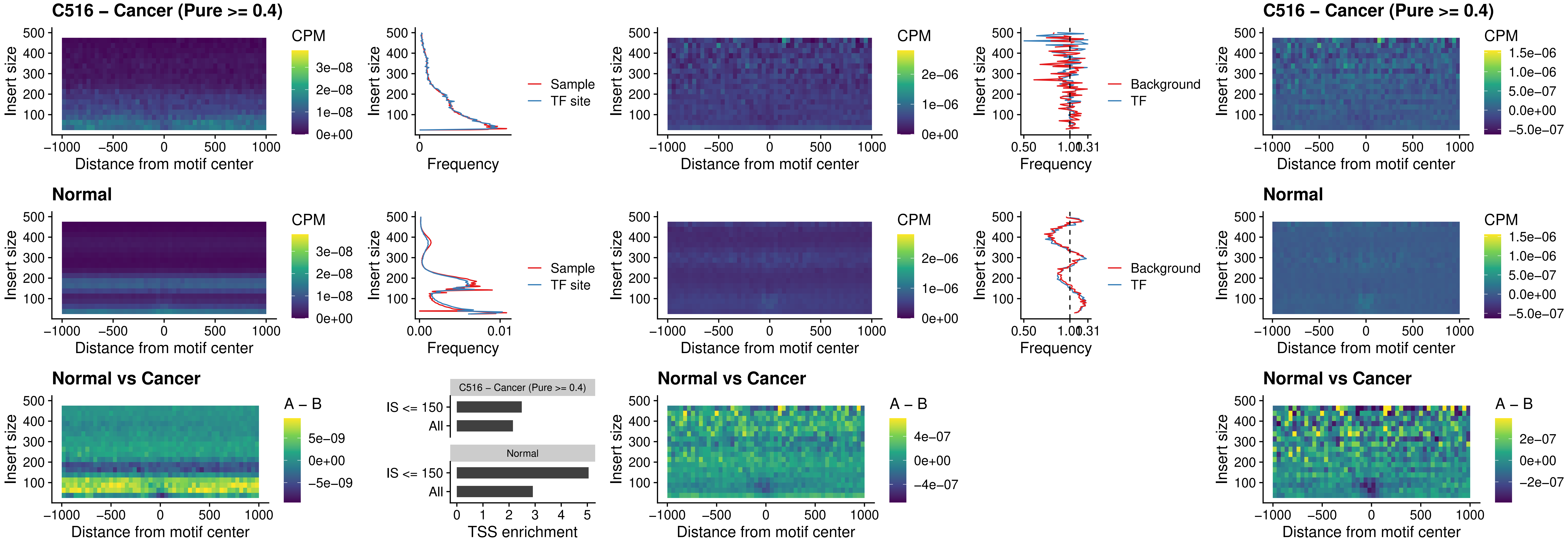

#### C516 – Cancer (Pure >= 0.4) – PRDM1_uc_cTSS_oPEAK

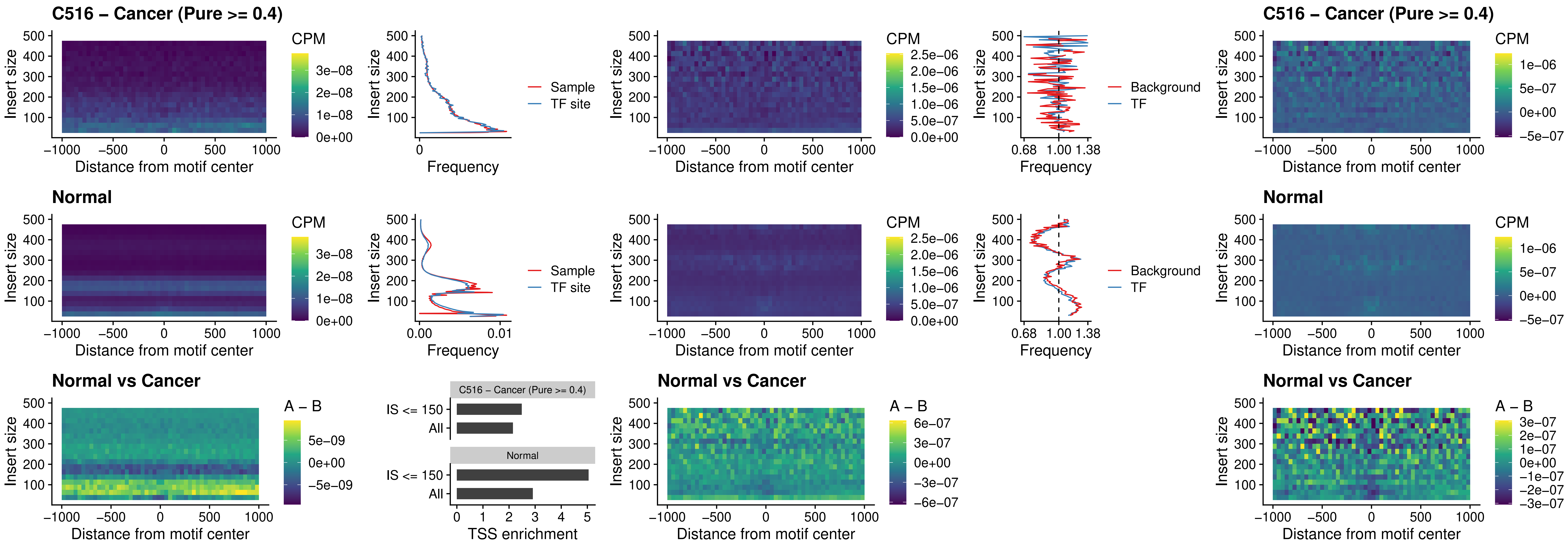

#### C516 – Cancer (Pure >= 0.4) – IRF3_uc_cTSS_oPEAK

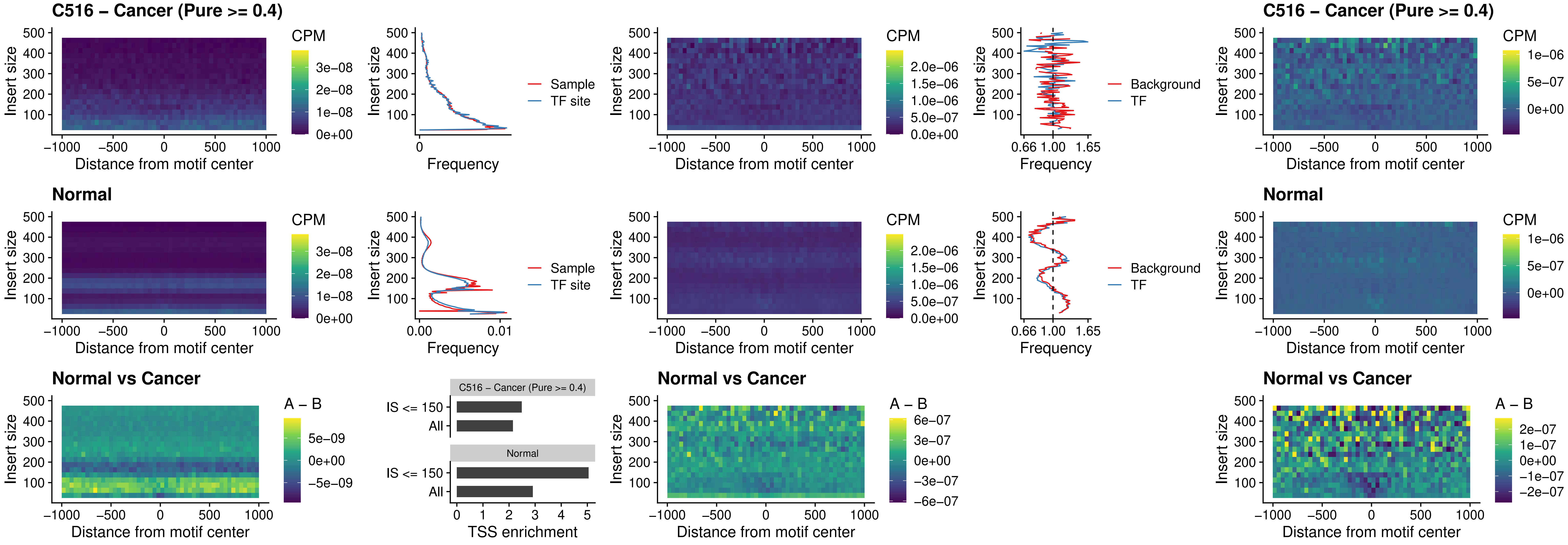

#### C516 – Cancer (Pure >= 0.4) – BCL11A_uc_dTSS_nPEAK

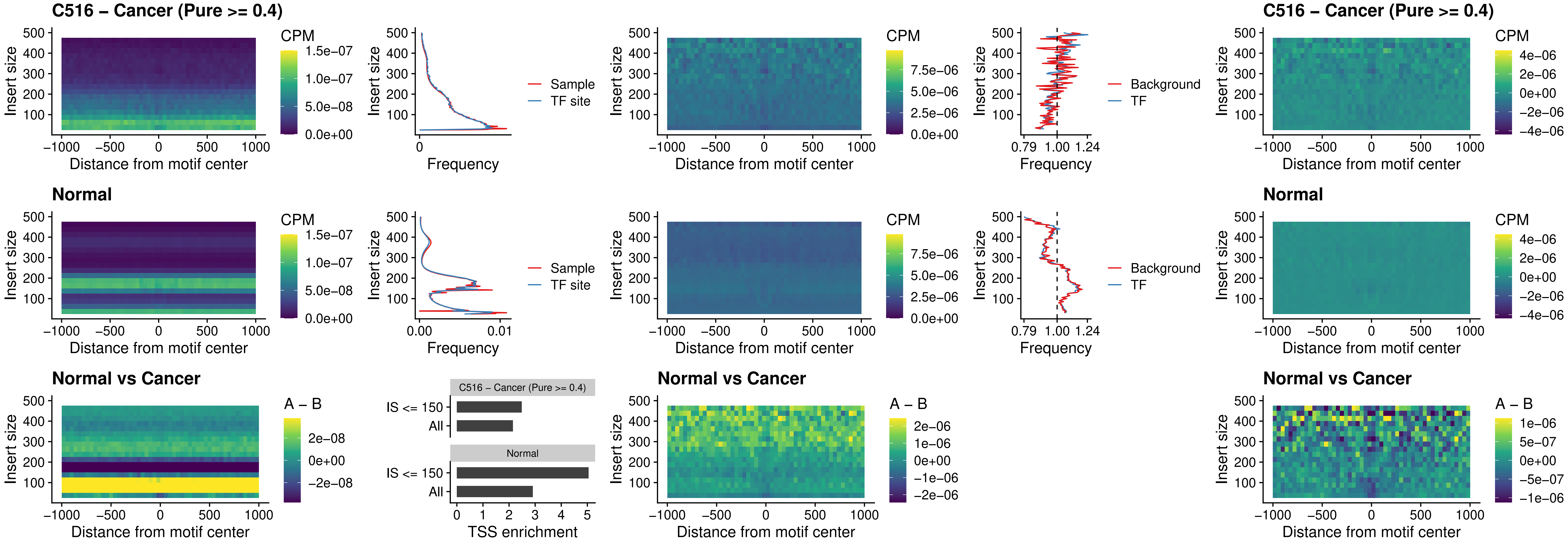

#### C516 – Cancer (Pure >= 0.4) – BCL11B_uc_dTSS_nPEAK

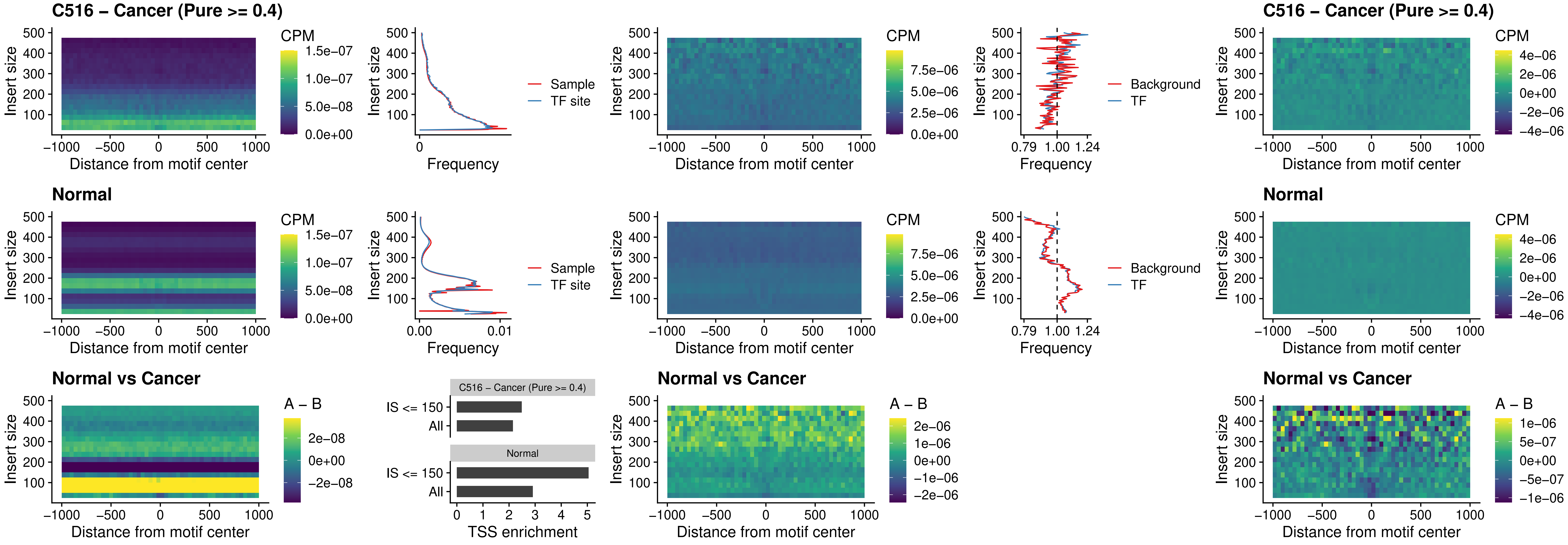

#### C516 – Cancer (Pure >= 0.4) – IRF1_uc_cTSS_oPEAK

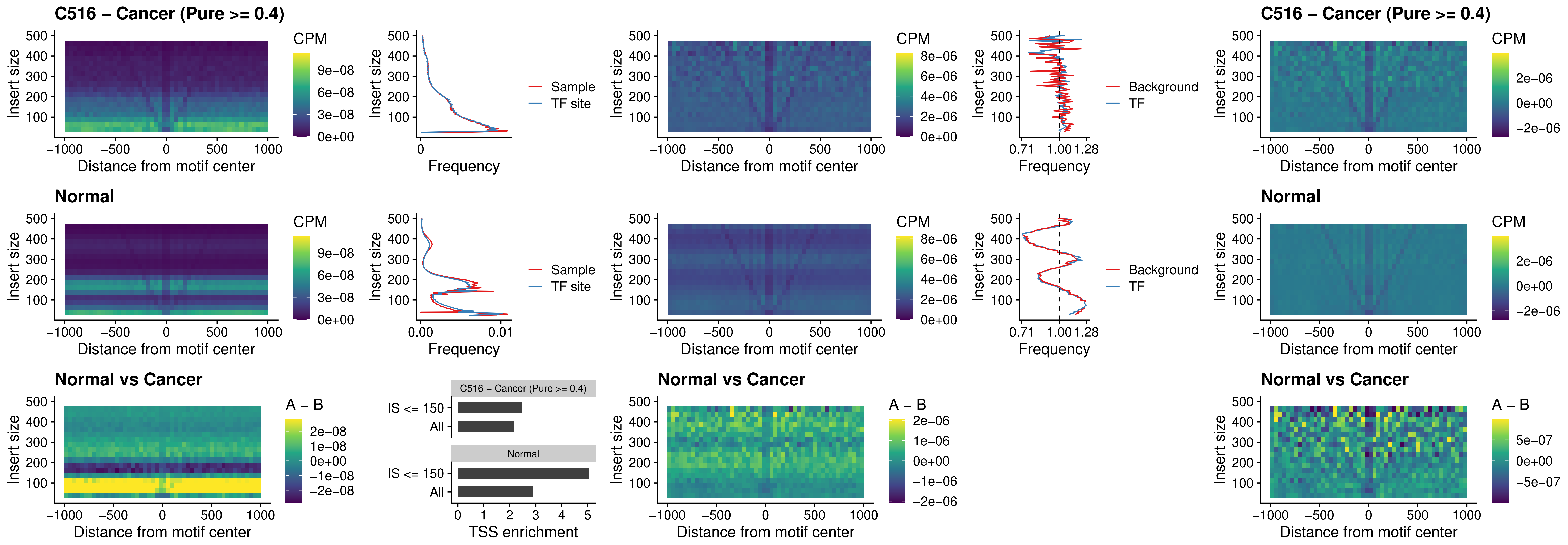

#### C516 – Cancer (Pure >= 0.4) – IRF1_uc_dTSS_oPEAK

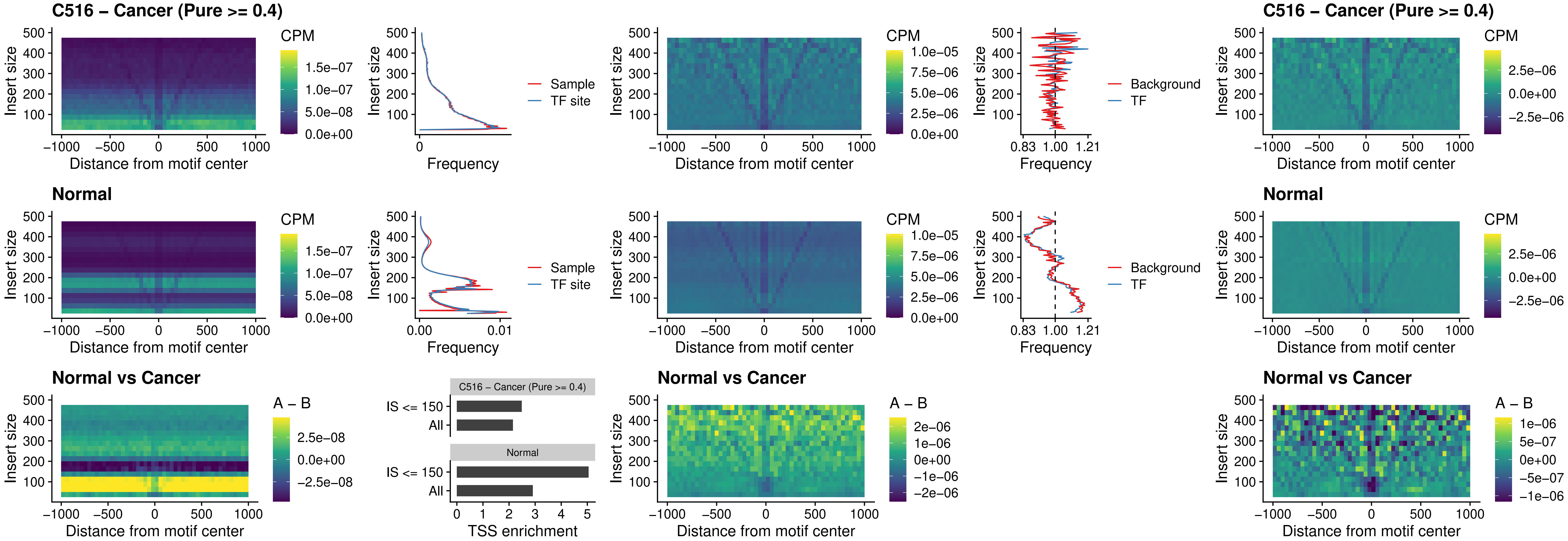

#### C516 – Cancer (Pure >= 0.4) – IRF1_uc_pTSS_oPEAK

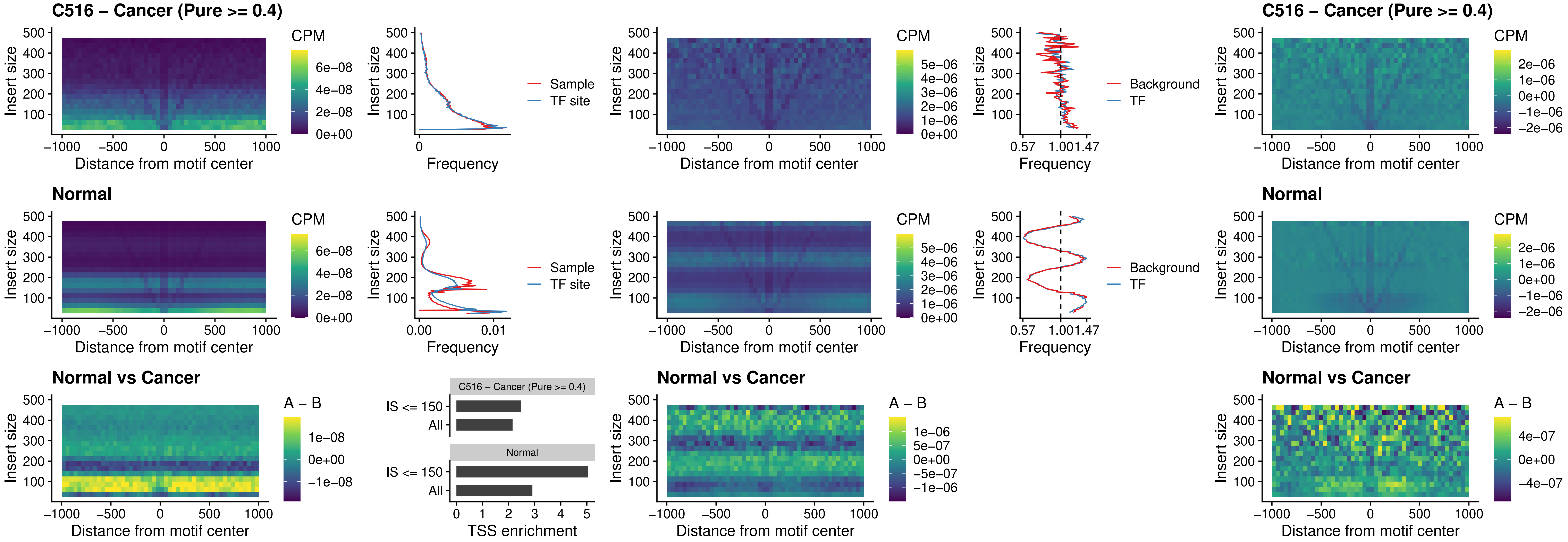

### C516 – Cancer (Pure >= 0.4) – SPIC_uc_dTSS_nPEAK

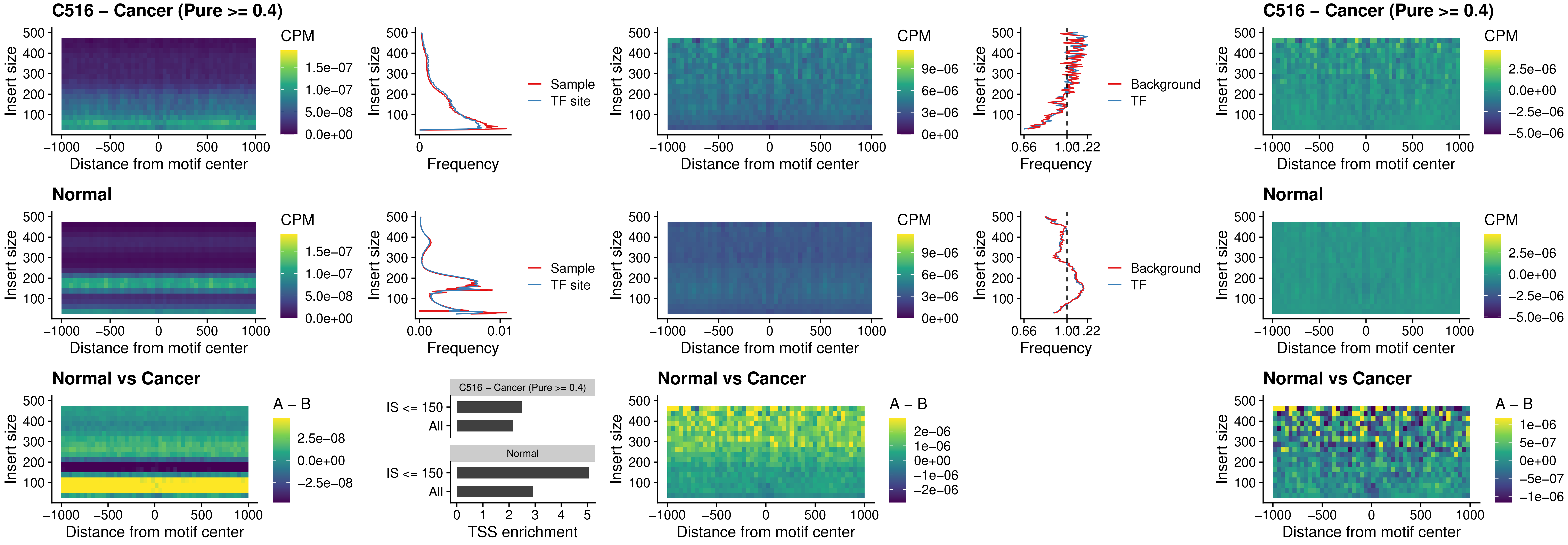

### C516 – Cancer (Pure >= 0.4) – SPIC_uc_dTSS_oPEAK

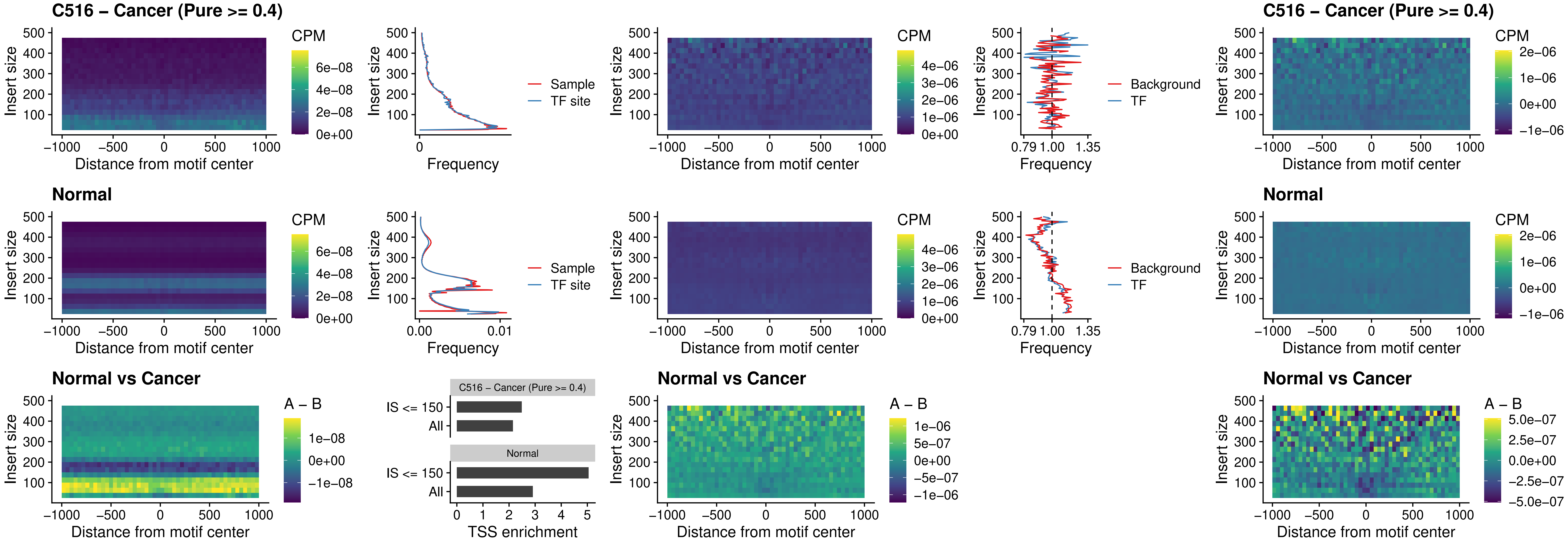

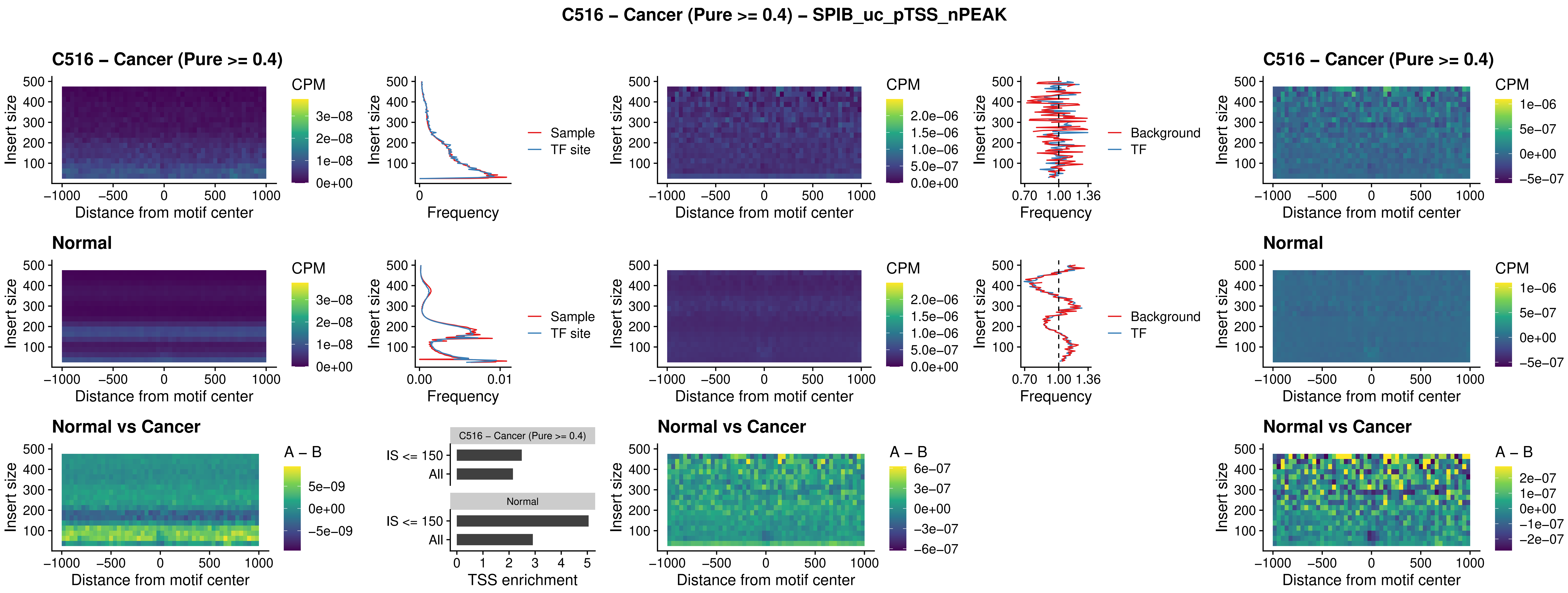
C516 – Cancer (Pure >= 0.4) – SPIB_uc_pTSS_nPEAK
C516 – Cancer (Pure >= 0.4)
Normal
Normal vs Cancer
Insert size
Distance from motif center
Frequency
CPM
A – B
Sample
TF site
Background
TF
C516 – Cancer (Pure >= 0.4)
Normal
IS <= 150
All
TSS enrichment

#### C516 – Cancer (Pure >= 0.4) – STAT2_uc_cTSS_oPEAK

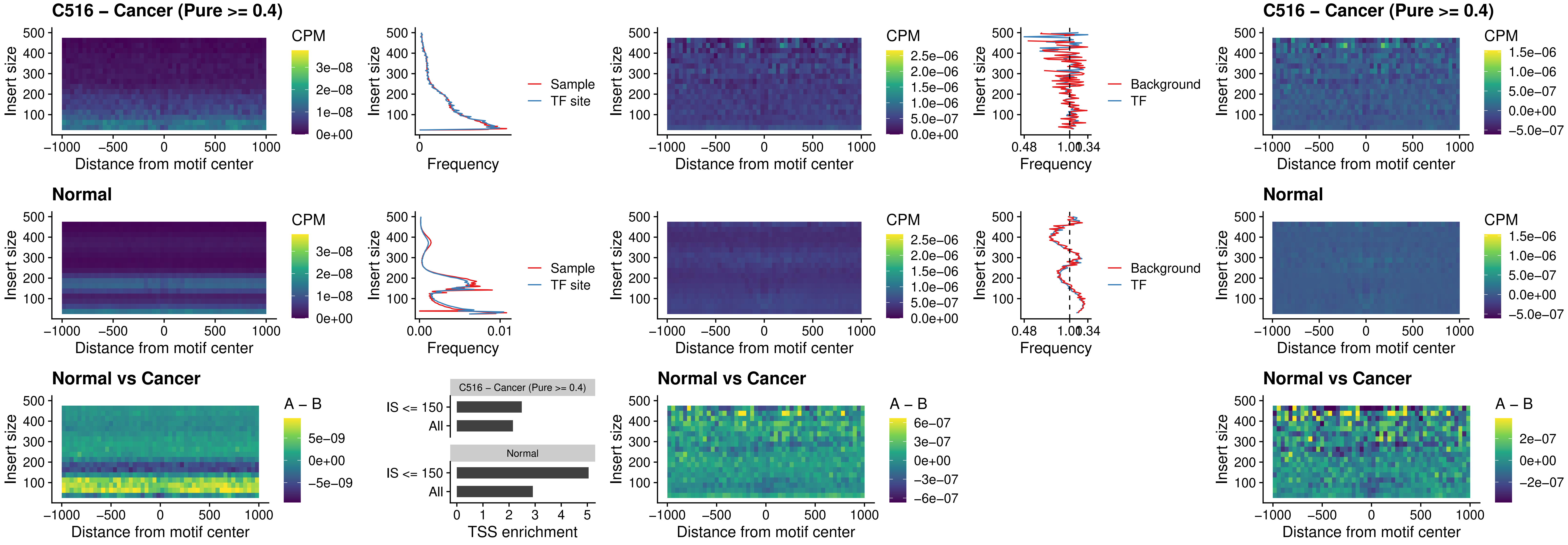

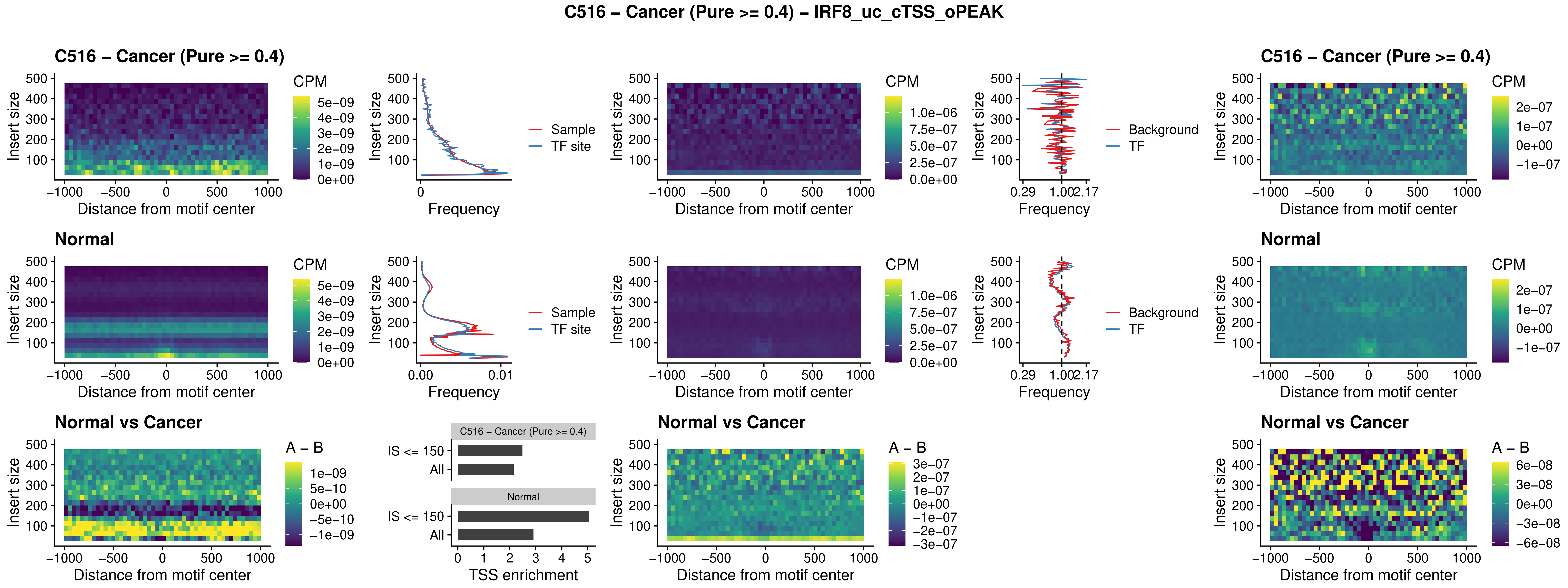
C516 – Cancer (Pure >= 0.4) – IRF8_uc_cTSS_oPEAK
C516 – Cancer (Pure >= 0.4)
Normal
Normal vs Cancer
Insert size
Distance from motif center
Frequency
CPM
A – B
Sample
TF site
Background
TF
C516 – Cancer (Pure >= 0.4)
Normal
IS <= 150
All
TSS enrichment

### C516 – Cancer (Pure >= 0.4) – IRF3_uc_dTSS_oPEAK

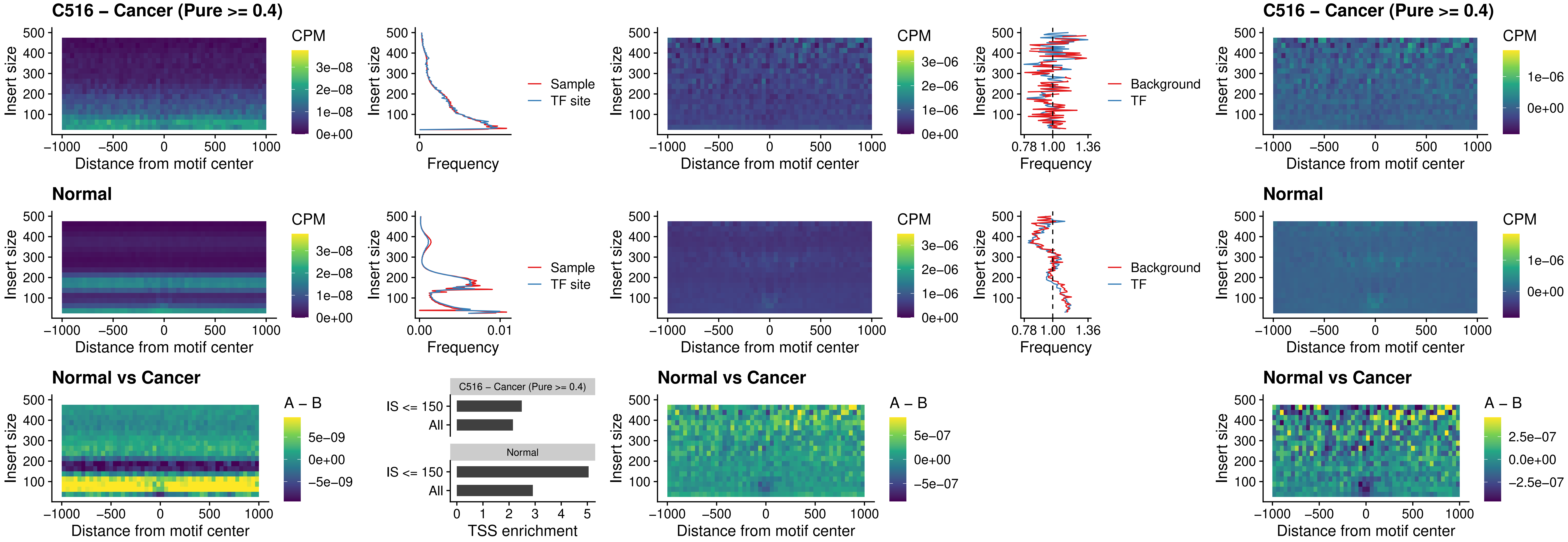

### C516 – Cancer (Pure >= 0.4) – IRF8_uc_dTSS_oPEAK

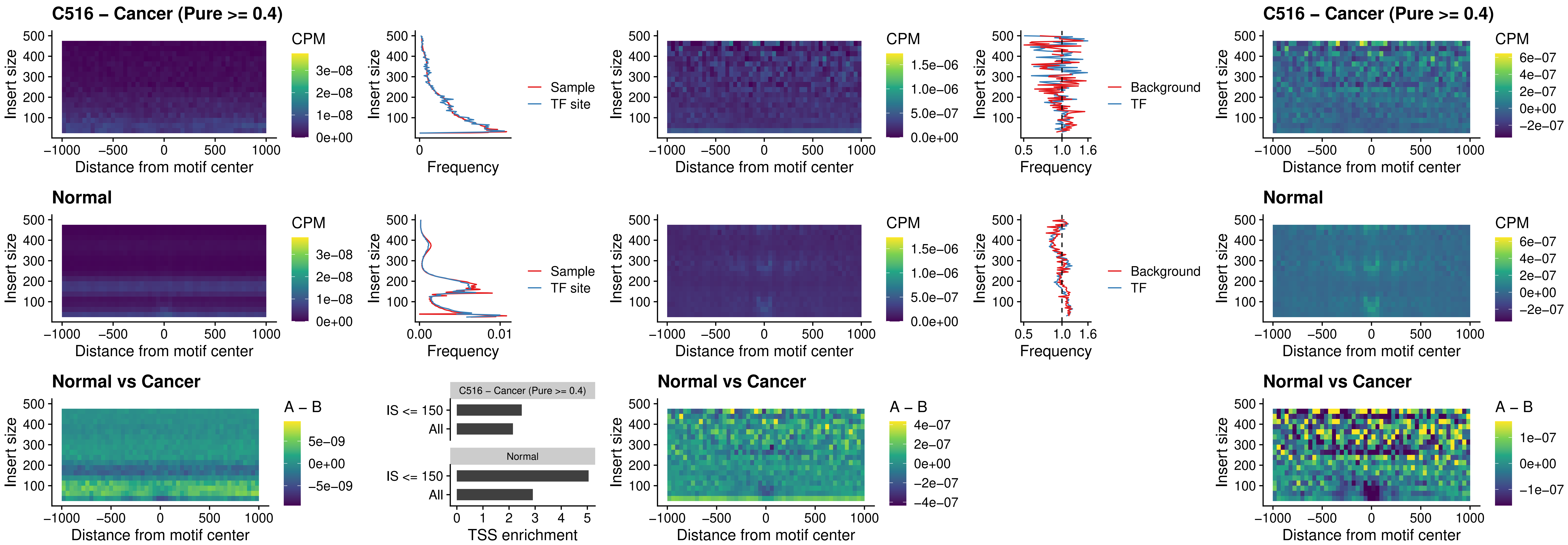

#### C516 – Cancer (Pure >= 0.4) – ETS2_uc_dTSS_oPEAK

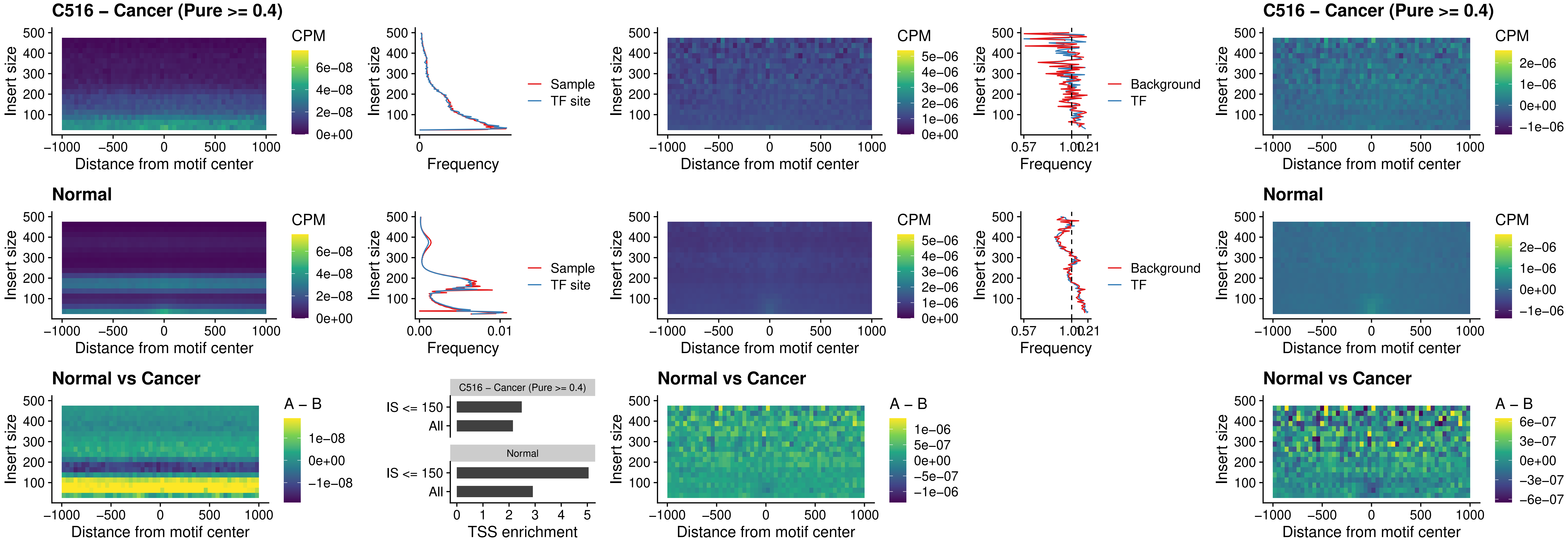

#### C516 – Cancer (Pure >= 0.4) – SMARCC2_uc_pTSS_oPEAK

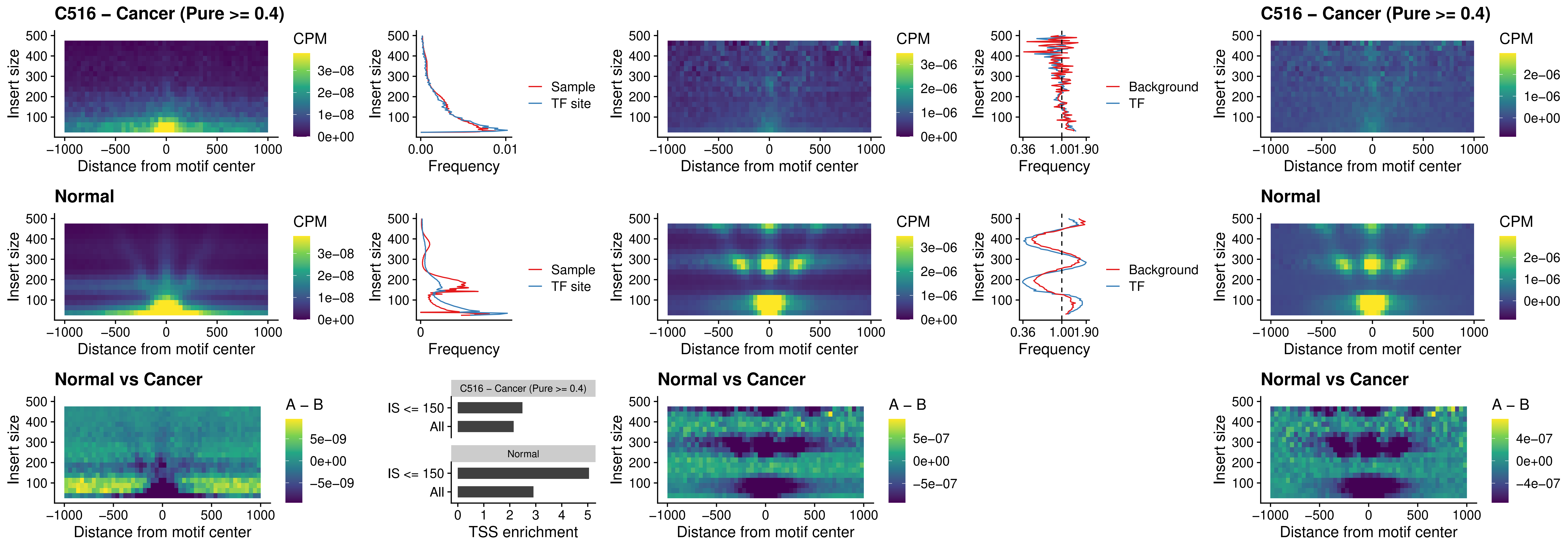

#### C516 – Cancer (Pure >= 0.4) – BCL11A_uc_cTSS_nPEAK

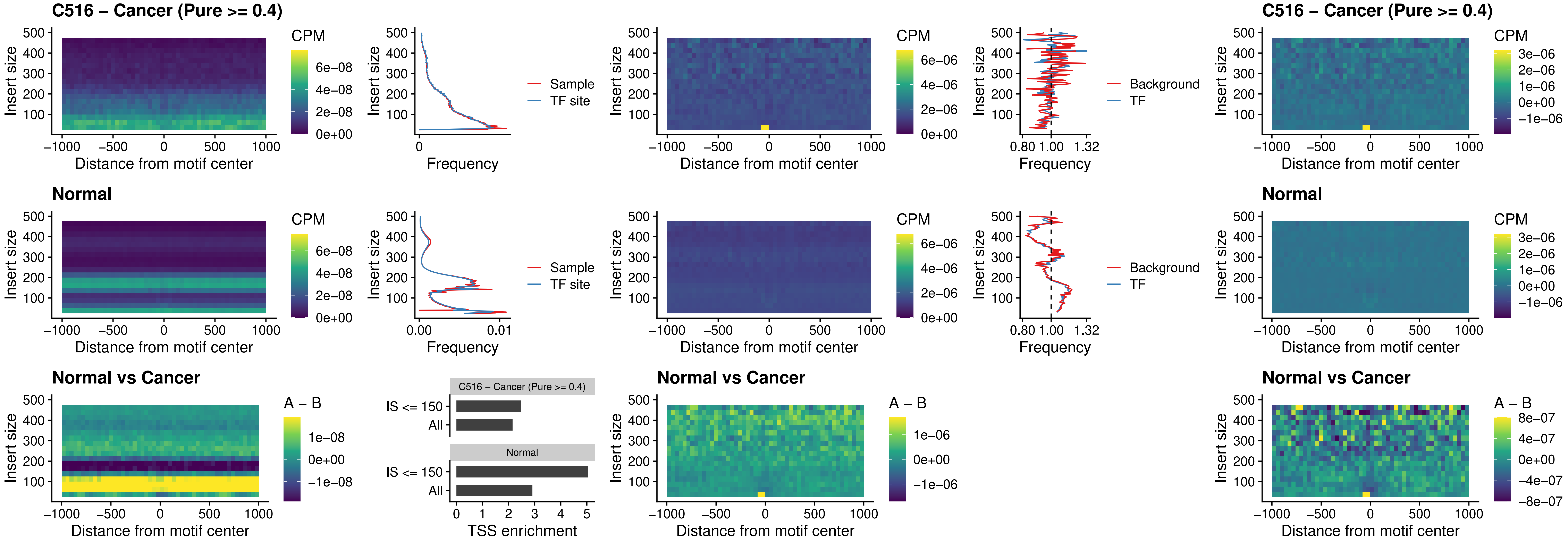

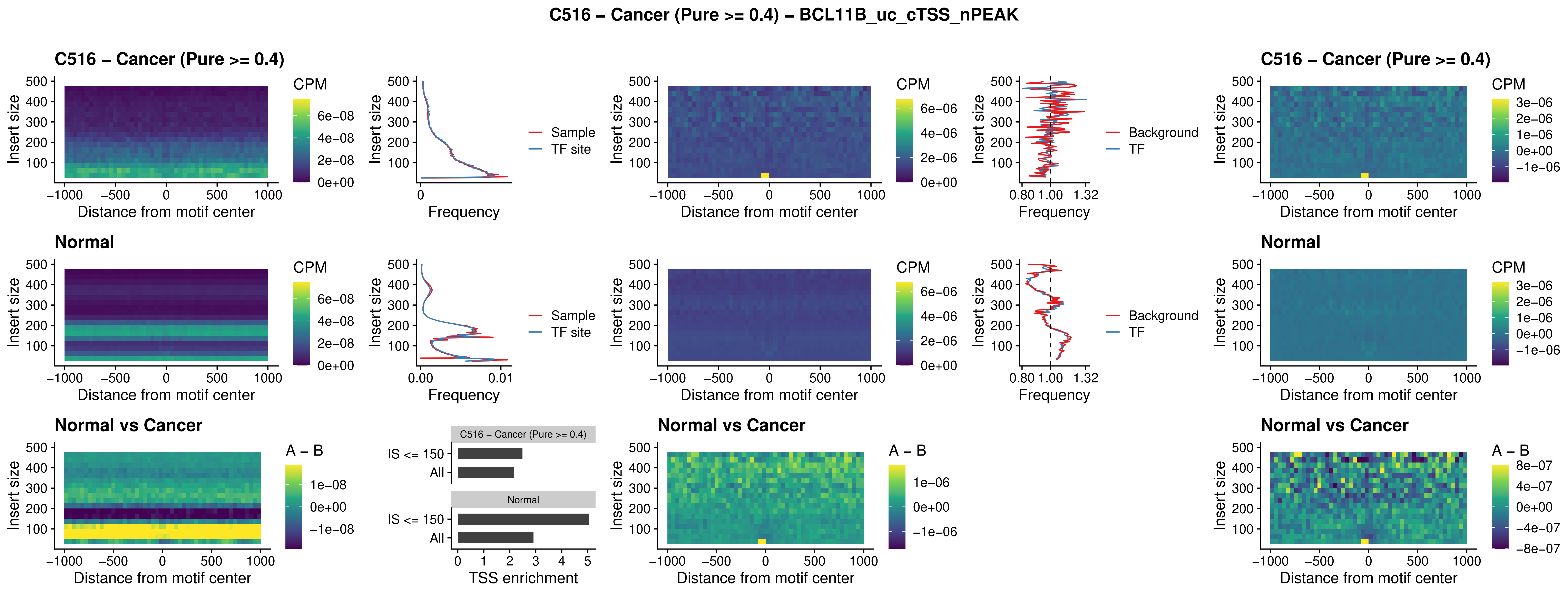
C516 – Cancer (Pure >= 0.4) – BCL11B_uc_cTSS_nPEAK
C516 – Cancer (Pure >= 0.4)
Normal
Normal vs Cancer
Insert size
Distance from motif center
Frequency
CPM
A – B
Sample
TF site
Background
TF
C516 – Cancer (Pure >= 0.4)
Normal
IS <= 150
All
TSS enrichment

#### C516 – Cancer (Pure >= 0.4) – STAT2_uc_dTSS_oPEAK

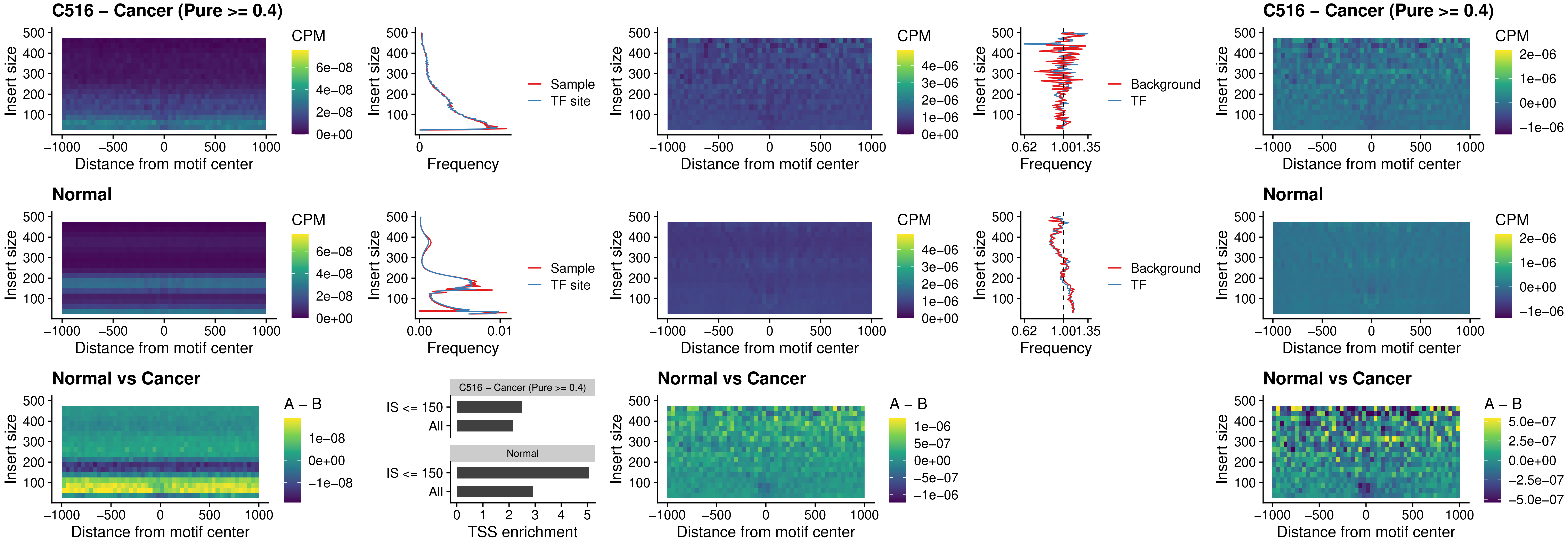

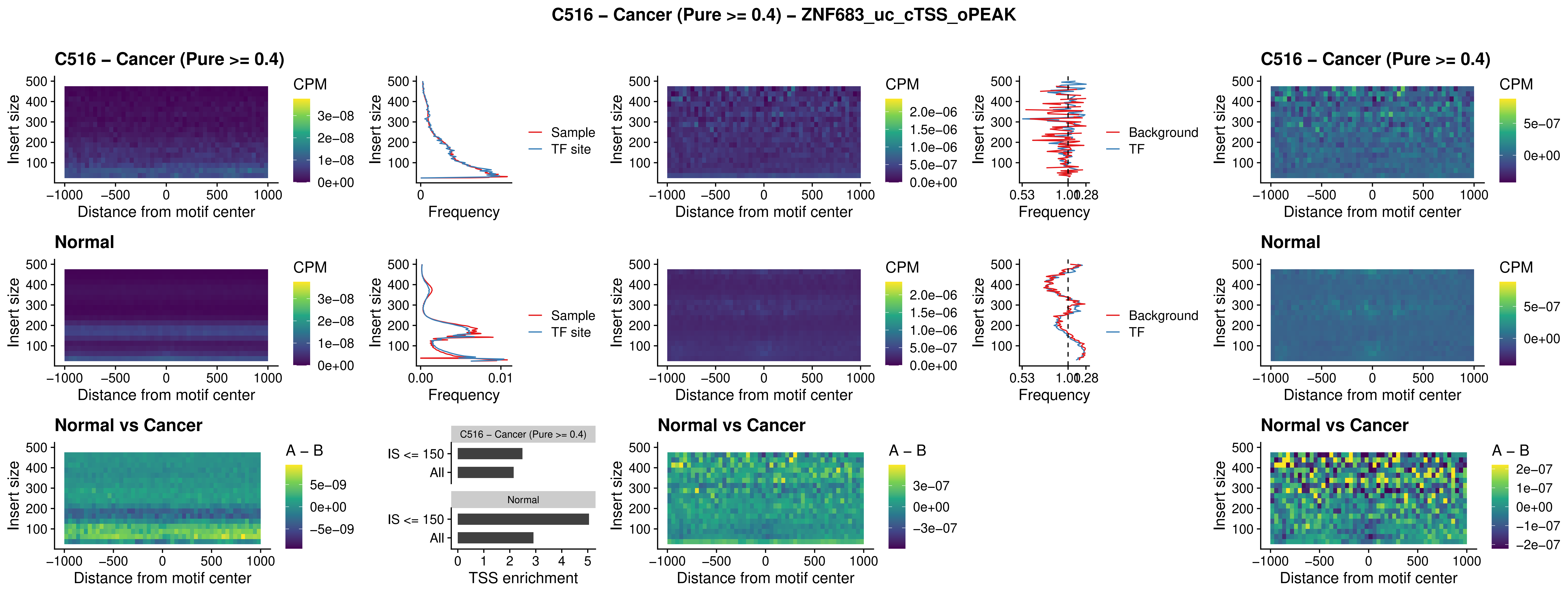
C516 – Cancer (Pure >= 0.4) – ZNF683_uc_cTSS_oPEAK
C516 – Cancer (Pure >= 0.4)
Normal
Normal vs Cancer
Insert size
Distance from motif center
Frequency
CPM
A – B
Sample
TF site
Background
TF
C516 – Cancer (Pure >= 0.4)
Normal
IS <= 150
All
TSS enrichment

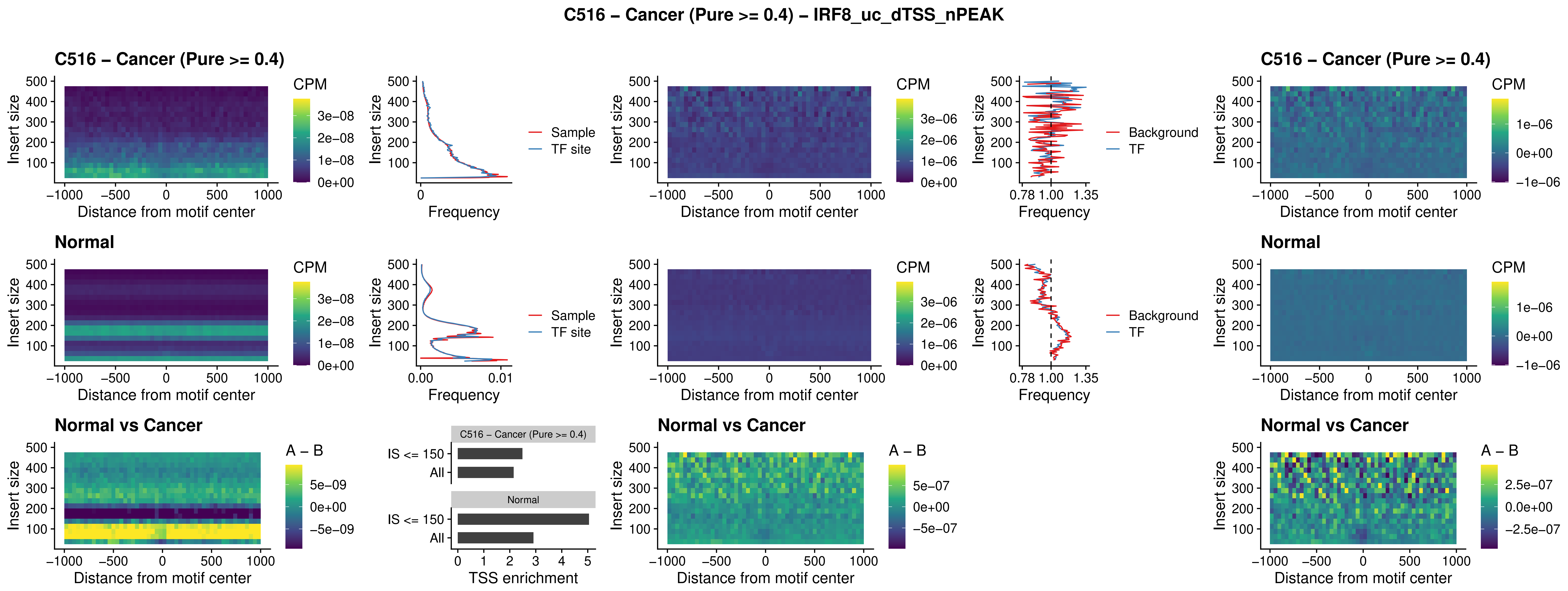
C516 – Cancer (Pure >= 0.4) – IRF8_uc_dTSS_nPEAK
C516 – Cancer (Pure >= 0.4)
Normal
Normal vs Cancer
Insert size
Distance from motif center
Frequency
CPM
A – B
Sample
TF site
Background
TF
C516 – Cancer (Pure >= 0.4)
Normal
IS <= 150
All
TSS enrichment

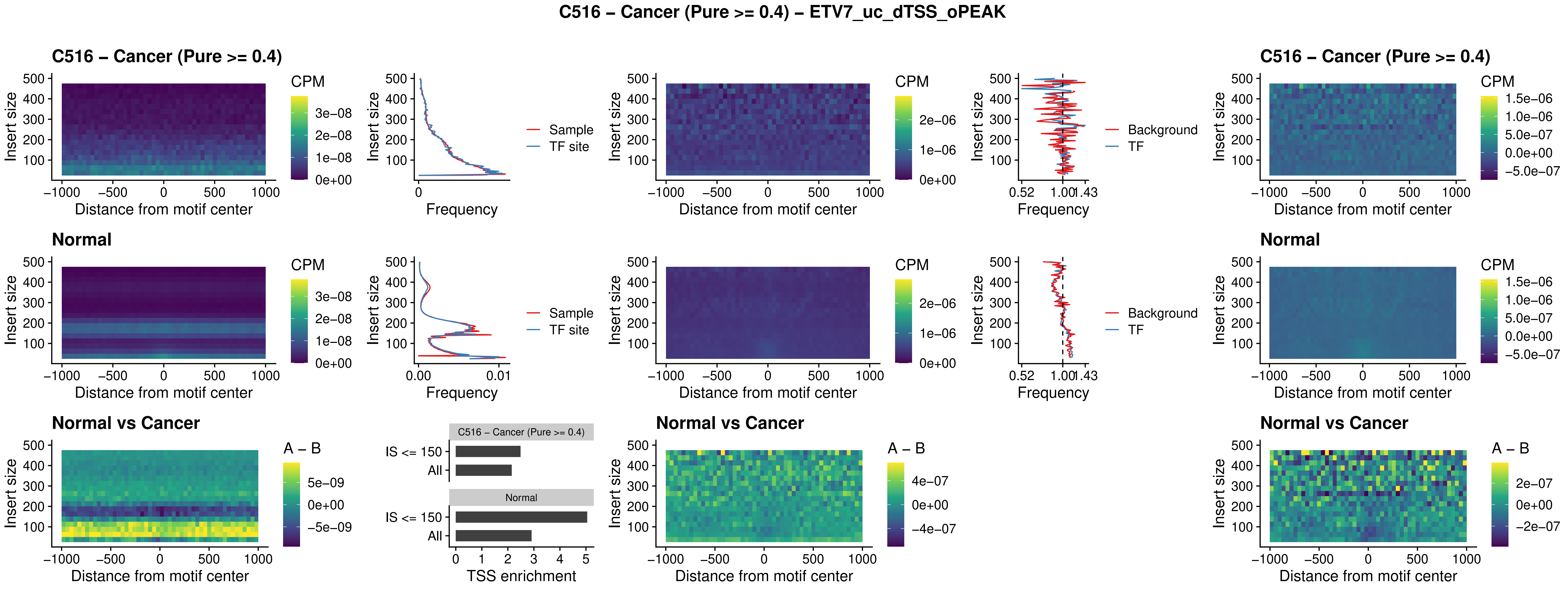
C516 – Cancer (Pure >= 0.4) – ETV7_uc_dTSS_oPEAK
C516 – Cancer (Pure >= 0.4)
Normal
Normal vs Cancer
Insert size
Distance from motif center
Frequency
CPM
A – B
Sample
TF site
Background
TF
C516 – Cancer (Pure >= 0.4)
Normal
IS <= 150
All
TSS enrichment

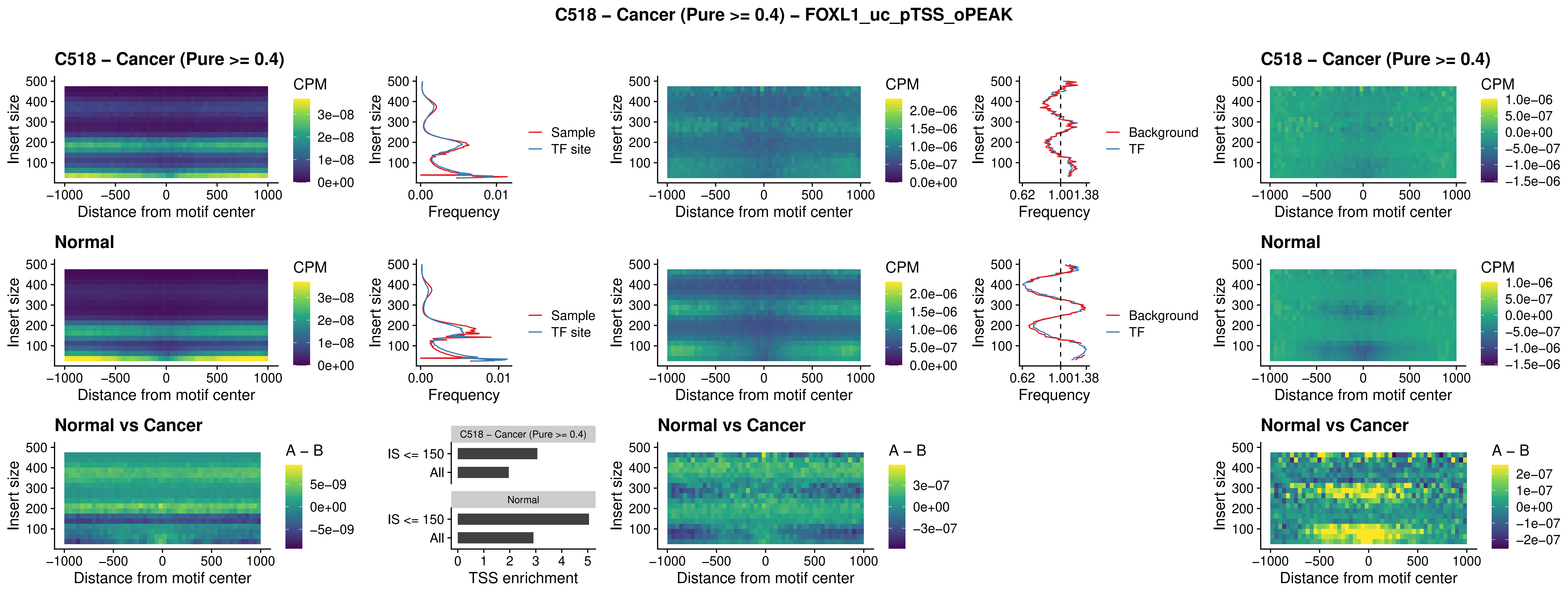
C518 – Cancer (Pure >= 0.4) – FOXL1_uc_pTSS_oPEAK
C518 – Cancer (Pure >= 0.4)
Normal
Normal vs Cancer
Insert size
Distance from motif center
CPM
A – B
Frequency
Sample
TF site
Background
TF
C518 – Cancer (Pure >= 0.4)
Normal
IS <= 150
All
TSS enrichment

#### C518 – Cancer (Pure >= 0.4) – CPEB1_uc_cTSS_oPEAK

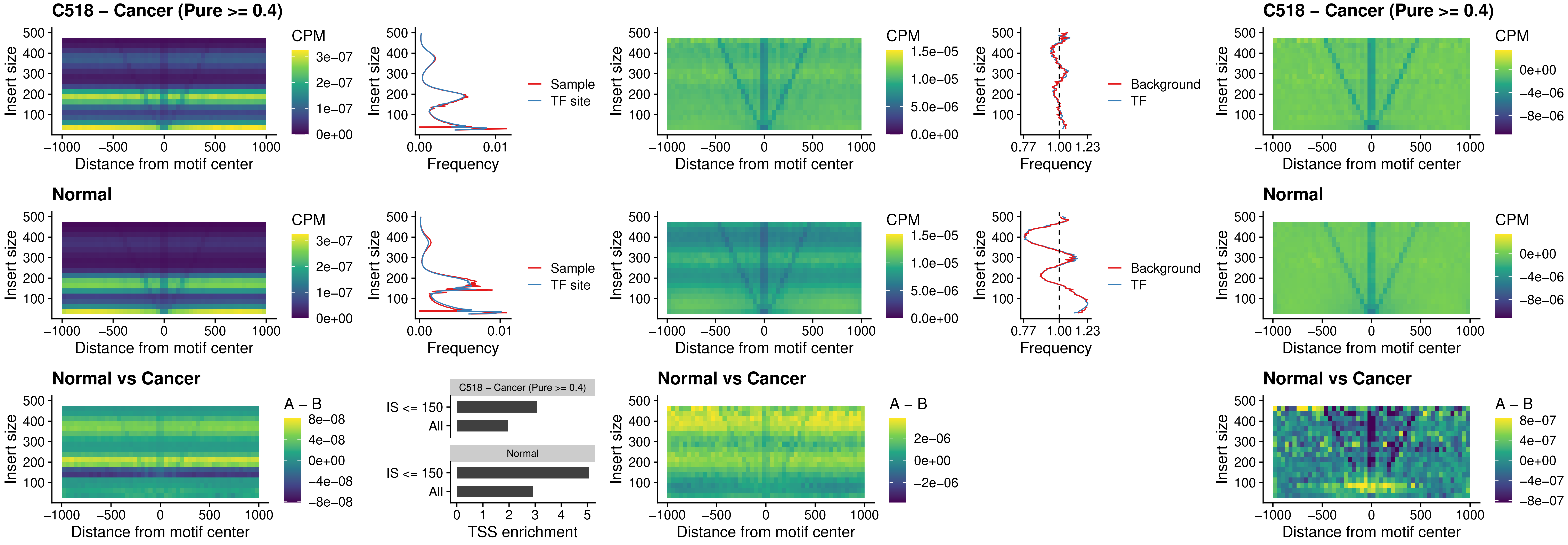

#### C518 – Cancer (Pure >= 0.4) – HMGA1_uc_pTSS_nPEAK

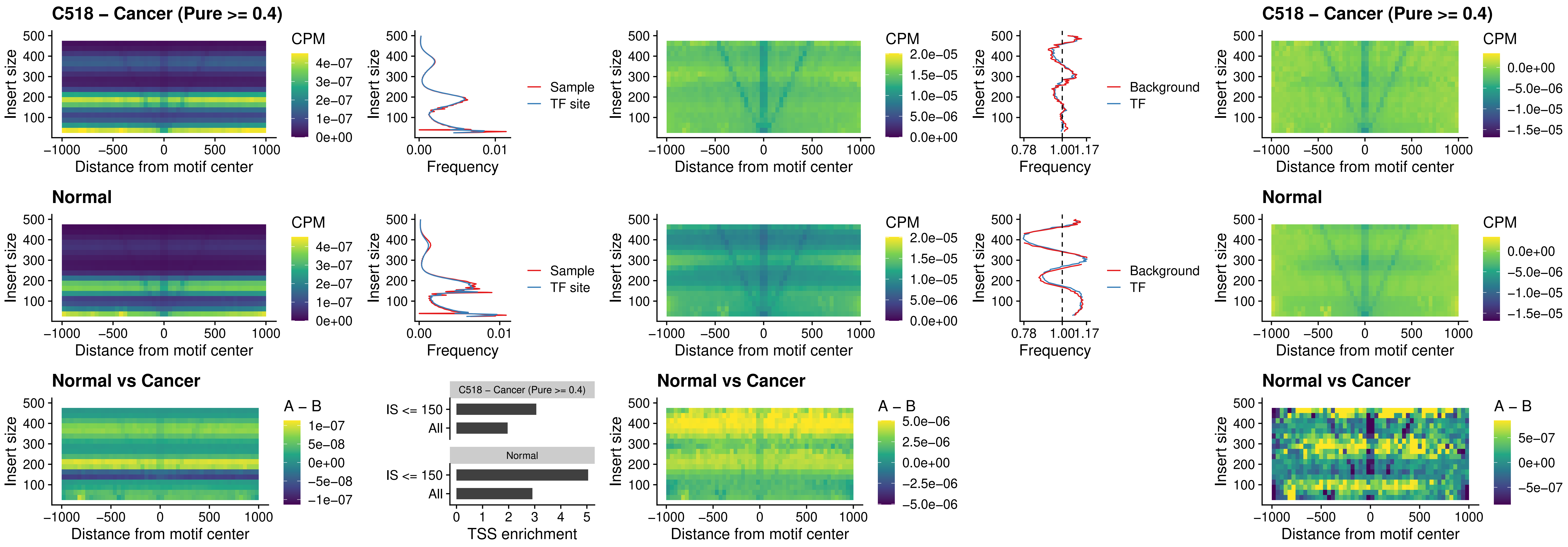

#### C518 – Cancer (Pure >= 0.4) – SOX13_uc_cTSS_nPEAK

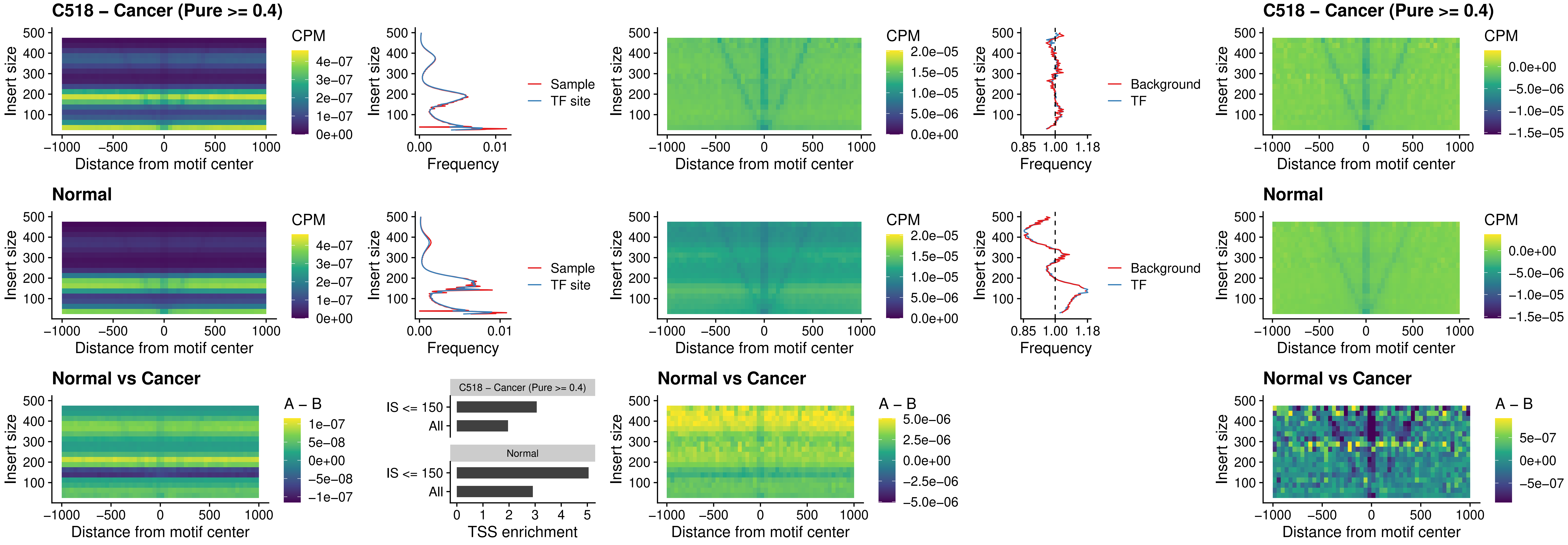

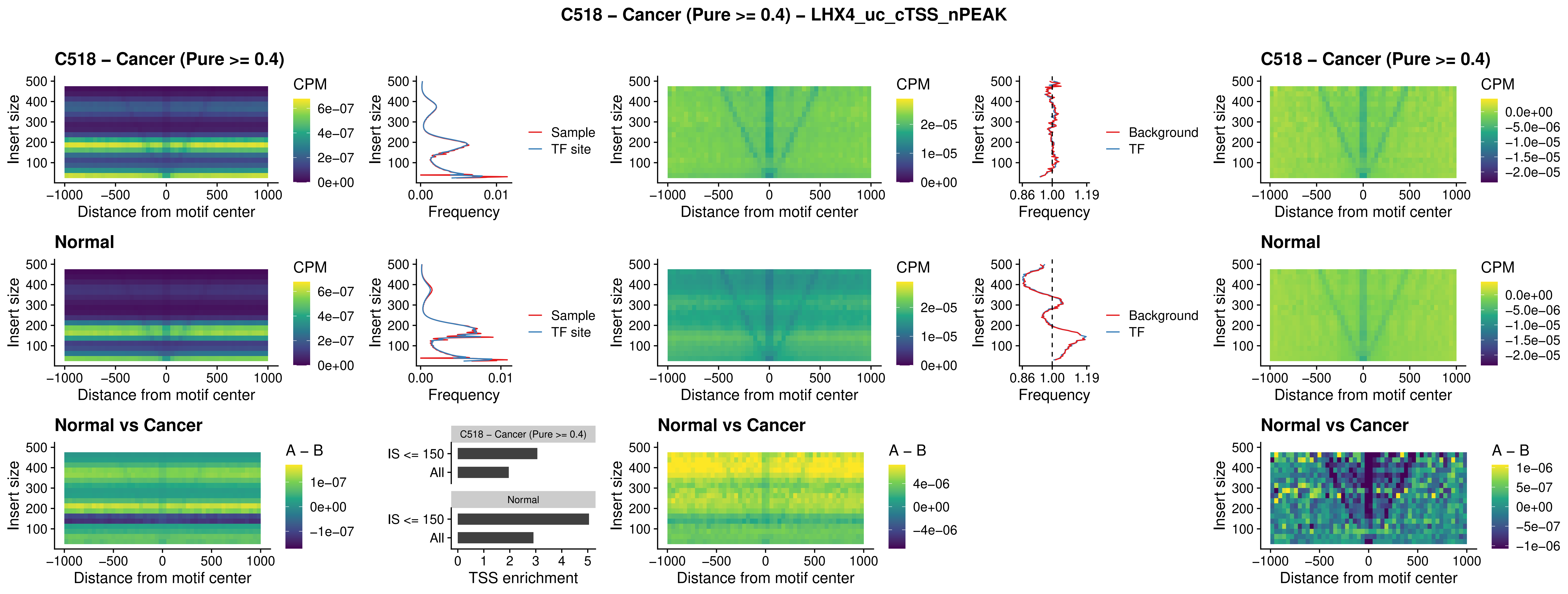
C518 – Cancer (Pure >= 0.4) – LHX4_uc_cTSS_nPEAK
C518 – Cancer (Pure >= 0.4)
Normal
Normal vs Cancer
Insert size
Distance from motif center
Frequency
CPM
A – B
Sample
TF site
Background
TF
C518 – Cancer (Pure >= 0.4)
Normal
IS <= 150
All
TSS enrichment

#### C518 – Cancer (Pure >= 0.4) – HOXA4_uc_cTSS_nPEAK

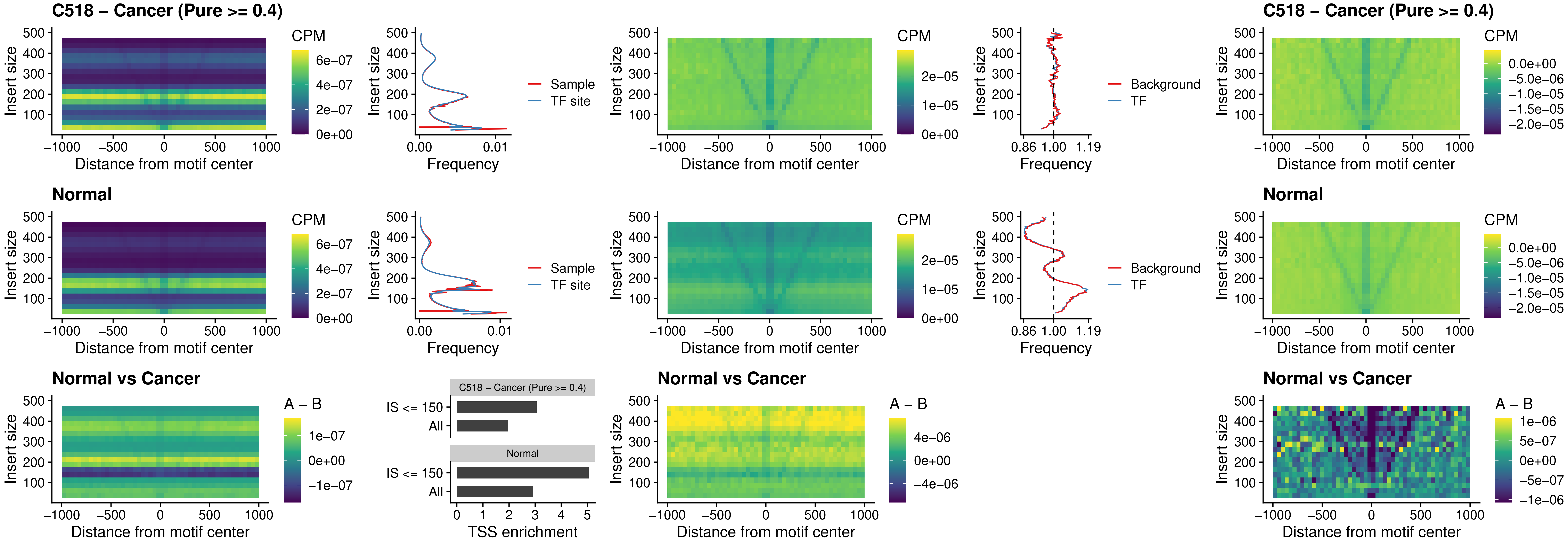

**C518 – Cancer (Pure >= 0.4) – HLX_uc_cTSS_nPEAK**

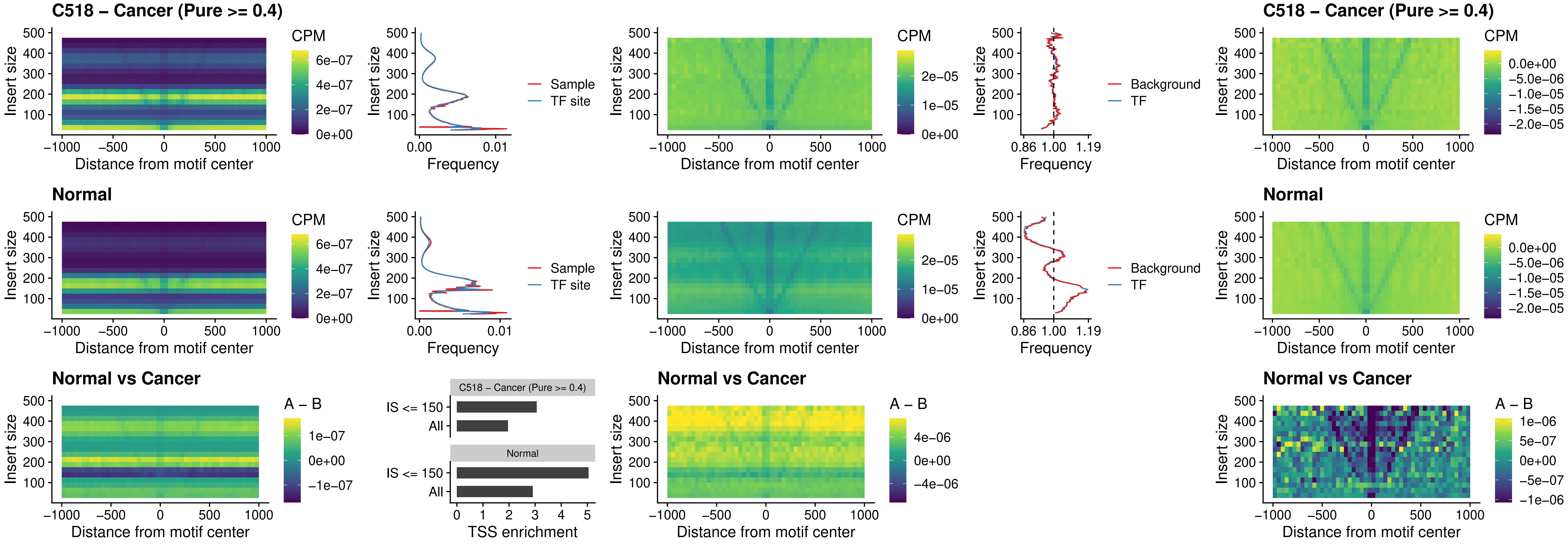

### C518 – Cancer (Pure >= 0.4) – AC0125311_uc_cTSS_nPEAK

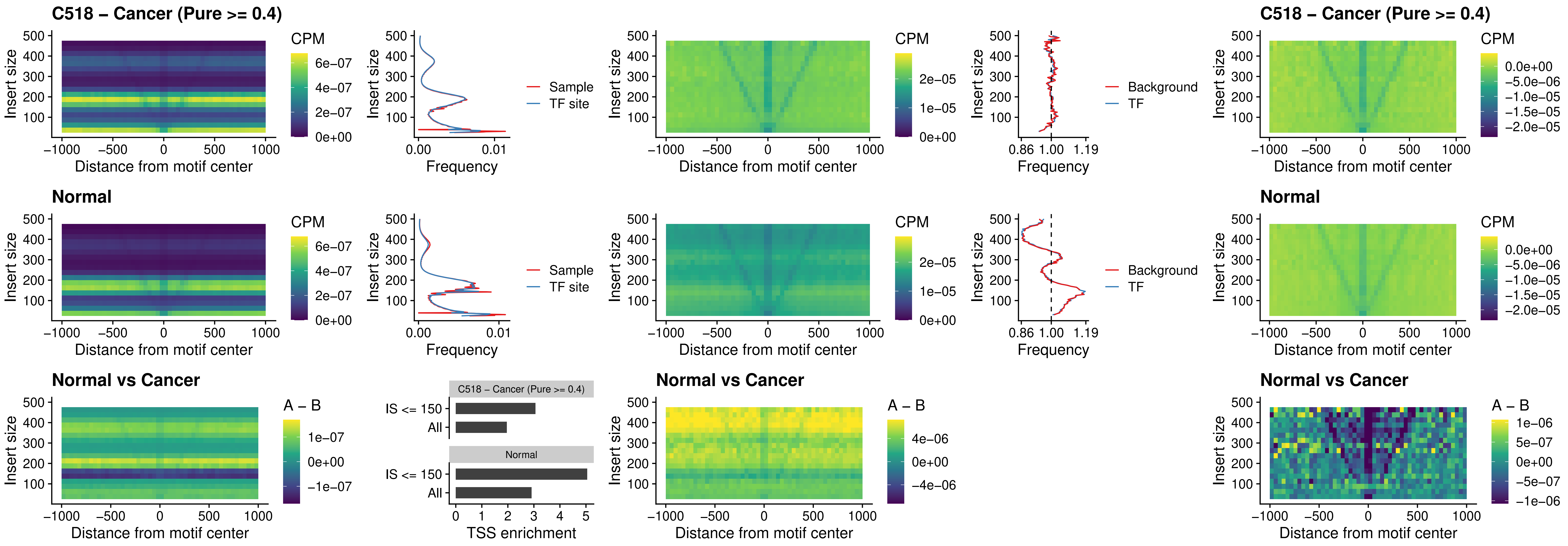

#### C518 – Cancer (Pure >= 0.4) – DBX2_uc_cTSS_nPEAK

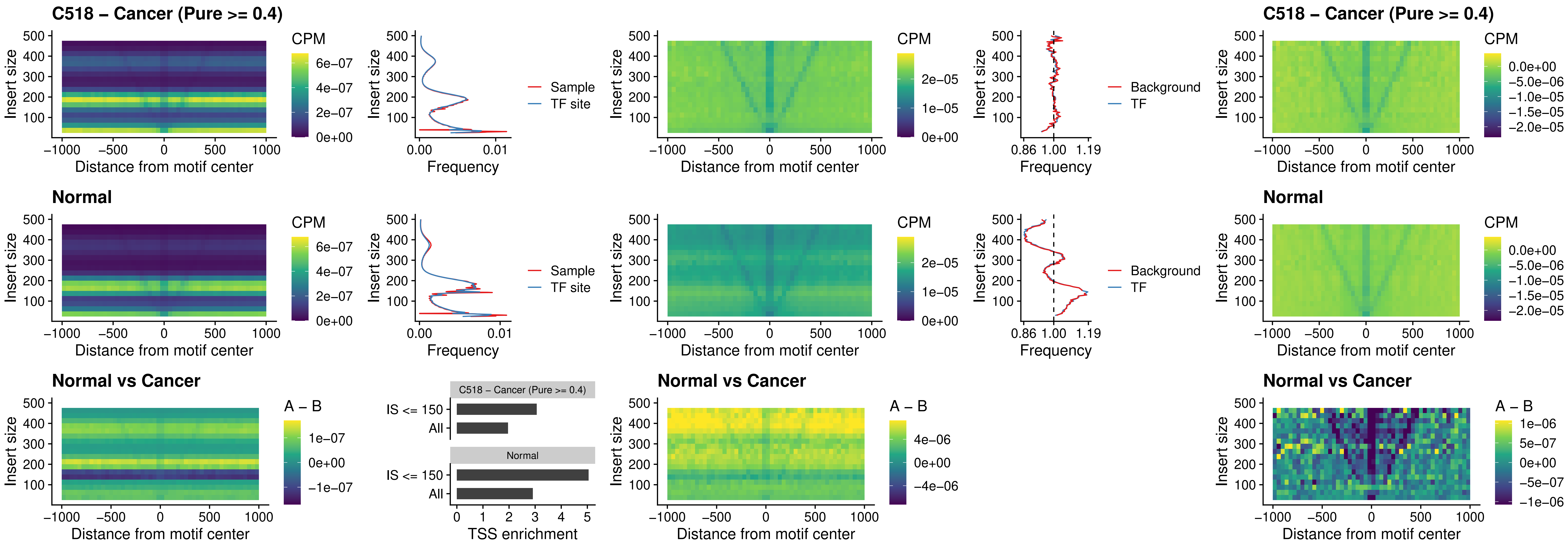

#### C518 – Cancer (Pure >= 0.4) – LHX6_uc_dTSS_nPEAK

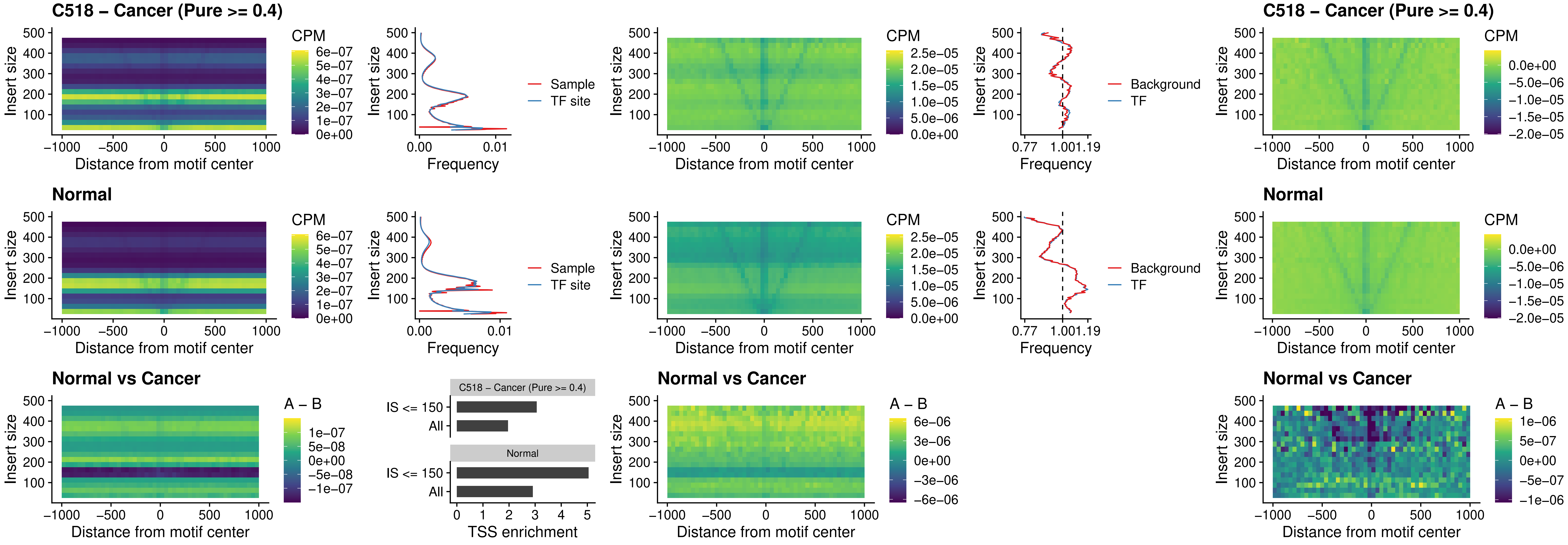

#### C518 – Cancer (Pure >= 0.4) – SOX1_uc_dTSS_nPEAK

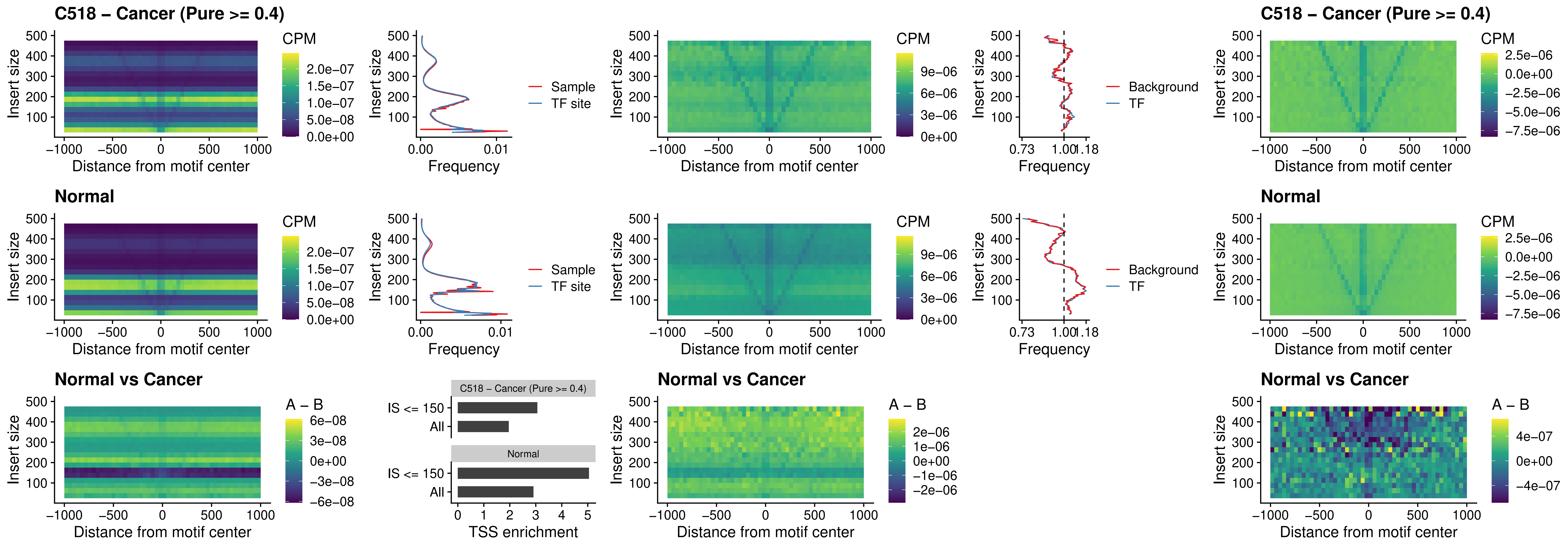

#### C518 – Cancer (Pure >= 0.4) – NKX24_uc_dTSS_nPEAK

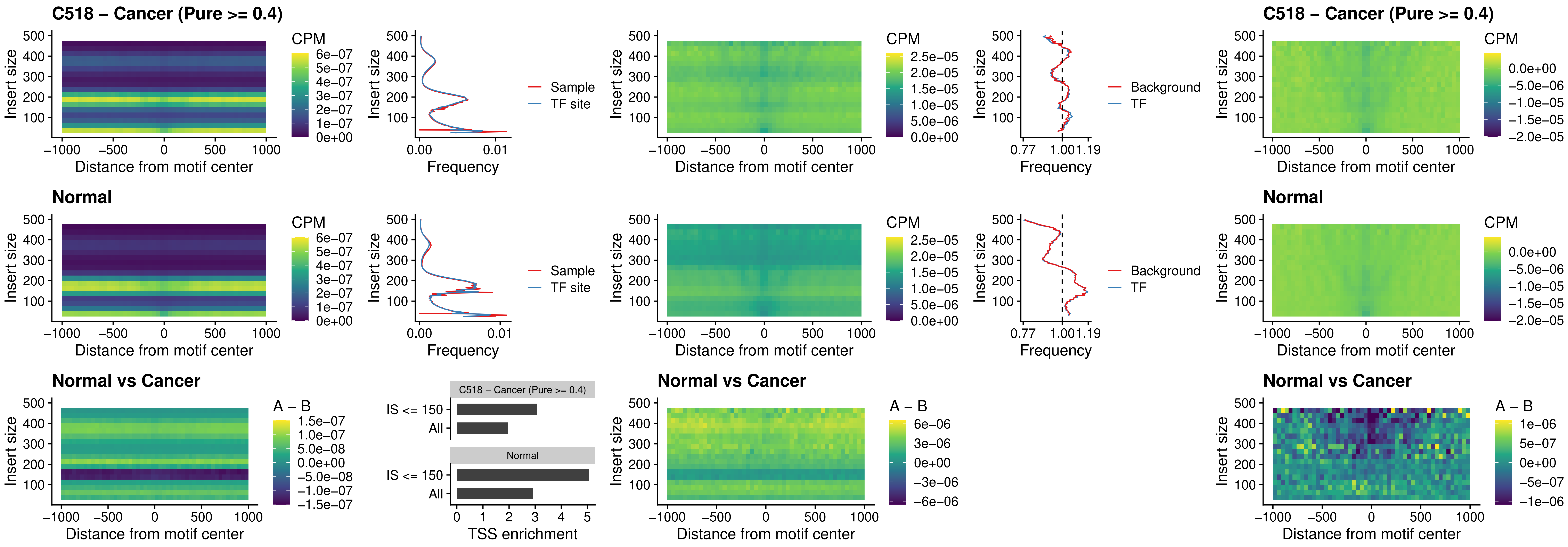

#### C518 – Cancer (Pure >= 0.4) – DLX3_uc_cTSS_nPEAK

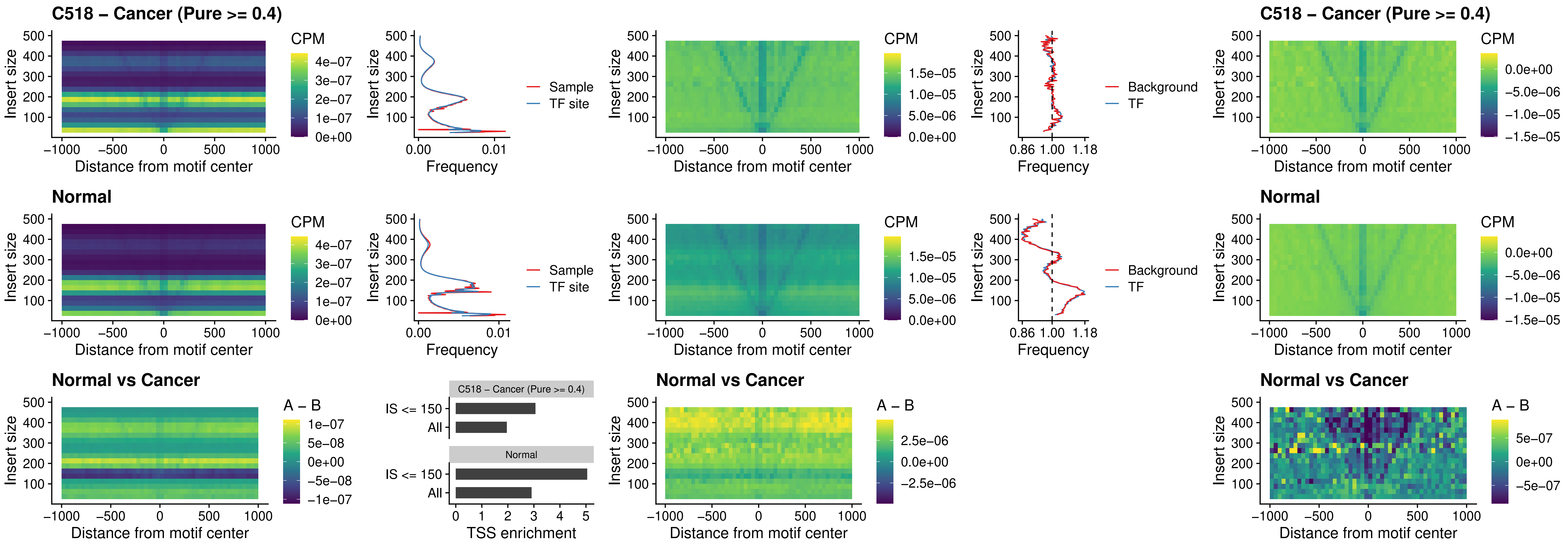

#### C518 – Cancer (Pure >= 0.4) – HOXD3_uc_cTSS_nPEAK

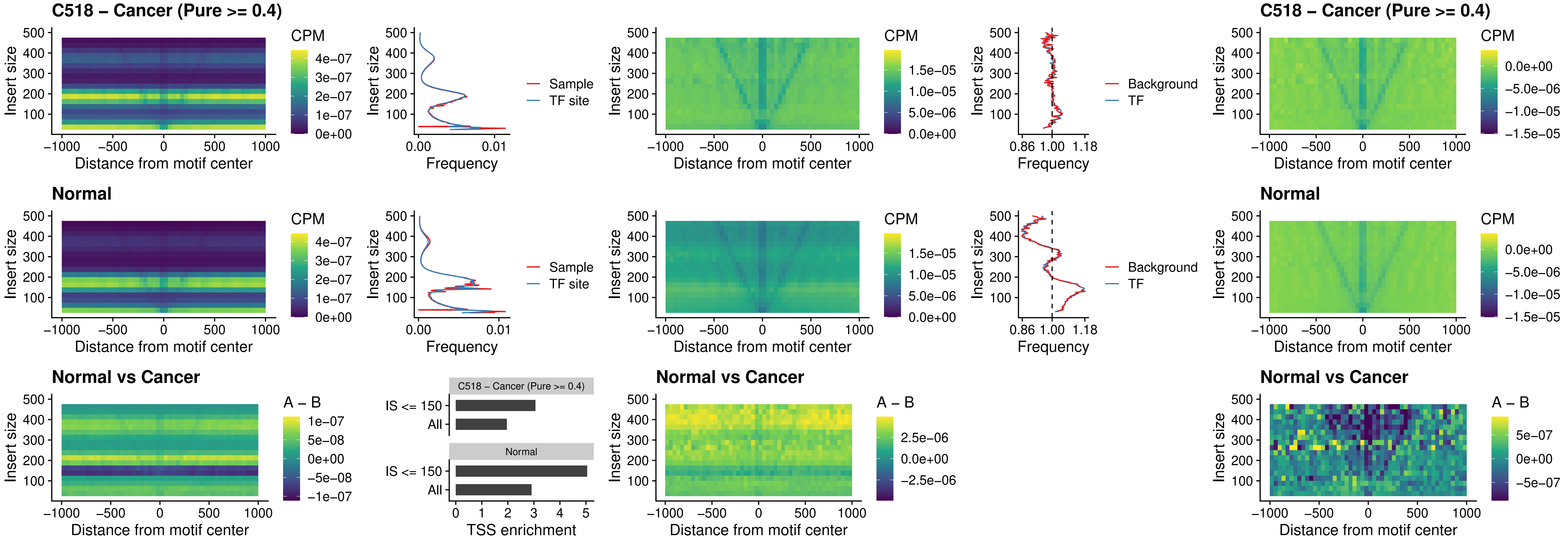

#### C518 – Cancer (Pure >= 0.4) – ISL1_uc_dTSS_oPEAK

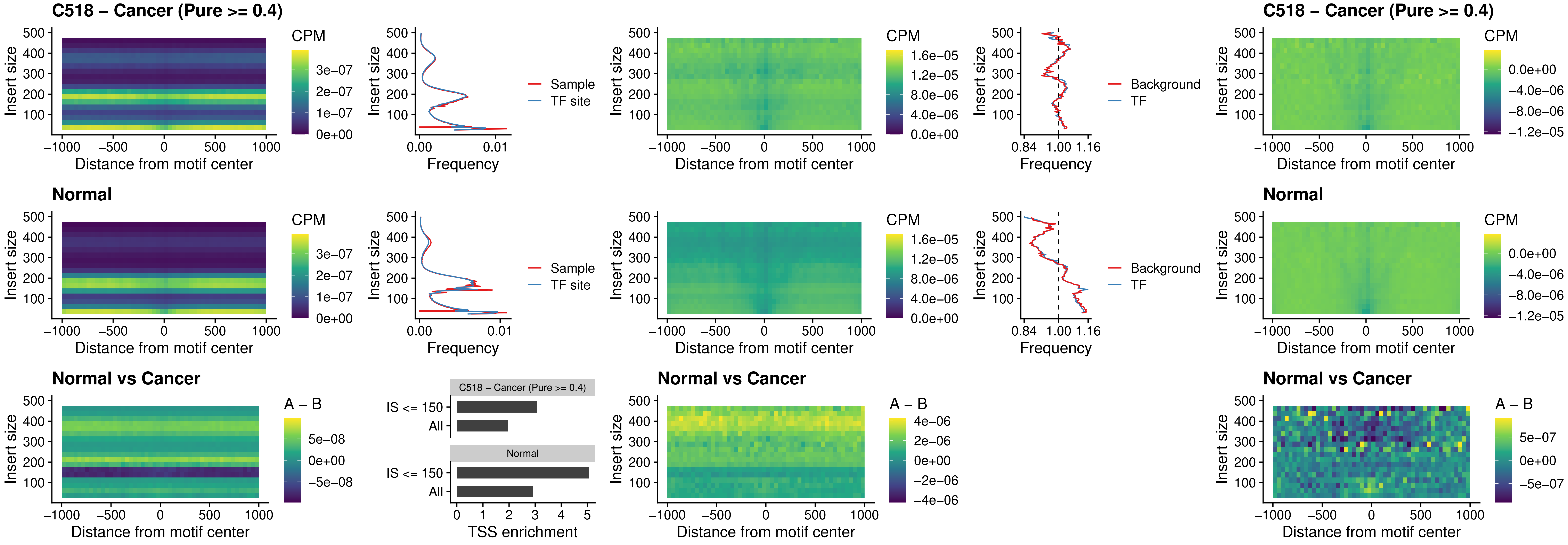

### C518 – Cancer (Pure >= 0.4) – HOXB4_uc_dTSS_oPEAK

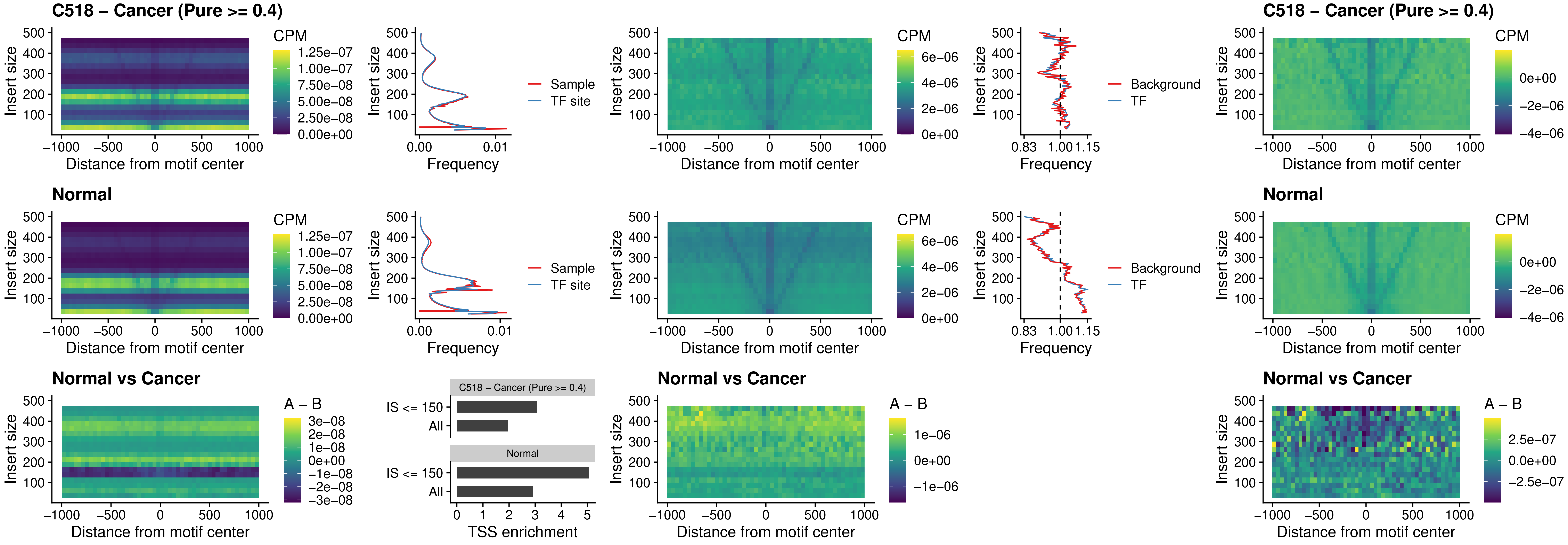

### C518 – Cancer (Pure >= 0.4) – VAX1_uc_dTSS_oPEAK

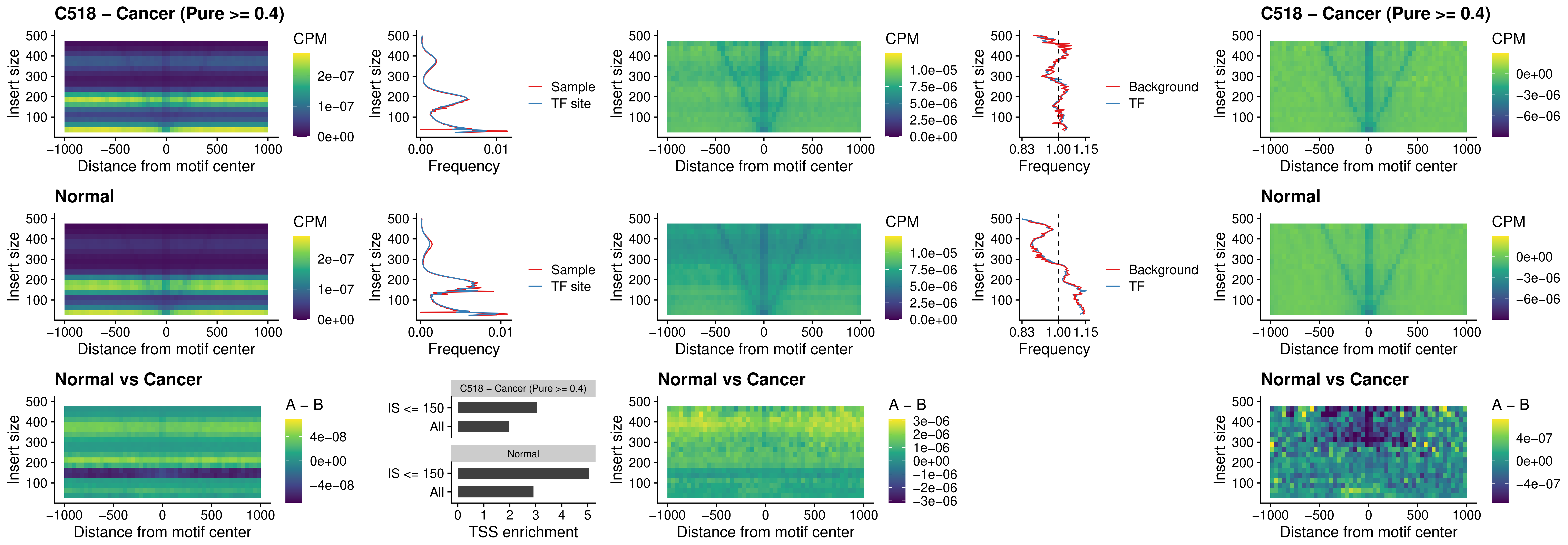

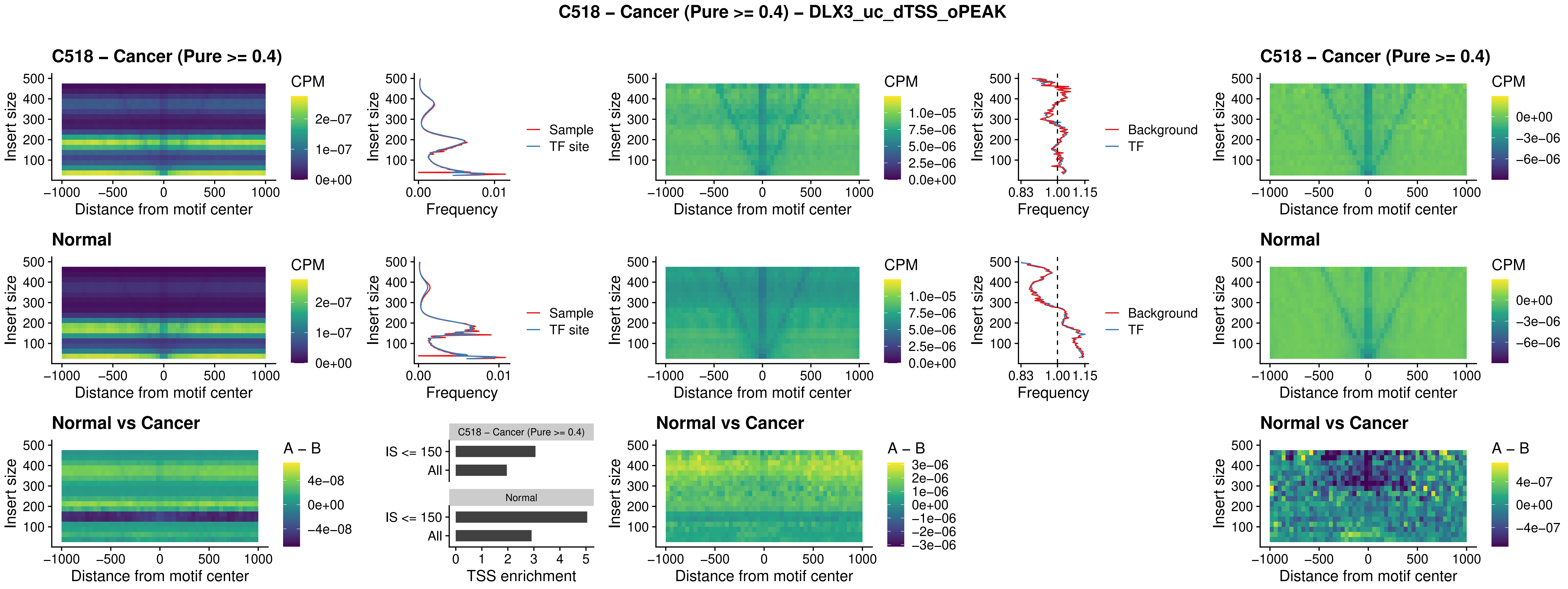
C518 – Cancer (Pure >= 0.4) – DLX3_uc_dTSS_oPEAK
C518 – Cancer (Pure >= 0.4)
Normal
Normal vs Cancer
Insert size
Distance from motif center
Frequency
CPM
A – B
Sample
TF site
Background
TF
C518 – Cancer (Pure >= 0.4)
Normal
IS <= 150
All
TSS enrichment

#### C518 – Cancer (Pure >= 0.4) – HOXD3_uc_dTSS_oPEAK

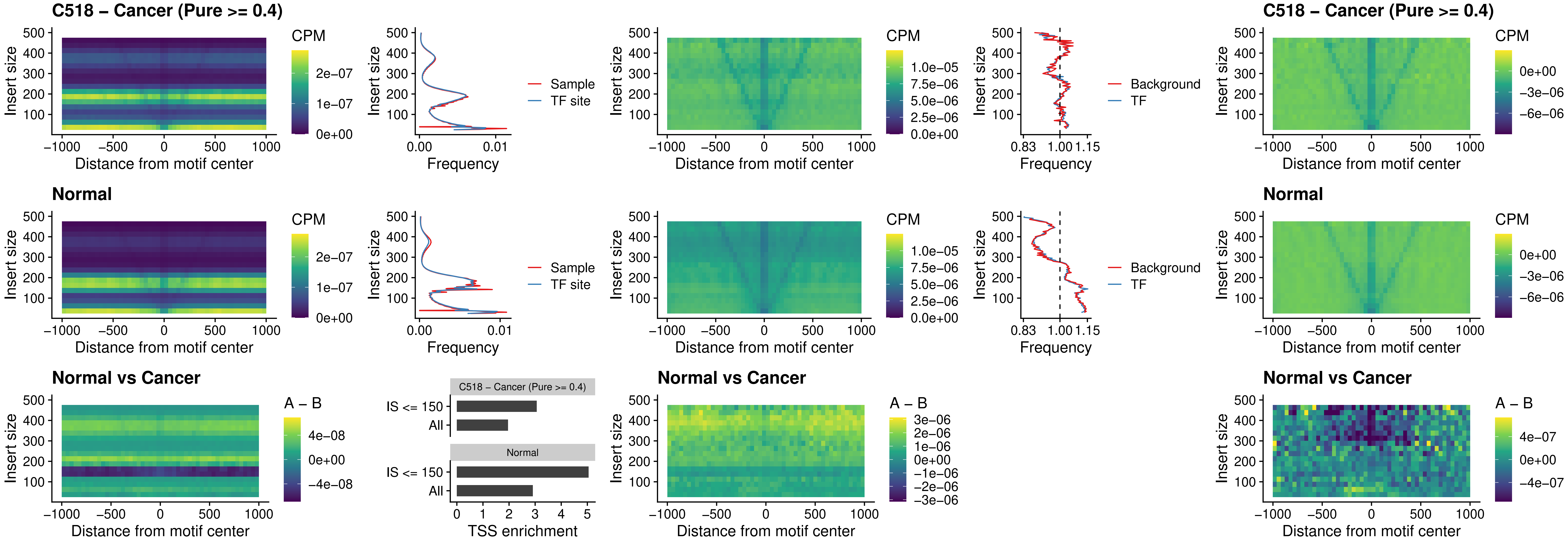

### C518 – Cancer (Pure >= 0.4) – BBX_uc_dTSS_nPEAK

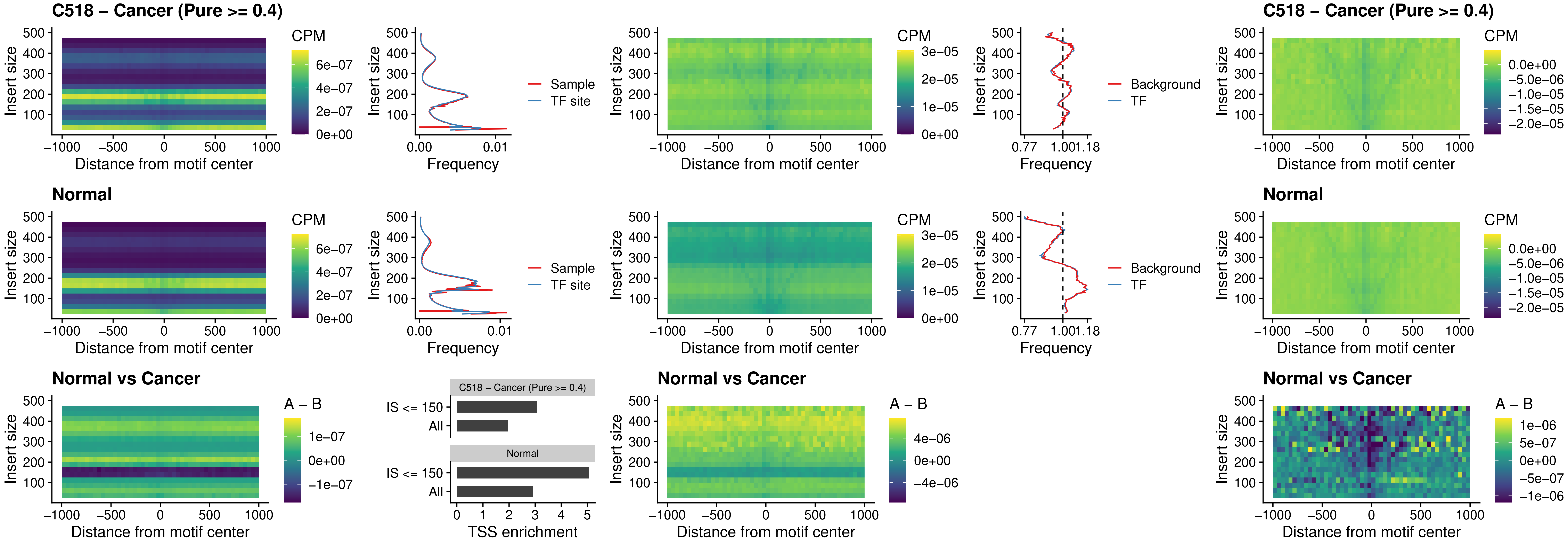

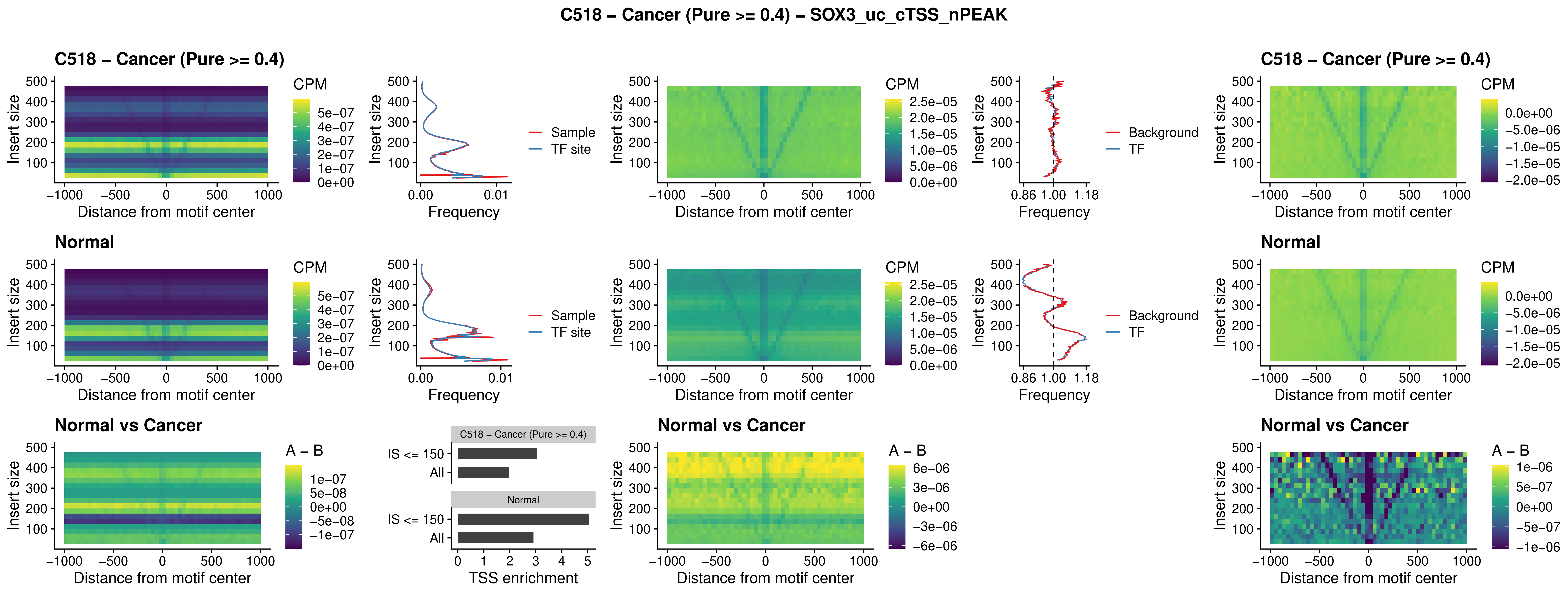
C518 – Cancer (Pure >= 0.4) – SOX3_uc_cTSS_nPEAK
C518 – Cancer (Pure >= 0.4)
Normal
Normal vs Cancer
Insert size
Distance from motif center
Frequency
CPM
A – B
Sample
TF site
Background
TF
C518 – Cancer (Pure >= 0.4)
Normal
IS <= 150
All
TSS enrichment

#### C518 – Cancer (Pure >= 0.4) – TBPL2_uc_cTSS_nPEAK

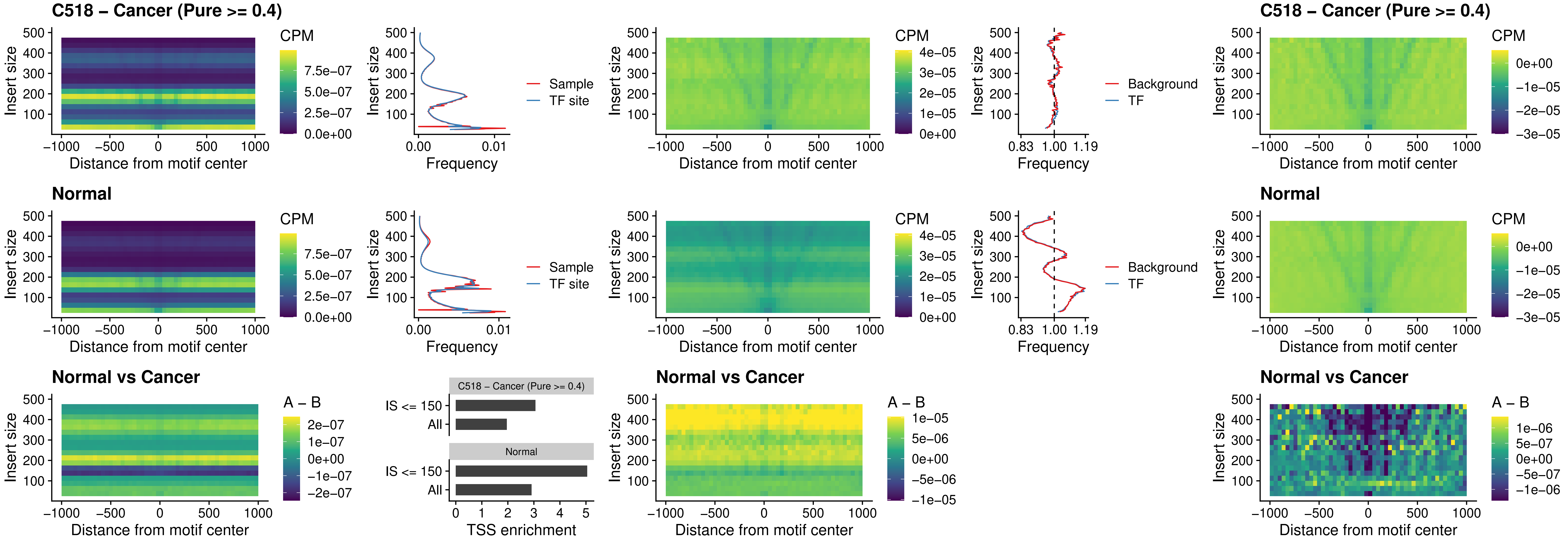

#### C518 – Cancer (Pure >= 0.4) – HOXA4_uc_cTSS_oPEAK

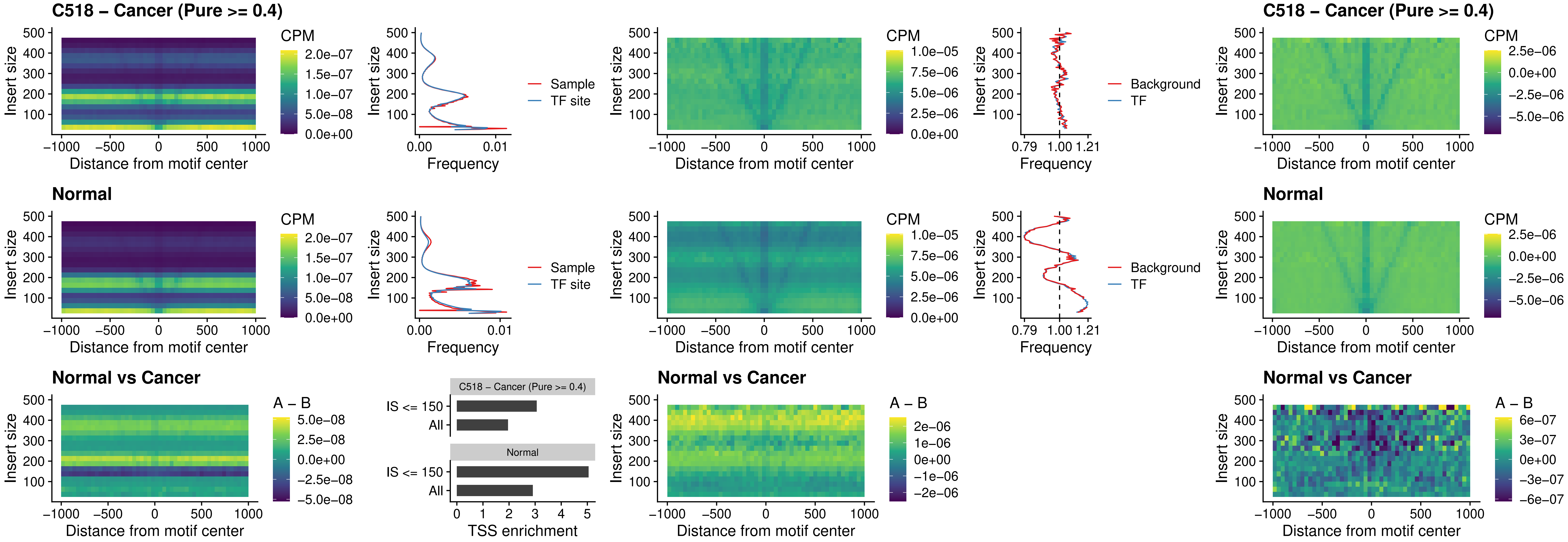

#### C518 – Cancer (Pure >= 0.4) – PDX1_uc_dTSS_oPEAK

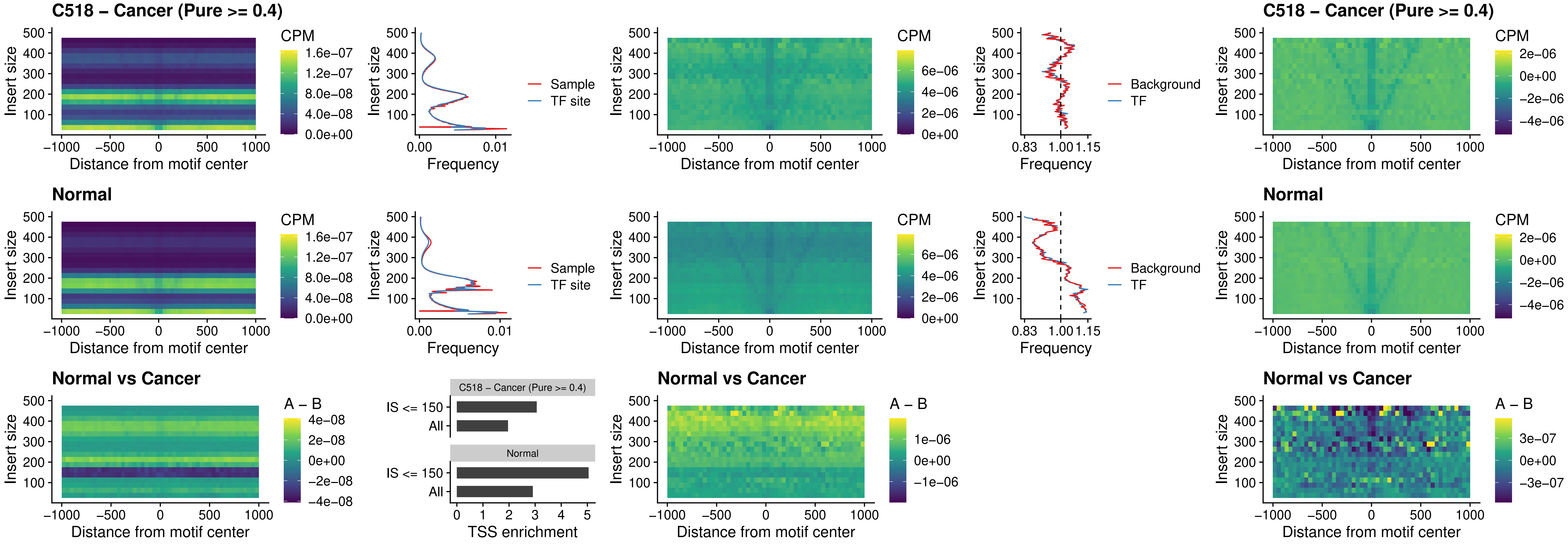

#### C518 – Cancer (Pure >= 0.4) – LHX4_uc_dTSS_oPEAK

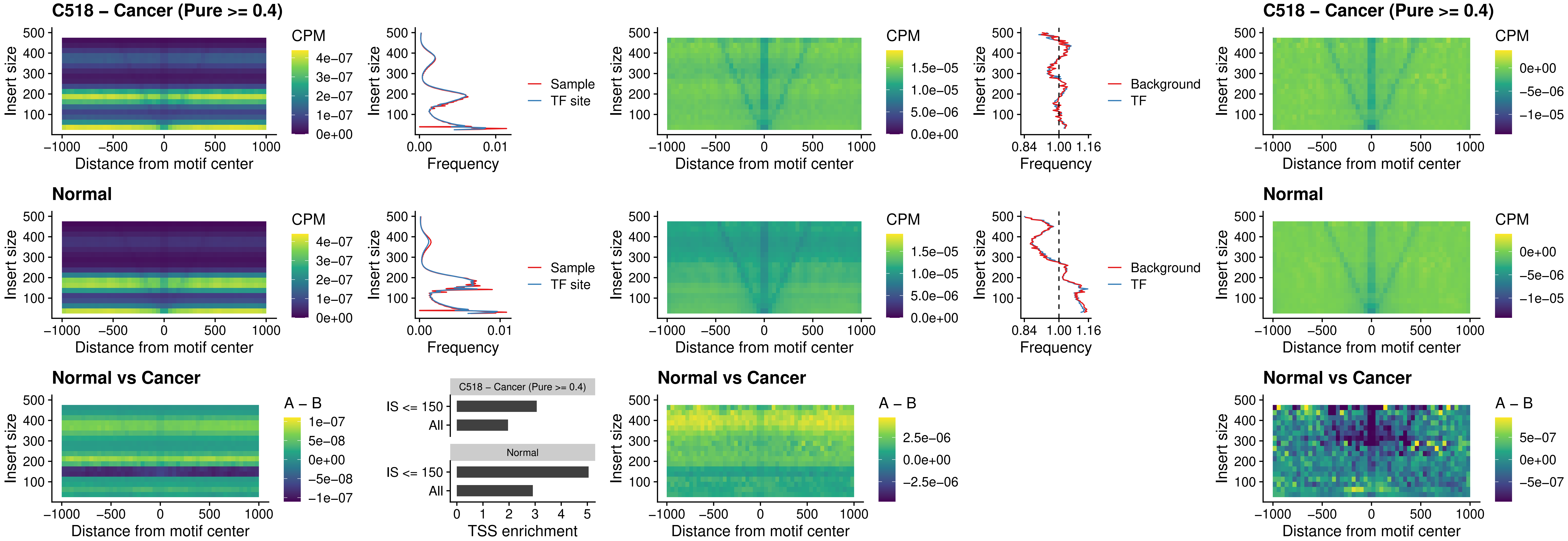

#### C518 – Cancer (Pure >= 0.4) – HOXA4_uc_dTSS_oPEAK

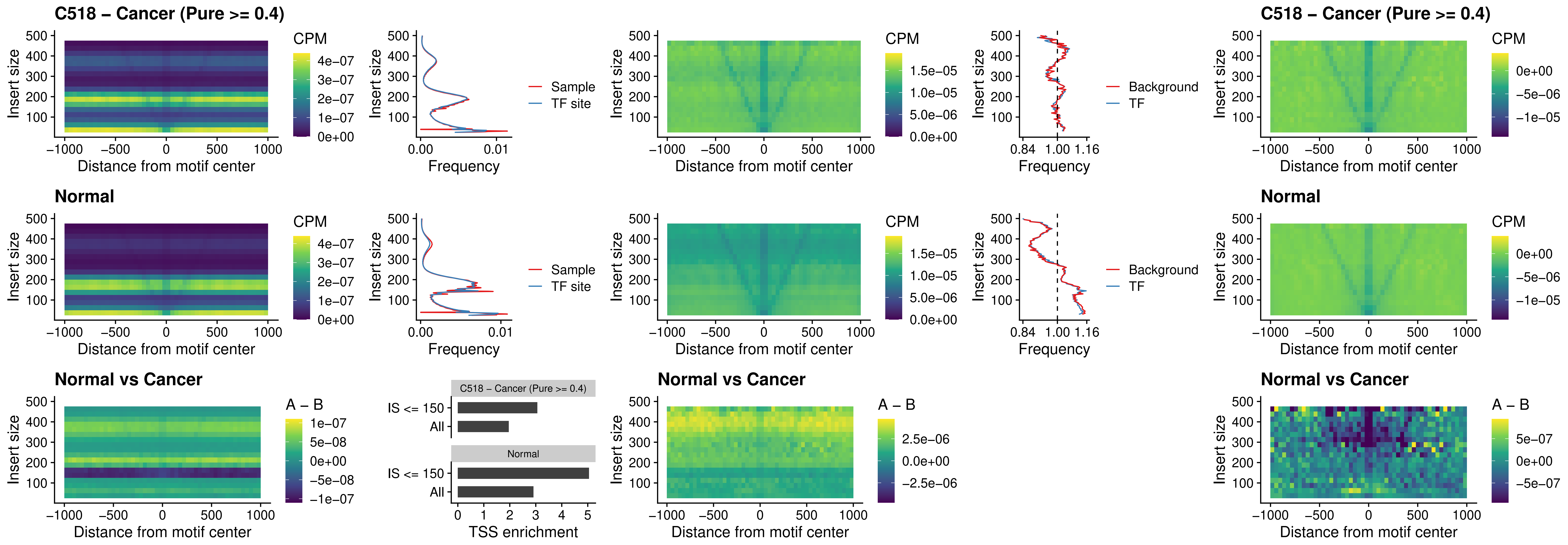

#### C518 – Cancer (Pure >= 0.4) – HLX_uc_dTSS_oPEAK

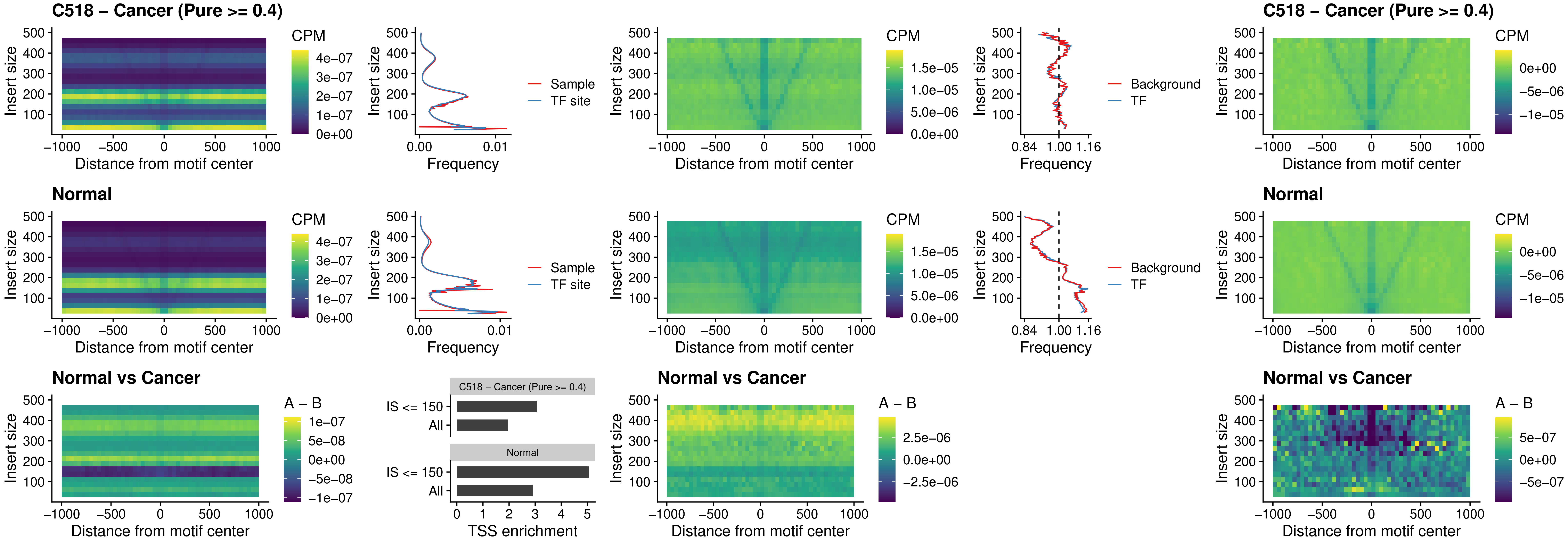

#### C518 – Cancer (Pure >= 0.4) – AC0125311_uc_dTSS_oPEAK

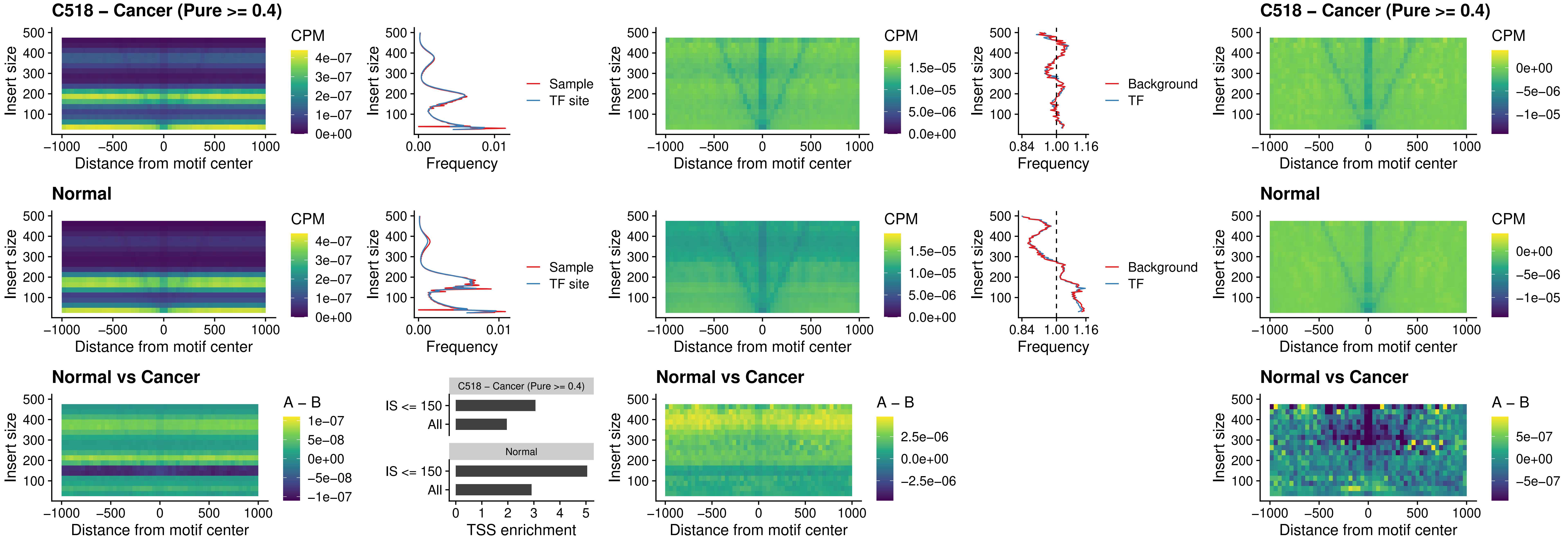

#### C518 – Cancer (Pure >= 0.4) – DBX2_uc_dTSS_oPEAK

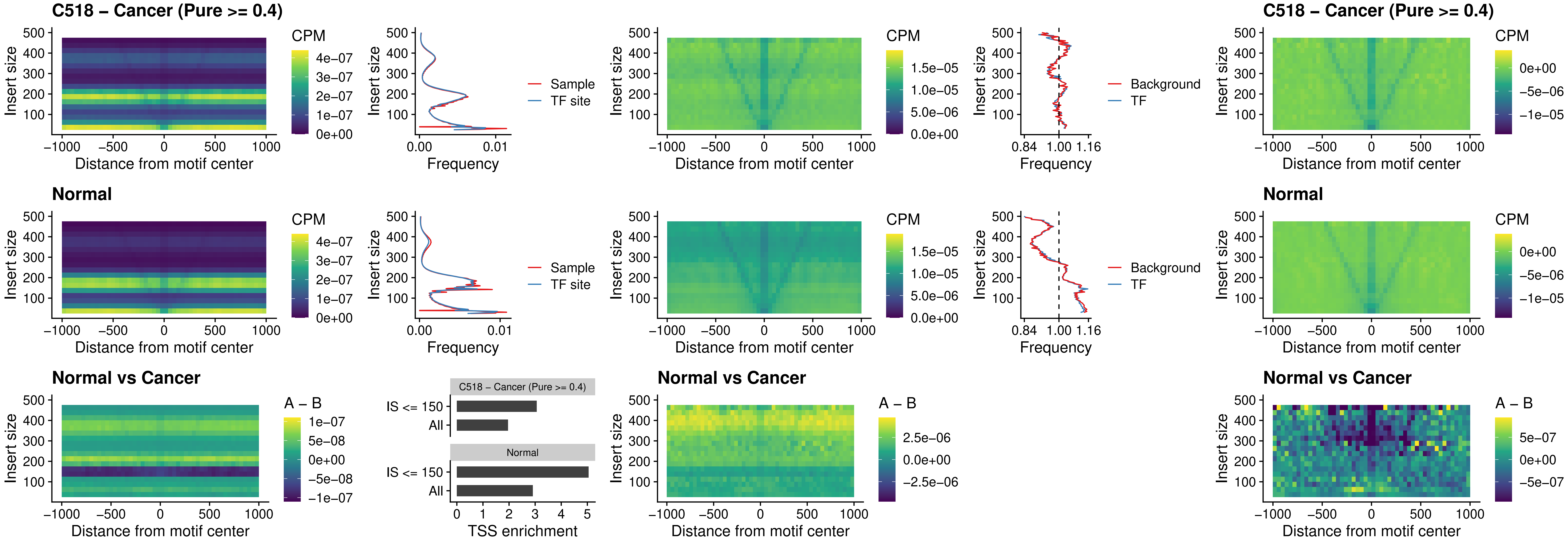

#### C518 – Cancer (Pure >= 0.4) – HOXA2_uc_dTSS_nPEAK

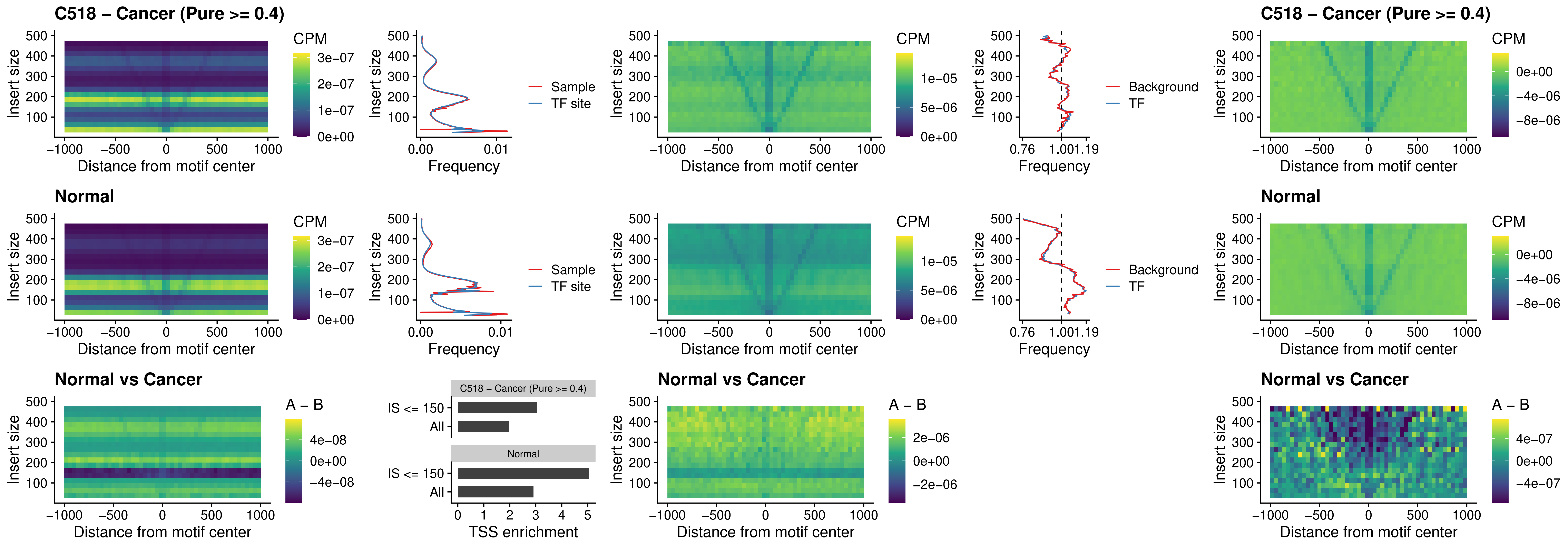

### C518 – Cancer (Pure >= 0.4) – SHOX_uc_dTSS_nPEAK

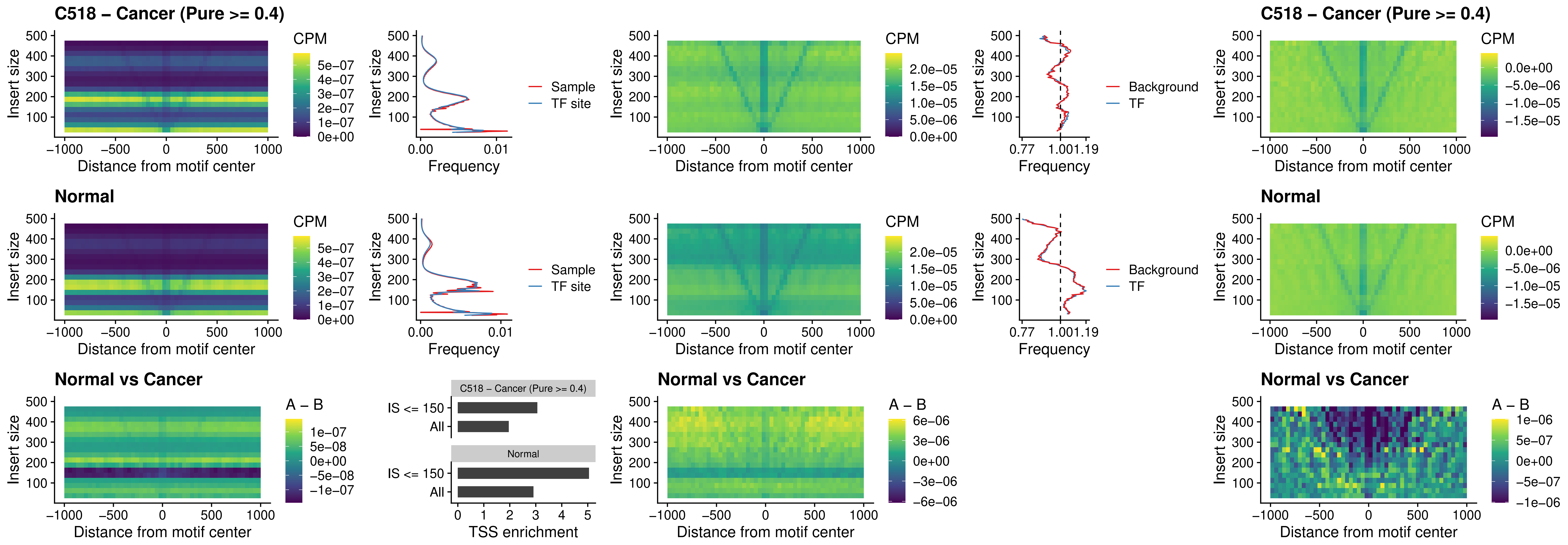

#### C518 – Cancer (Pure >= 0.4) – PRRX1_uc_dTSS_nPEAK

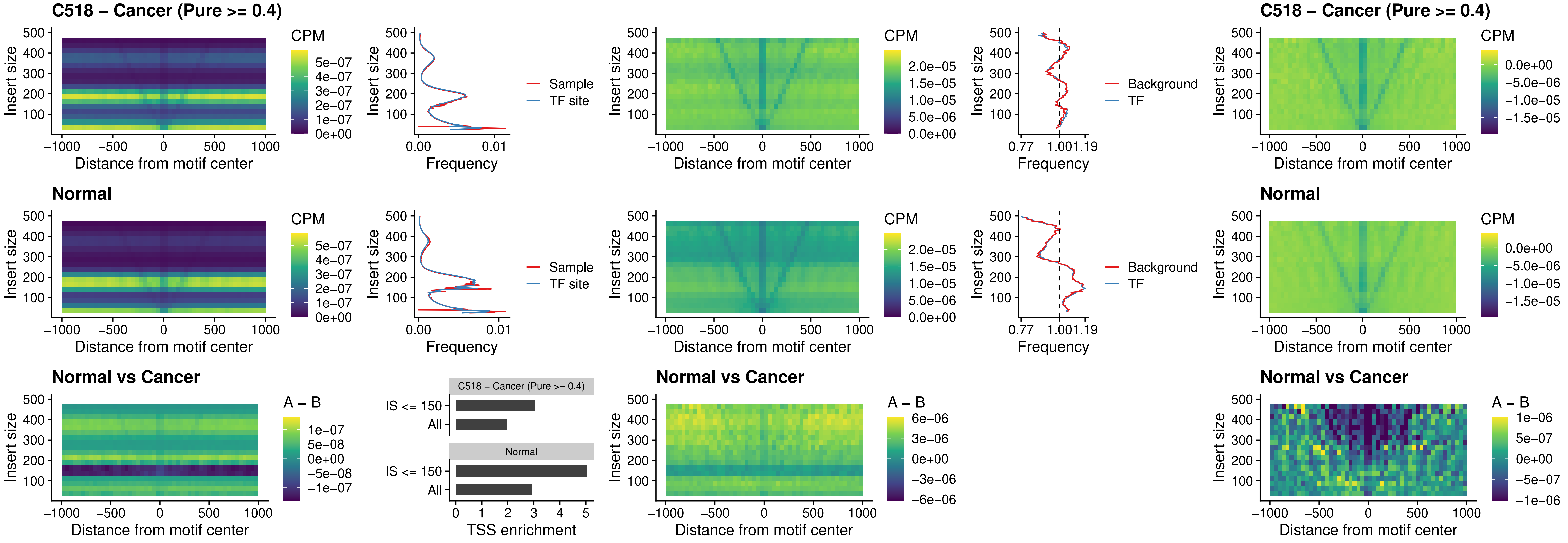

#### C518 – Cancer (Pure >= 0.4) – LMX1A_uc_dTSS_nPEAK

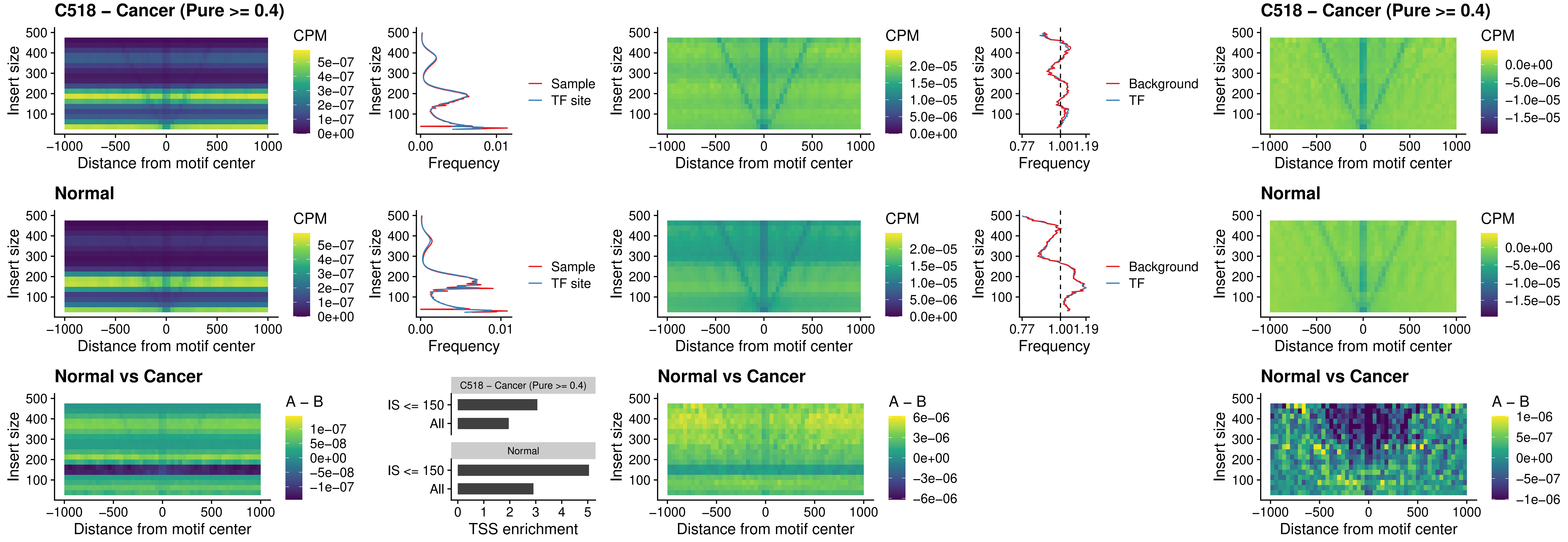

#### C518 – Cancer (Pure >= 0.4) – LMX1B_uc_dTSS_nPEAK

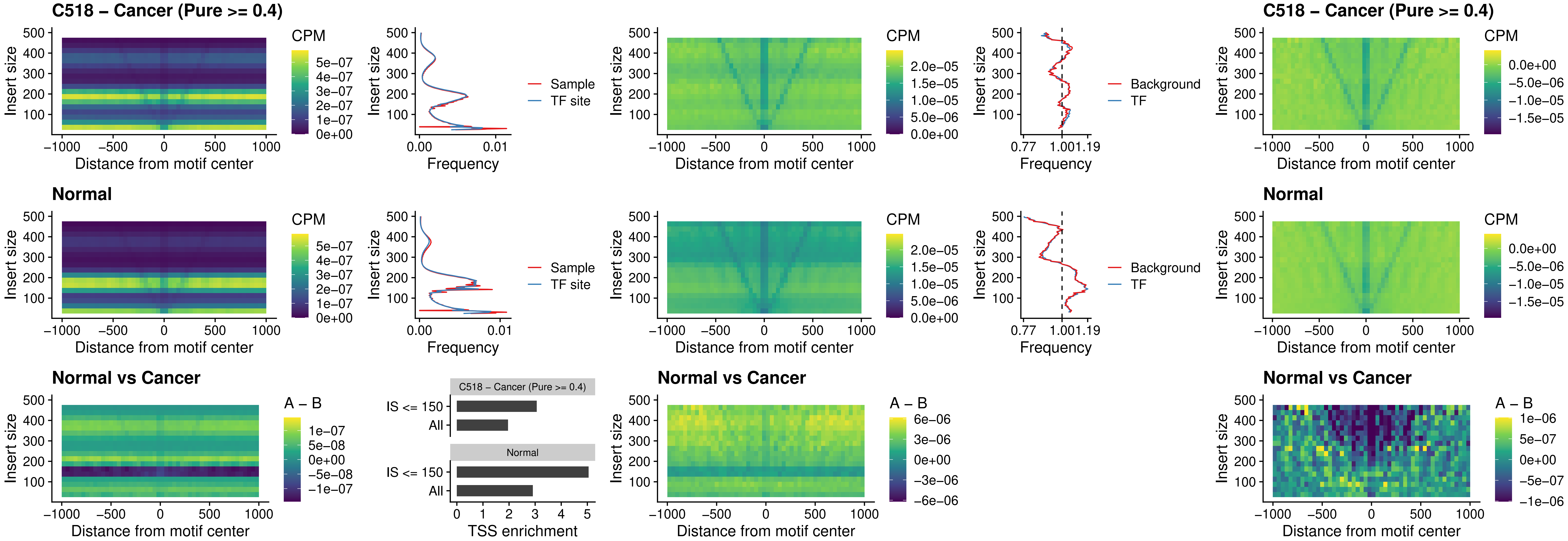

#### C518 – Cancer (Pure >= 0.4) – DLX5_uc_dTSS_oPEAK

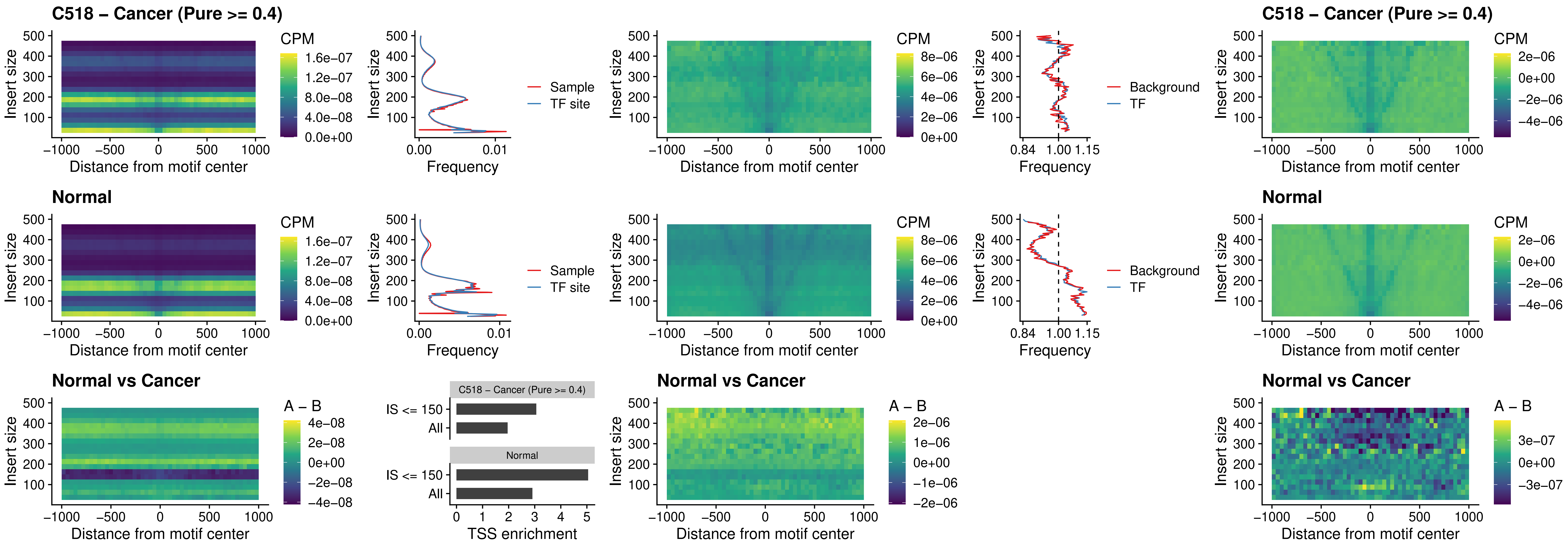

### C518 – Cancer (Pure >= 0.4) – ARID3C_uc_dTSS_nPEAK

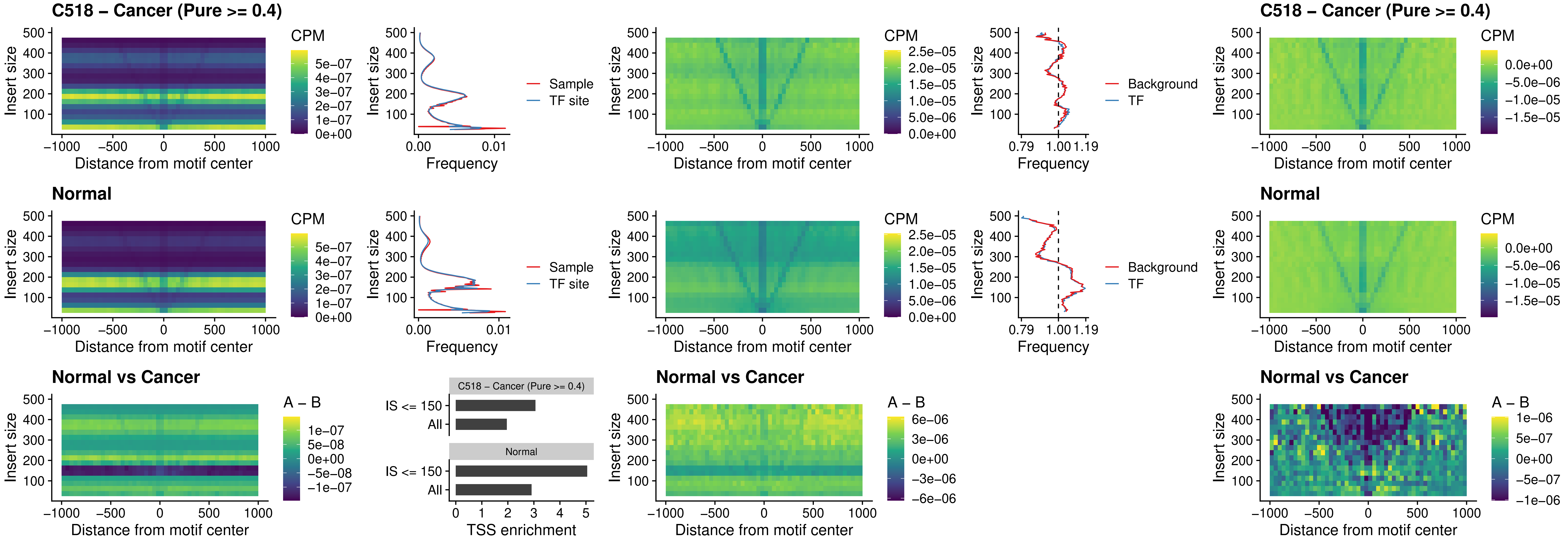

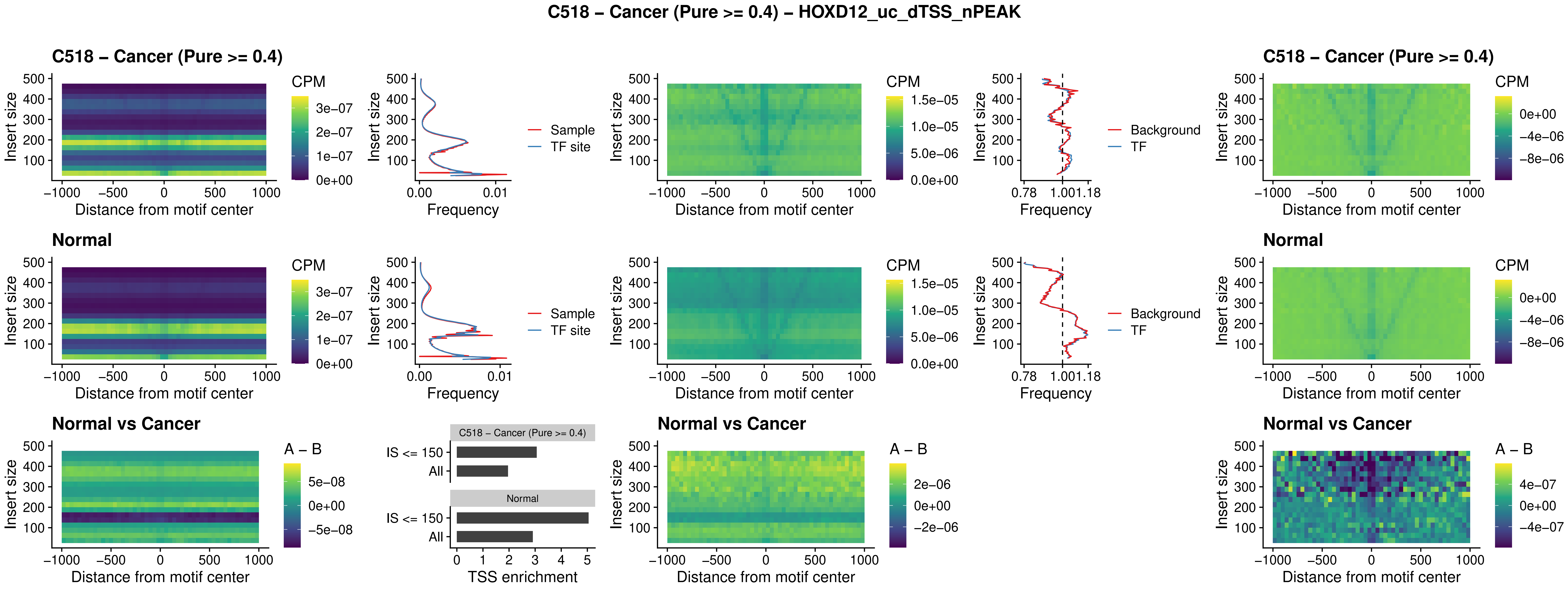
C518 – Cancer (Pure >= 0.4) – HOXD12_uc_dTSS_nPEAK
C518 – Cancer (Pure >= 0.4)
Normal
Normal vs Cancer
Insert size
Distance from motif center
Frequency
CPM
A – B
Sample
TF site
Background
TF
C518 – Cancer (Pure >= 0.4)
Normal
IS <= 150
All
TSS enrichment

### C518 – Cancer (Pure >= 0.4) – CDX2_uc_dTSS_nPEAK

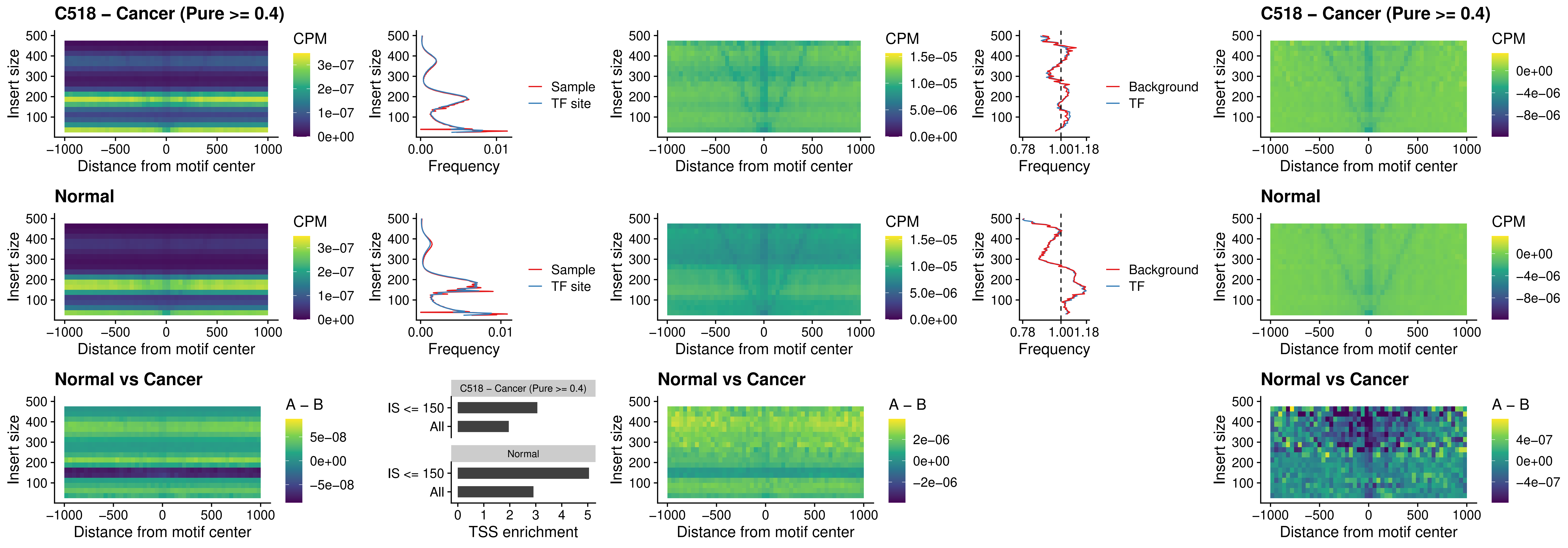

#### C518 – Cancer (Pure >= 0.4) – HOXC12_uc_dTSS_nPEAK

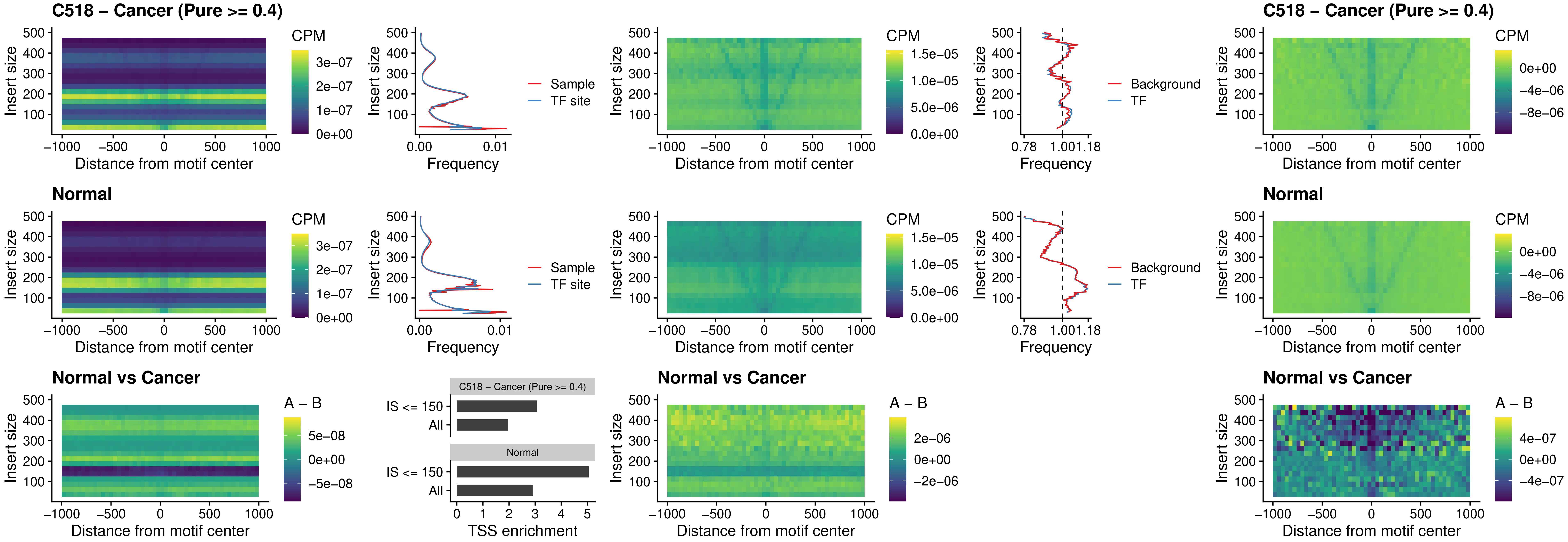

#### C518 – Cancer (Pure >= 0.4) – HHEX_uc_dTSS_oPEAK

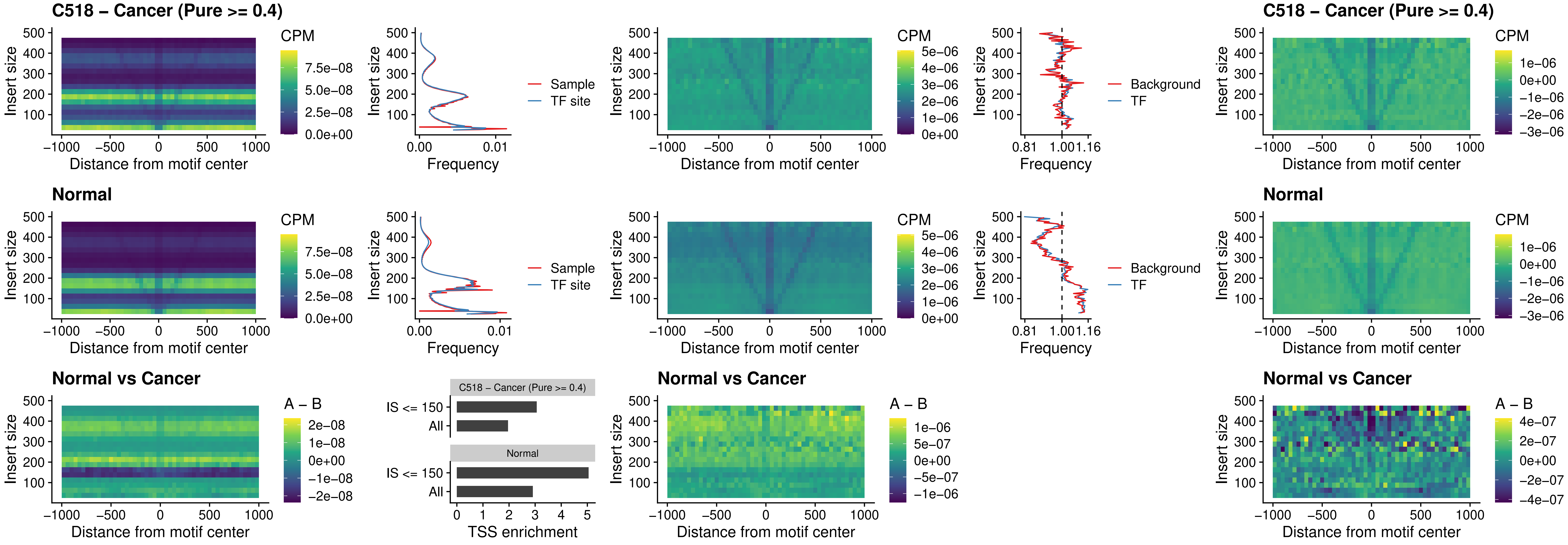

#### C518 – Cancer (Pure >= 0.4) – HOXA11_uc_dTSS_nPEAK

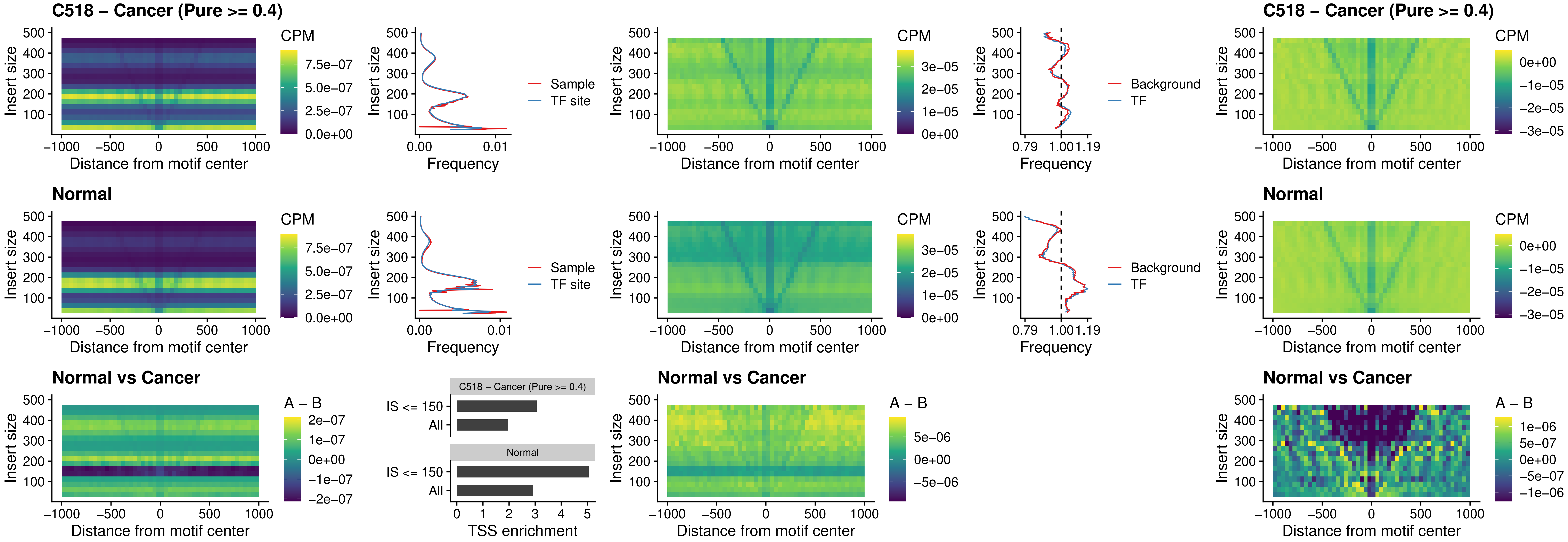

#### C518 – Cancer (Pure >= 0.4) – HOXA11_uc_cTSS_nPEAK

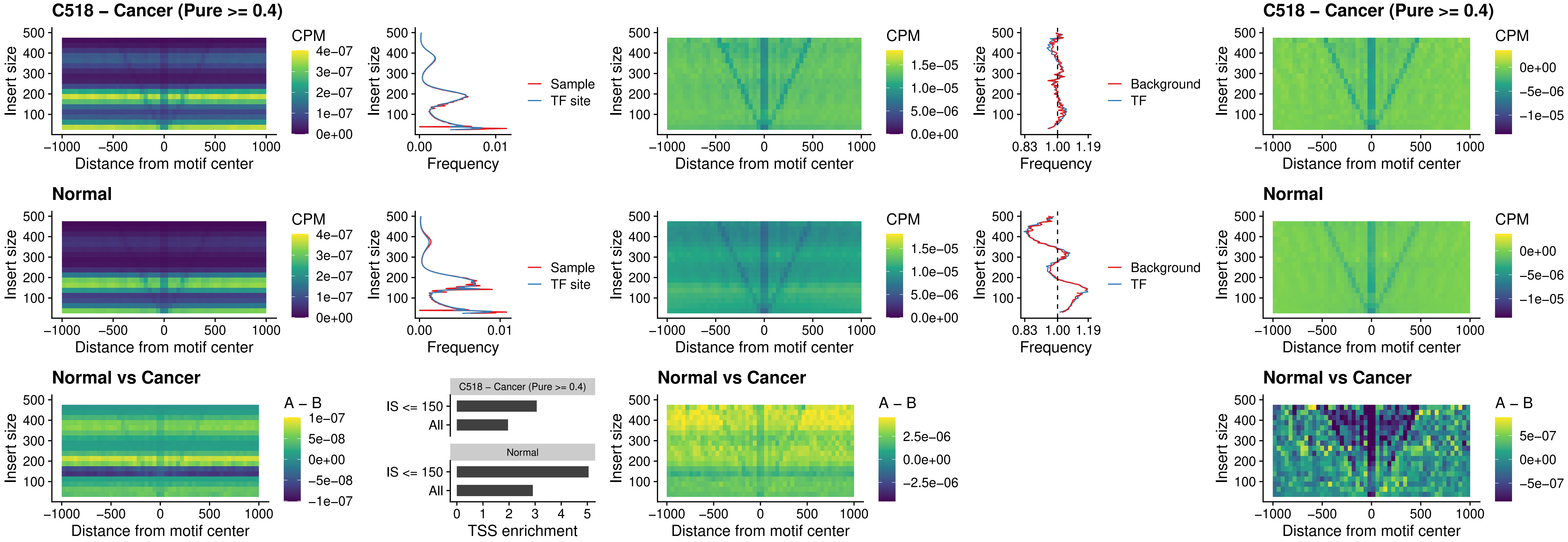

#### C518 – Cancer (Pure >= 0.4) – ZFHX3_uc_dTSS_nPEAK

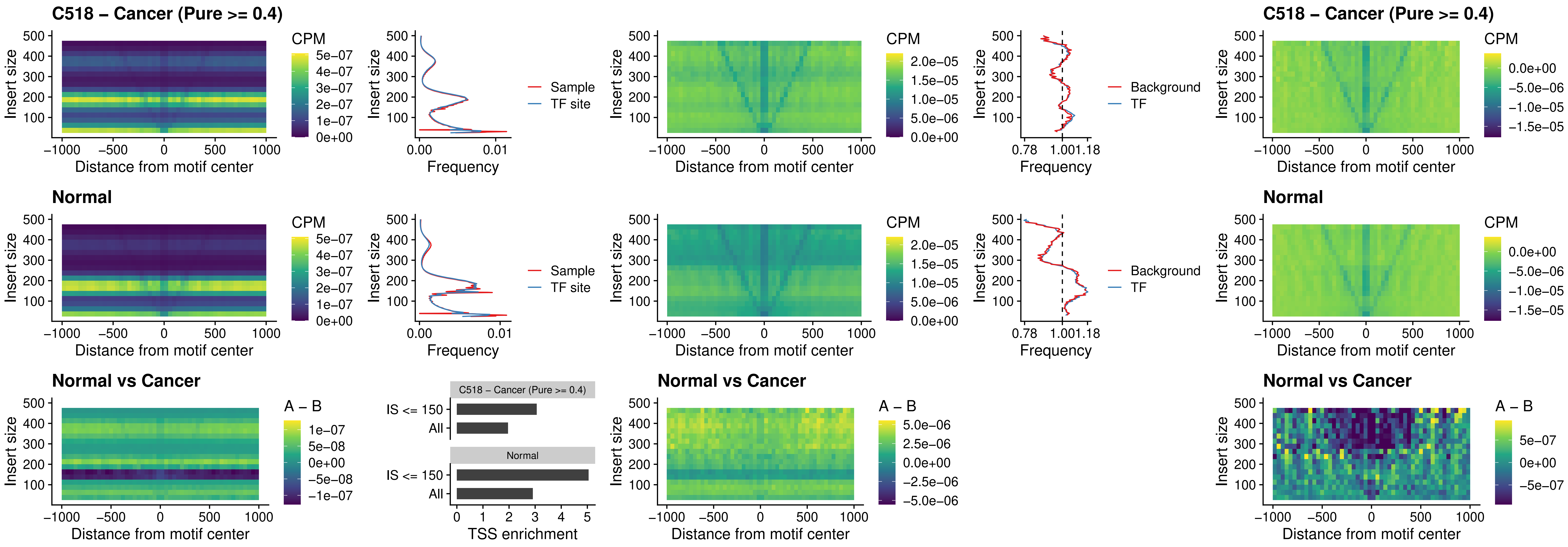

#### C518 – Cancer (Pure >= 0.4) – HOXB4_uc_dTSS_nPEAK

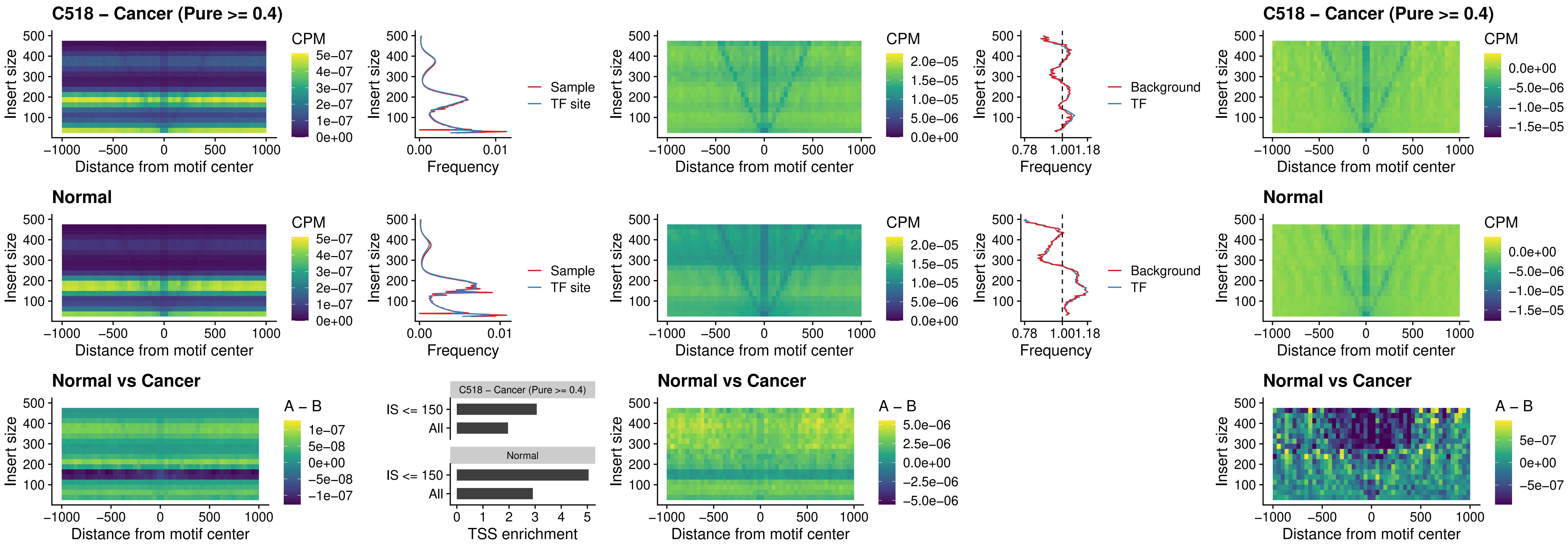

#### C518 – Cancer (Pure >= 0.4) – HOXC5_uc_dTSS_nPEAK

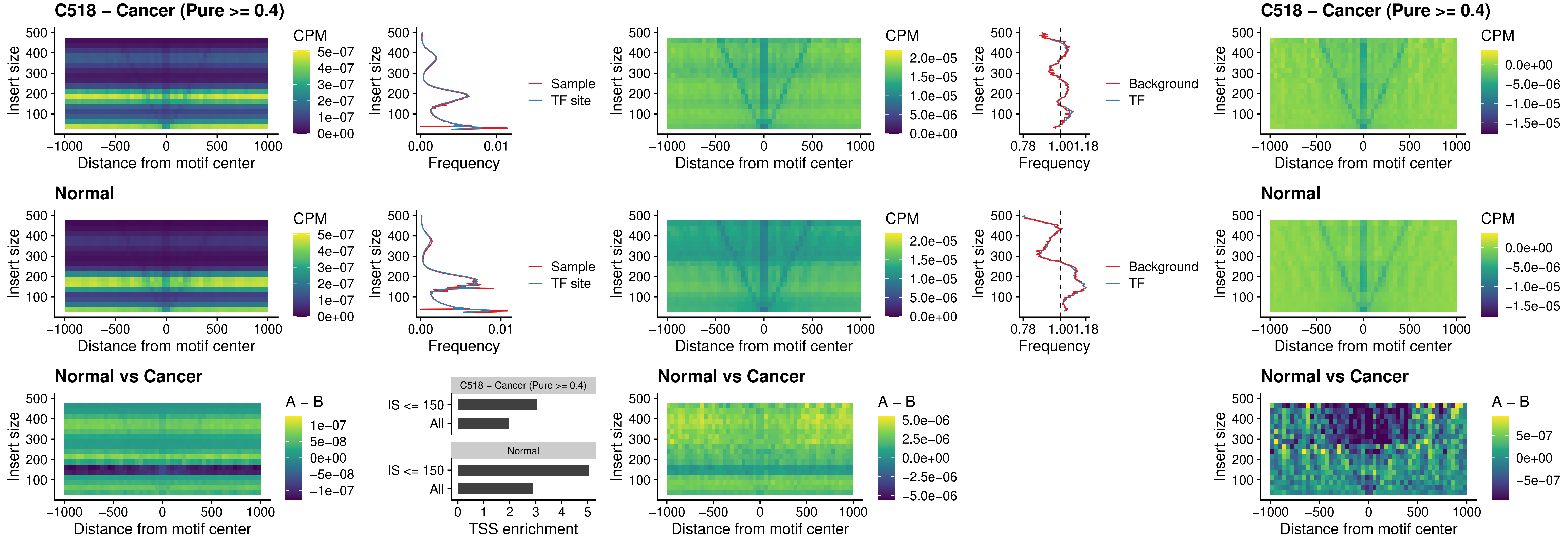

#### C518 – Cancer (Pure >= 0.4) – HOXD1_uc_dTSS_nPEAK

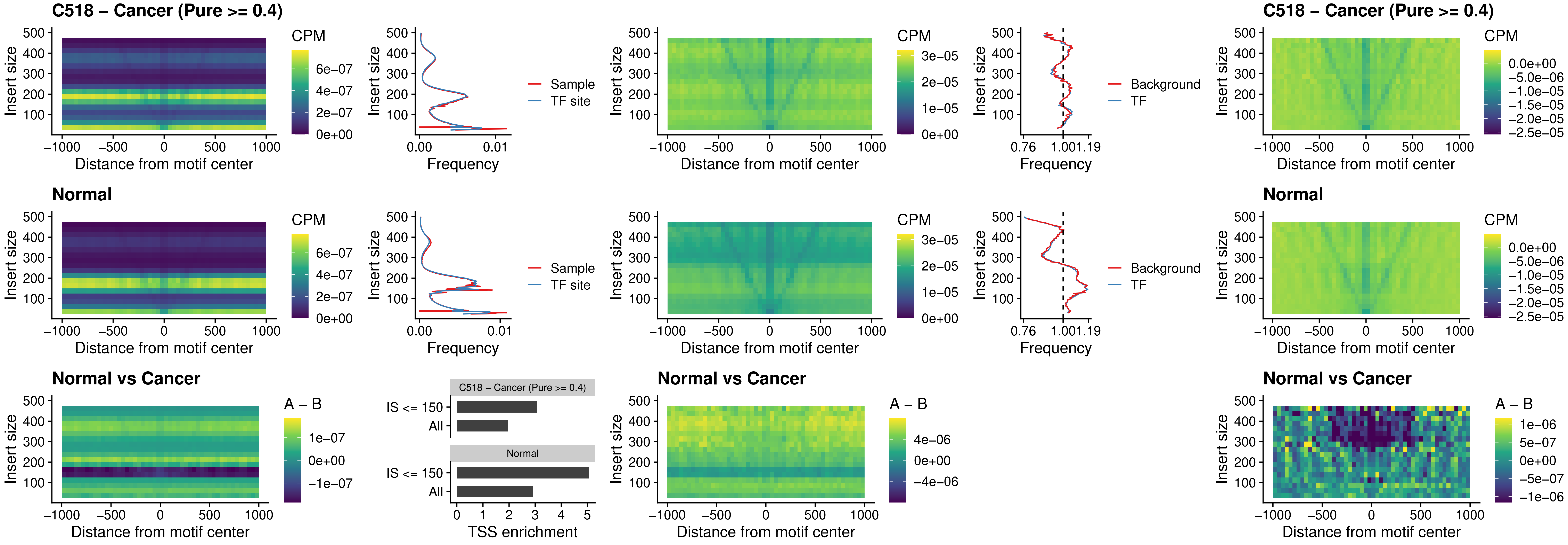

#### C518 – Cancer (Pure >= 0.4) – OTP_uc_dTSS_nPEAK

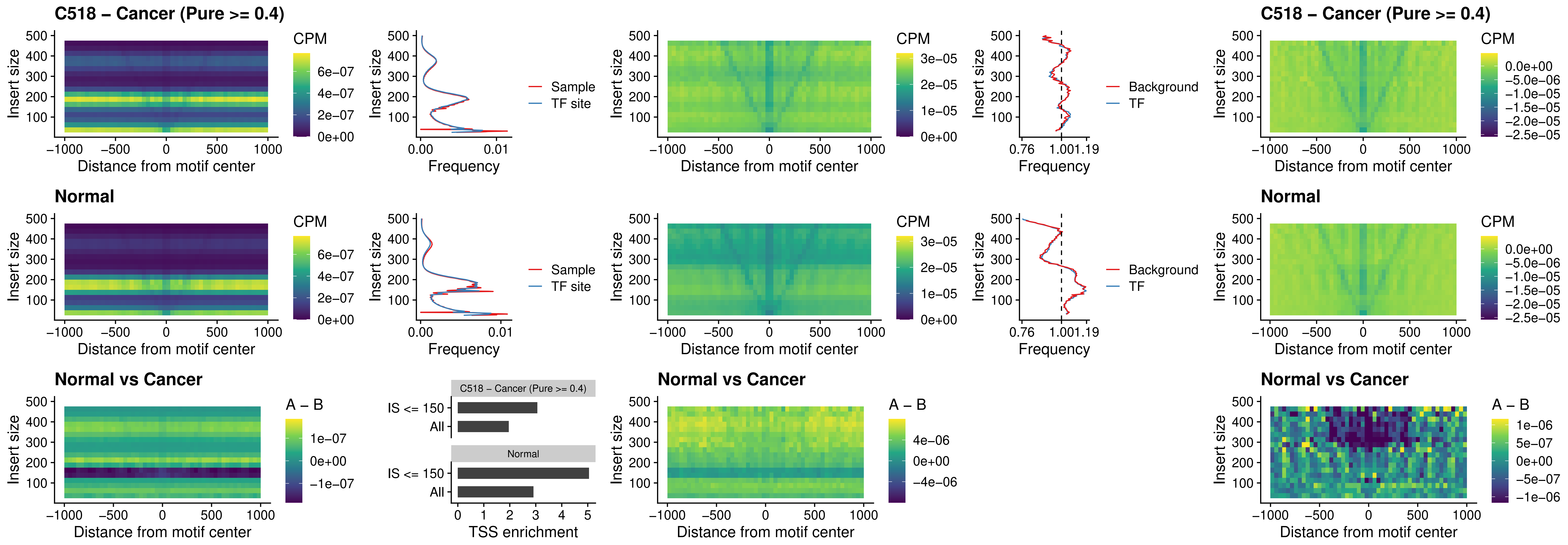

#### C518 – Cancer (Pure >= 0.4) – FOXL1_uc_dTSS_oPEAK

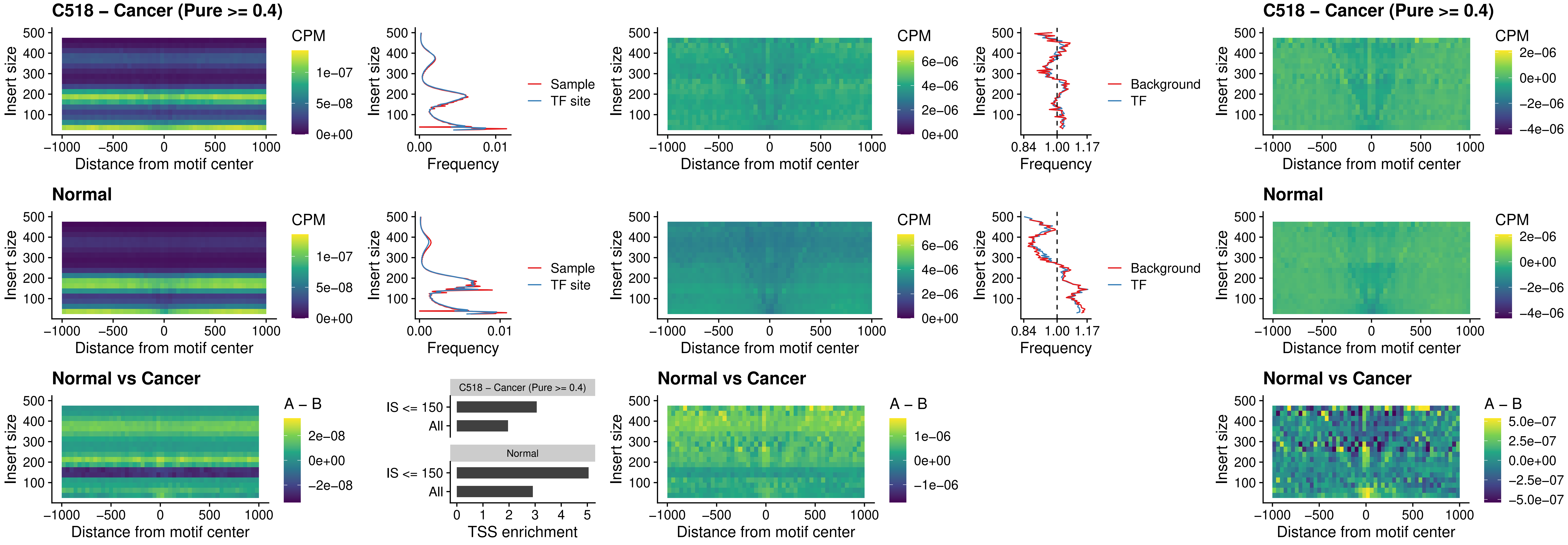

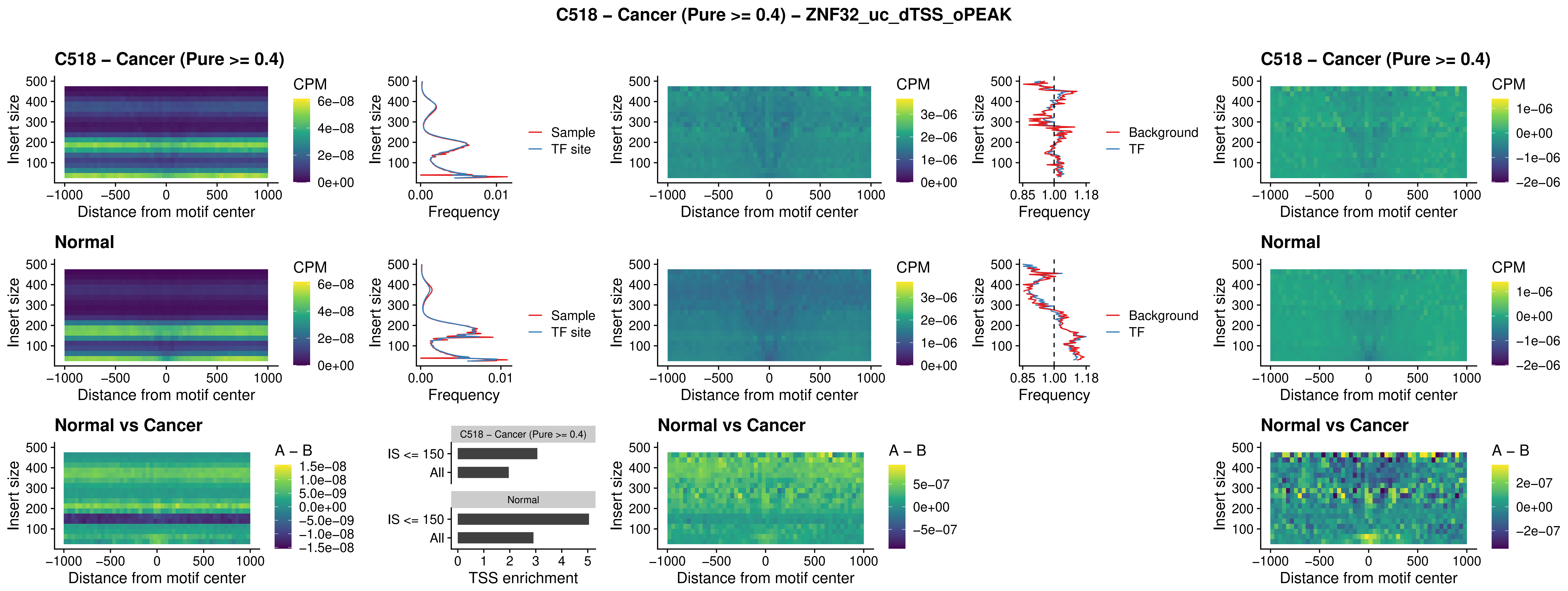
C518 – Cancer (Pure >= 0.4) – ZNF32_uc_dTSS_oPEAK
C518 – Cancer (Pure >= 0.4)
Normal
Normal vs Cancer
Insert size
Distance from motif center
Frequency
CPM
A – B
Sample
TF site
Background
TF
TSS enrichment
IS <= 150
All
C518 – Cancer (Pure >= 0.4)
Normal

#### C518 – Cancer (Pure >= 0.4) – FOXL1_uc_dTSS_nPEAK

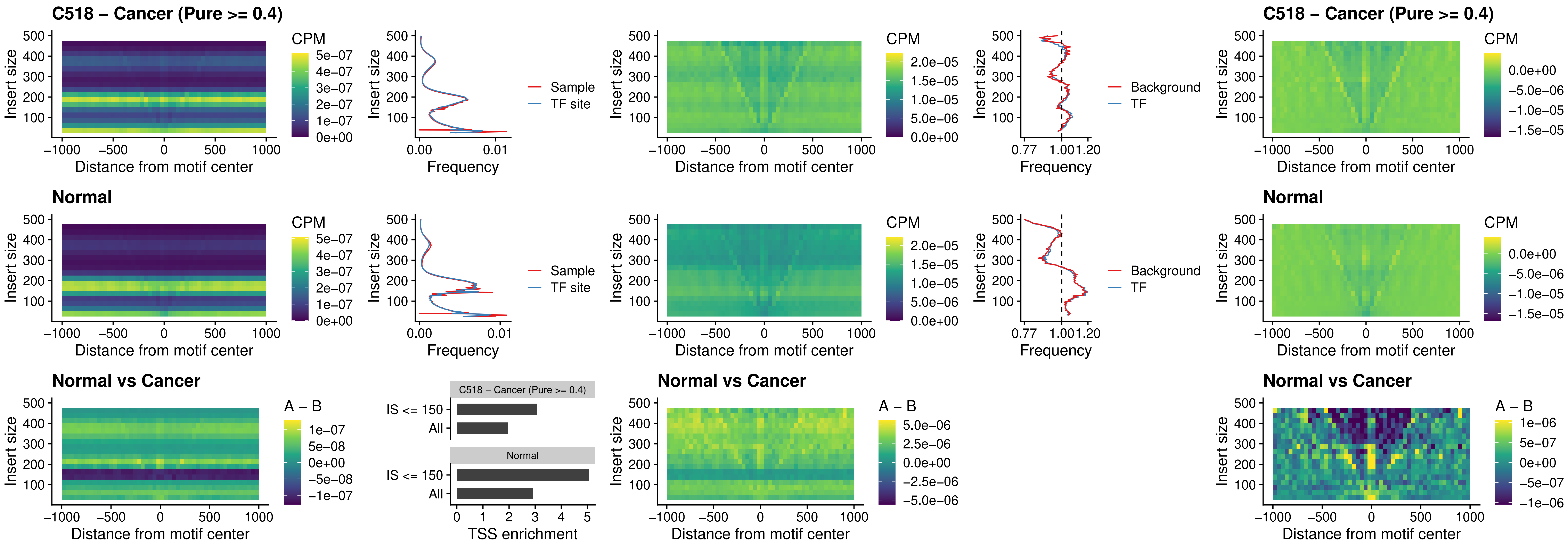

#### C518 – Cancer (Pure >= 0.4) – ZNF32_uc_dTSS_nPEAK

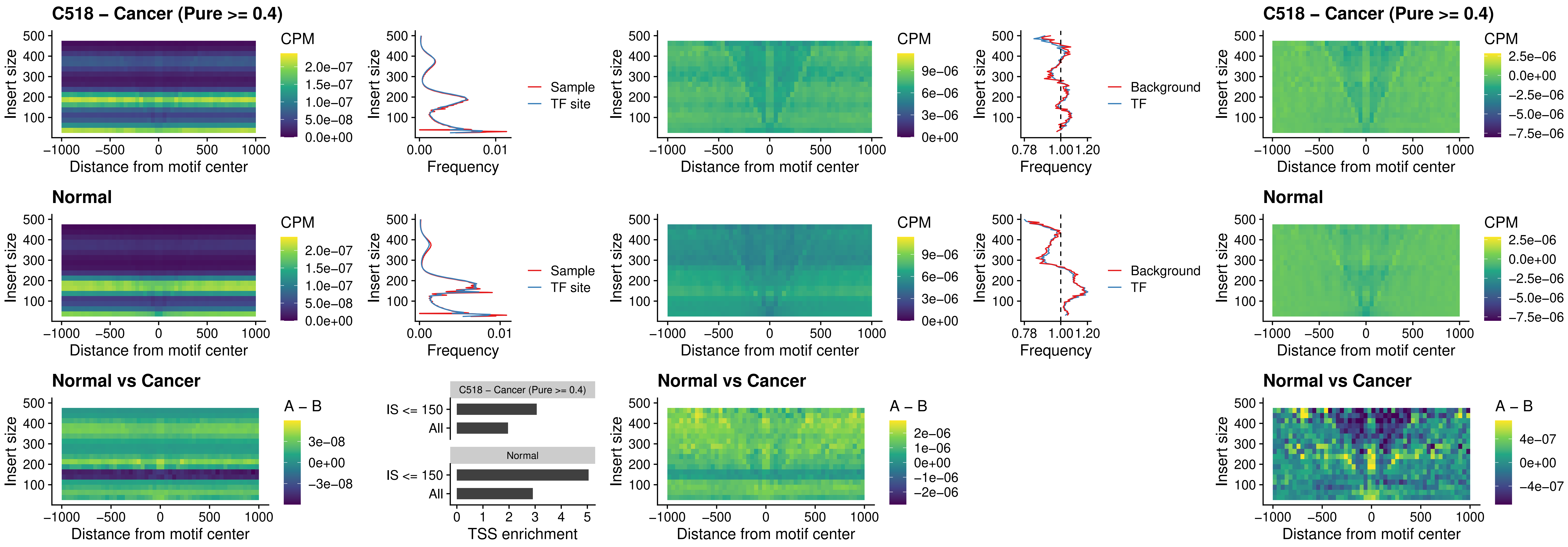

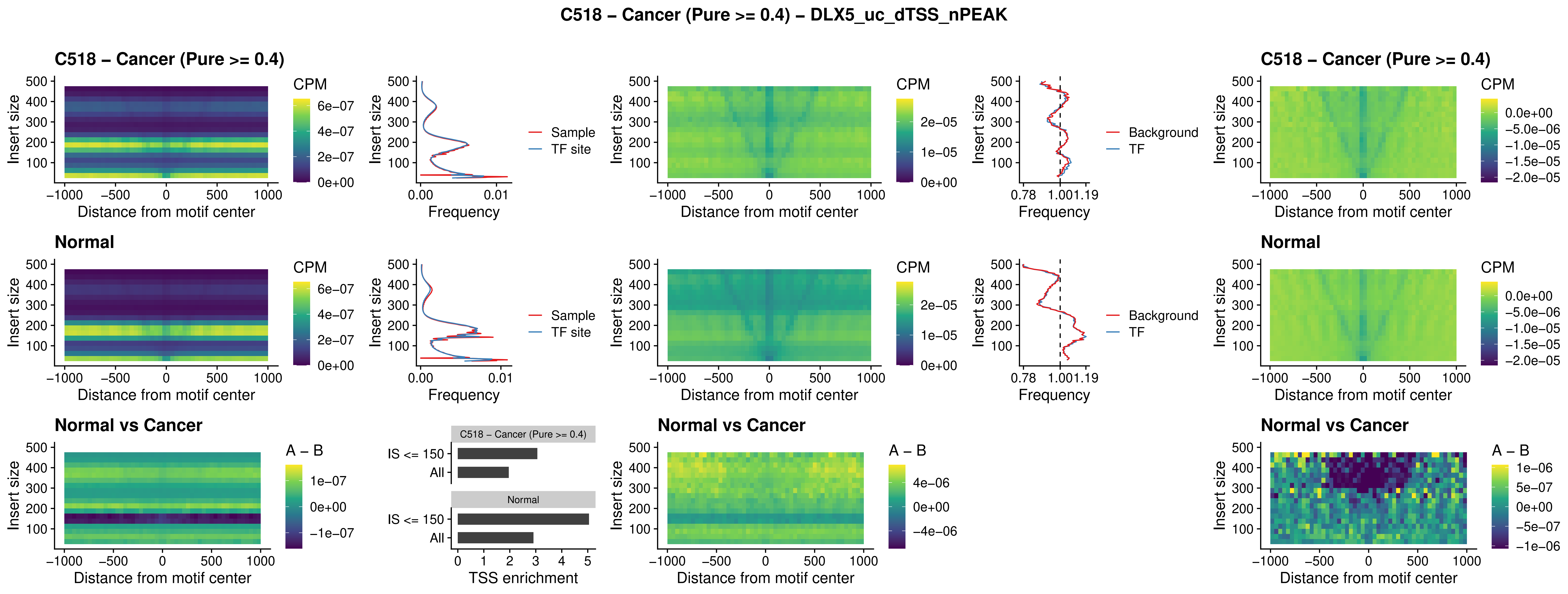
C518 – Cancer (Pure >= 0.4) – DLX5_uc_dTSS_nPEAK
C518 – Cancer (Pure >= 0.4)
Normal
Normal vs Cancer
Insert size
Distance from motif center
Frequency
CPM
A – B
Sample
TF site
Background
TF
C518 – Cancer (Pure >= 0.4)
Normal
IS <= 150
All
TSS enrichment

#### C518 – Cancer (Pure >= 0.4) – DMRT2_uc_dTSS_nPEAK

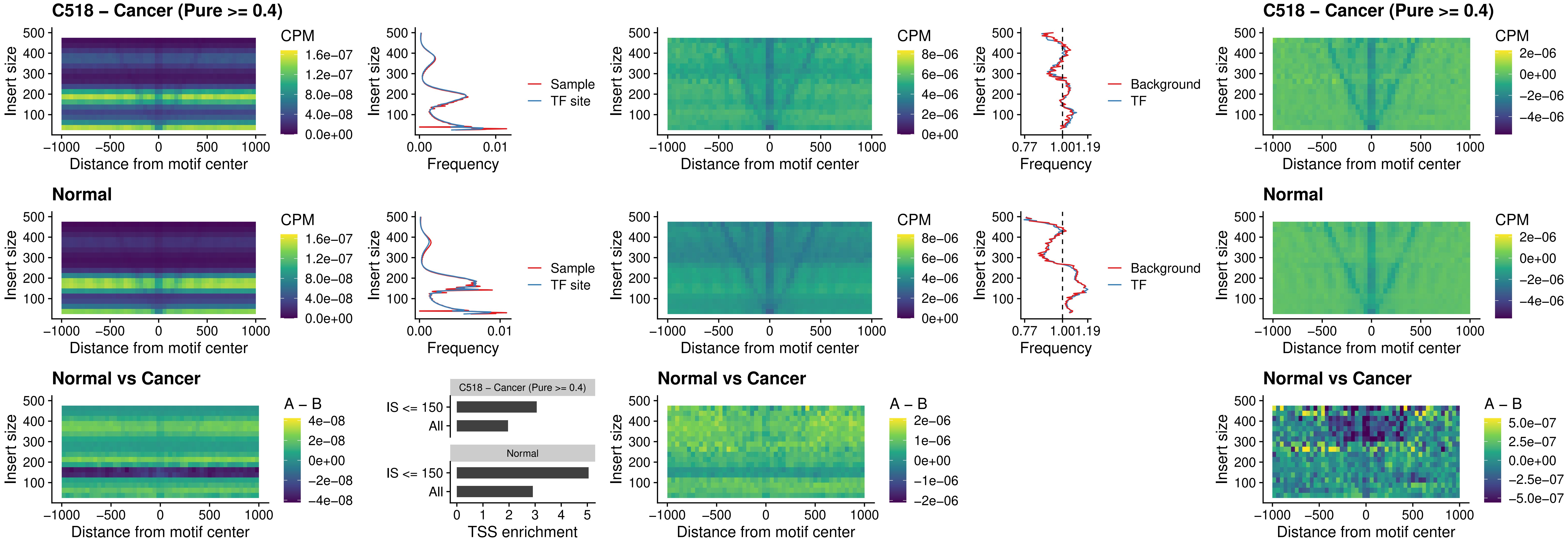

#### C518 – Cancer (Pure >= 0.4) – DMRTC2_uc_cTSS_nPEAK

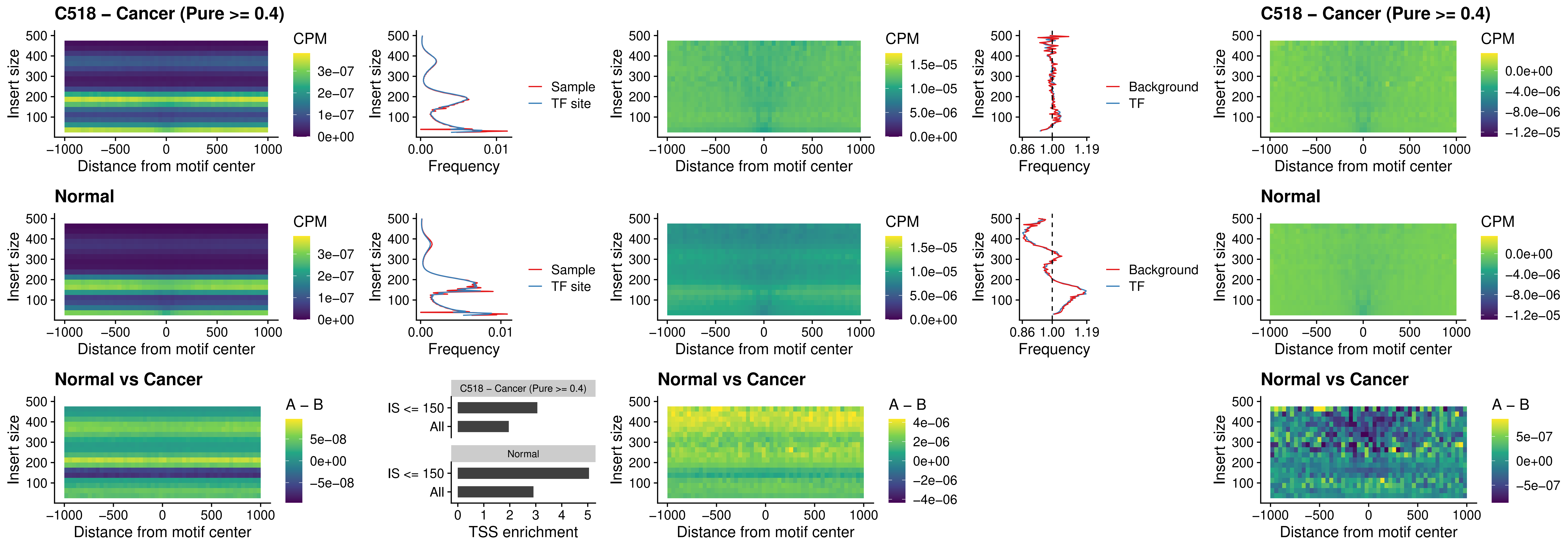

#### C518 – Cancer (Pure >= 0.4) – HOXD9_uc_dTSS_nPEAK

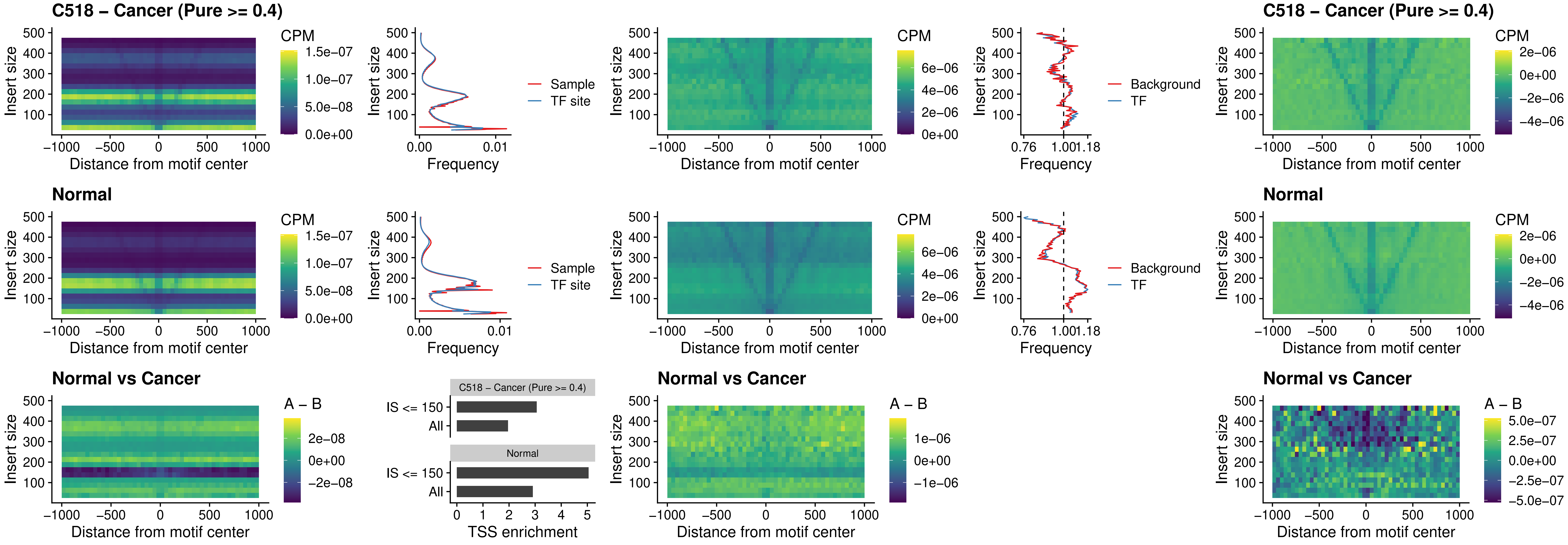

### C518 – Cancer (Pure >= 0.4) – HOXD4_uc_dTSS_nPEAK

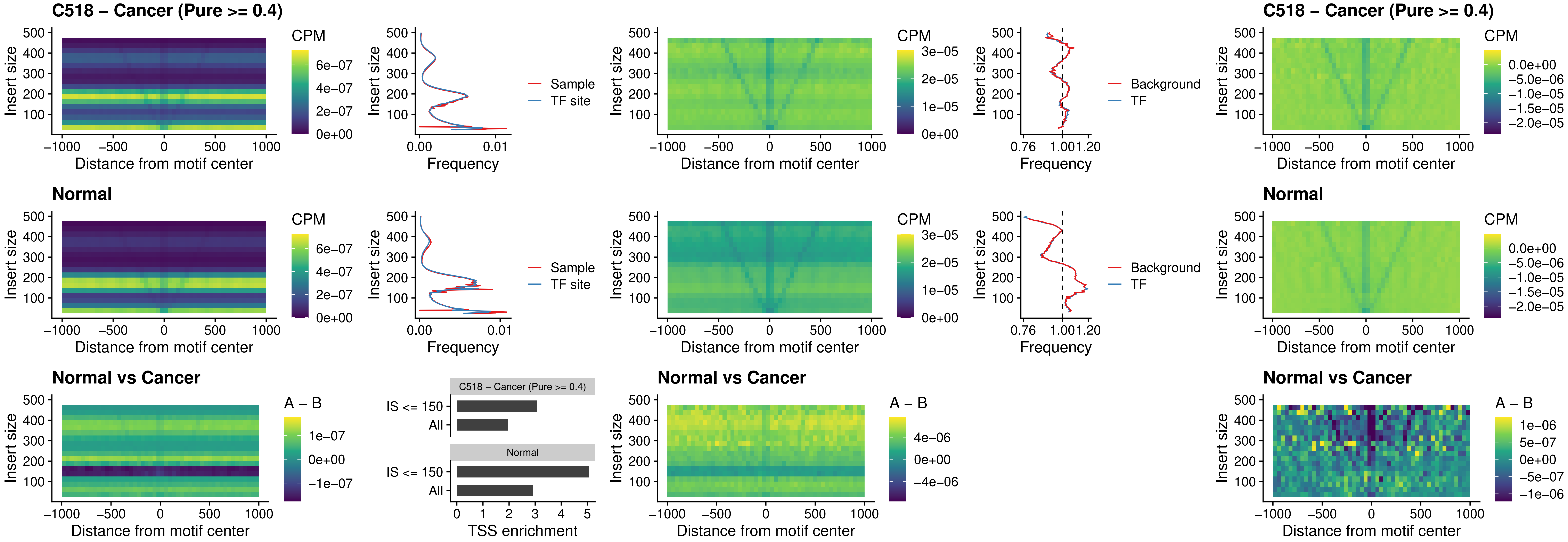

#### C518 – Cancer (Pure >= 0.4) – FOXP3_uc_cTSS_nPEAK

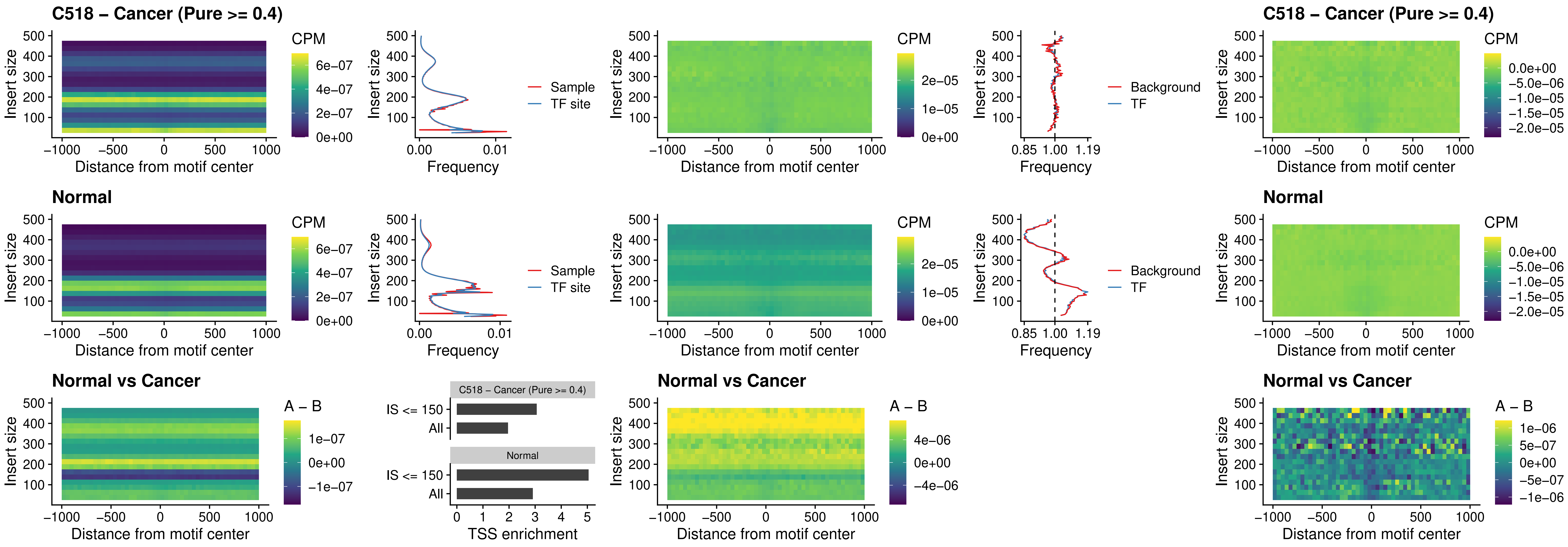

#### C518 – Cancer (Pure >= 0.4) – FOXD4L6_uc_cTSS_nPEAK

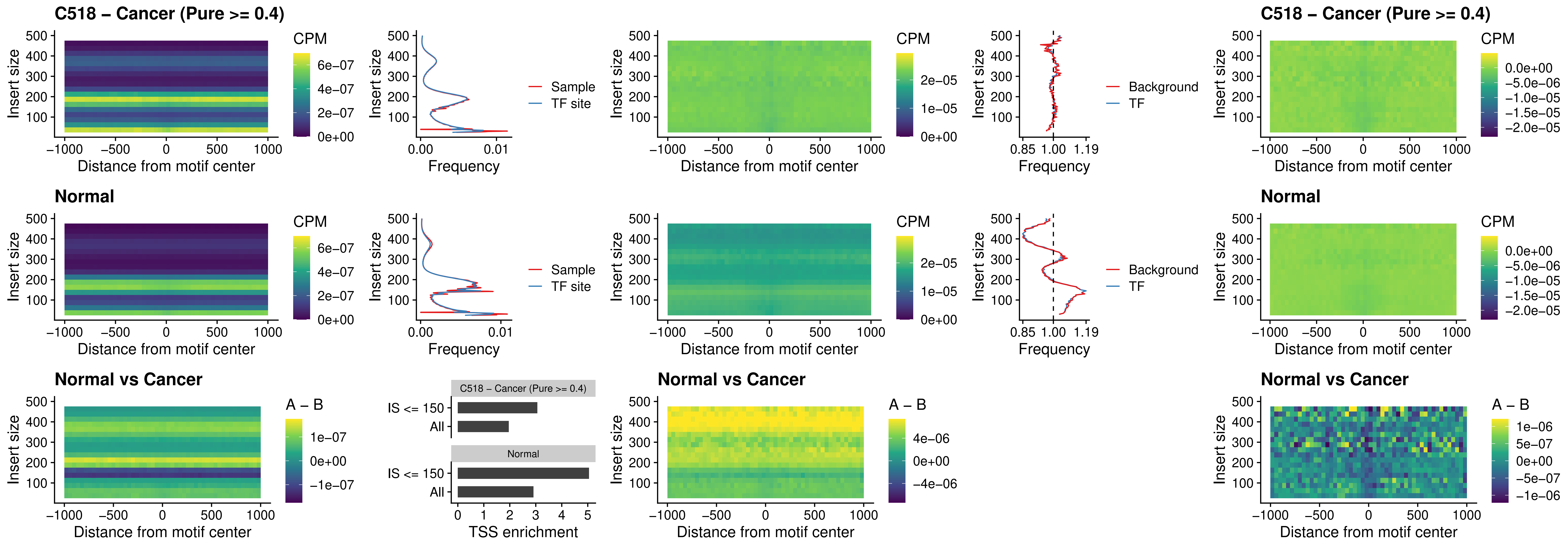

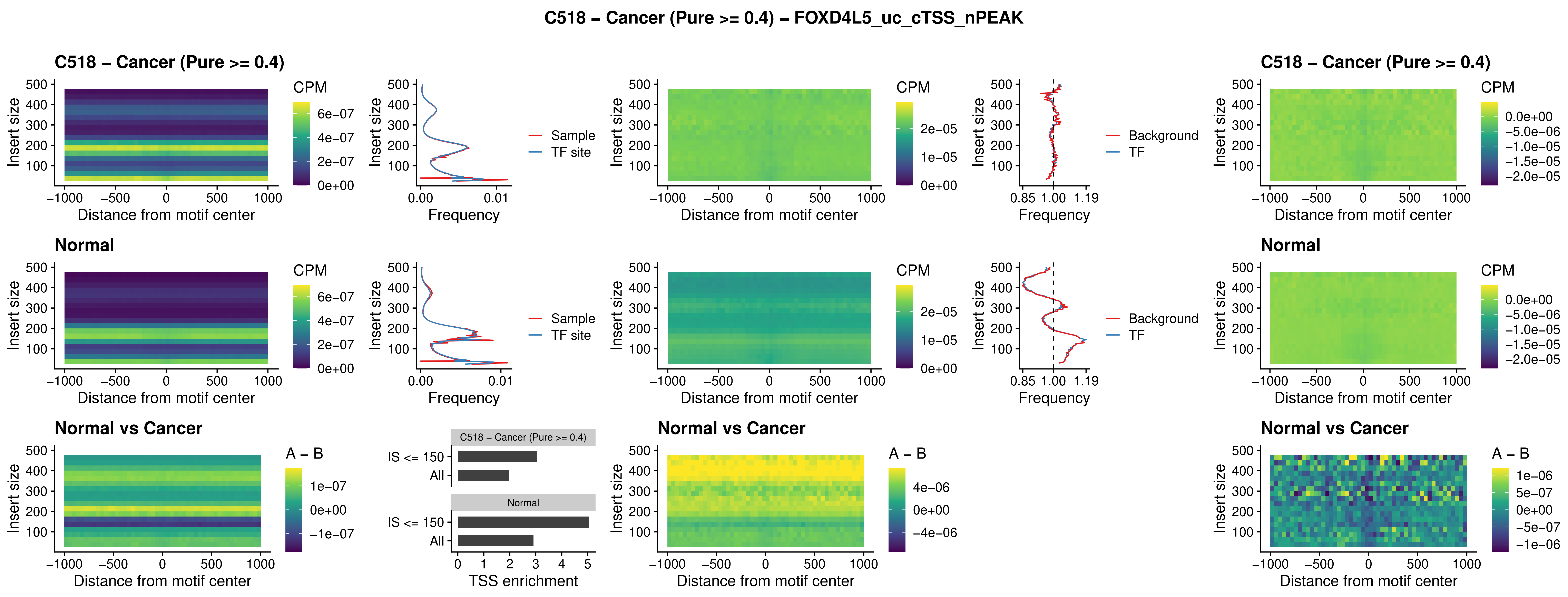
C518 – Cancer (Pure >= 0.4) – FOXD4L5_uc_cTSS_nPEAK
C518 – Cancer (Pure >= 0.4)
Normal
Normal vs Cancer
Insert size
Distance from motif center
Frequency
CPM
A – B
Sample
TF site
Background
TF
C518 – Cancer (Pure >= 0.4)
Normal
IS <= 150
All
TSS enrichment

#### C518 – Cancer (Pure >= 0.4) – FOXD4L4_uc_cTSS_nPEAK

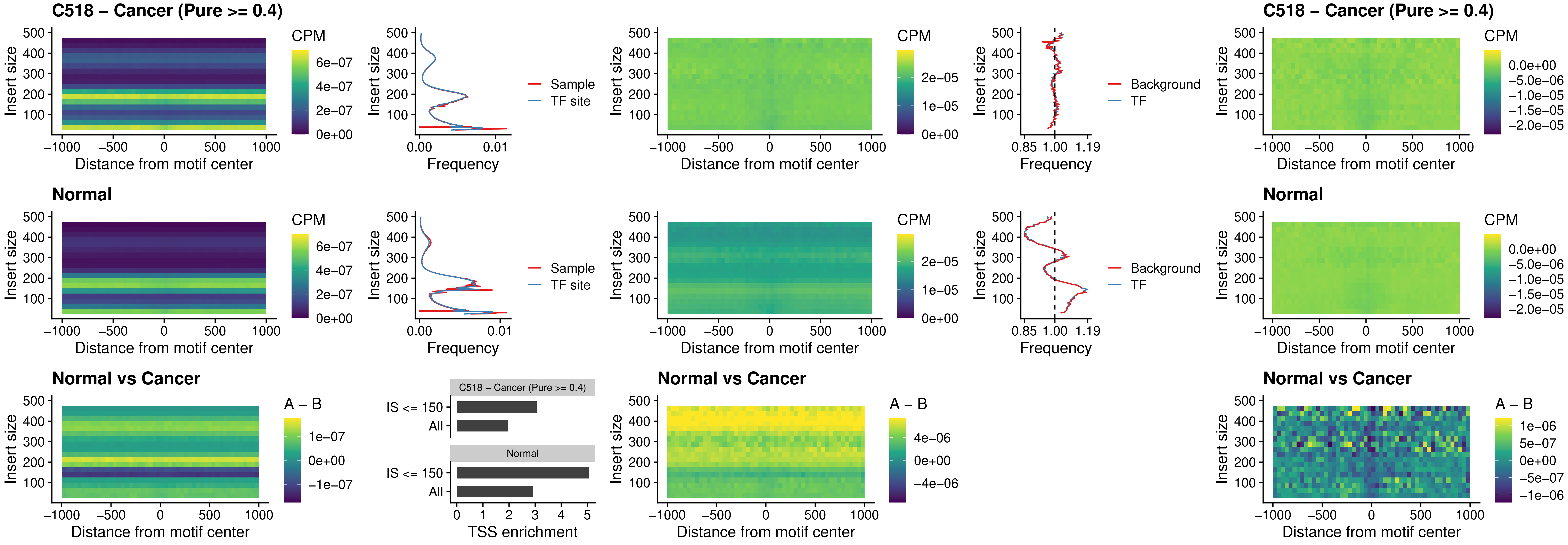

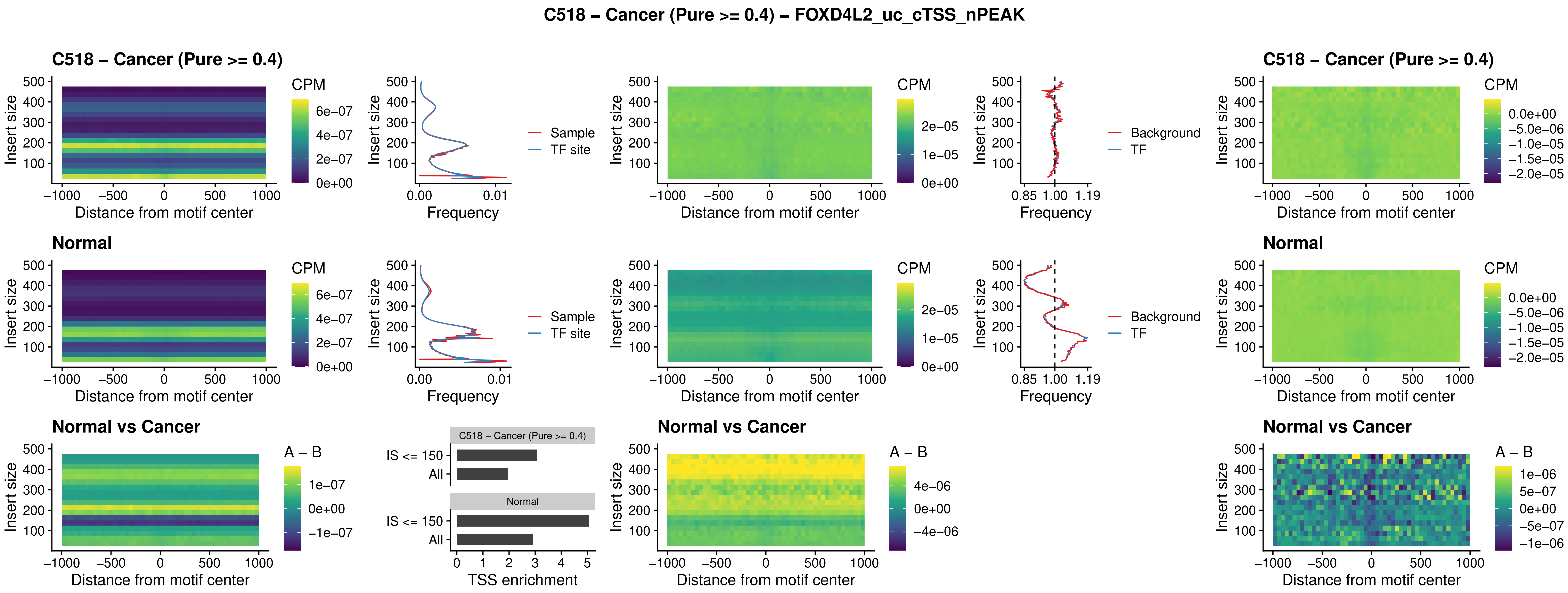
C518 – Cancer (Pure >= 0.4) – FOXD4L2_uc_cTSS_nPEAK
C518 – Cancer (Pure >= 0.4)
Normal
Normal vs Cancer
Insert size
Distance from motif center
Frequency
CPM
A – B
Sample
TF site
Background
TF
C518 – Cancer (Pure >= 0.4)
Normal
IS <= 150
All
TSS enrichment

#### C518 – Cancer (Pure >= 0.4) – FOXD4L3_uc_cTSS_nPEAK

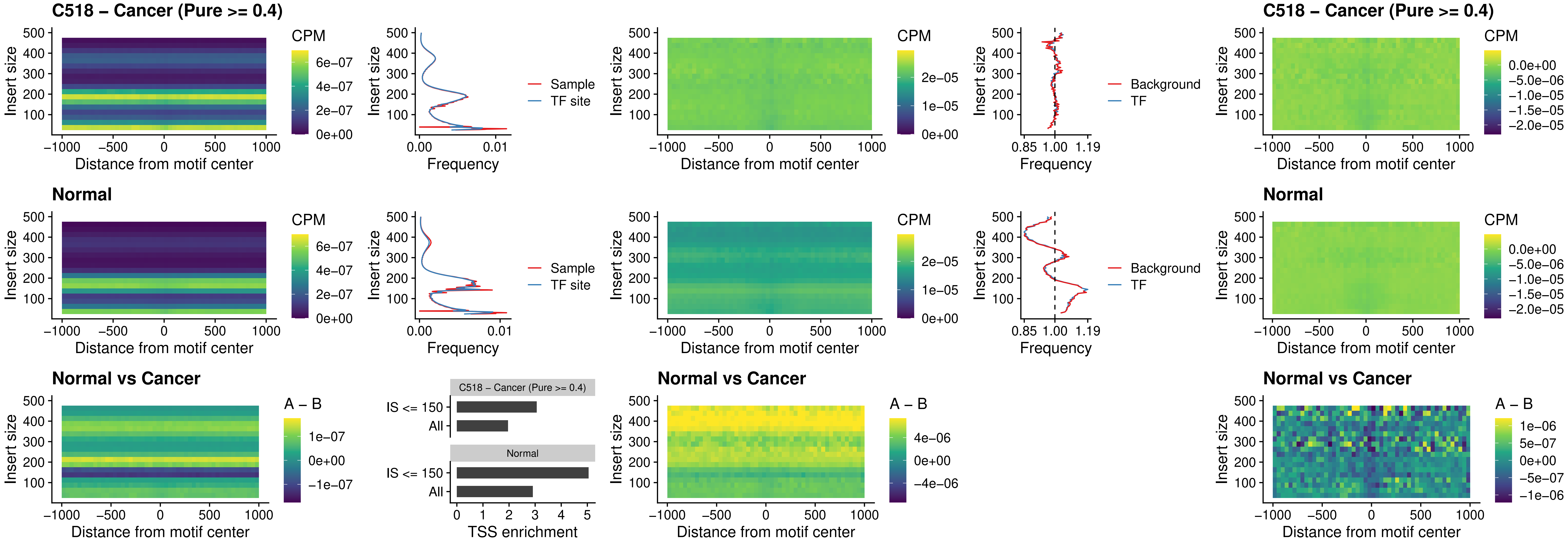

#### C518 – Cancer (Pure >= 0.4) – HOXD1_uc_dTSS_oPEAK

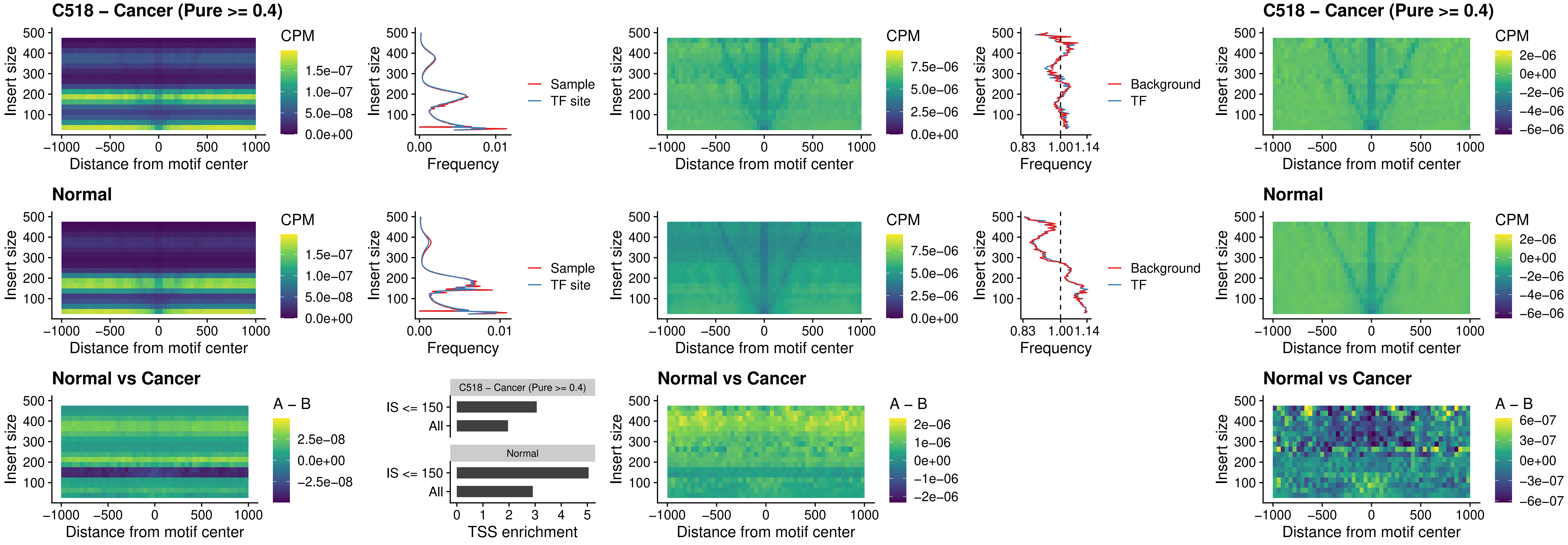

### C518 – Cancer (Pure >= 0.4) – VAX1_uc_pTSS_nPEAK

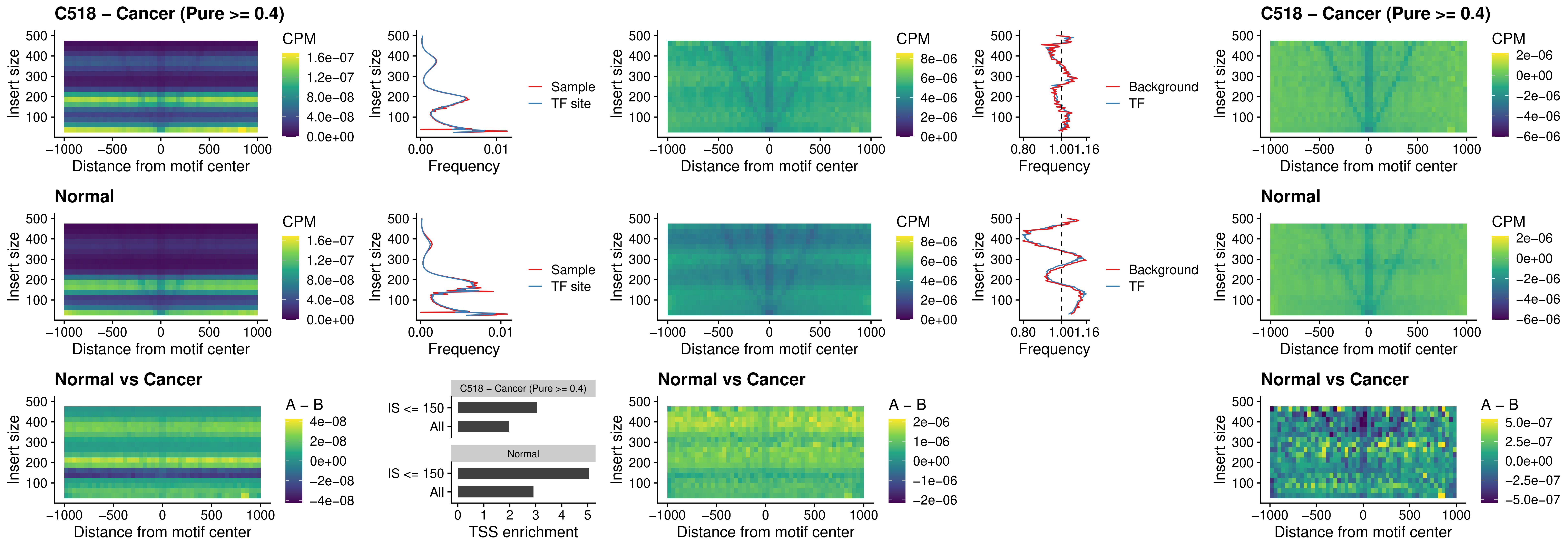

#### C518 – Cancer (Pure >= 0.4) – DLX3_uc_pTSS_nPEAK

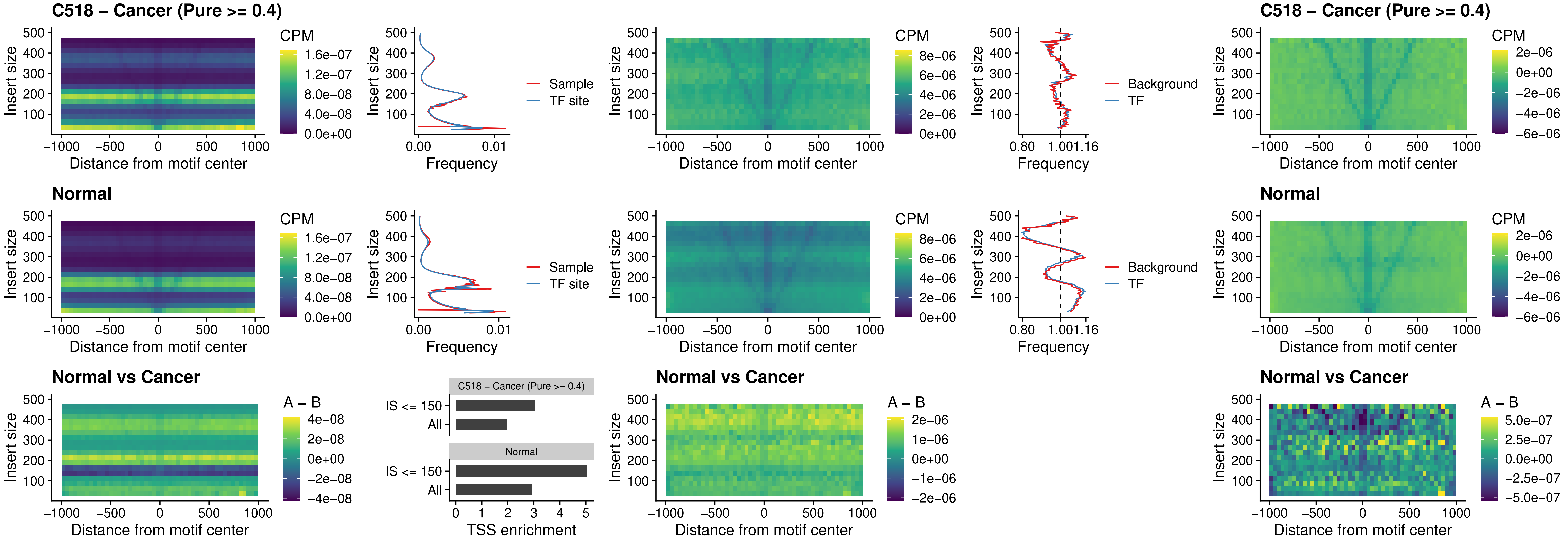

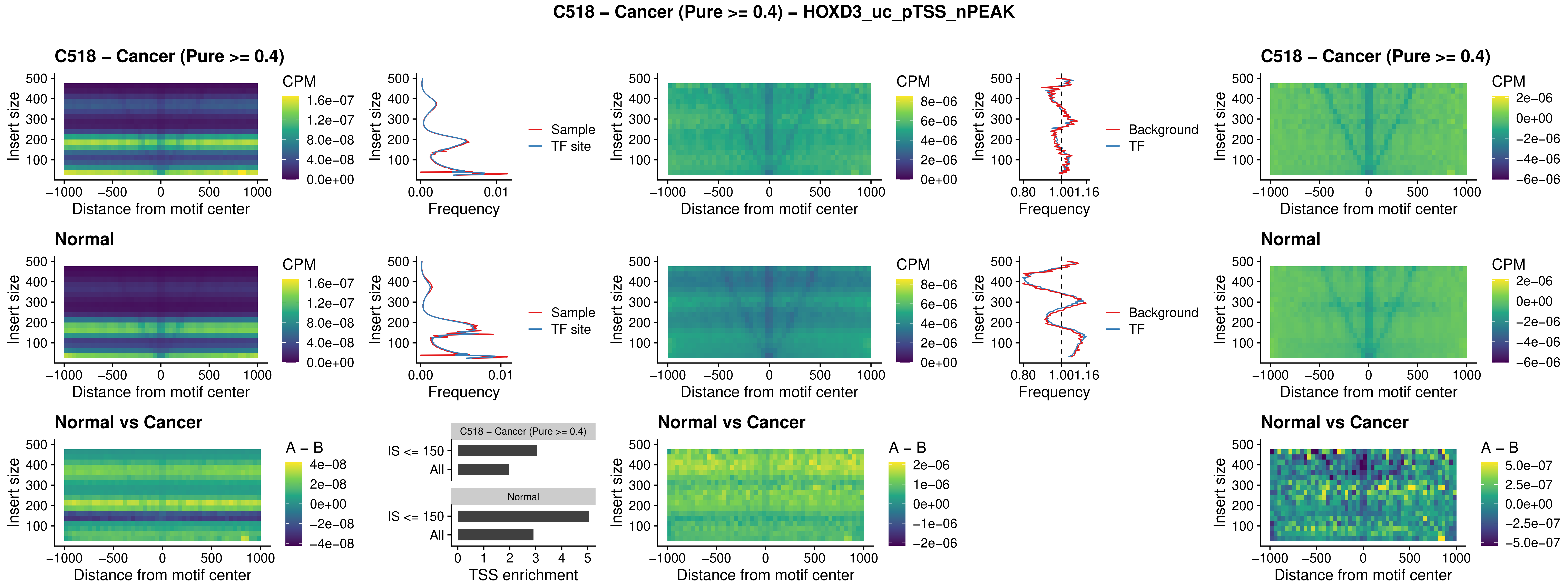
C518 – Cancer (Pure >= 0.4) – HOXD3_uc_pTSS_nPEAK
C518 – Cancer (Pure >= 0.4)
Normal
Normal vs Cancer
Insert size
Distance from motif center
Frequency
CPM
A – B
Sample
TF site
Background
TF
C518 – Cancer (Pure >= 0.4)
Normal
IS <= 150
All
TSS enrichment

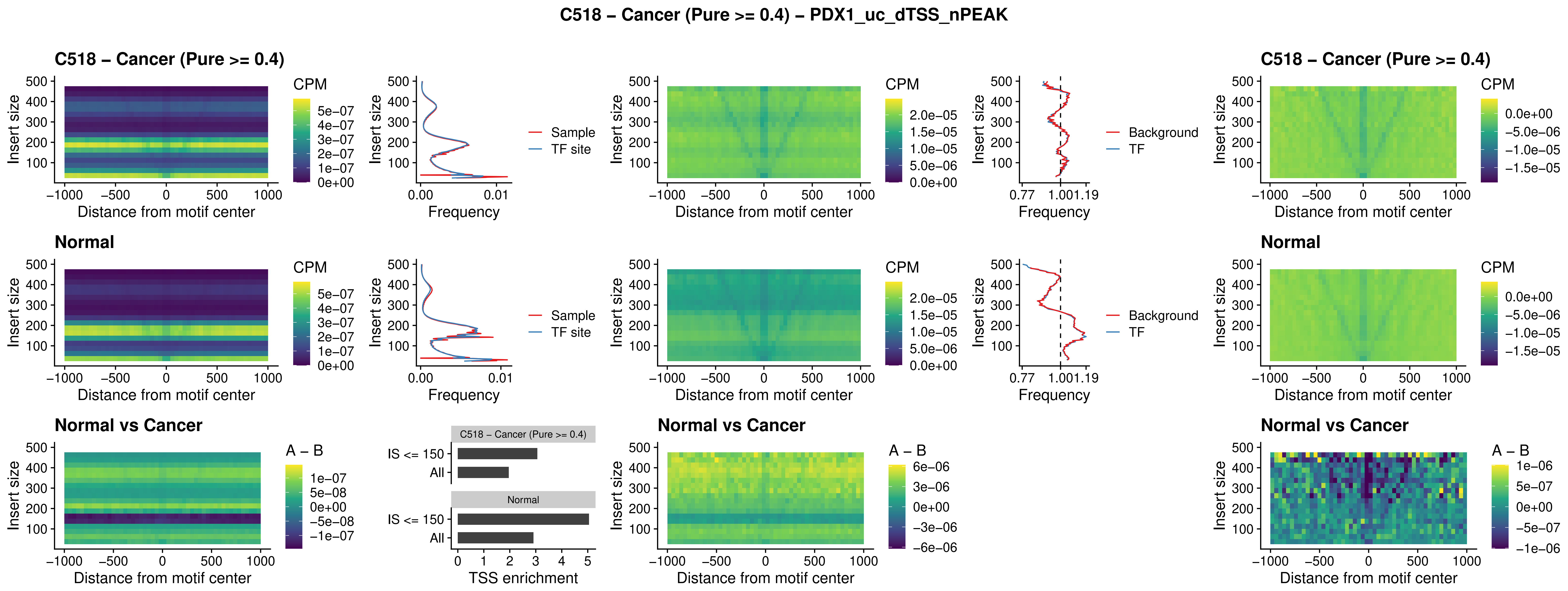
C518 – Cancer (Pure >= 0.4) – PDX1_uc_dTSS_nPEAK
C518 – Cancer (Pure >= 0.4)
Normal
Normal vs Cancer
Insert size
Distance from motif center
Frequency
CPM
A – B
Sample
TF site
Background
TF
C518 – Cancer (Pure >= 0.4)
Normal
IS <= 150
All
TSS enrichment

#### C518 – Cancer (Pure >= 0.4) – VAX2_uc_dTSS_nPEAK

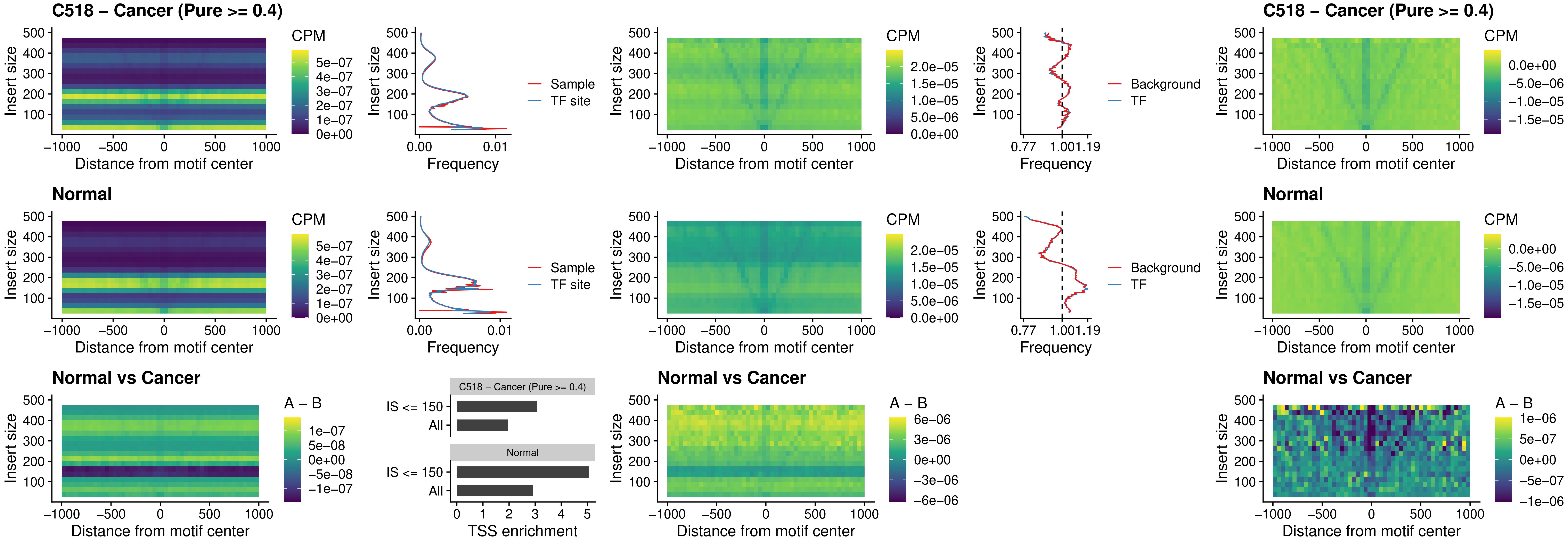

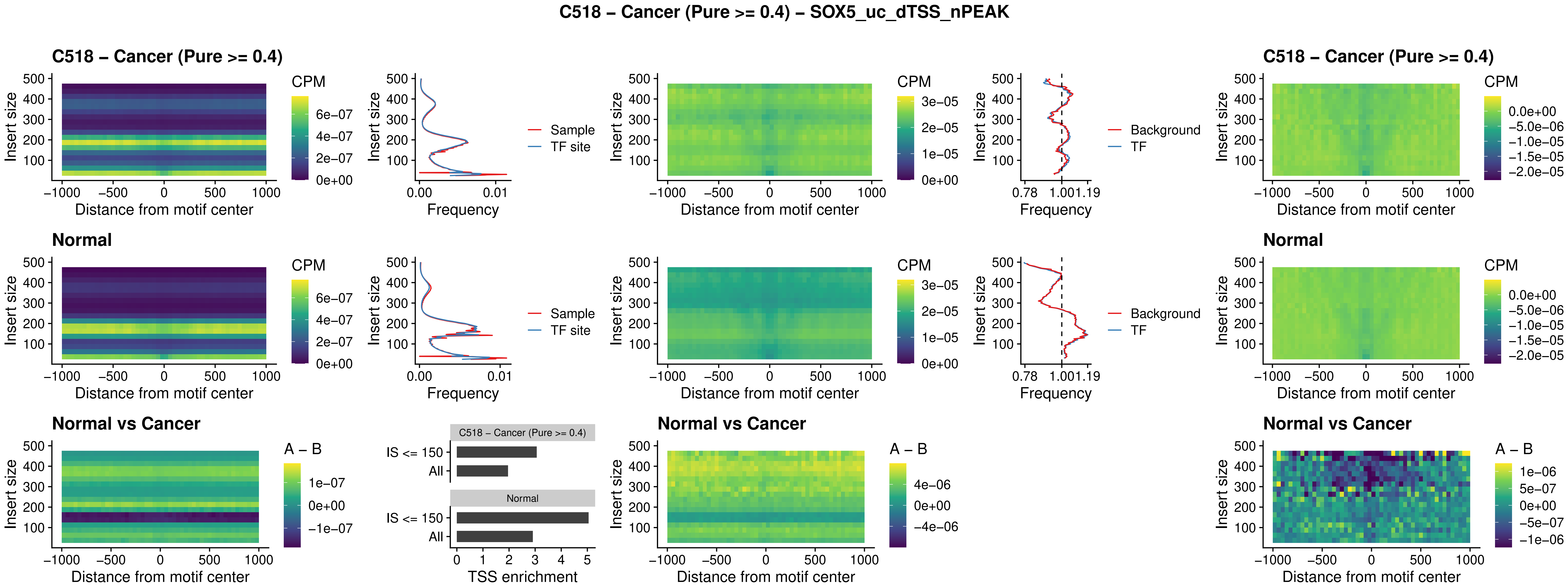
C518 – Cancer (Pure >= 0.4) – SOX5_uc_dTSS_nPEAK
C518 – Cancer (Pure >= 0.4)
Normal
Normal vs Cancer
Insert size
Distance from motif center
Frequency
CPM
A – B
Sample
TF site
Background
TF
C518 – Cancer (Pure >= 0.4)
Normal
IS <= 150
All
TSS enrichment

#### C518 – Cancer (Pure >= 0.4) – SOX6_uc_dTSS_nPEAK

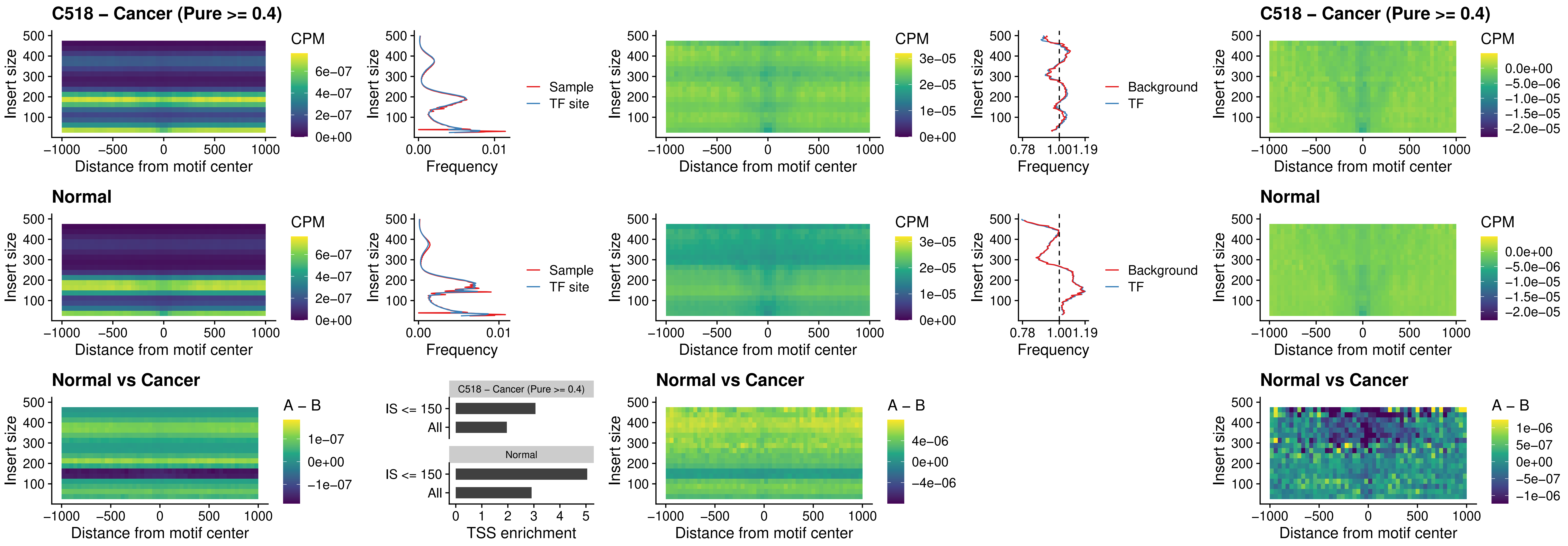

#### C518 – Cancer (Pure >= 0.4) – JDP2_uc_cTSS_nPEAK

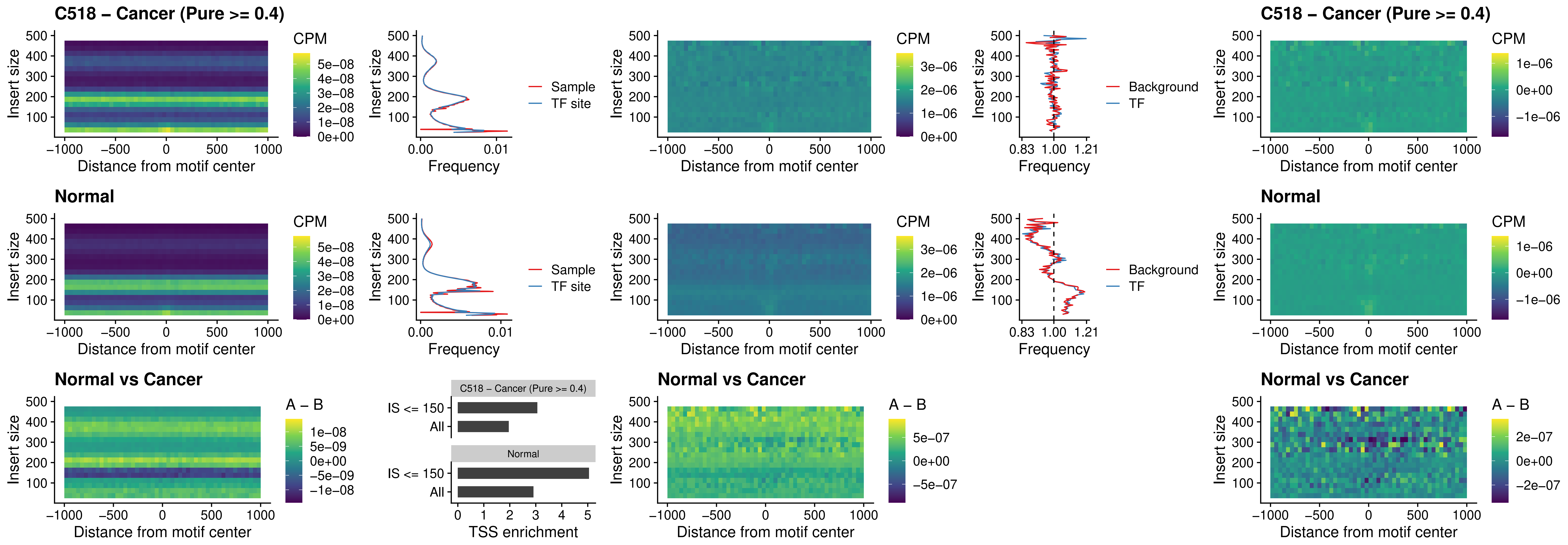

#### C518 – Cancer (Pure >= 0.4) – PURA_uc_dTSS_oPEAK

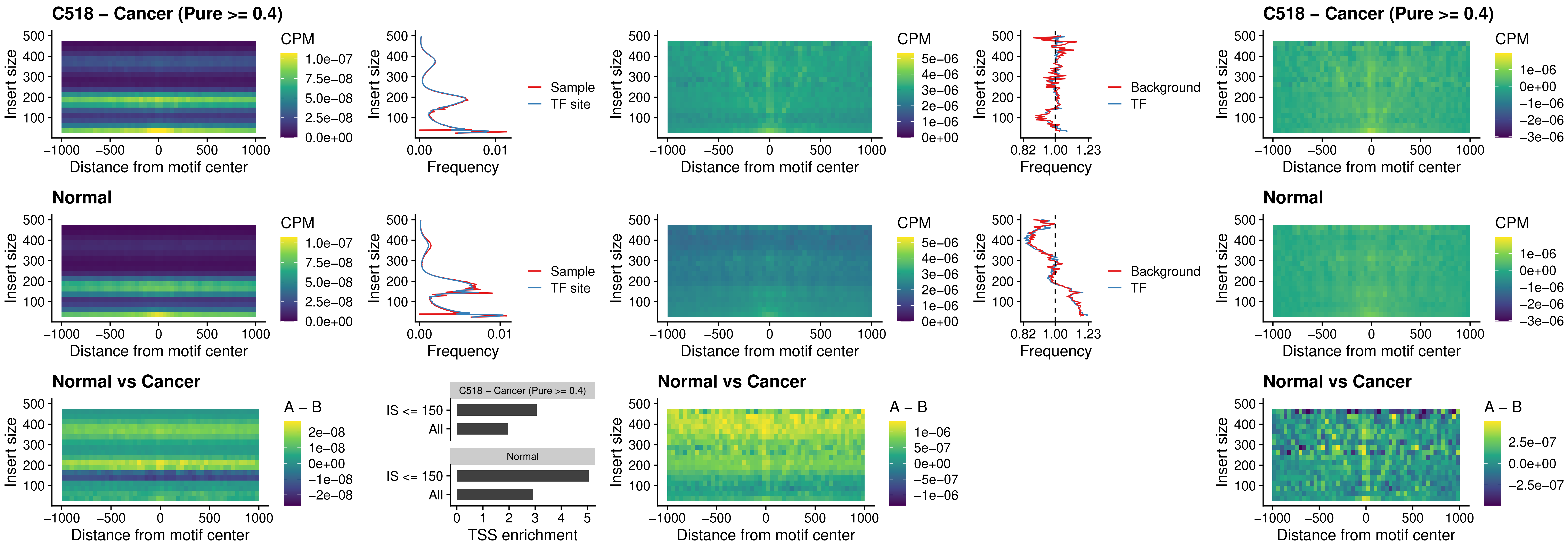

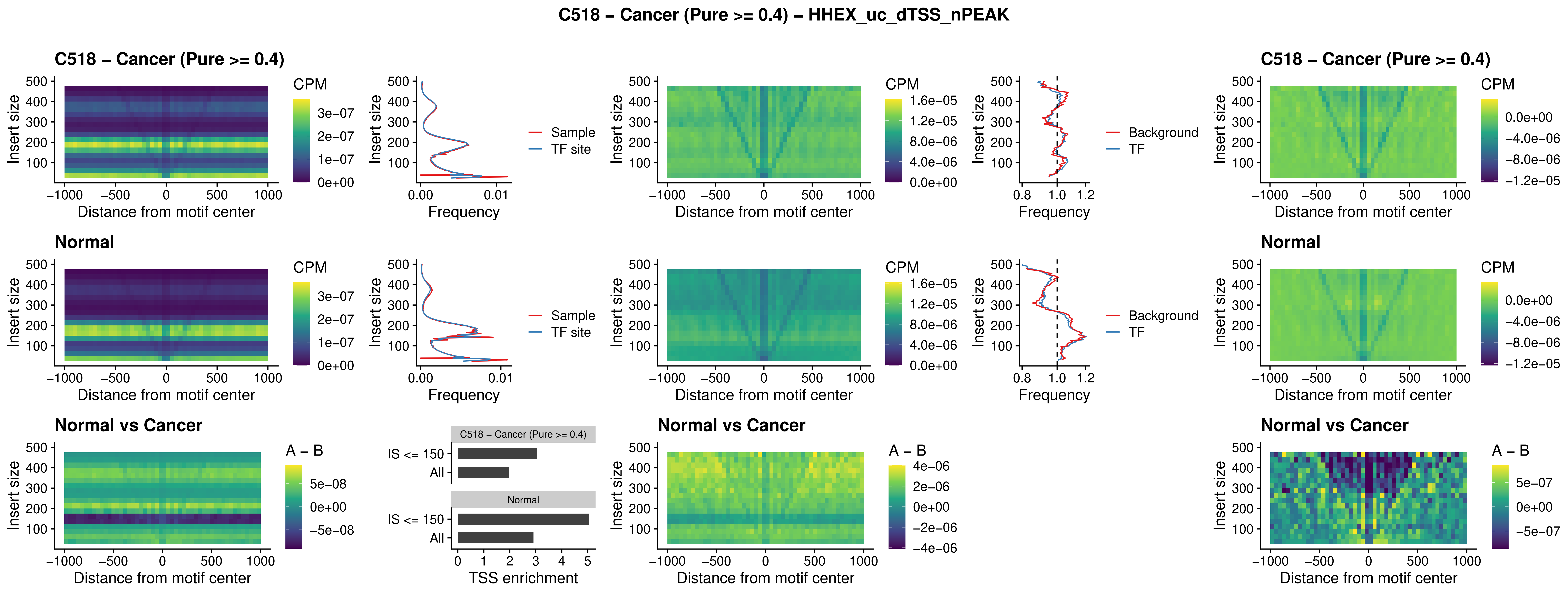
C518 – Cancer (Pure >= 0.4) – HHEX_uc_dTSS_nPEAK
C518 – Cancer (Pure >= 0.4)
Normal
Normal vs Cancer
Insert size
Distance from motif center
Frequency
CPM
A – B
Sample
TF site
Background
TF
C518 – Cancer (Pure >= 0.4)
Normal
IS <= 150
All
TSS enrichment

#### C518 – Cancer (Pure >= 0.4) – HMG20B_uc_dTSS_nPEAK

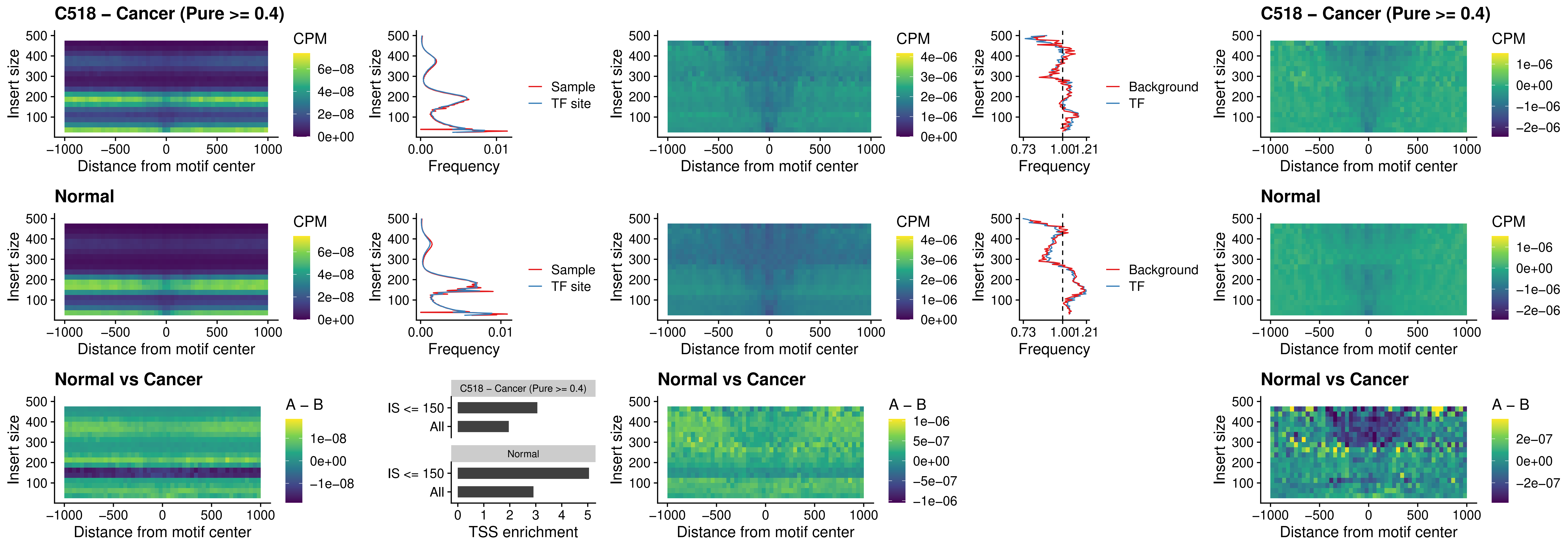

#### C518 – Cancer (Pure >= 0.4) – FOXB2_uc_dTSS_nPEAK

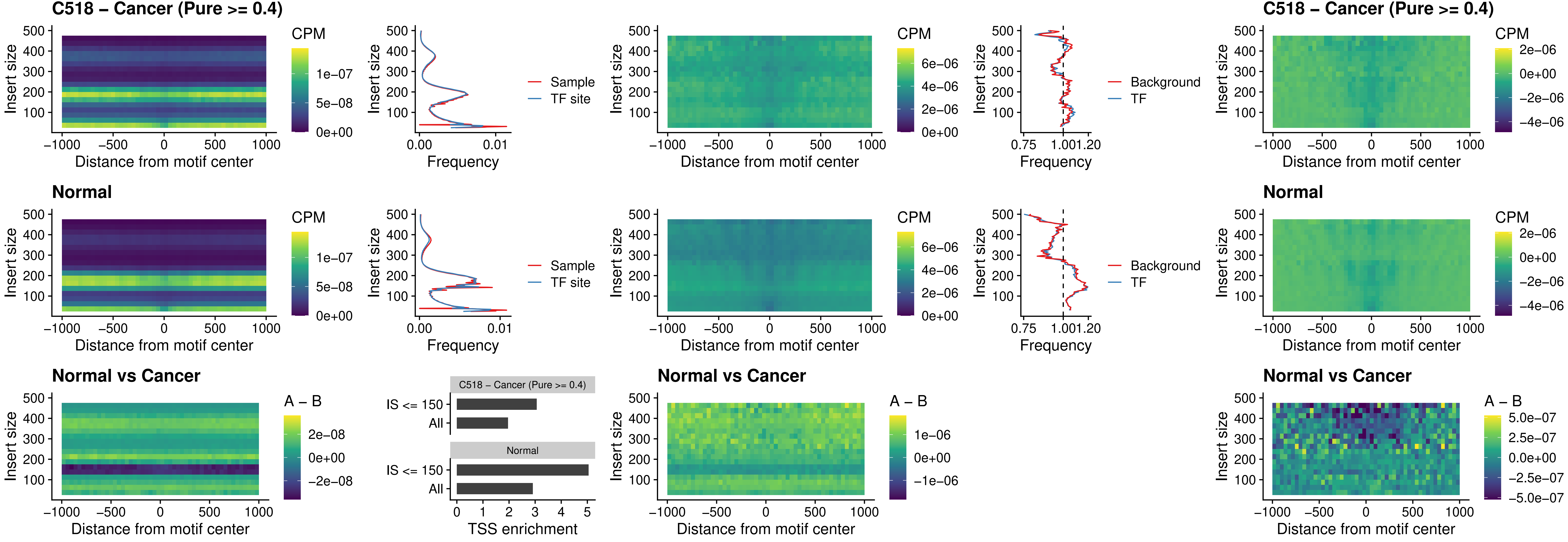

#### C518 – Cancer (Pure >= 0.4) – FOXL1_uc_cTSS_nPEAK

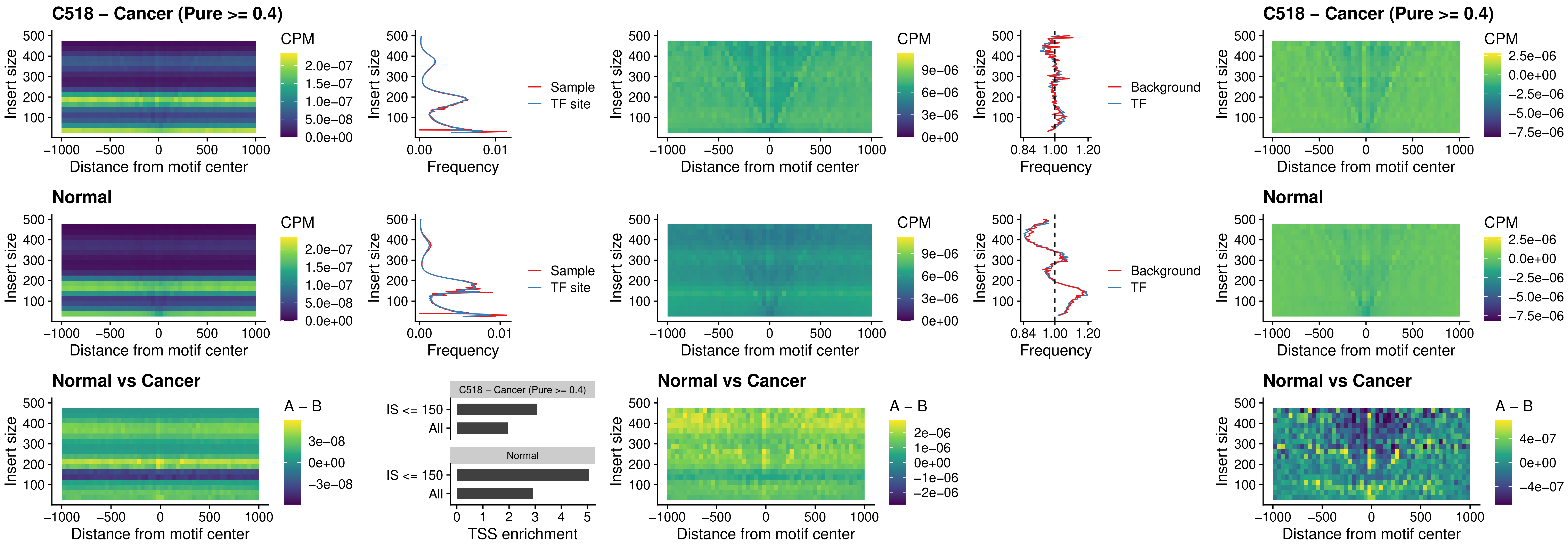

#### C518 – Cancer (Pure >= 0.4) – PBX3_uc_pTSS_oPEAK

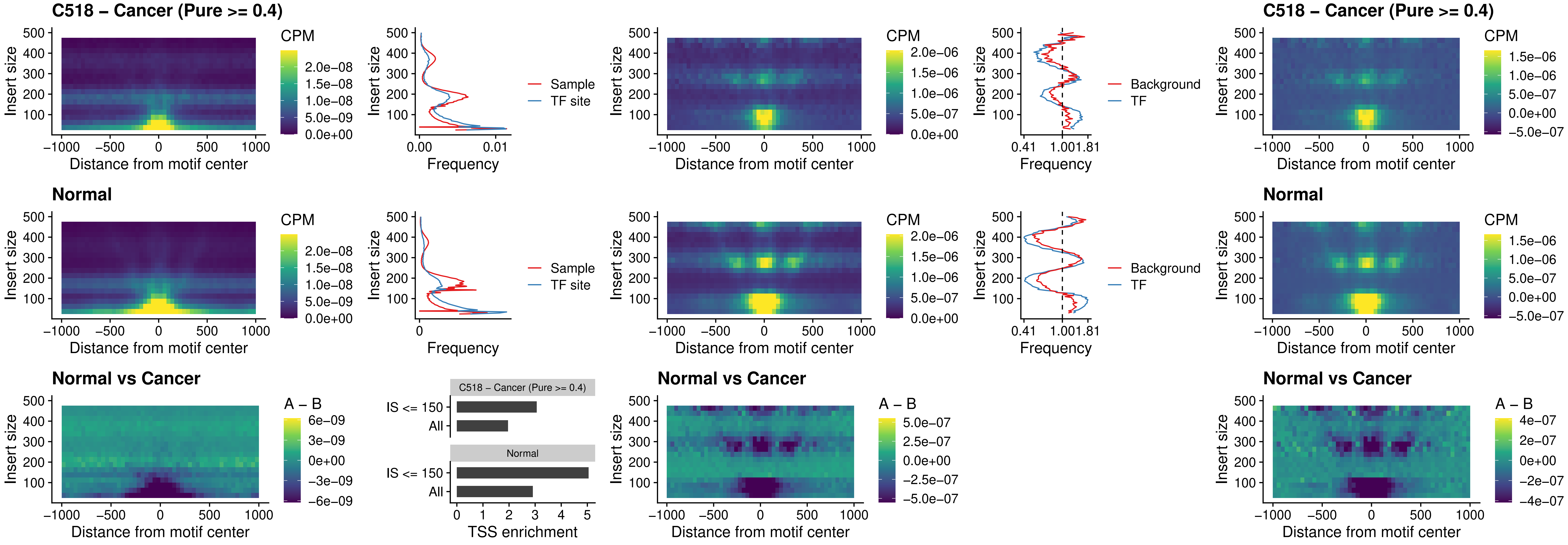

#### C518 – Cancer (Pure >= 0.4) – SNAI2_uc_cTSS_nPEAK

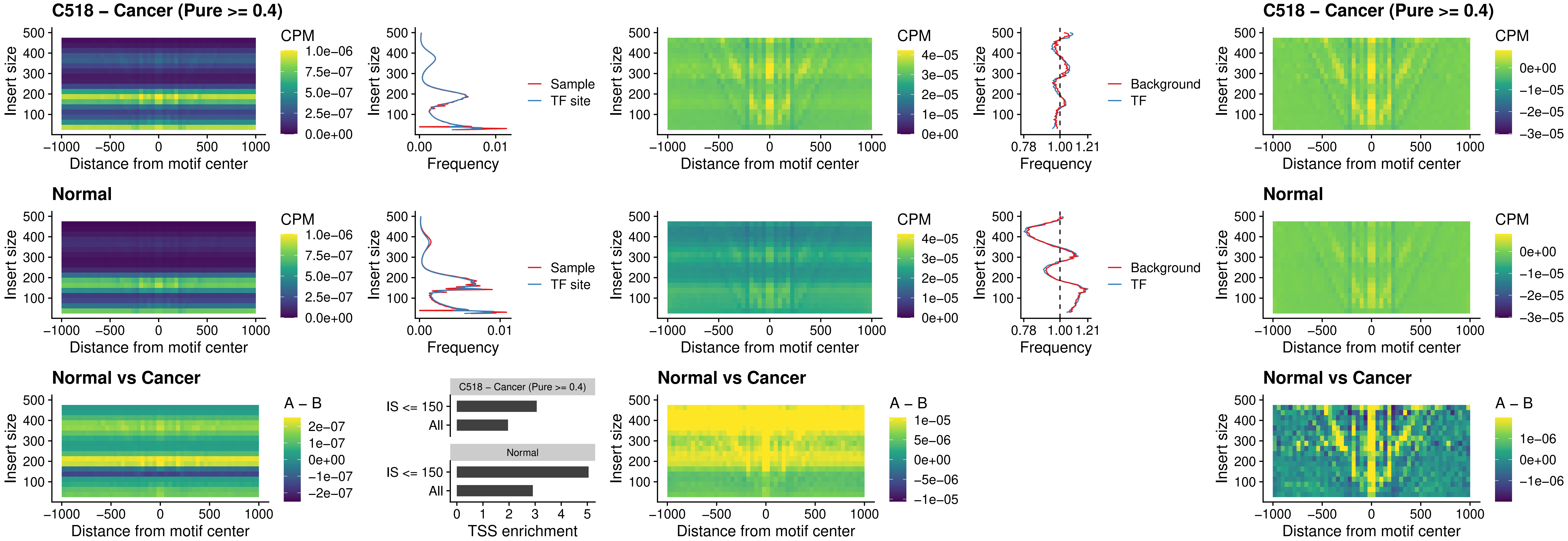

### C518 – Cancer (Pure >= 0.4) – SNAI2_uc_dTSS_oPEAK

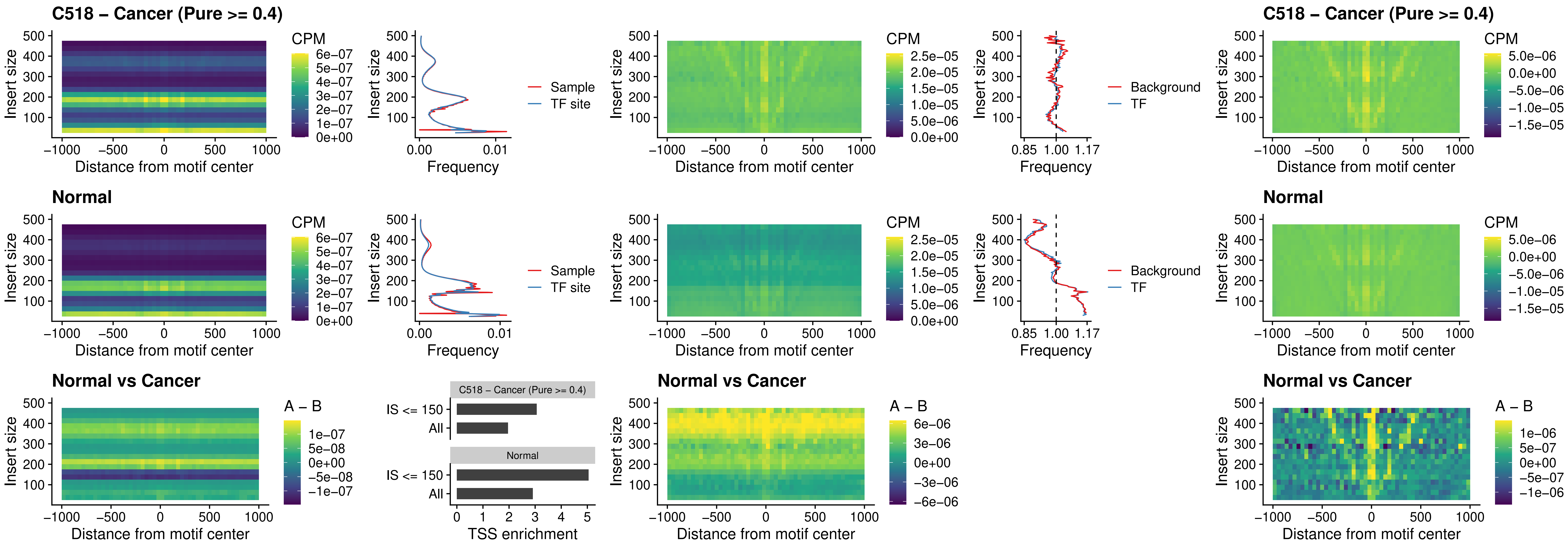

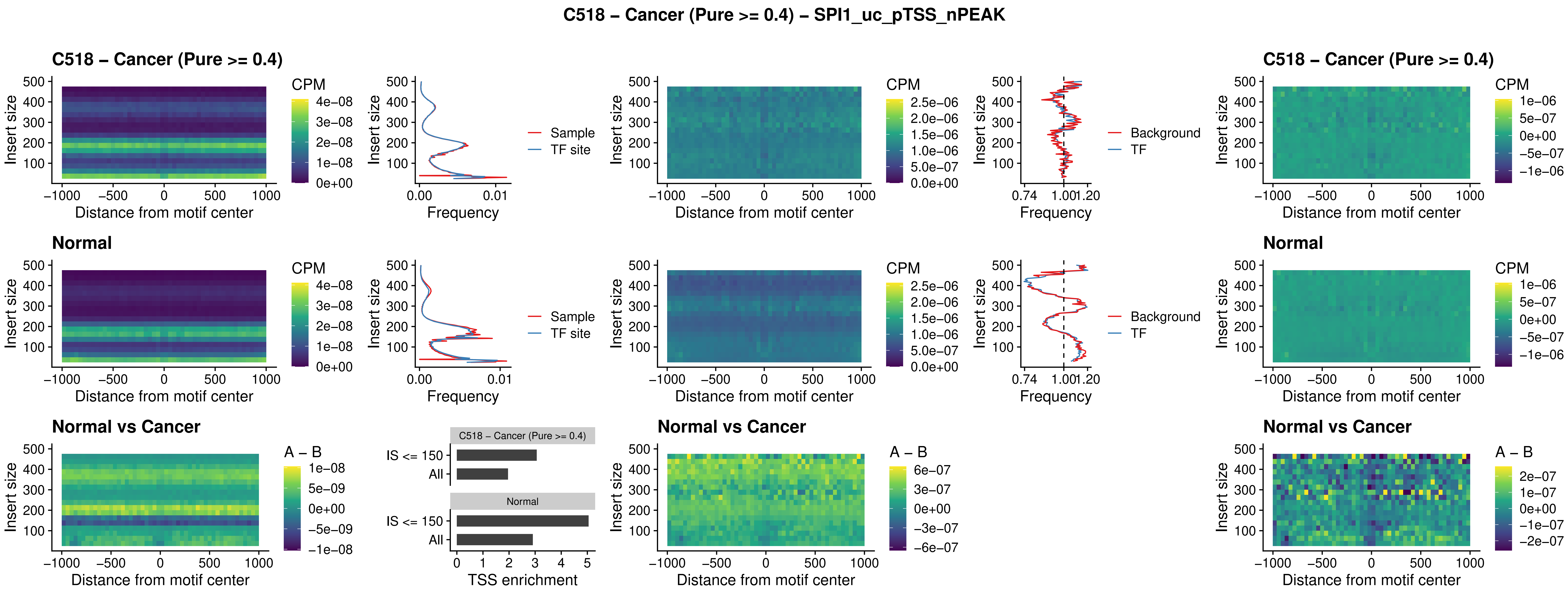
C518 – Cancer (Pure >= 0.4) – SPI1_uc_pTSS_nPEAK
C518 – Cancer (Pure >= 0.4)
Normal
Normal vs Cancer
Insert size
Distance from motif center
Frequency
CPM
A – B
Sample
TF site
Background
TF
C518 – Cancer (Pure >= 0.4)
Normal
IS <= 150
All
TSS enrichment

#### C518 – Cancer (Pure >= 0.4) – KLF16_uc_pTSS_nPEAK

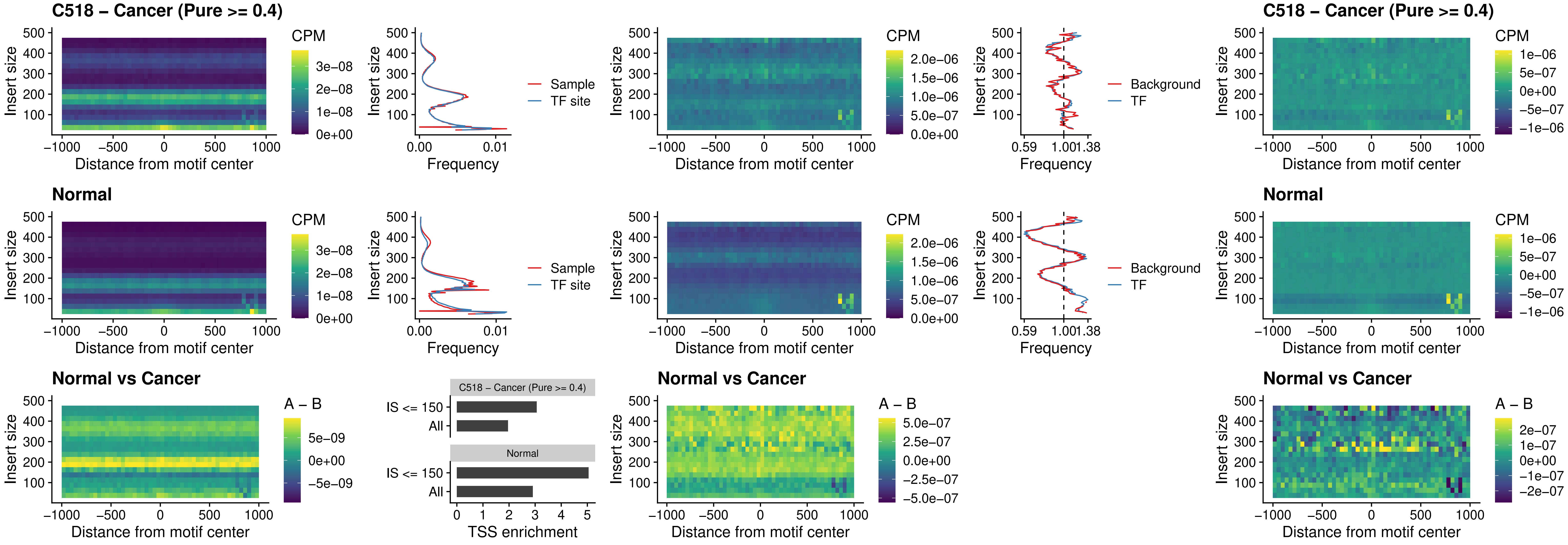

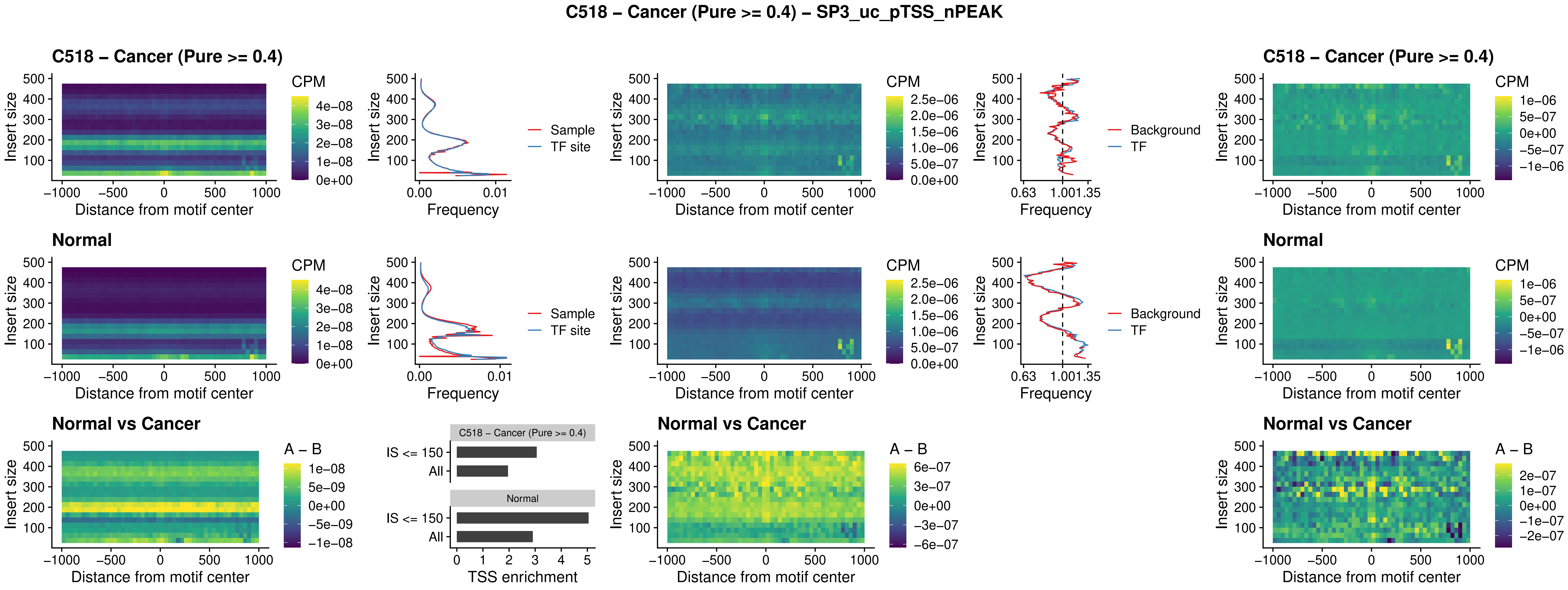
C518 – Cancer (Pure >= 0.4) – SP3_uc_pTSS_nPEAK
C518 – Cancer (Pure >= 0.4)
Normal
Normal vs Cancer
Insert size
Distance from motif center
CPM
A – B
Frequency
Sample
TF site
Background
TF
C518 – Cancer (Pure >= 0.4)
Normal
IS <= 150
All
TSS enrichment

#### C518 – Cancer (Pure >= 0.4) – NEUROD1_uc_dTSS_oPEAK

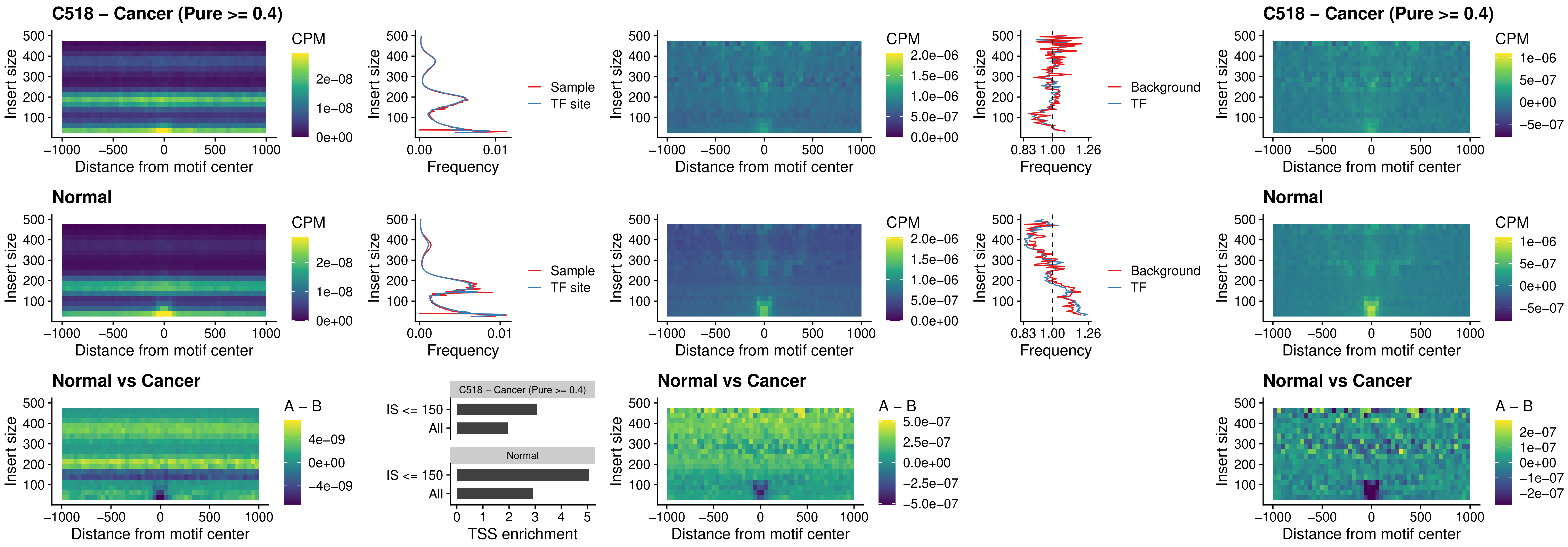

#### C518 – Cancer (Pure >= 0.4) – CTCF_uc_dTSS_oPEAK

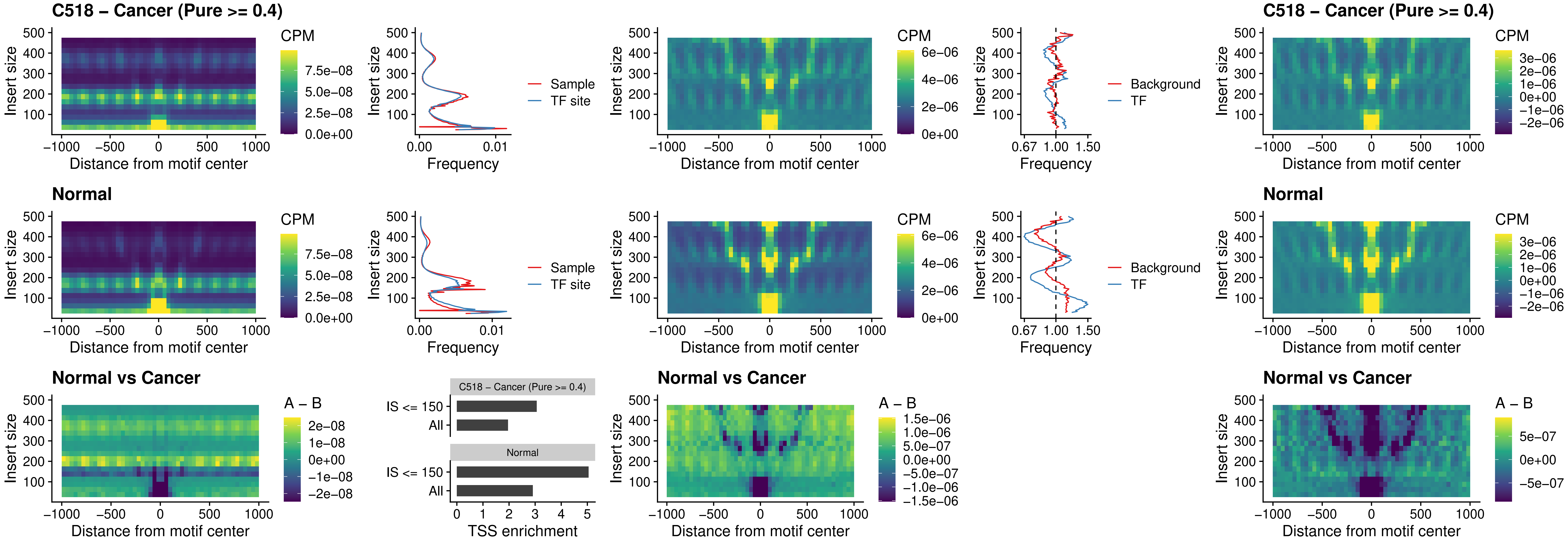

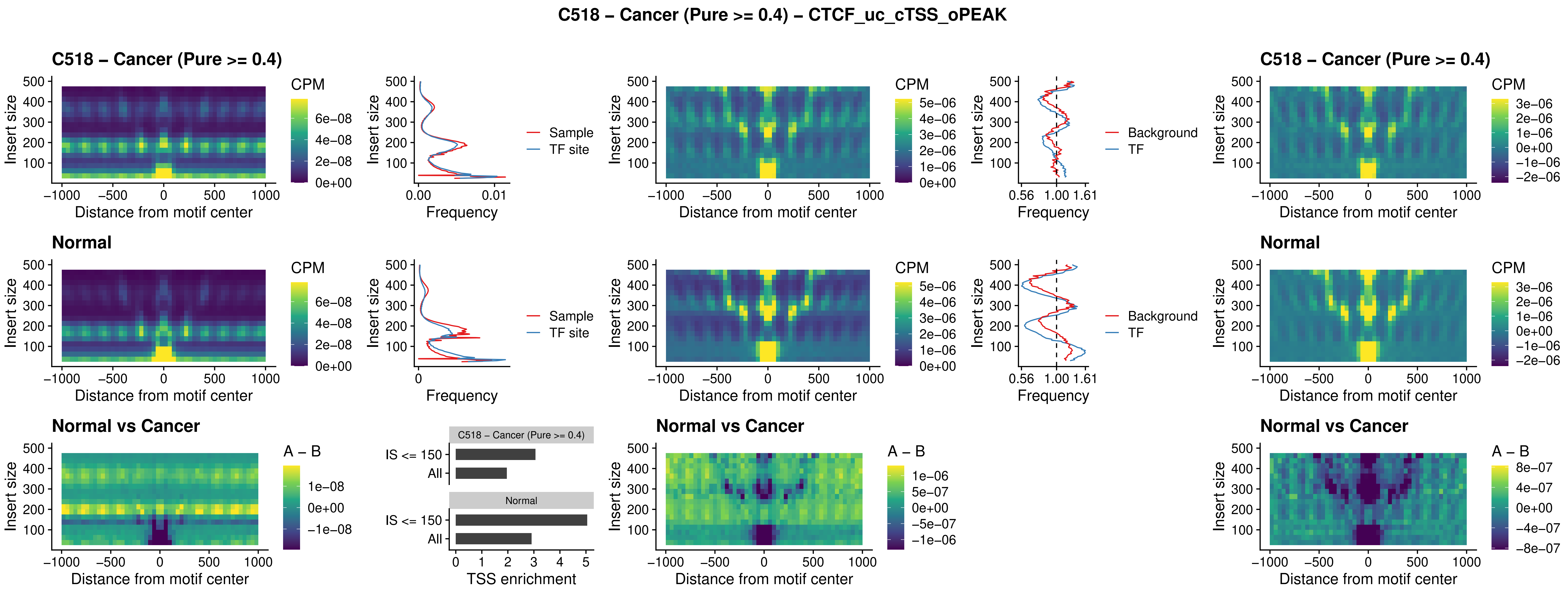
C518 – Cancer (Pure >= 0.4) – CTCF_uc_cTSS_oPEAK
C518 – Cancer (Pure >= 0.4)
Normal
Normal vs Cancer
Insert size
Distance from motif center
CPM
A – B
Frequency
Sample
TF site
Background
TF
C518 – Cancer (Pure >= 0.4)
Normal
IS <= 150
All
TSS enrichment

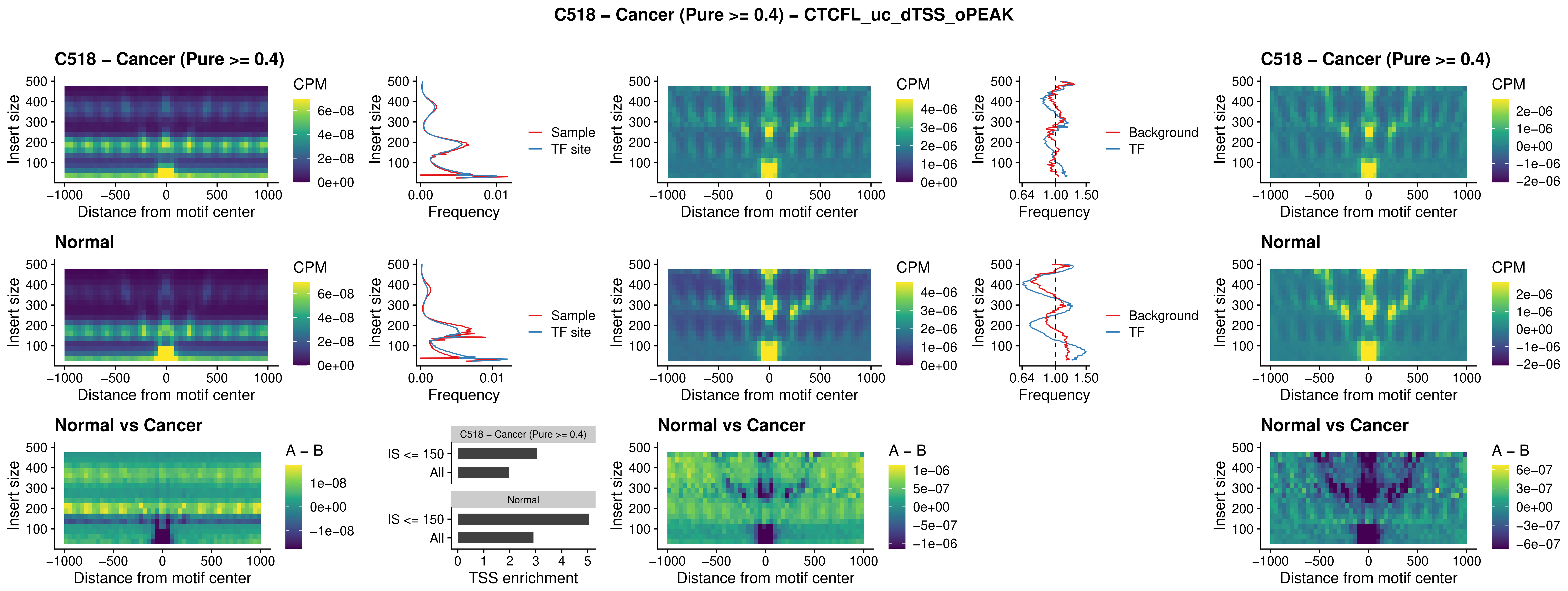
C518 – Cancer (Pure >= 0.4) – CTCFL_uc_dTSS_oPEAK
C518 – Cancer (Pure >= 0.4)
Normal
Normal vs Cancer
Insert size
Distance from motif center
Frequency
CPM
A – B
Sample
TF site
Background
TF
C518 – Cancer (Pure >= 0.4)
Normal
IS <= 150
All
TSS enrichment

#### C518 – Cancer (Pure >= 0.4) – ENCODE_CTCF_sites_COLON

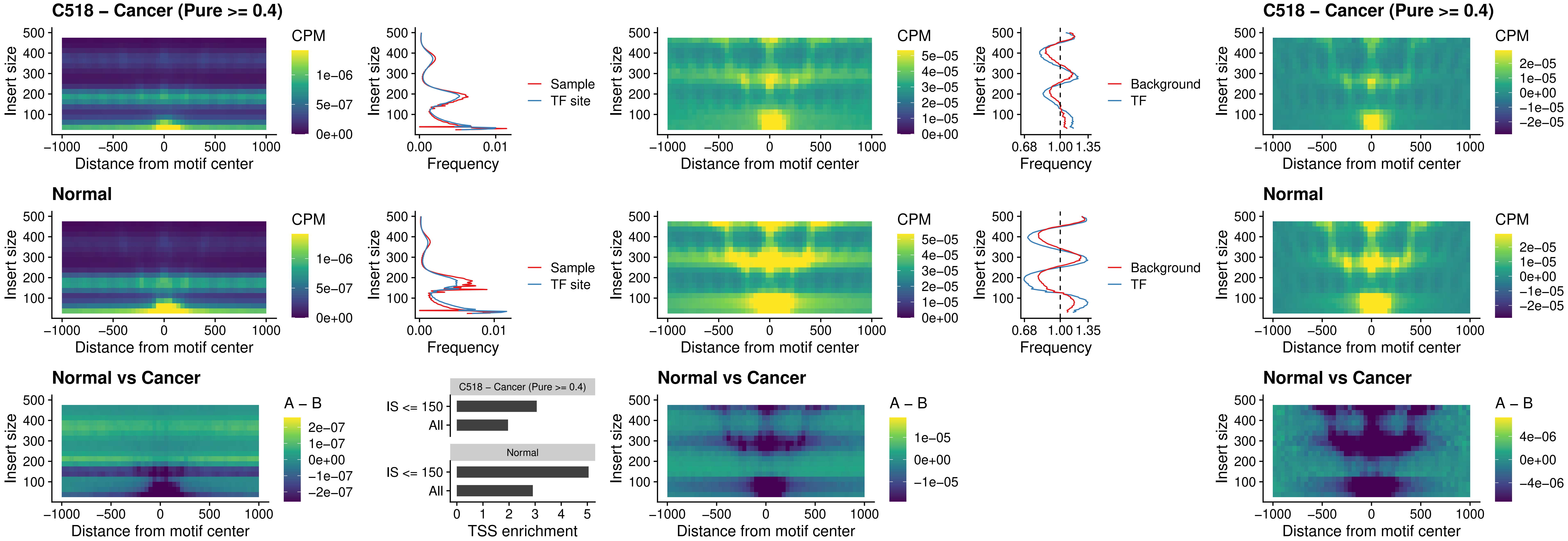

#### C518 – Cancer (Pure >= 0.4) – ENCODE_CTCF_sites_COLON_loop_ctcf

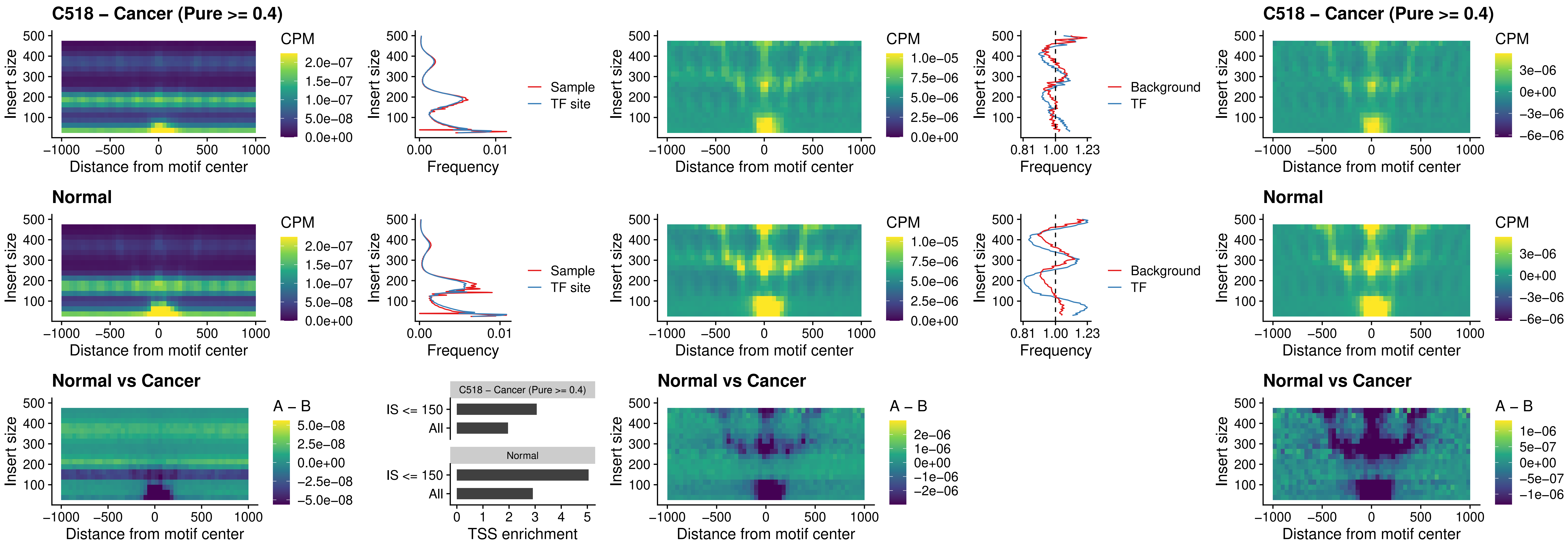

#### C518 – Cancer (Pure >= 0.4) – ENCODE_CTCF_sites_COLON_intergenic_ctcf

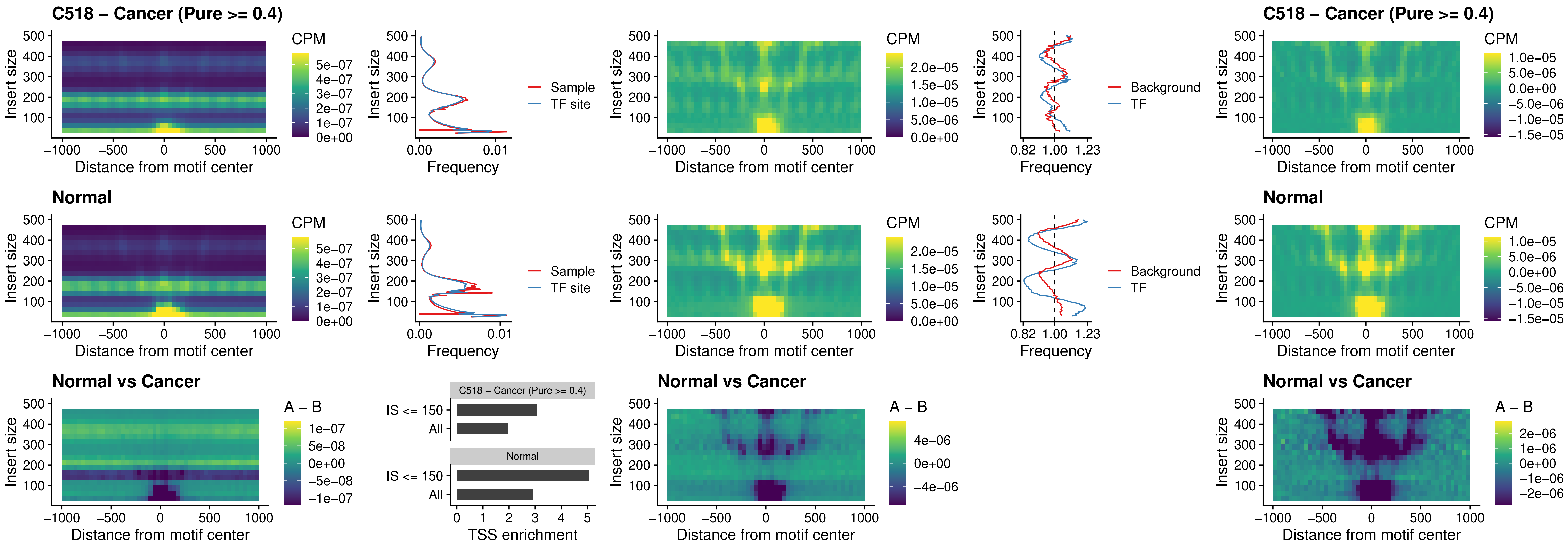

### C518 – Cancer (Pure >= 0.4) – ENCODE_CTCF_sites_COLON_nonpromotor

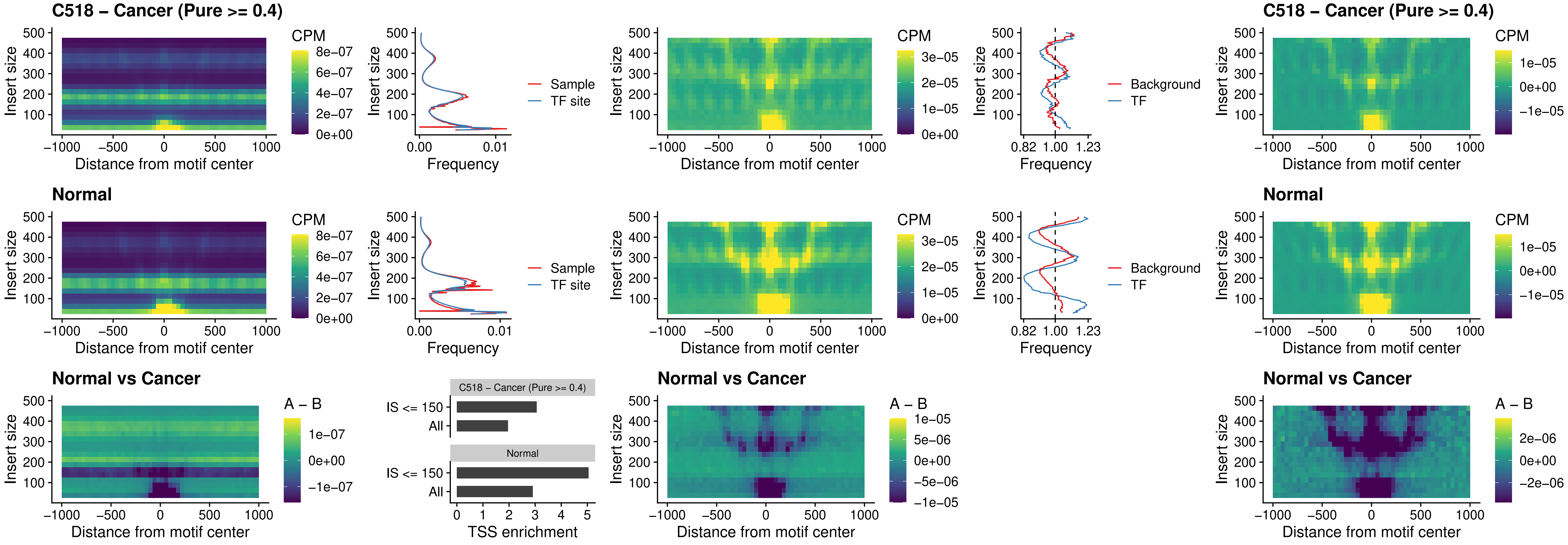

#### C518 – Cancer (Pure >= 0.4) – CTCFL_uc_cTSS_oPEAK

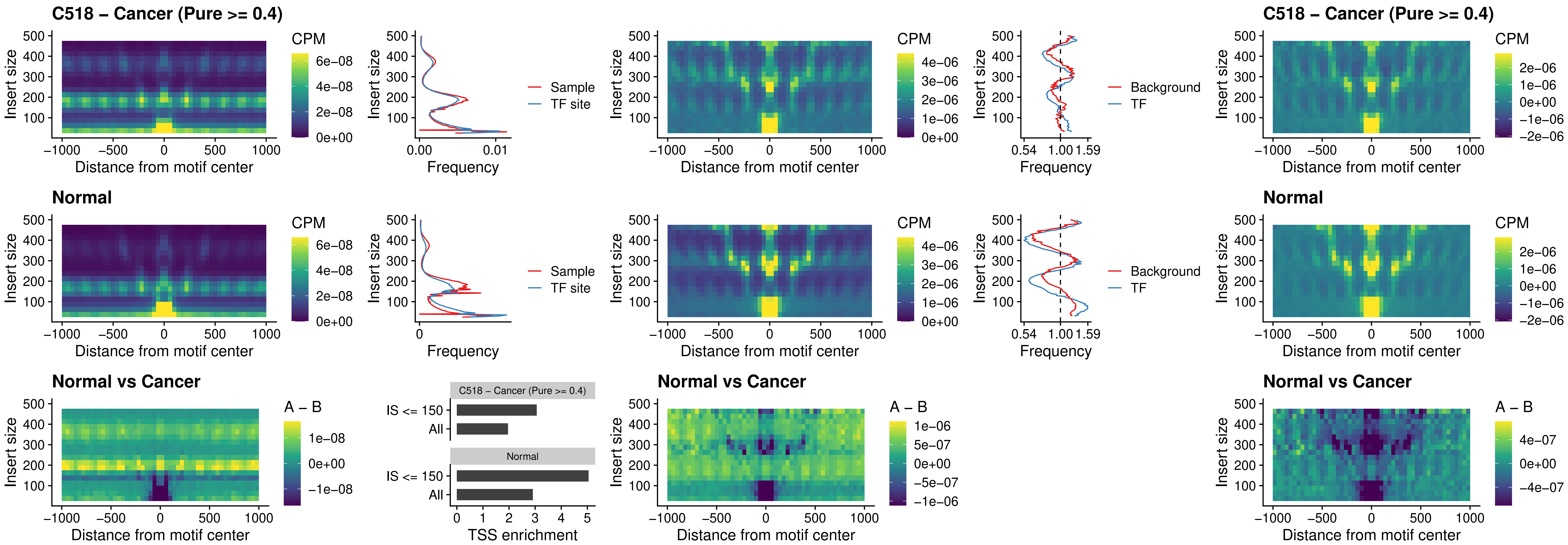

### C518 – Cancer (Pure >= 0.4) – NEUROD1_uc_cTSS_oPEAK

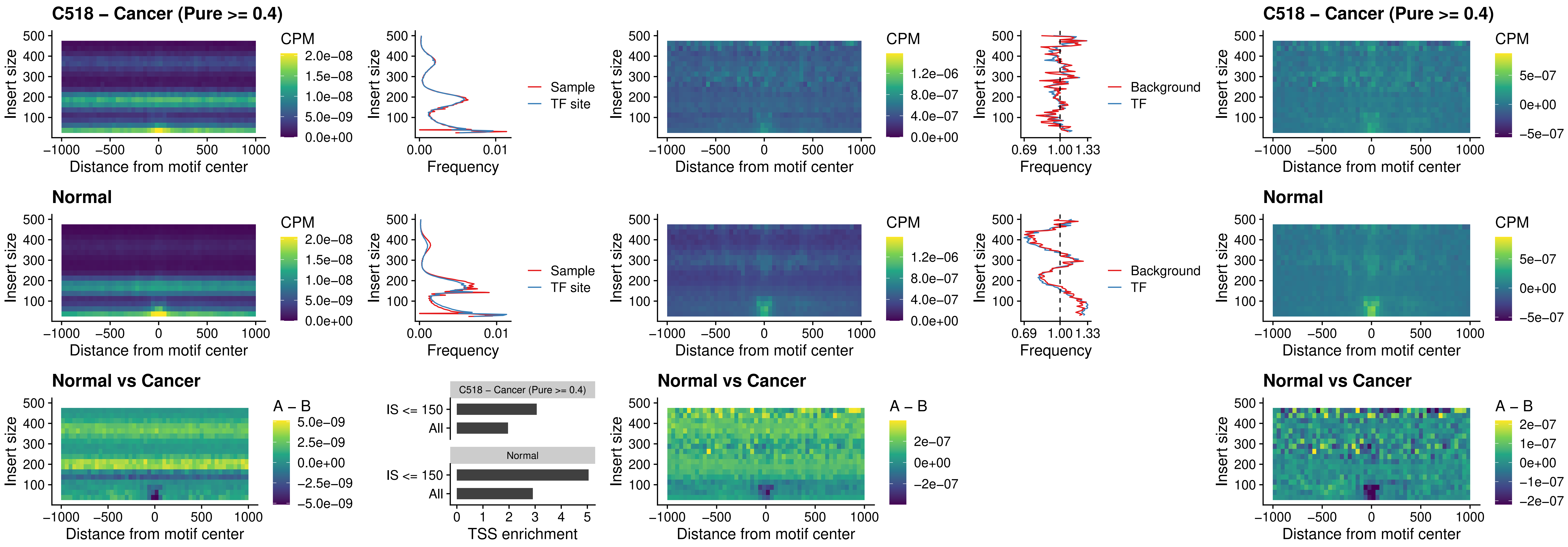

#### C518 – Cancer (Pure >= 0.4) – ENCODE_YY1_sites_COLON_nonpromotor

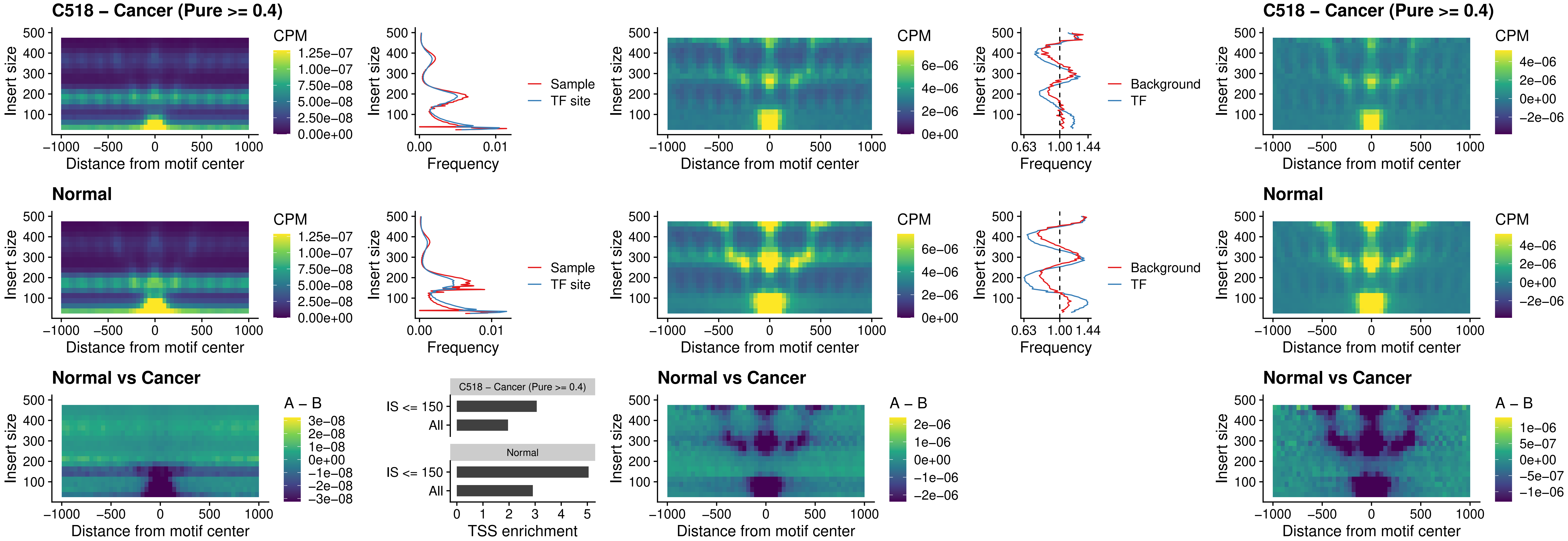

#### C518 – Cancer (Pure >= 0.4) – CTCF_uc_pTSS_oPEAK

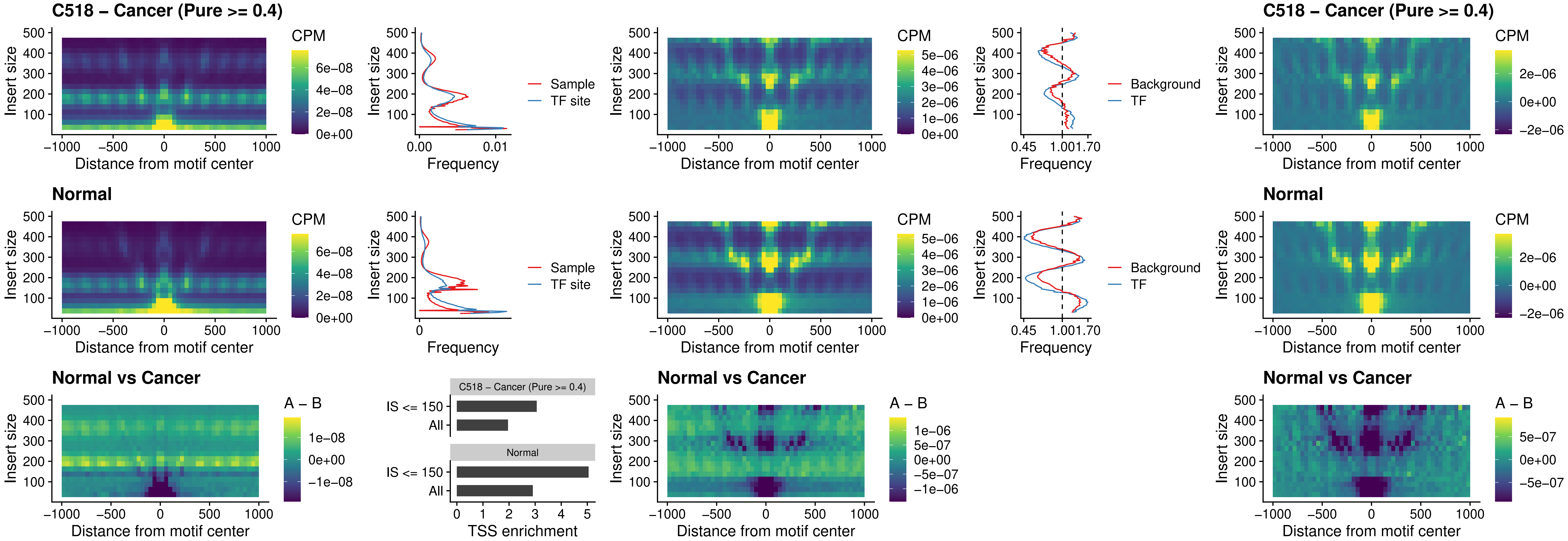

#### C518 – Cancer (Pure >= 0.4) – ENCODE_CTCF_sites_COLON_promotor

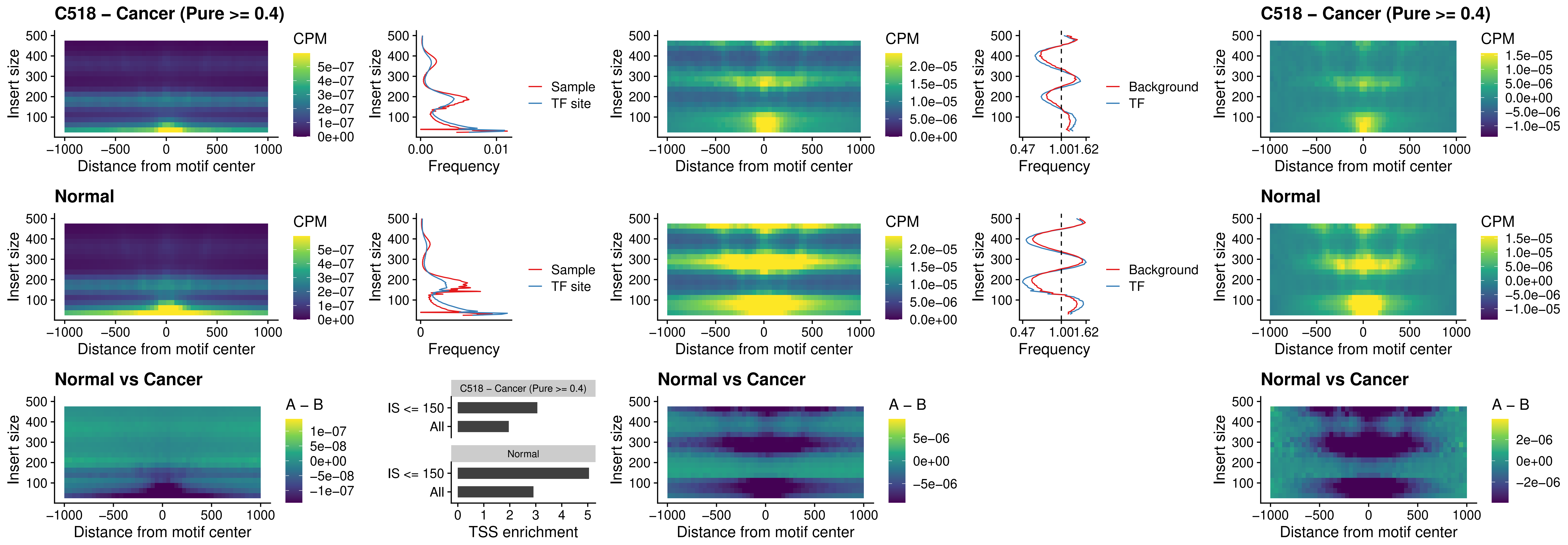

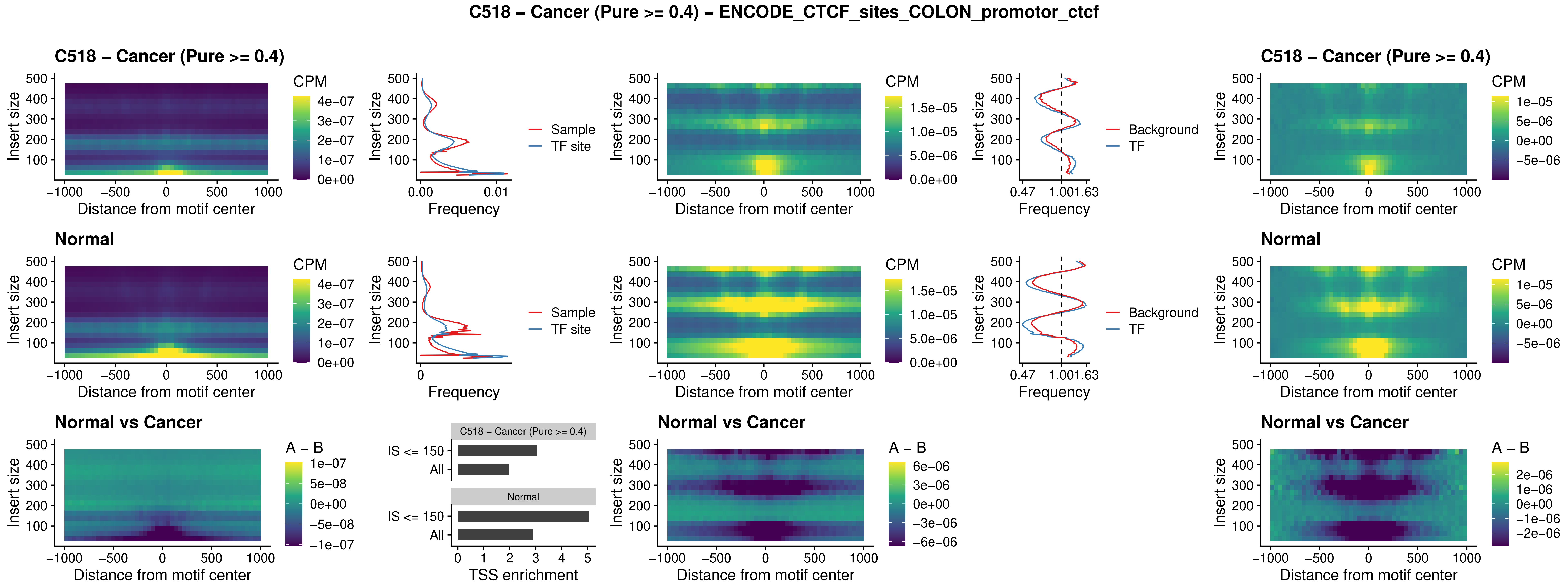
C518 – Cancer (Pure >= 0.4) – ENCODE_CTCF_sites_COLON_promotor_ctcf
C518 – Cancer (Pure >= 0.4)
Normal
Normal vs Cancer
Insert size
Distance from motif center
CPM
A – B
Frequency
Sample
TF site
Background
TF
C518 – Cancer (Pure >= 0.4)
Normal
IS <= 150
All
TSS enrichment

#### C518 – Cancer (Pure >= 0.4) – SPIC_uc_pTSS_oPEAK

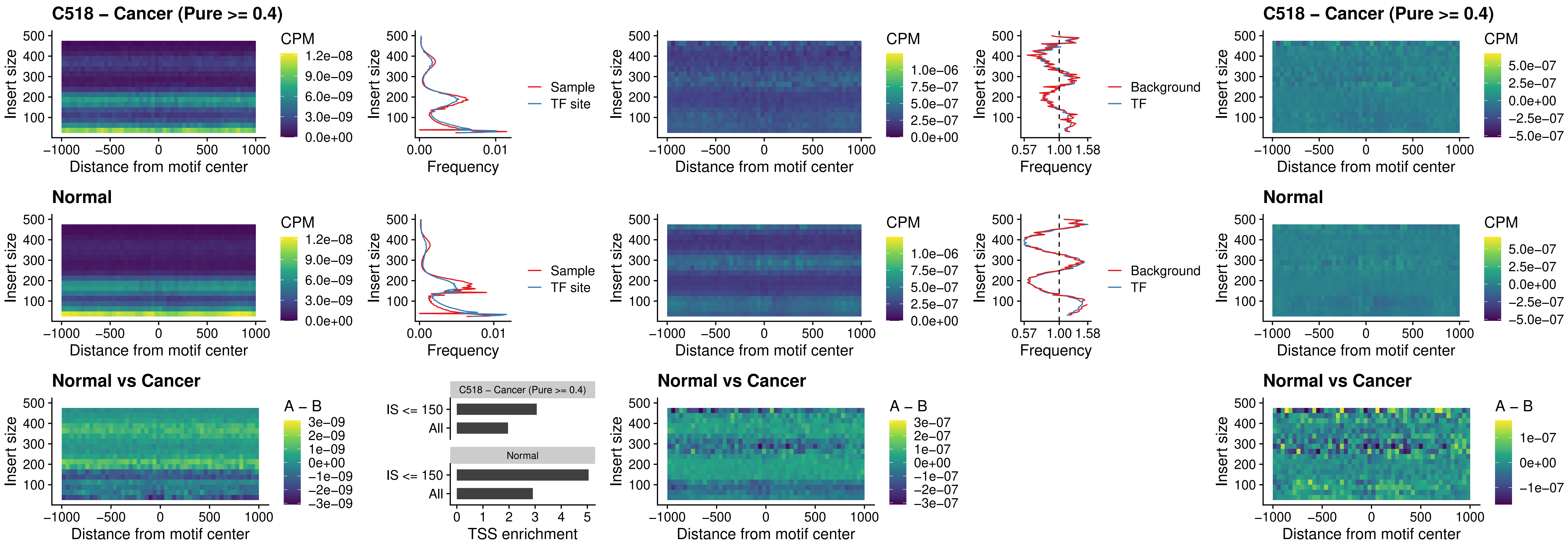

#### C518 – Cancer (Pure >= 0.4) – SPI1_uc_dTSS_nPEAK

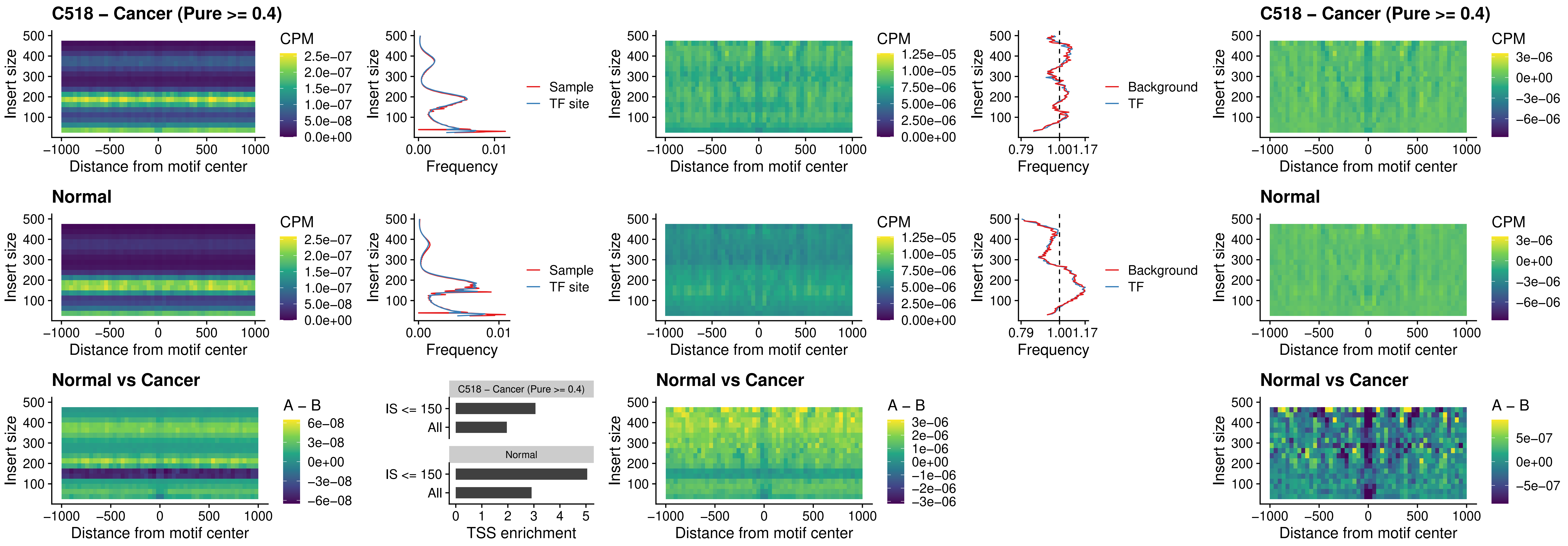

#### C518 – Cancer (Pure >= 0.4) – SPI1_uc_cTSS_nPEAK

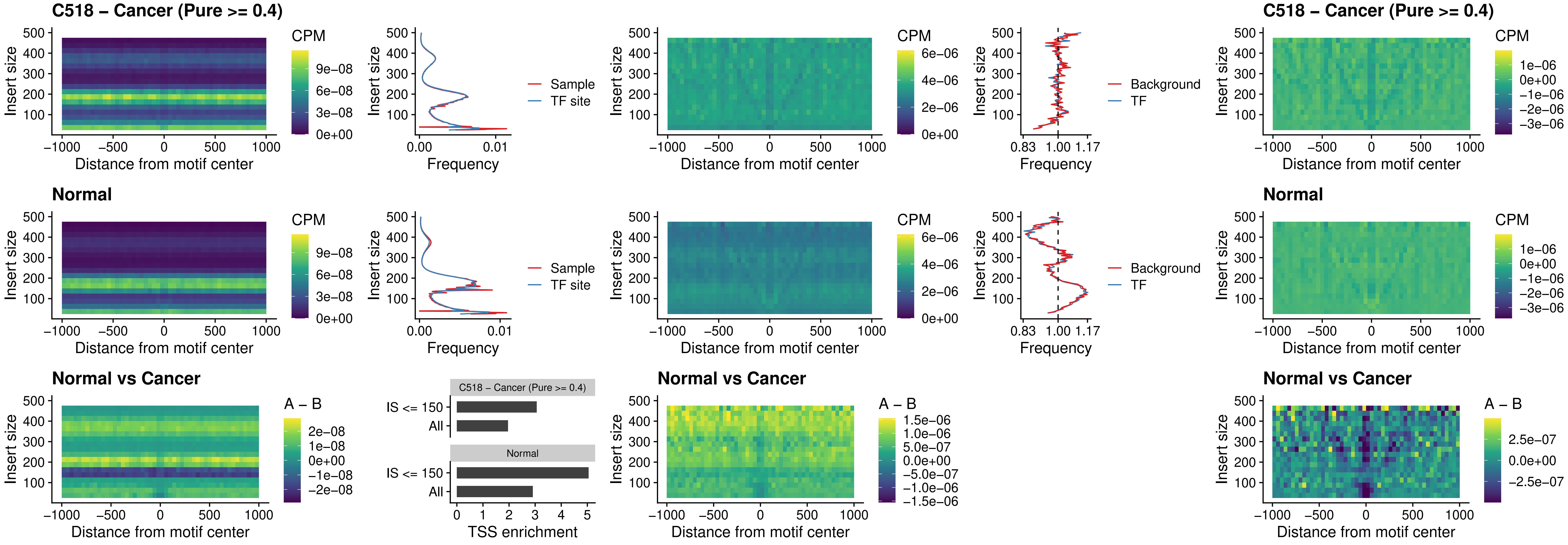

### C518 – Cancer (Pure >= 0.4) – BCL11A_uc_pTSS_nPEAK

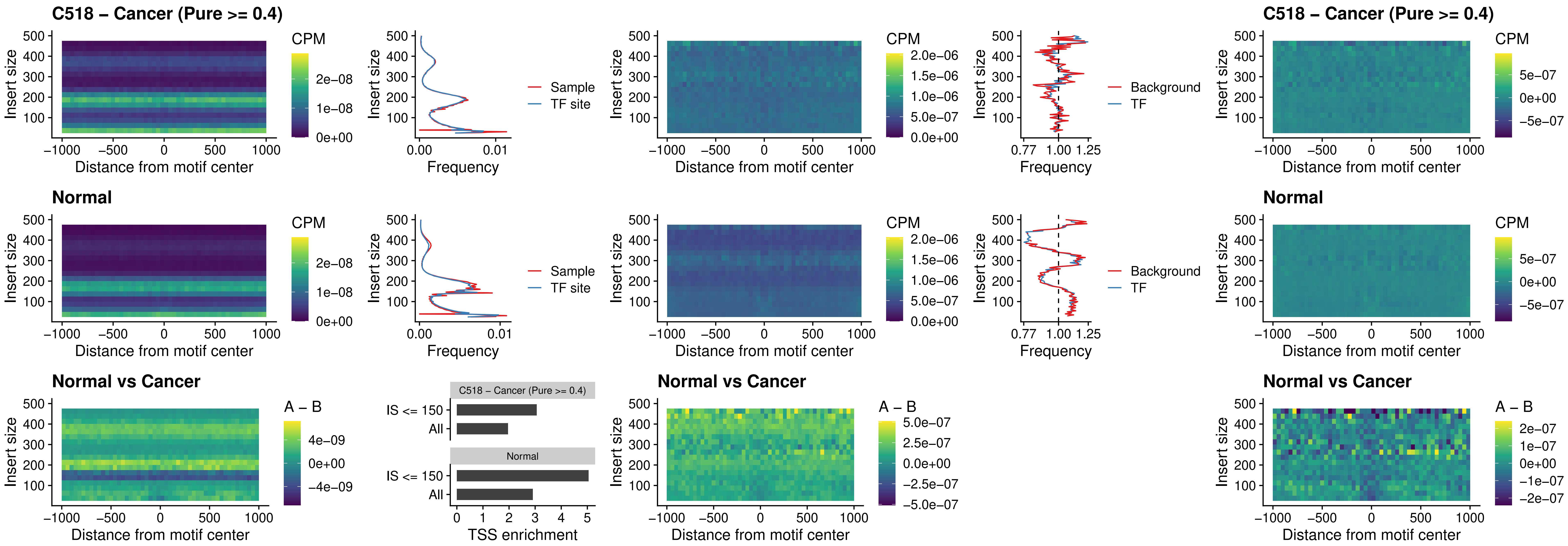

#### C518 – Cancer (Pure >= 0.4) – BCL11B_uc_pTSS_nPEAK

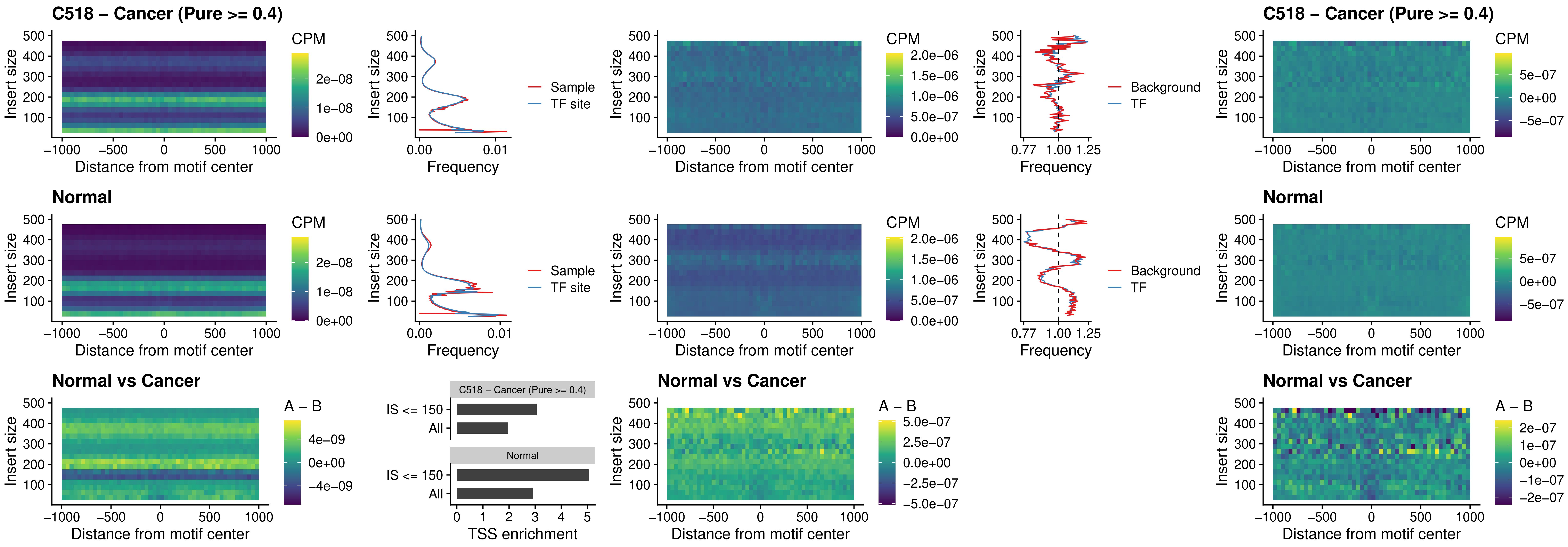

#### C518 – Cancer (Pure >= 0.4) – IRF9_uc_dTSS_oPEAK

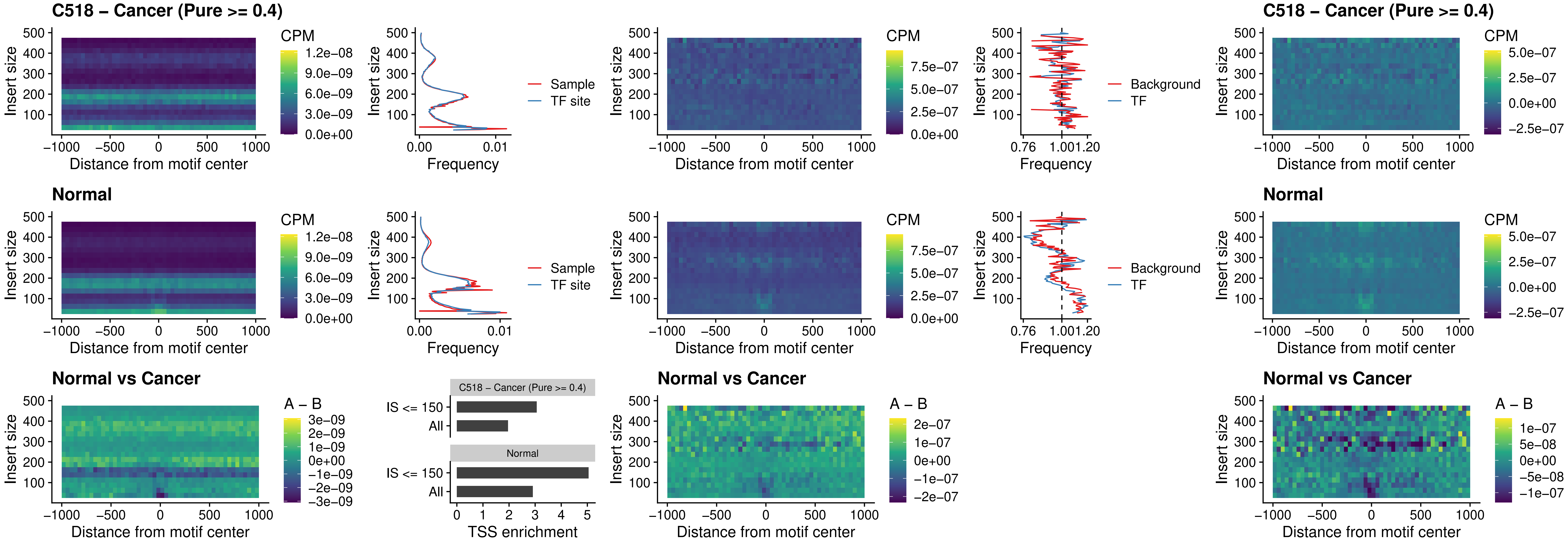

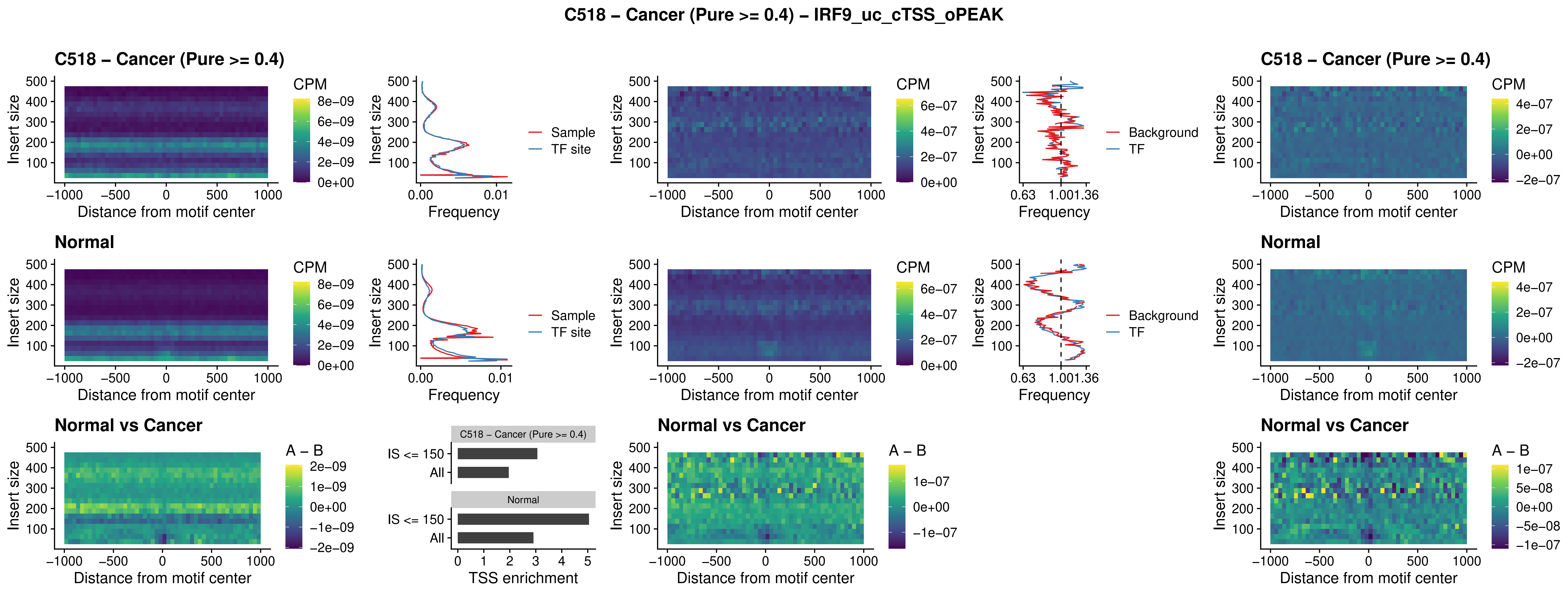
C518 – Cancer (Pure >= 0.4) – IRF9_uc_cTSS_oPEAK
C518 – Cancer (Pure >= 0.4)
Normal
Normal vs Cancer
Insert size
Distance from motif center
Frequency
CPM
A – B
Sample
TF site
Background
TF
C518 – Cancer (Pure >= 0.4)
Normal
IS <= 150
All
TSS enrichment

#### C518 – Cancer (Pure >= 0.4) – BCL11A_uc_cTSS_oPEAK

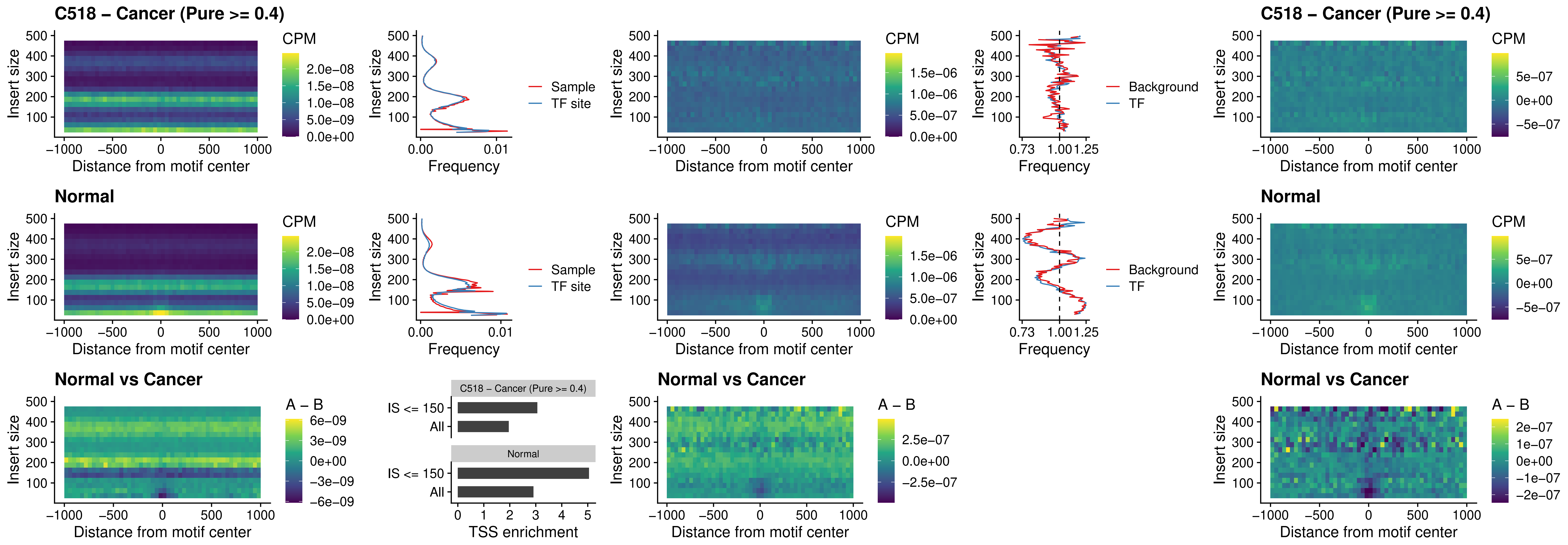

#### C518 – Cancer (Pure >= 0.4) – BCL11B_uc_cTSS_oPEAK

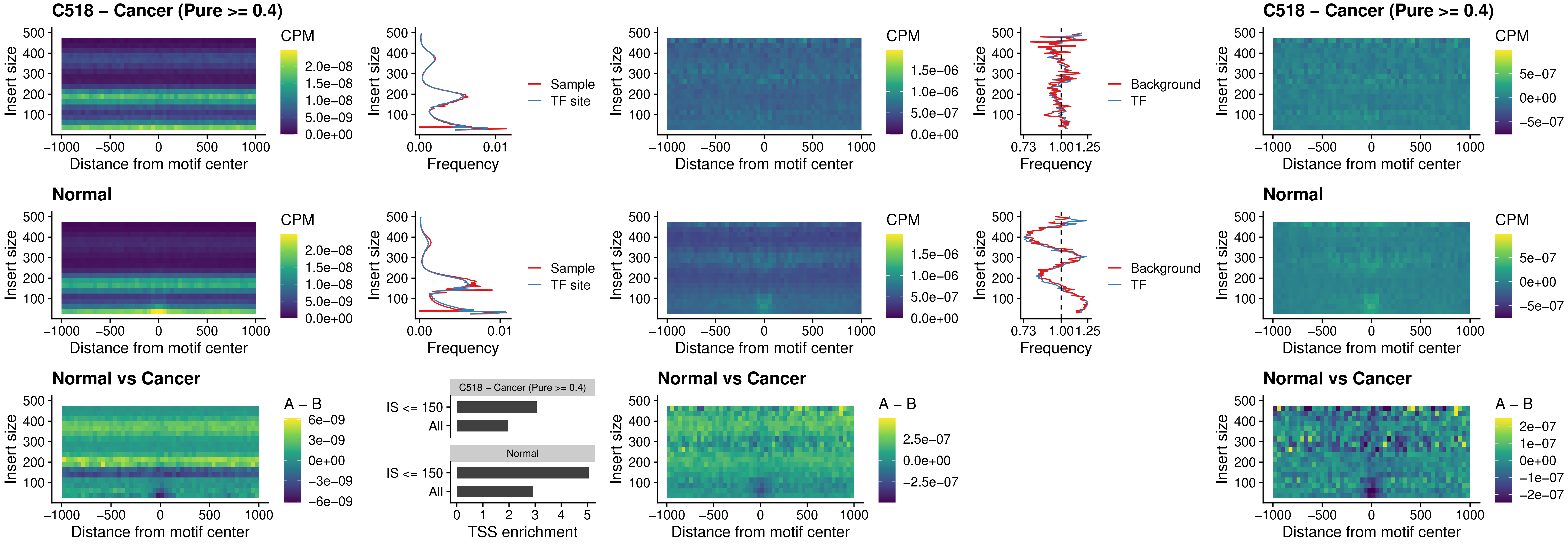

#### C518 – Cancer (Pure >= 0.4) – BCL11A_uc_dTSS_oPEAK

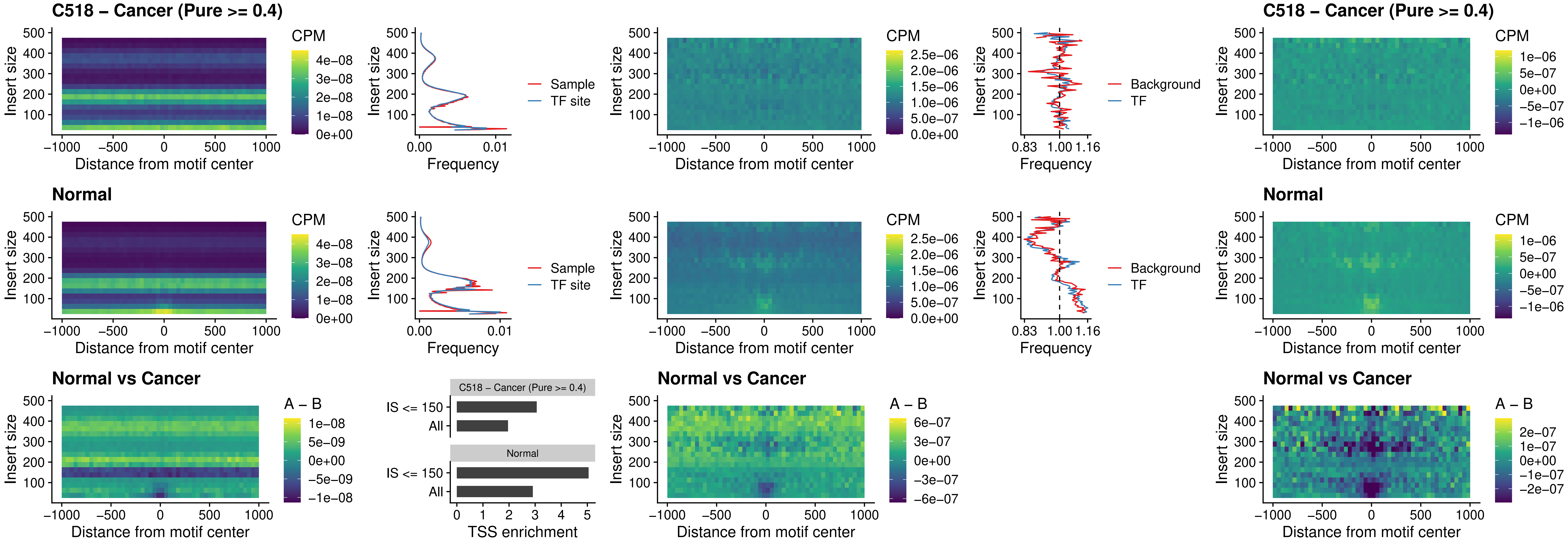

#### C518 – Cancer (Pure >= 0.4) – BCL11B_uc_dTSS_oPEAK

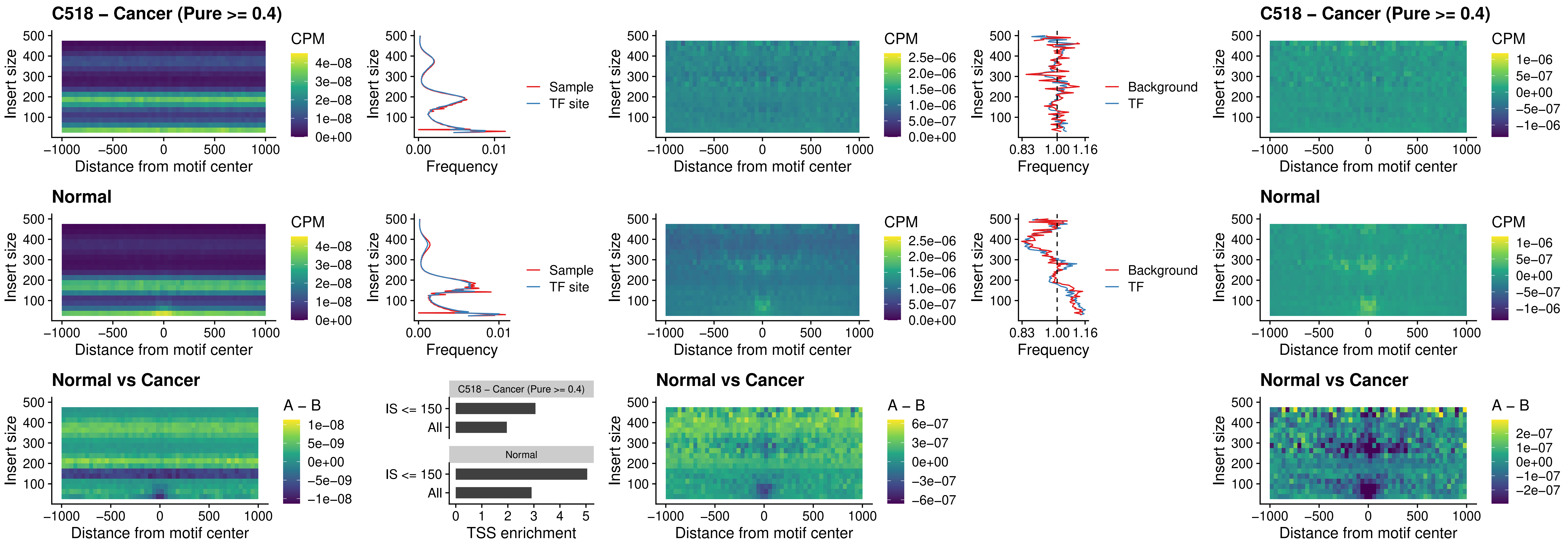

#### C518 – Cancer (Pure >= 0.4) – SPIB_uc_cTSS_nPEAK

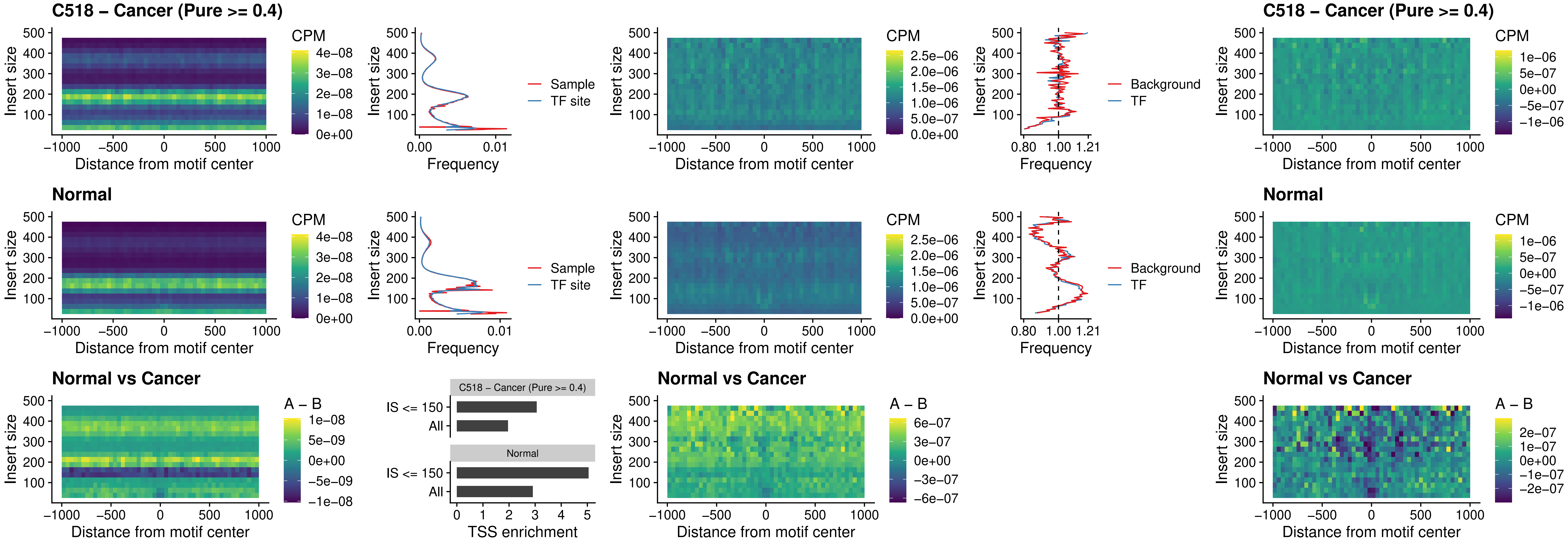

### C518 – Cancer (Pure >= 0.4) – SPIC_uc_cTSS_nPEAK

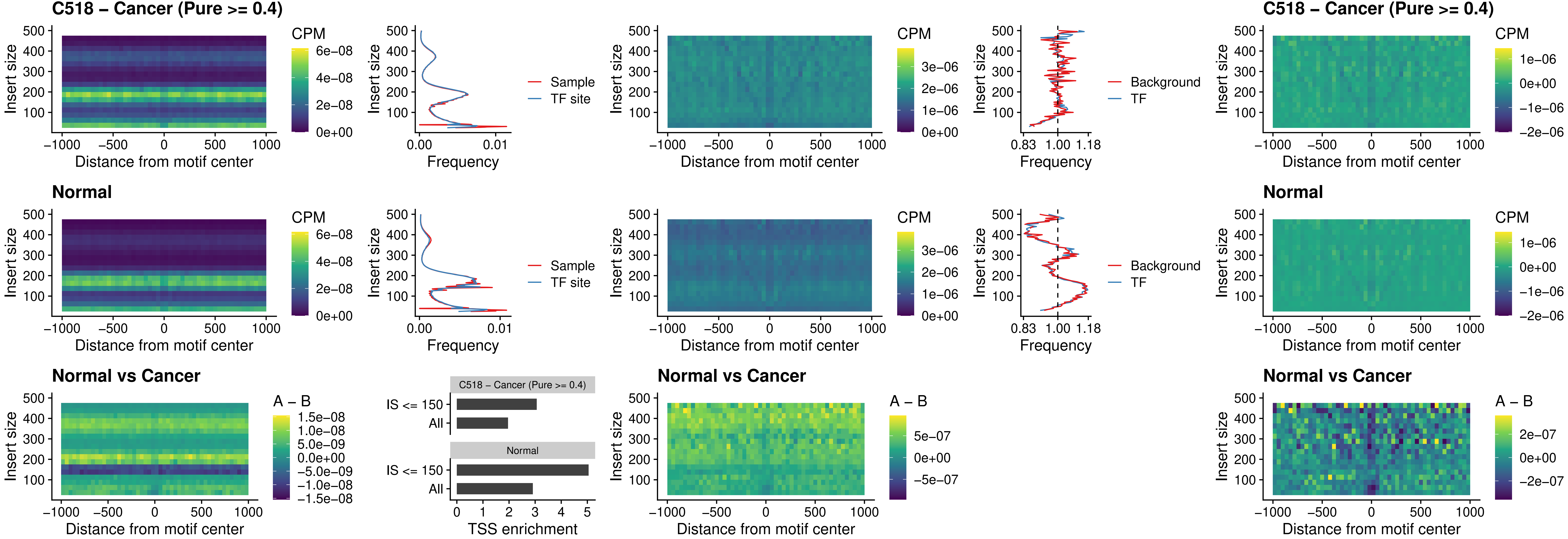

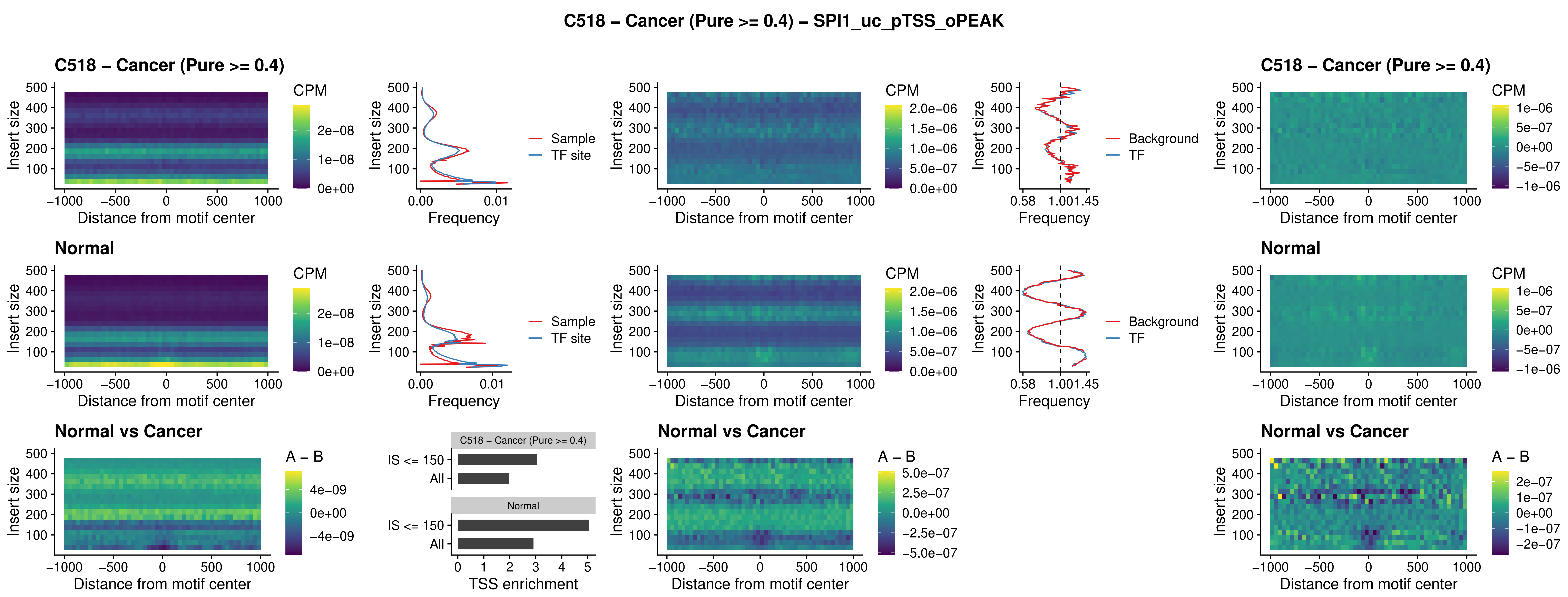
C518 – Cancer (Pure >= 0.4) – SPI1_uc_pTSS_oPEAK
C518 – Cancer (Pure >= 0.4)
Normal
Normal vs Cancer
Insert size
Distance from motif center
Frequency
CPM
A – B
Sample
TF site
Background
TF
C518 – Cancer (Pure >= 0.4)
Normal
IS <= 150
All
TSS enrichment

#### C518 – Cancer (Pure >= 0.4) – BCL11A_uc_pTSS_oPEAK

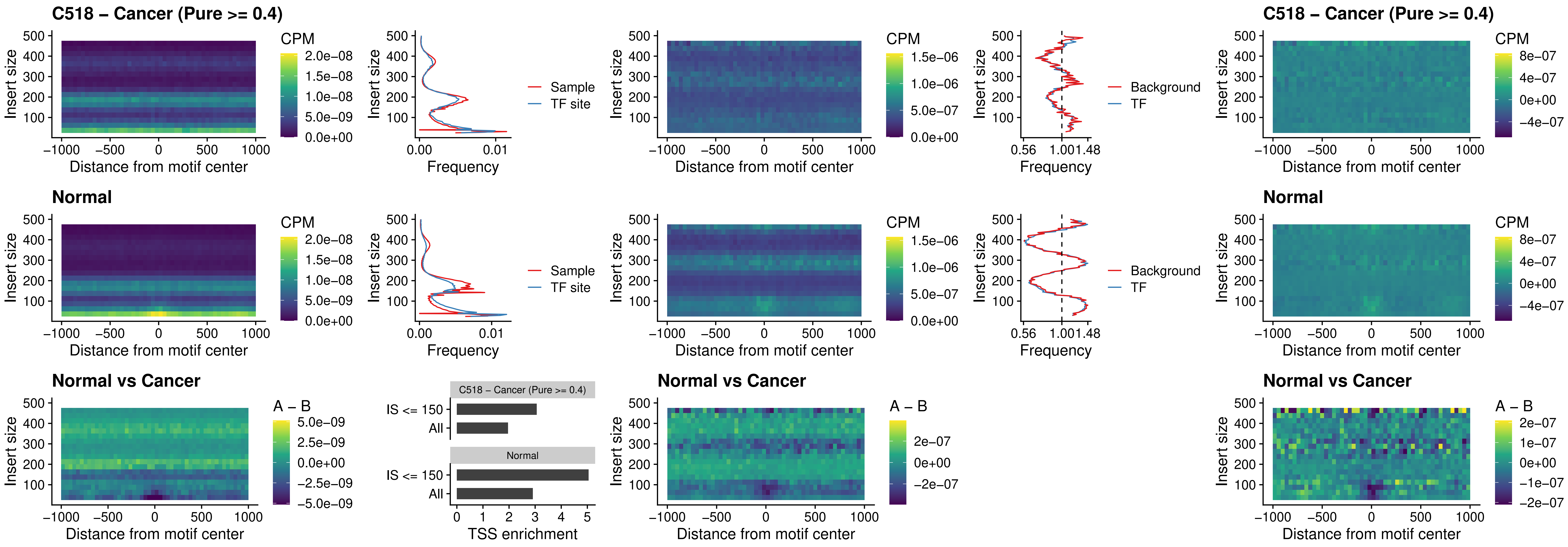

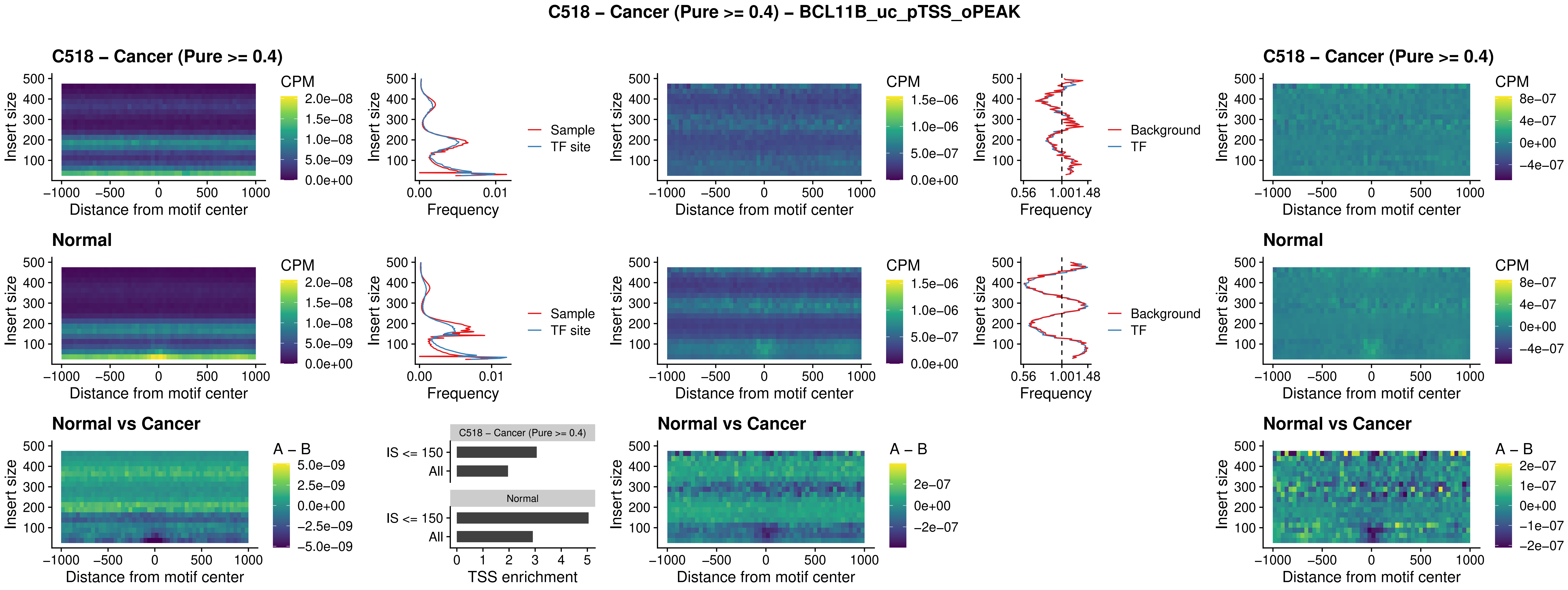
C518 – Cancer (Pure >= 0.4) – BCL11B_uc_pTSS_oPEAK
C518 – Cancer (Pure >= 0.4)
Normal
Normal vs Cancer
Insert size
Distance from motif center
Frequency
CPM
A – B
Sample
TF site
Background
TF
C518 – Cancer (Pure >= 0.4)
Normal
IS <= 150
All
TSS enrichment

#### C518 – Cancer (Pure >= 0.4) – SPI1_uc_cTSS_oPEAK

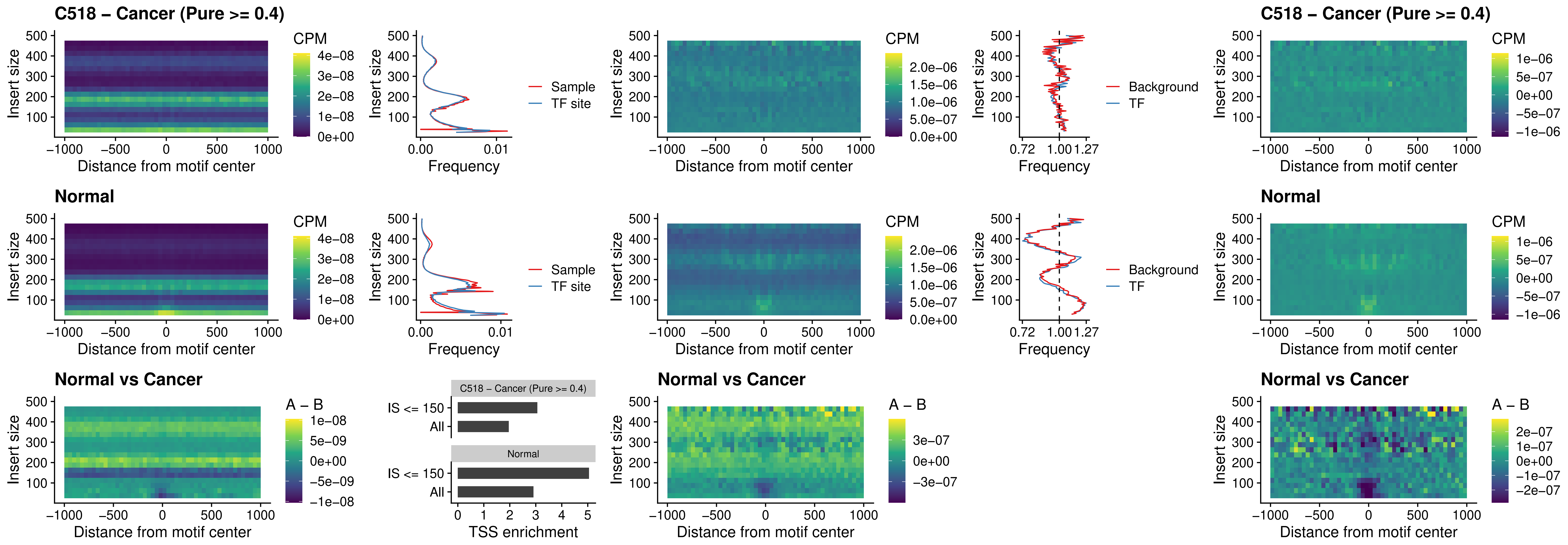

#### C518 – Cancer (Pure >= 0.4) – SPIB_uc_cTSS_oPEAK

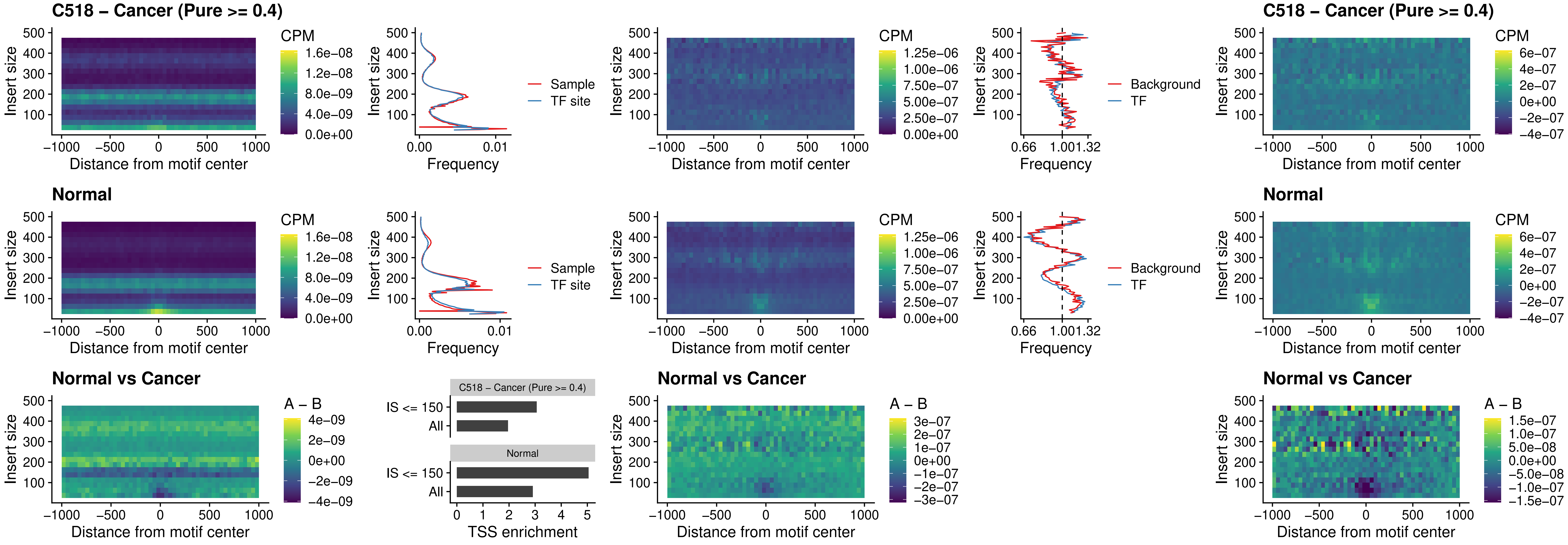

### C518 – Cancer (Pure >= 0.4) – SPIB_uc_pTSS_oPEAK

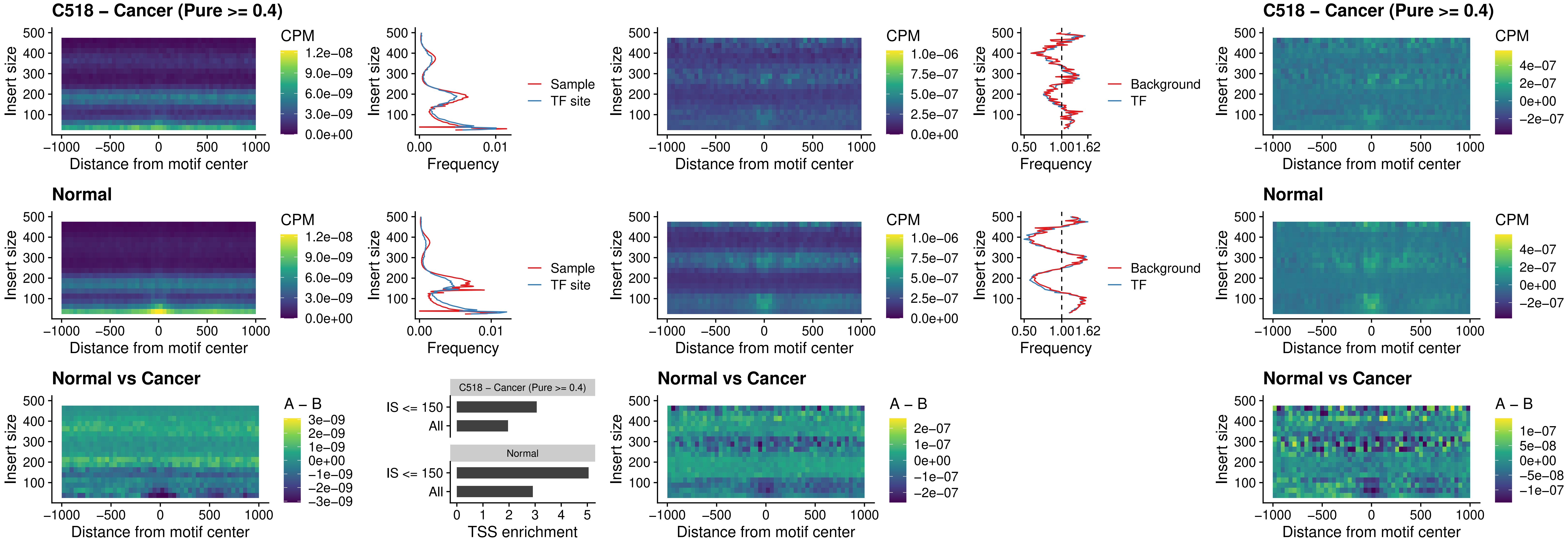

#### C518 – Cancer (Pure >= 0.4) – SPI1_uc_dTSS_oPEAK

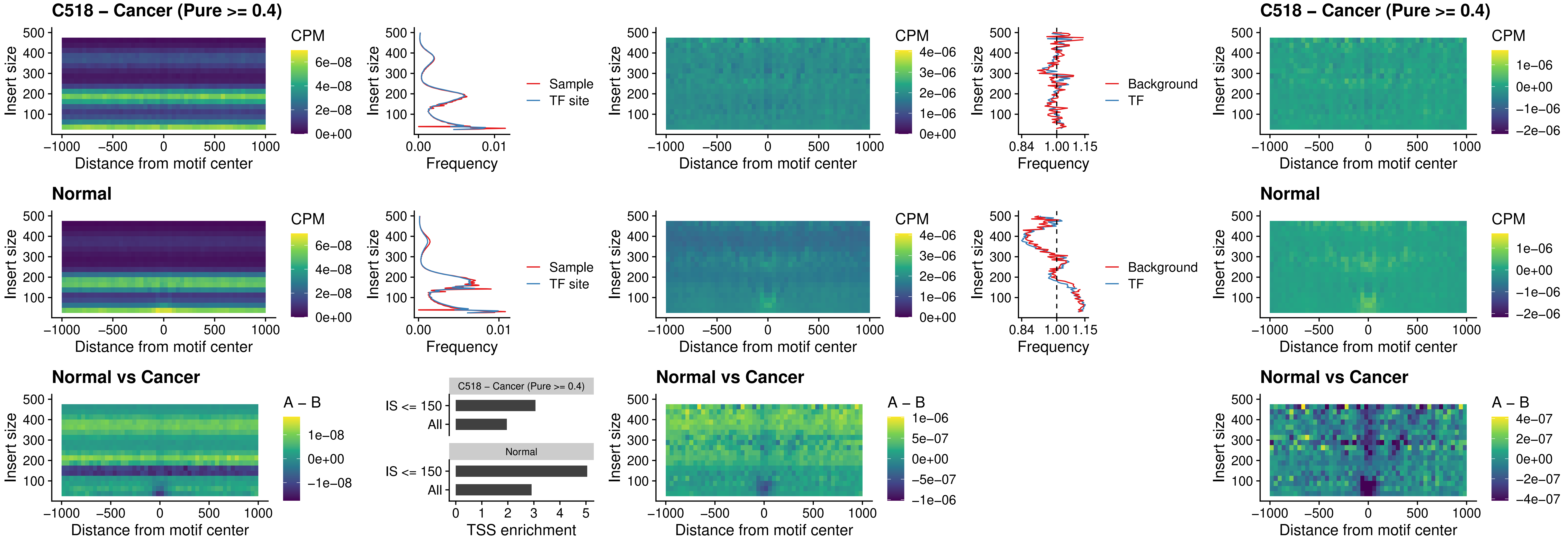

#### C518 – Cancer (Pure >= 0.4) – SPIB_uc_dTSS_oPEAK

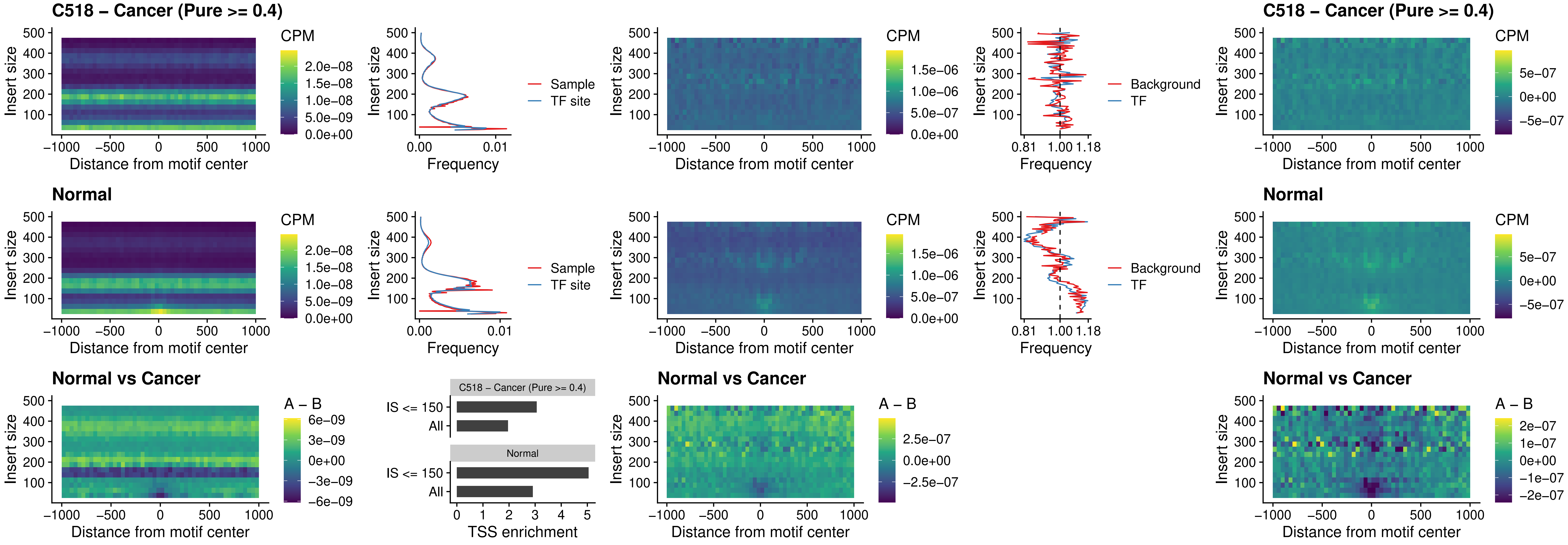

#### C518 – Cancer (Pure >= 0.4) – IRF3_uc_pTSS_oPEAK

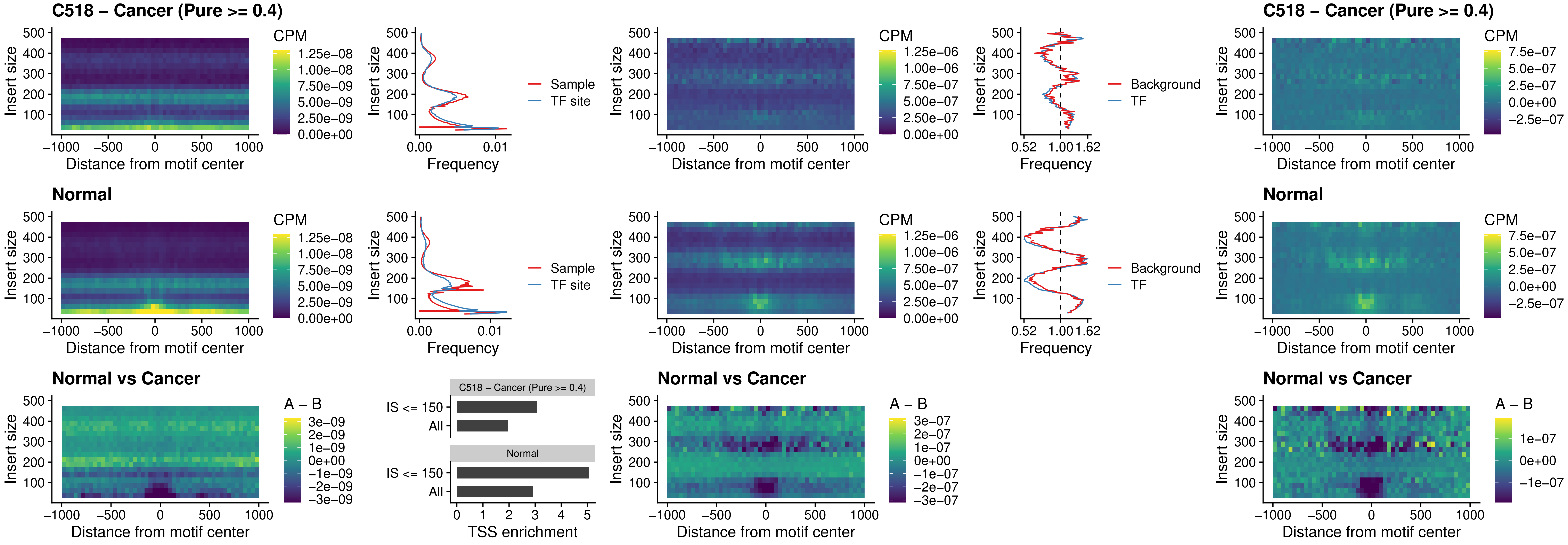

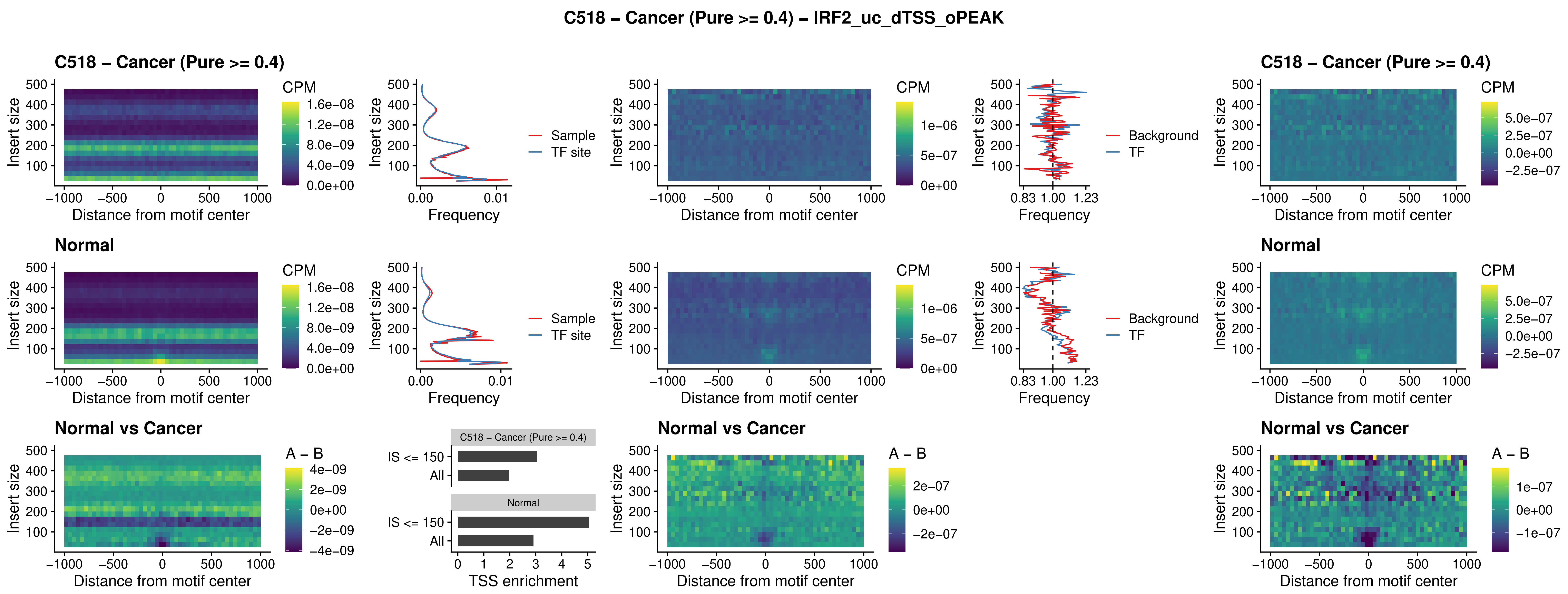
C518 – Cancer (Pure >= 0.4) – IRF2_uc_dTSS_oPEAK
C518 – Cancer (Pure >= 0.4)
Normal
Normal vs Cancer
Insert size
Distance from motif center
Frequency
CPM
A – B
Sample
TF site
Background
TF
C518 – Cancer (Pure >= 0.4)
Normal
IS <= 150
All
TSS enrichment

#### C518 – Cancer (Pure >= 0.4) – IRF2_uc_pTSS_oPEAK

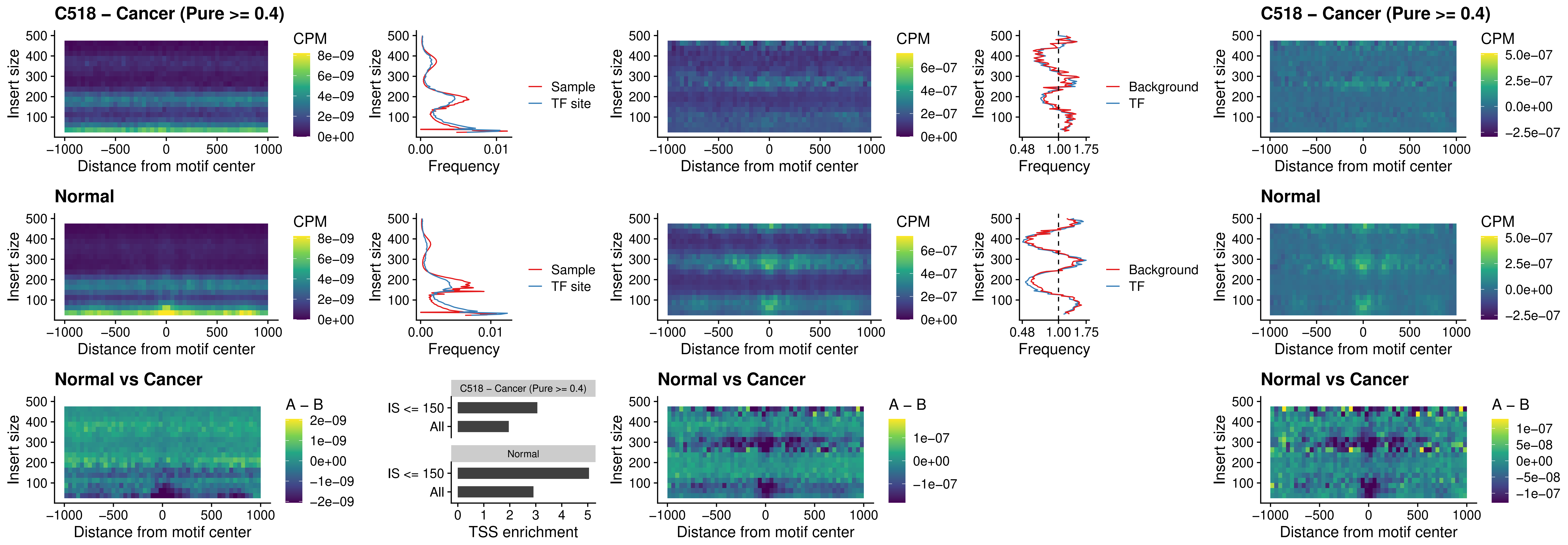

### C518 – Cancer (Pure >= 0.4) – STAT2_uc_pTSS_oPEAK

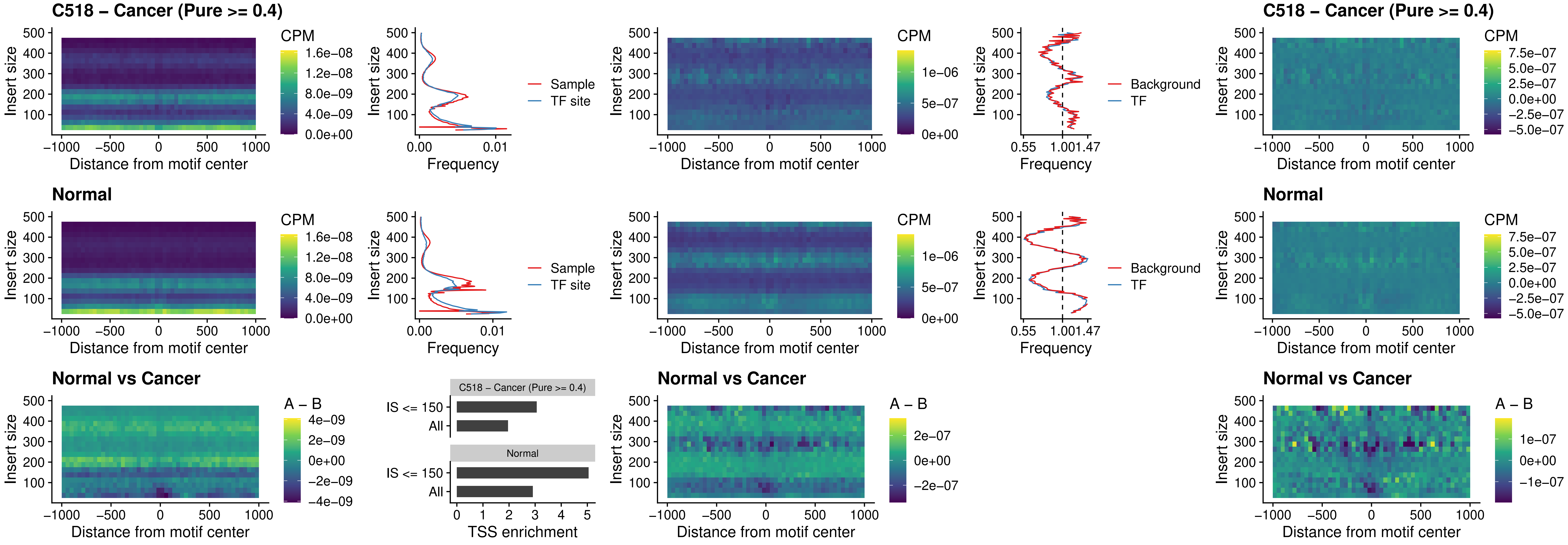

#### C518 – Cancer (Pure >= 0.4) – SPIB_uc_dTSS_nPEAK

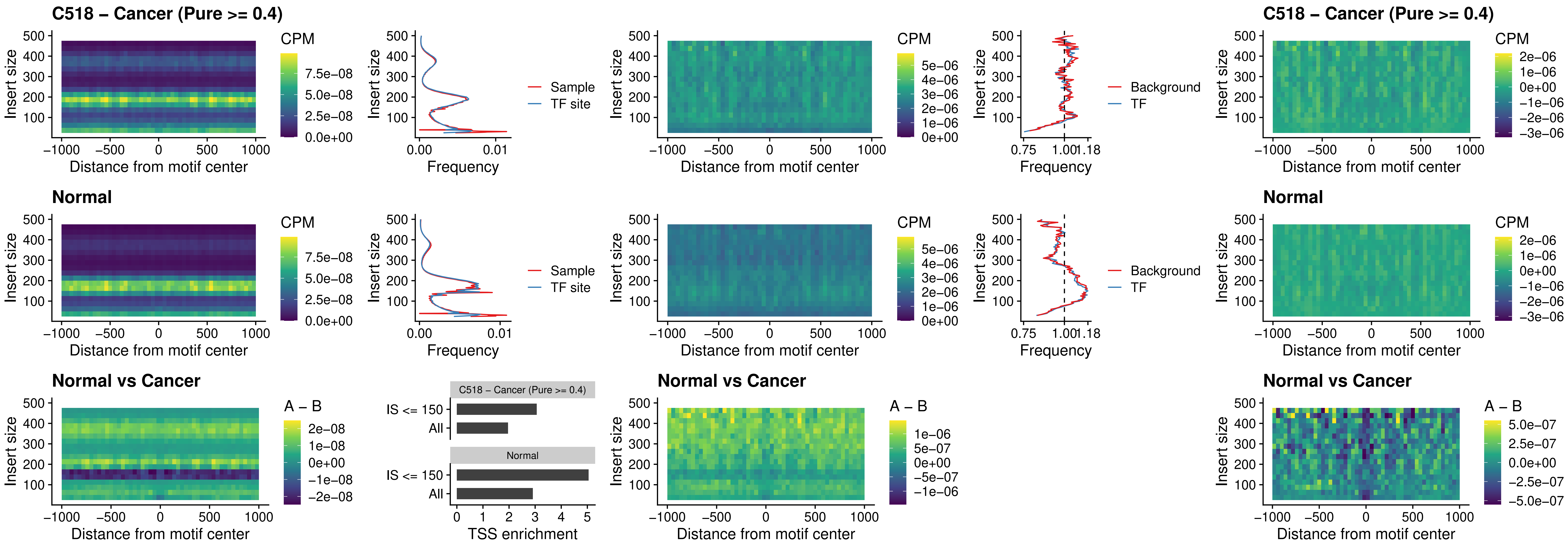

#### C518 – Cancer (Pure >= 0.4) – SPIC_uc_cTSS_oPEAK

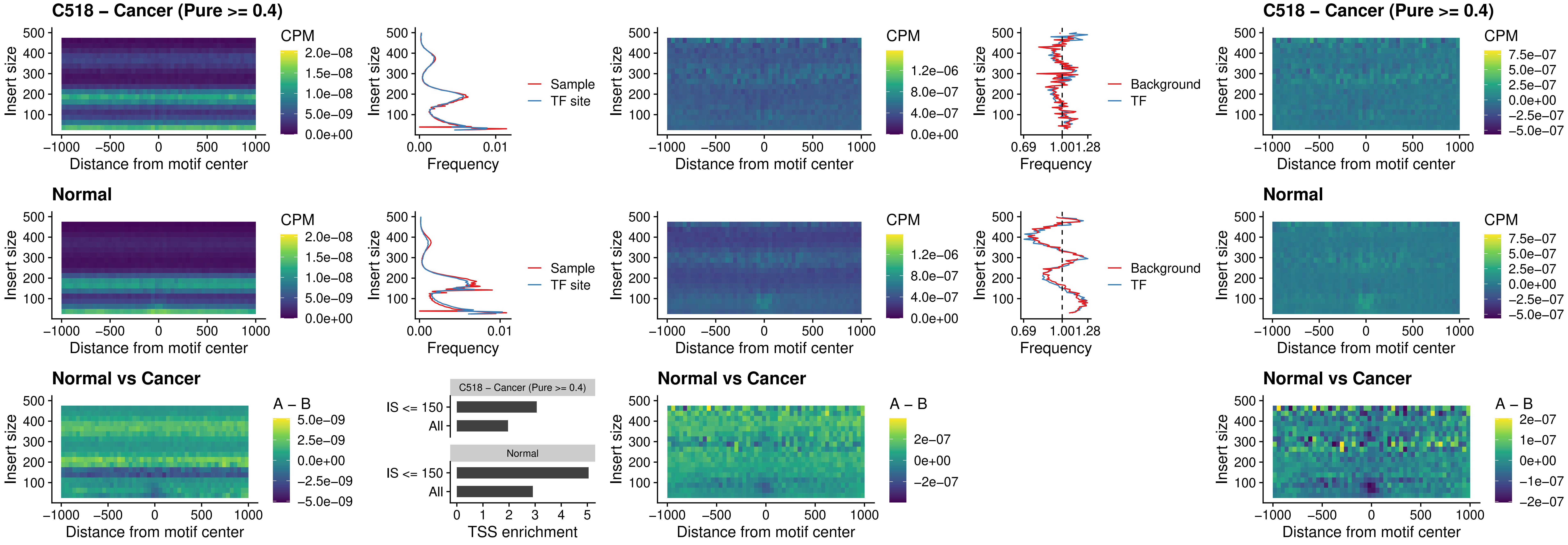

#### C518 – Cancer (Pure >= 0.4) – PRDM1_uc_cTSS_oPEAK

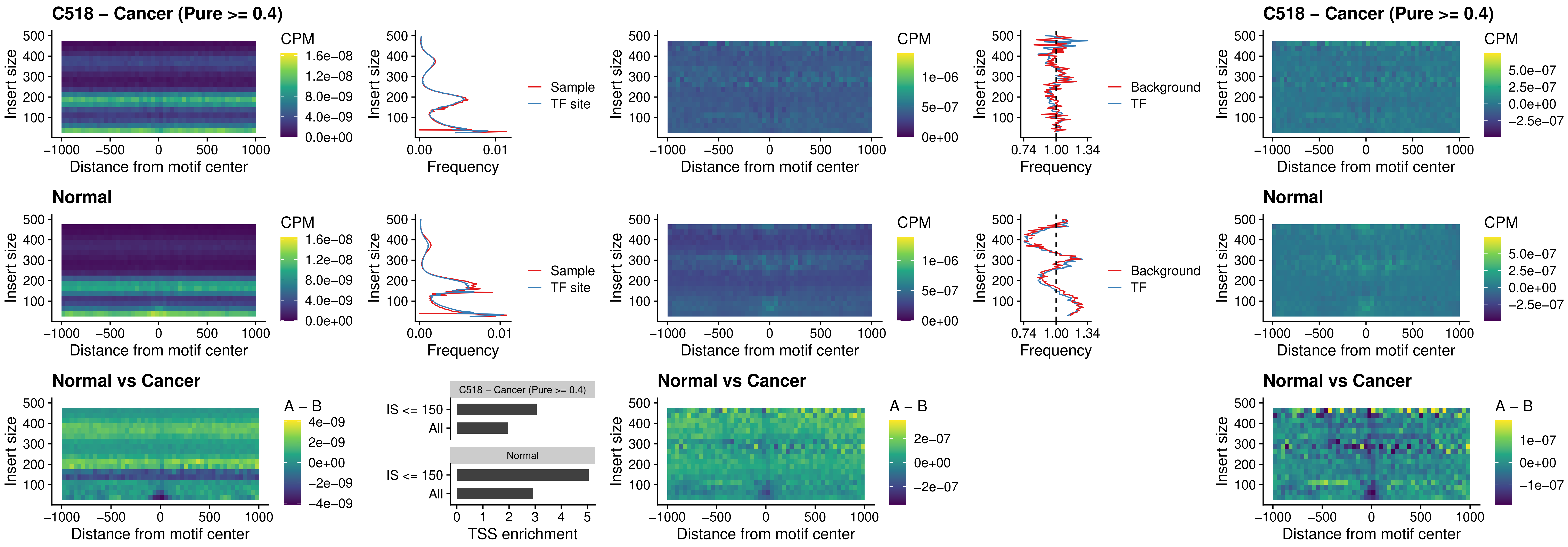

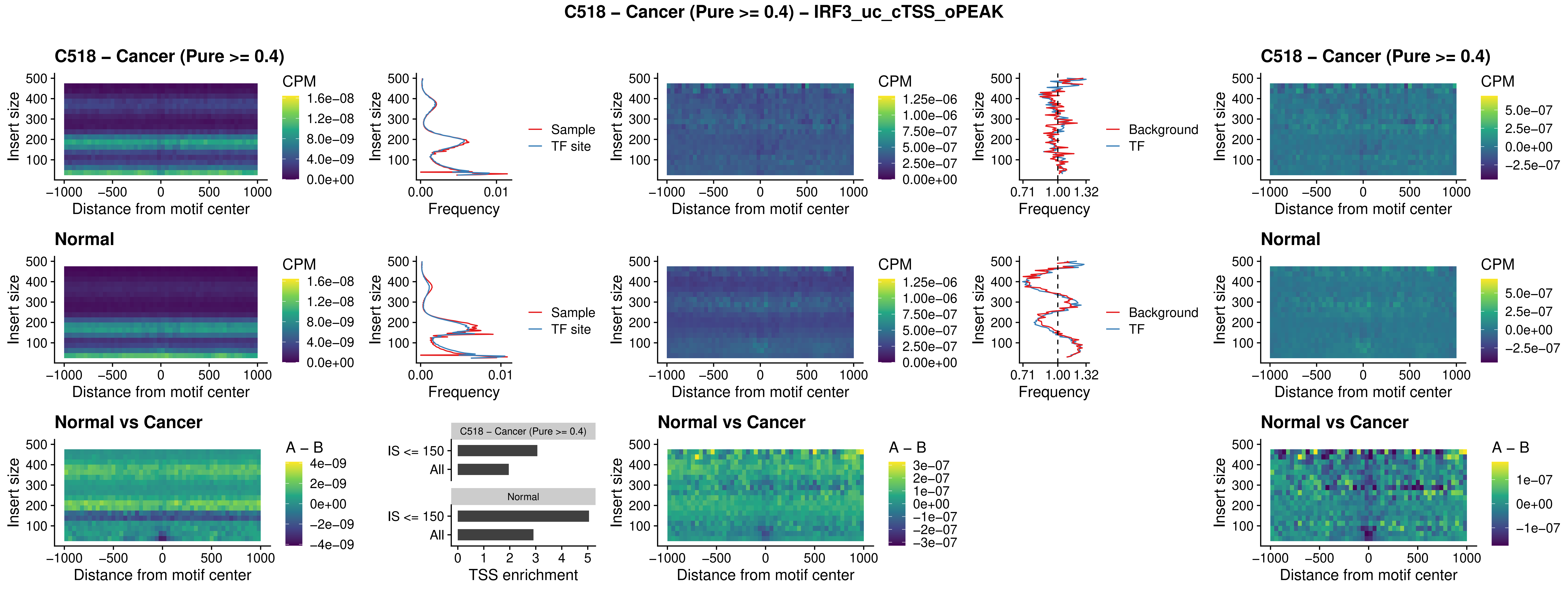

C518 – Cancer (Pure >= 0.4) – IRF3_uc_cTSS_oPEAK
C518 – Cancer (Pure >= 0.4)
Normal
Normal vs Cancer
Insert size
Distance from motif center
Frequency
CPM
A – B
Sample
TF site
Background
TF
C518 – Cancer (Pure >= 0.4)
Normal
IS <= 150
All
TSS enrichment

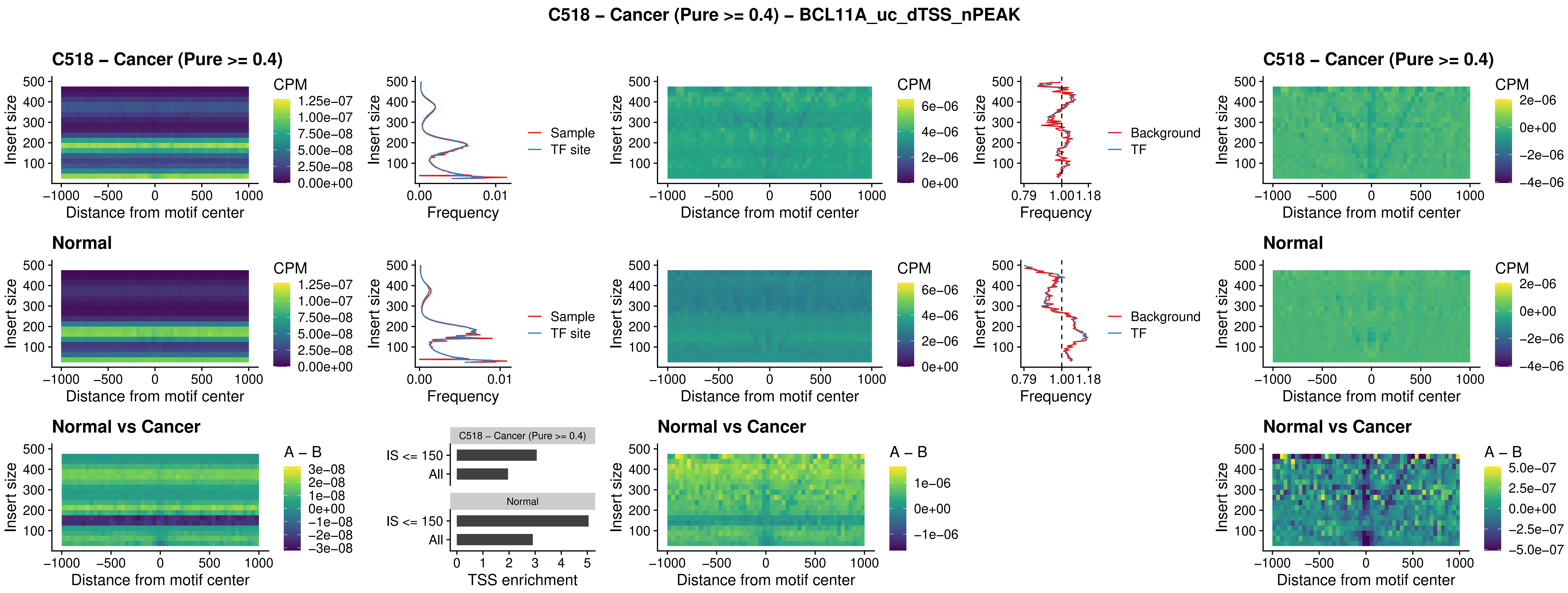
C518 – Cancer (Pure >= 0.4) – BCL11A_uc_dTSS_nPEAK
C518 – Cancer (Pure >= 0.4)
Normal
Normal vs Cancer
Insert size
Distance from motif center
CPM
A – B
Frequency
Sample
TF site
Background
TF
TSS enrichment
IS <= 150
All

#### C518 – Cancer (Pure >= 0.4) – BCL11B_uc_dTSS_nPEAK

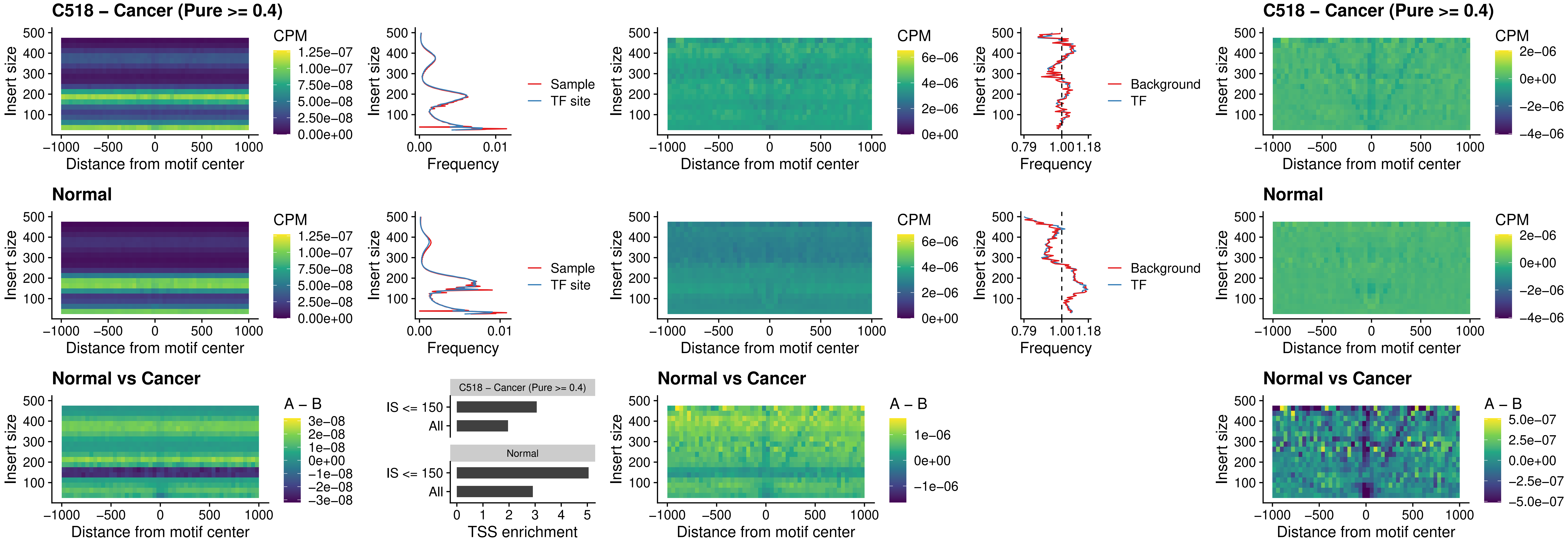

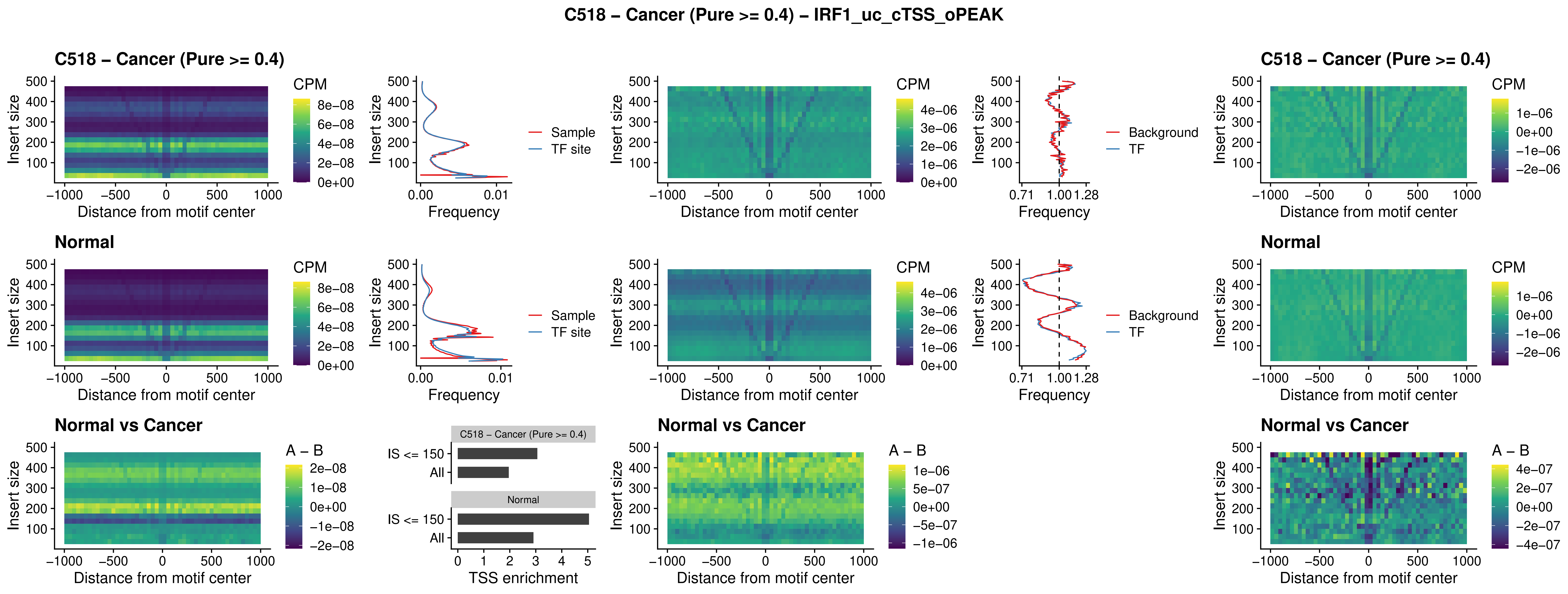
C518 – Cancer (Pure >= 0.4) – IRF1_uc_cTSS_oPEAK
C518 – Cancer (Pure >= 0.4)
Normal
Normal vs Cancer
Insert size
Distance from motif center
Frequency
CPM
A – B
Sample
TF site
Background
TF
C518 – Cancer (Pure >= 0.4)
Normal
IS <= 150
All
TSS enrichment

#### C518 – Cancer (Pure >= 0.4) – IRF1_uc_dTSS_oPEAK

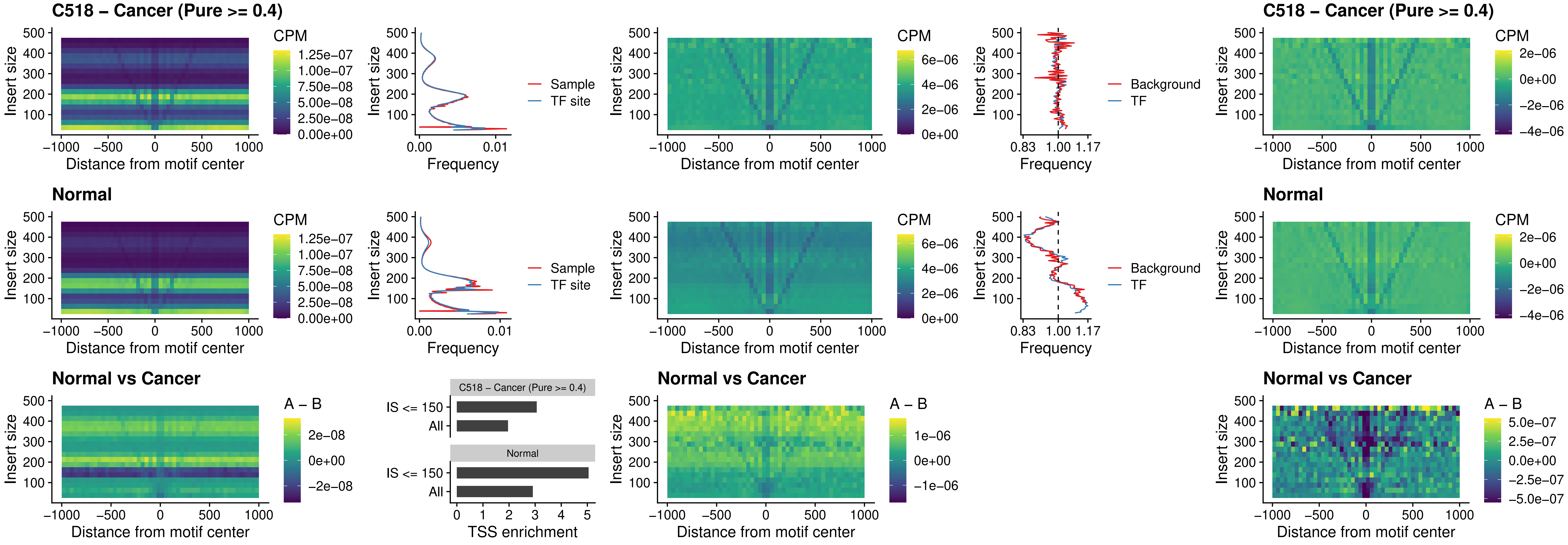

#### C518 – Cancer (Pure >= 0.4) – IRF1_uc_pTSS_oPEAK

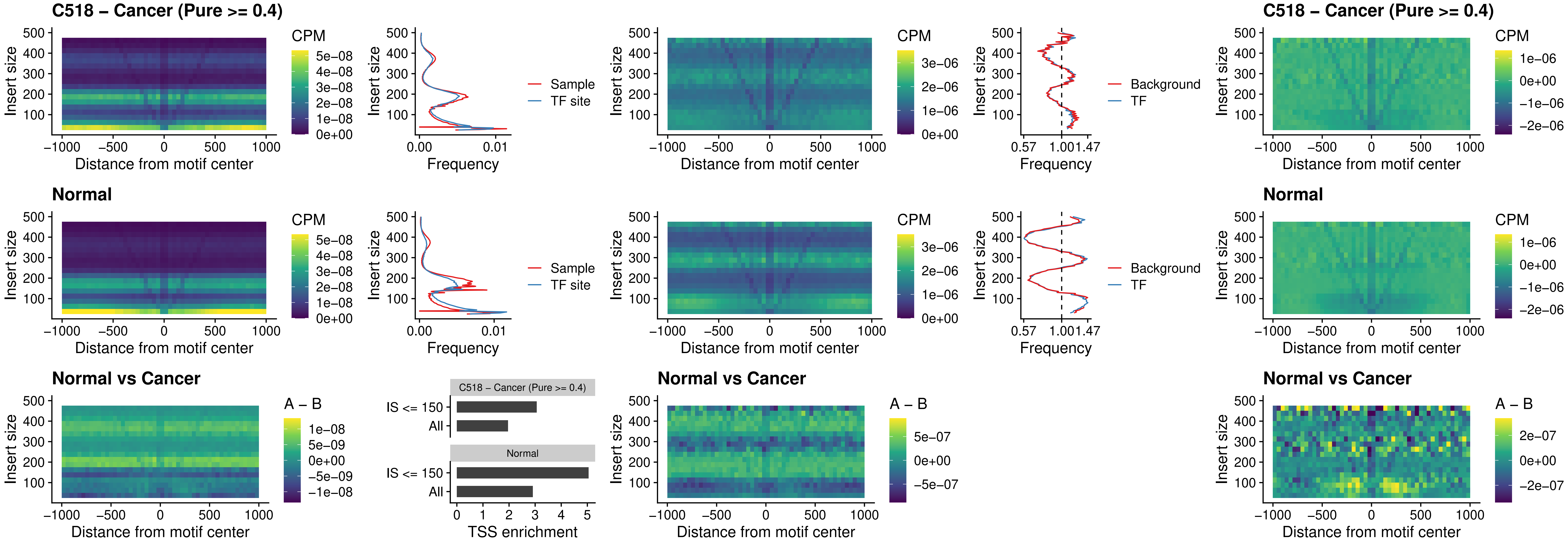

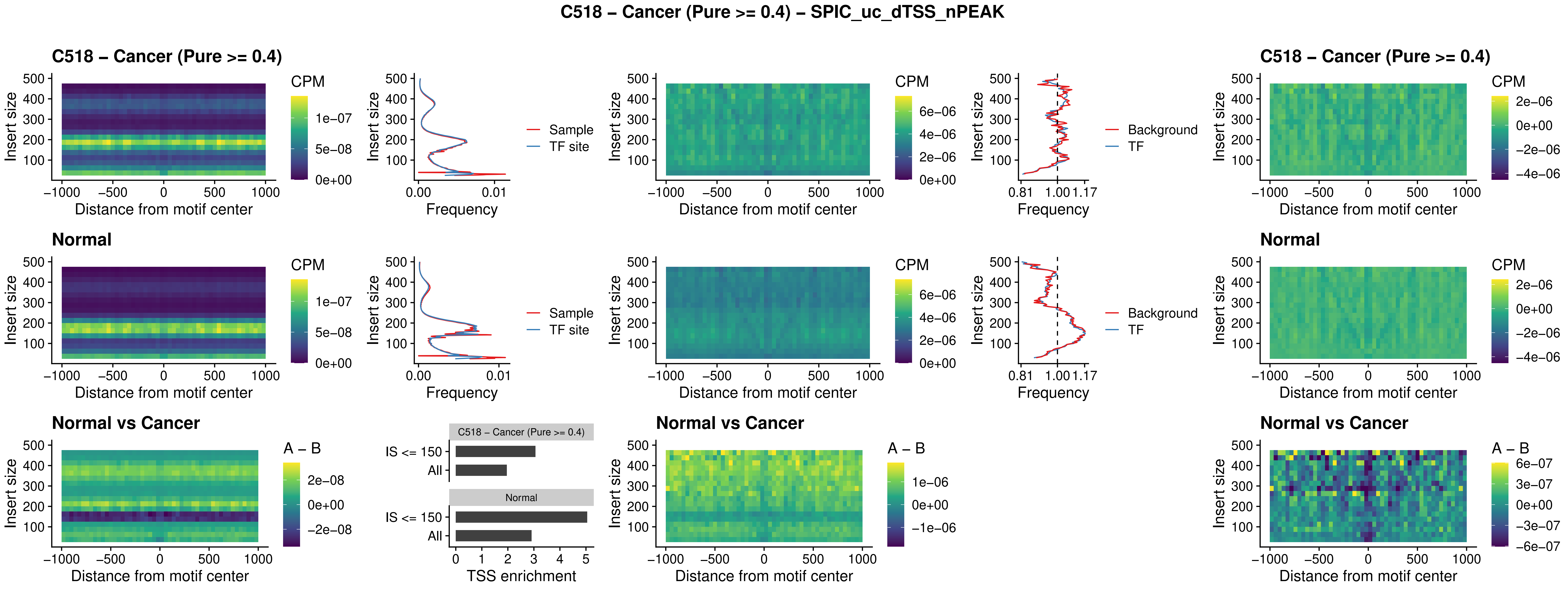
C518 – Cancer (Pure >= 0.4) – SPIC_uc_dTSS_nPEAK
C518 – Cancer (Pure >= 0.4)
Normal
Normal vs Cancer
Insert size
Distance from motif center
CPM
A – B
Frequency
Sample
TF site
Background
TF
C518 – Cancer (Pure >= 0.4)
Normal
IS <= 150
All
TSS enrichment

#### C518 – Cancer (Pure >= 0.4) – SPIC_uc_dTSS_oPEAK

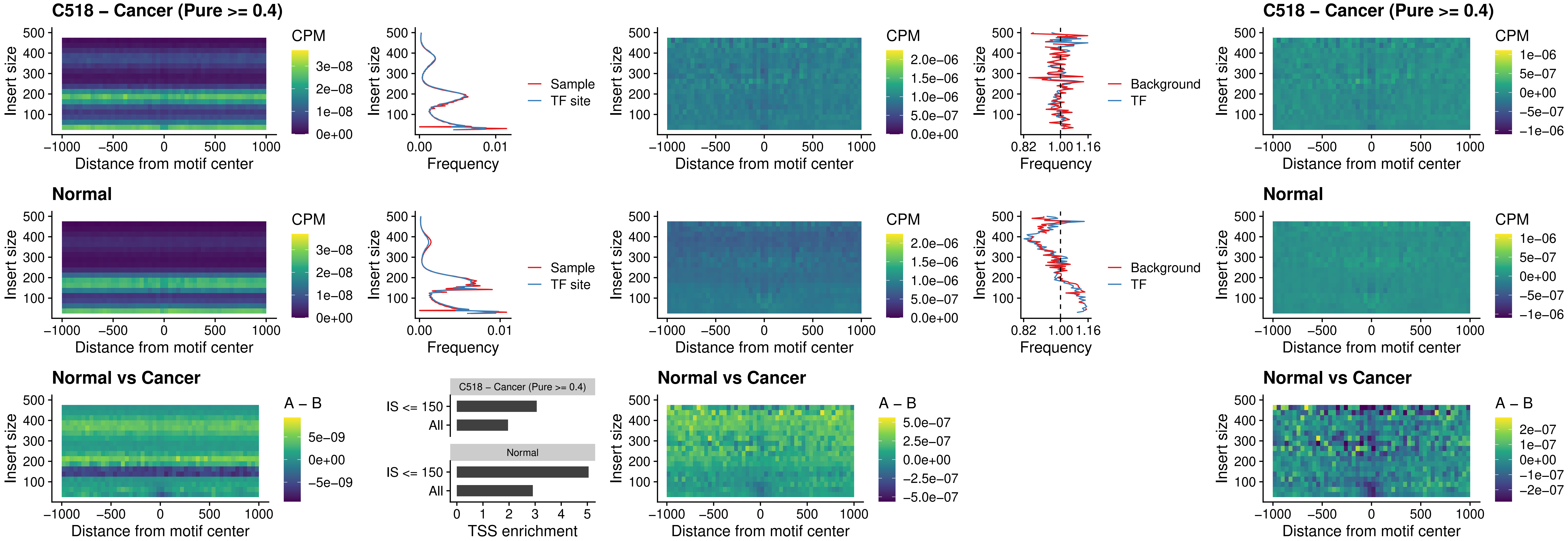

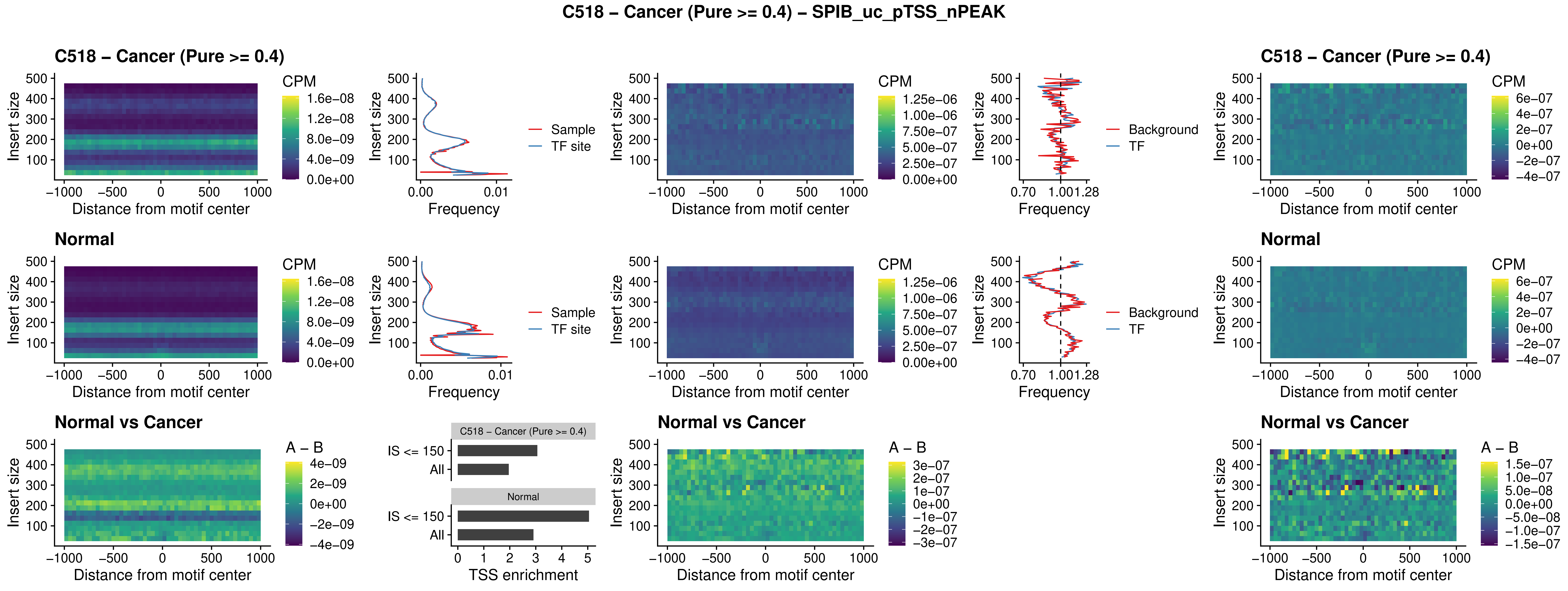
C518 – Cancer (Pure >= 0.4) – SPIB_uc_pTSS_nPEAK
C518 – Cancer (Pure >= 0.4)
Normal
Normal vs Cancer
Insert size
Distance from motif center
Frequency
CPM
A – B
Sample
TF site
Background
TF
C518 – Cancer (Pure >= 0.4)
Normal
IS <= 150
All
TSS enrichment

#### C518 – Cancer (Pure >= 0.4) – STAT2_uc_cTSS_oPEAK

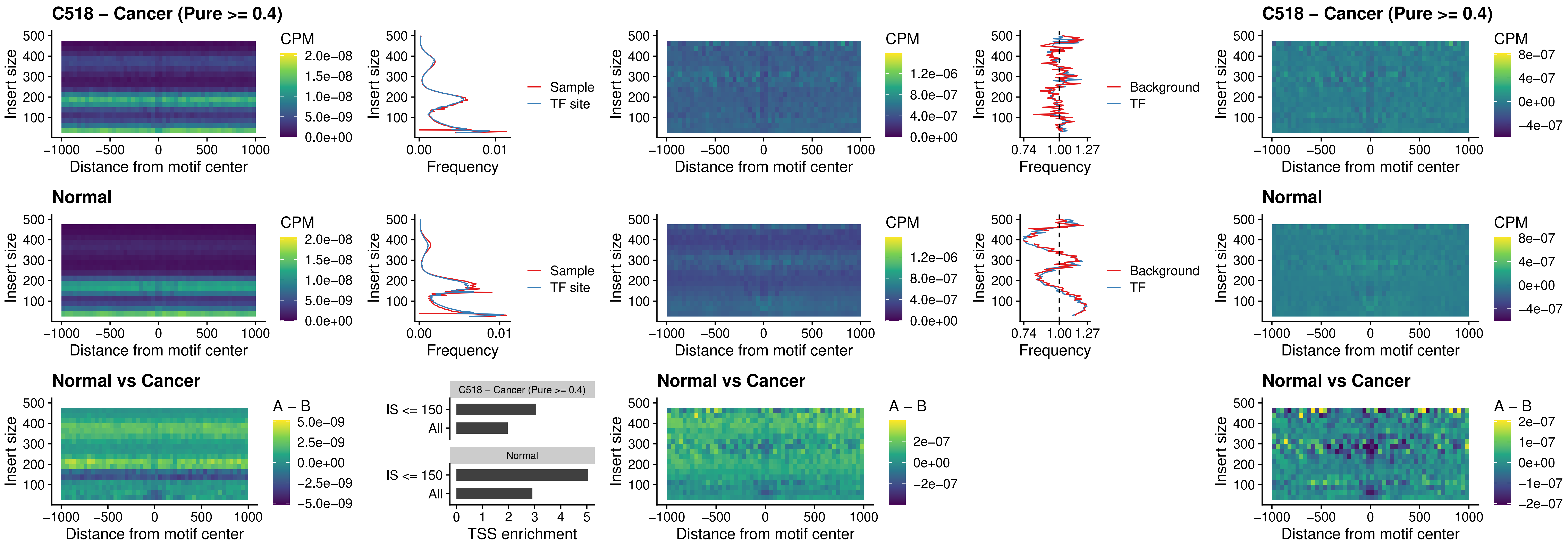

### C518 – Cancer (Pure >= 0.4) – IRF8_uc_cTSS_oPEAK

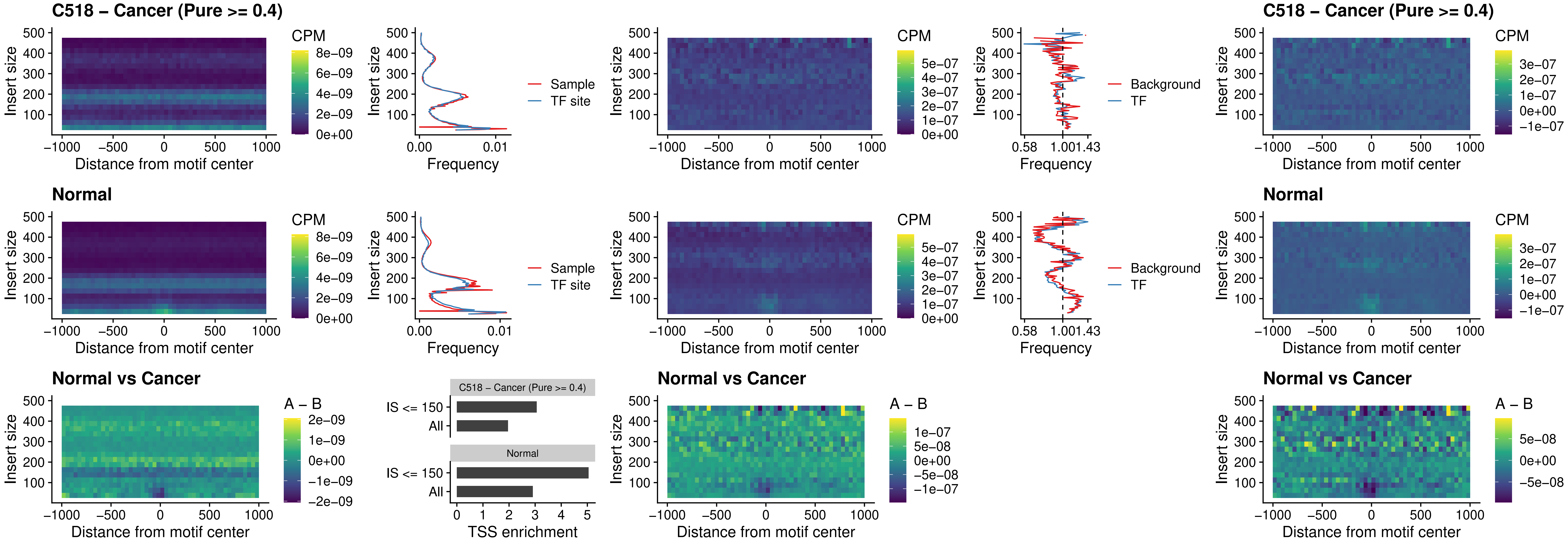

#### C518 – Cancer (Pure >= 0.4) – IRF3_uc_dTSS_oPEAK

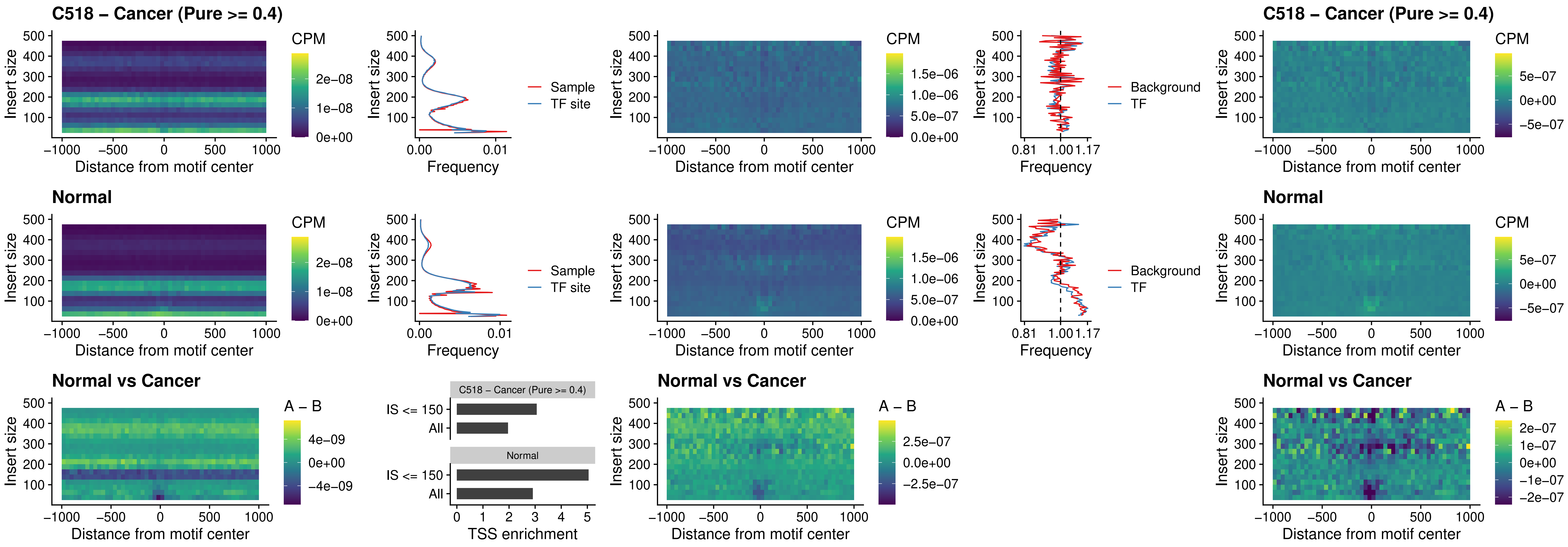

#### C518 – Cancer (Pure >= 0.4) – IRF8_uc_dTSS_oPEAK

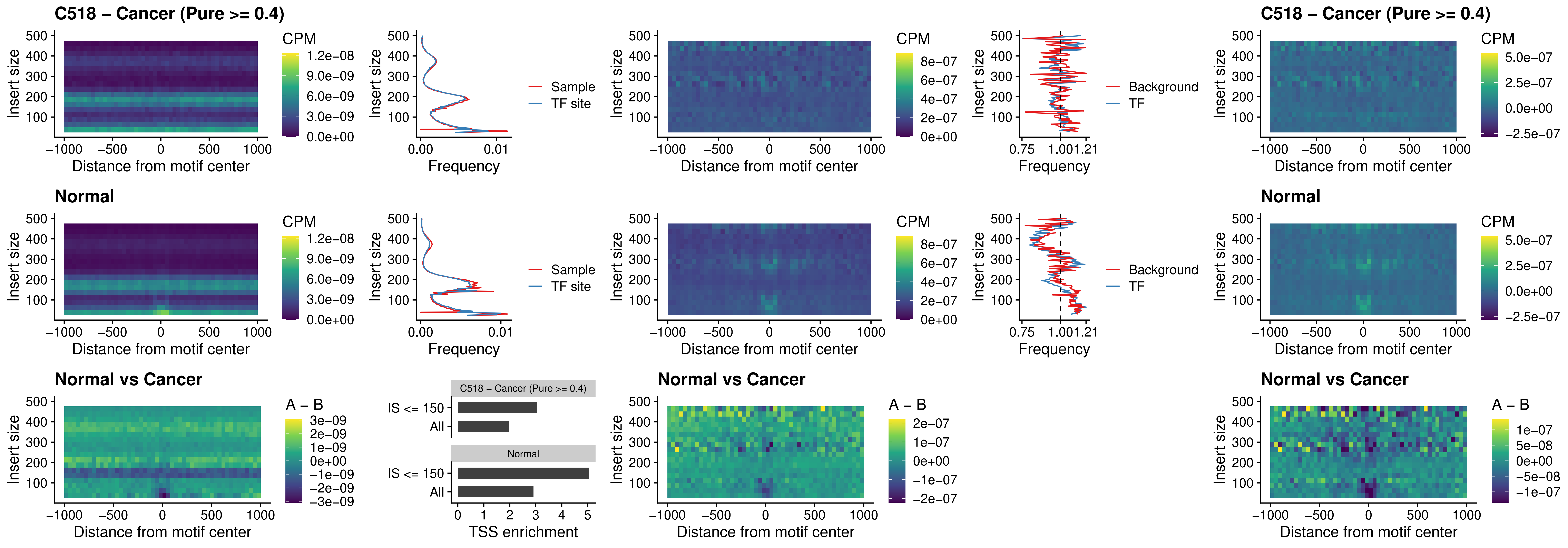

### C518 – Cancer (Pure >= 0.4) – ETS2_uc_dTSS_oPEAK

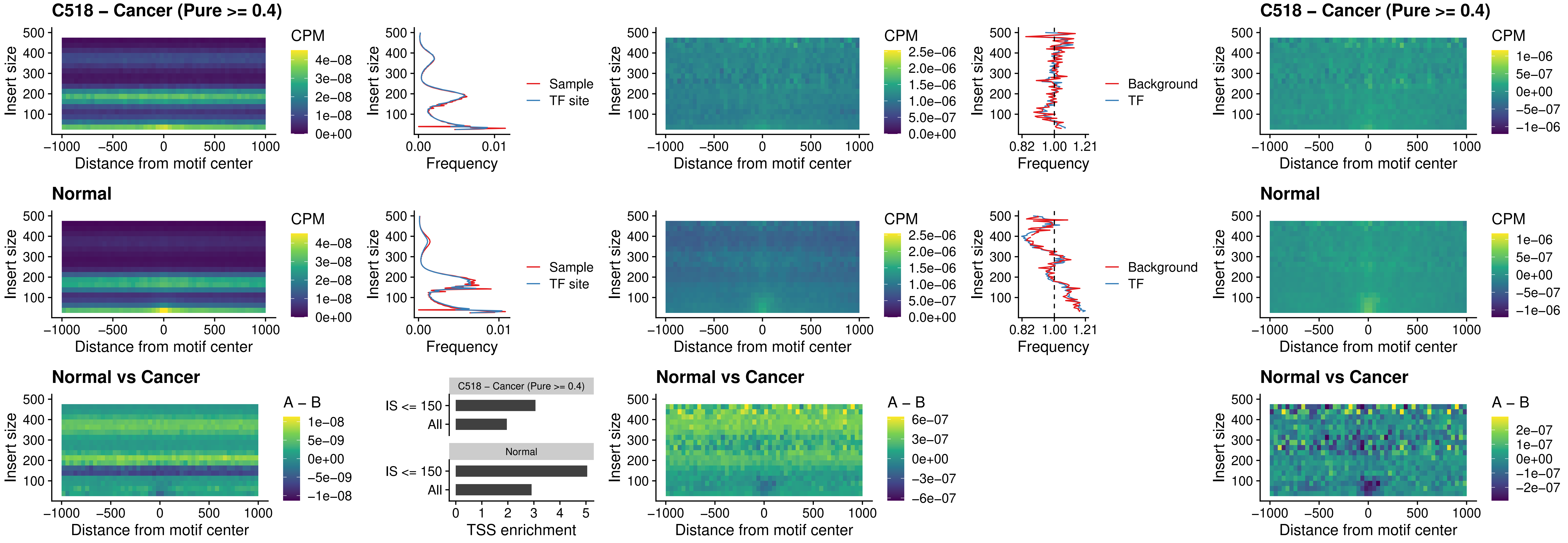

#### C518 – Cancer (Pure >= 0.4) – SMARCC2_uc_pTSS_oPEAK

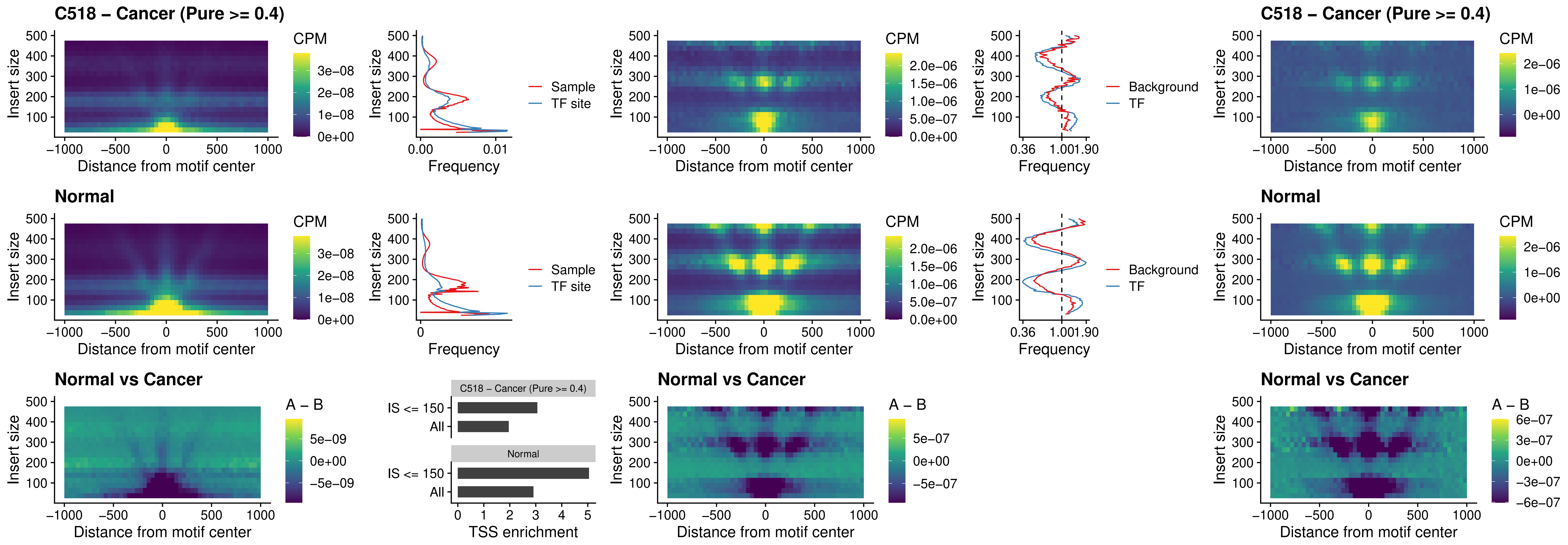

#### C518 – Cancer (Pure >= 0.4) – BCL11A_uc_cTSS_nPEAK

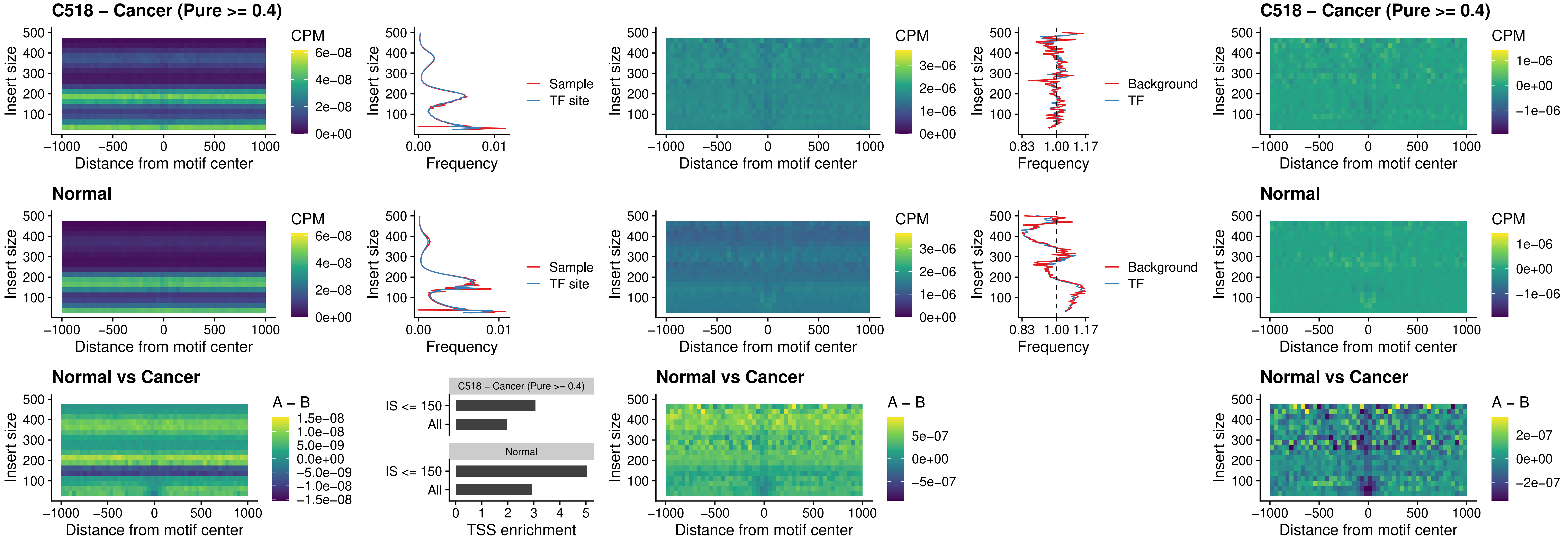

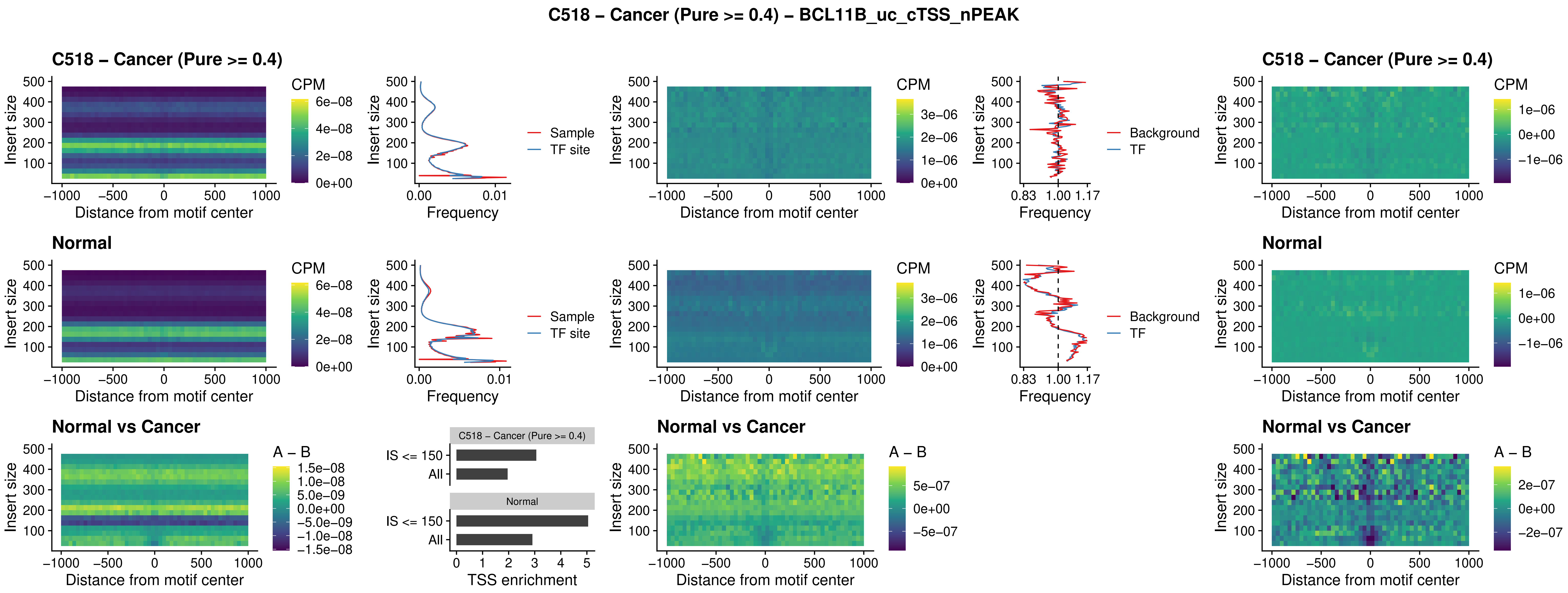

C518 – Cancer (Pure >= 0.4) – BCL11B_uc_cTSS_nPEAK
C518 – Cancer (Pure >= 0.4)
Normal
Normal vs Cancer
Insert size
Distance from motif center
Frequency
CPM
A – B
Sample
TF site
Background
TF
C518 – Cancer (Pure >= 0.4)
Normal
IS <= 150
All
TSS enrichment

#### C518 – Cancer (Pure >= 0.4) – STAT2_uc_dTSS_oPEAK

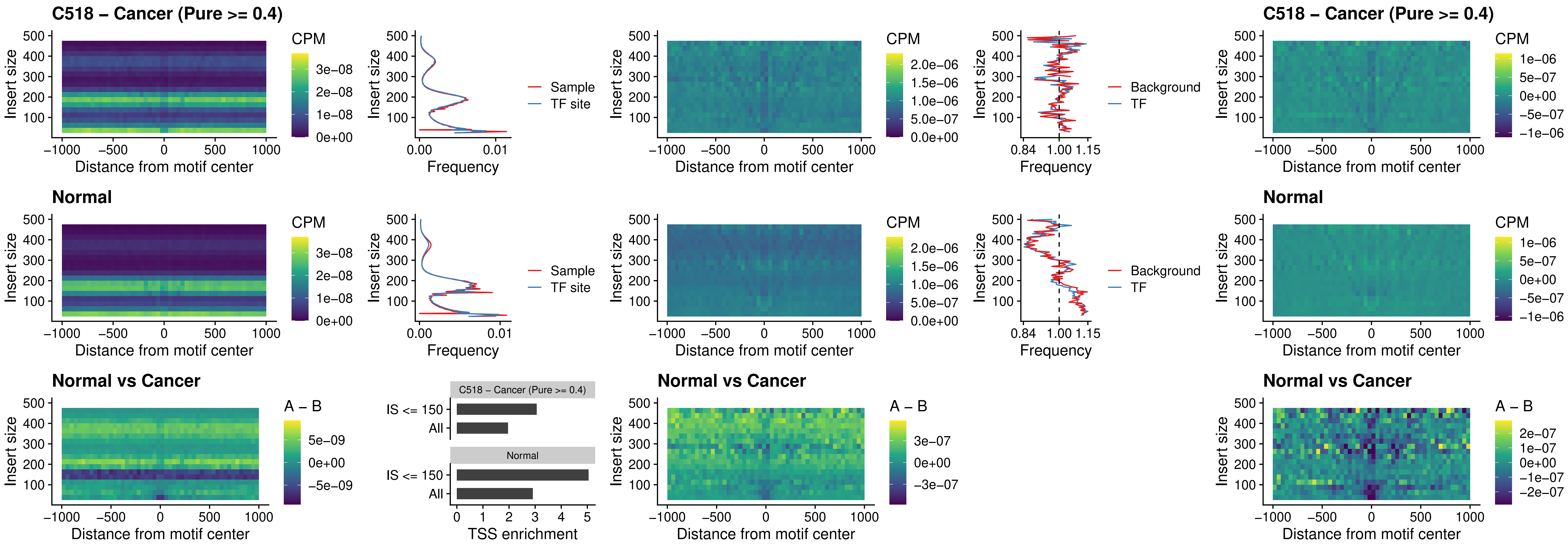

#### C518 – Cancer (Pure >= 0.4) – ZNF683_uc_cTSS_oPEAK

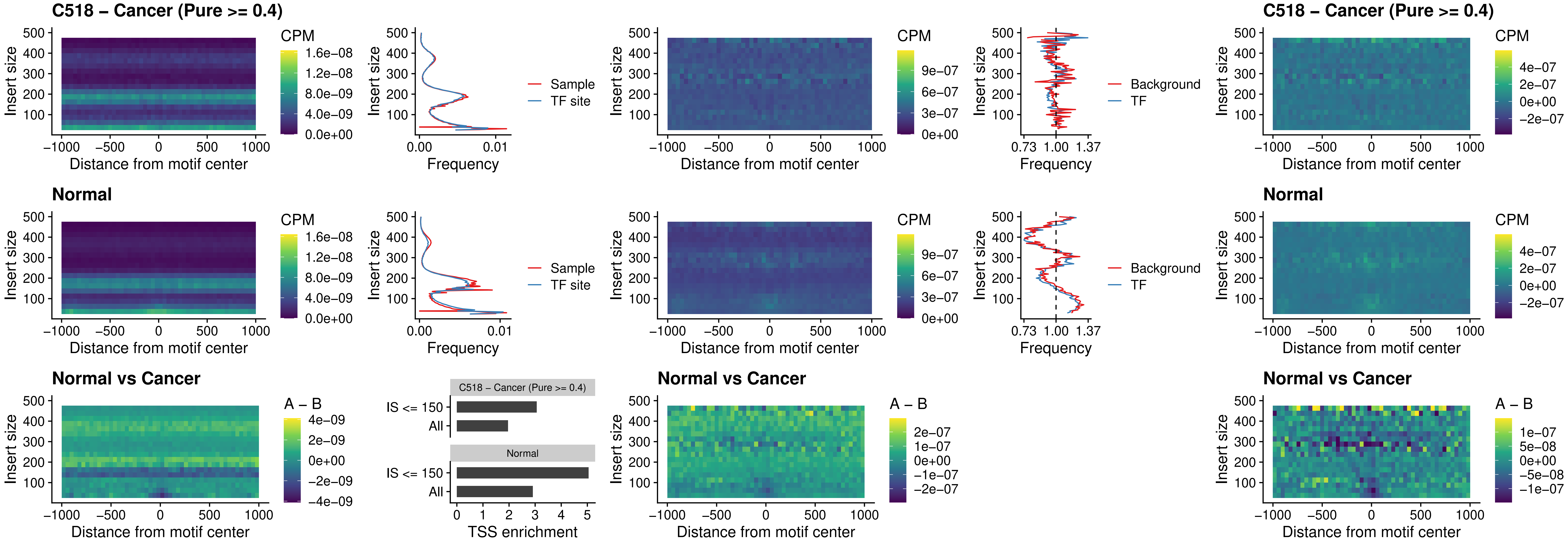

### C518 – Cancer (Pure >= 0.4) – IRF8_uc_dTSS_nPEAK

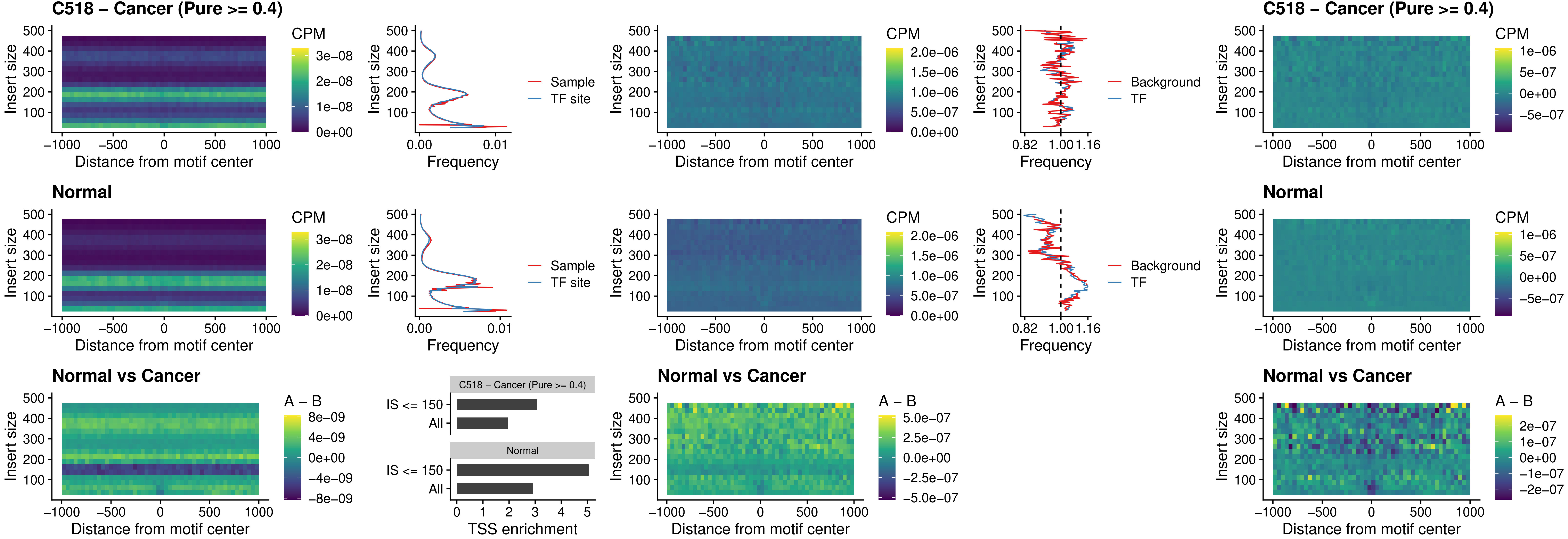

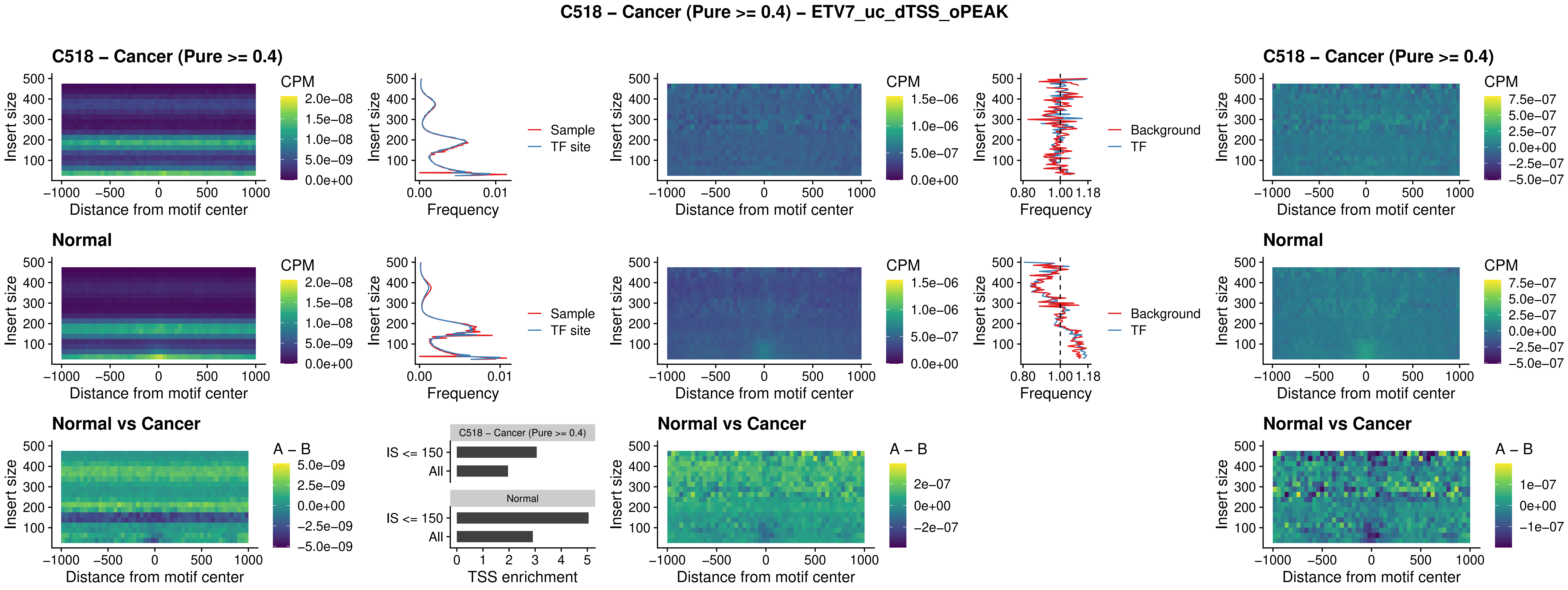

C518 – Cancer (Pure >= 0.4) – ETV7_uc_dTSS_oPEAK
C518 – Cancer (Pure >= 0.4)
Normal
Normal vs Cancer
Insert size
Distance from motif center
Frequency
CPM
A – B
Sample
TF site
Background
TF
C518 – Cancer (Pure >= 0.4)
Normal
IS <= 150
All
TSS enrichment

### C524 – Cancer (Pure >= 0.4) – FOXL1_uc_pTSS_oPEAK

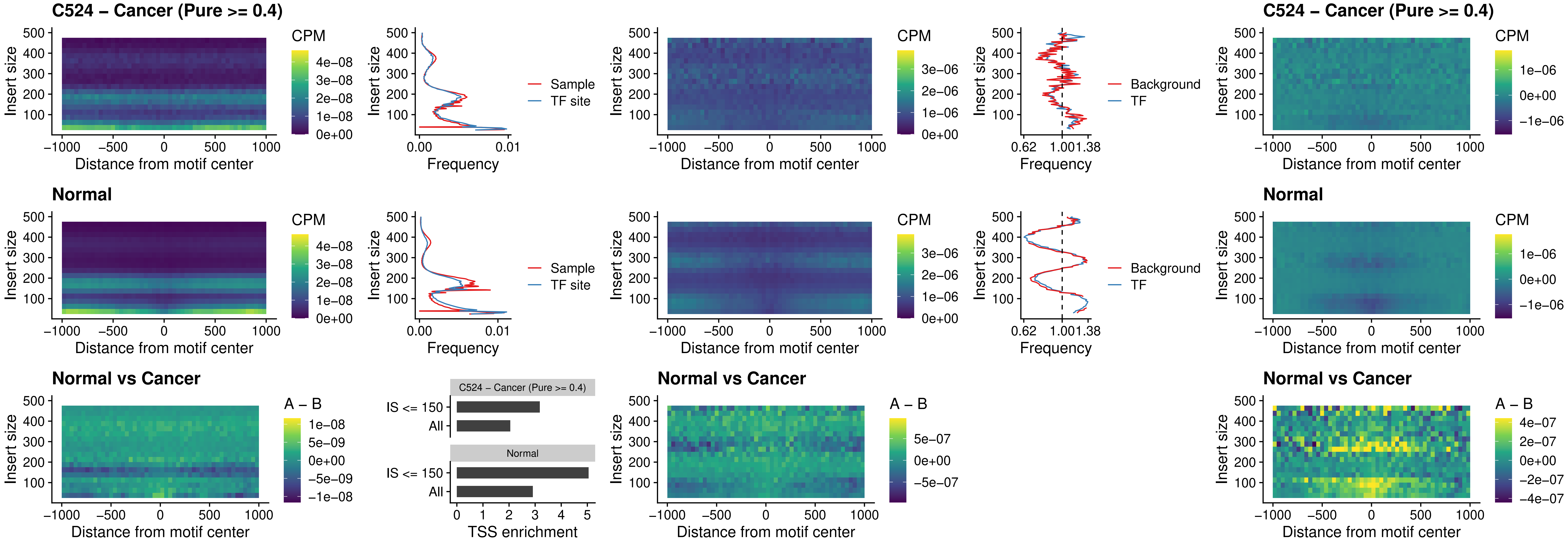

#### C524 – Cancer (Pure >= 0.4) – CPEB1_uc_cTSS_oPEAK

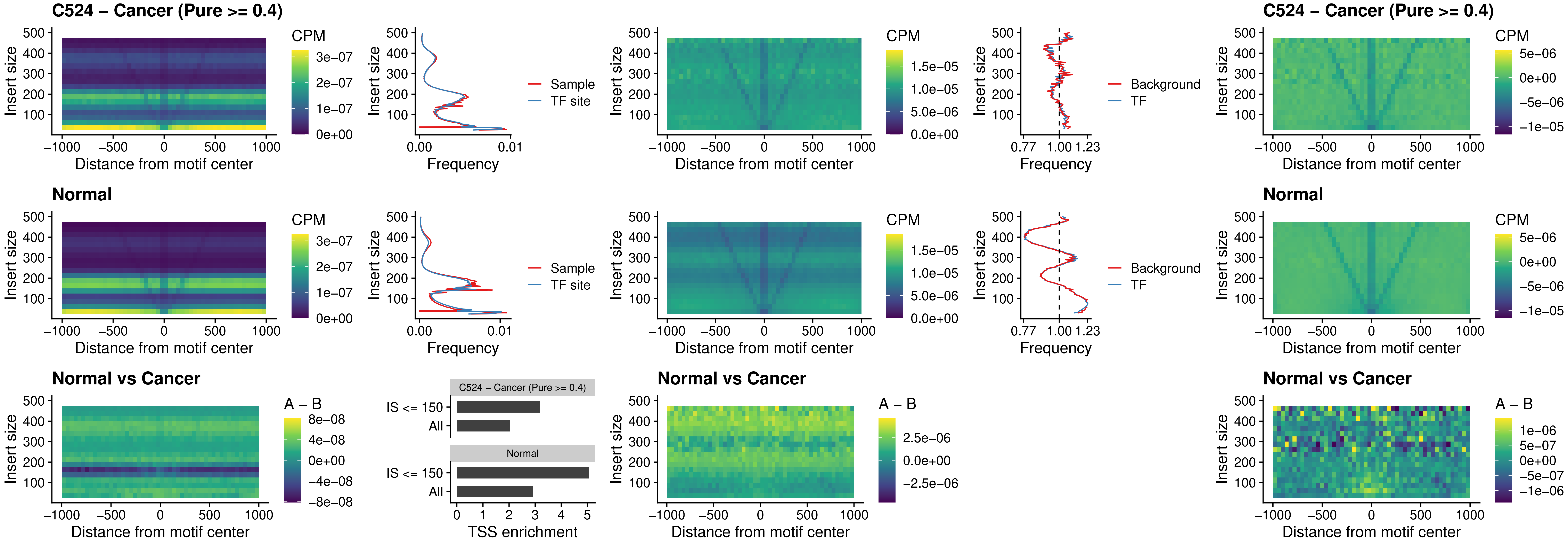

### C524 – Cancer (Pure >= 0.4) – HMGA1_uc_pTSS_nPEAK

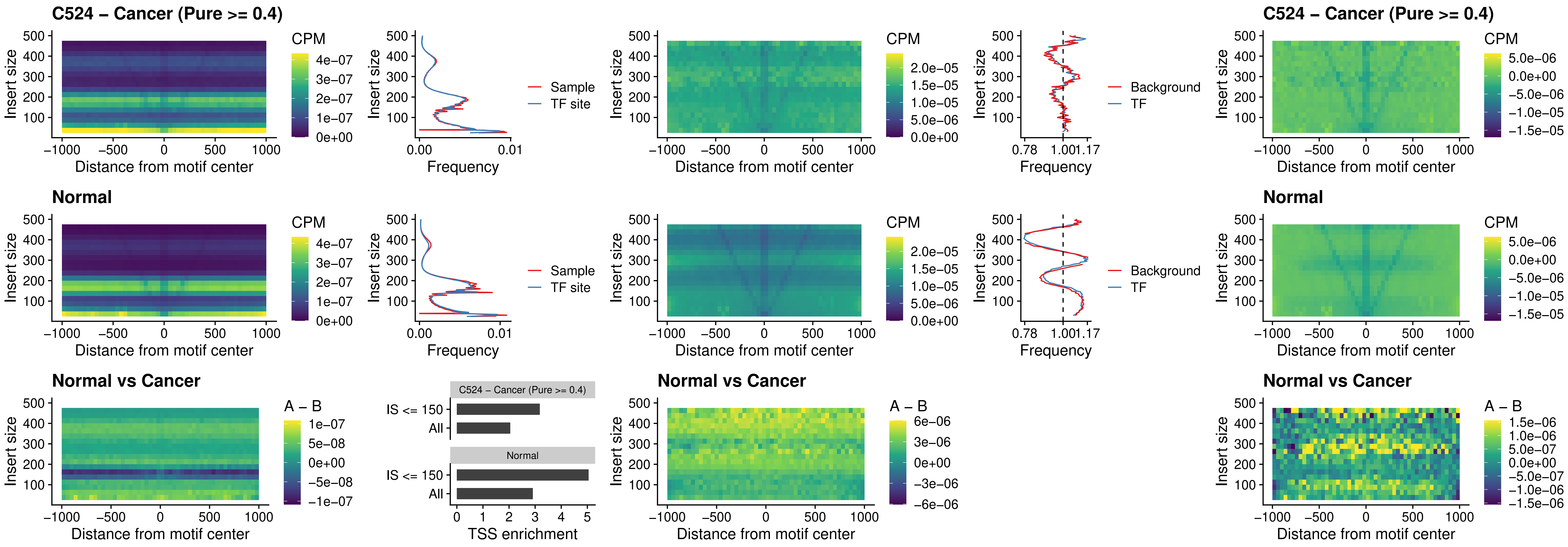

### C524 – Cancer (Pure >= 0.4) – SOX13_uc_cTSS_nPEAK

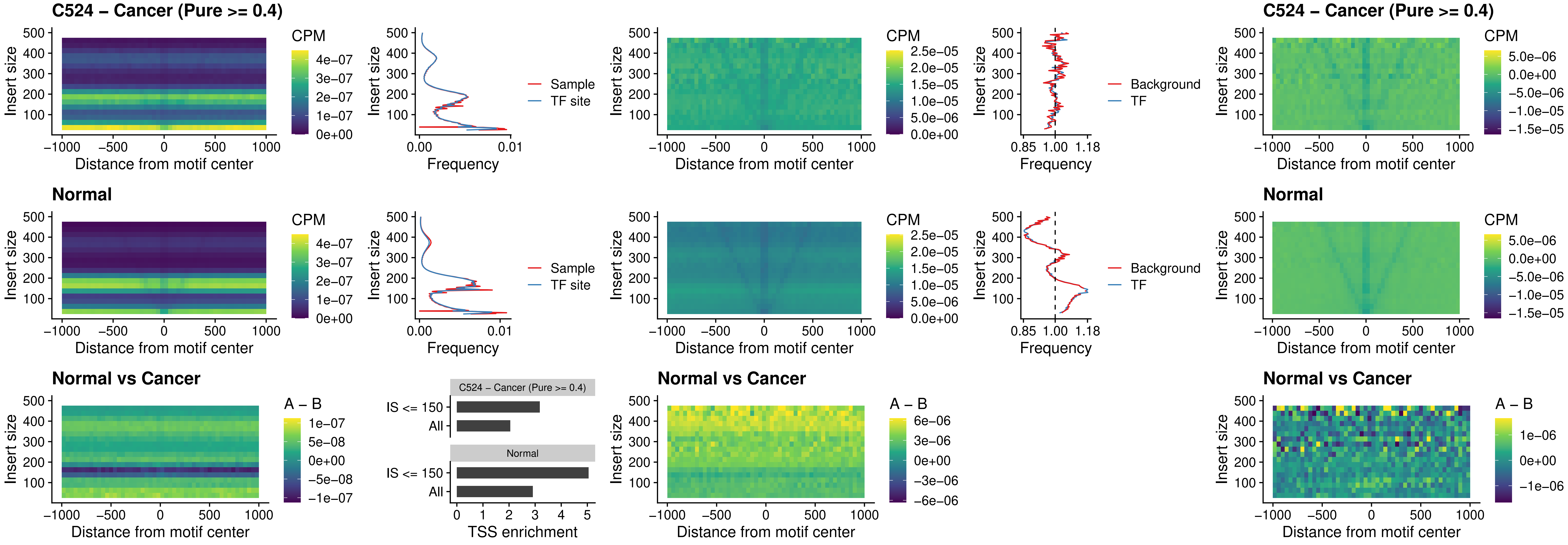

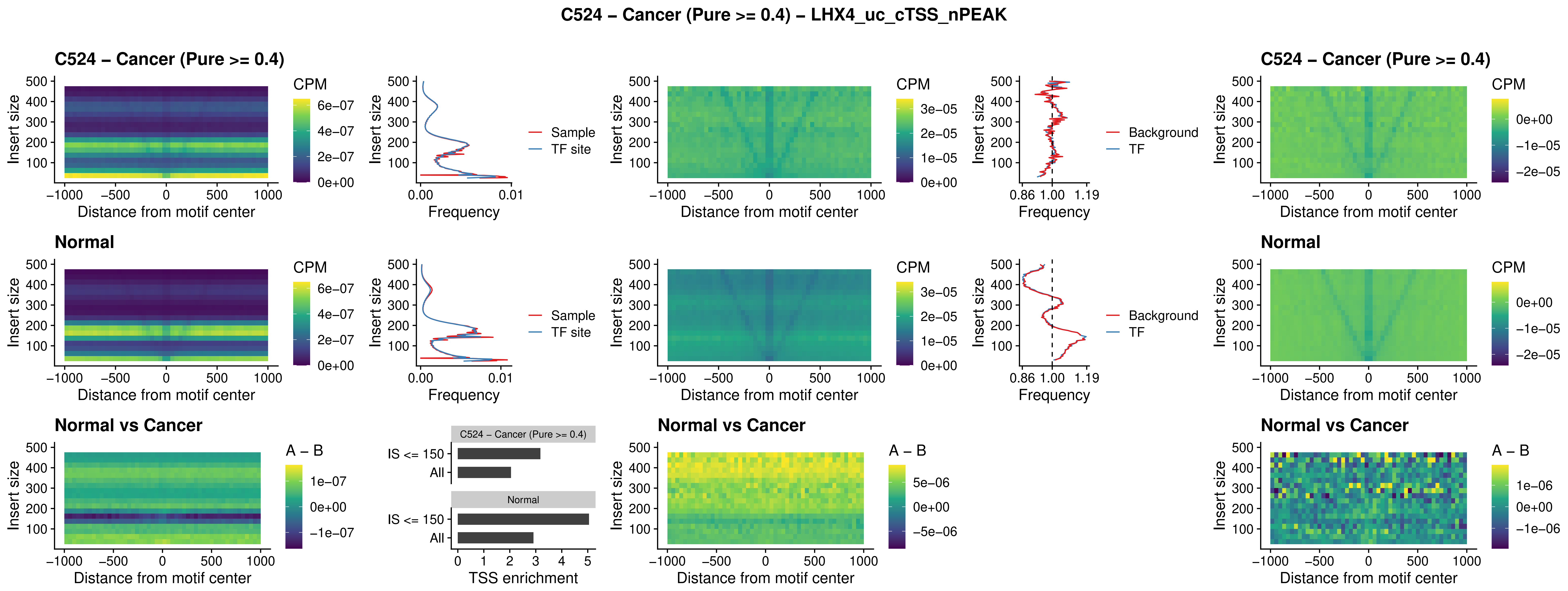
C524 – Cancer (Pure >= 0.4) – LHX4_uc_cTSS_nPEAK
C524 – Cancer (Pure >= 0.4)
Normal
Normal vs Cancer
Insert size
Distance from motif center
CPM
A – B
Frequency
Sample
TF site
Background
TF
C524 – Cancer (Pure >= 0.4)
Normal
IS <= 150
All
TSS enrichment

#### C524 – Cancer (Pure >= 0.4) – HOXA4_uc_cTSS_nPEAK

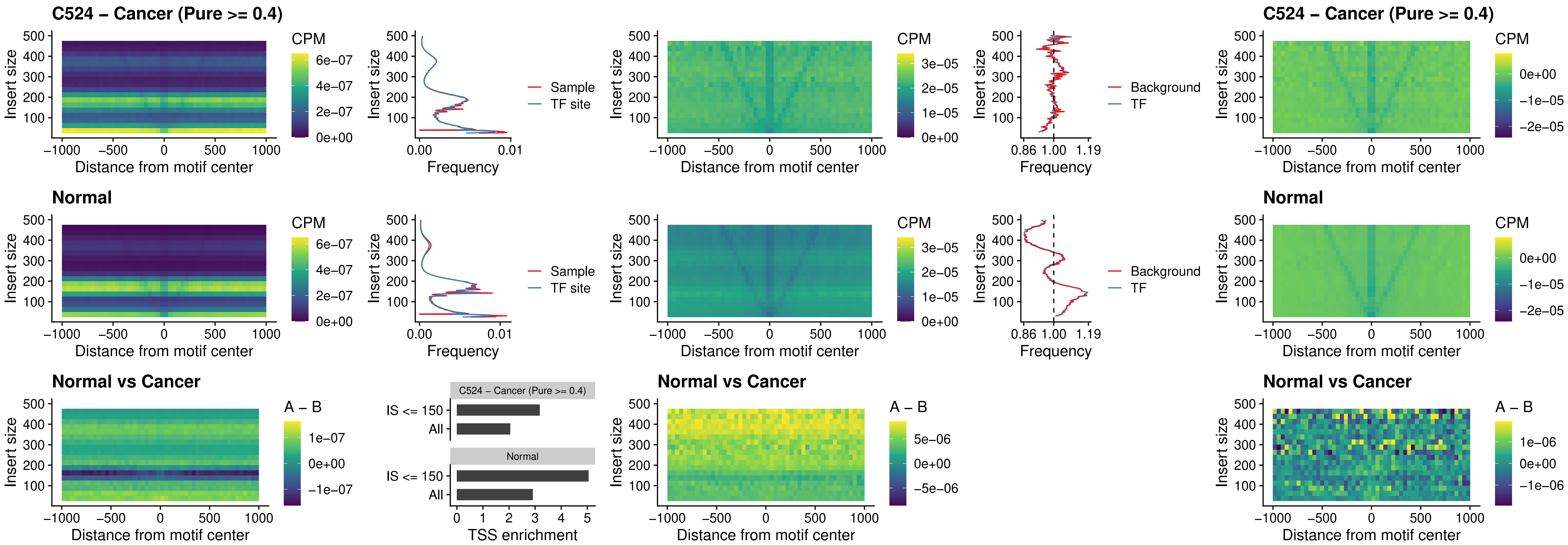

#### C524 – Cancer (Pure >= 0.4) – HLX_uc_cTSS_nPEAK

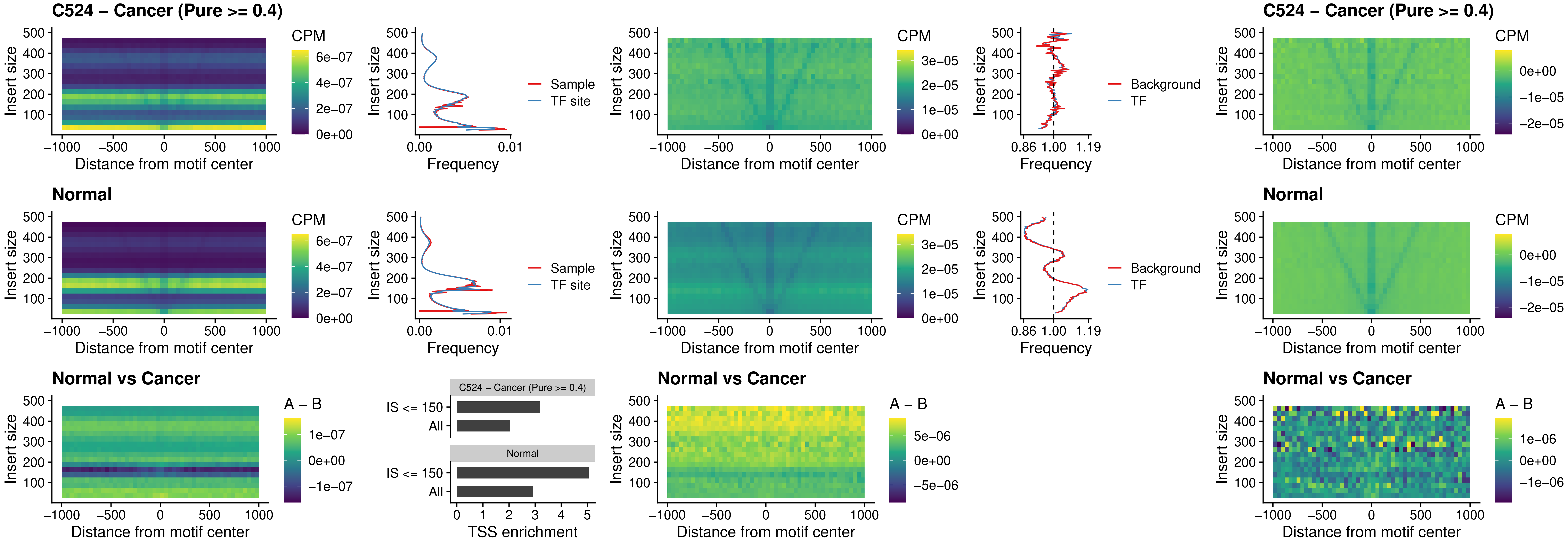

### C524 – Cancer (Pure >= 0.4) – AC0125311_uc_cTSS_nPEAK

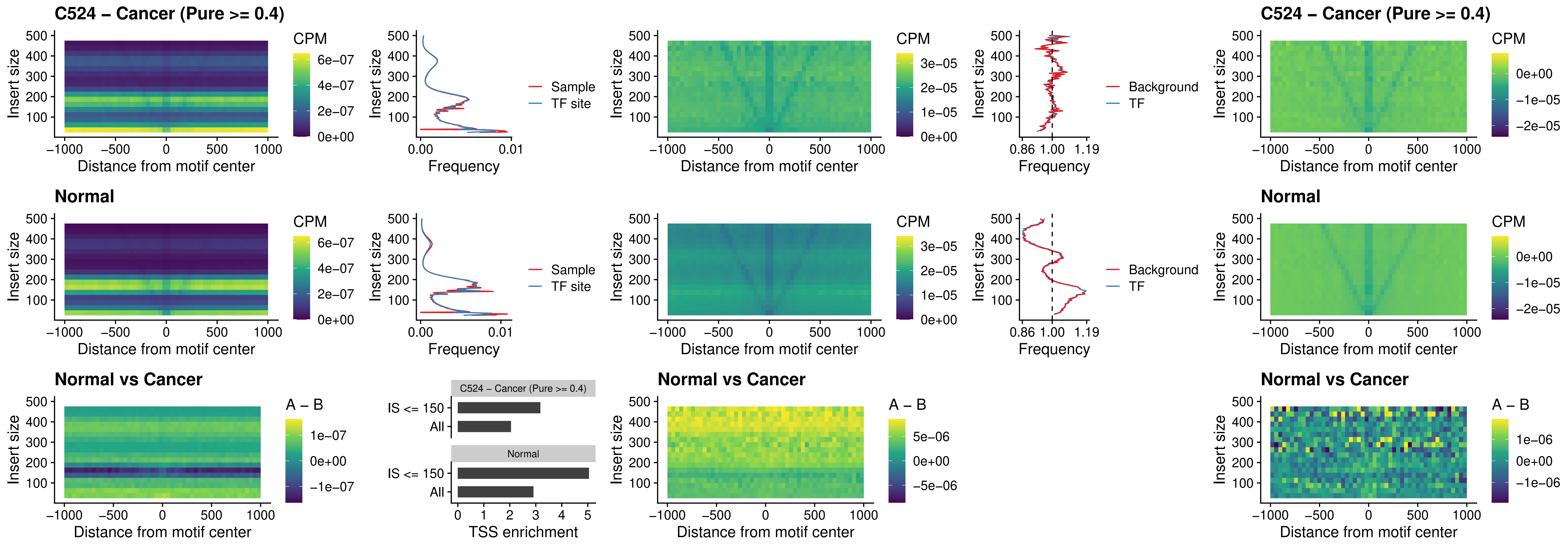

### C524 – Cancer (Pure >= 0.4) – DBX2_uc_cTSS_nPEAK

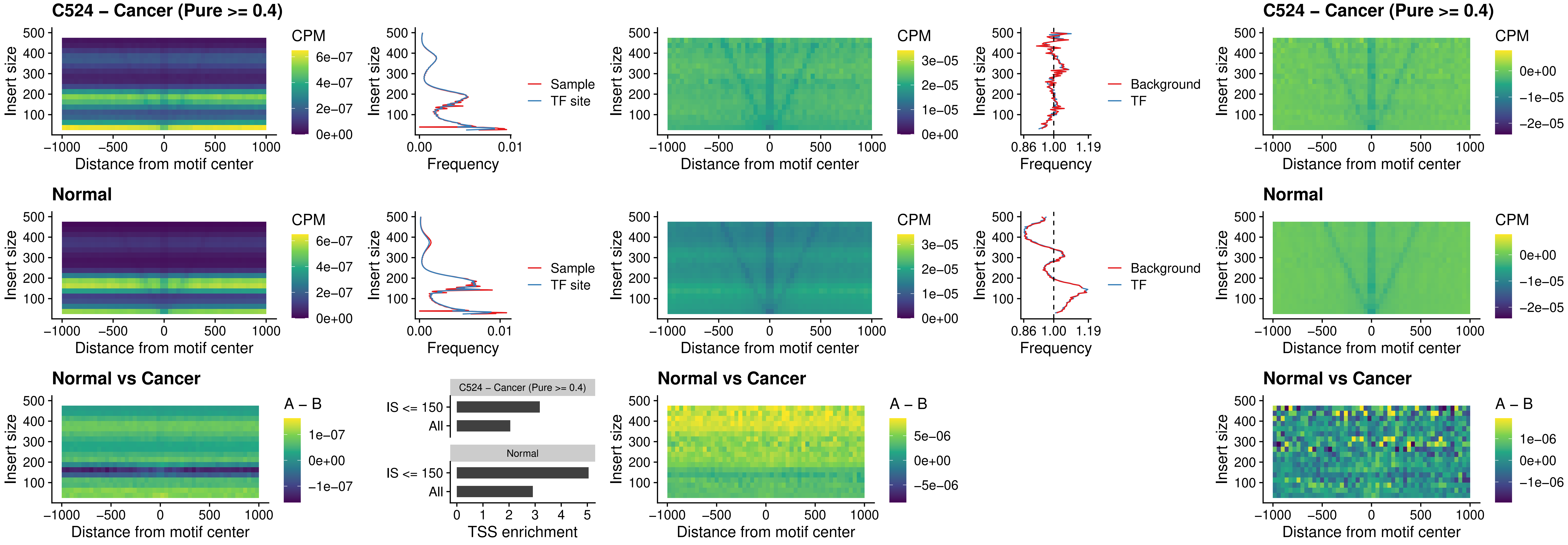

### C524 – Cancer (Pure >= 0.4) – LHX6_uc_dTSS_nPEAK

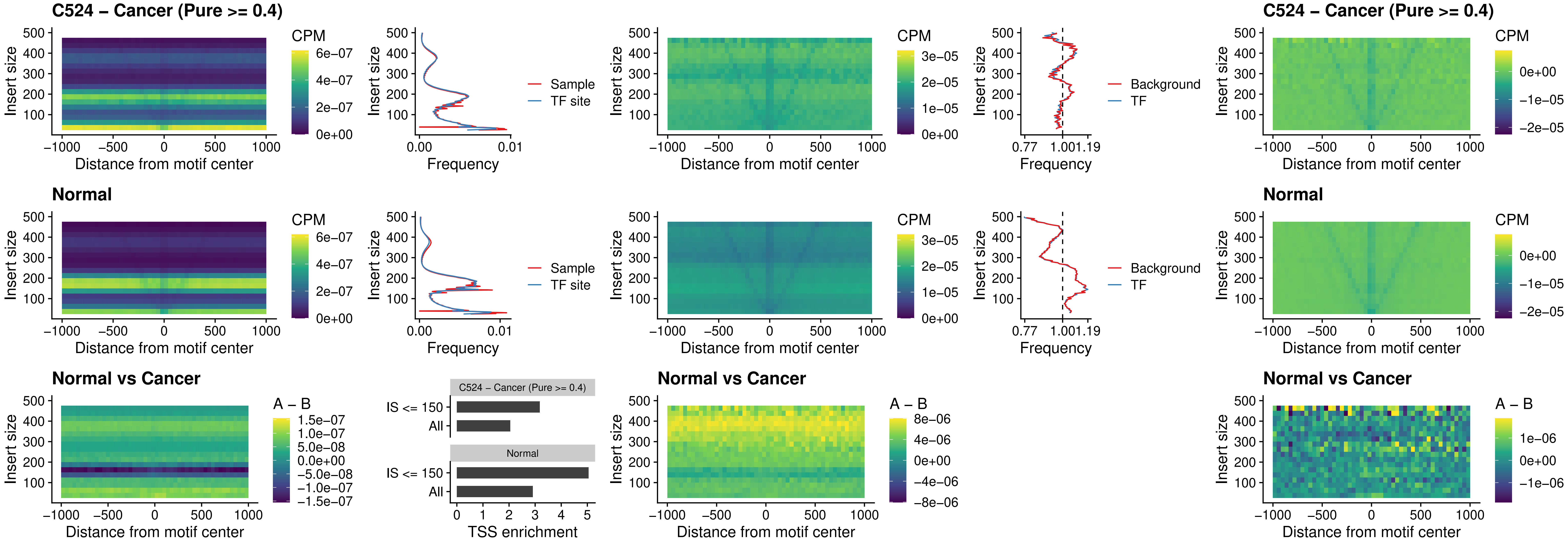

### C524 – Cancer (Pure >= 0.4) – SOX1_uc_dTSS_nPEAK

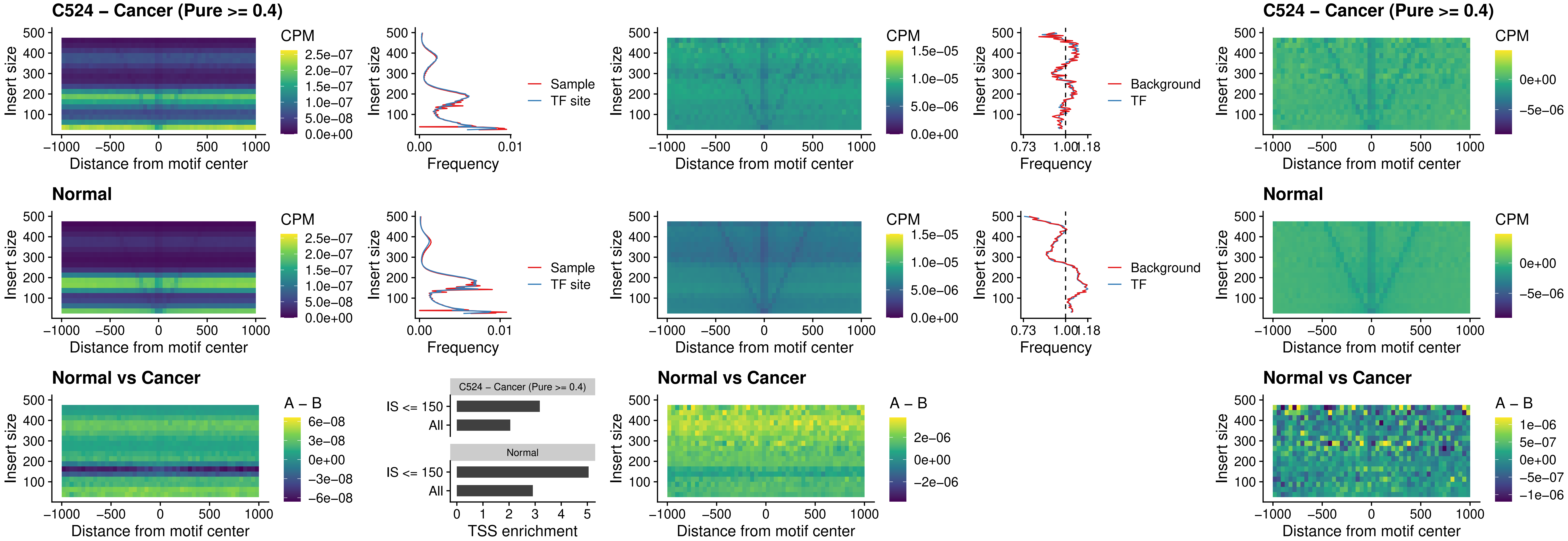

#### C524 – Cancer (Pure >= 0.4) – NKX24_uc_dTSS_nPEAK

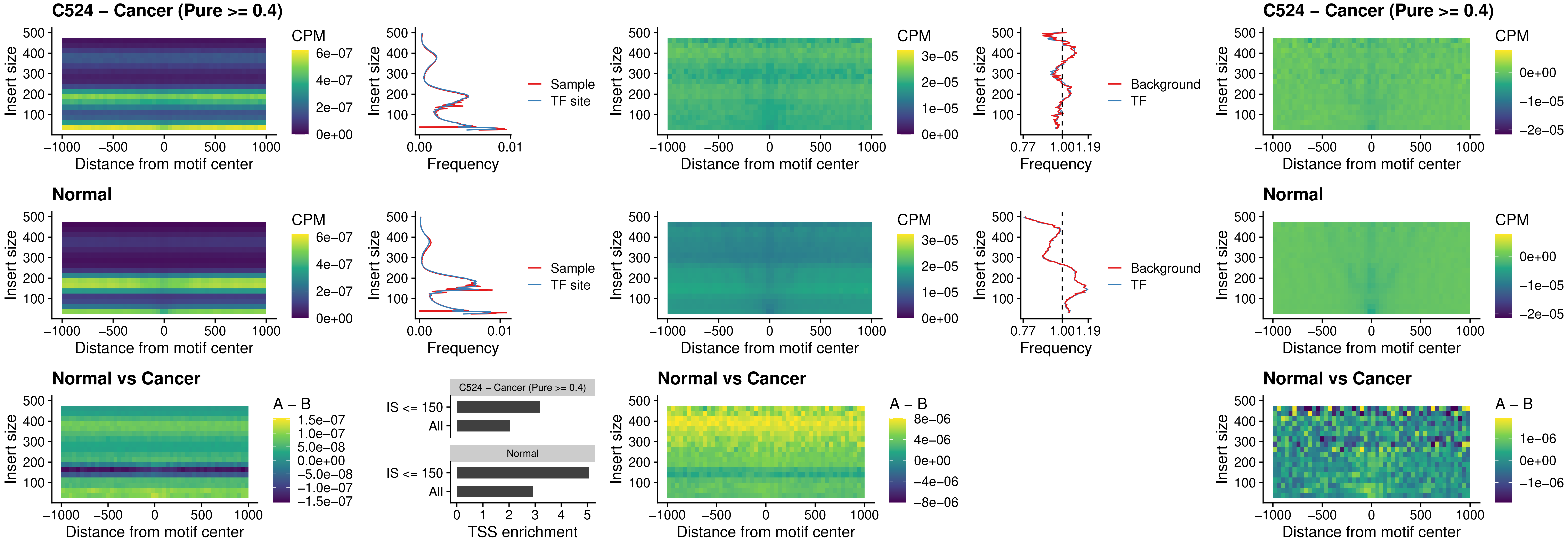

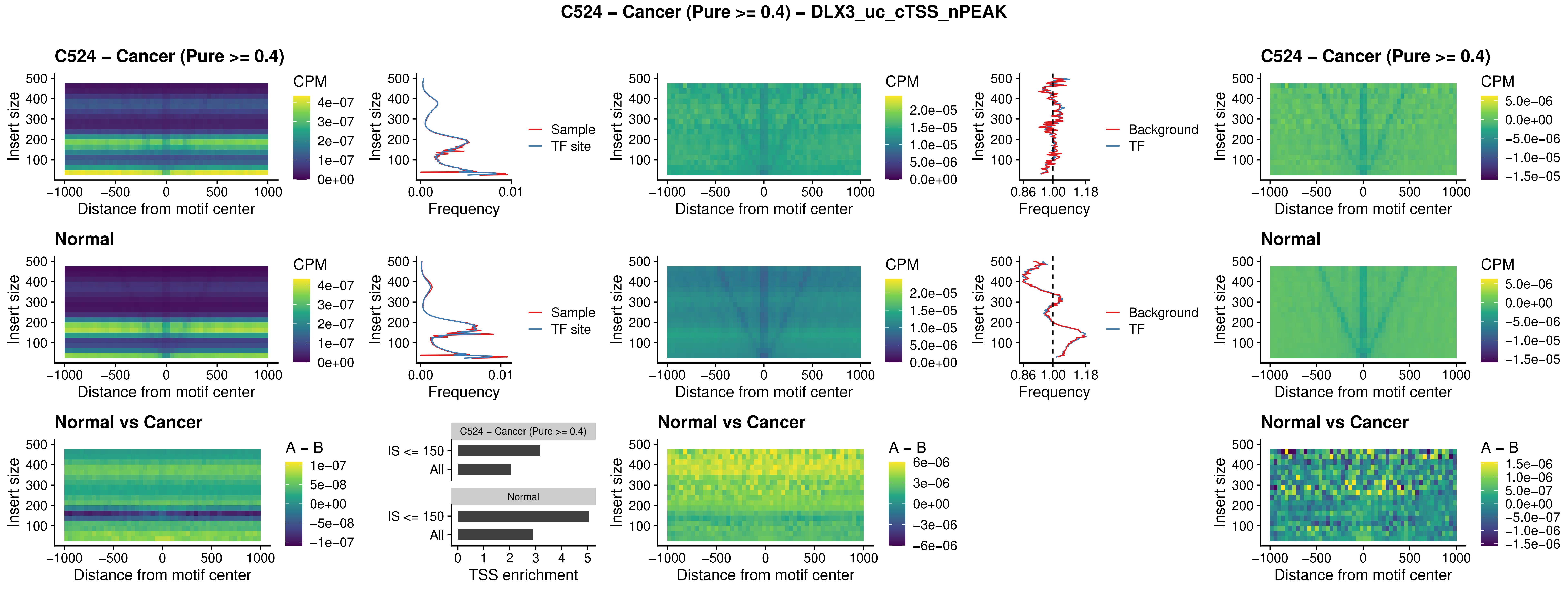
C524 – Cancer (Pure >= 0.4) – DLX3_uc_cTSS_nPEAK
C524 – Cancer (Pure >= 0.4)
Normal
Normal vs Cancer
Insert size
Distance from motif center
CPM
A – B
Frequency
Sample
TF site
Background
TF
C524 – Cancer (Pure >= 0.4)
Normal
IS <= 150
All
TSS enrichment

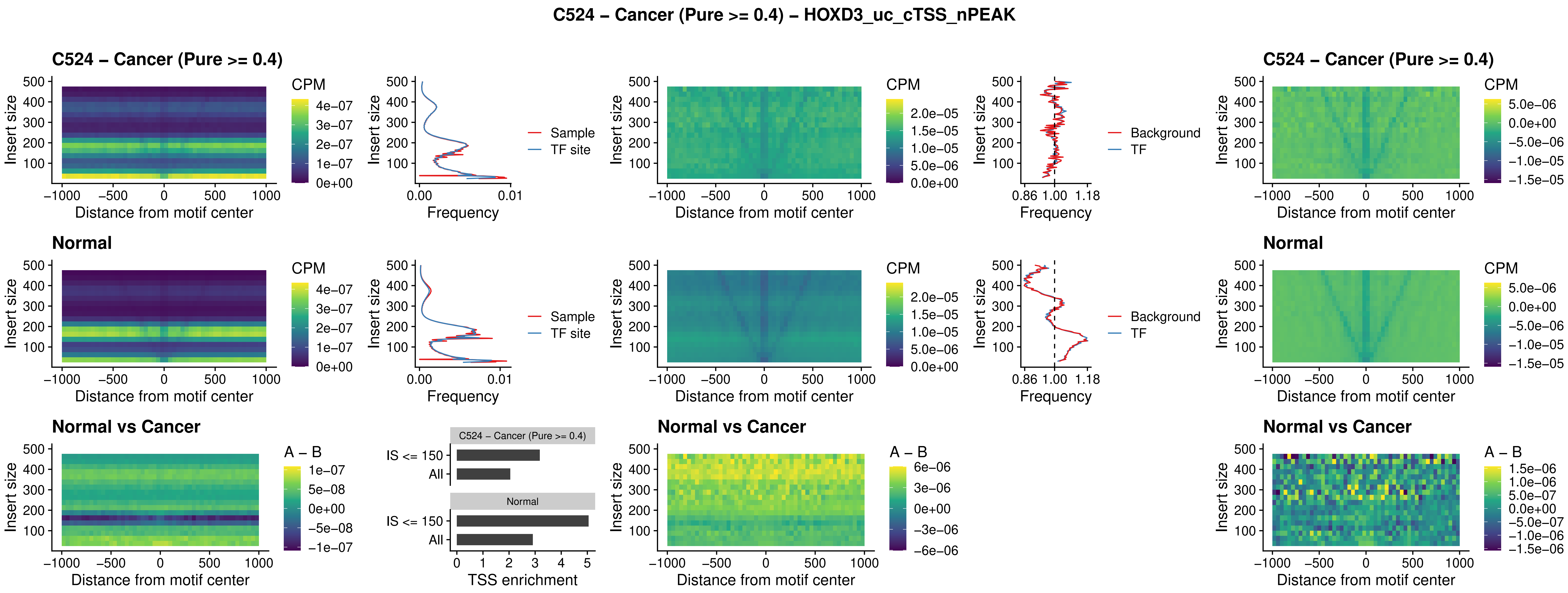
C524 – Cancer (Pure >= 0.4) – HOXD3_uc_cTSS_nPEAK
C524 – Cancer (Pure >= 0.4)
Normal
Normal vs Cancer
Insert size
Distance from motif center
Frequency
CPM
A – B
Sample
TF site
Background
TF
C524 – Cancer (Pure >= 0.4)
Normal
IS <= 150
All
TSS enrichment

#### C524 – Cancer (Pure >= 0.4) – ISL1_uc_dTSS_oPEAK

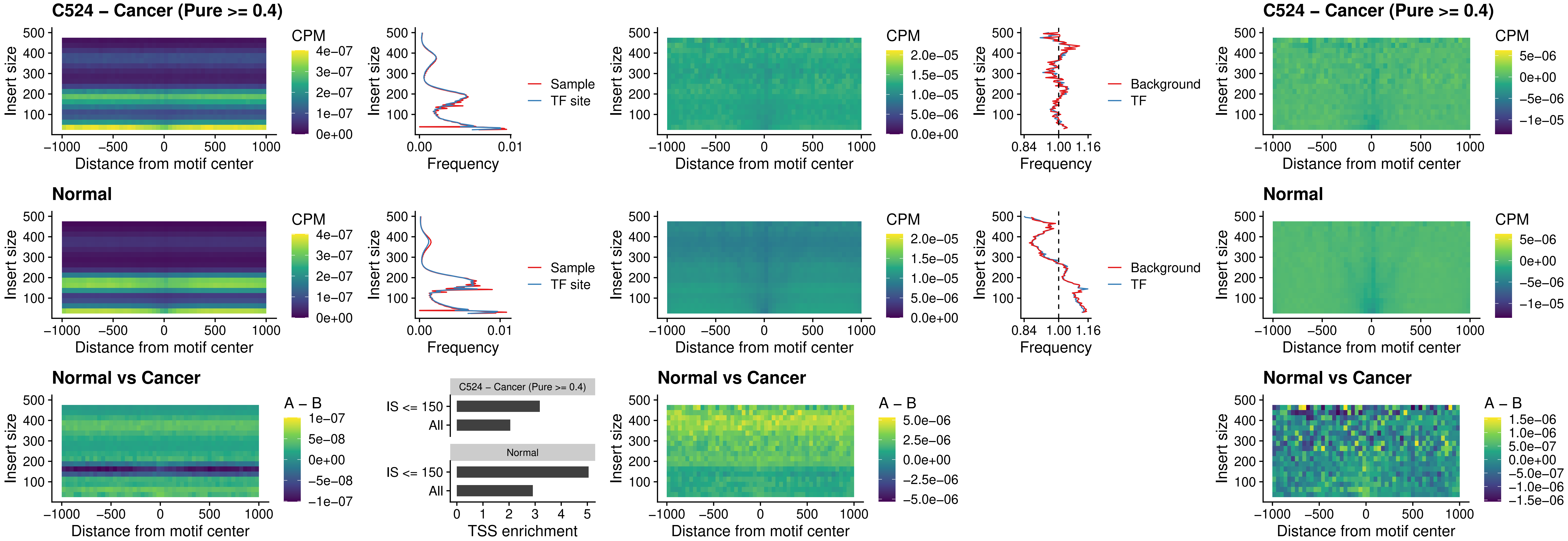

#### C524 – Cancer (Pure >= 0.4) – HOXB4_uc_dTSS_oPEAK

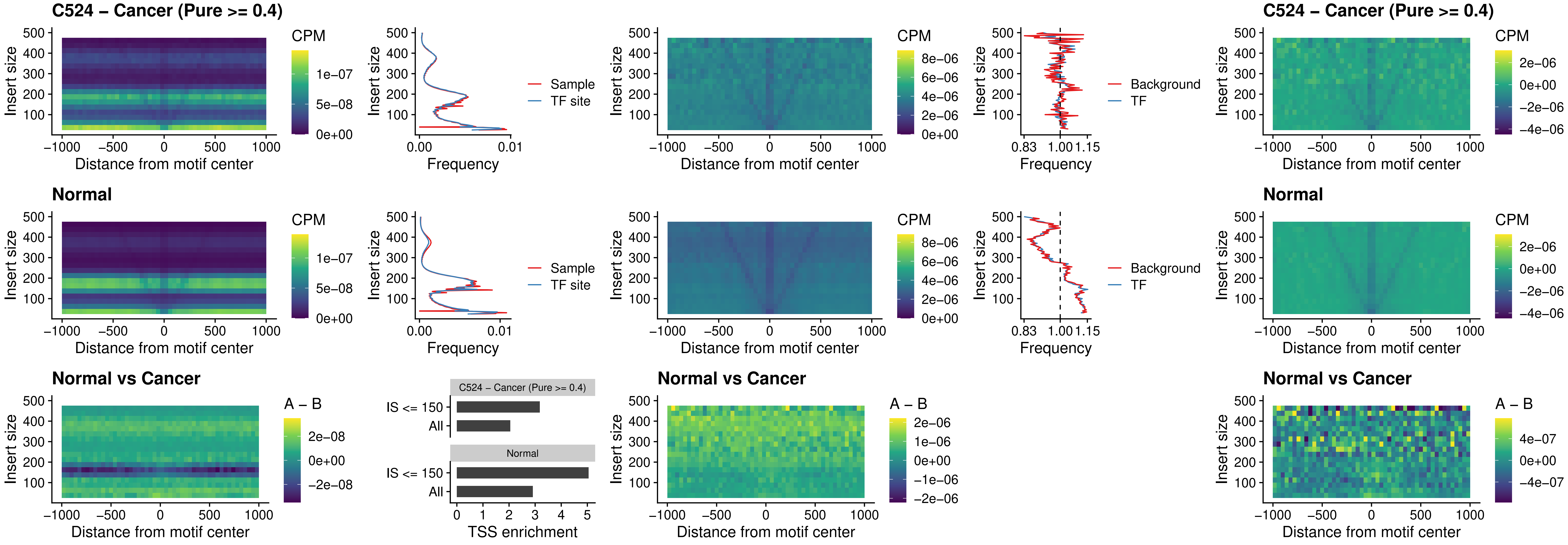

### C524 – Cancer (Pure >= 0.4) – VAX1_uc_dTSS_oPEAK

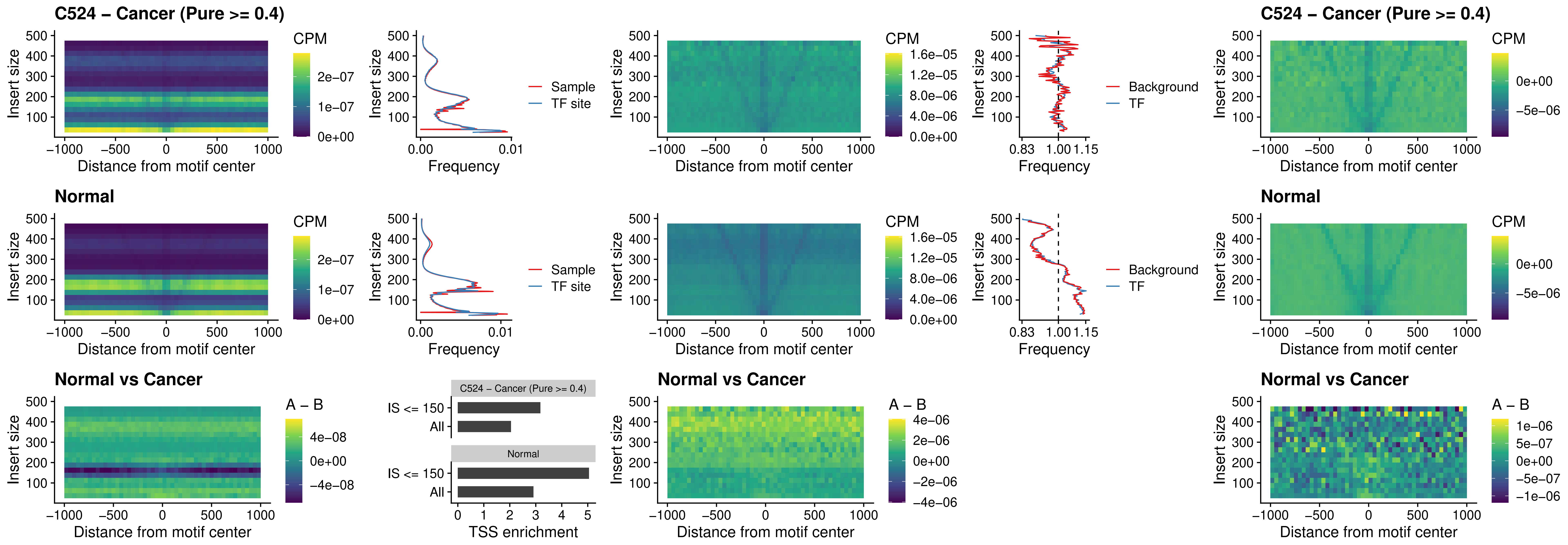

#### C524 – Cancer (Pure >= 0.4) – DLX3_uc_dTSS_oPEAK

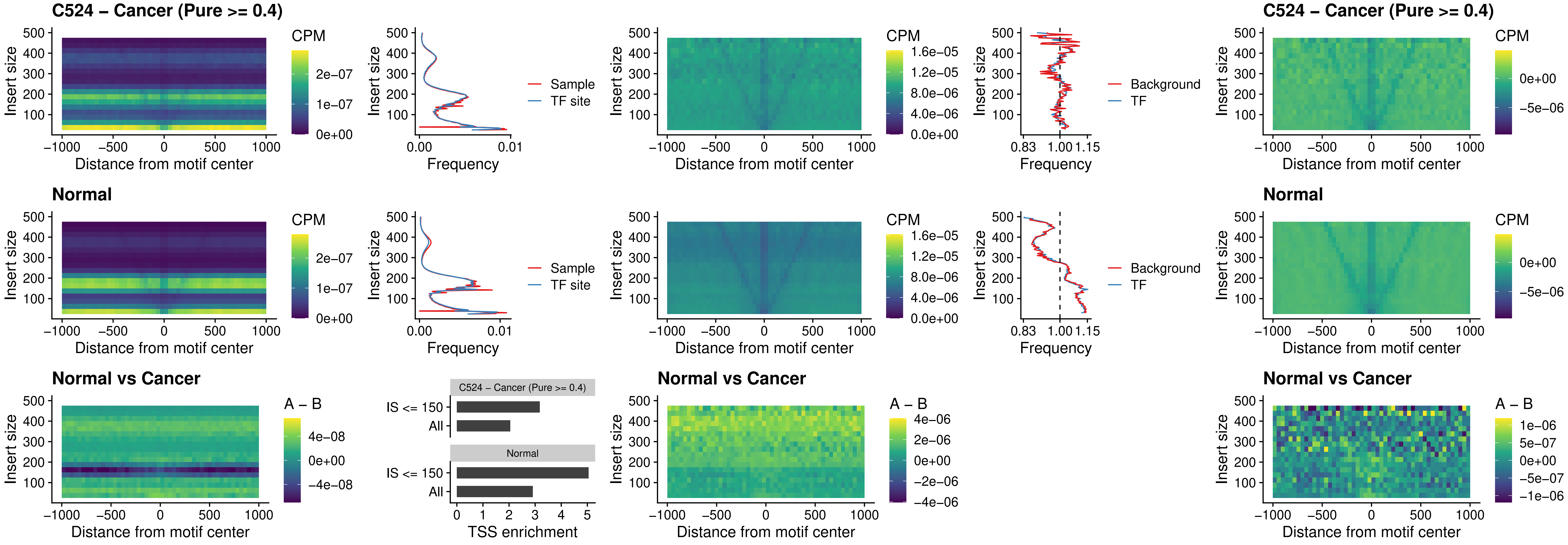

#### C524 – Cancer (Pure >= 0.4) – HOXD3_uc_dTSS_oPEAK

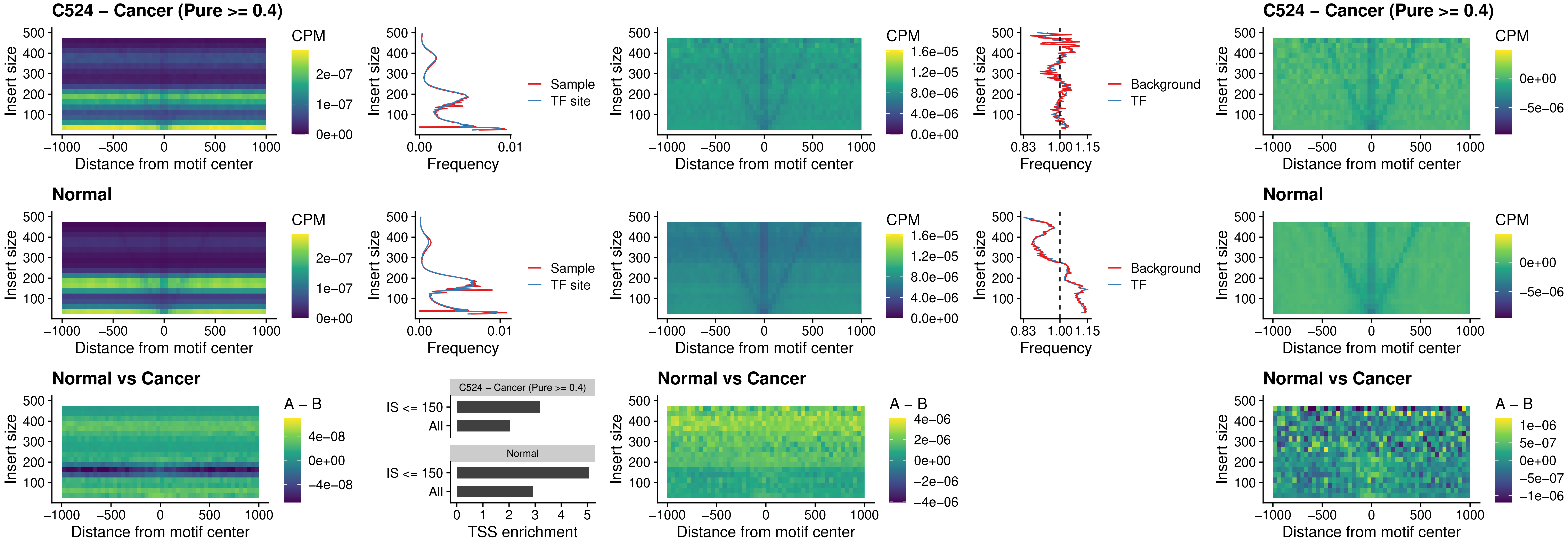

#### C524 – Cancer (Pure >= 0.4) – BBX_uc_dTSS_nPEAK

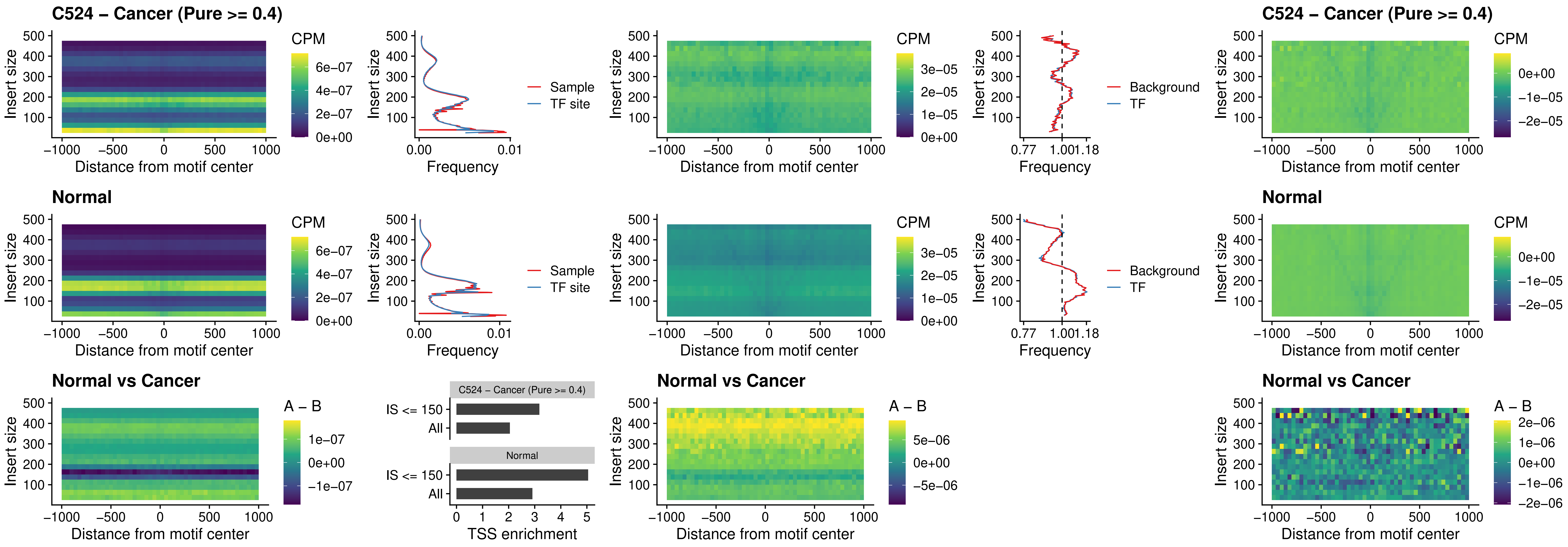

#### C524 – Cancer (Pure >= 0.4) – SOX3_uc_cTSS_nPEAK

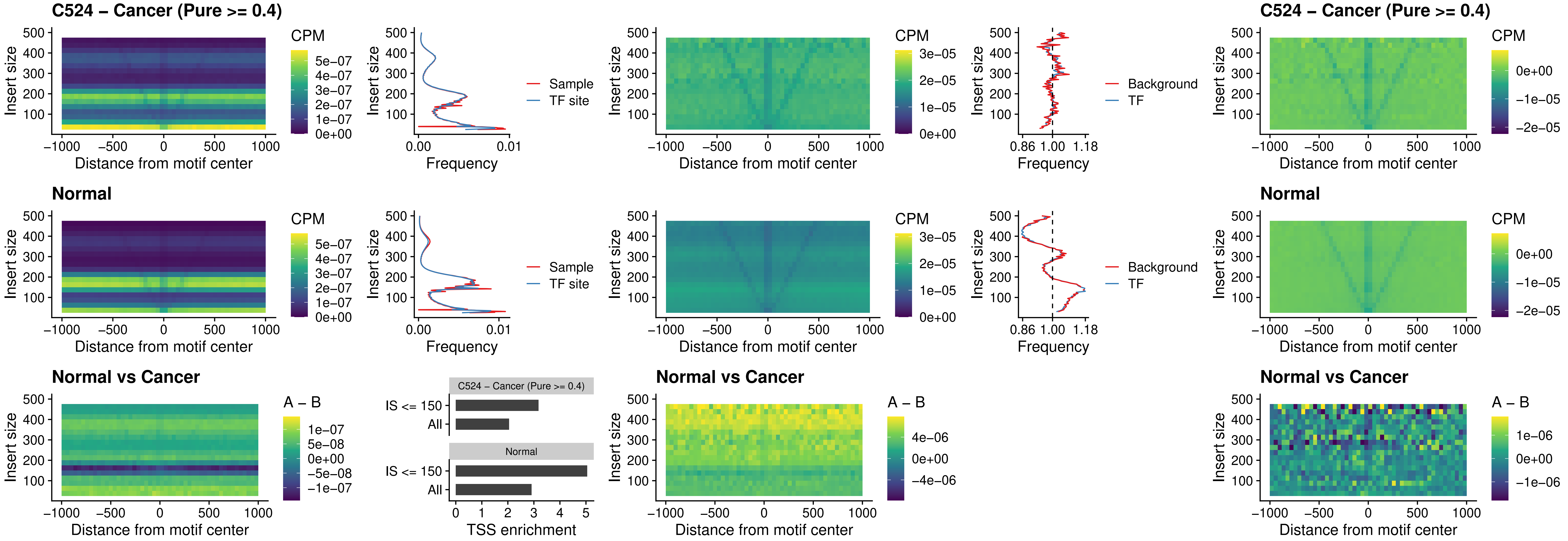

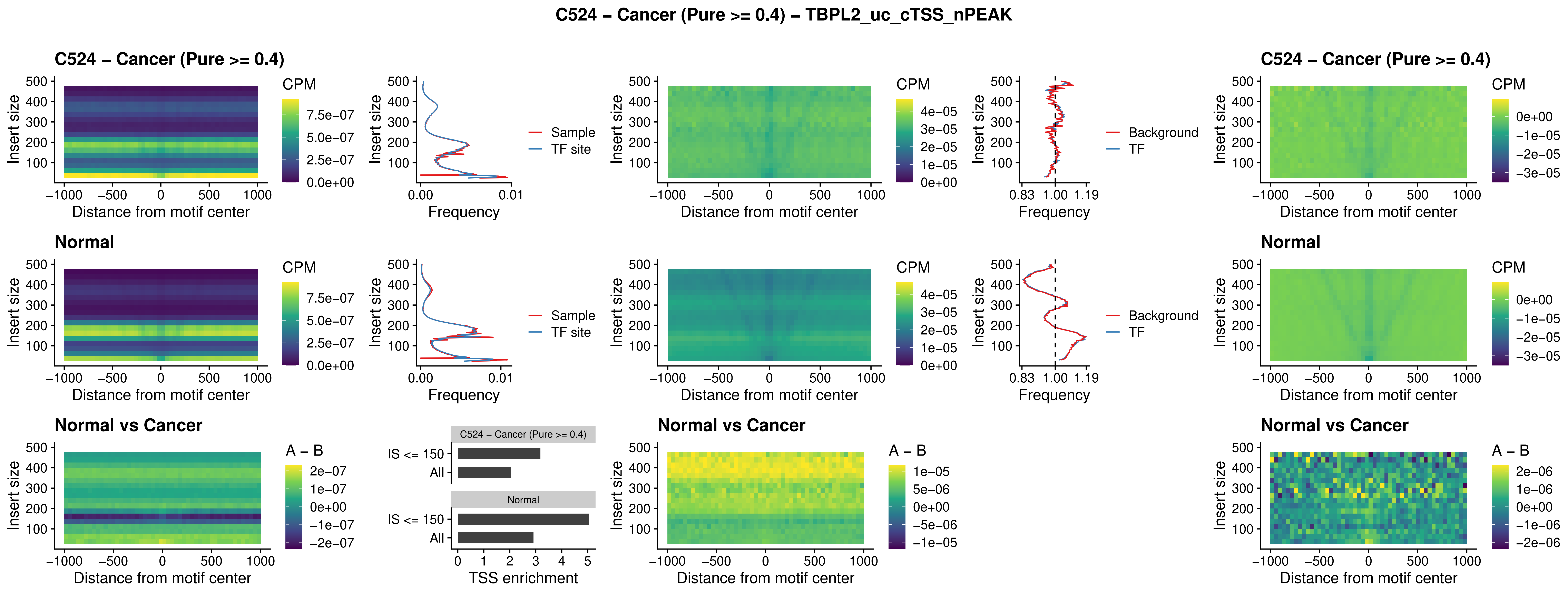
C524 – Cancer (Pure >= 0.4) – TBPL2_uc_cTSS_nPEAK
C524 – Cancer (Pure >= 0.4)
Normal
Normal vs Cancer
Insert size
Distance from motif center
Frequency
CPM
A – B
Sample
TF site
Background
TF
C524 – Cancer (Pure >= 0.4)
Normal
IS <= 150
All
TSS enrichment

#### C524 – Cancer (Pure >= 0.4) – HOXA4_uc_cTSS_oPEAK

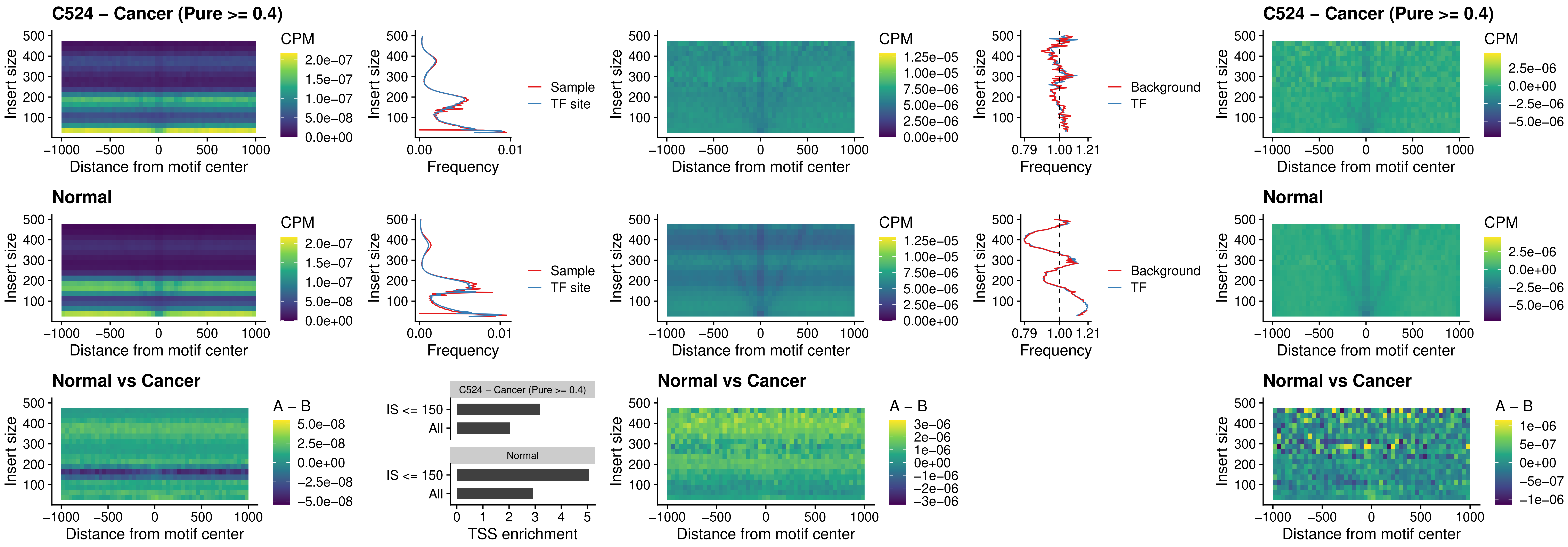

#### C524 – Cancer (Pure >= 0.4) – PDX1_uc_dTSS_oPEAK

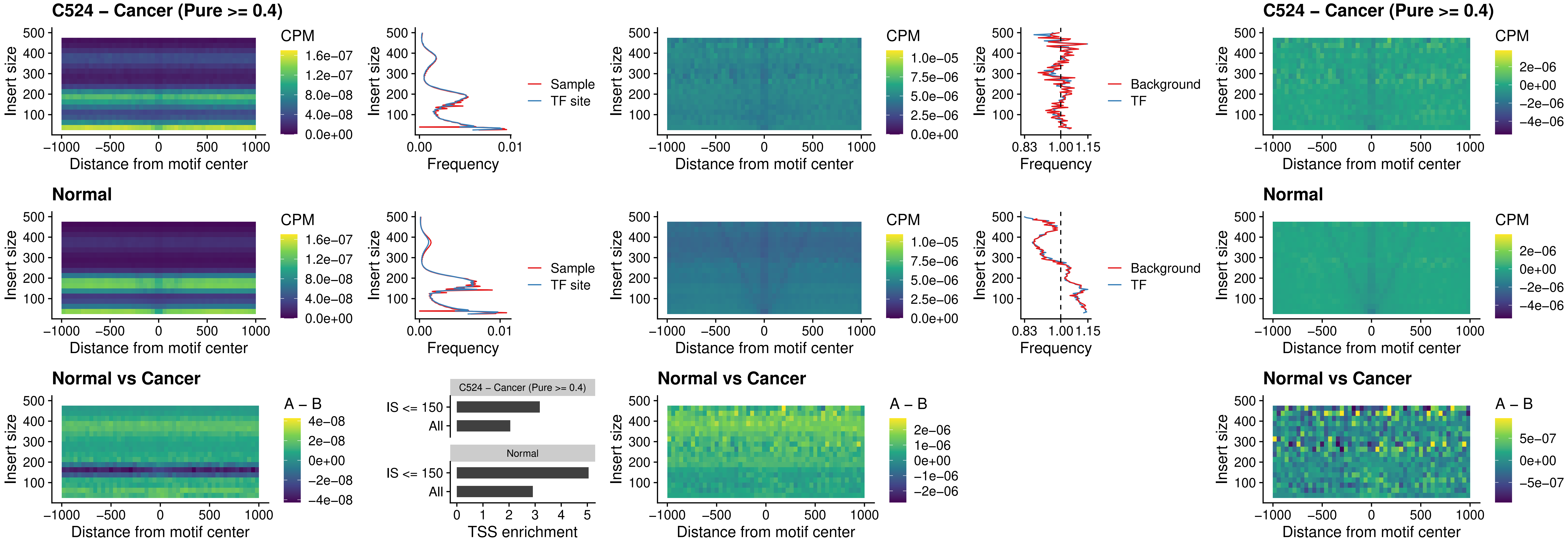

### C524 – Cancer (Pure >= 0.4) – LHX4_uc_dTSS_oPEAK

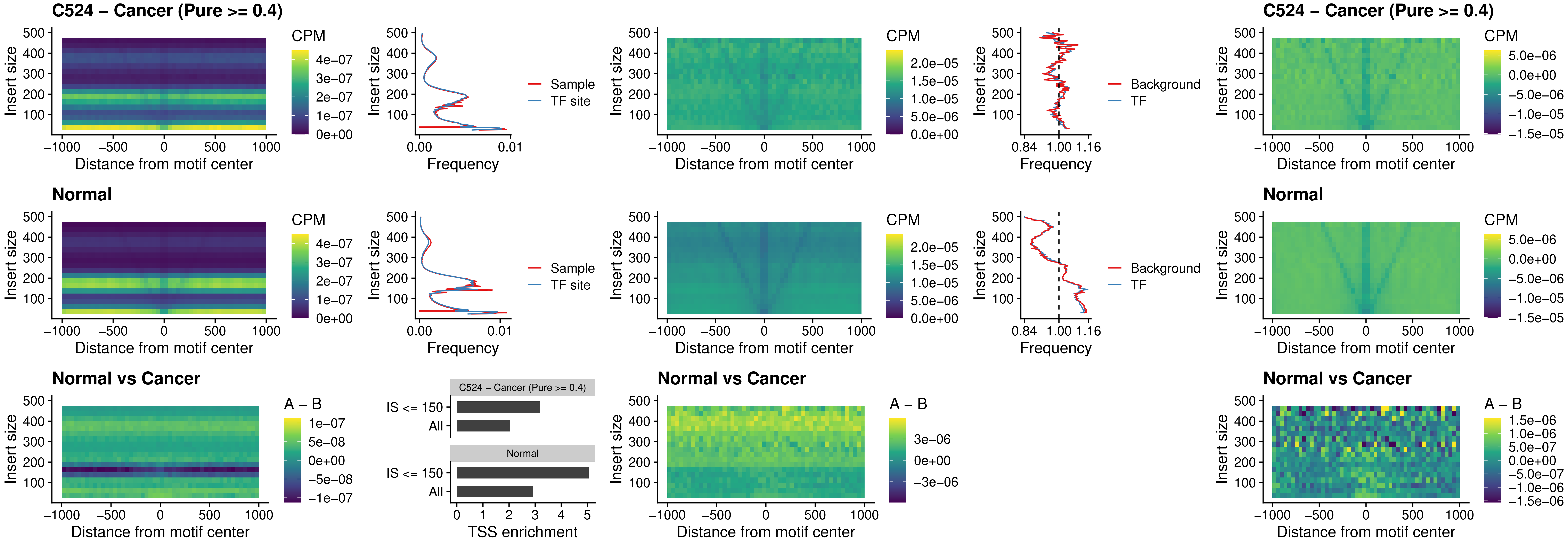

#### C524 – Cancer (Pure >= 0.4) – HOXA4_uc_dTSS_oPEAK

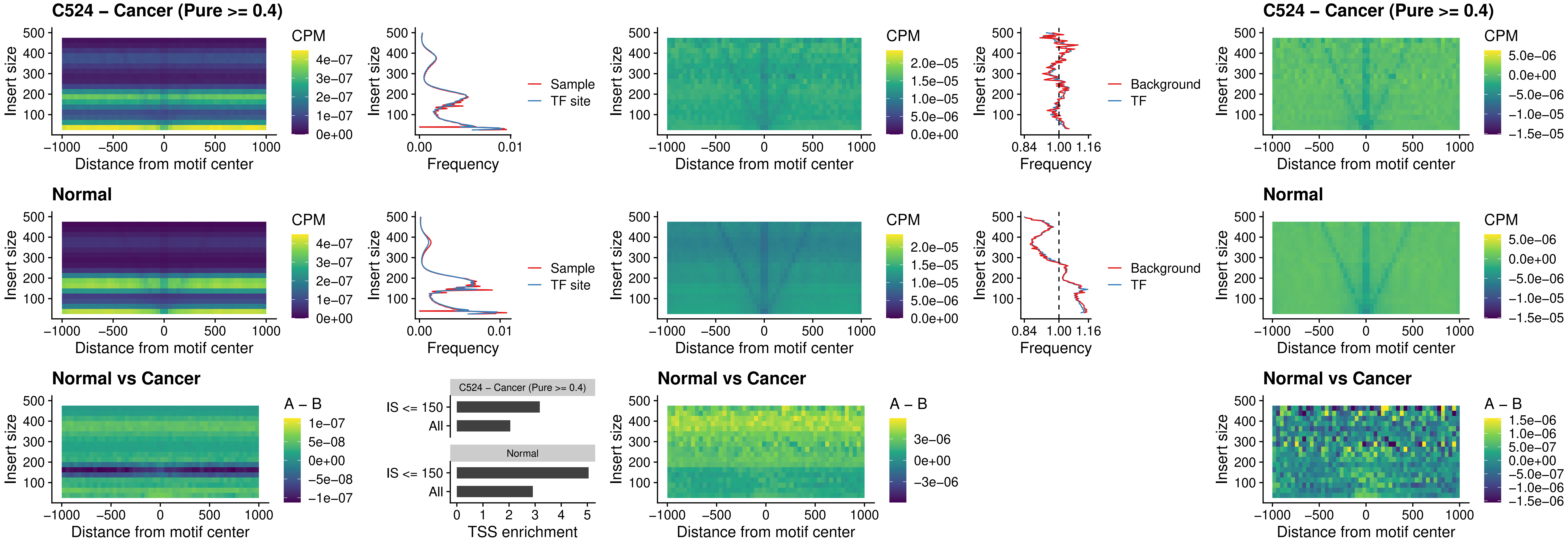

#### C524 – Cancer (Pure >= 0.4) – HLX_uc_dTSS_oPEAK

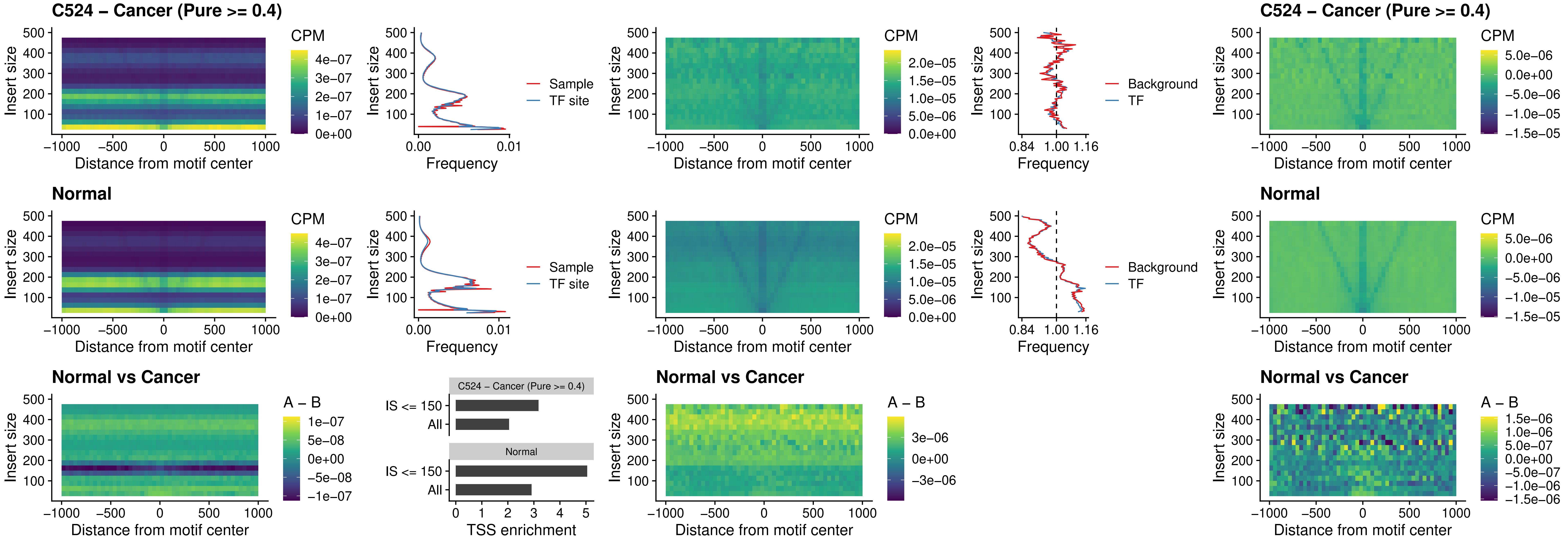

#### C524 – Cancer (Pure >= 0.4) – AC0125311_uc_dTSS_oPEAK

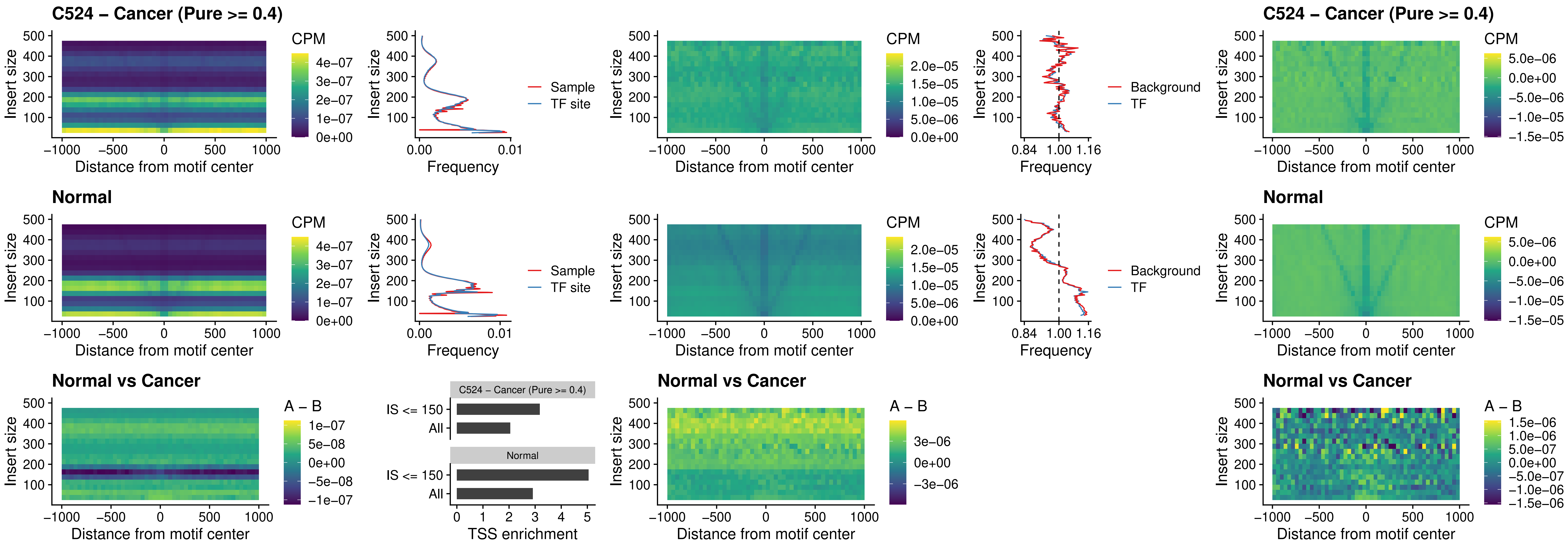

#### C524 – Cancer (Pure >= 0.4) – DBX2_uc_dTSS_oPEAK

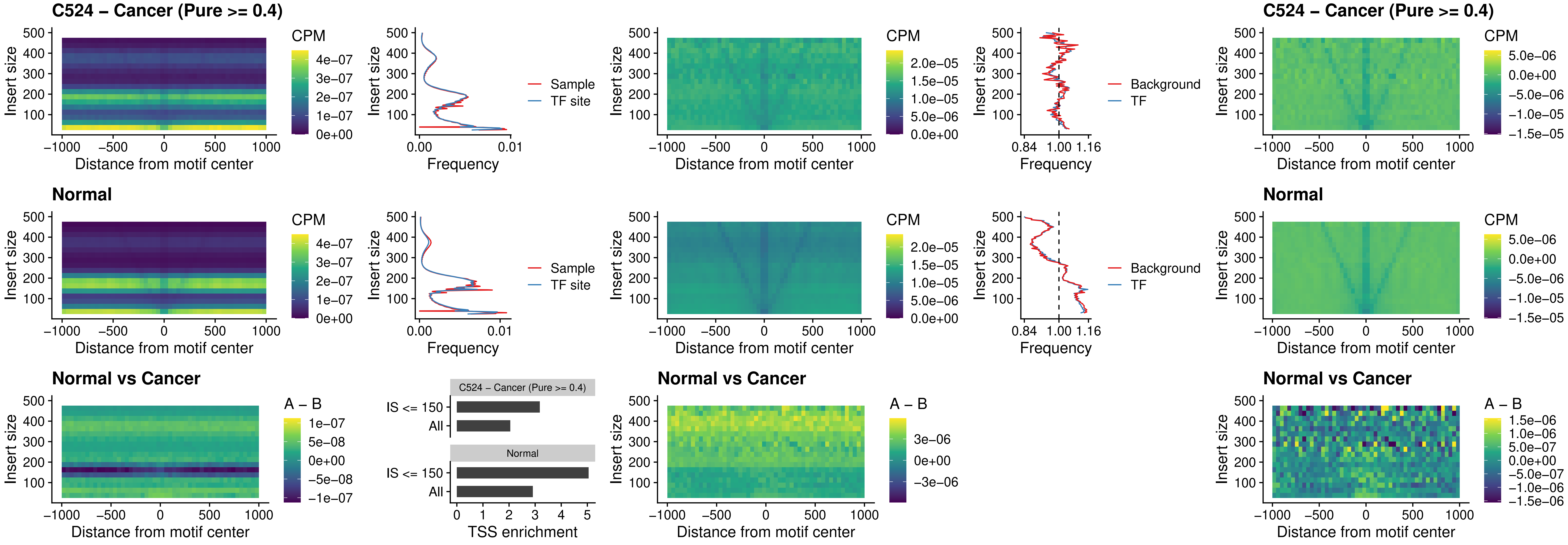

#### C524 – Cancer (Pure >= 0.4) – HOXA2_uc_dTSS_nPEAK

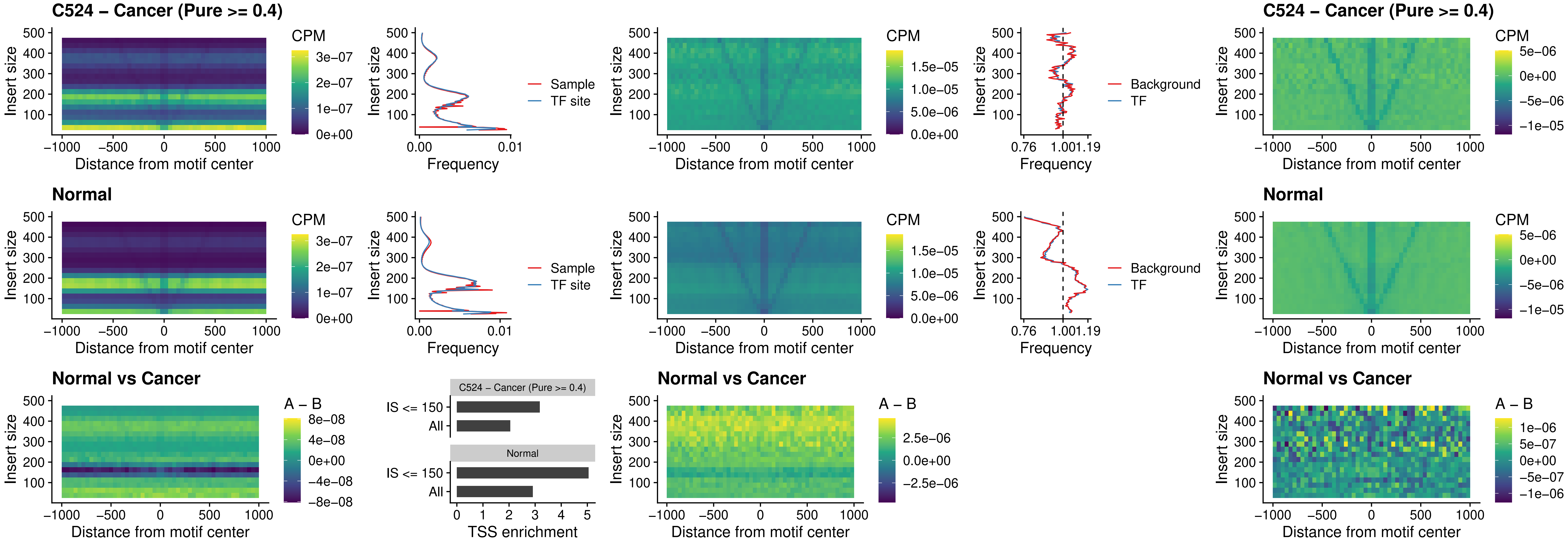

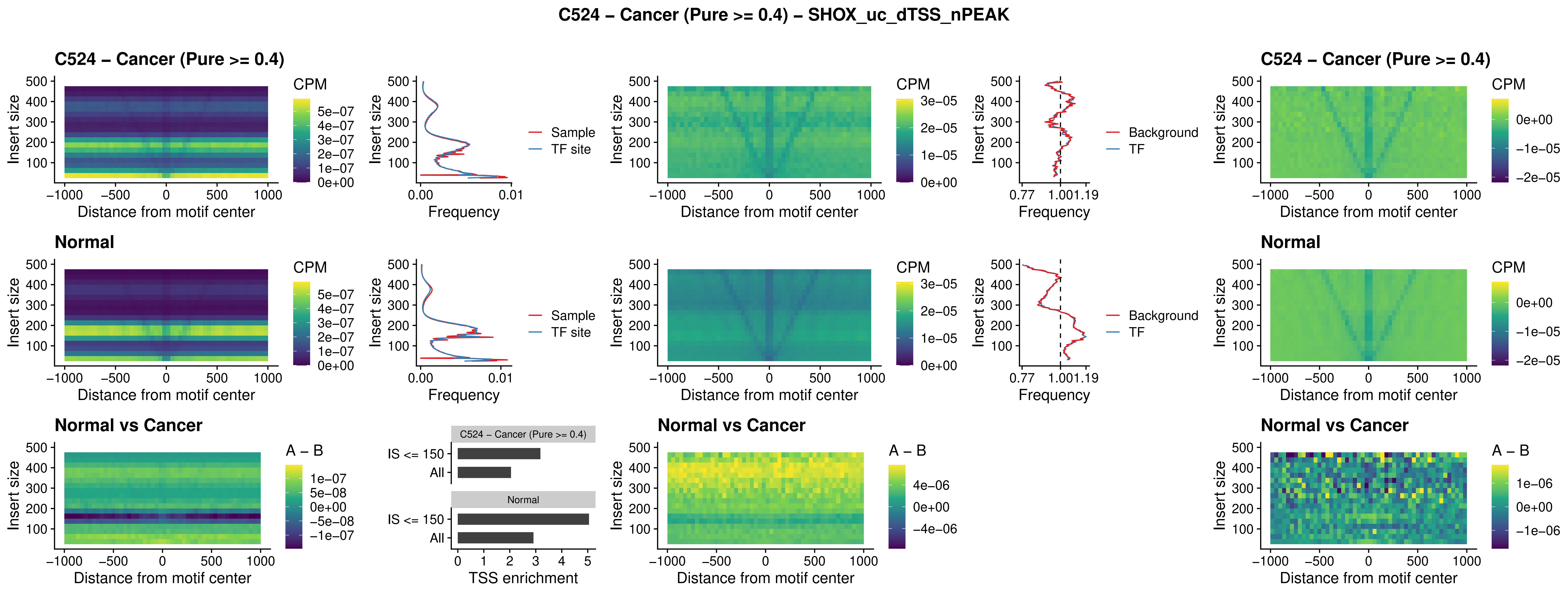
C524 – Cancer (Pure >= 0.4) – SHOX_uc_dTSS_nPEAK
C524 – Cancer (Pure >= 0.4)
Normal
Normal vs Cancer
Insert size
Distance from motif center
Frequency
CPM
A – B
Sample
TF site
Background
TF
C524 – Cancer (Pure >= 0.4)
Normal
IS <= 150
All
TSS enrichment

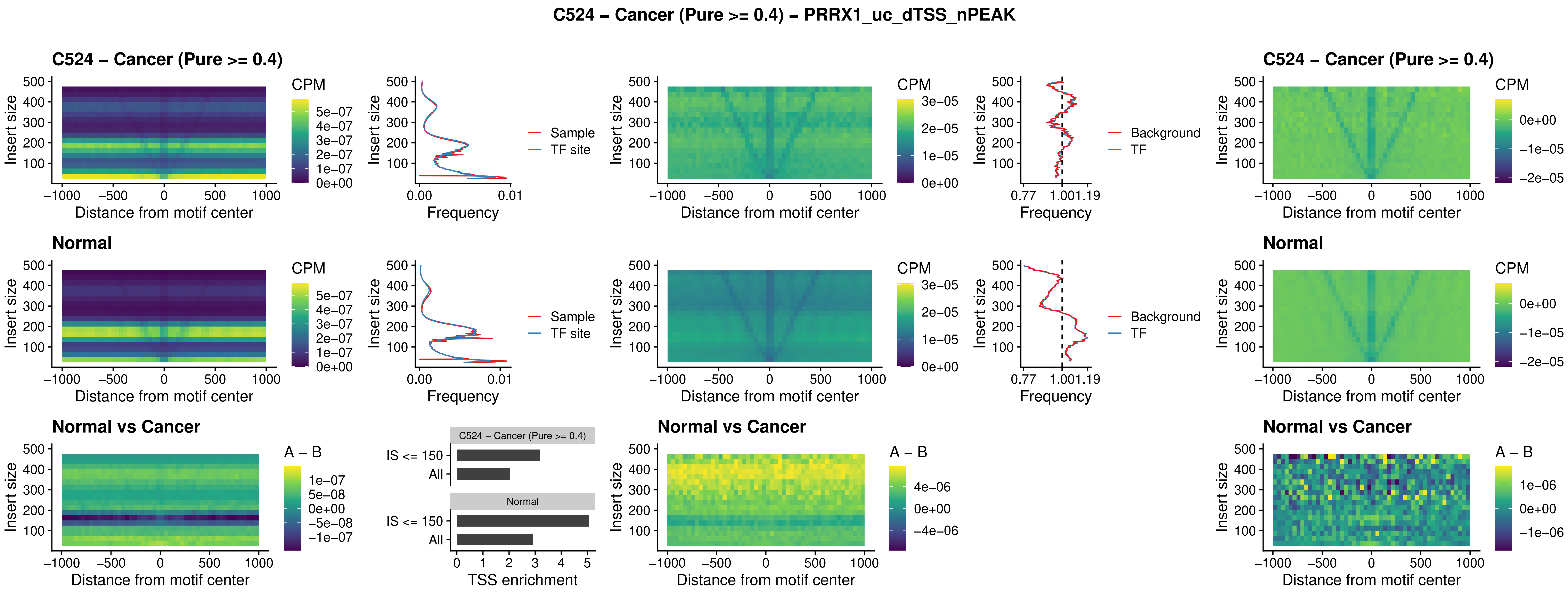

C524 – Cancer (Pure >= 0.4) – PRRX1_uc_dTSS_nPEAK
C524 – Cancer (Pure >= 0.4)
Normal
Normal vs Cancer
Insert size
Distance from motif center
Frequency
CPM
A – B
Sample
TF site
Background
TF
C524 – Cancer (Pure >= 0.4)
Normal
IS <= 150
All
TSS enrichment

#### C524 – Cancer (Pure >= 0.4) – LMX1A_uc_dTSS_nPEAK

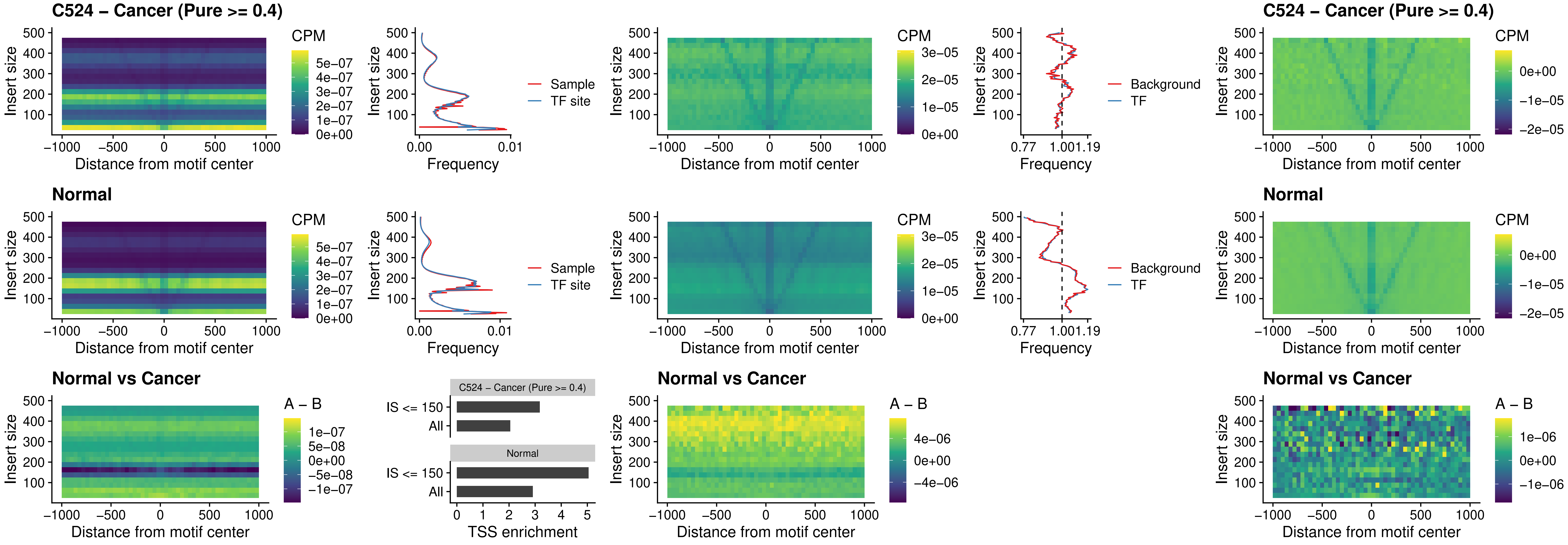

#### C524 – Cancer (Pure >= 0.4) – LMX1B_uc_dTSS_nPEAK

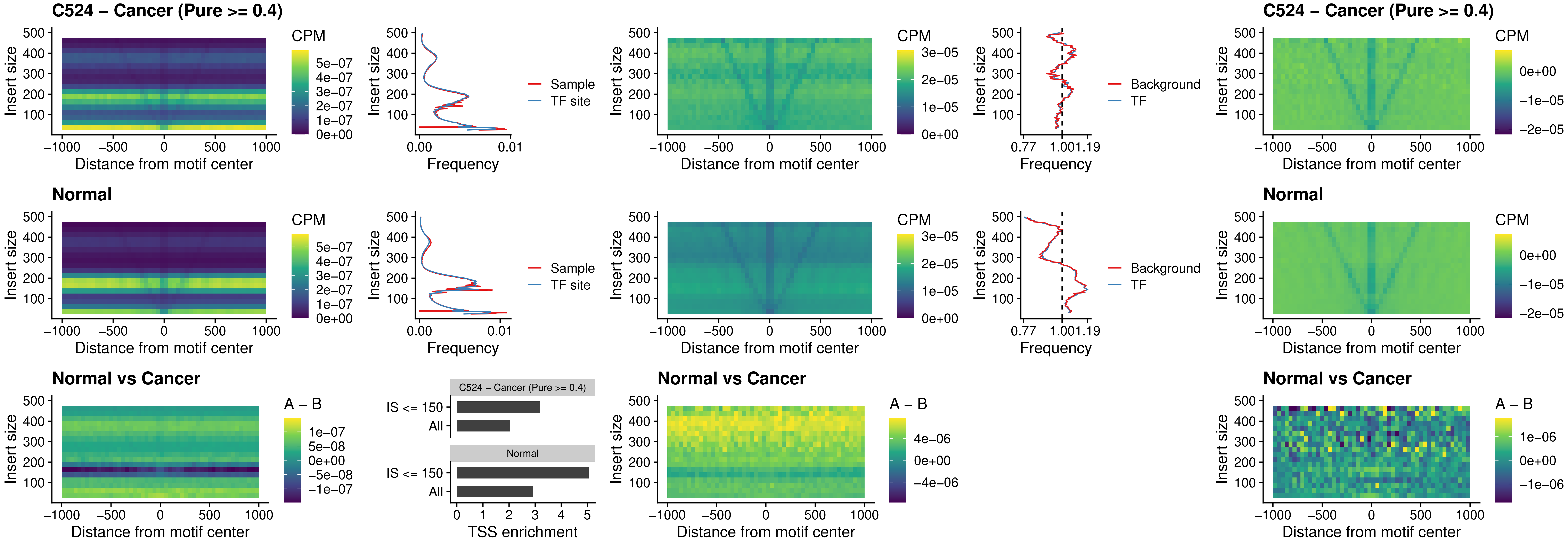

#### C524 – Cancer (Pure >= 0.4) – DLX5_uc_dTSS_oPEAK

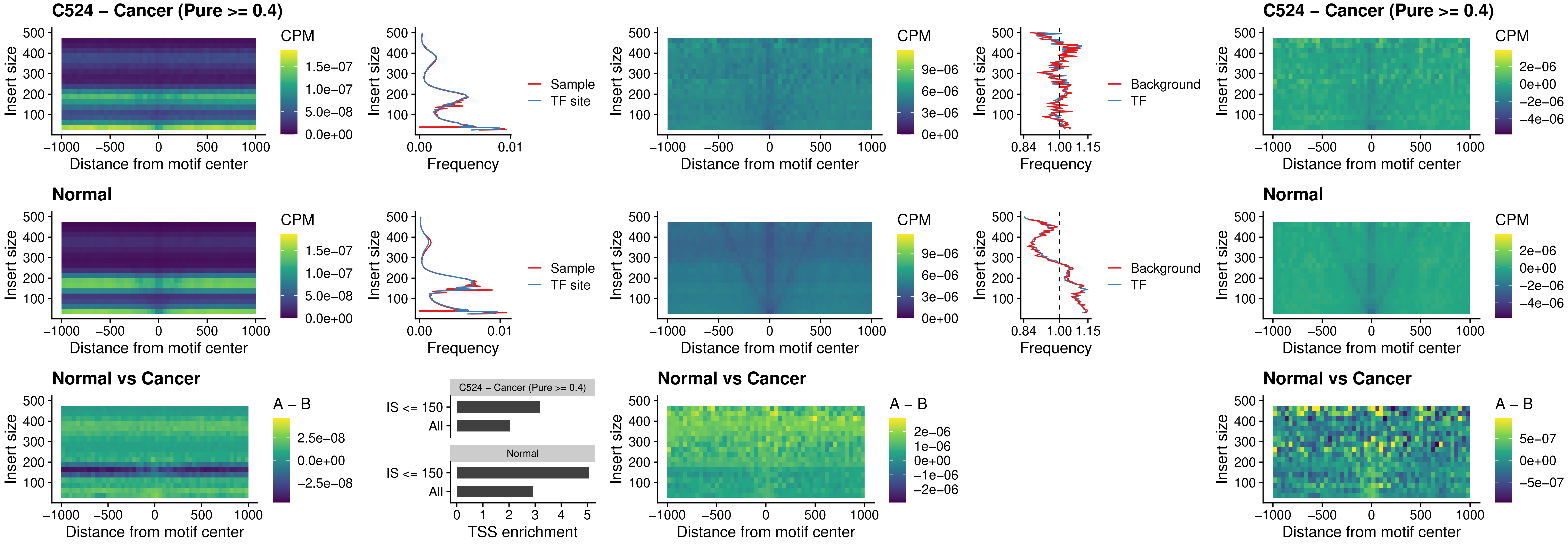

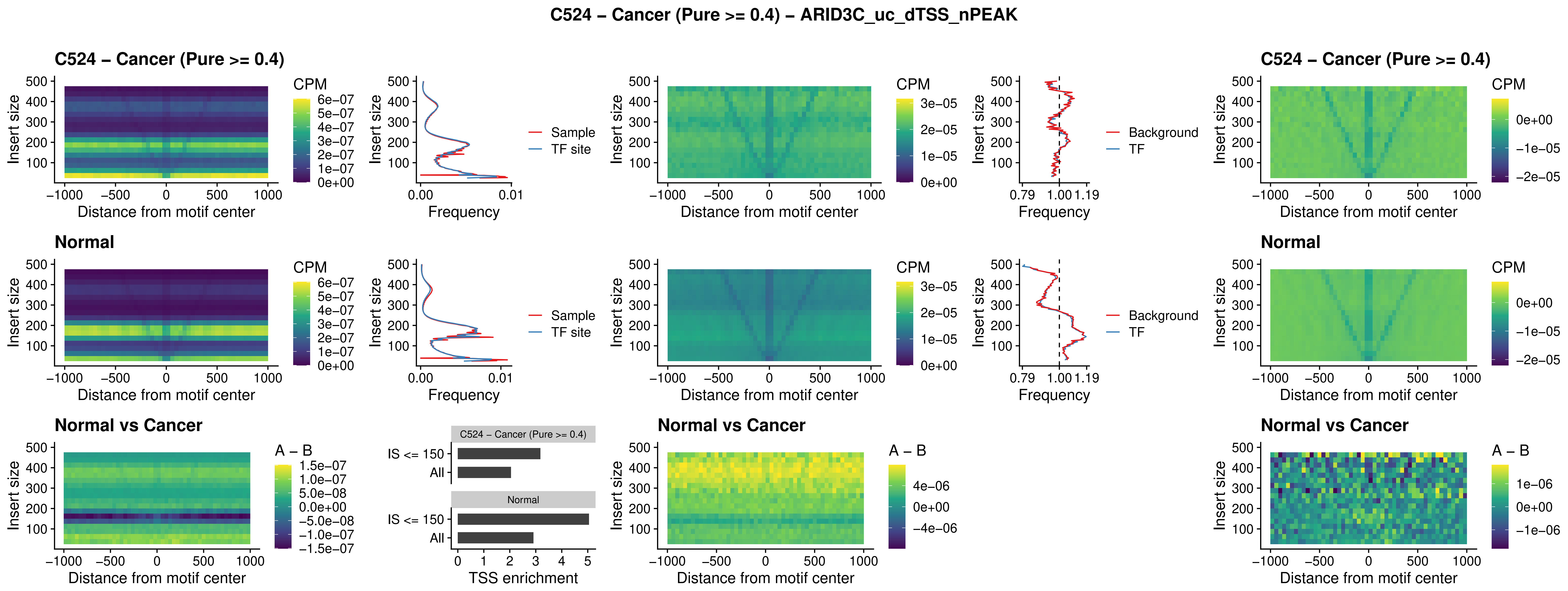
C524 – Cancer (Pure >= 0.4) – ARID3C_uc_dTSS_nPEAK
C524 – Cancer (Pure >= 0.4)
Normal
Normal vs Cancer
Insert size
Distance from motif center
Frequency
CPM
A – B
Sample
TF site
Background
TF
C524 – Cancer (Pure >= 0.4)
Normal
IS <= 150
All
TSS enrichment

#### C524 – Cancer (Pure >= 0.4) – HOXD12_uc_dTSS_nPEAK

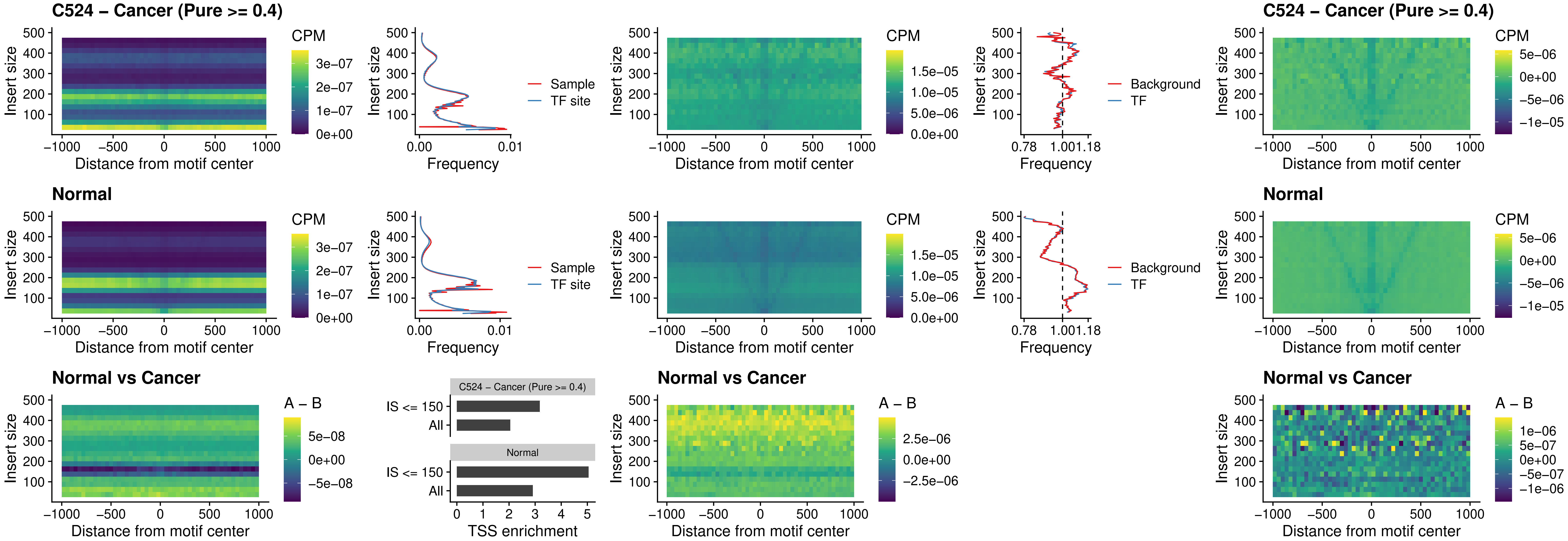

#### C524 – Cancer (Pure >= 0.4) – CDX2_uc_dTSS_nPEAK

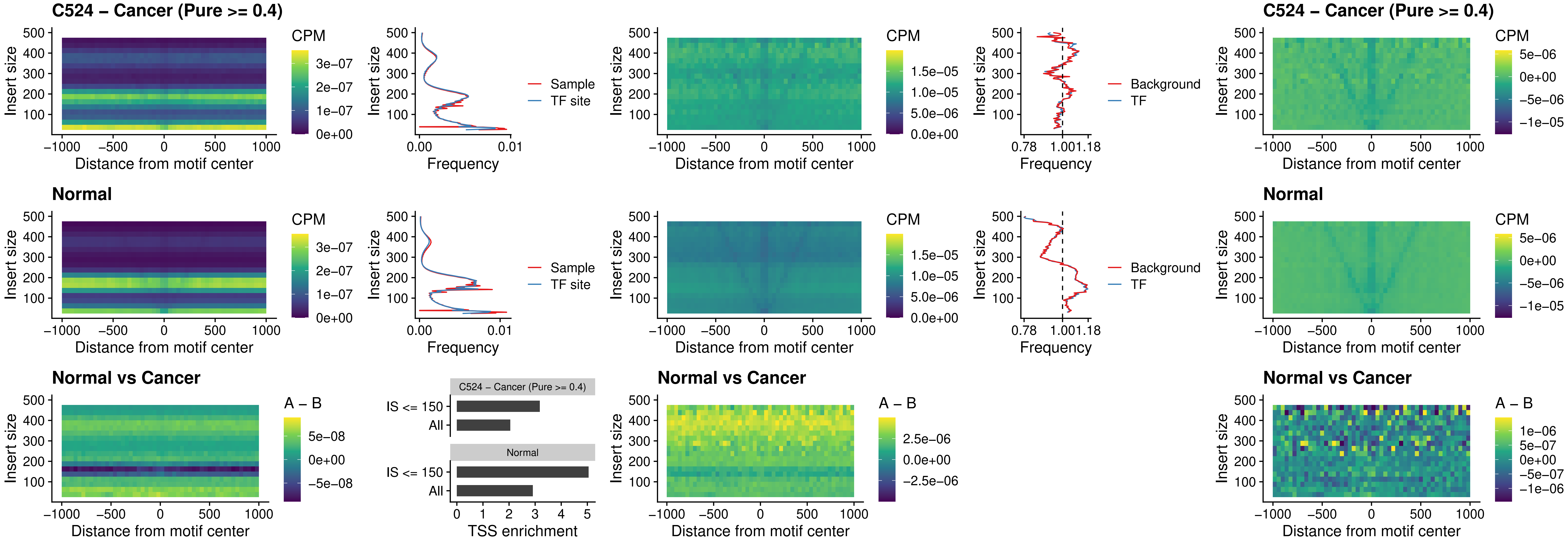

#### C524 – Cancer (Pure >= 0.4) – HOXC12_uc_dTSS_nPEAK

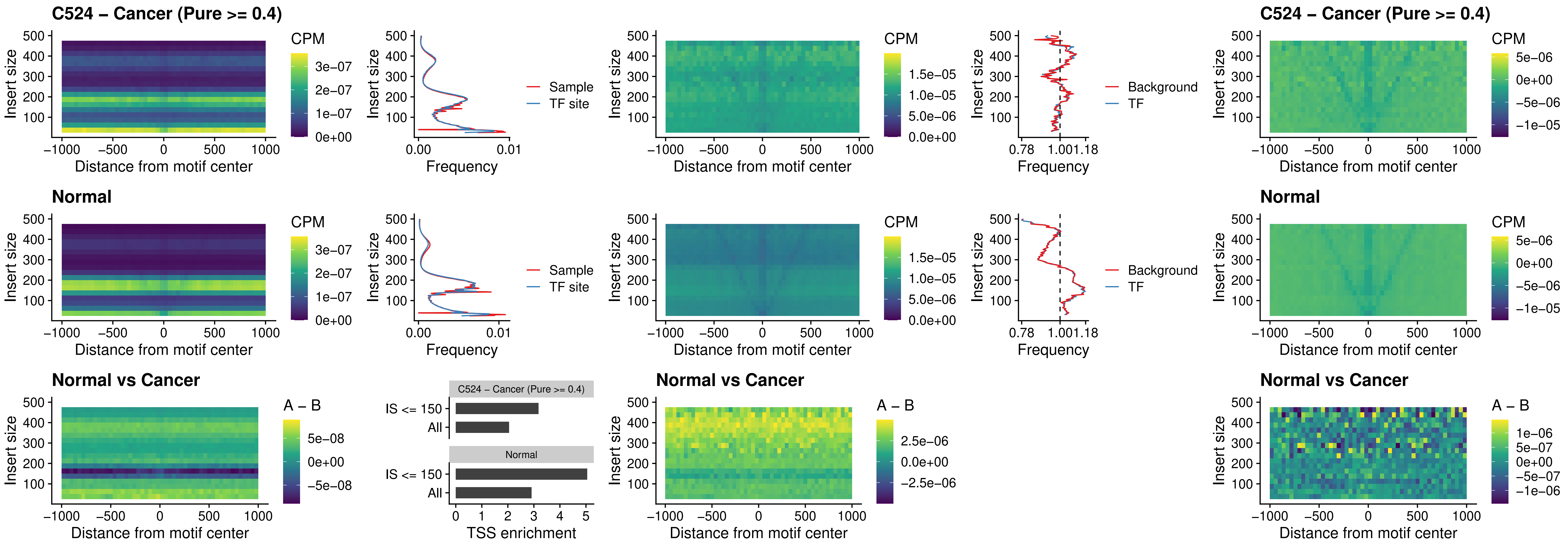

#### C524 – Cancer (Pure >= 0.4) – HHEX_uc_dTSS_oPEAK

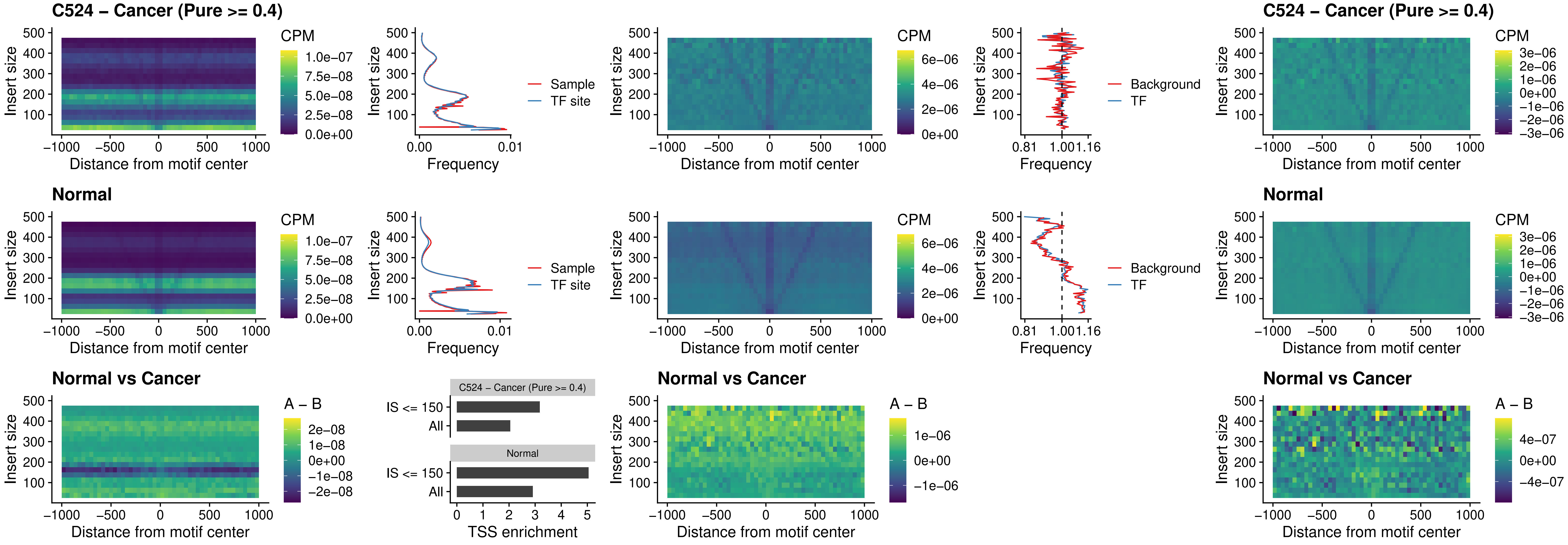

#### C524 – Cancer (Pure >= 0.4) – HOXA11_uc_dTSS_nPEAK

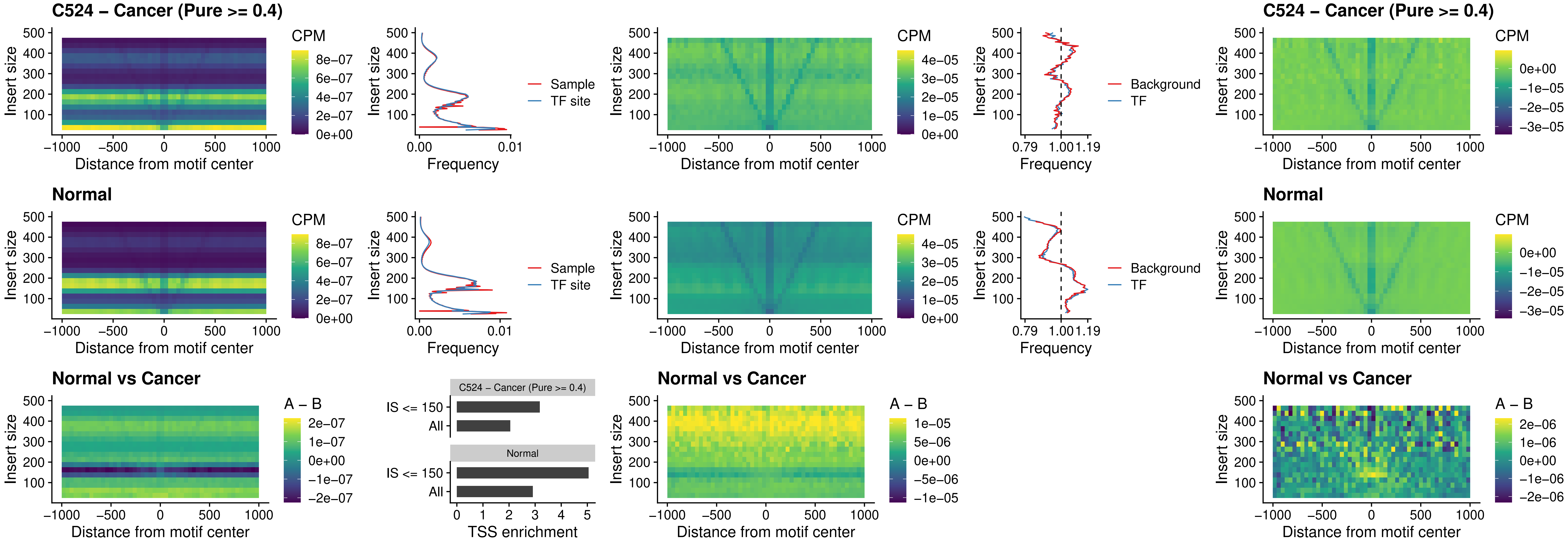

#### C524 – Cancer (Pure >= 0.4) – HOXA11_uc_cTSS_nPEAK

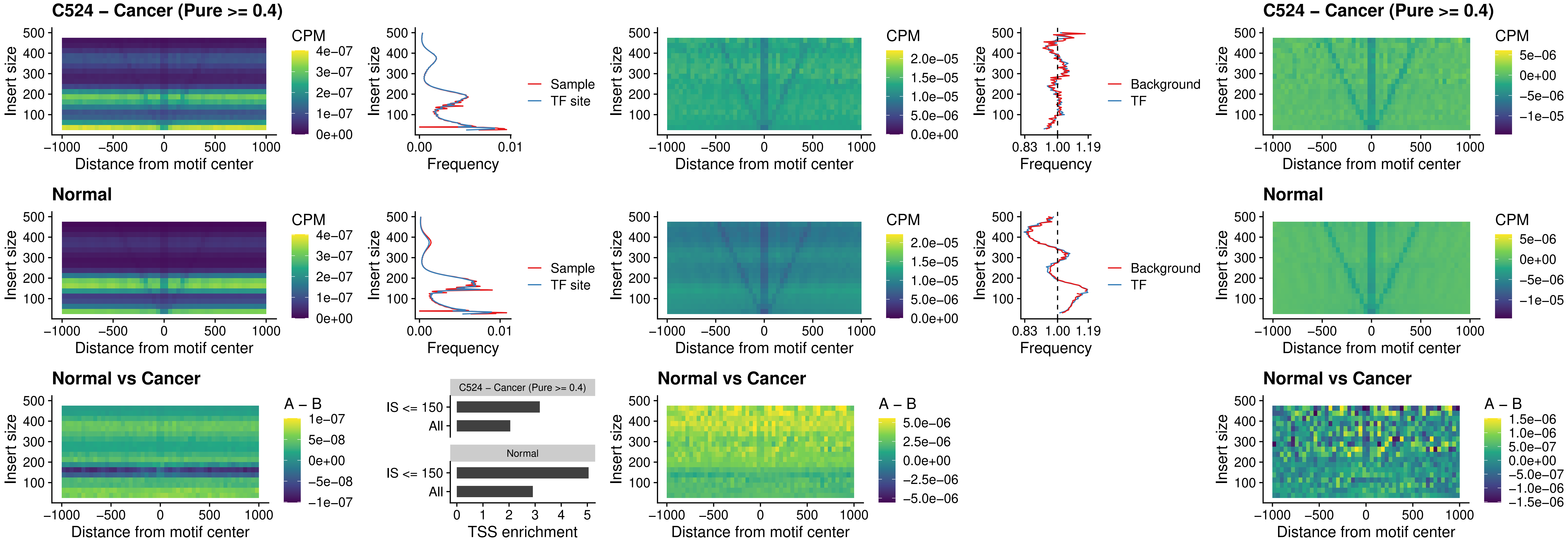

### C524 – Cancer (Pure >= 0.4) – ZFHX3_uc_dTSS_nPEAK

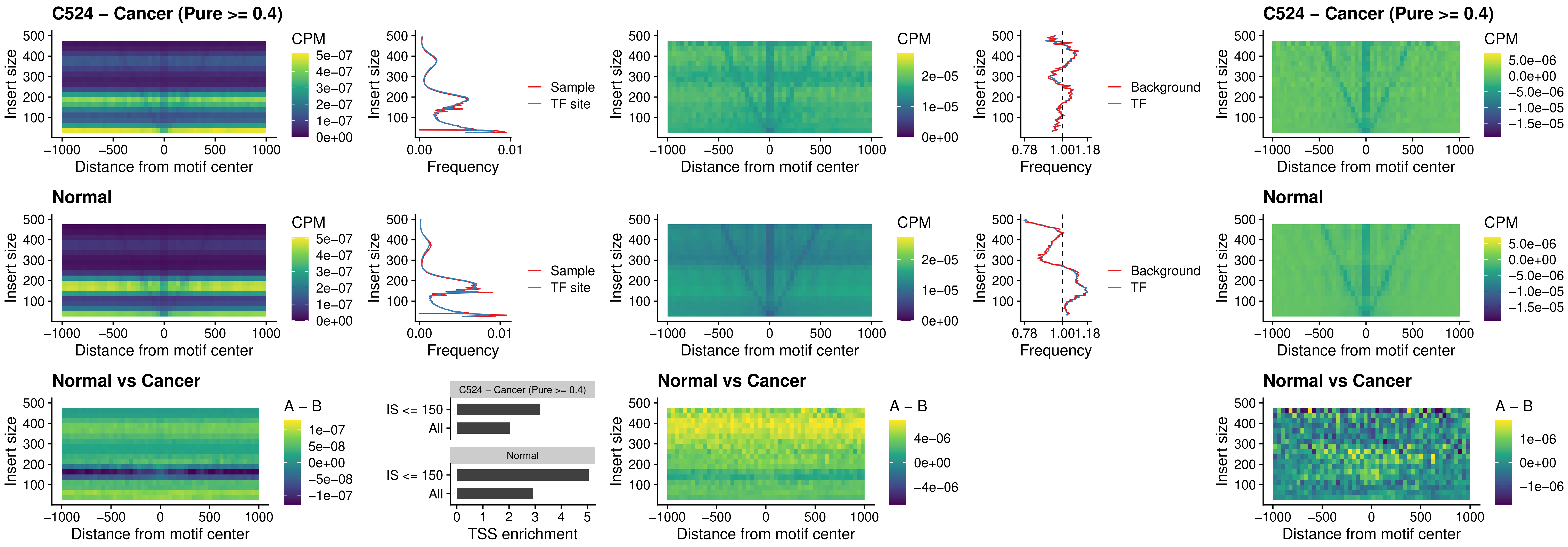

#### C524 – Cancer (Pure >= 0.4) – HOXB4_uc_dTSS_nPEAK

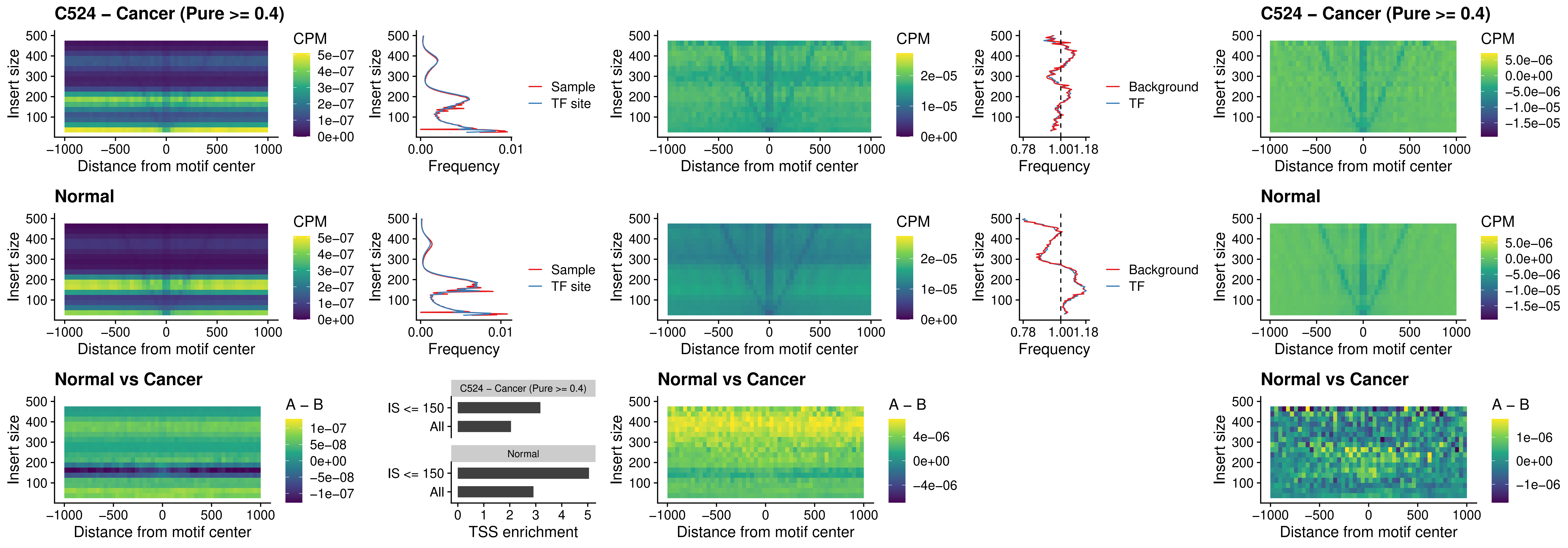

#### C524 – Cancer (Pure >= 0.4) – HOXC5_uc_dTSS_nPEAK

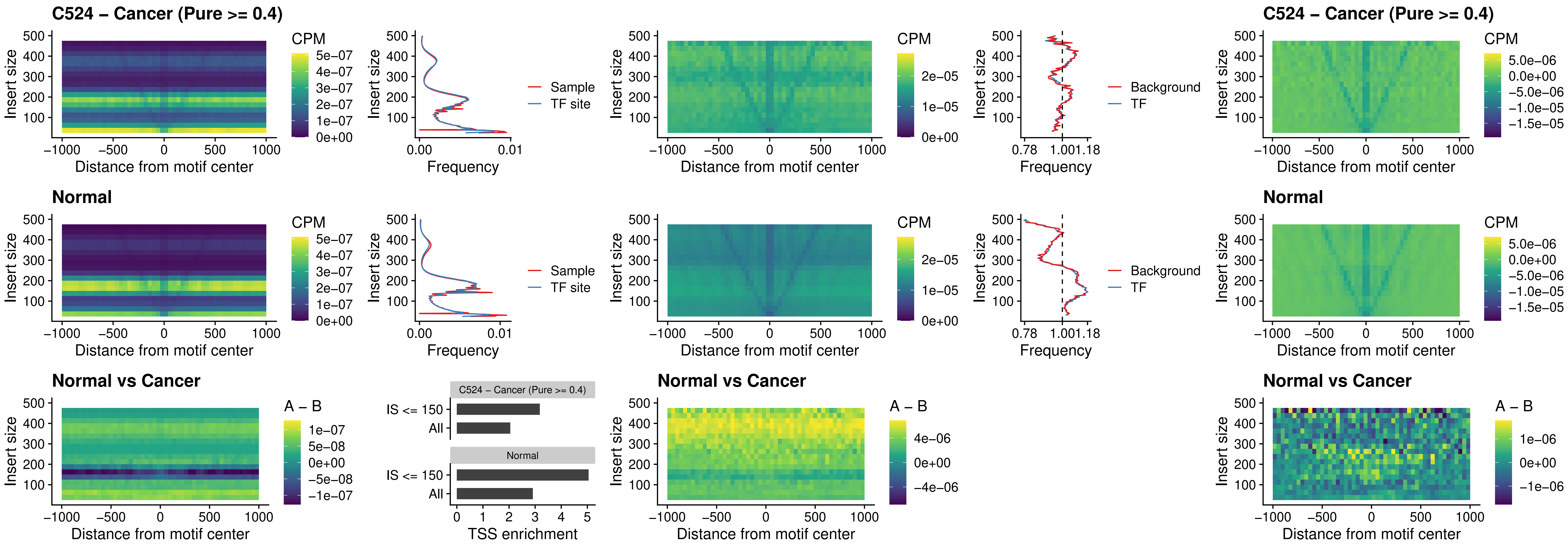

#### C524 – Cancer (Pure >= 0.4) – HOXD1_uc_dTSS_nPEAK

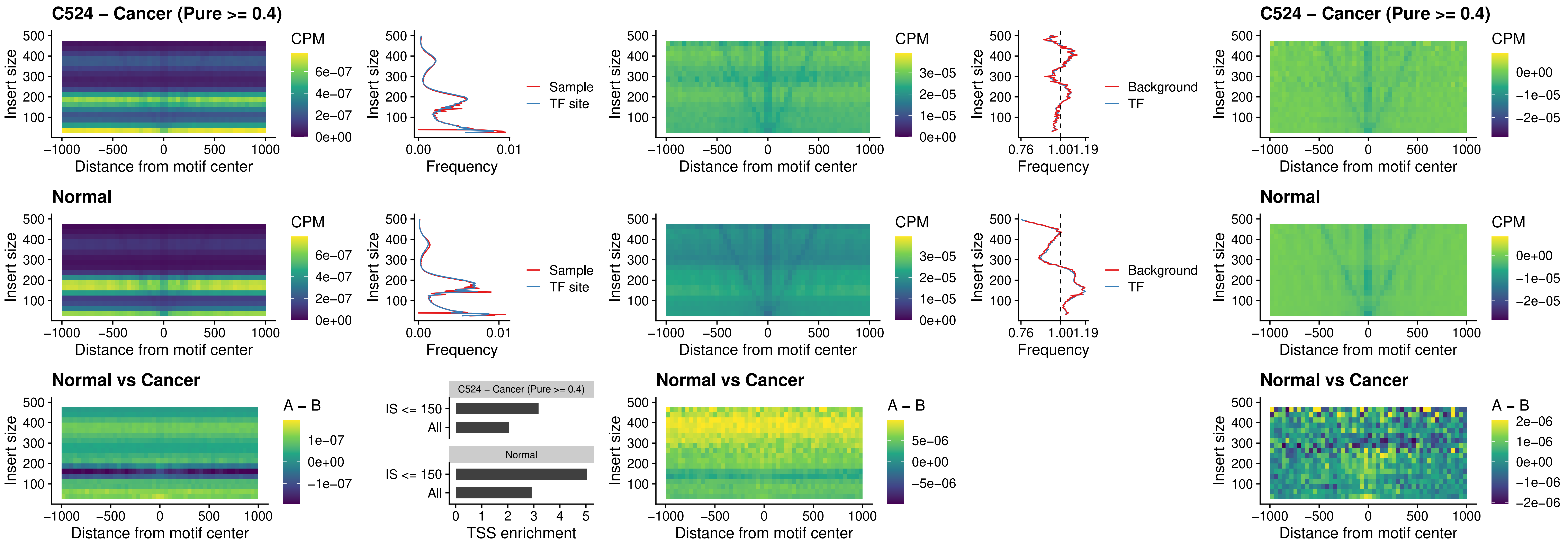

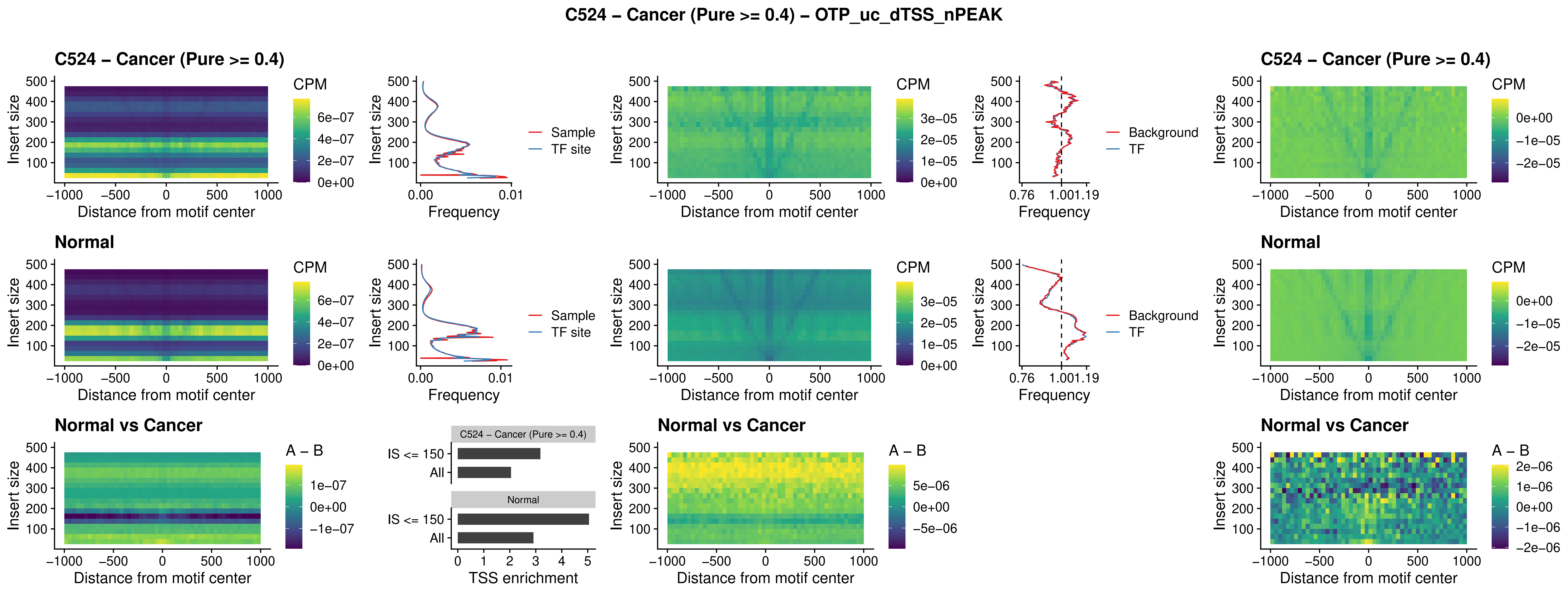
C524 – Cancer (Pure >= 0.4) – OTP_uc_dTSS_nPEAK
C524 – Cancer (Pure >= 0.4)
Normal
Normal vs Cancer
Insert size
Distance from motif center
Frequency
CPM
A – B
Sample
TF site
Background
TF
C524 – Cancer (Pure >= 0.4)
Normal
IS <= 150
All
TSS enrichment

#### C524 – Cancer (Pure >= 0.4) – FOXL1_uc_dTSS_oPEAK

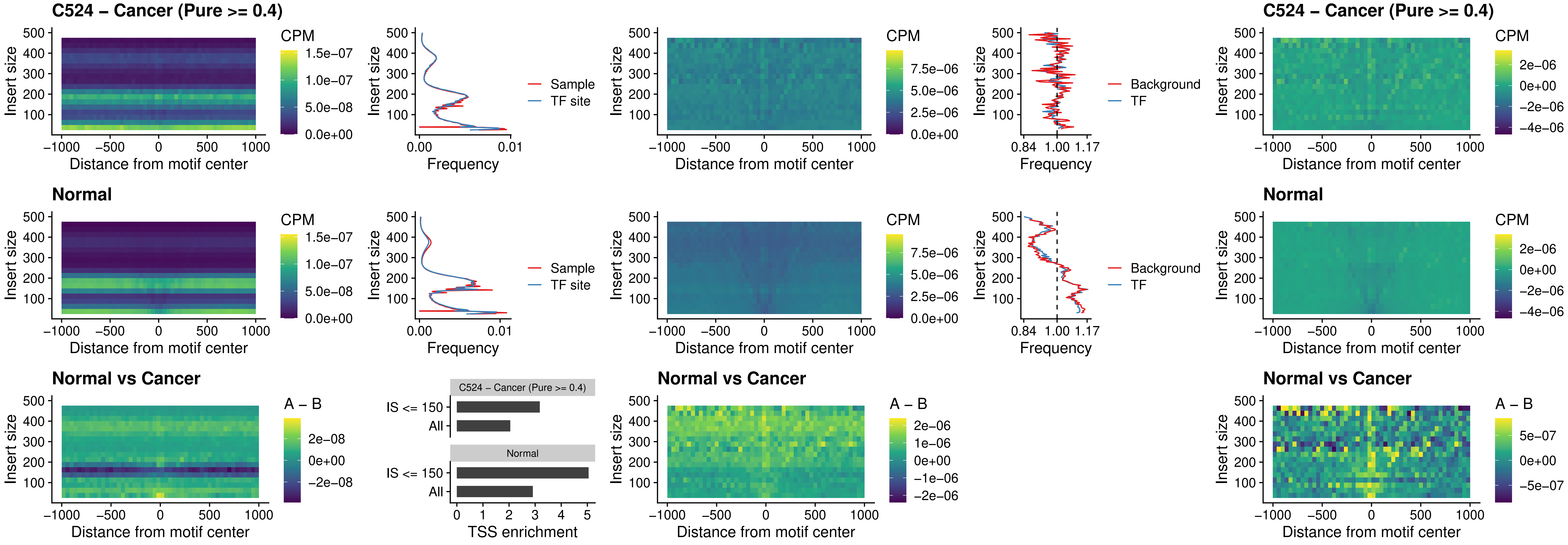

### C524 – Cancer (Pure >= 0.4) – ZNF32_uc_dTSS_oPEAK

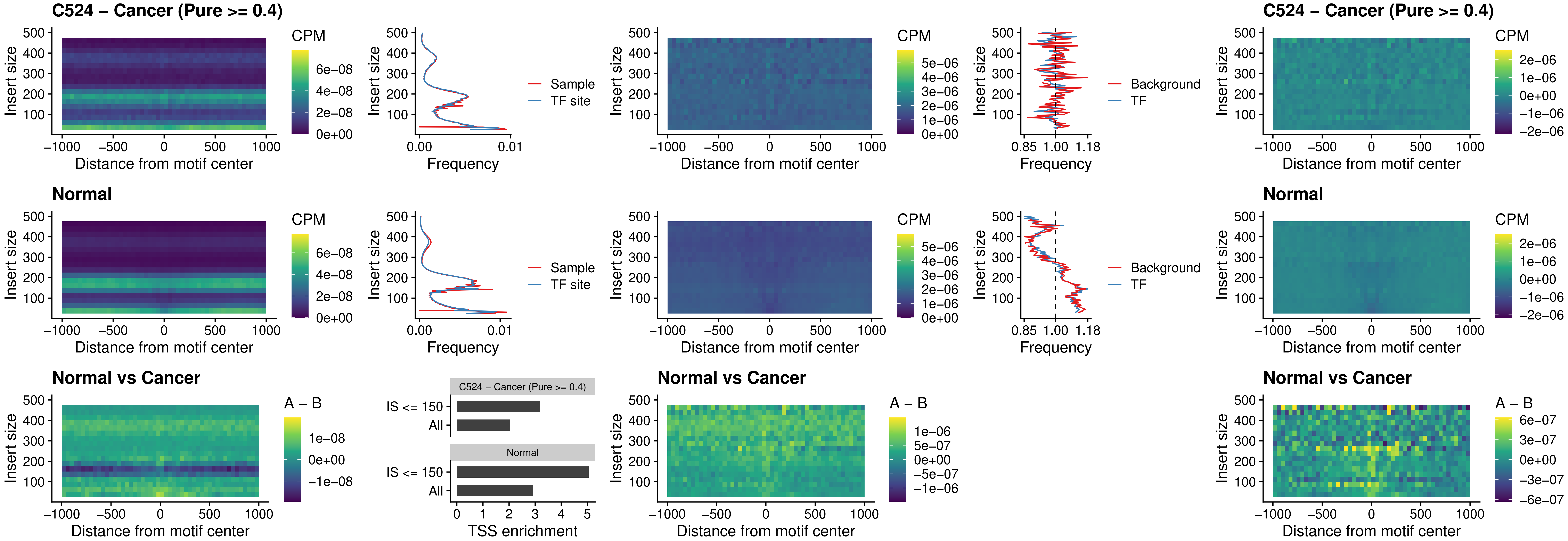

#### C524 – Cancer (Pure >= 0.4) – FOXL1_uc_dTSS_nPEAK

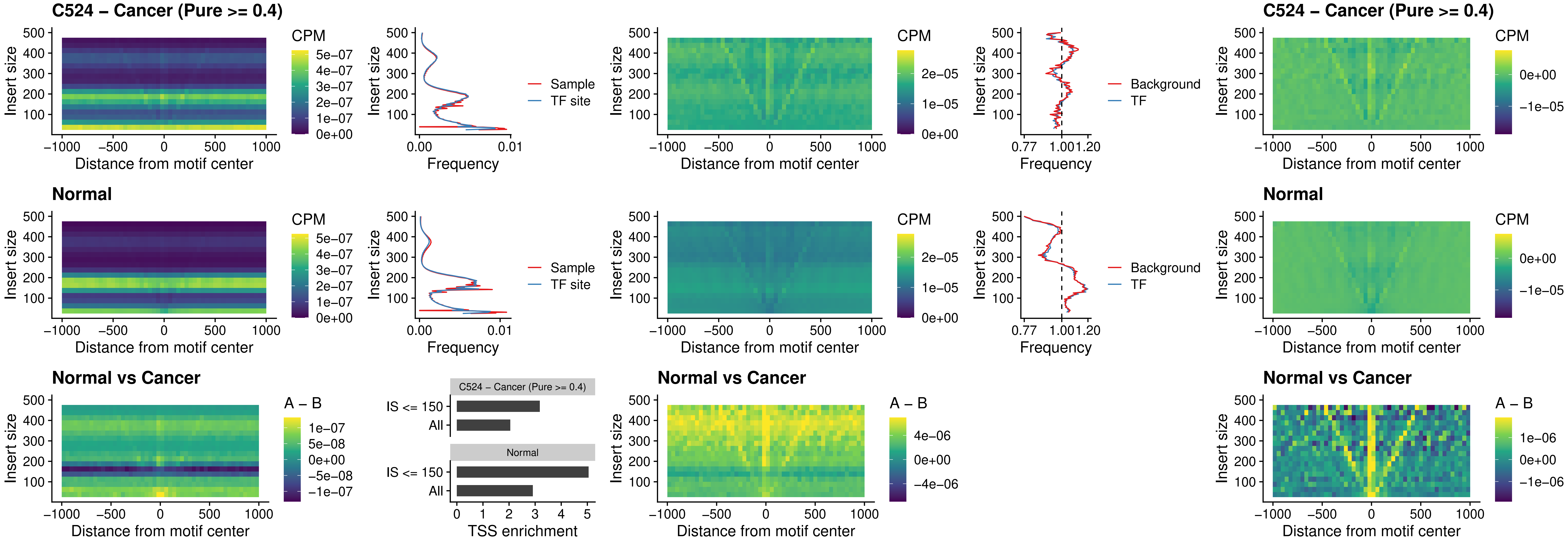

#### C524 – Cancer (Pure >= 0.4) – ZNF32_uc_dTSS_nPEAK

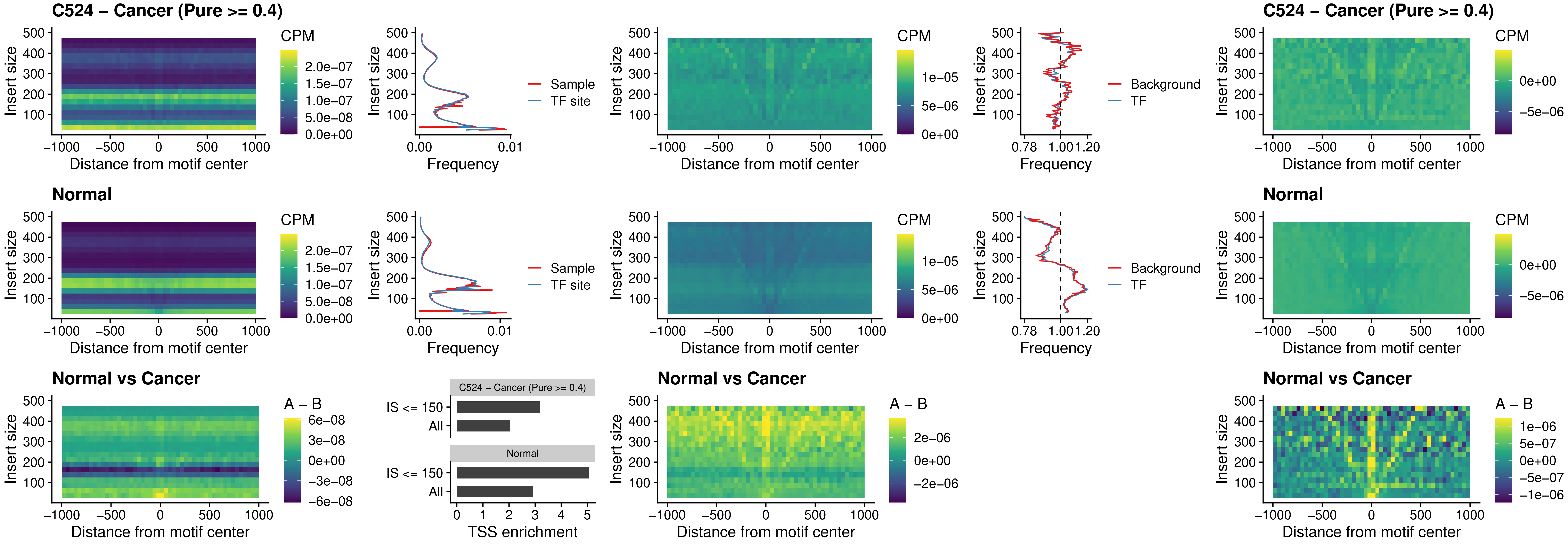

### C524 – Cancer (Pure >= 0.4) – DLX5_uc_dTSS_nPEAK

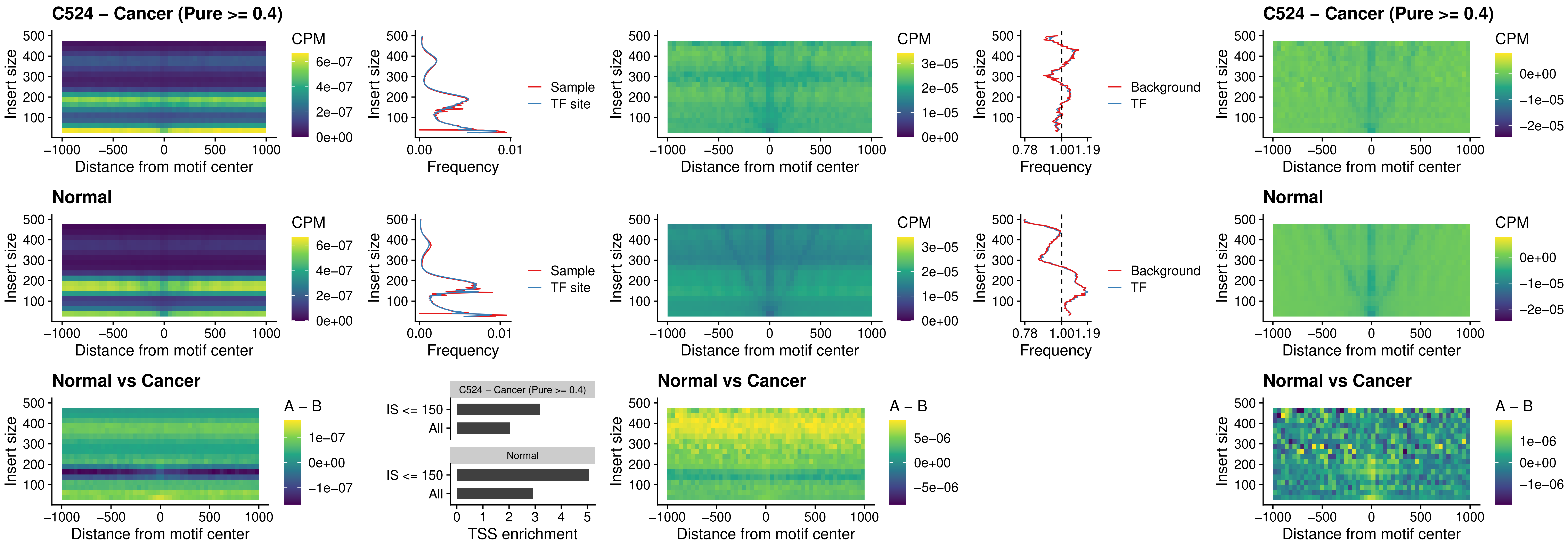

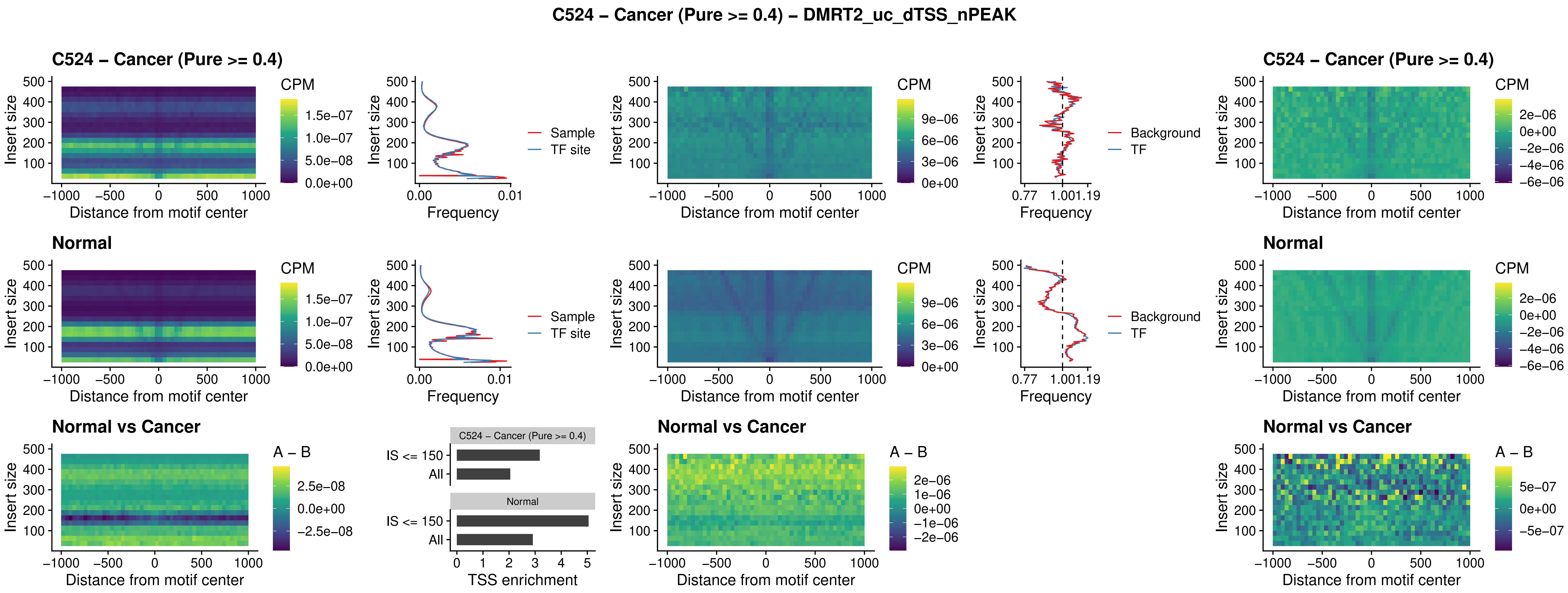
C524 – Cancer (Pure >= 0.4) – DMRT2_uc_dTSS_nPEAK
C524 – Cancer (Pure >= 0.4)
Normal
Normal vs Cancer
Insert size
Distance from motif center
Frequency
CPM
A – B
Sample
TF site
Background
TF
C524 – Cancer (Pure >= 0.4)
Normal
IS <= 150
All
TSS enrichment

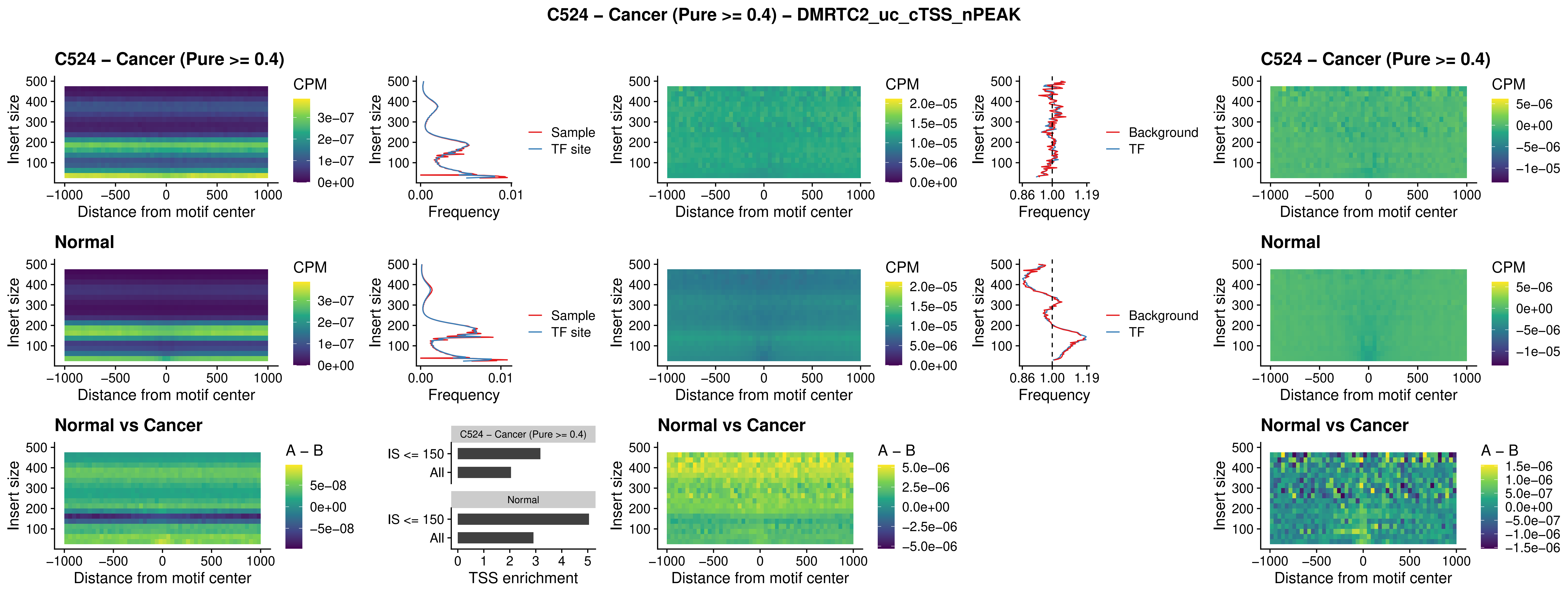
C524 – Cancer (Pure >= 0.4) – DMRTC2_uc_cTSS_nPEAK
C524 – Cancer (Pure >= 0.4)
Normal
Normal vs Cancer
Insert size
Distance from motif center
CPM
A – B
Frequency
Sample
TF site
Background
TF
C524 – Cancer (Pure >= 0.4)
Normal
IS <= 150
All
TSS enrichment

#### C524 – Cancer (Pure >= 0.4) – HOXD9_uc_dTSS_nPEAK

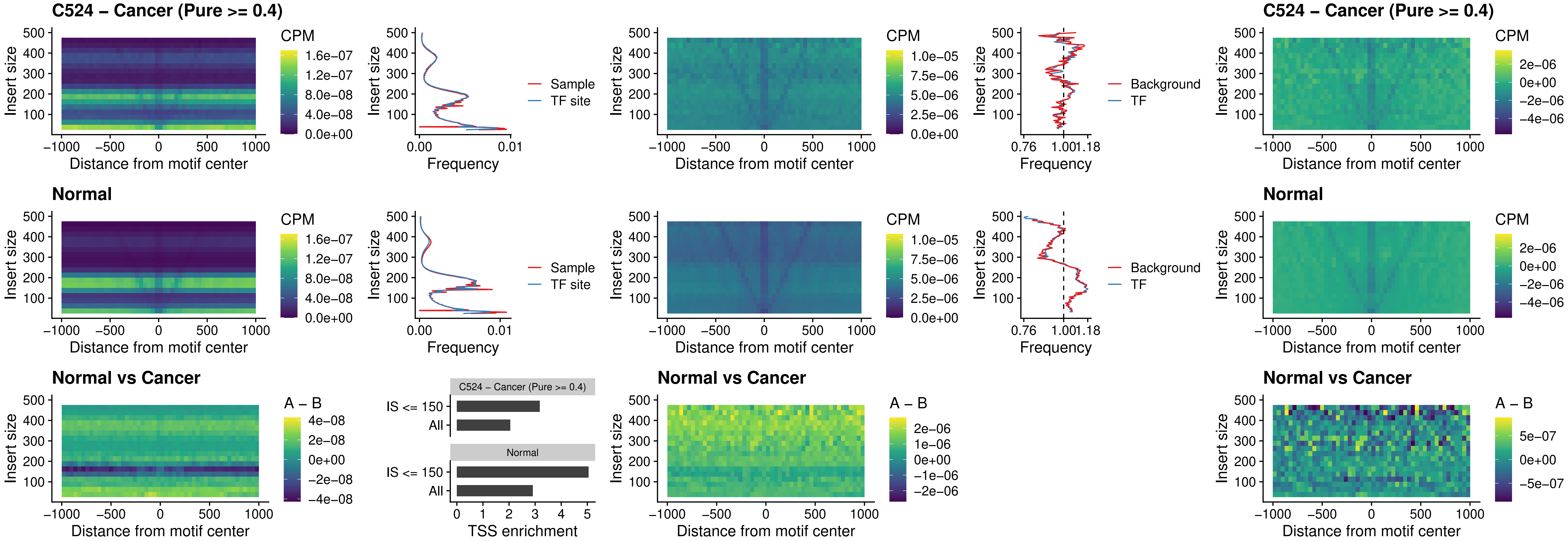

### C524 – Cancer (Pure >= 0.4) – HOXD4_uc_dTSS_nPEAK

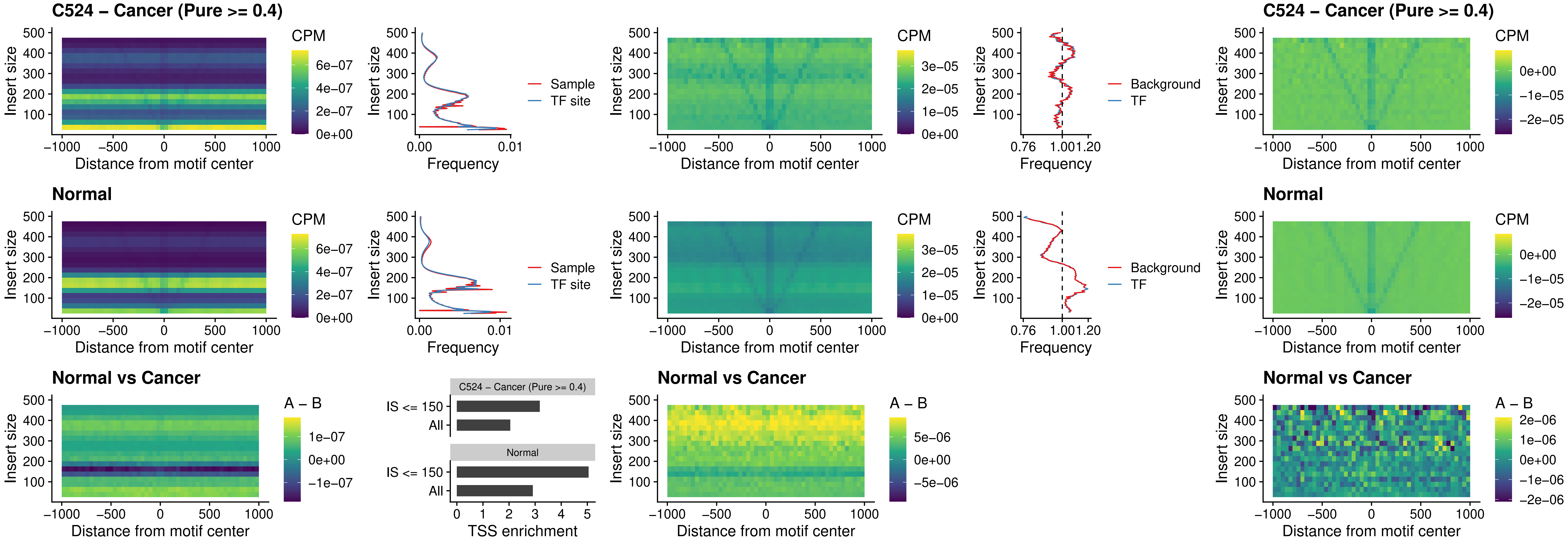

### C524 – Cancer (Pure >= 0.4) – FOXP3_uc_cTSS_nPEAK

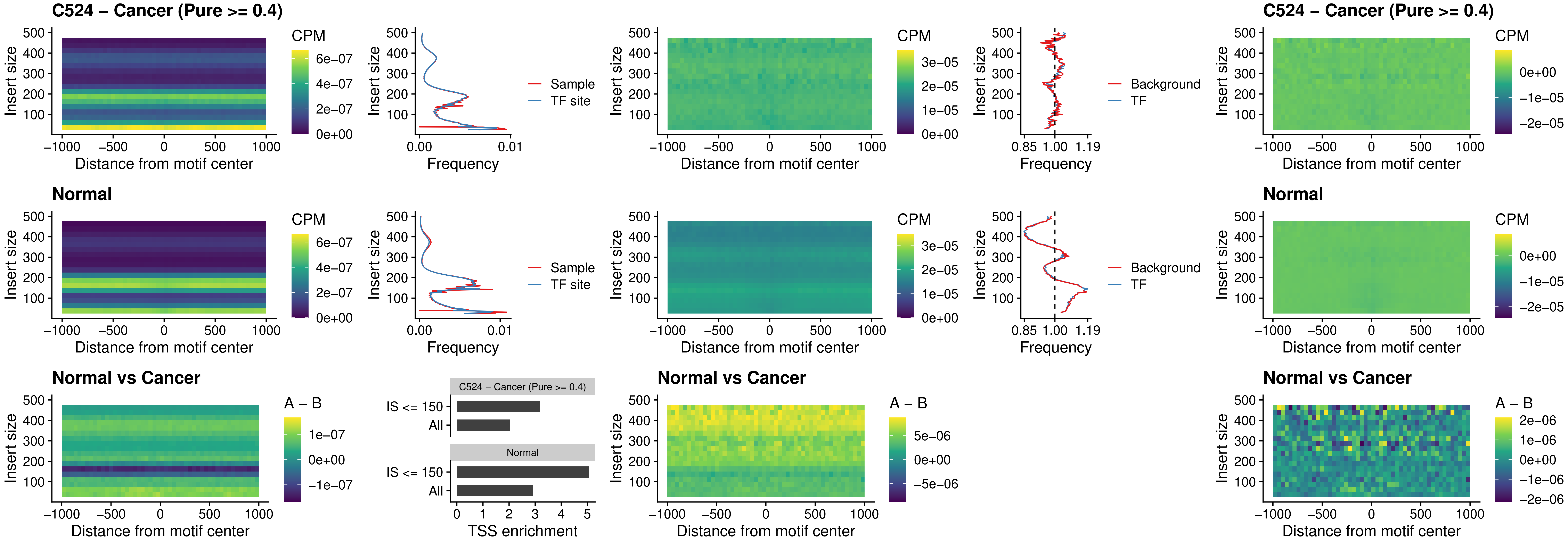

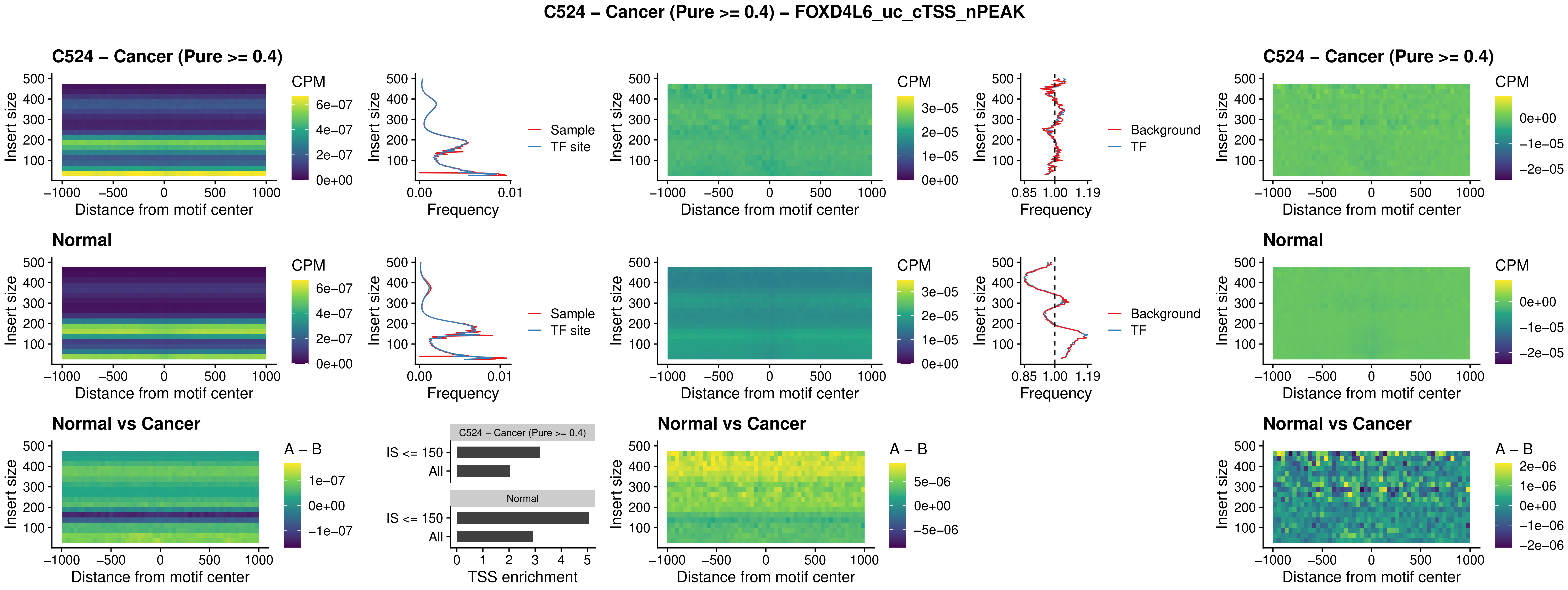
C524 – Cancer (Pure >= 0.4) – FOXD4L6_uc_cTSS_nPEAK
C524 – Cancer (Pure >= 0.4)
Normal
Normal vs Cancer
Insert size
Distance from motif center
CPM
A – B
Frequency
Sample
TF site
Background
TF
C524 – Cancer (Pure >= 0.4)
Normal
IS <= 150
All
TSS enrichment

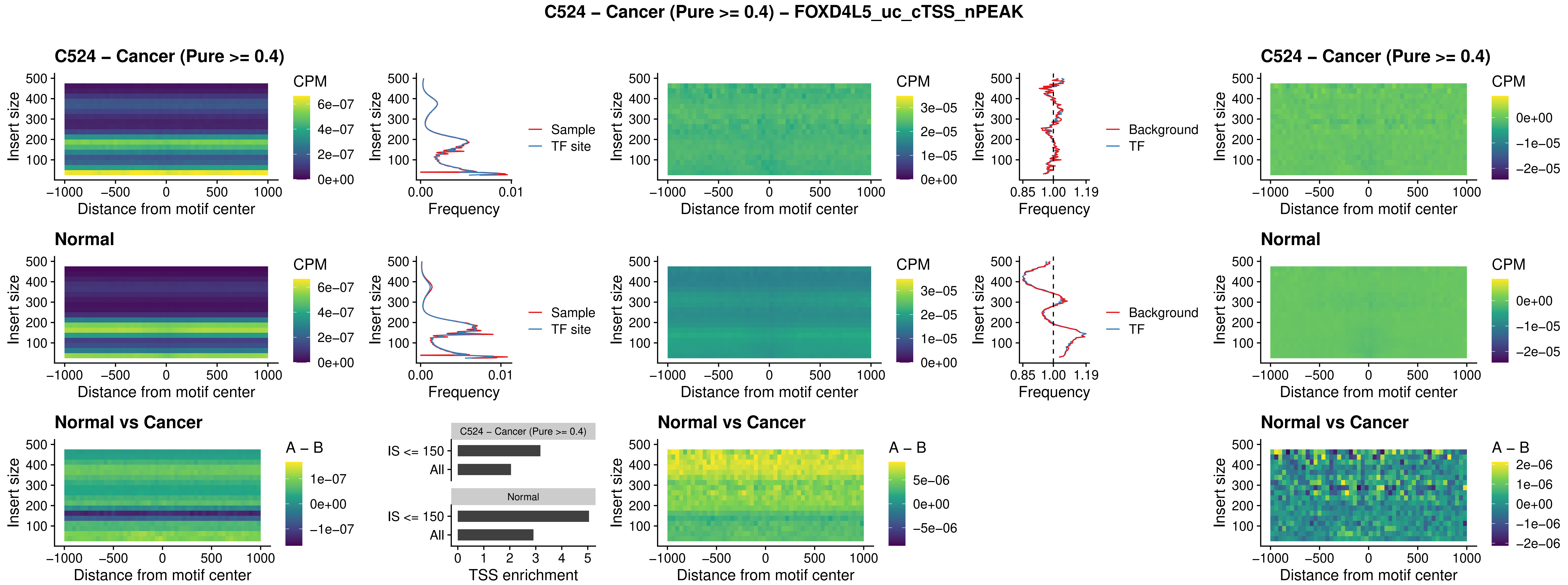

C524 – Cancer (Pure >= 0.4) – FOXD4L5_uc_cTSS_nPEAK
C524 – Cancer (Pure >= 0.4)
Normal
Normal vs Cancer
Insert size
Distance from motif center
Frequency
CPM
A – B
Sample
TF site
Background
TF
C524 – Cancer (Pure >= 0.4)
Normal
IS <= 150
All
TSS enrichment

#### C524 – Cancer (Pure >= 0.4) – FOXD4L4_uc_cTSS_nPEAK

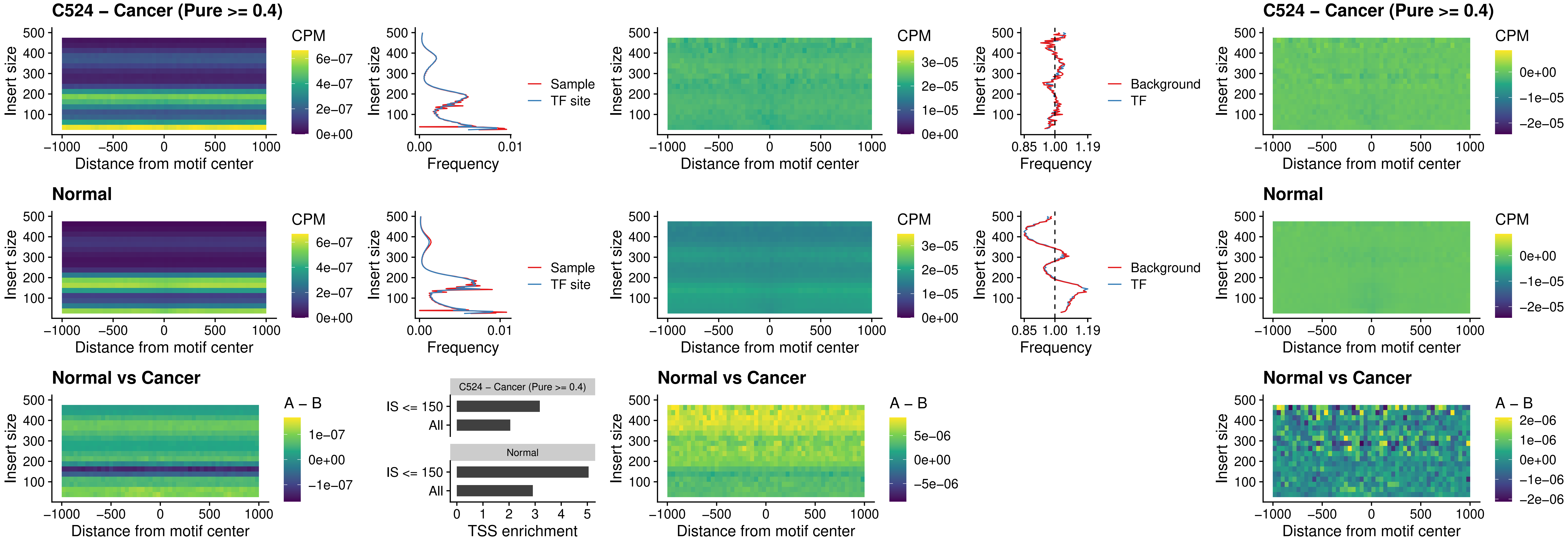

#### C524 – Cancer (Pure >= 0.4) – FOXD4L2_uc_cTSS_nPEAK

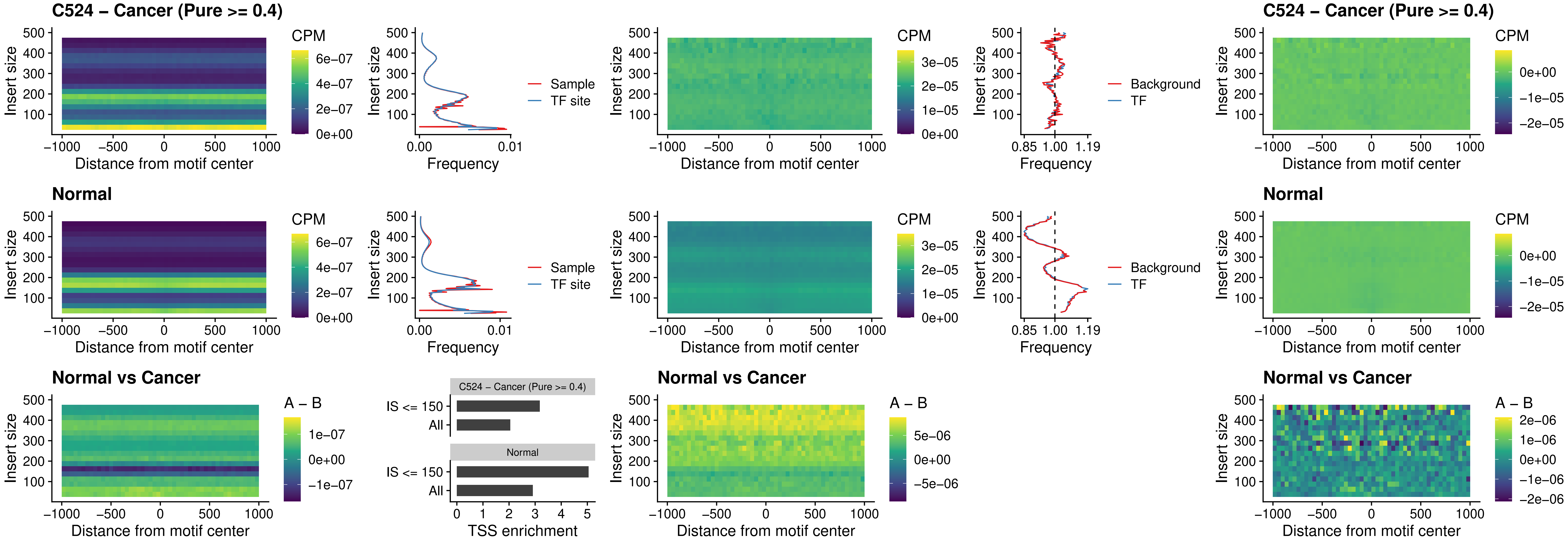

#### C524 – Cancer (Pure >= 0.4) – FOXD4L3_uc_cTSS_nPEAK

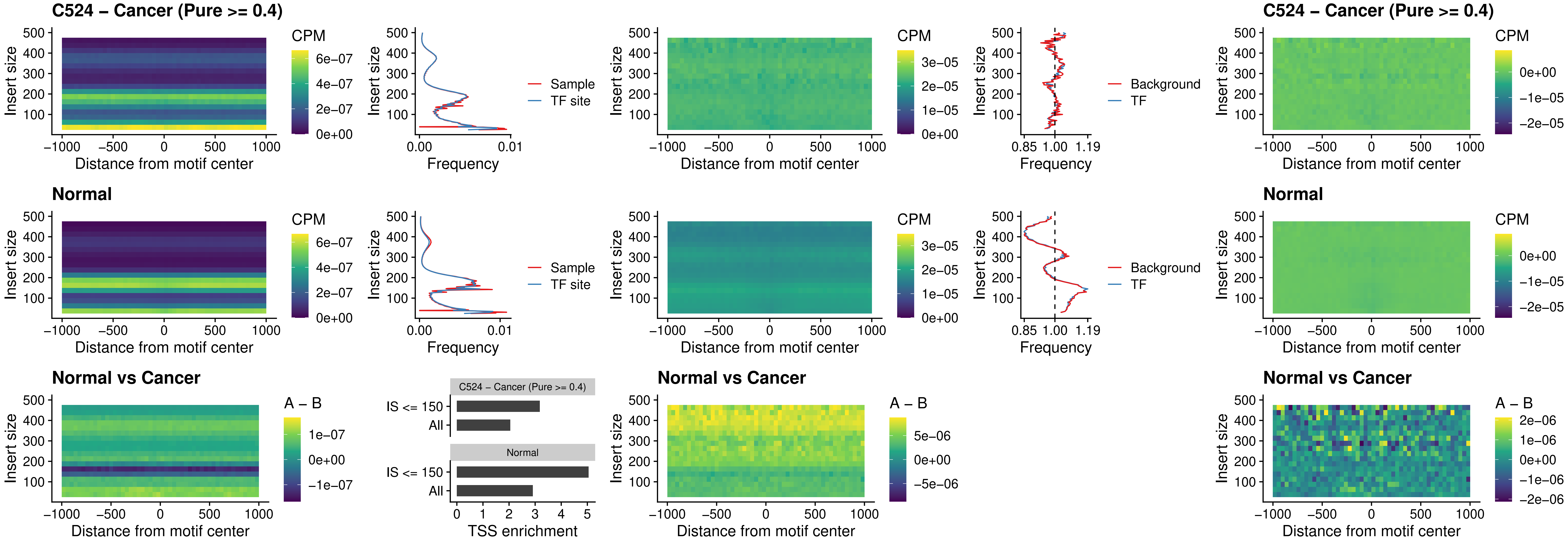

### C524 – Cancer (Pure >= 0.4) – HOXD1_uc_dTSS_oPEAK

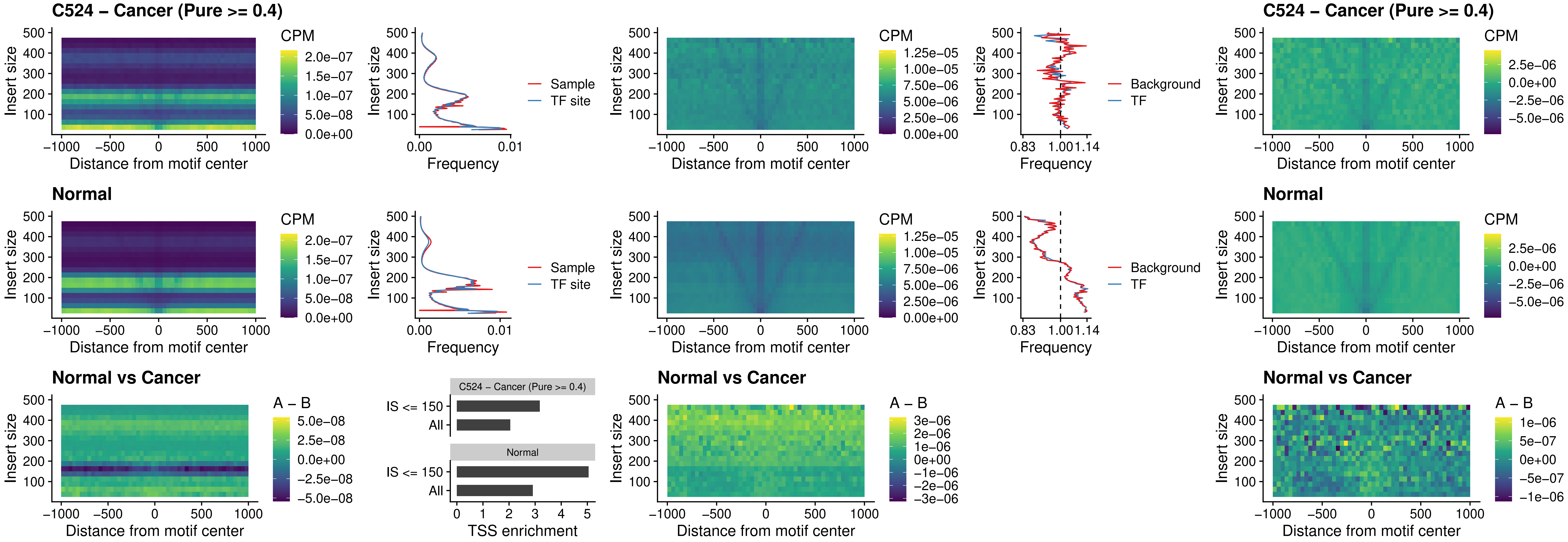

#### C524 – Cancer (Pure >= 0.4) – VAX1_uc_pTSS_nPEAK

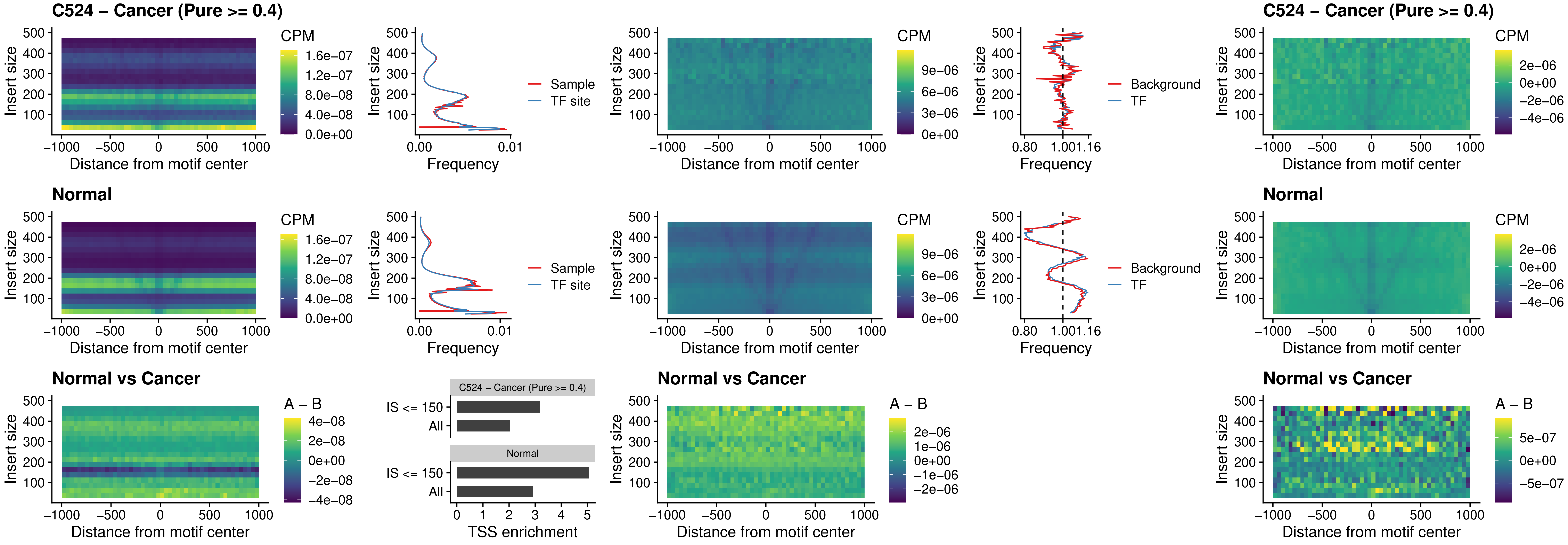

#### C524 – Cancer (Pure >= 0.4) – DLX3_uc_pTSS_nPEAK

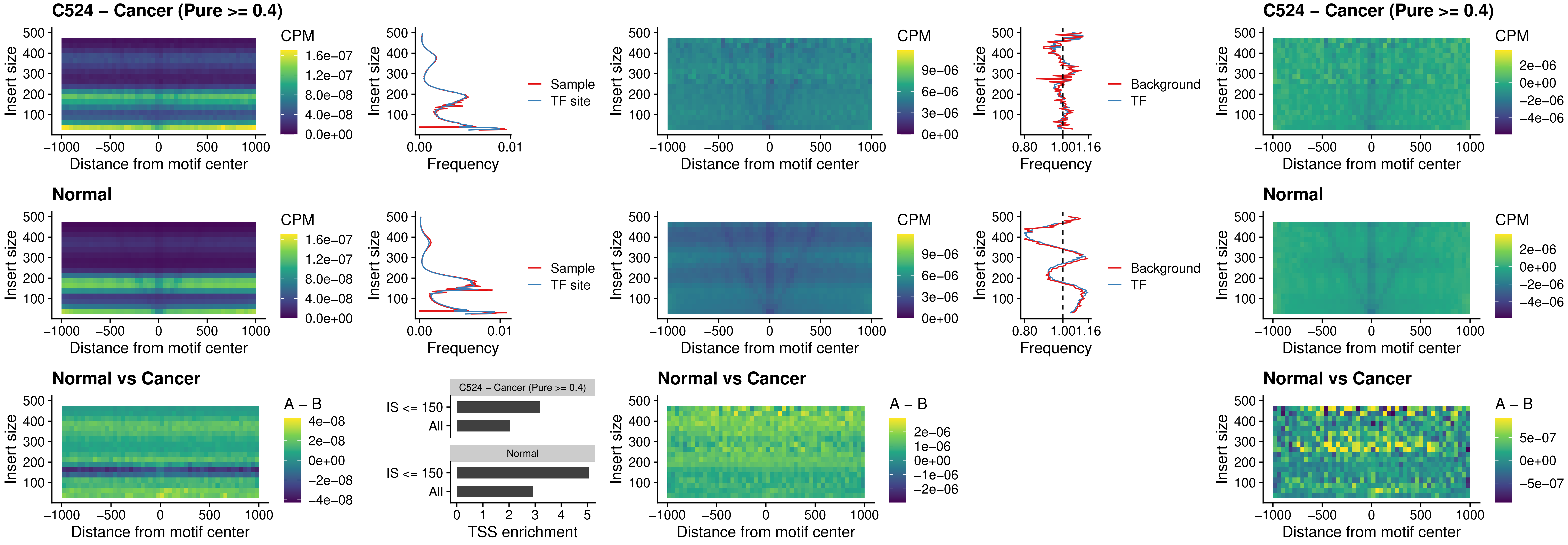

#### C524 – Cancer (Pure >= 0.4) – HOXD3_uc_pTSS_nPEAK

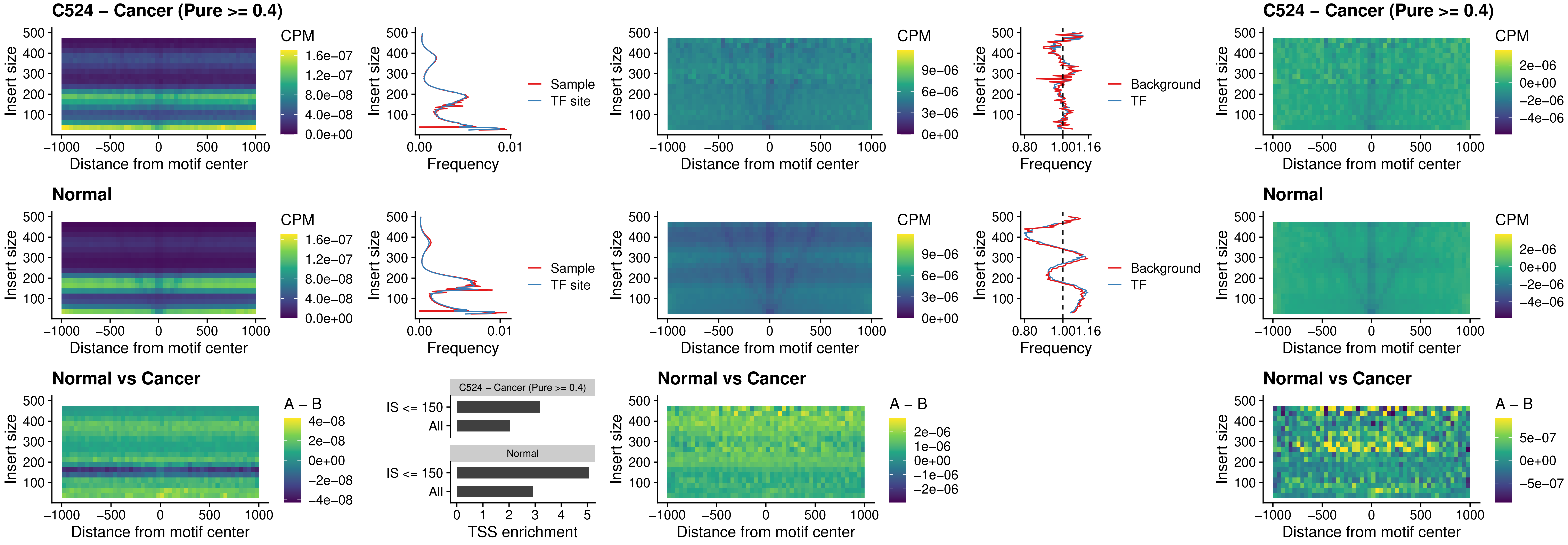

#### C524 – Cancer (Pure >= 0.4) – PDX1_uc_dTSS_nPEAK

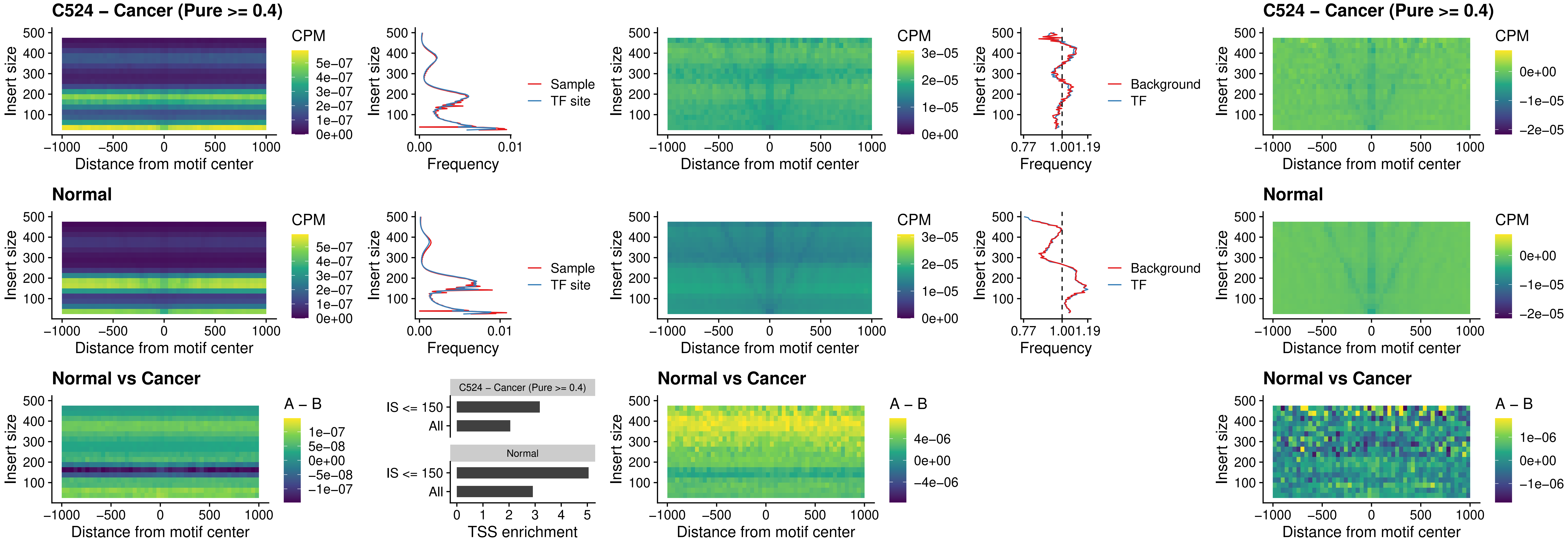

#### C524 – Cancer (Pure >= 0.4) – VAX2_uc_dTSS_nPEAK

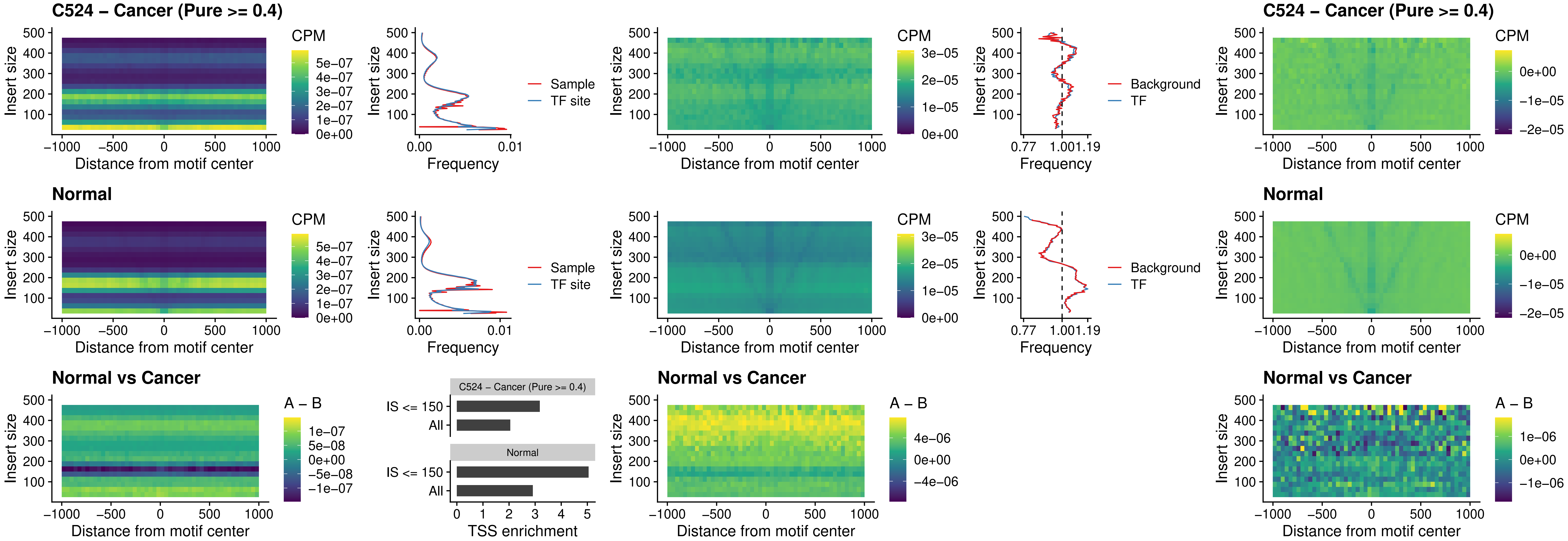

#### C524 – Cancer (Pure >= 0.4) – SOX5_uc_dTSS_nPEAK

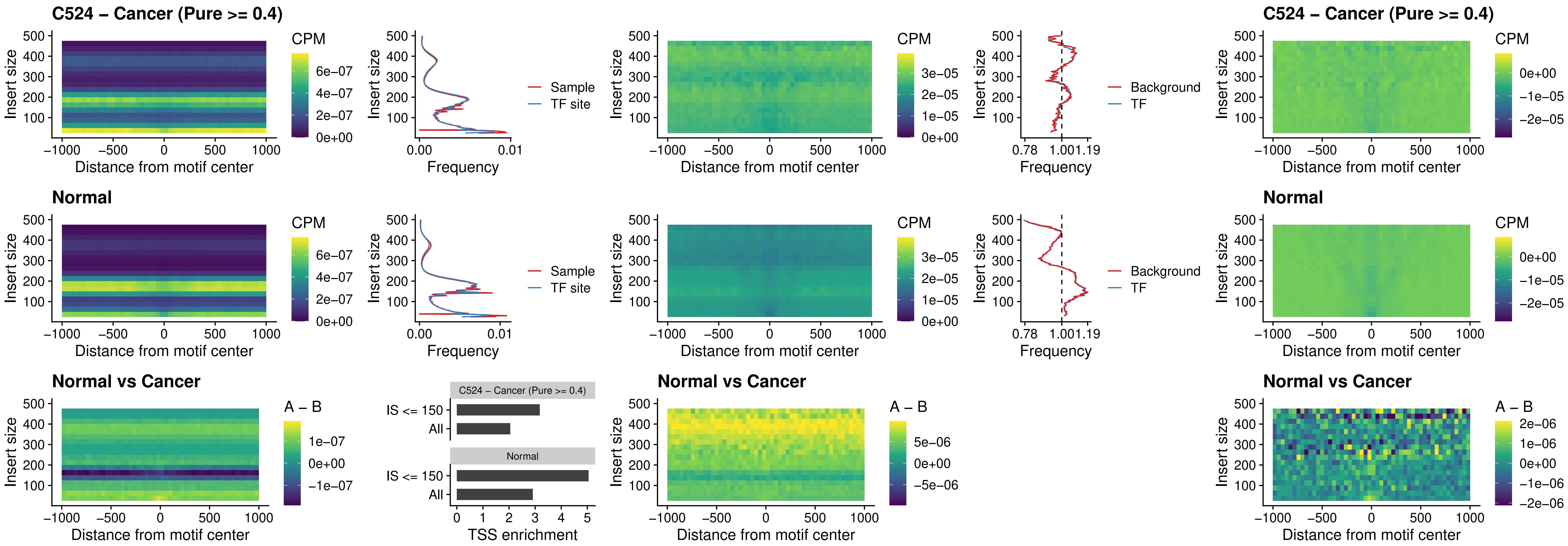

### C524 – Cancer (Pure >= 0.4) – SOX6_uc_dTSS_nPEAK

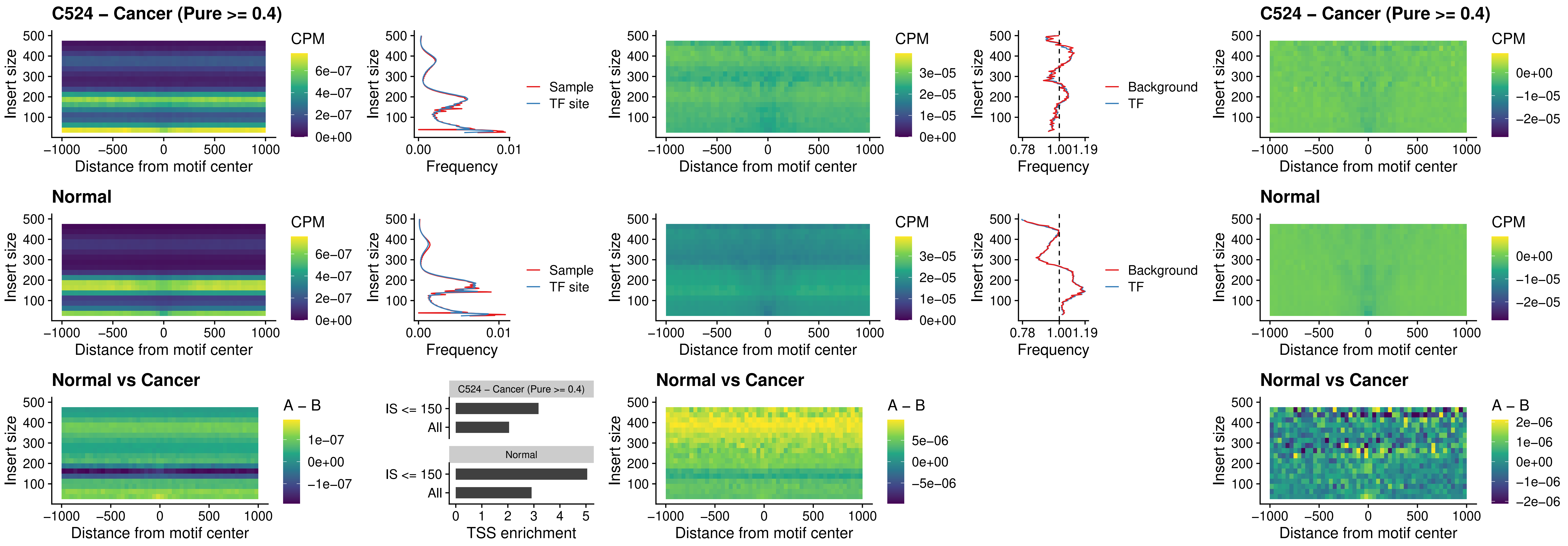

### C524 – Cancer (Pure >= 0.4) – JDP2_uc_cTSS_nPEAK

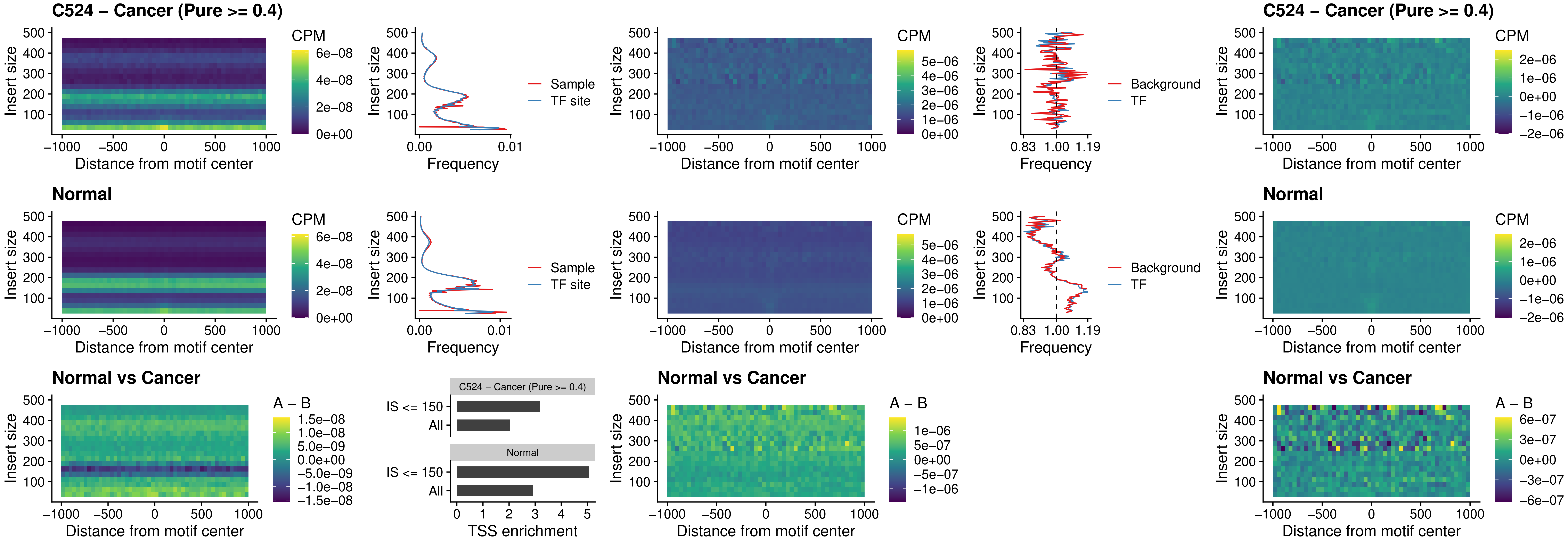

#### C524 – Cancer (Pure >= 0.4) – PURA_uc_dTSS_oPEAK

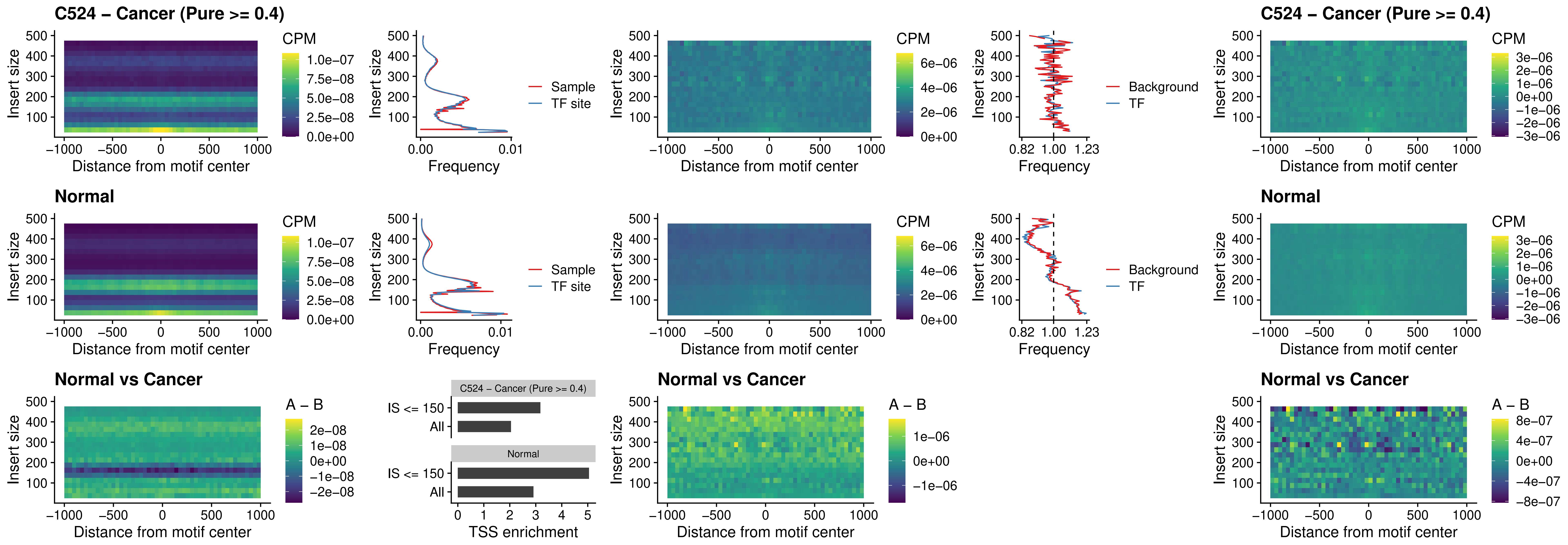

#### C524 – Cancer (Pure >= 0.4) – HHEX_uc_dTSS_nPEAK

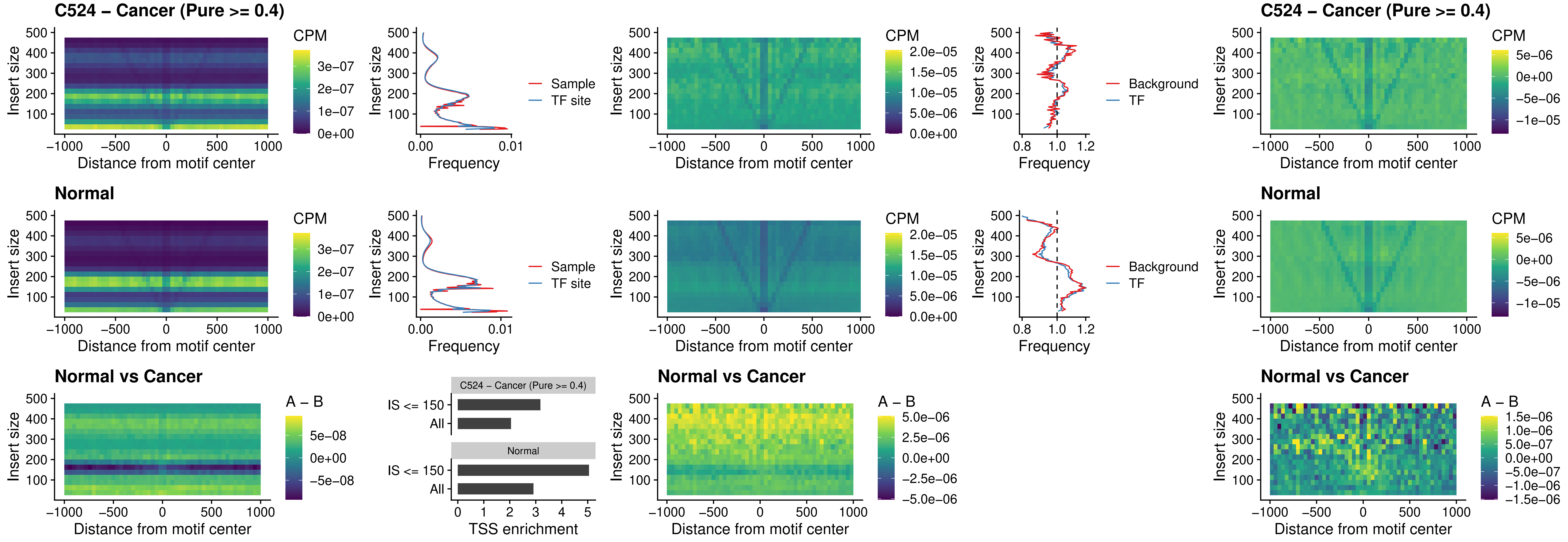

### C524 – Cancer (Pure >= 0.4) – HMG20B_uc_dTSS_nPEAK

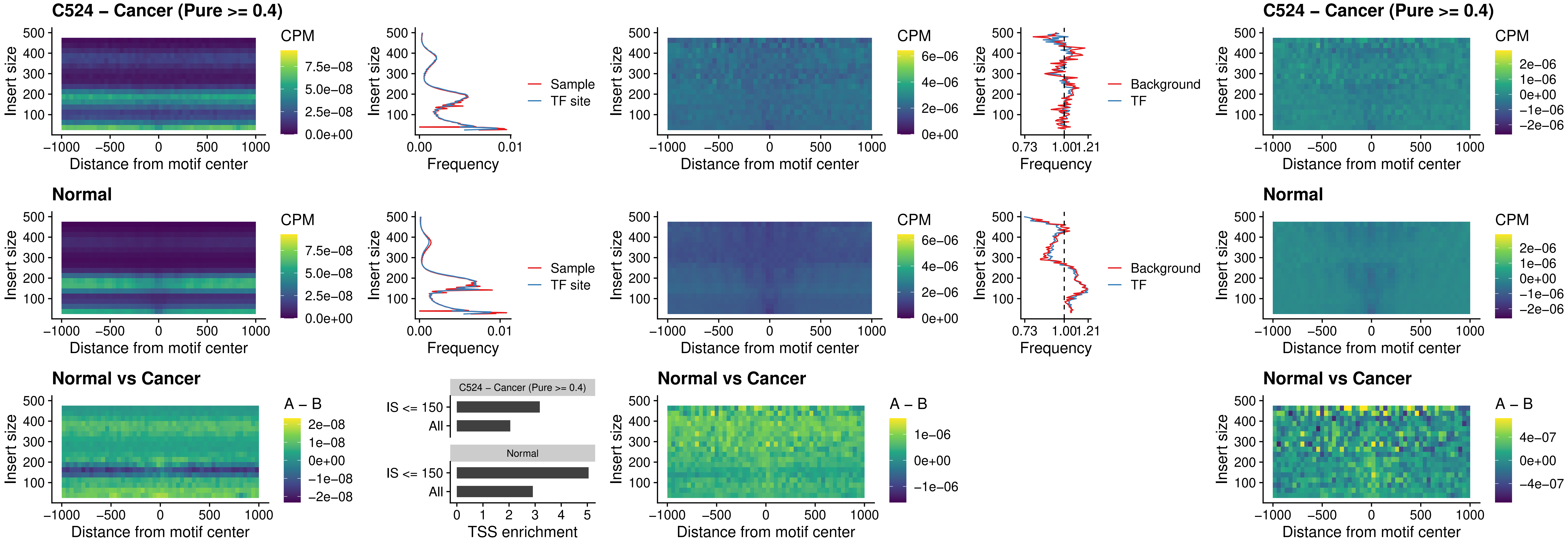

#### C524 – Cancer (Pure >= 0.4) – FOXB2_uc_dTSS_nPEAK

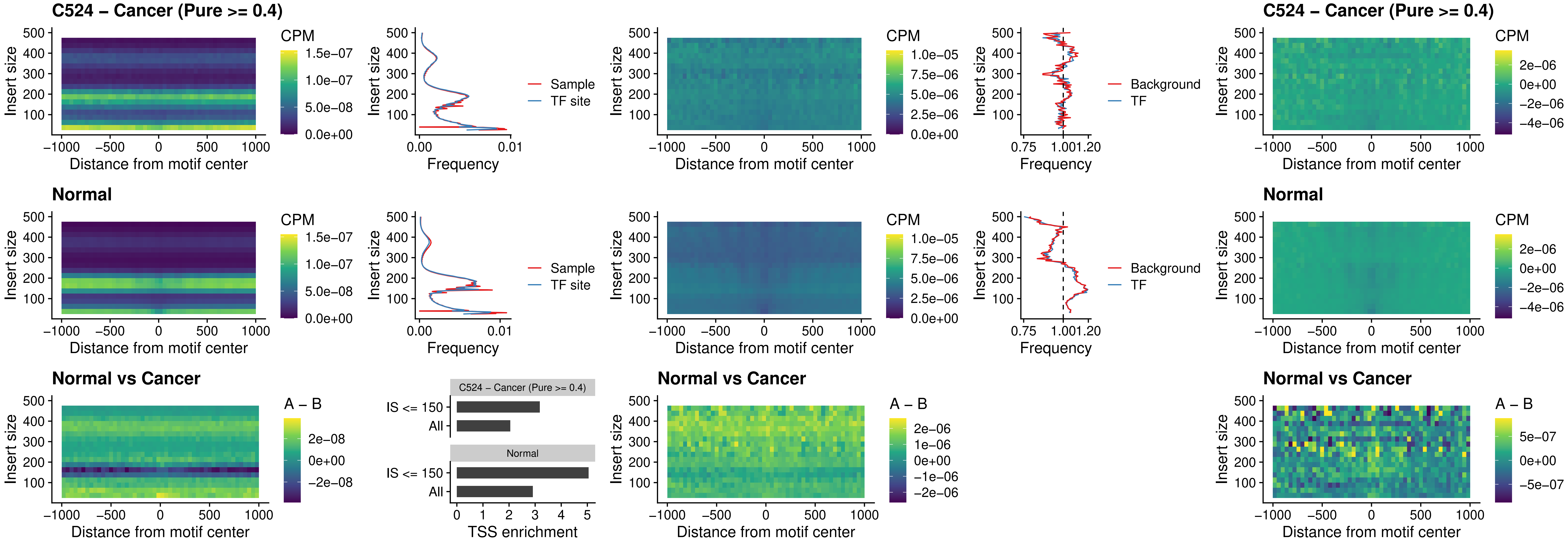

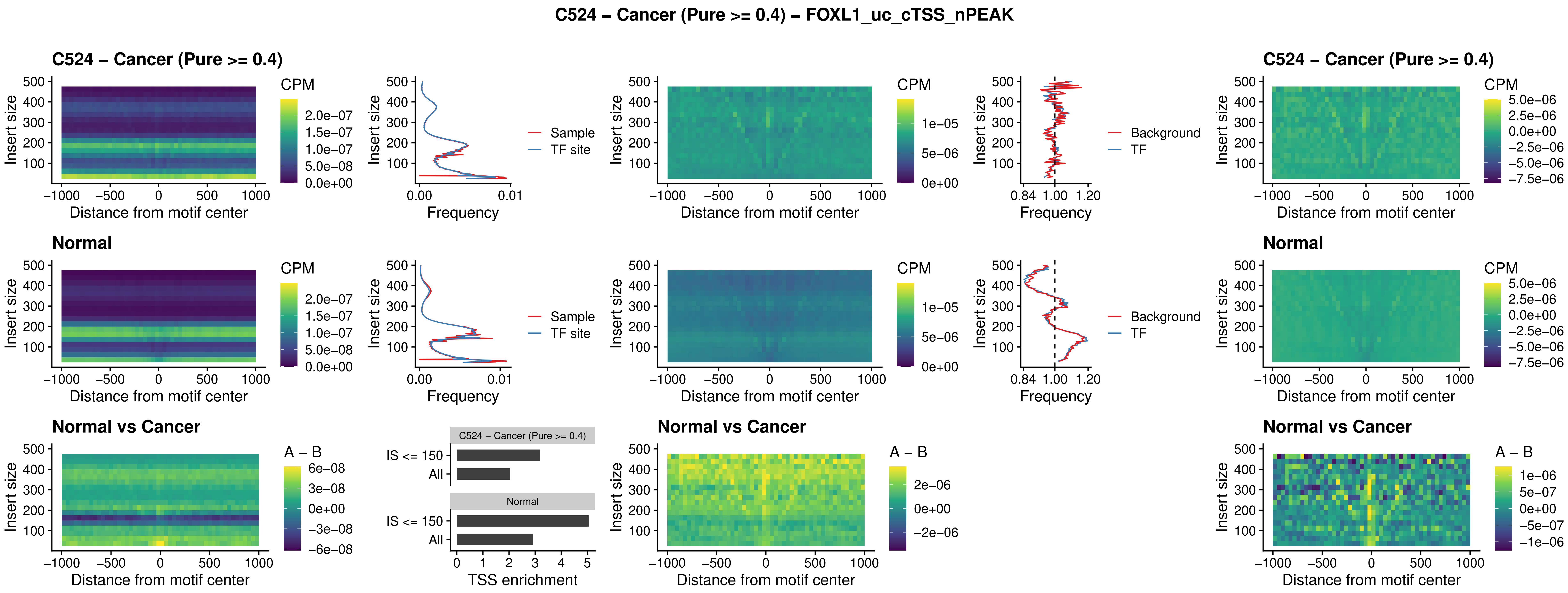
C524 – Cancer (Pure >= 0.4) – FOXL1_uc_cTSS_nPEAK
C524 – Cancer (Pure >= 0.4)
Normal
Normal vs Cancer
Insert size
Distance from motif center
Frequency
CPM
A – B
Sample
TF site
Background
TF
C524 – Cancer (Pure >= 0.4)
Normal
IS <= 150
All
TSS enrichment

#### C524 – Cancer (Pure >= 0.4) – PBX3_uc_pTSS_oPEAK

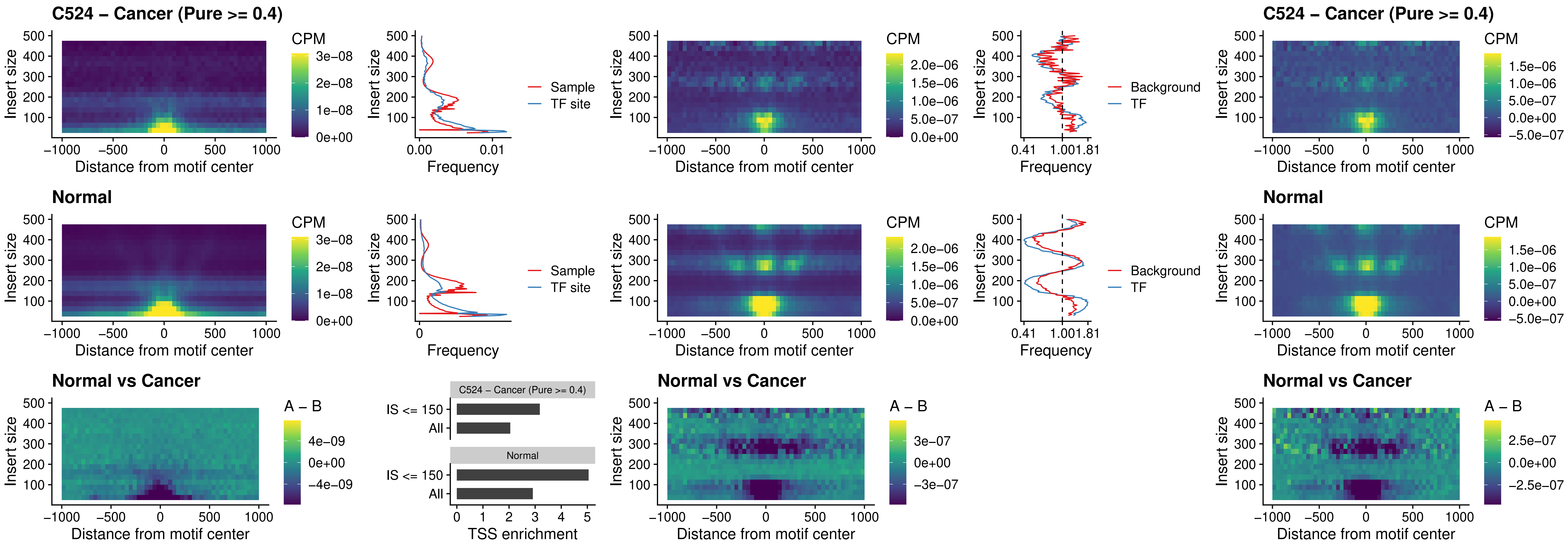

#### C524 – Cancer (Pure >= 0.4) – SNAI2_uc_cTSS_nPEAK

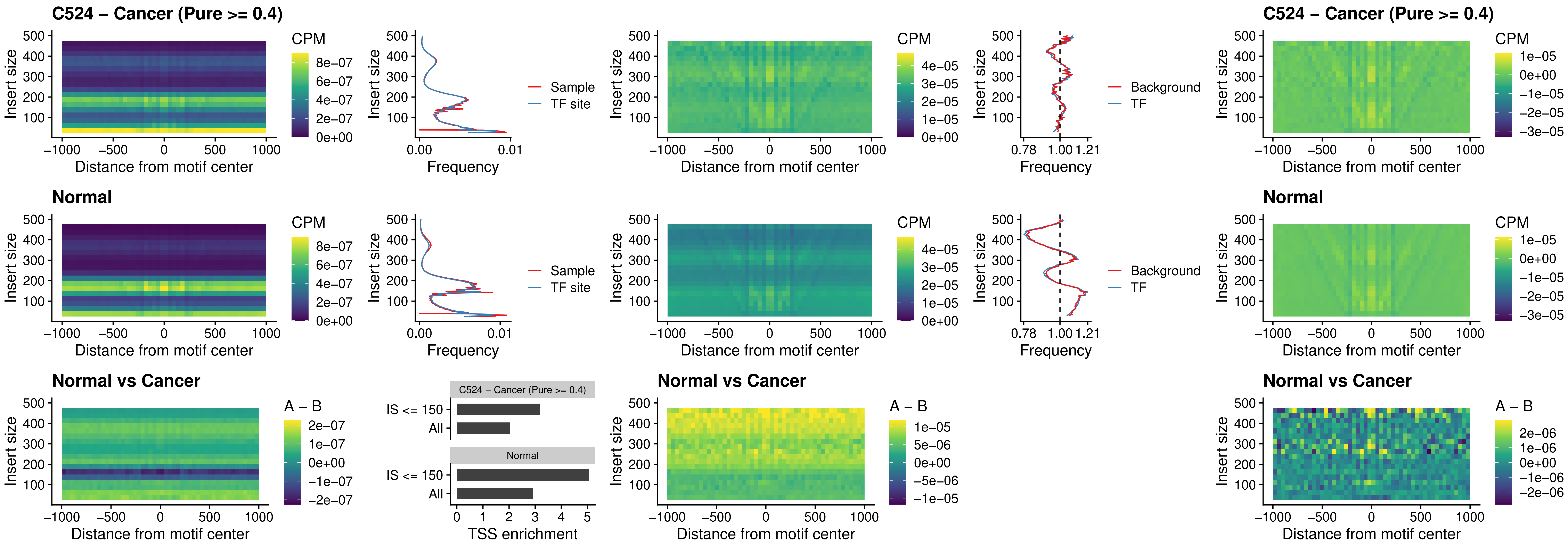

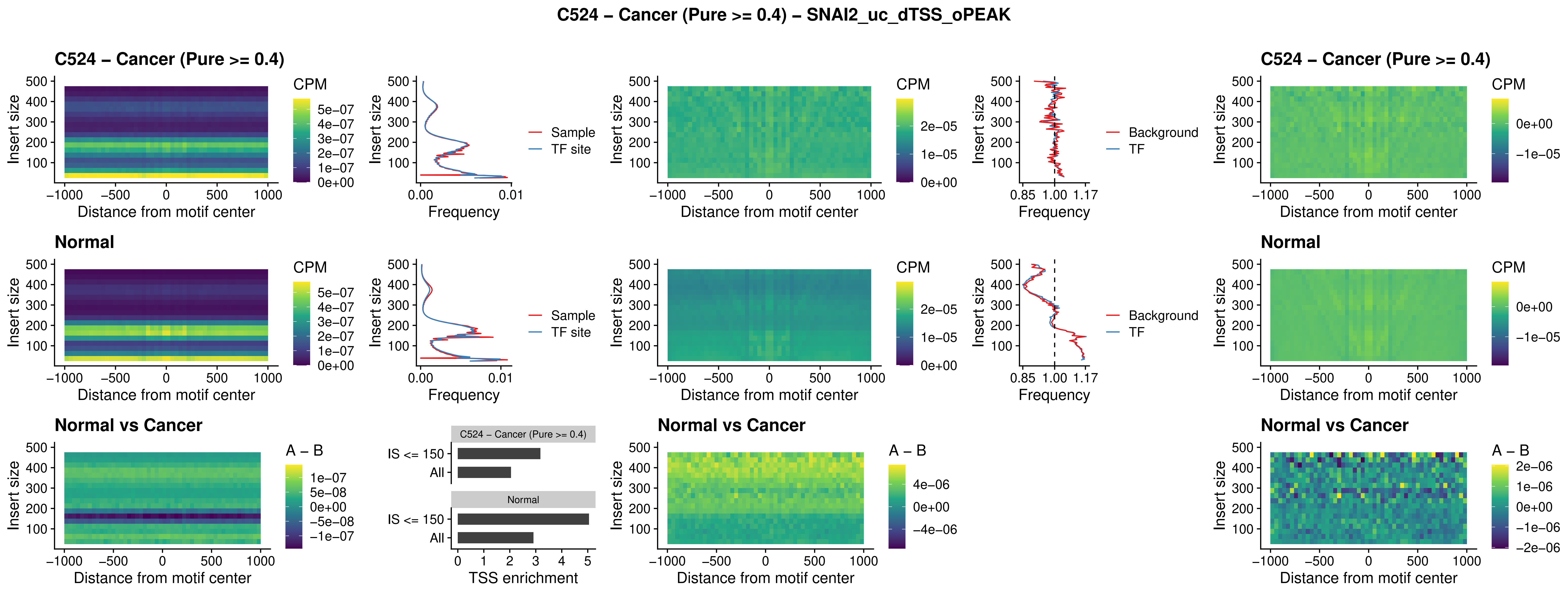
C524 – Cancer (Pure >= 0.4) – SNAI2_uc_dTSS_oPEAK
C524 – Cancer (Pure >= 0.4)
Normal
Normal vs Cancer
Insert size
Distance from motif center
Frequency
CPM
A – B
Sample
TF site
Background
TF
C524 – Cancer (Pure >= 0.4)
Normal
IS <= 150
All
TSS enrichment

#### C524 – Cancer (Pure >= 0.4) – SPI1_uc_pTSS_nPEAK

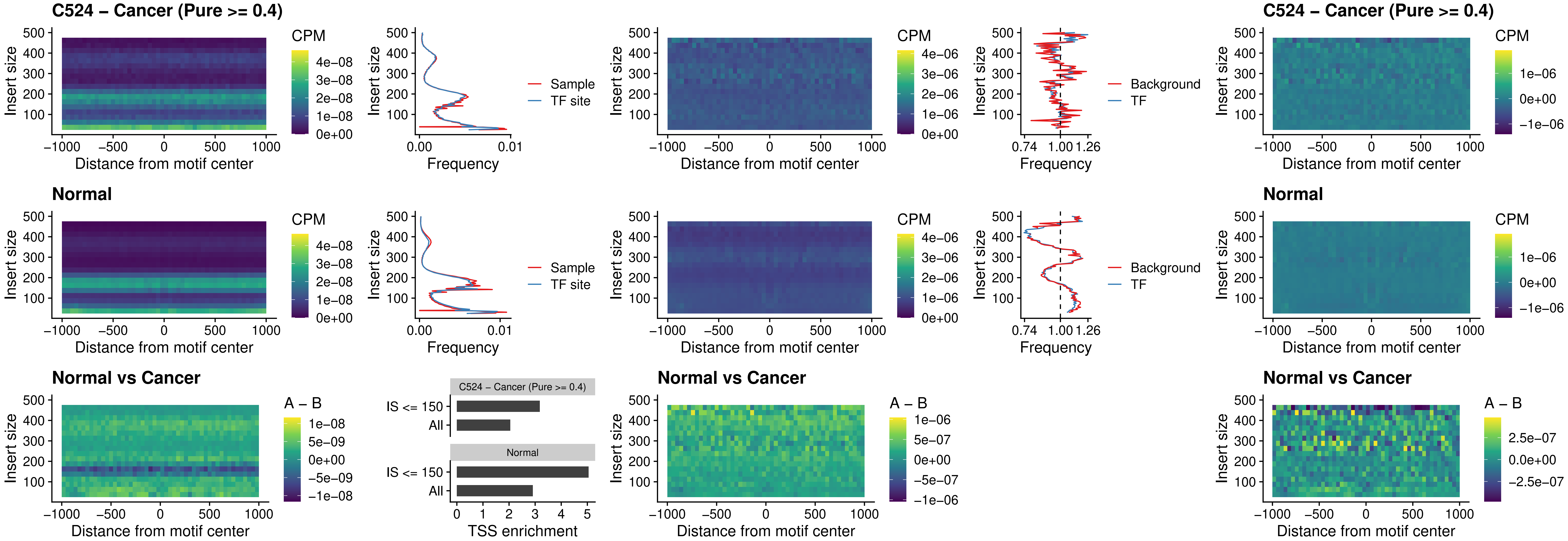

### C524 – Cancer (Pure >= 0.4) – KLF16_uc_pTSS_nPEAK

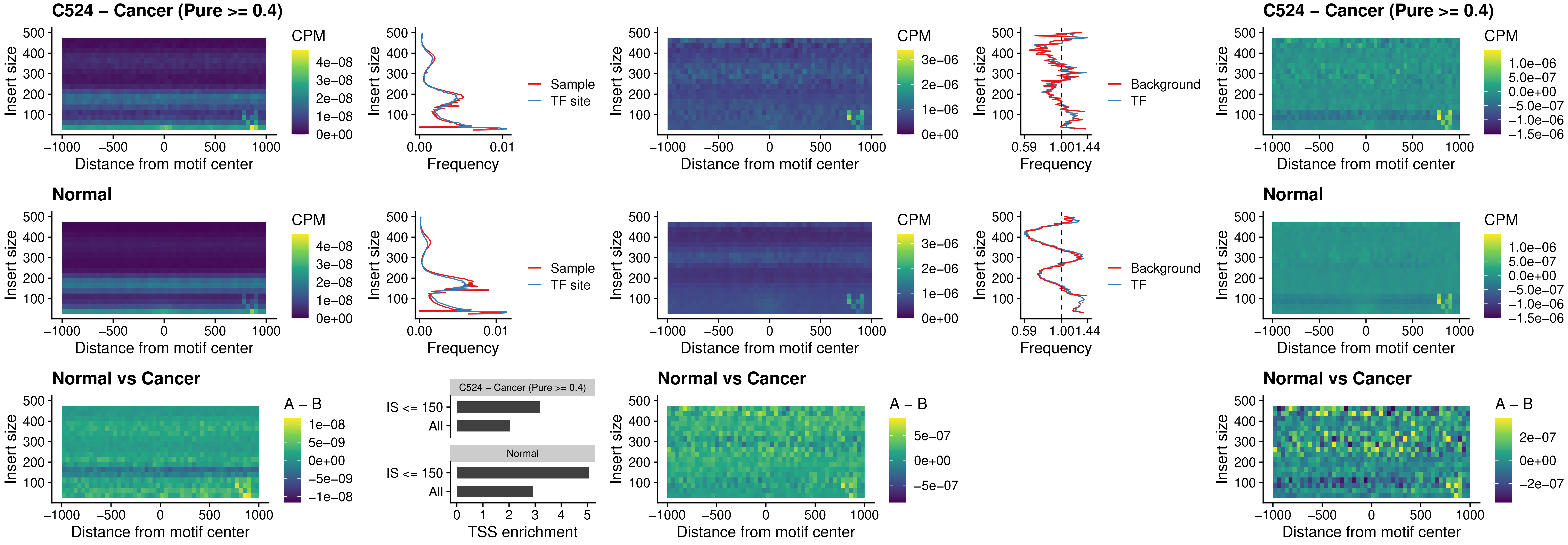

#### C524 – Cancer (Pure >= 0.4) – SP3_uc_pTSS_nPEAK

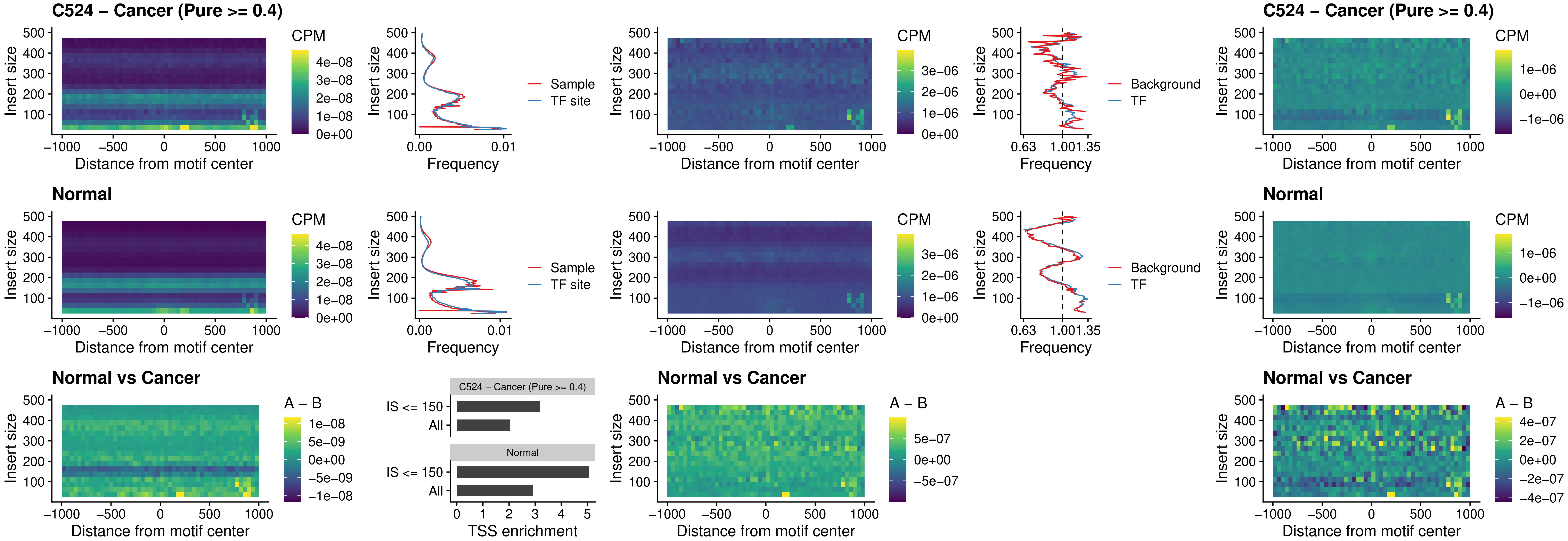

#### C524 – Cancer (Pure >= 0.4) – NEUROD1_uc_dTSS_oPEAK

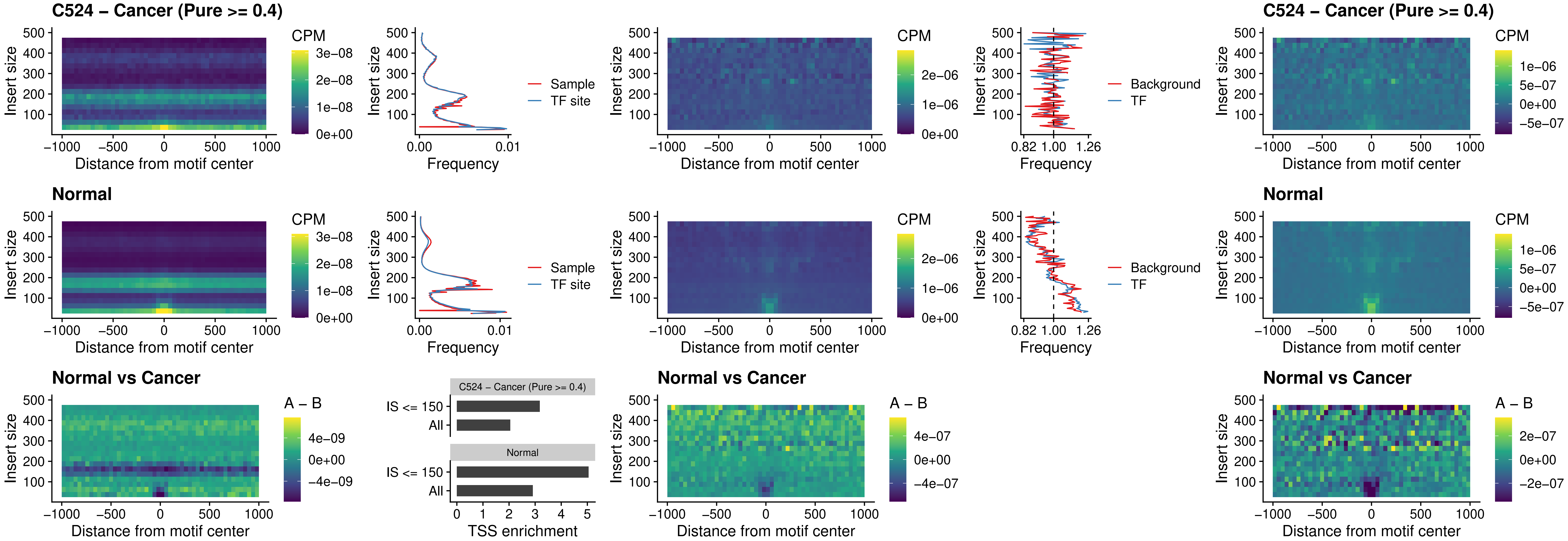

#### C524 – Cancer (Pure >= 0.4) – CTCF_uc_dTSS_oPEAK

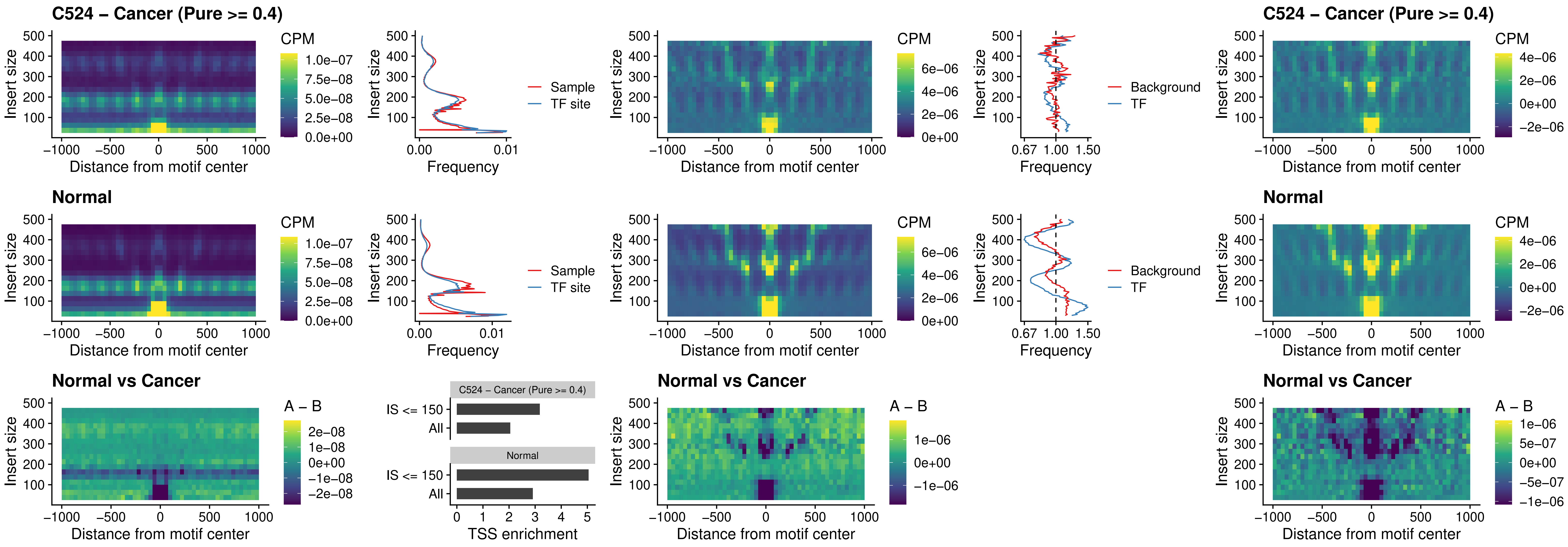

#### C524 – Cancer (Pure >= 0.4) – CTCF_uc_cTSS_oPEAK

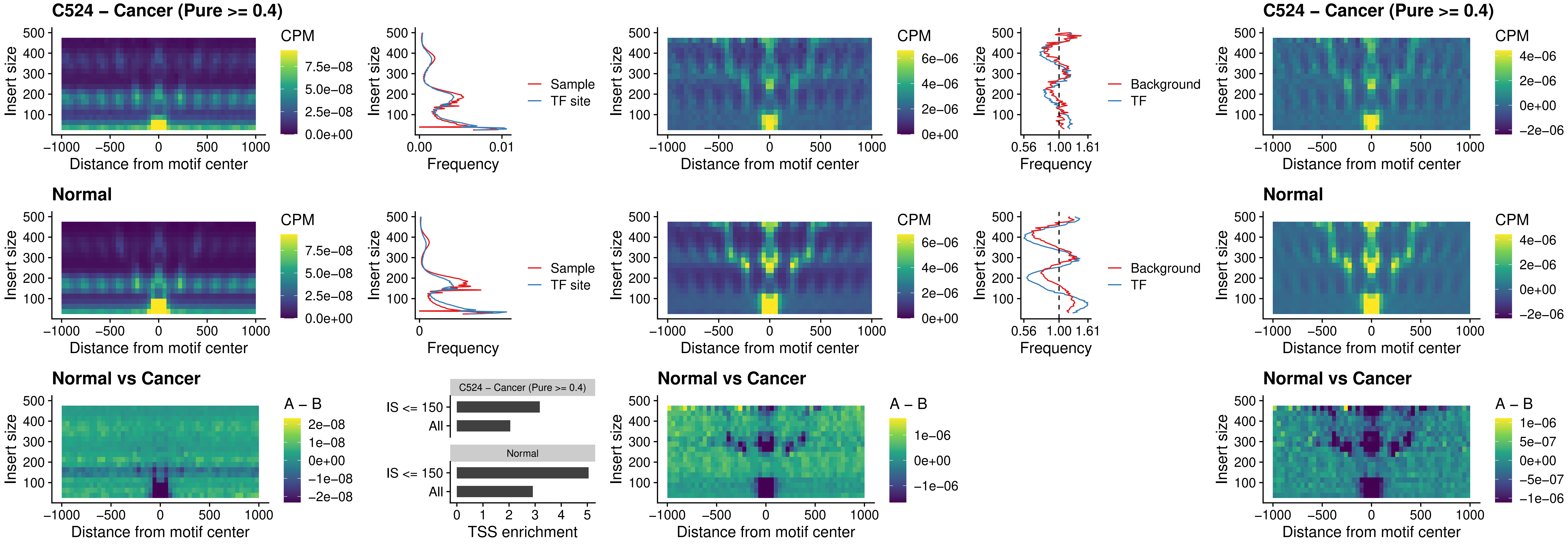

#### C524 – Cancer (Pure >= 0.4) – CTCFL_uc_dTSS_oPEAK

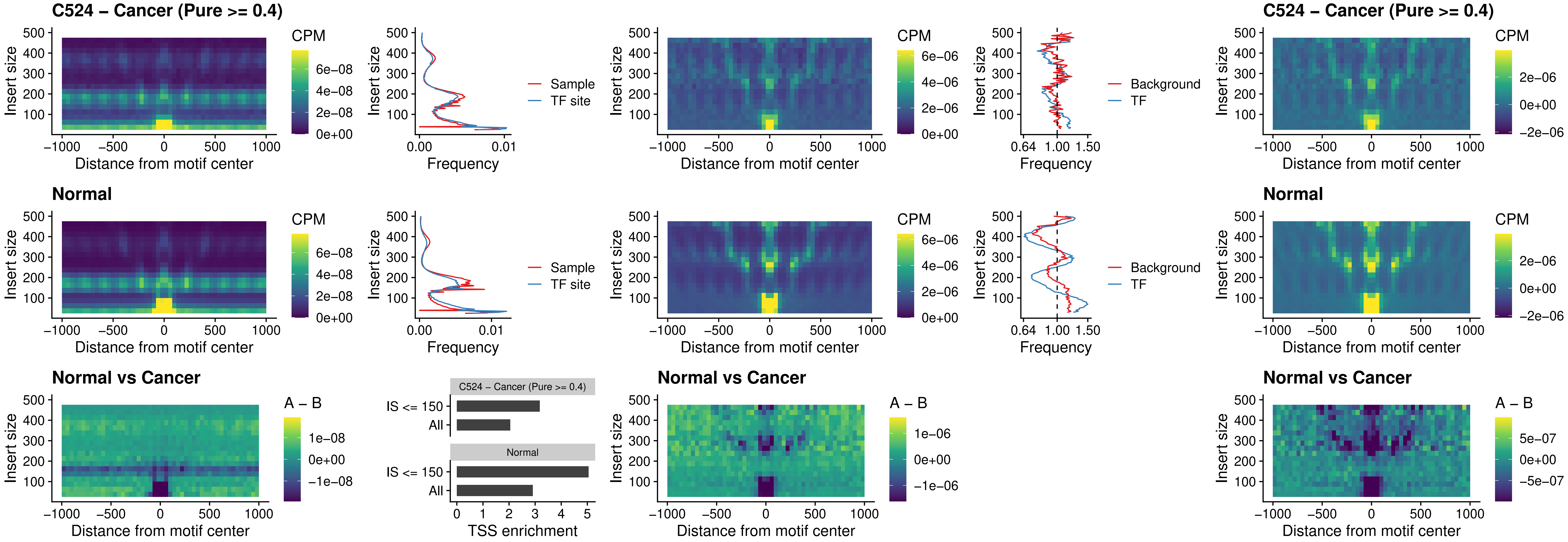

#### C524 – Cancer (Pure >= 0.4) – ENCODE_CTCF_sites_COLON

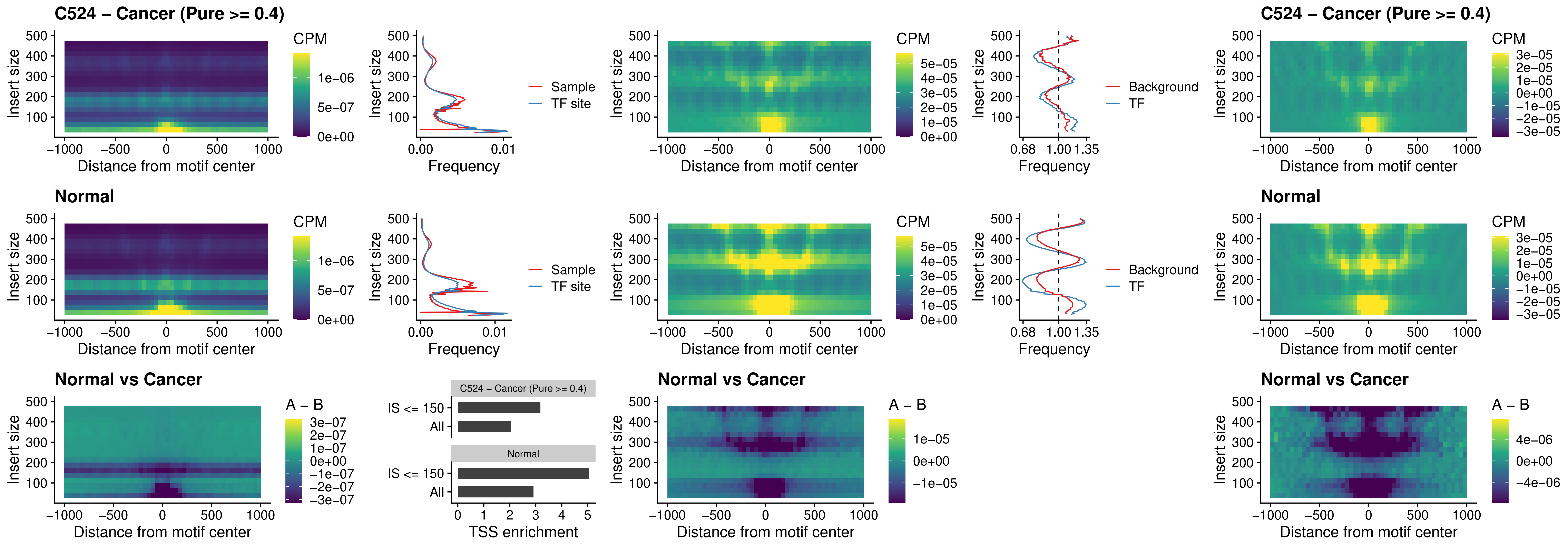

#### C524 – Cancer (Pure >= 0.4) – ENCODE_CTCF_sites_COLON_loop_ctcf

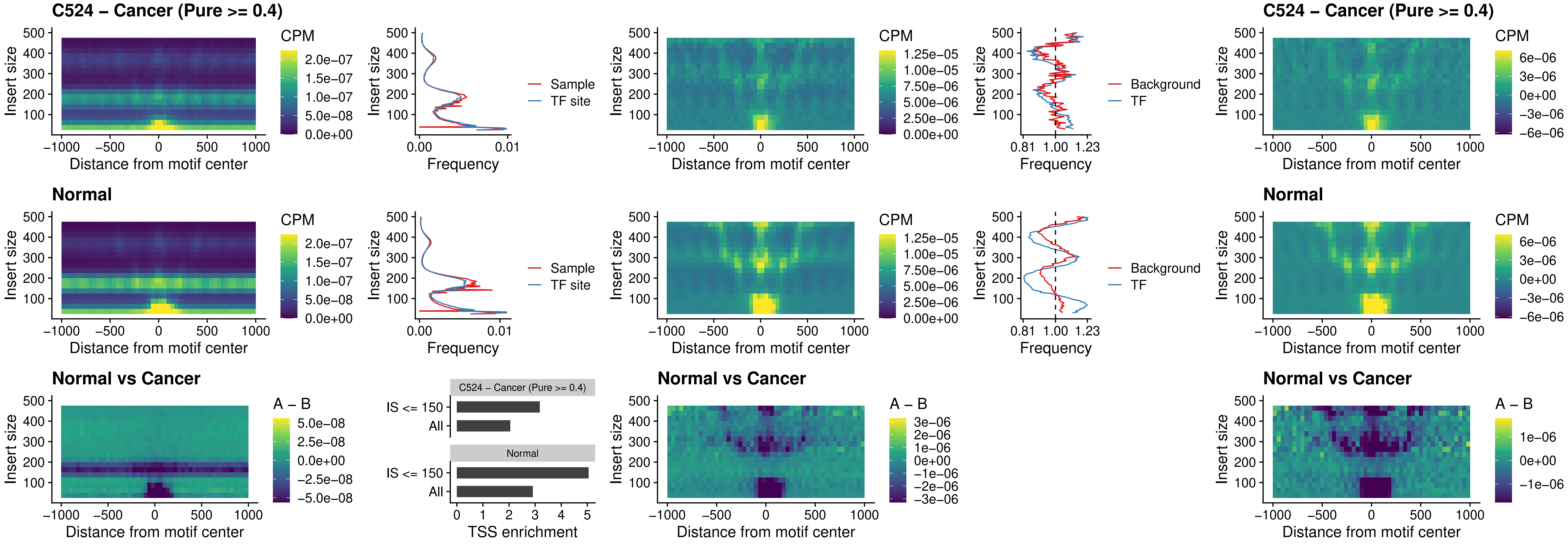

#### C524 – Cancer (Pure >= 0.4) – ENCODE_CTCF_sites_COLON_intergenic_ctcf

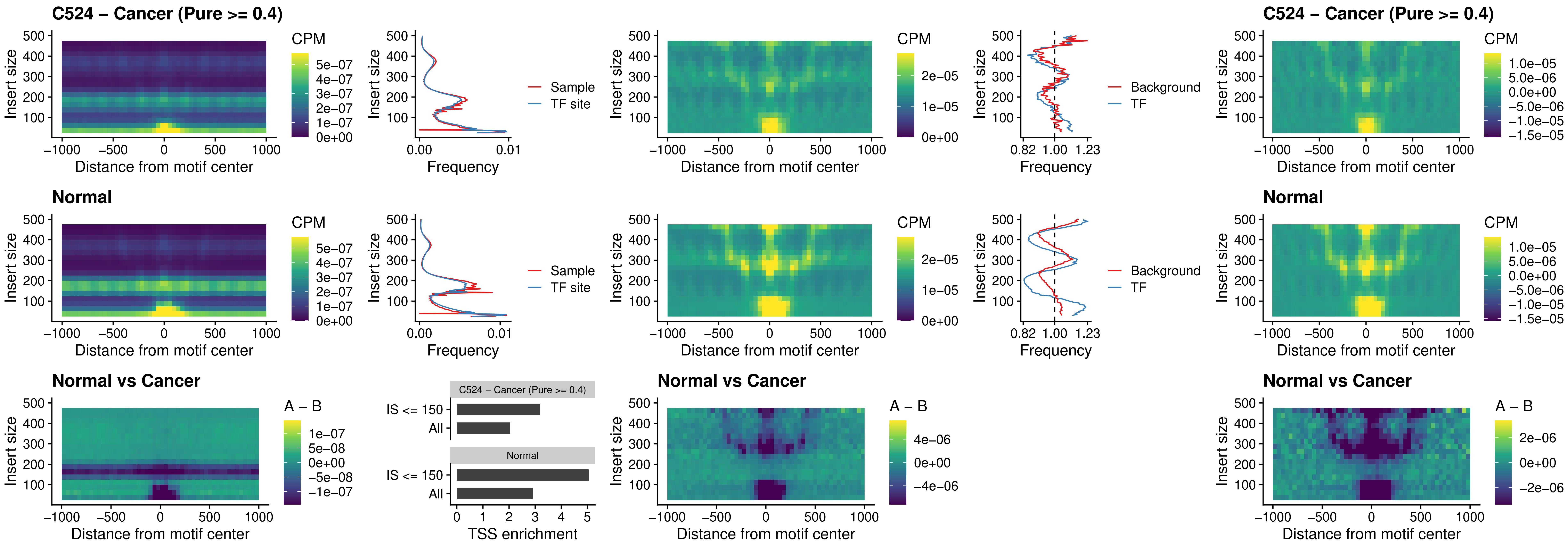

#### C524 – Cancer (Pure >= 0.4) – ENCODE_CTCF_sites_COLON_nonpromotor

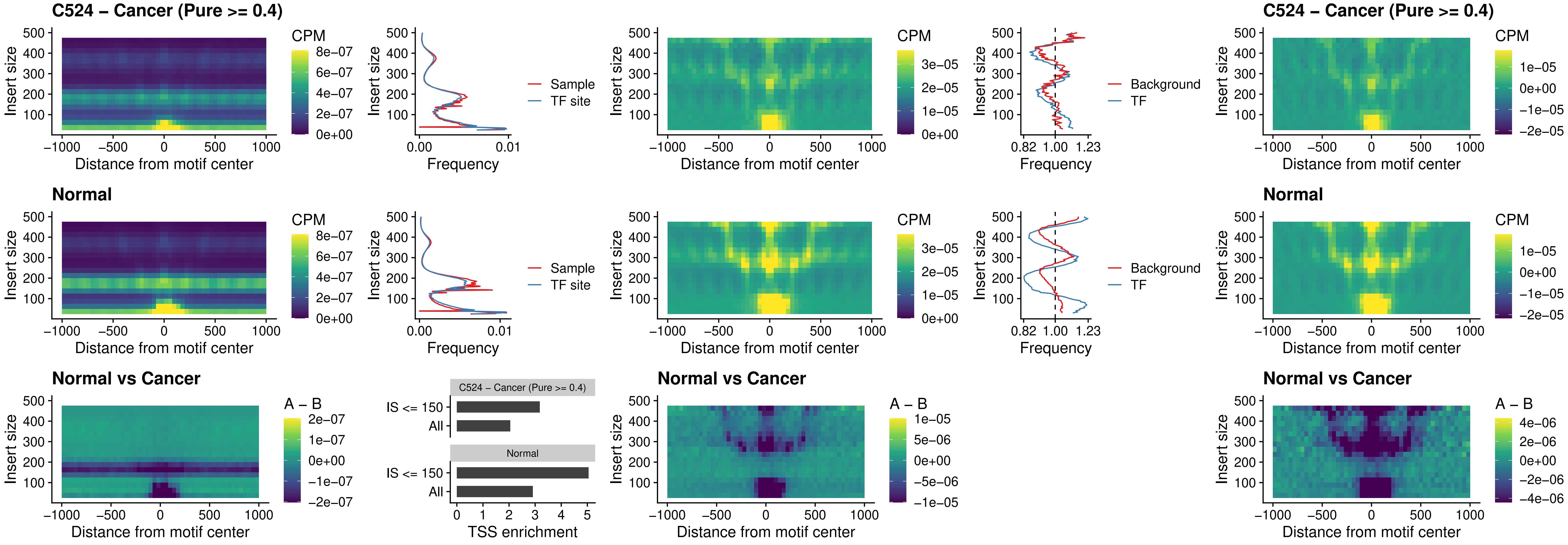

#### C524 – Cancer (Pure >= 0.4) – CTCFL_uc_cTSS_oPEAK

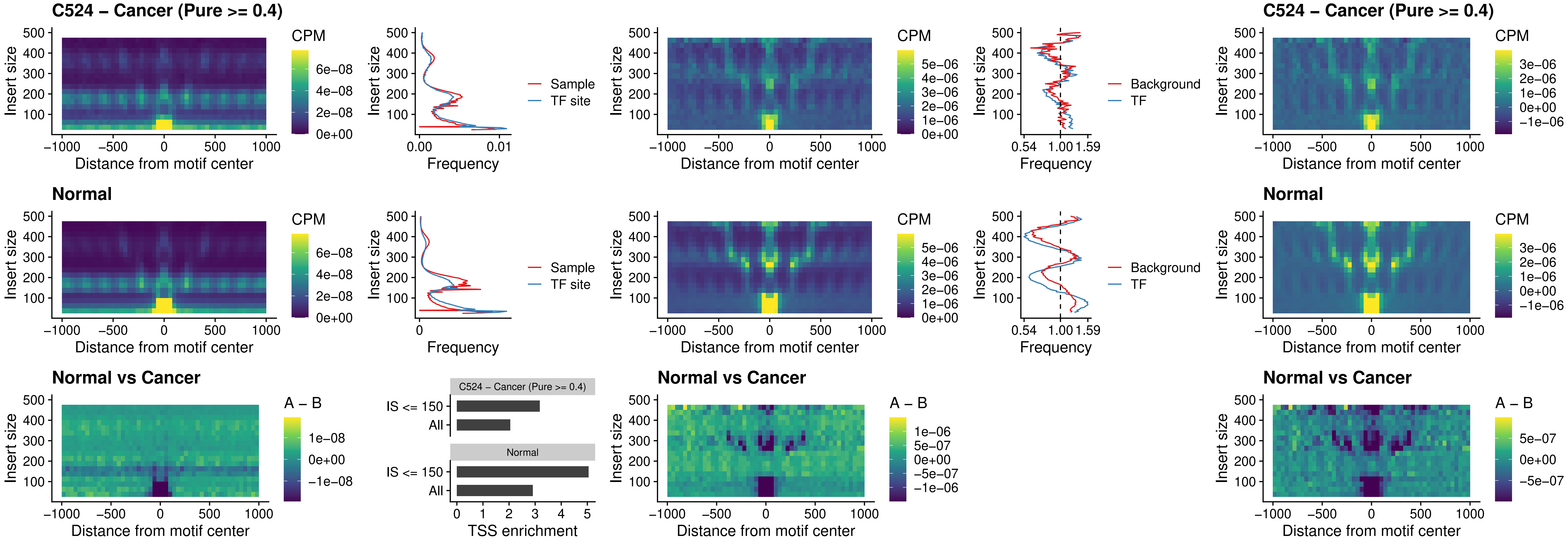

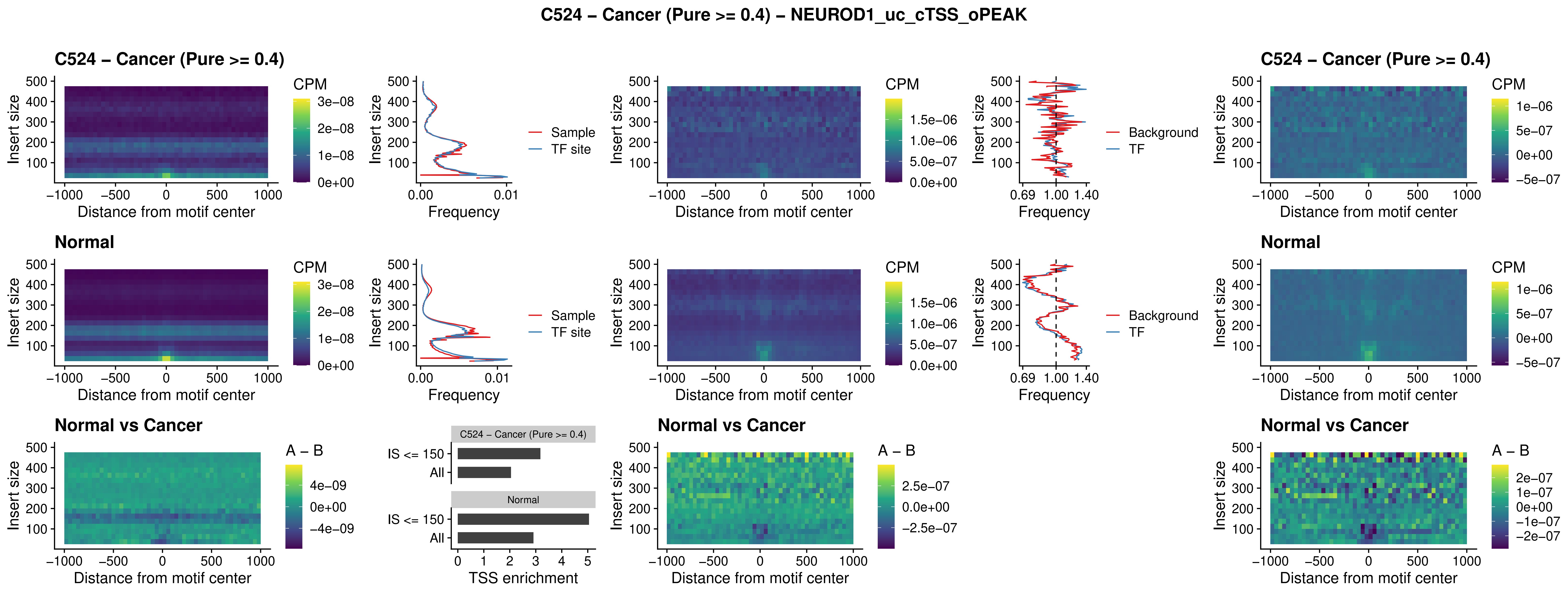
C524 – Cancer (Pure >= 0.4) – NEUROD1_uc_cTSS_oPEAK
C524 – Cancer (Pure >= 0.4)
Normal
Normal vs Cancer
Insert size
Distance from motif center
CPM
A – B
Frequency
Sample
TF site
Background
TF
TSS enrichment
C524 – Cancer (Pure >= 0.4)
Normal
IS <= 150
All

#### C524 – Cancer (Pure >= 0.4) – ENCODE_YY1_sites_COLON_nonpromotor

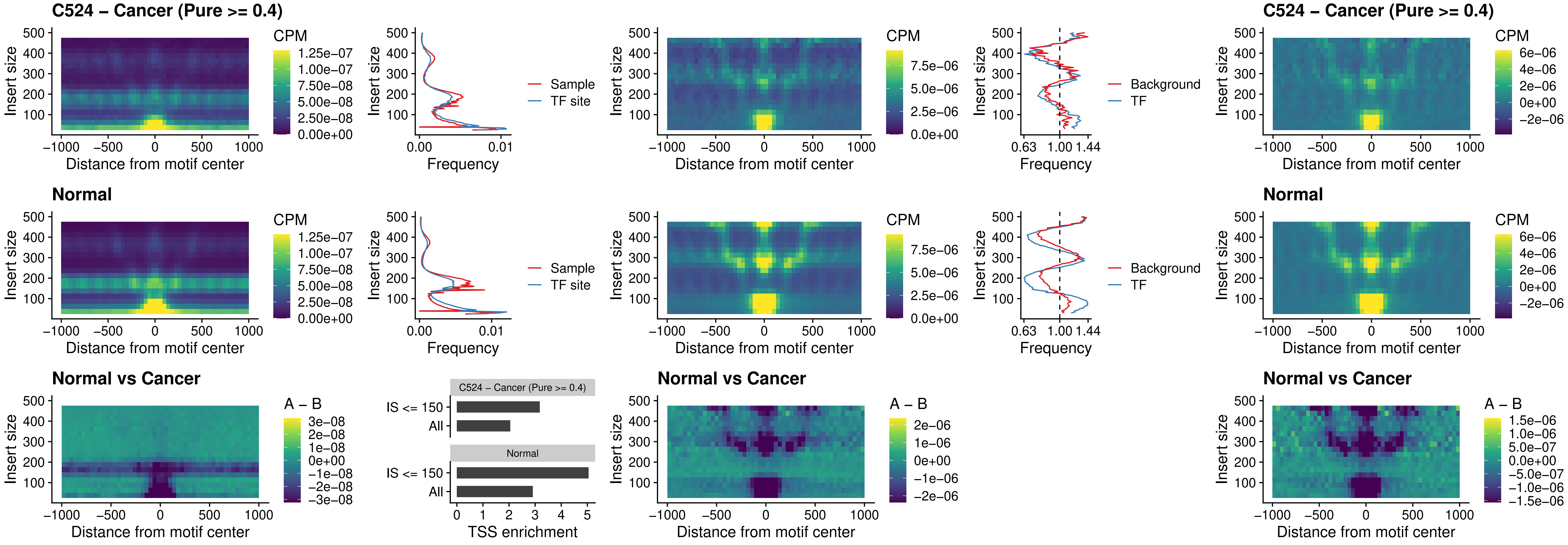

### C524 – Cancer (Pure >= 0.4) – CTCF_uc_pTSS_oPEAK

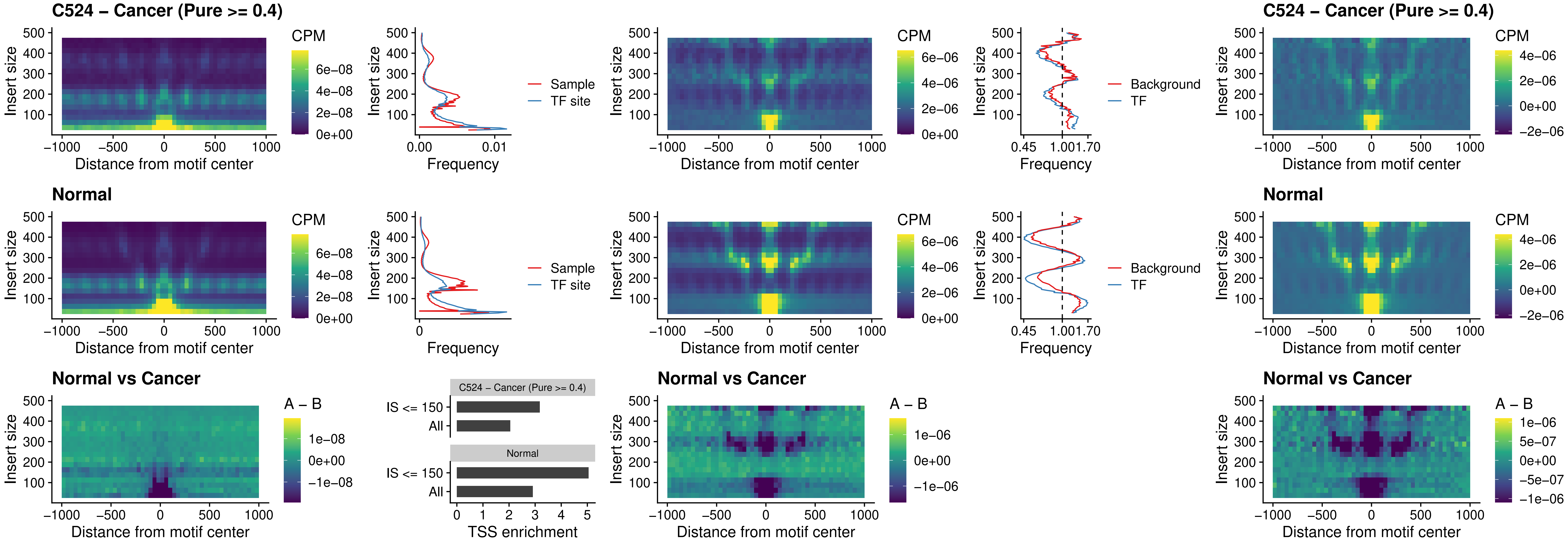

#### C524 – Cancer (Pure >= 0.4) – ENCODE_CTCF_sites_COLON_promotor

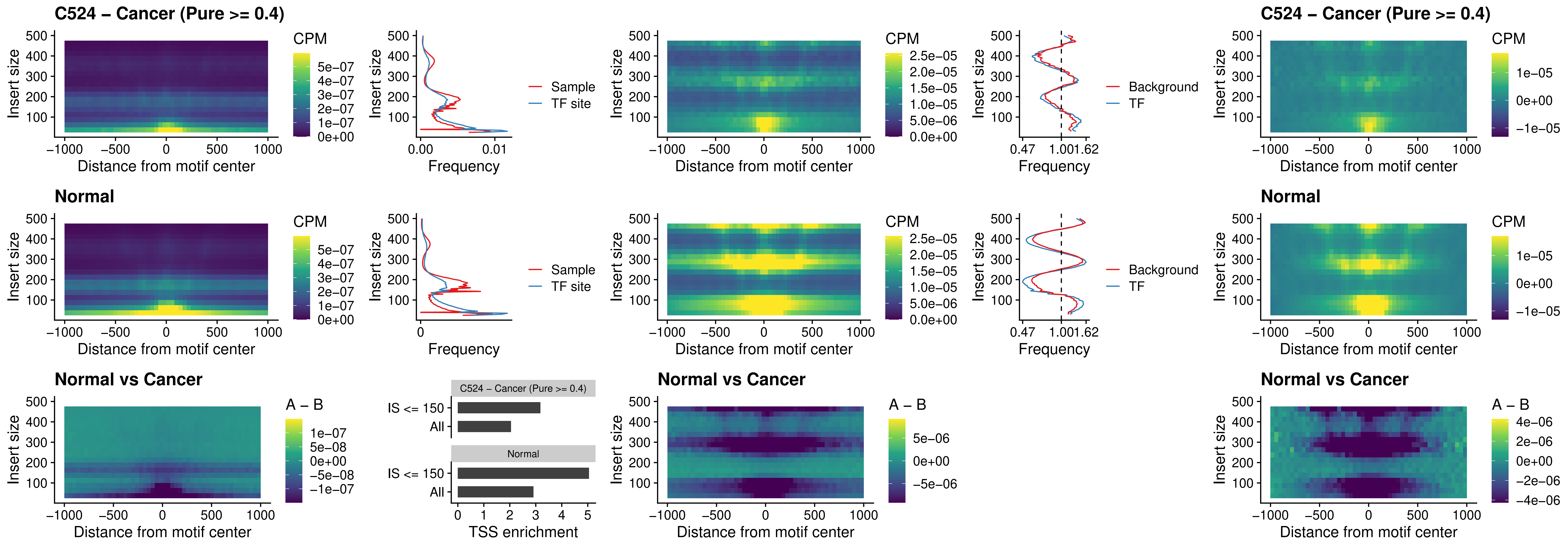

#### C524 – Cancer (Pure >= 0.4) – ENCODE_CTCF_sites_COLON_promotor_ctcf

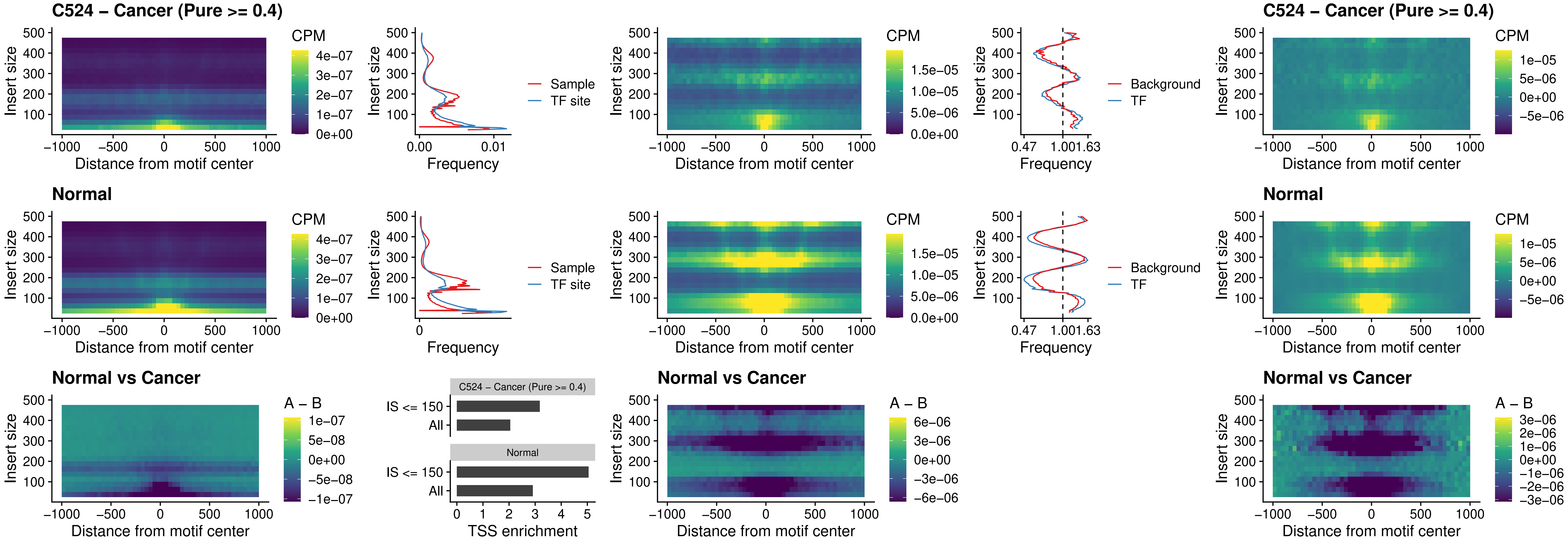

#### C524 – Cancer (Pure >= 0.4) – SPIC_uc_pTSS_oPEAK

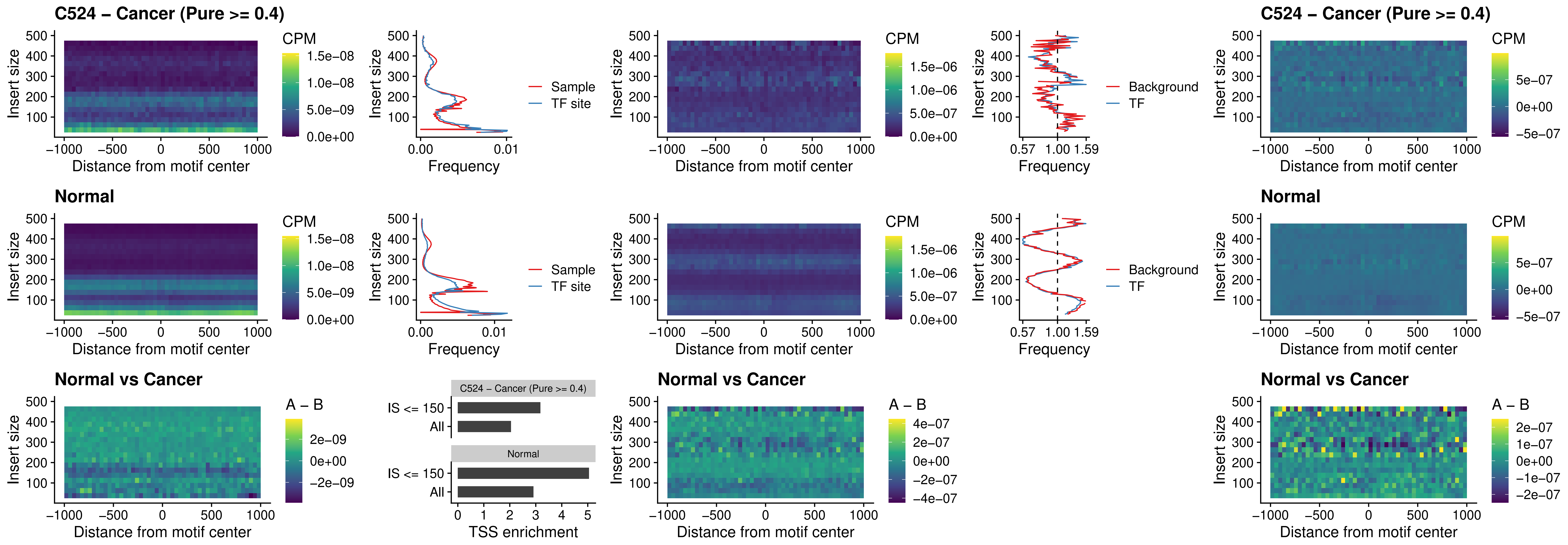

#### C524 – Cancer (Pure >= 0.4) – SPI1_uc_dTSS_nPEAK

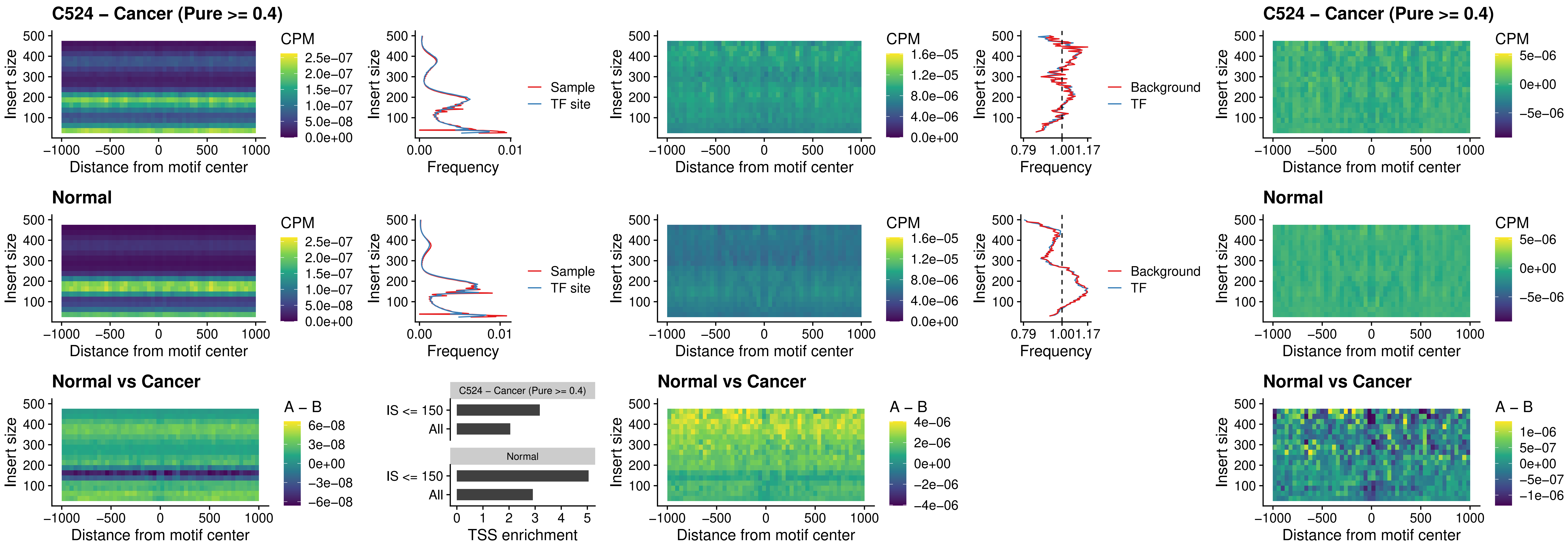

#### C524 – Cancer (Pure >= 0.4) – SPI1_uc_cTSS_nPEAK

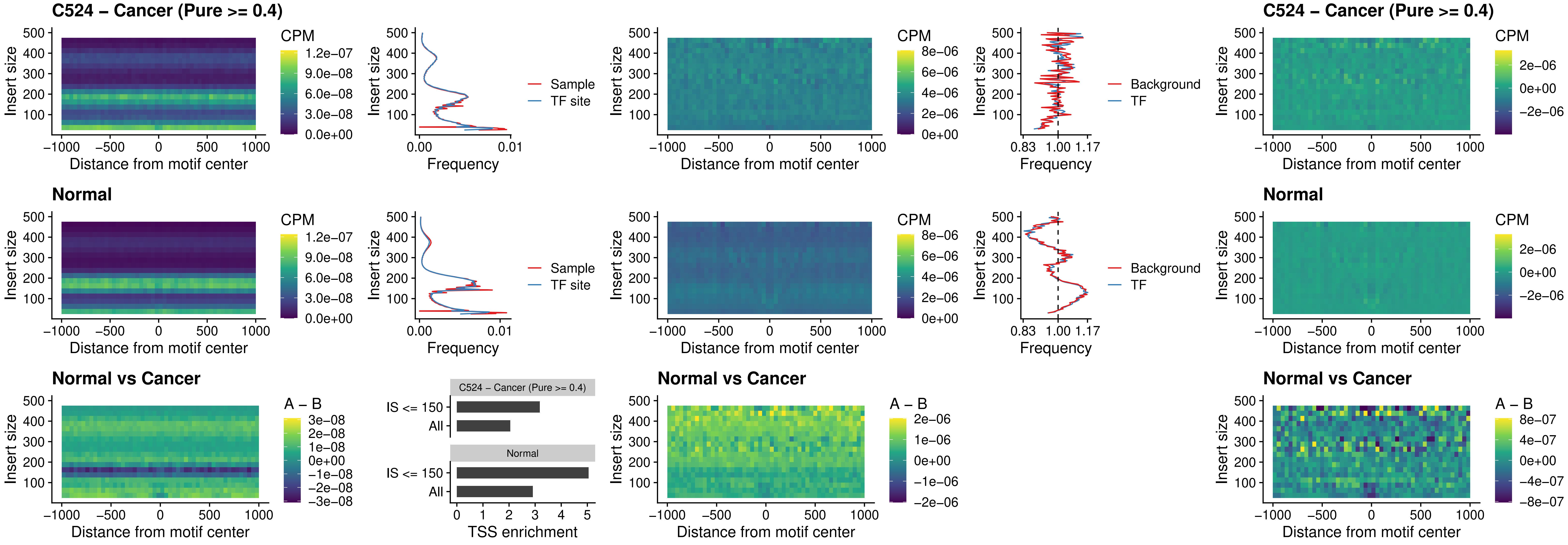

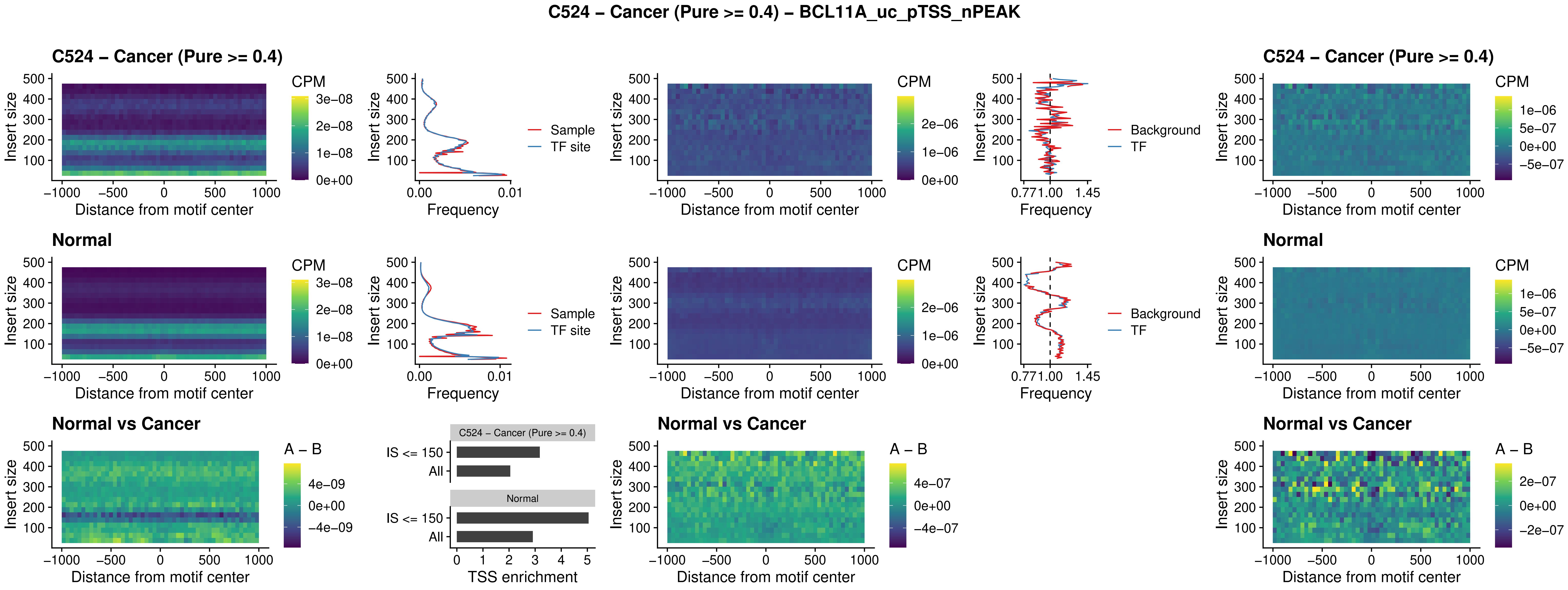
C524 – Cancer (Pure >= 0.4) – BCL11A_uc_pTSS_nPEAK
C524 – Cancer (Pure >= 0.4)
Normal
Normal vs Cancer
Insert size
Distance from motif center
Frequency
CPM
A – B
Sample
TF site
Background
TF
C524 – Cancer (Pure >= 0.4)
Normal
IS <= 150
All
TSS enrichment

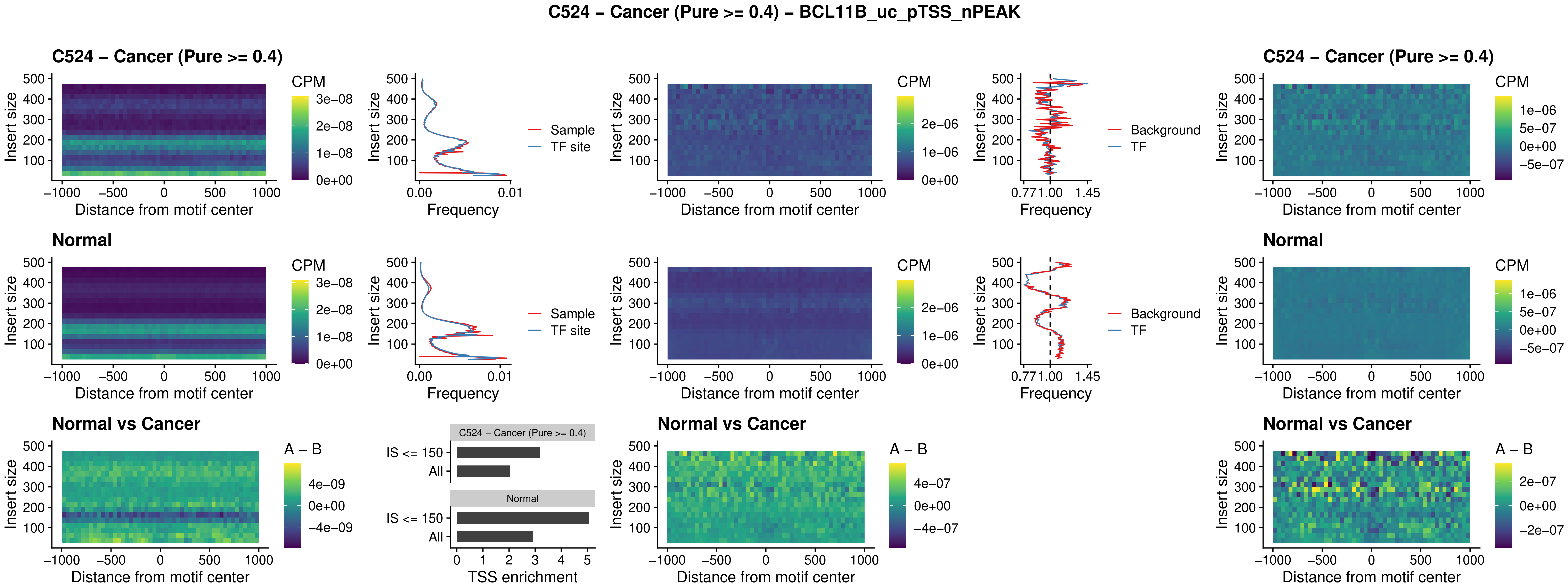
C524 – Cancer (Pure >= 0.4) – BCL11B_uc_pTSS_nPEAK
C524 – Cancer (Pure >= 0.4)
Normal
Normal vs Cancer
Insert size
Distance from motif center
Frequency
CPM
A – B
Sample
TF site
Background
TF
C524 – Cancer (Pure >= 0.4)
Normal
IS <= 150
All
TSS enrichment

### C524 – Cancer (Pure >= 0.4) – IRF9_uc_dTSS_oPEAK

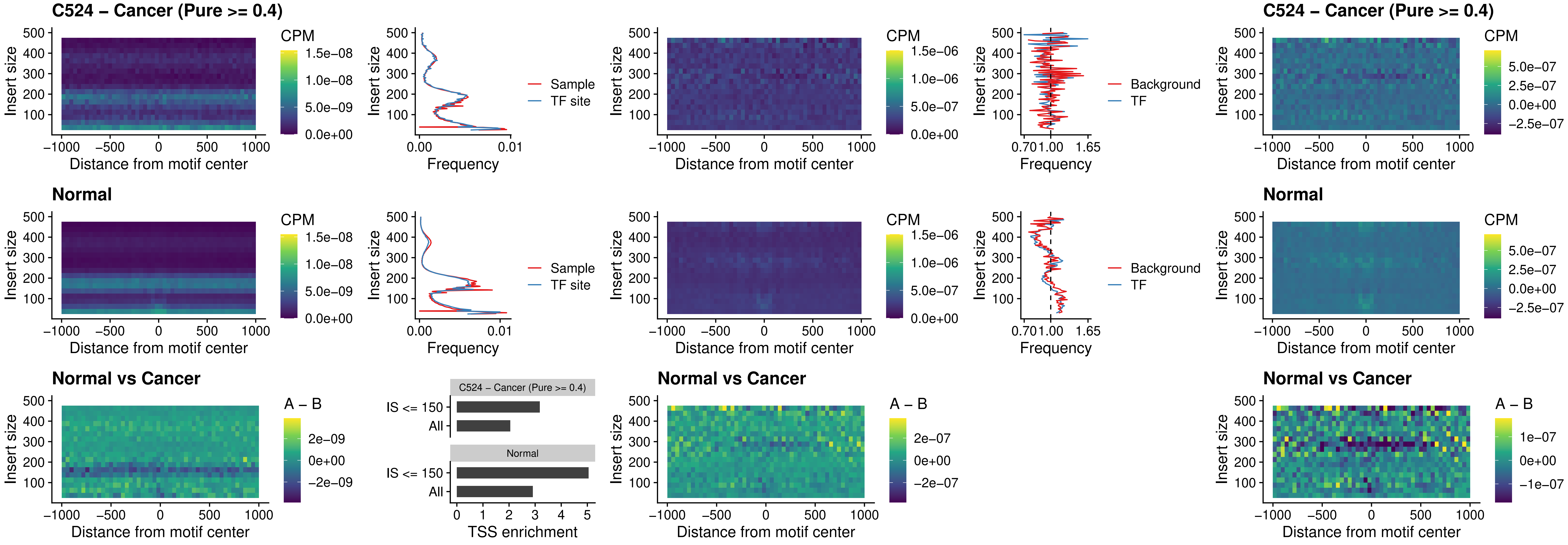

#### C524 – Cancer (Pure >= 0.4) – IRF9_uc_cTSS_oPEAK

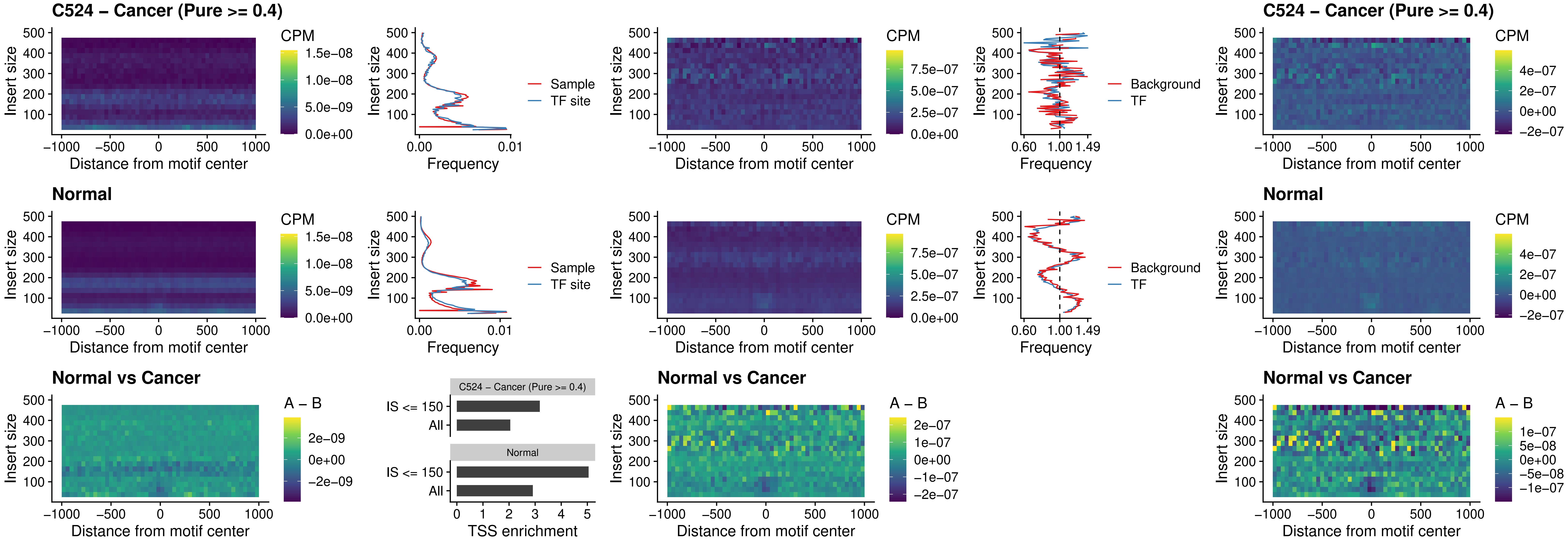

### C524 – Cancer (Pure >= 0.4) – BCL11A_uc_cTSS_oPEAK

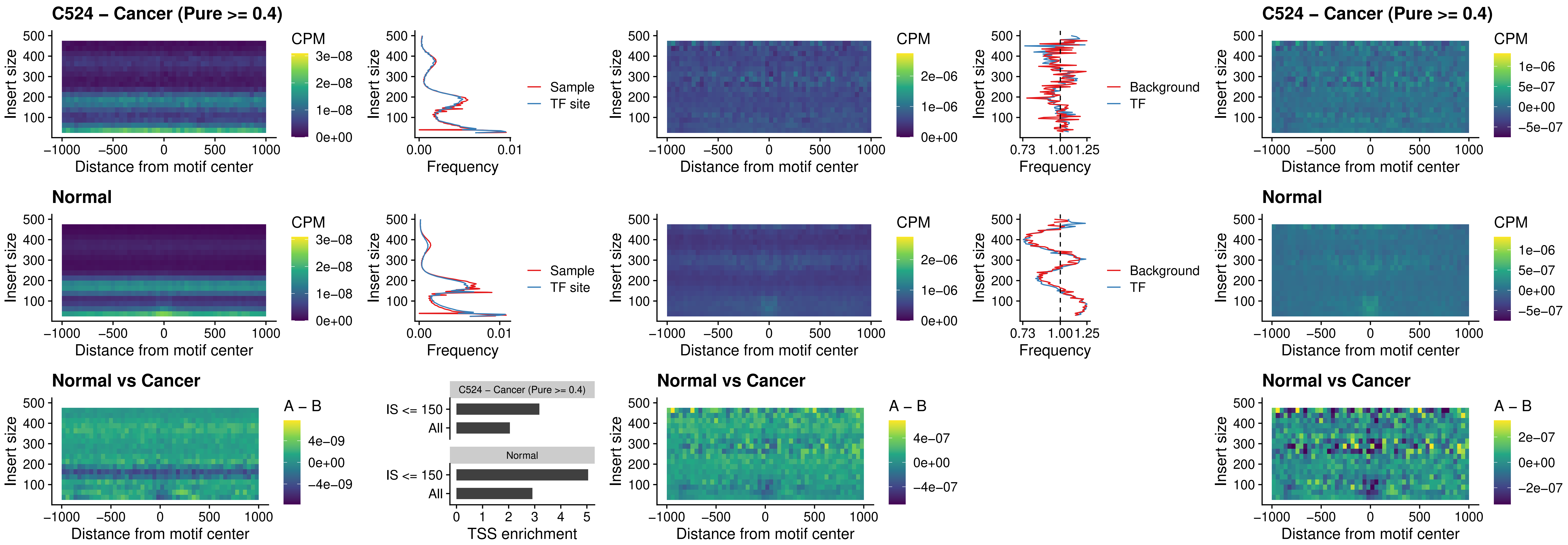

#### C524 – Cancer (Pure >= 0.4) – BCL11B_uc_cTSS_oPEAK

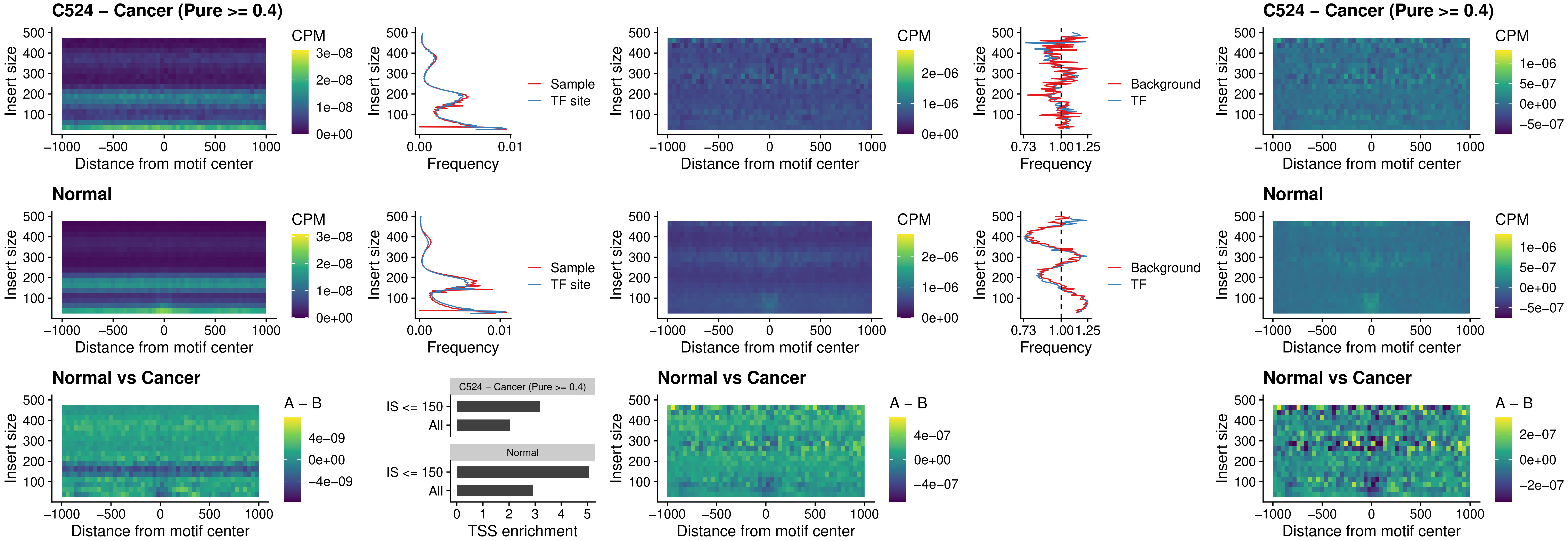

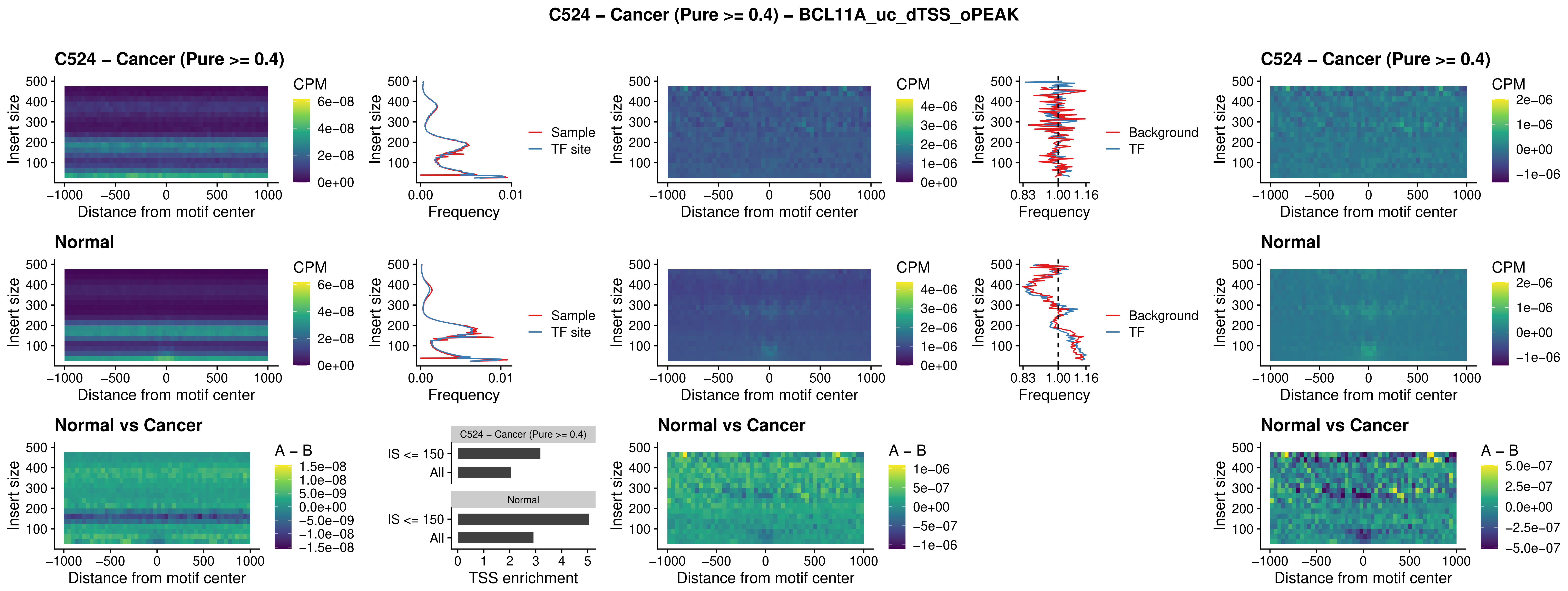
C524 – Cancer (Pure >= 0.4) – BCL11A_uc_dTSS_oPEAK
C524 – Cancer (Pure >= 0.4)
Normal
Normal vs Cancer
Insert size
Distance from motif center
Frequency
CPM
A – B
Sample
TF site
Background
TF
C524 – Cancer (Pure >= 0.4)
Normal
IS <= 150
All
TSS enrichment

#### C524 – Cancer (Pure >= 0.4) – BCL11B_uc_dTSS_oPEAK

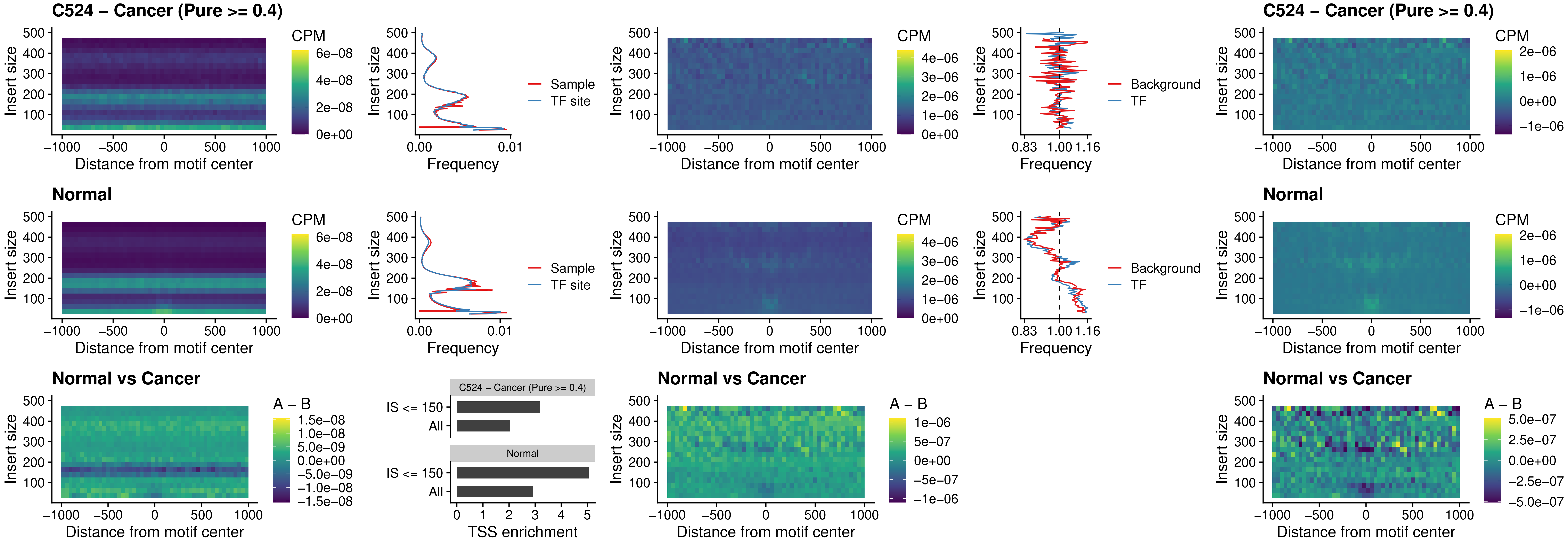

#### C524 – Cancer (Pure >= 0.4) – SPIB_uc_cTSS_nPEAK

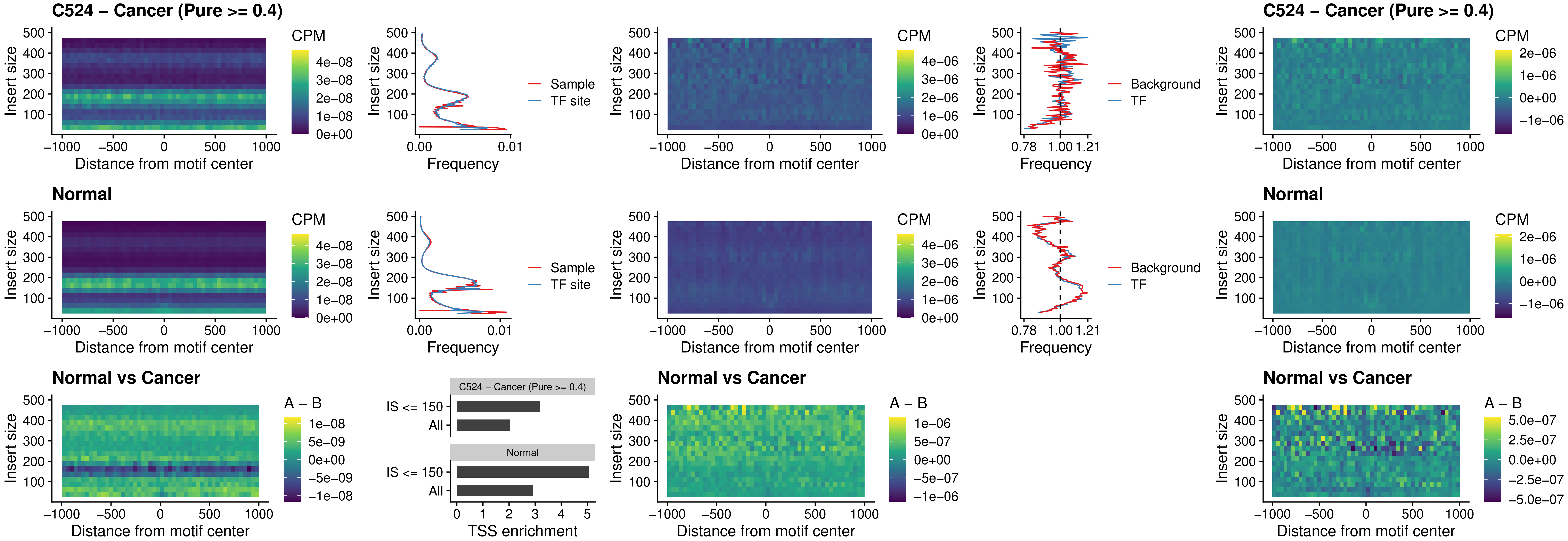

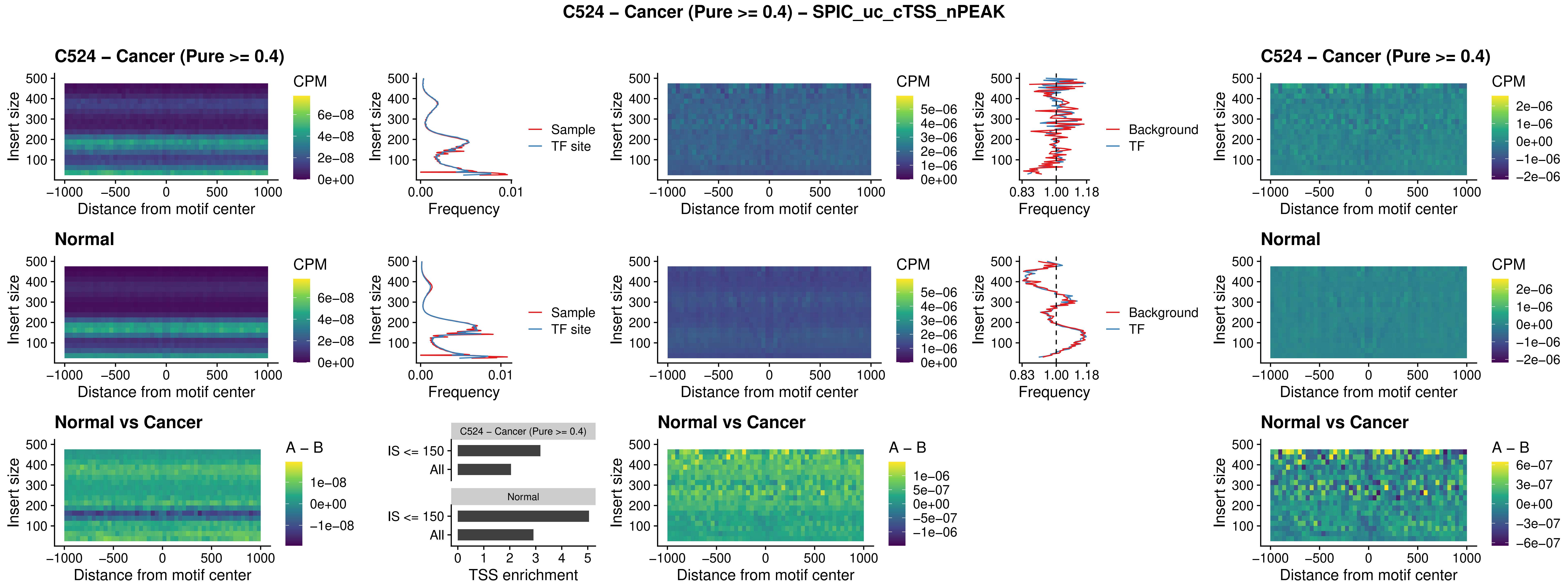
C524 – Cancer (Pure >= 0.4) – SPIC_uc_cTSS_nPEAK
C524 – Cancer (Pure >= 0.4)
Normal
Normal vs Cancer
Insert size
Distance from motif center
Frequency
CPM
A – B
Sample
TF site
Background
TF
C524 – Cancer (Pure >= 0.4)
Normal
IS <= 150
All
TSS enrichment

#### C524 – Cancer (Pure >= 0.4) – SPI1_uc_pTSS_oPEAK

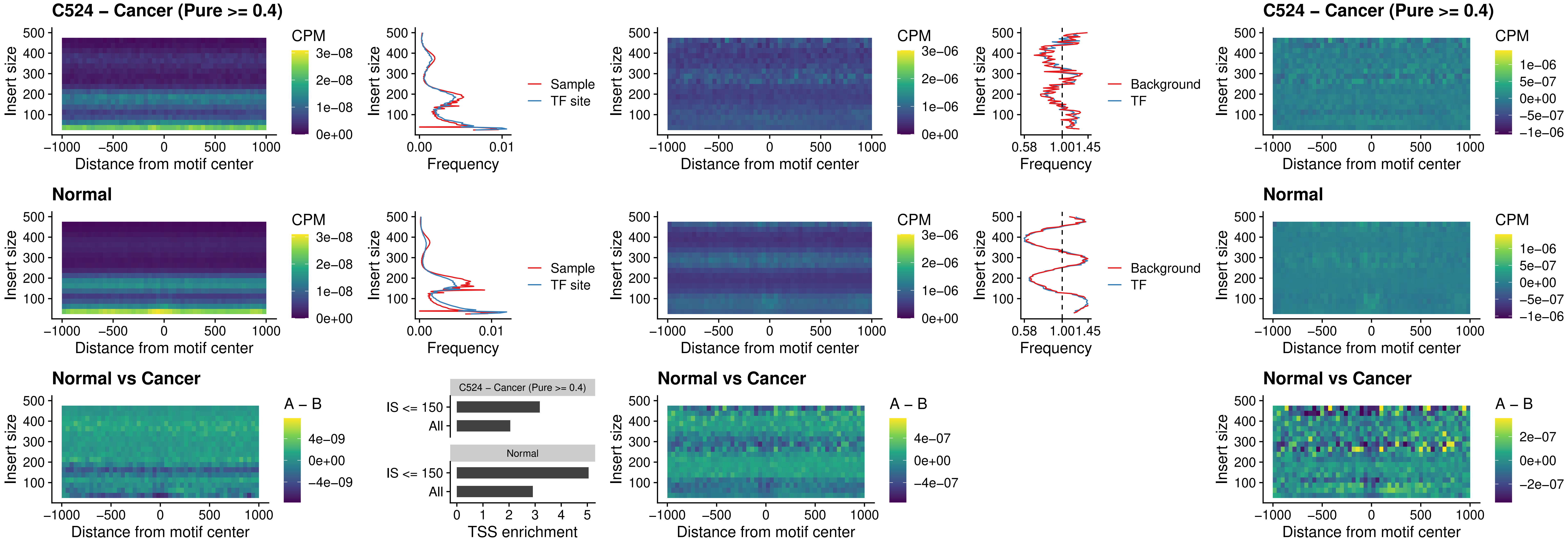

#### C524 – Cancer (Pure >= 0.4) – BCL11A_uc_pTSS_oPEAK

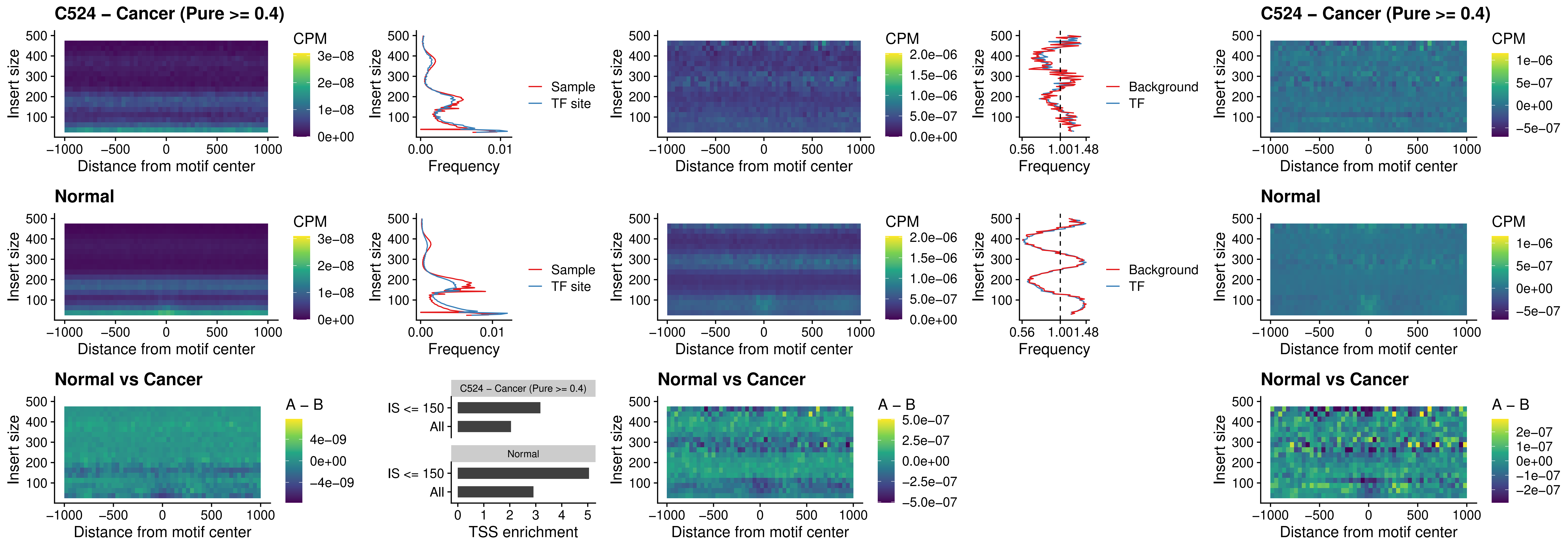

#### C524 – Cancer (Pure >= 0.4) – BCL11B_uc_pTSS_oPEAK

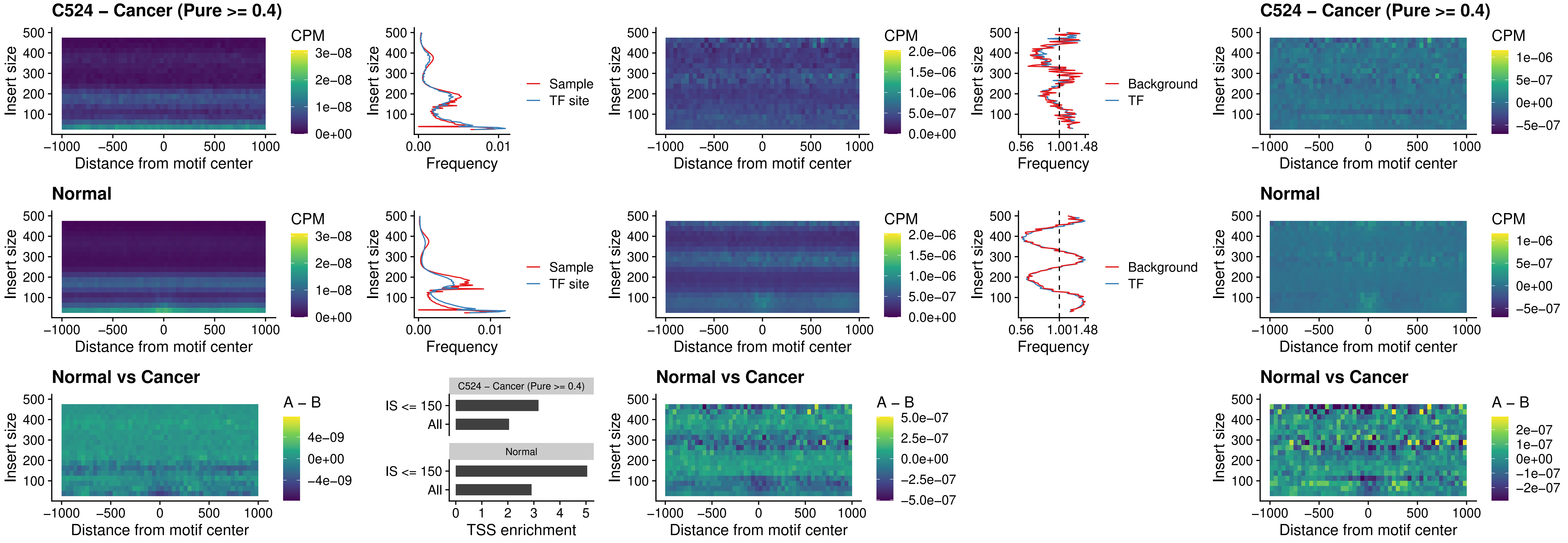

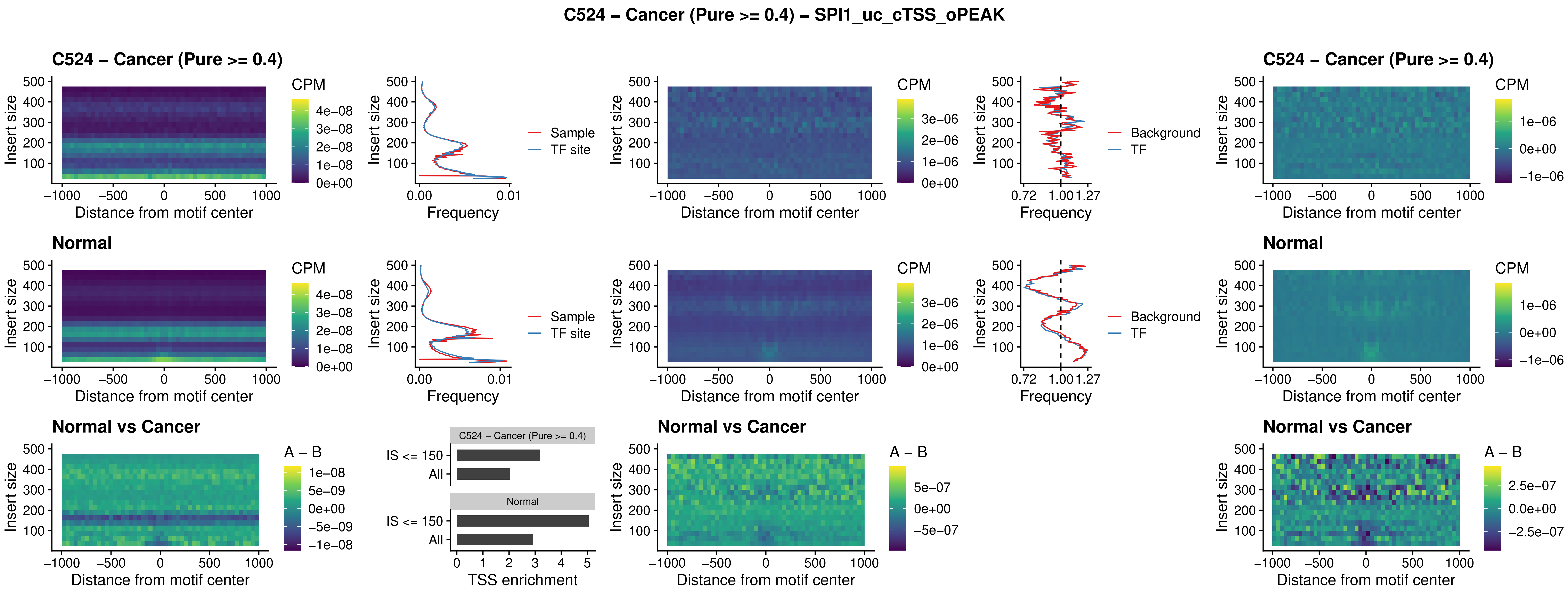
C524 – Cancer (Pure >= 0.4) – SPI1_uc_cTSS_oPEAK
C524 – Cancer (Pure >= 0.4)
Normal
Normal vs Cancer
Insert size
Distance from motif center
Frequency
CPM
A – B
Sample
TF site
Background
TF
TSS enrichment
IS <= 150
All
C524 – Cancer (Pure >= 0.4)
Normal

#### C524 – Cancer (Pure >= 0.4) – SPIB_uc_cTSS_oPEAK

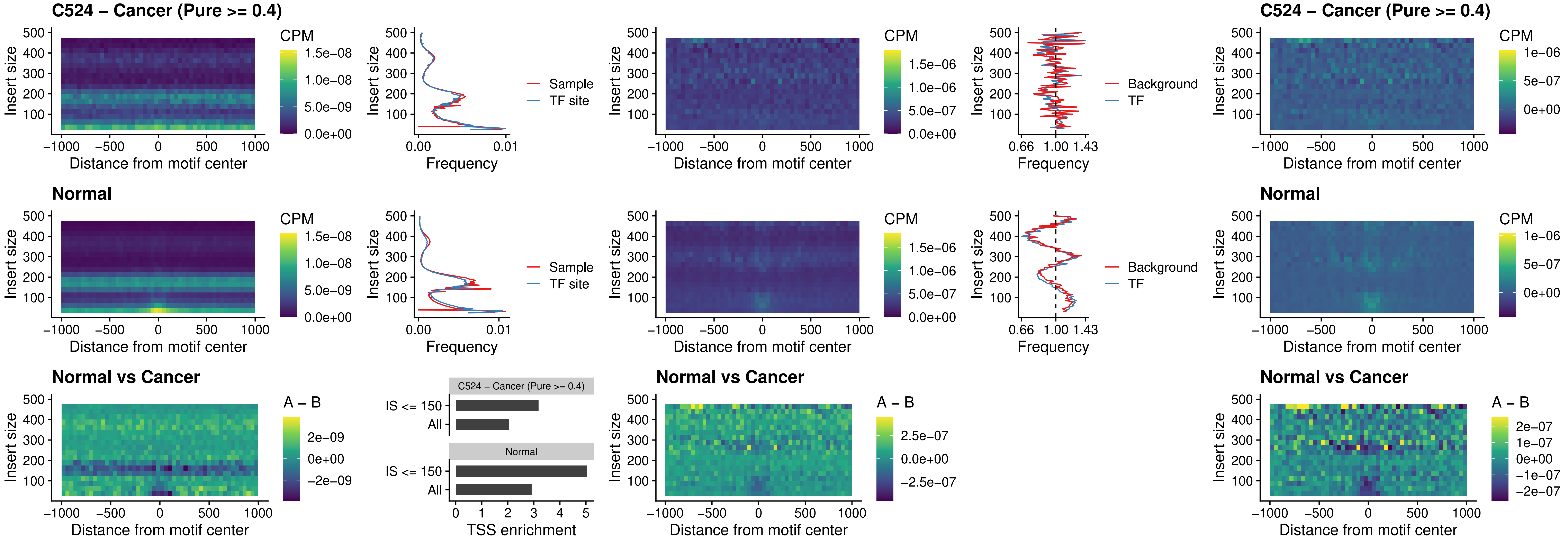

### C524 – Cancer (Pure >= 0.4) – SPIB_uc_pTSS_oPEAK

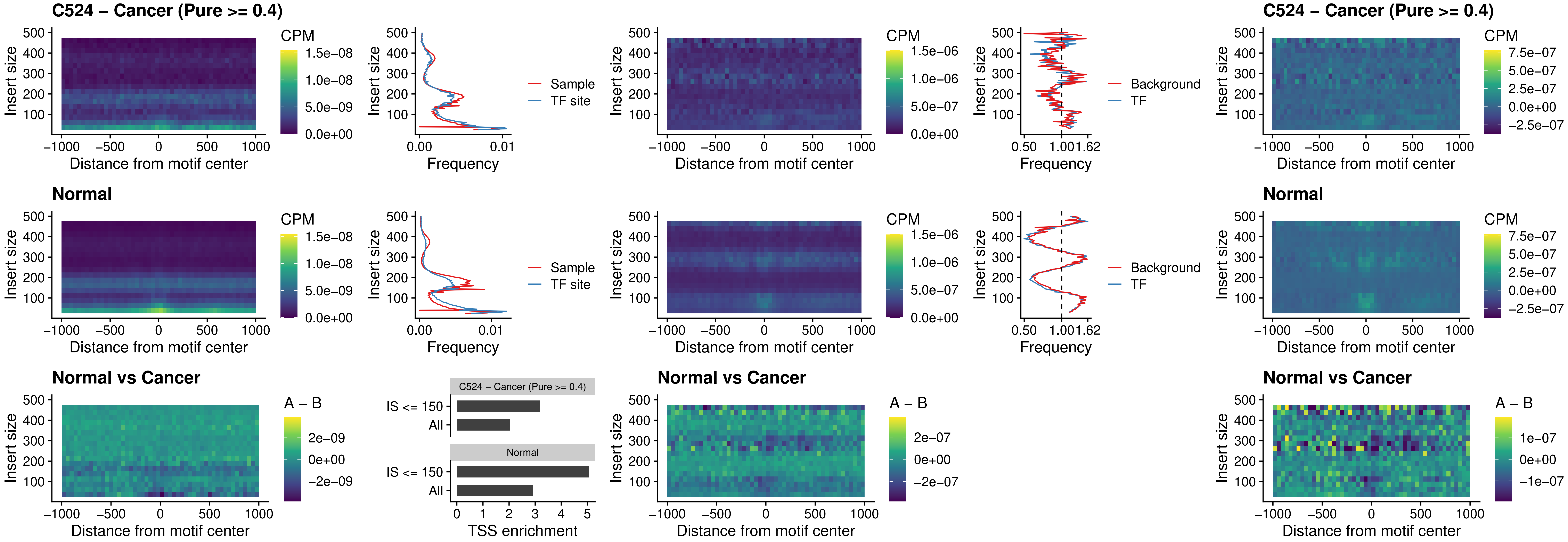

#### C524 – Cancer (Pure >= 0.4) – SPI1_uc_dTSS_oPEAK

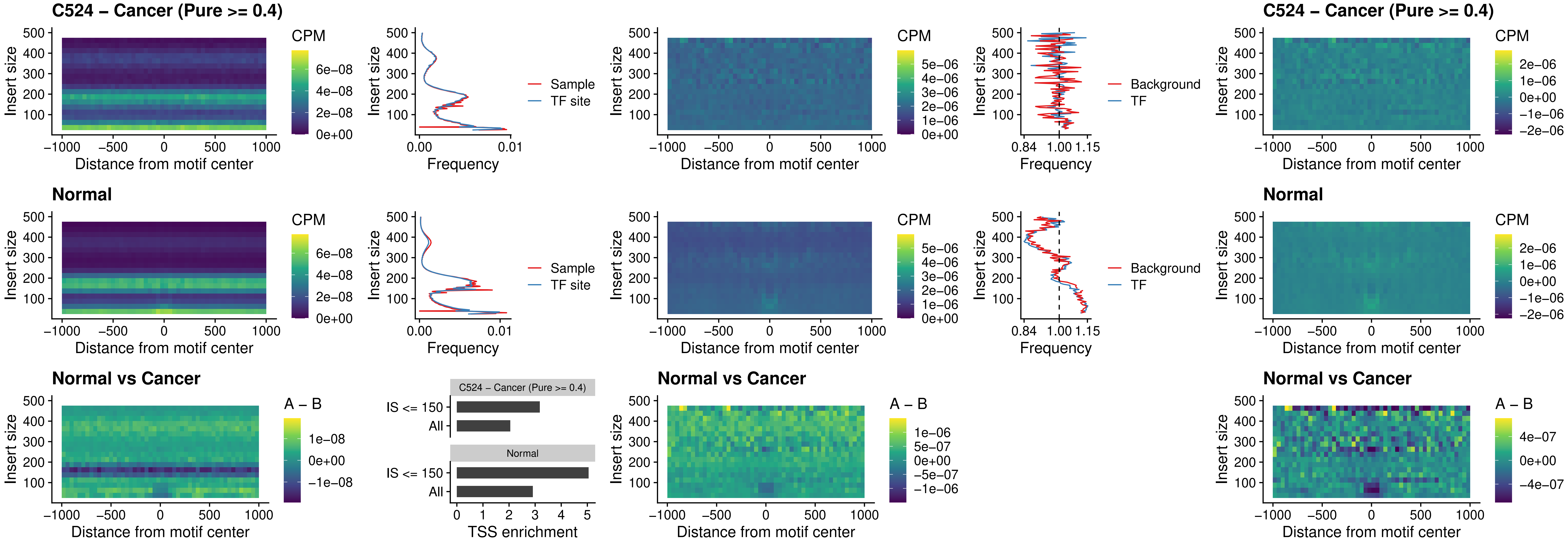

#### C524 – Cancer (Pure >= 0.4) – SPIB_uc_dTSS_oPEAK

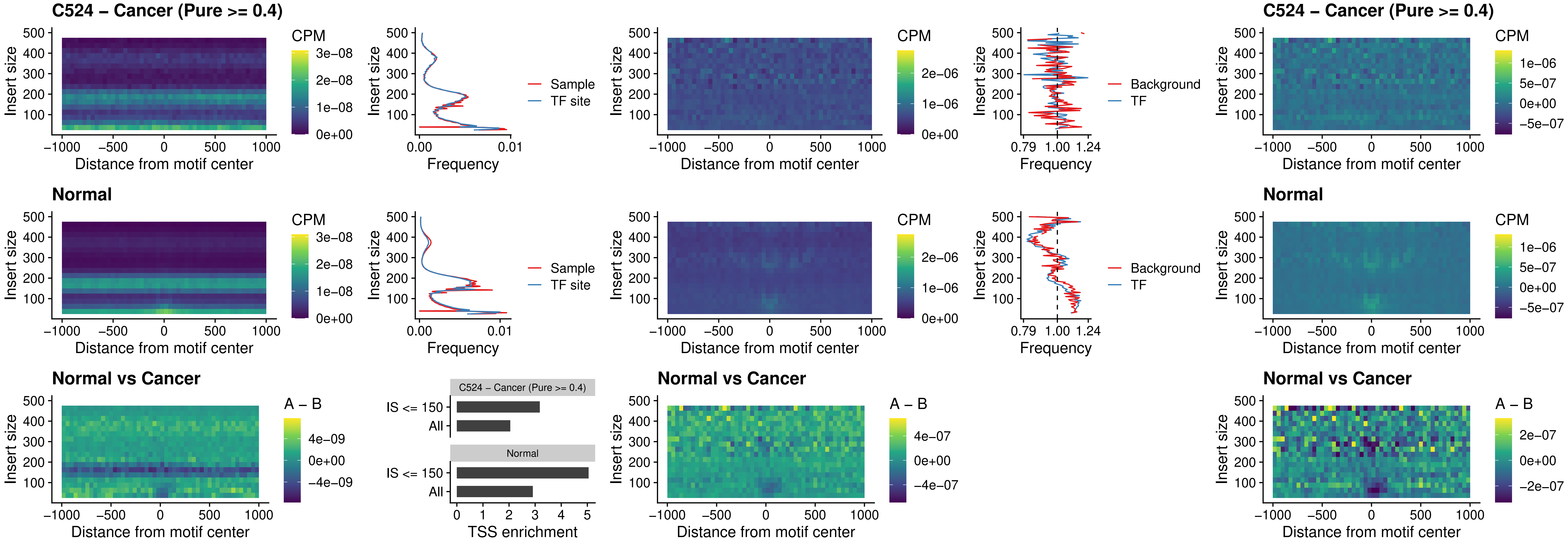

#### C524 – Cancer (Pure >= 0.4) – IRF3_uc_pTSS_oPEAK

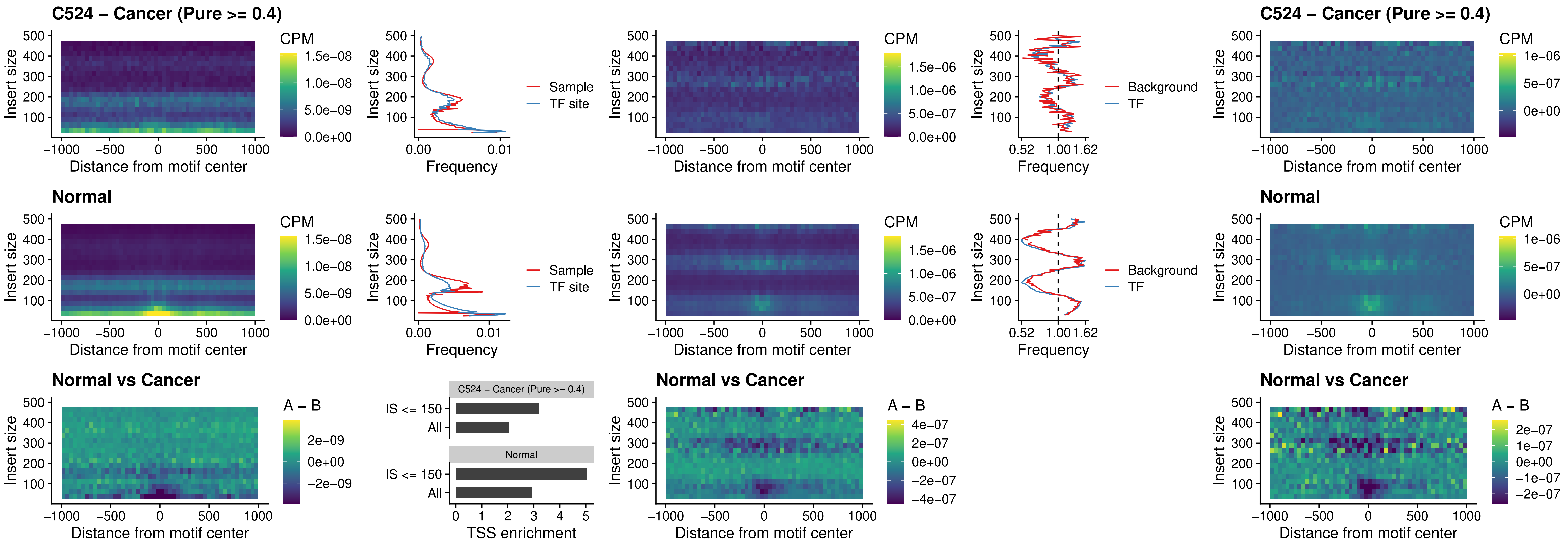

### C524 – Cancer (Pure >= 0.4) – IRF2_uc_dTSS_oPEAK

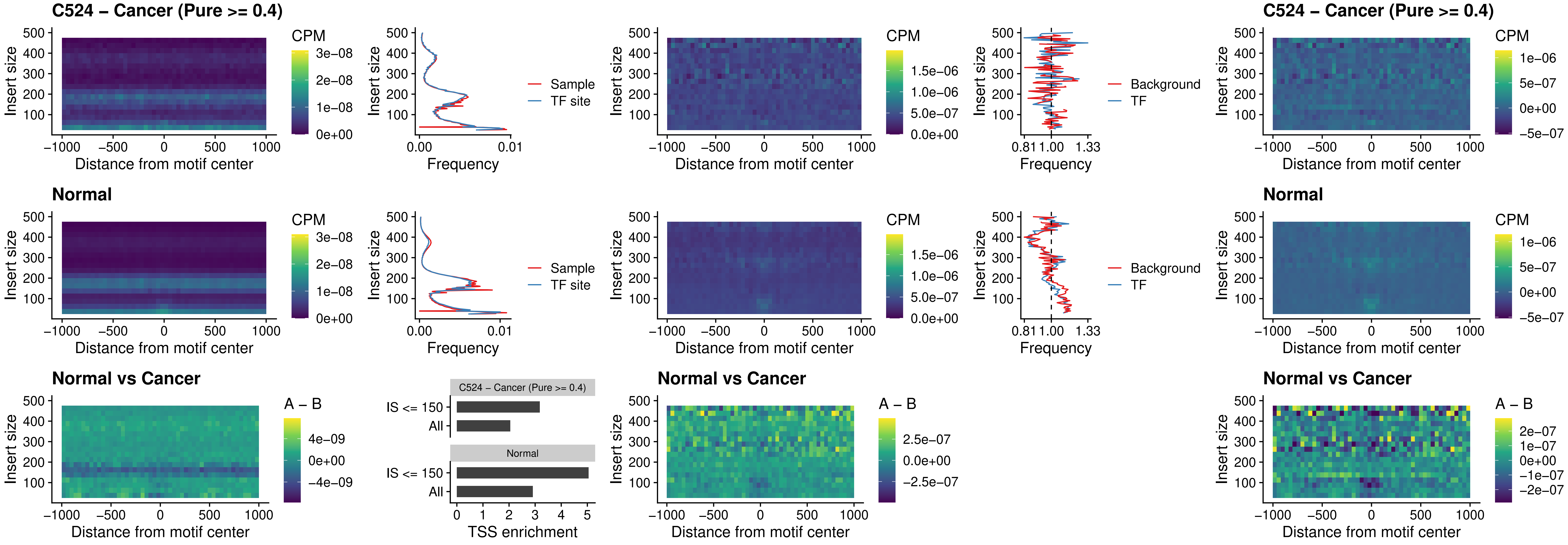

### C524 – Cancer (Pure >= 0.4) – IRF2_uc_pTSS_oPEAK

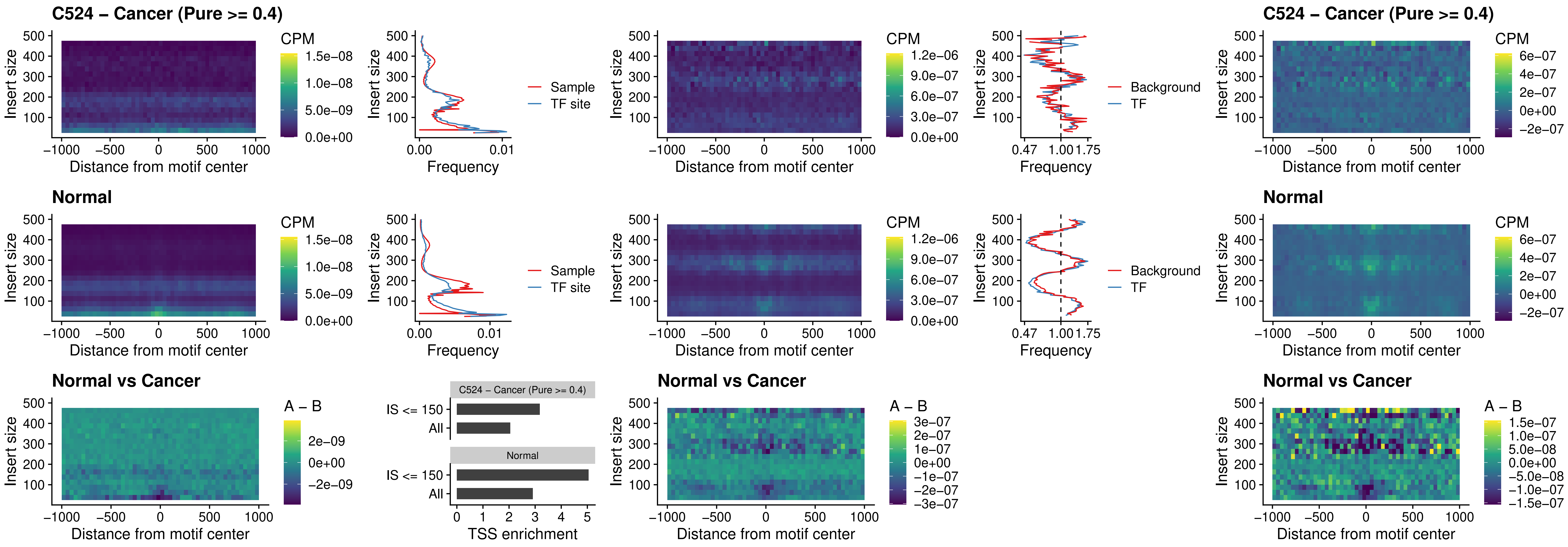

### C524 – Cancer (Pure >= 0.4) – STAT2_uc_pTSS_oPEAK

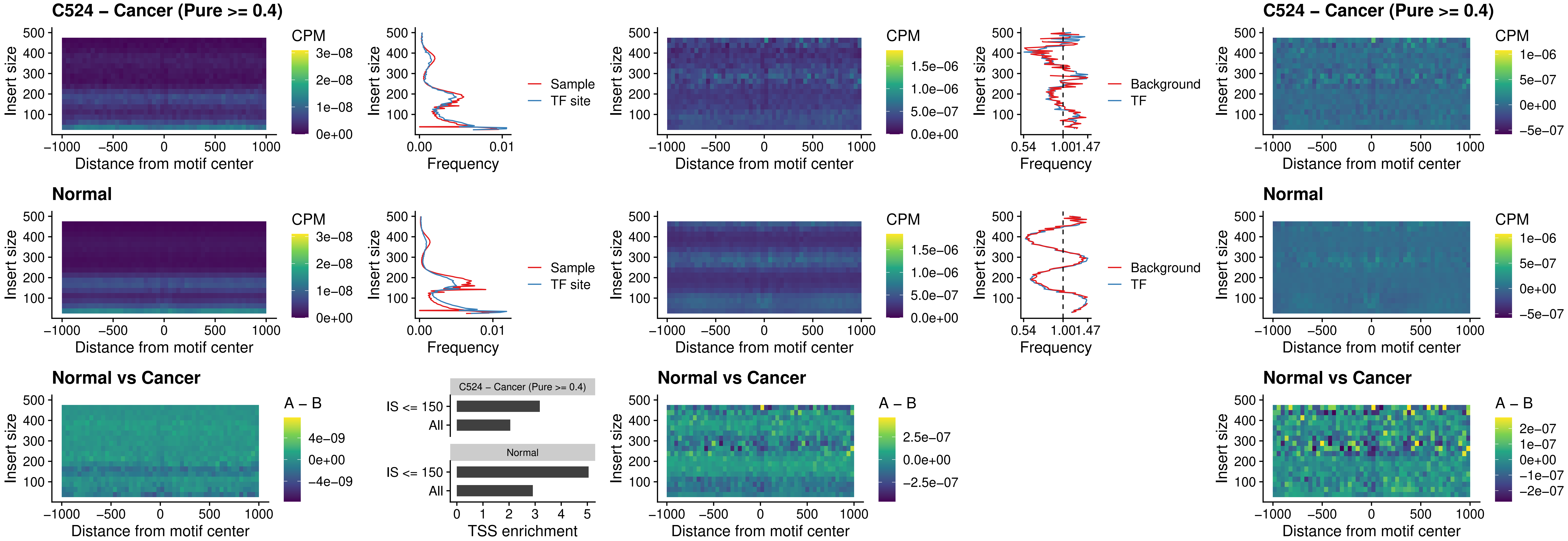

### C524 – Cancer (Pure >= 0.4) – SPIB_uc_dTSS_nPEAK

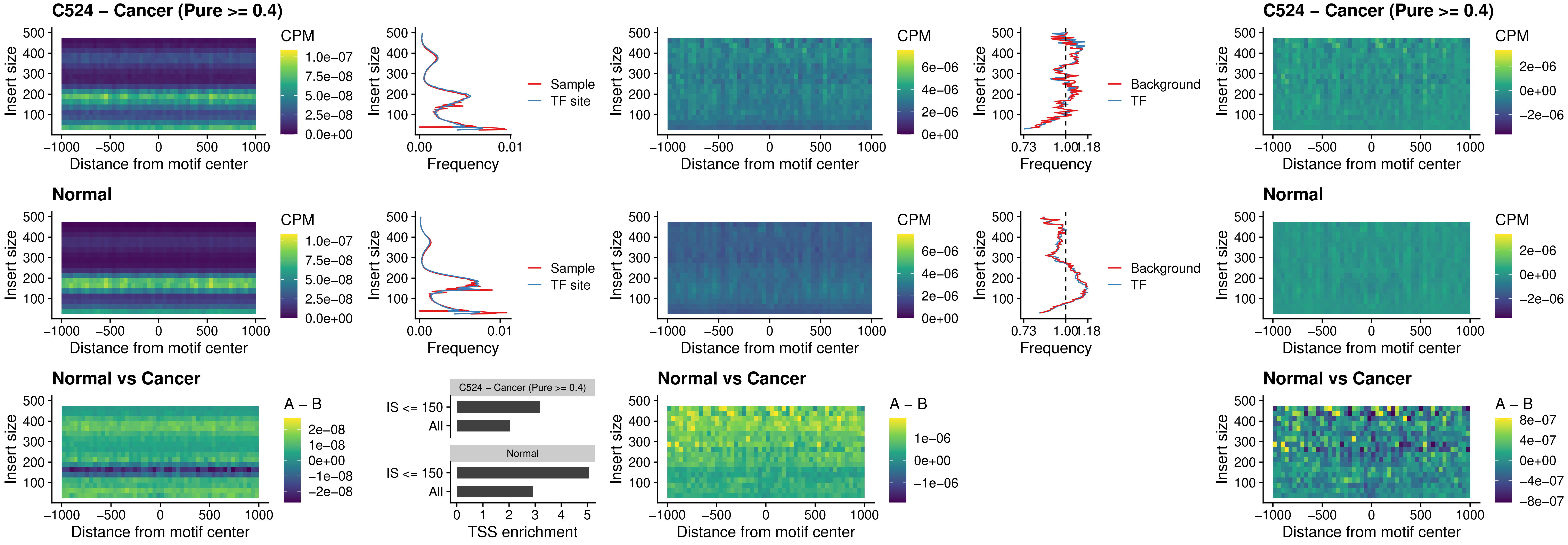

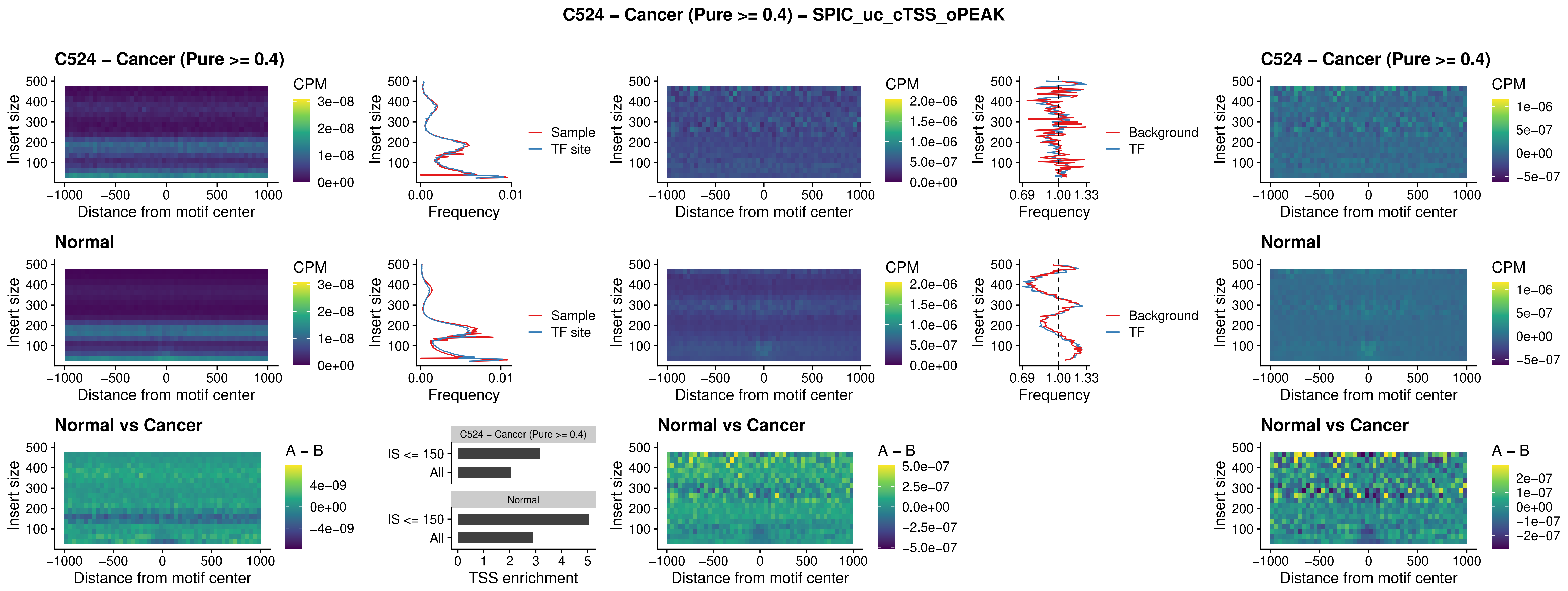
C524 – Cancer (Pure >= 0.4) – SPIC_uc_cTSS_oPEAK
C524 – Cancer (Pure >= 0.4)
Normal
Normal vs Cancer
Insert size
Distance from motif center
Frequency
CPM
A – B
Sample
TF site
Background
TF
C524 – Cancer (Pure >= 0.4)
Normal
IS <= 150
All
TSS enrichment

#### C524 – Cancer (Pure >= 0.4) – PRDM1_uc_cTSS_oPEAK

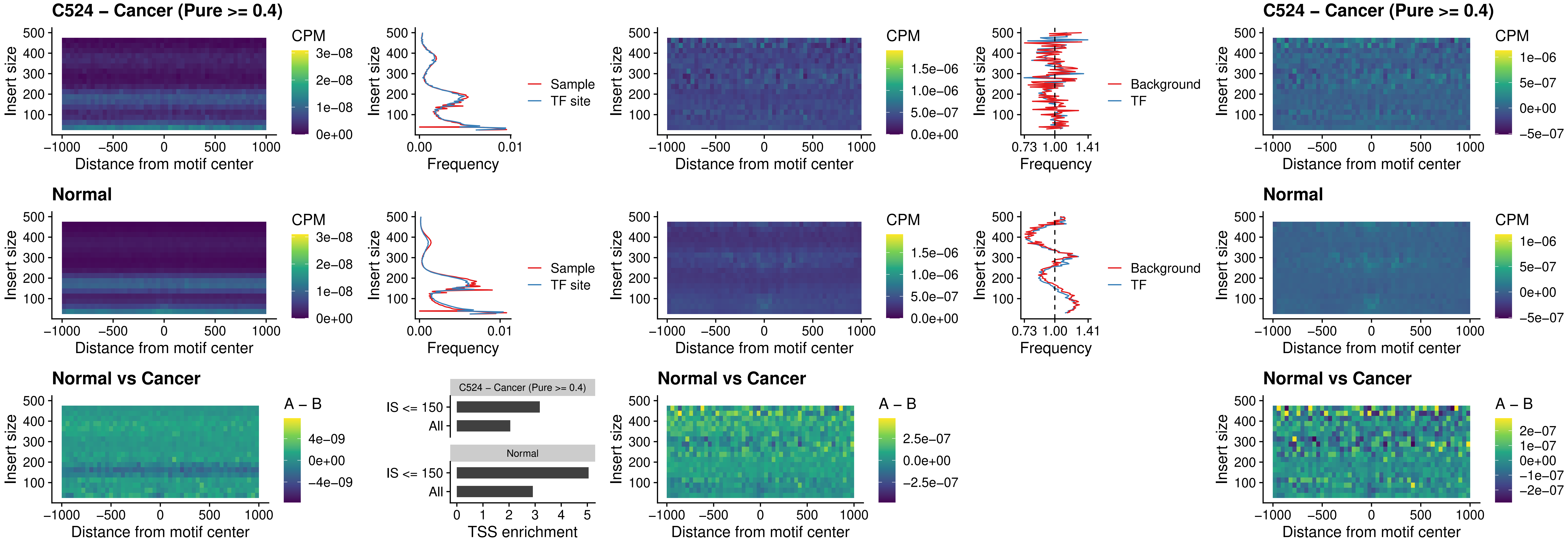

#### C524 – Cancer (Pure >= 0.4) – IRF3_uc_cTSS_oPEAK

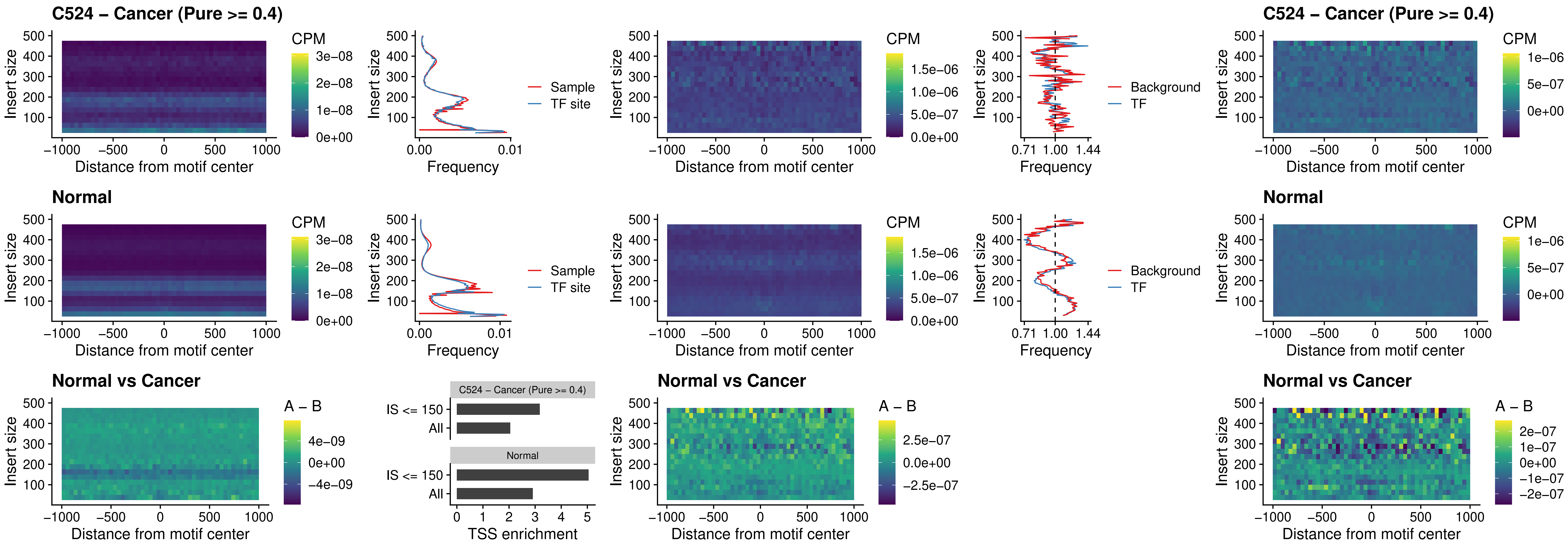

### C524 – Cancer (Pure >= 0.4) – BCL11A_uc_dTSS_nPEAK

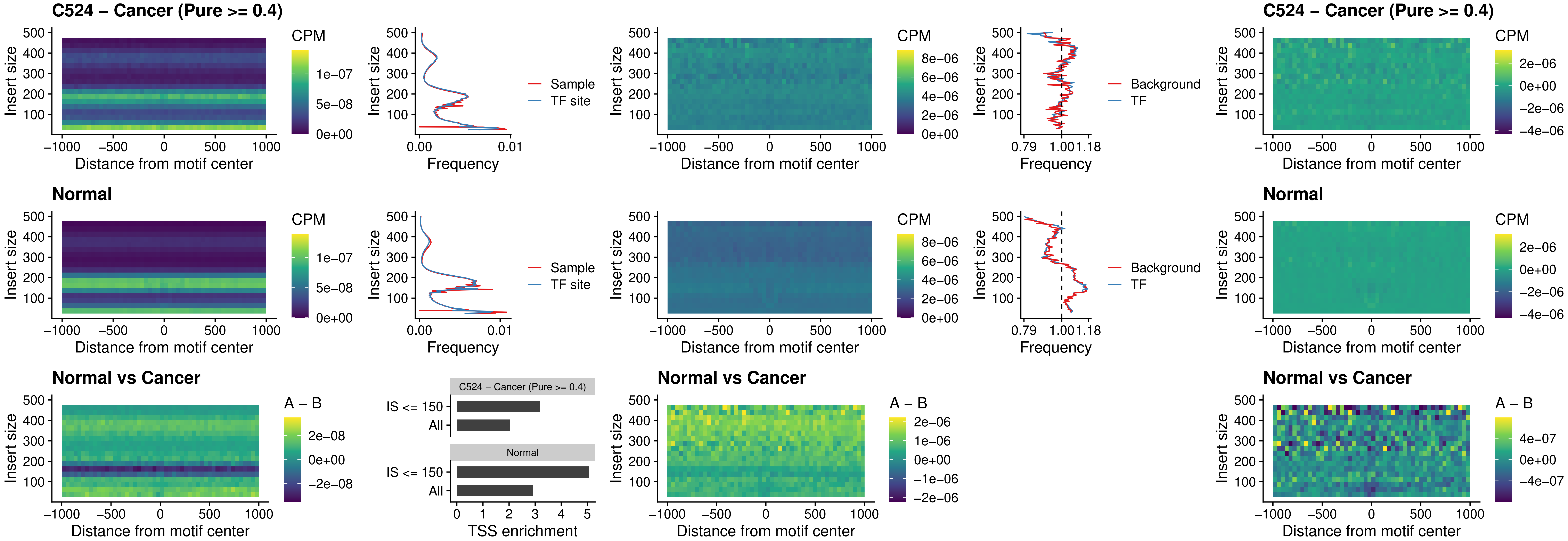

#### C524 – Cancer (Pure >= 0.4) – BCL11B_uc_dTSS_nPEAK

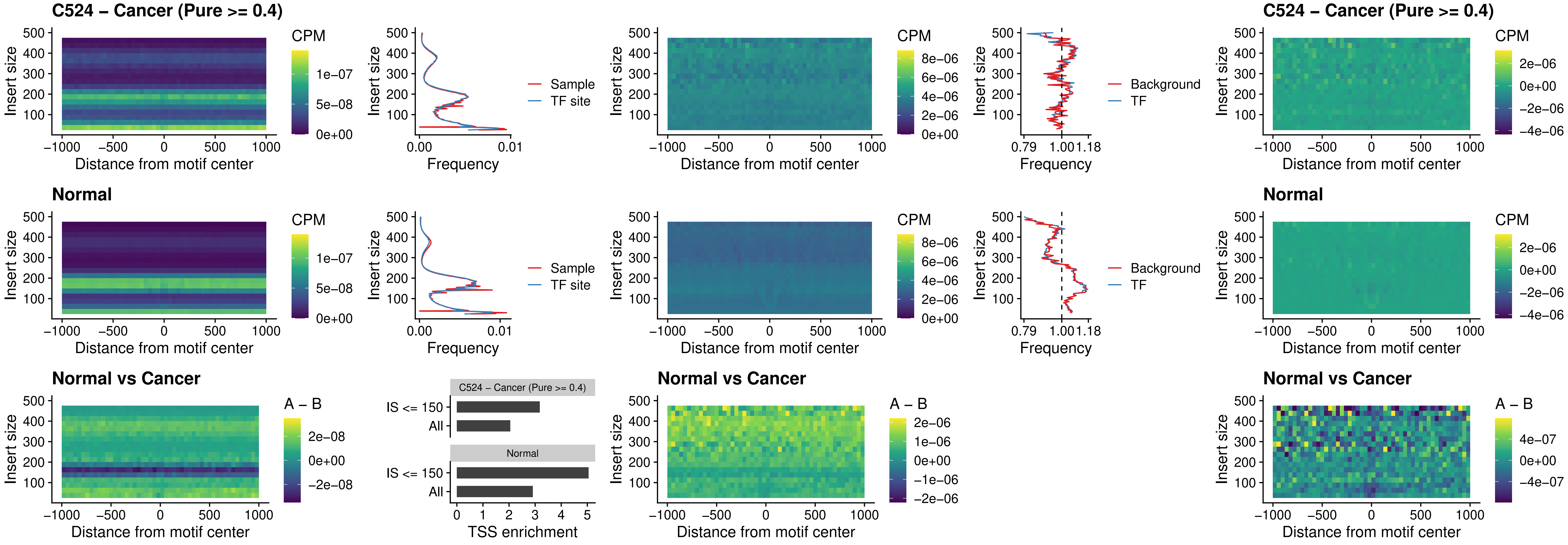

#### C524 – Cancer (Pure >= 0.4) – IRF1_uc_cTSS_oPEAK

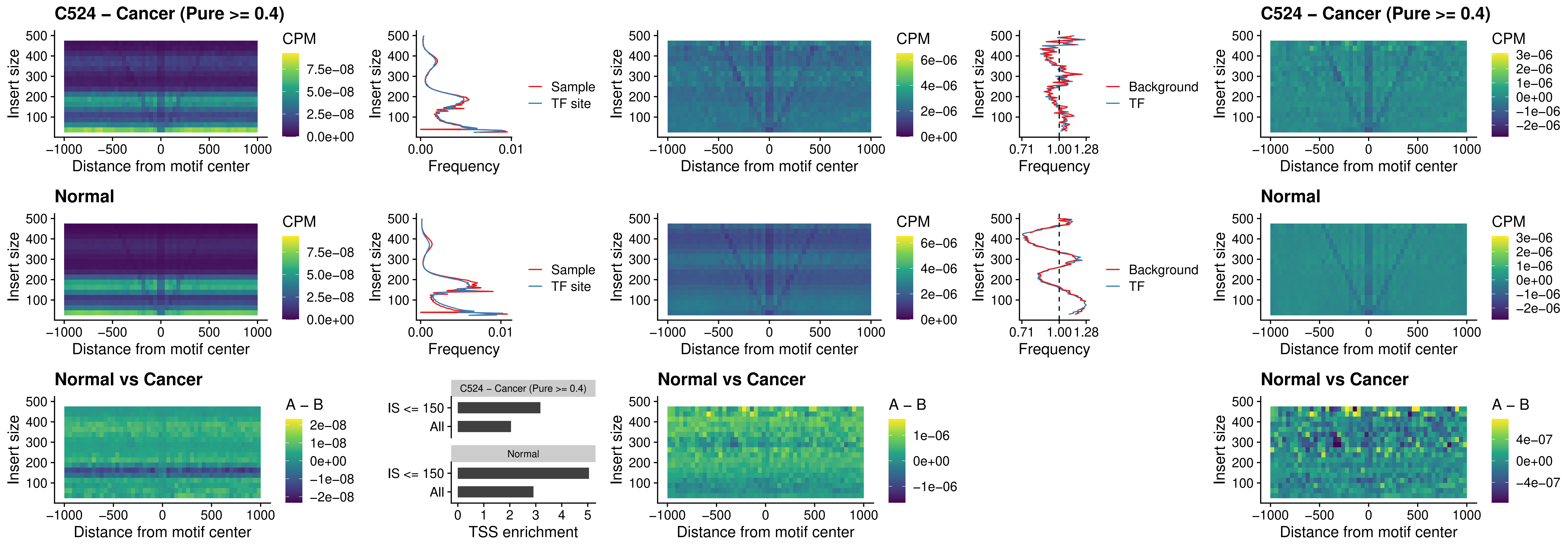

#### C524 – Cancer (Pure >= 0.4) – IRF1_uc_dTSS_oPEAK

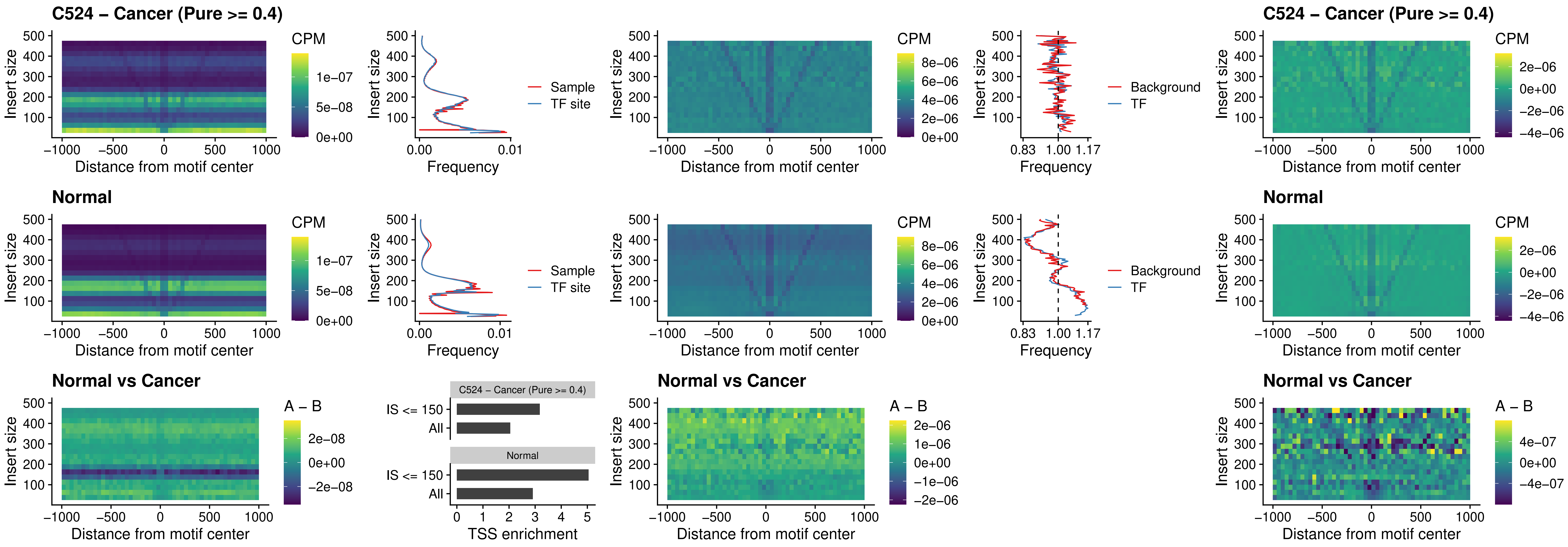

### C524 – Cancer (Pure >= 0.4) – IRF1_uc_pTSS_oPEAK

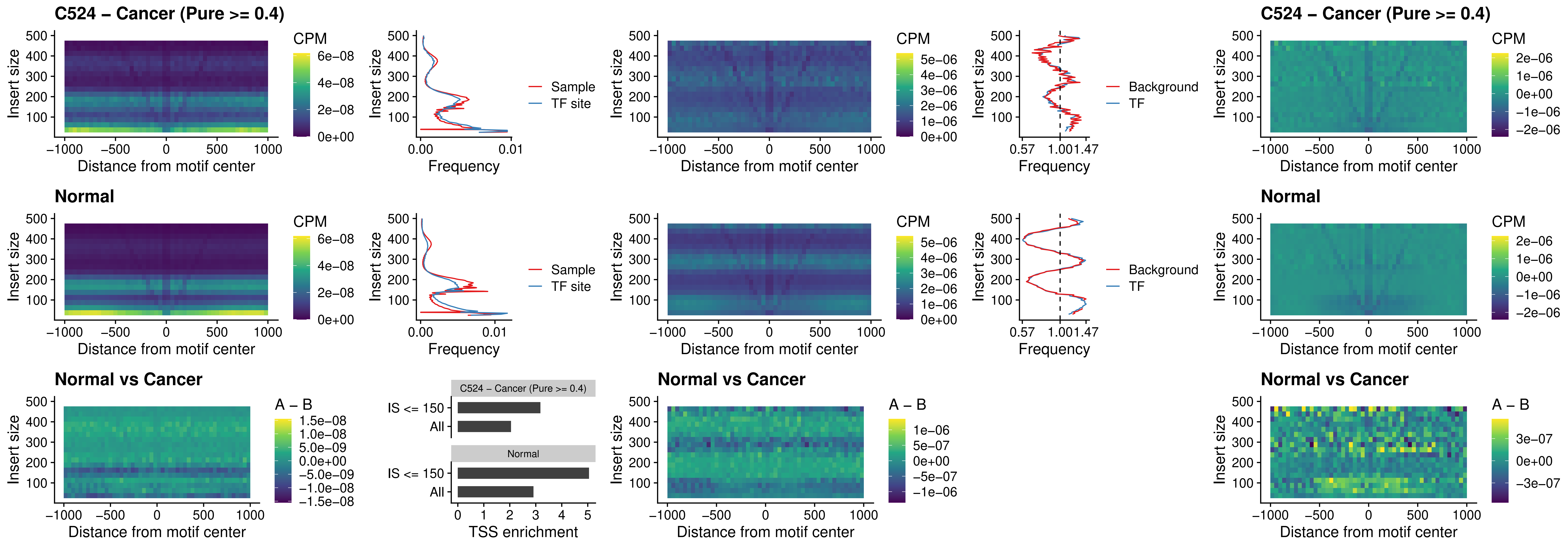

#### C524 – Cancer (Pure >= 0.4) – SPIC_uc_dTSS_nPEAK

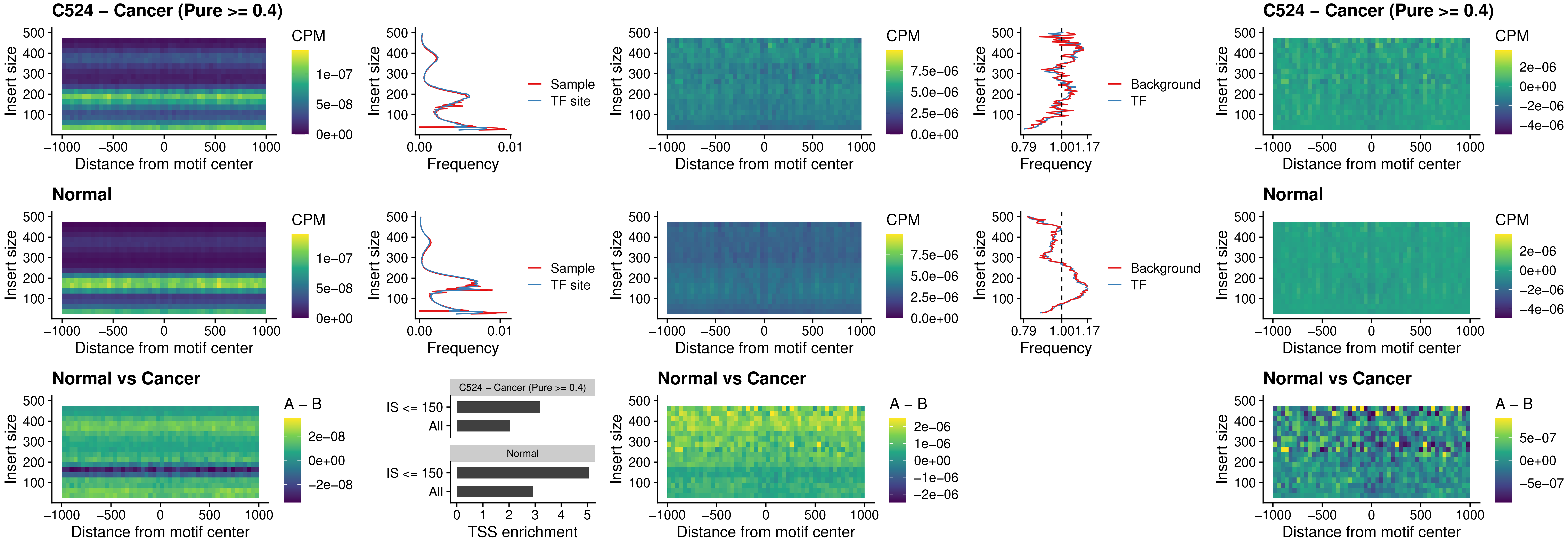

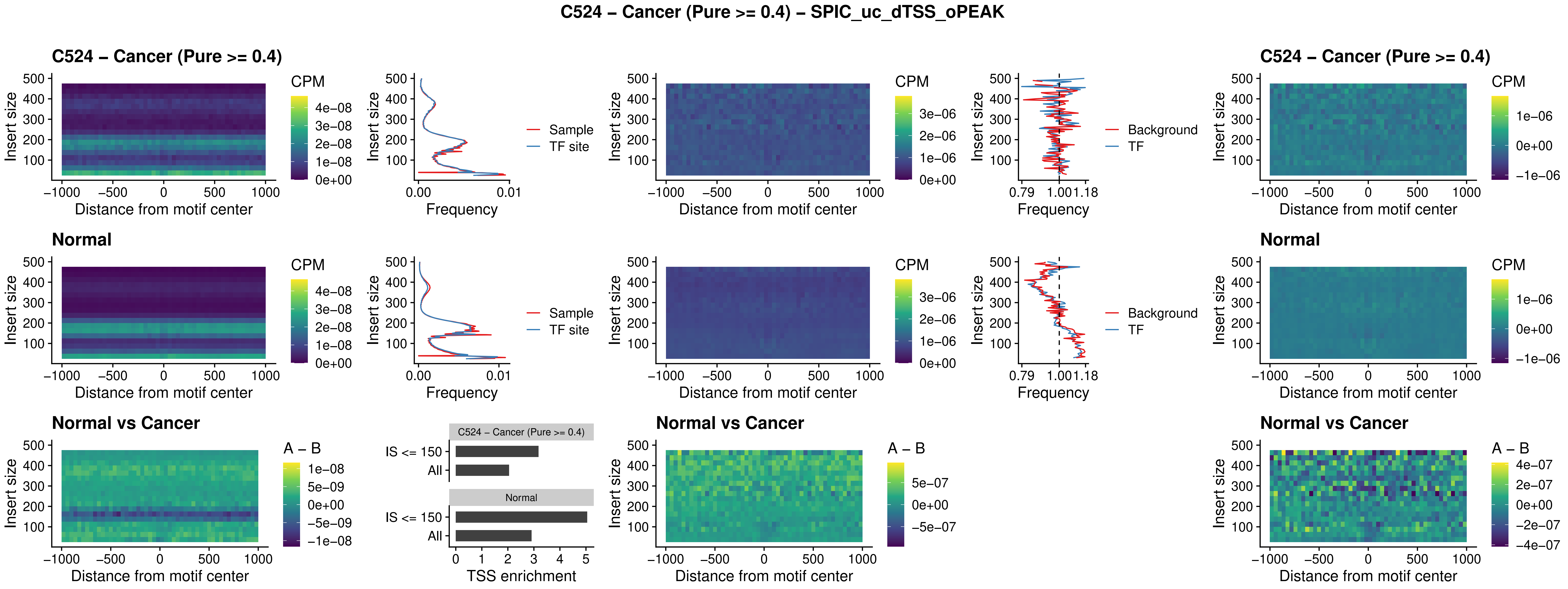

C524 – Cancer (Pure >= 0.4) – SPIC_uc_dTSS_oPEAK
C524 – Cancer (Pure >= 0.4)
Normal
Normal vs Cancer
Insert size
Distance from motif center
Frequency
CPM
A – B
Sample
TF site
Background
TF
C524 – Cancer (Pure >= 0.4)
Normal
IS <= 150
All
TSS enrichment

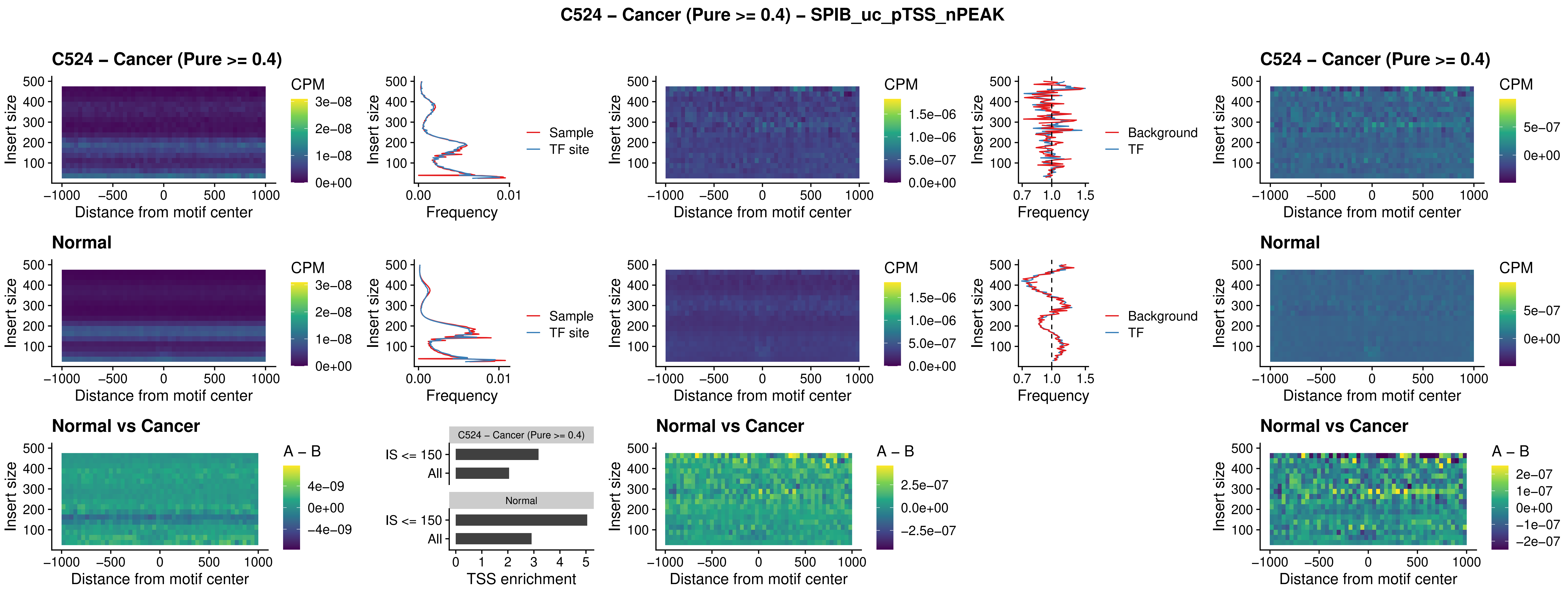
C524 – Cancer (Pure >= 0.4) – SPIB_uc_pTSS_nPEAK
C524 – Cancer (Pure >= 0.4)
Normal
Normal vs Cancer
Insert size
Distance from motif center
CPM
A – B
Frequency
Sample
TF site
Background
TF
C524 – Cancer (Pure >= 0.4)
Normal
IS <= 150
All
TSS enrichment

#### C524 – Cancer (Pure >= 0.4) – STAT2_uc_cTSS_oPEAK

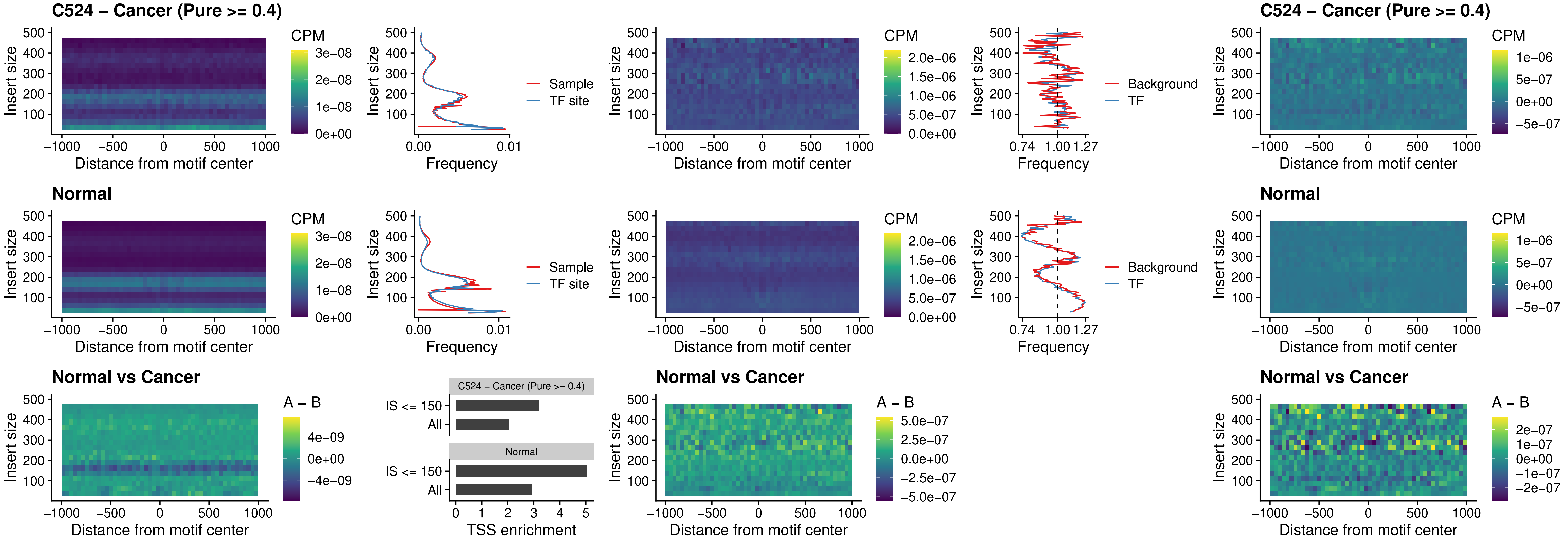

#### C524 – Cancer (Pure >= 0.4) – IRF8_uc_cTSS_oPEAK

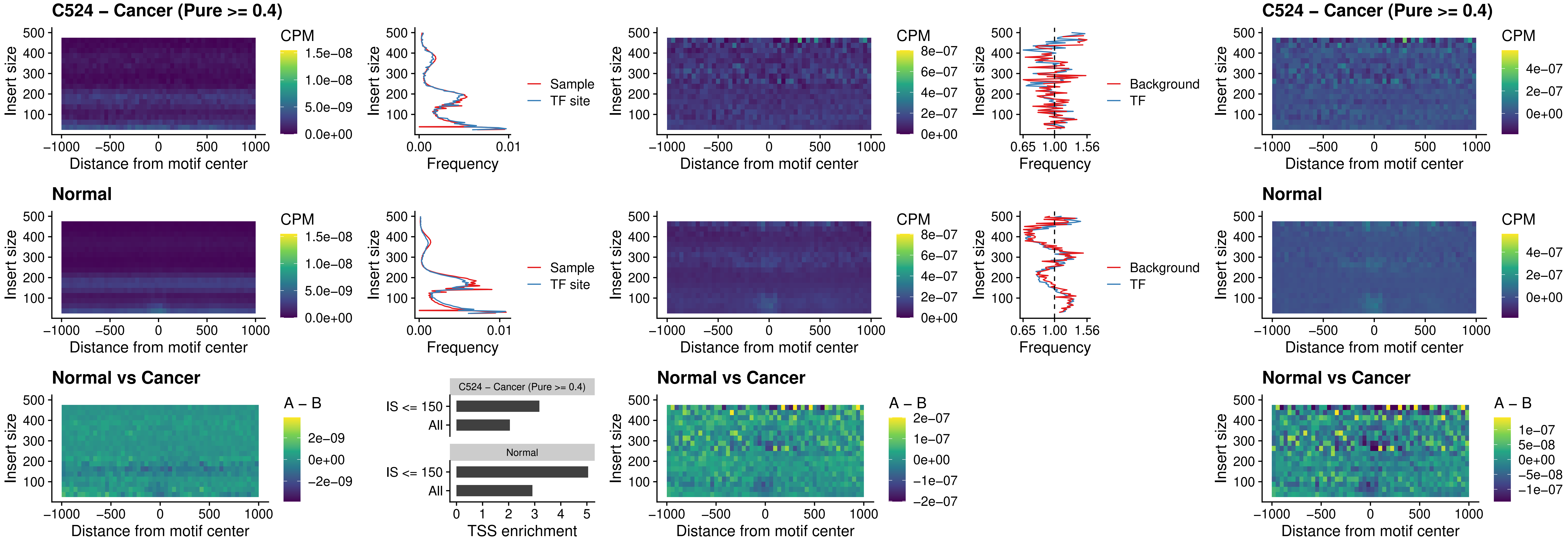

#### C524 – Cancer (Pure >= 0.4) – IRF3_uc_dTSS_oPEAK

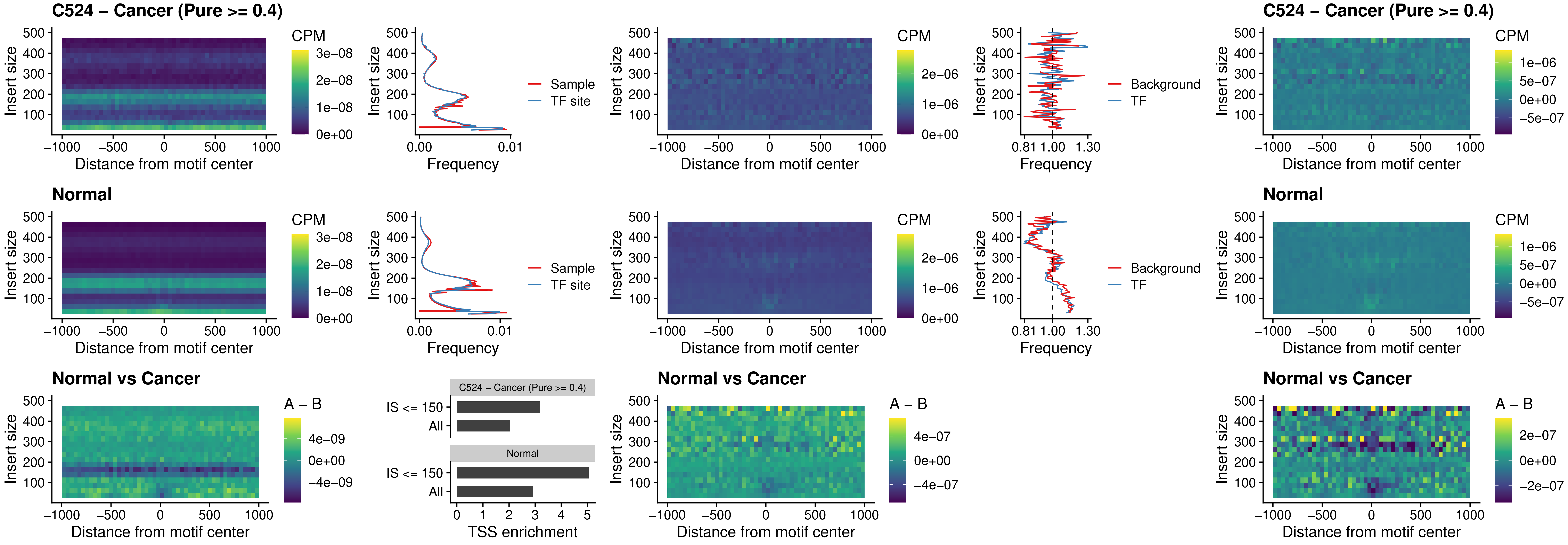

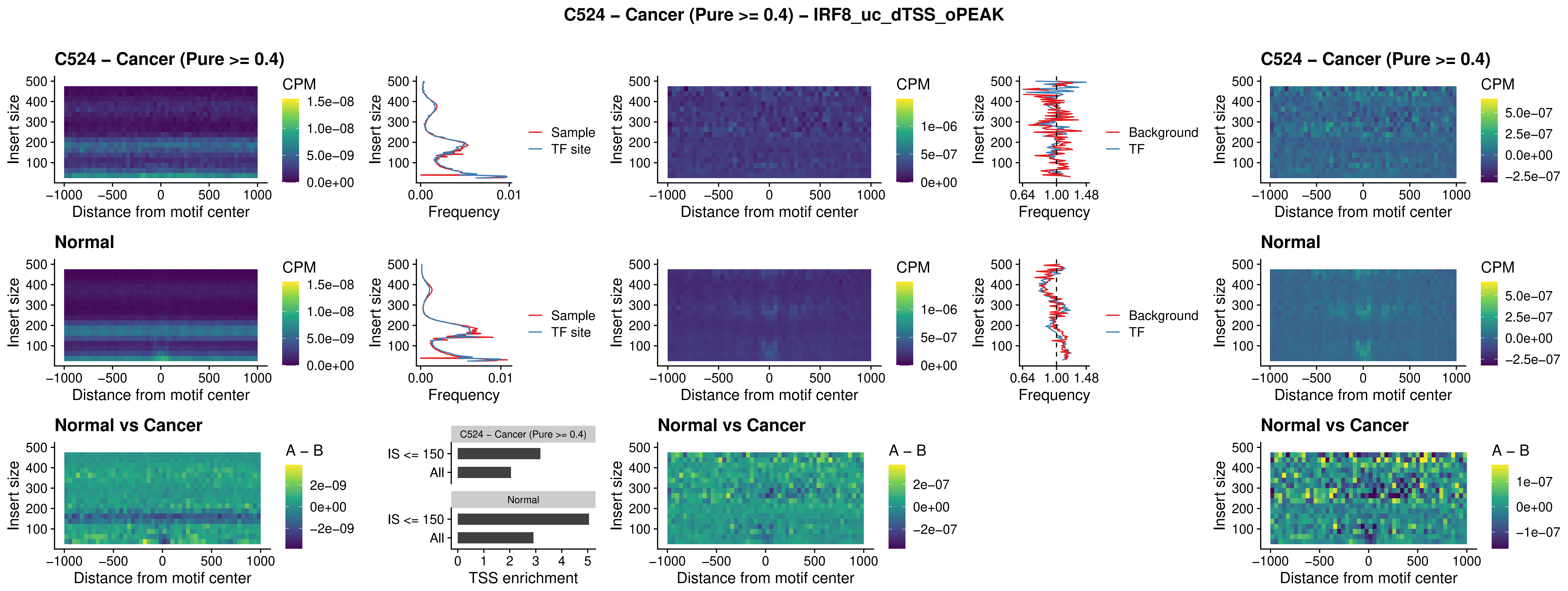
C524 – Cancer (Pure >= 0.4) – IRF8_uc_dTSS_oPEAK
C524 – Cancer (Pure >= 0.4)
Normal
Normal vs Cancer
Insert size
Distance from motif center
Frequency
CPM
A – B
Sample
TF site
Background
TF
C524 – Cancer (Pure >= 0.4)
Normal
IS <= 150
All
TSS enrichment

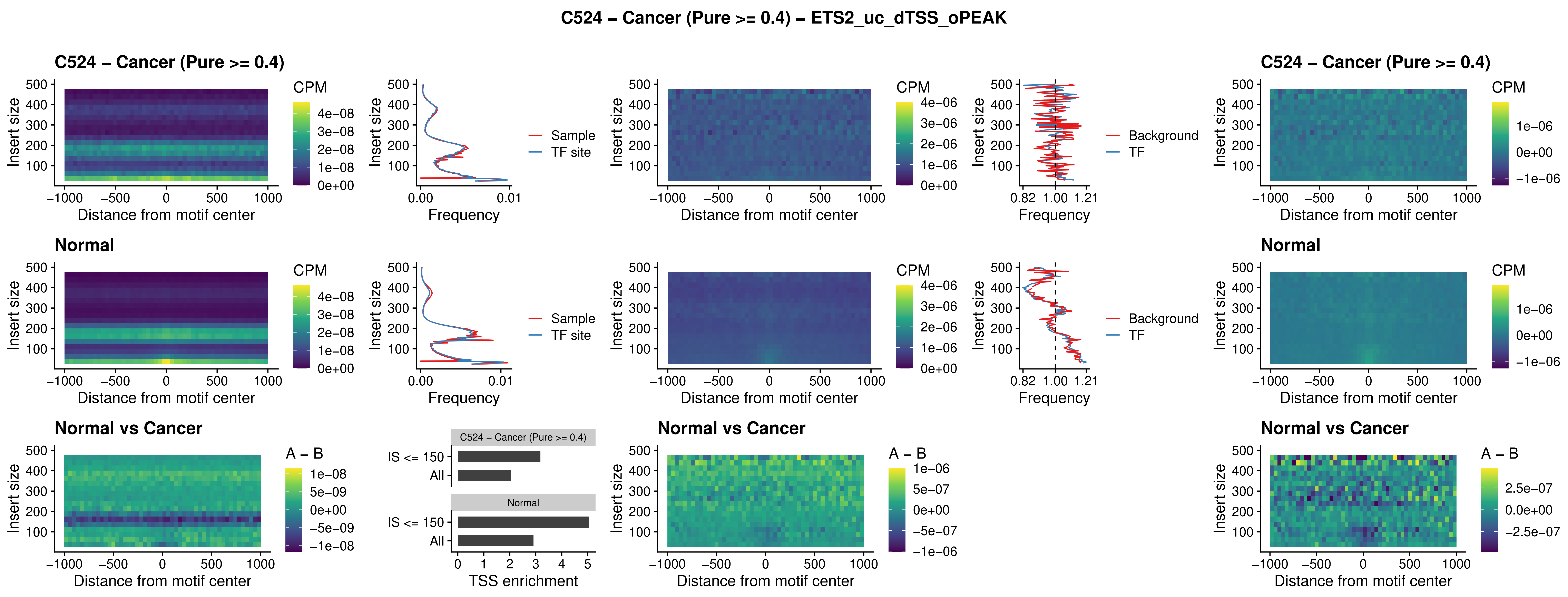
C524 – Cancer (Pure >= 0.4) – ETS2_uc_dTSS_oPEAK
C524 – Cancer (Pure >= 0.4)
Normal
Normal vs Cancer
Insert size
Distance from motif center
Frequency
CPM
A – B
Sample
TF site
Background
TF
C524 – Cancer (Pure >= 0.4)
IS <= 150
All
TSS enrichment

### C524 – Cancer (Pure >= 0.4) – SMARCC2_uc_pTSS_oPEAK

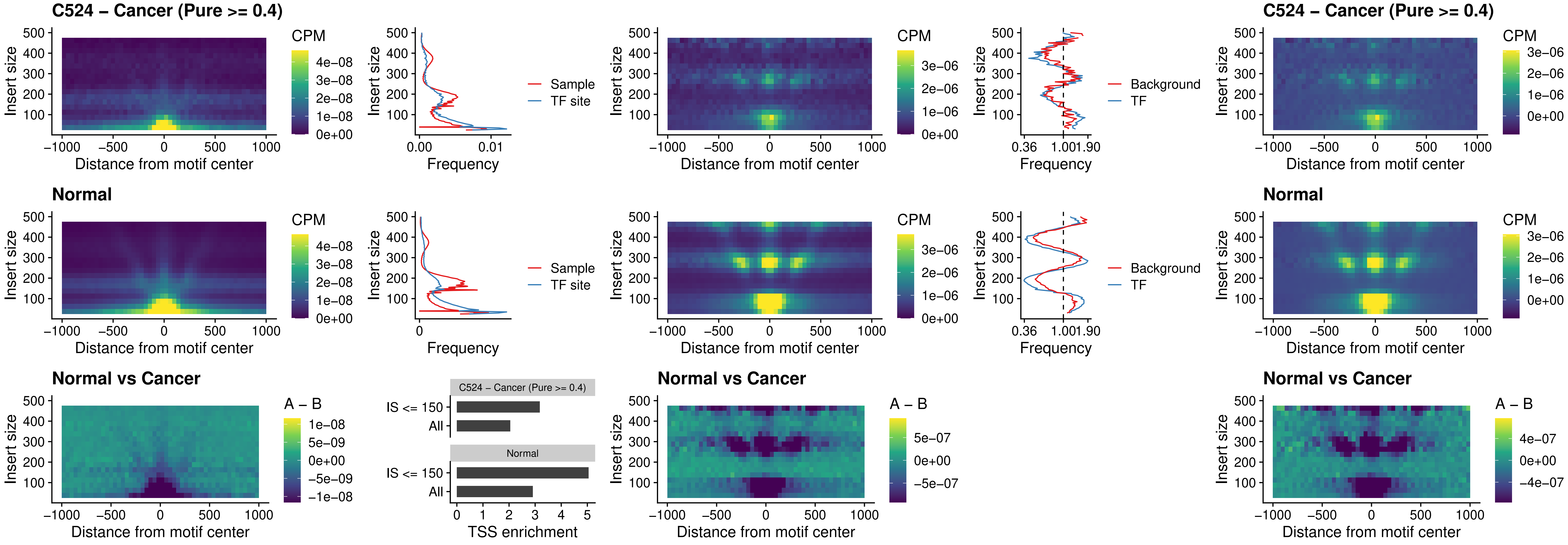

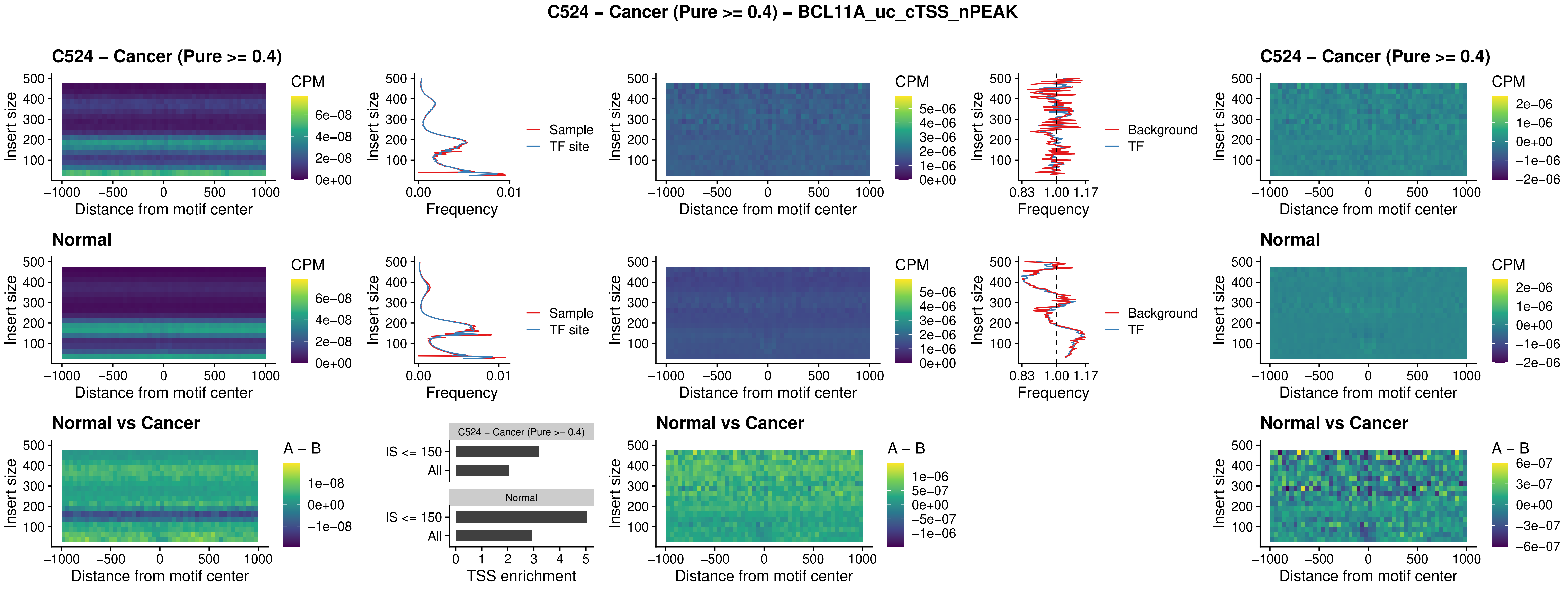
C524 – Cancer (Pure >= 0.4) – BCL11A_uc_cTSS_nPEAK
C524 – Cancer (Pure >= 0.4)
Normal
Normal vs Cancer
Insert size
Distance from motif center
Frequency
CPM
A – B
Sample
TF site
Background
TF
TSS enrichment
IS <= 150
All
C524 – Cancer (Pure >= 0.4)
Normal

#### C524 – Cancer (Pure >= 0.4) – BCL11B_uc_cTSS_nPEAK

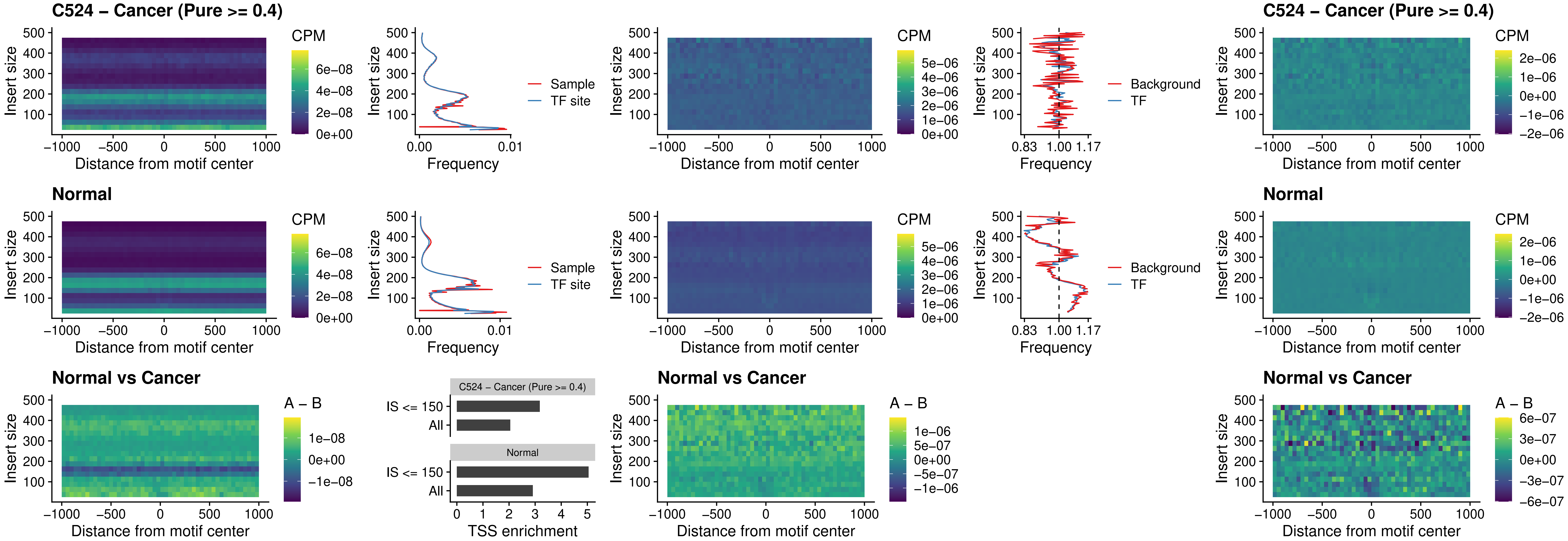

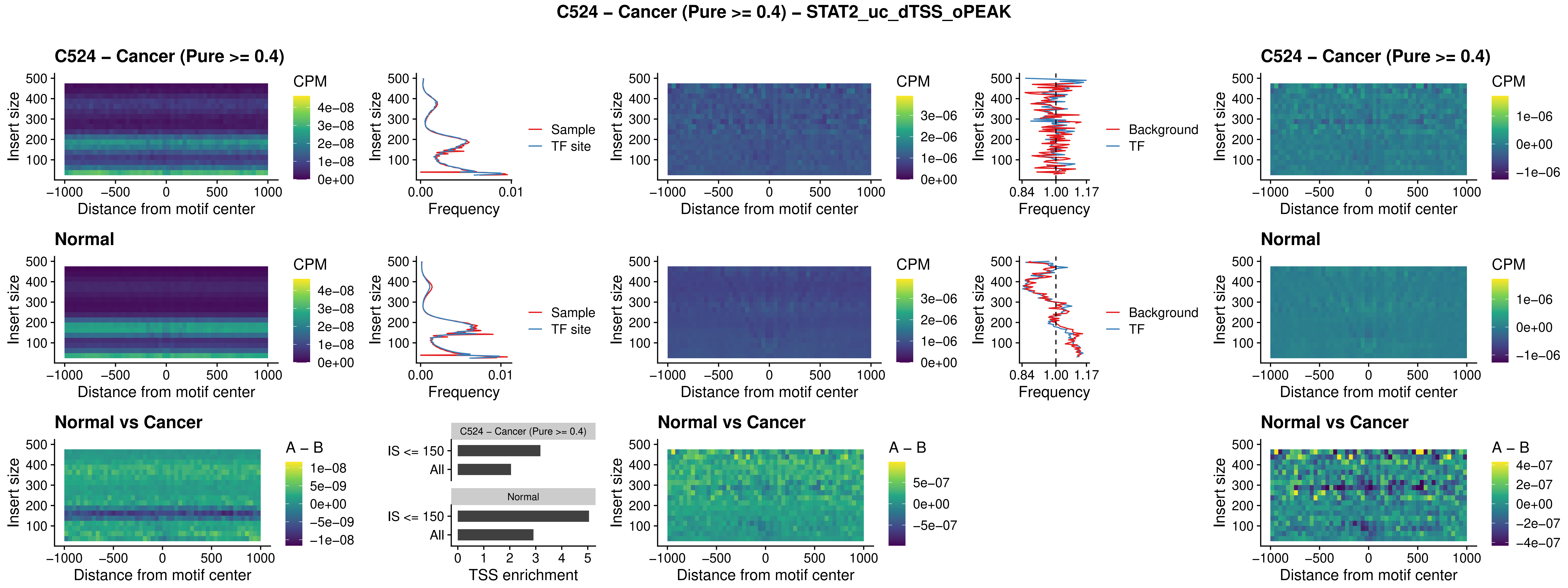
C524 – Cancer (Pure >= 0.4) – STAT2_uc_dTSS_oPEAK
C524 – Cancer (Pure >= 0.4)
Normal
Normal vs Cancer
Insert size
Distance from motif center
Frequency
CPM
A – B
Sample
TF site
Background
TF
C524 – Cancer (Pure >= 0.4)
Normal
IS <= 150
All
TSS enrichment

#### C524 – Cancer (Pure >= 0.4) – ZNF683_uc_cTSS_oPEAK

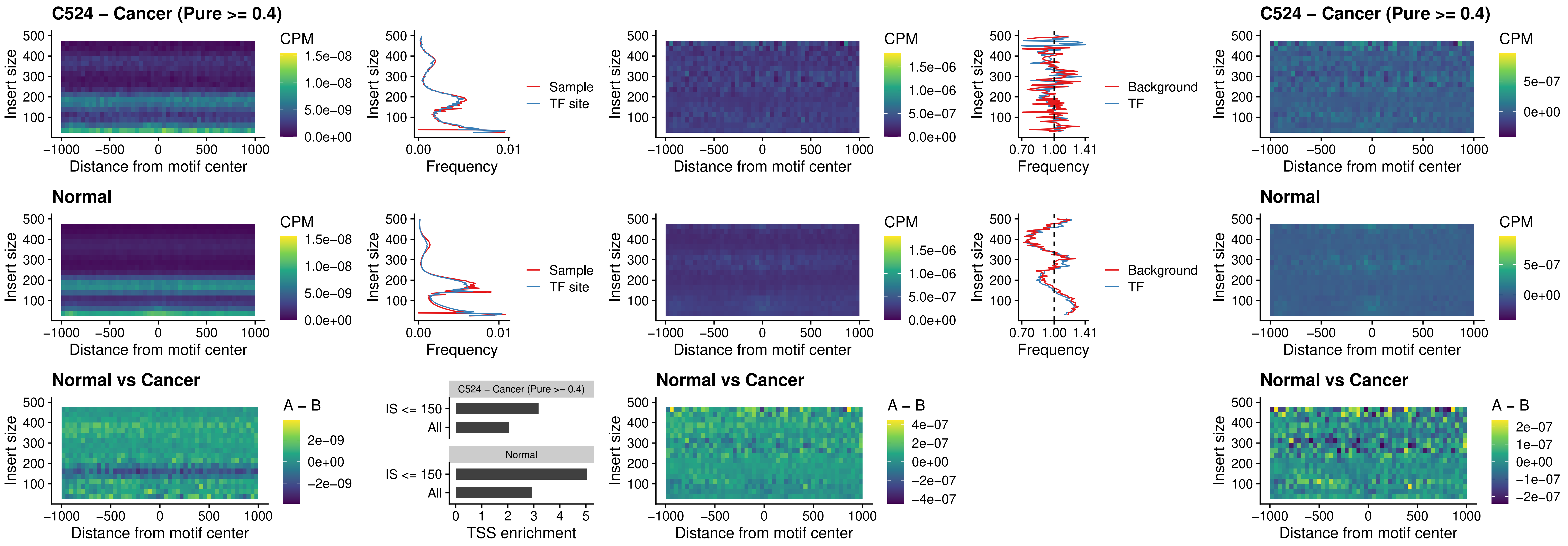

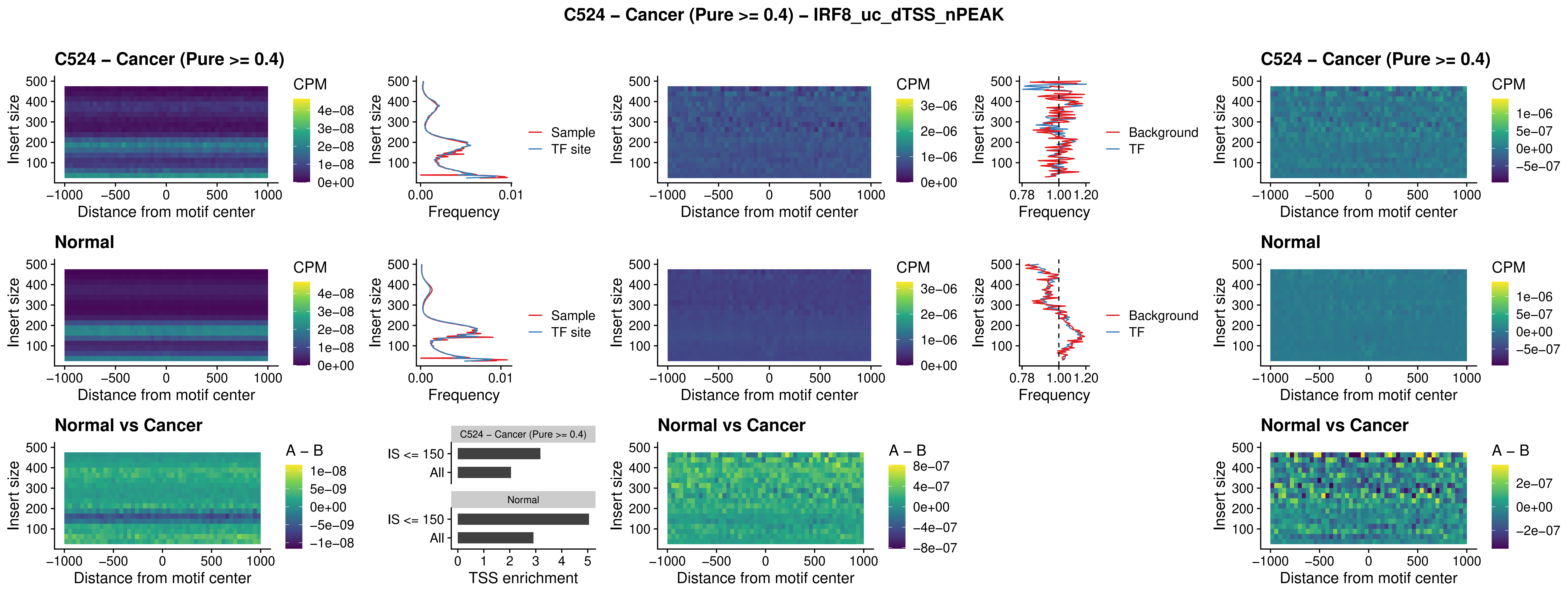
C524 – Cancer (Pure >= 0.4) – IRF8_uc_dTSS_nPEAK
C524 – Cancer (Pure >= 0.4)
Normal
Normal vs Cancer
Insert size
Distance from motif center
Frequency
CPM
A – B
Sample
TF site
Background
TF
TSS enrichment
C524 – Cancer (Pure >= 0.4)
Normal
IS <= 150
All

### C524 – Cancer (Pure >= 0.4) – ETV7_uc_dTSS_oPEAK

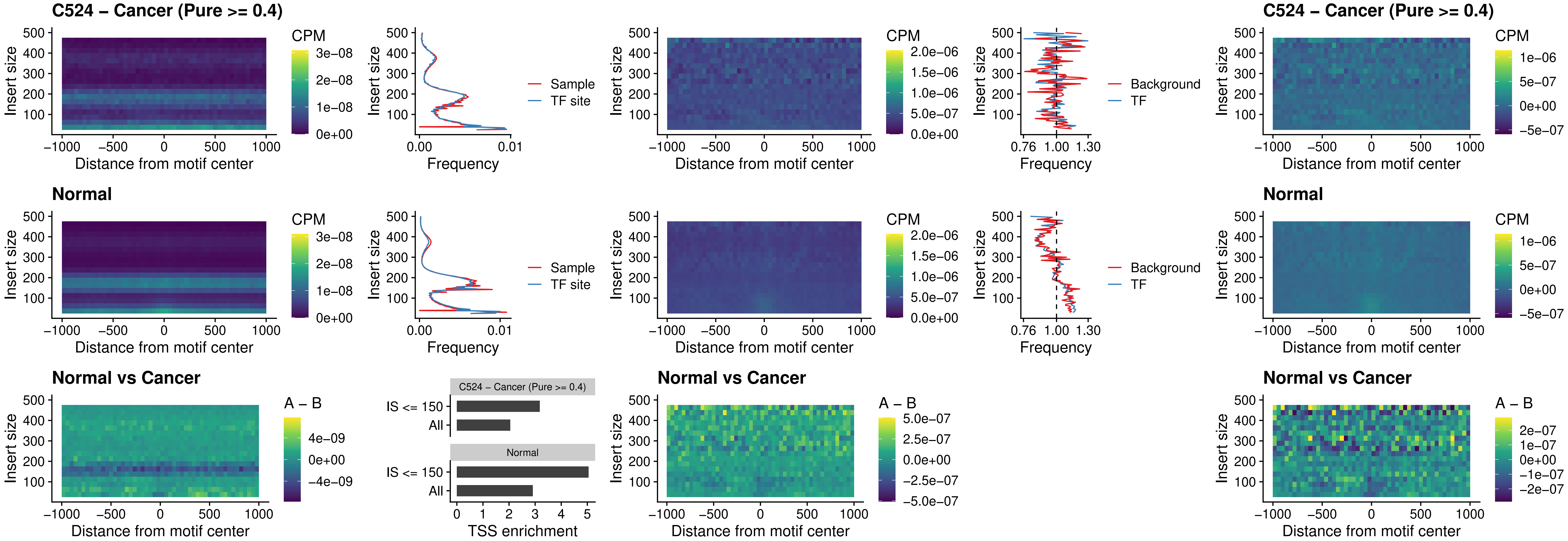

#### C525 – Cancer (Pure >= 0.4) – FOXL1_uc_pTSS_oPEAK

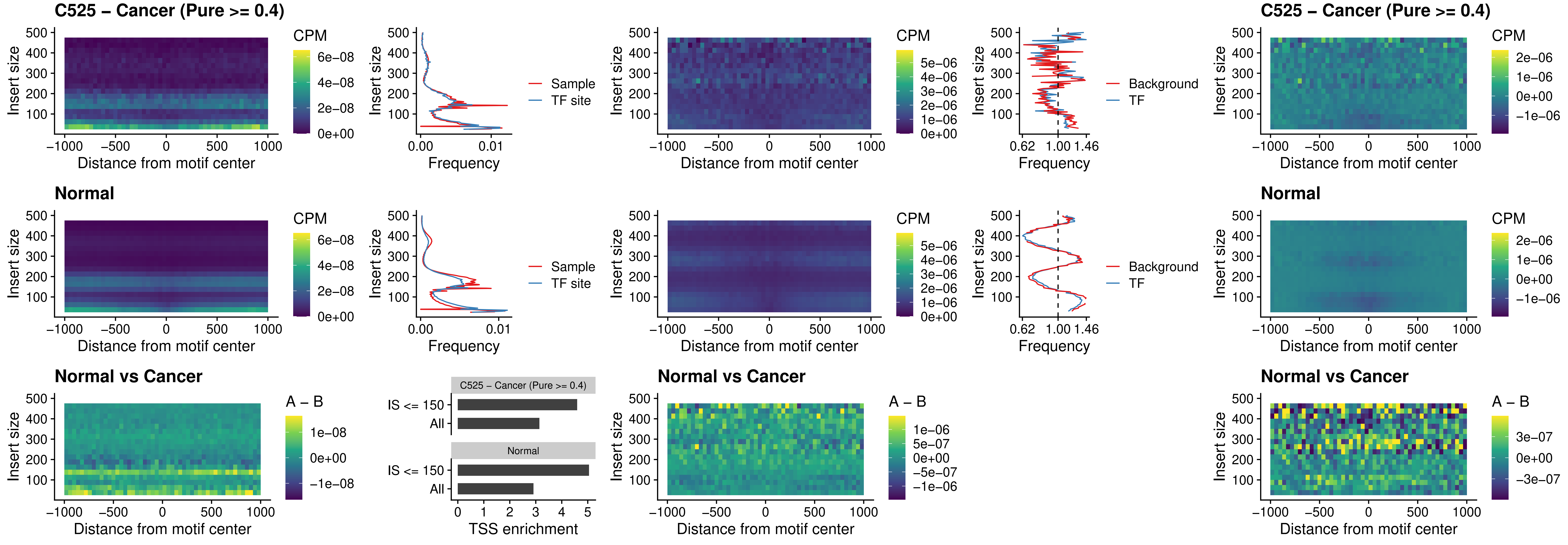

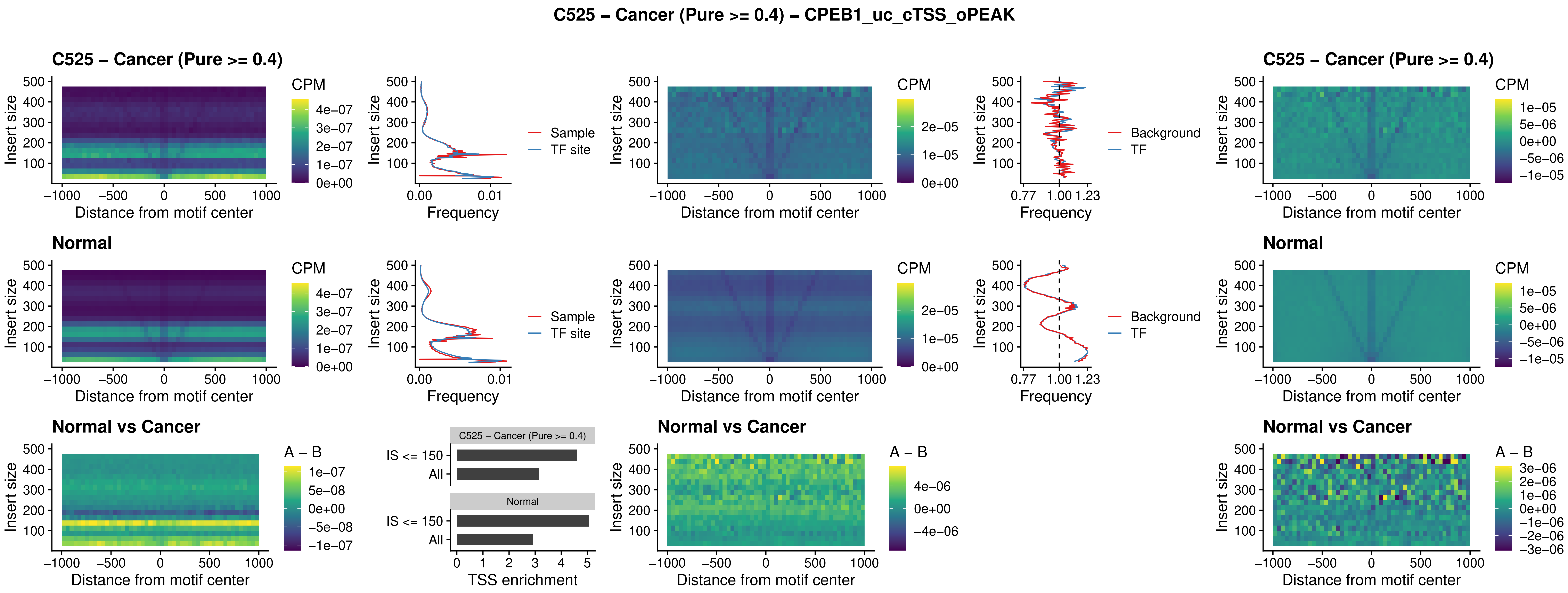
C525 – Cancer (Pure >= 0.4) – CPEB1_uc_cTSS_oPEAK
C525 – Cancer (Pure >= 0.4)
Normal
Normal vs Cancer
Insert size
Distance from motif center
Frequency
CPM
A – B
Sample
TF site
Background
TF
C525 – Cancer (Pure >= 0.4)
Normal
IS <= 150
All
TSS enrichment

### C525 – Cancer (Pure >= 0.4) – HMGA1_uc_pTSS_nPEAK

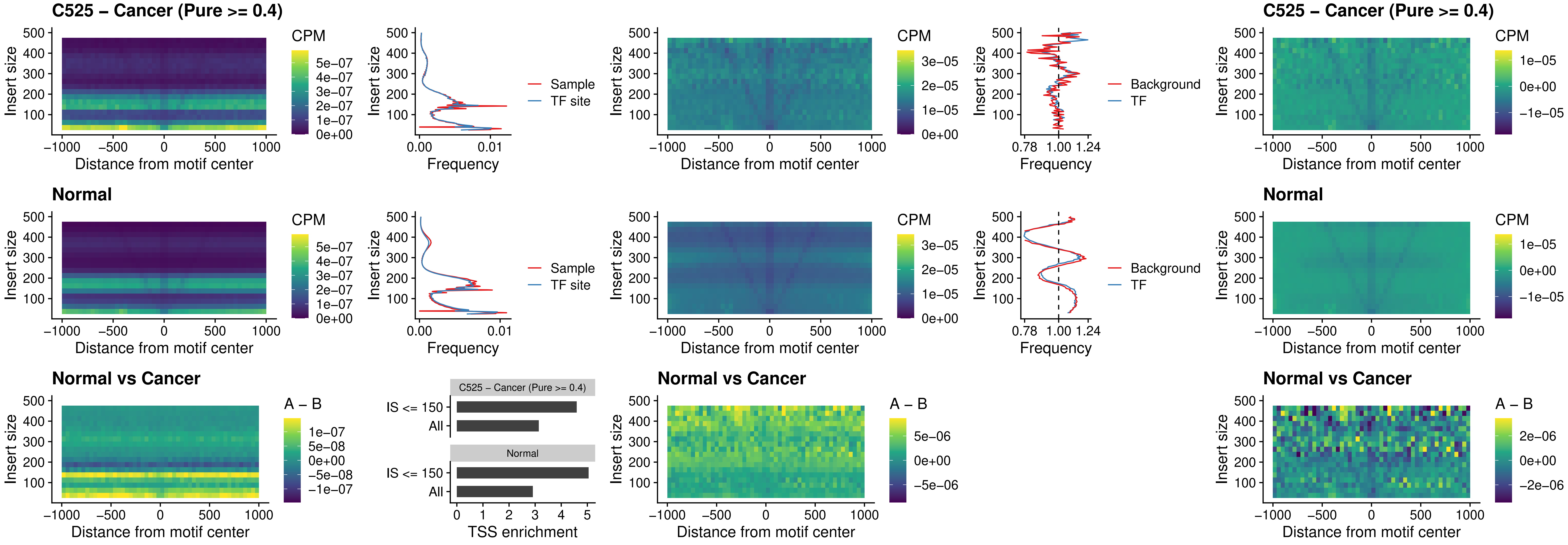

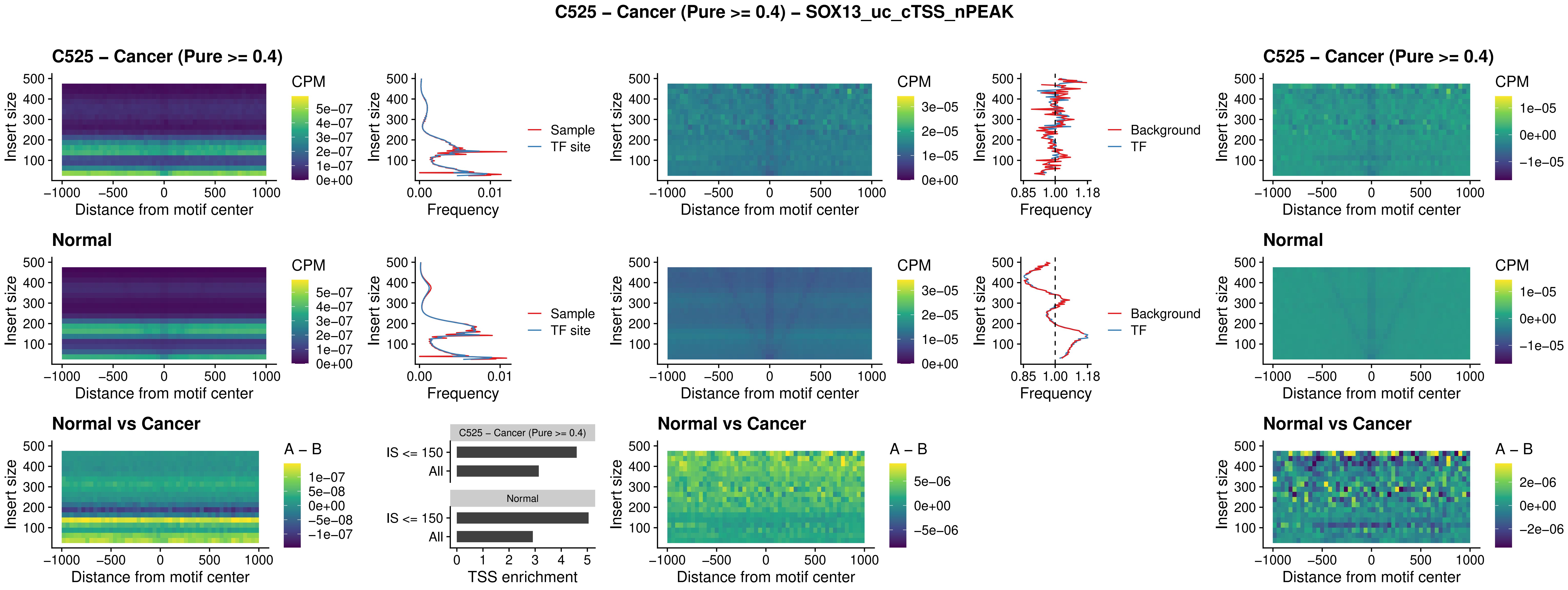
C525 – Cancer (Pure >= 0.4) – SOX13_uc_cTSS_nPEAK
C525 – Cancer (Pure >= 0.4)
Normal
Normal vs Cancer
Insert size
Distance from motif center
Frequency
CPM
A – B
Sample
TF site
Background
TF
C525 – Cancer (Pure >= 0.4)
IS <= 150
All
TSS enrichment

#### C525 – Cancer (Pure >= 0.4) – LHX4_uc_cTSS_nPEAK

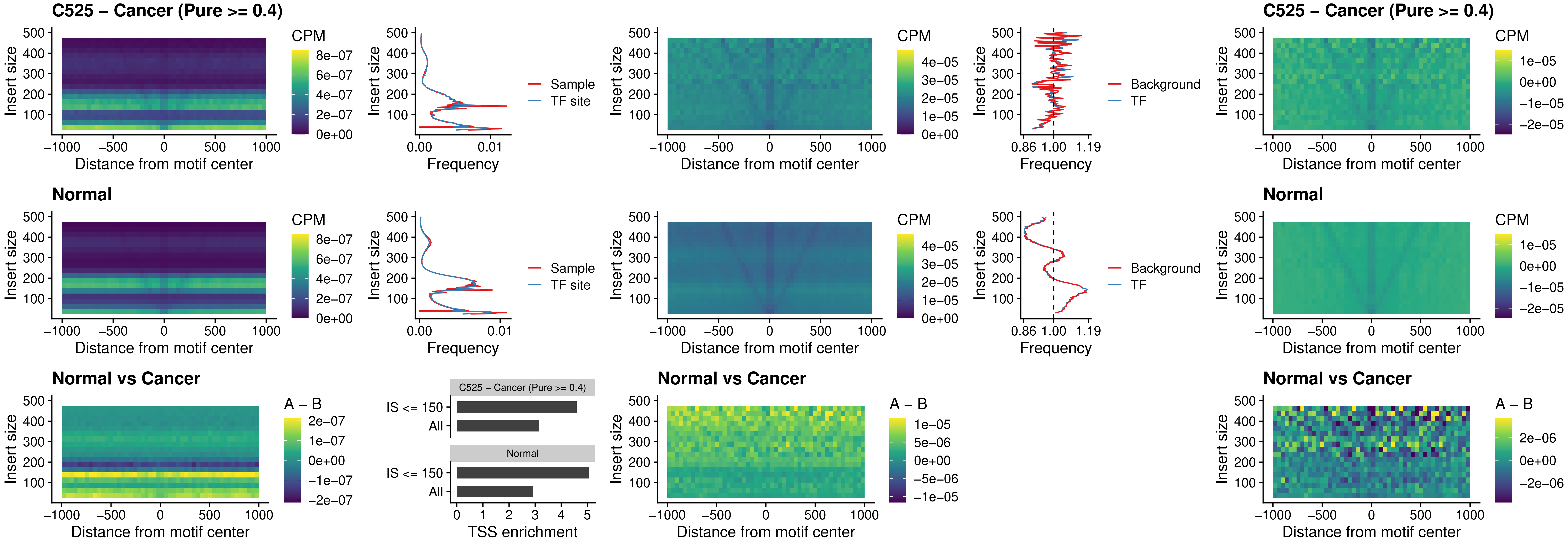

#### C525 – Cancer (Pure >= 0.4) – HOXA4_uc_cTSS_nPEAK

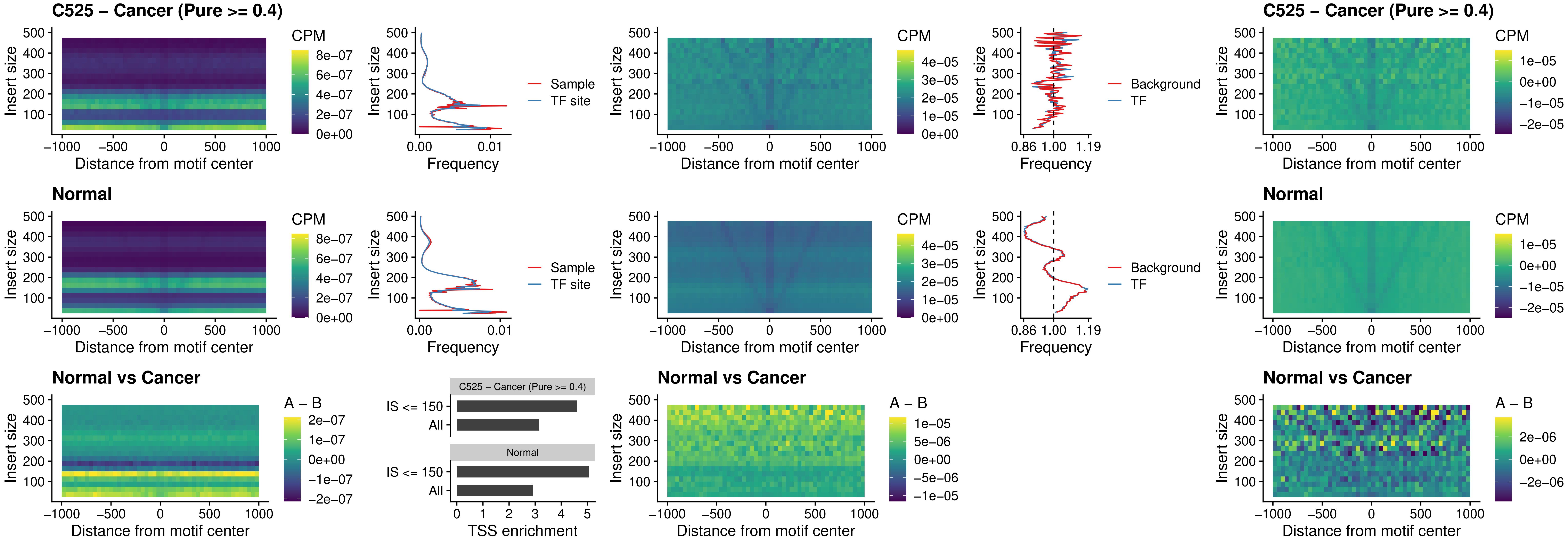

#### C525 – Cancer (Pure >= 0.4) – HLX_uc_cTSS_nPEAK

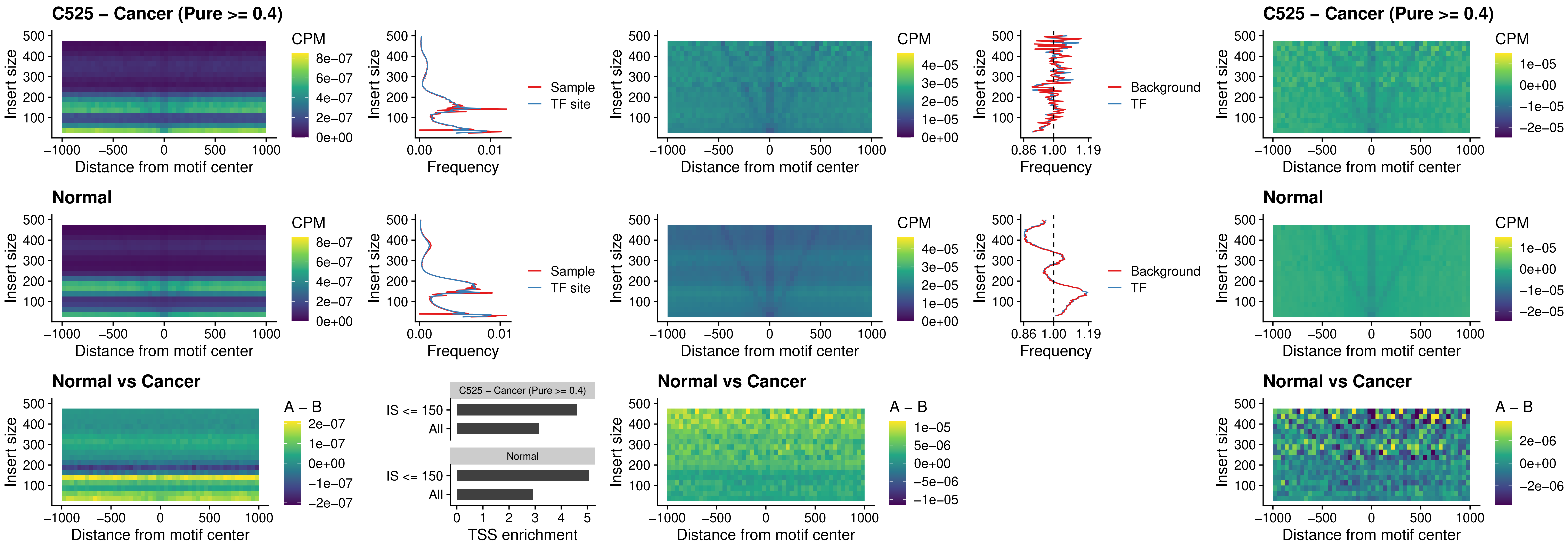

### C525 – Cancer (Pure >= 0.4) – AC0125311_uc_cTSS_nPEAK

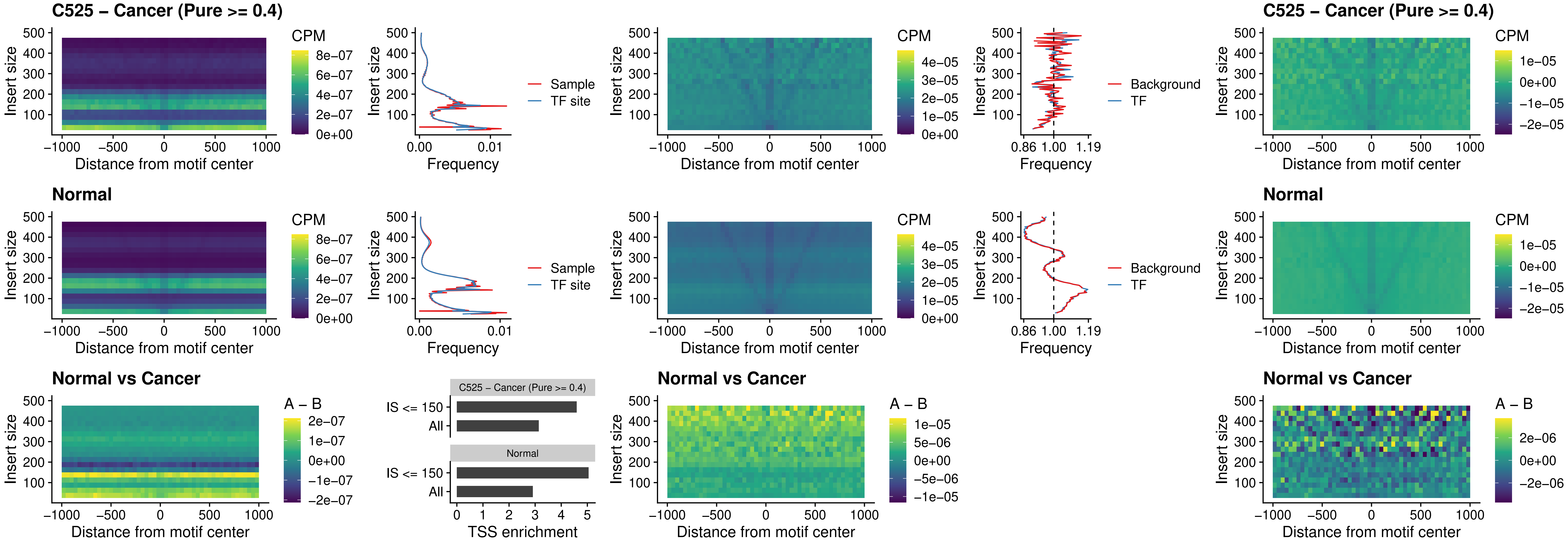

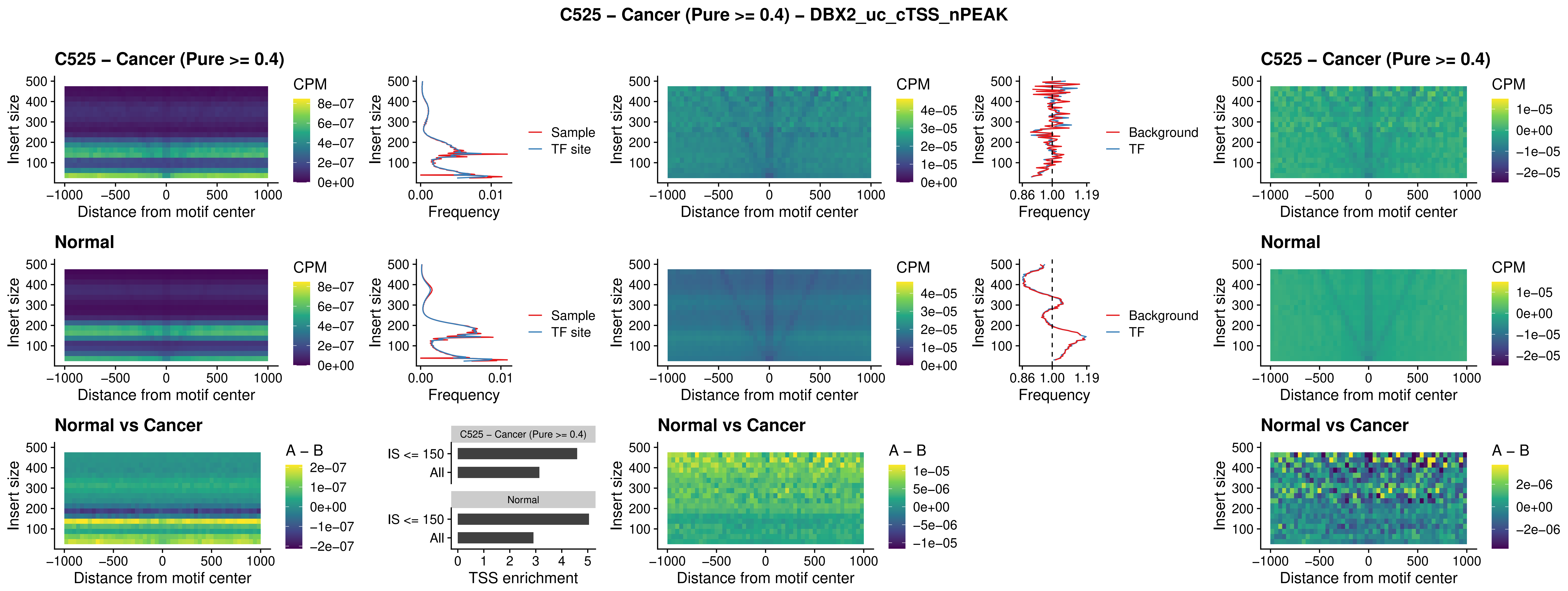
C525 – Cancer (Pure >= 0.4) – DBX2_uc_cTSS_nPEAK
C525 – Cancer (Pure >= 0.4)
Normal
Normal vs Cancer
Insert size
Distance from motif center
Frequency
CPM
A – B
Sample
TF site
Background
TF
C525 – Cancer (Pure >= 0.4)
Normal
IS <= 150
All
TSS enrichment

### C525 – Cancer (Pure >= 0.4) – LHX6_uc_dTSS_nPEAK

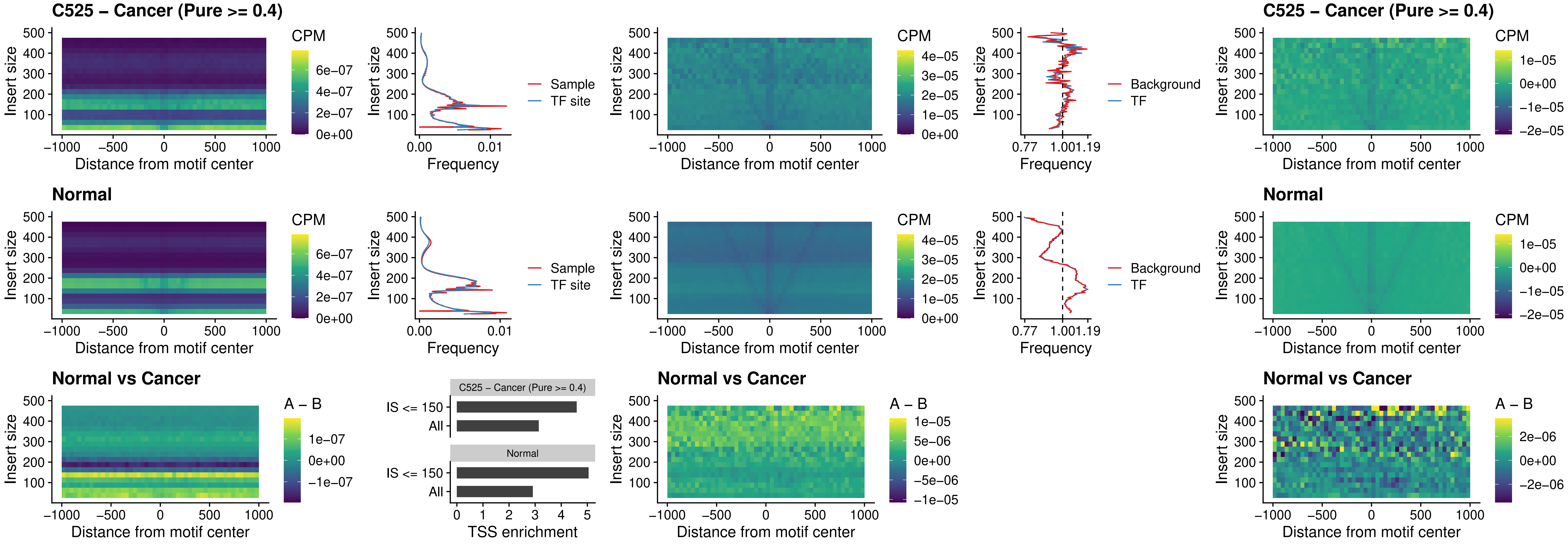

#### C525 – Cancer (Pure >= 0.4) – SOX1_uc_dTSS_nPEAK

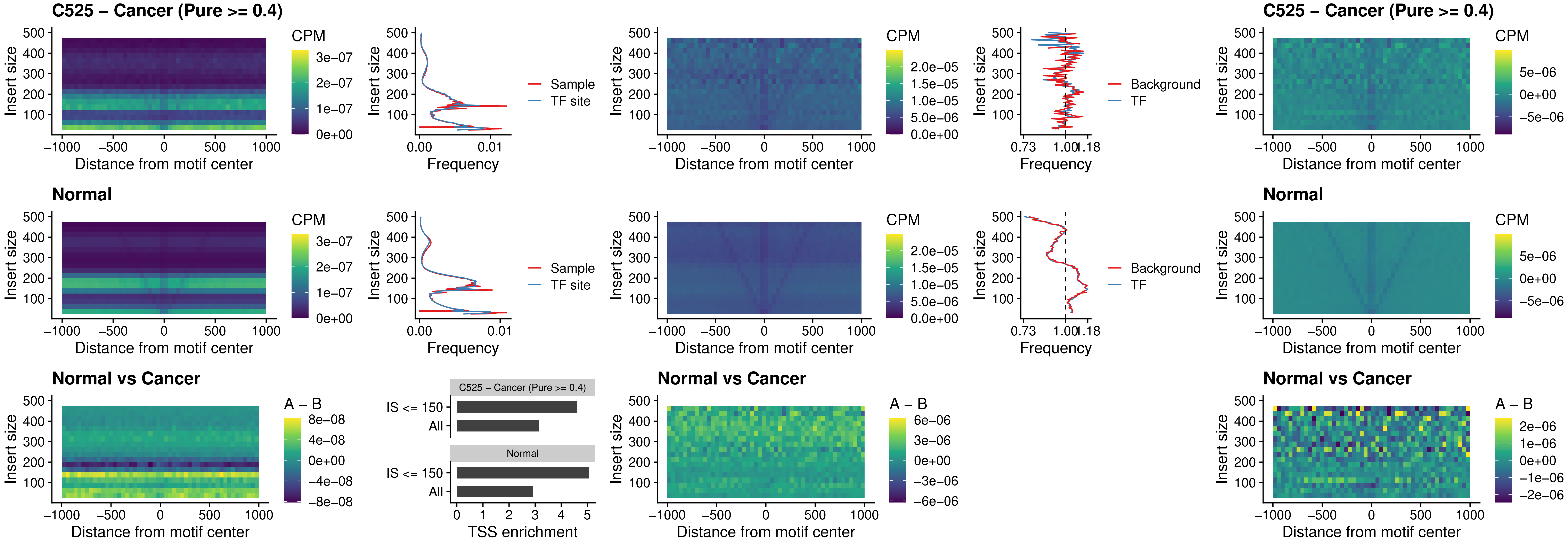

#### C525 – Cancer (Pure >= 0.4) – NKX24_uc_dTSS_nPEAK

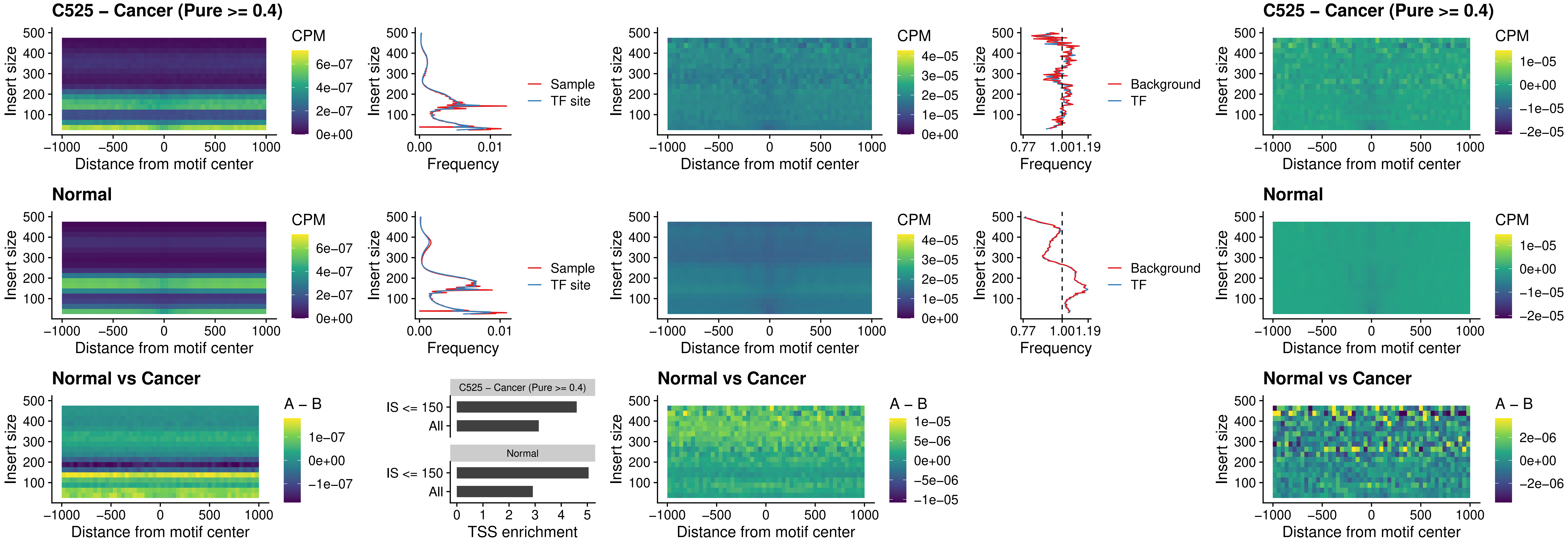

#### C525 – Cancer (Pure >= 0.4) – DLX3_uc_cTSS_nPEAK

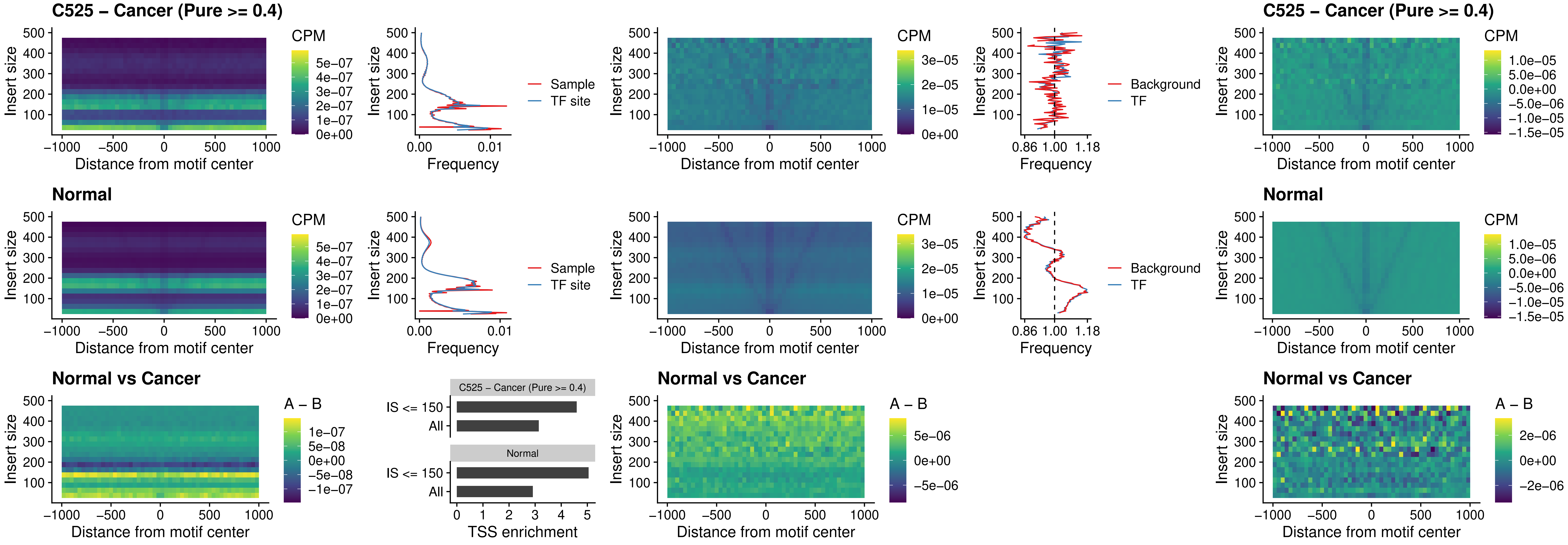

### C525 – Cancer (Pure >= 0.4) – HOXD3_uc_cTSS_nPEAK

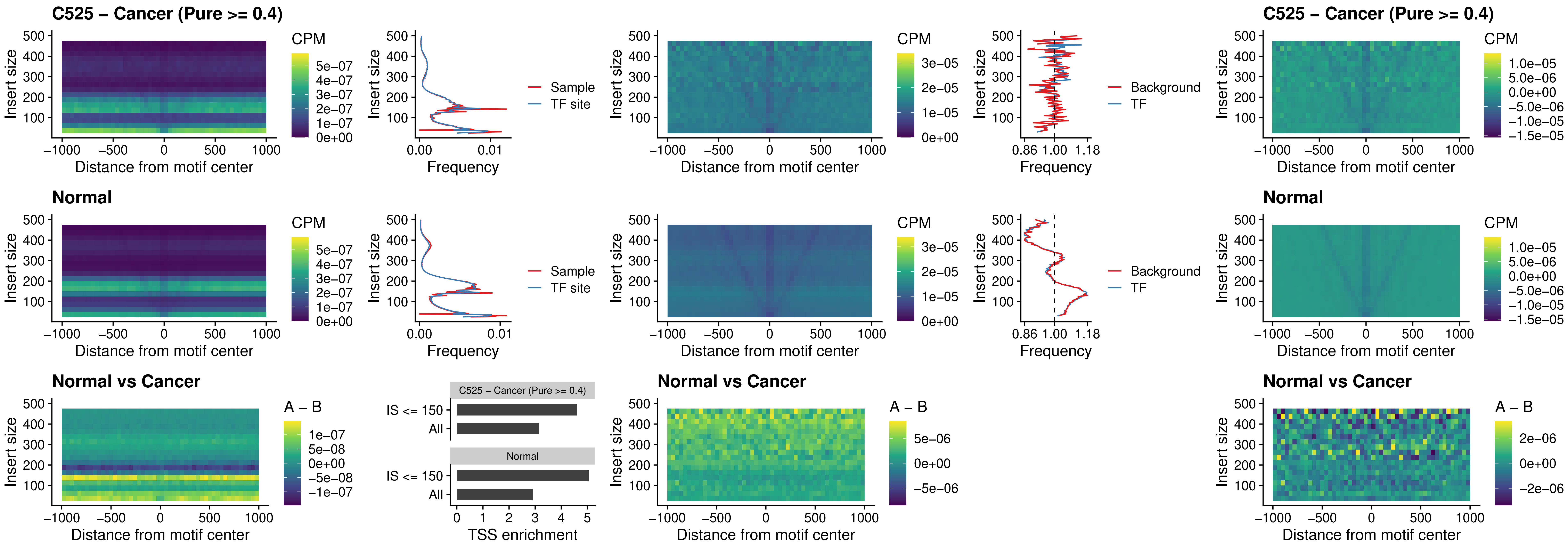

### C525 – Cancer (Pure >= 0.4) – ISL1_uc_dTSS_oPEAK

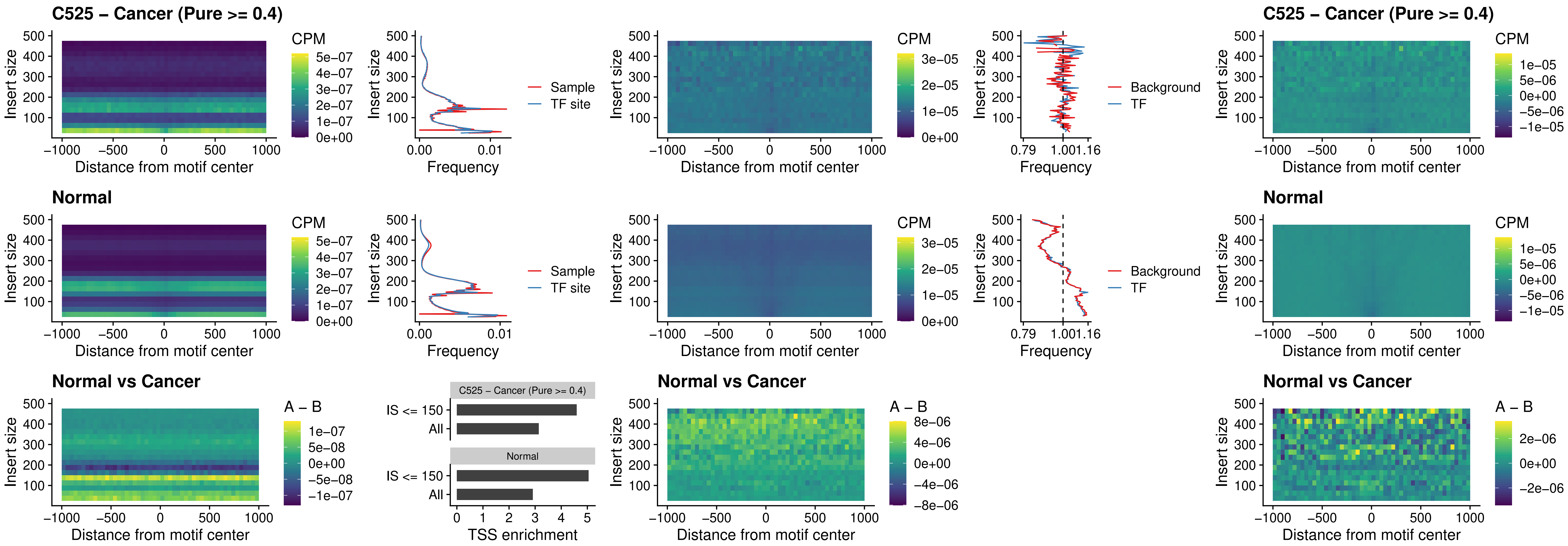

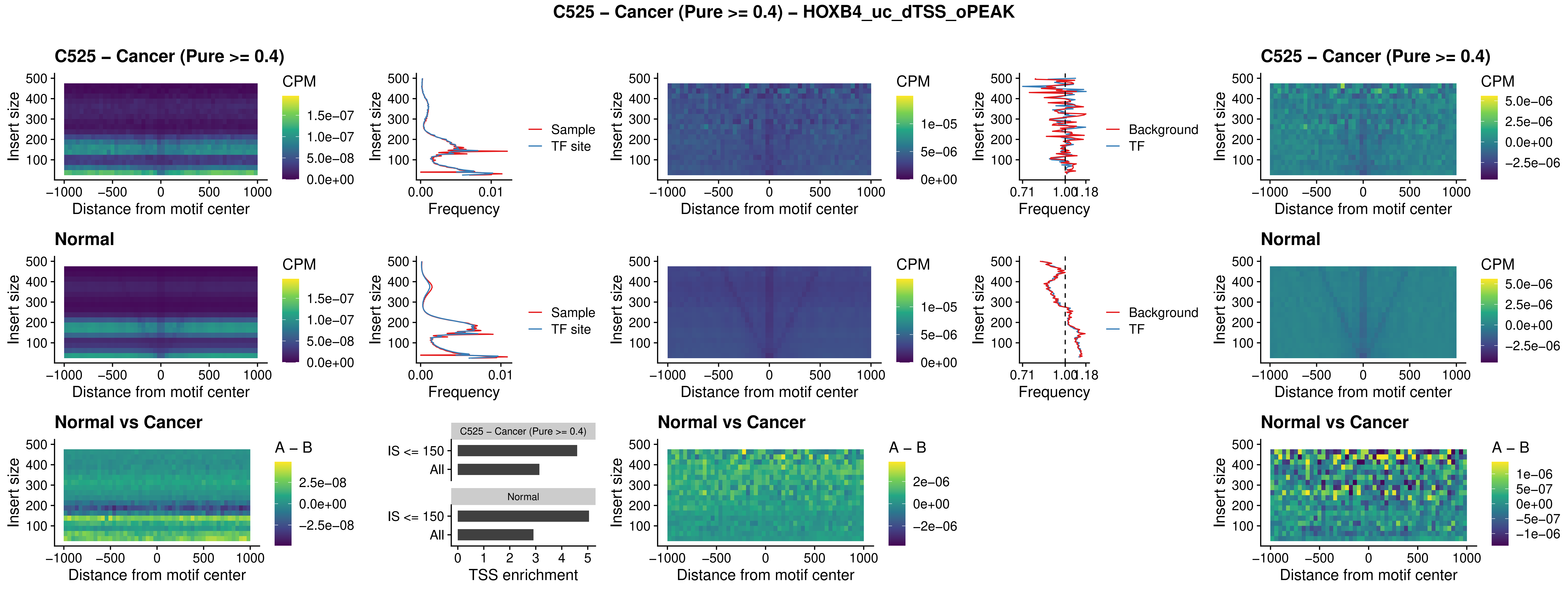

C525 – Cancer (Pure >= 0.4) – HOXB4_uc_dTSS_oPEAK
C525 – Cancer (Pure >= 0.4)
Normal
Normal vs Cancer
Insert size
Distance from motif center
Frequency
CPM
A – B
Sample
TF site
Background
TF
C525 – Cancer (Pure >= 0.4)
Normal
IS <= 150
All
TSS enrichment

#### C525 – Cancer (Pure >= 0.4) – VAX1_uc_dTSS_oPEAK

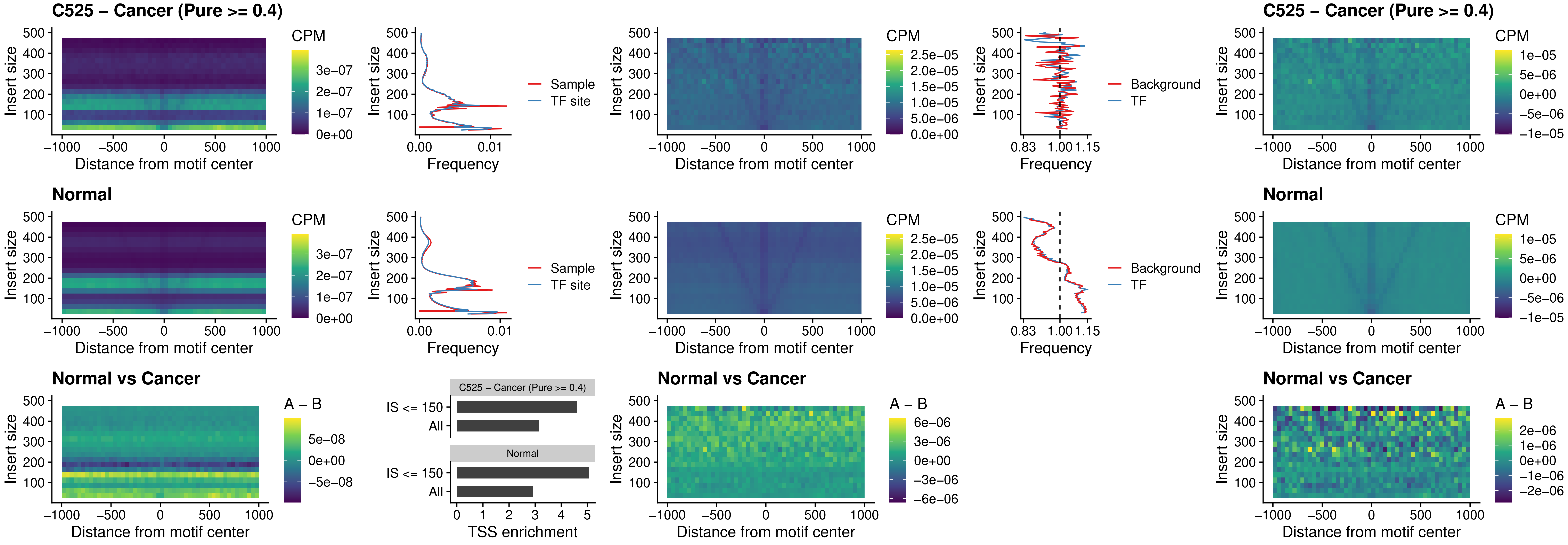

#### C525 – Cancer (Pure >= 0.4) – DLX3_uc_dTSS_oPEAK

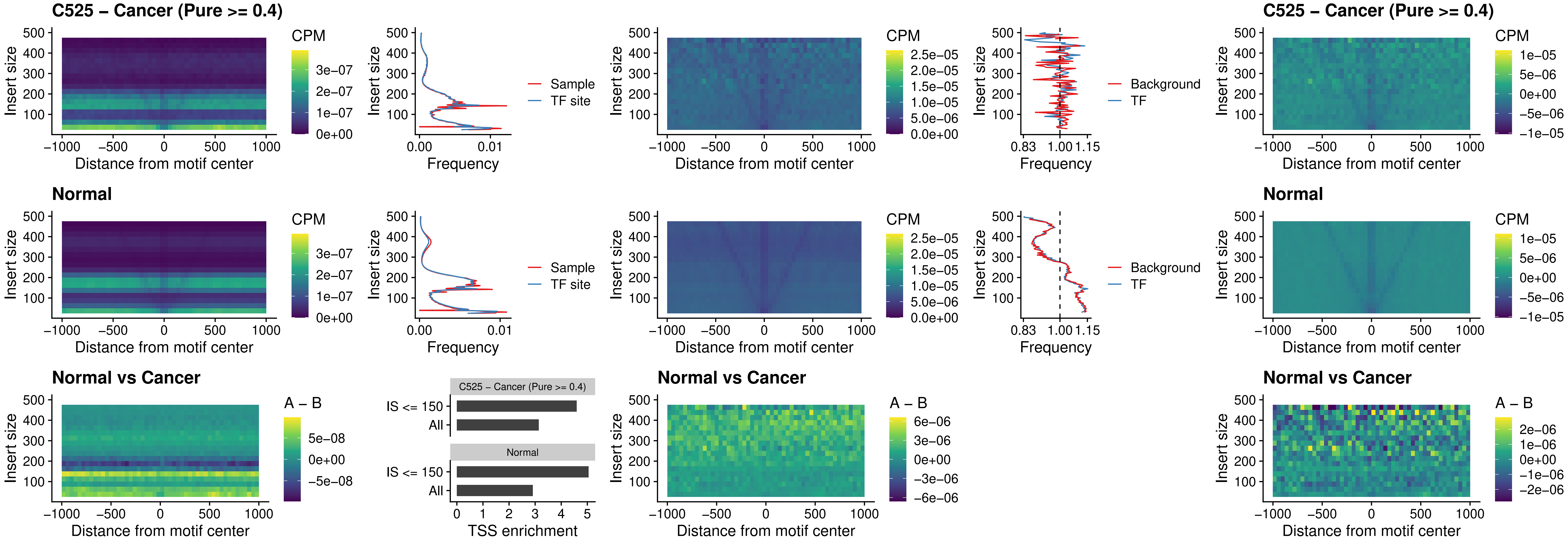

#### C525 – Cancer (Pure >= 0.4) – HOXD3_uc_dTSS_oPEAK

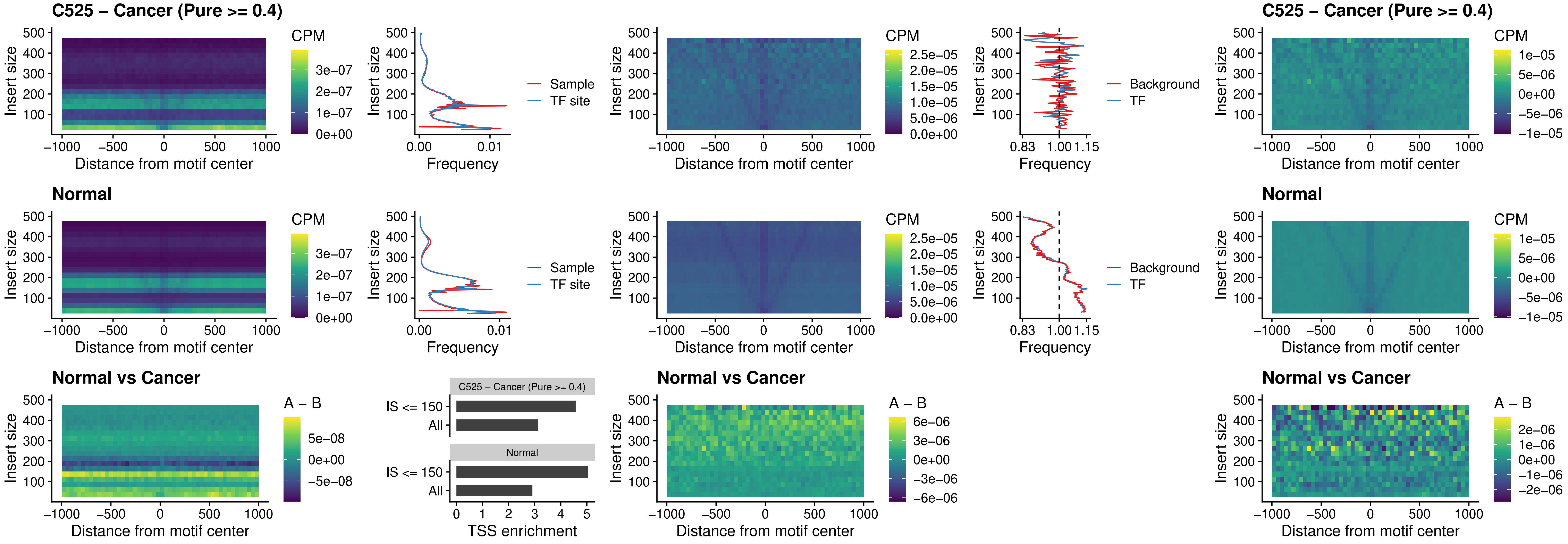

#### C525 – Cancer (Pure >= 0.4) – BBX_uc_dTSS_nPEAK

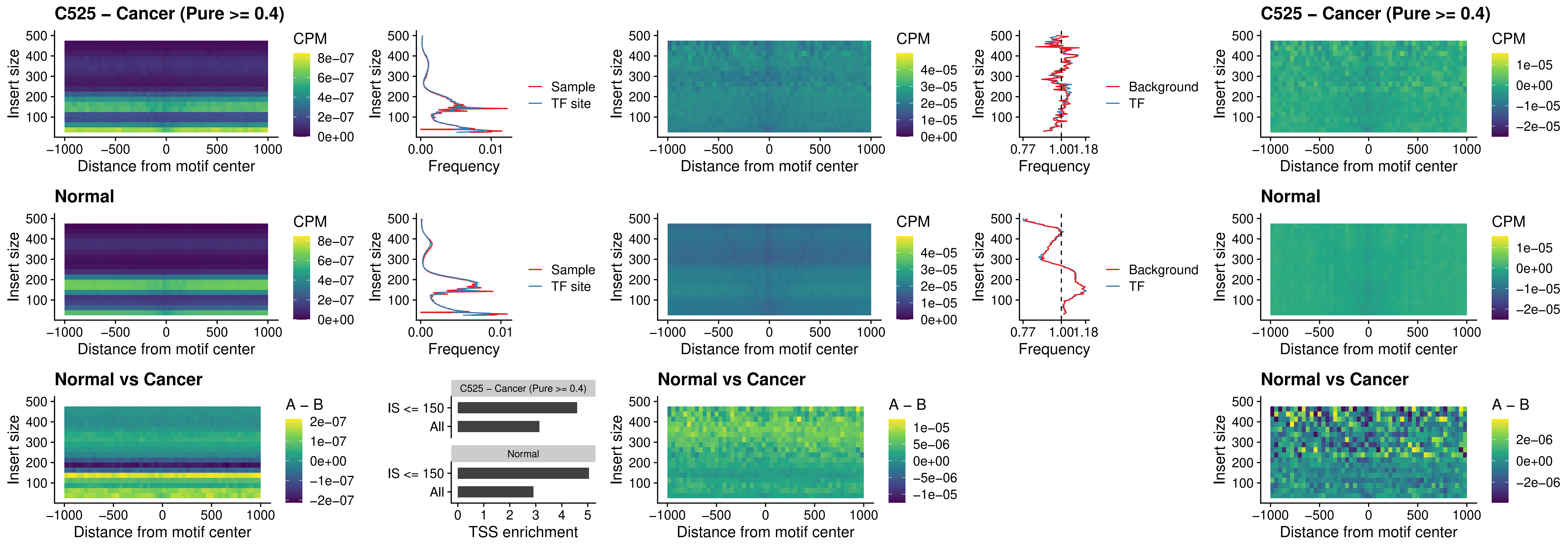

#### C525 – Cancer (Pure >= 0.4) – SOX3_uc_cTSS_nPEAK

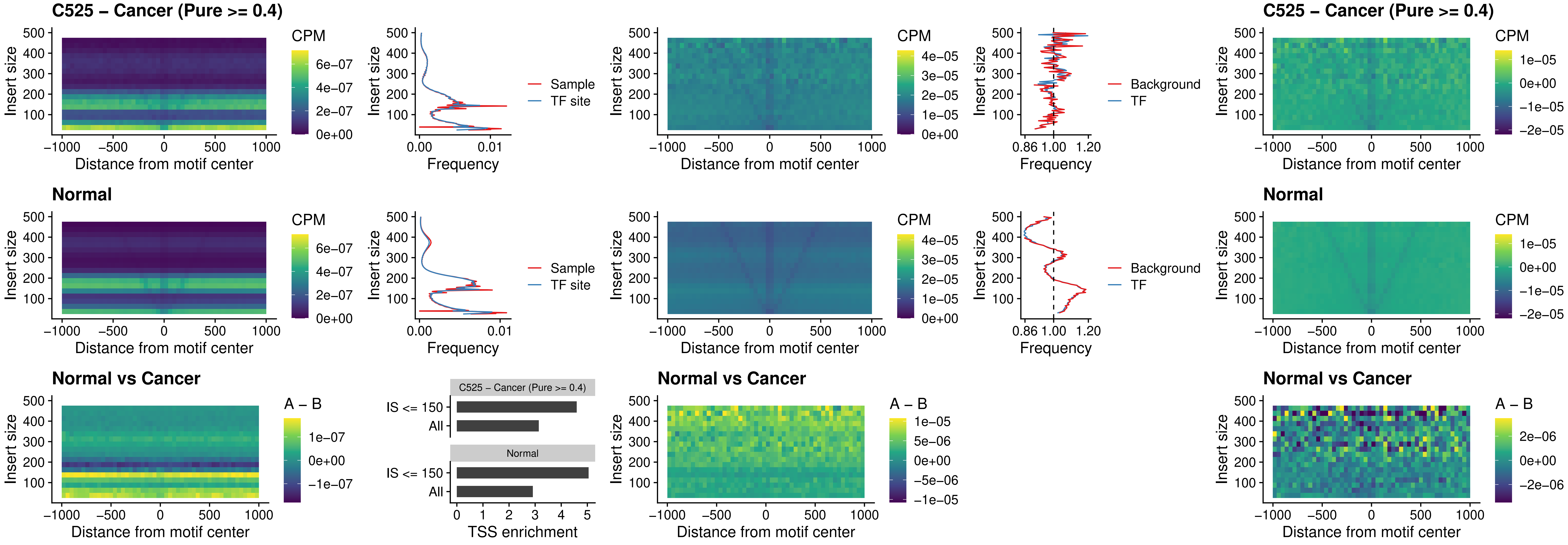

#### C525 – Cancer (Pure >= 0.4) – TBPL2_uc_cTSS_nPEAK

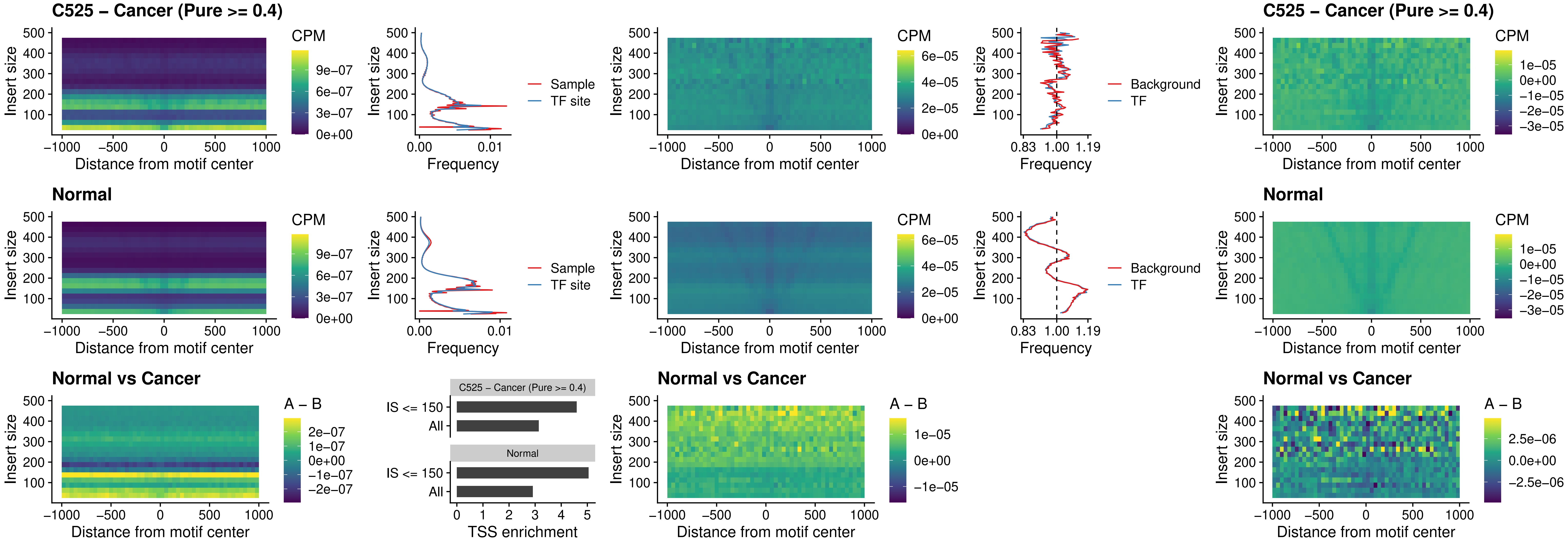

#### C525 – Cancer (Pure >= 0.4) – HOXA4_uc_cTSS_oPEAK

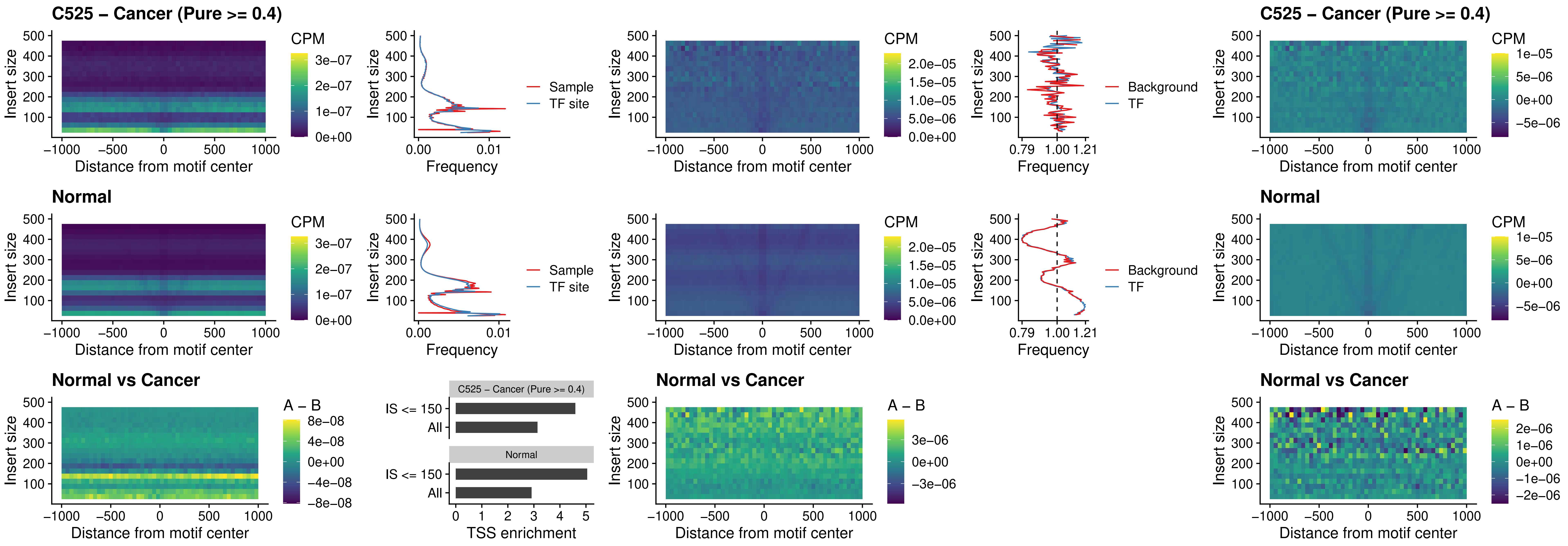

#### C525 – Cancer (Pure >= 0.4) – PDX1_uc_dTSS_oPEAK

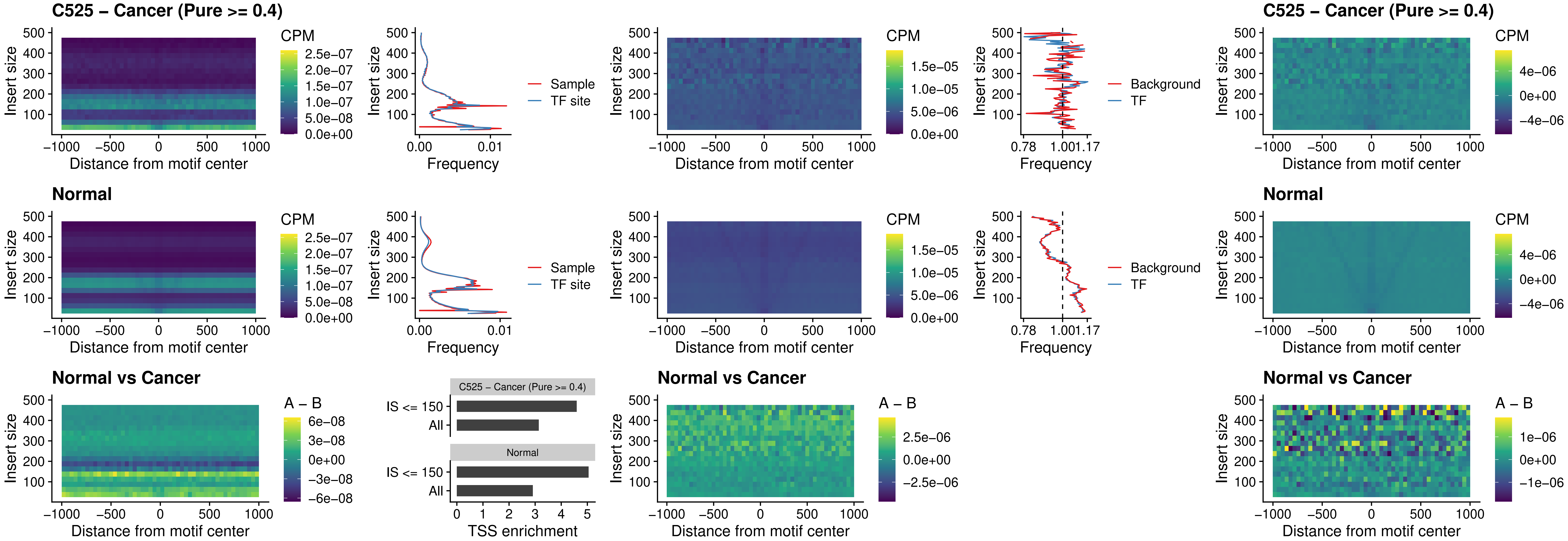

#### C525 – Cancer (Pure >= 0.4) – LHX4_uc_dTSS_oPEAK

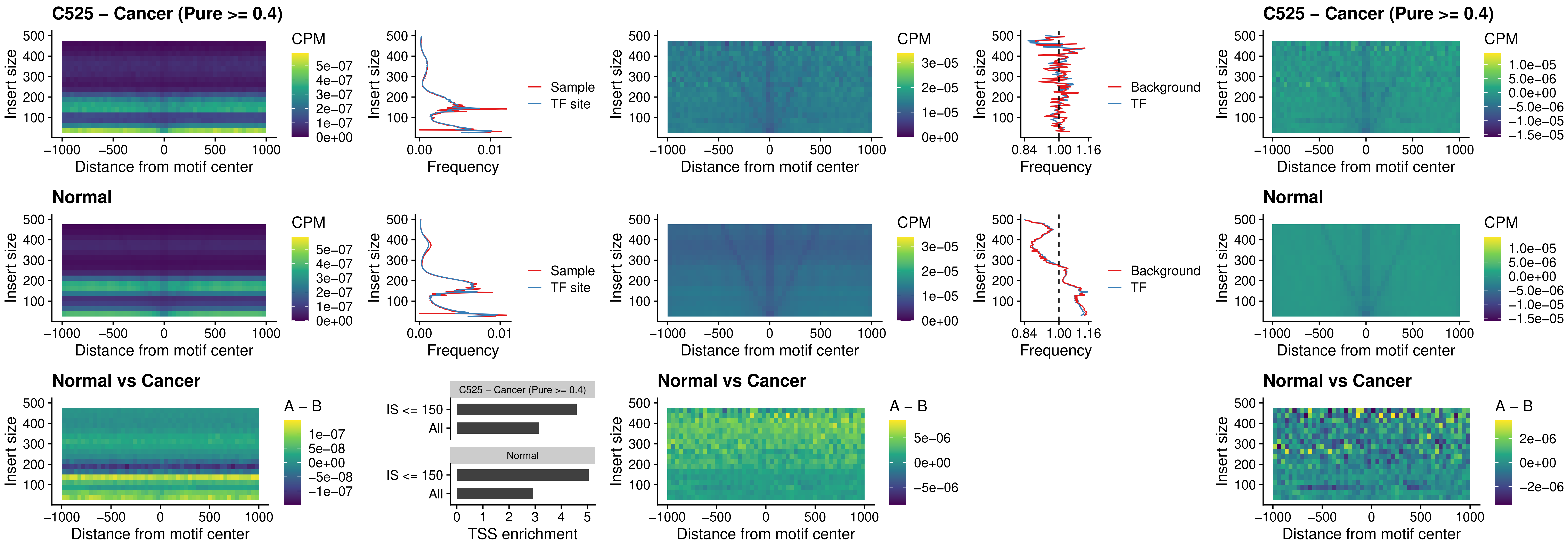

#### C525 – Cancer (Pure >= 0.4) – HOXA4_uc_dTSS_oPEAK

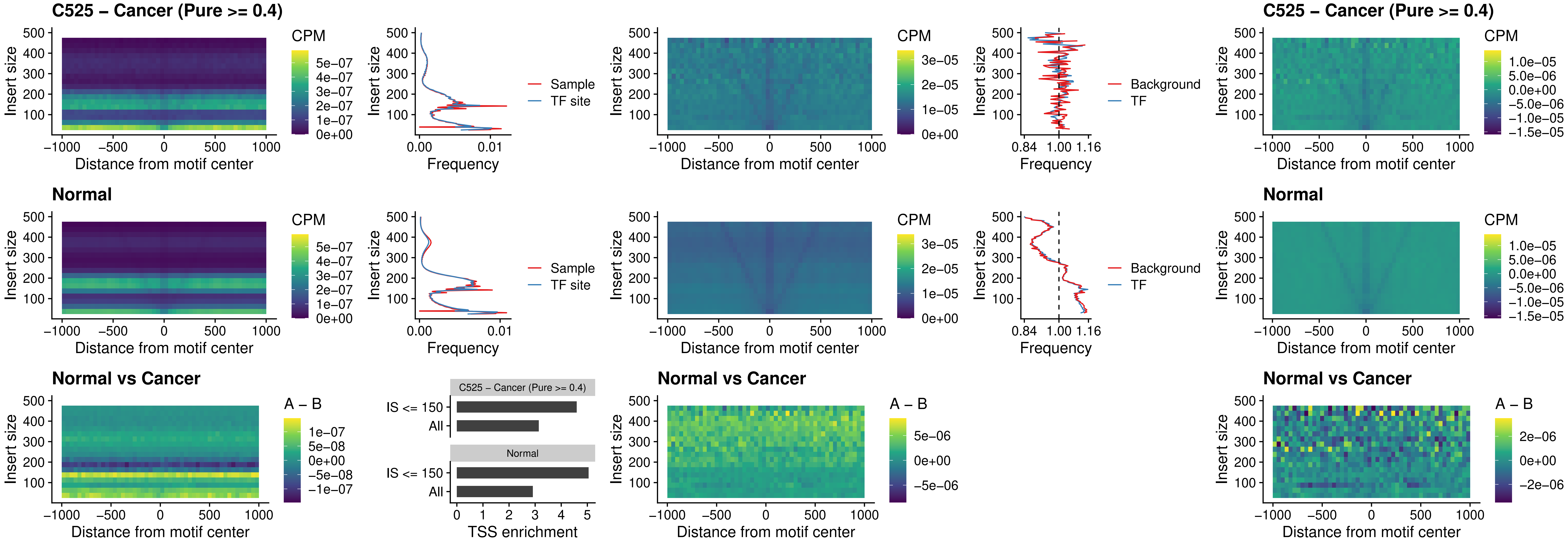

#### C525 – Cancer (Pure >= 0.4) – HLX_uc_dTSS_oPEAK

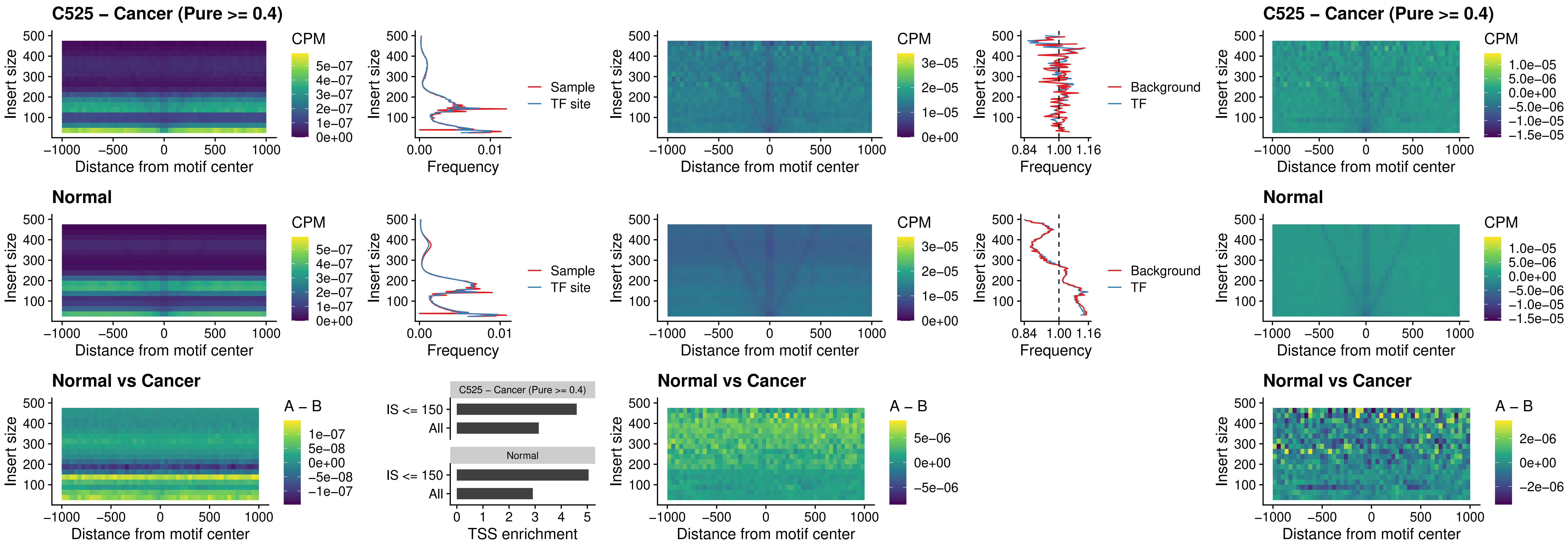

#### C525 – Cancer (Pure >= 0.4) – AC0125311_uc_dTSS_oPEAK

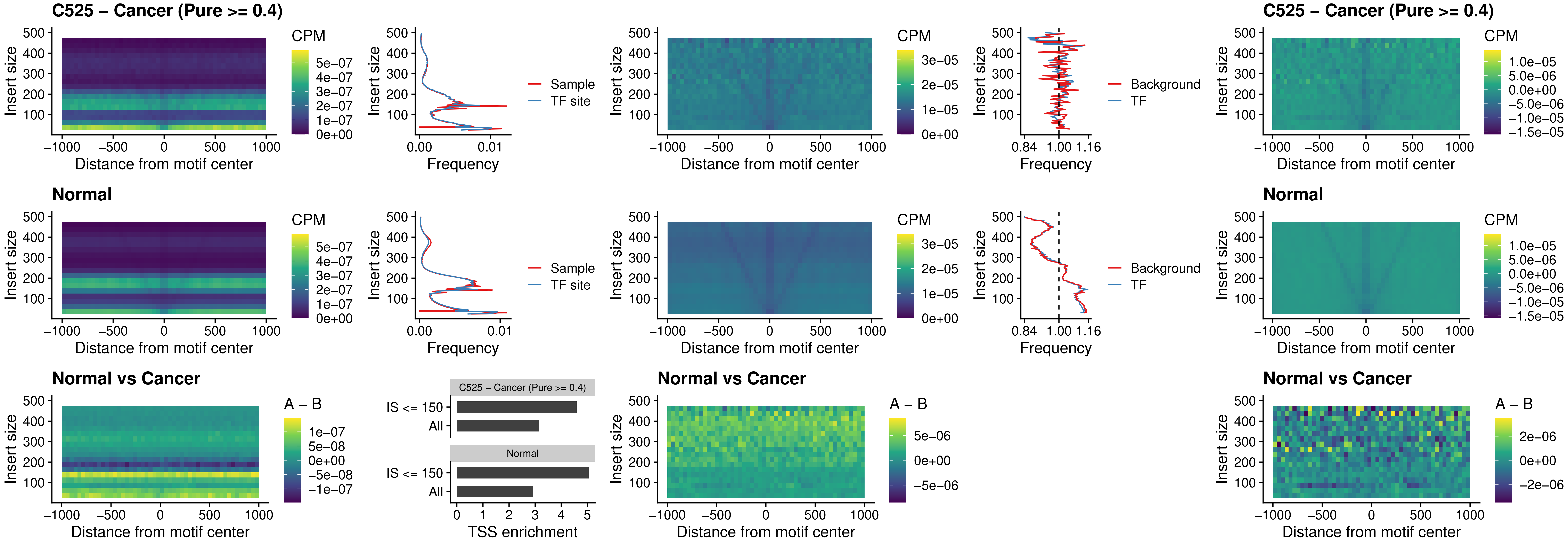

#### C525 – Cancer (Pure >= 0.4) – DBX2_uc_dTSS_oPEAK

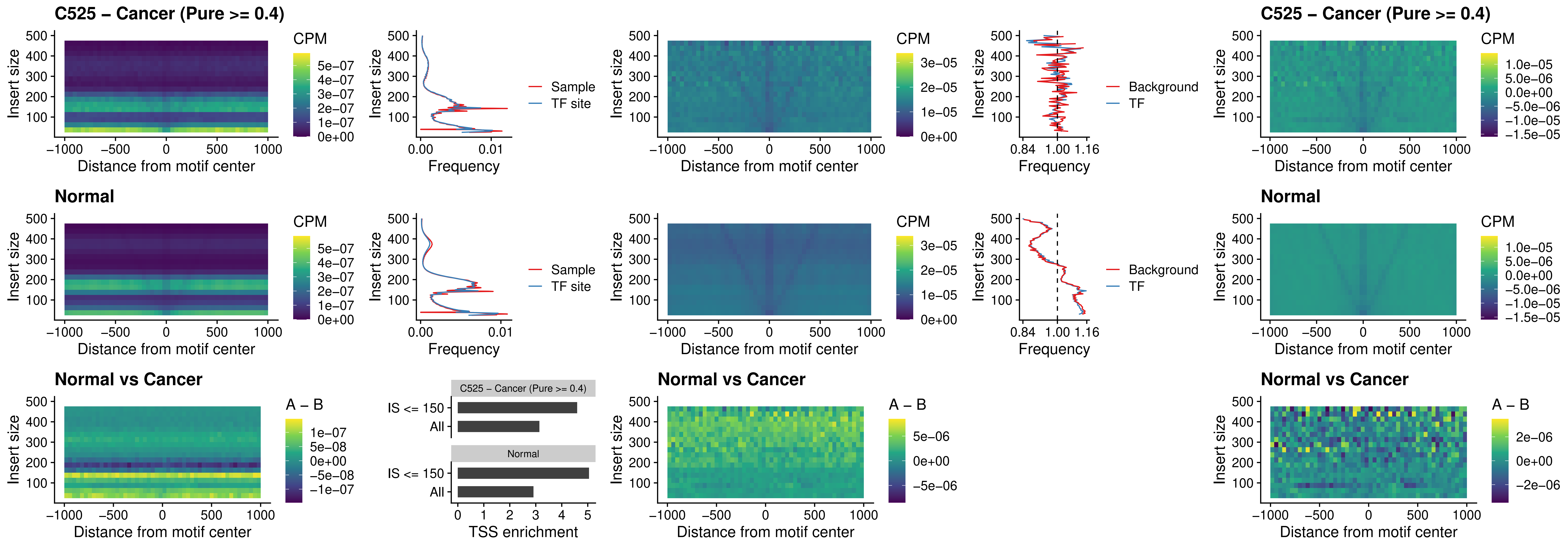

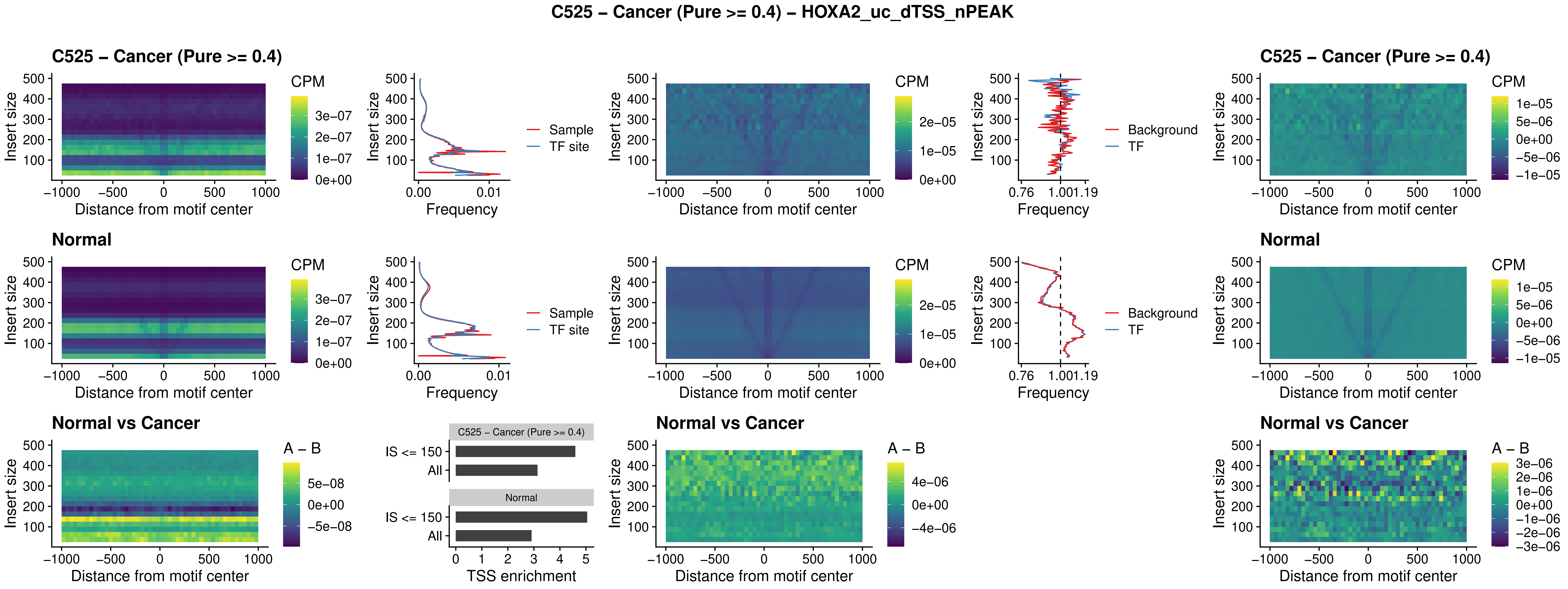
C525 – Cancer (Pure >= 0.4) – HOXA2_uc_dTSS_nPEAK
C525 – Cancer (Pure >= 0.4)
Normal
Normal vs Cancer
Insert size
Distance from motif center
Frequency
CPM
A – B
Sample
TF site
Background
TF
C525 – Cancer (Pure >= 0.4)
Normal
IS <= 150
All
TSS enrichment

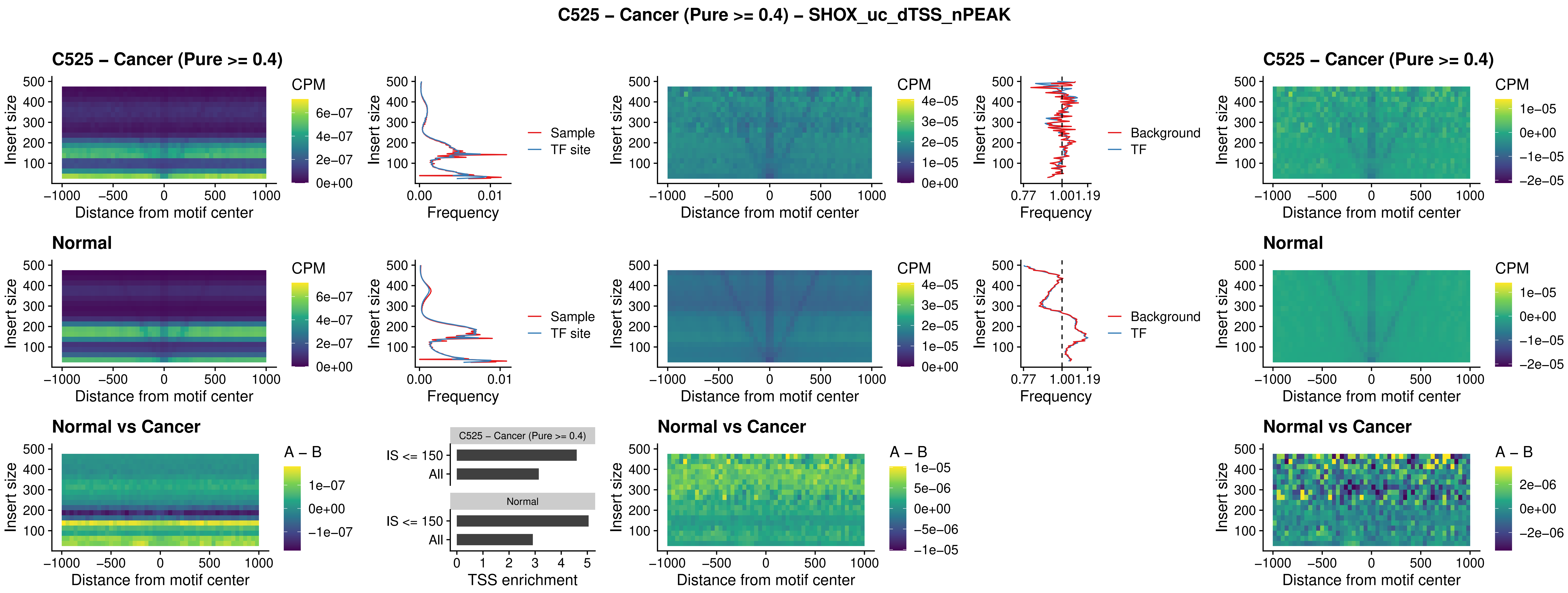

C525 – Cancer (Pure >= 0.4) – SHOX_uc_dTSS_nPEAK
C525 – Cancer (Pure >= 0.4)
Normal
Normal vs Cancer
Insert size
Distance from motif center
CPM
A – B
Frequency
Sample
TF site
Background
TF
C525 – Cancer (Pure >= 0.4)
Normal
IS <= 150
All
TSS enrichment

### C525 – Cancer (Pure >= 0.4) – PRRX1_uc_dTSS_nPEAK

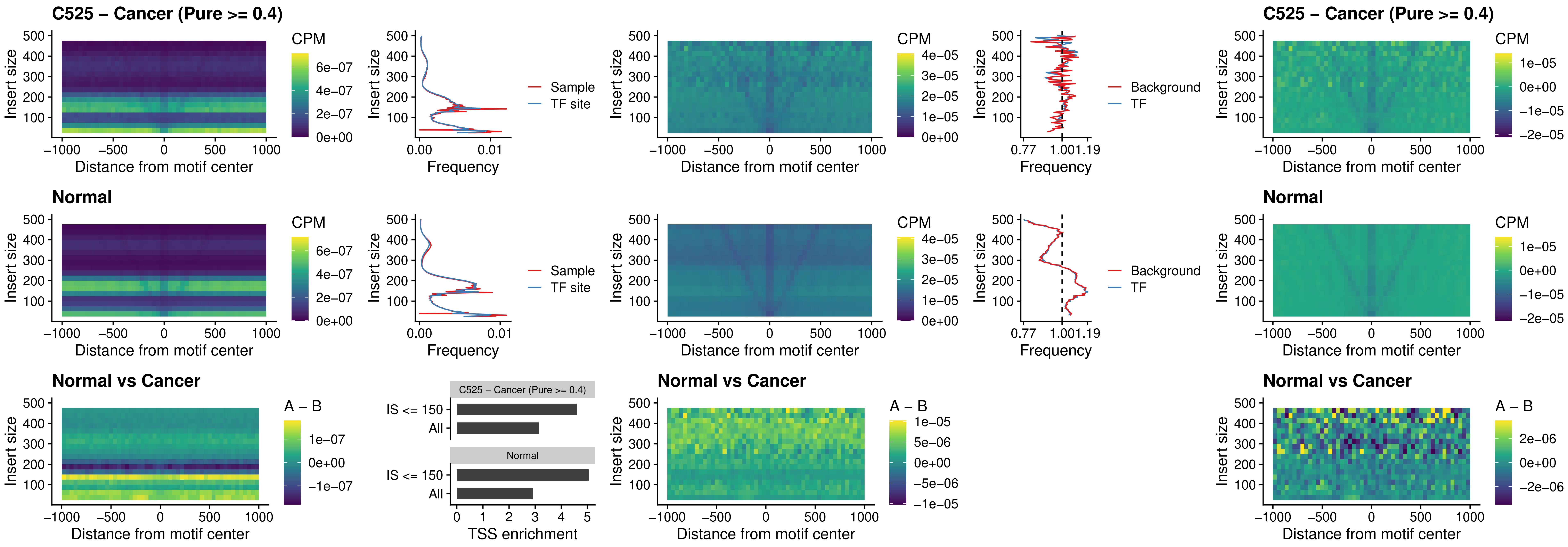

#### C525 – Cancer (Pure >= 0.4) – LMX1A_uc_dTSS_nPEAK

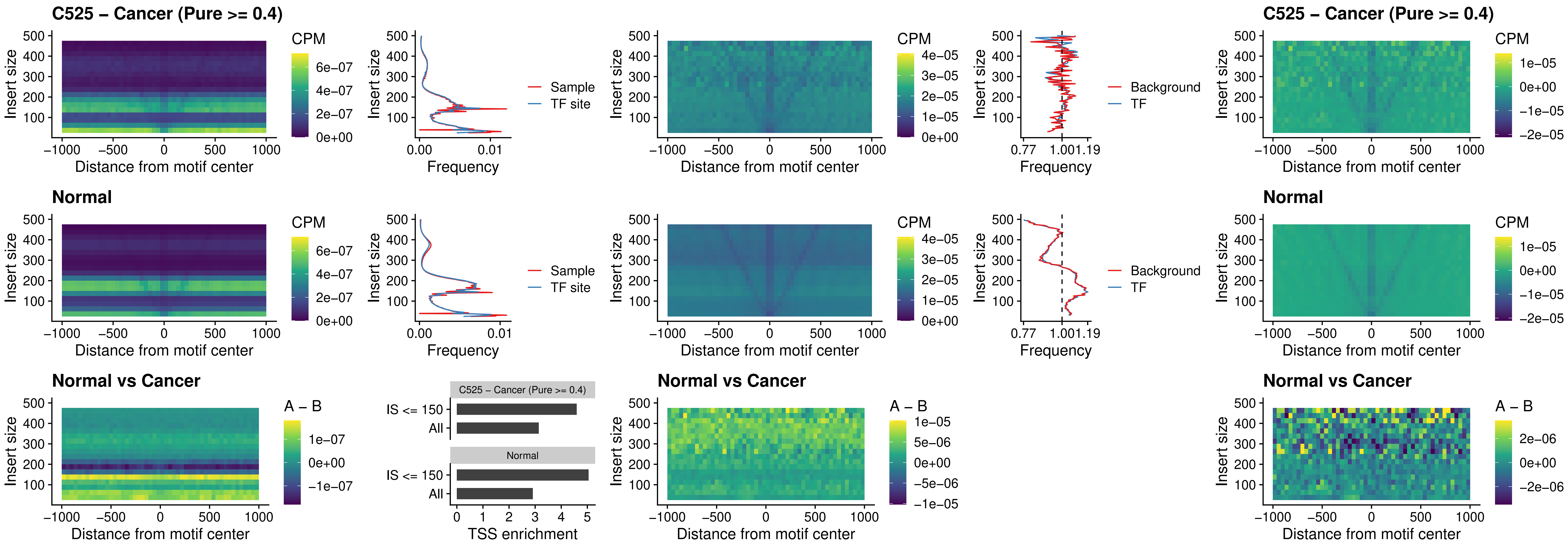

#### C525 – Cancer (Pure >= 0.4) – LMX1B_uc_dTSS_nPEAK

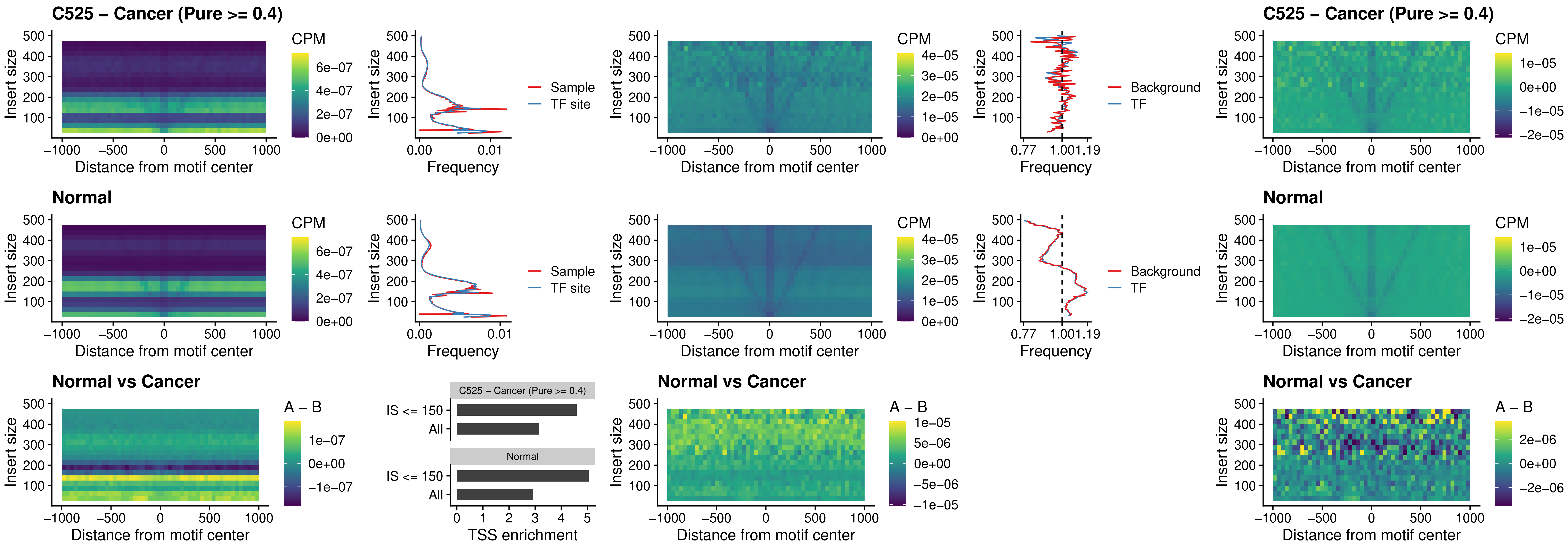

#### C525 – Cancer (Pure >= 0.4) – DLX5_uc_dTSS_oPEAK

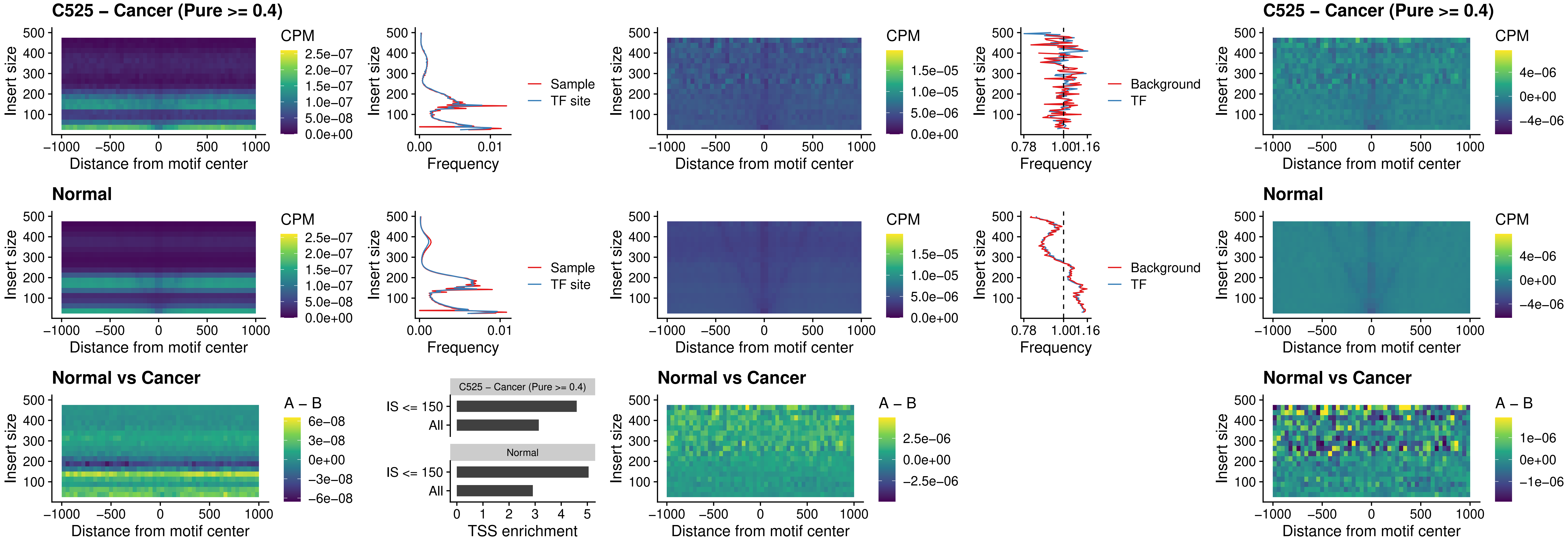

#### C525 – Cancer (Pure >= 0.4) – ARID3C_uc_dTSS_nPEAK

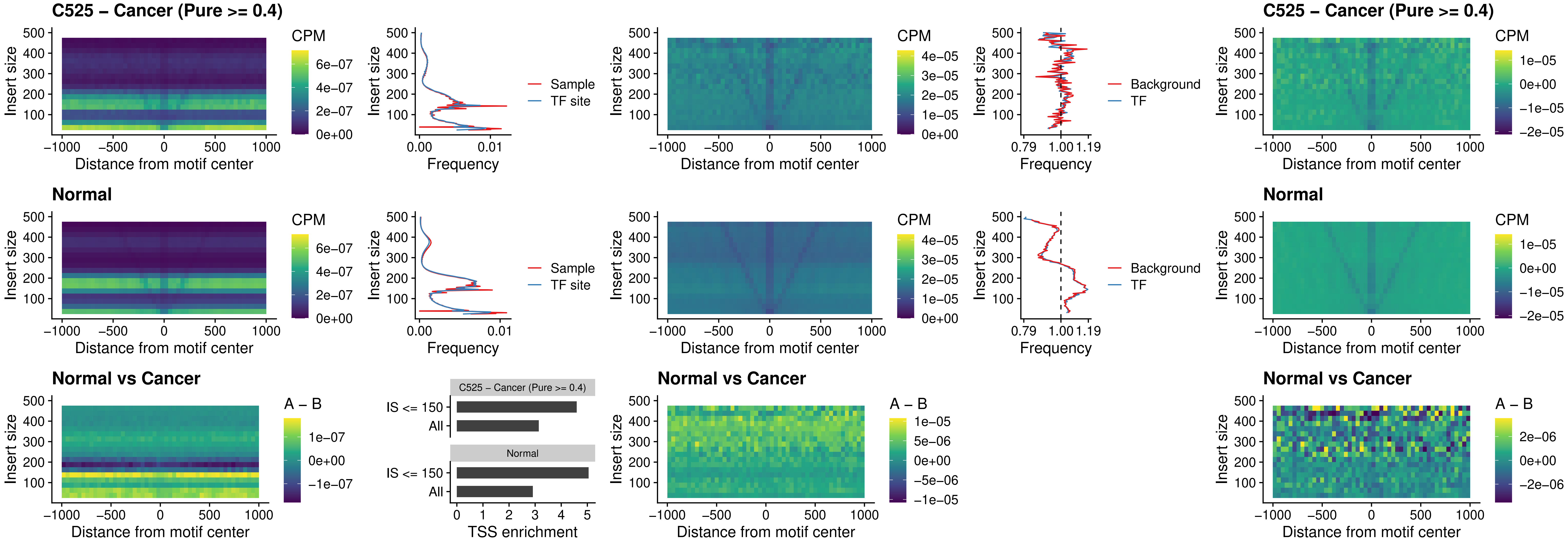

#### C525 – Cancer (Pure >= 0.4) – HOXD12_uc_dTSS_nPEAK

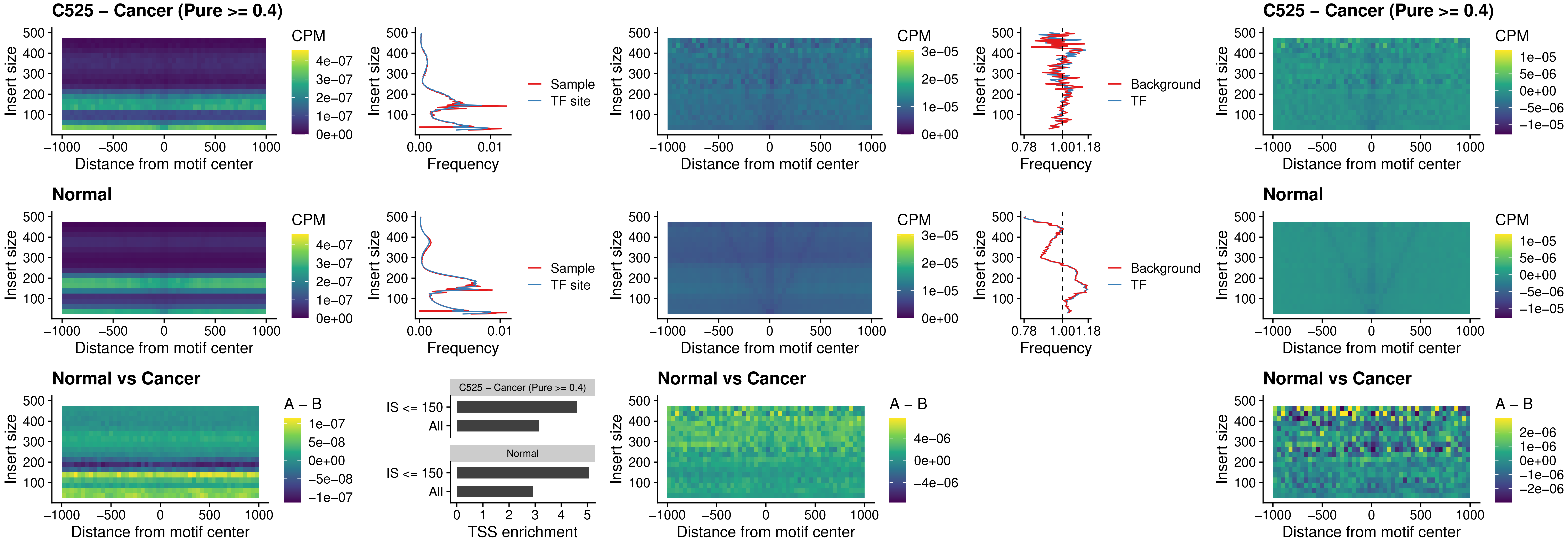

#### C525 – Cancer (Pure >= 0.4) – CDX2_uc_dTSS_nPEAK

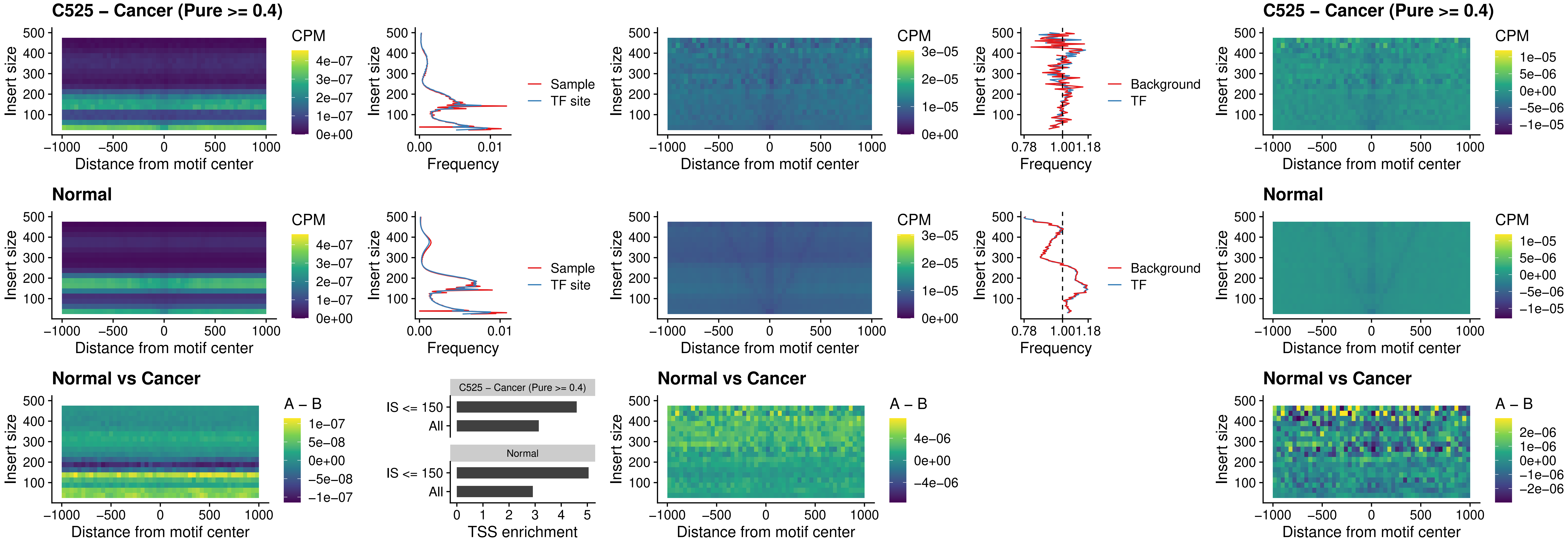

#### C525 – Cancer (Pure >= 0.4) – HOXC12_uc_dTSS_nPEAK

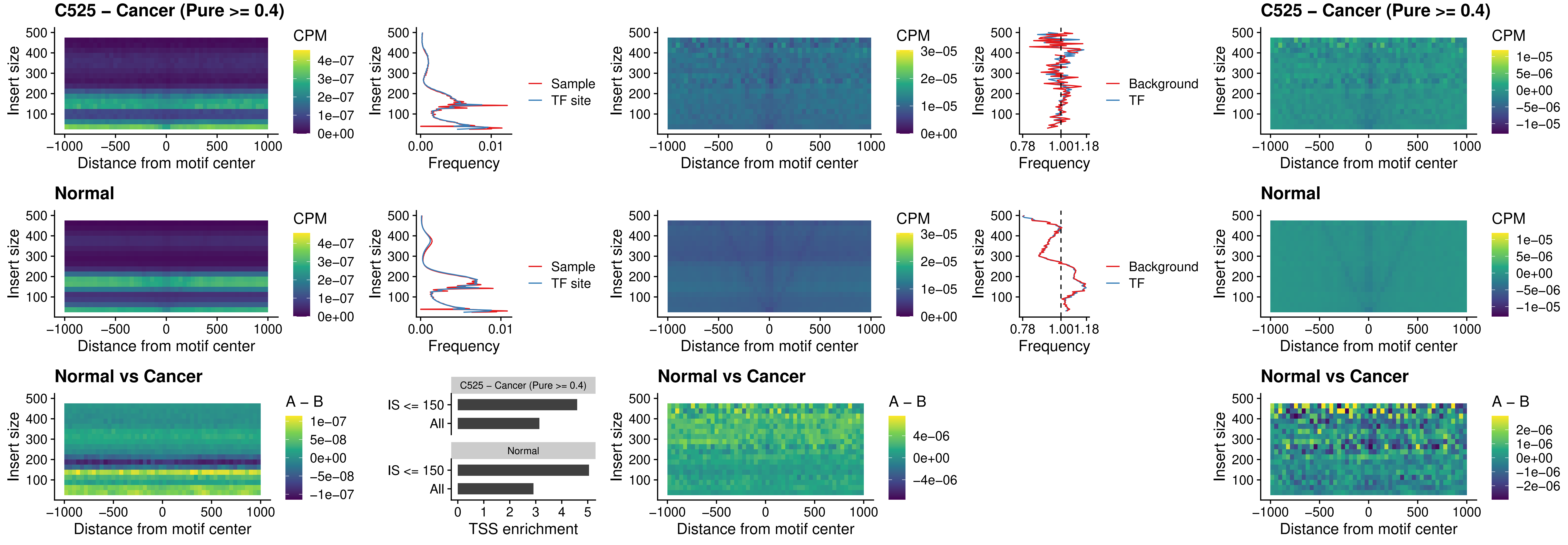

### C525 – Cancer (Pure >= 0.4) – HHEX_uc_dTSS_oPEAK

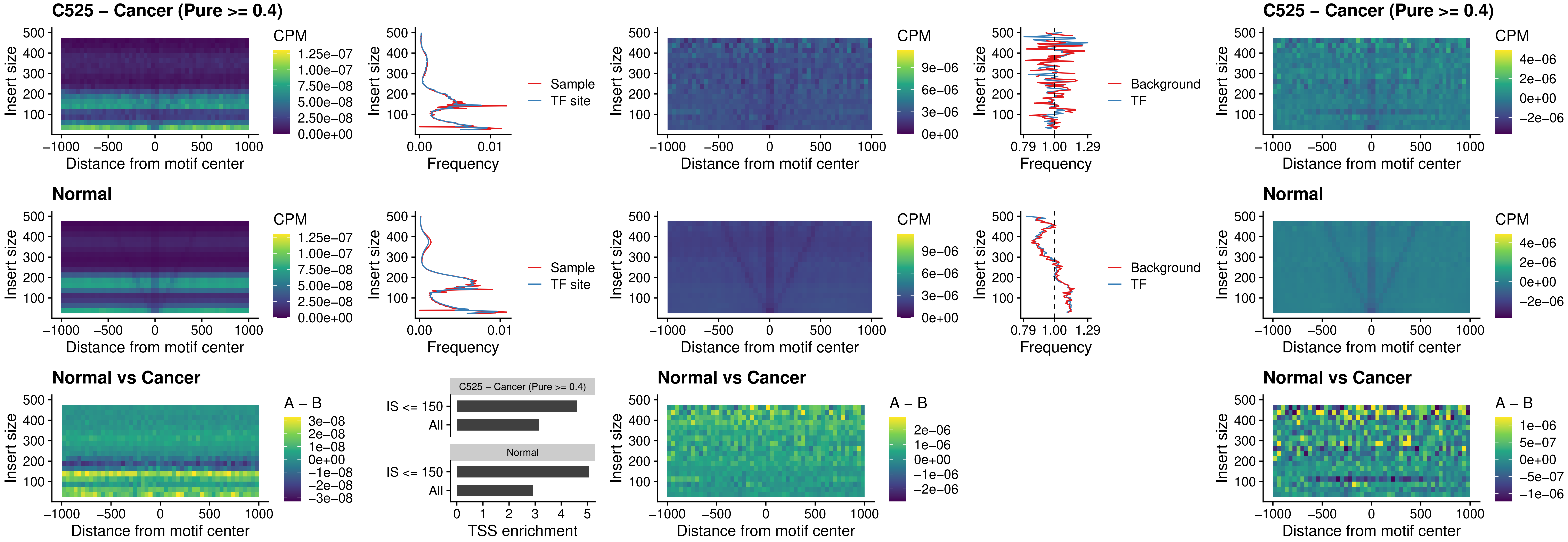

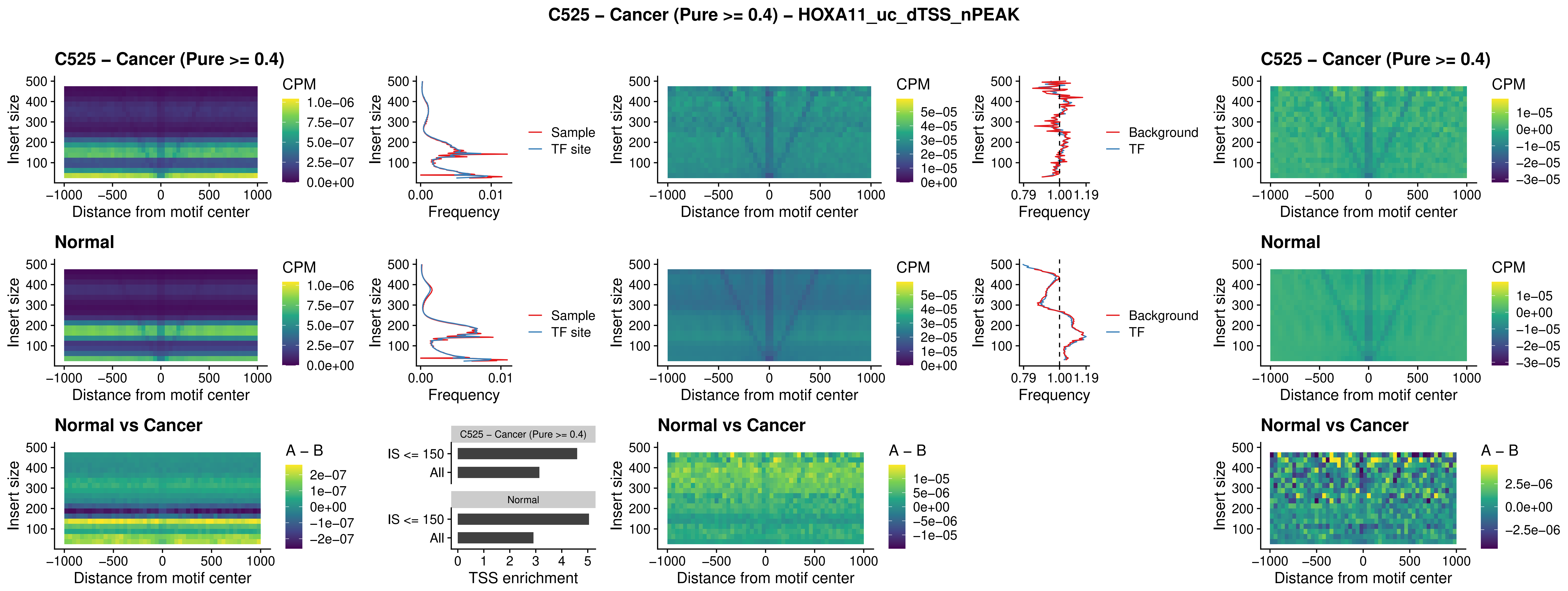
C525 – Cancer (Pure >= 0.4) – HOXA11_uc_dTSS_nPEAK
C525 – Cancer (Pure >= 0.4)
Normal
Normal vs Cancer
CPM
A – B
Insert size
Distance from motif center
Frequency
Sample
TF site
Background
TF
C525 – Cancer (Pure >= 0.4)
Normal
IS <= 150
All
TSS enrichment

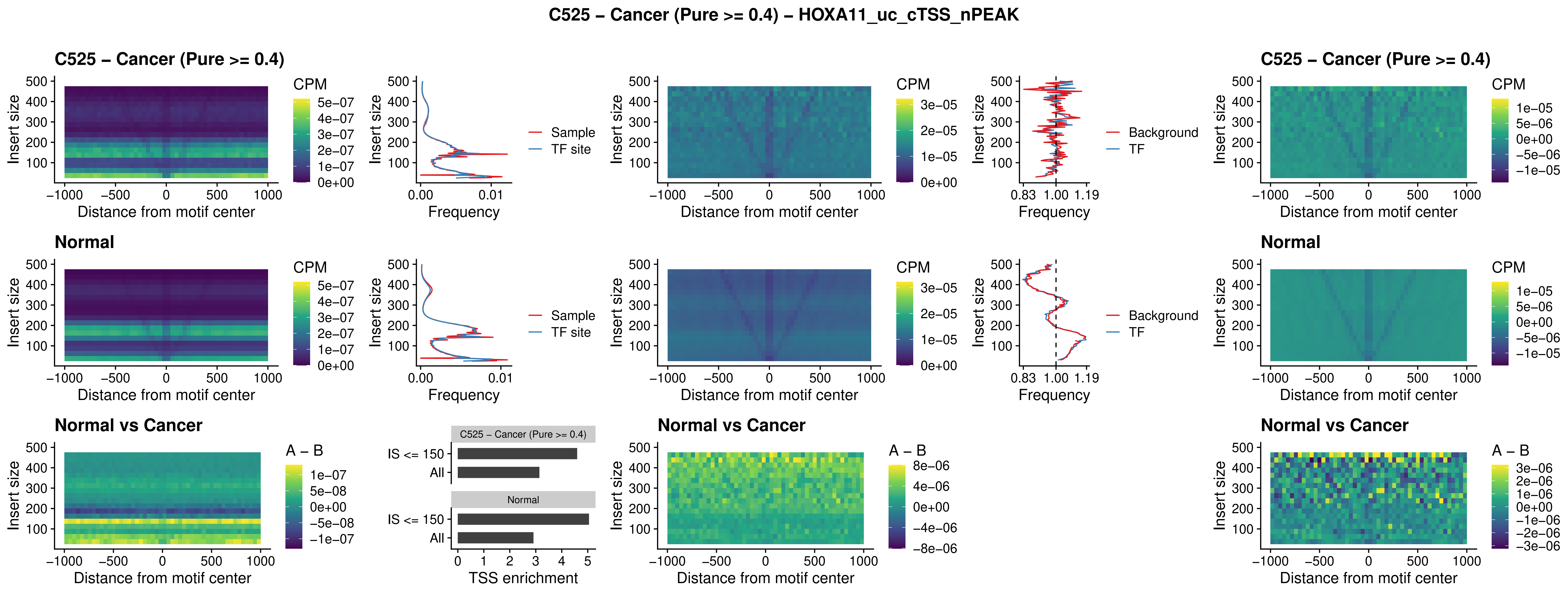

C525 – Cancer (Pure >= 0.4) – HOXA11_uc_cTSS_nPEAK
C525 – Cancer (Pure >= 0.4)
Normal
Normal vs Cancer
Insert size
Distance from motif center
Frequency
CPM
A – B
Sample
TF site
Background
TF
C525 – Cancer (Pure >= 0.4)
Normal
IS <= 150
All
TSS enrichment

#### C525 – Cancer (Pure >= 0.4) – ZFHX3_uc_dTSS_nPEAK

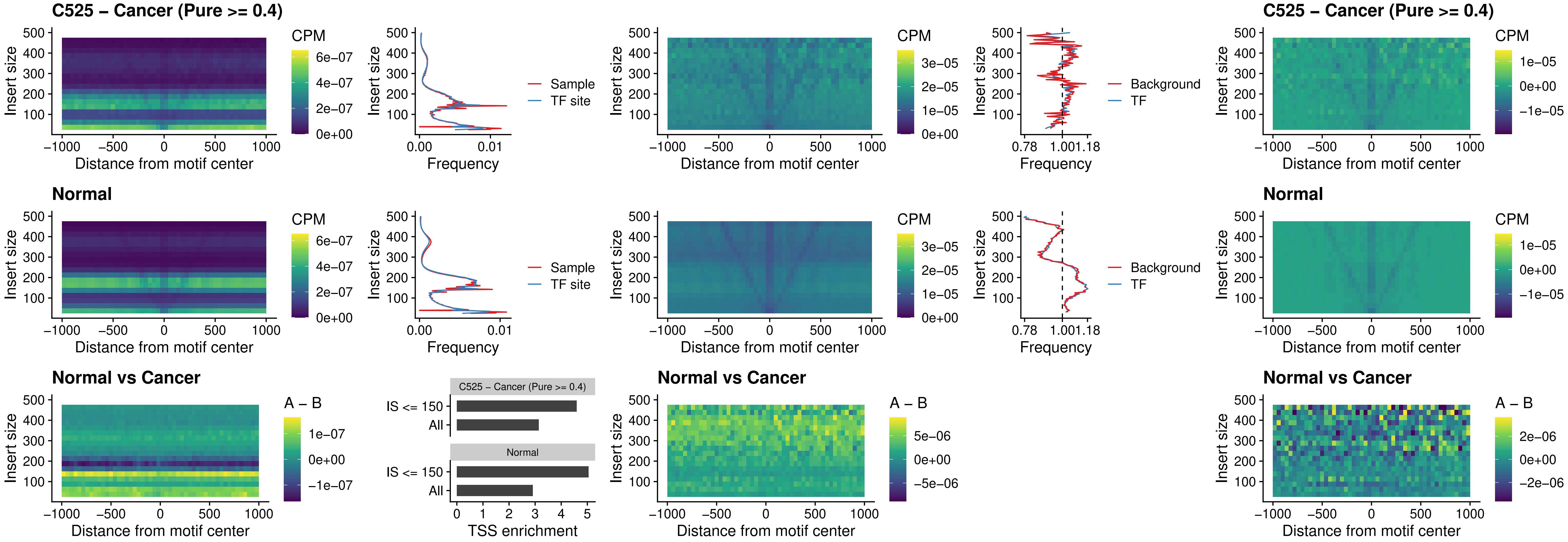

#### C525 – Cancer (Pure >= 0.4) – HOXB4_uc_dTSS_nPEAK

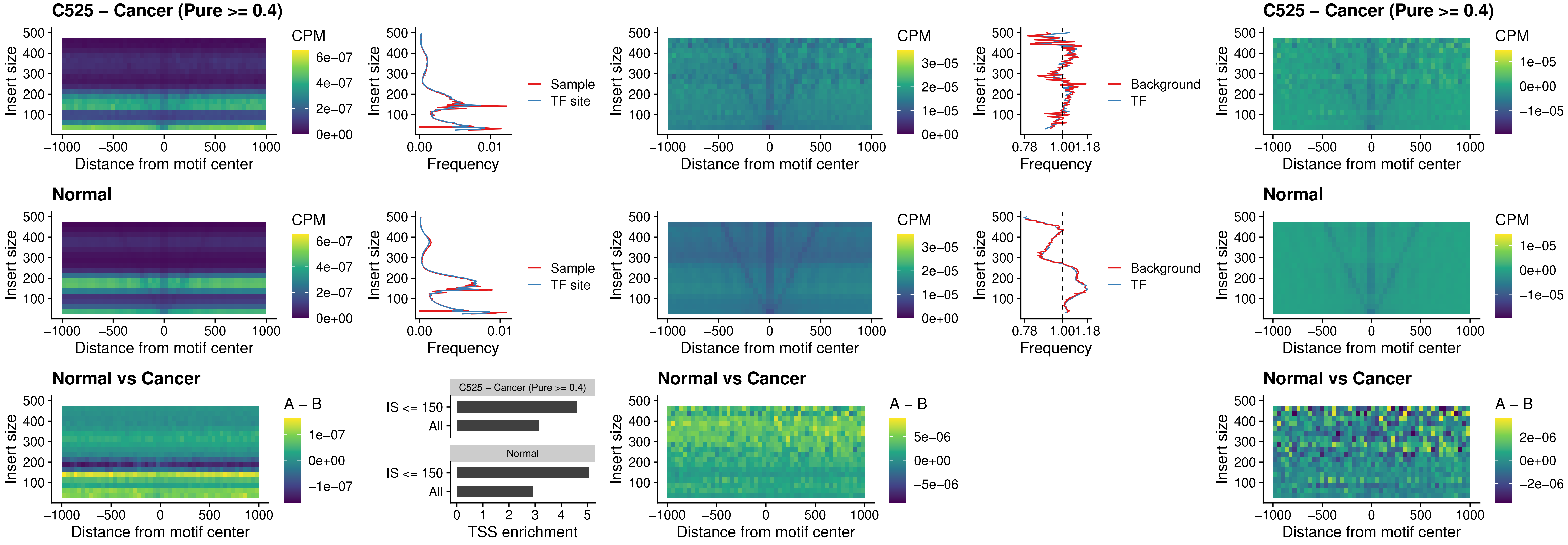

#### C525 – Cancer (Pure >= 0.4) – HOXC5_uc_dTSS_nPEAK

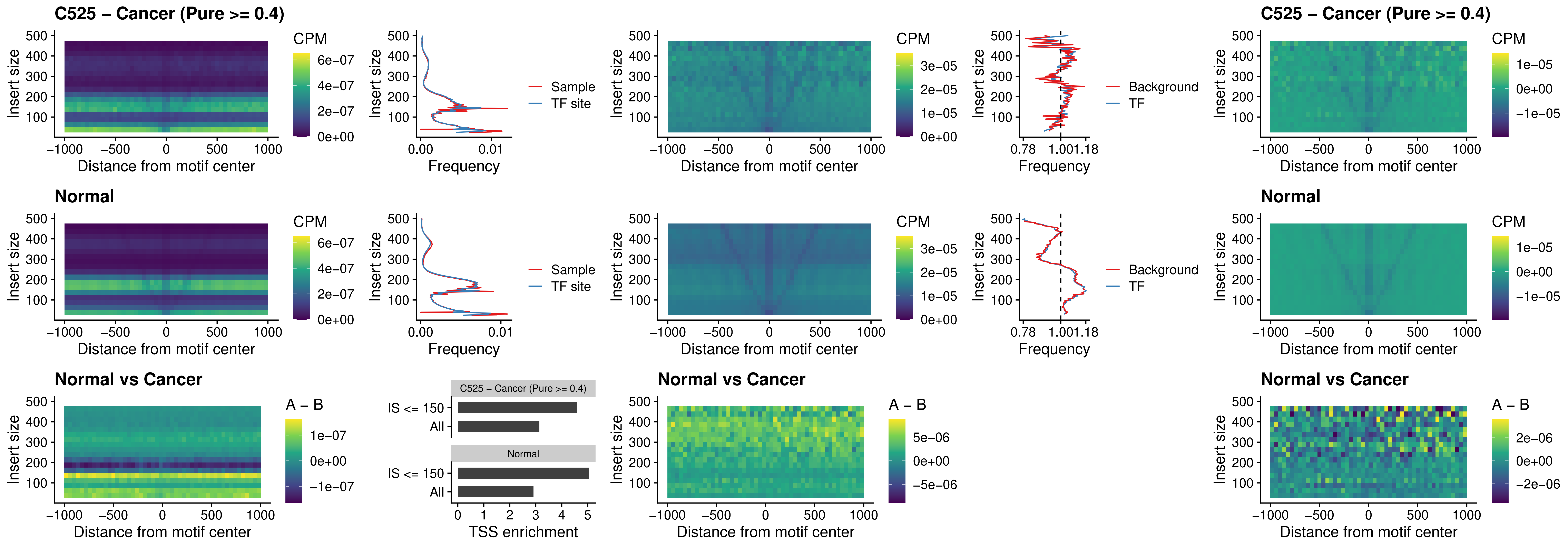

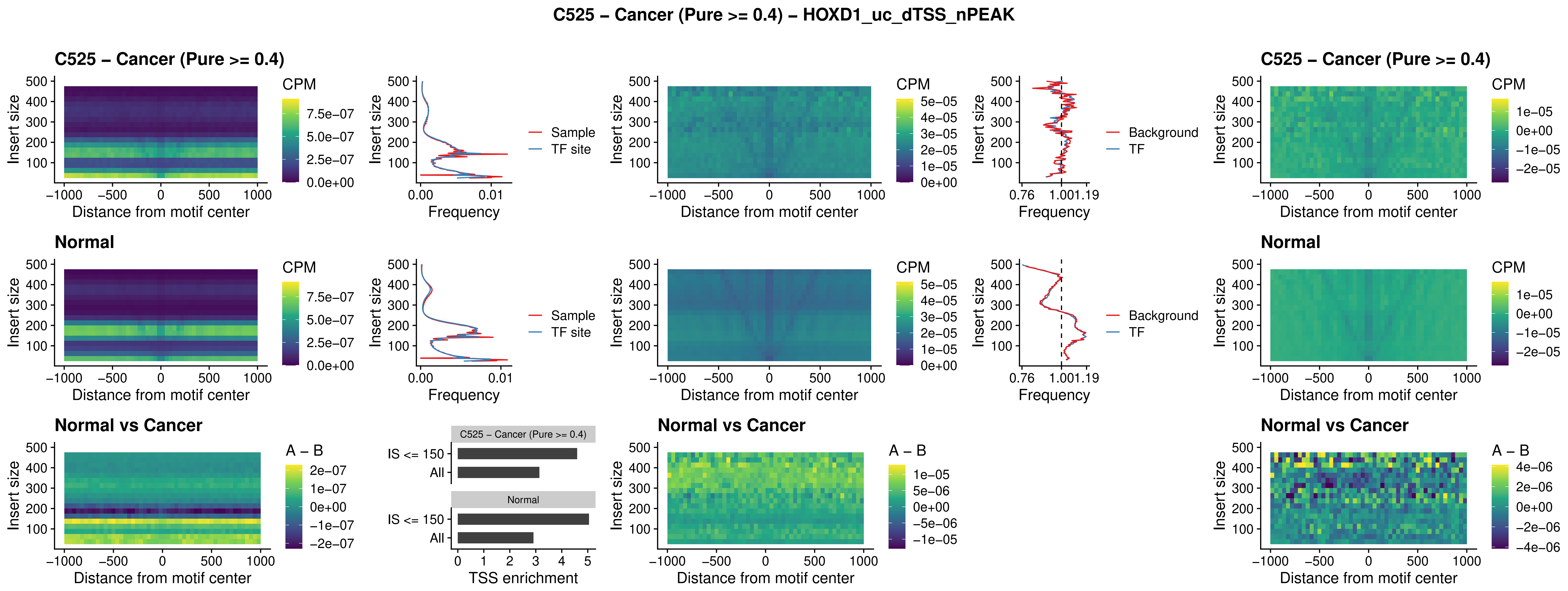
C525 – Cancer (Pure >= 0.4) – HOXD1_uc_dTSS_nPEAK
C525 – Cancer (Pure >= 0.4)
Normal
Normal vs Cancer
Insert size
Distance from motif center
Frequency
CPM
A – B
Sample
TF site
Background
TF
C525 – Cancer (Pure >= 0.4)
Normal
IS <= 150
All
TSS enrichment

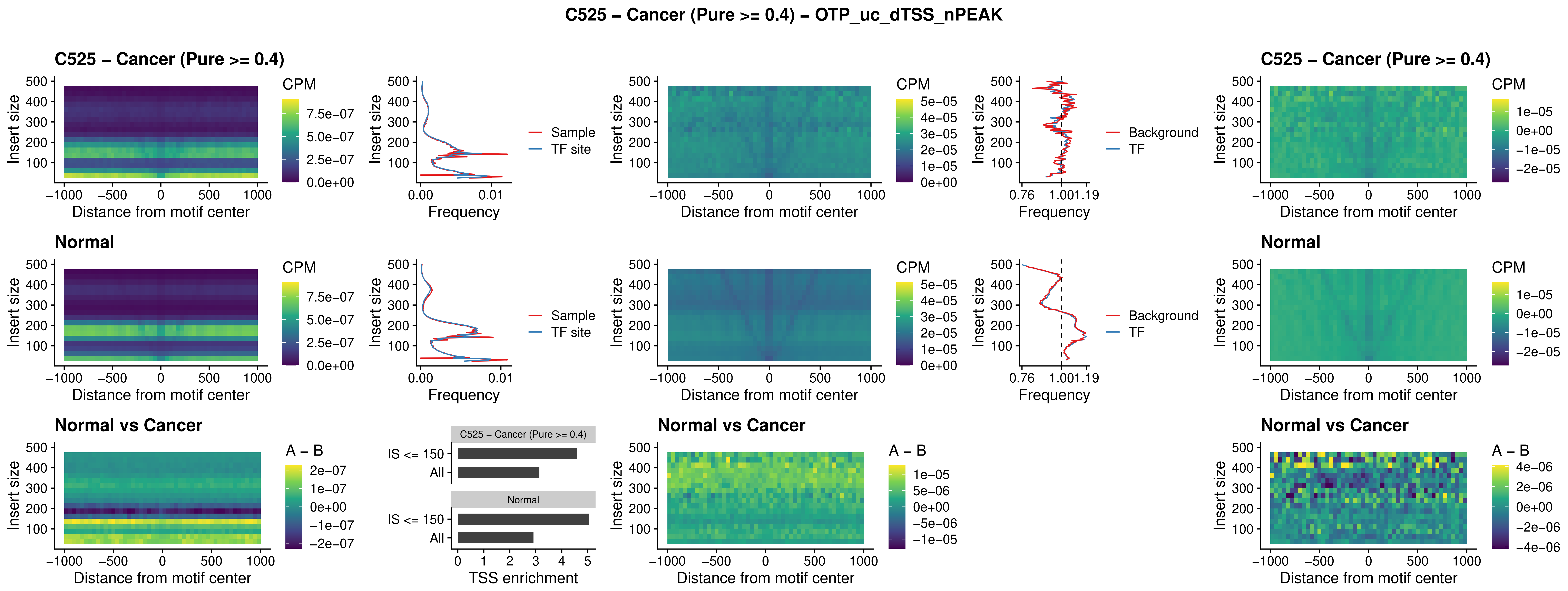
C525 – Cancer (Pure >= 0.4) – OTP_uc_dTSS_nPEAK
C525 – Cancer (Pure >= 0.4)
Normal
Normal vs Cancer
Insert size
Distance from motif center
Frequency
CPM
A – B
Sample
TF site
Background
TF
C525 – Cancer (Pure >= 0.4)
Normal
IS <= 150
All
TSS enrichment

#### C525 – Cancer (Pure >= 0.4) – FOXL1_uc_dTSS_oPEAK

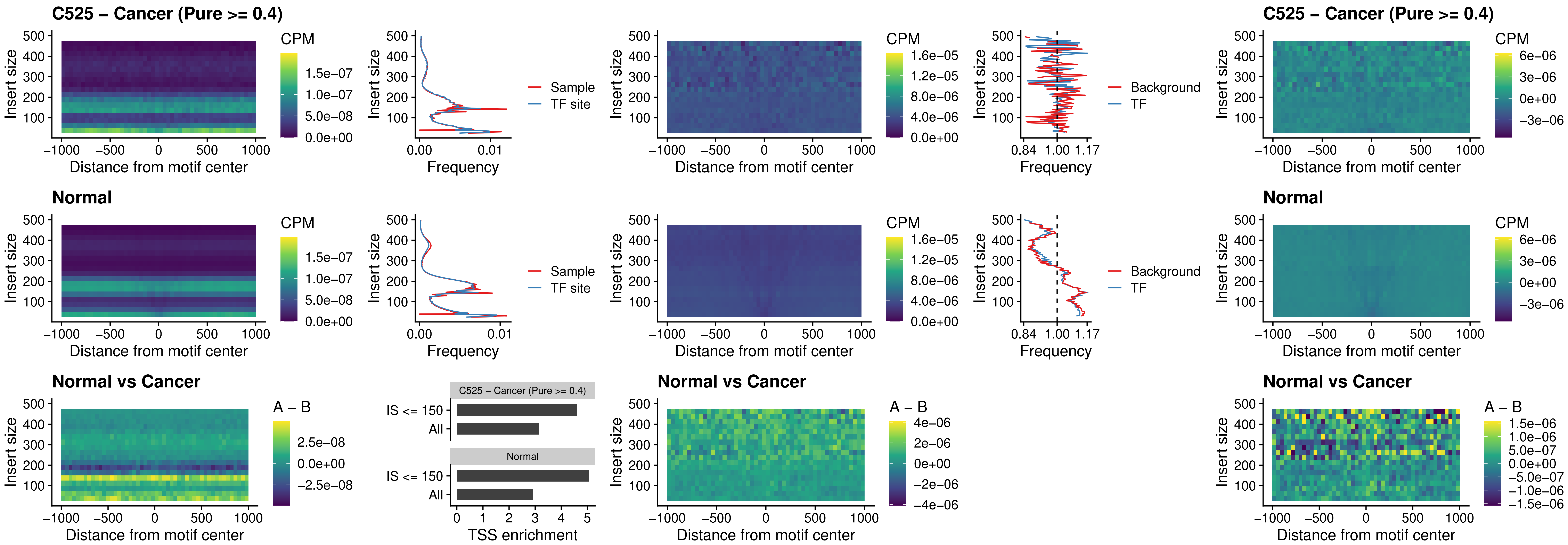

### C525 – Cancer (Pure >= 0.4) – ZNF32_uc_dTSS_oPEAK

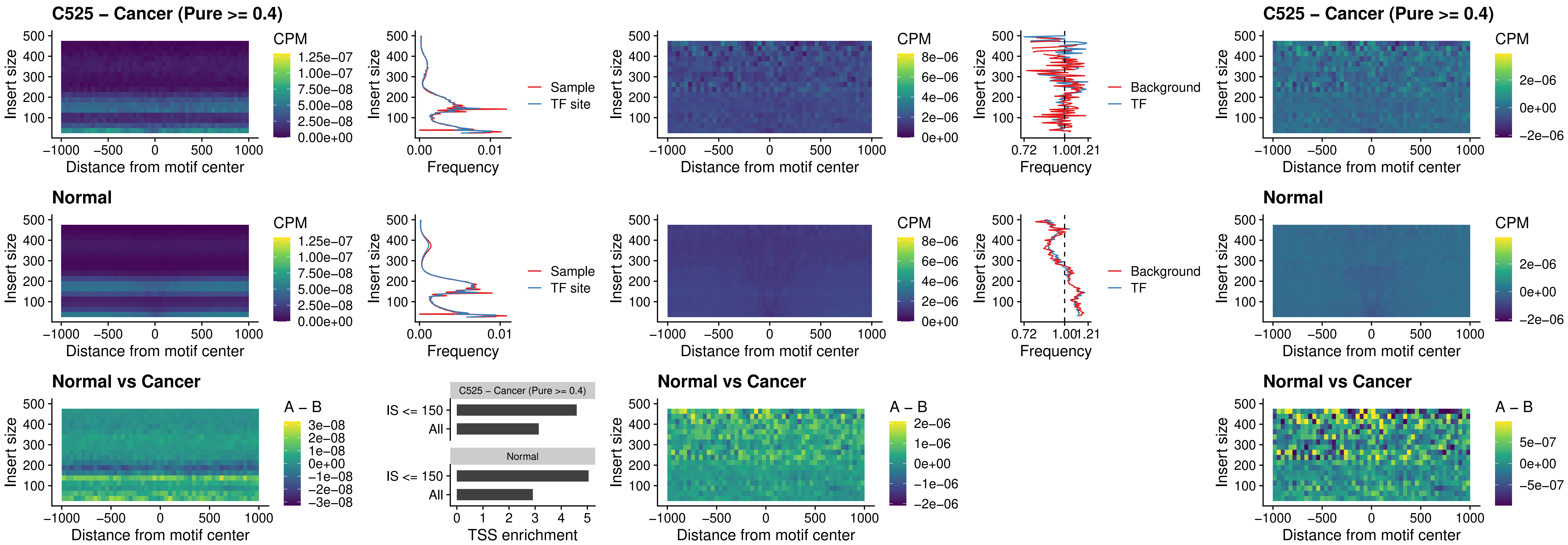

#### C525 – Cancer (Pure >= 0.4) – FOXL1_uc_dTSS_nPEAK

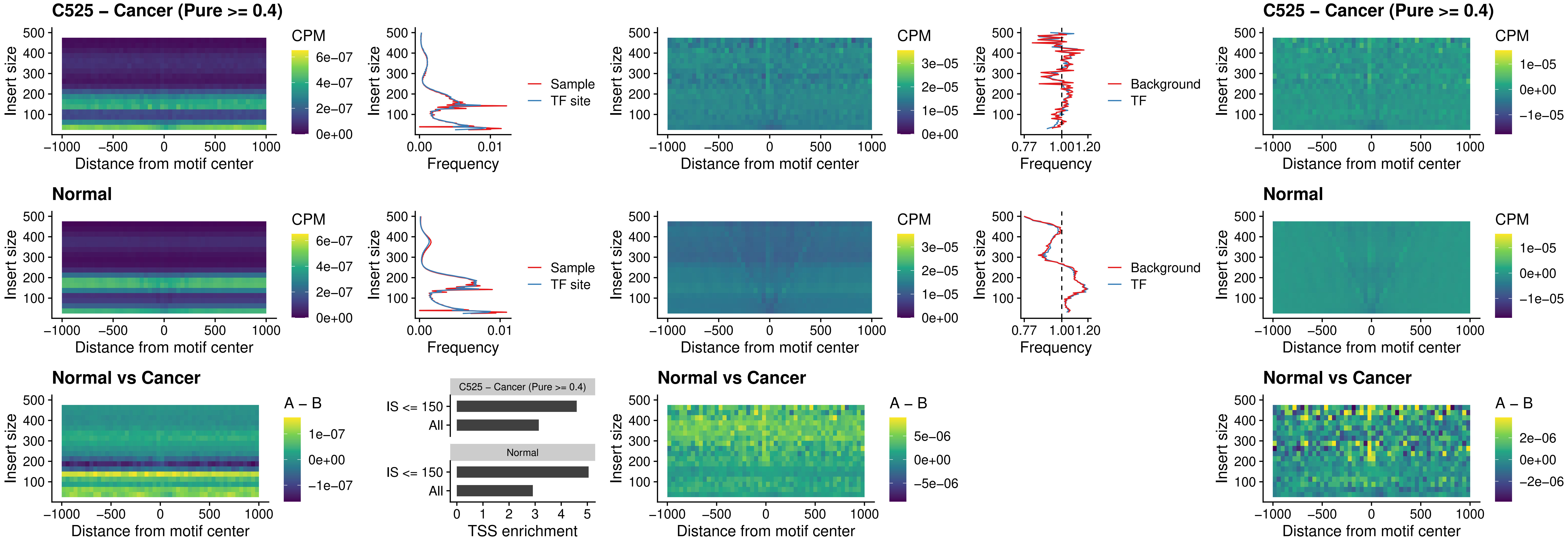

#### C525 – Cancer (Pure >= 0.4) – ZNF32_uc_dTSS_nPEAK

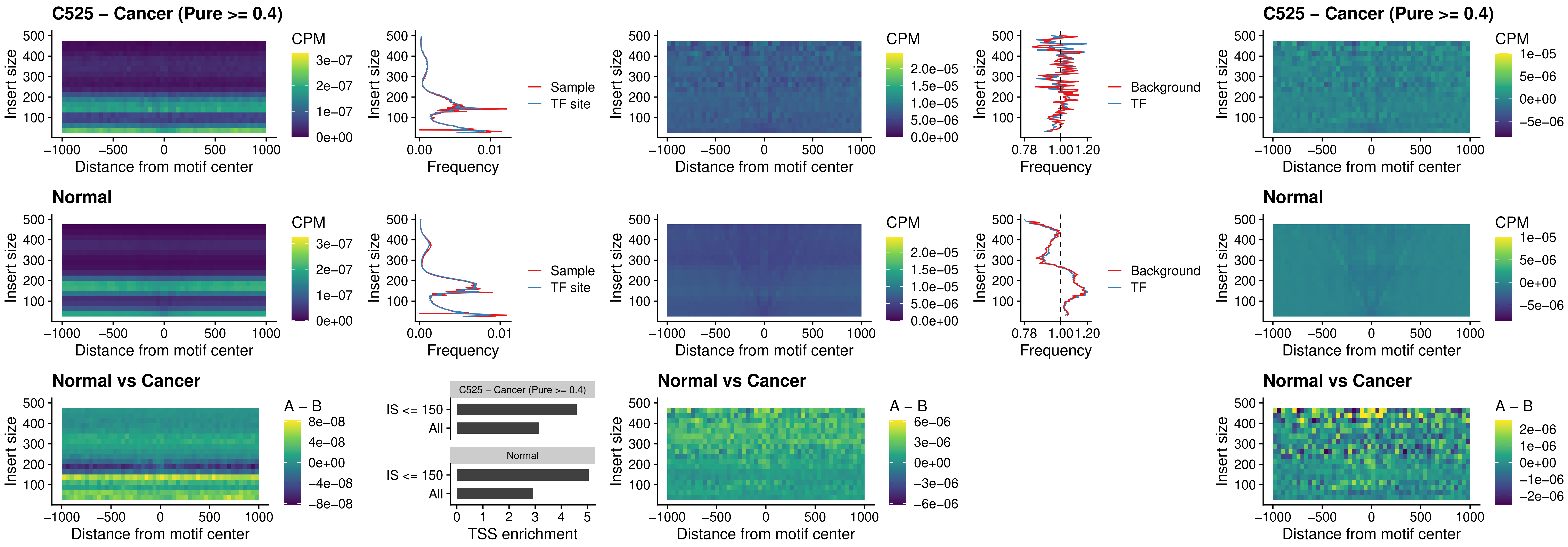

#### C525 – Cancer (Pure >= 0.4) – DLX5_uc_dTSS_nPEAK

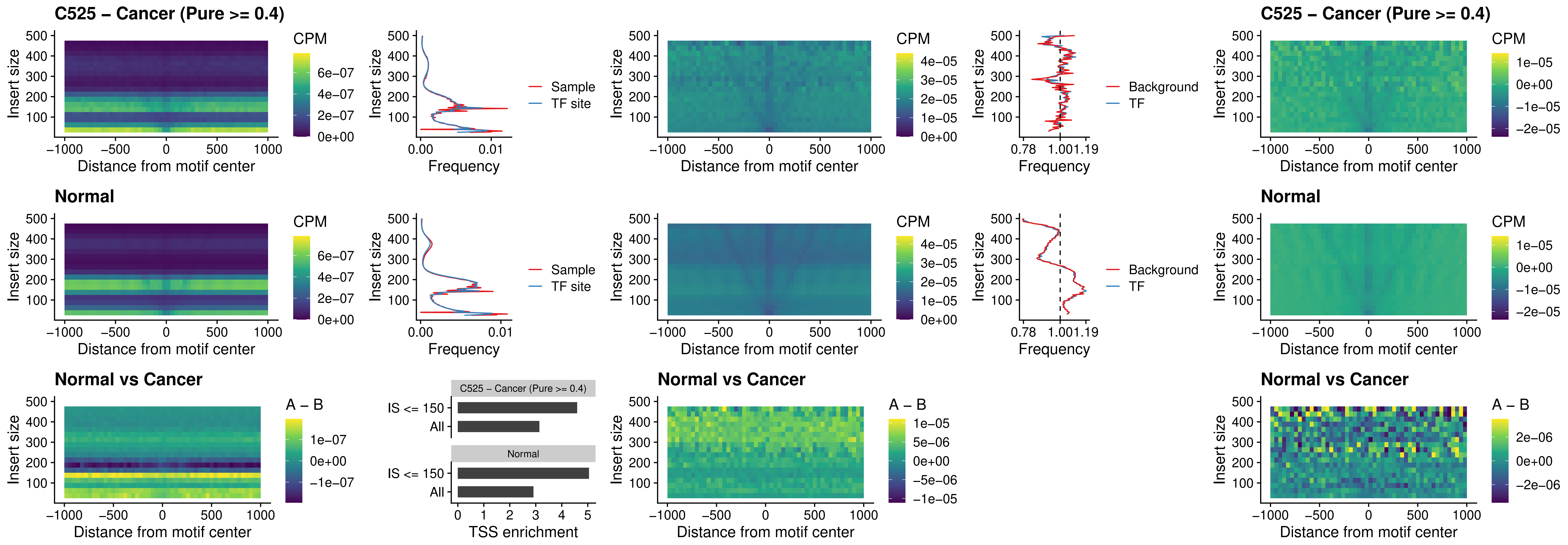

#### C525 – Cancer (Pure >= 0.4) – DMRT2_uc_dTSS_nPEAK

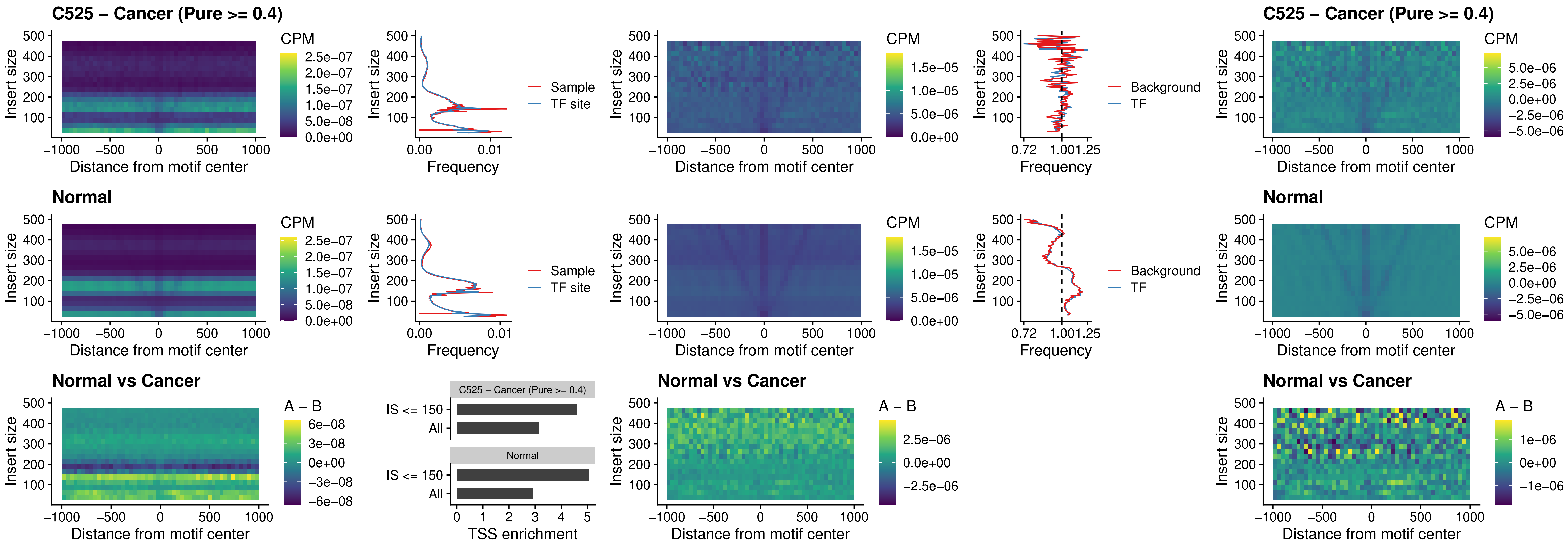

### C525 – Cancer (Pure >= 0.4) – DMRTC2_uc_cTSS_nPEAK

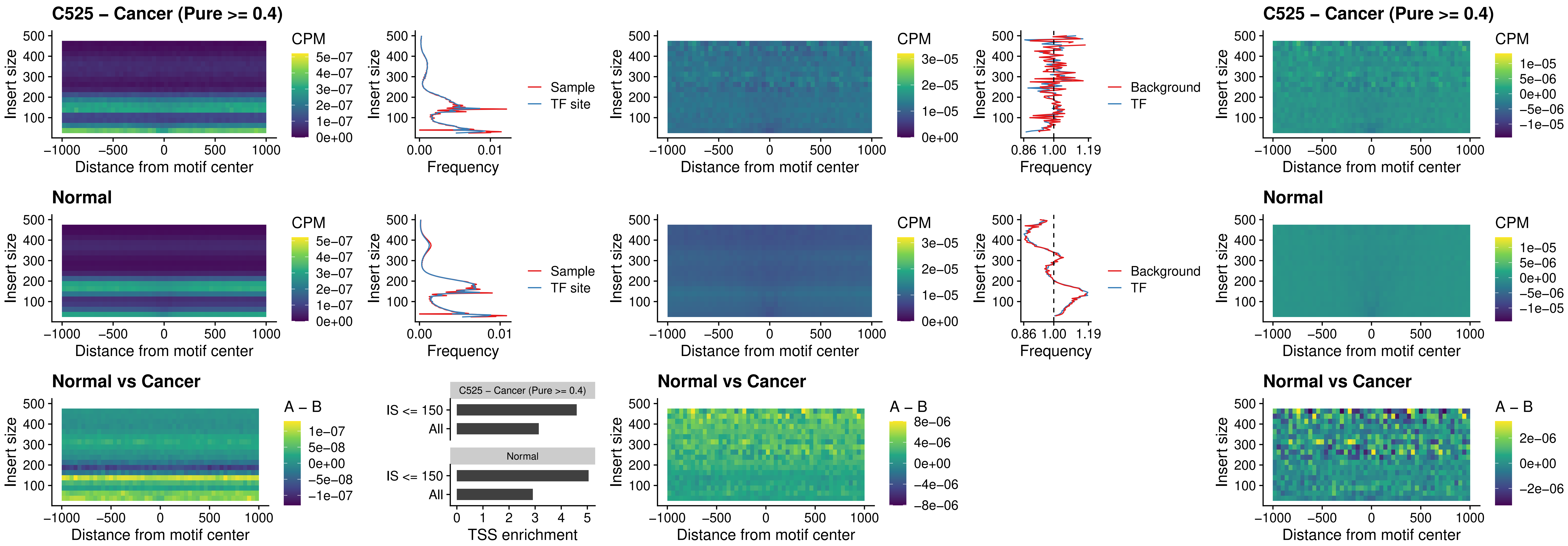

**C525 – Cancer (Pure >= 0.4) – HOXD9_uc_dTSS_nPEAK**

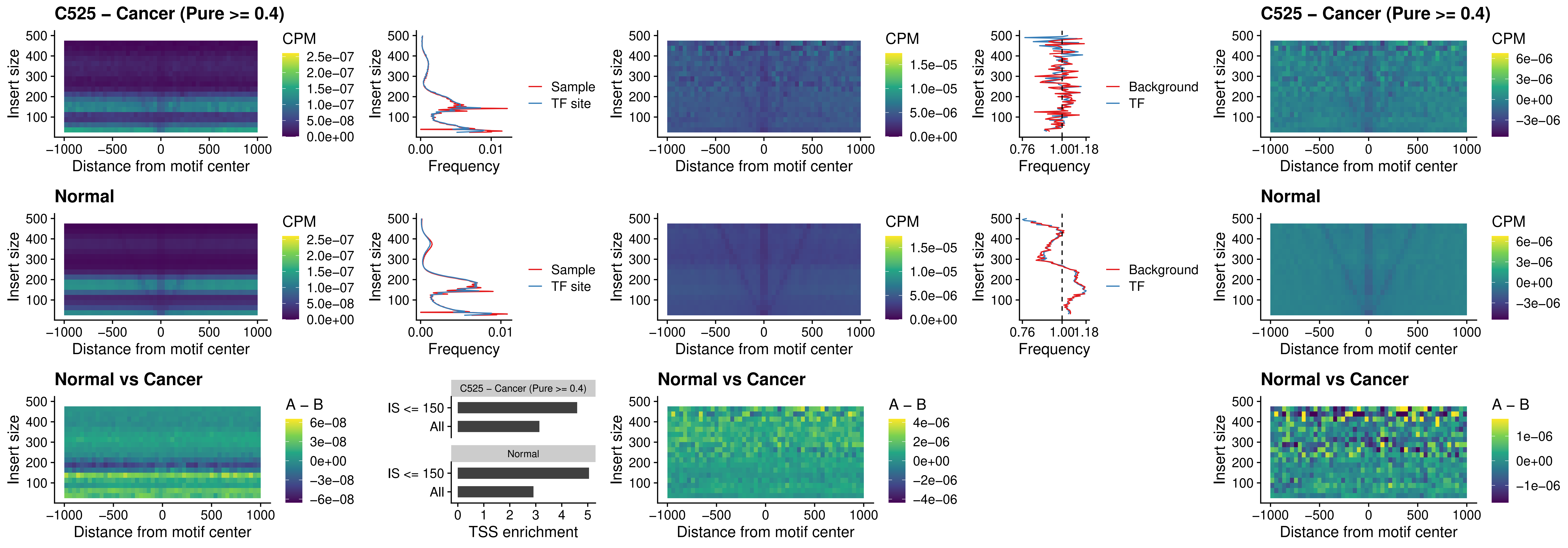

#### C525 – Cancer (Pure >= 0.4) – HOXD4_uc_dTSS_nPEAK

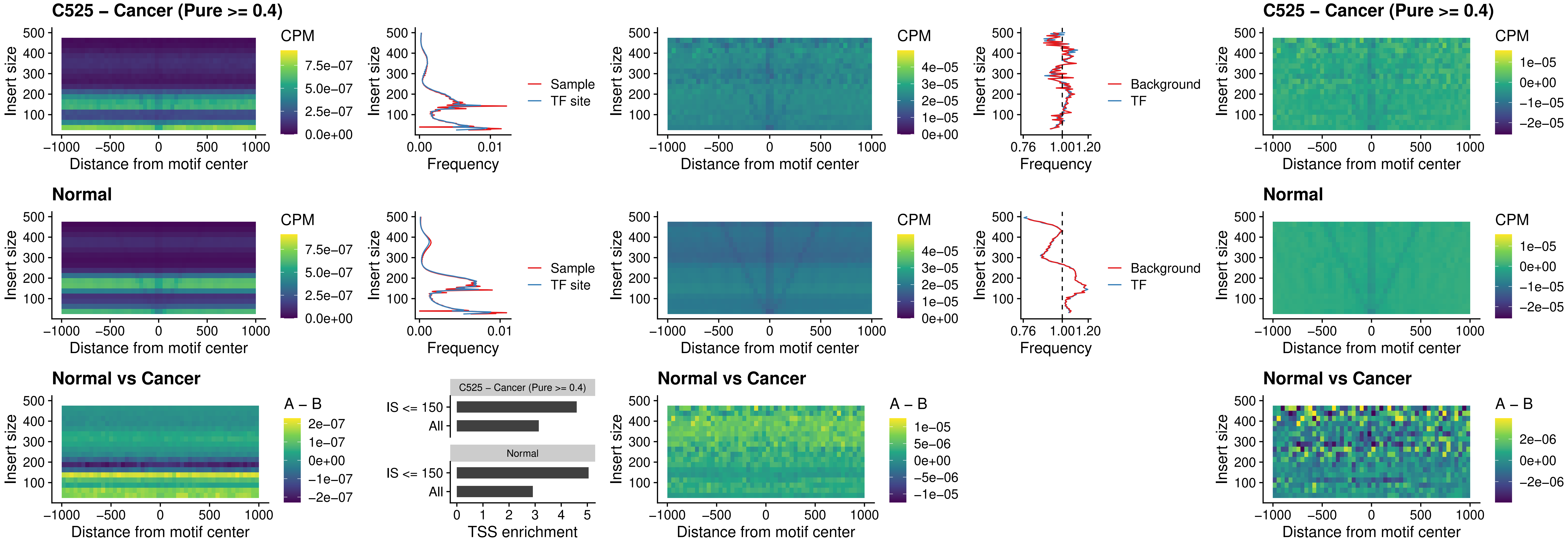

#### C525 – Cancer (Pure >= 0.4) – FOXP3_uc_cTSS_nPEAK

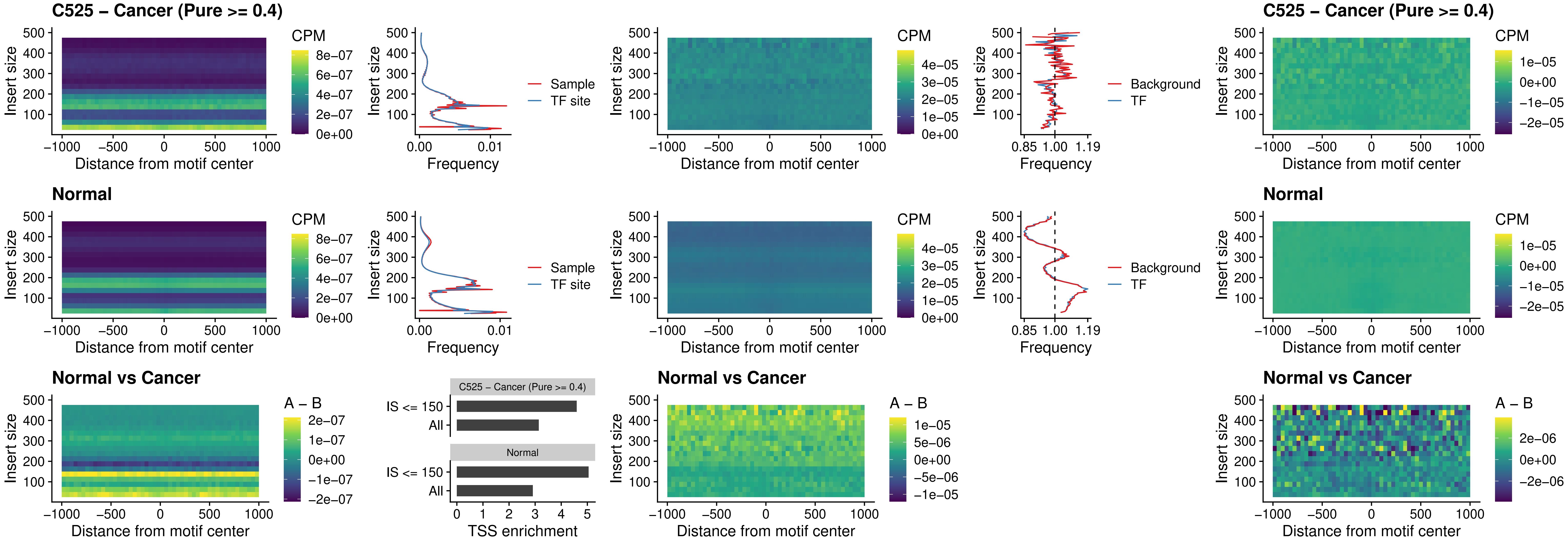

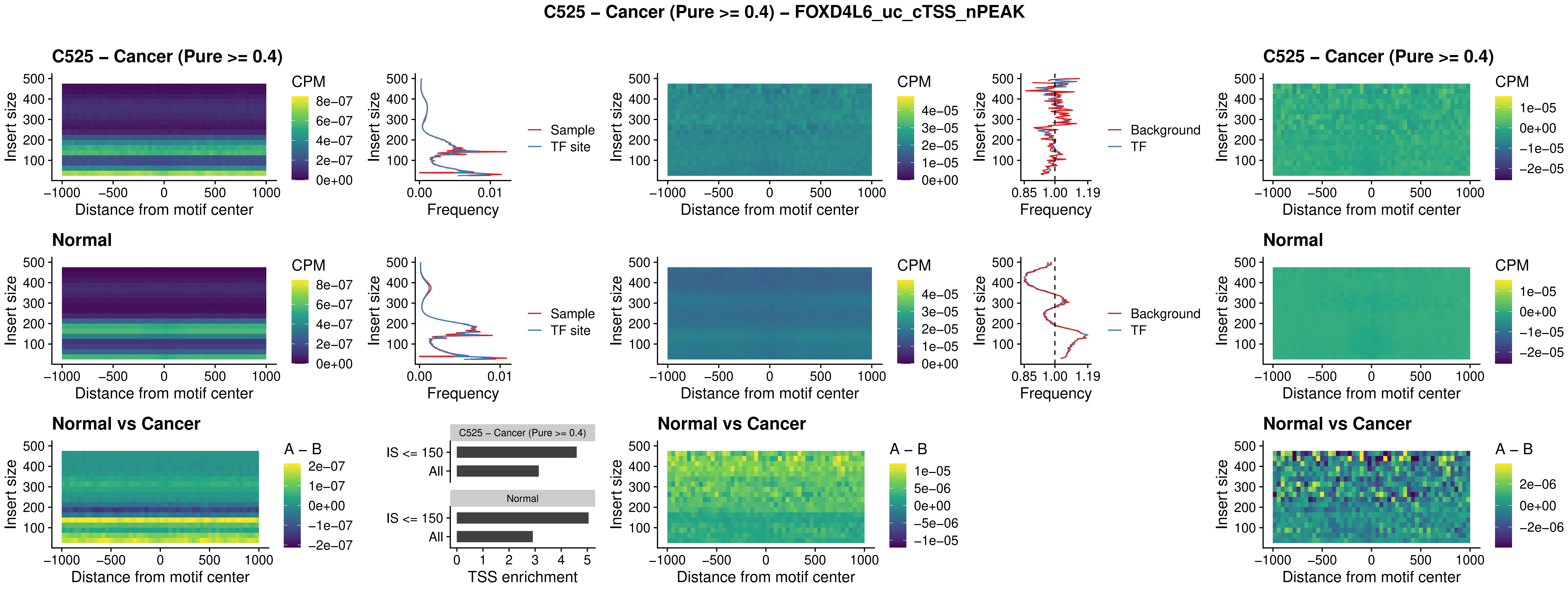
C525 – Cancer (Pure >= 0.4) – FOXD4L6_uc_cTSS_nPEAK
C525 – Cancer (Pure >= 0.4)
Normal
Normal vs Cancer
Insert size
Distance from motif center
Frequency
CPM
A – B
Sample
TF site
Background
TF
C525 – Cancer (Pure >= 0.4)
Normal
IS <= 150
All
TSS enrichment

#### C525 – Cancer (Pure >= 0.4) – FOXD4L5_uc_cTSS_nPEAK

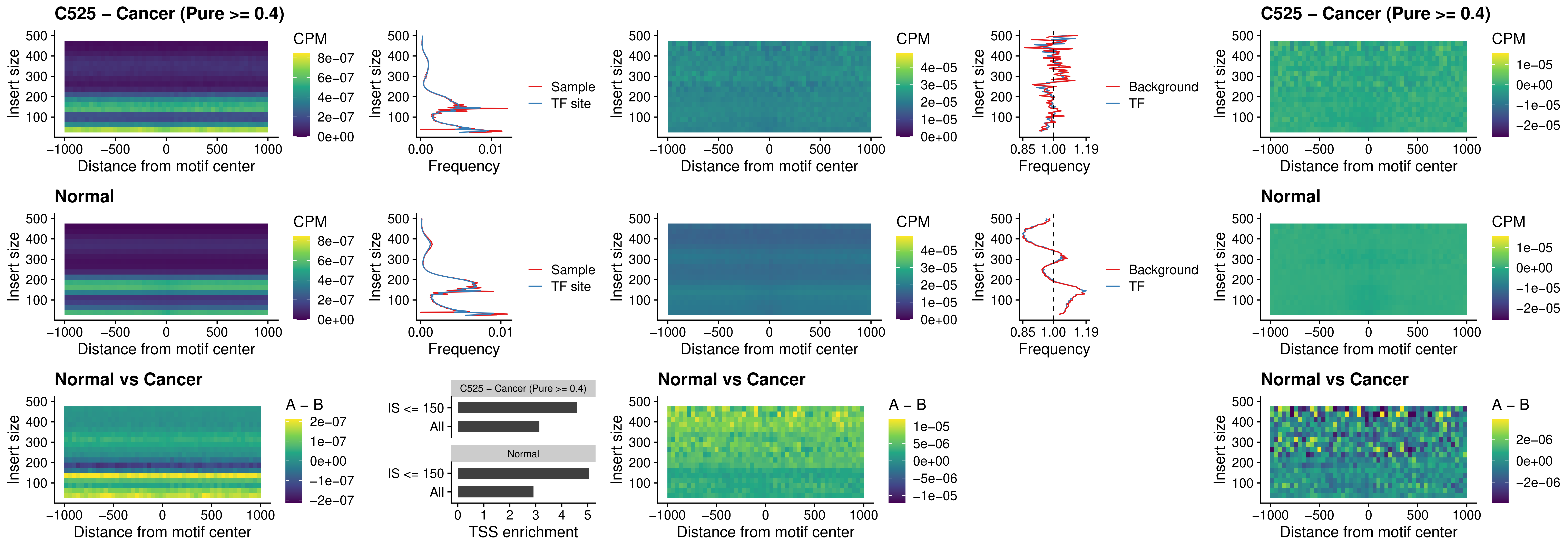

### C525 – Cancer (Pure >= 0.4) – FOXD4L4_uc_cTSS_nPEAK

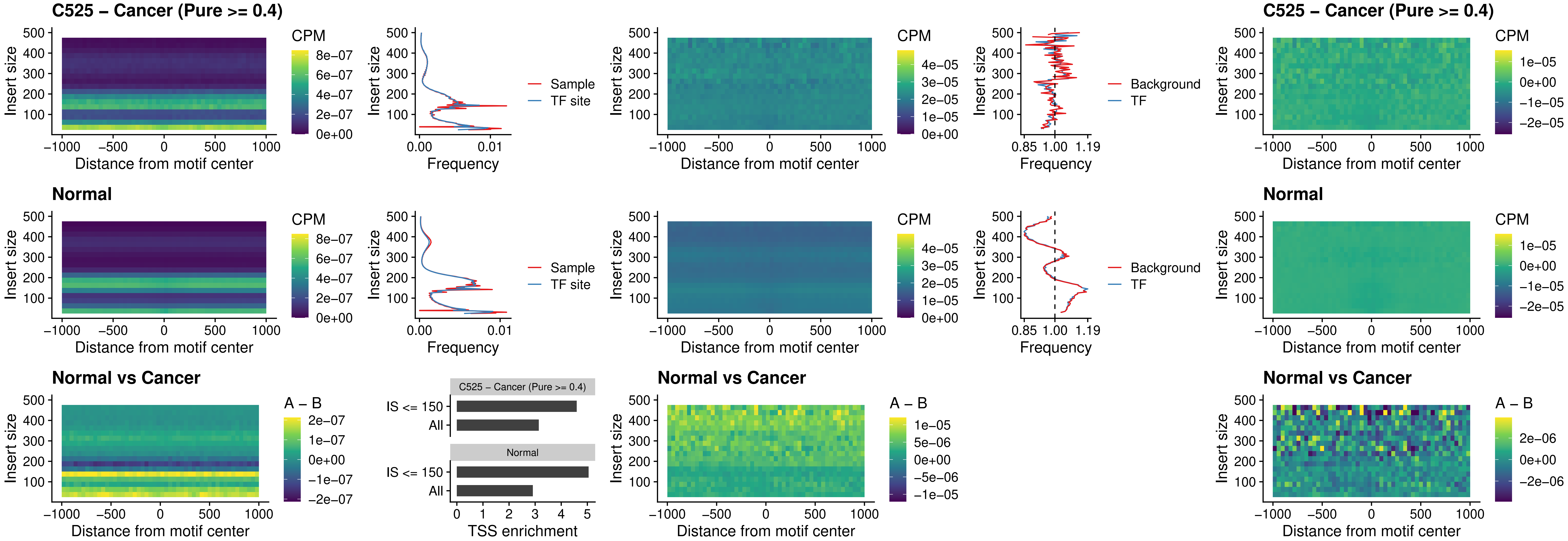

#### C525 – Cancer (Pure >= 0.4) – FOXD4L2_uc_cTSS_nPEAK

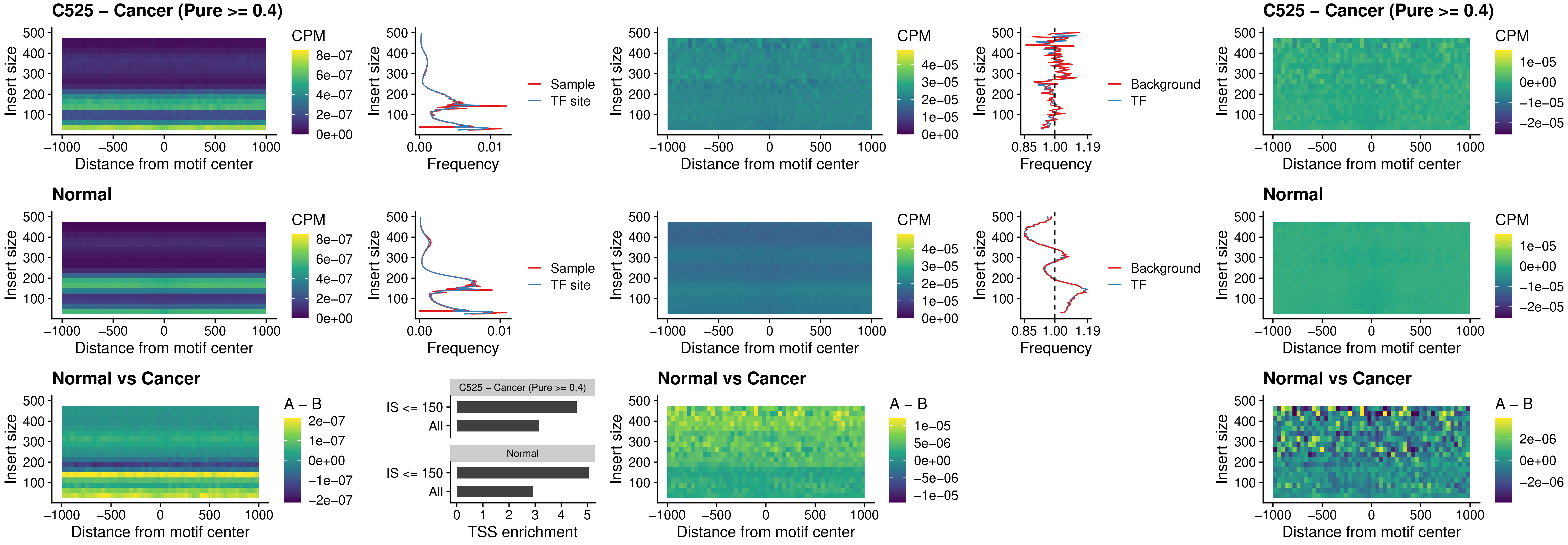

#### C525 – Cancer (Pure >= 0.4) – FOXD4L3_uc_cTSS_nPEAK

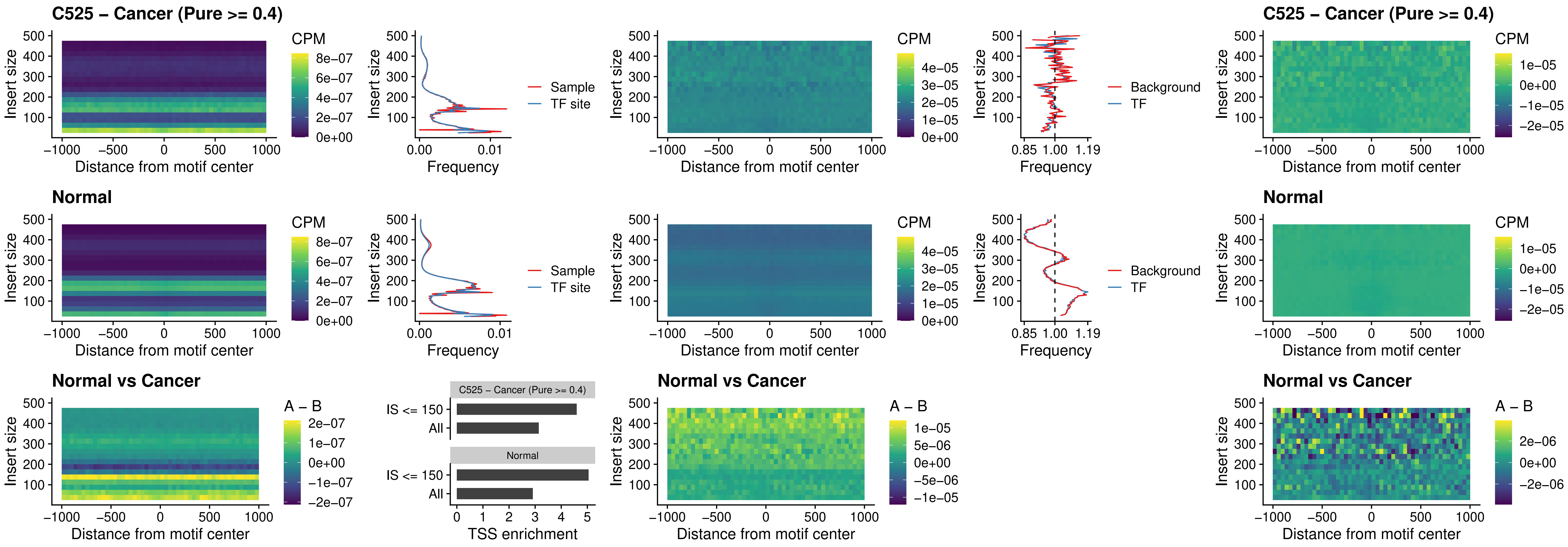

### C525 – Cancer (Pure >= 0.4) – HOXD1_uc_dTSS_oPEAK

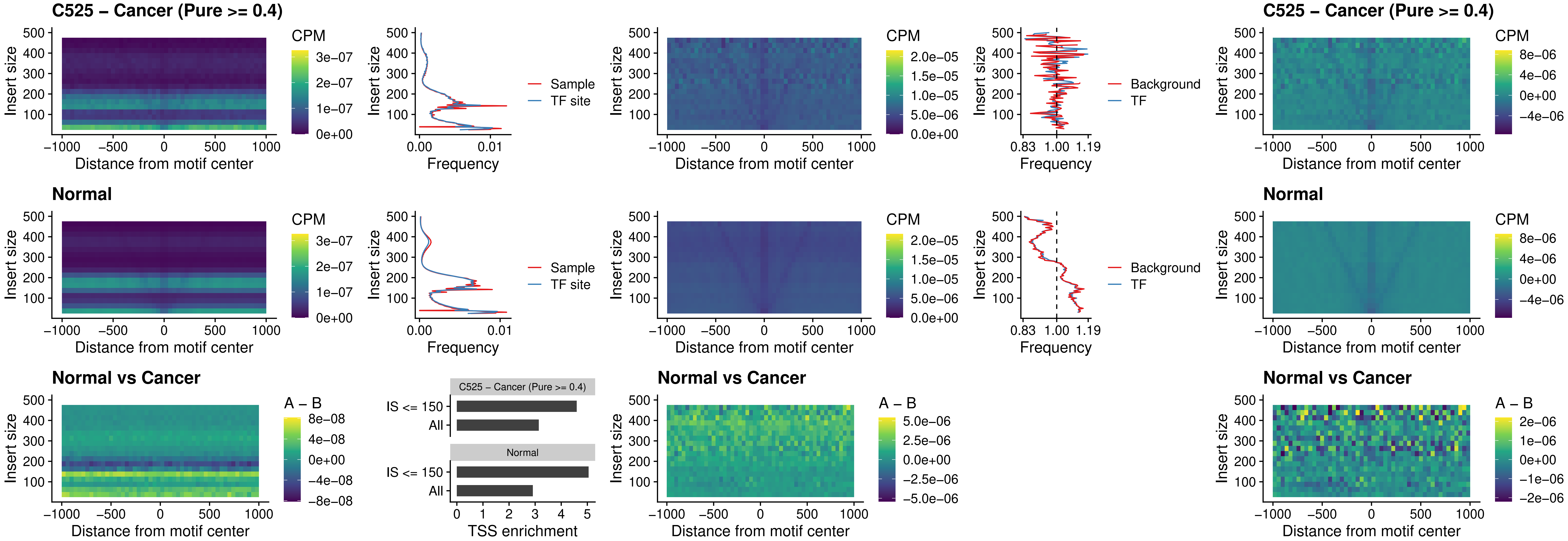

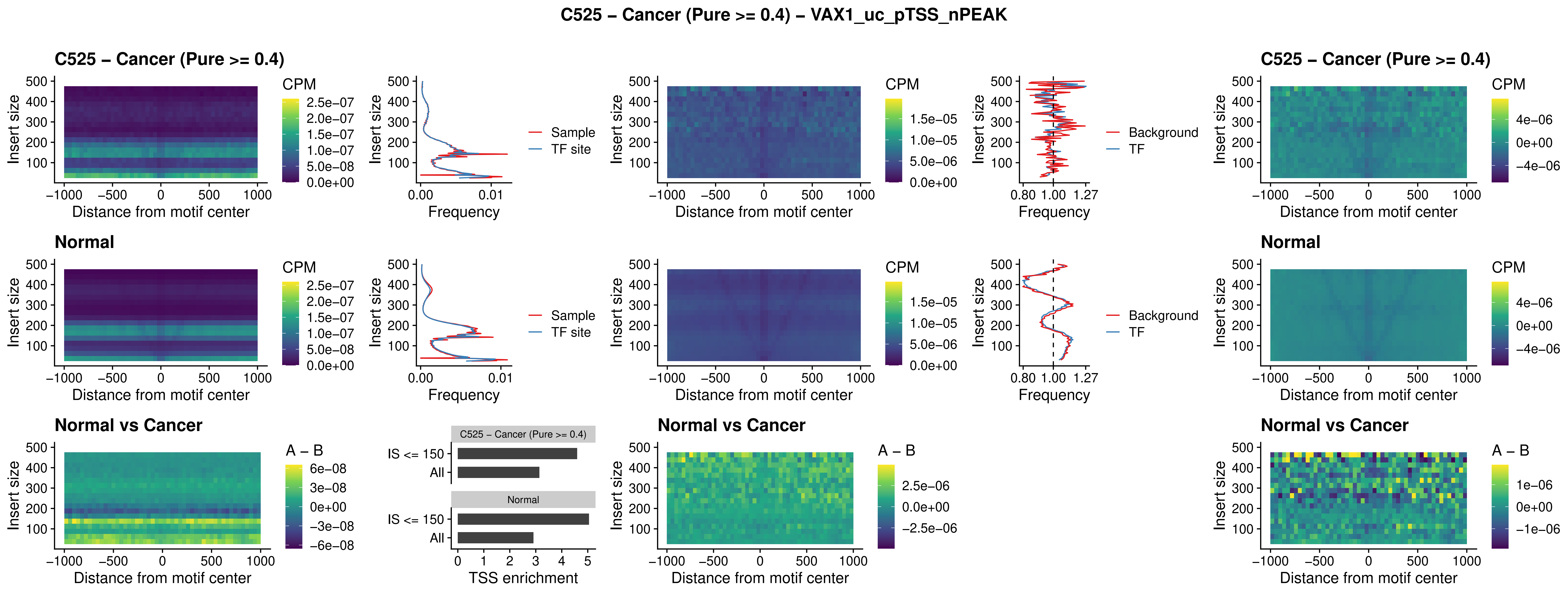
C525 – Cancer (Pure >= 0.4) – VAX1_uc_pTSS_nPEAK
C525 – Cancer (Pure >= 0.4)
Normal
Normal vs Cancer
Insert size
Distance from motif center
Frequency
CPM
A – B
Sample
TF site
Background
TF
C525 – Cancer (Pure >= 0.4)
Normal
IS <= 150
All
TSS enrichment

### C525 – Cancer (Pure >= 0.4) – DLX3_uc_pTSS_nPEAK

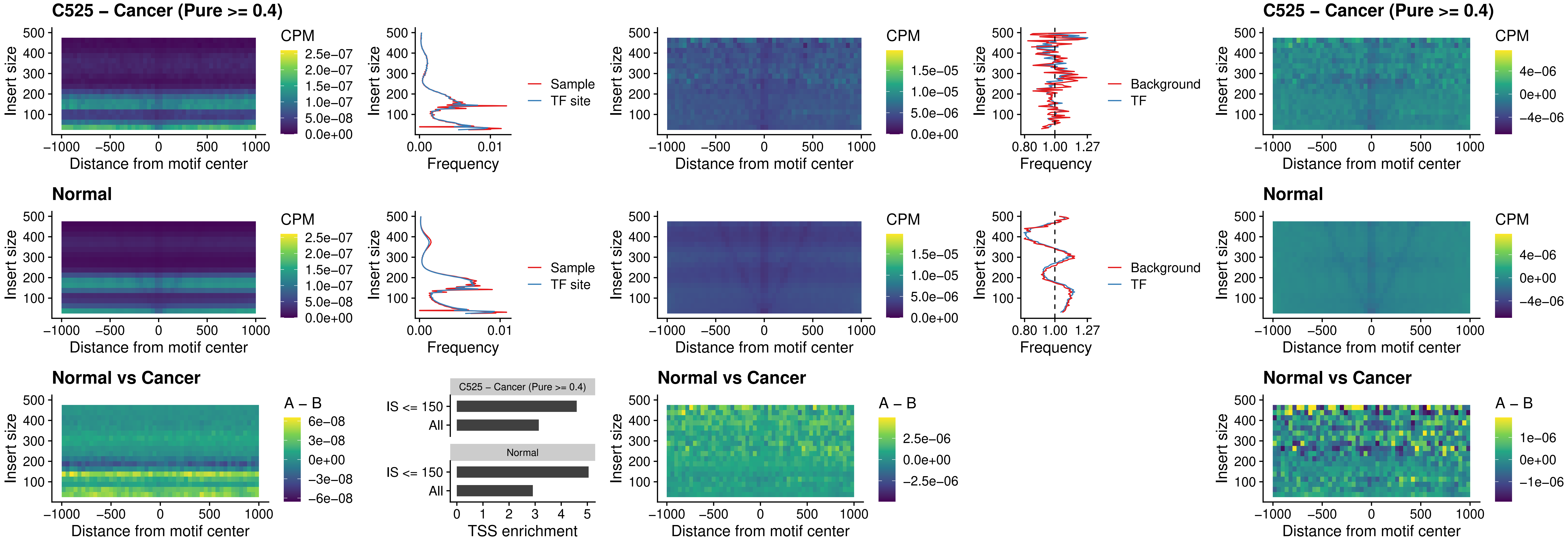

### C525 – Cancer (Pure >= 0.4) – HOXD3_uc_pTSS_nPEAK

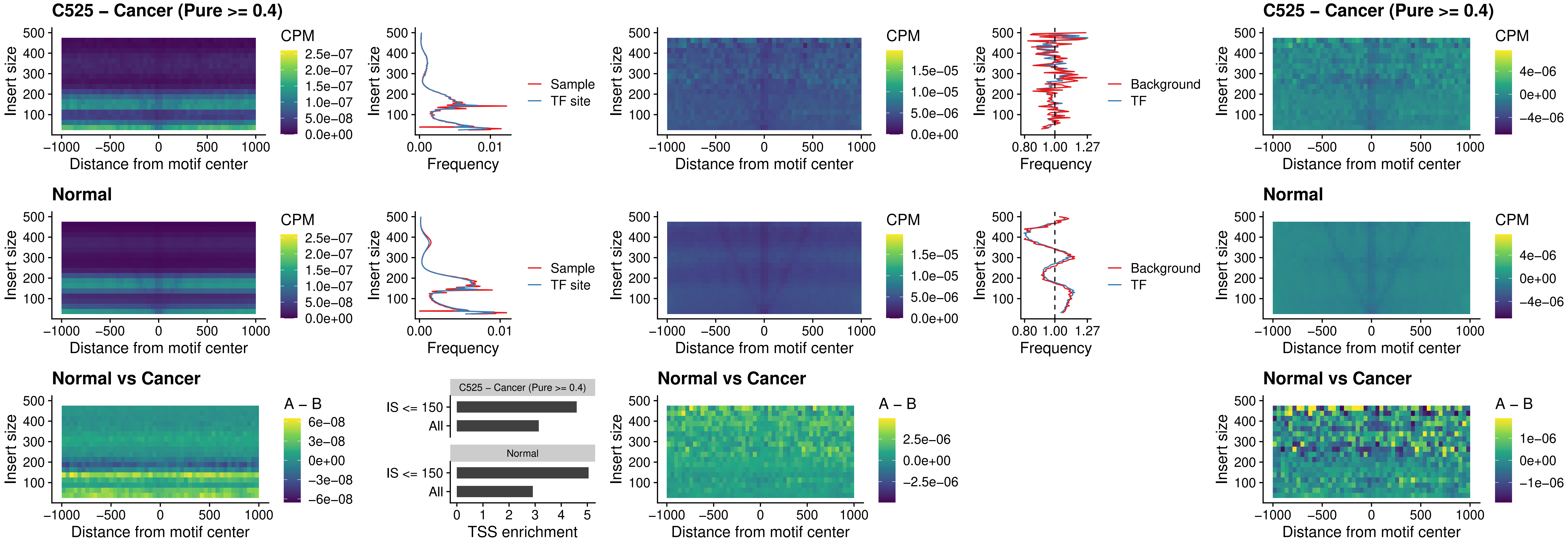

### C525 – Cancer (Pure >= 0.4) – PDX1_uc_dTSS_nPEAK

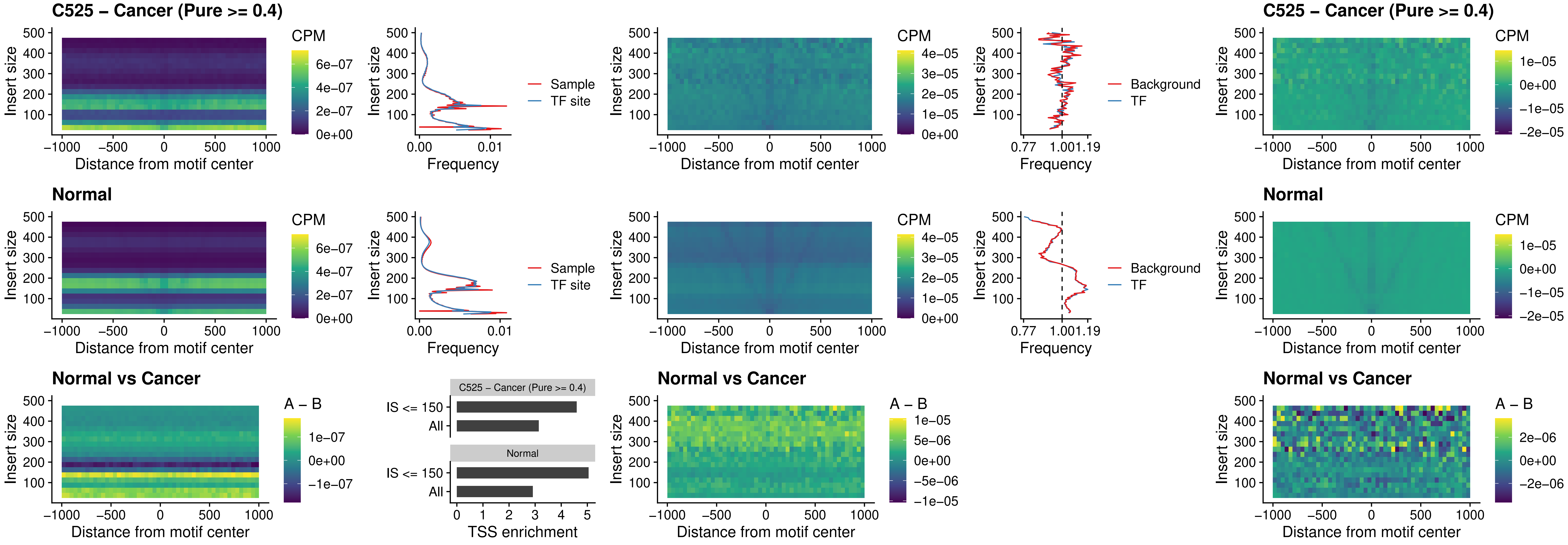

### C525 – Cancer (Pure >= 0.4) – VAX2_uc_dTSS_nPEAK

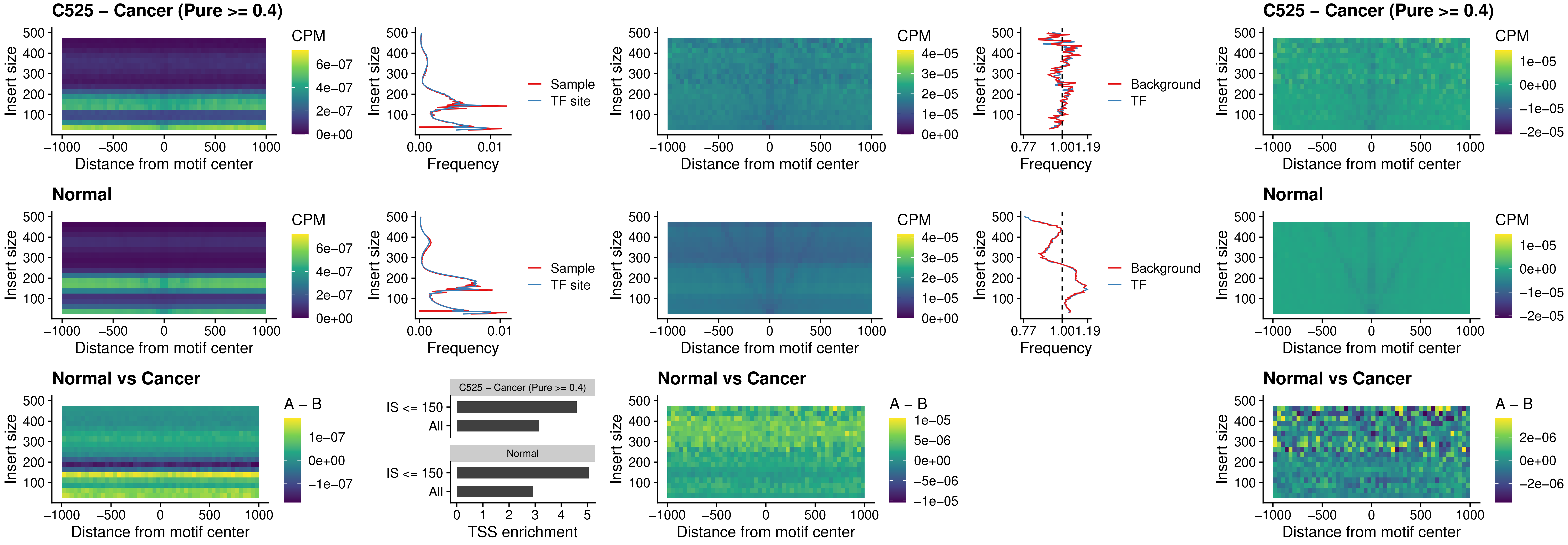

#### C525 – Cancer (Pure >= 0.4) – SOX5_uc_dTSS_nPEAK

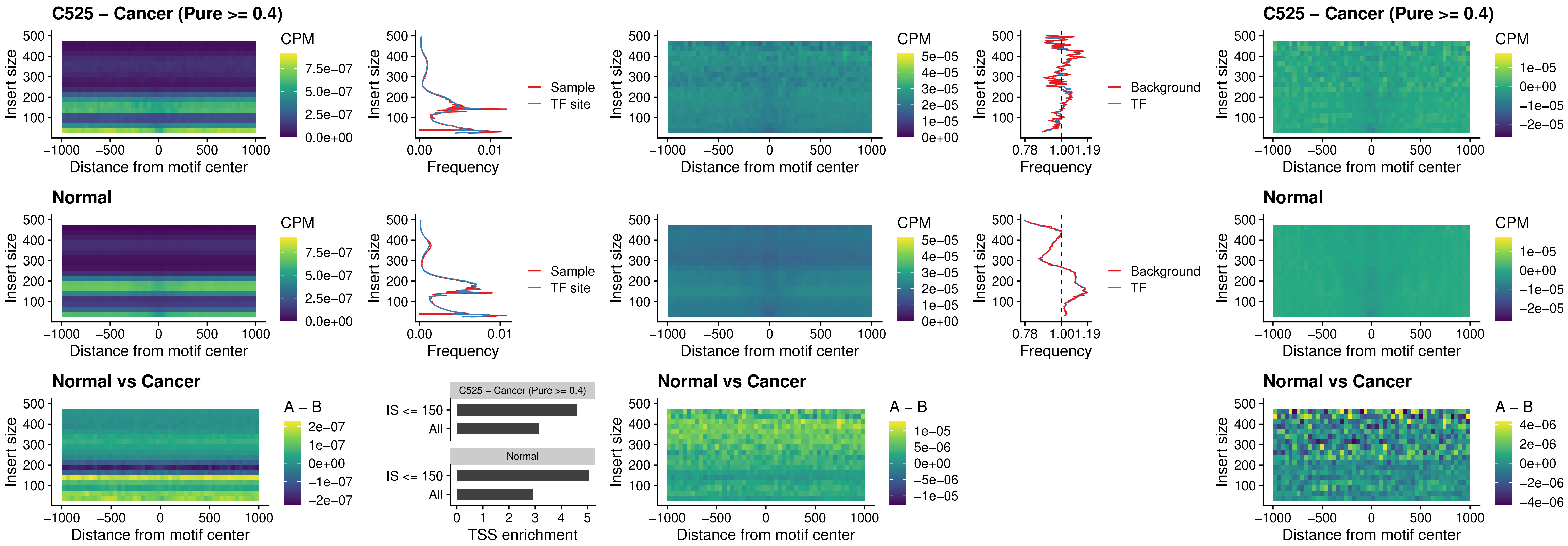

#### C525 – Cancer (Pure >= 0.4) – SOX6_uc_dTSS_nPEAK

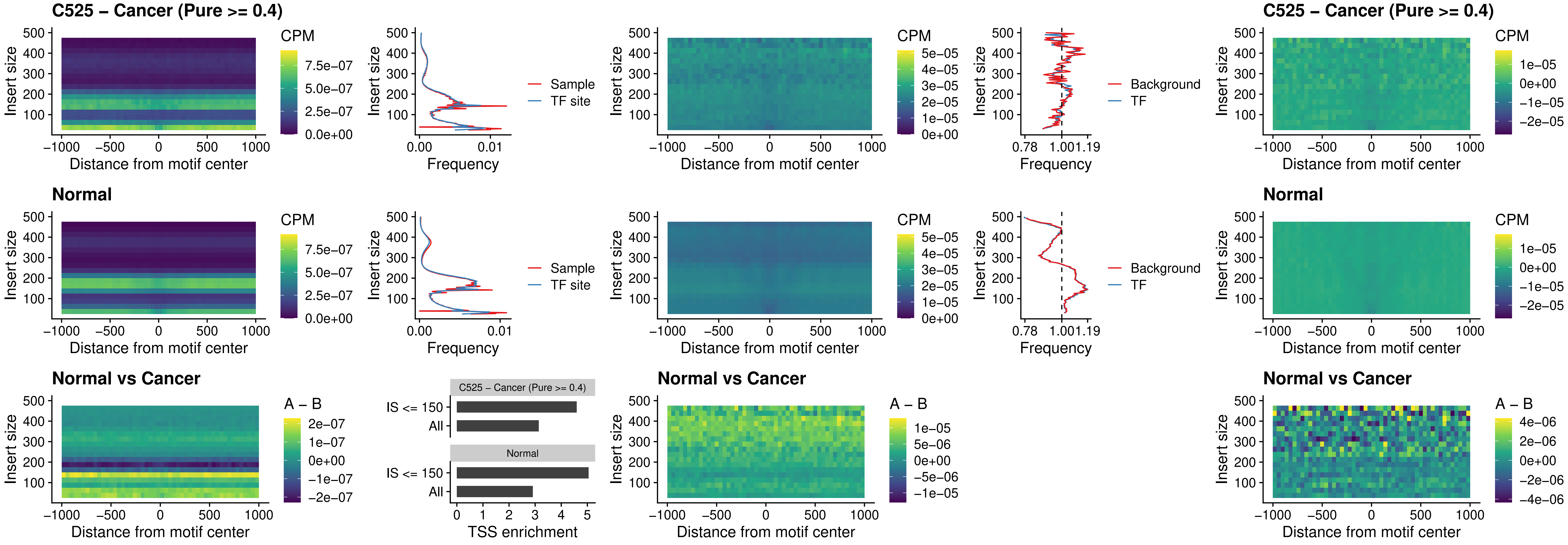

### C525 – Cancer (Pure >= 0.4) – JDP2_uc_cTSS_nPEAK

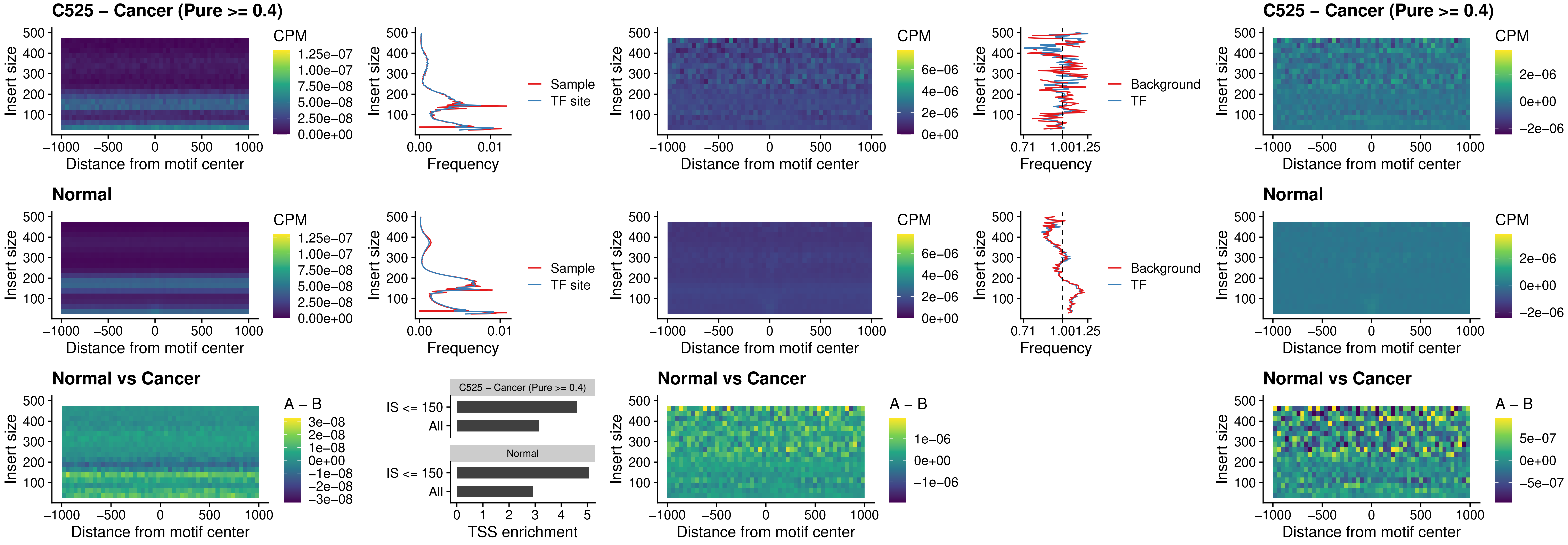

### C525 – Cancer (Pure >= 0.4) – PURA_uc_dTSS_oPEAK

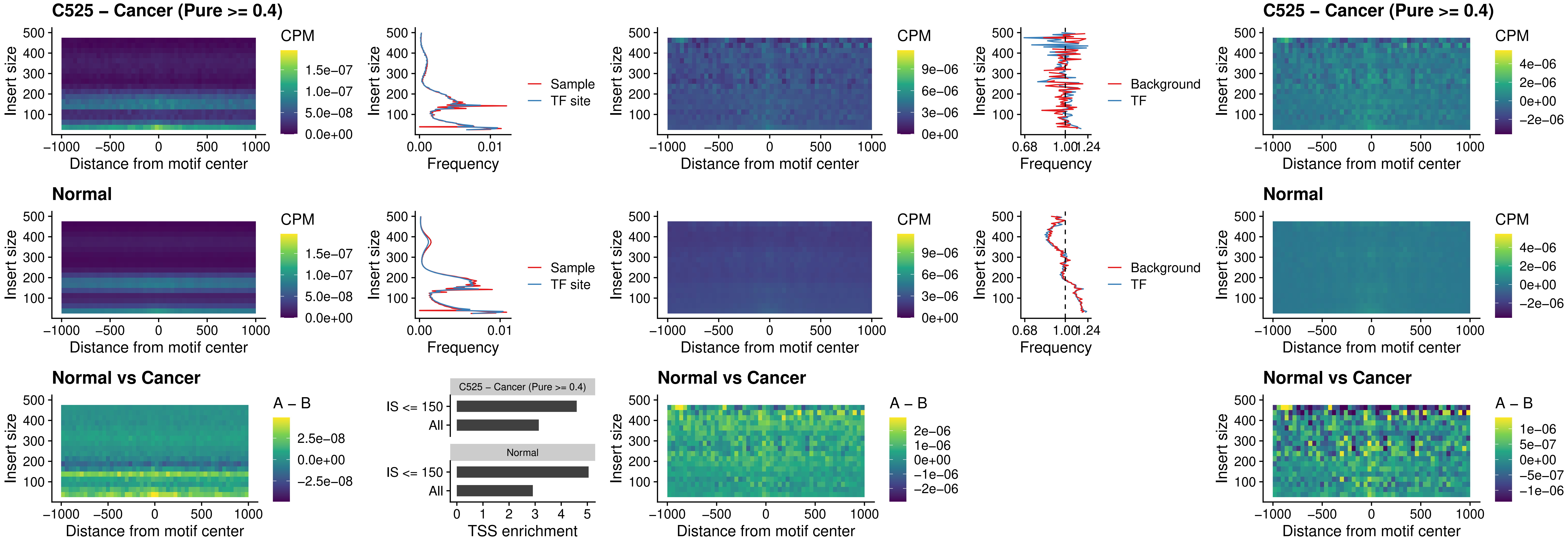

#### C525 – Cancer (Pure >= 0.4) – HHEX_uc_dTSS_nPEAK

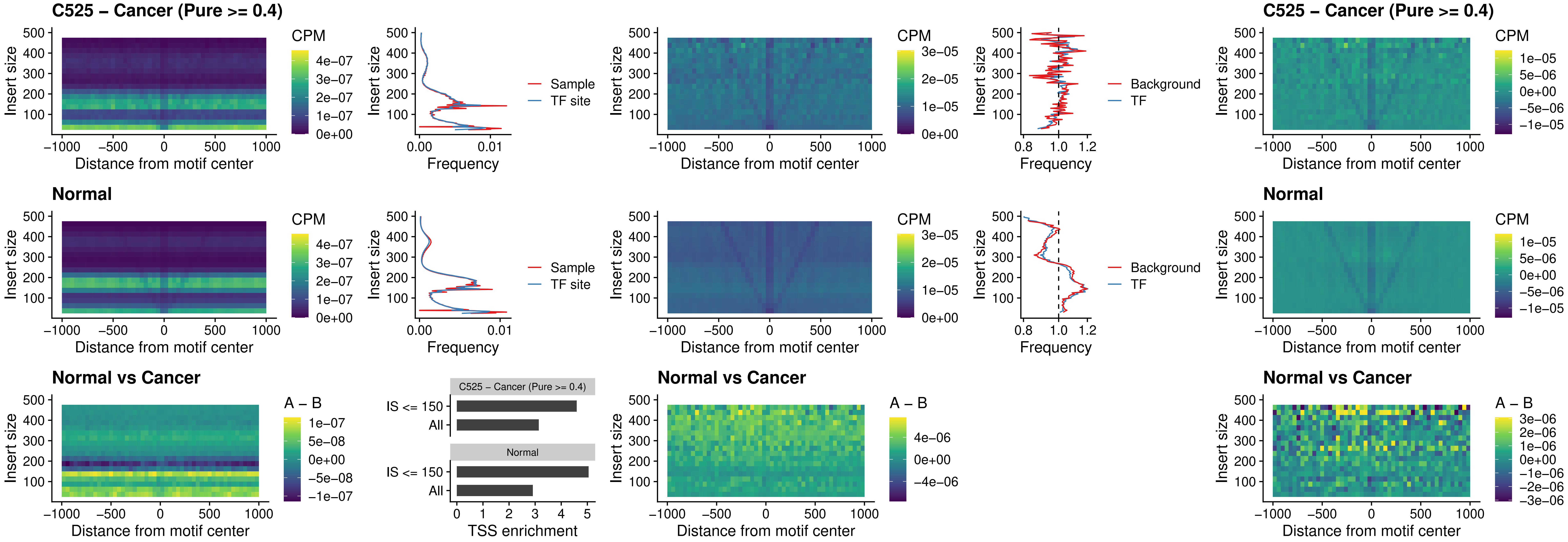

#### C525 – Cancer (Pure >= 0.4) – HMG20B_uc_dTSS_nPEAK

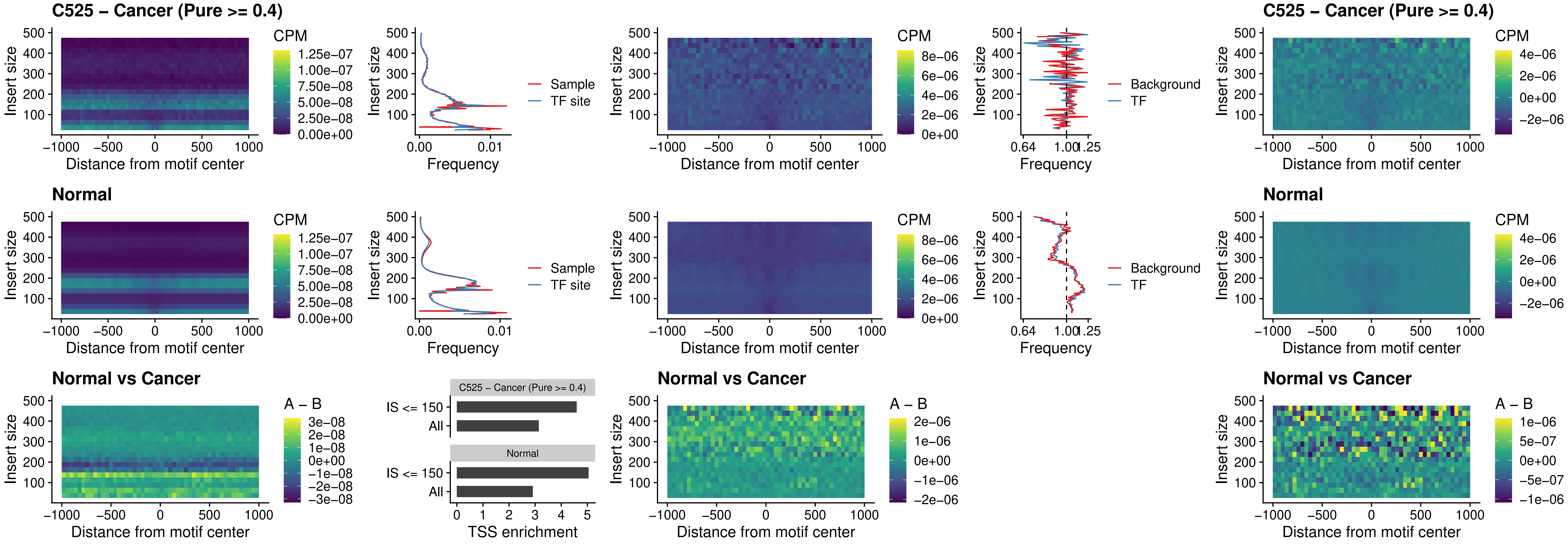

#### C525 – Cancer (Pure >= 0.4) – FOXB2_uc_dTSS_nPEAK

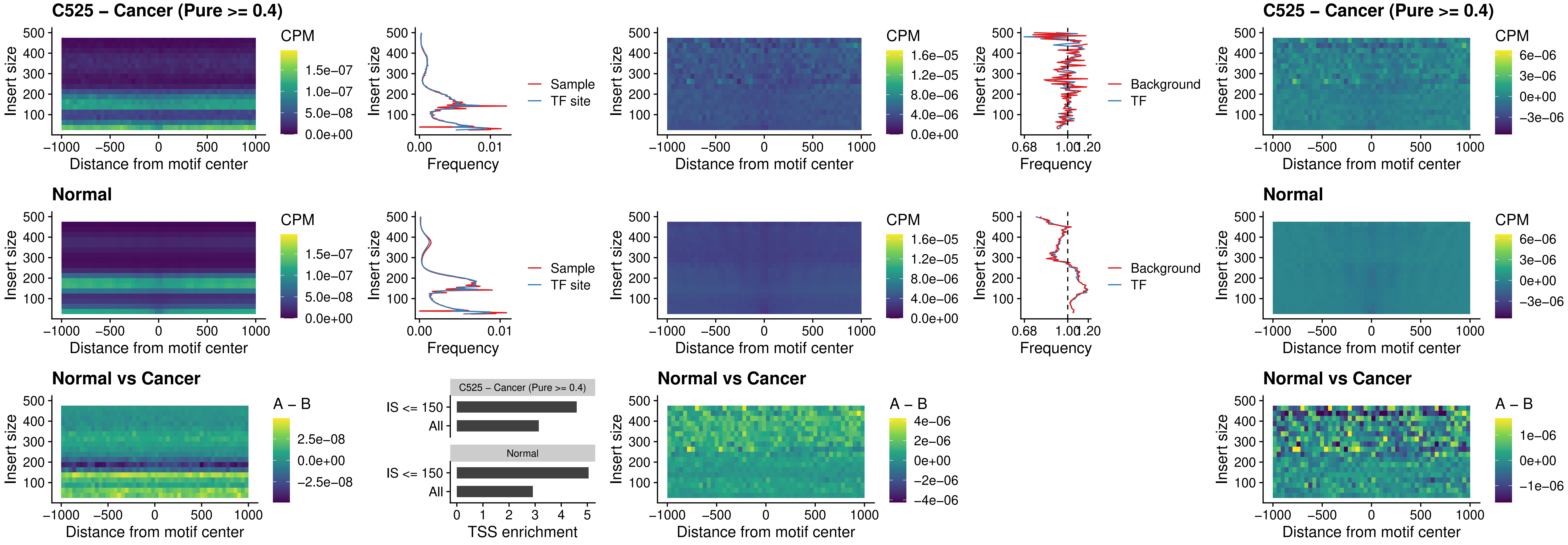

#### C525 – Cancer (Pure >= 0.4) – FOXL1_uc_cTSS_nPEAK

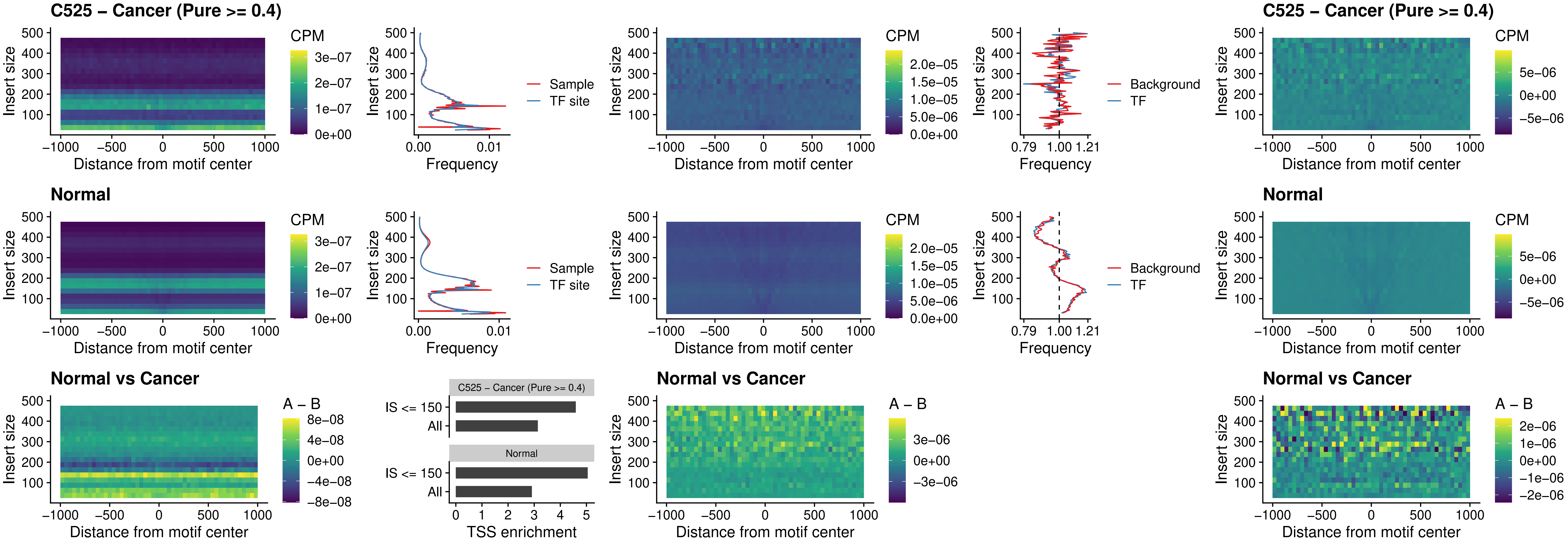

### C525 – Cancer (Pure >= 0.4) – PBX3_uc_pTSS_oPEAK

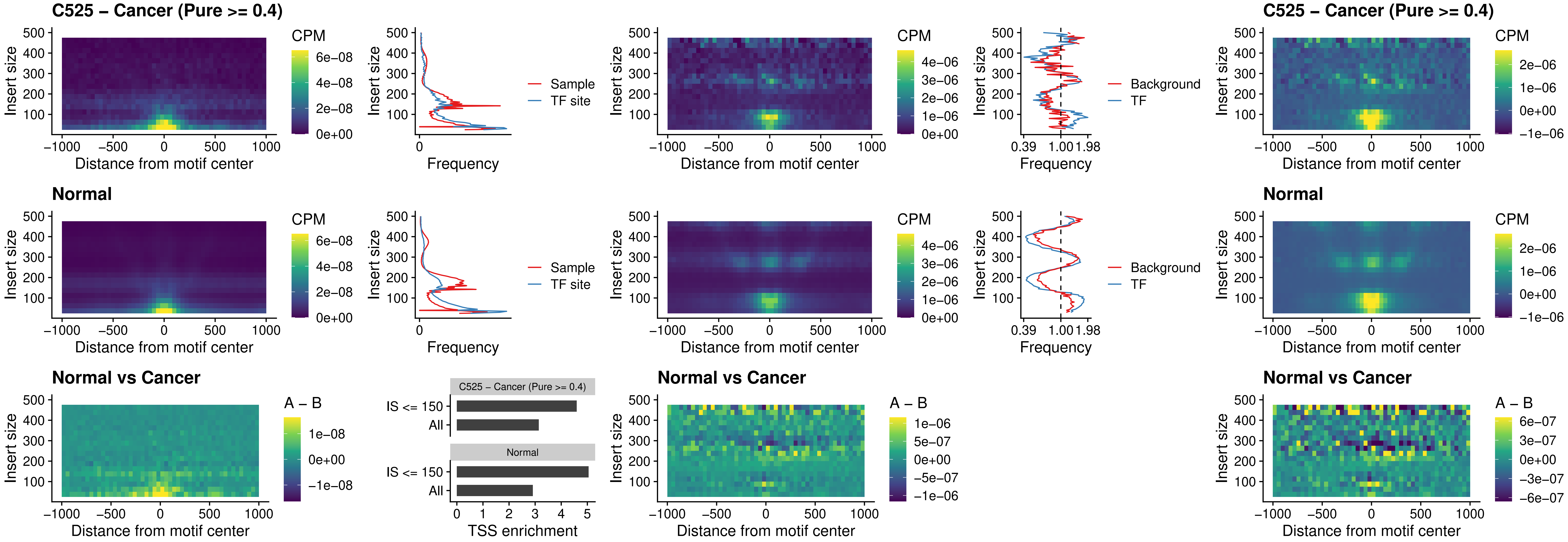

### C525 – Cancer (Pure >= 0.4) – SNAI2_uc_cTSS_nPEAK

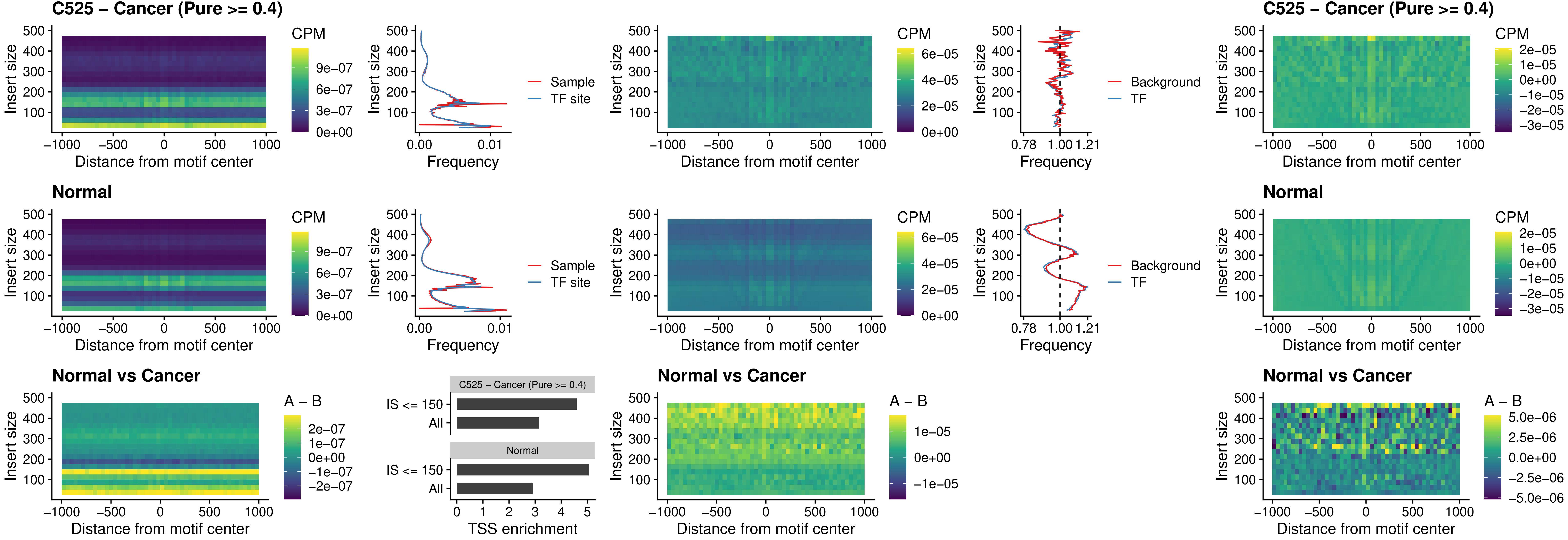

### C525 – Cancer (Pure >= 0.4) – SNAI2_uc_dTSS_oPEAK

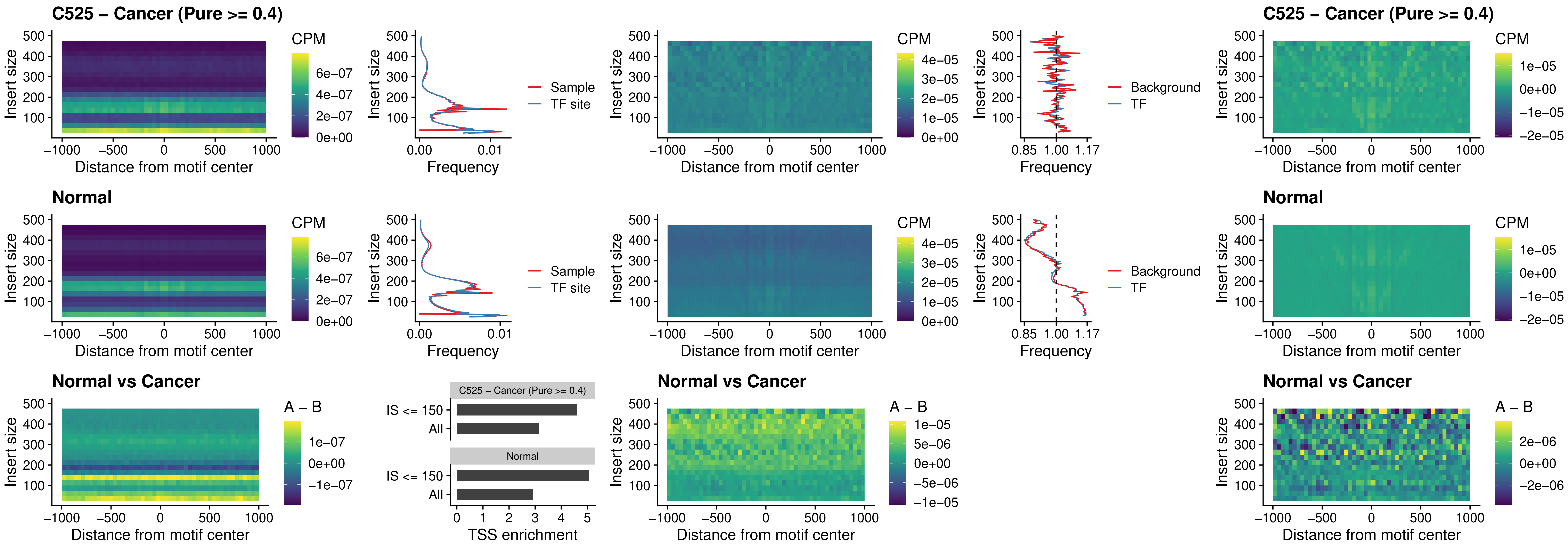

#### C525 – Cancer (Pure >= 0.4) – SPI1_uc_pTSS_nPEAK

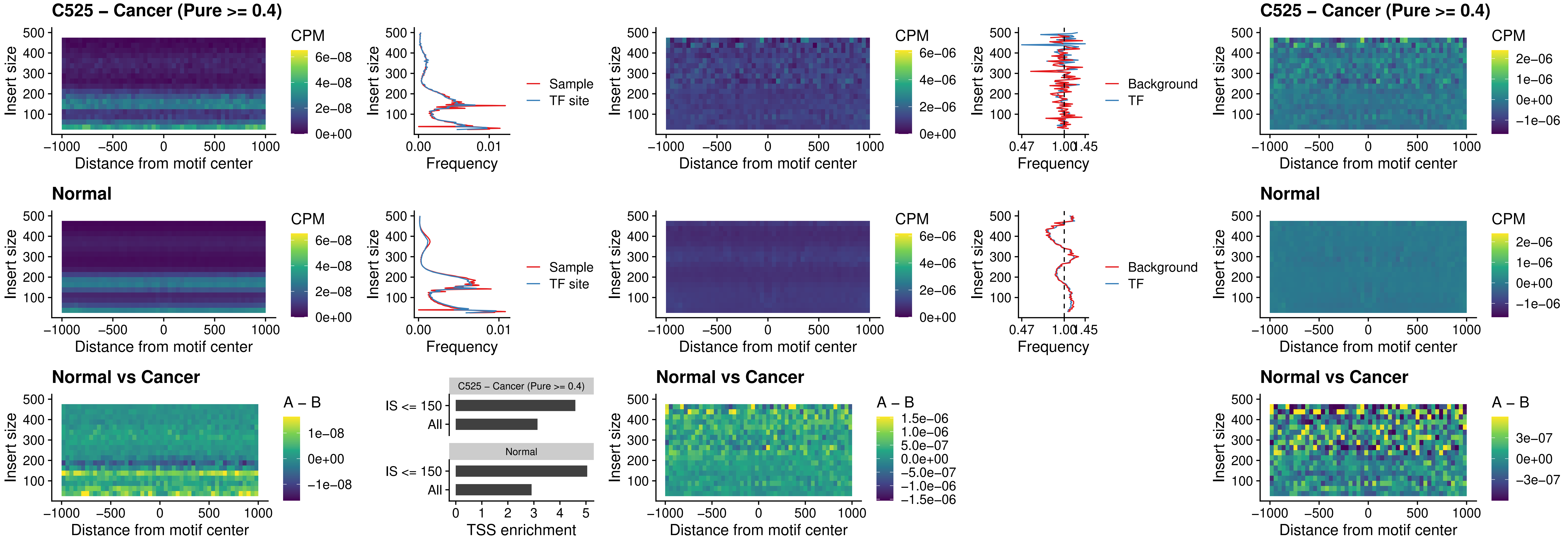

#### C525 – Cancer (Pure >= 0.4) – KLF16_uc_pTSS_nPEAK

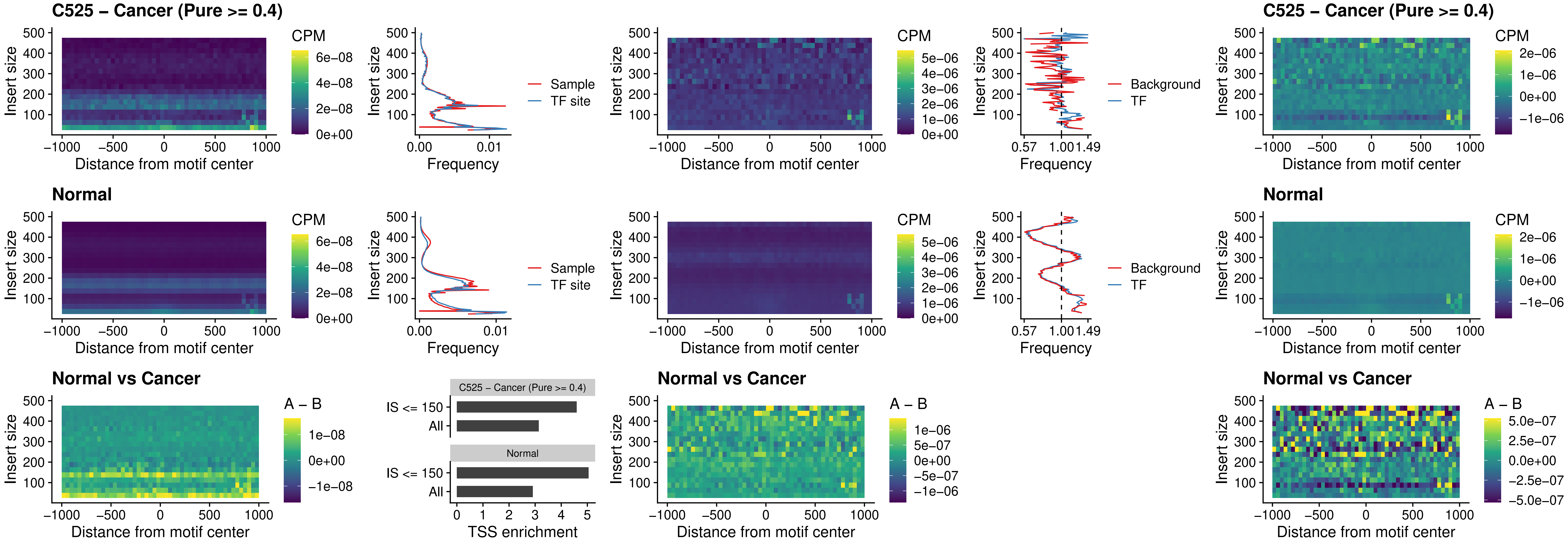

#### C525 – Cancer (Pure >= 0.4) – SP3_uc_pTSS_nPEAK

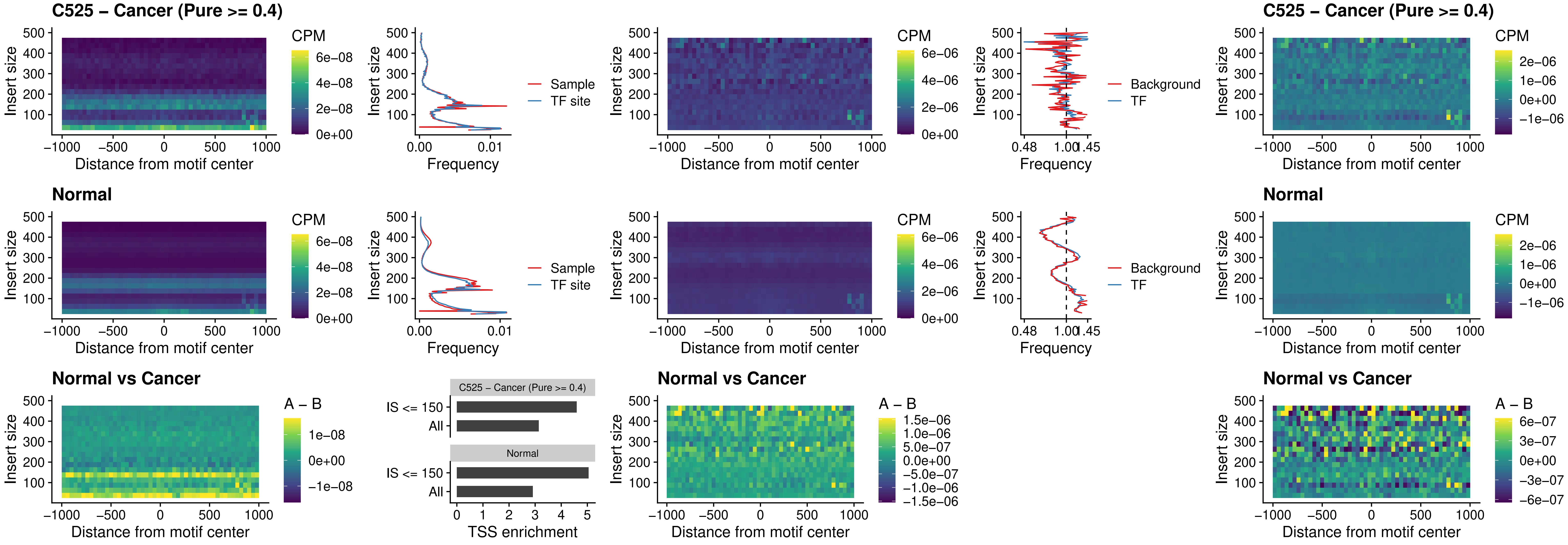

### C525 – Cancer (Pure >= 0.4) – NEUROD1_uc_dTSS_oPEAK

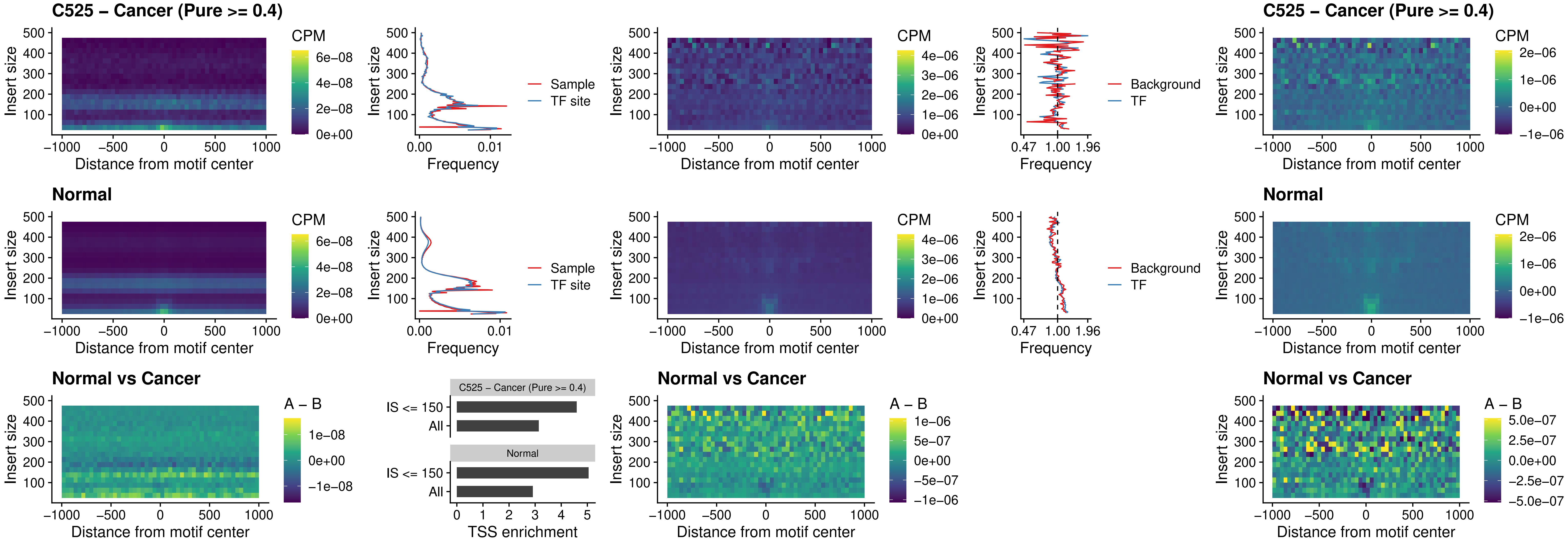

### C525 – Cancer (Pure >= 0.4) – CTCF_uc_dTSS_oPEAK

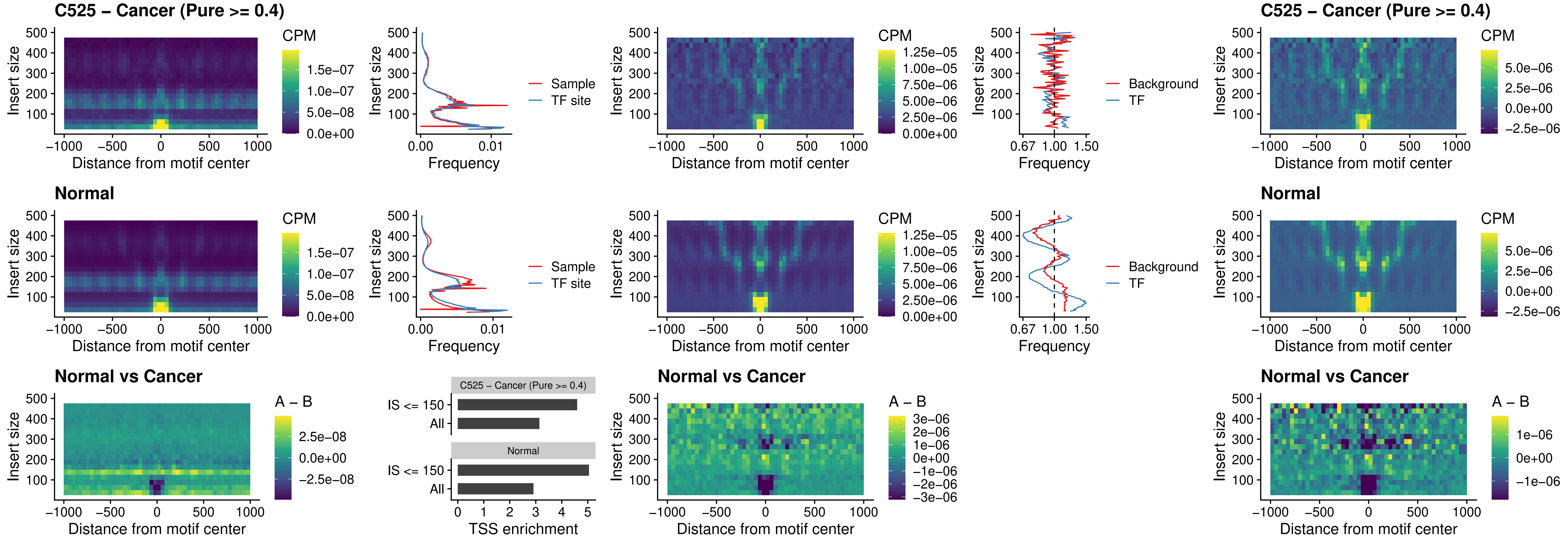

#### C525 – Cancer (Pure >= 0.4) – CTCF_uc_cTSS_oPEAK

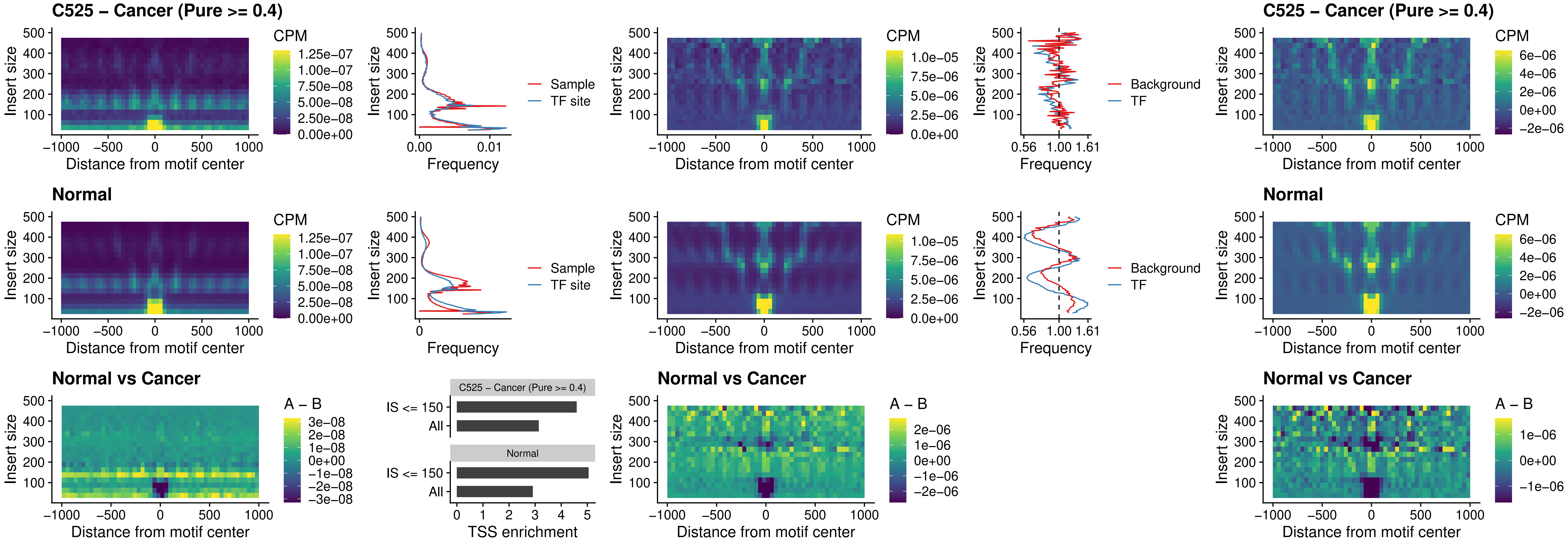

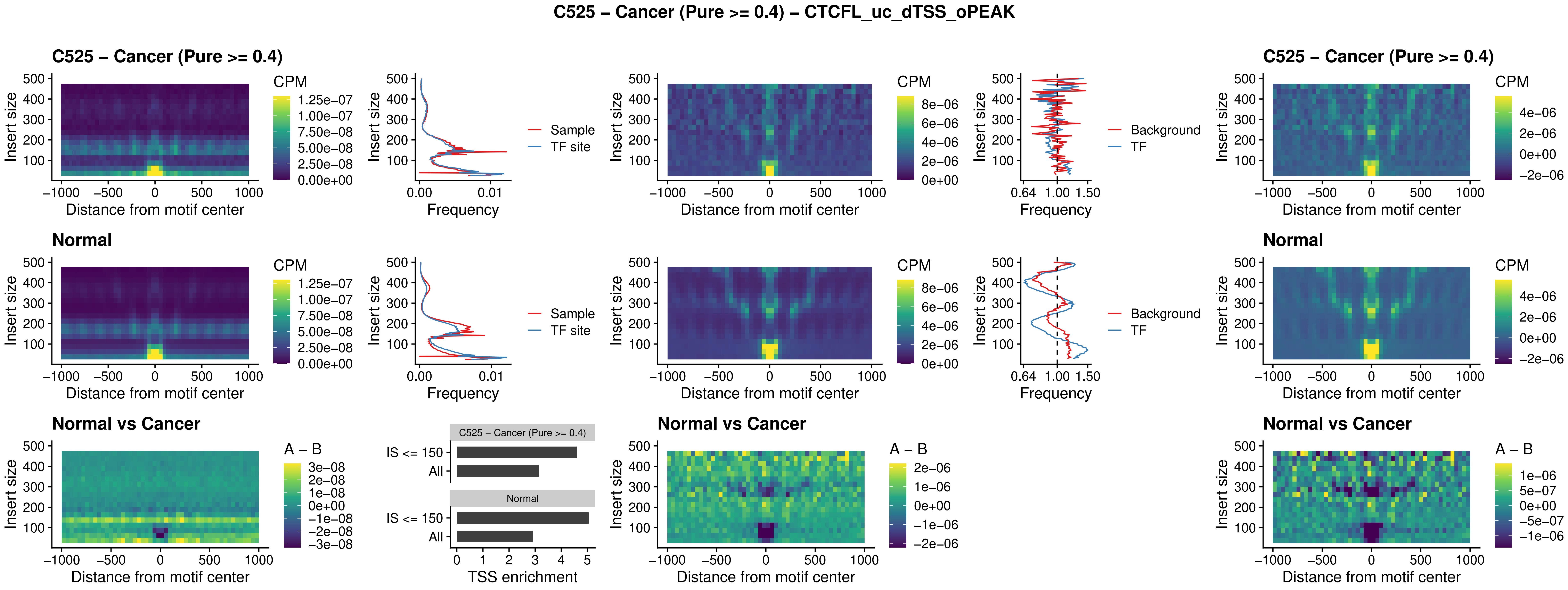
C525 – Cancer (Pure >= 0.4) – CTCFL_uc_dTSS_oPEAK
C525 – Cancer (Pure >= 0.4)
Normal
Normal vs Cancer
Insert size
Distance from motif center
CPM
A – B
Frequency
Sample
TF site
Background
TF
C525 – Cancer (Pure >= 0.4)
Normal
IS <= 150
All
TSS enrichment

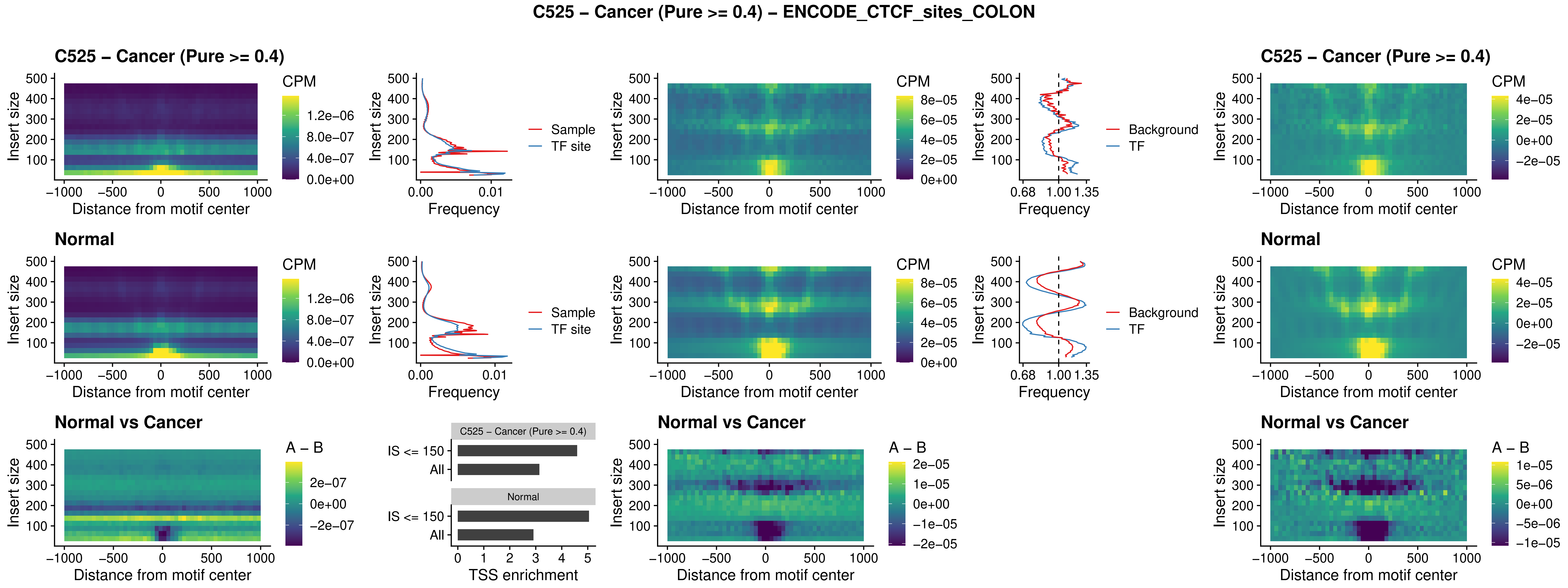

C525 – Cancer (Pure >= 0.4) – ENCODE_CTCF_sites_COLON
C525 – Cancer (Pure >= 0.4)
Normal
Normal vs Cancer
Insert size
Distance from motif center
Frequency
CPM
A – B
Sample
TF site
Background
TF
C525 – Cancer (Pure >= 0.4)
Normal
IS <= 150
All
TSS enrichment

#### C525 – Cancer (Pure >= 0.4) – ENCODE_CTCF_sites_COLON_loop_ctcf

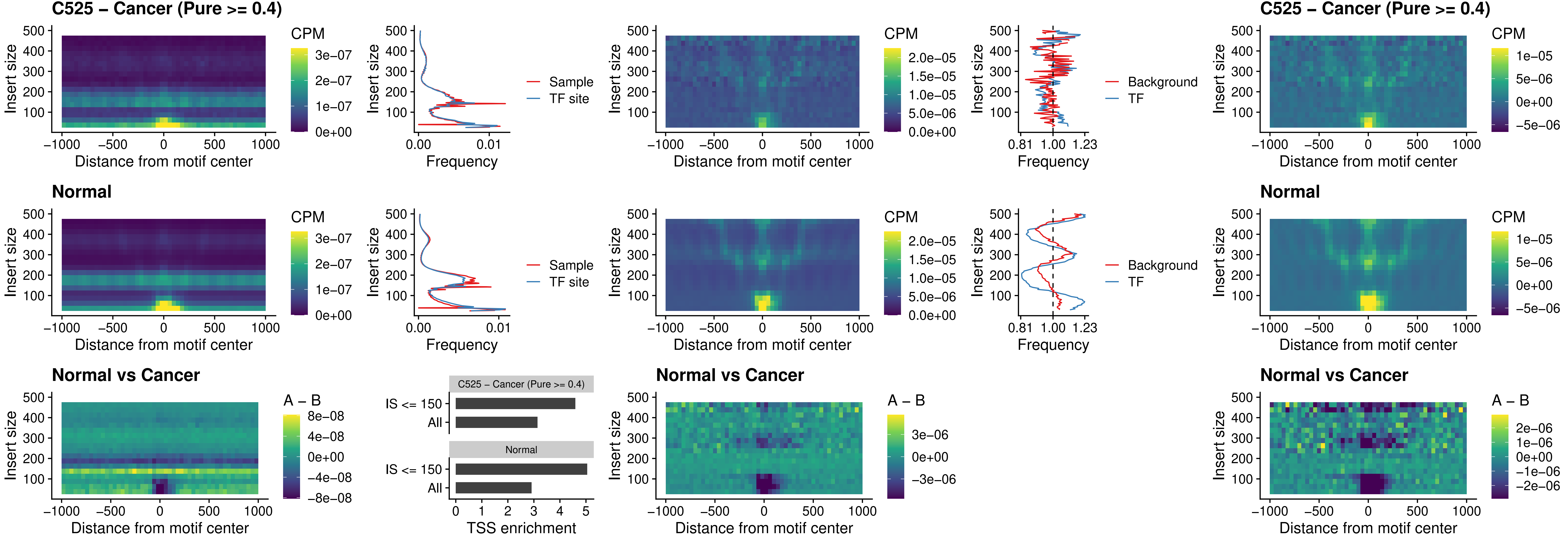

#### C525 – Cancer (Pure >= 0.4) – ENCODE_CTCF_sites_COLON_intergenic_ctcf

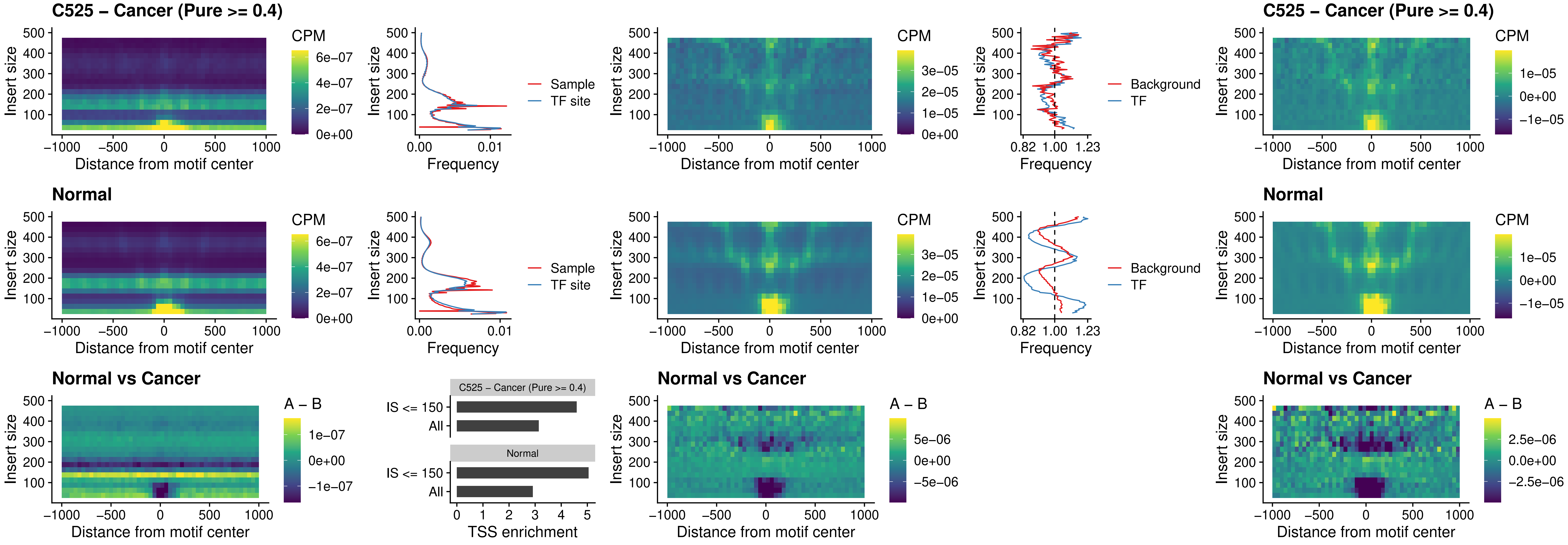

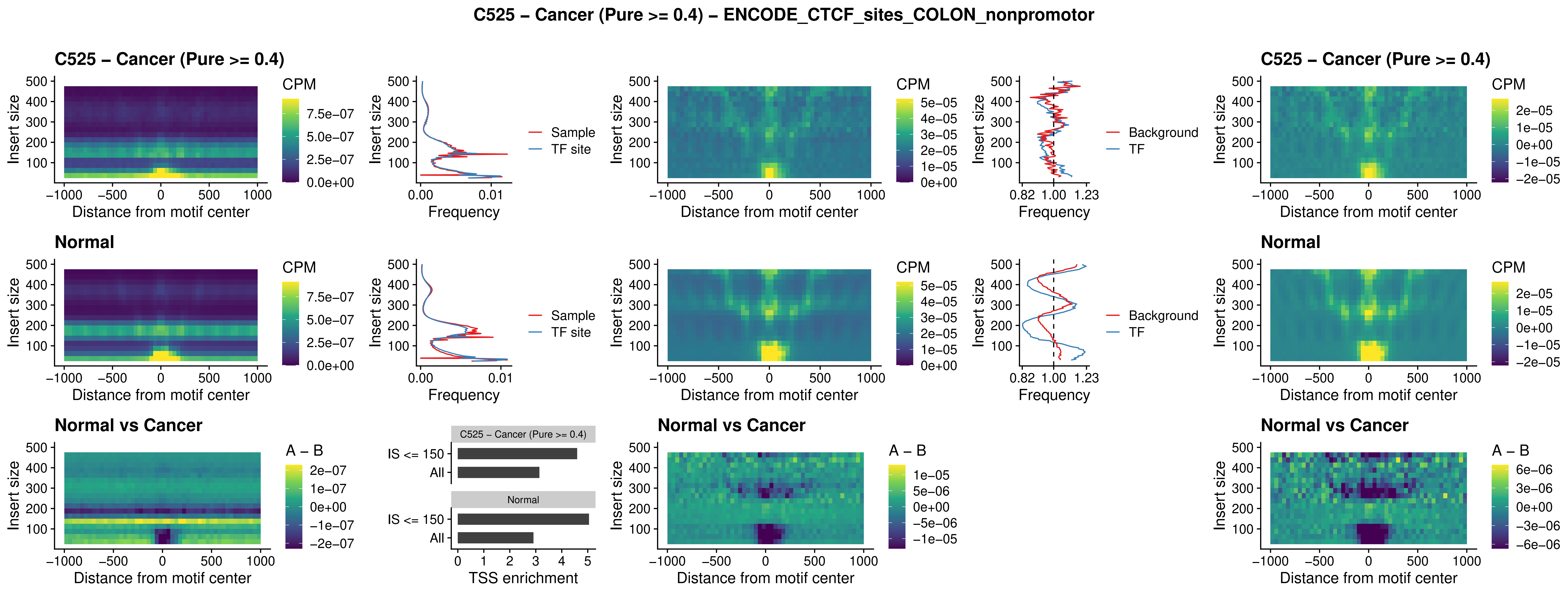
C525 – Cancer (Pure >= 0.4) – ENCODE_CTCF_sites_COLON_nonpromotor
C525 – Cancer (Pure >= 0.4)
Normal
Normal vs Cancer
Insert size
Distance from motif center
Frequency
CPM
A – B
Sample
TF site
Background
TF
C525 – Cancer (Pure >= 0.4)
IS <= 150
All
TSS enrichment

#### C525 – Cancer (Pure >= 0.4) – CTCFL_uc_cTSS_oPEAK

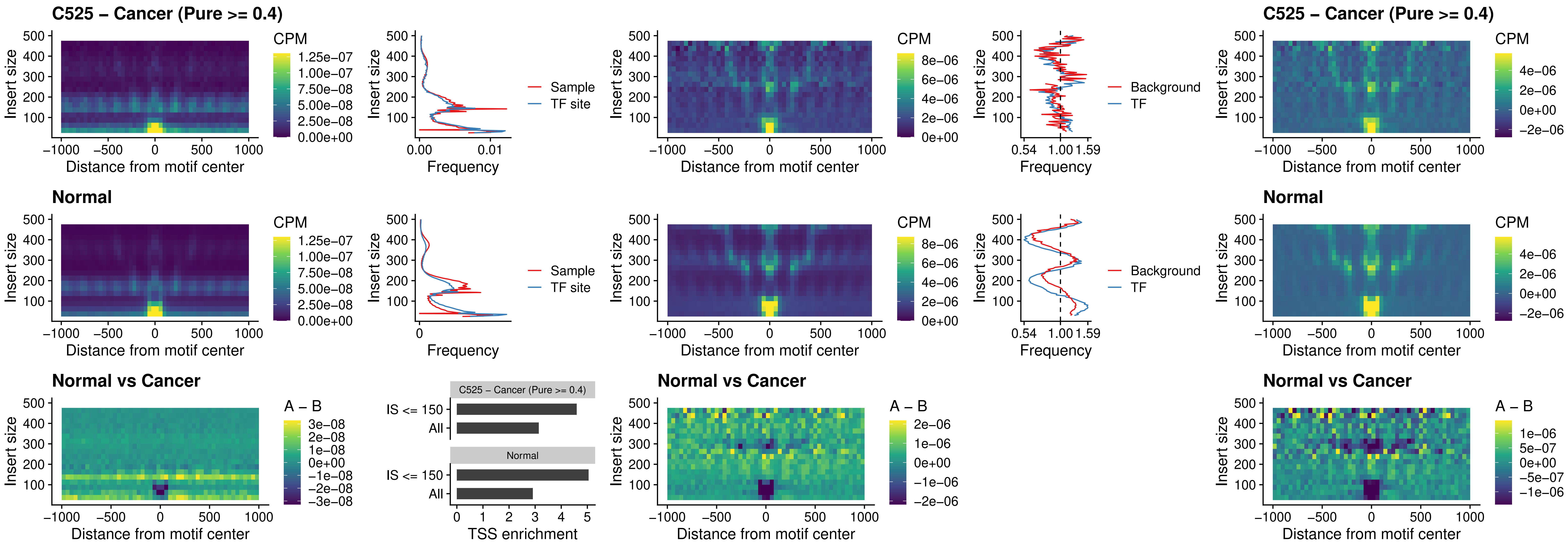

### C525 – Cancer (Pure >= 0.4) – NEUROD1_uc_cTSS_oPEAK

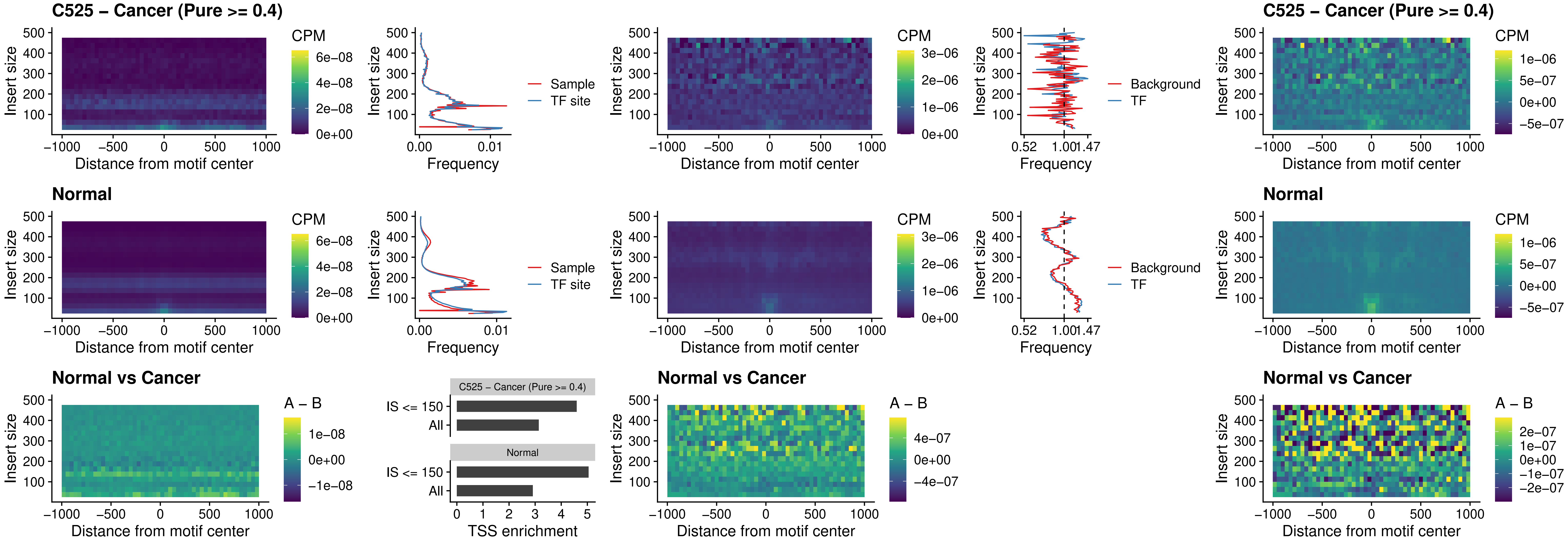

#### C525 – Cancer (Pure >= 0.4) – ENCODE_YY1_sites_COLON_nonpromotor

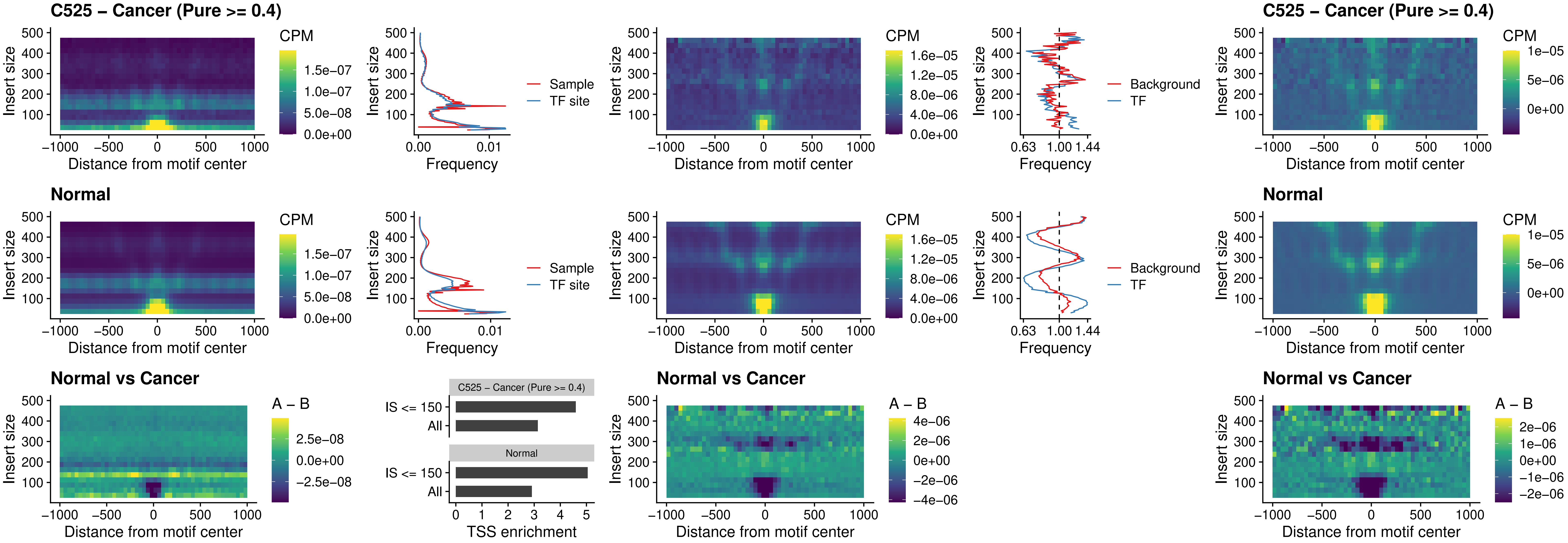

#### C525 – Cancer (Pure >= 0.4) – CTCF_uc_pTSS_oPEAK

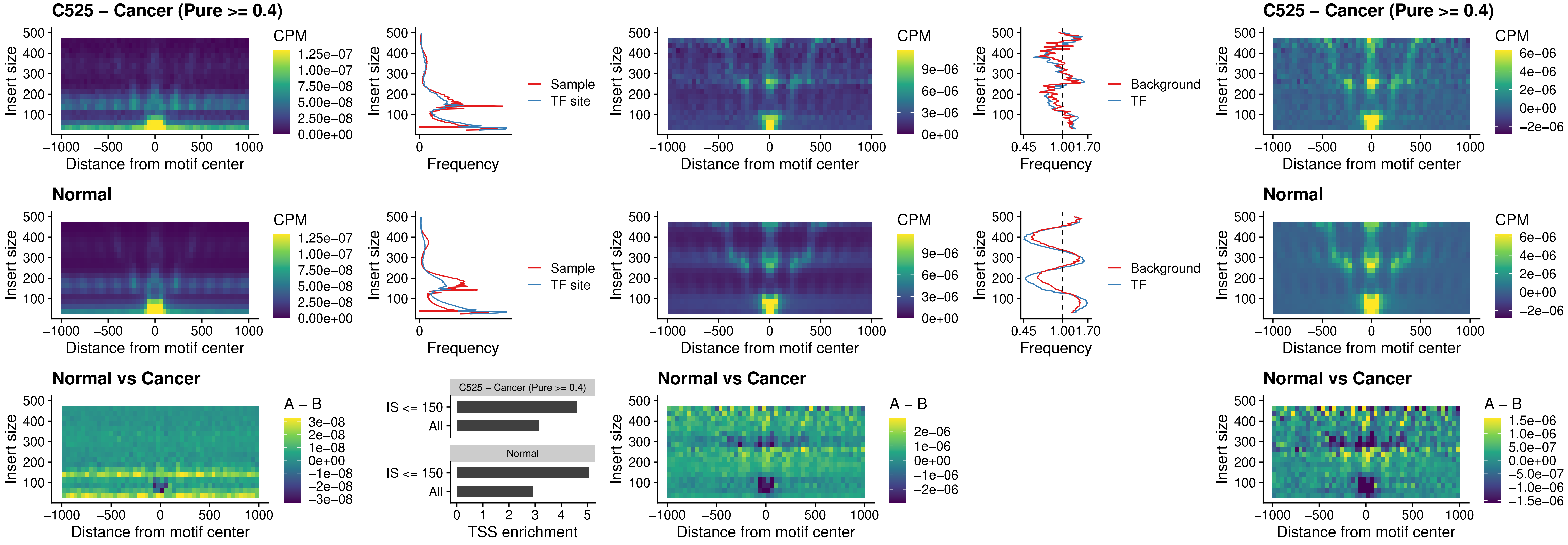

#### C525 – Cancer (Pure >= 0.4) – ENCODE_CTCF_sites_COLON_promotor

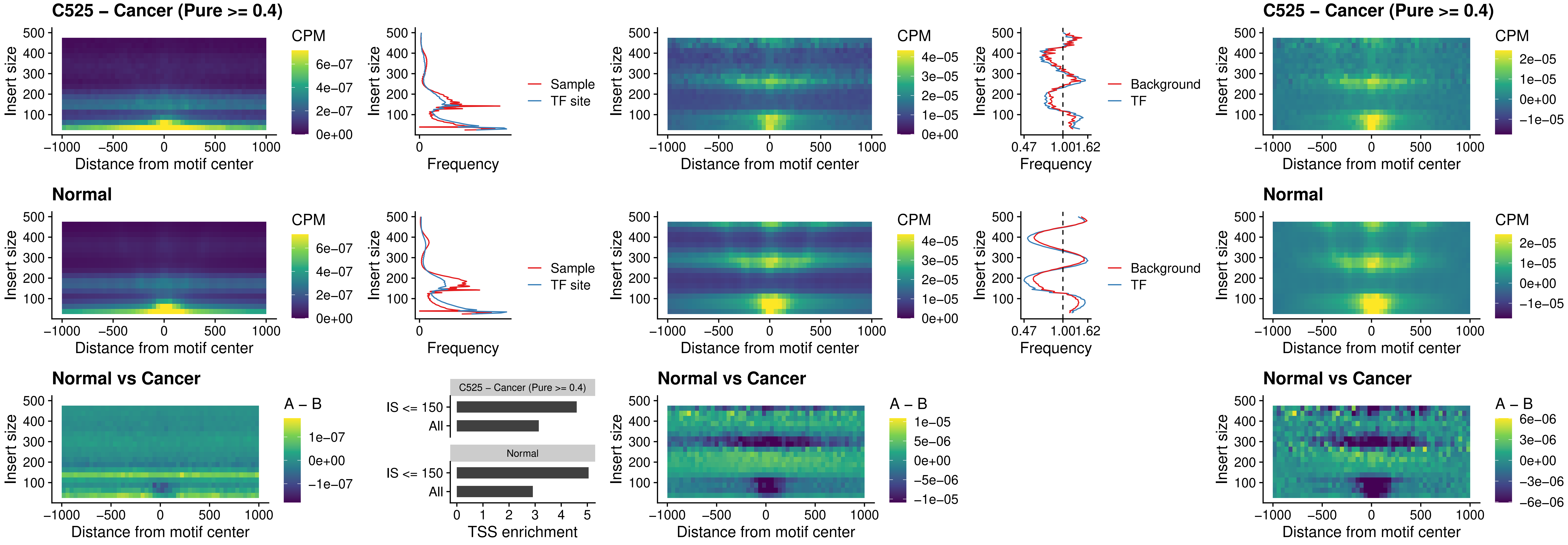

#### C525 – Cancer (Pure >= 0.4) – ENCODE_CTCF_sites_COLON_promotor_ctcf

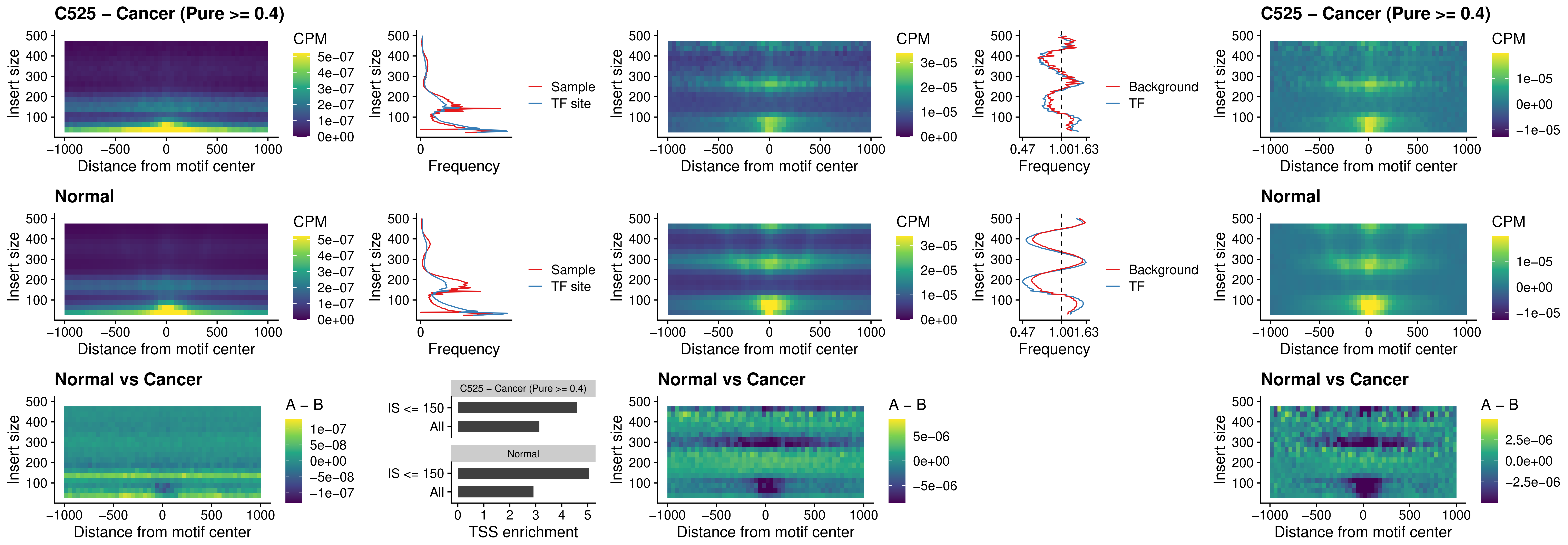

#### C525 – Cancer (Pure >= 0.4) – SPIC_uc_pTSS_oPEAK

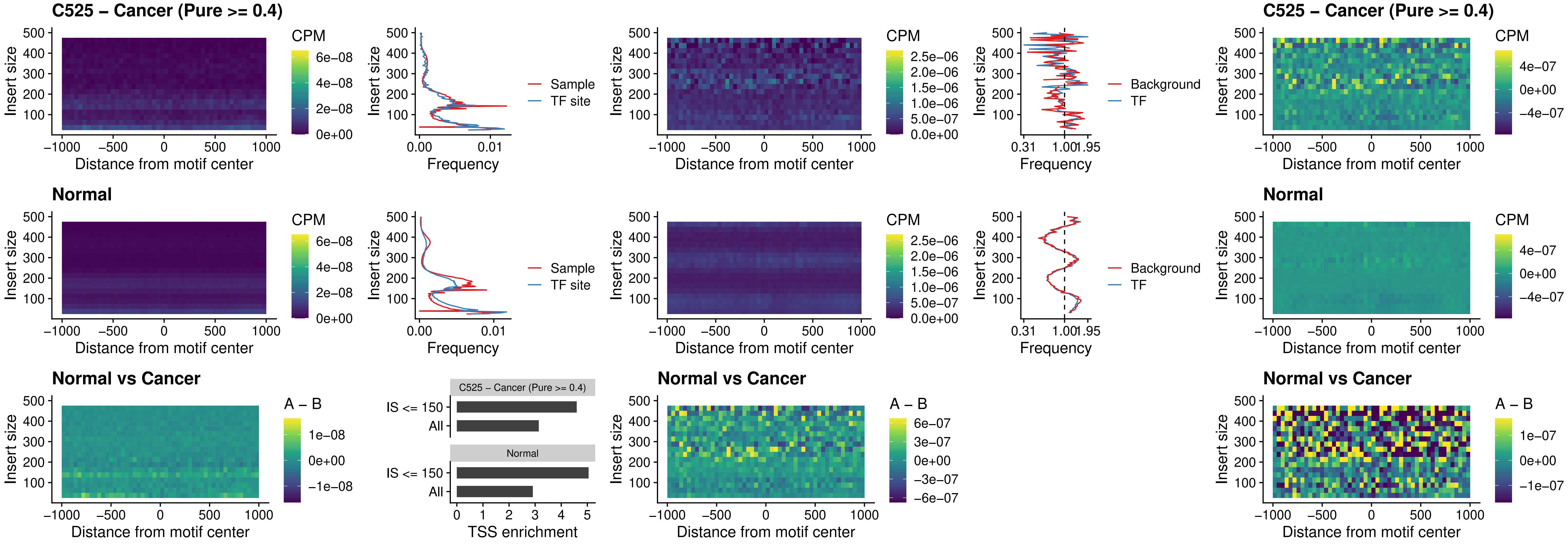

### C525 – Cancer (Pure >= 0.4) – SPI1_uc_dTSS_nPEAK

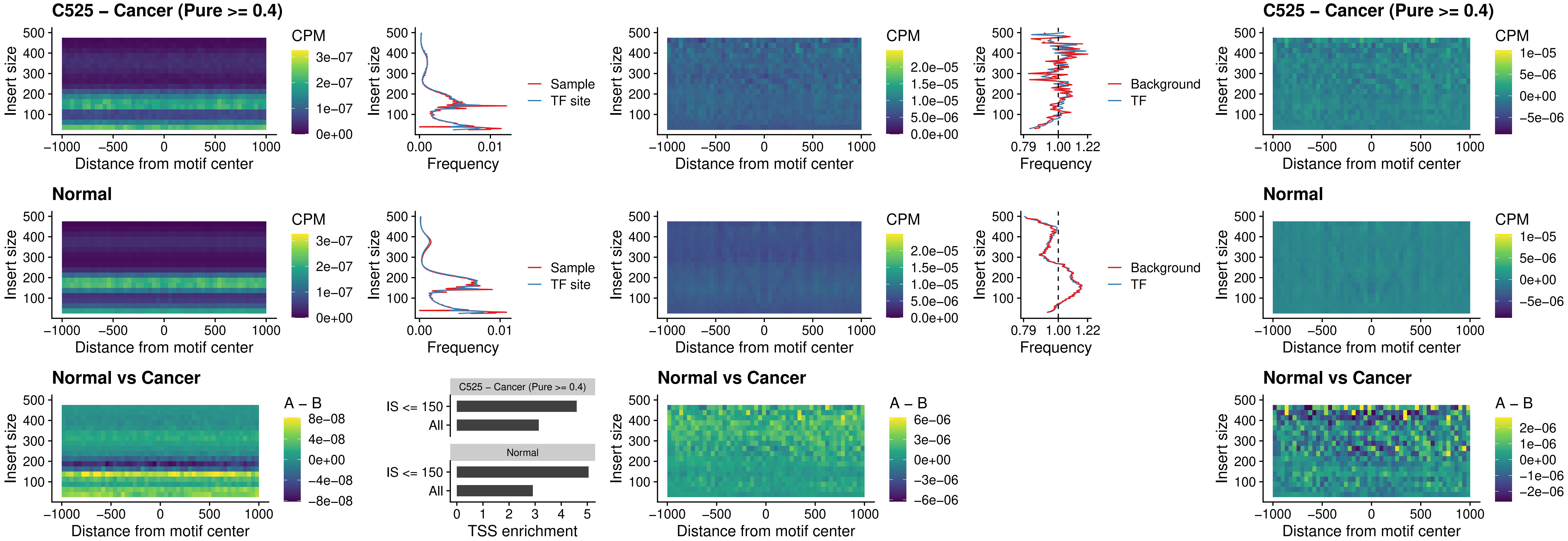

### C525 – Cancer (Pure >= 0.4) – SPI1_uc_cTSS_nPEAK

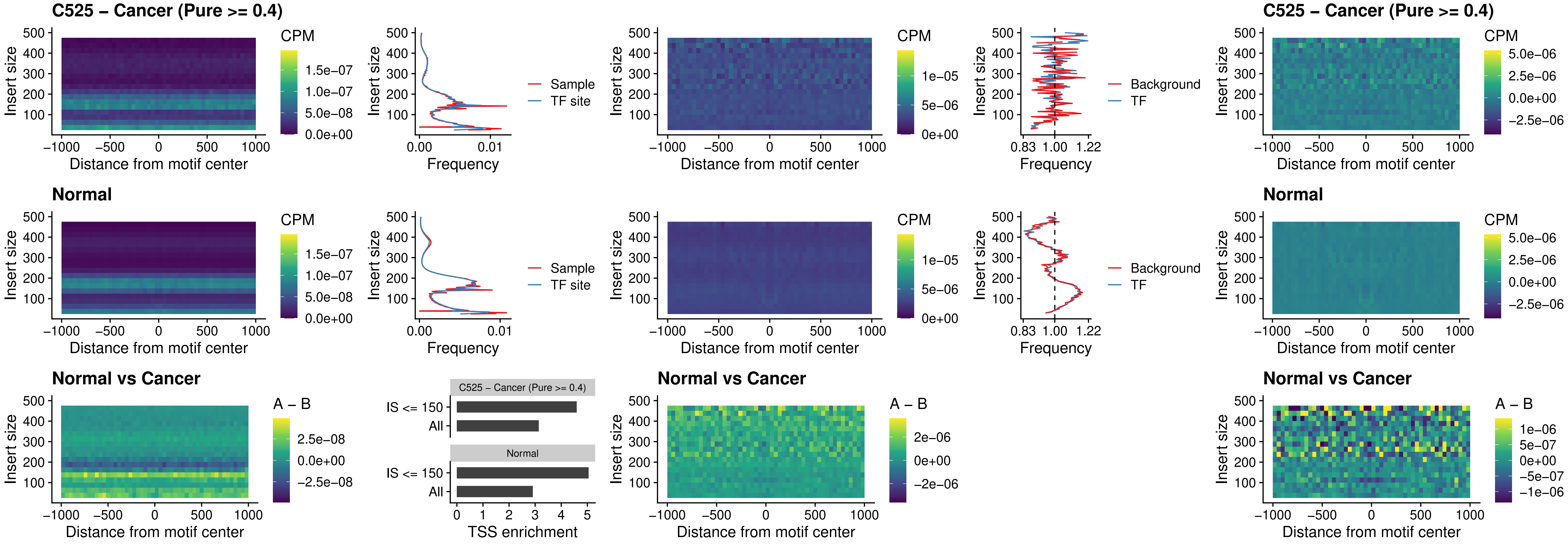

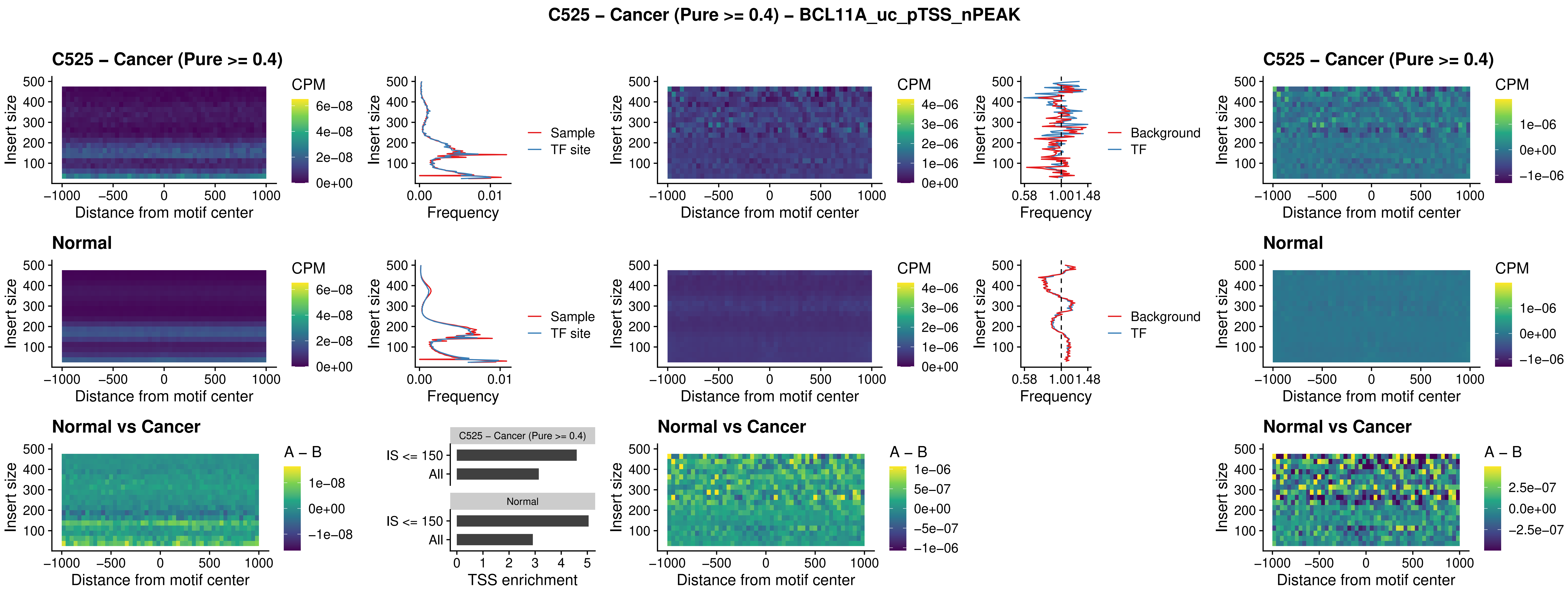
C525 – Cancer (Pure >= 0.4) – BCL11A_uc_pTSS_nPEAK
C525 – Cancer (Pure >= 0.4)
Normal
Normal vs Cancer
Insert size
Distance from motif center
Frequency
CPM
A – B
Sample
TF site
Background
TF
C525 – Cancer (Pure >= 0.4)
Normal
IS <= 150
All
TSS enrichment

#### C525 – Cancer (Pure >= 0.4) – BCL11B_uc_pTSS_nPEAK

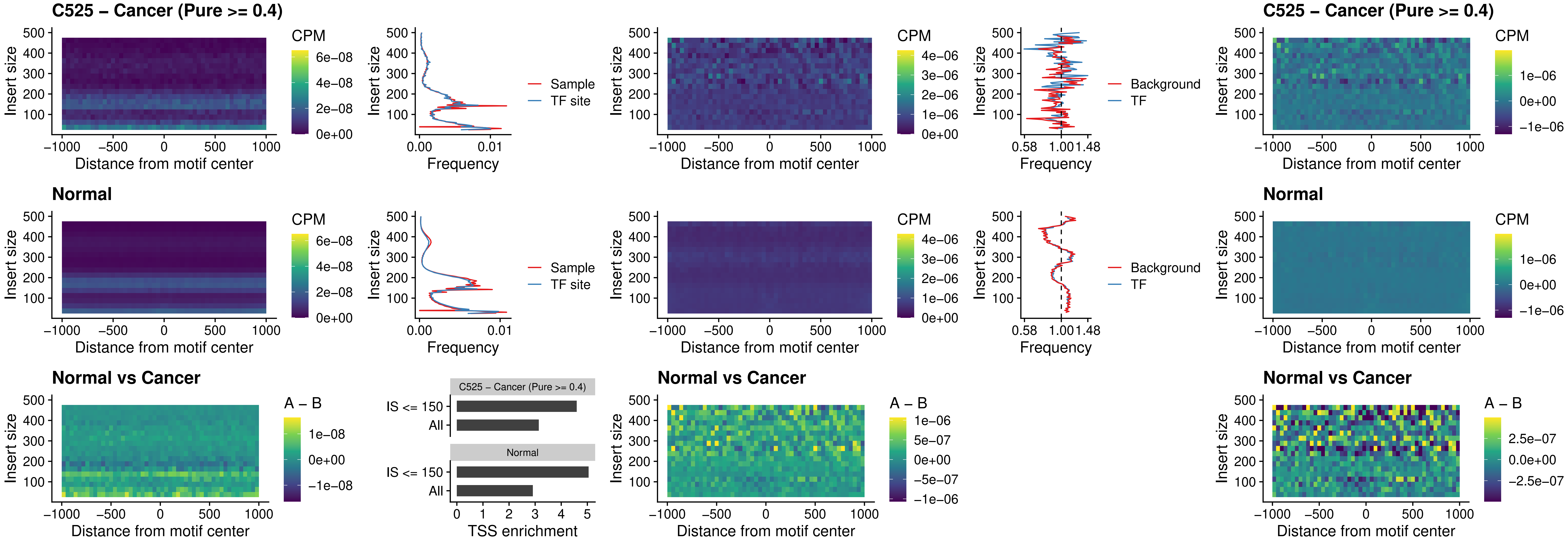

#### C525 – Cancer (Pure >= 0.4) – IRF9_uc_dTSS_oPEAK

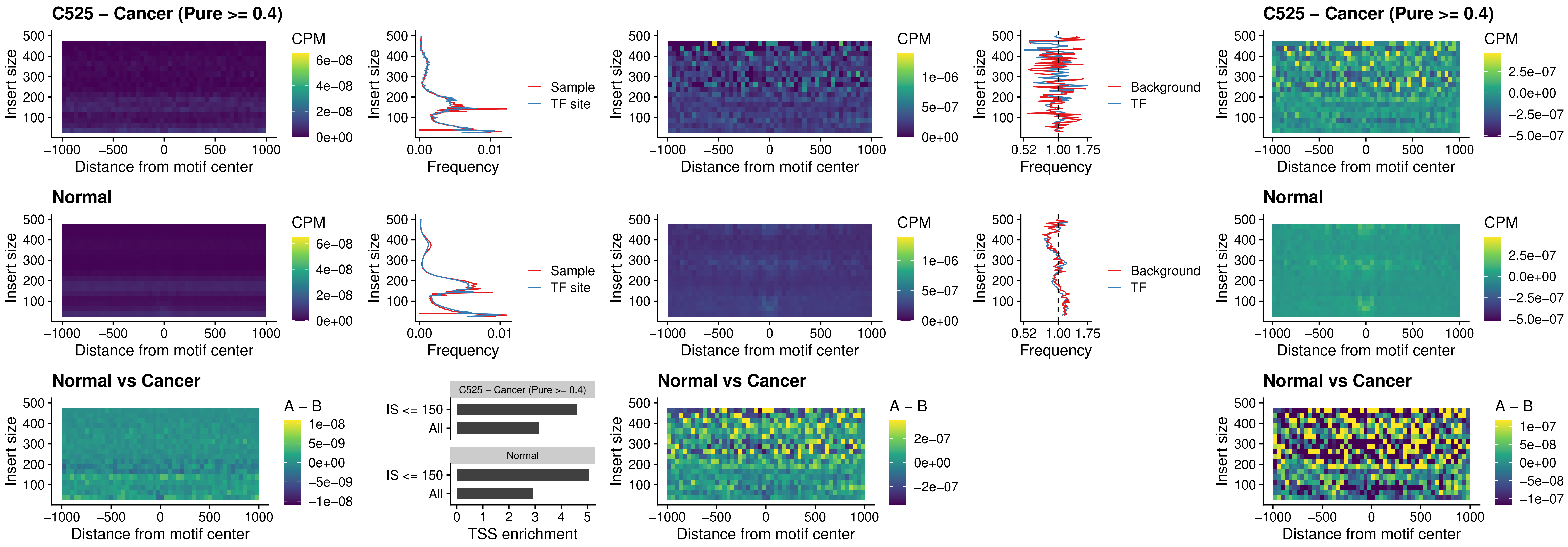

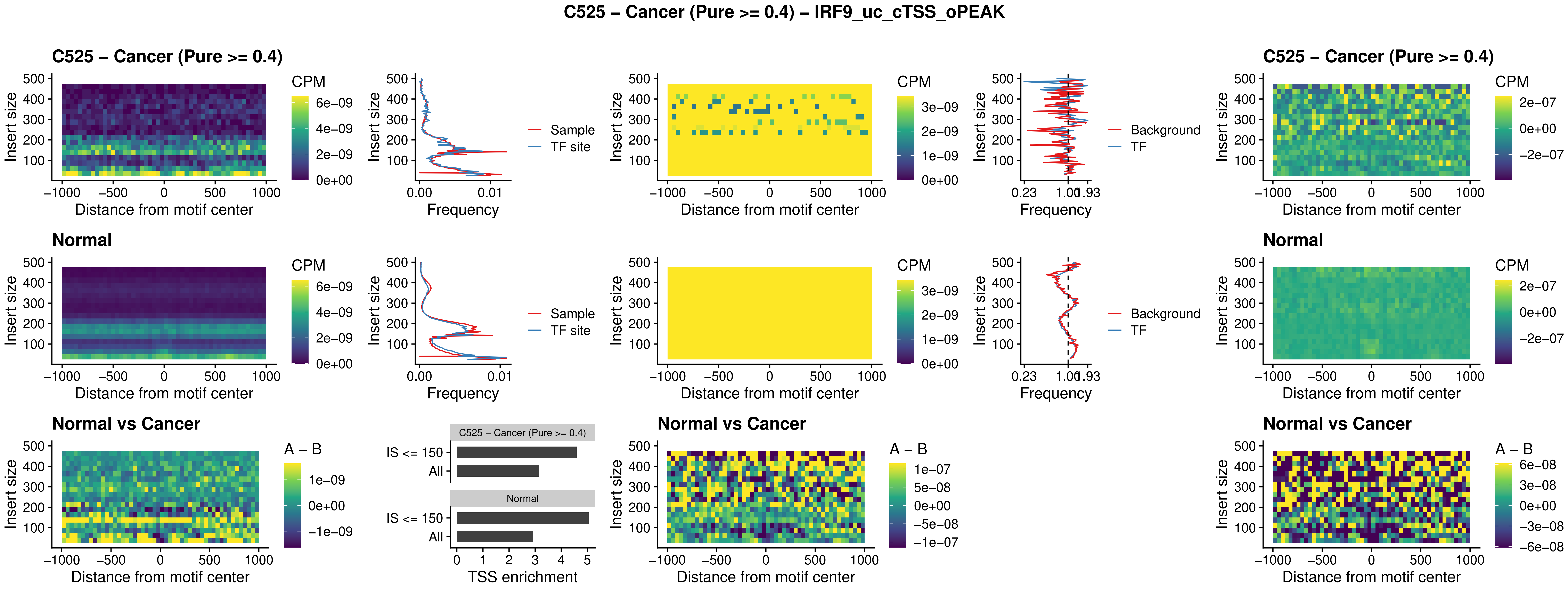
C525 – Cancer (Pure >= 0.4) – IRF9_uc_cTSS_oPEAK
C525 – Cancer (Pure >= 0.4)
Normal
Normal vs Cancer
Insert size
Distance from motif center
Frequency
CPM
A – B
Sample
TF site
Background
TF
C525 – Cancer (Pure >= 0.4)
Normal
IS <= 150
All
TSS enrichment

#### C525 – Cancer (Pure >= 0.4) – BCL11A_uc_cTSS_oPEAK

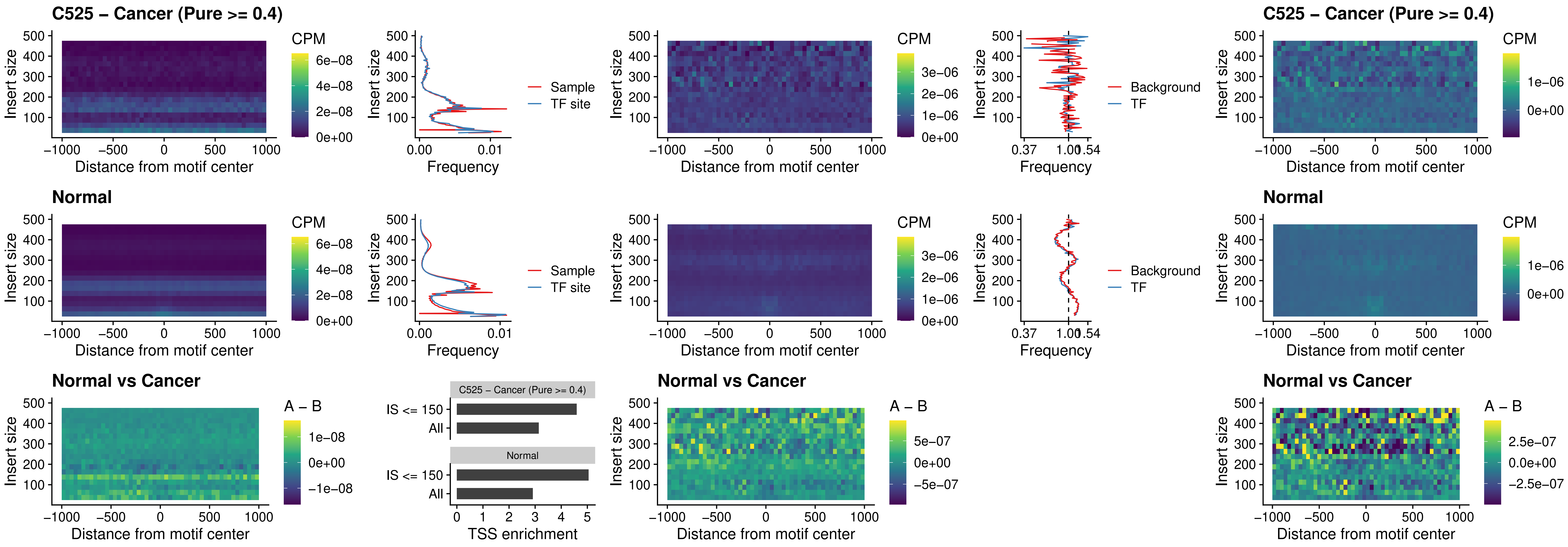

#### C525 – Cancer (Pure >= 0.4) – BCL11B_uc_cTSS_oPEAK

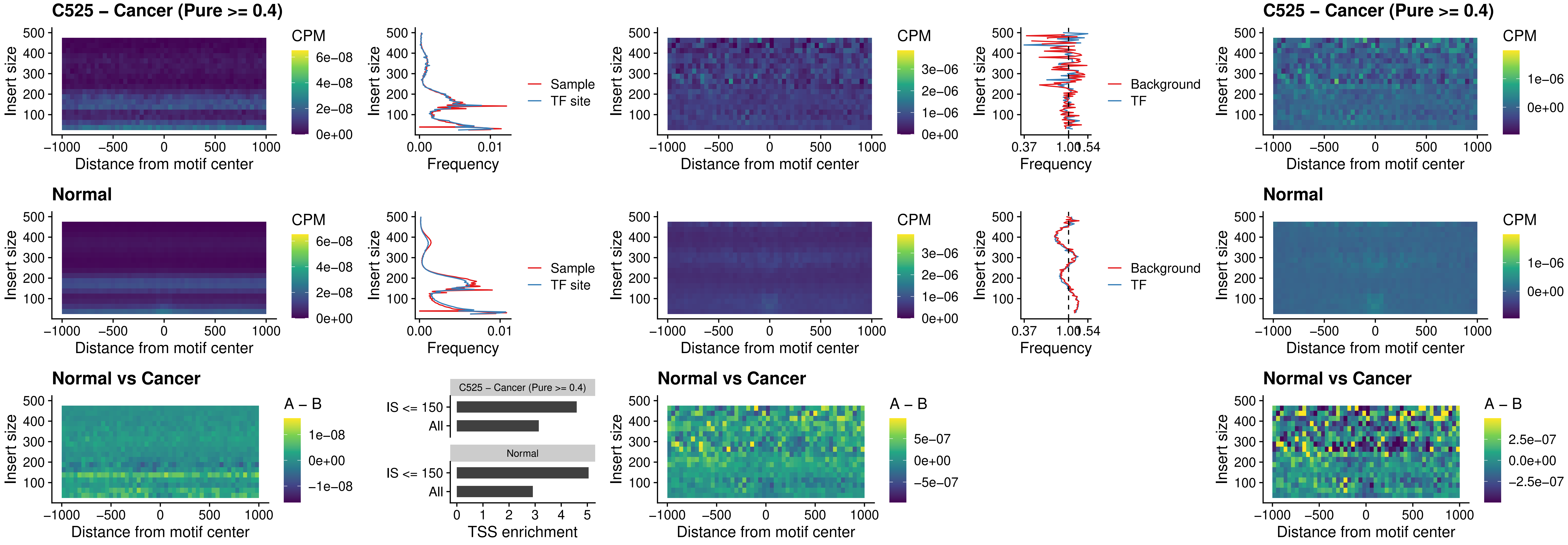

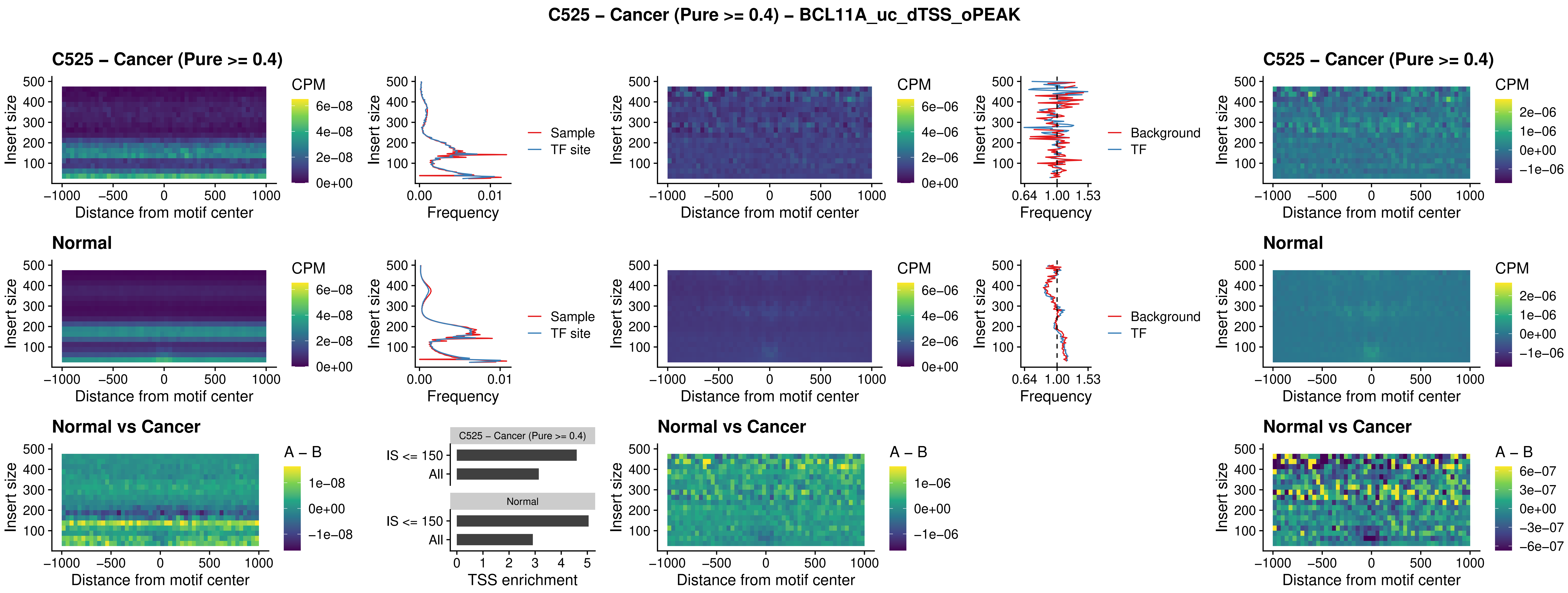
C525 – Cancer (Pure >= 0.4) – BCL11A_uc_dTSS_oPEAK
C525 – Cancer (Pure >= 0.4)
Normal
Normal vs Cancer
Insert size
Distance from motif center
Frequency
CPM
A – B
Sample
TF site
Background
TF
C525 – Cancer (Pure >= 0.4)
Normal
IS <= 150
All
TSS enrichment

#### C525 – Cancer (Pure >= 0.4) – BCL11B_uc_dTSS_oPEAK

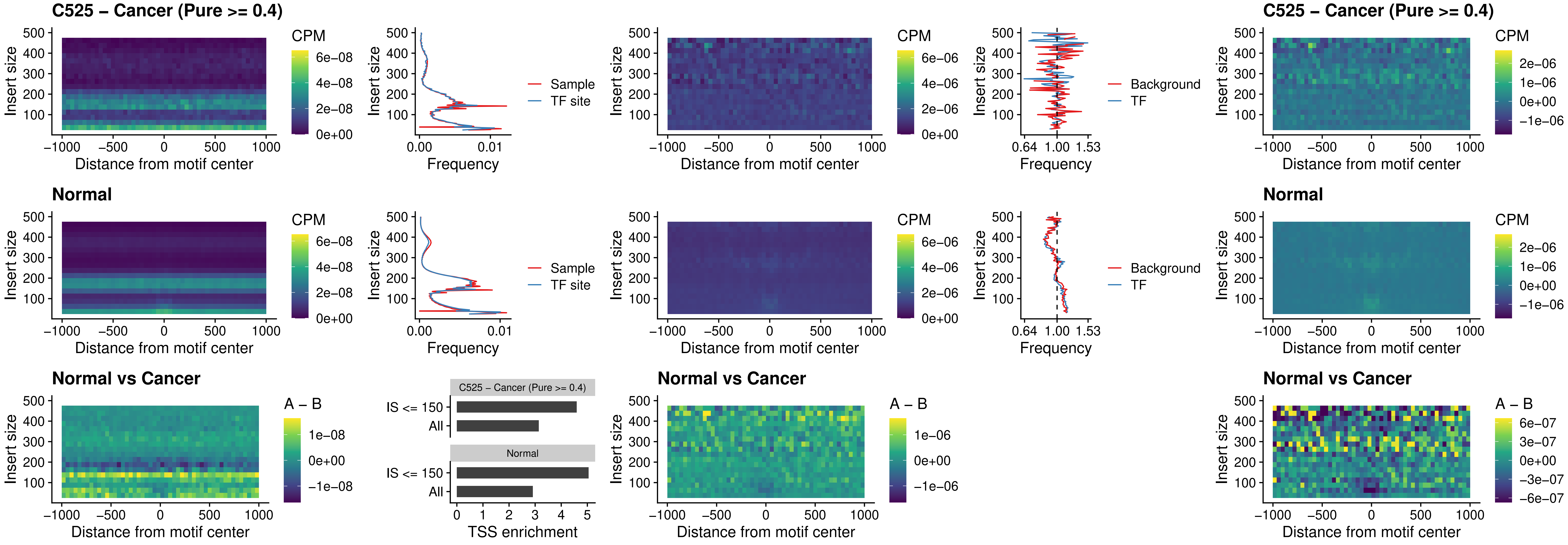

#### C525 – Cancer (Pure >= 0.4) – SPIB_uc_cTSS_nPEAK

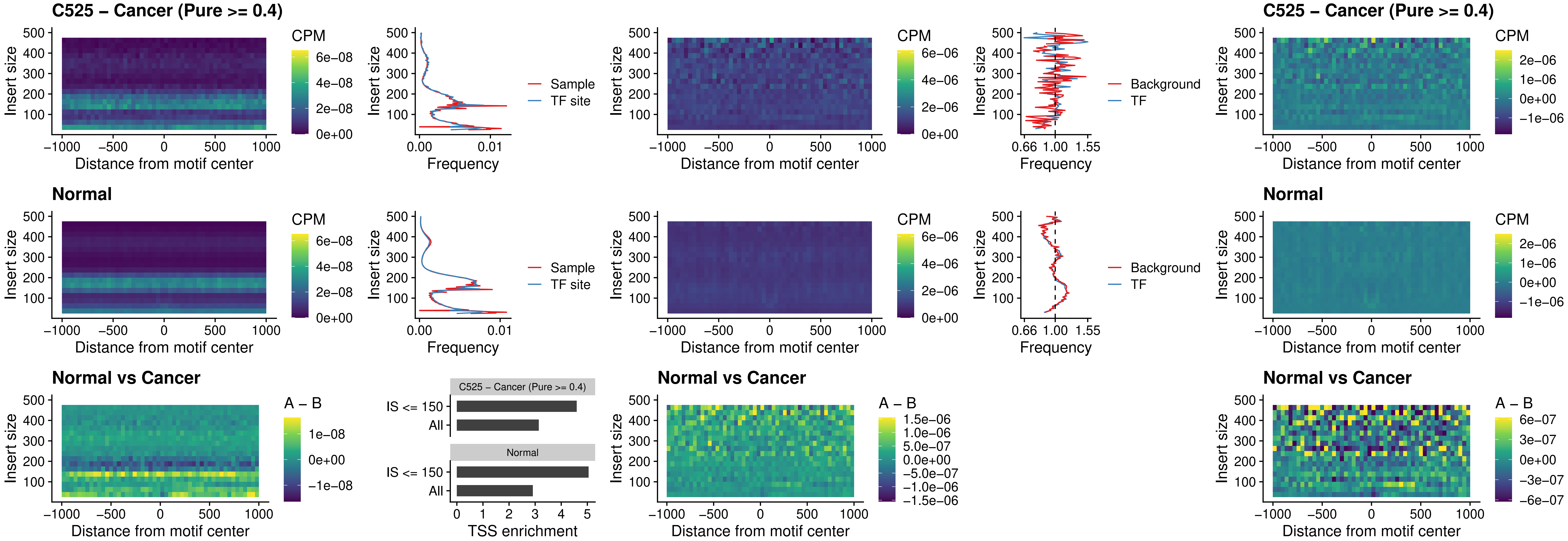

#### C525 – Cancer (Pure >= 0.4) – SPIC_uc_cTSS_nPEAK

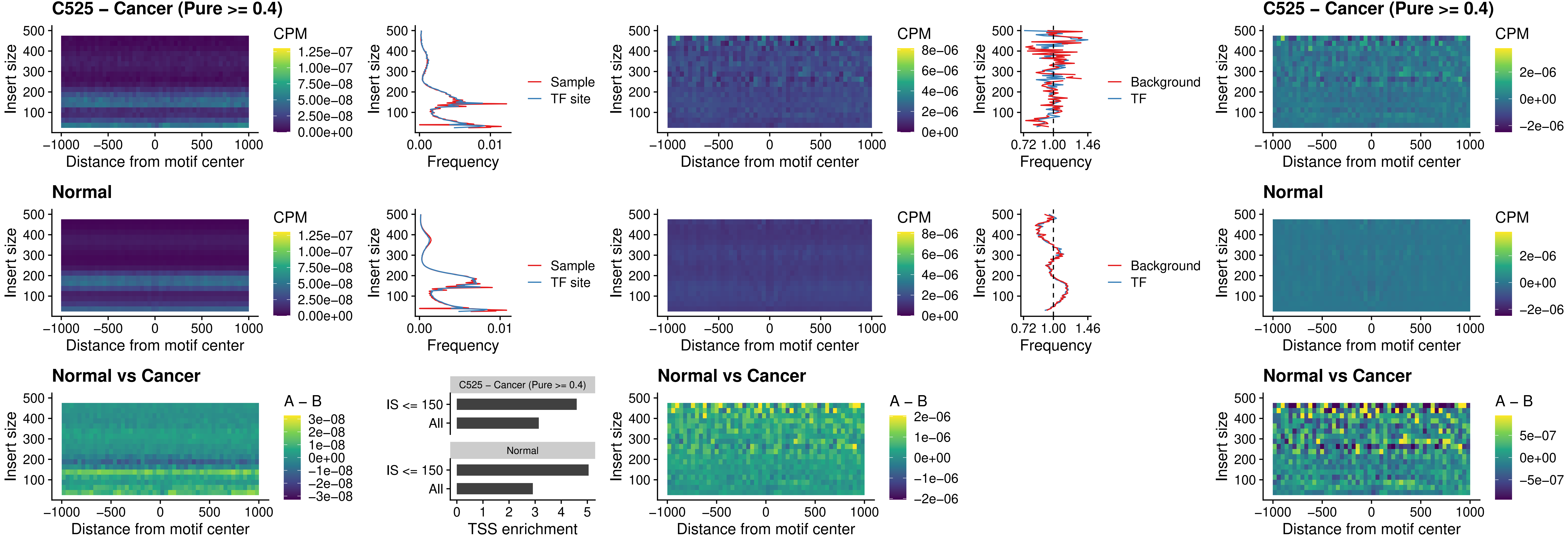

#### C525 – Cancer (Pure >= 0.4) – SPI1_uc_pTSS_oPEAK

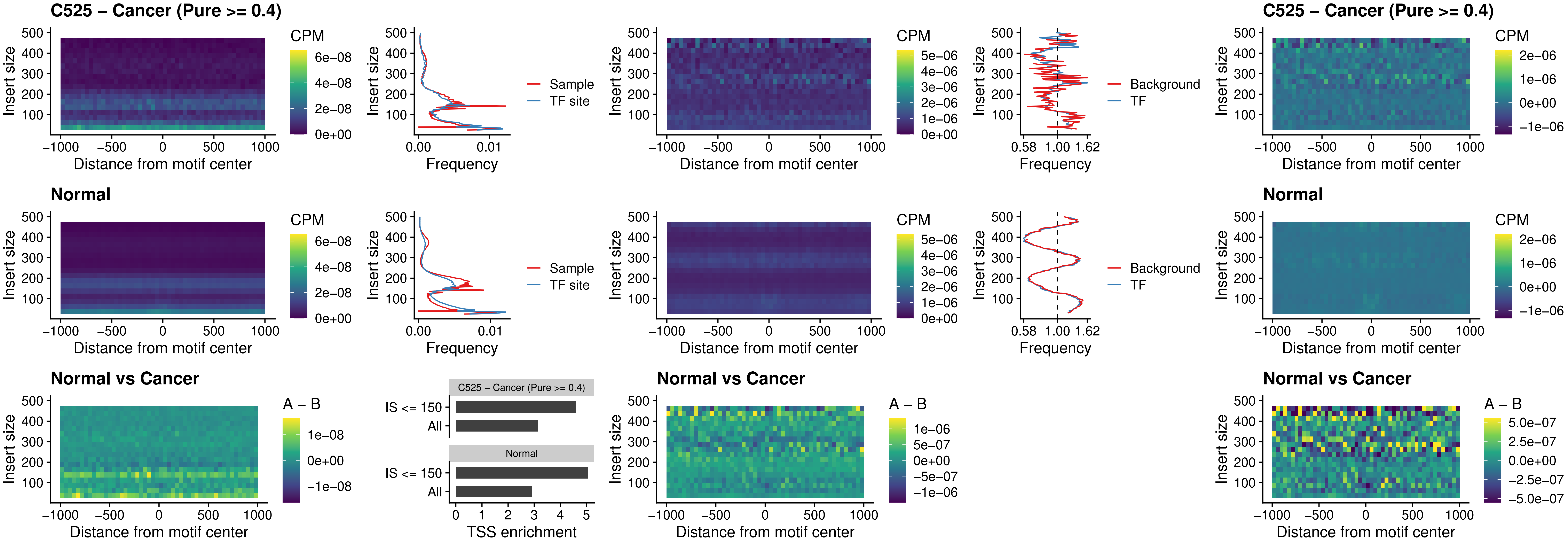

### C525 – Cancer (Pure >= 0.4) – BCL11A_uc_pTSS_oPEAK

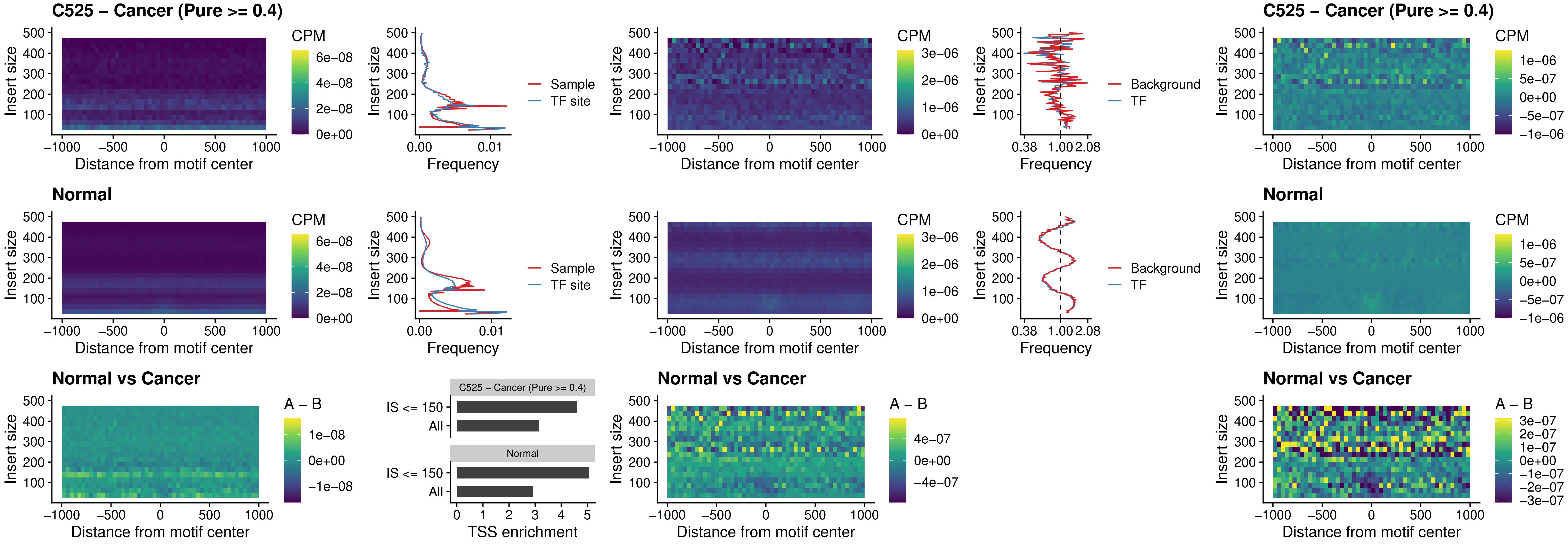

#### C525 – Cancer (Pure >= 0.4) – BCL11B_uc_pTSS_oPEAK

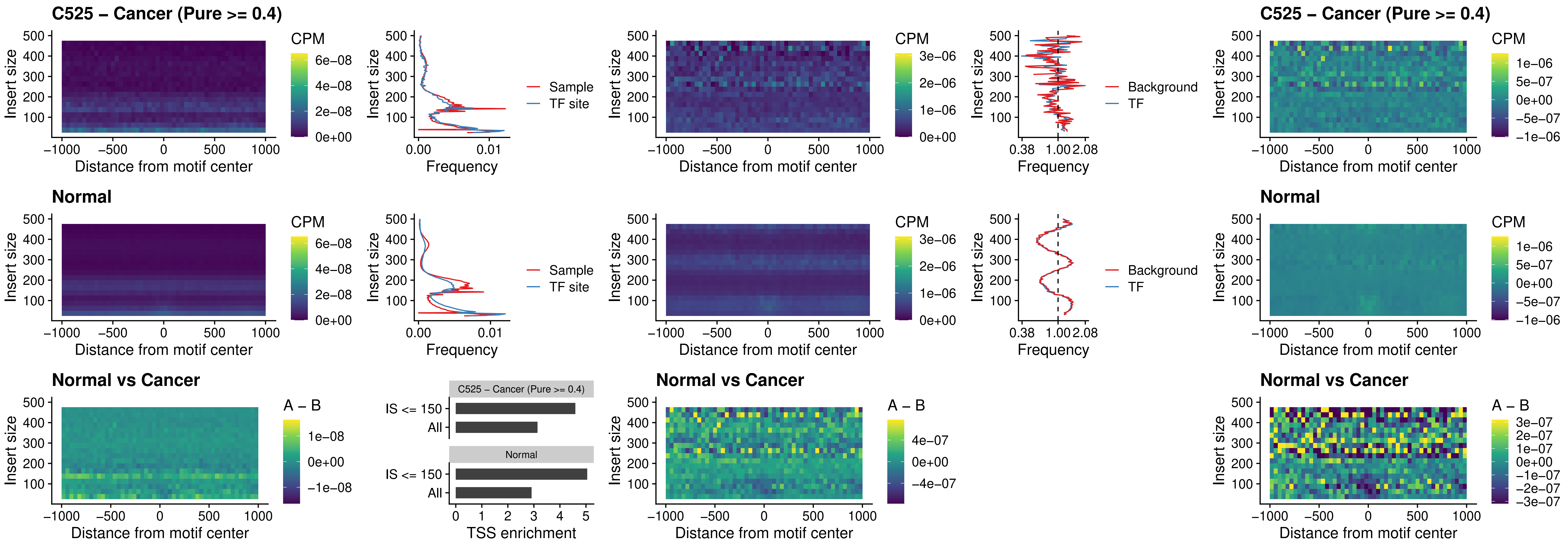

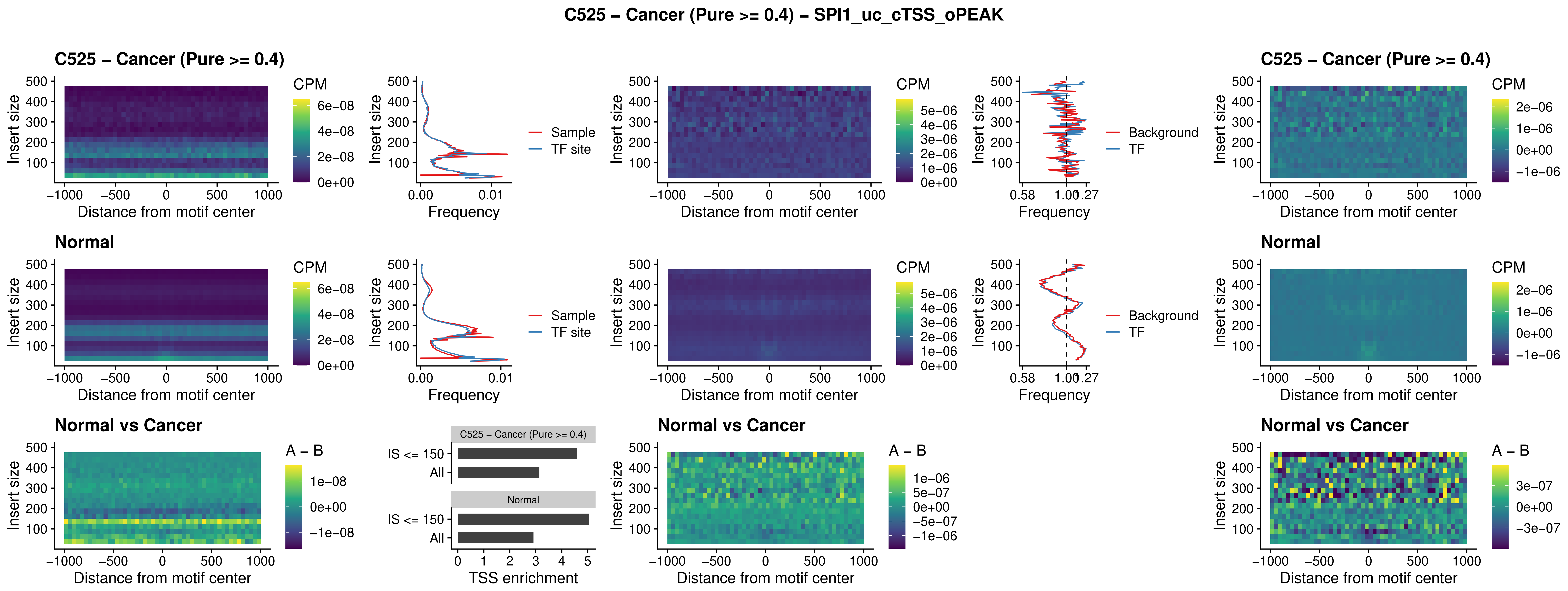
C525 – Cancer (Pure >= 0.4) – SPI1_uc_cTSS_oPEAK
C525 – Cancer (Pure >= 0.4)
Normal
Normal vs Cancer
Insert size
Distance from motif center
CPM
A – B
Frequency
Sample
TF site
Background
TF
TSS enrichment
C525 – Cancer (Pure >= 0.4)
IS <= 150
All

#### C525 – Cancer (Pure >= 0.4) – SPIB_uc_cTSS_oPEAK

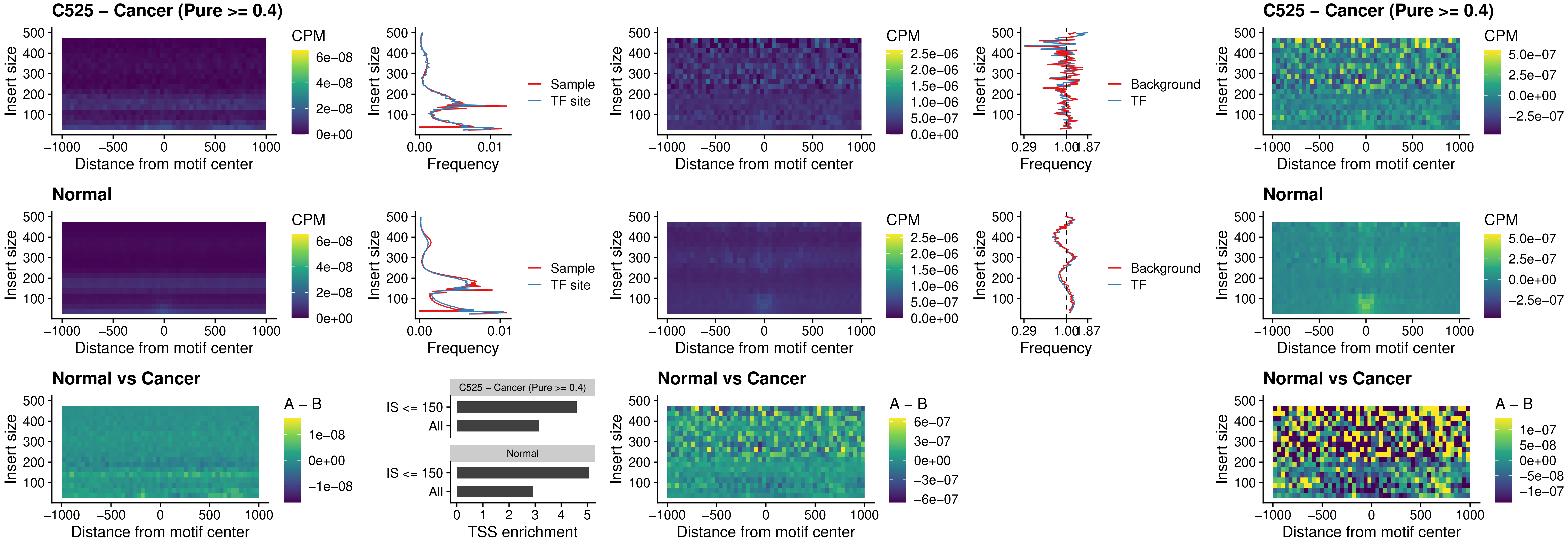

#### C525 – Cancer (Pure >= 0.4) – SPIB_uc_pTSS_oPEAK

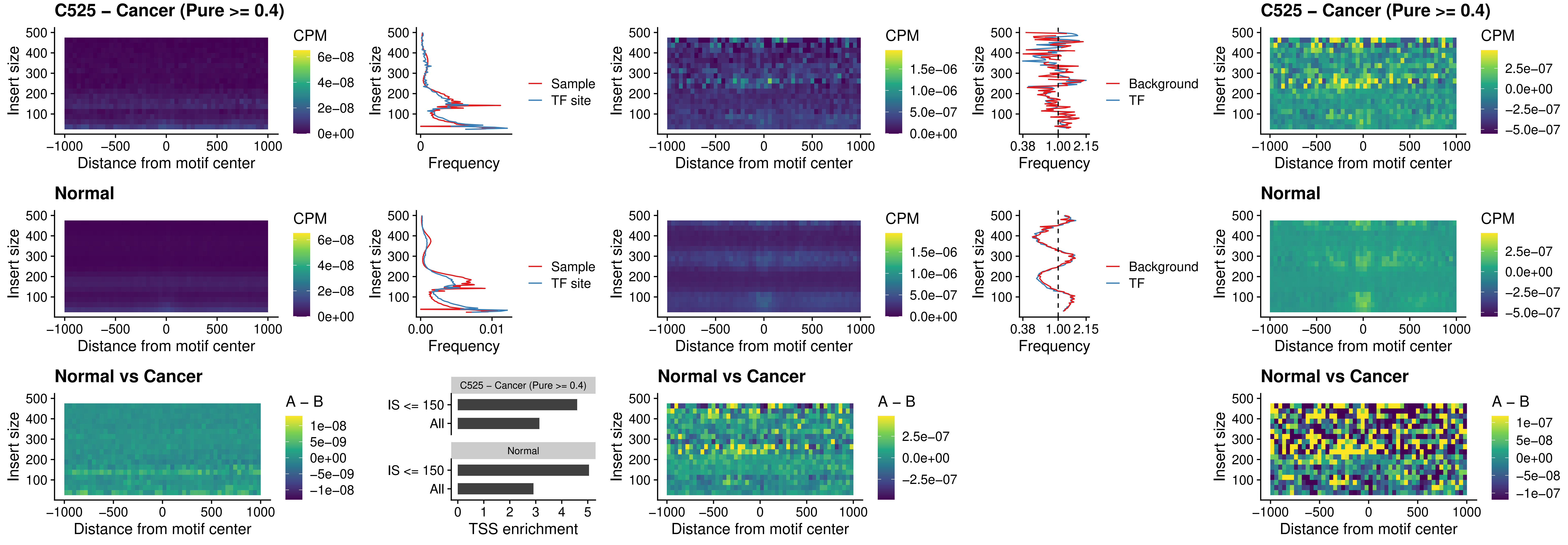

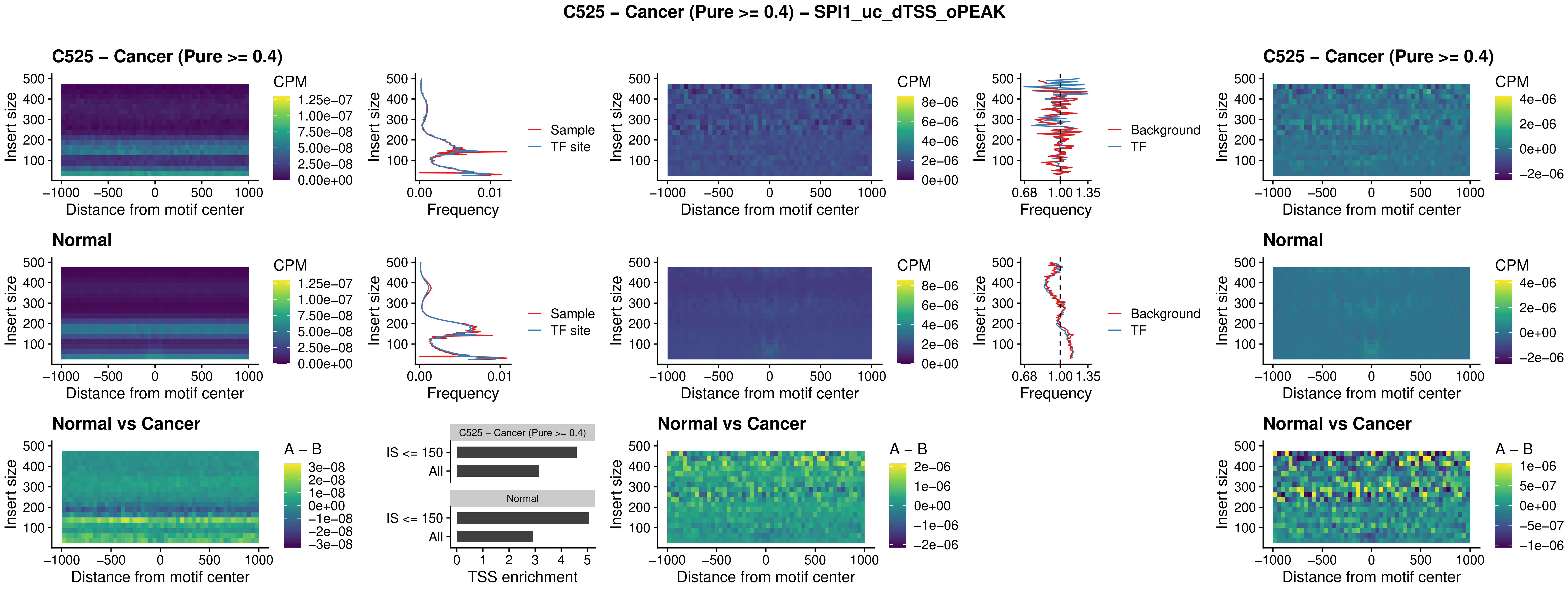
C525 – Cancer (Pure >= 0.4) – SPI1_uc_dTSS_oPEAK
C525 – Cancer (Pure >= 0.4)
Normal
Normal vs Cancer
Insert size
Distance from motif center
Frequency
CPM
A – B
Sample
TF site
Background
TF
TSS enrichment
IS <= 150
All
C525 – Cancer (Pure >= 0.4)
Normal

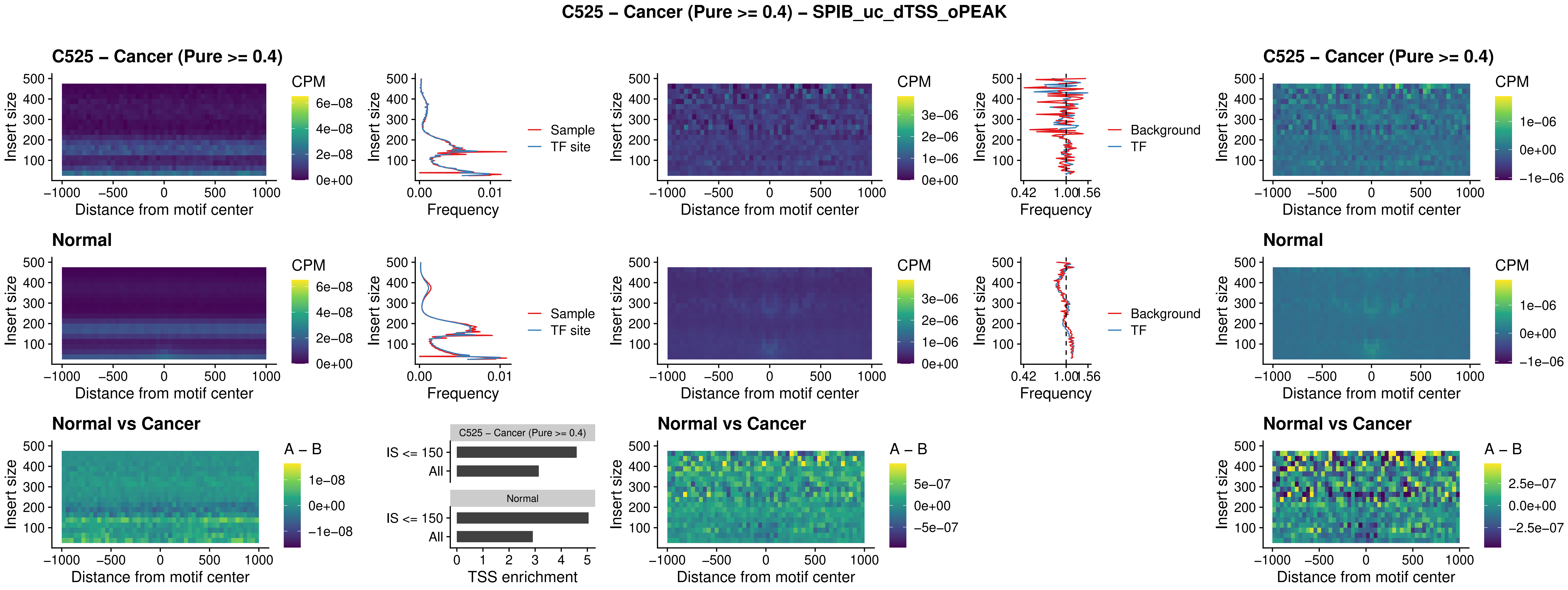
C525 – Cancer (Pure >= 0.4) – SPIB_uc_dTSS_oPEAK
C525 – Cancer (Pure >= 0.4)
Normal
Normal vs Cancer
Insert size
Distance from motif center
Frequency
CPM
A – B
Sample
TF site
Background
TF
C525 – Cancer (Pure >= 0.4)
Normal
IS <= 150
All
TSS enrichment

#### C525 – Cancer (Pure >= 0.4) – IRF3_uc_pTSS_oPEAK

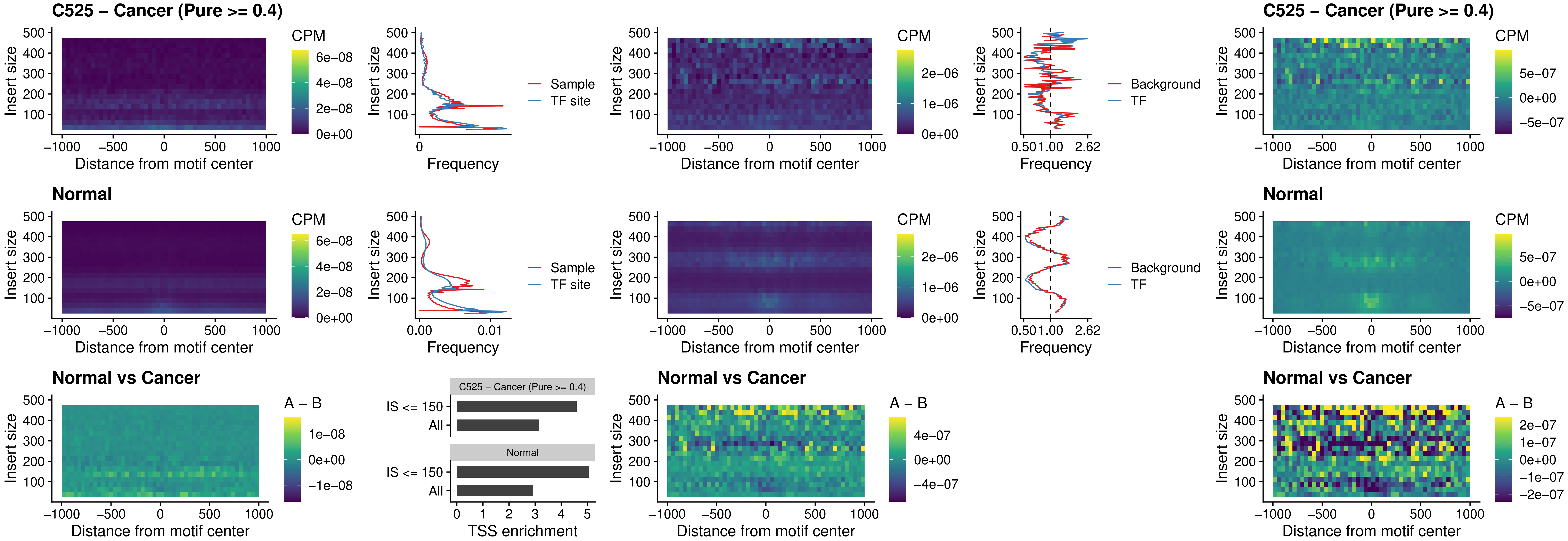

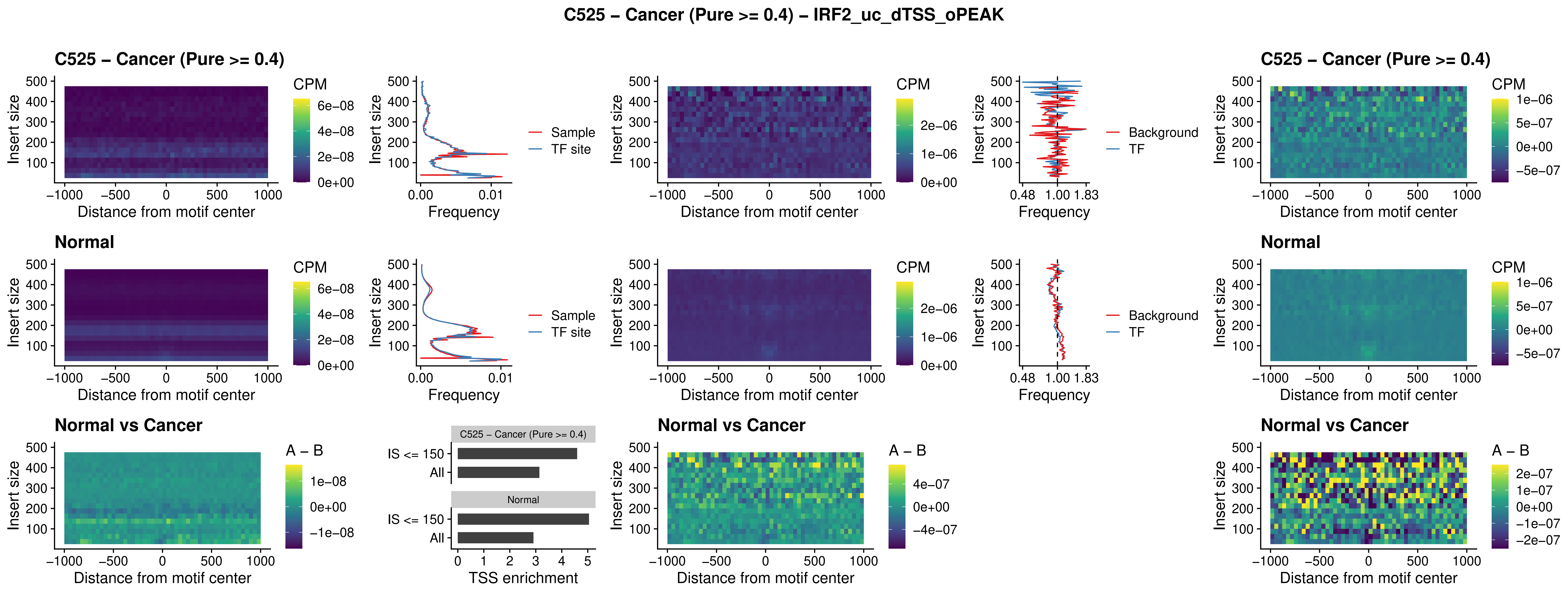
C525 – Cancer (Pure >= 0.4) – IRF2_uc_dTSS_oPEAK
C525 – Cancer (Pure >= 0.4)
Normal
Normal vs Cancer
Insert size
Distance from motif center
CPM
A – B
Frequency
Sample
TF site
Background
TF
C525 – Cancer (Pure >= 0.4)
Normal
IS <= 150
All
TSS enrichment

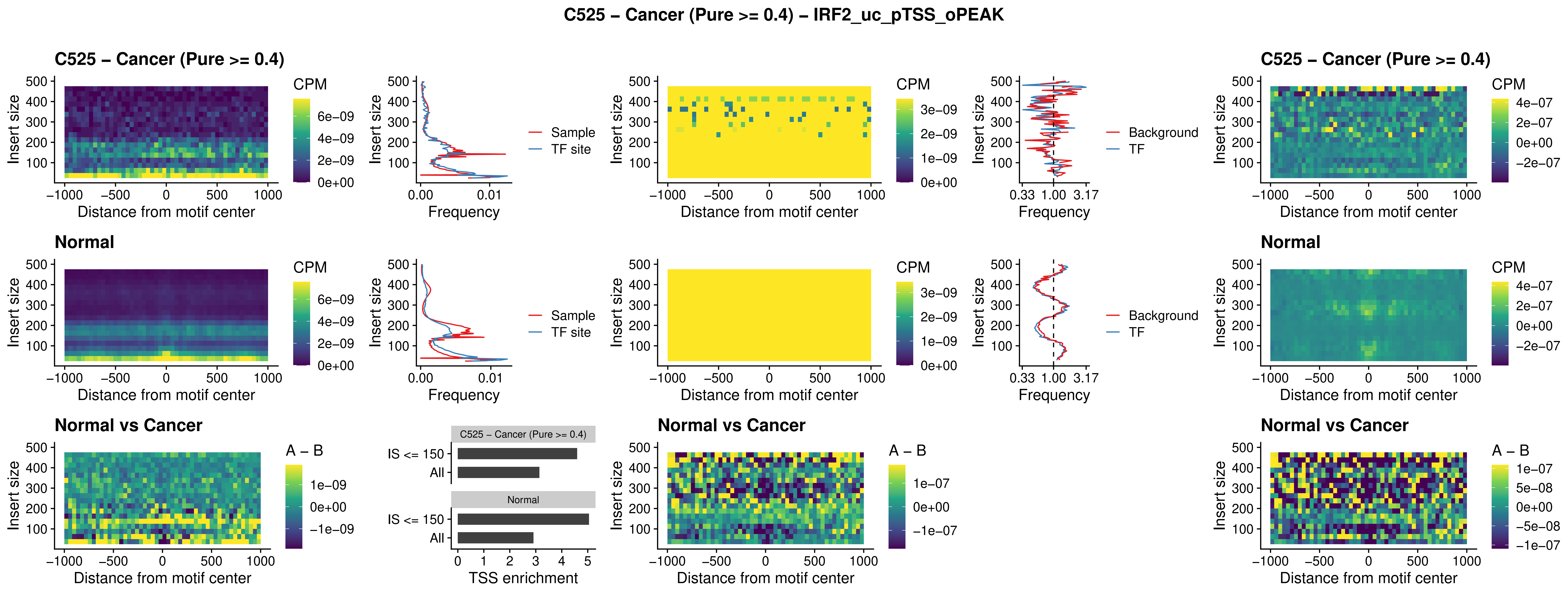

C525 – Cancer (Pure >= 0.4) – IRF2_uc_pTSS_oPEAK
C525 – Cancer (Pure >= 0.4)
Normal
Normal vs Cancer
Insert size
Distance from motif center
Frequency
CPM
A – B
Sample
TF site
Background
TF
C525 – Cancer (Pure >= 0.4)
Normal
IS <= 150
All
TSS enrichment

### C525 – Cancer (Pure >= 0.4) – STAT2_uc_pTSS_oPEAK

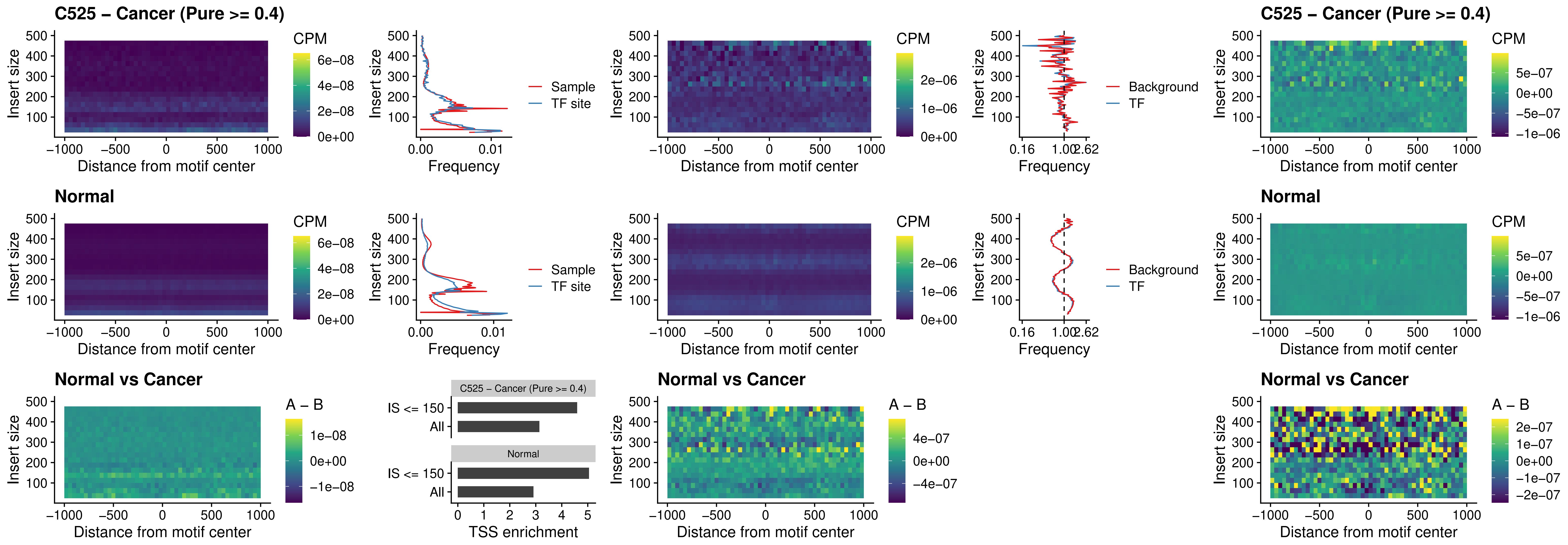

### C525 – Cancer (Pure >= 0.4) – SPIB_uc_dTSS_nPEAK

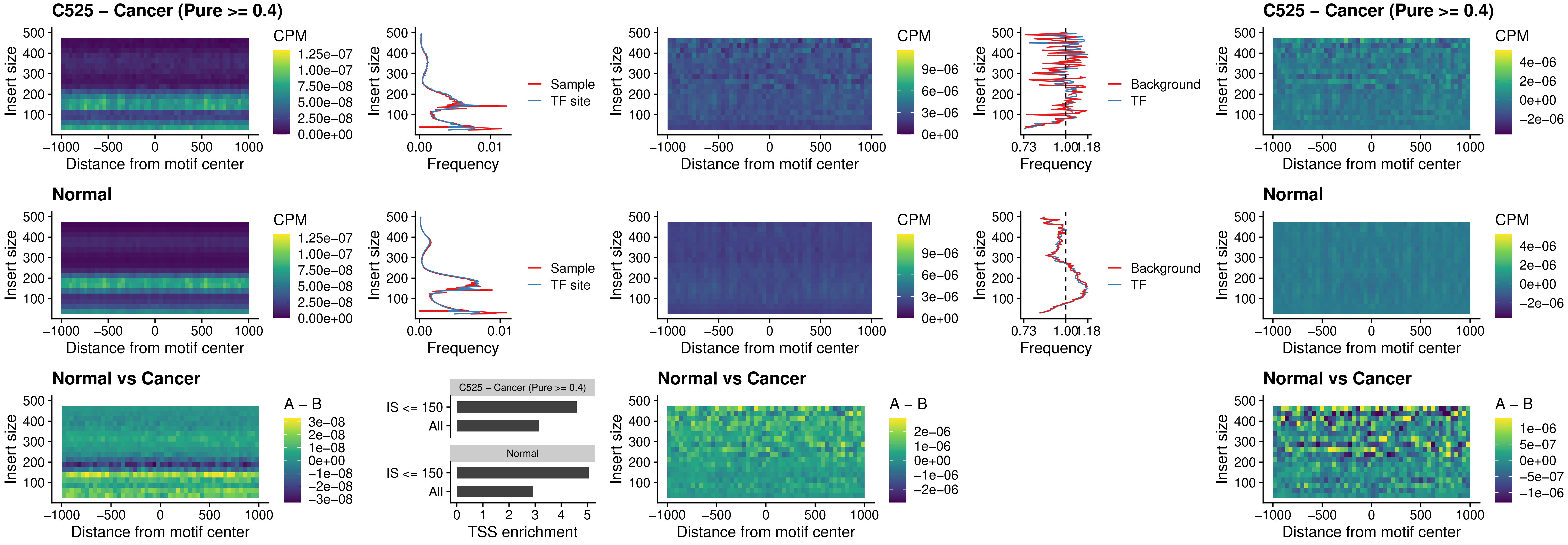

#### C525 – Cancer (Pure >= 0.4) – SPIC_uc_cTSS_oPEAK

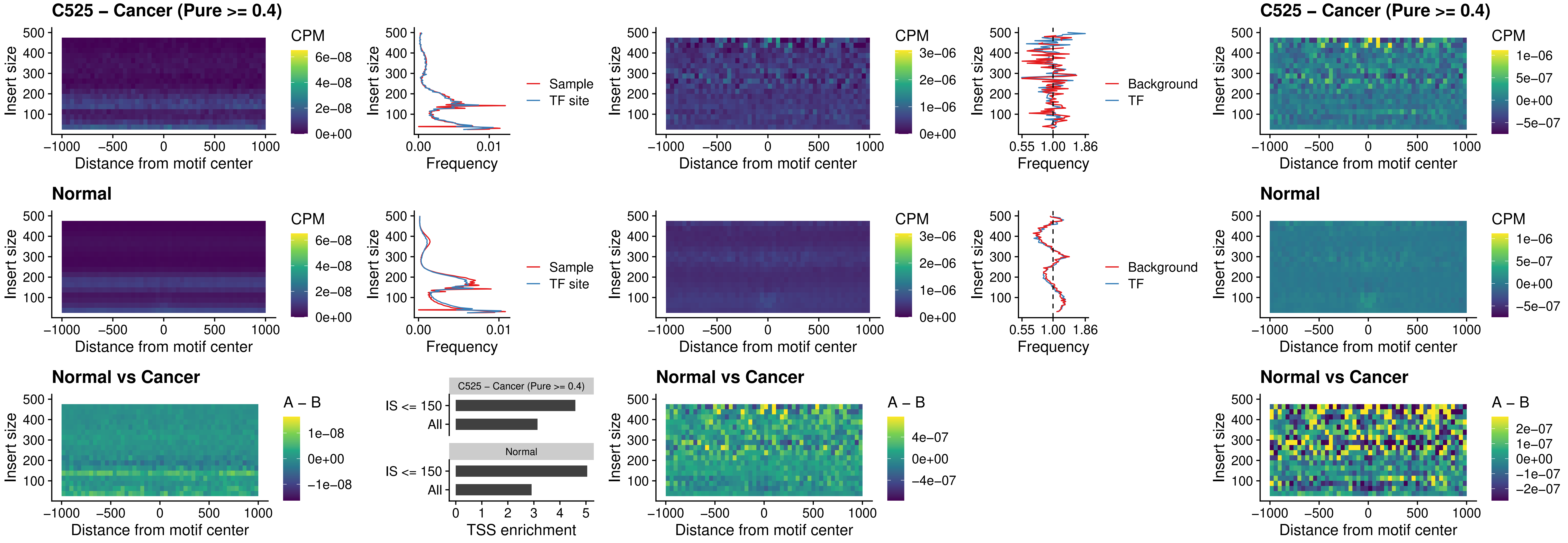

#### C525 – Cancer (Pure >= 0.4) – PRDM1_uc_cTSS_oPEAK

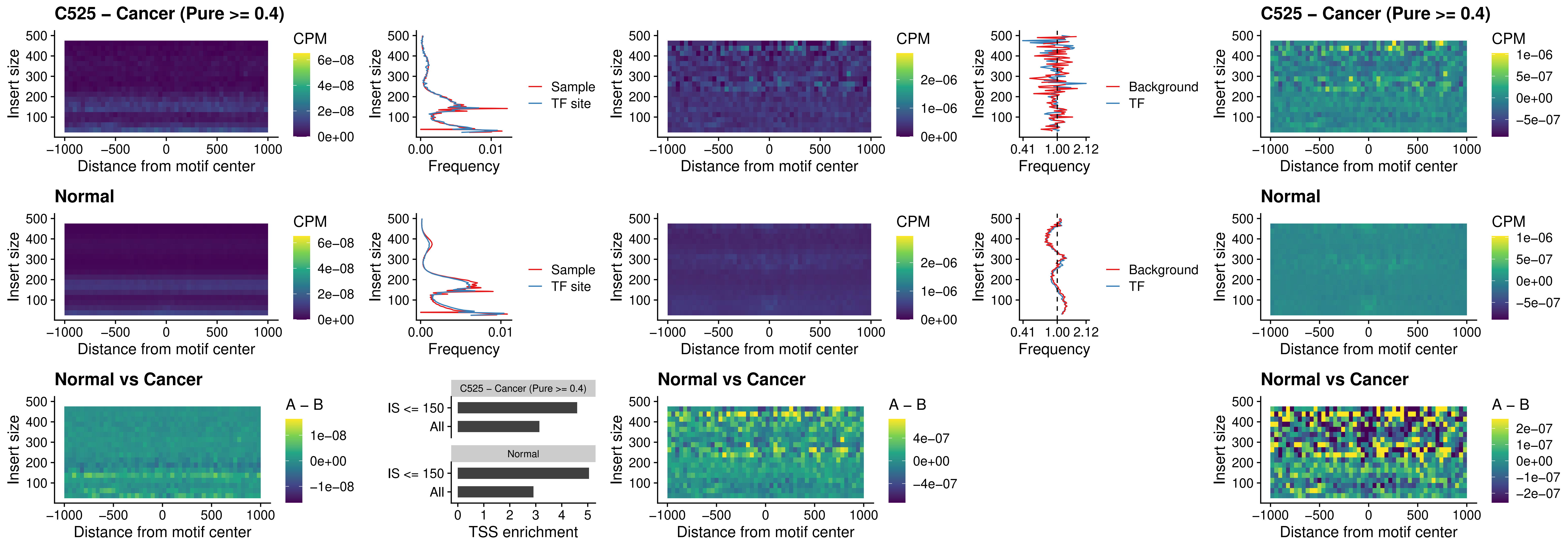

#### C525 – Cancer (Pure >= 0.4) – IRF3_uc_cTSS_oPEAK

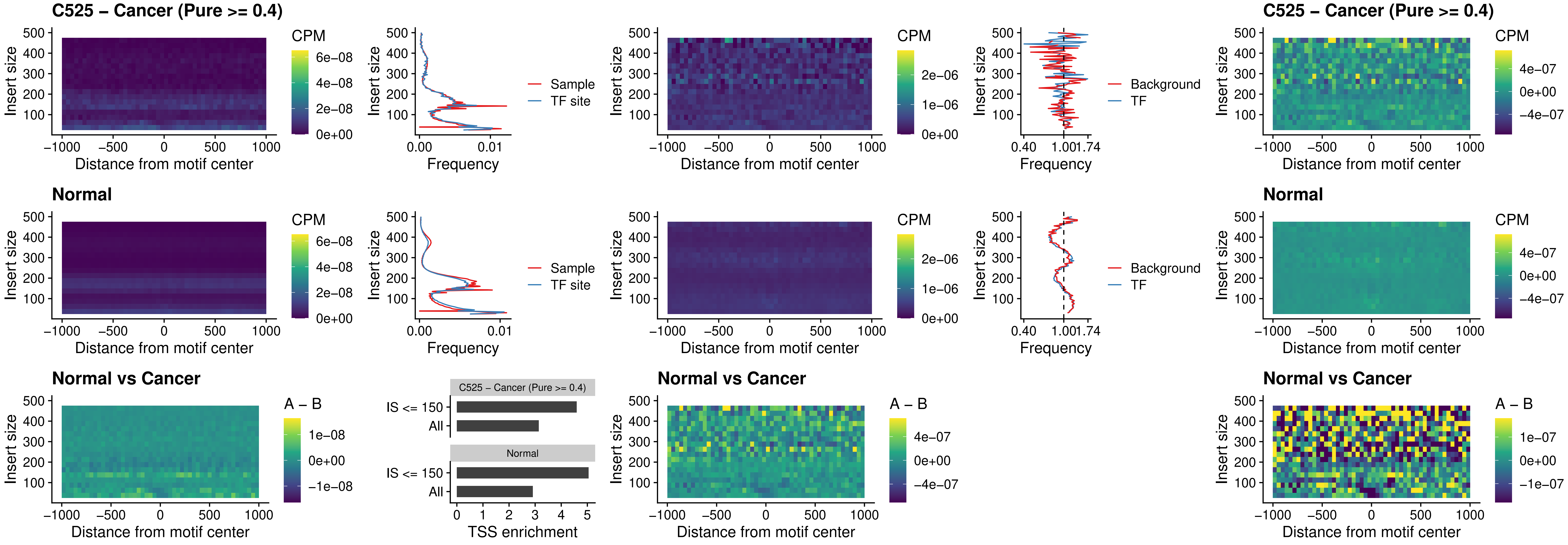

#### C525 – Cancer (Pure >= 0.4) – BCL11A_uc_dTSS_nPEAK

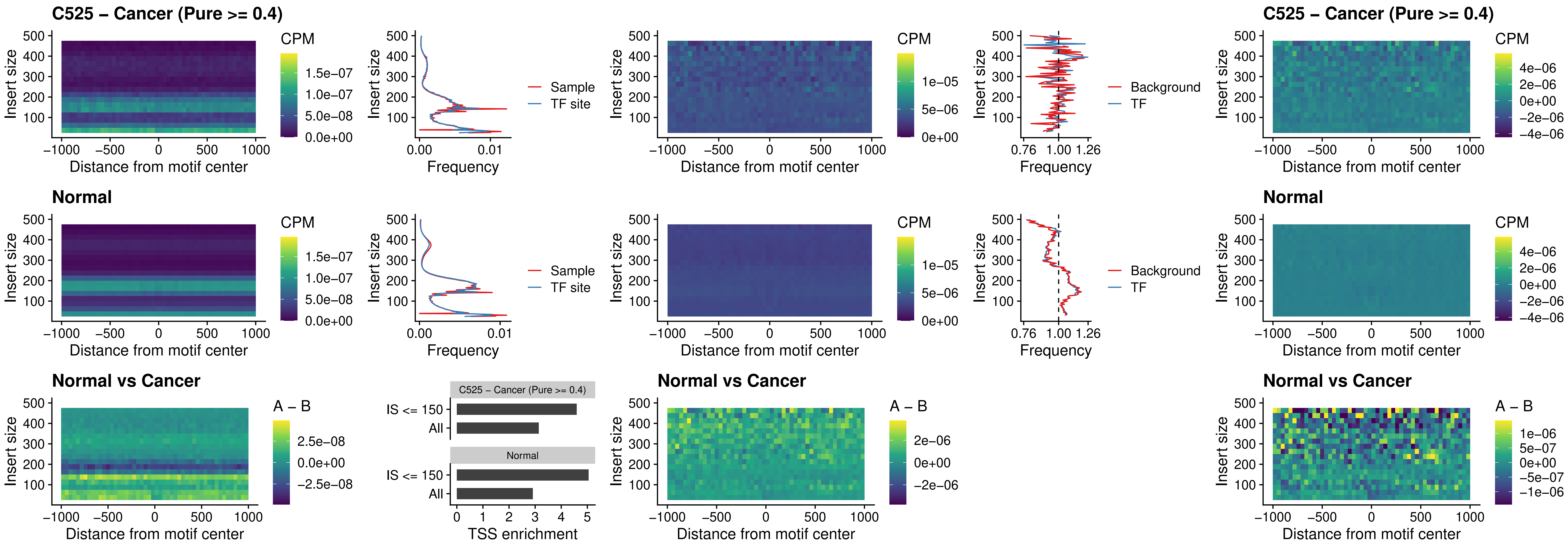

#### C525 – Cancer (Pure >= 0.4) – BCL11B_uc_dTSS_nPEAK

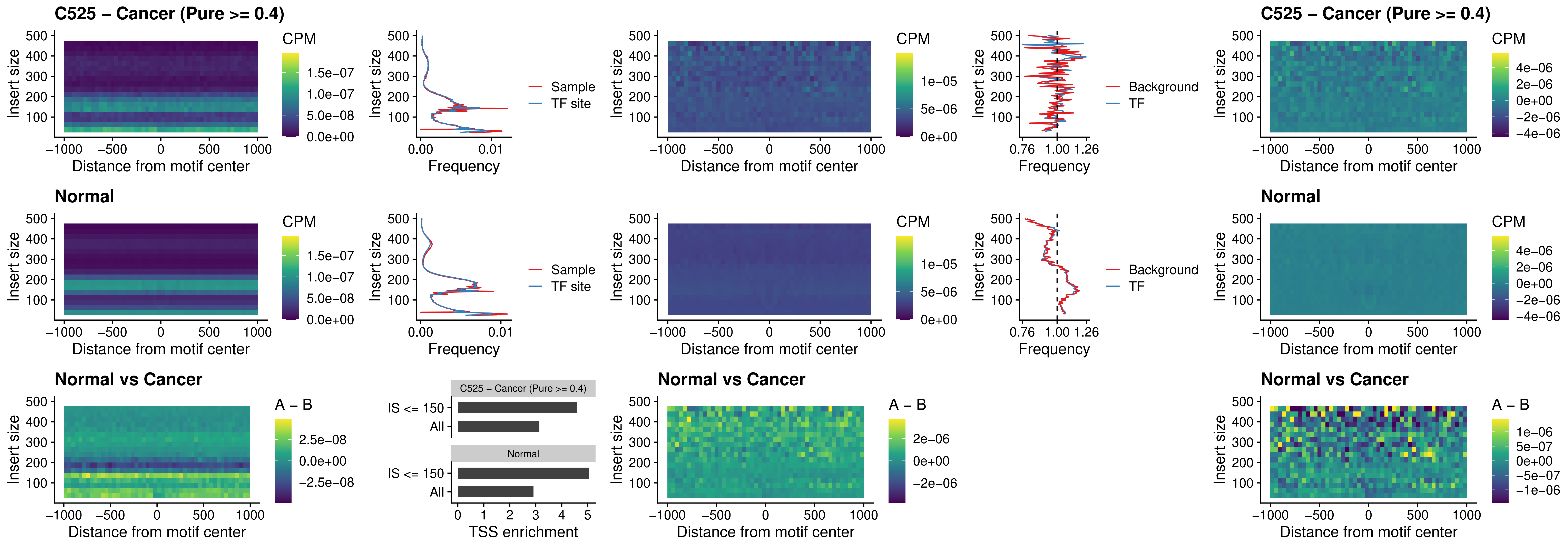

### C525 – Cancer (Pure >= 0.4) – IRF1_uc_cTSS_oPEAK

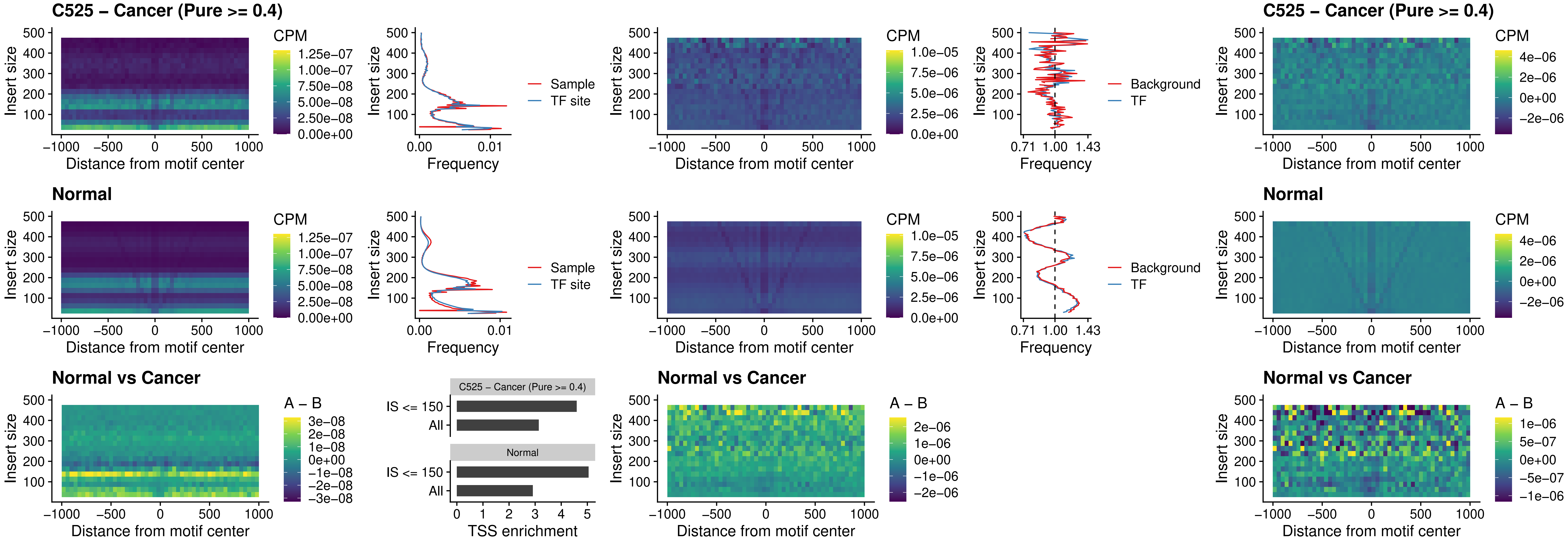

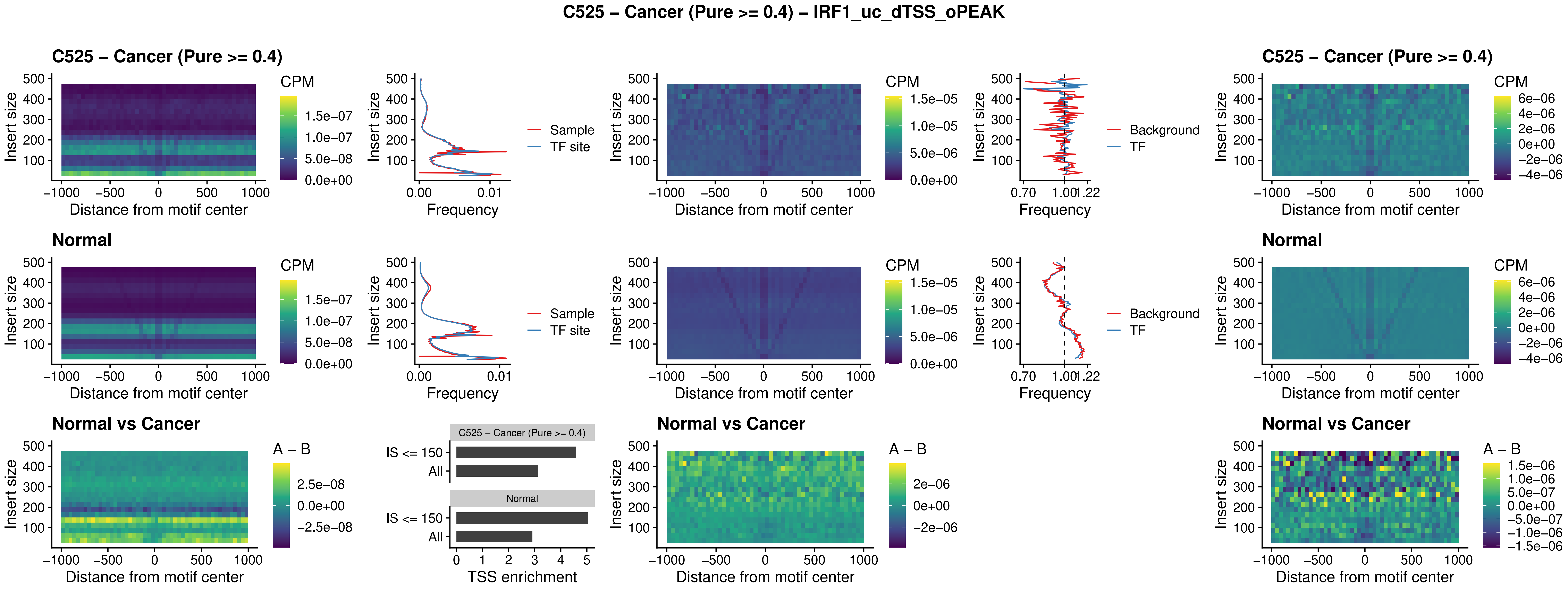
C525 – Cancer (Pure >= 0.4) – IRF1_uc_dTSS_oPEAK
C525 – Cancer (Pure >= 0.4)
Normal
Normal vs Cancer
Insert size
Distance from motif center
Frequency
CPM
A – B
Sample
TF site
Background
TF
C525 – Cancer (Pure >= 0.4)
Normal
IS <= 150
All
TSS enrichment

#### C525 – Cancer (Pure >= 0.4) – IRF1_uc_pTSS_oPEAK

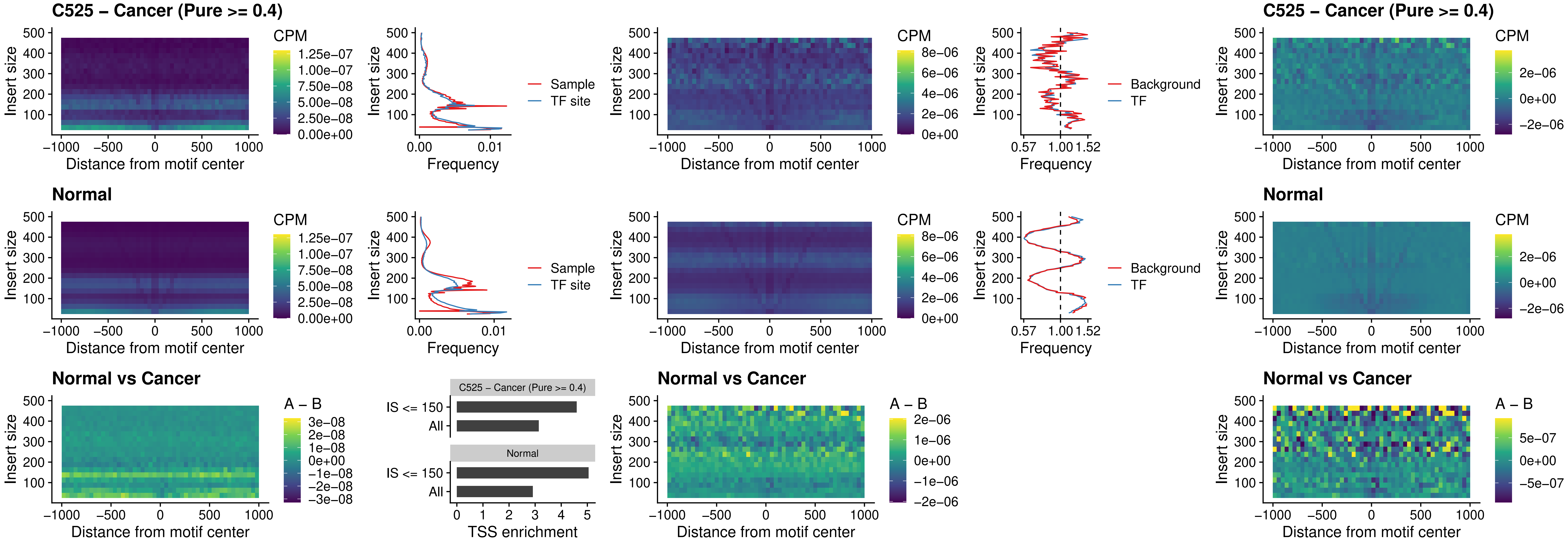

### C525 – Cancer (Pure >= 0.4) – SPIC_uc_dTSS_nPEAK

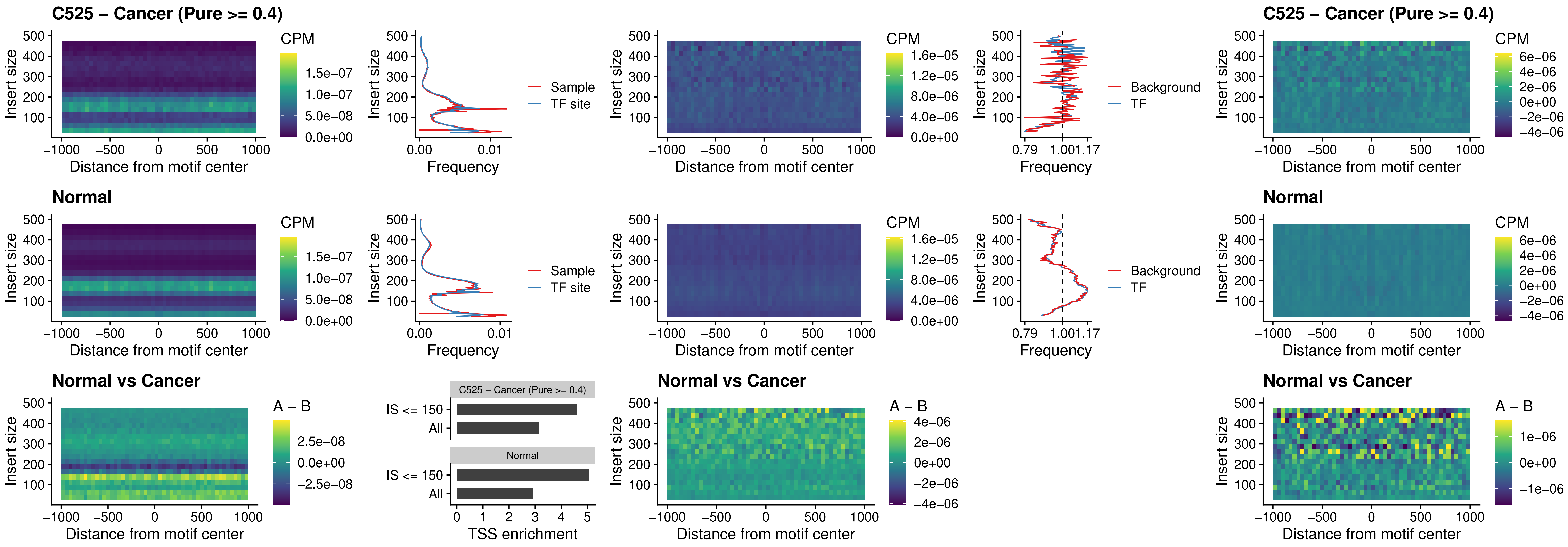

#### C525 – Cancer (Pure >= 0.4) – SPIC_uc_dTSS_oPEAK

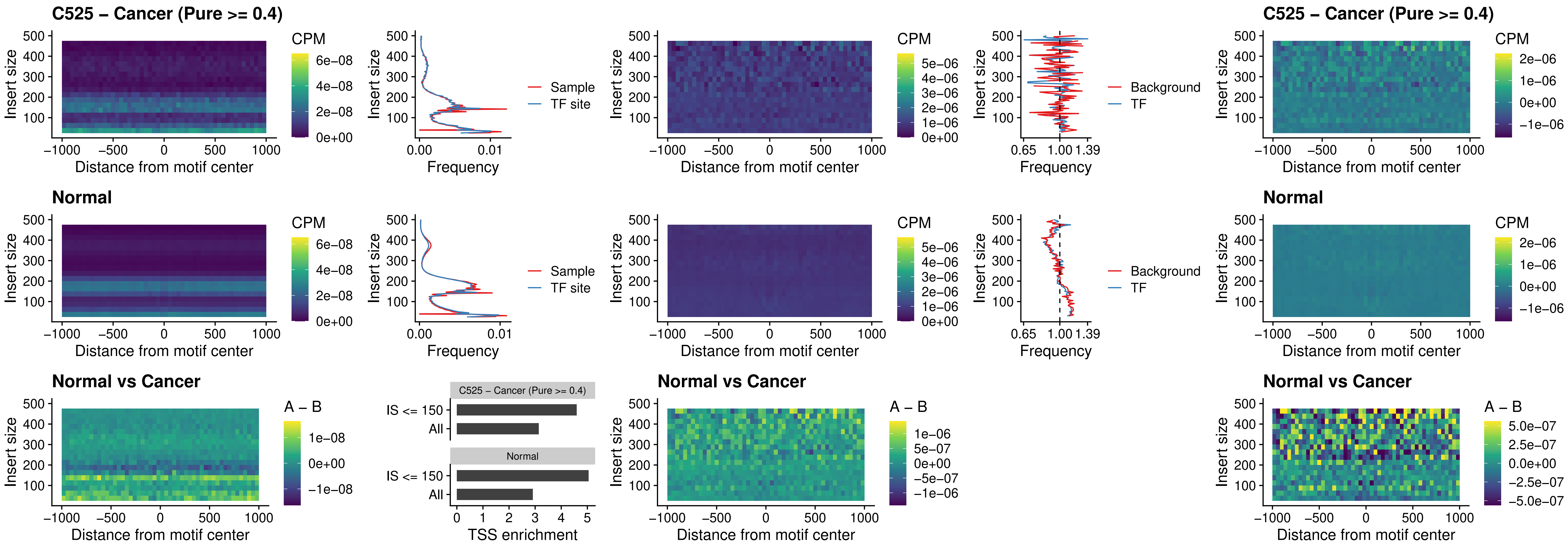

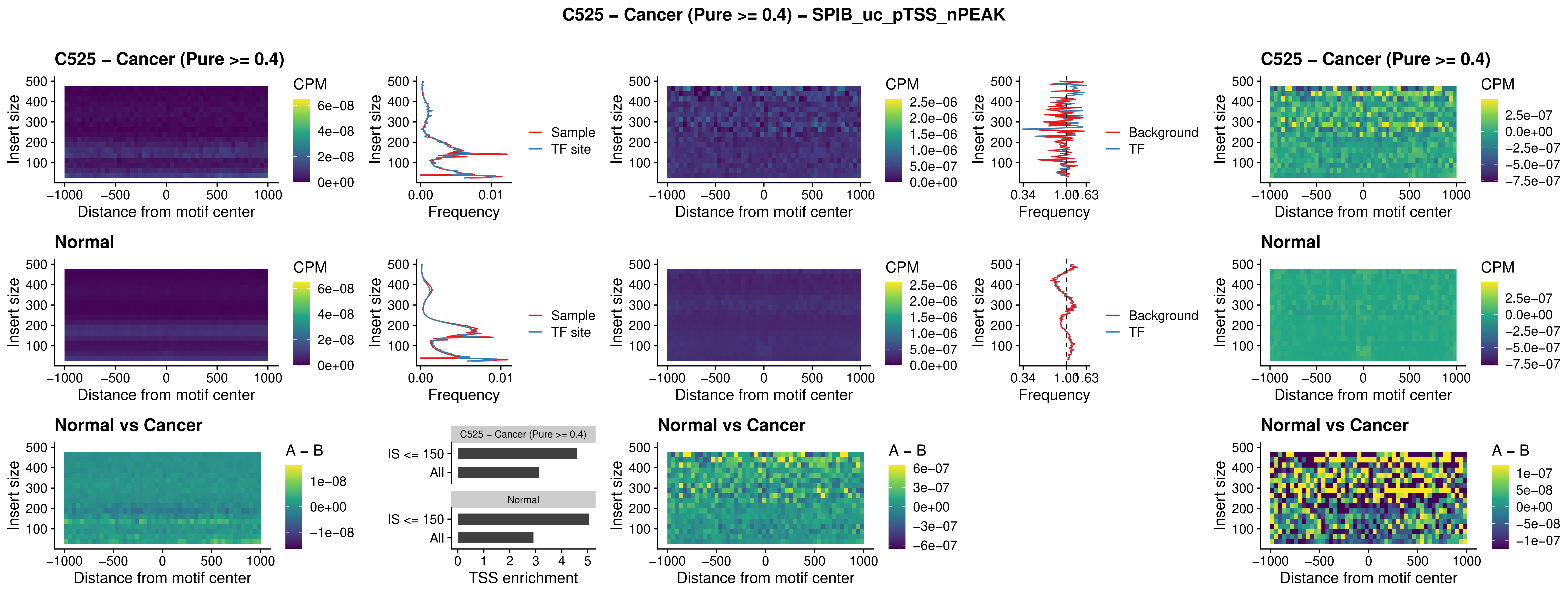
C525 – Cancer (Pure >= 0.4) – SPIB_uc_pTSS_nPEAK
C525 – Cancer (Pure >= 0.4)
Normal
Normal vs Cancer
Insert size
Distance from motif center
Frequency
CPM
A – B
Sample
TF site
Background
TF
TSS enrichment
IS <= 150
All

#### C525 – Cancer (Pure >= 0.4) – STAT2_uc_cTSS_oPEAK

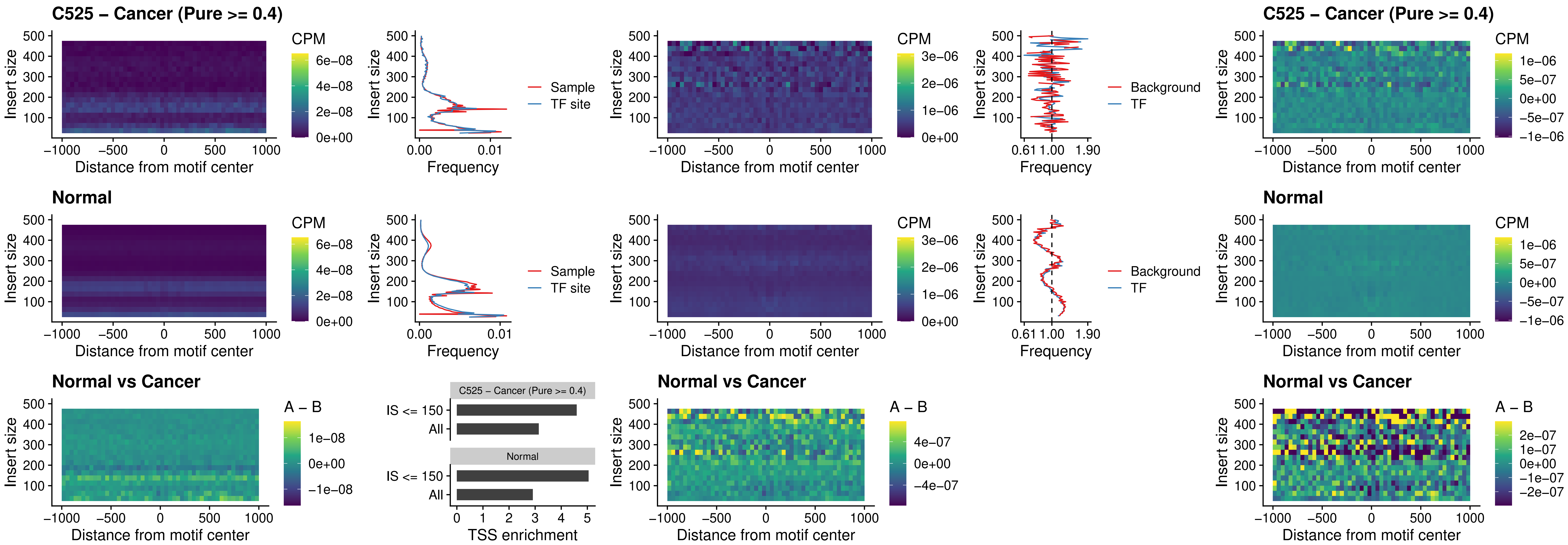

#### C525 – Cancer (Pure >= 0.4) – IRF8_uc_cTSS_oPEAK

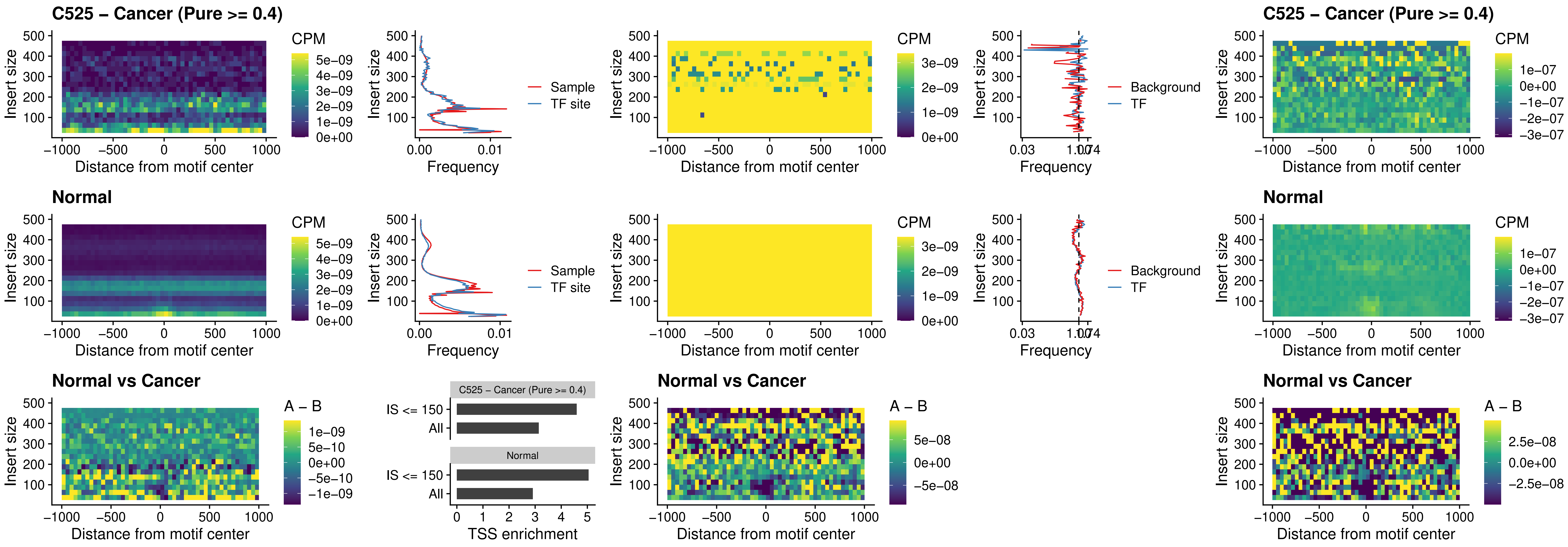

#### C525 – Cancer (Pure >= 0.4) – IRF3_uc_dTSS_oPEAK

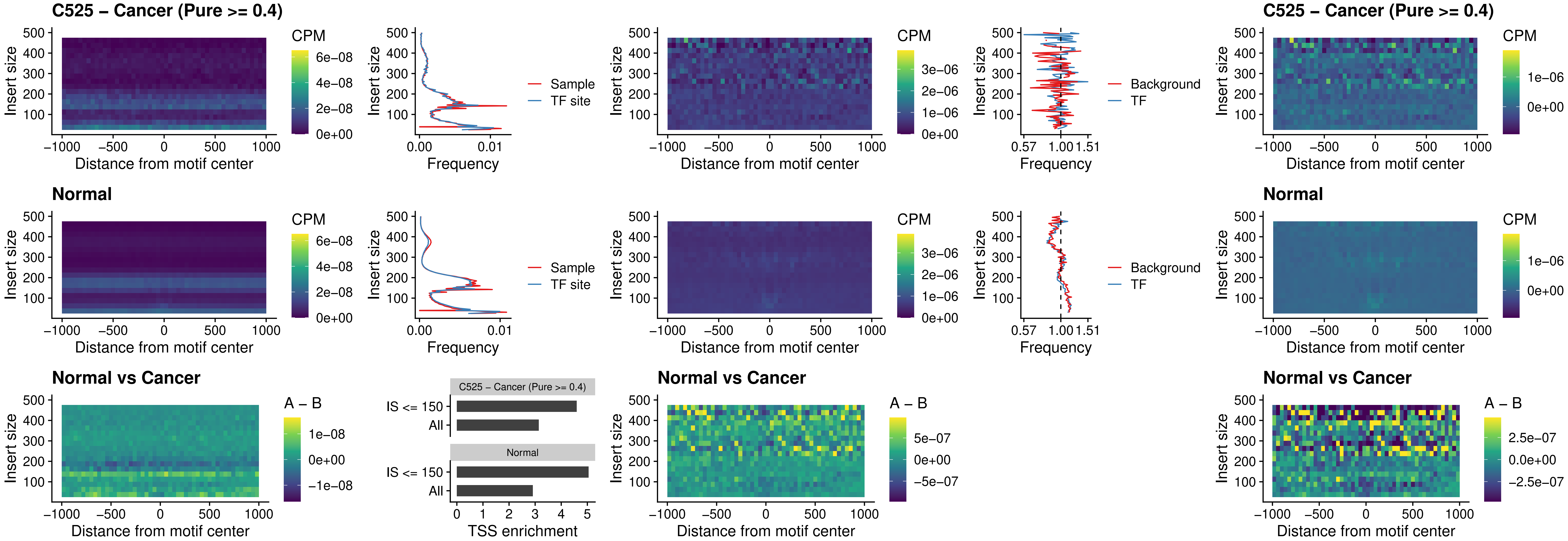

#### C525 – Cancer (Pure >= 0.4) – IRF8_uc_dTSS_oPEAK

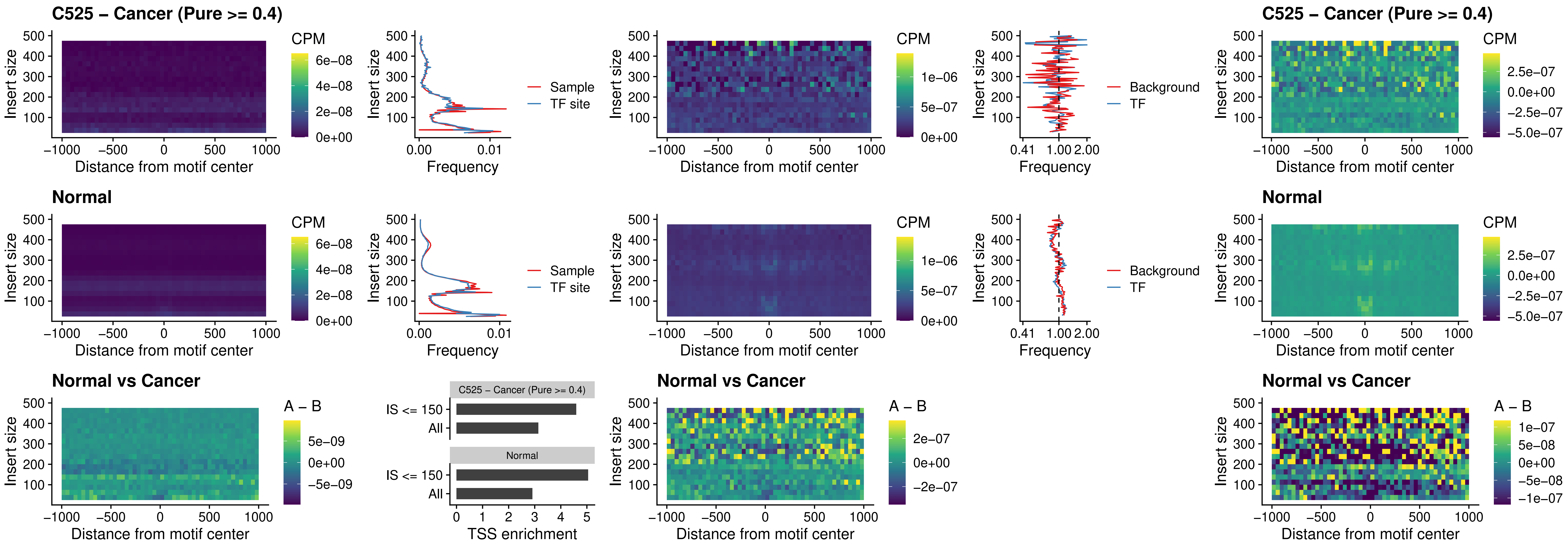

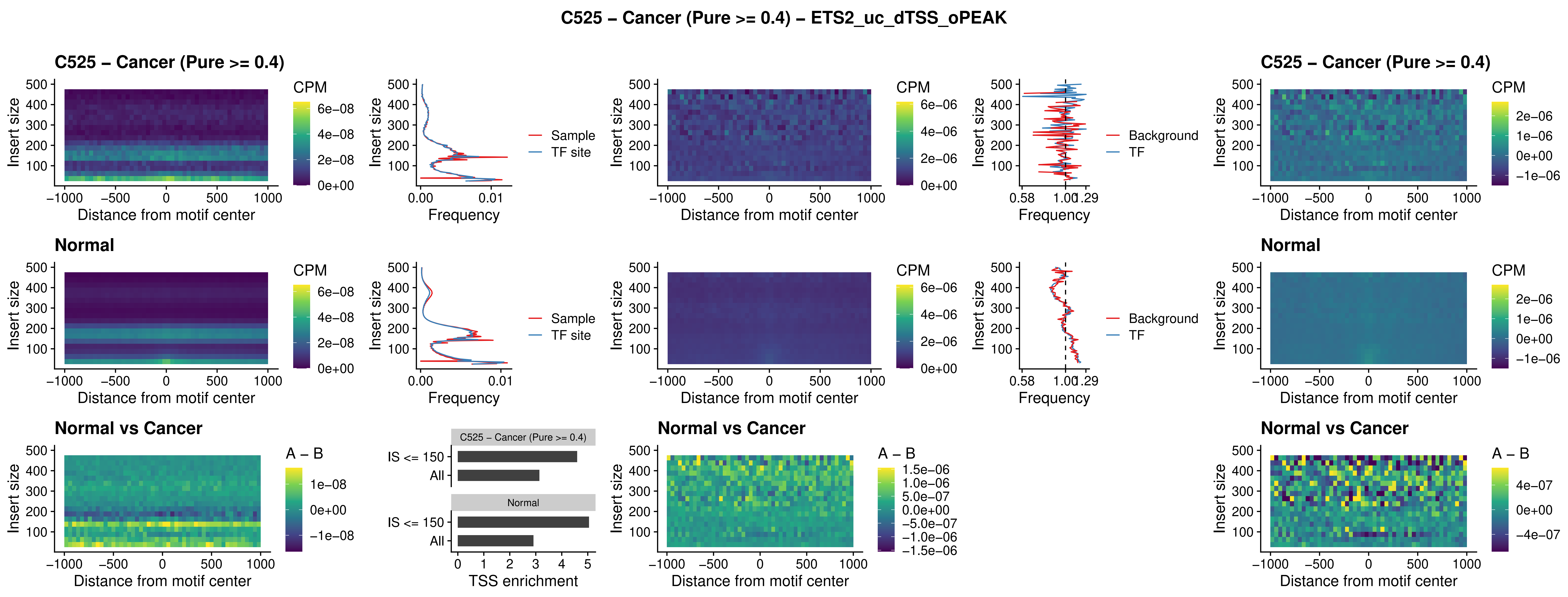
C525 – Cancer (Pure >= 0.4) – ETS2_uc_dTSS_oPEAK
C525 – Cancer (Pure >= 0.4)
Normal
Normal vs Cancer
Insert size
Distance from motif center
Frequency
CPM
A – B
Sample
TF site
Background
TF
C525 – Cancer (Pure >= 0.4)
Normal
IS <= 150
All
TSS enrichment

#### C525 – Cancer (Pure >= 0.4) – SMARCC2_uc_pTSS_oPEAK

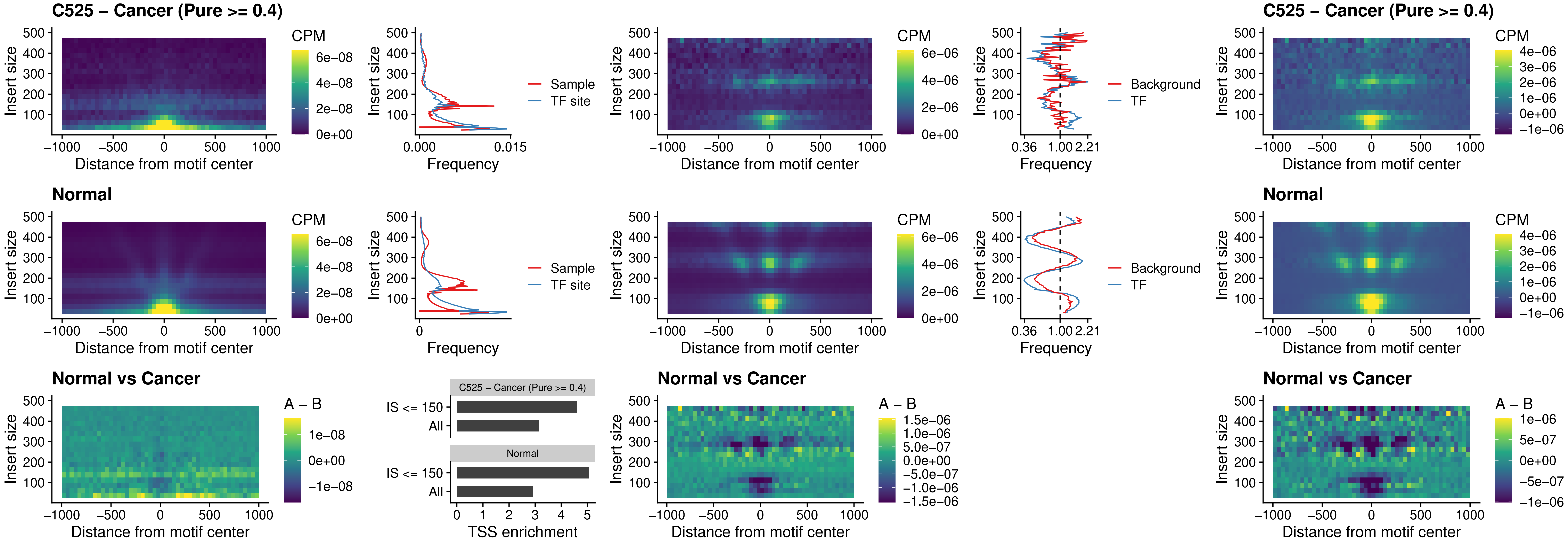

#### C525 – Cancer (Pure >= 0.4) – BCL11A_uc_cTSS_nPEAK

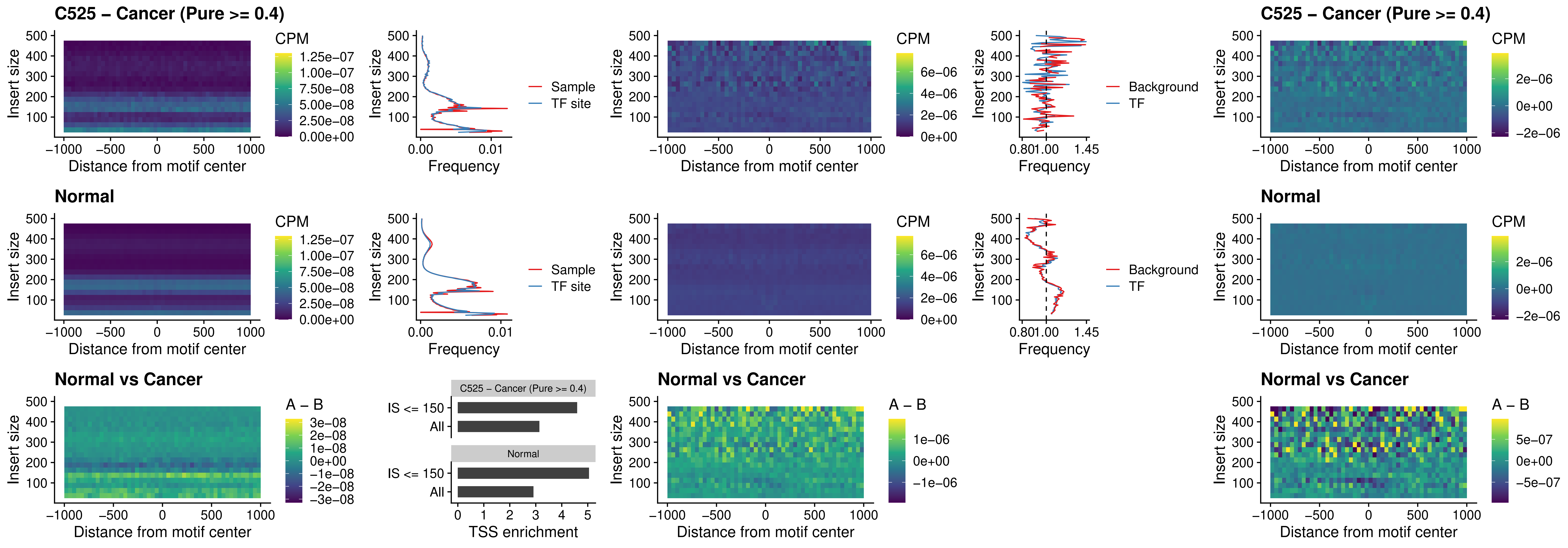

#### C525 – Cancer (Pure >= 0.4) – BCL11B_uc_cTSS_nPEAK

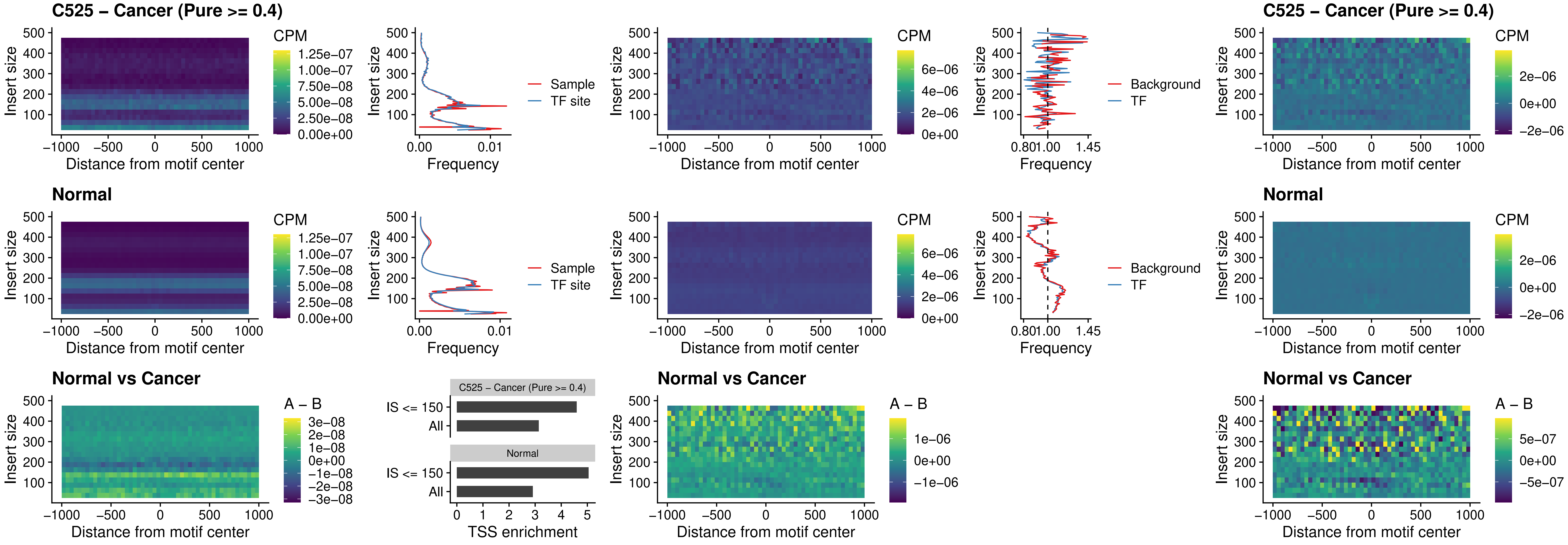

#### C525 – Cancer (Pure >= 0.4) – STAT2_uc_dTSS_oPEAK

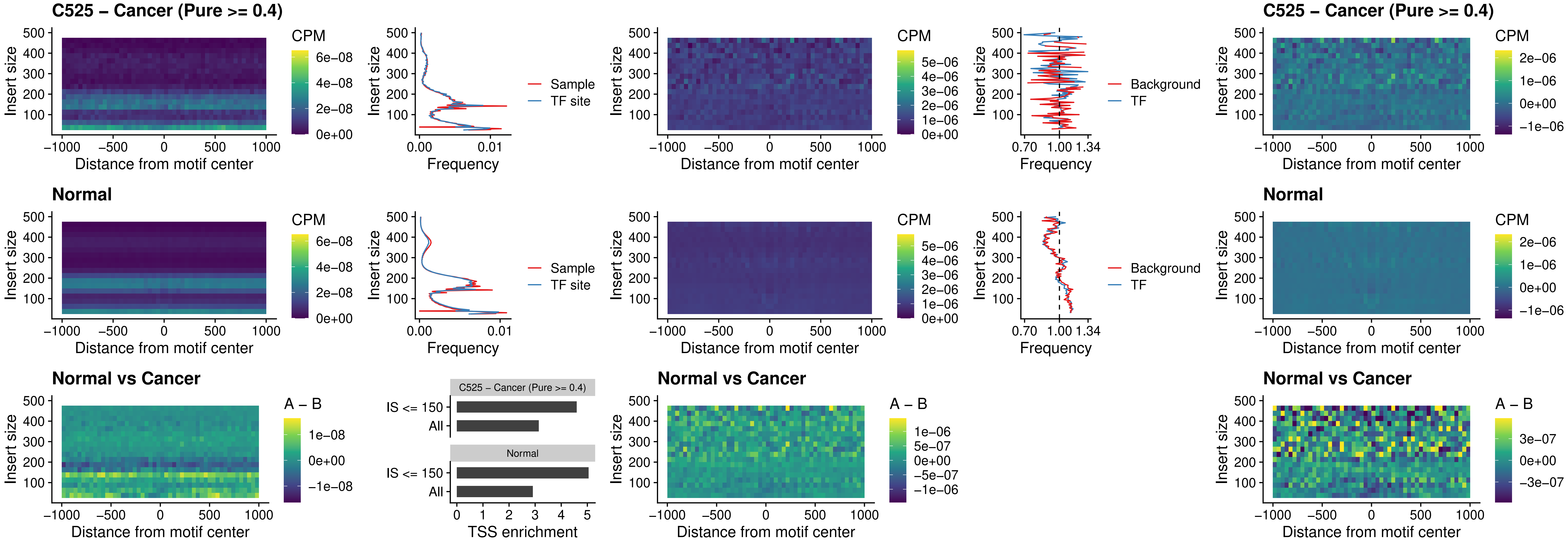

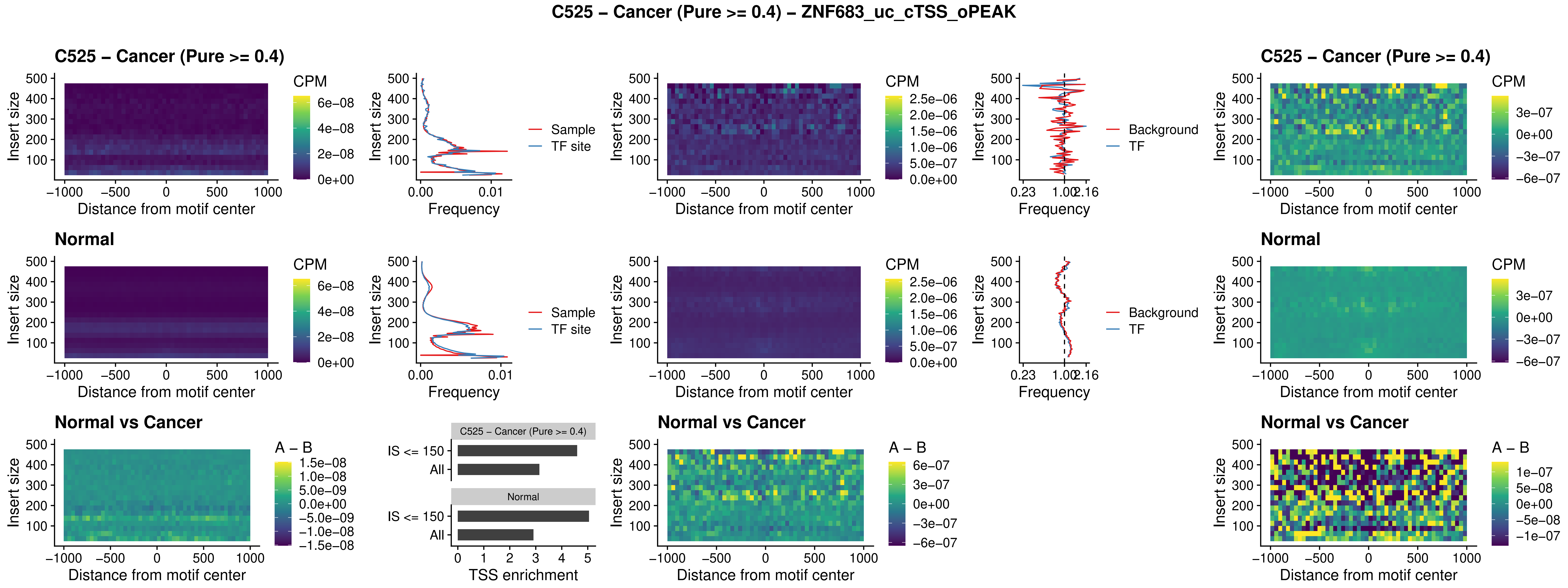
C525 – Cancer (Pure >= 0.4) – ZNF683_uc_cTSS_oPEAK
C525 – Cancer (Pure >= 0.4)
Normal
Normal vs Cancer
Insert size
Distance from motif center
CPM
Frequency
Sample
TF site
Background
TF
A – B
C525 – Cancer (Pure >= 0.4)
Normal
IS <= 150
All
TSS enrichment

#### C525 – Cancer (Pure >= 0.4) – IRF8_uc_dTSS_nPEAK

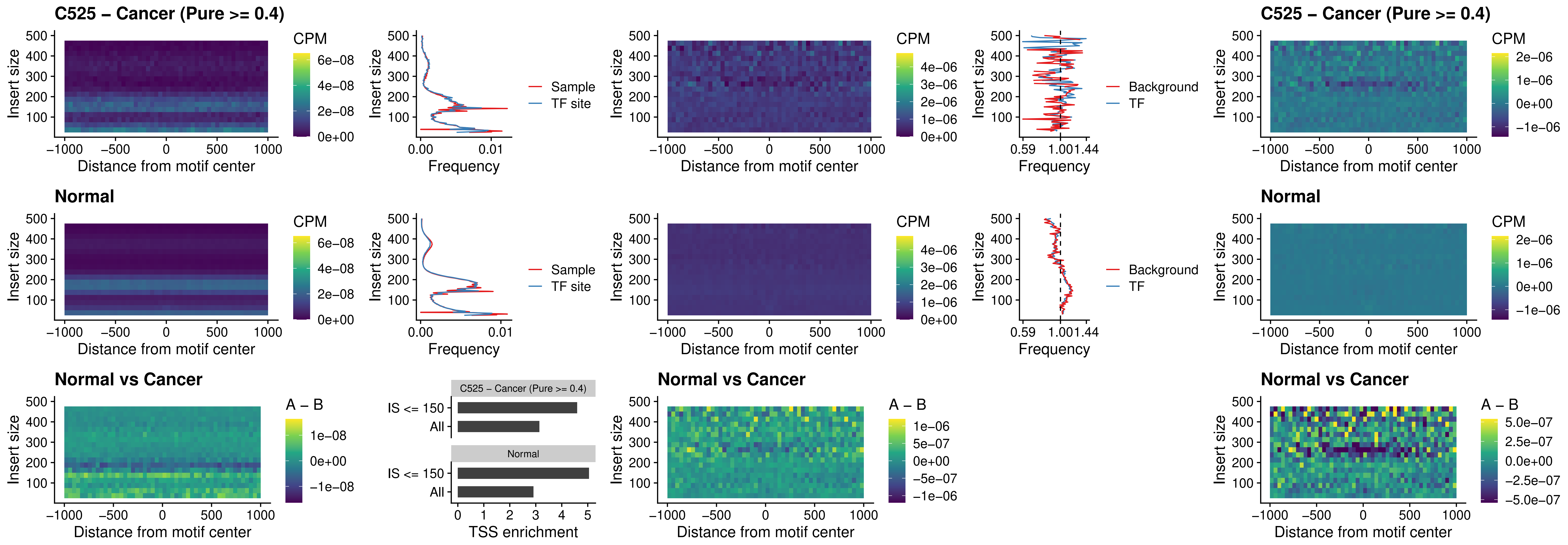

#### C525 – Cancer (Pure >= 0.4) – ETV7_uc_dTSS_oPEAK

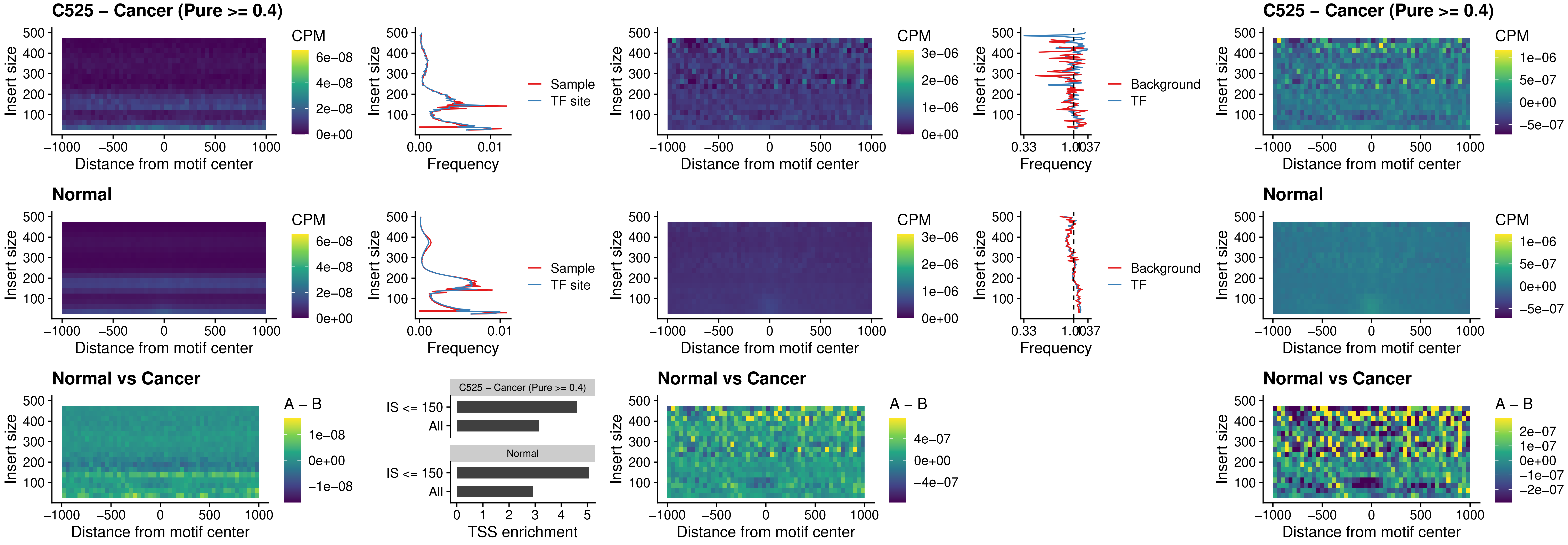

#### C528 – Cancer (Pure >= 0.4) – FOXL1_uc_pTSS_oPEAK

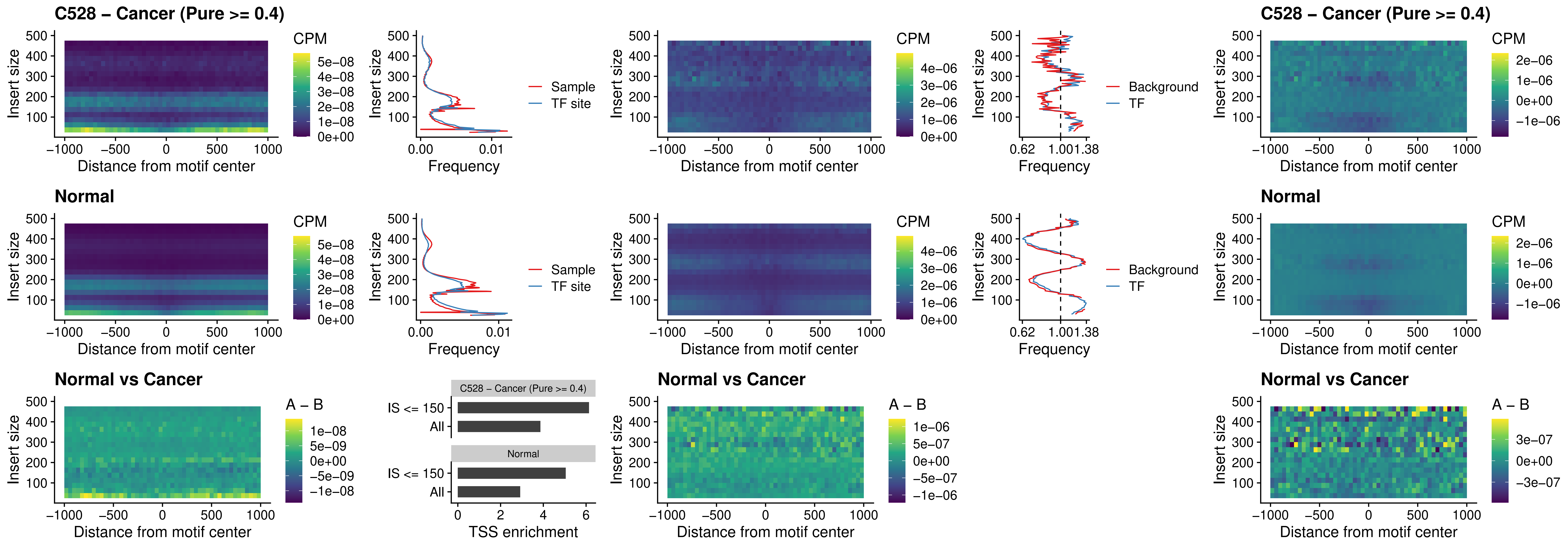

#### C528 – Cancer (Pure >= 0.4) – CPEB1_uc_cTSS_oPEAK

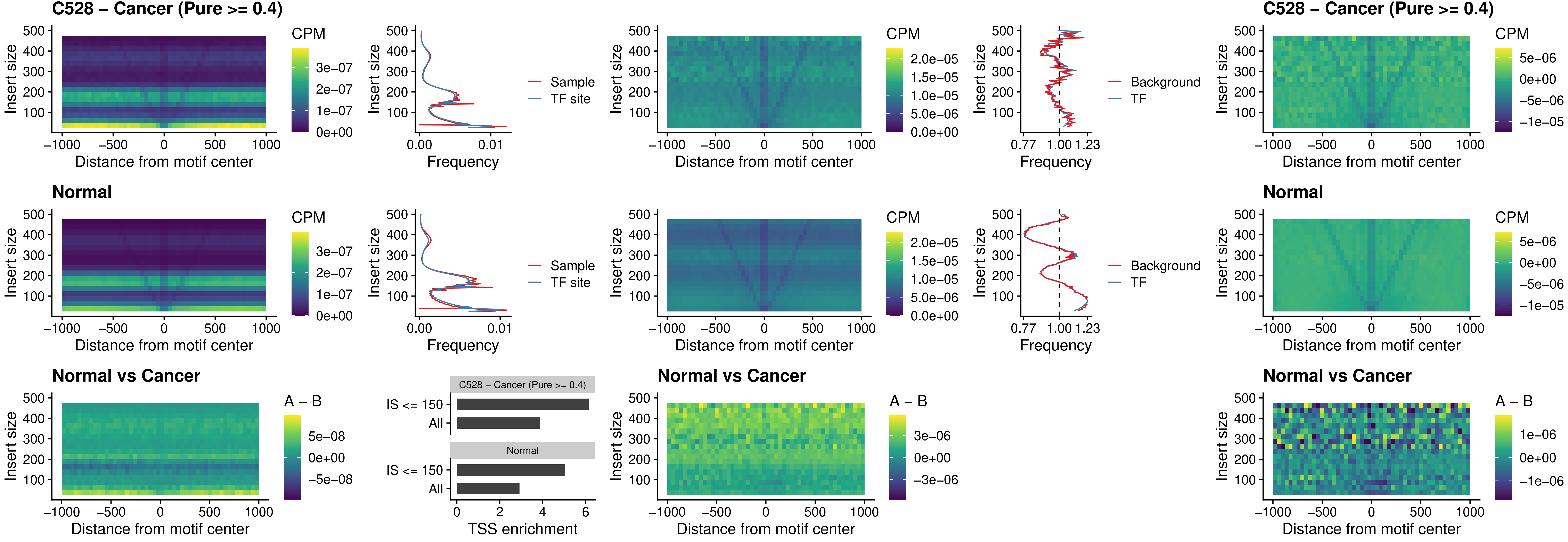

#### C528 – Cancer (Pure >= 0.4) – HMGA1_uc_pTSS_nPEAK

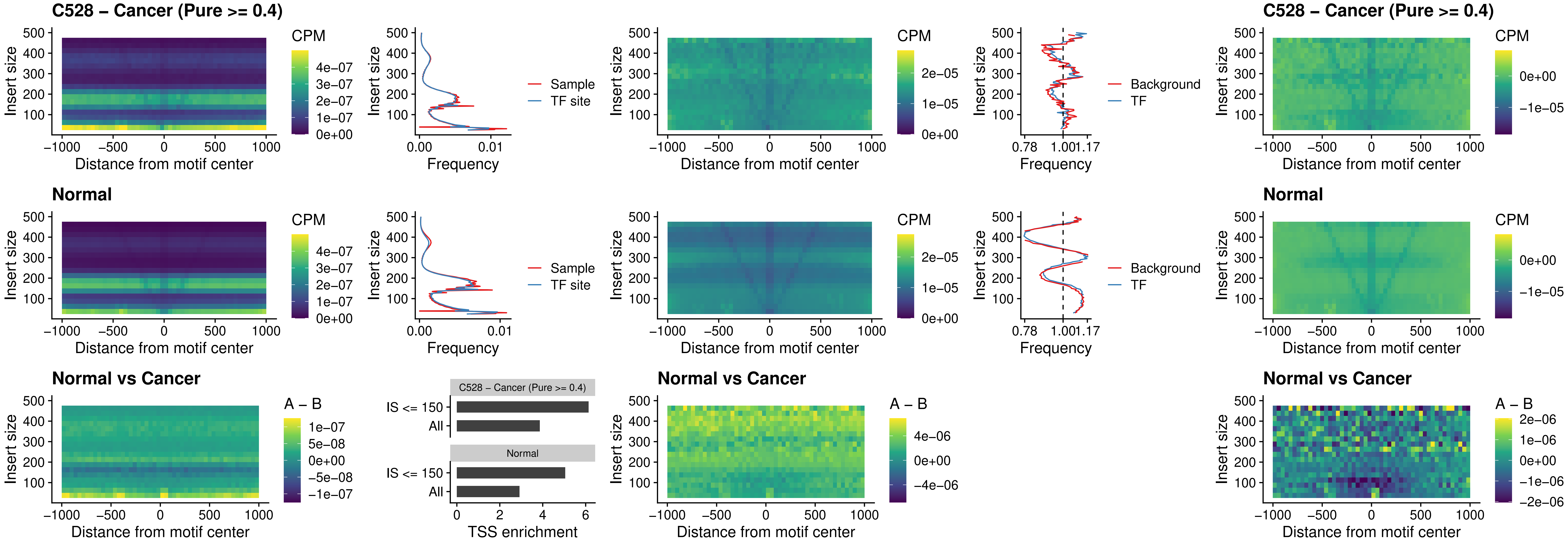

### C528 – Cancer (Pure >= 0.4) – SOX13_uc_cTSS_nPEAK

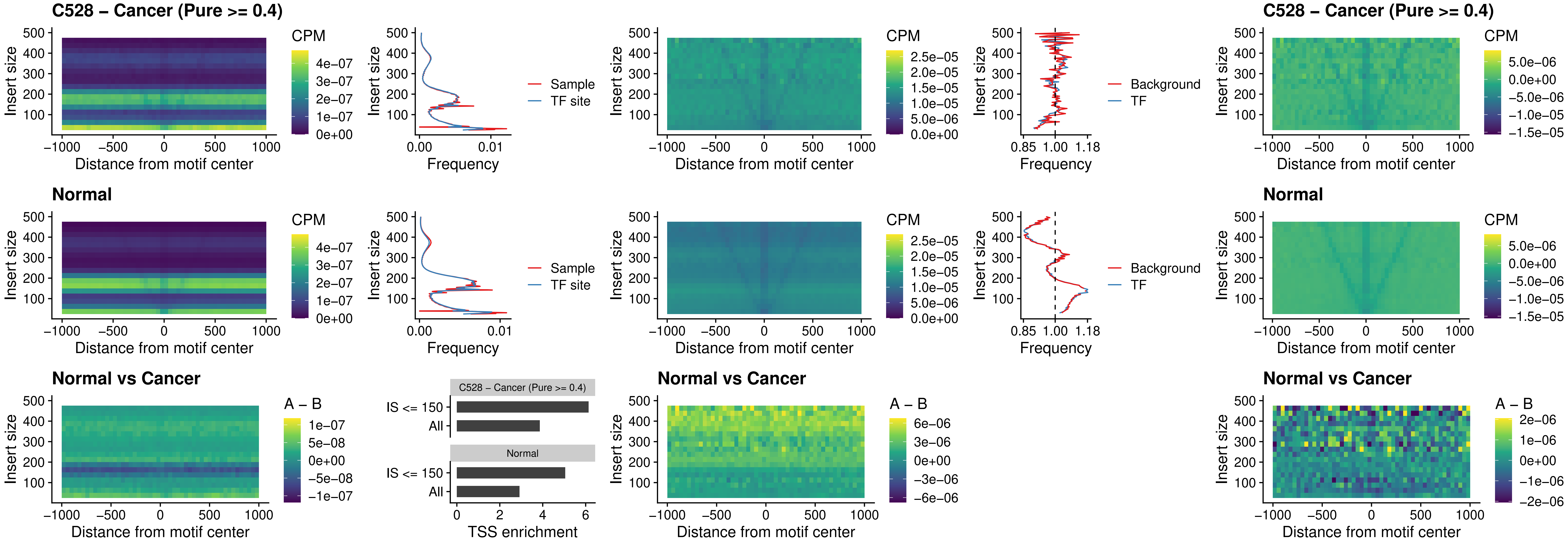

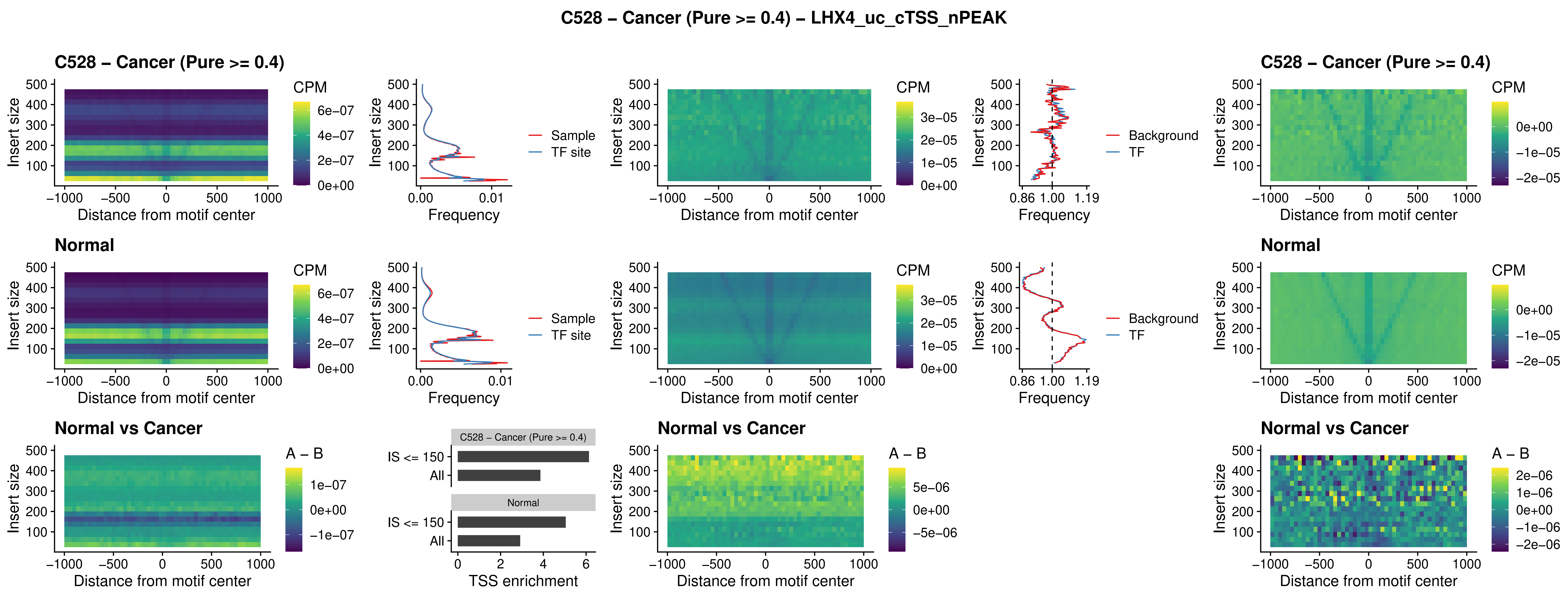
C528 – Cancer (Pure >= 0.4) – LHX4_uc_cTSS_nPEAK
C528 – Cancer (Pure >= 0.4)
Normal
Normal vs Cancer
Insert size
Distance from motif center
Frequency
CPM
A – B
Sample
TF site
Background
TF
C528 – Cancer (Pure >= 0.4)
Normal
IS <= 150
All
TSS enrichment

#### C528 – Cancer (Pure >= 0.4) – HOXA4_uc_cTSS_nPEAK

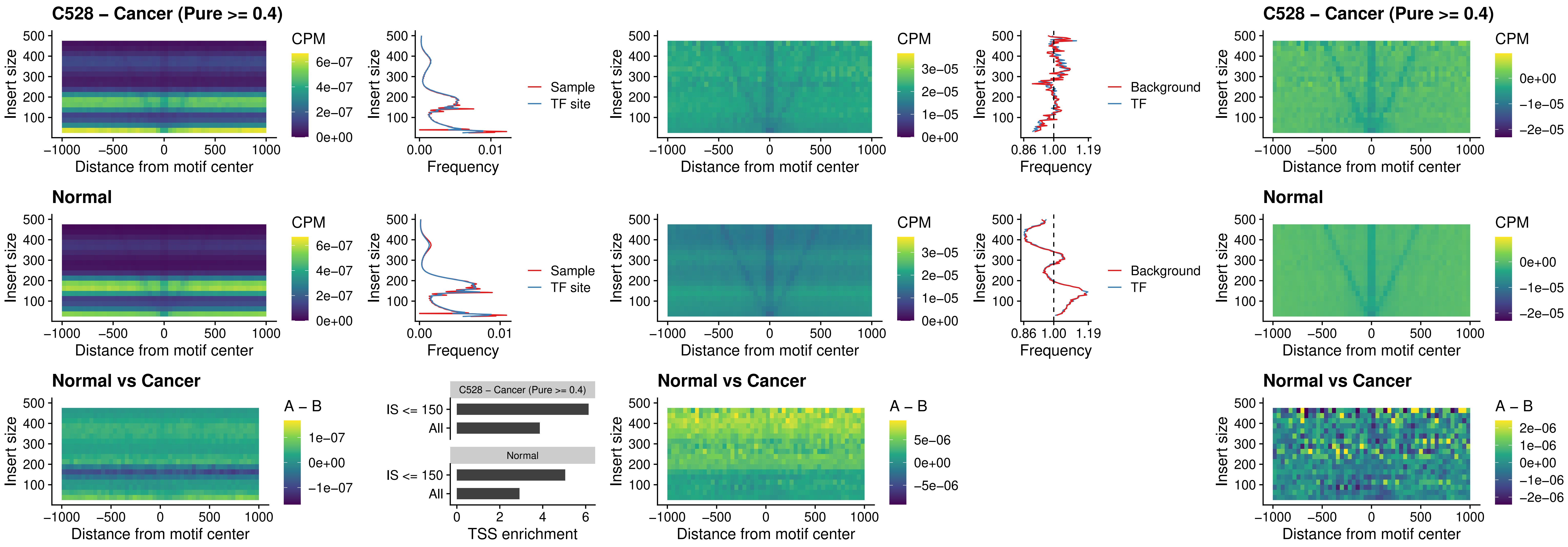

#### C528 – Cancer (Pure >= 0.4) – HLX_uc_cTSS_nPEAK

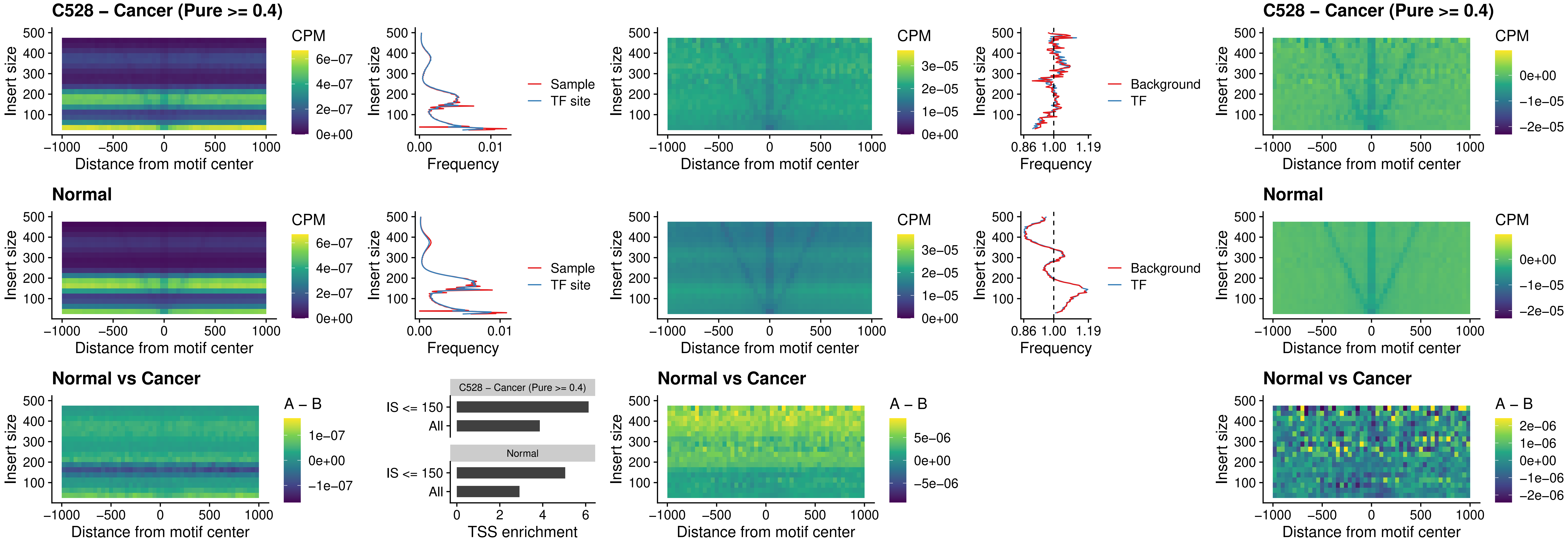

### C528 – Cancer (Pure >= 0.4) – AC0125311_uc_cTSS_nPEAK

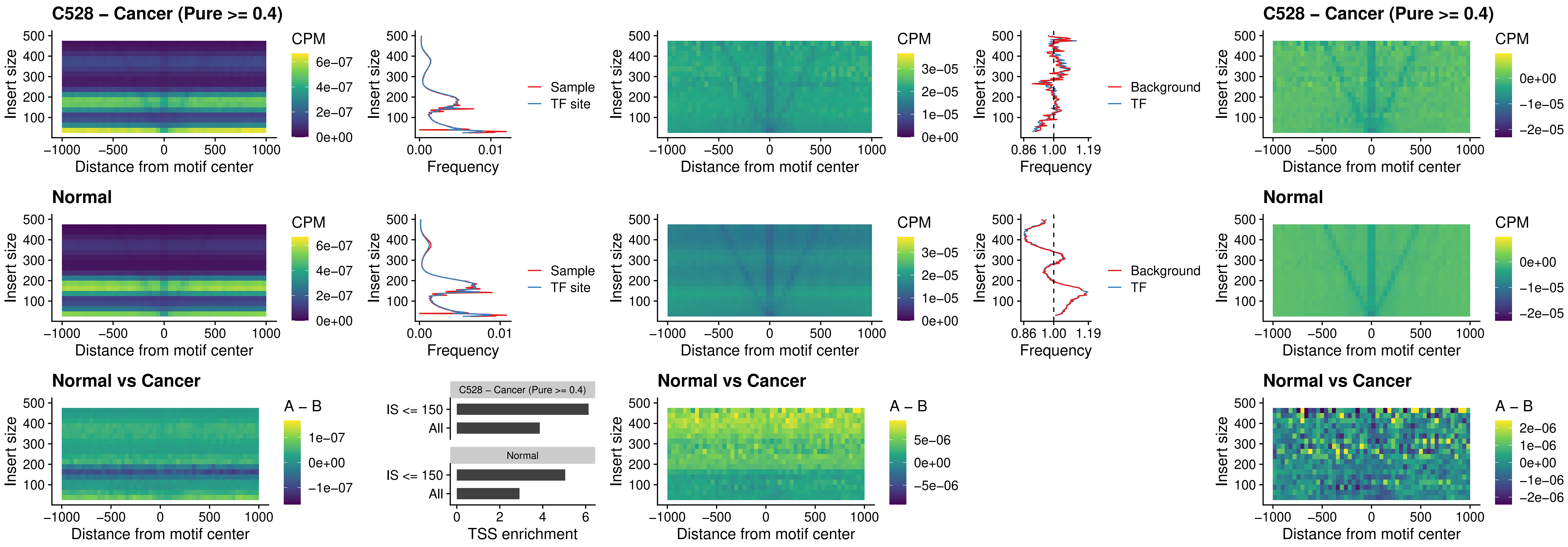

#### C528 – Cancer (Pure >= 0.4) – DBX2_uc_cTSS_nPEAK

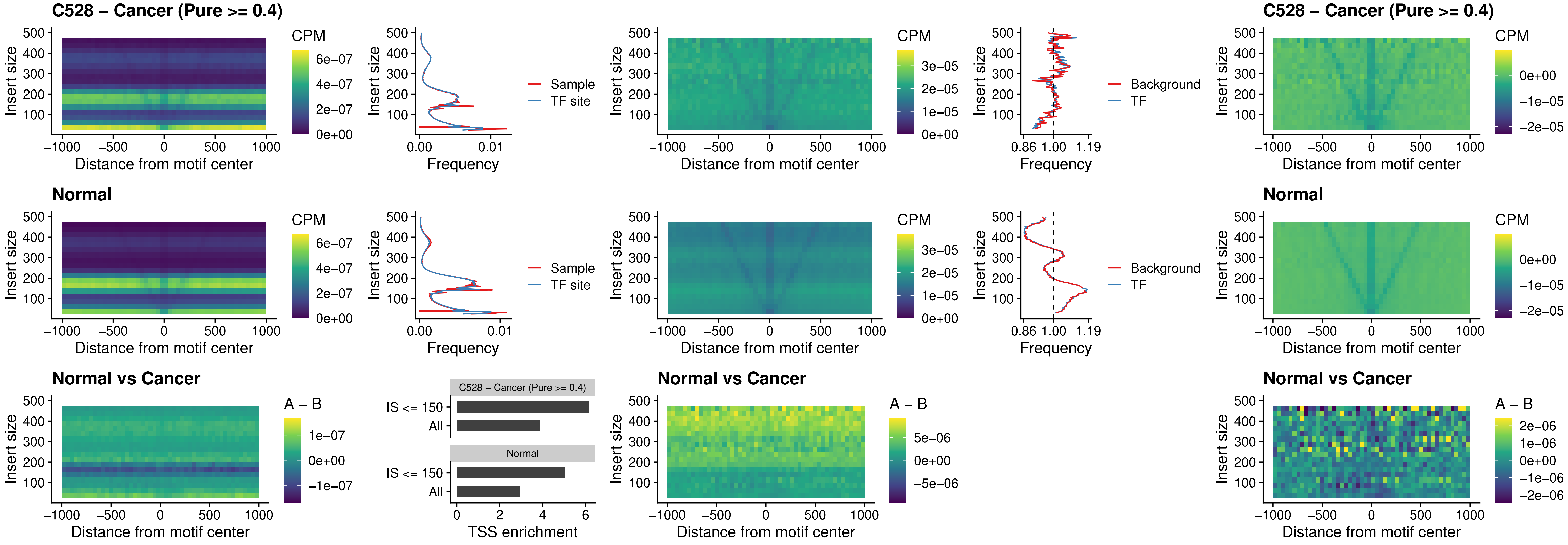

#### C528 – Cancer (Pure >= 0.4) – LHX6_uc_dTSS_nPEAK

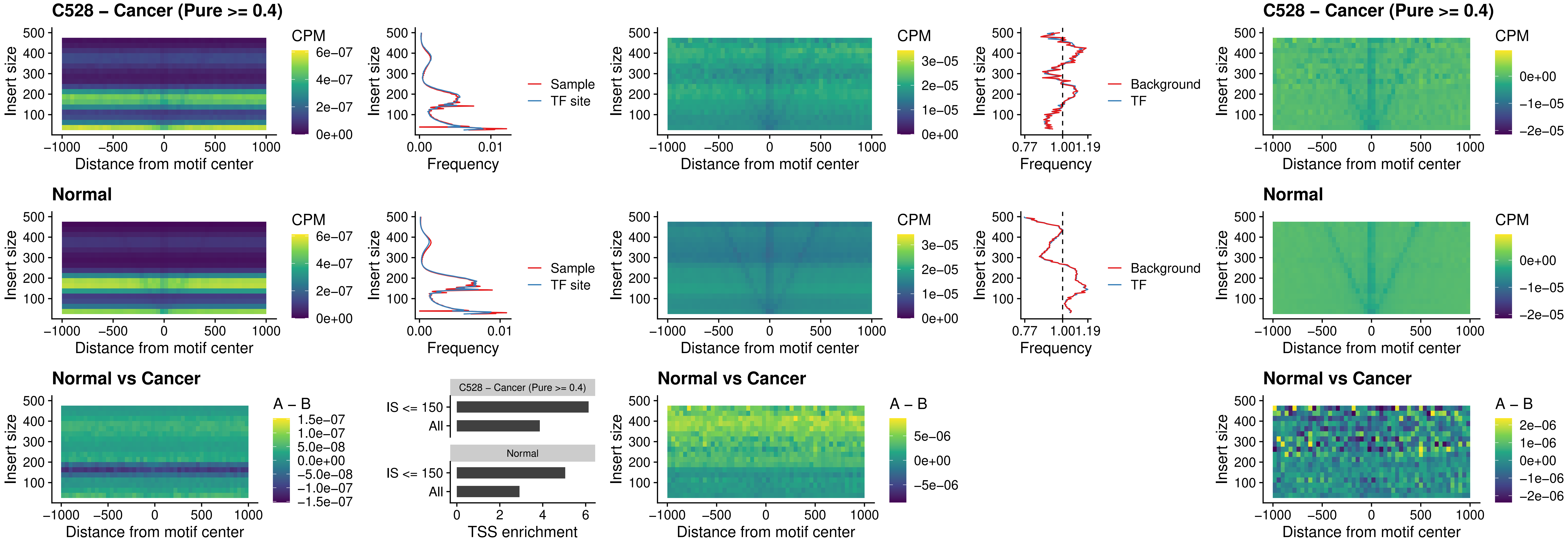

#### C528 – Cancer (Pure >= 0.4) – SOX1_uc_dTSS_nPEAK

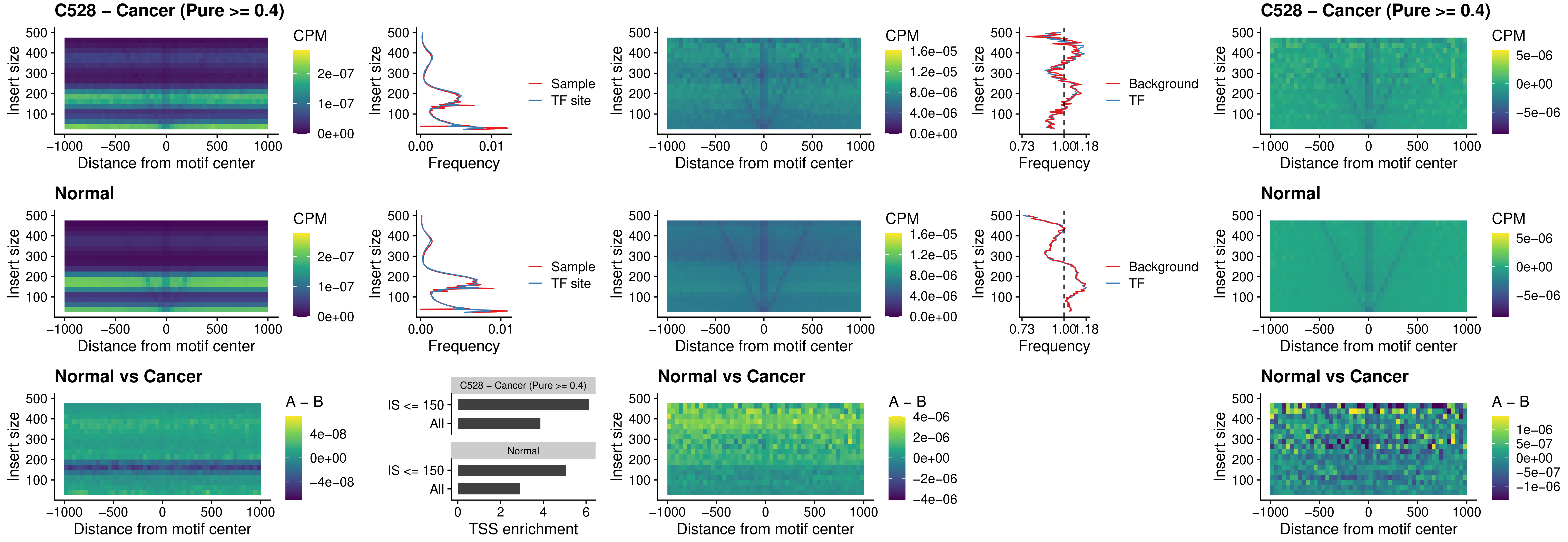

#### C528 – Cancer (Pure >= 0.4) – NKX24_uc_dTSS_nPEAK

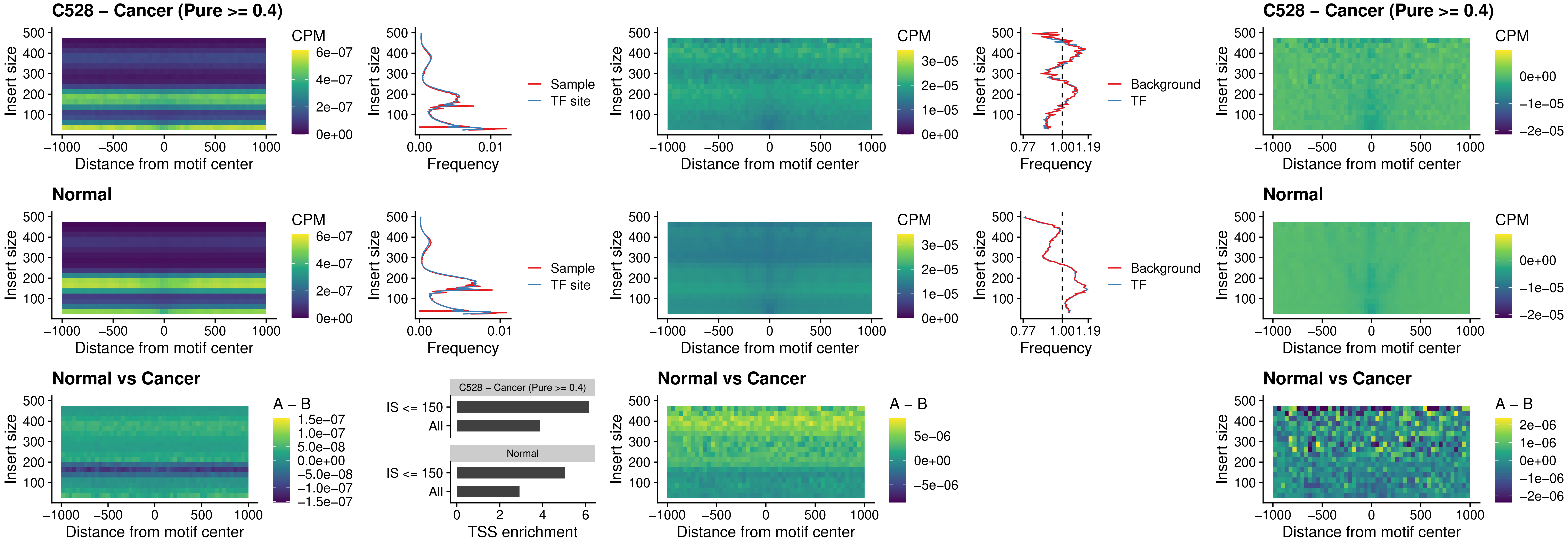

#### C528 – Cancer (Pure >= 0.4) – DLX3_uc_cTSS_nPEAK

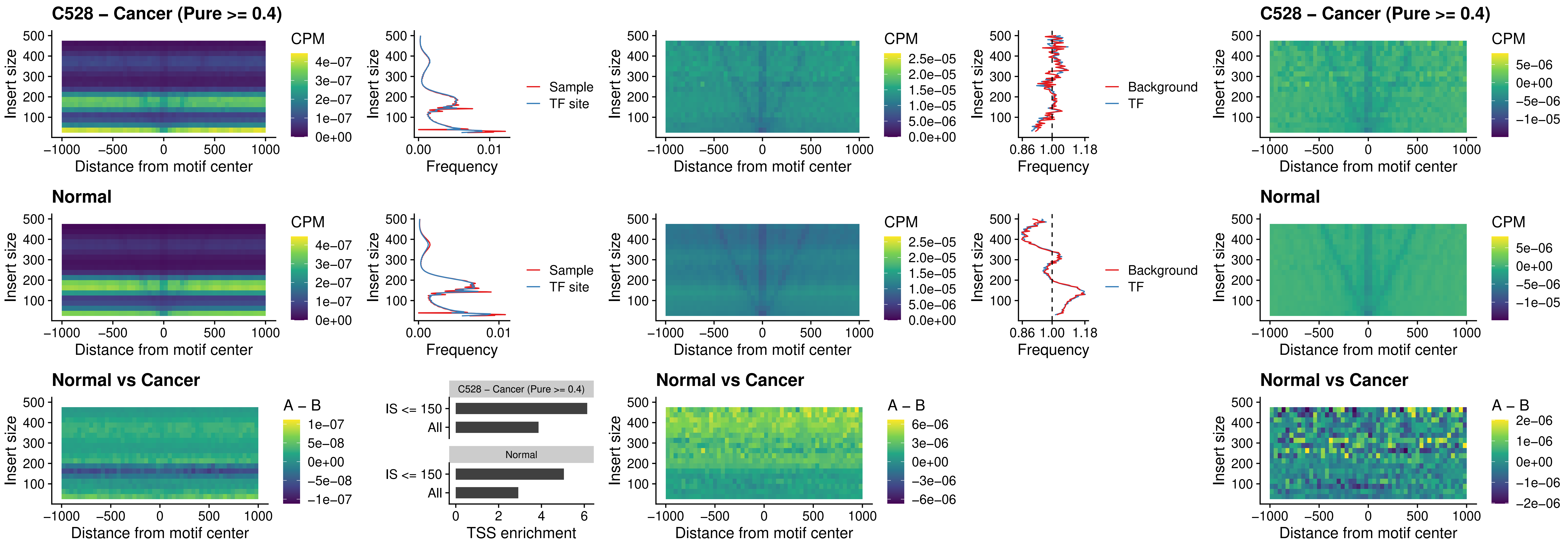

#### C528 – Cancer (Pure >= 0.4) – HOXD3_uc_cTSS_nPEAK

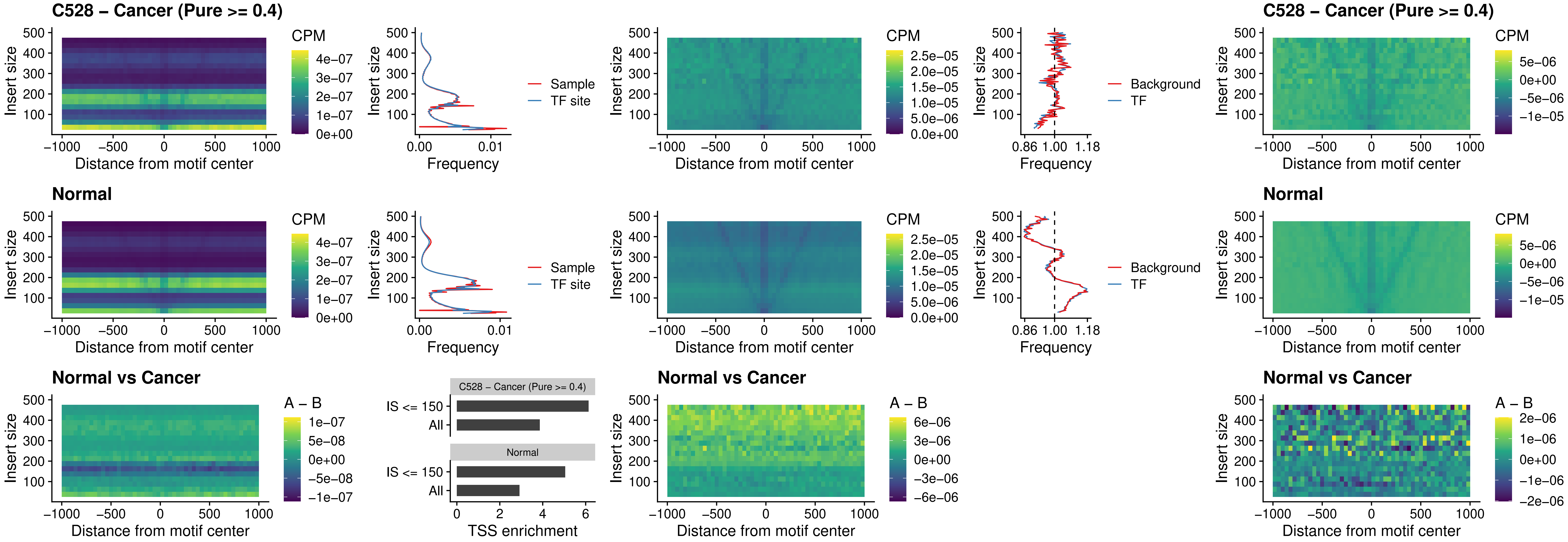

#### C528 – Cancer (Pure >= 0.4) – ISL1_uc_dTSS_oPEAK

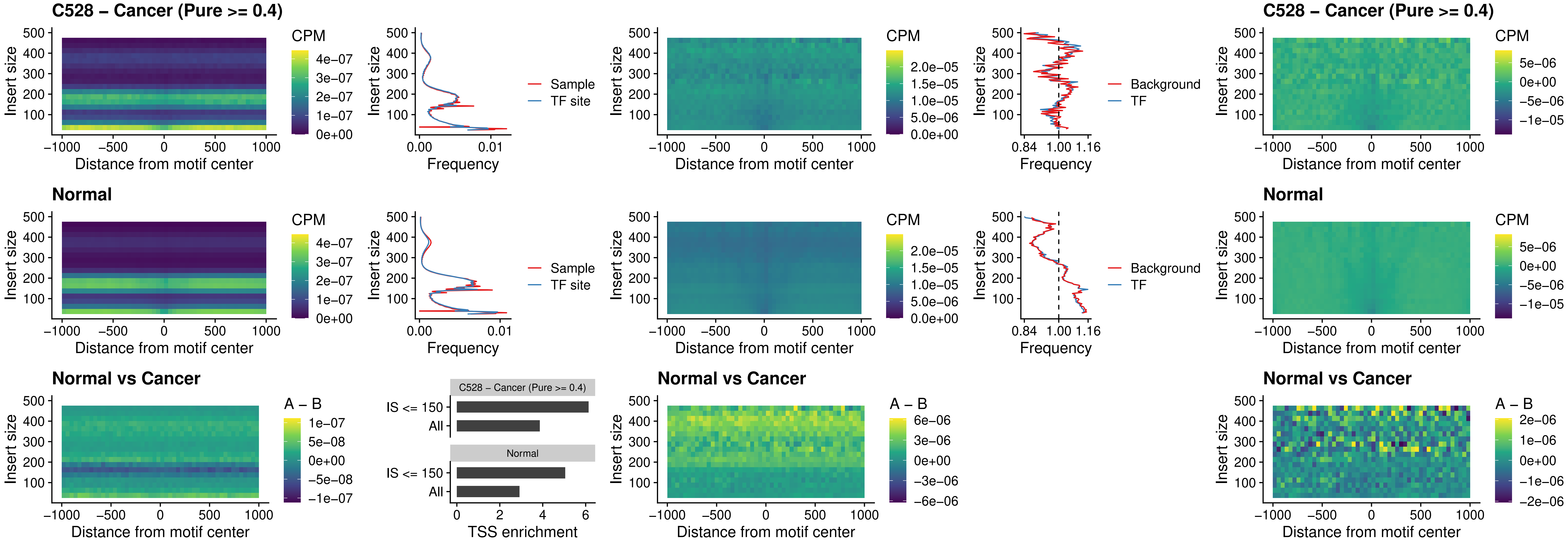

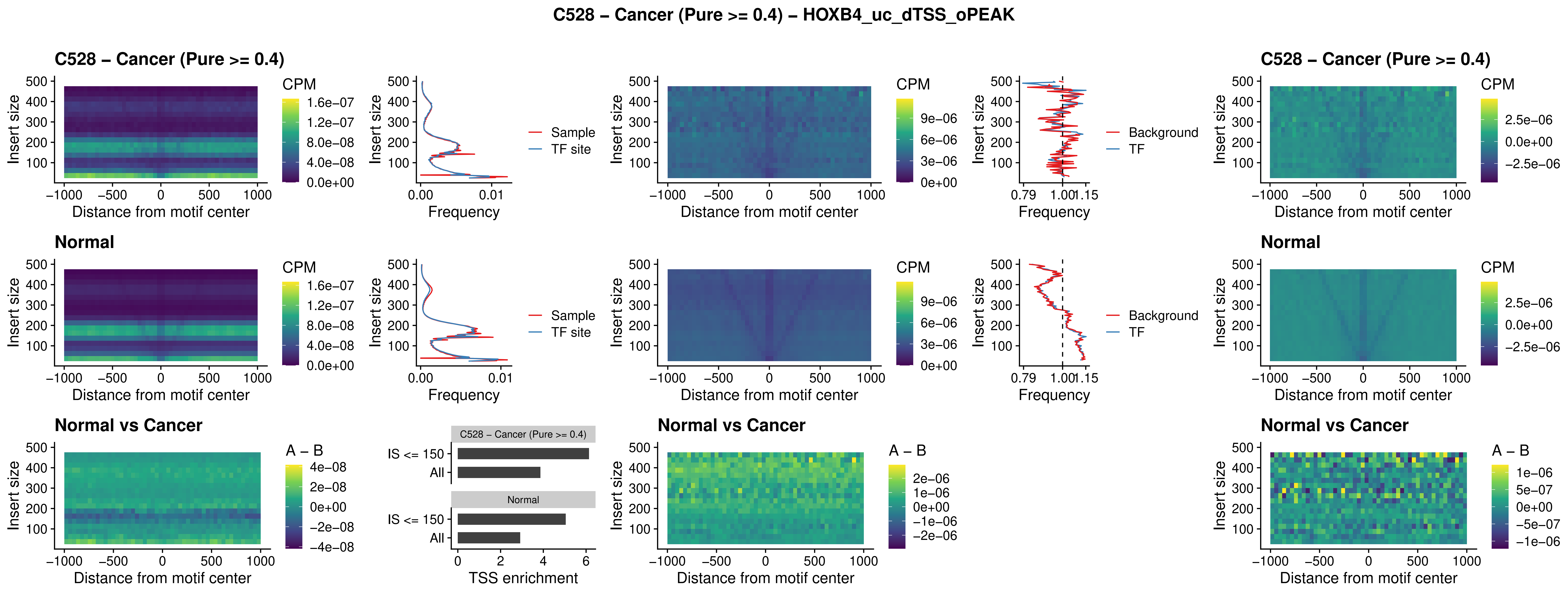
C528 – Cancer (Pure >= 0.4) – HOXB4_uc_dTSS_oPEAK
C528 – Cancer (Pure >= 0.4)
Normal
Normal vs Cancer
Insert size
Distance from motif center
Frequency
CPM
A – B
Sample
TF site
Background
TF
C528 – Cancer (Pure >= 0.4)
Normal
IS <= 150
All
TSS enrichment

### C528 – Cancer (Pure >= 0.4) – VAX1_uc_dTSS_oPEAK

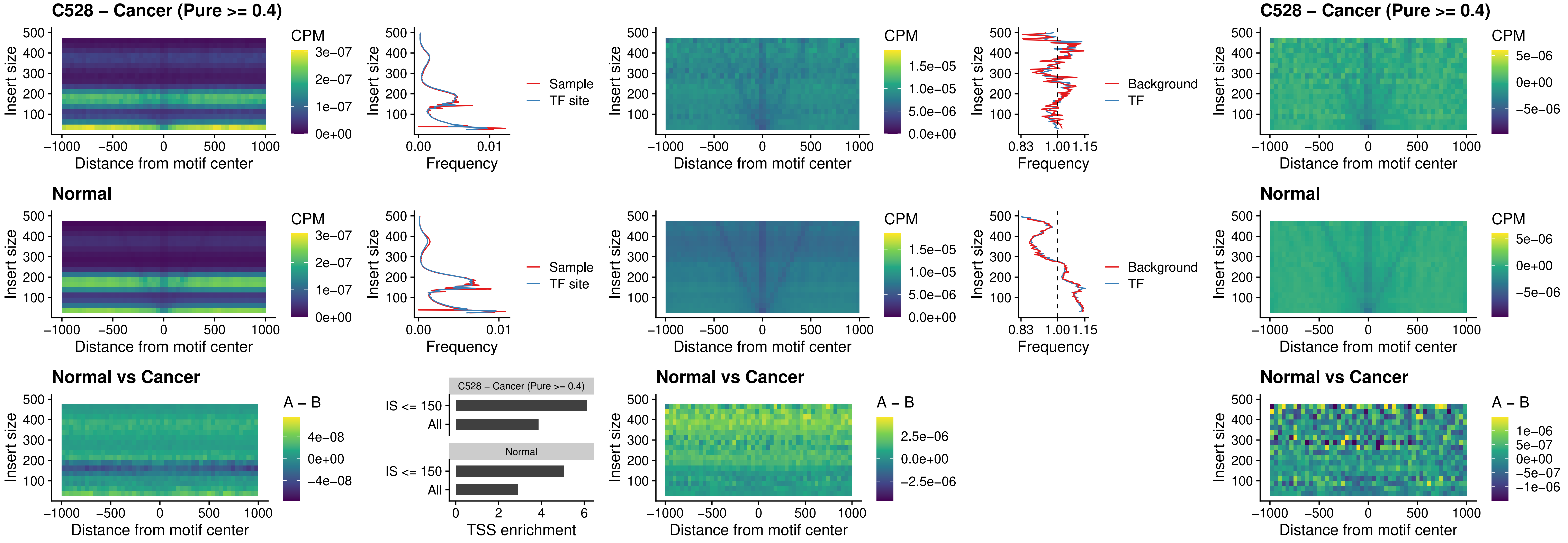

#### C528 – Cancer (Pure >= 0.4) – DLX3_uc_dTSS_oPEAK

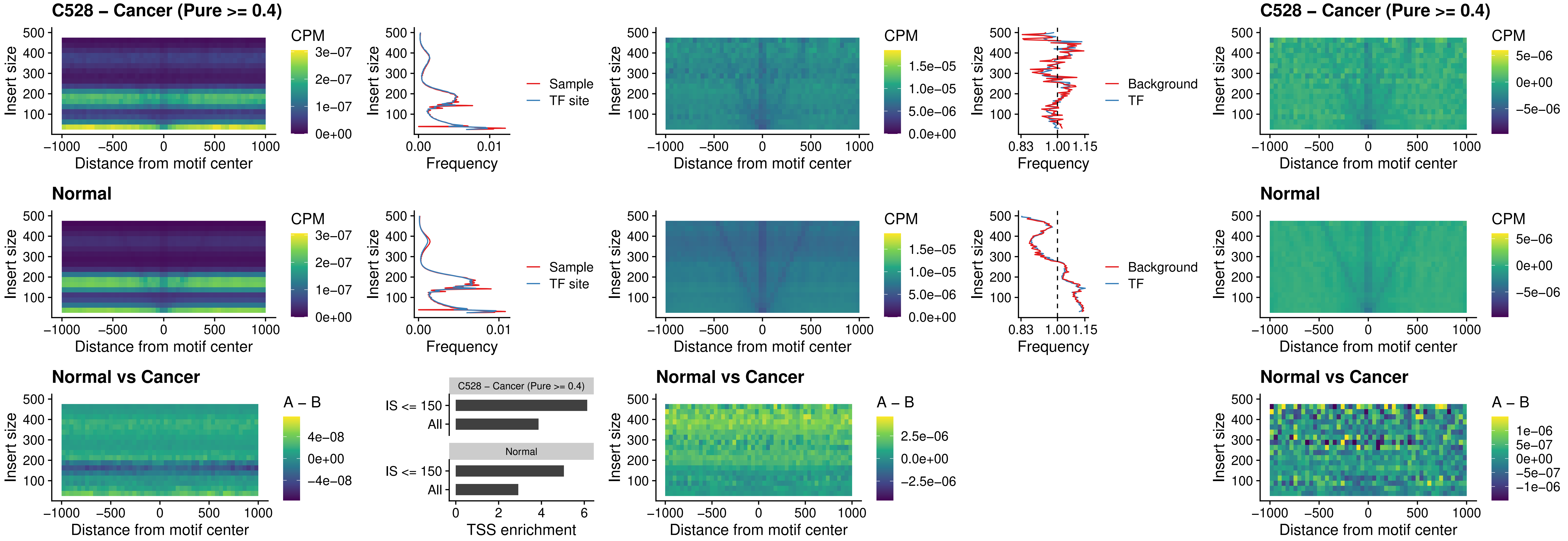

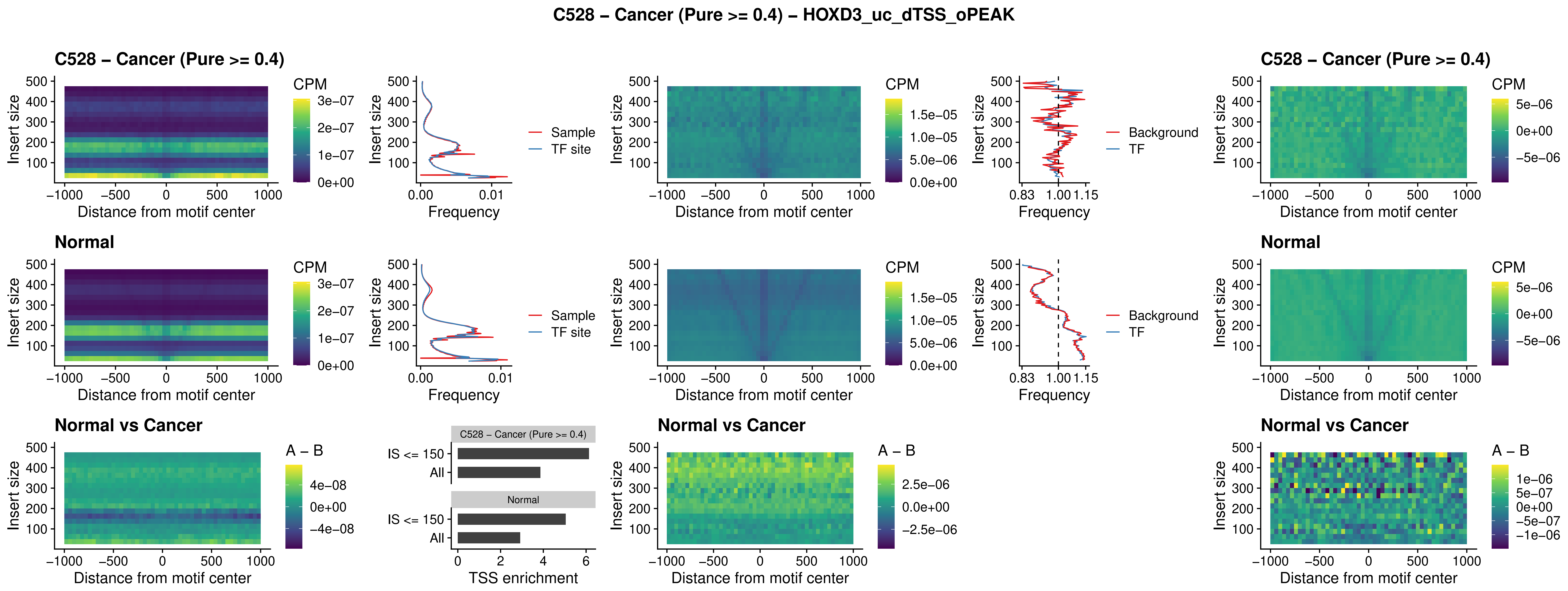
C528 – Cancer (Pure >= 0.4) – HOXD3_uc_dTSS_oPEAK
C528 – Cancer (Pure >= 0.4)
Normal
Normal vs Cancer
Insert size
Distance from motif center
Frequency
CPM
A – B
Sample
TF site
Background
TF
C528 – Cancer (Pure >= 0.4)
Normal
IS <= 150
All
TSS enrichment

#### C528 – Cancer (Pure >= 0.4) – BBX_uc_dTSS_nPEAK

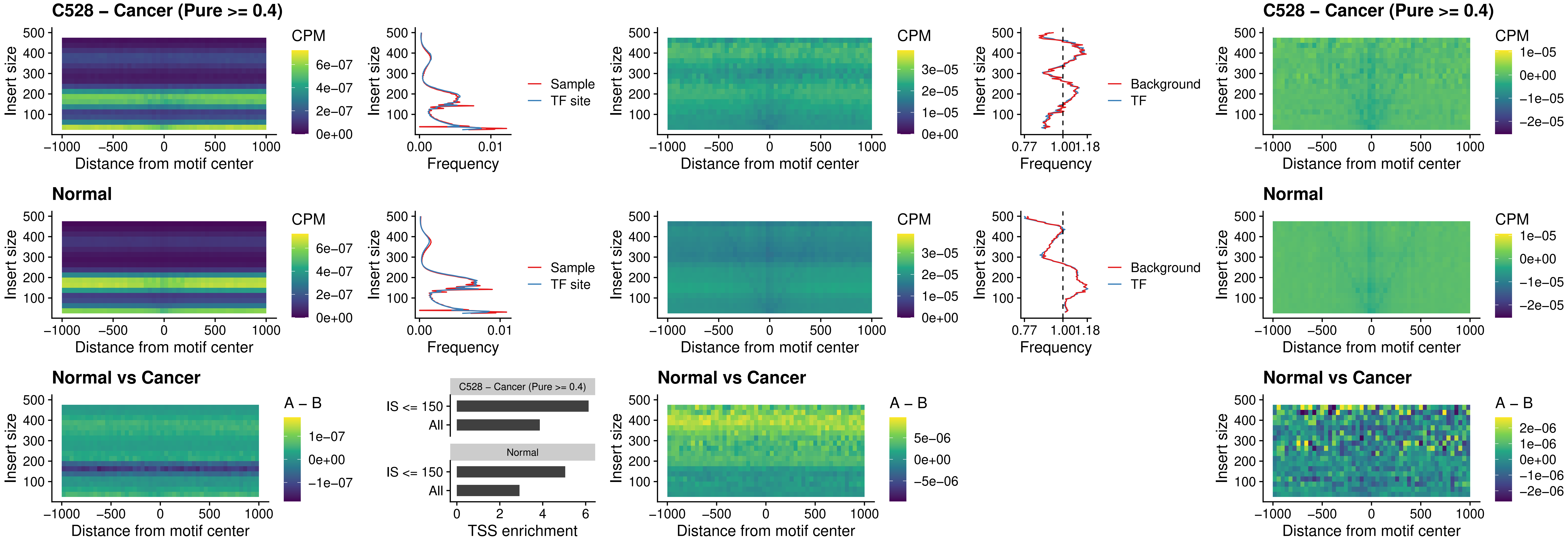

#### C528 – Cancer (Pure >= 0.4) – SOX3_uc_cTSS_nPEAK

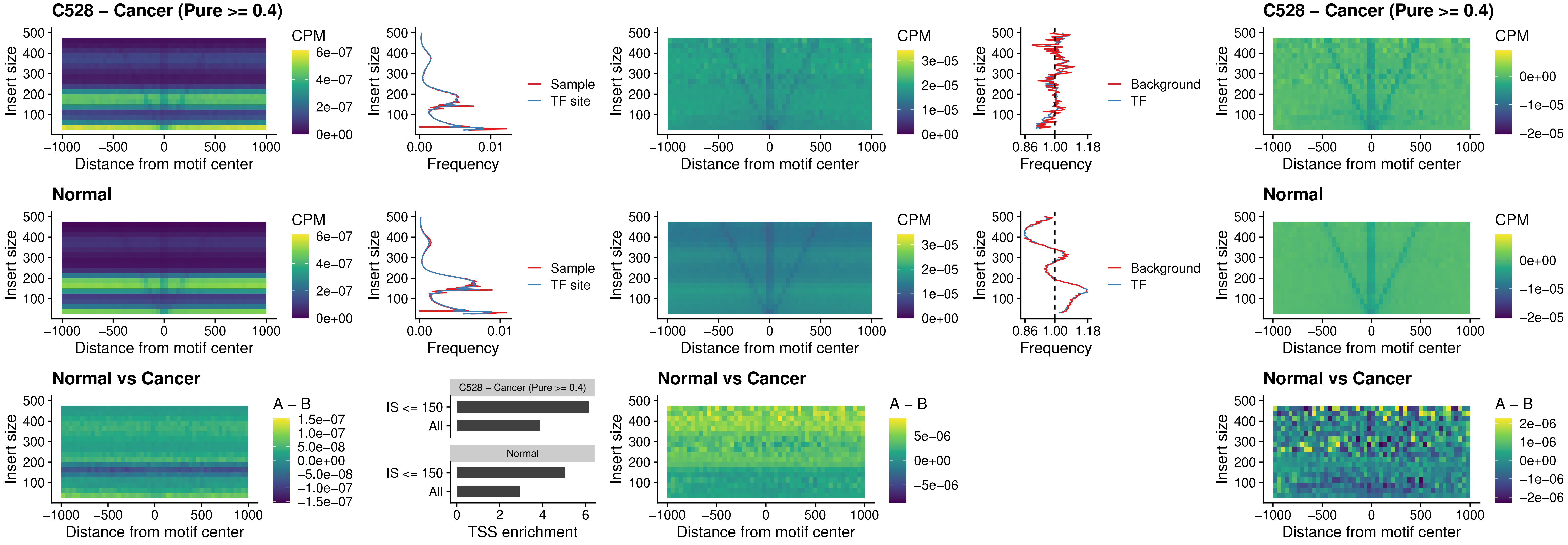

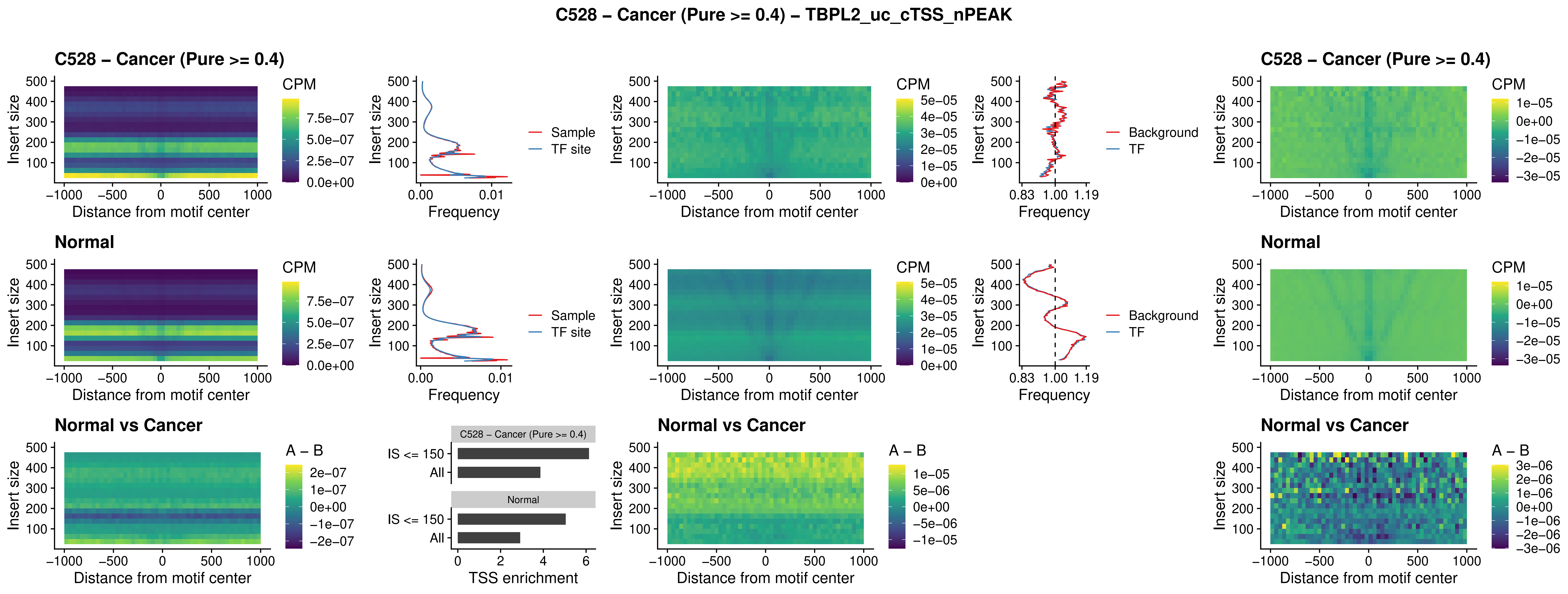
C528 – Cancer (Pure >= 0.4) – TBPL2_uc_cTSS_nPEAK
C528 – Cancer (Pure >= 0.4)
Normal
Normal vs Cancer
Insert size
Distance from motif center
Frequency
CPM
A – B
Sample
TF site
Background
TF
C528 – Cancer (Pure >= 0.4)
Normal
IS <= 150
All
TSS enrichment

#### C528 – Cancer (Pure >= 0.4) – HOXA4_uc_cTSS_oPEAK

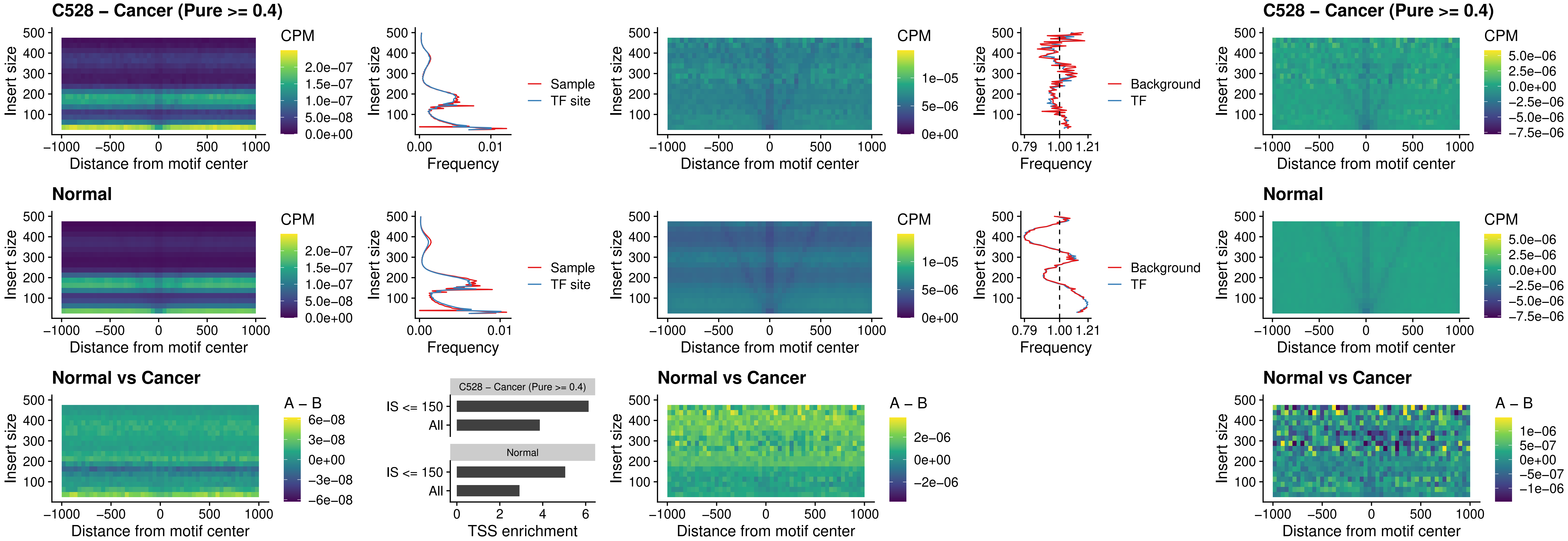

#### C528 – Cancer (Pure >= 0.4) – PDX1_uc_dTSS_oPEAK

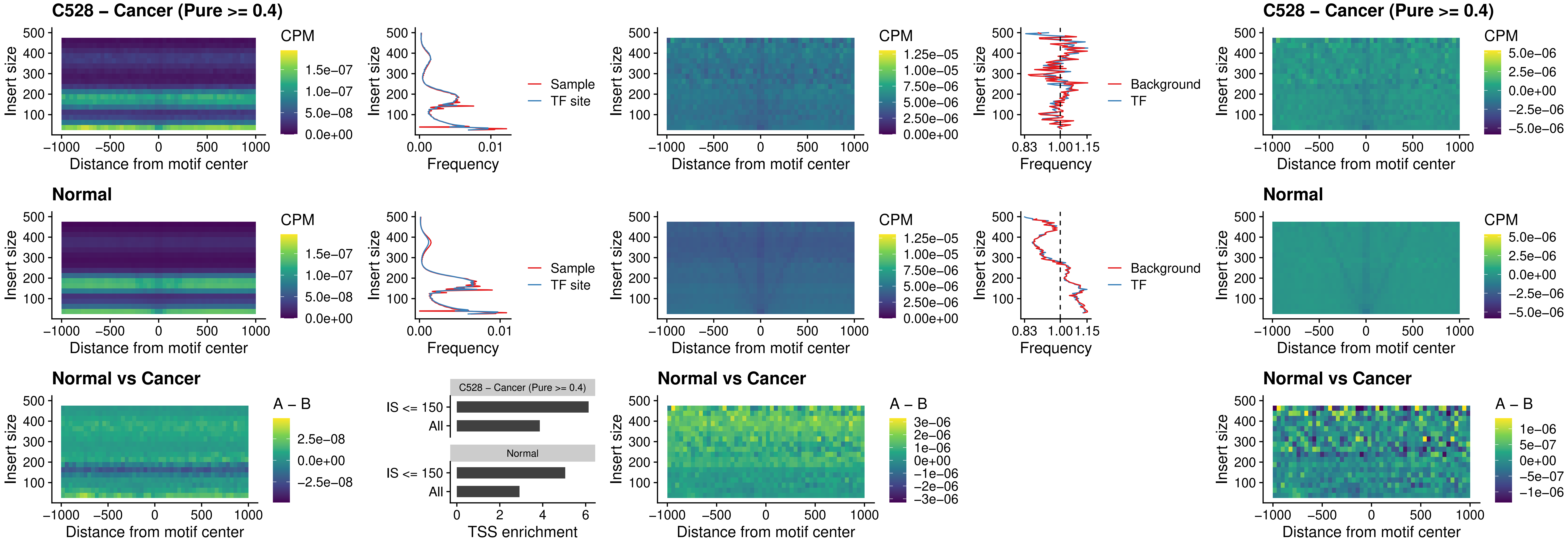

#### C528 – Cancer (Pure >= 0.4) – LHX4_uc_dTSS_oPEAK

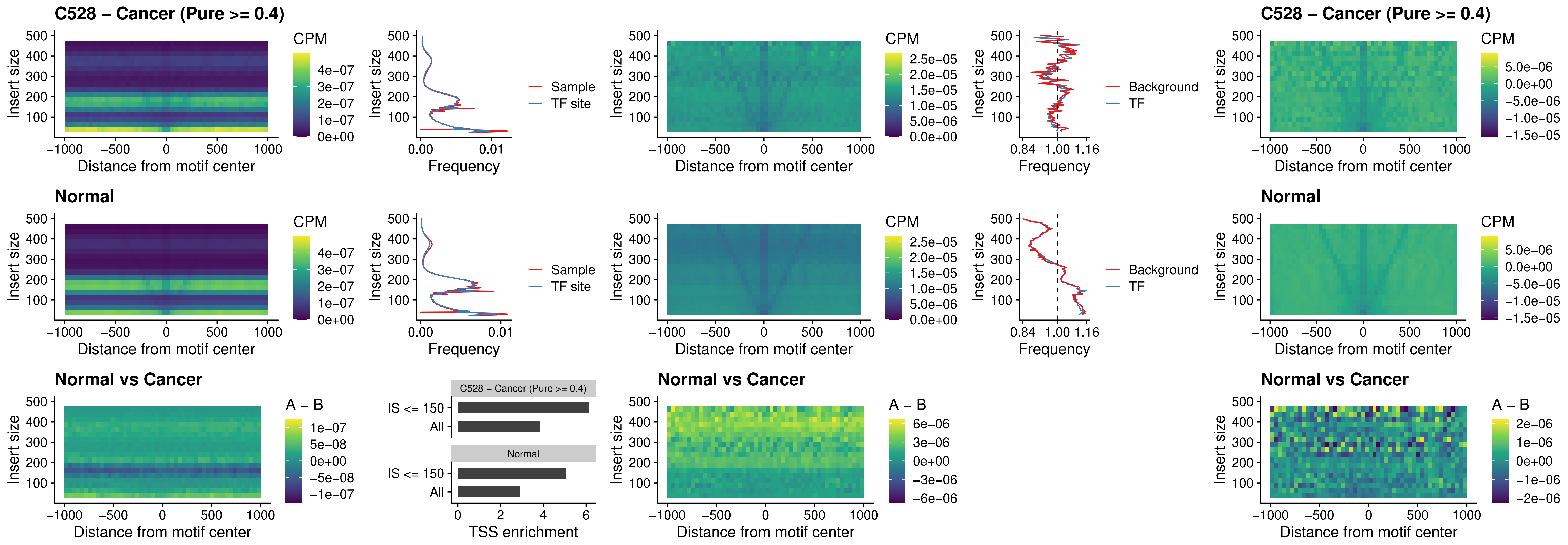

#### C528 – Cancer (Pure >= 0.4) – HOXA4_uc_dTSS_oPEAK

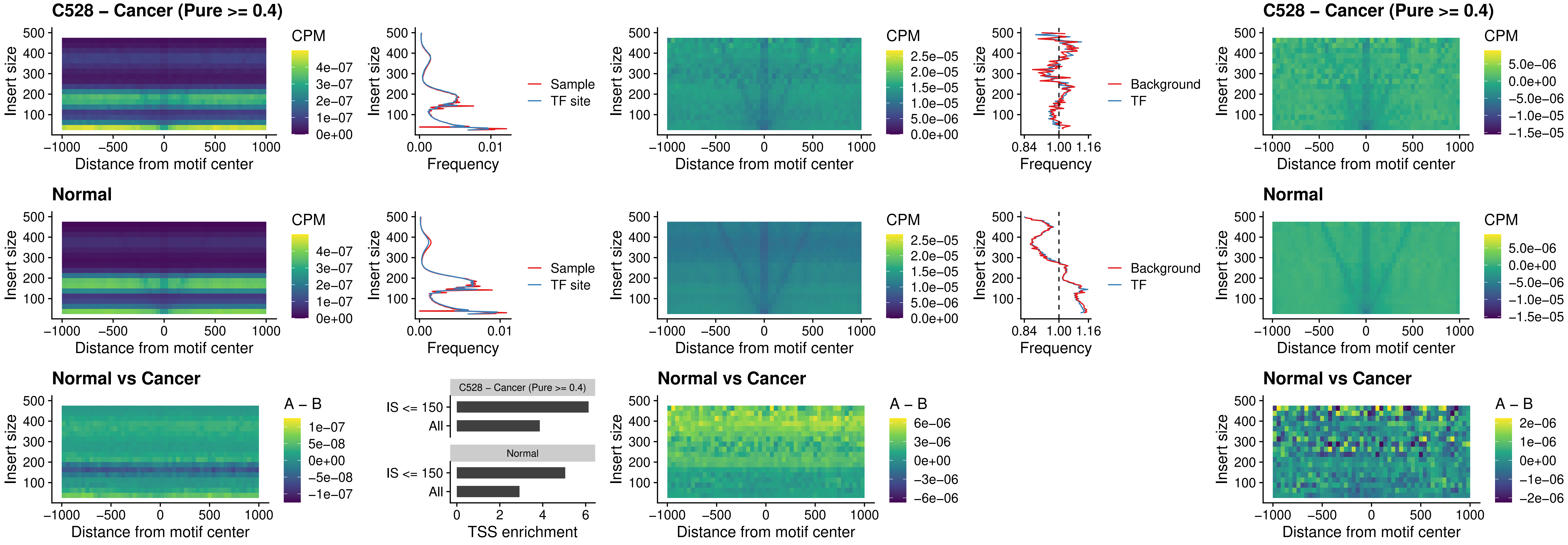

#### C528 – Cancer (Pure >= 0.4) – HLX_uc_dTSS_oPEAK

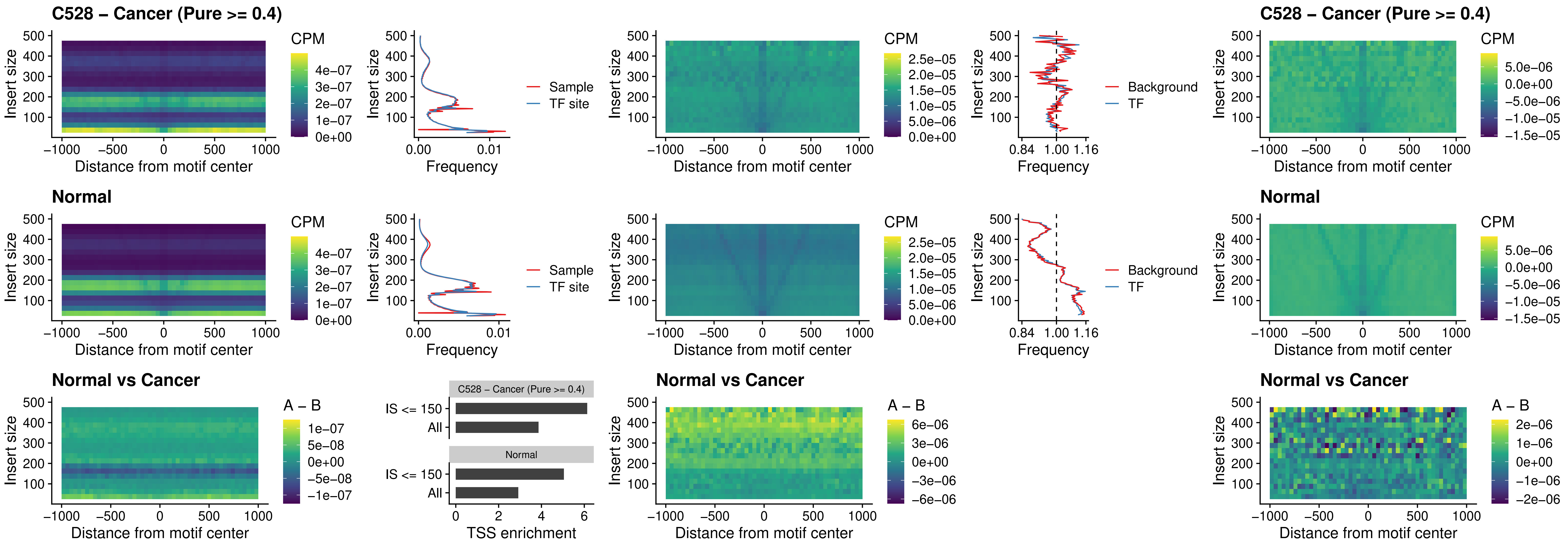

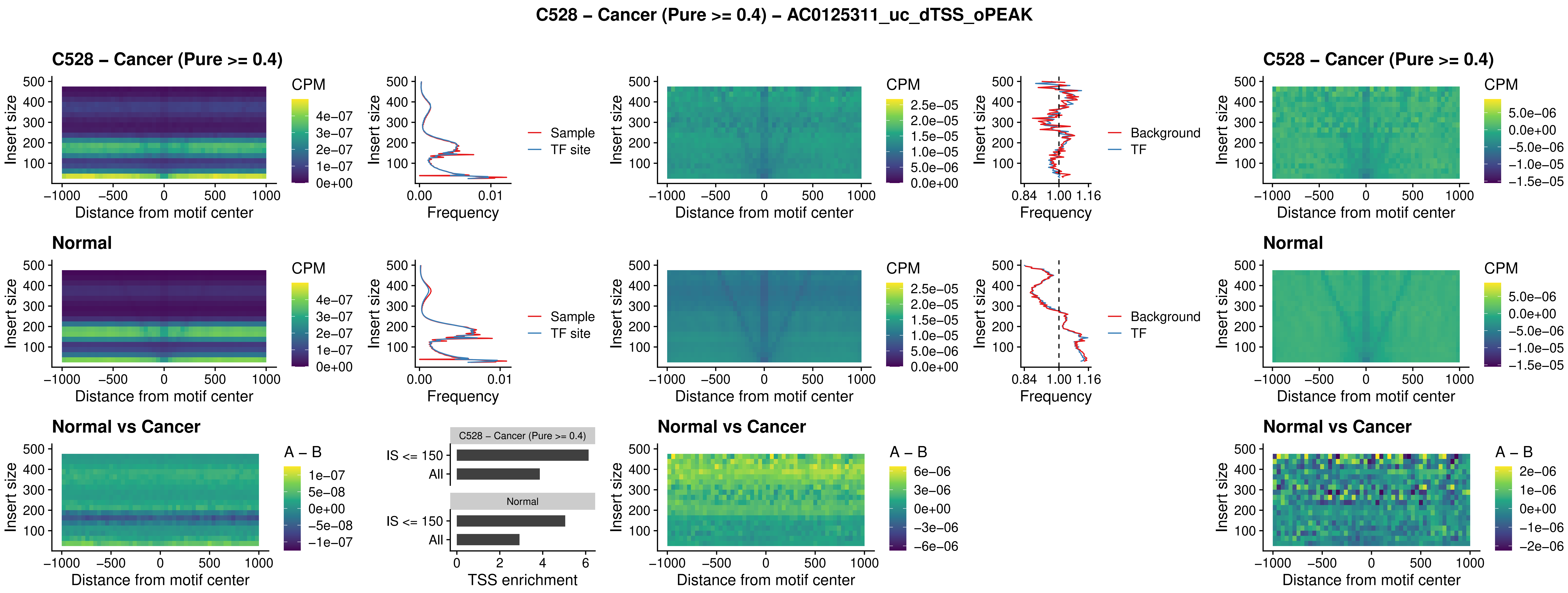
C528 – Cancer (Pure >= 0.4) – AC0125311_uc_dTSS_oPEAK
C528 – Cancer (Pure >= 0.4)
Normal
Normal vs Cancer
Insert size
Distance from motif center
CPM
A – B
Frequency
Sample
TF site
Background
TF
C528 – Cancer (Pure >= 0.4)
Normal
IS <= 150
All
TSS enrichment

### C528 – Cancer (Pure >= 0.4) – DBX2_uc_dTSS_oPEAK

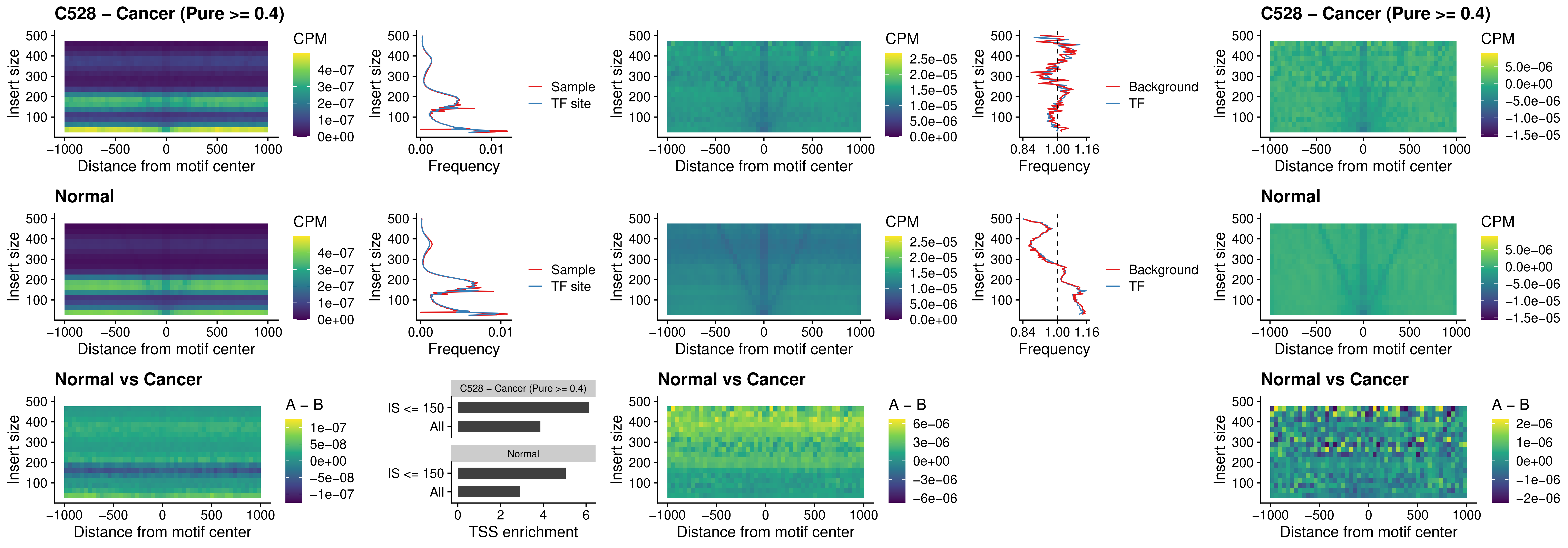

#### C528 – Cancer (Pure >= 0.4) – HOXA2_uc_dTSS_nPEAK

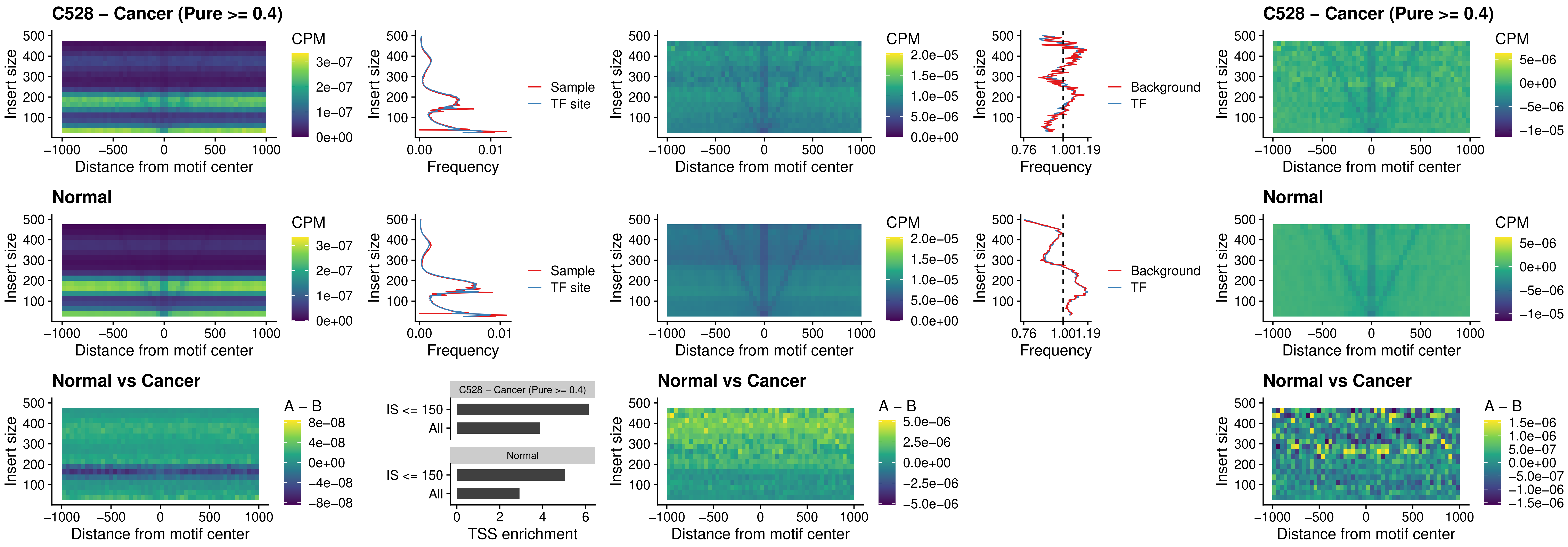

#### C528 – Cancer (Pure >= 0.4) – SHOX_uc_dTSS_nPEAK

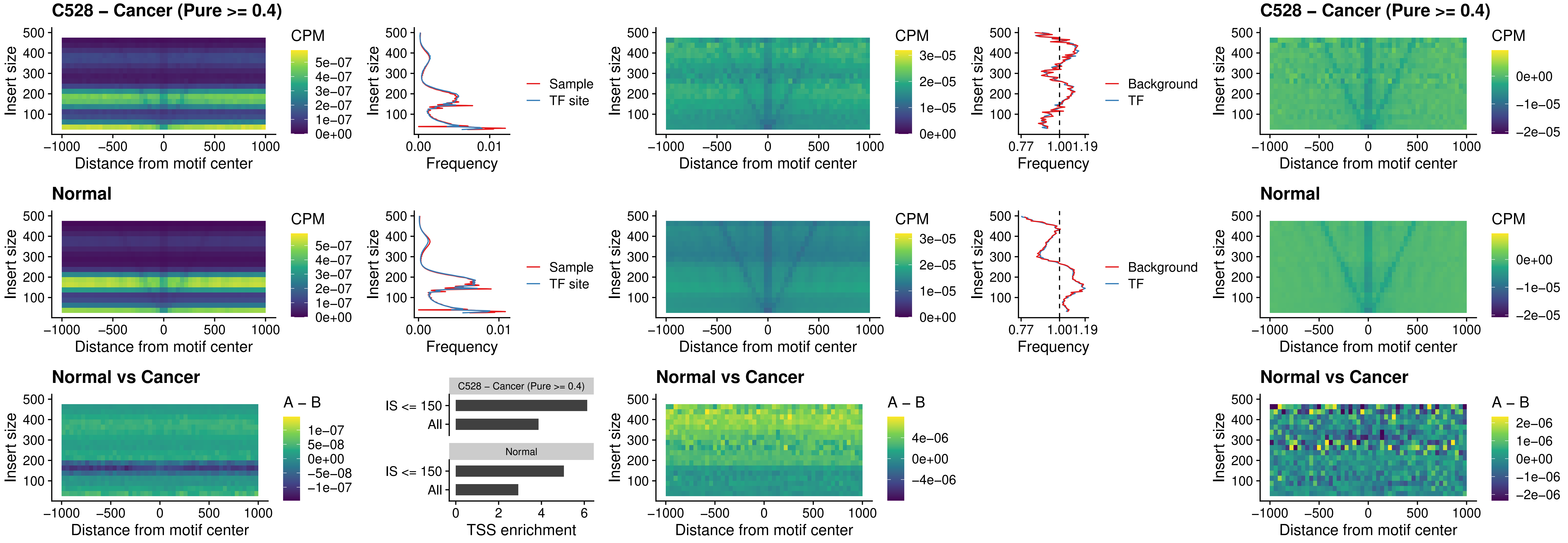

#### C528 – Cancer (Pure >= 0.4) – PRRX1_uc_dTSS_nPEAK

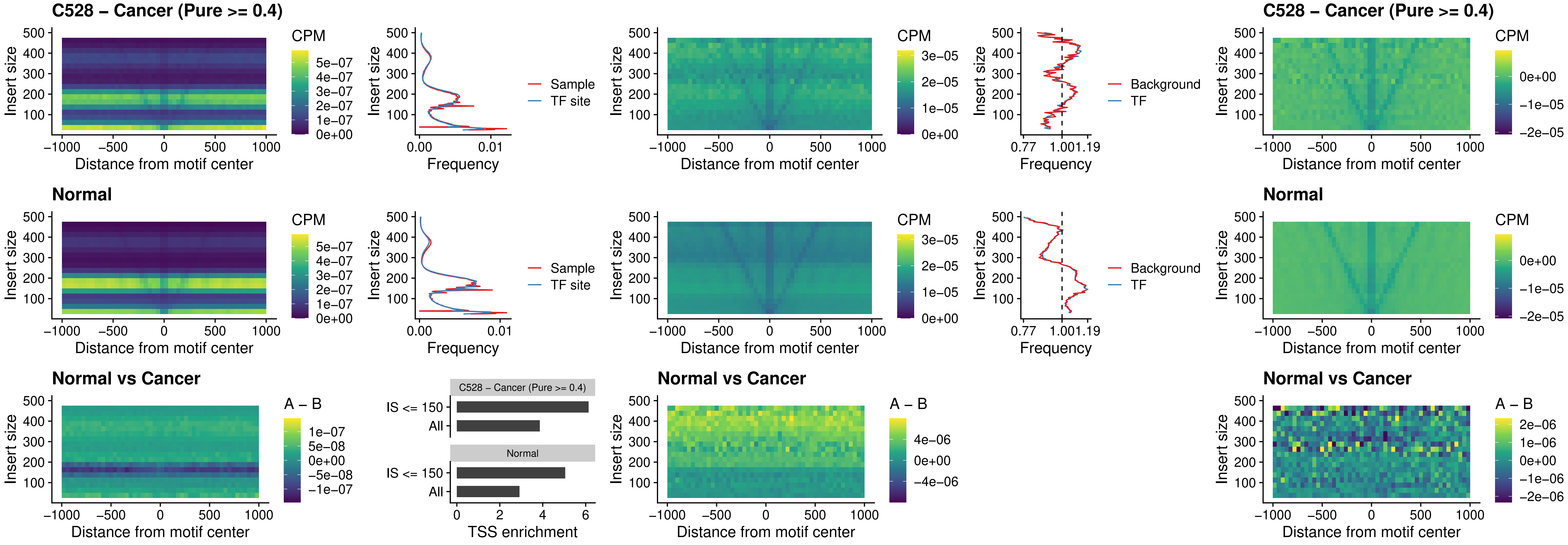

#### C528 – Cancer (Pure >= 0.4) – LMX1A_uc_dTSS_nPEAK

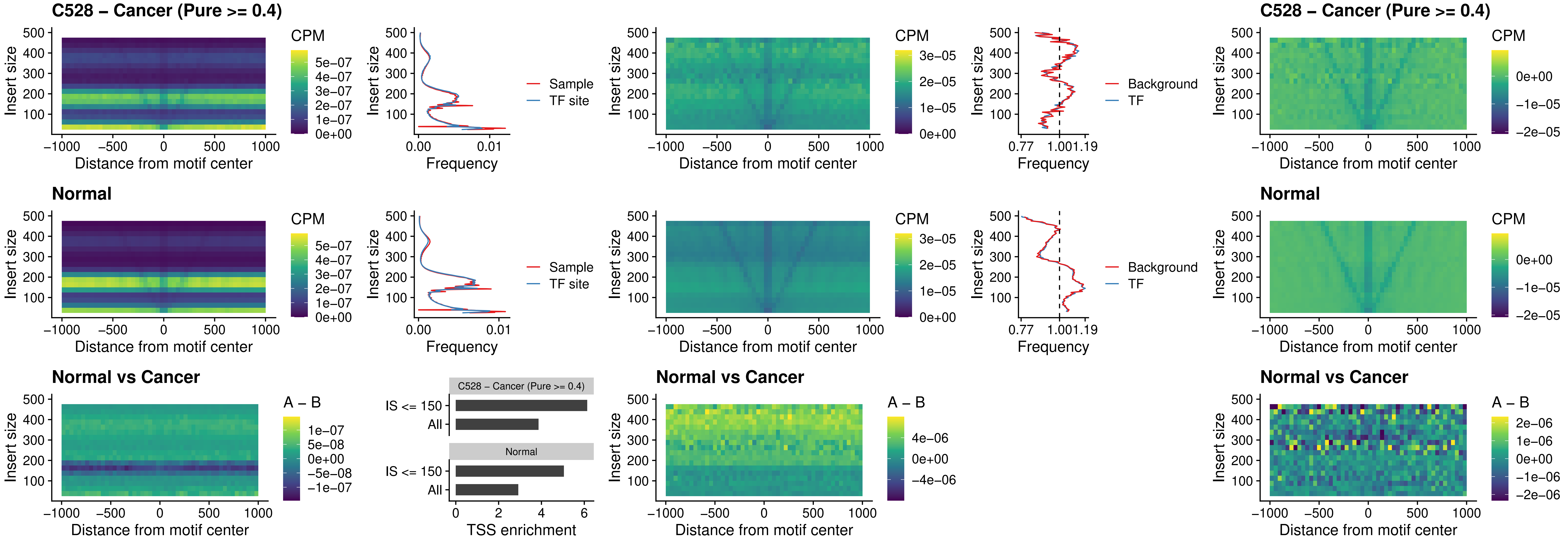

#### C528 – Cancer (Pure >= 0.4) – LMX1B_uc_dTSS_nPEAK

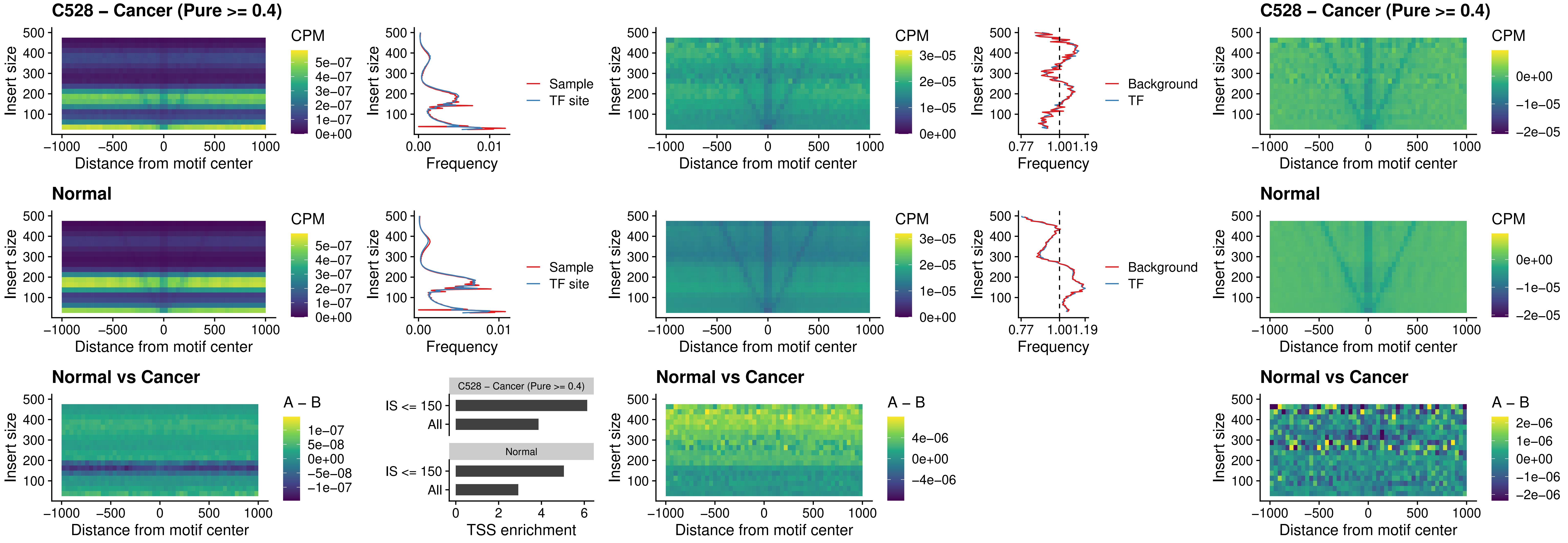

#### C528 – Cancer (Pure >= 0.4) – DLX5_uc_dTSS_oPEAK

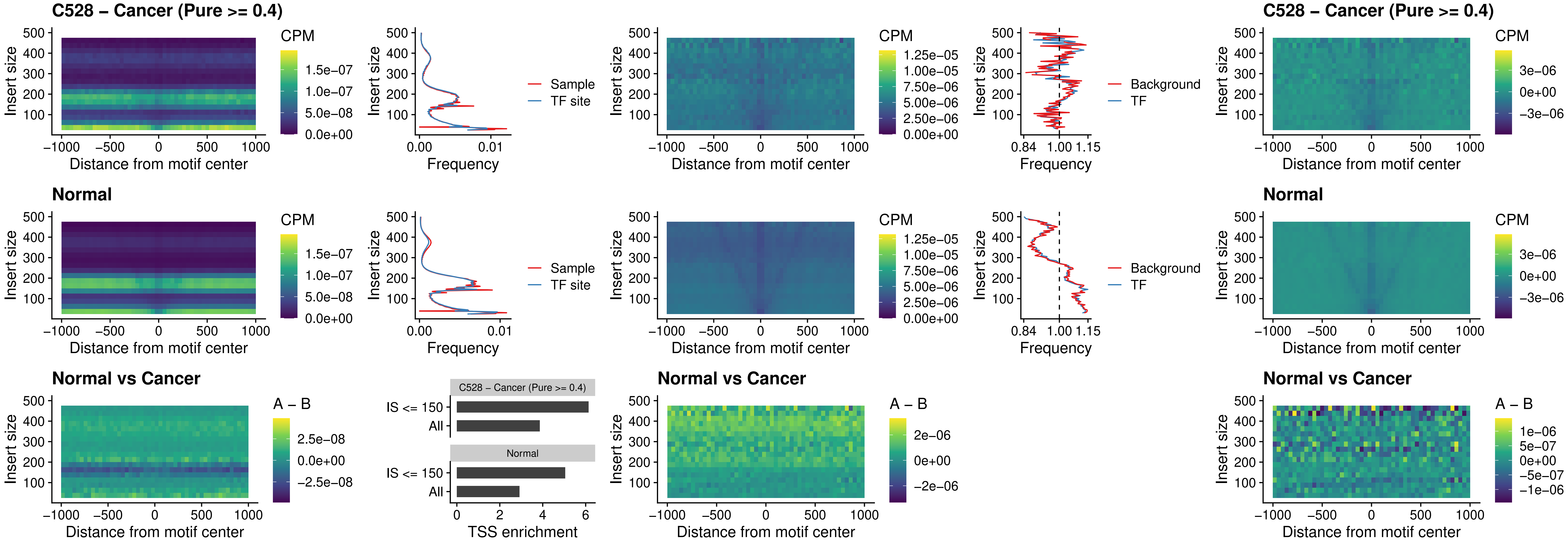

#### C528 – Cancer (Pure >= 0.4) – ARID3C_uc_dTSS_nPEAK

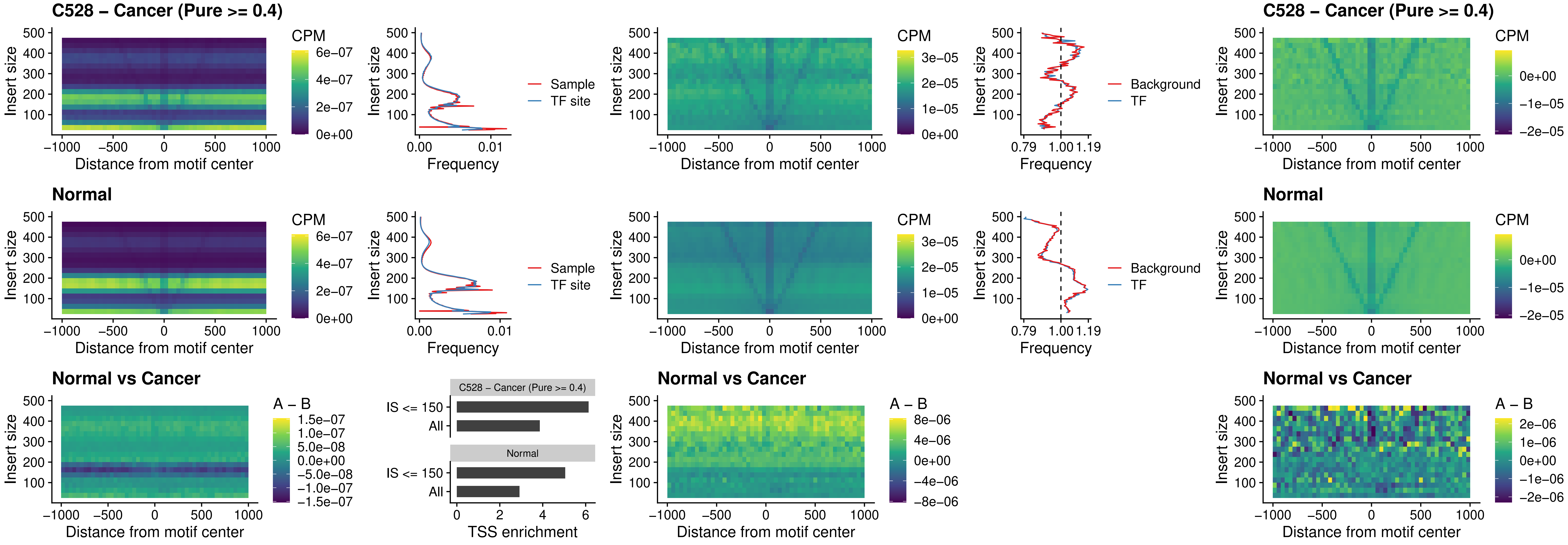

#### C528 – Cancer (Pure >= 0.4) – HOXD12_uc_dTSS_nPEAK

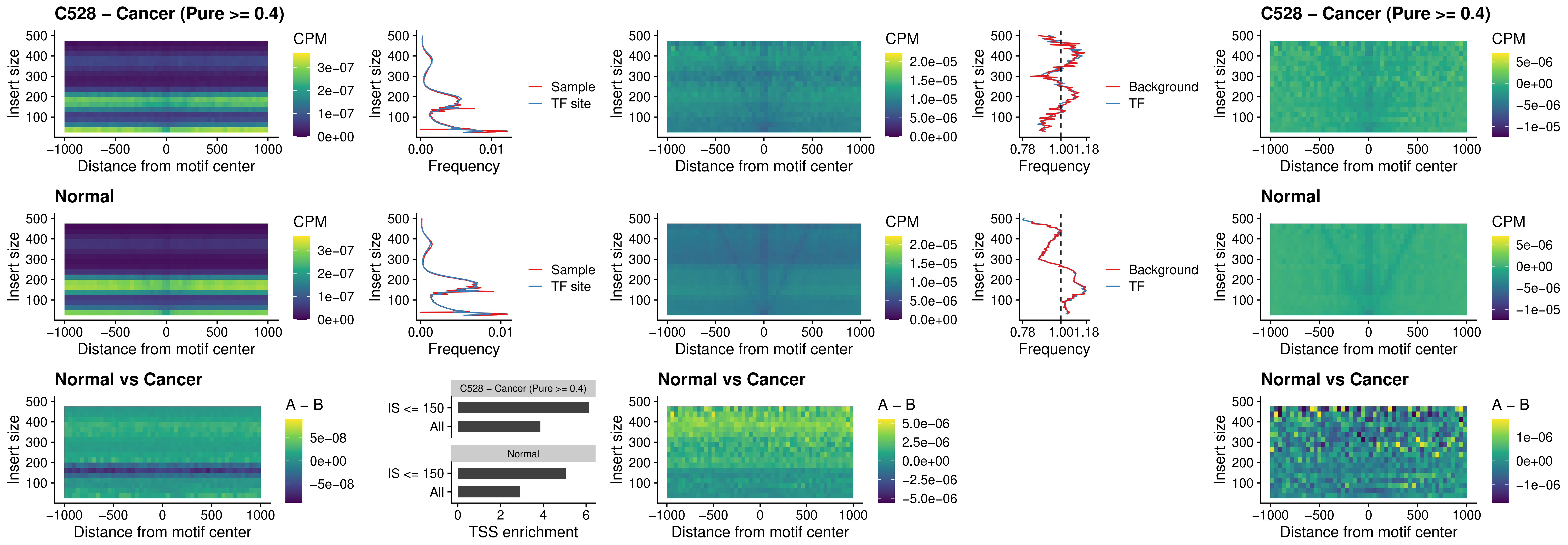

#### C528 – Cancer (Pure >= 0.4) – CDX2_uc_dTSS_nPEAK

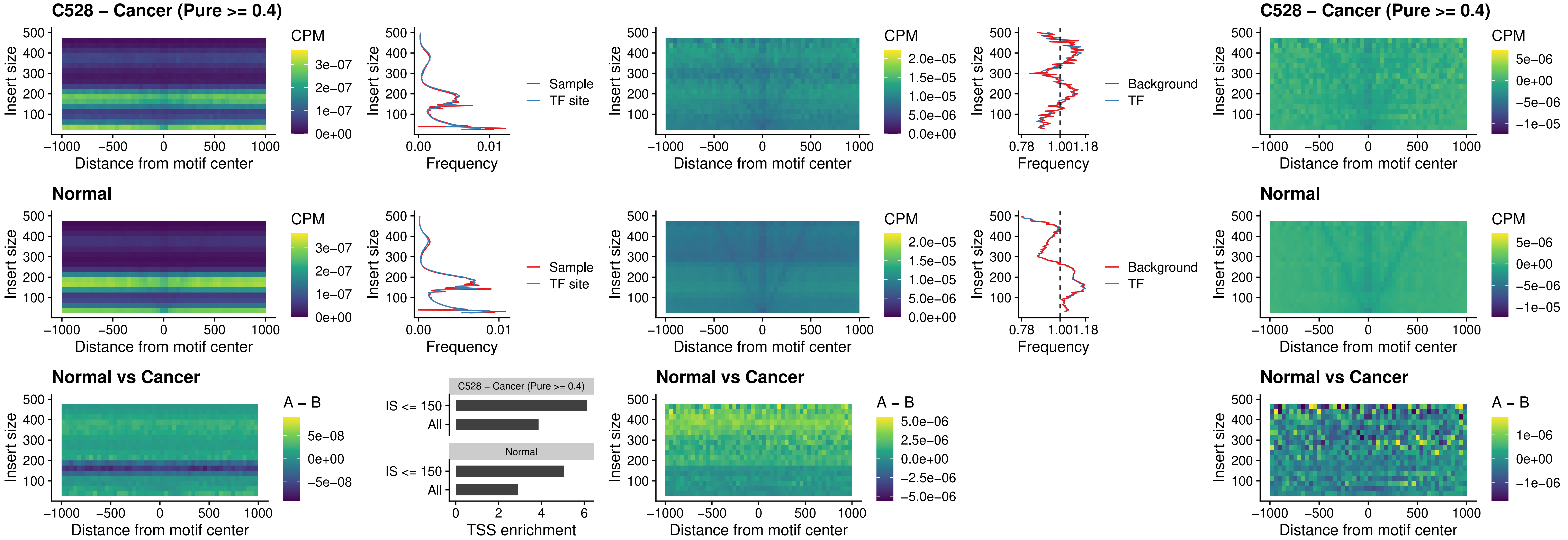

#### C528 – Cancer (Pure >= 0.4) – HOXC12_uc_dTSS_nPEAK

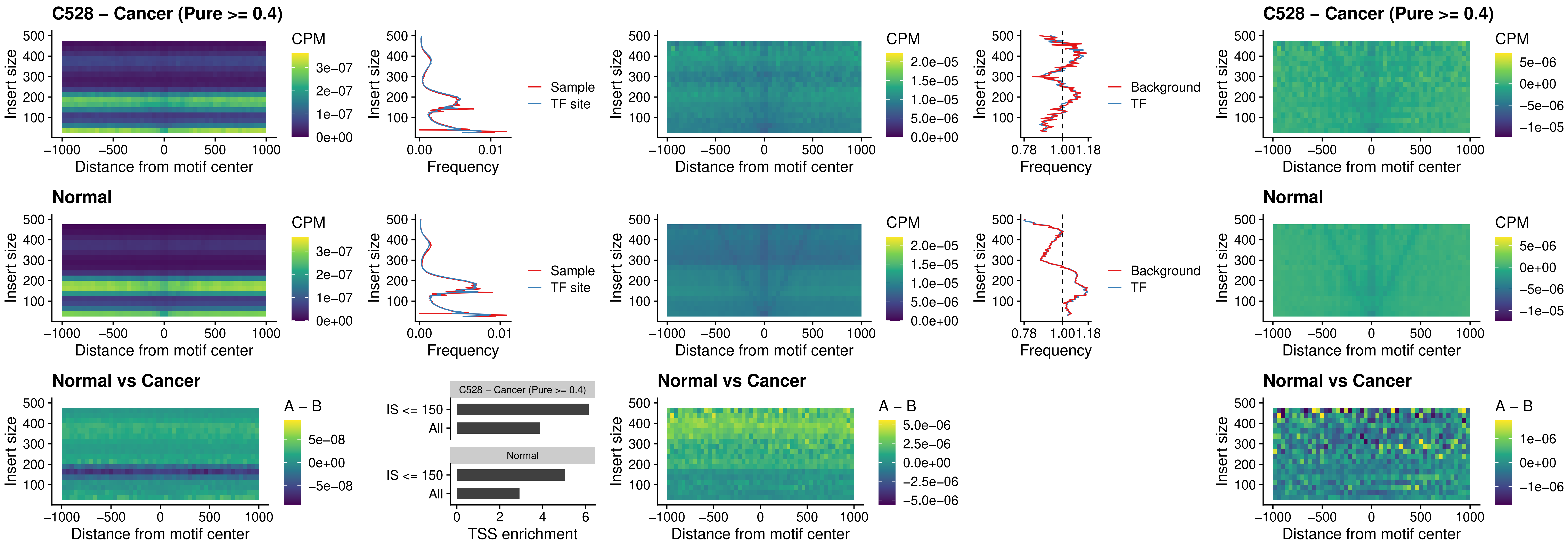

### C528 – Cancer (Pure >= 0.4) – HHEX_uc_dTSS_oPEAK

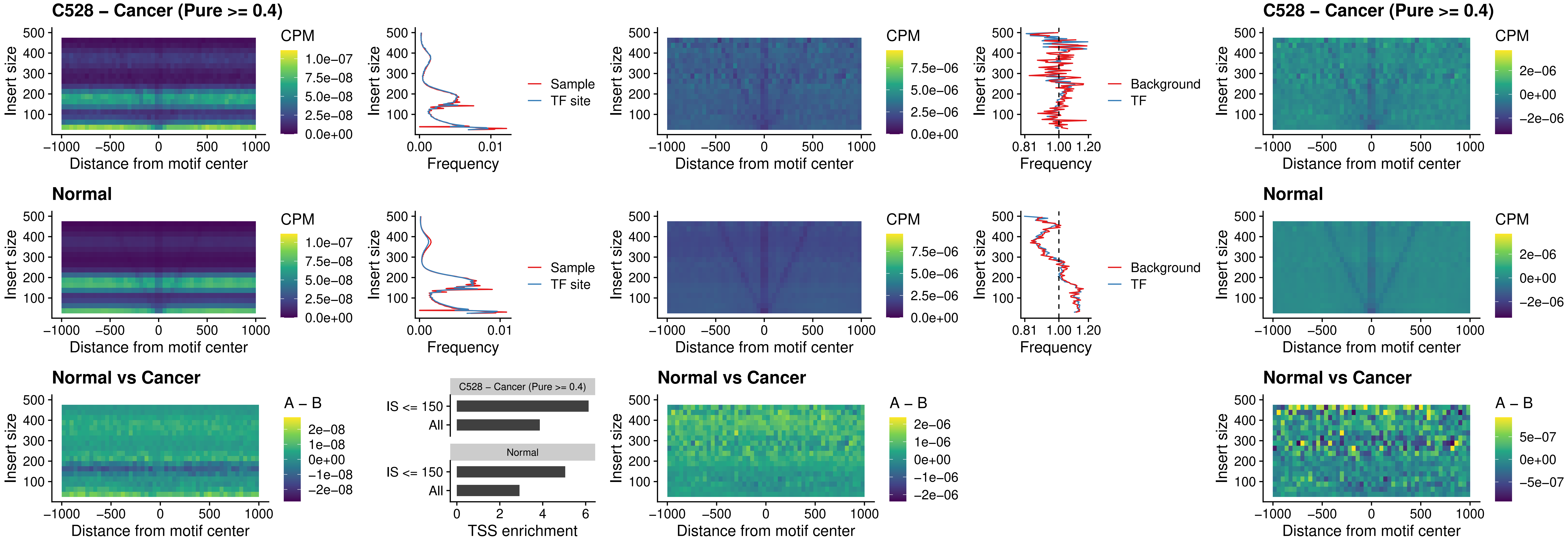

#### C528 – Cancer (Pure >= 0.4) – HOXA11_uc_dTSS_nPEAK

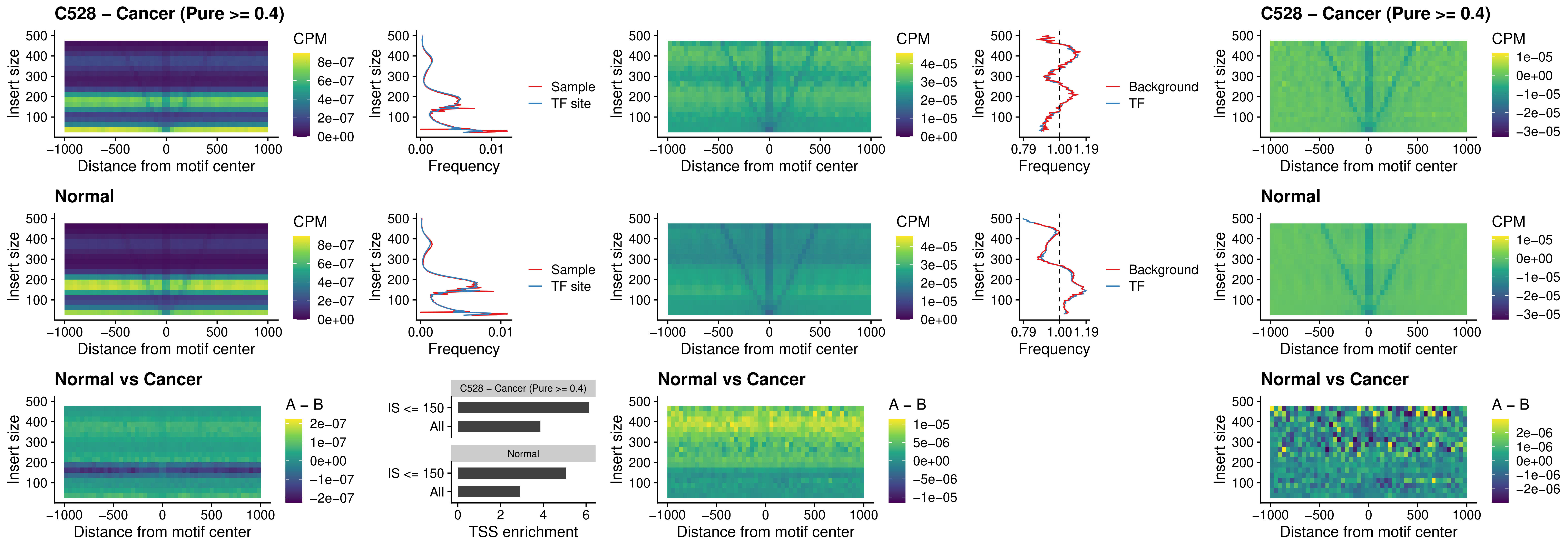

#### C528 – Cancer (Pure >= 0.4) – HOXA11_uc_cTSS_nPEAK

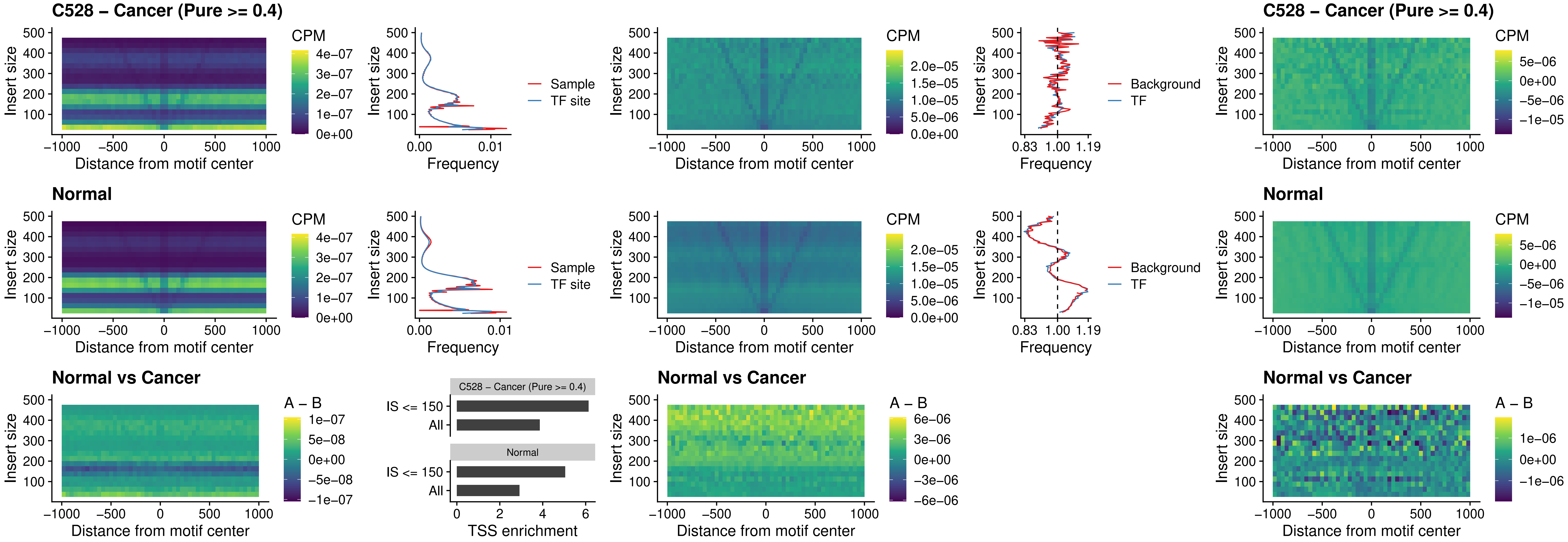

#### C528 – Cancer (Pure >= 0.4) – ZFHX3_uc_dTSS_nPEAK

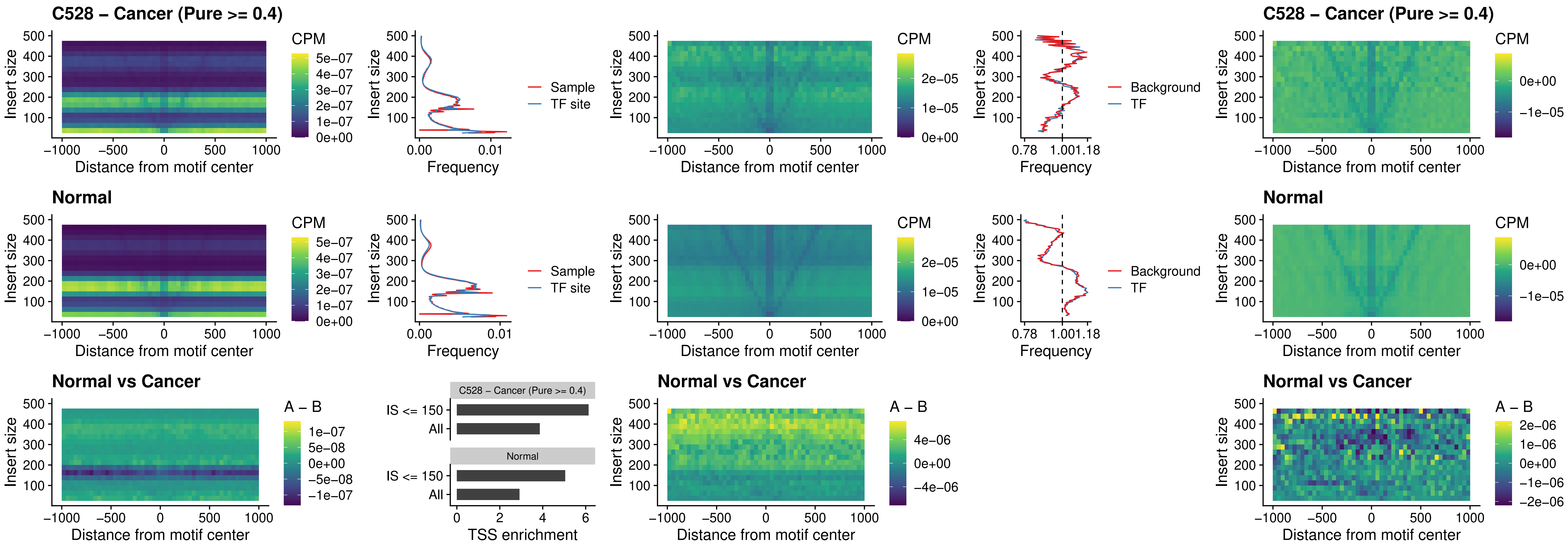

#### C528 – Cancer (Pure >= 0.4) – HOXB4_uc_dTSS_nPEAK

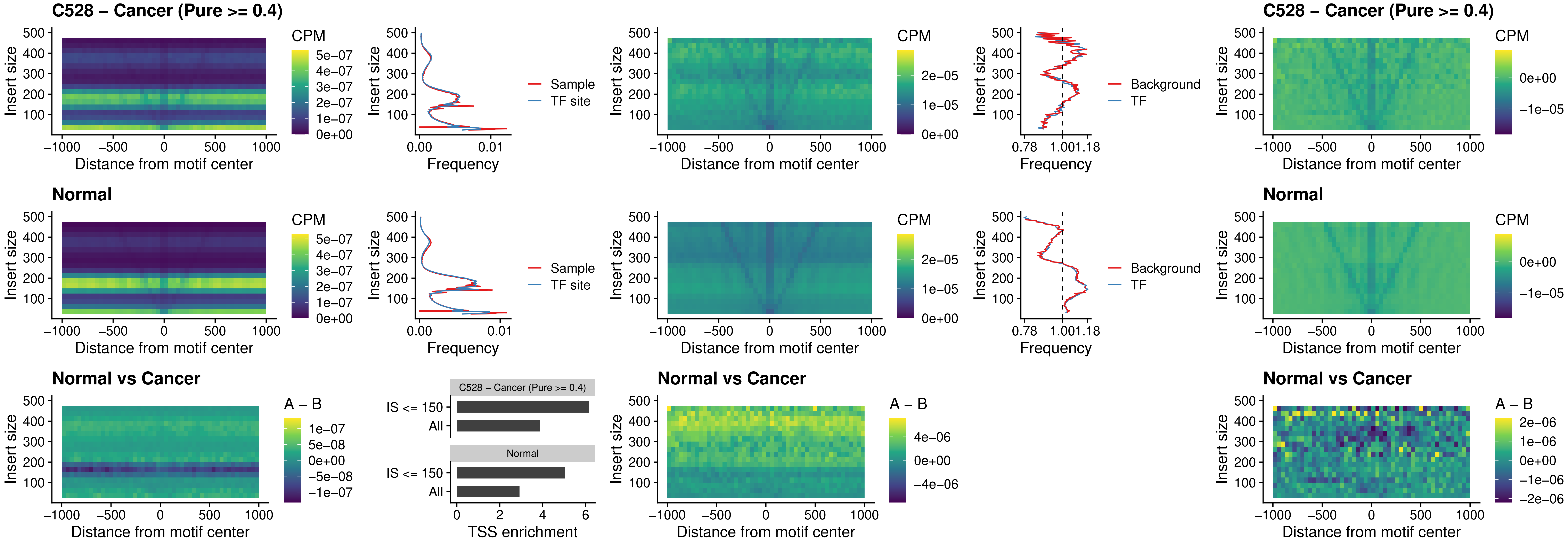

#### C528 – Cancer (Pure >= 0.4) – HOXC5_uc_dTSS_nPEAK

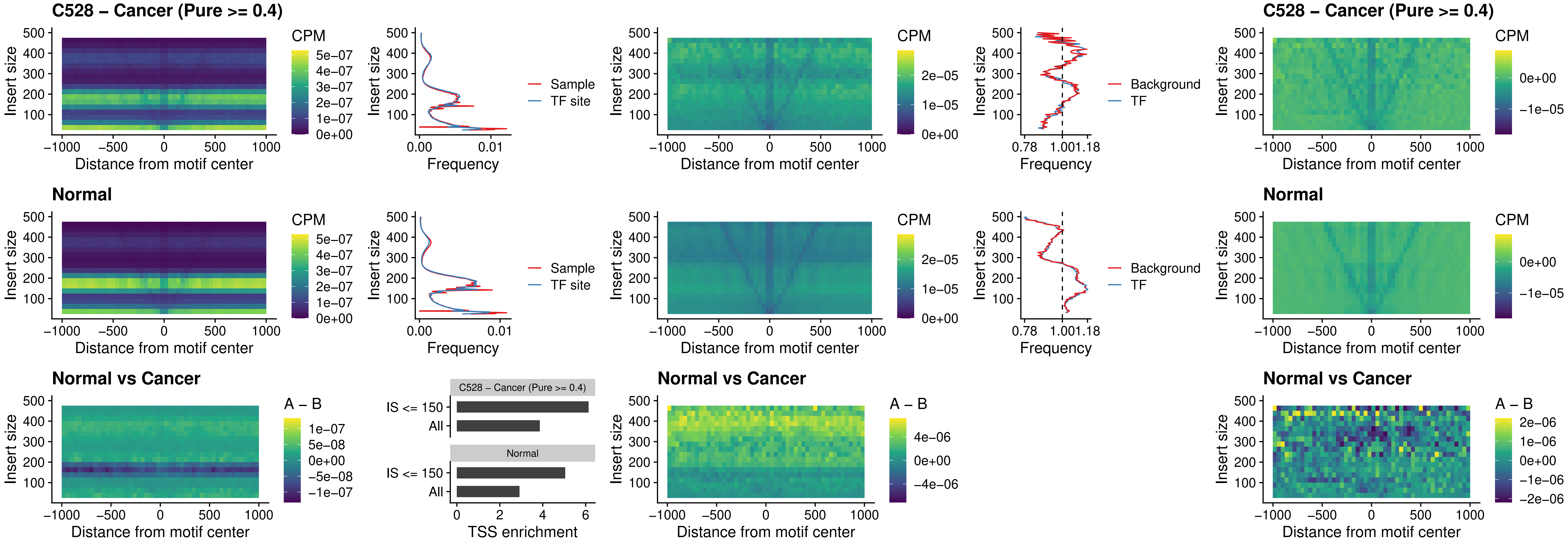

#### C528 – Cancer (Pure >= 0.4) – HOXD1_uc_dTSS_nPEAK

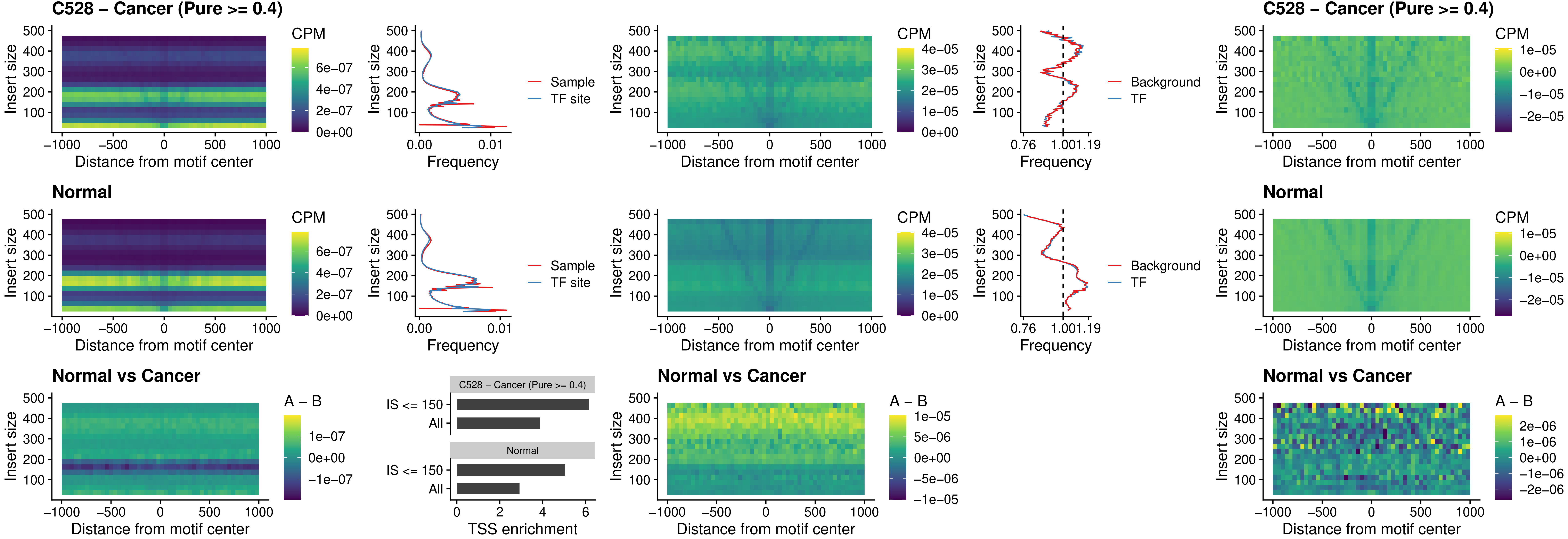

#### C528 – Cancer (Pure >= 0.4) – OTP_uc_dTSS_nPEAK

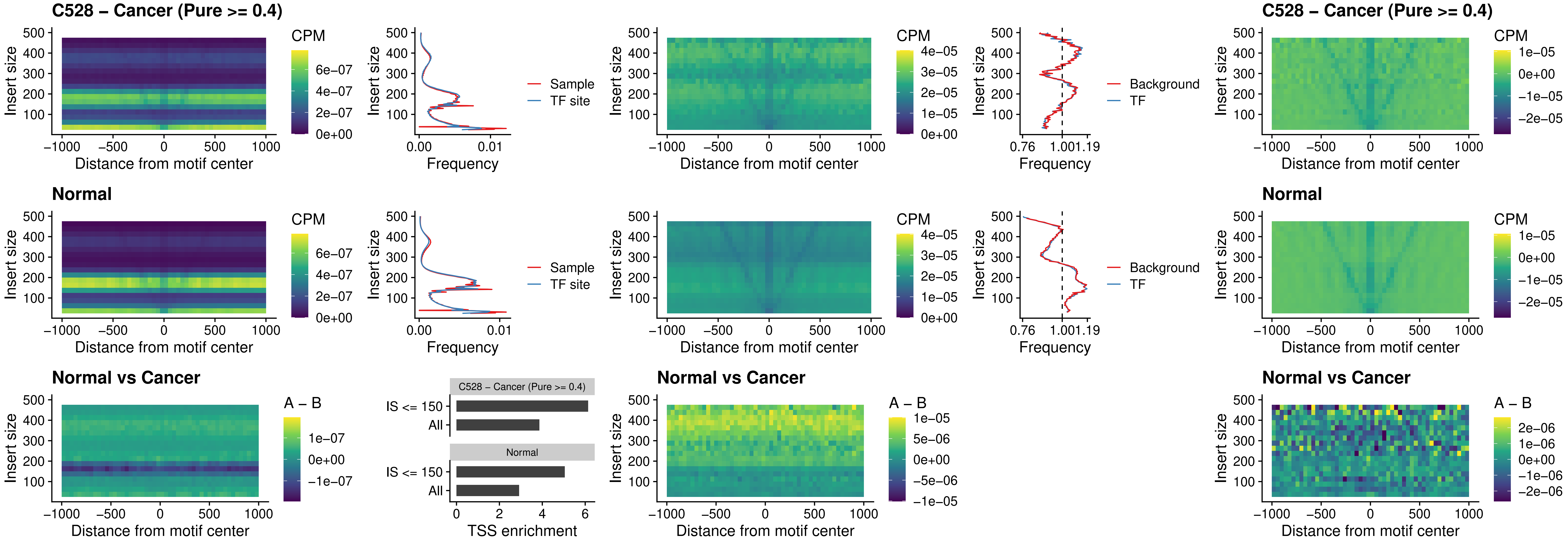

### C528 – Cancer (Pure >= 0.4) – FOXL1_uc_dTSS_oPEAK

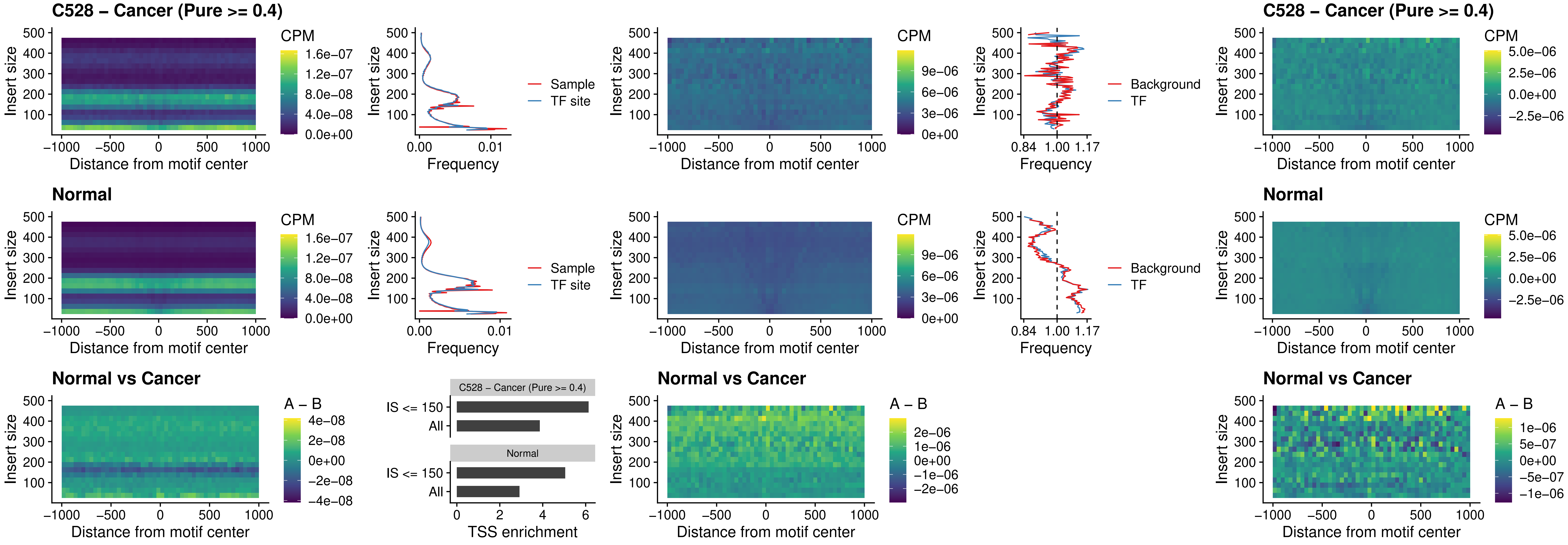

#### C528 – Cancer (Pure >= 0.4) – ZNF32_uc_dTSS_oPEAK

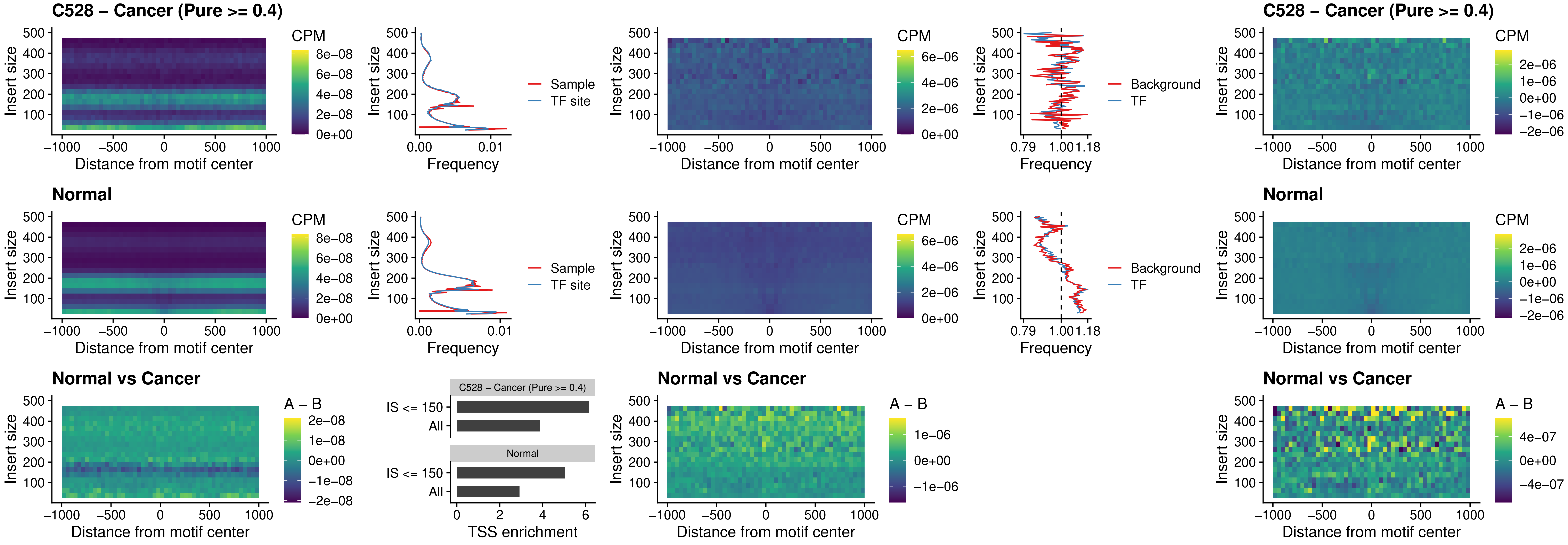

#### C528 – Cancer (Pure >= 0.4) – FOXL1_uc_dTSS_nPEAK

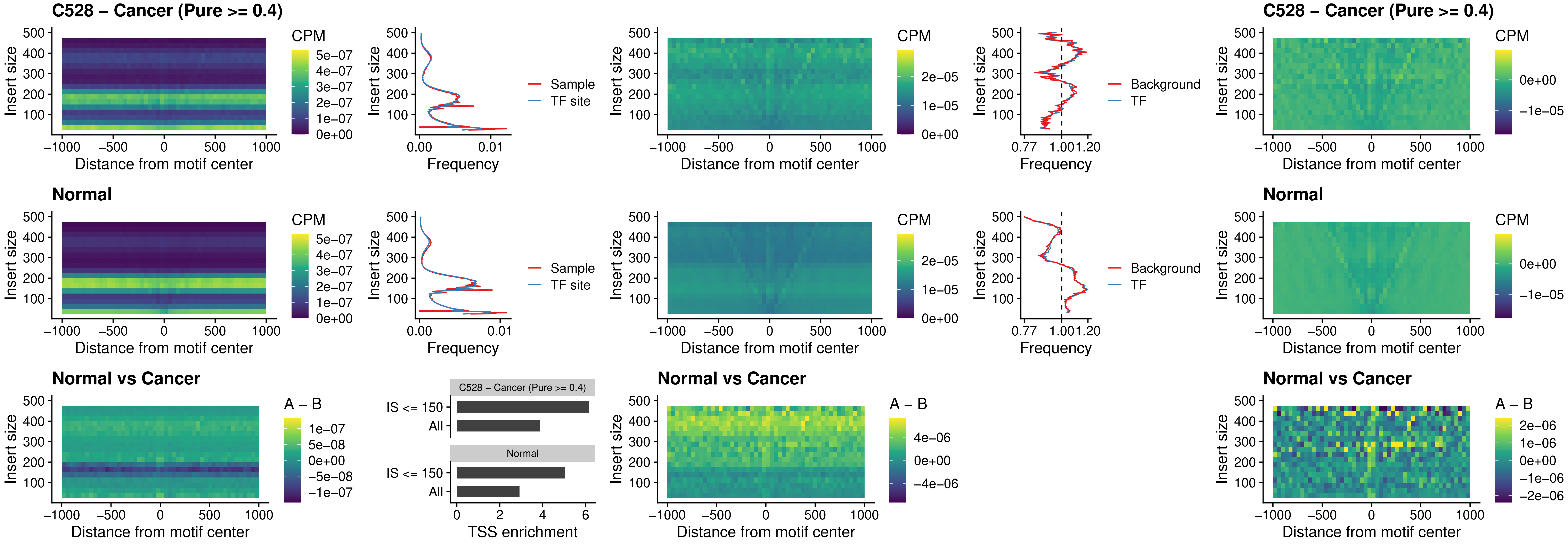

#### C528 – Cancer (Pure >= 0.4) – ZNF32_uc_dTSS_nPEAK

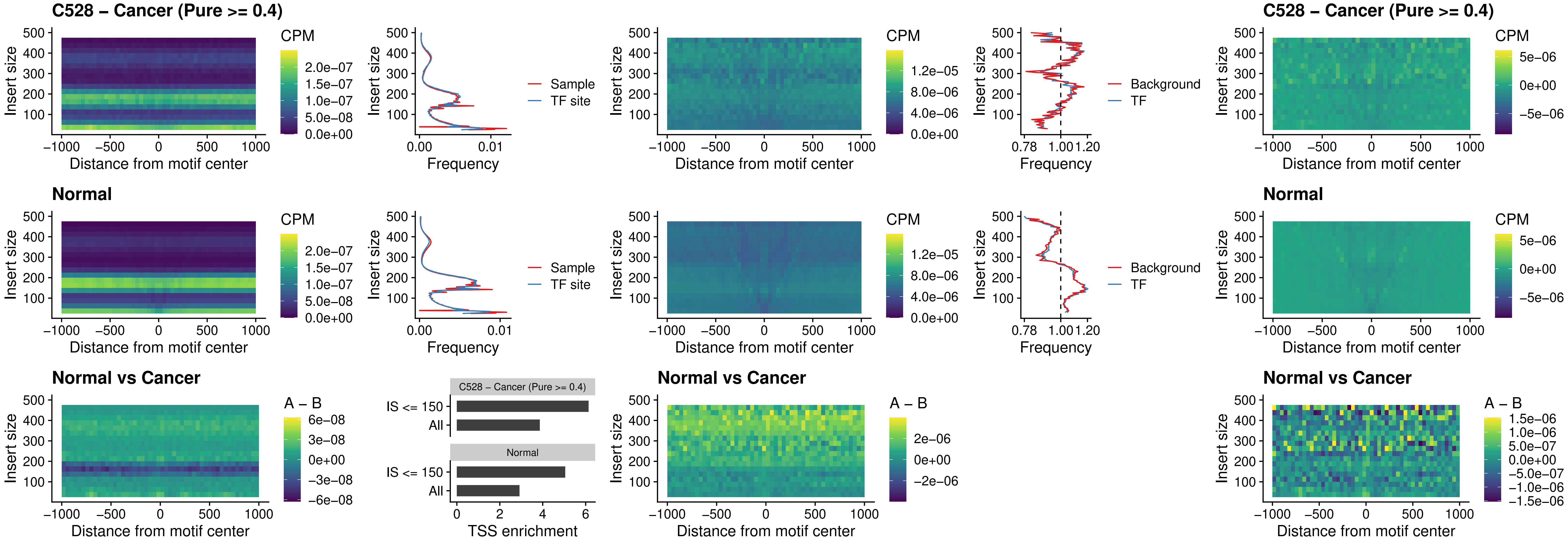

#### C528 – Cancer (Pure >= 0.4) – DLX5_uc_dTSS_nPEAK

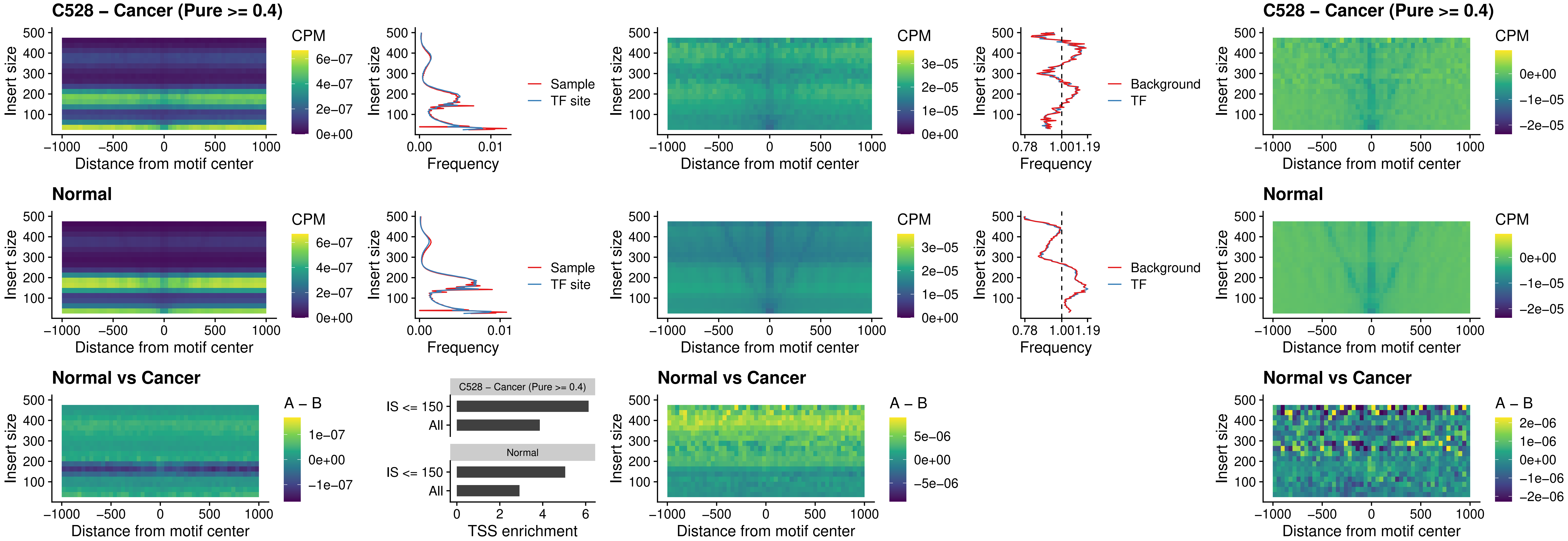

#### C528 – Cancer (Pure >= 0.4) – DMRT2_uc_dTSS_nPEAK

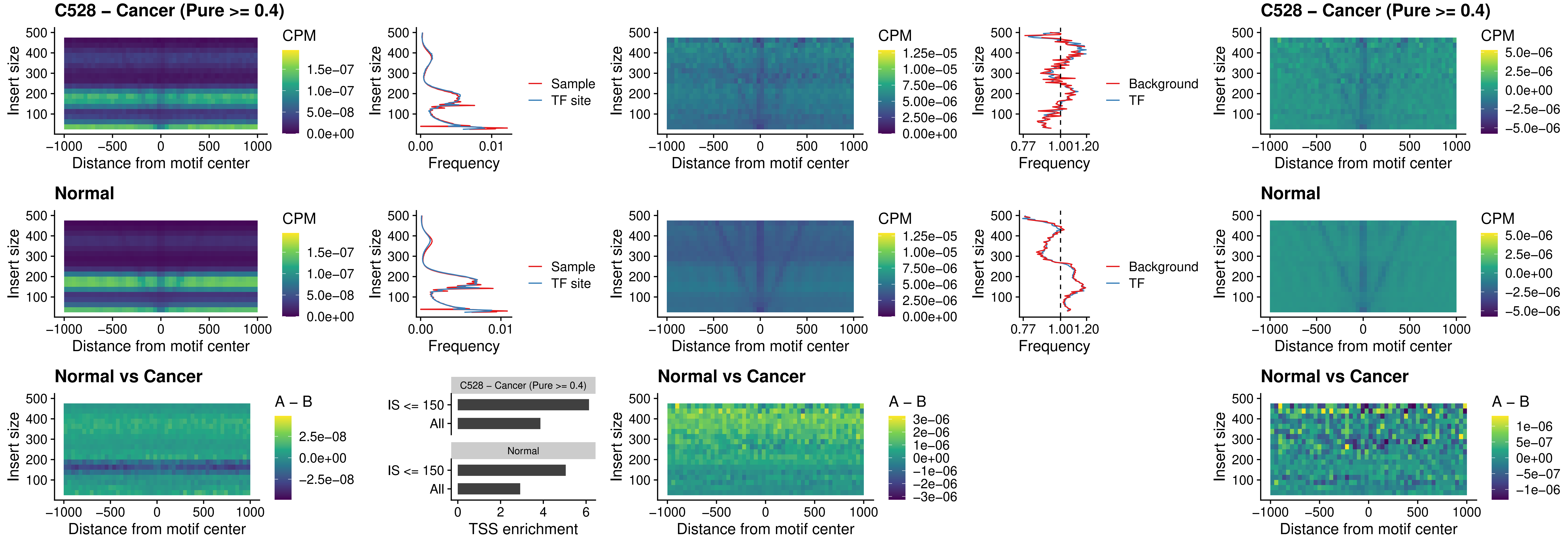

#### C528 – Cancer (Pure >= 0.4) – DMRTC2_uc_cTSS_nPEAK

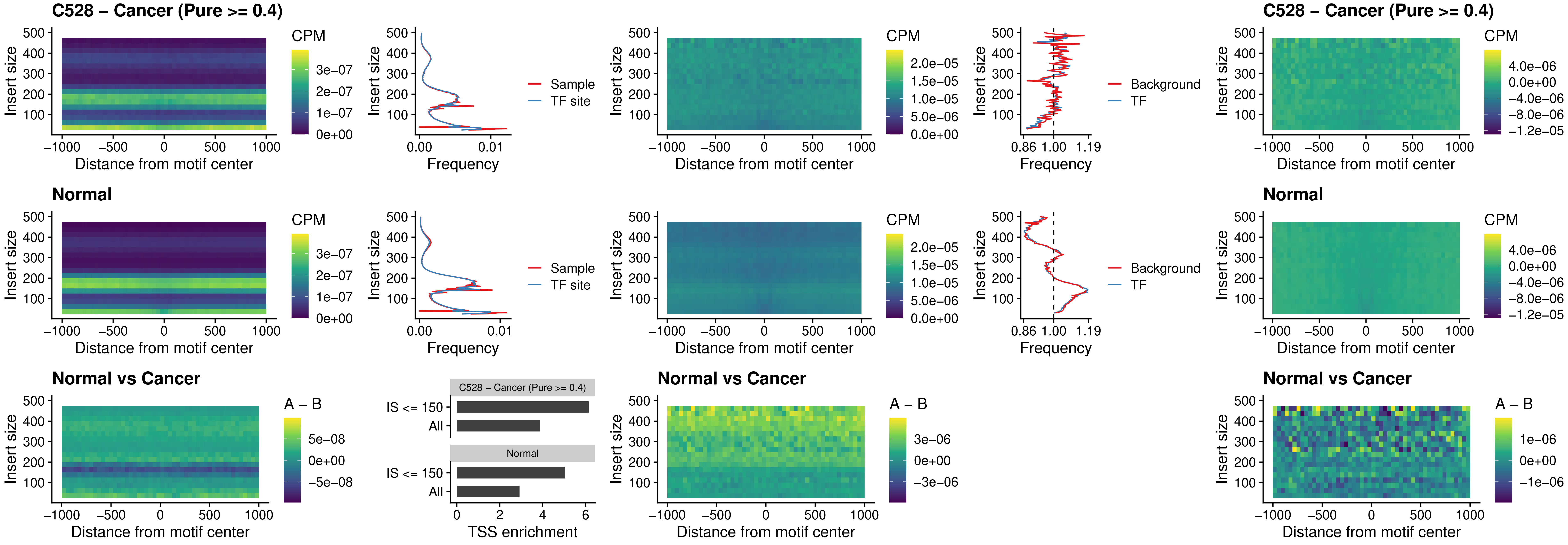

#### C528 – Cancer (Pure >= 0.4) – HOXD9_uc_dTSS_nPEAK

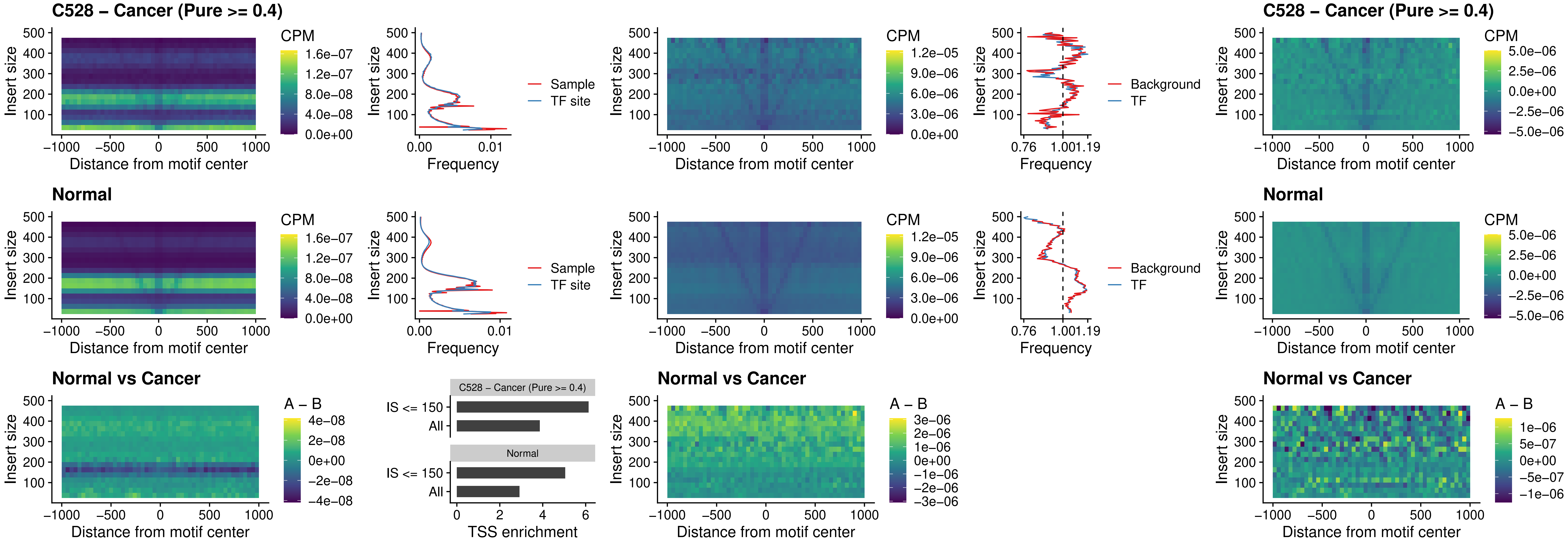

#### C528 – Cancer (Pure >= 0.4) – HOXD4_uc_dTSS_nPEAK

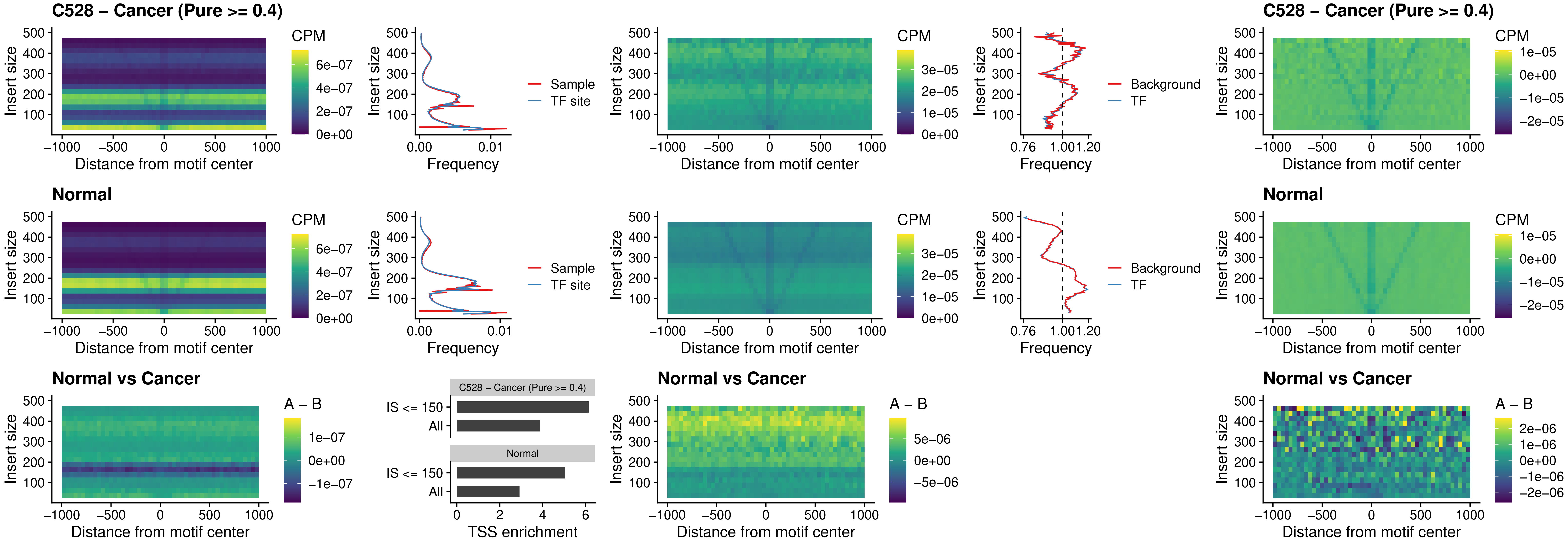

#### C528 – Cancer (Pure >= 0.4) – FOXP3_uc_cTSS_nPEAK

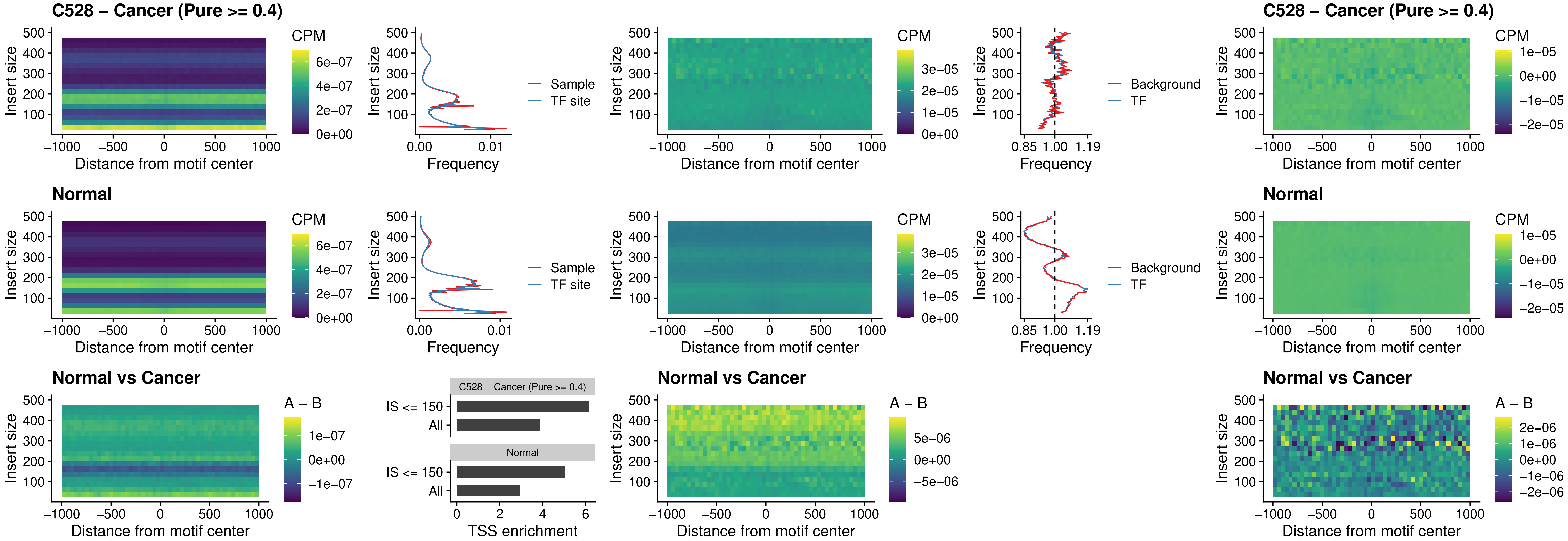

### C528 – Cancer (Pure >= 0.4) – FOXD4L6_uc_cTSS_nPEAK

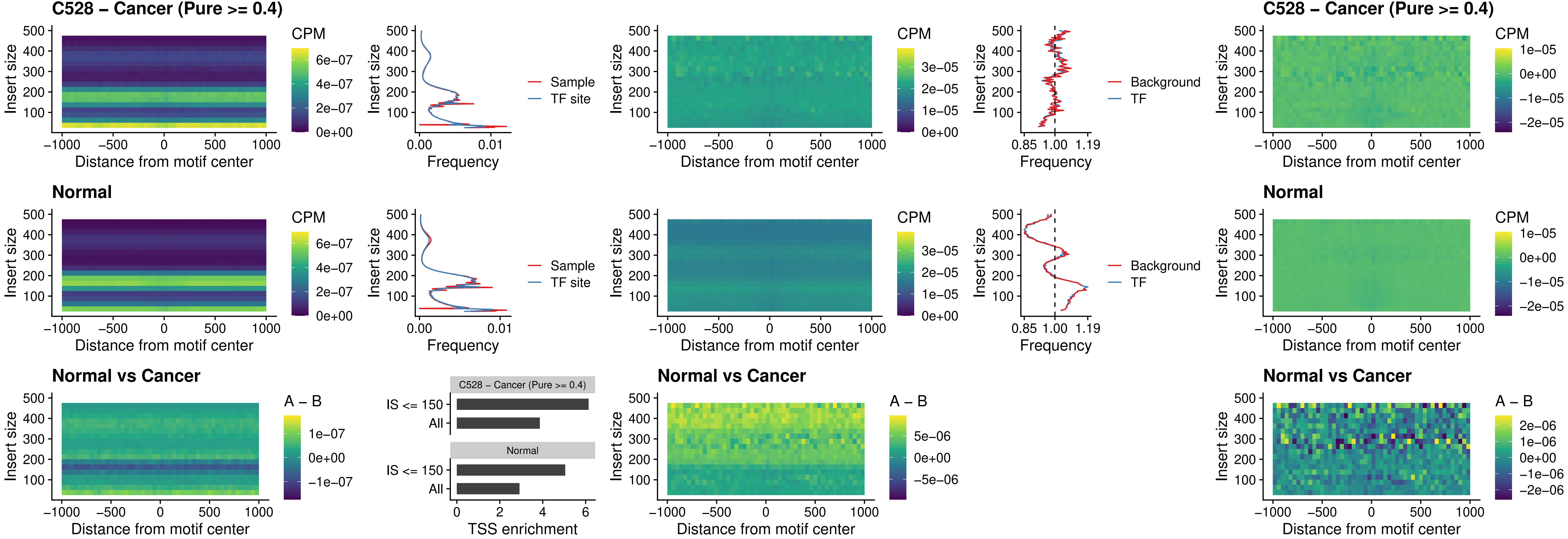

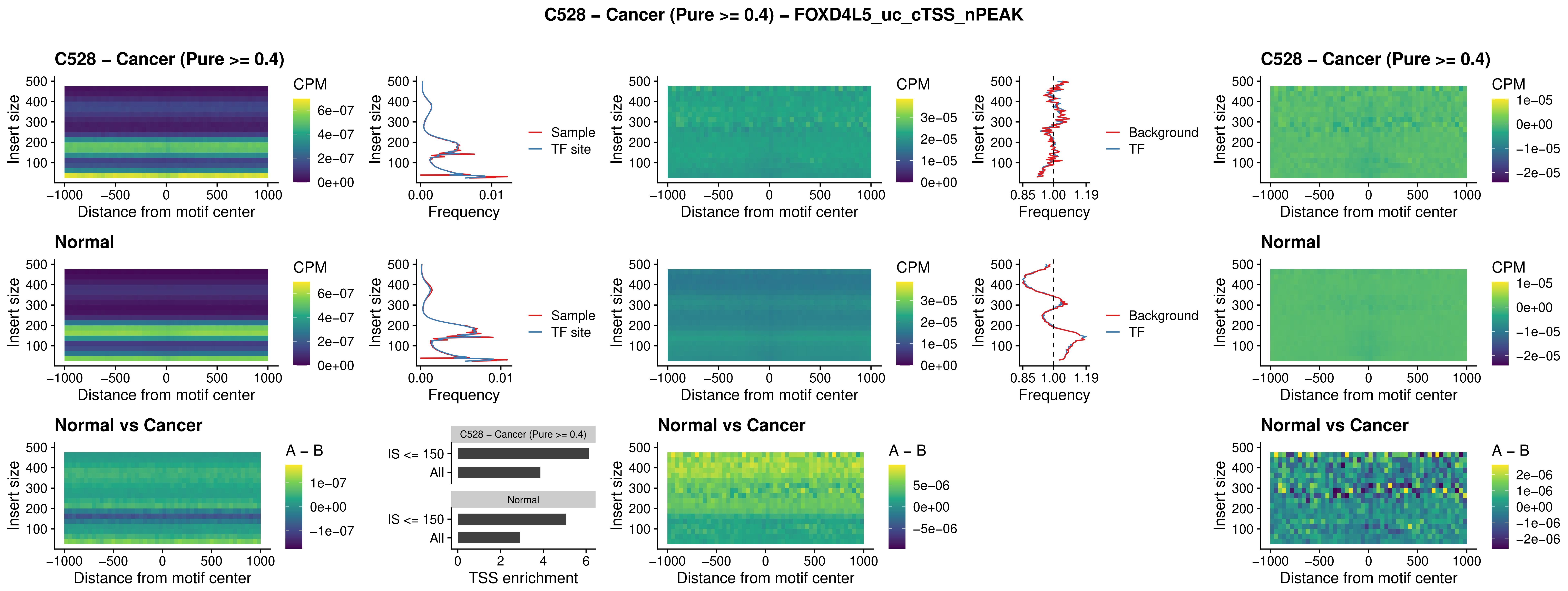
C528 – Cancer (Pure >= 0.4) – FOXD4L5_uc_cTSS_nPEAK
C528 – Cancer (Pure >= 0.4)
Normal
Normal vs Cancer
Insert size
Distance from motif center
Frequency
CPM
A – B
Sample
TF site
Background
TF
C528 – Cancer (Pure >= 0.4)
Normal
IS <= 150
All
TSS enrichment

#### C528 – Cancer (Pure >= 0.4) – FOXD4L4_uc_cTSS_nPEAK

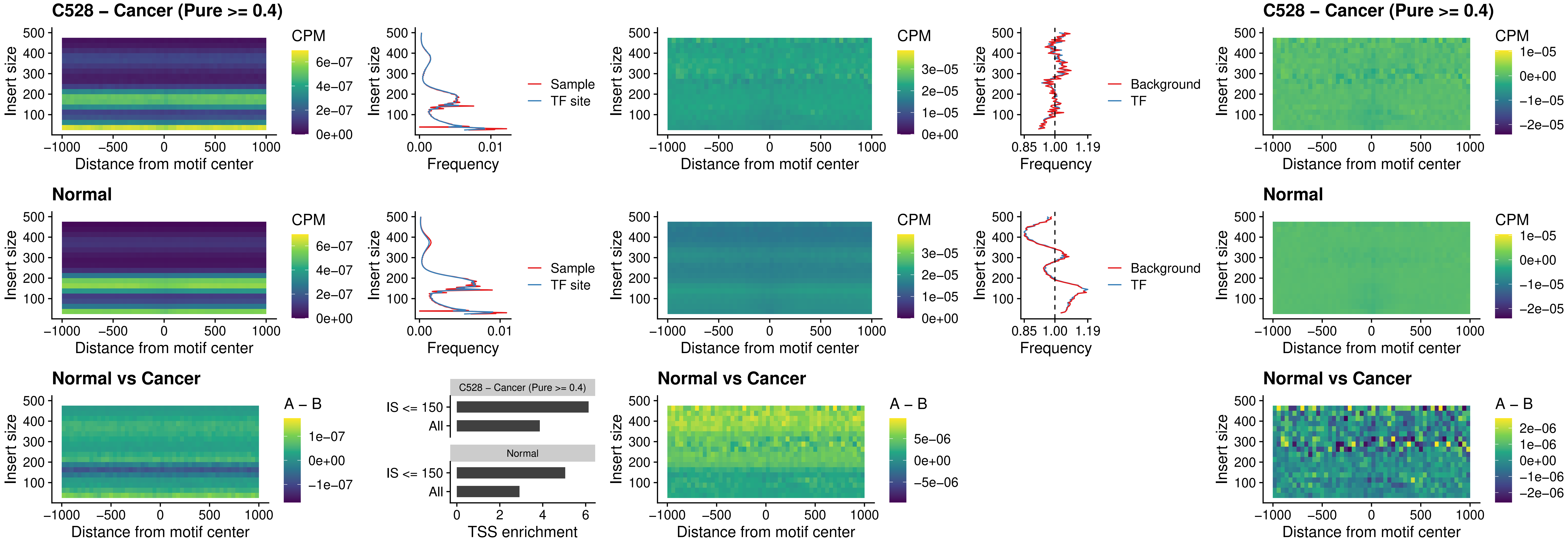

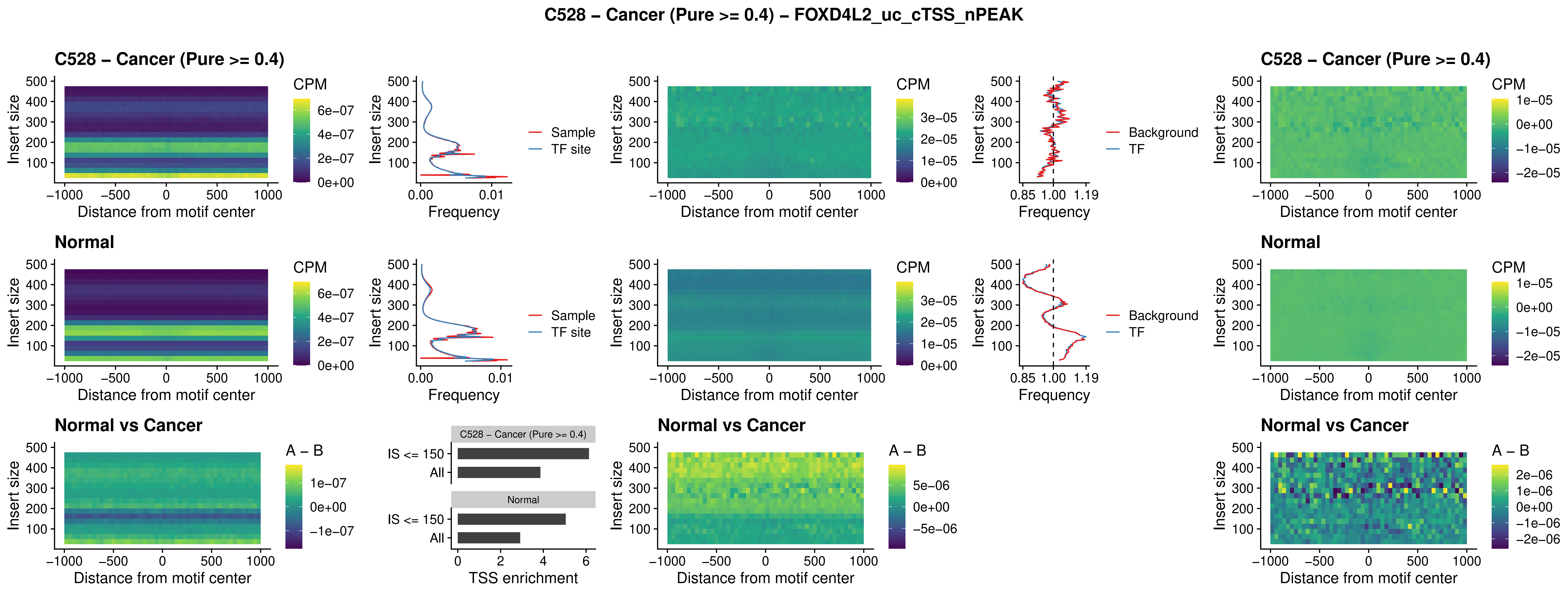

C528 – Cancer (Pure >= 0.4) – FOXD4L2_uc_cTSS_nPEAK
C528 – Cancer (Pure >= 0.4)
Normal
Normal vs Cancer
Insert size
Distance from motif center
Frequency
CPM
A – B
Sample
TF site
Background
TF
C528 – Cancer (Pure >= 0.4)
Normal
IS <= 150
All
TSS enrichment

#### C528 – Cancer (Pure >= 0.4) – FOXD4L3_uc_cTSS_nPEAK

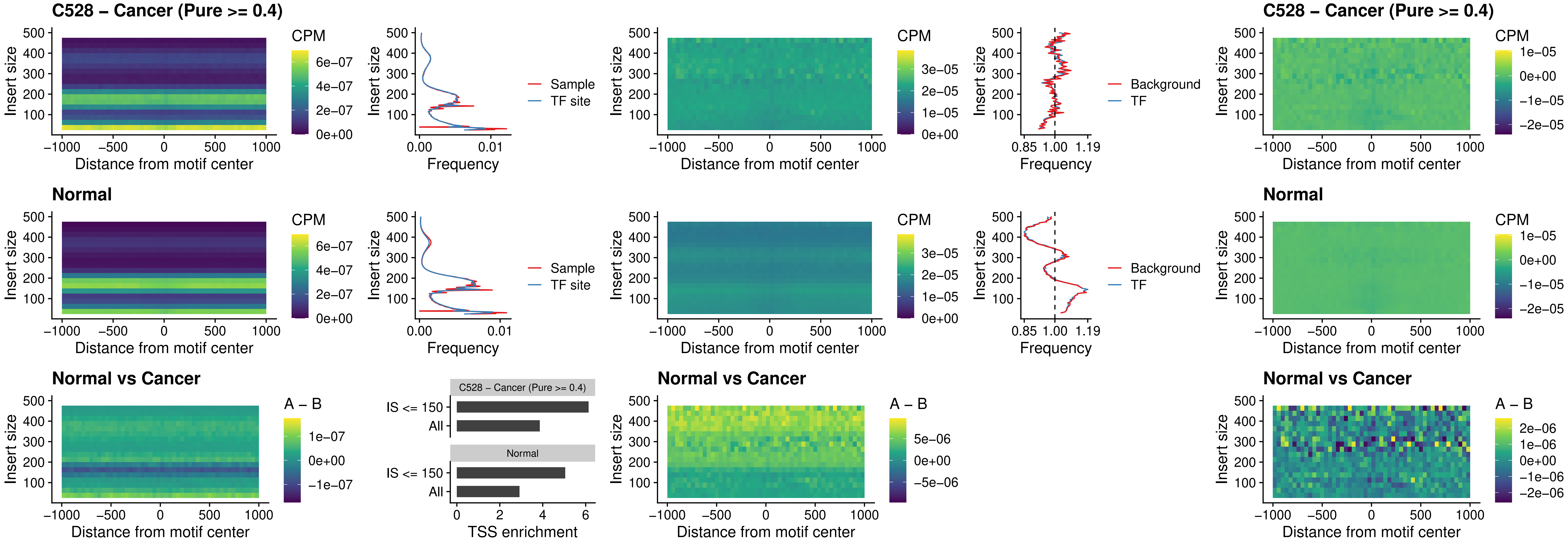

#### C528 – Cancer (Pure >= 0.4) – HOXD1_uc_dTSS_oPEAK

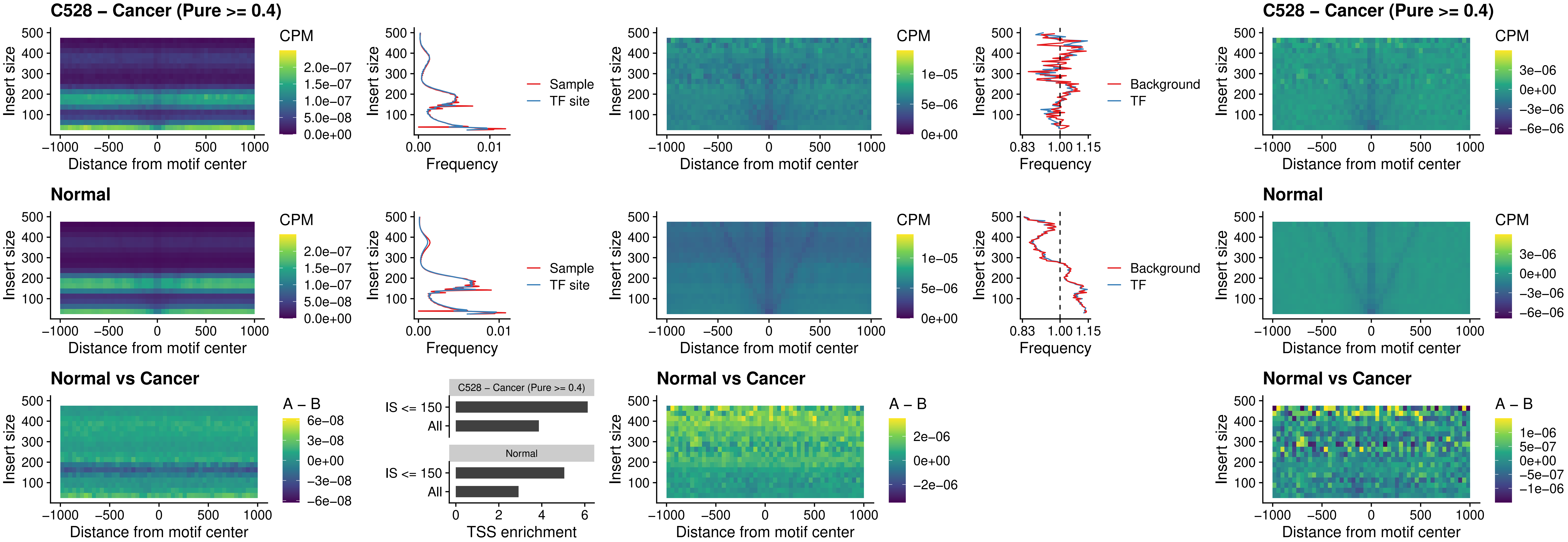

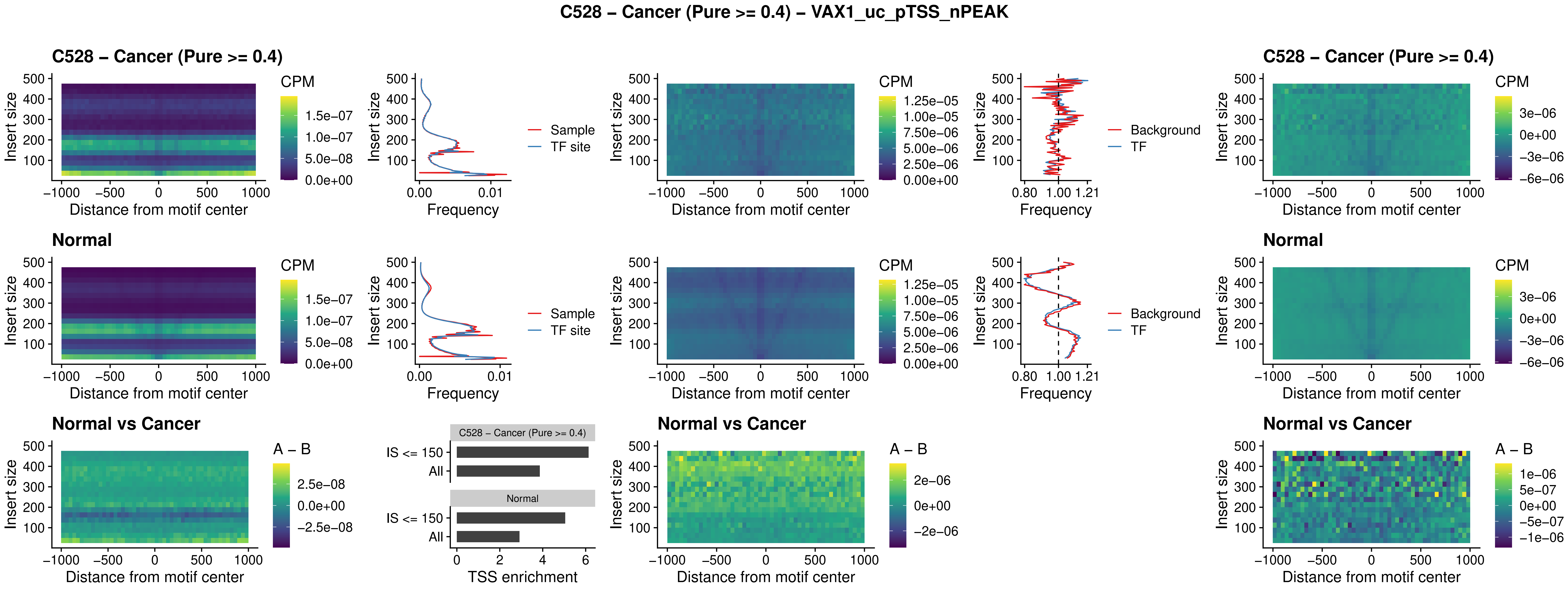
C528 – Cancer (Pure >= 0.4) – VAX1_uc_pTSS_nPEAK
C528 – Cancer (Pure >= 0.4)
Normal
Normal vs Cancer
Insert size
Distance from motif center
Frequency
CPM
A – B
Sample
TF site
Background
TF
C528 – Cancer (Pure >= 0.4)
Normal
IS <= 150
All
TSS enrichment

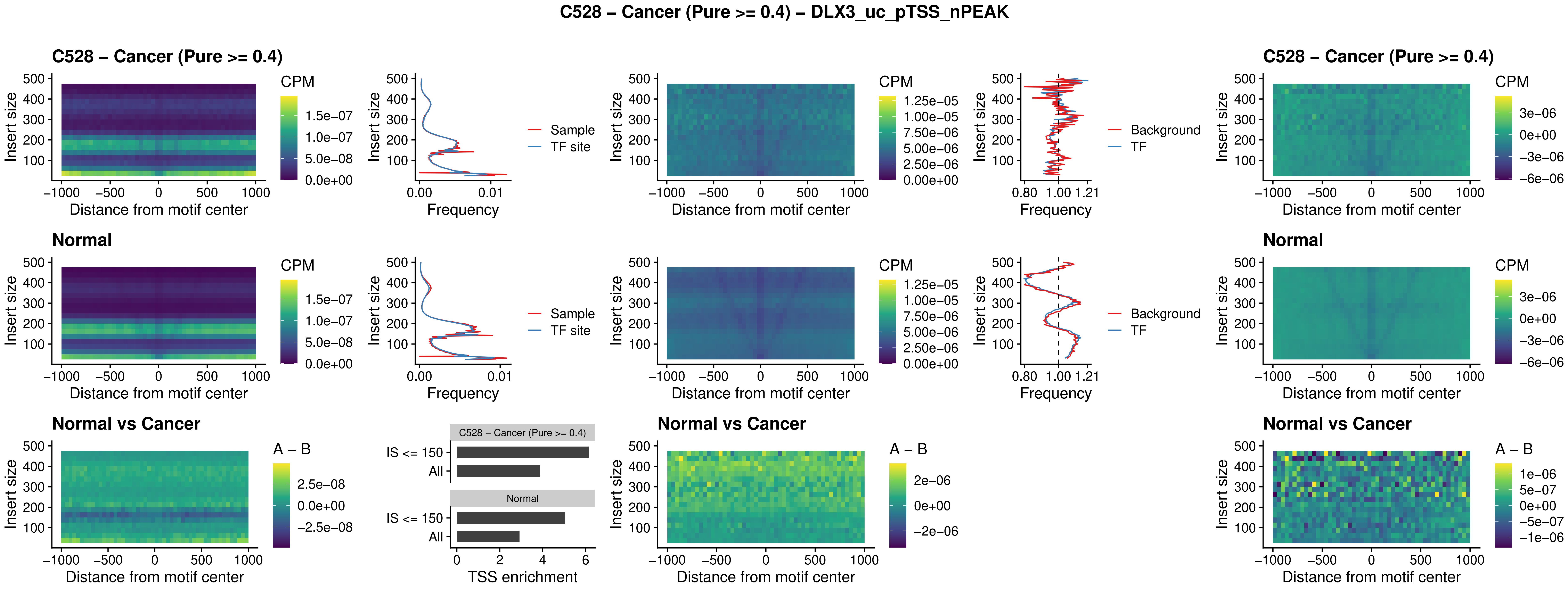

C528 – Cancer (Pure >= 0.4) – DLX3_uc_pTSS_nPEAK
C528 – Cancer (Pure >= 0.4)
Normal
Normal vs Cancer
Insert size
Distance from motif center
Frequency
CPM
A – B
Sample
TF site
Background
TF
C528 – Cancer (Pure >= 0.4)
Normal
IS <= 150
All
TSS enrichment

#### C528 – Cancer (Pure >= 0.4) – HOXD3_uc_pTSS_nPEAK

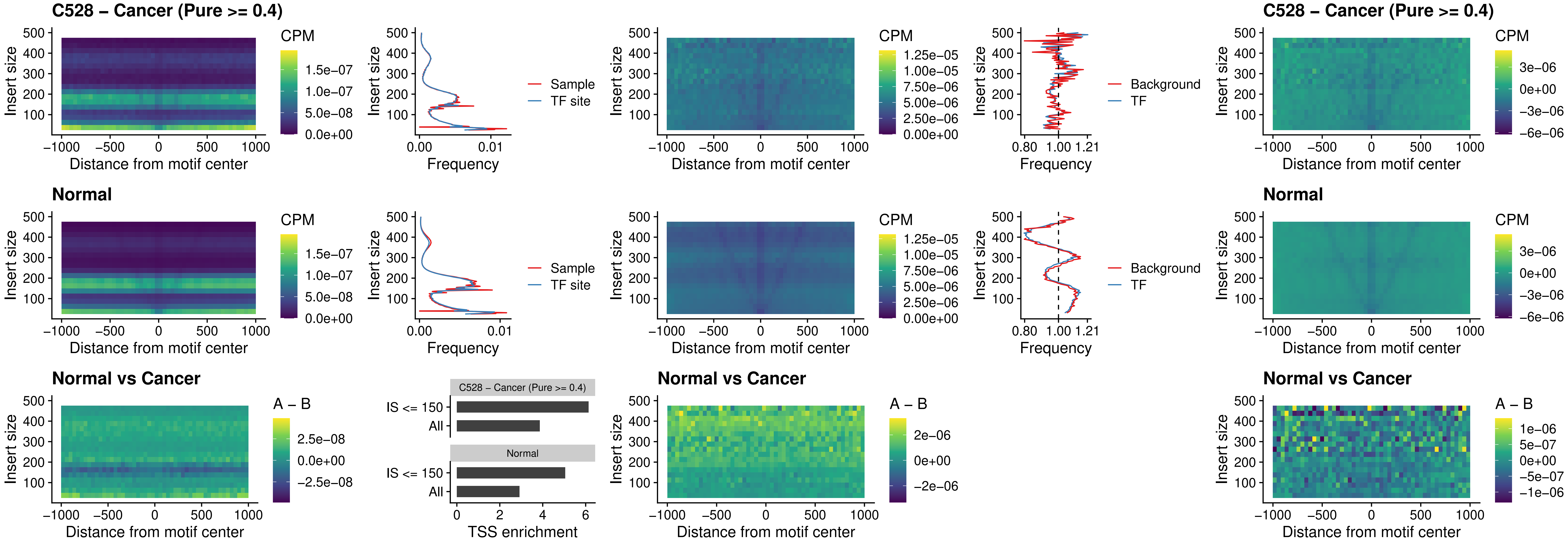

#### C528 – Cancer (Pure >= 0.4) – PDX1_uc_dTSS_nPEAK

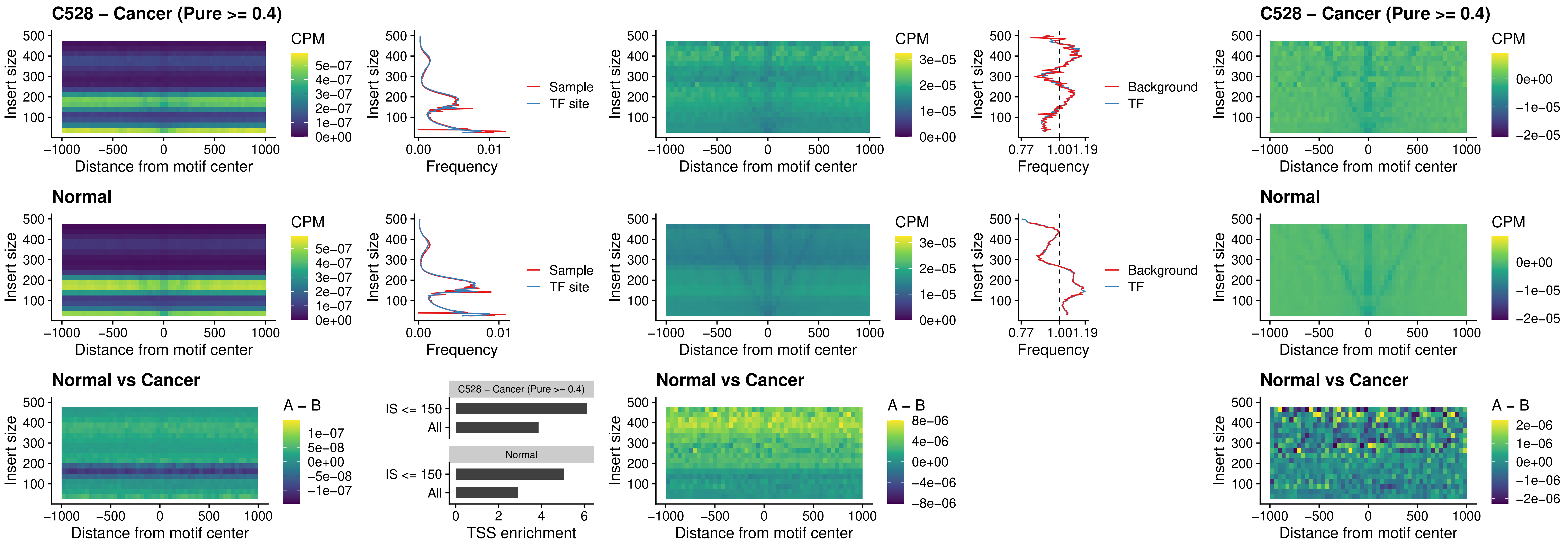

### C528 – Cancer (Pure >= 0.4) – VAX2_uc_dTSS_nPEAK

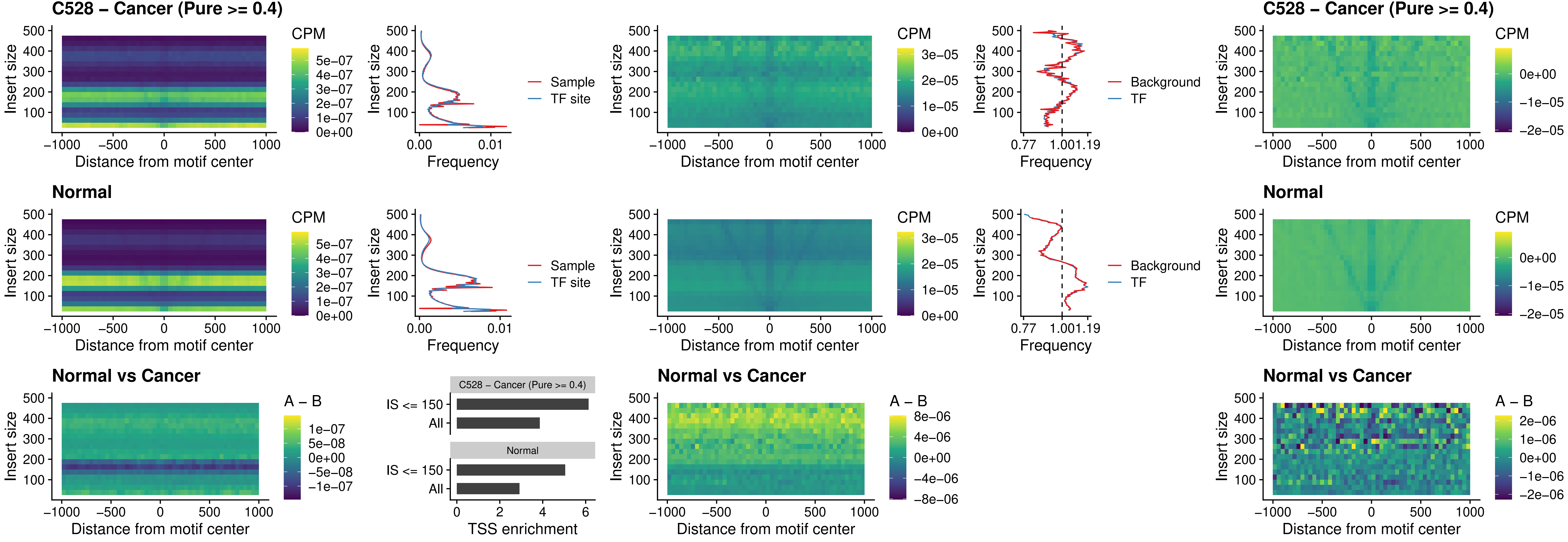

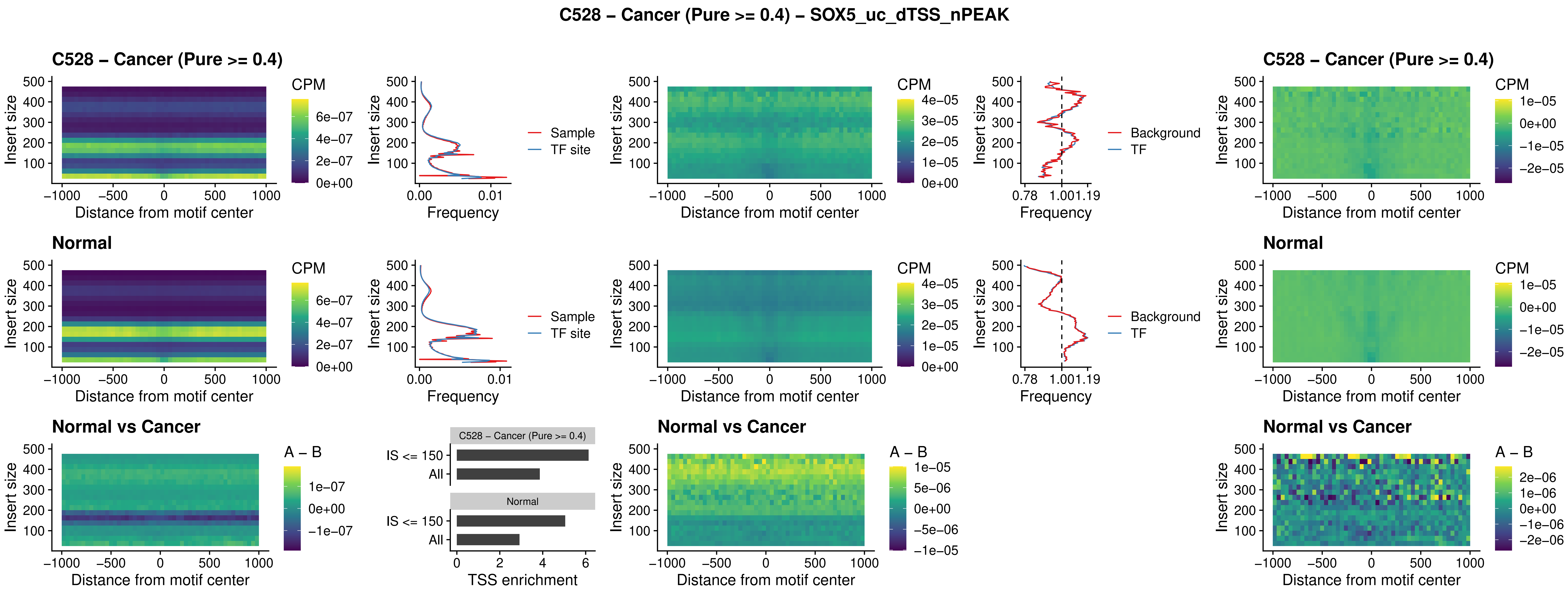
C528 – Cancer (Pure >= 0.4) – SOX5_uc_dTSS_nPEAK
C528 – Cancer (Pure >= 0.4)
Normal
Normal vs Cancer
Insert size
Distance from motif center
Frequency
CPM
A – B
Sample
TF site
Background
TF
C528 – Cancer (Pure >= 0.4)
Normal
IS <= 150
All
TSS enrichment

#### C528 – Cancer (Pure >= 0.4) – SOX6_uc_dTSS_nPEAK

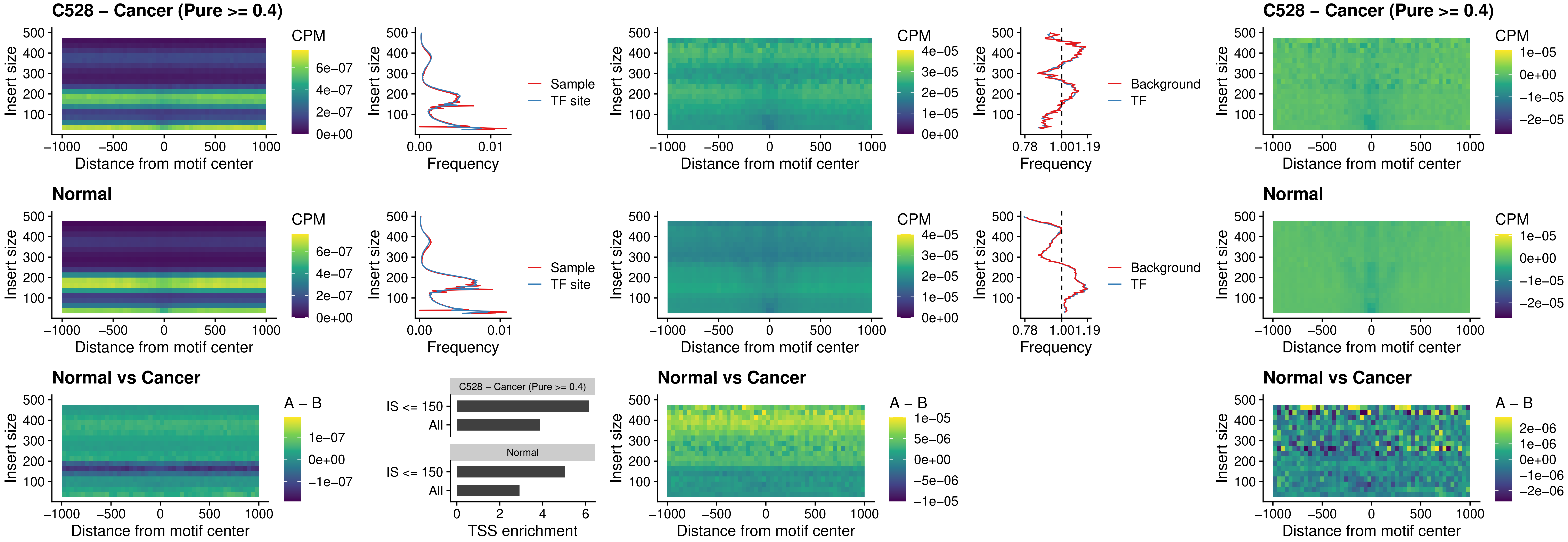

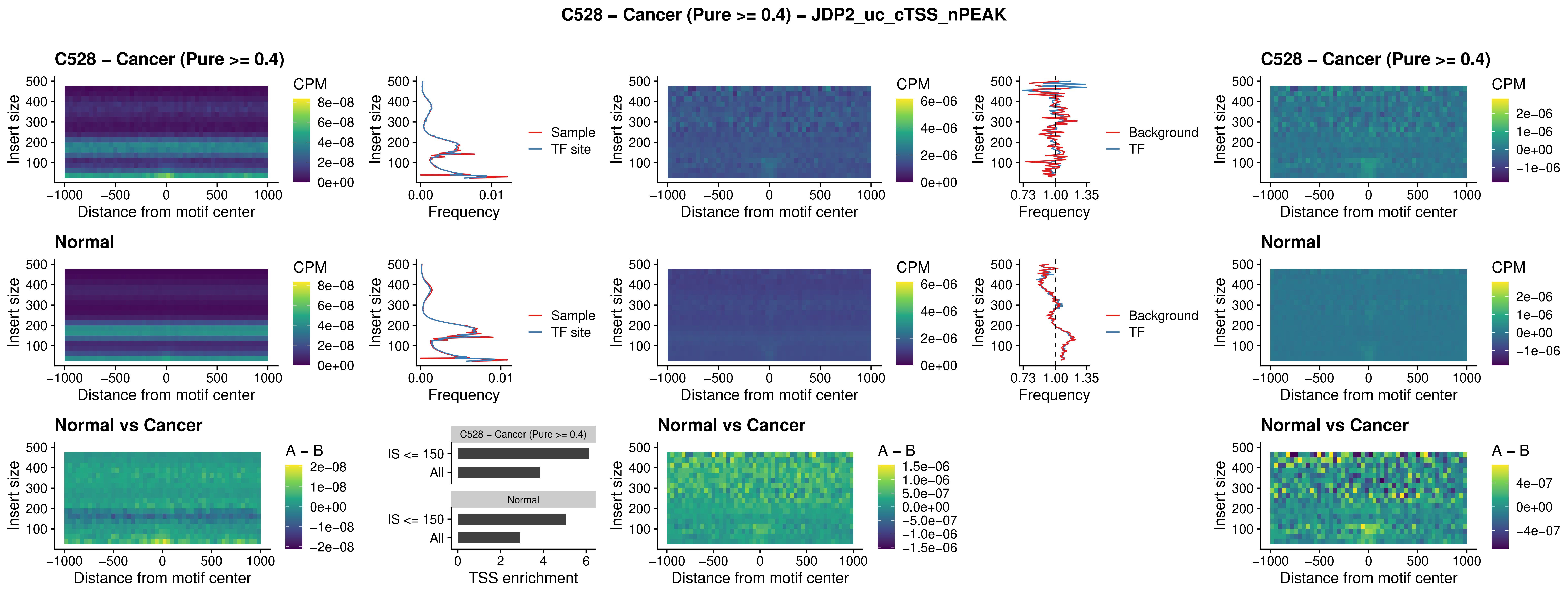
C528 – Cancer (Pure >= 0.4) – JDP2_uc_cTSS_nPEAK
C528 – Cancer (Pure >= 0.4)
Normal
Normal vs Cancer
Insert size
Distance from motif center
Frequency
CPM
A – B
Sample
TF site
Background
TF
C528 – Cancer (Pure >= 0.4)
Normal
IS <= 150
All
TSS enrichment

#### C528 – Cancer (Pure >= 0.4) – PURA_uc_dTSS_oPEAK

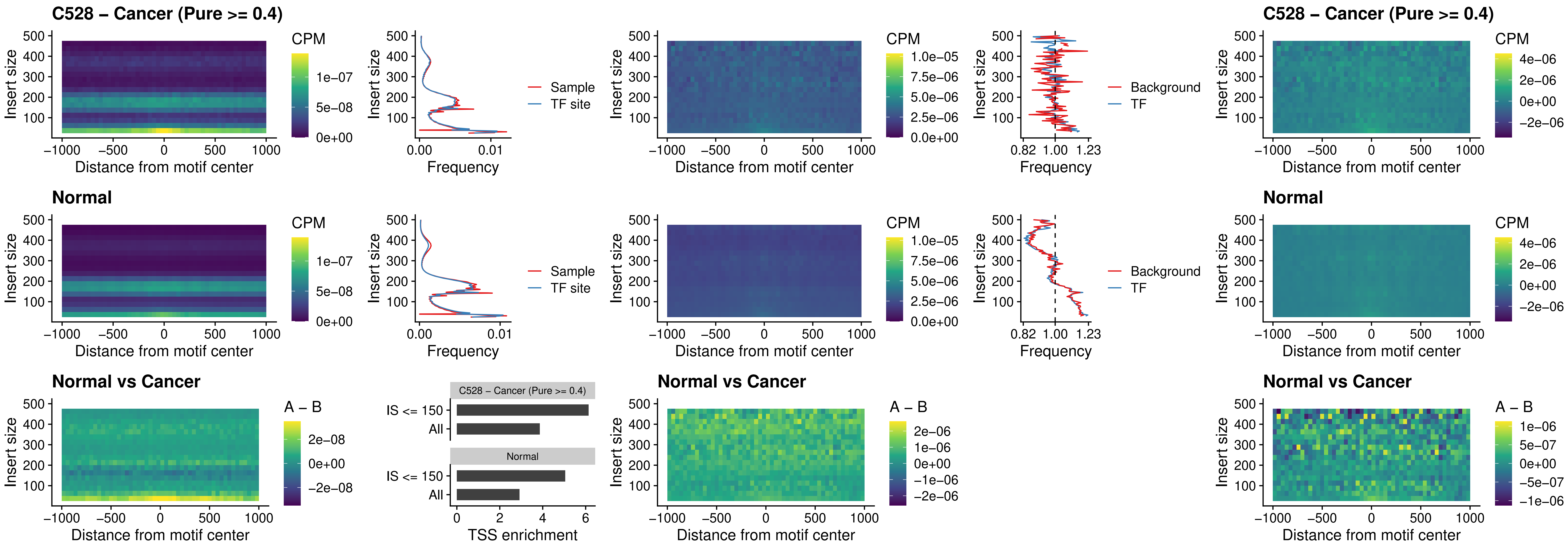

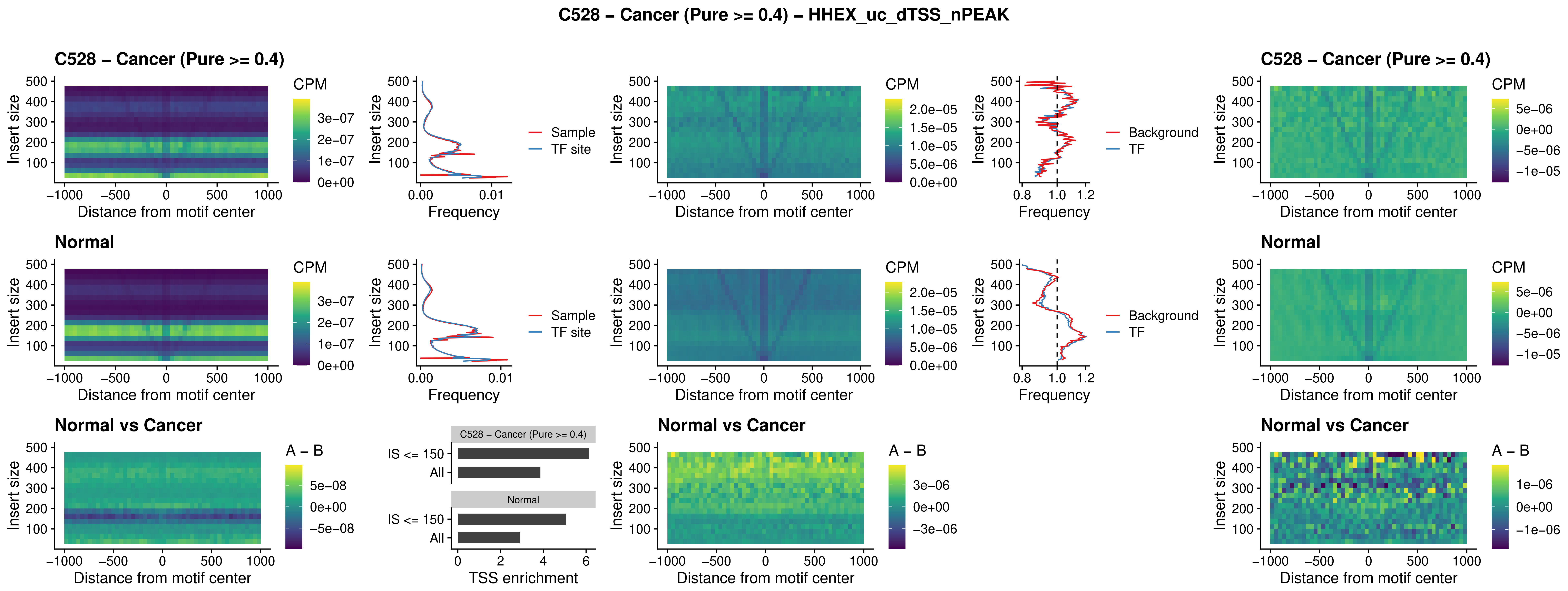
C528 – Cancer (Pure >= 0.4) – HHEX_uc_dTSS_nPEAK
C528 – Cancer (Pure >= 0.4)
Normal
Normal vs Cancer
Insert size
Distance from motif center
Frequency
CPM
A – B
Sample
TF site
Background
TF
C528 – Cancer (Pure >= 0.4)
Normal
IS <= 150
All
TSS enrichment

#### C528 – Cancer (Pure >= 0.4) – HMG20B_uc_dTSS_nPEAK

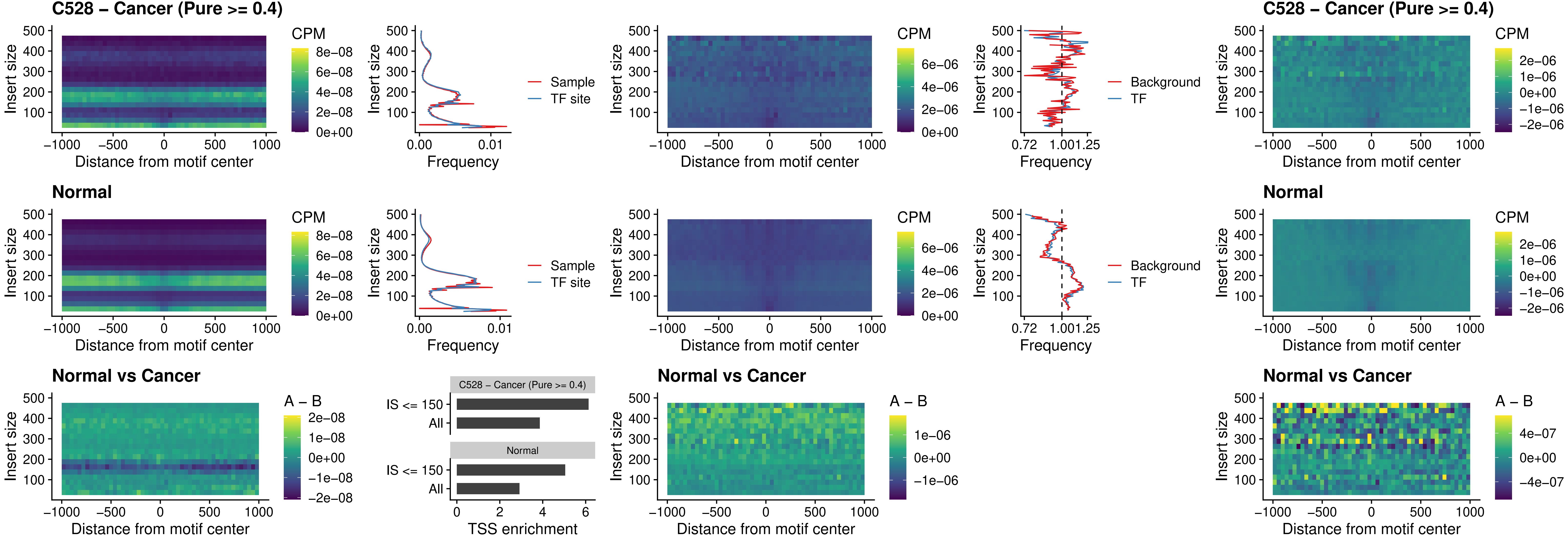

#### C528 – Cancer (Pure >= 0.4) – FOXB2_uc_dTSS_nPEAK

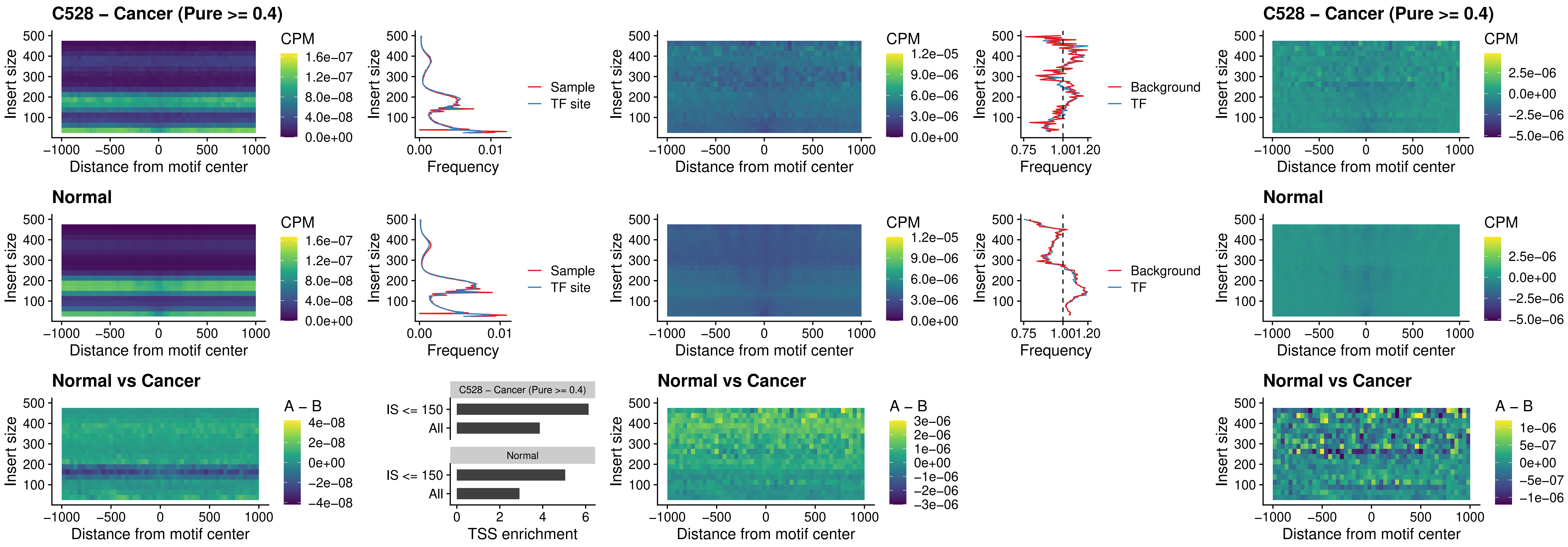

### C528 – Cancer (Pure >= 0.4) – FOXL1_uc_cTSS_nPEAK

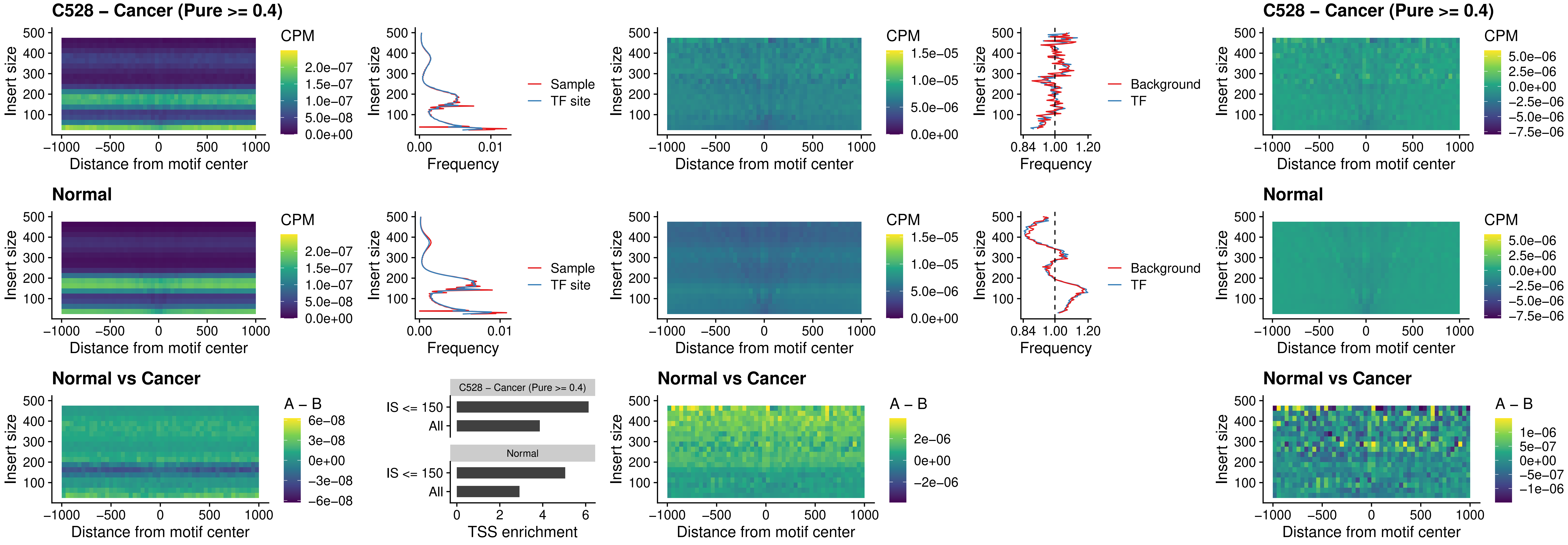

#### C528 – Cancer (Pure >= 0.4) – PBX3_uc_pTSS_oPEAK

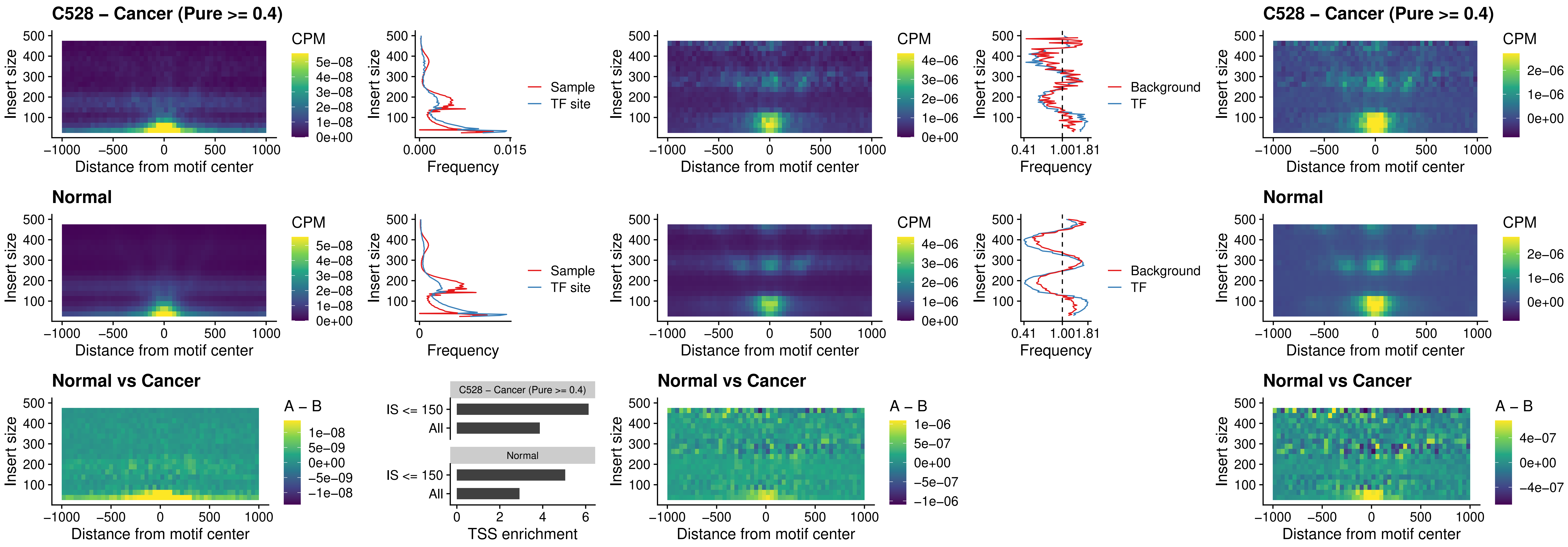

#### C528 – Cancer (Pure >= 0.4) – SNAI2_uc_cTSS_nPEAK

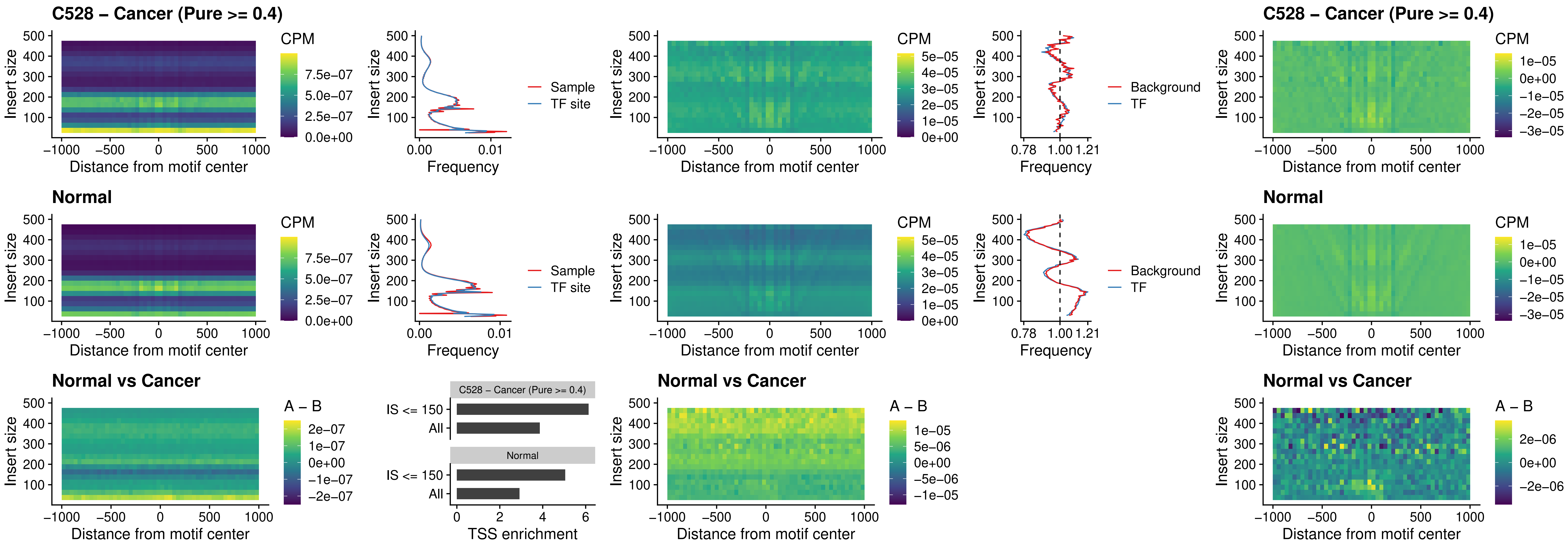

#### C528 – Cancer (Pure >= 0.4) – SNAI2_uc_dTSS_oPEAK

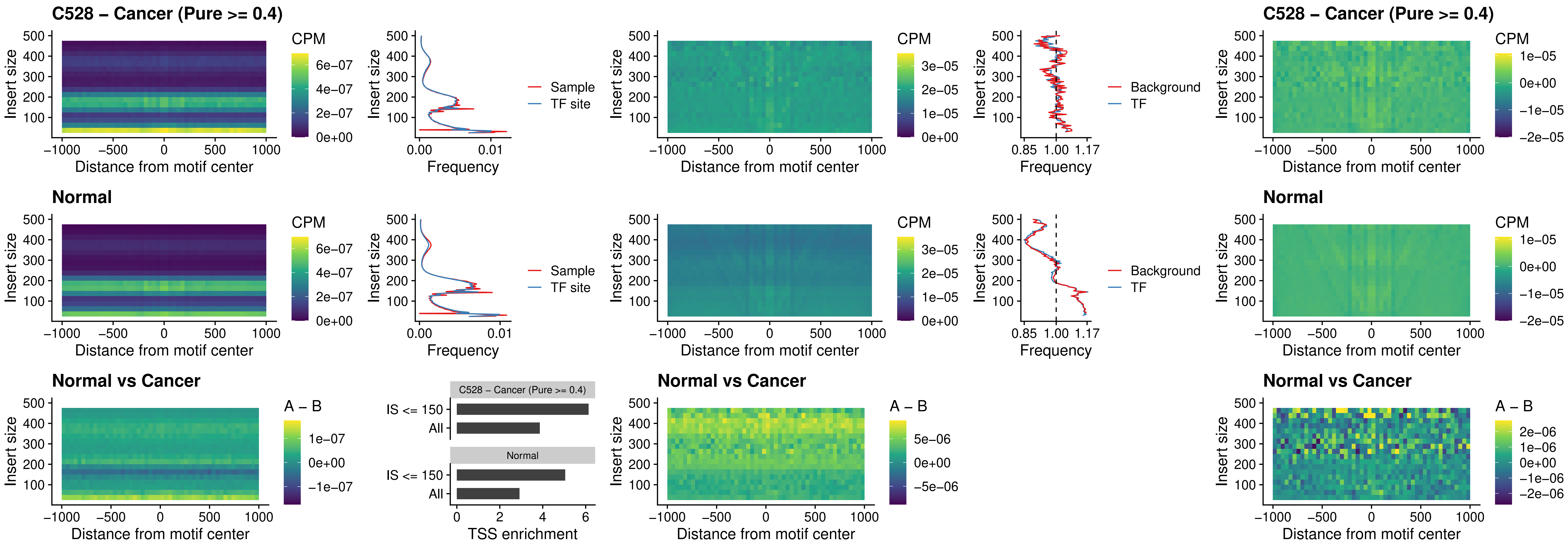

#### C528 – Cancer (Pure >= 0.4) – SPI1_uc_pTSS_nPEAK

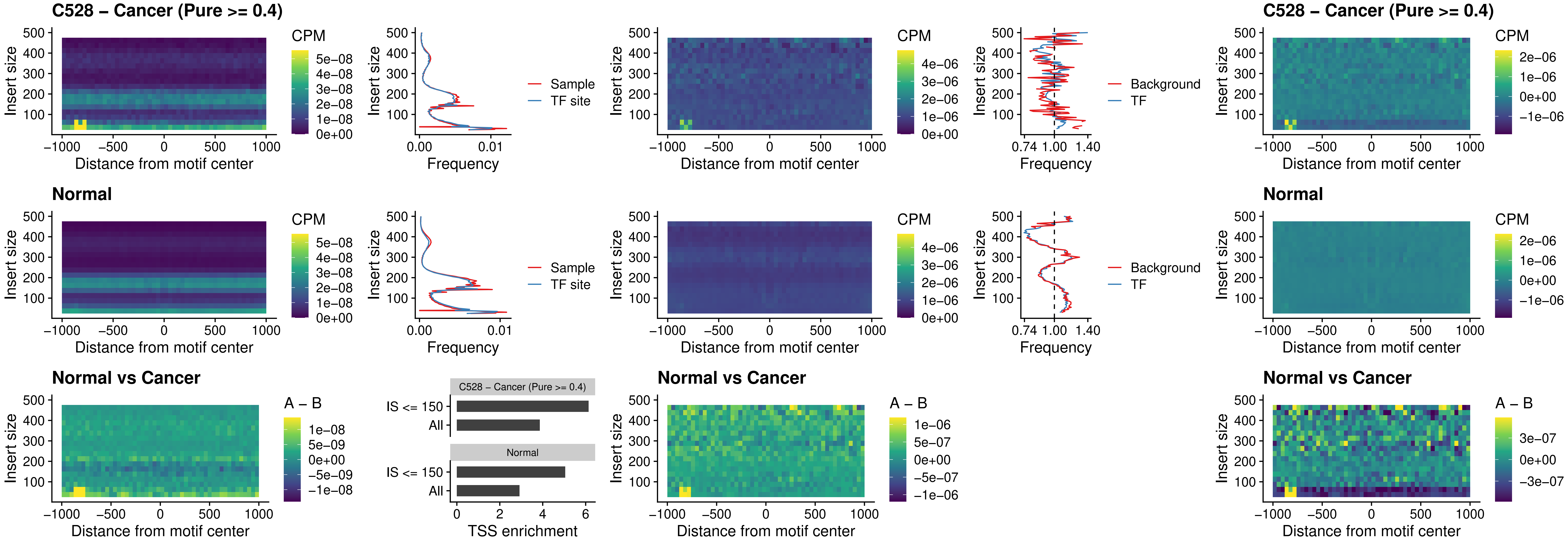

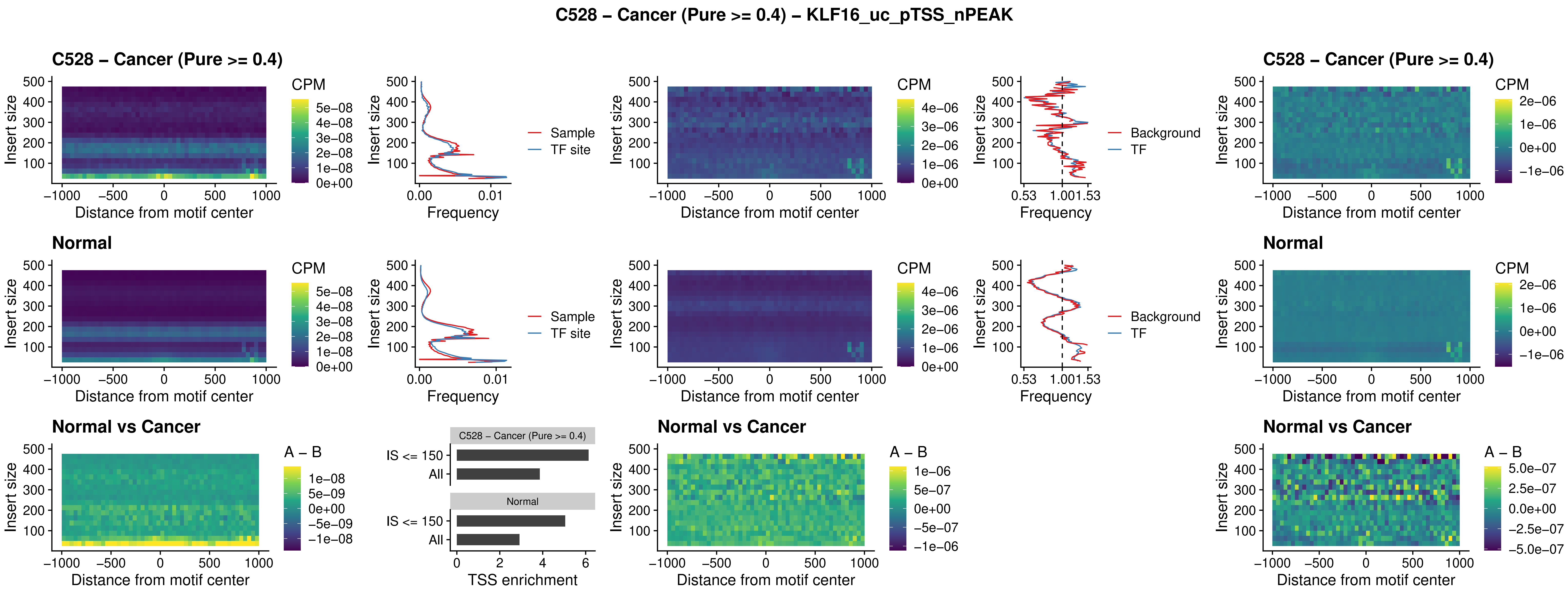
C528 – Cancer (Pure >= 0.4) – KLF16_uc_pTSS_nPEAK
C528 – Cancer (Pure >= 0.4)
Normal
Normal vs Cancer
Insert size
Distance from motif center
Frequency
CPM
A – B
Sample
TF site
Background
TF
TSS enrichment
IS <= 150
All
C528 – Cancer (Pure >= 0.4)
Normal

#### C528 – Cancer (Pure >= 0.4) – SP3_uc_pTSS_nPEAK

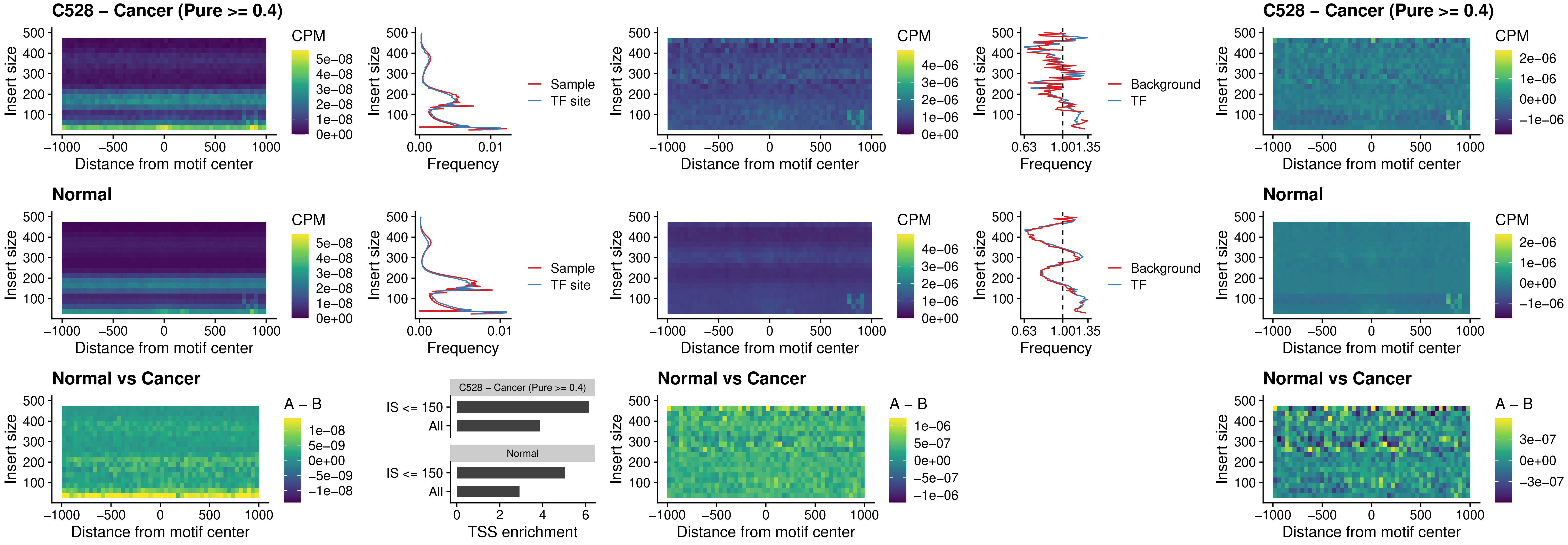

#### C528 – Cancer (Pure >= 0.4) – NEUROD1_uc_dTSS_oPEAK

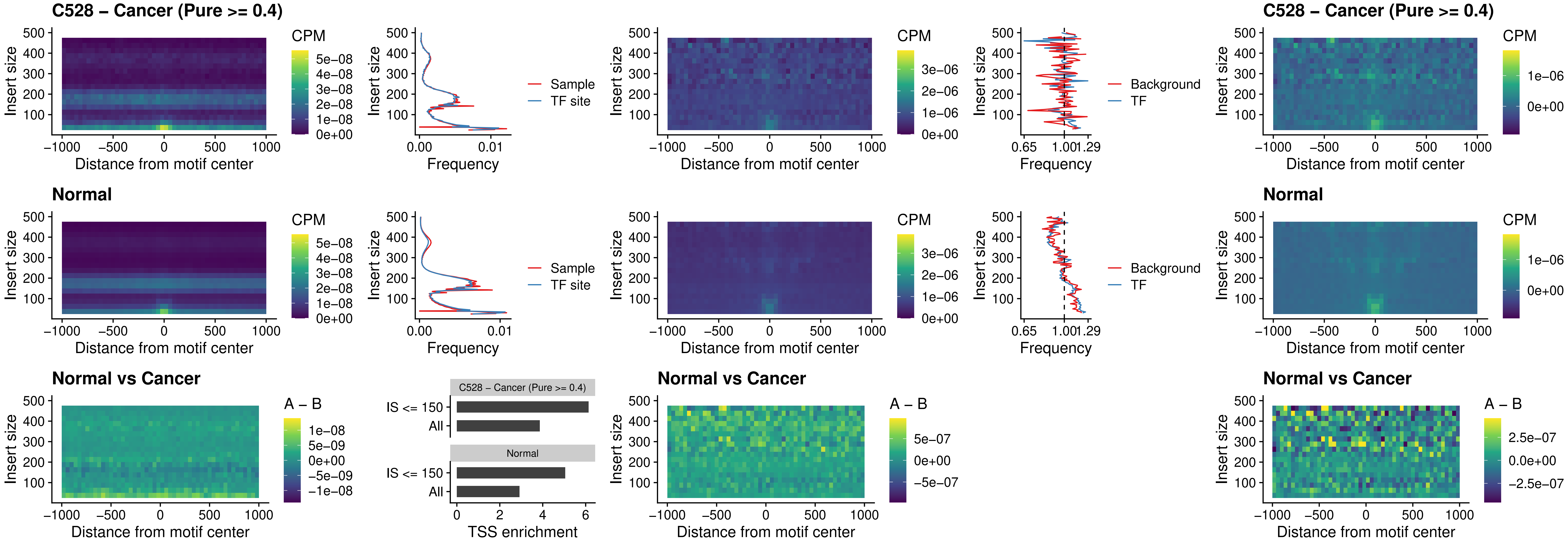

### C528 – Cancer (Pure >= 0.4) – CTCF_uc_dTSS_oPEAK

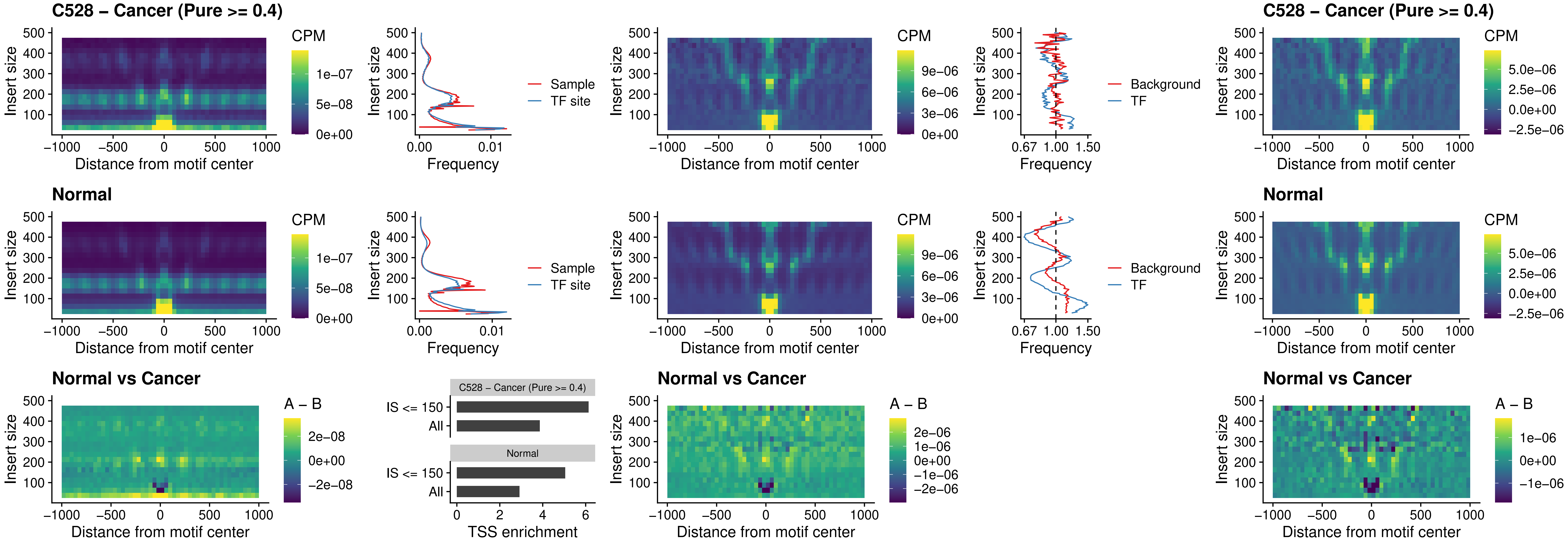

#### C528 – Cancer (Pure >= 0.4) – CTCF_uc_cTSS_oPEAK

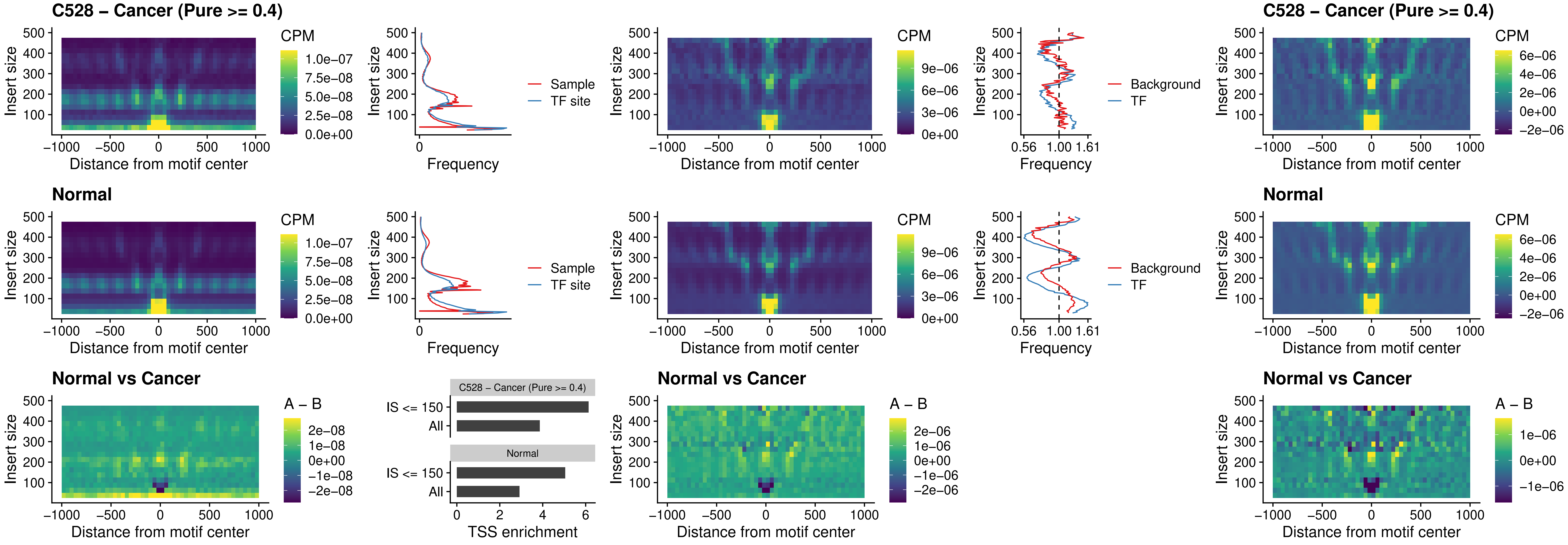

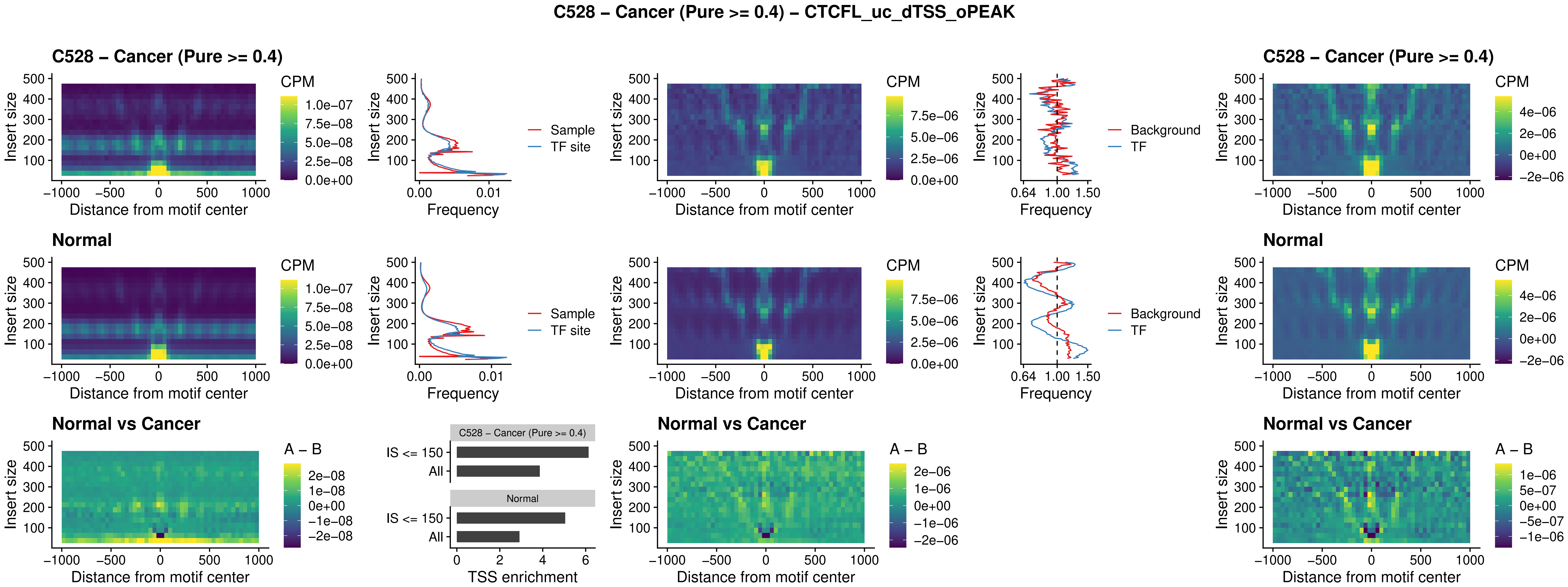

C528 – Cancer (Pure >= 0.4) – CTCFL_uc_dTSS_oPEAK
C528 – Cancer (Pure >= 0.4)
Normal
Normal vs Cancer
Insert size
Distance from motif center
Frequency
CPM
A – B
Sample
TF site
Background
TF
C528 – Cancer (Pure >= 0.4)
Normal
IS <= 150
All
TSS enrichment

#### C528 – Cancer (Pure >= 0.4) – ENCODE_CTCF_sites_COLON

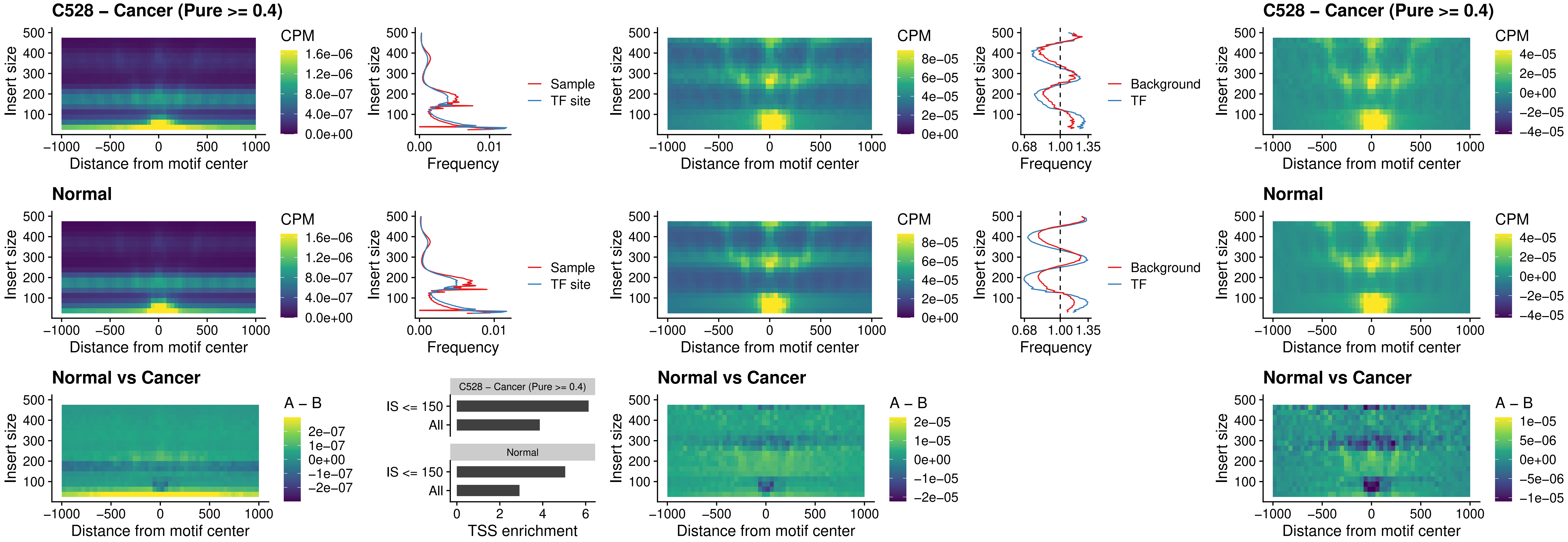

#### C528 – Cancer (Pure >= 0.4) – ENCODE_CTCF_sites_COLON_loop_ctcf

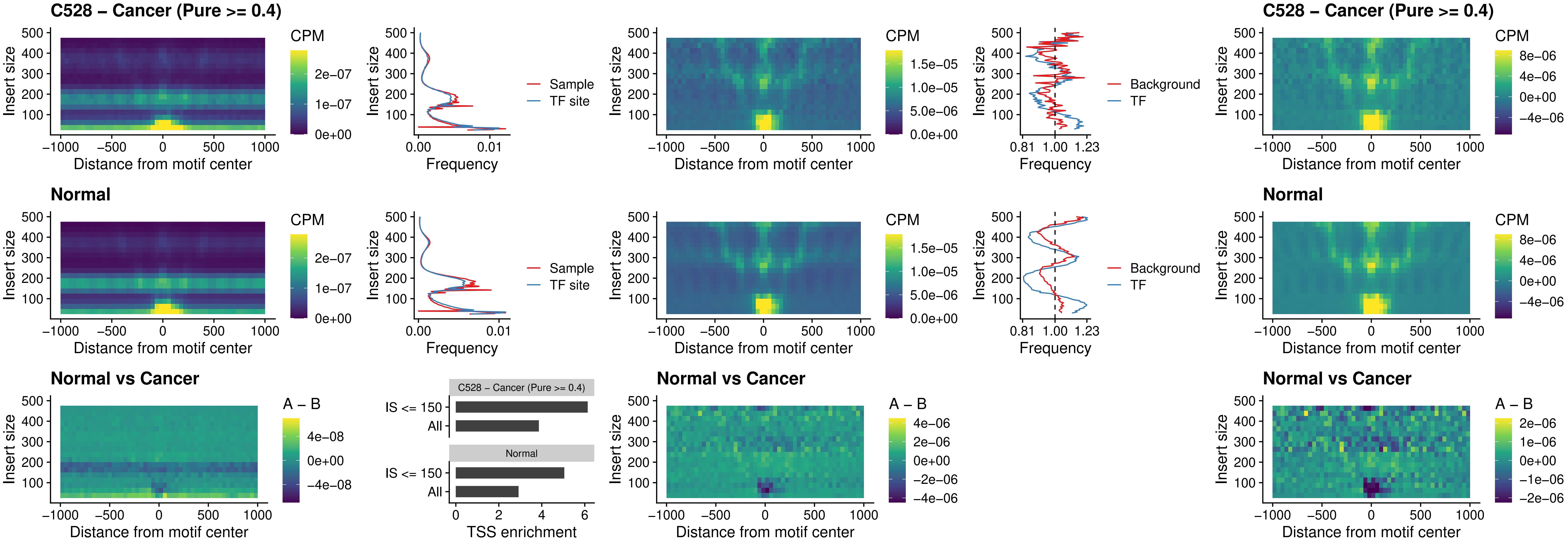

#### C528 – Cancer (Pure >= 0.4) – ENCODE_CTCF_sites_COLON_intergenic_ctcf

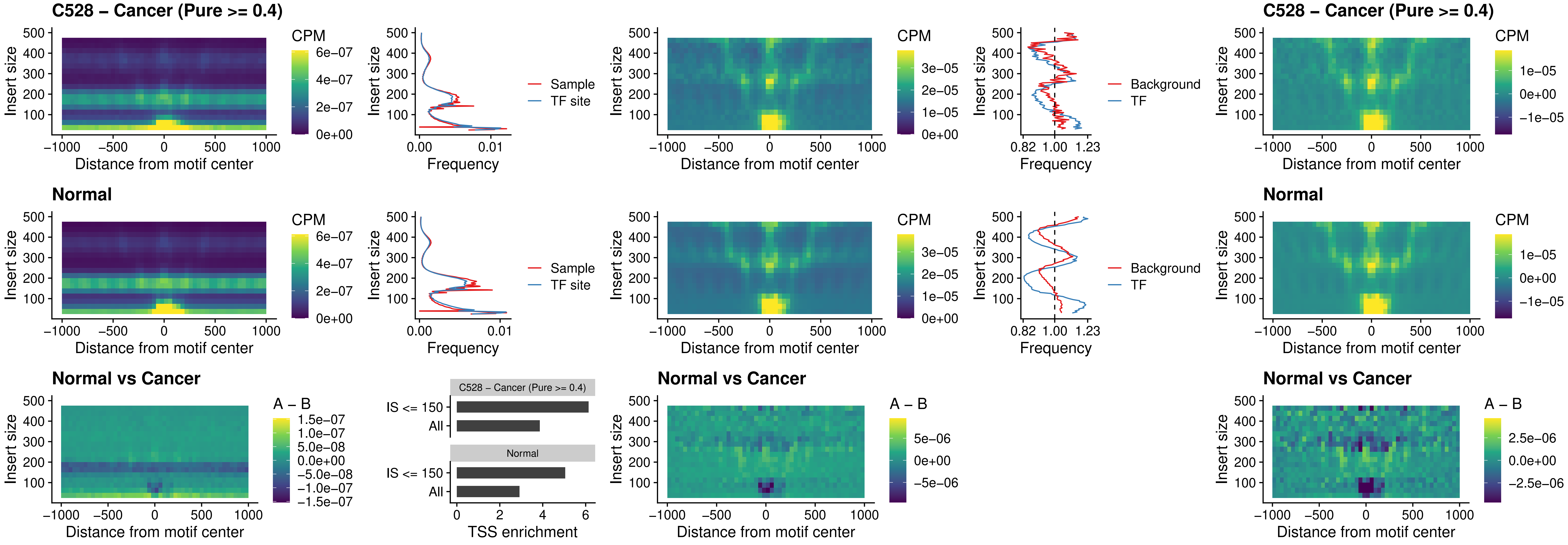

#### C528 – Cancer (Pure >= 0.4) – ENCODE_CTCF_sites_COLON_nonpromotor

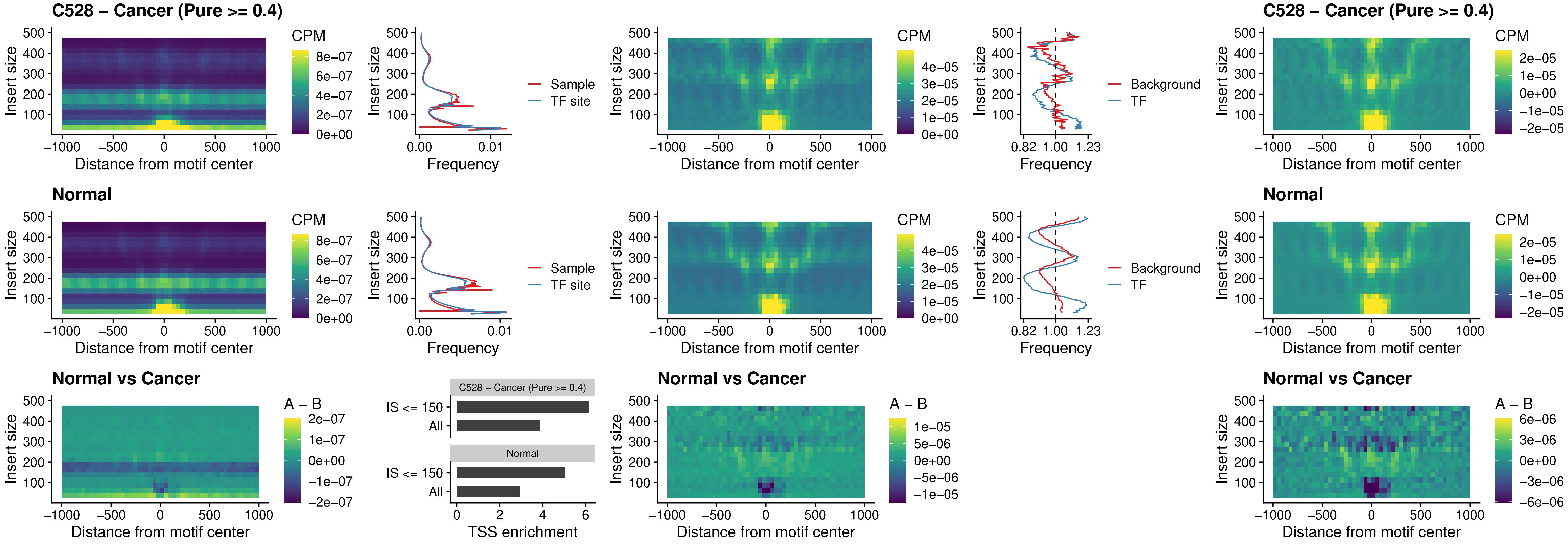

#### C528 – Cancer (Pure >= 0.4) – CTCFL_uc_cTSS_oPEAK

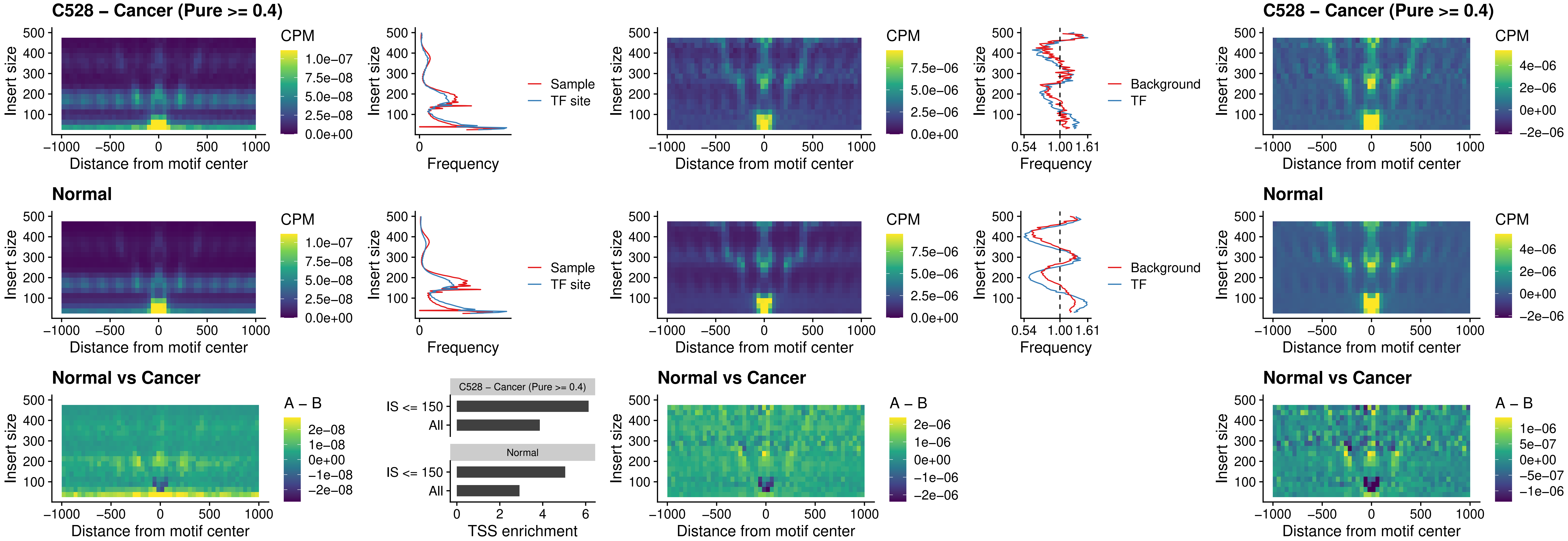

### C528 – Cancer (Pure >= 0.4) – NEUROD1_uc_cTSS_oPEAK

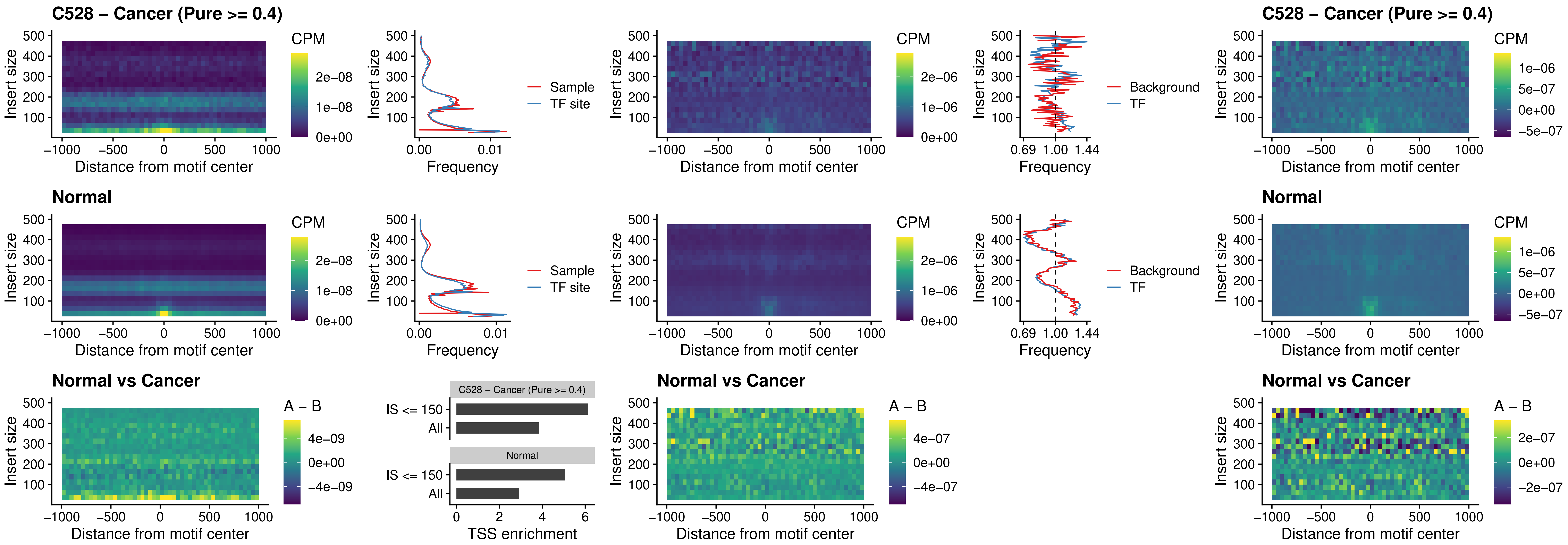

#### C528 – Cancer (Pure >= 0.4) – ENCODE_YY1_sites_COLON_nonpromotor

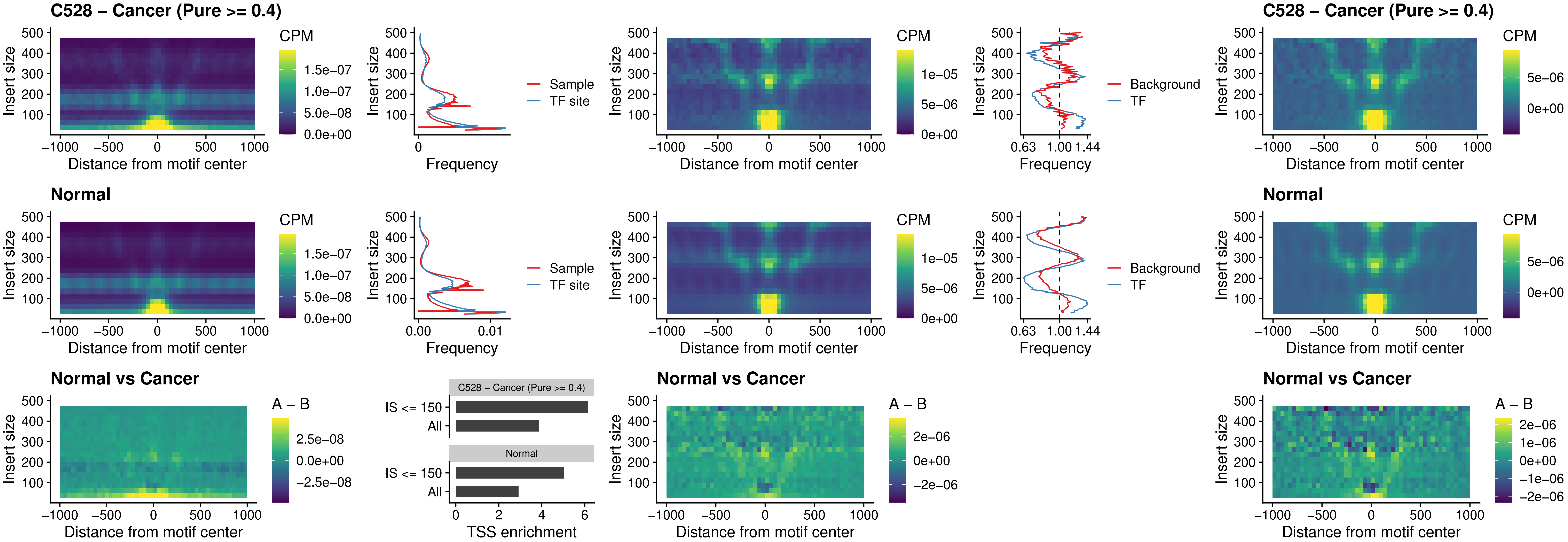

#### C528 – Cancer (Pure >= 0.4) – CTCF_uc_pTSS_oPEAK

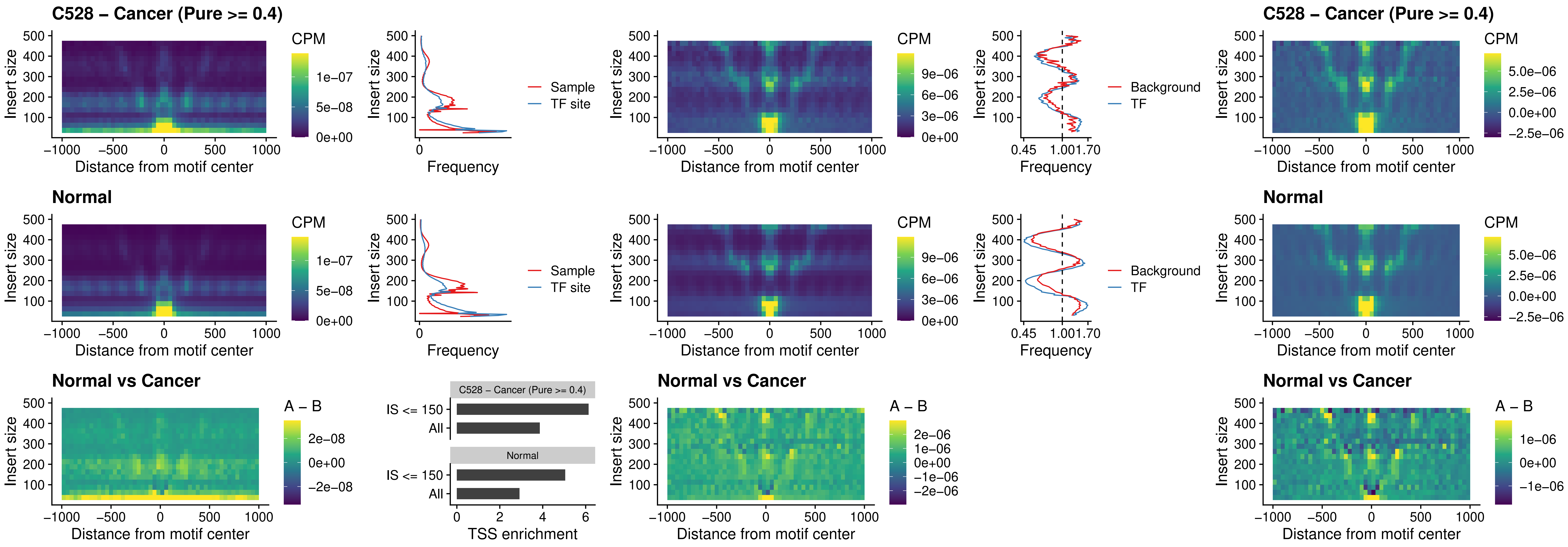

#### C528 – Cancer (Pure >= 0.4) – ENCODE_CTCF_sites_COLON_promotor

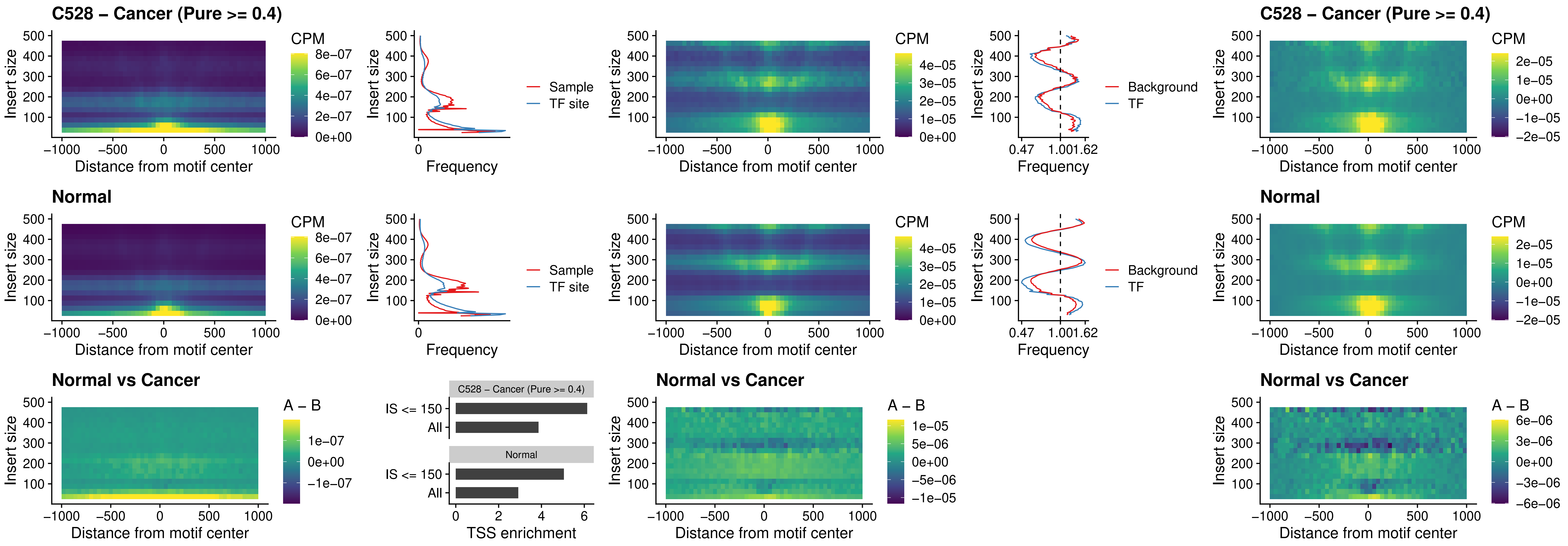

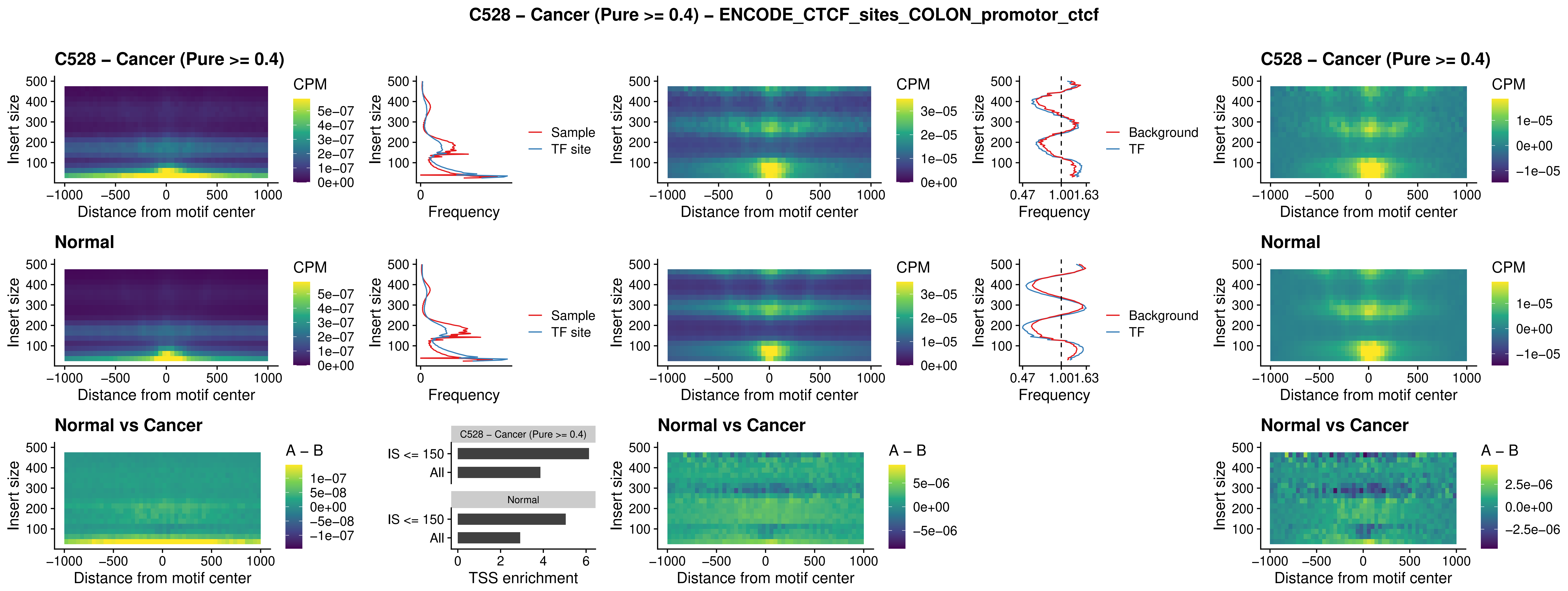
C528 – Cancer (Pure >= 0.4) – ENCODE_CTCF_sites_COLON_promotor_ctcf
C528 – Cancer (Pure >= 0.4)
Normal
Normal vs Cancer
Insert size
Distance from motif center
Frequency
CPM
A – B
Sample
TF site
Background
TF
C528 – Cancer (Pure >= 0.4)
IS <= 150
All
TSS enrichment

#### C528 – Cancer (Pure >= 0.4) – SPIC_uc_pTSS_oPEAK

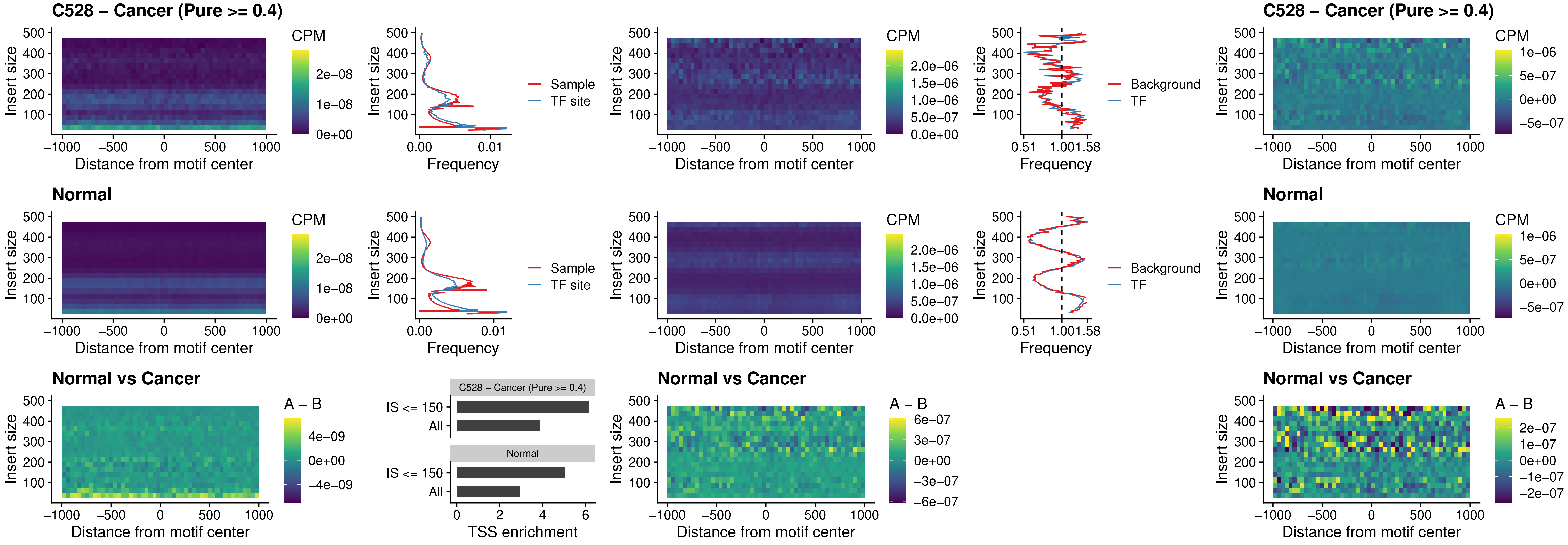

#### C528 – Cancer (Pure >= 0.4) – SPI1_uc_dTSS_nPEAK

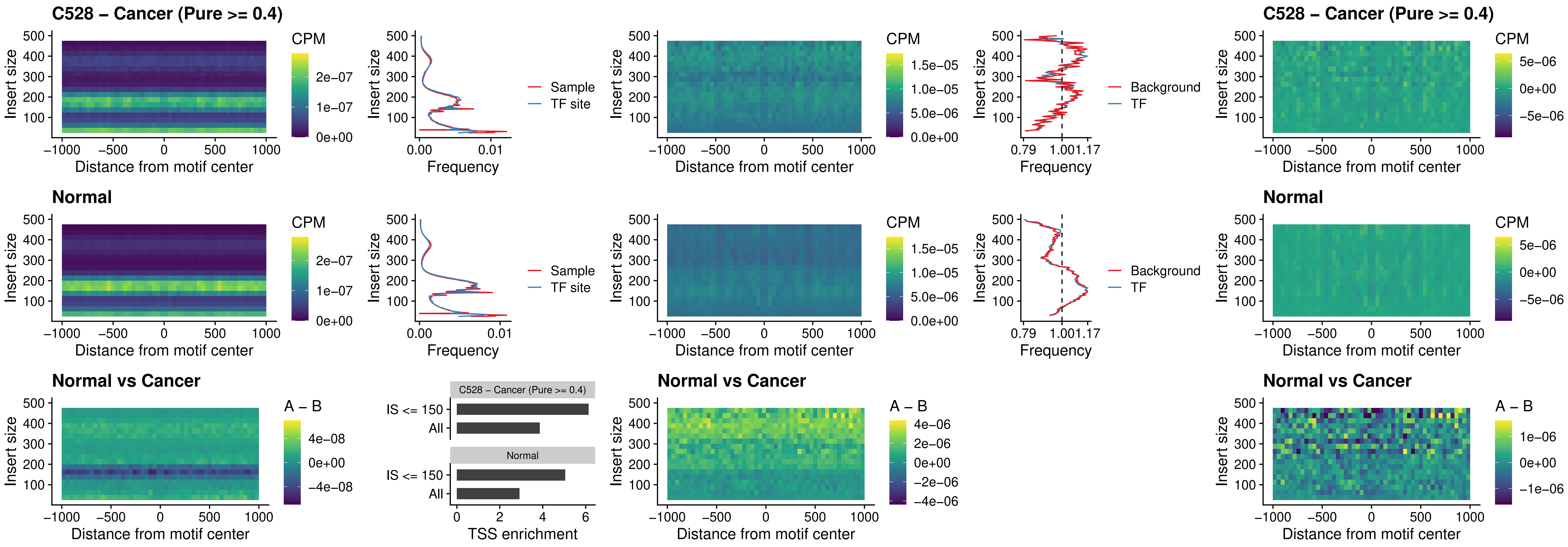

#### C528 – Cancer (Pure >= 0.4) – SPI1_uc_cTSS_nPEAK

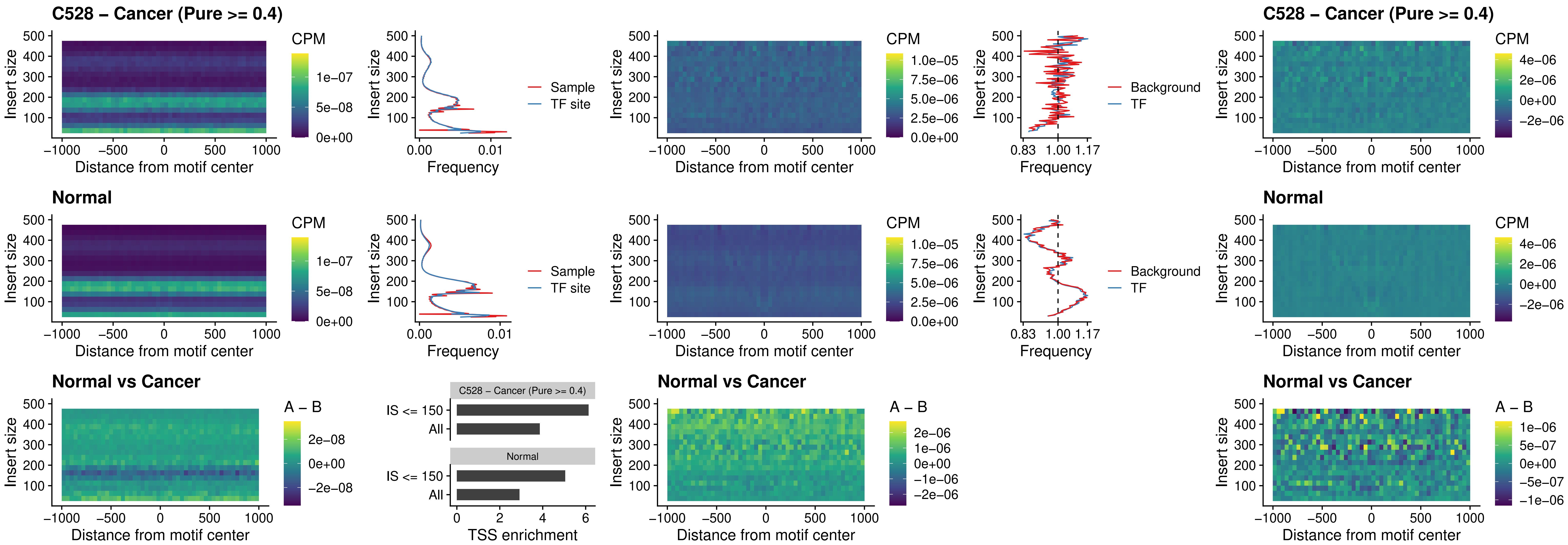

#### C528 – Cancer (Pure >= 0.4) – BCL11A_uc_pTSS_nPEAK

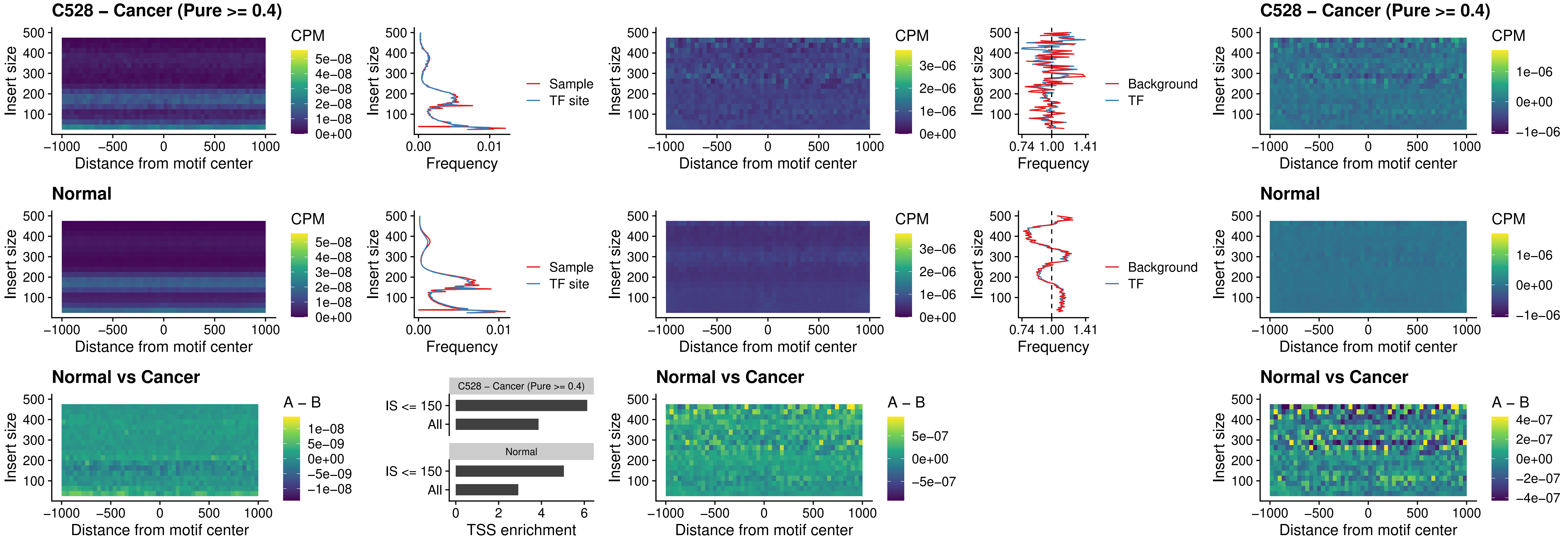

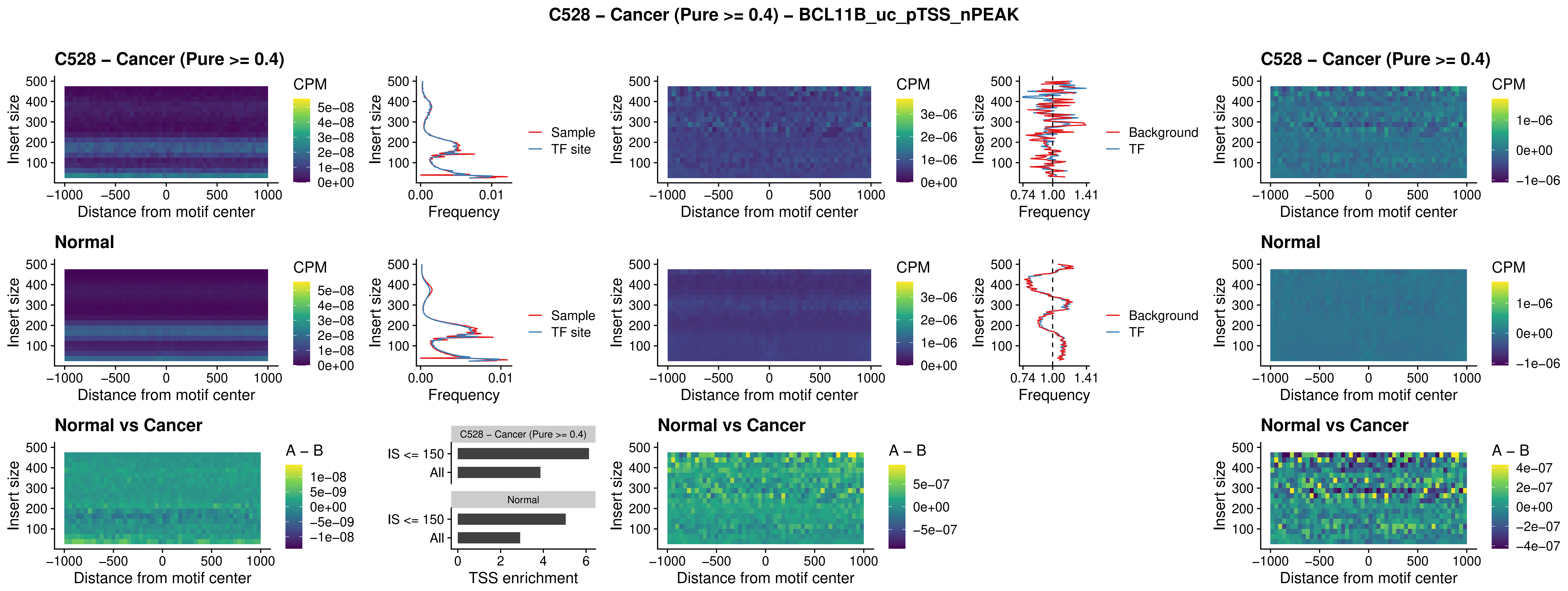
C528 – Cancer (Pure >= 0.4) – BCL11B_uc_pTSS_nPEAK
C528 – Cancer (Pure >= 0.4)
Normal
Normal vs Cancer
Insert size
Distance from motif center
Frequency
CPM
A – B
Sample
TF site
Background
TF
C528 – Cancer (Pure >= 0.4)
Normal
IS <= 150
All
TSS enrichment

#### C528 – Cancer (Pure >= 0.4) – IRF9_uc_dTSS_oPEAK

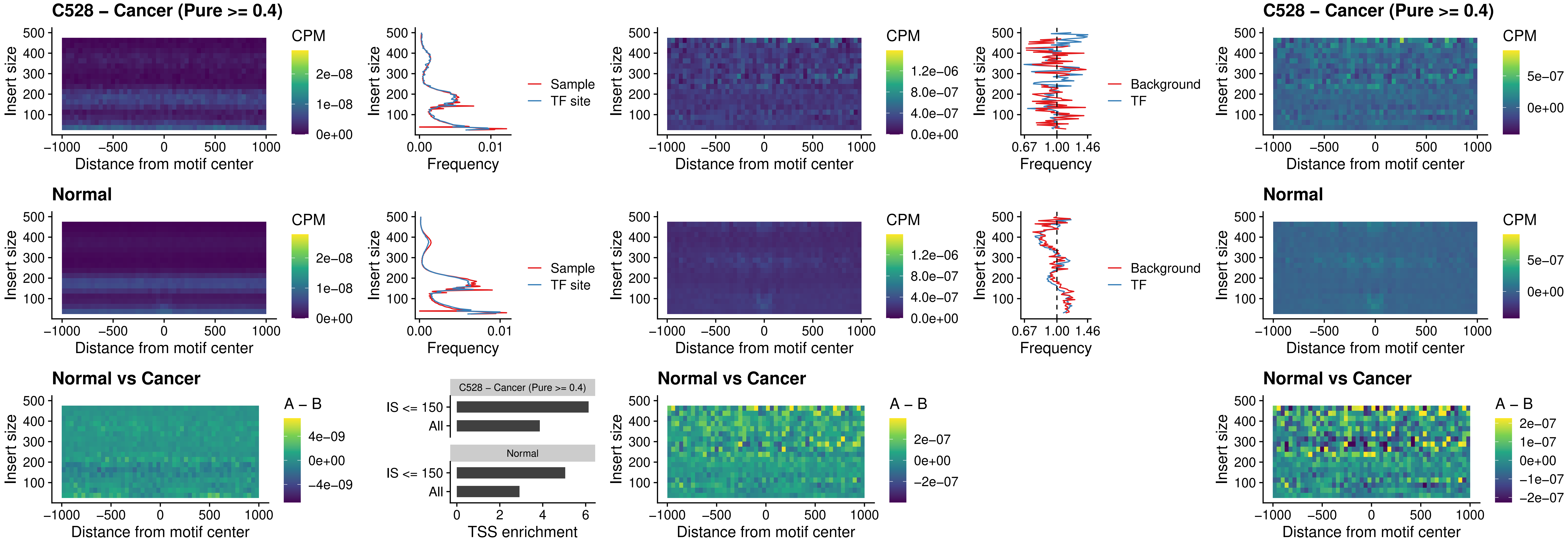

#### C528 – Cancer (Pure >= 0.4) – IRF9_uc_cTSS_oPEAK

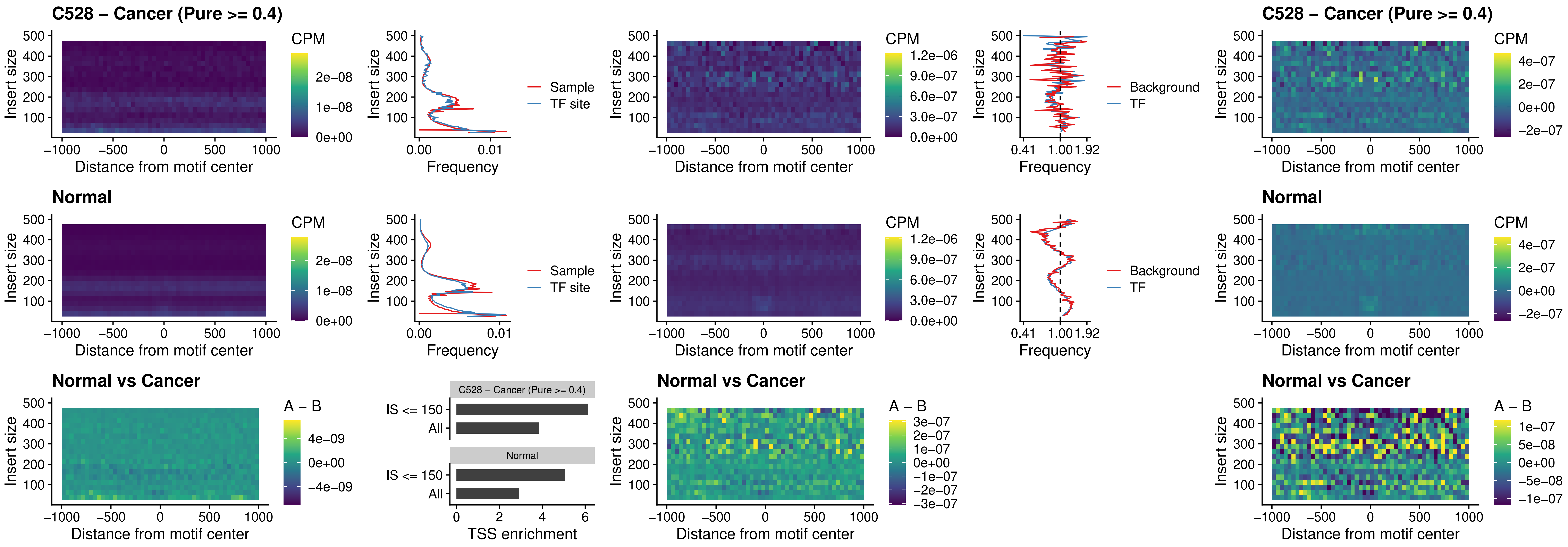

#### C528 – Cancer (Pure >= 0.4) – BCL11A_uc_cTSS_oPEAK

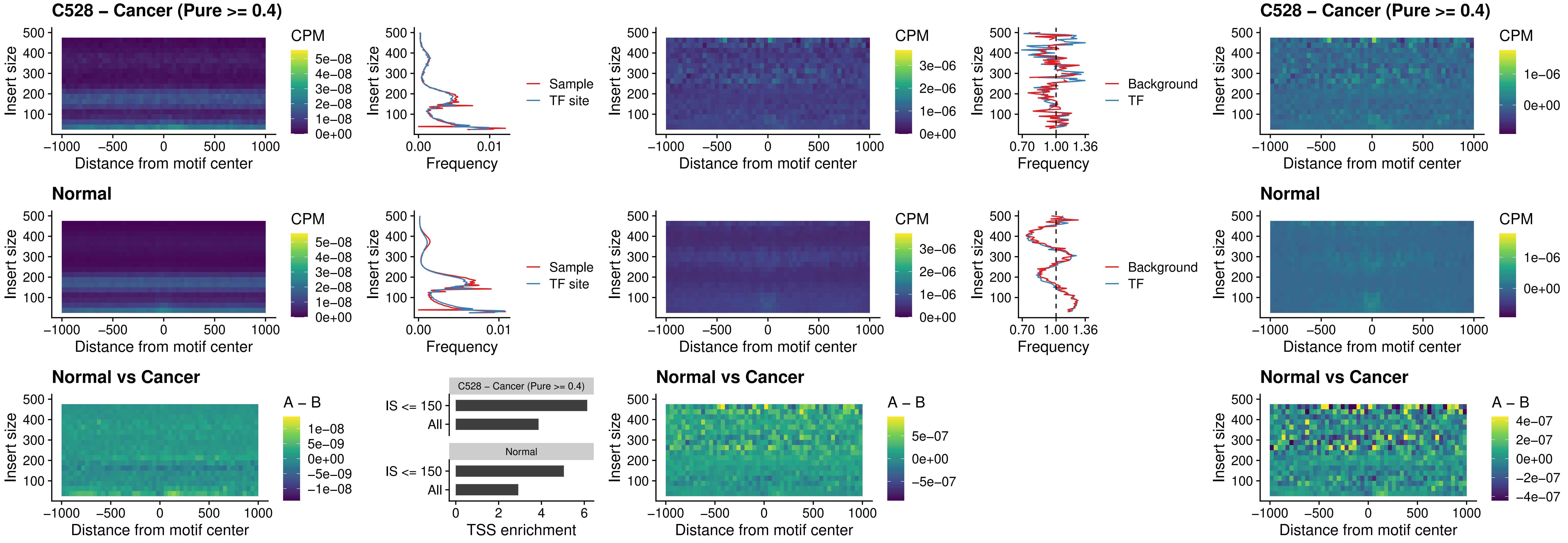

#### C528 – Cancer (Pure >= 0.4) – BCL11B_uc_cTSS_oPEAK

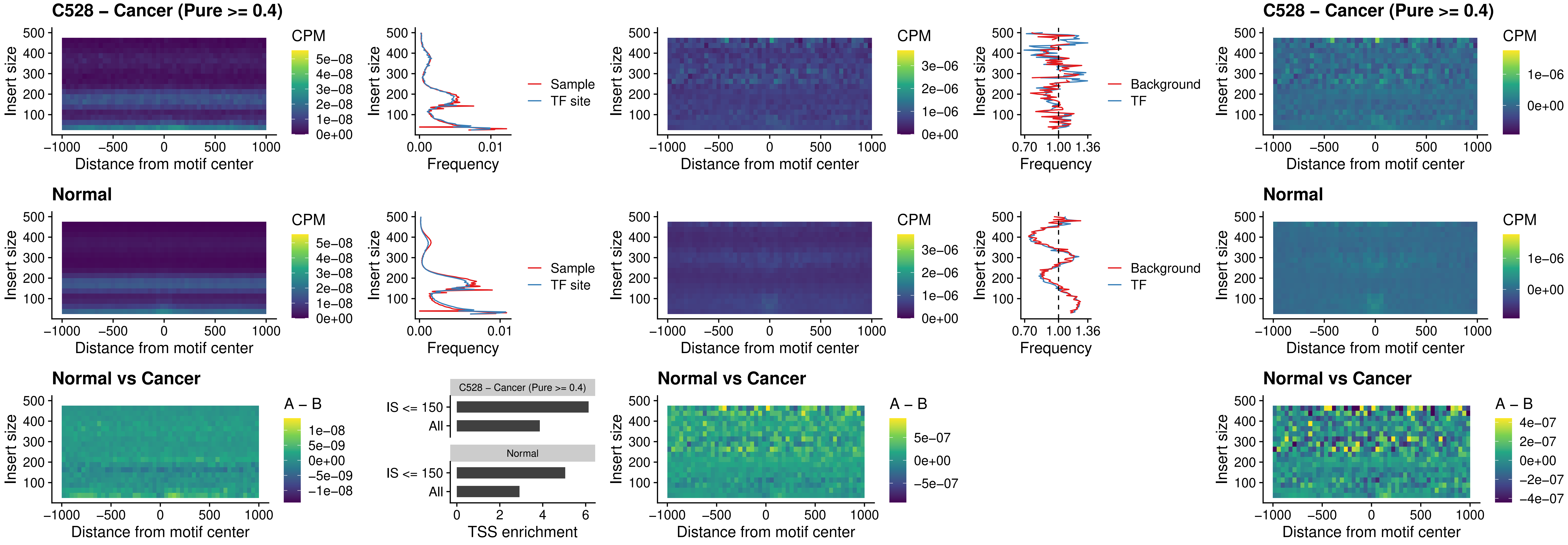

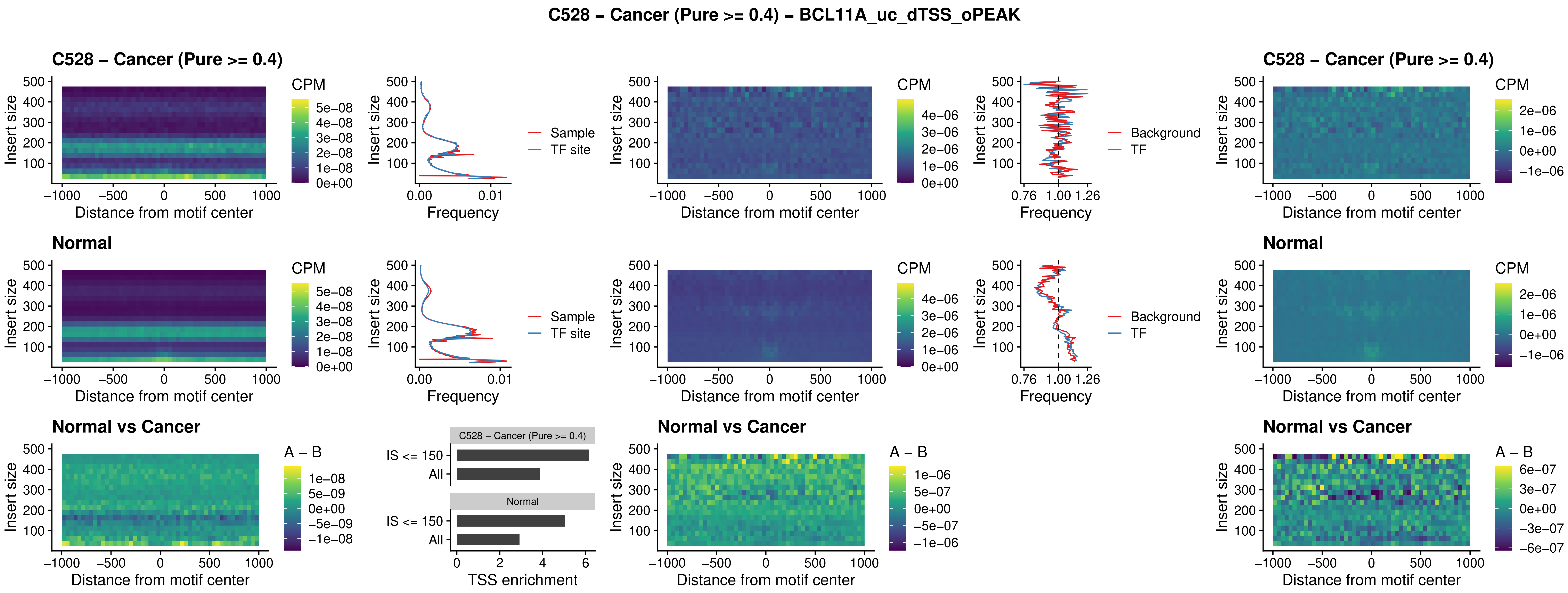
C528 – Cancer (Pure >= 0.4) – BCL11A_uc_dTSS_oPEAK
C528 – Cancer (Pure >= 0.4)
Normal
Normal vs Cancer
Insert size
Distance from motif center
Frequency
CPM
A – B
Sample
TF site
Background
TF
C528 – Cancer (Pure >= 0.4)
Normal
IS <= 150
All
TSS enrichment

#### C528 – Cancer (Pure >= 0.4) – BCL11B_uc_dTSS_oPEAK

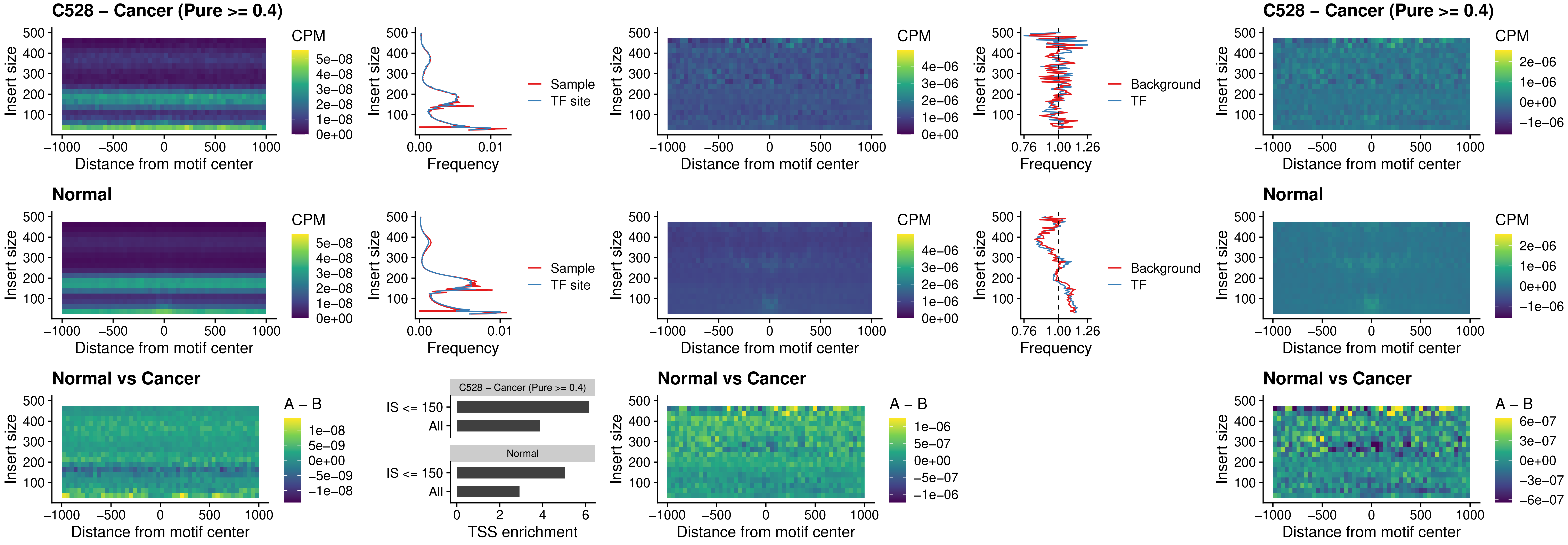

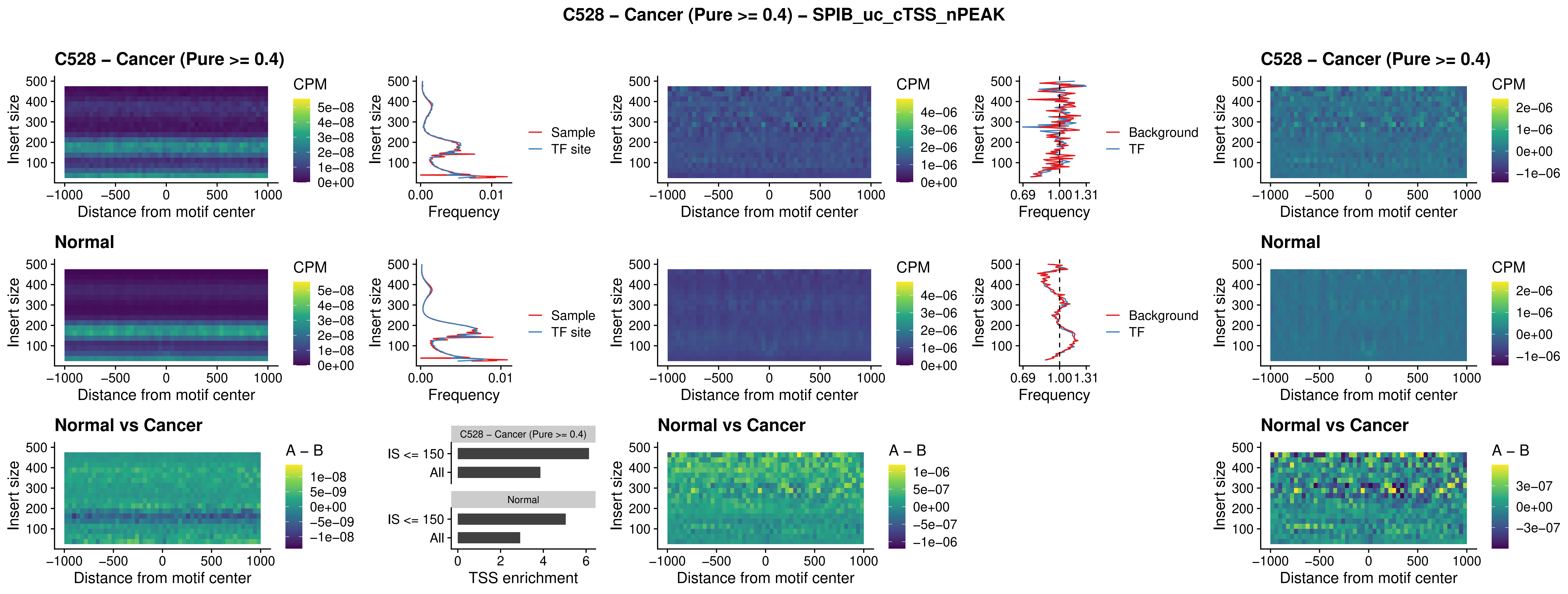
C528 – Cancer (Pure >= 0.4) – SPIB_uc_cTSS_nPEAK
C528 – Cancer (Pure >= 0.4)
Normal
Normal vs Cancer
Insert size
Distance from motif center
CPM
A – B
Frequency
Sample
TF site
Background
TF
C528 – Cancer (Pure >= 0.4)
Normal
IS <= 150
All
TSS enrichment

#### C528 – Cancer (Pure >= 0.4) – SPIC_uc_cTSS_nPEAK

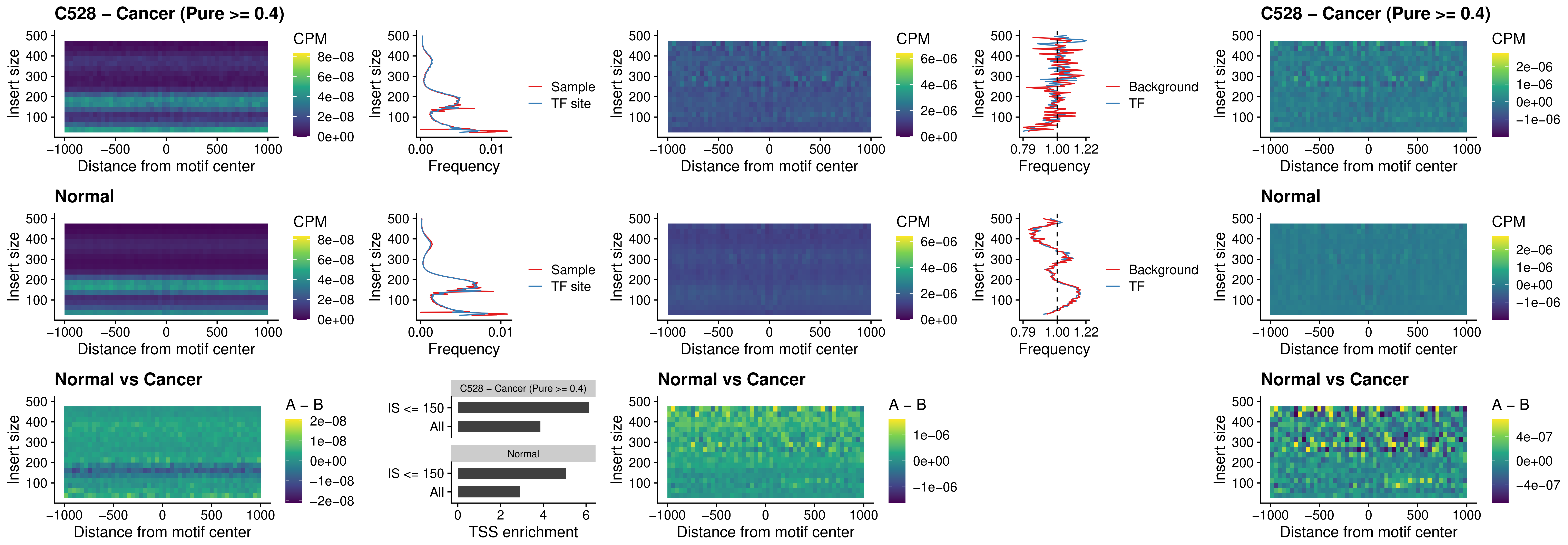

#### C528 – Cancer (Pure >= 0.4) – SPI1_uc_pTSS_oPEAK

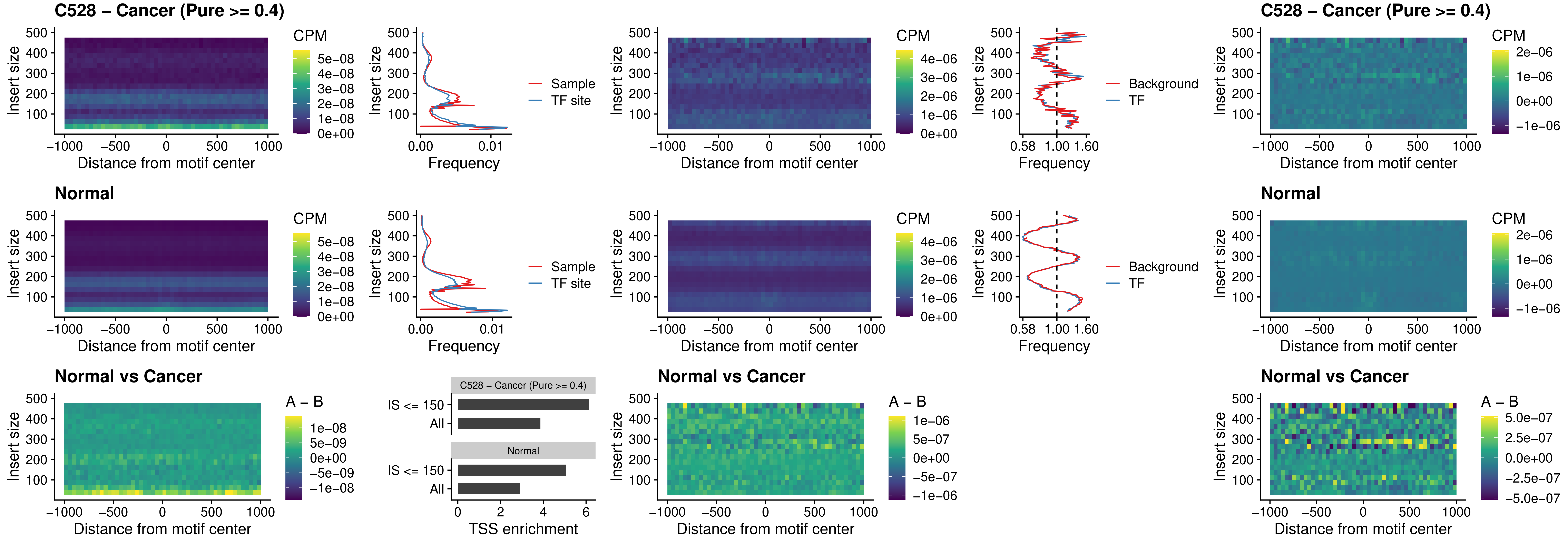

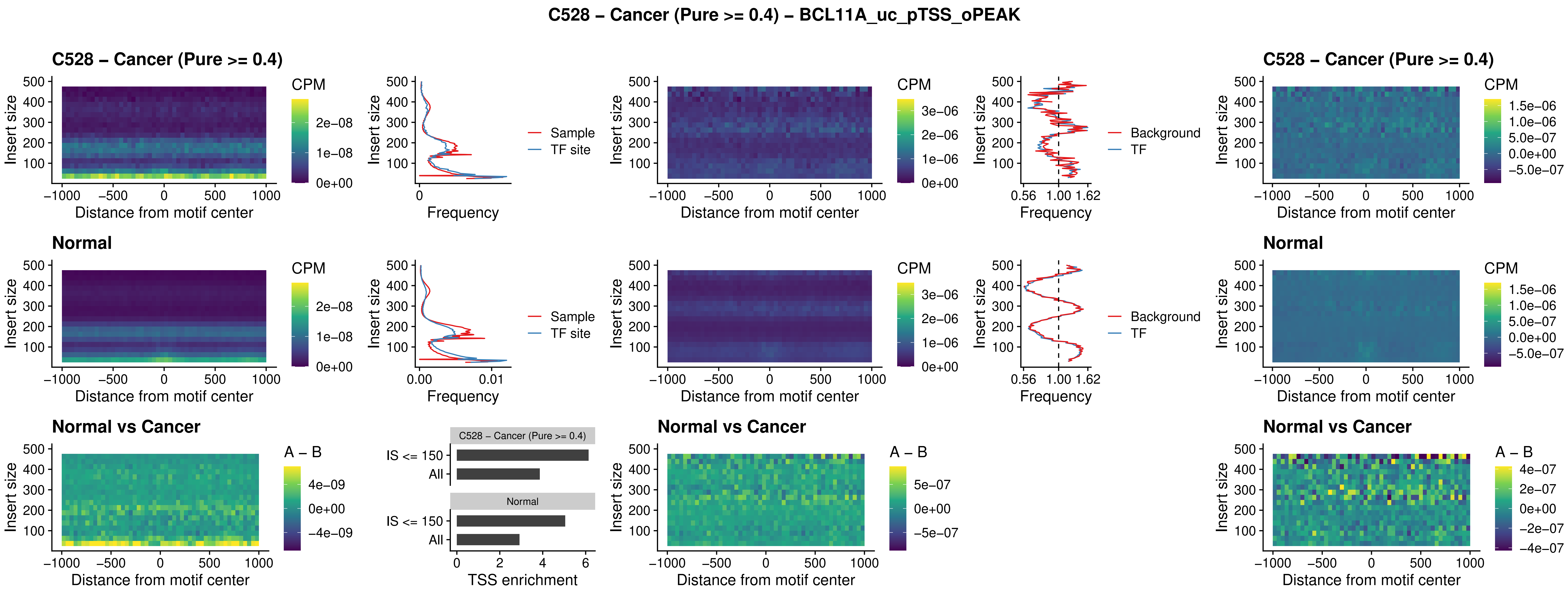
C528 – Cancer (Pure >= 0.4) – BCL11A_uc_pTSS_oPEAK
C528 – Cancer (Pure >= 0.4)
Normal
Normal vs Cancer
Insert size
Distance from motif center
Frequency
CPM
A – B
Sample
TF site
Background
TF
C528 – Cancer (Pure >= 0.4)
Normal
IS <= 150
All
TSS enrichment

#### C528 – Cancer (Pure >= 0.4) – BCL11B_uc_pTSS_oPEAK

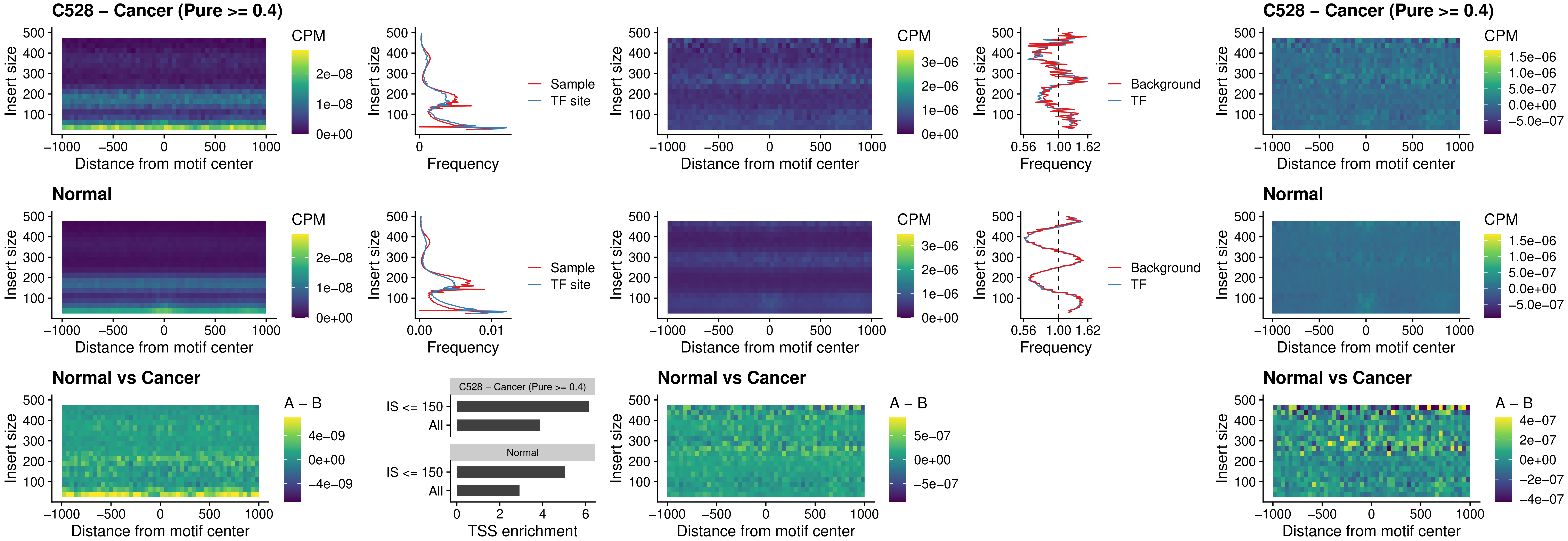

#### C528 – Cancer (Pure >= 0.4) – SPI1_uc_cTSS_oPEAK

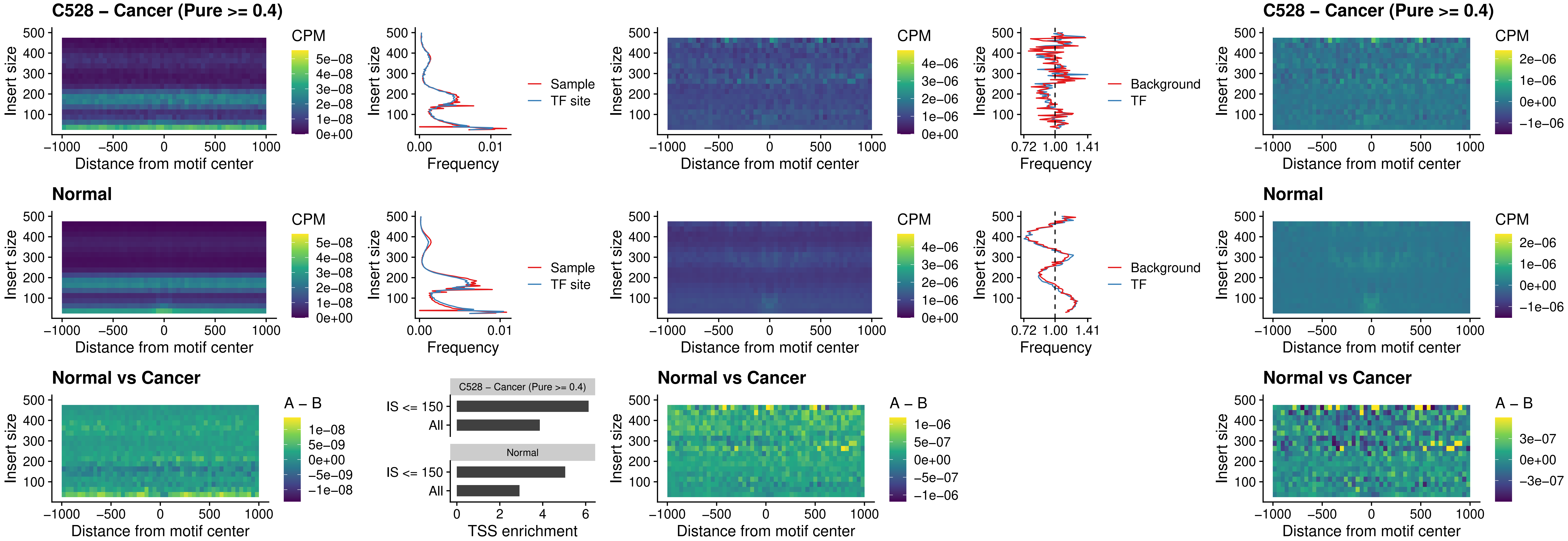

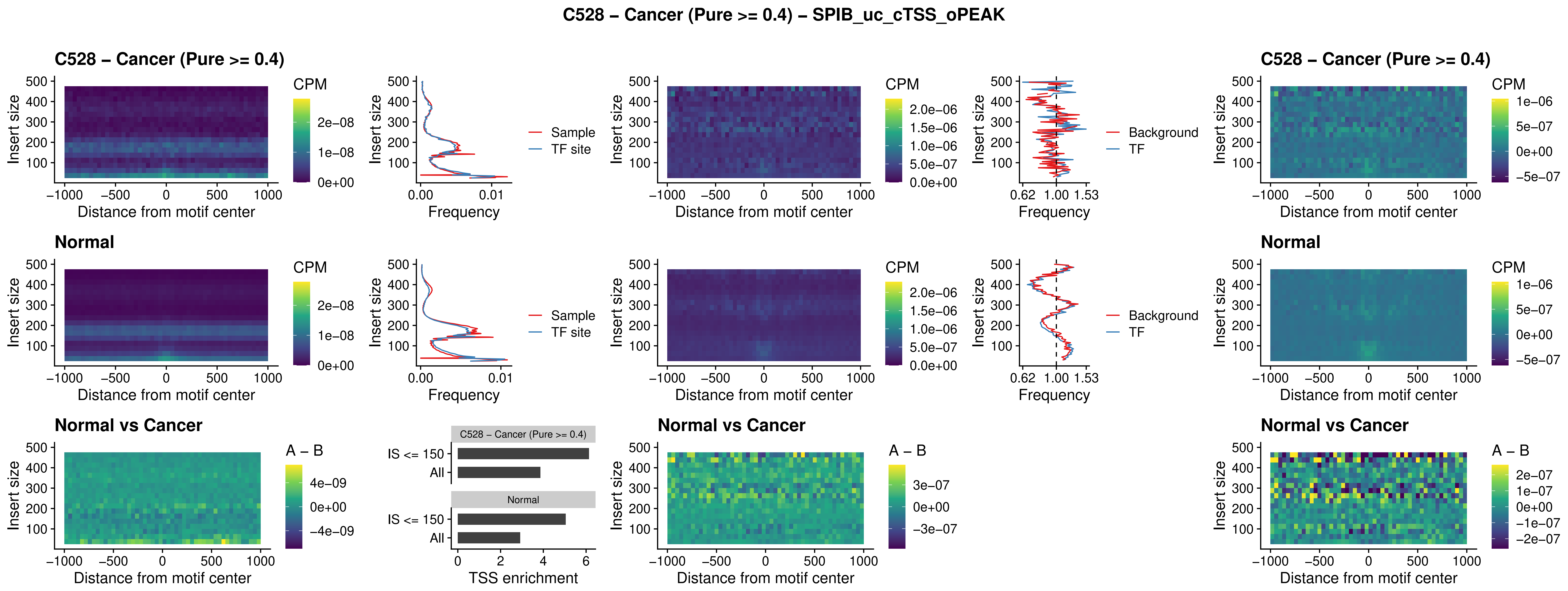
C528 – Cancer (Pure >= 0.4) – SPIB_uc_cTSS_oPEAK
C528 – Cancer (Pure >= 0.4)
Normal
Normal vs Cancer
Insert size
Distance from motif center
Frequency
CPM
A – B
Sample
TF site
Background
TF
C528 – Cancer (Pure >= 0.4)
Normal
IS <= 150
All
TSS enrichment

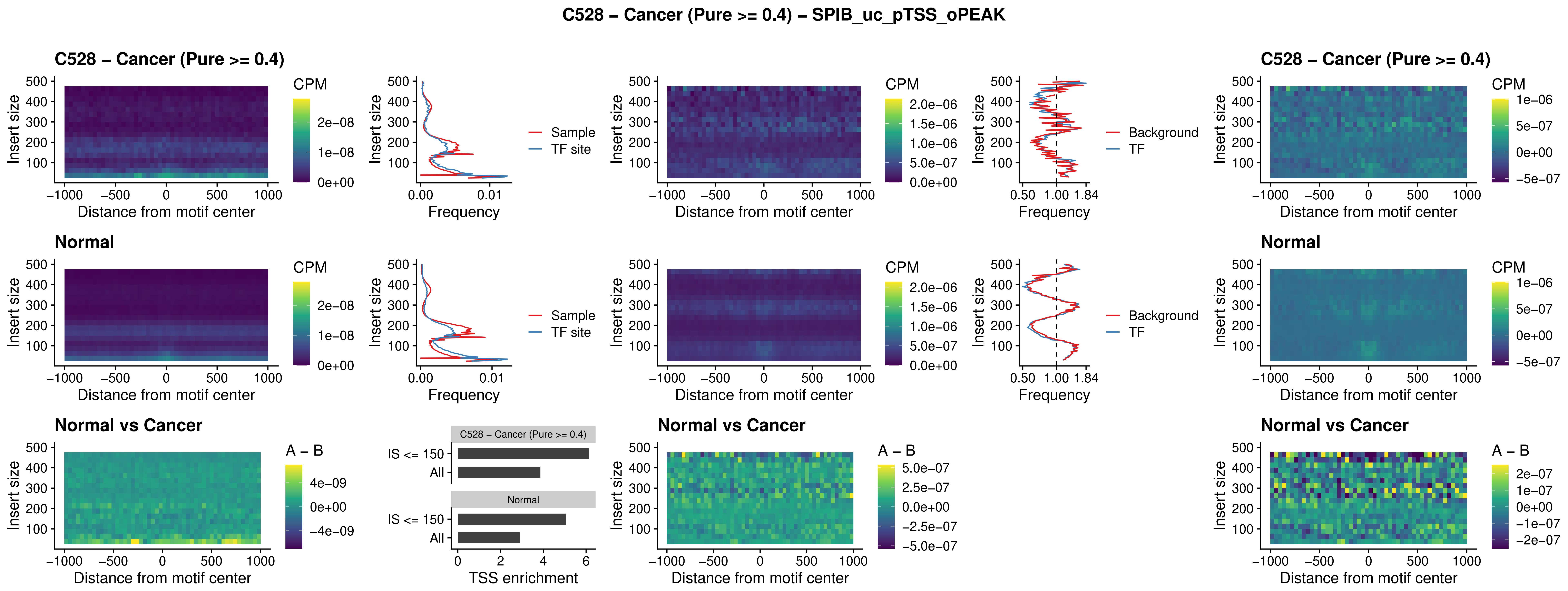
C528 – Cancer (Pure >= 0.4) – SPIB_uc_pTSS_oPEAK
C528 – Cancer (Pure >= 0.4)
Normal
Normal vs Cancer
Insert size
Distance from motif center
Frequency
CPM
A – B
Sample
TF site
Background
TF
C528 – Cancer (Pure >= 0.4)
Normal
IS <= 150
All
TSS enrichment

#### C528 – Cancer (Pure >= 0.4) – SPI1_uc_dTSS_oPEAK

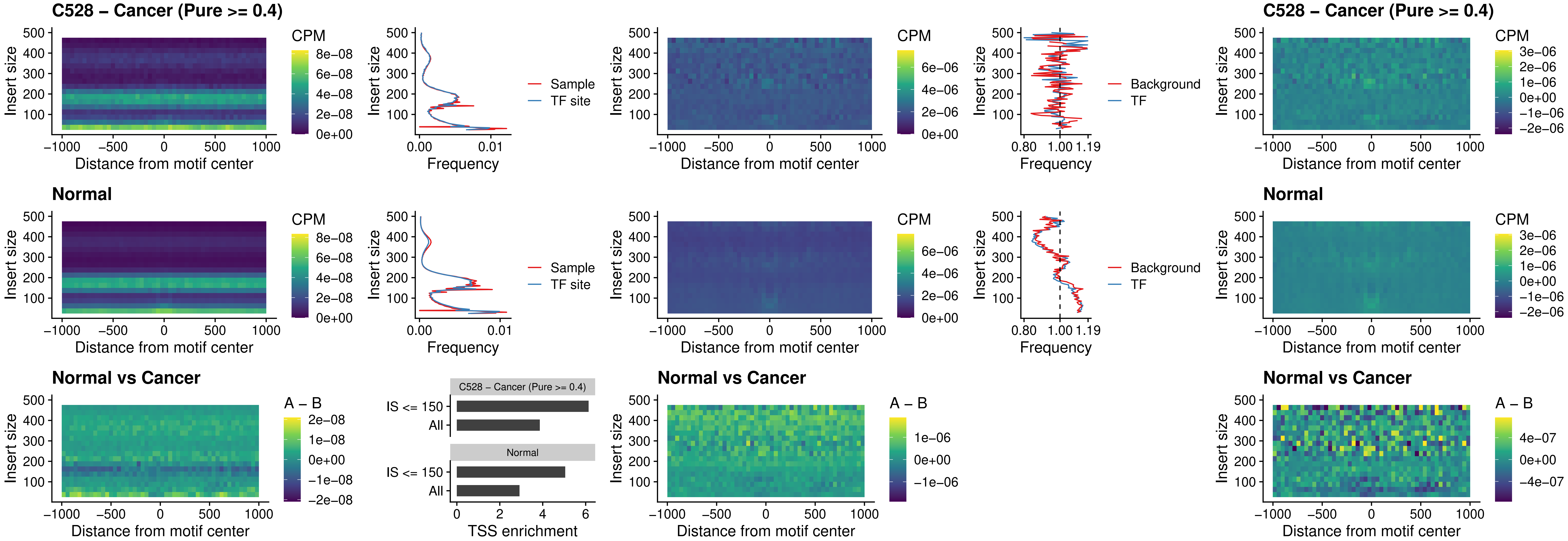

#### C528 – Cancer (Pure >= 0.4) – SPIB_uc_dTSS_oPEAK

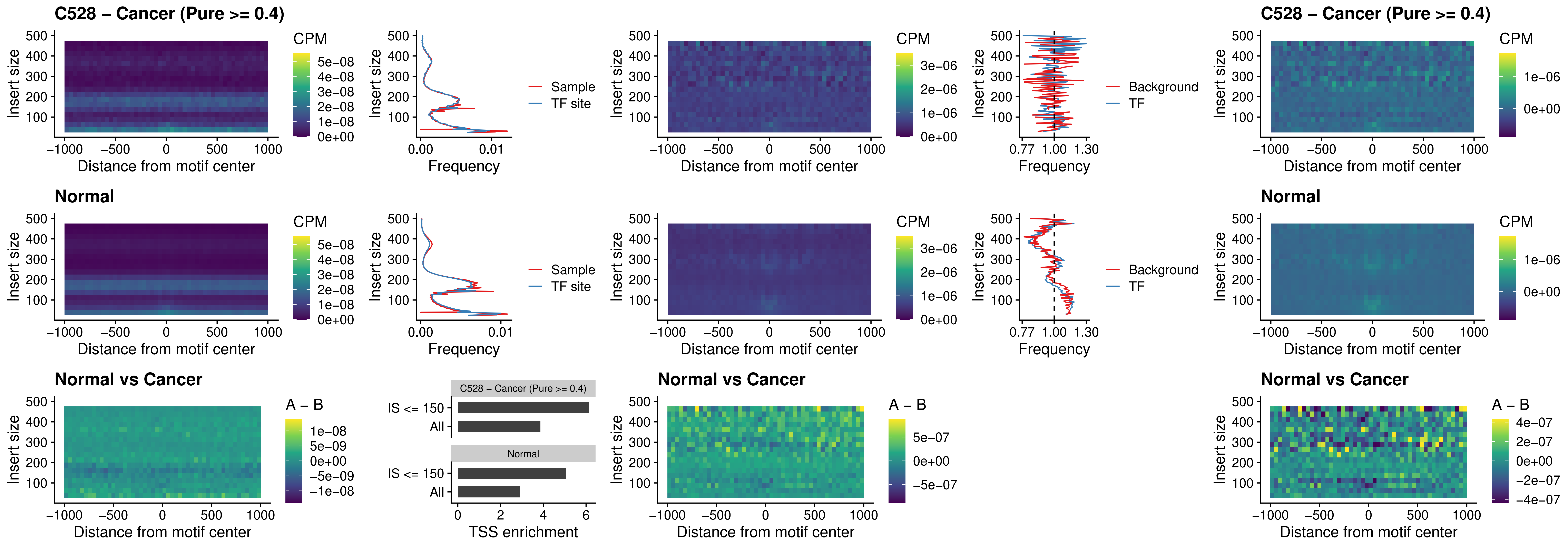

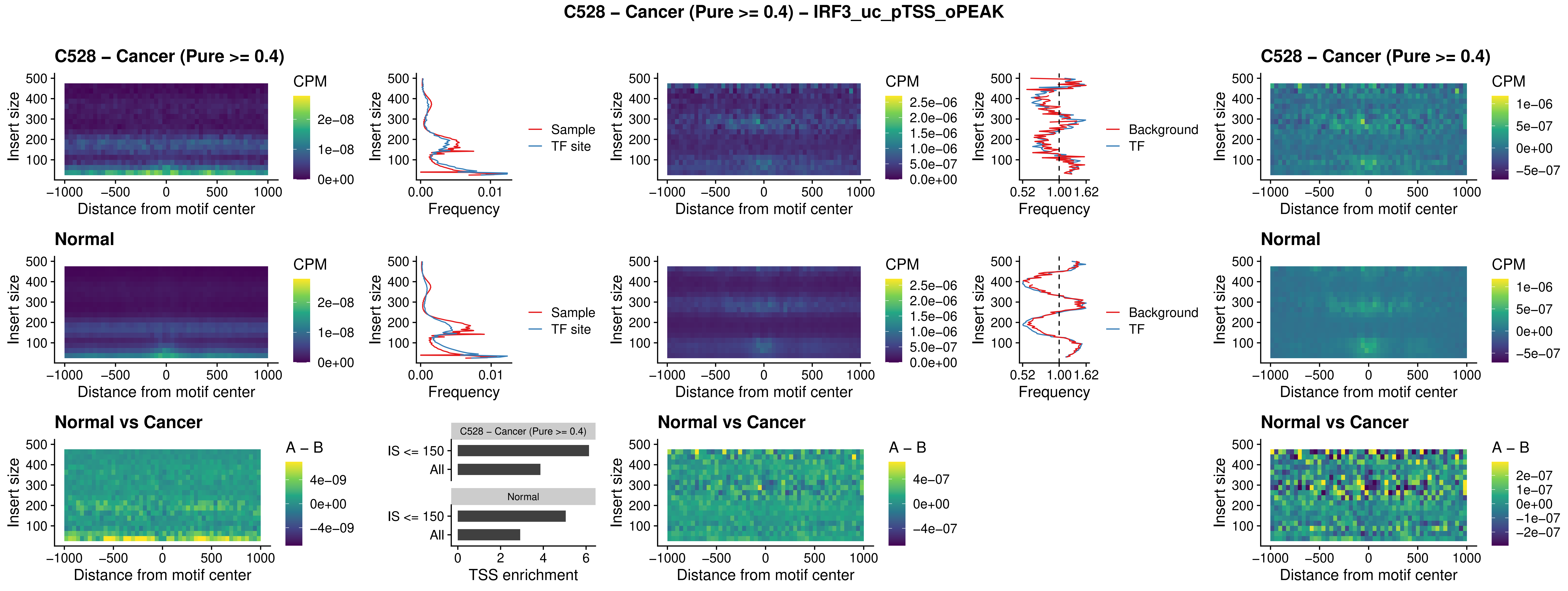
C528 – Cancer (Pure >= 0.4) – IRF3_uc_pTSS_oPEAK
C528 – Cancer (Pure >= 0.4)
Normal
Normal vs Cancer
Insert size
Distance from motif center
Frequency
CPM
A – B
Sample
TF site
Background
TF
C528 – Cancer (Pure >= 0.4)
Normal
IS <= 150
All
TSS enrichment

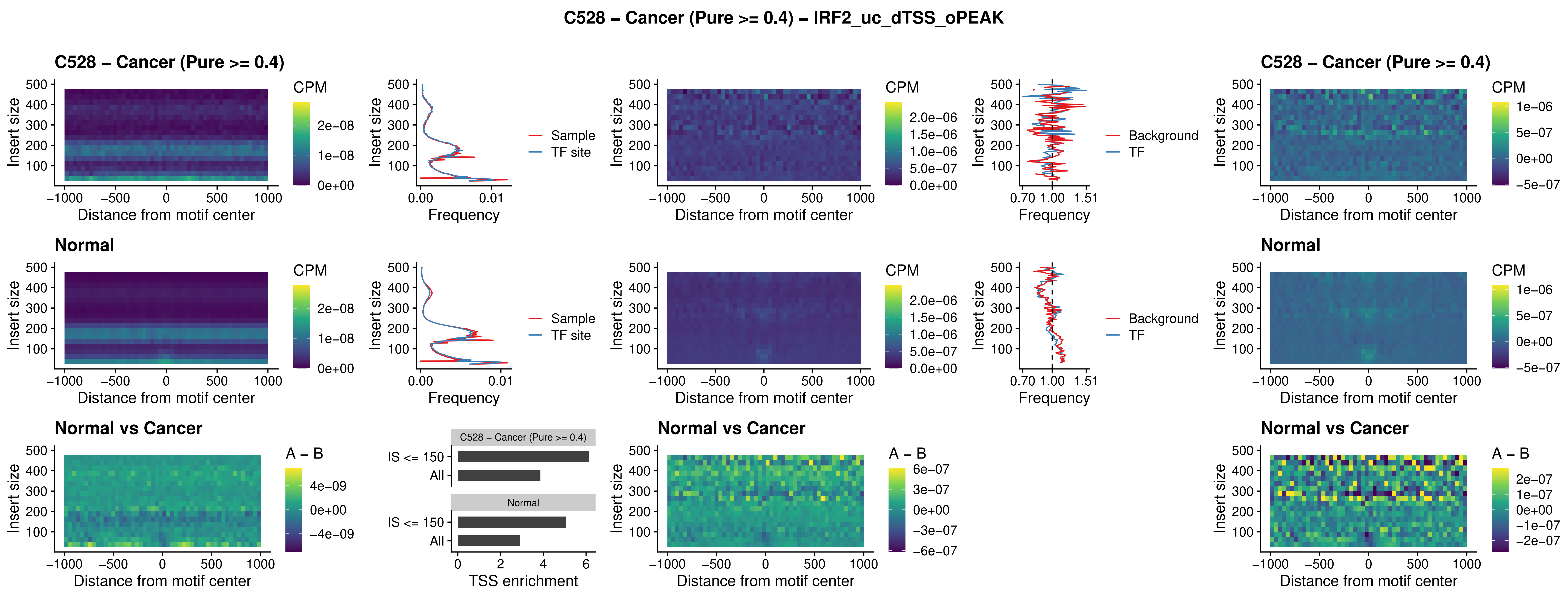

C528 – Cancer (Pure >= 0.4) – IRF2_uc_dTSS_oPEAK
C528 – Cancer (Pure >= 0.4)
Normal
Normal vs Cancer
Insert size
Distance from motif center
Frequency
CPM
A – B
Sample
TF site
Background
TF
C528 – Cancer (Pure >= 0.4)
Normal
IS <= 150
All
TSS enrichment

#### C528 – Cancer (Pure >= 0.4) – IRF2_uc_pTSS_oPEAK

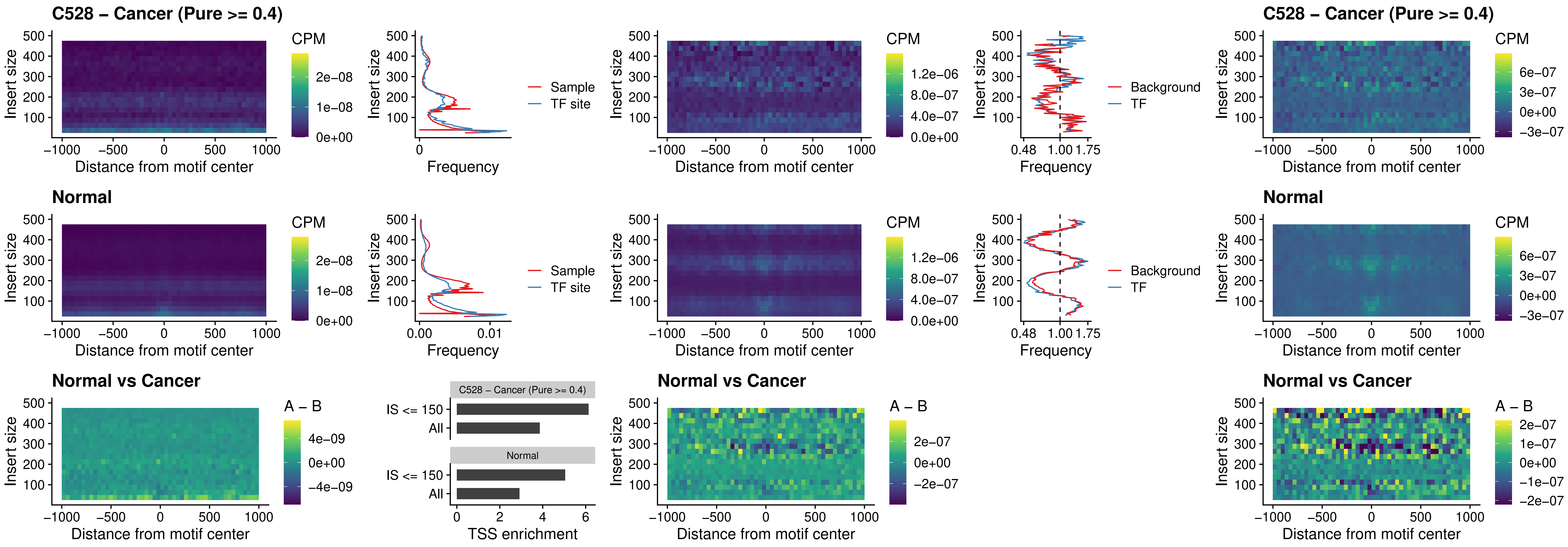

### C528 – Cancer (Pure >= 0.4) – STAT2_uc_pTSS_oPEAK

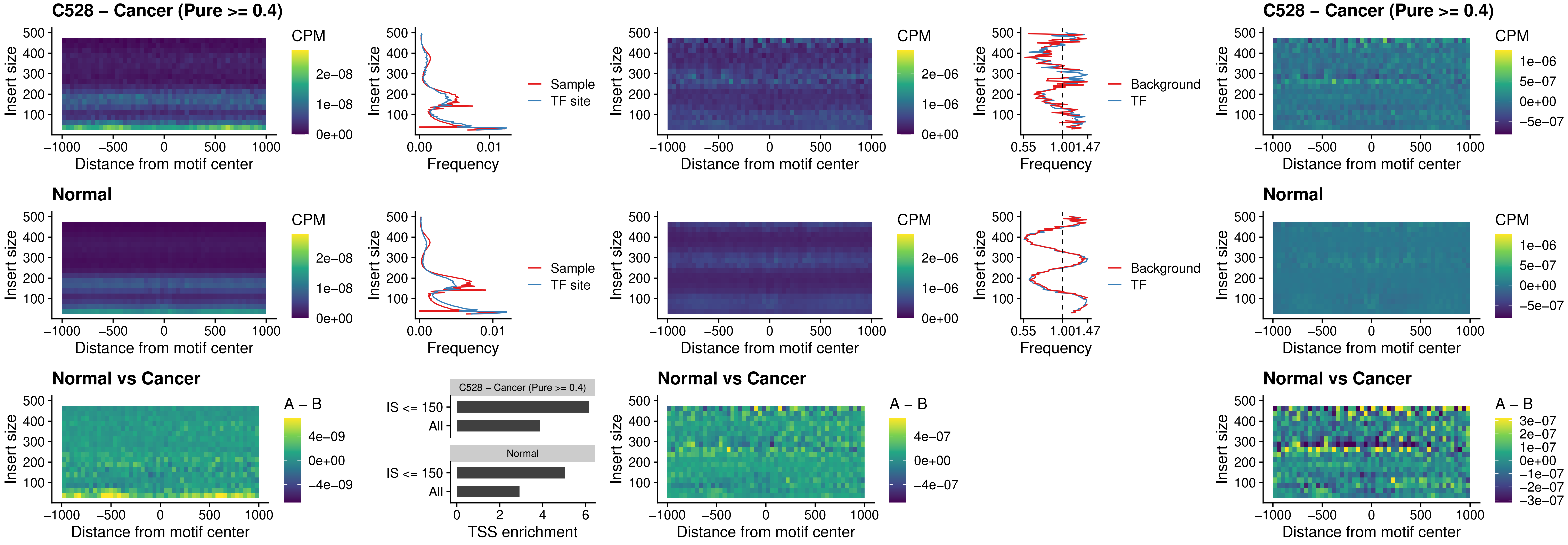

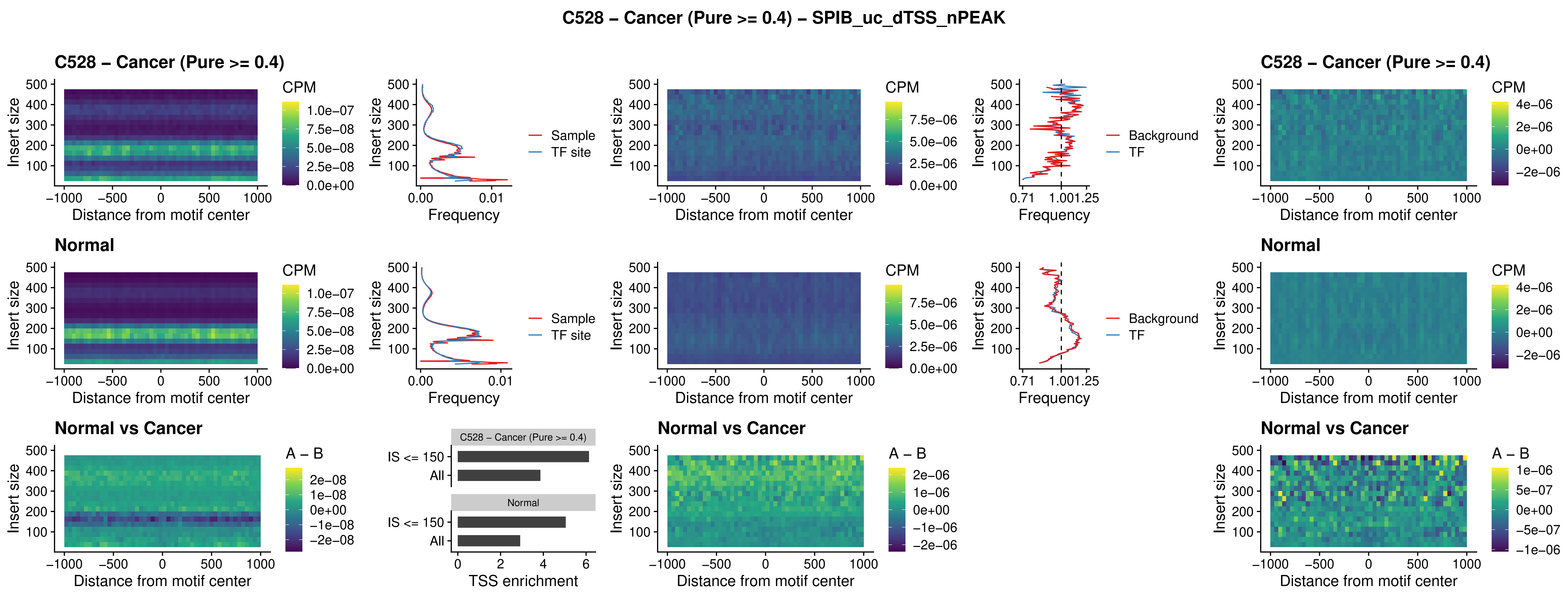
C528 – Cancer (Pure >= 0.4) – SPIB_uc_dTSS_nPEAK
C528 – Cancer (Pure >= 0.4)
Normal
Normal vs Cancer
Insert size
Distance from motif center
Frequency
CPM
A – B
Sample
TF site
Background
TF
C528 – Cancer (Pure >= 0.4)
Normal
IS <= 150
All
TSS enrichment

### C528 – Cancer (Pure >= 0.4) – SPIC_uc_cTSS_oPEAK

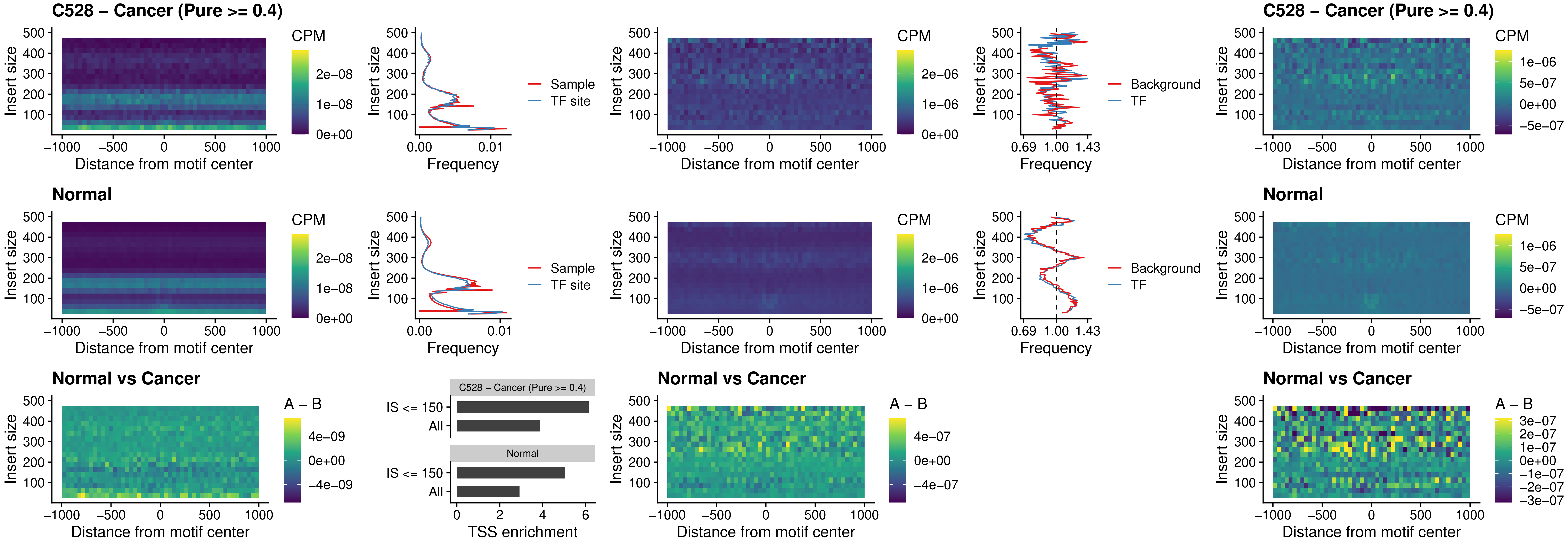

**C528 – Cancer (Pure >= 0.4) – PRDM1_uc_cTSS_oPEAK**

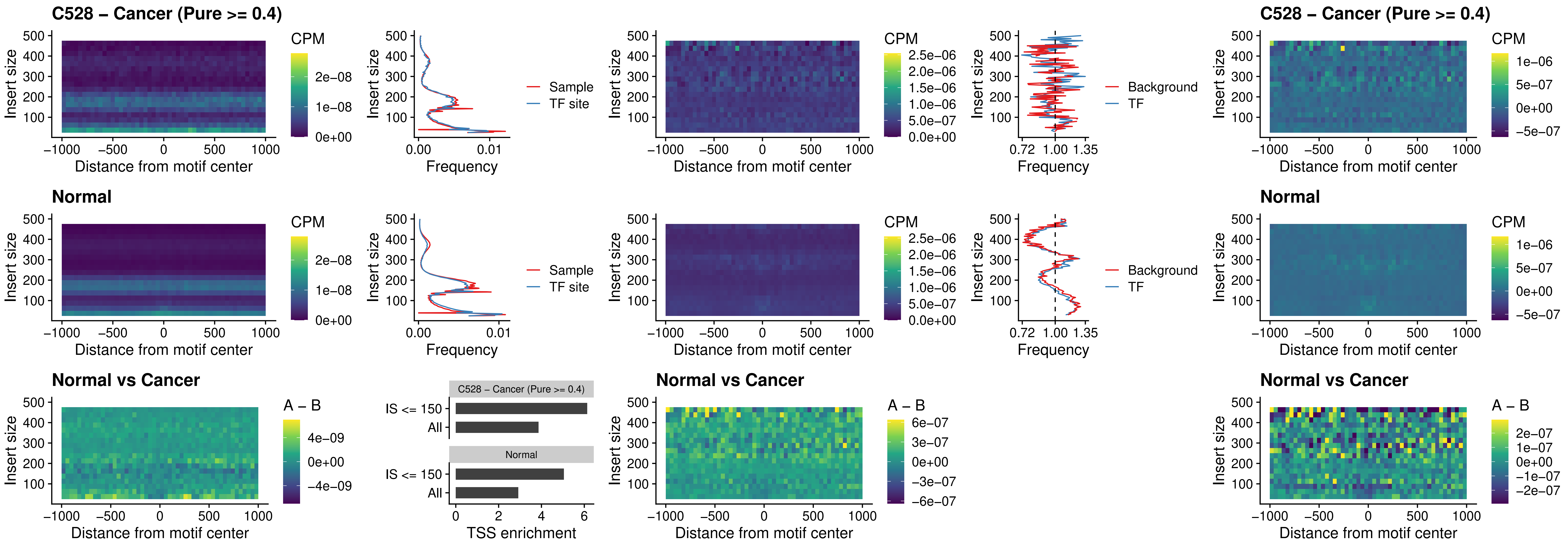

#### C528 – Cancer (Pure >= 0.4) – IRF3_uc_cTSS_oPEAK

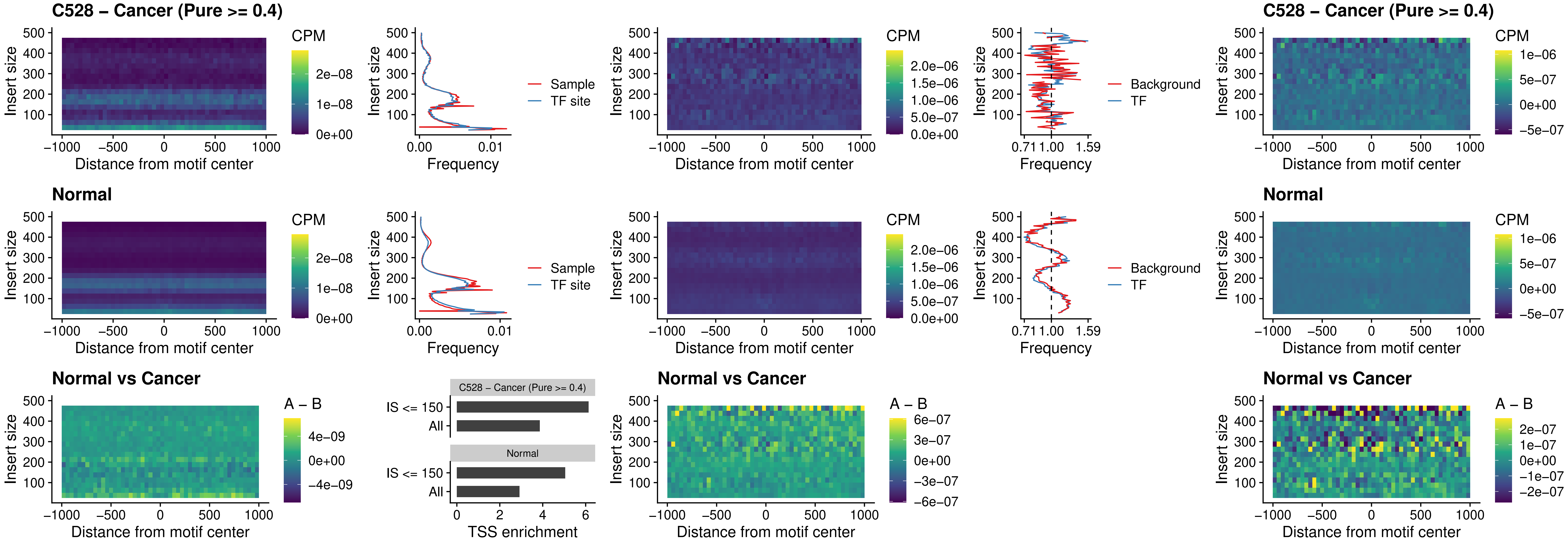

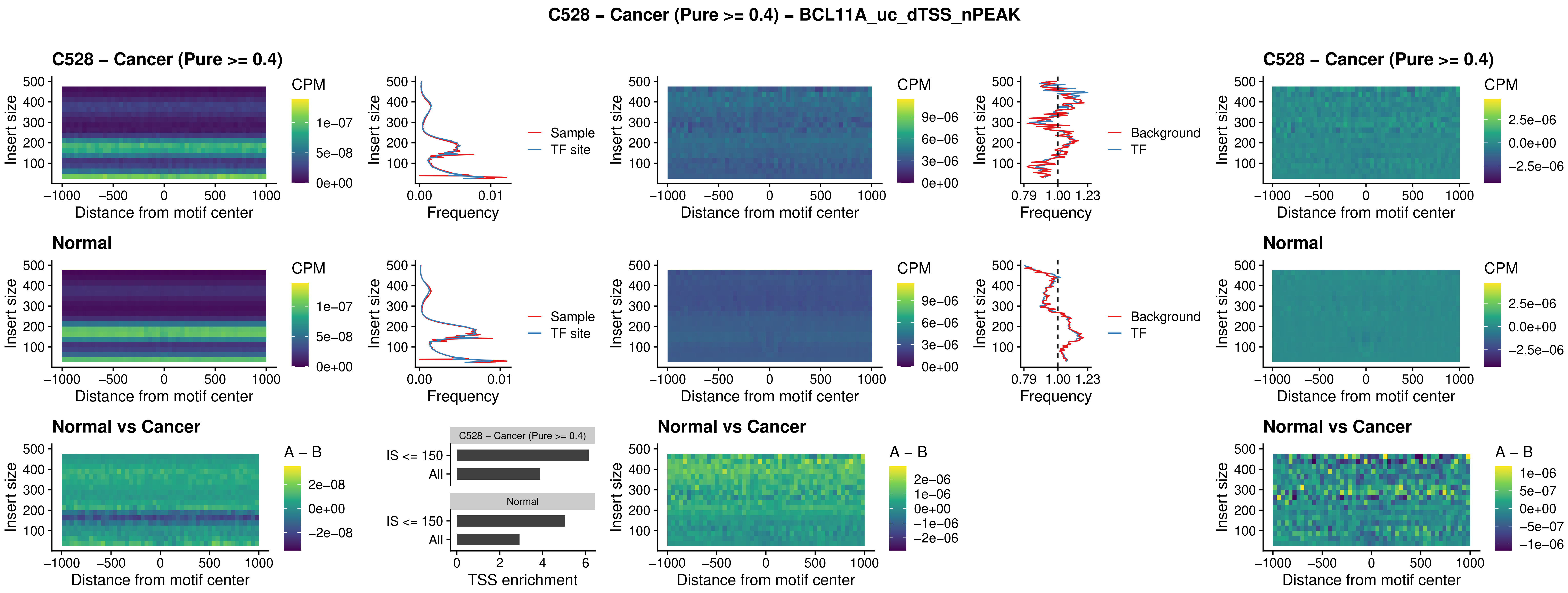
C528 – Cancer (Pure >= 0.4) – BCL11A_uc_dTSS_nPEAK
C528 – Cancer (Pure >= 0.4)
Normal
Normal vs Cancer
Insert size
Distance from motif center
Frequency
CPM
A – B
Sample
TF site
Background
TF
C528 – Cancer (Pure >= 0.4)
Normal
IS <= 150
All
TSS enrichment

#### C528 – Cancer (Pure >= 0.4) – BCL11B_uc_dTSS_nPEAK

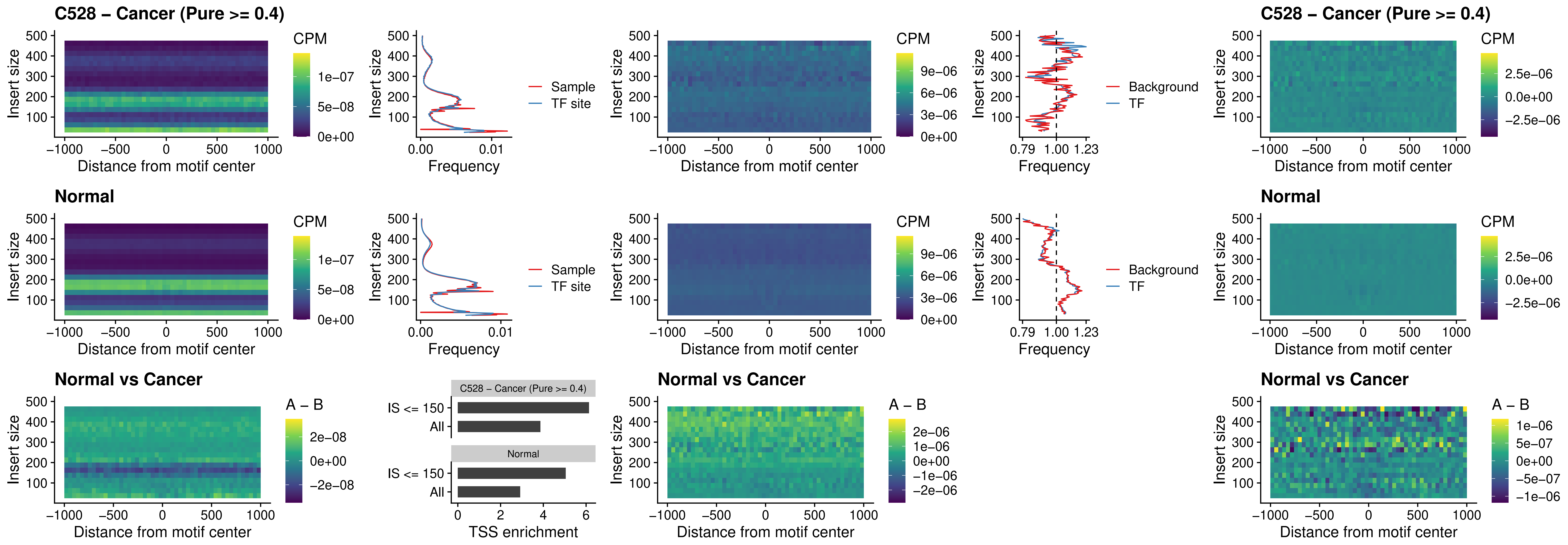

### C528 – Cancer (Pure >= 0.4) – IRF1_uc_cTSS_oPEAK

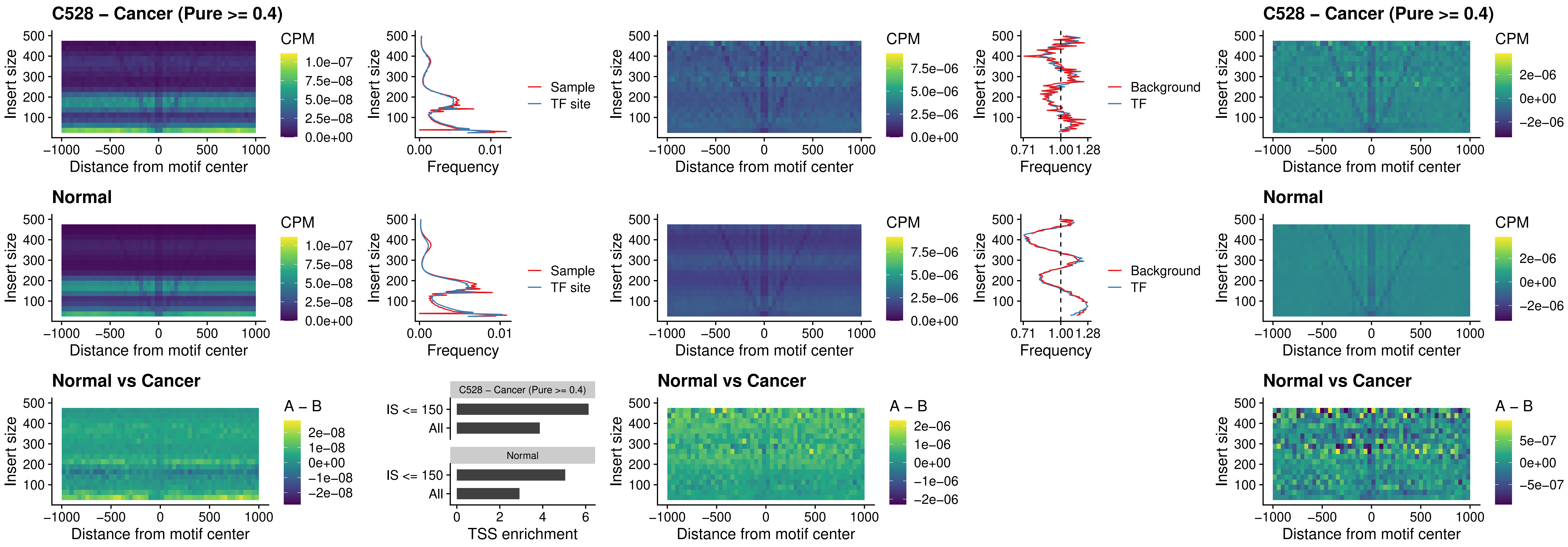

#### C528 – Cancer (Pure >= 0.4) – IRF1_uc_dTSS_oPEAK

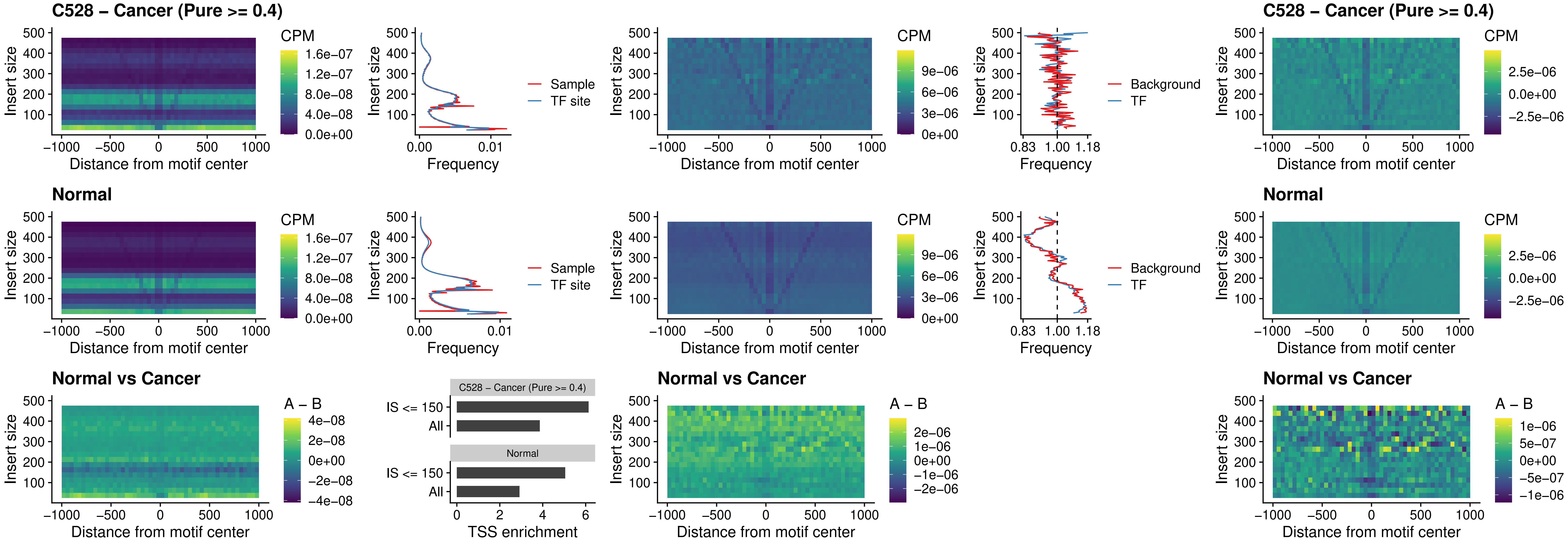

#### C528 – Cancer (Pure >= 0.4) – IRF1_uc_pTSS_oPEAK

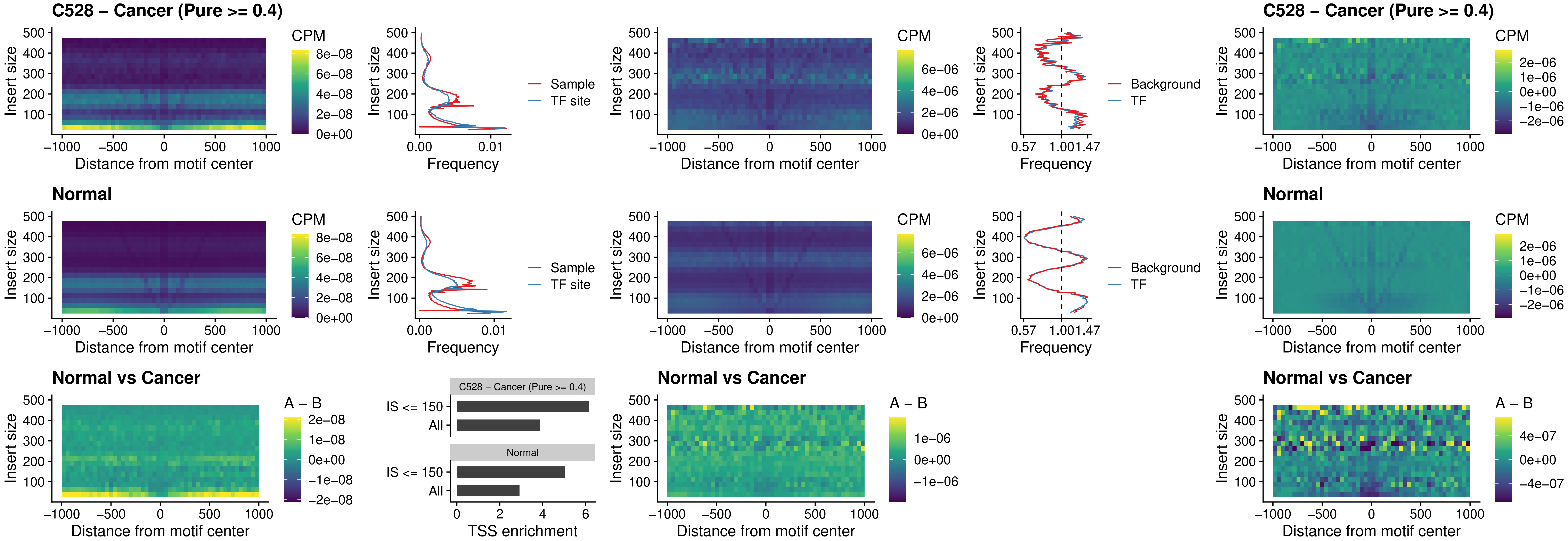

### C528 – Cancer (Pure >= 0.4) – SPIC_uc_dTSS_nPEAK

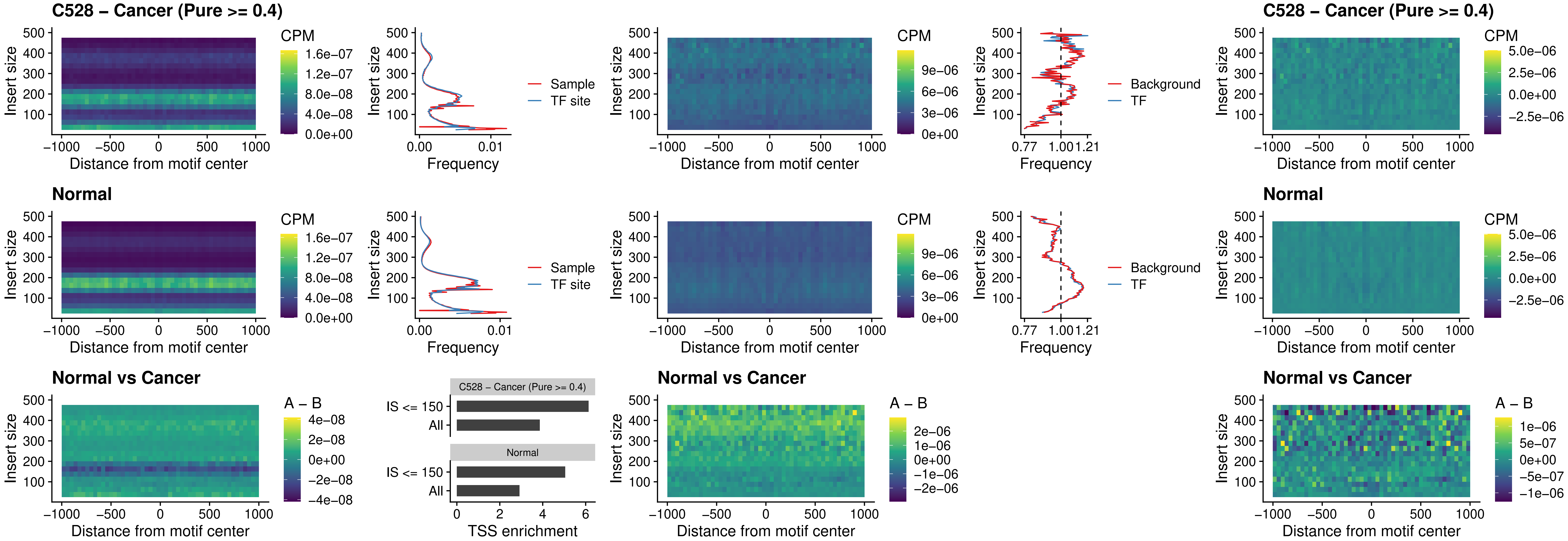

#### C528 – Cancer (Pure >= 0.4) – SPIC_uc_dTSS_oPEAK

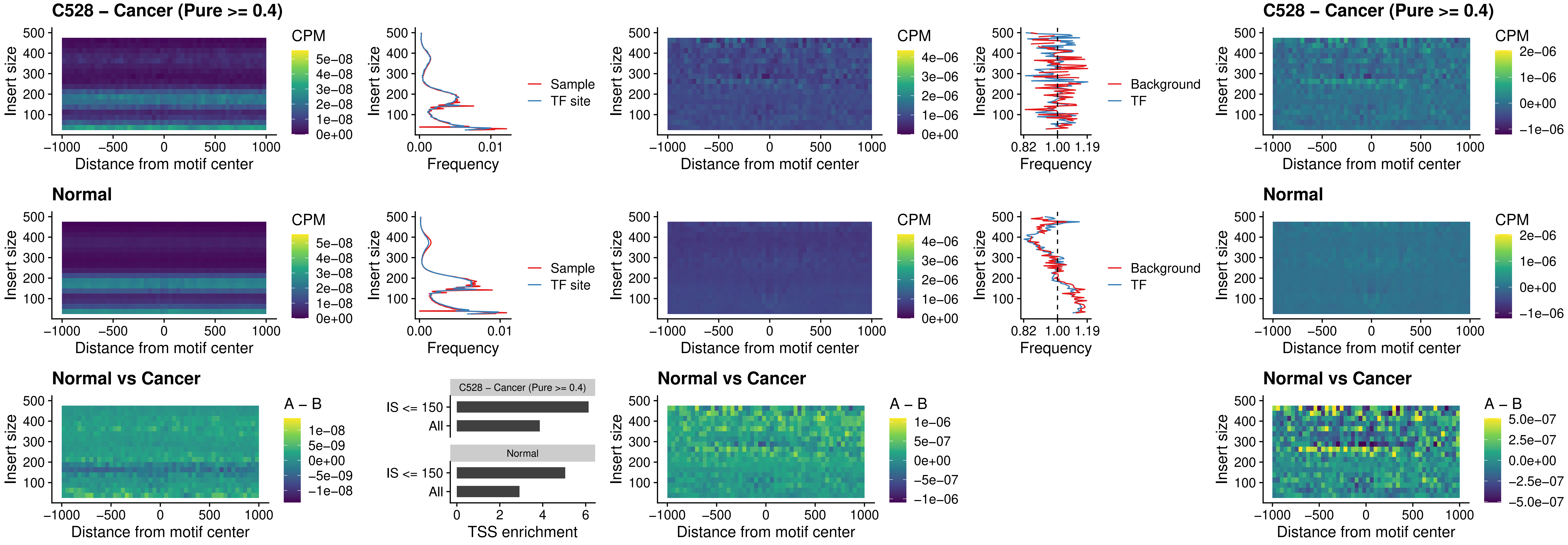

### C528 – Cancer (Pure >= 0.4) – SPIB_uc_pTSS_nPEAK

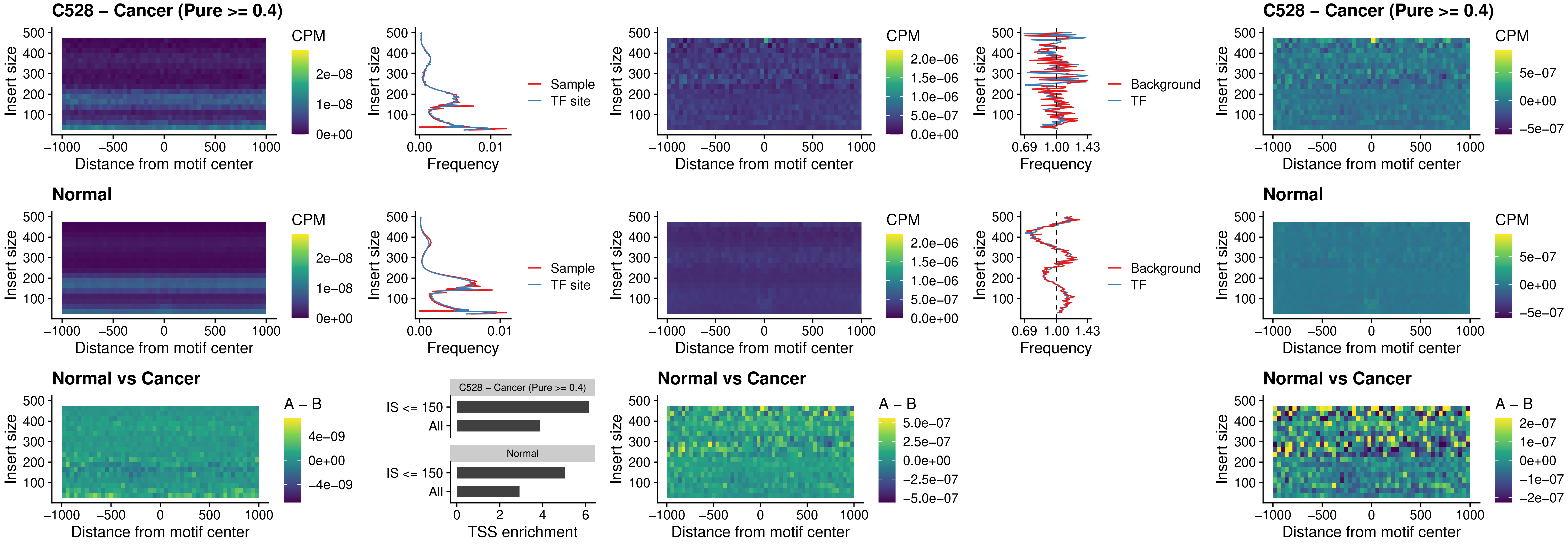

### C528 – Cancer (Pure >= 0.4) – STAT2_uc_cTSS_oPEAK

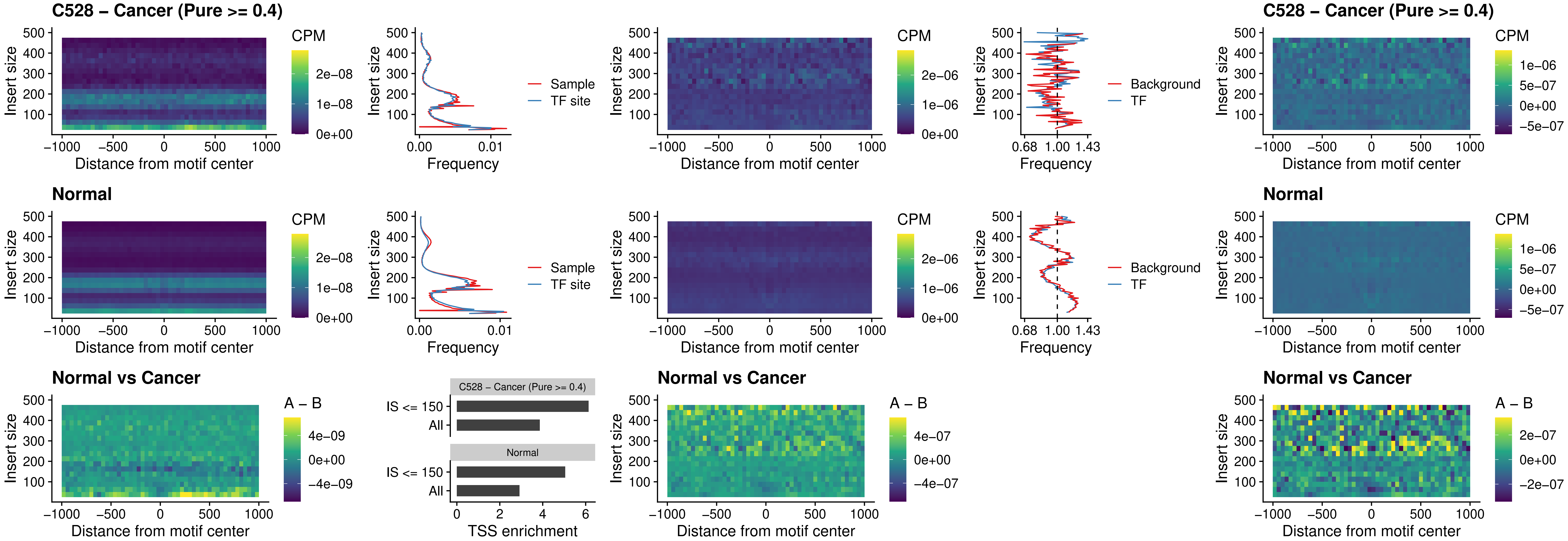

#### C528 – Cancer (Pure >= 0.4) – IRF8_uc_cTSS_oPEAK

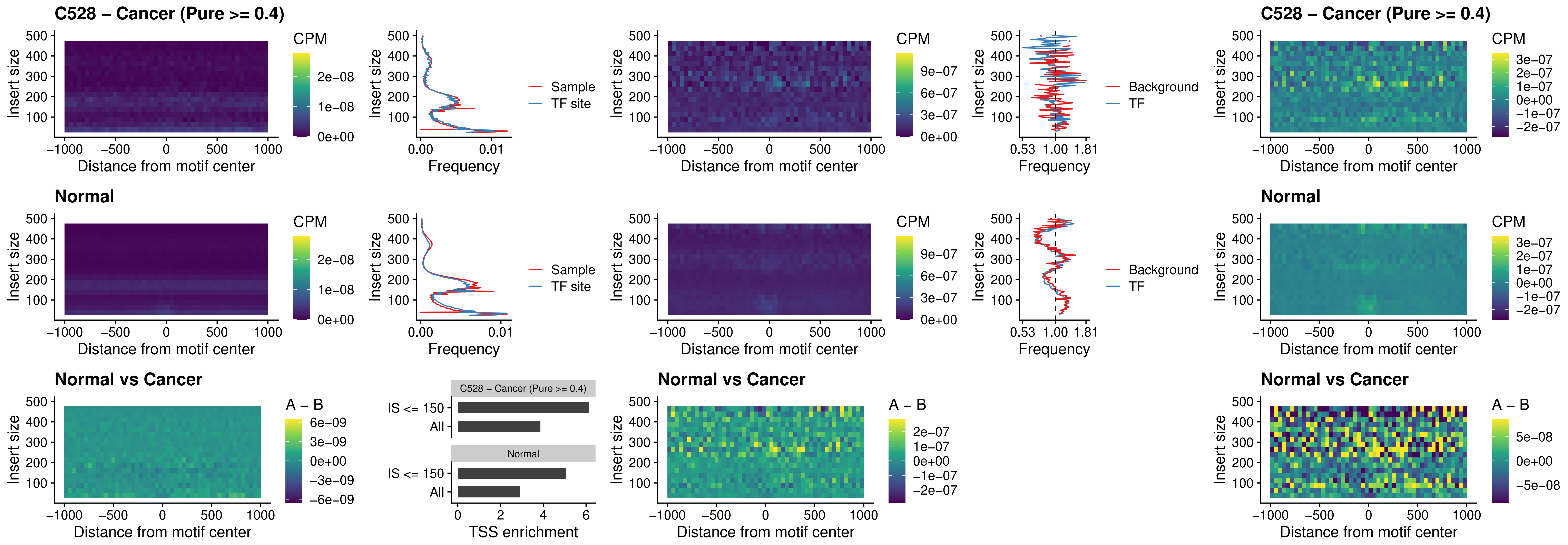

#### C528 – Cancer (Pure >= 0.4) – IRF3_uc_dTSS_oPEAK

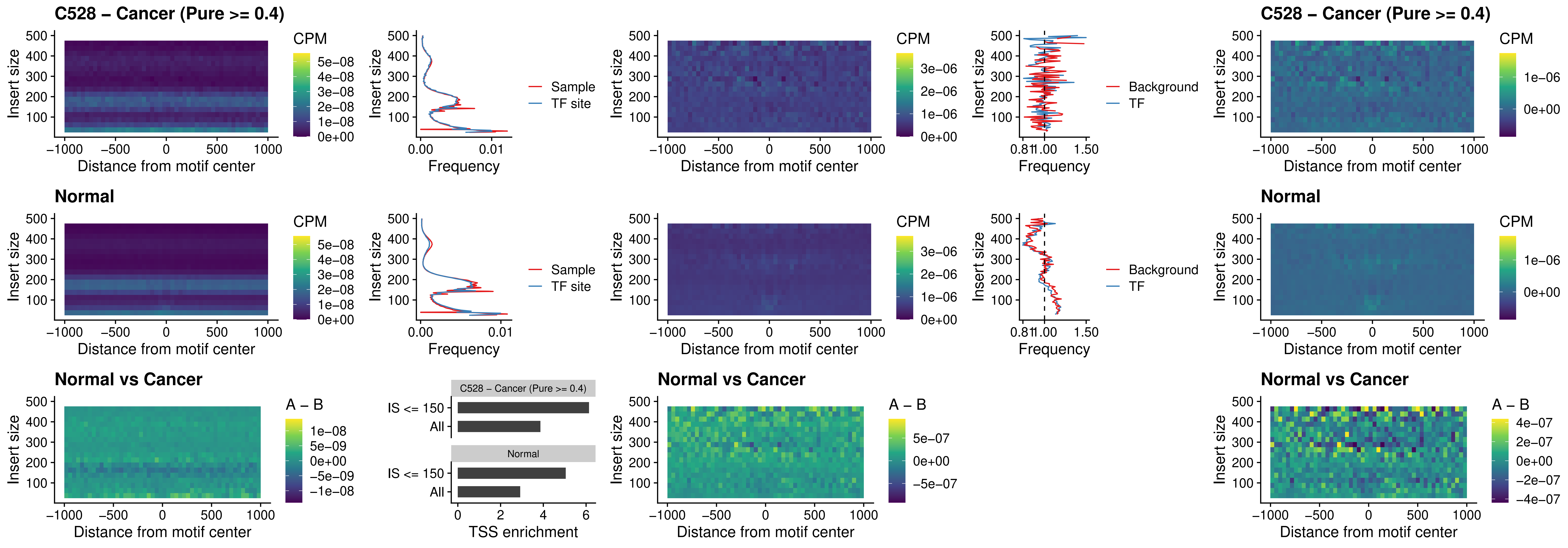

### C528 – Cancer (Pure >= 0.4) – IRF8_uc_dTSS_oPEAK

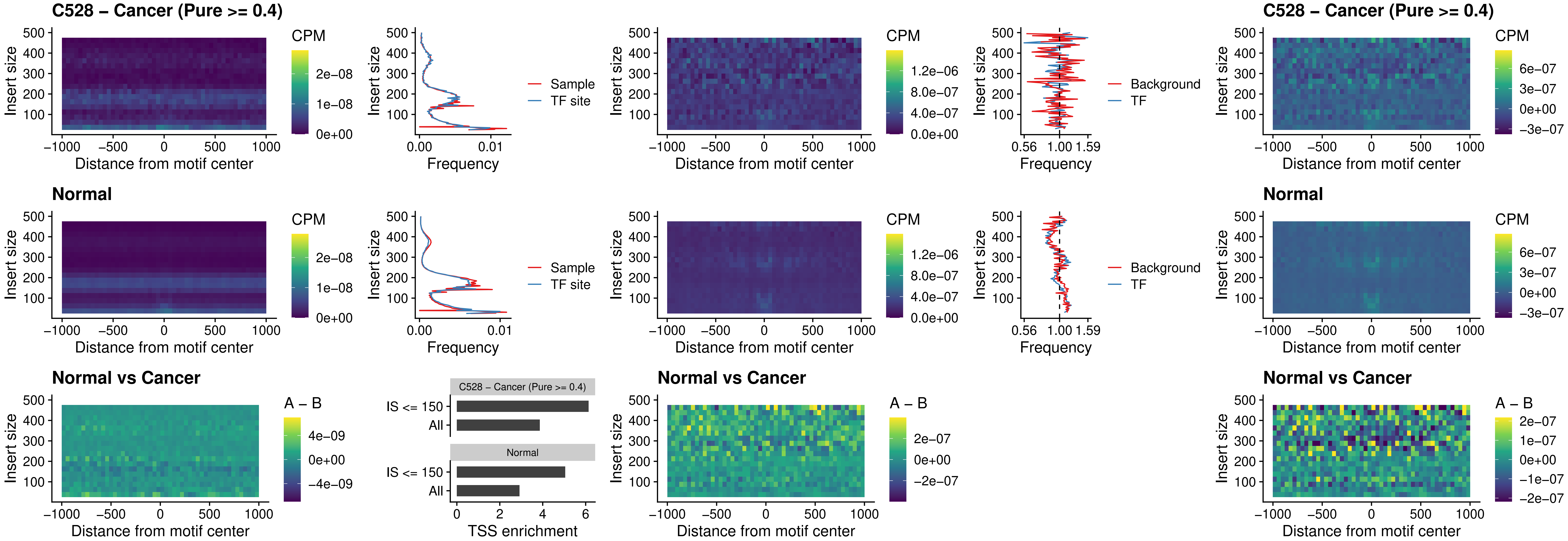

### C528 – Cancer (Pure >= 0.4) – ETS2_uc_dTSS_oPEAK

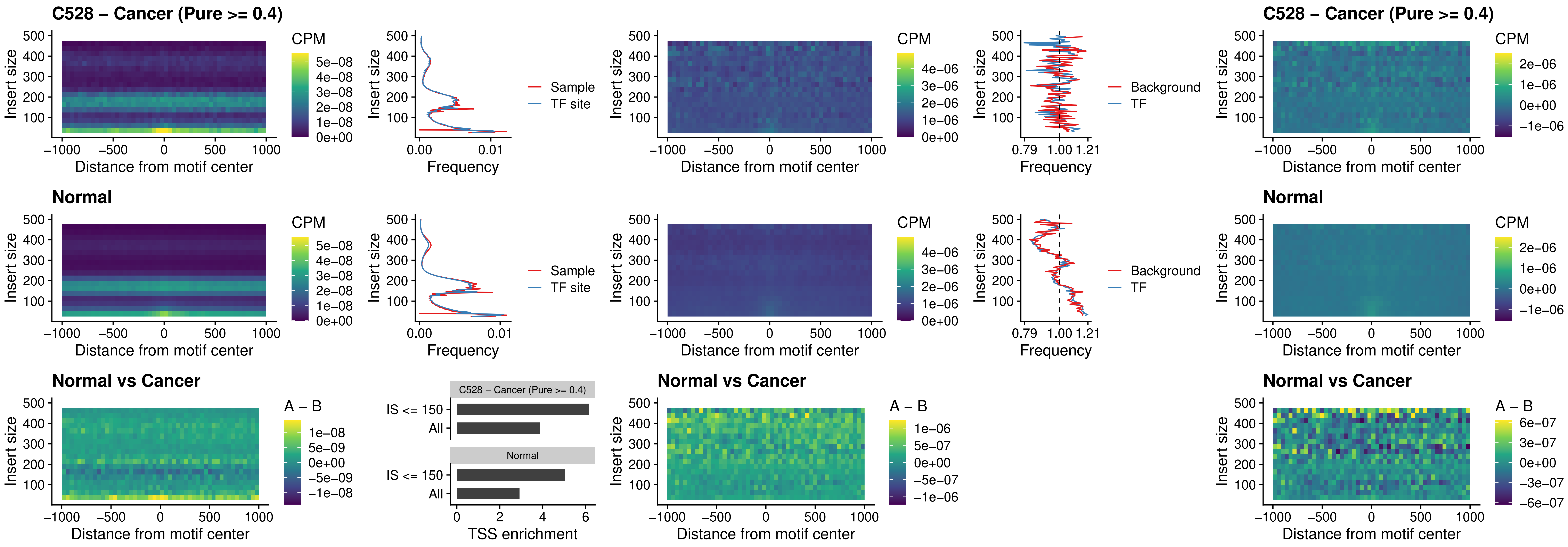

#### C528 – Cancer (Pure >= 0.4) – SMARCC2_uc_pTSS_oPEAK

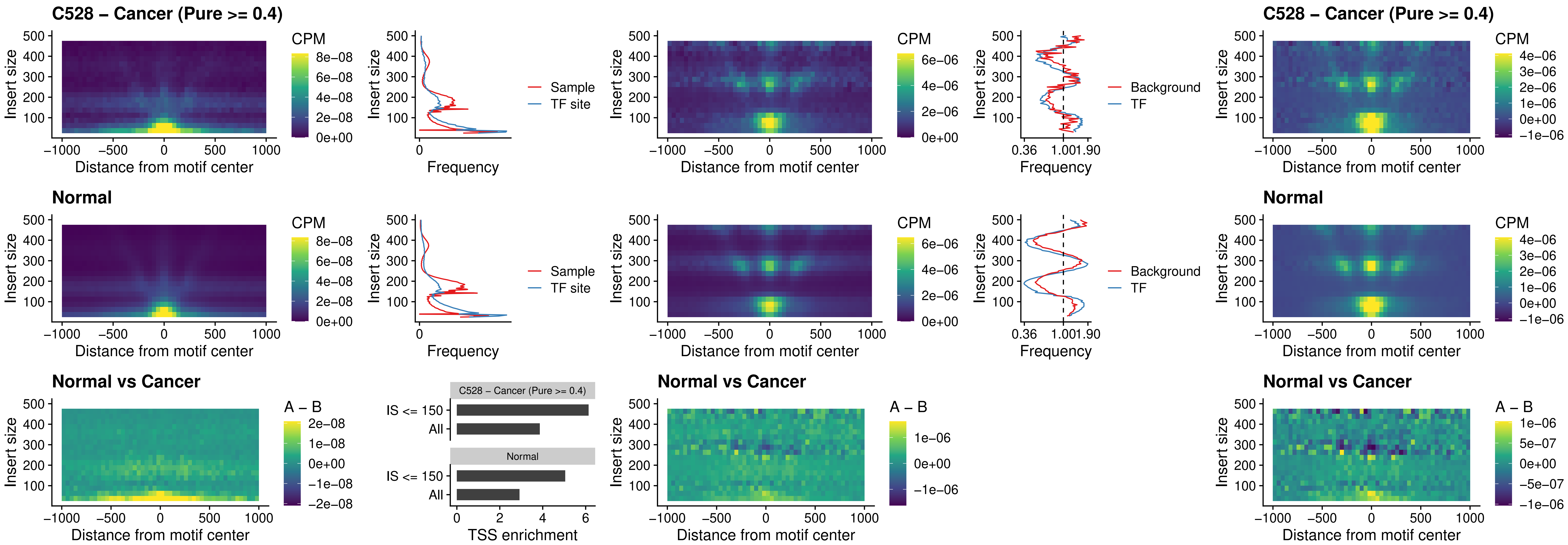

#### C528 – Cancer (Pure >= 0.4) – BCL11A_uc_cTSS_nPEAK

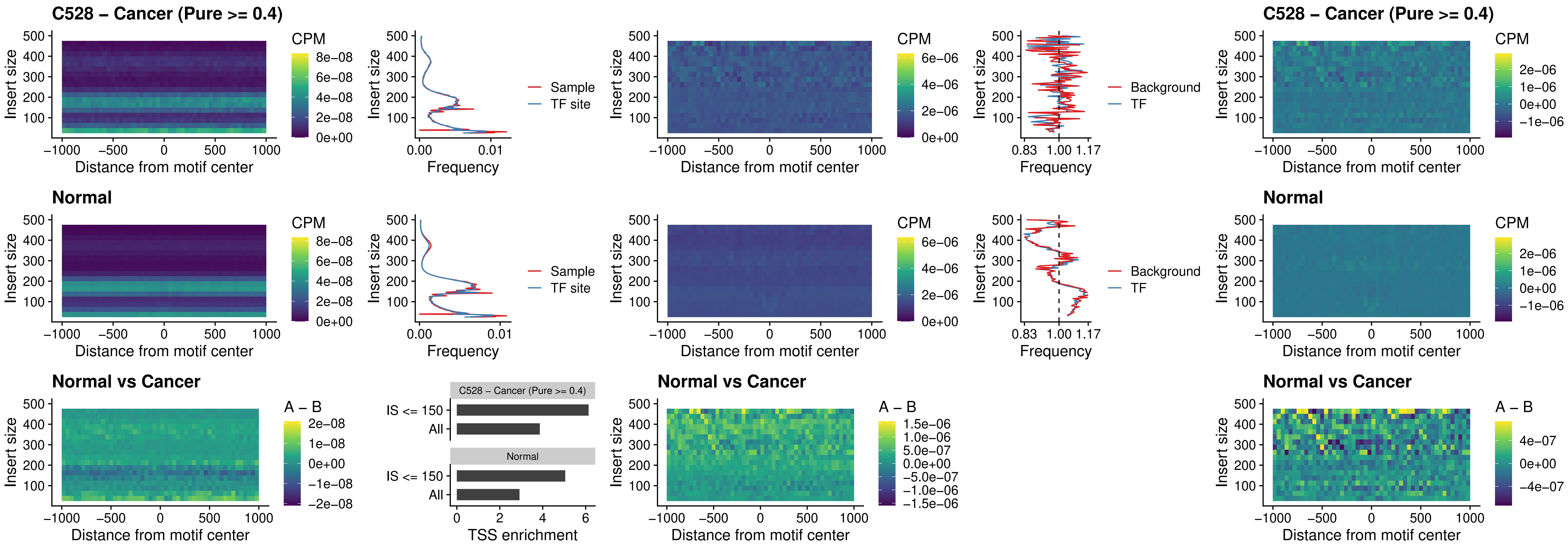

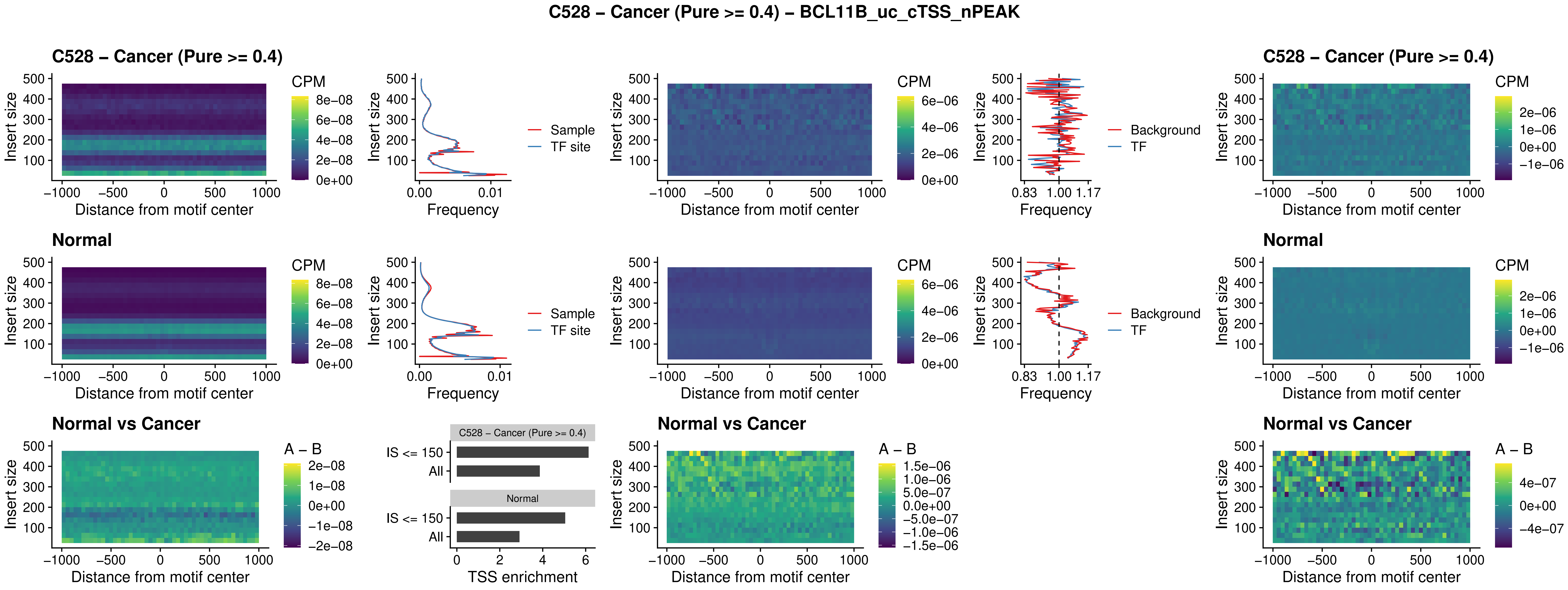
C528 – Cancer (Pure >= 0.4) – BCL11B_uc_cTSS_nPEAK
C528 – Cancer (Pure >= 0.4)
Normal
Normal vs Cancer
Insert size
Distance from motif center
Frequency
CPM
A – B
Sample
TF site
Background
TF
C528 – Cancer (Pure >= 0.4)
Normal
IS <= 150
All
TSS enrichment

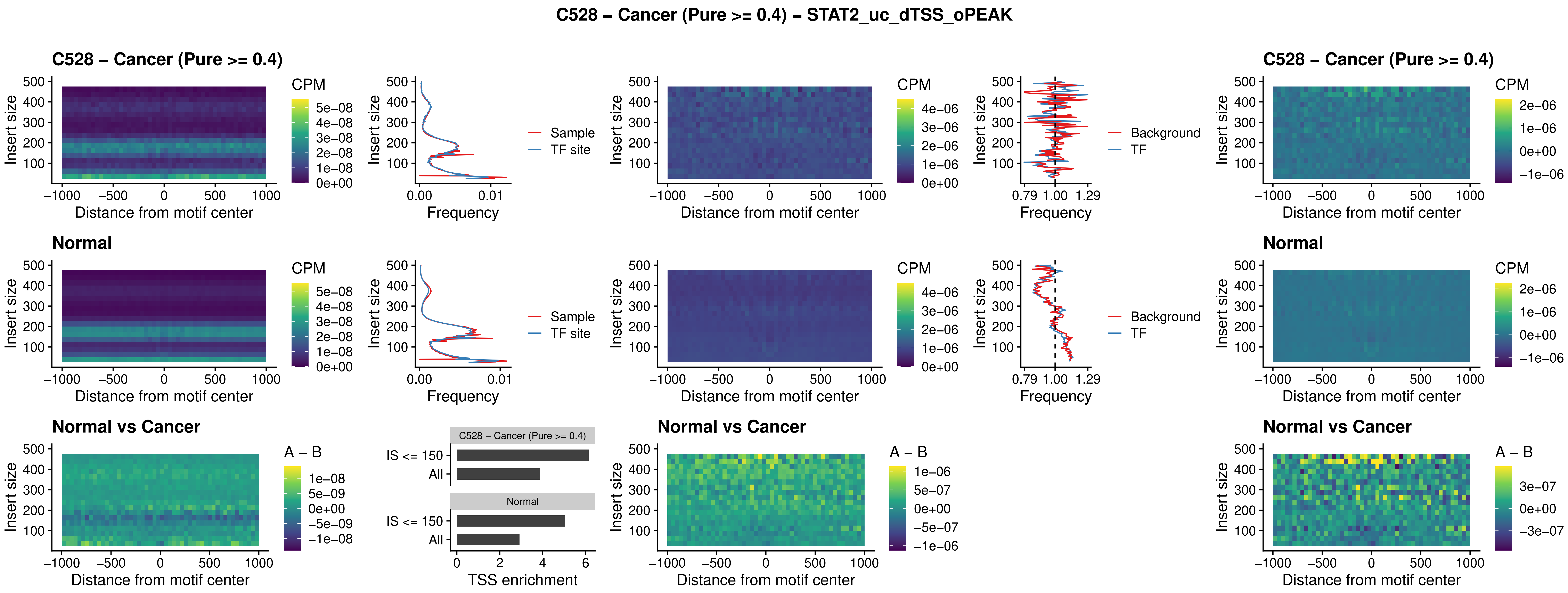

C528 – Cancer (Pure >= 0.4) – STAT2_uc_dTSS_oPEAK
C528 – Cancer (Pure >= 0.4)
Normal
Normal vs Cancer
Insert size
Distance from motif center
Frequency
CPM
A – B
Sample
TF site
Background
TF
C528 – Cancer (Pure >= 0.4)
Normal
IS <= 150
All
TSS enrichment

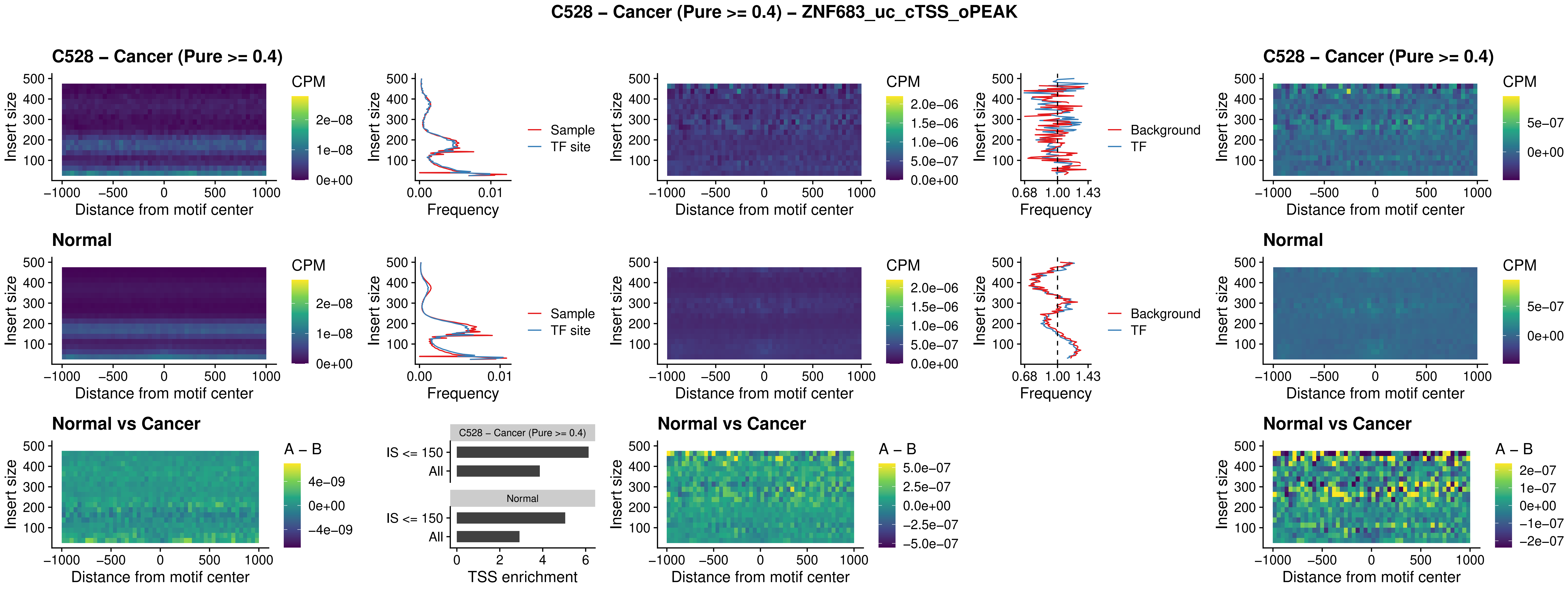
C528 – Cancer (Pure >= 0.4) – ZNF683_uc_cTSS_oPEAK
C528 – Cancer (Pure >= 0.4)
Normal
Normal vs Cancer
Insert size
Distance from motif center
CPM
A – B
Frequency
Sample
TF site
Background
TF
C528 – Cancer (Pure >= 0.4)
Normal
IS <= 150
All
TSS enrichment

#### C528 – Cancer (Pure >= 0.4) – IRF8_uc_dTSS_nPEAK

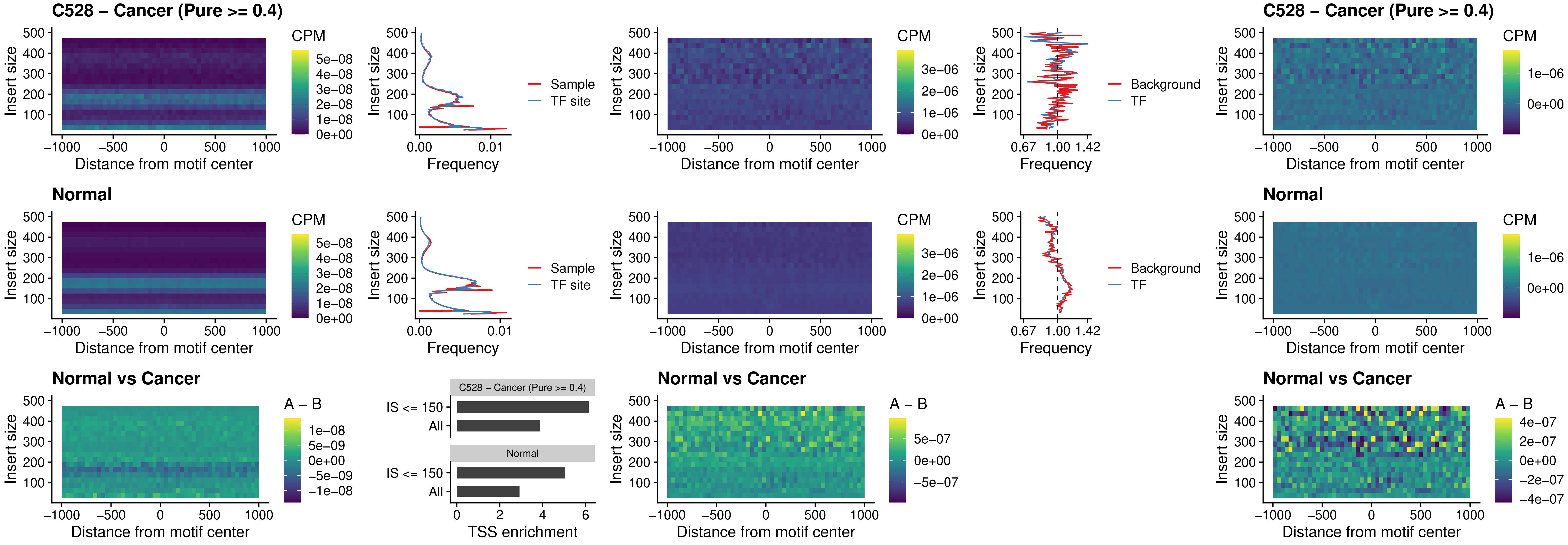

### C528 – Cancer (Pure >= 0.4) – ETV7_uc_dTSS_oPEAK

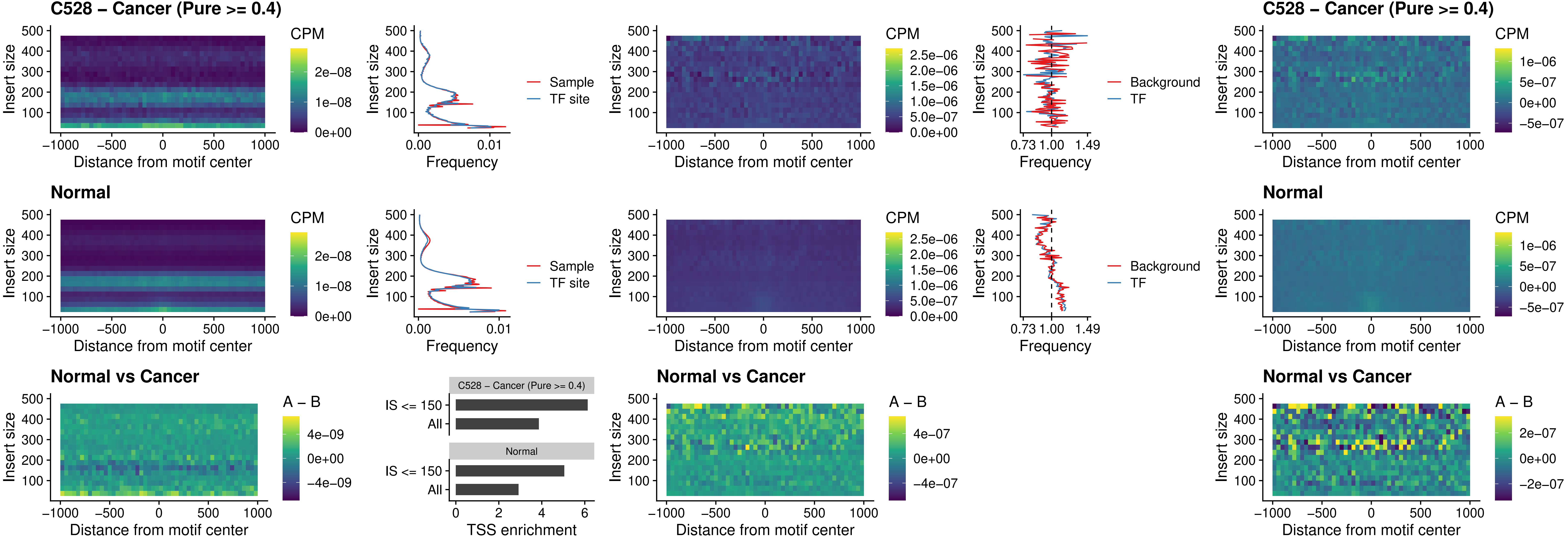

### C530 – Cancer (Pure >= 0.4) – FOXL1_uc_pTSS_oPEAK

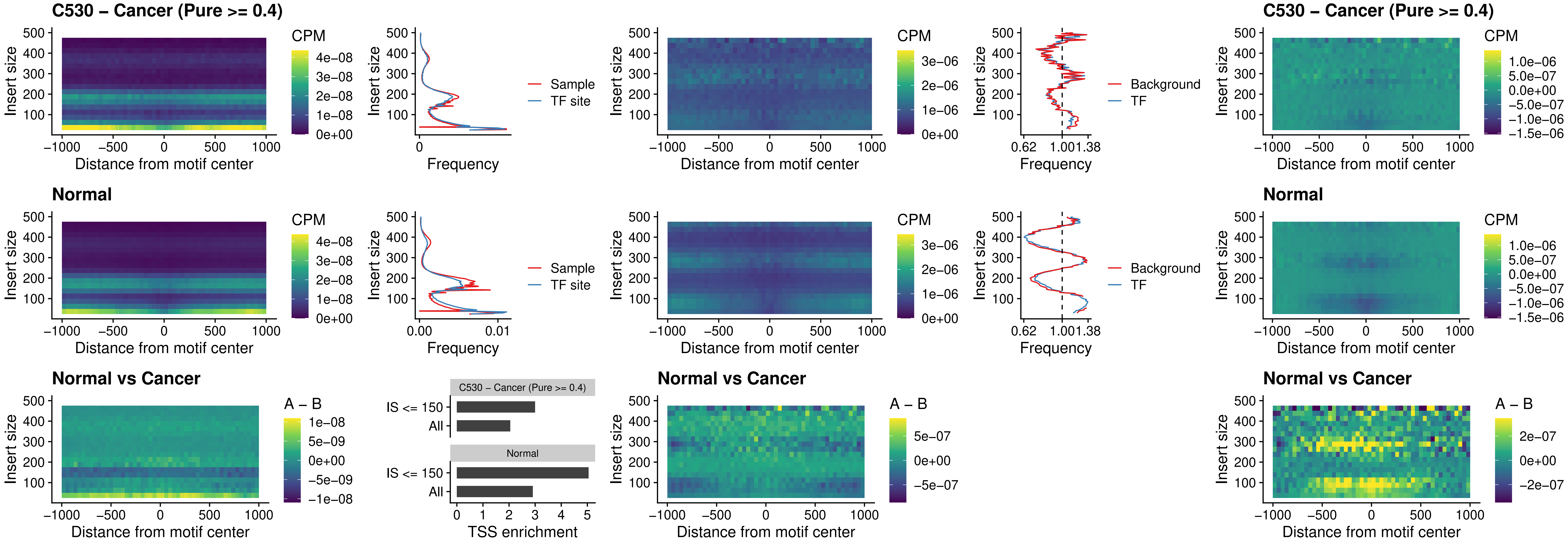

#### C530 – Cancer (Pure >= 0.4) – CPEB1_uc_cTSS_oPEAK

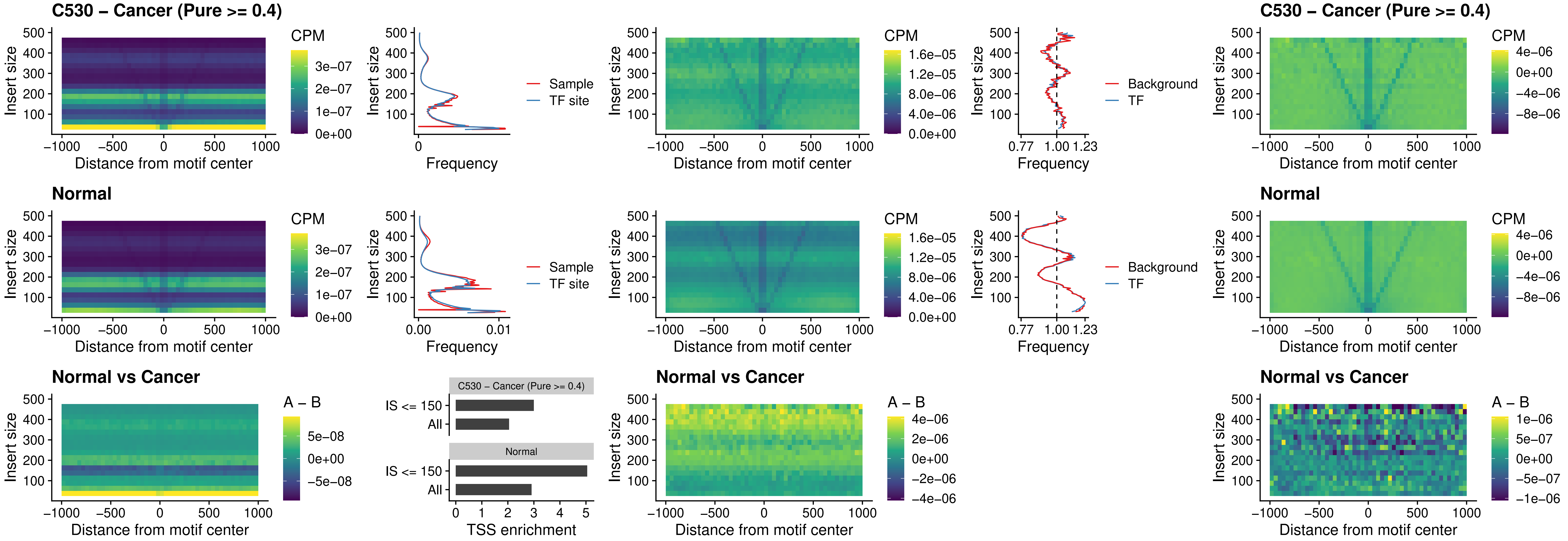

**C530 – Cancer (Pure >= 0.4) – HMGA1_uc_pTSS_nPEAK**

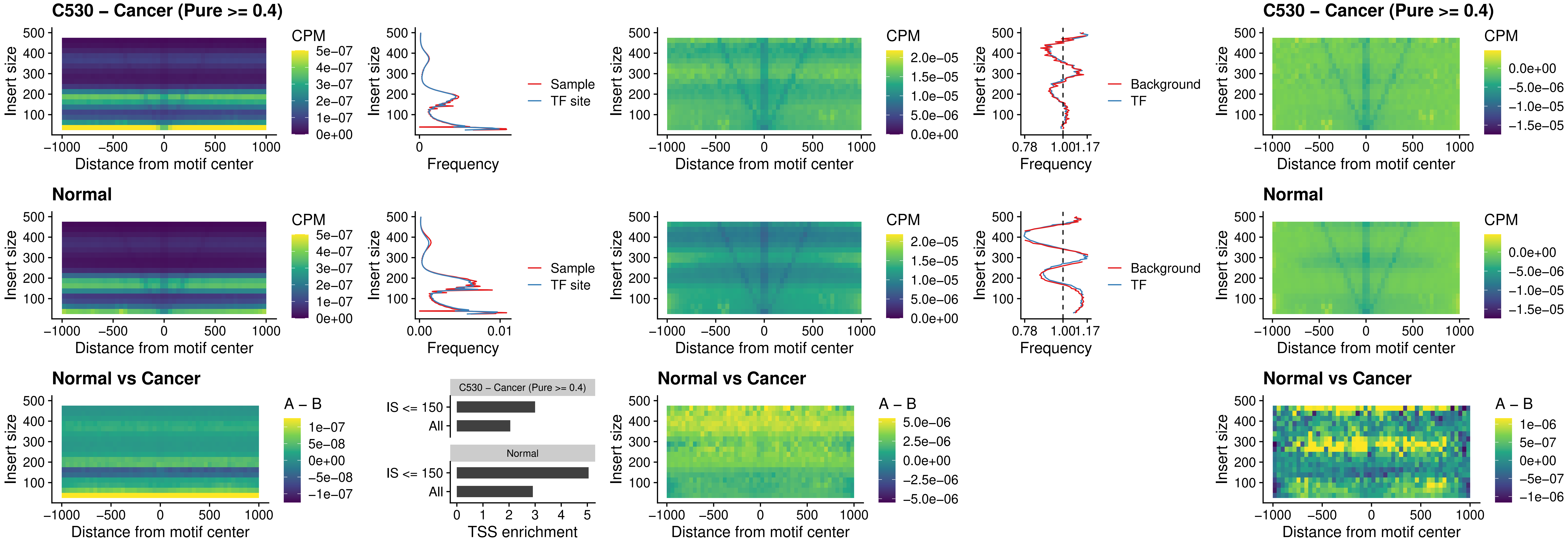

### C530 – Cancer (Pure >= 0.4) – SOX13_uc_cTSS_nPEAK

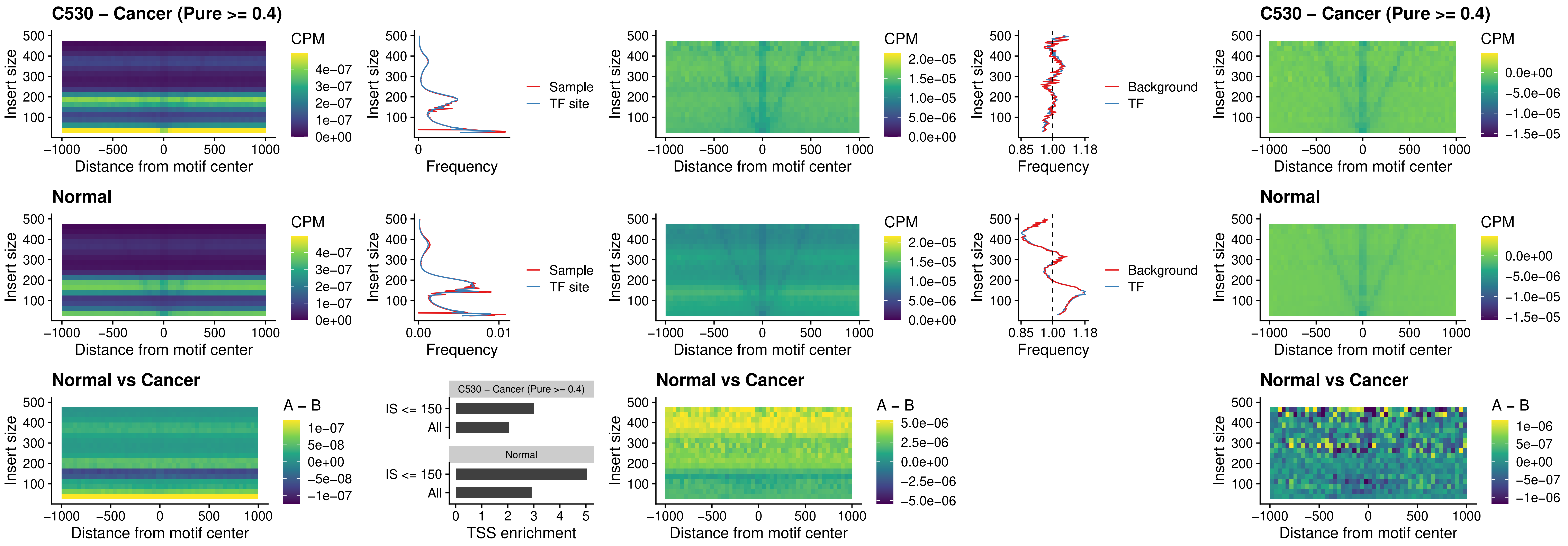

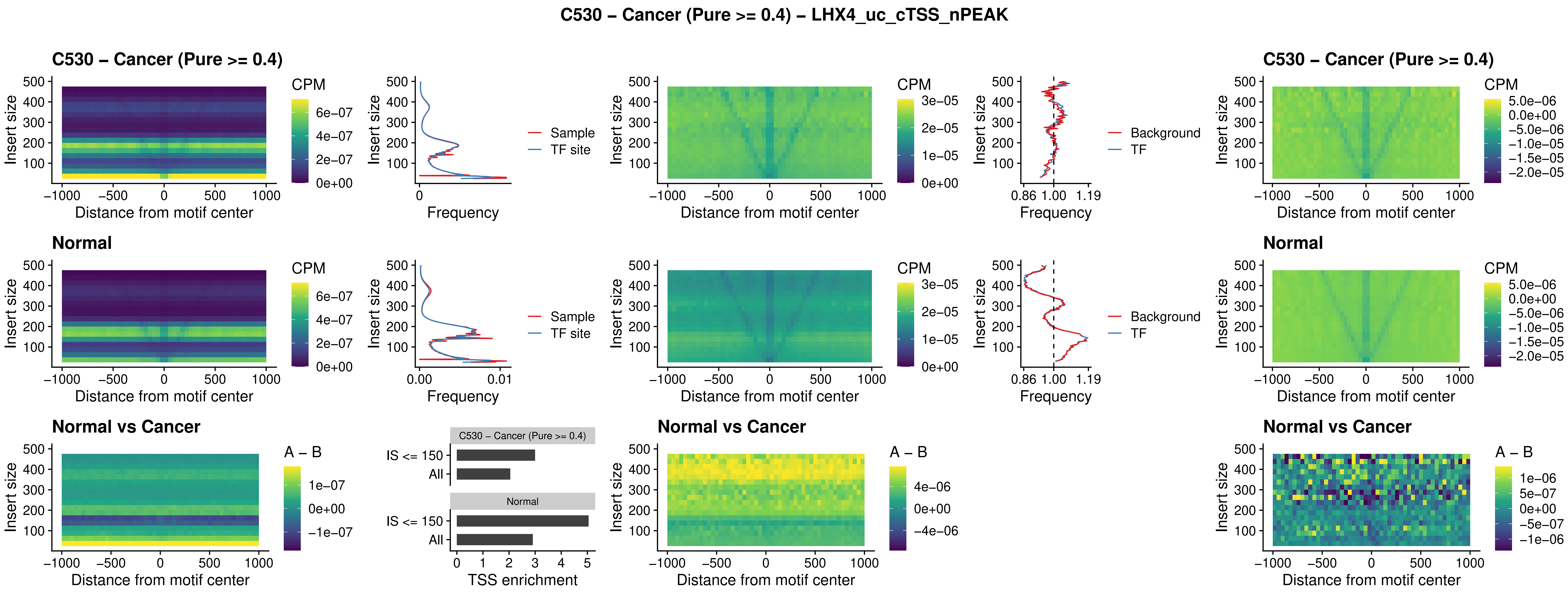
C530 – Cancer (Pure >= 0.4) – LHX4_uc_cTSS_nPEAK
C530 – Cancer (Pure >= 0.4)
Normal
Normal vs Cancer
Insert size
Distance from motif center
Frequency
CPM
A – B
Sample
TF site
Background
TF
C530 – Cancer (Pure >= 0.4)
Normal
IS <= 150
All
TSS enrichment

#### C530 – Cancer (Pure >= 0.4) – HOXA4_uc_cTSS_nPEAK

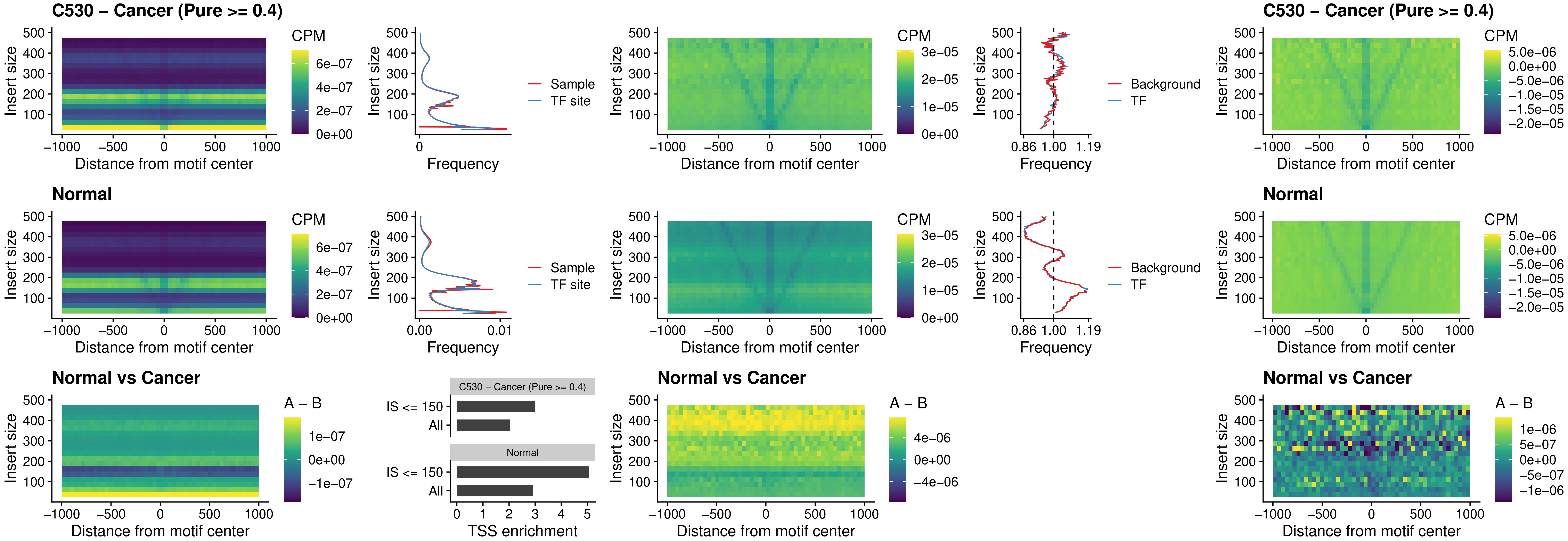

#### C530 – Cancer (Pure >= 0.4) – HLX_uc_cTSS_nPEAK

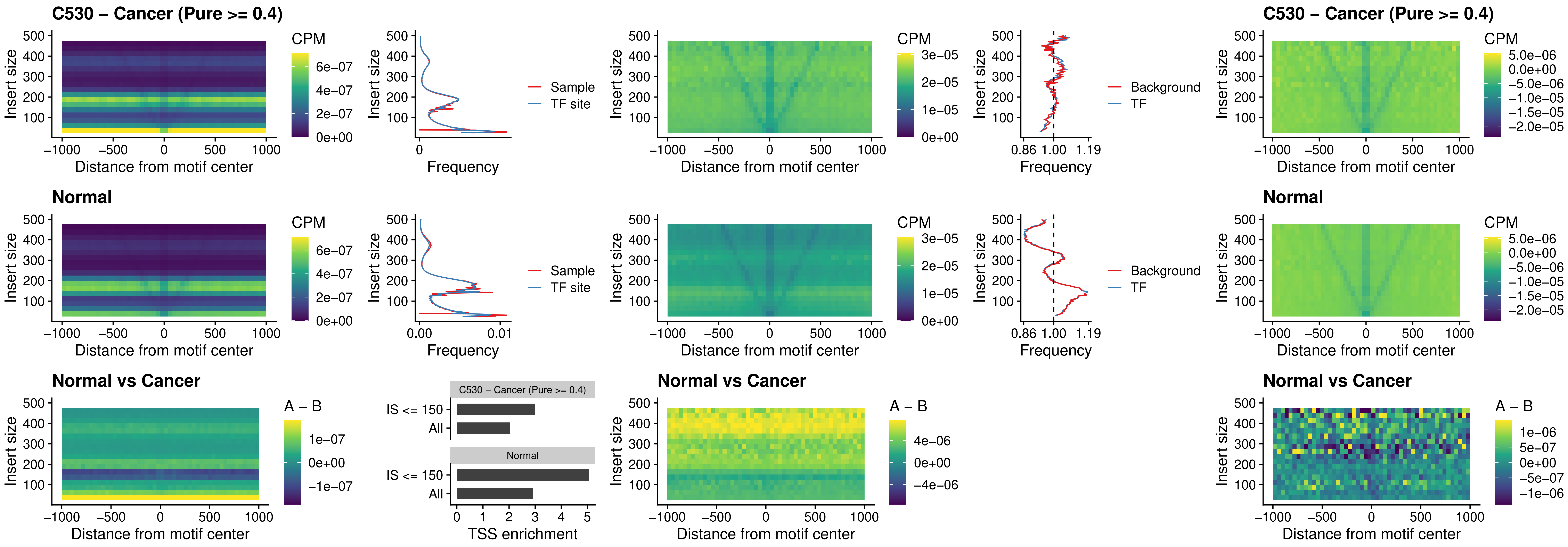

#### C530 – Cancer (Pure >= 0.4) – AC0125311_uc_cTSS_nPEAK

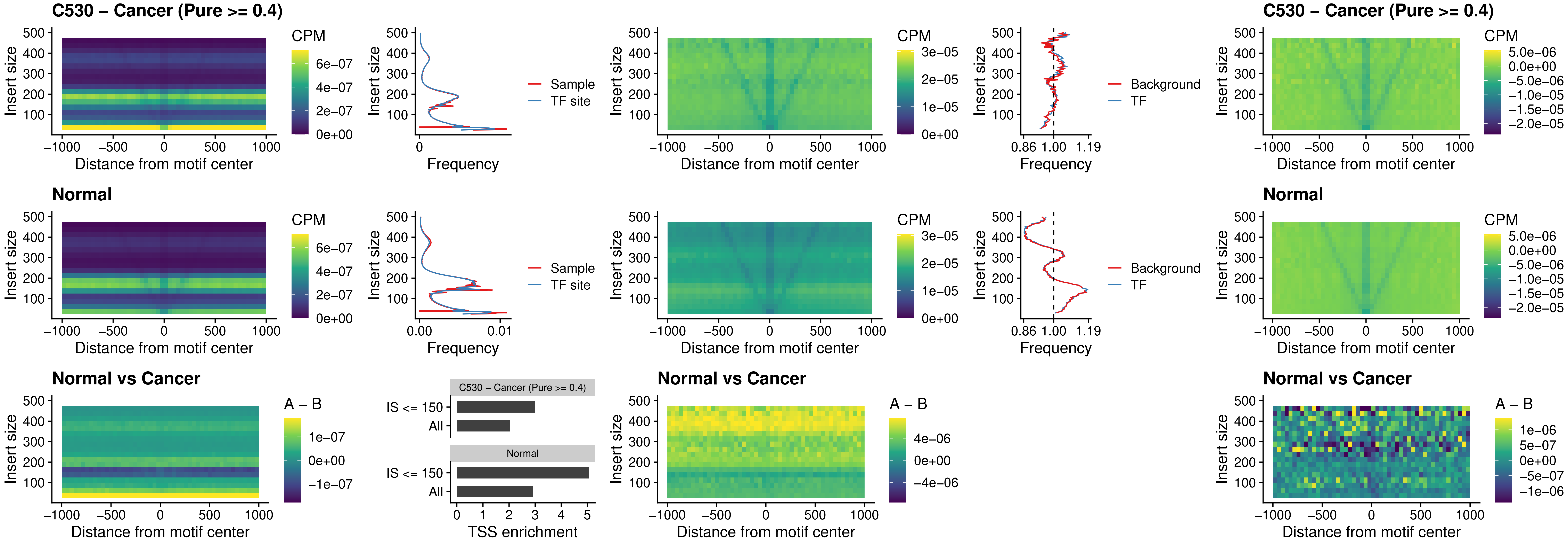

#### C530 – Cancer (Pure >= 0.4) – DBX2_uc_cTSS_nPEAK

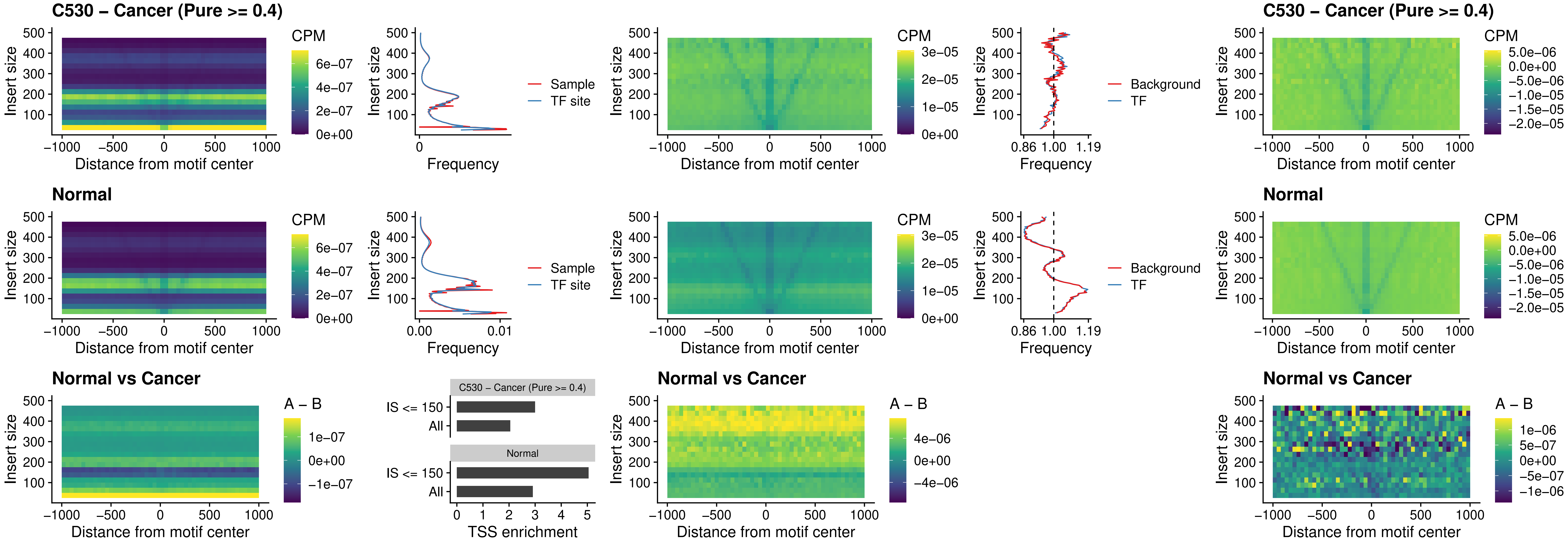

#### C530 – Cancer (Pure >= 0.4) – LHX6_uc_dTSS_nPEAK

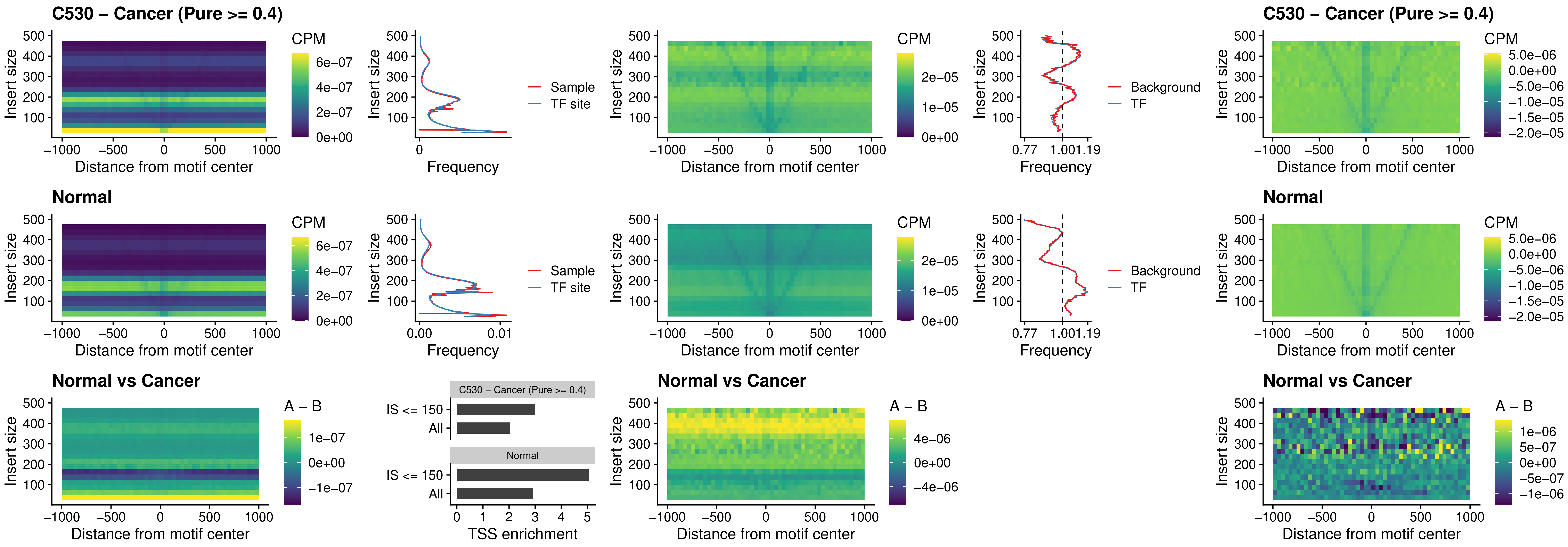

#### C530 – Cancer (Pure >= 0.4) – SOX1_uc_dTSS_nPEAK

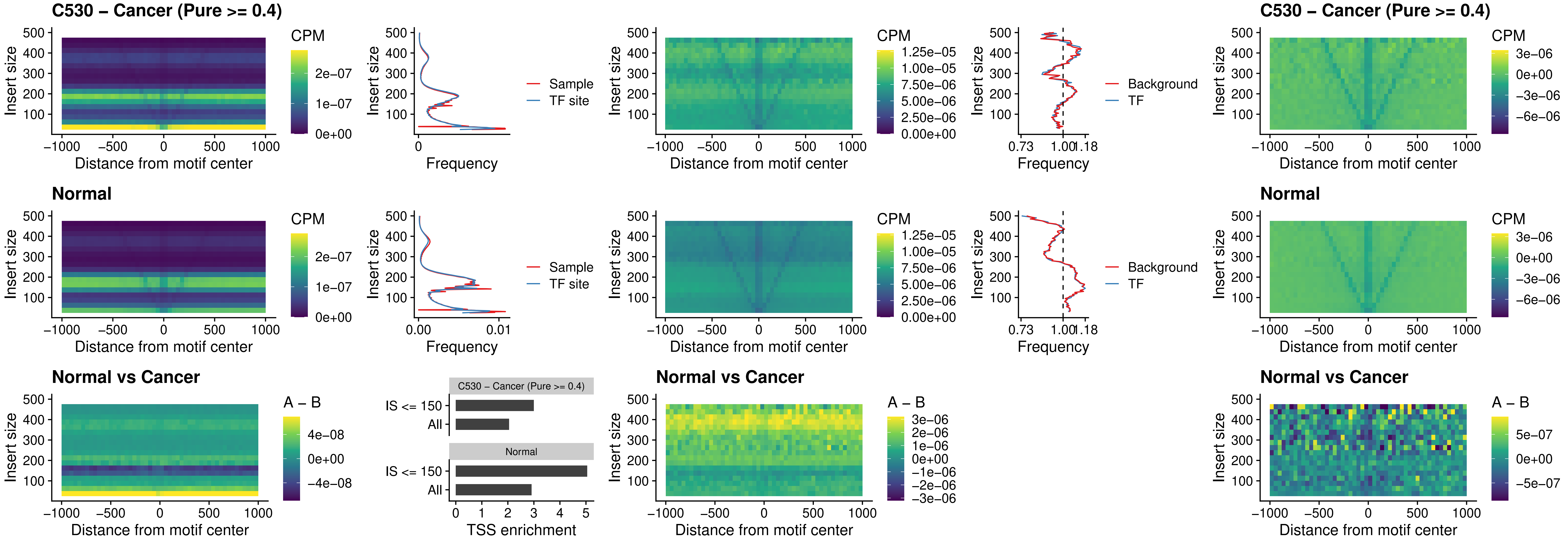

### C530 – Cancer (Pure >= 0.4) – NKX24_uc_dTSS_nPEAK

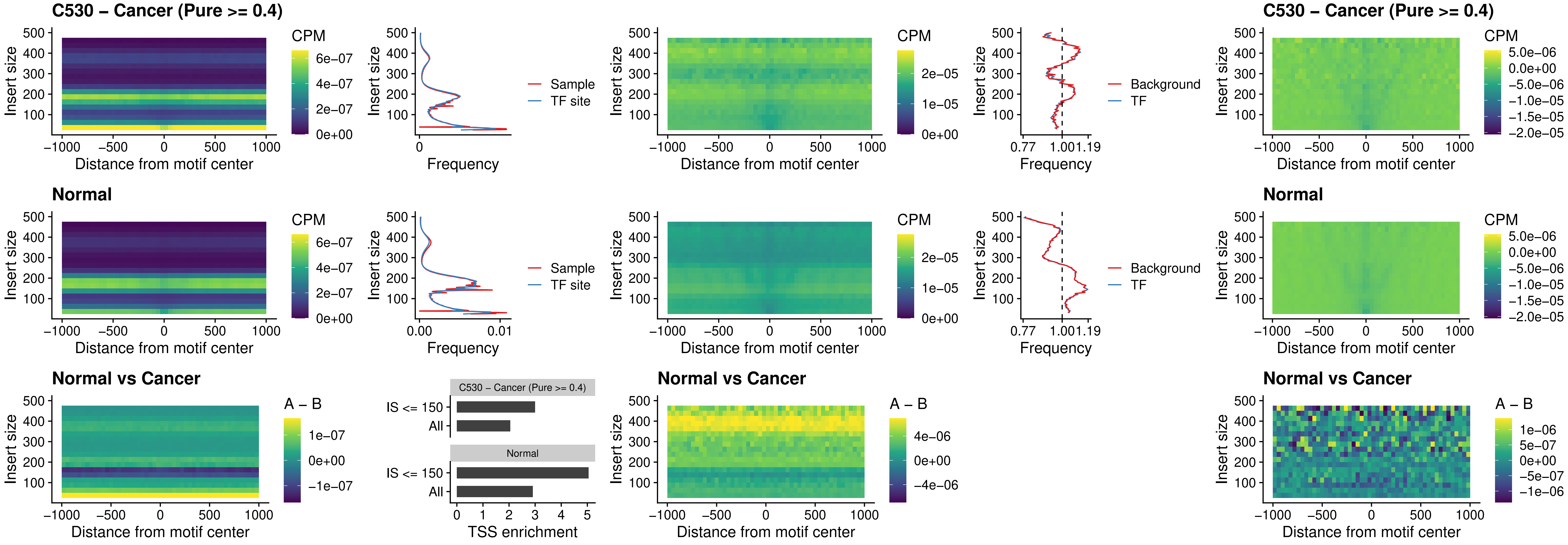

#### C530 – Cancer (Pure >= 0.4) – DLX3_uc_cTSS_nPEAK

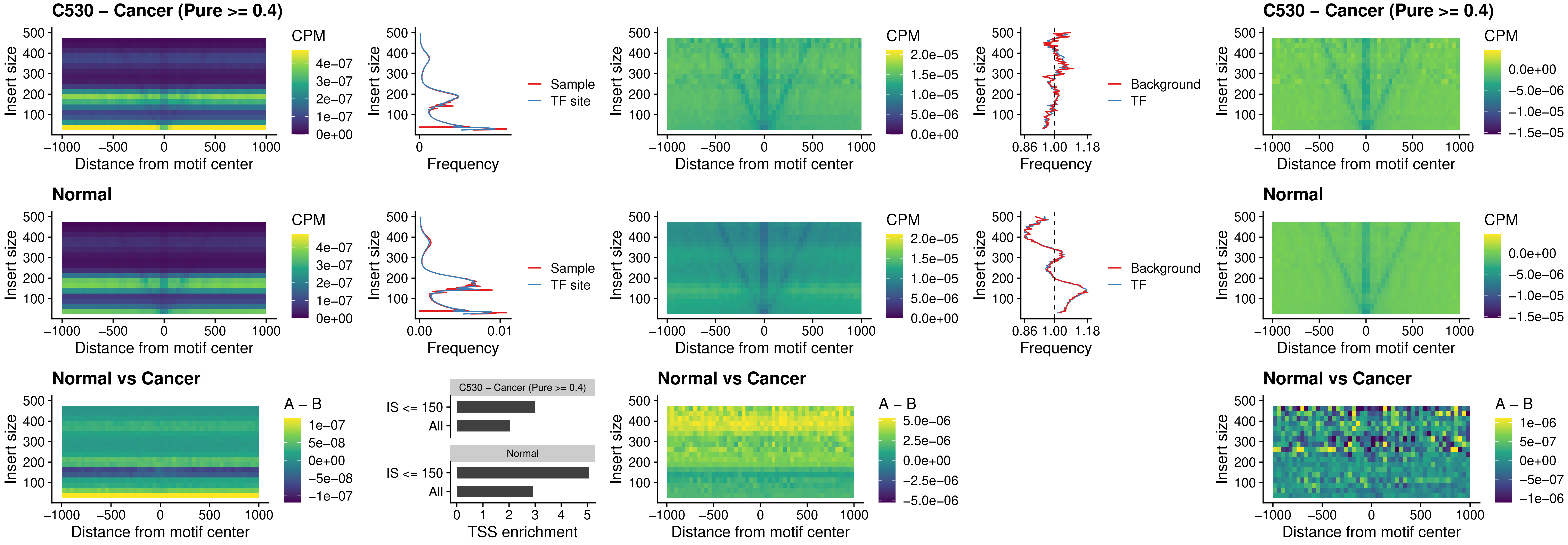

#### C530 – Cancer (Pure >= 0.4) – HOXD3_uc_cTSS_nPEAK

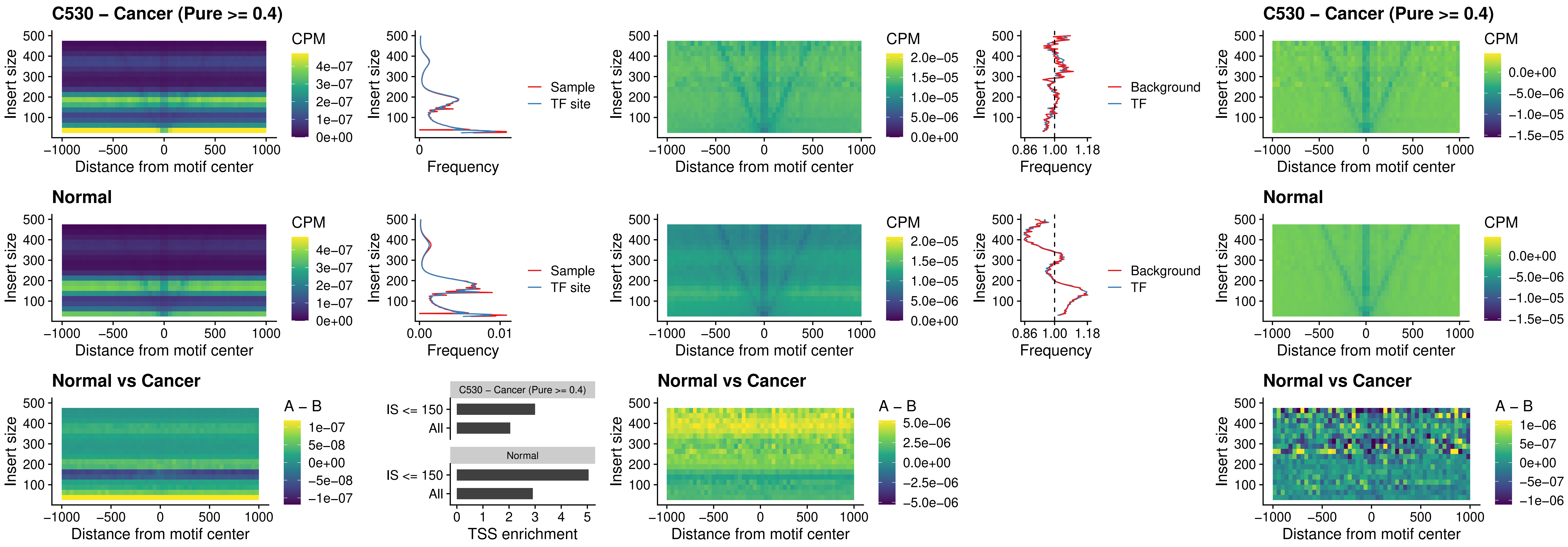

### C530 – Cancer (Pure >= 0.4) – ISL1_uc_dTSS_oPEAK

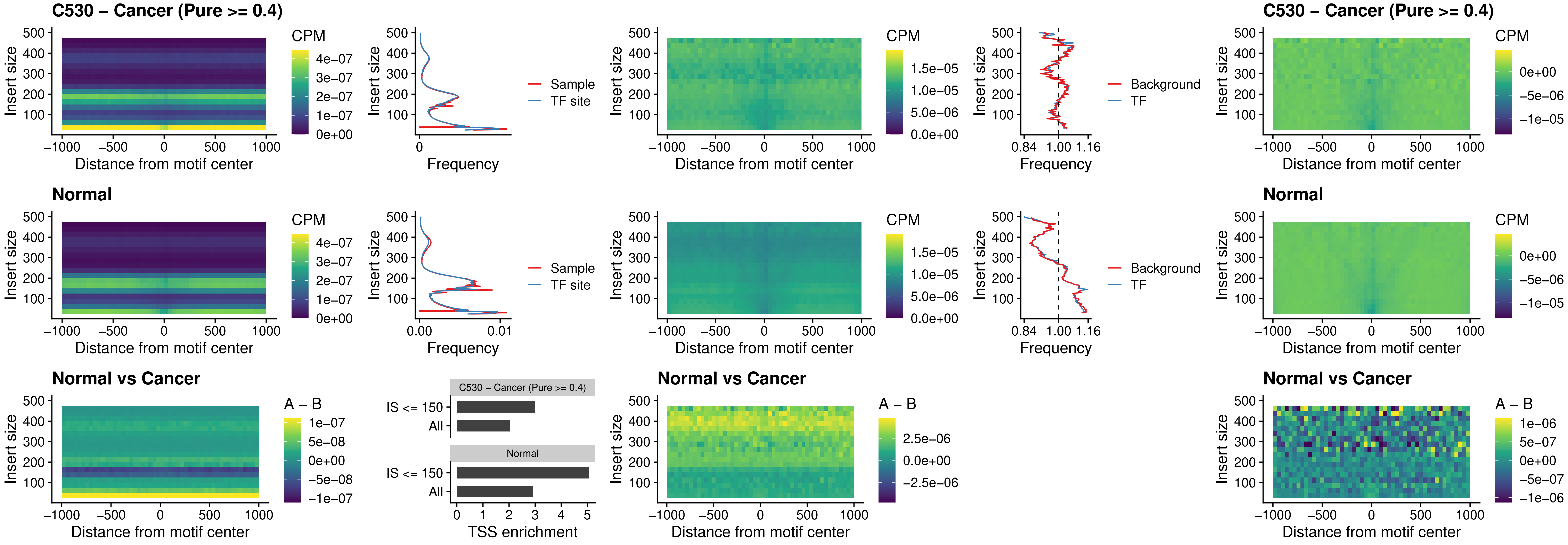

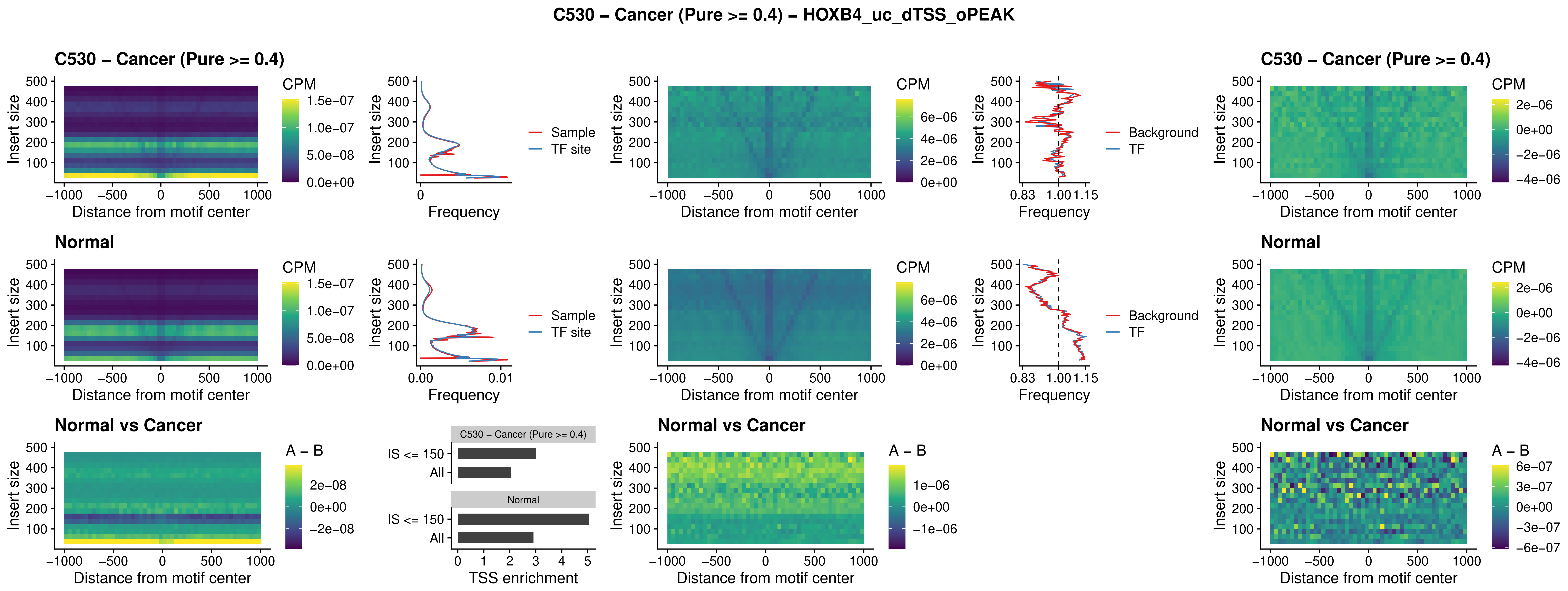
C530 – Cancer (Pure >= 0.4) – HOXB4_uc_dTSS_oPEAK
C530 – Cancer (Pure >= 0.4)
Normal
Normal vs Cancer
Insert size
Distance from motif center
Frequency
CPM
A – B
Sample
TF site
Background
TF
C530 – Cancer (Pure >= 0.4)
Normal
IS <= 150
All
TSS enrichment

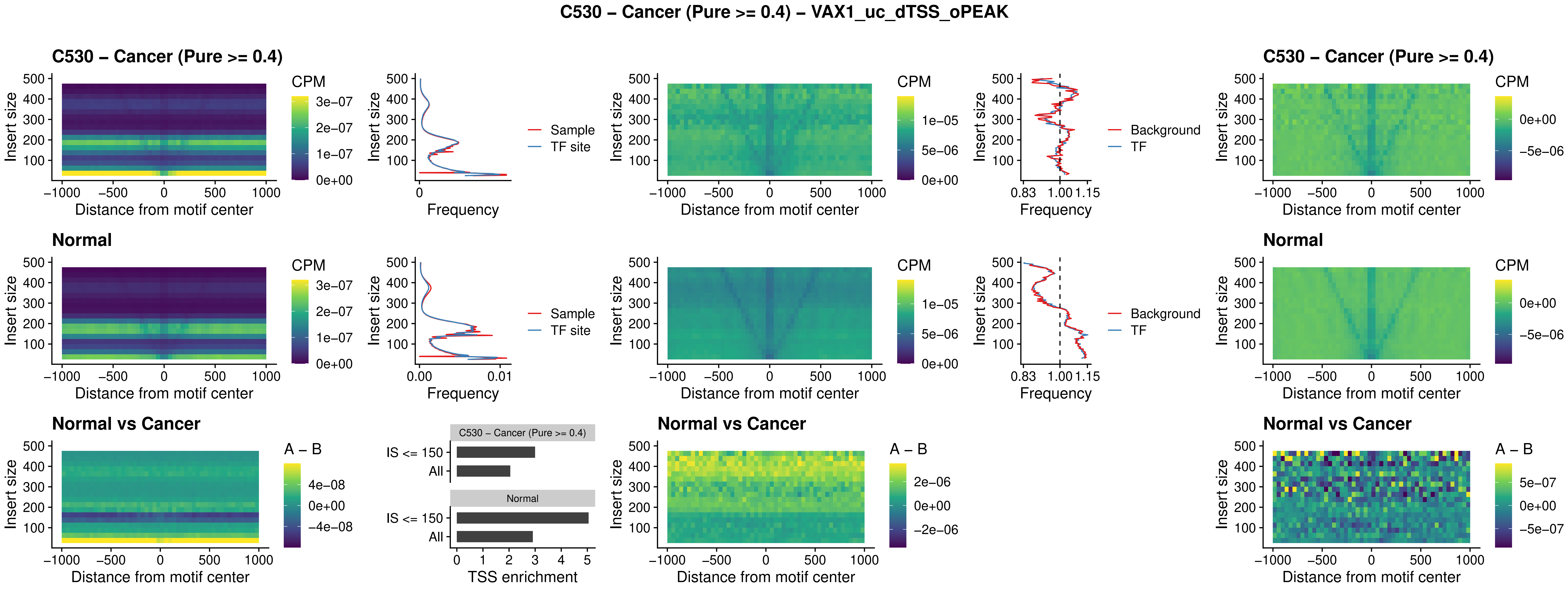
C530 – Cancer (Pure >= 0.4) – VAX1_uc_dTSS_oPEAK
C530 – Cancer (Pure >= 0.4)
Normal
Normal vs Cancer
Insert size
Distance from motif center
Frequency
CPM
A – B
Sample
TF site
Background
TF
C530 – Cancer (Pure >= 0.4)
Normal
IS <= 150
All
TSS enrichment

#### C530 – Cancer (Pure >= 0.4) – DLX3_uc_dTSS_oPEAK

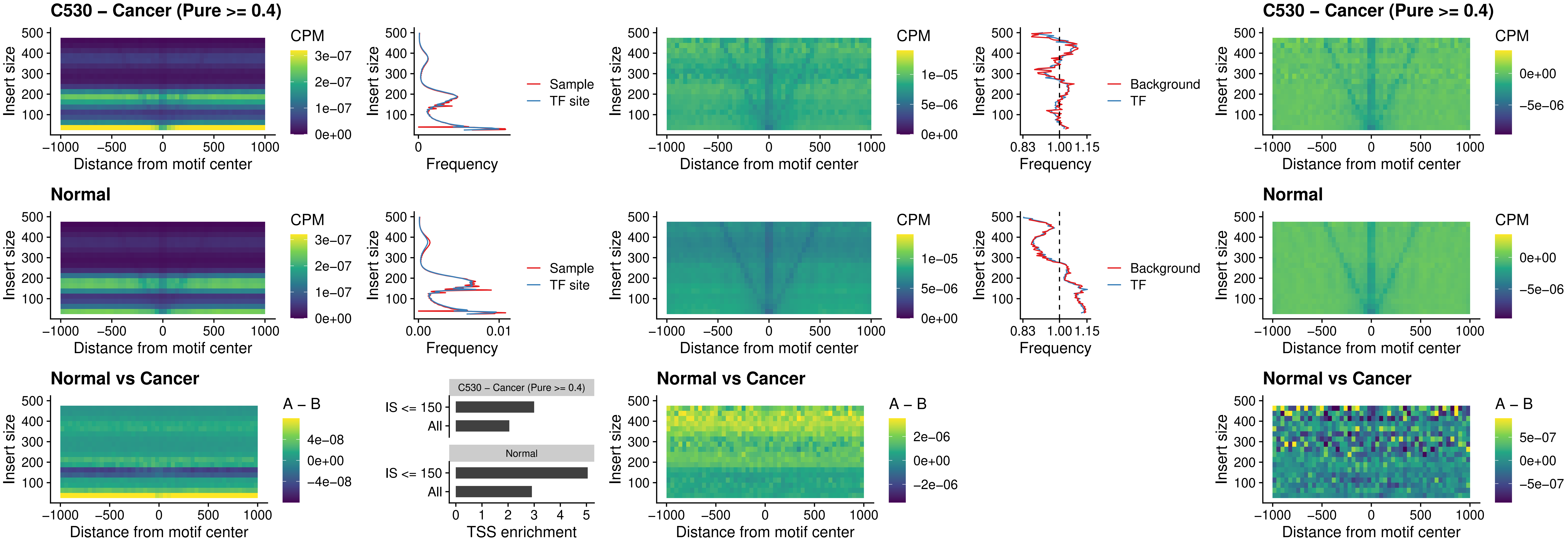

#### C530 – Cancer (Pure >= 0.4) – HOXD3_uc_dTSS_oPEAK

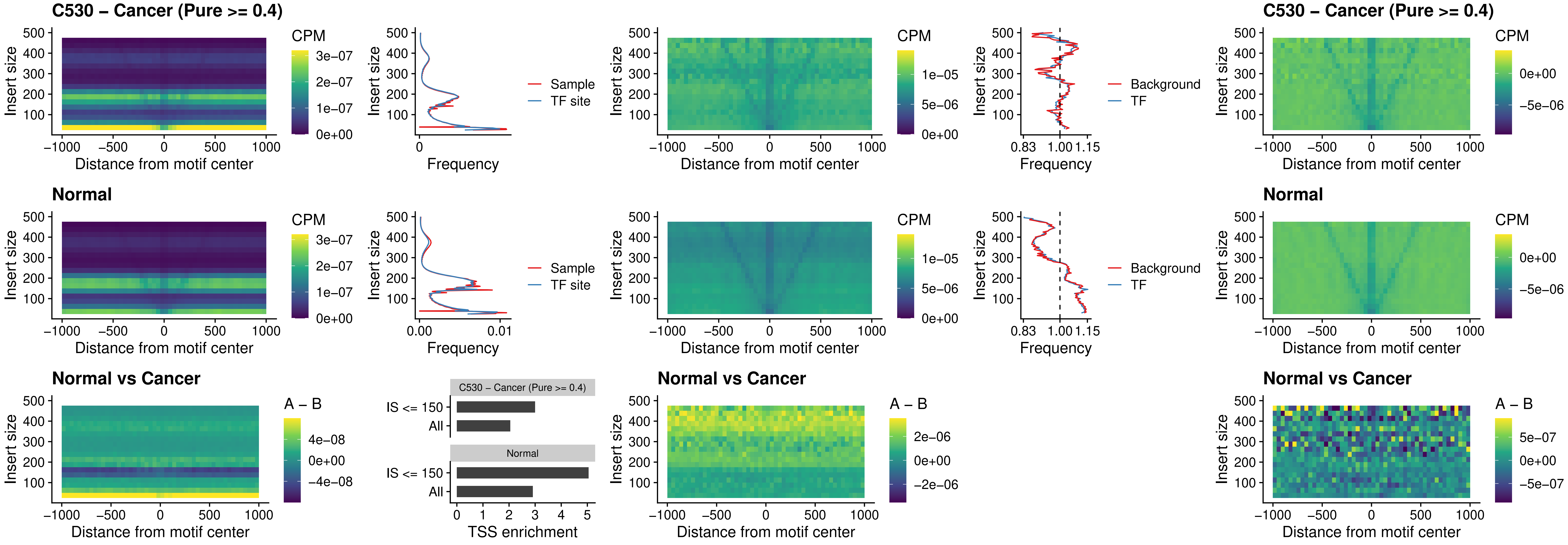

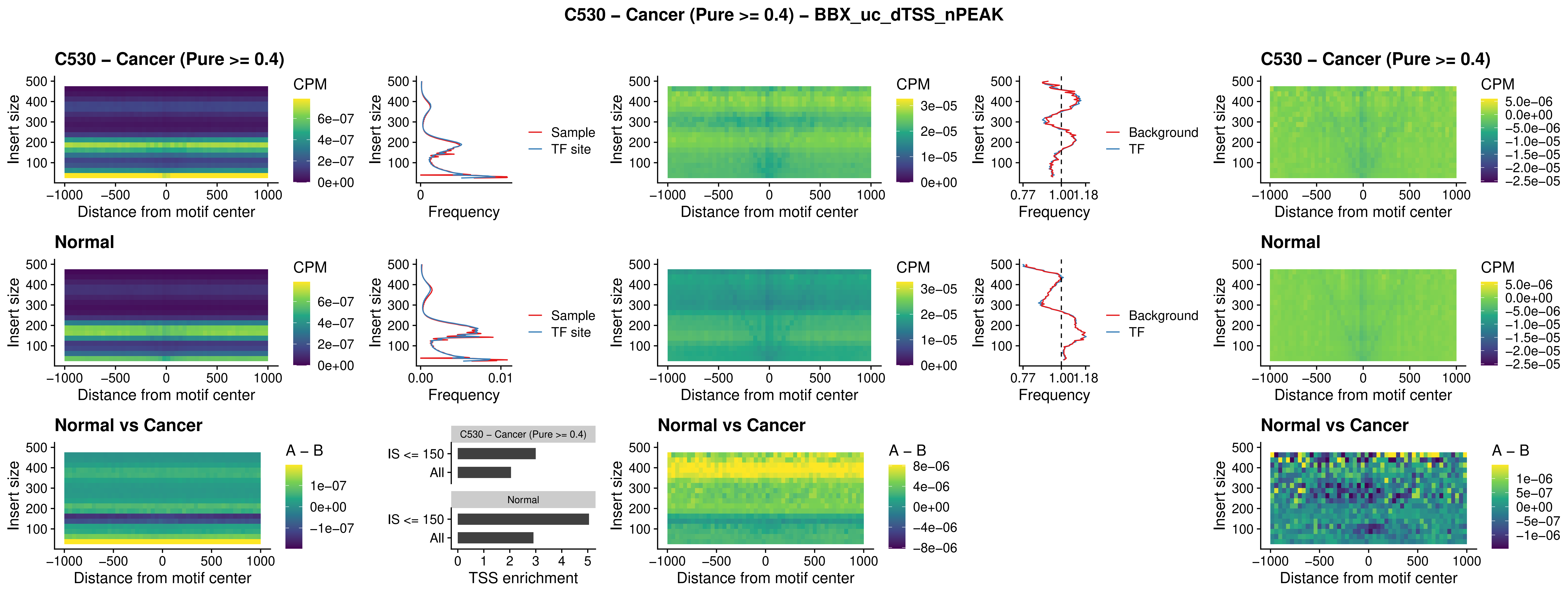
C530 – Cancer (Pure >= 0.4) – BBX_uc_dTSS_nPEAK
C530 – Cancer (Pure >= 0.4)
Normal
Normal vs Cancer
Insert size
Distance from motif center
Frequency
CPM
A – B
Sample
TF site
Background
TF
TSS enrichment
IS <= 150
All
C530 – Cancer (Pure >= 0.4)
Normal

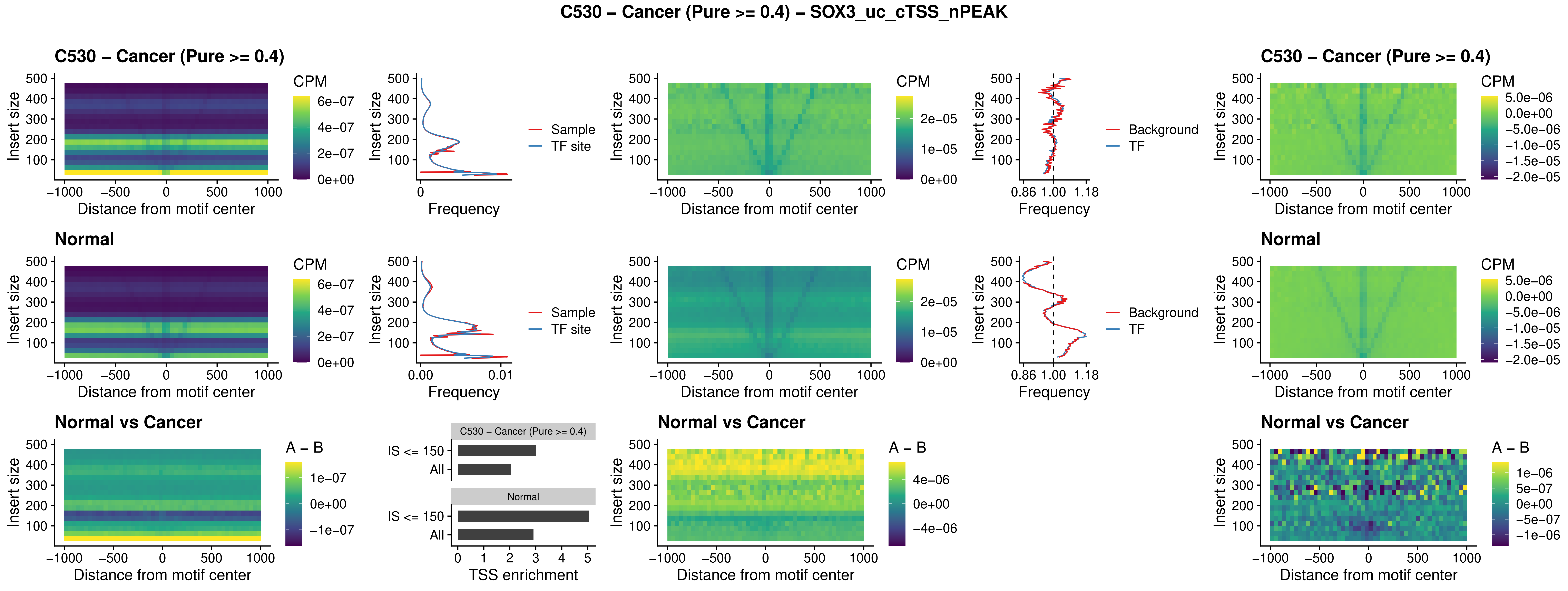
C530 – Cancer (Pure >= 0.4) – SOX3_uc_cTSS_nPEAK
C530 – Cancer (Pure >= 0.4)
Normal
Normal vs Cancer
Insert size
Distance from motif center
CPM
A – B
Frequency
Sample
TF site
Background
TF
C530 – Cancer (Pure >= 0.4)
IS <= 150
All
TSS enrichment

### C530 – Cancer (Pure >= 0.4) – TBPL2_uc_cTSS_nPEAK

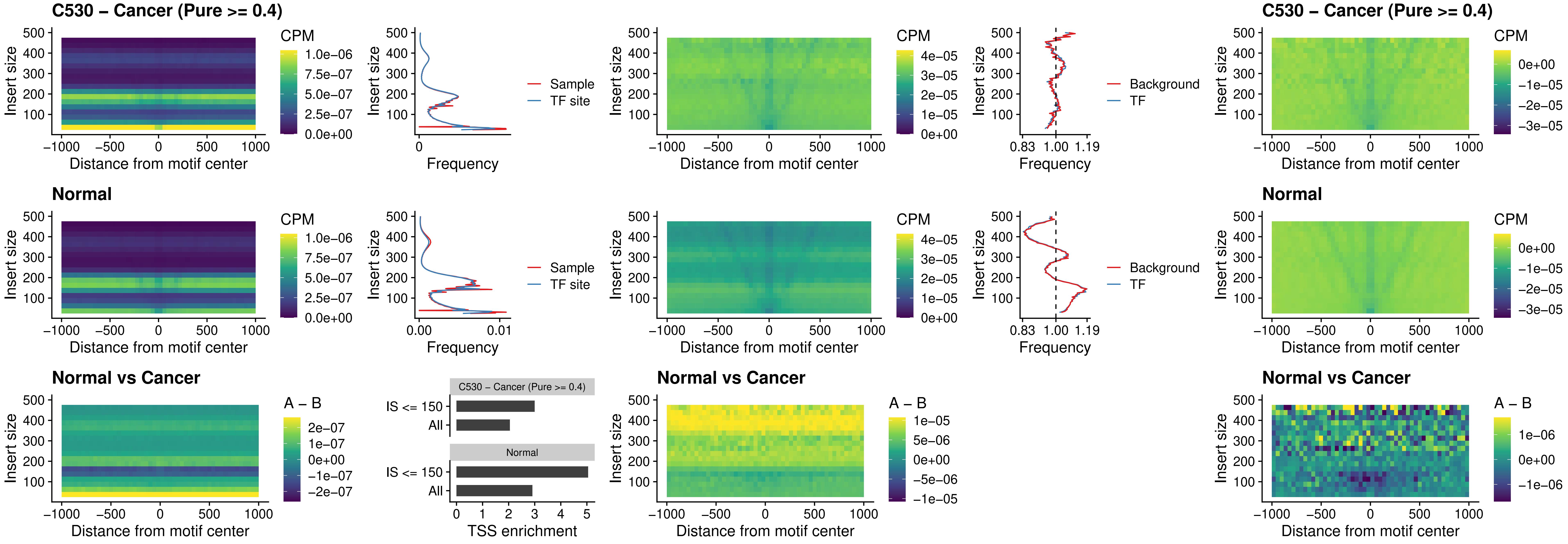

#### C530 – Cancer (Pure >= 0.4) – HOXA4_uc_cTSS_oPEAK

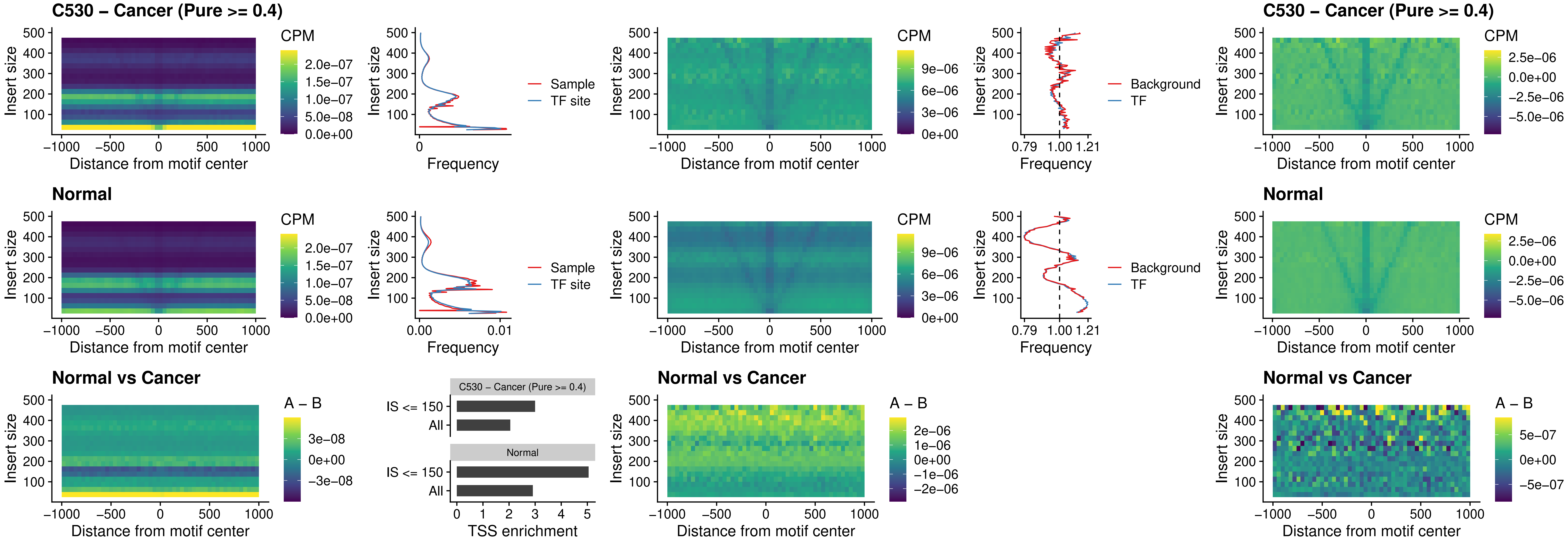

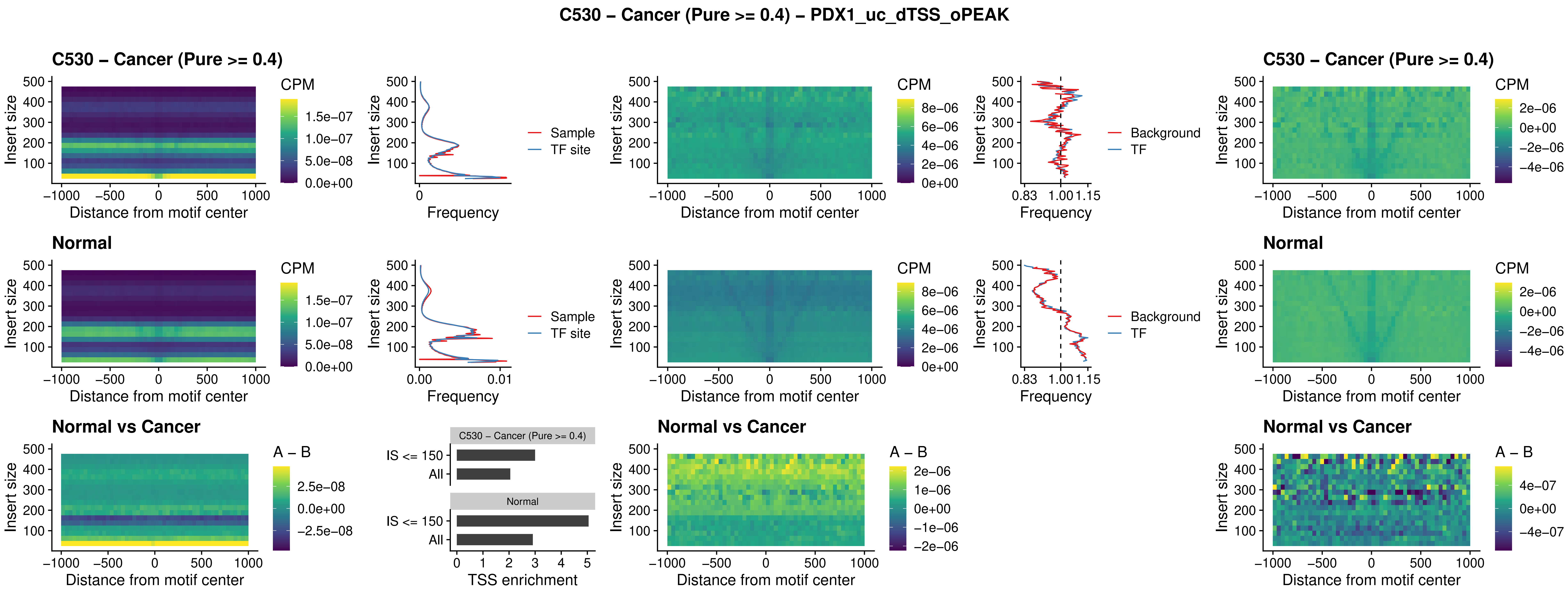

C530 – Cancer (Pure >= 0.4) – PDX1_uc_dTSS_oPEAK
C530 – Cancer (Pure >= 0.4)
Normal
Normal vs Cancer
Insert size
Distance from motif center
Frequency
CPM
A – B
Sample
TF site
Background
TF
C530 – Cancer (Pure >= 0.4)
Normal
IS <= 150
All
TSS enrichment

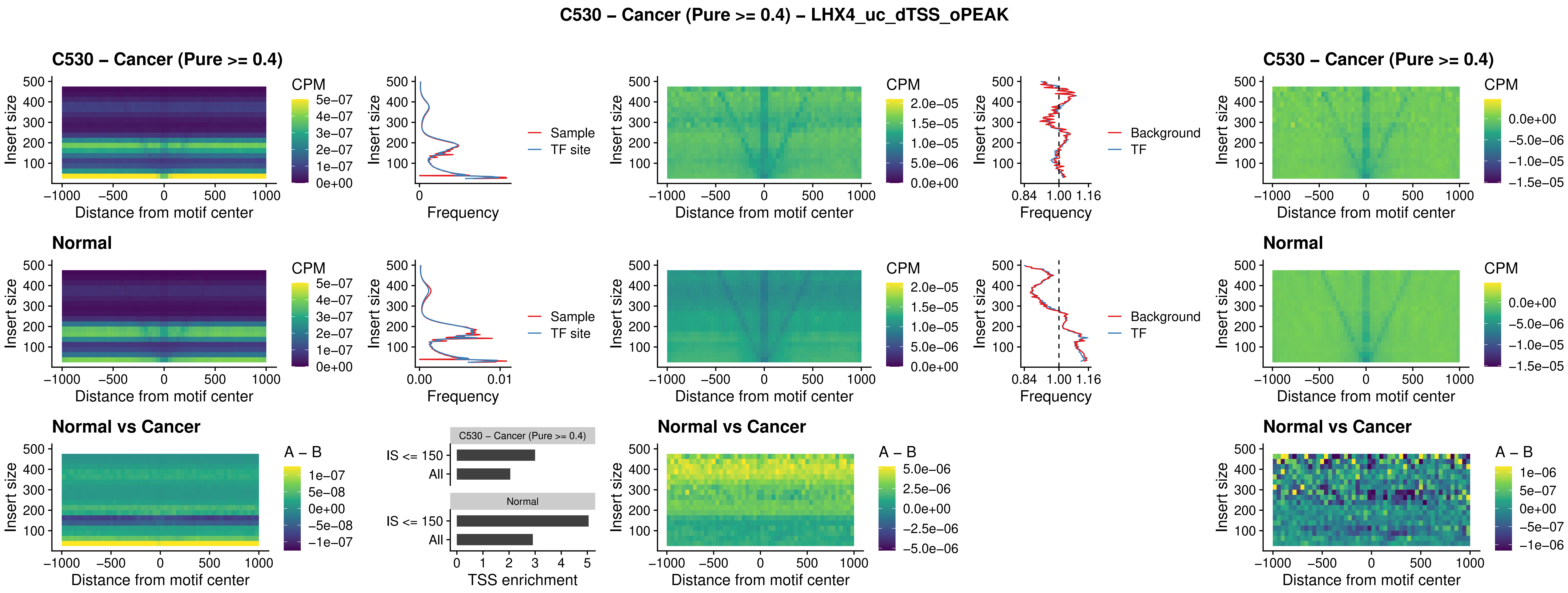
C530 – Cancer (Pure >= 0.4) – LHX4_uc_dTSS_oPEAK
C530 – Cancer (Pure >= 0.4)
Normal
Normal vs Cancer
Insert size
Distance from motif center
Frequency
CPM
A – B
Sample
TF site
Background
TF
C530 – Cancer (Pure >= 0.4)
Normal
IS <= 150
All
TSS enrichment

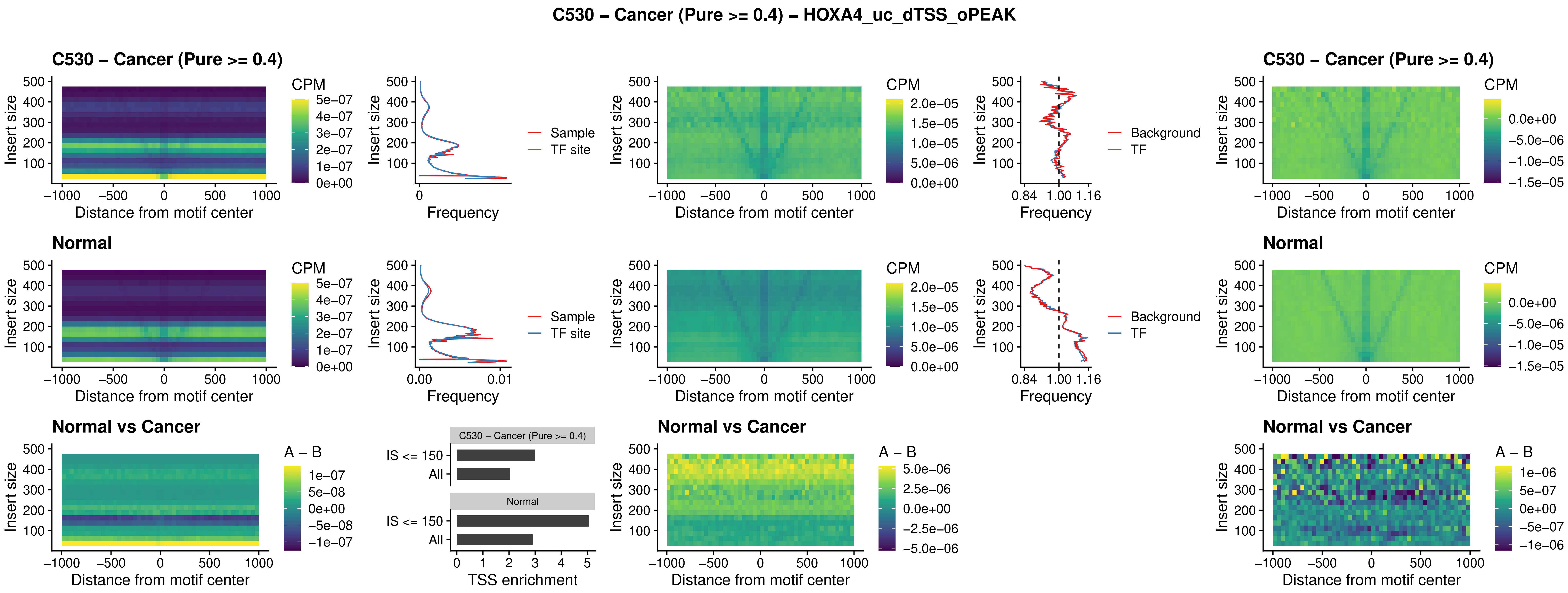
C530 – Cancer (Pure >= 0.4) – HOXA4_uc_dTSS_oPEAK
C530 – Cancer (Pure >= 0.4)
Normal
Normal vs Cancer
Insert size
Distance from motif center
Frequency
CPM
A – B
Sample
TF site
Background
TF
C530 – Cancer (Pure >= 0.4)
Normal
IS <= 150
All
TSS enrichment

### C530 – Cancer (Pure >= 0.4) – HLX_uc_dTSS_oPEAK

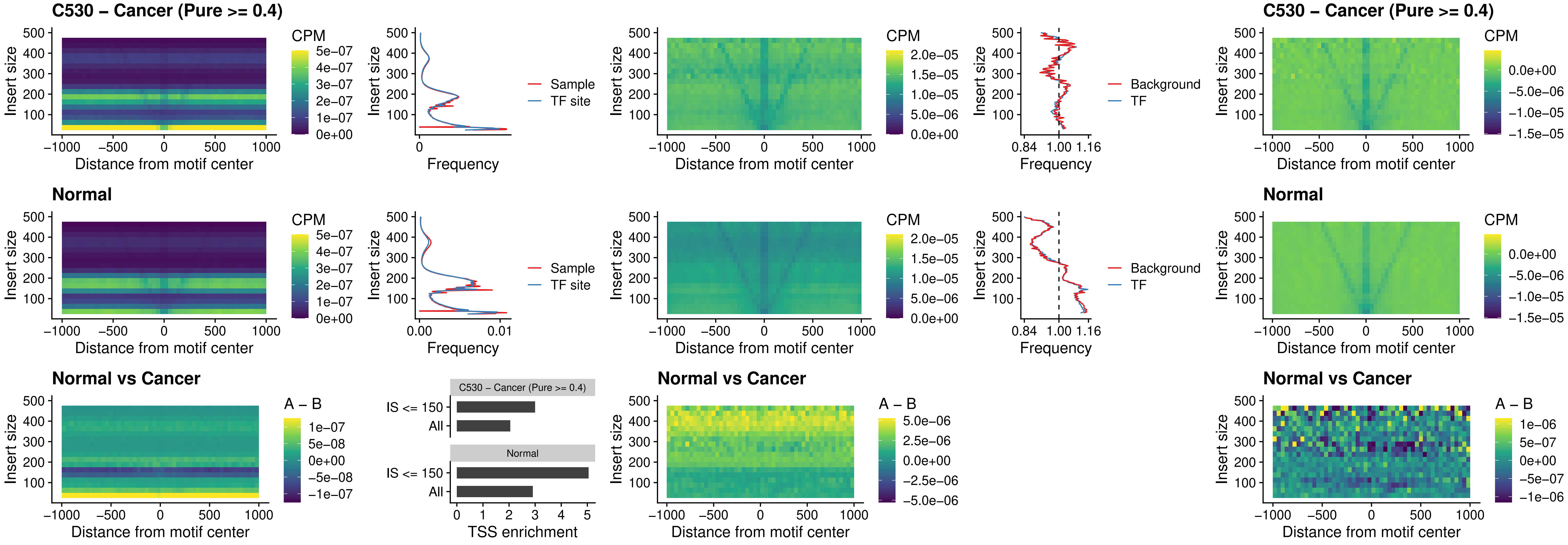

#### C530 – Cancer (Pure >= 0.4) – AC0125311_uc_dTSS_oPEAK

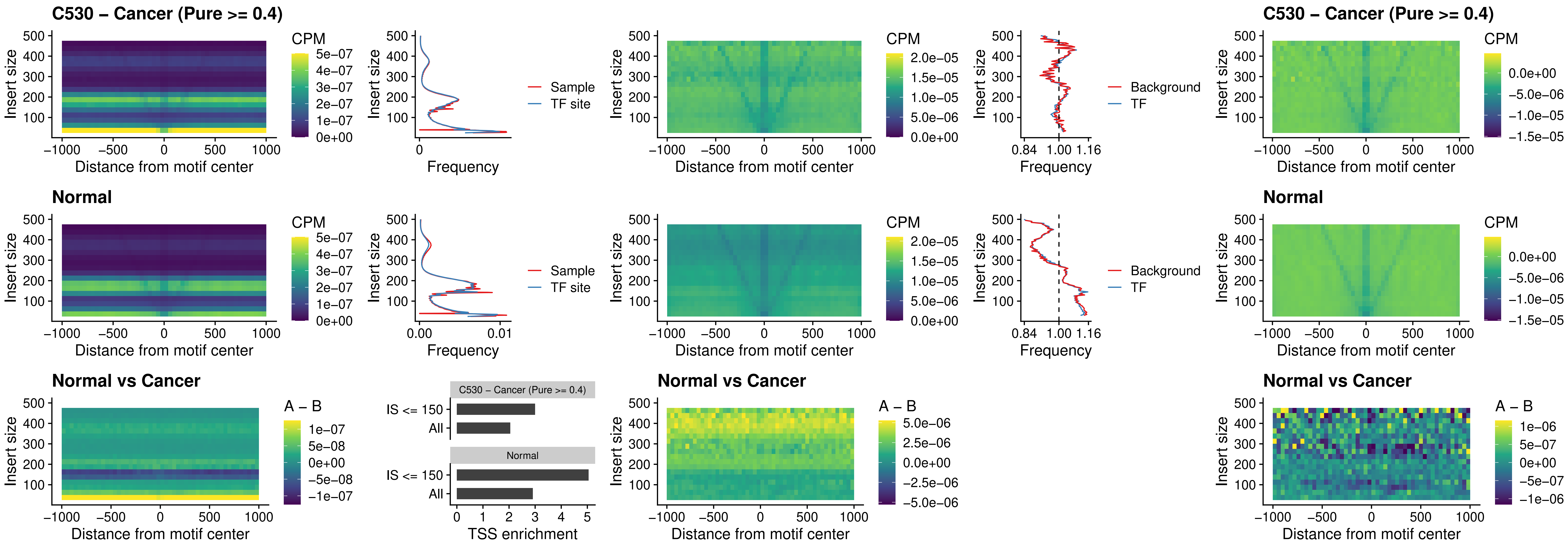

#### C530 – Cancer (Pure >= 0.4) – DBX2_uc_dTSS_oPEAK

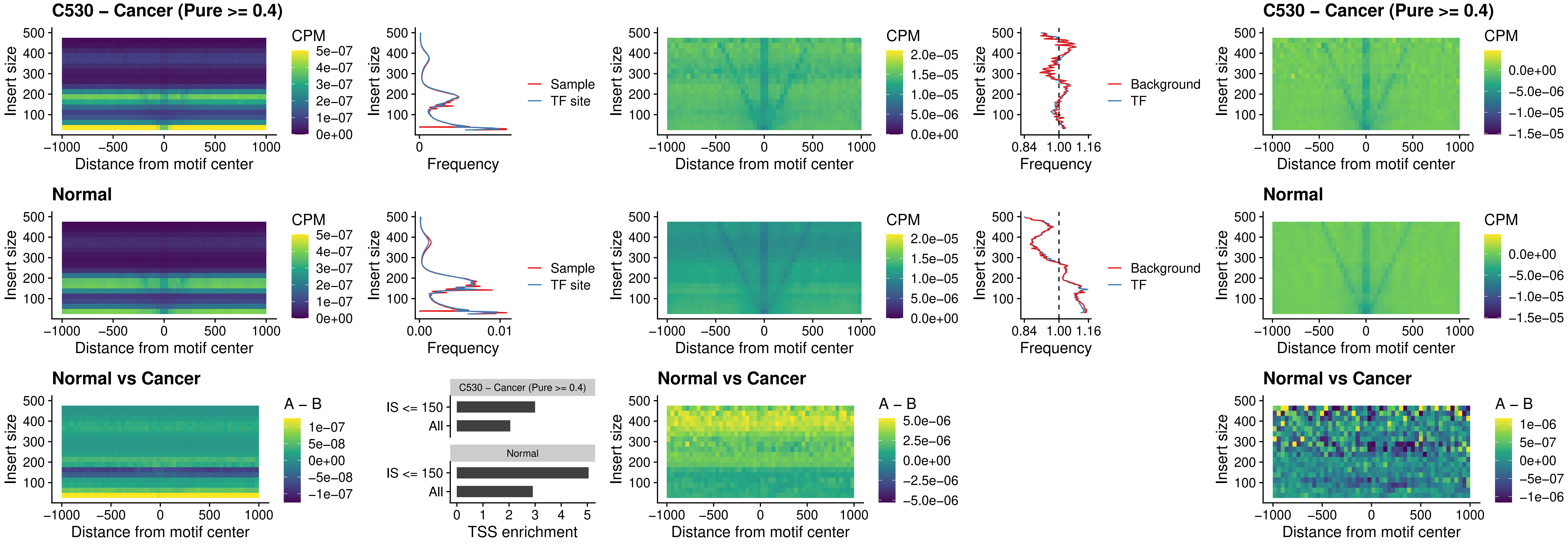

#### C530 – Cancer (Pure >= 0.4) – HOXA2_uc_dTSS_nPEAK

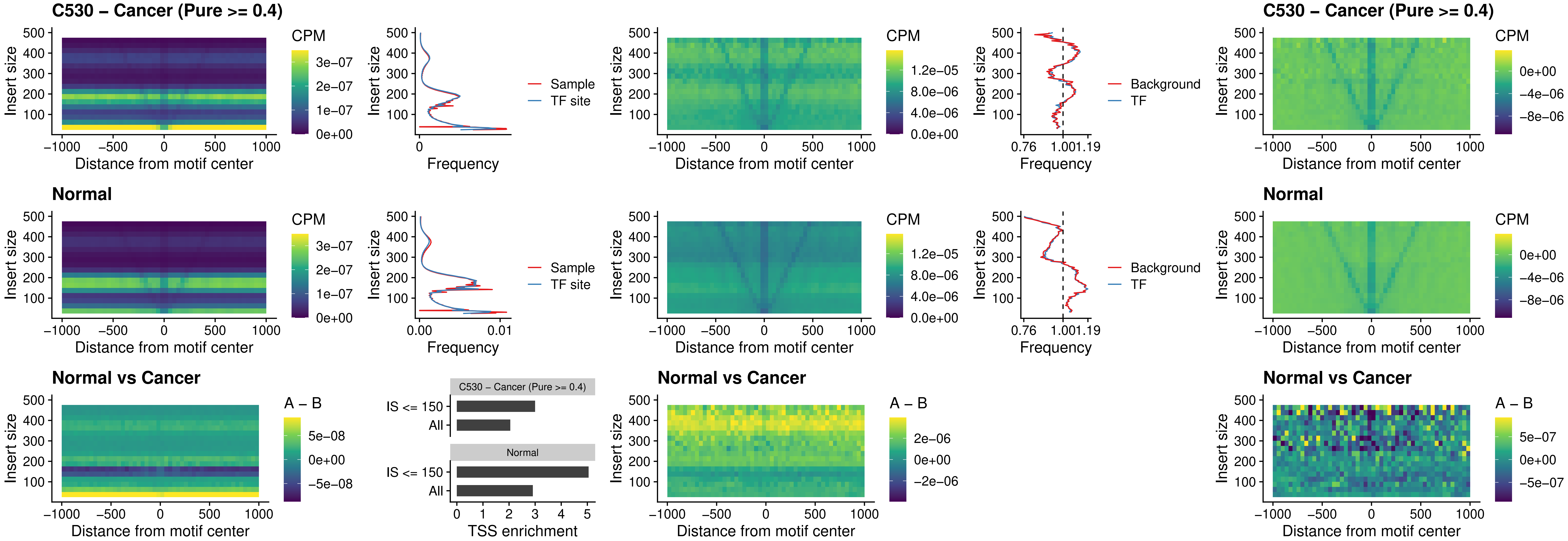

#### C530 – Cancer (Pure >= 0.4) – SHOX_uc_dTSS_nPEAK

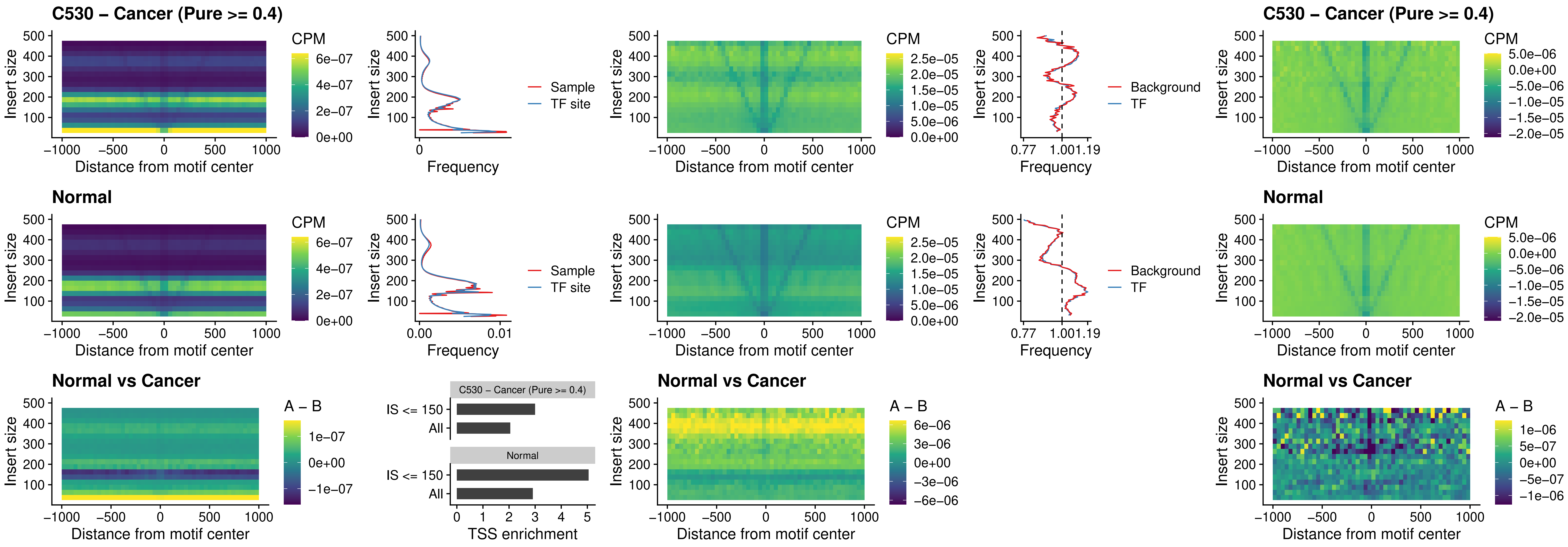

#### C530 – Cancer (Pure >= 0.4) – PRRX1_uc_dTSS_nPEAK

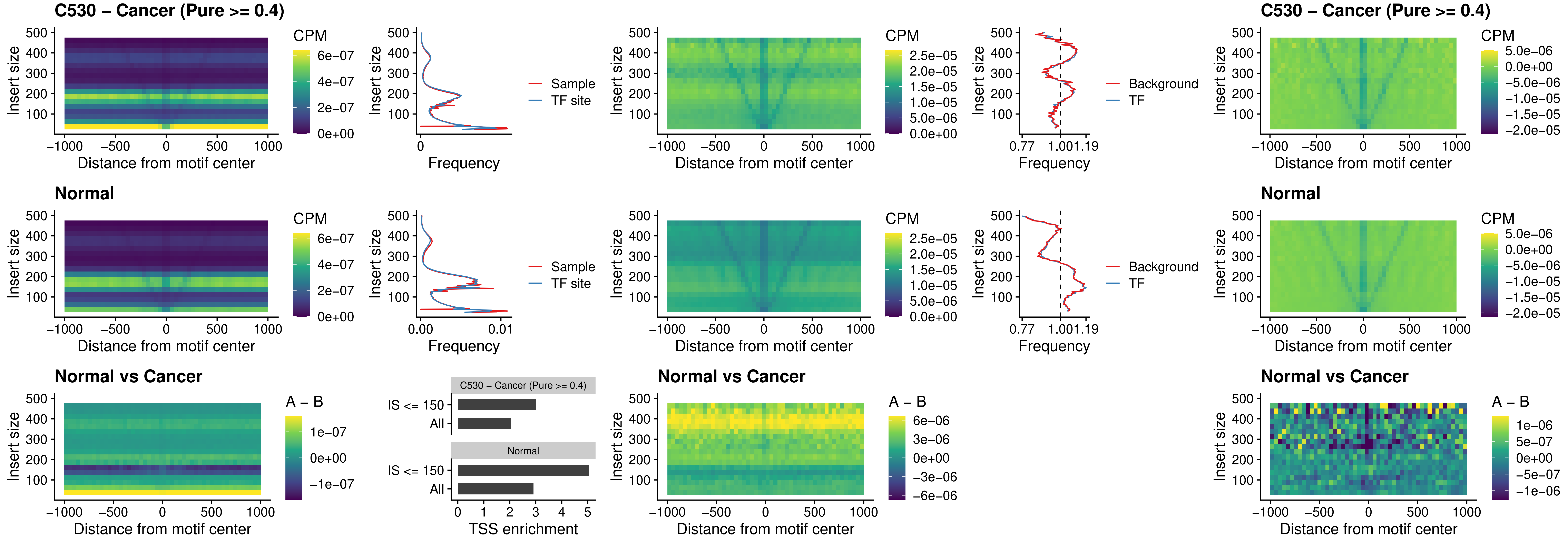

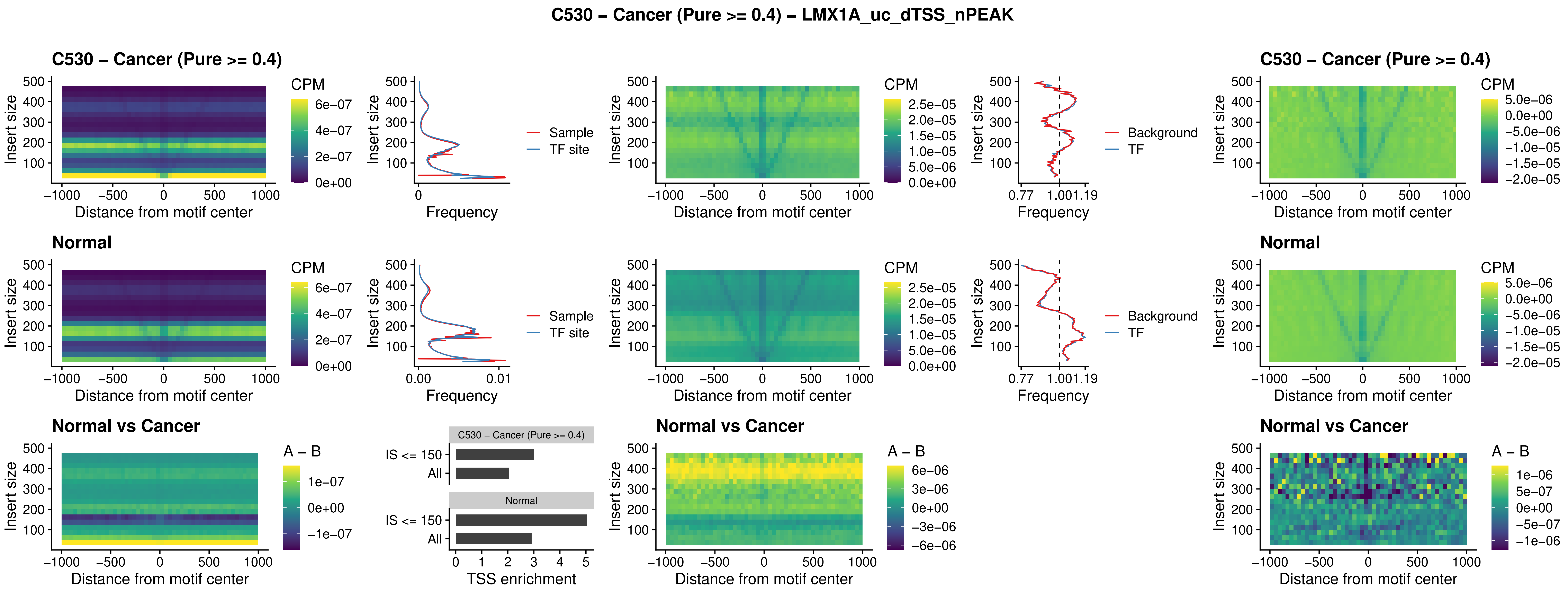
C530 – Cancer (Pure >= 0.4) – LMX1A_uc_dTSS_nPEAK
C530 – Cancer (Pure >= 0.4)
Normal
Normal vs Cancer
Insert size
Distance from motif center
Frequency
CPM
A – B
Sample
TF site
Background
TF
C530 – Cancer (Pure >= 0.4)
Normal
IS <= 150
All
TSS enrichment

#### C530 – Cancer (Pure >= 0.4) – LMX1B_uc_dTSS_nPEAK

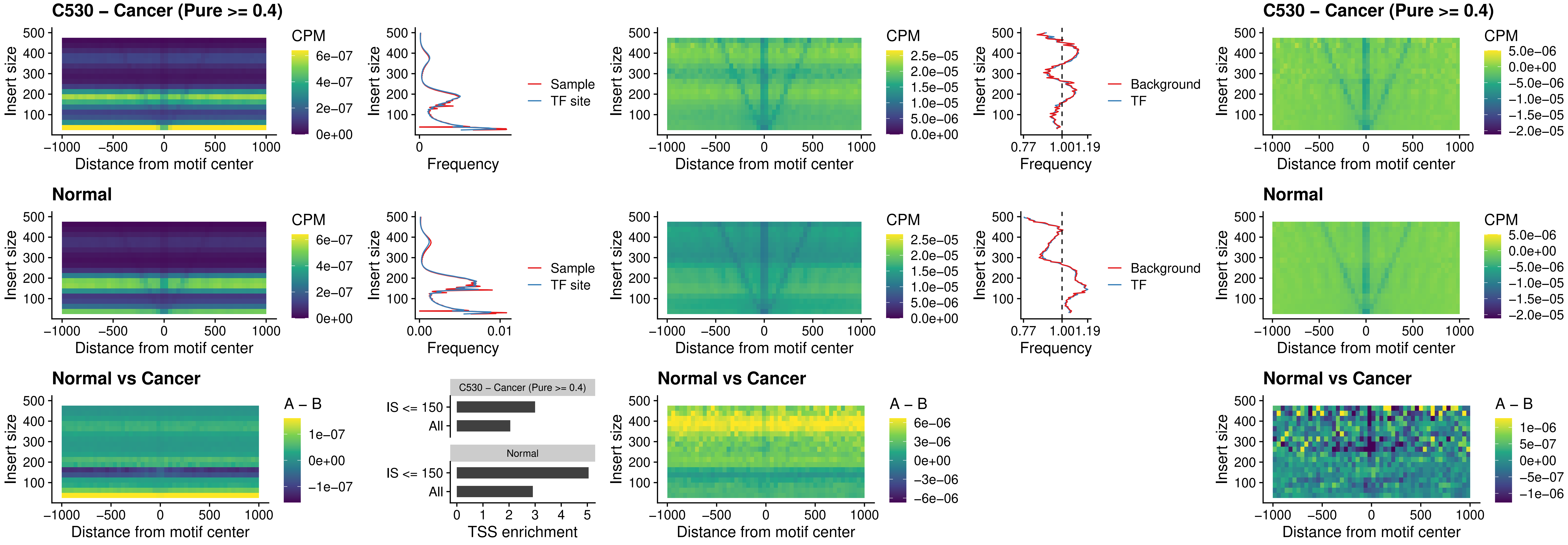

#### C530 – Cancer (Pure >= 0.4) – DLX5_uc_dTSS_oPEAK

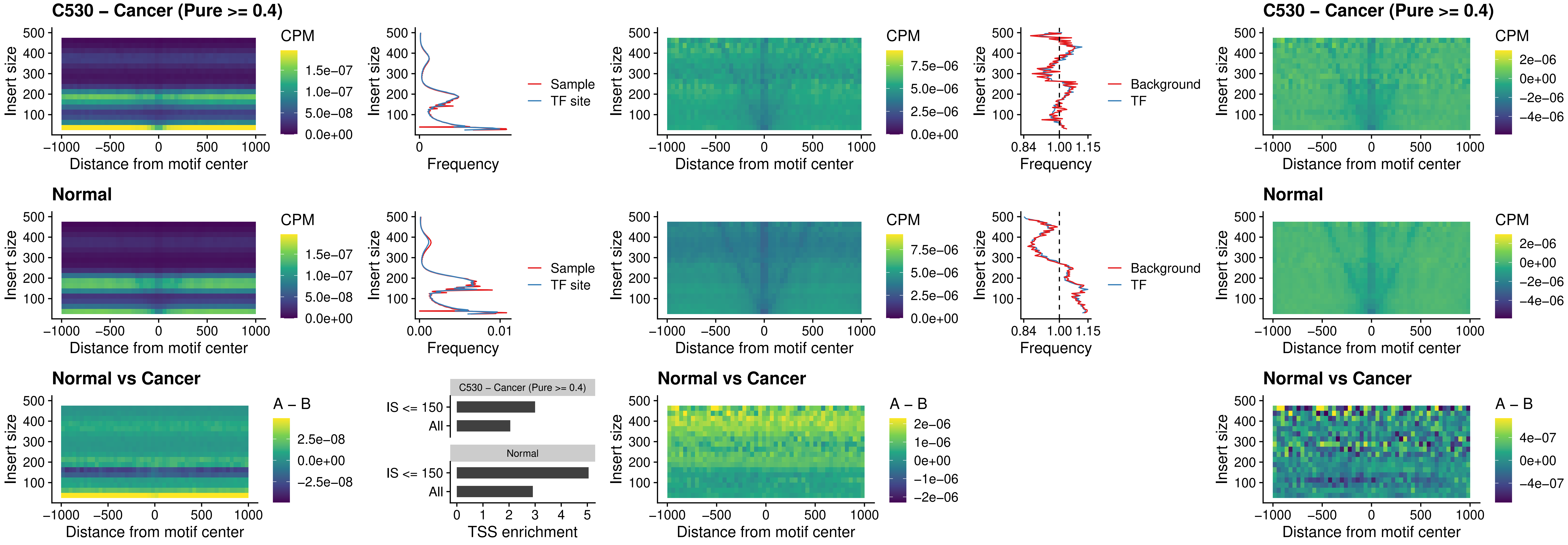

#### C530 – Cancer (Pure >= 0.4) – ARID3C_uc_dTSS_nPEAK

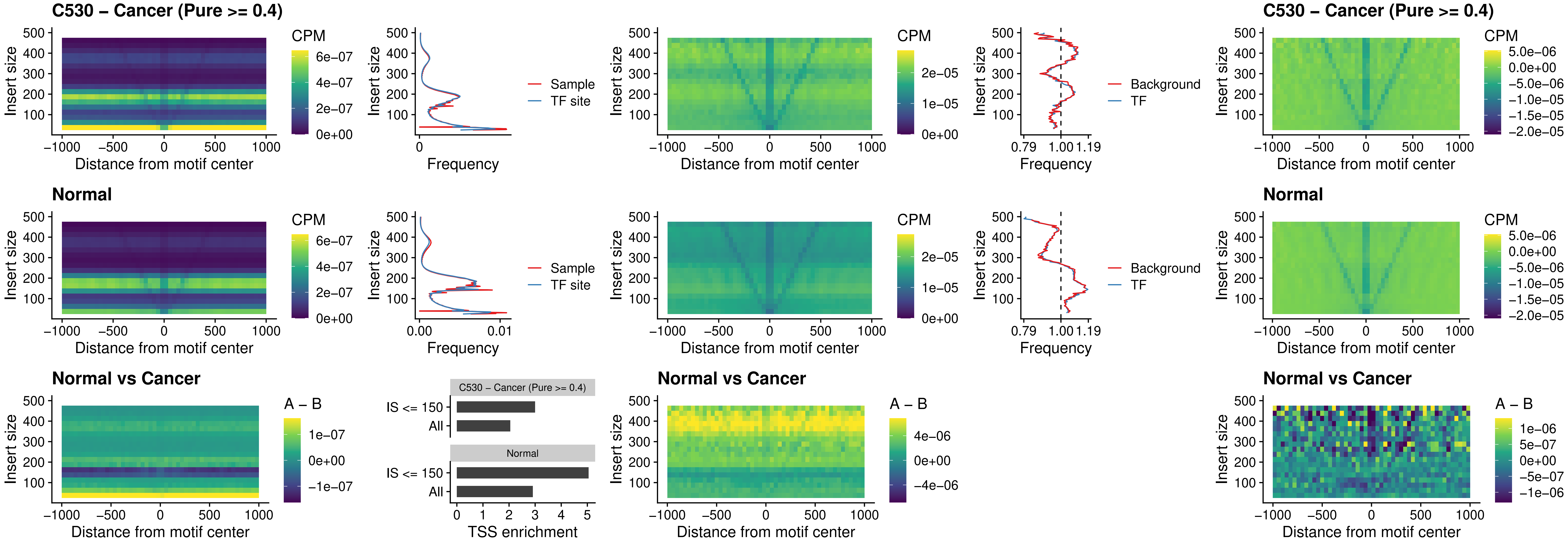

#### C530 – Cancer (Pure >= 0.4) – HOXD12_uc_dTSS_nPEAK

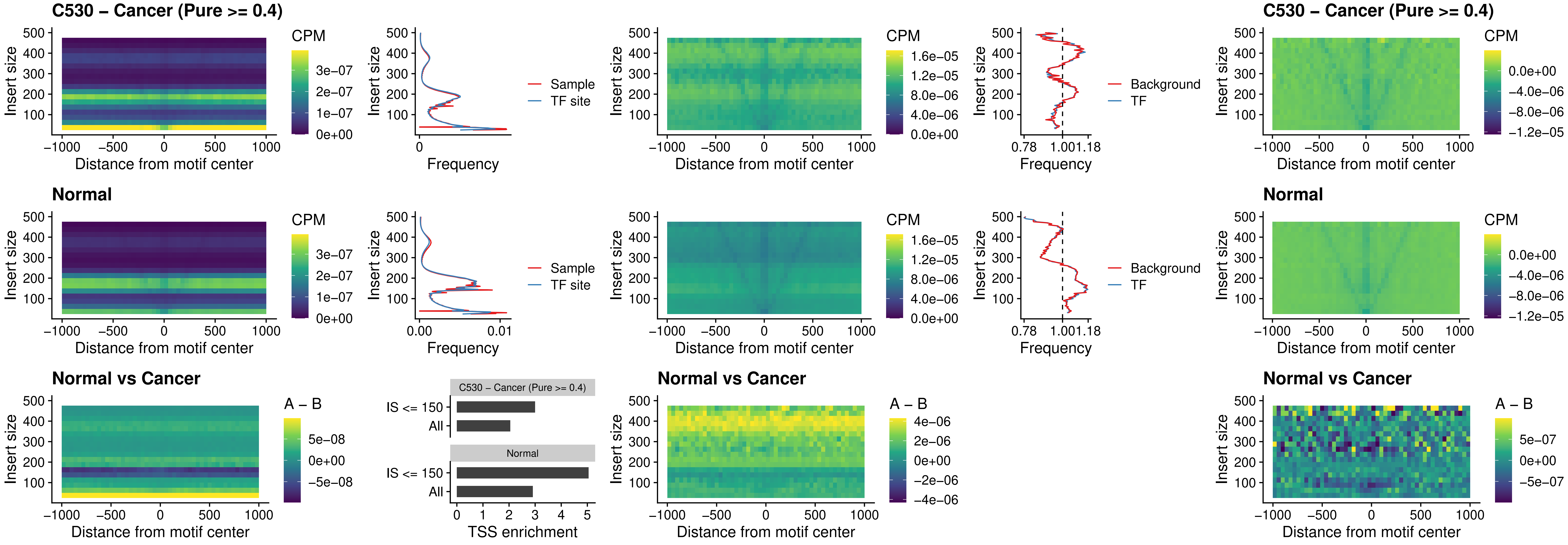

### C530 – Cancer (Pure >= 0.4) – CDX2_uc_dTSS_nPEAK

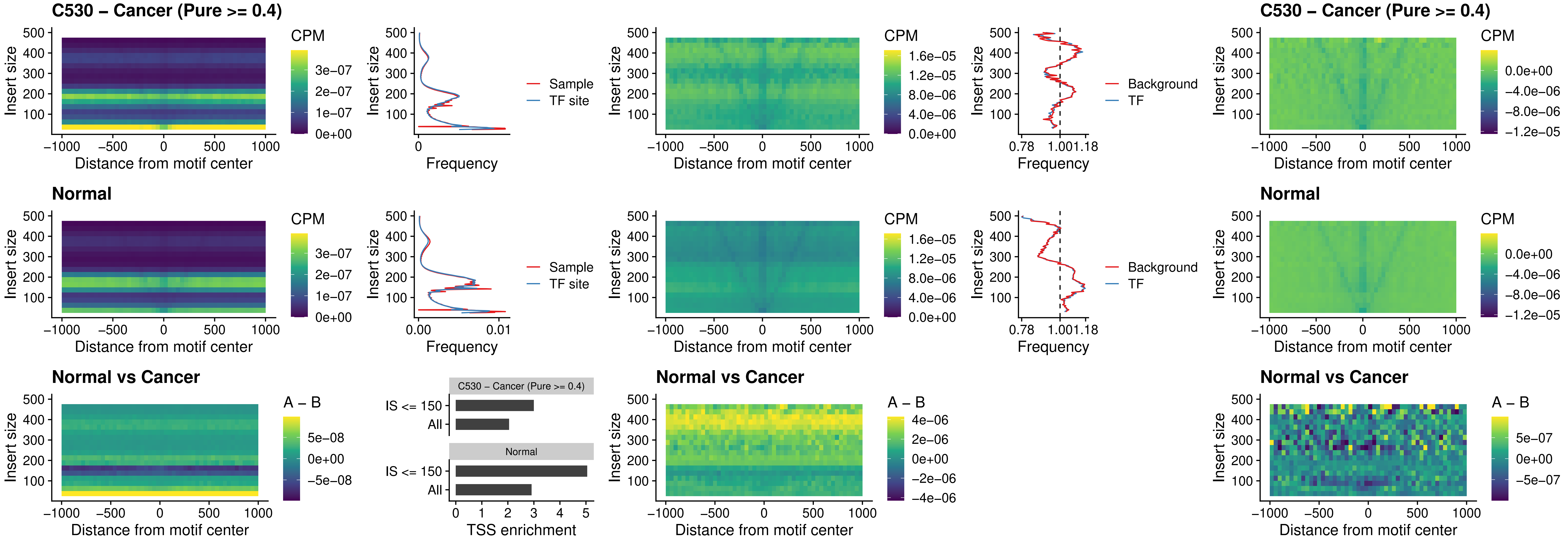

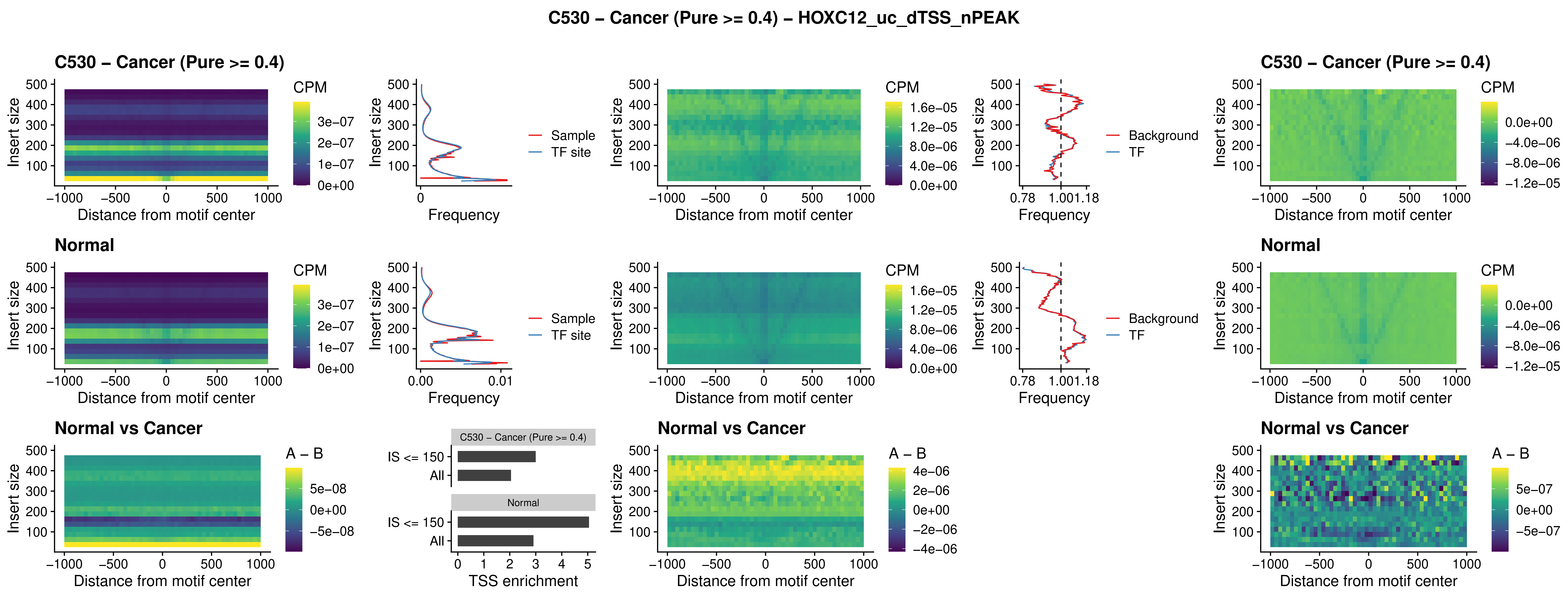
C530 – Cancer (Pure >= 0.4) – HOXC12_uc_dTSS_nPEAK
C530 – Cancer (Pure >= 0.4)
Normal
Normal vs Cancer
Insert size
Distance from motif center
Frequency
CPM
A – B
Sample
TF site
Background
TF
C530 – Cancer (Pure >= 0.4)
Normal
IS <= 150
All
TSS enrichment

### C530 – Cancer (Pure >= 0.4) – HHEX_uc_dTSS_oPEAK

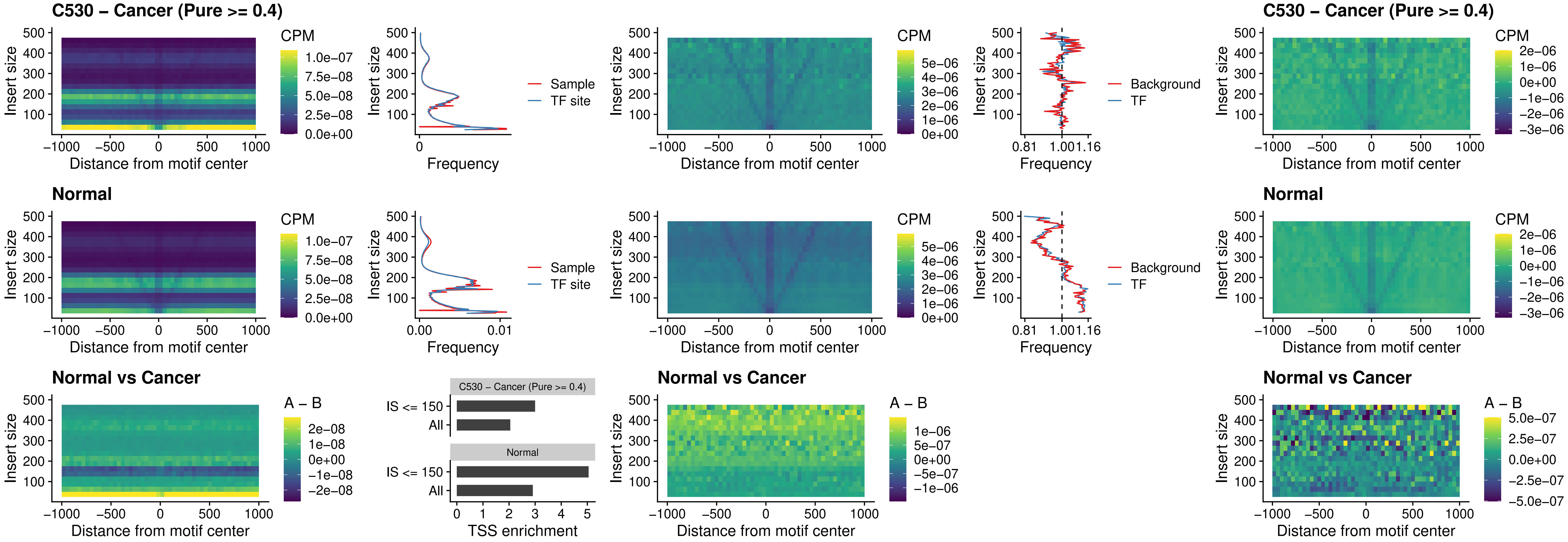

#### C530 – Cancer (Pure >= 0.4) – HOXA11_uc_dTSS_nPEAK

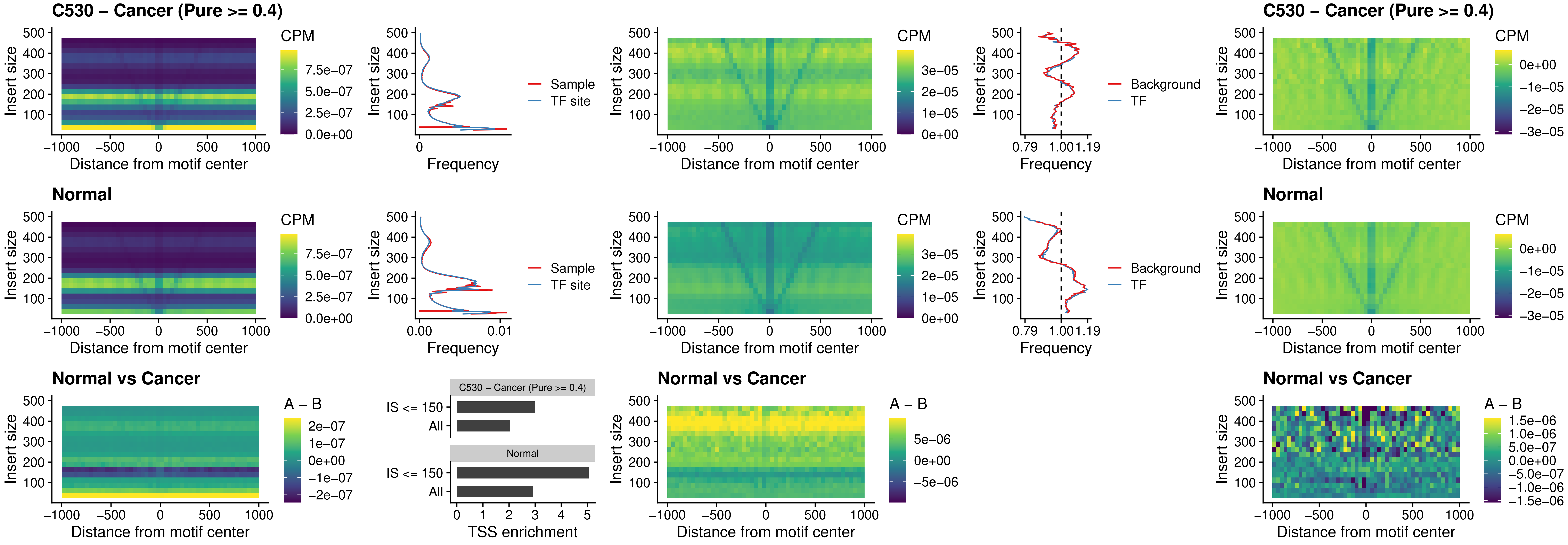

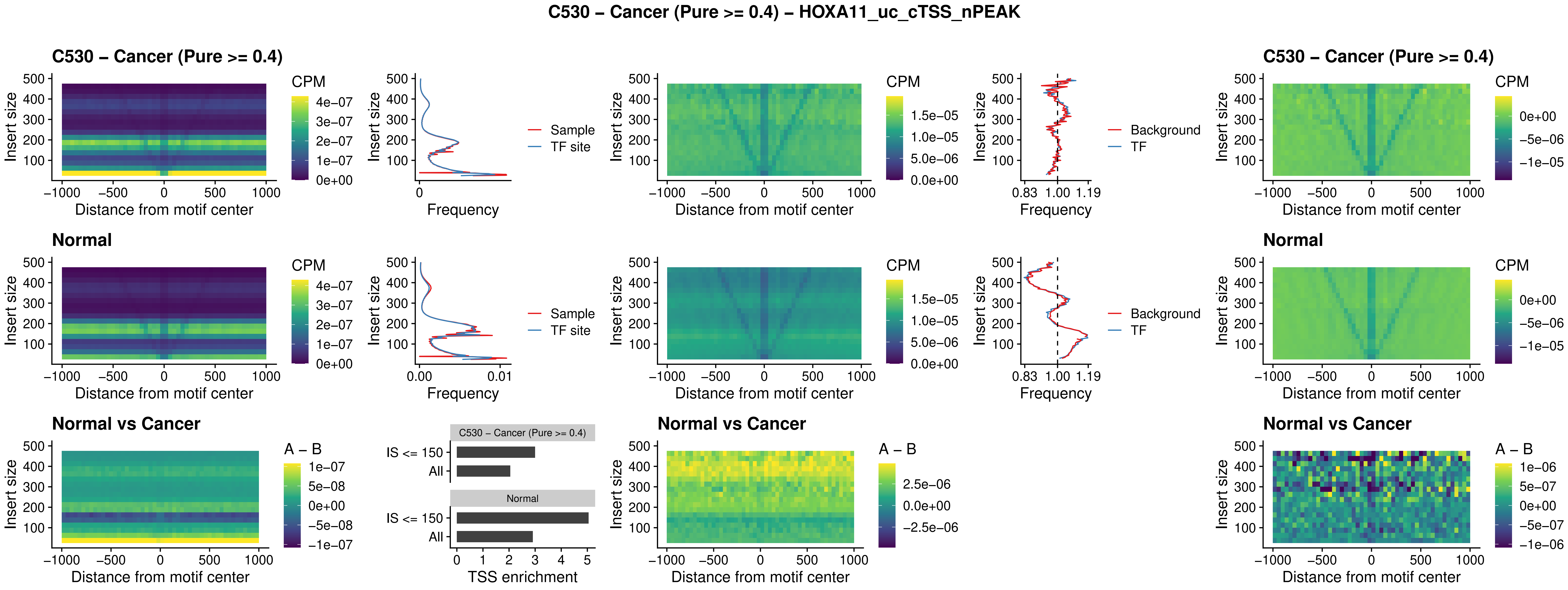
C530 – Cancer (Pure >= 0.4) – HOXA11_uc_cTSS_nPEAK
C530 – Cancer (Pure >= 0.4)
Normal
Normal vs Cancer
Insert size
Distance from motif center
Frequency
CPM
A – B
Sample
TF site
Background
TF
TSS enrichment
IS <= 150
All

#### C530 – Cancer (Pure >= 0.4) – ZFHX3_uc_dTSS_nPEAK

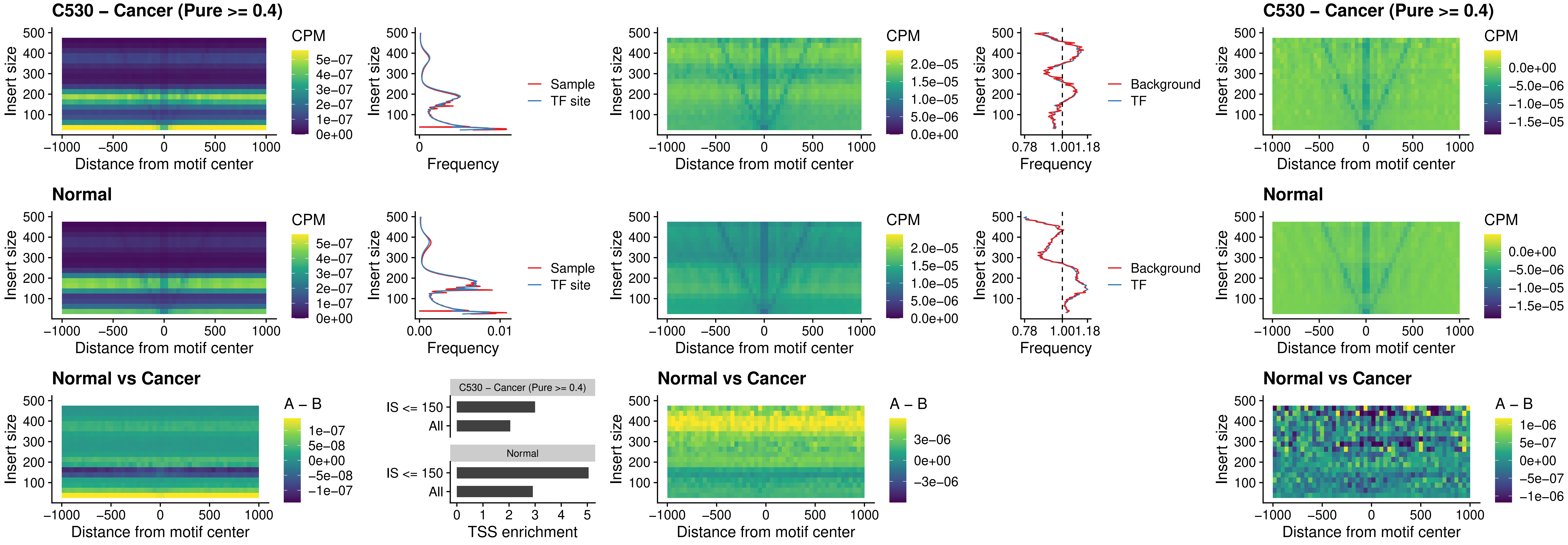

#### C530 – Cancer (Pure >= 0.4) – HOXB4_uc_dTSS_nPEAK

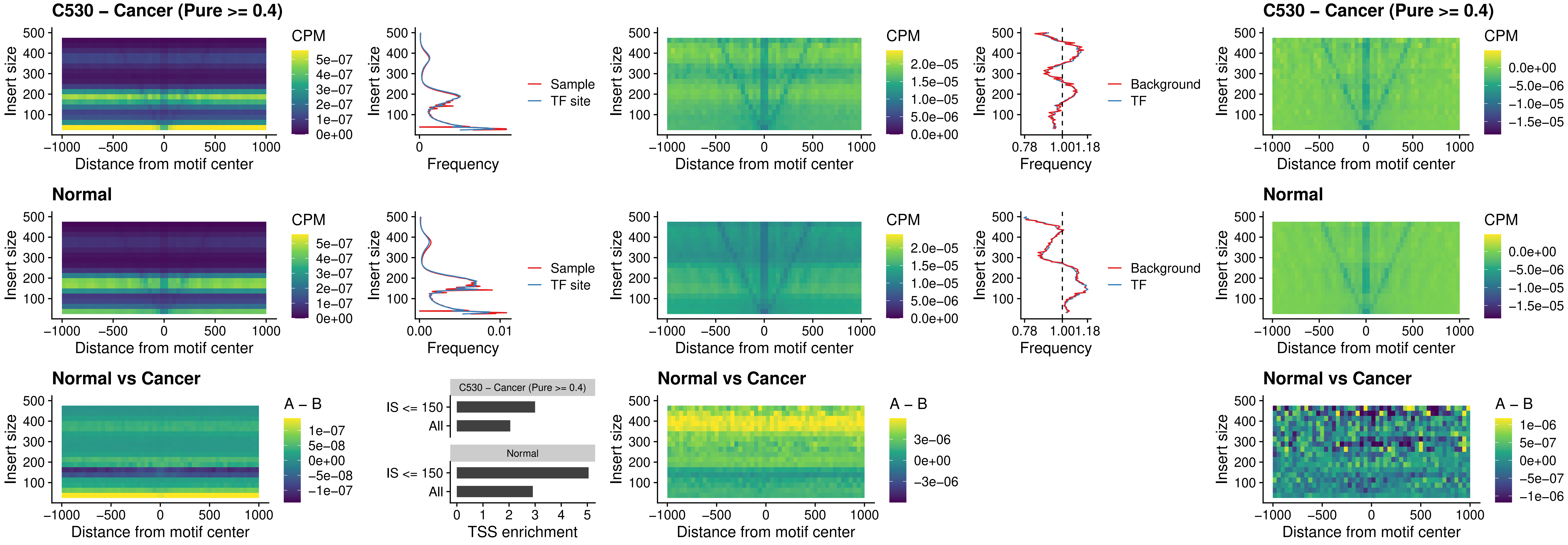

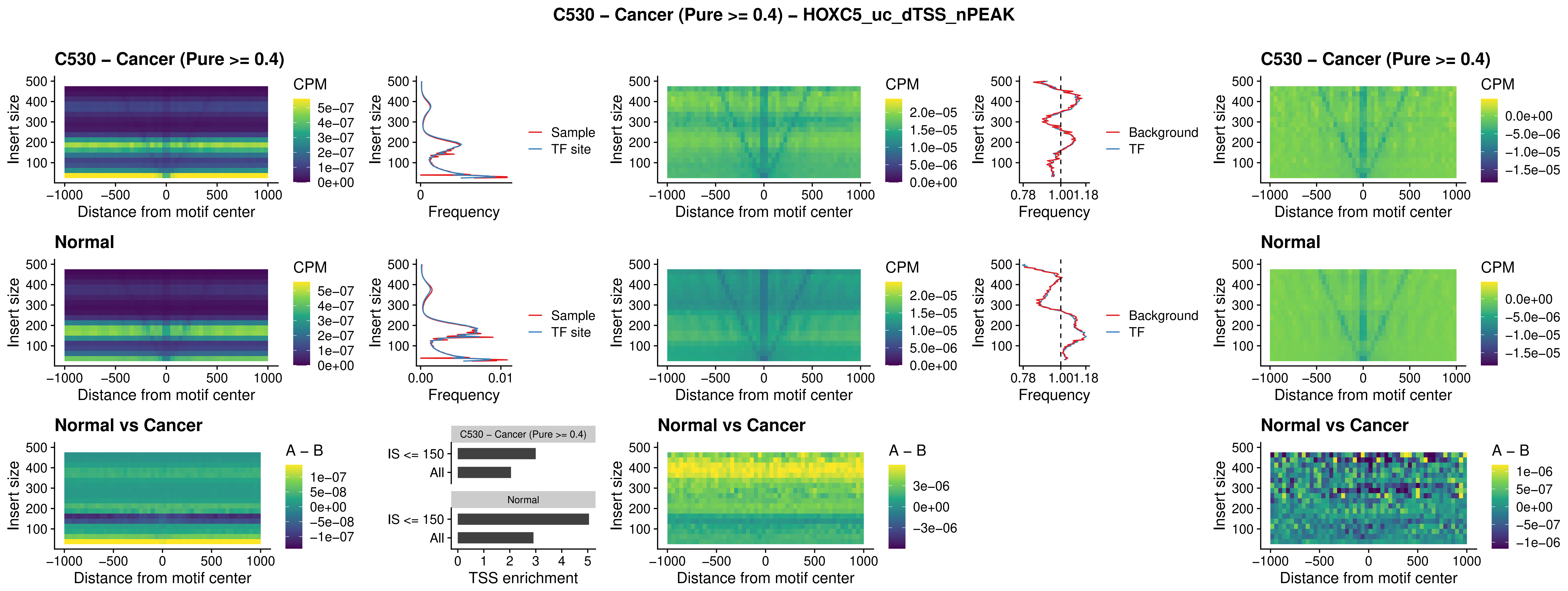
C530 – Cancer (Pure >= 0.4) – HOXC5_uc_dTSS_nPEAK
C530 – Cancer (Pure >= 0.4)
Normal
Normal vs Cancer
Insert size
Distance from motif center
Frequency
CPM
A – B
Sample
TF site
Background
TF
C530 – Cancer (Pure >= 0.4)
Normal
IS <= 150
All
TSS enrichment

### C530 – Cancer (Pure >= 0.4) – HOXD1_uc_dTSS_nPEAK

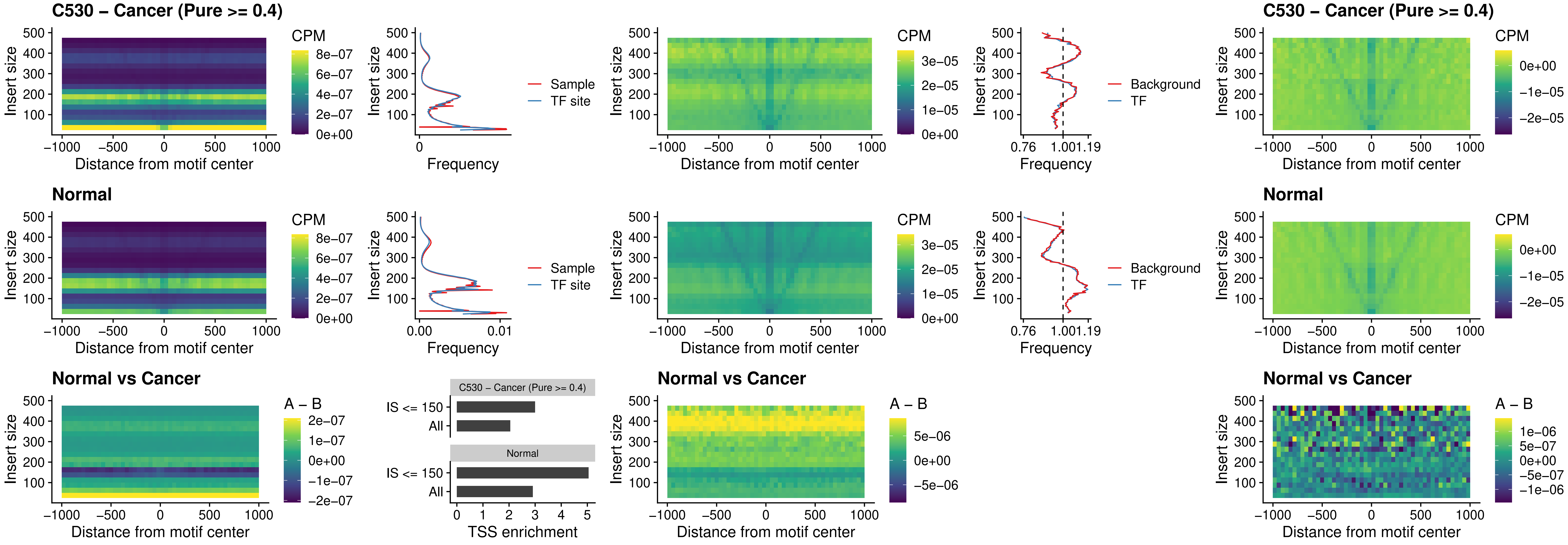

#### C530 – Cancer (Pure >= 0.4) – OTP_uc_dTSS_nPEAK

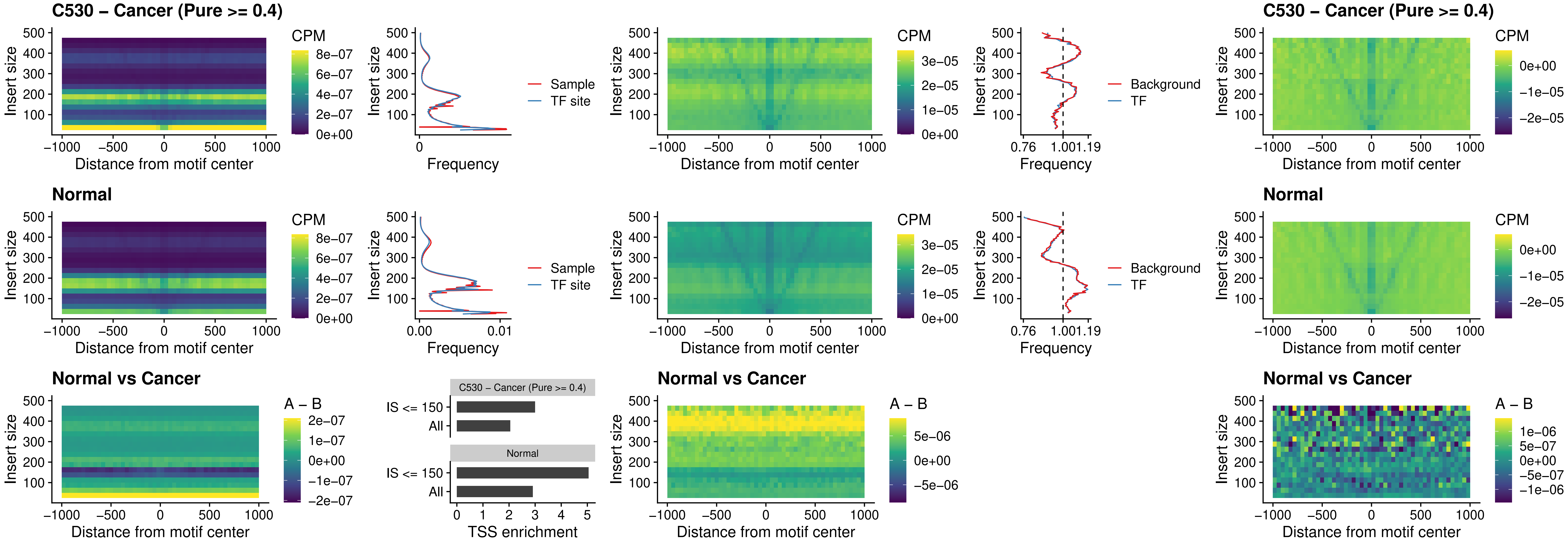

#### C530 – Cancer (Pure >= 0.4) – FOXL1_uc_dTSS_oPEAK

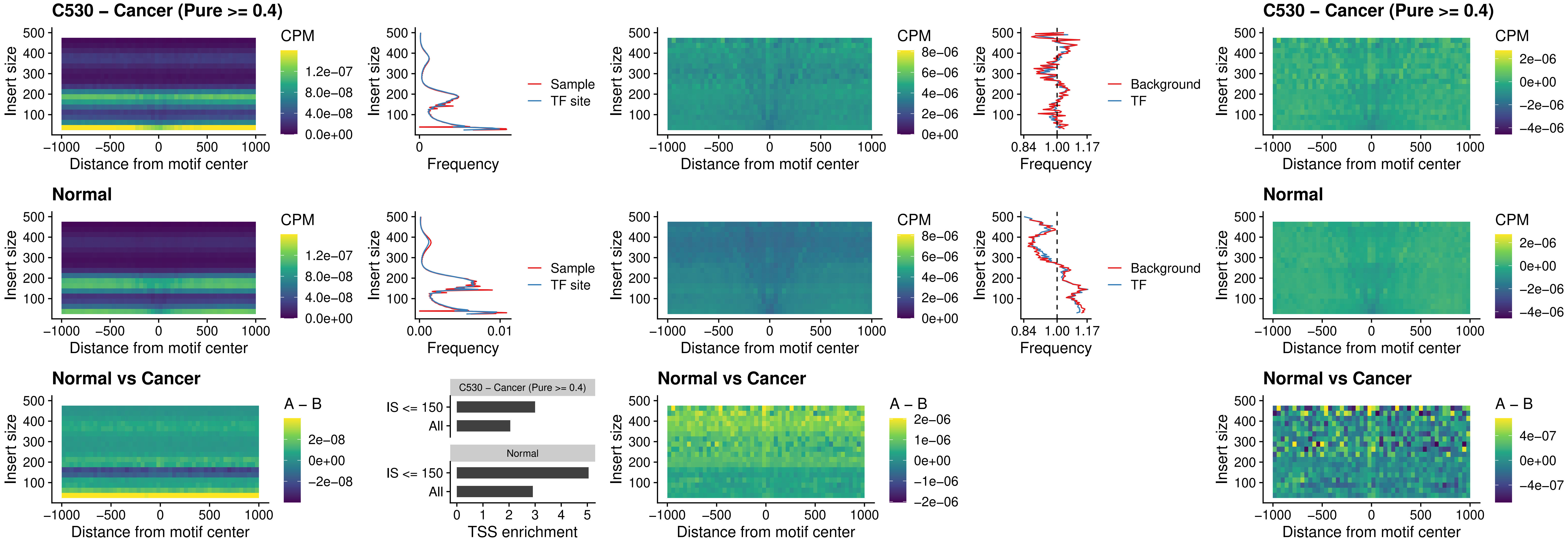

#### C530 – Cancer (Pure >= 0.4) – ZNF32_uc_dTSS_oPEAK

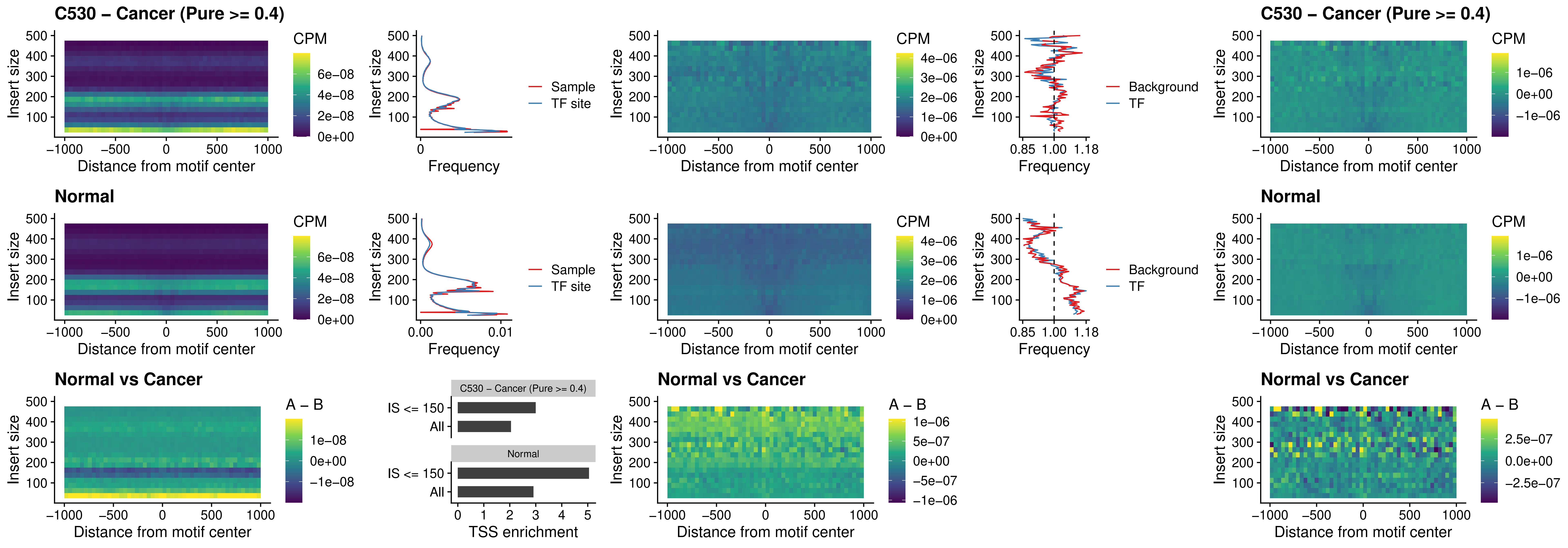

#### C530 – Cancer (Pure >= 0.4) – FOXL1_uc_dTSS_nPEAK

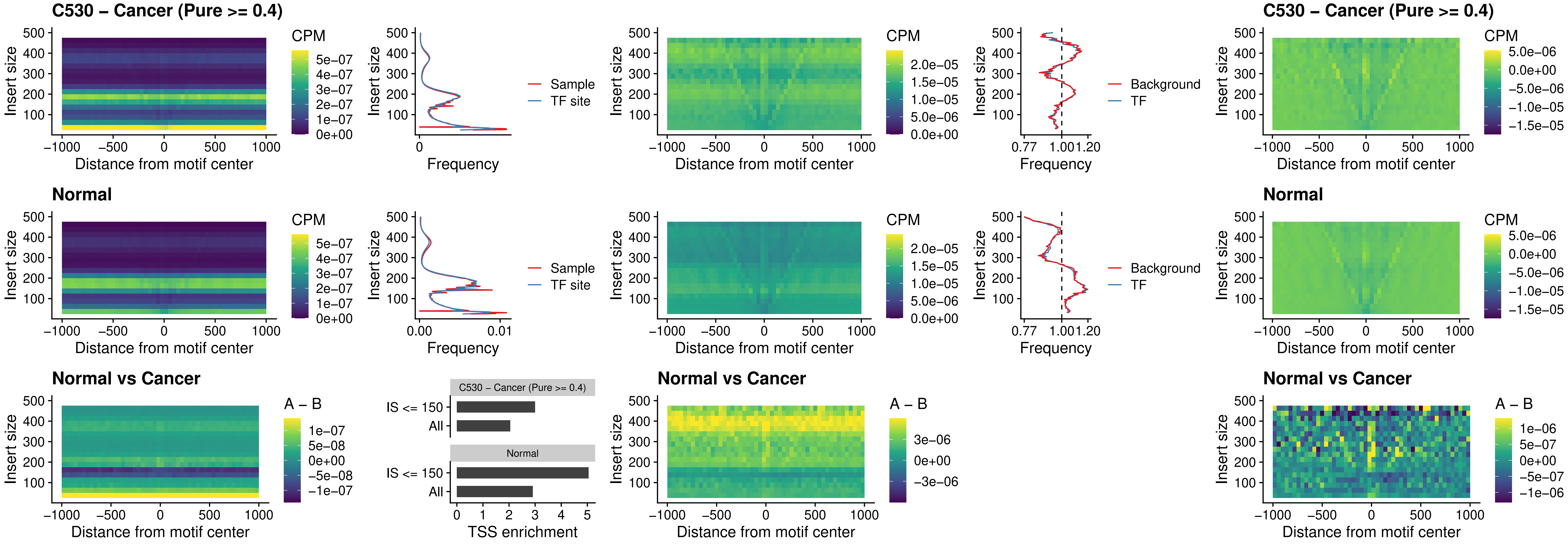

### C530 – Cancer (Pure >= 0.4) – ZNF32_uc_dTSS_nPEAK

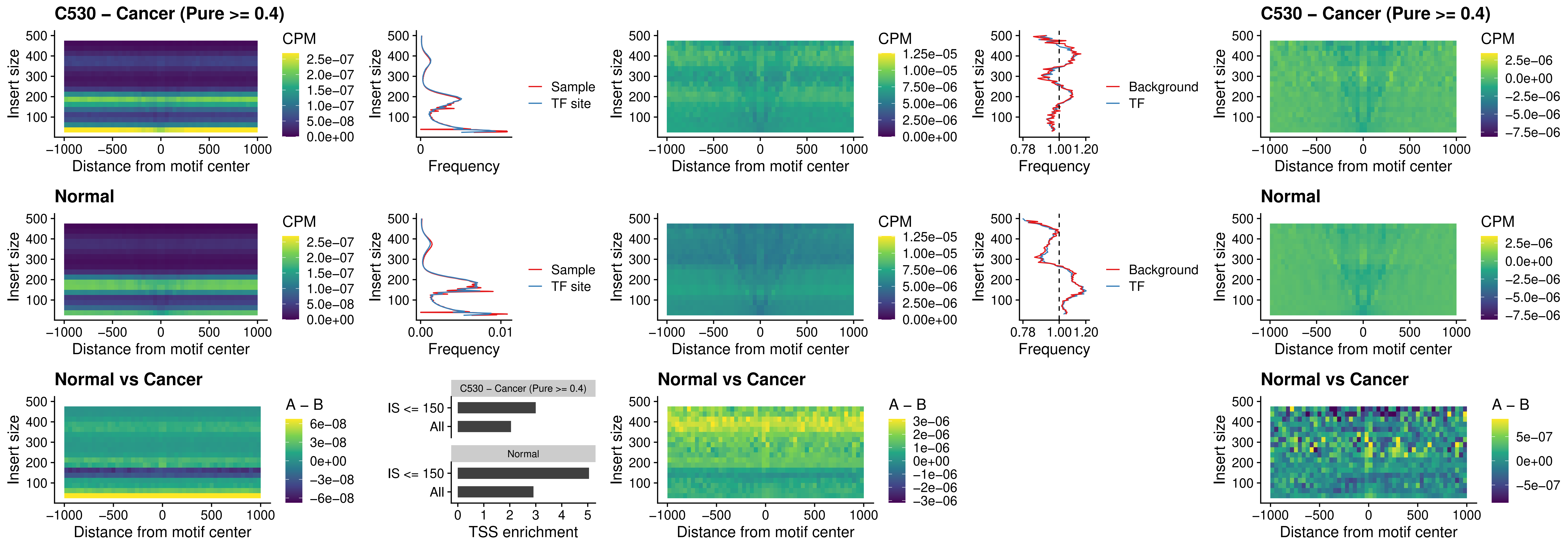

#### C530 – Cancer (Pure >= 0.4) – DLX5_uc_dTSS_nPEAK

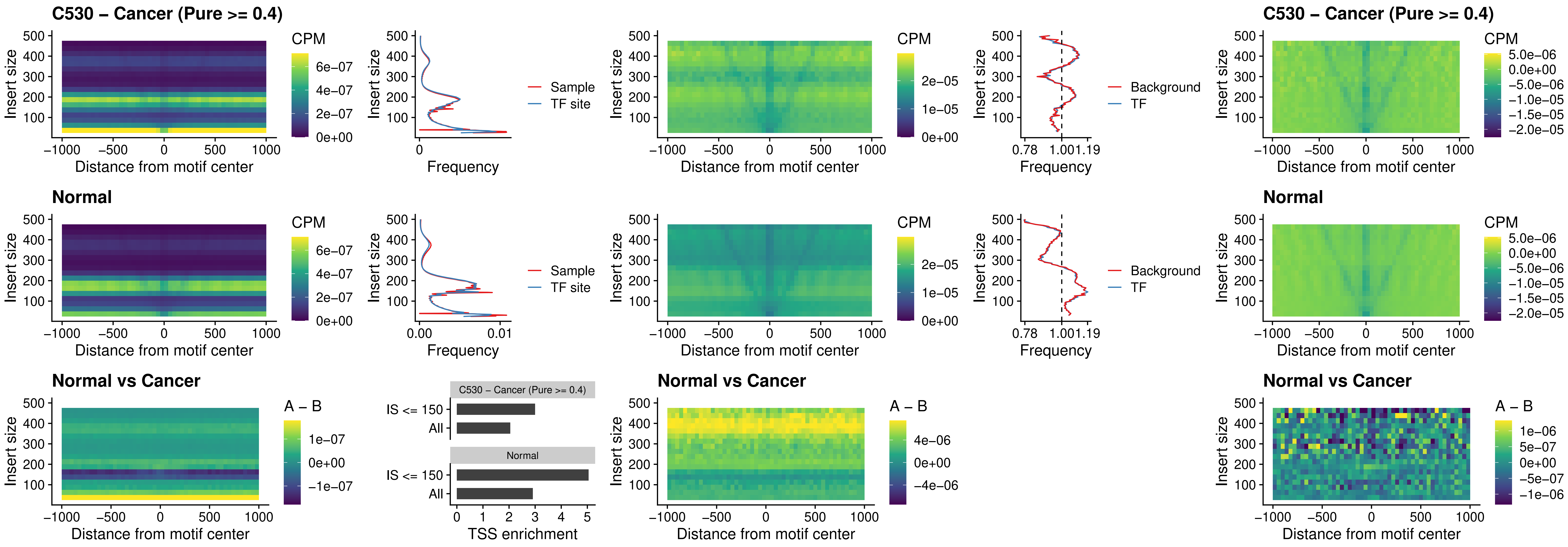

### C530 – Cancer (Pure >= 0.4) – DMRT2_uc_dTSS_nPEAK

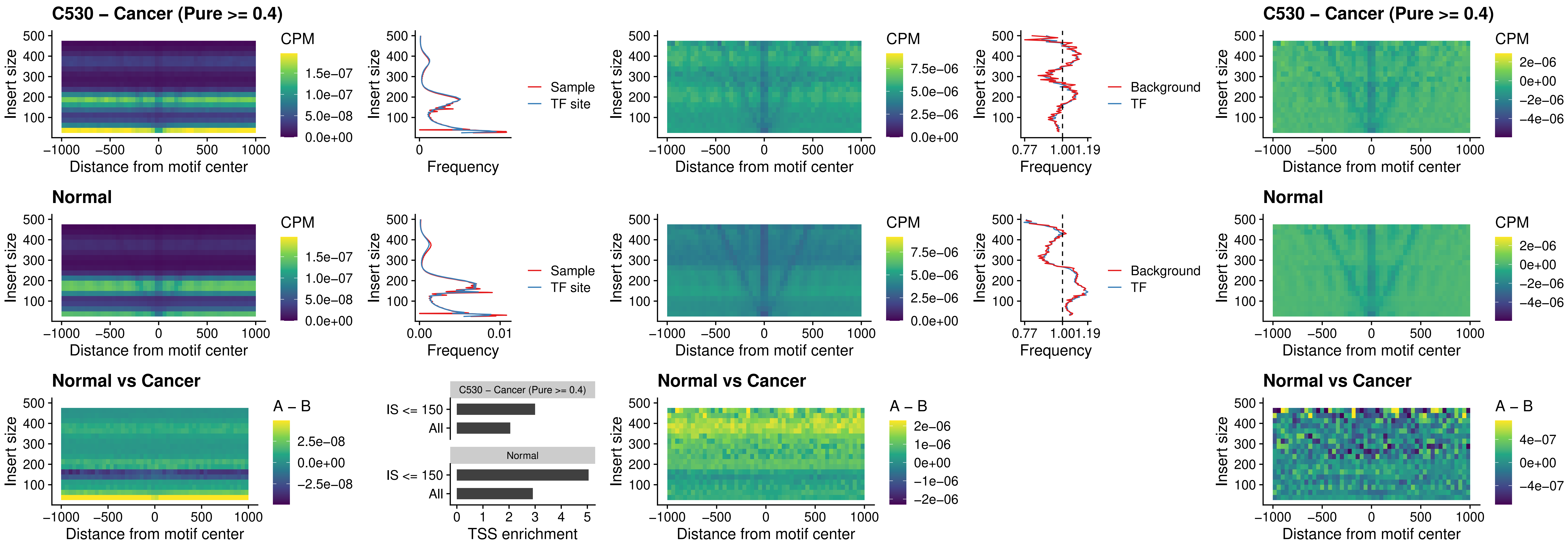

#### C530 – Cancer (Pure >= 0.4) – DMRTC2_uc_cTSS_nPEAK

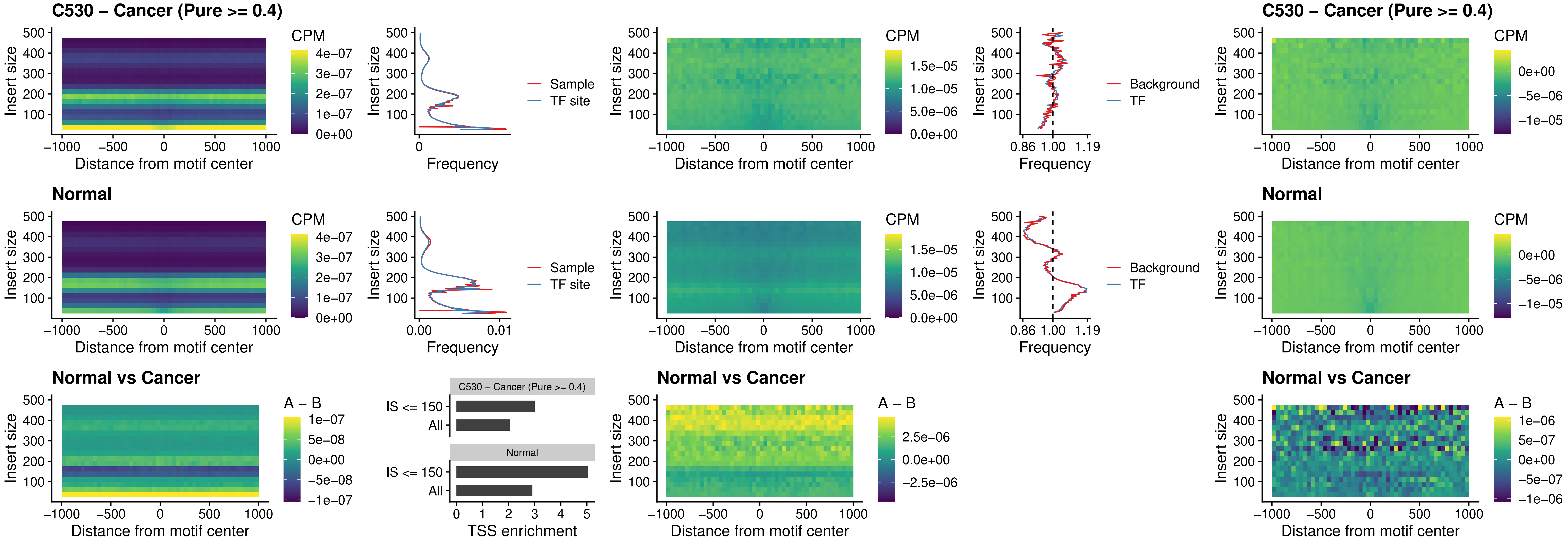

#### C530 – Cancer (Pure >= 0.4) – HOXD9_uc_dTSS_nPEAK

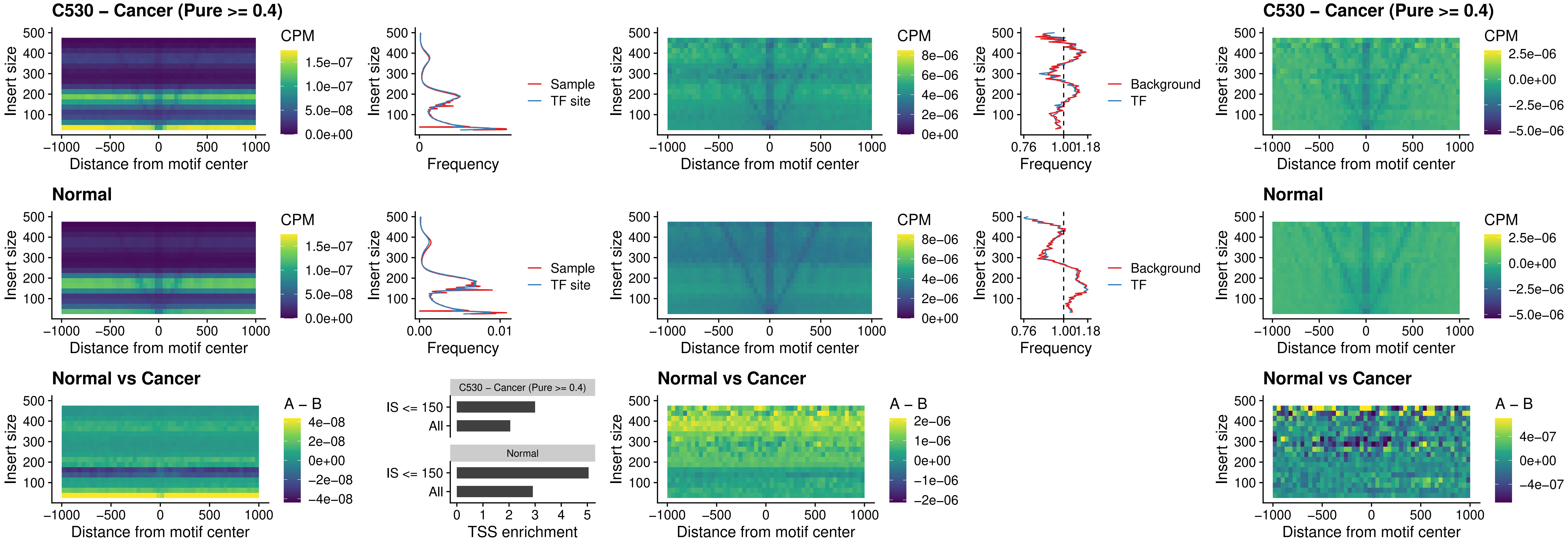

#### C530 – Cancer (Pure >= 0.4) – HOXD4_uc_dTSS_nPEAK

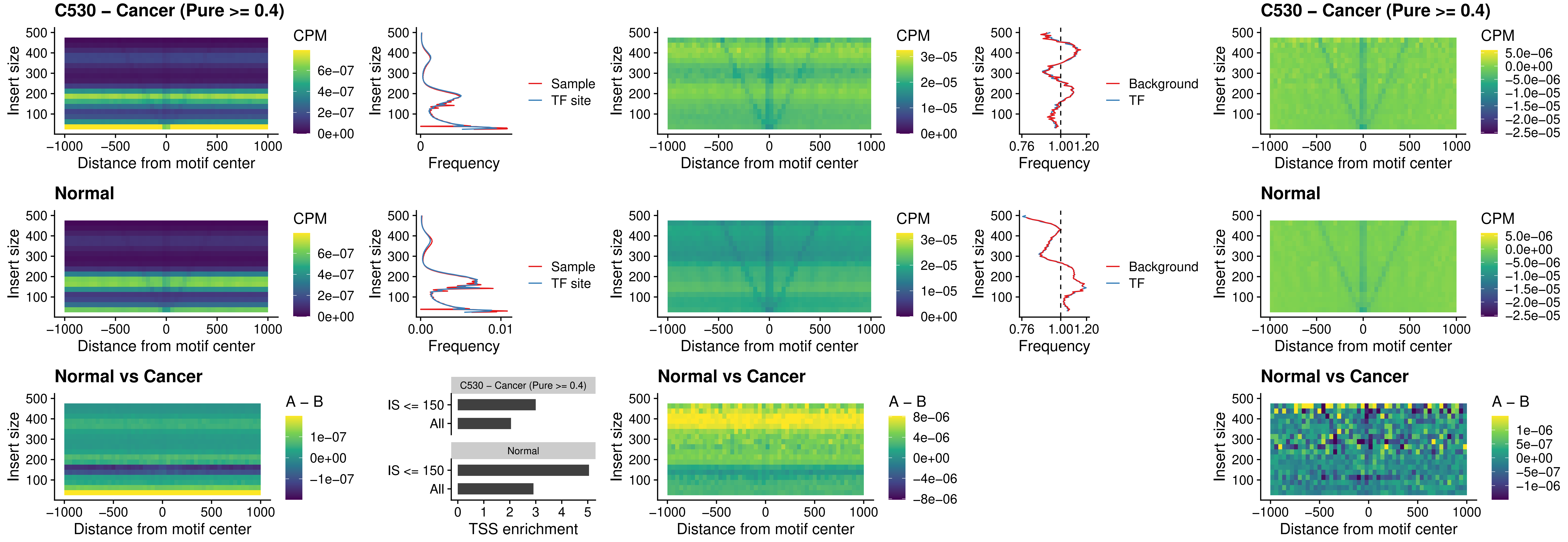

### C530 – Cancer (Pure >= 0.4) – FOXP3_uc_cTSS_nPEAK

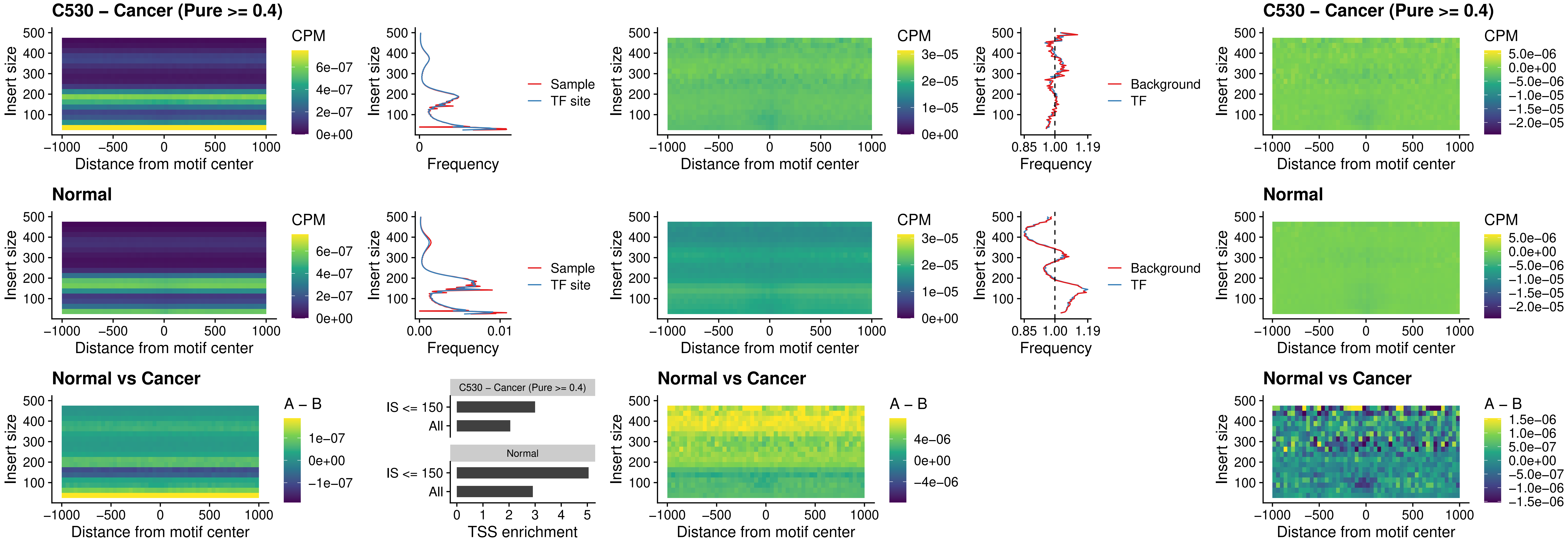

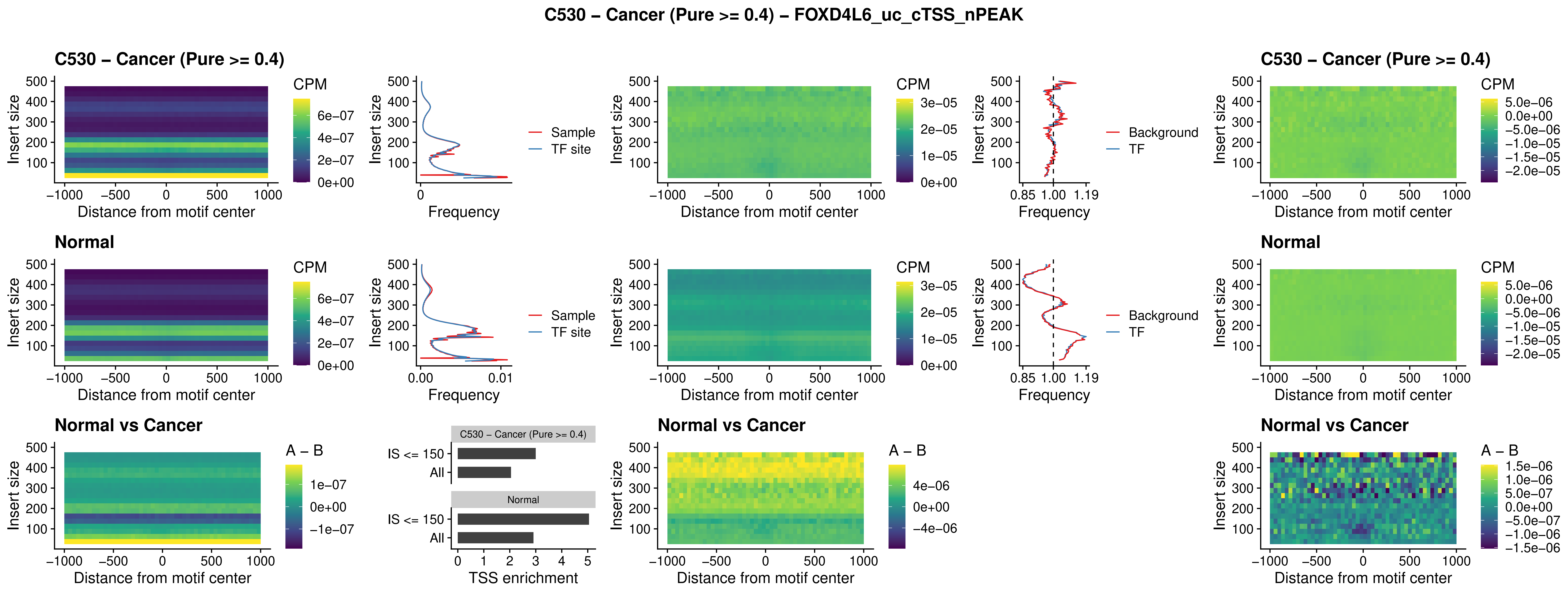
C530 – Cancer (Pure >= 0.4) – FOXD4L6_uc_cTSS_nPEAK
C530 – Cancer (Pure >= 0.4)
Normal
Normal vs Cancer
Insert size
Distance from motif center
Frequency
CPM
A – B
Sample
TF site
Background
TF
TSS enrichment
IS <= 150
All
C530 – Cancer (Pure >= 0.4)
Normal

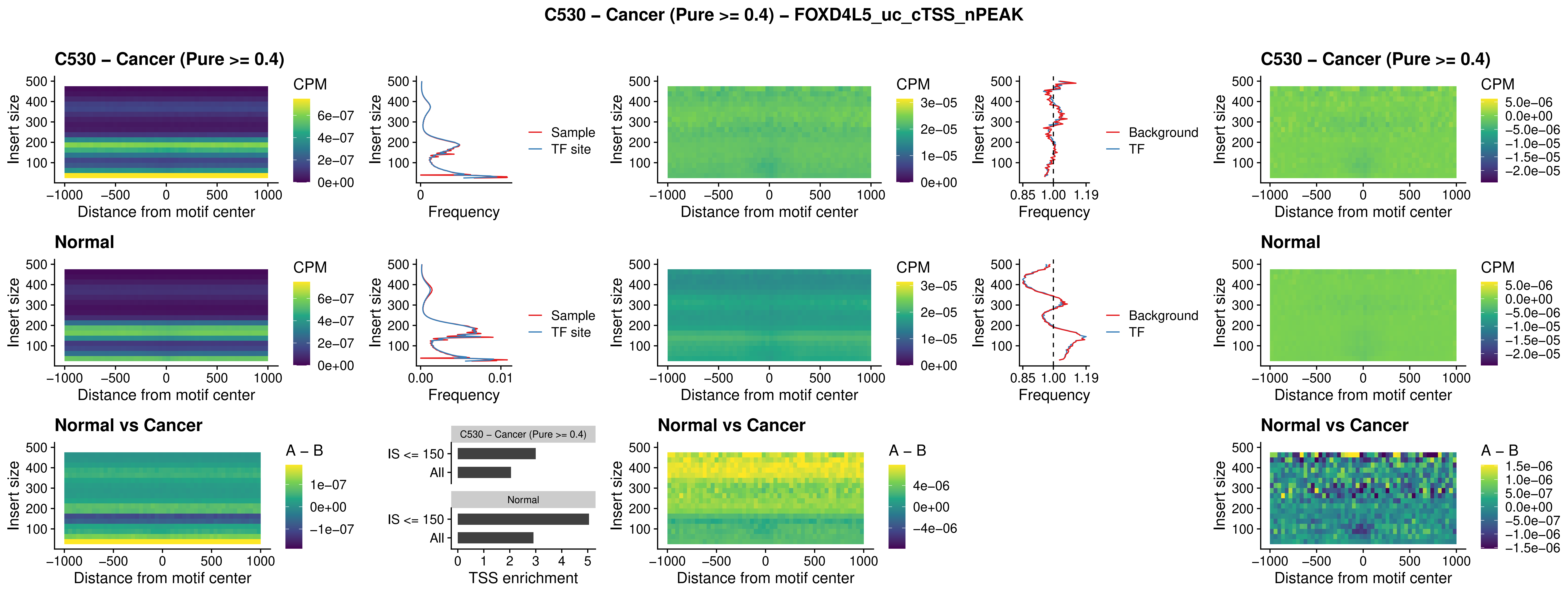
C530 – Cancer (Pure >= 0.4) – FOXD4L5_uc_cTSS_nPEAK
C530 – Cancer (Pure >= 0.4)
Normal
Normal vs Cancer
Insert size
Distance from motif center
Frequency
CPM
A – B
Sample
TF site
Background
TF
C530 – Cancer (Pure >= 0.4)
Normal
IS <= 150
All
TSS enrichment

#### C530 – Cancer (Pure >= 0.4) – FOXD4L4_uc_cTSS_nPEAK

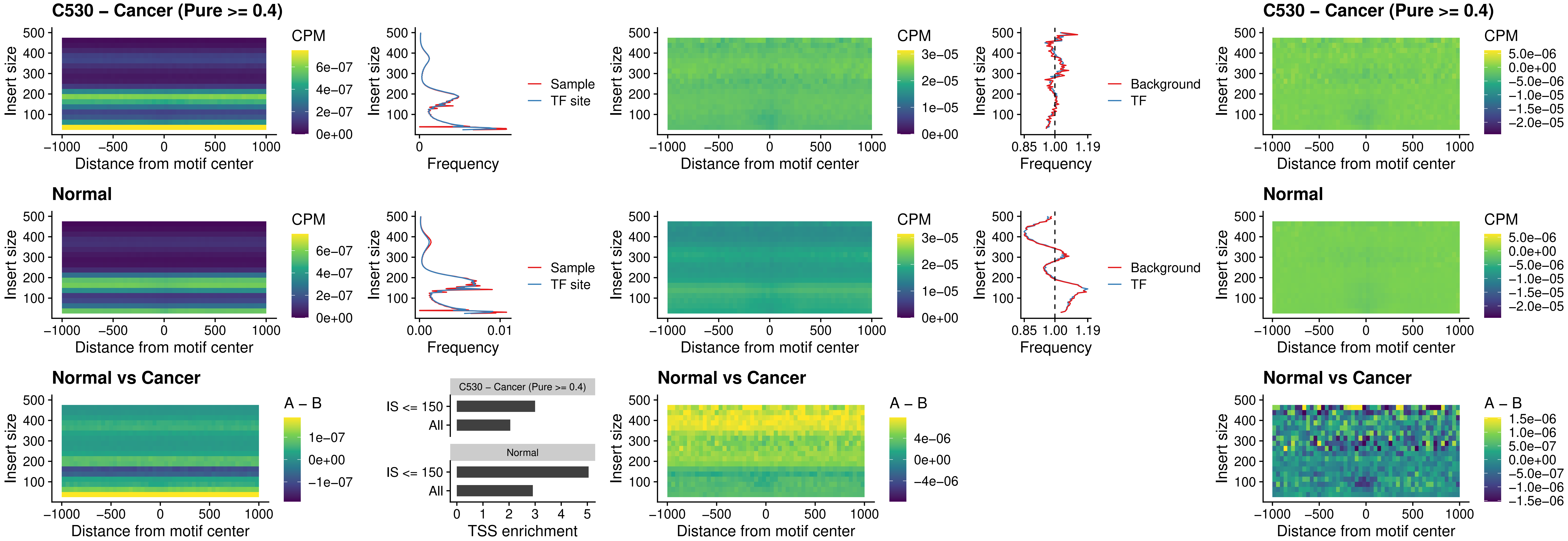

#### C530 – Cancer (Pure >= 0.4) – FOXD4L2_uc_cTSS_nPEAK

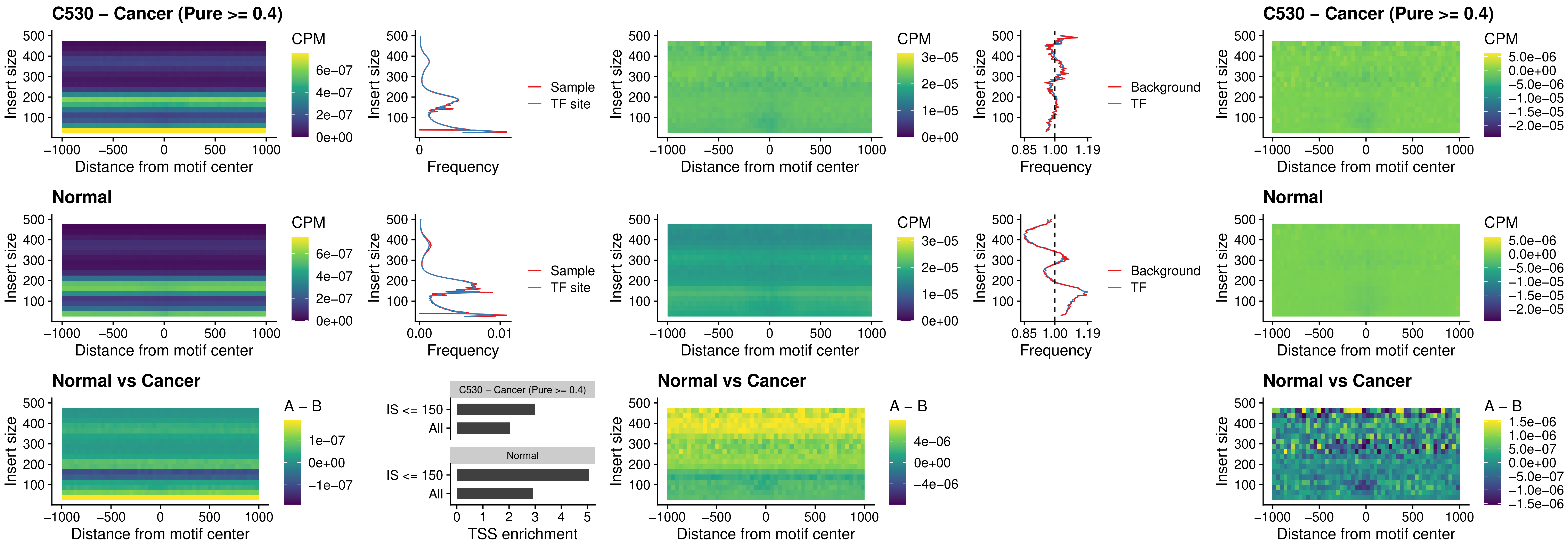

### C530 – Cancer (Pure >= 0.4) – FOXD4L3_uc_cTSS_nPEAK

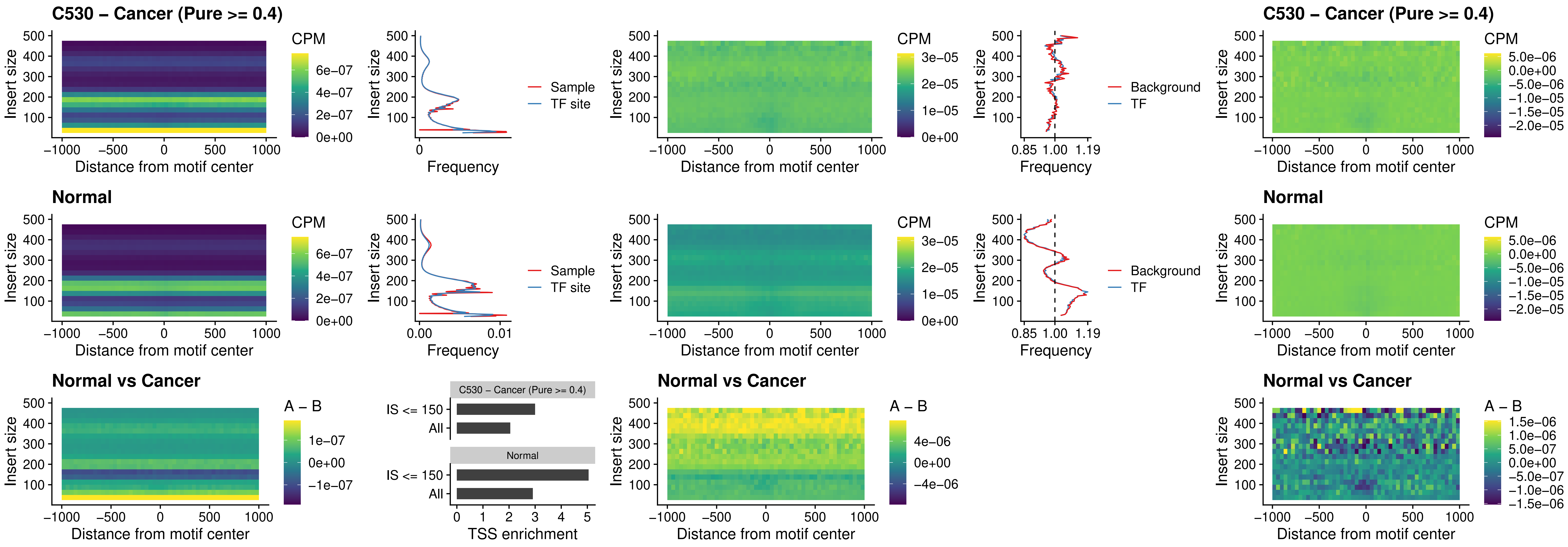

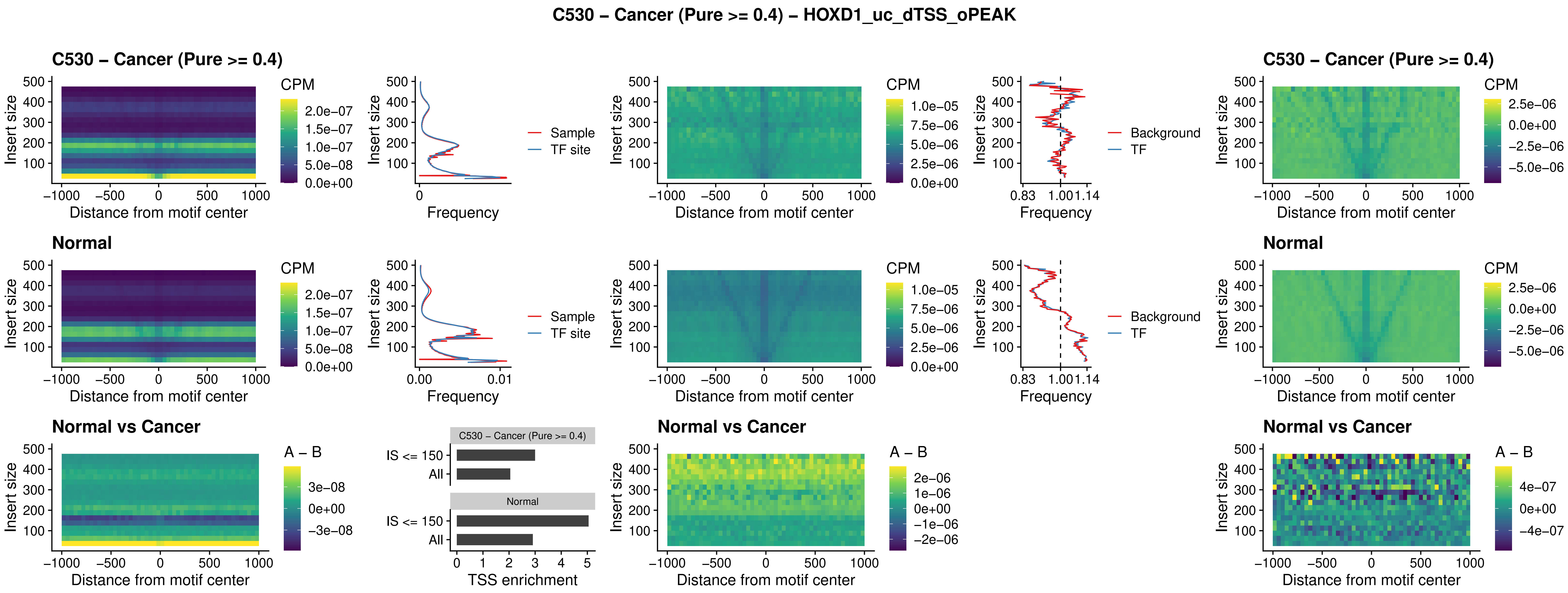

C530 – Cancer (Pure >= 0.4) – HOXD1_uc_dTSS_oPEAK
C530 – Cancer (Pure >= 0.4)
Normal
Normal vs Cancer
Insert size
Distance from motif center
Frequency
CPM
A – B
Sample
TF site
Background
TF
C530 – Cancer (Pure >= 0.4)
Normal
IS <= 150
All
TSS enrichment

#### C530 – Cancer (Pure >= 0.4) – VAX1_uc_pTSS_nPEAK

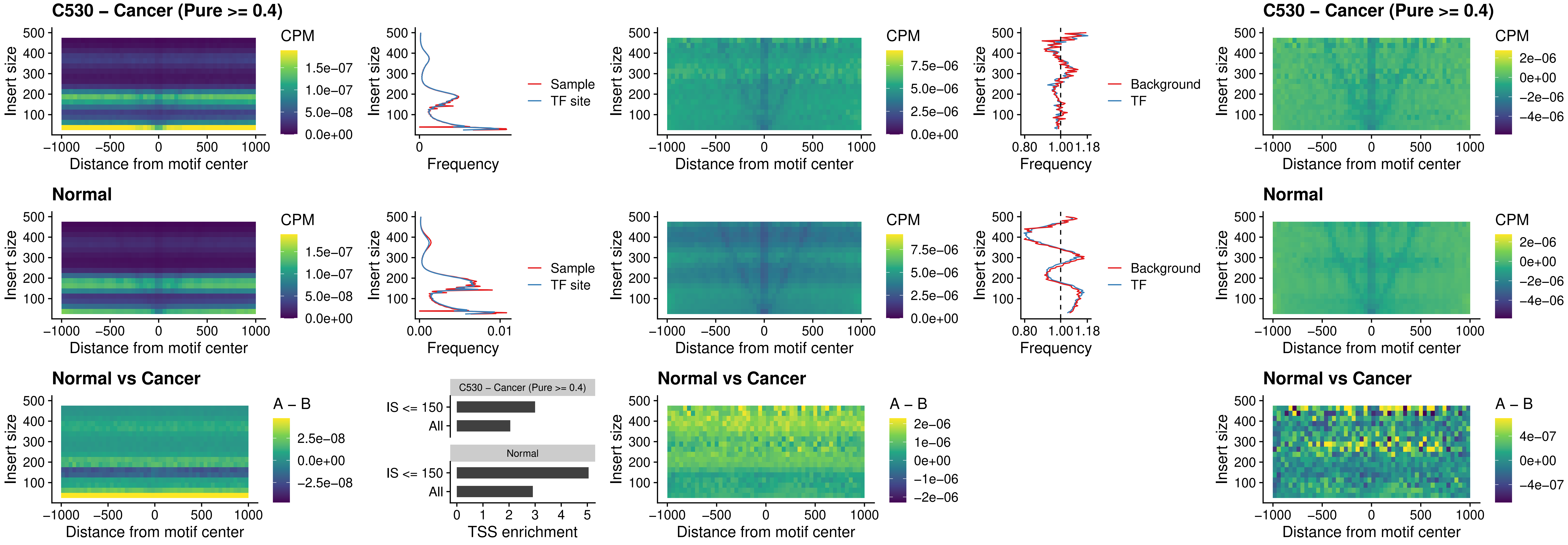

#### C530 – Cancer (Pure >= 0.4) – DLX3_uc_pTSS_nPEAK

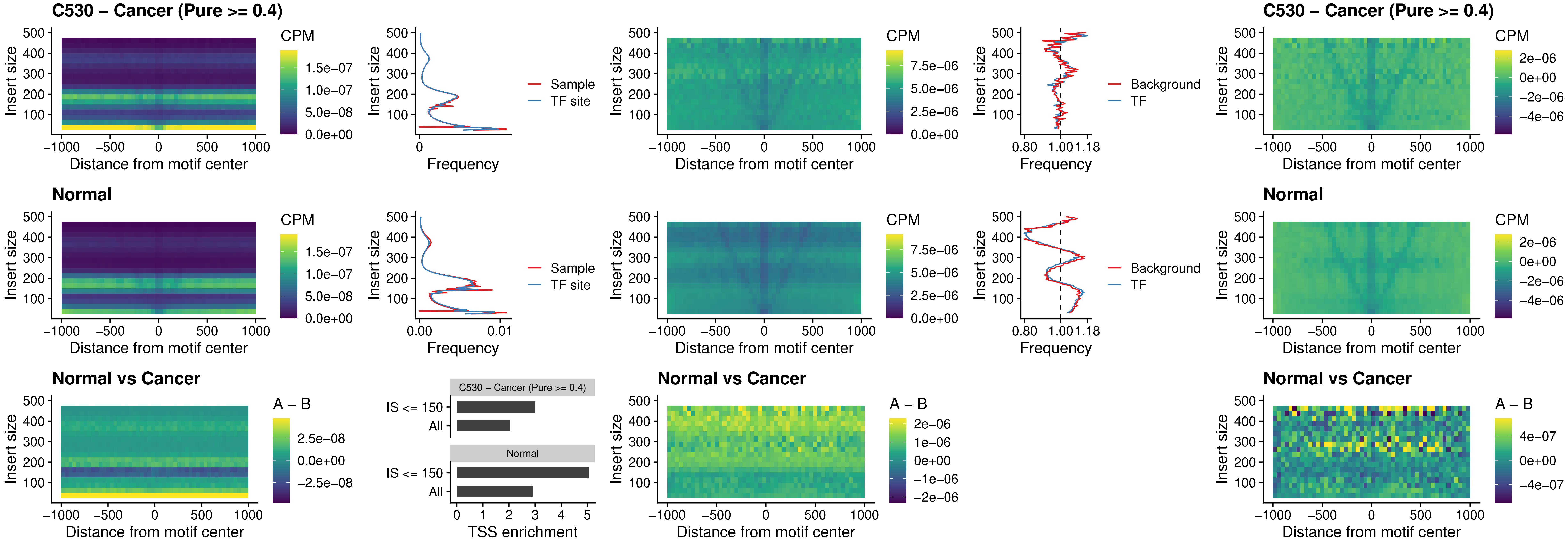

#### C530 – Cancer (Pure >= 0.4) – HOXD3_uc_pTSS_nPEAK

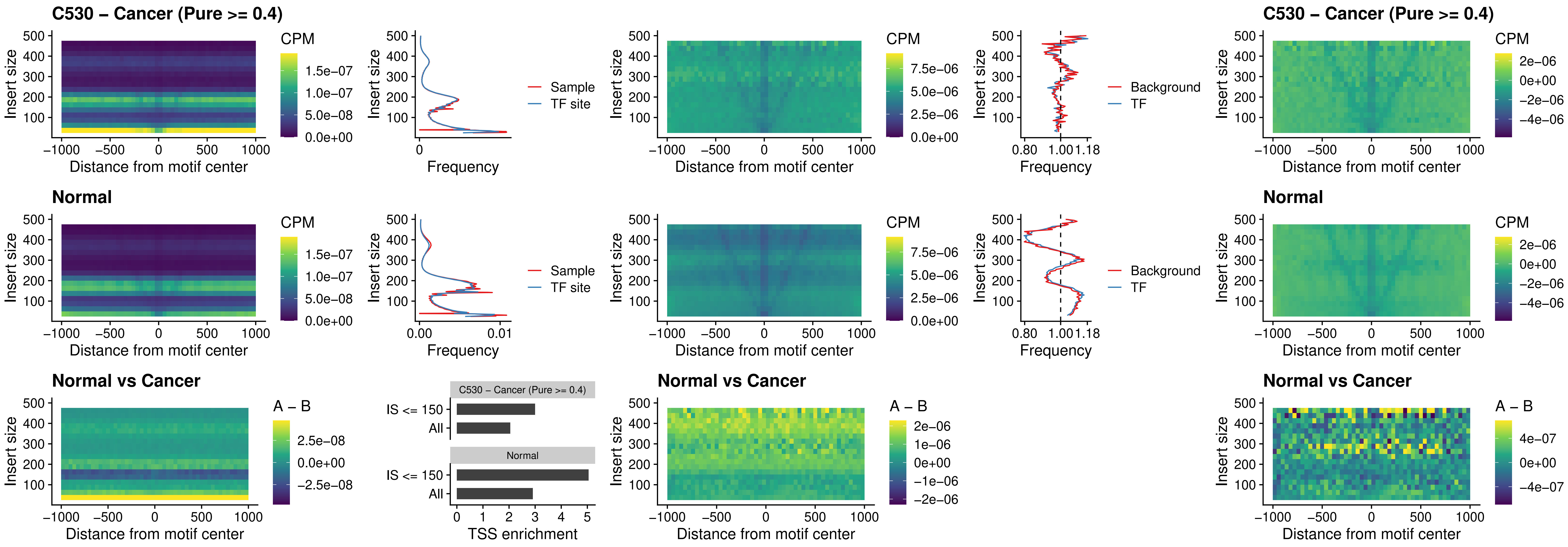

#### C530 – Cancer (Pure >= 0.4) – PDX1_uc_dTSS_nPEAK

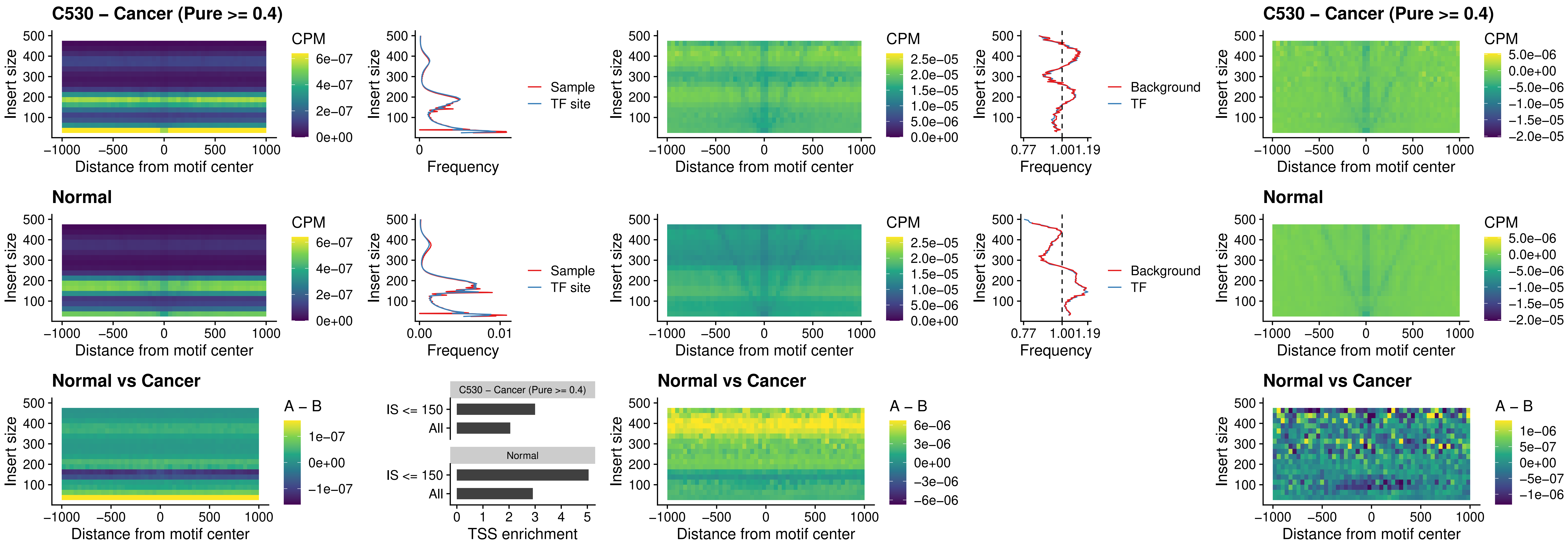

### C530 – Cancer (Pure >= 0.4) – VAX2_uc_dTSS_nPEAK

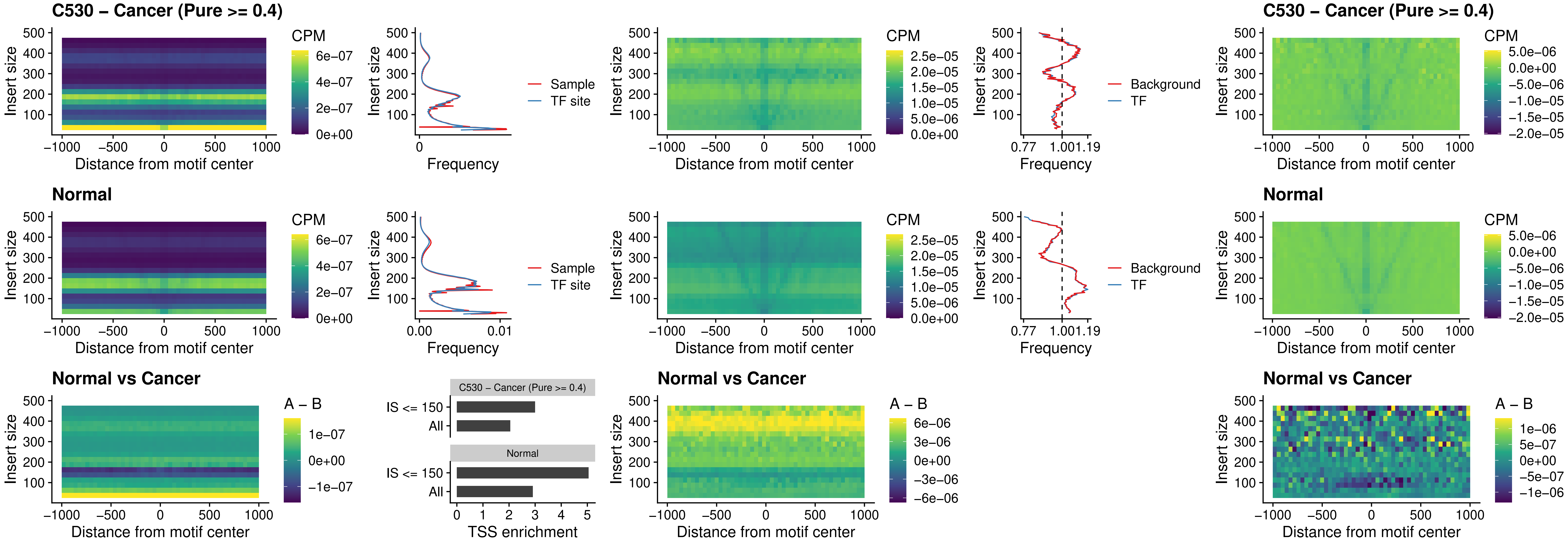

#### C530 – Cancer (Pure >= 0.4) – SOX5_uc_dTSS_nPEAK

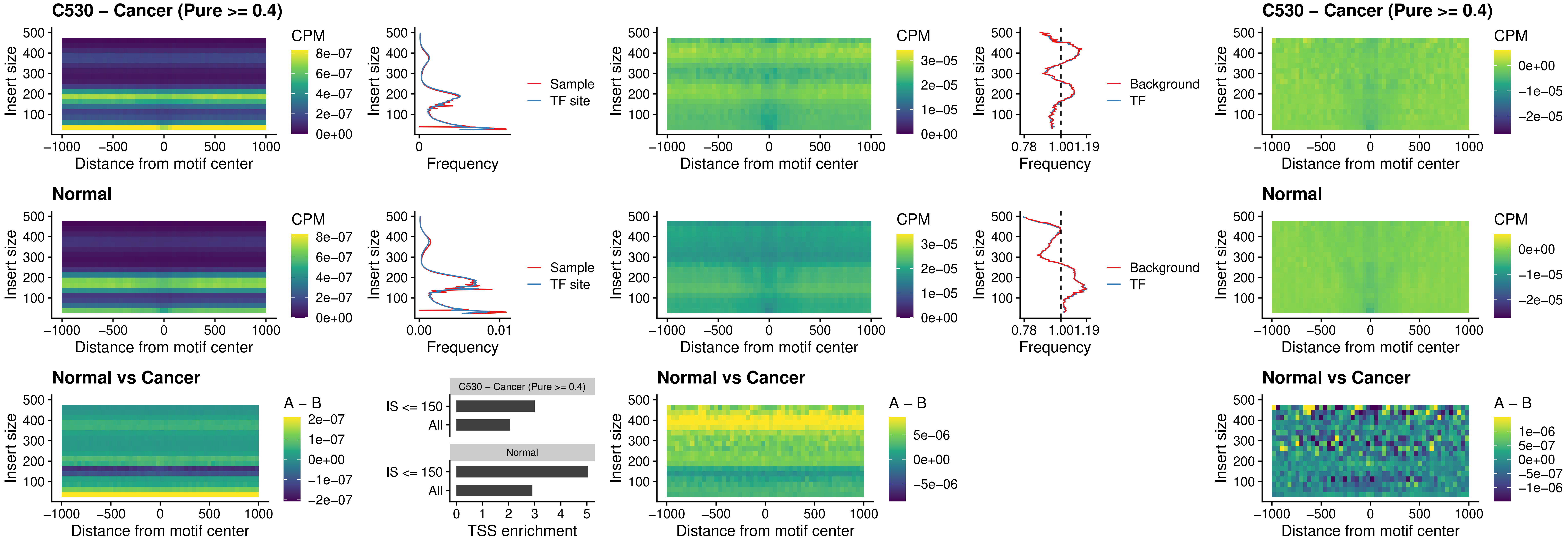

### C530 – Cancer (Pure >= 0.4) – SOX6_uc_dTSS_nPEAK

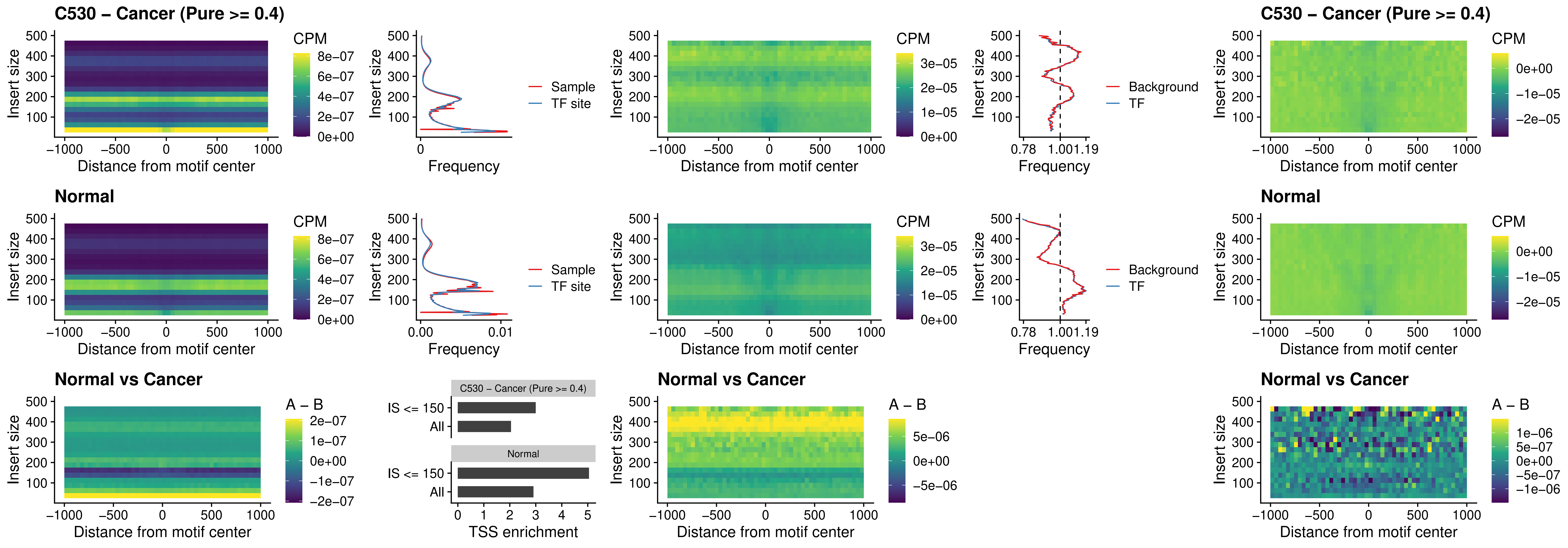

### C530 – Cancer (Pure >= 0.4) – JDP2_uc_cTSS_nPEAK

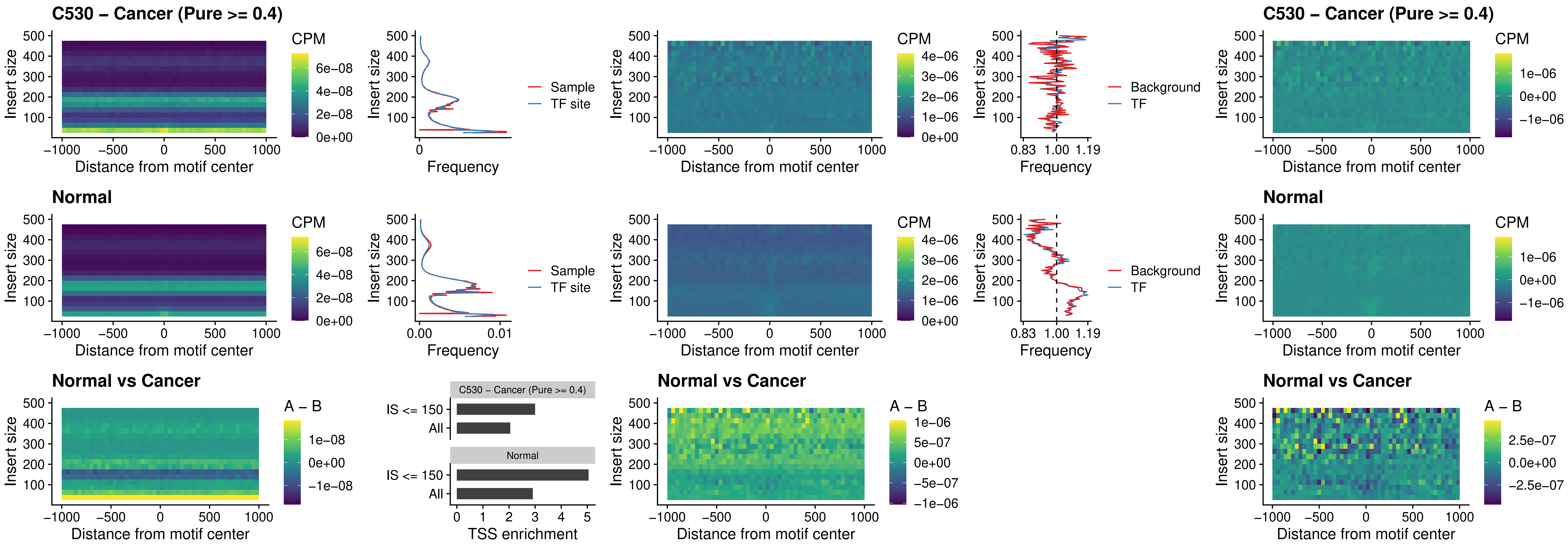

### C530 – Cancer (Pure >= 0.4) – PURA_uc_dTSS_oPEAK

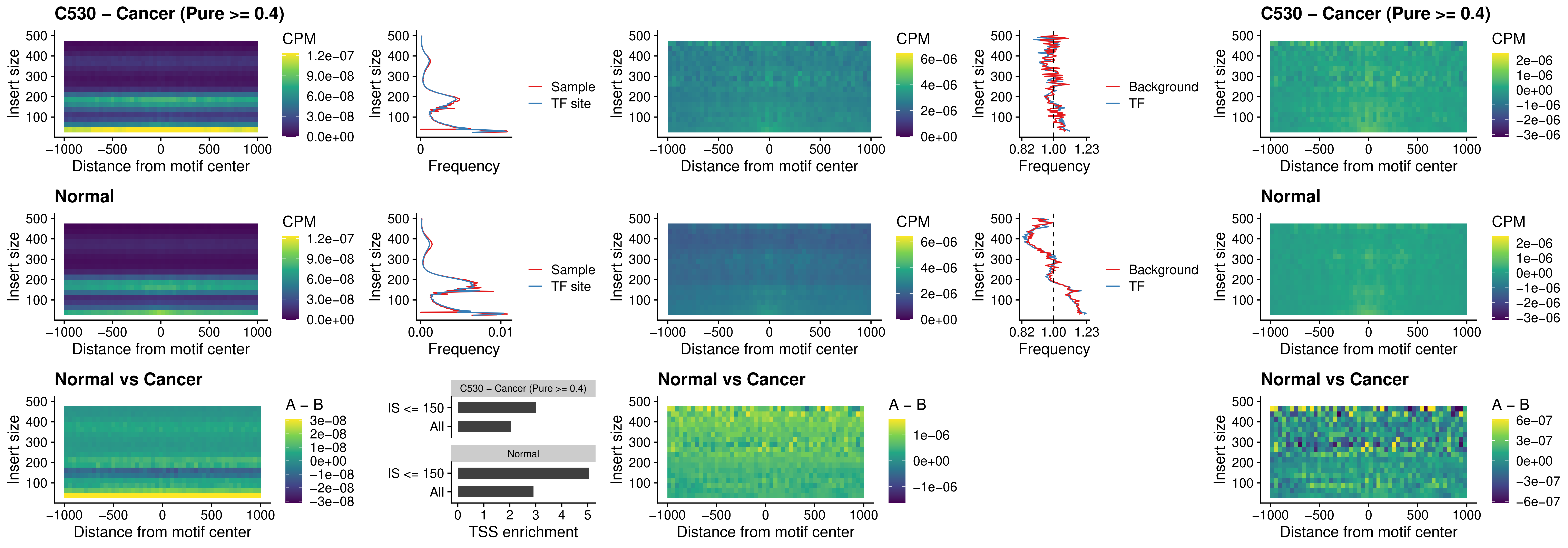

#### C530 – Cancer (Pure >= 0.4) – HHEX_uc_dTSS_nPEAK

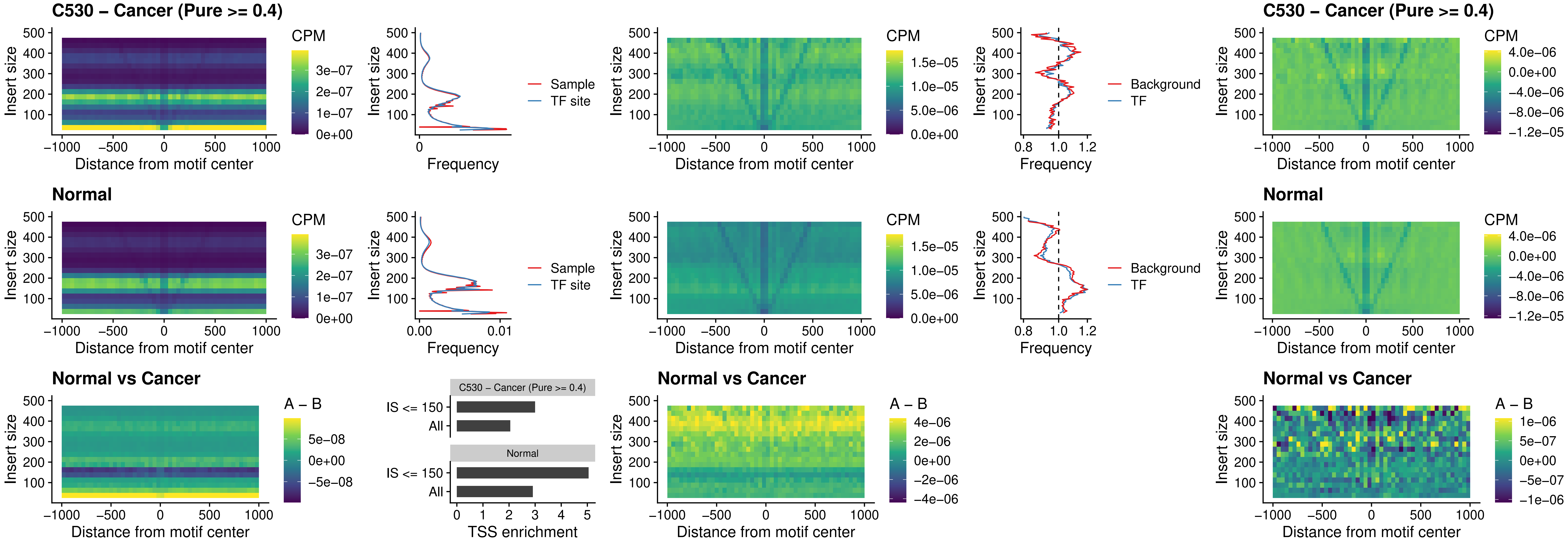

#### C530 – Cancer (Pure >= 0.4) – HMG20B_uc_dTSS_nPEAK

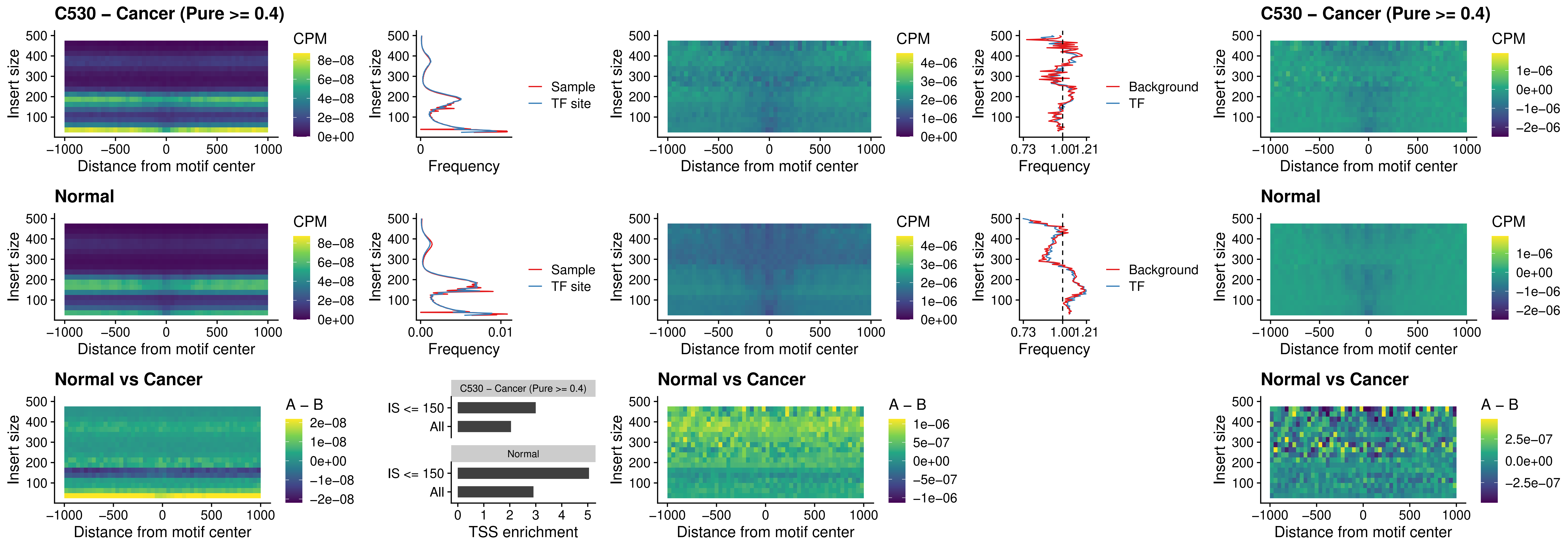

#### C530 – Cancer (Pure >= 0.4) – FOXB2_uc_dTSS_nPEAK

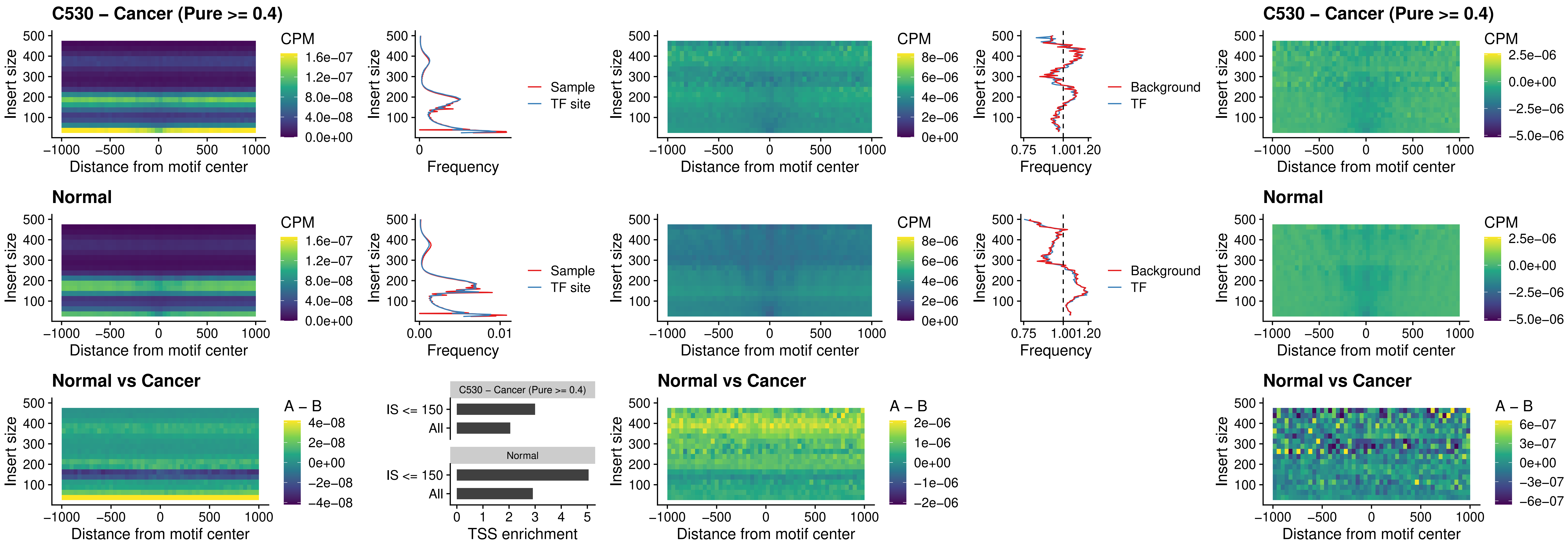

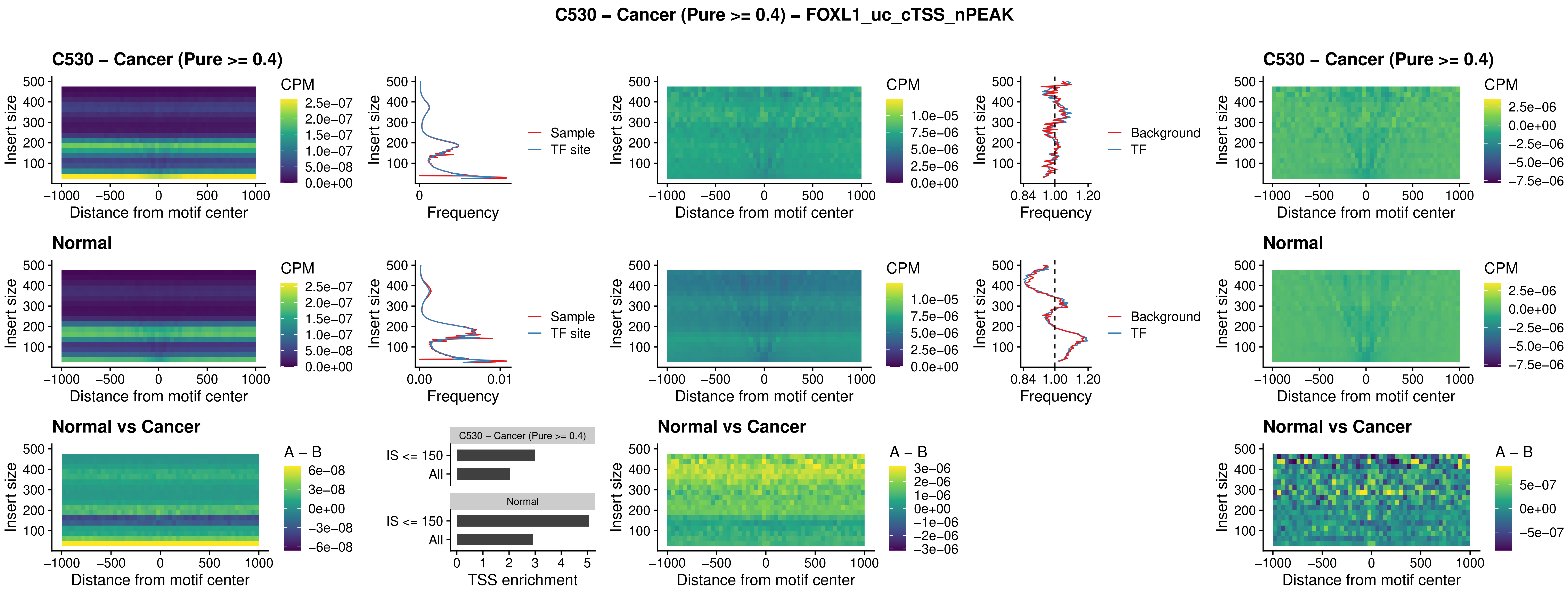
C530 – Cancer (Pure >= 0.4) – FOXL1_uc_cTSS_nPEAK
C530 – Cancer (Pure >= 0.4)
Normal
Normal vs Cancer
Insert size
Distance from motif center
Frequency
CPM
A – B
Sample
TF site
Background
TF
C530 – Cancer (Pure >= 0.4)
Normal
IS <= 150
All
TSS enrichment

#### C530 – Cancer (Pure >= 0.4) – PBX3_uc_pTSS_oPEAK

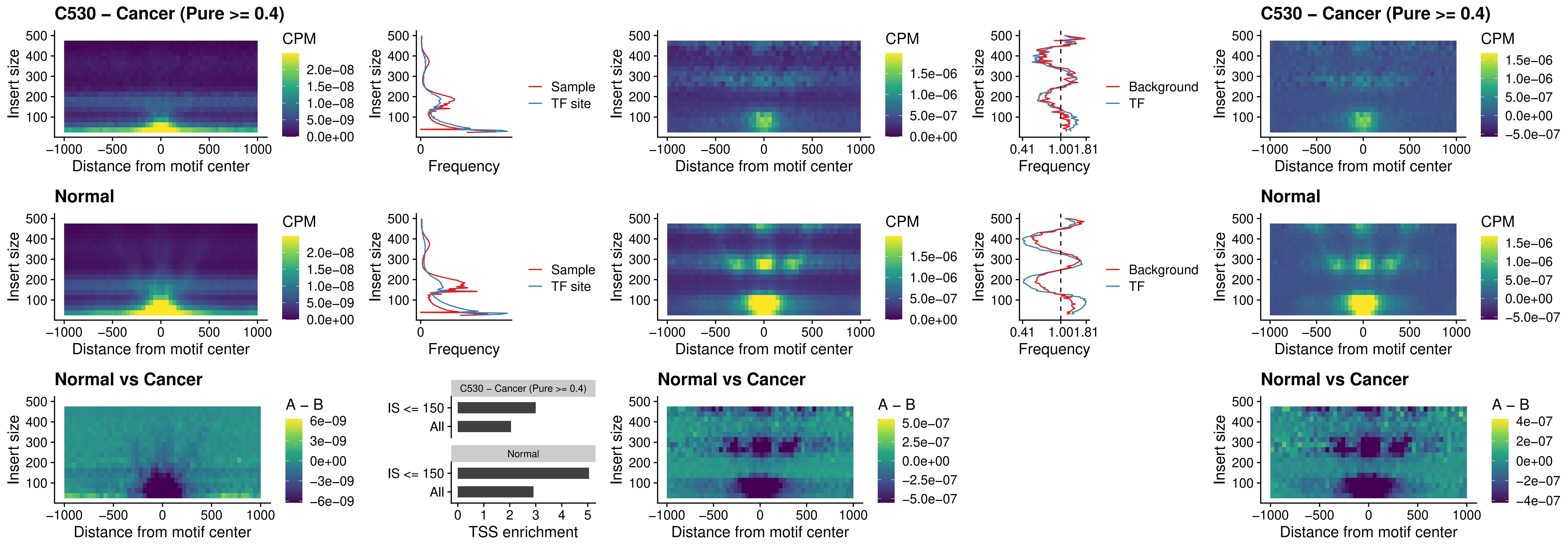

#### C530 – Cancer (Pure >= 0.4) – SNAI2_uc_cTSS_nPEAK

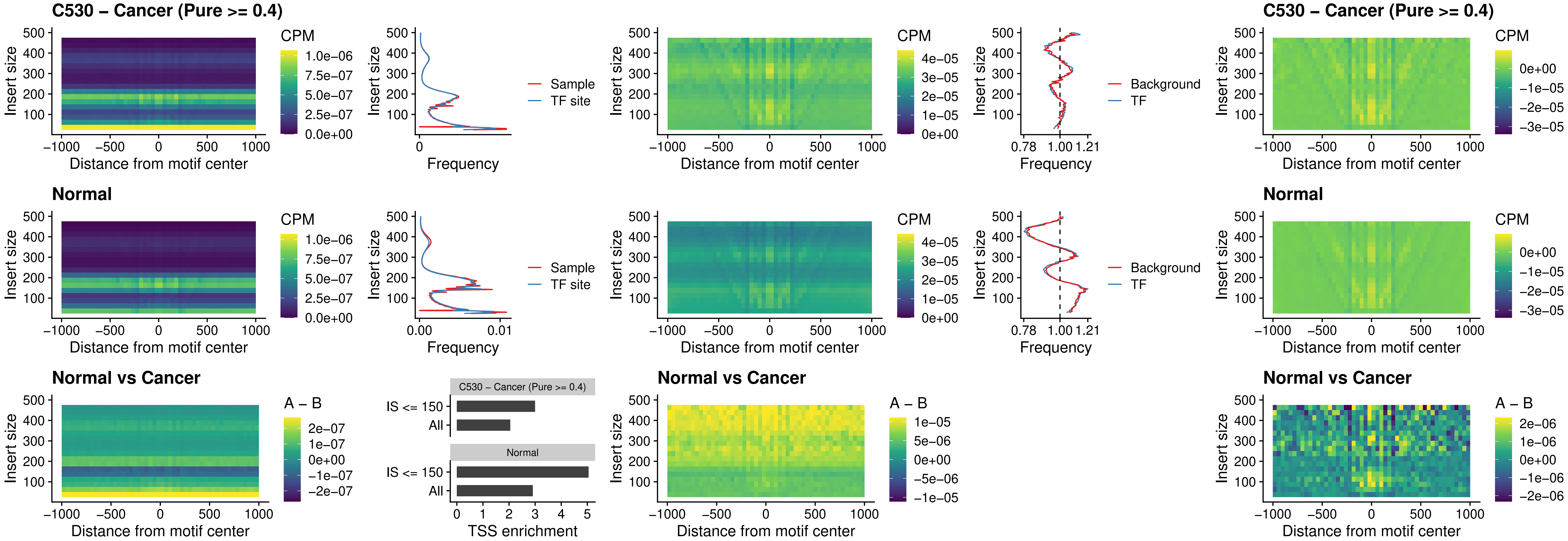

### C530 – Cancer (Pure >= 0.4) – SNAI2_uc_dTSS_oPEAK

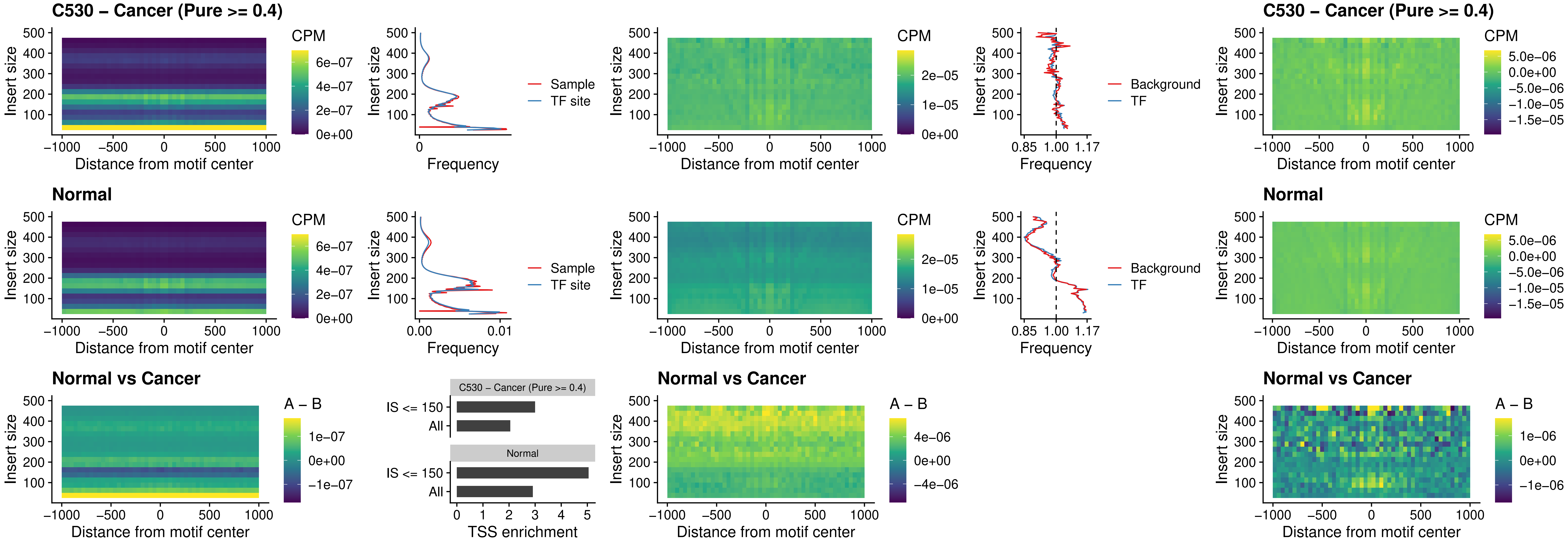

### C530 – Cancer (Pure >= 0.4) – SPI1_uc_pTSS_nPEAK

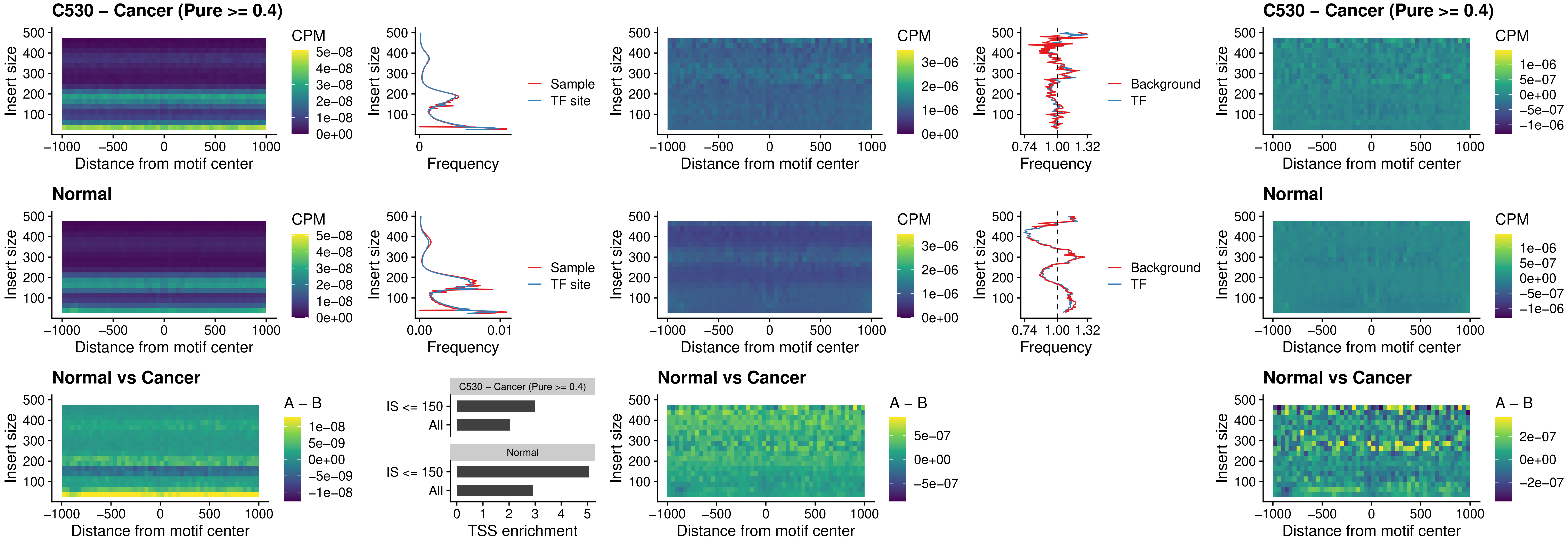

### C530 – Cancer (Pure >= 0.4) – KLF16_uc_pTSS_nPEAK

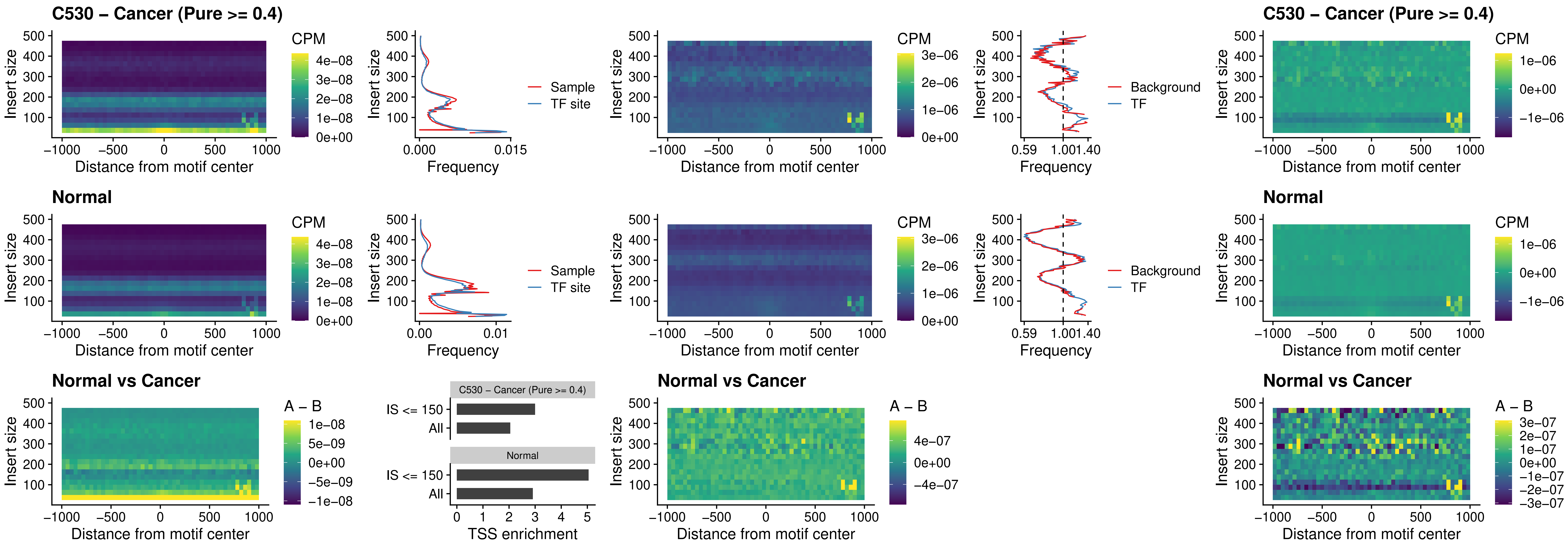

#### C530 – Cancer (Pure >= 0.4) – SP3_uc_pTSS_nPEAK

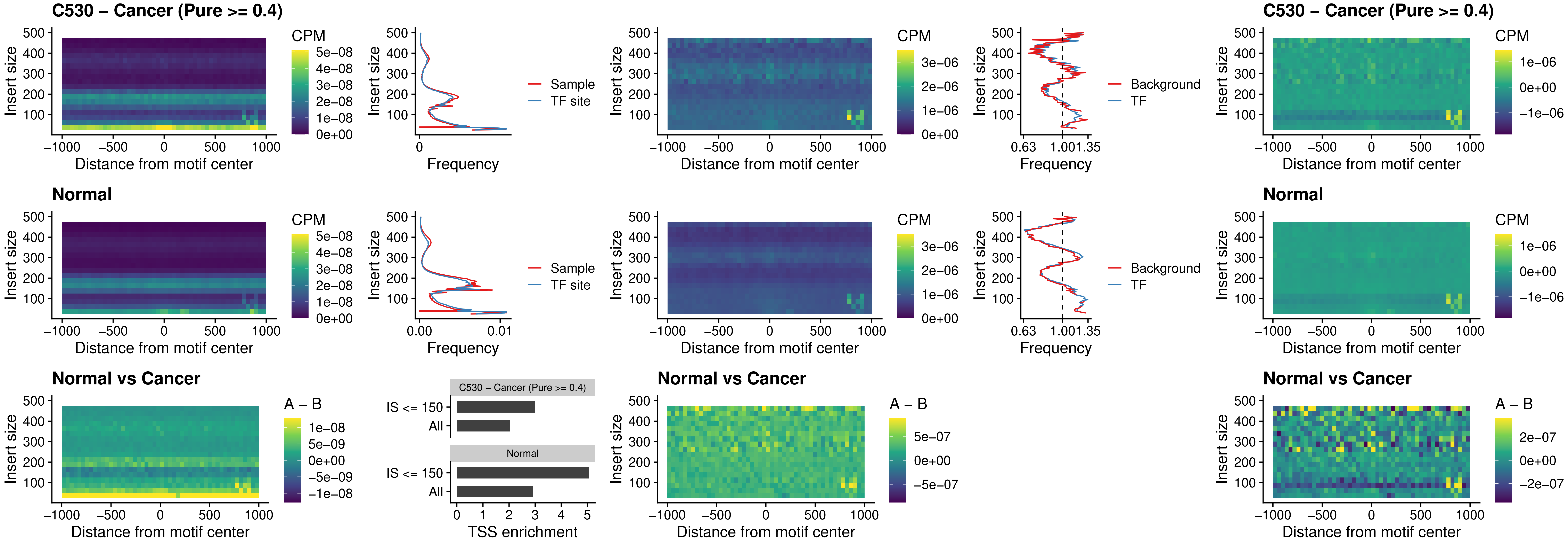

#### C530 – Cancer (Pure >= 0.4) – NEUROD1_uc_dTSS_oPEAK

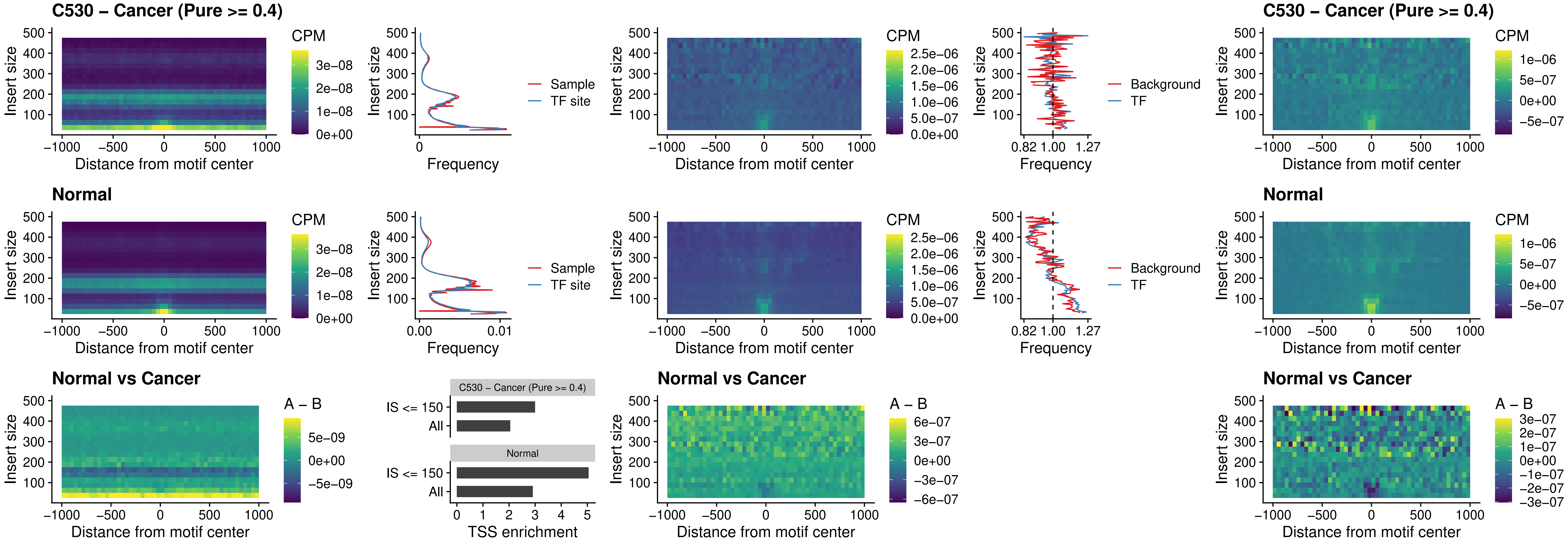

#### C530 – Cancer (Pure >= 0.4) – CTCF_uc_dTSS_oPEAK

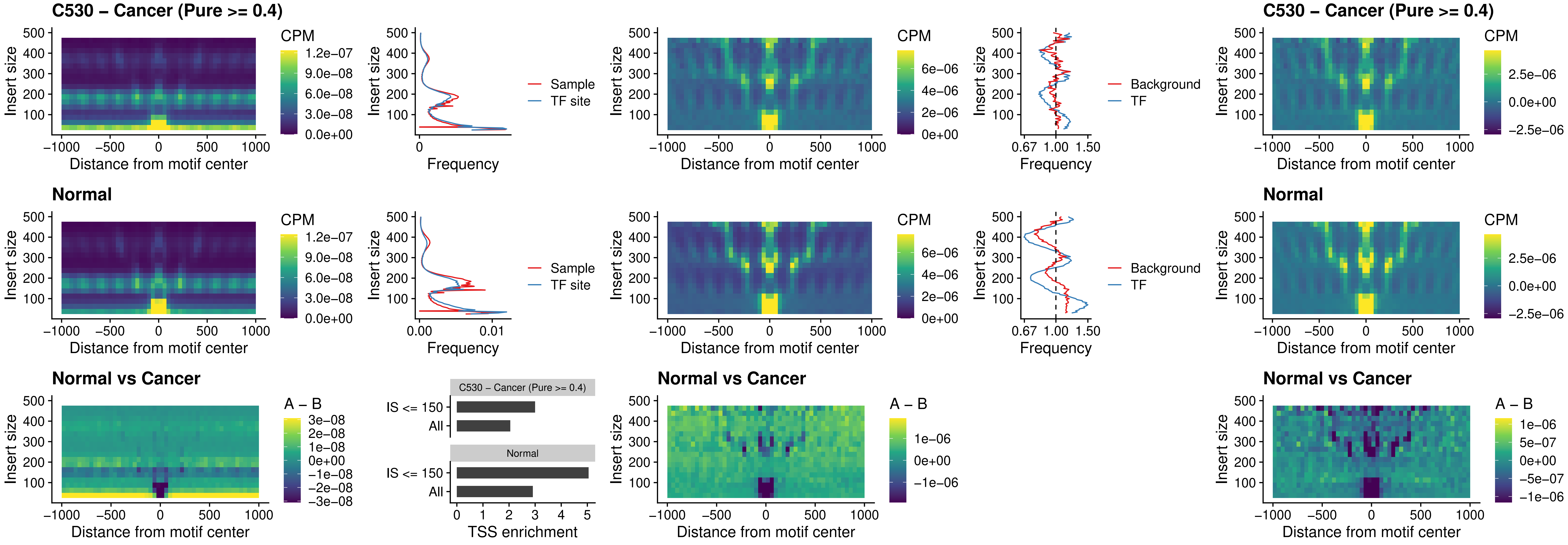

### C530 – Cancer (Pure >= 0.4) – CTCF_uc_cTSS_oPEAK

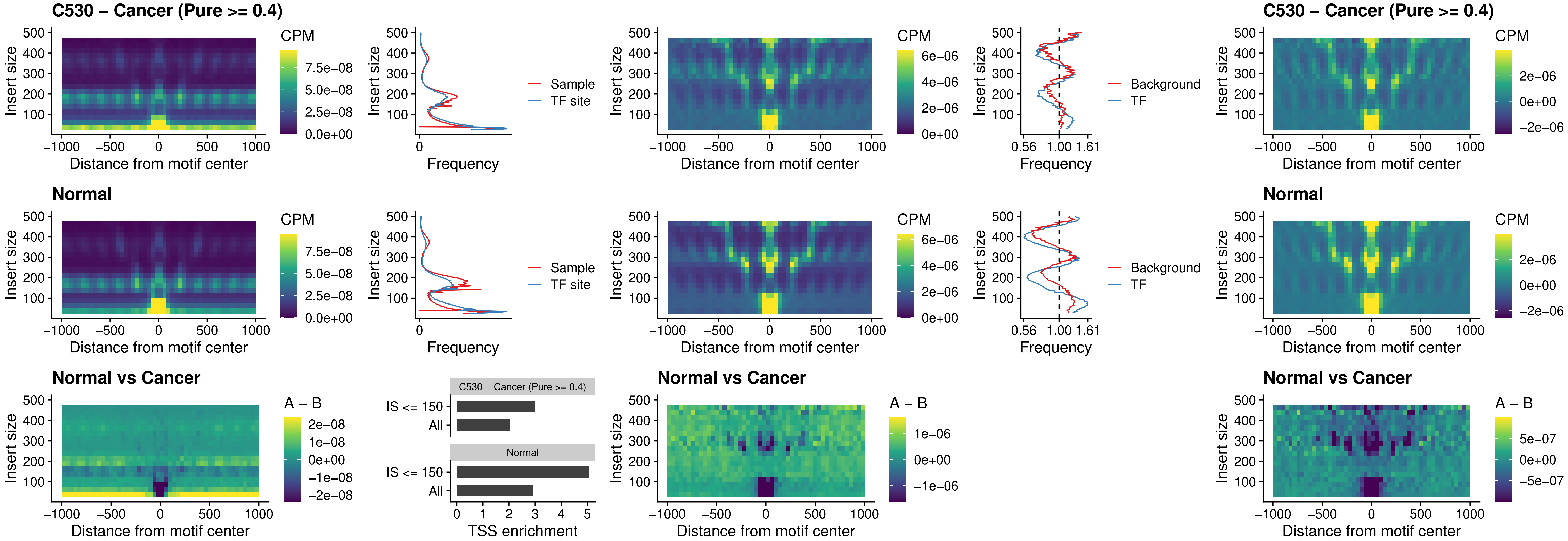

#### C530 – Cancer (Pure >= 0.4) – CTCFL_uc_dTSS_oPEAK

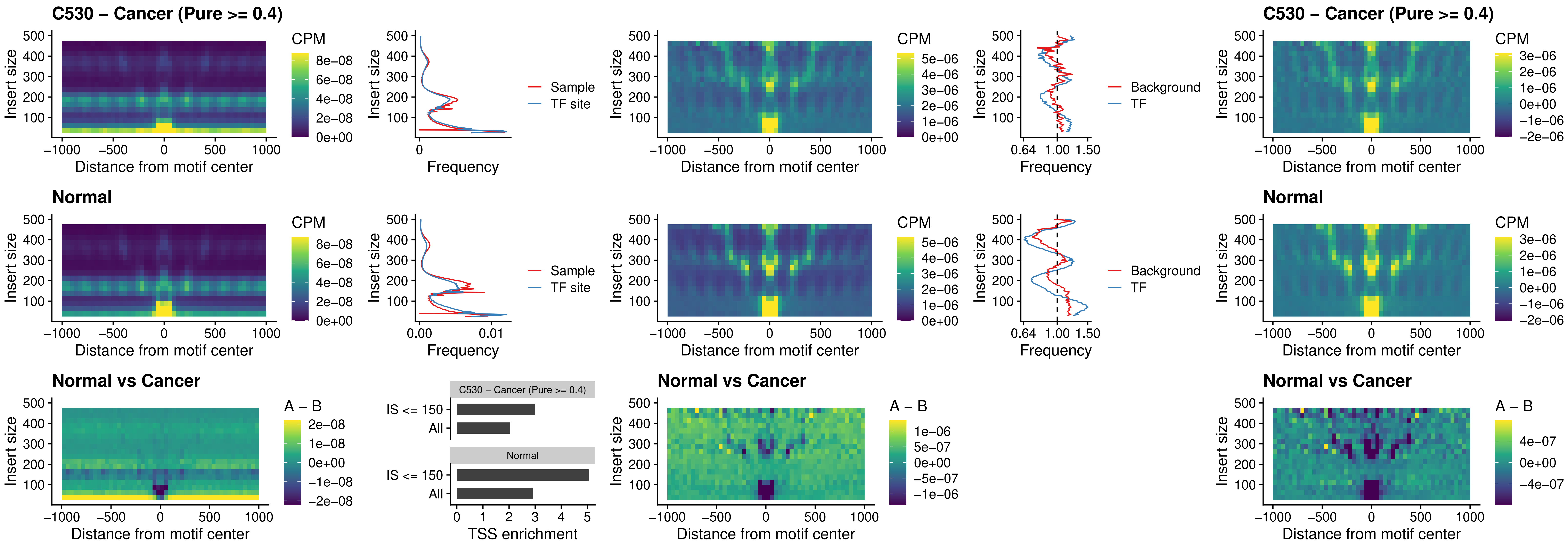

#### C530 – Cancer (Pure >= 0.4) – ENCODE_CTCF_sites_COLON

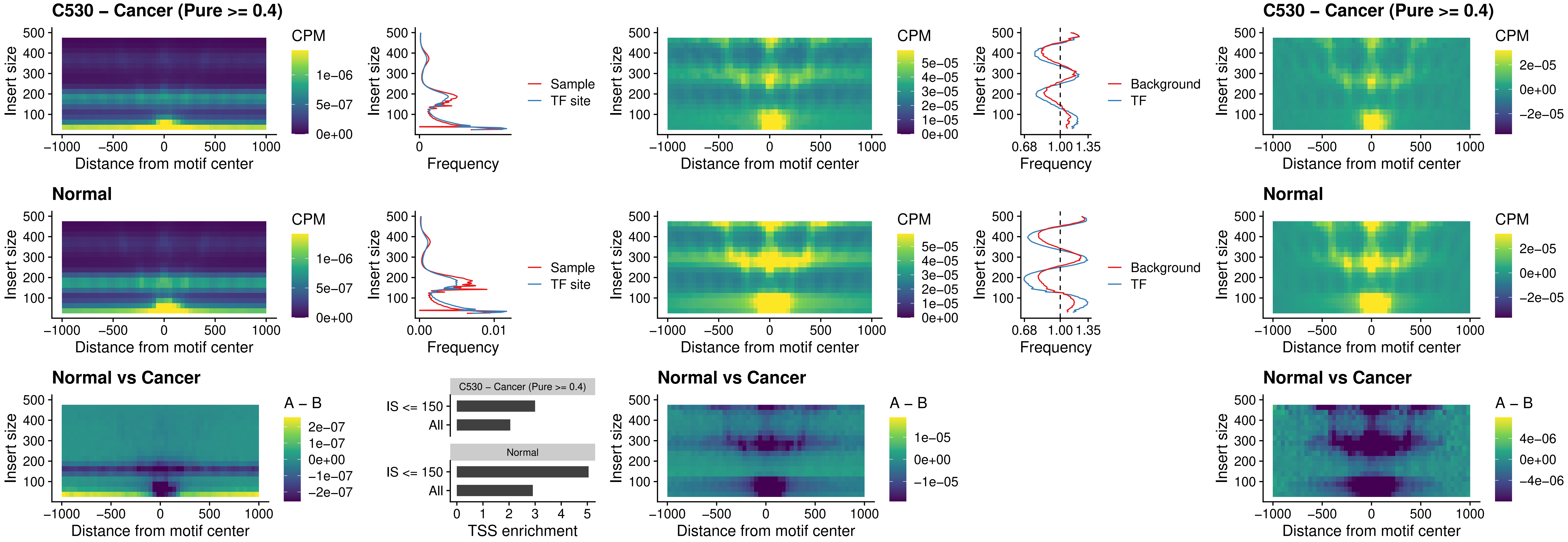

#### C530 – Cancer (Pure >= 0.4) – ENCODE_CTCF_sites_COLON_loop_ctcf

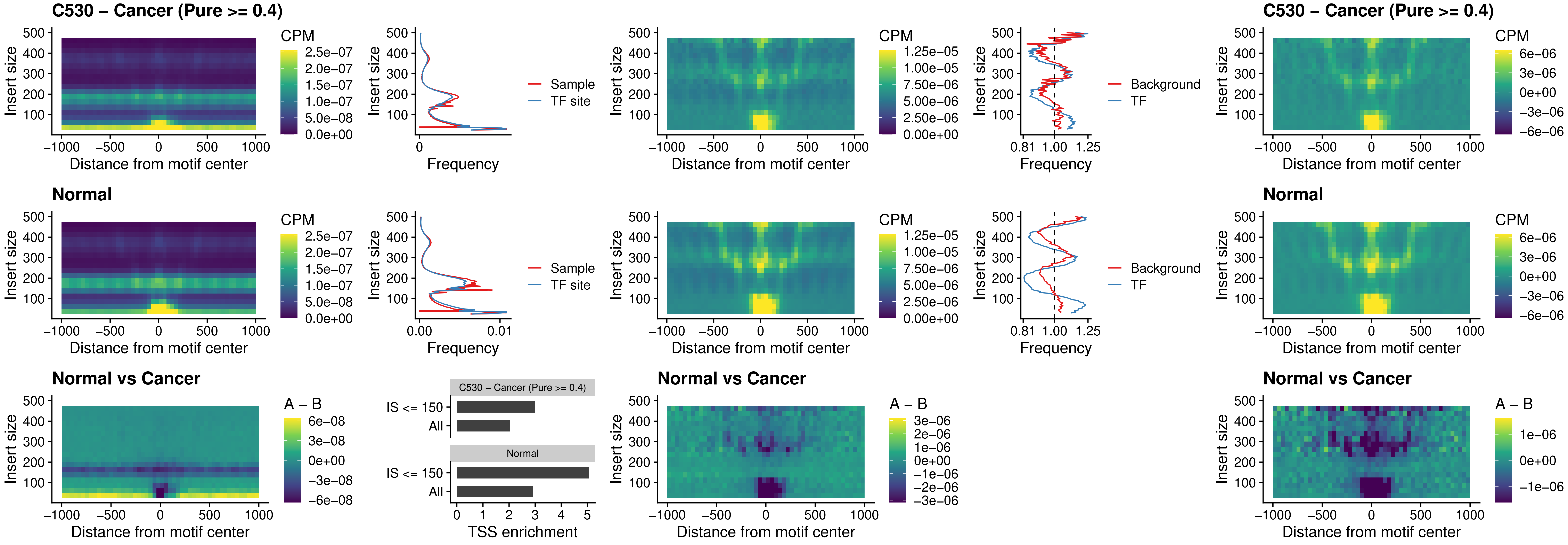

#### C530 – Cancer (Pure >= 0.4) – ENCODE_CTCF_sites_COLON_intergenic_ctcf

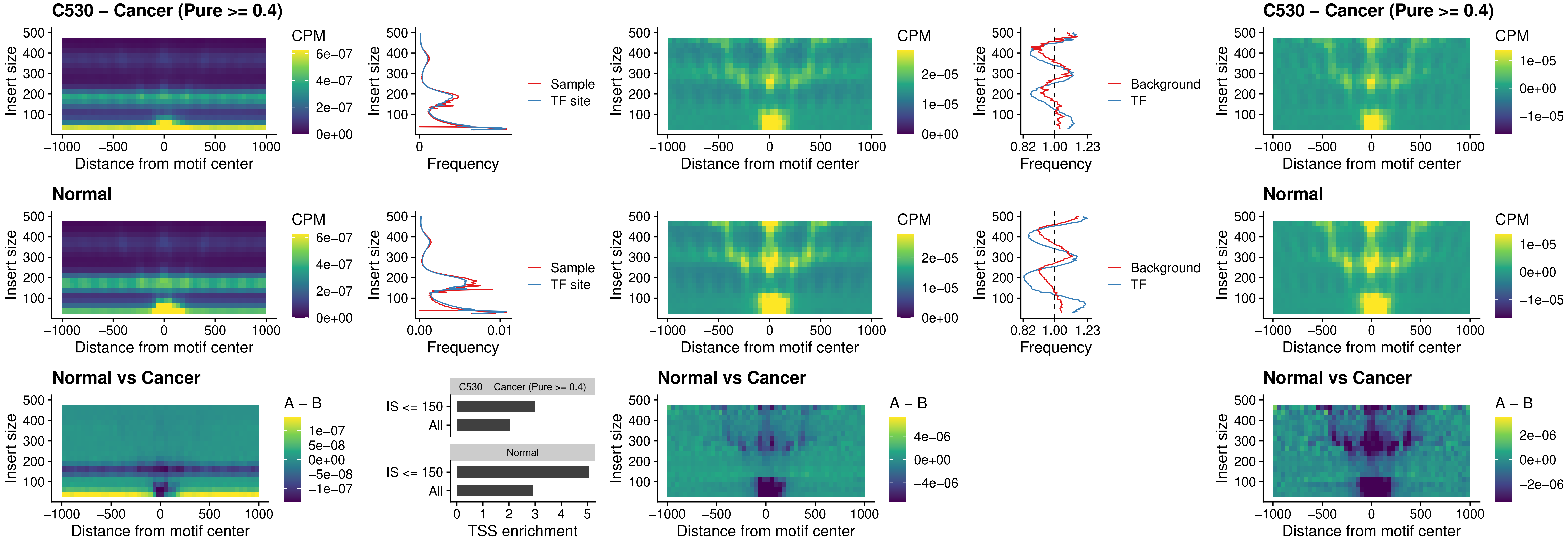

#### C530 – Cancer (Pure >= 0.4) – ENCODE_CTCF_sites_COLON_nonpromotor

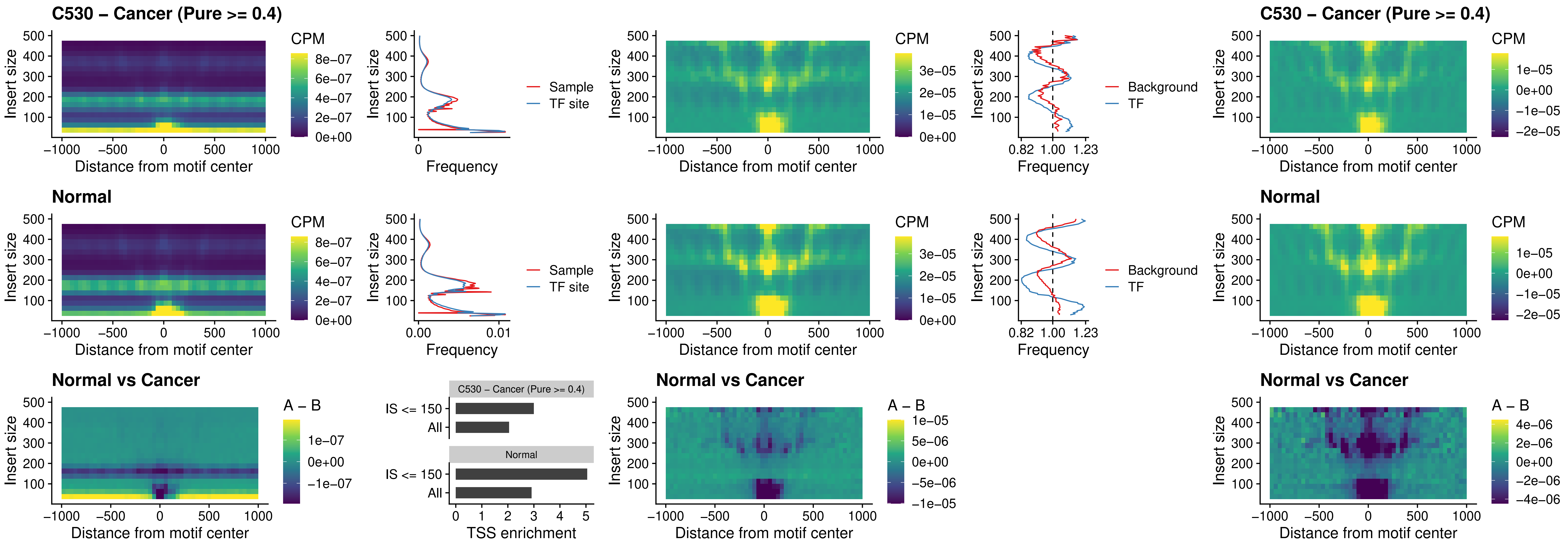

#### C530 – Cancer (Pure >= 0.4) – CTCFL_uc_cTSS_oPEAK

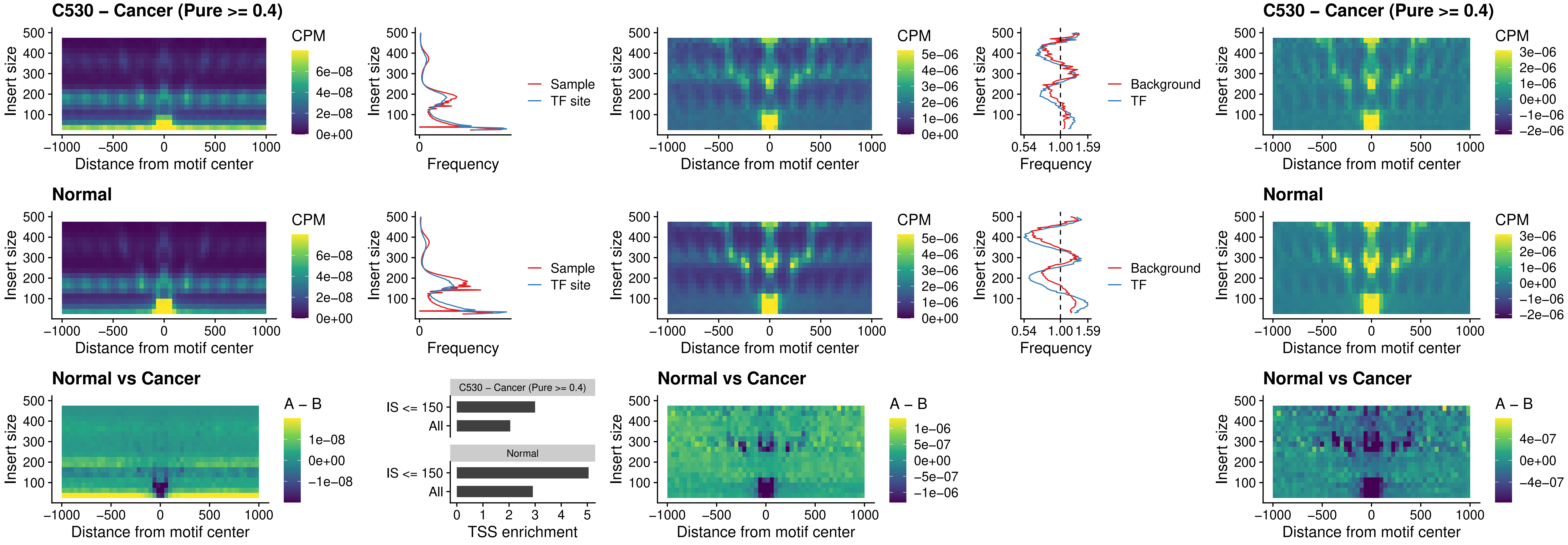

#### C530 – Cancer (Pure >= 0.4) – NEUROD1_uc_cTSS_oPEAK

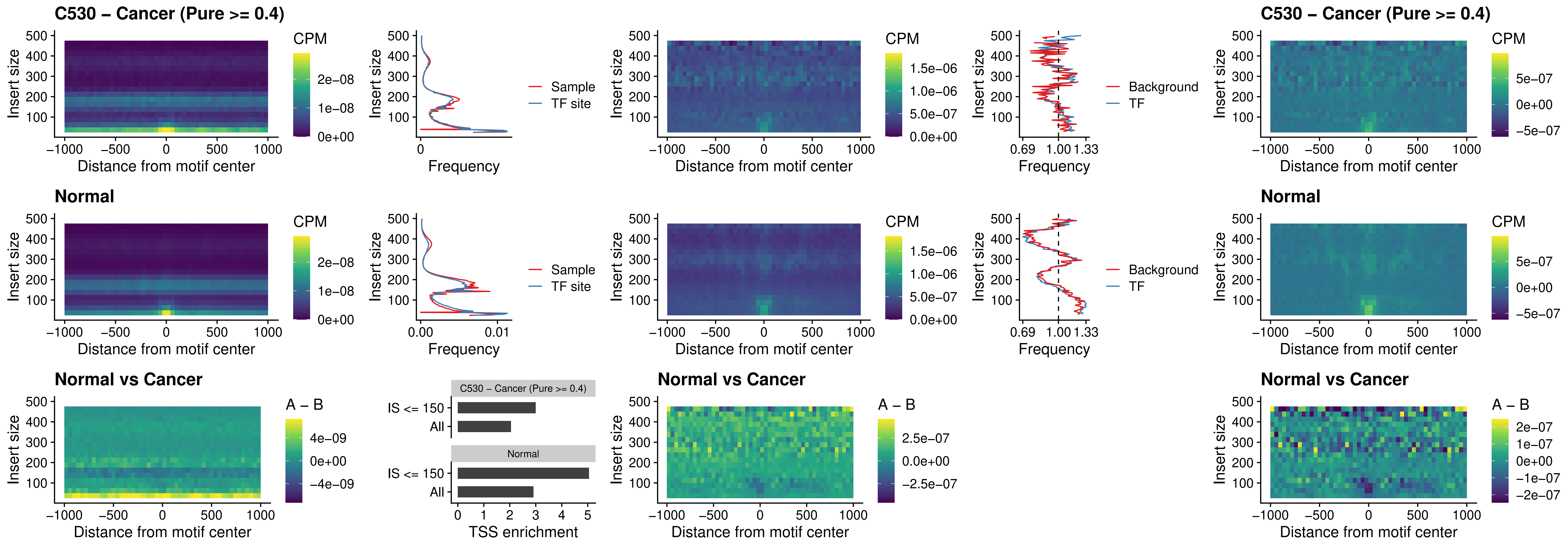

### C530 – Cancer (Pure >= 0.4) – ENCODE_YY1_sites_COLON_nonpromotor

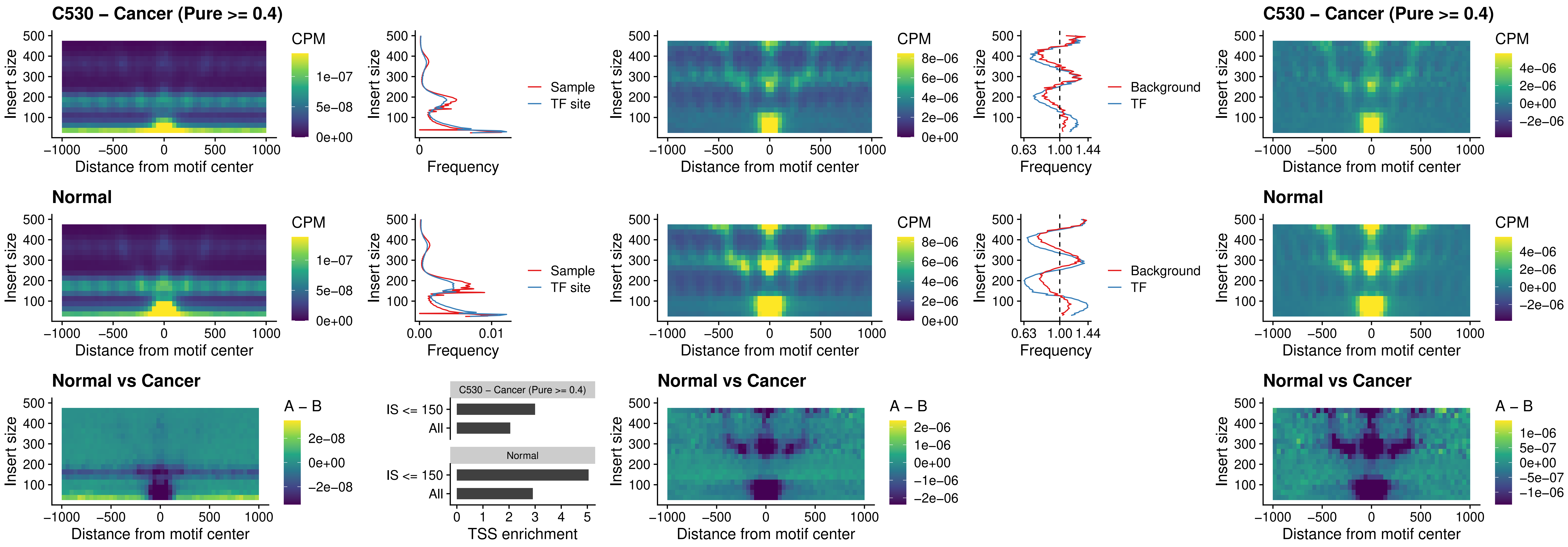

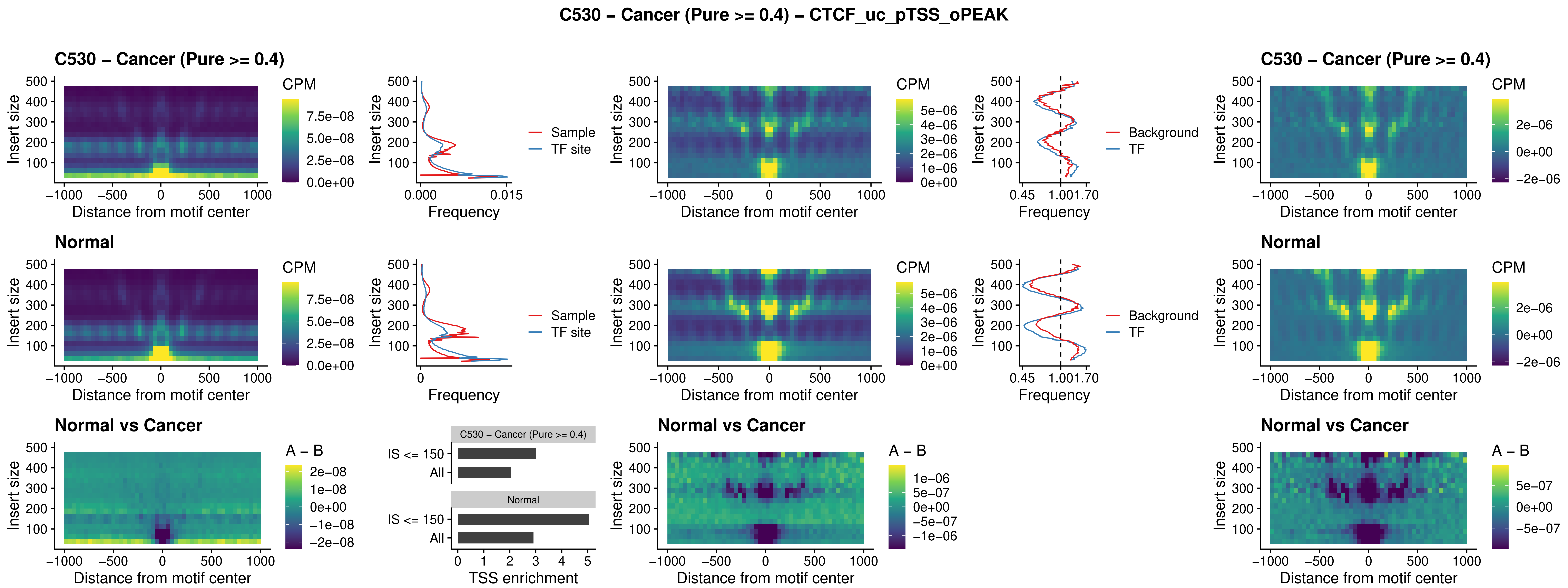
C530 – Cancer (Pure >= 0.4) – CTCF_uc_pTSS_oPEAK
C530 – Cancer (Pure >= 0.4)
Normal
Normal vs Cancer
Insert size
Distance from motif center
Frequency
CPM
A – B
Sample
TF site
Background
TF
C530 – Cancer (Pure >= 0.4)
IS <= 150
All
TSS enrichment

#### C530 – Cancer (Pure >= 0.4) – ENCODE_CTCF_sites_COLON_promotor

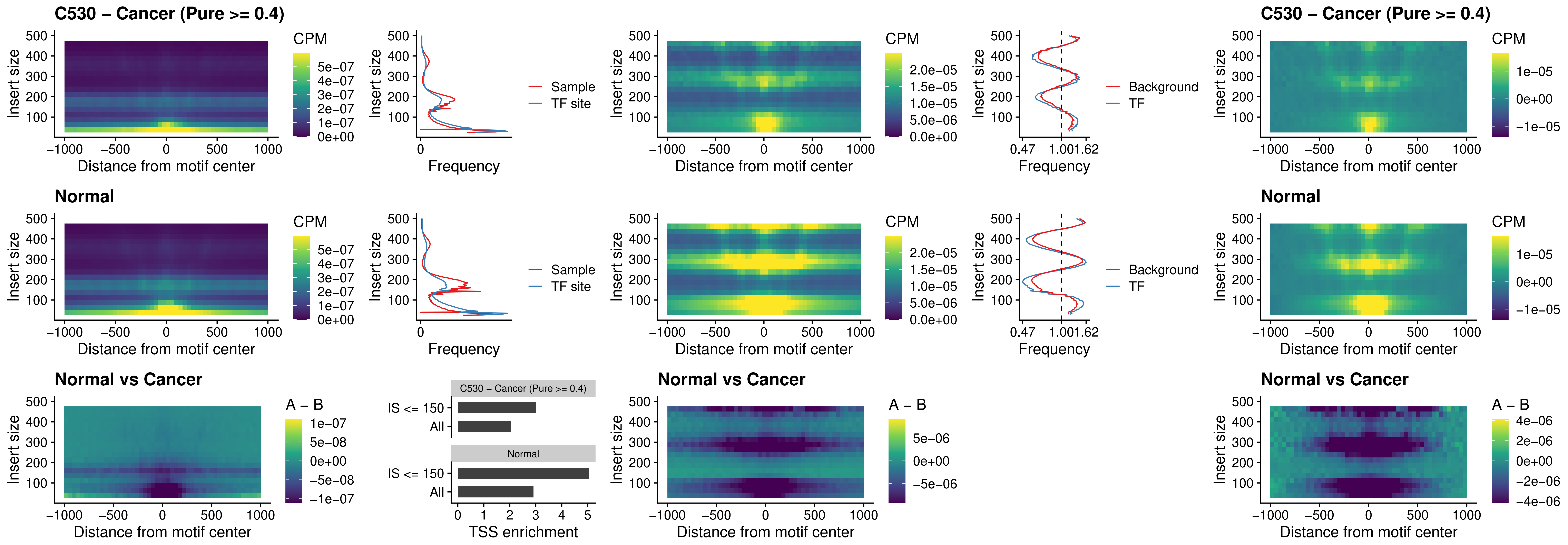

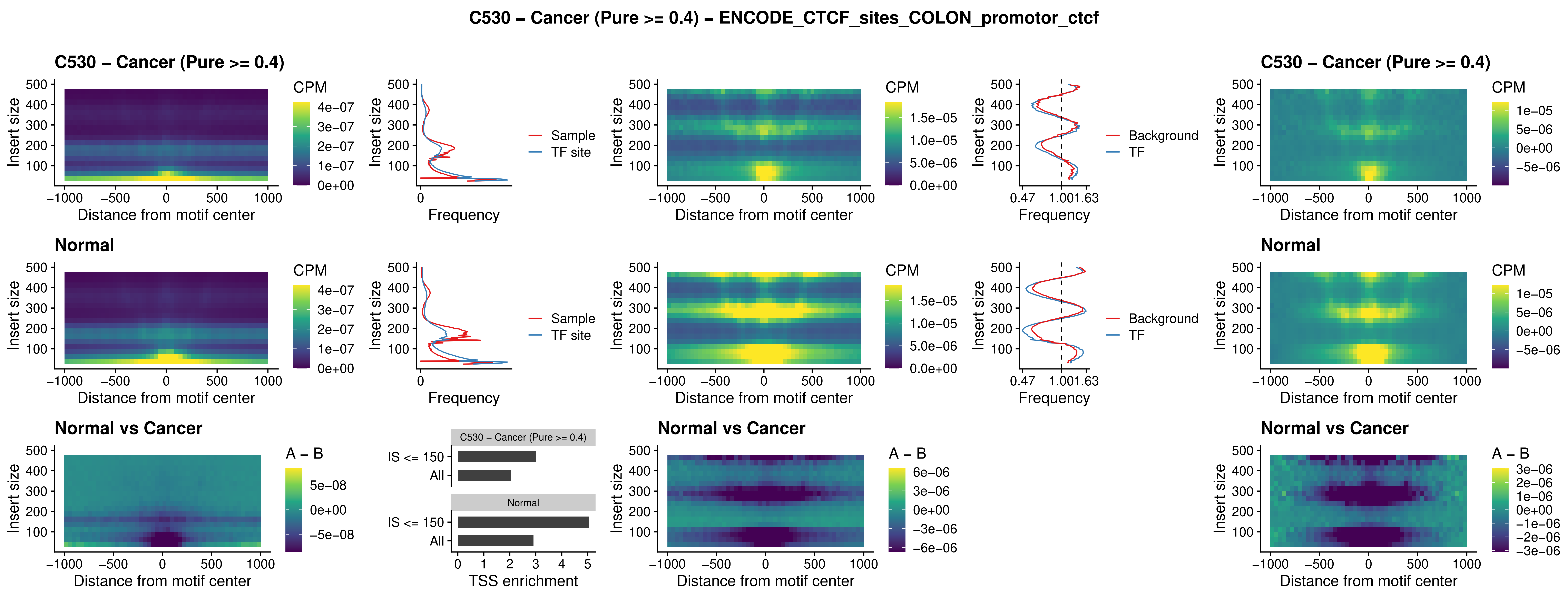
C530 – Cancer (Pure >= 0.4) – ENCODE_CTCF_sites_COLON_promotor_ctcf
C530 – Cancer (Pure >= 0.4)
Normal
Normal vs Cancer
Insert size
Distance from motif center
Frequency
CPM
A – B
Sample
TF site
Background
TF
TSS enrichment
IS <= 150
All
C530 – Cancer (Pure >= 0.4)
Normal

#### C530 – Cancer (Pure >= 0.4) – SPIC_uc_pTSS_oPEAK

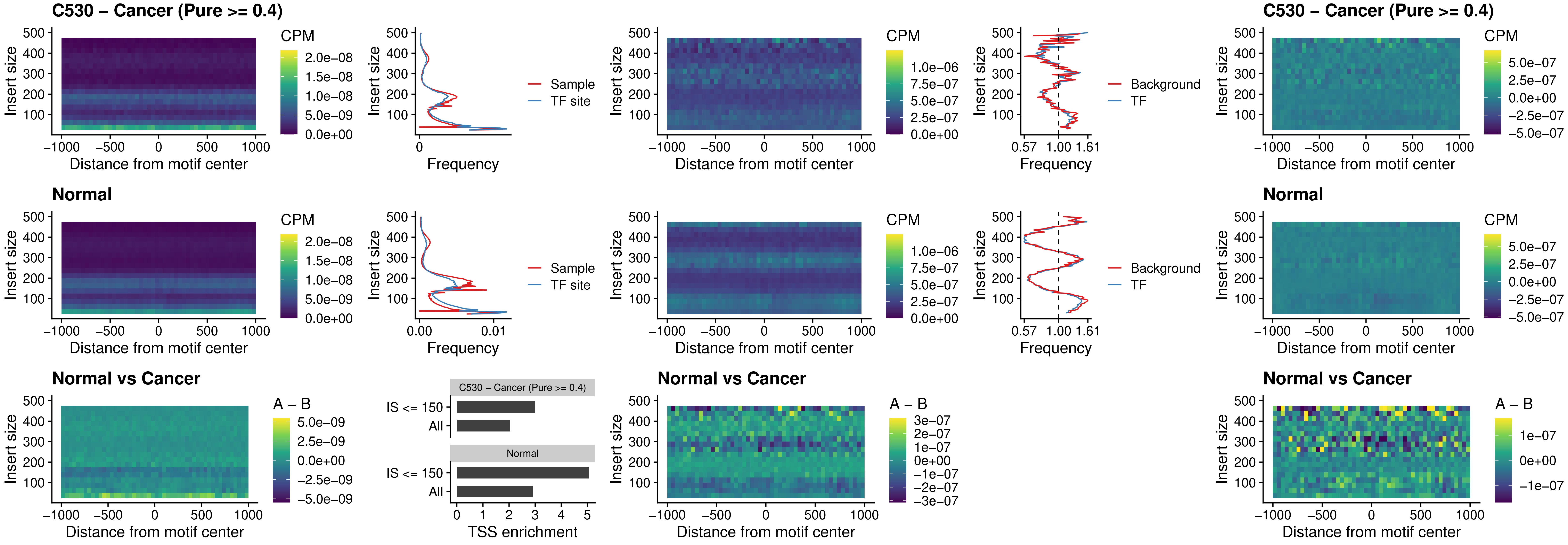

#### C530 – Cancer (Pure >= 0.4) – SPI1_uc_dTSS_nPEAK

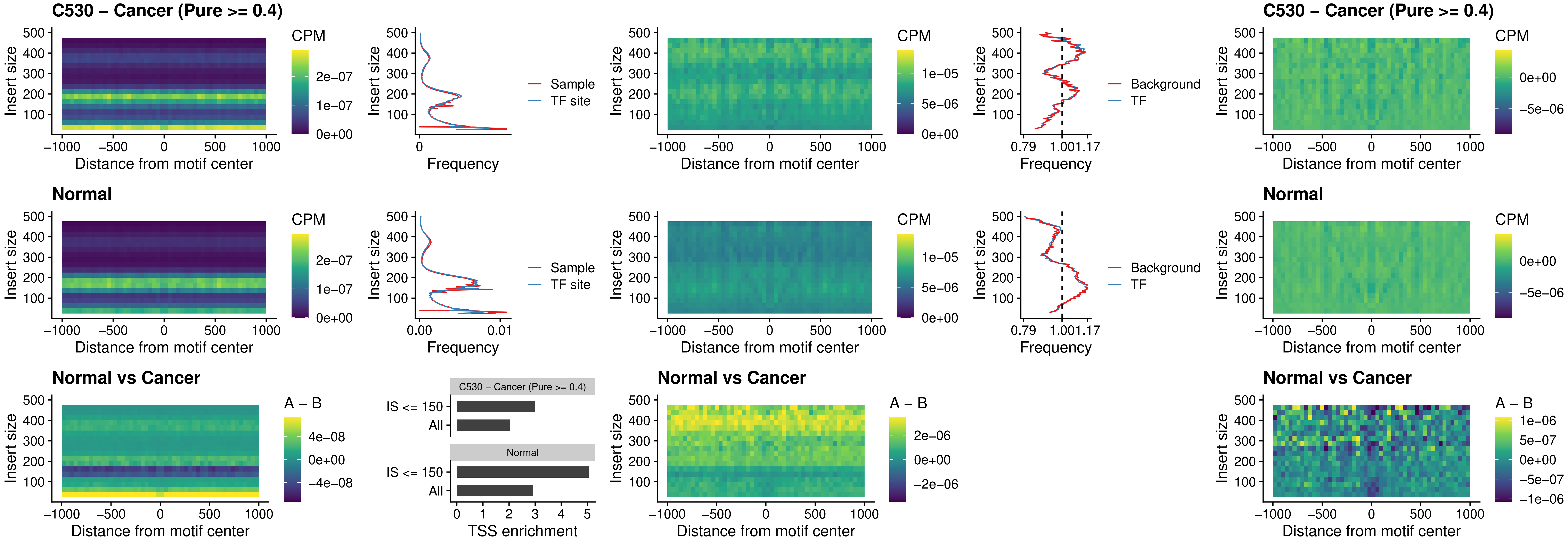

#### C530 – Cancer (Pure >= 0.4) – SPI1_uc_cTSS_nPEAK

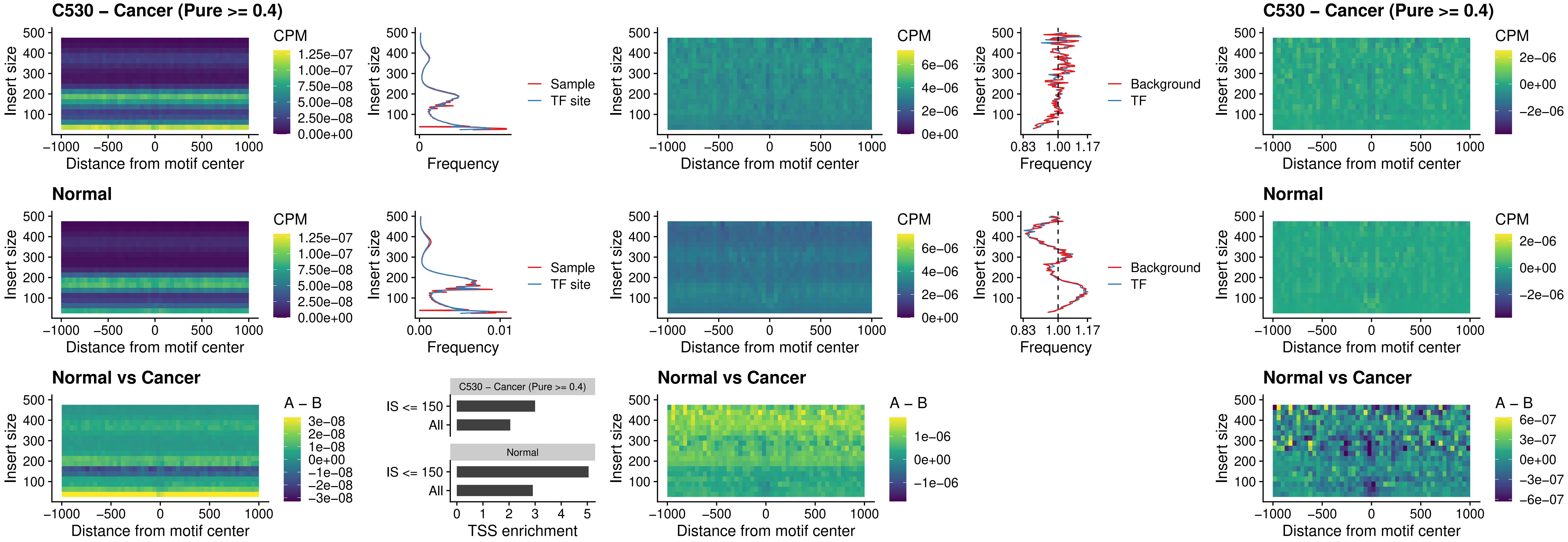

#### C530 – Cancer (Pure >= 0.4) – BCL11A_uc_pTSS_nPEAK

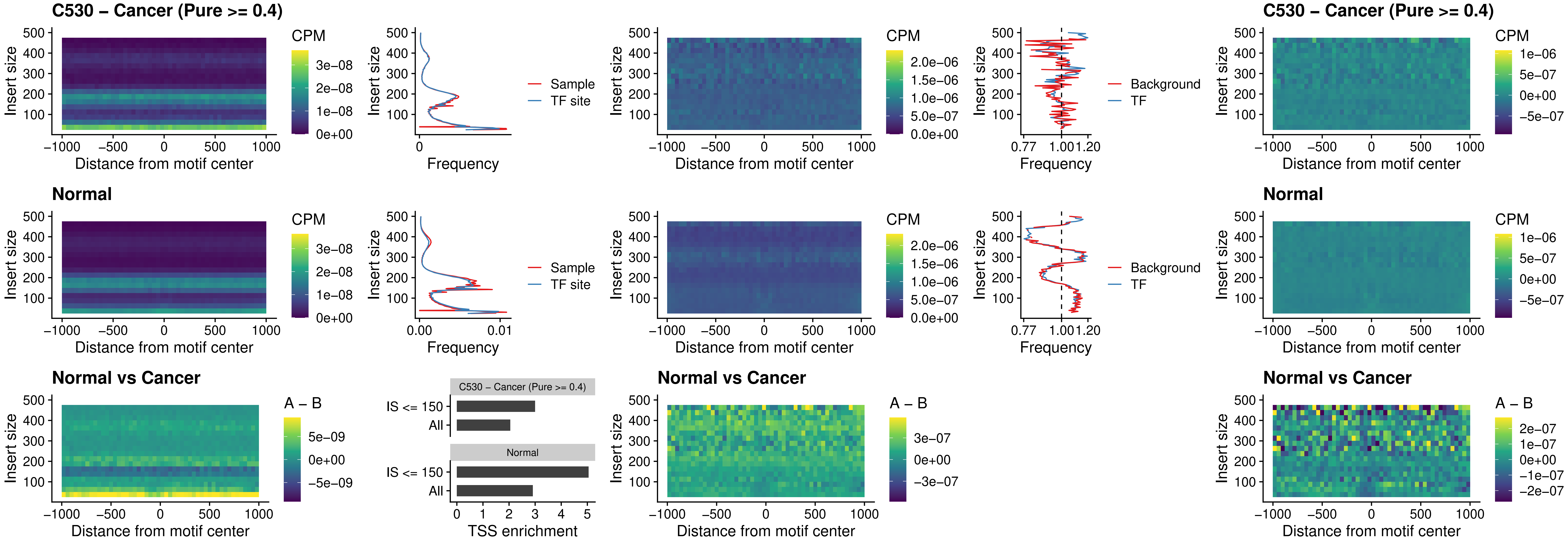

#### C530 – Cancer (Pure >= 0.4) – BCL11B_uc_pTSS_nPEAK

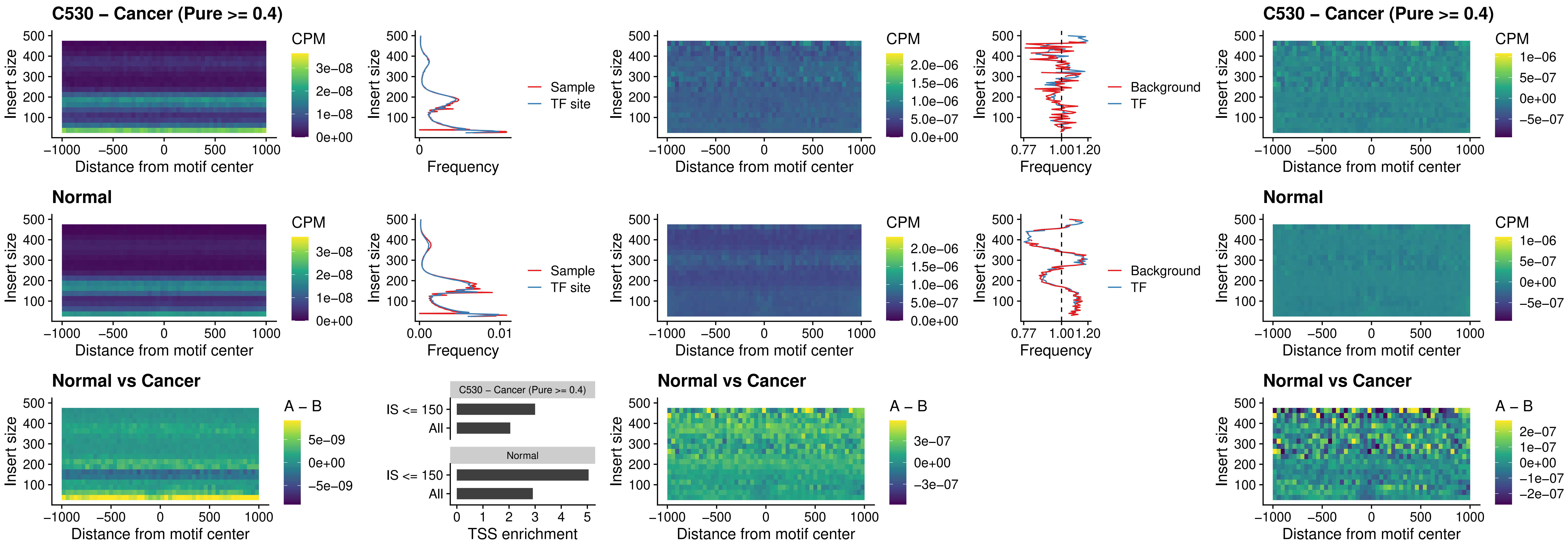

#### C530 – Cancer (Pure >= 0.4) – IRF9_uc_dTSS_oPEAK

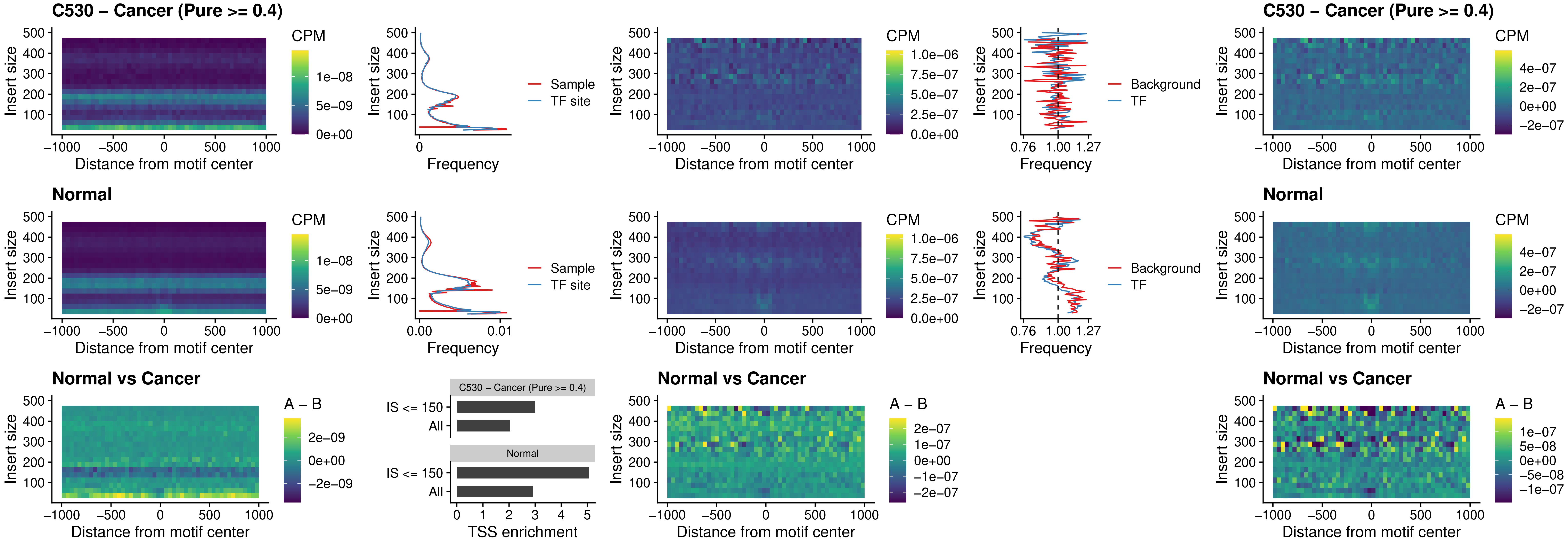

#### C530 – Cancer (Pure >= 0.4) – IRF9_uc_cTSS_oPEAK

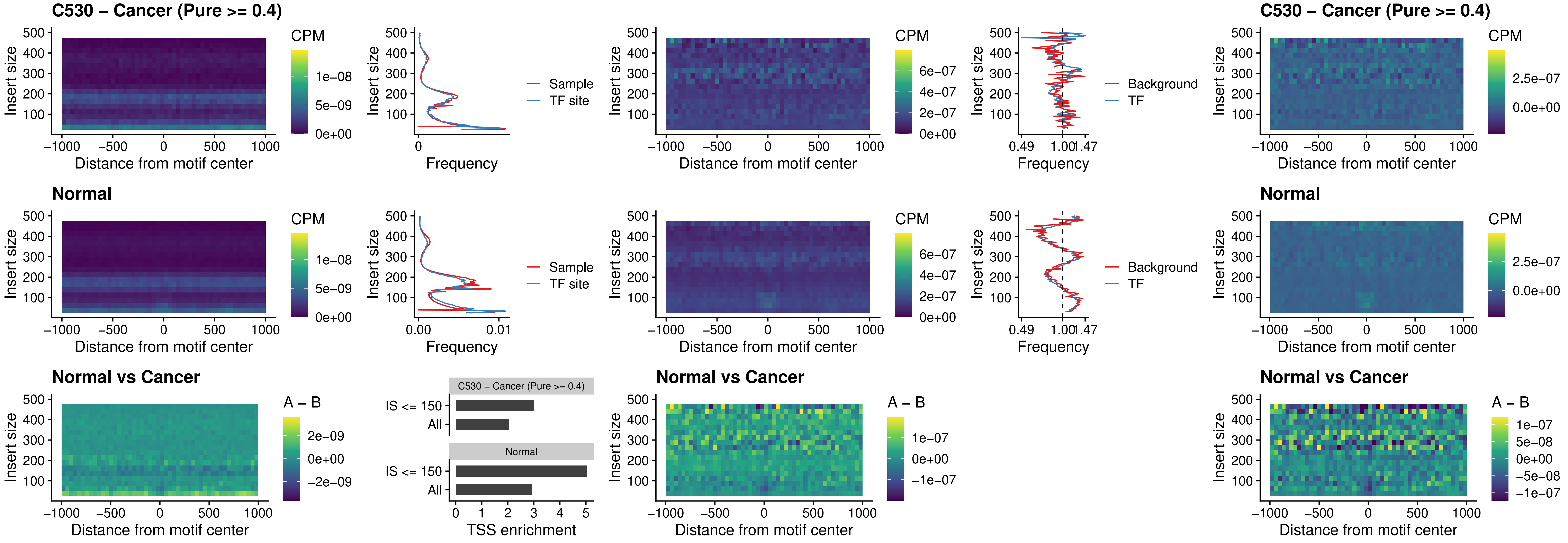

#### C530 – Cancer (Pure >= 0.4) – BCL11A_uc_cTSS_oPEAK

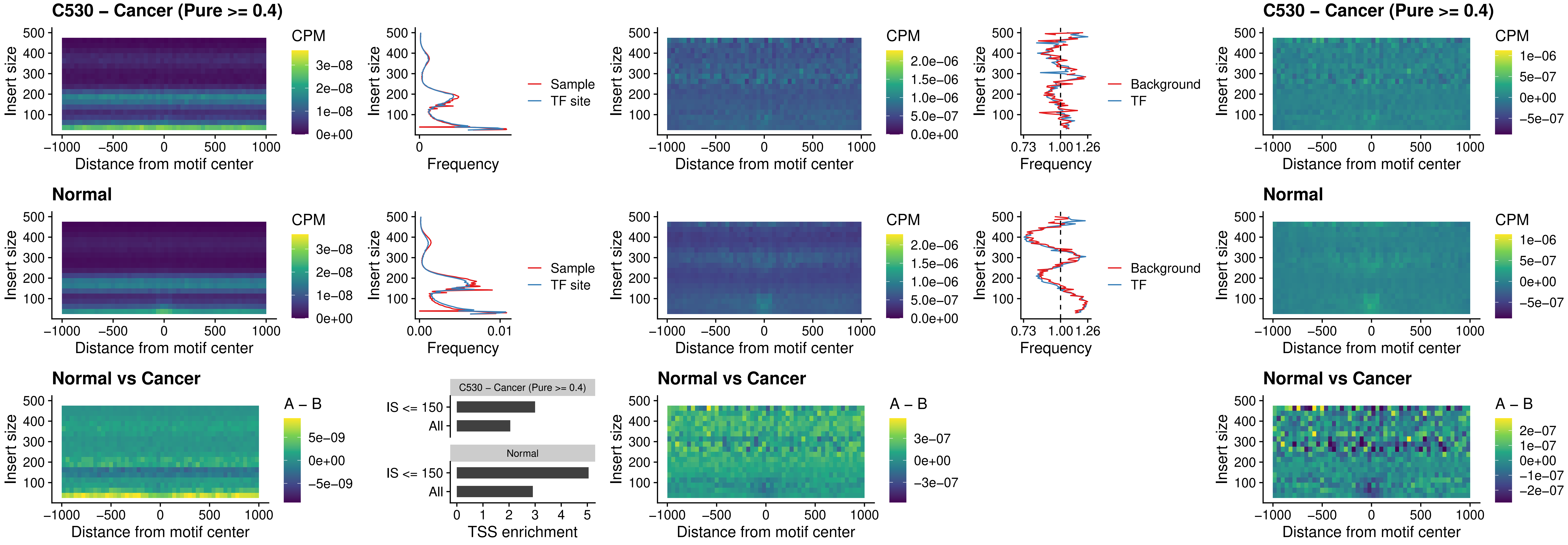

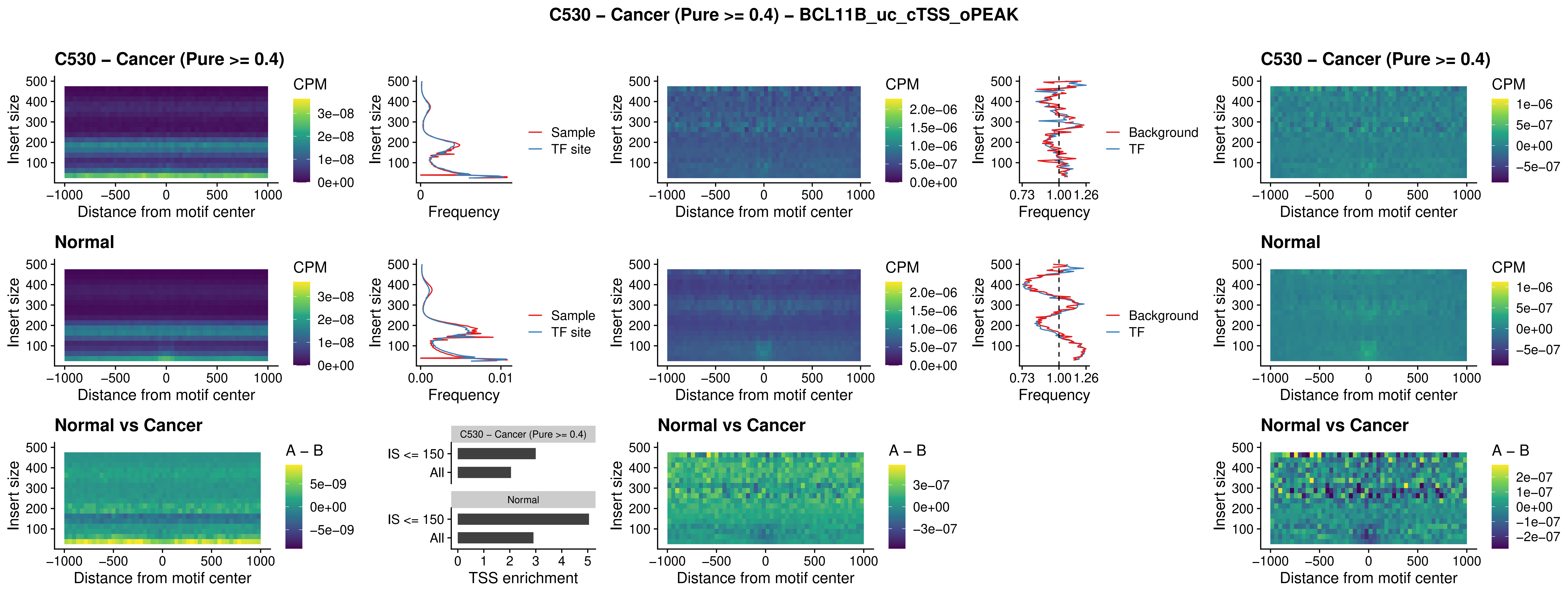
C530 – Cancer (Pure >= 0.4) – BCL11B_uc_cTSS_oPEAK
C530 – Cancer (Pure >= 0.4)
Normal
Normal vs Cancer
Insert size
Distance from motif center
Frequency
CPM
A – B
Sample
TF site
Background
TF
C530 – Cancer (Pure >= 0.4)
Normal
IS <= 150
All
TSS enrichment

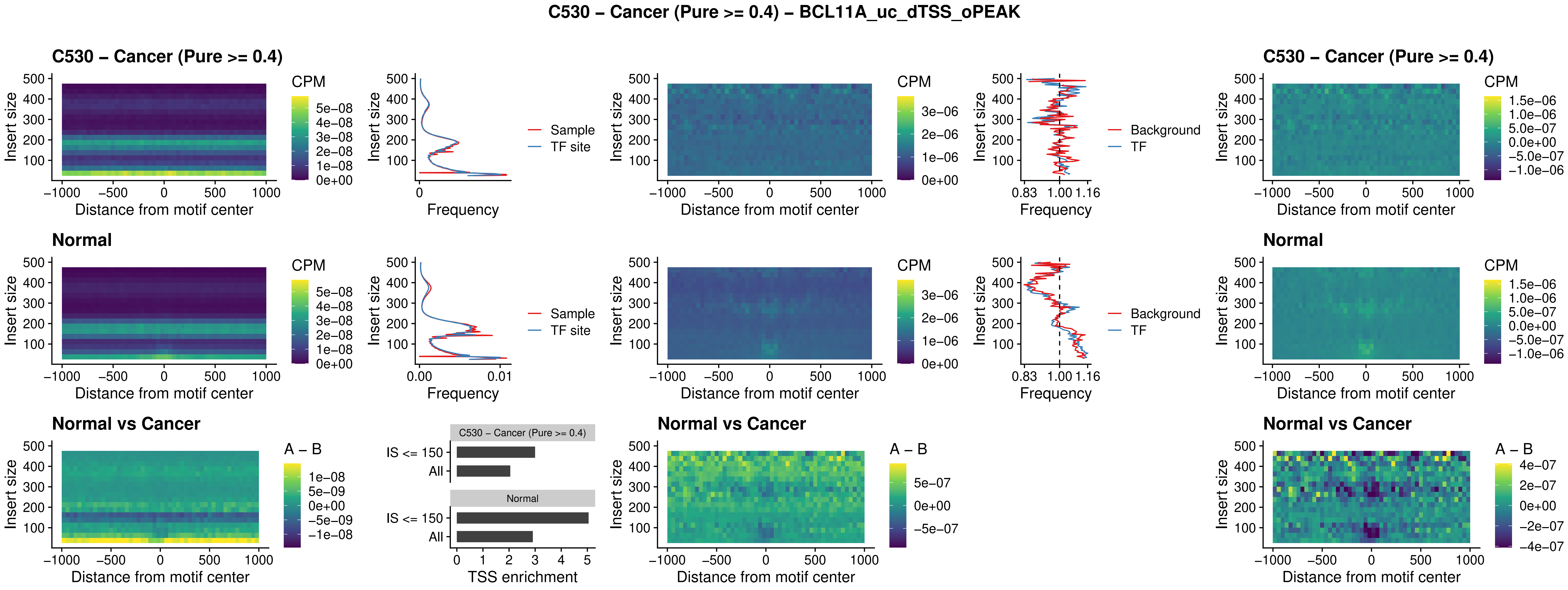
C530 – Cancer (Pure >= 0.4) – BCL11A_uc_dTSS_oPEAK
C530 – Cancer (Pure >= 0.4)
Normal
Normal vs Cancer
Insert size
Distance from motif center
Frequency
CPM
A – B
Sample
TF site
Background
TF
C530 – Cancer (Pure >= 0.4)
Normal
IS <= 150
All
TSS enrichment

#### C530 – Cancer (Pure >= 0.4) – BCL11B_uc_dTSS_oPEAK

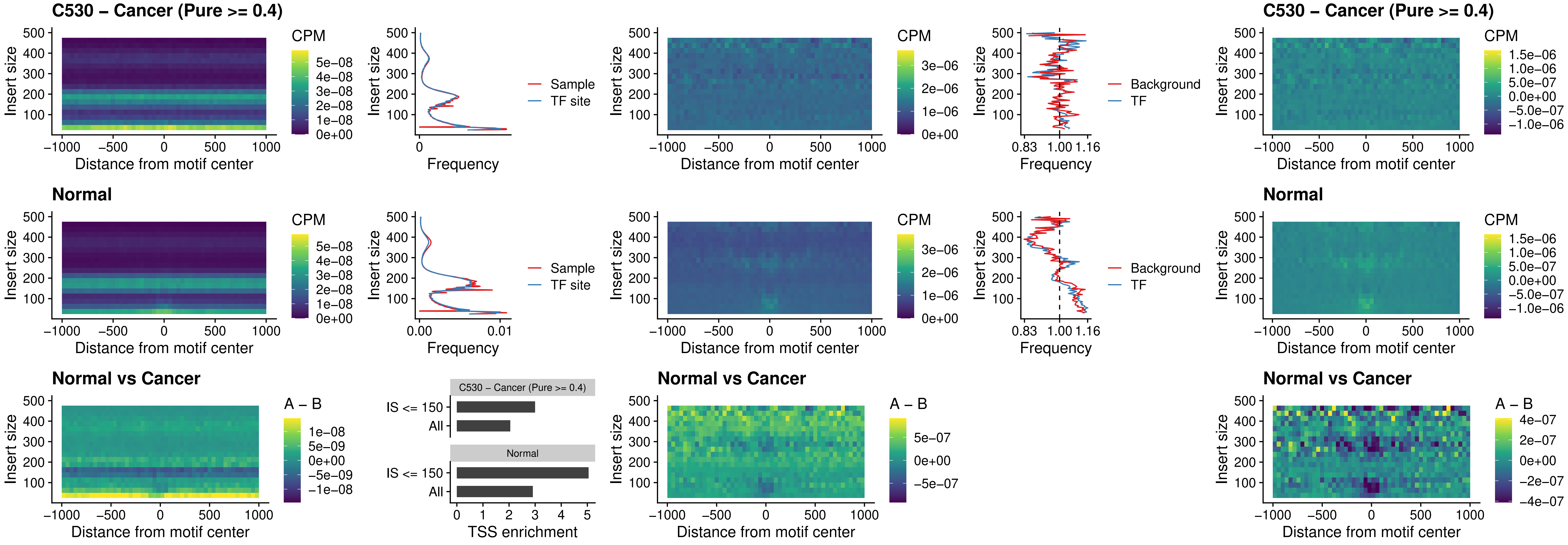

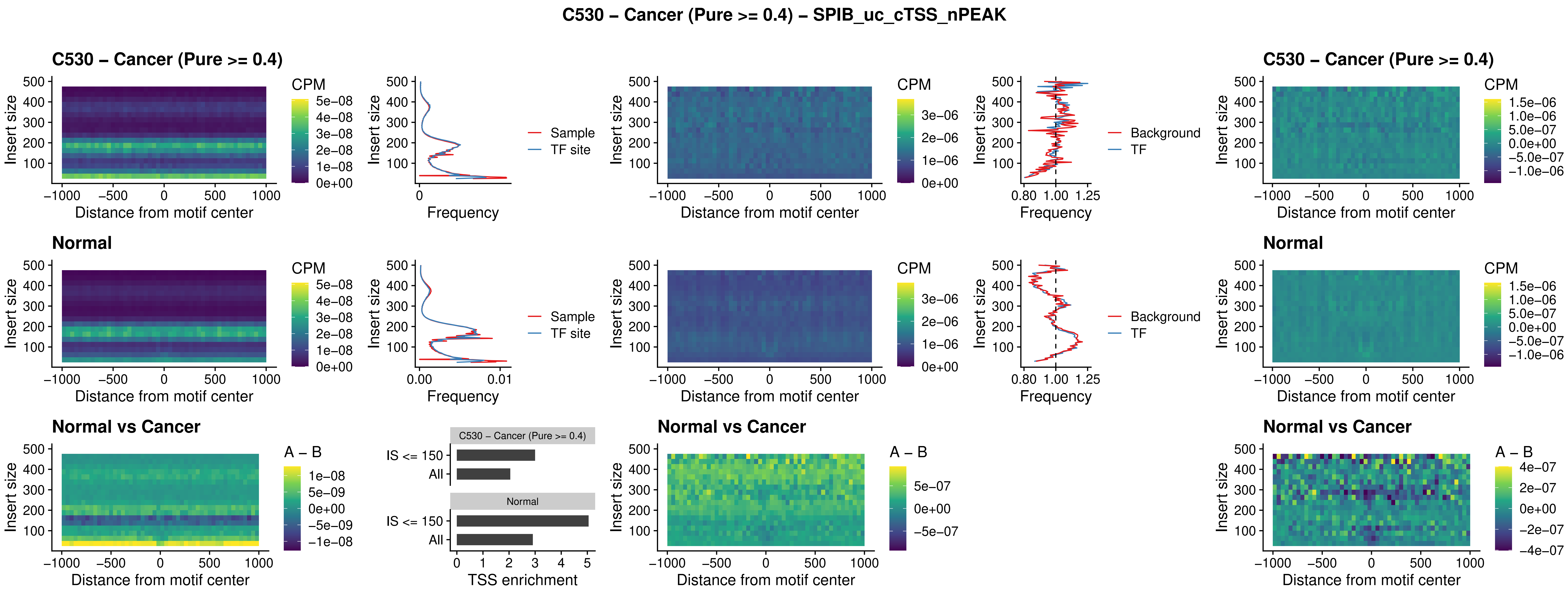
C530 – Cancer (Pure >= 0.4) – SPIB_uc_cTSS_nPEAK
C530 – Cancer (Pure >= 0.4)
Normal
Normal vs Cancer
Insert size
Distance from motif center
Frequency
CPM
A – B
Sample
TF site
Background
TF
C530 – Cancer (Pure >= 0.4)
Normal
IS <= 150
All
TSS enrichment

#### C530 – Cancer (Pure >= 0.4) – SPIC_uc_cTSS_nPEAK

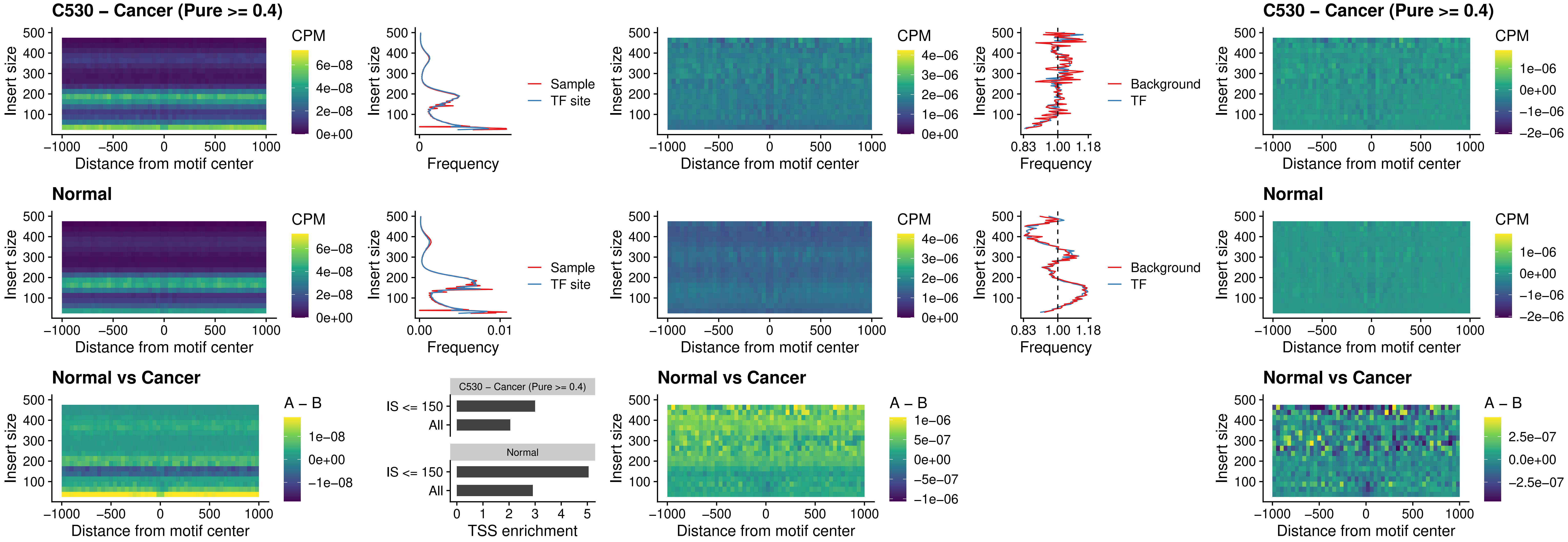

#### C530 – Cancer (Pure >= 0.4) – SPI1_uc_pTSS_oPEAK

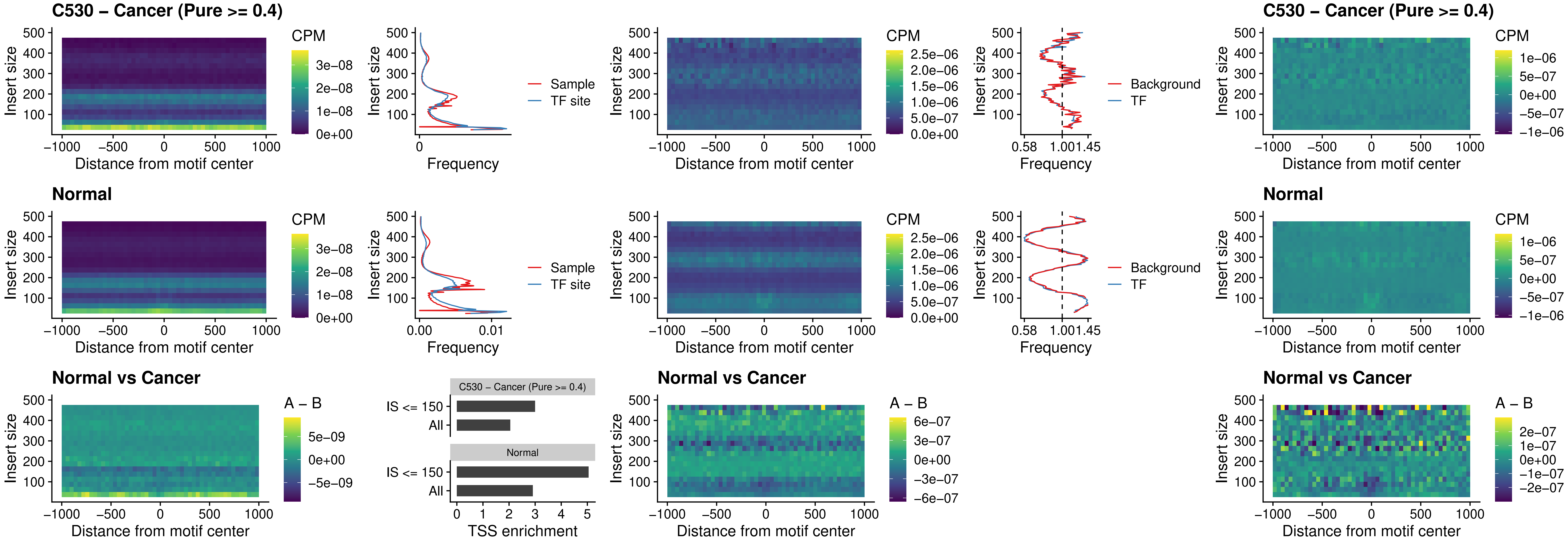

#### C530 – Cancer (Pure >= 0.4) – BCL11A_uc_pTSS_oPEAK

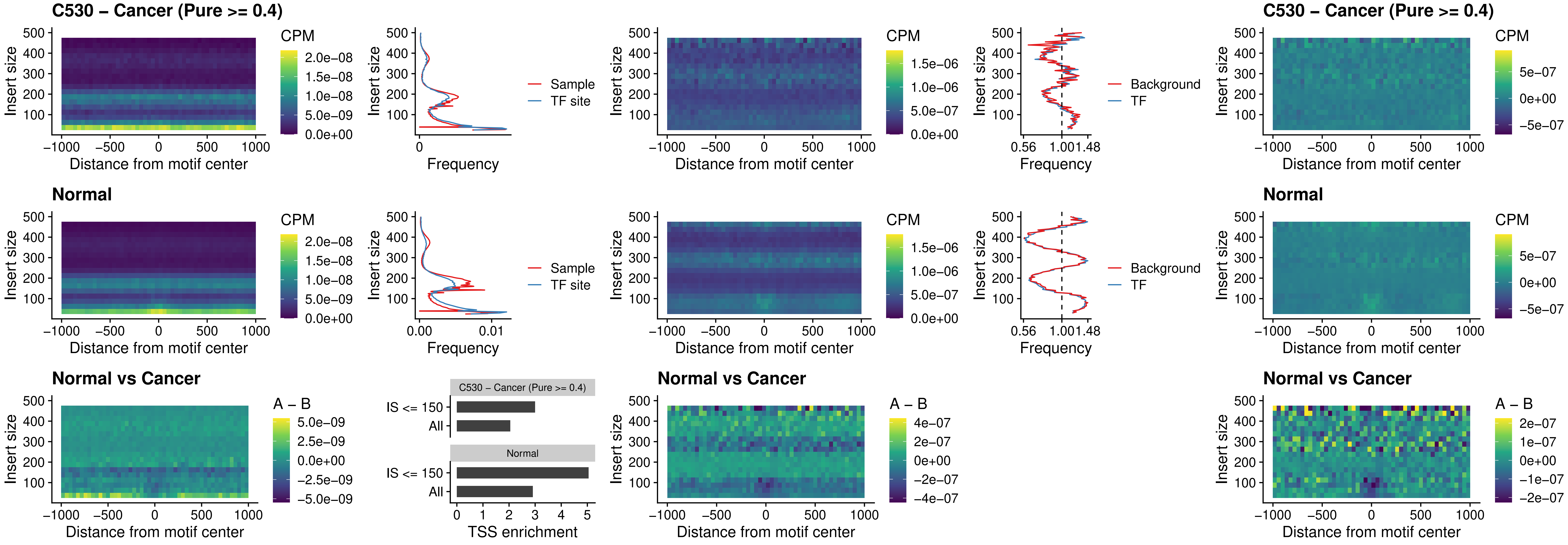

#### C530 – Cancer (Pure >= 0.4) – BCL11B_uc_pTSS_oPEAK

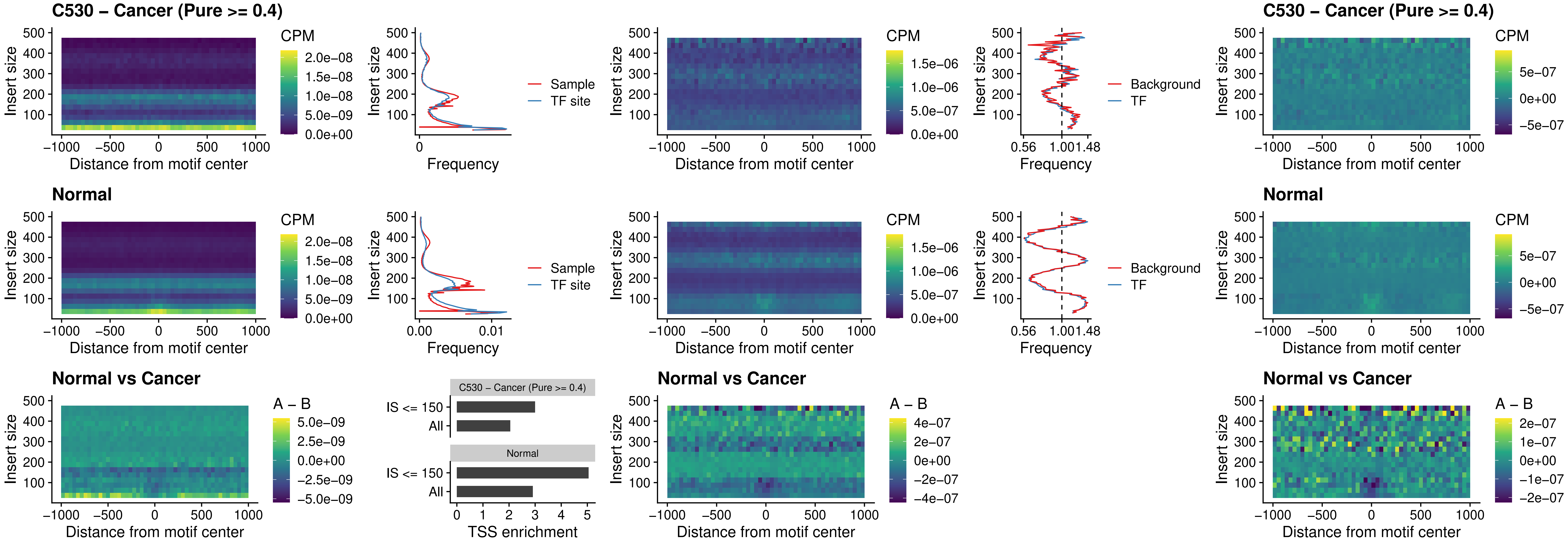

#### C530 – Cancer (Pure >= 0.4) – SPI1_uc_cTSS_oPEAK

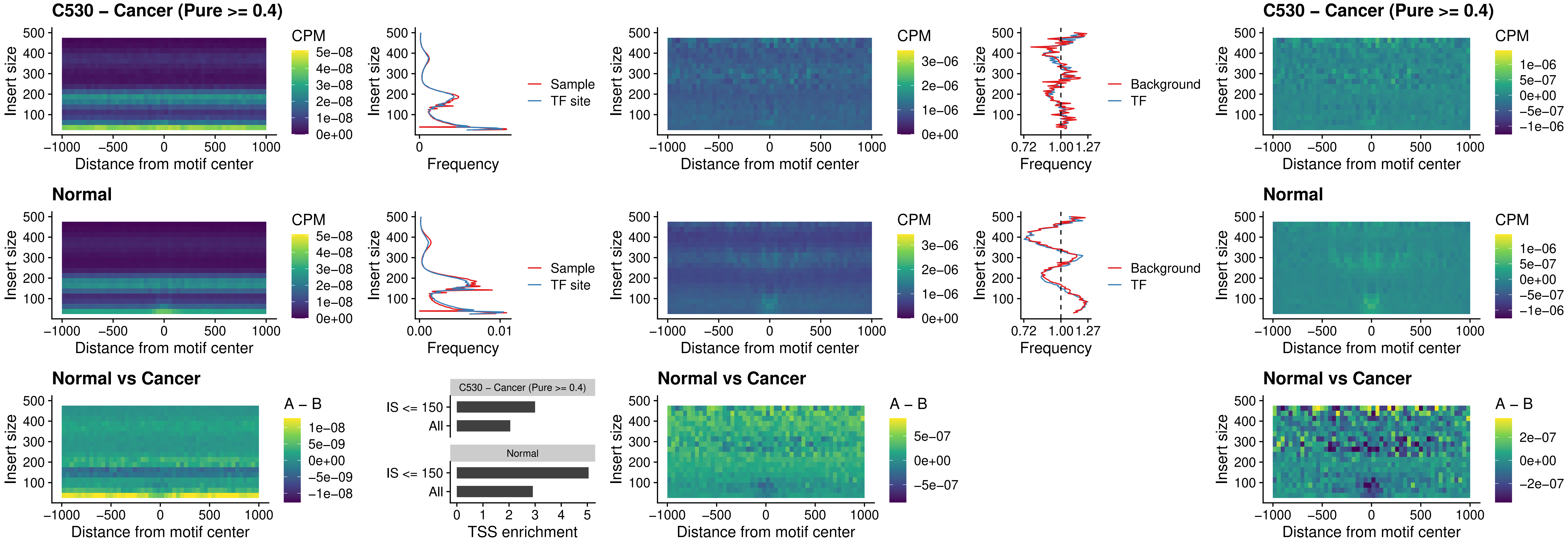

#### C530 – Cancer (Pure >= 0.4) – SPIB_uc_cTSS_oPEAK

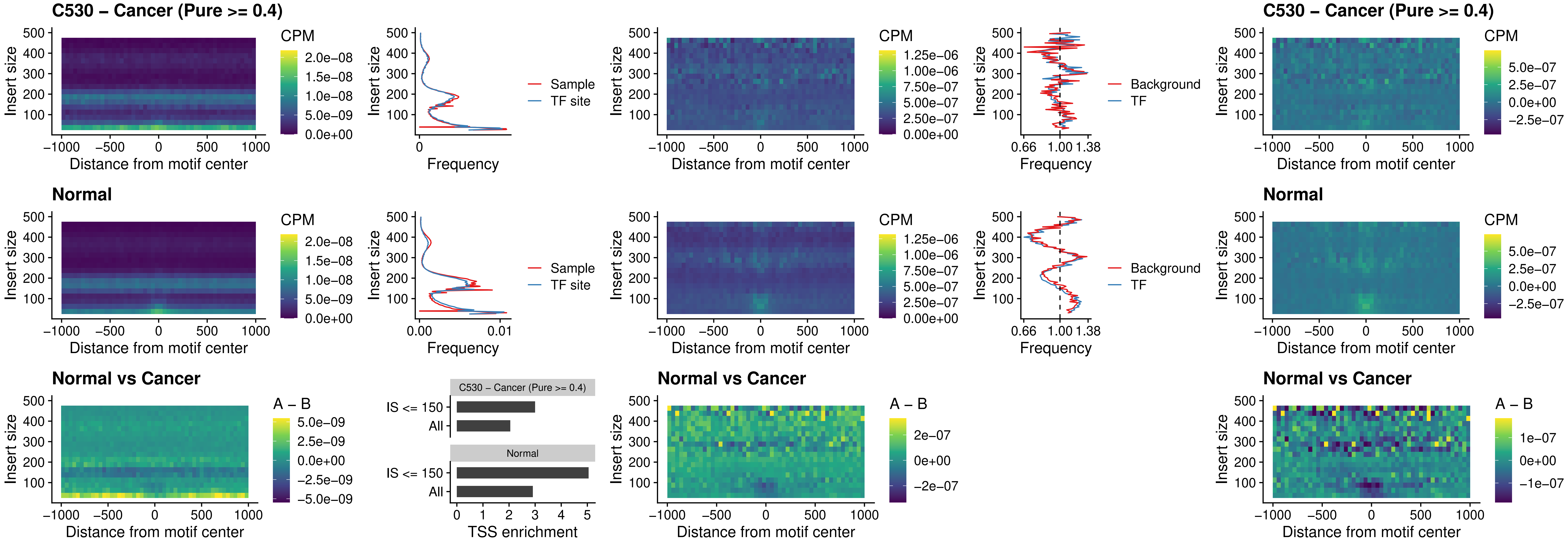

#### C530 – Cancer (Pure >= 0.4) – SPIB_uc_pTSS_oPEAK

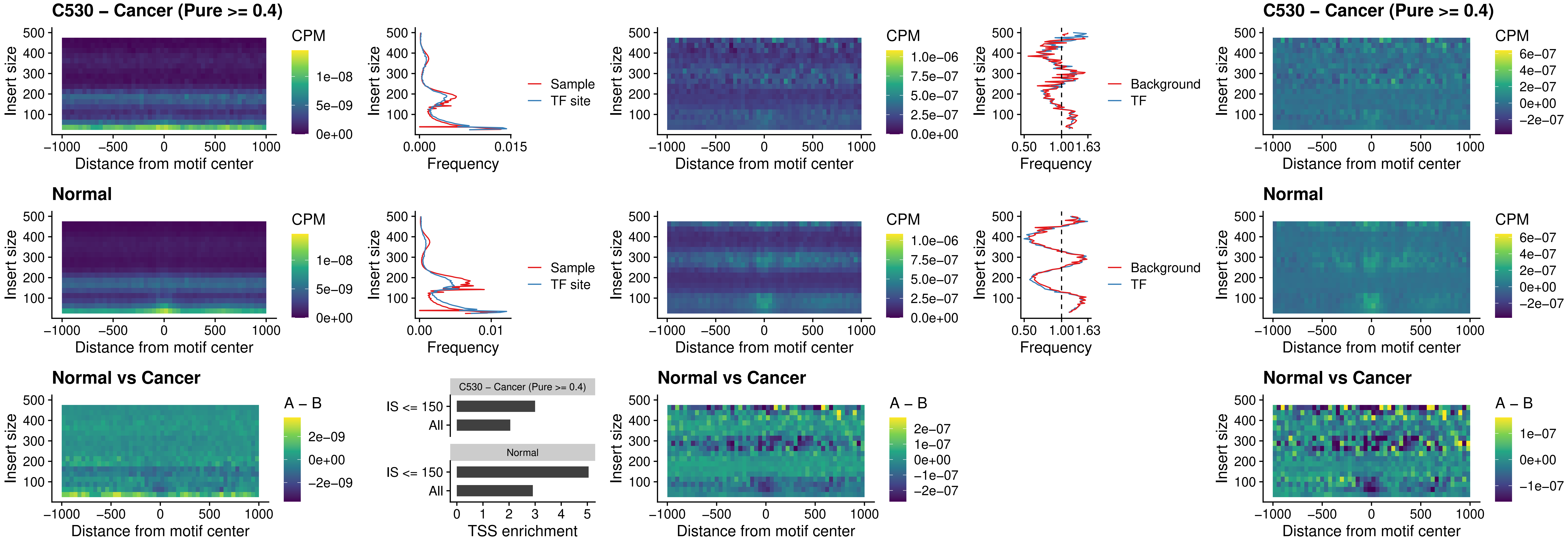

#### C530 – Cancer (Pure >= 0.4) – SPI1_uc_dTSS_oPEAK

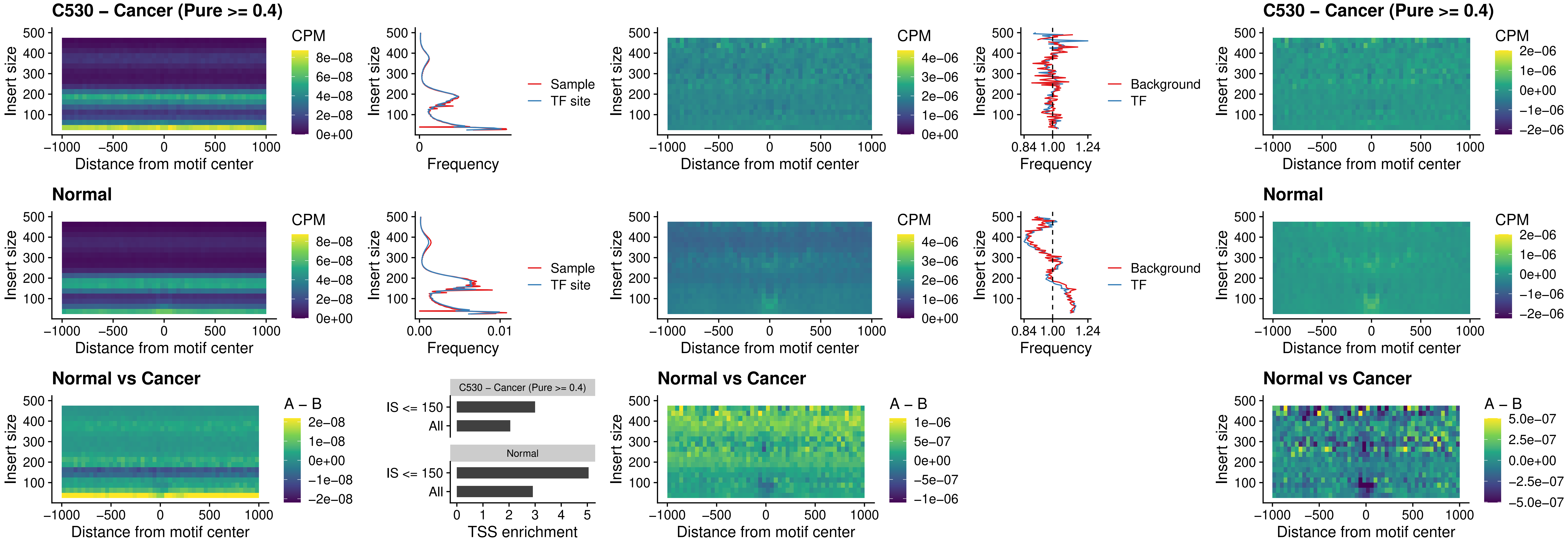

### C530 – Cancer (Pure >= 0.4) – SPIB_uc_dTSS_oPEAK

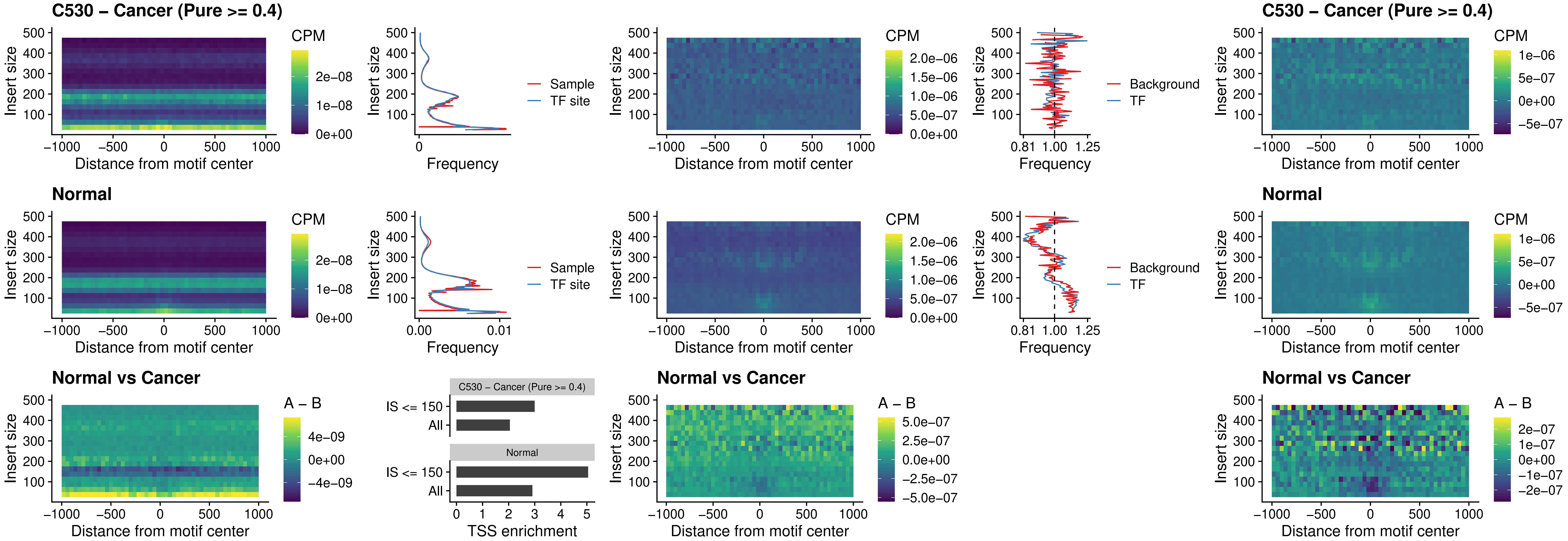

#### C530 – Cancer (Pure >= 0.4) – IRF3_uc_pTSS_oPEAK

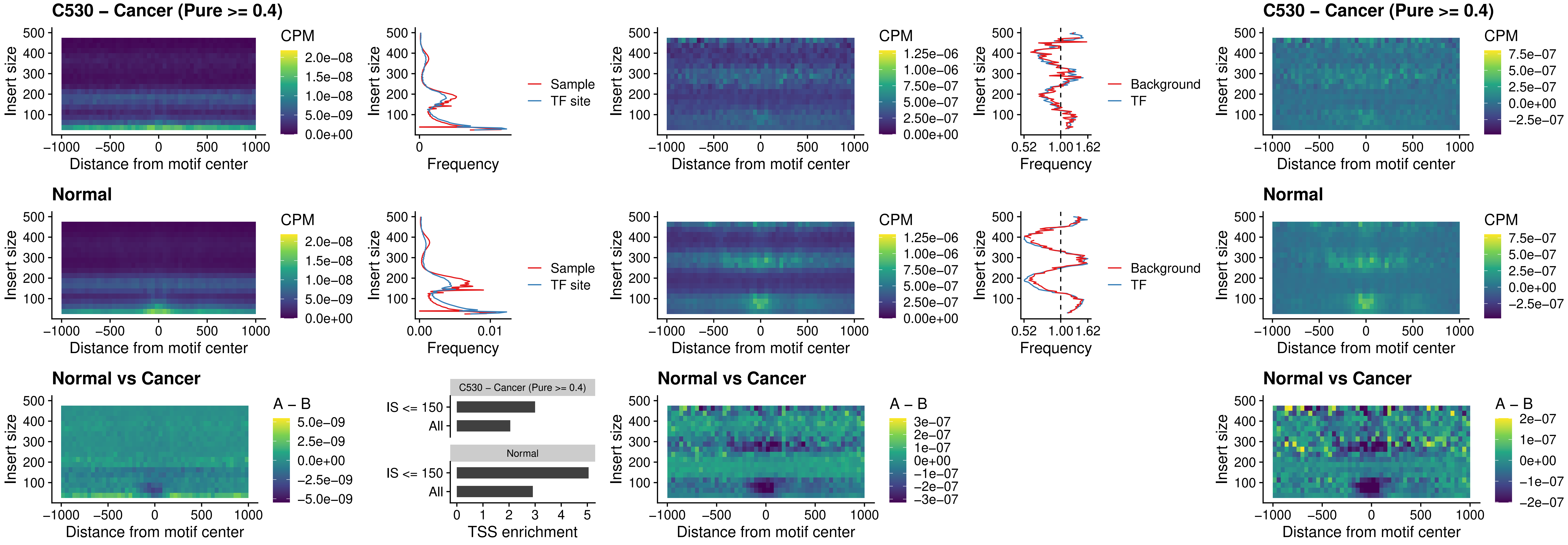

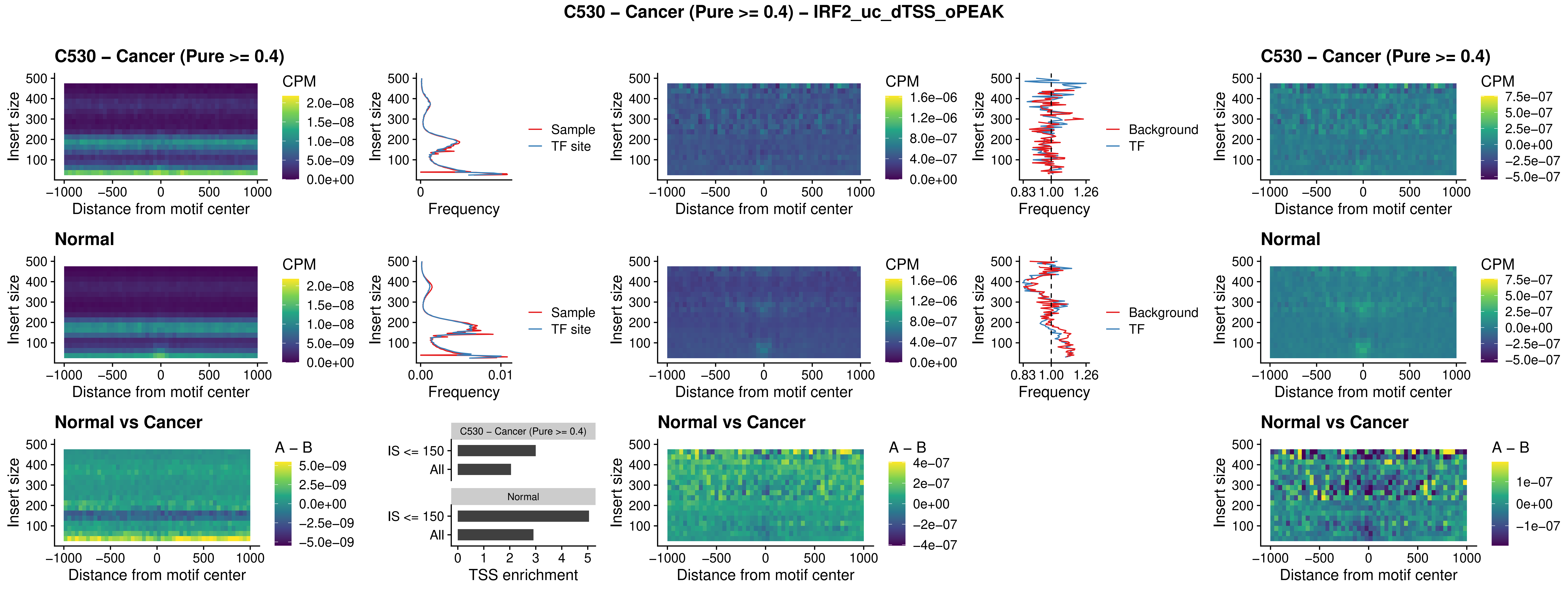

C530 – Cancer (Pure >= 0.4) – IRF2_uc_dTSS_oPEAK
C530 – Cancer (Pure >= 0.4)
Normal
Normal vs Cancer
Insert size
Distance from motif center
Frequency
CPM
A – B
Sample
TF site
Background
TF
C530 – Cancer (Pure >= 0.4)
Normal
IS <= 150
All
TSS enrichment

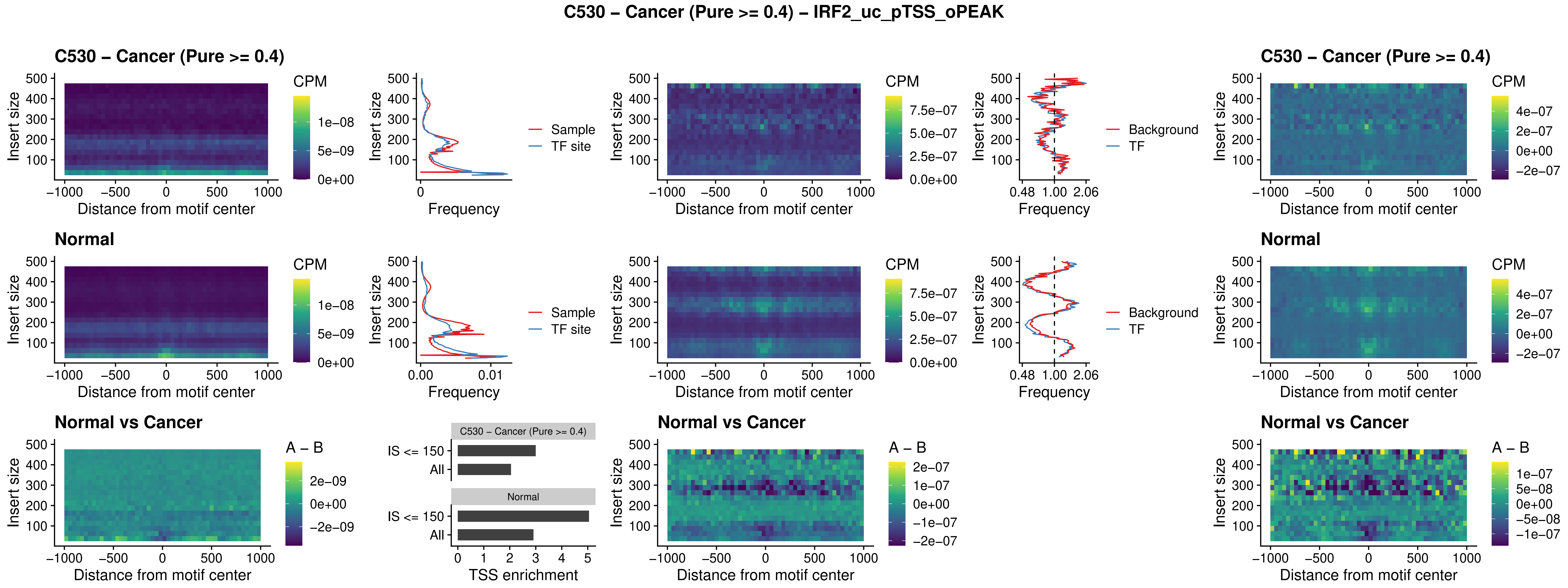
C530 – Cancer (Pure >= 0.4) – IRF2_uc_pTSS_oPEAK
C530 – Cancer (Pure >= 0.4)
Normal
Normal vs Cancer
Insert size
Distance from motif center
Frequency
CPM
A – B
Sample
TF site
Background
TF
C530 – Cancer (Pure >= 0.4)
Normal
IS <= 150
All
TSS enrichment

#### C530 – Cancer (Pure >= 0.4) – STAT2_uc_pTSS_oPEAK

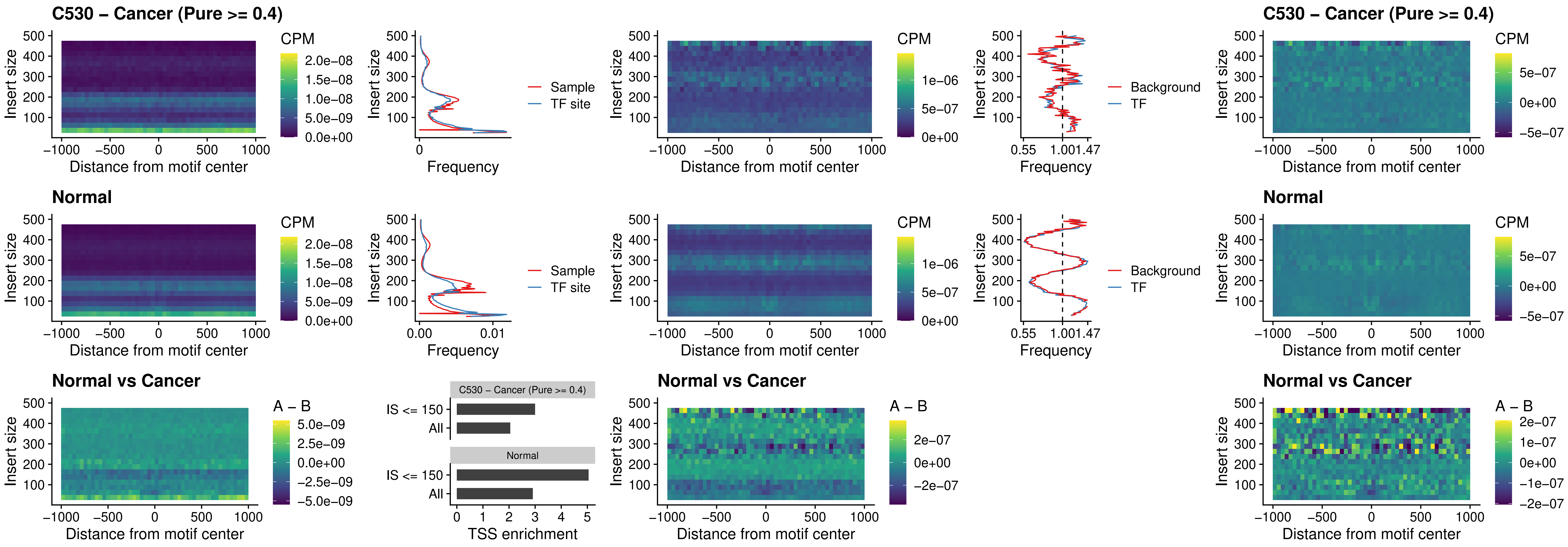

#### C530 – Cancer (Pure >= 0.4) – SPIB_uc_dTSS_nPEAK

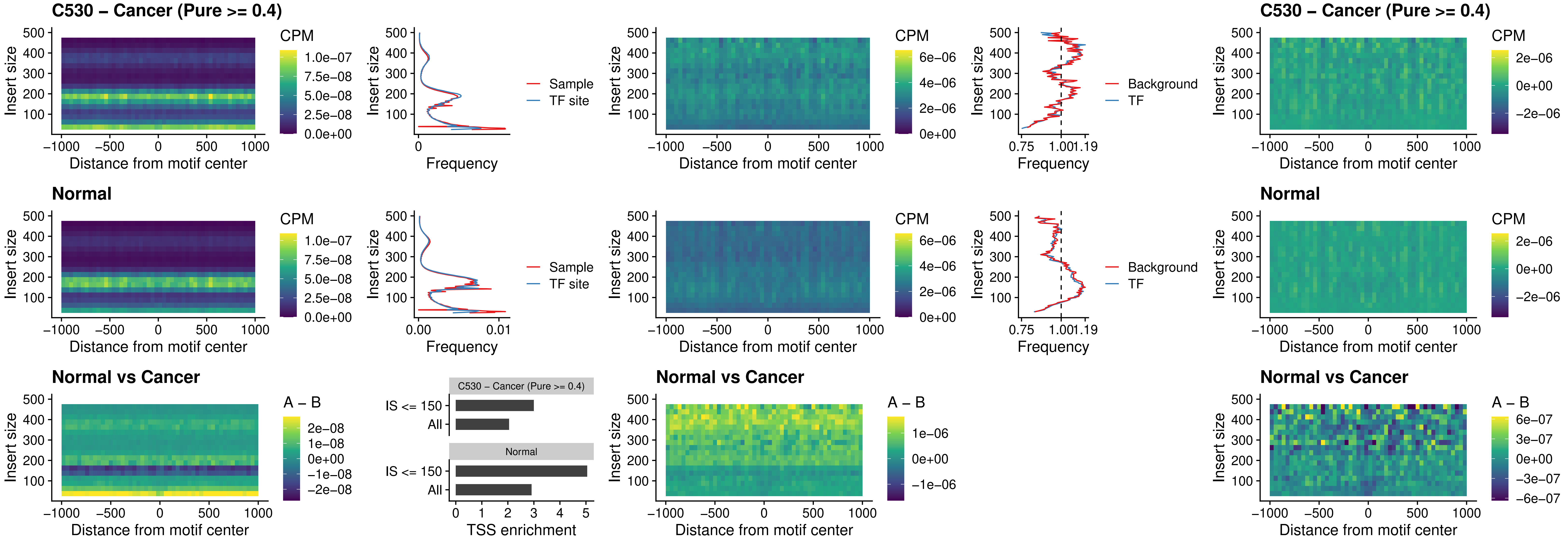

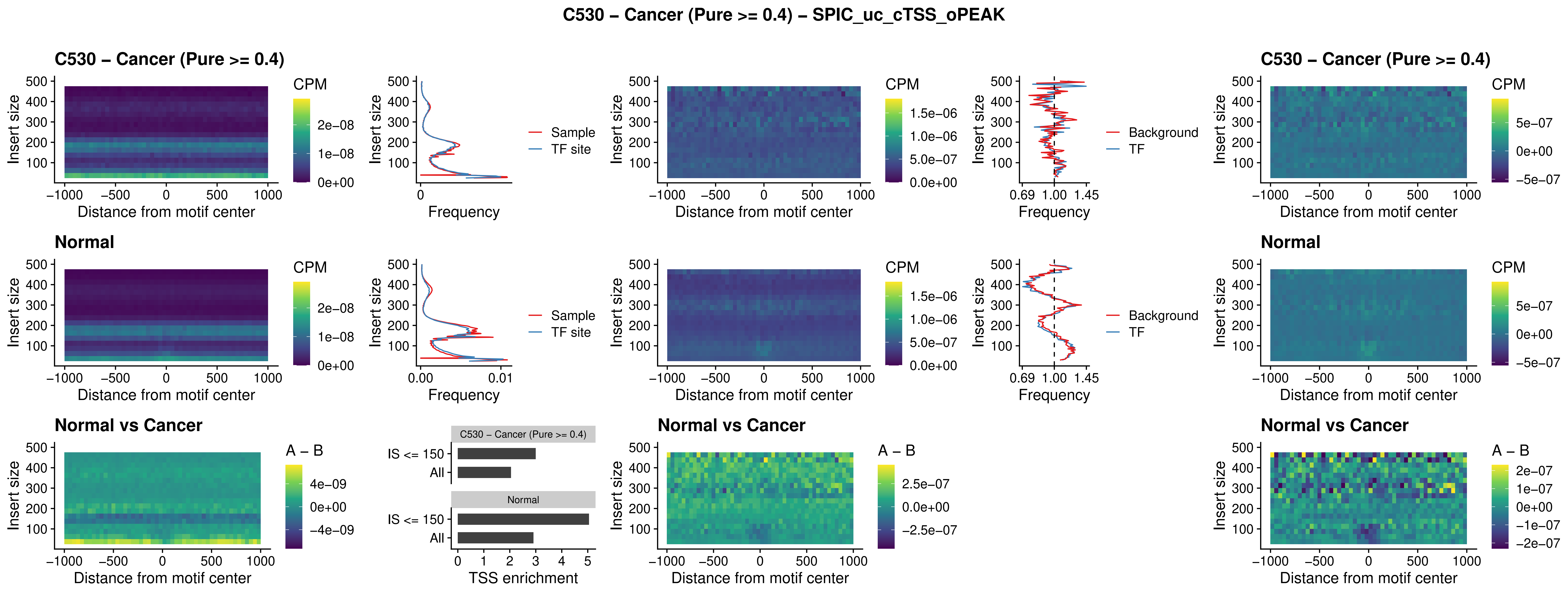

C530 – Cancer (Pure >= 0.4) – SPIC_uc_cTSS_oPEAK
C530 – Cancer (Pure >= 0.4)
Normal
Normal vs Cancer
Insert size
Distance from motif center
Frequency
CPM
A – B
Sample
TF site
Background
TF
C530 – Cancer (Pure >= 0.4)
Normal
IS <= 150
All
TSS enrichment

#### C530 – Cancer (Pure >= 0.4) – PRDM1_uc_cTSS_oPEAK

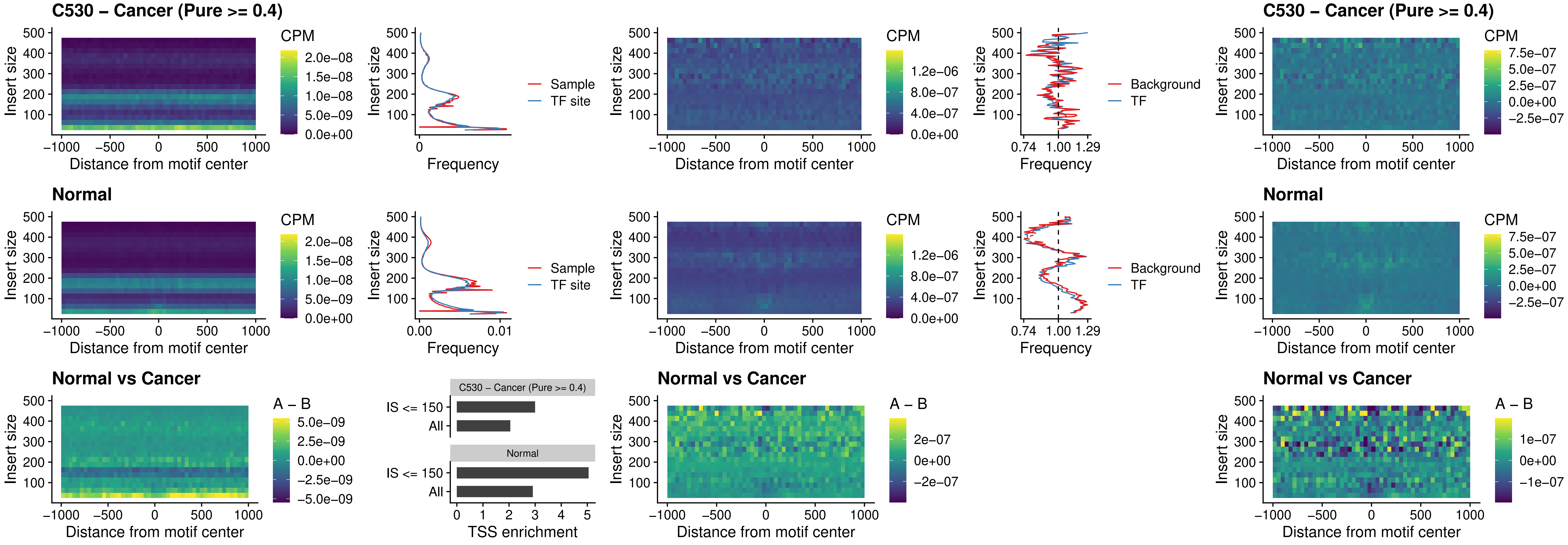

#### C530 – Cancer (Pure >= 0.4) – IRF3_uc_cTSS_oPEAK

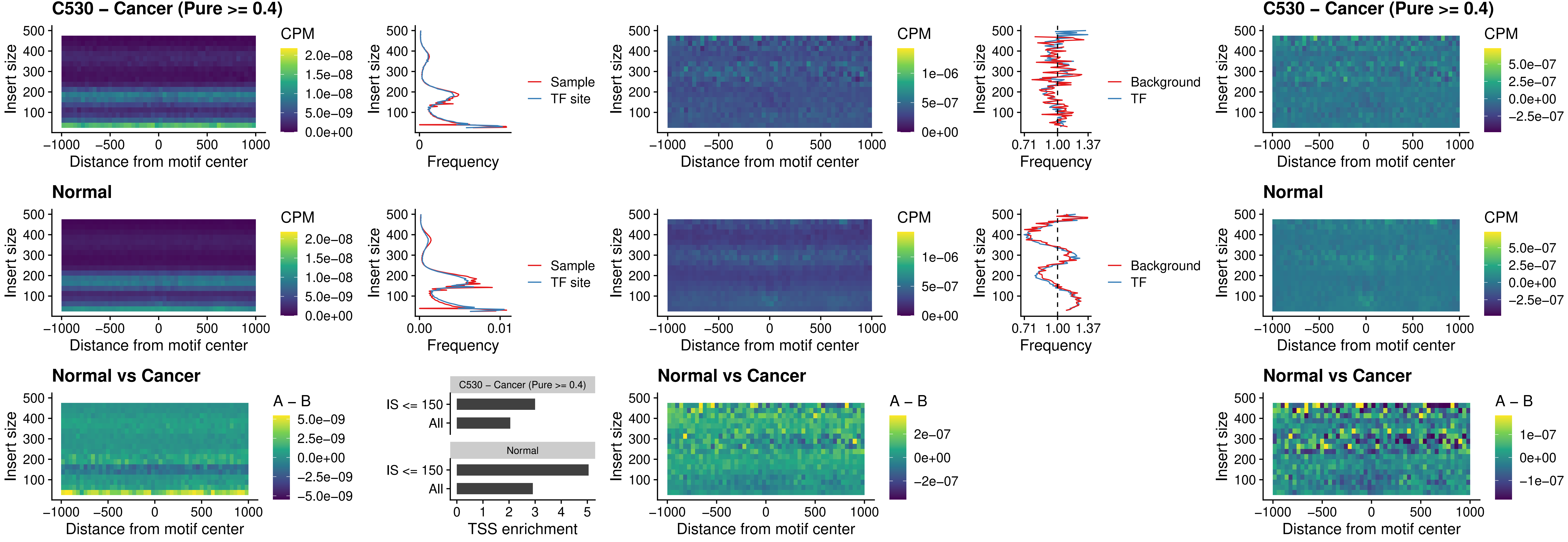

#### C530 – Cancer (Pure >= 0.4) – BCL11A_uc_dTSS_nPEAK

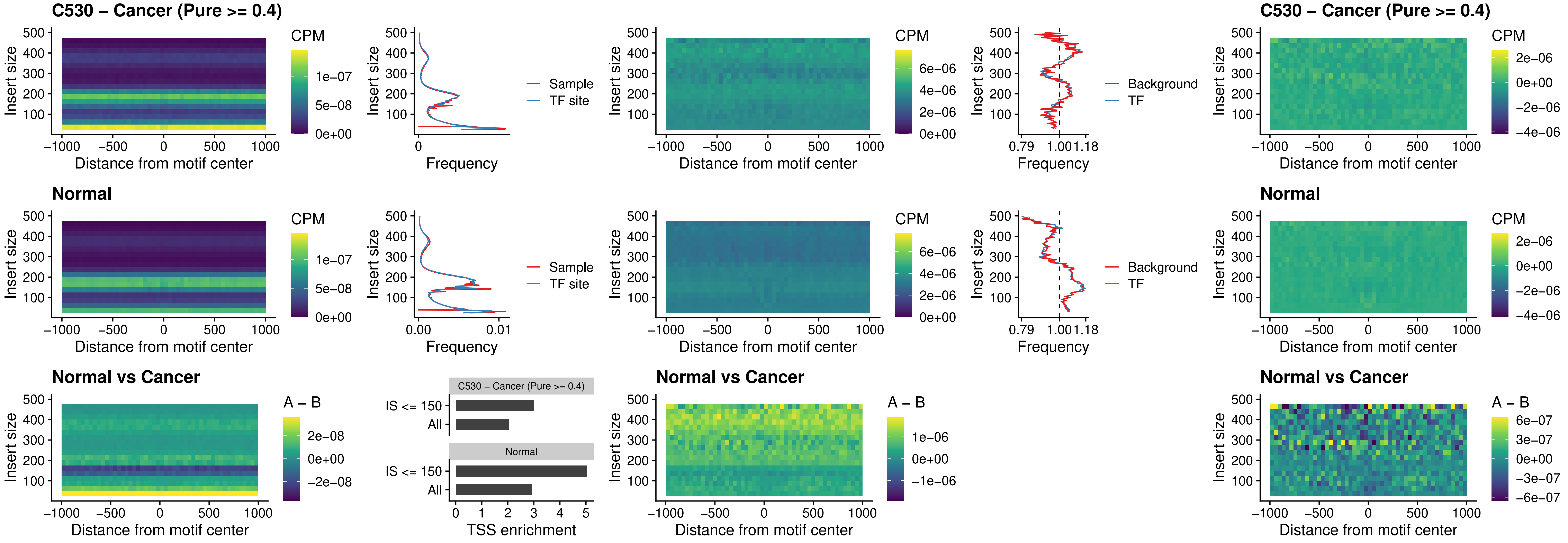

#### C530 – Cancer (Pure >= 0.4) – BCL11B_uc_dTSS_nPEAK

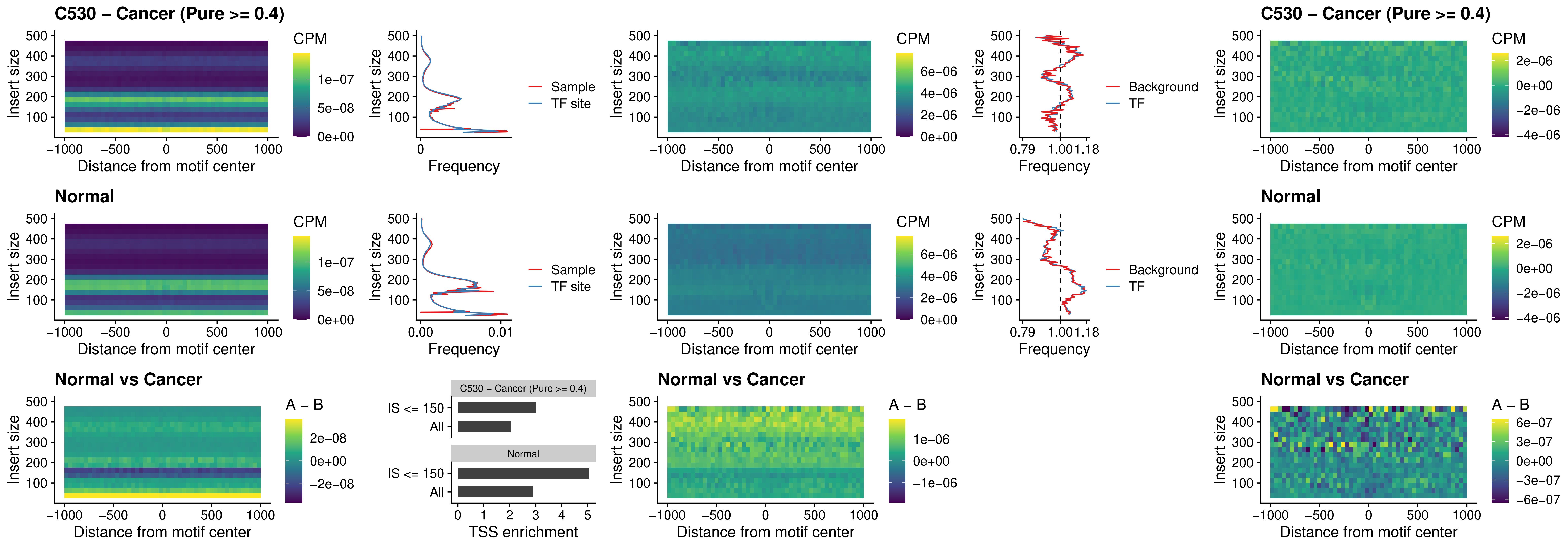

#### C530 – Cancer (Pure >= 0.4) – IRF1_uc_cTSS_oPEAK

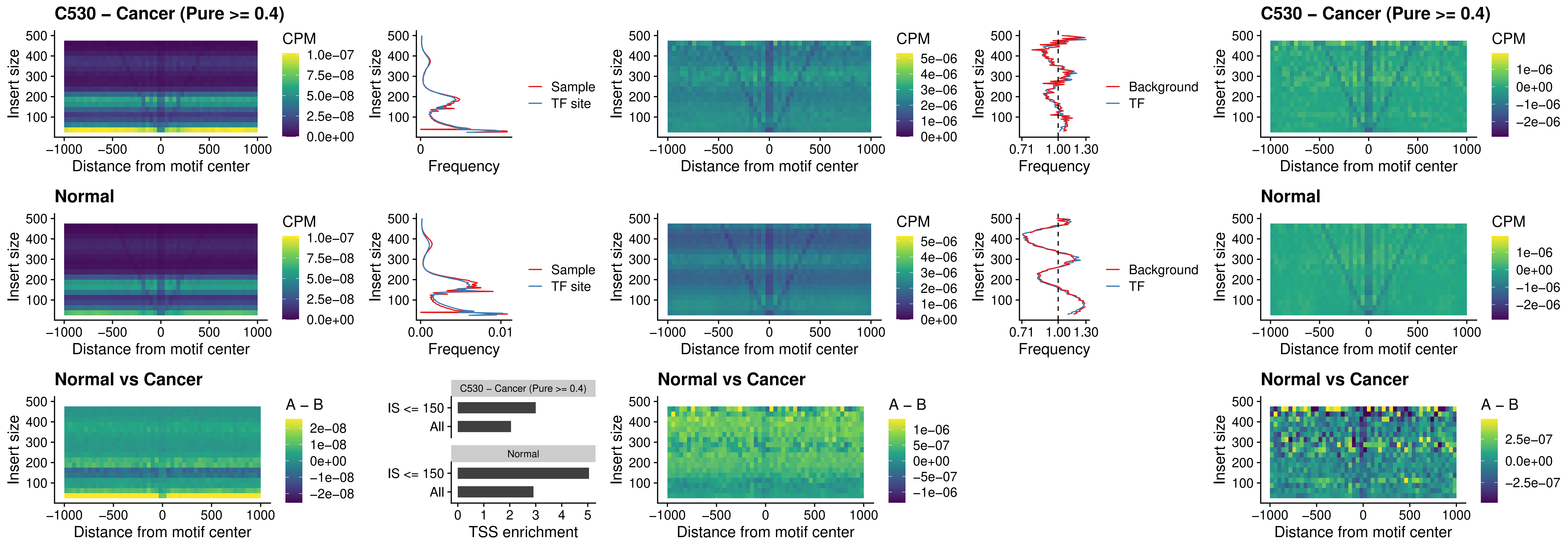

#### C530 – Cancer (Pure >= 0.4) – IRF1_uc_dTSS_oPEAK

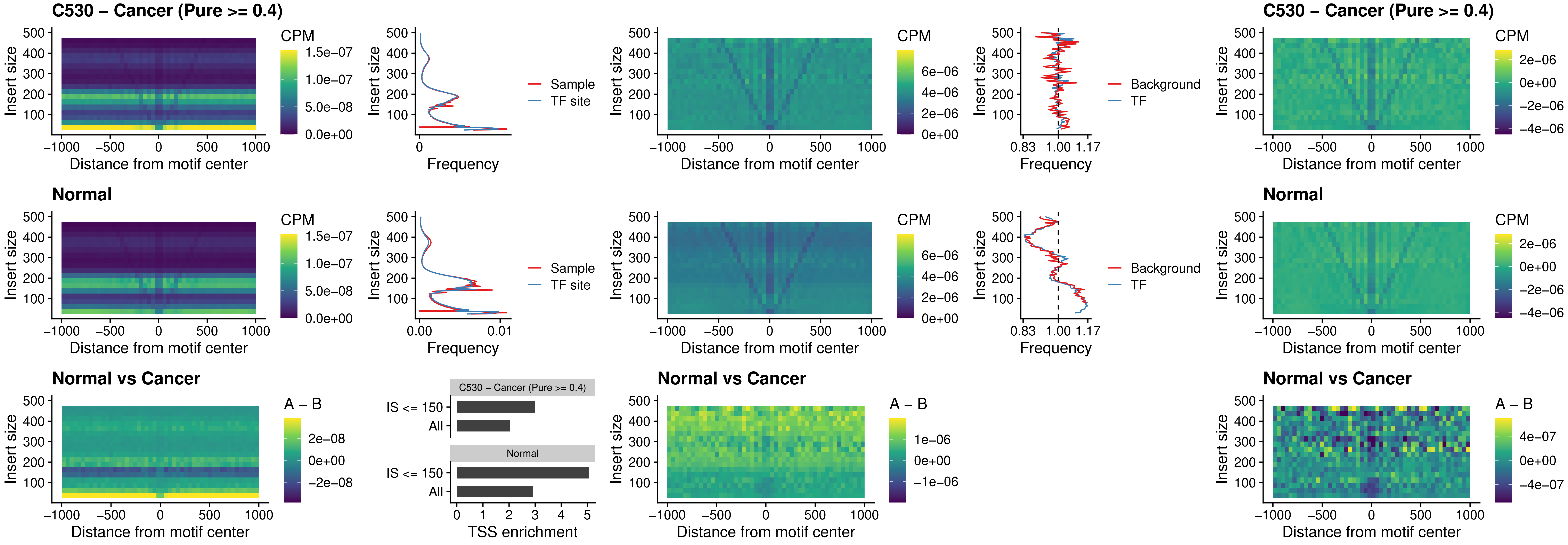

#### C530 – Cancer (Pure >= 0.4) – IRF1_uc_pTSS_oPEAK

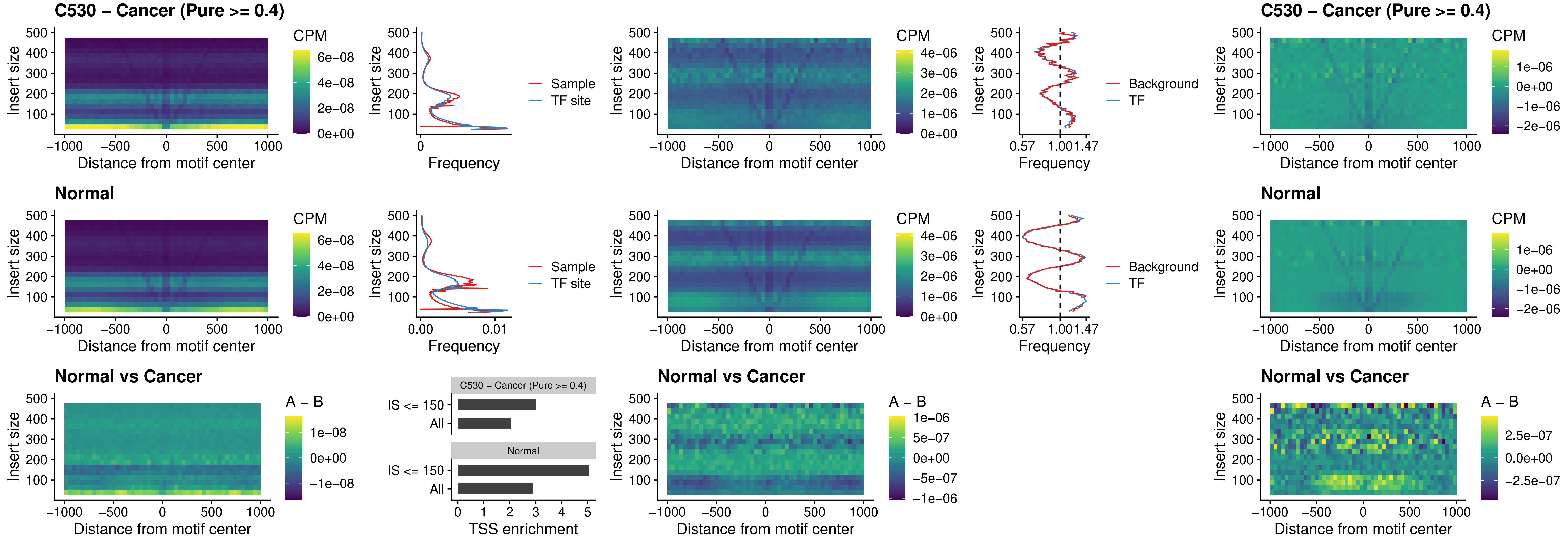

#### C530 – Cancer (Pure >= 0.4) – SPIC_uc_dTSS_nPEAK

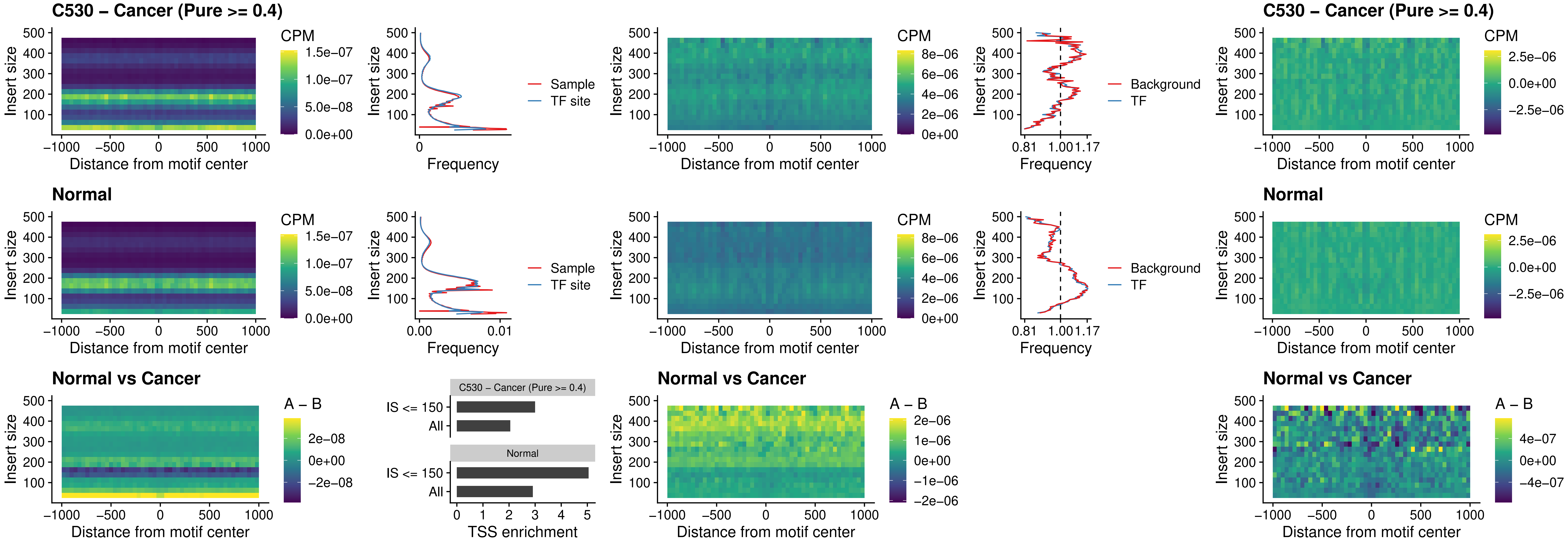

#### C530 – Cancer (Pure >= 0.4) – SPIC_uc_dTSS_oPEAK

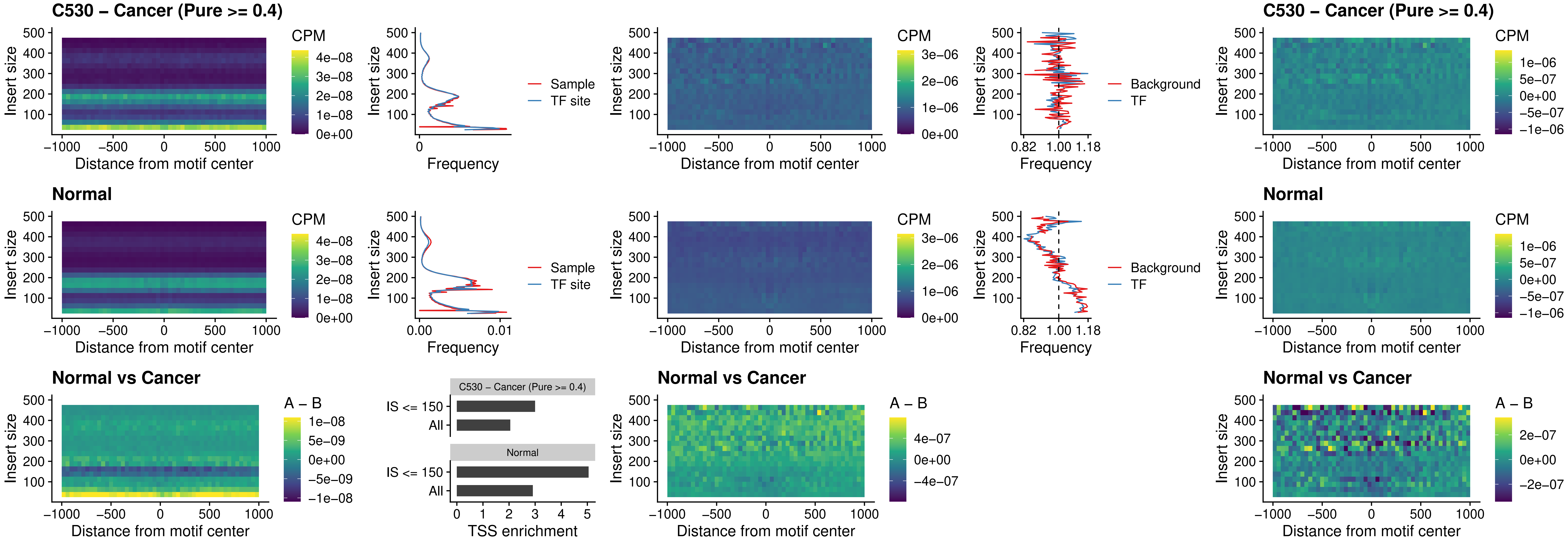

### C530 – Cancer (Pure >= 0.4) – SPIB_uc_pTSS_nPEAK

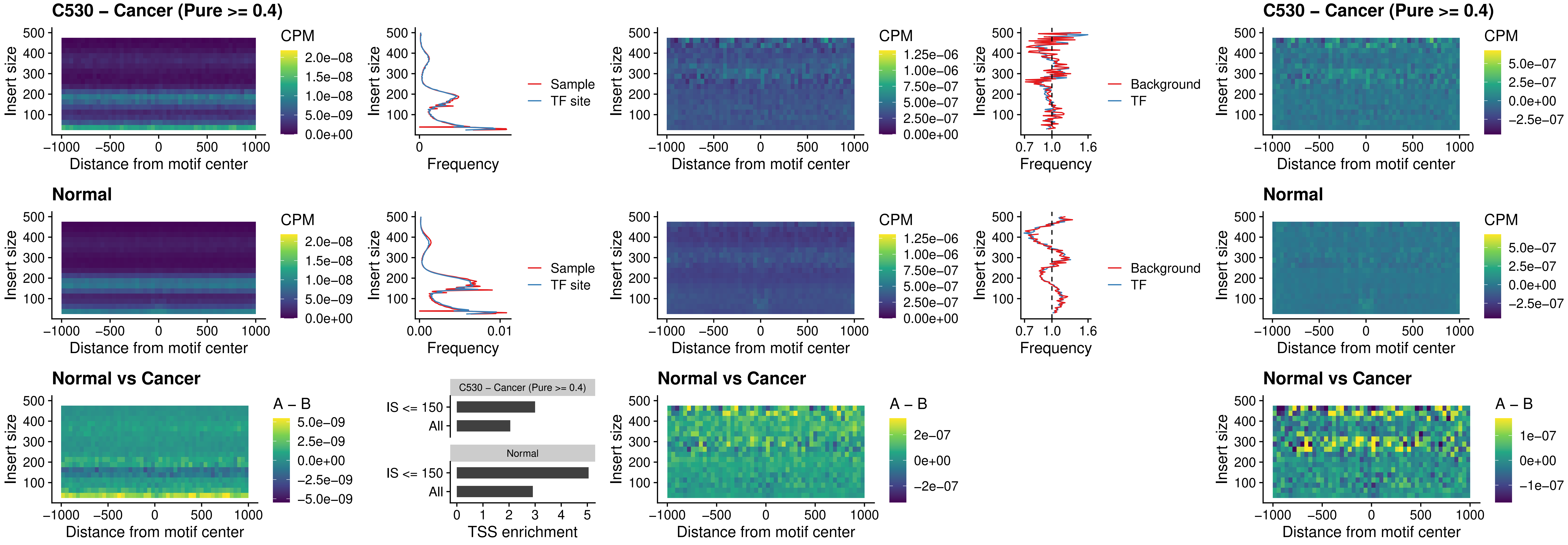

### C530 – Cancer (Pure >= 0.4) – STAT2_uc_cTSS_oPEAK

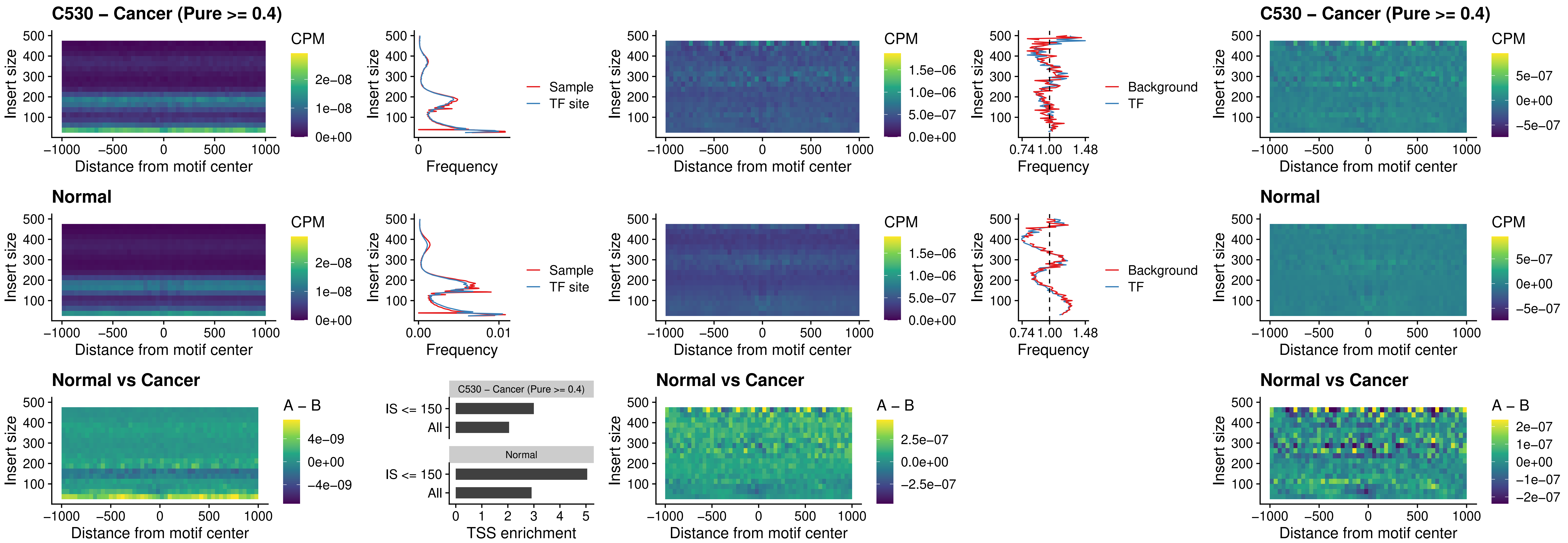

#### C530 – Cancer (Pure >= 0.4) – IRF8_uc_cTSS_oPEAK

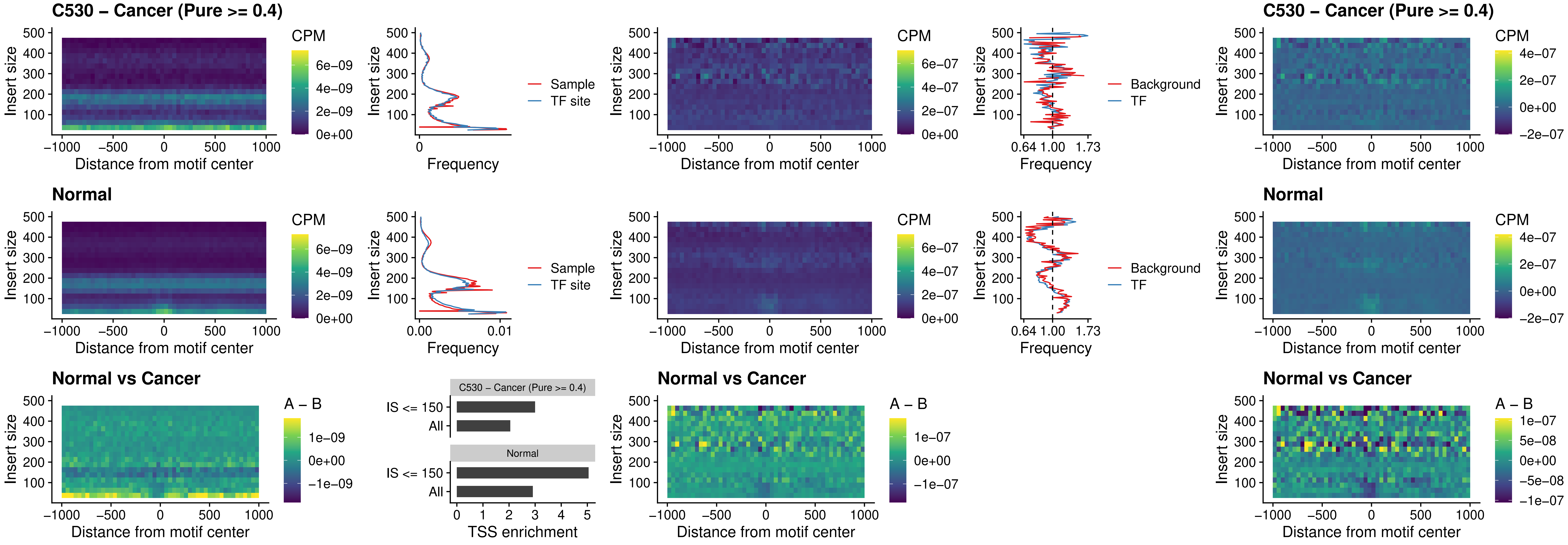

### C530 – Cancer (Pure >= 0.4) – IRF3_uc_dTSS_oPEAK

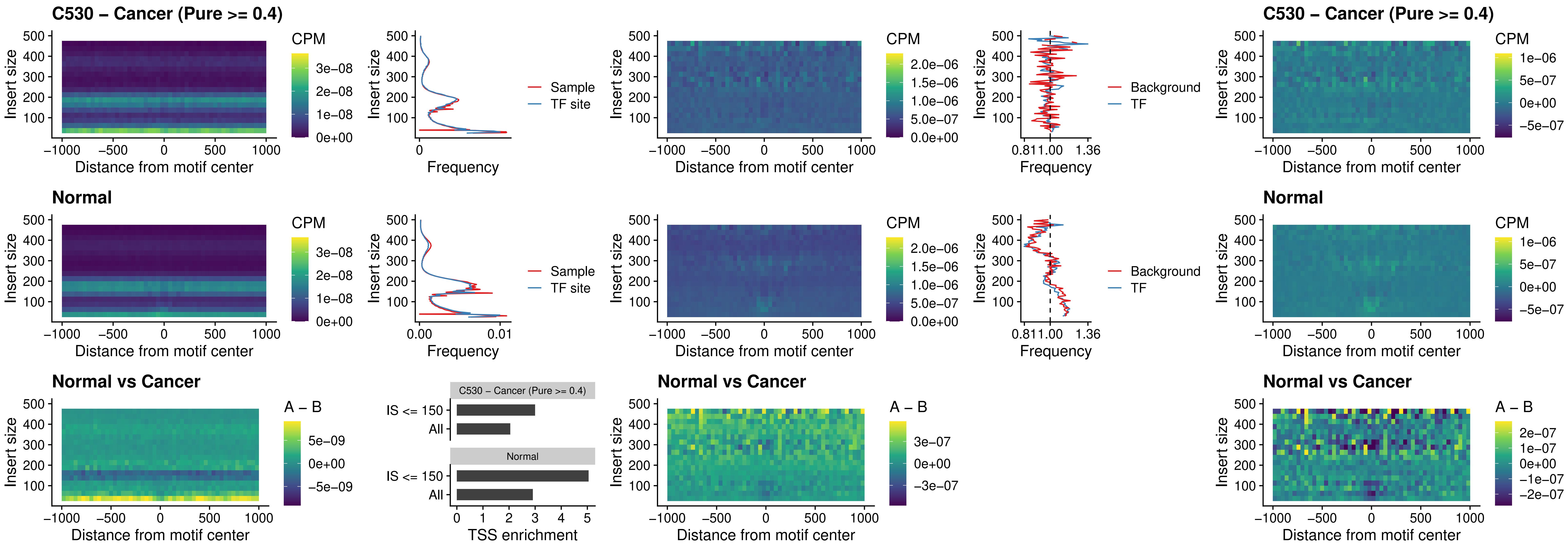

#### C530 – Cancer (Pure >= 0.4) – IRF8_uc_dTSS_oPEAK

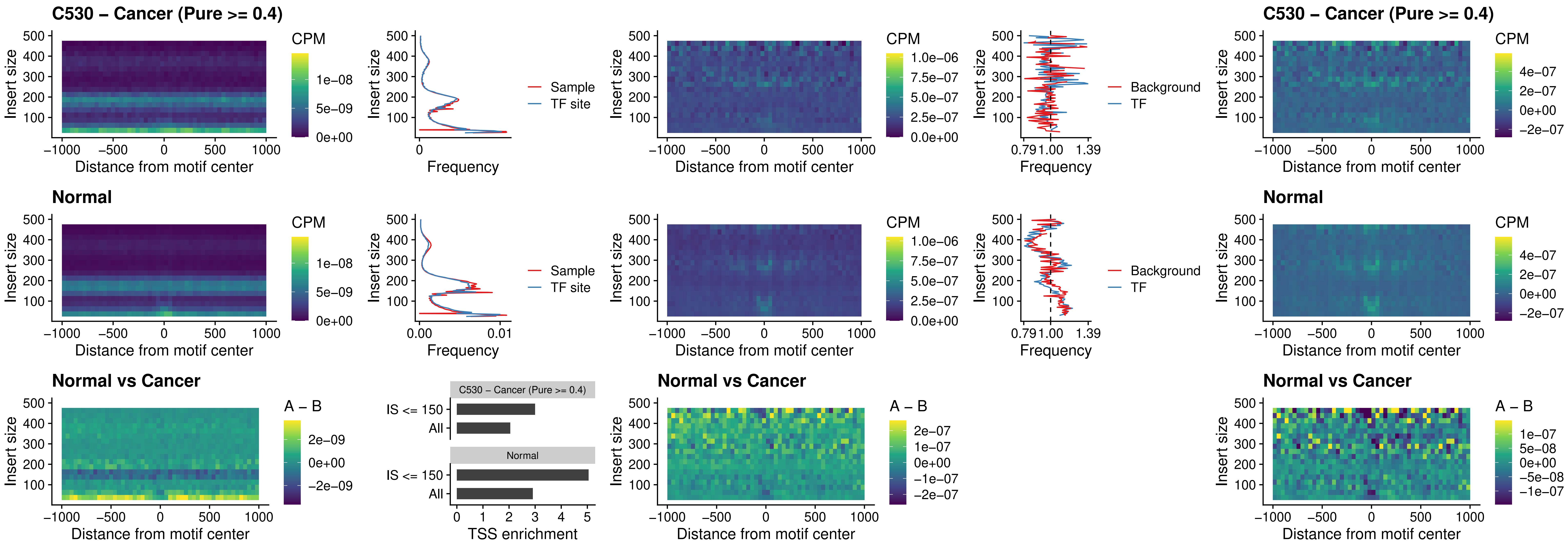

### C530 – Cancer (Pure >= 0.4) – ETS2_uc_dTSS_oPEAK

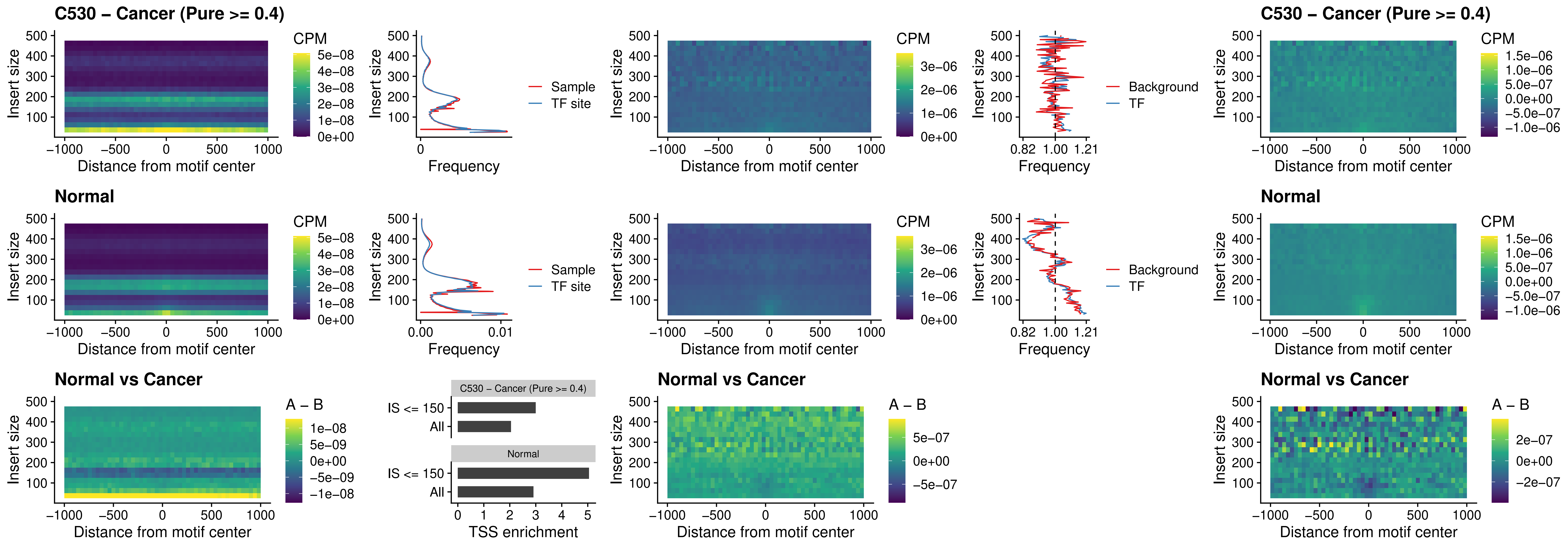

#### C530 – Cancer (Pure >= 0.4) – SMARCC2_uc_pTSS_oPEAK

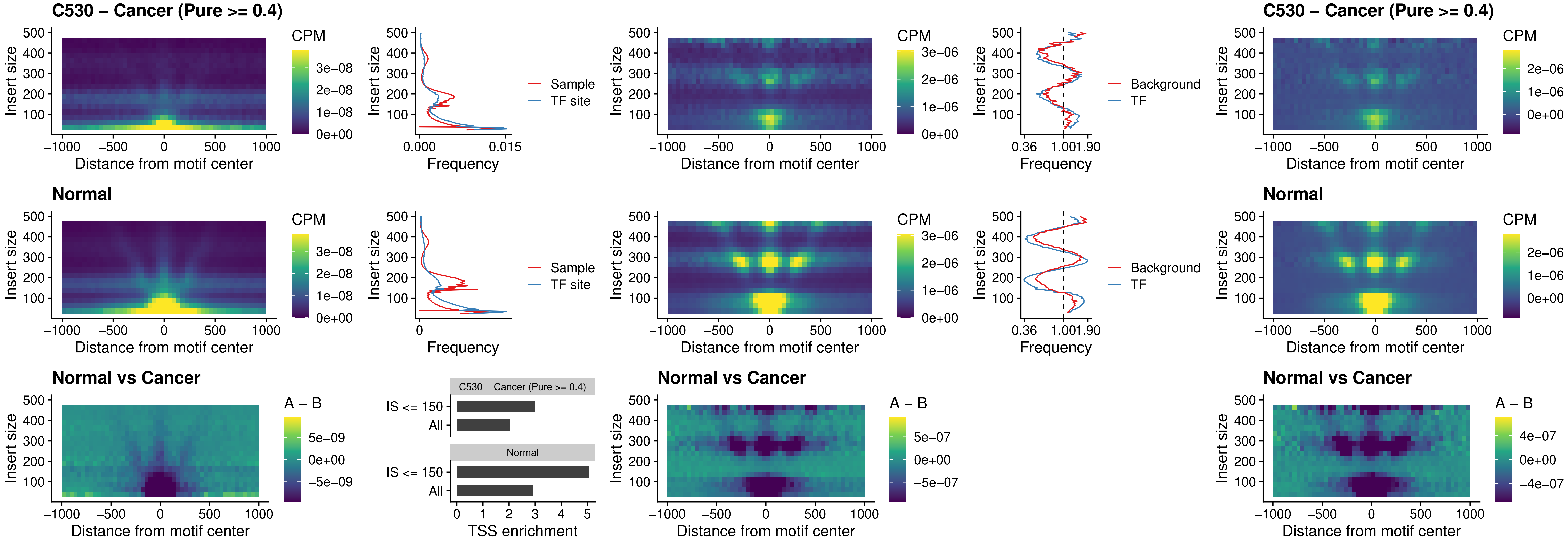

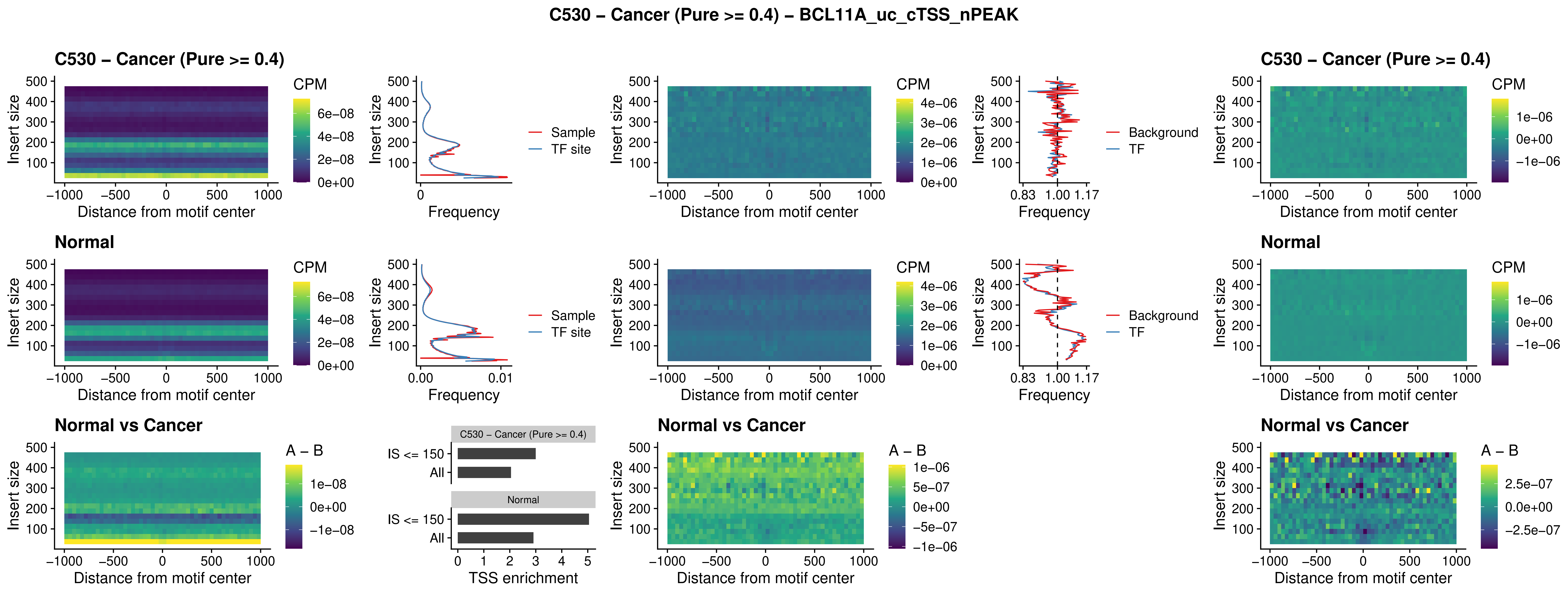
C530 – Cancer (Pure >= 0.4) – BCL11A_uc_cTSS_nPEAK
C530 – Cancer (Pure >= 0.4)
Normal
Normal vs Cancer
Insert size
Distance from motif center
Frequency
CPM
A – B
Sample
TF site
Background
TF
C530 – Cancer (Pure >= 0.4)
Normal
IS <= 150
All
TSS enrichment

### C530 – Cancer (Pure >= 0.4) – BCL11B_uc_cTSS_nPEAK

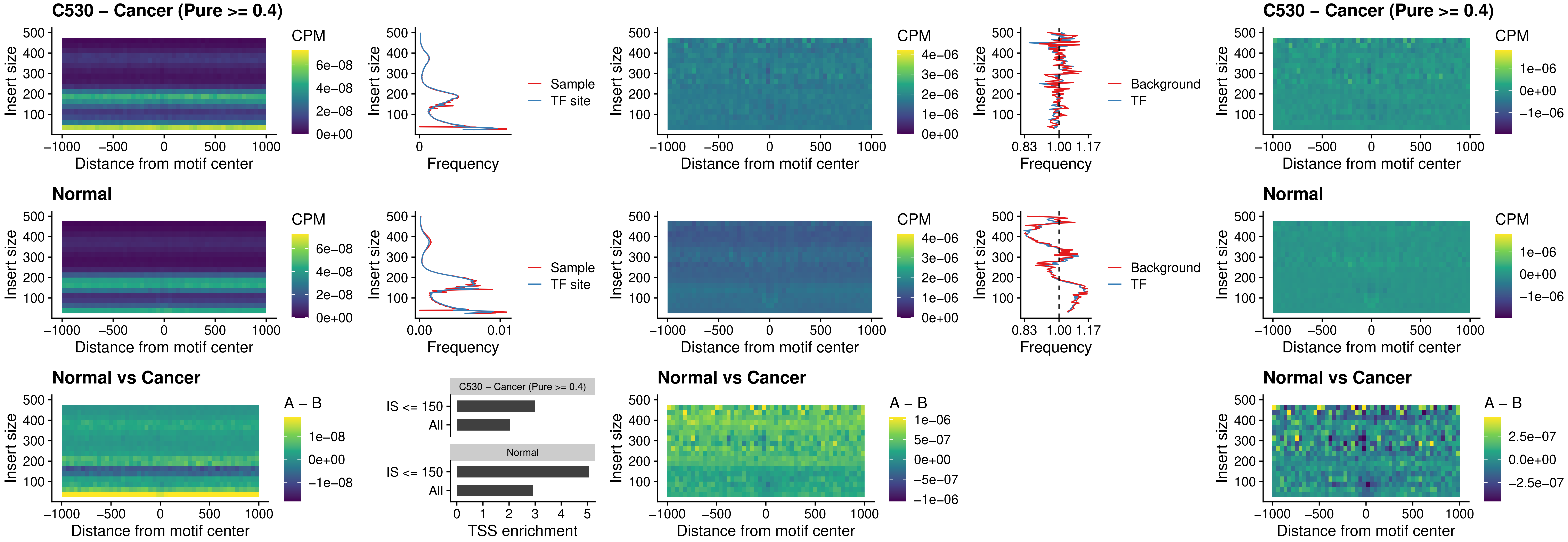

#### C530 – Cancer (Pure >= 0.4) – STAT2_uc_dTSS_oPEAK

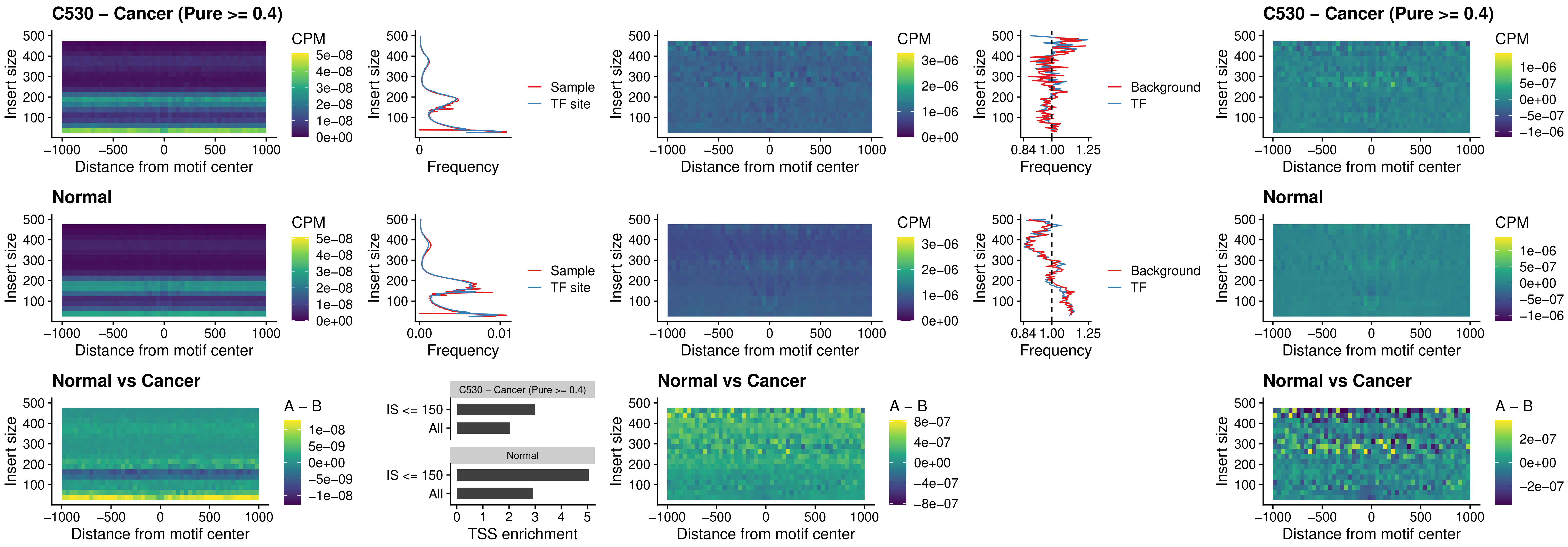

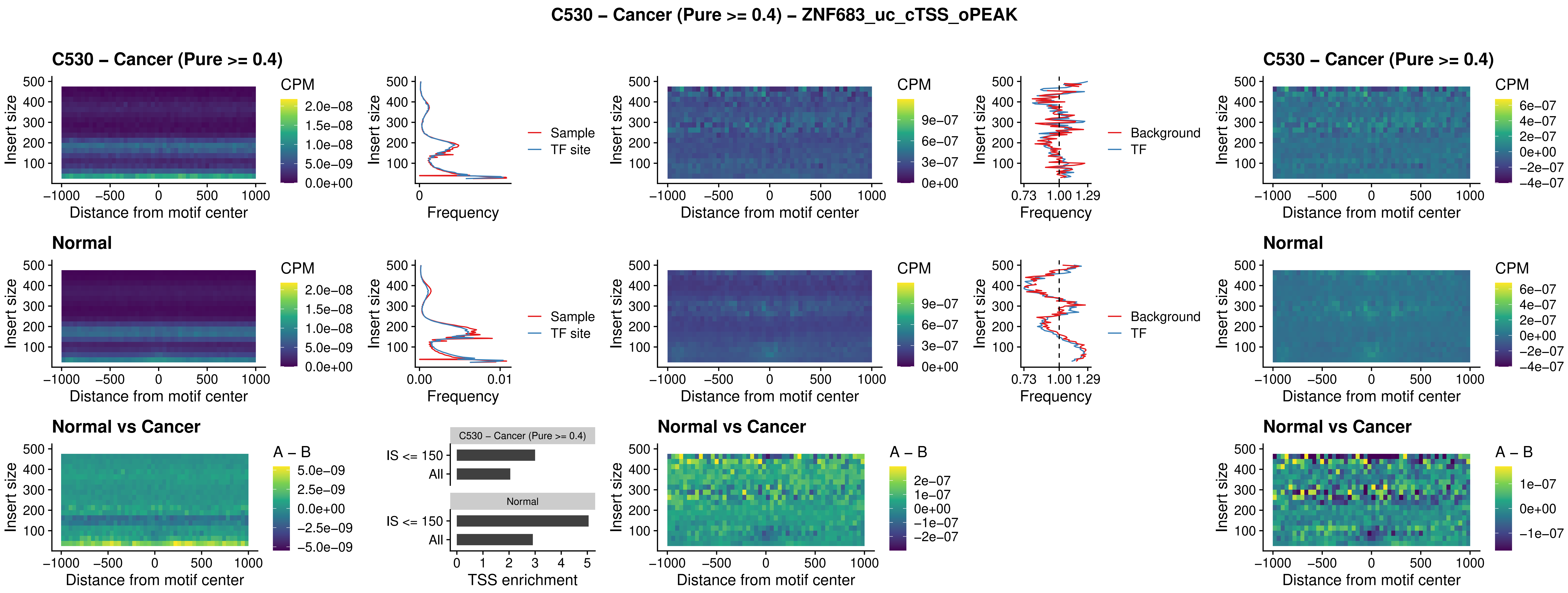
C530 – Cancer (Pure >= 0.4) – ZNF683_uc_cTSS_oPEAK
C530 – Cancer (Pure >= 0.4)
Normal
Normal vs Cancer
Insert size
Distance from motif center
Frequency
CPM
A – B
Sample
TF site
Background
TF
IS <= 150
All
Normal
TSS enrichment

### C530 – Cancer (Pure >= 0.4) – IRF8_uc_dTSS_nPEAK

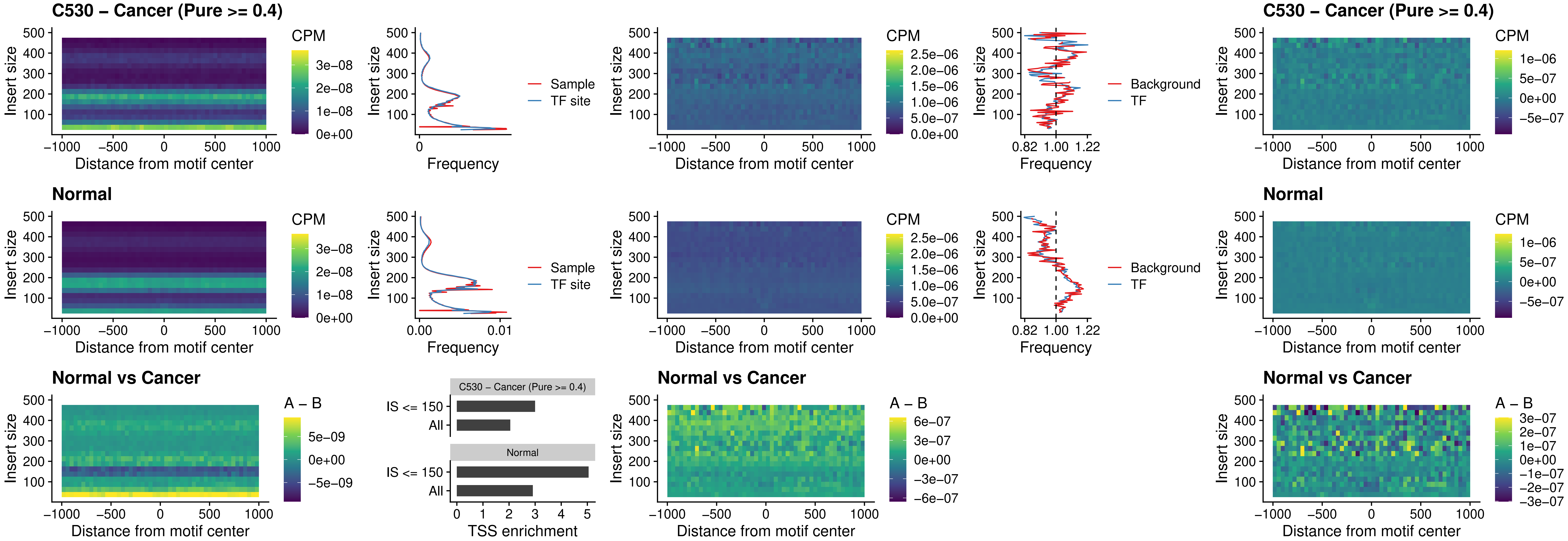

### C530 – Cancer (Pure >= 0.4) – ETV7_uc_dTSS_oPEAK

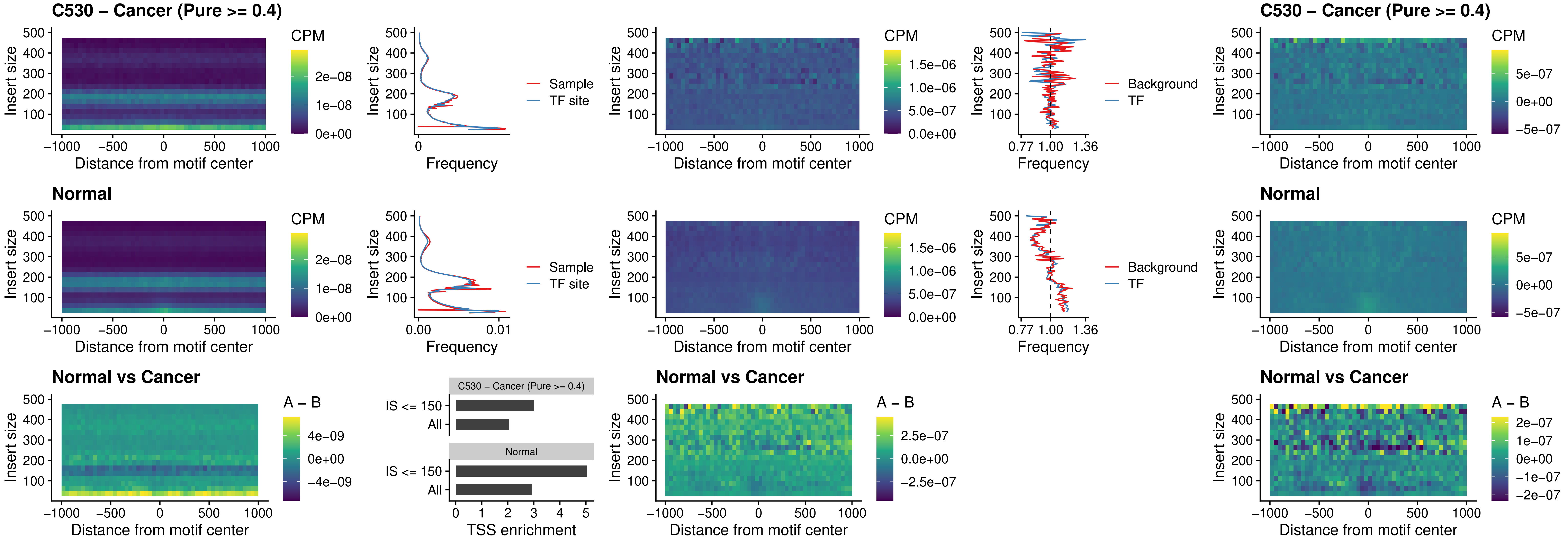

### C531 – Cancer (Pure >= 0.4) – FOXL1_uc_pTSS_oPEAK

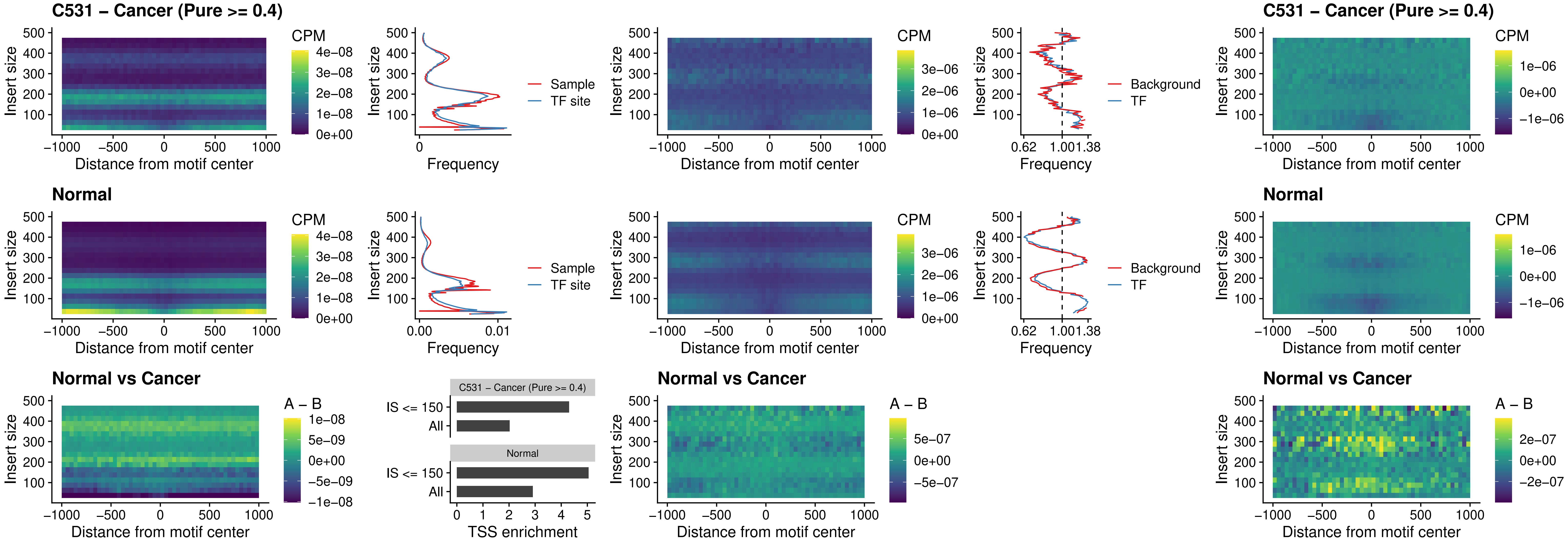

#### C531 – Cancer (Pure >= 0.4) – CPEB1_uc_cTSS_oPEAK

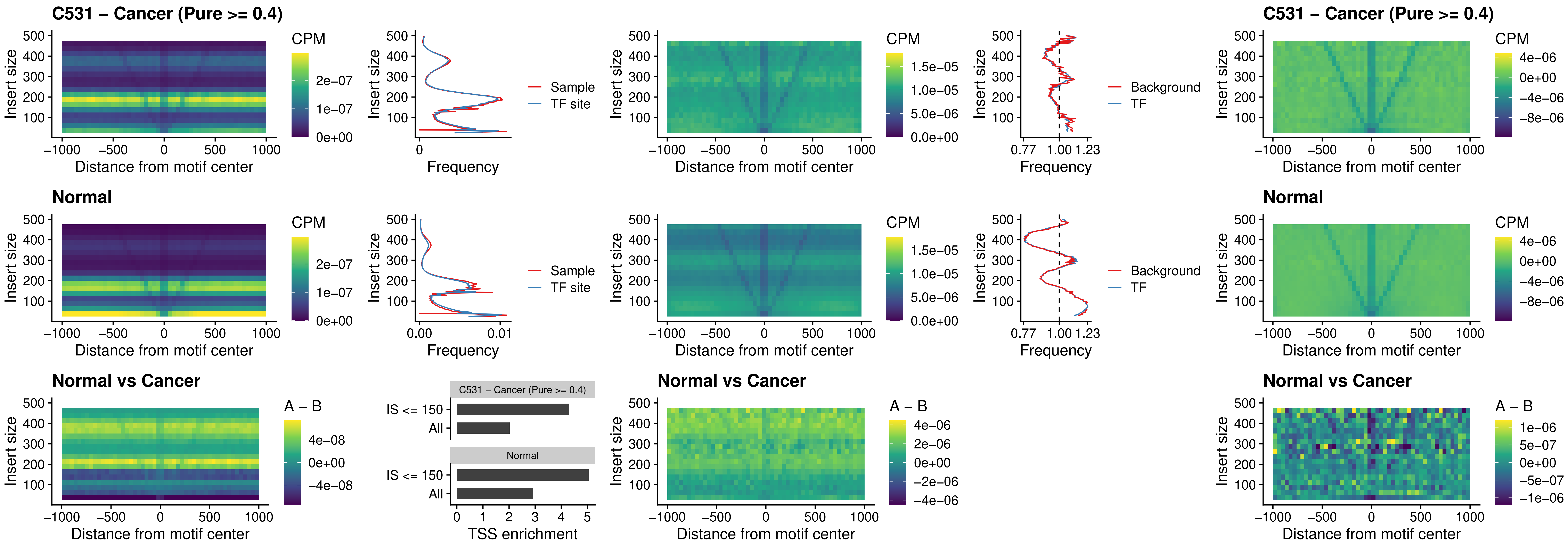

### C531 – Cancer (Pure >= 0.4) – HMGA1_uc_pTSS_nPEAK

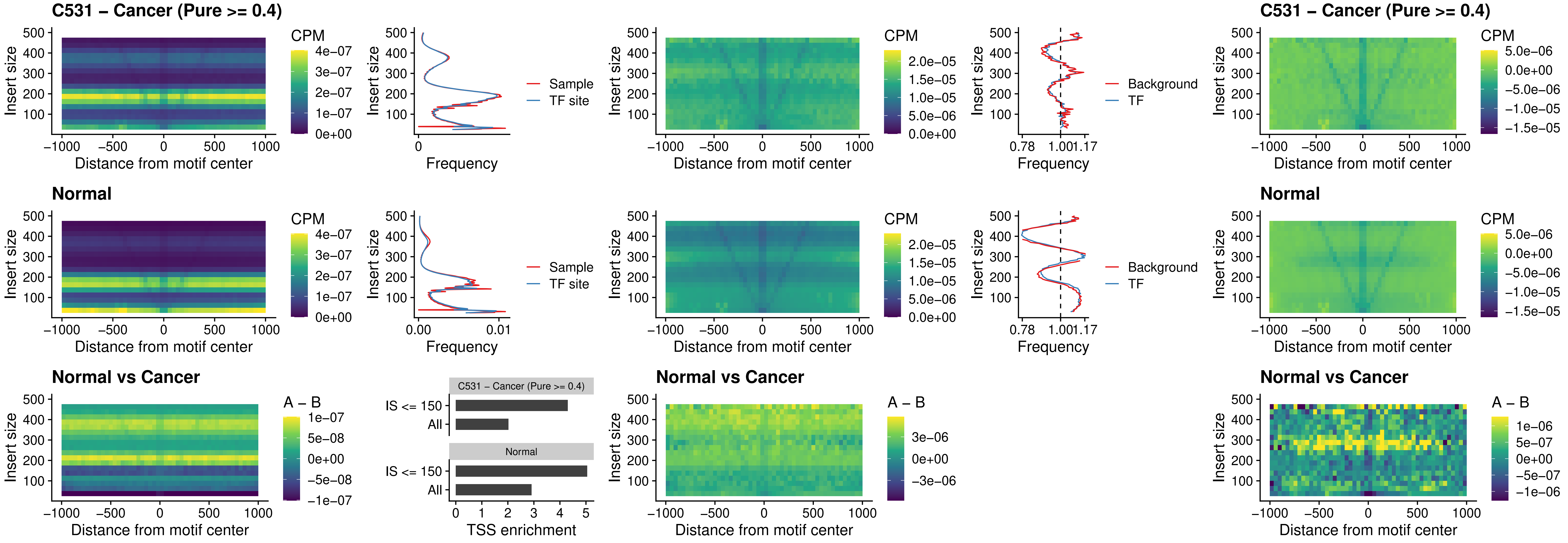

### C531 – Cancer (Pure >= 0.4) – SOX13_uc_cTSS_nPEAK

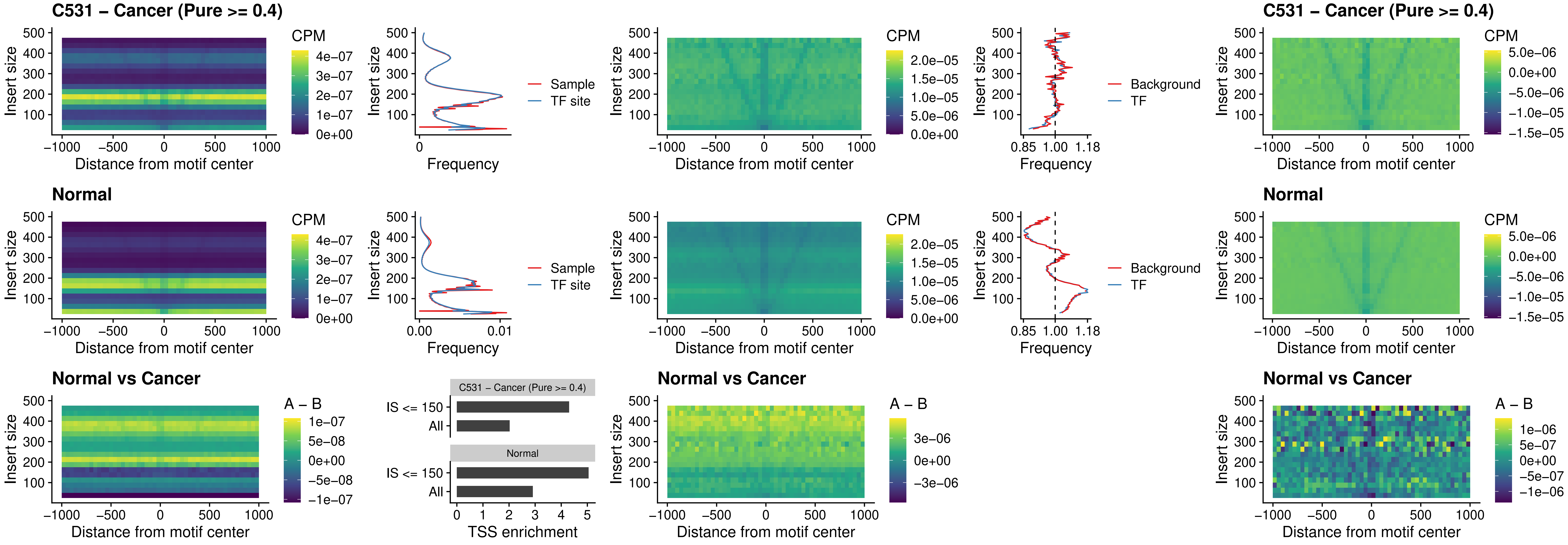

### C531 – Cancer (Pure >= 0.4) – LHX4_uc_cTSS_nPEAK

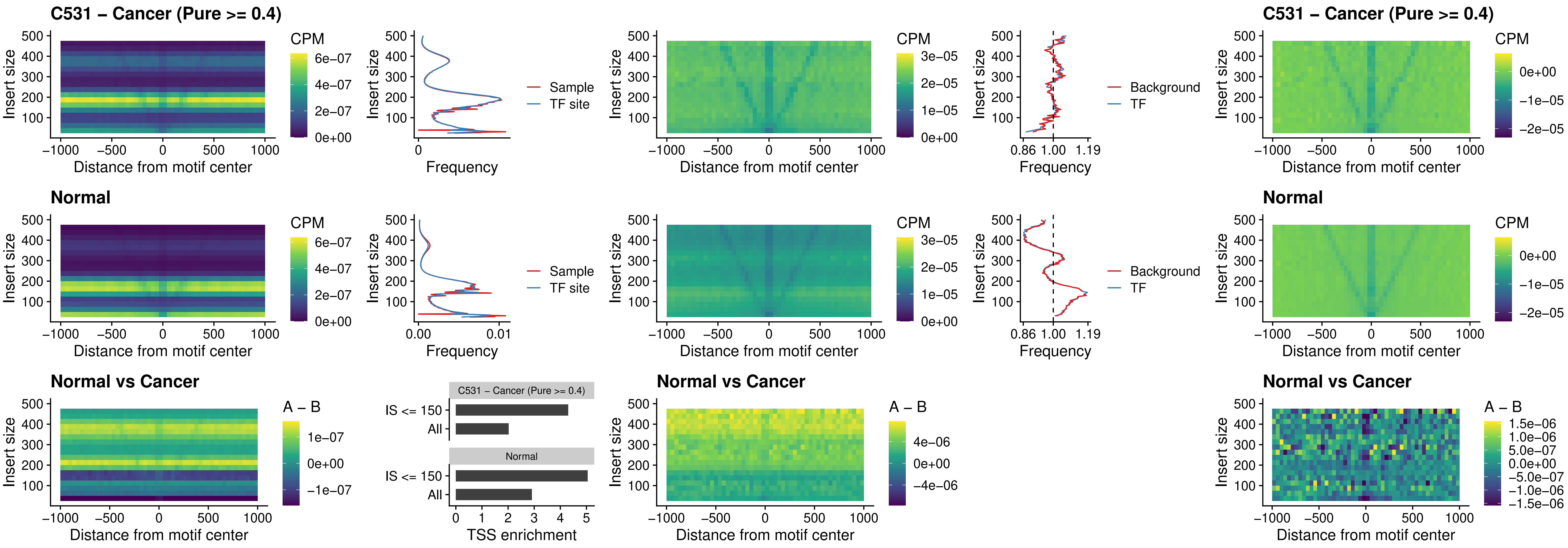

#### C531 – Cancer (Pure >= 0.4) – HOXA4_uc_cTSS_nPEAK

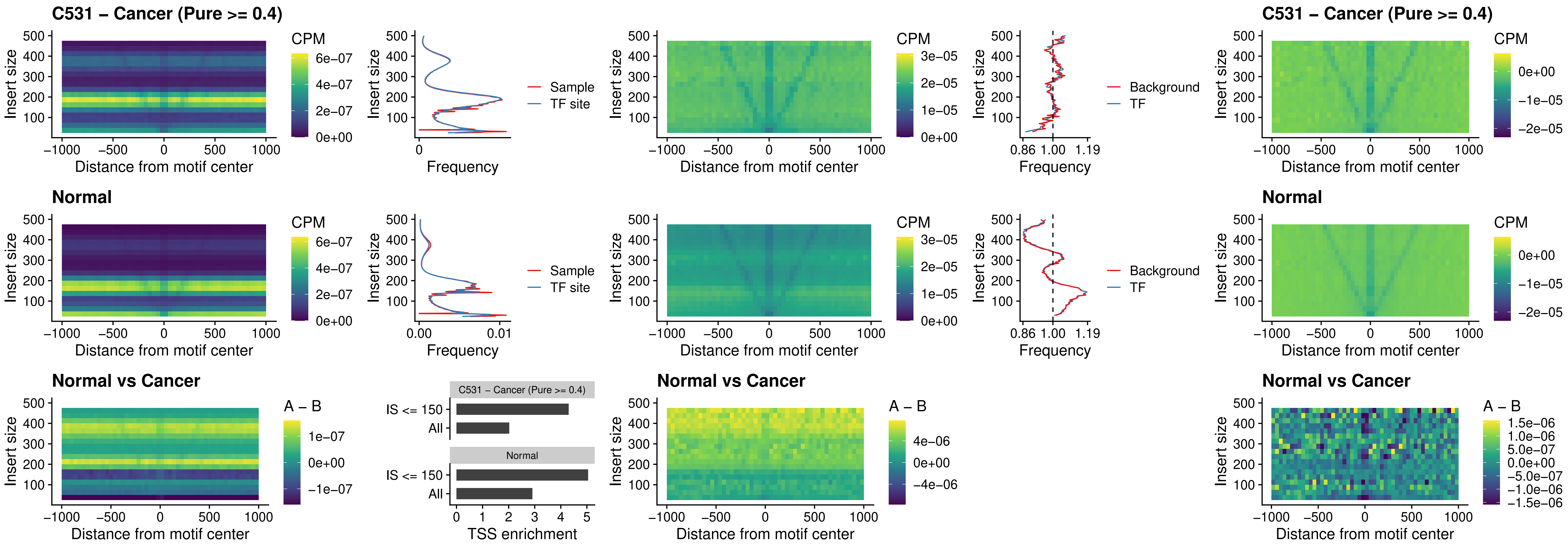

#### C531 – Cancer (Pure >= 0.4) – HLX_uc_cTSS_nPEAK

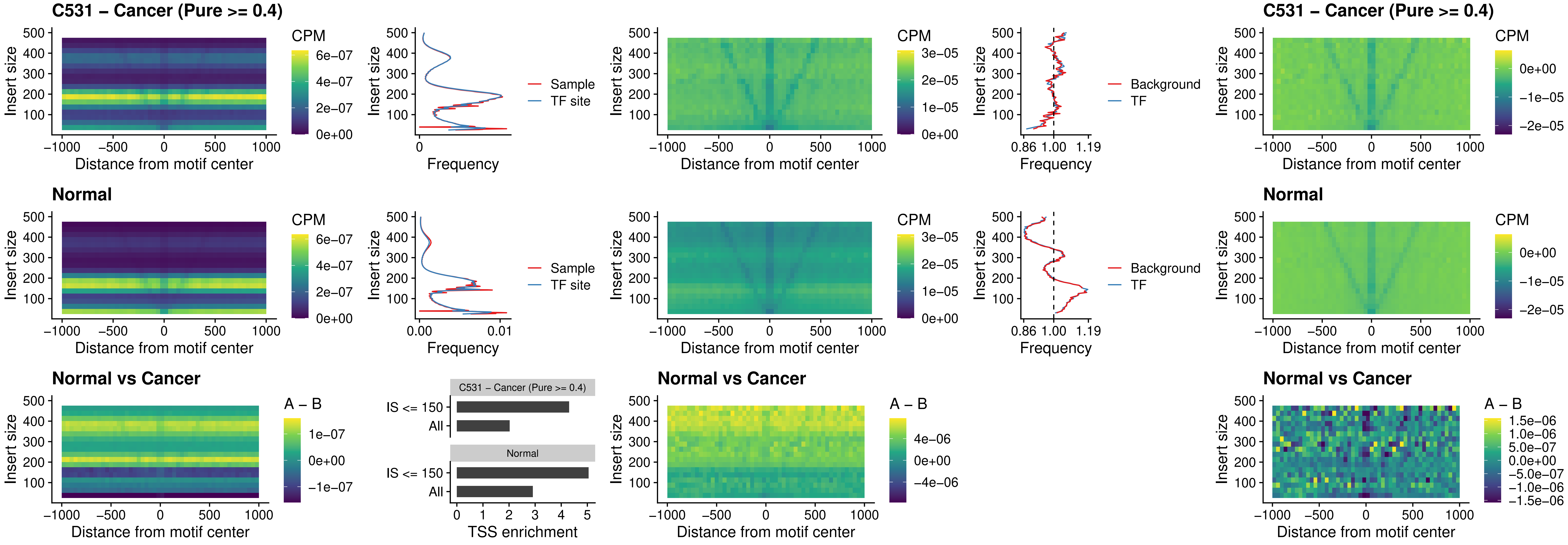

#### C531 – Cancer (Pure >= 0.4) – AC0125311_uc_cTSS_nPEAK

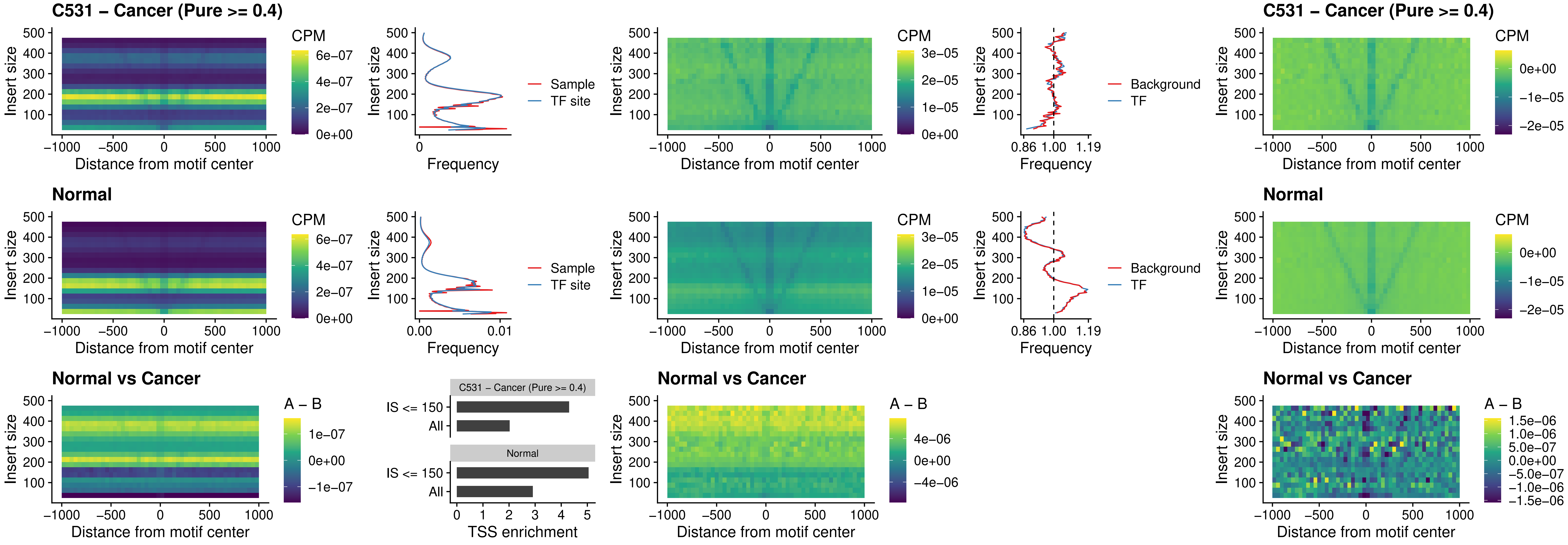

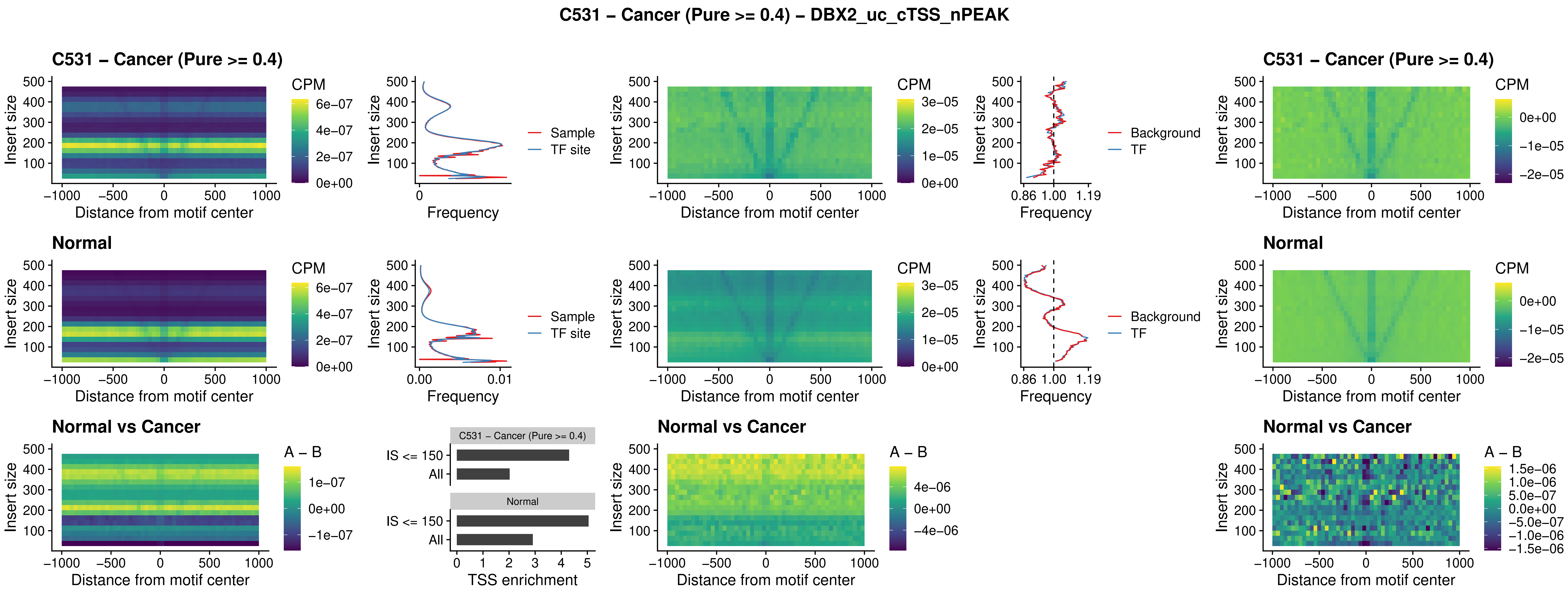
C531 – Cancer (Pure >= 0.4) – DBX2_uc_cTSS_nPEAK
C531 – Cancer (Pure >= 0.4)
Normal
Normal vs Cancer
Insert size
Distance from motif center
Frequency
CPM
A – B
Sample
TF site
Background
TF
C531 – Cancer (Pure >= 0.4)
Normal
IS <= 150
All
TSS enrichment

#### C531 – Cancer (Pure >= 0.4) – LHX6_uc_dTSS_nPEAK

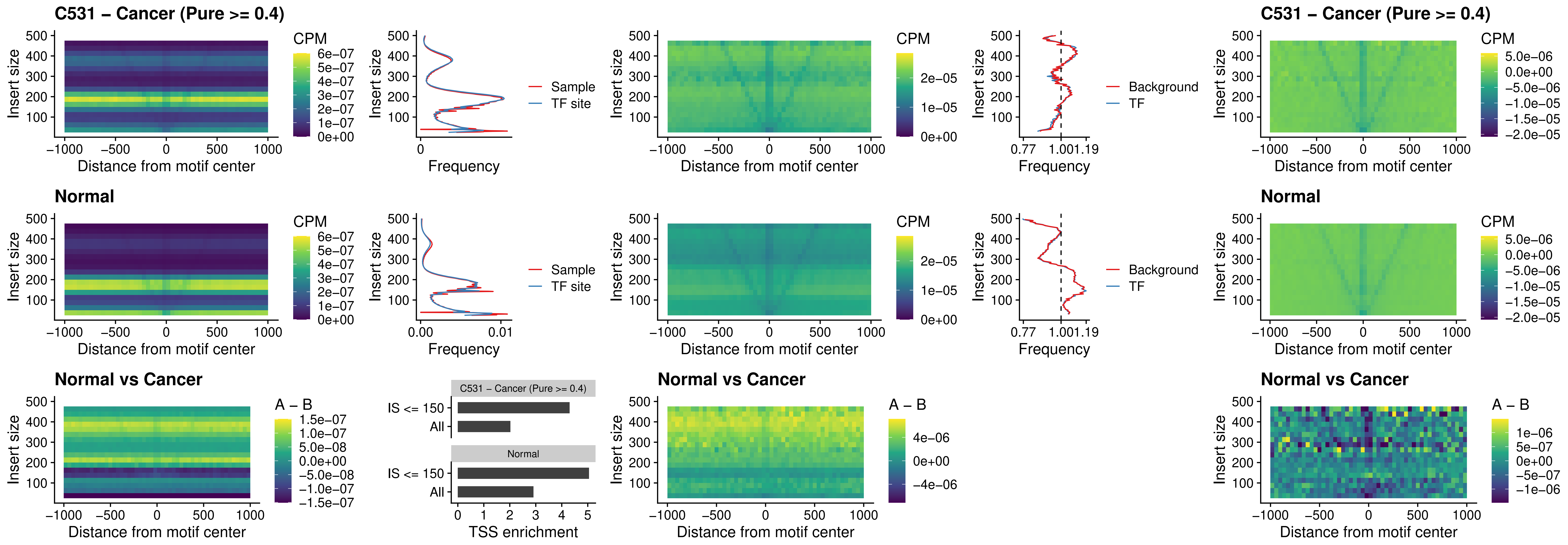

#### C531 – Cancer (Pure >= 0.4) – SOX1_uc_dTSS_nPEAK

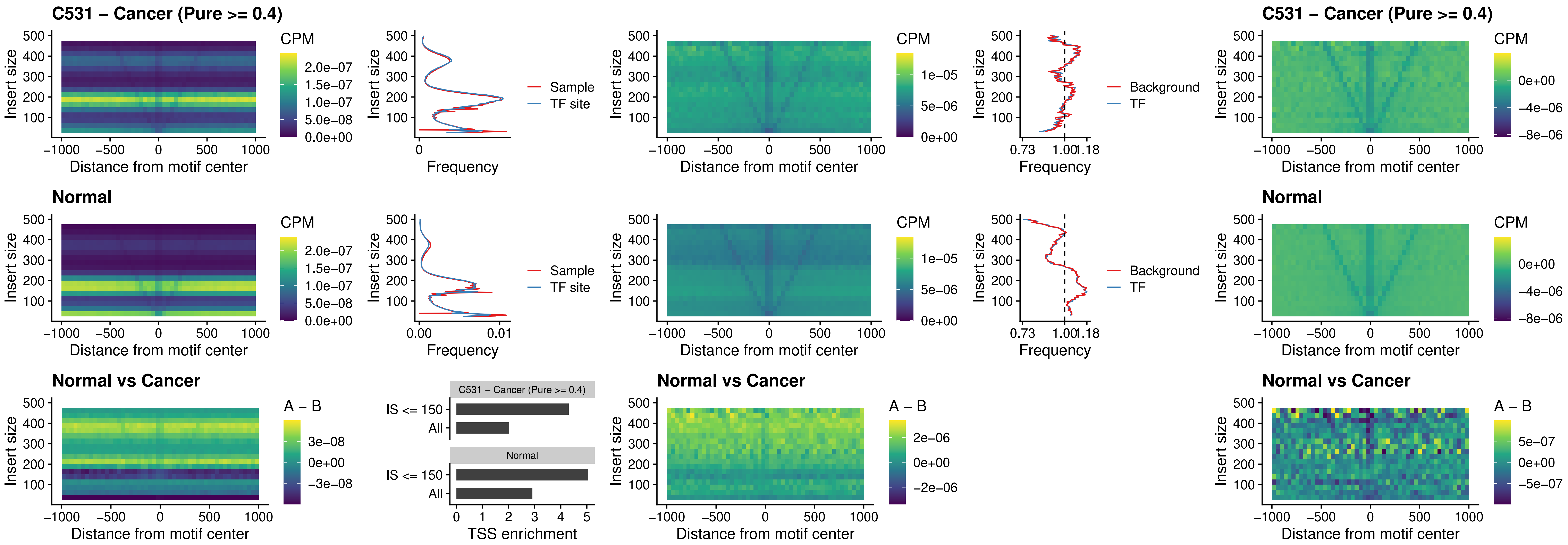

### C531 – Cancer (Pure >= 0.4) – NKX24_uc_dTSS_nPEAK

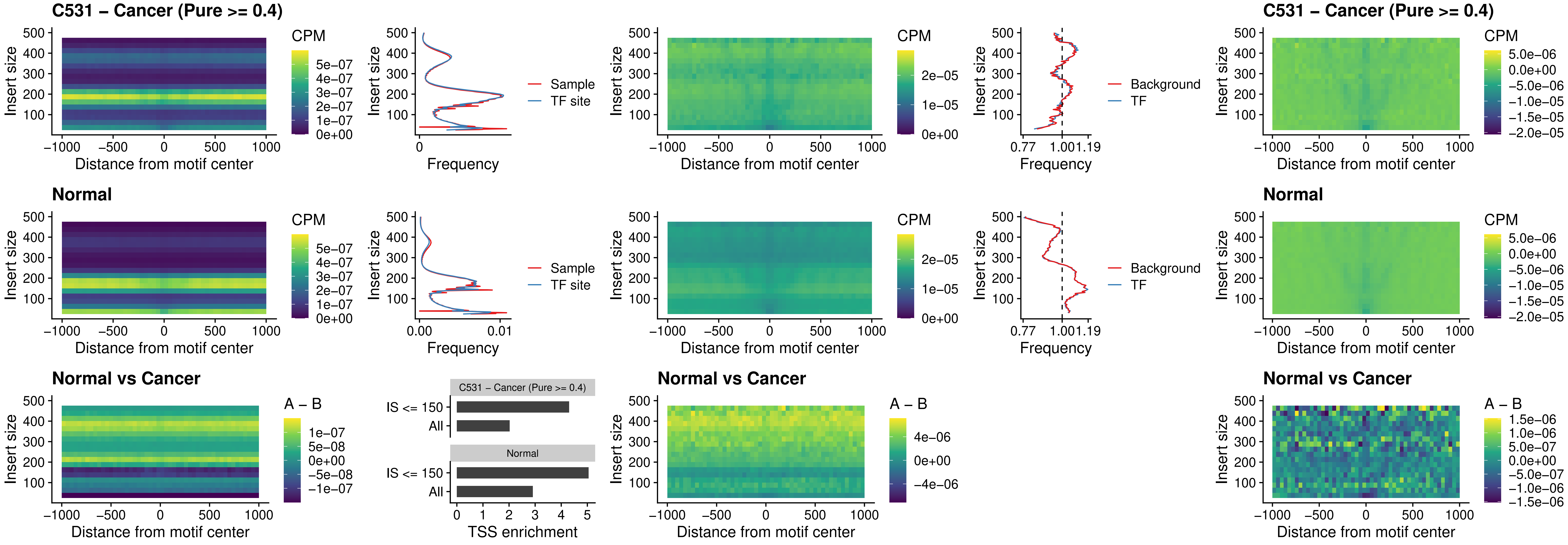

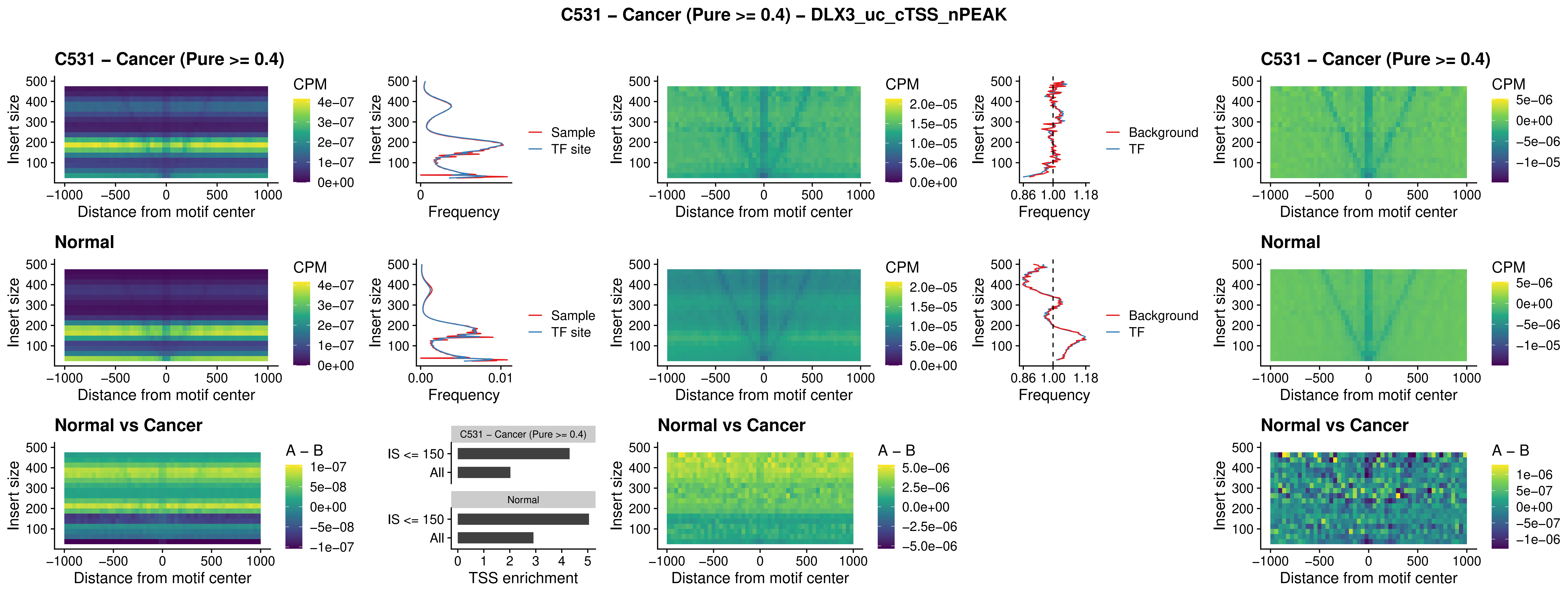
C531 – Cancer (Pure >= 0.4) – DLX3_uc_cTSS_nPEAK
C531 – Cancer (Pure >= 0.4)
Normal
Normal vs Cancer
Insert size
Distance from motif center
Frequency
CPM
A – B
Sample
TF site
Background
TF
C531 – Cancer (Pure >= 0.4)
Normal
IS <= 150
All
TSS enrichment

### C531 – Cancer (Pure >= 0.4) – HOXD3_uc_cTSS_nPEAK

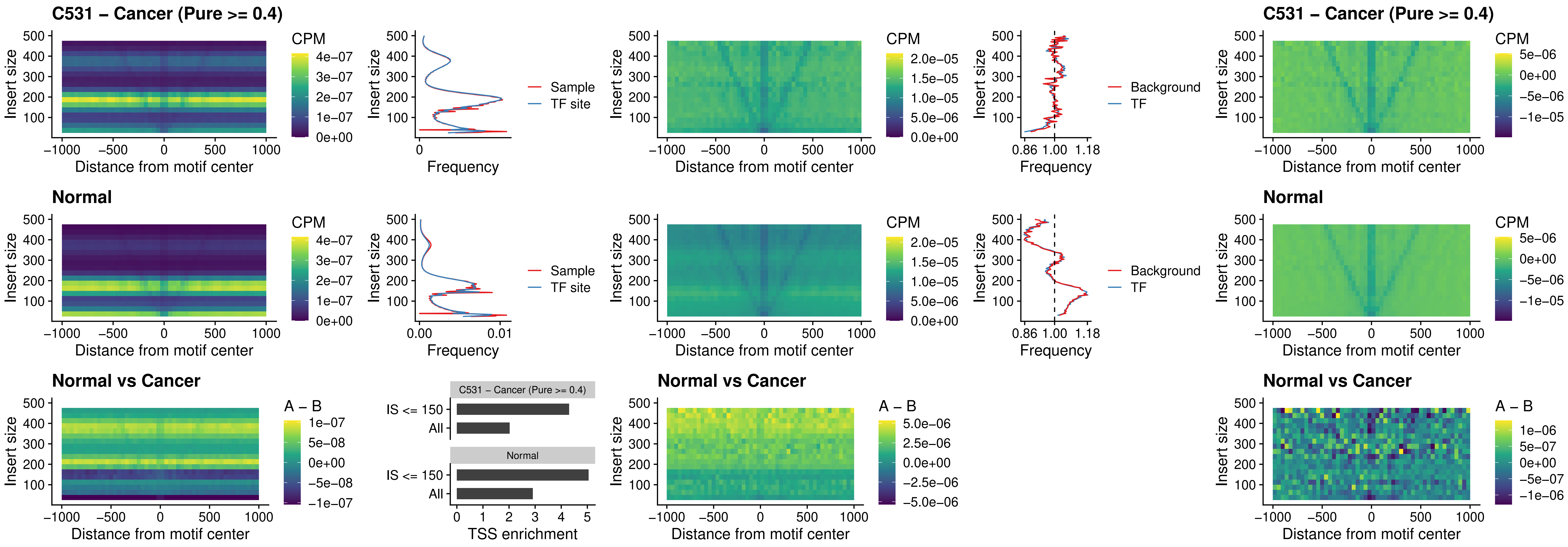

#### C531 – Cancer (Pure >= 0.4) – ISL1_uc_dTSS_oPEAK

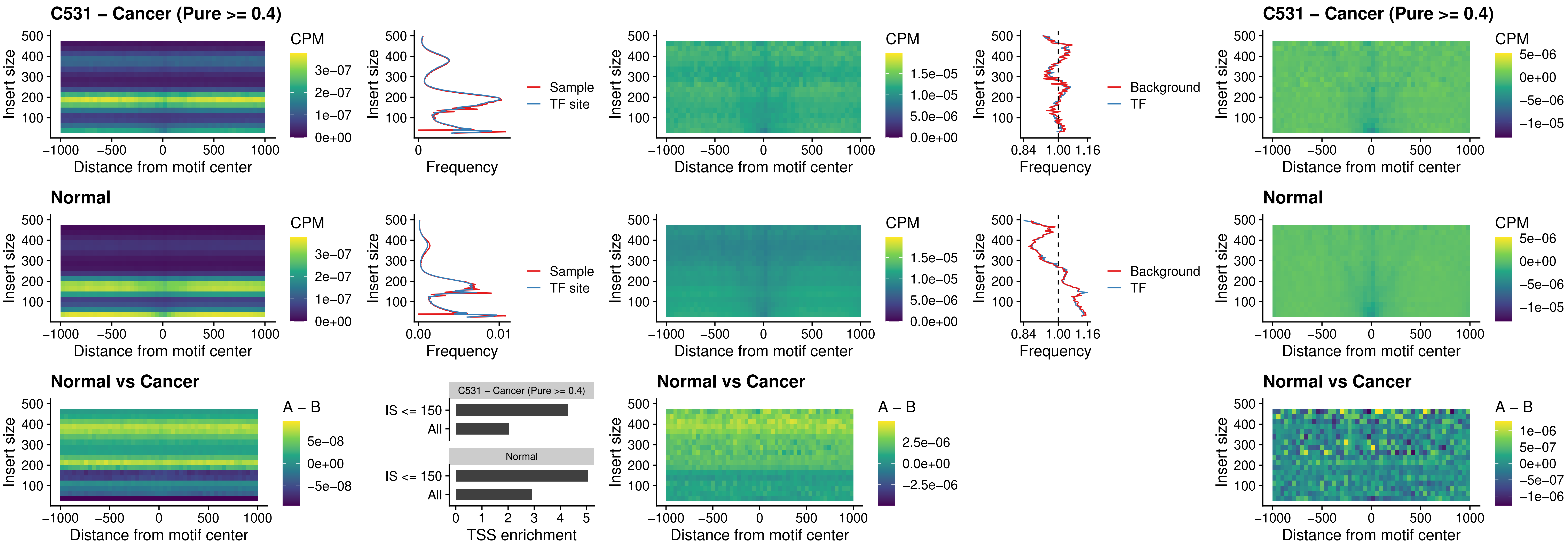

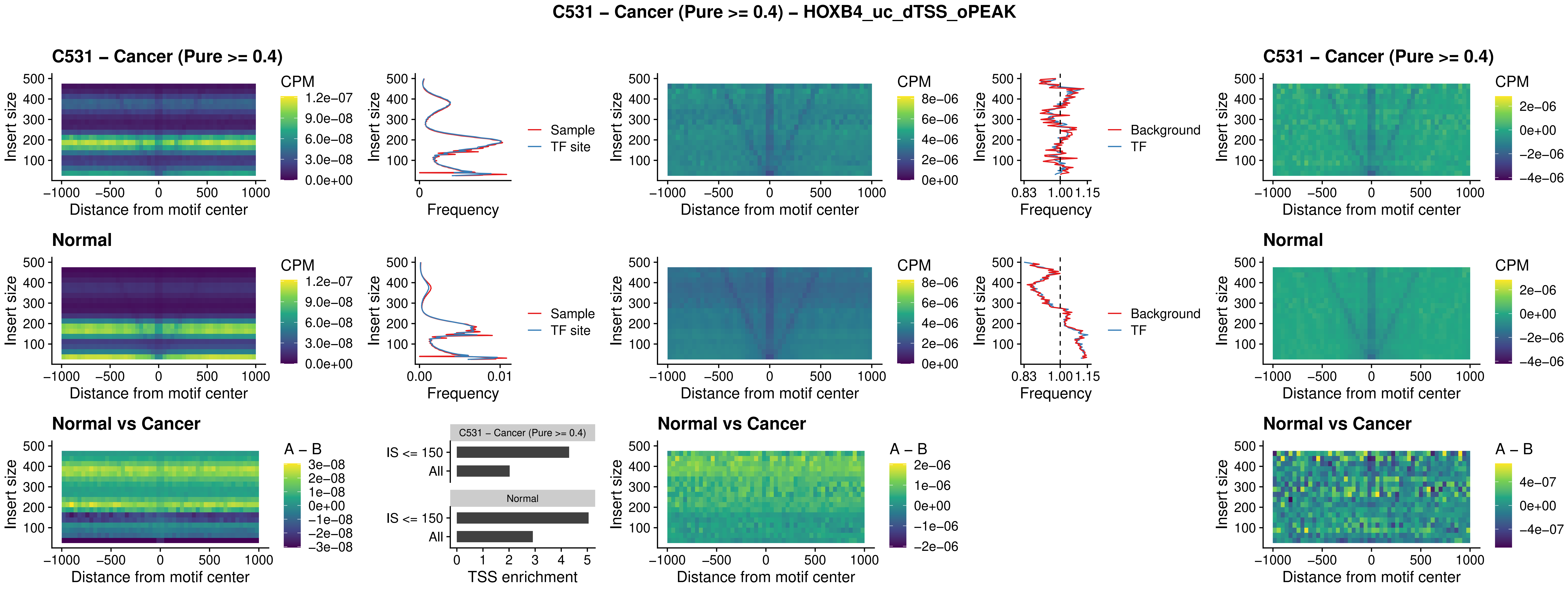
C531 – Cancer (Pure >= 0.4) – HOXB4_uc_dTSS_oPEAK
C531 – Cancer (Pure >= 0.4)
Normal
Normal vs Cancer
Insert size
Distance from motif center
Frequency
CPM
A – B
Sample
TF site
Background
TF
C531 – Cancer (Pure >= 0.4)
Normal
IS <= 150
All
TSS enrichment

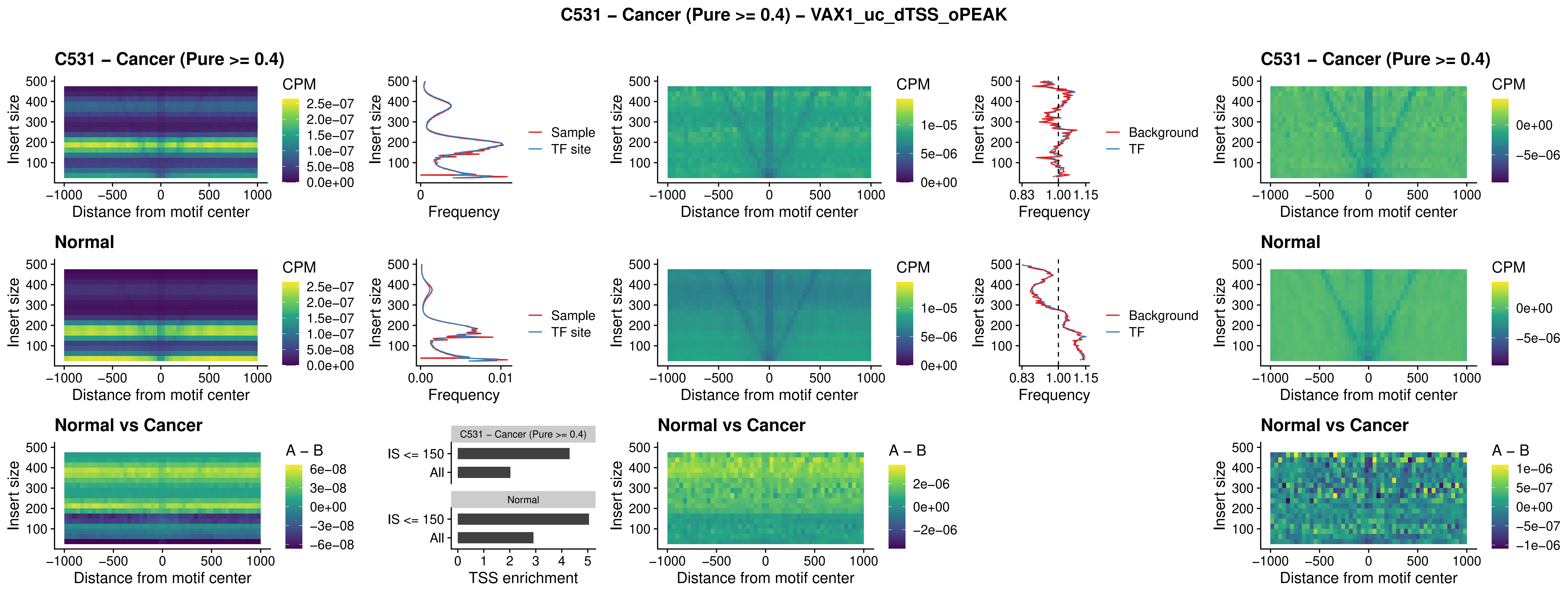
C531 – Cancer (Pure >= 0.4) – VAX1_uc_dTSS_oPEAK
C531 – Cancer (Pure >= 0.4)
Normal
Normal vs Cancer
Insert size
Distance from motif center
Frequency
CPM
A – B
Sample
TF site
Background
TF
C531 – Cancer (Pure >= 0.4)
Normal
IS <= 150
All
TSS enrichment

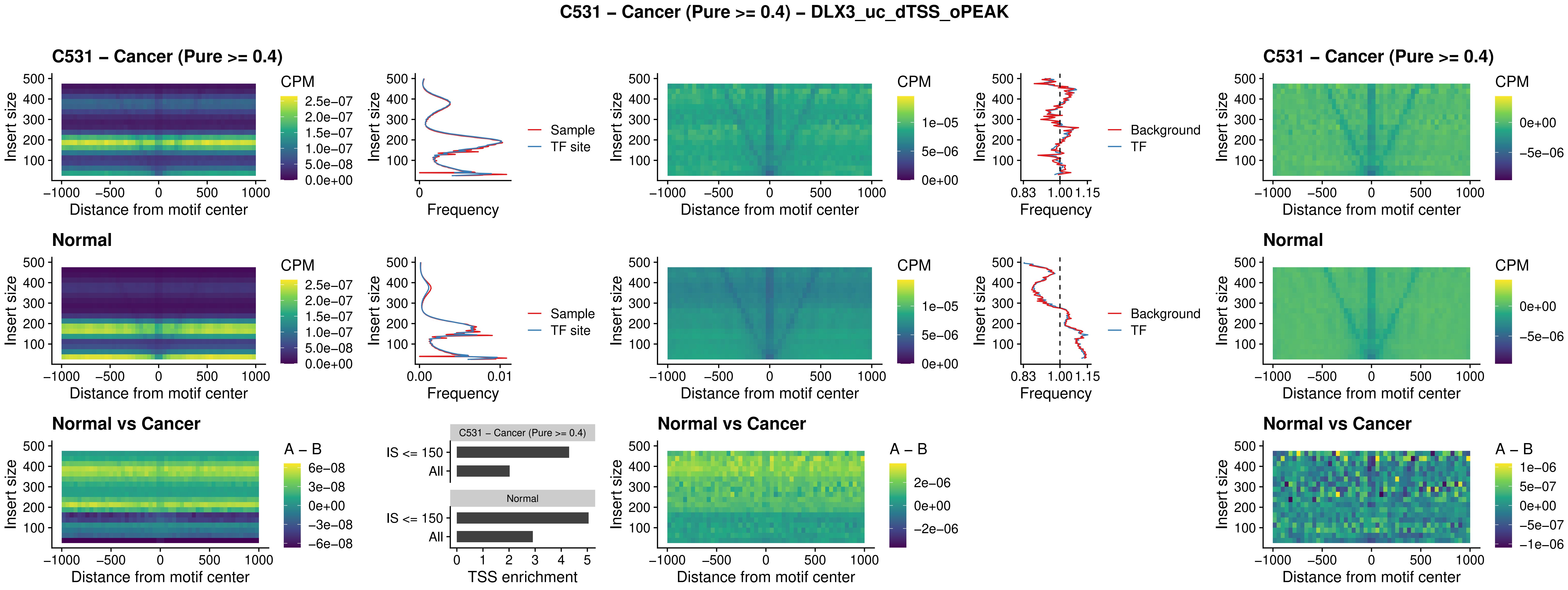
C531 – Cancer (Pure >= 0.4) – DLX3_uc_dTSS_oPEAK
C531 – Cancer (Pure >= 0.4)
Normal
Normal vs Cancer
Insert size
Distance from motif center
Frequency
CPM
A – B
Sample
TF site
Background
TF
C531 – Cancer (Pure >= 0.4)
Normal
IS <= 150
All
TSS enrichment

#### C531 – Cancer (Pure >= 0.4) – HOXD3_uc_dTSS_oPEAK

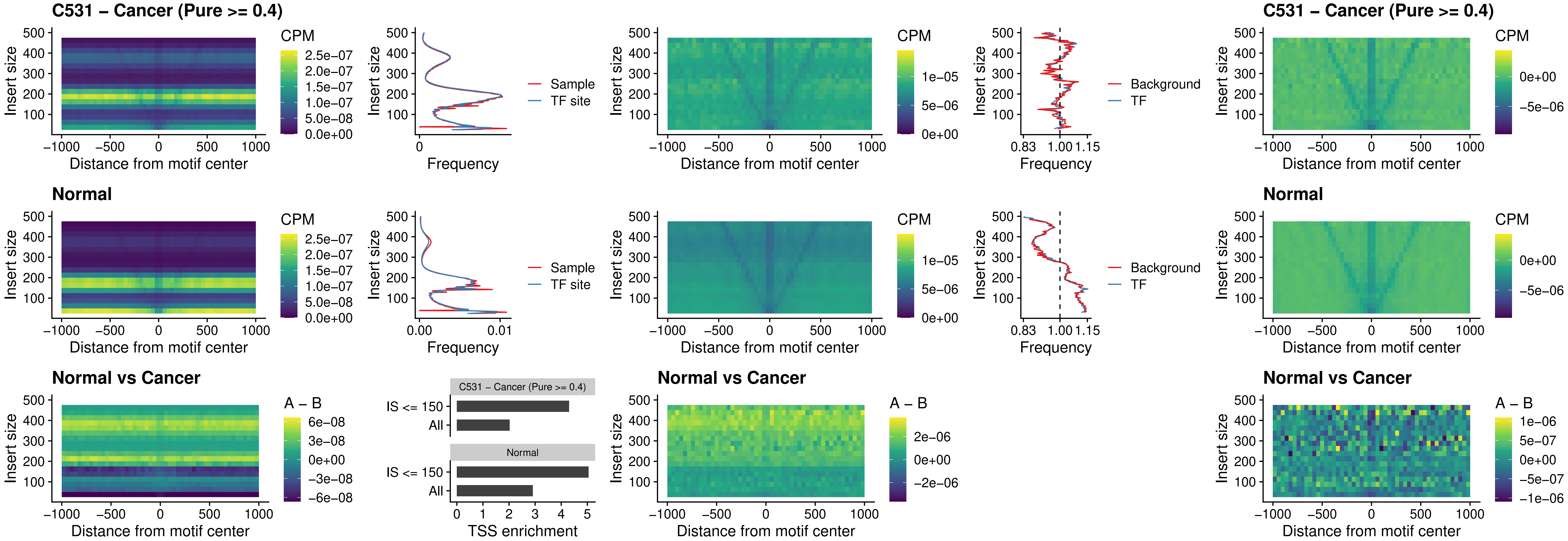

#### C531 – Cancer (Pure >= 0.4) – BBX_uc_dTSS_nPEAK

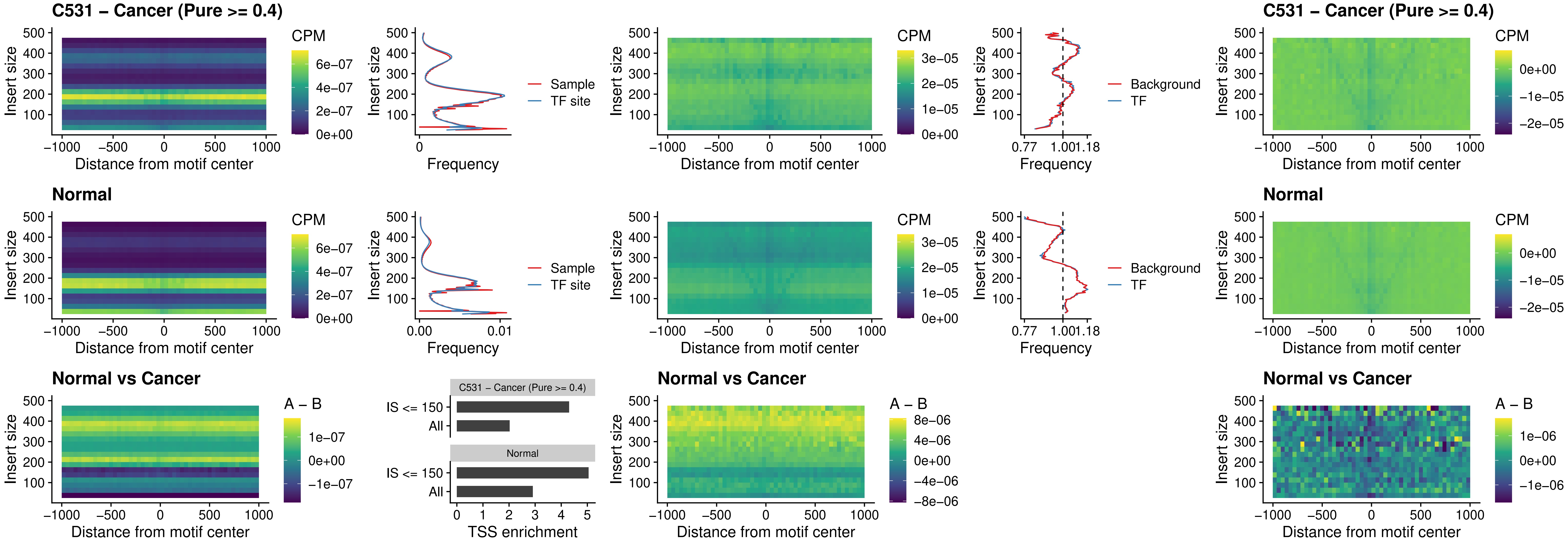

#### C531 – Cancer (Pure >= 0.4) – SOX3_uc_cTSS_nPEAK

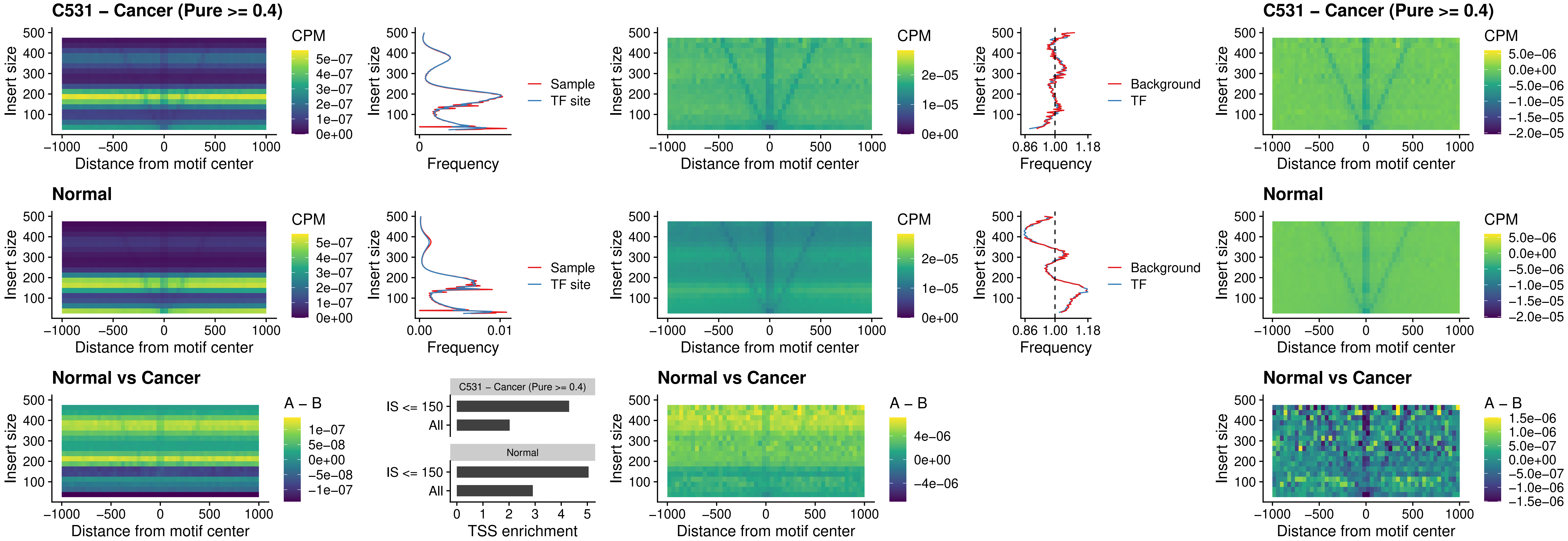

### C531 – Cancer (Pure >= 0.4) – TBPL2_uc_cTSS_nPEAK

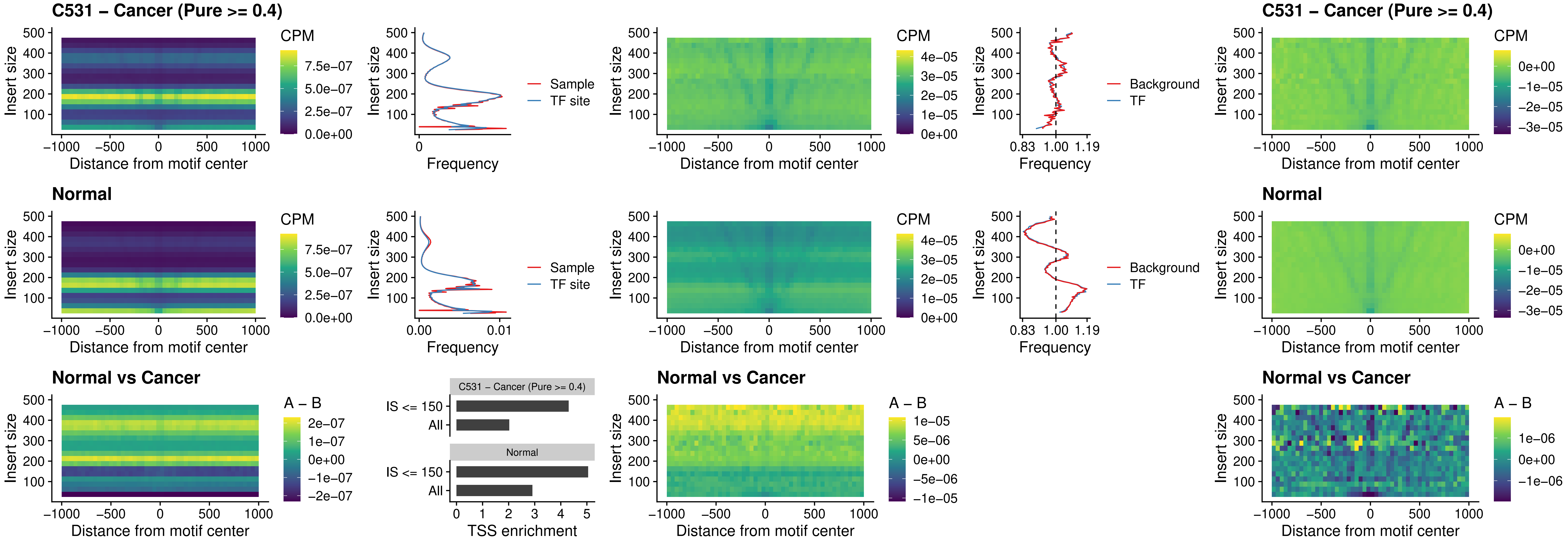

#### C531 – Cancer (Pure >= 0.4) – HOXA4_uc_cTSS_oPEAK

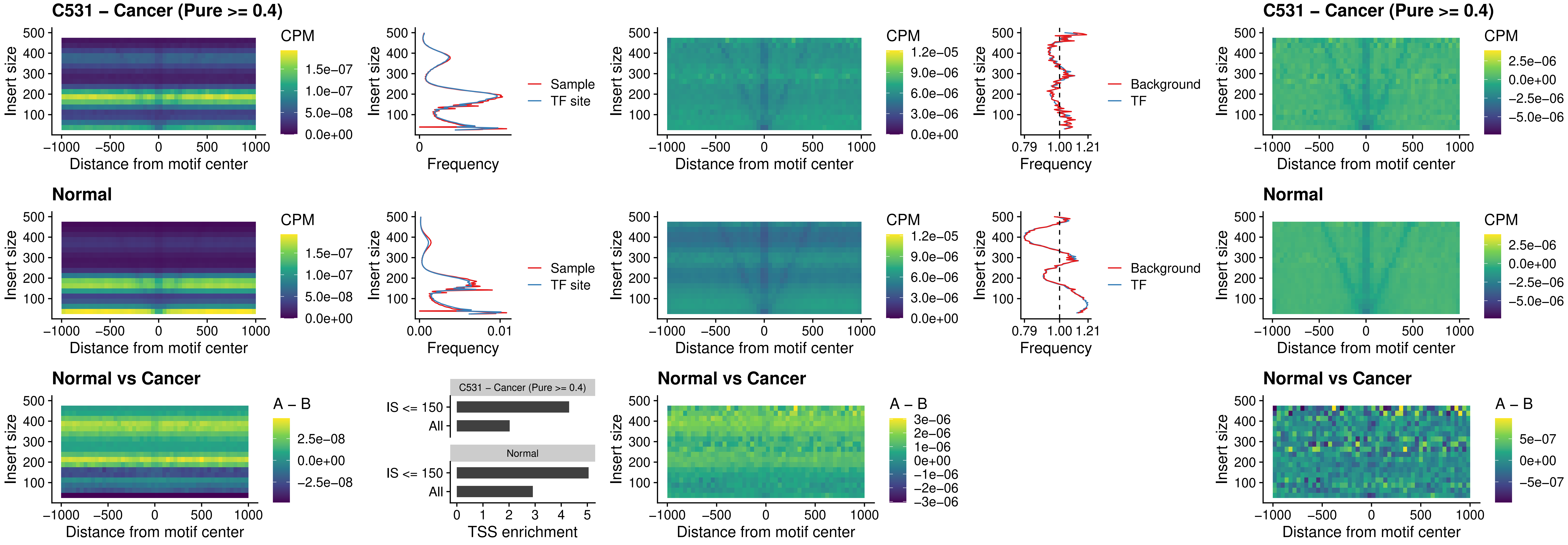

### C531 – Cancer (Pure >= 0.4) – PDX1_uc_dTSS_oPEAK

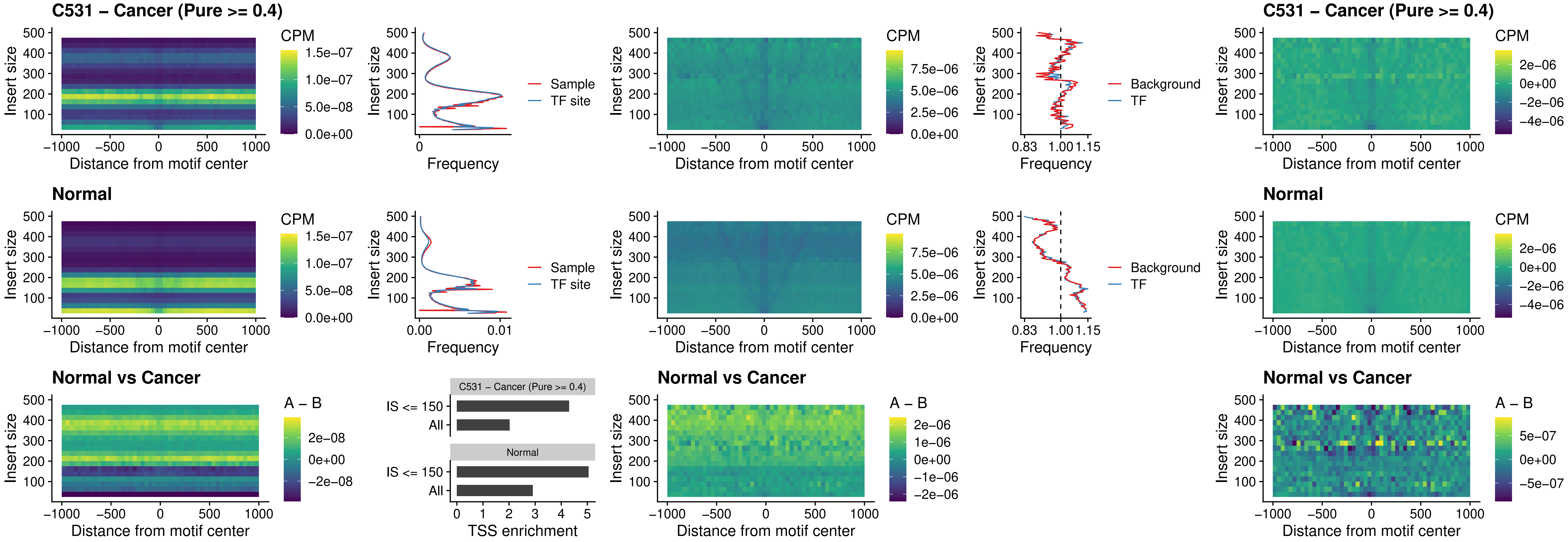

#### C531 – Cancer (Pure >= 0.4) – LHX4_uc_dTSS_oPEAK

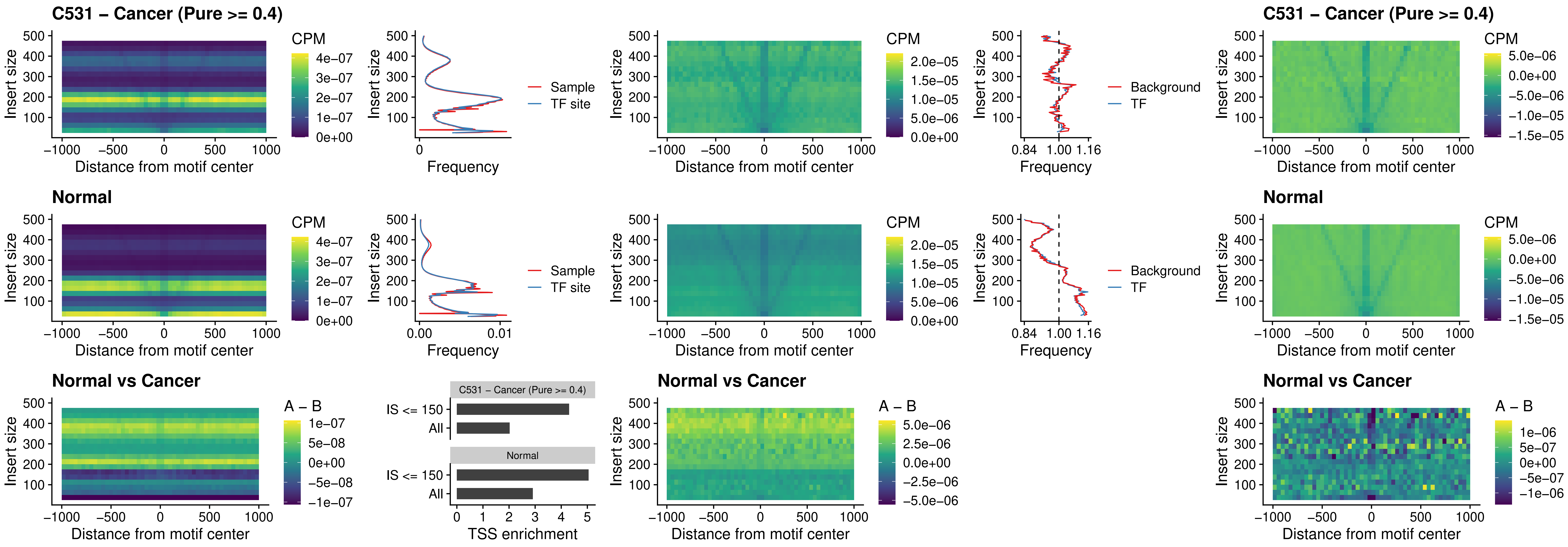

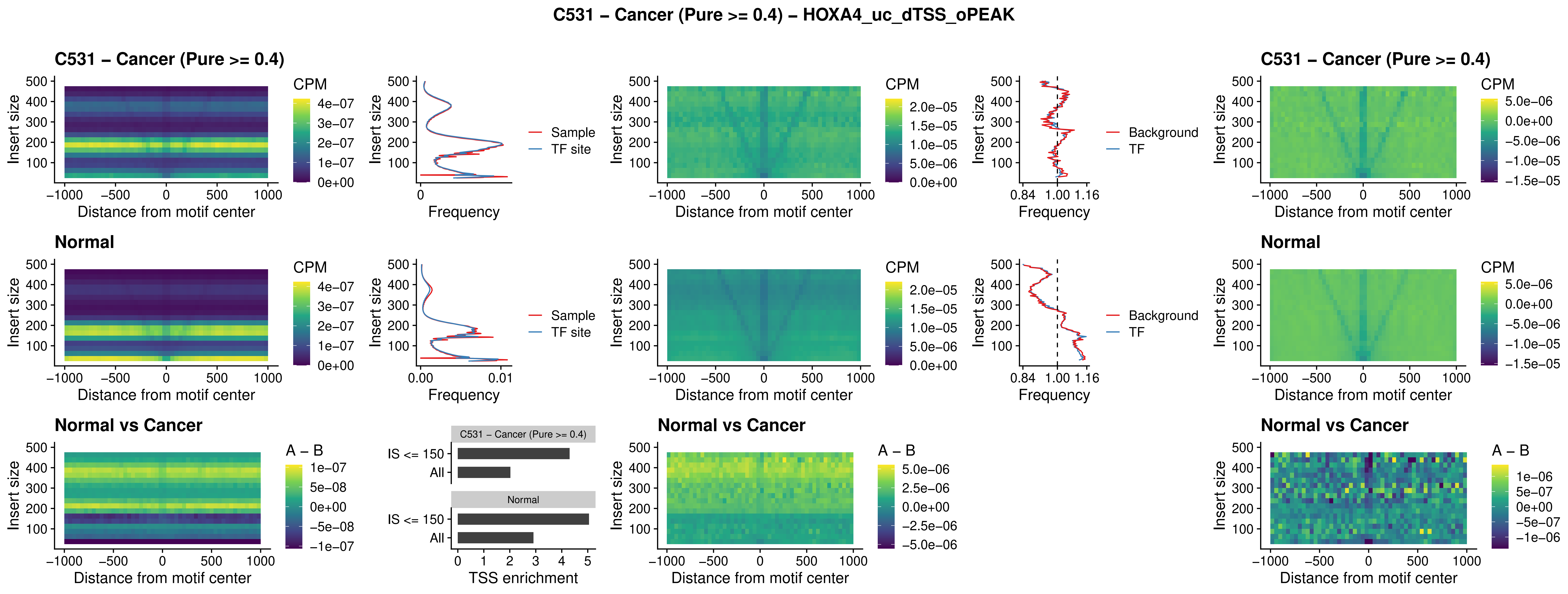
C531 – Cancer (Pure >= 0.4) – HOXA4_uc_dTSS_oPEAK
C531 – Cancer (Pure >= 0.4)
Normal
Normal vs Cancer
Insert size
Distance from motif center
Frequency
CPM
A – B
Sample
TF site
Background
TF
C531 – Cancer (Pure >= 0.4)
Normal
IS <= 150
All
TSS enrichment

#### C531 – Cancer (Pure >= 0.4) – HLX_uc_dTSS_oPEAK

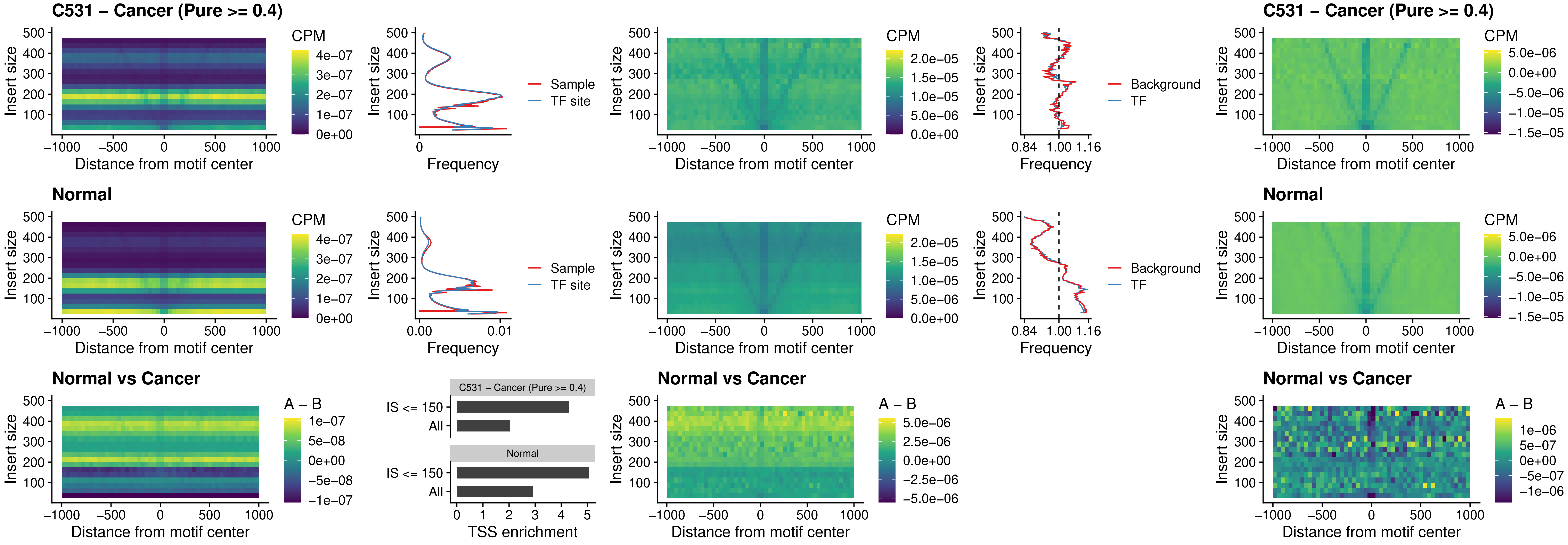

#### C531 – Cancer (Pure >= 0.4) – AC0125311_uc_dTSS_oPEAK

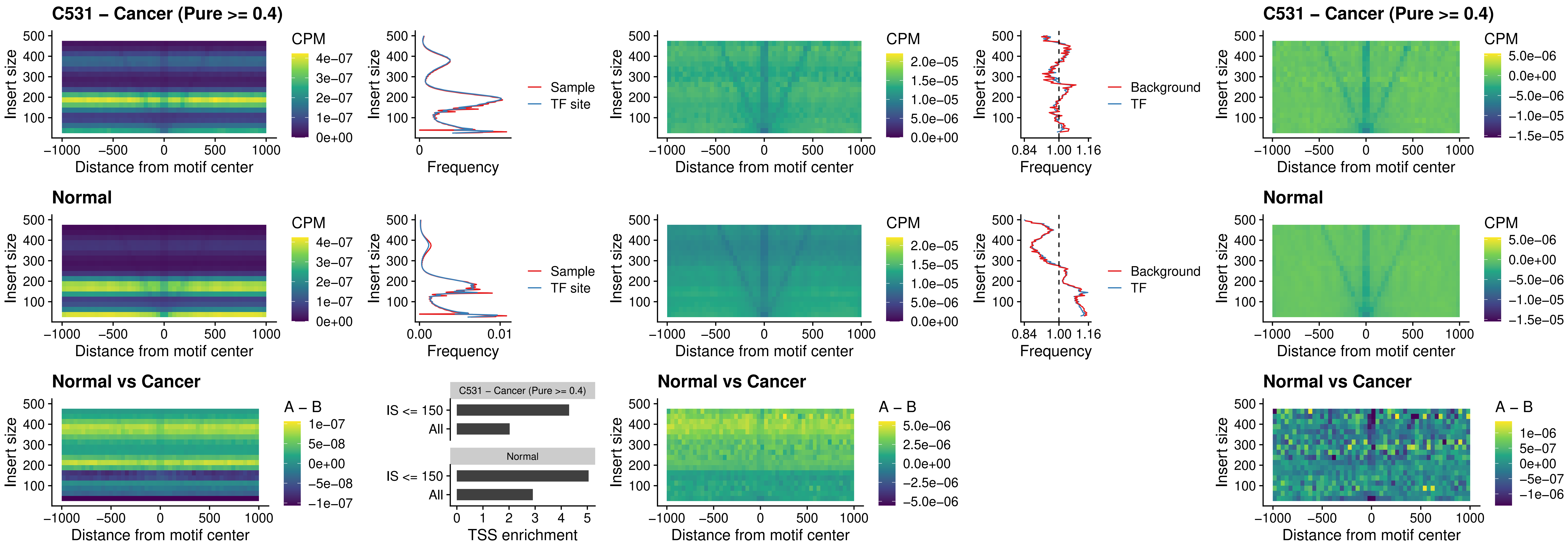

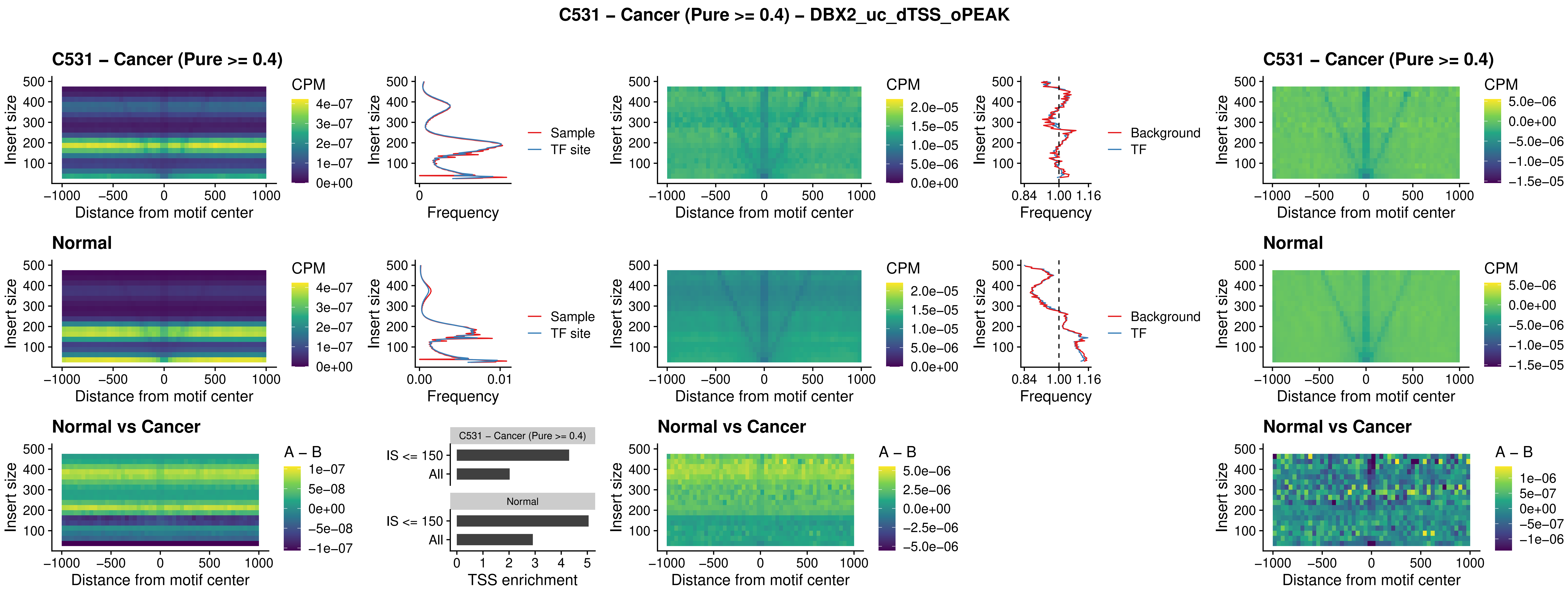
C531 – Cancer (Pure >= 0.4) – DBX2_uc_dTSS_oPEAK
C531 – Cancer (Pure >= 0.4)
Normal
Normal vs Cancer
Insert size
Distance from motif center
Frequency
CPM
A – B
Sample
TF site
Background
TF
C531 – Cancer (Pure >= 0.4)
Normal
IS <= 150
All
TSS enrichment

#### C531 – Cancer (Pure >= 0.4) – HOXA2_uc_dTSS_nPEAK

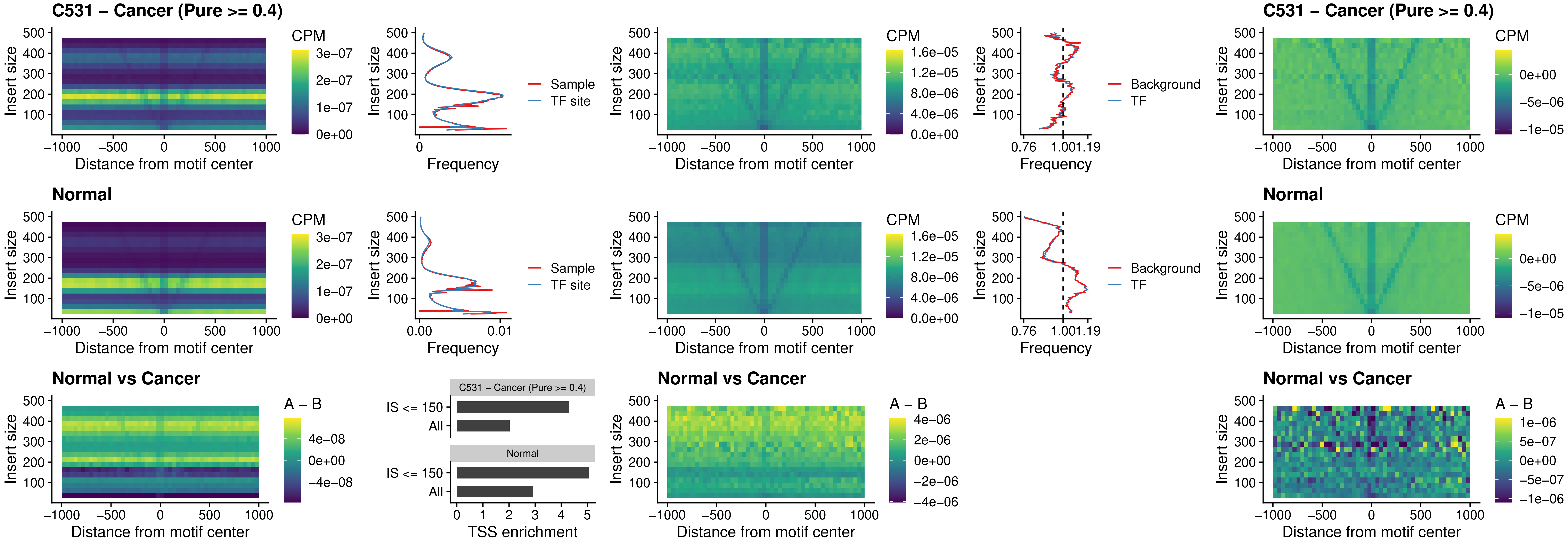

### C531 – Cancer (Pure >= 0.4) – SHOX_uc_dTSS_nPEAK

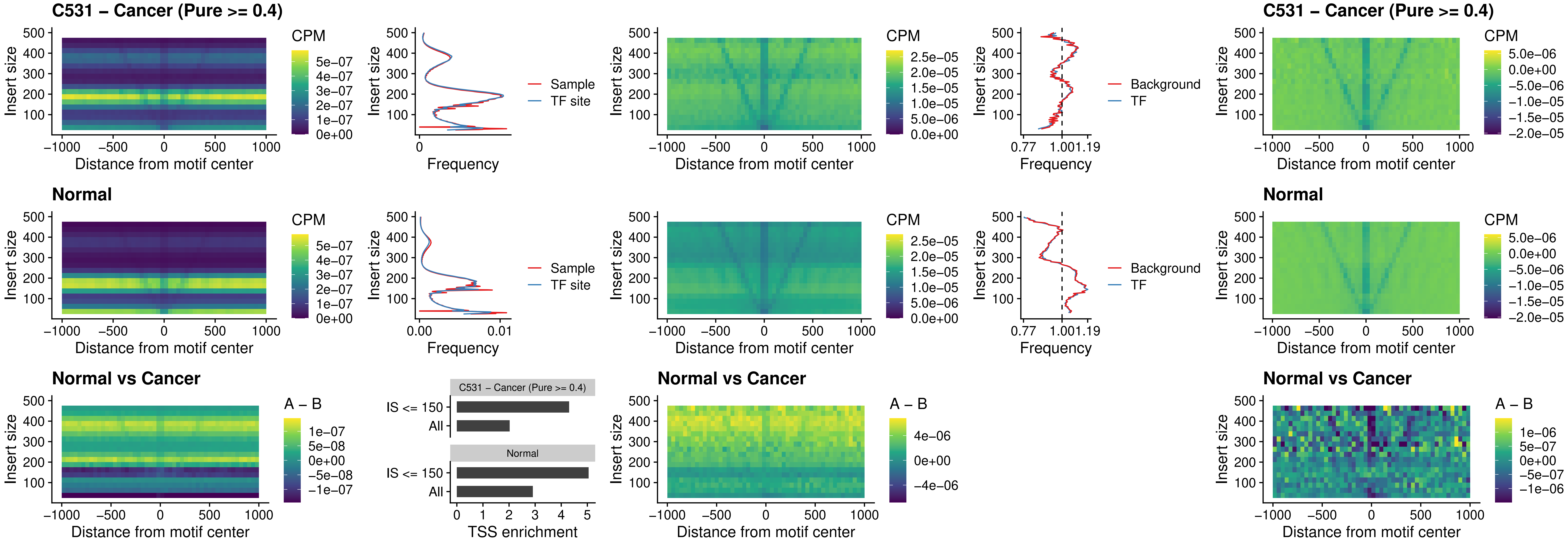

### C531 – Cancer (Pure >= 0.4) – PRRX1_uc_dTSS_nPEAK

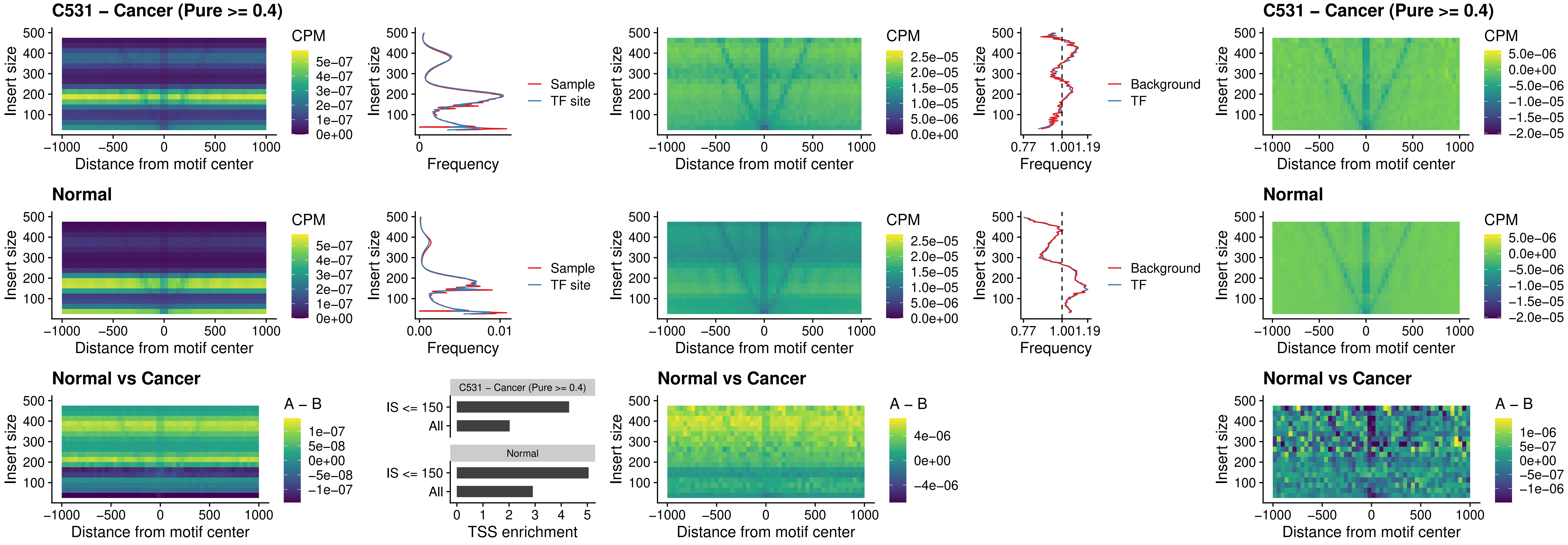

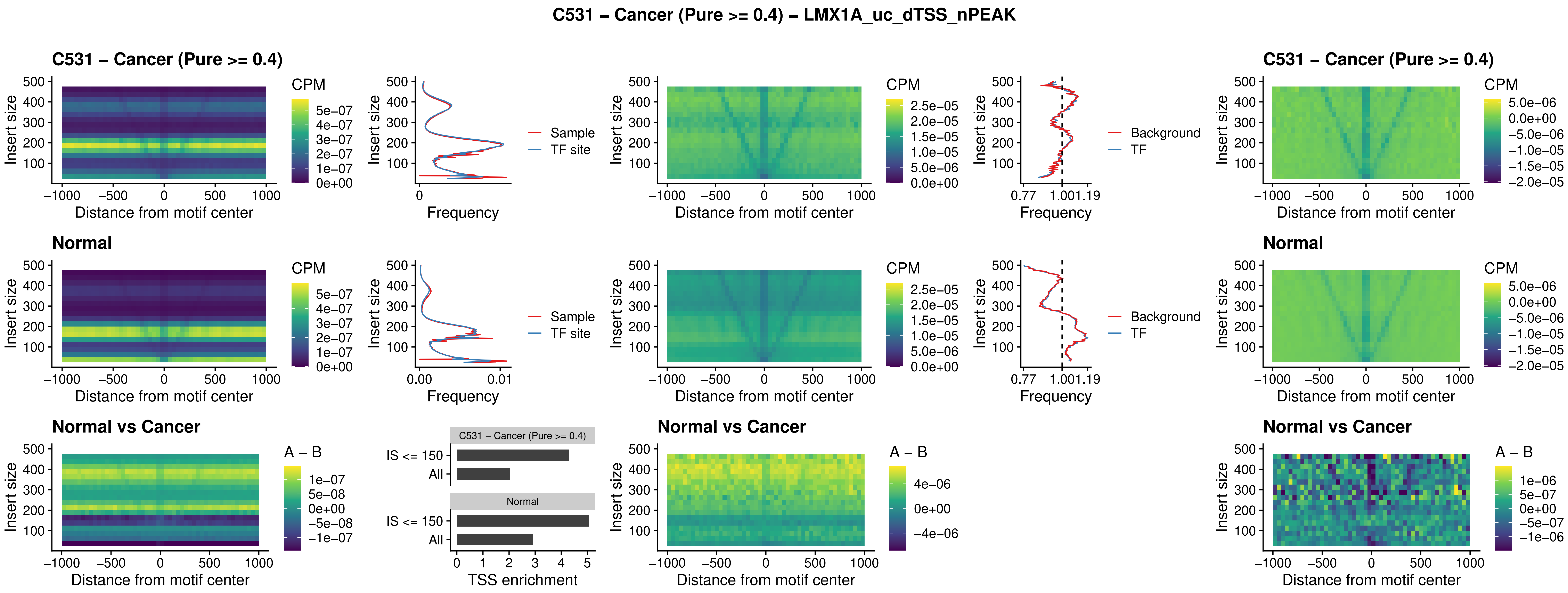

C531 – Cancer (Pure >= 0.4) – LMX1A_uc_dTSS_nPEAK
C531 – Cancer (Pure >= 0.4)
Normal
Normal vs Cancer
Insert size
Distance from motif center
CPM
A – B
Frequency
Sample
TF site
Background
TF
C531 – Cancer (Pure >= 0.4)
Normal
IS <= 150
All
TSS enrichment

#### C531 – Cancer (Pure >= 0.4) – LMX1B_uc_dTSS_nPEAK

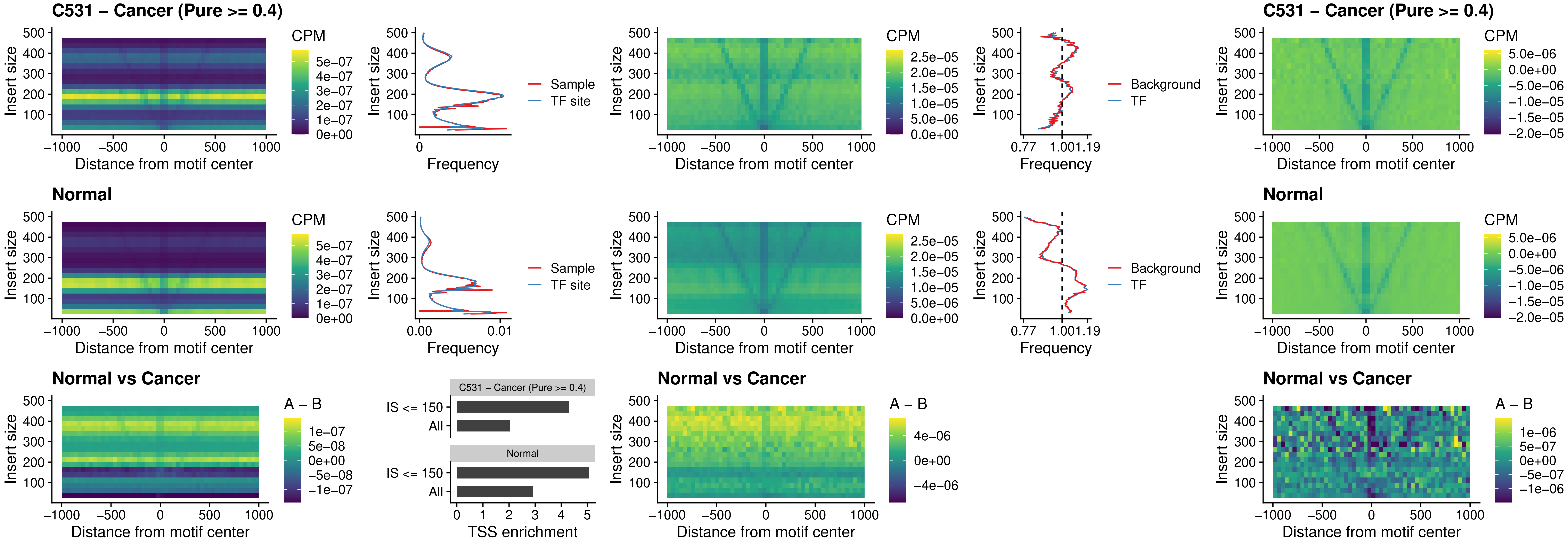

#### C531 – Cancer (Pure >= 0.4) – DLX5_uc_dTSS_oPEAK

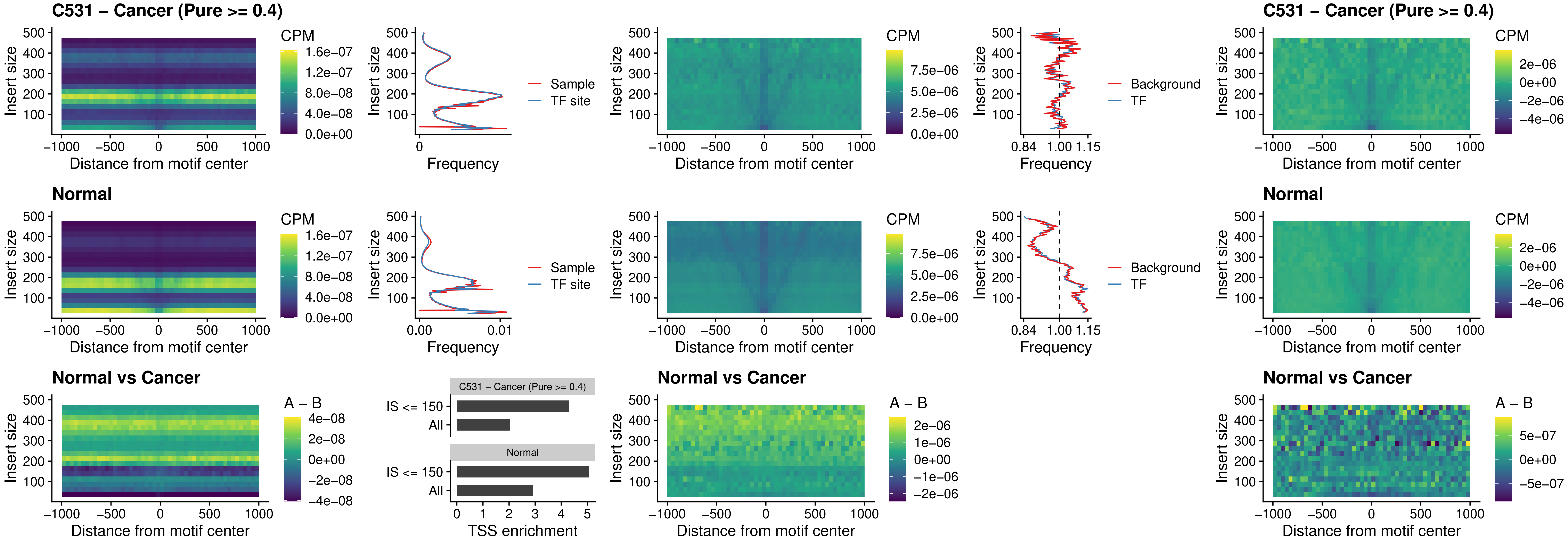

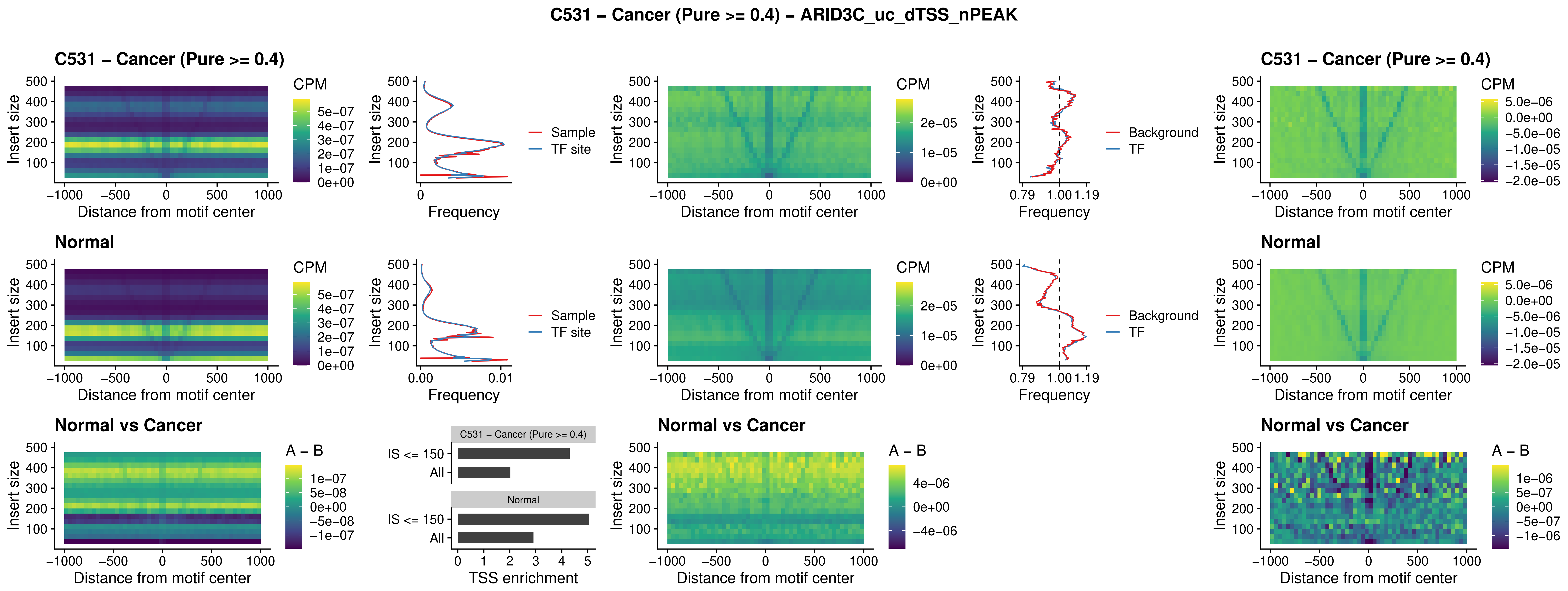

C531 – Cancer (Pure >= 0.4) – ARID3C_uc_dTSS_nPEAK
C531 – Cancer (Pure >= 0.4)
Normal
Normal vs Cancer
Insert size
Distance from motif center
CPM
A – B
Frequency
Sample
TF site
Background
TF
C531 – Cancer (Pure >= 0.4)
Normal
IS <= 150
All
TSS enrichment

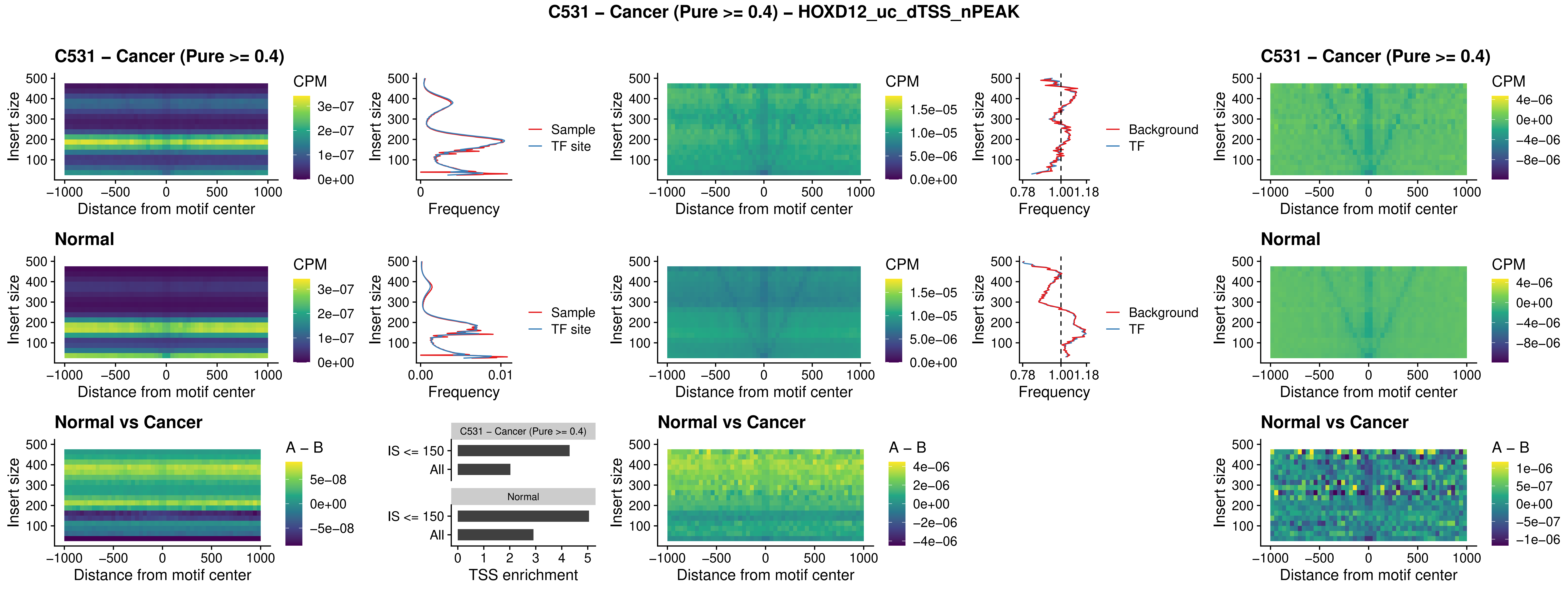
C531 – Cancer (Pure >= 0.4) – HOXD12_uc_dTSS_nPEAK
C531 – Cancer (Pure >= 0.4)
Normal
Normal vs Cancer
Insert size
Distance from motif center
Frequency
CPM
A – B
Sample
TF site
Background
TF
C531 – Cancer (Pure >= 0.4)
Normal
IS <= 150
All
TSS enrichment

#### C531 – Cancer (Pure >= 0.4) – CDX2_uc_dTSS_nPEAK

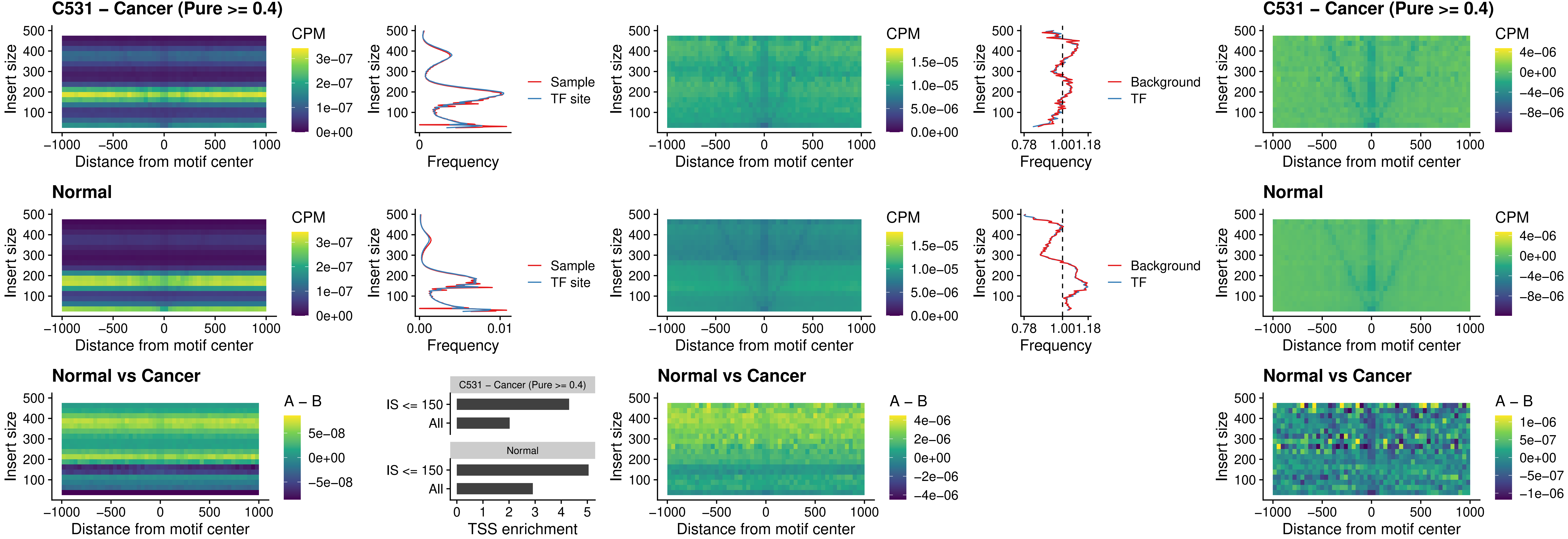

#### C531 – Cancer (Pure >= 0.4) – HOXC12_uc_dTSS_nPEAK

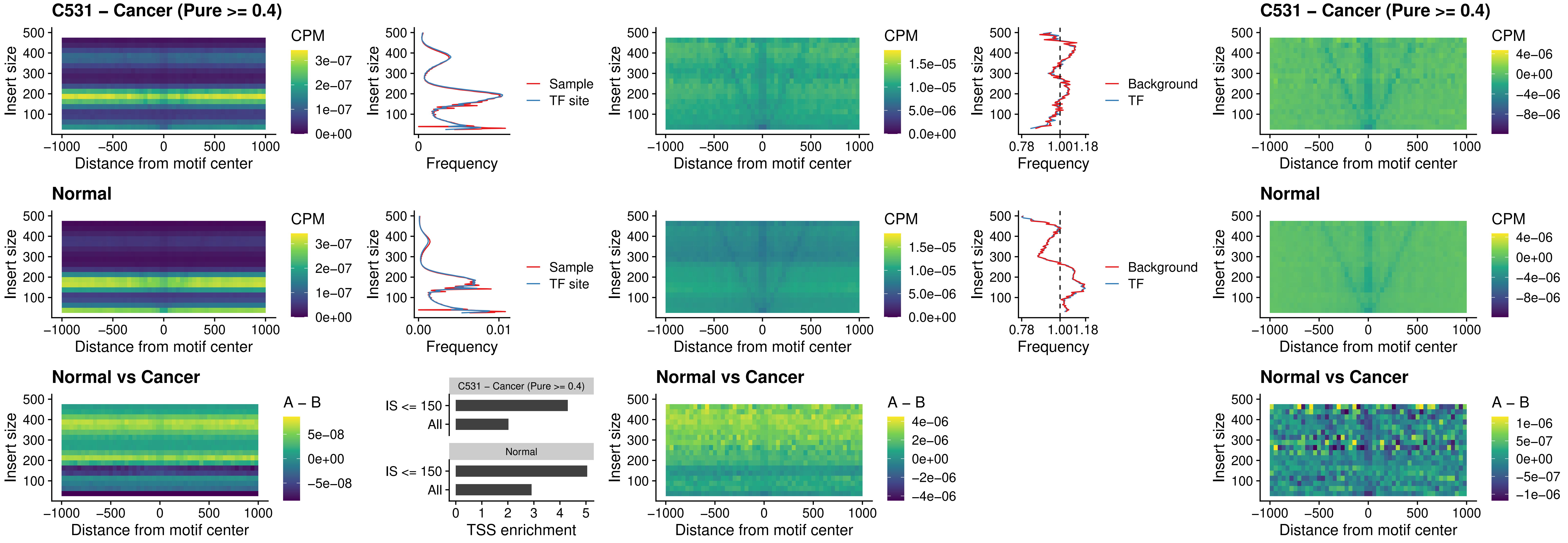

#### C531 – Cancer (Pure >= 0.4) – HHEX_uc_dTSS_oPEAK

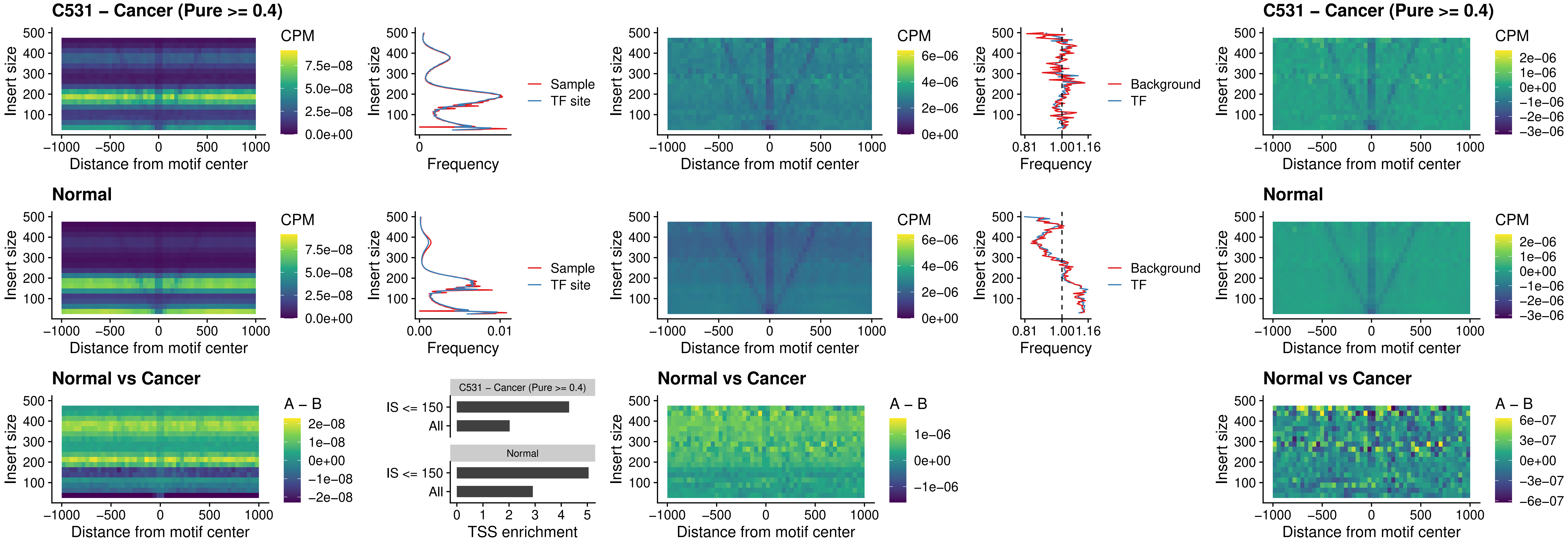

#### C531 – Cancer (Pure >= 0.4) – HOXA11_uc_dTSS_nPEAK

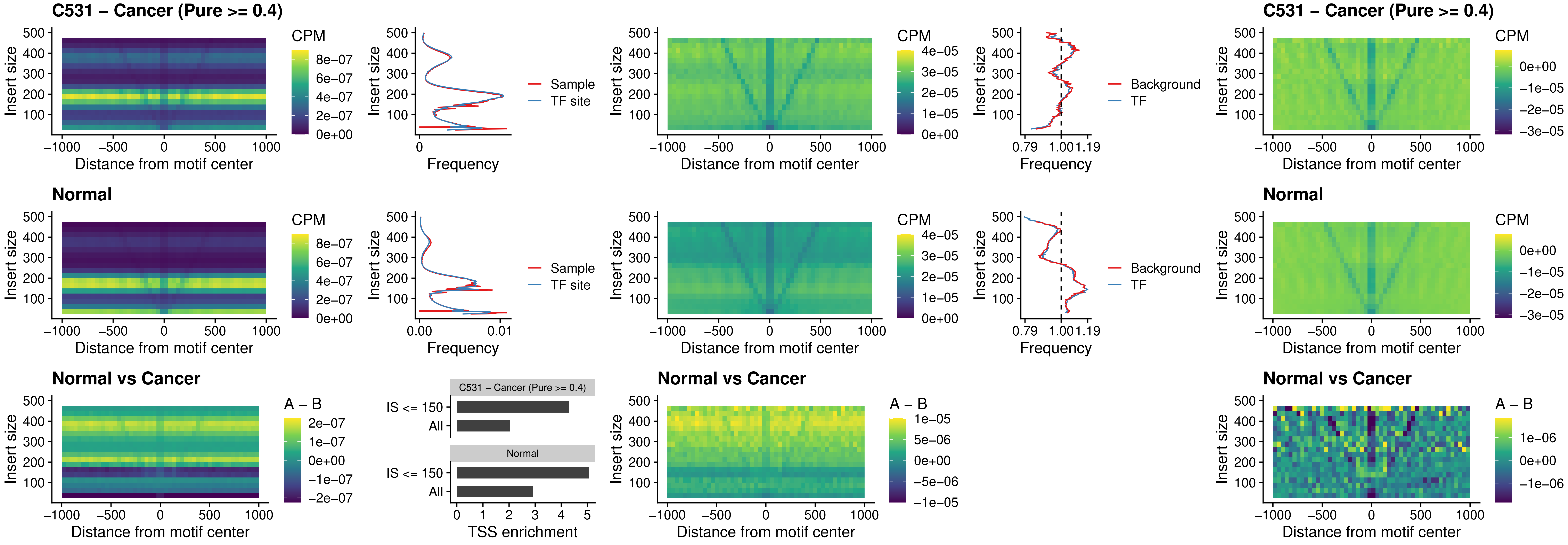

#### C531 – Cancer (Pure >= 0.4) – HOXA11_uc_cTSS_nPEAK

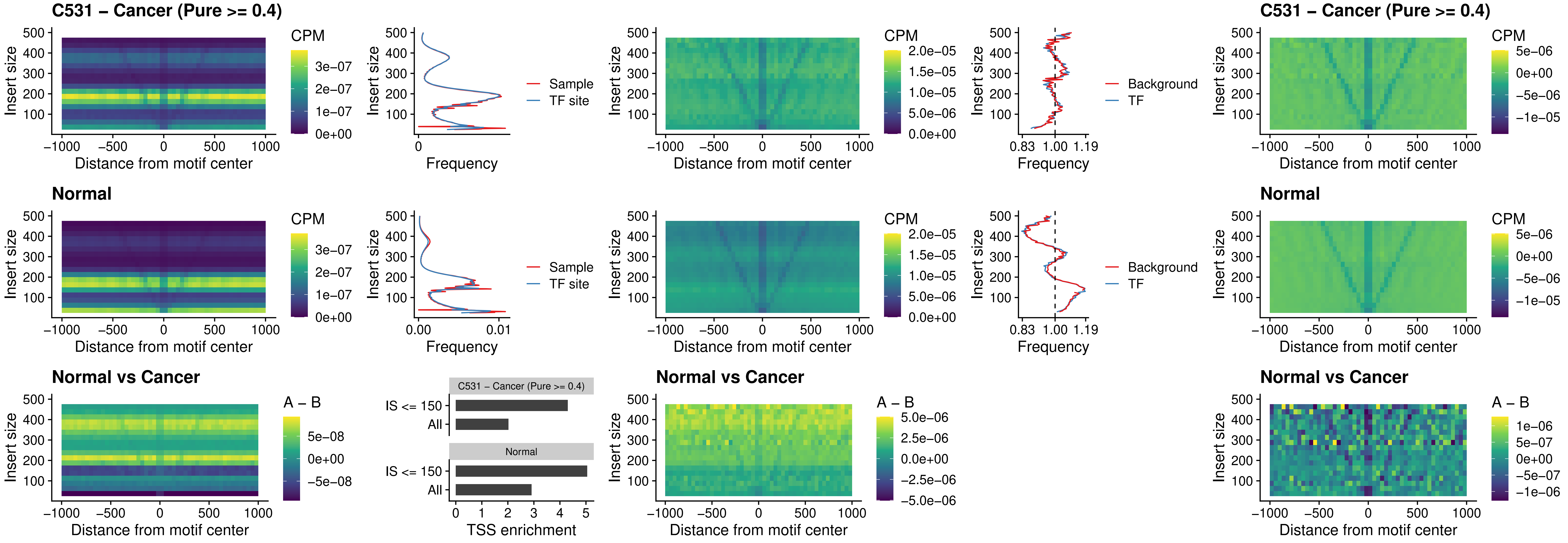

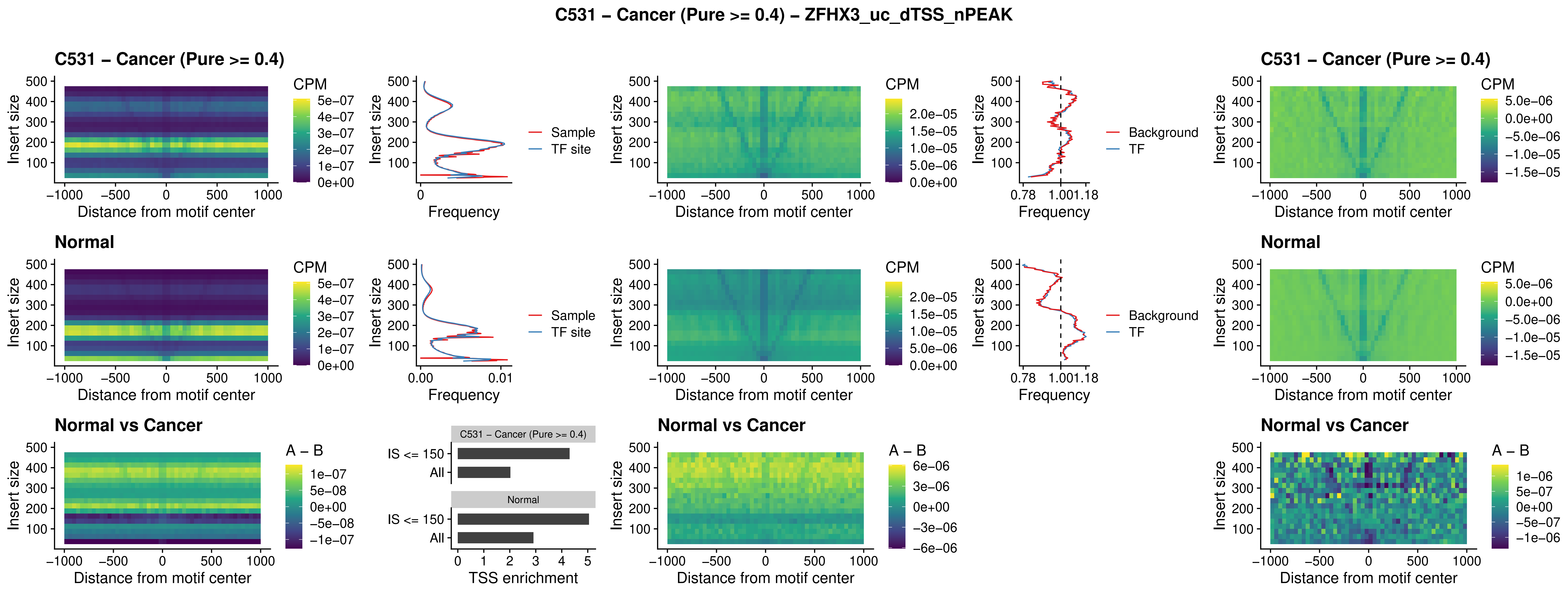
C531 – Cancer (Pure >= 0.4) – ZFHX3_uc_dTSS_nPEAK
C531 – Cancer (Pure >= 0.4)
Normal
Normal vs Cancer
Insert size
Distance from motif center
Frequency
CPM
A – B
Sample
TF site
Background
TF
TSS enrichment
IS <= 150
All
C531 – Cancer (Pure >= 0.4)
Normal

#### C531 – Cancer (Pure >= 0.4) – HOXB4_uc_dTSS_nPEAK

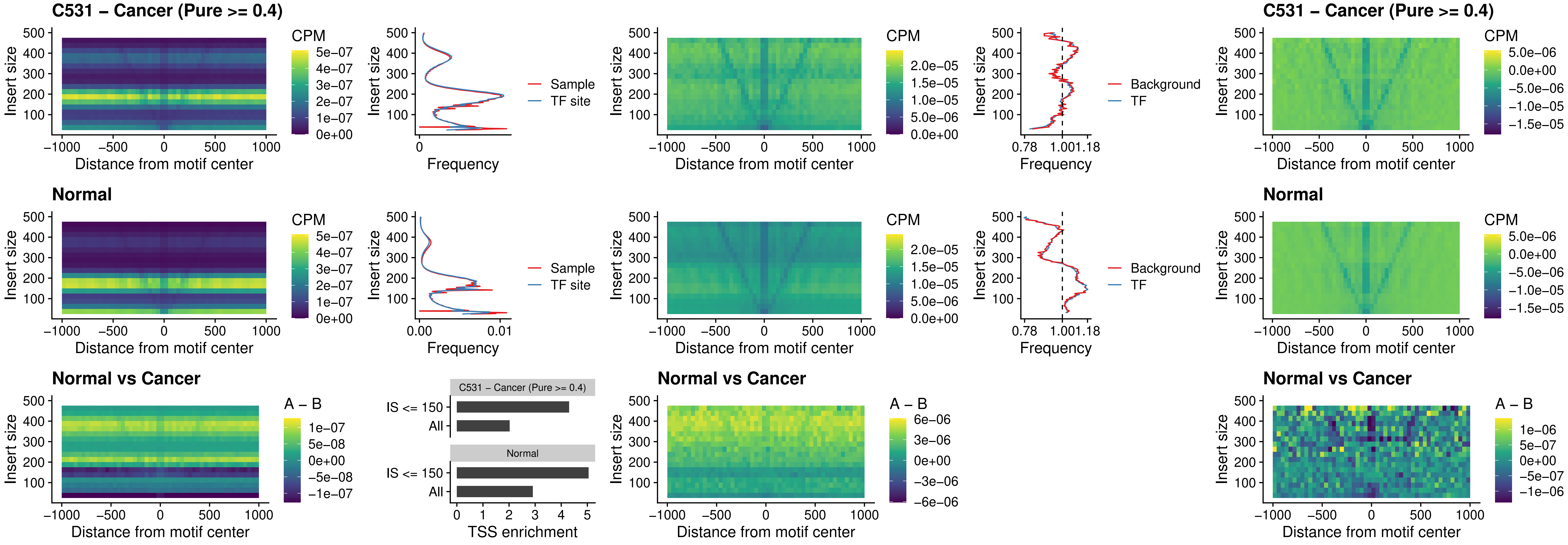

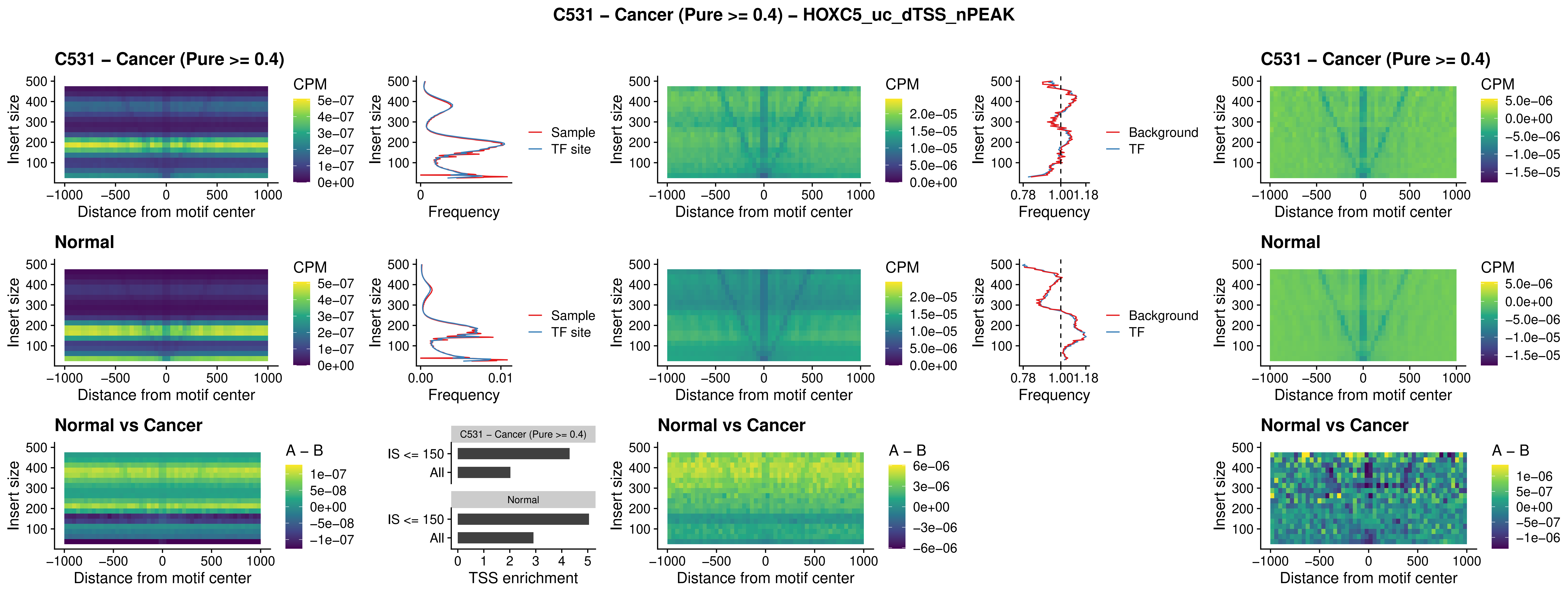
C531 – Cancer (Pure >= 0.4) – HOXC5_uc_dTSS_nPEAK
C531 – Cancer (Pure >= 0.4)
Normal
Normal vs Cancer
Insert size
Distance from motif center
Frequency
CPM
A – B
Sample
TF site
Background
TF
C531 – Cancer (Pure >= 0.4)
Normal
IS <= 150
All
TSS enrichment

#### C531 – Cancer (Pure >= 0.4) – HOXD1_uc_dTSS_nPEAK

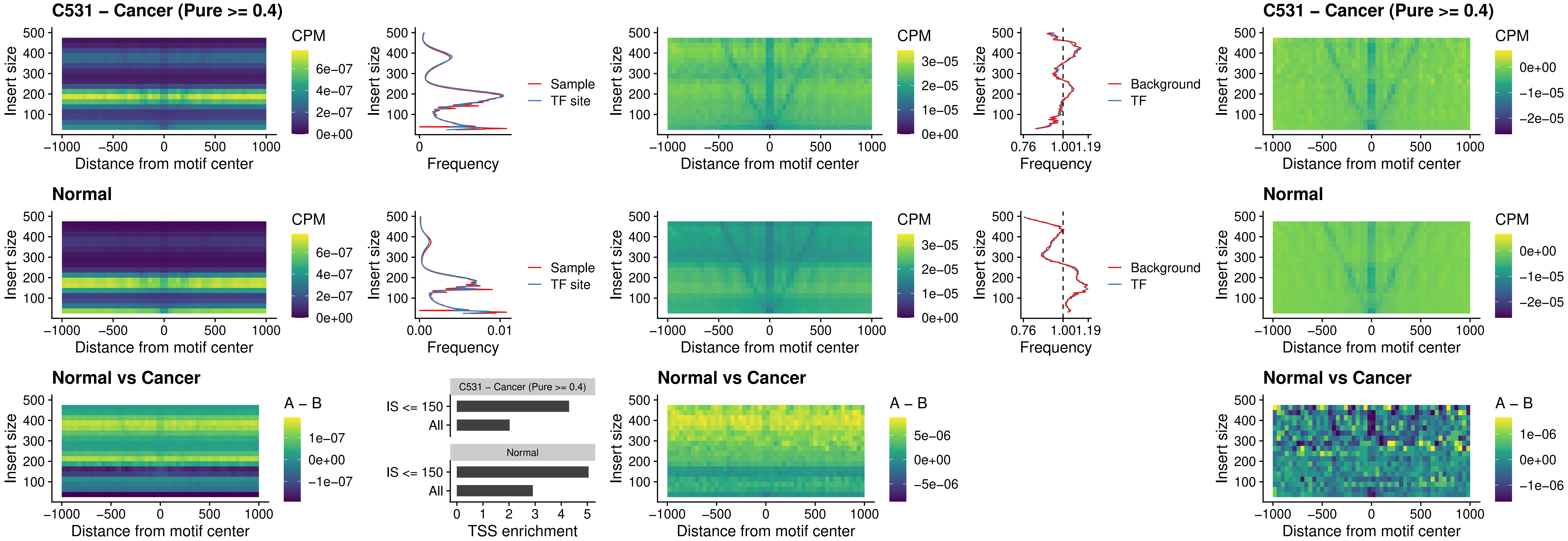

#### C531 – Cancer (Pure >= 0.4) – OTP_uc_dTSS_nPEAK

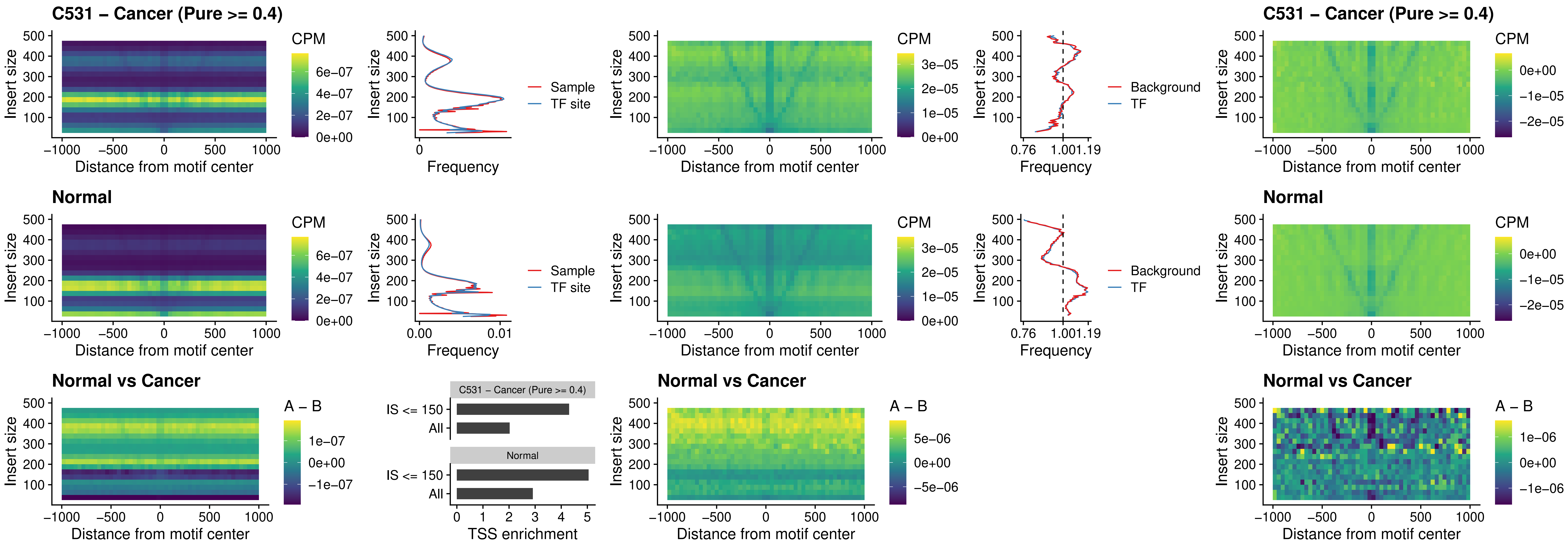

#### C531 – Cancer (Pure >= 0.4) – FOXL1_uc_dTSS_oPEAK

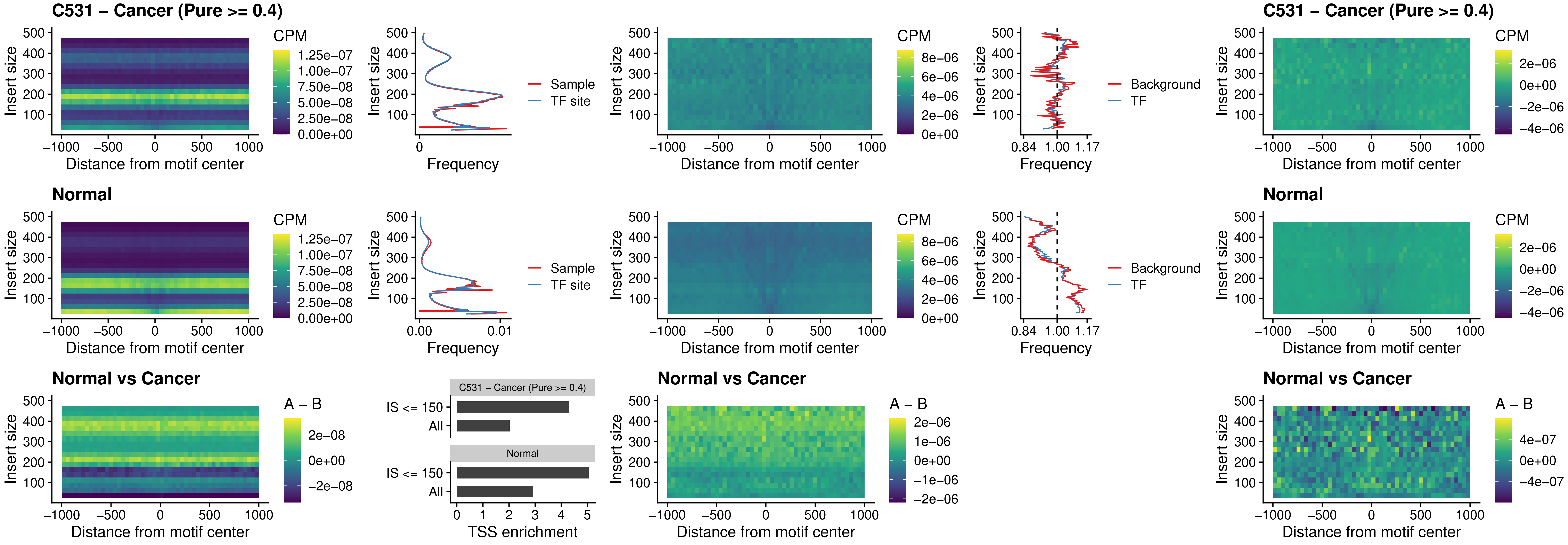

#### C531 – Cancer (Pure >= 0.4) – ZNF32_uc_dTSS_oPEAK

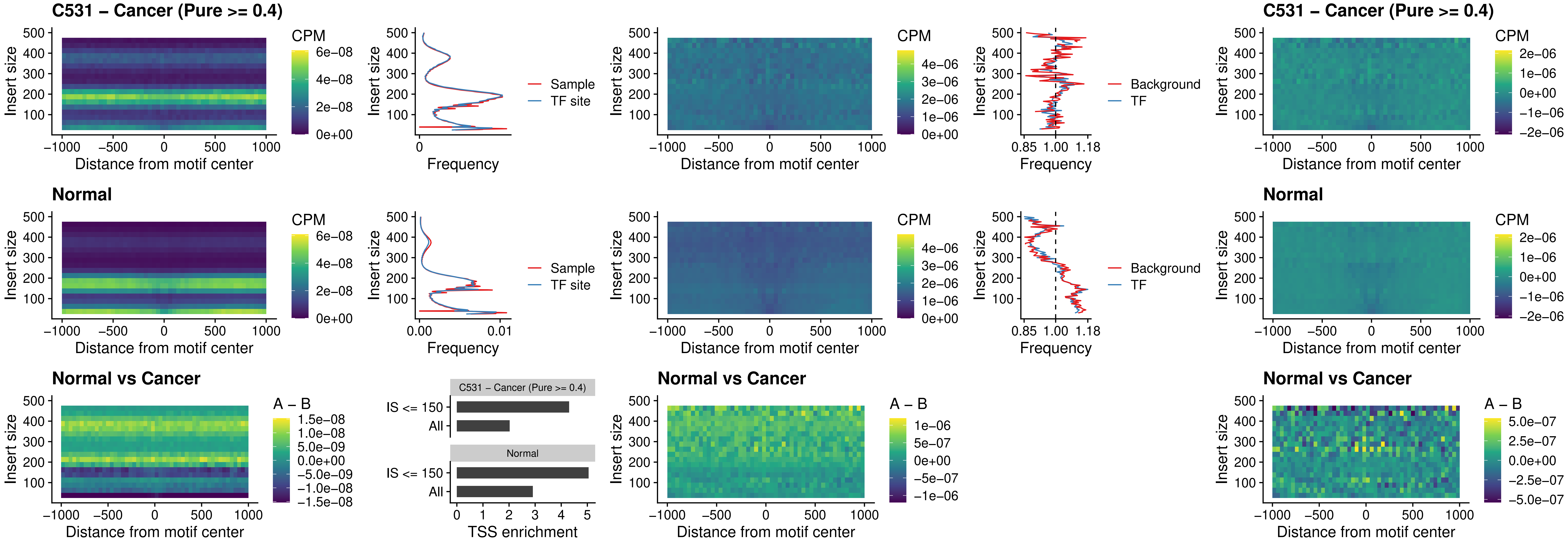

### C531 – Cancer (Pure >= 0.4) – FOXL1_uc_dTSS_nPEAK

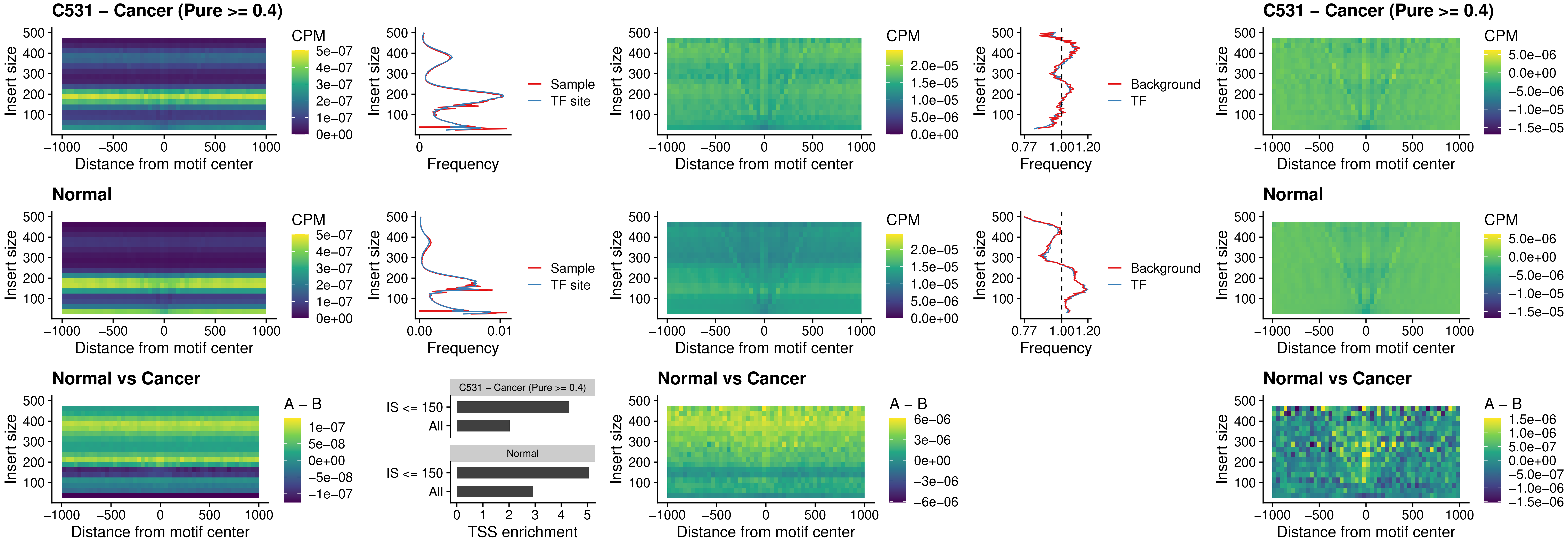

#### C531 – Cancer (Pure >= 0.4) – ZNF32_uc_dTSS_nPEAK

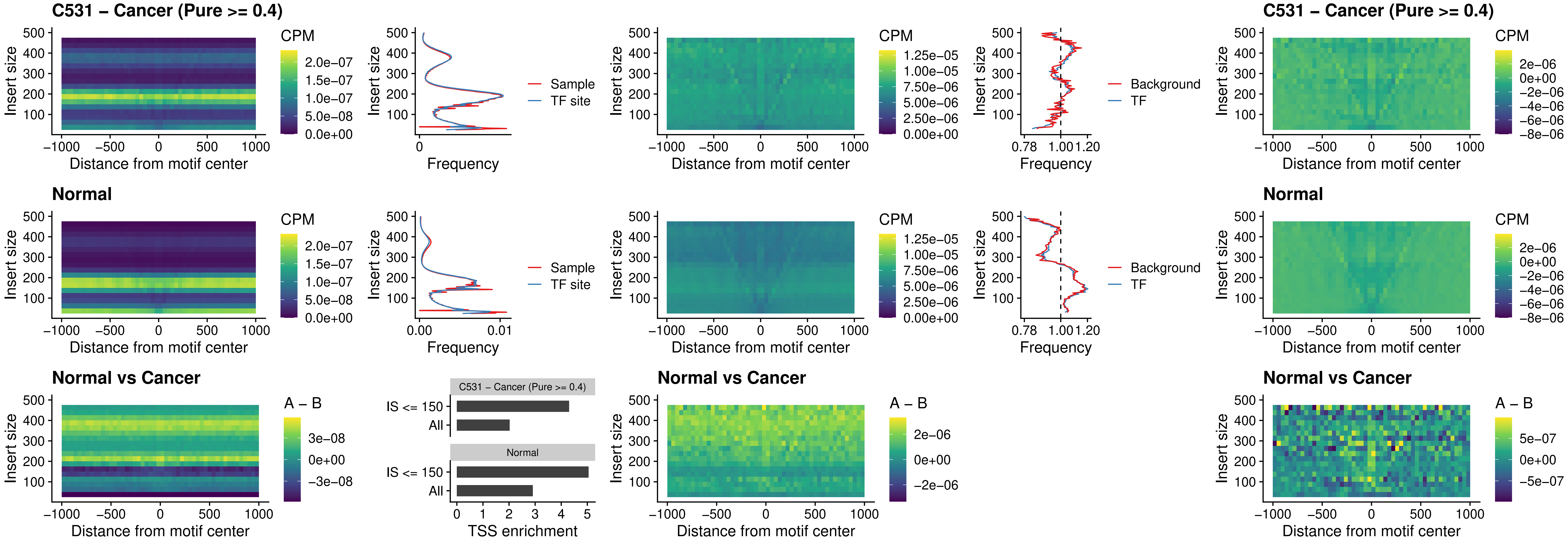

#### C531 – Cancer (Pure >= 0.4) – DLX5_uc_dTSS_nPEAK

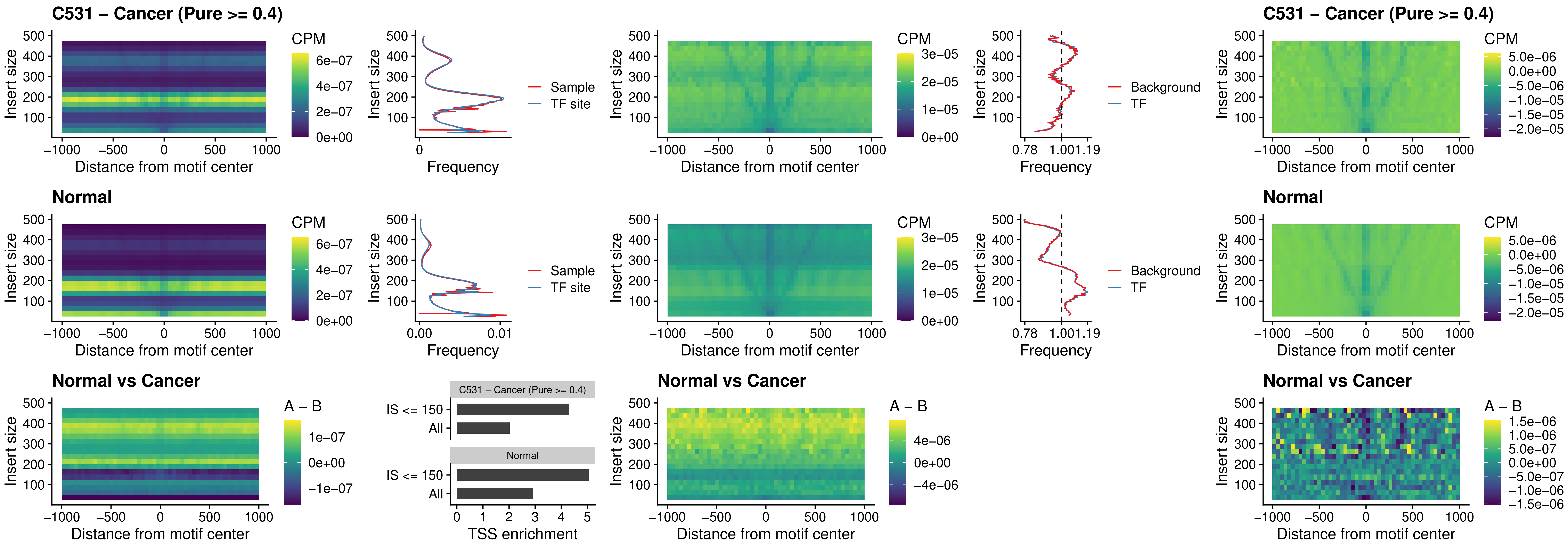

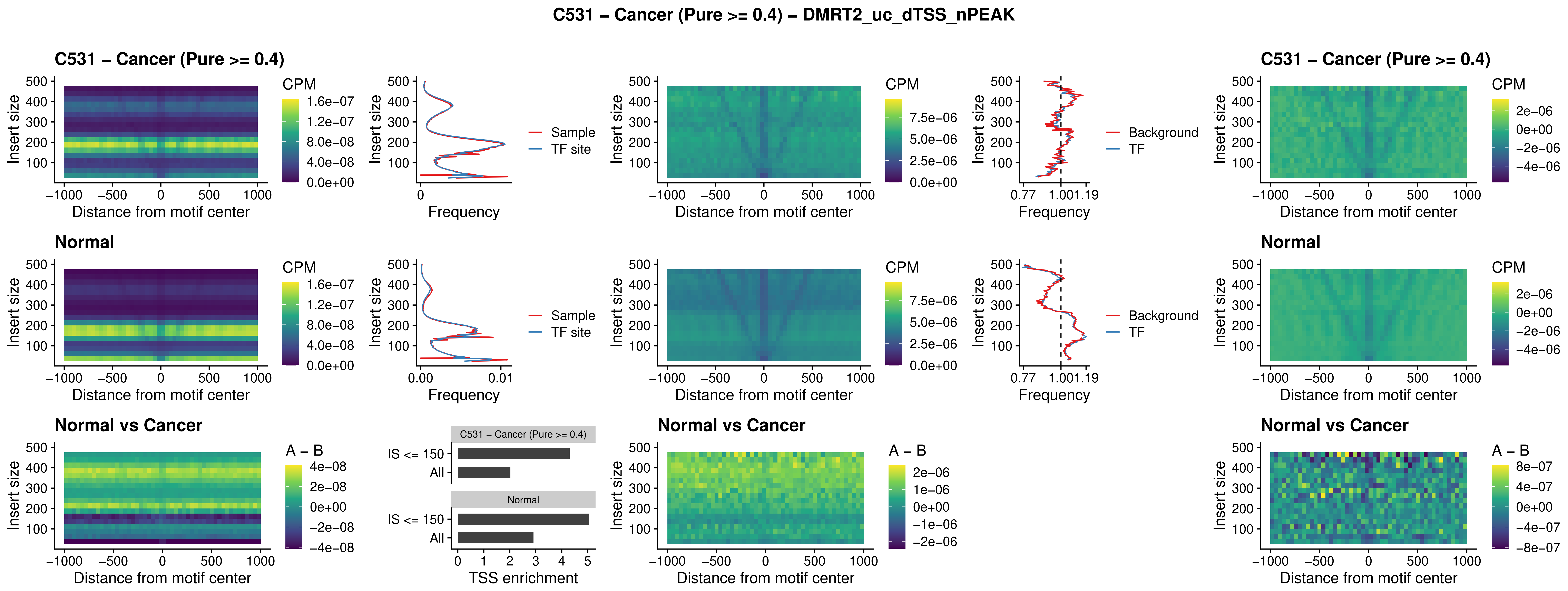
C531 – Cancer (Pure >= 0.4) – DMRT2_uc_dTSS_nPEAK
C531 – Cancer (Pure >= 0.4)
Normal
Normal vs Cancer
Insert size
Distance from motif center
Frequency
CPM
A – B
Sample
TF site
Background
TF
C531 – Cancer (Pure >= 0.4)
Normal
IS <= 150
All
TSS enrichment

#### C531 – Cancer (Pure >= 0.4) – DMRTC2_uc_cTSS_nPEAK

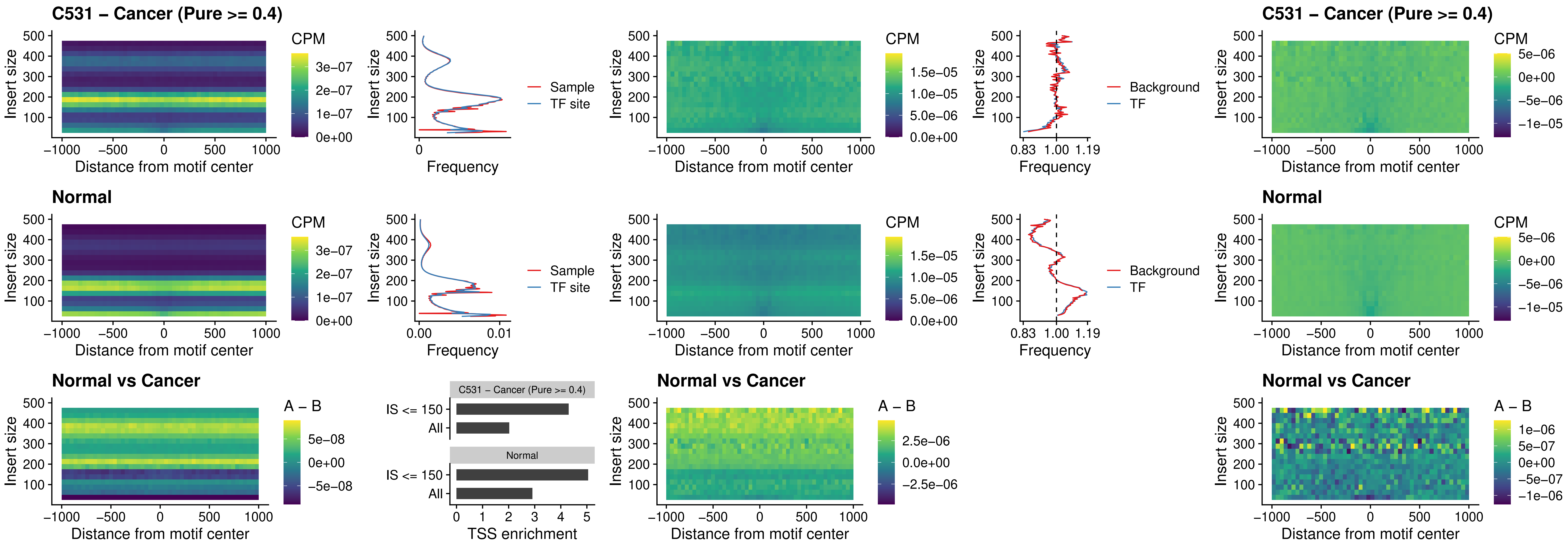

#### C531 – Cancer (Pure >= 0.4) – HOXD9_uc_dTSS_nPEAK

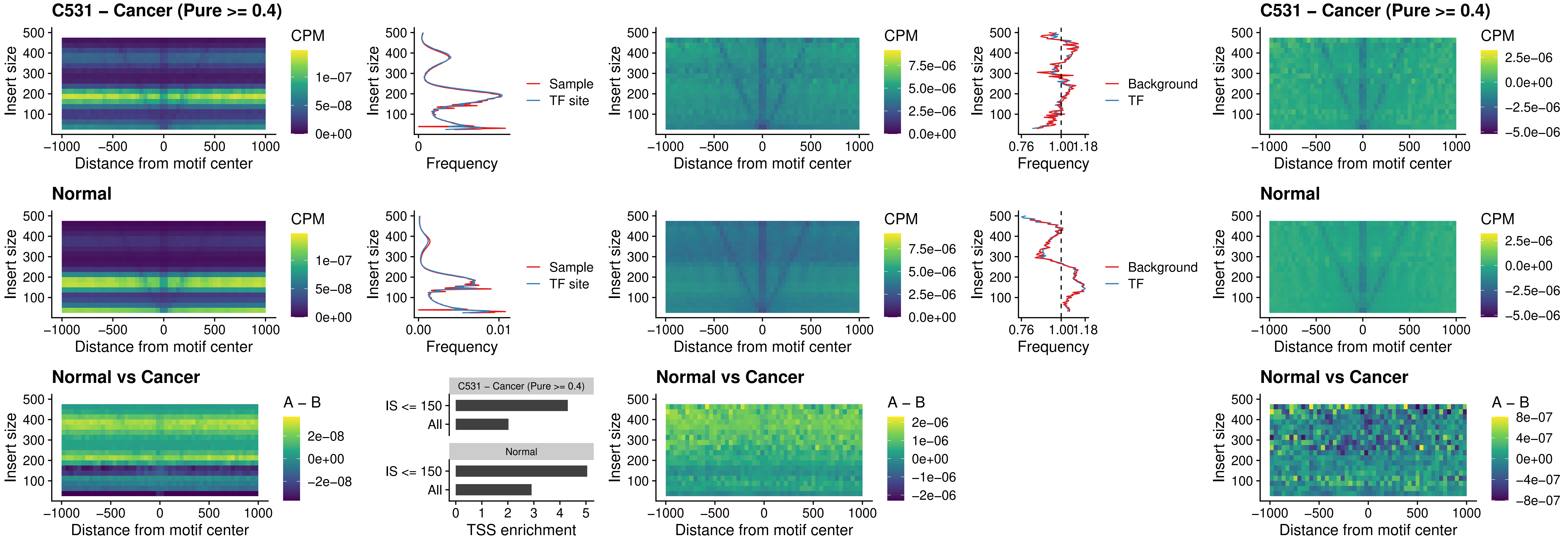

#### C531 – Cancer (Pure >= 0.4) – HOXD4_uc_dTSS_nPEAK

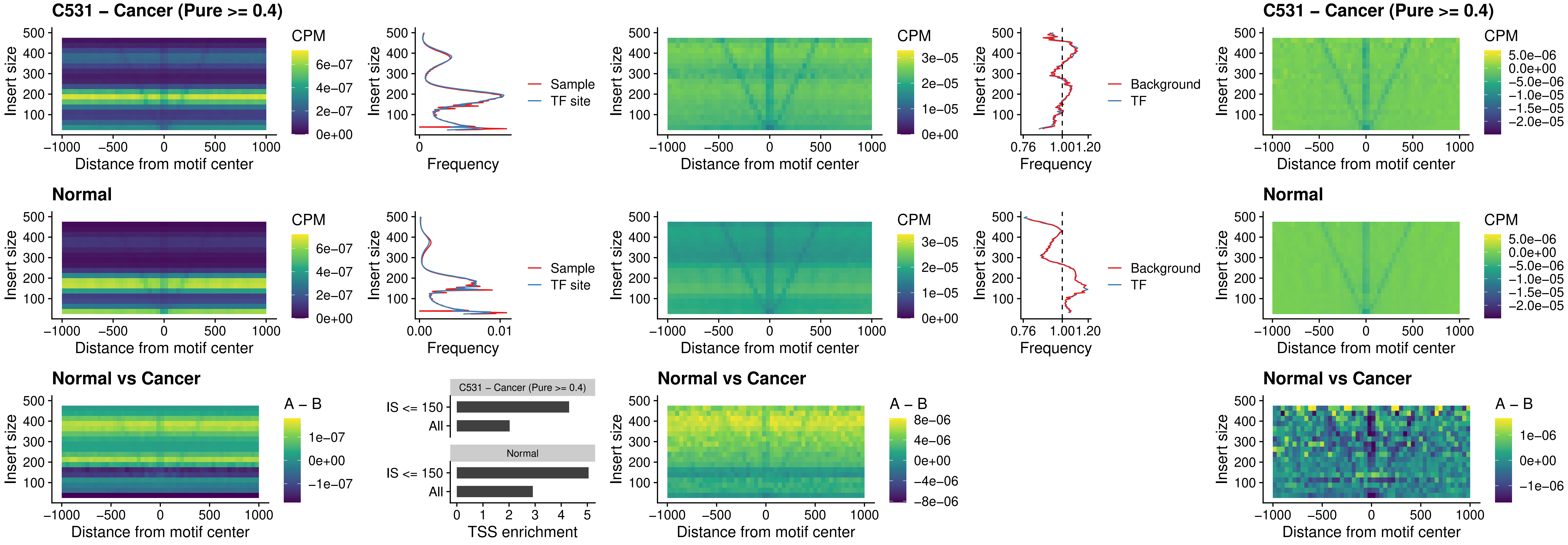

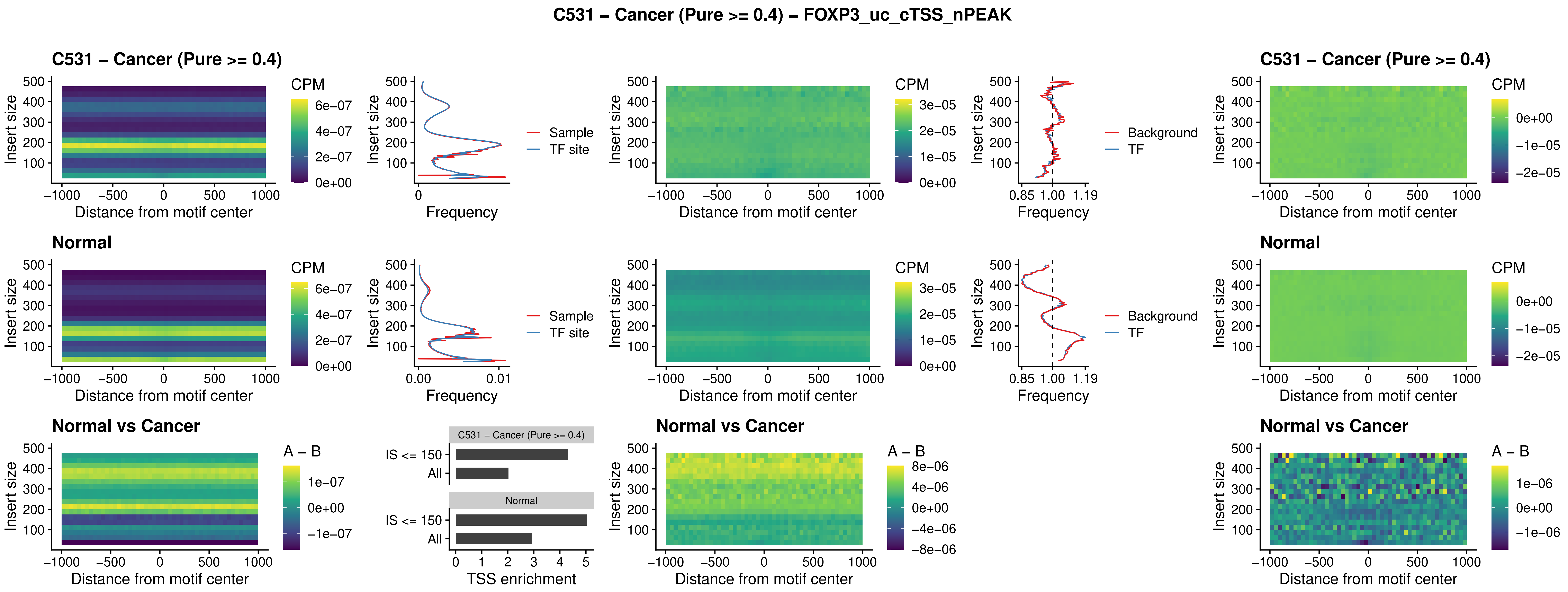
C531 – Cancer (Pure >= 0.4) – FOXP3_uc_cTSS_nPEAK
C531 – Cancer (Pure >= 0.4)
Normal
Normal vs Cancer
Insert size
Distance from motif center
CPM
A – B
Frequency
Sample
TF site
Background
TF
C531 – Cancer (Pure >= 0.4)
Normal
IS <= 150
All
TSS enrichment

#### C531 – Cancer (Pure >= 0.4) – FOXD4L6_uc_cTSS_nPEAK

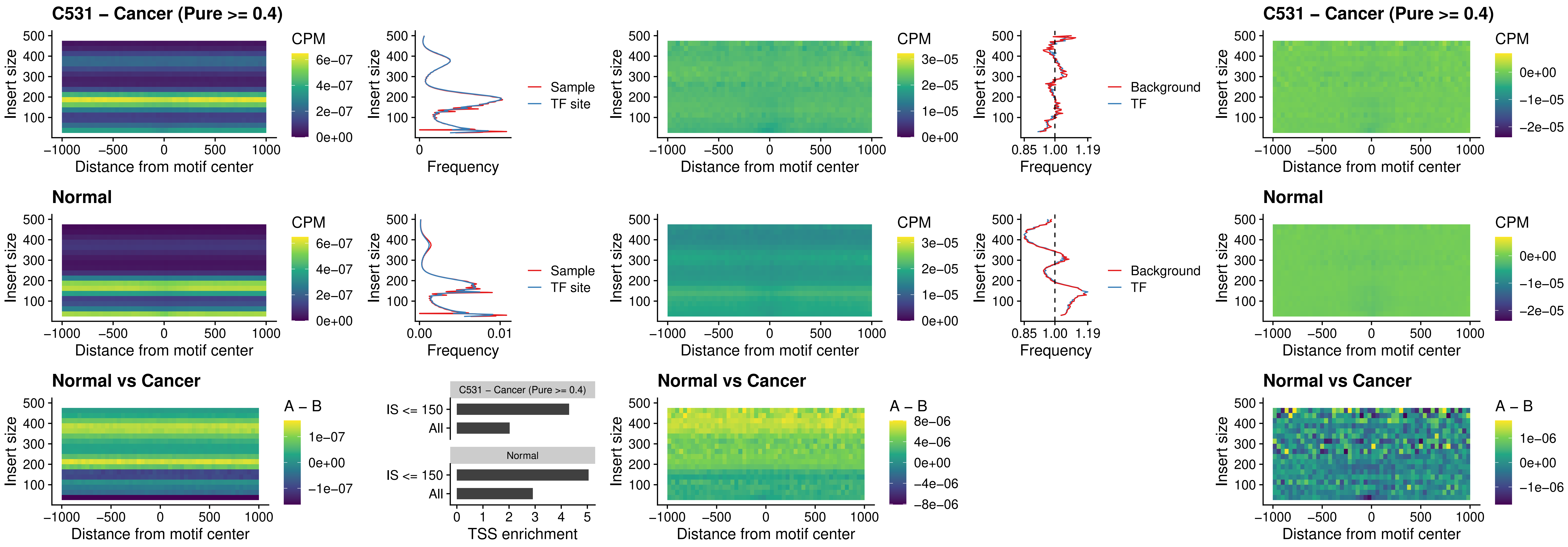

#### C531 – Cancer (Pure >= 0.4) – FOXD4L5_uc_cTSS_nPEAK

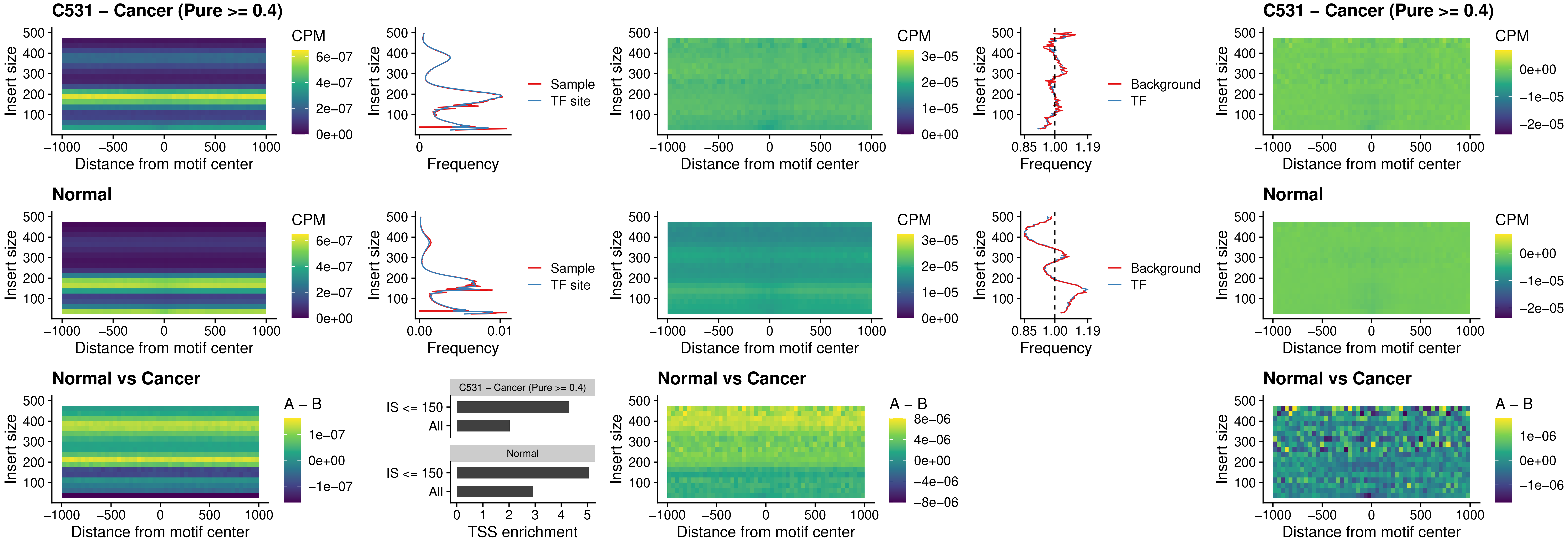

### C531 – Cancer (Pure >= 0.4) – FOXD4L4_uc_cTSS_nPEAK

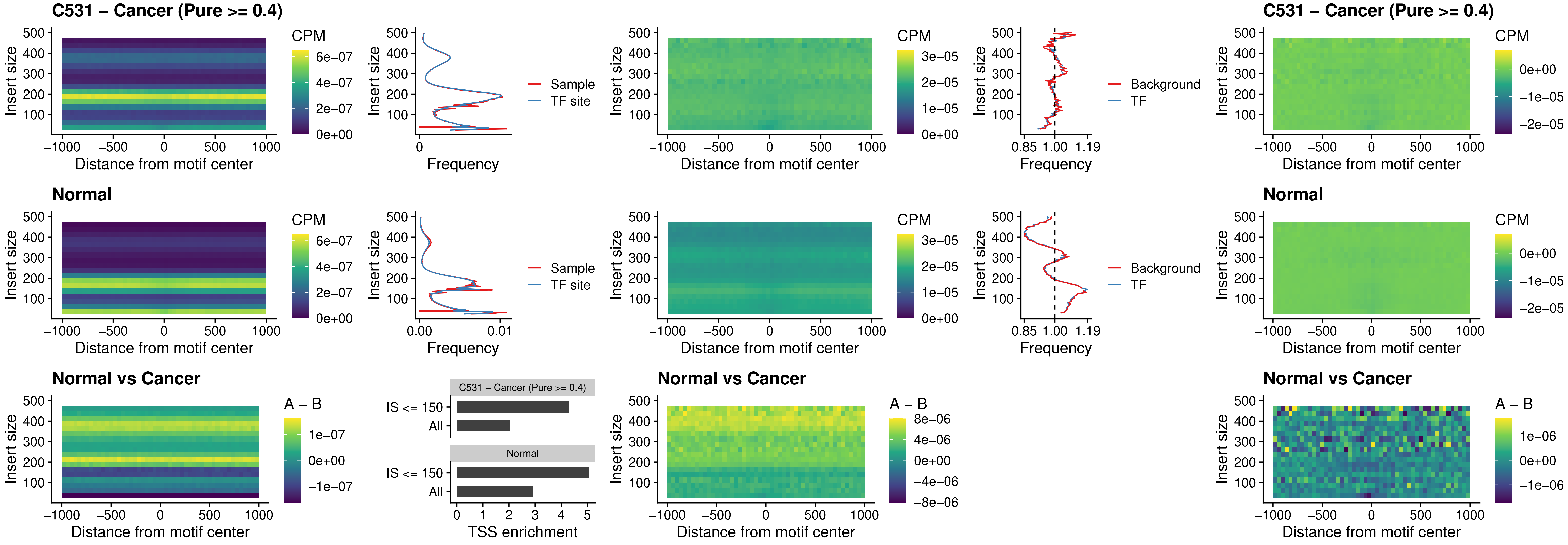

#### C531 – Cancer (Pure >= 0.4) – FOXD4L2_uc_cTSS_nPEAK

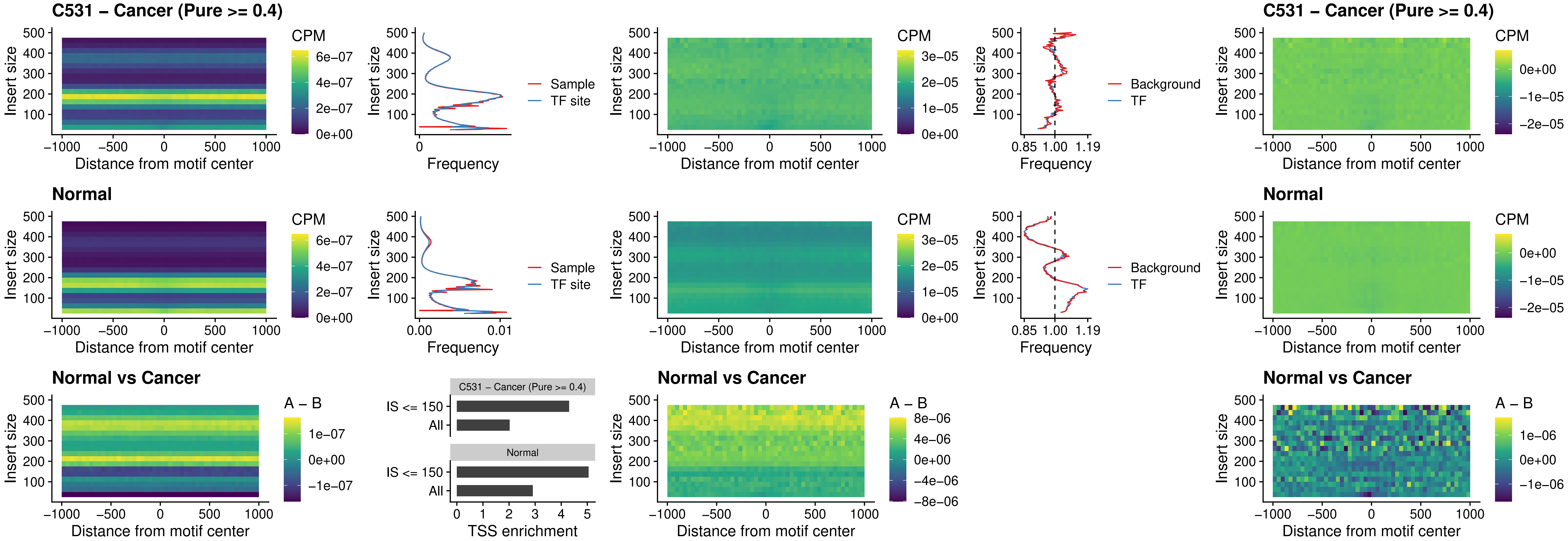

#### C531 – Cancer (Pure >= 0.4) – FOXD4L3_uc_cTSS_nPEAK

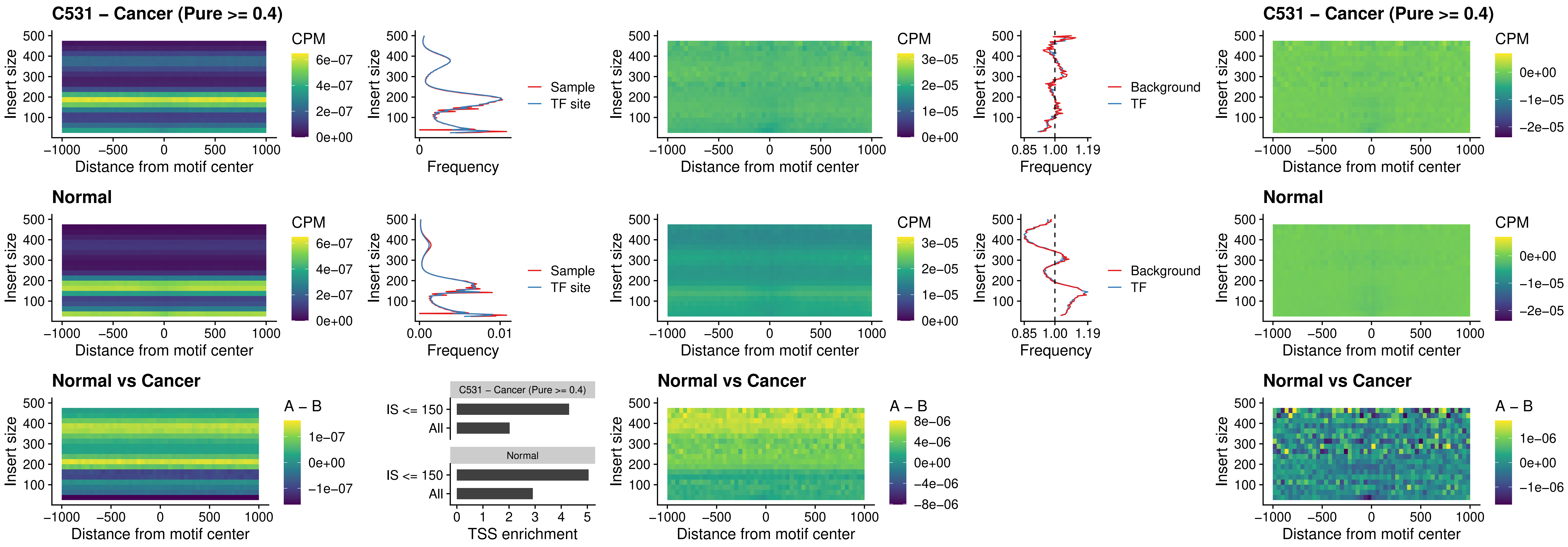

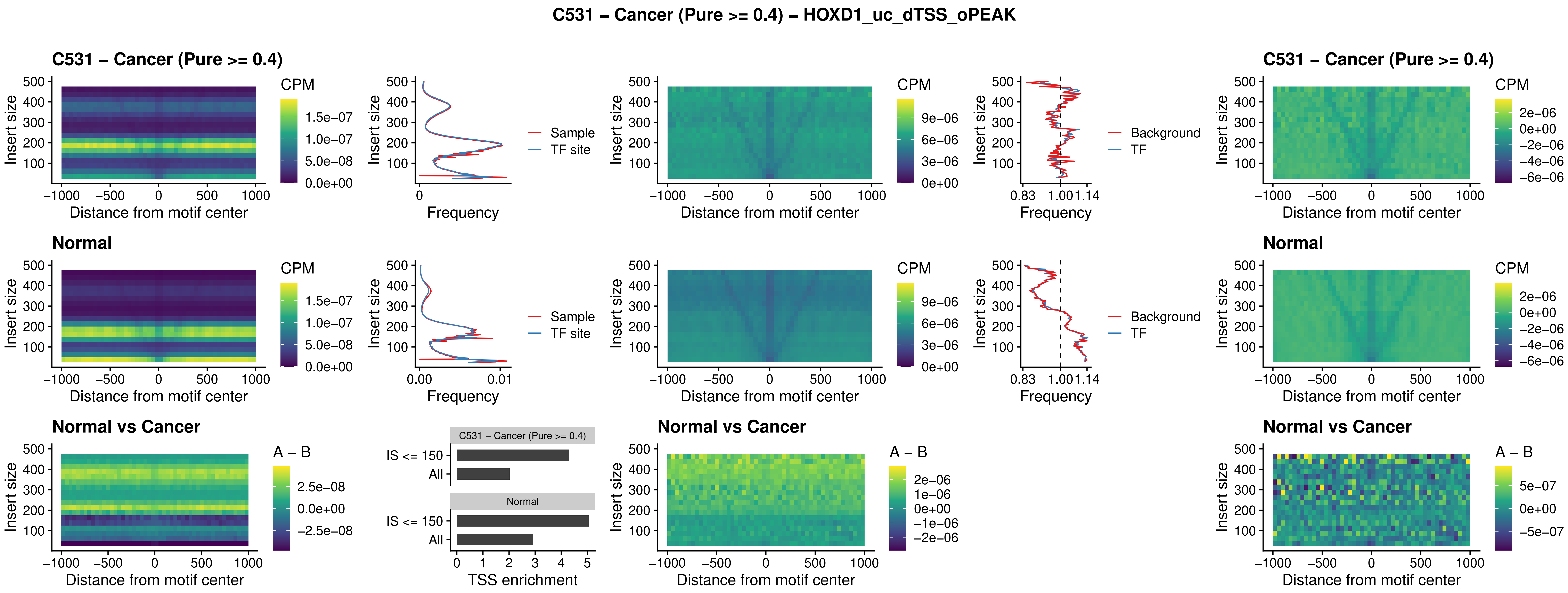
C531 – Cancer (Pure >= 0.4) – HOXD1_uc_dTSS_oPEAK
C531 – Cancer (Pure >= 0.4)
Normal
Normal vs Cancer
Insert size
Distance from motif center
Frequency
CPM
A – B
Sample
TF site
Background
TF
C531 – Cancer (Pure >= 0.4)
Normal
IS <= 150
All
TSS enrichment

### C531 – Cancer (Pure >= 0.4) – VAX1_uc_pTSS_nPEAK

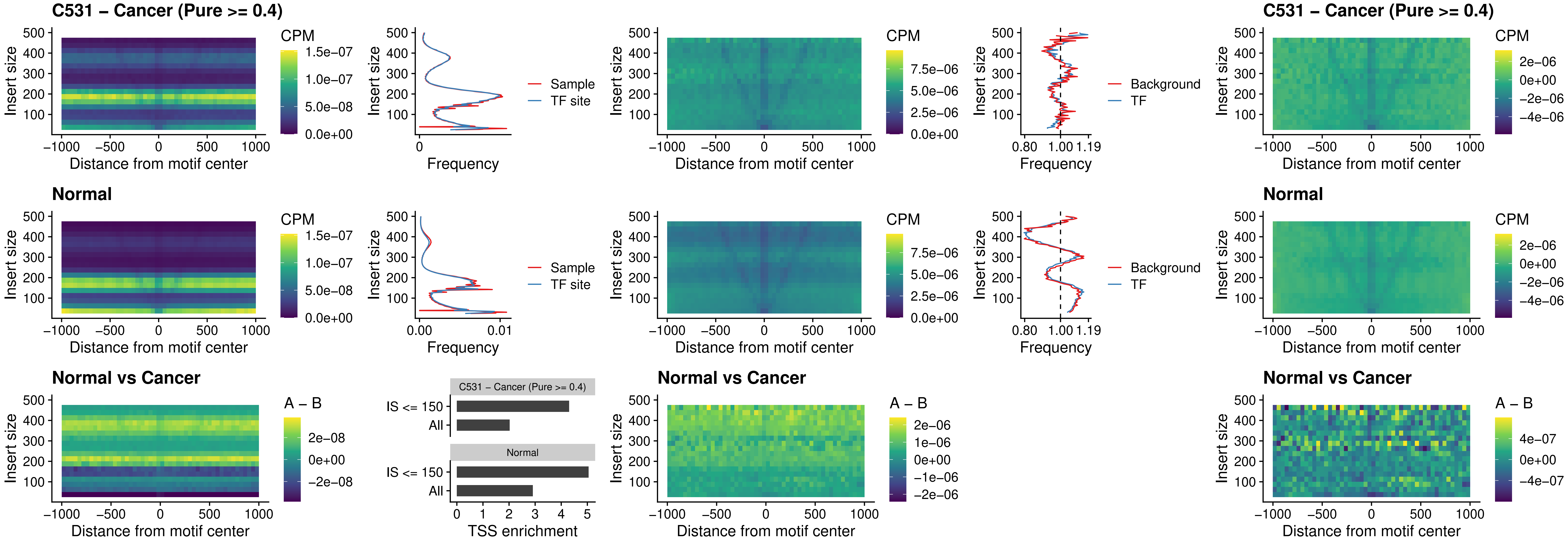

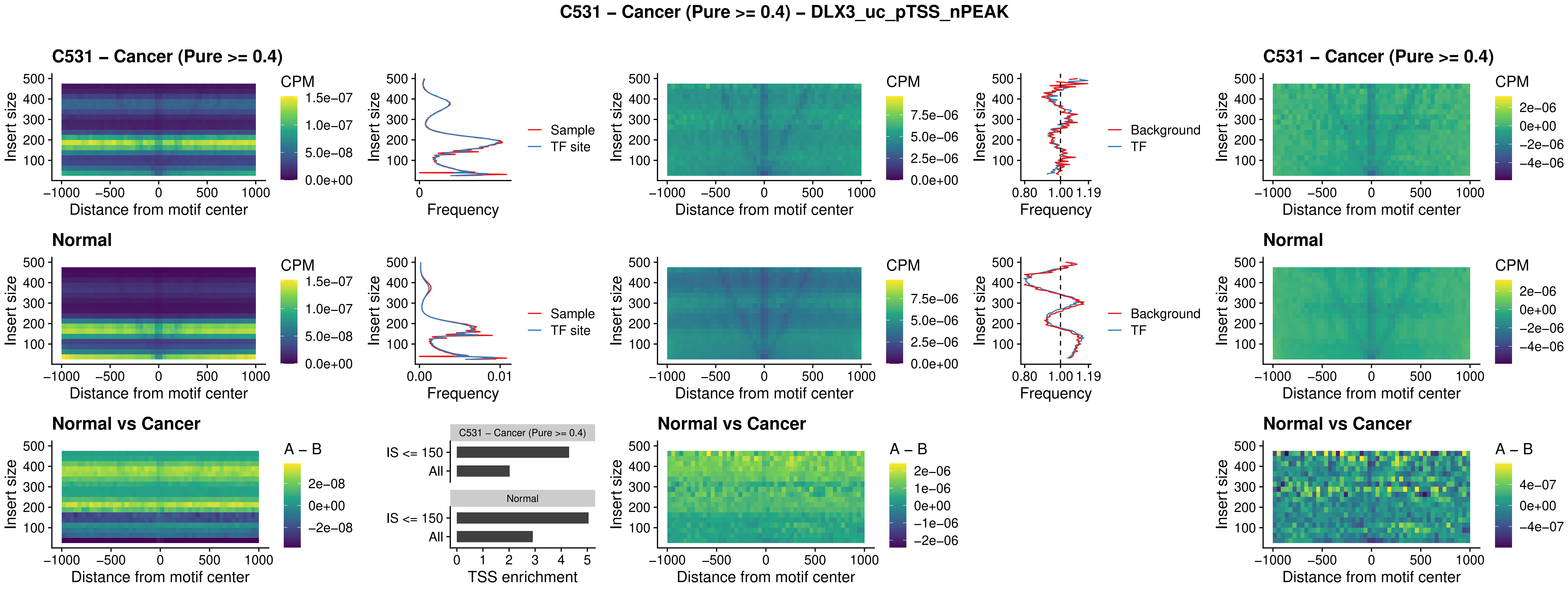
C531 – Cancer (Pure >= 0.4) – DLX3_uc_pTSS_nPEAK
C531 – Cancer (Pure >= 0.4)
Normal
Normal vs Cancer
Insert size
Distance from motif center
Frequency
CPM
A – B
Sample
TF site
Background
TF
C531 – Cancer (Pure >= 0.4)
Normal
IS <= 150
All
TSS enrichment

#### C531 – Cancer (Pure >= 0.4) – HOXD3_uc_pTSS_nPEAK

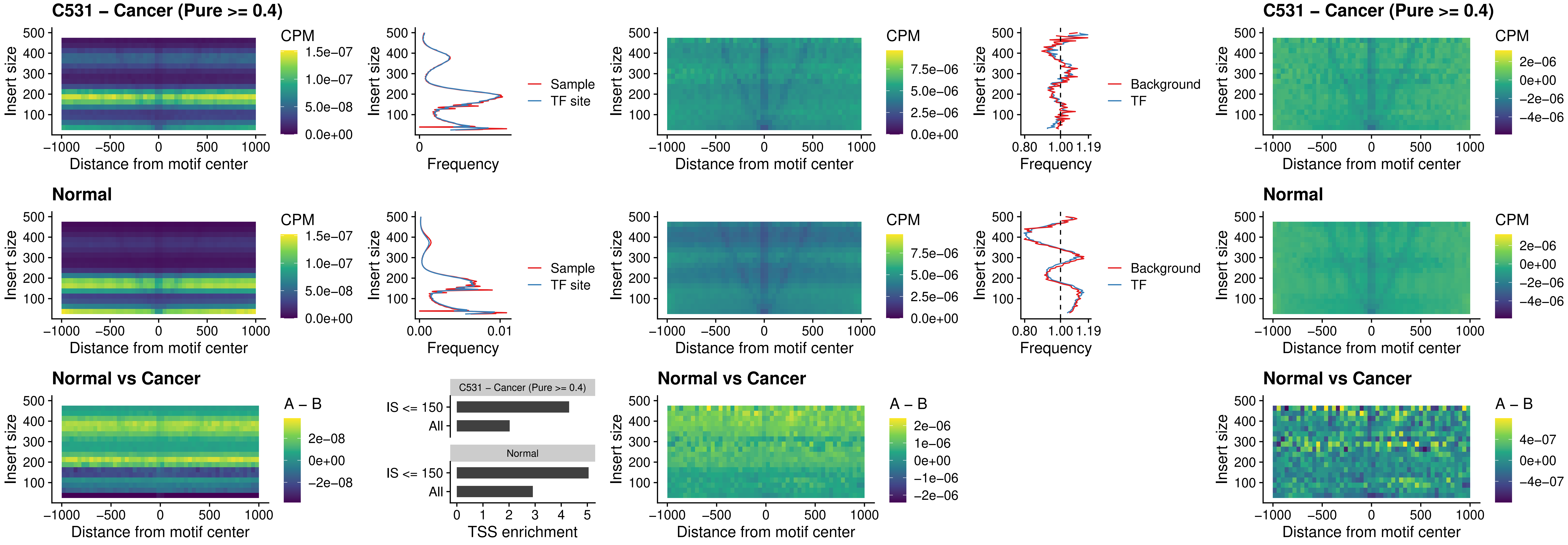

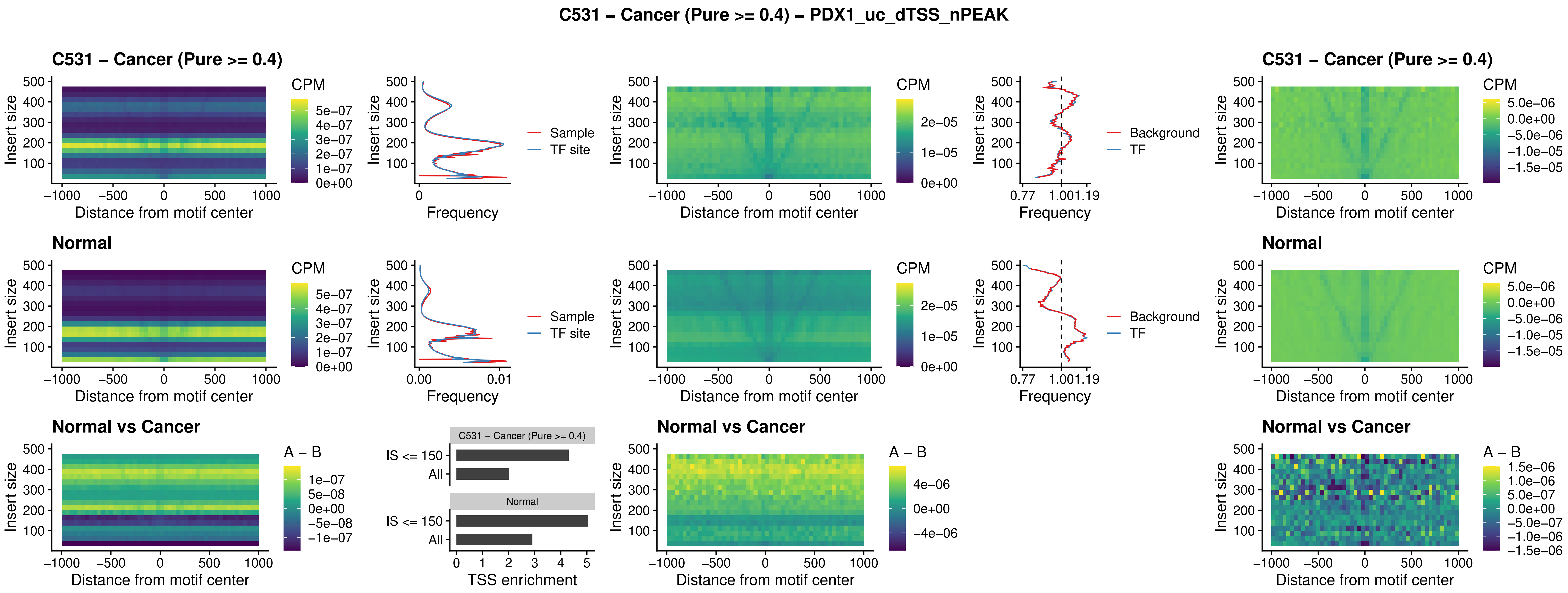

C531 – Cancer (Pure >= 0.4) – PDX1_uc_dTSS_nPEAK
C531 – Cancer (Pure >= 0.4)
Normal
Normal vs Cancer
Insert size
Distance from motif center
Frequency
CPM
A – B
Sample
TF site
Background
TF
TSS enrichment
IS <= 150
All
C531 – Cancer (Pure >= 0.4)
Normal

### C531 – Cancer (Pure >= 0.4) – VAX2_uc_dTSS_nPEAK

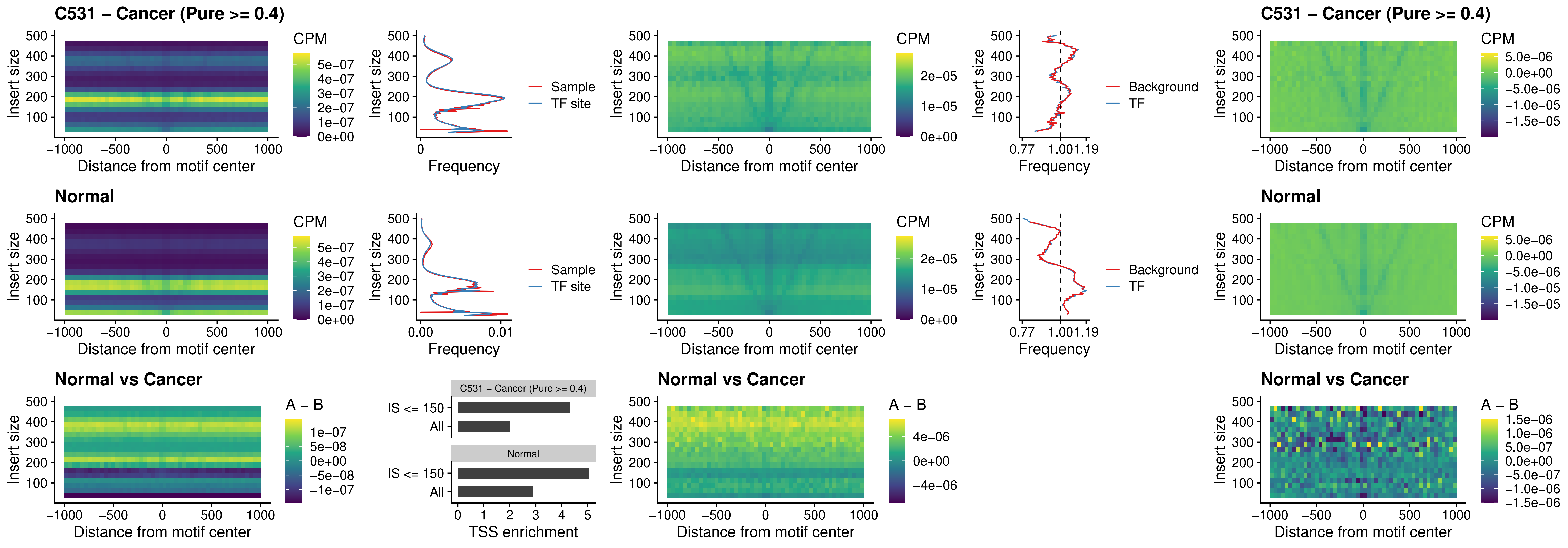

#### C531 – Cancer (Pure >= 0.4) – SOX5_uc_dTSS_nPEAK

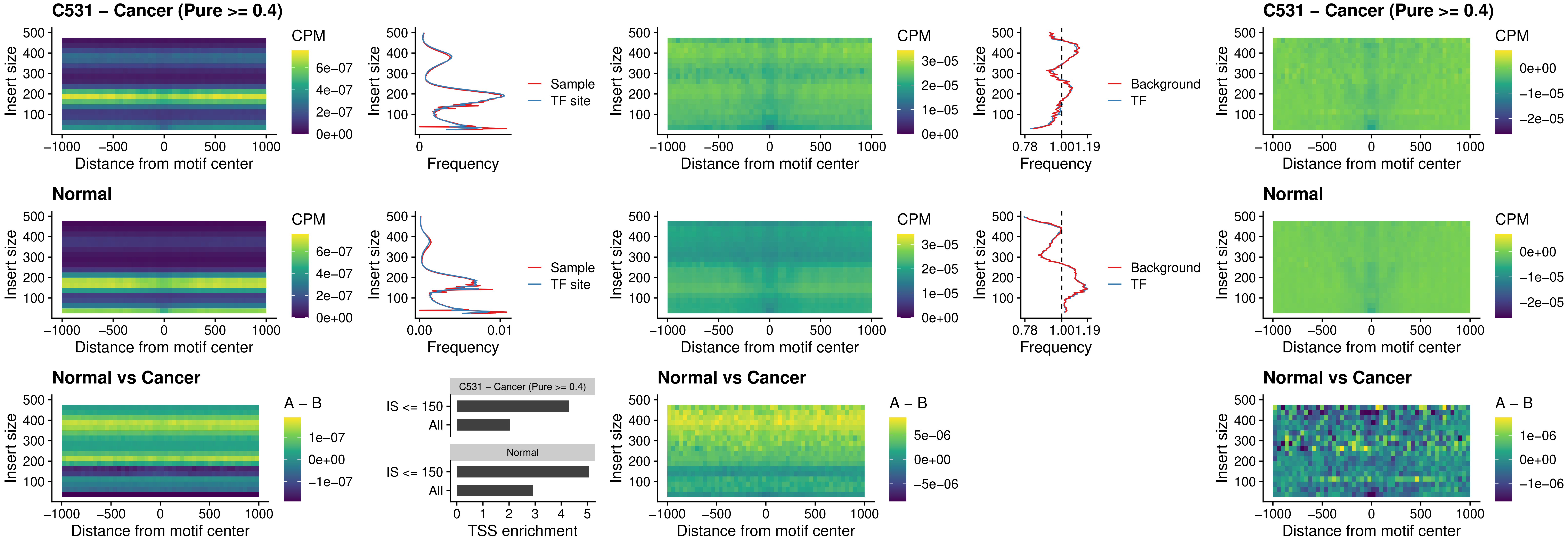

#### C531 – Cancer (Pure >= 0.4) – SOX6_uc_dTSS_nPEAK

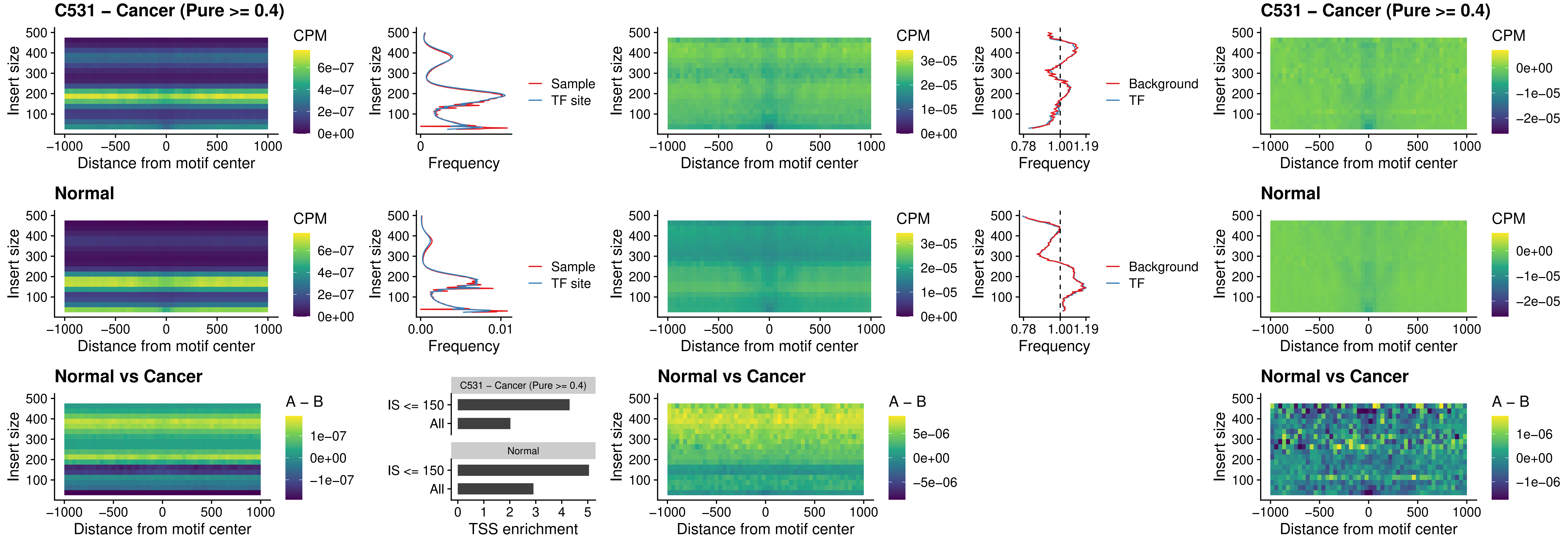

#### C531 – Cancer (Pure >= 0.4) – JDP2_uc_cTSS_nPEAK

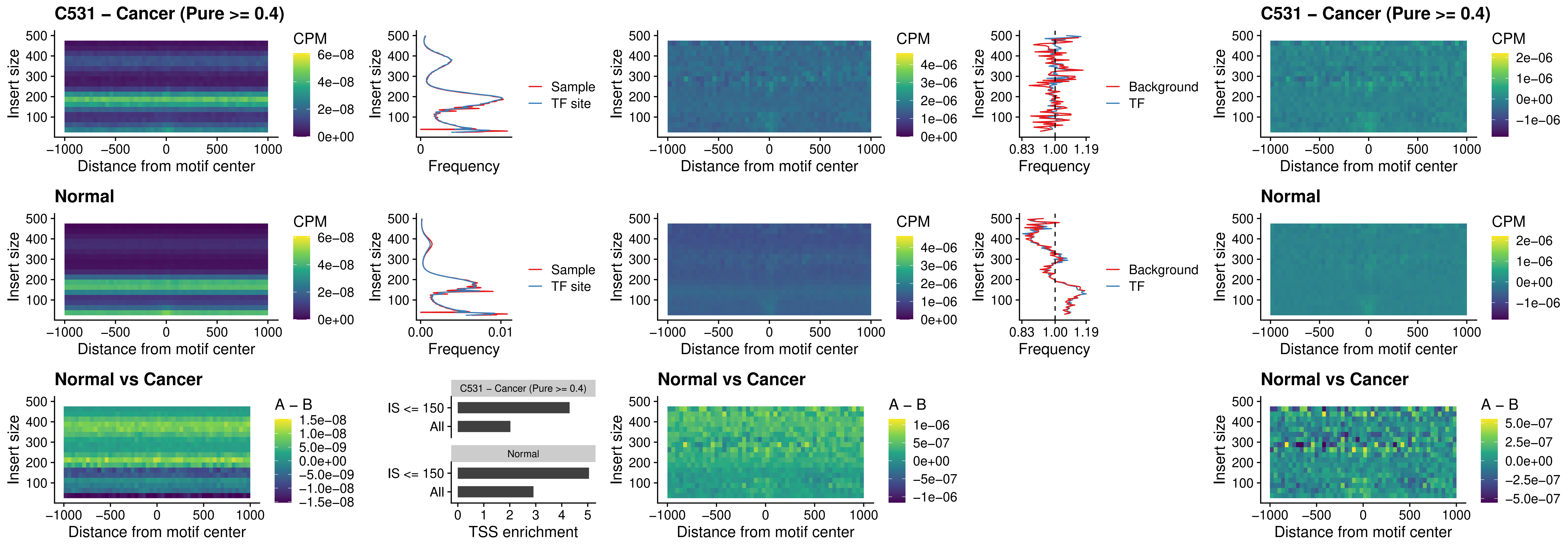

#### C531 – Cancer (Pure >= 0.4) – PURA_uc_dTSS_oPEAK

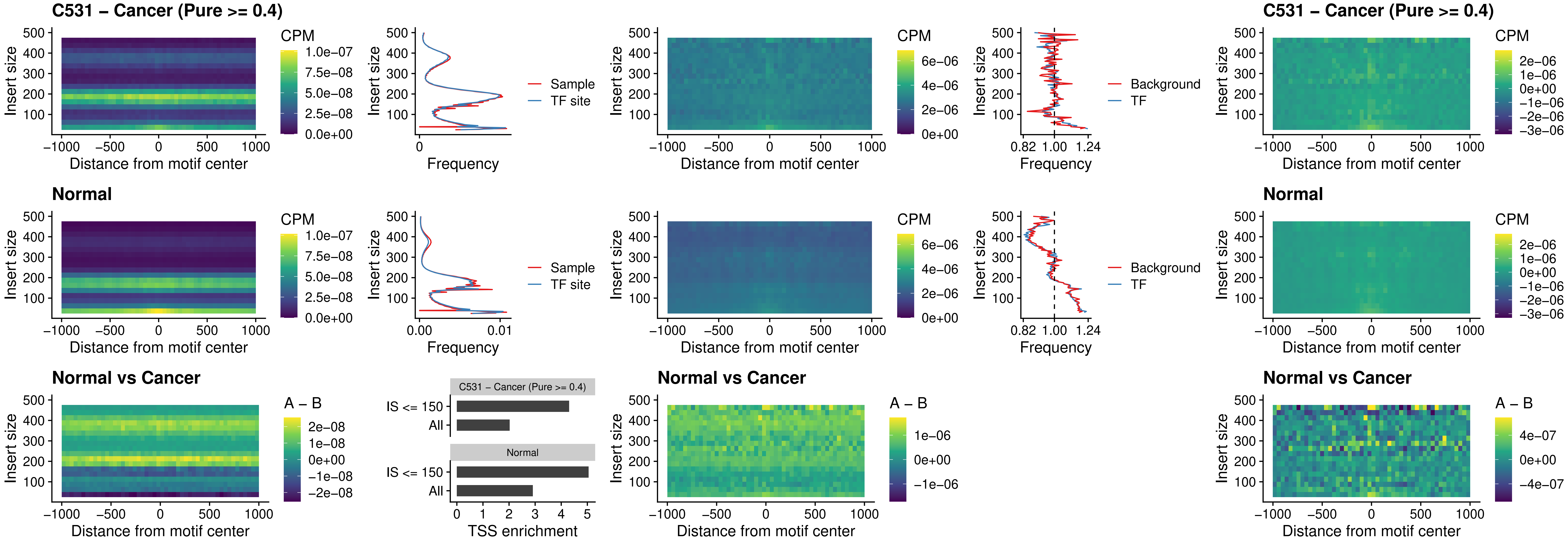

#### C531 – Cancer (Pure >= 0.4) – HHEX_uc_dTSS_nPEAK

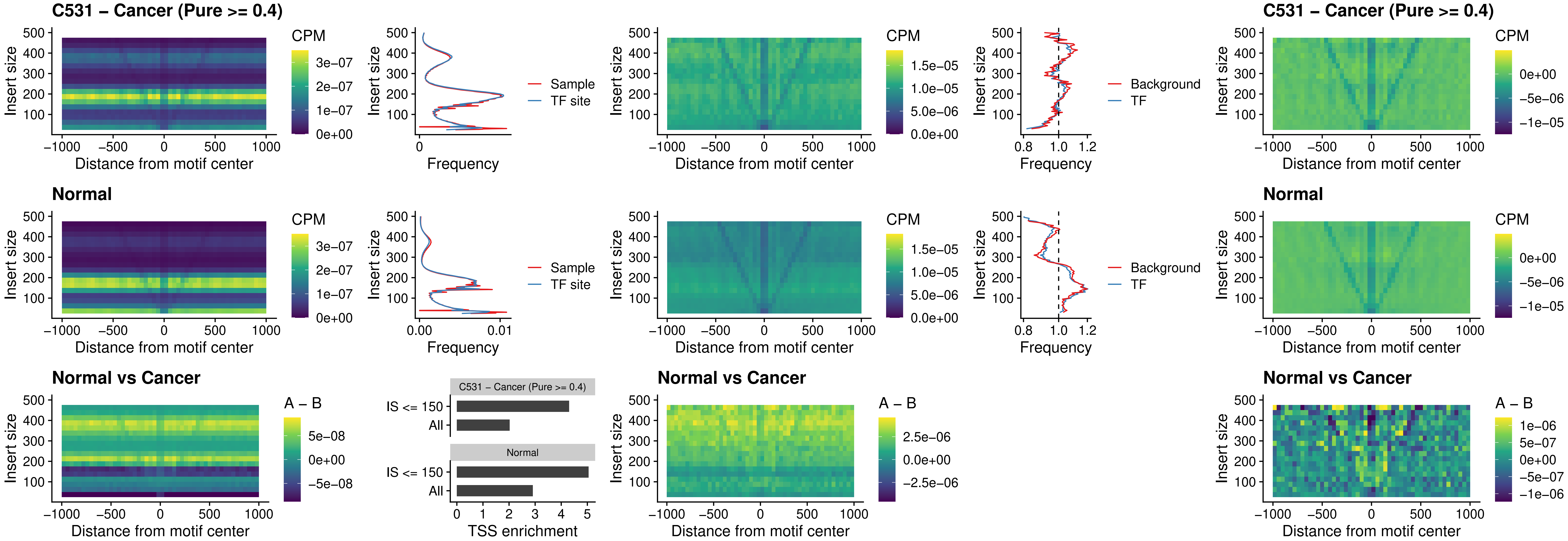

#### C531 – Cancer (Pure >= 0.4) – HMG20B_uc_dTSS_nPEAK

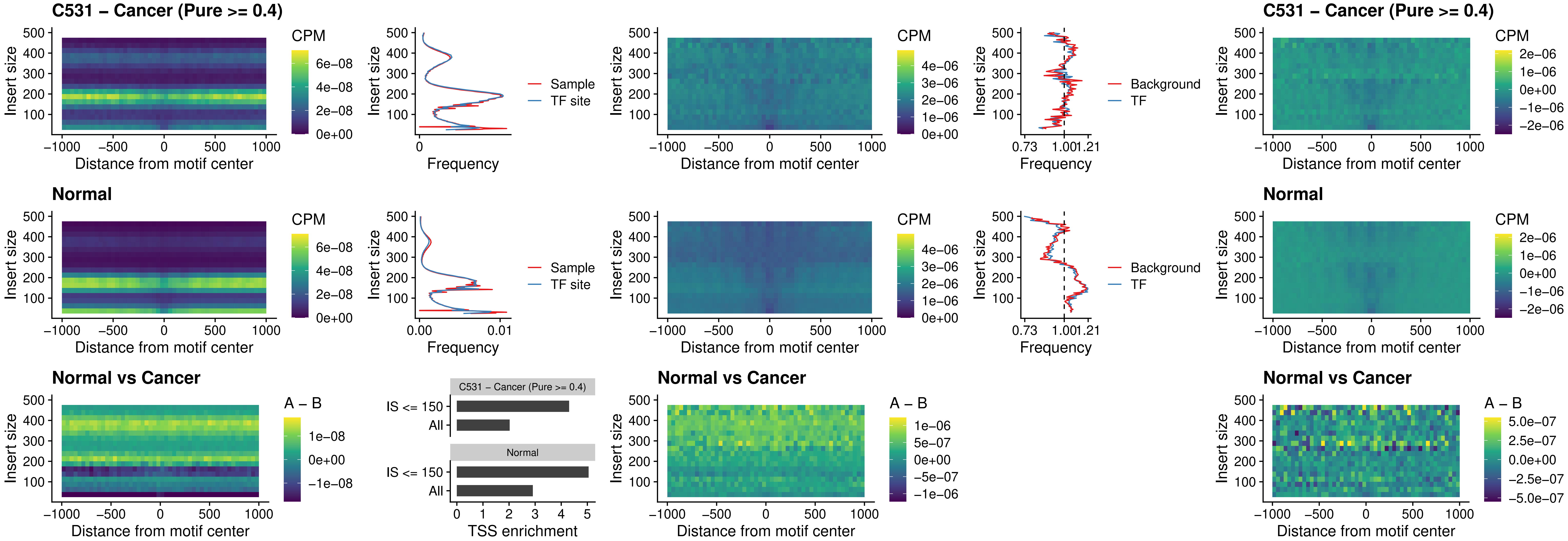

#### C531 – Cancer (Pure >= 0.4) – FOXB2_uc_dTSS_nPEAK

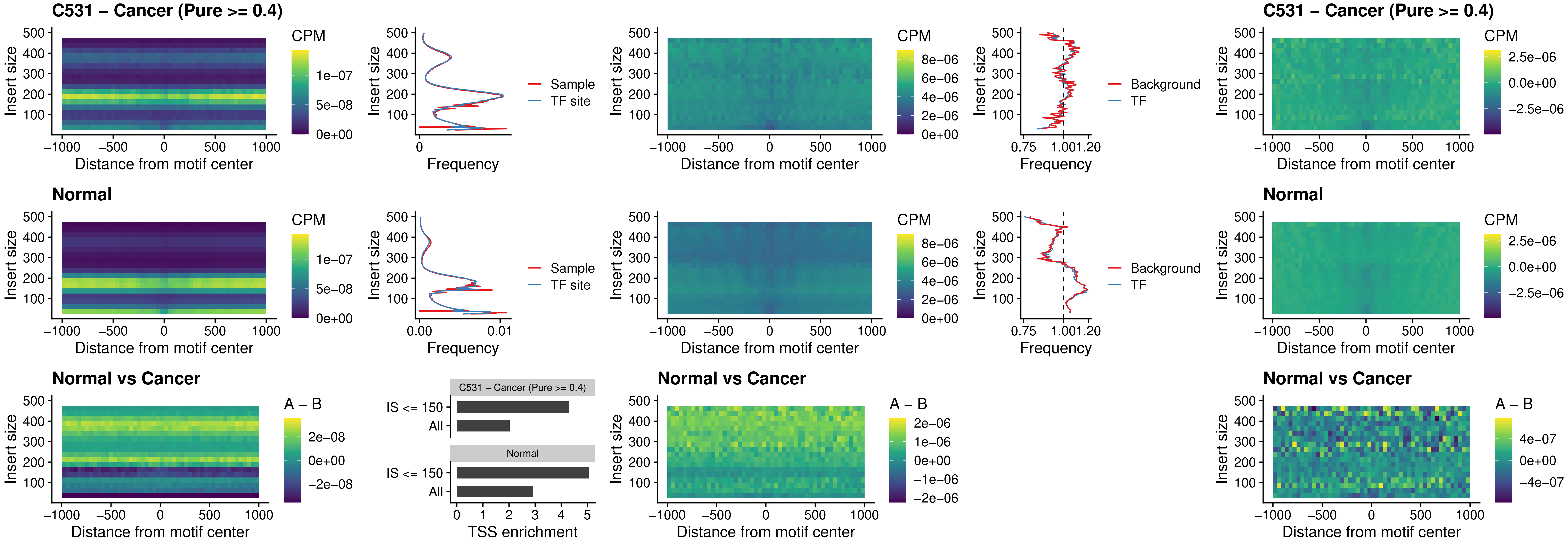

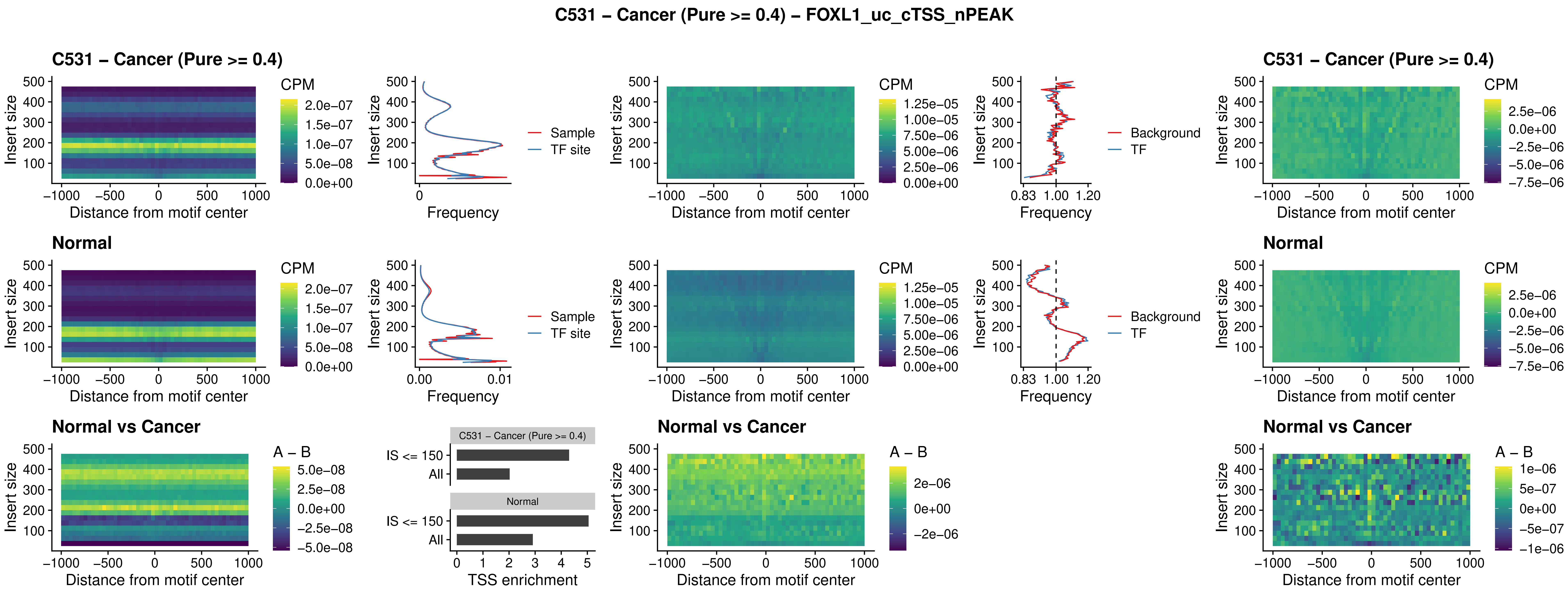
C531 – Cancer (Pure >= 0.4) – FOXL1_uc_cTSS_nPEAK
C531 – Cancer (Pure >= 0.4)
Normal
Normal vs Cancer
Insert size
Distance from motif center
CPM
A – B
Frequency
Sample
TF site
Background
TF
C531 – Cancer (Pure >= 0.4)
Normal
IS <= 150
All
TSS enrichment

#### C531 – Cancer (Pure >= 0.4) – PBX3_uc_pTSS_oPEAK

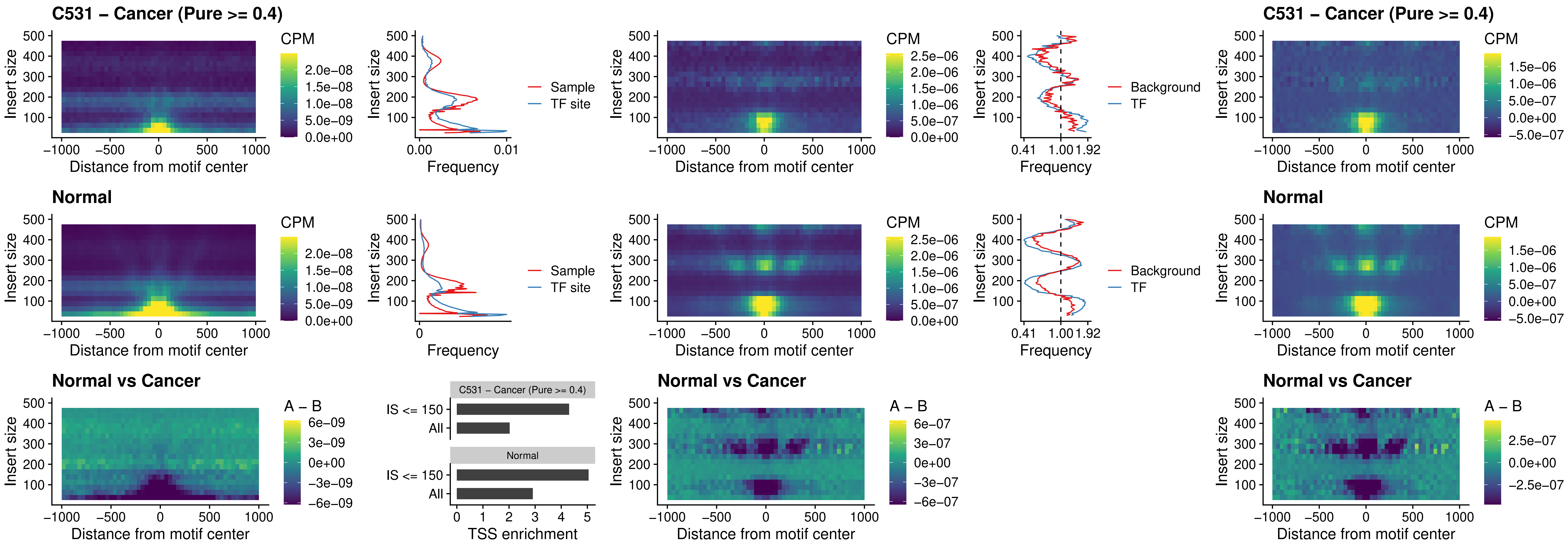

#### C531 – Cancer (Pure >= 0.4) – SNAI2_uc_cTSS_nPEAK

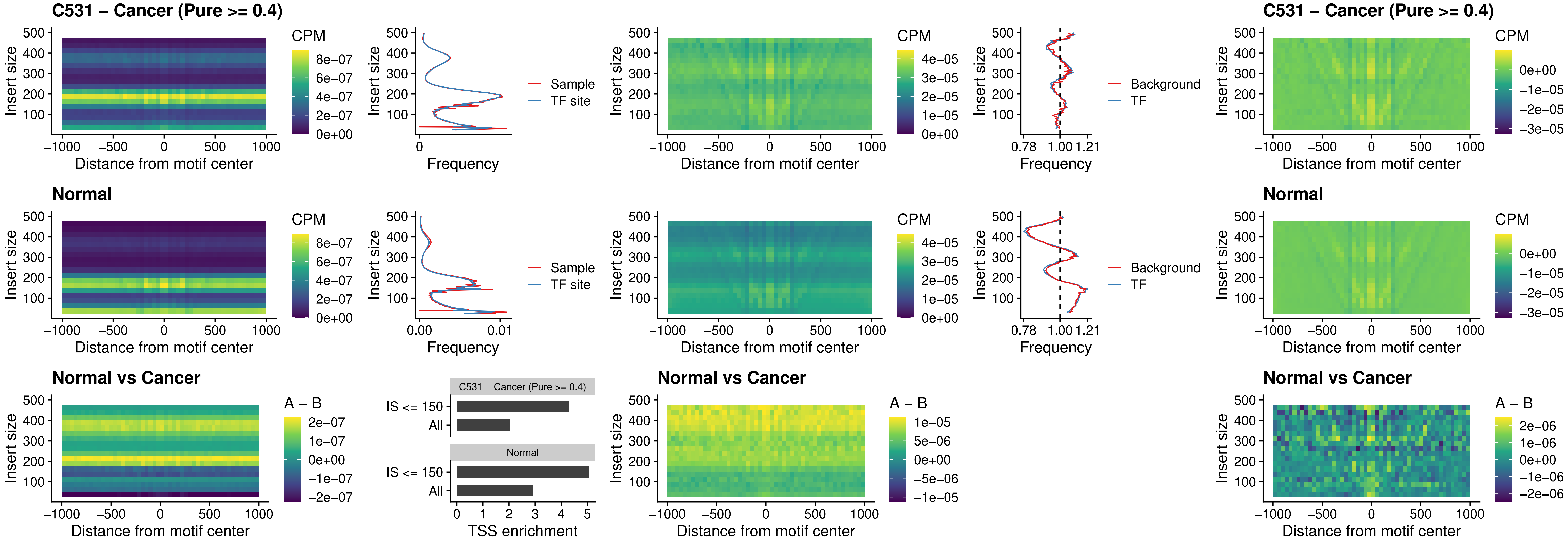

#### C531 – Cancer (Pure >= 0.4) – SNAI2_uc_dTSS_oPEAK

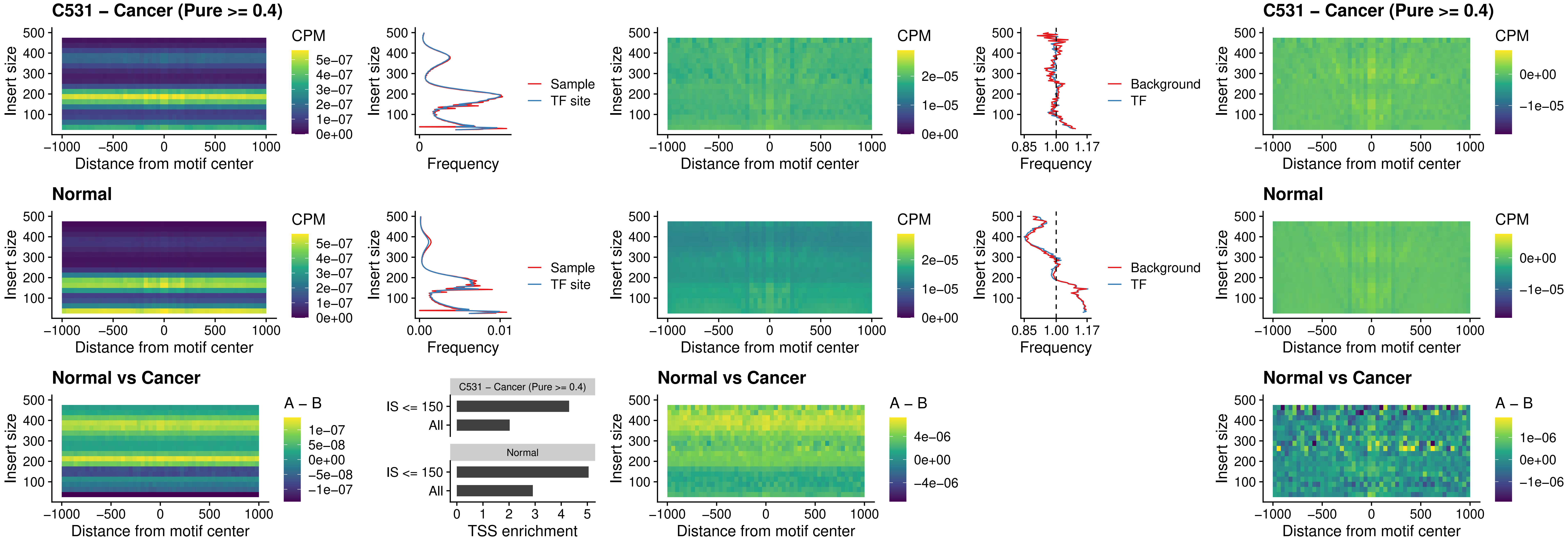

### C531 – Cancer (Pure >= 0.4) – SPI1_uc_pTSS_nPEAK

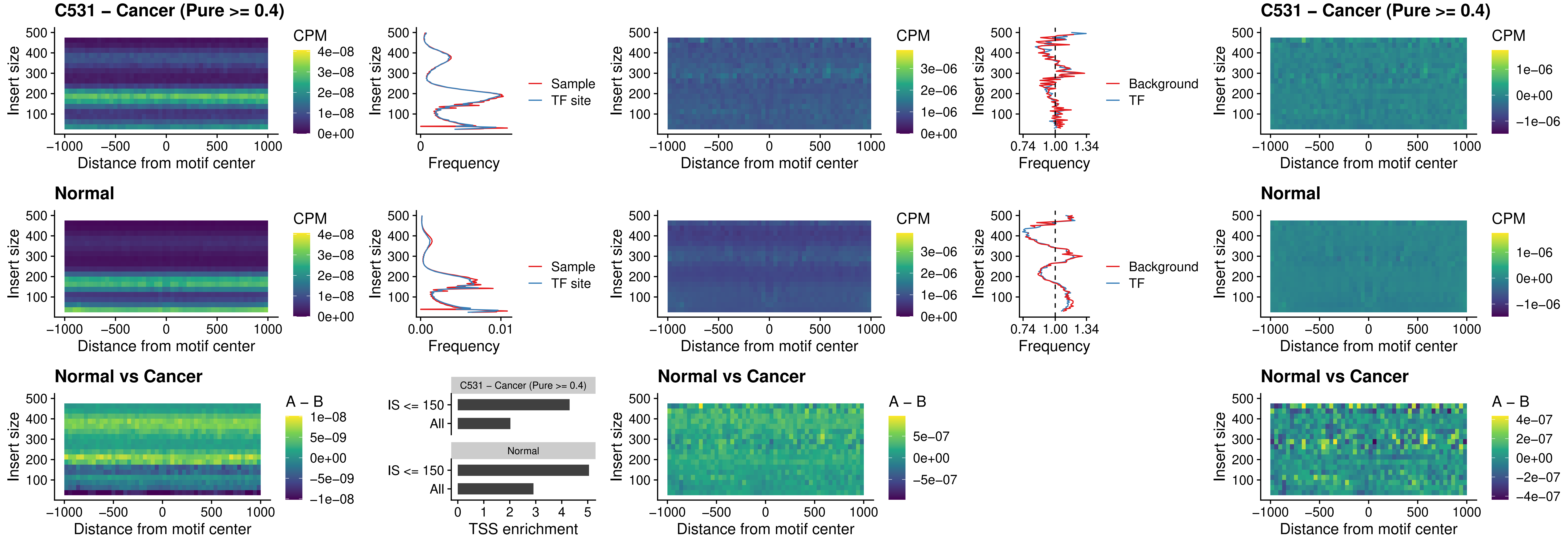

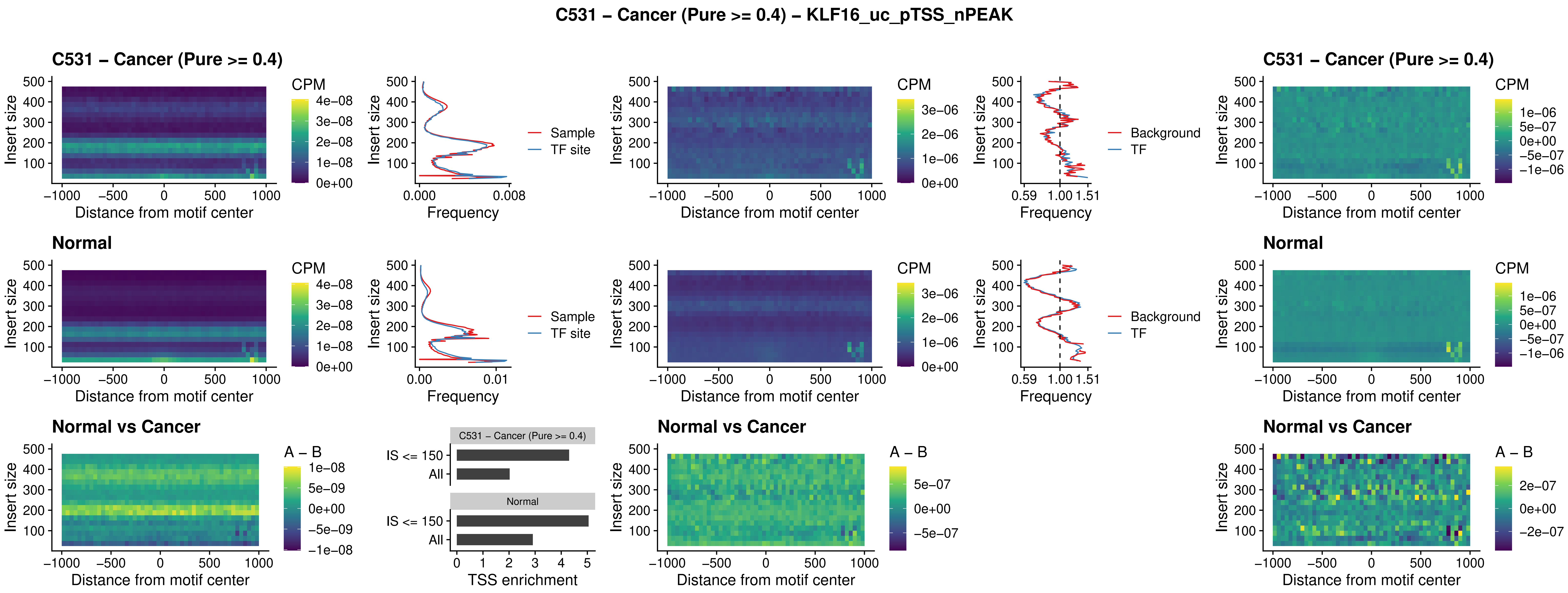

C531 – Cancer (Pure >= 0.4) – KLF16_uc_pTSS_nPEAK
C531 – Cancer (Pure >= 0.4)
Normal
Normal vs Cancer
Insert size
Distance from motif center
Frequency
CPM
A – B
Sample
TF site
Background
TF
C531 – Cancer (Pure >= 0.4)
Normal
IS <= 150
All
TSS enrichment

#### C531 – Cancer (Pure >= 0.4) – SP3_uc_pTSS_nPEAK

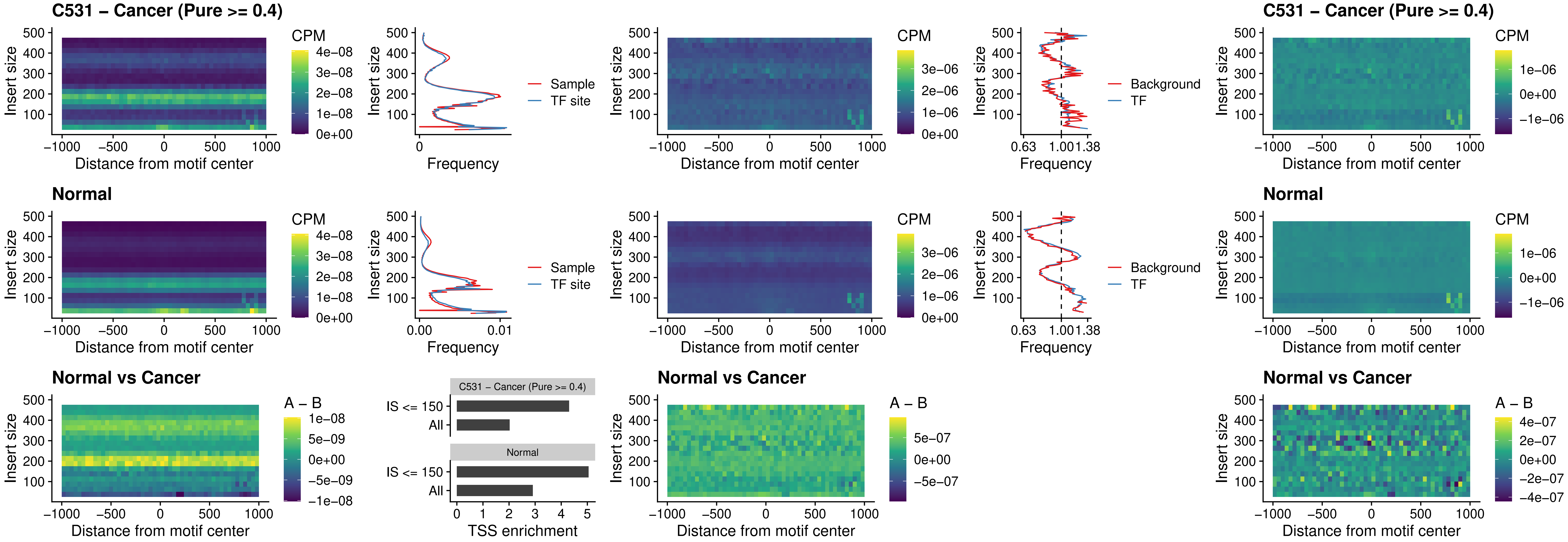

#### C531 – Cancer (Pure >= 0.4) – NEUROD1_uc_dTSS_oPEAK

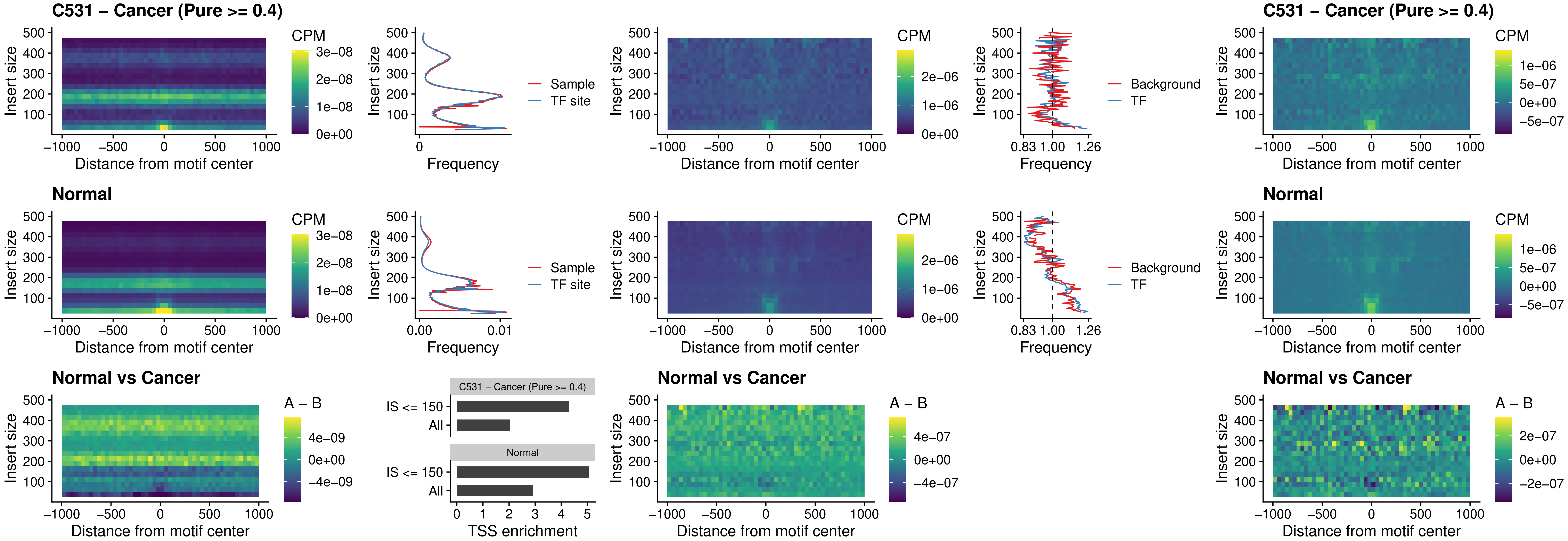

#### C531 – Cancer (Pure >= 0.4) – CTCF_uc_dTSS_oPEAK

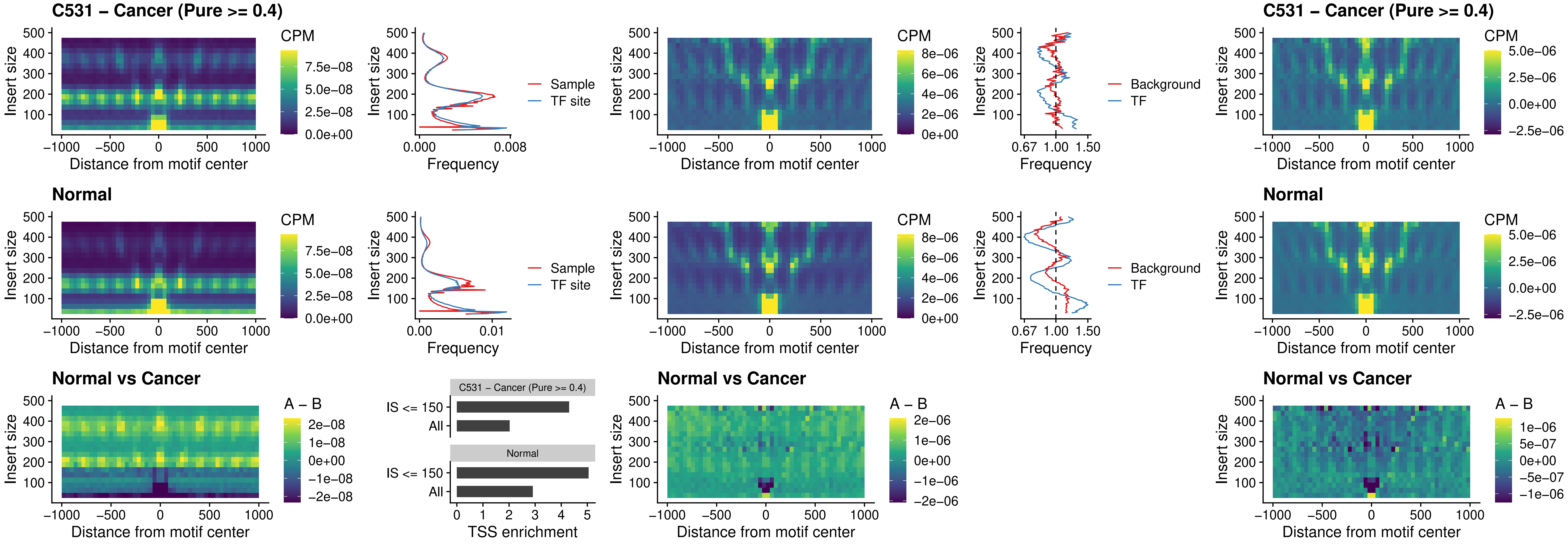

#### C531 – Cancer (Pure >= 0.4) – CTCF_uc_cTSS_oPEAK

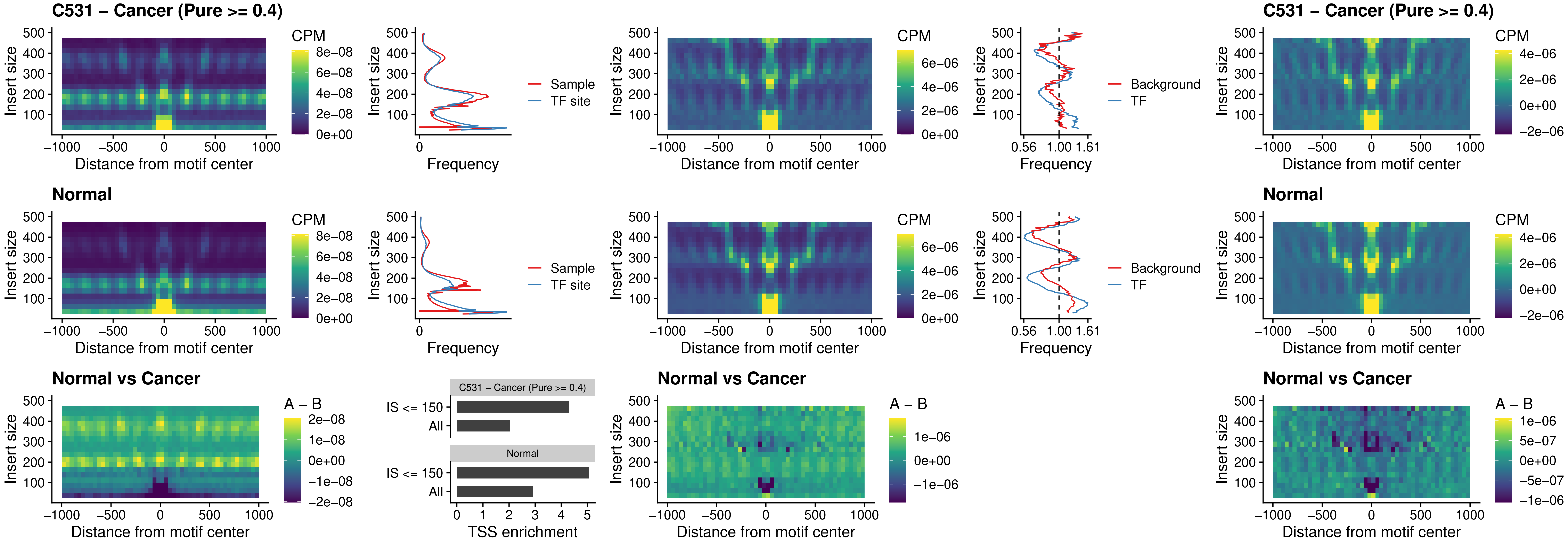

#### C531 – Cancer (Pure >= 0.4) – CTCFL_uc_dTSS_oPEAK

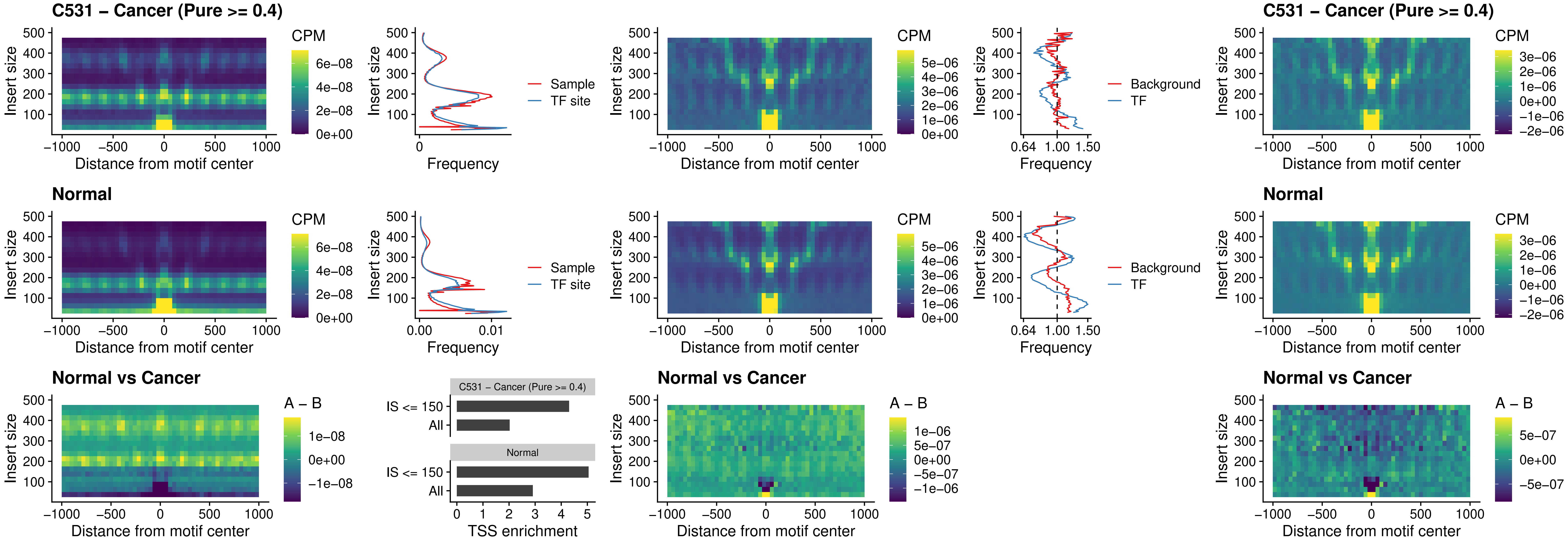

#### C531 – Cancer (Pure >= 0.4) – ENCODE_CTCF_sites_COLON

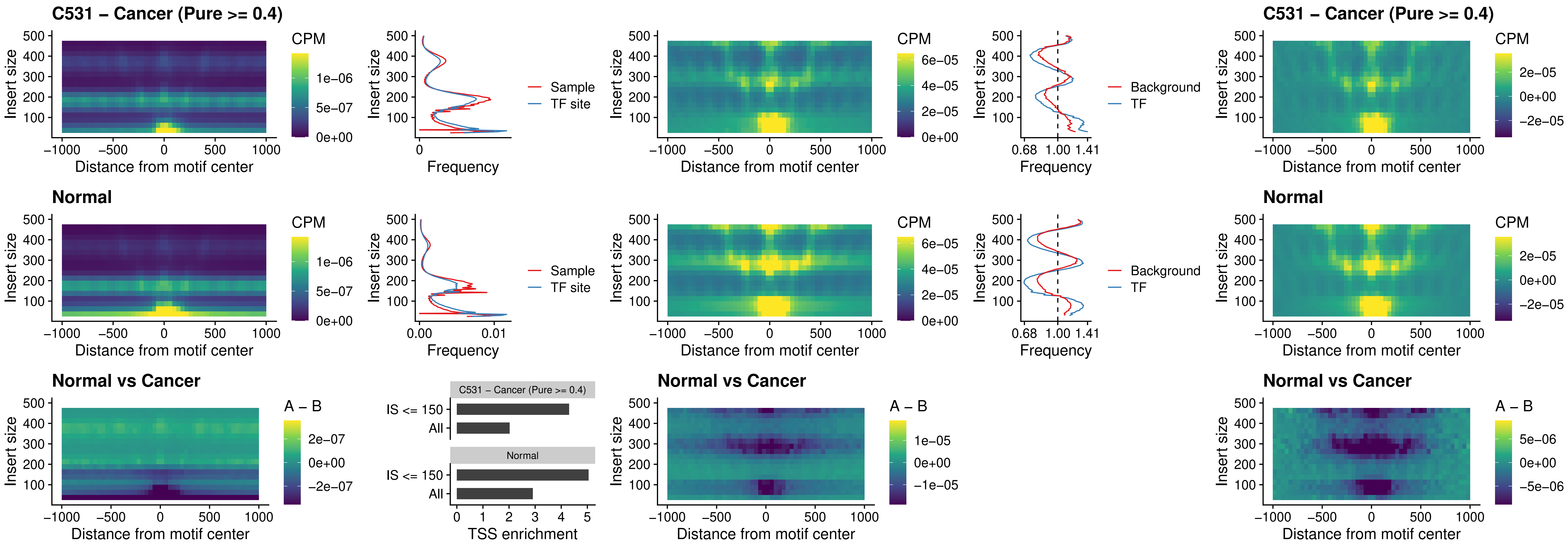

#### C531 – Cancer (Pure >= 0.4) – ENCODE_CTCF_sites_COLON_loop_ctcf

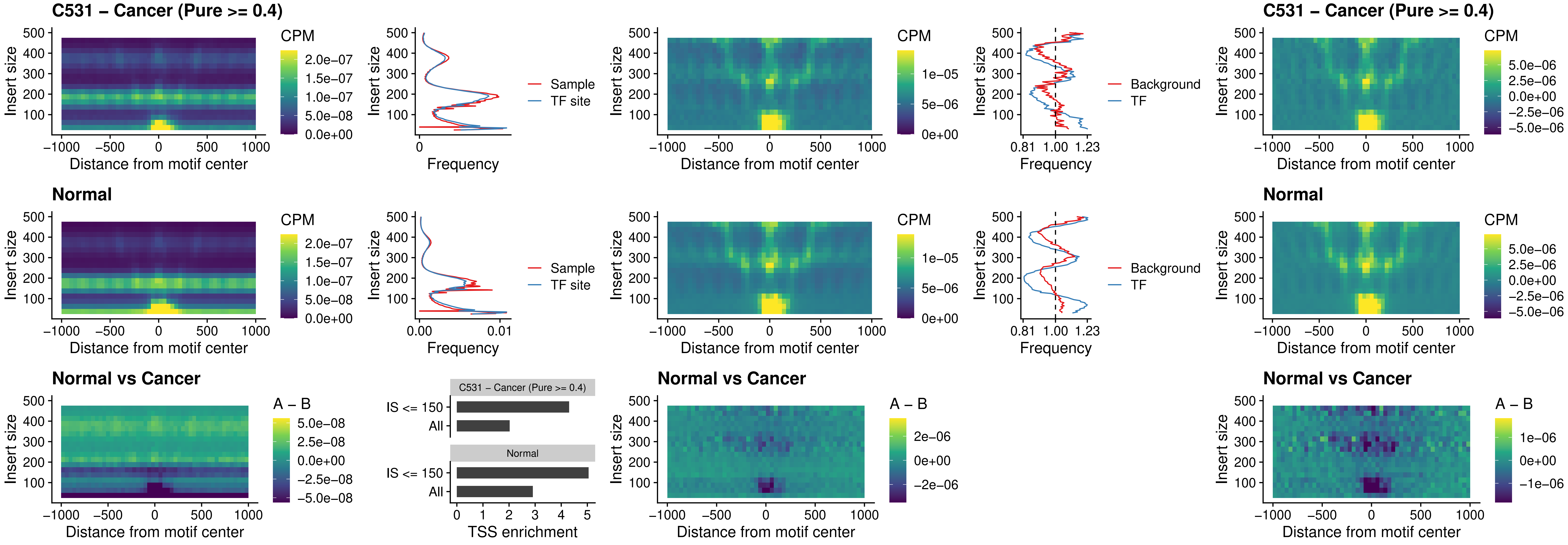

#### C531 – Cancer (Pure >= 0.4) – ENCODE_CTCF_sites_COLON_intergenic_ctcf

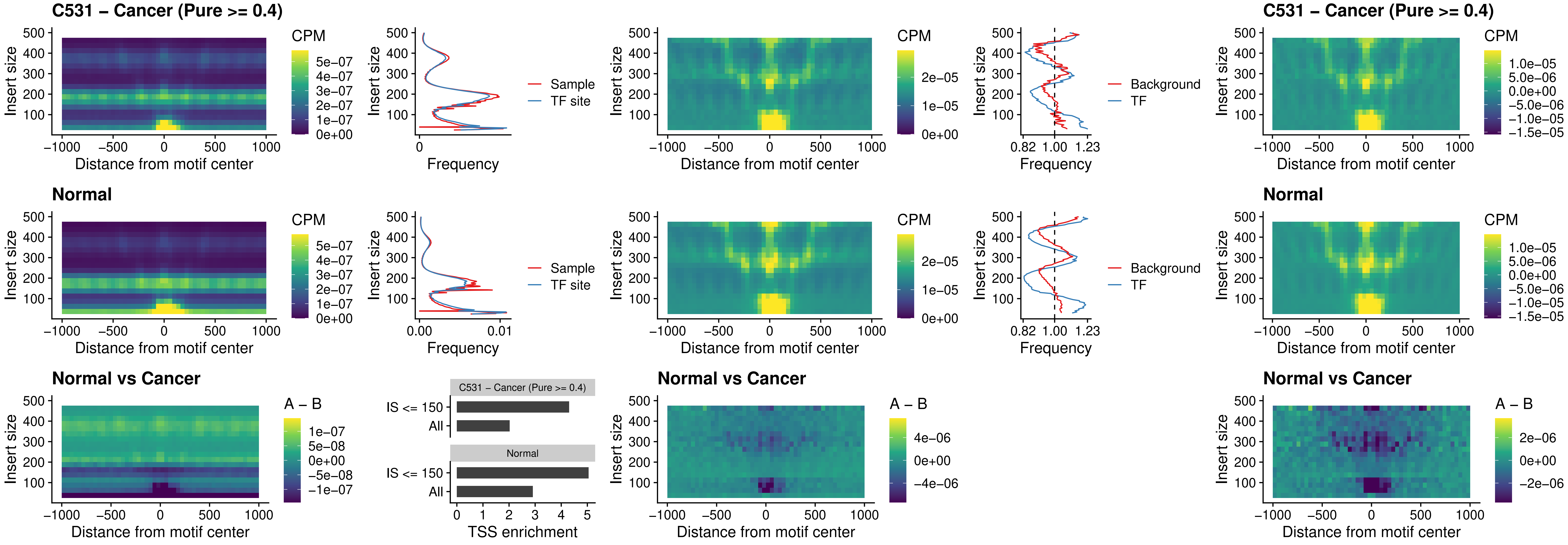

#### C531 – Cancer (Pure >= 0.4) – ENCODE_CTCF_sites_COLON_nonpromotor

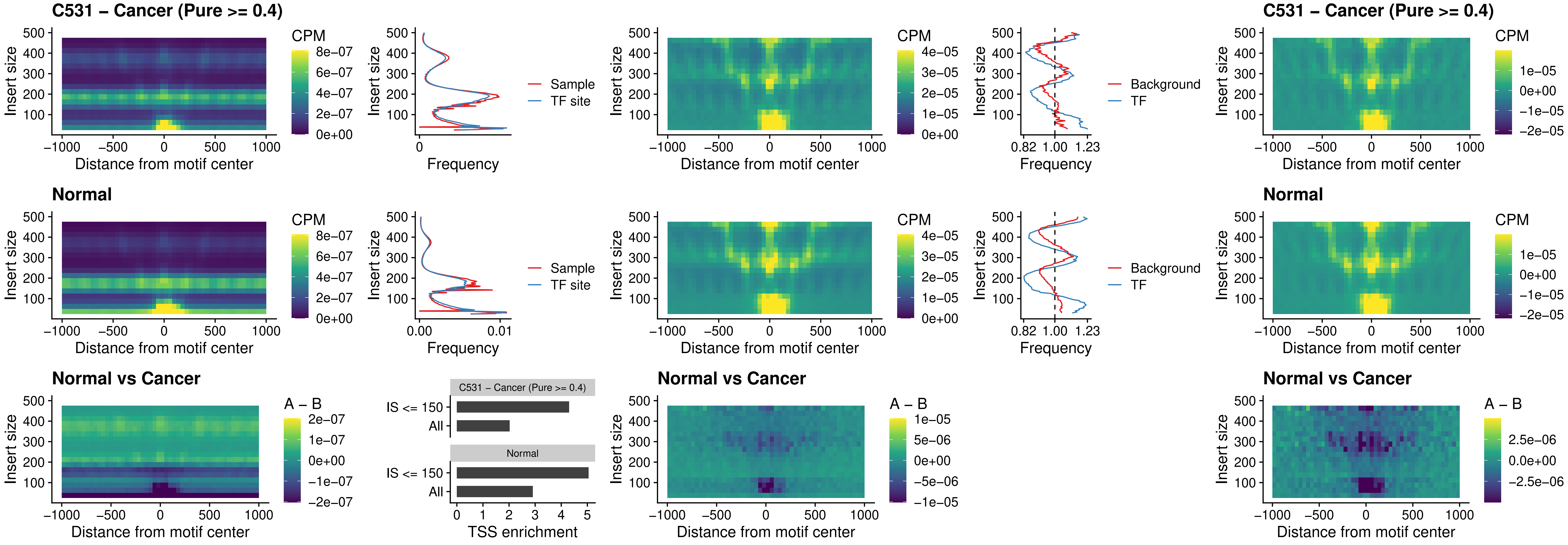

#### C531 – Cancer (Pure >= 0.4) – CTCFL_uc_cTSS_oPEAK

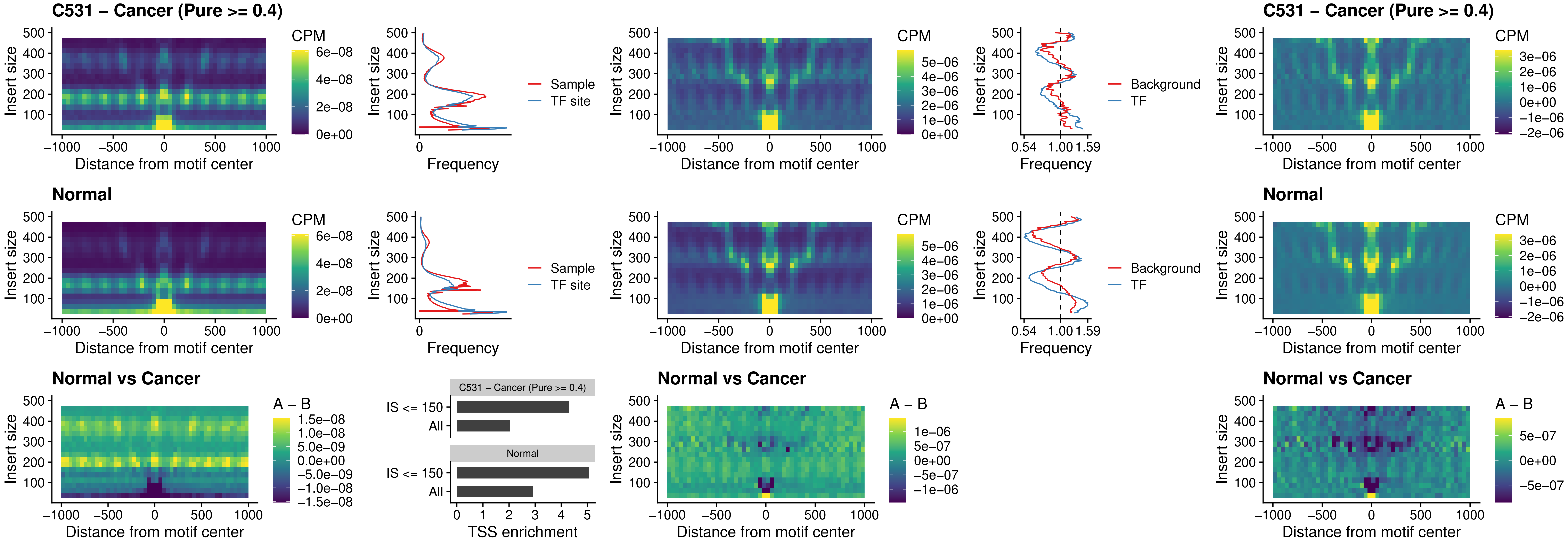

### C531 – Cancer (Pure >= 0.4) – NEUROD1_uc_cTSS_oPEAK

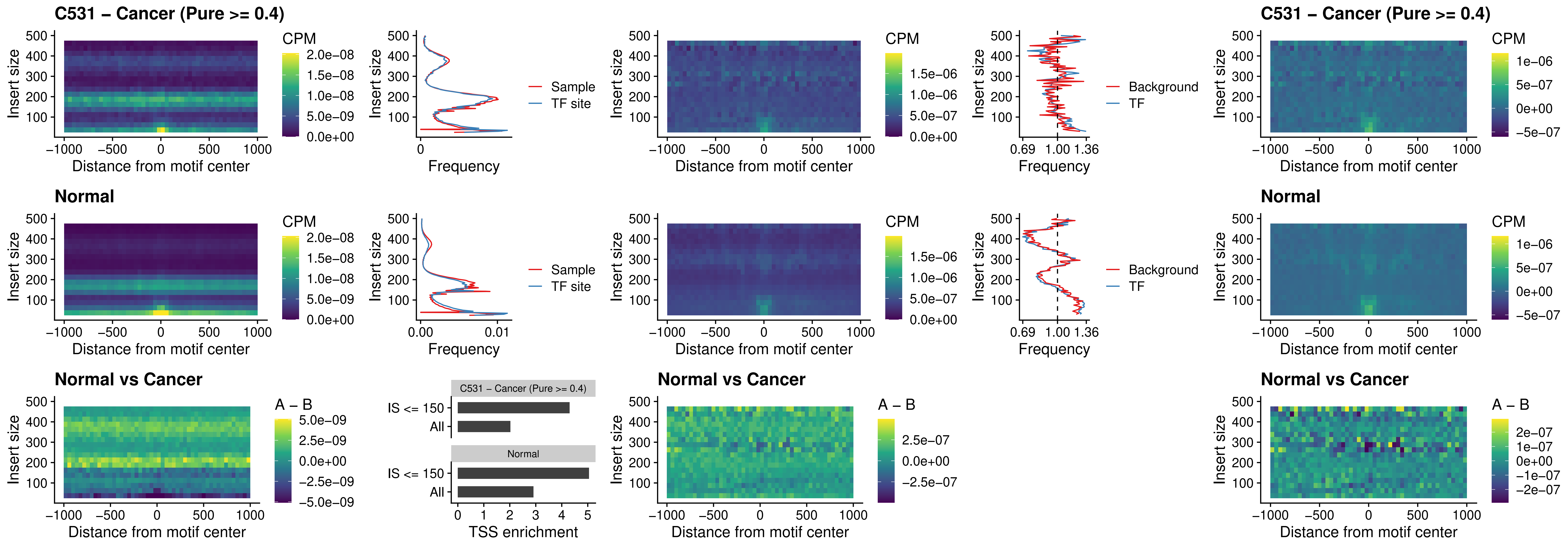

#### C531 – Cancer (Pure >= 0.4) – ENCODE_YY1_sites_COLON_nonpromotor

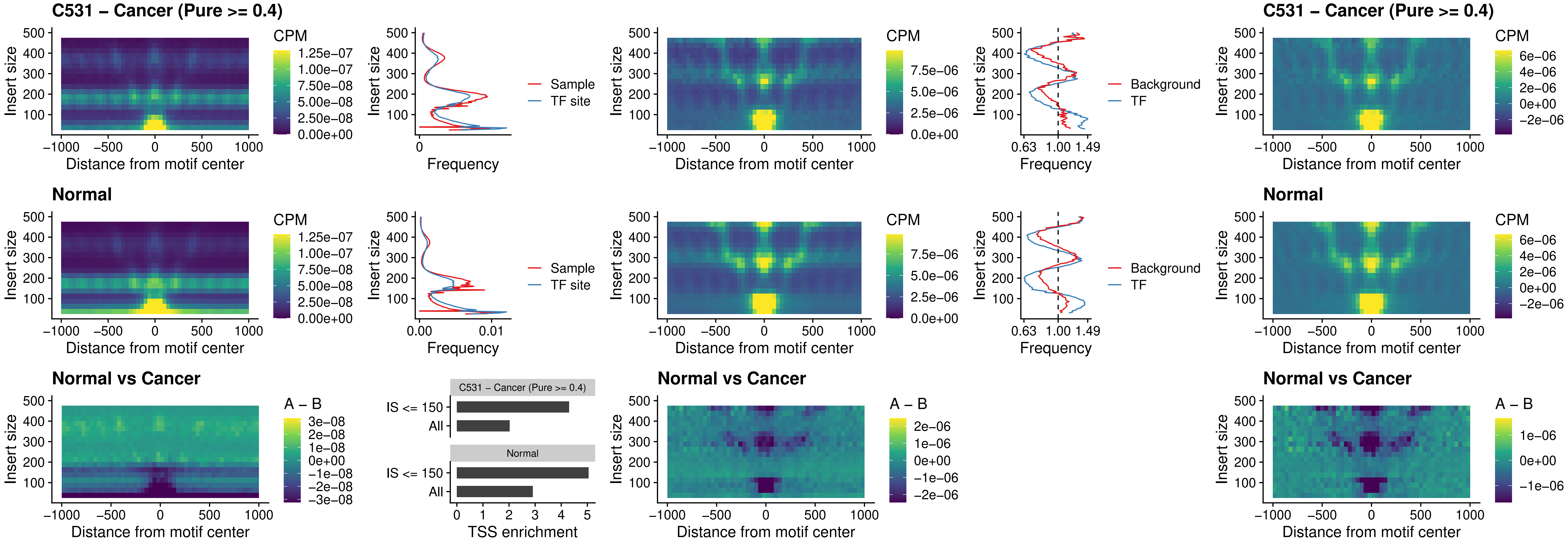

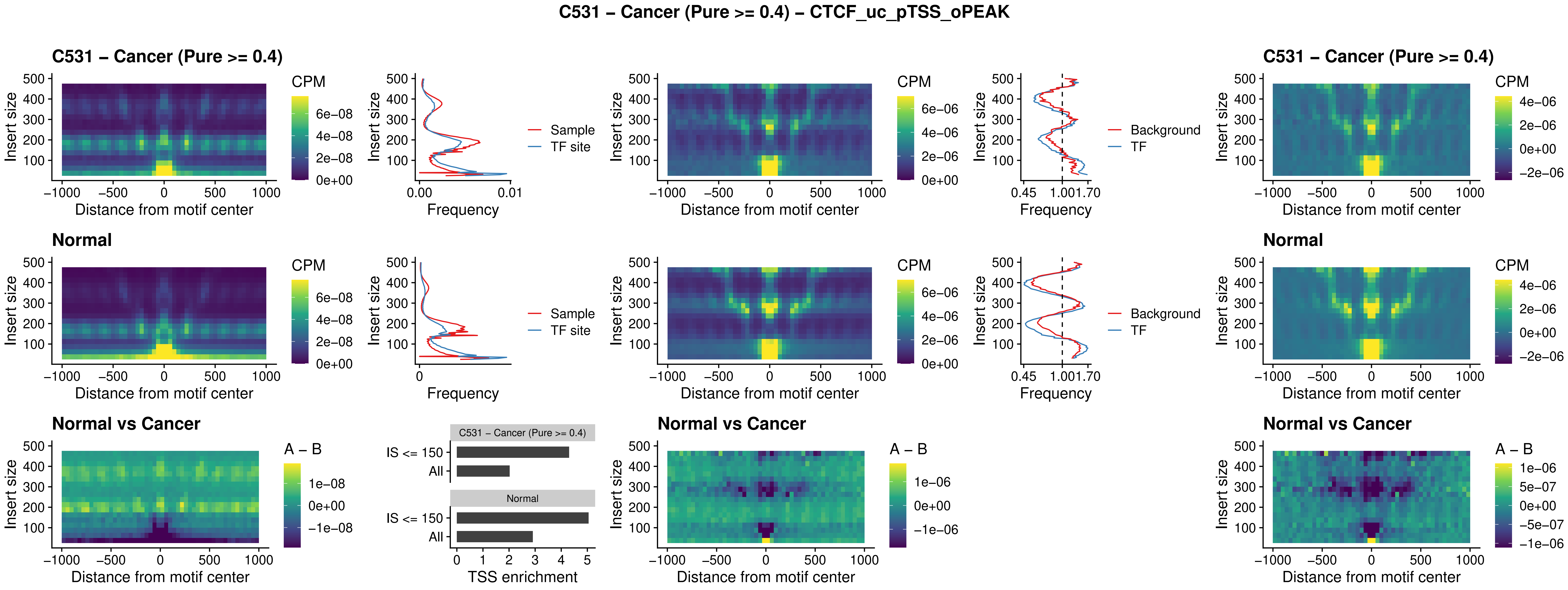
C531 – Cancer (Pure >= 0.4) – CTCF_uc_pTSS_oPEAK
C531 – Cancer (Pure >= 0.4)
Normal
Normal vs Cancer
Insert size
Distance from motif center
Frequency
CPM
A – B
Sample
TF site
Background
TF
C531 – Cancer (Pure >= 0.4)
Normal
IS <= 150
All
TSS enrichment

#### C531 – Cancer (Pure >= 0.4) – ENCODE_CTCF_sites_COLON_promotor

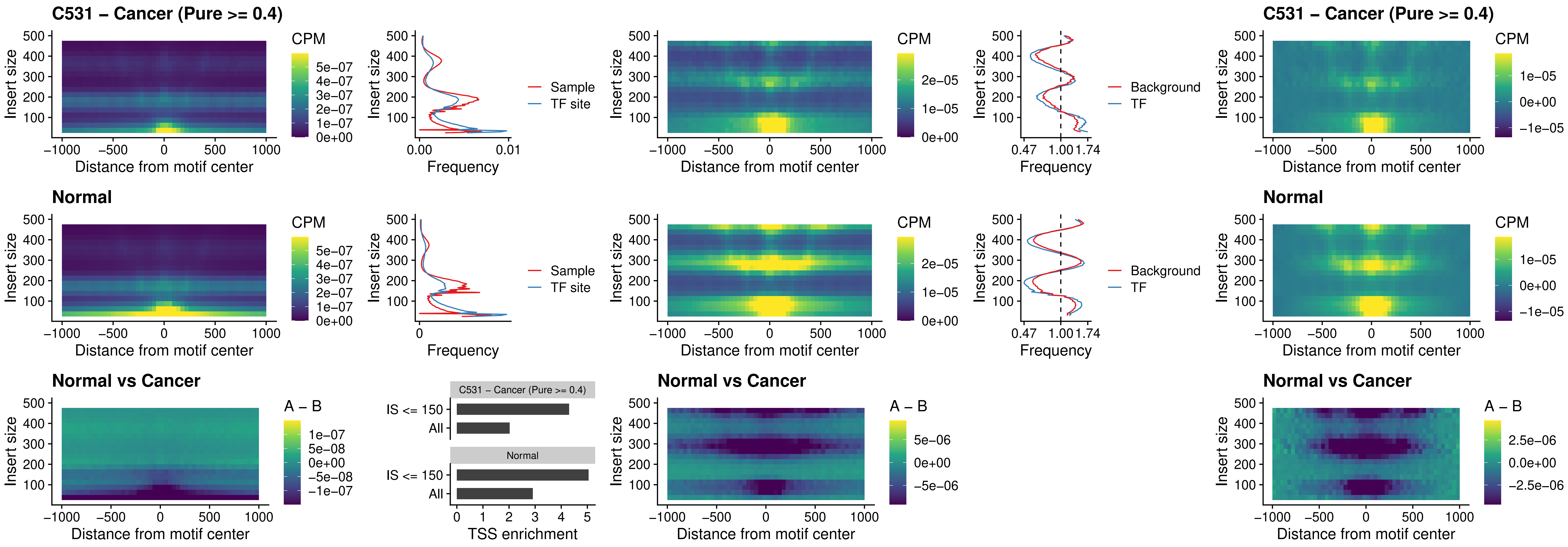

### C531 – Cancer (Pure >= 0.4) – ENCODE_CTCF_sites_COLON_promotor_ctcf

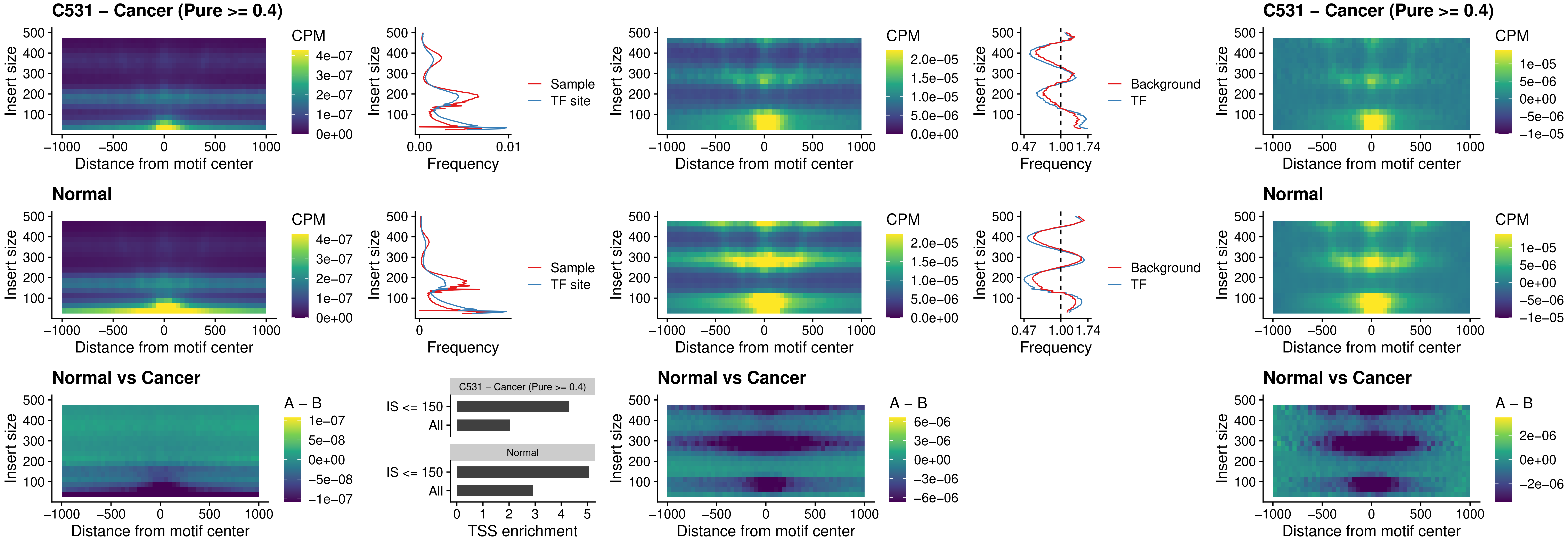

#### C531 – Cancer (Pure >= 0.4) – SPIC_uc_pTSS_oPEAK

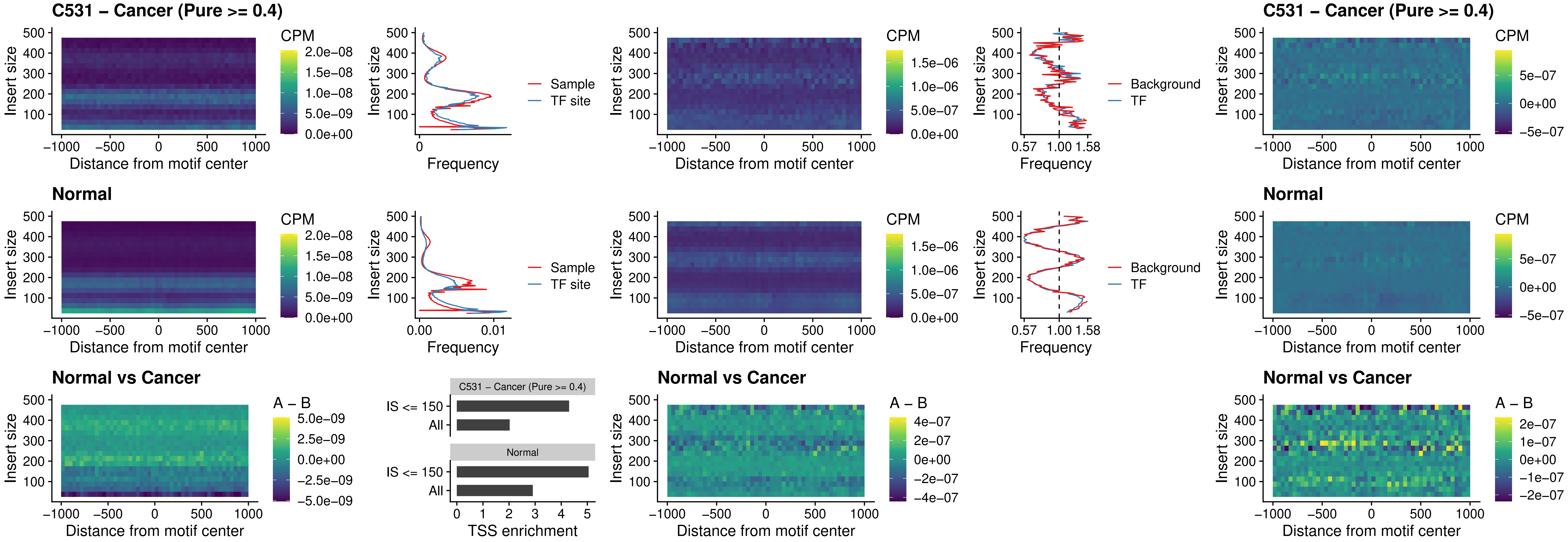

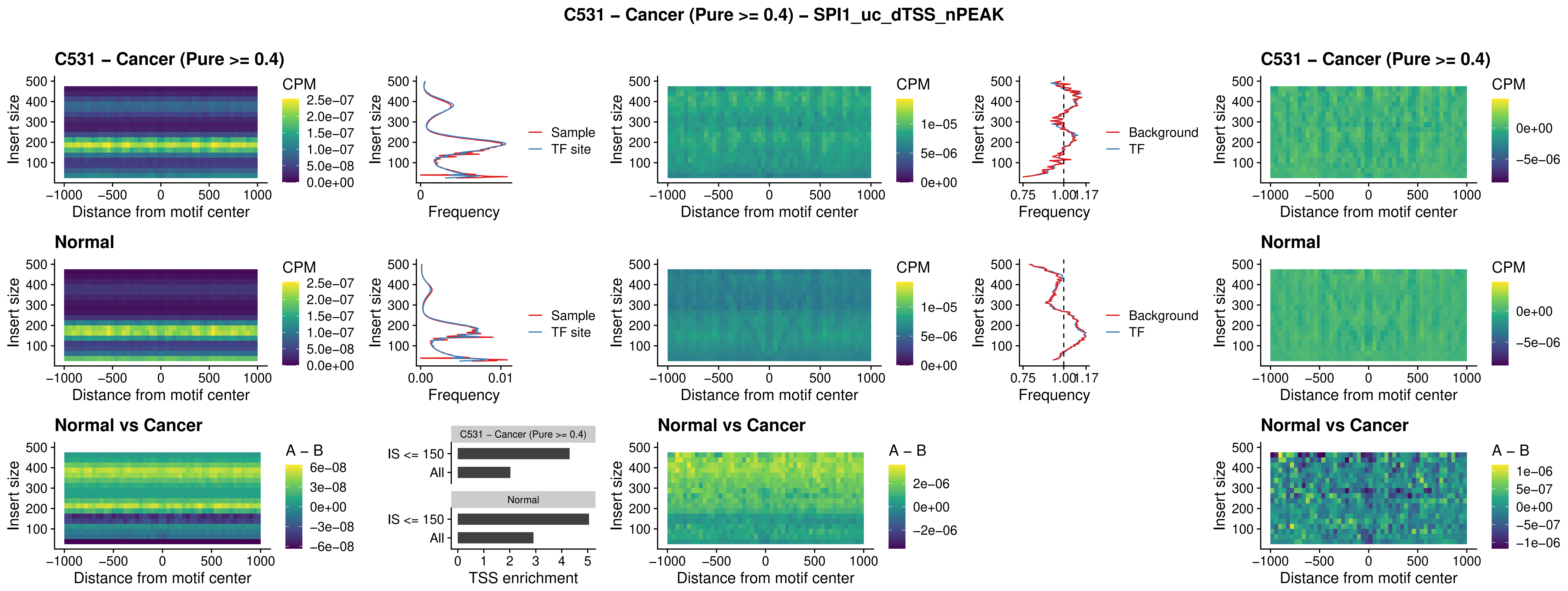
C531 – Cancer (Pure >= 0.4) – SPI1_uc_dTSS_nPEAK
C531 – Cancer (Pure >= 0.4)
Normal
Normal vs Cancer
CPM
A – B
Insert size
Distance from motif center
Frequency
Sample
TF site
Background
TF
TSS enrichment
C531 – Cancer (Pure >= 0.4)
Normal
IS <= 150
All

#### C531 – Cancer (Pure >= 0.4) – SPI1_uc_cTSS_nPEAK

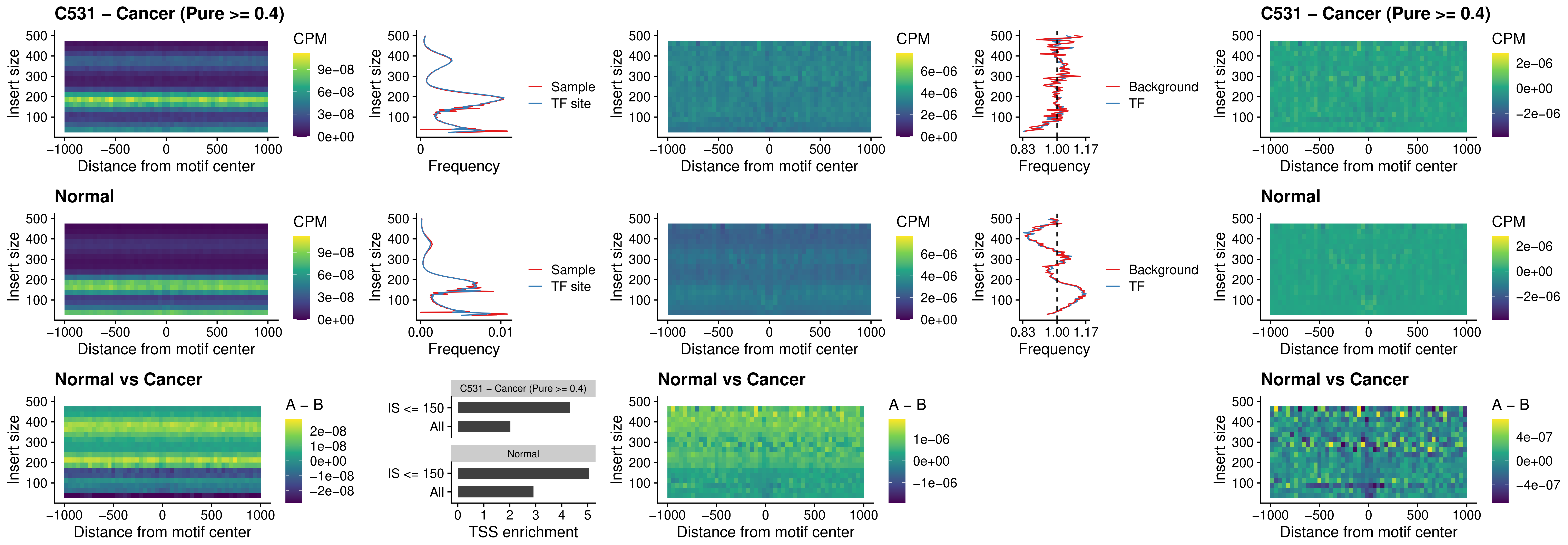

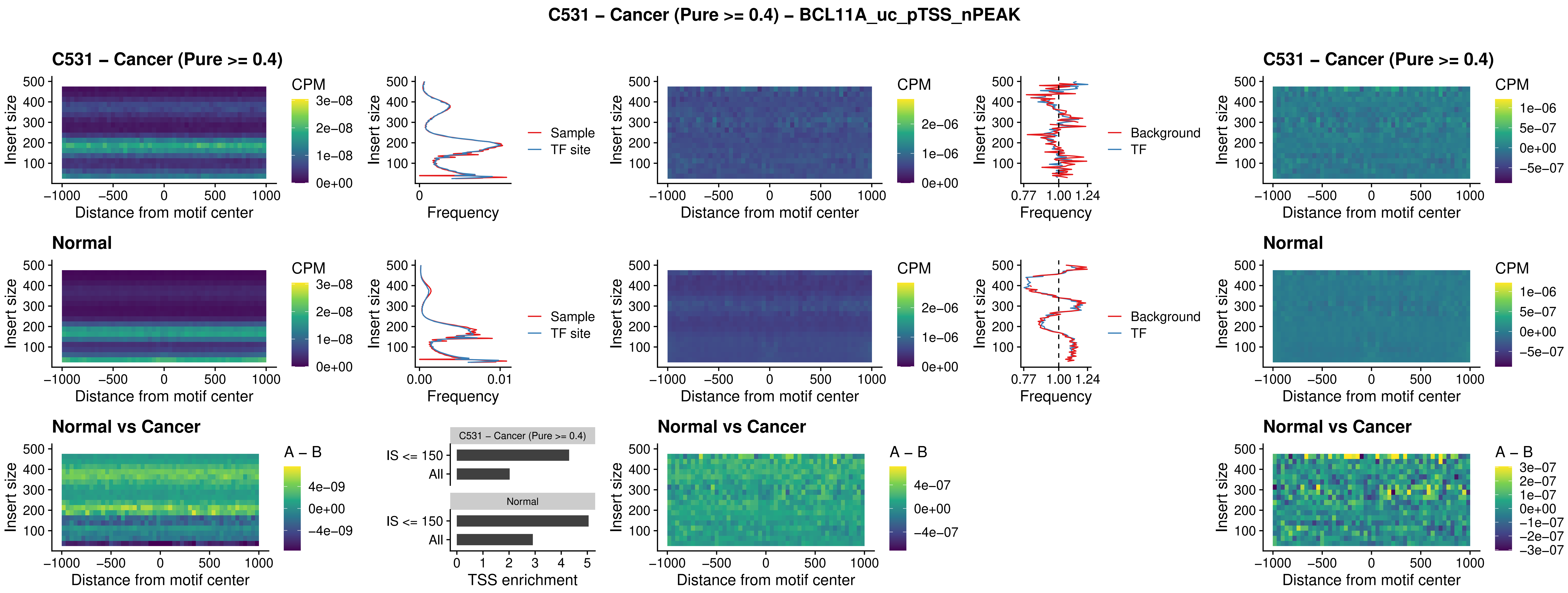

C531 – Cancer (Pure >= 0.4) – BCL11A_uc_pTSS_nPEAK
C531 – Cancer (Pure >= 0.4)
Normal
Normal vs Cancer
Insert size
Distance from motif center
CPM
A – B
Frequency
Sample
TF site
Background
TF
C531 – Cancer (Pure >= 0.4)
Normal
IS <= 150
All
TSS enrichment

#### C531 – Cancer (Pure >= 0.4) – BCL11B_uc_pTSS_nPEAK

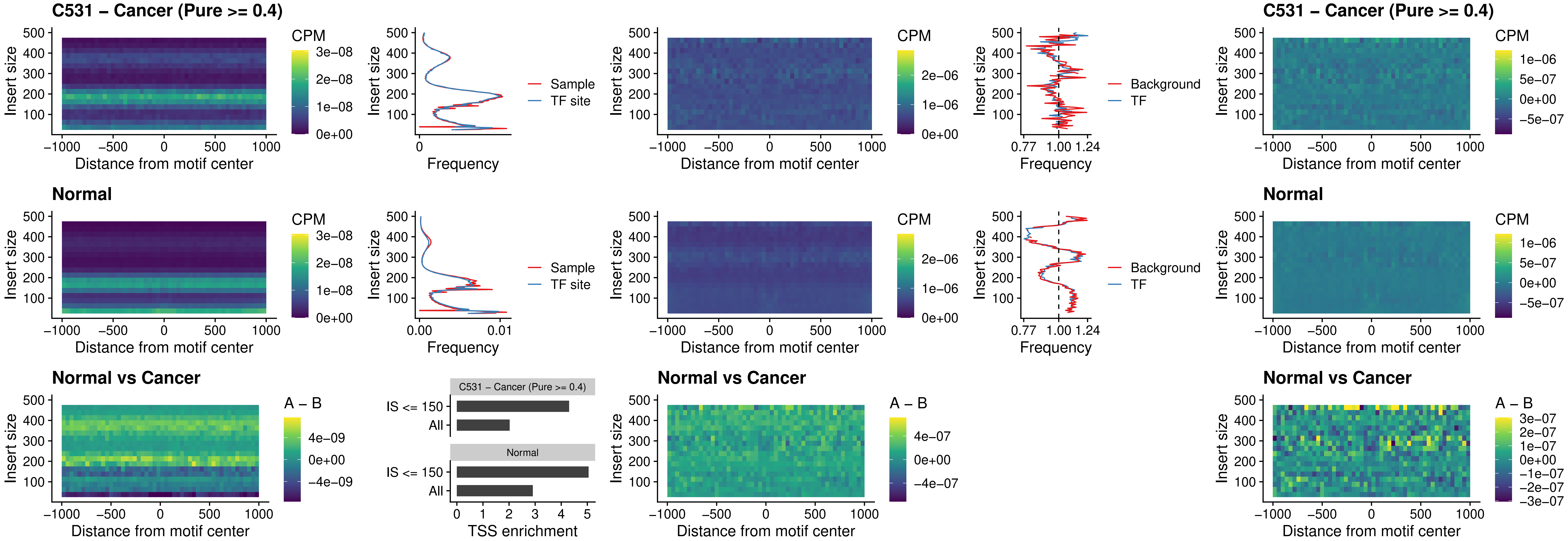

#### C531 – Cancer (Pure >= 0.4) – IRF9_uc_dTSS_oPEAK

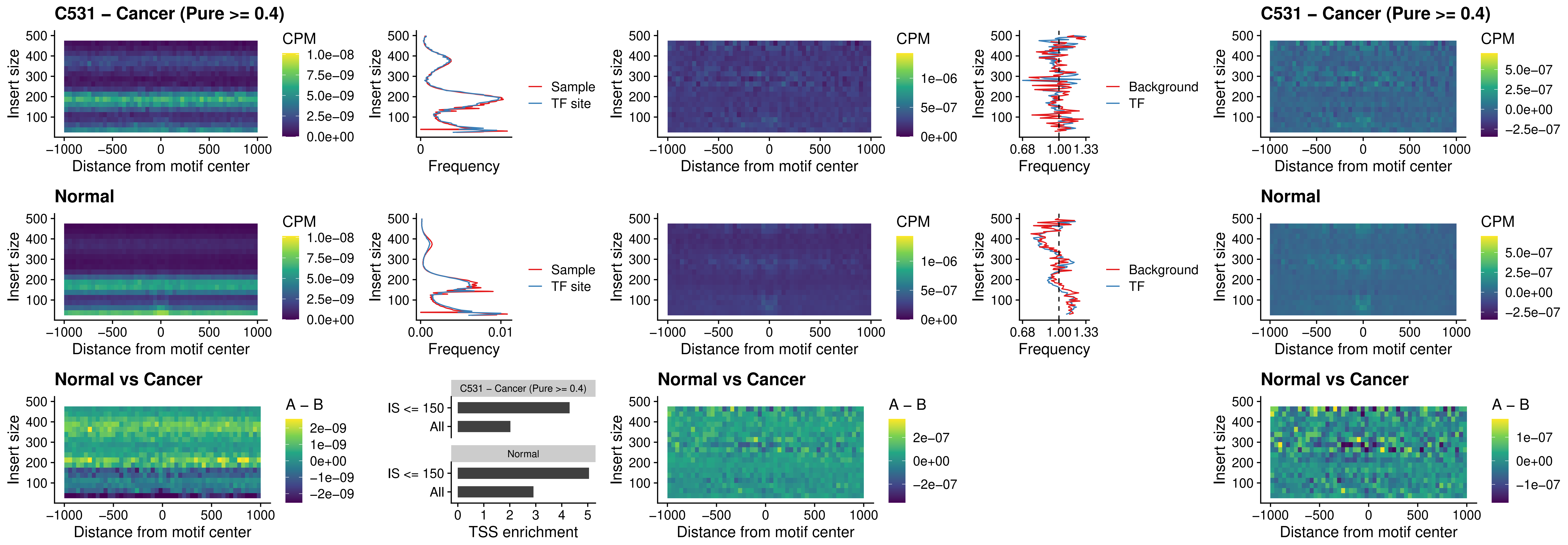

#### C531 – Cancer (Pure >= 0.4) – IRF9_uc_cTSS_oPEAK

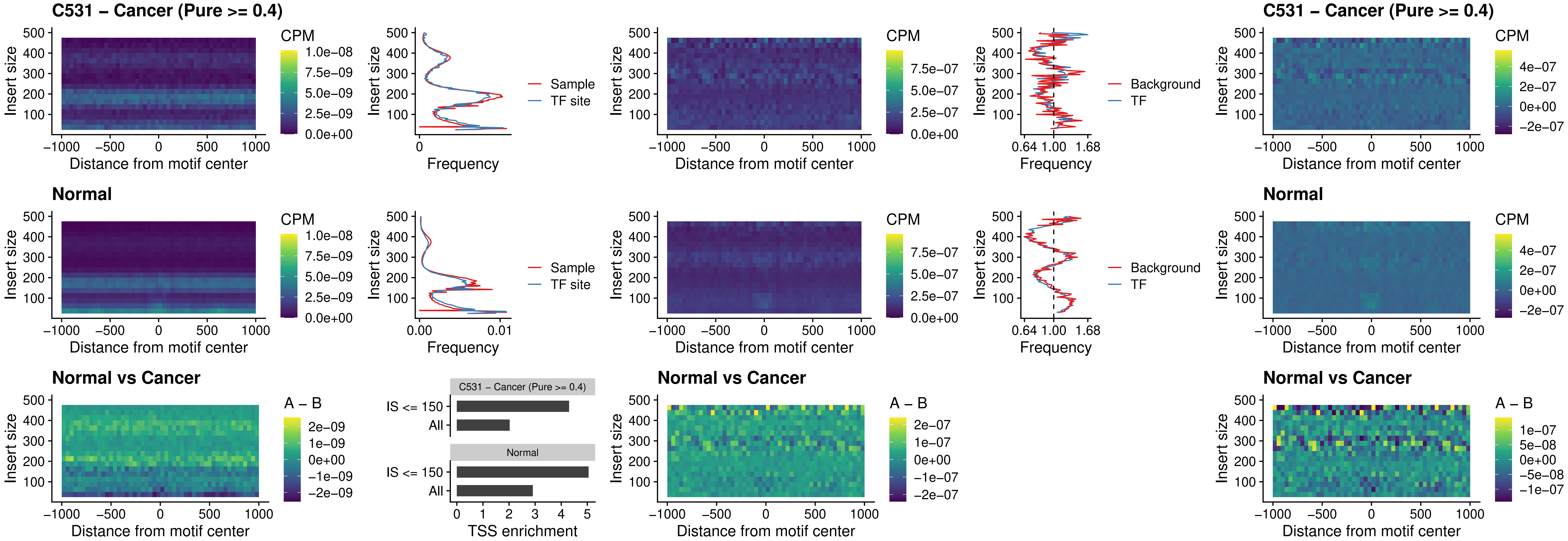

#### C531 – Cancer (Pure >= 0.4) – BCL11A_uc_cTSS_oPEAK

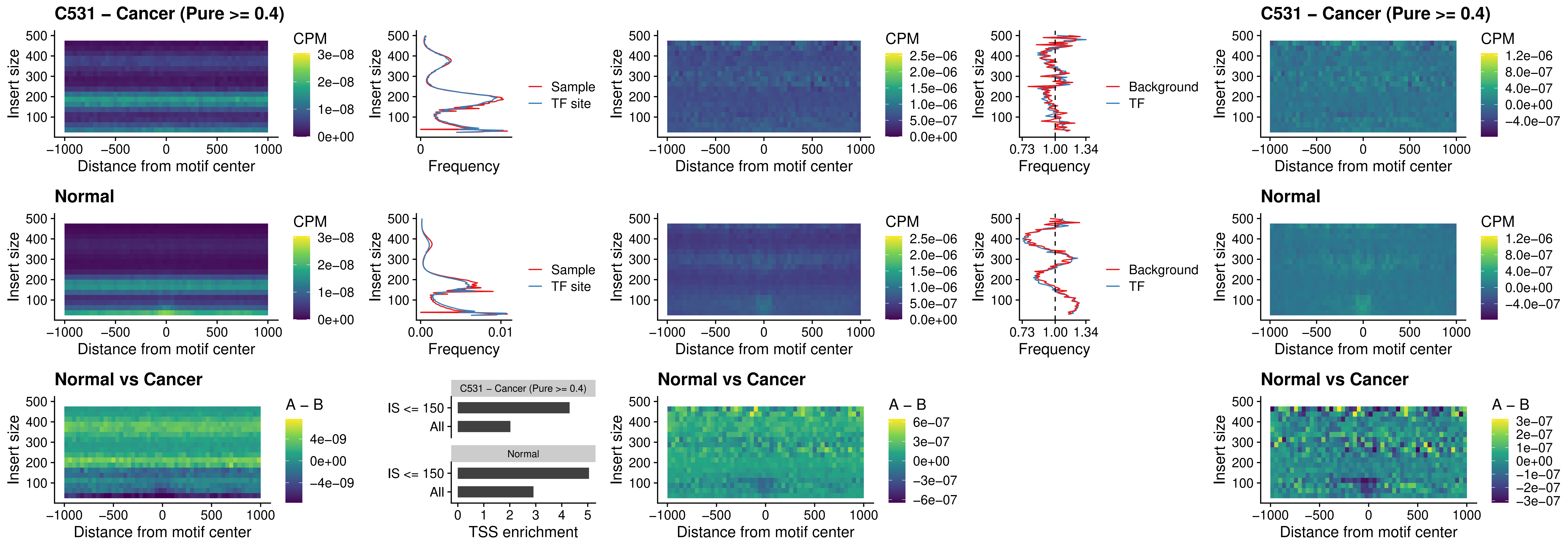

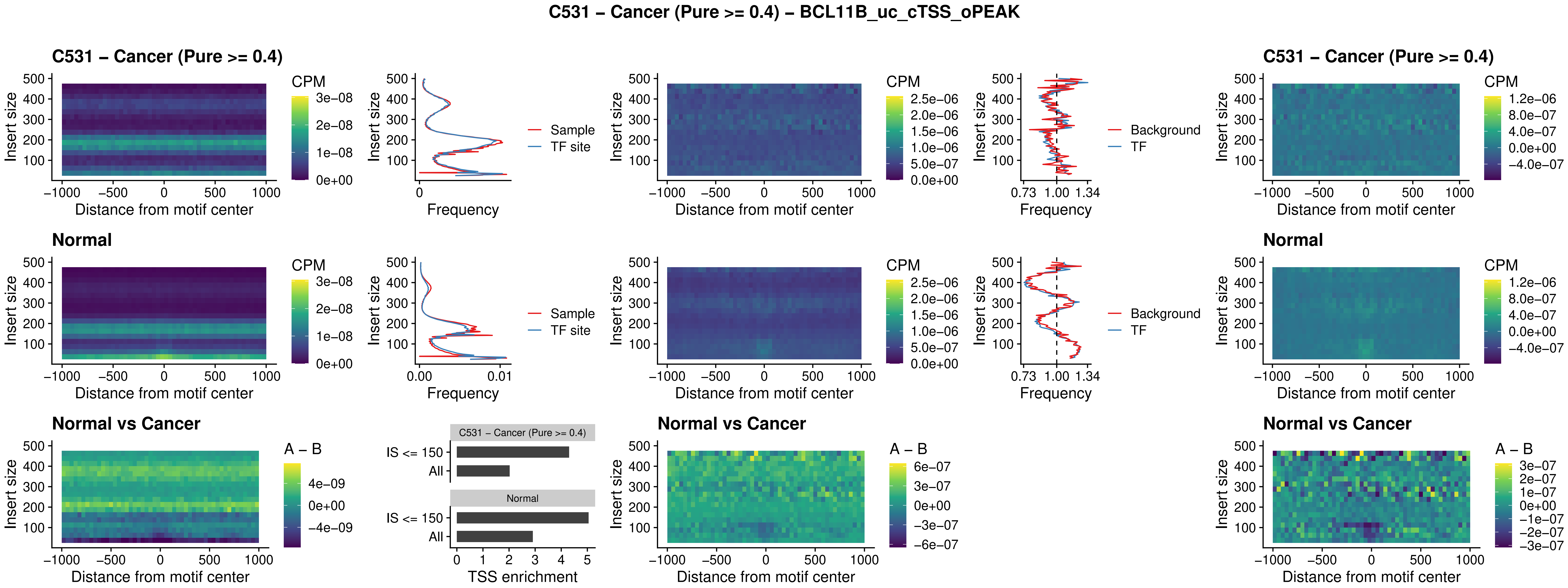
C531 – Cancer (Pure >= 0.4) – BCL11B_uc_cTSS_oPEAK
C531 – Cancer (Pure >= 0.4)
Normal
Normal vs Cancer
Insert size
Distance from motif center
CPM
A – B
Frequency
Sample
TF site
Background
TF
TSS enrichment
IS <= 150
All
C531 – Cancer (Pure >= 0.4)
Normal

#### C531 – Cancer (Pure >= 0.4) – BCL11A_uc_dTSS_oPEAK

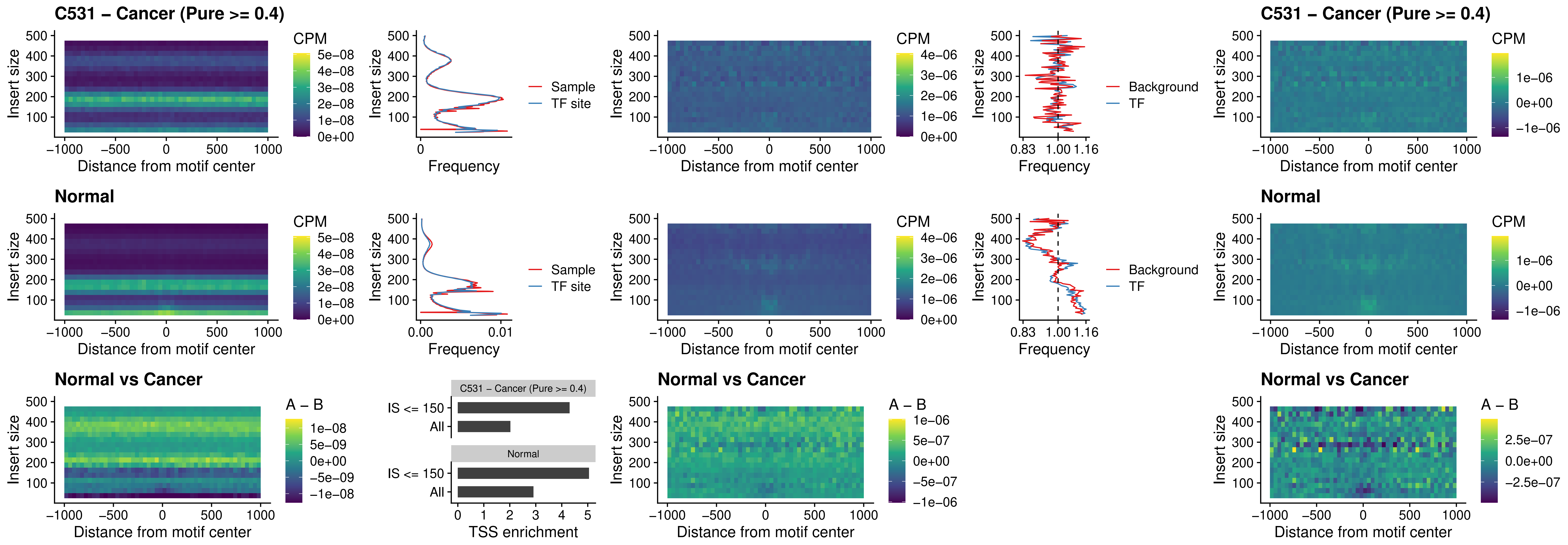

#### C531 – Cancer (Pure >= 0.4) – BCL11B_uc_dTSS_oPEAK

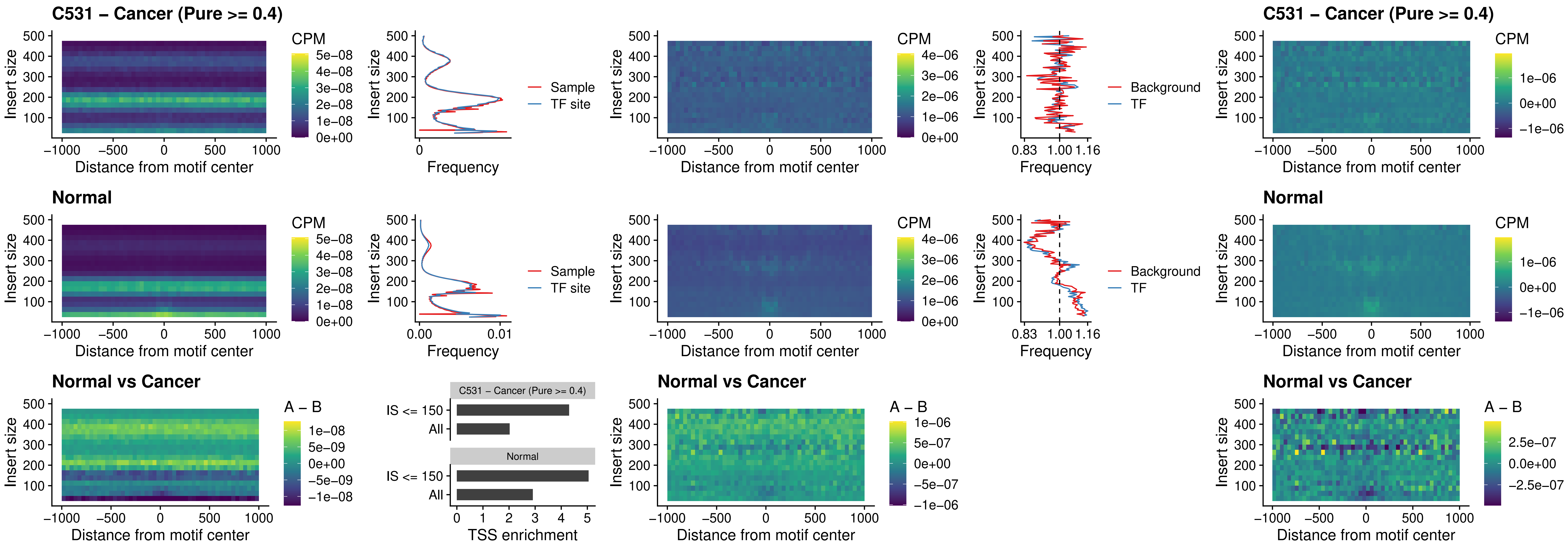

#### C531 – Cancer (Pure >= 0.4) – SPIB_uc_cTSS_nPEAK

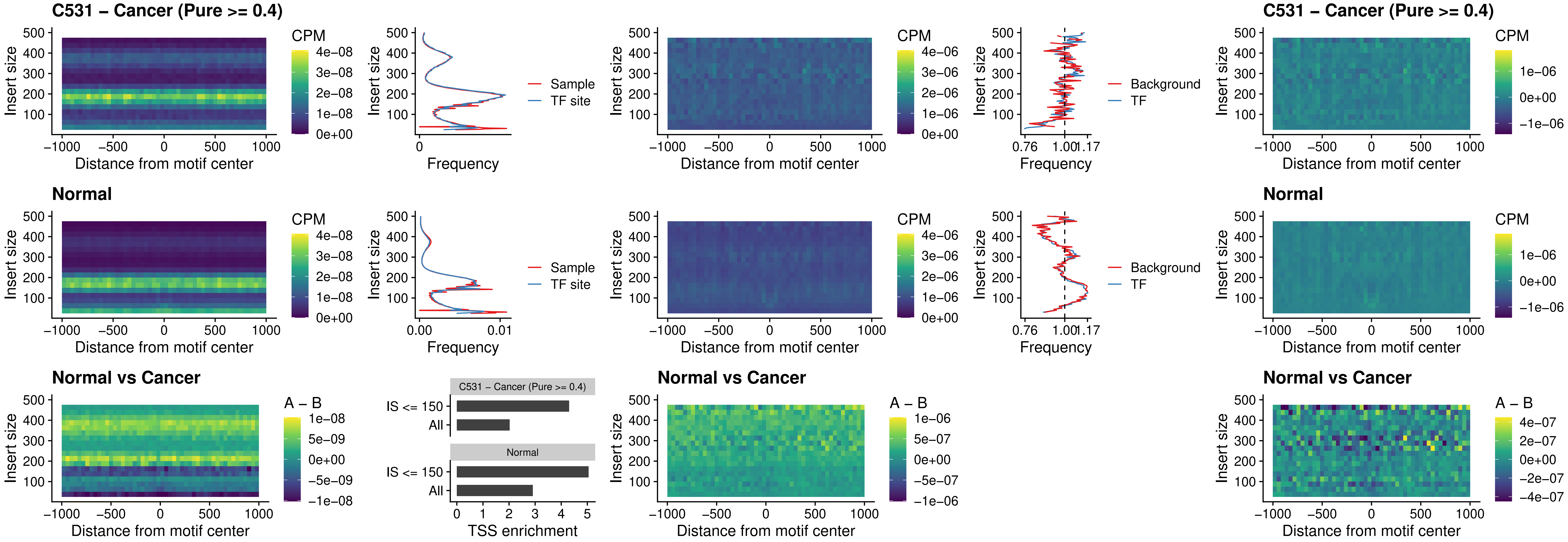

#### C531 – Cancer (Pure >= 0.4) – SPIC_uc_cTSS_nPEAK

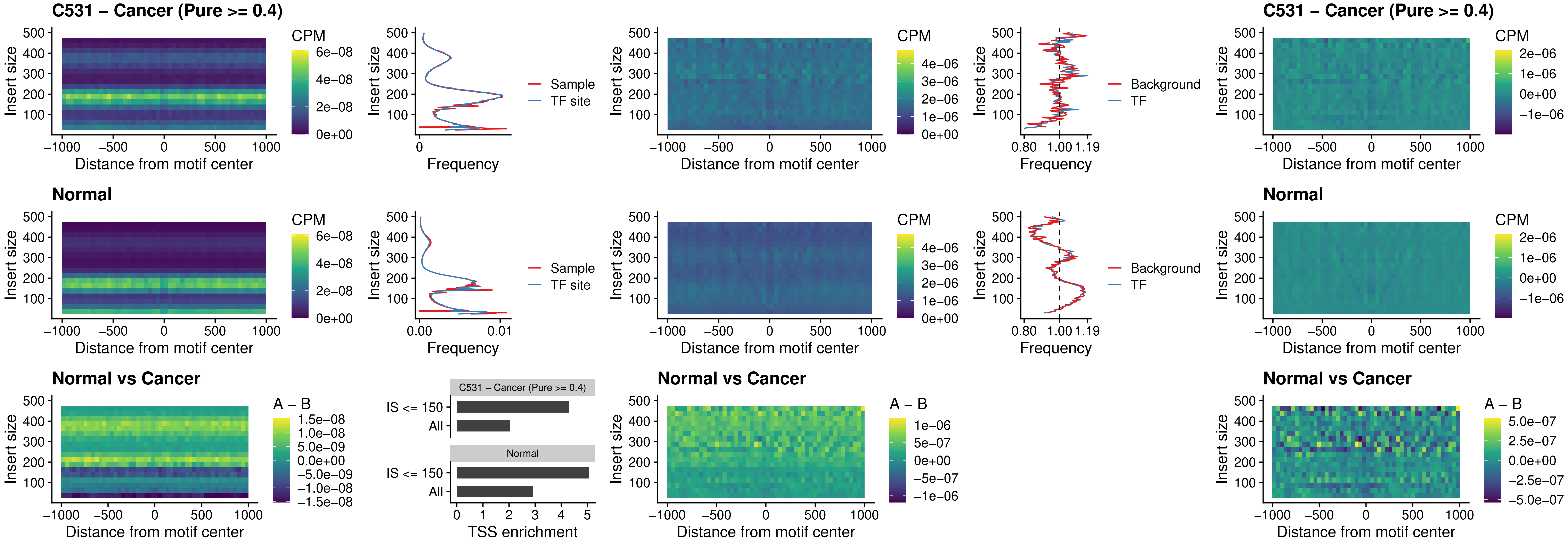

#### C531 – Cancer (Pure >= 0.4) – SPI1_uc_pTSS_oPEAK

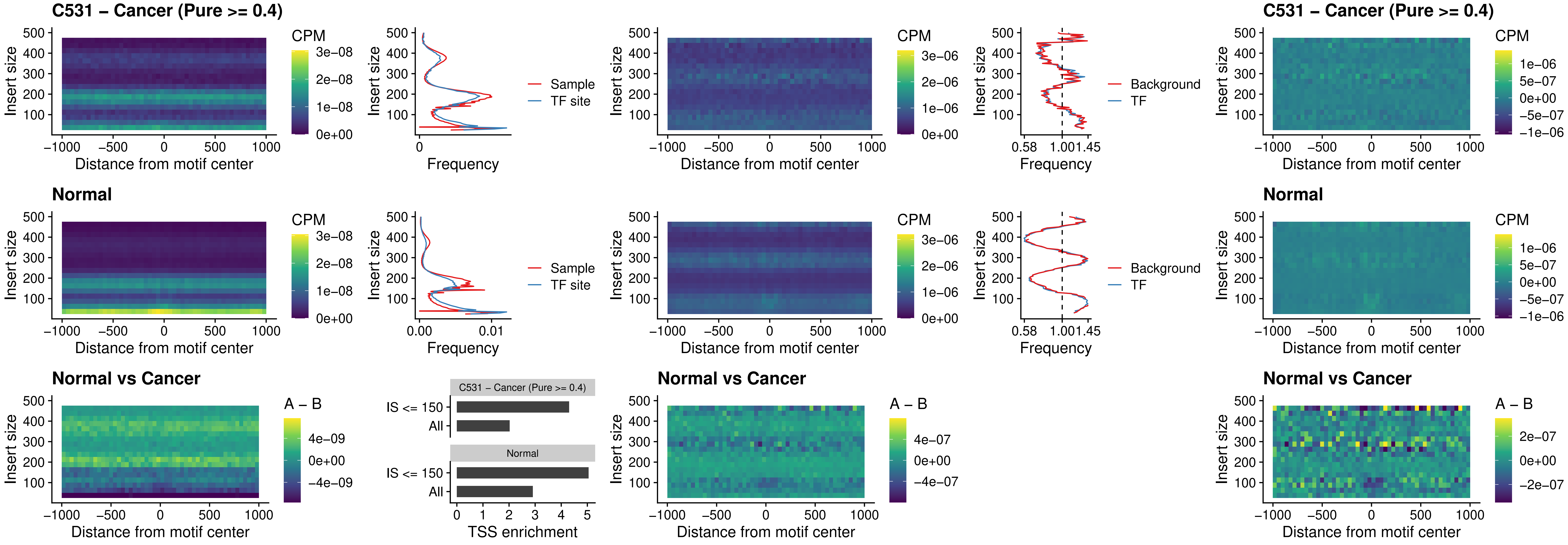

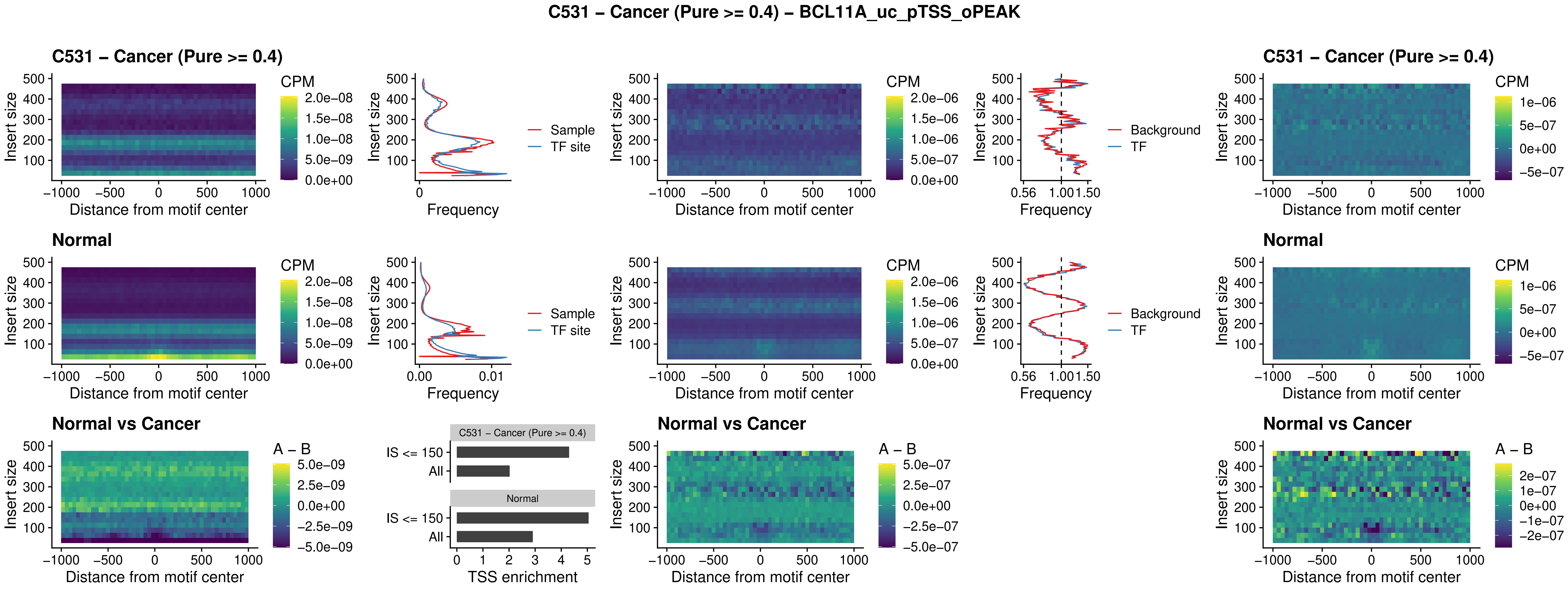

C531 – Cancer (Pure >= 0.4) – BCL11A_uc_pTSS_oPEAK
C531 – Cancer (Pure >= 0.4)
Normal
Normal vs Cancer
Insert size
Distance from motif center
Frequency
CPM
A – B
Sample
TF site
Background
TF
C531 – Cancer (Pure >= 0.4)
Normal
IS <= 150
All
TSS enrichment

### C531 – Cancer (Pure >= 0.4) – BCL11B_uc_pTSS_oPEAK

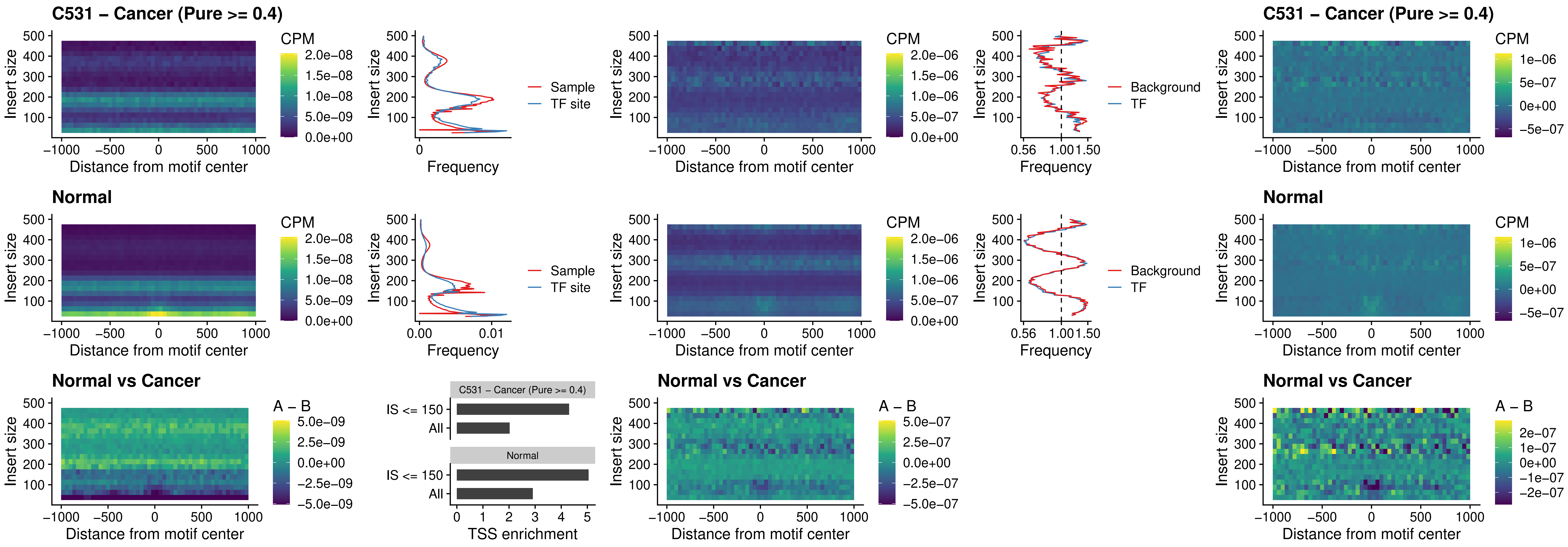

#### C531 – Cancer (Pure >= 0.4) – SPI1_uc_cTSS_oPEAK

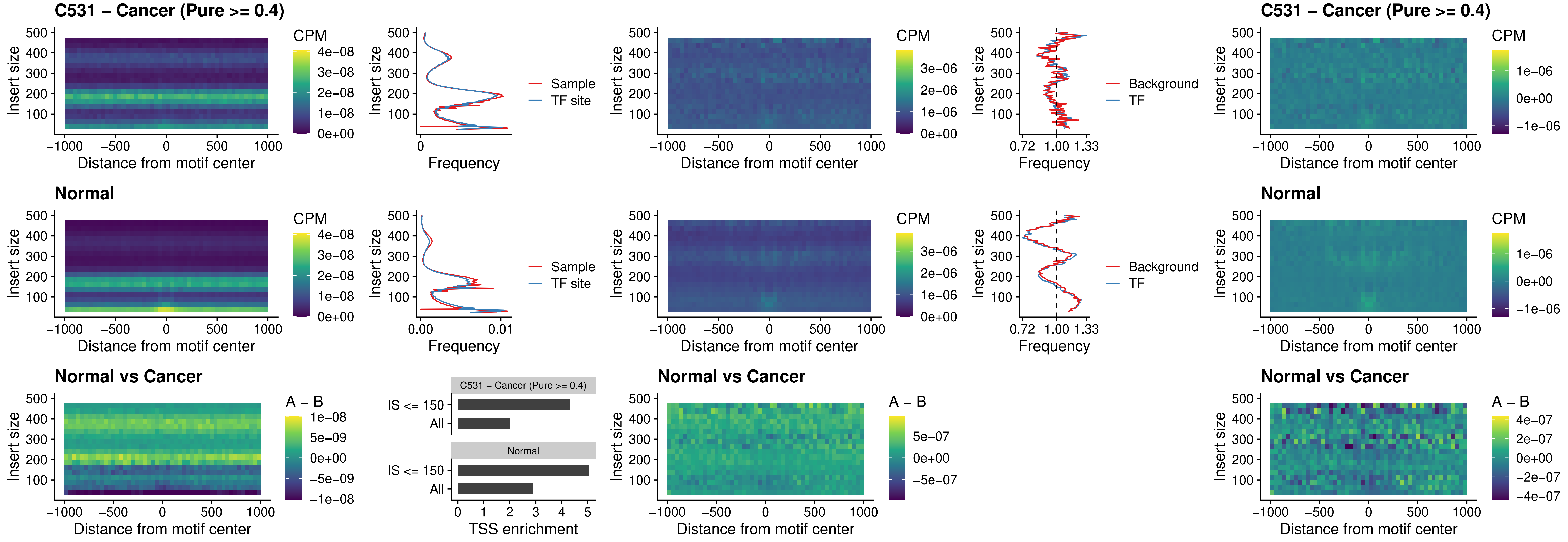

#### C531 – Cancer (Pure >= 0.4) – SPIB_uc_cTSS_oPEAK

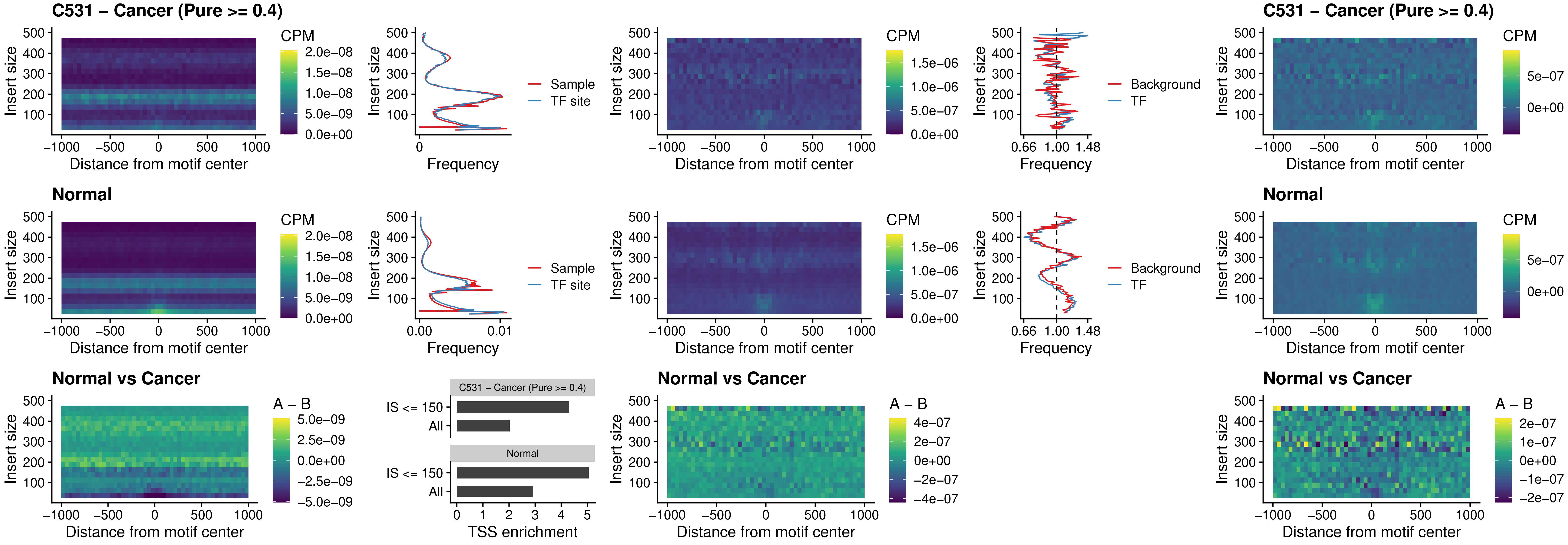

#### C531 – Cancer (Pure >= 0.4) – SPIB_uc_pTSS_oPEAK

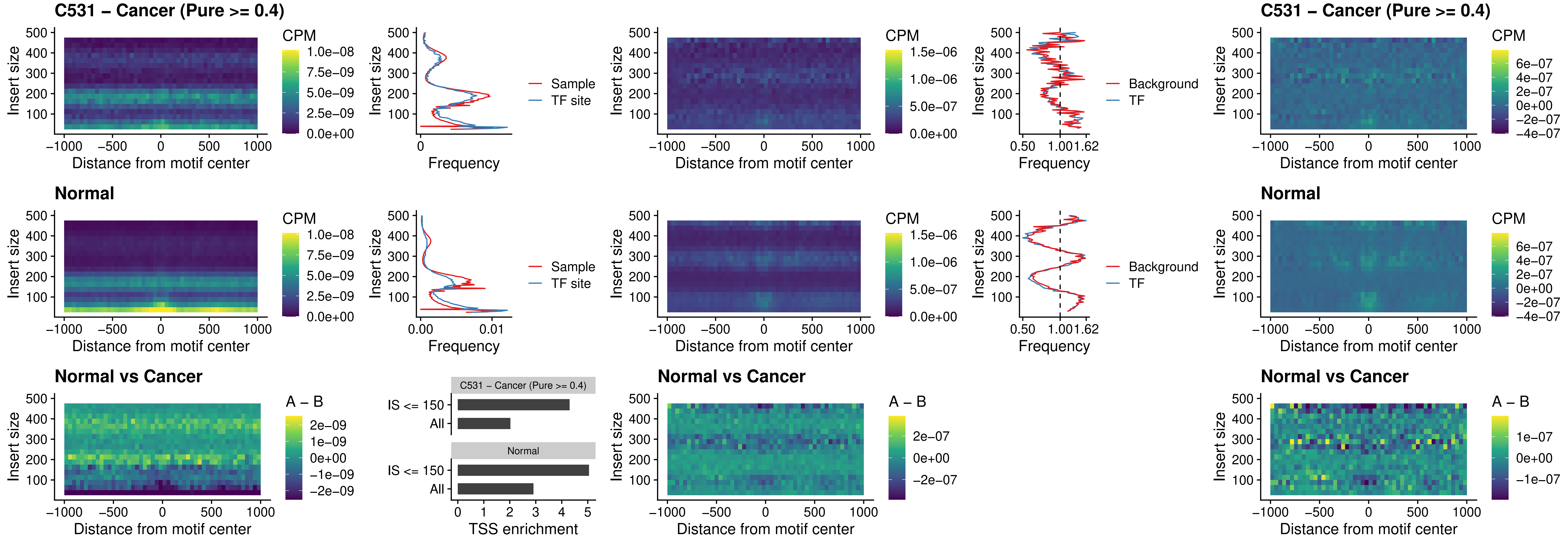

### C531 – Cancer (Pure >= 0.4) – SPI1_uc_dTSS_oPEAK

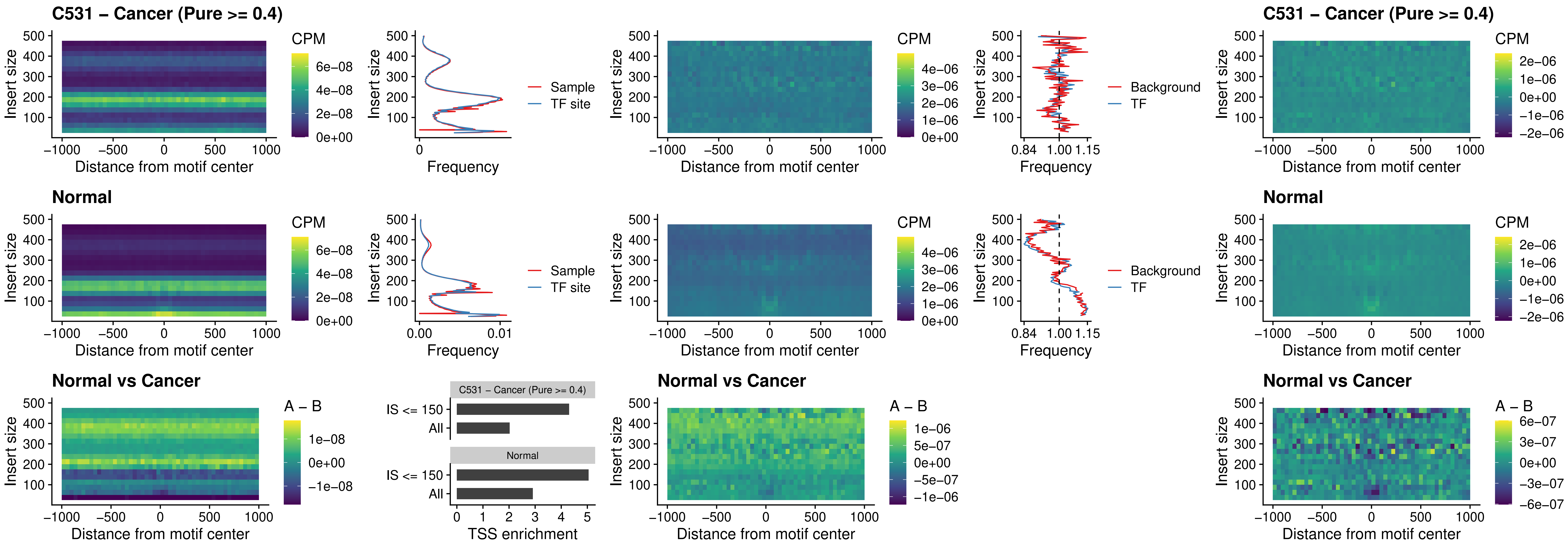

#### C531 – Cancer (Pure >= 0.4) – SPIB_uc_dTSS_oPEAK

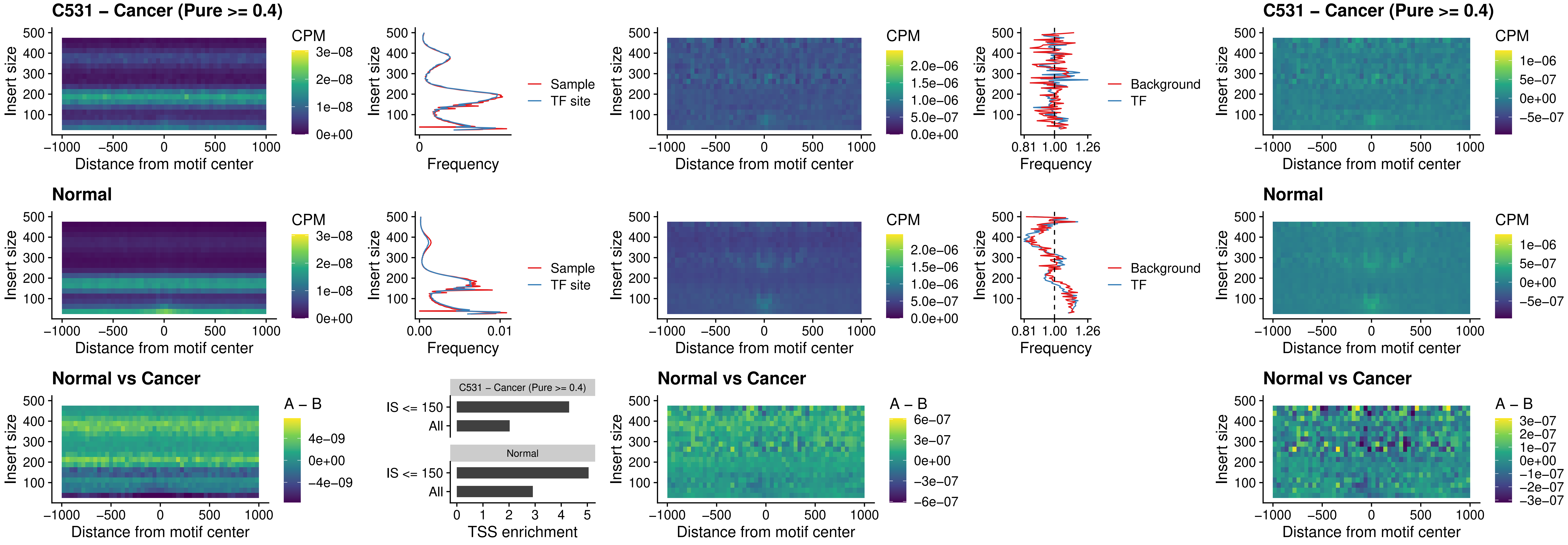

#### C531 – Cancer (Pure >= 0.4) – IRF3_uc_pTSS_oPEAK

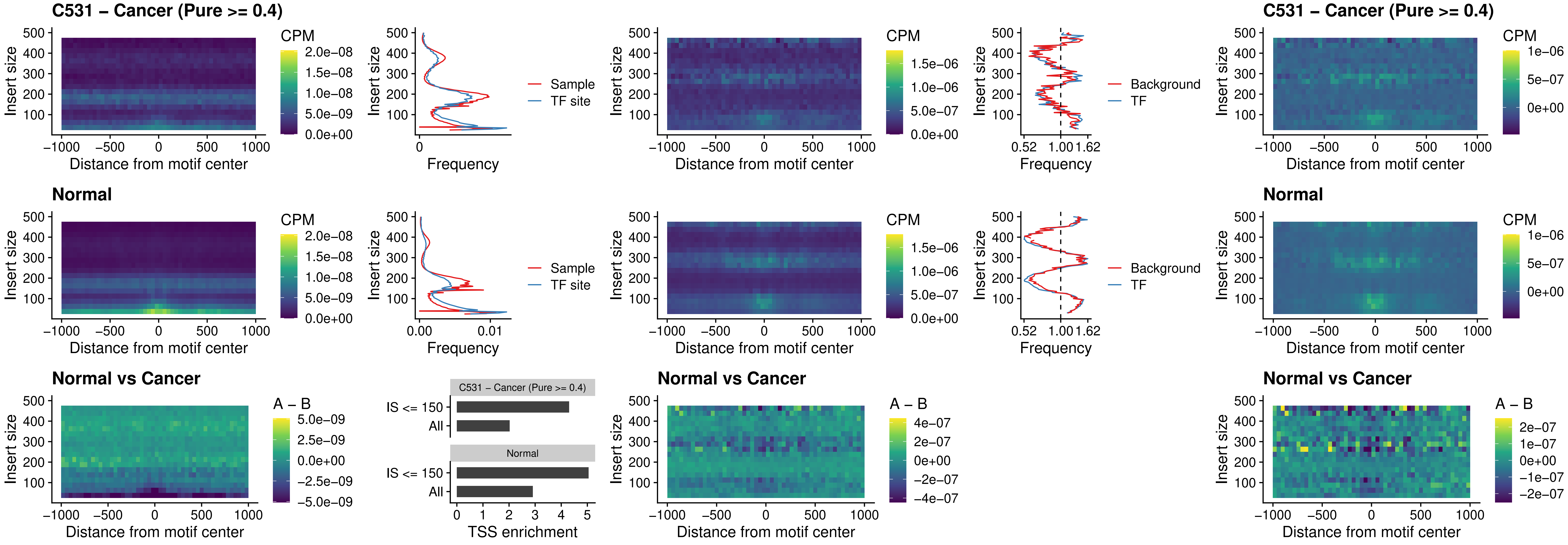

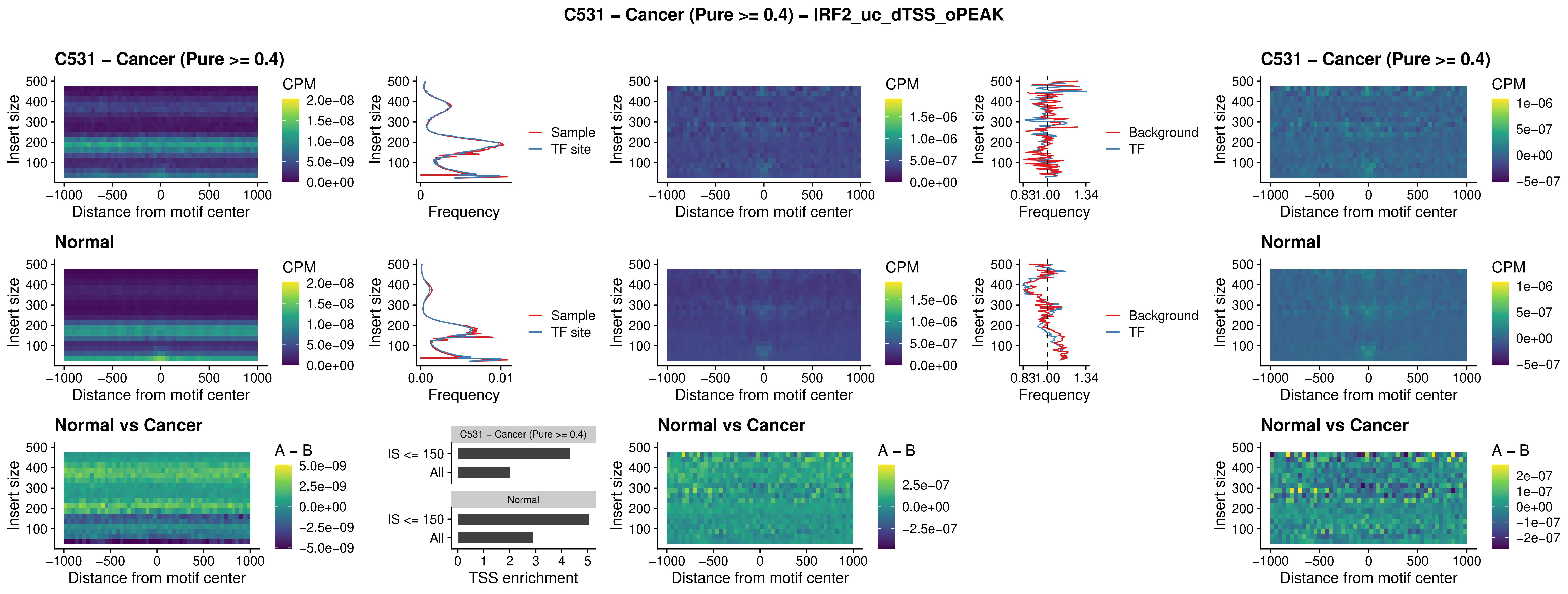
C531 – Cancer (Pure >= 0.4) – IRF2_uc_dTSS_oPEAK
C531 – Cancer (Pure >= 0.4)
Normal
Normal vs Cancer
Insert size
Distance from motif center
Frequency
CPM
A – B
Sample
TF site
Background
TF
C531 – Cancer (Pure >= 0.4)
Normal
IS <= 150
All
TSS enrichment

### C531 – Cancer (Pure >= 0.4) – IRF2_uc_pTSS_oPEAK

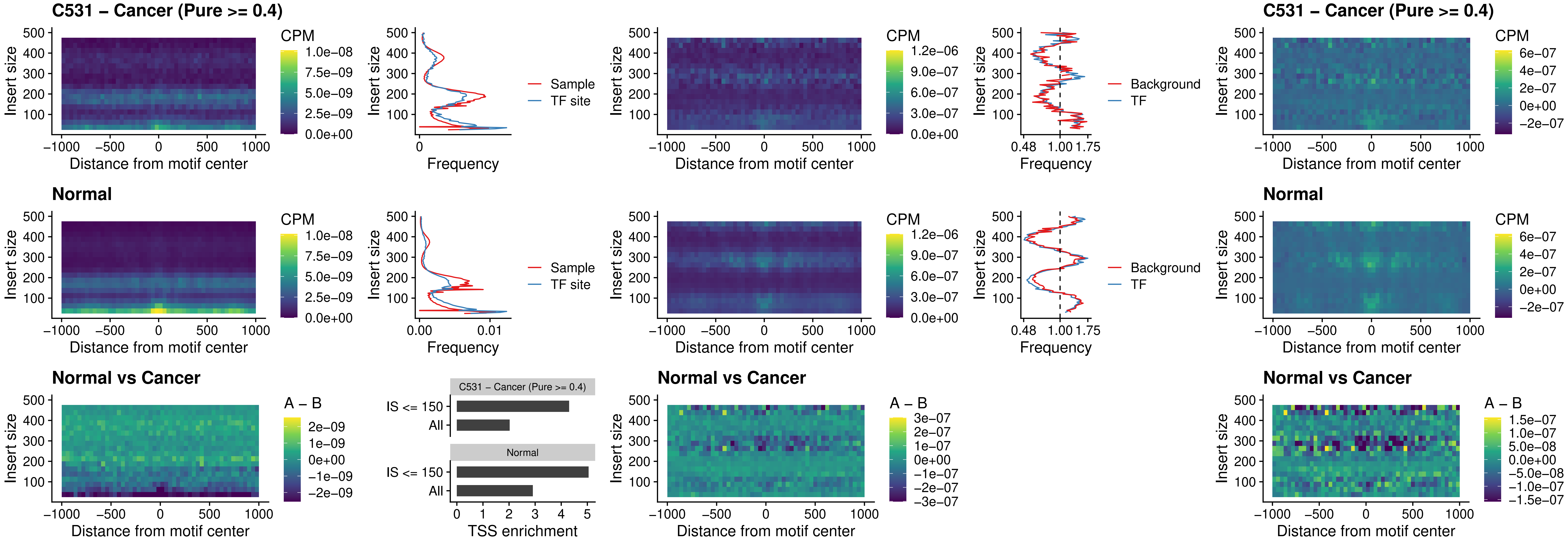

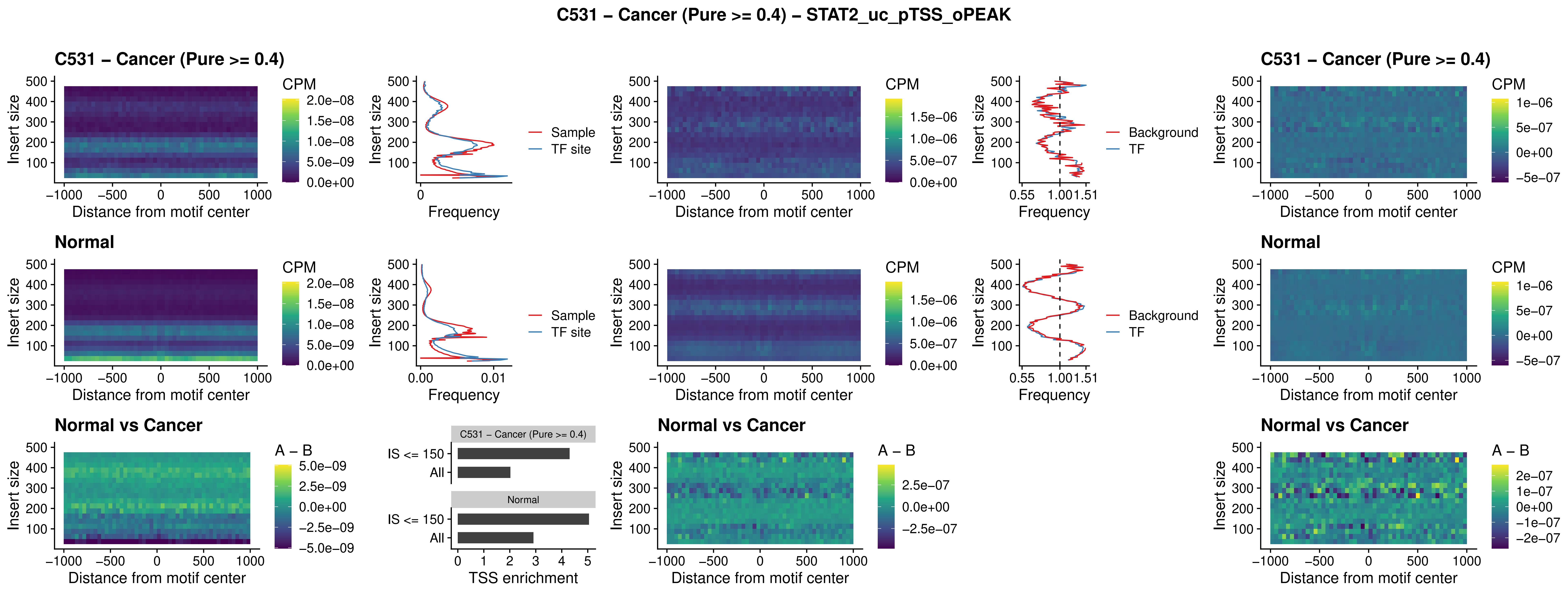
C531 – Cancer (Pure >= 0.4) – STAT2_uc_pTSS_oPEAK
C531 – Cancer (Pure >= 0.4)
Normal
Normal vs Cancer
Insert size
Distance from motif center
Frequency
CPM
A – B
Sample
TF site
Background
TF
C531 – Cancer (Pure >= 0.4)
Normal
IS <= 150
All
TSS enrichment

### C531 – Cancer (Pure >= 0.4) – SPIB_uc_dTSS_nPEAK

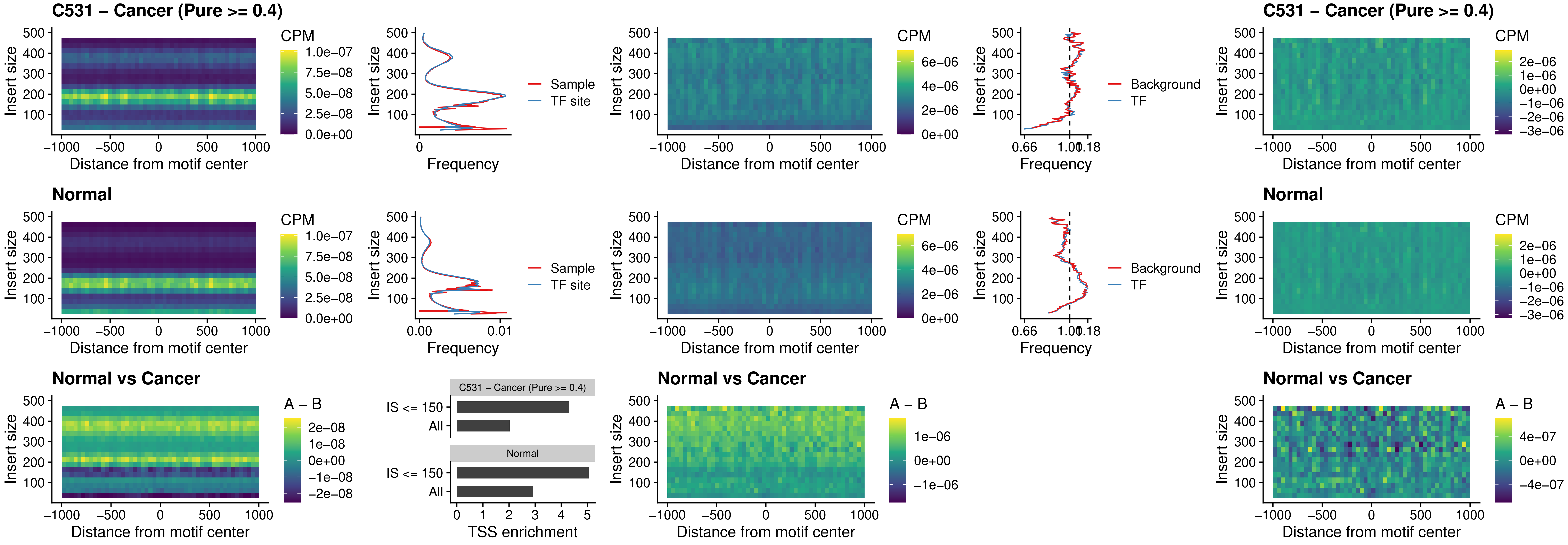

#### C531 – Cancer (Pure >= 0.4) – SPIC_uc_cTSS_oPEAK

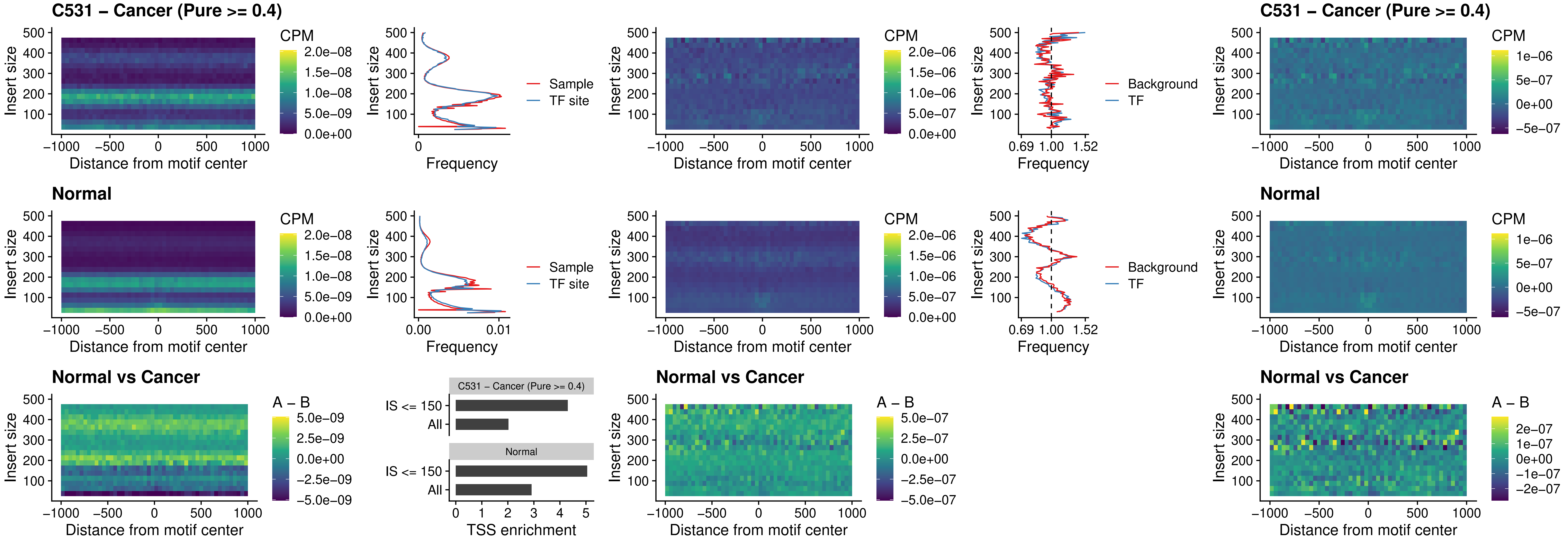

#### C531 – Cancer (Pure >= 0.4) – PRDM1_uc_cTSS_oPEAK

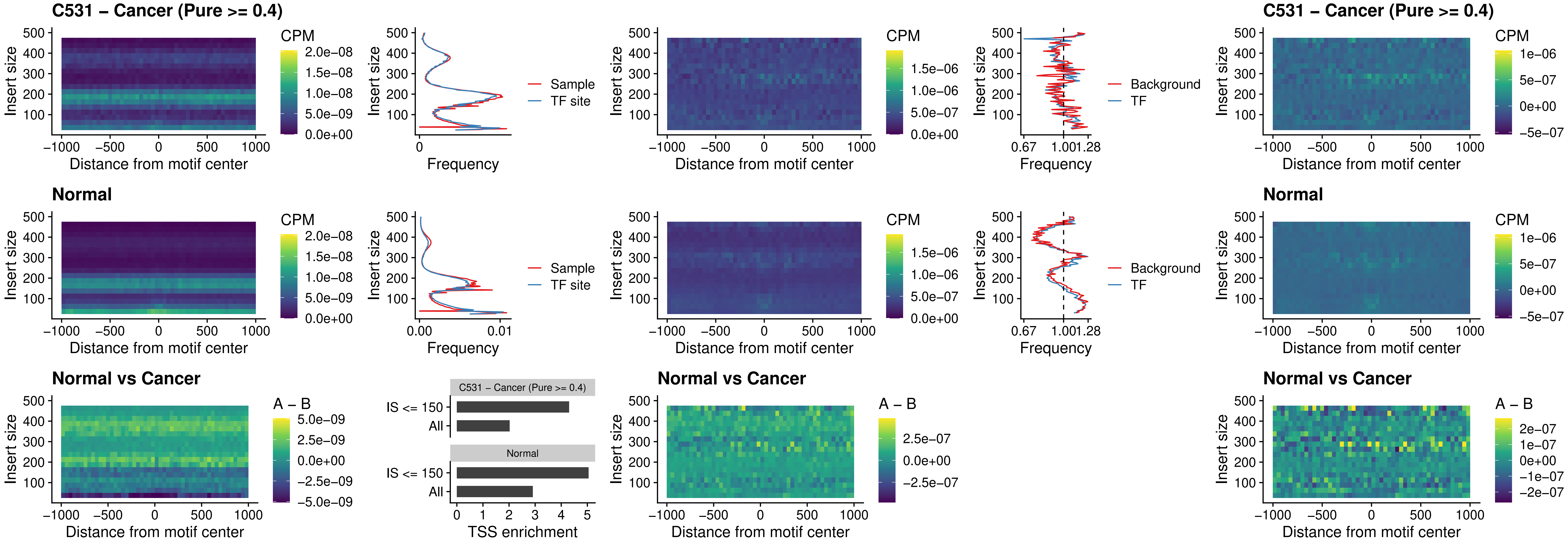

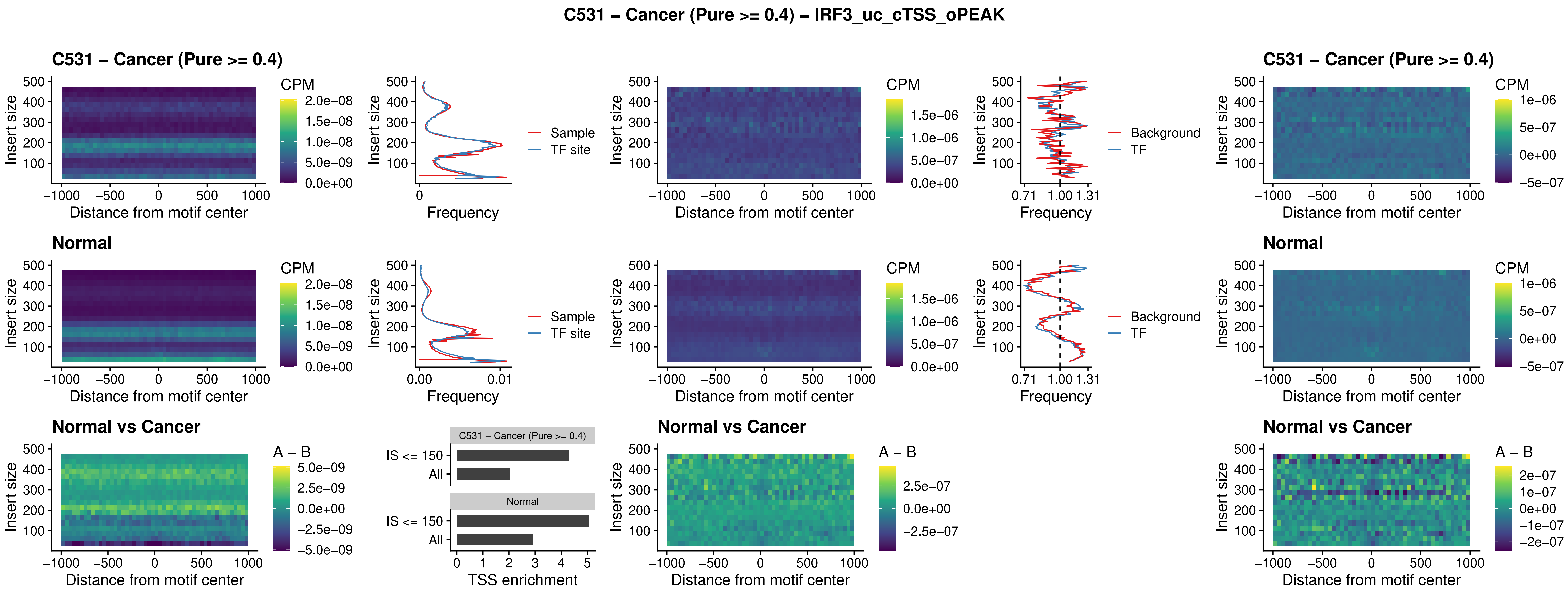
C531 – Cancer (Pure >= 0.4) – IRF3_uc_cTSS_oPEAK
C531 – Cancer (Pure >= 0.4)
Normal
Normal vs Cancer
Insert size
Distance from motif center
Frequency
CPM
A – B
Sample
TF site
Background
TF
C531 – Cancer (Pure >= 0.4)
Normal
IS <= 150
All
TSS enrichment

#### C531 – Cancer (Pure >= 0.4) – BCL11A_uc_dTSS_nPEAK

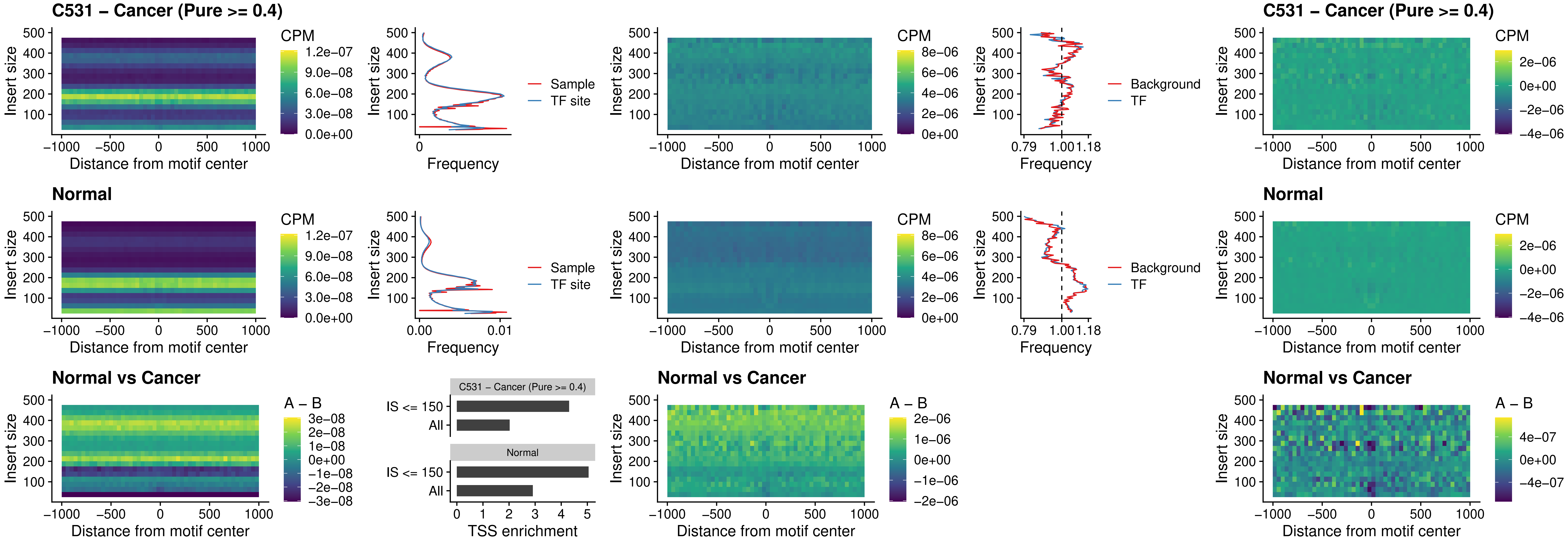

#### C531 – Cancer (Pure >= 0.4) – BCL11B_uc_dTSS_nPEAK

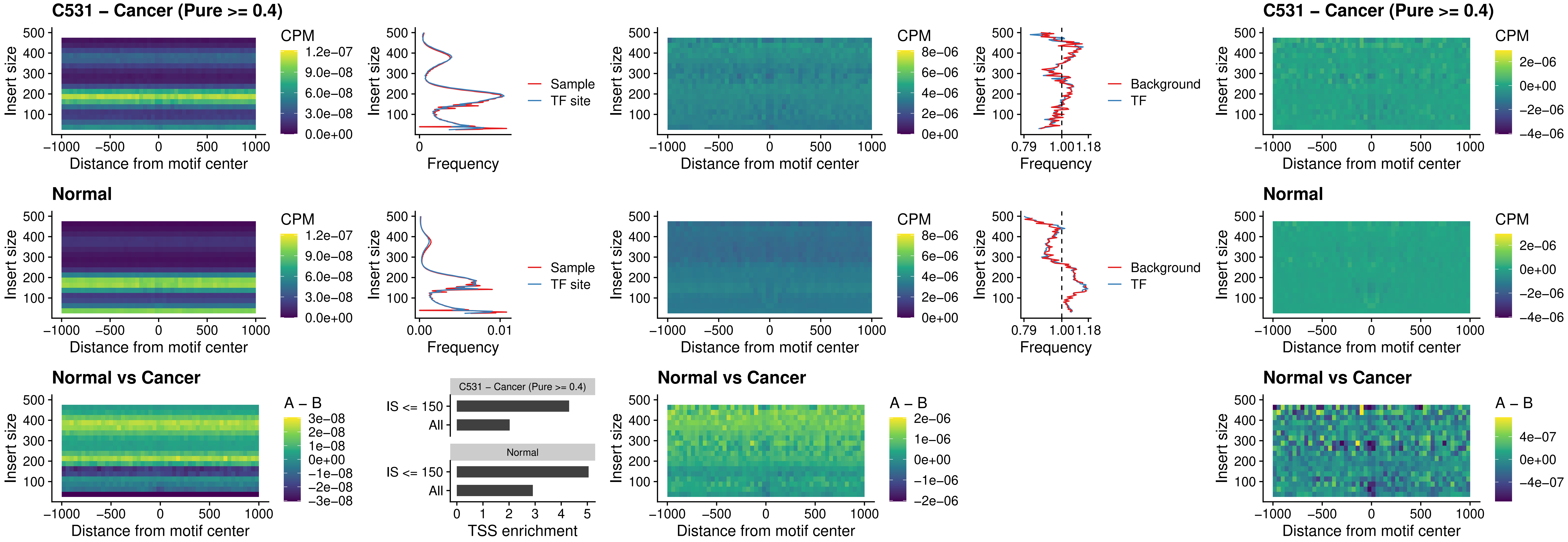

#### C531 – Cancer (Pure >= 0.4) – IRF1_uc_cTSS_oPEAK

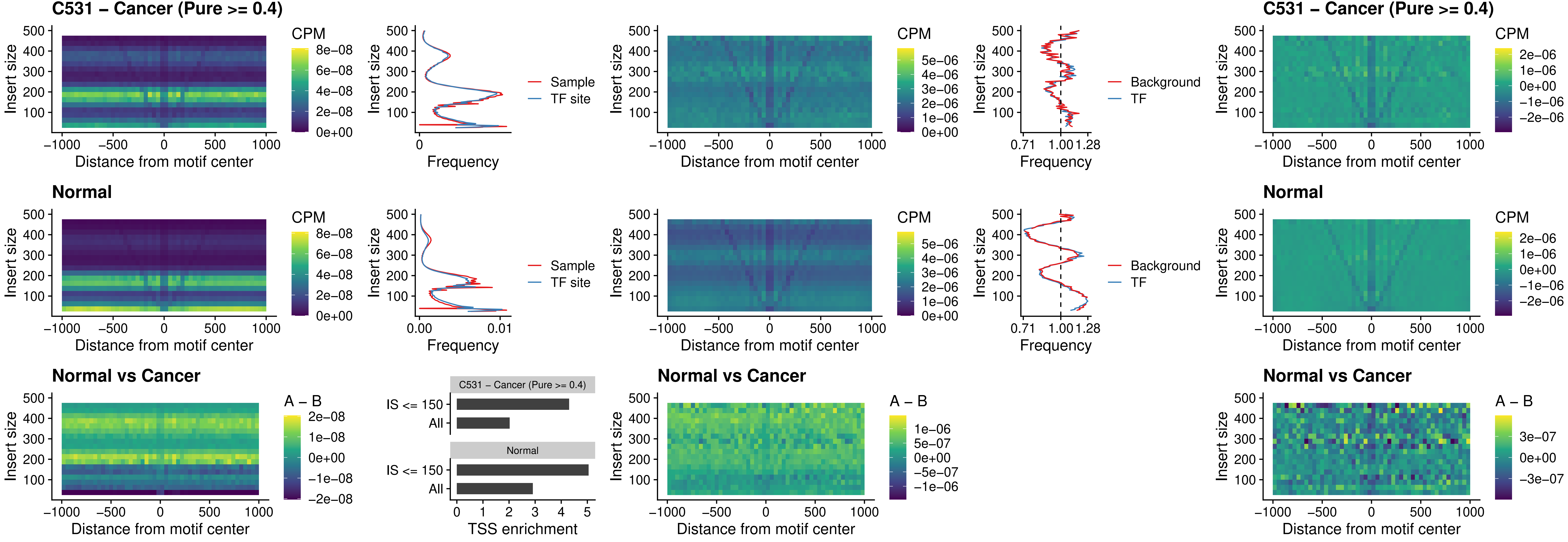

#### C531 – Cancer (Pure >= 0.4) – IRF1_uc_dTSS_oPEAK

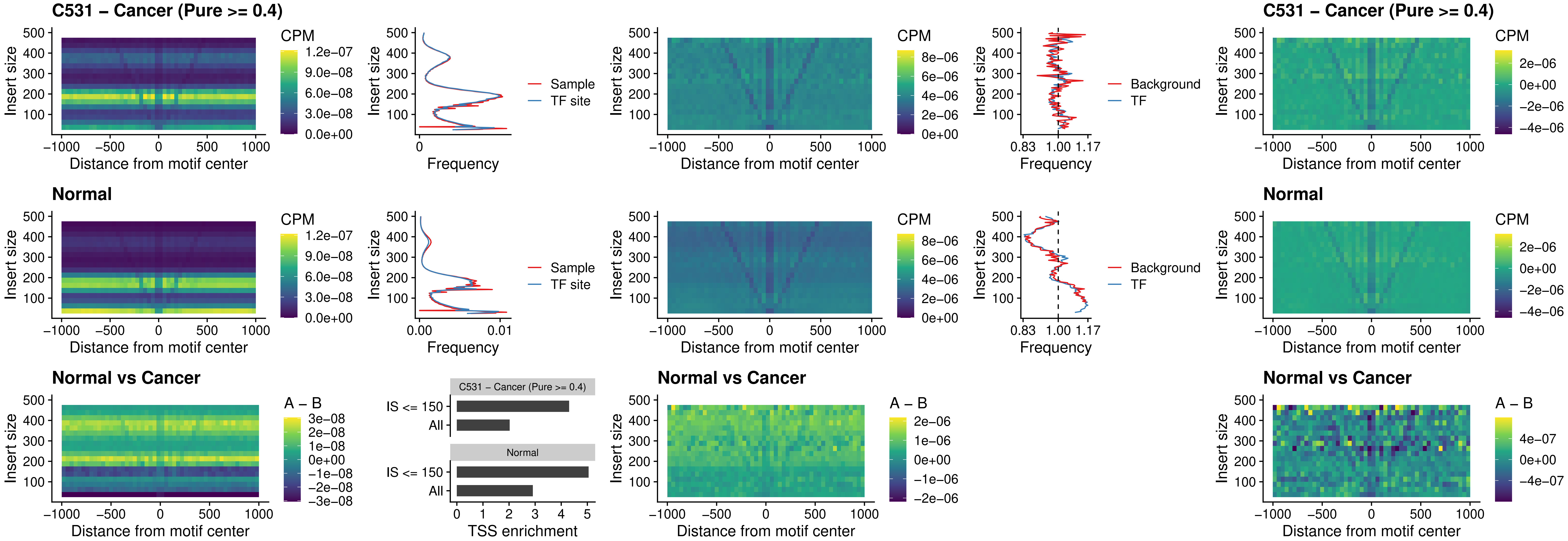

#### C531 – Cancer (Pure >= 0.4) – IRF1_uc_pTSS_oPEAK

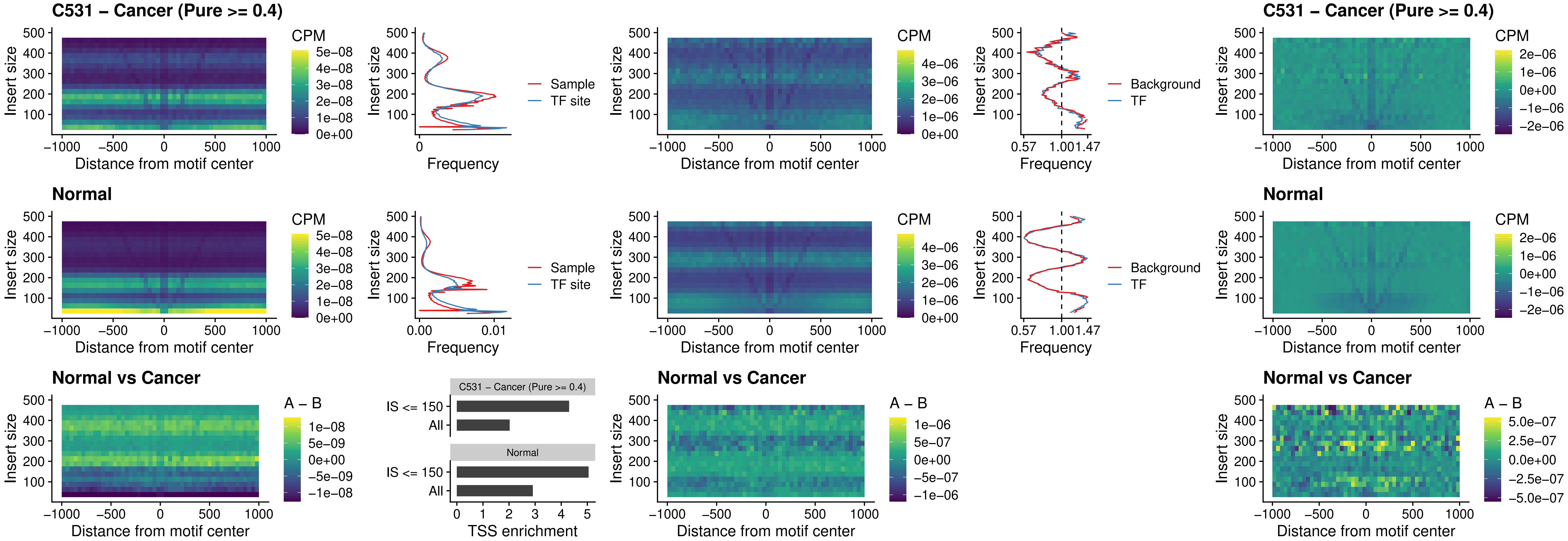

#### C531 – Cancer (Pure >= 0.4) – SPIC_uc_dTSS_nPEAK

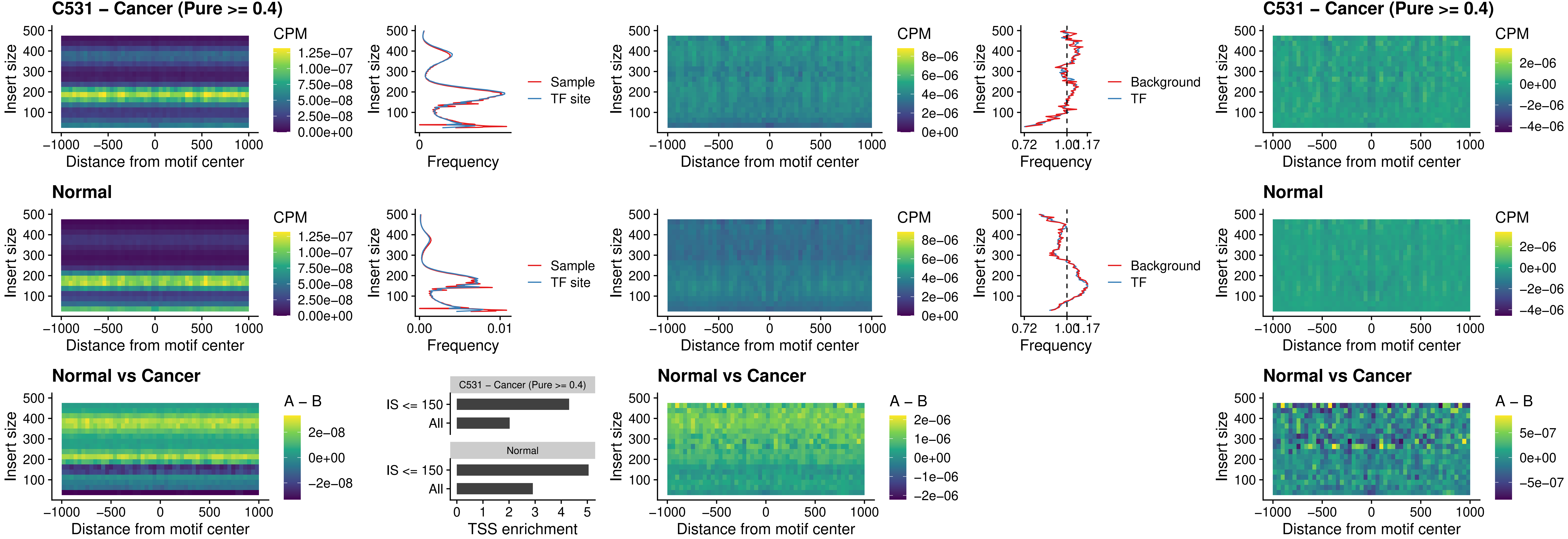

#### C531 – Cancer (Pure >= 0.4) – SPIC_uc_dTSS_oPEAK

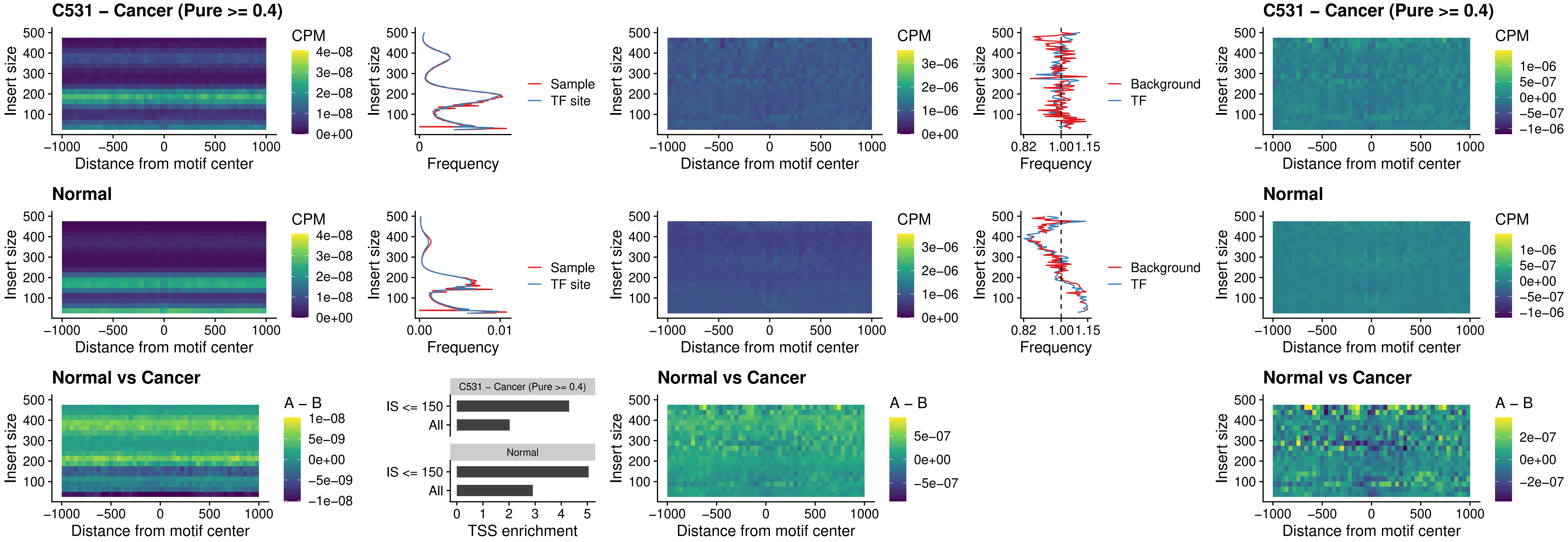

### C531 – Cancer (Pure >= 0.4) – SPIB_uc_pTSS_nPEAK

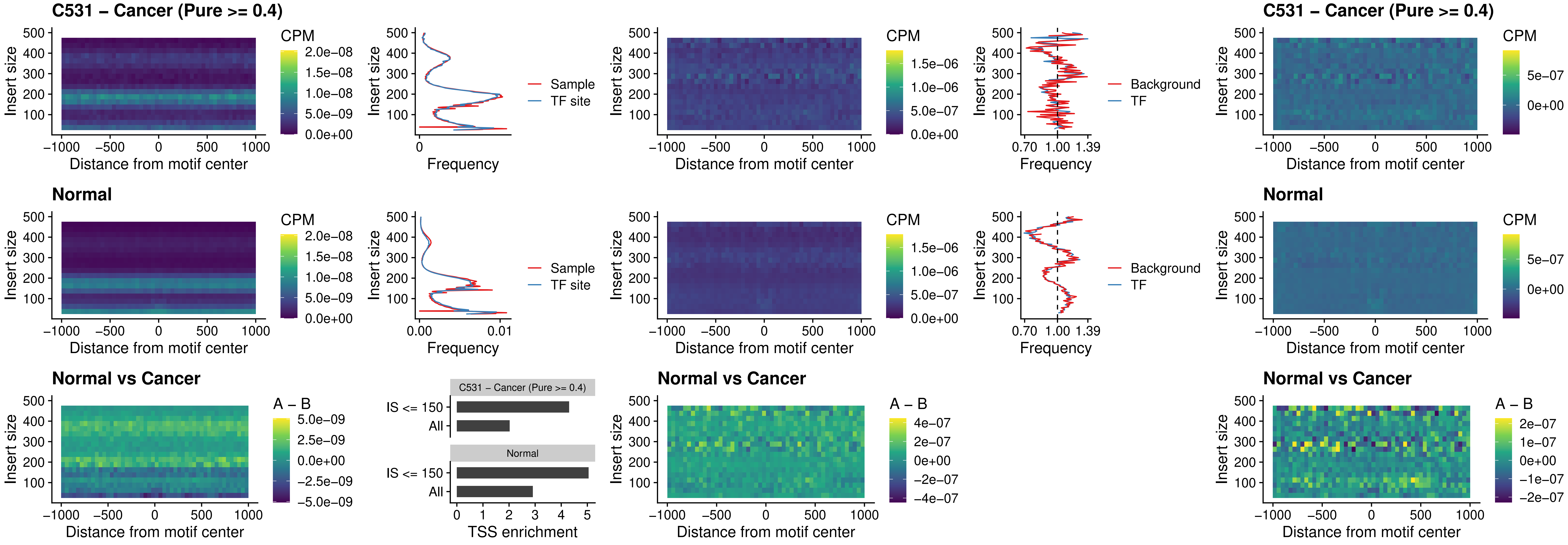

### C531 – Cancer (Pure >= 0.4) – STAT2_uc_cTSS_oPEAK

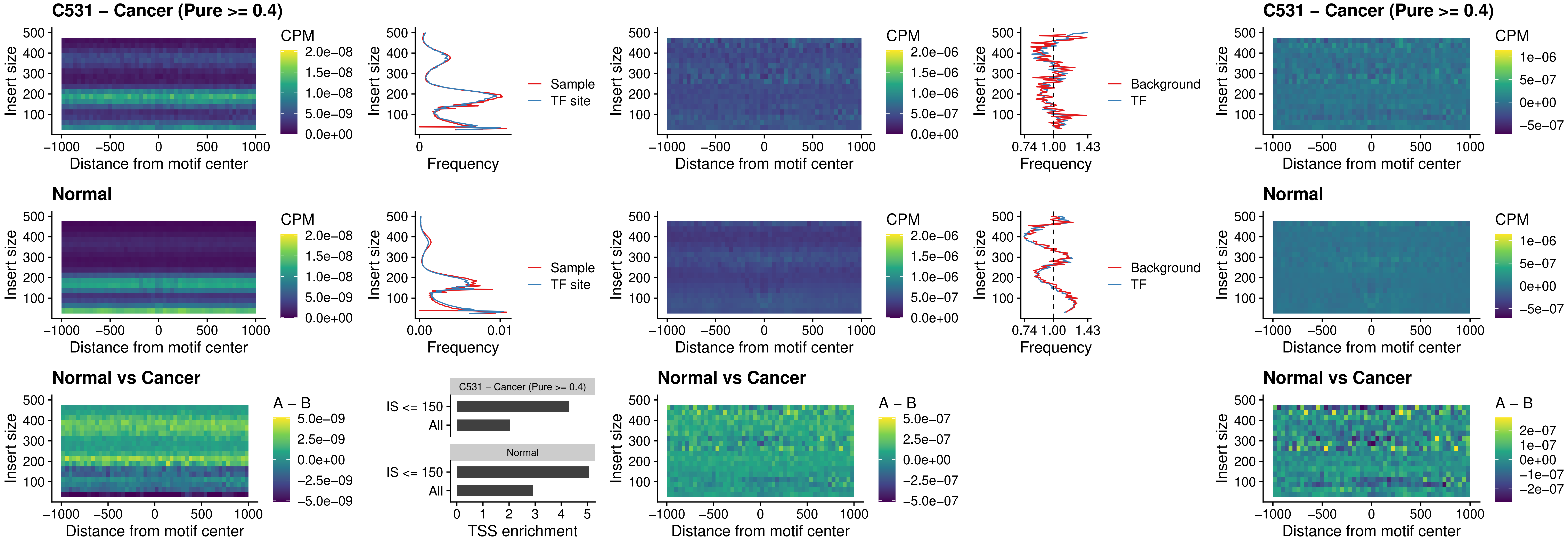

#### C531 – Cancer (Pure >= 0.4) – IRF8_uc_cTSS_oPEAK

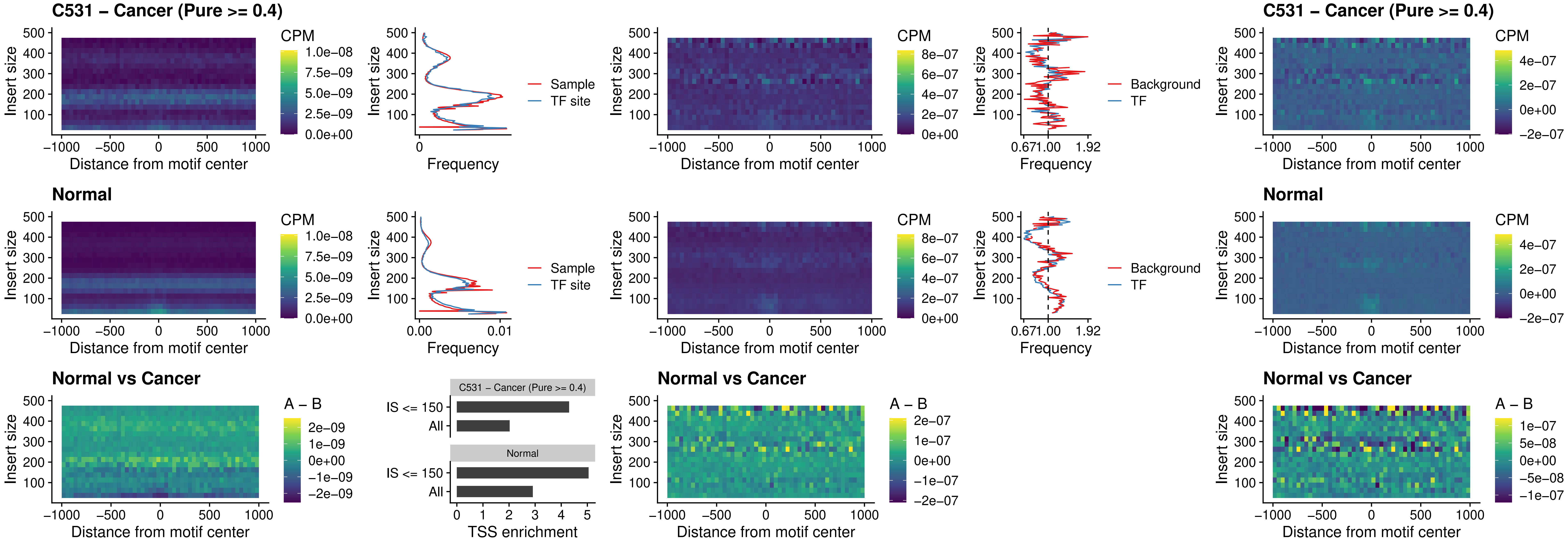

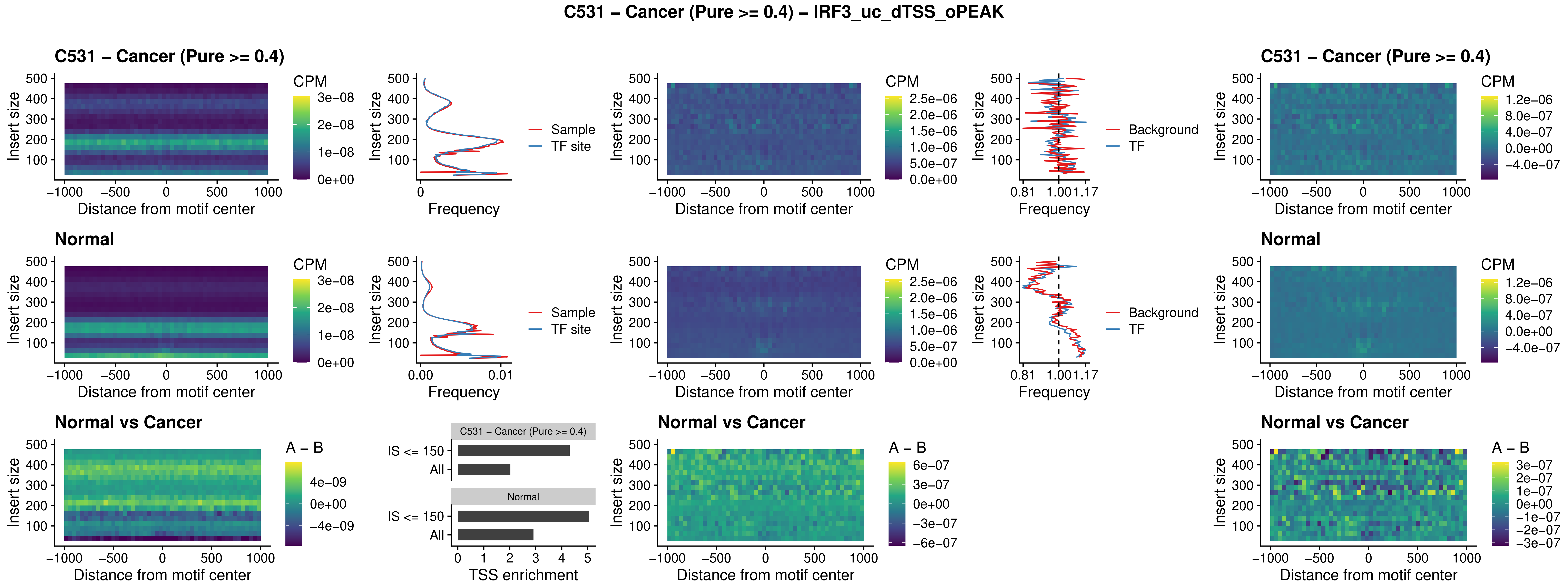
C531 – Cancer (Pure >= 0.4) – IRF3_uc_dTSS_oPEAK
C531 – Cancer (Pure >= 0.4)
Normal
Normal vs Cancer
Insert size
Distance from motif center
Frequency
CPM
A – B
Sample
TF site
Background
TF
C531 – Cancer (Pure >= 0.4)
Normal
IS <= 150
All
TSS enrichment

#### C531 – Cancer (Pure >= 0.4) – IRF8_uc_dTSS_oPEAK

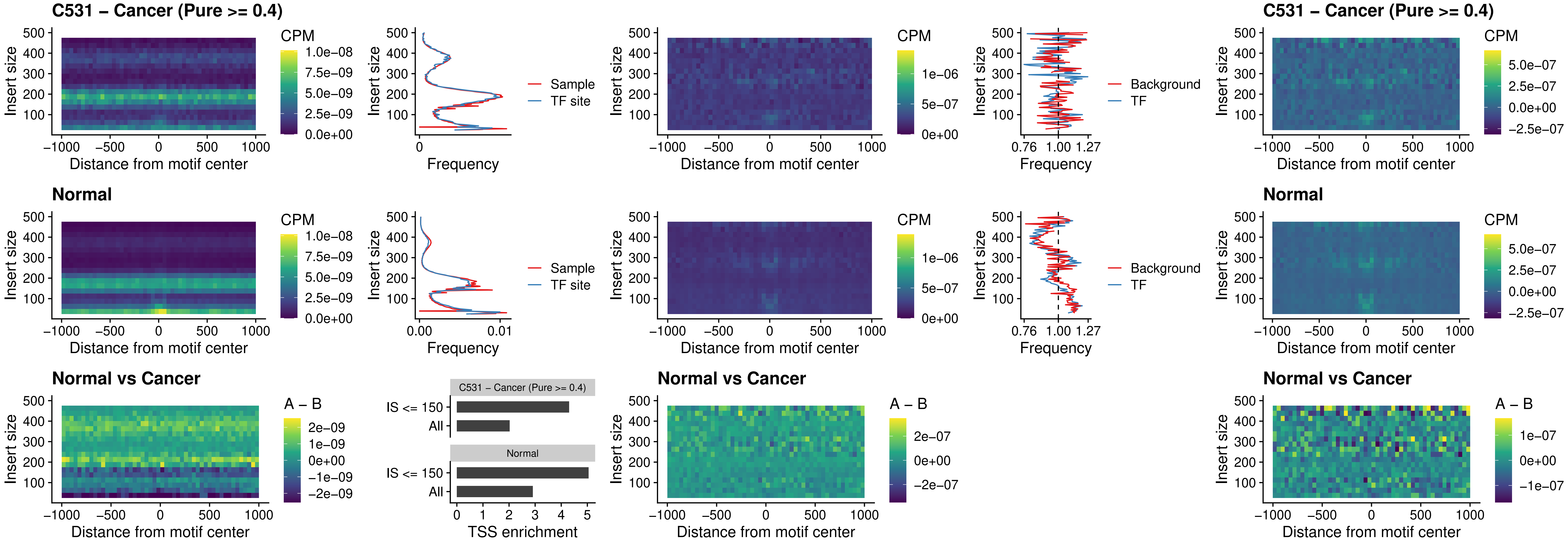

#### C531 – Cancer (Pure >= 0.4) – ETS2_uc_dTSS_oPEAK

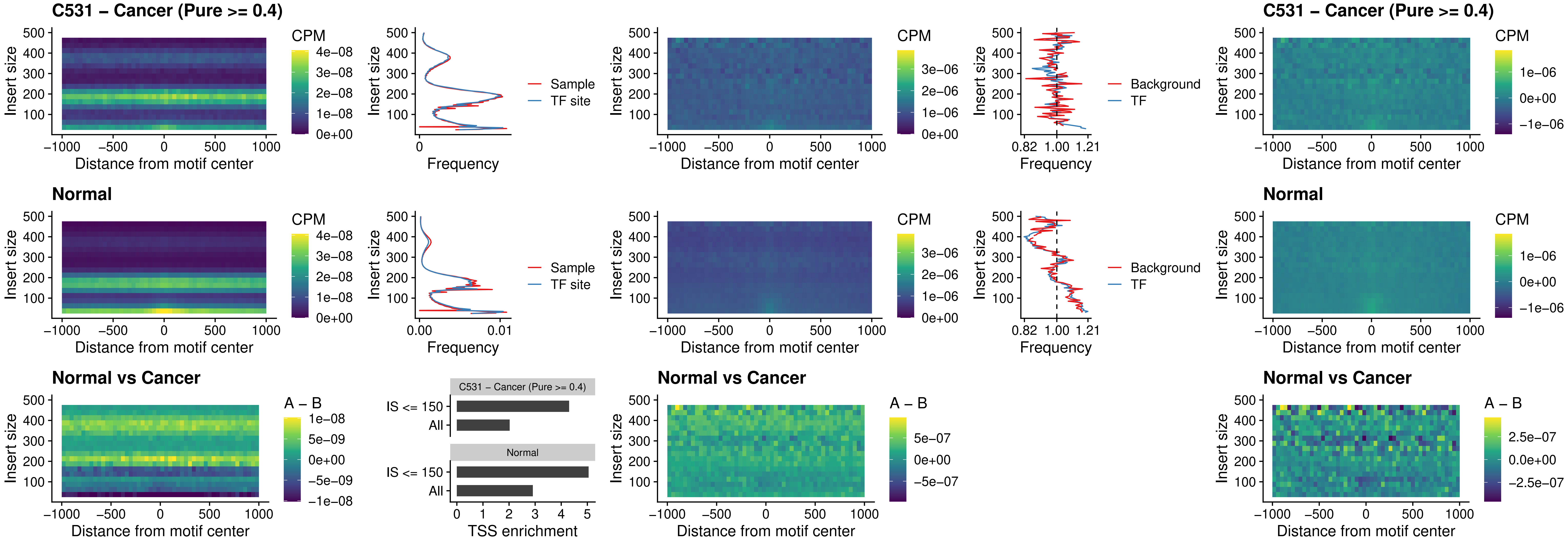

#### C531 – Cancer (Pure >= 0.4) – SMARCC2_uc_pTSS_oPEAK

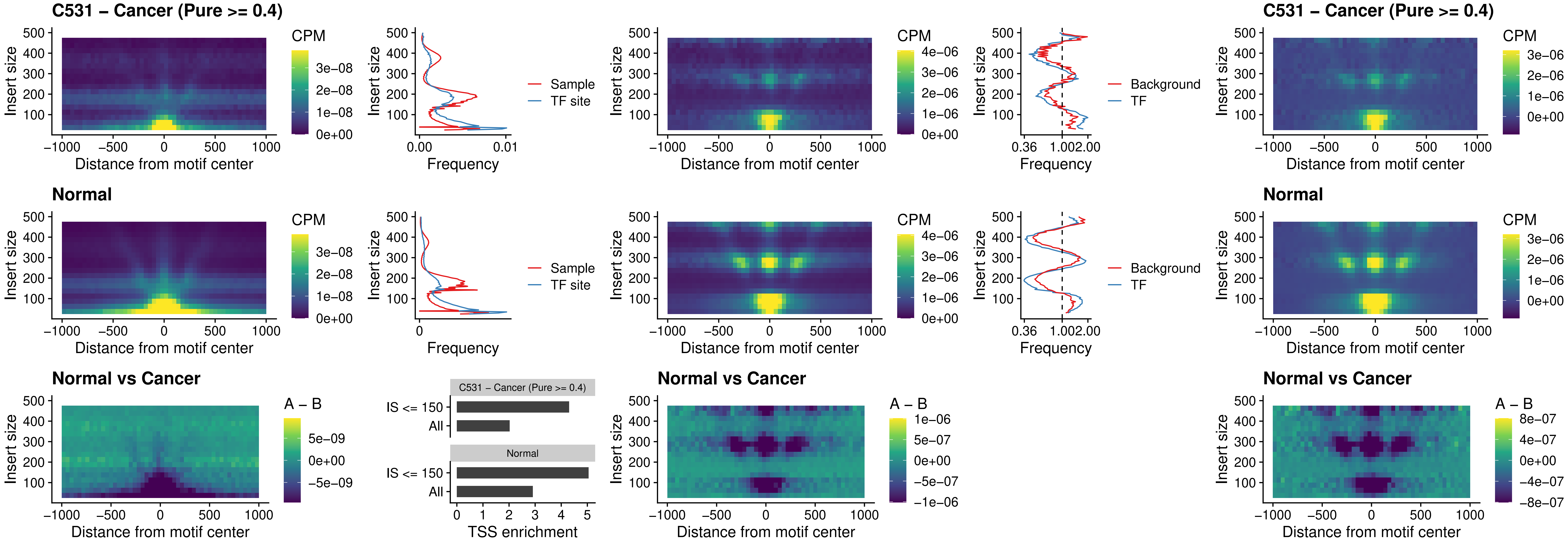

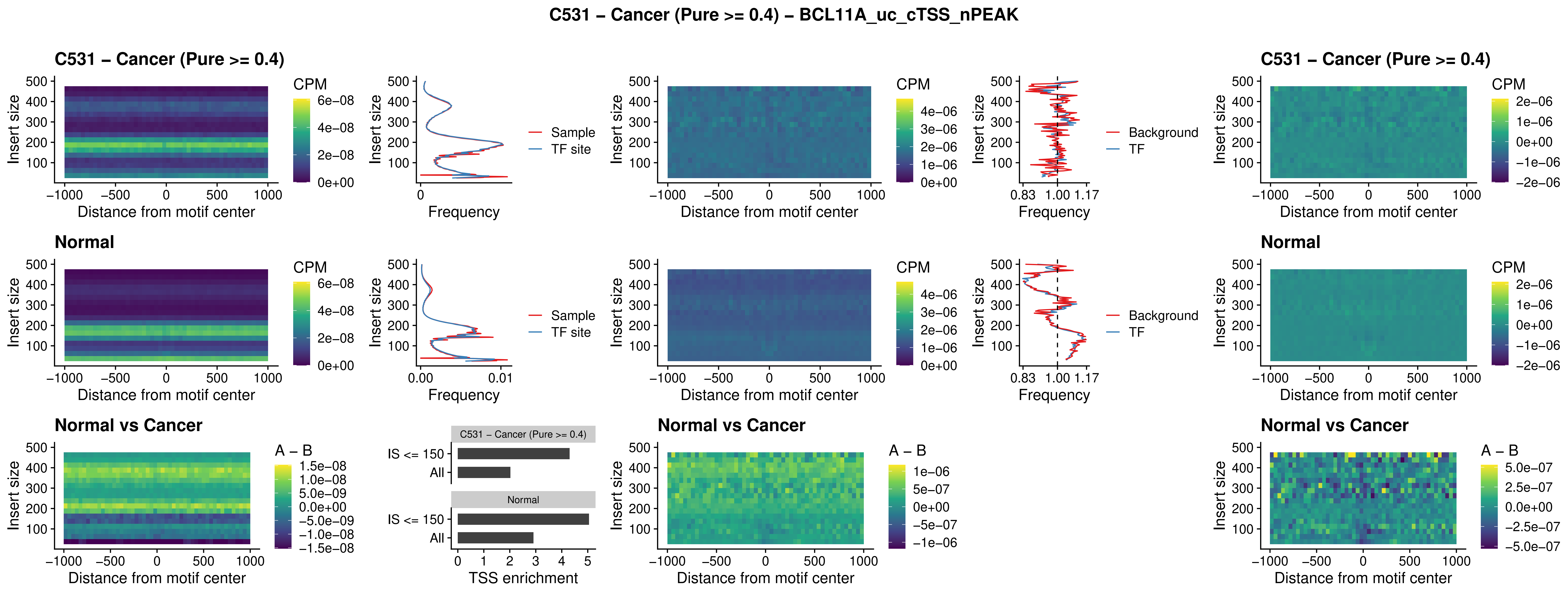
C531 – Cancer (Pure >= 0.4) – BCL11A_uc_cTSS_nPEAK
C531 – Cancer (Pure >= 0.4)
Normal
Normal vs Cancer
Insert size
Distance from motif center
CPM
A – B
Frequency
Sample
TF site
Background
TF
TSS enrichment
IS <= 150
All
C531 – Cancer (Pure >= 0.4)
Normal

#### C531 – Cancer (Pure >= 0.4) – BCL11B_uc_cTSS_nPEAK

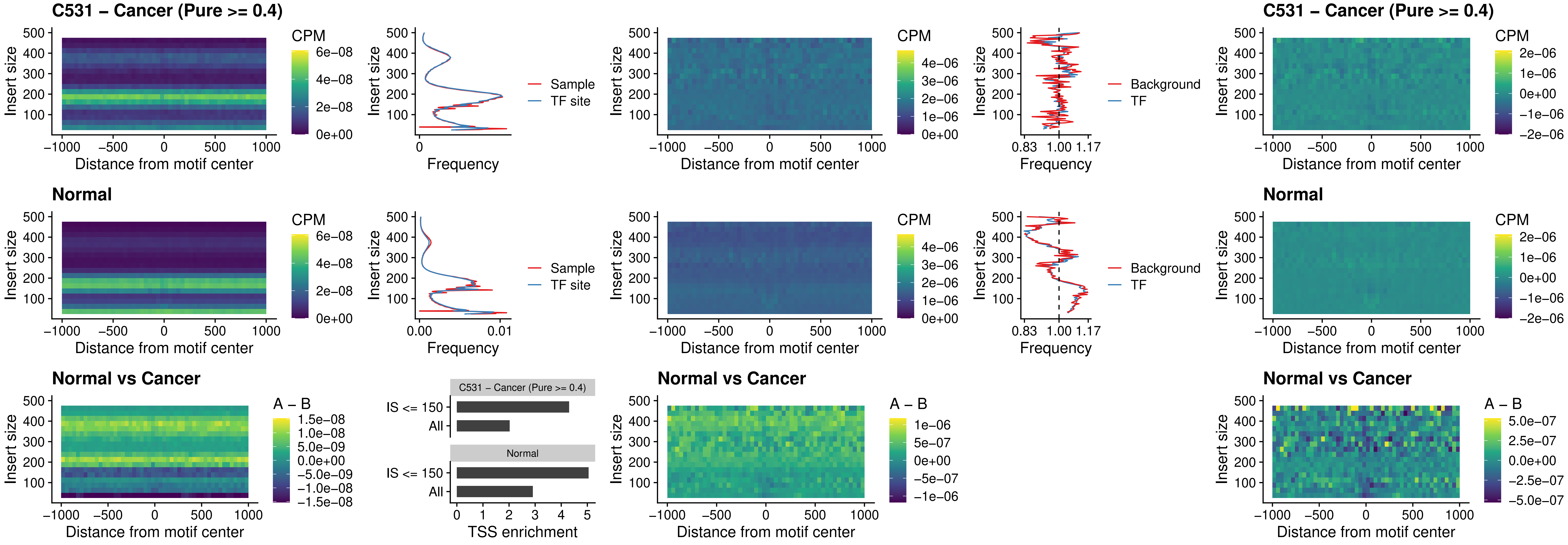

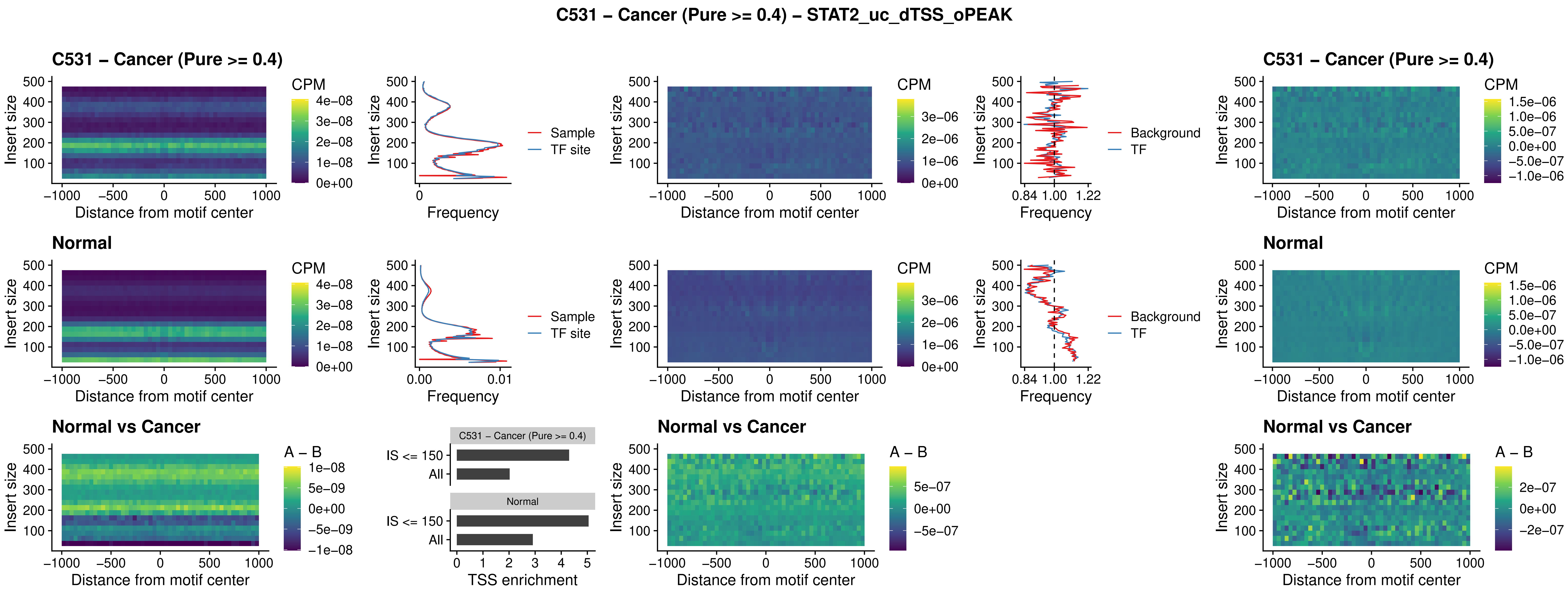
C531 – Cancer (Pure >= 0.4) – STAT2_uc_dTSS_oPEAK
C531 – Cancer (Pure >= 0.4)
Normal
Normal vs Cancer
Insert size
Distance from motif center
Frequency
CPM
A – B
Sample
TF site
Background
TF
TSS enrichment
IS <= 150
All

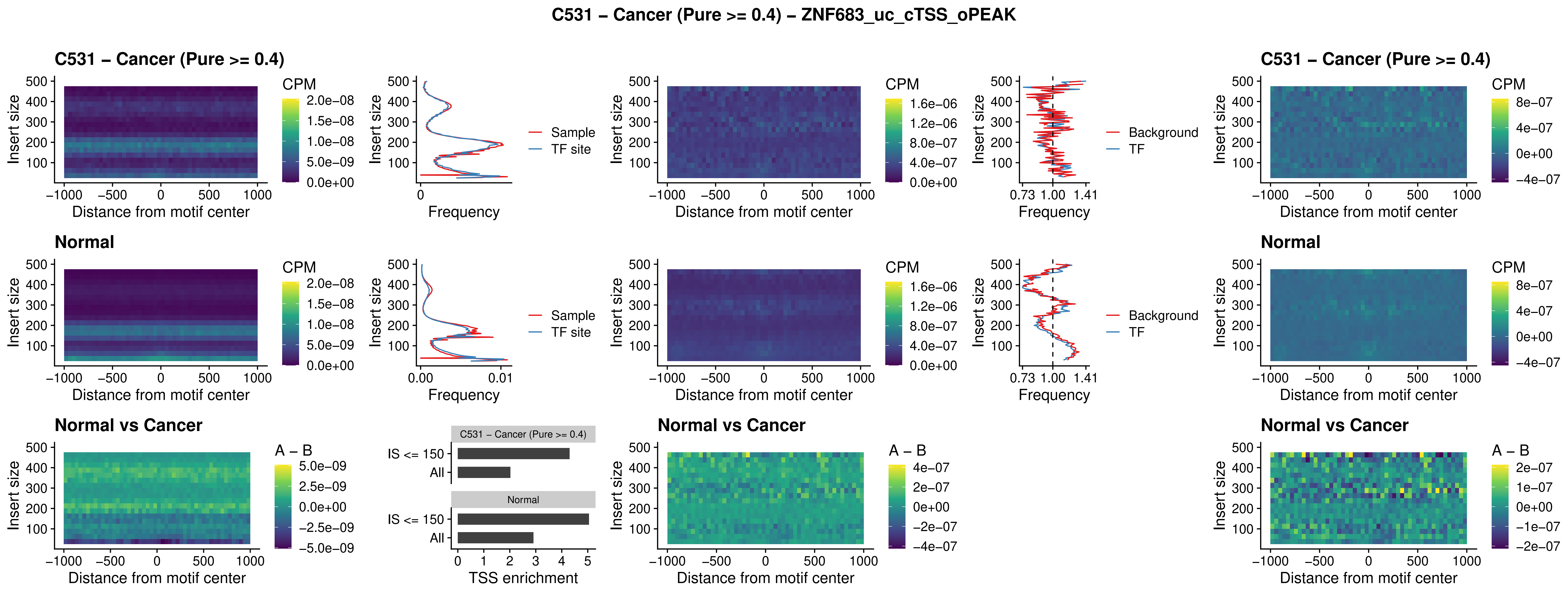

C531 – Cancer (Pure >= 0.4) – ZNF683_uc_cTSS_oPEAK
C531 – Cancer (Pure >= 0.4)
Normal
Normal vs Cancer
Insert size
Distance from motif center
CPM
A – B
Frequency
Sample
TF site
Background
TF
C531 – Cancer (Pure >= 0.4)
Normal
IS <= 150
All
TSS enrichment

#### C531 – Cancer (Pure >= 0.4) – IRF8_uc_dTSS_nPEAK

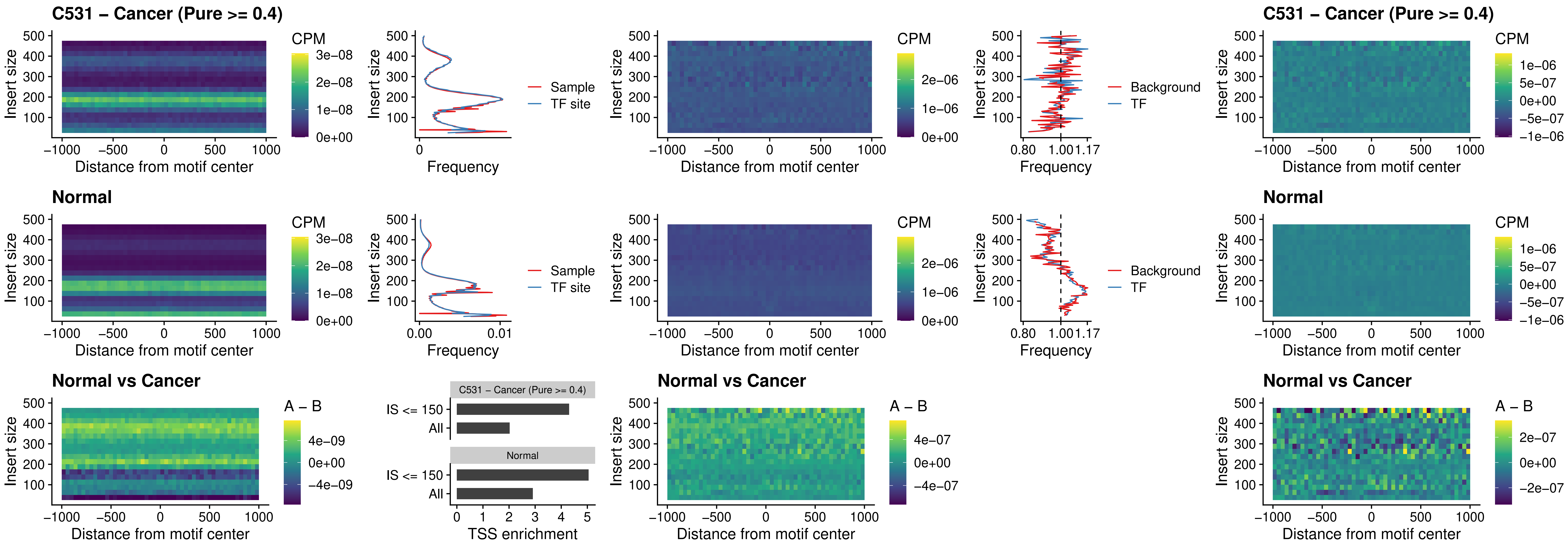

#### C531 – Cancer (Pure >= 0.4) – ETV7_uc_dTSS_oPEAK

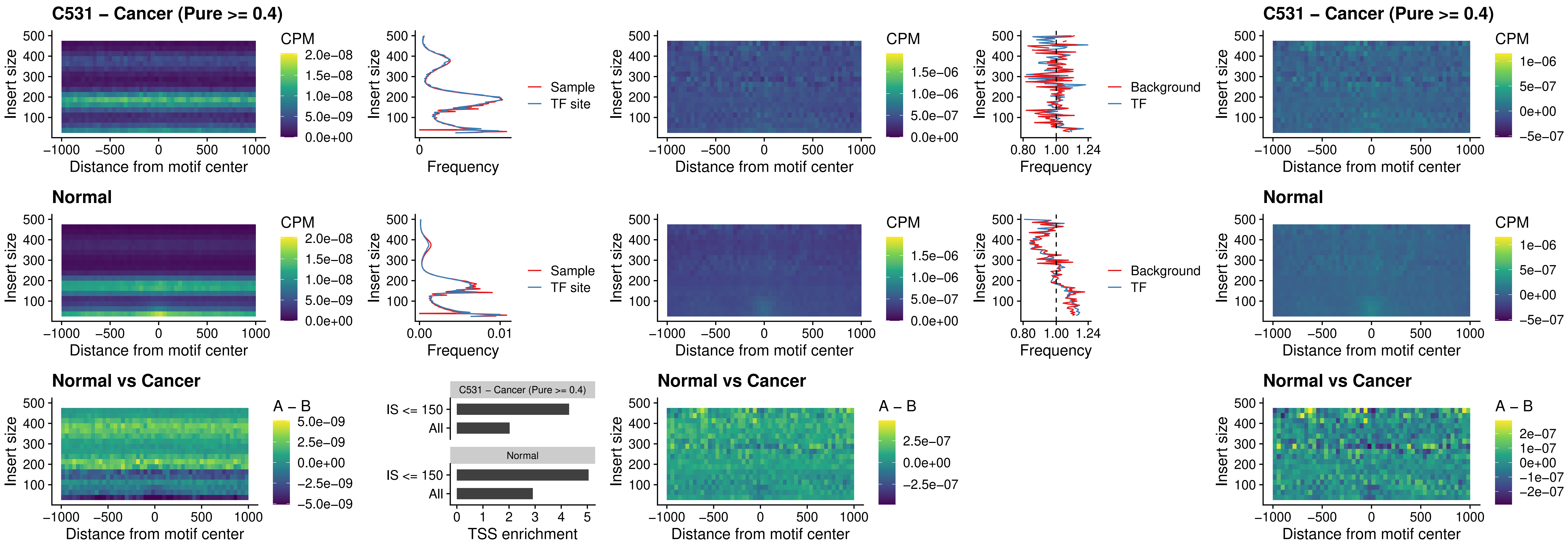

#### C532 – Cancer (Pure >= 0.4) – FOXL1_uc_pTSS_oPEAK

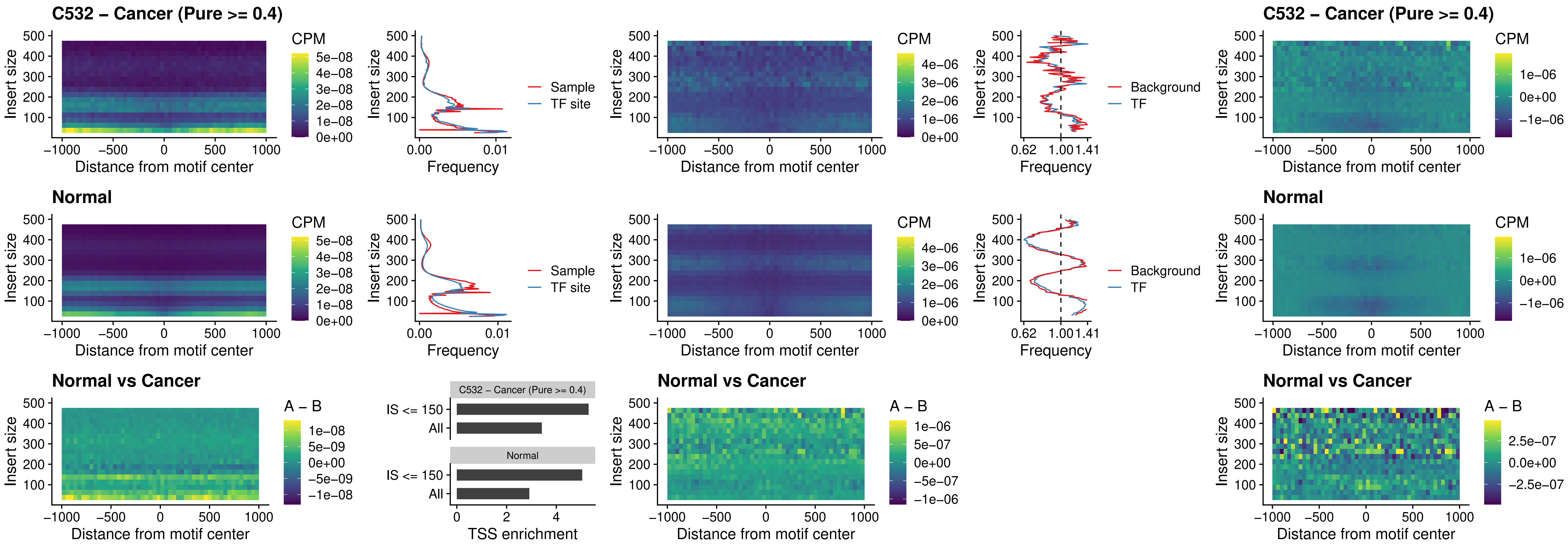

#### C532 – Cancer (Pure >= 0.4) – CPEB1_uc_cTSS_oPEAK

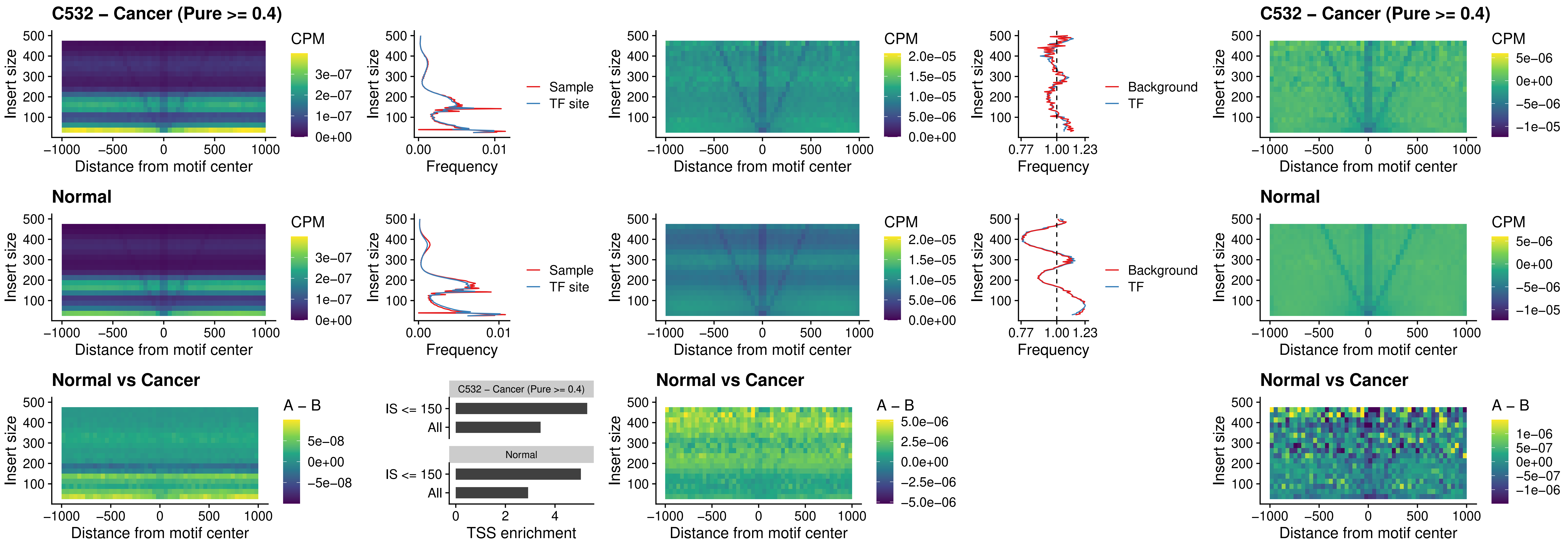

#### C532 – Cancer (Pure >= 0.4) – HMGA1_uc_pTSS_nPEAK

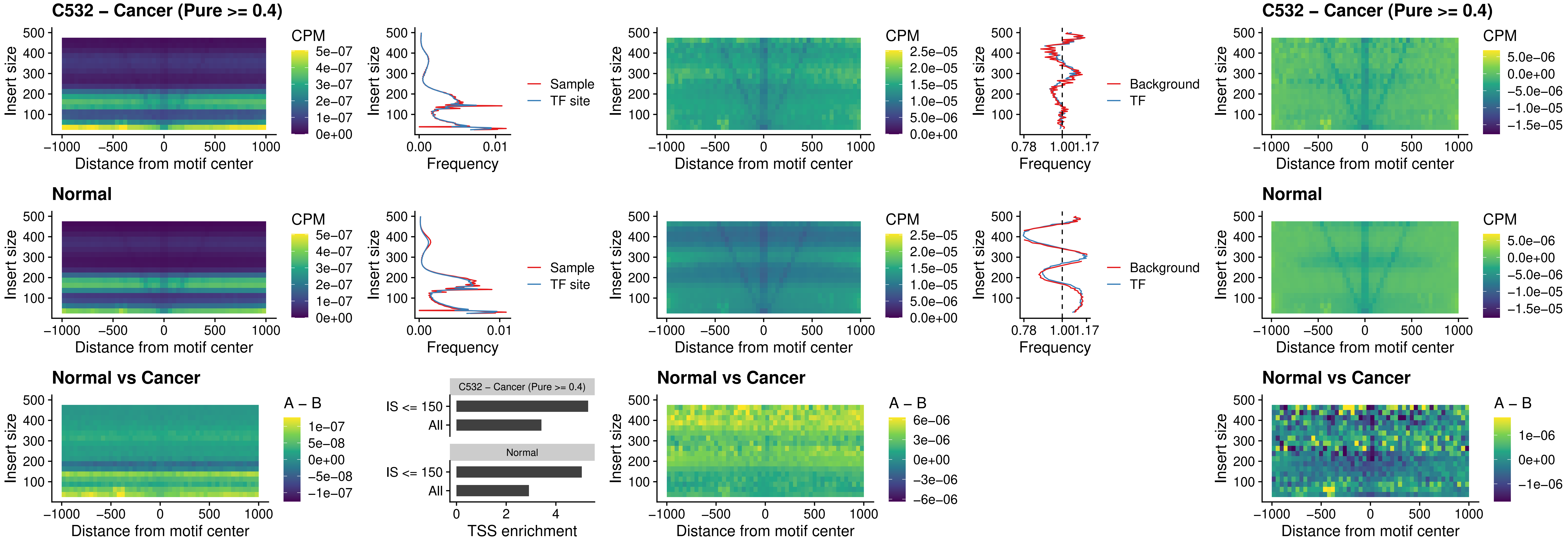

#### C532 – Cancer (Pure >= 0.4) – SOX13_uc_cTSS_nPEAK

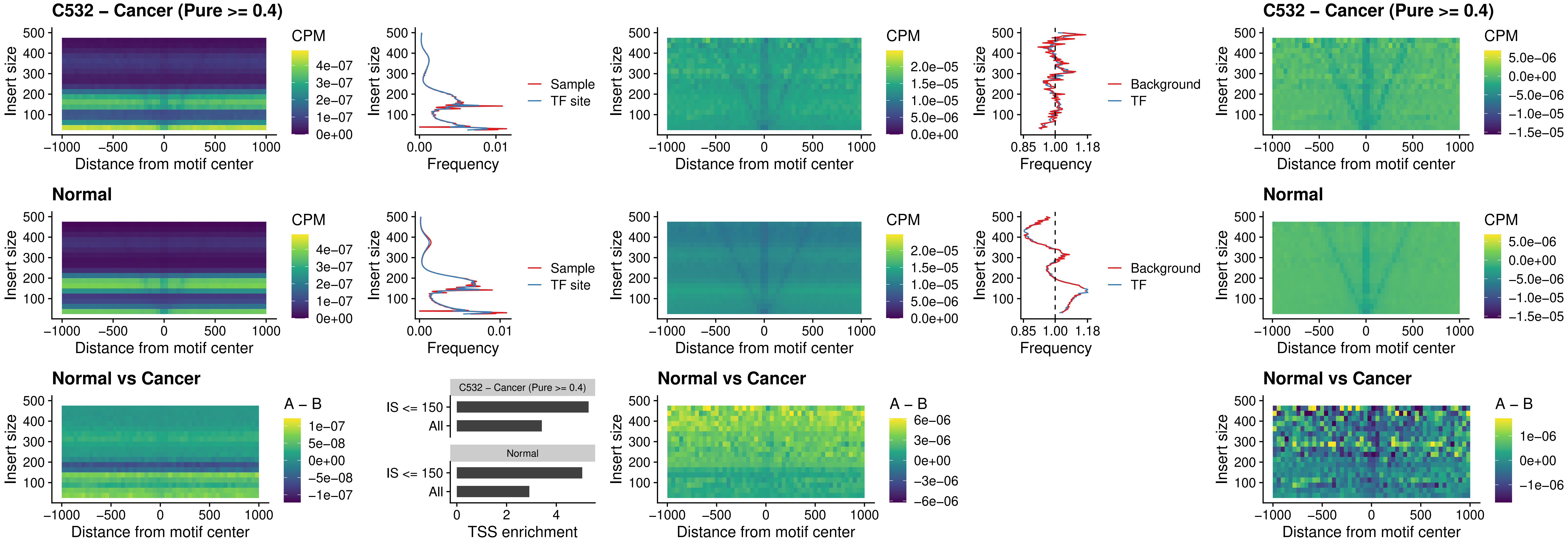

#### C532 – Cancer (Pure >= 0.4) – LHX4_uc_cTSS_nPEAK

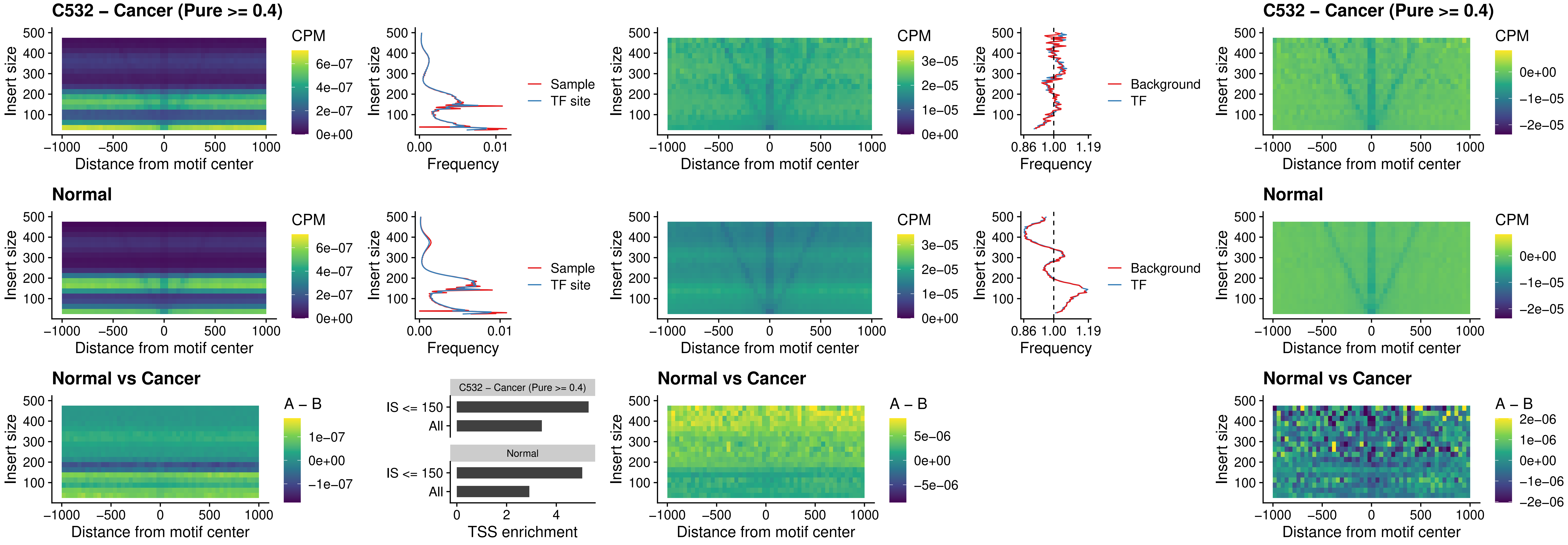

#### C532 – Cancer (Pure >= 0.4) – HOXA4_uc_cTSS_nPEAK

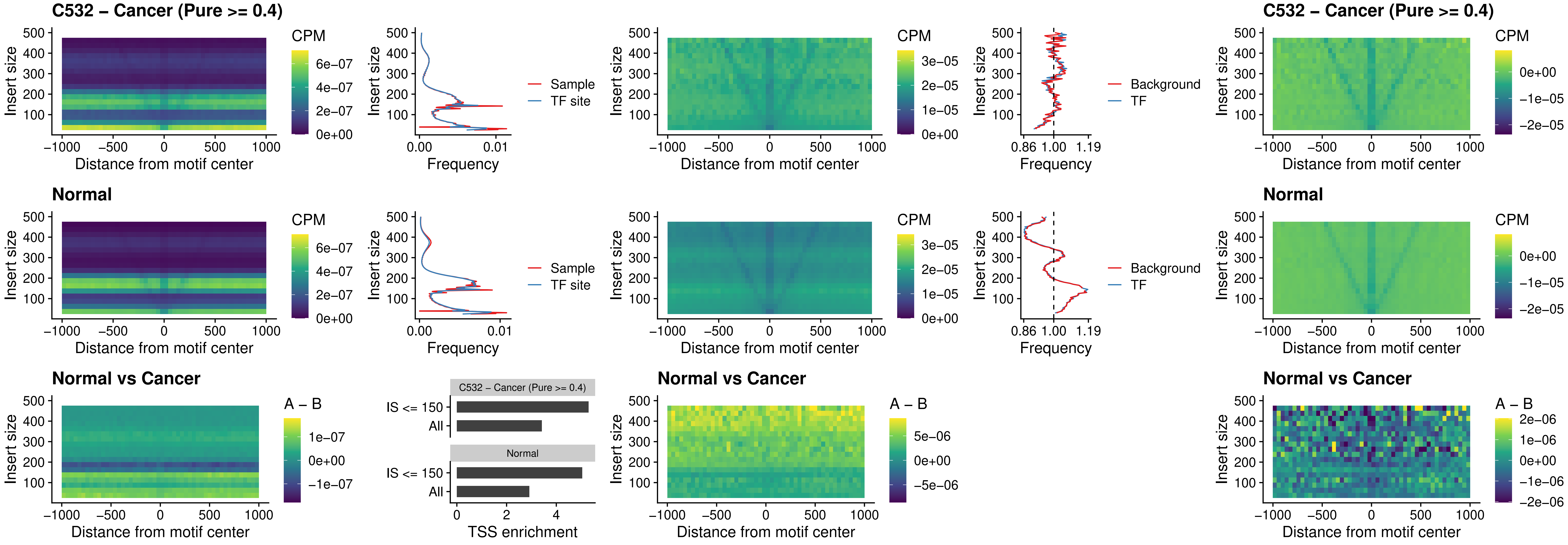

### C532 – Cancer (Pure >= 0.4) – HLX_uc_cTSS_nPEAK

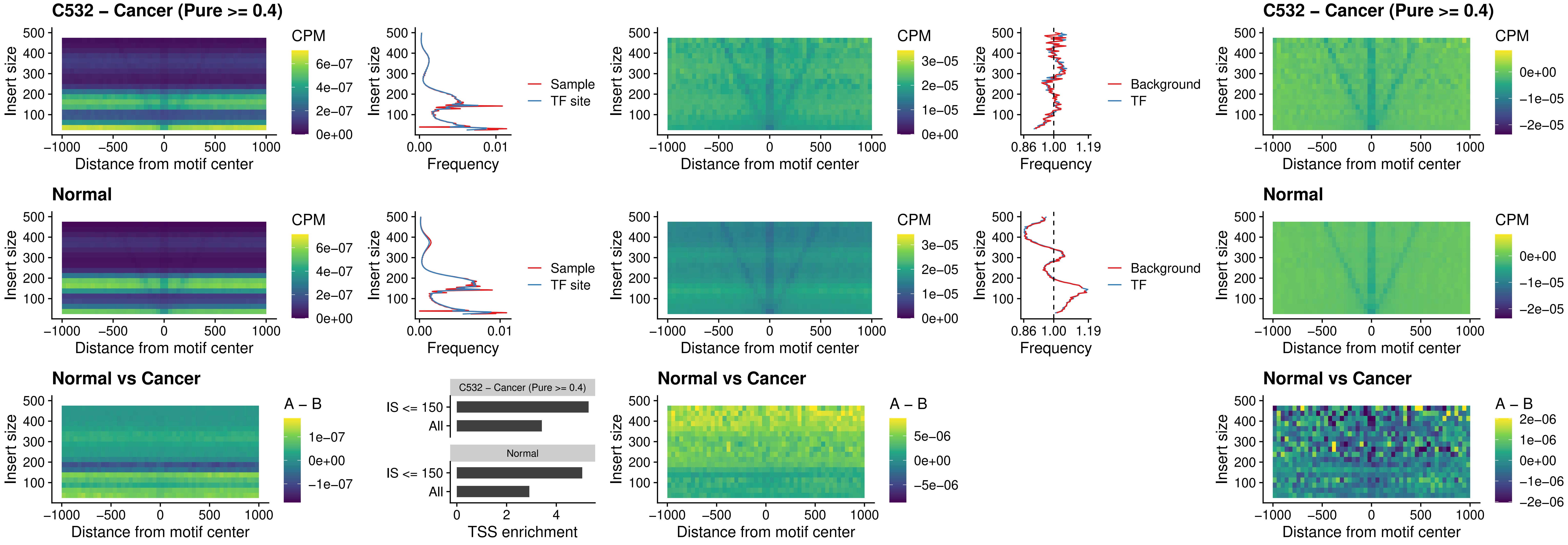

#### C532 – Cancer (Pure >= 0.4) – AC0125311_uc_cTSS_nPEAK

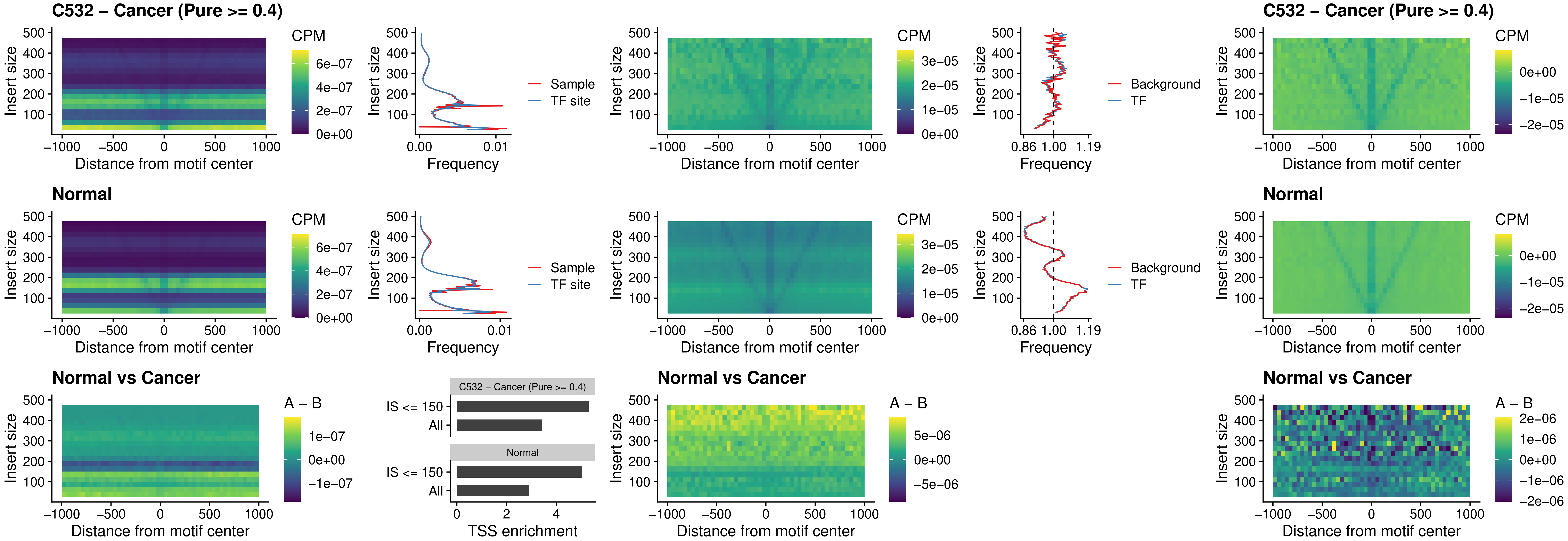

### C532 – Cancer (Pure >= 0.4) – DBX2_uc_cTSS_nPEAK

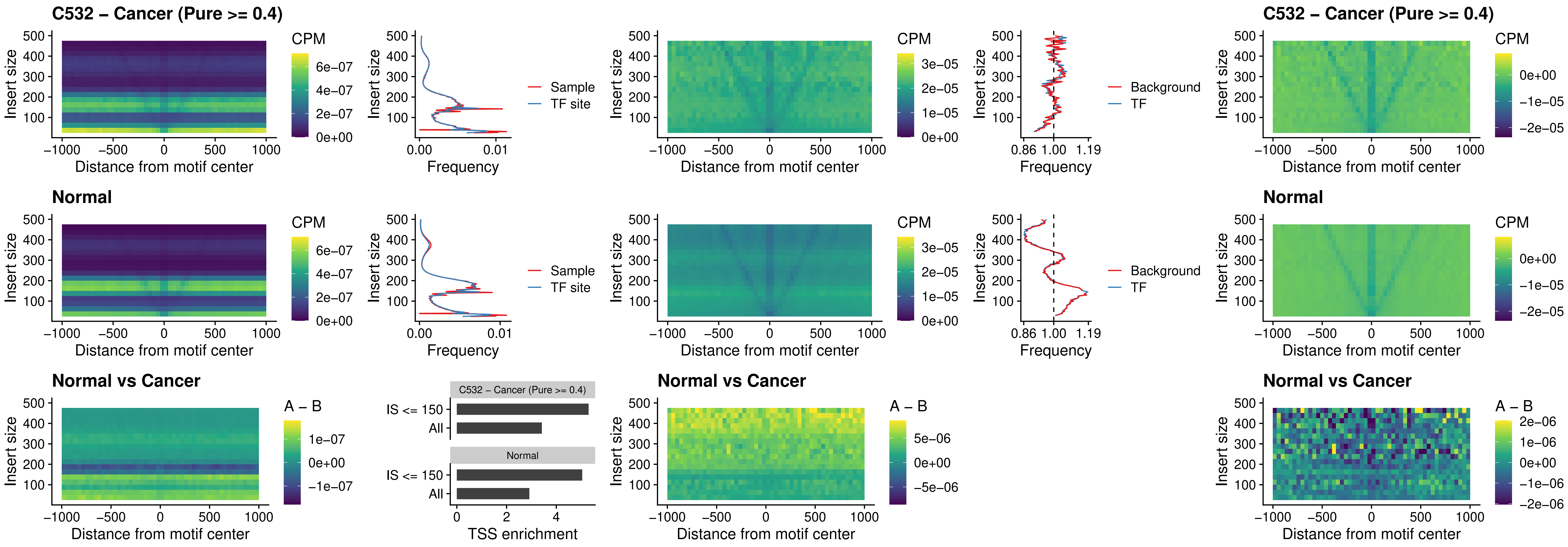

#### C532 – Cancer (Pure >= 0.4) – LHX6_uc_dTSS_nPEAK

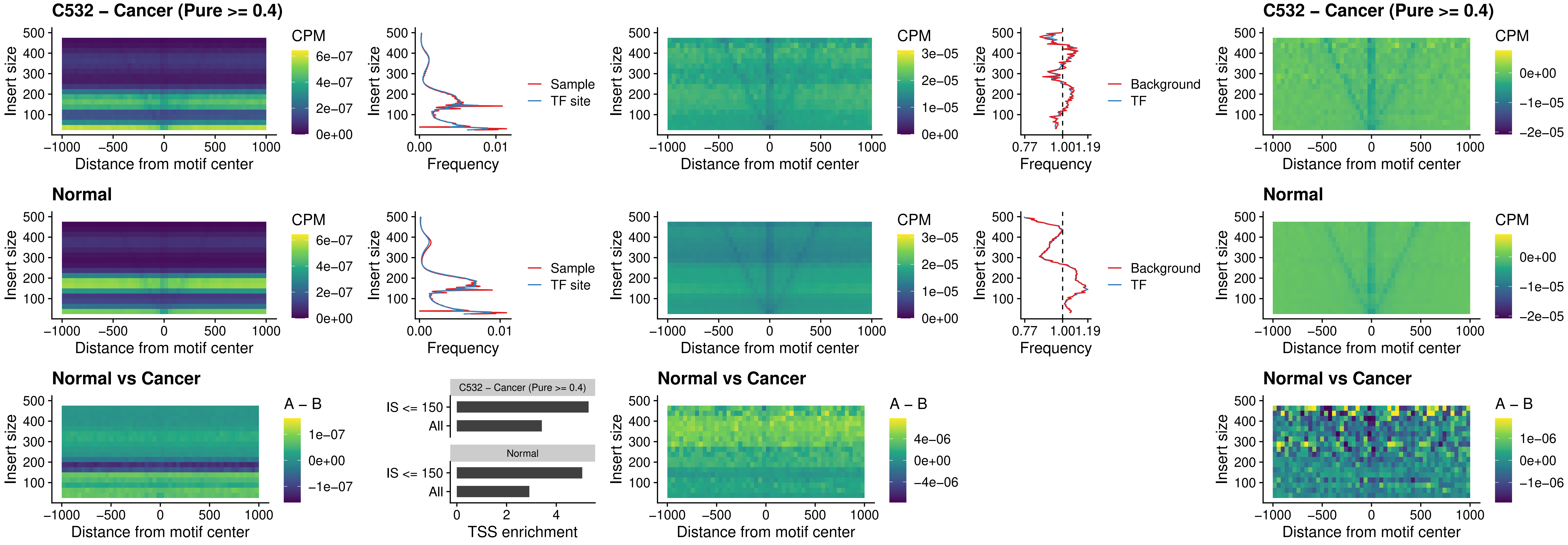

#### C532 – Cancer (Pure >= 0.4) – SOX1_uc_dTSS_nPEAK

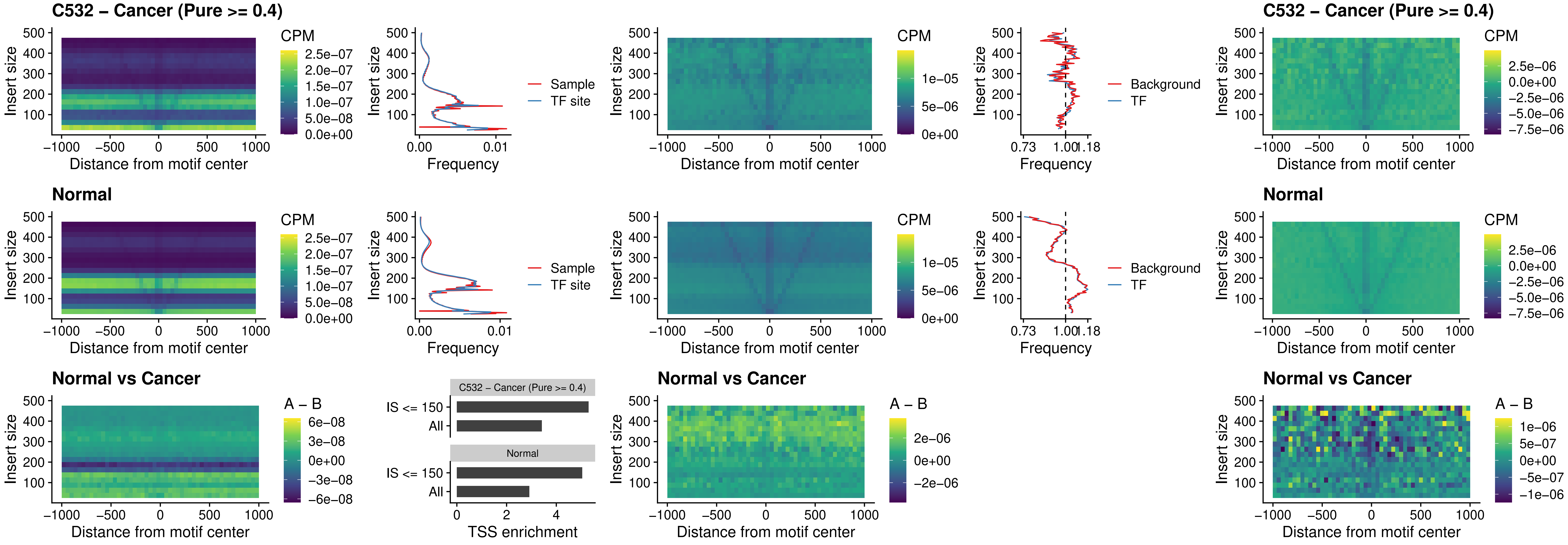

#### C532 – Cancer (Pure >= 0.4) – NKX24_uc_dTSS_nPEAK

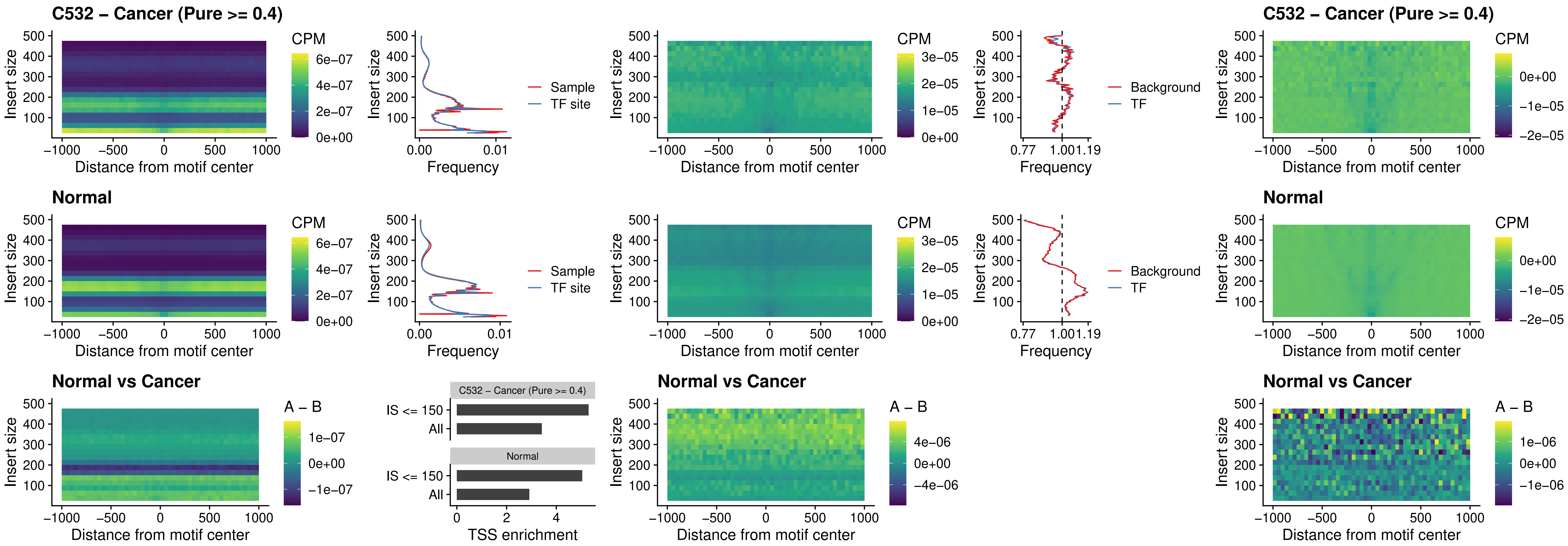

### C532 – Cancer (Pure >= 0.4) – DLX3_uc_cTSS_nPEAK

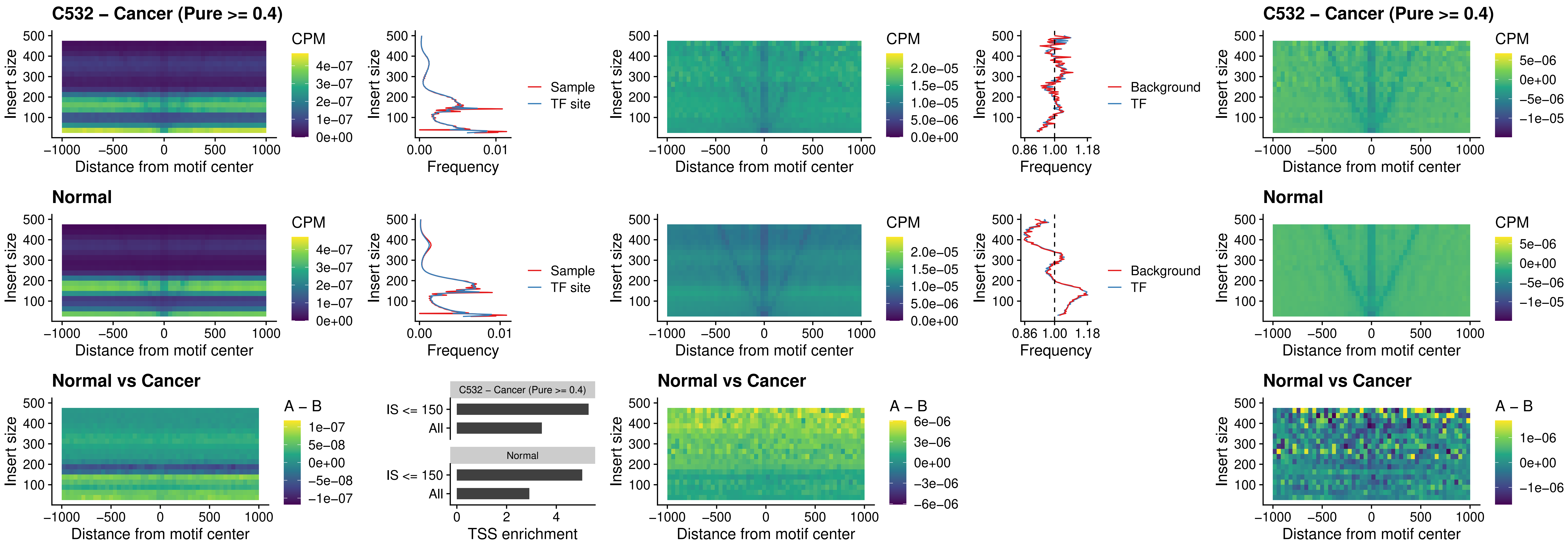

#### C532 – Cancer (Pure >= 0.4) – HOXD3_uc_cTSS_nPEAK

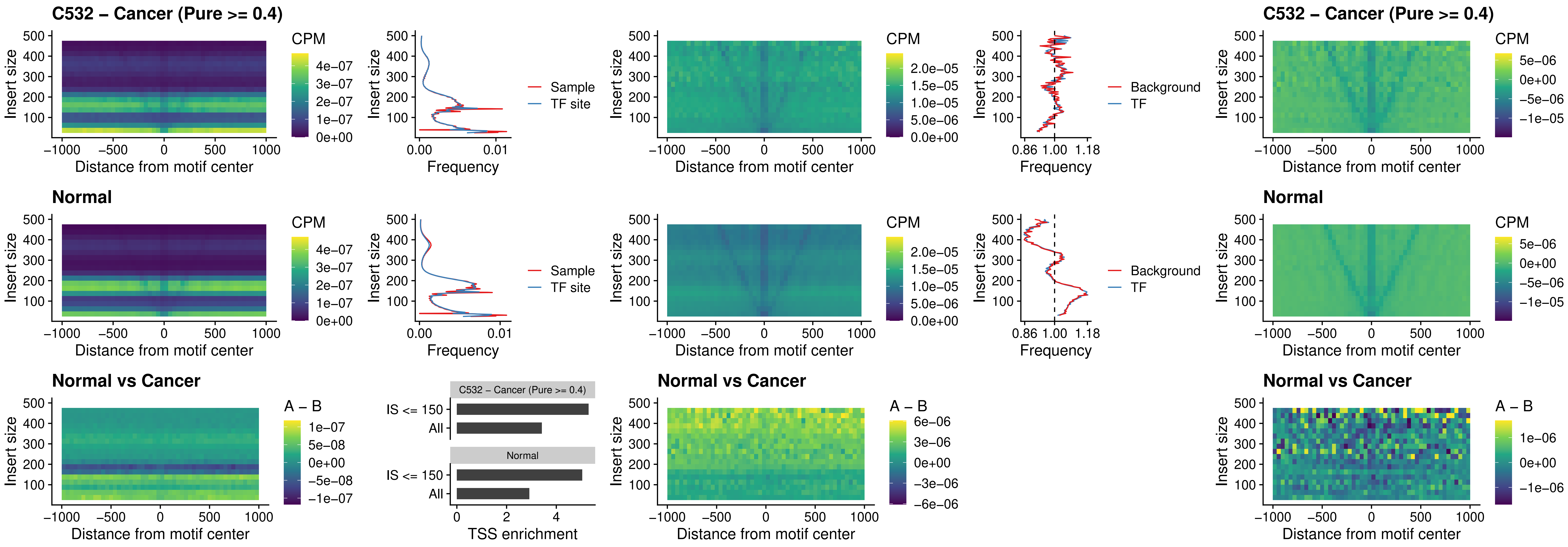

#### C532 – Cancer (Pure >= 0.4) – ISL1_uc_dTSS_oPEAK

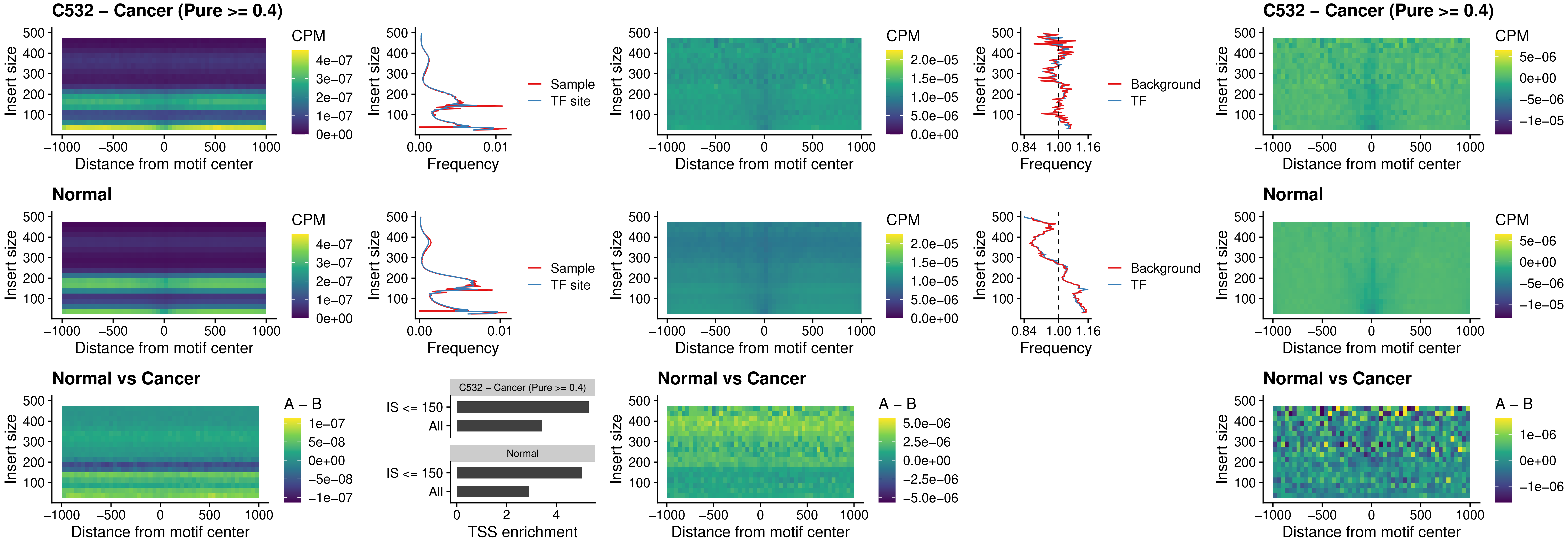

#### C532 – Cancer (Pure >= 0.4) – HOXB4_uc_dTSS_oPEAK

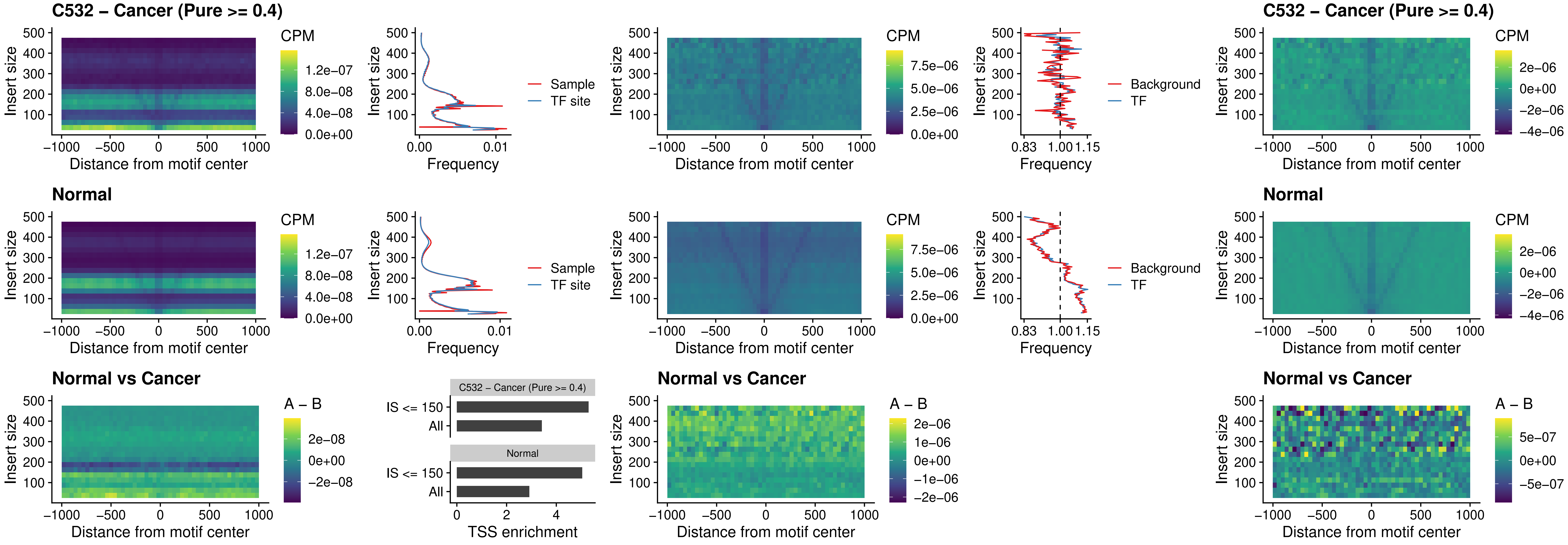

#### C532 – Cancer (Pure >= 0.4) – VAX1_uc_dTSS_oPEAK

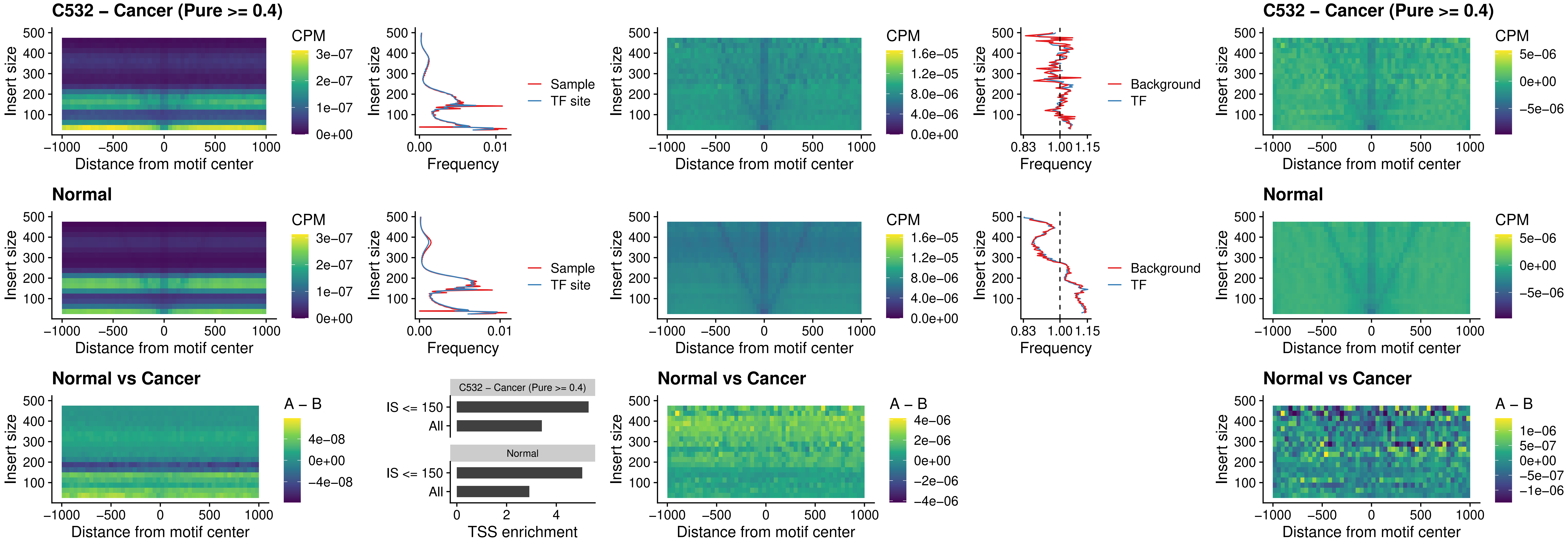

#### C532 – Cancer (Pure >= 0.4) – DLX3_uc_dTSS_oPEAK

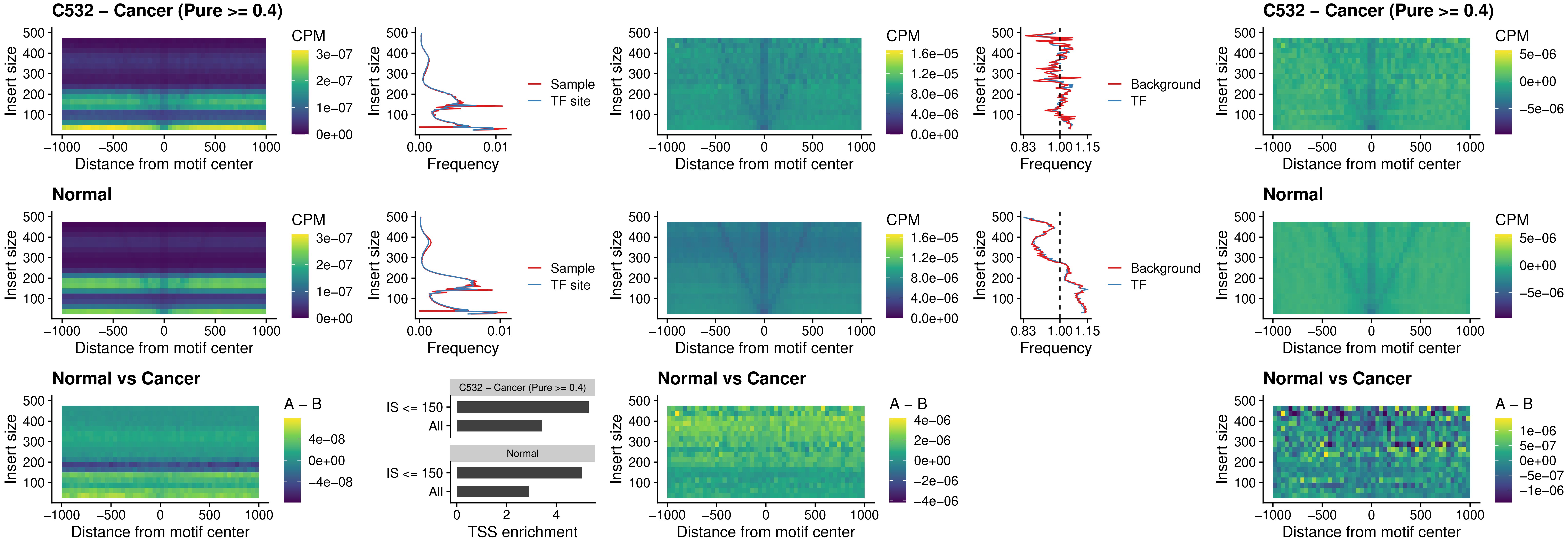

#### C532 – Cancer (Pure >= 0.4) – HOXD3_uc_dTSS_oPEAK

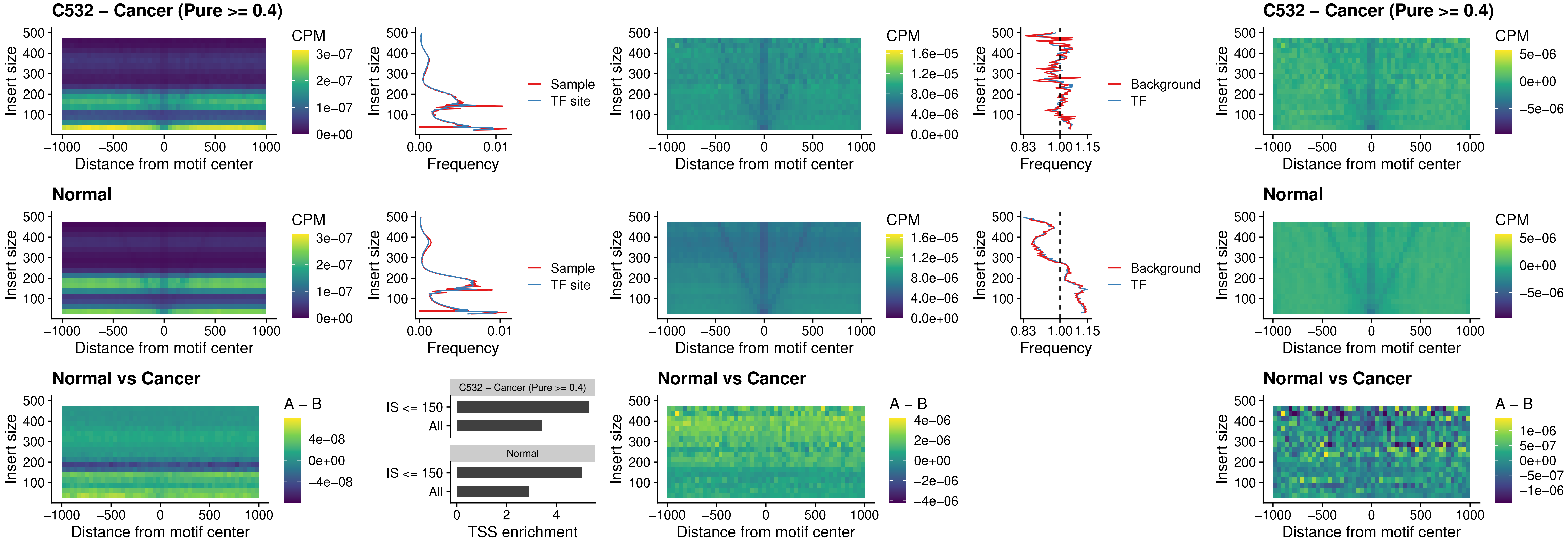

### C532 – Cancer (Pure >= 0.4) – BBX_uc_dTSS_nPEAK

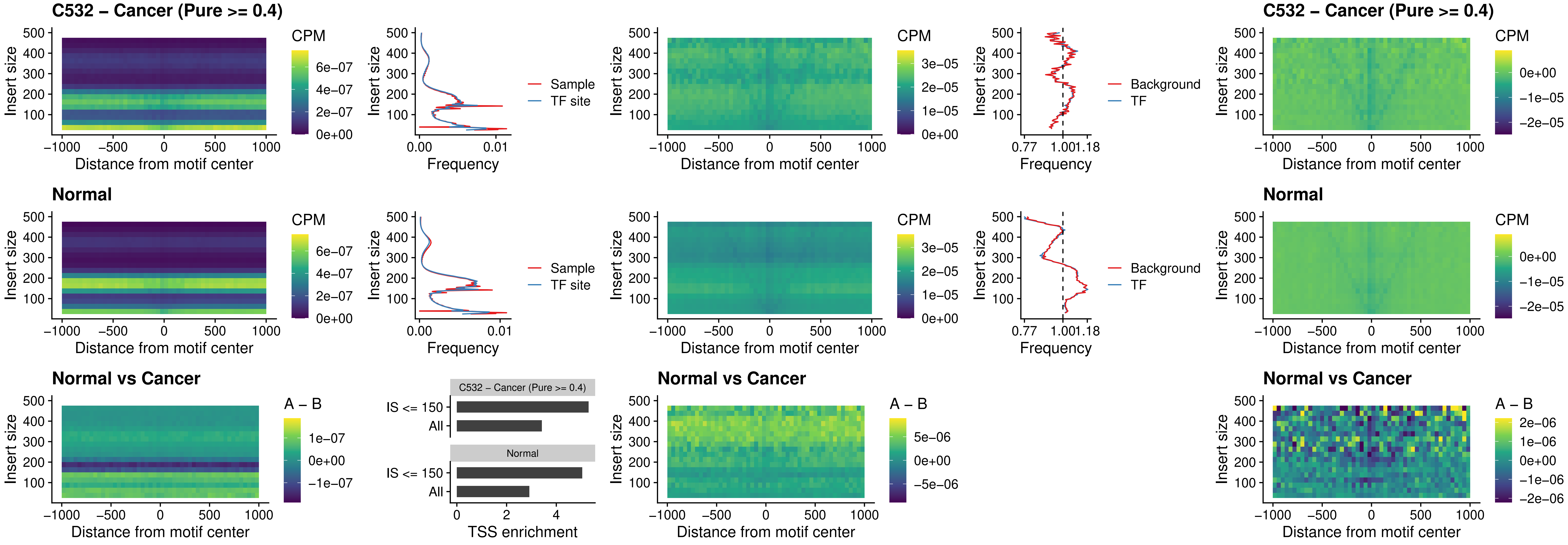

#### C532 – Cancer (Pure >= 0.4) – SOX3_uc_cTSS_nPEAK

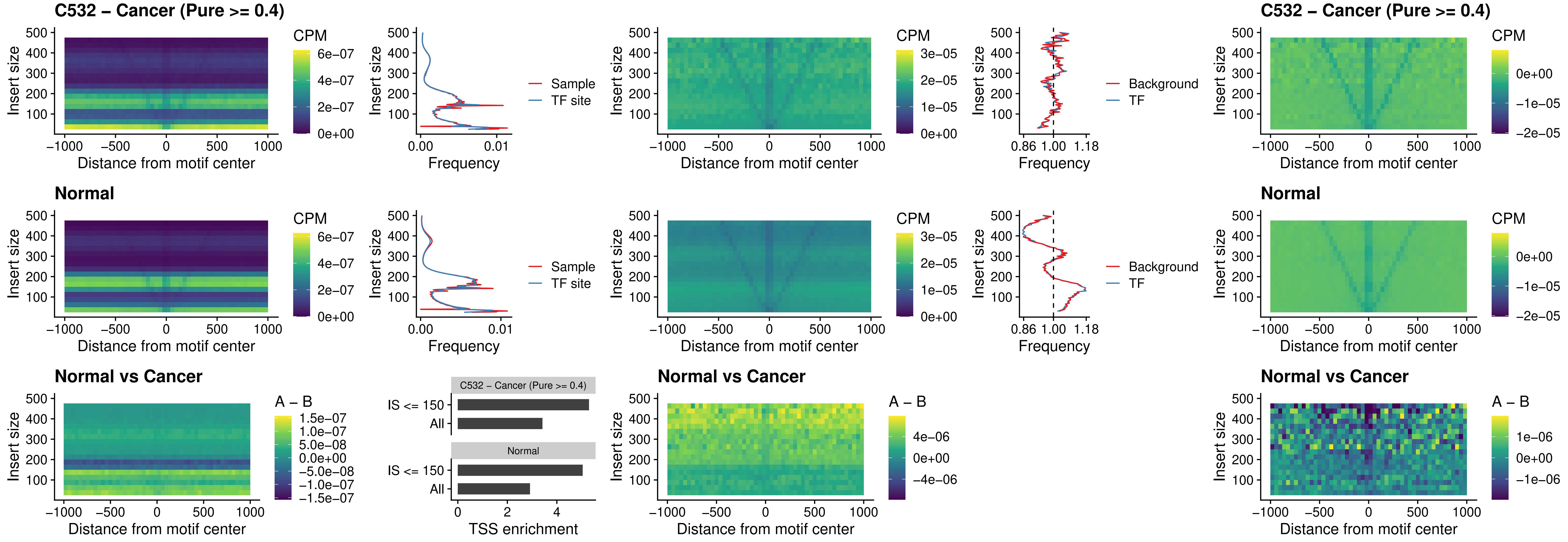

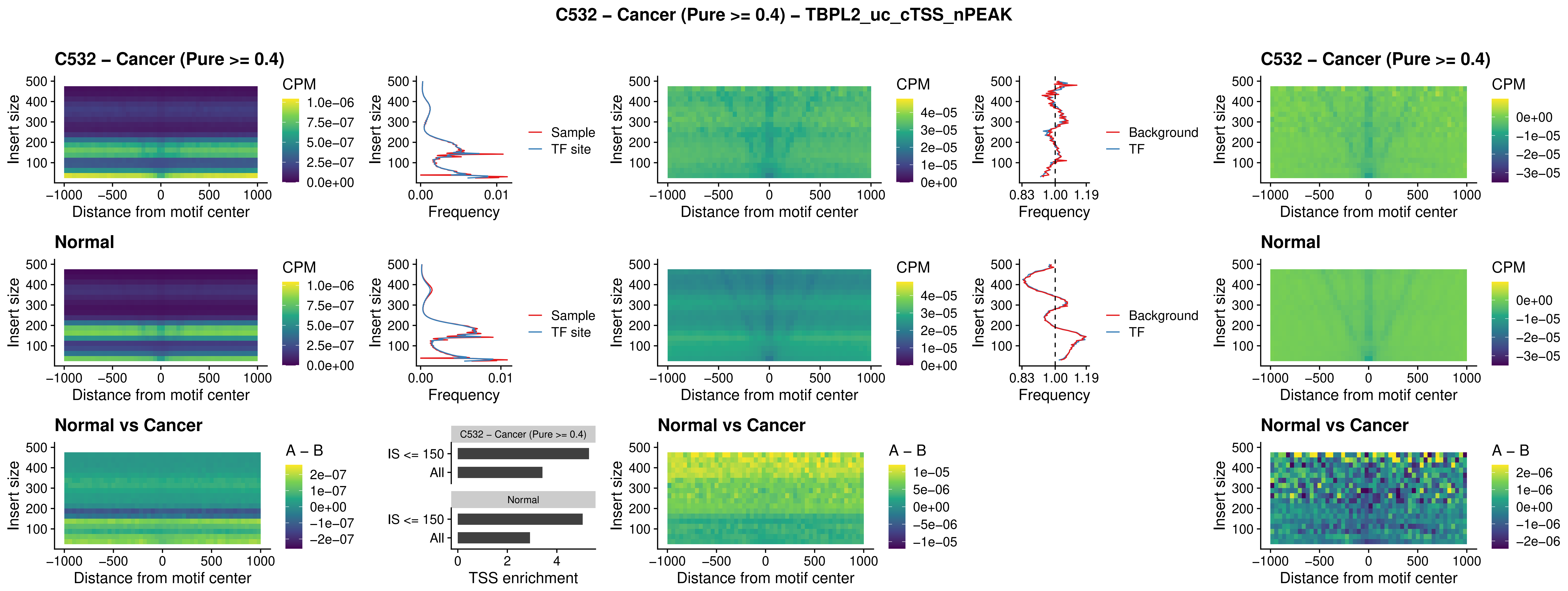

C532 – Cancer (Pure >= 0.4) – TBPL2_uc_cTSS_nPEAK
C532 – Cancer (Pure >= 0.4)
Normal
Normal vs Cancer
Insert size
Distance from motif center
Frequency
CPM
A – B
Sample
TF site
Background
TF
C532 – Cancer (Pure >= 0.4)
Normal
IS <= 150
All
TSS enrichment

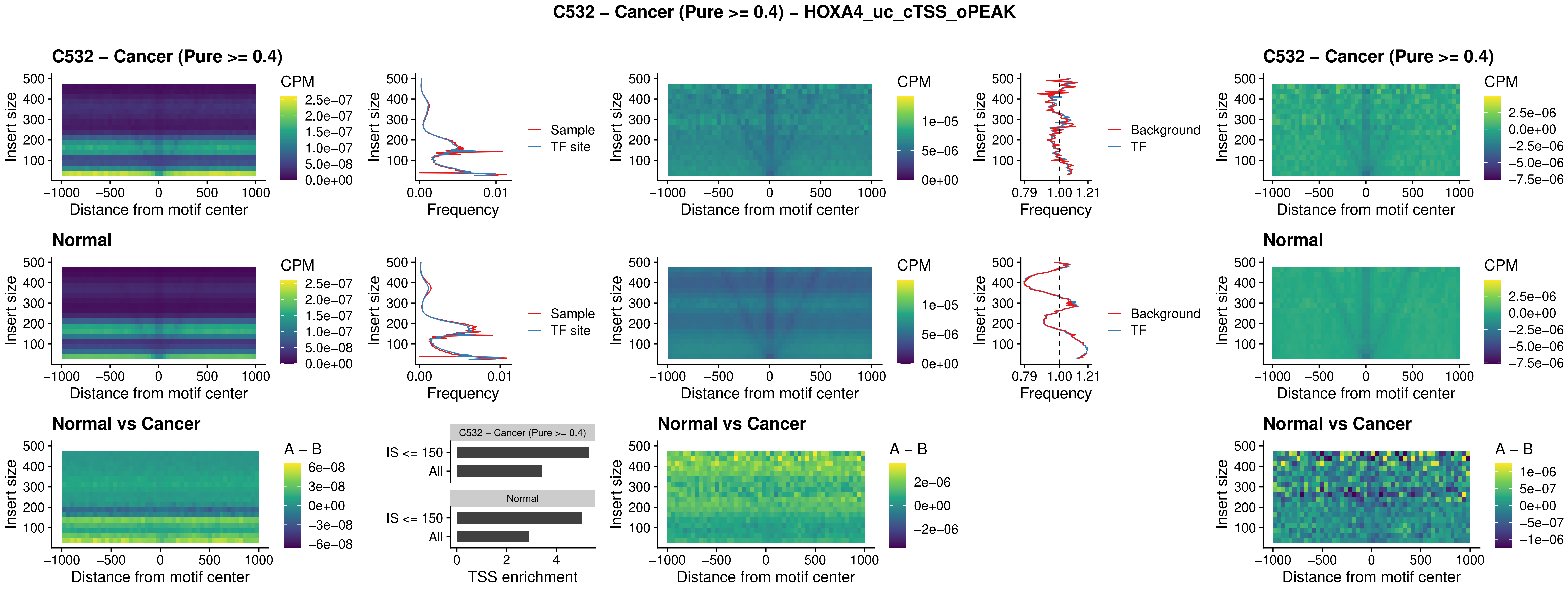
C532 – Cancer (Pure >= 0.4) – HOXA4_uc_cTSS_oPEAK
C532 – Cancer (Pure >= 0.4)
Normal
Normal vs Cancer
Insert size
Distance from motif center
Frequency
CPM
A – B
Sample
TF site
Background
TF
C532 – Cancer (Pure >= 0.4)
Normal
IS <= 150
All
TSS enrichment

#### C532 – Cancer (Pure >= 0.4) – PDX1_uc_dTSS_oPEAK

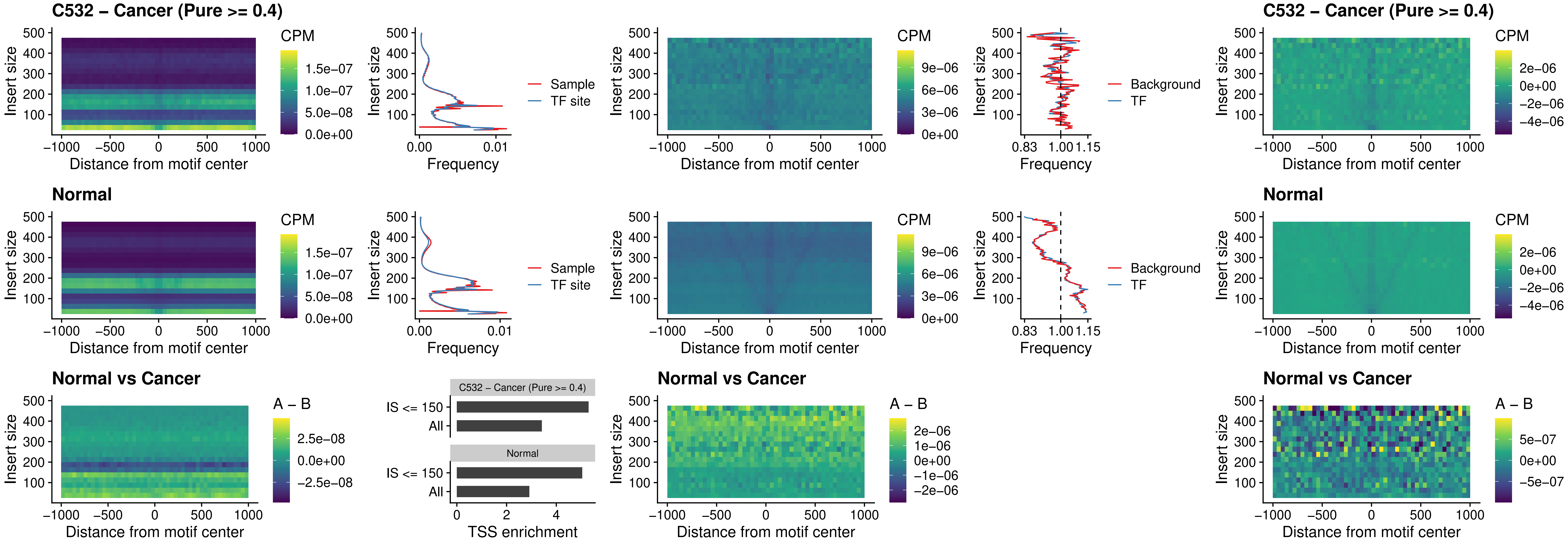

#### C532 – Cancer (Pure >= 0.4) – LHX4_uc_dTSS_oPEAK

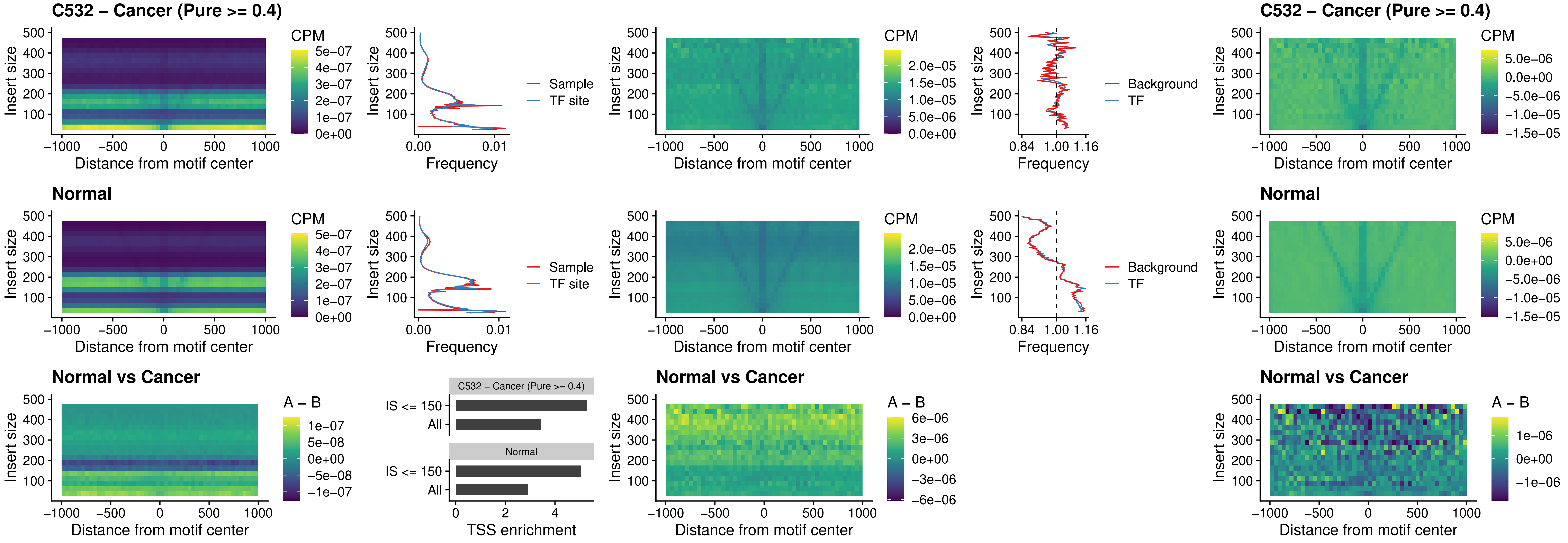

### C532 – Cancer (Pure >= 0.4) – HOXA4_uc_dTSS_oPEAK

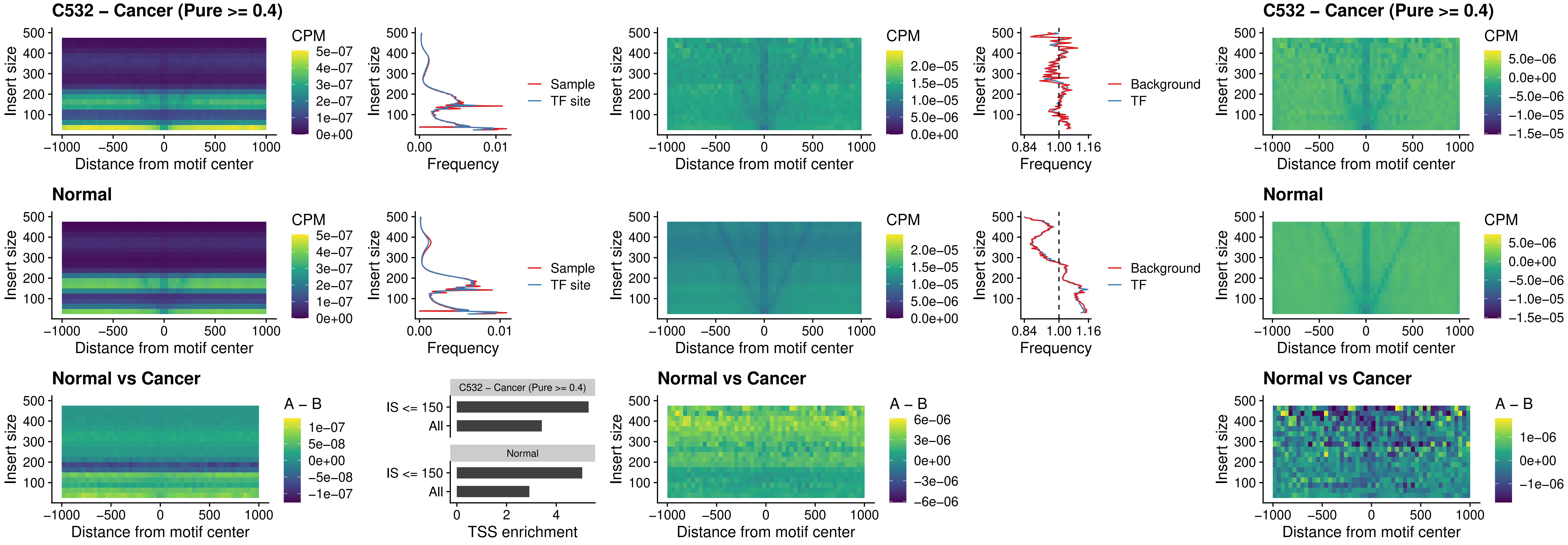

#### C532 – Cancer (Pure >= 0.4) – HLX_uc_dTSS_oPEAK

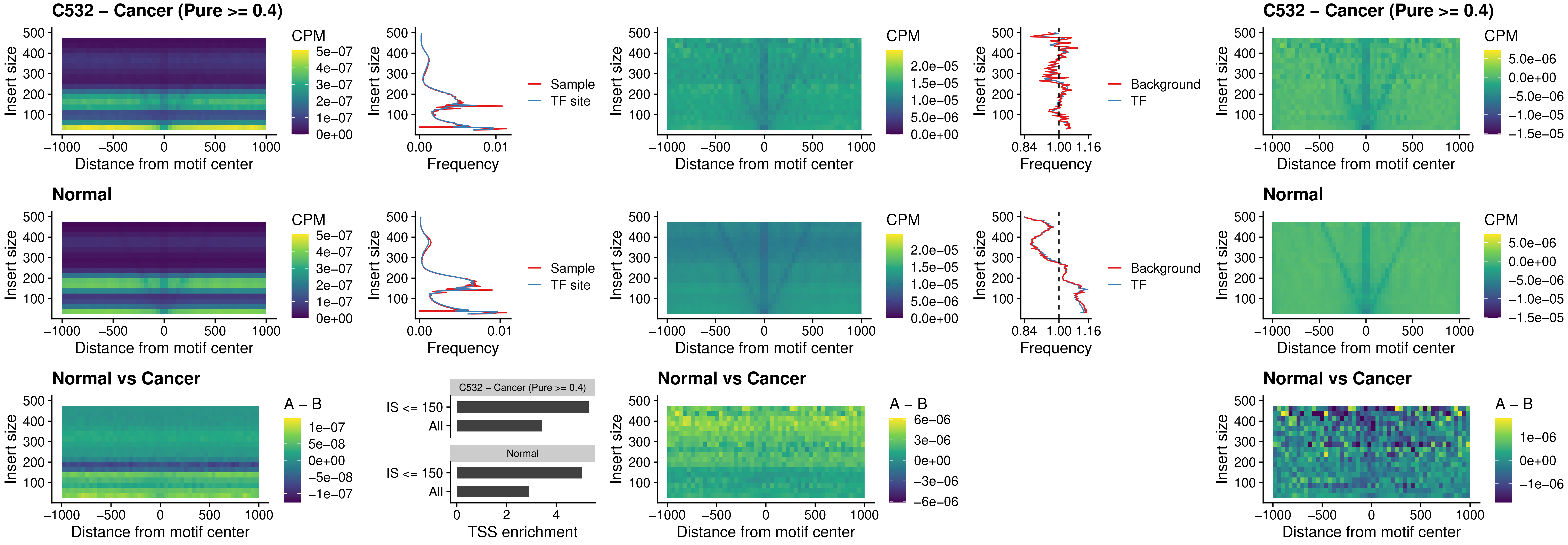

#### C532 – Cancer (Pure >= 0.4) – AC0125311_uc_dTSS_oPEAK

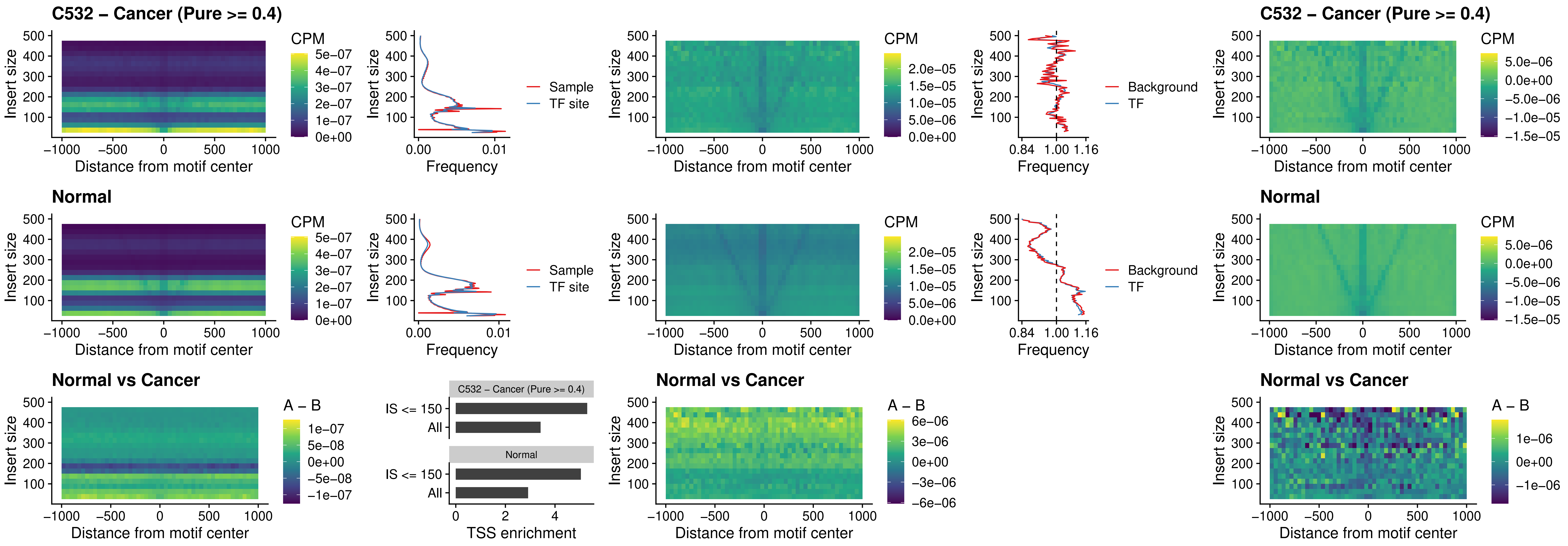

#### C532 – Cancer (Pure >= 0.4) – DBX2_uc_dTSS_oPEAK

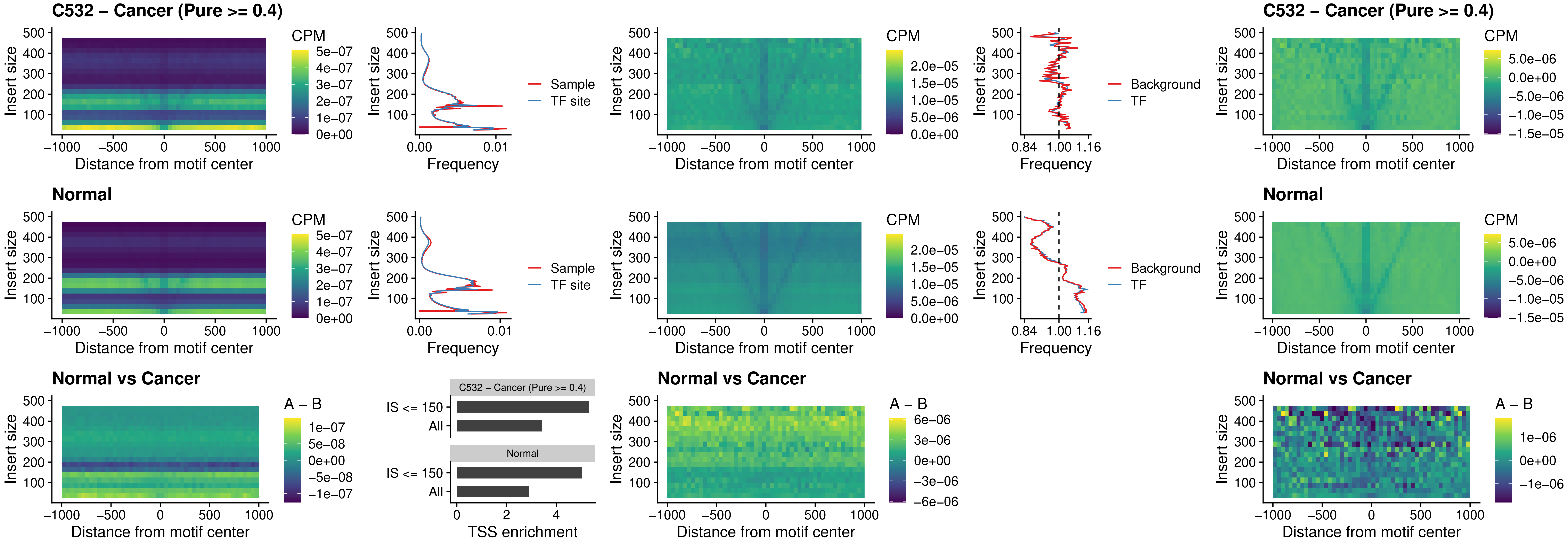

#### C532 – Cancer (Pure >= 0.4) – HOXA2_uc_dTSS_nPEAK

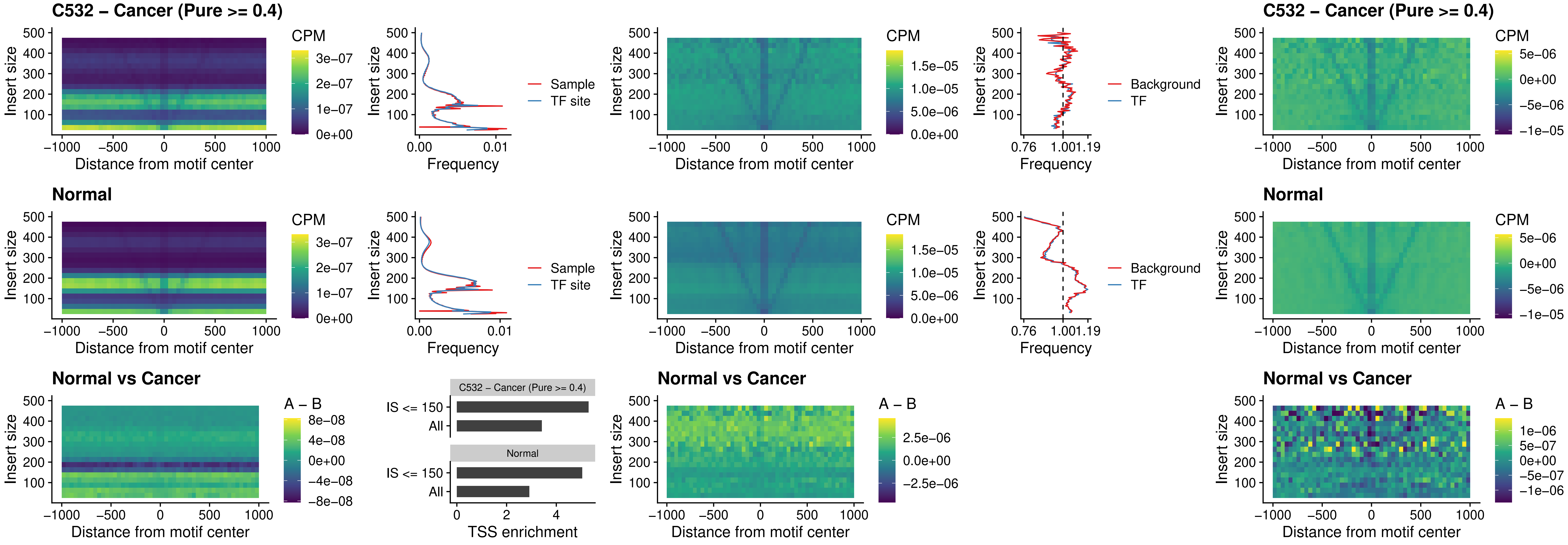

#### C532 – Cancer (Pure >= 0.4) – SHOX_uc_dTSS_nPEAK

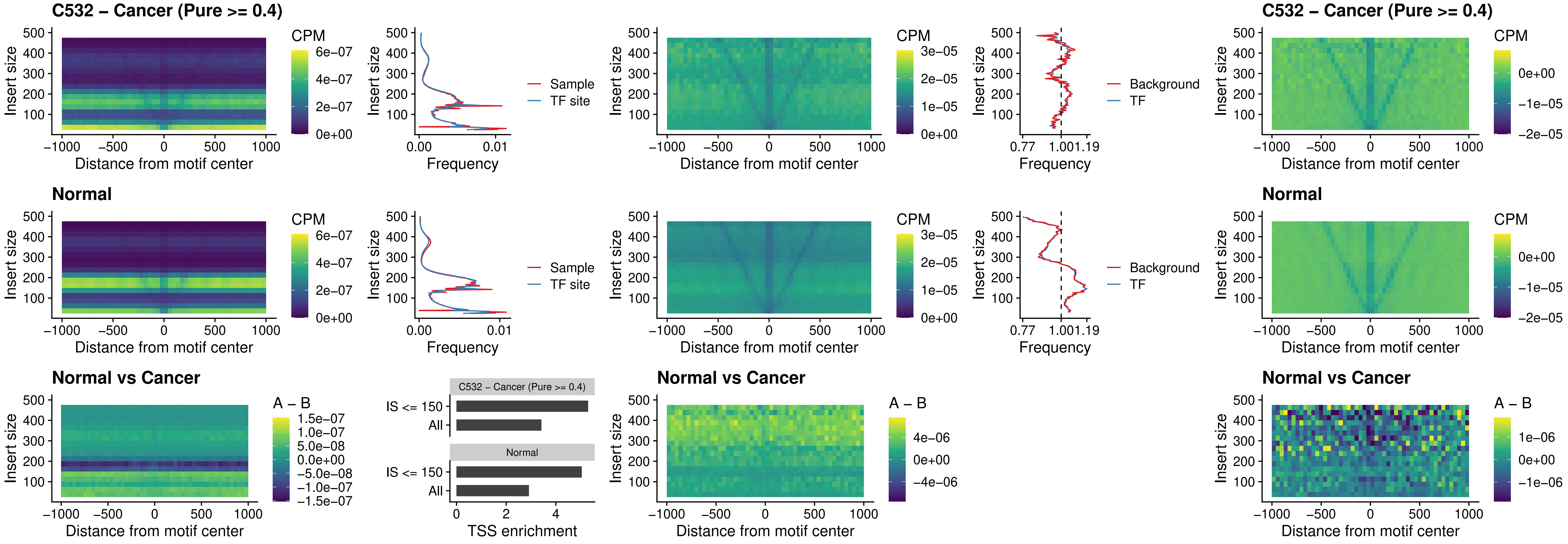

#### C532 – Cancer (Pure >= 0.4) – PRRX1_uc_dTSS_nPEAK

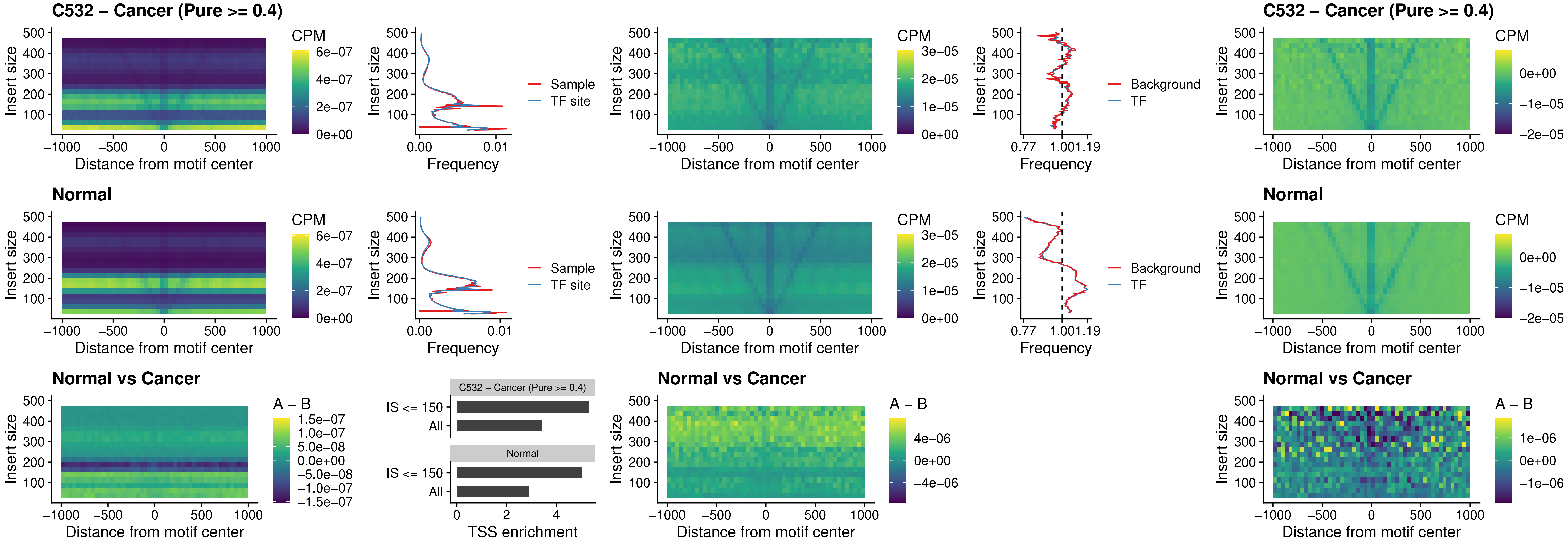

#### C532 – Cancer (Pure >= 0.4) – LMX1A_uc_dTSS_nPEAK

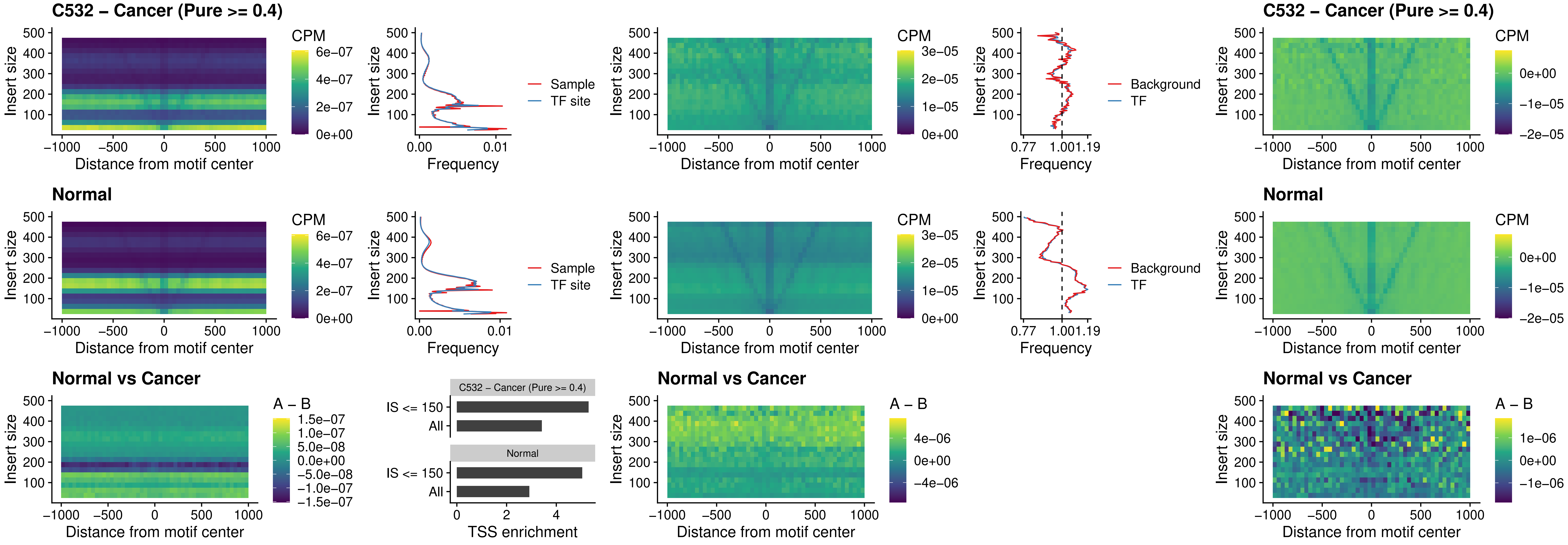

#### C532 – Cancer (Pure >= 0.4) – LMX1B_uc_dTSS_nPEAK

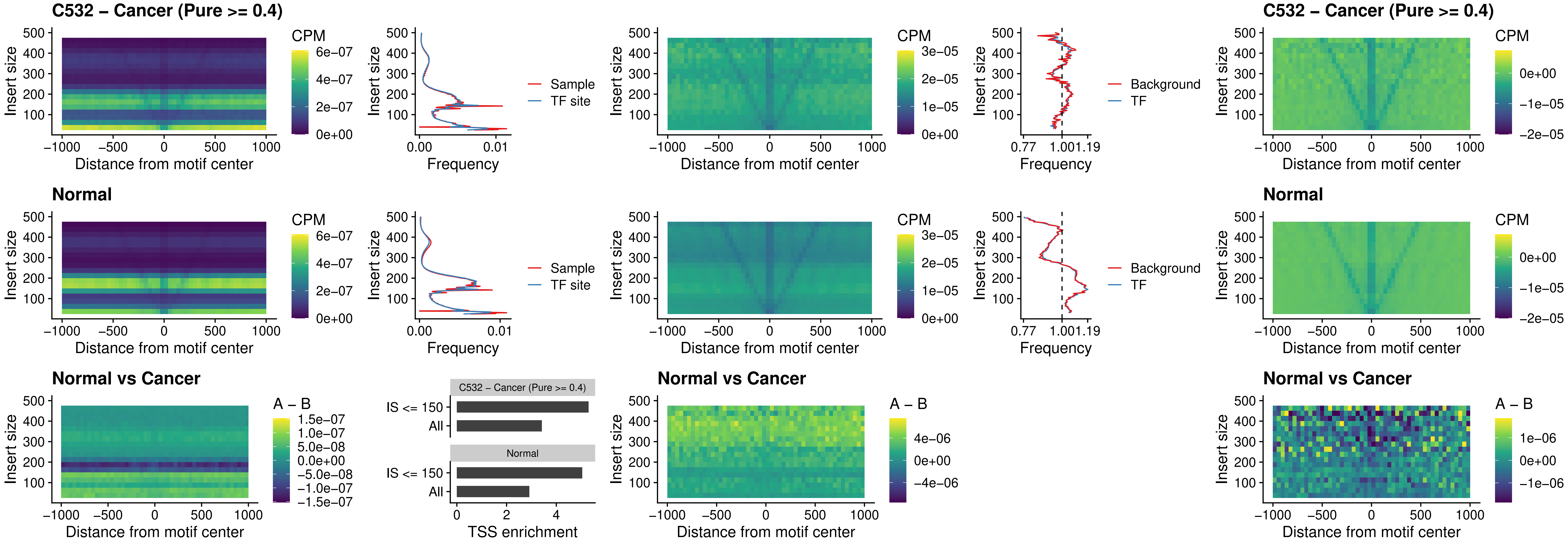

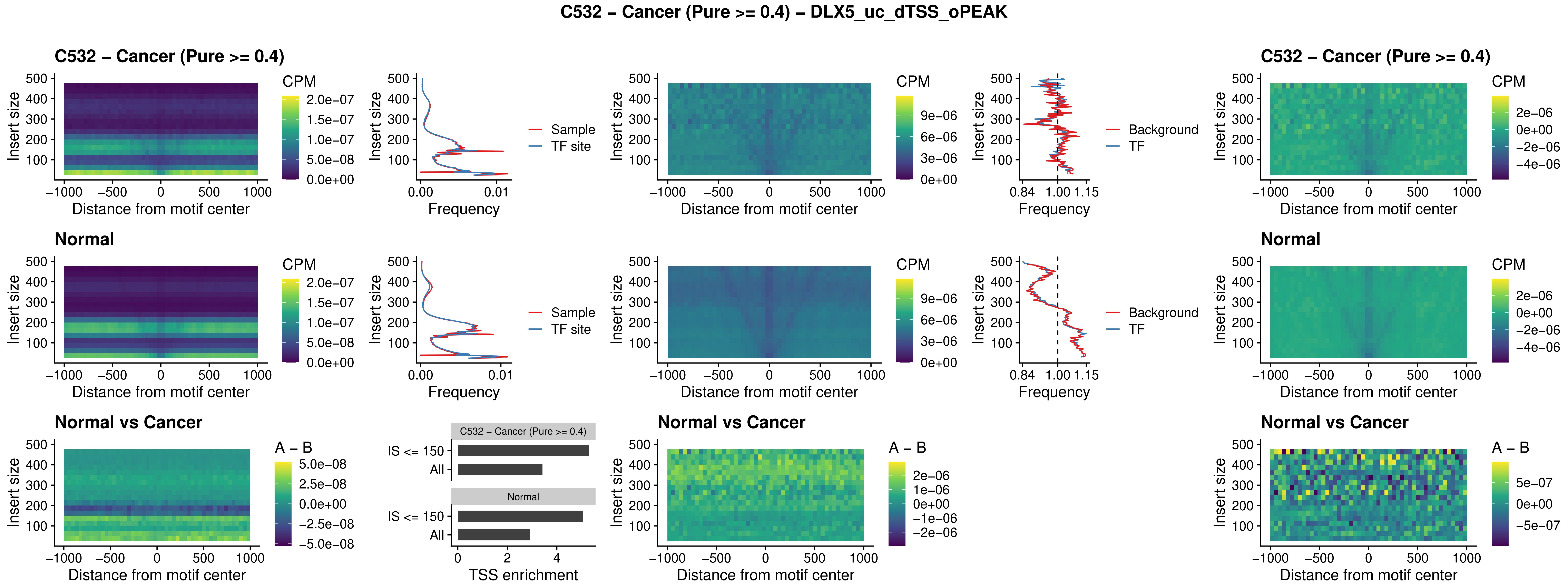
C532 – Cancer (Pure >= 0.4) – DLX5_uc_dTSS_oPEAK
C532 – Cancer (Pure >= 0.4)
Normal
Normal vs Cancer
Insert size
Distance from motif center
Frequency
CPM
A – B
Sample
TF site
Background
TF
C532 – Cancer (Pure >= 0.4)
Normal
IS <= 150
All
TSS enrichment

#### C532 – Cancer (Pure >= 0.4) – ARID3C_uc_dTSS_nPEAK

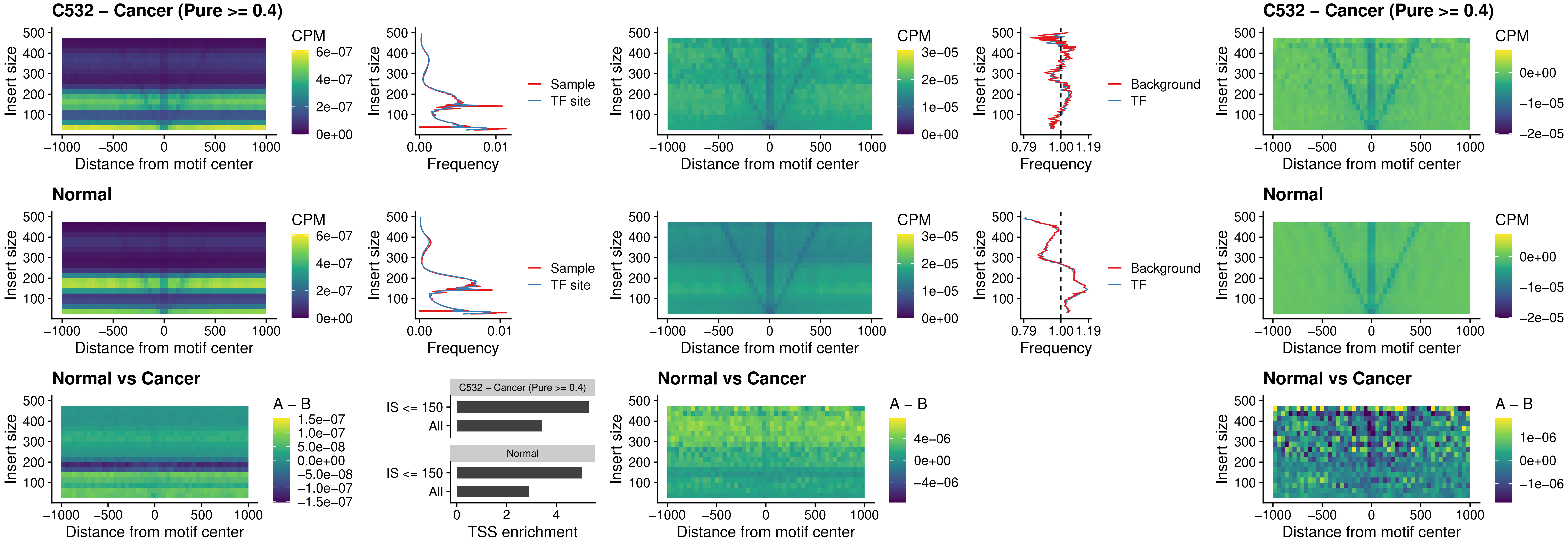

#### C532 – Cancer (Pure >= 0.4) – HOXD12_uc_dTSS_nPEAK

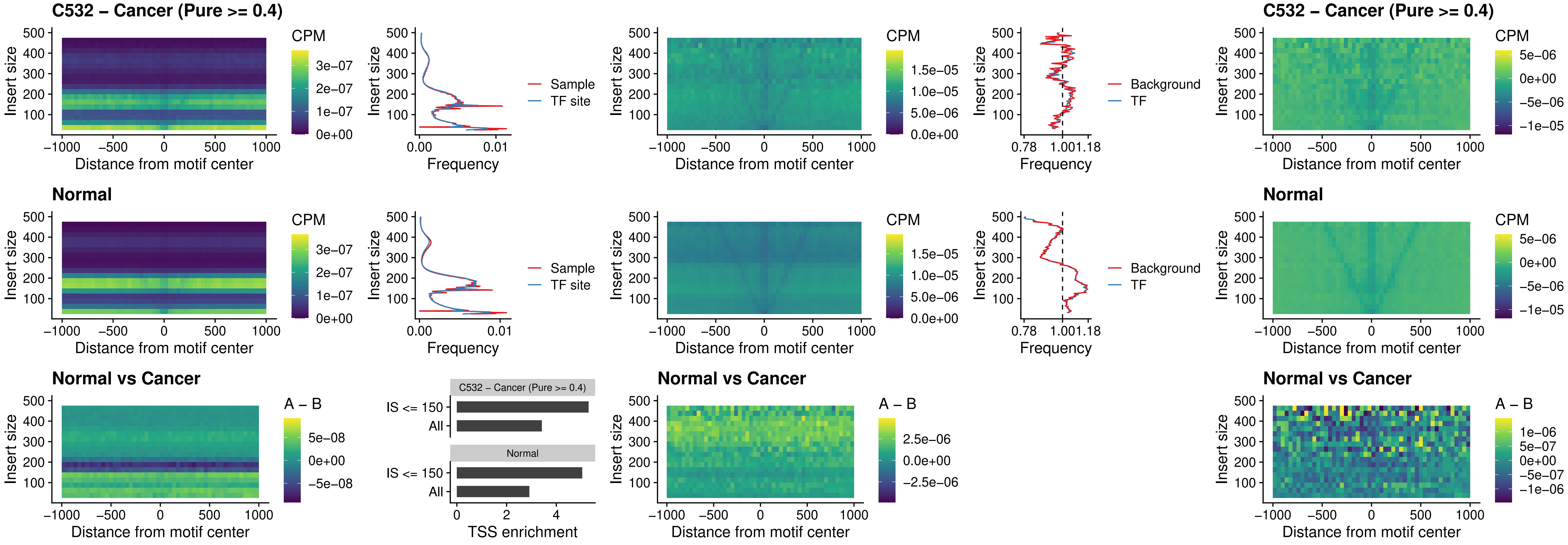

#### C532 – Cancer (Pure >= 0.4) – CDX2_uc_dTSS_nPEAK

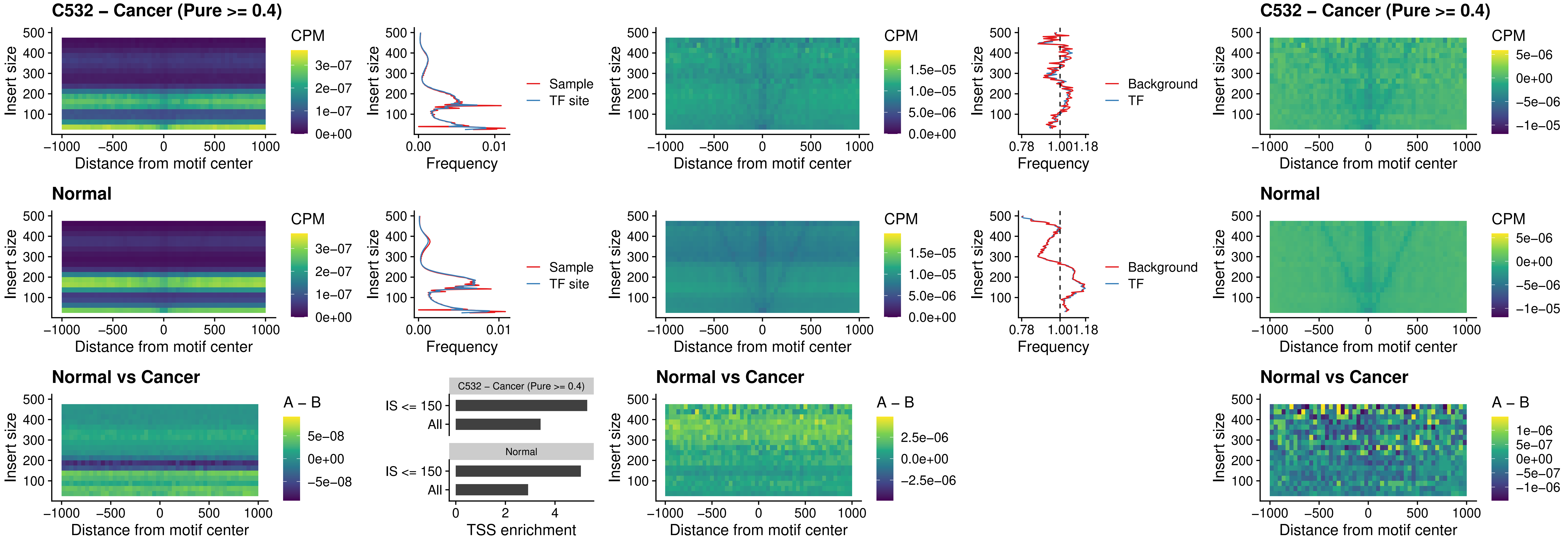

#### C532 – Cancer (Pure >= 0.4) – HOXC12_uc_dTSS_nPEAK

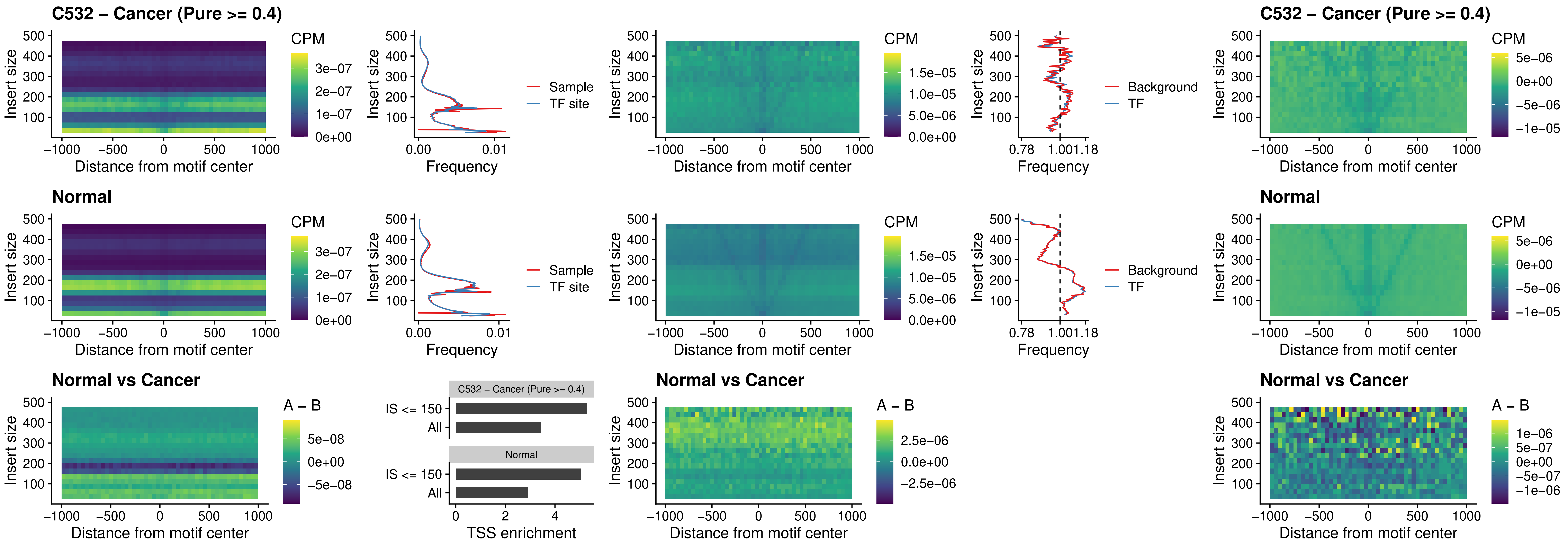

#### C532 – Cancer (Pure >= 0.4) – HHEX_uc_dTSS_oPEAK

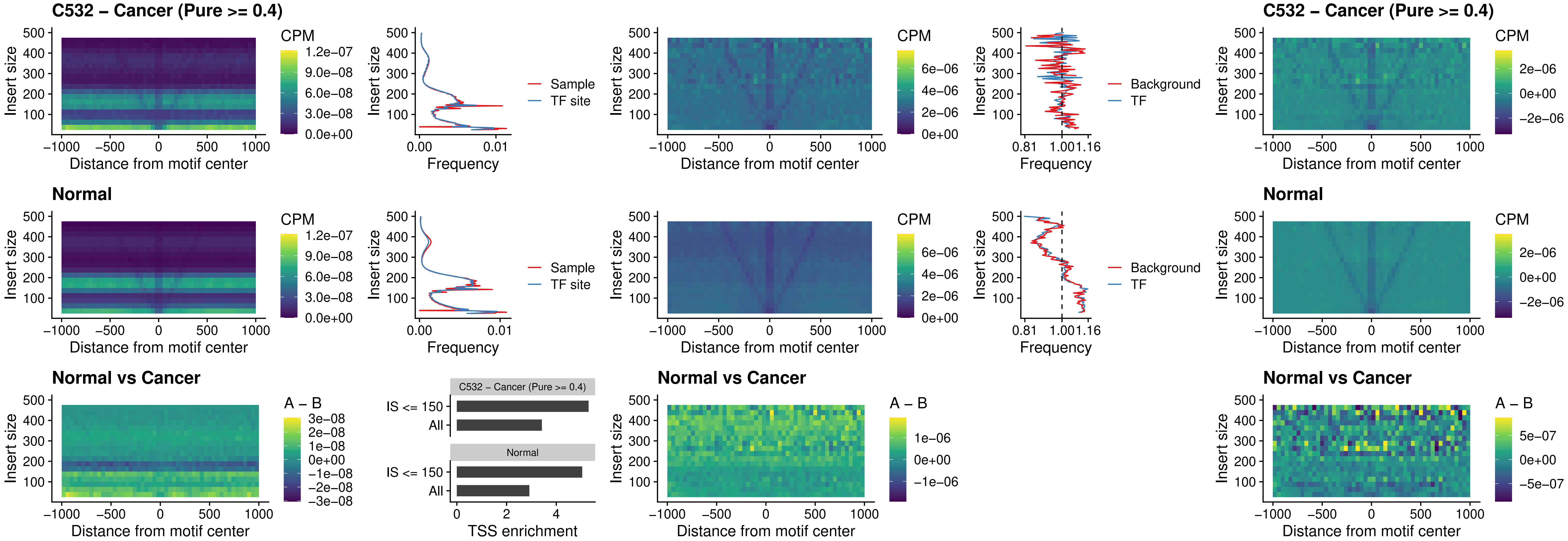

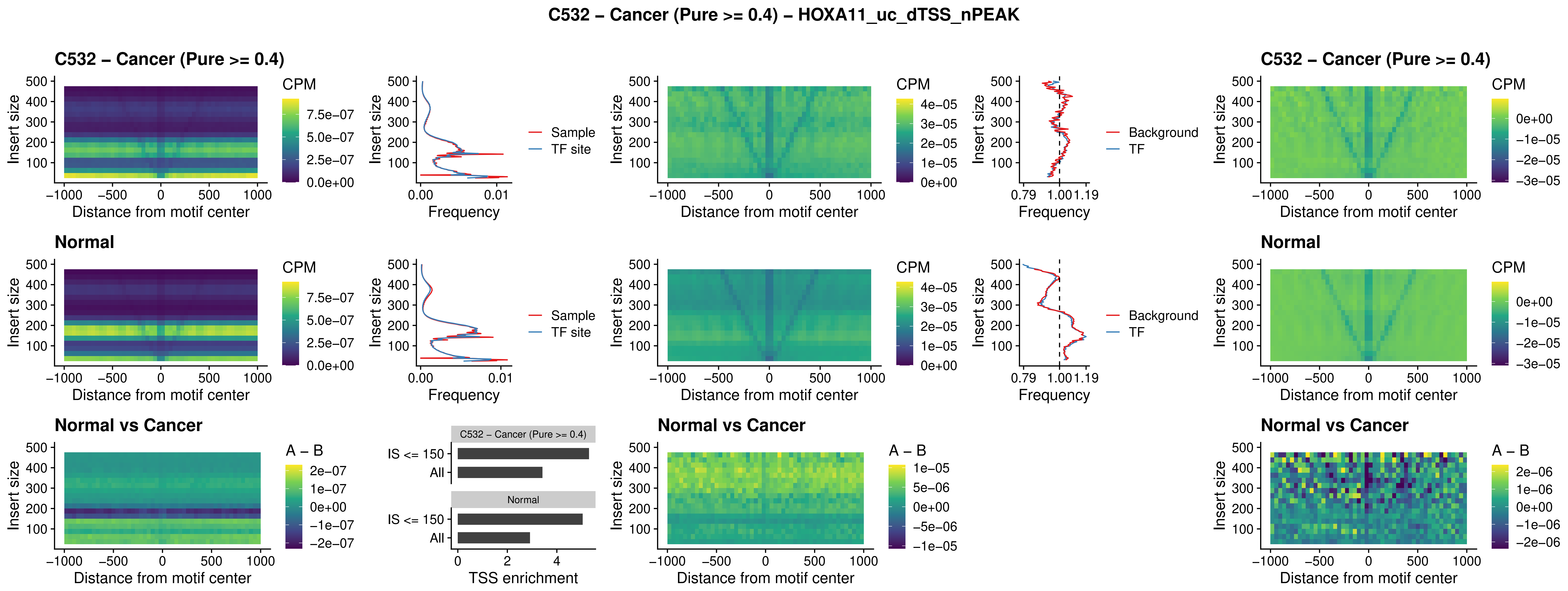
C532 – Cancer (Pure >= 0.4) – HOXA11_uc_dTSS_nPEAK
C532 – Cancer (Pure >= 0.4)
Normal
Normal vs Cancer
Insert size
Distance from motif center
Frequency
CPM
A – B
Sample
TF site
Background
TF
C532 – Cancer (Pure >= 0.4)
Normal
IS <= 150
All
TSS enrichment

#### C532 – Cancer (Pure >= 0.4) – HOXA11_uc_cTSS_nPEAK

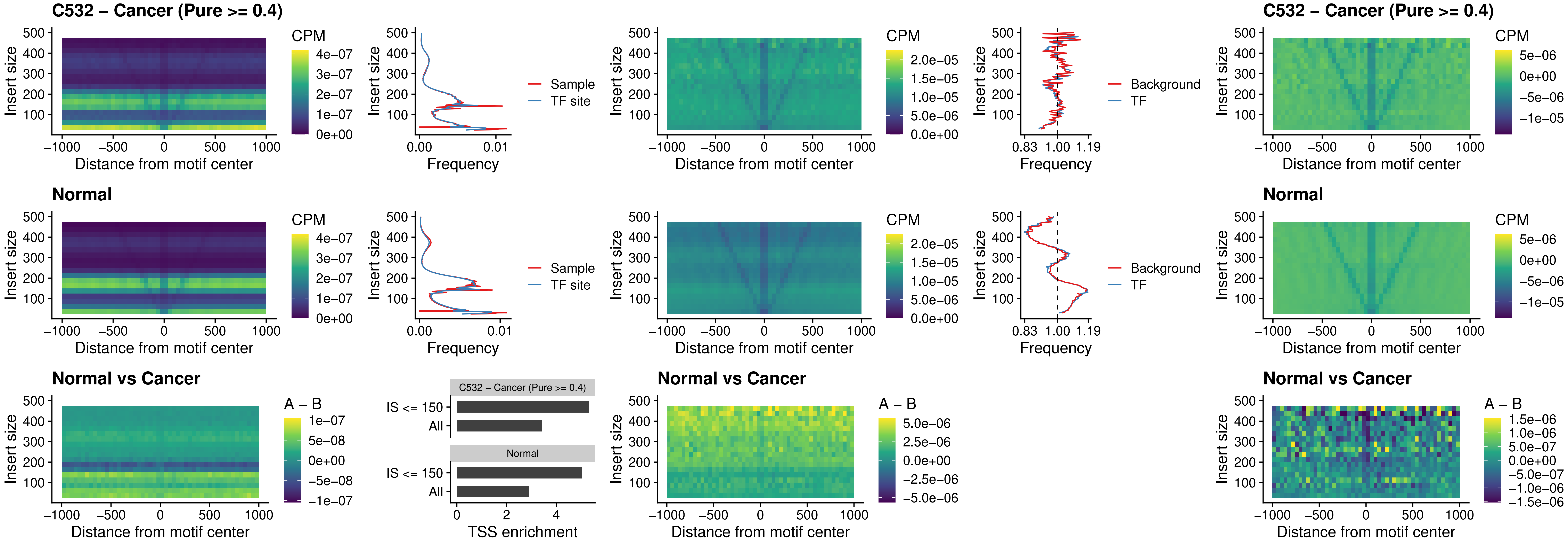

### C532 – Cancer (Pure >= 0.4) – ZFHX3_uc_dTSS_nPEAK

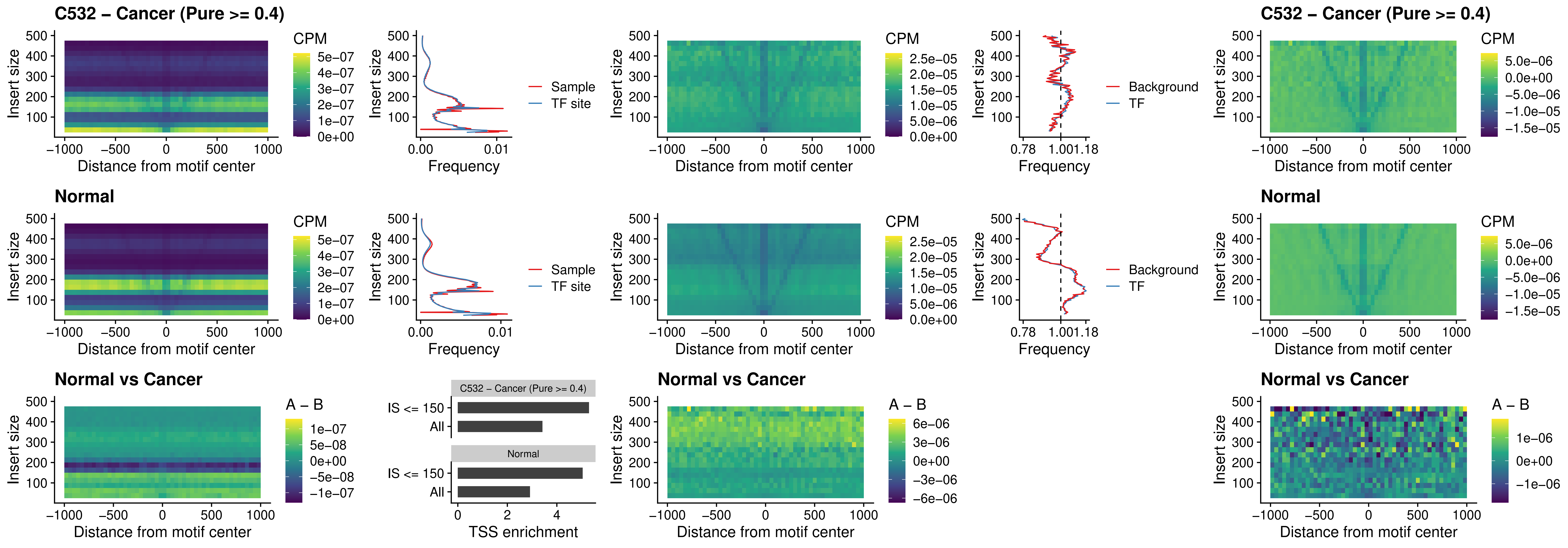

### C532 – Cancer (Pure >= 0.4) – HOXB4_uc_dTSS_nPEAK

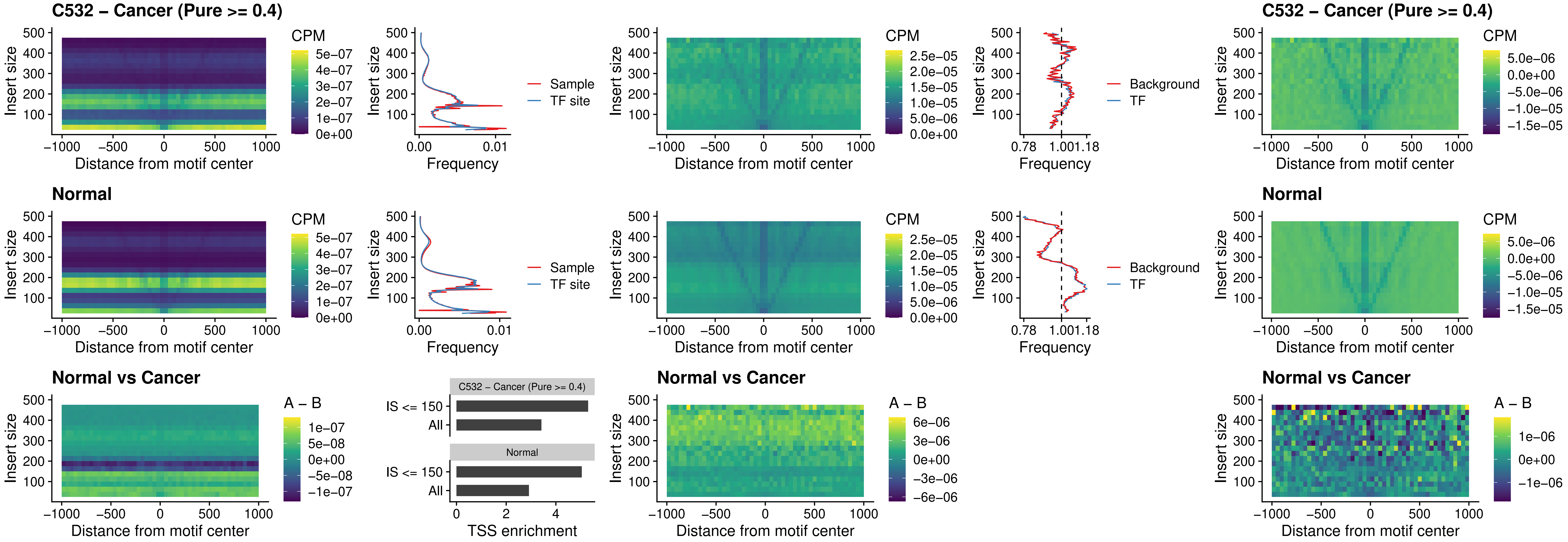

#### C532 – Cancer (Pure >= 0.4) – HOXC5_uc_dTSS_nPEAK

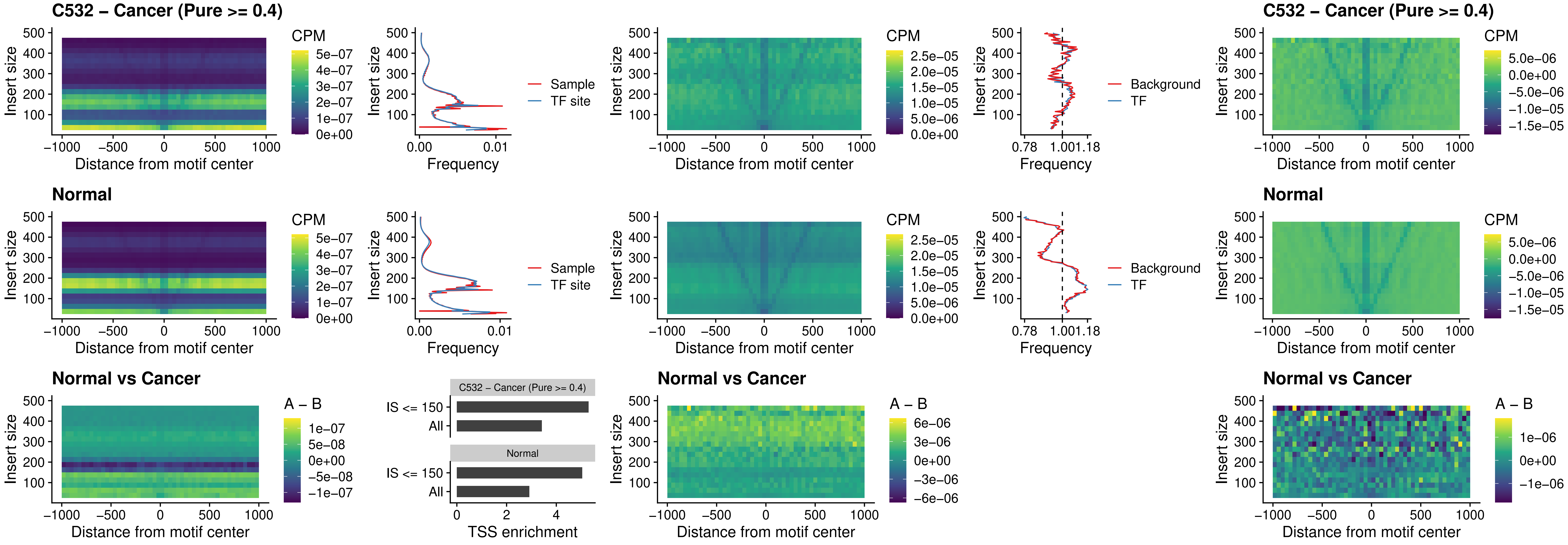

#### C532 – Cancer (Pure >= 0.4) – HOXD1_uc_dTSS_nPEAK

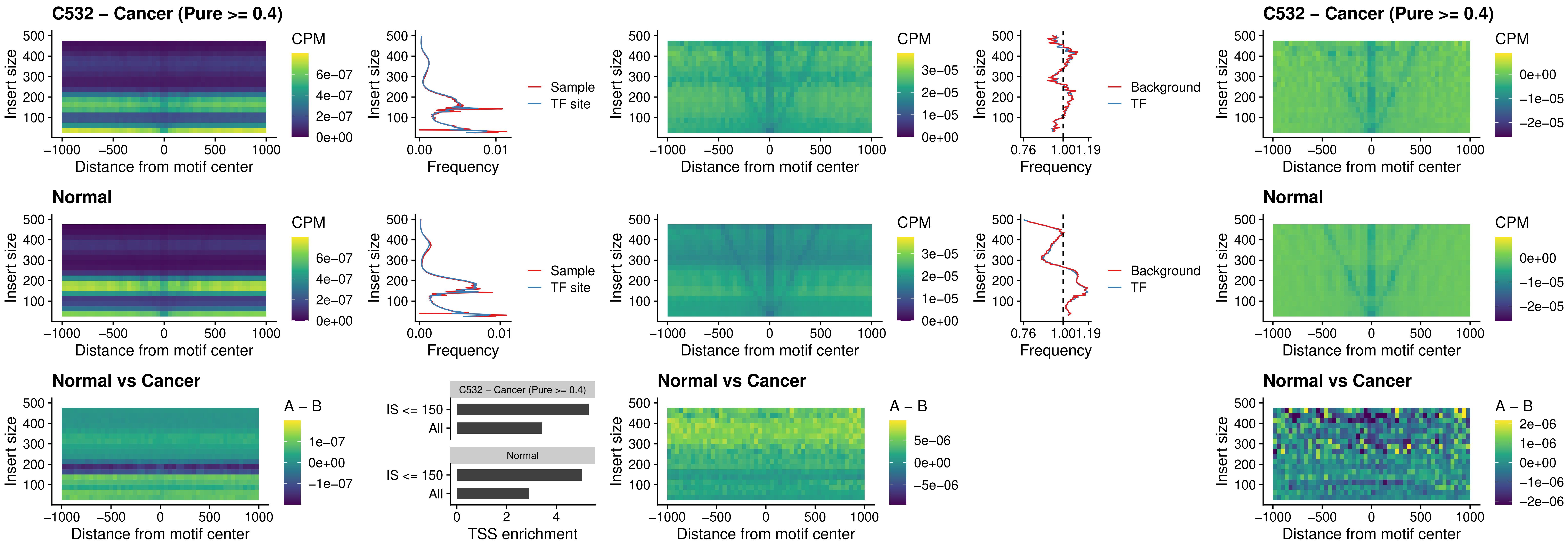

#### C532 – Cancer (Pure >= 0.4) – OTP_uc_dTSS_nPEAK

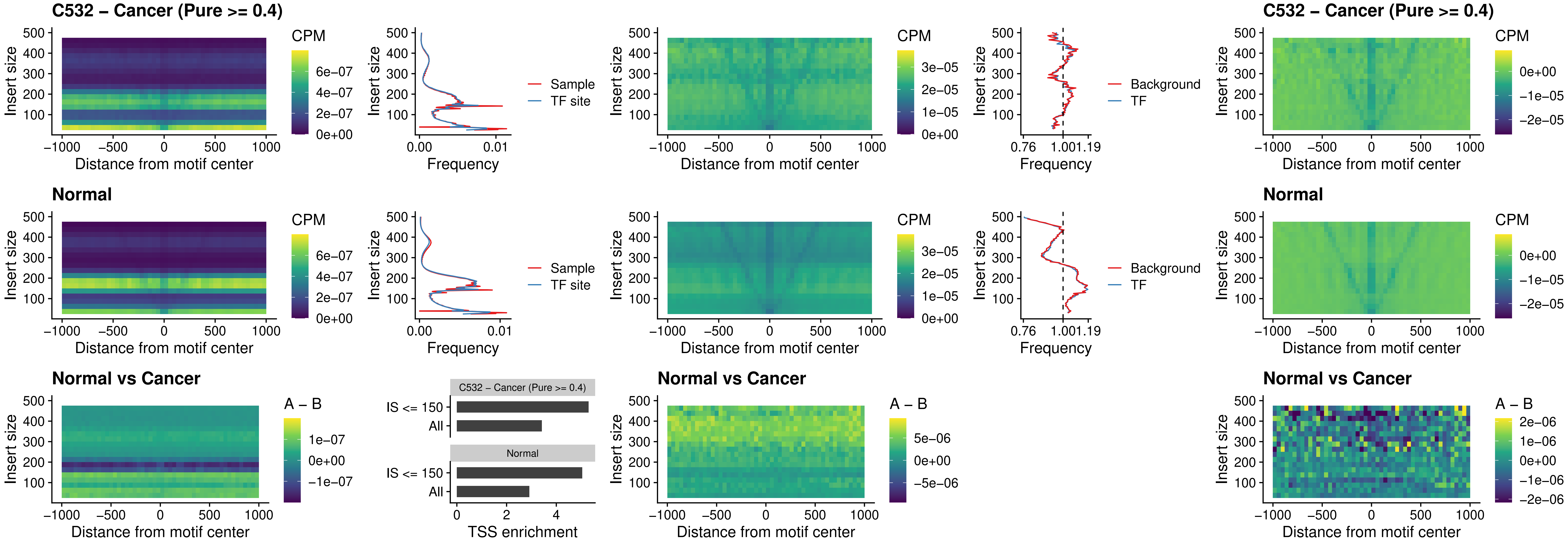

### C532 – Cancer (Pure >= 0.4) – FOXL1_uc_dTSS_oPEAK

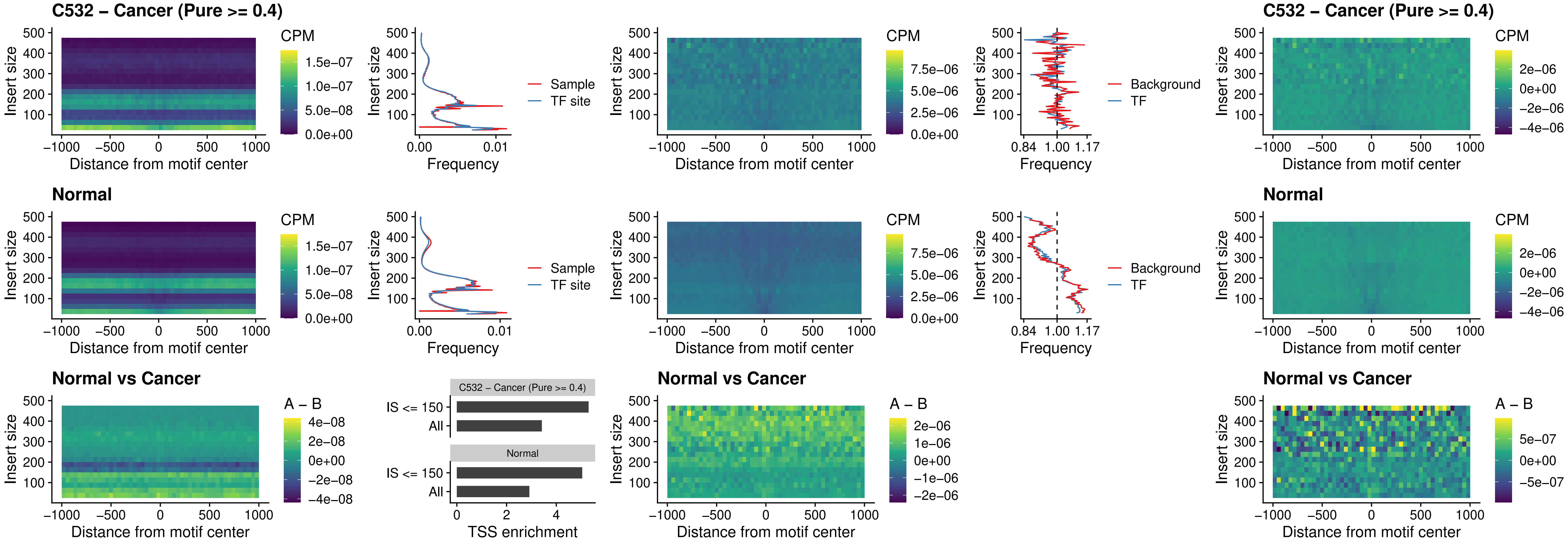

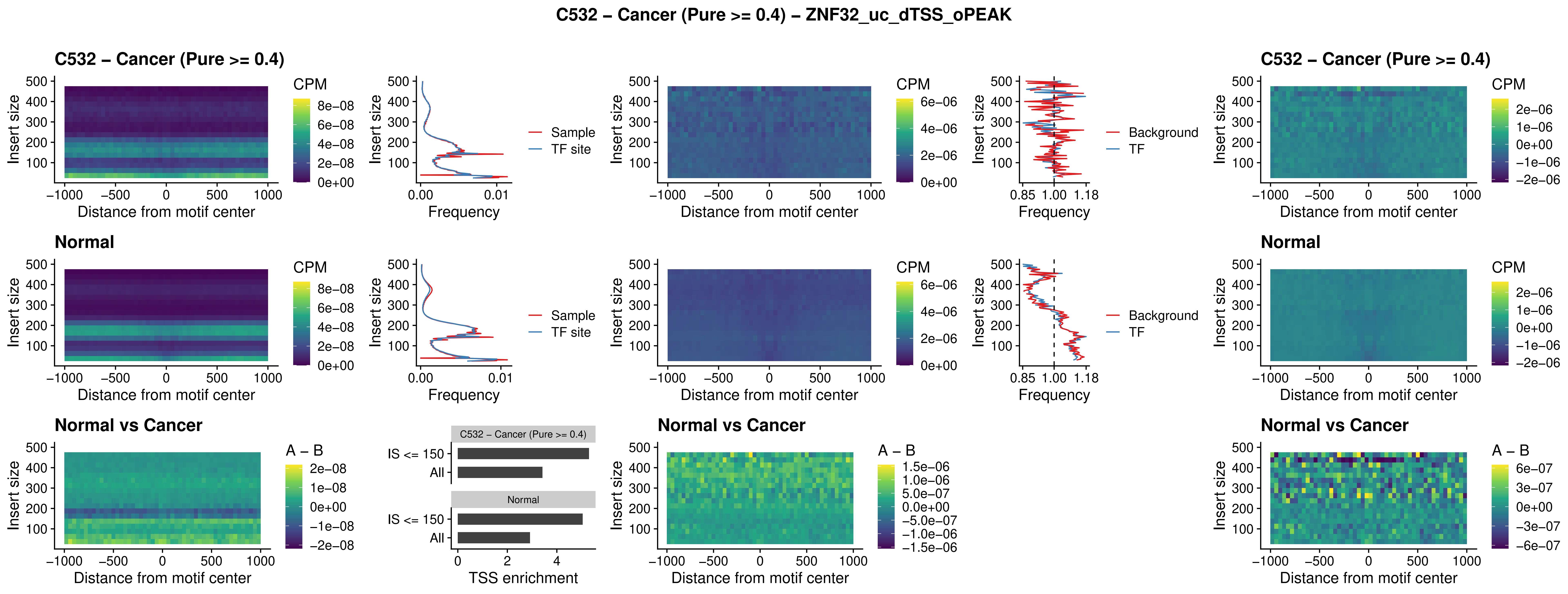
C532 – Cancer (Pure >= 0.4) – ZNF32_uc_dTSS_oPEAK
C532 – Cancer (Pure >= 0.4)
Normal
Normal vs Cancer
Insert size
Distance from motif center
CPM
A – B
Frequency
Sample
TF site
Background
TF
C532 – Cancer (Pure >= 0.4)
Normal
IS <= 150
All
TSS enrichment

#### C532 – Cancer (Pure >= 0.4) – FOXL1_uc_dTSS_nPEAK

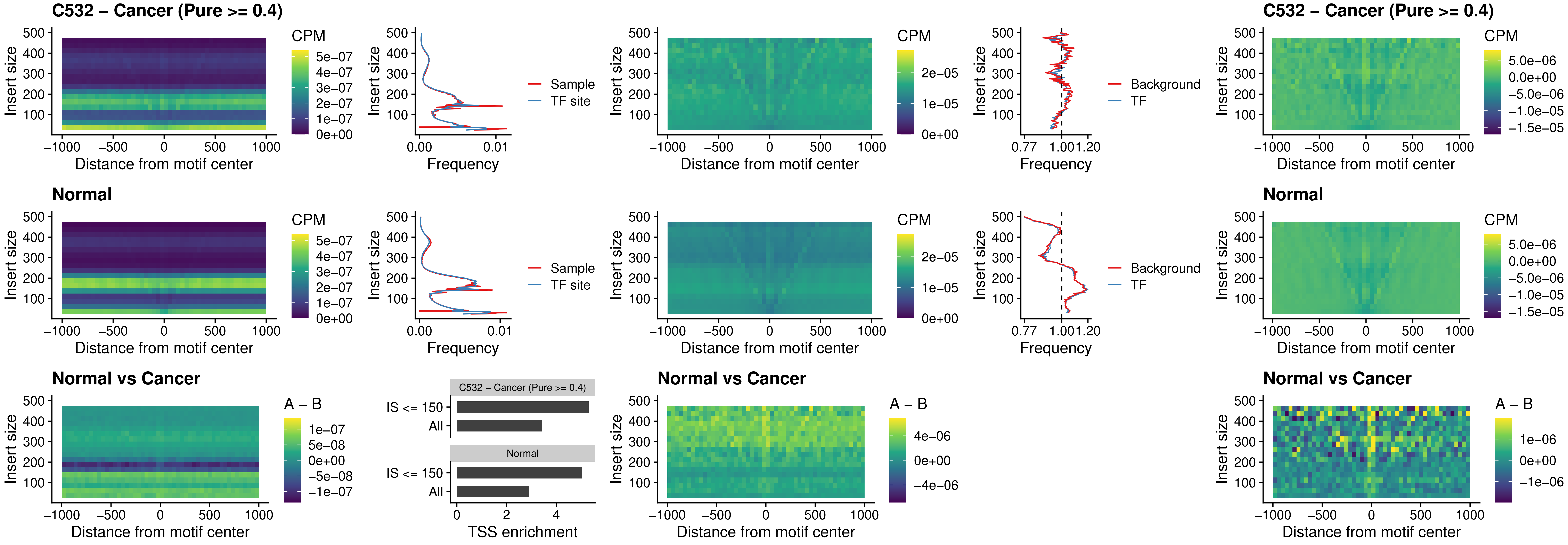

### C532 – Cancer (Pure >= 0.4) – ZNF32_uc_dTSS_nPEAK

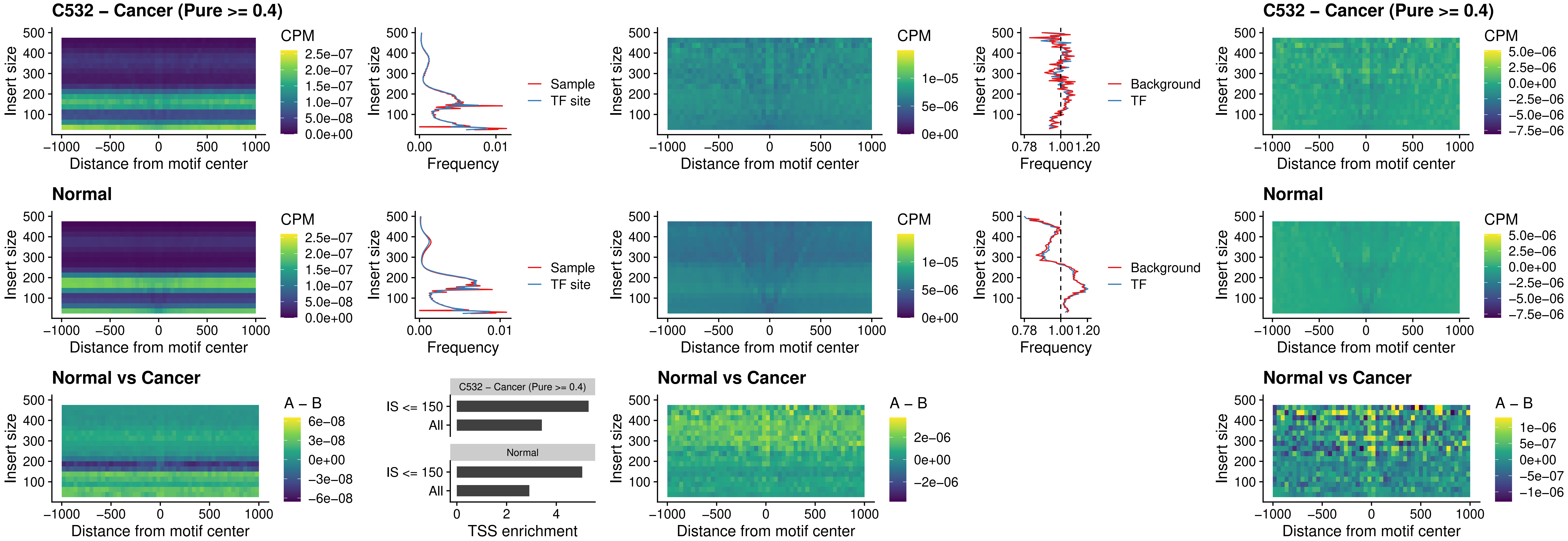

#### C532 – Cancer (Pure >= 0.4) – DLX5_uc_dTSS_nPEAK

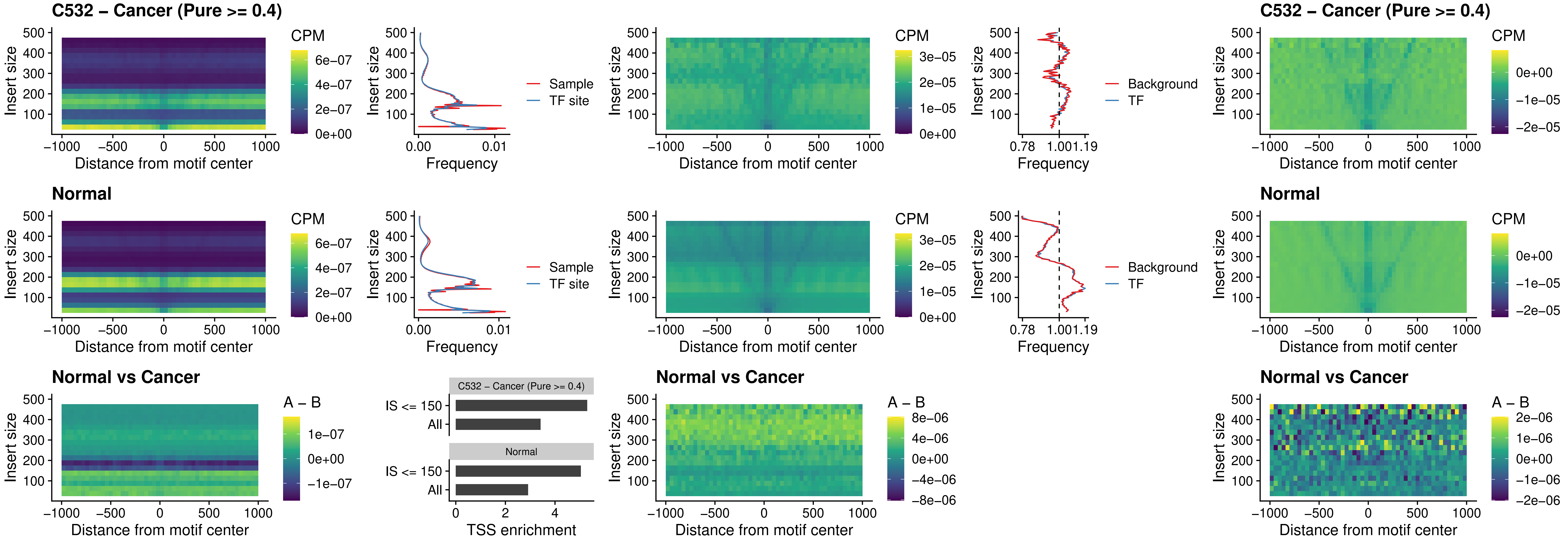

#### C532 – Cancer (Pure >= 0.4) – DMRT2_uc_dTSS_nPEAK

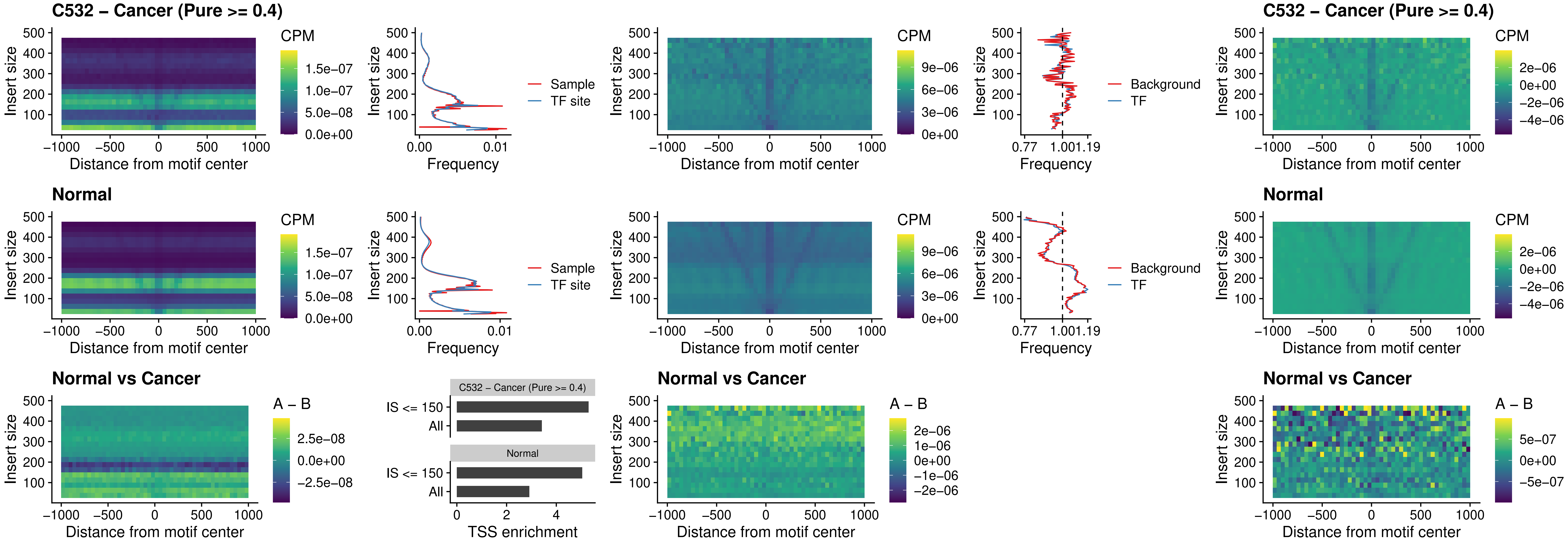

#### C532 – Cancer (Pure >= 0.4) – DMRTC2_uc_cTSS_nPEAK

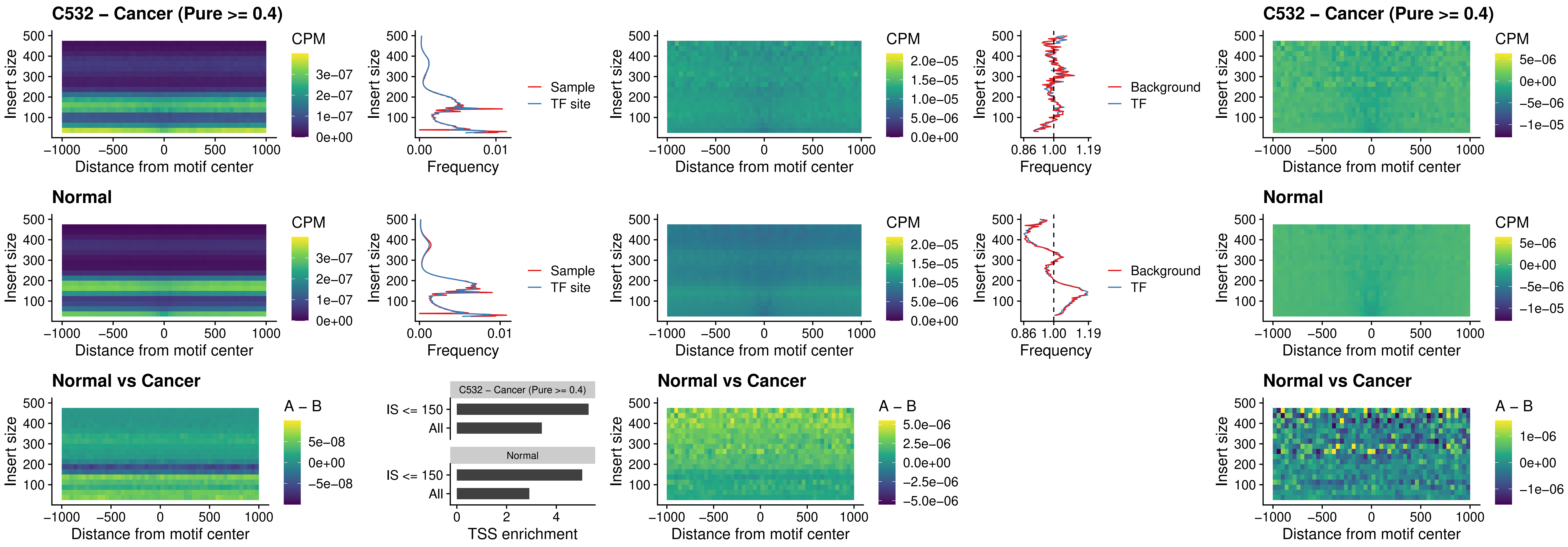

#### C532 – Cancer (Pure >= 0.4) – HOXD9_uc_dTSS_nPEAK

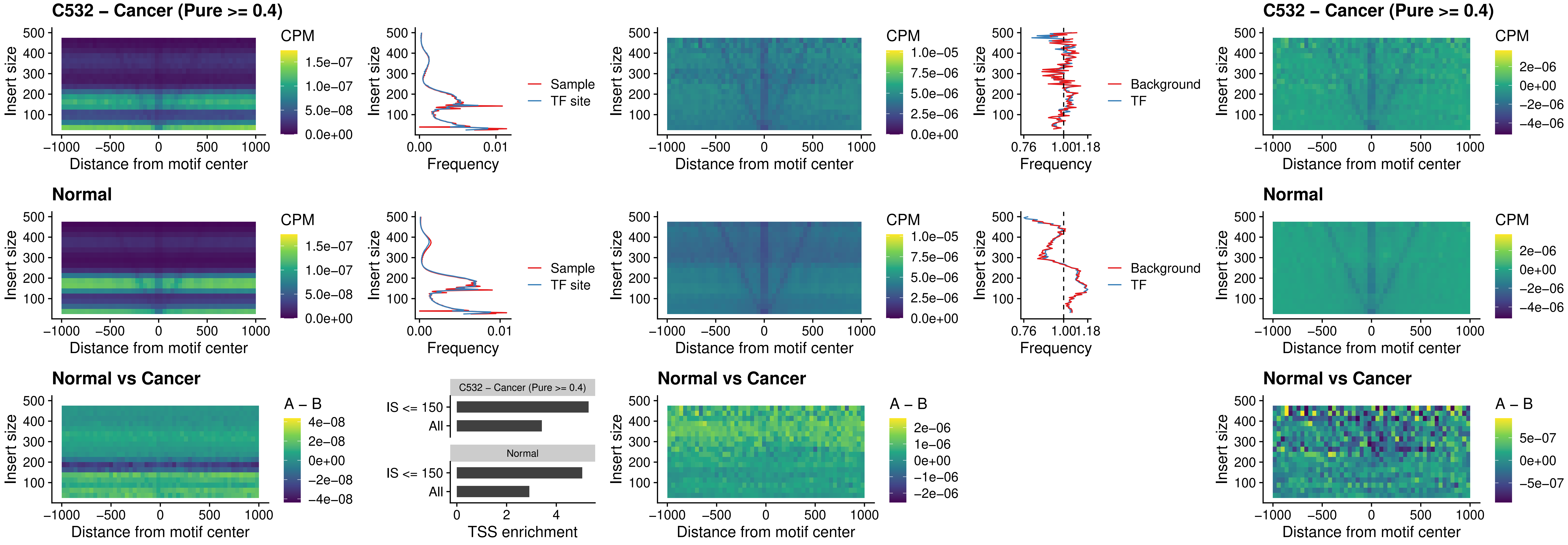

#### C532 – Cancer (Pure >= 0.4) – HOXD4_uc_dTSS_nPEAK

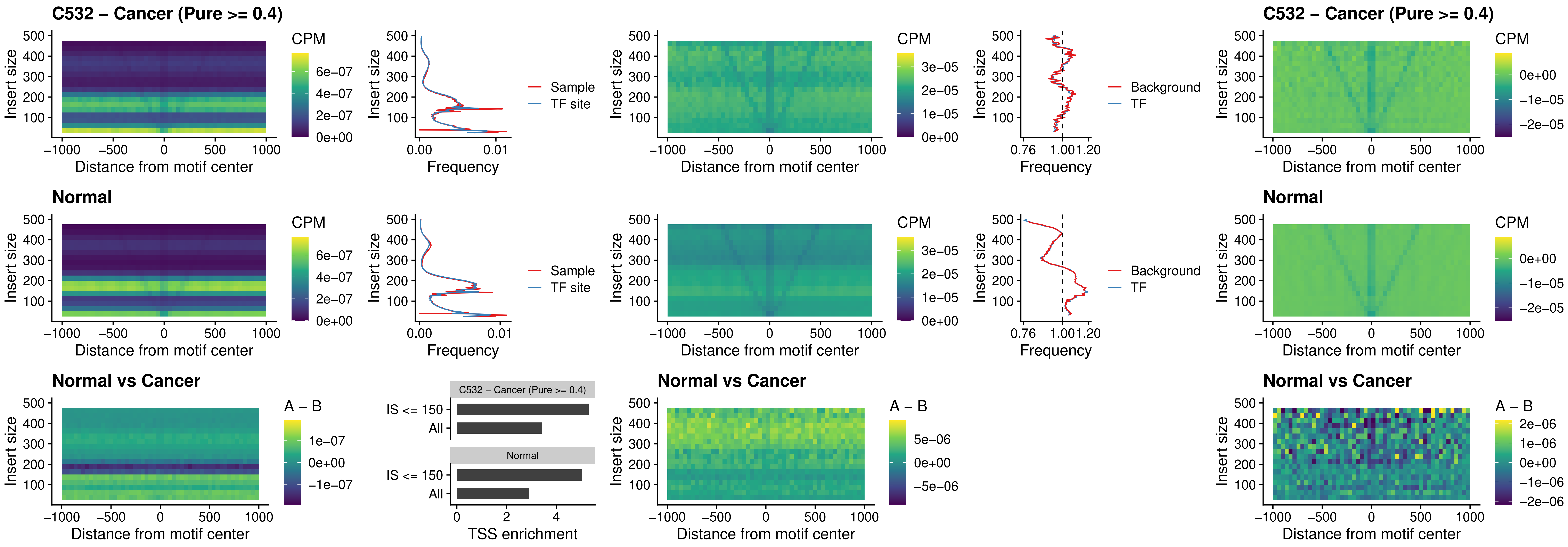

#### C532 – Cancer (Pure >= 0.4) – FOXP3_uc_cTSS_nPEAK

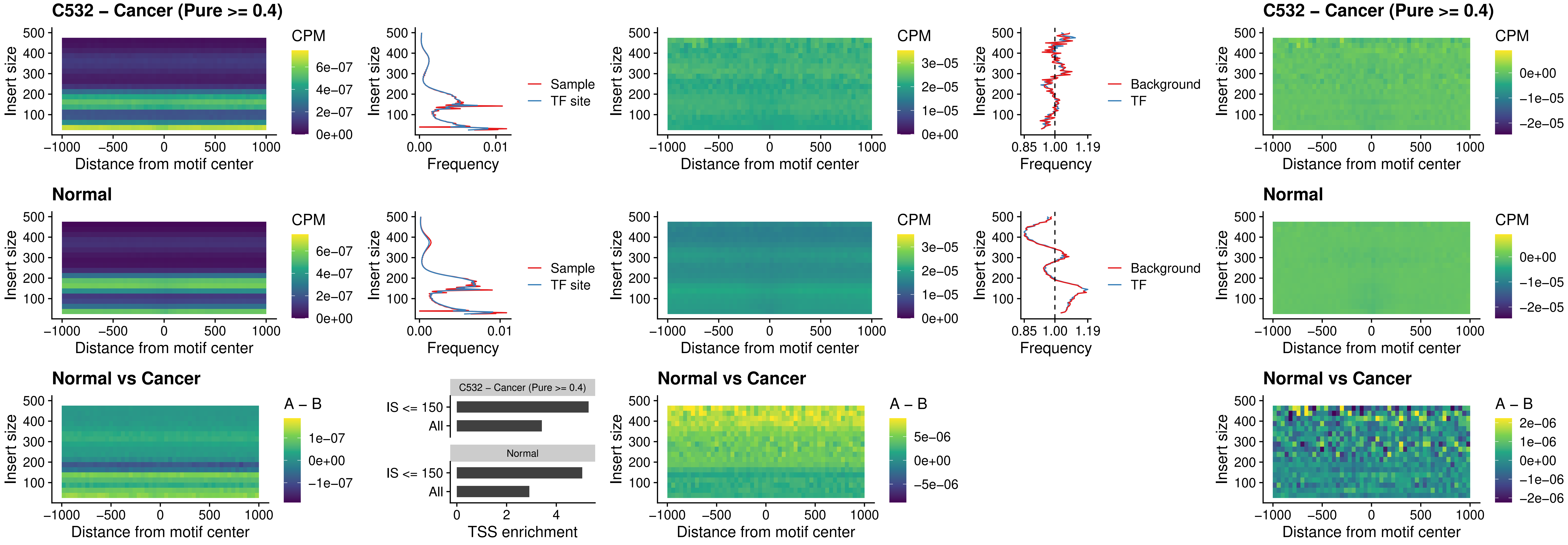

### C532 – Cancer (Pure >= 0.4) – FOXD4L6_uc_cTSS_nPEAK

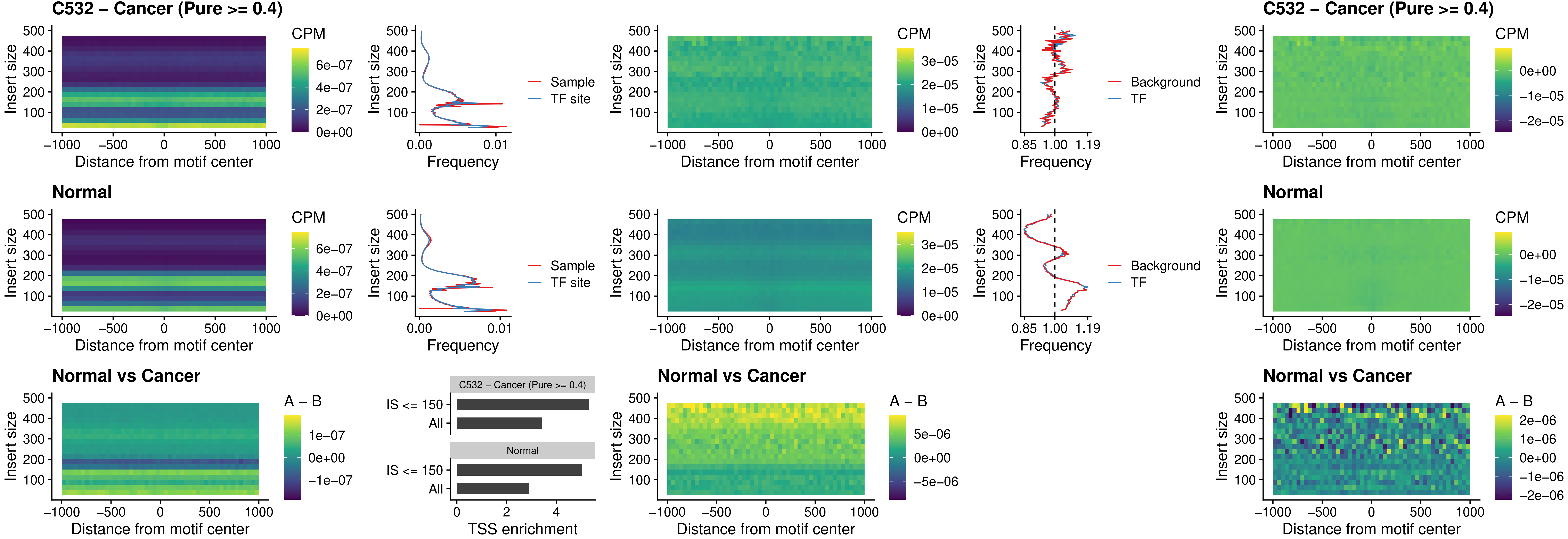

#### C532 – Cancer (Pure >= 0.4) – FOXD4L5_uc_cTSS_nPEAK

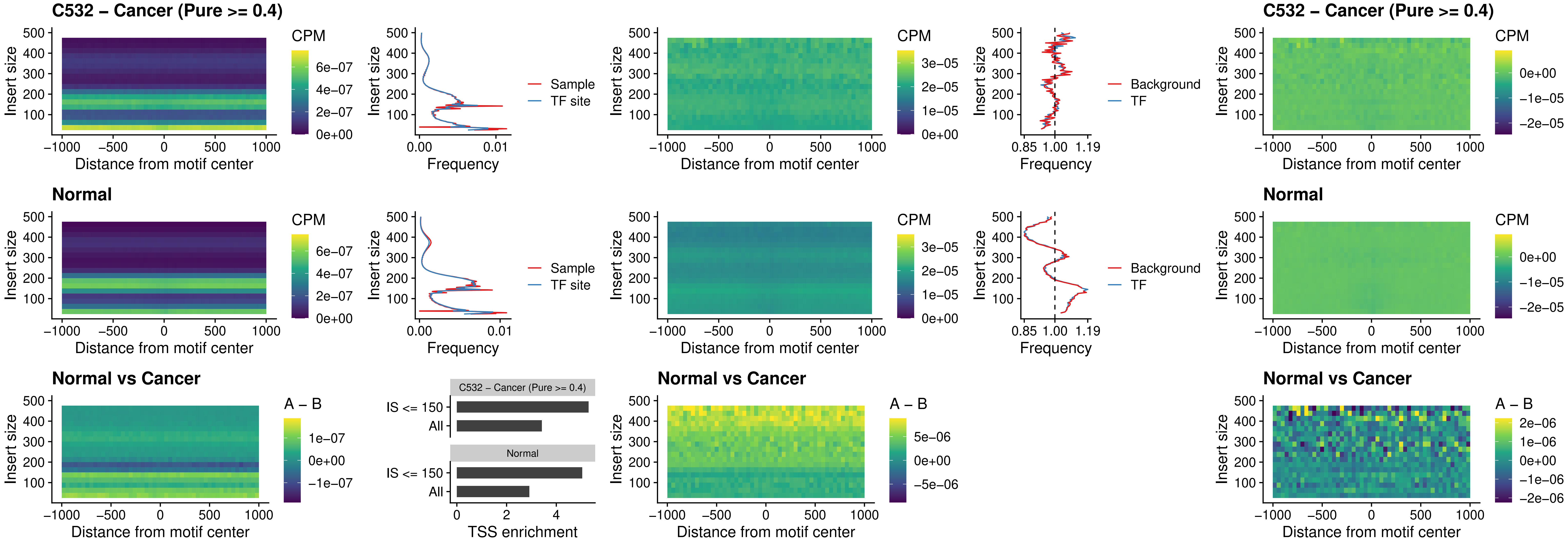

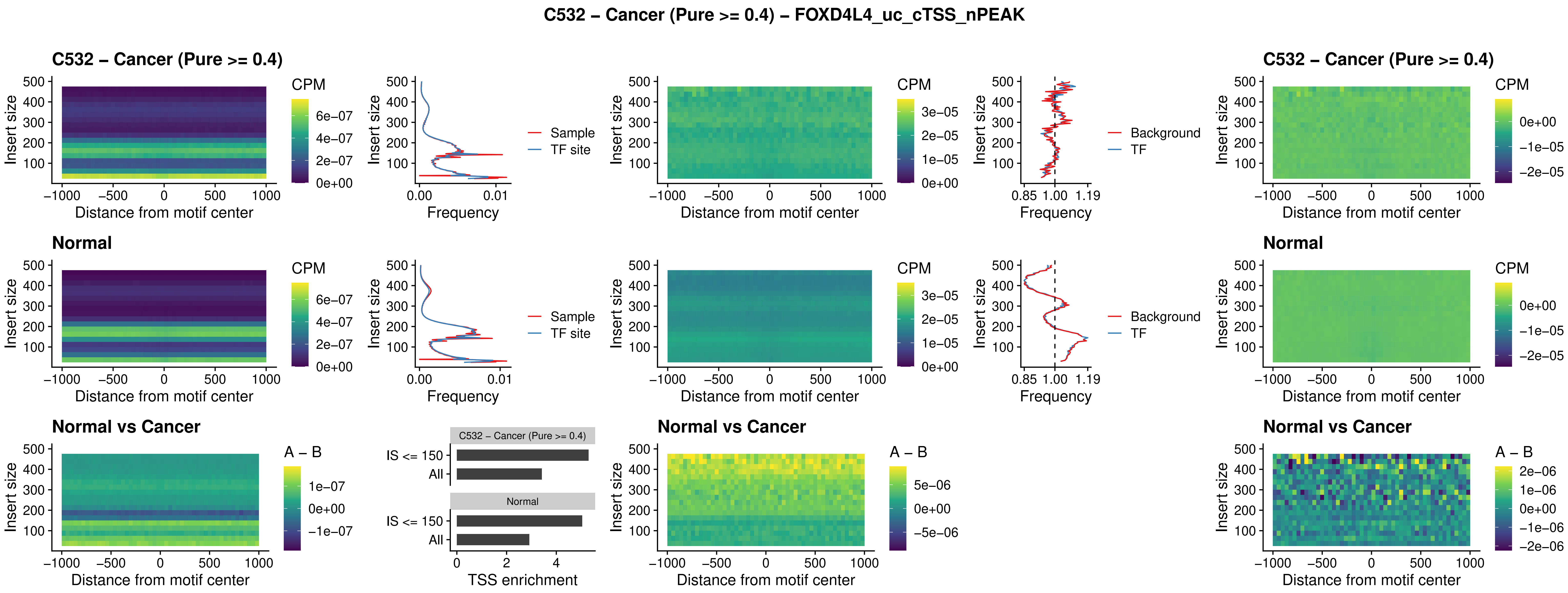
C532 – Cancer (Pure >= 0.4) – FOXD4L4_uc_cTSS_nPEAK
C532 – Cancer (Pure >= 0.4)
Normal
Normal vs Cancer
Insert size
Distance from motif center
Frequency
CPM
A – B
Sample
TF site
Background
TF
C532 – Cancer (Pure >= 0.4)
Normal
IS <= 150
All
TSS enrichment

#### C532 – Cancer (Pure >= 0.4) – FOXD4L2_uc_cTSS_nPEAK

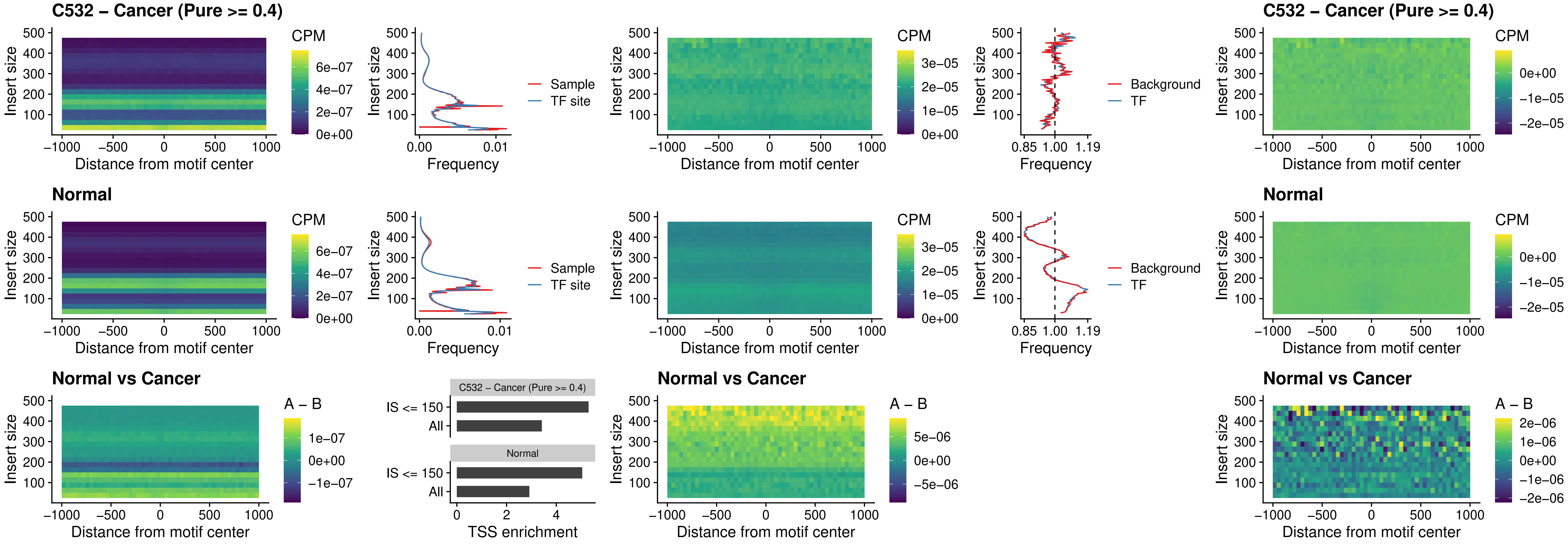

### C532 – Cancer (Pure >= 0.4) – FOXD4L3_uc_cTSS_nPEAK

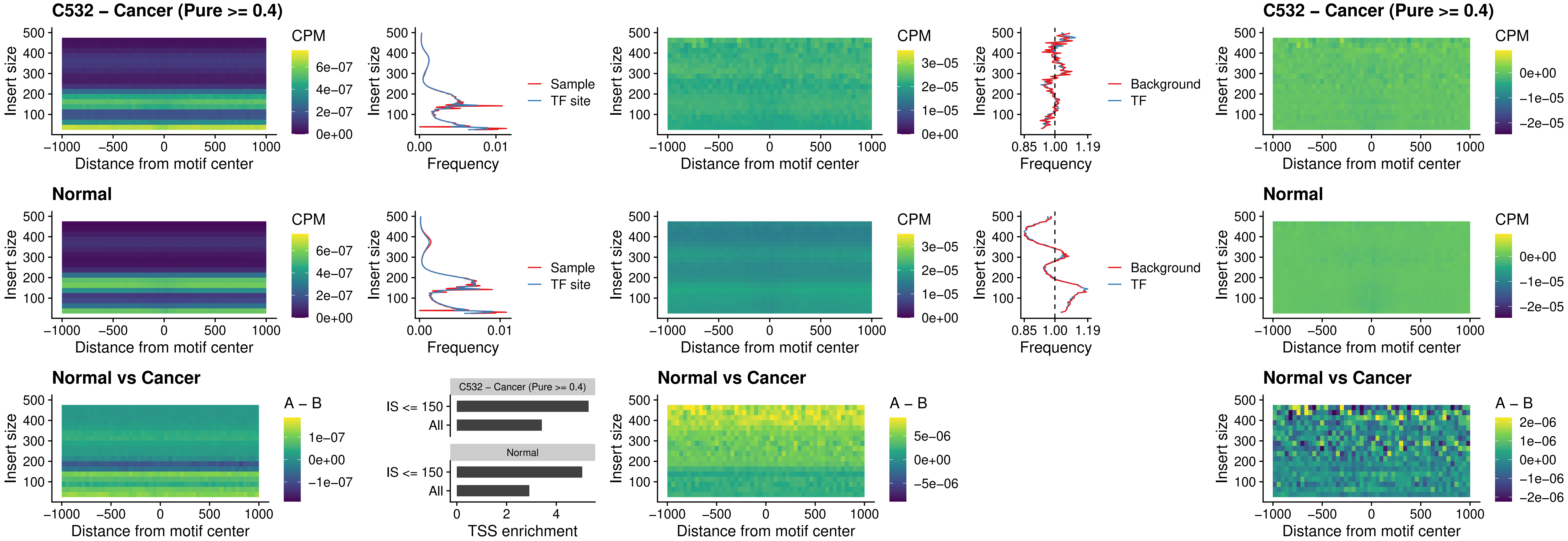

#### C532 – Cancer (Pure >= 0.4) – HOXD1_uc_dTSS_oPEAK

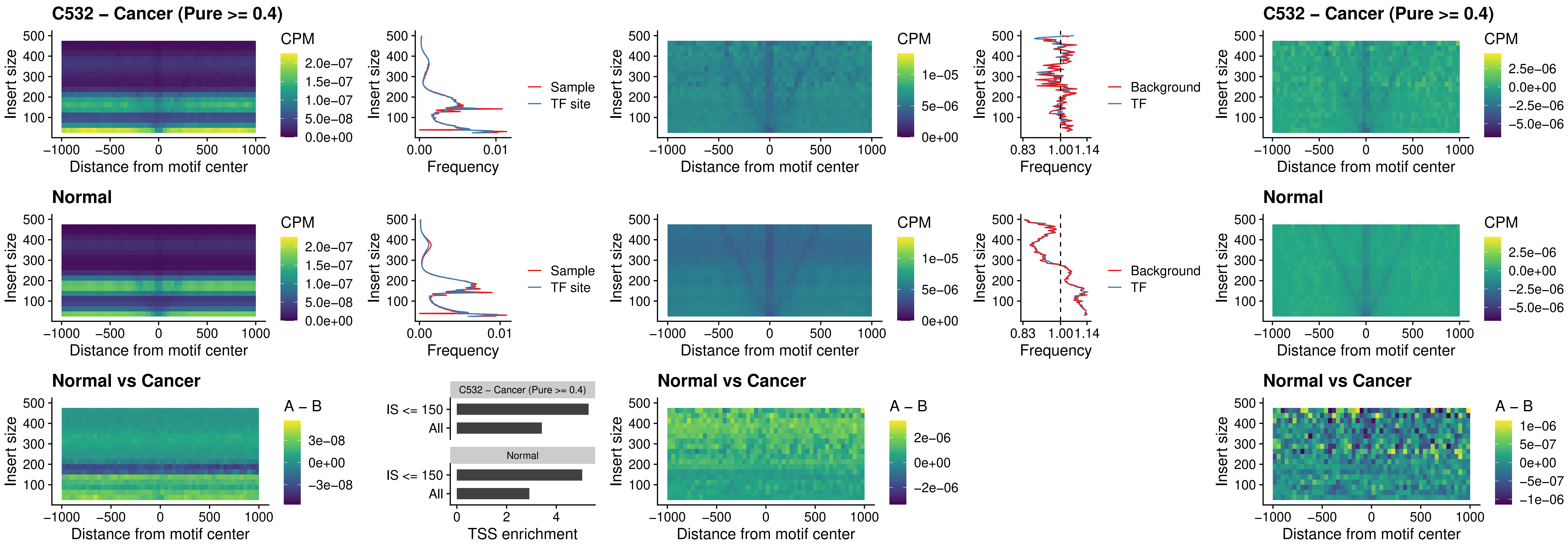

#### C532 – Cancer (Pure >= 0.4) – VAX1_uc_pTSS_nPEAK

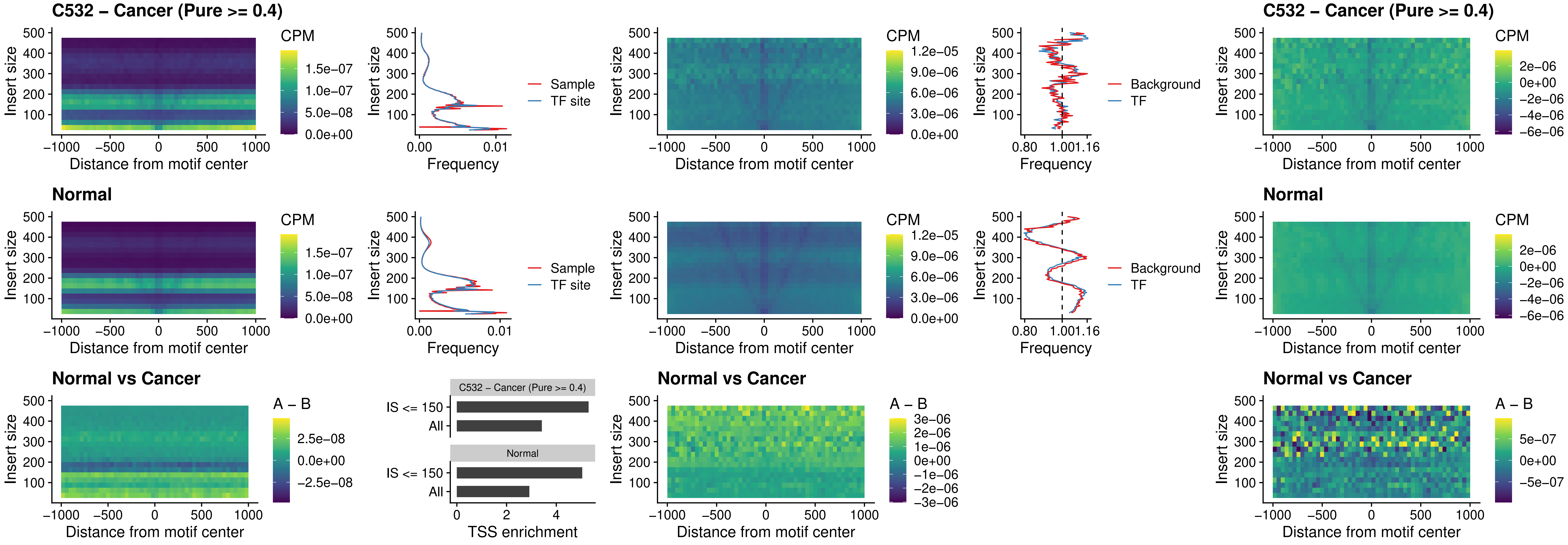

#### C532 – Cancer (Pure >= 0.4) – DLX3_uc_pTSS_nPEAK

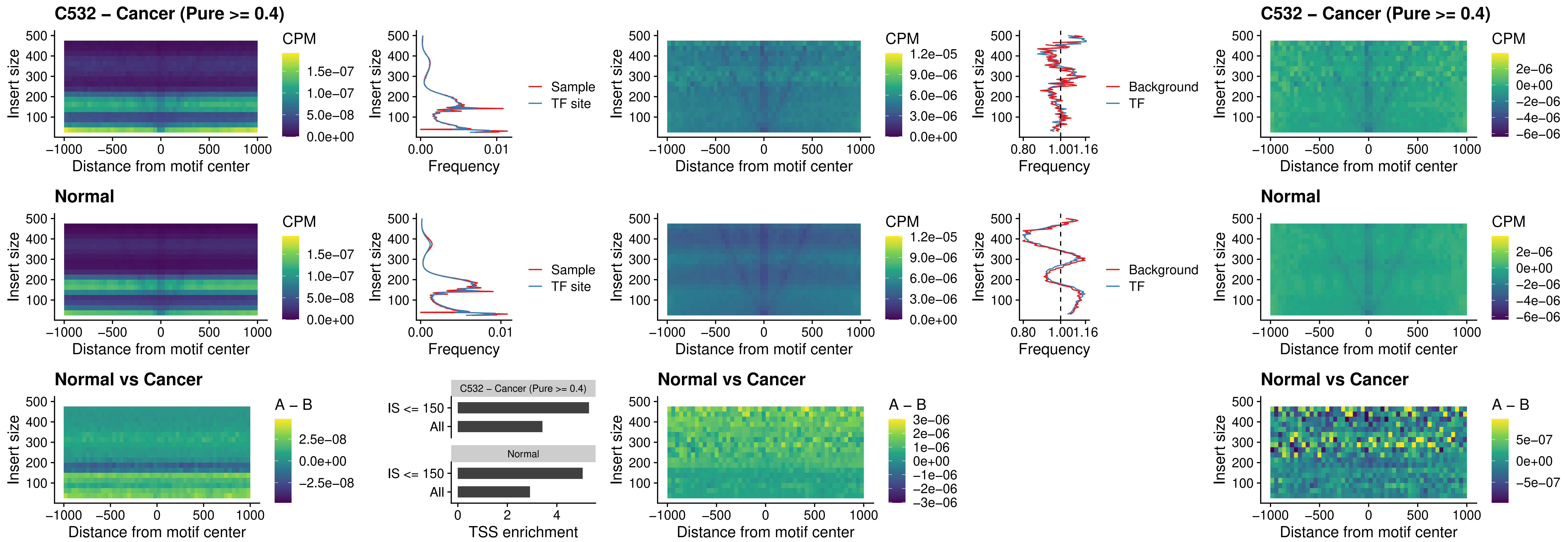

#### C532 – Cancer (Pure >= 0.4) – HOXD3_uc_pTSS_nPEAK

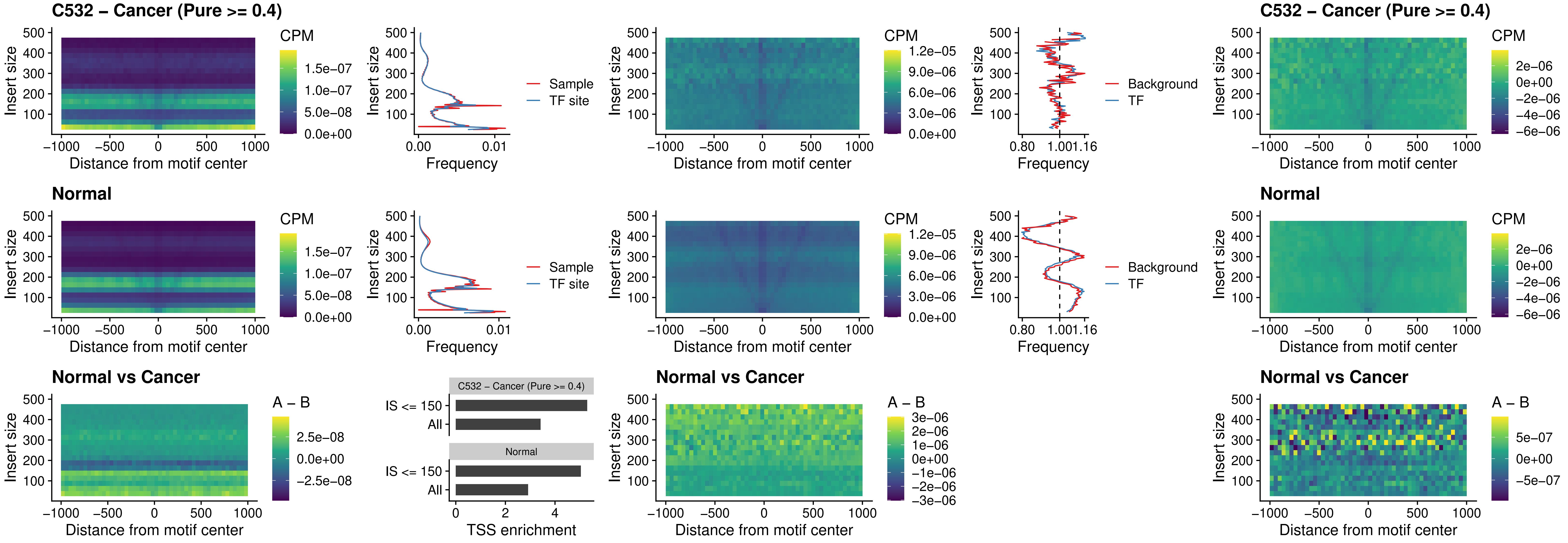

### C532 – Cancer (Pure >= 0.4) – PDX1_uc_dTSS_nPEAK

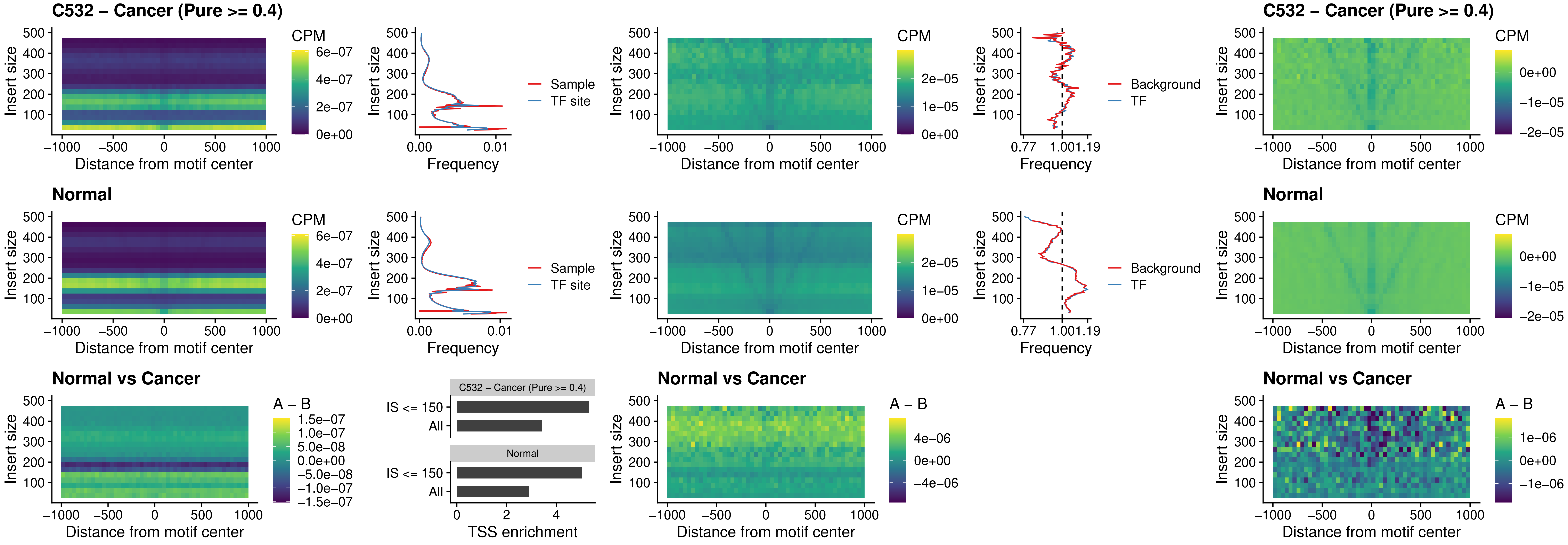

#### C532 – Cancer (Pure >= 0.4) – VAX2_uc_dTSS_nPEAK

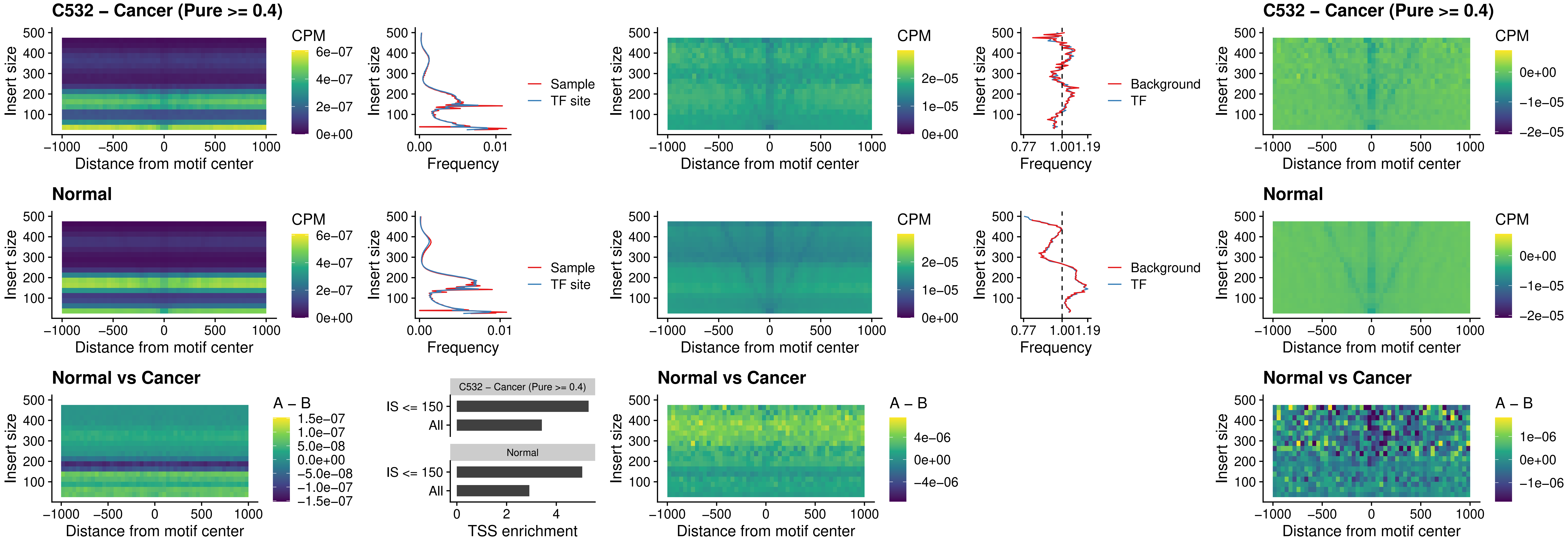

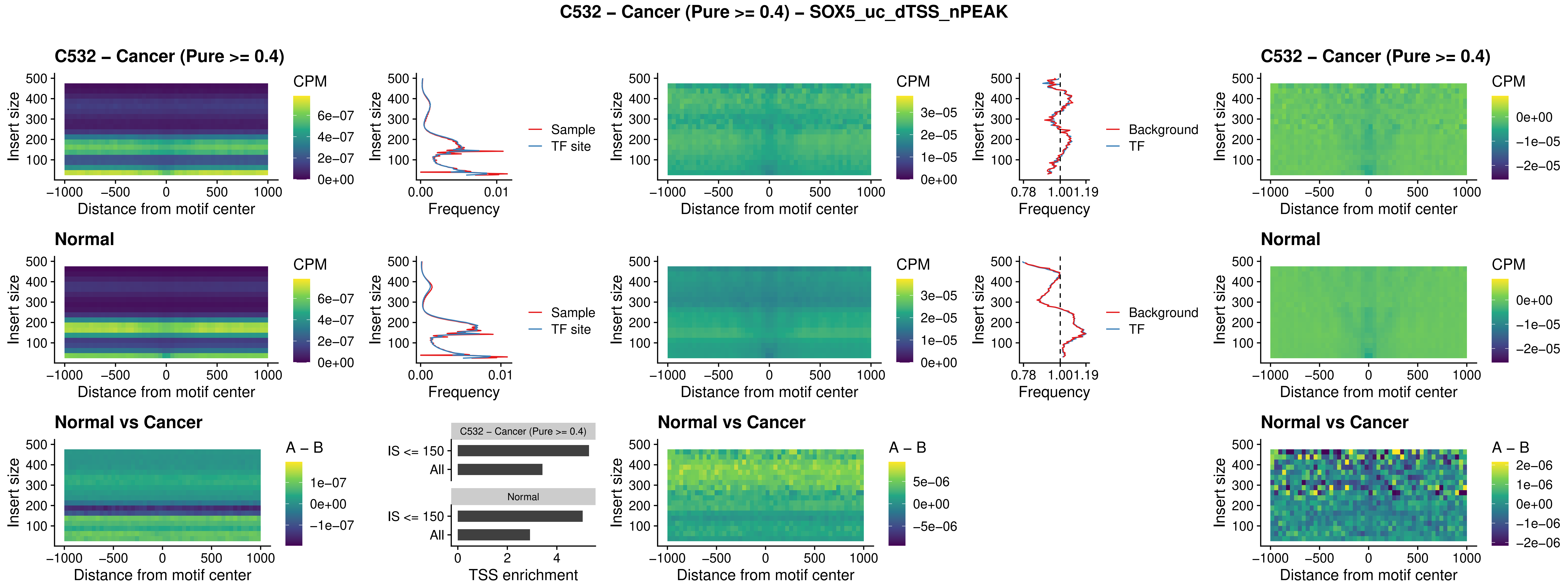
C532 – Cancer (Pure >= 0.4) – SOX5_uc_dTSS_nPEAK
C532 – Cancer (Pure >= 0.4)
Normal
Normal vs Cancer
Insert size
Distance from motif center
CPM
A – B
Frequency
Sample
TF site
Background
TF
TSS enrichment
C532 – Cancer (Pure >= 0.4)
Normal
IS <= 150
All

#### C532 – Cancer (Pure >= 0.4) – SOX6_uc_dTSS_nPEAK

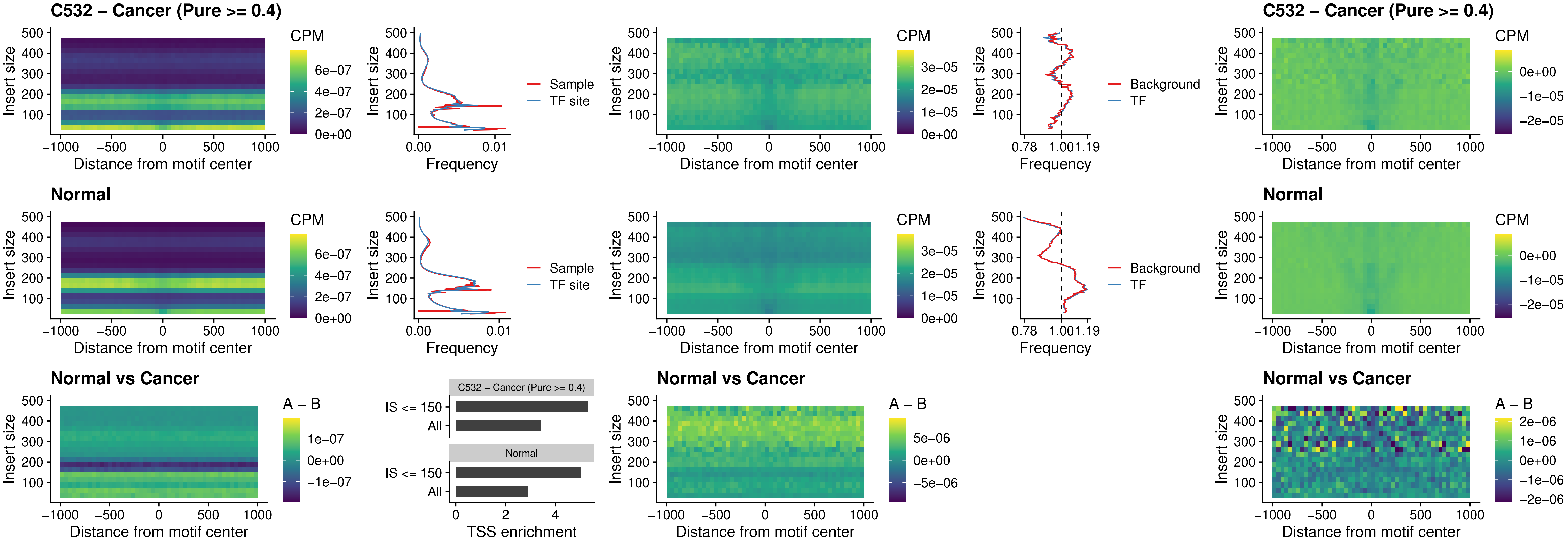

#### C532 – Cancer (Pure >= 0.4) – JDP2_uc_cTSS_nPEAK

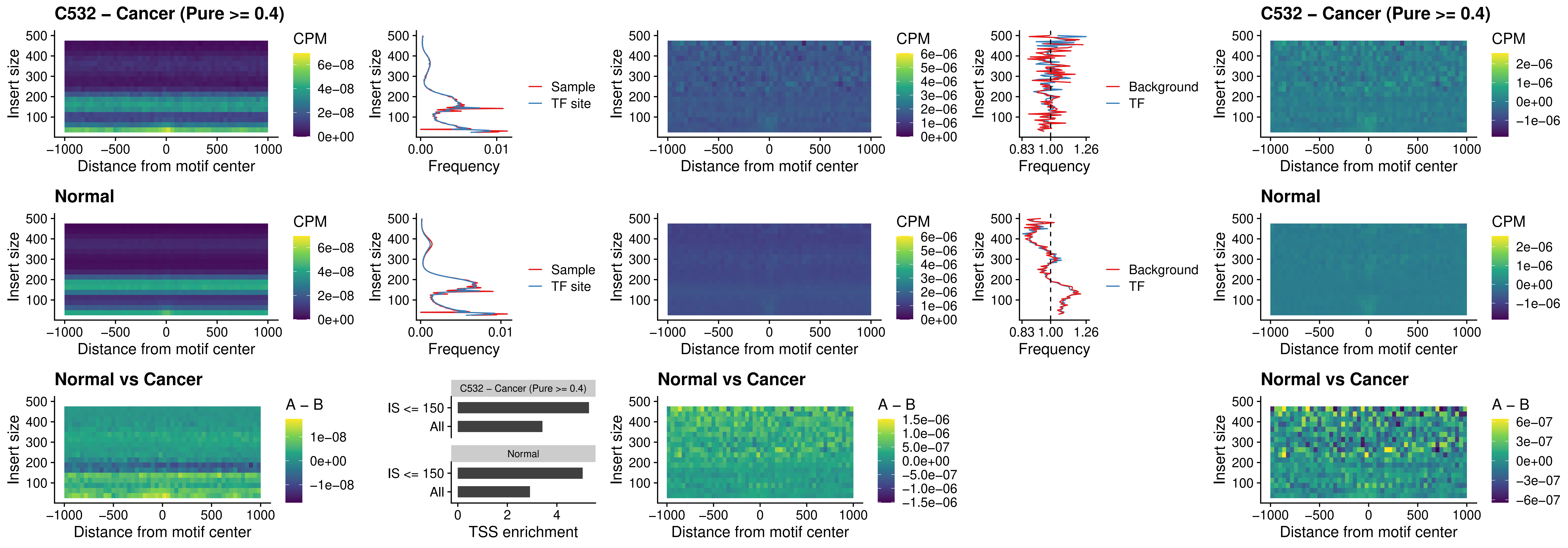

#### C532 – Cancer (Pure >= 0.4) – PURA_uc_dTSS_oPEAK

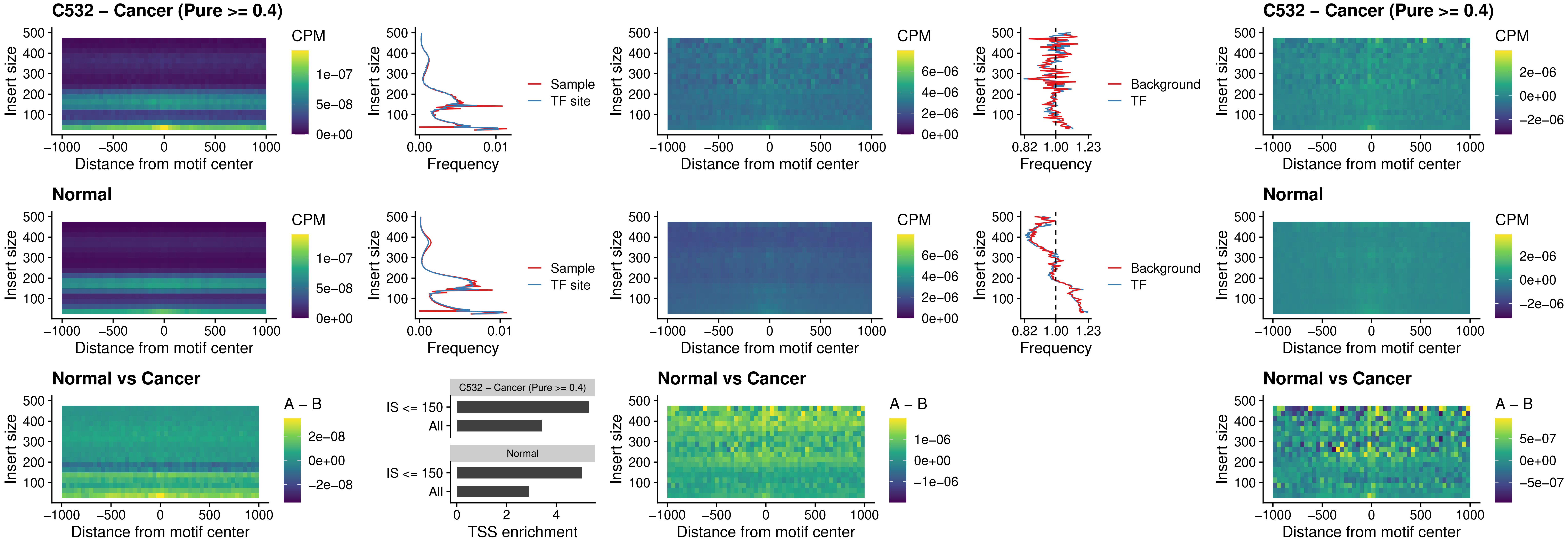

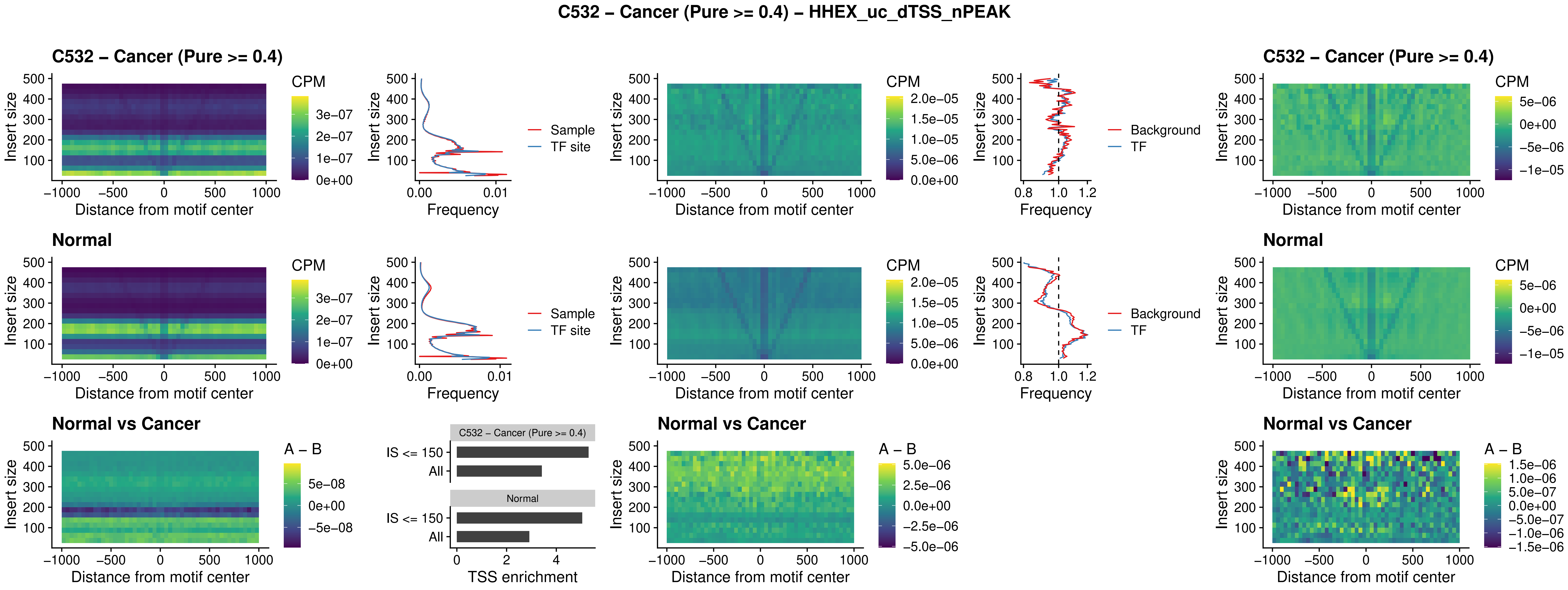
C532 – Cancer (Pure >= 0.4) – HHEX_uc_dTSS_nPEAK
C532 – Cancer (Pure >= 0.4)
Normal
Normal vs Cancer
Insert size
Distance from motif center
Frequency
CPM
A – B
Sample
TF site
Background
TF
C532 – Cancer (Pure >= 0.4)
Normal
IS <= 150
All
TSS enrichment

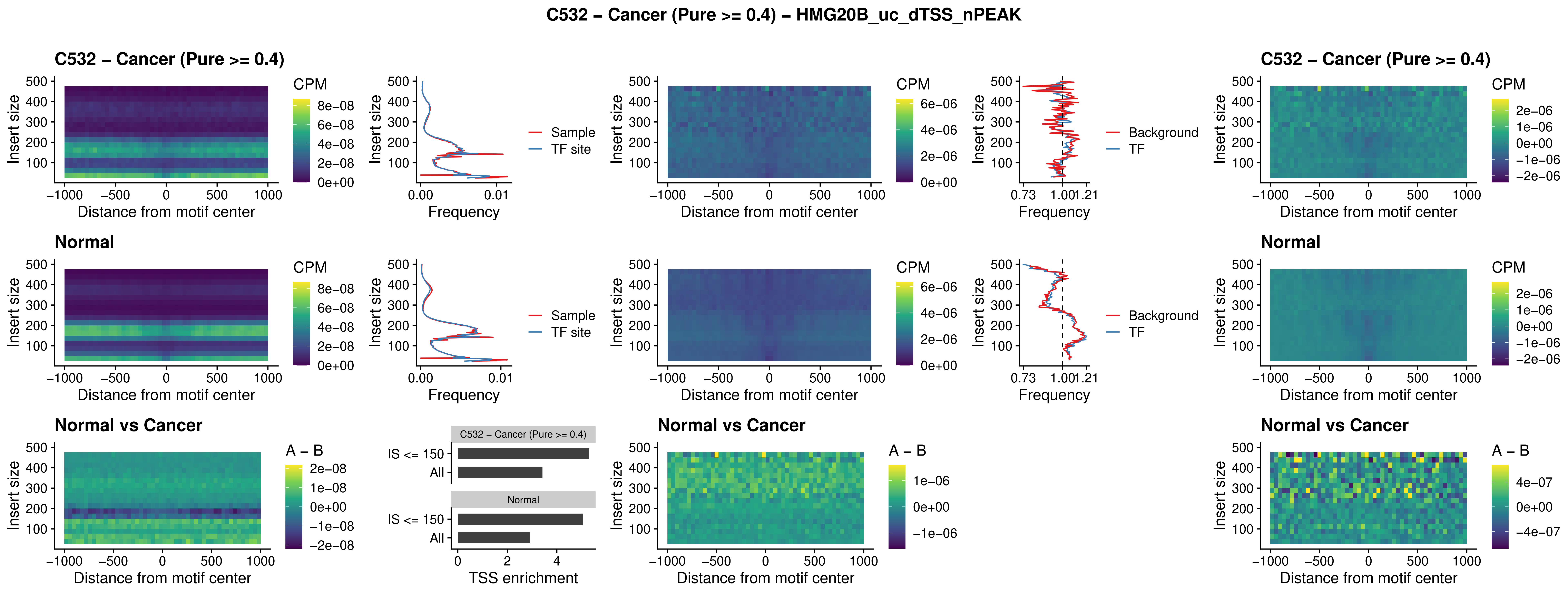

C532 – Cancer (Pure >= 0.4) – HMG20B_uc_dTSS_nPEAK
C532 – Cancer (Pure >= 0.4)
Normal
Normal vs Cancer
Insert size
Distance from motif center
CPM
A – B
Frequency
Sample
TF site
Background
TF
C532 – Cancer (Pure >= 0.4)
Normal
IS <= 150
All
TSS enrichment

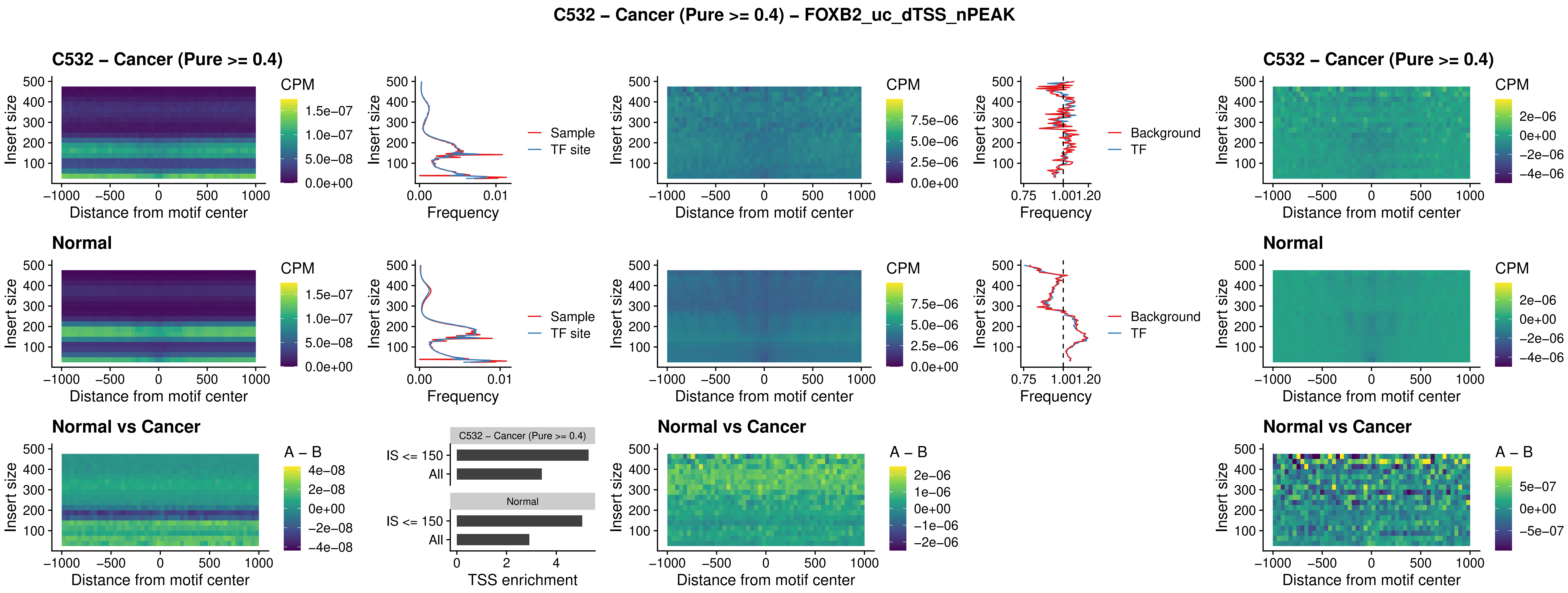
C532 – Cancer (Pure >= 0.4) – FOXB2_uc_dTSS_nPEAK
C532 – Cancer (Pure >= 0.4)
Normal
Normal vs Cancer
Insert size
Distance from motif center
Frequency
CPM
A – B
Sample
TF site
Background
TF
C532 – Cancer (Pure >= 0.4)
Normal
IS <= 150
All
TSS enrichment

### C532 – Cancer (Pure >= 0.4) – FOXL1_uc_cTSS_nPEAK

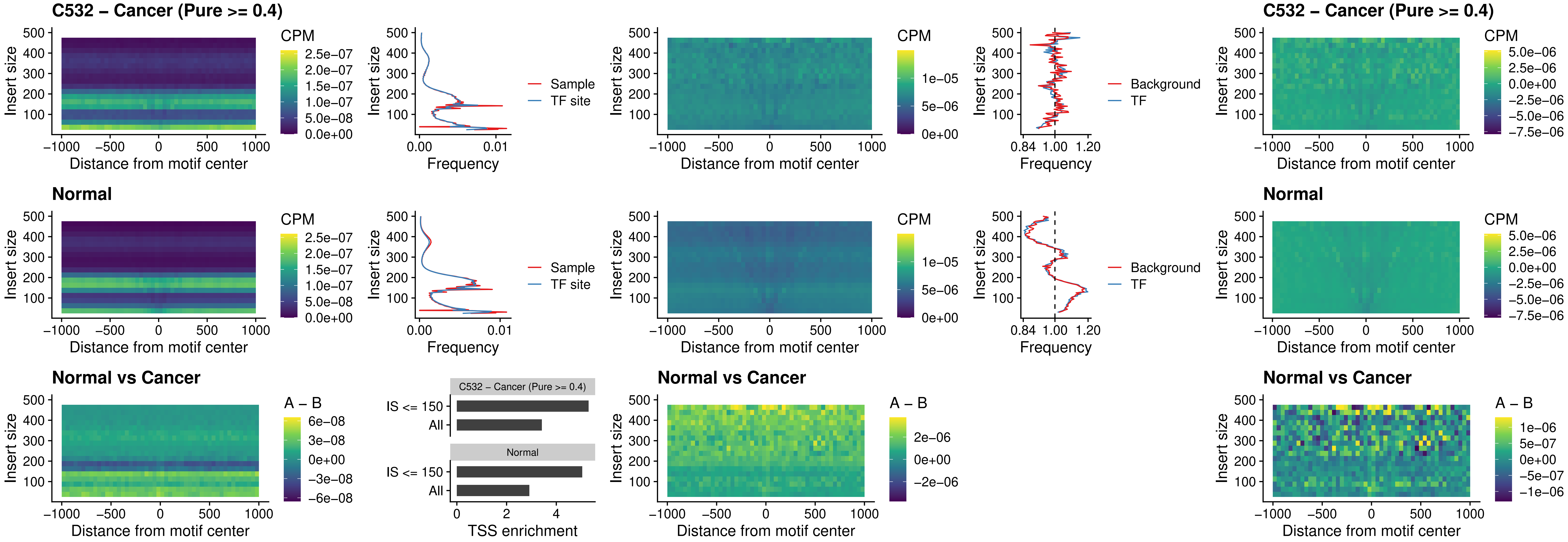

#### C532 – Cancer (Pure >= 0.4) – PBX3_uc_pTSS_oPEAK

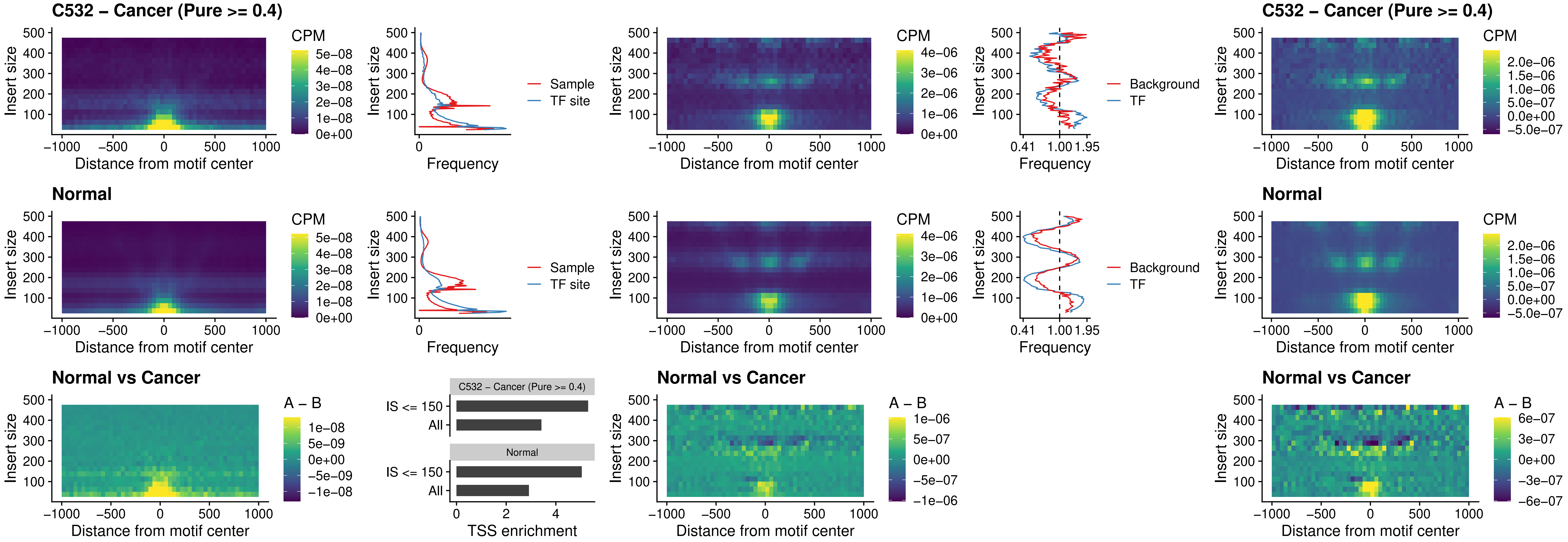

#### C532 – Cancer (Pure >= 0.4) – SNAI2_uc_cTSS_nPEAK

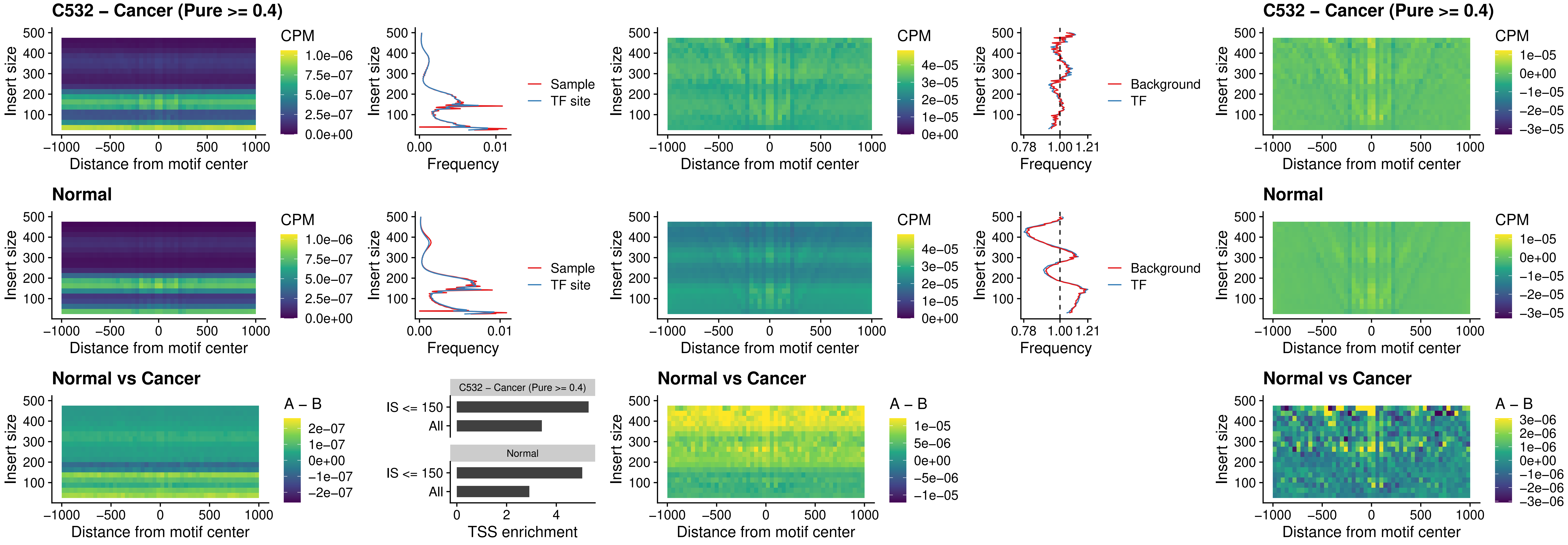

#### C532 – Cancer (Pure >= 0.4) – SNAI2_uc_dTSS_oPEAK

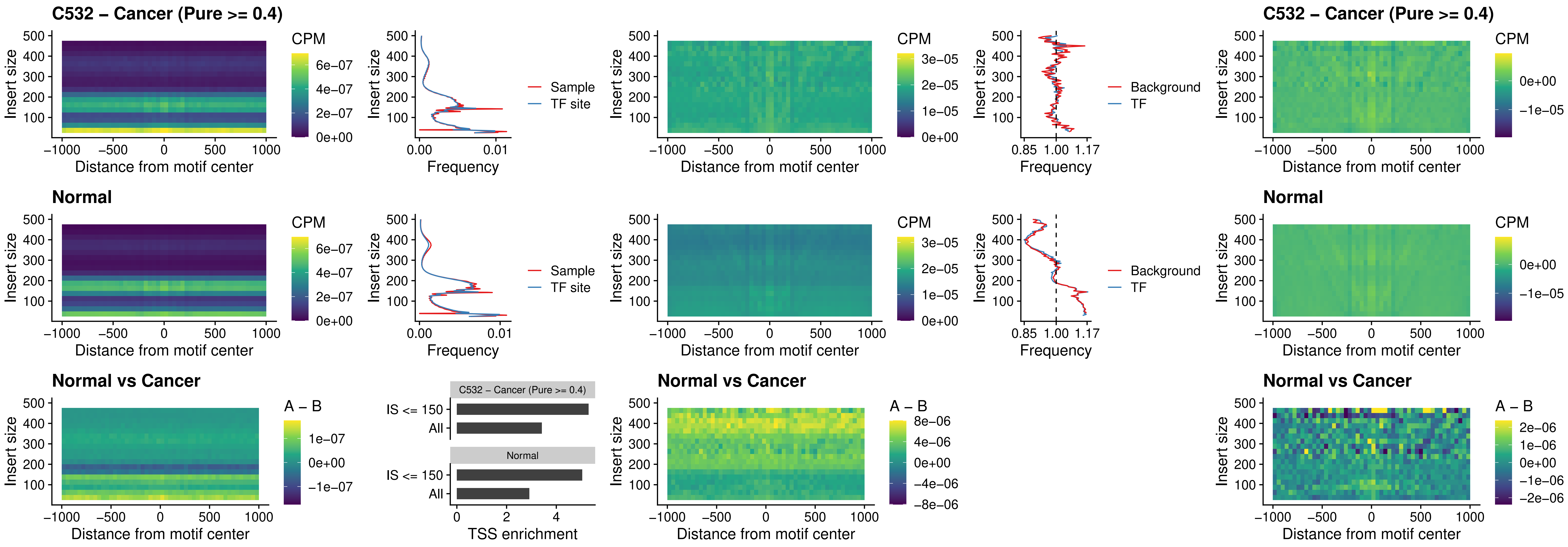

#### C532 – Cancer (Pure >= 0.4) – SPI1_uc_pTSS_nPEAK

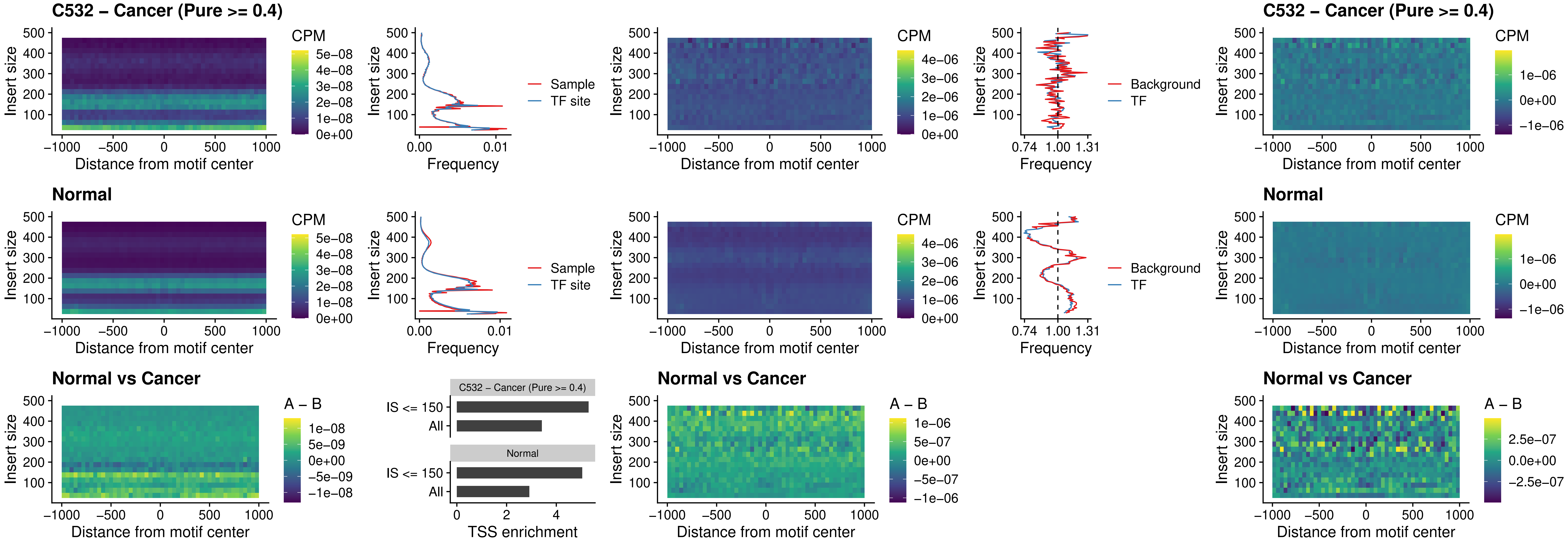

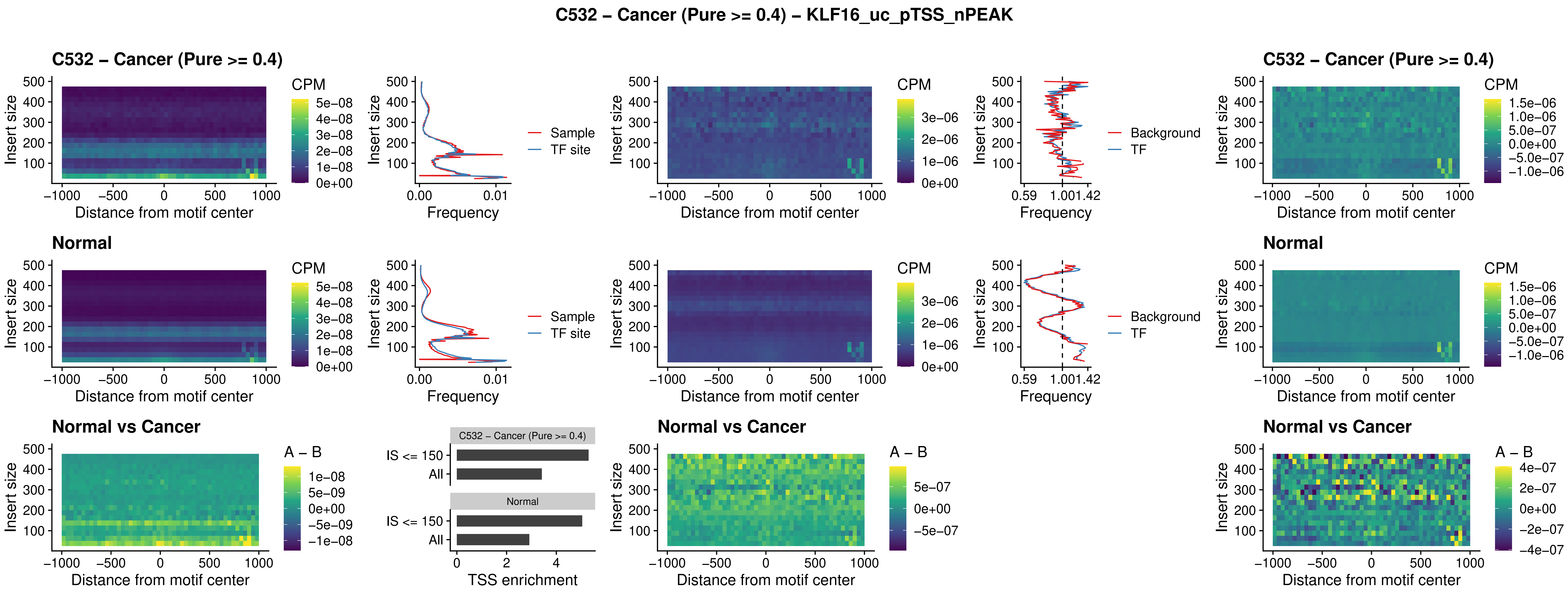
C532 – Cancer (Pure >= 0.4) – KLF16_uc_pTSS_nPEAK
C532 – Cancer (Pure >= 0.4)
Normal
Normal vs Cancer
Insert size
Distance from motif center
CPM
A – B
Frequency
Sample
TF site
Background
TF
C532 – Cancer (Pure >= 0.4)
Normal
IS <= 150
All
TSS enrichment

#### C532 – Cancer (Pure >= 0.4) – SP3_uc_pTSS_nPEAK

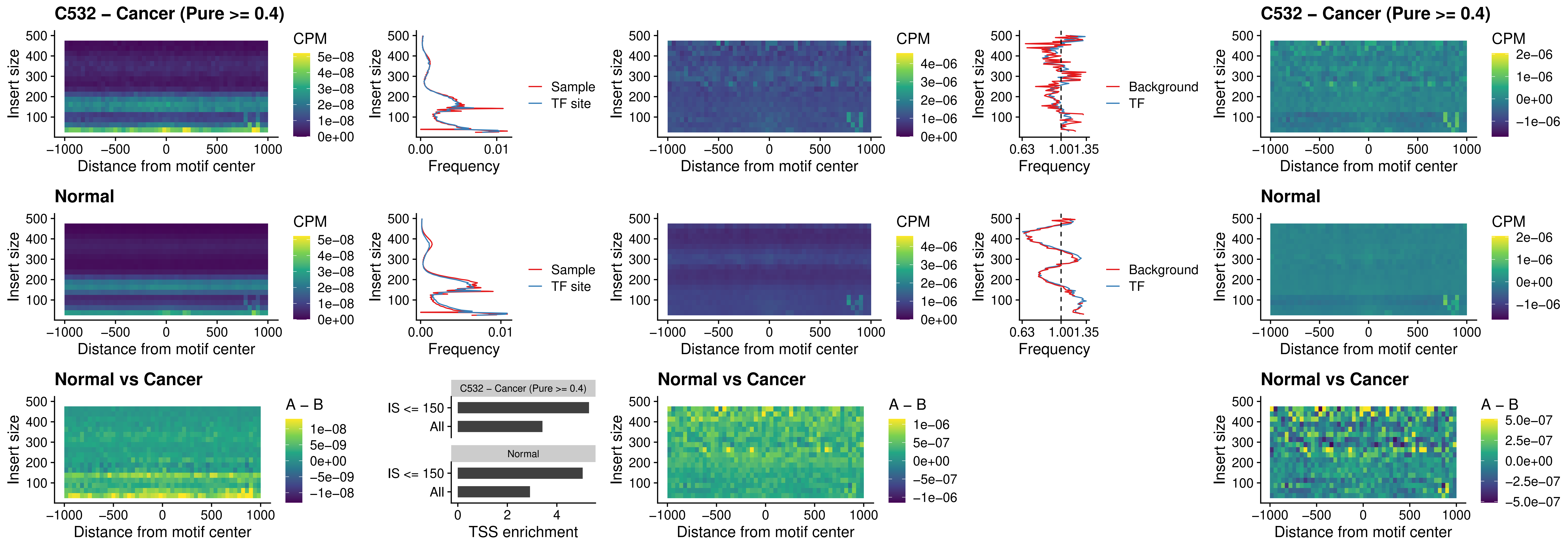

#### C532 – Cancer (Pure >= 0.4) – NEUROD1_uc_dTSS_oPEAK

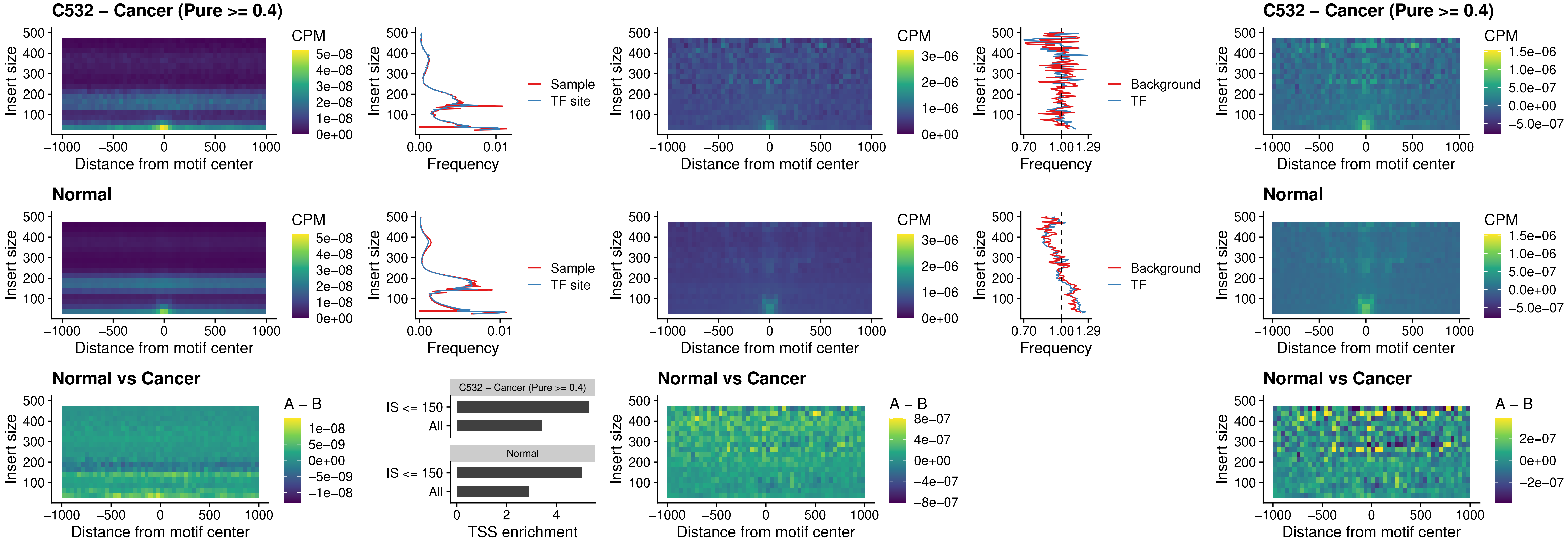

### C532 – Cancer (Pure >= 0.4) – CTCF_uc_dTSS_oPEAK

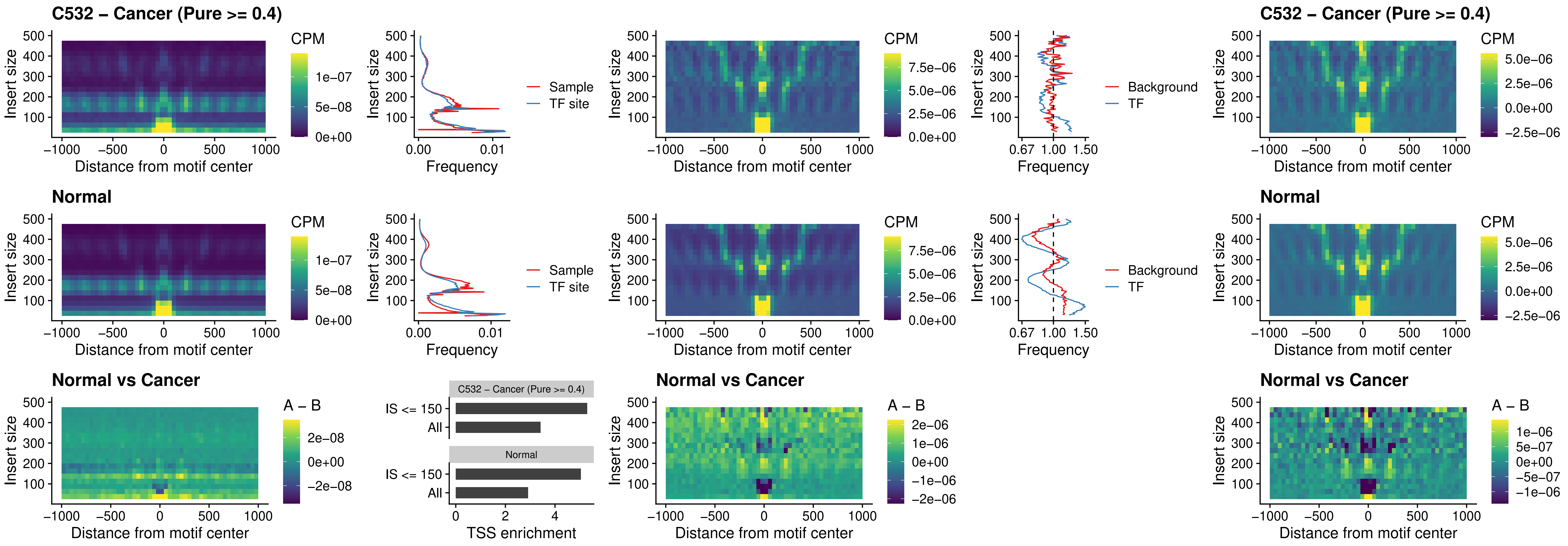

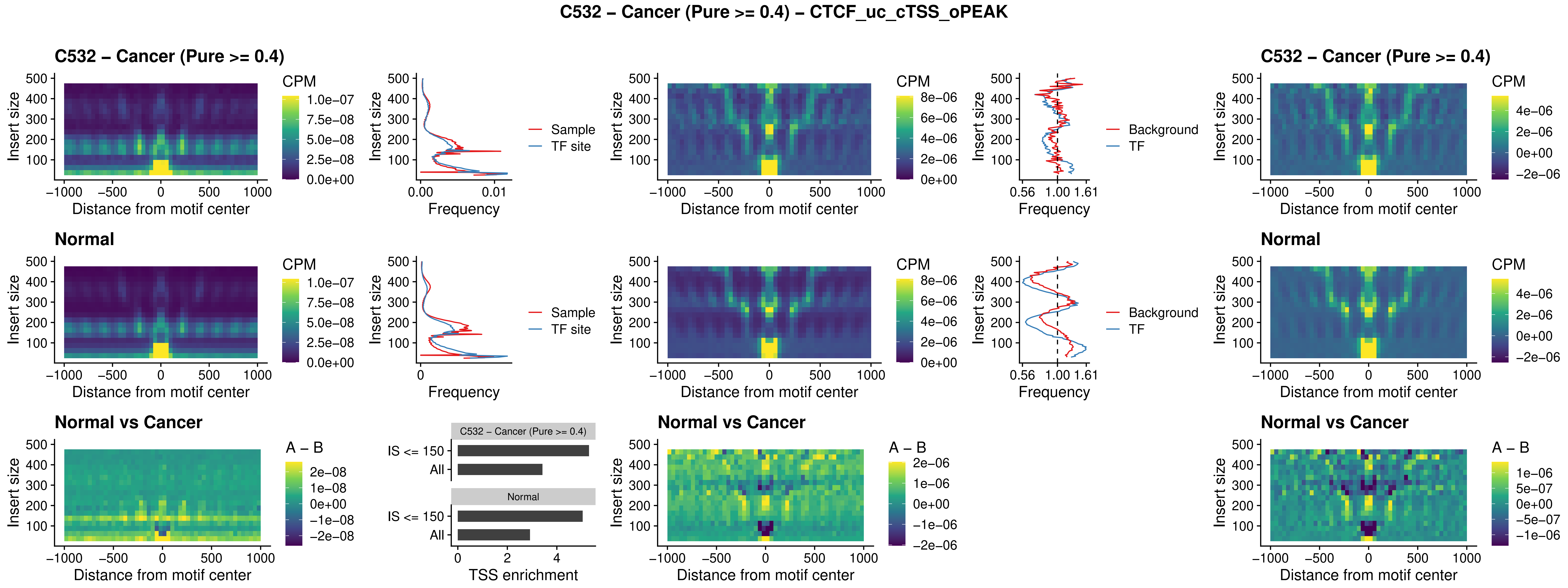

C532 – Cancer (Pure >= 0.4) – CTCF_uc_cTSS_oPEAK
C532 – Cancer (Pure >= 0.4)
Normal
Normal vs Cancer
Insert size
Distance from motif center
Frequency
CPM
A – B
Sample
TF site
Background
TF
C532 – Cancer (Pure >= 0.4)
Normal
IS <= 150
All
TSS enrichment

#### C532 – Cancer (Pure >= 0.4) – CTCFL_uc_dTSS_oPEAK

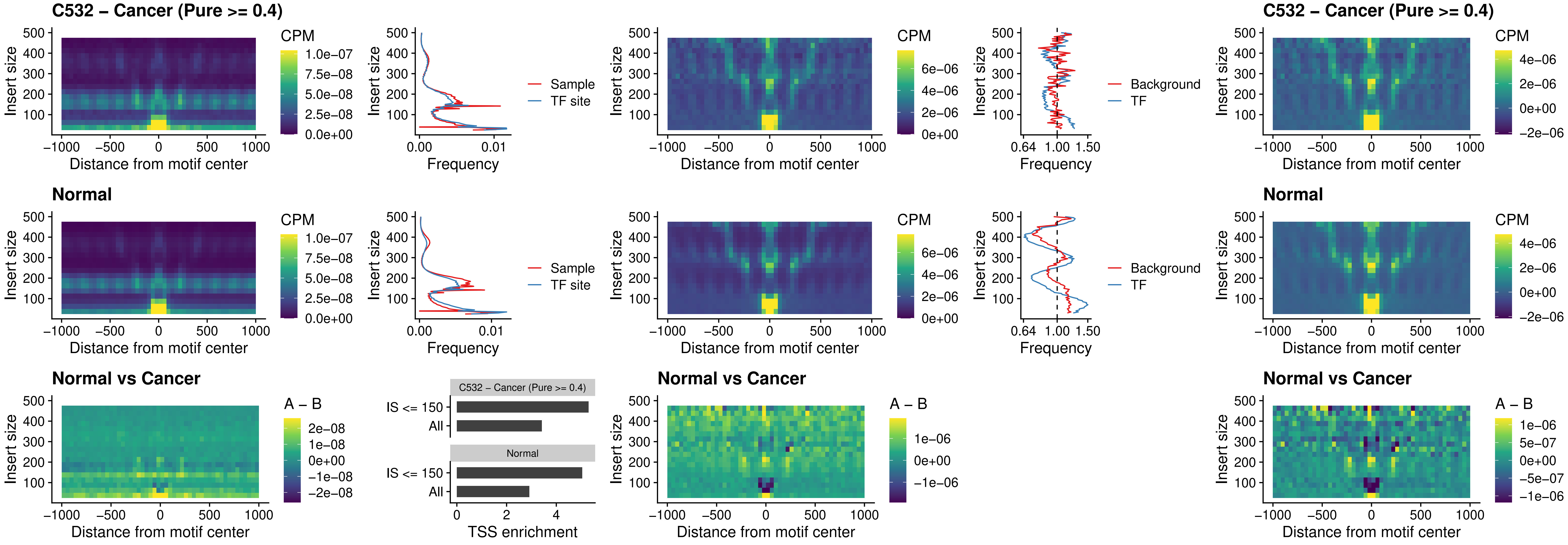

#### C532 – Cancer (Pure >= 0.4) – ENCODE_CTCF_sites_COLON

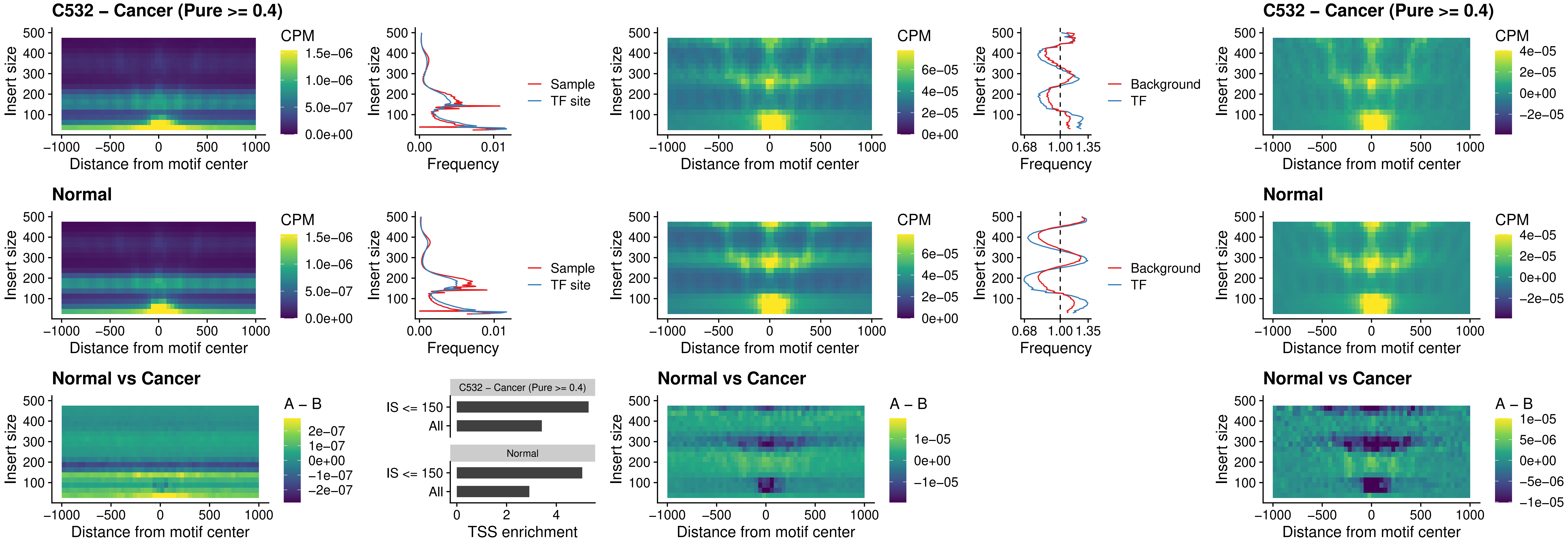

### C532 – Cancer (Pure >= 0.4) – ENCODE_CTCF_sites_COLON_loop_ctcf

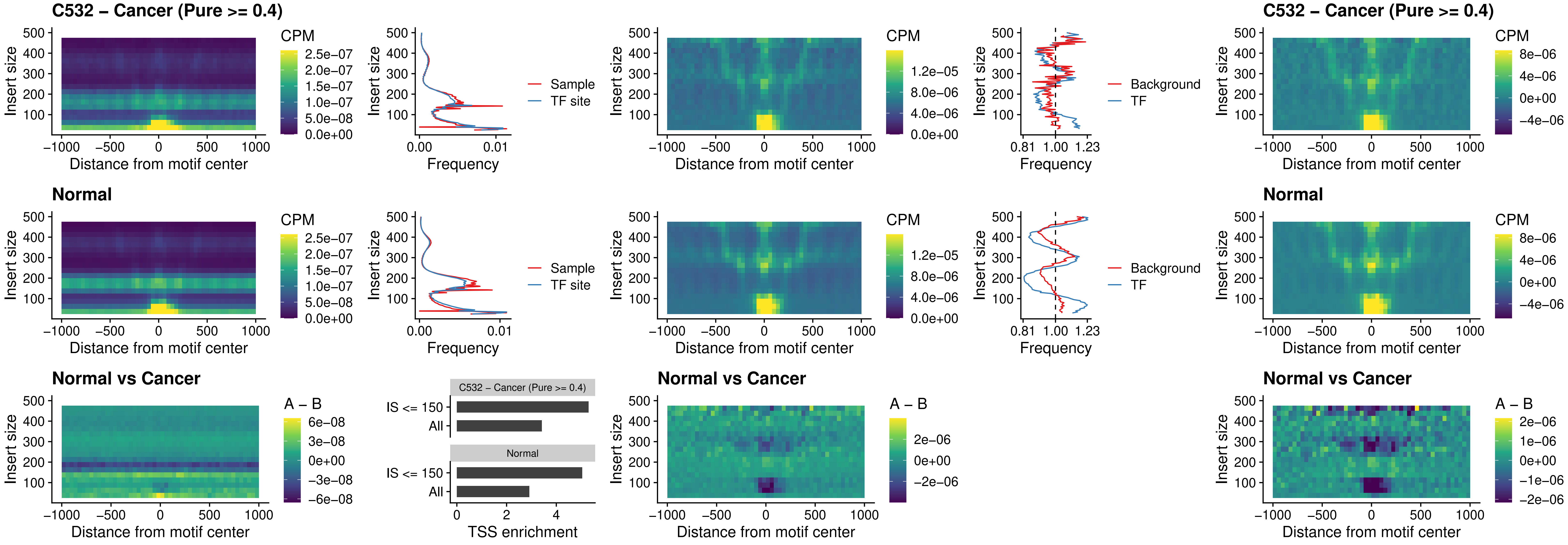

#### C532 – Cancer (Pure >= 0.4) – ENCODE_CTCF_sites_COLON_intergenic_ctcf

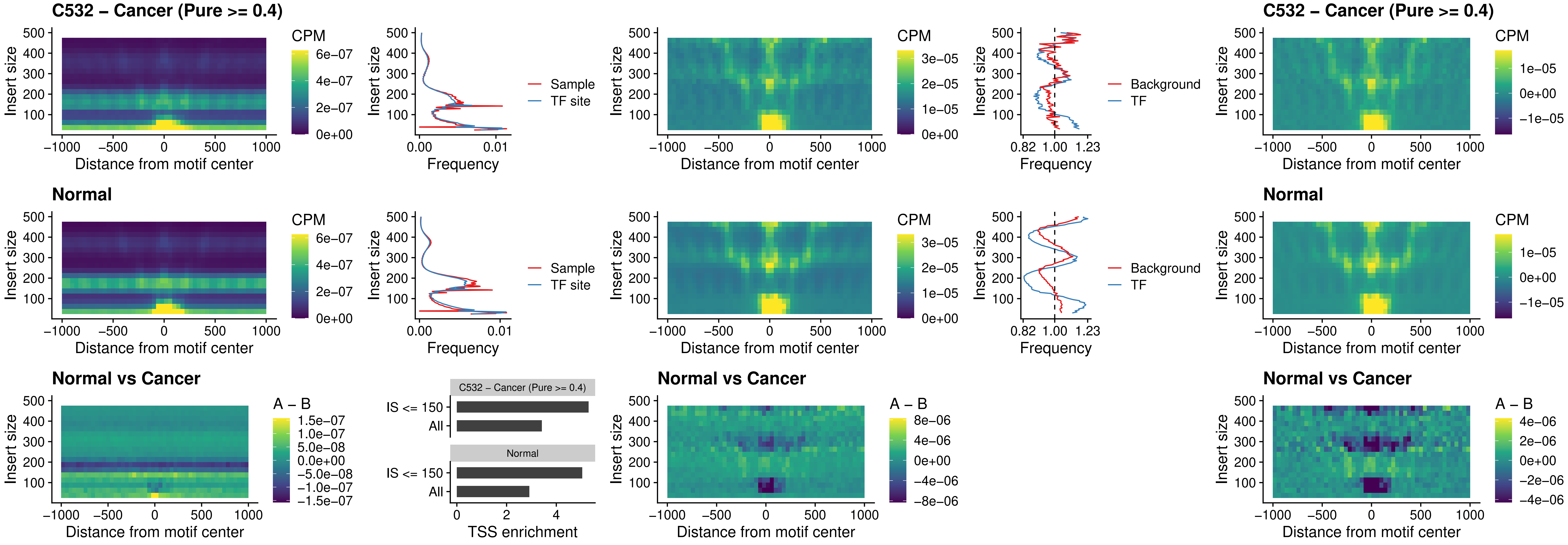

#### C532 – Cancer (Pure >= 0.4) – ENCODE_CTCF_sites_COLON_nonpromotor

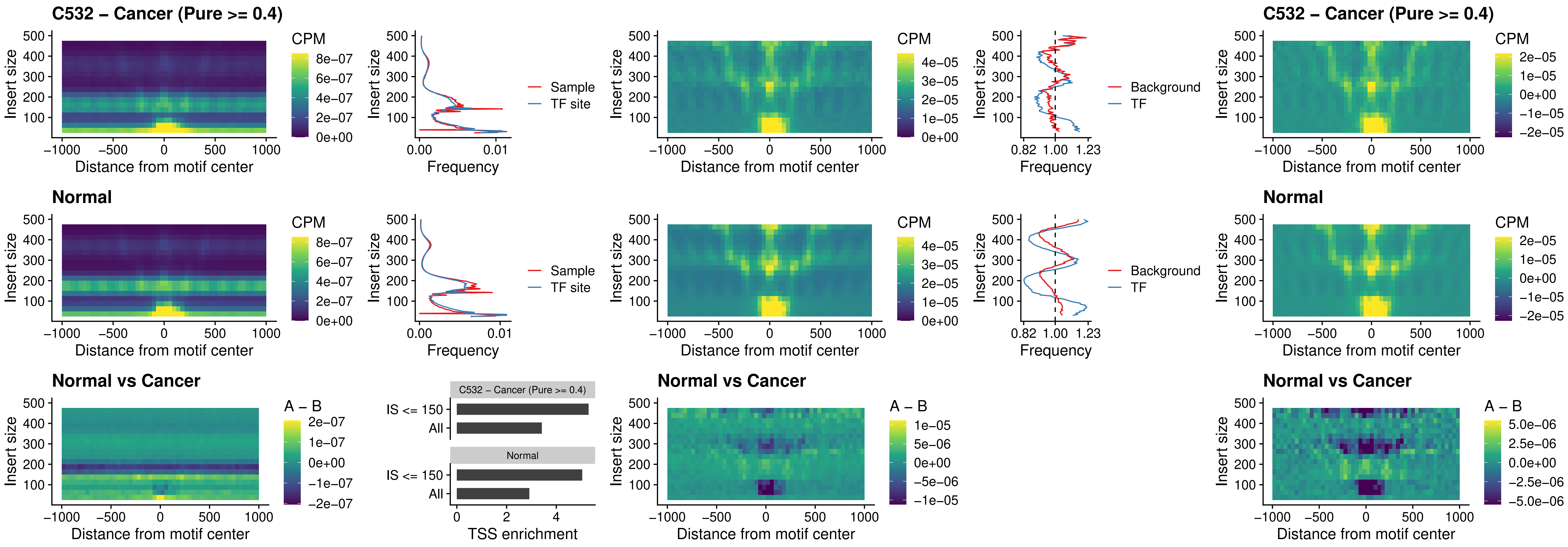

#### C532 – Cancer (Pure >= 0.4) – CTCFL_uc_cTSS_oPEAK

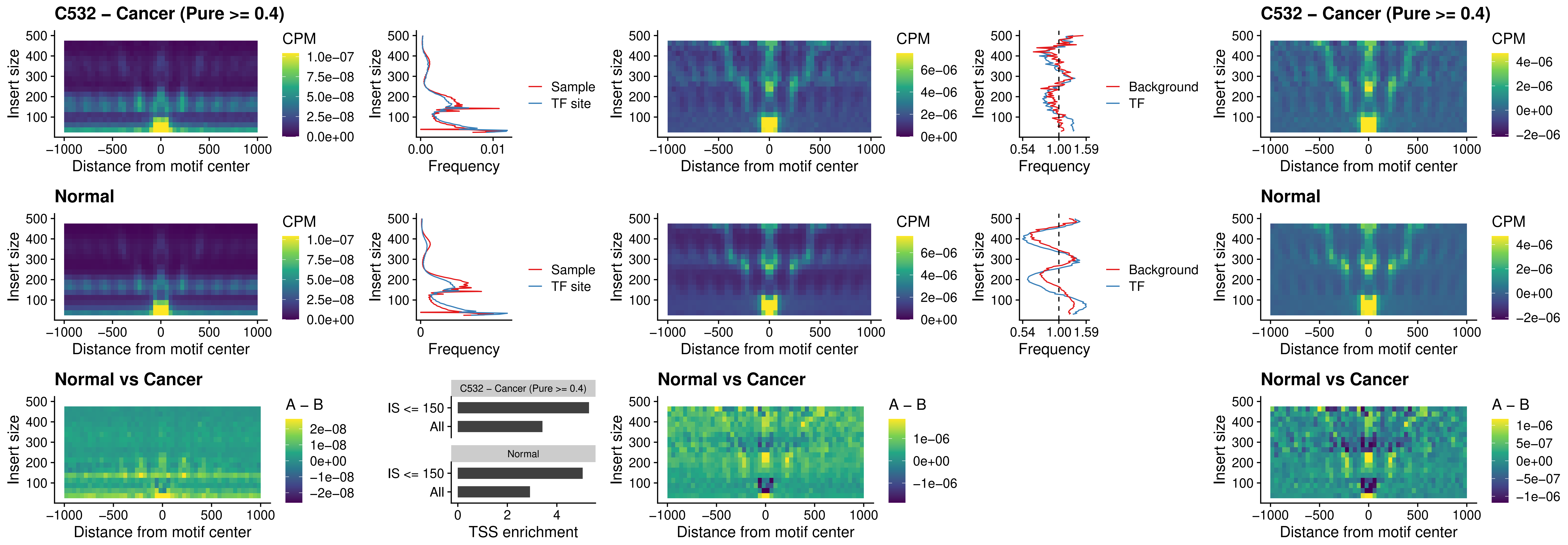

### C532 – Cancer (Pure >= 0.4) – NEUROD1_uc_cTSS_oPEAK

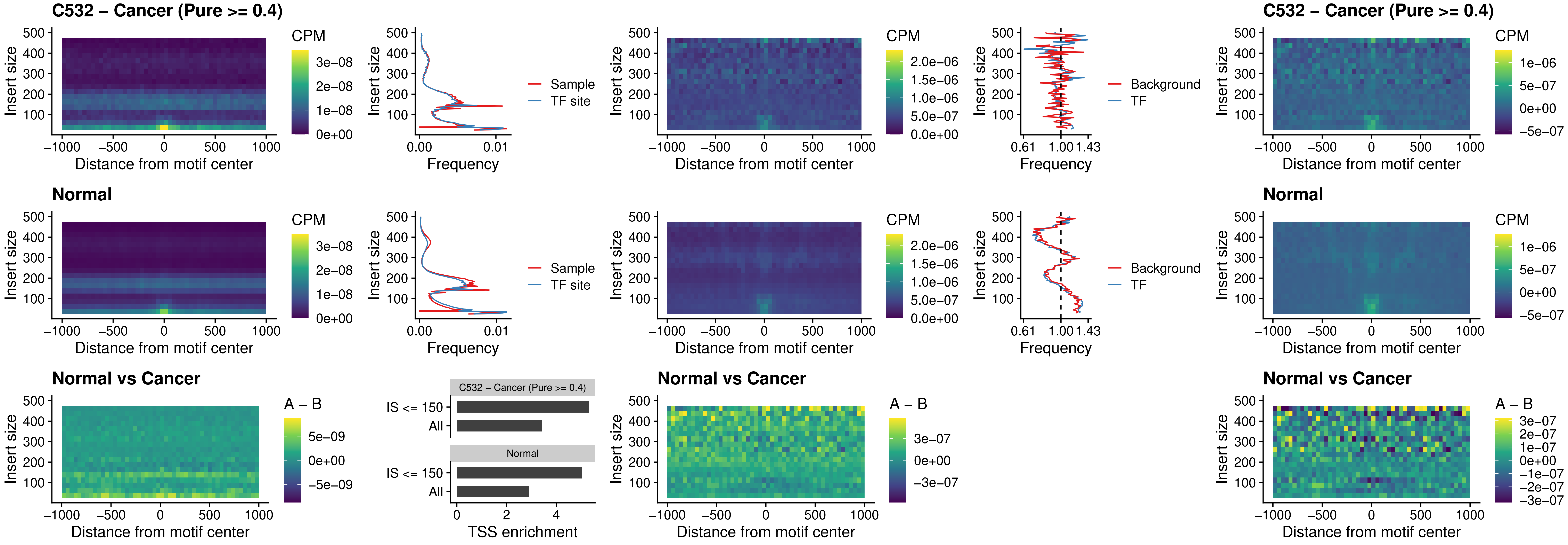

### C532 – Cancer (Pure >= 0.4) – ENCODE_YY1_sites_COLON_nonpromotor

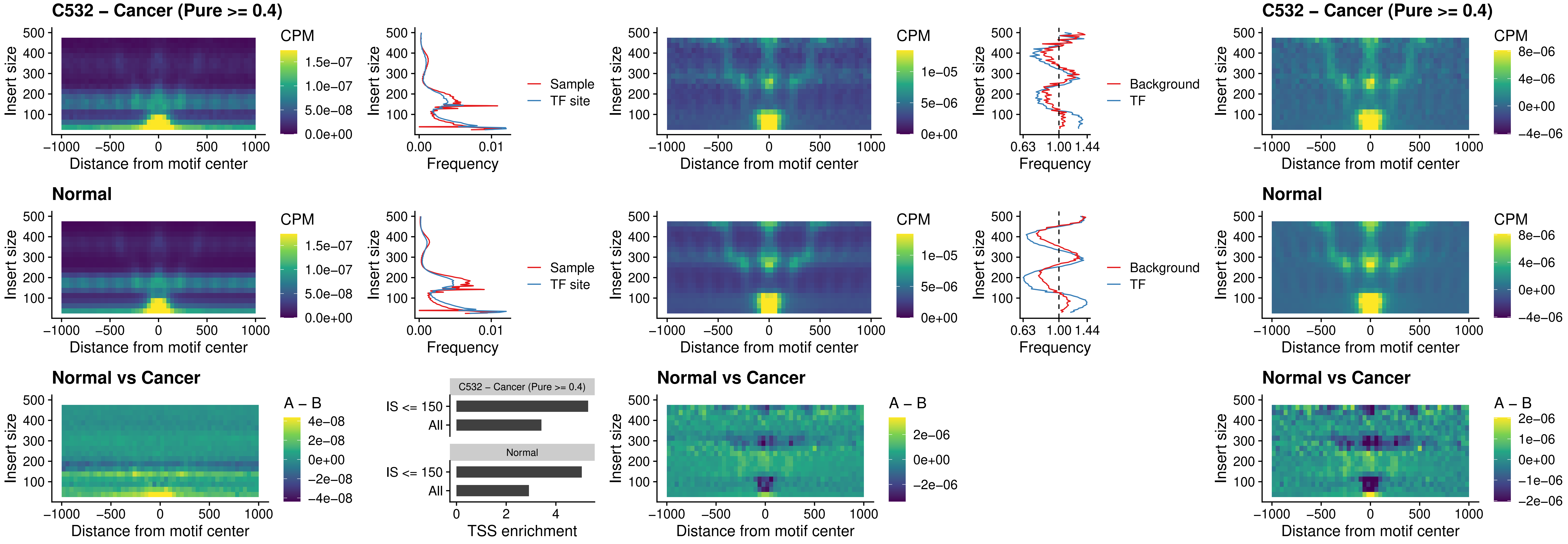

#### C532 – Cancer (Pure >= 0.4) – CTCF_uc_pTSS_oPEAK

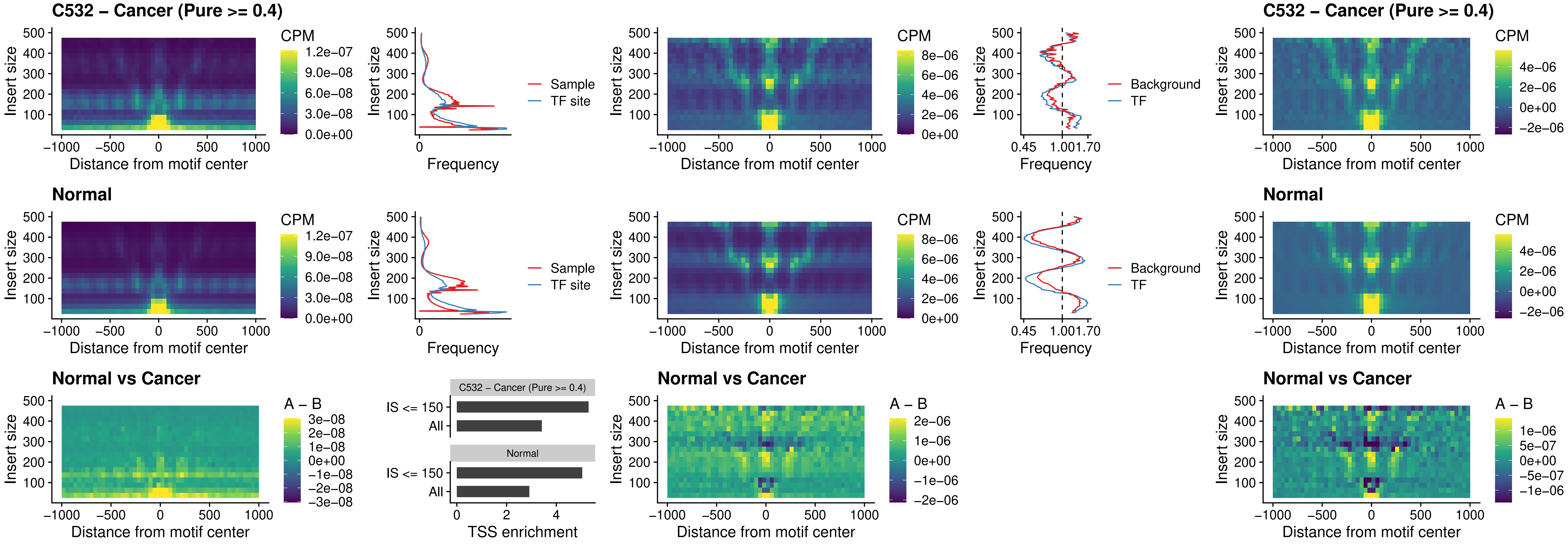

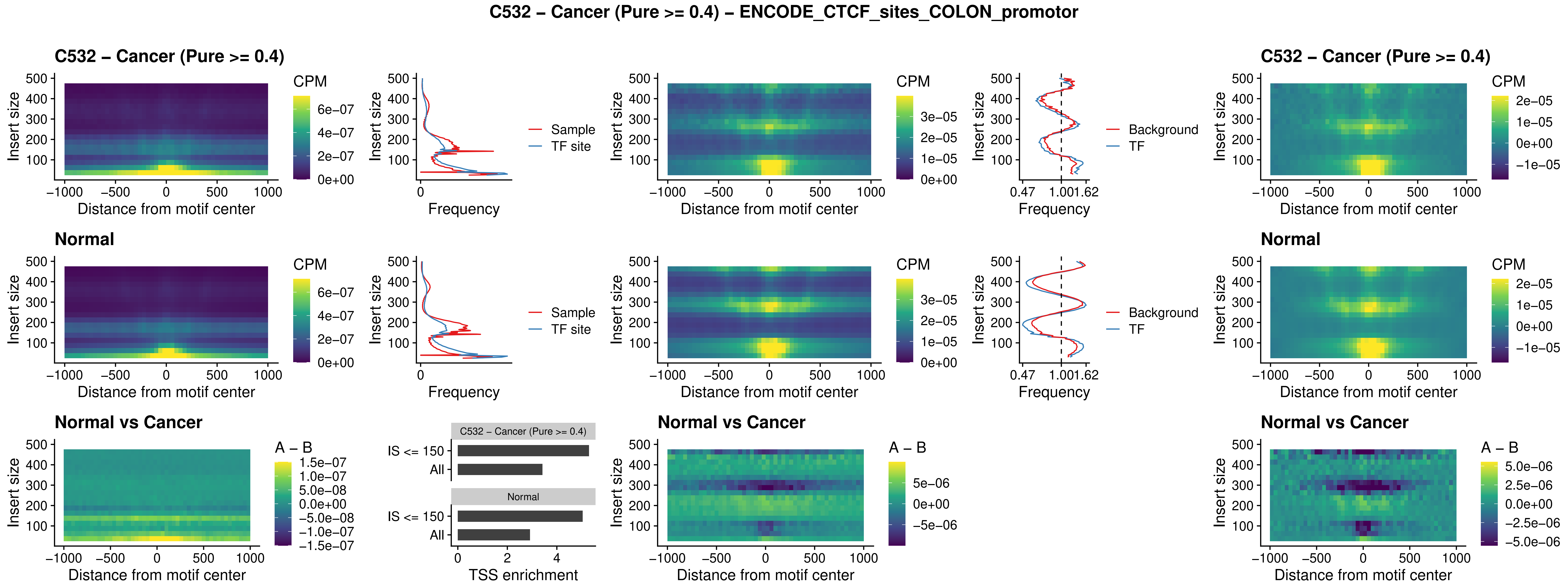
C532 – Cancer (Pure >= 0.4) – ENCODE_CTCF_sites_COLON_promotor
C532 – Cancer (Pure >= 0.4)
Normal
Normal vs Cancer
Insert size
Distance from motif center
CPM
A – B
Frequency
Sample
TF site
Background
TF
0.47
1.00
1.62
C532 – Cancer (Pure >= 0.4)
Normal
IS <= 150
All
TSS enrichment

#### C532 – Cancer (Pure >= 0.4) – ENCODE_CTCF_sites_COLON_promotor_ctcf

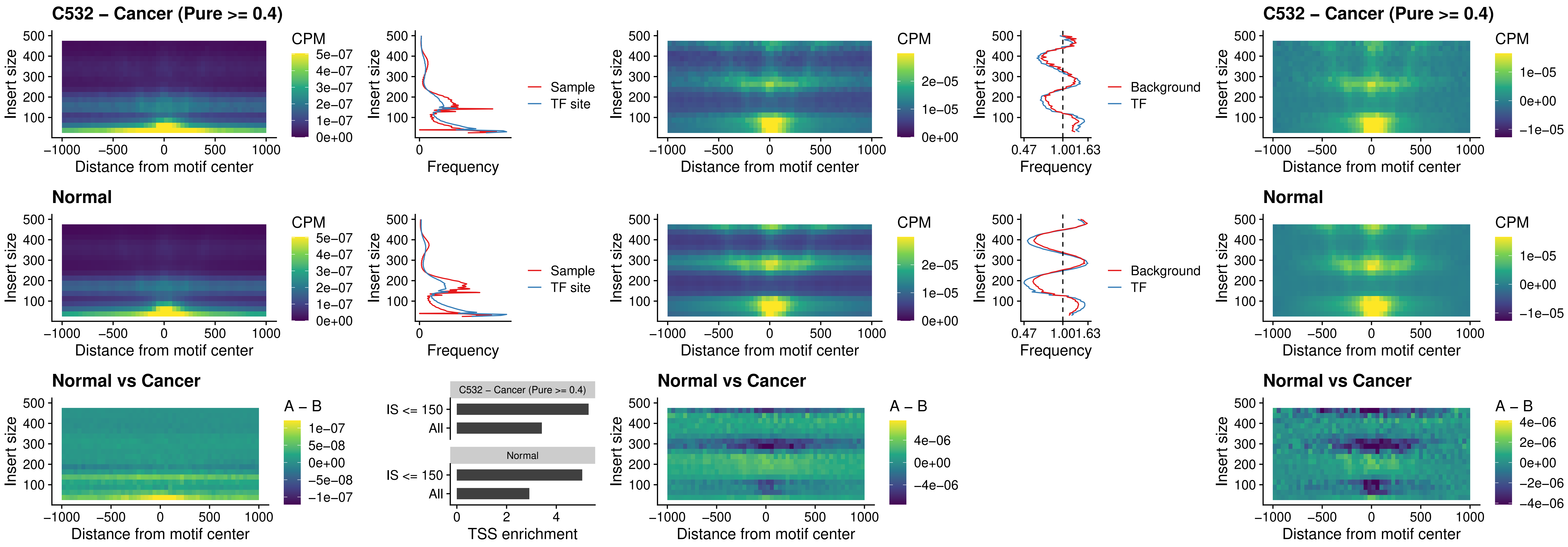

### C532 – Cancer (Pure >= 0.4) – SPIC_uc_pTSS_oPEAK

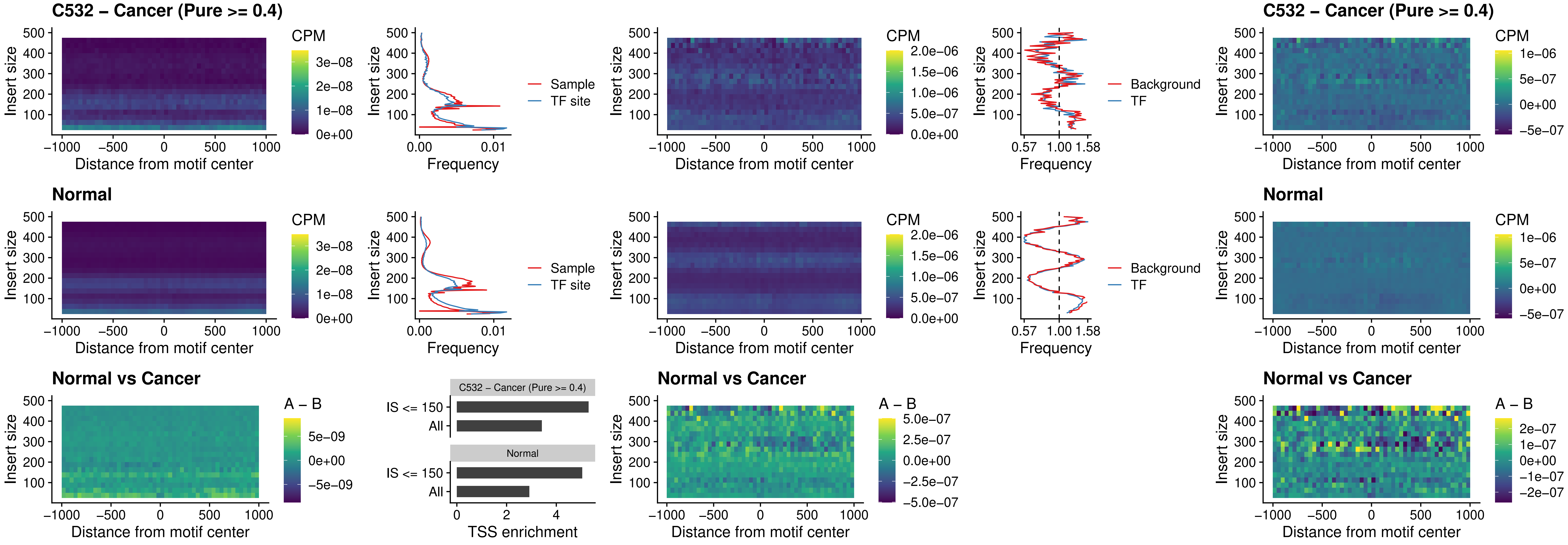

#### C532 – Cancer (Pure >= 0.4) – SPI1_uc_dTSS_nPEAK

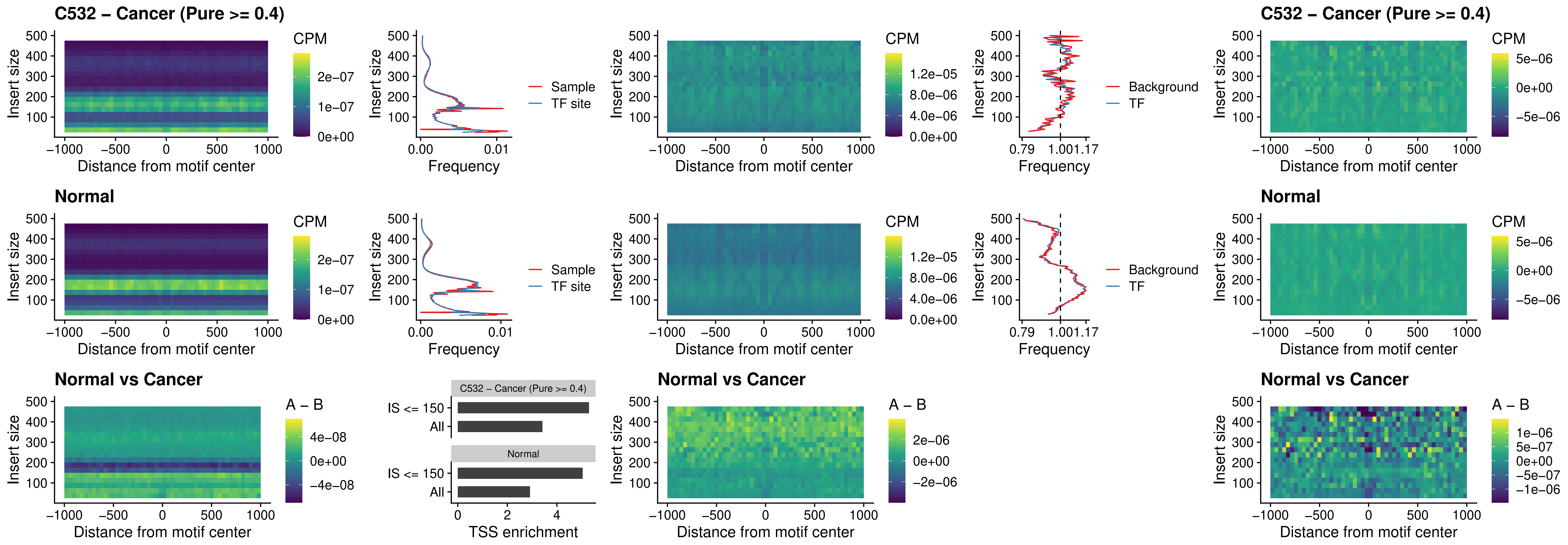

#### C532 – Cancer (Pure >= 0.4) – SPI1_uc_cTSS_nPEAK

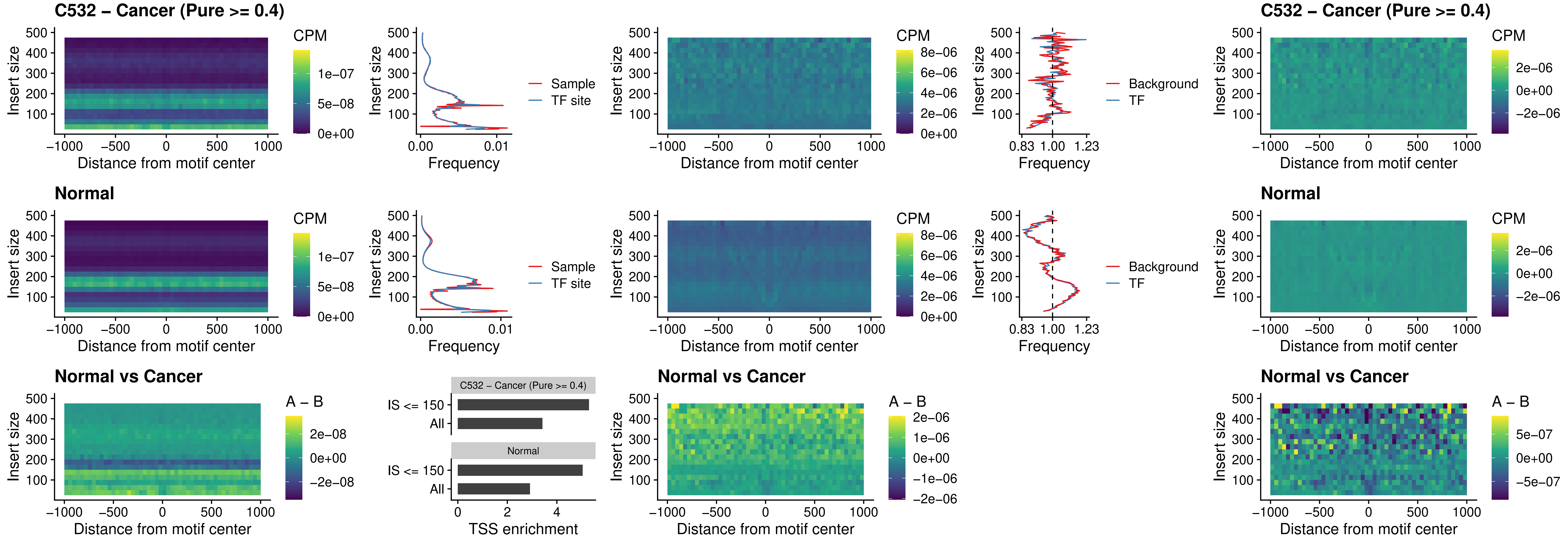

#### C532 – Cancer (Pure >= 0.4) – BCL11A_uc_pTSS_nPEAK

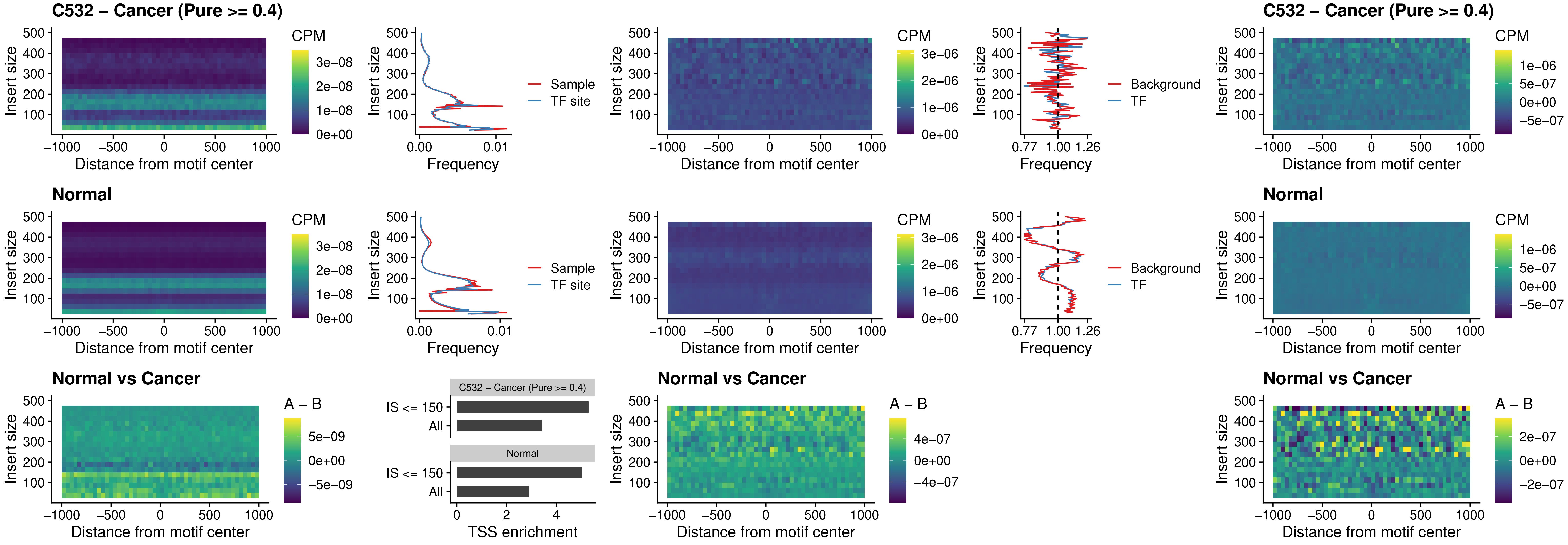

#### C532 – Cancer (Pure >= 0.4) – BCL11B_uc_pTSS_nPEAK

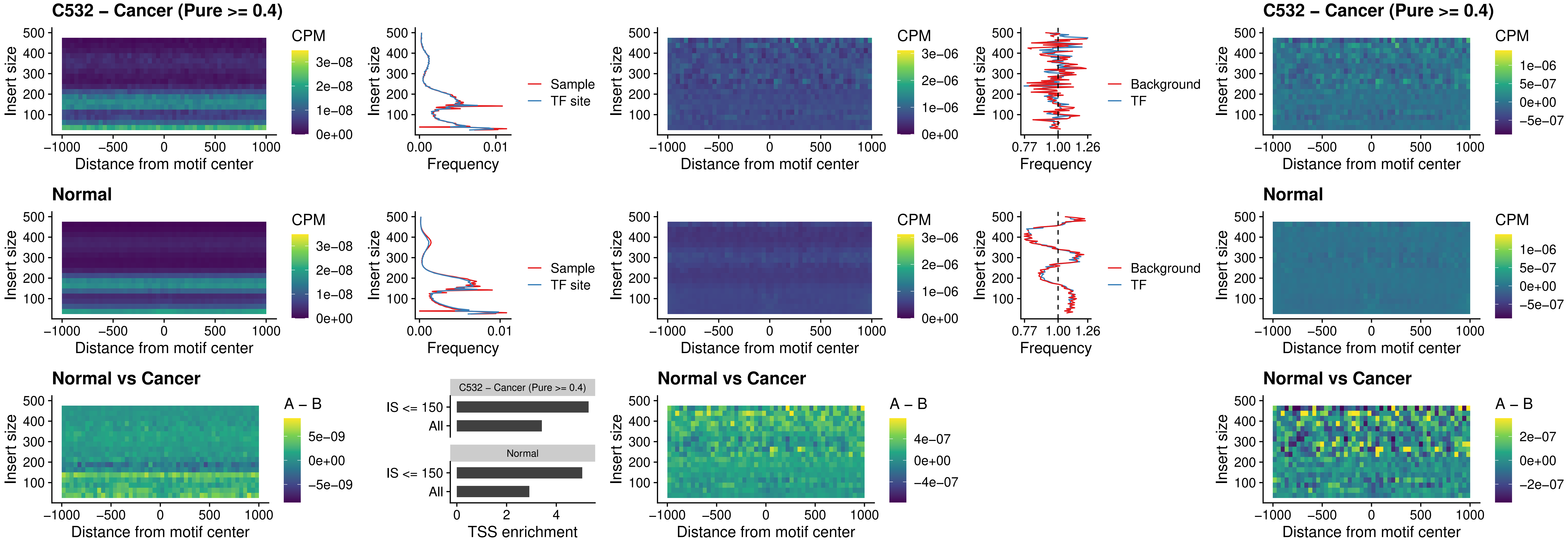

#### C532 – Cancer (Pure >= 0.4) – IRF9_uc_dTSS_oPEAK

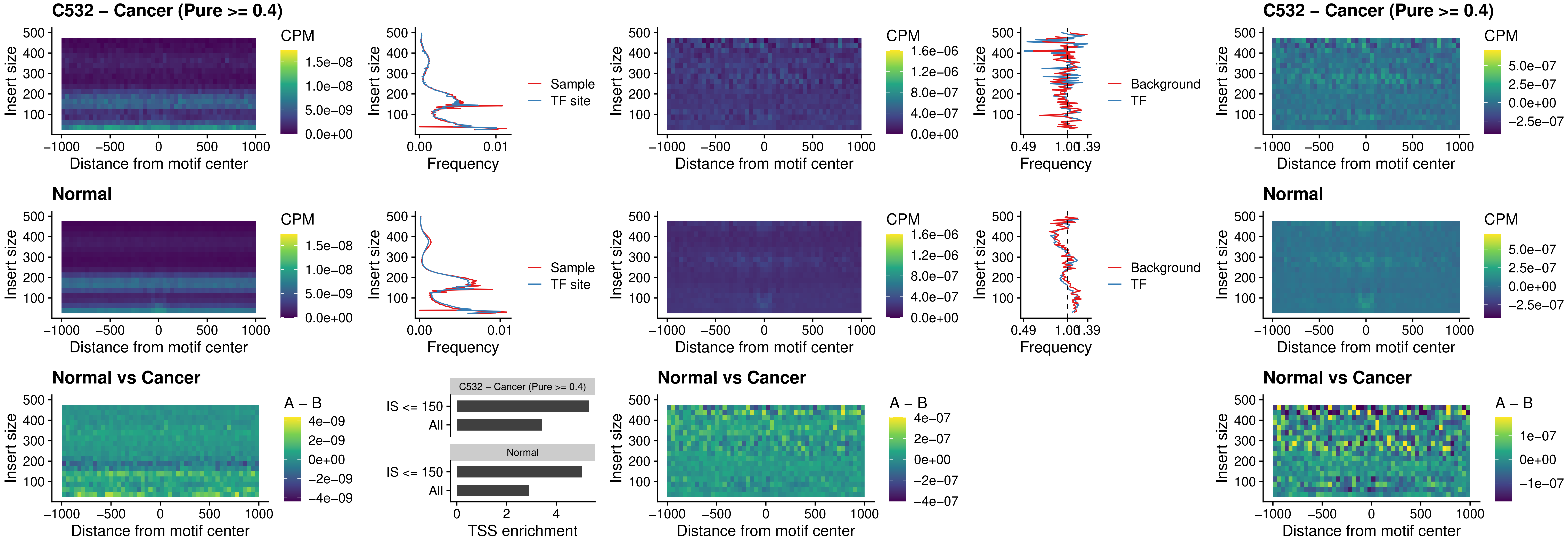

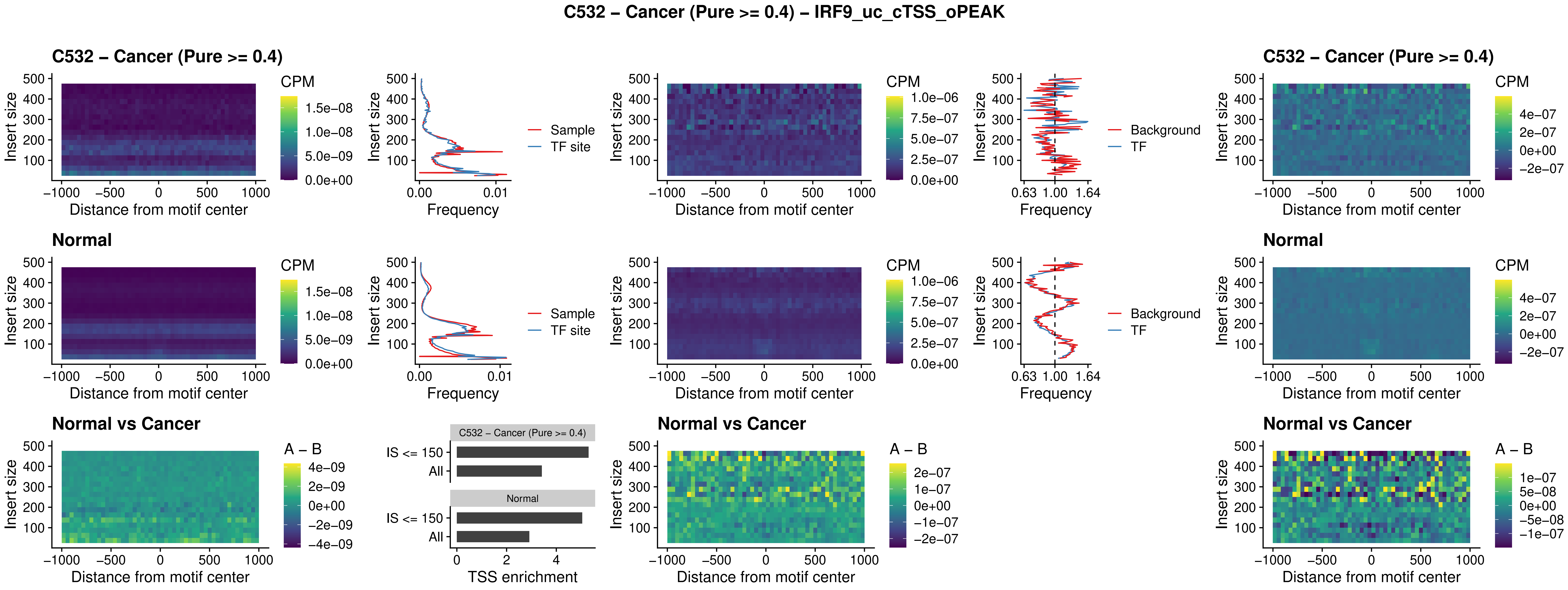
C532 – Cancer (Pure >= 0.4) – IRF9_uc_cTSS_oPEAK
C532 – Cancer (Pure >= 0.4)
Normal
Normal vs Cancer
Insert size
Distance from motif center
Frequency
CPM
A – B
Sample
TF site
Background
TF
C532 – Cancer (Pure >= 0.4)
Normal
IS <= 150
All
TSS enrichment

#### C532 – Cancer (Pure >= 0.4) – BCL11A_uc_cTSS_oPEAK

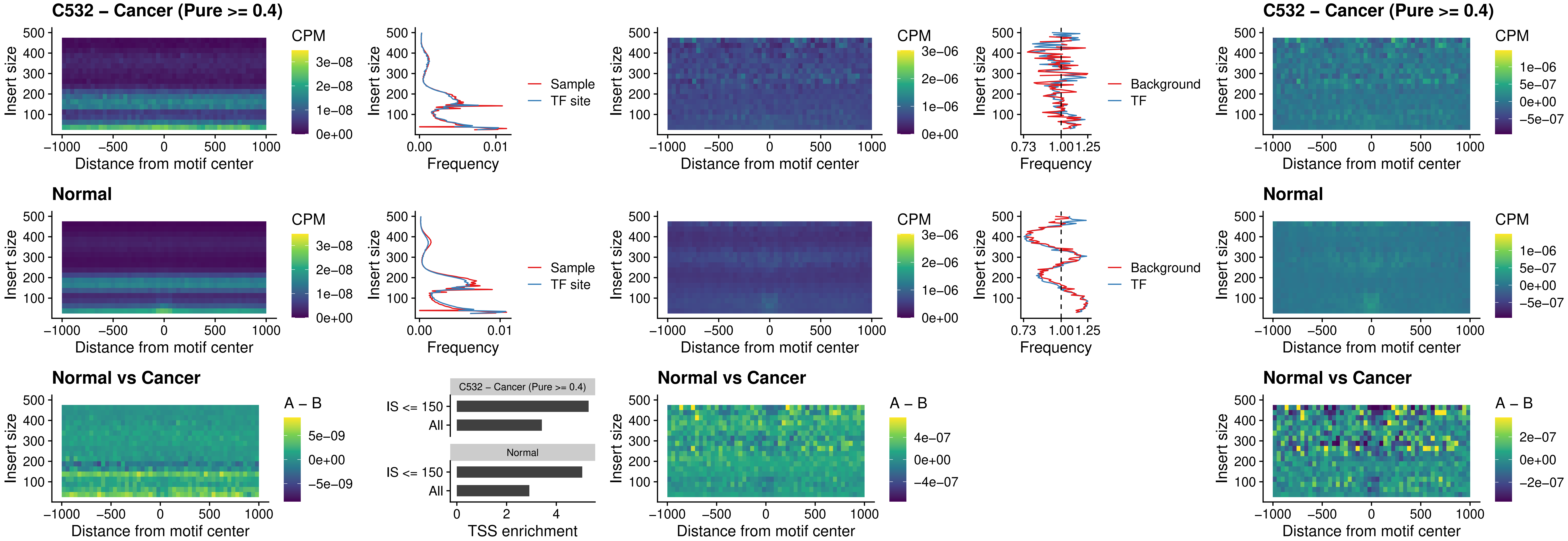

#### C532 – Cancer (Pure >= 0.4) – BCL11B_uc_cTSS_oPEAK

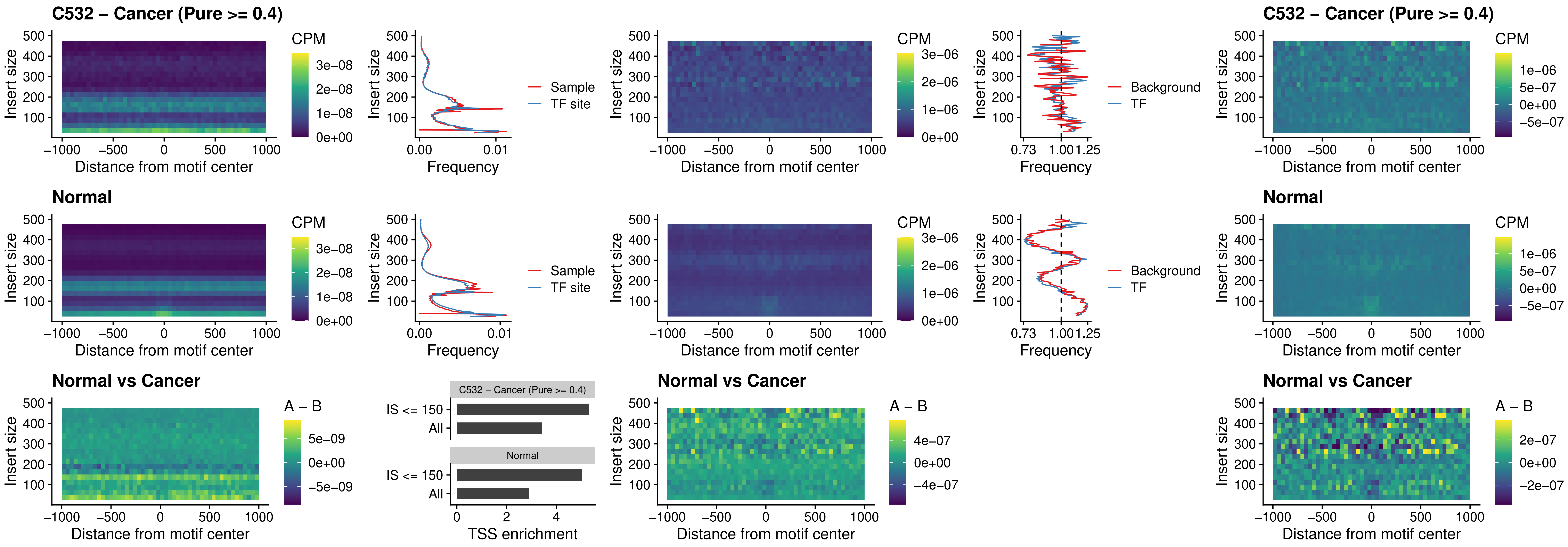

#### C532 – Cancer (Pure >= 0.4) – BCL11A_uc_dTSS_oPEAK

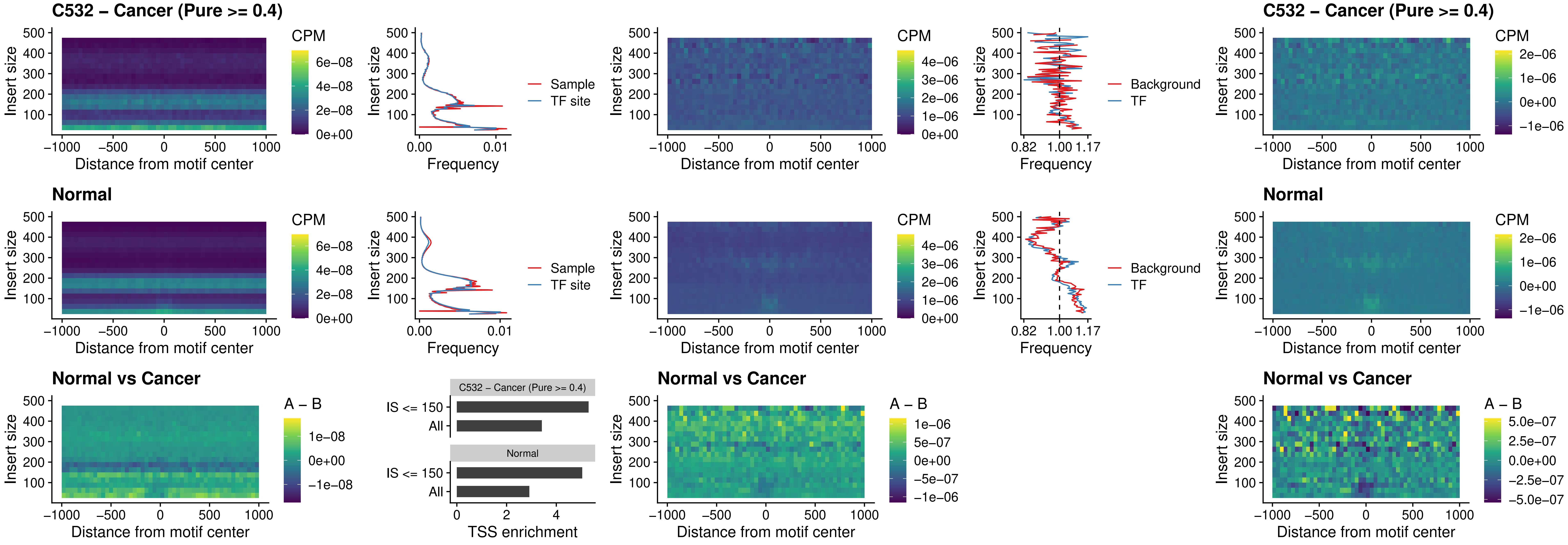

#### C532 – Cancer (Pure >= 0.4) – BCL11B_uc_dTSS_oPEAK

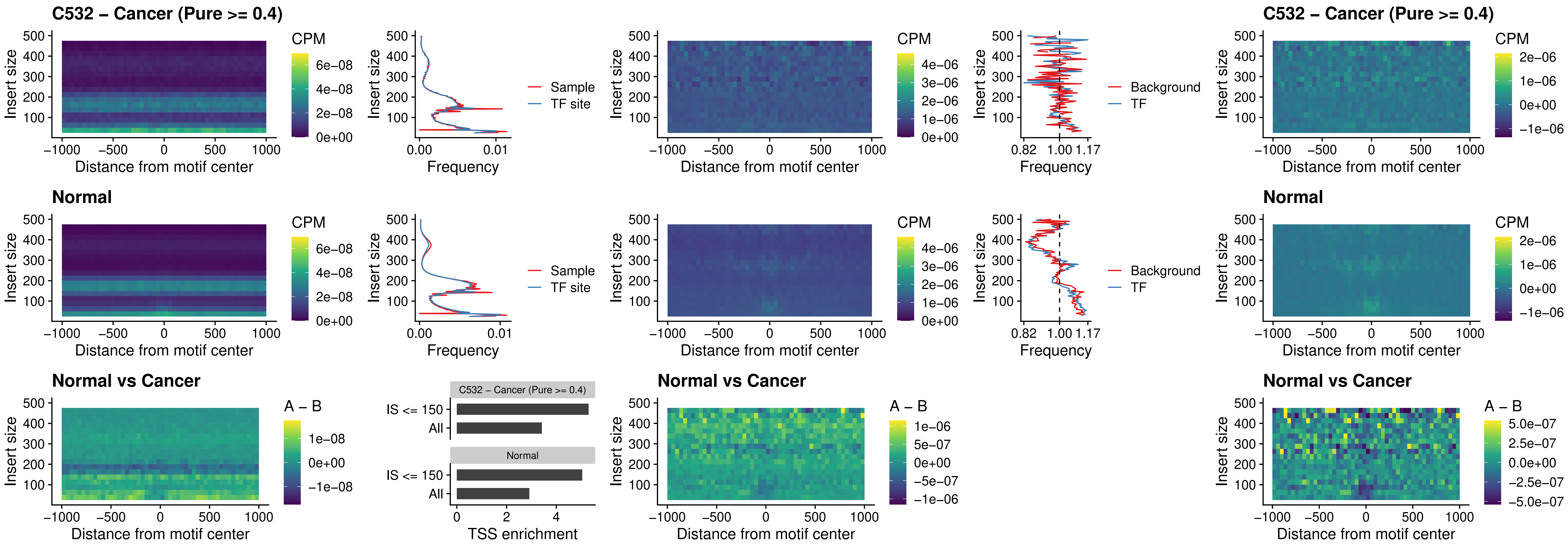

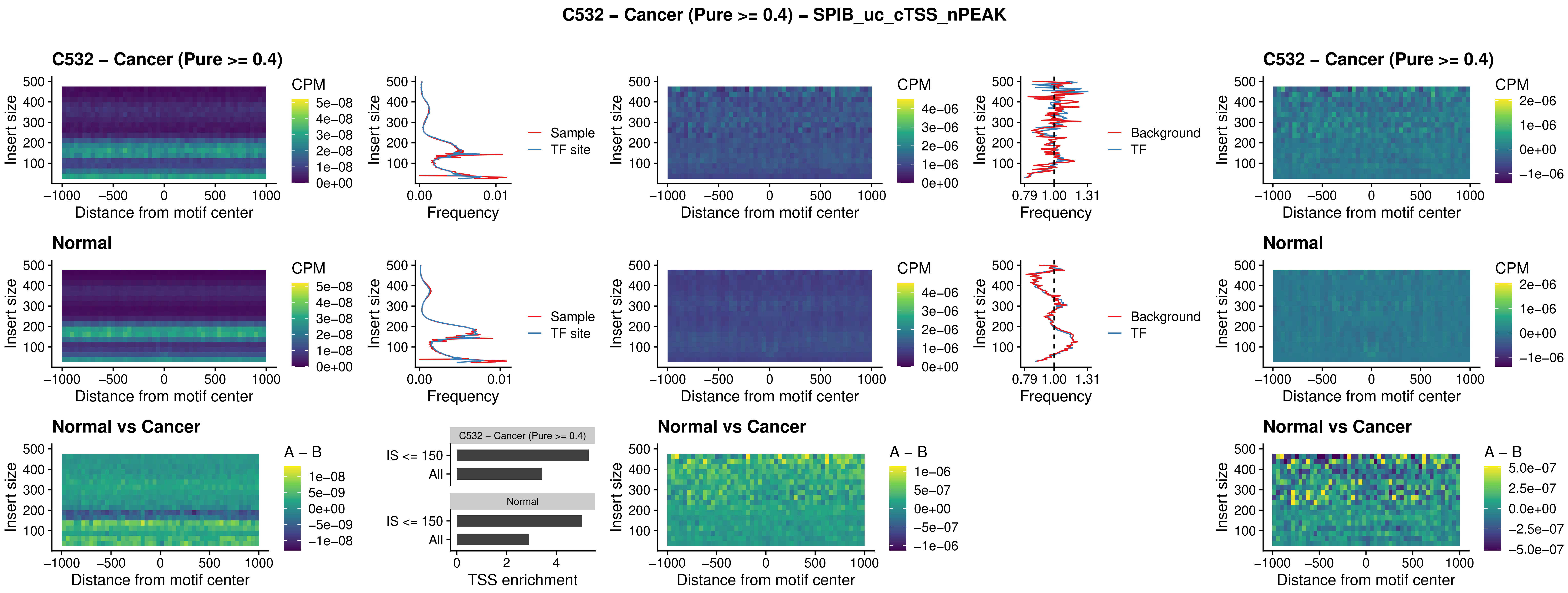
C532 – Cancer (Pure >= 0.4) – SPIB_uc_cTSS_nPEAK
C532 – Cancer (Pure >= 0.4)
Normal
Normal vs Cancer
Insert size
Distance from motif center
CPM
A – B
Frequency
Sample
TF site
Background
TF
C532 – Cancer (Pure >= 0.4)
IS <= 150
All
TSS enrichment

#### C532 – Cancer (Pure >= 0.4) – SPIC_uc_cTSS_nPEAK

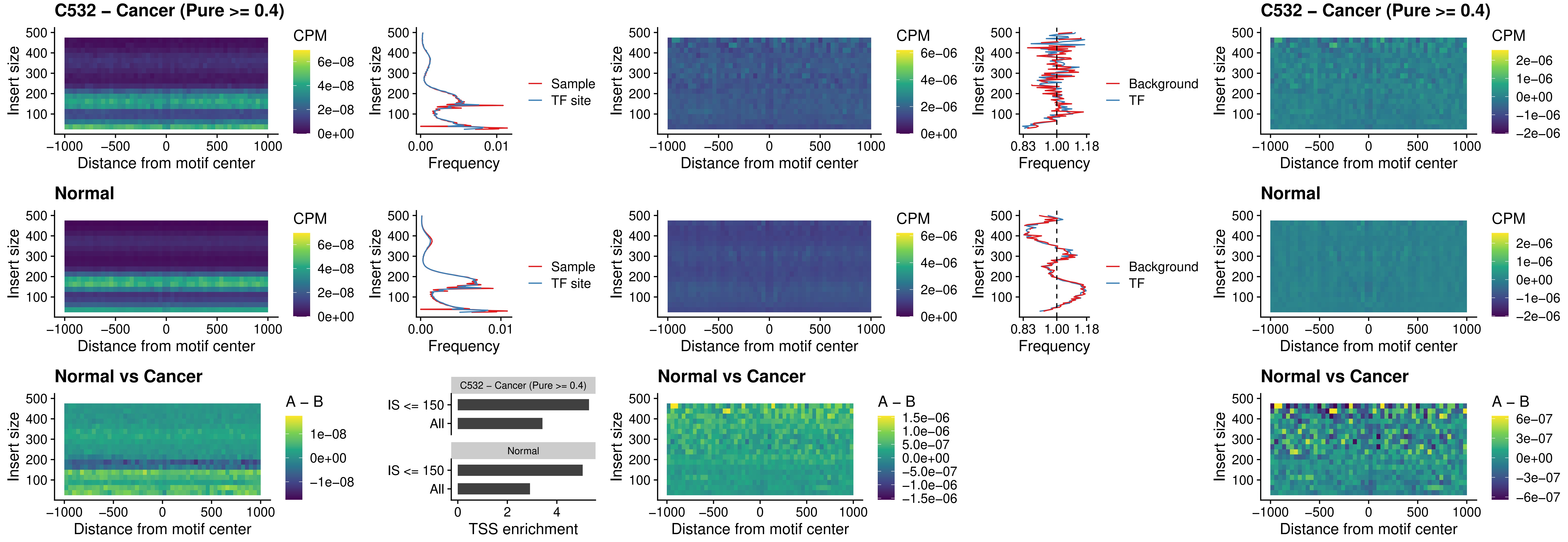

#### C532 – Cancer (Pure >= 0.4) – SPI1_uc_pTSS_oPEAK

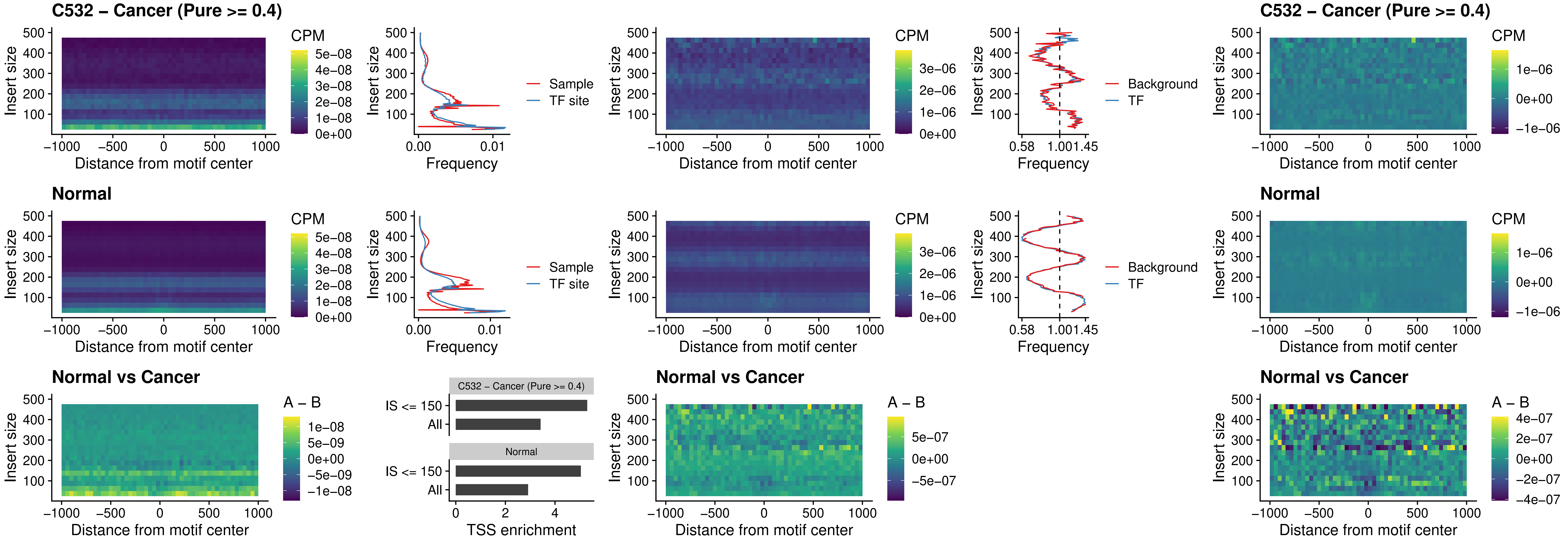

#### C532 – Cancer (Pure >= 0.4) – BCL11A_uc_pTSS_oPEAK

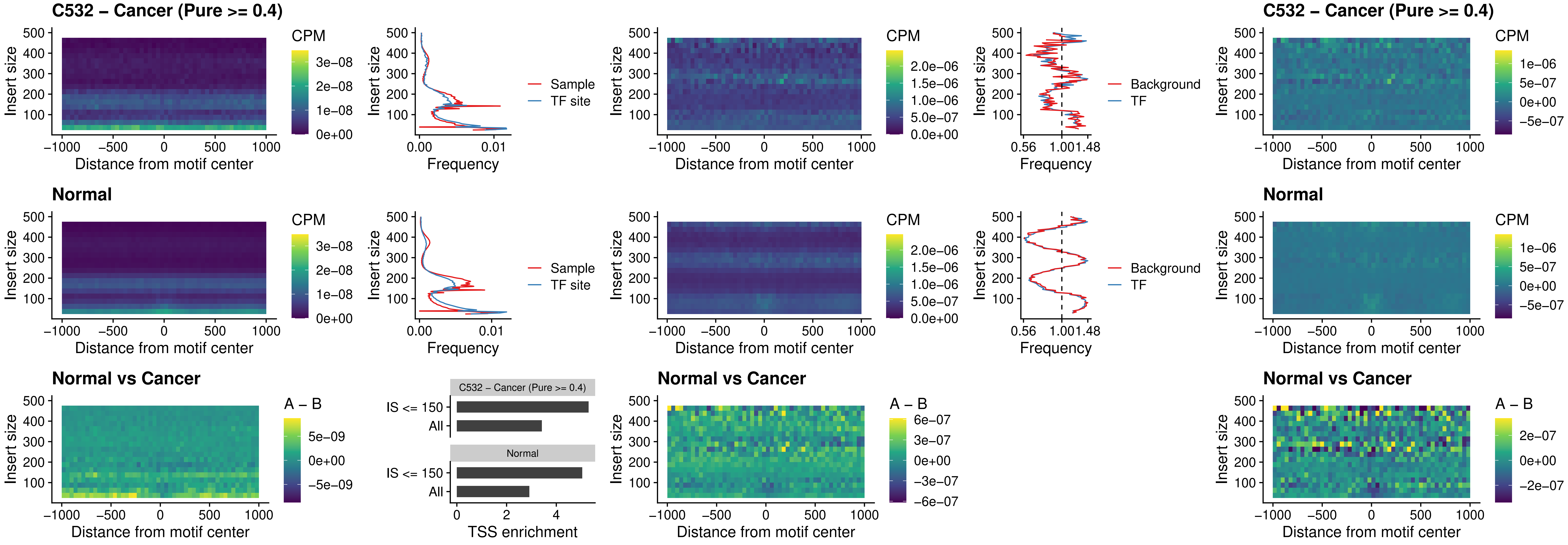

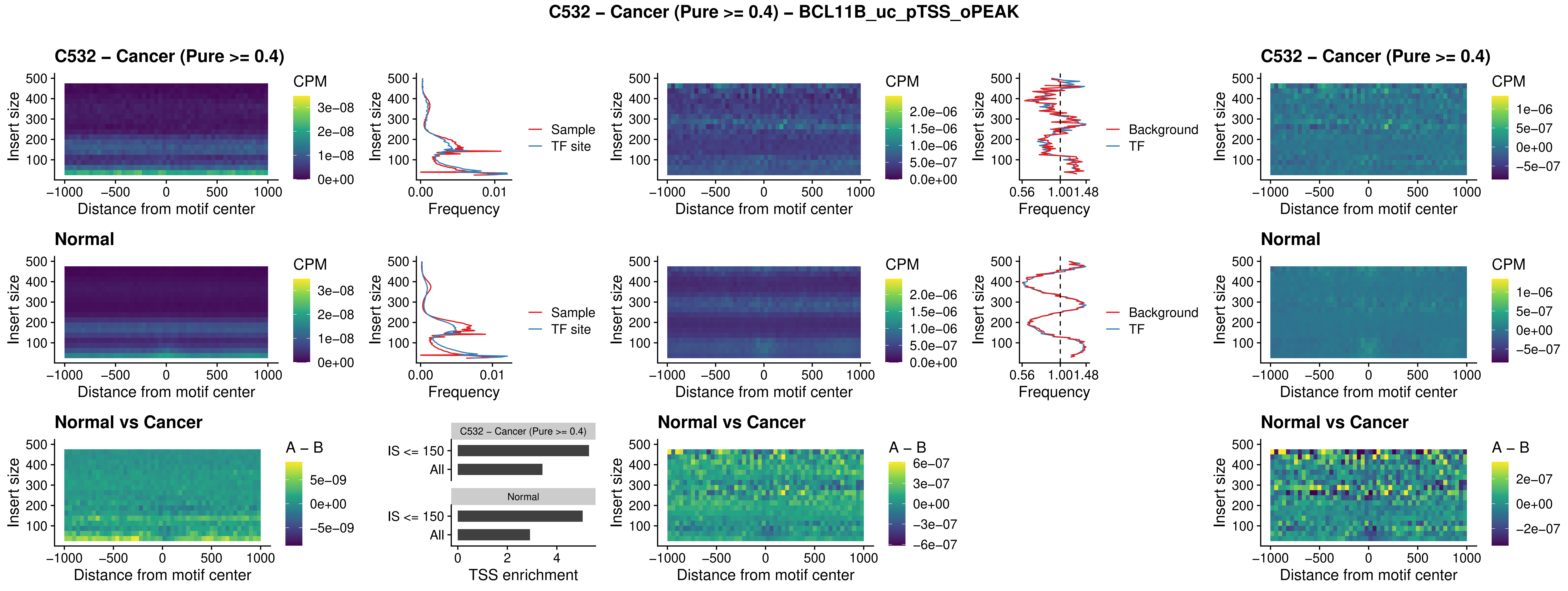
C532 – Cancer (Pure >= 0.4) – BCL11B_uc_pTSS_oPEAK
C532 – Cancer (Pure >= 0.4)
Normal
Normal vs Cancer
Insert size
Distance from motif center
CPM
A – B
Frequency
Sample
TF site
Background
TF
C532 – Cancer (Pure >= 0.4)
Normal
IS <= 150
All
TSS enrichment

#### C532 – Cancer (Pure >= 0.4) – SPI1_uc_cTSS_oPEAK

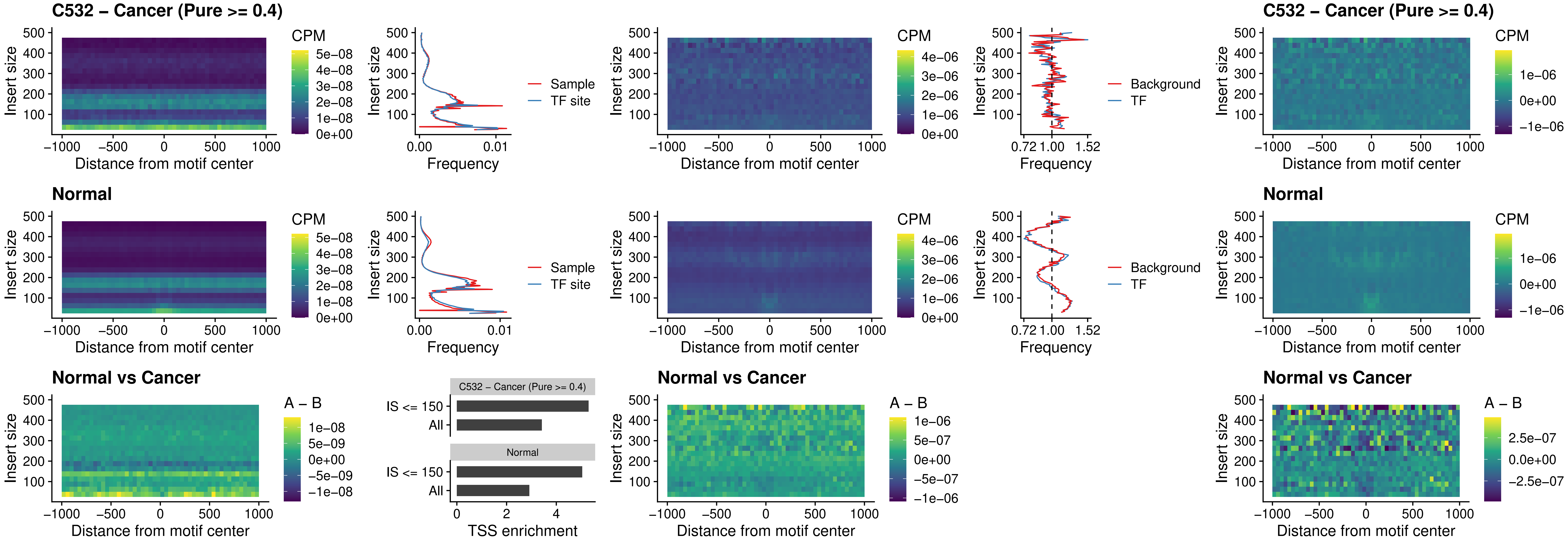

#### C532 – Cancer (Pure >= 0.4) – SPIB_uc_cTSS_oPEAK

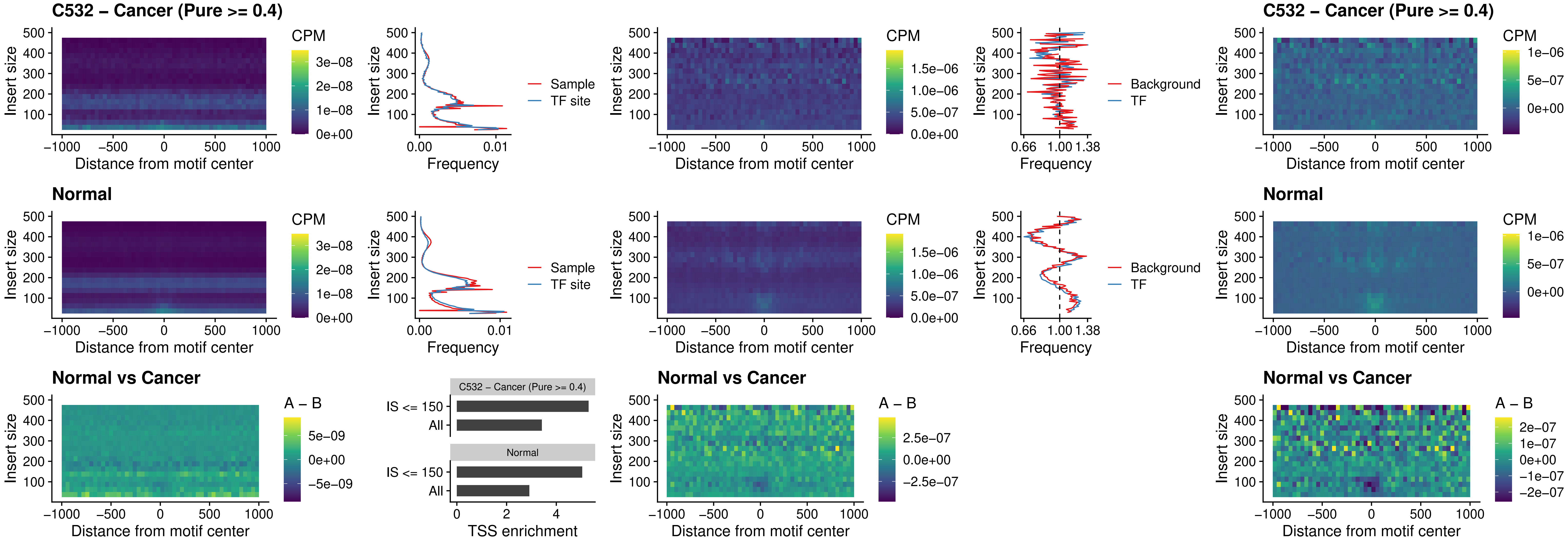

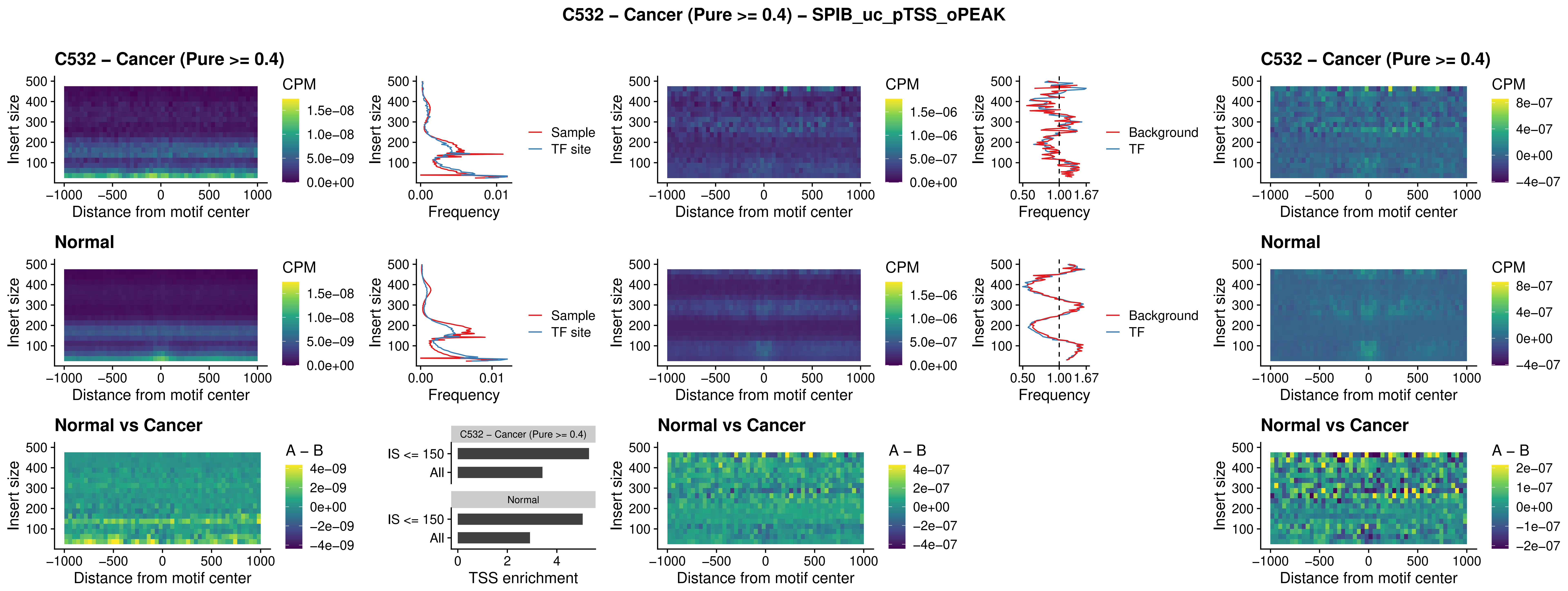

C532 – Cancer (Pure >= 0.4) – SPIB_uc_pTSS_oPEAK
C532 – Cancer (Pure >= 0.4)
Normal
Normal vs Cancer
Insert size
Distance from motif center
Frequency
CPM
A – B
Sample
TF site
Background
TF
C532 – Cancer (Pure >= 0.4)
Normal
IS <= 150
All
TSS enrichment

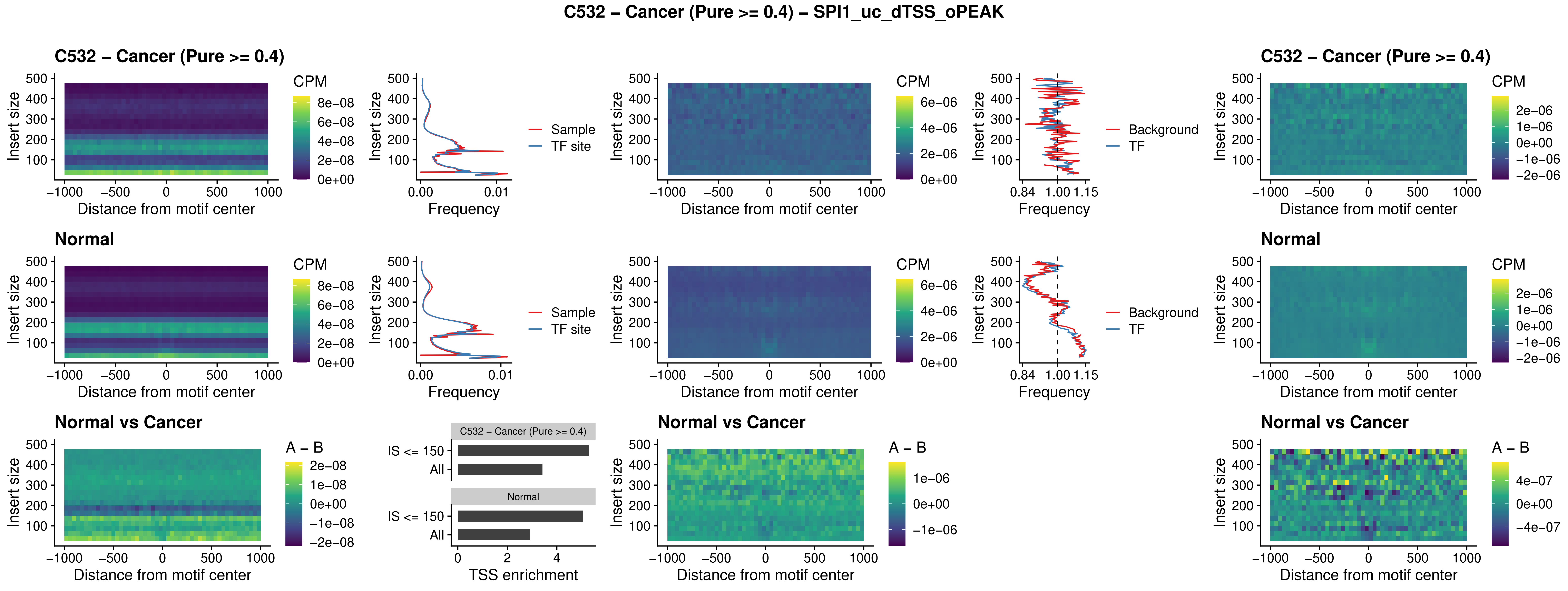
C532 – Cancer (Pure >= 0.4) – SPI1_uc_dTSS_oPEAK
C532 – Cancer (Pure >= 0.4)
Normal
Normal vs Cancer
Insert size
Distance from motif center
Frequency
CPM
A – B
Sample
TF site
Background
TF
C532 – Cancer (Pure >= 0.4)
Normal
IS <= 150
All
TSS enrichment

#### C532 – Cancer (Pure >= 0.4) – SPIB_uc_dTSS_oPEAK

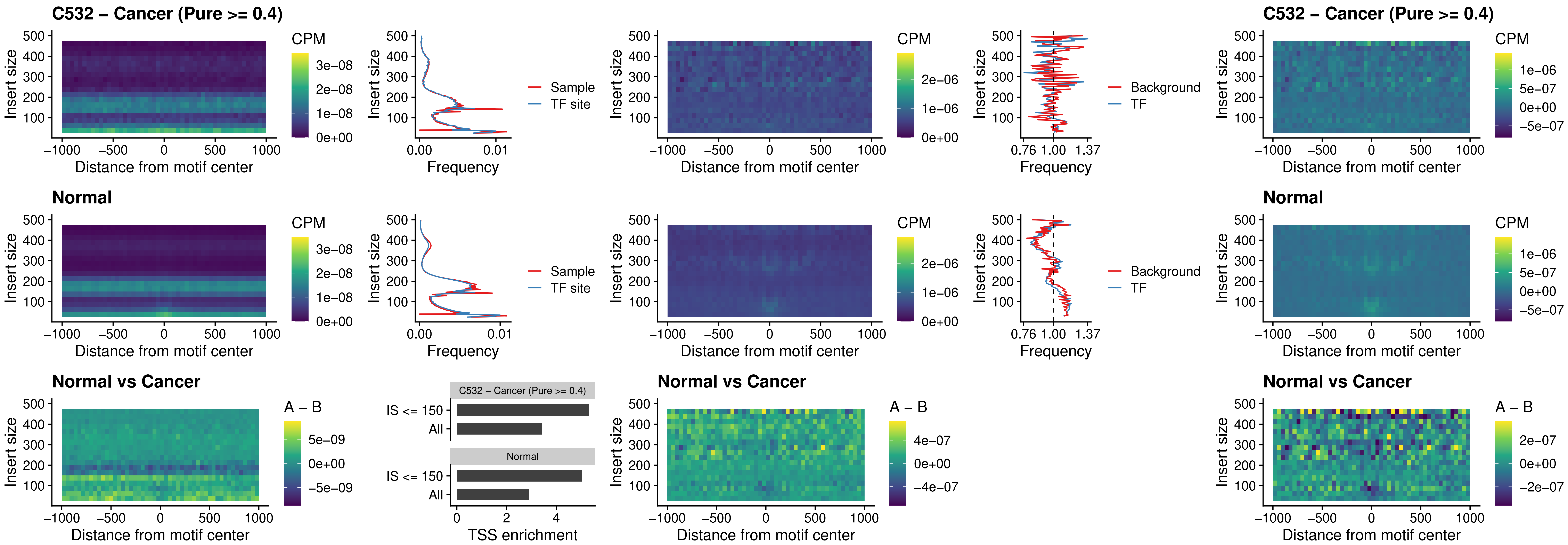

#### C532 – Cancer (Pure >= 0.4) – IRF3_uc_pTSS_oPEAK

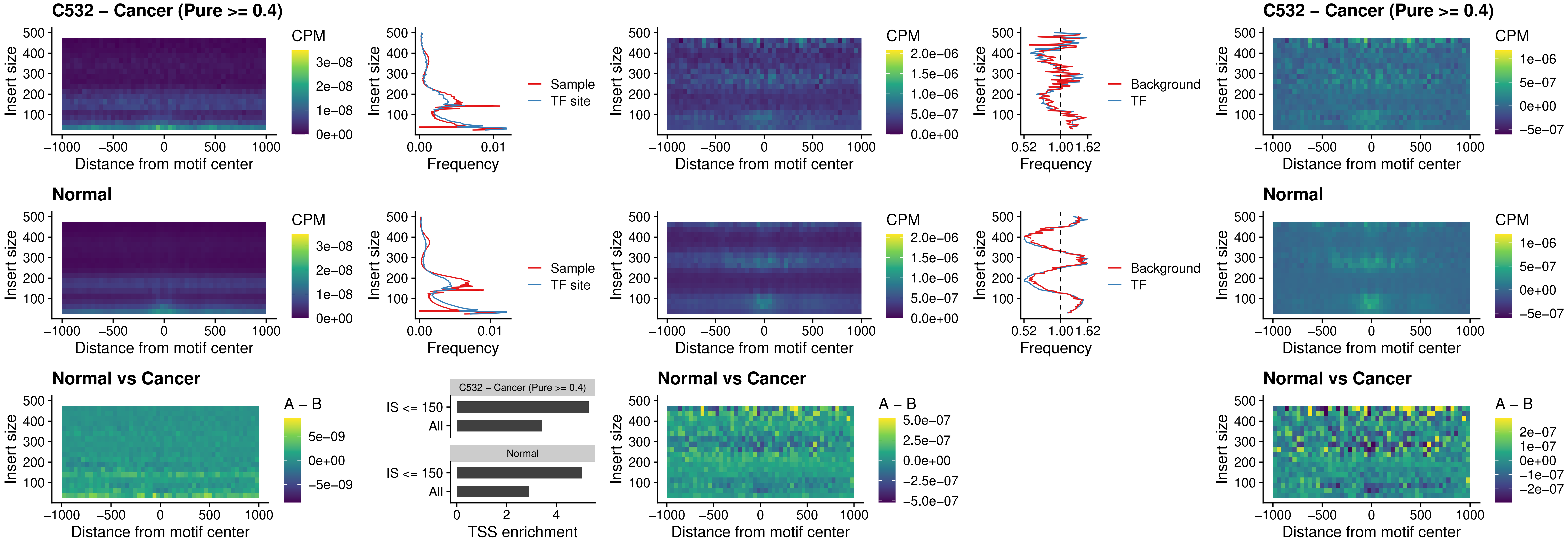

### C532 – Cancer (Pure >= 0.4) – IRF2_uc_dTSS_oPEAK

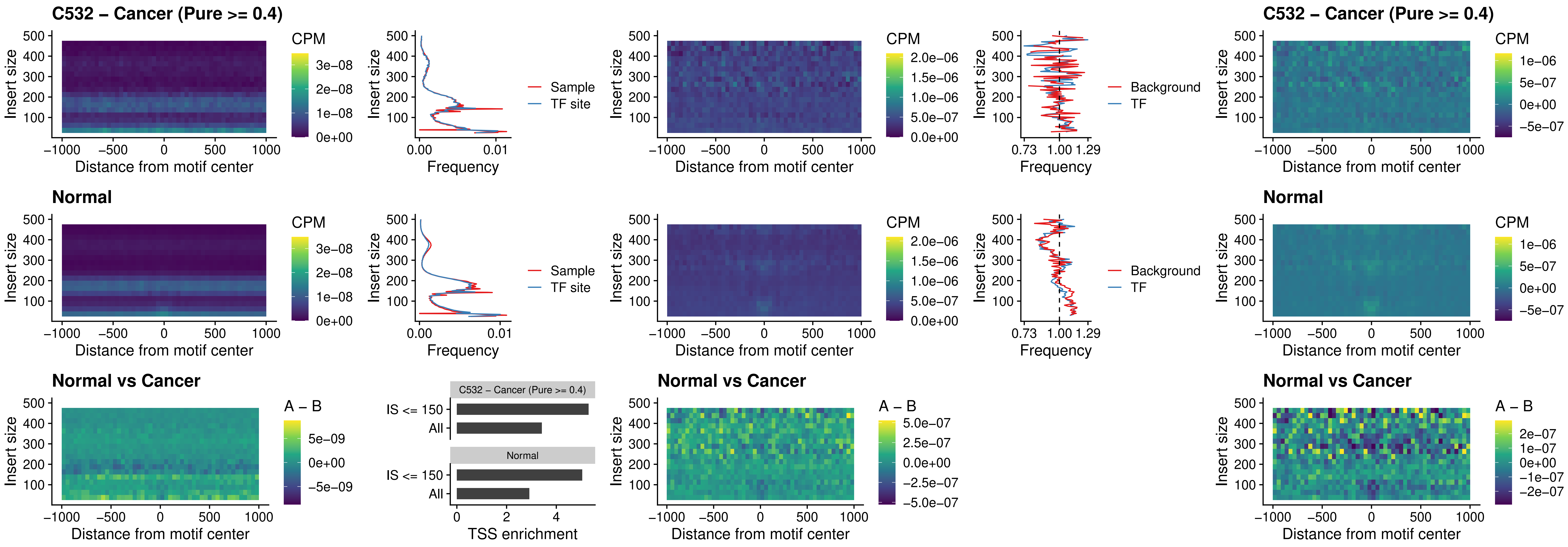

#### C532 – Cancer (Pure >= 0.4) – IRF2_uc_pTSS_oPEAK

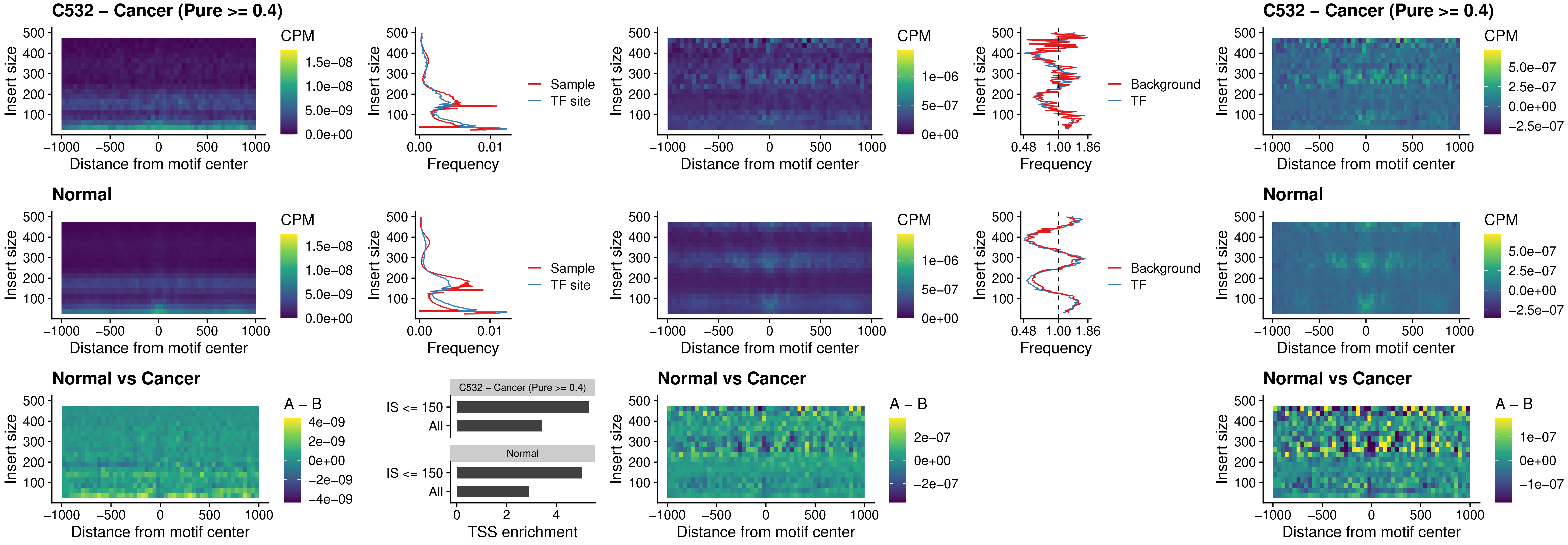

### C532 – Cancer (Pure >= 0.4) – STAT2_uc_pTSS_oPEAK

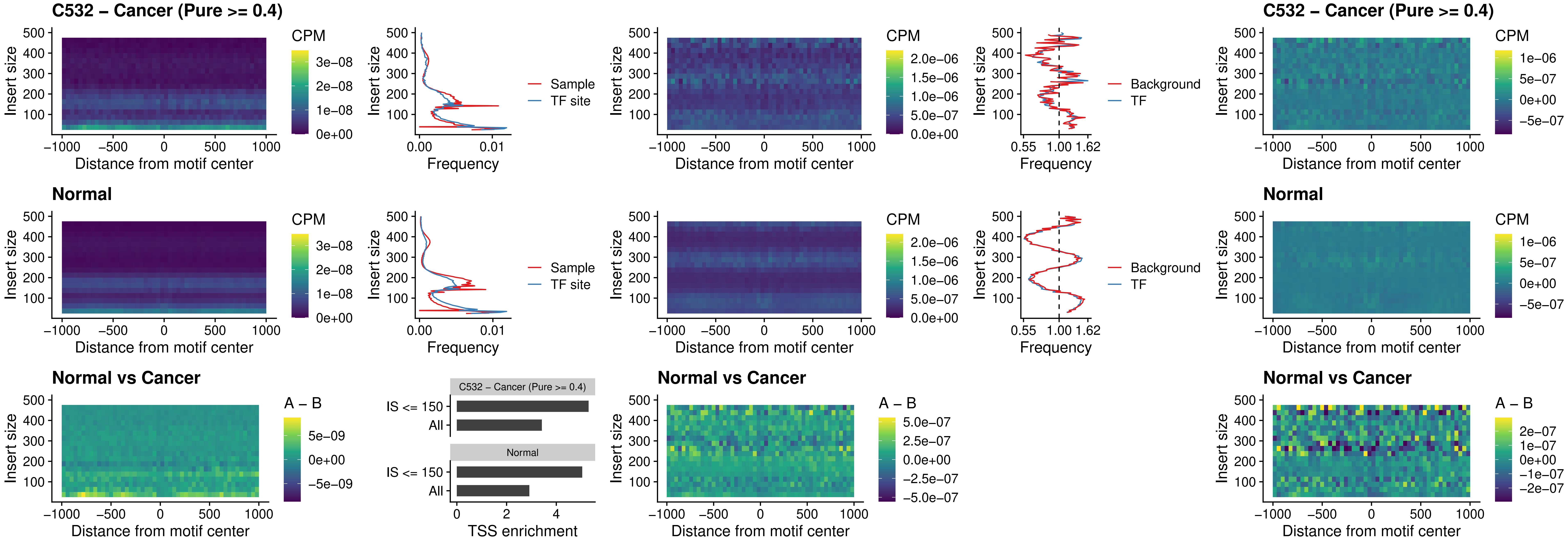

### C532 – Cancer (Pure >= 0.4) – SPIB_uc_dTSS_nPEAK

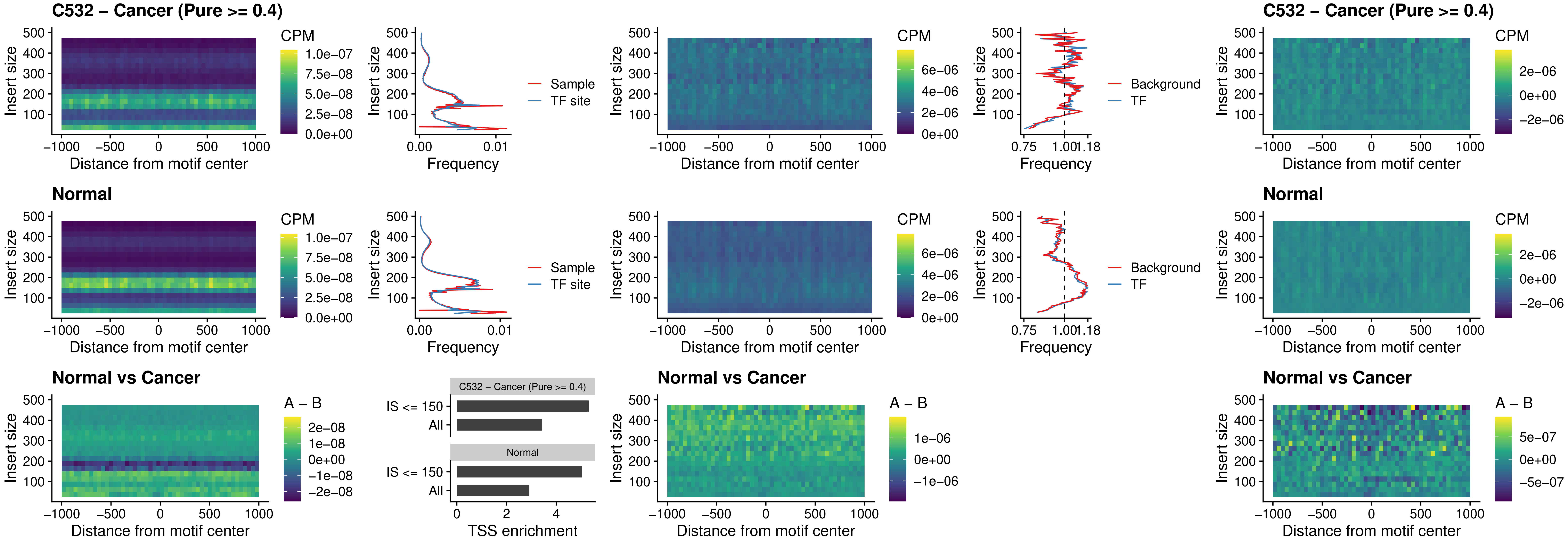

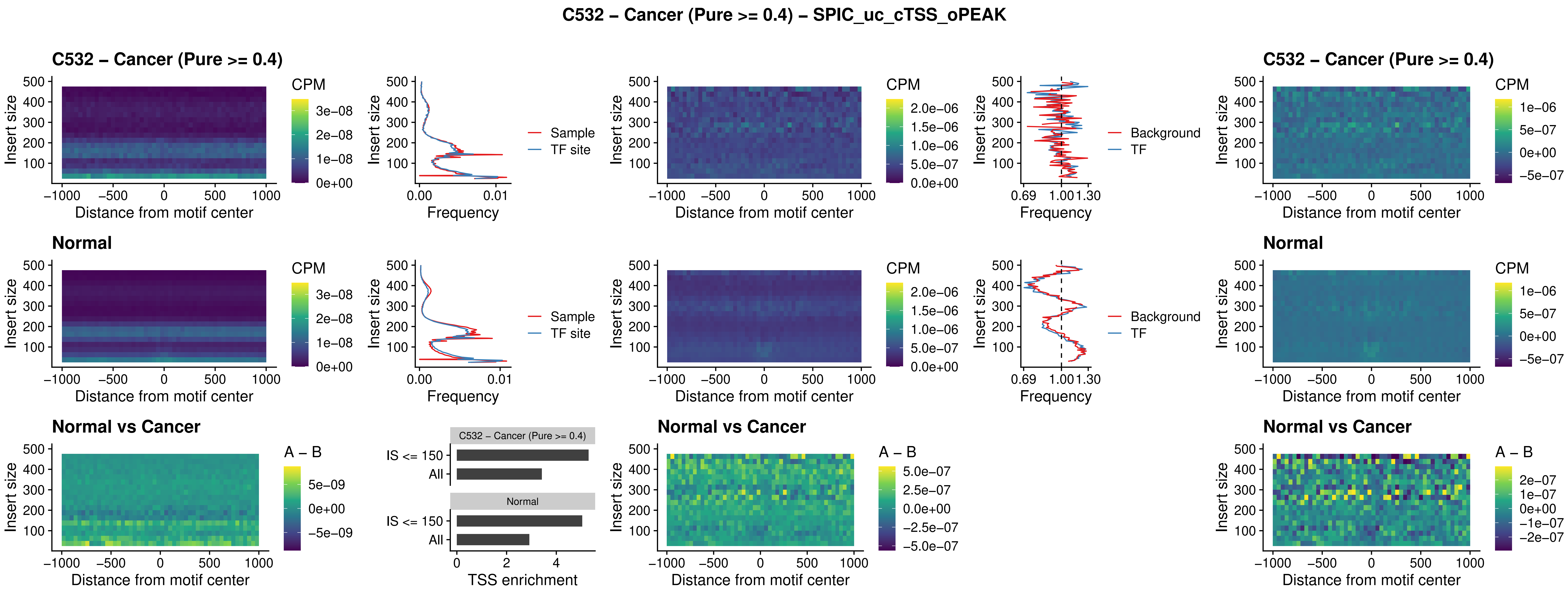
C532 – Cancer (Pure >= 0.4) – SPIC_uc_cTSS_oPEAK
C532 – Cancer (Pure >= 0.4)
Normal
Normal vs Cancer
Insert size
Distance from motif center
Frequency
CPM
A – B
Sample
TF site
Background
TF
C532 – Cancer (Pure >= 0.4)
Normal
IS <= 150
All
TSS enrichment

#### C532 – Cancer (Pure >= 0.4) – PRDM1_uc_cTSS_oPEAK

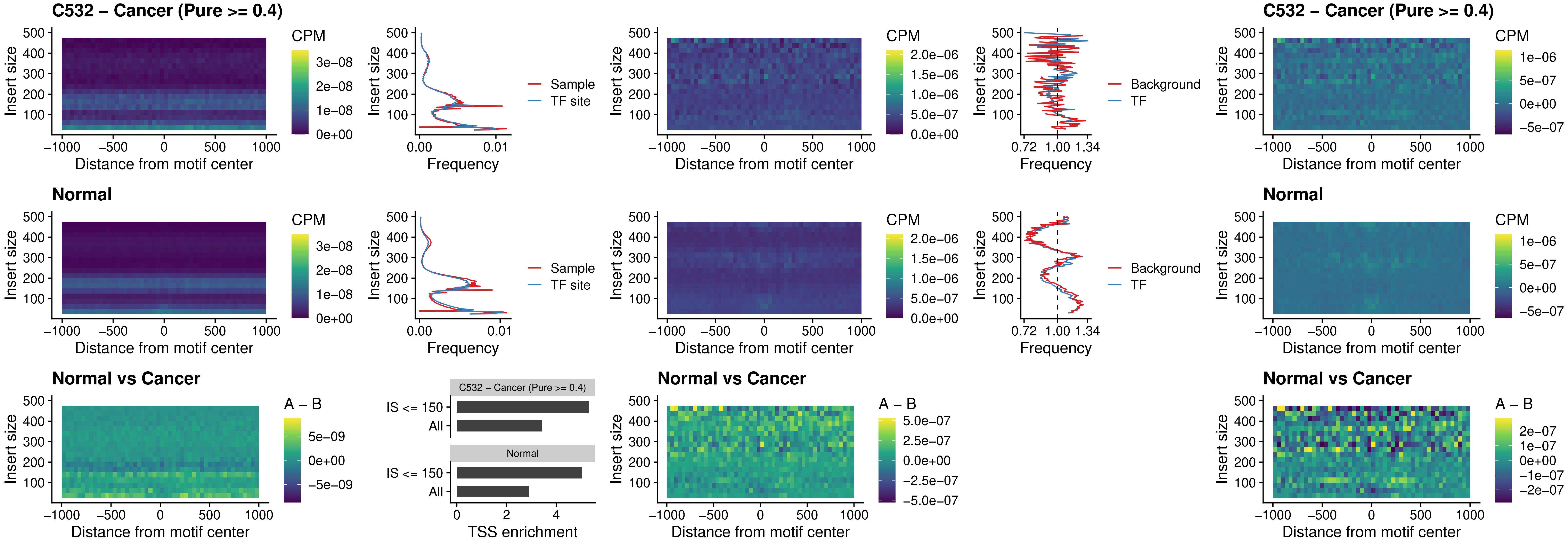

#### C532 – Cancer (Pure >= 0.4) – IRF3_uc_cTSS_oPEAK

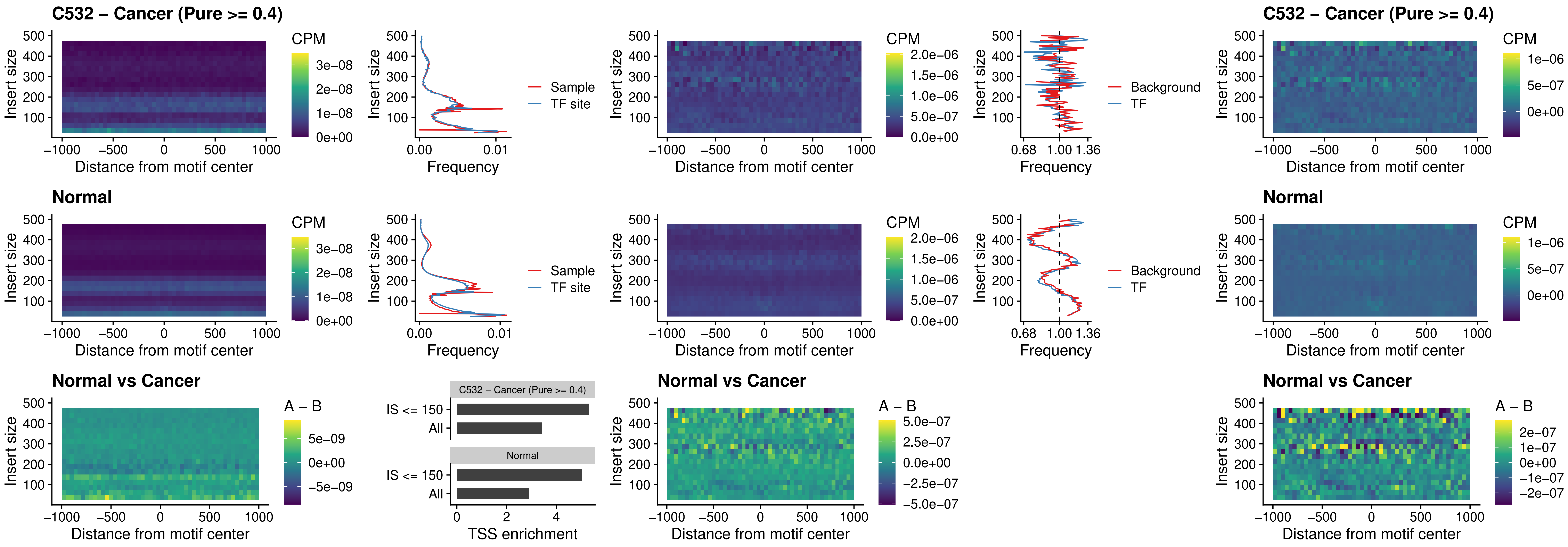

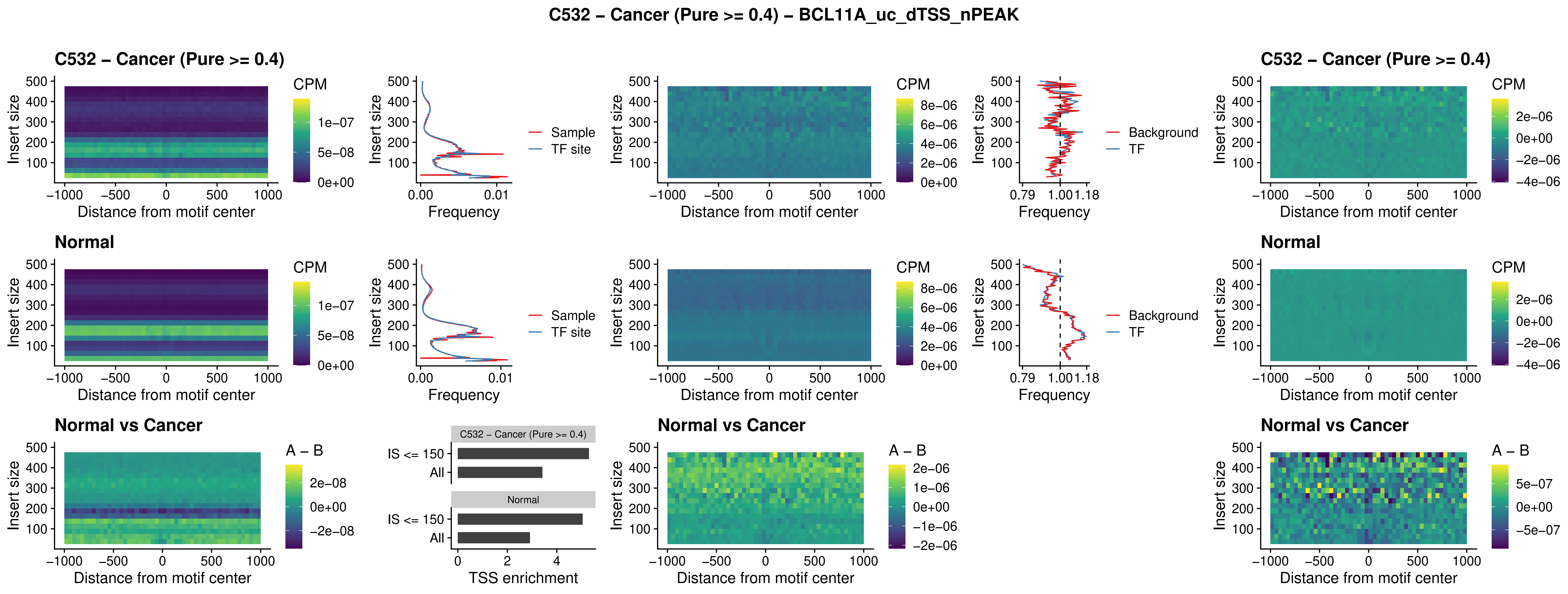

C532 – Cancer (Pure >= 0.4) – BCL11A_uc_dTSS_nPEAK
C532 – Cancer (Pure >= 0.4)
Normal
Normal vs Cancer
Insert size
Distance from motif center
CPM
A – B
Frequency
Sample
TF site
Background
TF
C532 – Cancer (Pure >= 0.4)
Normal
IS <= 150
All
TSS enrichment

#### C532 – Cancer (Pure >= 0.4) – BCL11B_uc_dTSS_nPEAK

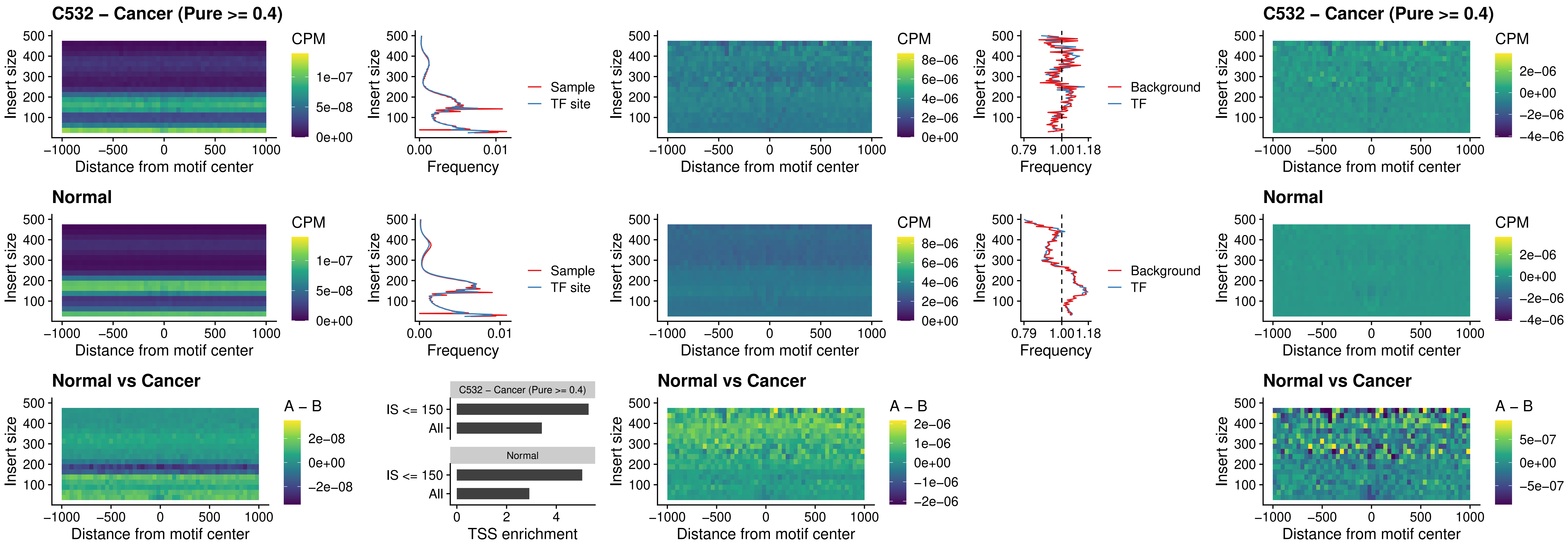

#### C532 – Cancer (Pure >= 0.4) – IRF1_uc_cTSS_oPEAK

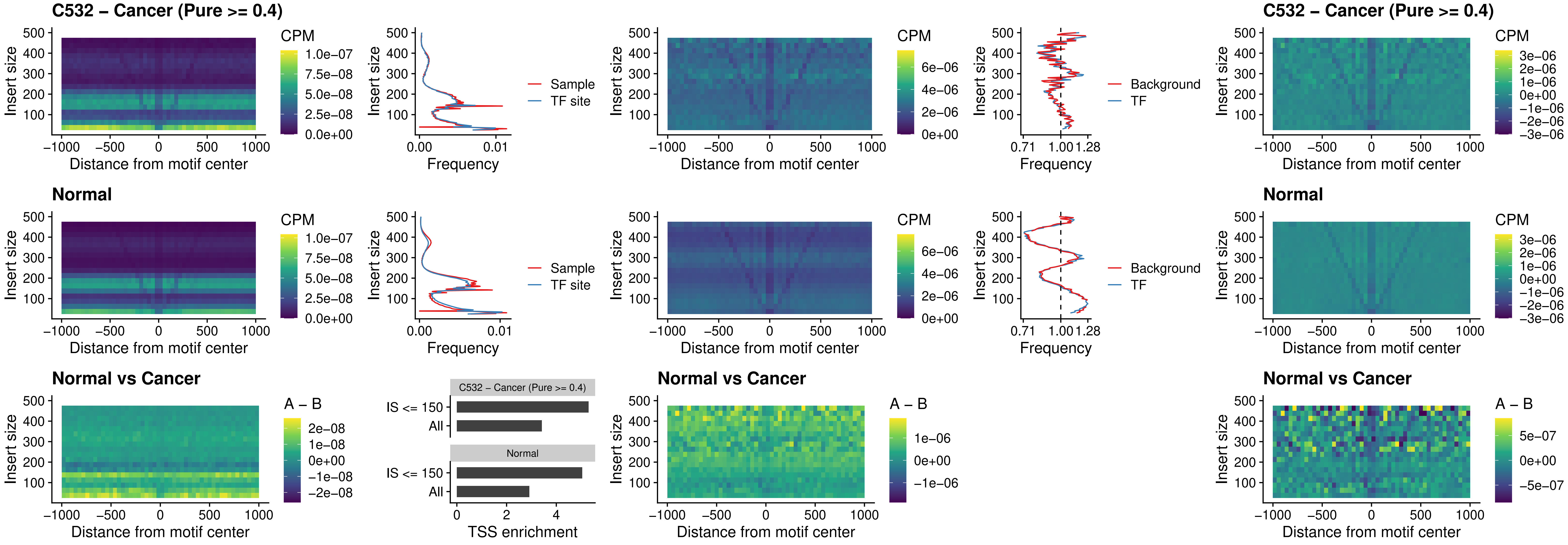

### C532 – Cancer (Pure >= 0.4) – IRF1_uc_dTSS_oPEAK

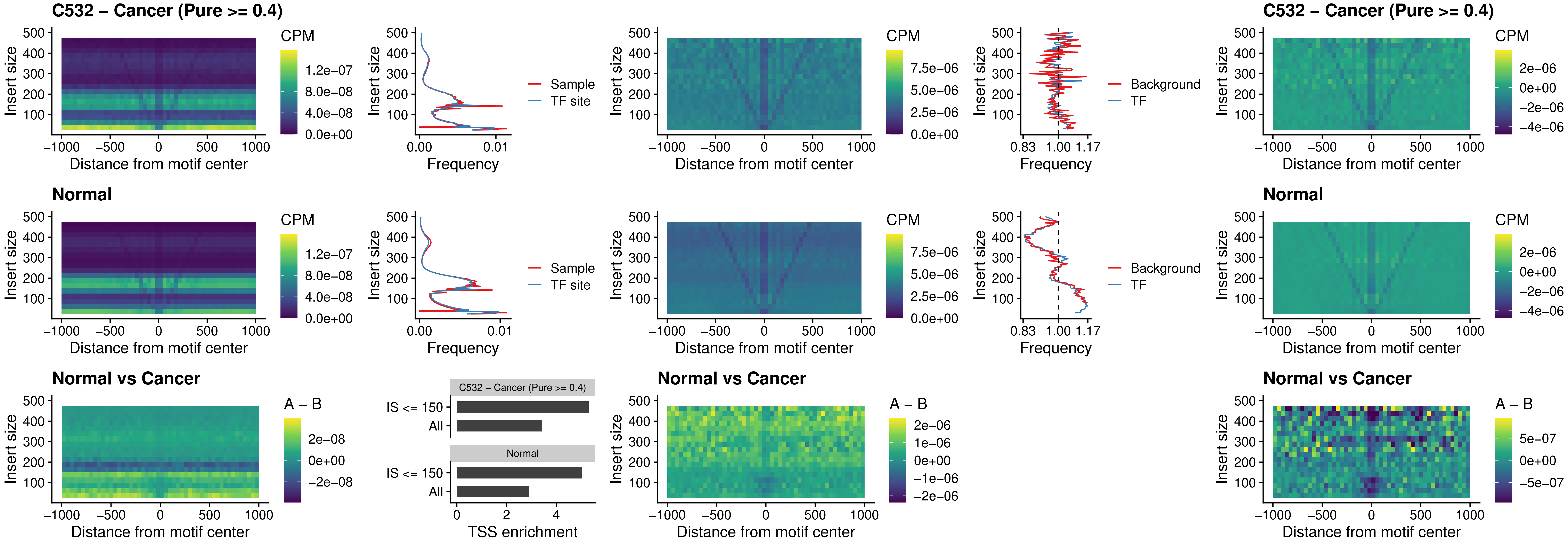

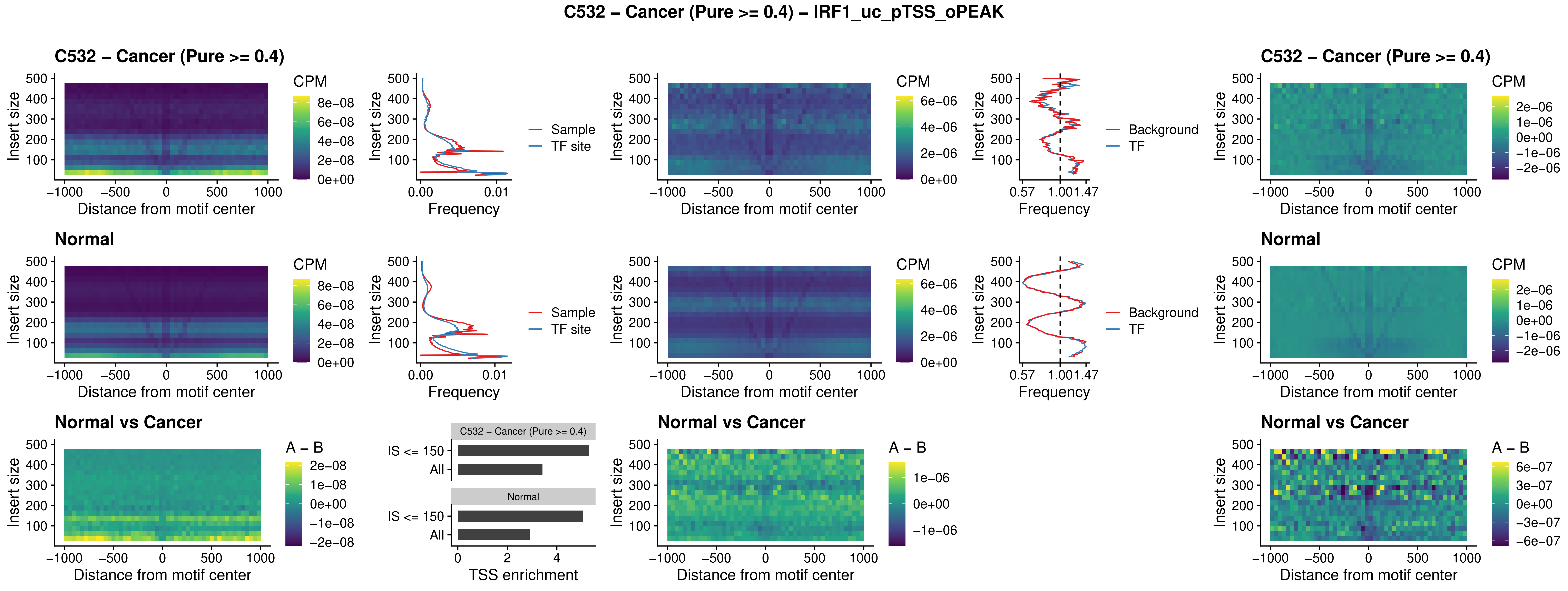

C532 – Cancer (Pure >= 0.4) – IRF1_uc_pTSS_oPEAK
C532 – Cancer (Pure >= 0.4)
Normal
Normal vs Cancer
Insert size
Distance from motif center
Frequency
CPM
A – B
Sample
TF site
Background
TF
C532 – Cancer (Pure >= 0.4)
Normal
IS <= 150
All
TSS enrichment

#### C532 – Cancer (Pure >= 0.4) – SPIC_uc_dTSS_nPEAK

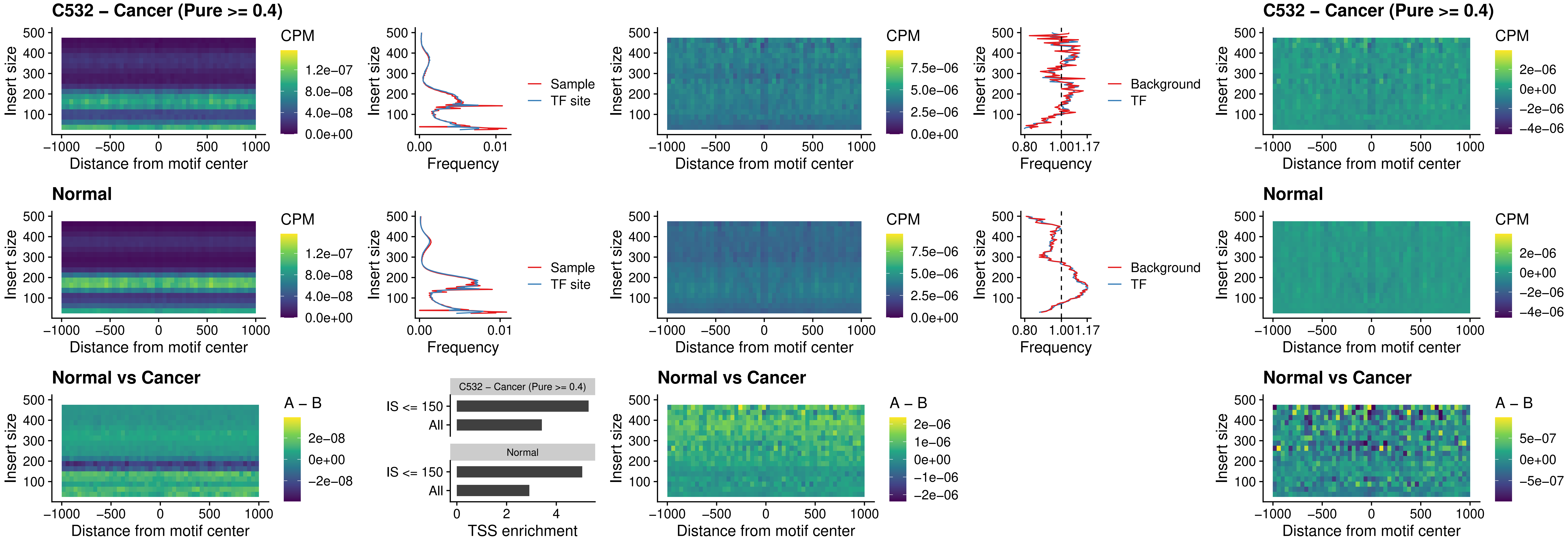

#### C532 – Cancer (Pure >= 0.4) – SPIC_uc_dTSS_oPEAK

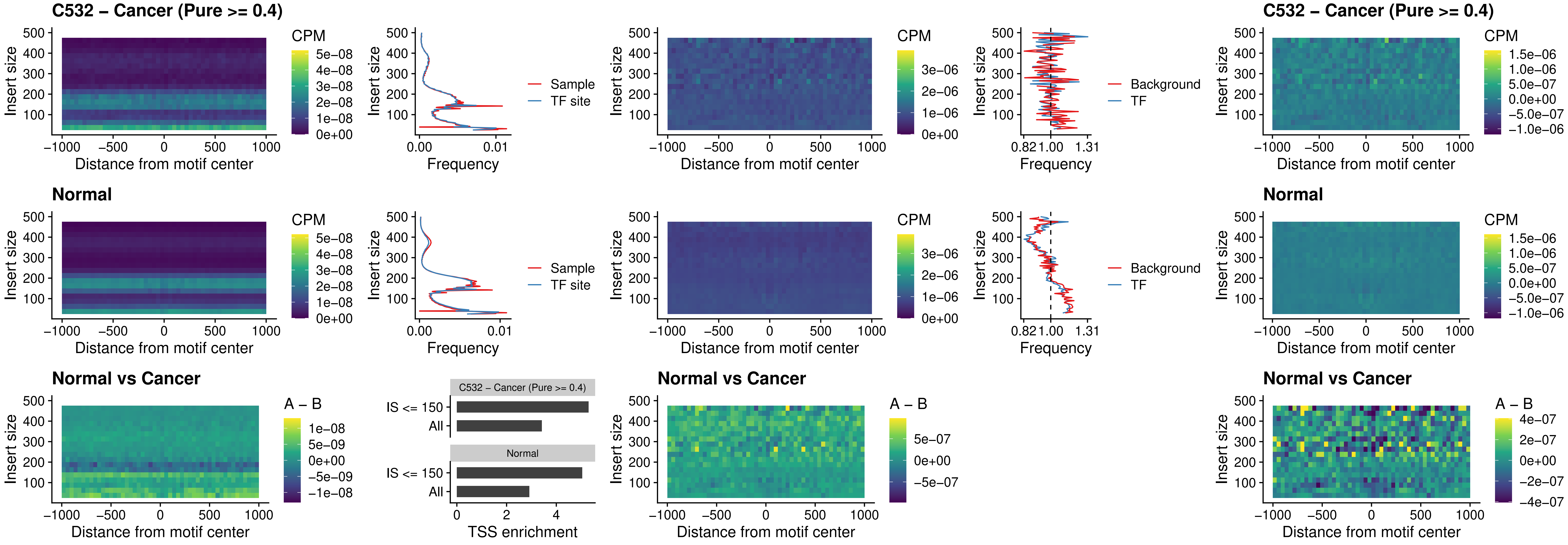

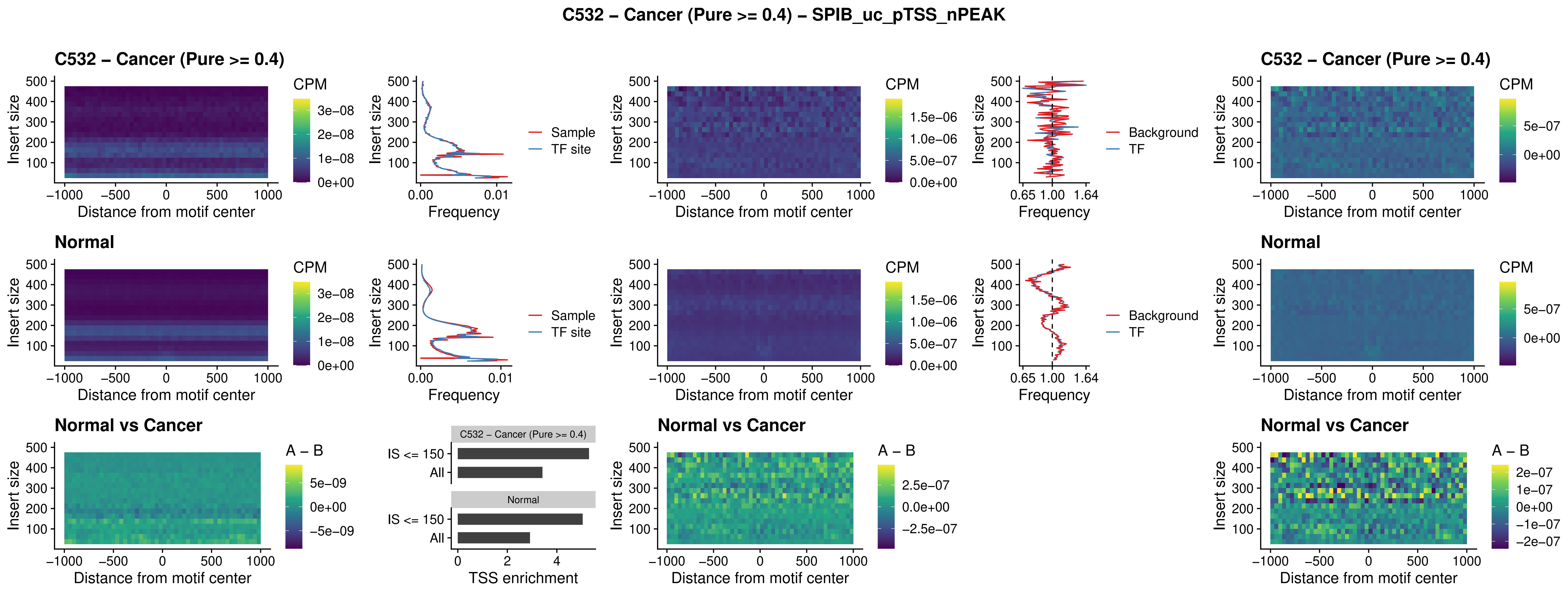

C532 – Cancer (Pure >= 0.4) – SPIB_uc_pTSS_nPEAK
C532 – Cancer (Pure >= 0.4)
Normal
Normal vs Cancer
Insert size
Distance from motif center
CPM
A – B
Frequency
Sample
TF site
Background
TF
C532 – Cancer (Pure >= 0.4)
Normal
IS <= 150
All
TSS enrichment

#### C532 – Cancer (Pure >= 0.4) – STAT2_uc_cTSS_oPEAK

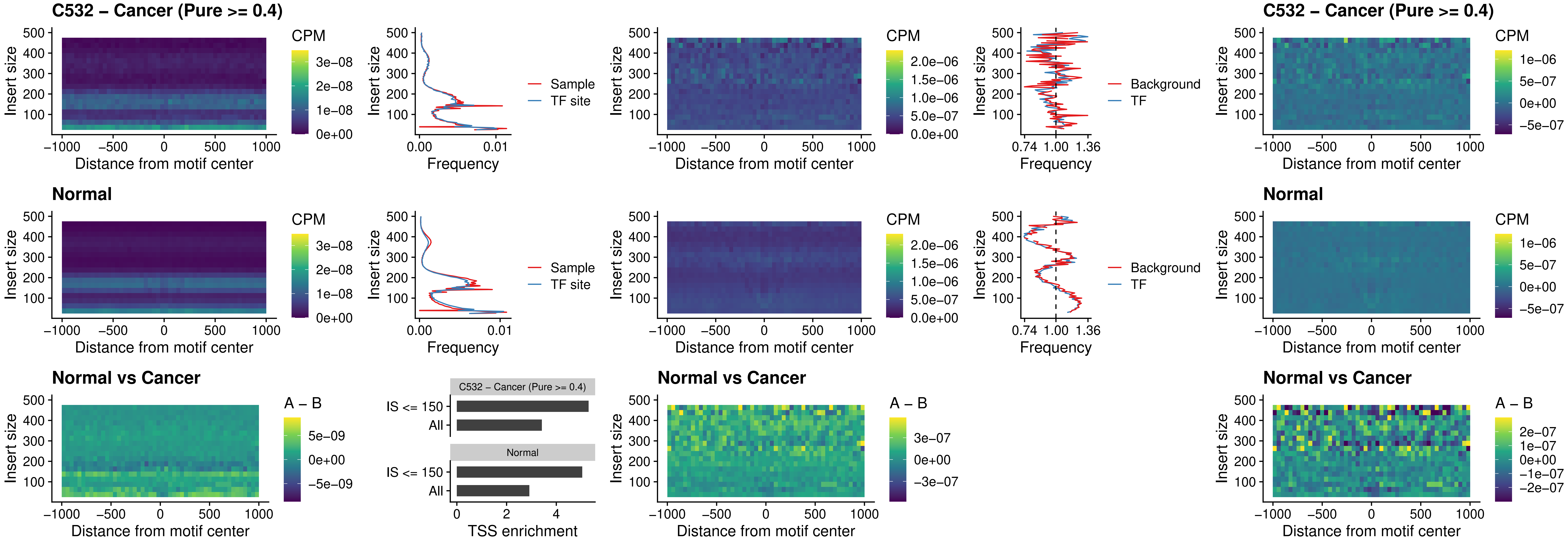

#### C532 – Cancer (Pure >= 0.4) – IRF8_uc_cTSS_oPEAK

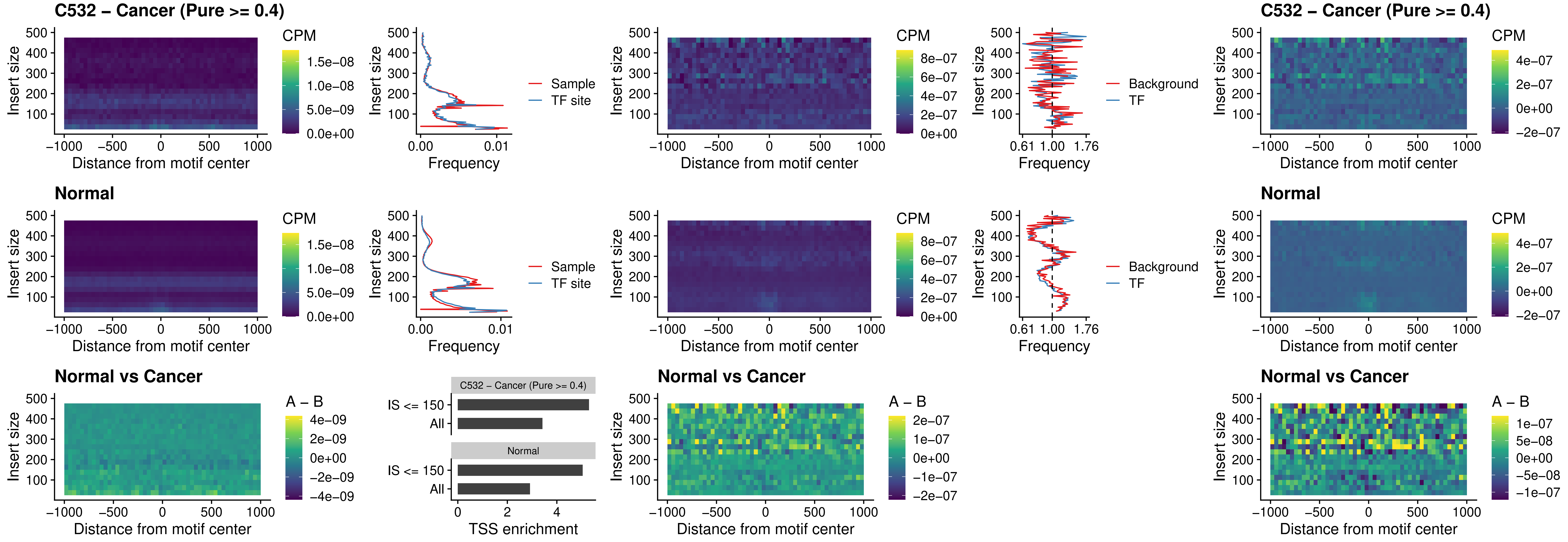

#### C532 – Cancer (Pure >= 0.4) – IRF3_uc_dTSS_oPEAK

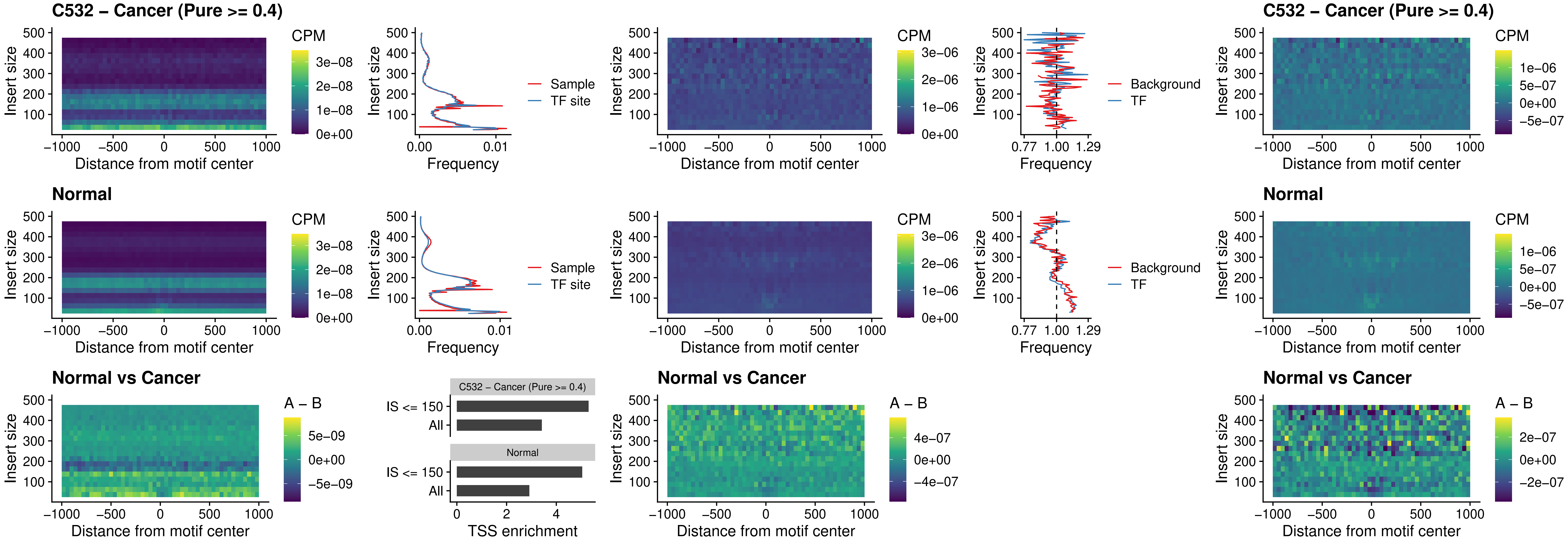

### C532 – Cancer (Pure >= 0.4) – IRF8_uc_dTSS_oPEAK

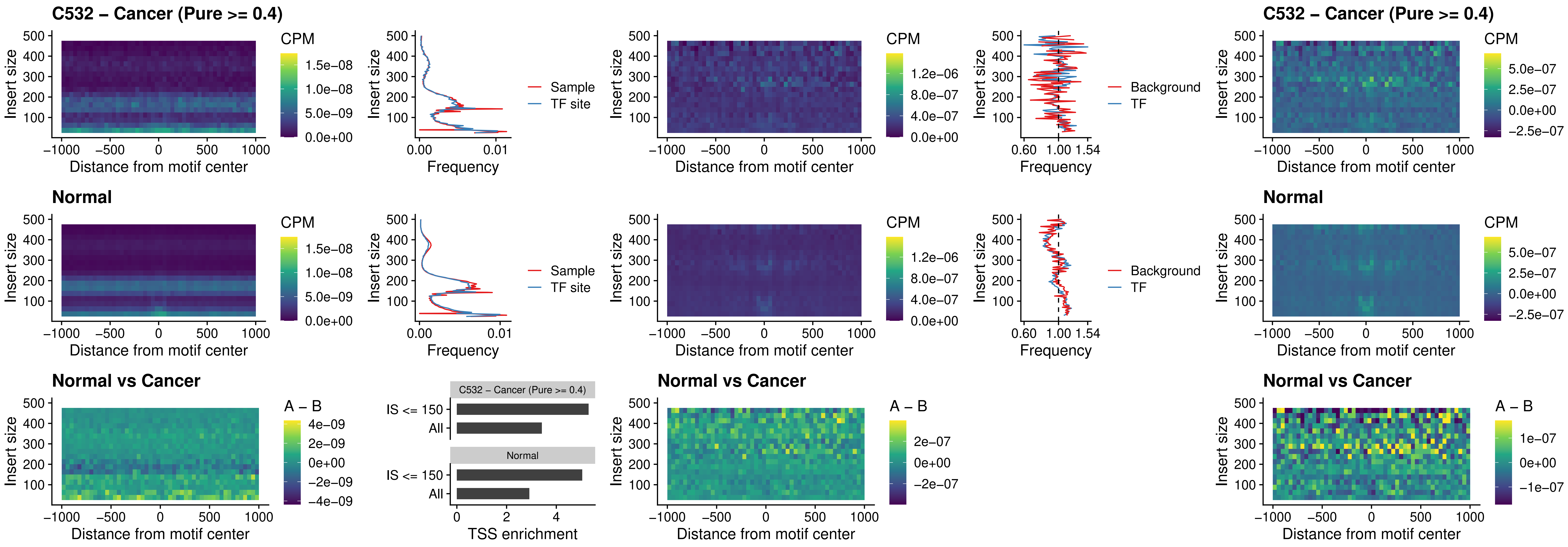

### C532 – Cancer (Pure >= 0.4) – ETS2_uc_dTSS_oPEAK

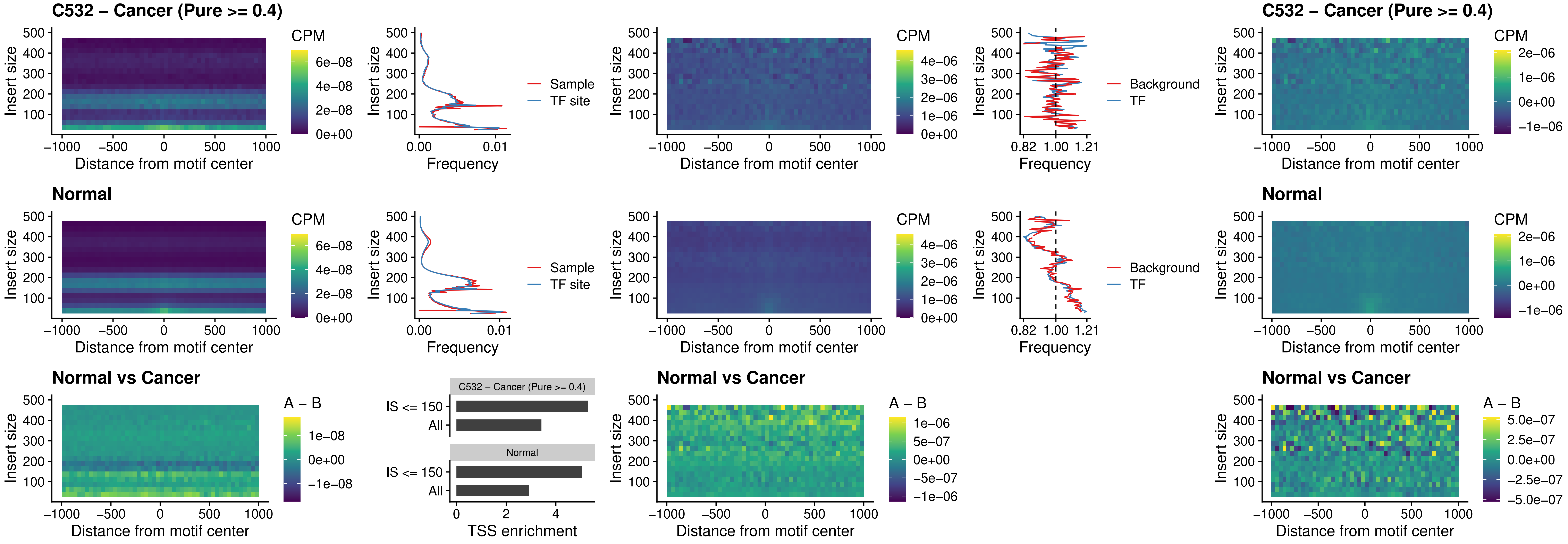

### C532 – Cancer (Pure >= 0.4) – SMARCC2_uc_pTSS_oPEAK

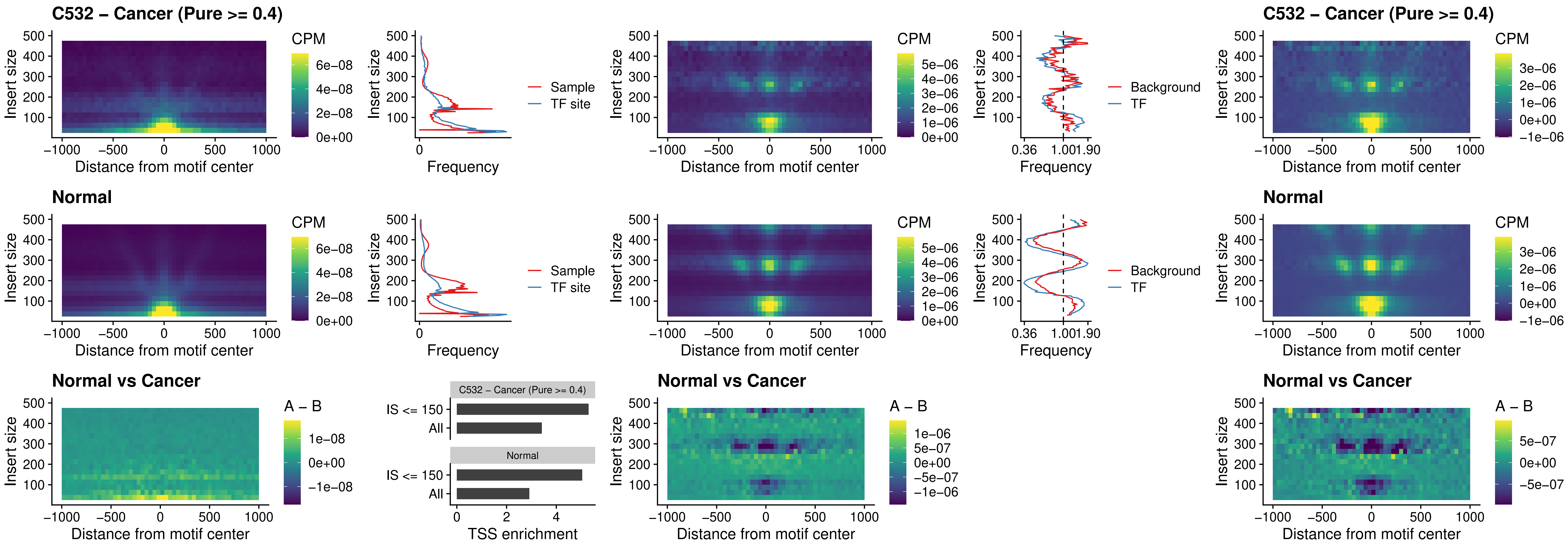

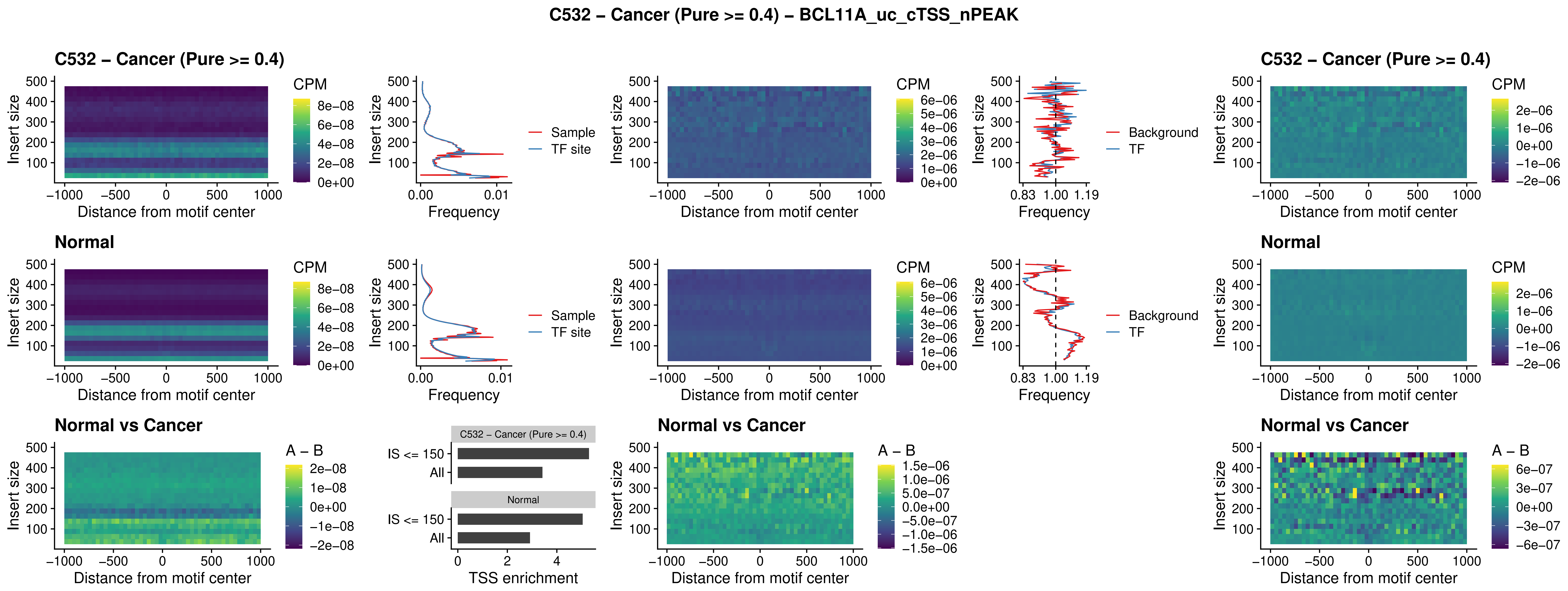
C532 – Cancer (Pure >= 0.4) – BCL11A_uc_cTSS_nPEAK
C532 – Cancer (Pure >= 0.4)
Normal
Normal vs Cancer
Insert size
Distance from motif center
Frequency
CPM
A – B
Sample
TF site
Background
TF
C532 – Cancer (Pure >= 0.4)
Normal
IS <= 150
All
TSS enrichment

#### C532 – Cancer (Pure >= 0.4) – BCL11B_uc_cTSS_nPEAK

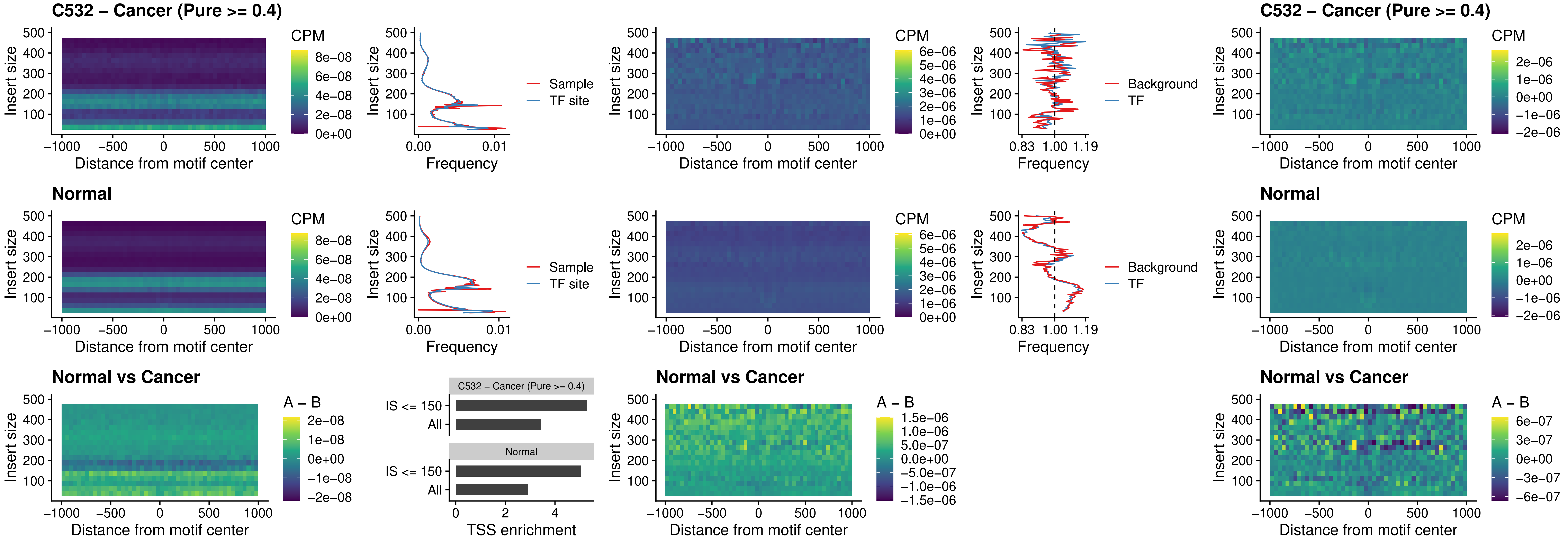

#### C532 – Cancer (Pure >= 0.4) – STAT2_uc_dTSS_oPEAK

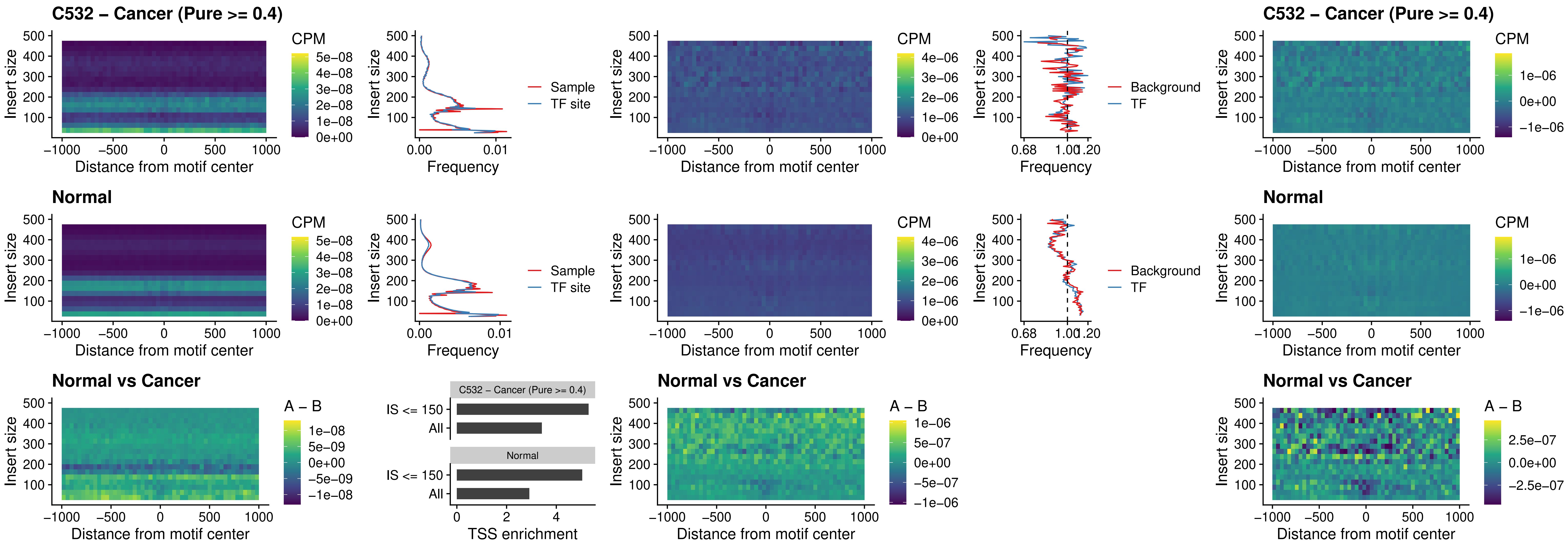

#### C532 – Cancer (Pure >= 0.4) – ZNF683_uc_cTSS_oPEAK

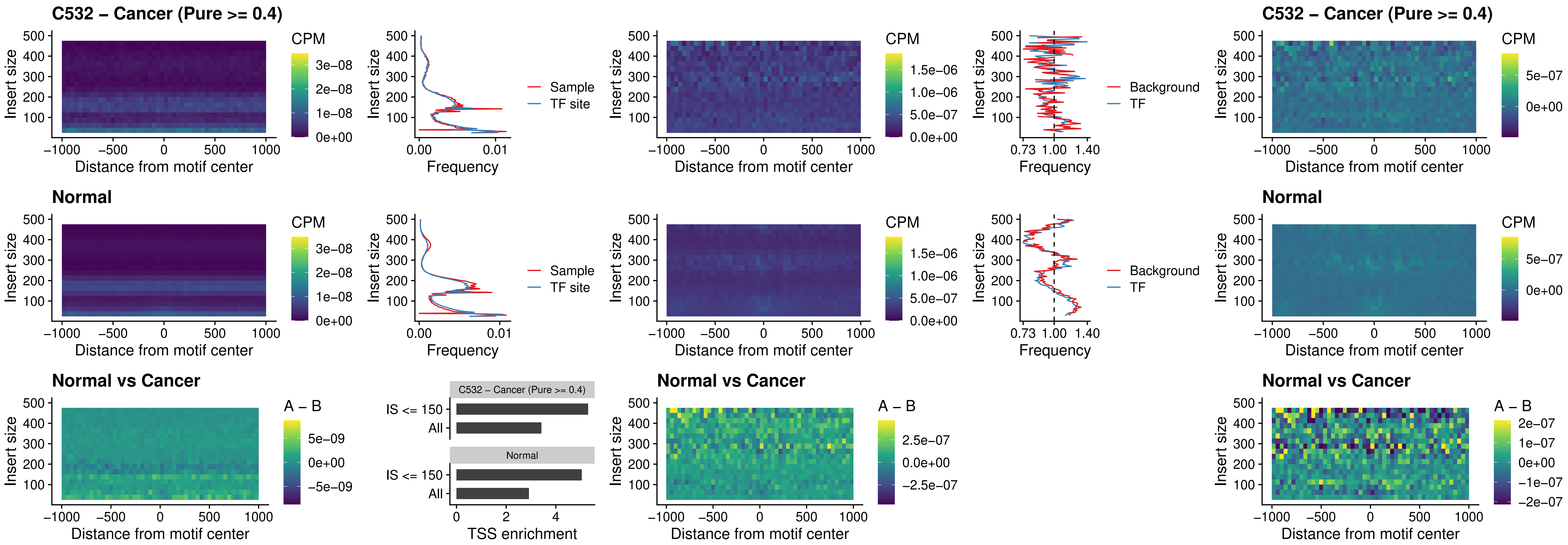

#### C532 – Cancer (Pure >= 0.4) – IRF8_uc_dTSS_nPEAK

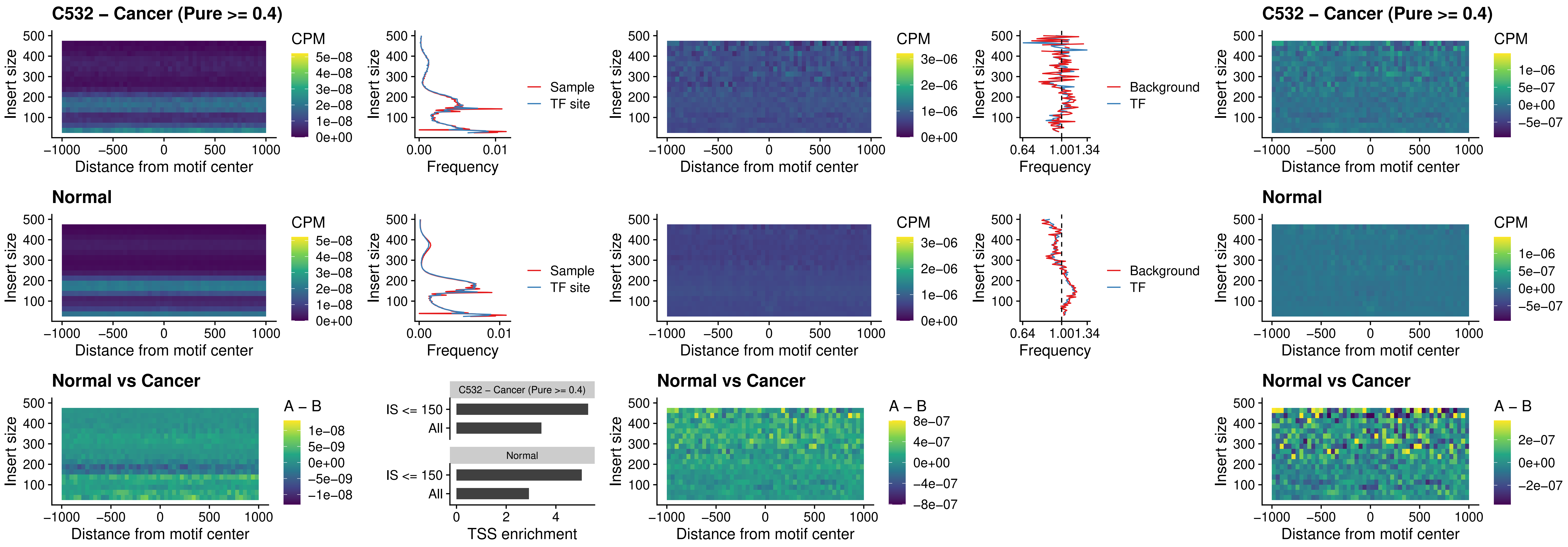

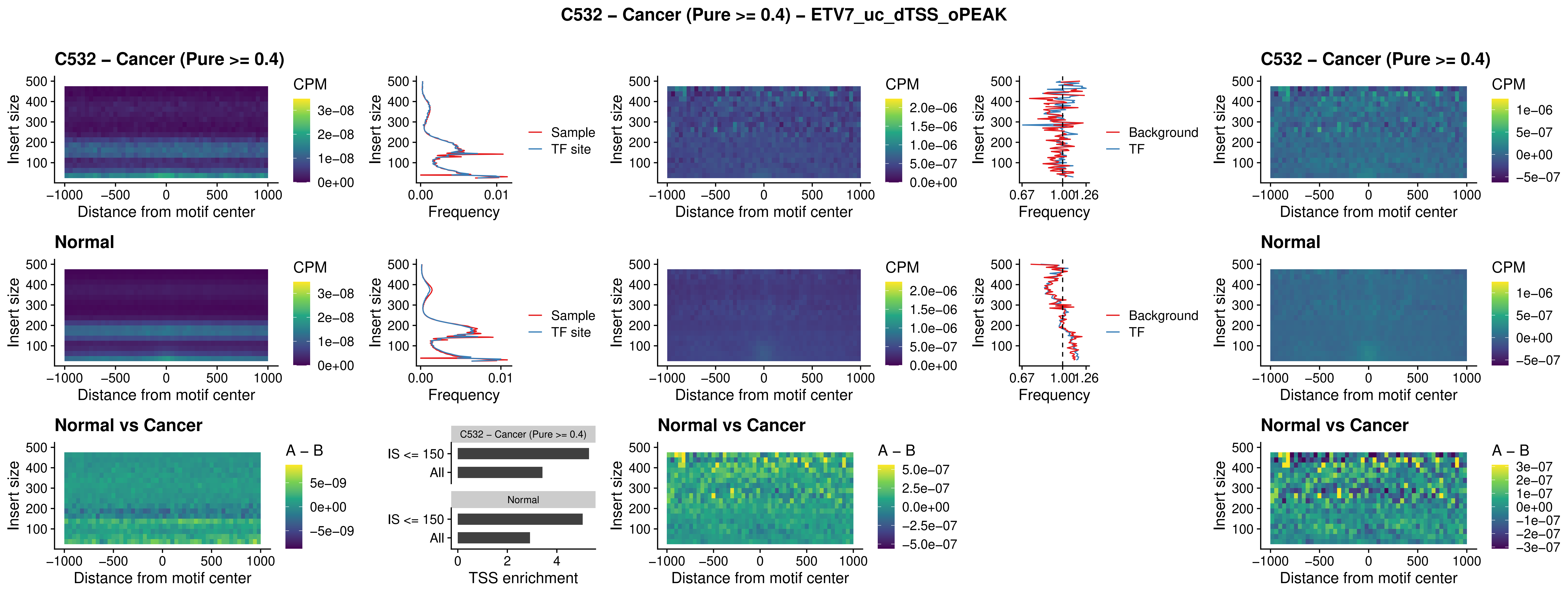
C532 – Cancer (Pure >= 0.4) – ETV7_uc_dTSS_oPEAK
C532 – Cancer (Pure >= 0.4)
Normal
Normal vs Cancer
Insert size
Distance from motif center
Frequency
CPM
A – B
Sample
TF site
Background
TF
C532 – Cancer (Pure >= 0.4)
Normal
IS <= 150
All
TSS enrichment

#### C536 – Cancer (Pure >= 0.4) – FOXL1_uc_pTSS_oPEAK

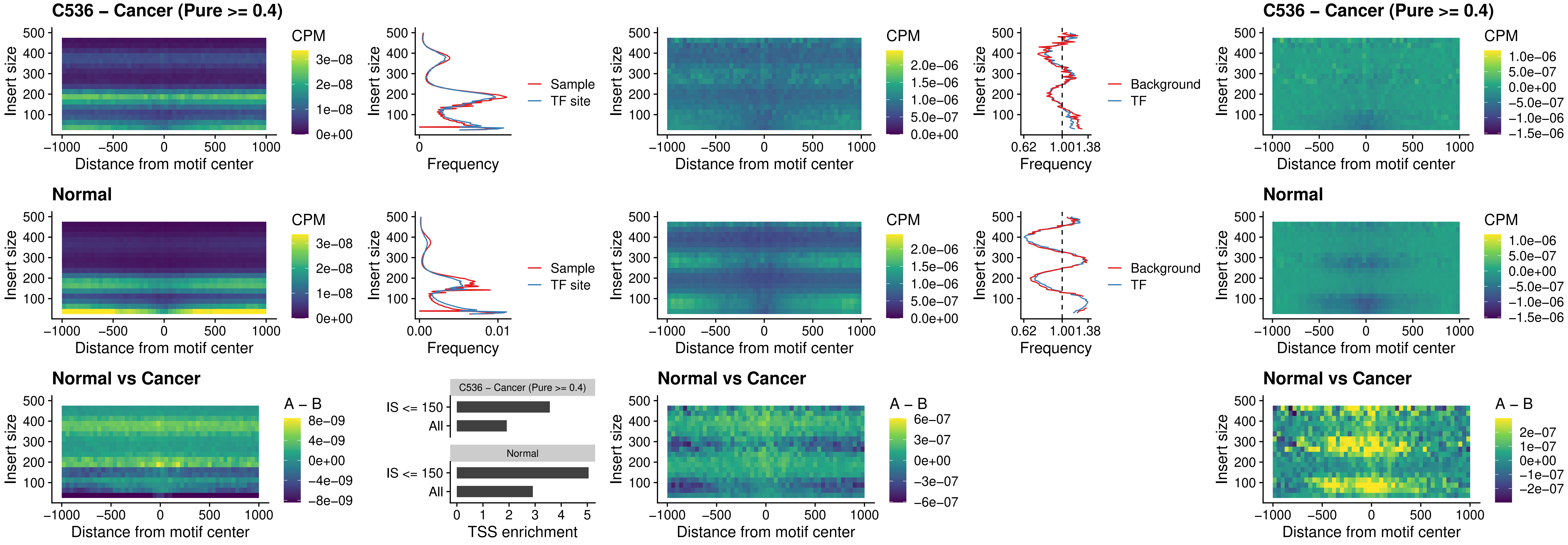

### C536 – Cancer (Pure >= 0.4) – CPEB1_uc_cTSS_oPEAK

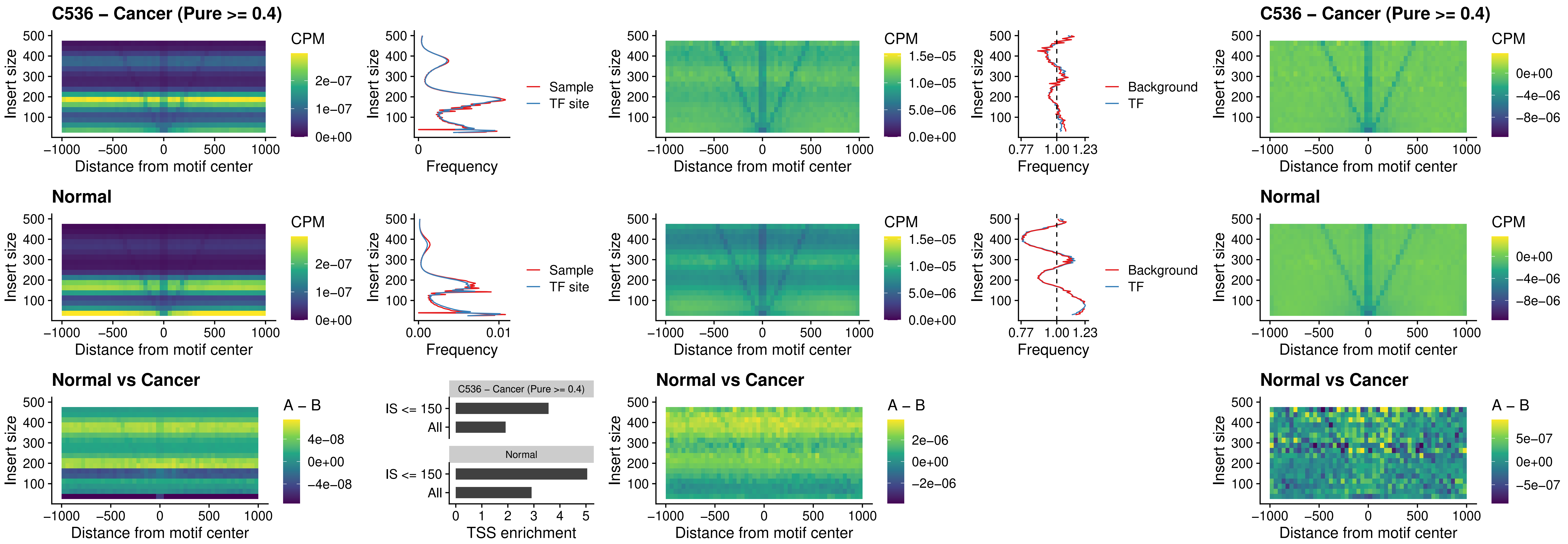

### C536 – Cancer (Pure >= 0.4) – HMGA1_uc_pTSS_nPEAK

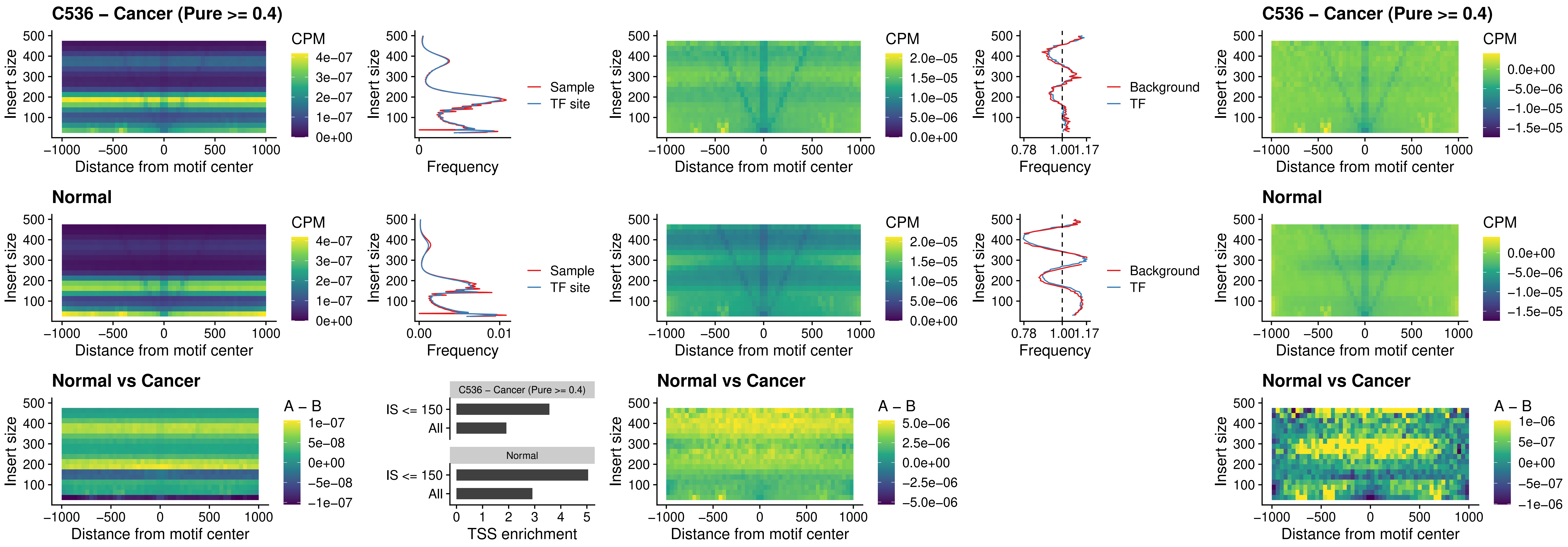

### C536 – Cancer (Pure >= 0.4) – SOX13_uc_cTSS_nPEAK

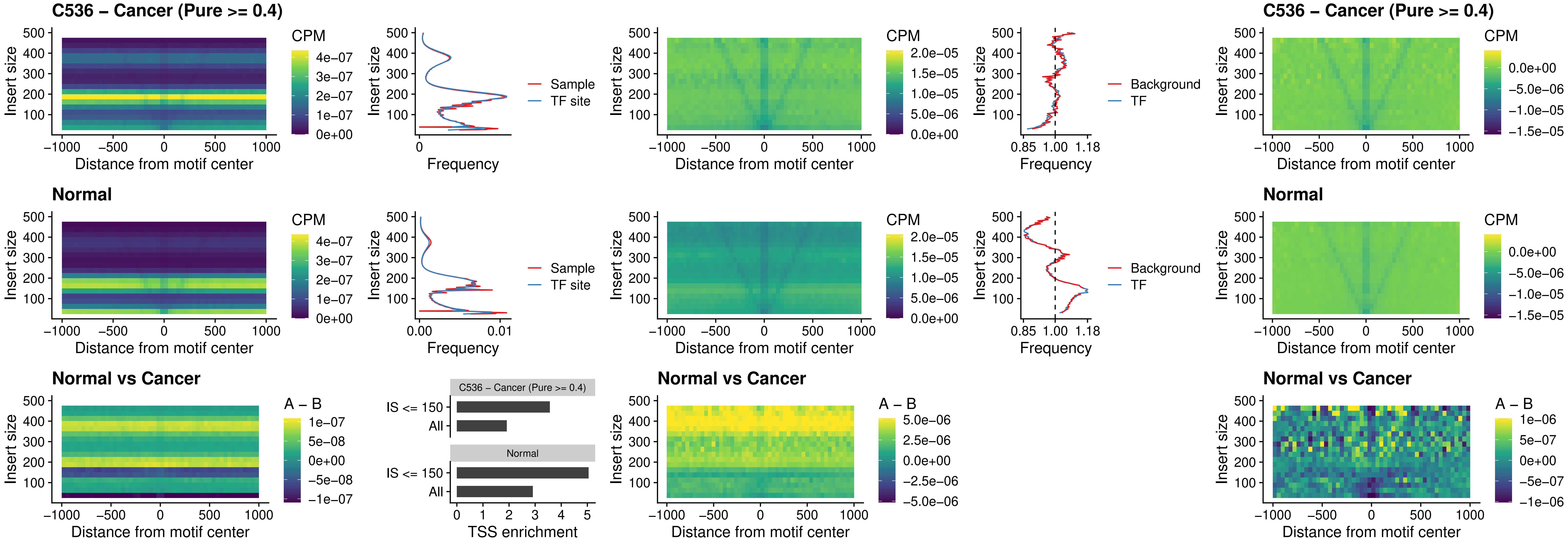

#### C536 – Cancer (Pure >= 0.4) – LHX4_uc_cTSS_nPEAK

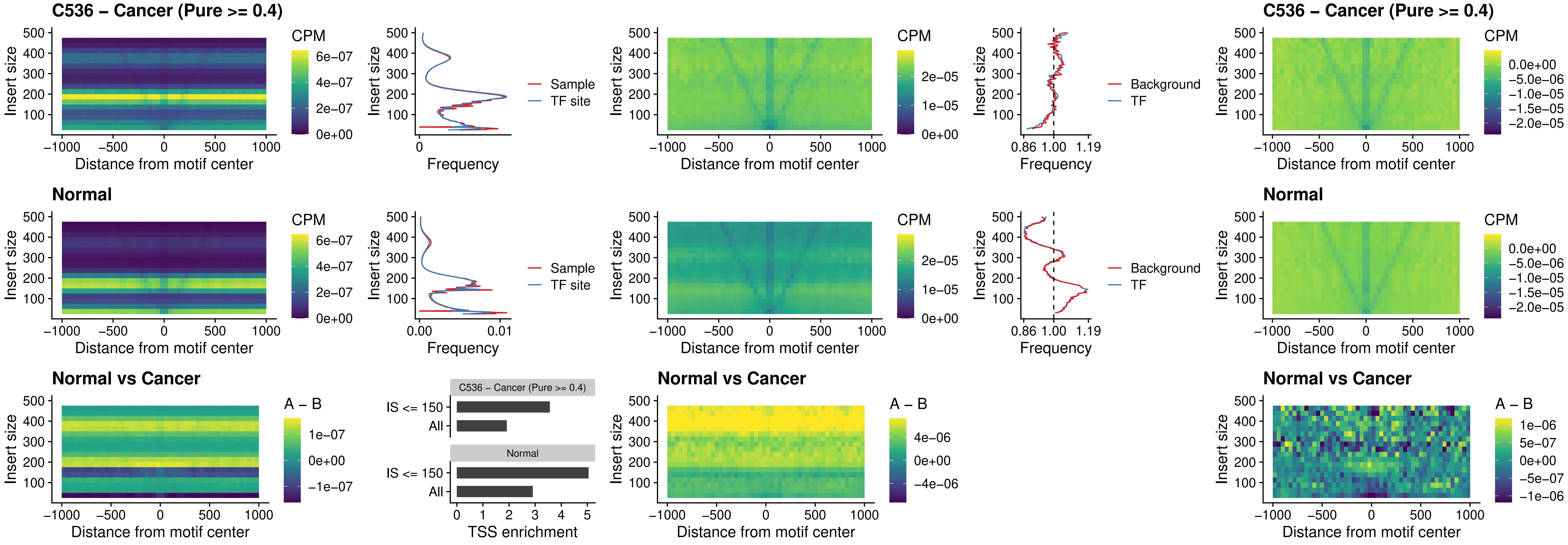

#### C536 – Cancer (Pure >= 0.4) – HOXA4_uc_cTSS_nPEAK

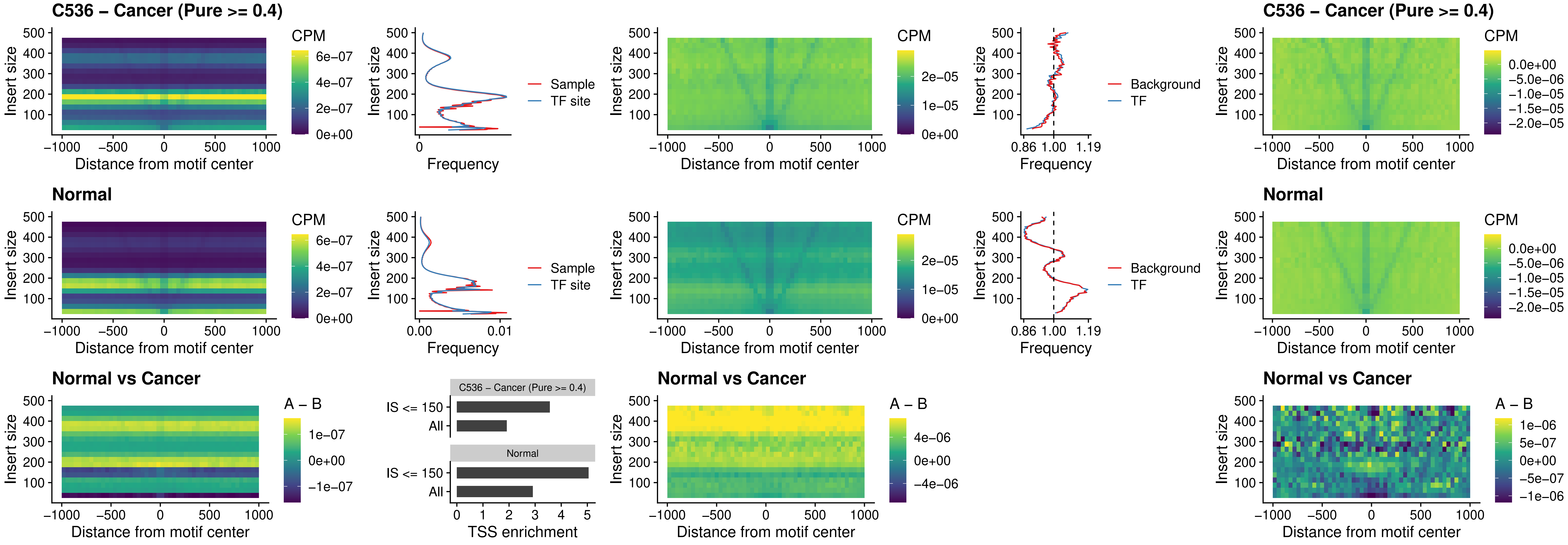

#### C536 – Cancer (Pure >= 0.4) – HLX_uc_cTSS_nPEAK

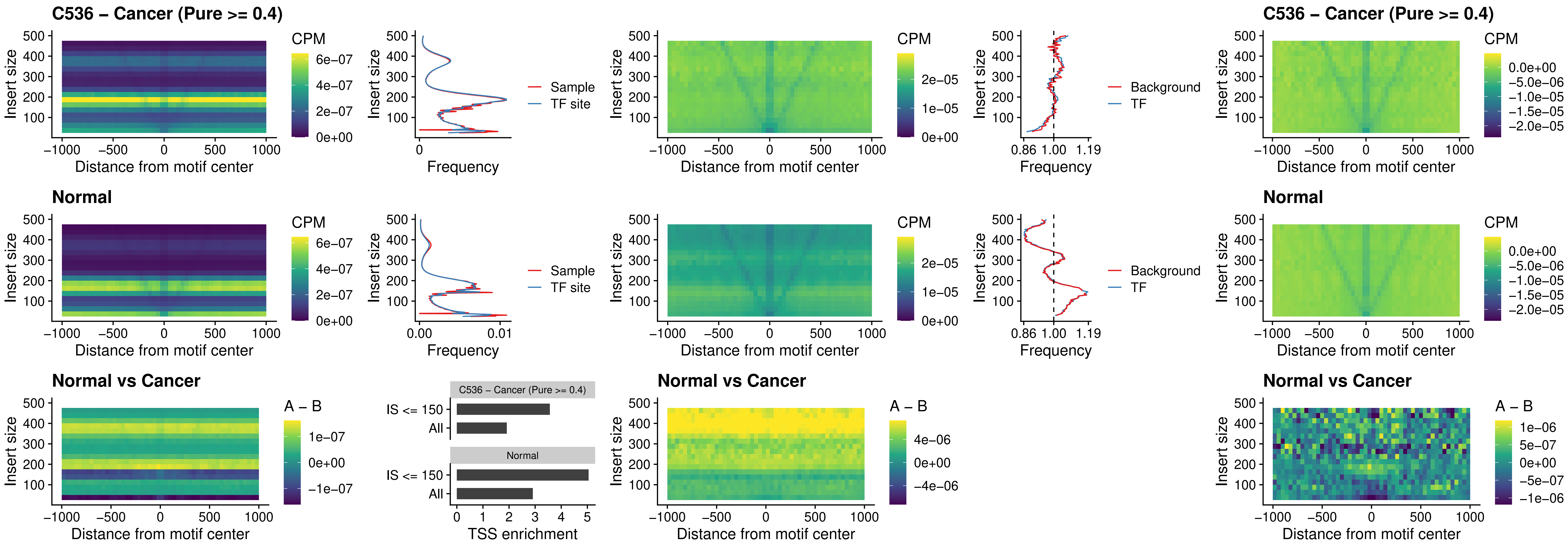

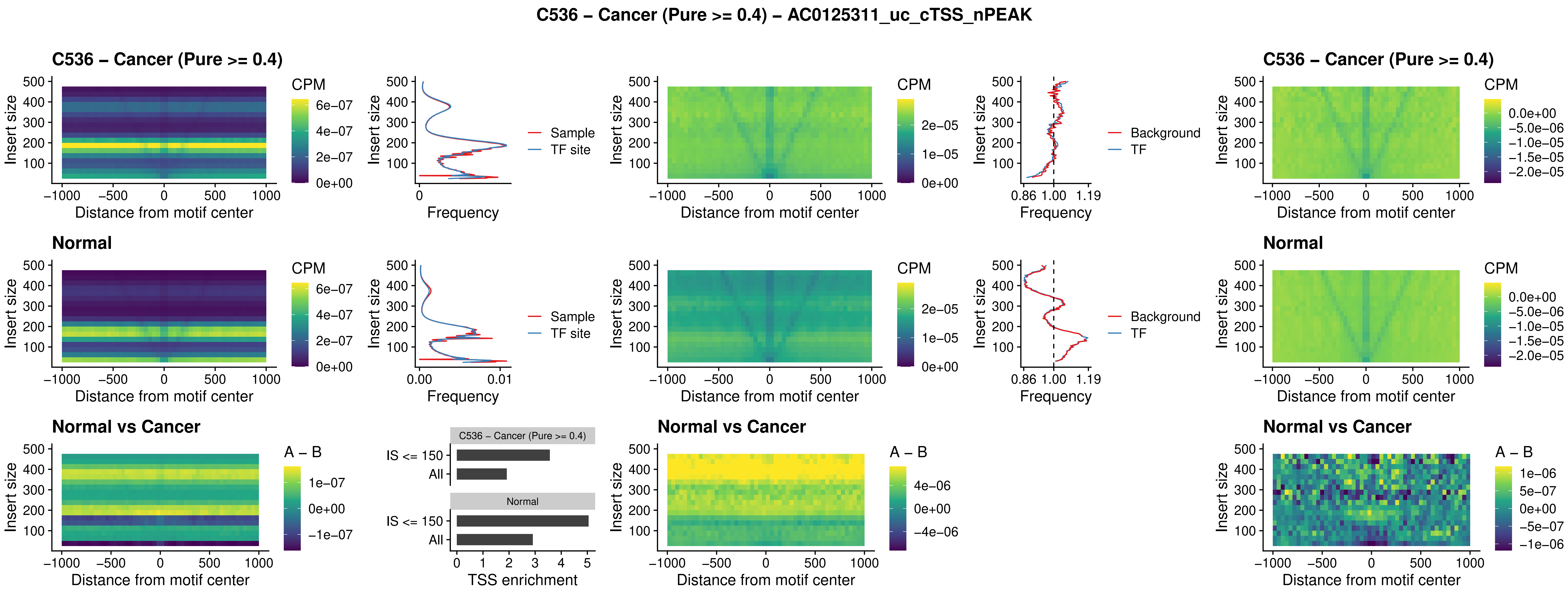
C536 – Cancer (Pure >= 0.4) – AC0125311_uc_cTSS_nPEAK
C536 – Cancer (Pure >= 0.4)
Normal
Normal vs Cancer
Insert size
Distance from motif center
Frequency
CPM
A – B
Sample
TF site
Background
TF
C536 – Cancer (Pure >= 0.4)
Normal
IS <= 150
All
TSS enrichment

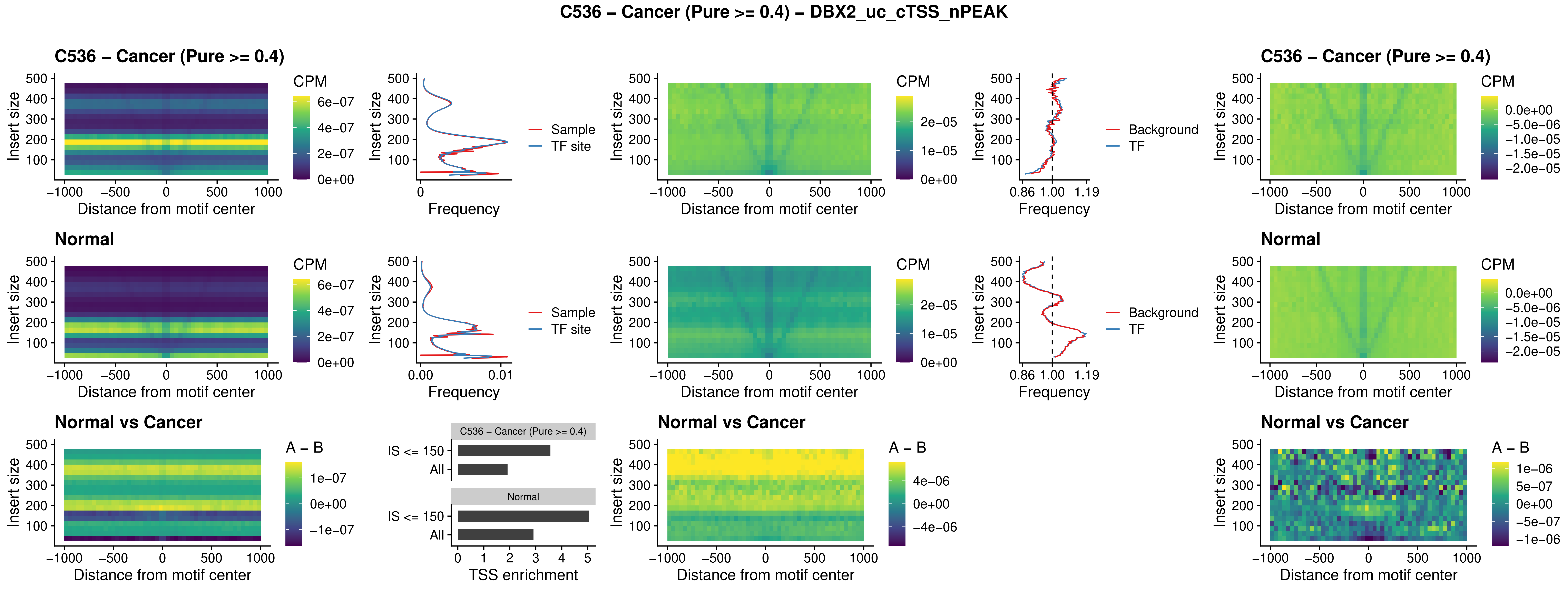
C536 – Cancer (Pure >= 0.4) – DBX2_uc_cTSS_nPEAK
C536 – Cancer (Pure >= 0.4)
Normal
Normal vs Cancer
Insert size
Distance from motif center
Frequency
CPM
A – B
Sample
TF site
Background
TF
C536 – Cancer (Pure >= 0.4)
Normal
IS <= 150
All
TSS enrichment

#### C536 – Cancer (Pure >= 0.4) – LHX6_uc_dTSS_nPEAK

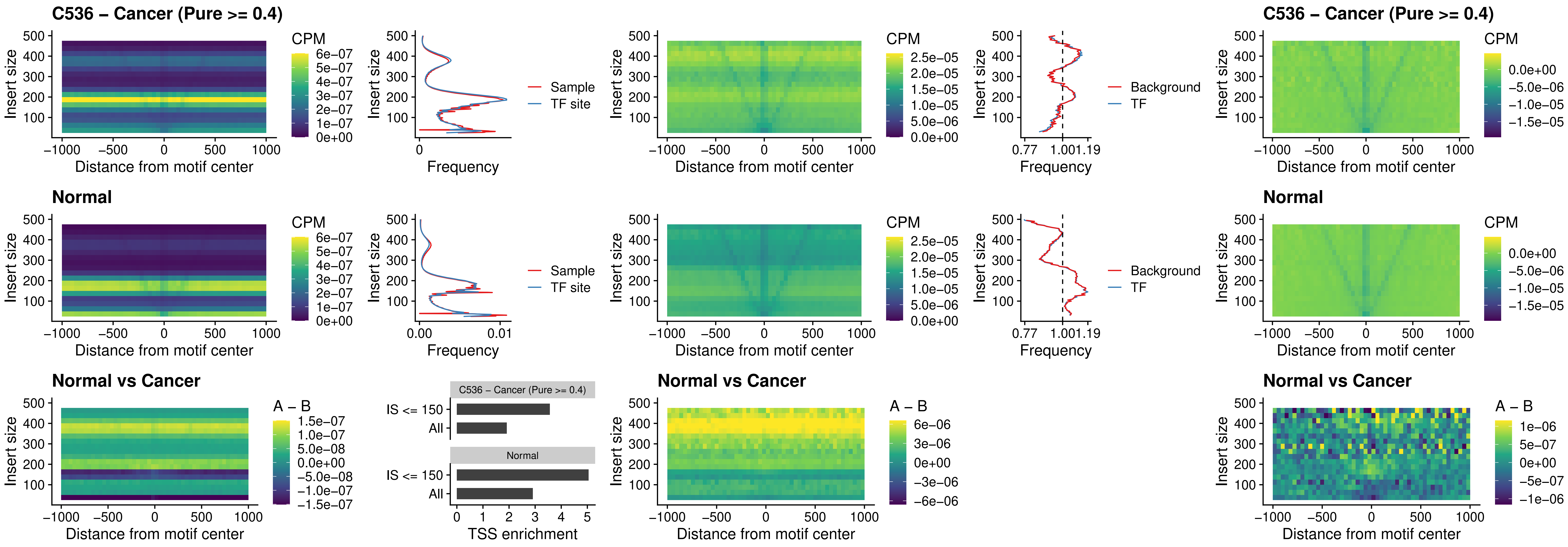

#### C536 – Cancer (Pure >= 0.4) – SOX1_uc_dTSS_nPEAK

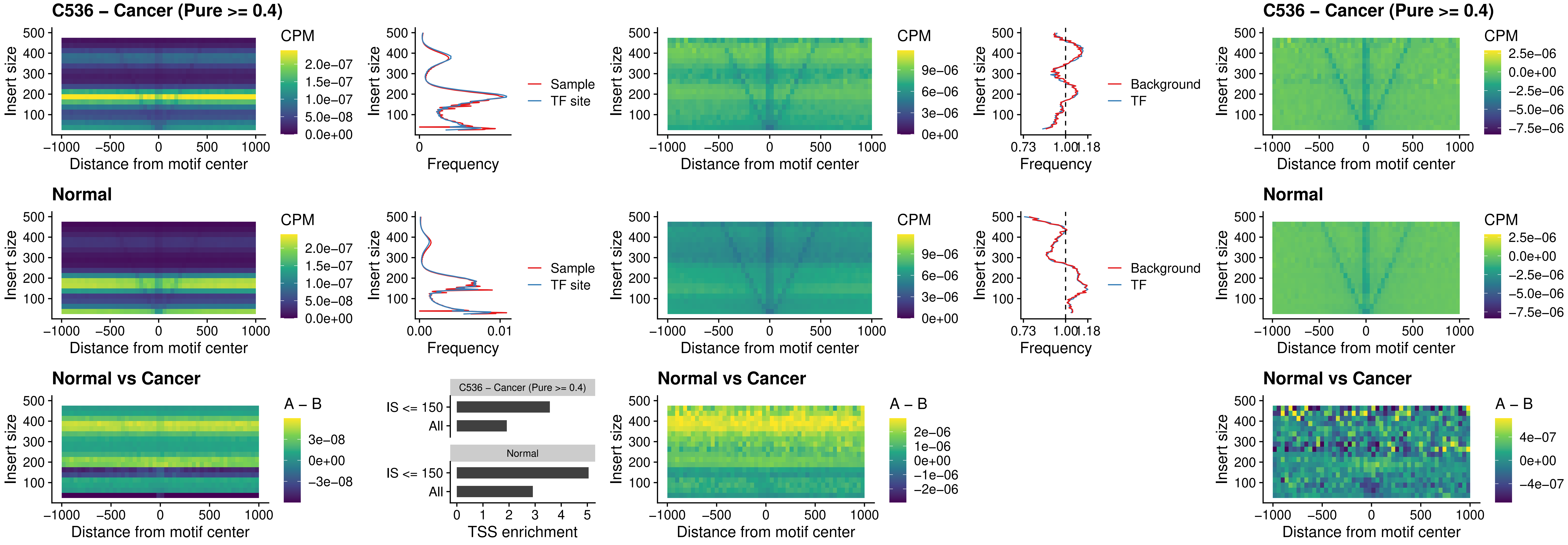

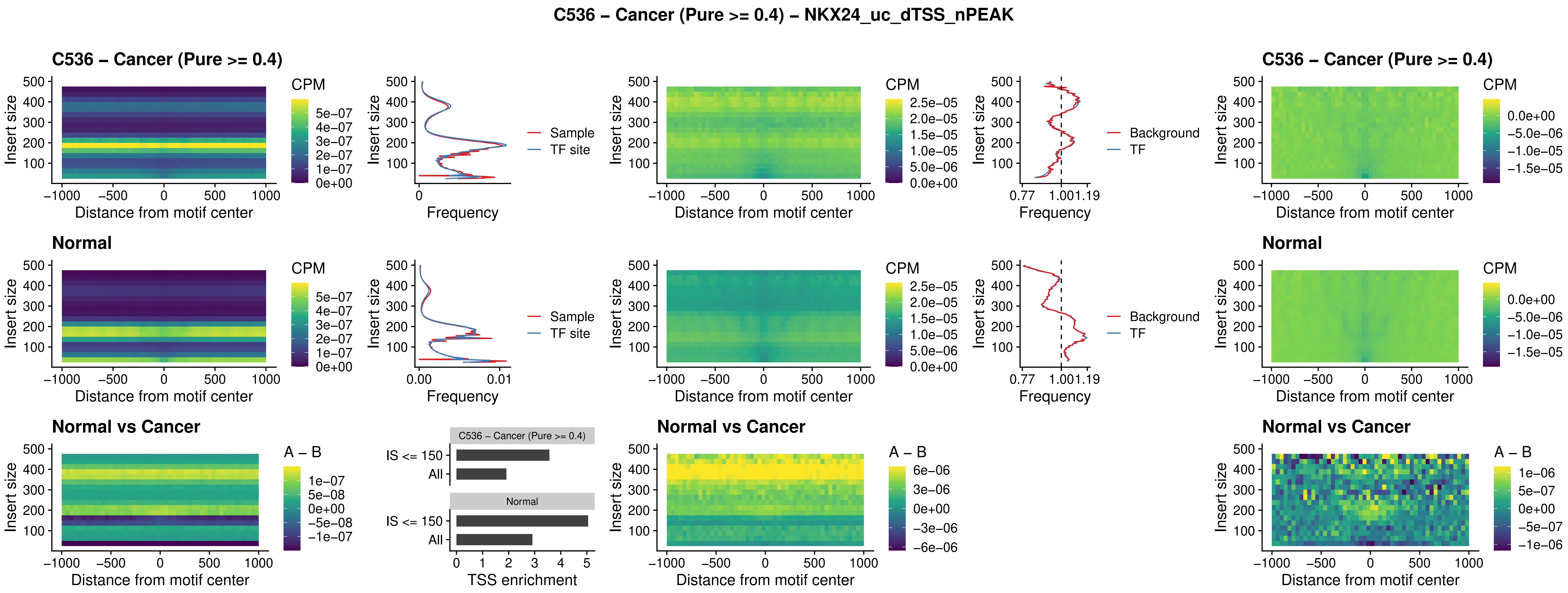

C536 – Cancer (Pure >= 0.4) – NKX24_uc_dTSS_nPEAK
C536 – Cancer (Pure >= 0.4)
Normal
Normal vs Cancer
Insert size
Distance from motif center
Frequency
CPM
A – B
Sample
TF site
Background
TF
C536 – Cancer (Pure >= 0.4)
Normal
IS <= 150
All
TSS enrichment

### C536 – Cancer (Pure >= 0.4) – DLX3_uc_cTSS_nPEAK

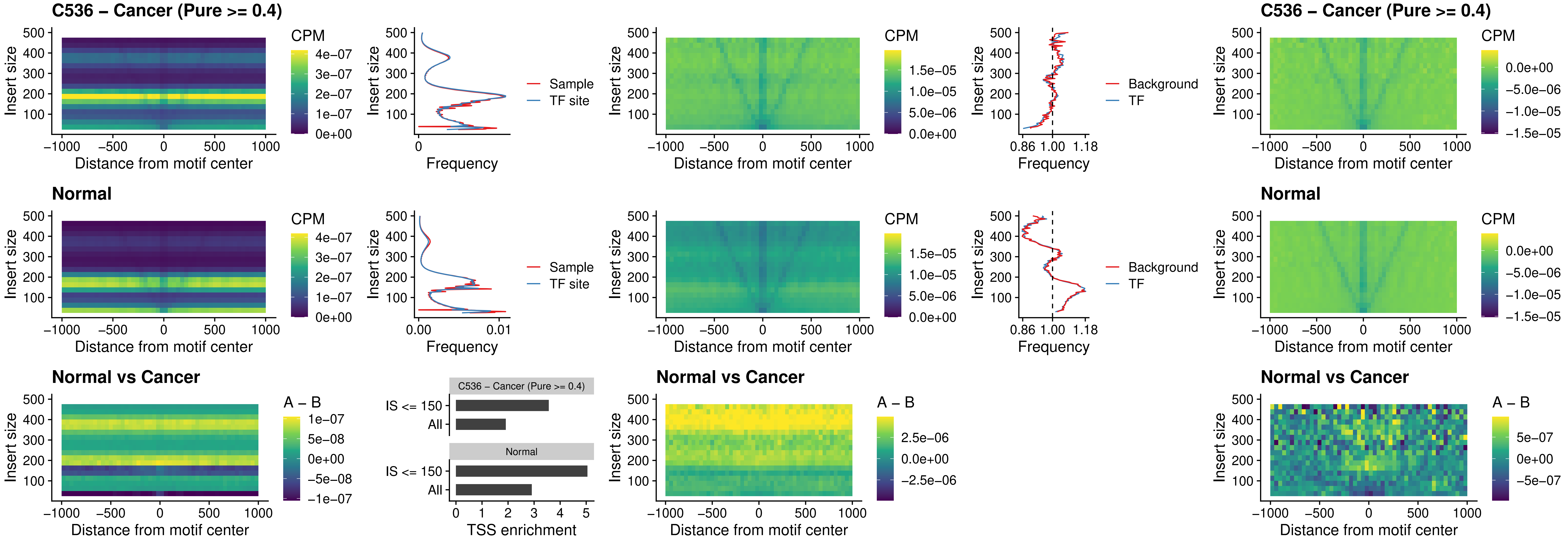

#### C536 – Cancer (Pure >= 0.4) – HOXD3_uc_cTSS_nPEAK

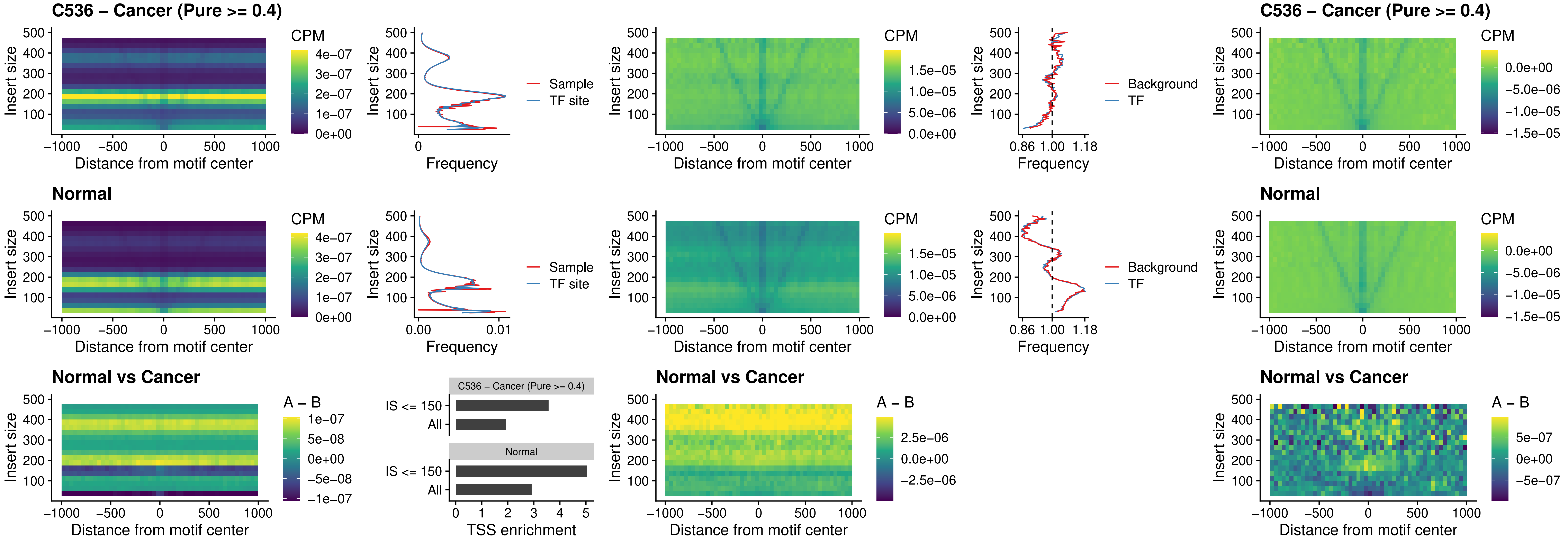

#### C536 – Cancer (Pure >= 0.4) – ISL1_uc_dTSS_oPEAK

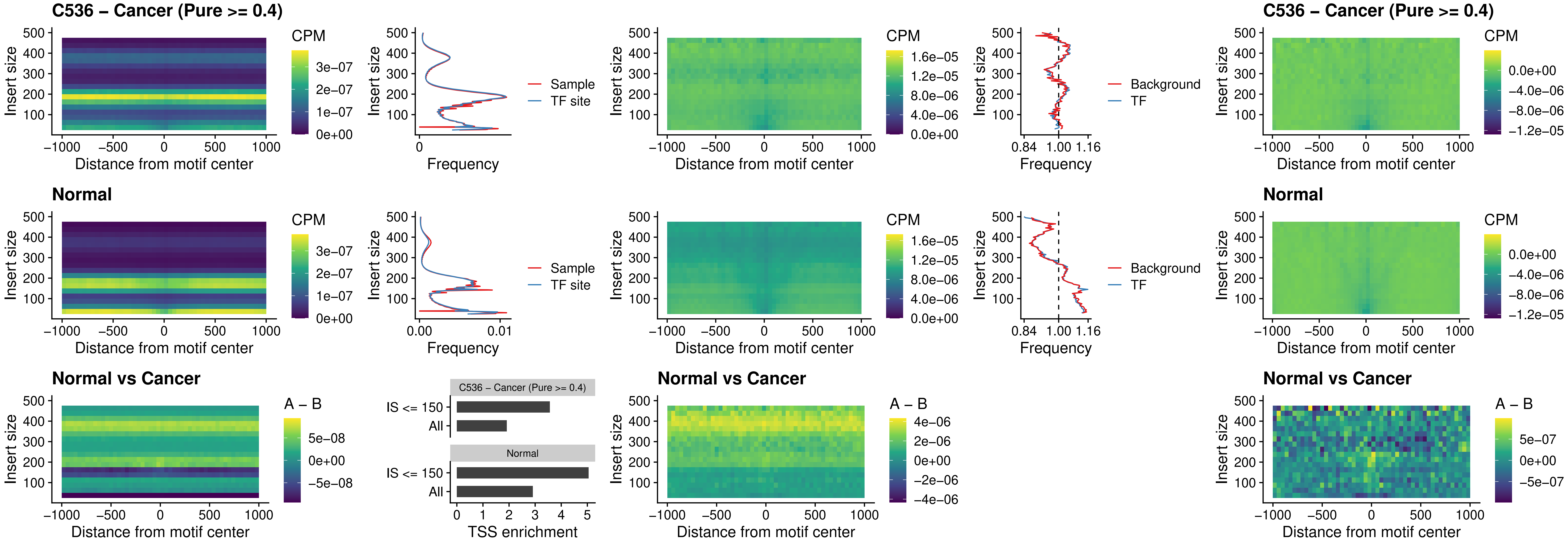

#### C536 – Cancer (Pure >= 0.4) – HOXB4_uc_dTSS_oPEAK

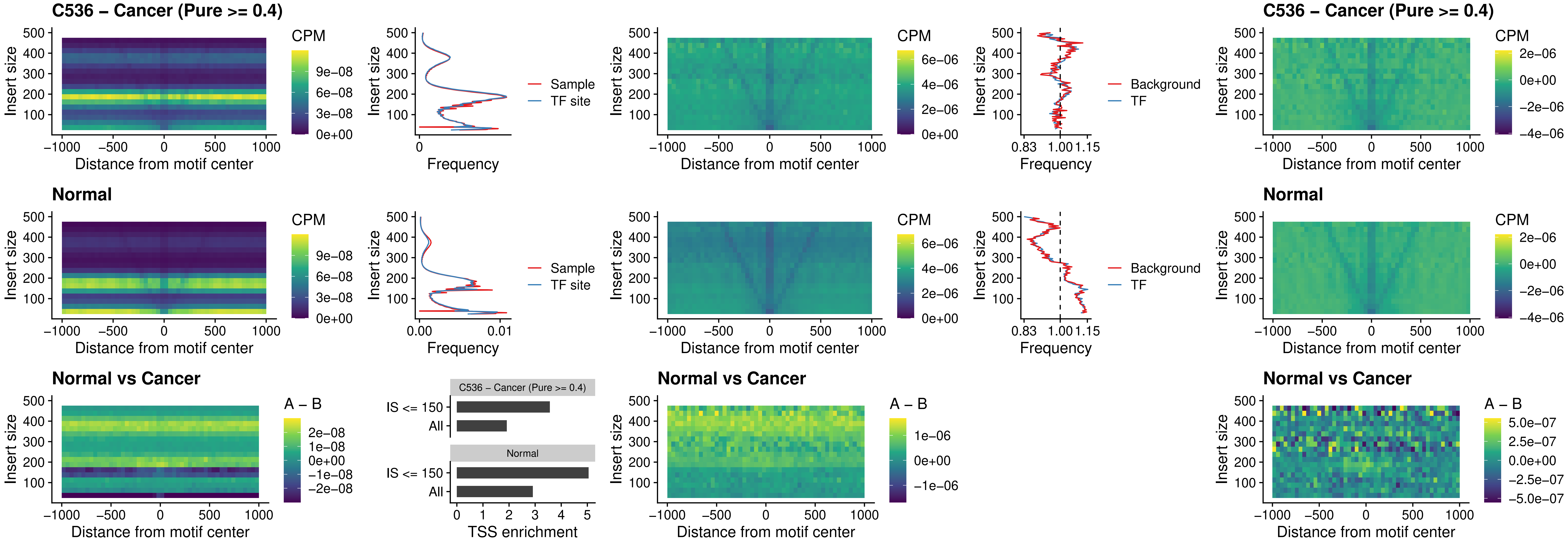

#### C536 – Cancer (Pure >= 0.4) – VAX1_uc_dTSS_oPEAK

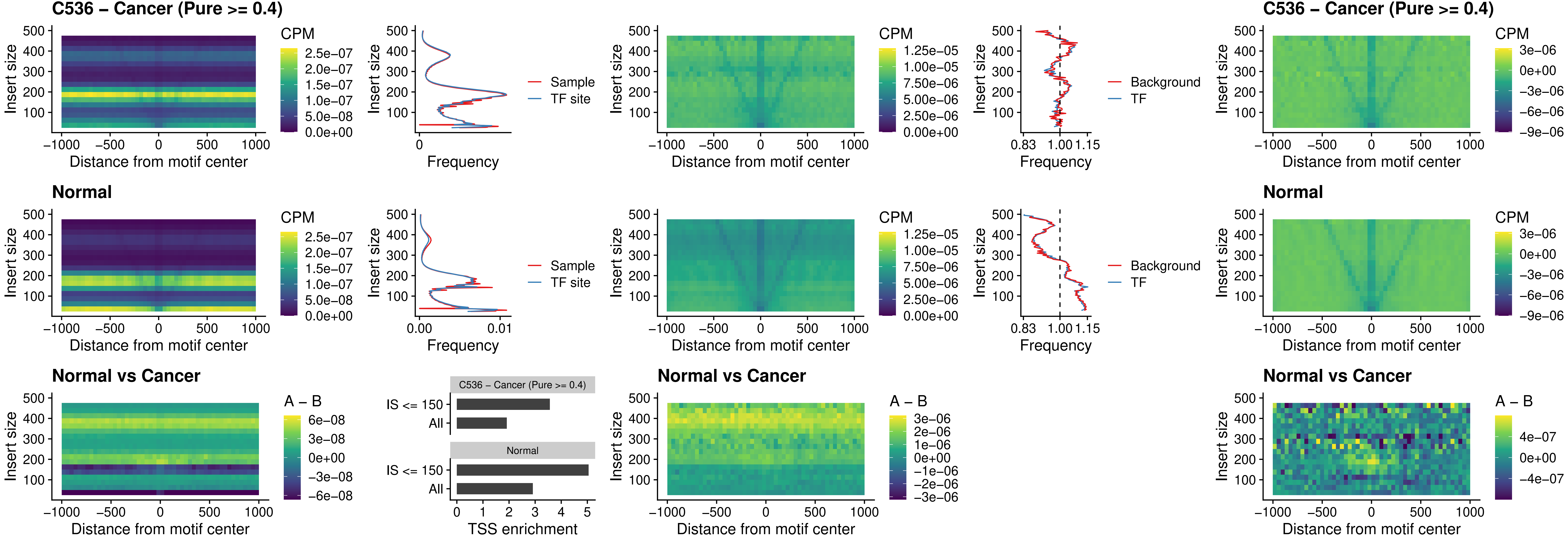

#### C536 – Cancer (Pure >= 0.4) – DLX3_uc_dTSS_oPEAK

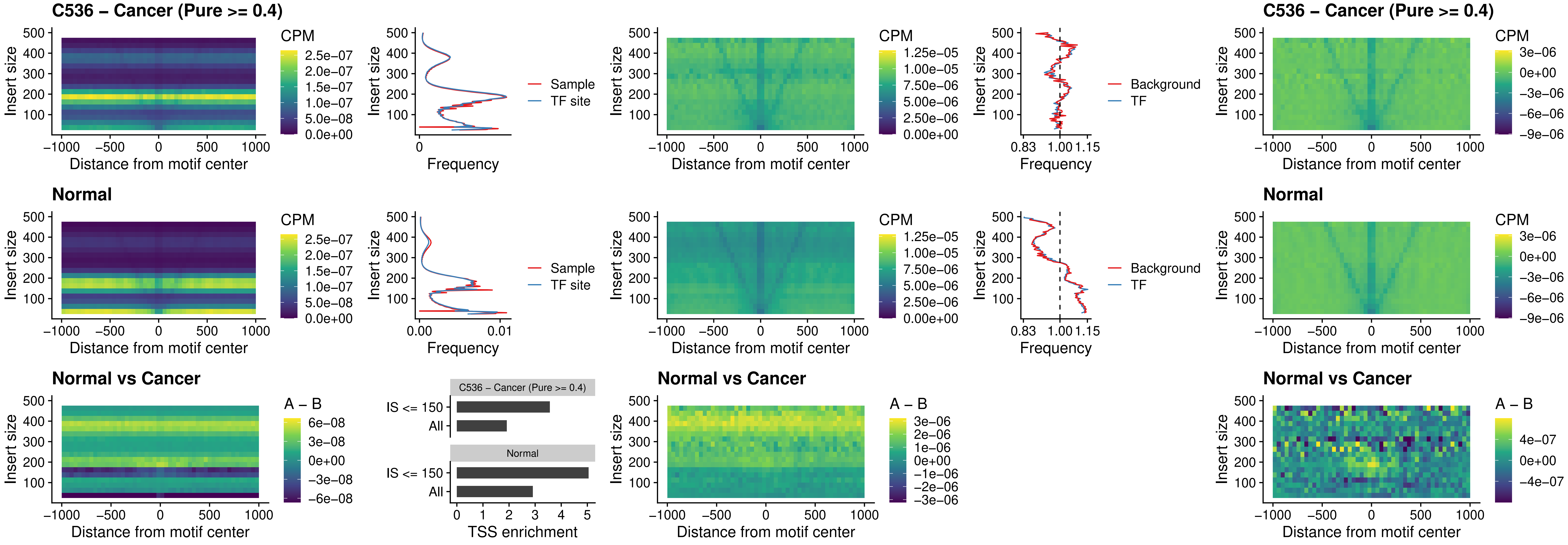

#### C536 – Cancer (Pure >= 0.4) – HOXD3_uc_dTSS_oPEAK

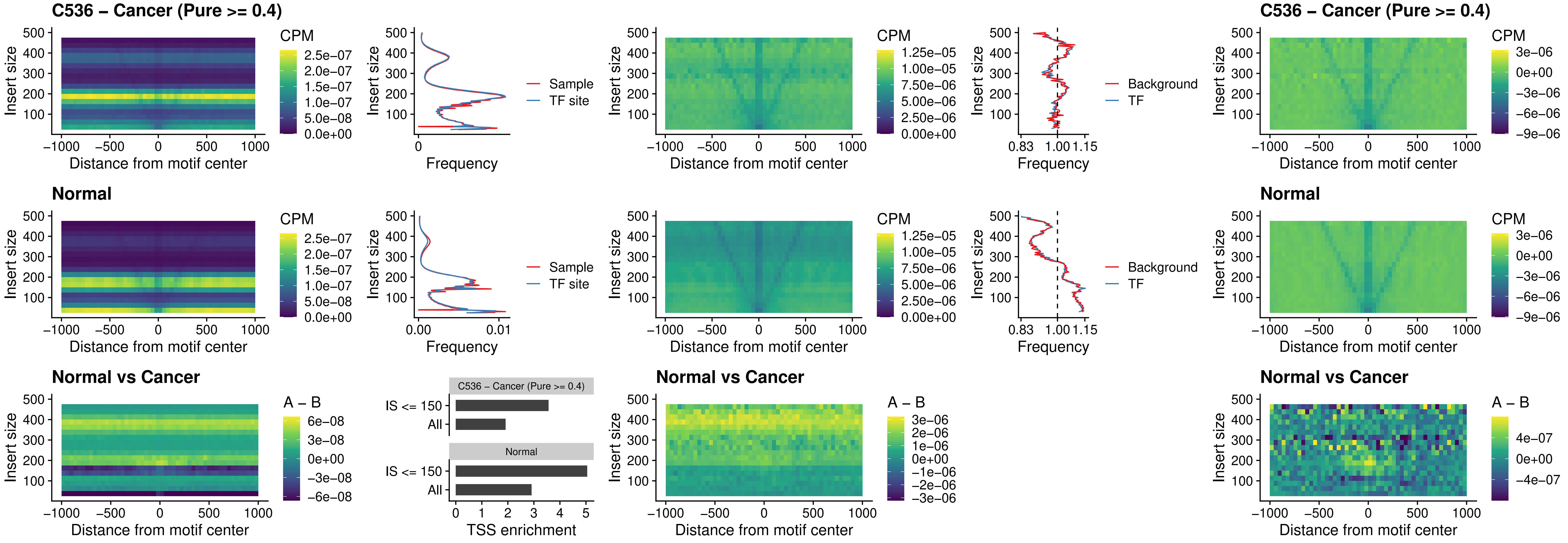

#### C536 – Cancer (Pure >= 0.4) – BBX_uc_dTSS_nPEAK

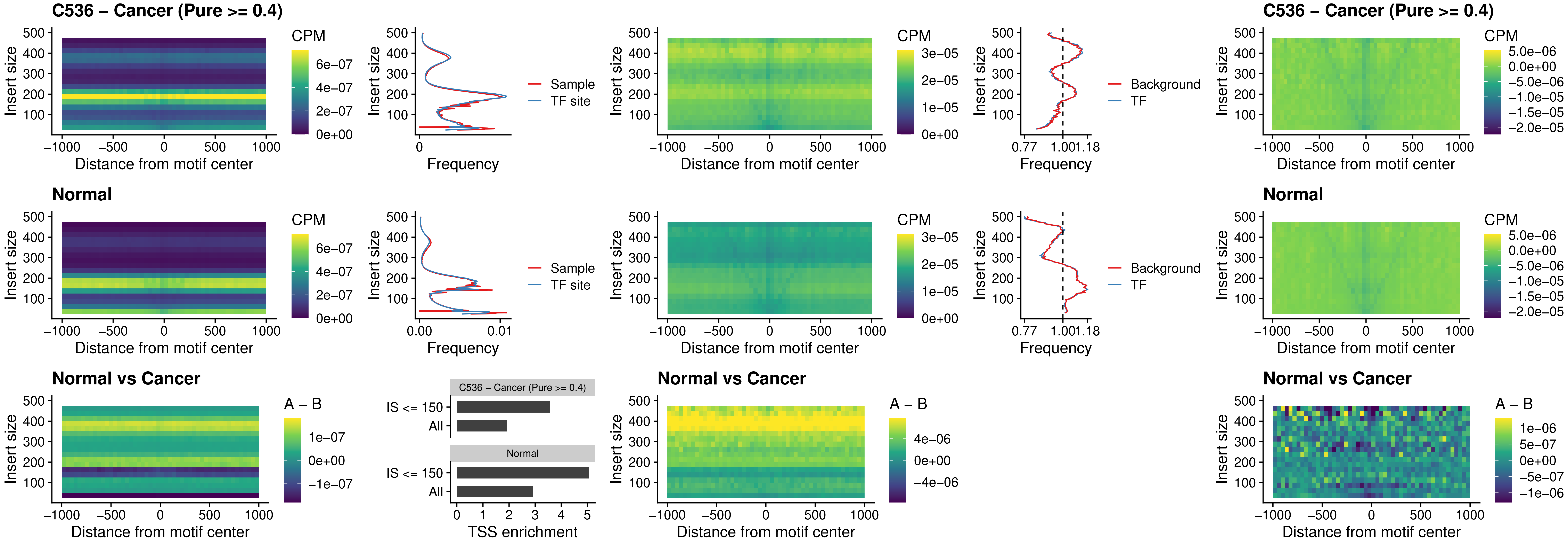

### C536 – Cancer (Pure >= 0.4) – SOX3_uc_cTSS_nPEAK

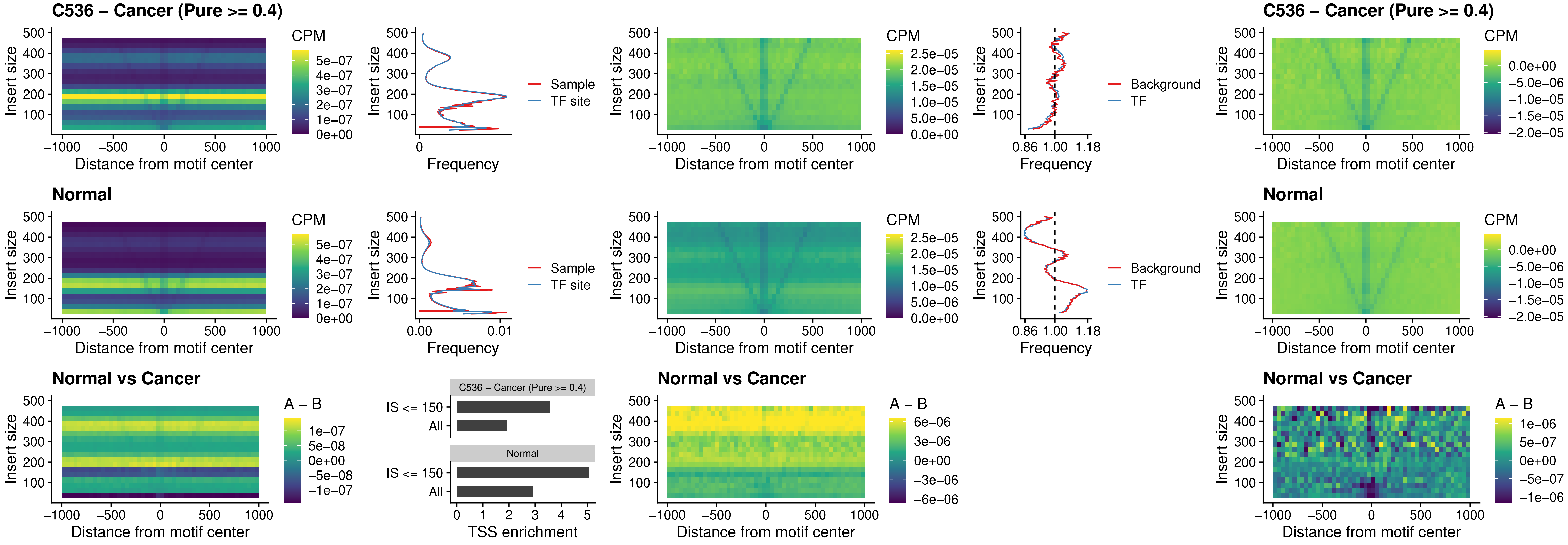

#### C536 – Cancer (Pure >= 0.4) – TBPL2_uc_cTSS_nPEAK

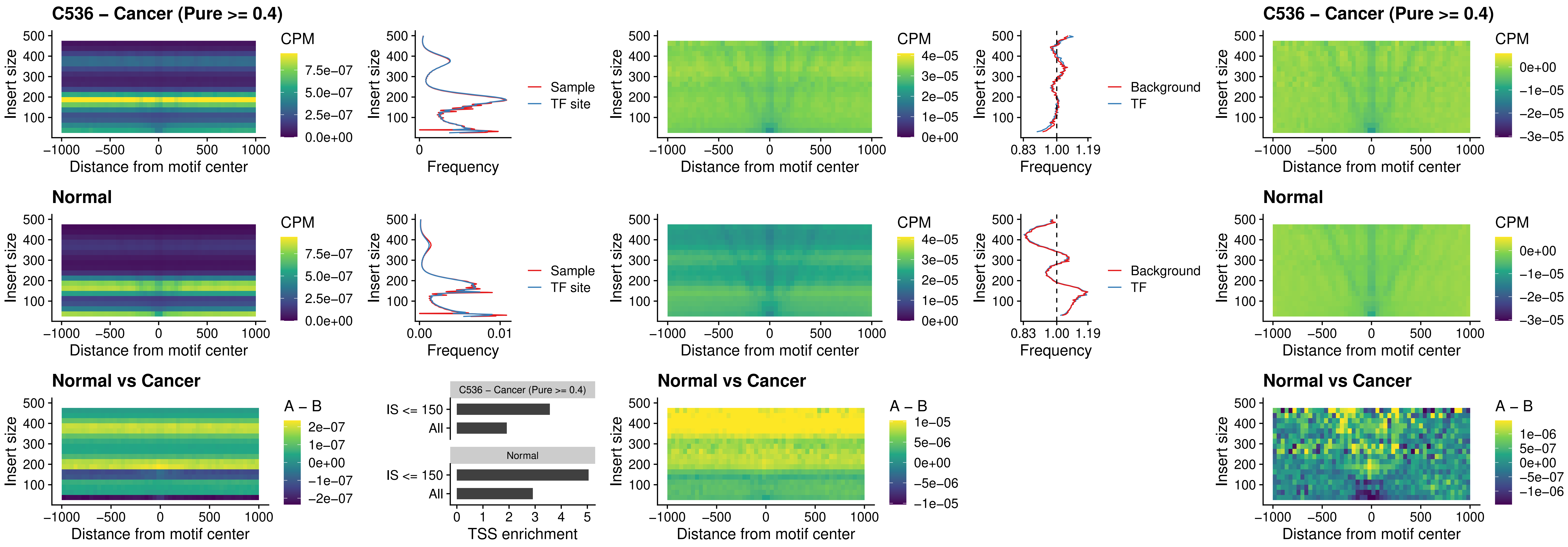

#### C536 – Cancer (Pure >= 0.4) – HOXA4_uc_cTSS_oPEAK

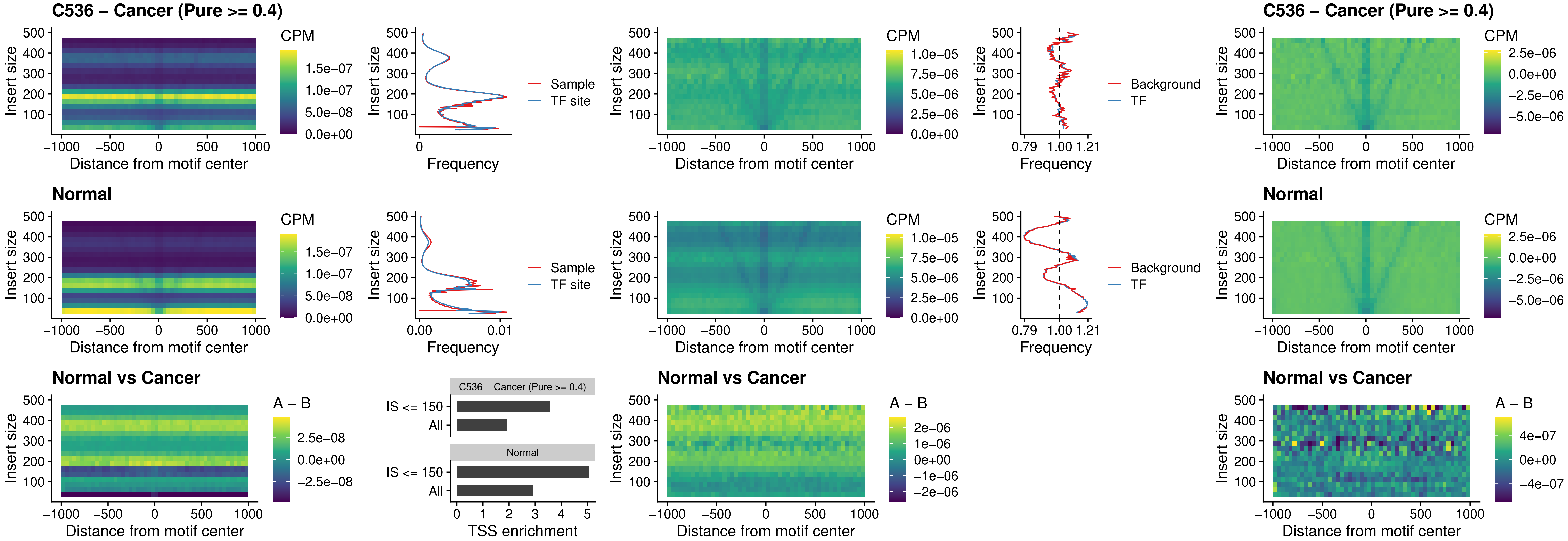

#### C536 – Cancer (Pure >= 0.4) – PDX1_uc_dTSS_oPEAK

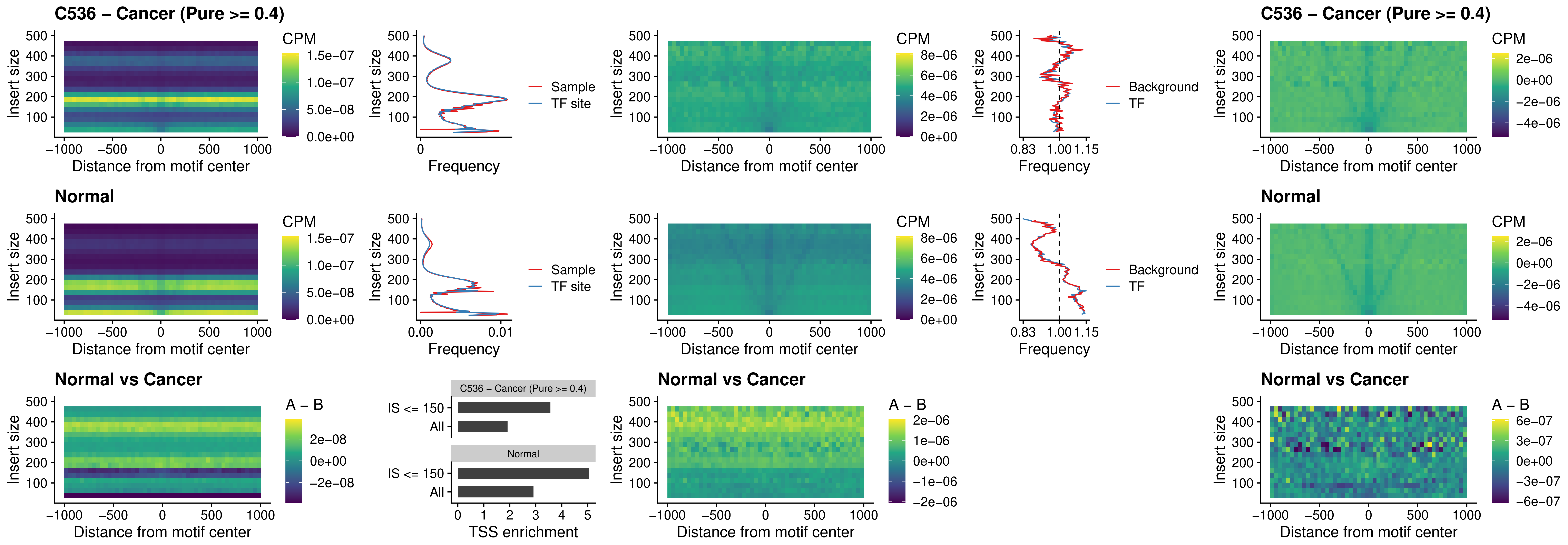

#### C536 – Cancer (Pure >= 0.4) – LHX4_uc_dTSS_oPEAK

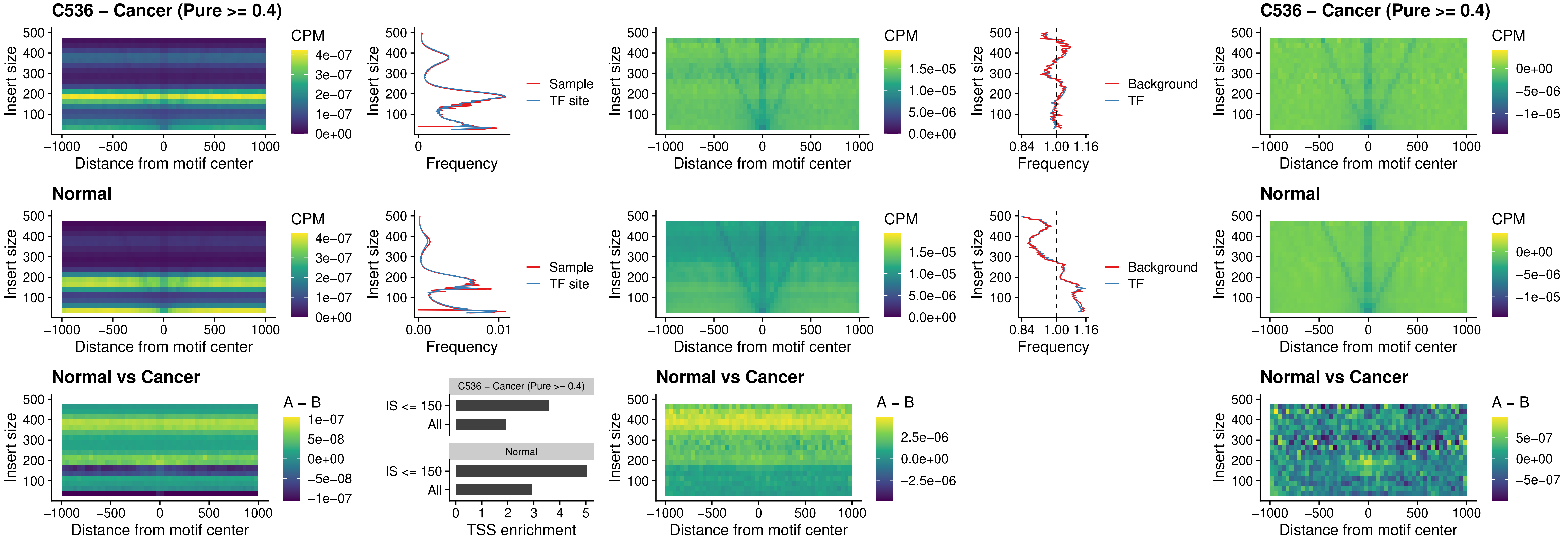

#### C536 – Cancer (Pure >= 0.4) – HOXA4_uc_dTSS_oPEAK

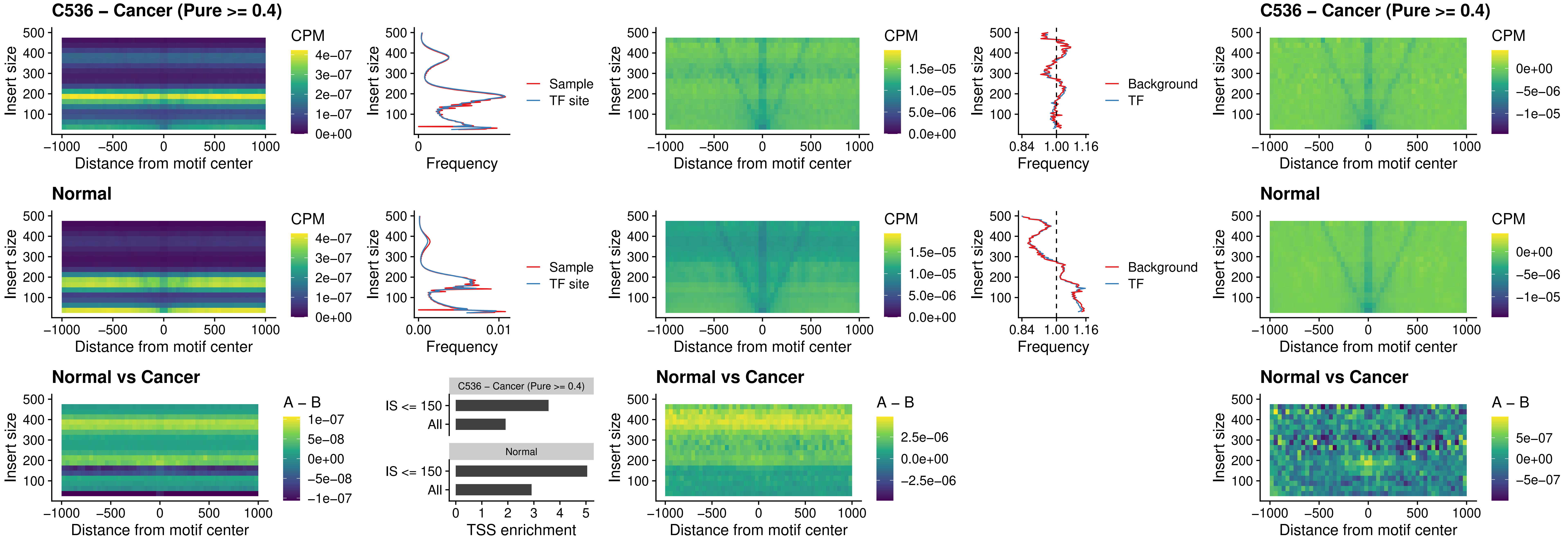

#### C536 – Cancer (Pure >= 0.4) – HLX_uc_dTSS_oPEAK

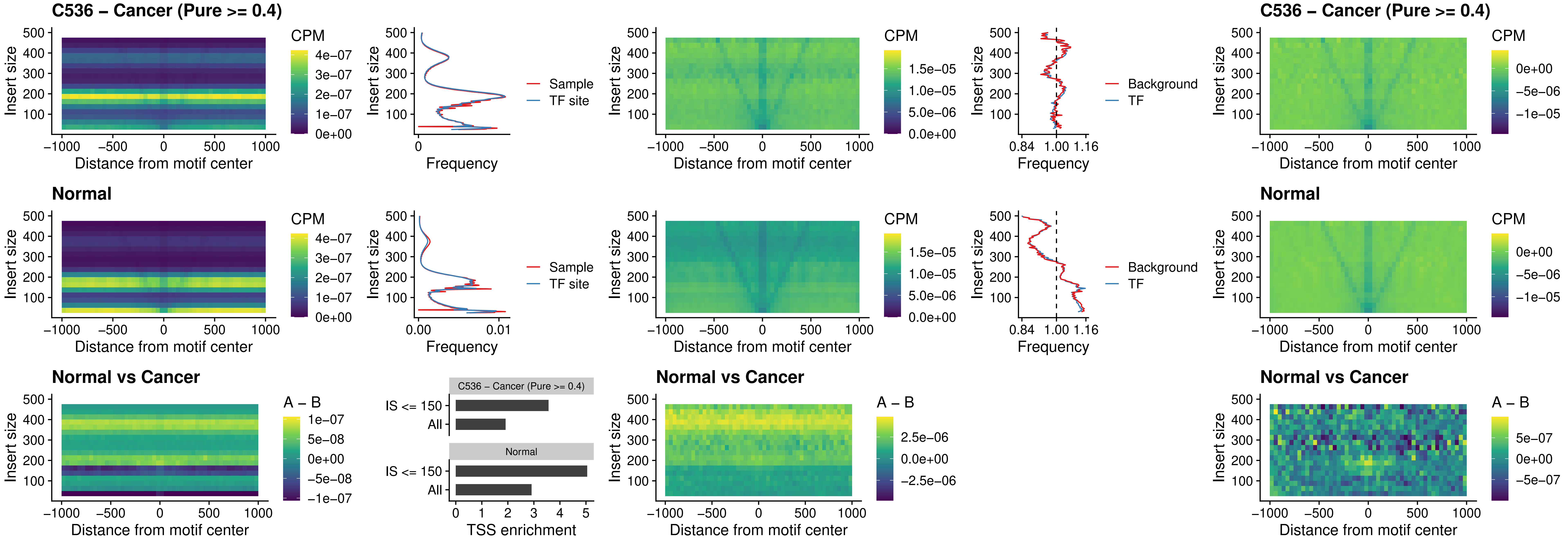

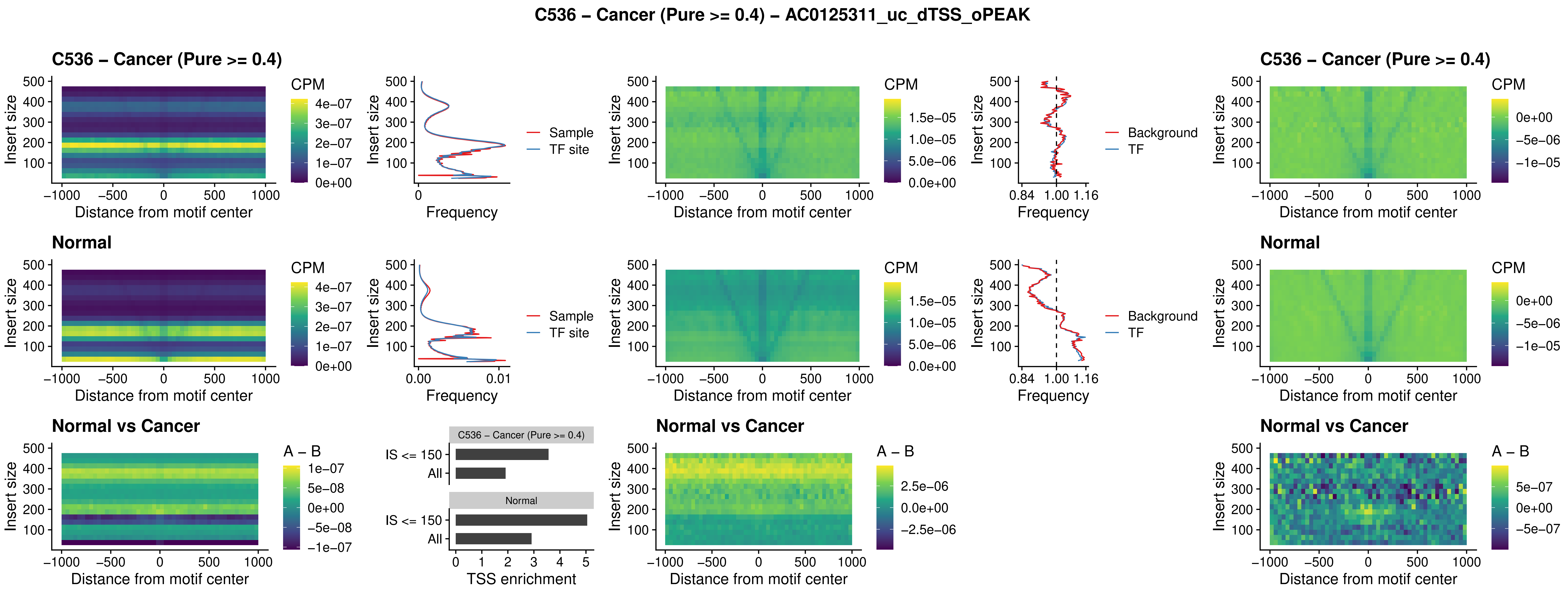

C536 – Cancer (Pure >= 0.4) – AC0125311_uc_dTSS_oPEAK
C536 – Cancer (Pure >= 0.4)
Normal
Normal vs Cancer
Insert size
Distance from motif center
Frequency
CPM
A – B
Sample
TF site
Background
TF
C536 – Cancer (Pure >= 0.4)
Normal
IS <= 150
All
TSS enrichment

### C536 – Cancer (Pure >= 0.4) – DBX2_uc_dTSS_oPEAK

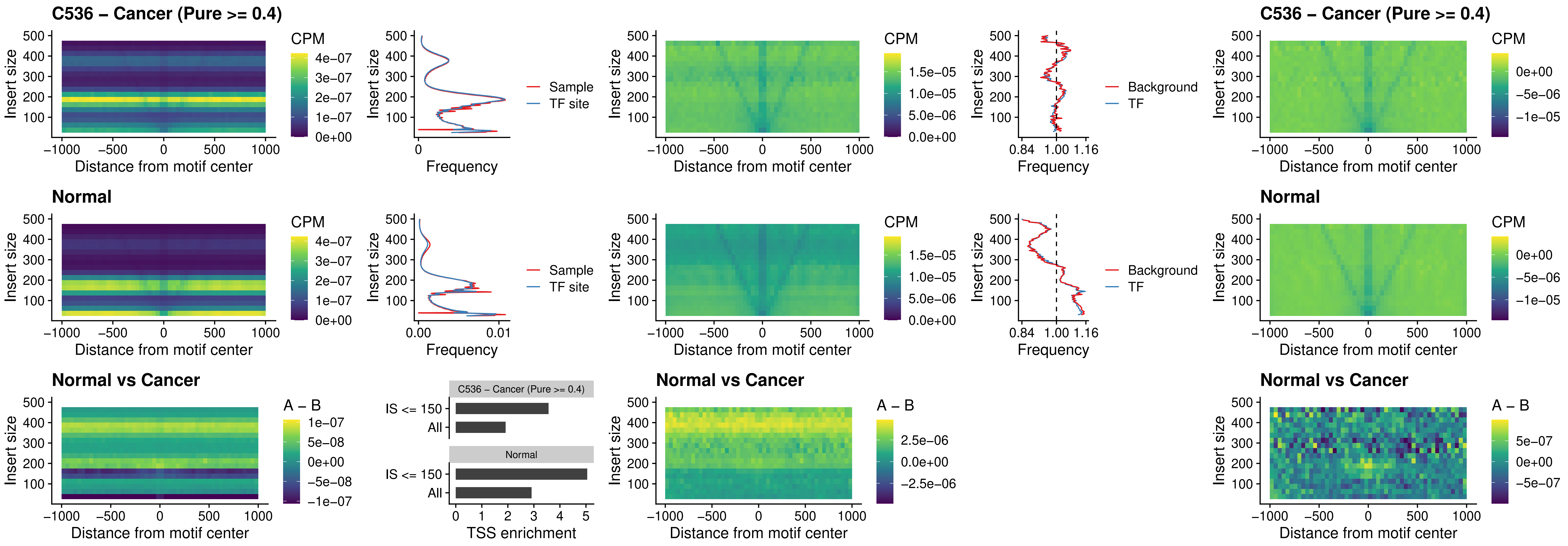

#### C536 – Cancer (Pure >= 0.4) – HOXA2_uc_dTSS_nPEAK

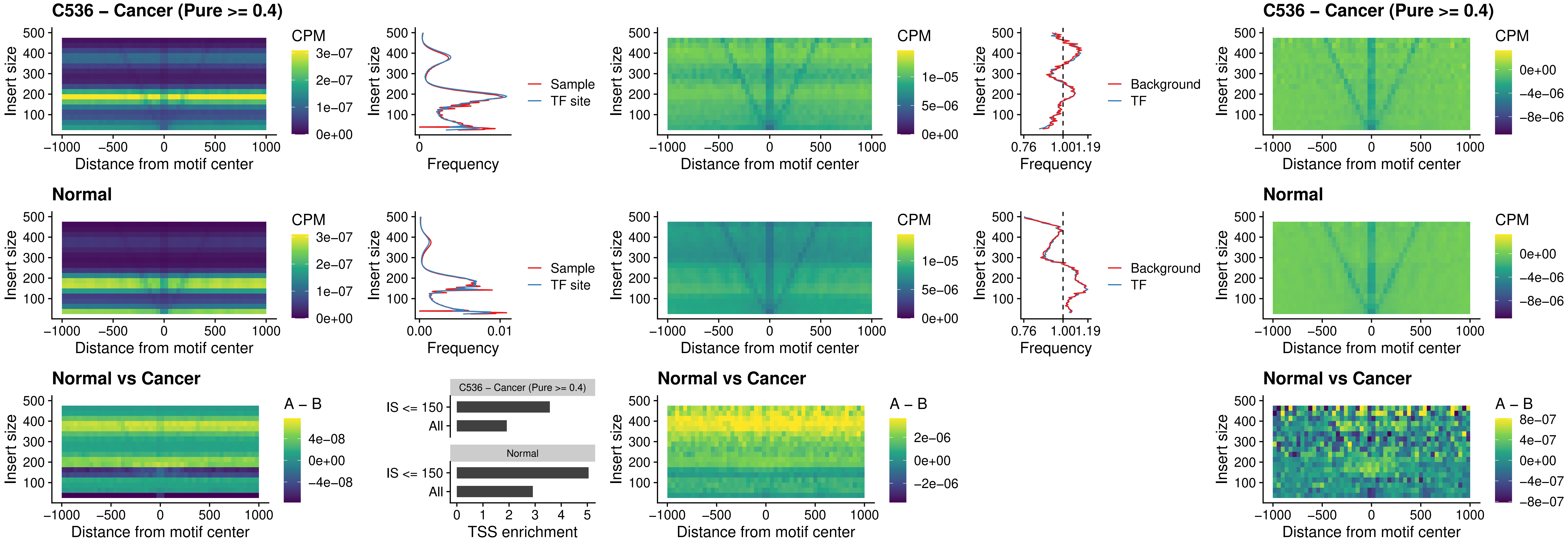

### C536 – Cancer (Pure >= 0.4) – SHOX_uc_dTSS_nPEAK

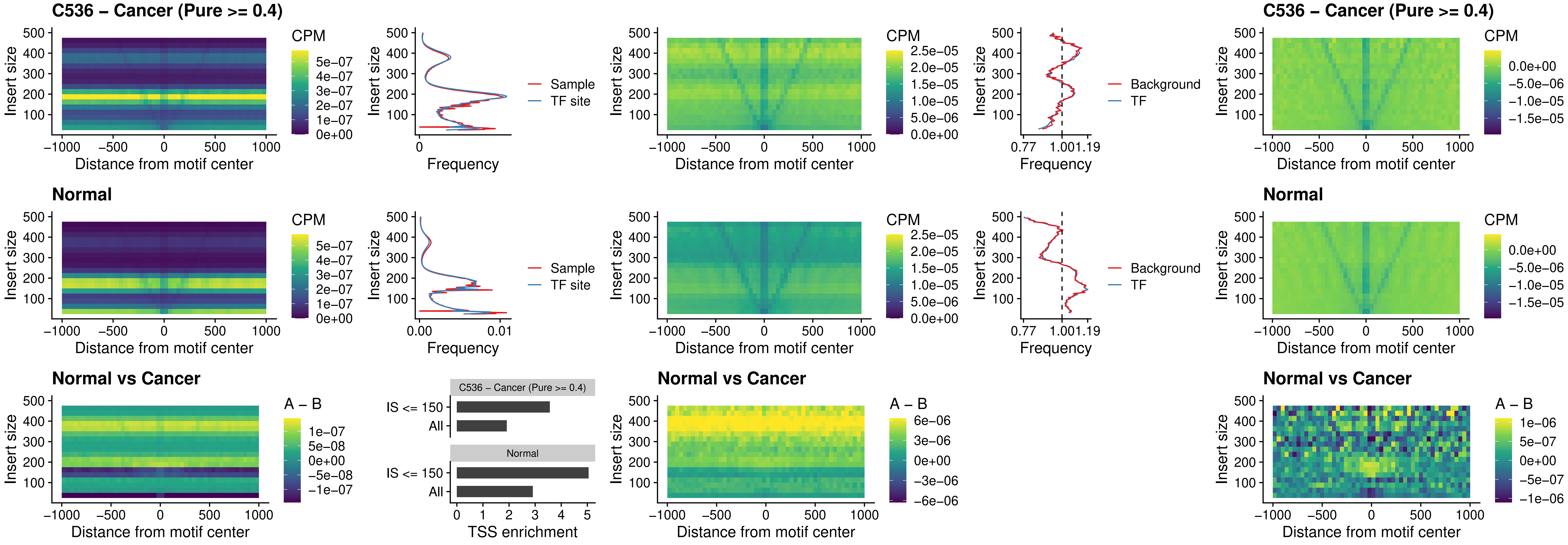

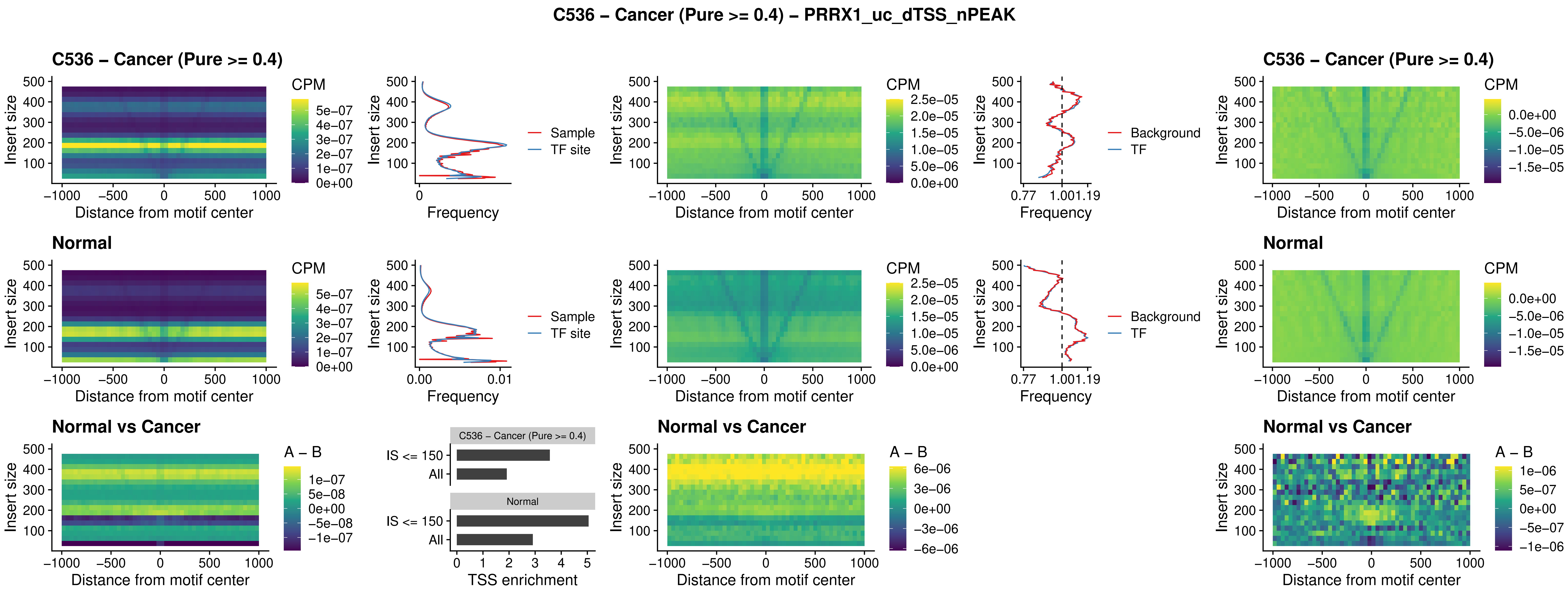
C536 – Cancer (Pure >= 0.4) – PRRX1_uc_dTSS_nPEAK
C536 – Cancer (Pure >= 0.4)
Normal
Normal vs Cancer
Insert size
Distance from motif center
Frequency
CPM
A – B
Sample
TF site
Background
TF
C536 – Cancer (Pure >= 0.4)
Normal
IS <= 150
All
TSS enrichment

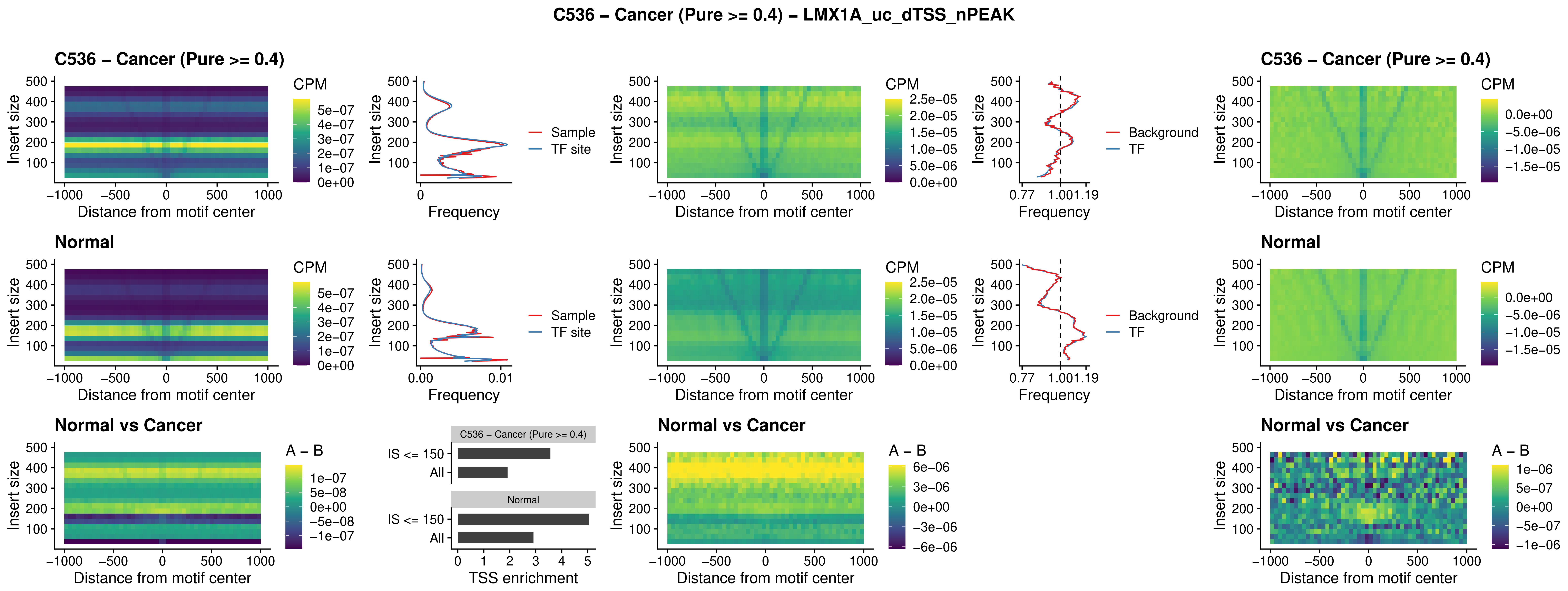
C536 – Cancer (Pure >= 0.4) – LMX1A_uc_dTSS_nPEAK
C536 – Cancer (Pure >= 0.4)
Normal
Normal vs Cancer
Insert size
Distance from motif center
Frequency
CPM
A – B
Sample
TF site
Background
TF
C536 – Cancer (Pure >= 0.4)
Normal
IS <= 150
All
TSS enrichment

#### C536 – Cancer (Pure >= 0.4) – LMX1B_uc_dTSS_nPEAK

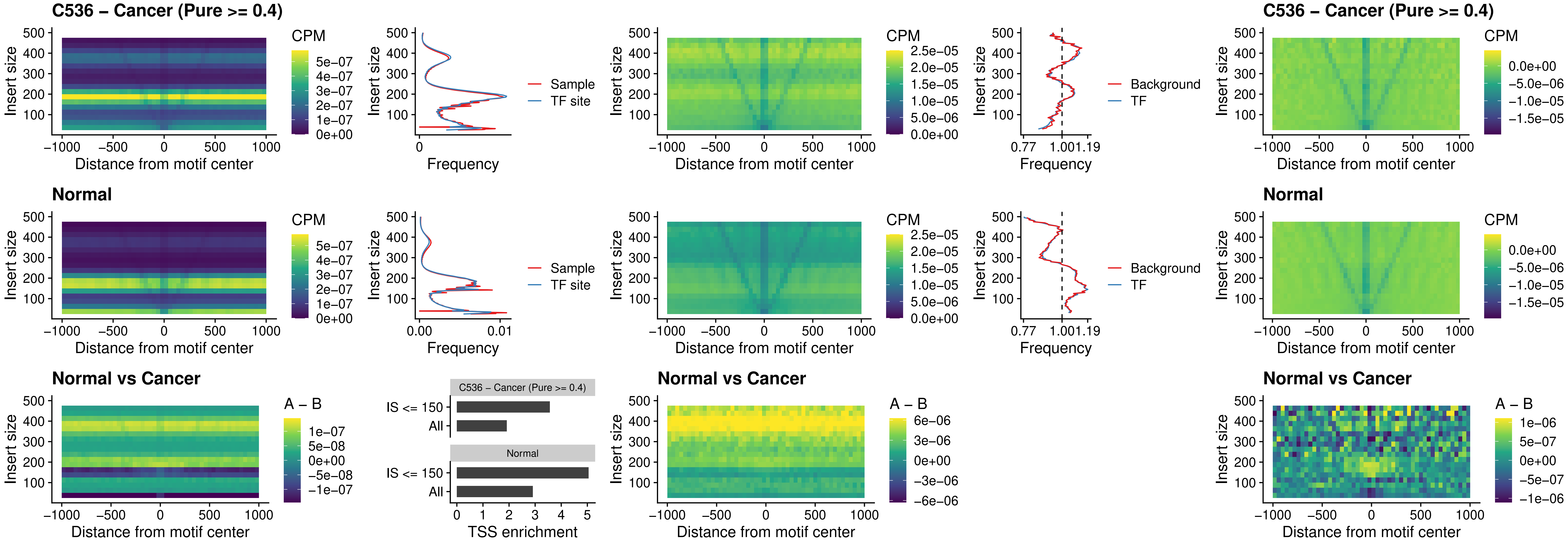

#### C536 – Cancer (Pure >= 0.4) – DLX5_uc_dTSS_oPEAK

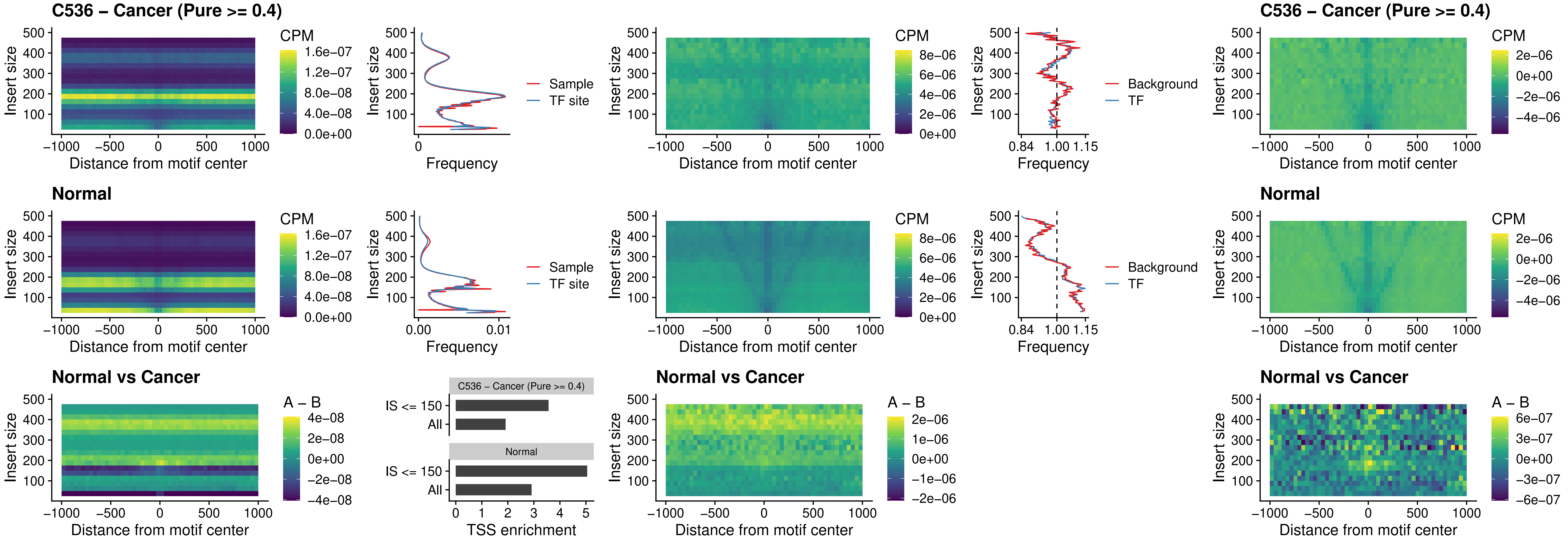

#### C536 – Cancer (Pure >= 0.4) – ARID3C_uc_dTSS_nPEAK

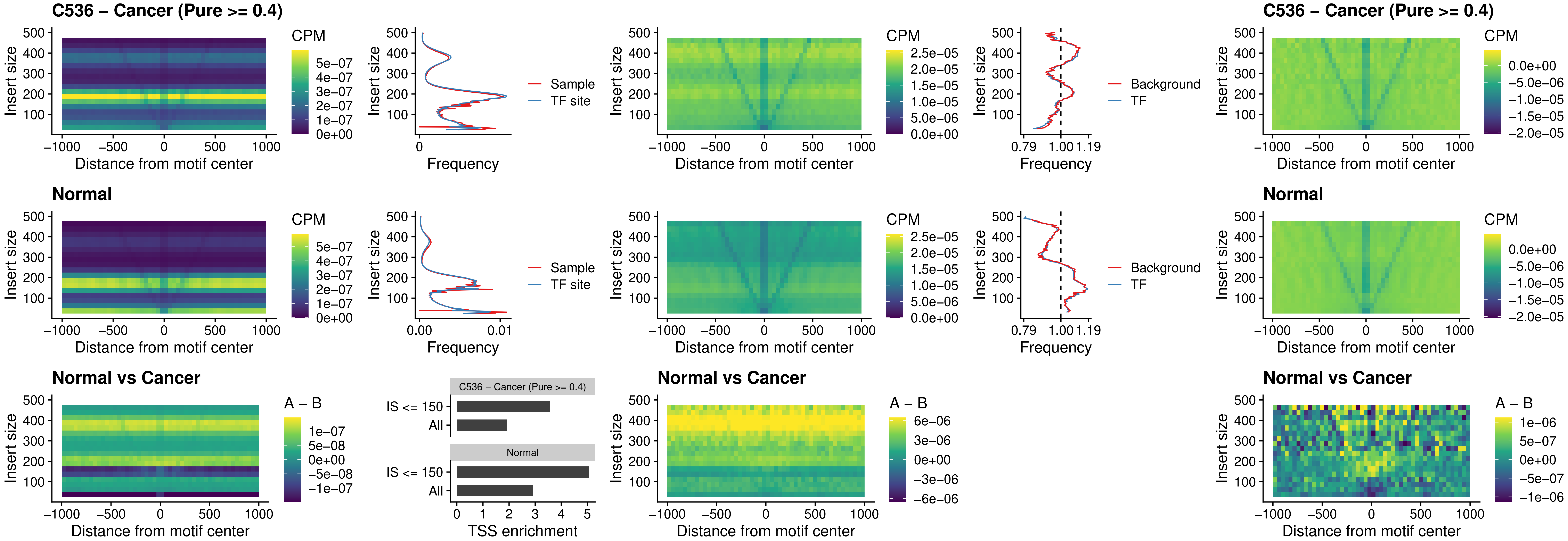

#### C536 – Cancer (Pure >= 0.4) – HOXD12_uc_dTSS_nPEAK

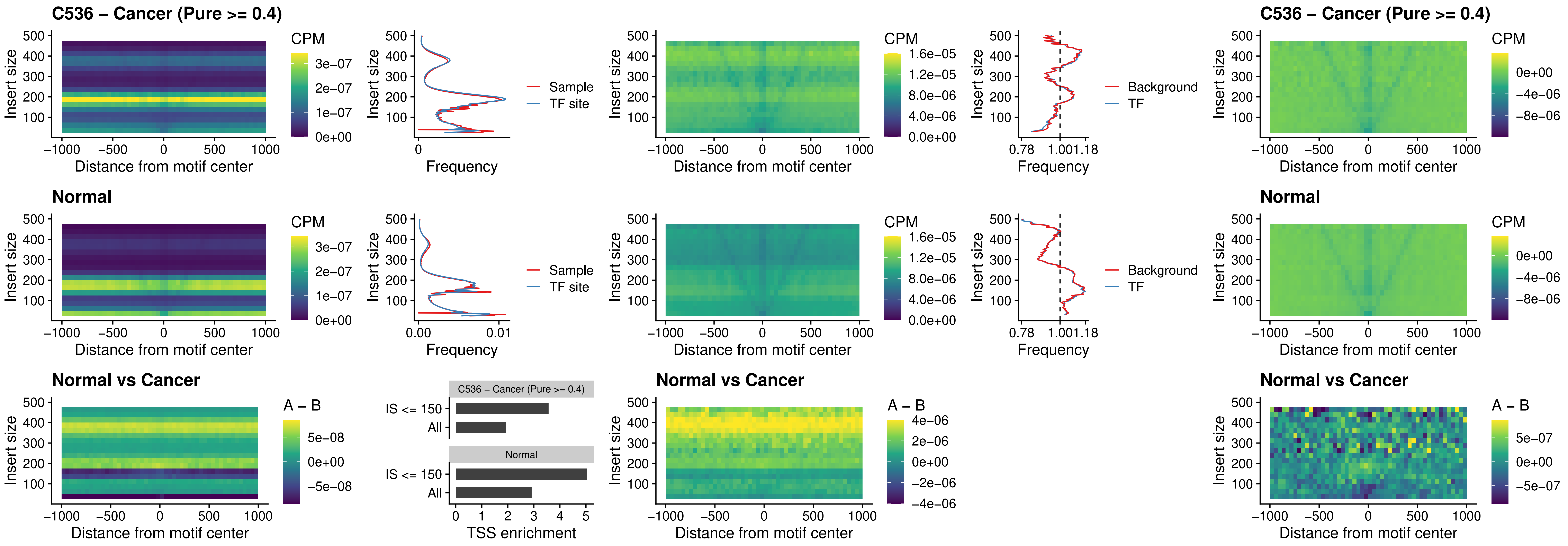

#### C536 – Cancer (Pure >= 0.4) – CDX2_uc_dTSS_nPEAK

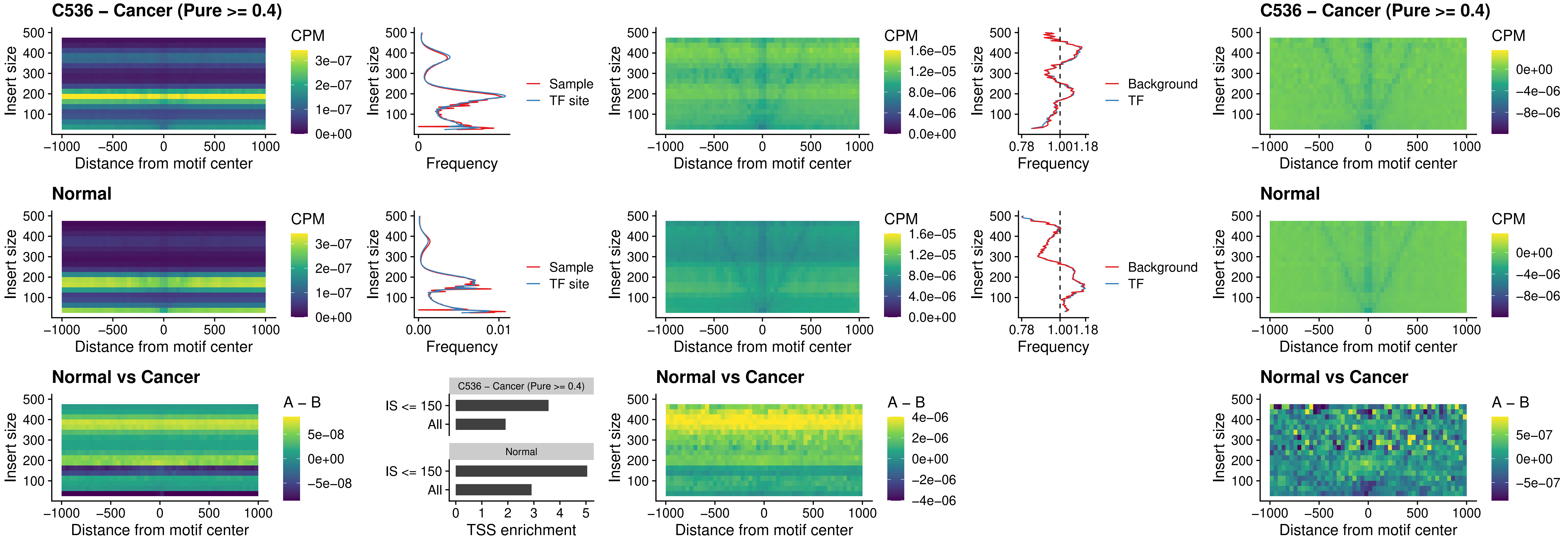

#### C536 – Cancer (Pure >= 0.4) – HOXC12_uc_dTSS_nPEAK

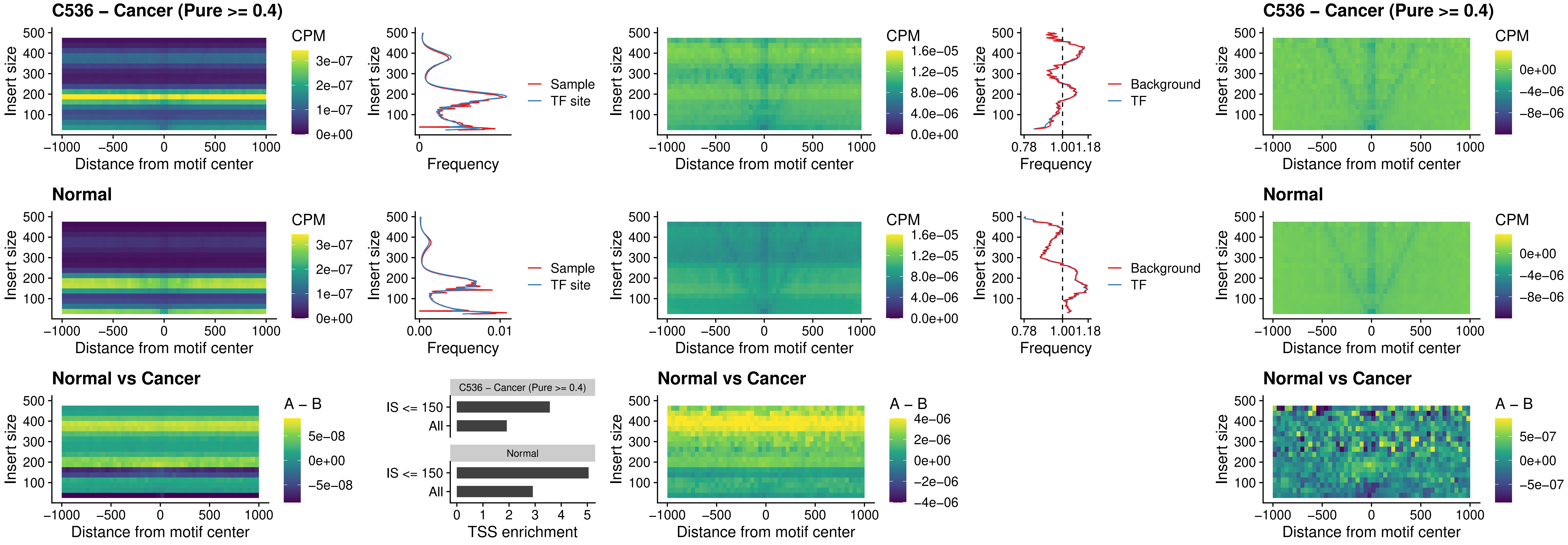

#### C536 – Cancer (Pure >= 0.4) – HHEX_uc_dTSS_oPEAK

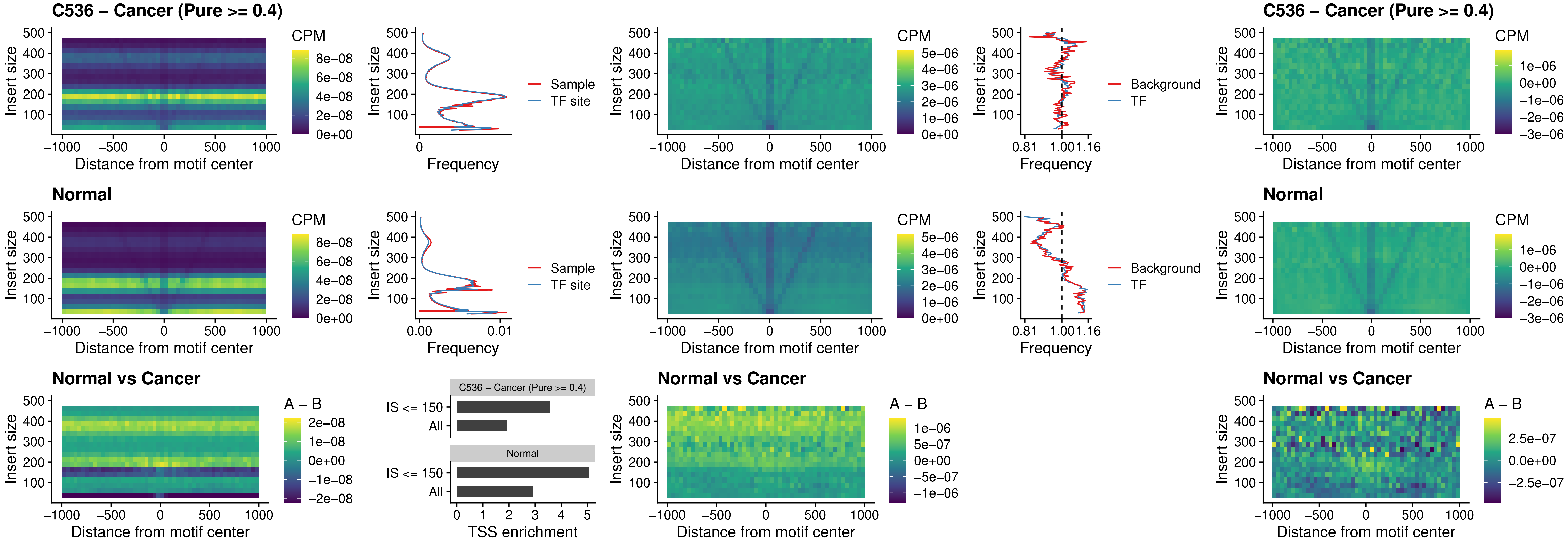

### C536 – Cancer (Pure >= 0.4) – HOXA11_uc_dTSS_nPEAK

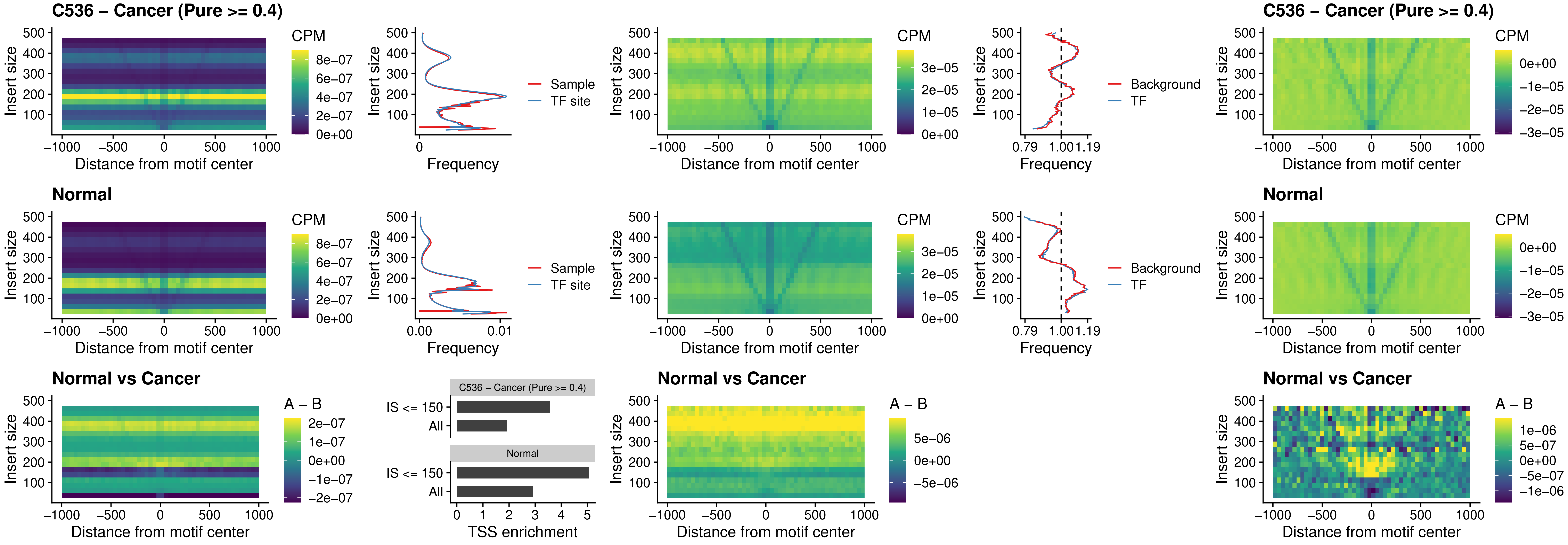

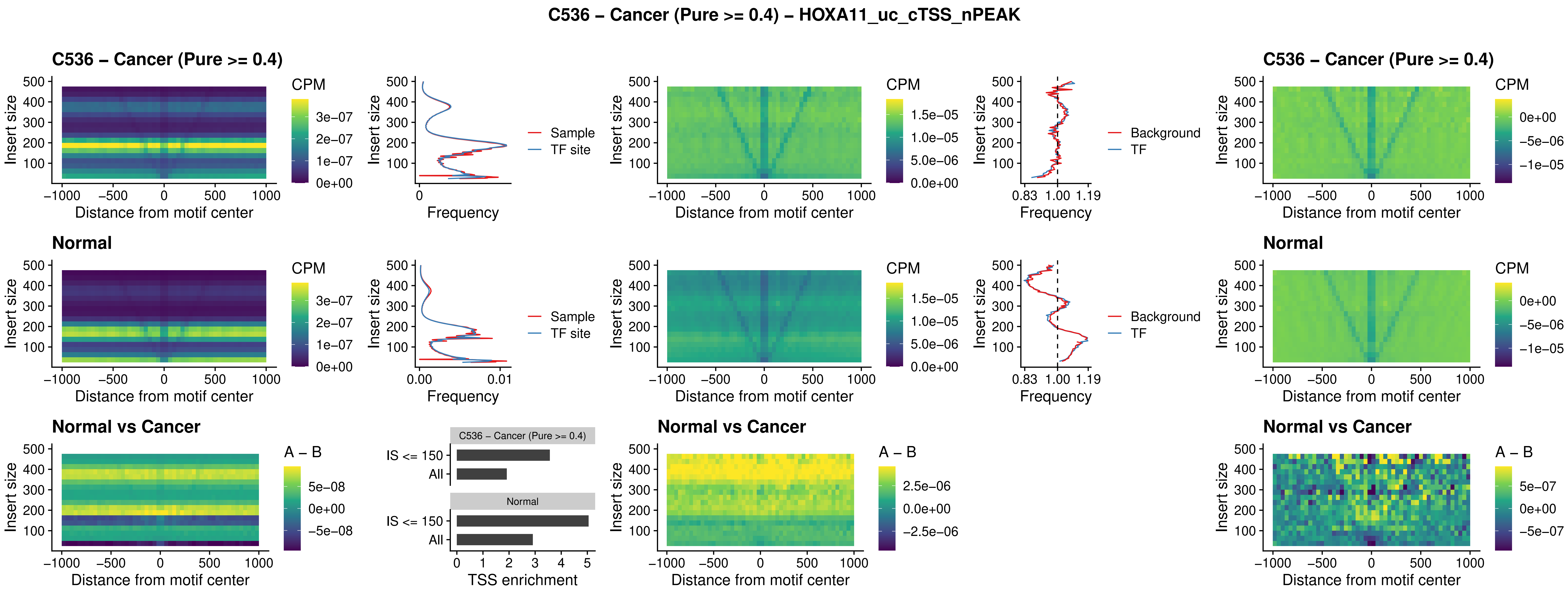
C536 – Cancer (Pure >= 0.4) – HOXA11_uc_cTSS_nPEAK
C536 – Cancer (Pure >= 0.4)
Normal
Normal vs Cancer
Insert size
Distance from motif center
CPM
A – B
Frequency
Sample
TF site
Background
TF
C536 – Cancer (Pure >= 0.4)
IS <= 150
All
TSS enrichment

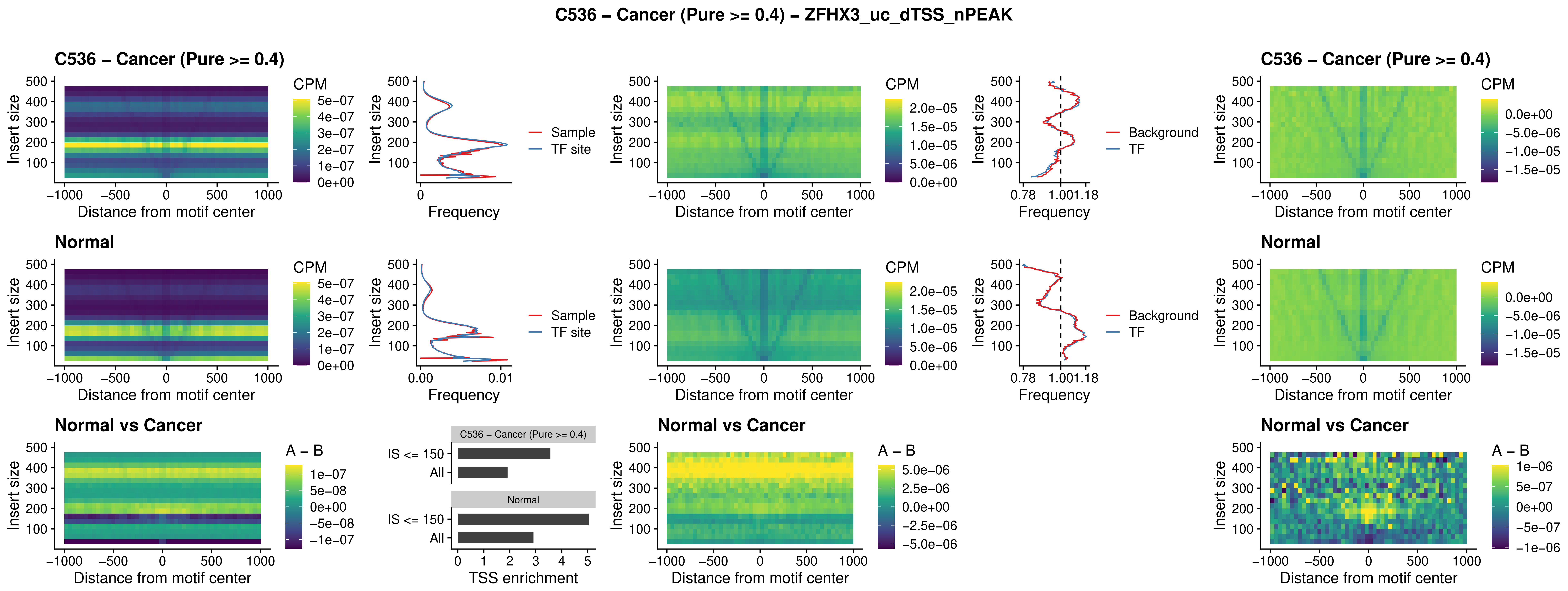
C536 – Cancer (Pure >= 0.4) – ZFHX3_uc_dTSS_nPEAK
C536 – Cancer (Pure >= 0.4)
Normal
Normal vs Cancer
Insert size
Distance from motif center
Frequency
CPM
A – B
Sample
TF site
Background
TF
C536 – Cancer (Pure >= 0.4)
Normal
IS <= 150
All
TSS enrichment

#### C536 – Cancer (Pure >= 0.4) – HOXB4_uc_dTSS_nPEAK

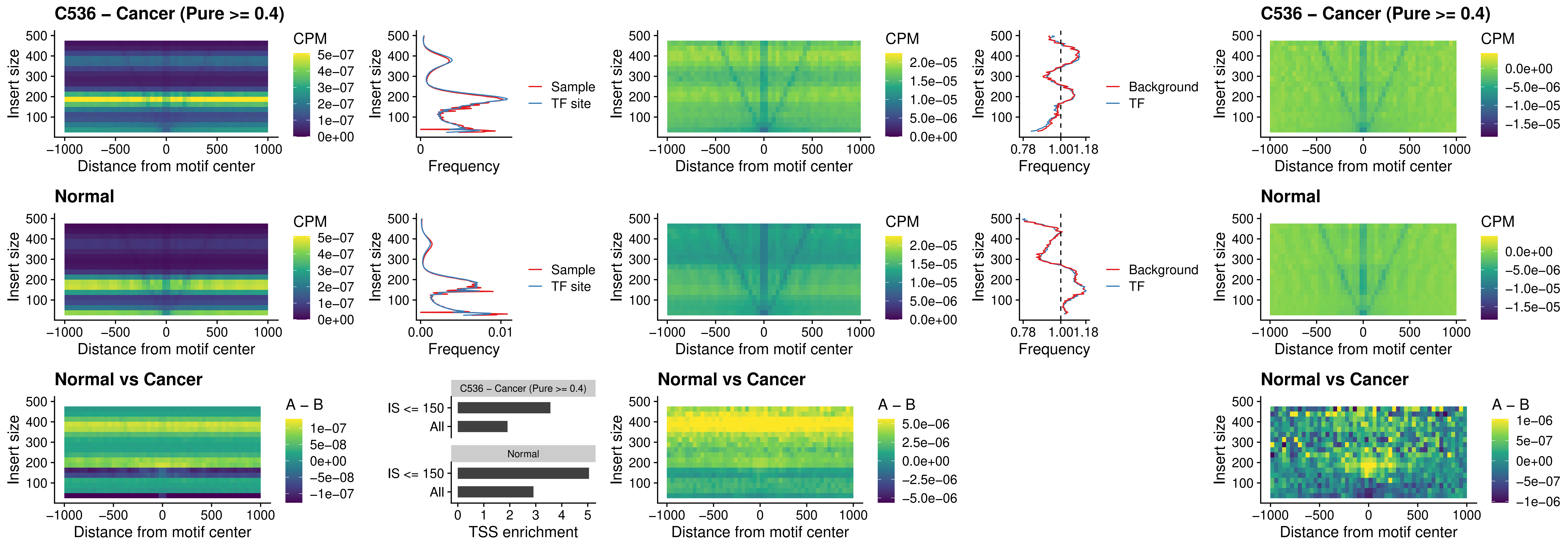

#### C536 – Cancer (Pure >= 0.4) – HOXC5_uc_dTSS_nPEAK

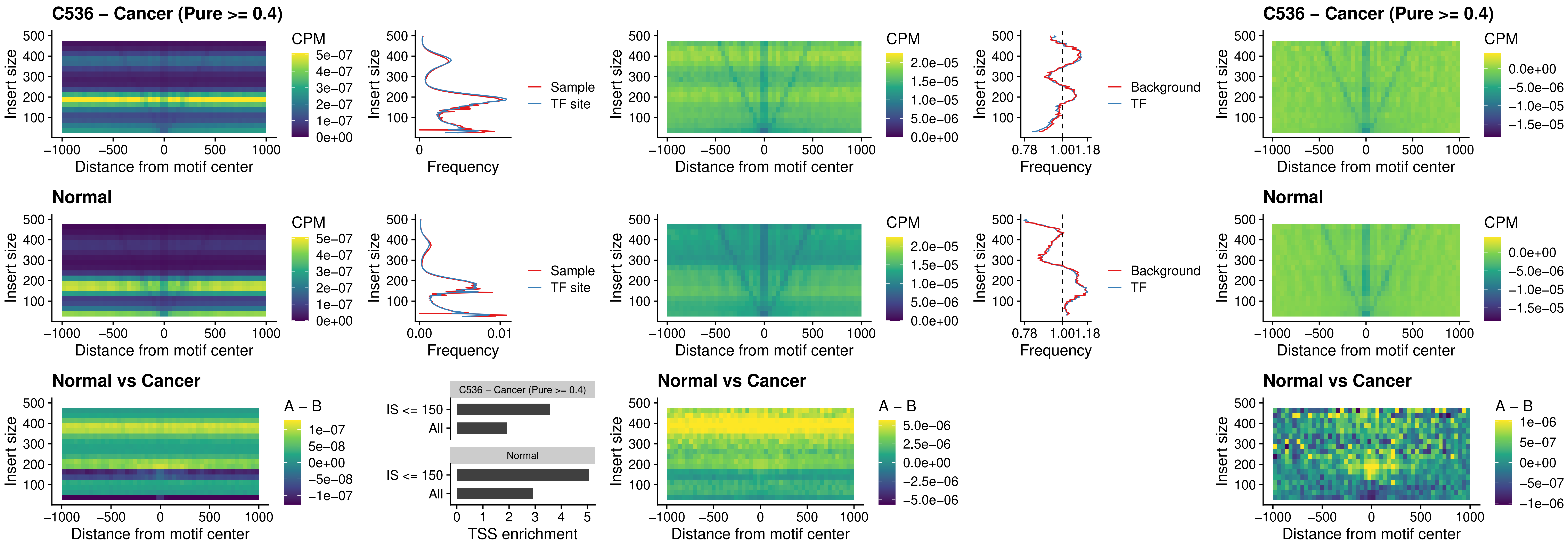

#### C536 – Cancer (Pure >= 0.4) – HOXD1_uc_dTSS_nPEAK

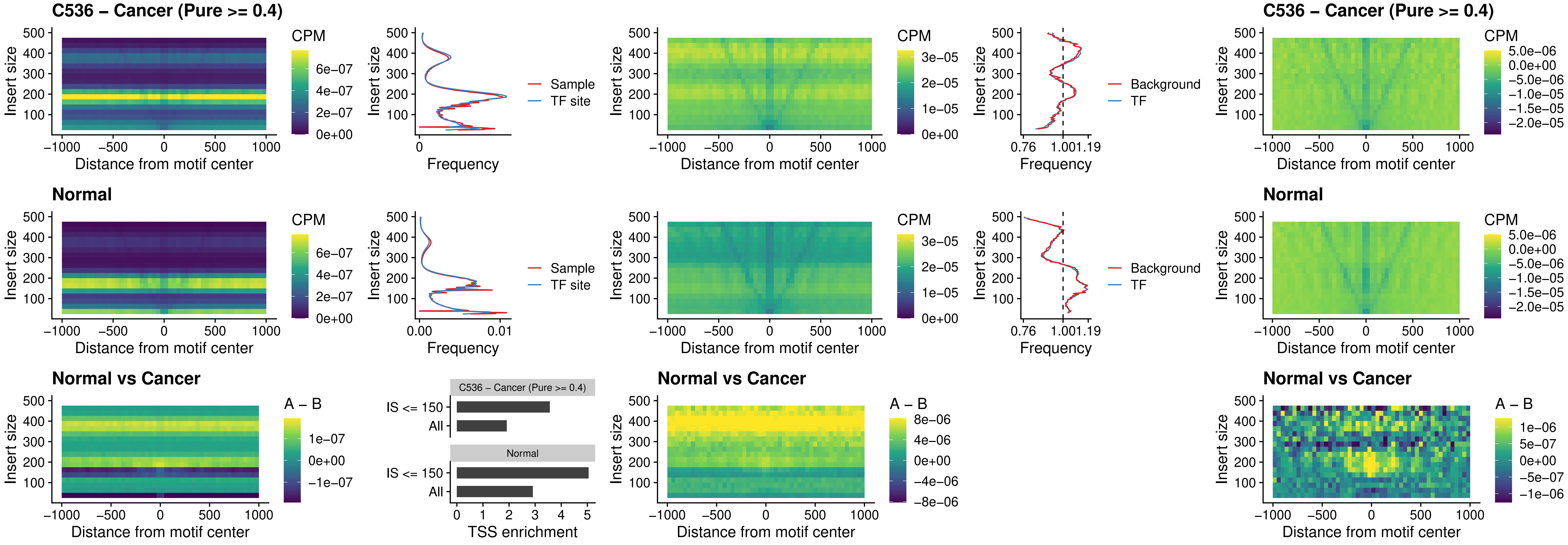

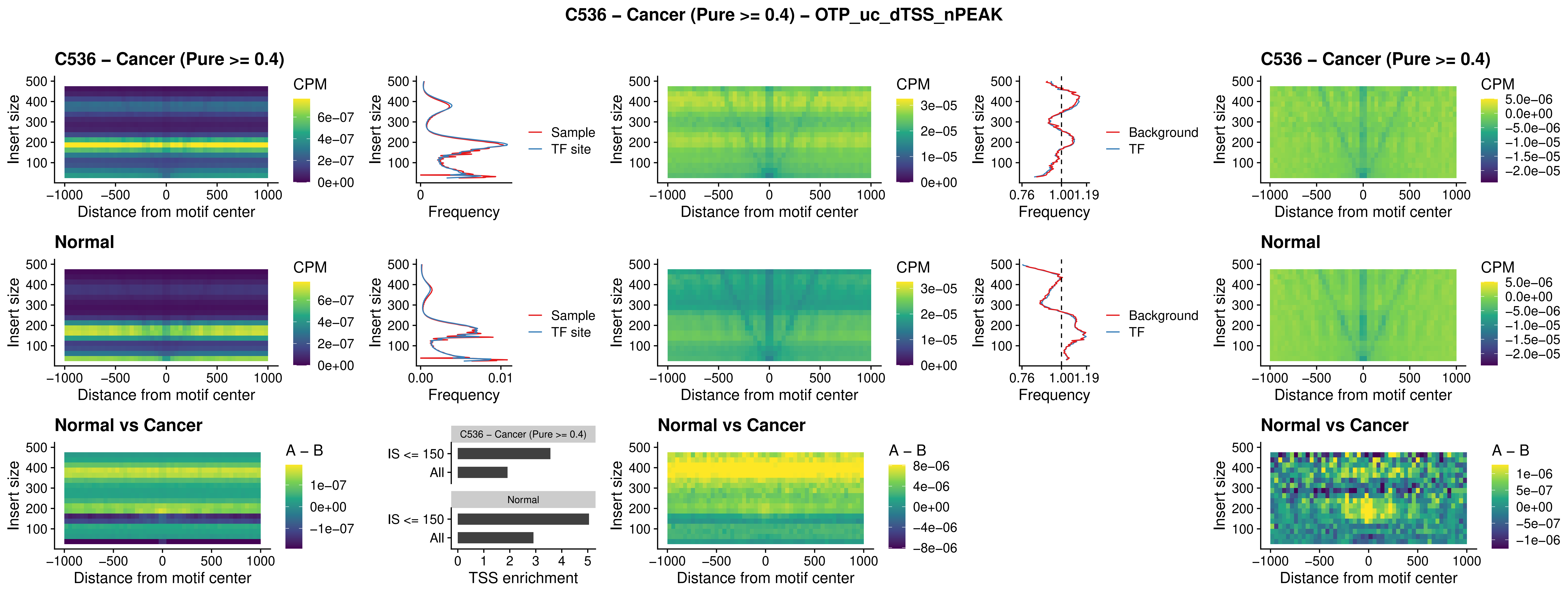
C536 – Cancer (Pure >= 0.4) – OTP_uc_dTSS_nPEAK
C536 – Cancer (Pure >= 0.4)
Normal
Normal vs Cancer
Insert size
Distance from motif center
Frequency
CPM
A – B
Sample
TF site
Background
TF
C536 – Cancer (Pure >= 0.4)
Normal
IS <= 150
All
TSS enrichment

#### C536 – Cancer (Pure >= 0.4) – FOXL1_uc_dTSS_oPEAK

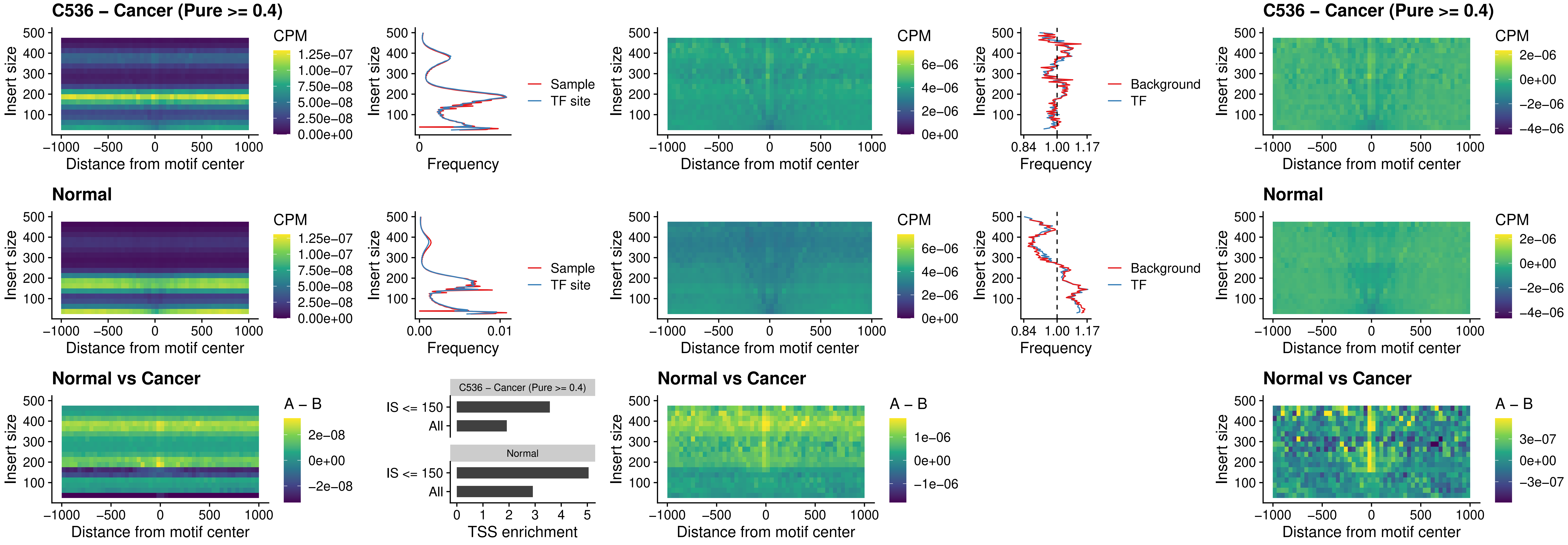

#### C536 – Cancer (Pure >= 0.4) – ZNF32_uc_dTSS_oPEAK

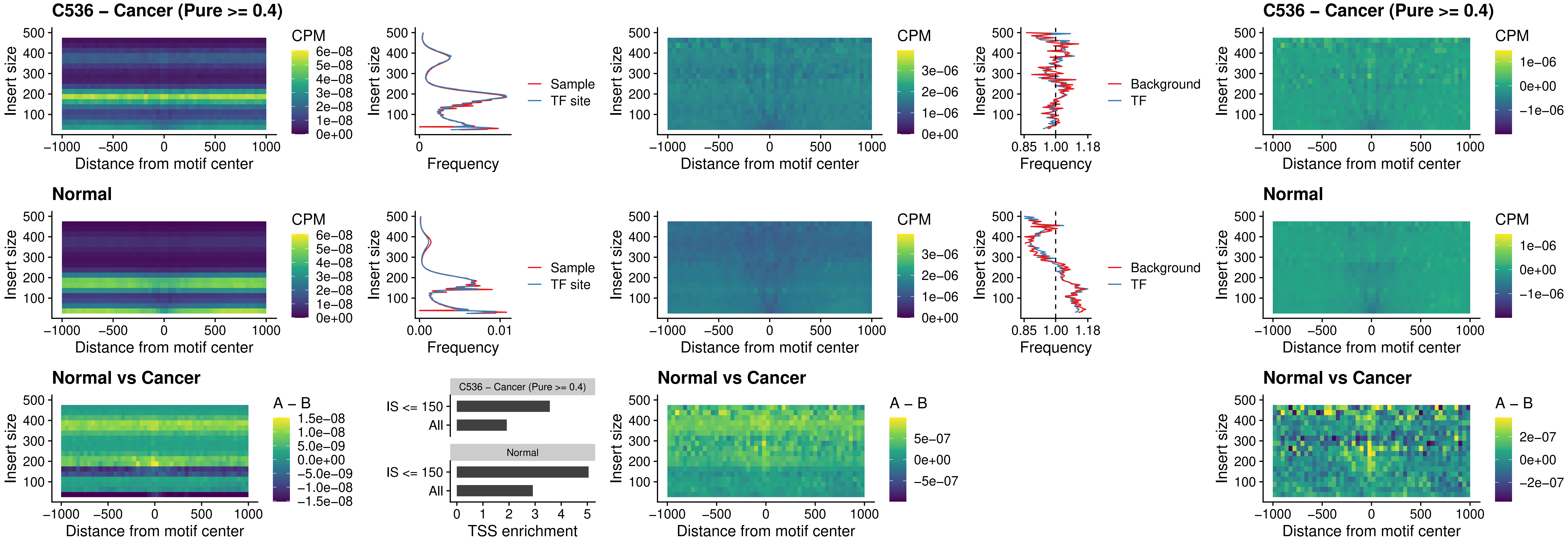

#### C536 – Cancer (Pure >= 0.4) – FOXL1_uc_dTSS_nPEAK

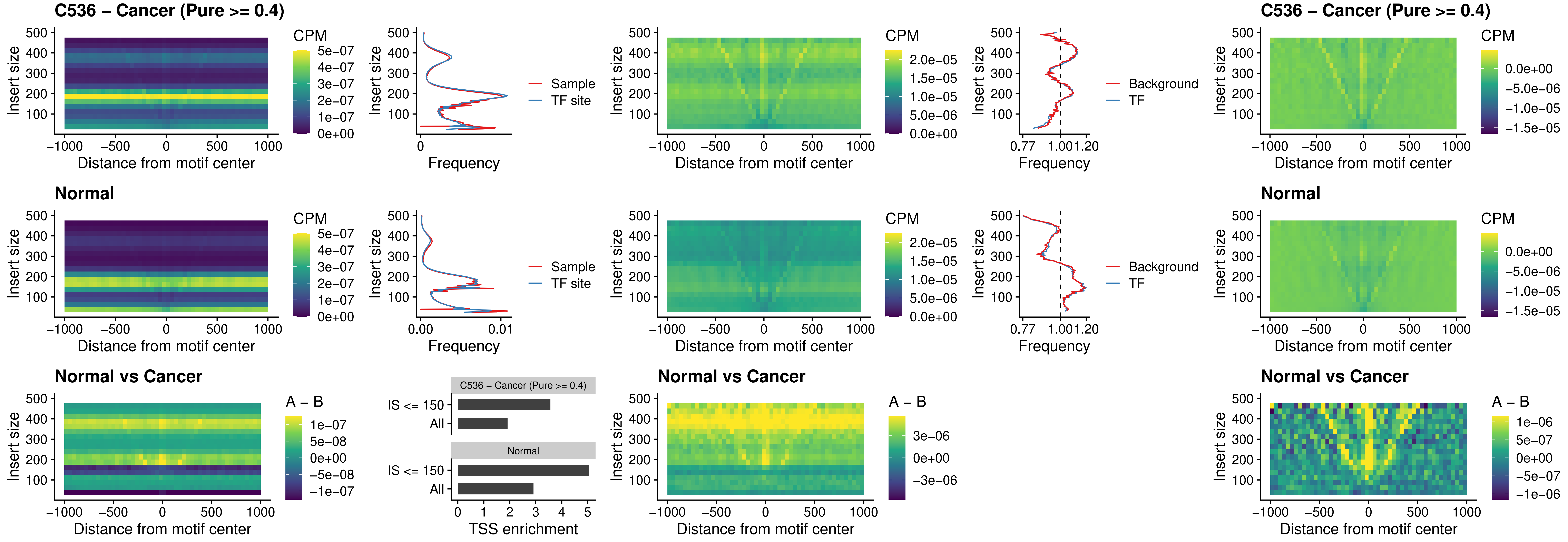

#### C536 – Cancer (Pure >= 0.4) – ZNF32_uc_dTSS_nPEAK

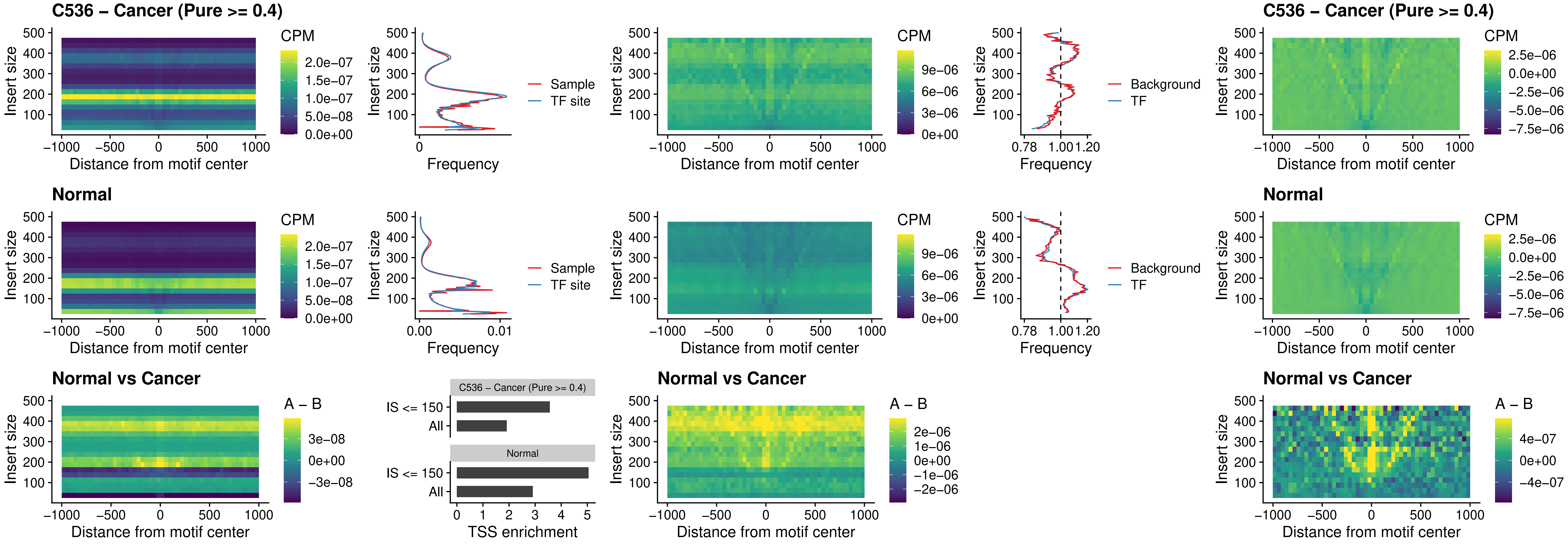

#### C536 – Cancer (Pure >= 0.4) – DLX5_uc_dTSS_nPEAK

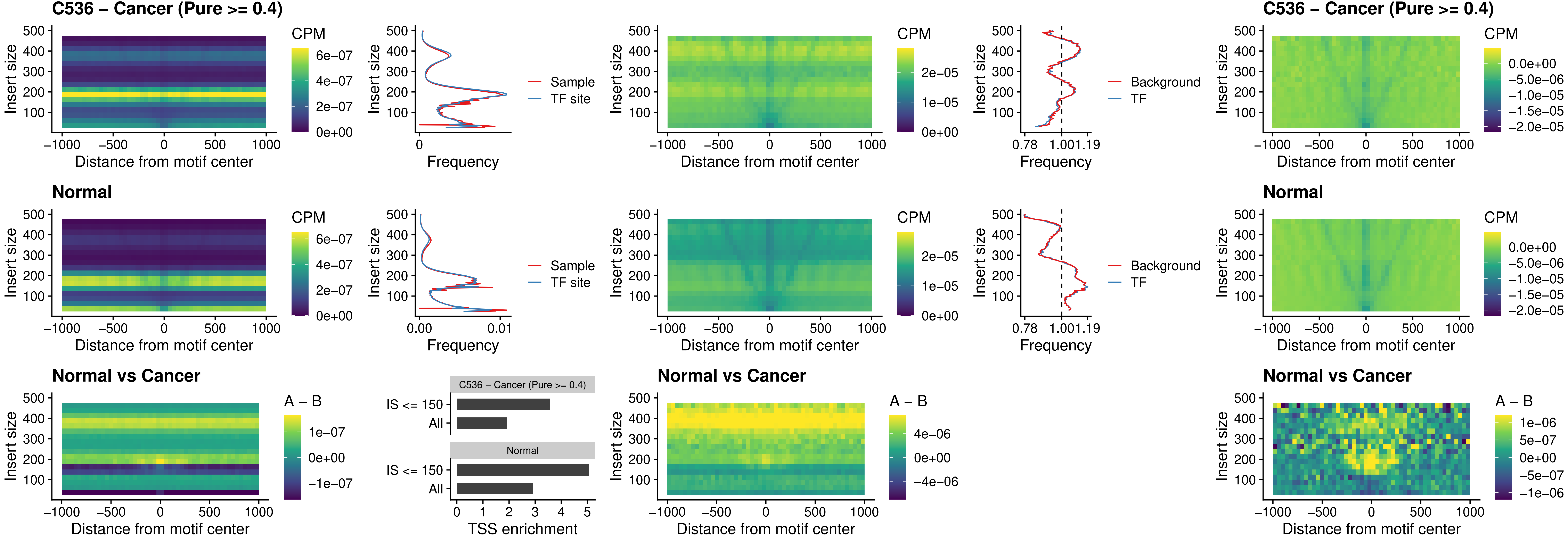

#### C536 – Cancer (Pure >= 0.4) – DMRT2_uc_dTSS_nPEAK

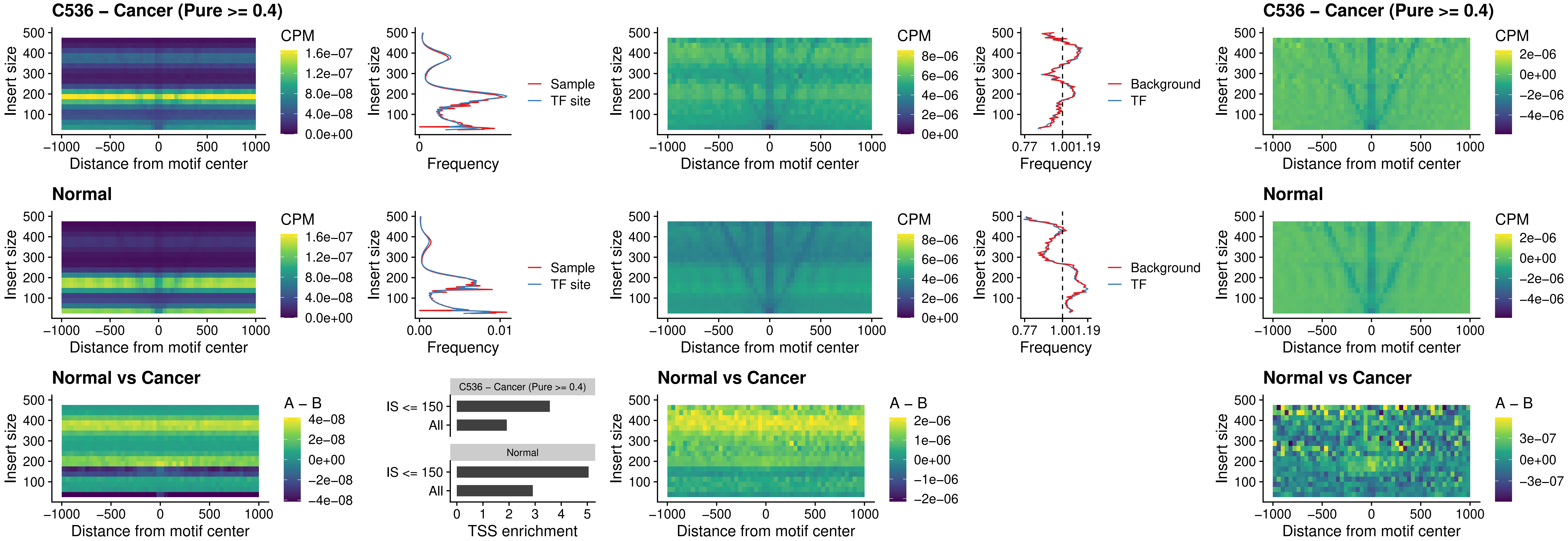

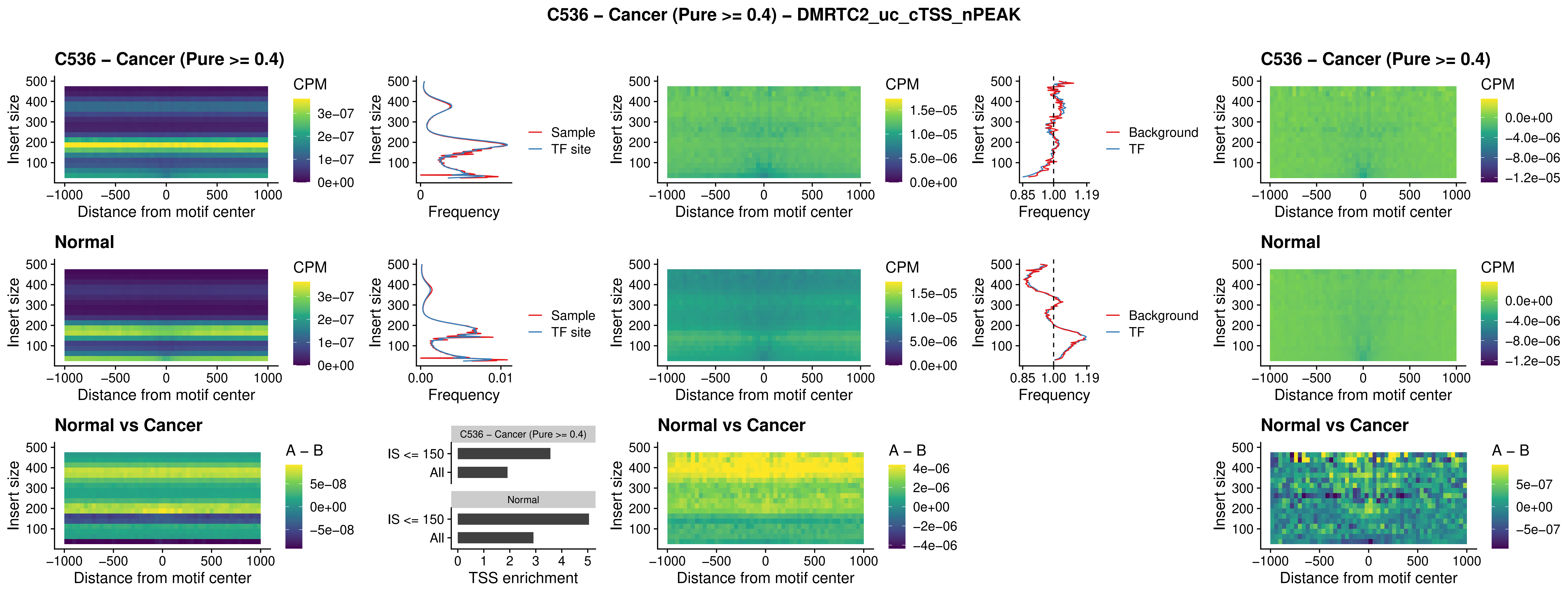
C536 – Cancer (Pure >= 0.4) – DMRTC2_uc_cTSS_nPEAK
C536 – Cancer (Pure >= 0.4)
Normal
Normal vs Cancer
Insert size
Distance from motif center
CPM
A – B
Frequency
Sample
TF site
Background
TF
C536 – Cancer (Pure >= 0.4)
Normal
IS <= 150
All
TSS enrichment

#### C536 – Cancer (Pure >= 0.4) – HOXD9_uc_dTSS_nPEAK

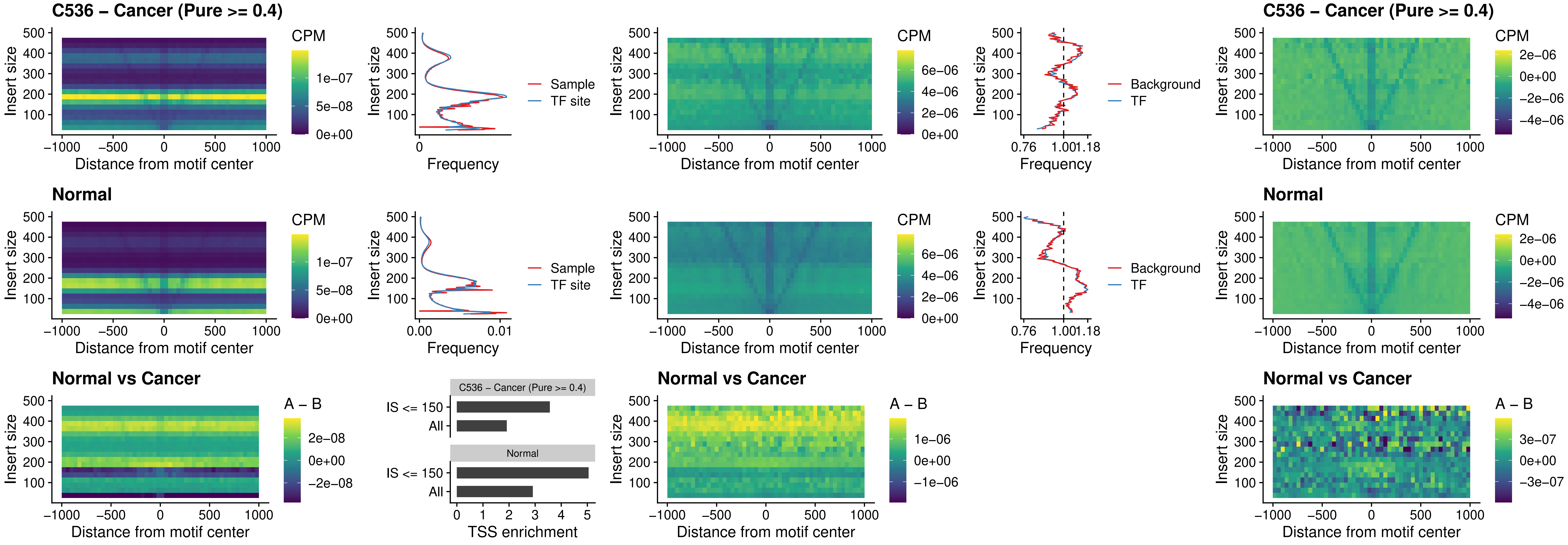

#### C536 – Cancer (Pure >= 0.4) – HOXD4_uc_dTSS_nPEAK

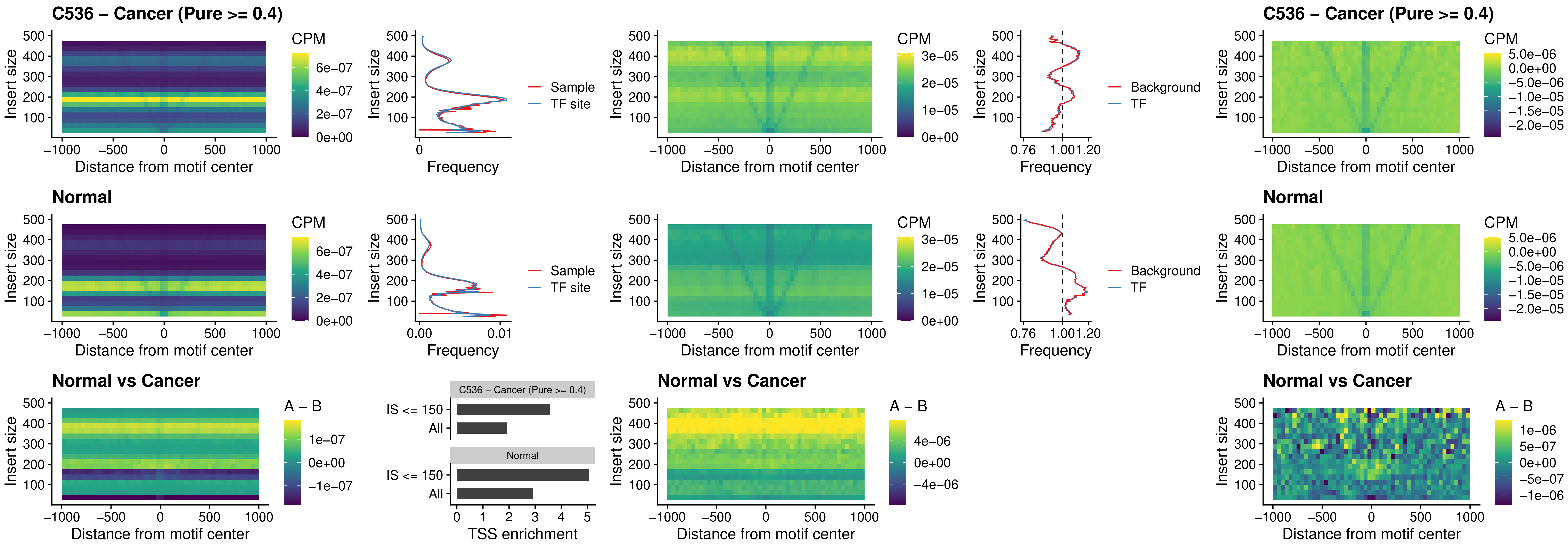

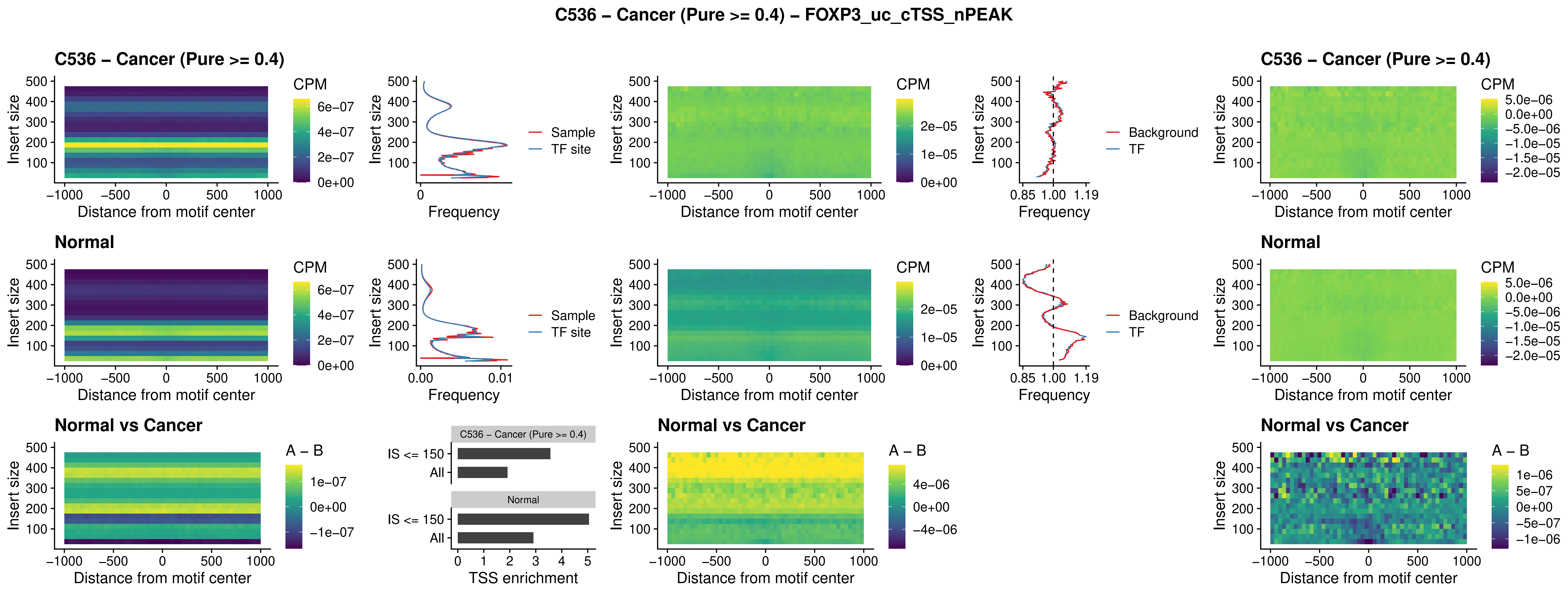
C536 – Cancer (Pure >= 0.4) – FOXP3_uc_cTSS_nPEAK
C536 – Cancer (Pure >= 0.4)
Normal
Normal vs Cancer
Insert size
Distance from motif center
Frequency
CPM
A – B
Sample
TF site
Background
TF
C536 – Cancer (Pure >= 0.4)
Normal
IS <= 150
All
TSS enrichment

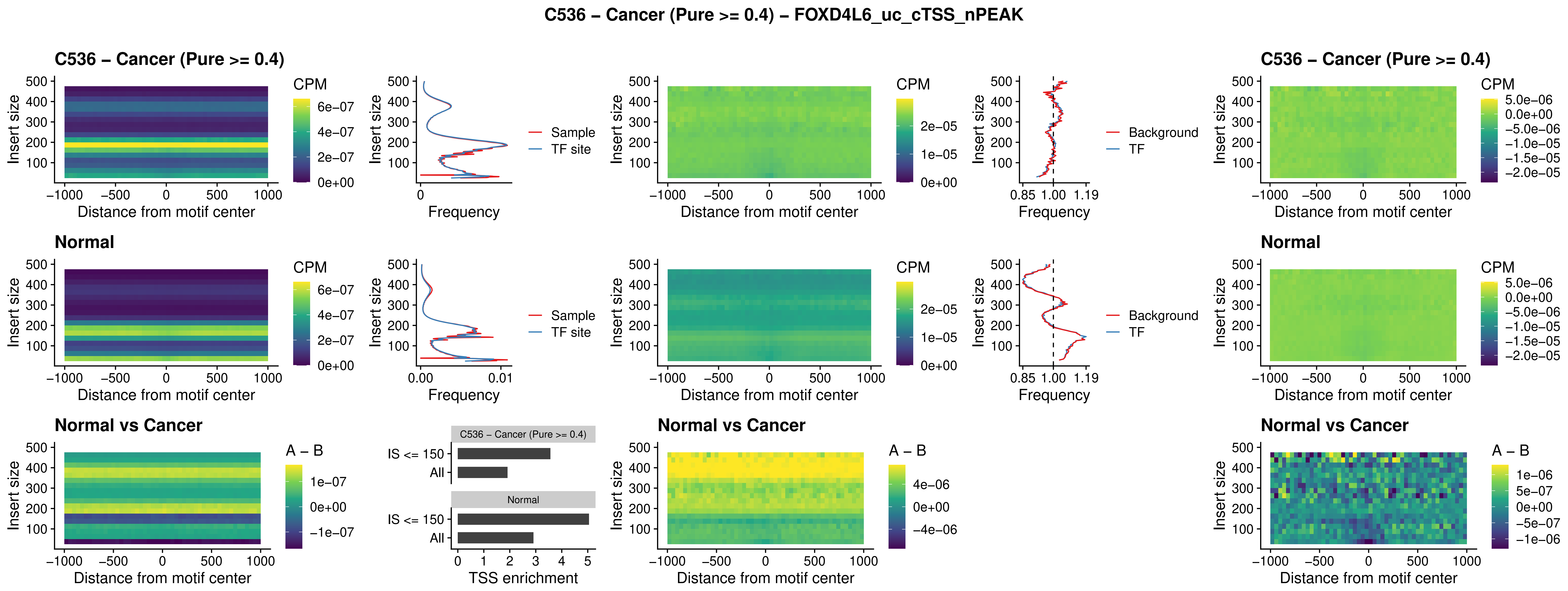
C536 – Cancer (Pure >= 0.4) – FOXD4L6_uc_cTSS_nPEAK
C536 – Cancer (Pure >= 0.4)
Normal
Normal vs Cancer
Insert size
Distance from motif center
Frequency
CPM
A – B
Sample
TF site
Background
TF
C536 – Cancer (Pure >= 0.4)
Normal
IS <= 150
All
TSS enrichment

### C536 – Cancer (Pure >= 0.4) – FOXD4L5_uc_cTSS_nPEAK

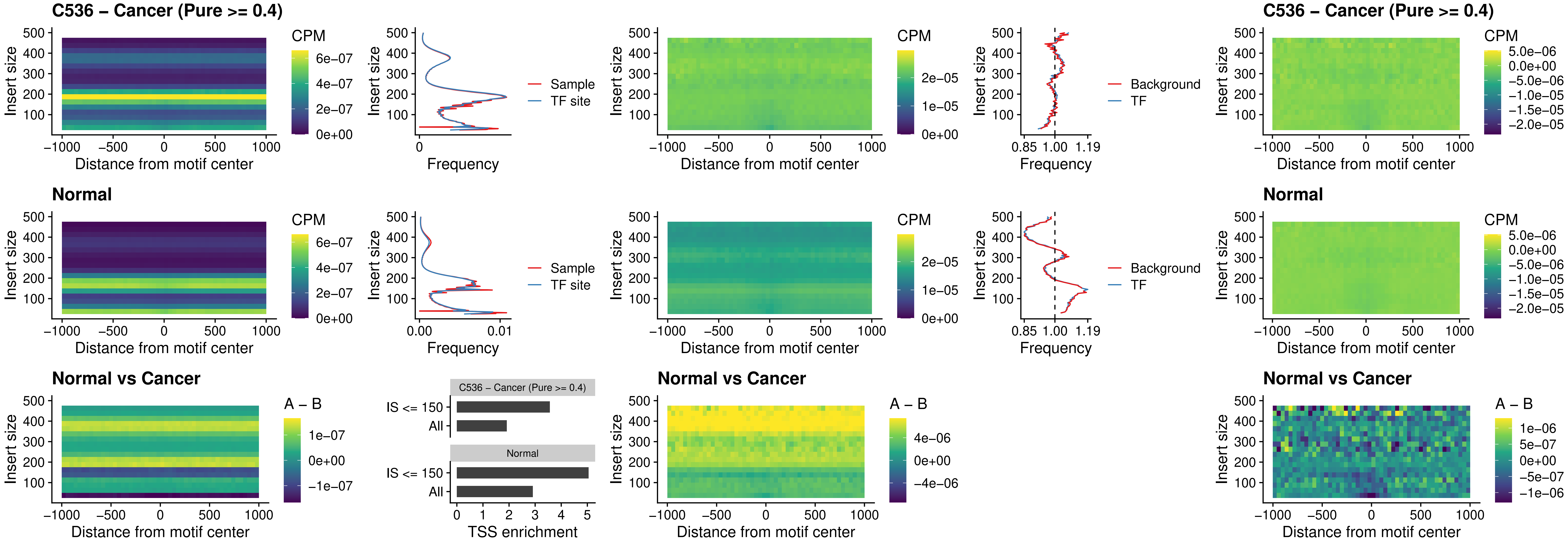

### C536 – Cancer (Pure >= 0.4) – FOXD4L4_uc_cTSS_nPEAK

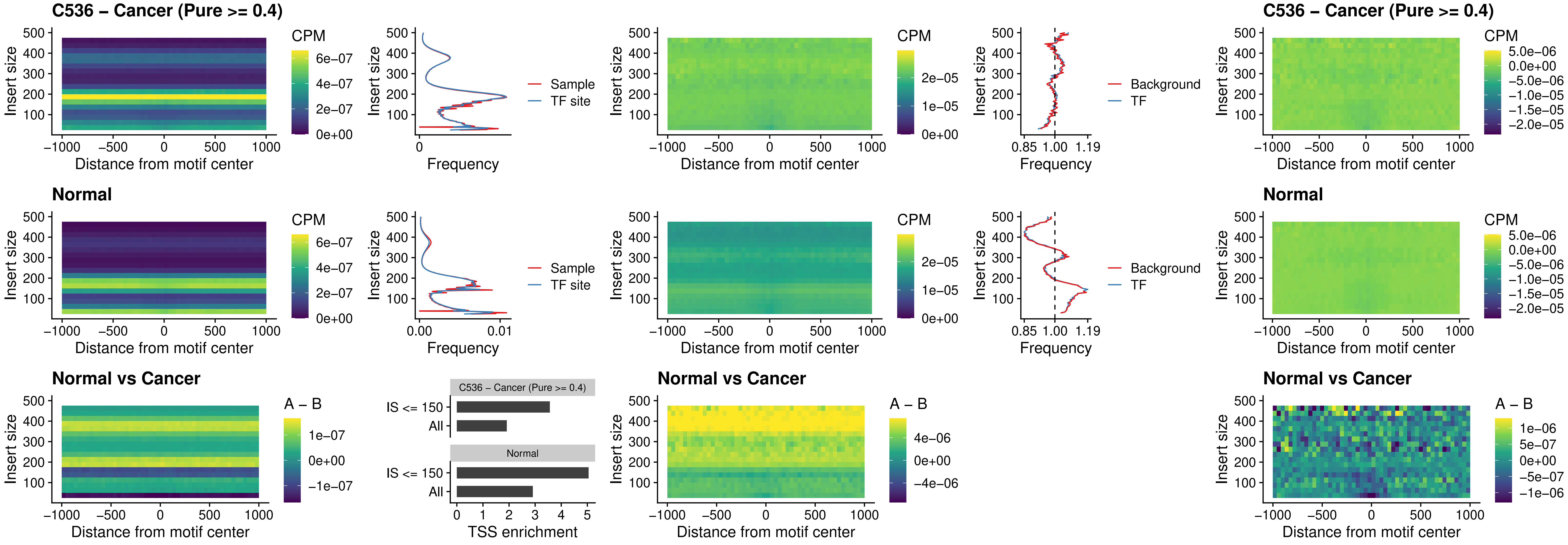

### C536 – Cancer (Pure >= 0.4) – FOXD4L2_uc_cTSS_nPEAK

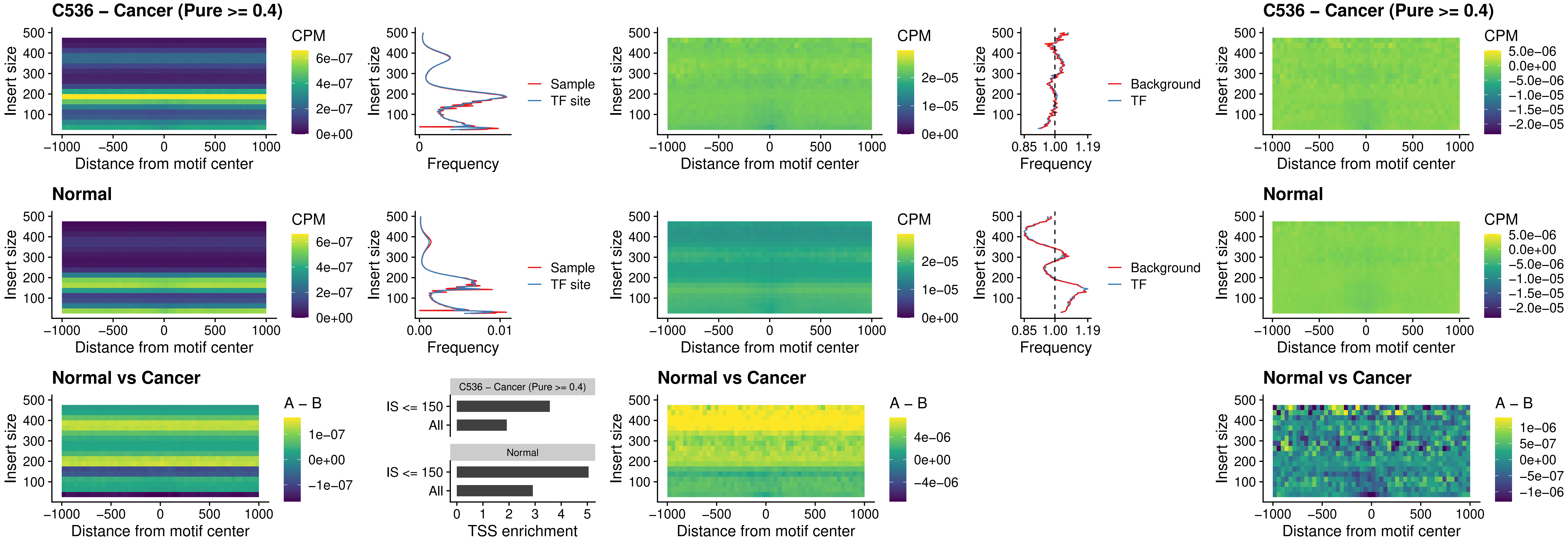

#### C536 – Cancer (Pure >= 0.4) – FOXD4L3_uc_cTSS_nPEAK

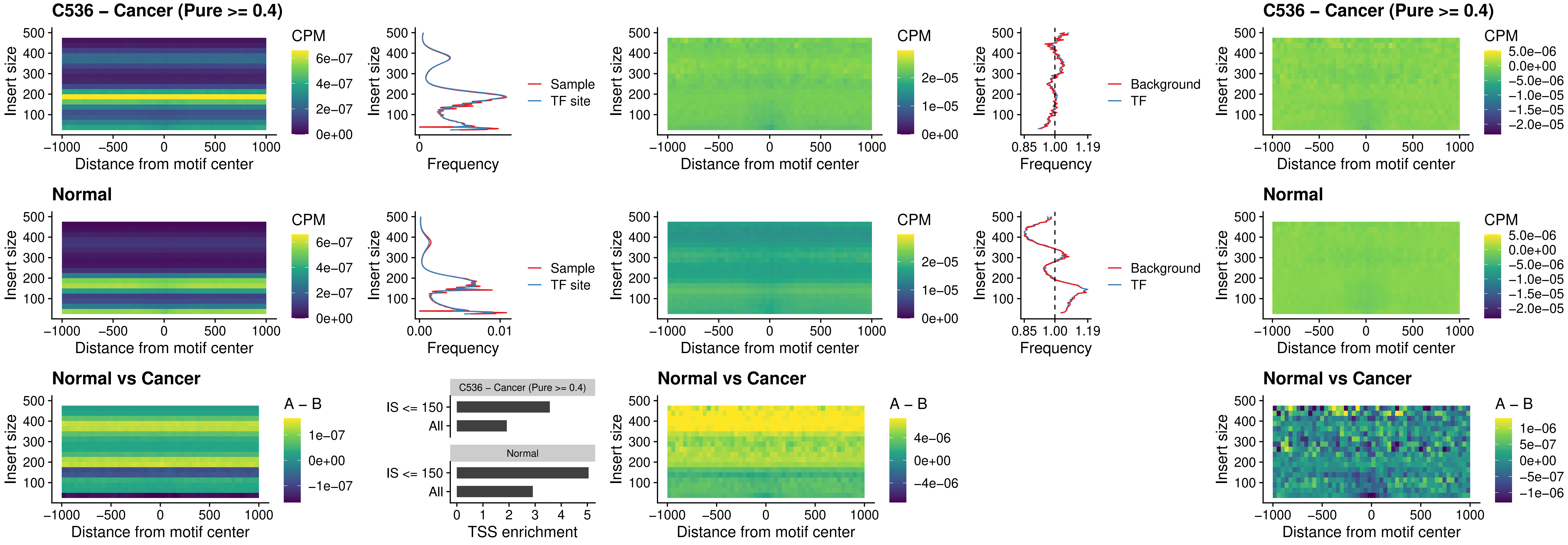

#### C536 – Cancer (Pure >= 0.4) – HOXD1_uc_dTSS_oPEAK

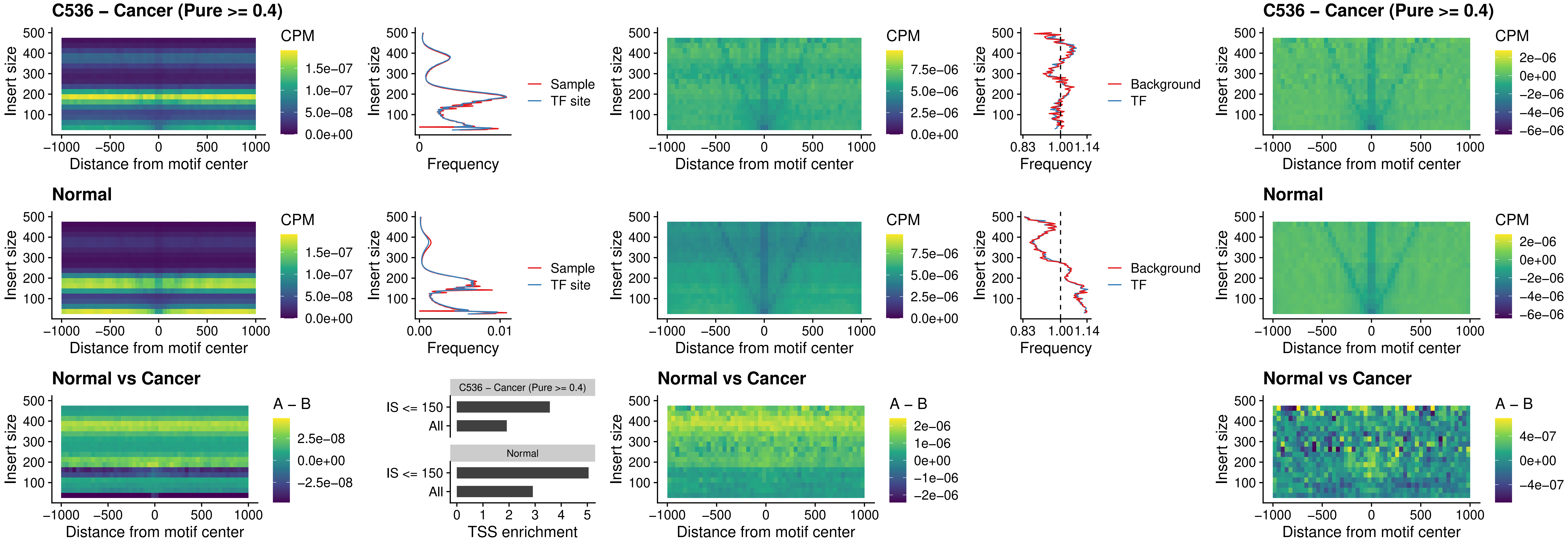

### C536 – Cancer (Pure >= 0.4) – VAX1_uc_pTSS_nPEAK

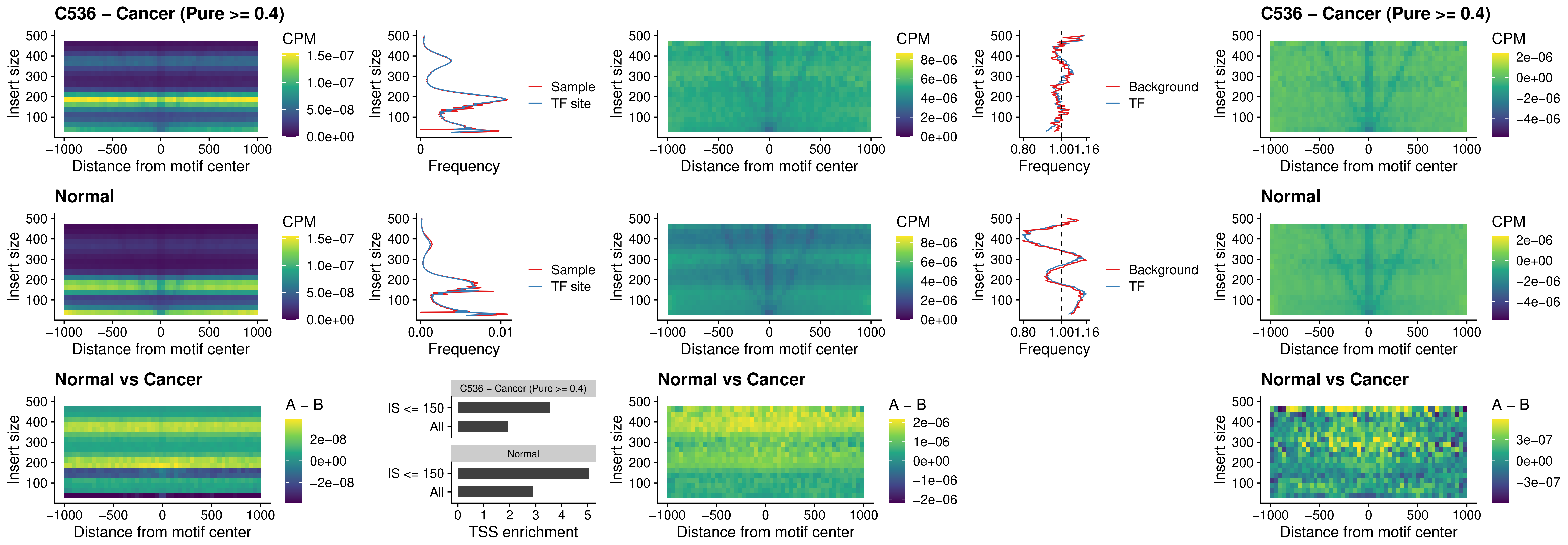

#### C536 – Cancer (Pure >= 0.4) – DLX3_uc_pTSS_nPEAK

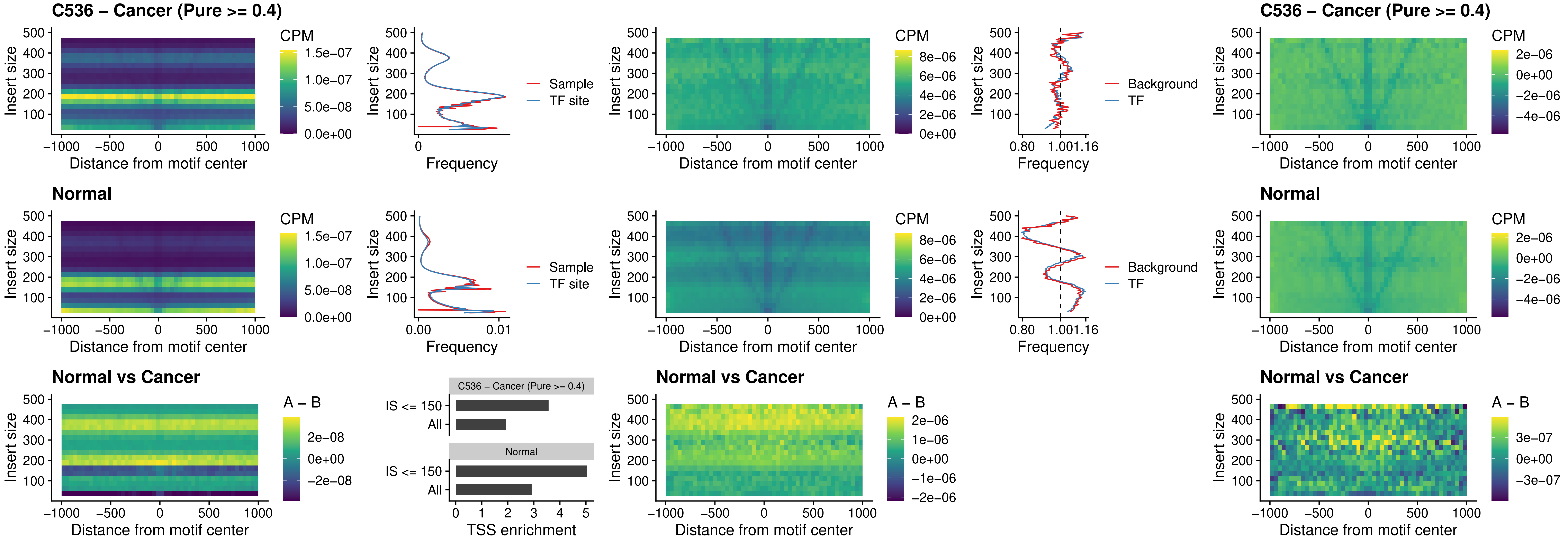

#### C536 – Cancer (Pure >= 0.4) – HOXD3_uc_pTSS_nPEAK

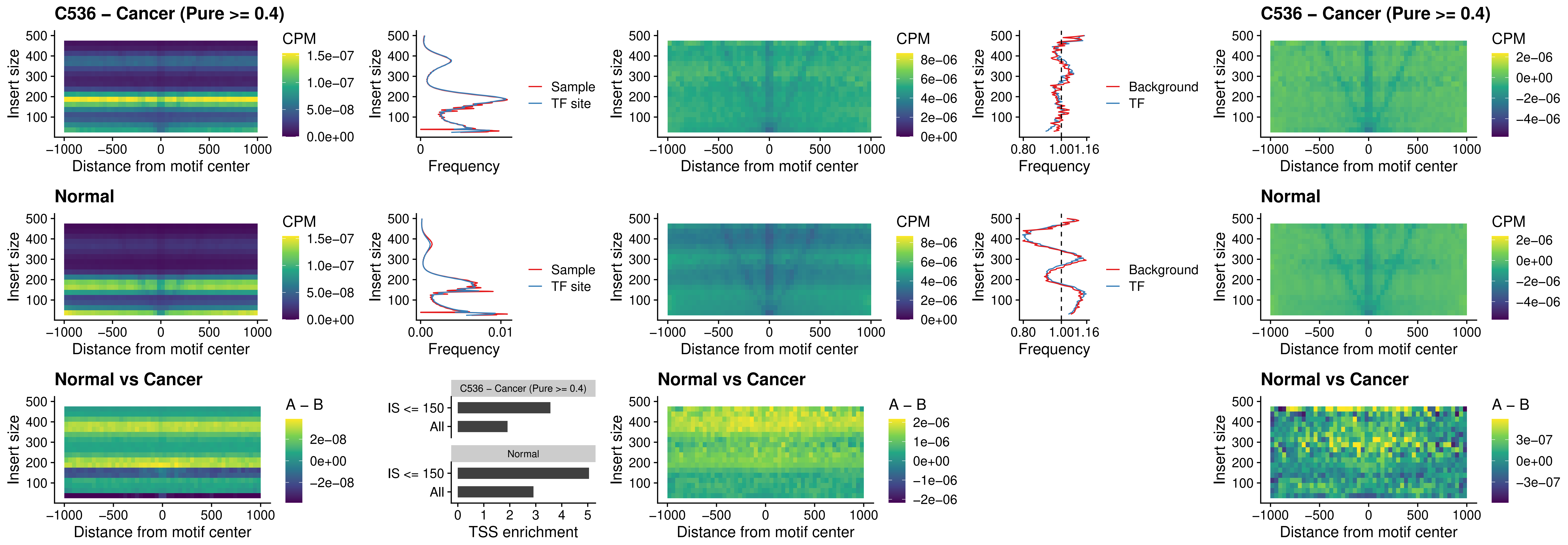

#### C536 – Cancer (Pure >= 0.4) – PDX1_uc_dTSS_nPEAK

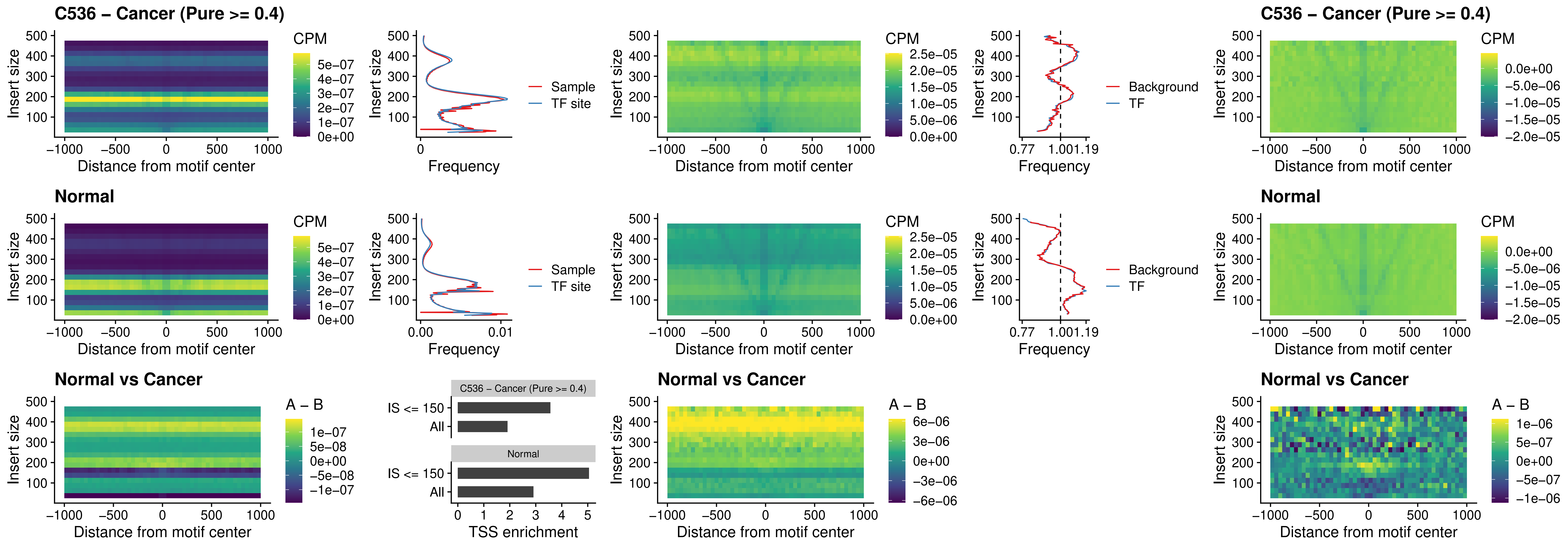

#### C536 – Cancer (Pure >= 0.4) – VAX2_uc_dTSS_nPEAK

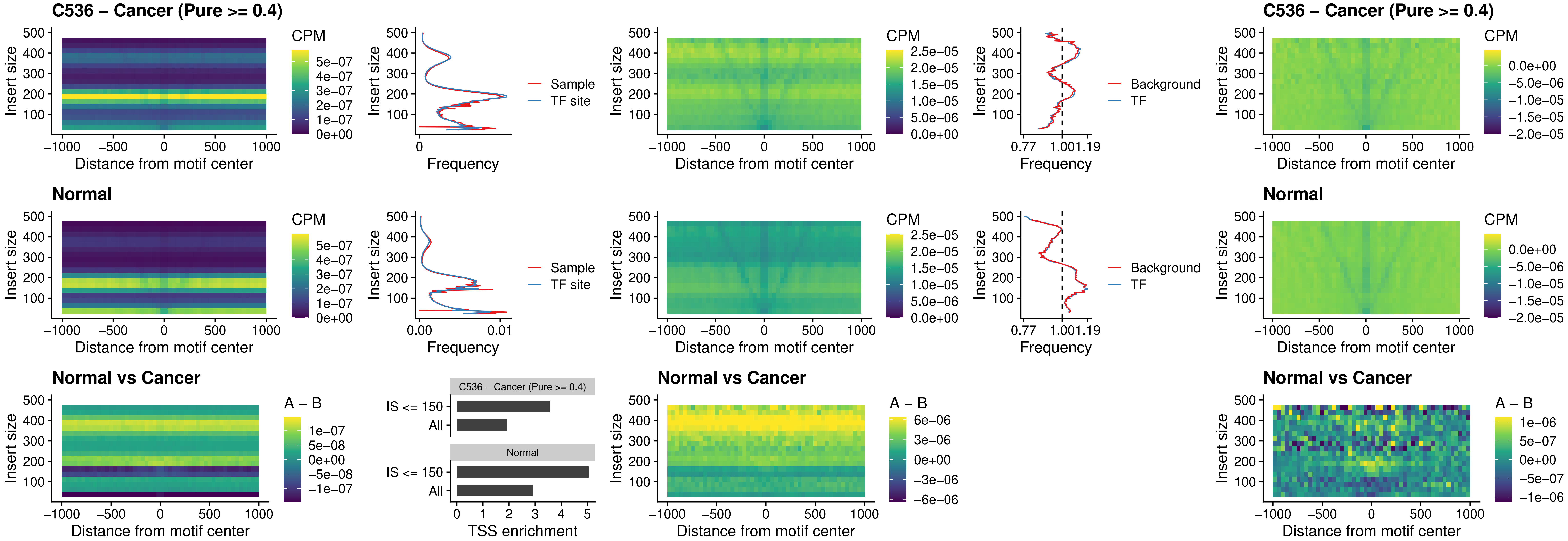

#### C536 – Cancer (Pure >= 0.4) – SOX5_uc_dTSS_nPEAK

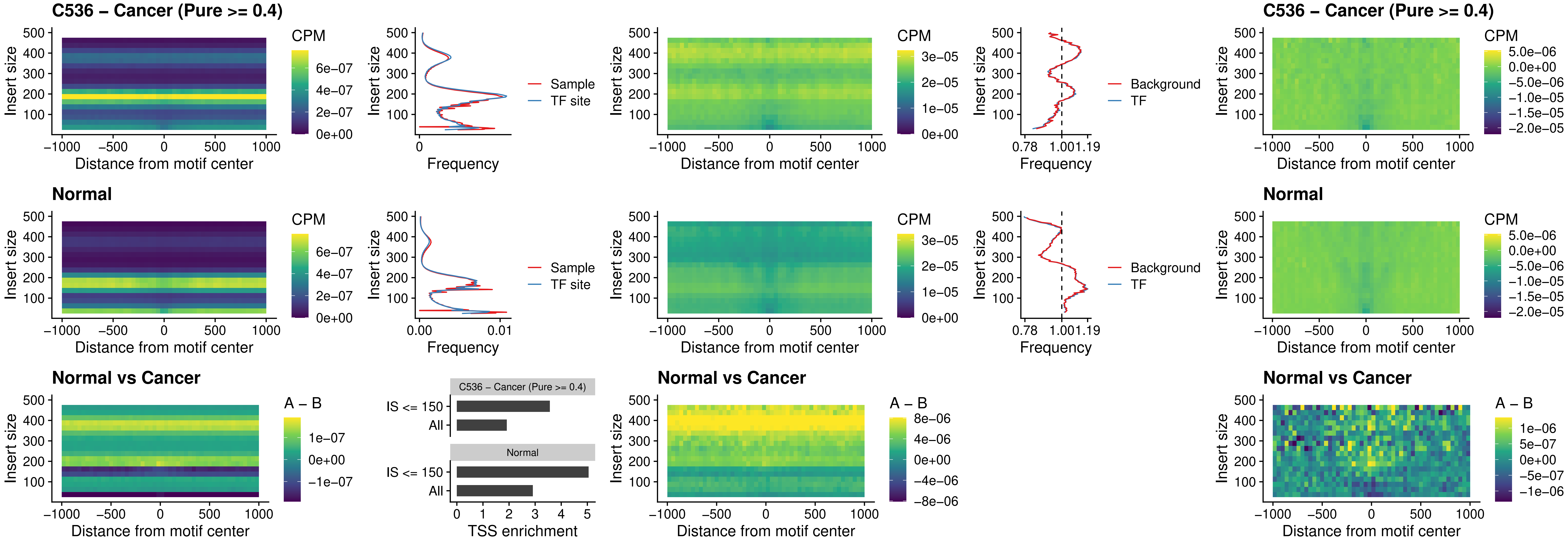

### C536 – Cancer (Pure >= 0.4) – SOX6_uc_dTSS_nPEAK

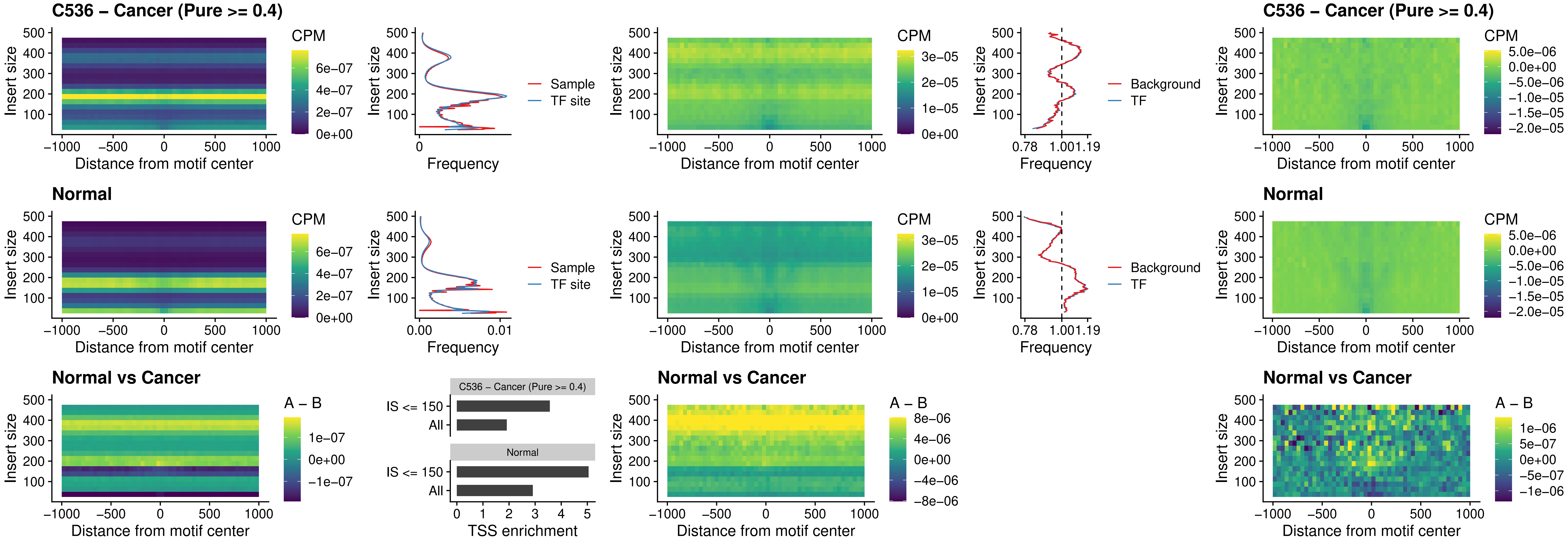

#### C536 – Cancer (Pure >= 0.4) – JDP2_uc_cTSS_nPEAK

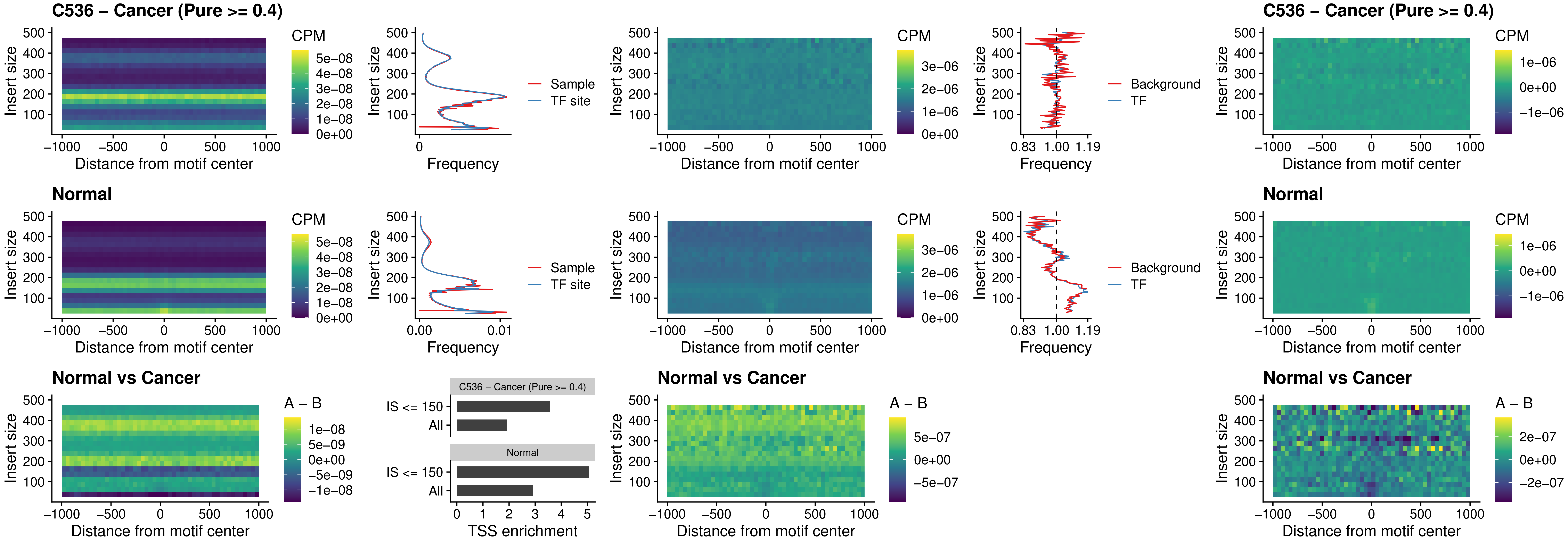

#### C536 – Cancer (Pure >= 0.4) – PURA_uc_dTSS_oPEAK

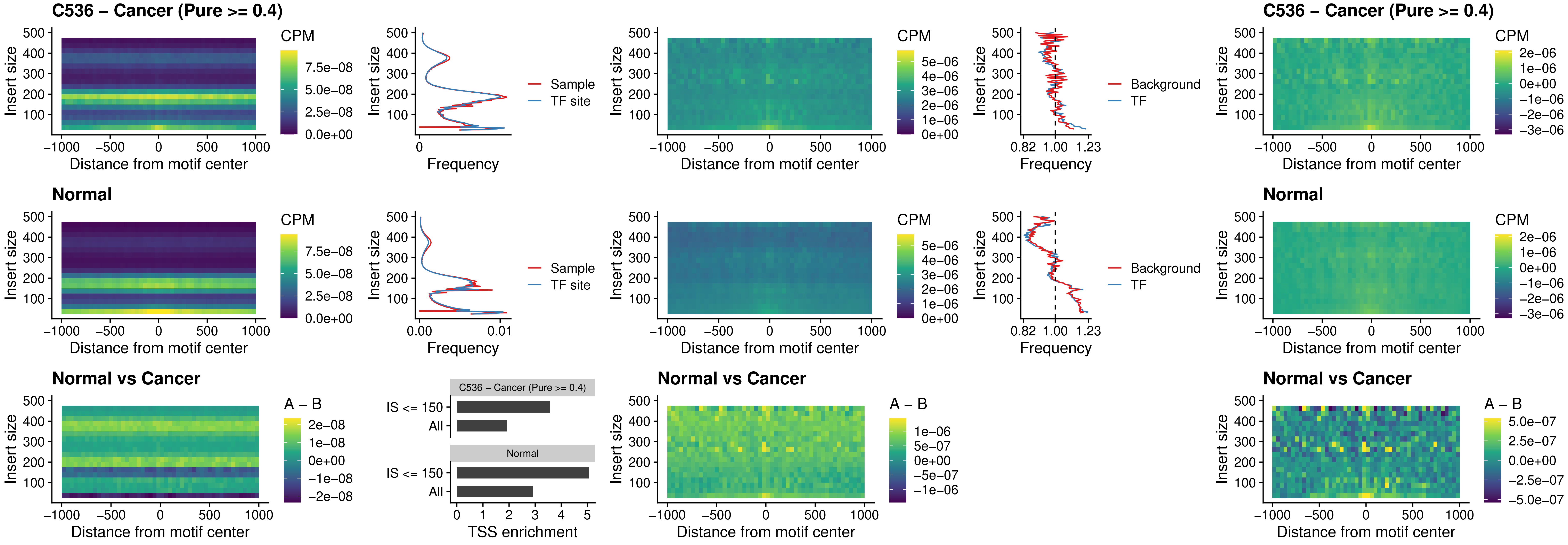

#### C536 – Cancer (Pure >= 0.4) – HHEX_uc_dTSS_nPEAK

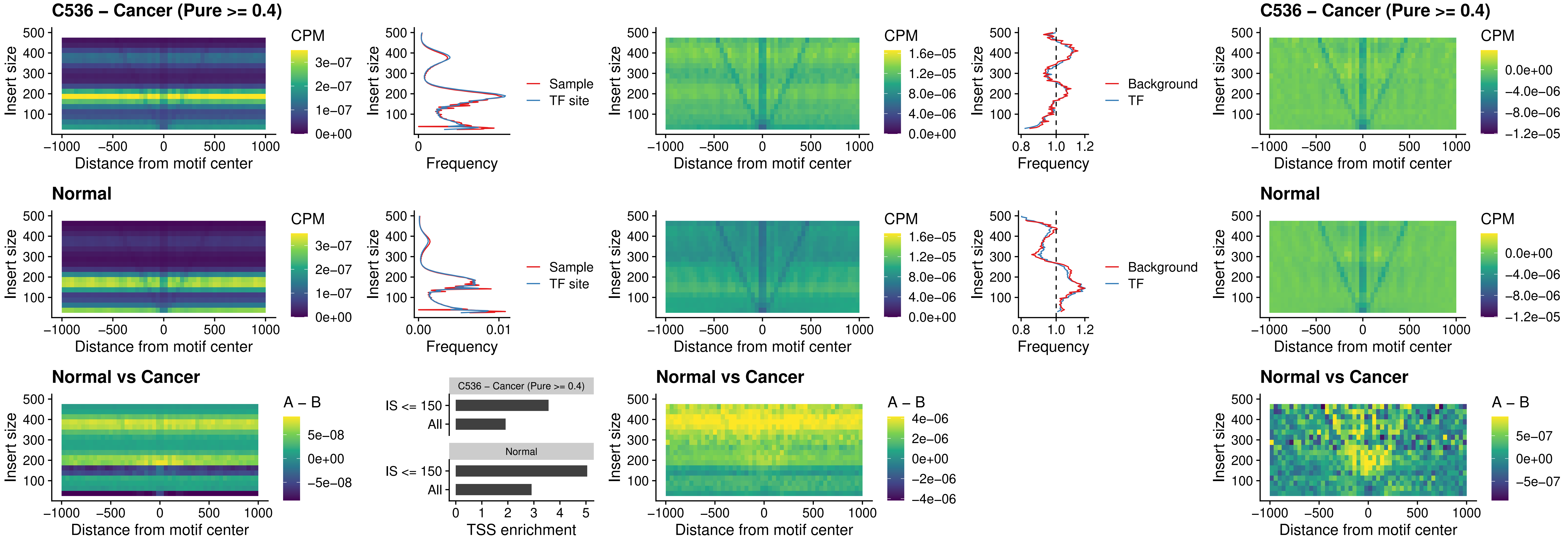

#### C536 – Cancer (Pure >= 0.4) – HMG20B_uc_dTSS_nPEAK

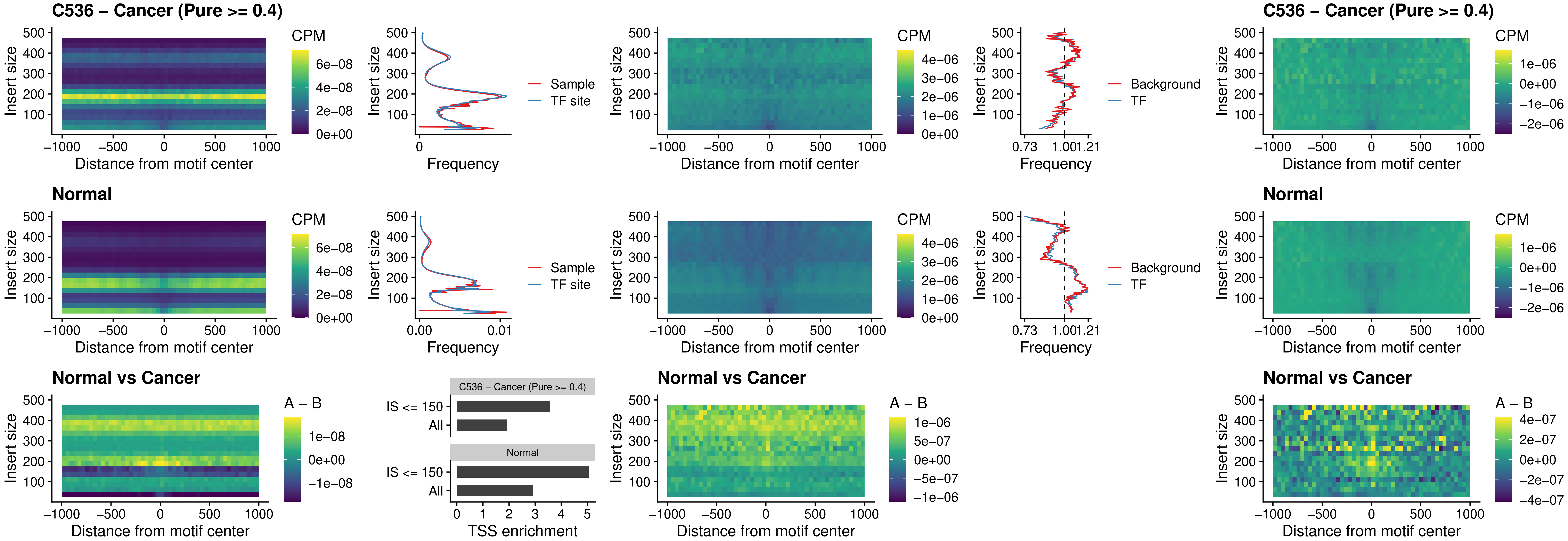

#### C536 – Cancer (Pure >= 0.4) – FOXB2_uc_dTSS_nPEAK

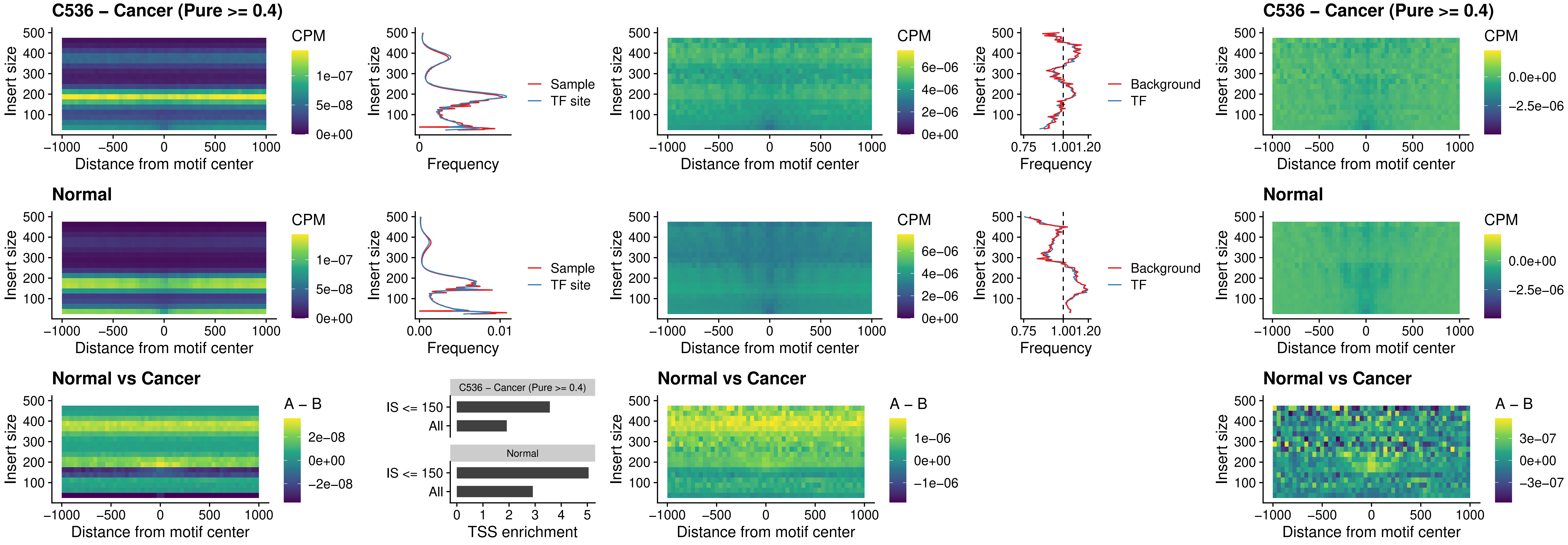

#### C536 – Cancer (Pure >= 0.4) – FOXL1_uc_cTSS_nPEAK

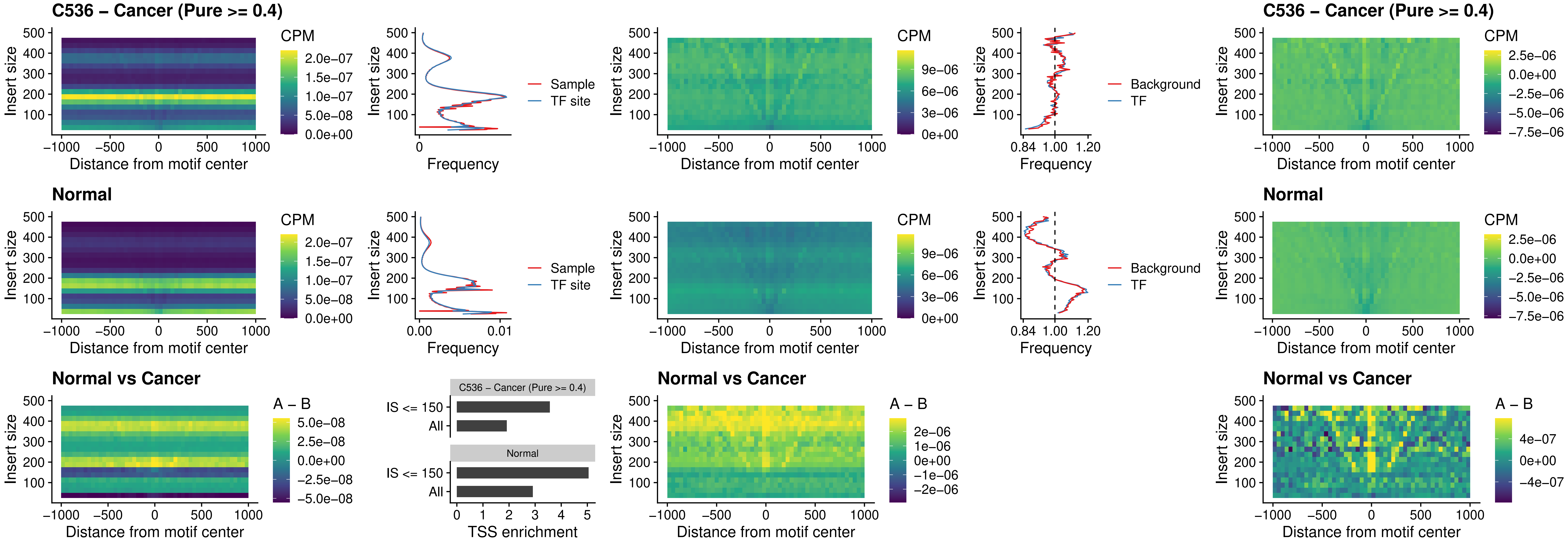

#### C536 – Cancer (Pure >= 0.4) – PBX3_uc_pTSS_oPEAK

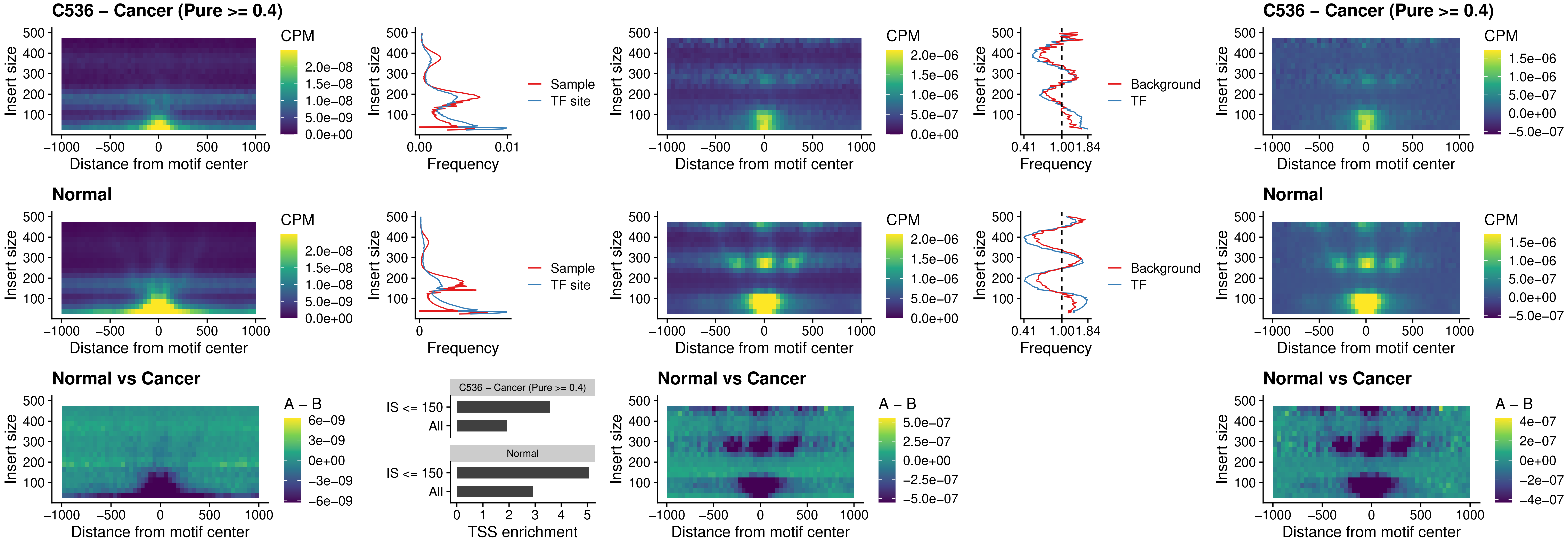

#### C536 – Cancer (Pure >= 0.4) – SNAI2_uc_cTSS_nPEAK

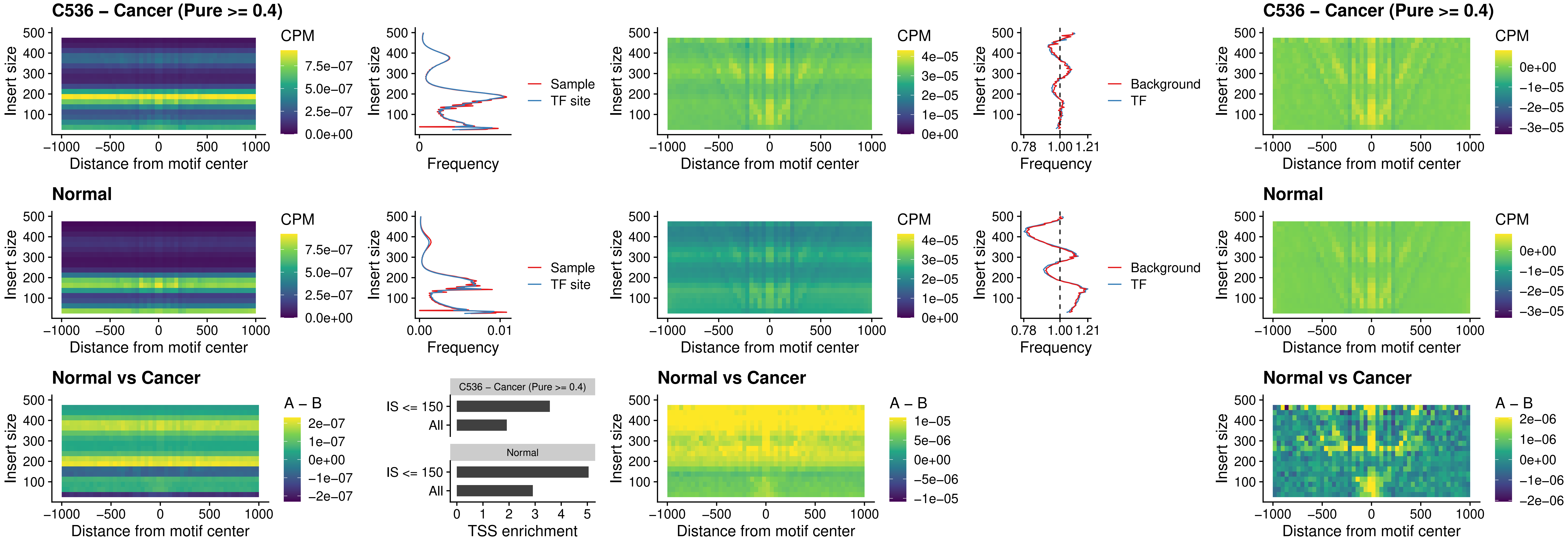

#### C536 – Cancer (Pure >= 0.4) – SNAI2_uc_dTSS_oPEAK

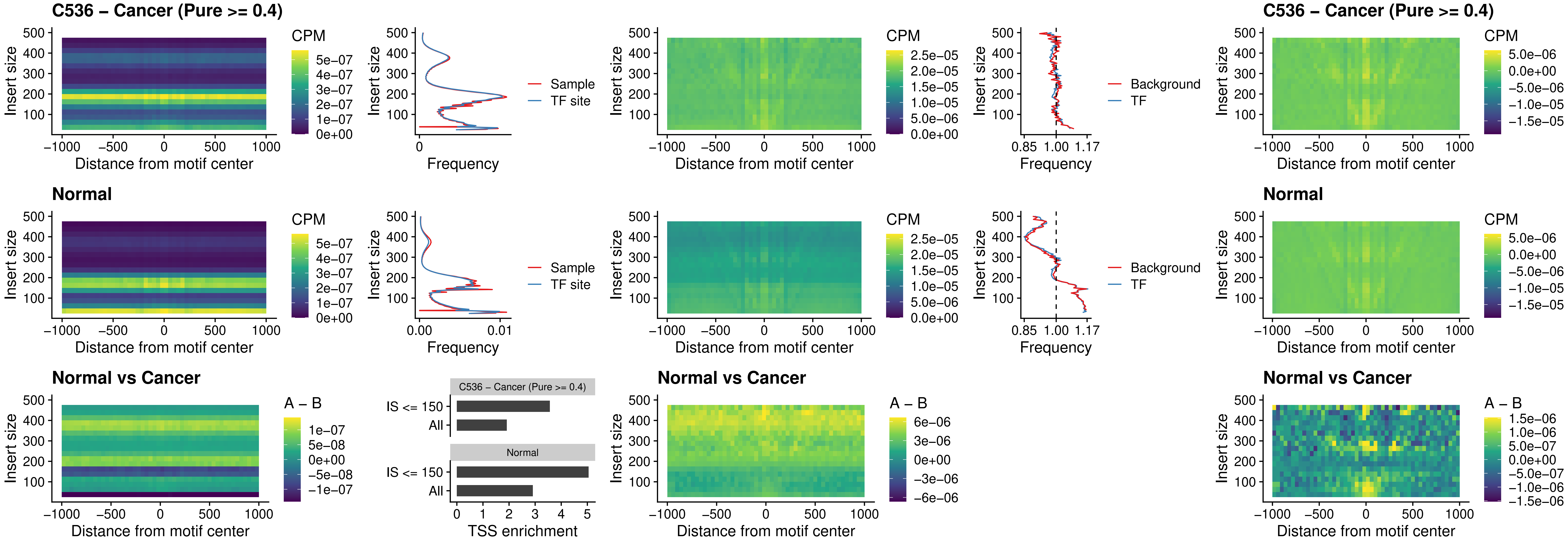

### C536 – Cancer (Pure >= 0.4) – SPI1_uc_pTSS_nPEAK

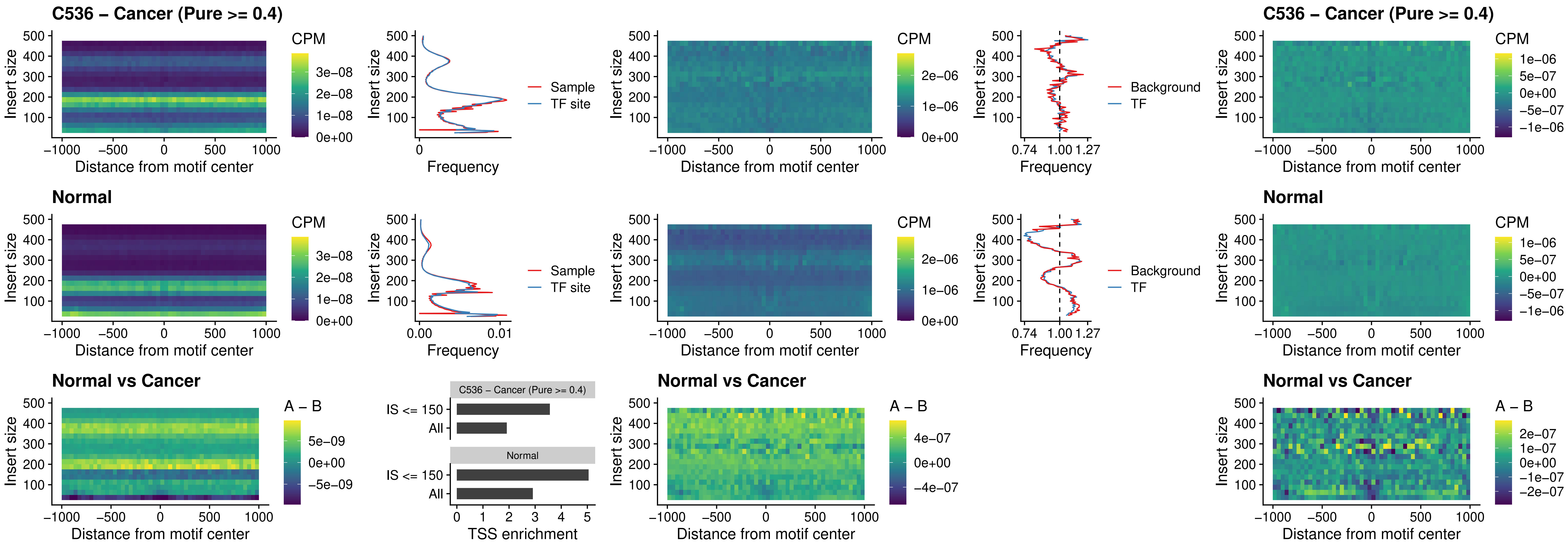

### C536 – Cancer (Pure >= 0.4) – KLF16_uc_pTSS_nPEAK

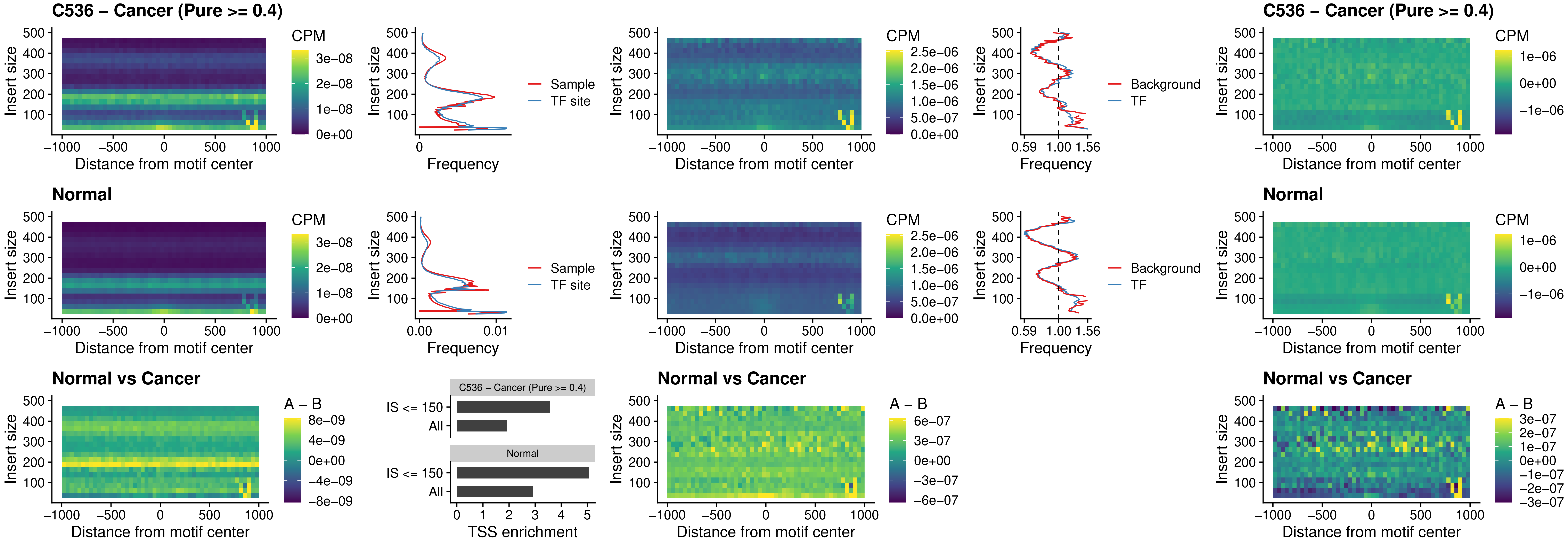

#### C536 – Cancer (Pure >= 0.4) – SP3_uc_pTSS_nPEAK

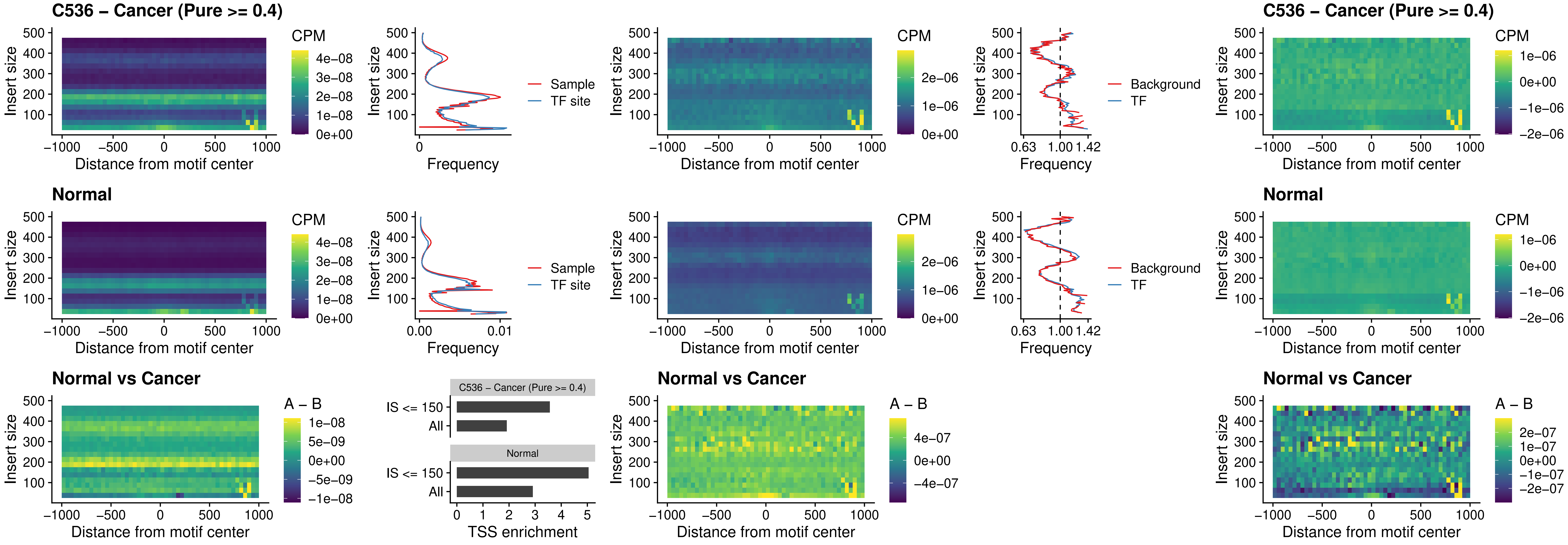

#### C536 – Cancer (Pure >= 0.4) – NEUROD1_uc_dTSS_oPEAK

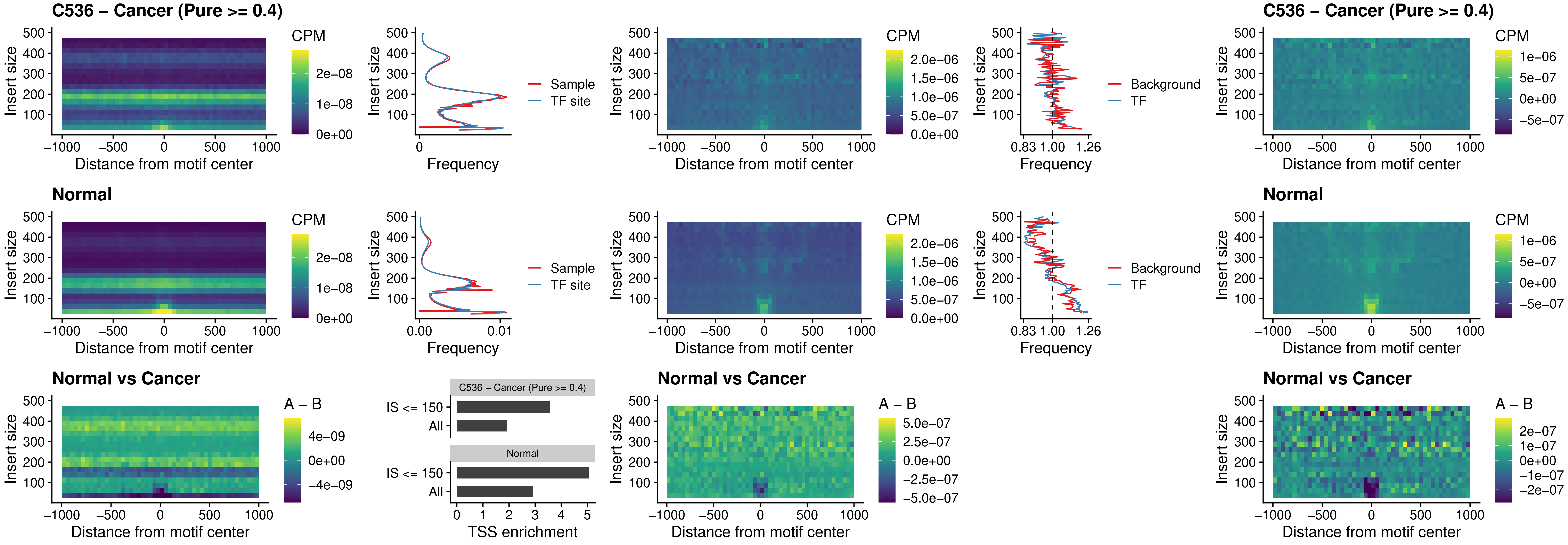

#### C536 – Cancer (Pure >= 0.4) – CTCF_uc_dTSS_oPEAK

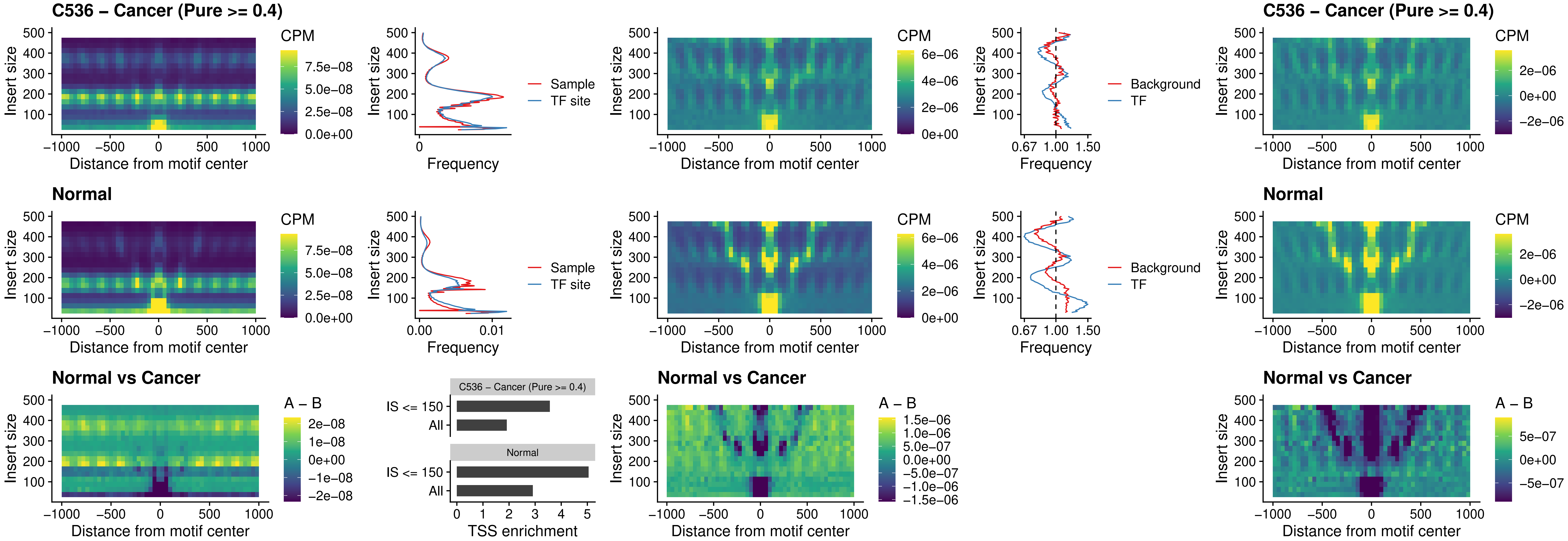

#### C536 – Cancer (Pure >= 0.4) – CTCF_uc_cTSS_oPEAK

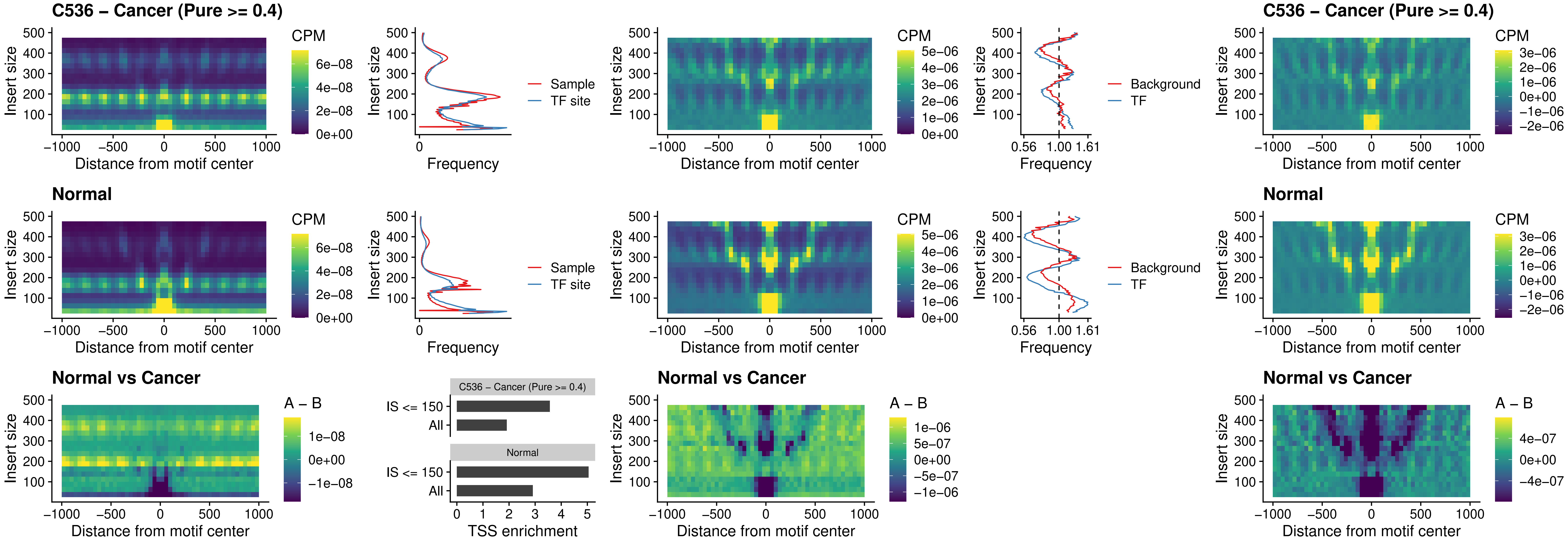

#### C536 – Cancer (Pure >= 0.4) – CTCFL_uc_dTSS_oPEAK

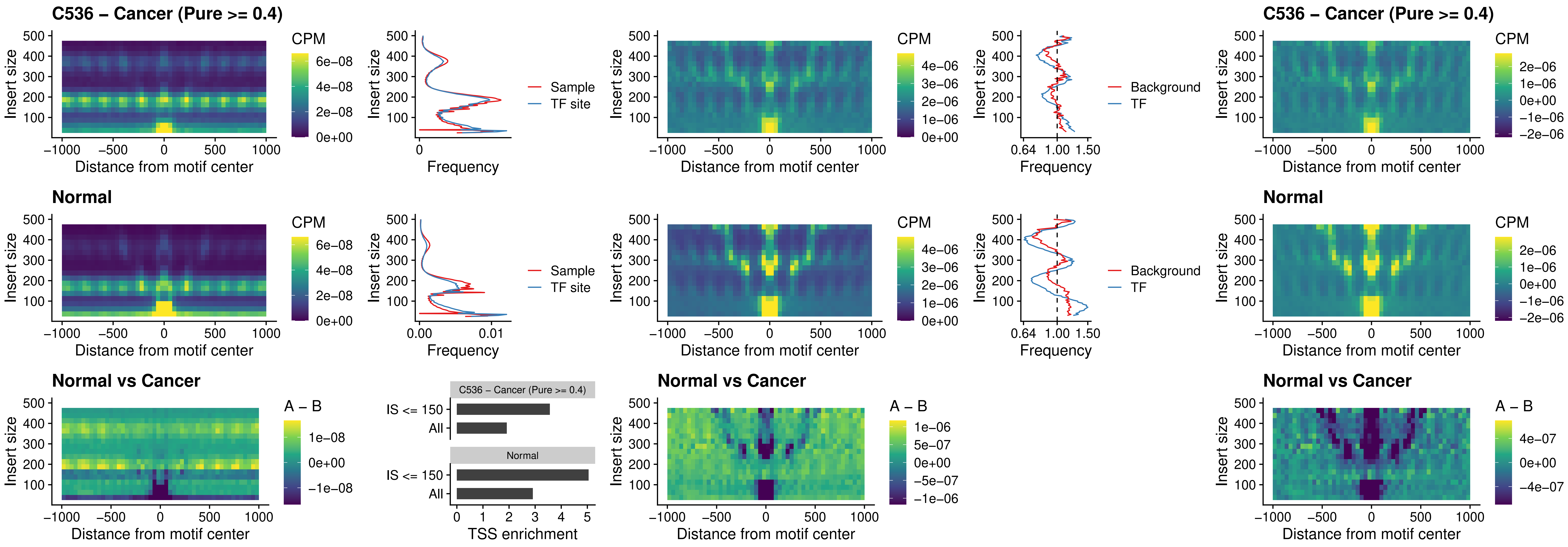

#### C536 – Cancer (Pure >= 0.4) – ENCODE_CTCF_sites_COLON

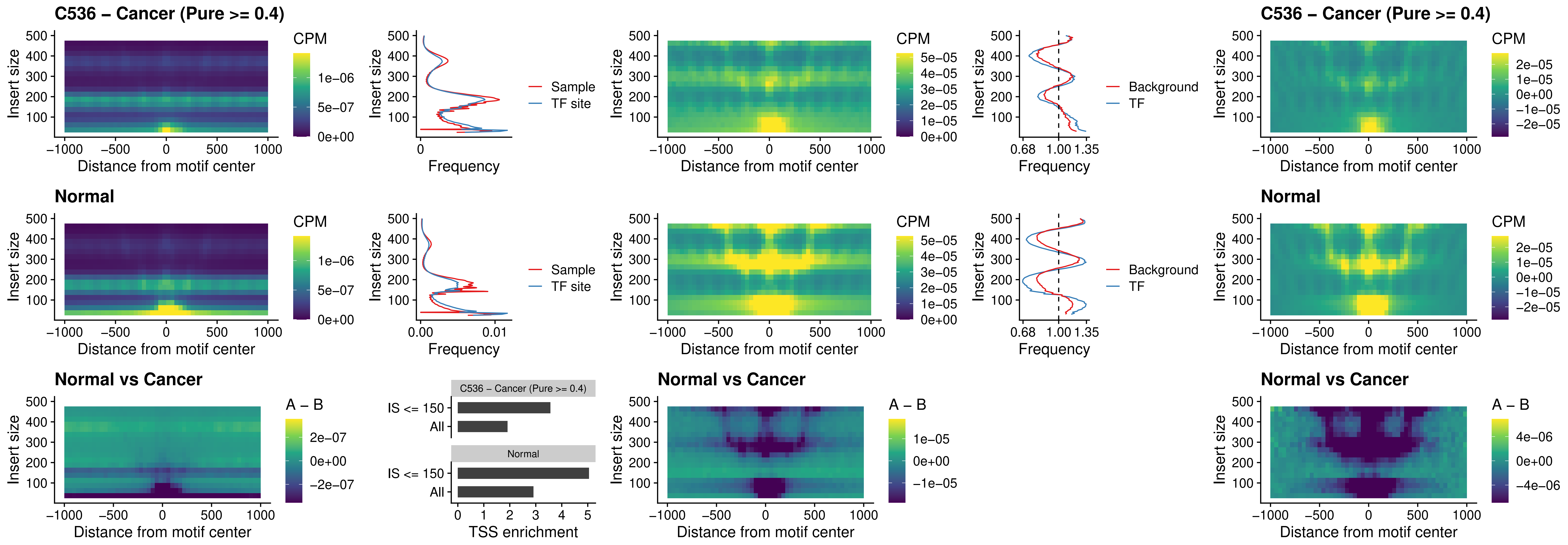

#### C536 – Cancer (Pure >= 0.4) – ENCODE_CTCF_sites_COLON_loop_ctcf

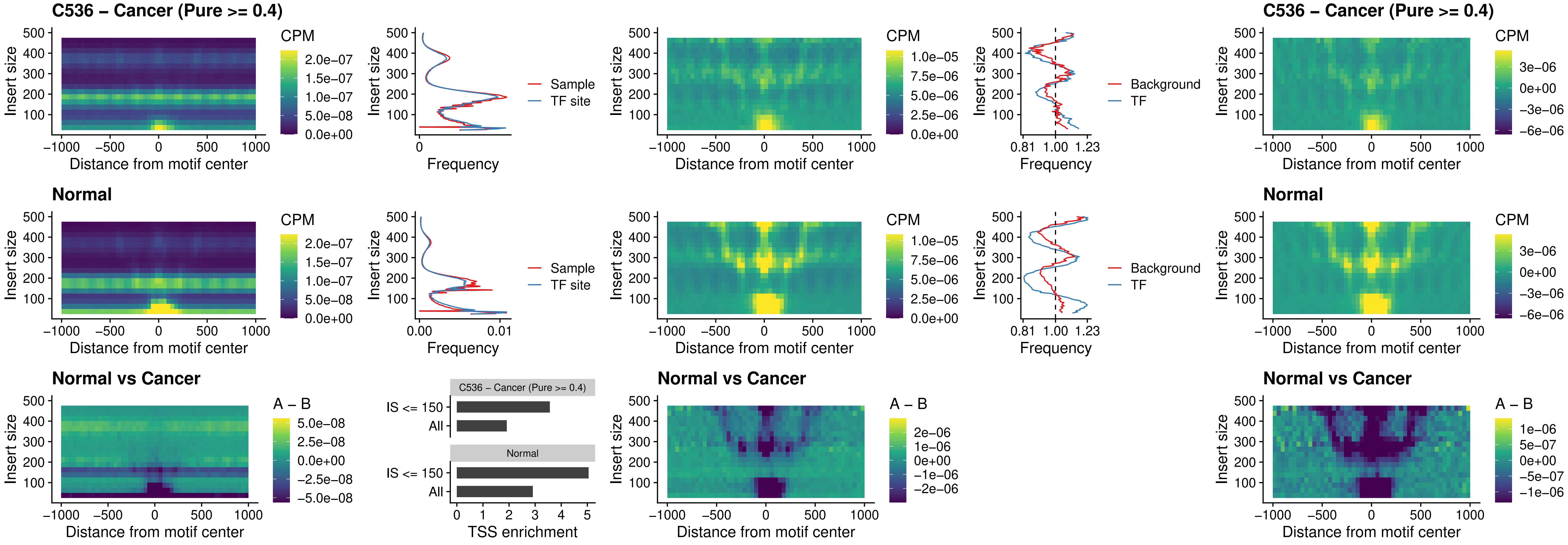

#### C536 – Cancer (Pure >= 0.4) – ENCODE_CTCF_sites_COLON_intergenic_ctcf

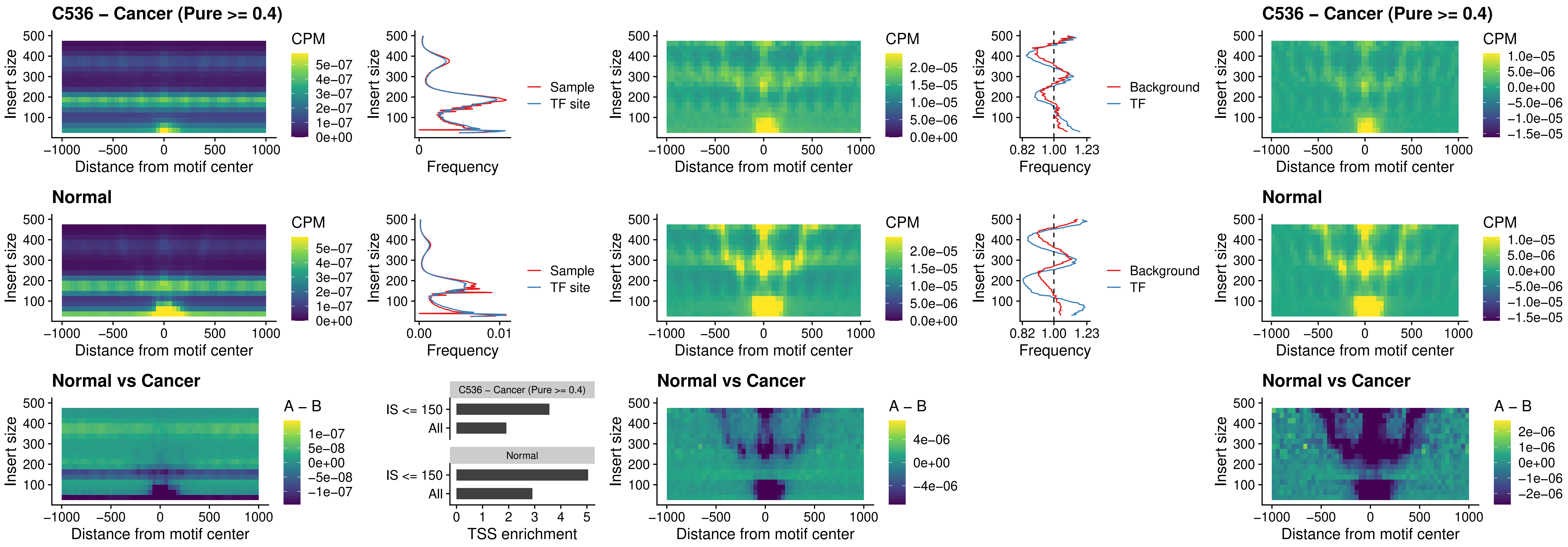

#### C536 – Cancer (Pure >= 0.4) – ENCODE_CTCF_sites_COLON_nonpromotor

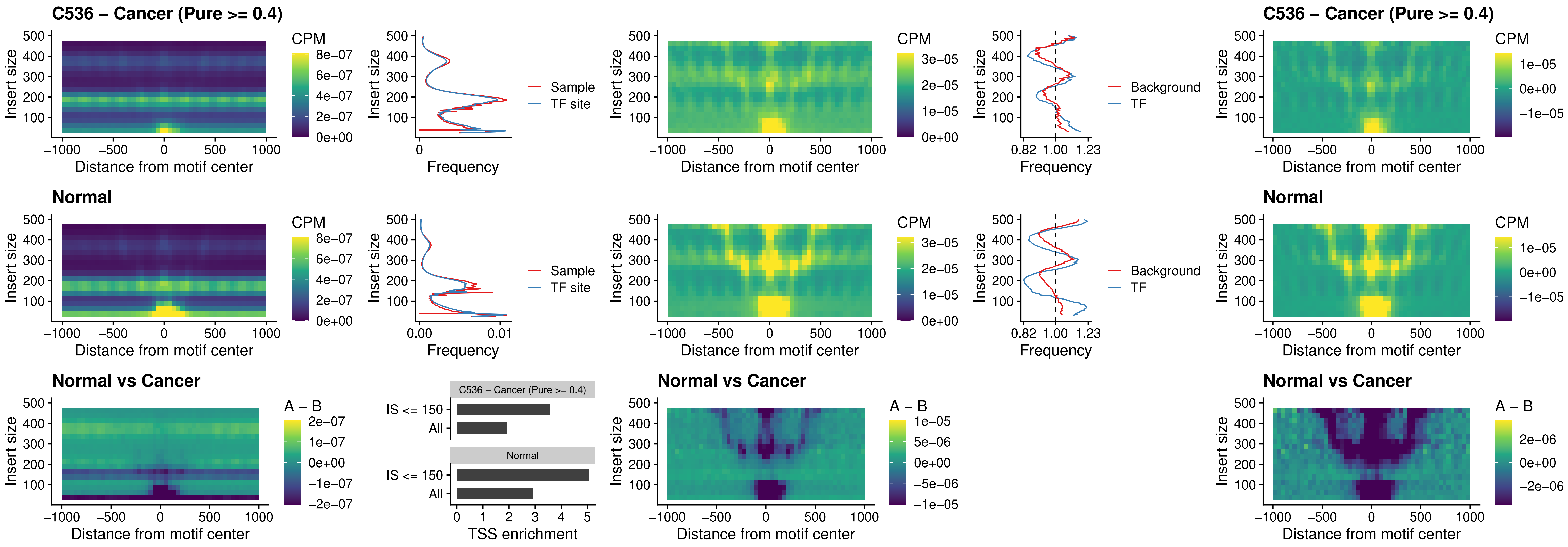

#### C536 – Cancer (Pure >= 0.4) – CTCFL_uc_cTSS_oPEAK

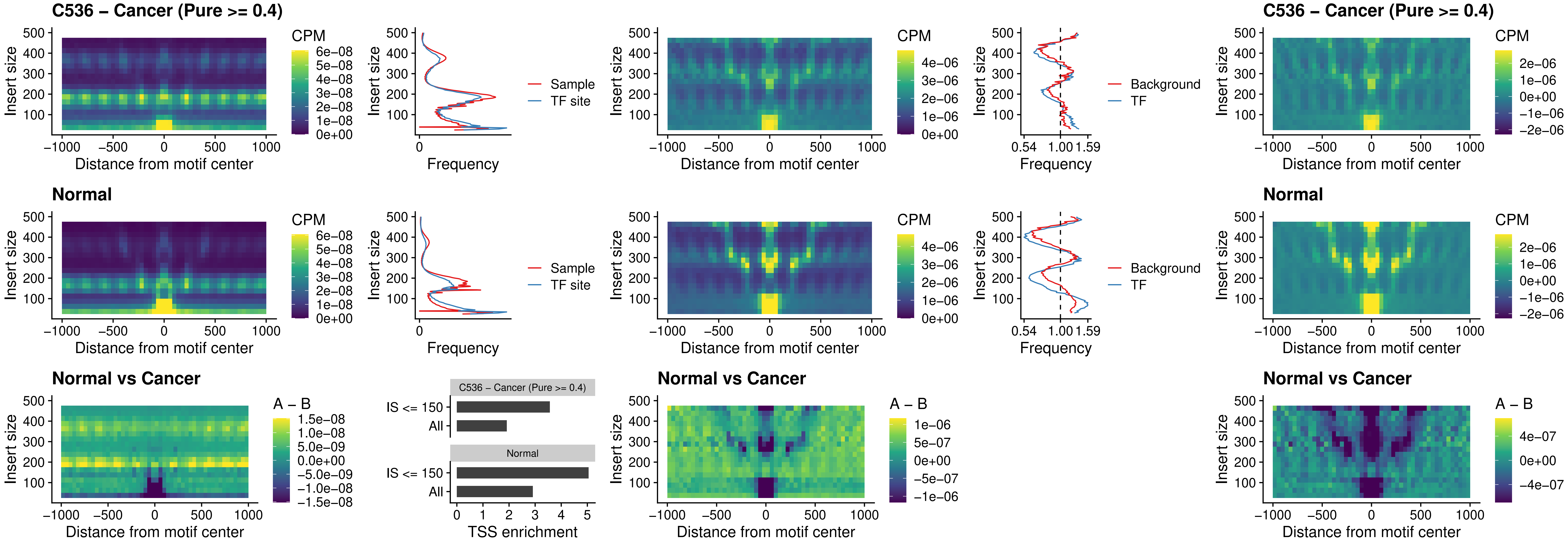

#### C536 – Cancer (Pure >= 0.4) – NEUROD1_uc_cTSS_oPEAK

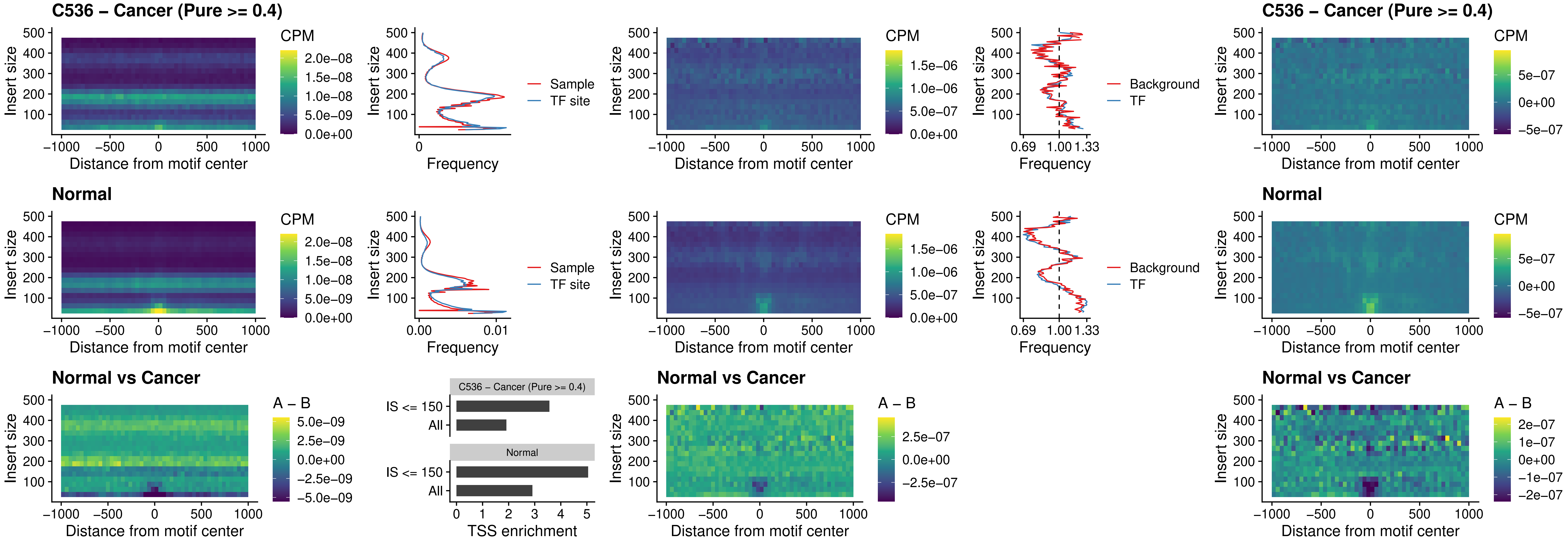

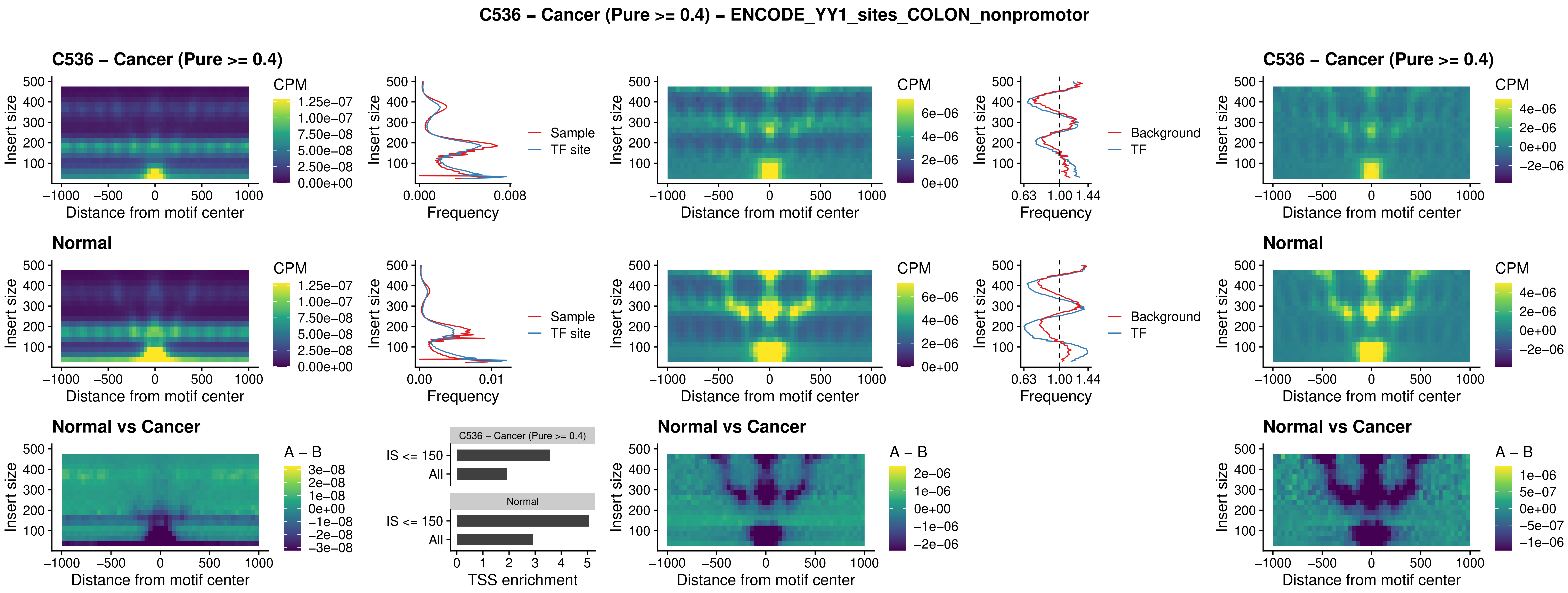
C536 – Cancer (Pure >= 0.4) – ENCODE_YY1_sites_COLON_nonpromotor
C536 – Cancer (Pure >= 0.4)
Normal
Normal vs Cancer
Insert size
Distance from motif center
Frequency
CPM
A – B
Sample
TF site
Background
TF
C536 – Cancer (Pure >= 0.4)
Normal
IS <= 150
All
TSS enrichment

#### C536 – Cancer (Pure >= 0.4) – CTCF_uc_pTSS_oPEAK

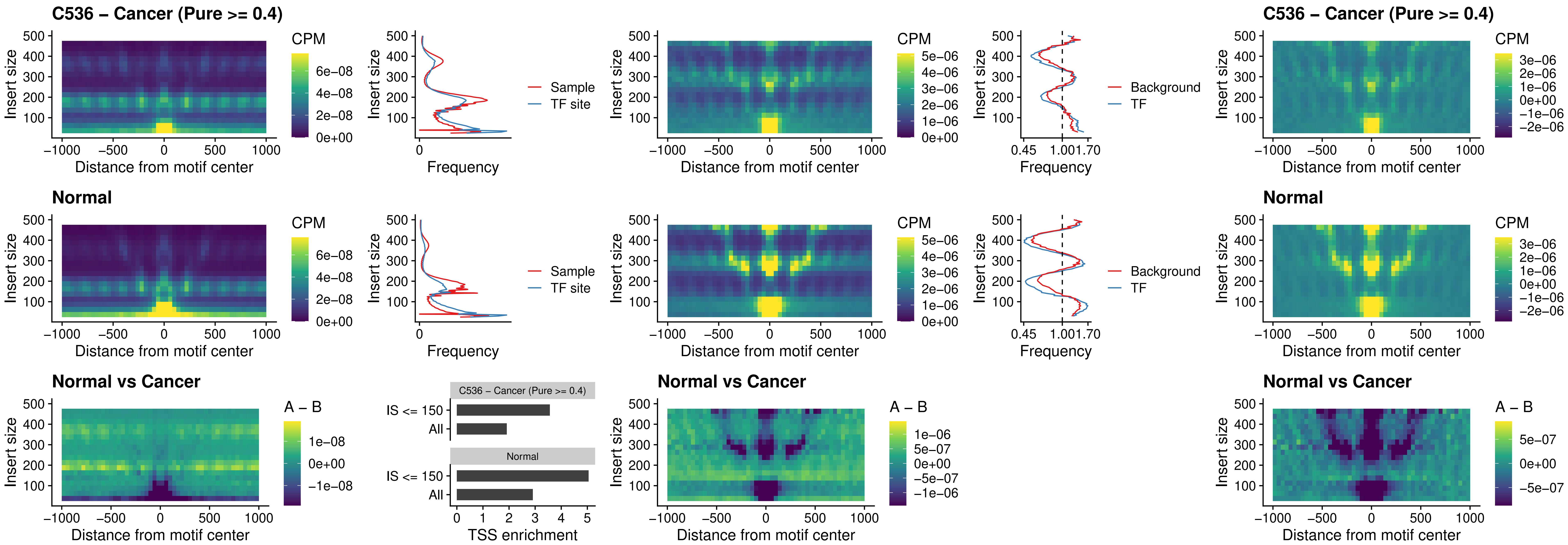

#### C536 – Cancer (Pure >= 0.4) – ENCODE_CTCF_sites_COLON_promotor

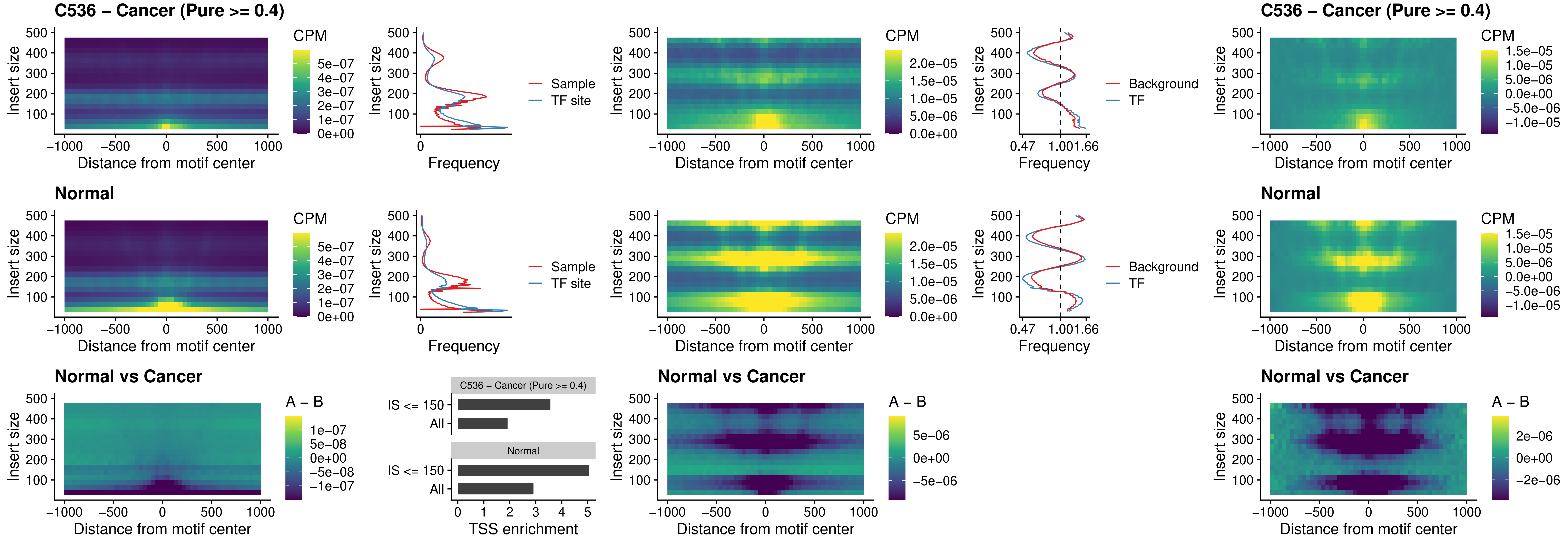

#### C536 – Cancer (Pure >= 0.4) – ENCODE_CTCF_sites_COLON_promotor_ctcf

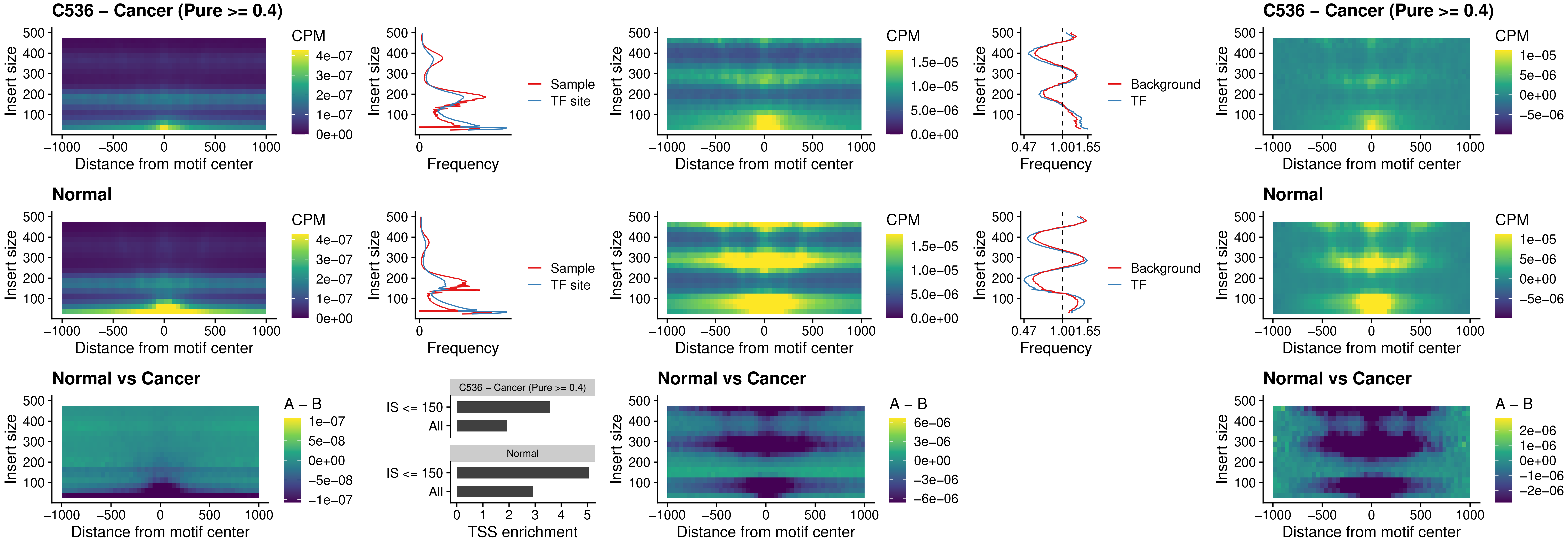

#### C536 – Cancer (Pure >= 0.4) – SPIC_uc_pTSS_oPEAK

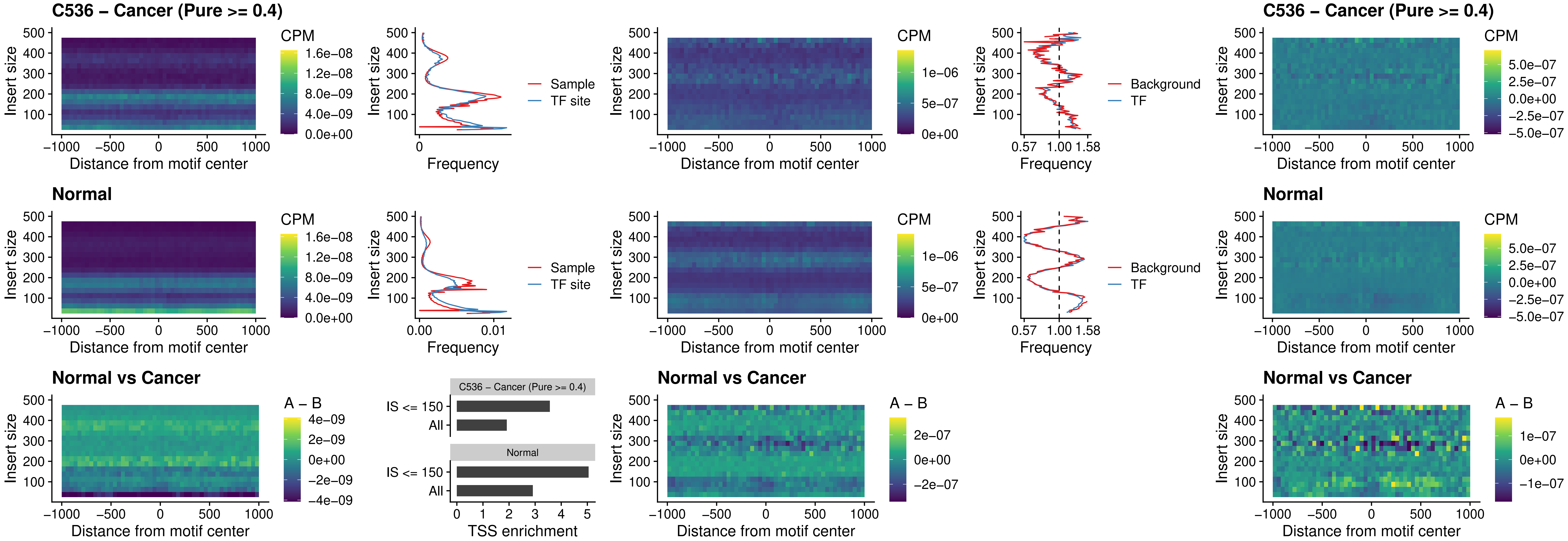

#### C536 – Cancer (Pure >= 0.4) – SPI1_uc_dTSS_nPEAK

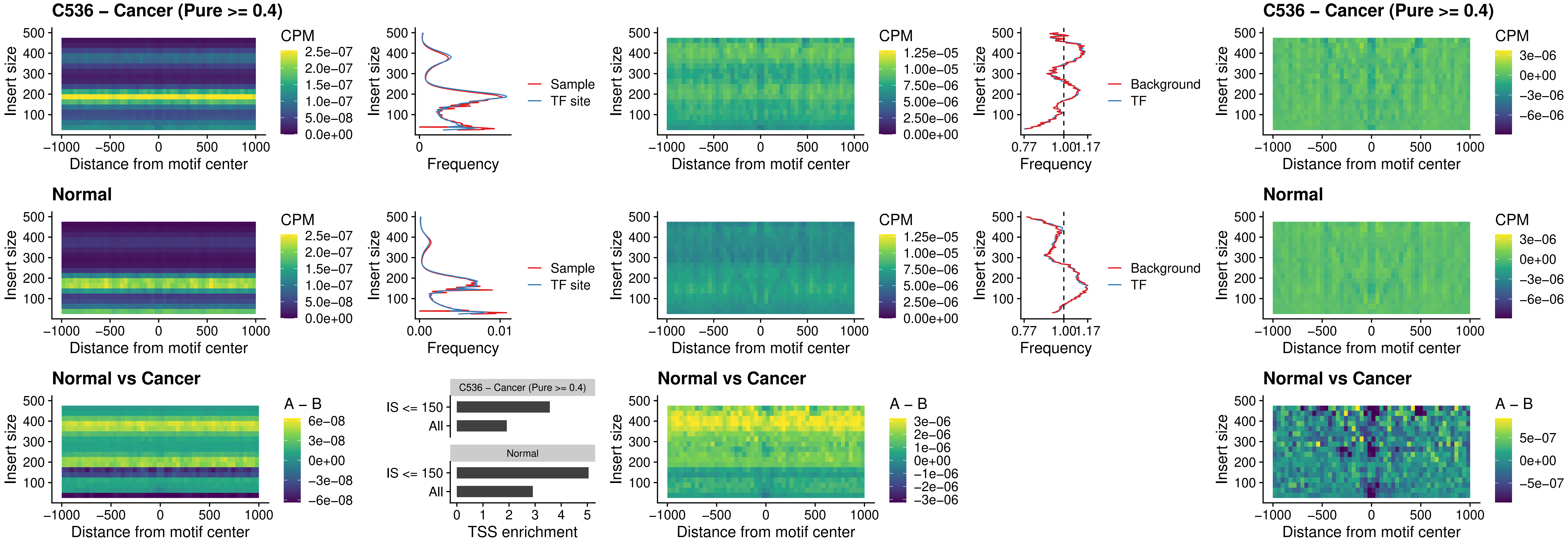

#### C536 – Cancer (Pure >= 0.4) – SPI1_uc_cTSS_nPEAK

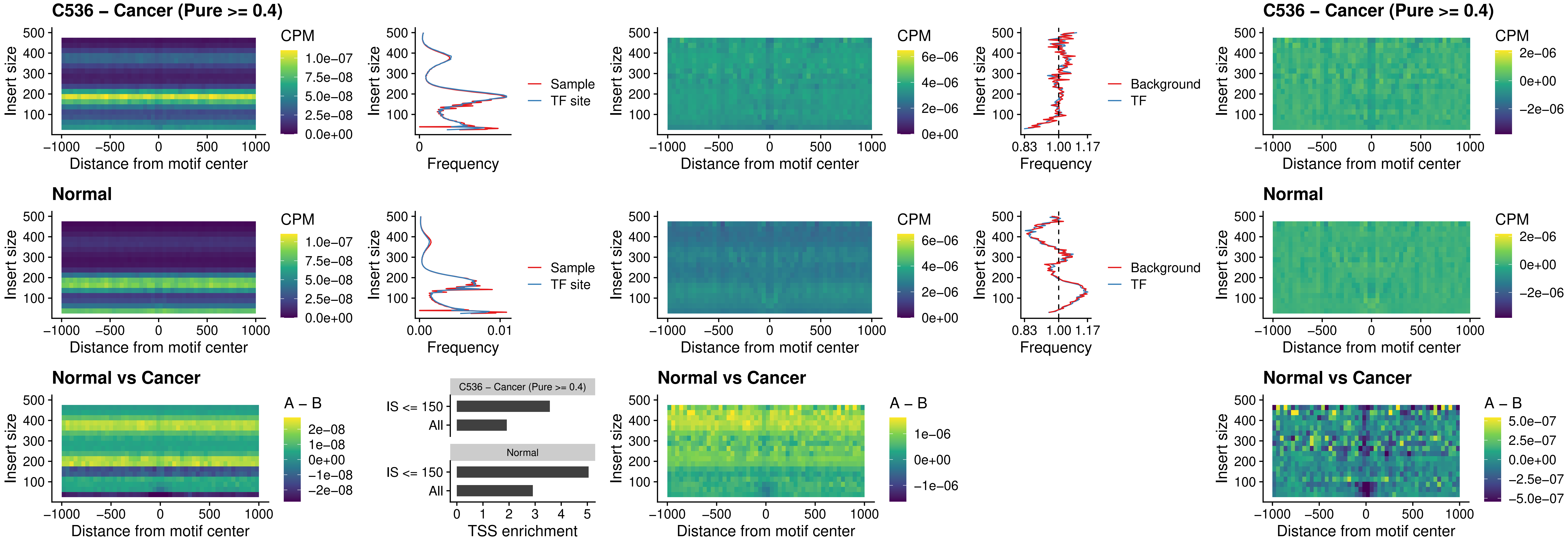

#### C536 – Cancer (Pure >= 0.4) – BCL11A_uc_pTSS_nPEAK

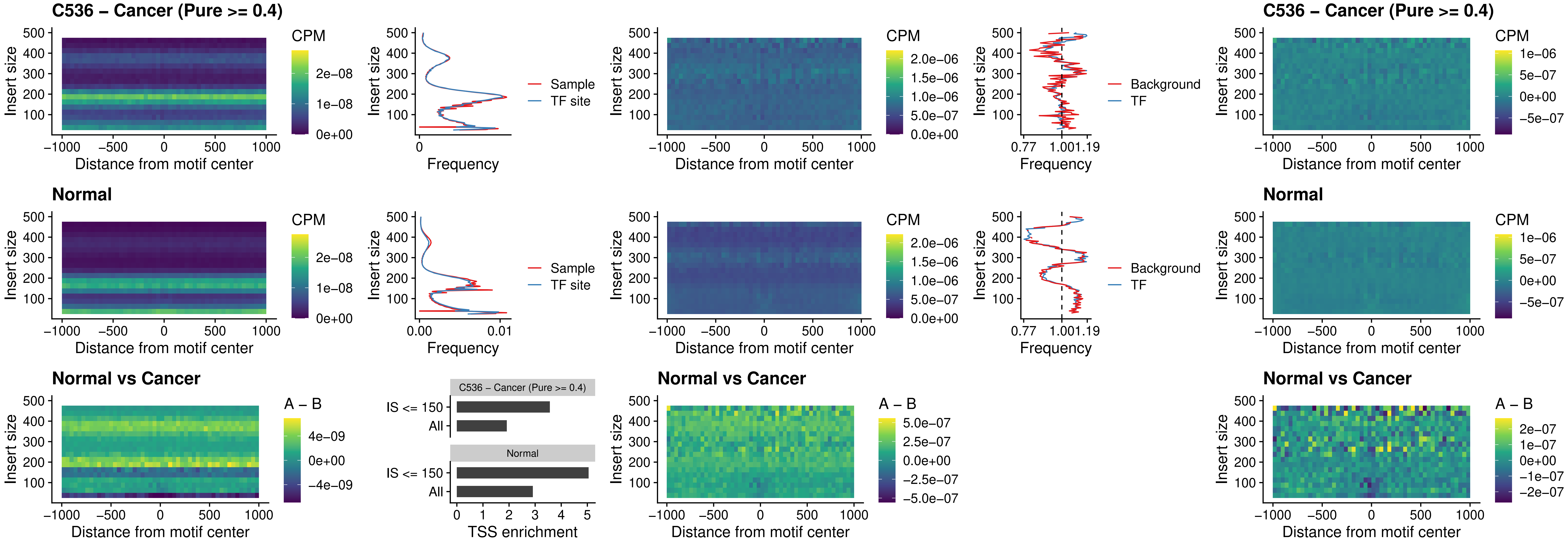

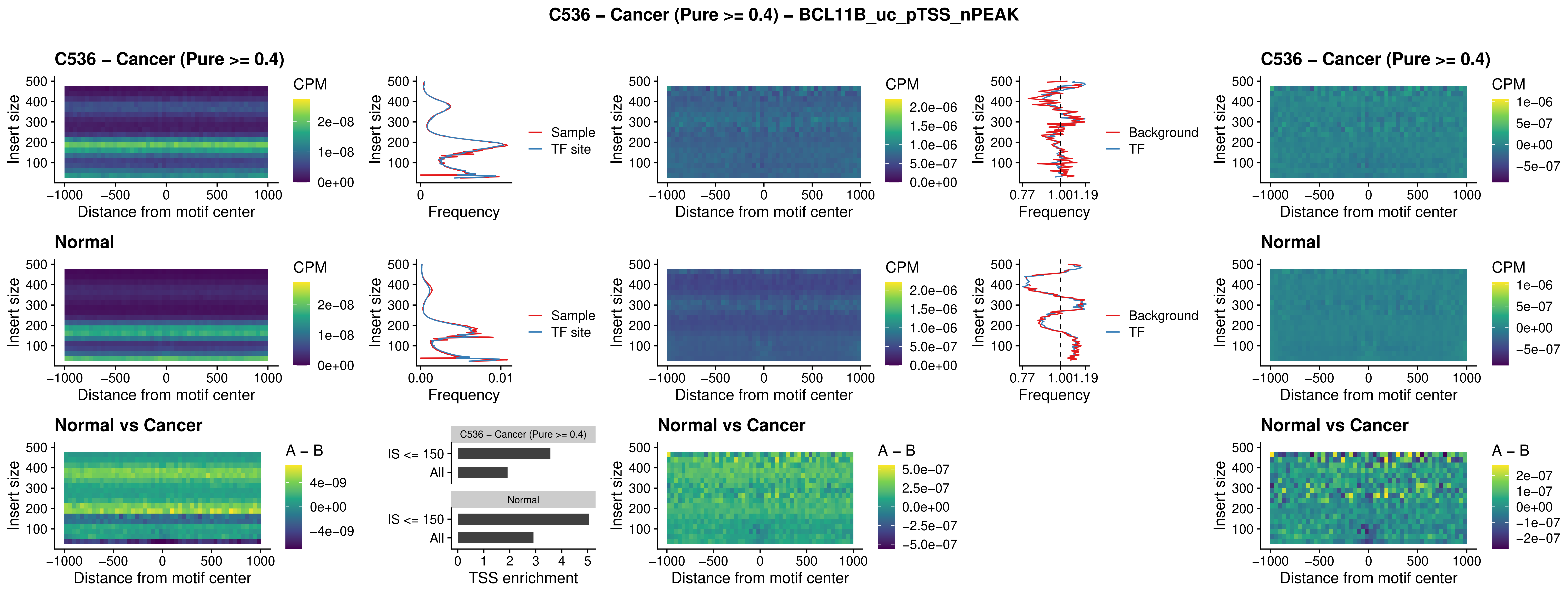
C536 – Cancer (Pure >= 0.4) – BCL11B_uc_pTSS_nPEAK
C536 – Cancer (Pure >= 0.4)
Normal
Normal vs Cancer
Insert size
Distance from motif center
Frequency
CPM
A – B
Sample
TF site
Background
TF
TSS enrichment
IS <= 150
All
C536 – Cancer (Pure >= 0.4)
Normal

### C536 – Cancer (Pure >= 0.4) – IRF9_uc_dTSS_oPEAK

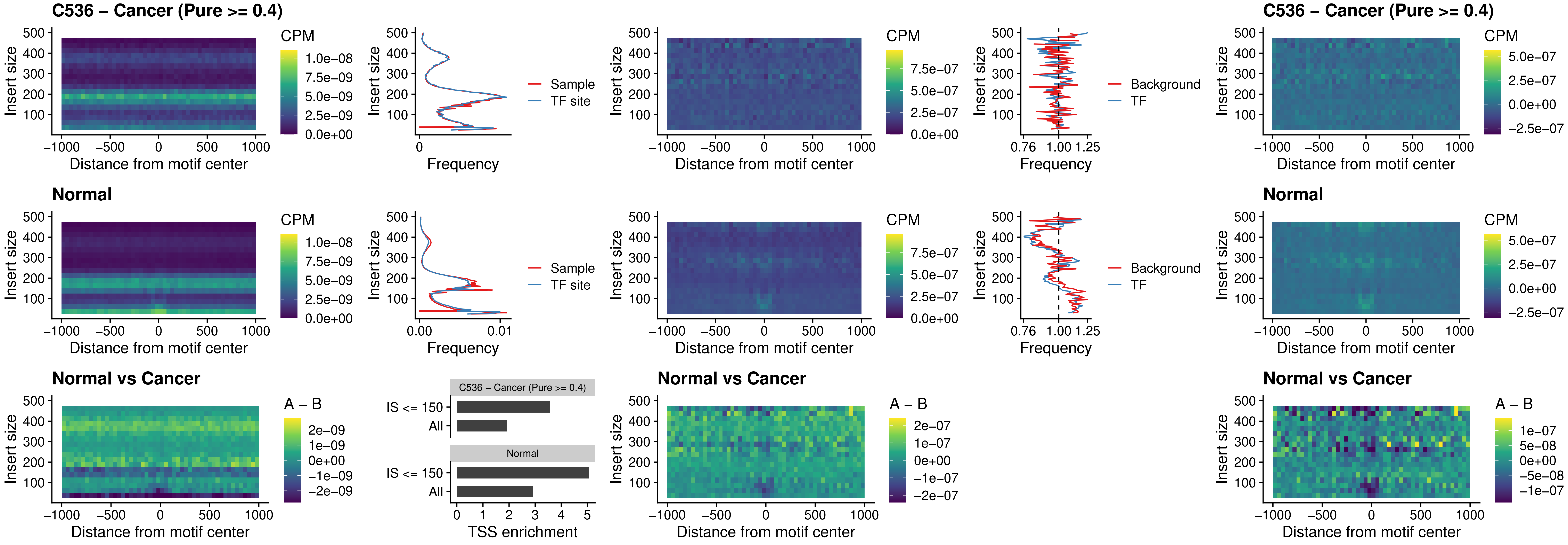

#### C536 – Cancer (Pure >= 0.4) – IRF9_uc_cTSS_oPEAK

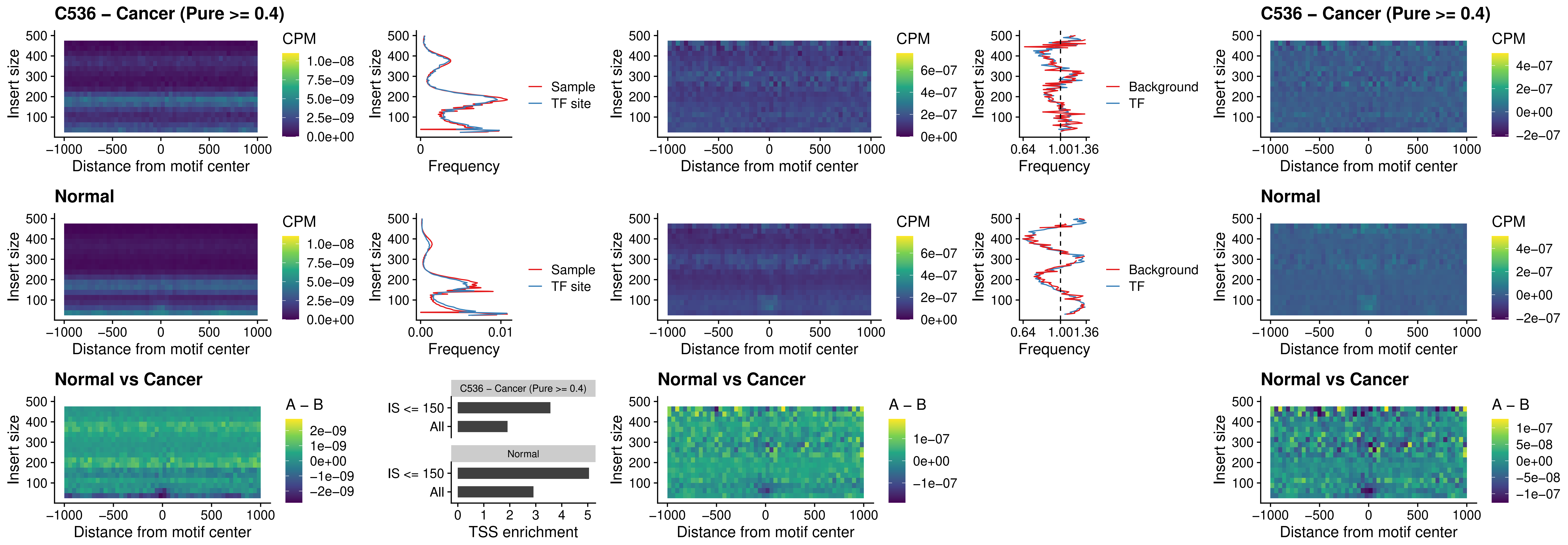

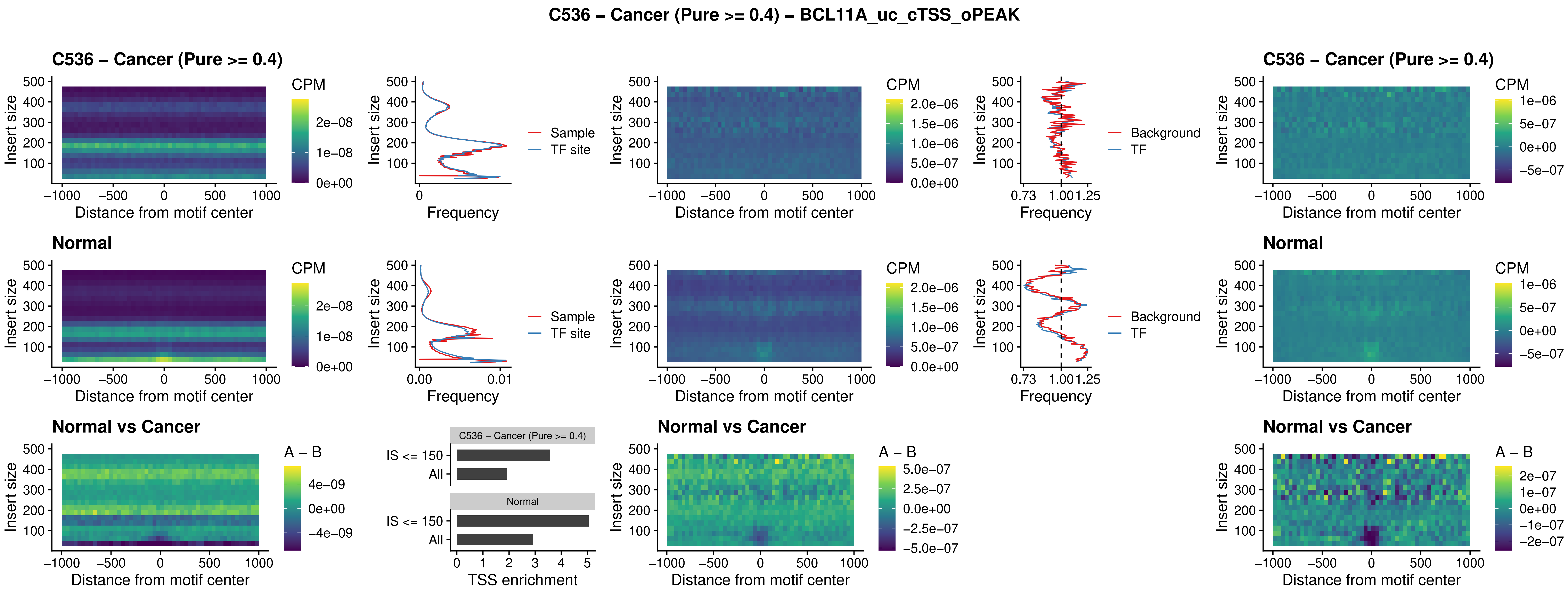
C536 – Cancer (Pure >= 0.4) – BCL11A_uc_cTSS_oPEAK
C536 – Cancer (Pure >= 0.4)
Normal
Normal vs Cancer
Insert size
Distance from motif center
Frequency
CPM
A – B
Sample
TF site
Background
TF
C536 – Cancer (Pure >= 0.4)
Normal
IS <= 150
All
TSS enrichment

#### C536 – Cancer (Pure >= 0.4) – BCL11B_uc_cTSS_oPEAK

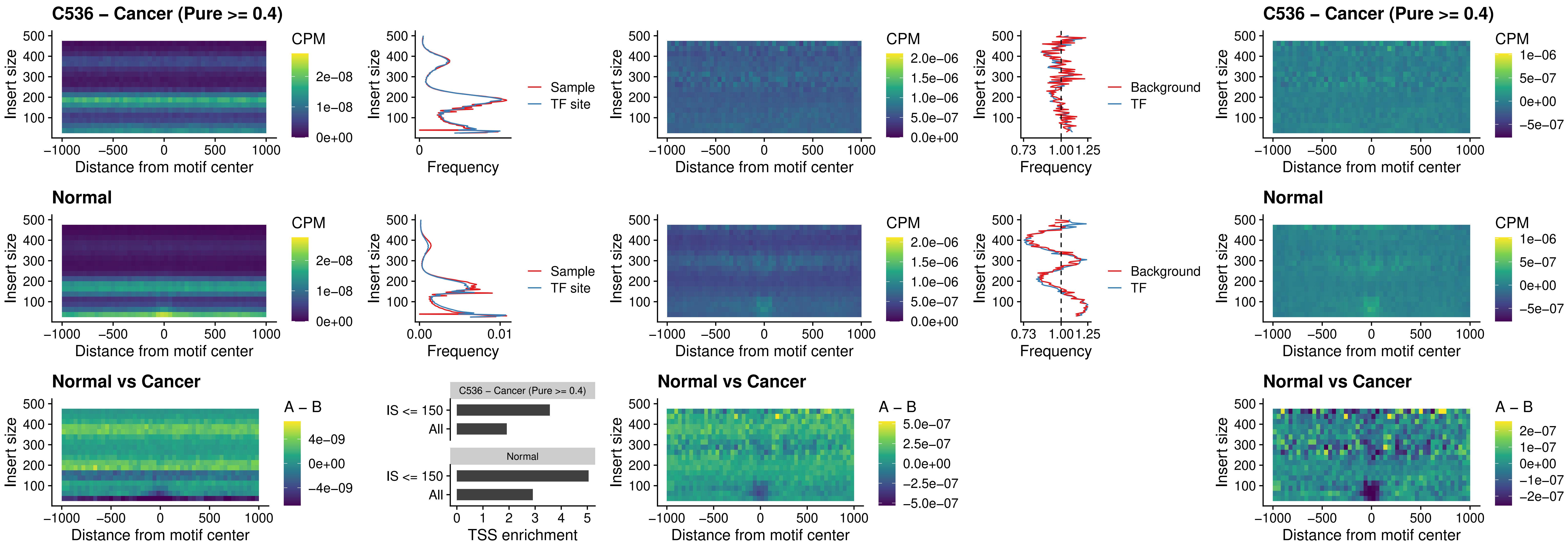

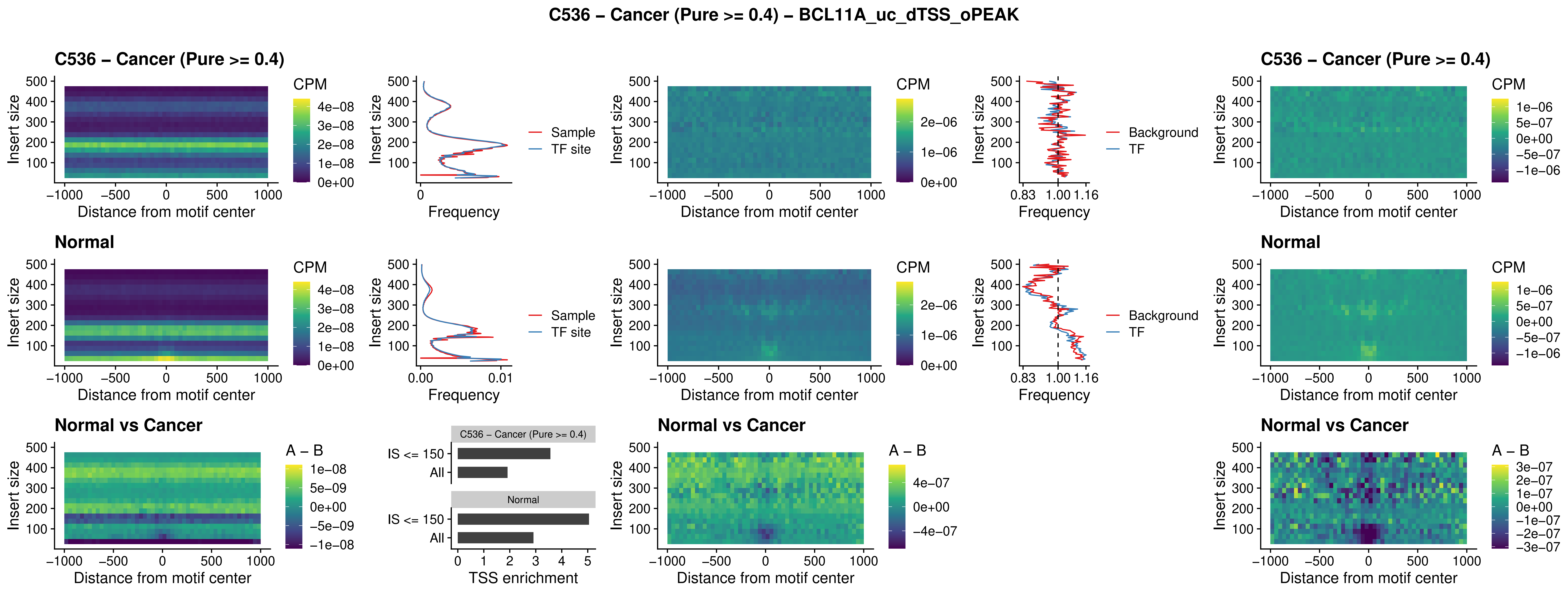
C536 – Cancer (Pure >= 0.4) – BCL11A_uc_dTSS_oPEAK
C536 – Cancer (Pure >= 0.4)
Normal
Normal vs Cancer
Insert size
Distance from motif center
Frequency
CPM
A – B
Sample
TF site
Background
TF
IS <= 150
All
TSS enrichment

#### C536 – Cancer (Pure >= 0.4) – BCL11B_uc_dTSS_oPEAK

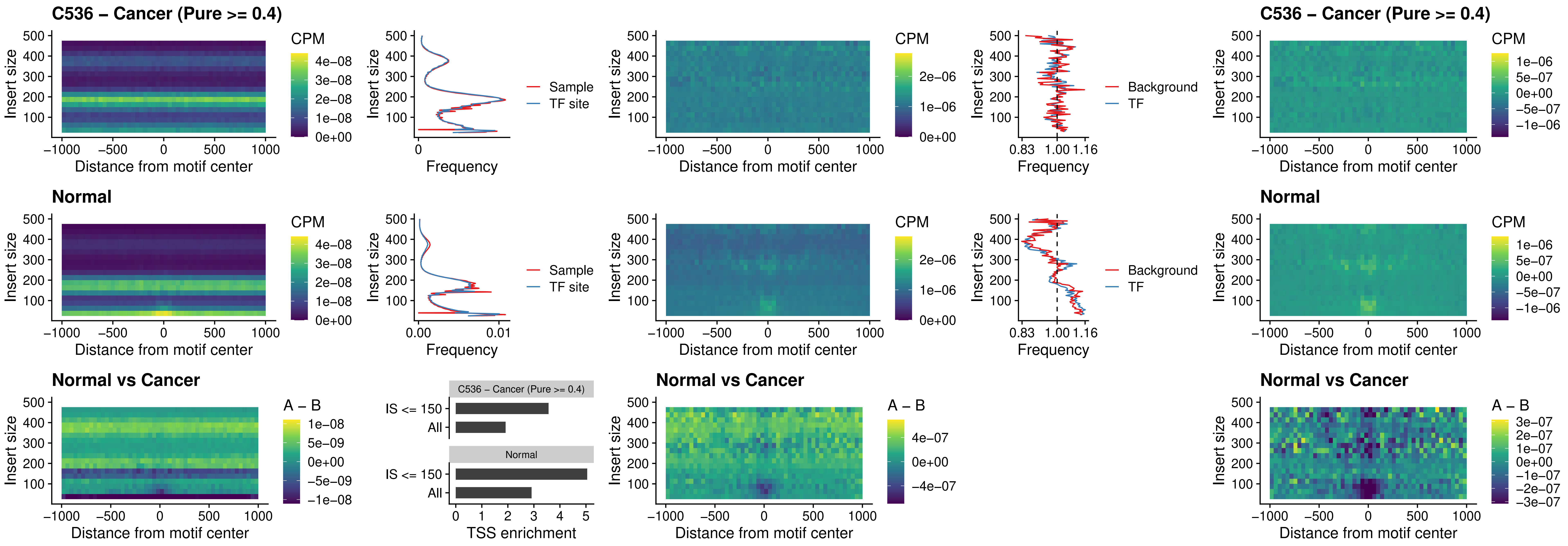

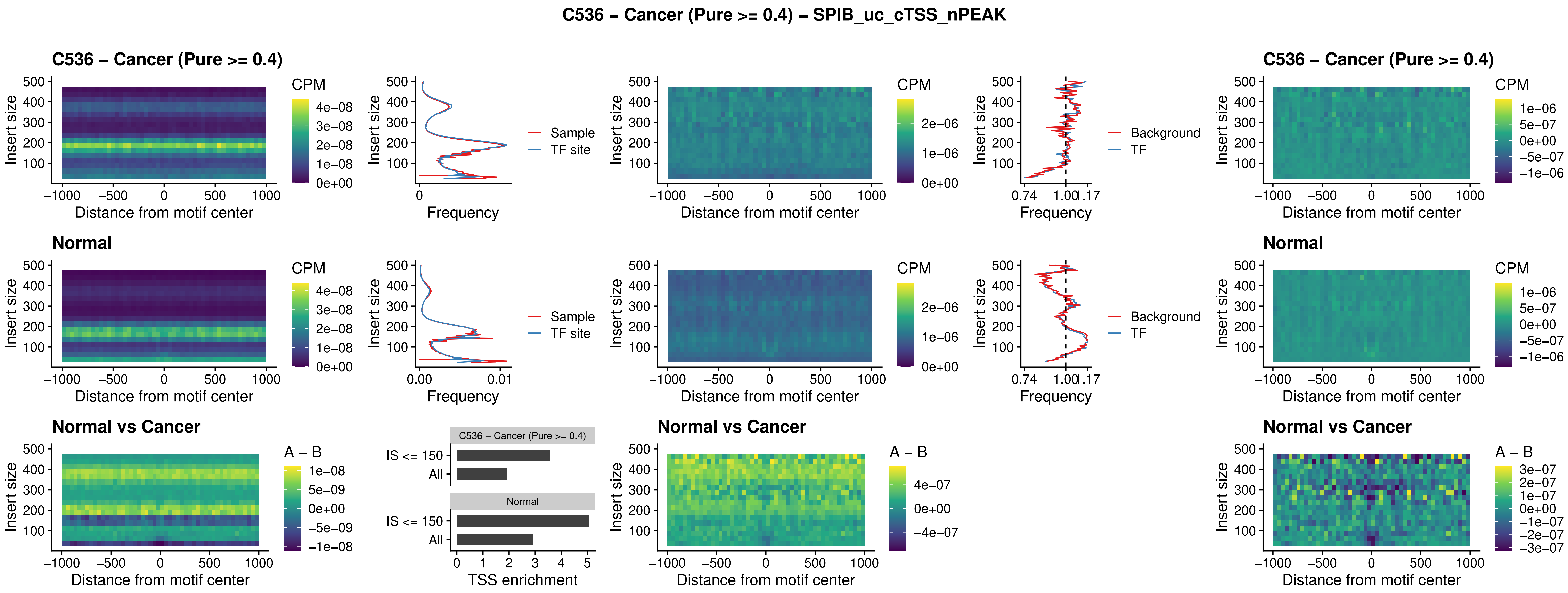
C536 – Cancer (Pure >= 0.4) – SPIB_uc_cTSS_nPEAK
C536 – Cancer (Pure >= 0.4)
Normal
Normal vs Cancer
Insert size
Distance from motif center
CPM
A – B
Frequency
Sample
TF site
Background
TF
C536 – Cancer (Pure >= 0.4)
Normal
IS <= 150
All
TSS enrichment

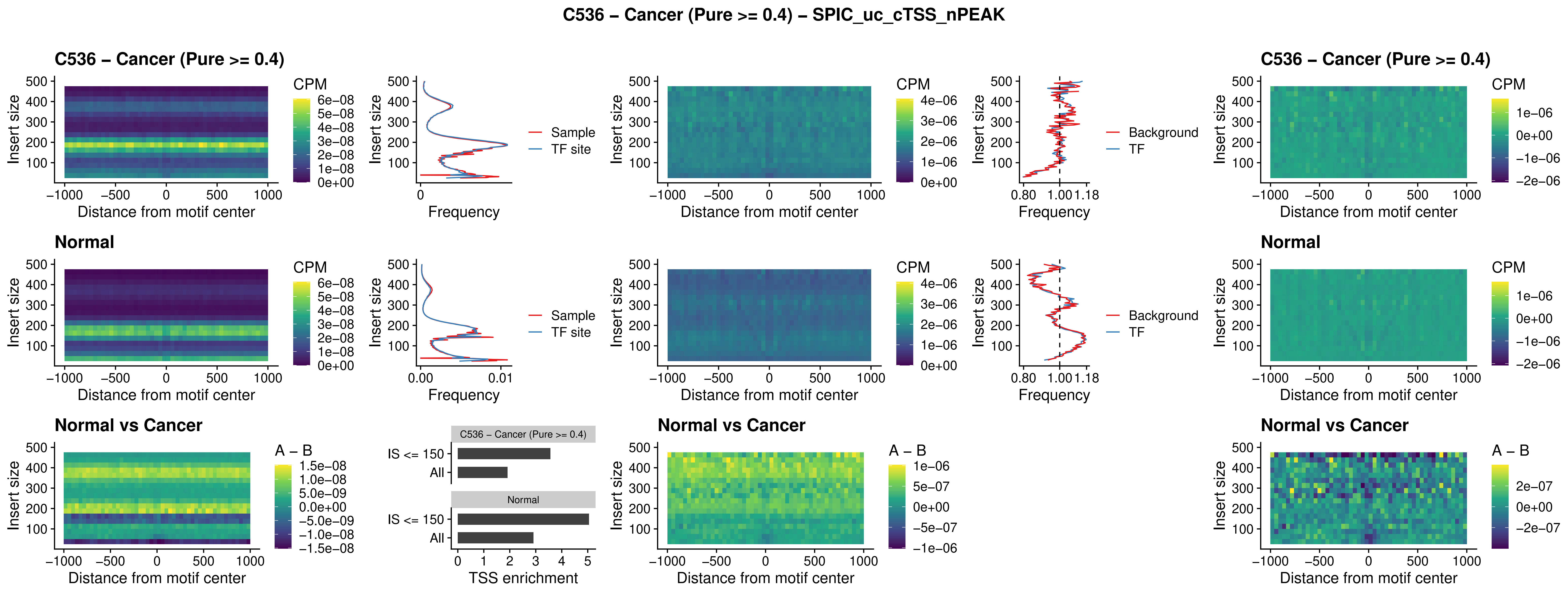

C536 – Cancer (Pure >= 0.4) – SPIC_uc_cTSS_nPEAK
C536 – Cancer (Pure >= 0.4)
Normal
Normal vs Cancer
Insert size
Distance from motif center
Frequency
CPM
A – B
Sample
TF site
Background
TF
TSS enrichment
IS <= 150
All
C536 – Cancer (Pure >= 0.4)
Normal

#### C536 – Cancer (Pure >= 0.4) – SPI1_uc_pTSS_oPEAK

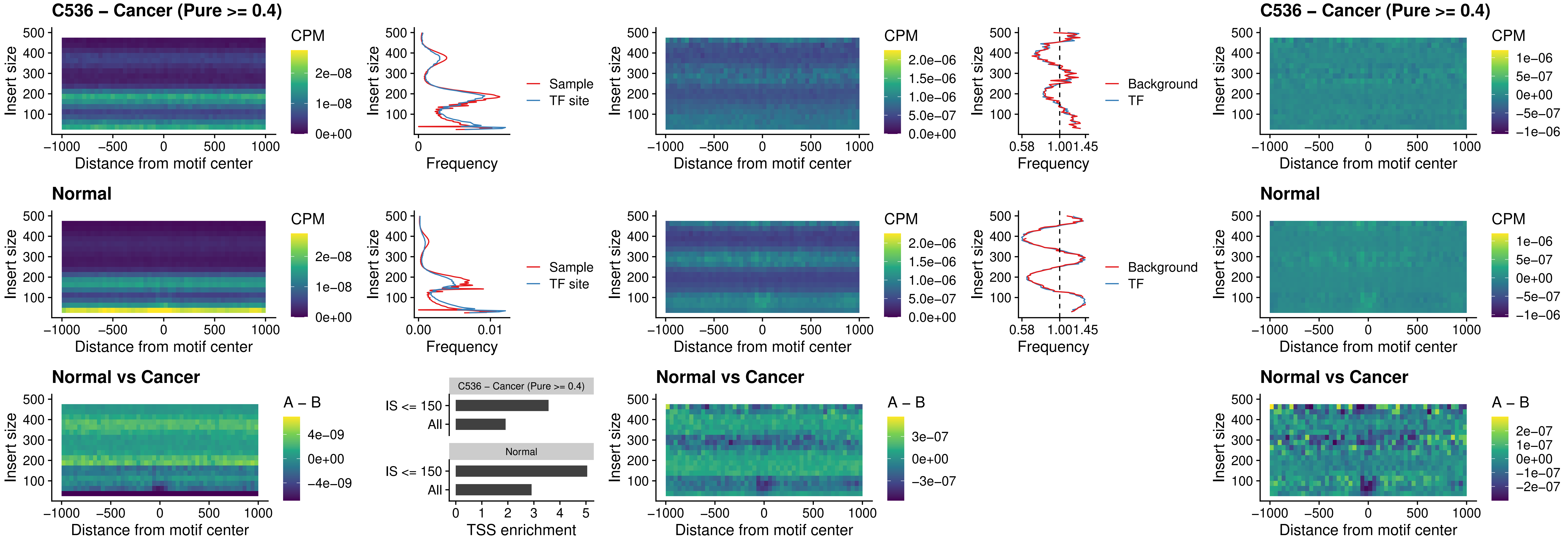

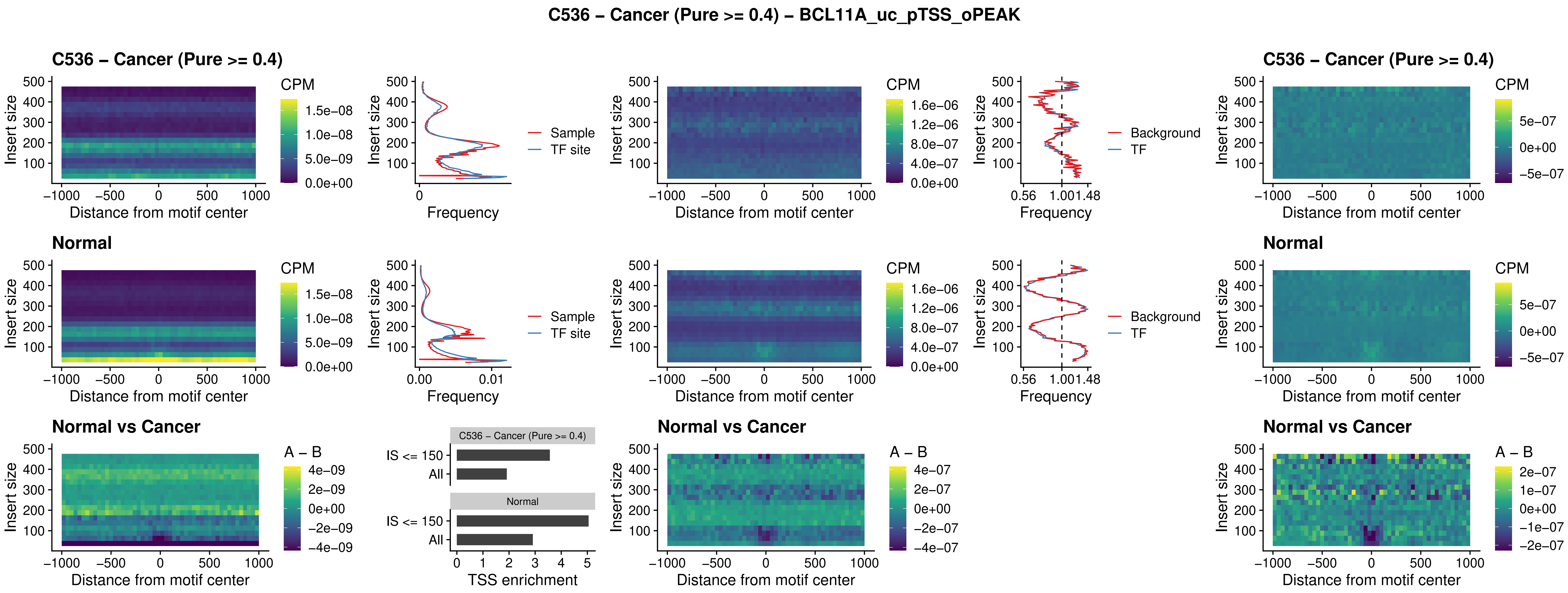
C536 – Cancer (Pure >= 0.4) – BCL11A_uc_pTSS_oPEAK
C536 – Cancer (Pure >= 0.4)
Normal
Normal vs Cancer
Insert size
Distance from motif center
Frequency
CPM
A – B
Sample
TF site
Background
TF
TSS enrichment
IS <= 150
All
C536 – Cancer (Pure >= 0.4)
Normal

#### C536 – Cancer (Pure >= 0.4) – BCL11B_uc_pTSS_oPEAK

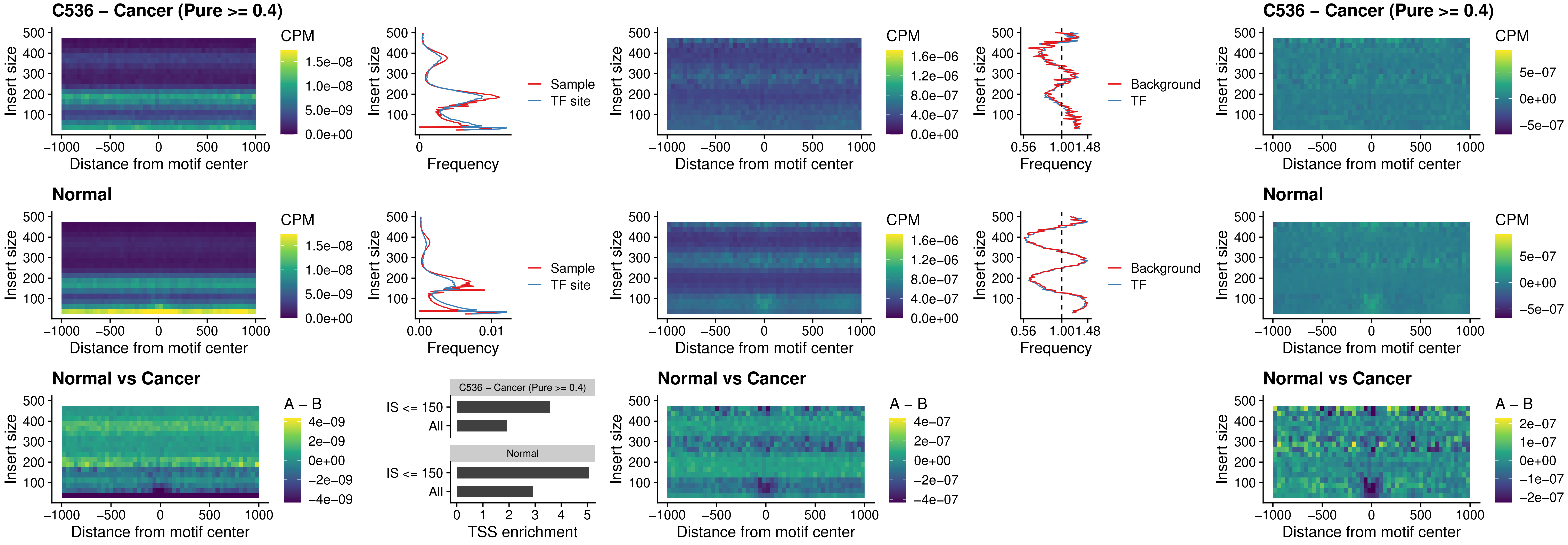

### C536 – Cancer (Pure >= 0.4) – SPI1_uc_cTSS_oPEAK

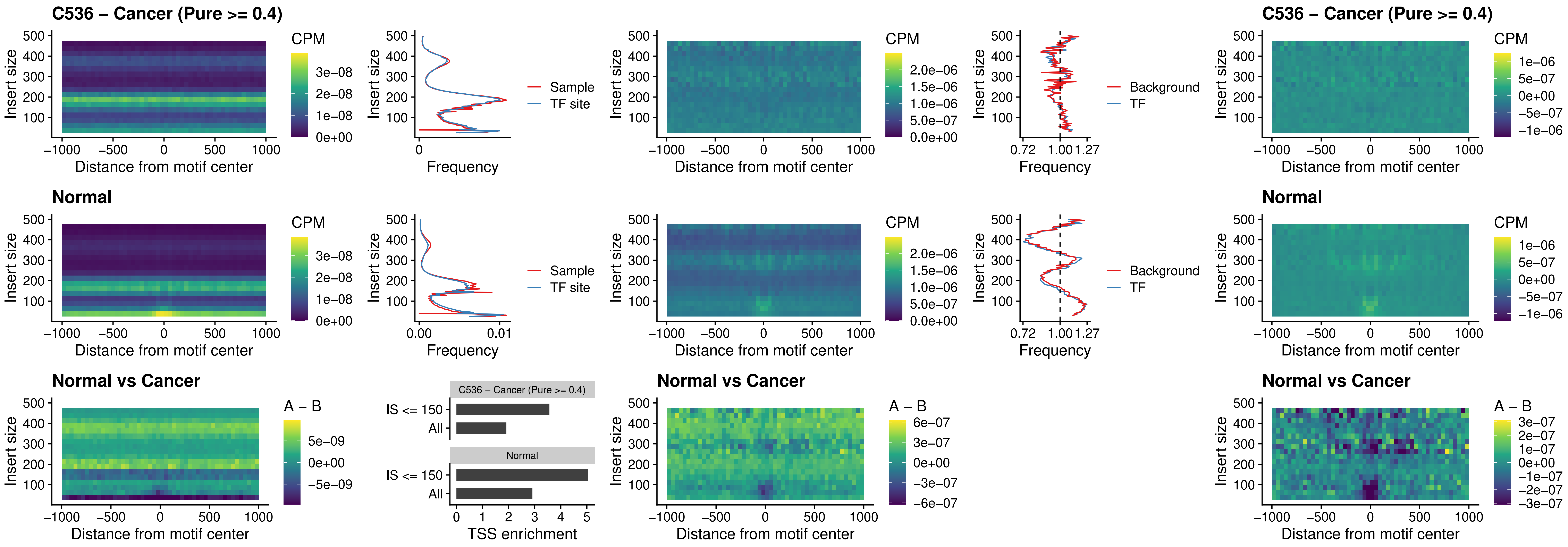

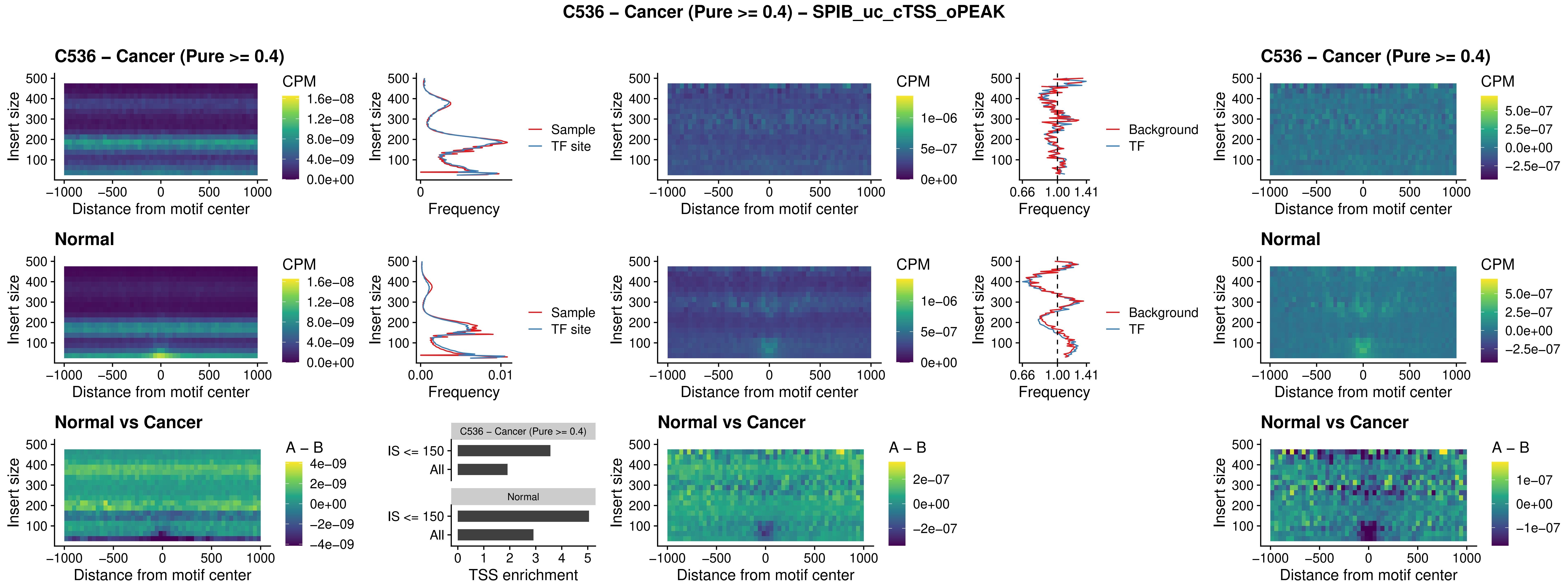
C536 – Cancer (Pure >= 0.4) – SPIB_uc_cTSS_oPEAK
C536 – Cancer (Pure >= 0.4)
Normal
Normal vs Cancer
Insert size
Distance from motif center
Frequency
CPM
A – B
Sample
TF site
Background
TF
TSS enrichment
C536 – Cancer (Pure >= 0.4)
Normal
IS <= 150
All

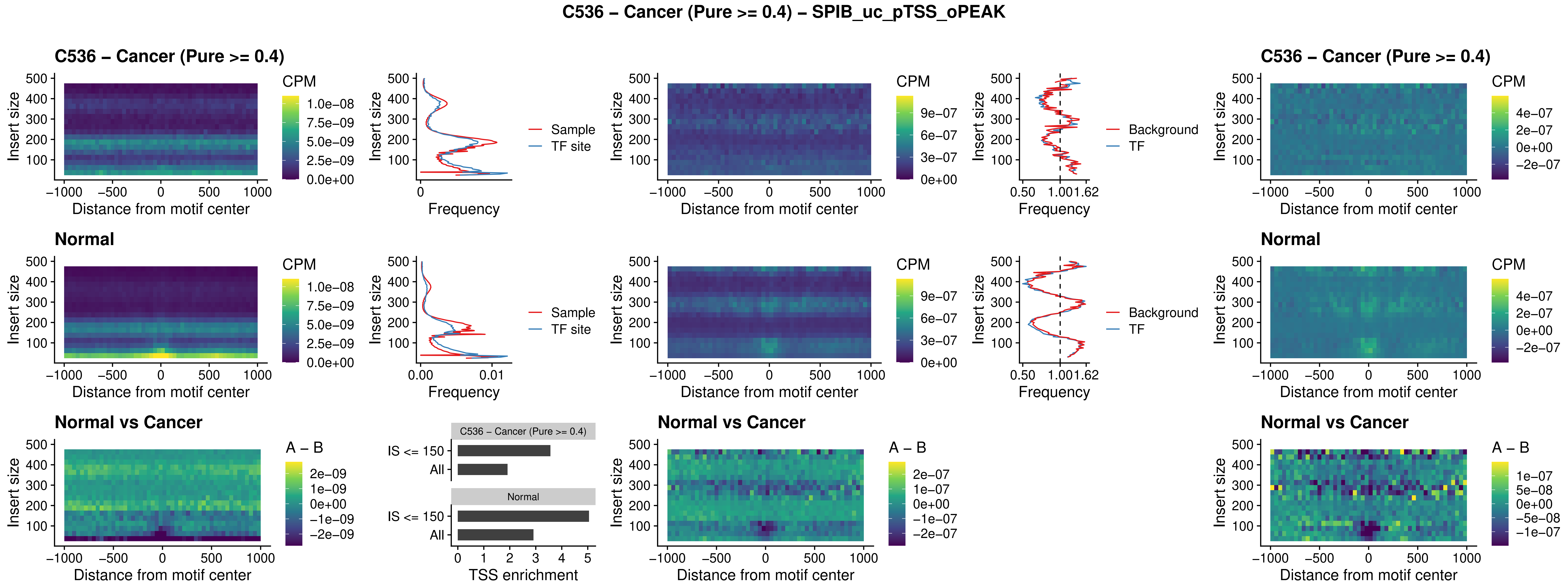

C536 – Cancer (Pure >= 0.4) – SPIB_uc_pTSS_oPEAK
C536 – Cancer (Pure >= 0.4)
Normal
Normal vs Cancer
Insert size
Distance from motif center
Frequency
CPM
A – B
Sample
TF site
Background
TF
TSS enrichment
C536 – Cancer (Pure >= 0.4)
Normal
IS <= 150
All

#### C536 – Cancer (Pure >= 0.4) – SPI1_uc_dTSS_oPEAK

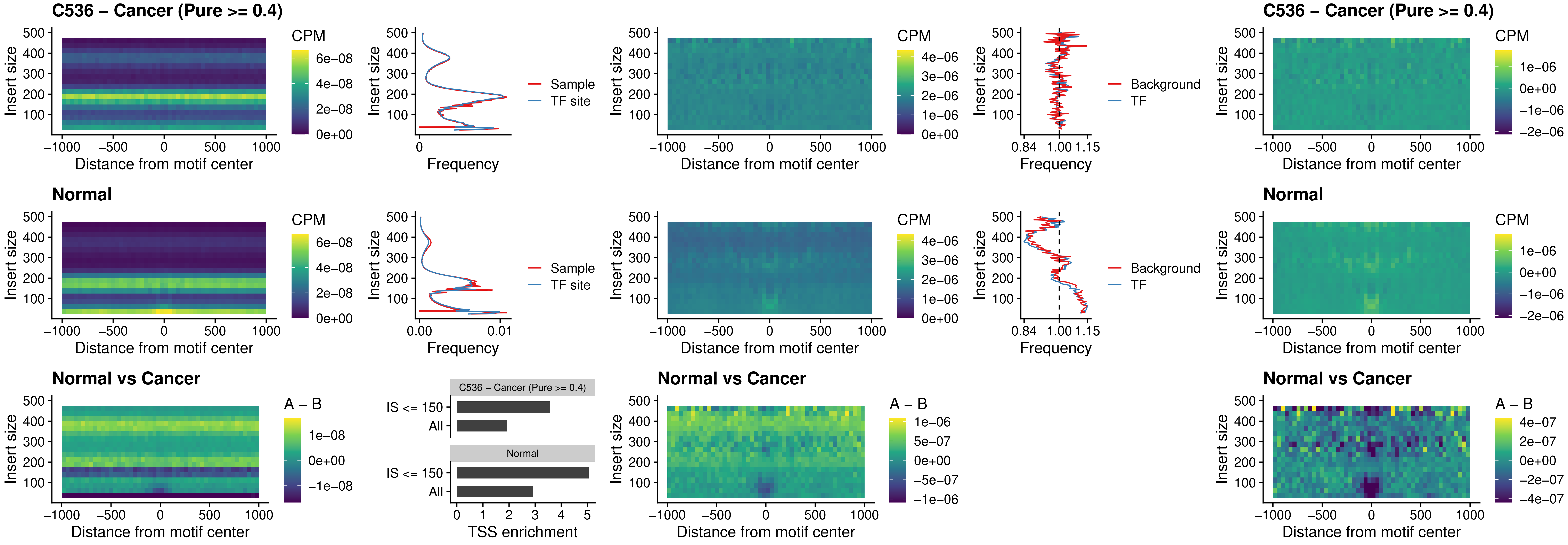

#### C536 – Cancer (Pure >= 0.4) – SPIB_uc_dTSS_oPEAK

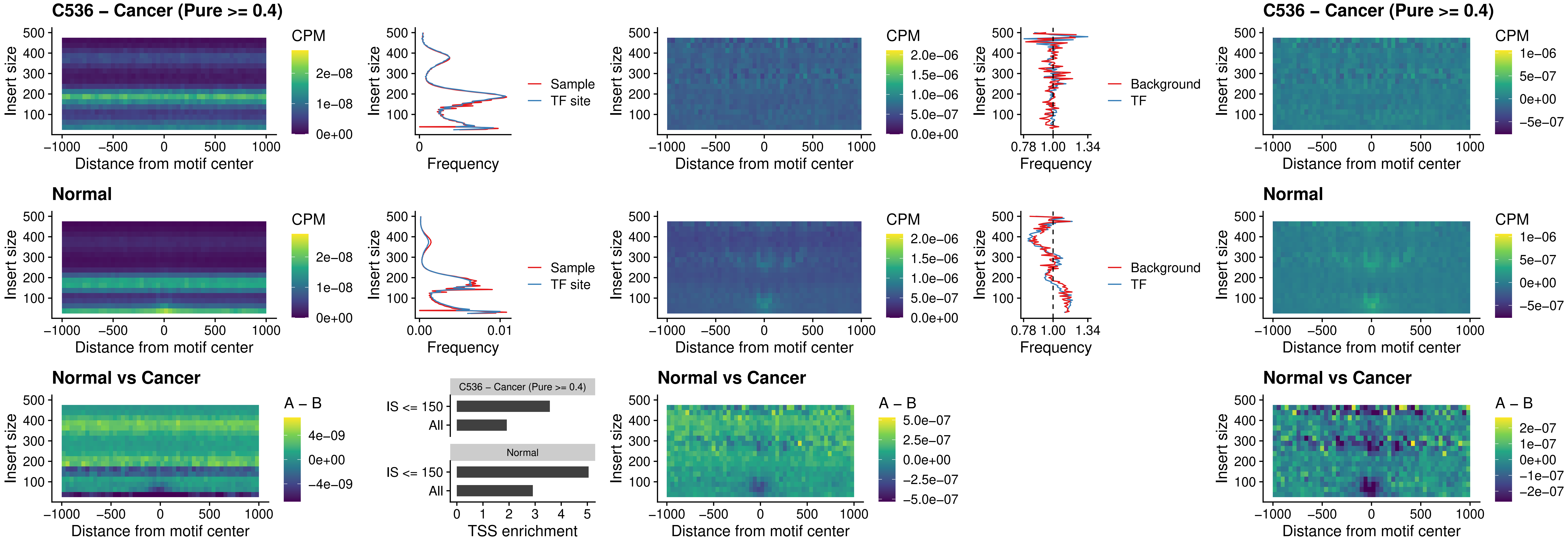

#### C536 – Cancer (Pure >= 0.4) – IRF3_uc_pTSS_oPEAK

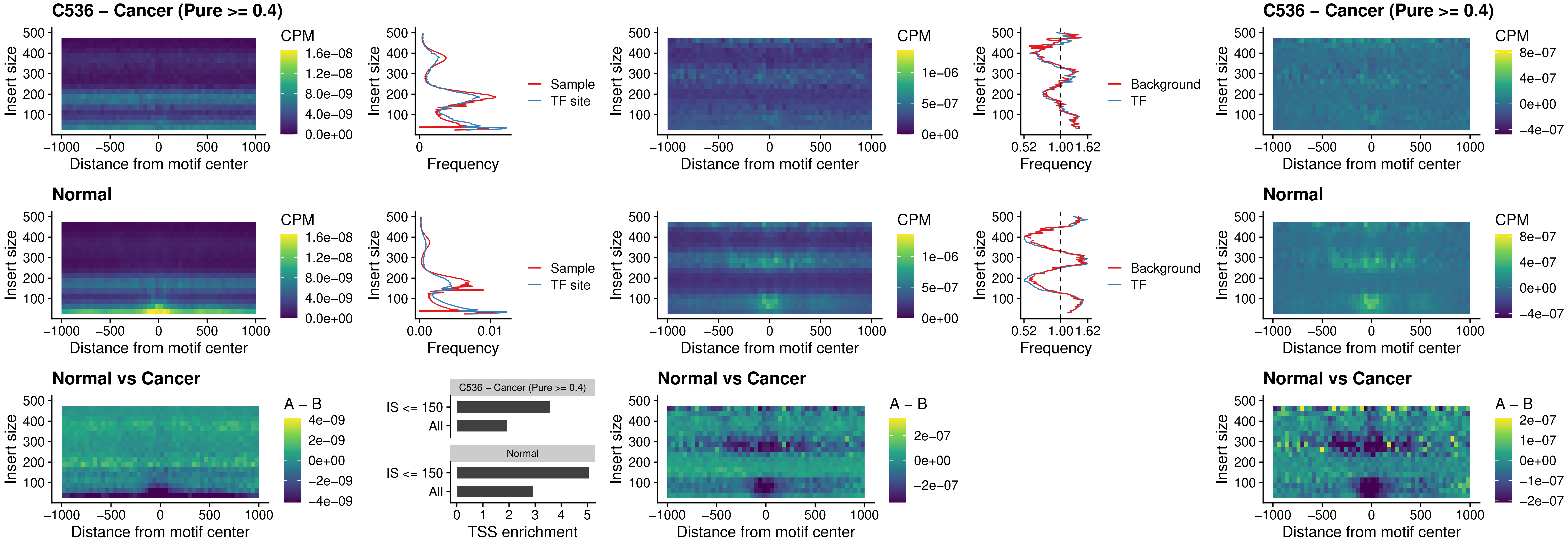

### C536 – Cancer (Pure >= 0.4) – IRF2_uc_dTSS_oPEAK

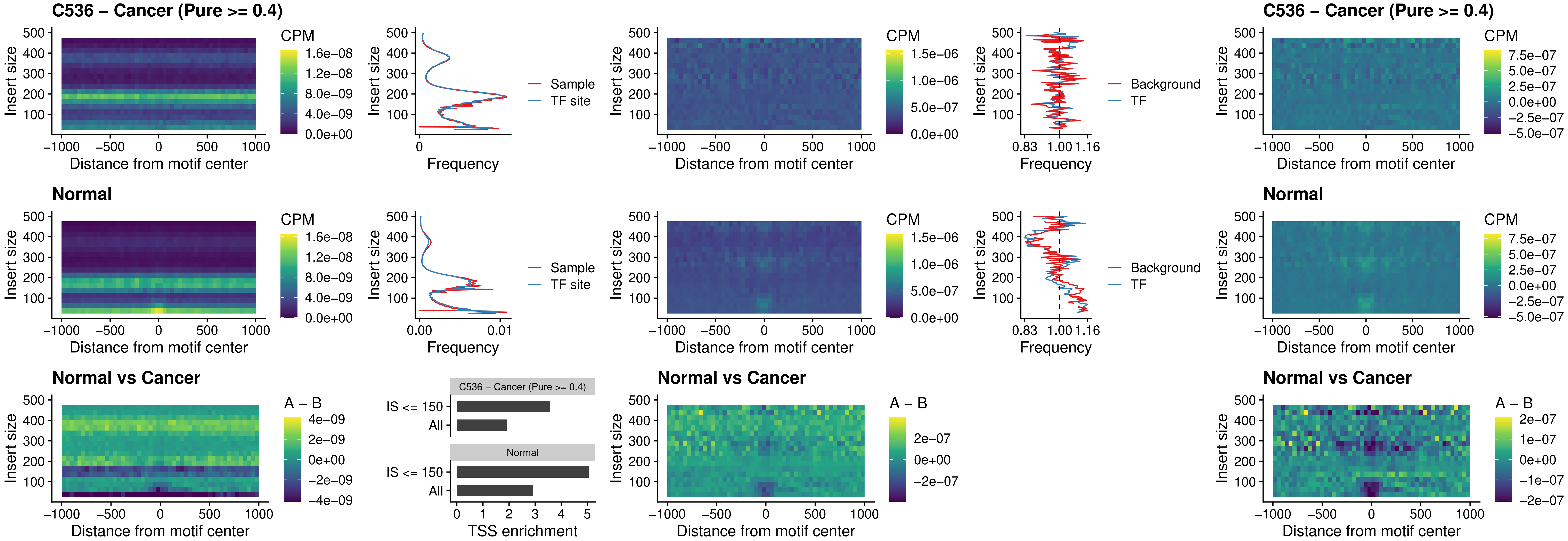

#### C536 – Cancer (Pure >= 0.4) – IRF2_uc_pTSS_oPEAK

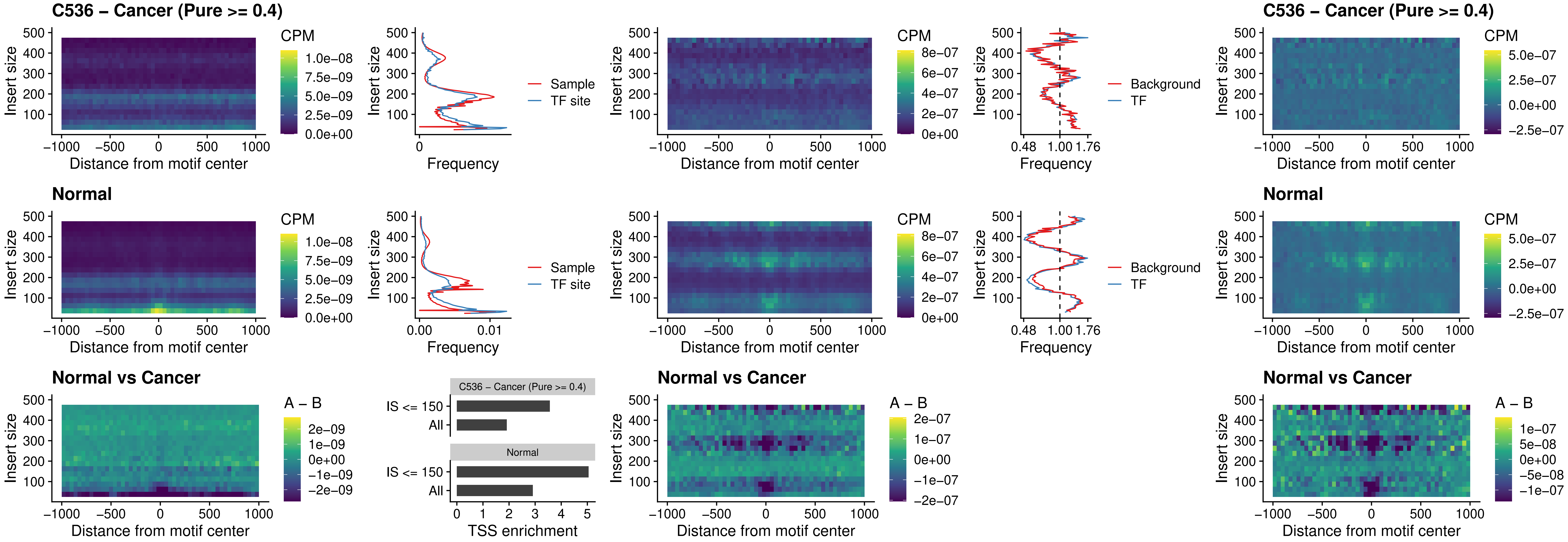

#### C536 – Cancer (Pure >= 0.4) – STAT2_uc_pTSS_oPEAK

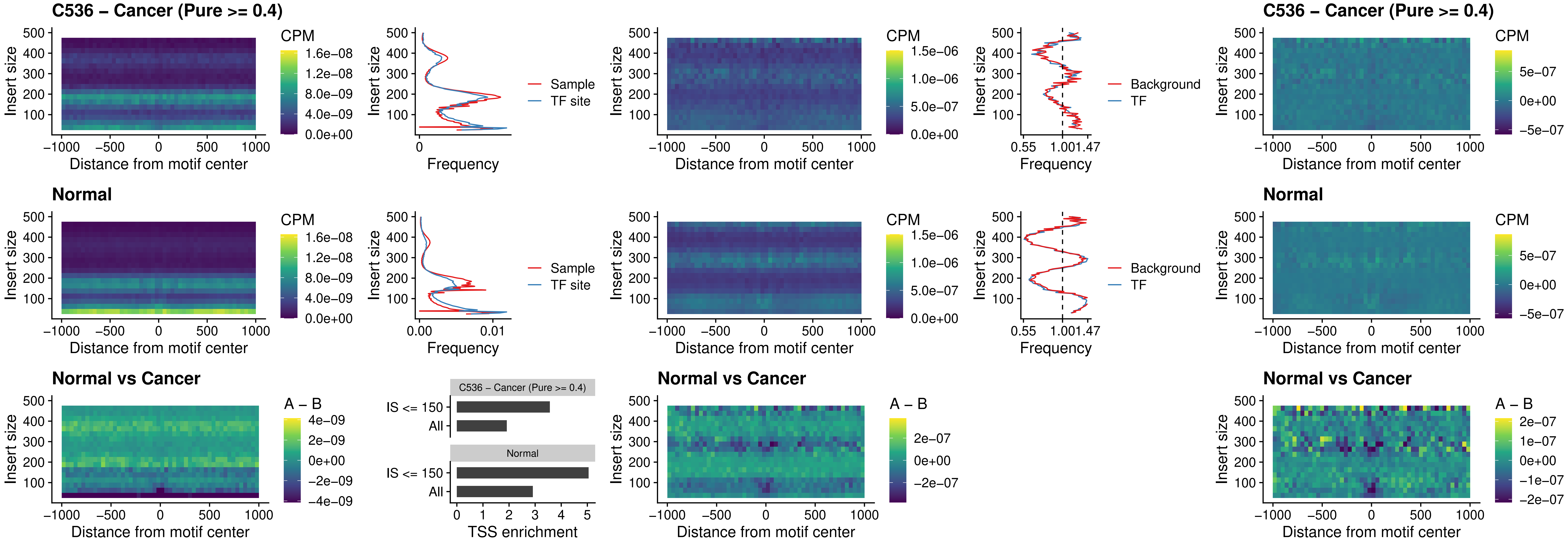

#### C536 – Cancer (Pure >= 0.4) – SPIB_uc_dTSS_nPEAK

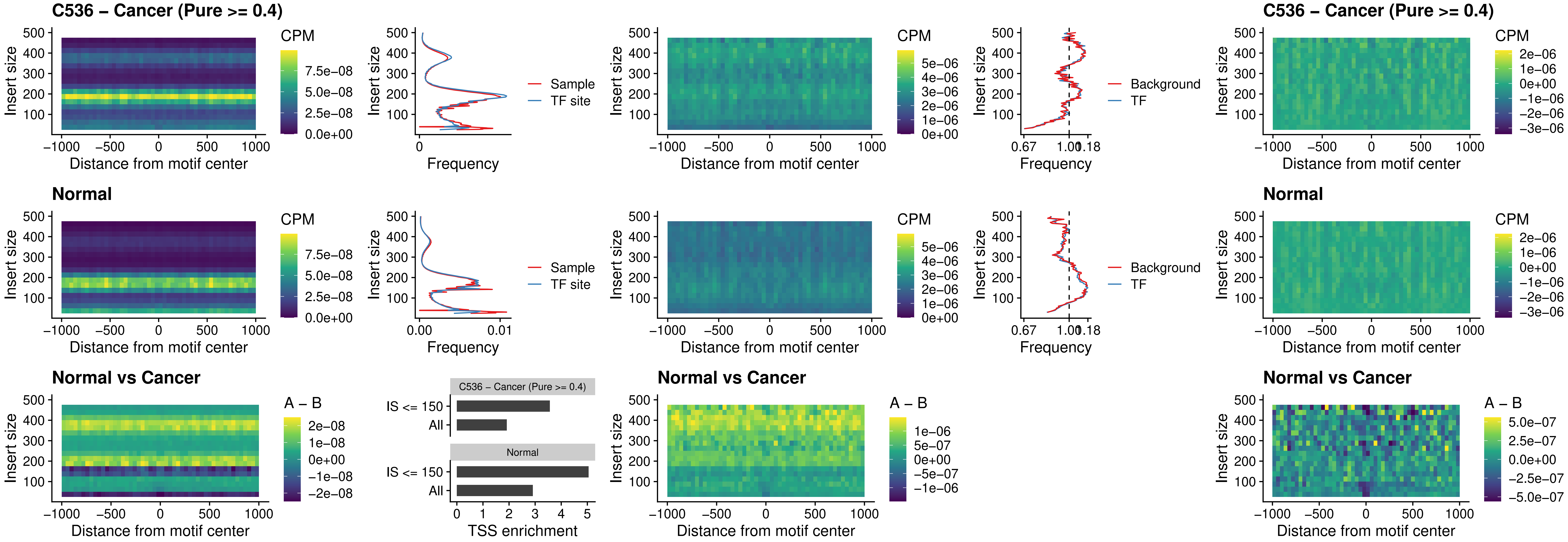

#### C536 – Cancer (Pure >= 0.4) – SPIC_uc_cTSS_oPEAK

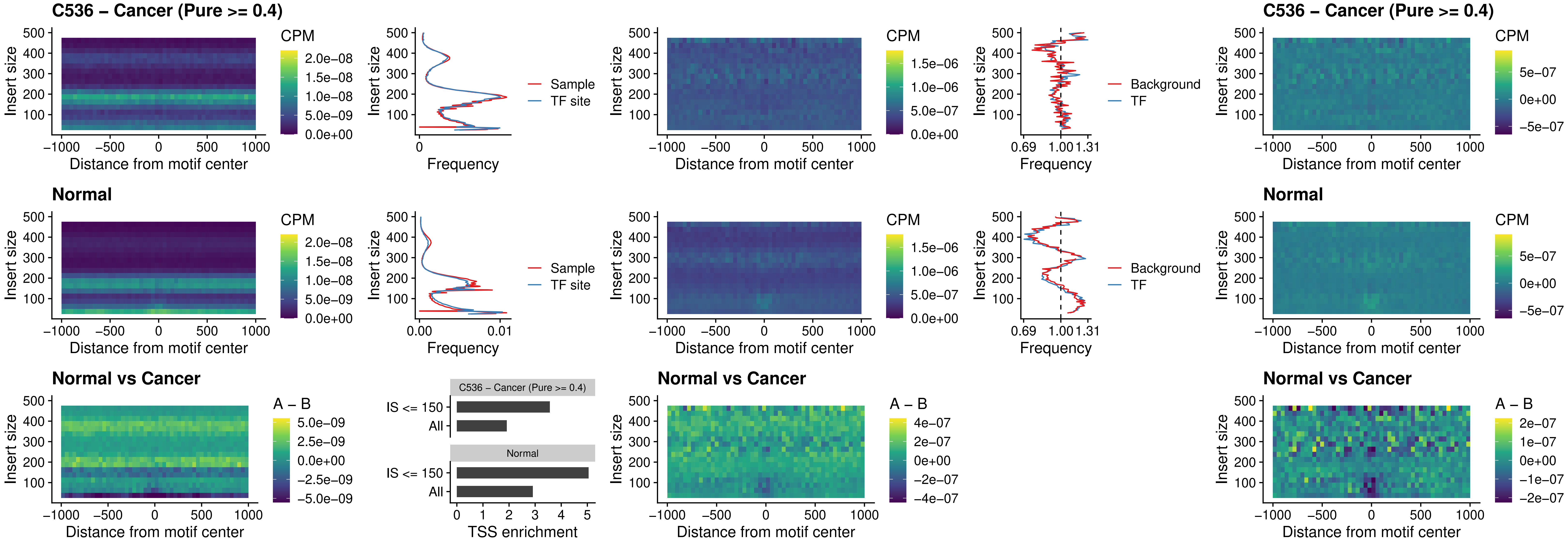

#### C536 – Cancer (Pure >= 0.4) – PRDM1_uc_cTSS_oPEAK

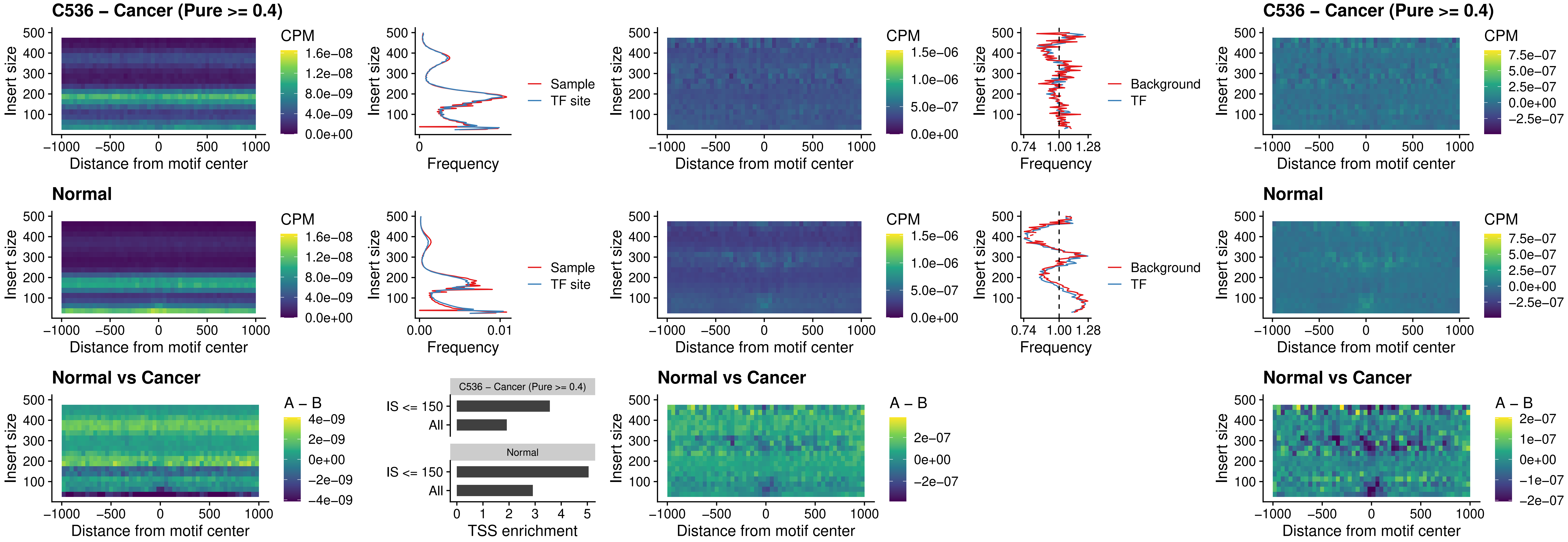

#### C536 – Cancer (Pure >= 0.4) – IRF3_uc_cTSS_oPEAK

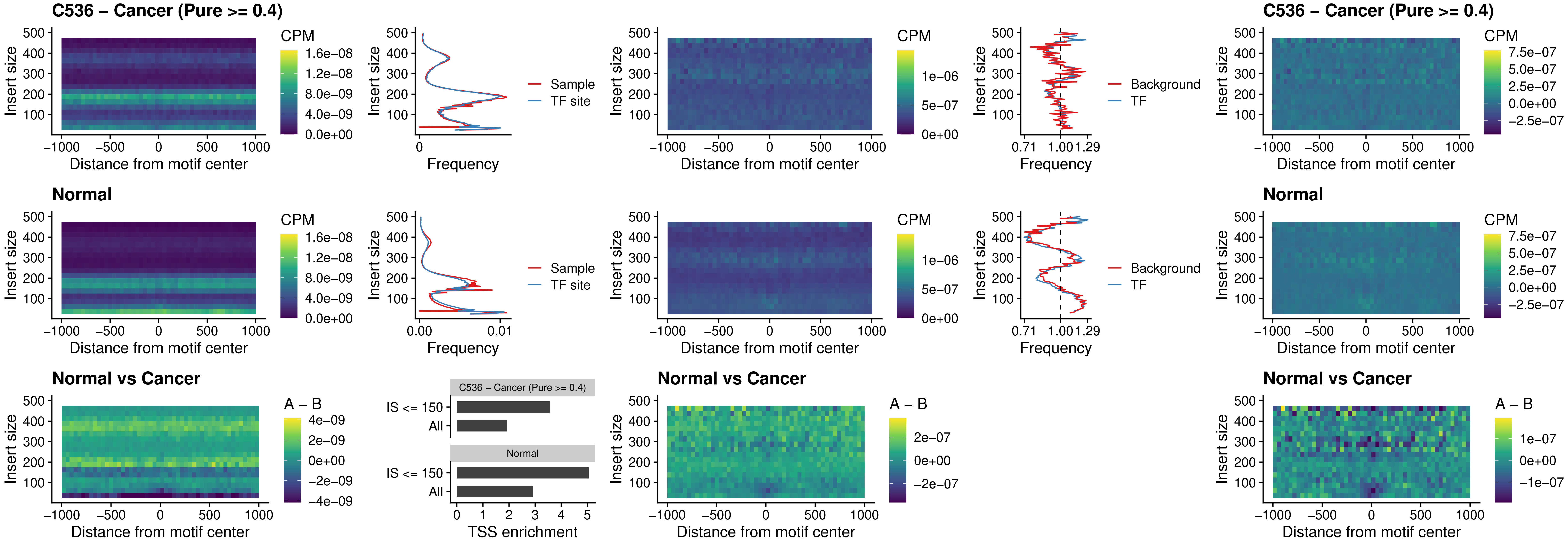

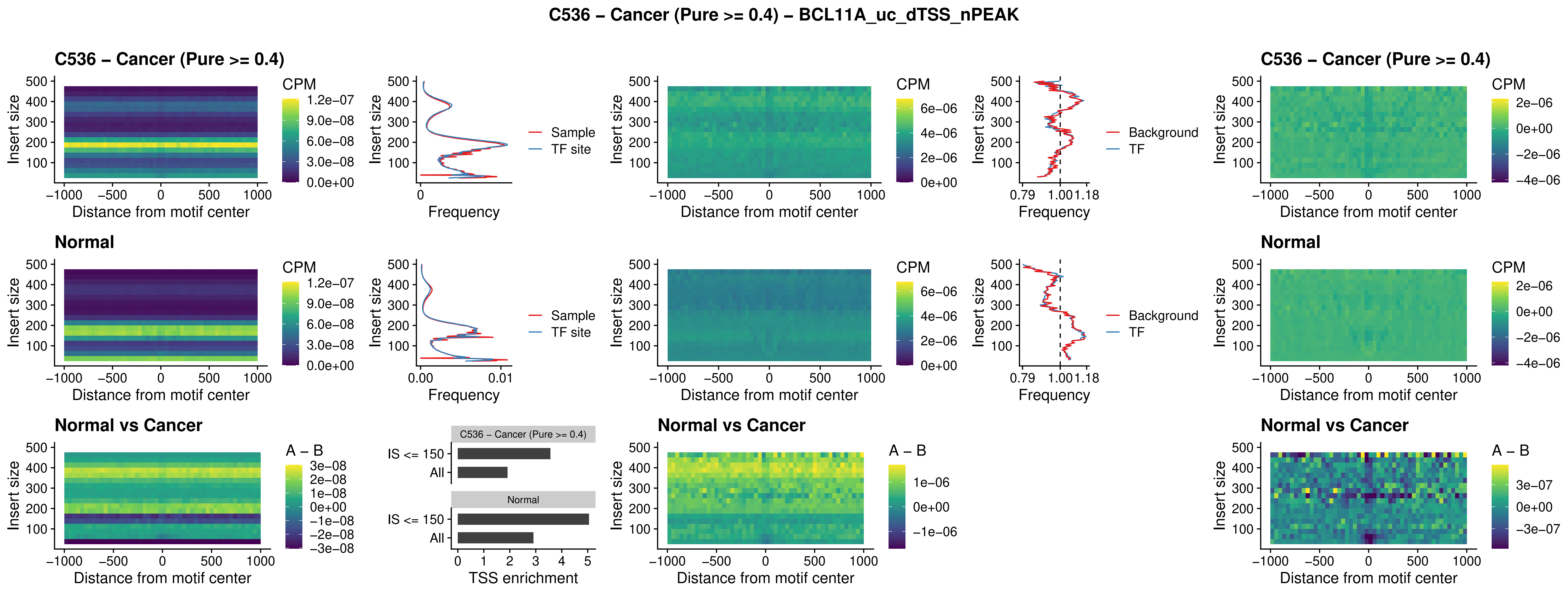
C536 – Cancer (Pure >= 0.4) – BCL11A_uc_dTSS_nPEAK
C536 – Cancer (Pure >= 0.4)
Normal
Normal vs Cancer
Insert size
Distance from motif center
Frequency
CPM
A – B
Sample
TF site
Background
TF
C536 – Cancer (Pure >= 0.4)
Normal
IS <= 150
All
TSS enrichment

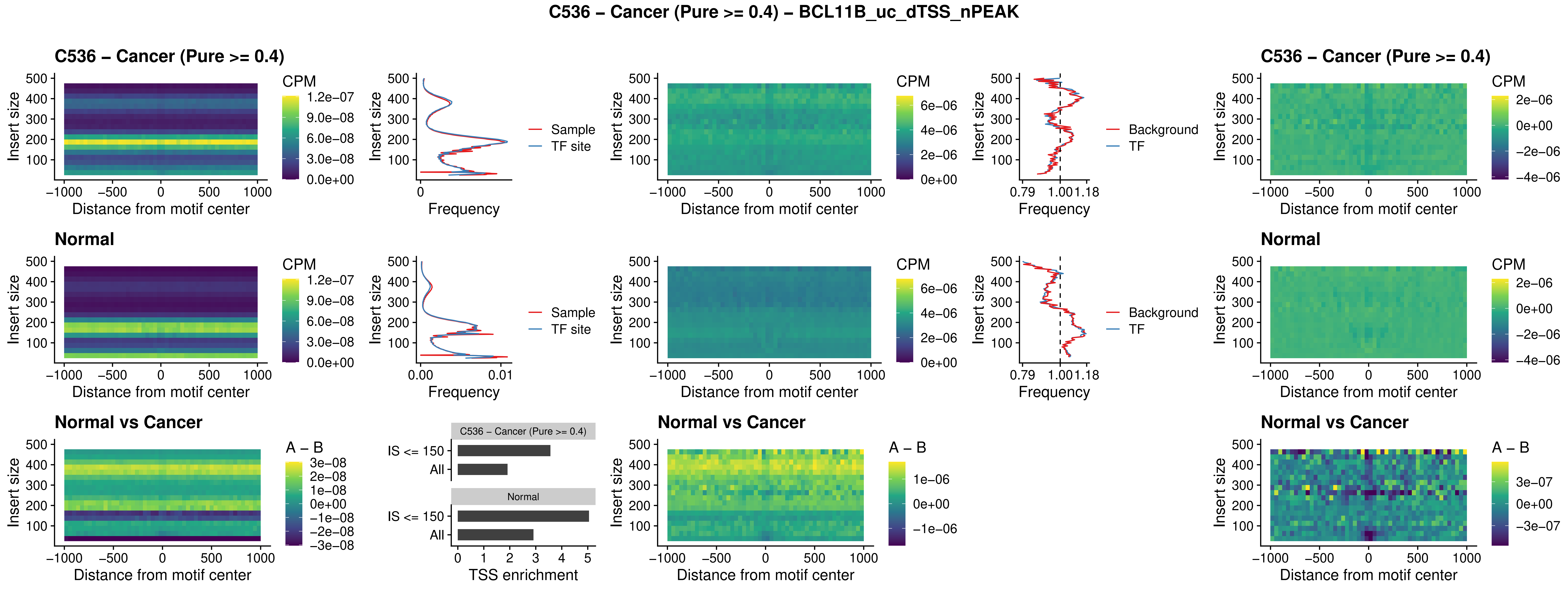
C536 – Cancer (Pure >= 0.4) – BCL11B_uc_dTSS_nPEAK
C536 – Cancer (Pure >= 0.4)
Normal
Normal vs Cancer
Insert size
Distance from motif center
Frequency
CPM
A – B
Sample
TF site
Background
TF
C536 – Cancer (Pure >= 0.4)
Normal
IS <= 150
All
TSS enrichment

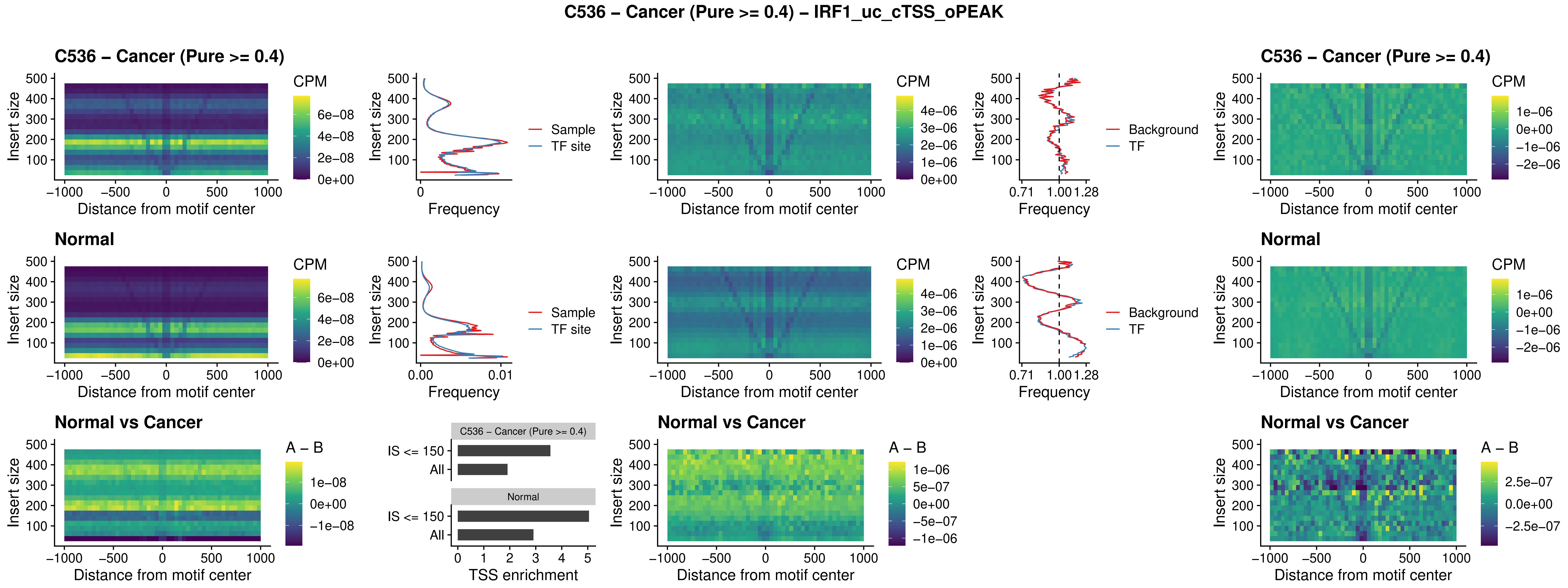
C536 – Cancer (Pure >= 0.4) – IRF1_uc_cTSS_oPEAK
C536 – Cancer (Pure >= 0.4)
Normal
Normal vs Cancer
Insert size
Distance from motif center
Frequency
CPM
A – B
Sample
TF site
Background
TF
C536 – Cancer (Pure >= 0.4)
Normal
IS <= 150
All
TSS enrichment

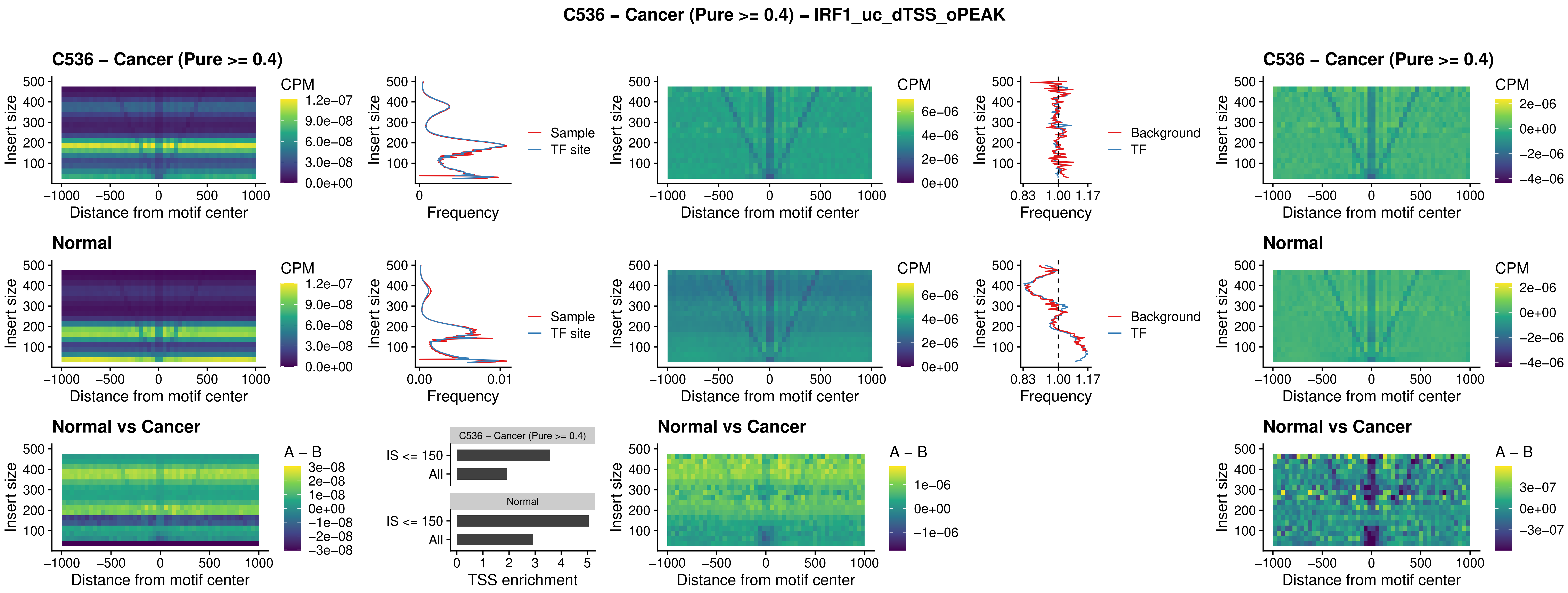
C536 – Cancer (Pure >= 0.4) – IRF1_uc_dTSS_oPEAK
C536 – Cancer (Pure >= 0.4)
Normal
Normal vs Cancer
Insert size
Distance from motif center
Frequency
CPM
A – B
Sample
TF site
Background
TF
C536 – Cancer (Pure >= 0.4)
Normal
IS <= 150
All
TSS enrichment

#### C536 – Cancer (Pure >= 0.4) – IRF1_uc_pTSS_oPEAK

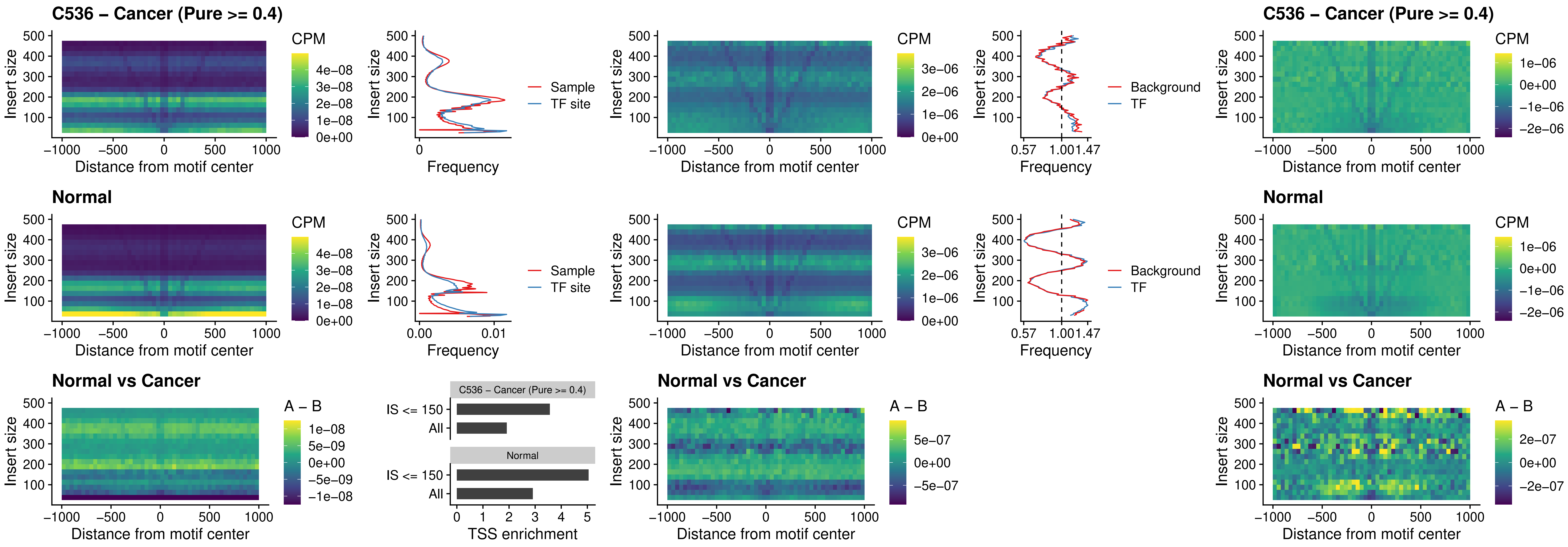

#### C536 – Cancer (Pure >= 0.4) – SPIC_uc_dTSS_nPEAK

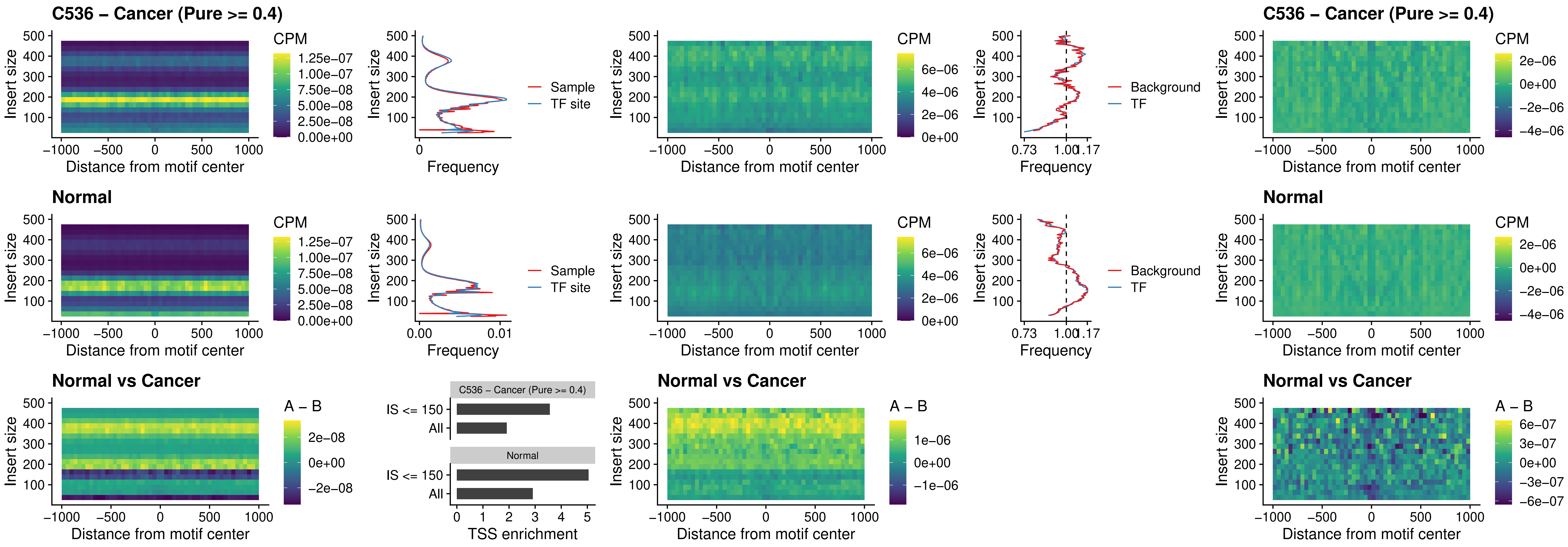

### C536 – Cancer (Pure >= 0.4) – SPIC_uc_dTSS_oPEAK

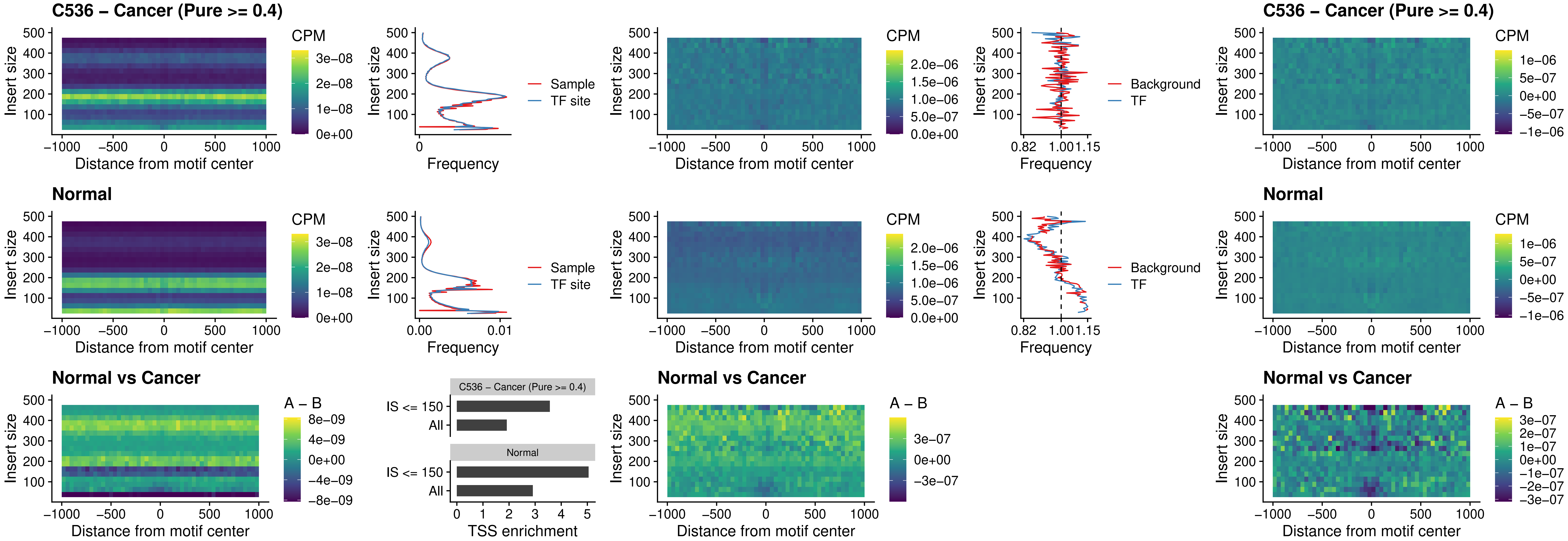

#### C536 – Cancer (Pure >= 0.4) – SPIB_uc_pTSS_nPEAK

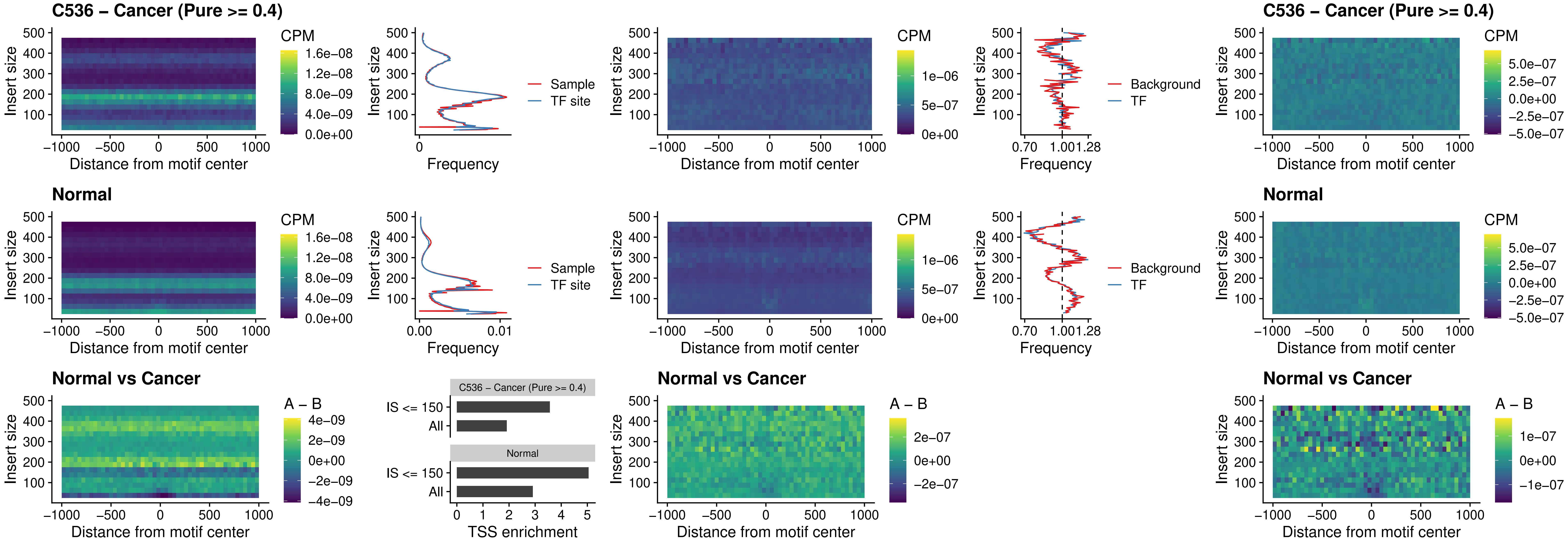

### C536 – Cancer (Pure >= 0.4) – STAT2_uc_cTSS_oPEAK

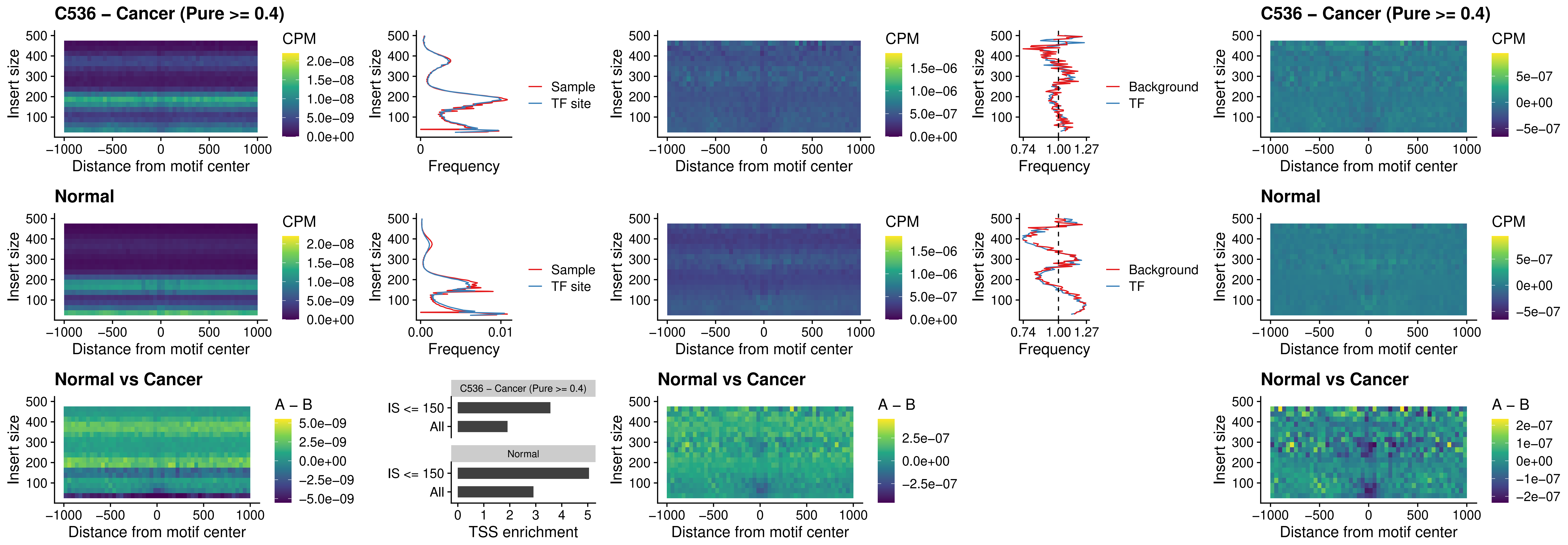

#### C536 – Cancer (Pure >= 0.4) – IRF8_uc_cTSS_oPEAK

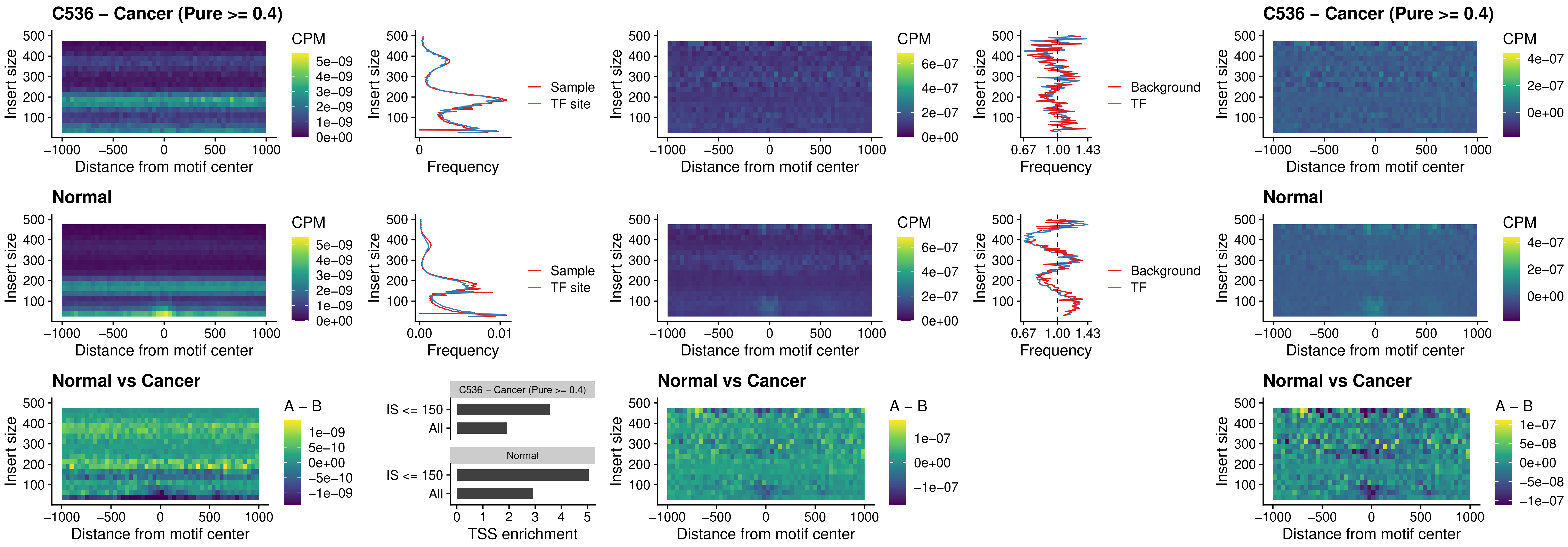

#### C536 – Cancer (Pure >= 0.4) – IRF3_uc_dTSS_oPEAK

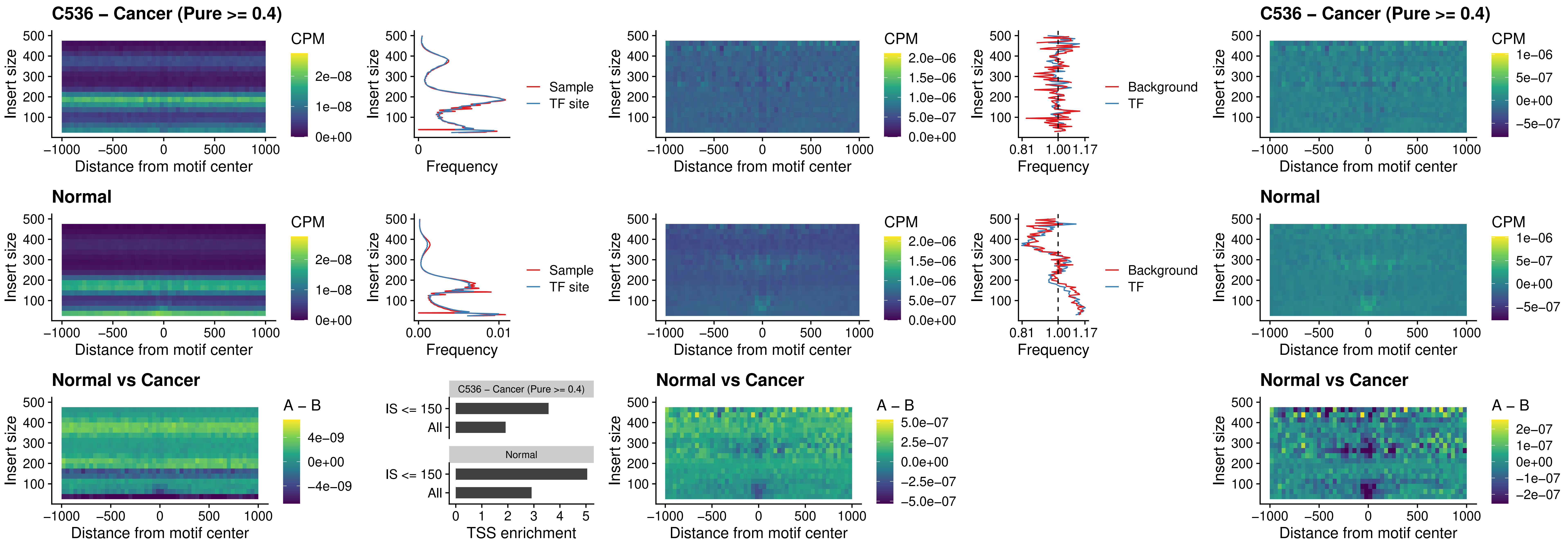

### C536 – Cancer (Pure >= 0.4) – IRF8_uc_dTSS_oPEAK

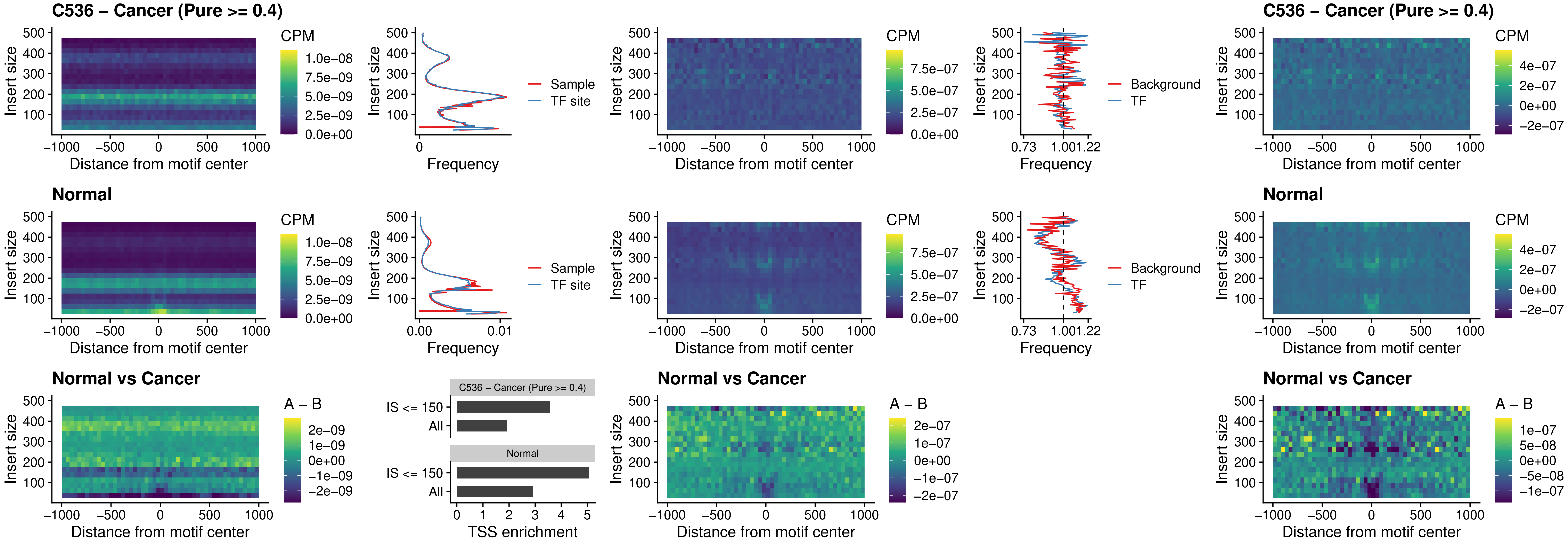

### C536 – Cancer (Pure >= 0.4) – ETS2_uc_dTSS_oPEAK

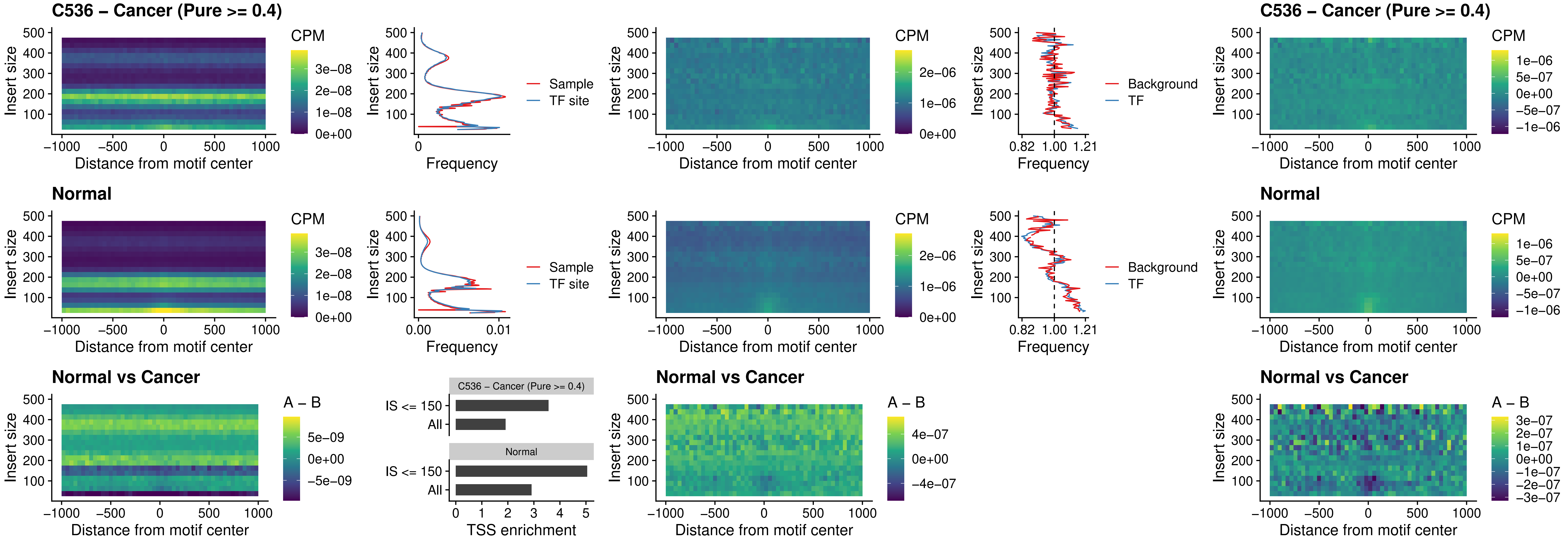

#### C536 – Cancer (Pure >= 0.4) – SMARCC2_uc_pTSS_oPEAK

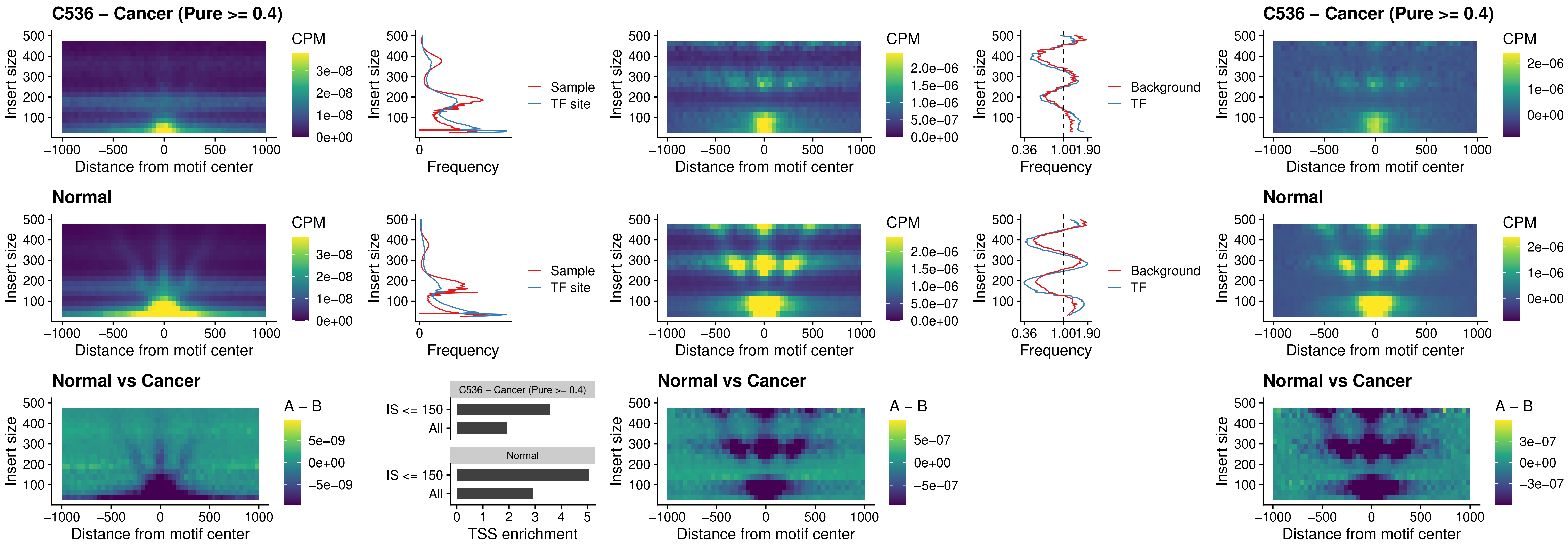

#### C536 – Cancer (Pure >= 0.4) – BCL11A_uc_cTSS_nPEAK

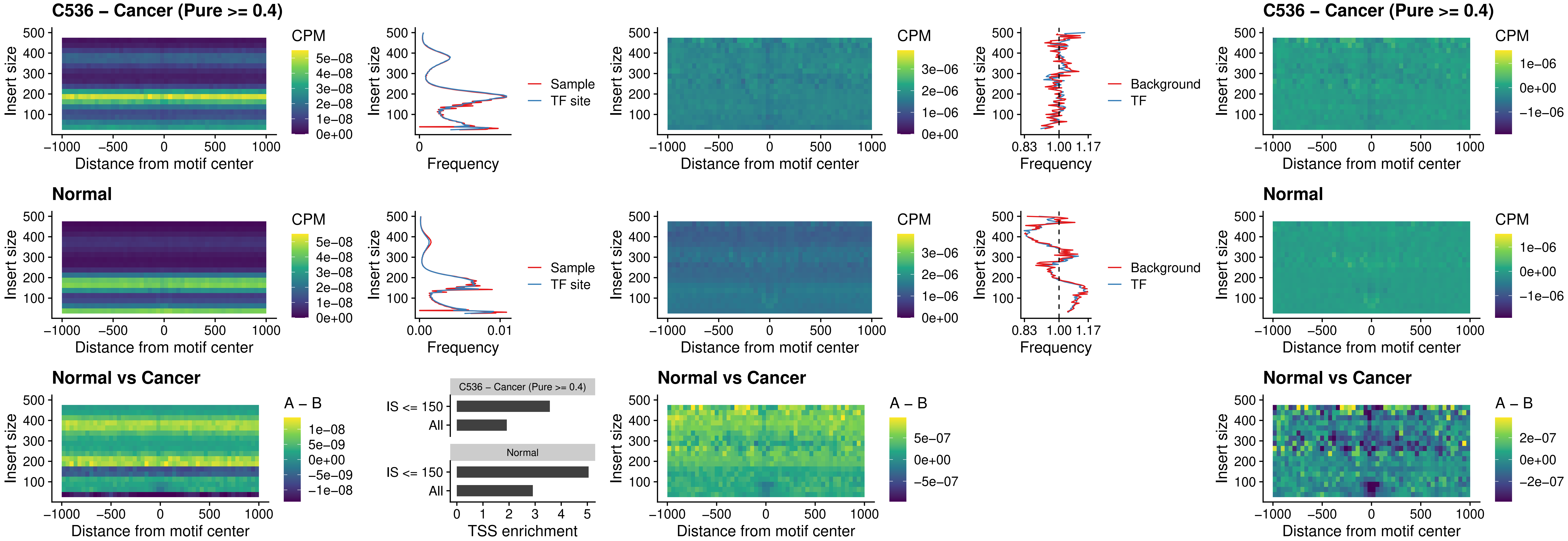

#### C536 – Cancer (Pure >= 0.4) – BCL11B_uc_cTSS_nPEAK

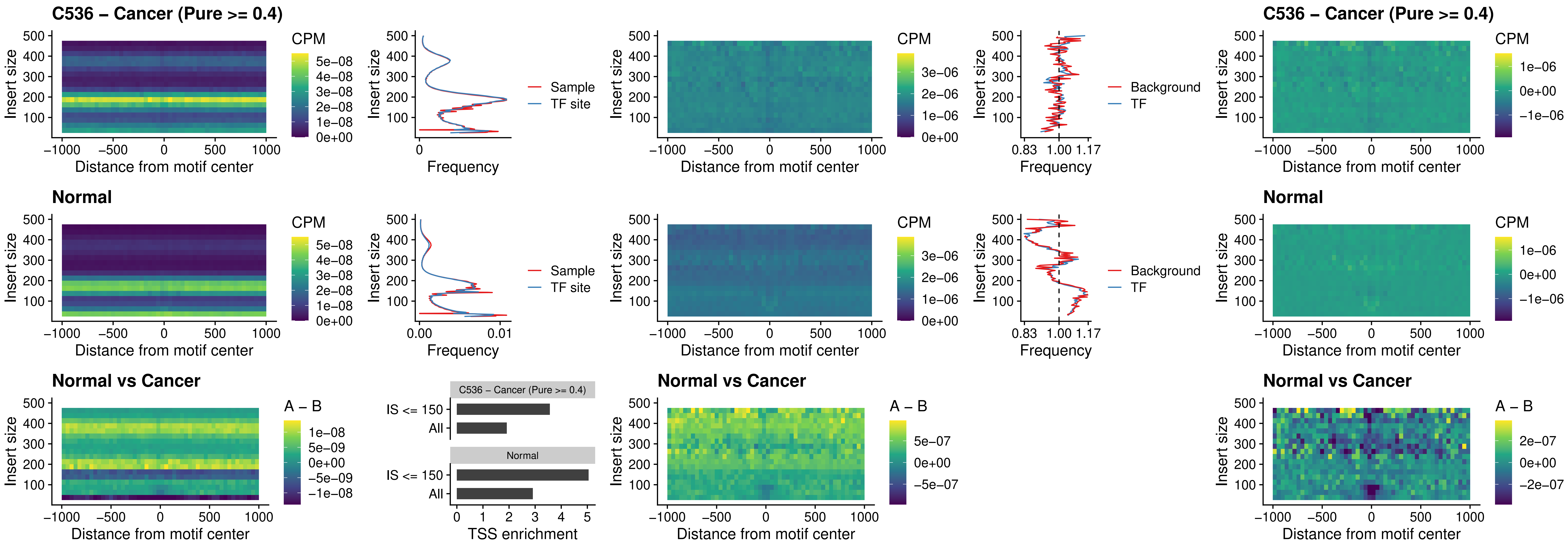

#### C536 – Cancer (Pure >= 0.4) – STAT2_uc_dTSS_oPEAK

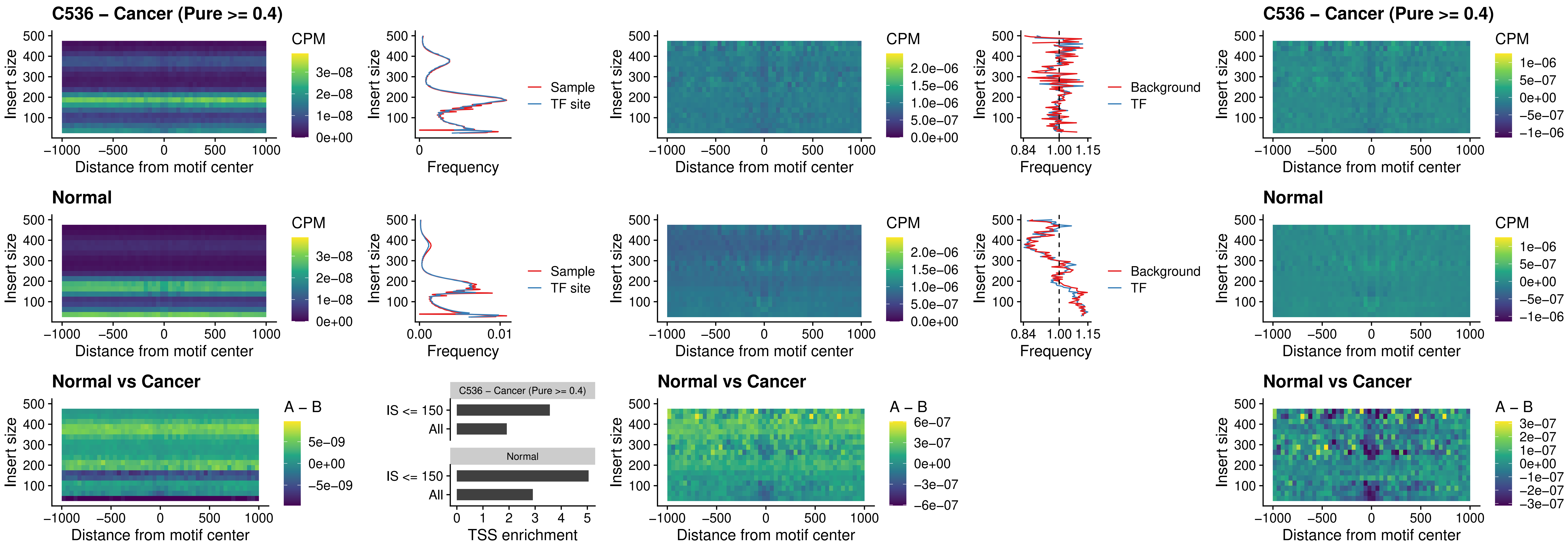

#### C536 – Cancer (Pure >= 0.4) – ZNF683_uc_cTSS_oPEAK

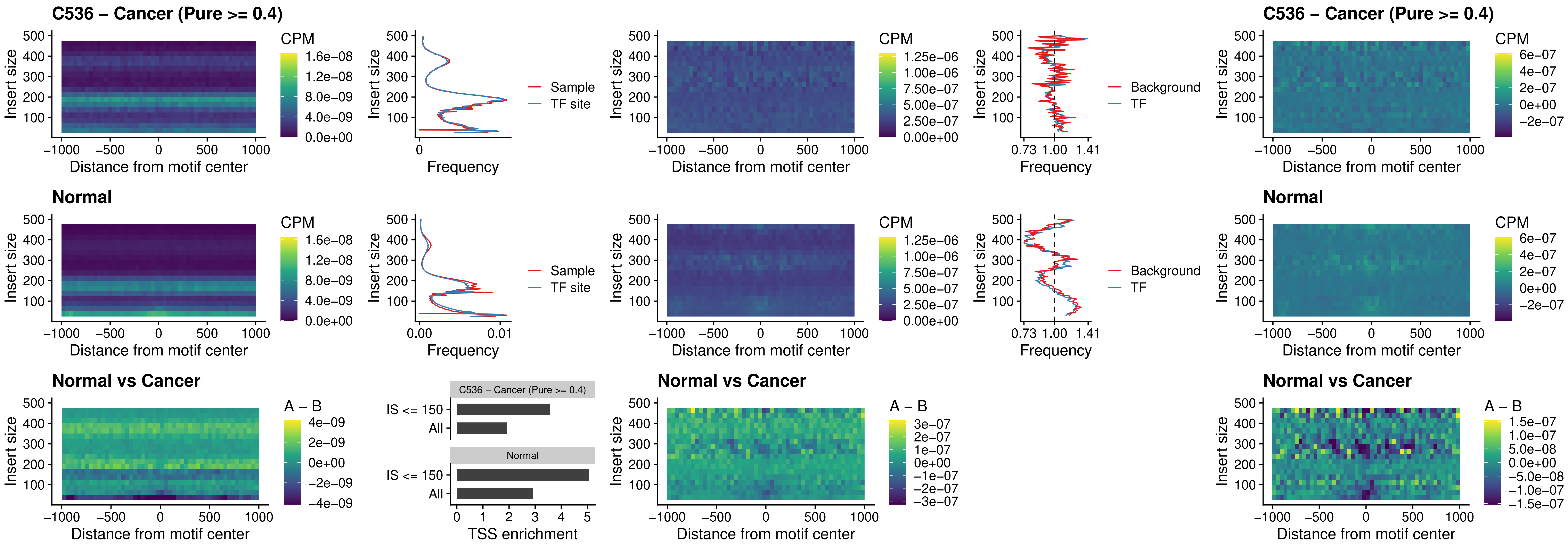

#### C536 – Cancer (Pure >= 0.4) – IRF8_uc_dTSS_nPEAK

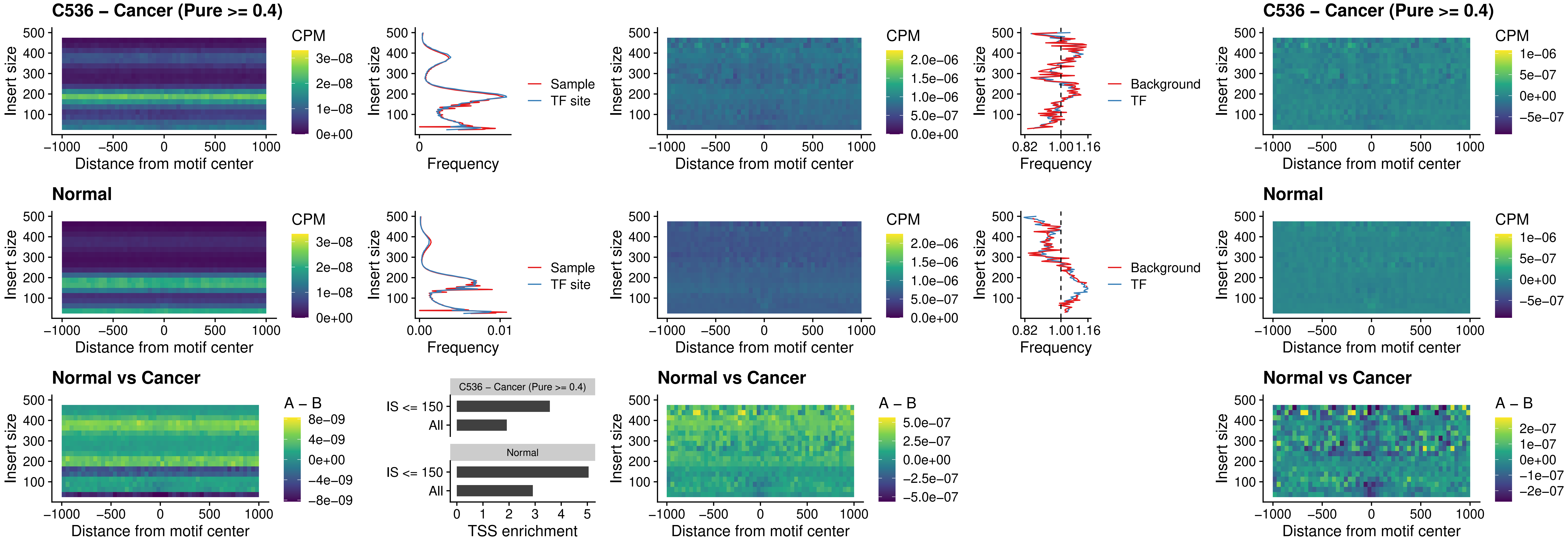

#### C536 – Cancer (Pure >= 0.4) – ETV7_uc_dTSS_oPEAK

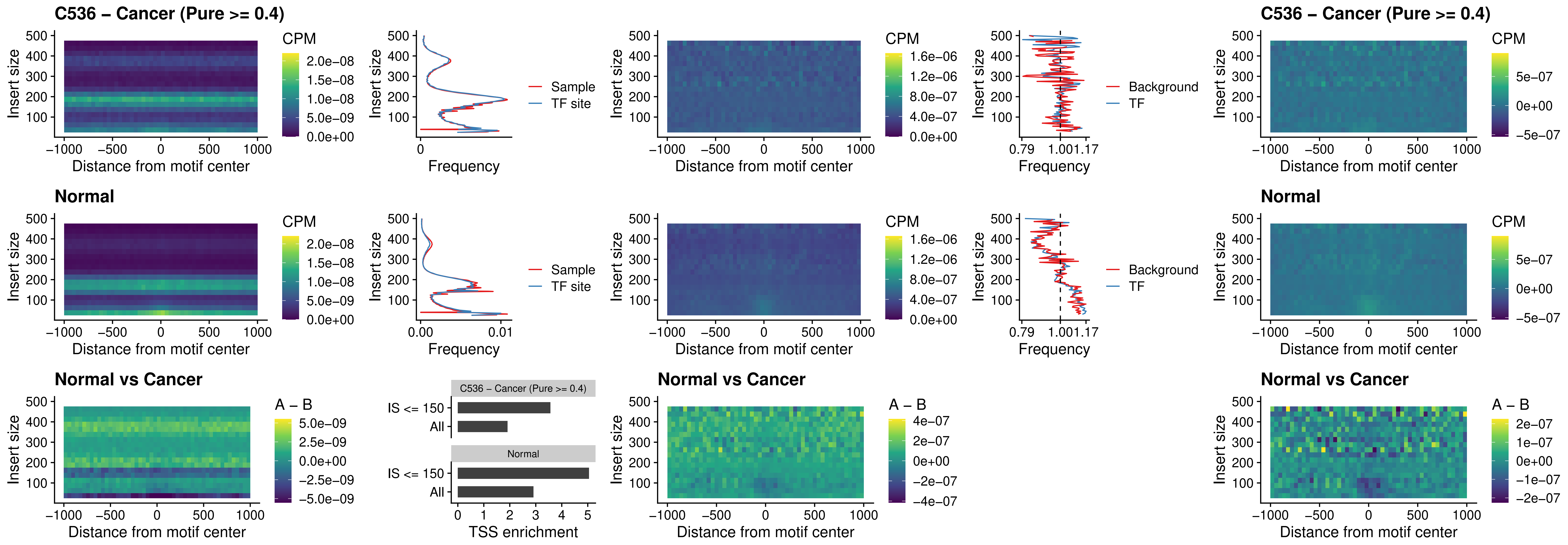

#### C537 – Cancer (Pure >= 0.4) – FOXL1_uc_pTSS_oPEAK

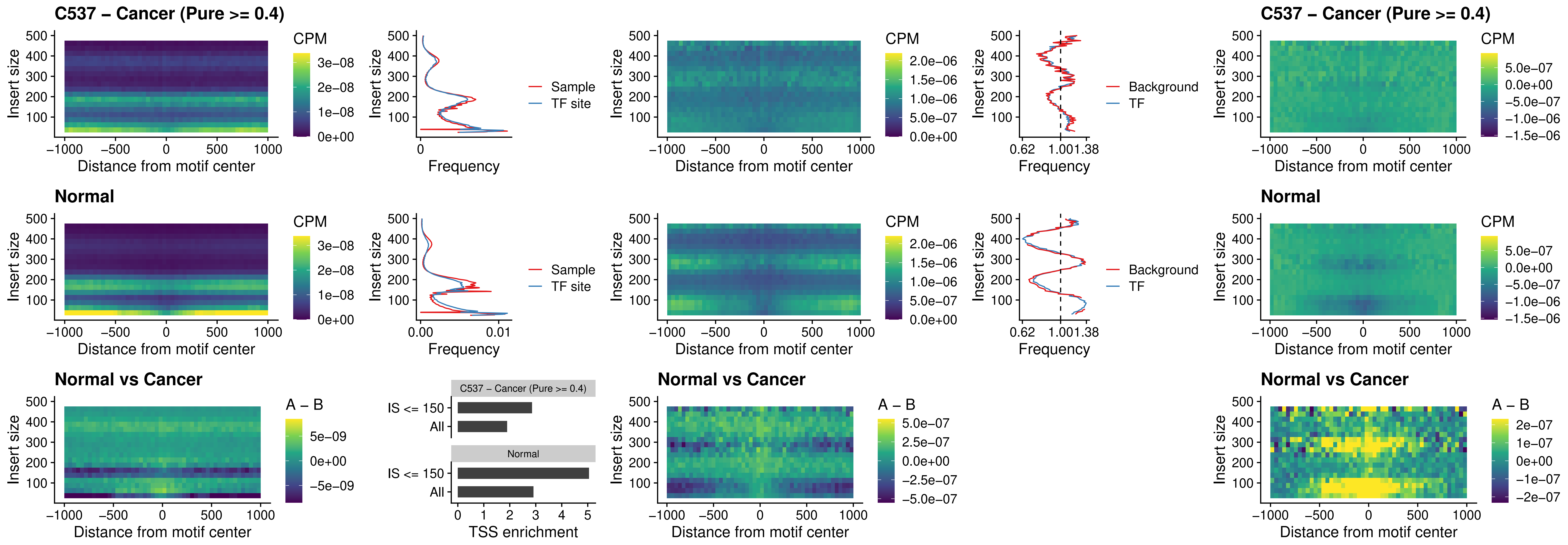

#### C537 – Cancer (Pure >= 0.4) – CPEB1_uc_cTSS_oPEAK

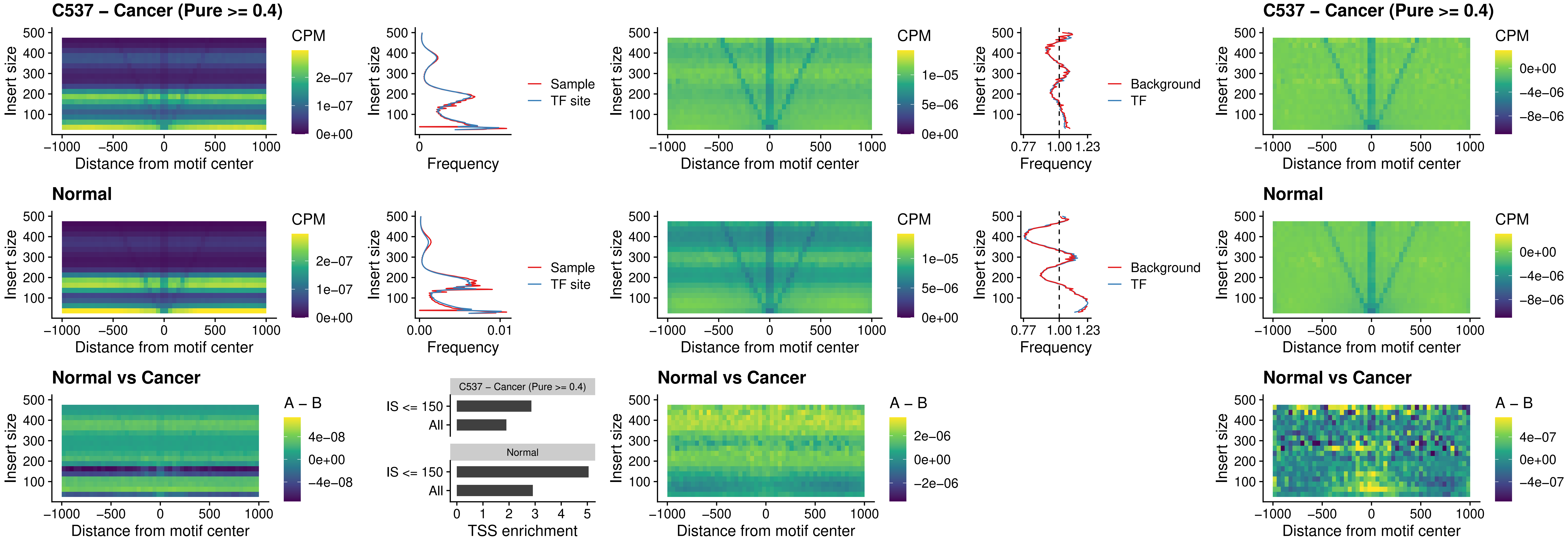

#### C537 – Cancer (Pure >= 0.4) – HMGA1_uc_pTSS_nPEAK

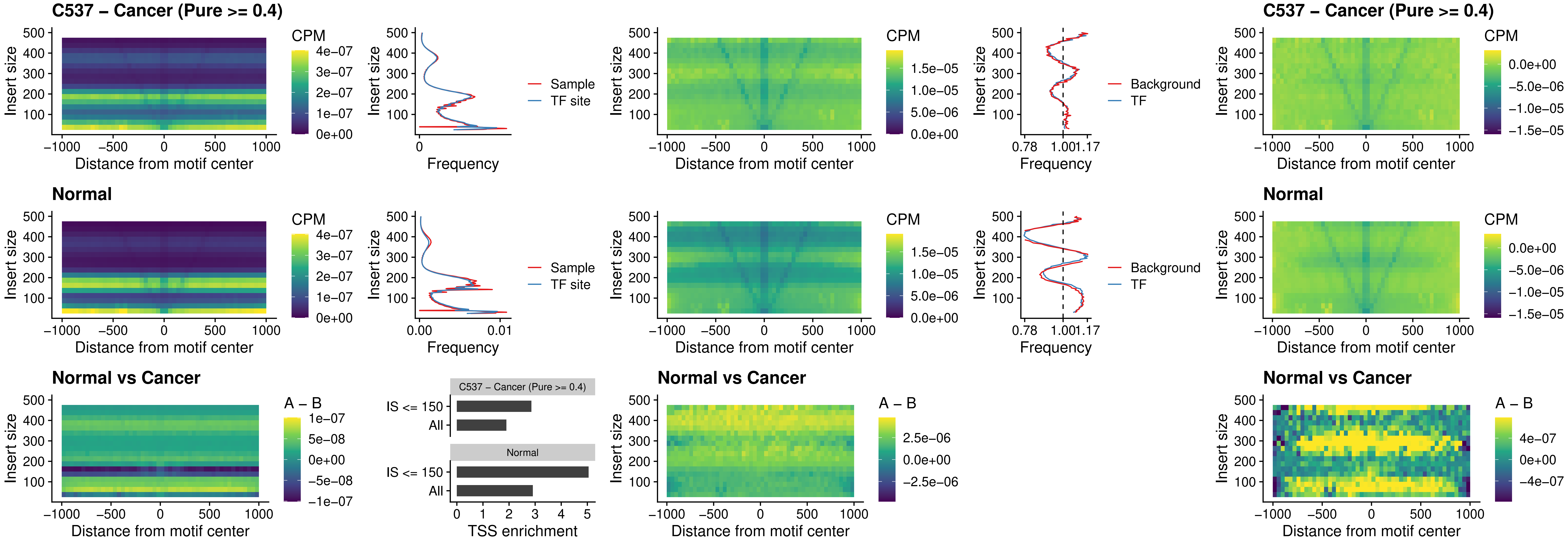

#### C537 – Cancer (Pure >= 0.4) – SOX13_uc_cTSS_nPEAK

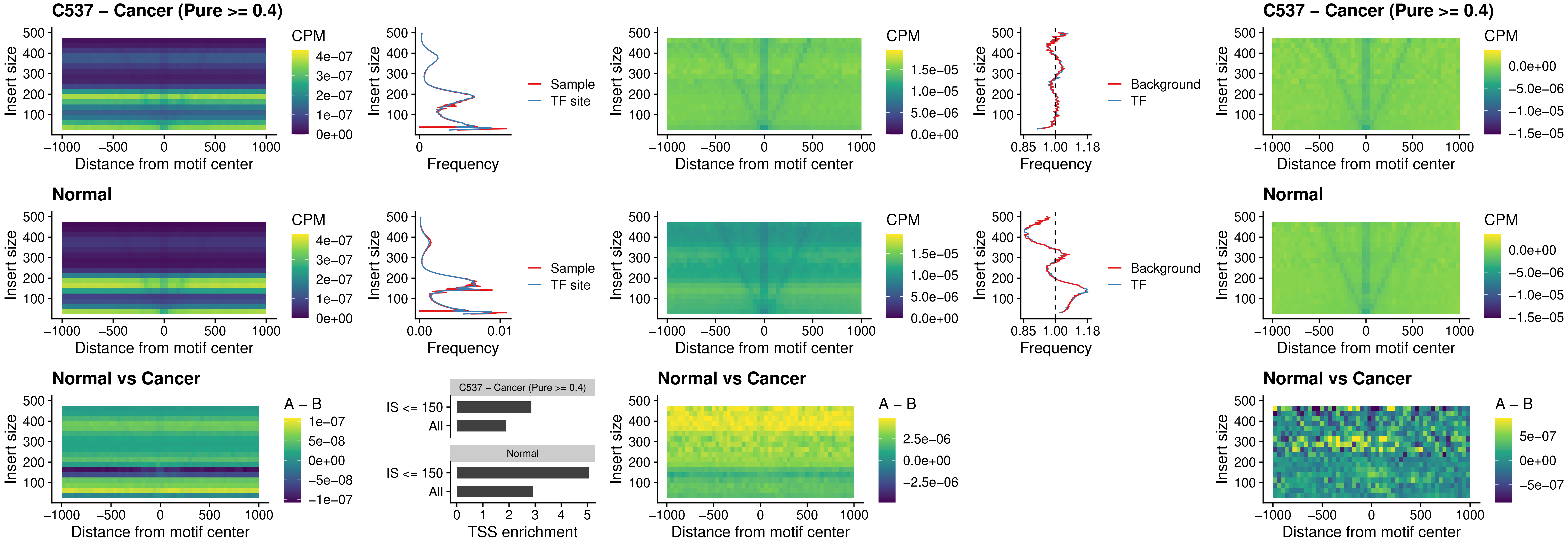

### C537 – Cancer (Pure >= 0.4) – LHX4_uc_cTSS_nPEAK

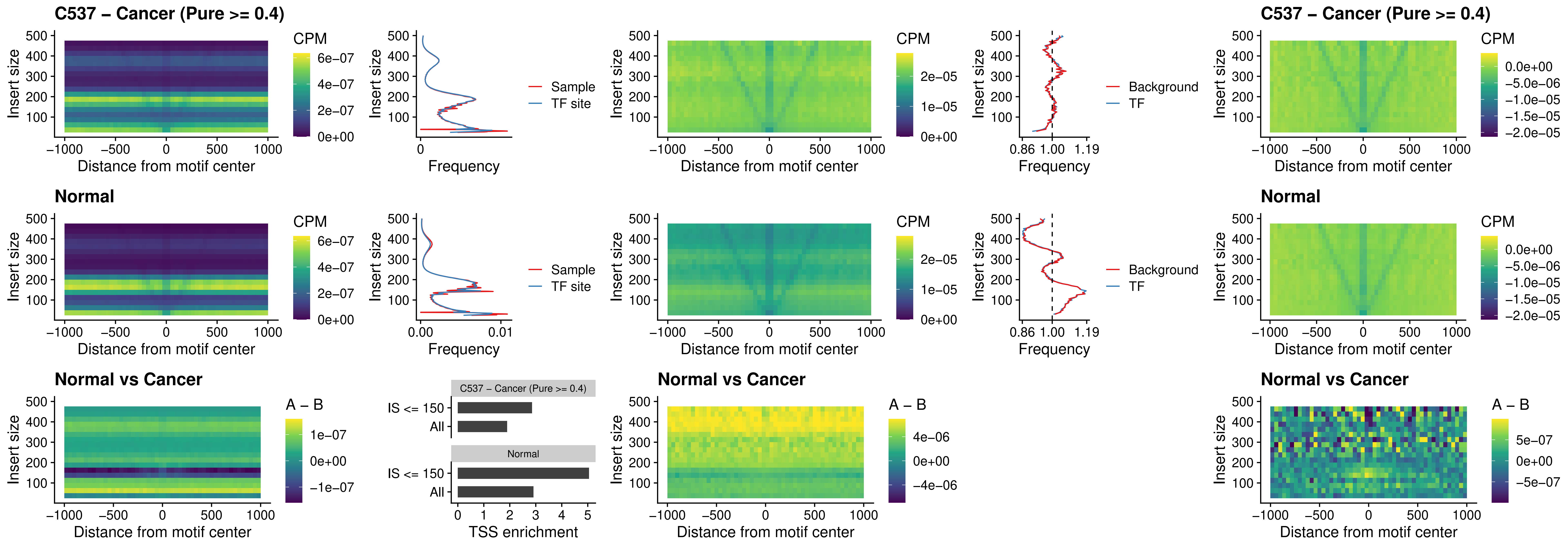

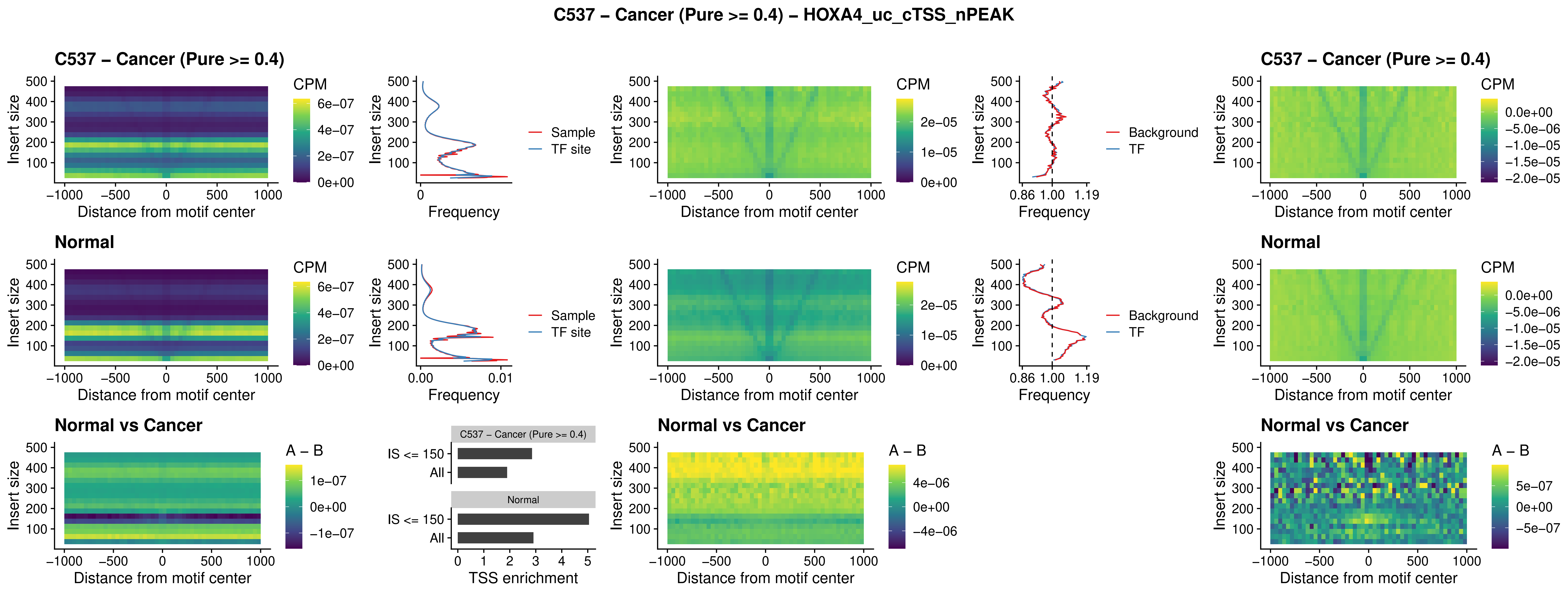
C537 – Cancer (Pure >= 0.4) – HOXA4_uc_cTSS_nPEAK
C537 – Cancer (Pure >= 0.4)
Normal
Normal vs Cancer
Insert size
Distance from motif center
Frequency
CPM
A – B
Sample
TF site
Background
TF
C537 – Cancer (Pure >= 0.4)
Normal
IS <= 150
All
TSS enrichment

**C537 – Cancer (Pure >= 0.4) – HLX_uc_cTSS_nPEAK**

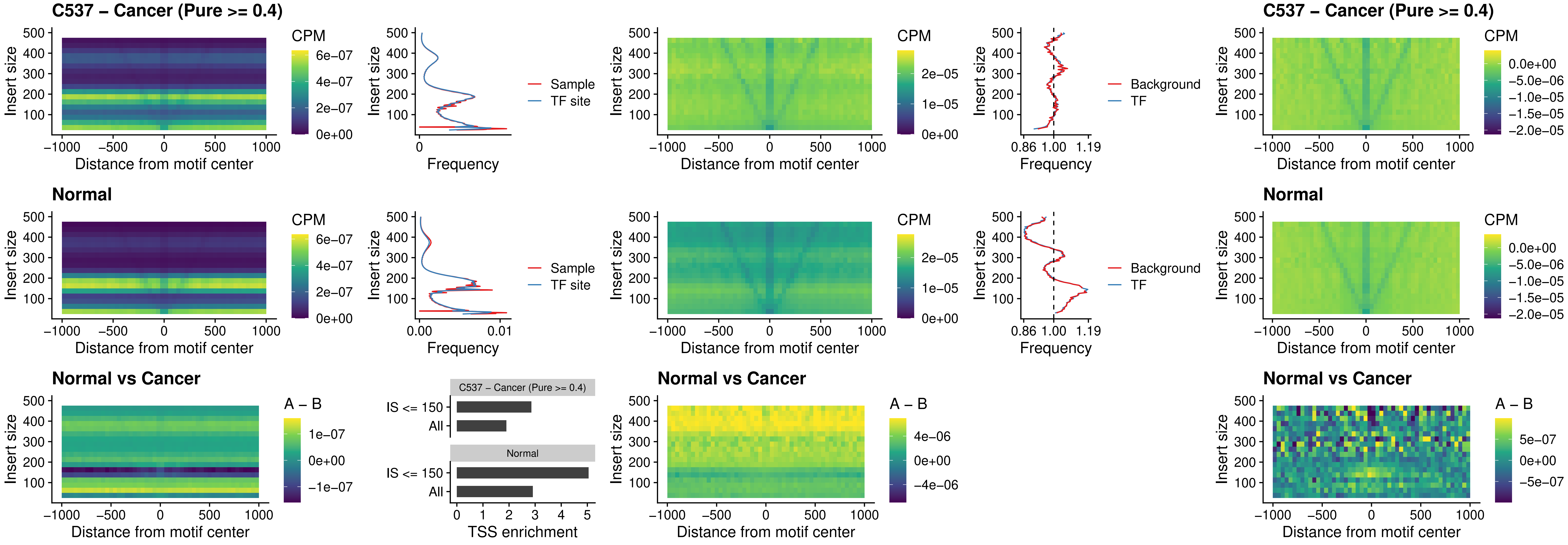

### C537 – Cancer (Pure >= 0.4) – AC0125311_uc_cTSS_nPEAK

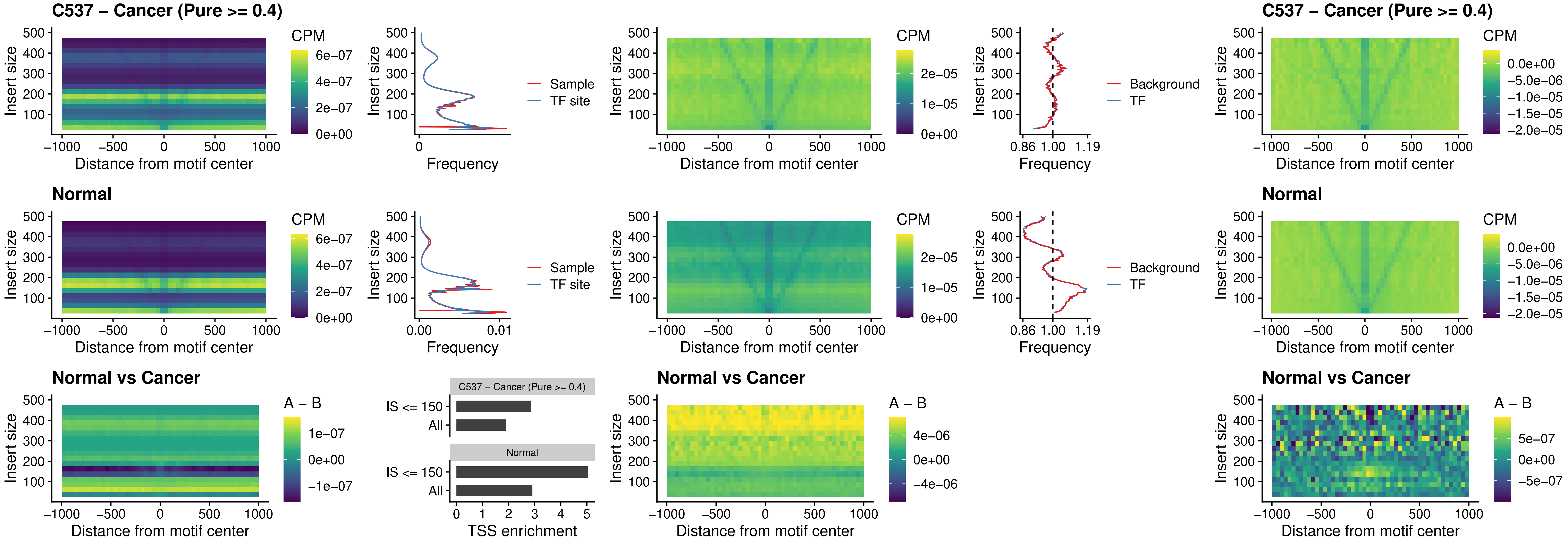

#### C537 – Cancer (Pure >= 0.4) – DBX2_uc_cTSS_nPEAK

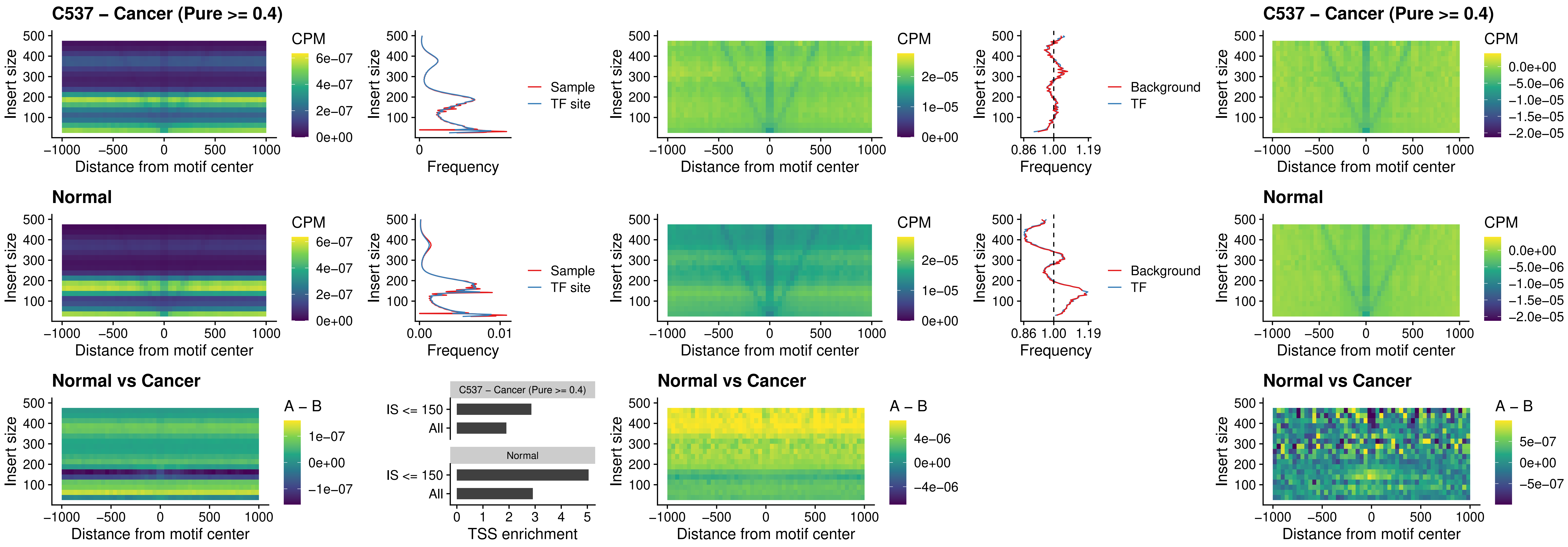

#### C537 – Cancer (Pure >= 0.4) – LHX6_uc_dTSS_nPEAK

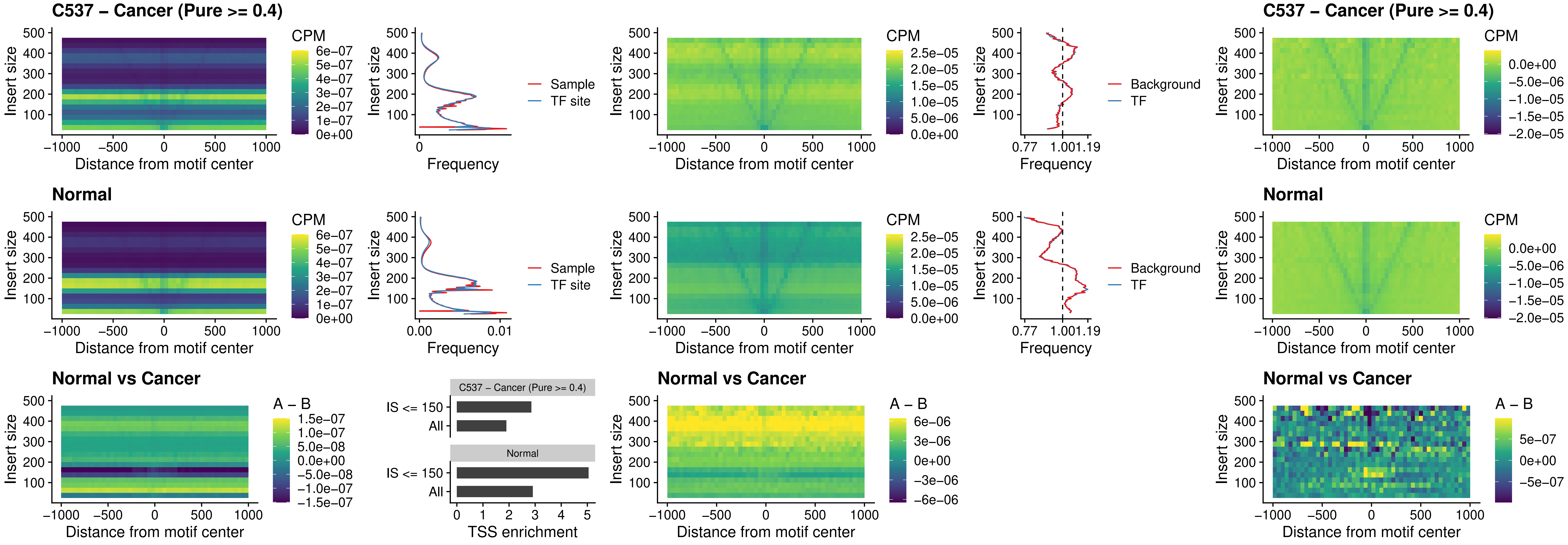

#### C537 – Cancer (Pure >= 0.4) – SOX1_uc_dTSS_nPEAK

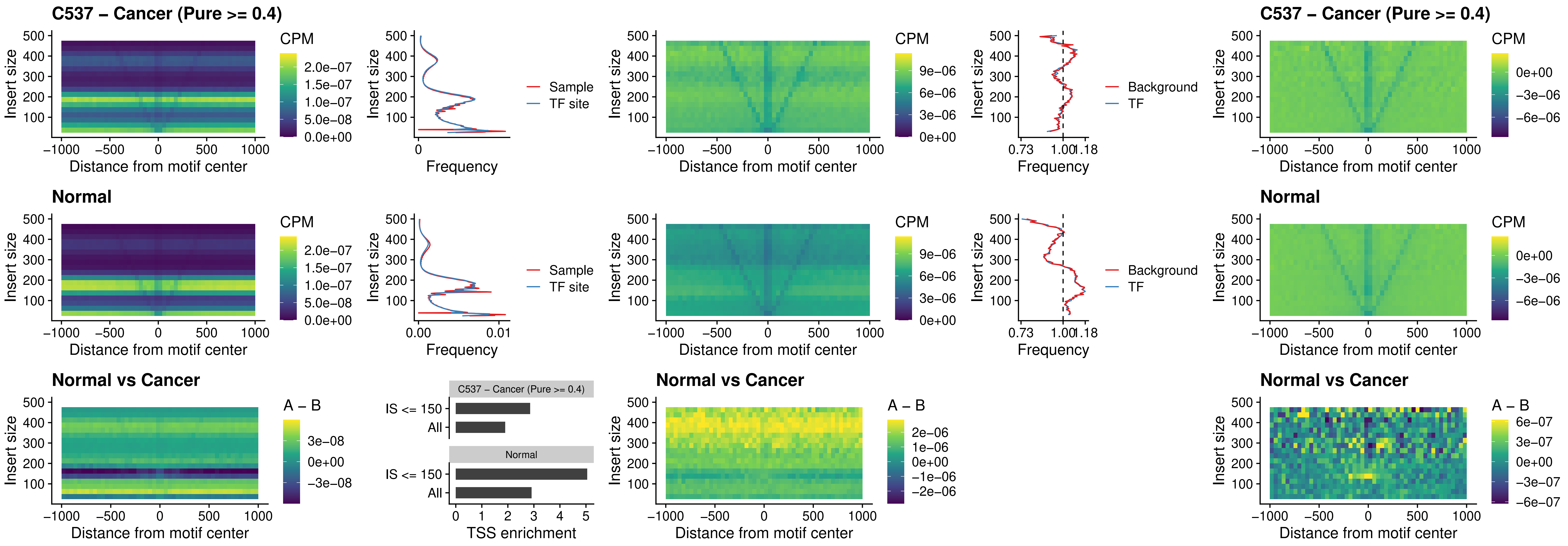

#### C537 – Cancer (Pure >= 0.4) – NKX24_uc_dTSS_nPEAK

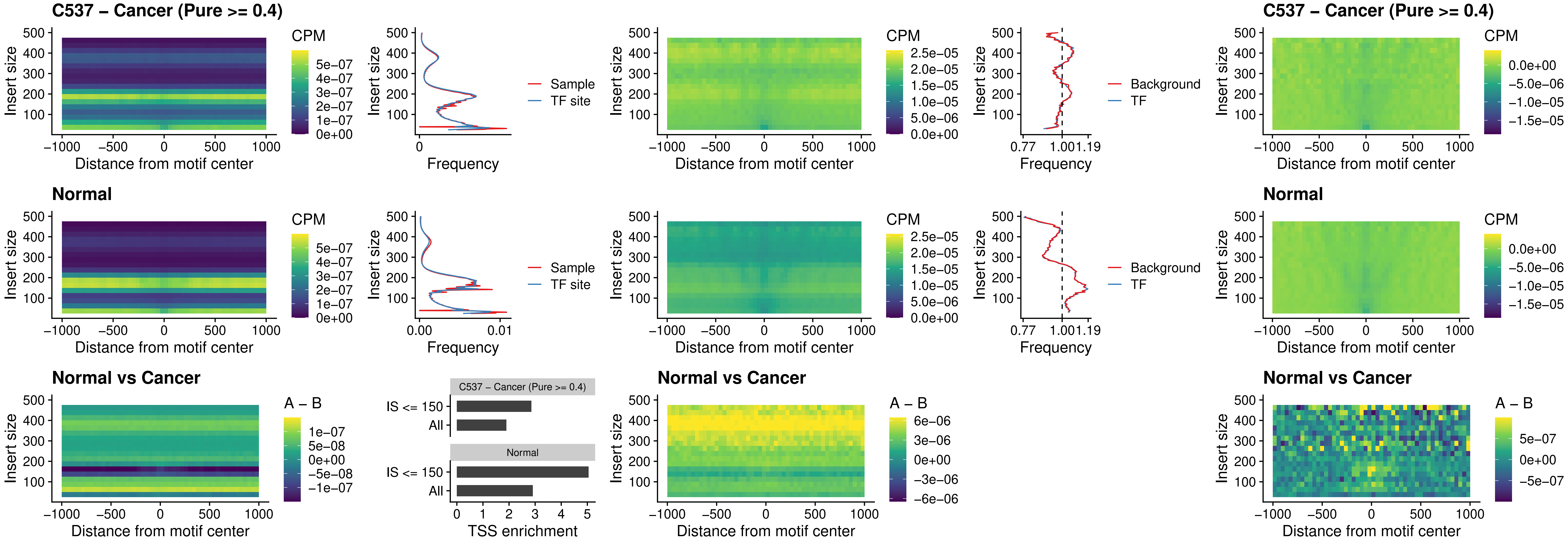

#### C537 – Cancer (Pure >= 0.4) – DLX3_uc_cTSS_nPEAK

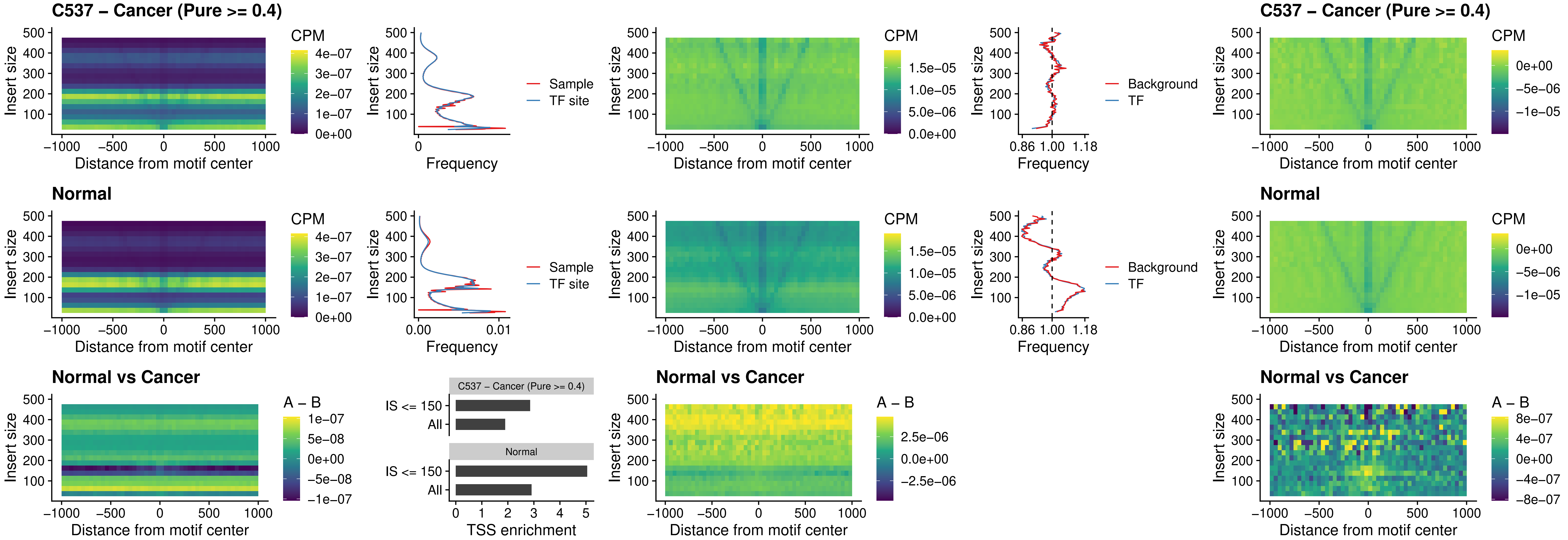

#### C537 – Cancer (Pure >= 0.4) – HOXD3_uc_cTSS_nPEAK

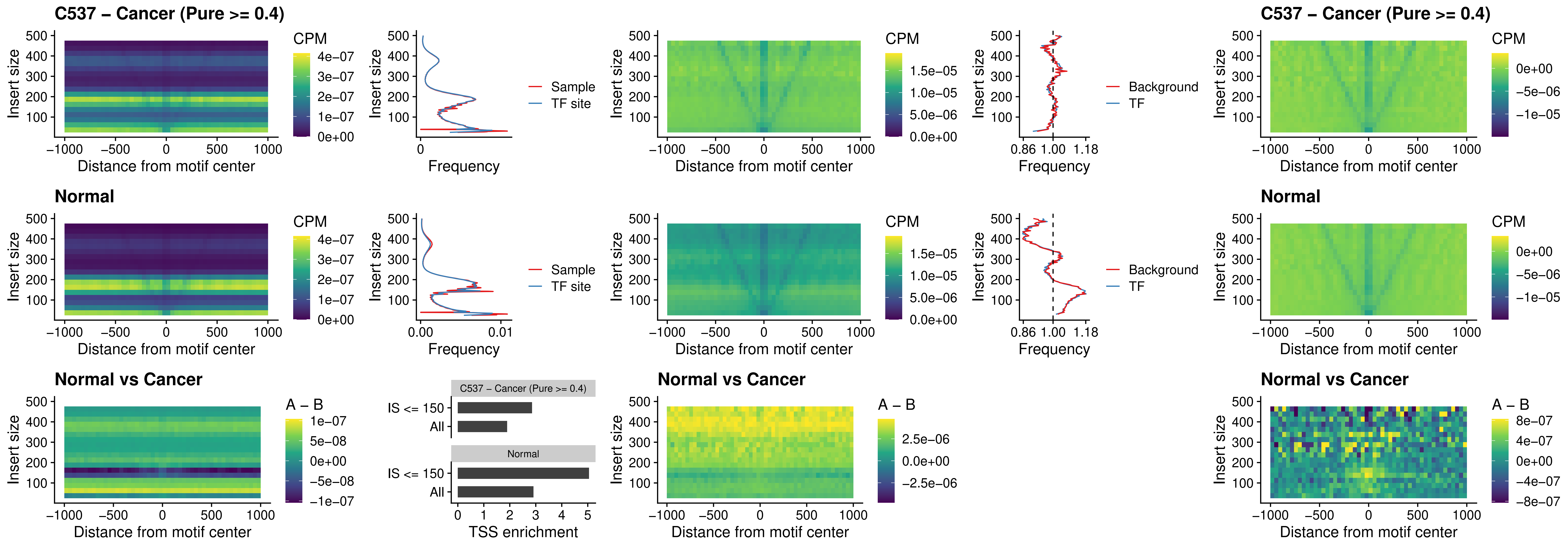

#### C537 – Cancer (Pure >= 0.4) – ISL1_uc_dTSS_oPEAK

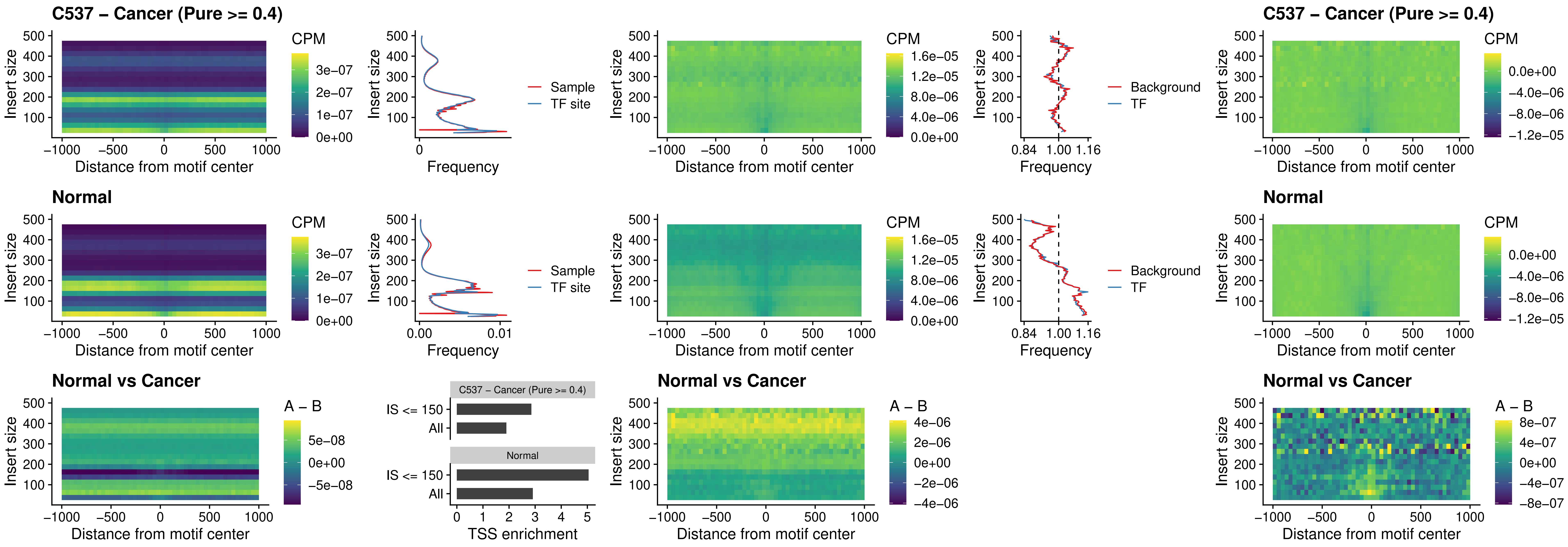

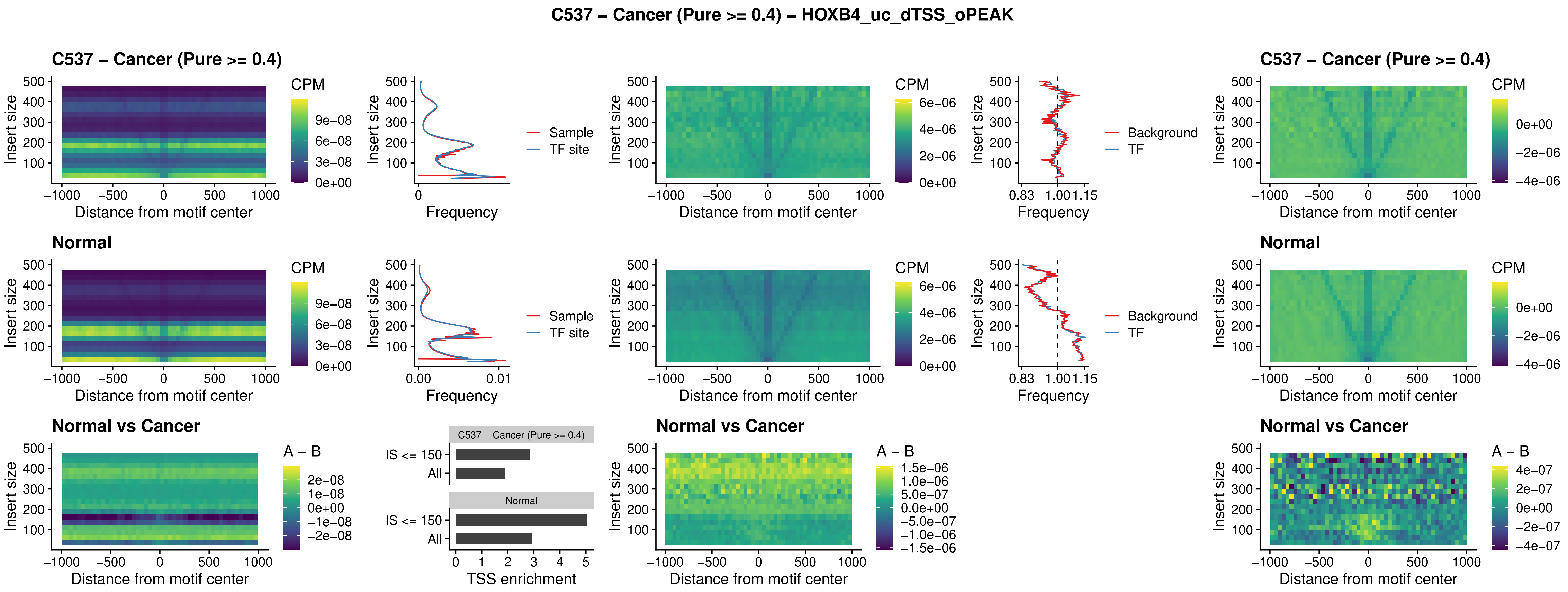
C537 – Cancer (Pure >= 0.4) – HOXB4_uc_dTSS_oPEAK
C537 – Cancer (Pure >= 0.4)
Normal
Normal vs Cancer
Insert size
Distance from motif center
Frequency
CPM
A – B
Sample
TF site
Background
TF
C537 – Cancer (Pure >= 0.4)
IS <= 150
All
TSS enrichment

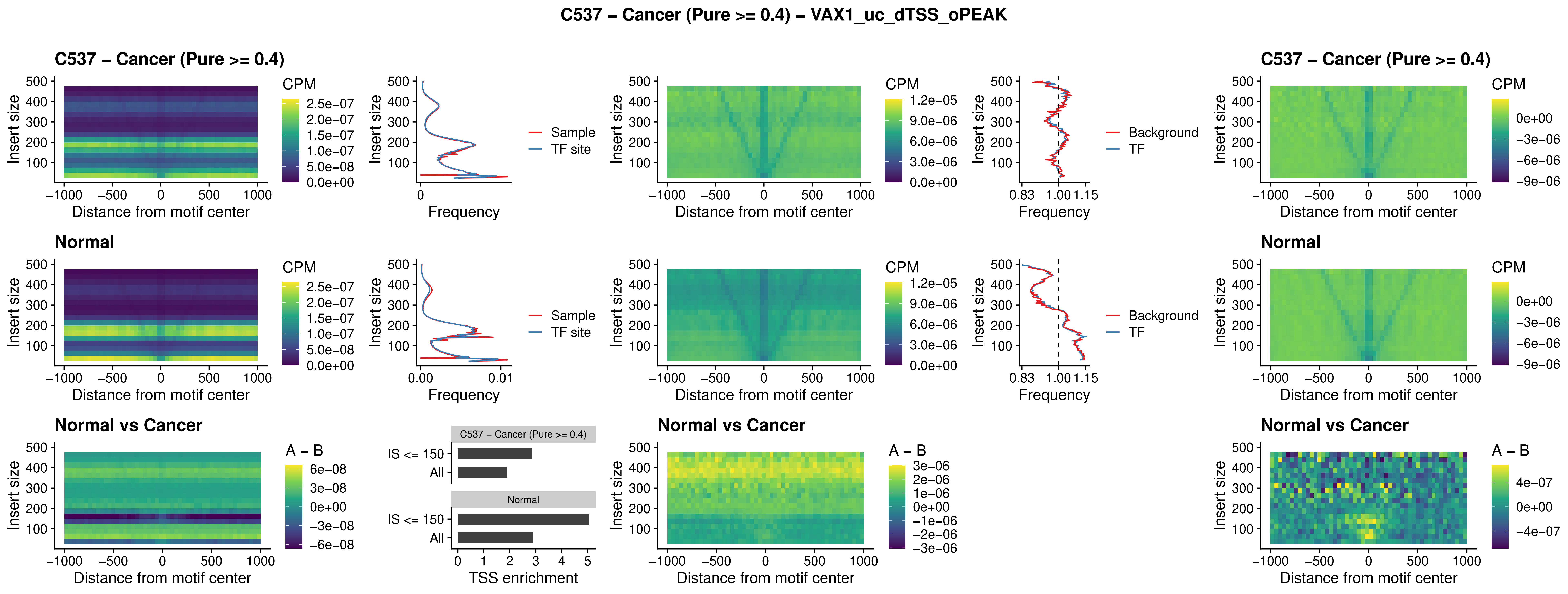
C537 – Cancer (Pure >= 0.4) – VAX1_uc_dTSS_oPEAK
C537 – Cancer (Pure >= 0.4)
Normal
Normal vs Cancer
Insert size
Distance from motif center
Frequency
CPM
A – B
Sample
TF site
Background
TF
C537 – Cancer (Pure >= 0.4)
Normal
IS <= 150
All
TSS enrichment

#### C537 – Cancer (Pure >= 0.4) – DLX3_uc_dTSS_oPEAK

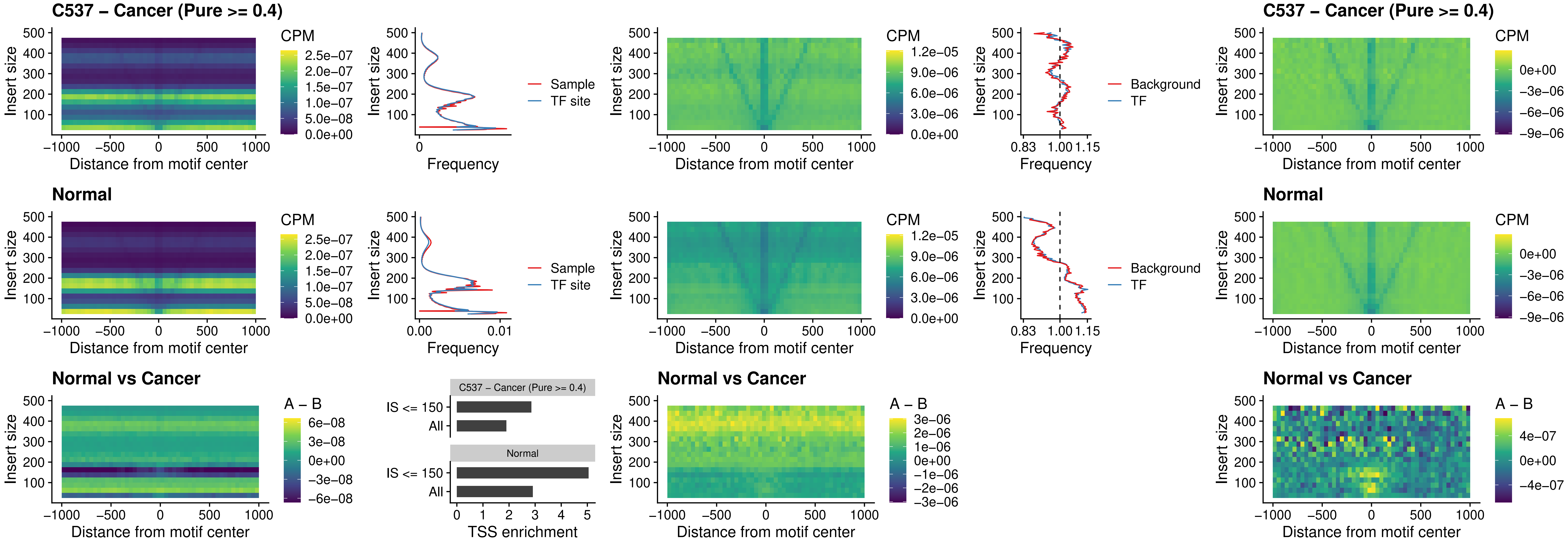

#### C537 – Cancer (Pure >= 0.4) – HOXD3_uc_dTSS_oPEAK

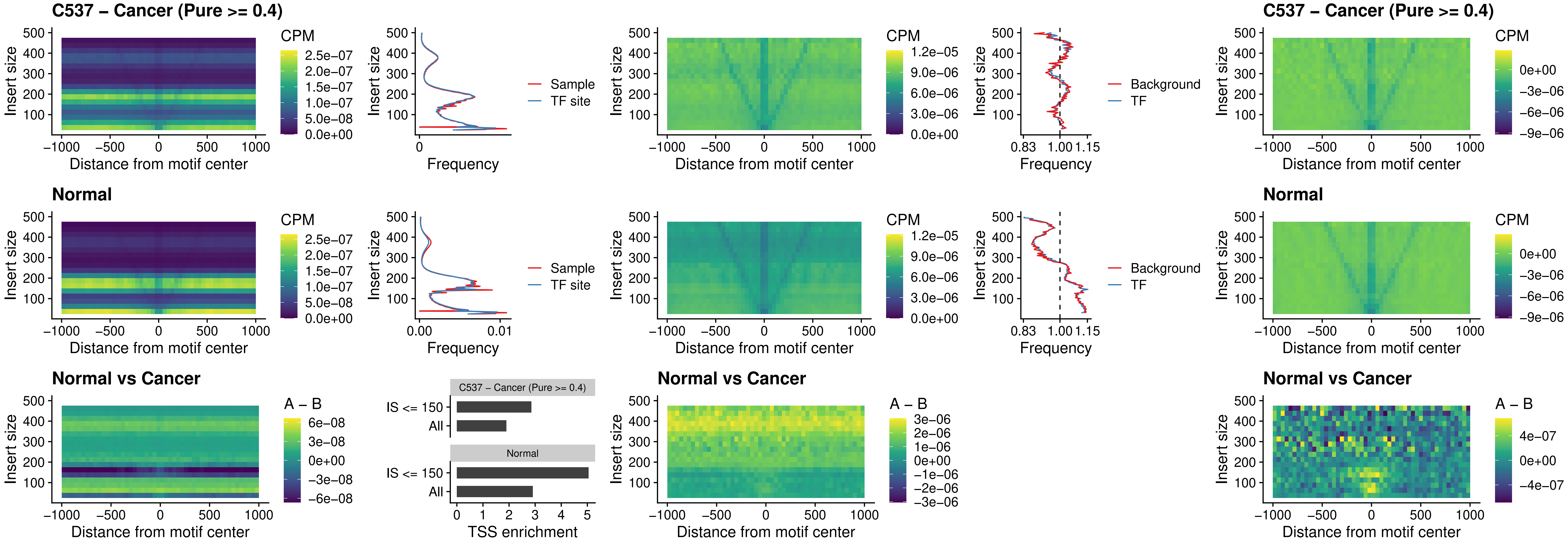

#### C537 – Cancer (Pure >= 0.4) – BBX_uc_dTSS_nPEAK

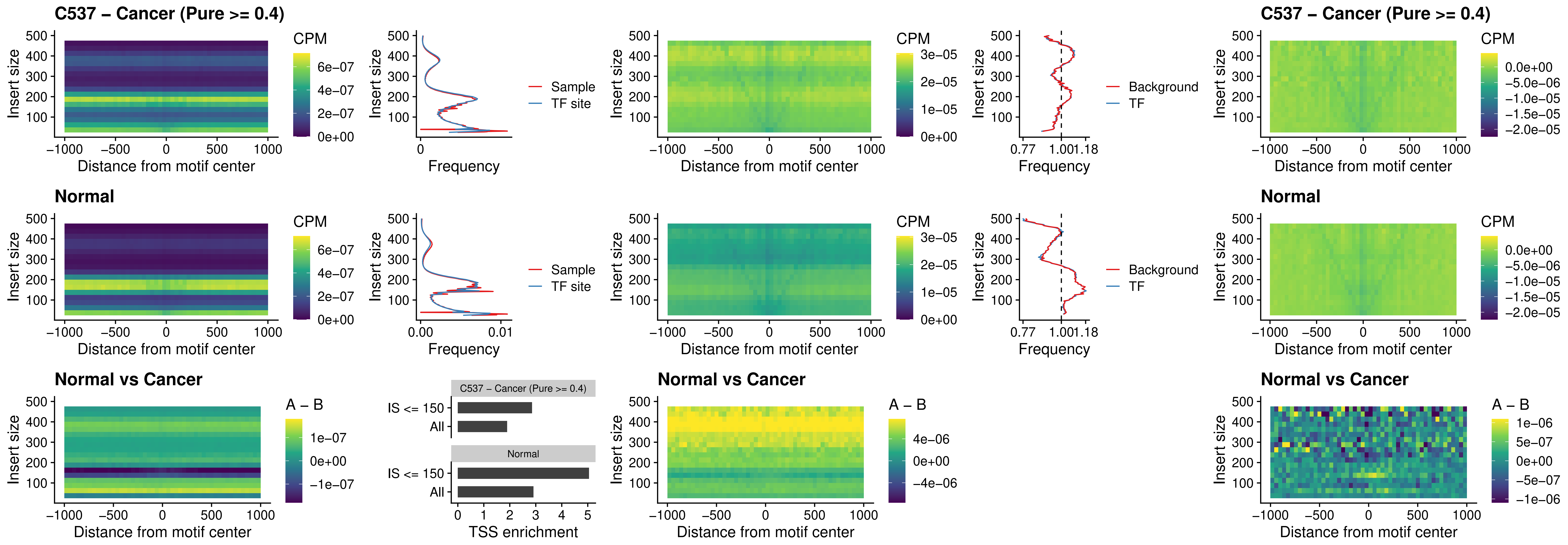

#### C537 – Cancer (Pure >= 0.4) – SOX3_uc_cTSS_nPEAK

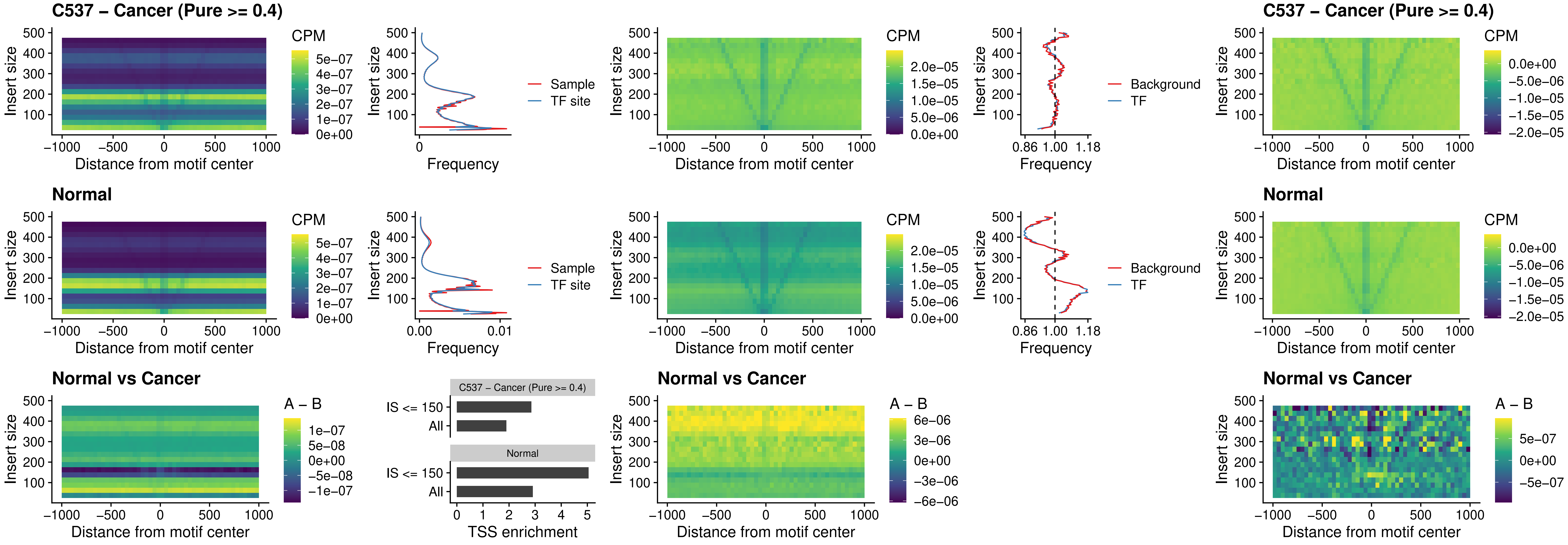

#### C537 – Cancer (Pure >= 0.4) – TBPL2_uc_cTSS_nPEAK

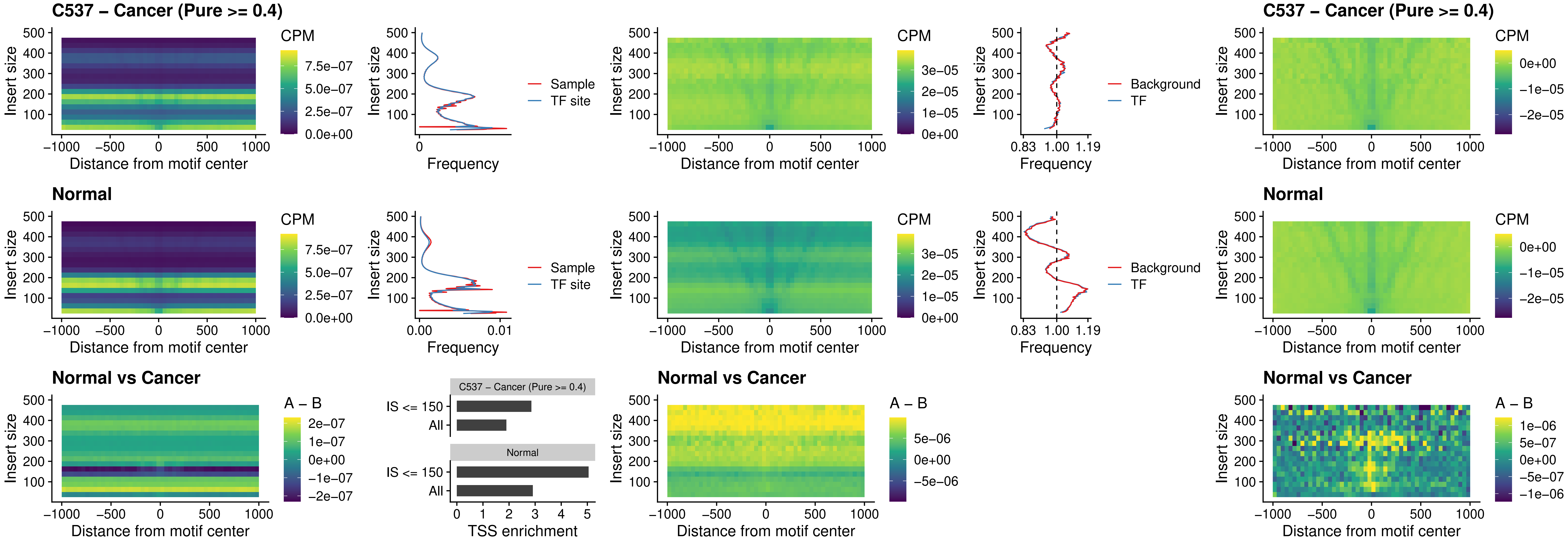

#### C537 – Cancer (Pure >= 0.4) – HOXA4_uc_cTSS_oPEAK

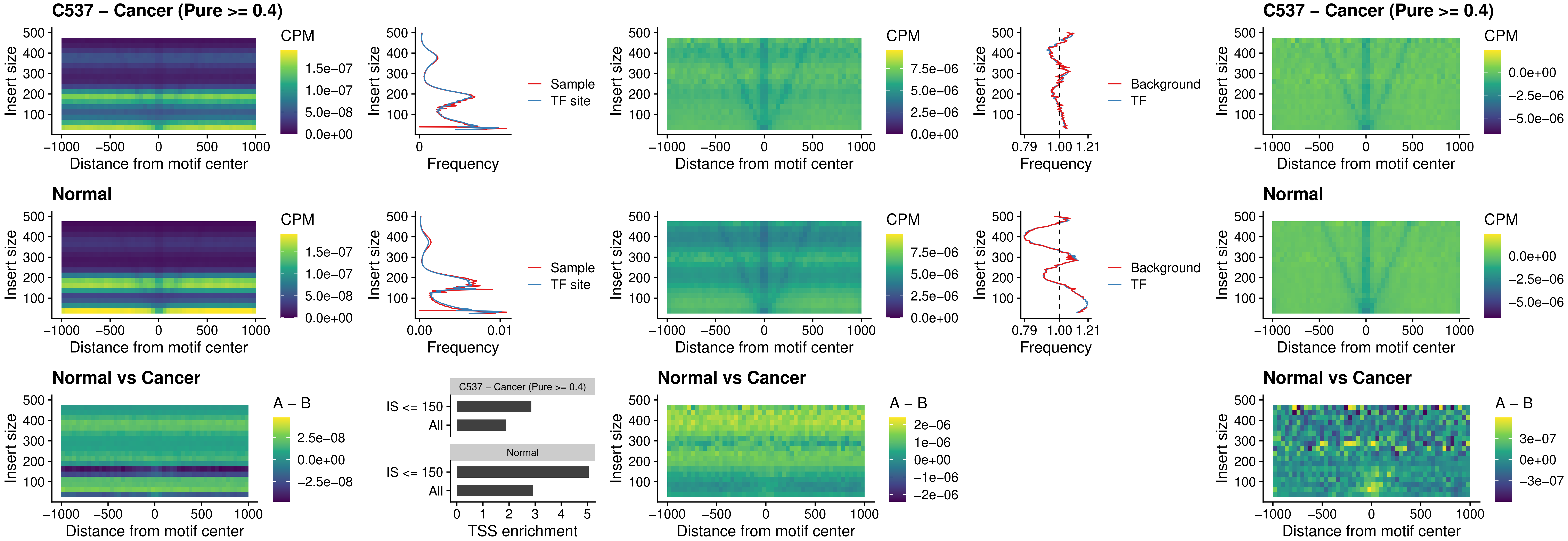

### C537 – Cancer (Pure >= 0.4) – PDX1_uc_dTSS_oPEAK

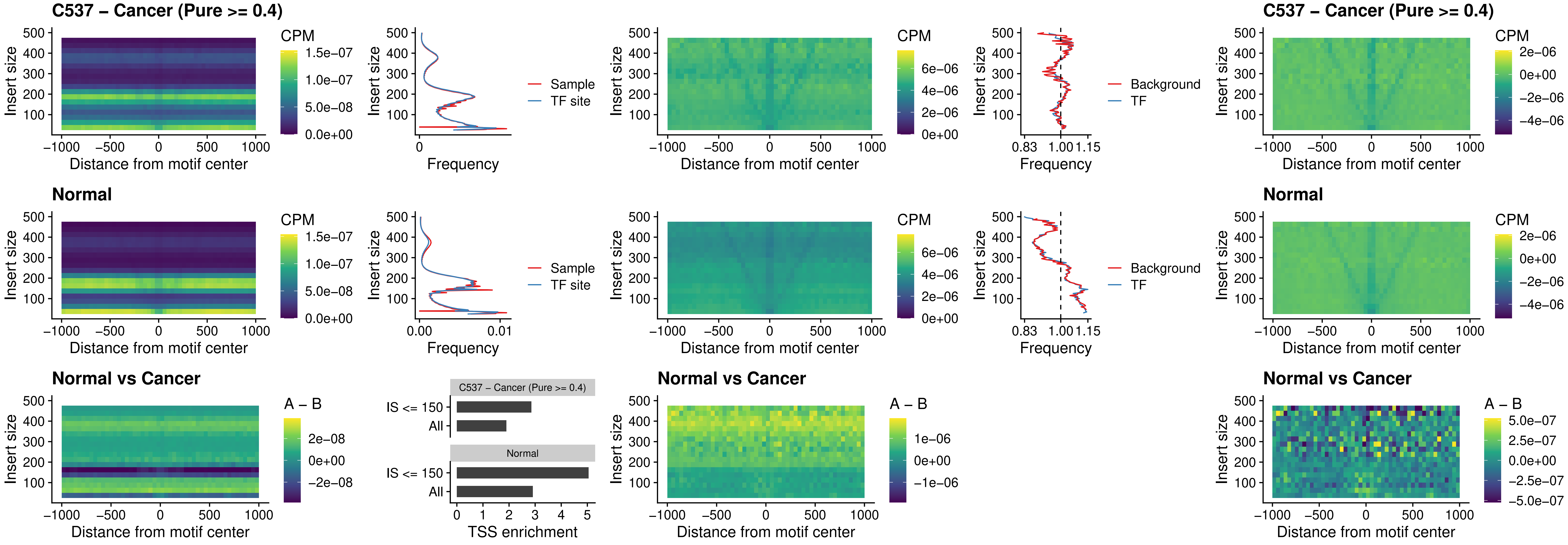

### C537 – Cancer (Pure >= 0.4) – LHX4_uc_dTSS_oPEAK

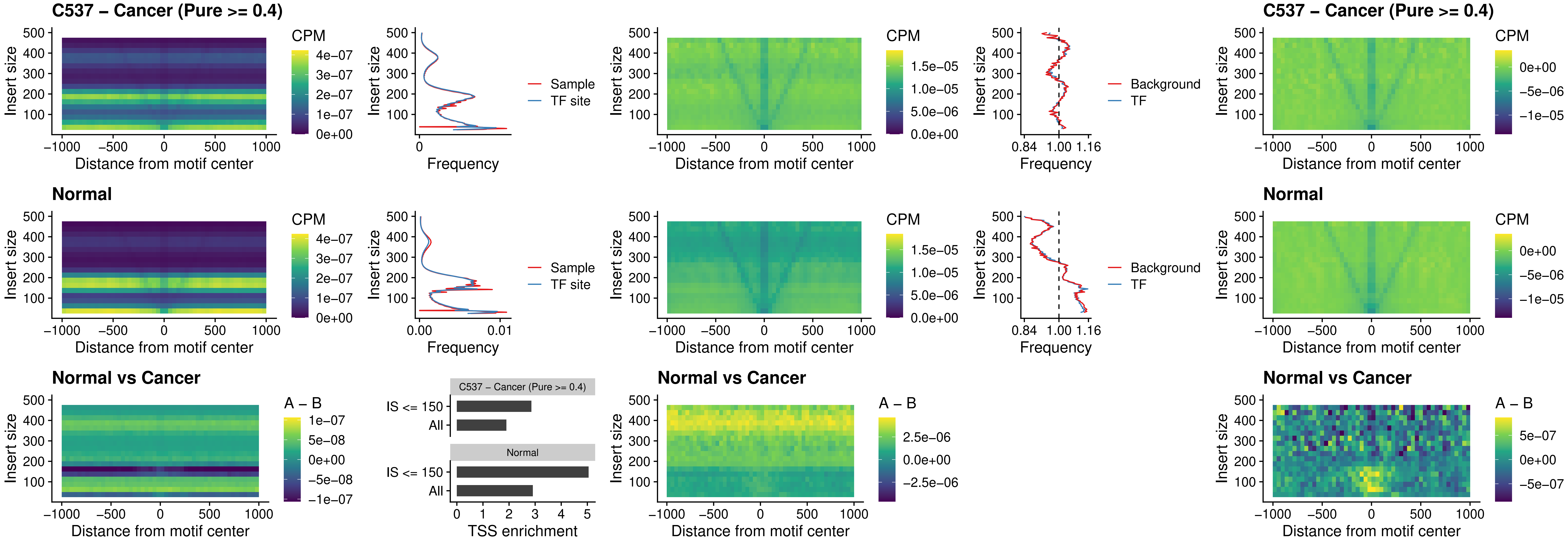

### C537 – Cancer (Pure >= 0.4) – HOXA4_uc_dTSS_oPEAK

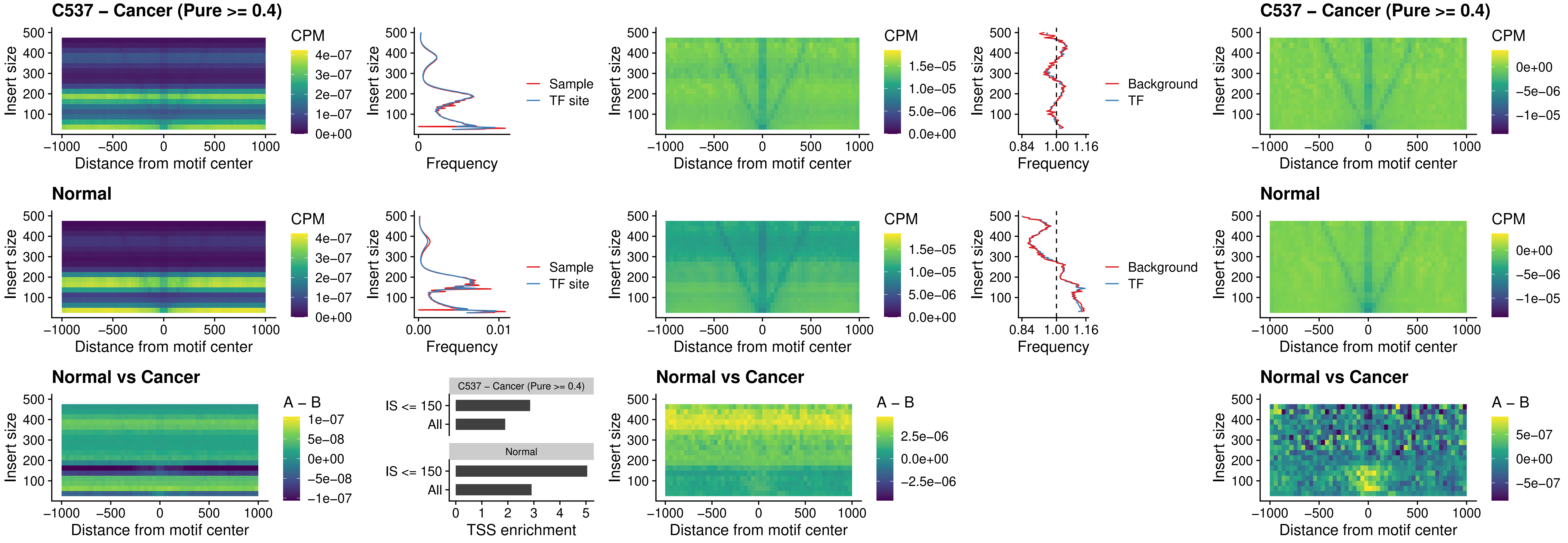

#### C537 – Cancer (Pure >= 0.4) – HLX_uc_dTSS_oPEAK

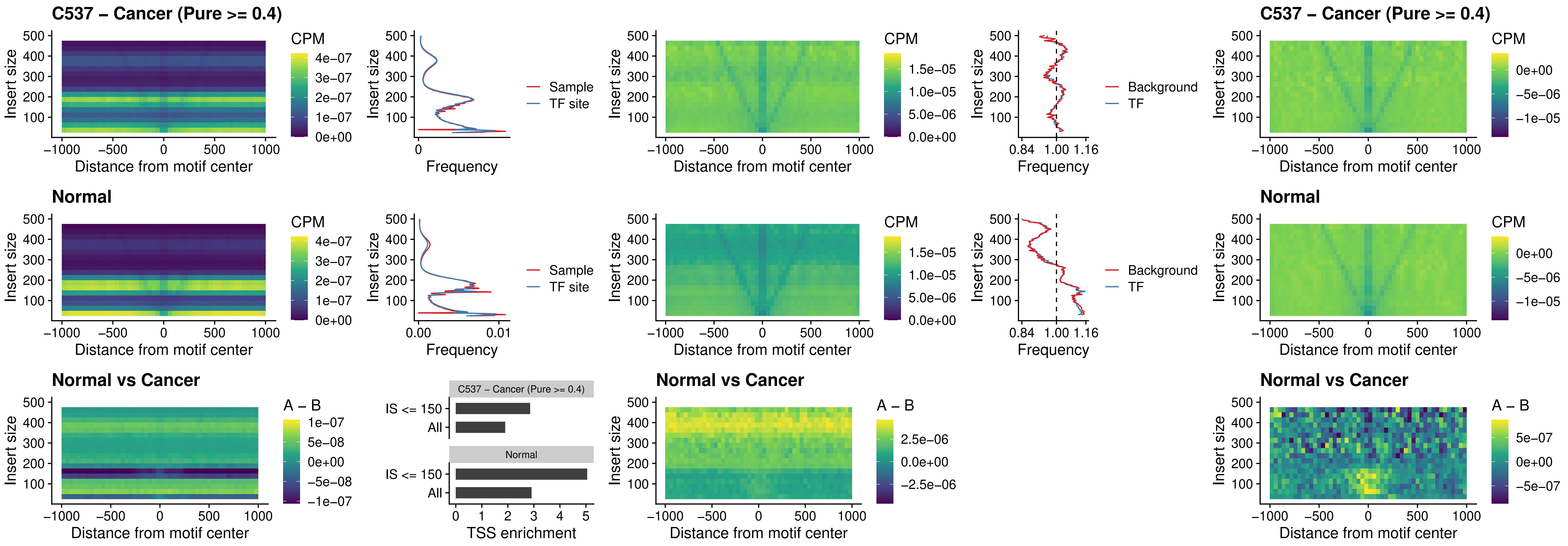

#### C537 – Cancer (Pure >= 0.4) – AC0125311_uc_dTSS_oPEAK

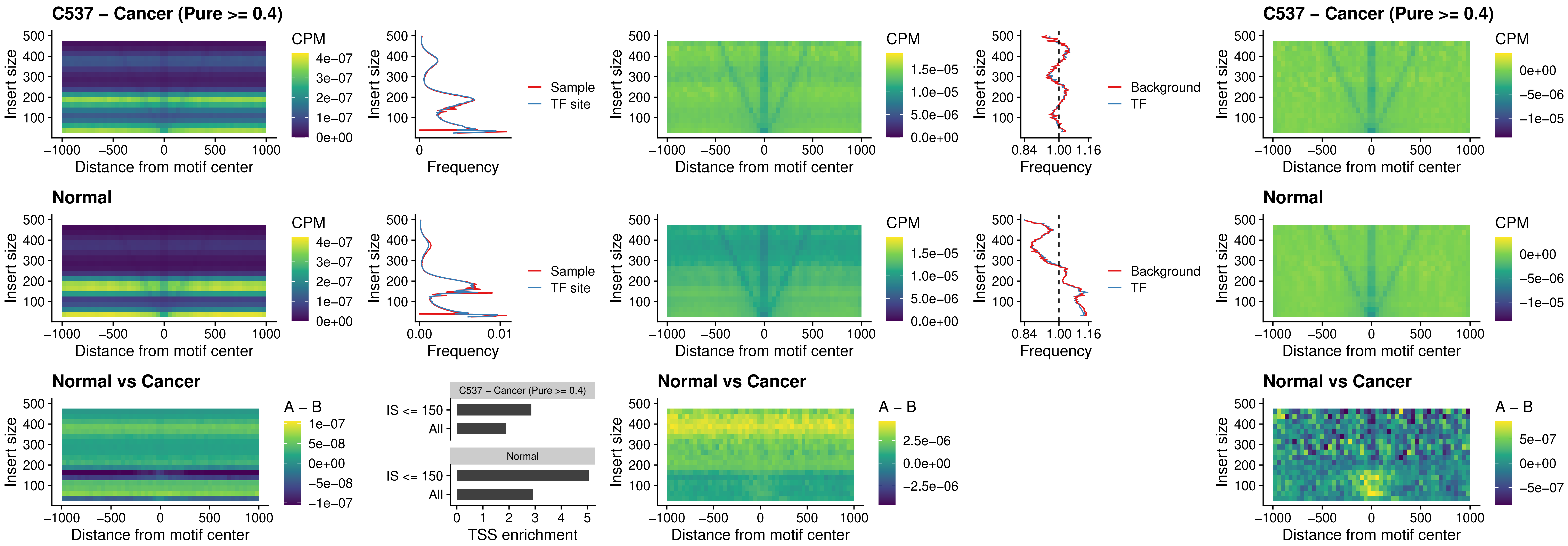

### C537 – Cancer (Pure >= 0.4) – DBX2_uc_dTSS_oPEAK

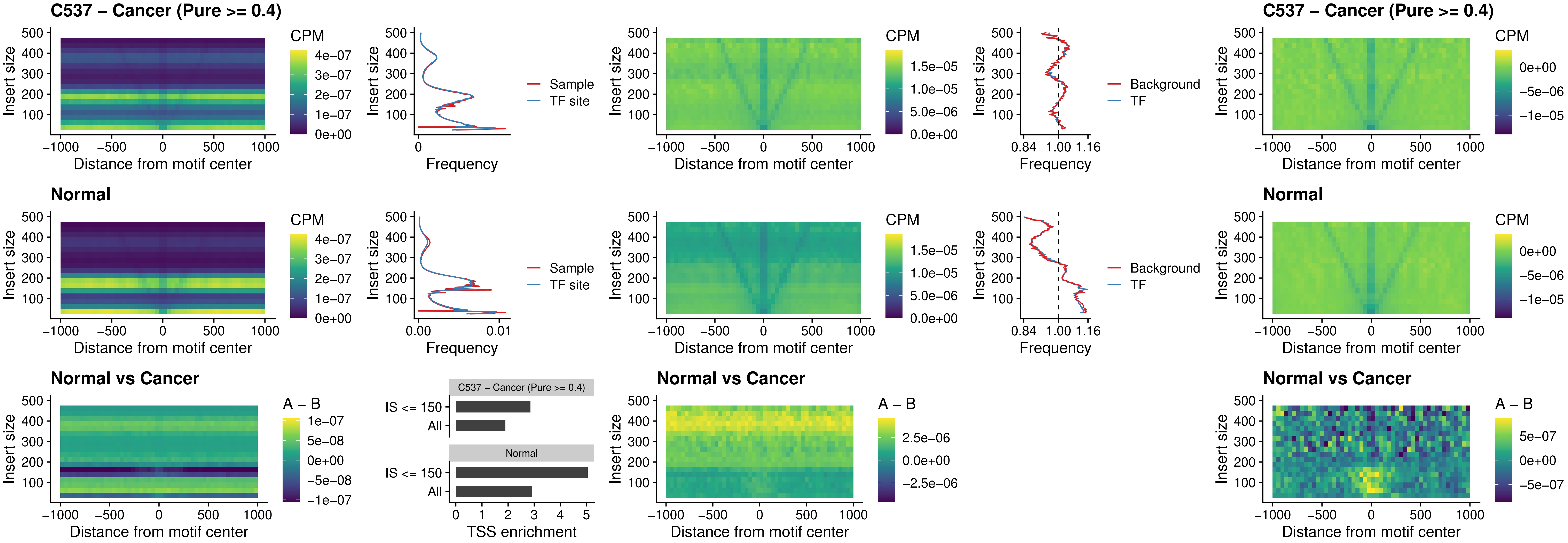

#### C537 – Cancer (Pure >= 0.4) – HOXA2_uc_dTSS_nPEAK

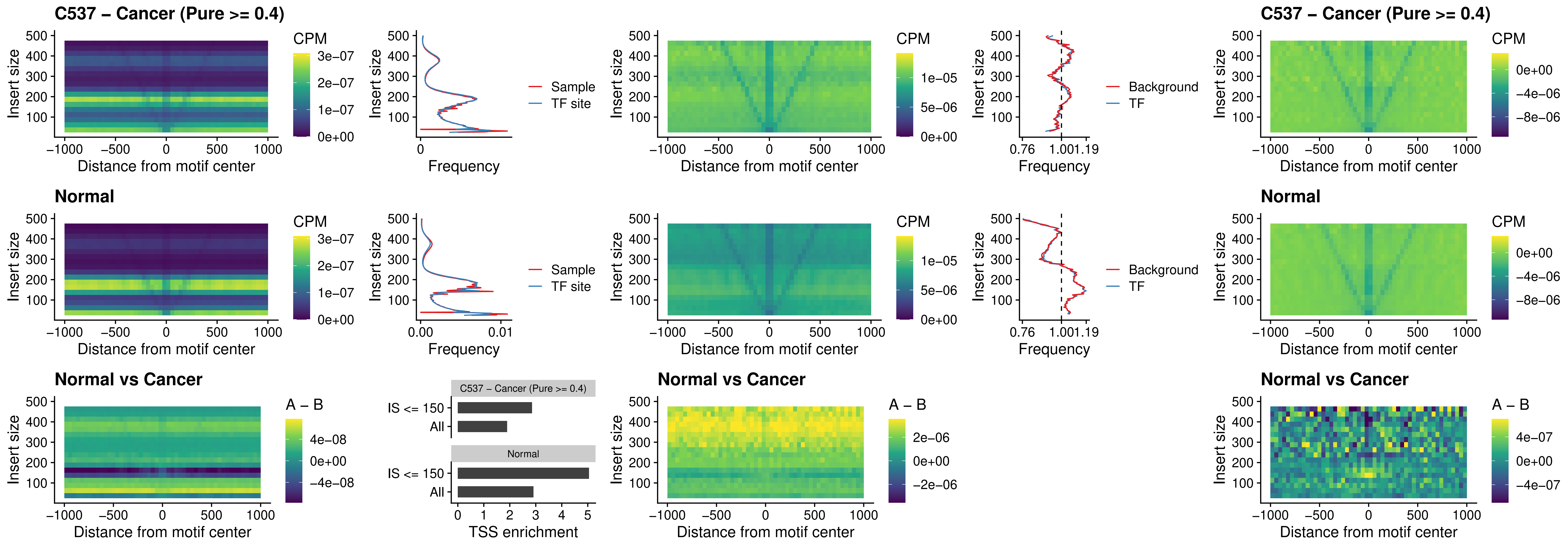

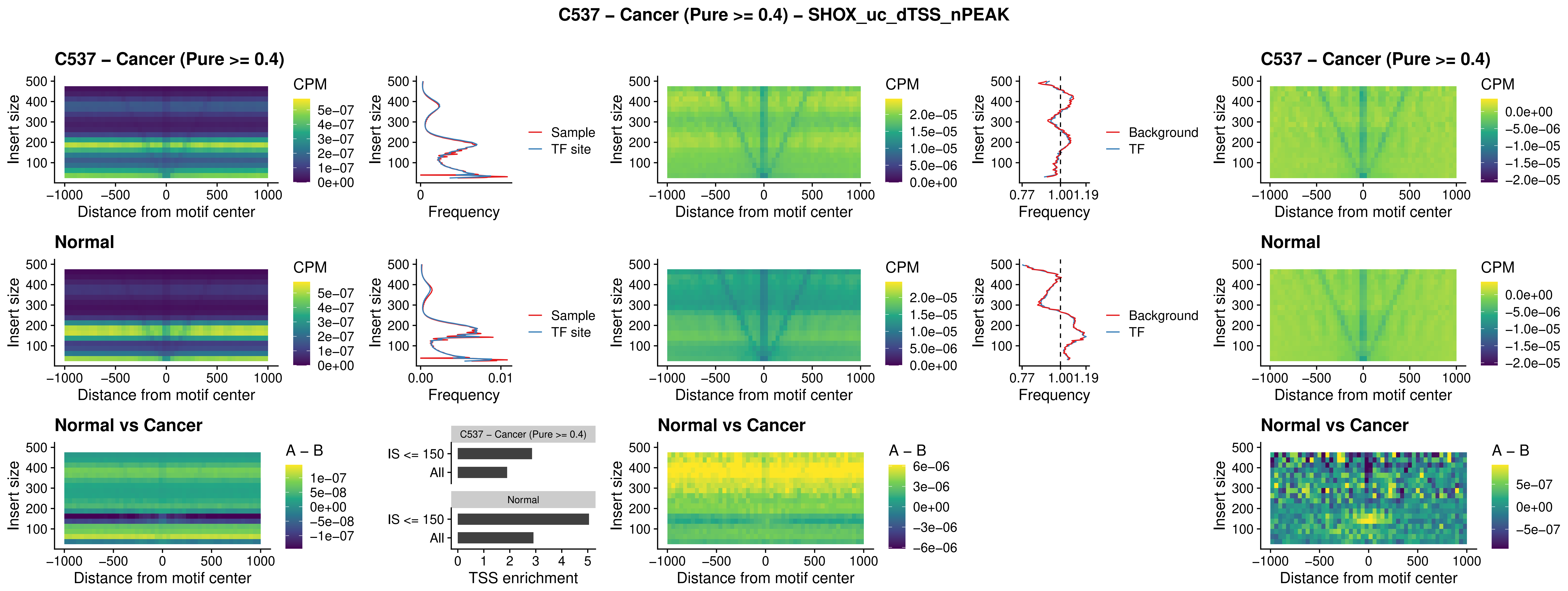
C537 – Cancer (Pure >= 0.4) – SHOX_uc_dTSS_nPEAK
C537 – Cancer (Pure >= 0.4)
Normal
Normal vs Cancer
Insert size
Distance from motif center
Frequency
CPM
A – B
Sample
TF site
Background
TF
TSS enrichment
IS <= 150
All
C537 – Cancer (Pure >= 0.4)
Normal

#### C537 – Cancer (Pure >= 0.4) – PRRX1_uc_dTSS_nPEAK

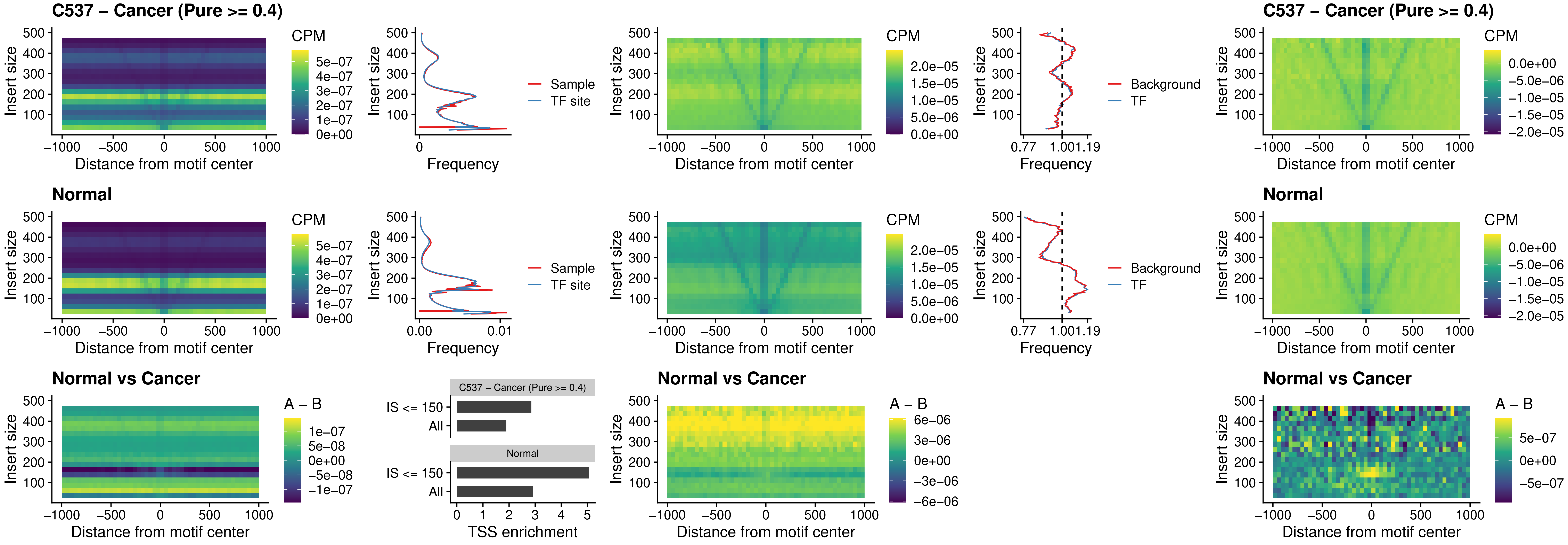

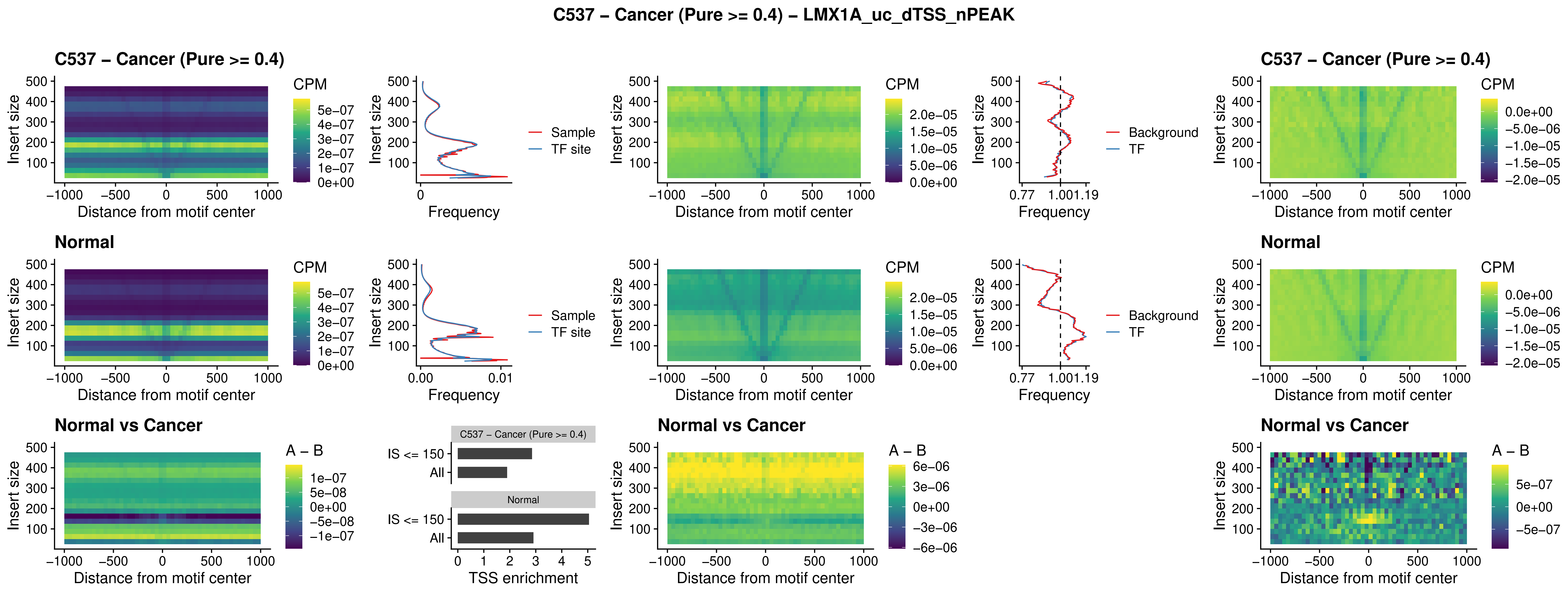

C537 – Cancer (Pure >= 0.4) – LMX1A_uc_dTSS_nPEAK
C537 – Cancer (Pure >= 0.4)
Normal
Normal vs Cancer
Insert size
Distance from motif center
Frequency
CPM
A – B
Sample
TF site
Background
TF
TSS enrichment
C537 – Cancer (Pure >= 0.4)
Normal
IS <= 150
All

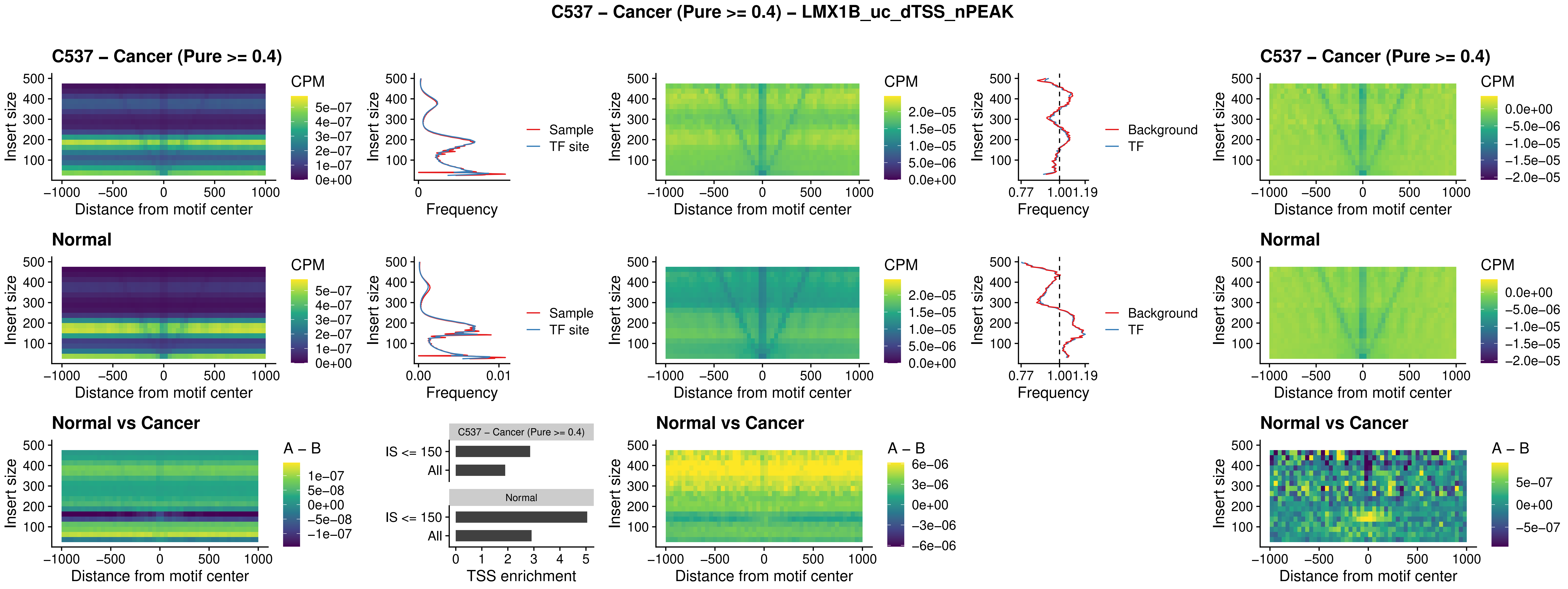
C537 – Cancer (Pure >= 0.4) – LMX1B_uc_dTSS_nPEAK
C537 – Cancer (Pure >= 0.4)
Normal
Normal vs Cancer
Insert size
Distance from motif center
Frequency
CPM
A – B
Sample
TF site
Background
TF
C537 – Cancer (Pure >= 0.4)
Normal
IS <= 150
All
TSS enrichment

#### C537 – Cancer (Pure >= 0.4) – DLX5_uc_dTSS_oPEAK

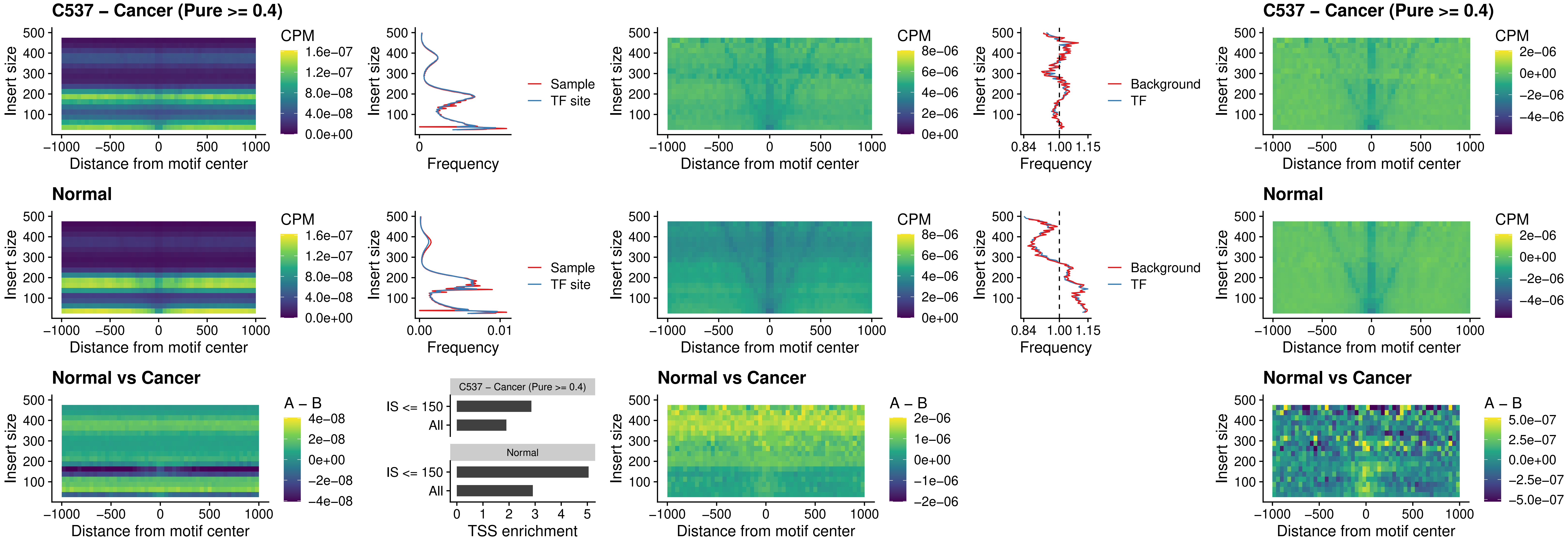

#### C537 – Cancer (Pure >= 0.4) – ARID3C_uc_dTSS_nPEAK

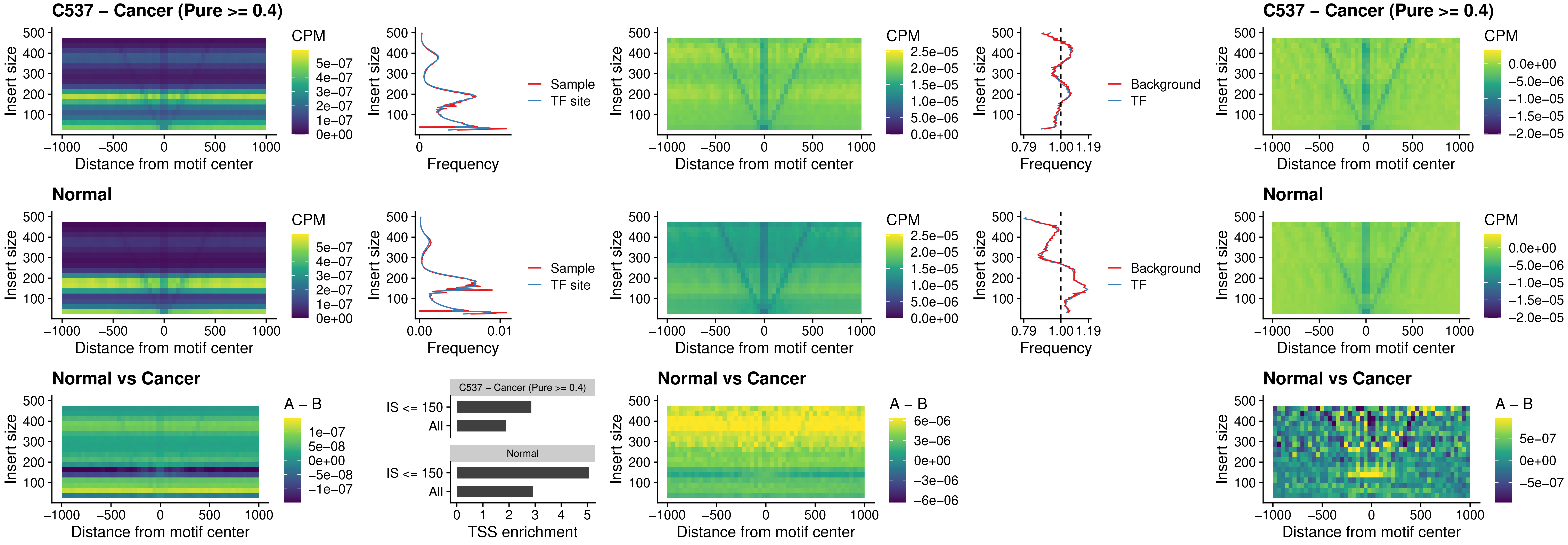

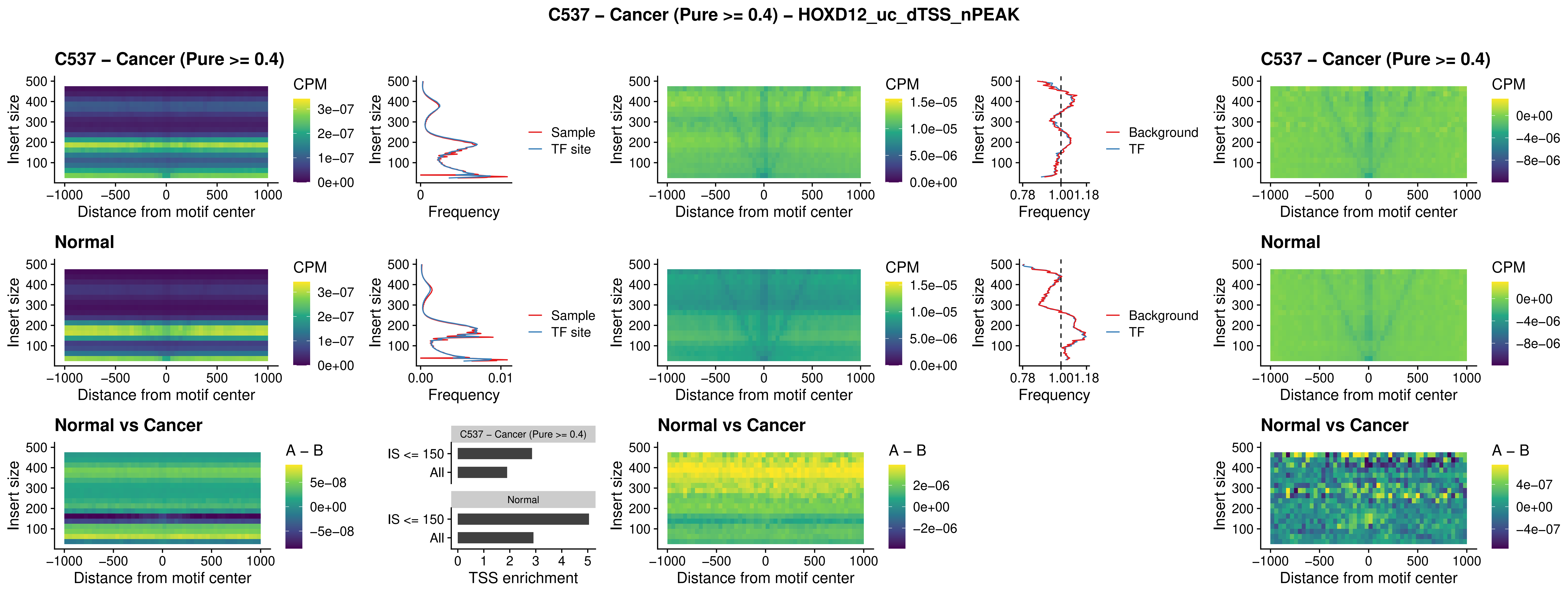
C537 – Cancer (Pure >= 0.4) – HOXD12_uc_dTSS_nPEAK
C537 – Cancer (Pure >= 0.4)
Normal
Normal vs Cancer
Insert size
Distance from motif center
Frequency
CPM
A – B
Sample
TF site
Background
TF
C537 – Cancer (Pure >= 0.4)
Normal
IS <= 150
All
TSS enrichment

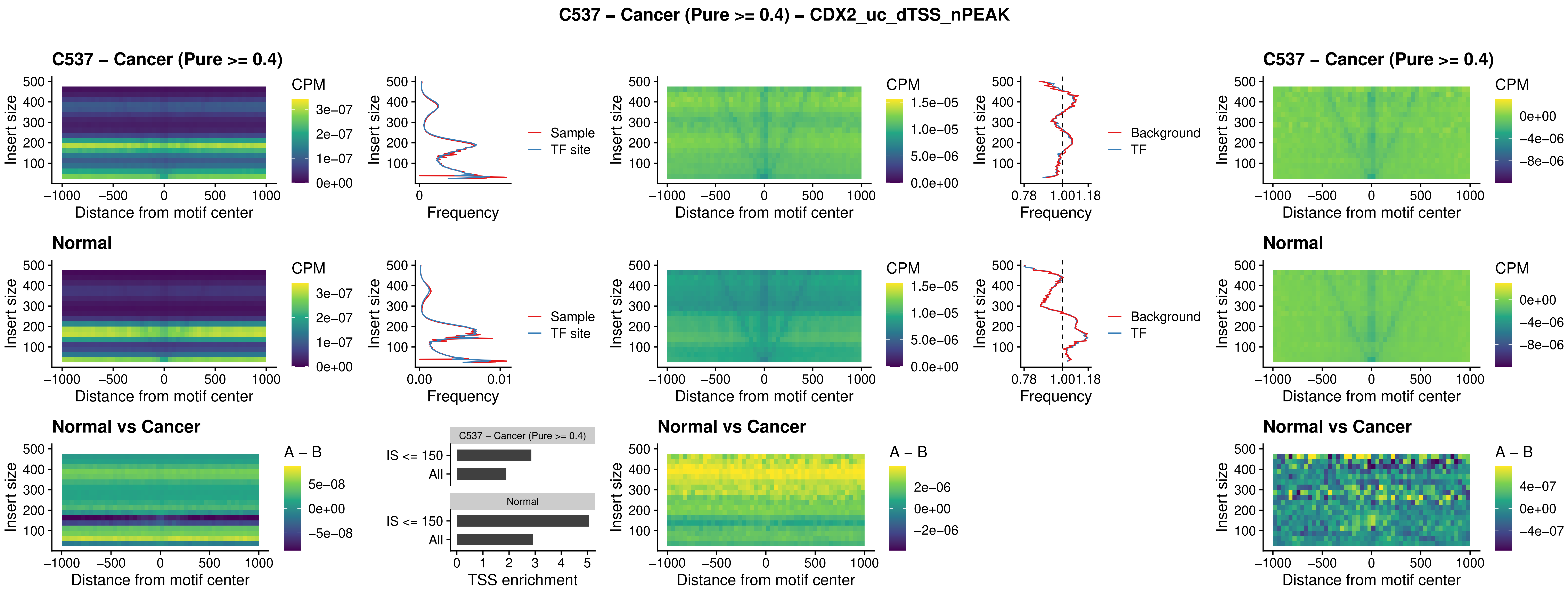
C537 – Cancer (Pure >= 0.4) – CDX2_uc_dTSS_nPEAK
C537 – Cancer (Pure >= 0.4)
Normal
Normal vs Cancer
Insert size
Distance from motif center
Frequency
CPM
A – B
Sample
TF site
Background
TF
C537 – Cancer (Pure >= 0.4)
Normal
IS <= 150
All
TSS enrichment

#### C537 – Cancer (Pure >= 0.4) – HOXC12_uc_dTSS_nPEAK

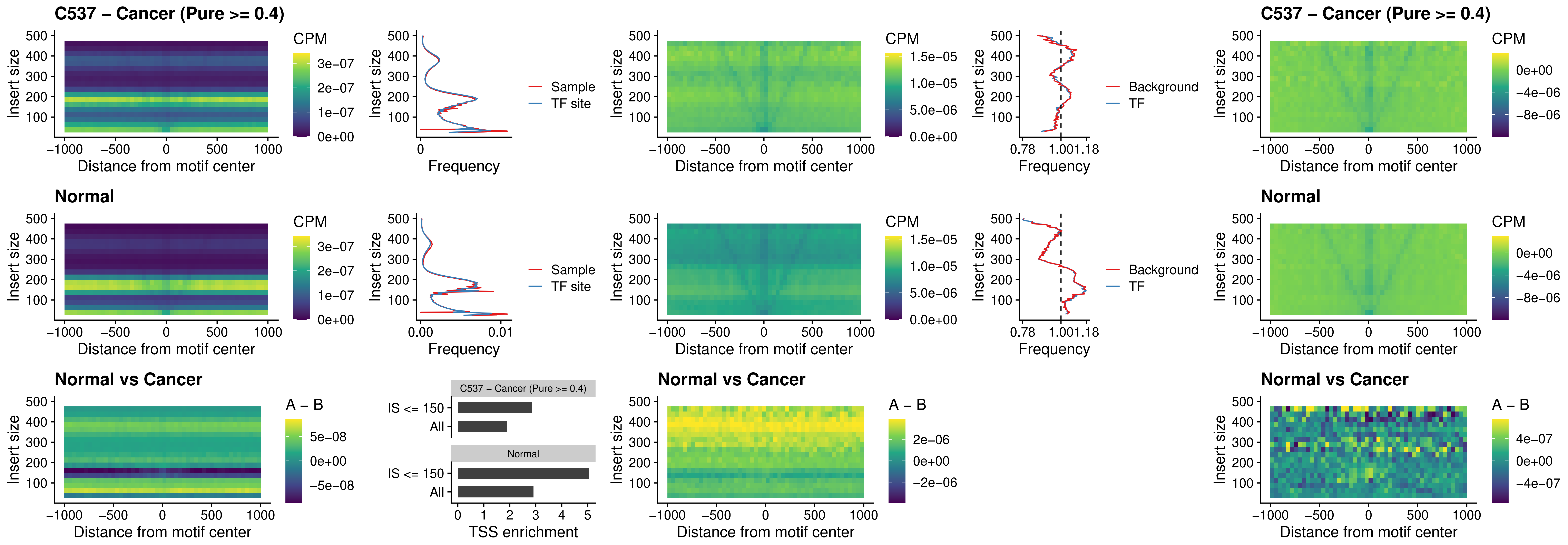

#### C537 – Cancer (Pure >= 0.4) – HHEX_uc_dTSS_oPEAK

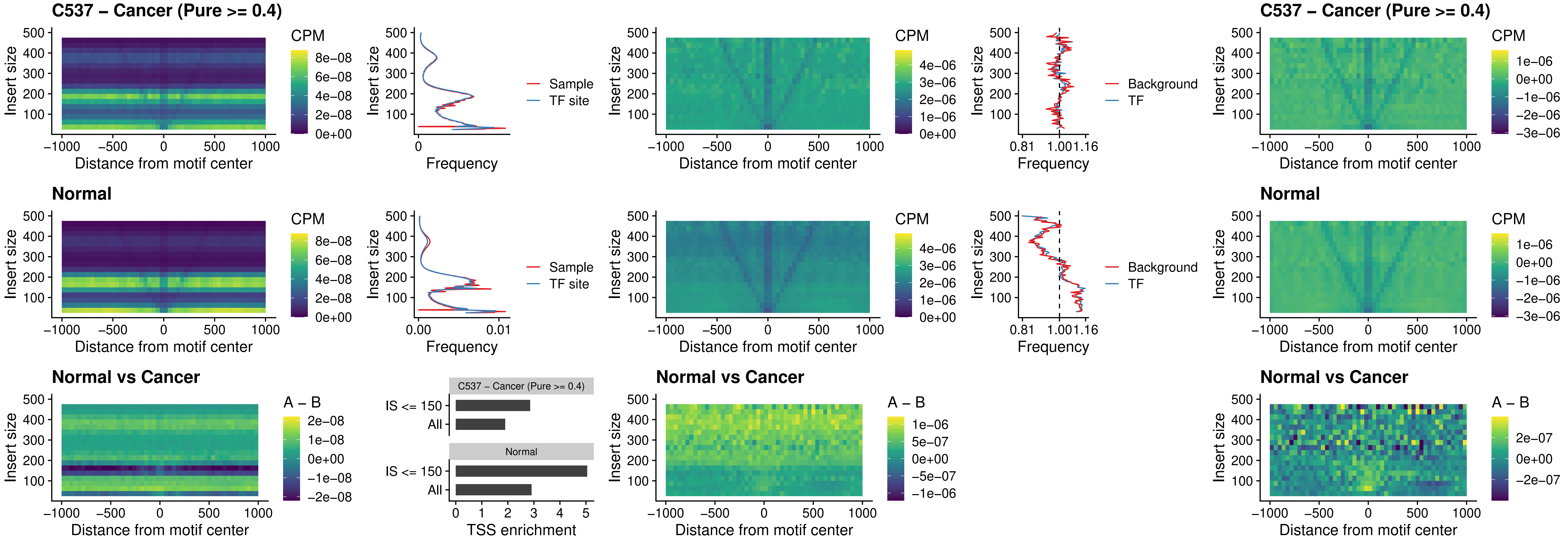

#### C537 – Cancer (Pure >= 0.4) – HOXA11_uc_dTSS_nPEAK

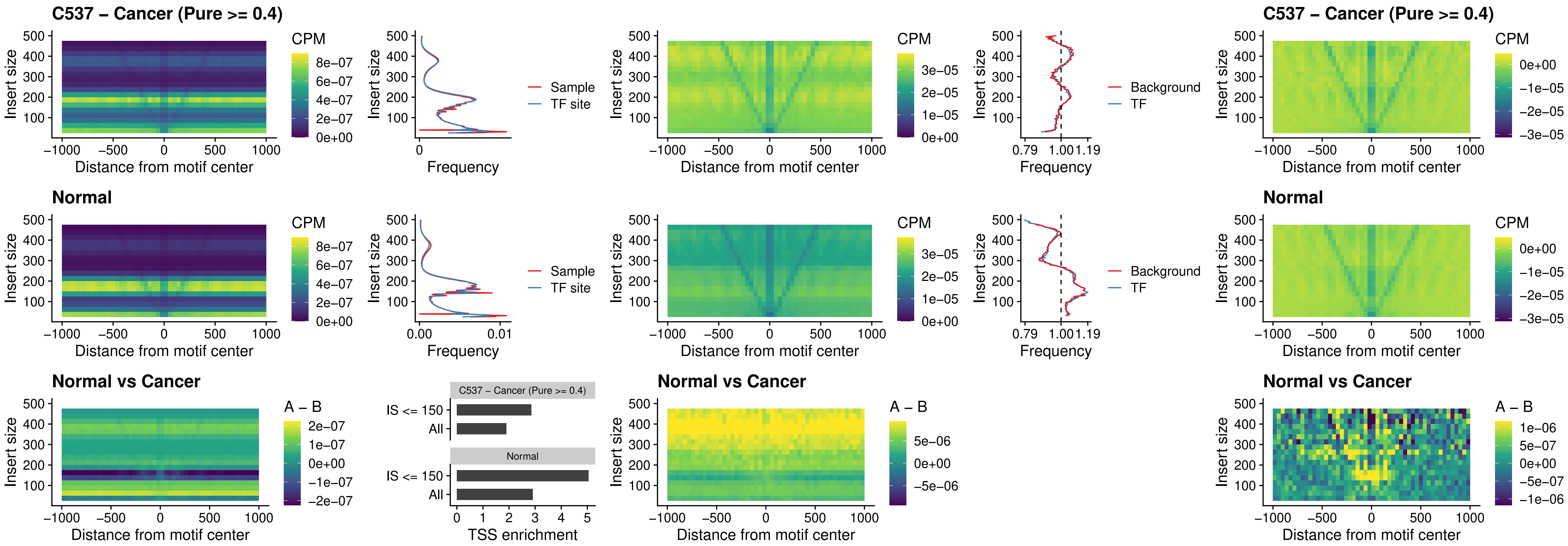

#### C537 – Cancer (Pure >= 0.4) – HOXA11_uc_cTSS_nPEAK

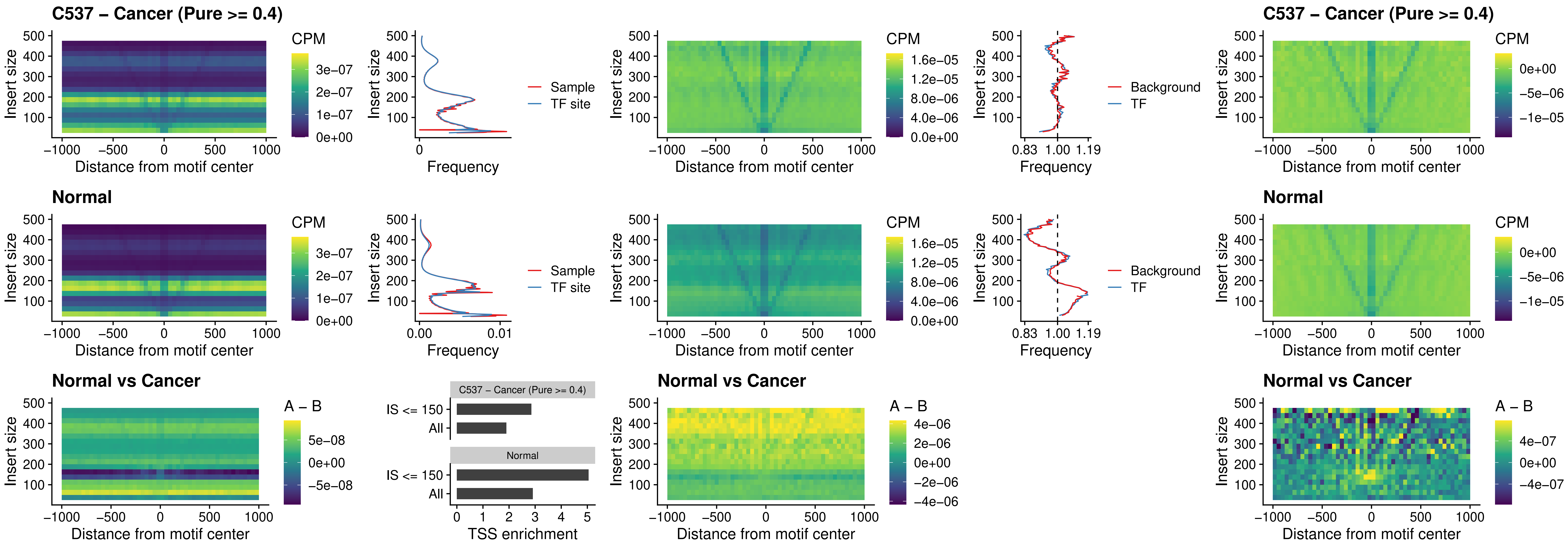

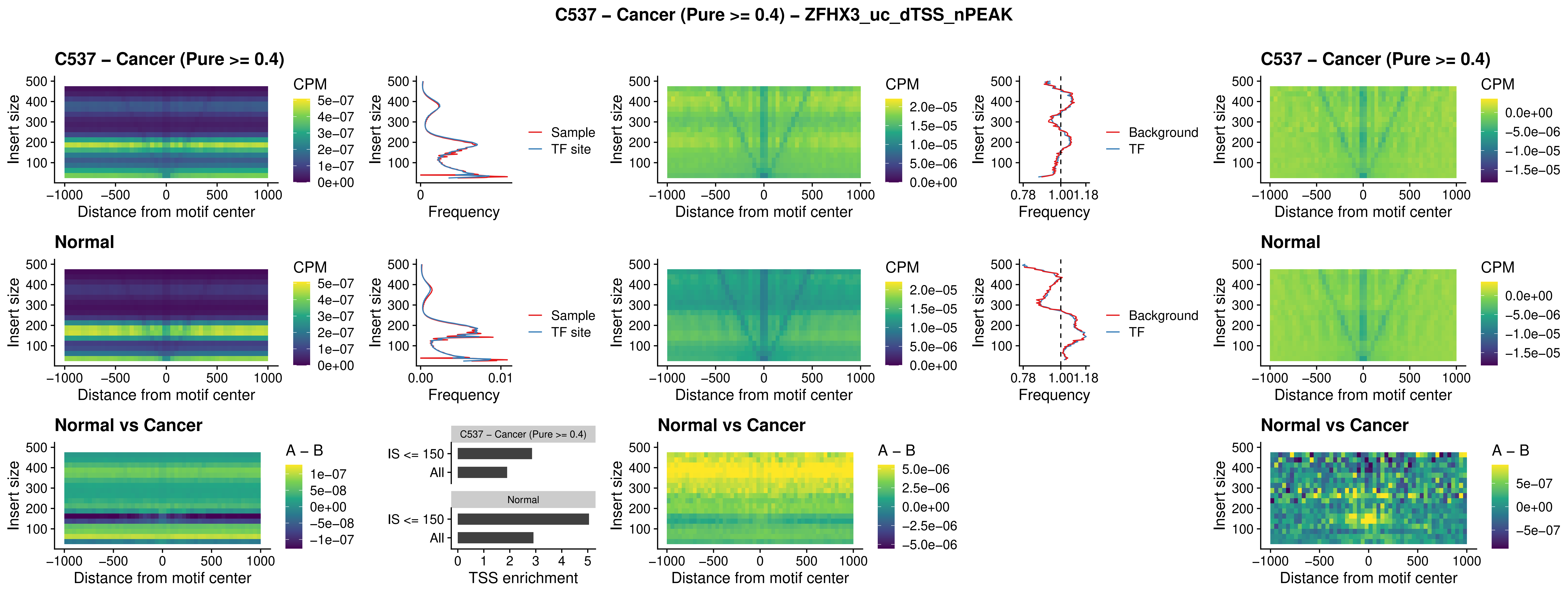
C537 – Cancer (Pure >= 0.4) – ZFHX3_uc_dTSS_nPEAK
C537 – Cancer (Pure >= 0.4)
Normal
Normal vs Cancer
Insert size
Distance from motif center
Frequency
CPM
A – B
Sample
TF site
Background
TF
C537 – Cancer (Pure >= 0.4)
Normal
IS <= 150
All
TSS enrichment

#### C537 – Cancer (Pure >= 0.4) – HOXB4_uc_dTSS_nPEAK

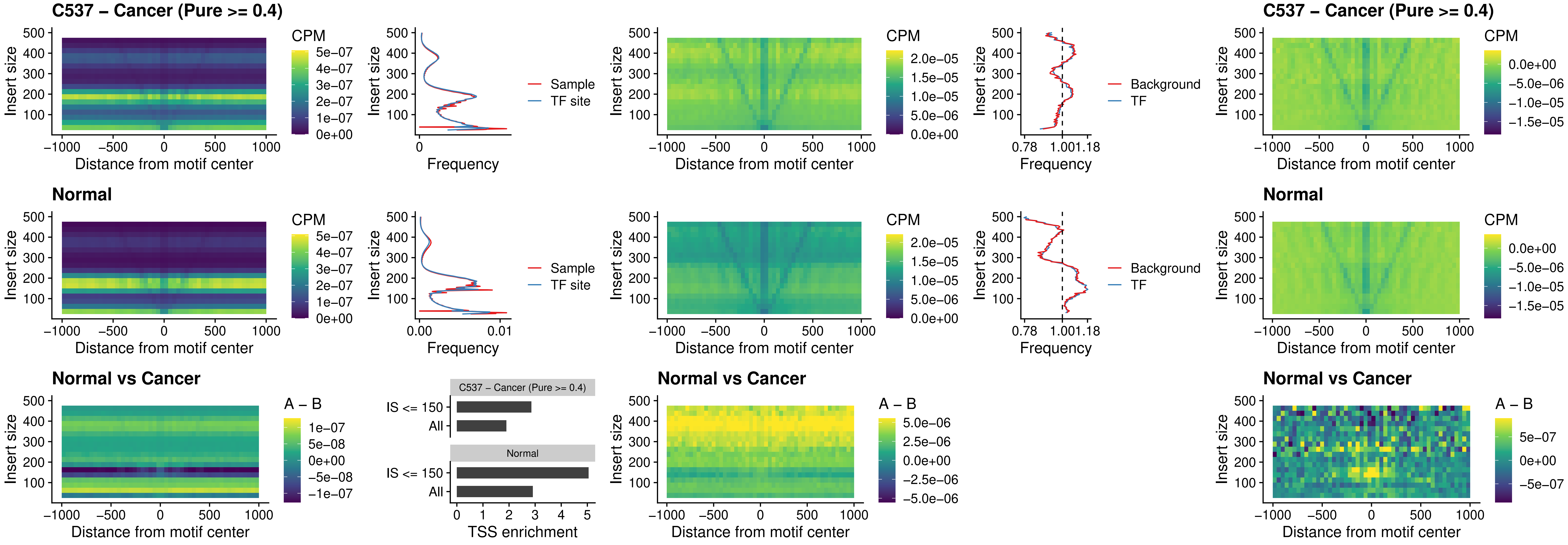

#### C537 – Cancer (Pure >= 0.4) – HOXC5_uc_dTSS_nPEAK

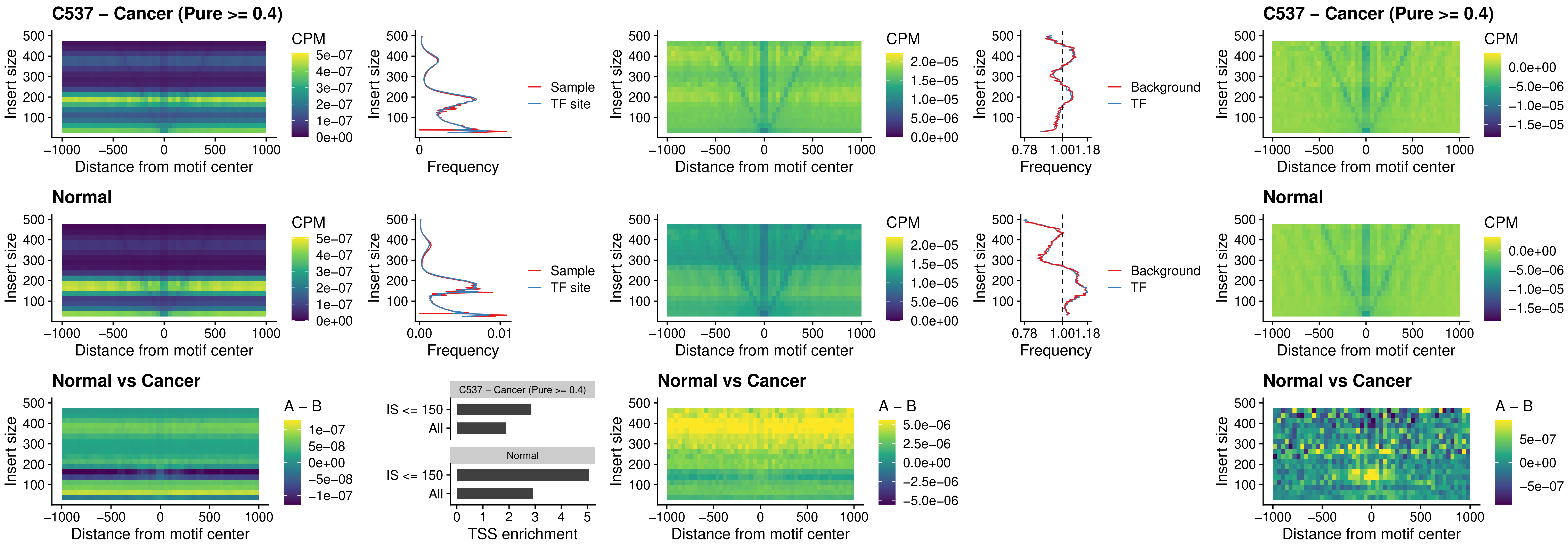

#### C537 – Cancer (Pure >= 0.4) – HOXD1_uc_dTSS_nPEAK

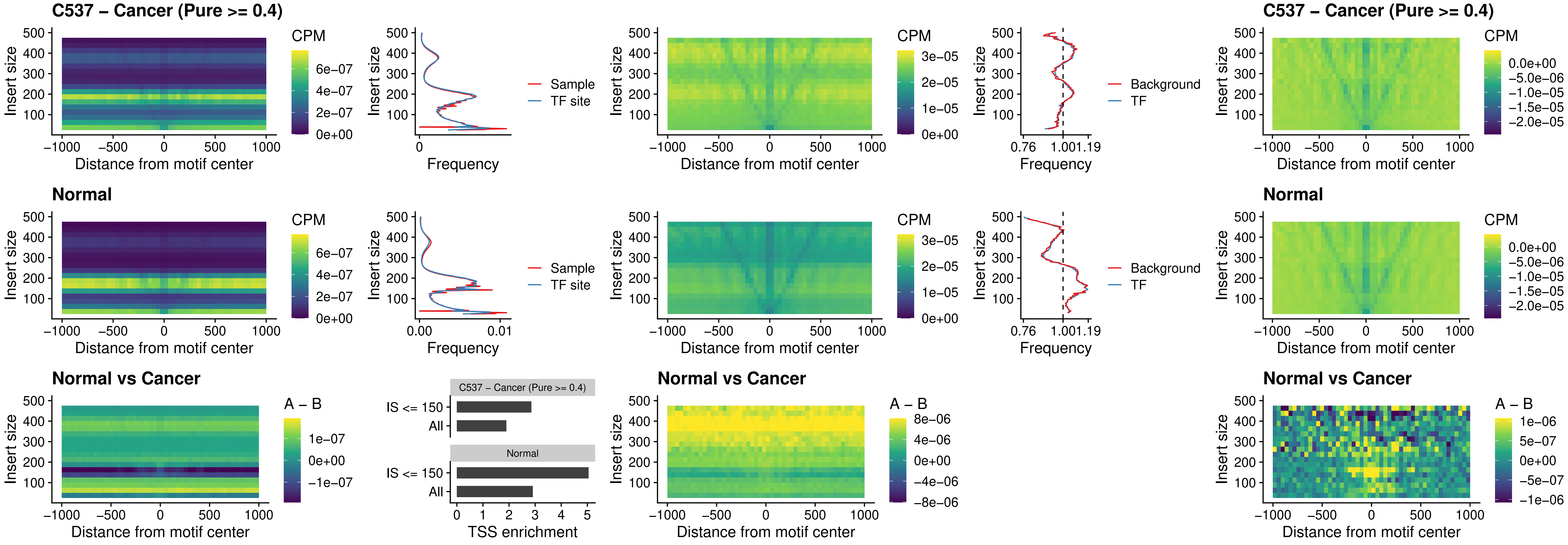

#### C537 – Cancer (Pure >= 0.4) – OTP_uc_dTSS_nPEAK

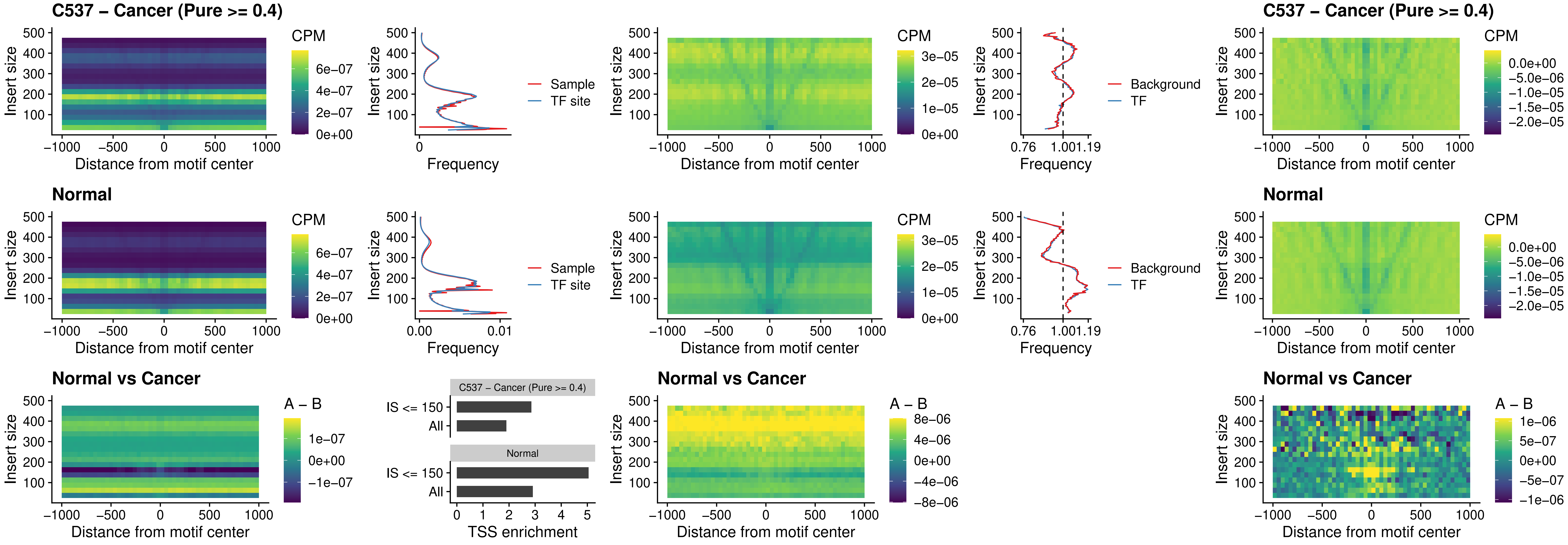

#### C537 – Cancer (Pure >= 0.4) – FOXL1_uc_dTSS_oPEAK

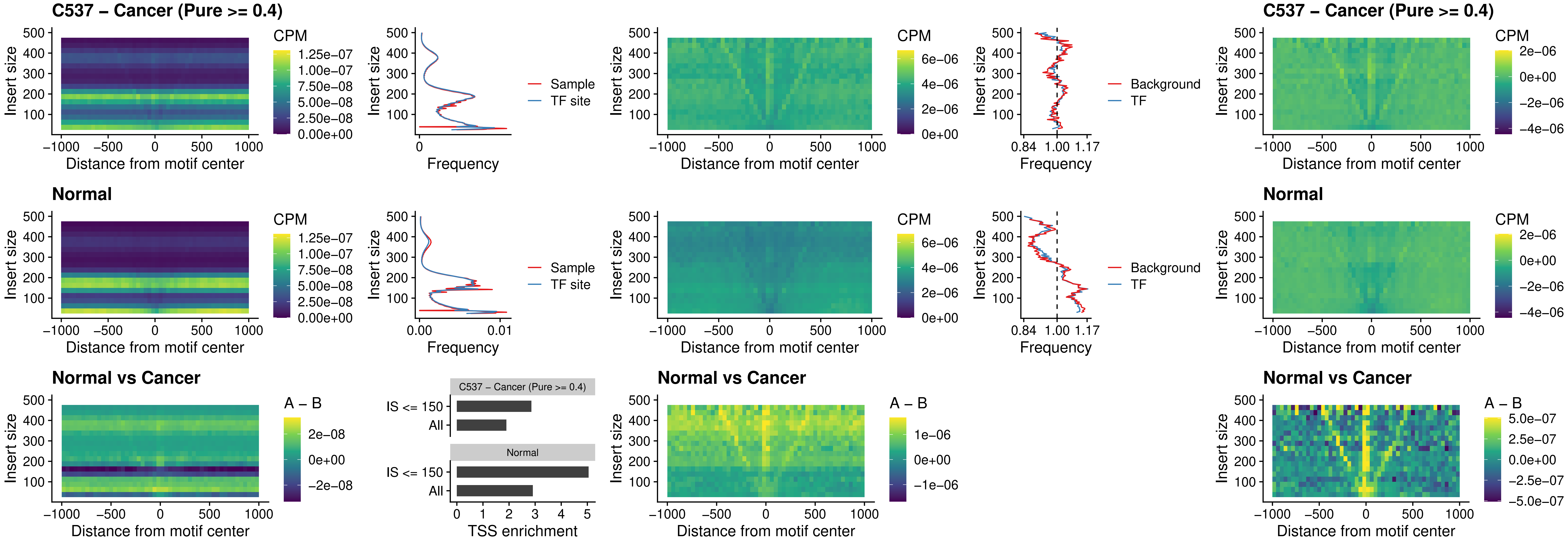

#### C537 – Cancer (Pure >= 0.4) – ZNF32_uc_dTSS_oPEAK

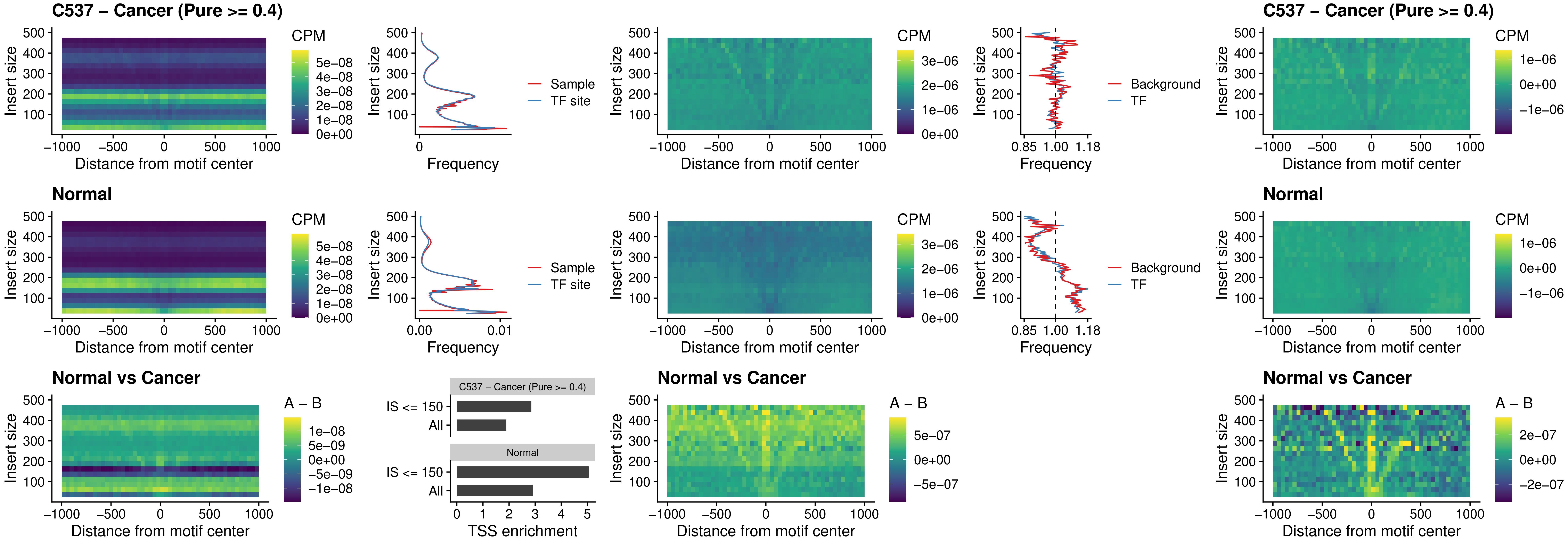

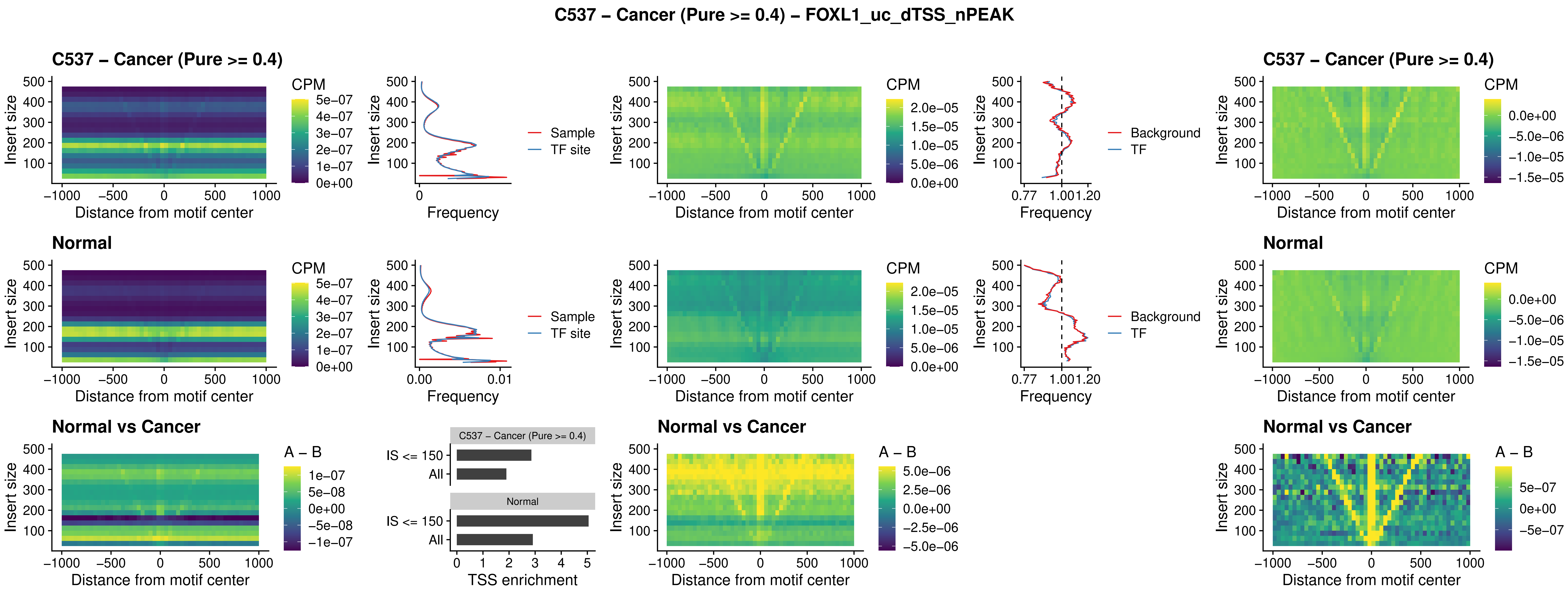

C537 – Cancer (Pure >= 0.4) – FOXL1_uc_dTSS_nPEAK
C537 – Cancer (Pure >= 0.4)
Normal
Normal vs Cancer
Insert size
Distance from motif center
Frequency
CPM
A – B
Sample
TF site
Background
TF
C537 – Cancer (Pure >= 0.4)
Normal
IS <= 150
All
TSS enrichment

#### C537 – Cancer (Pure >= 0.4) – ZNF32_uc_dTSS_nPEAK

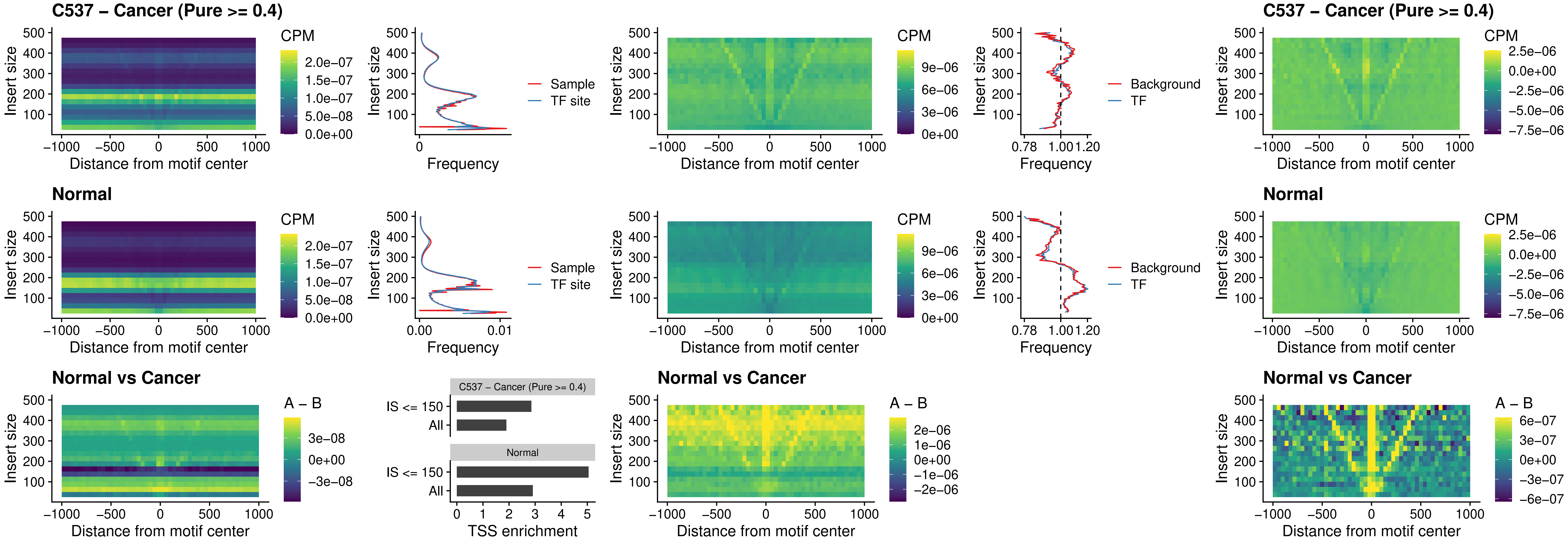

#### C537 – Cancer (Pure >= 0.4) – DLX5_uc_dTSS_nPEAK

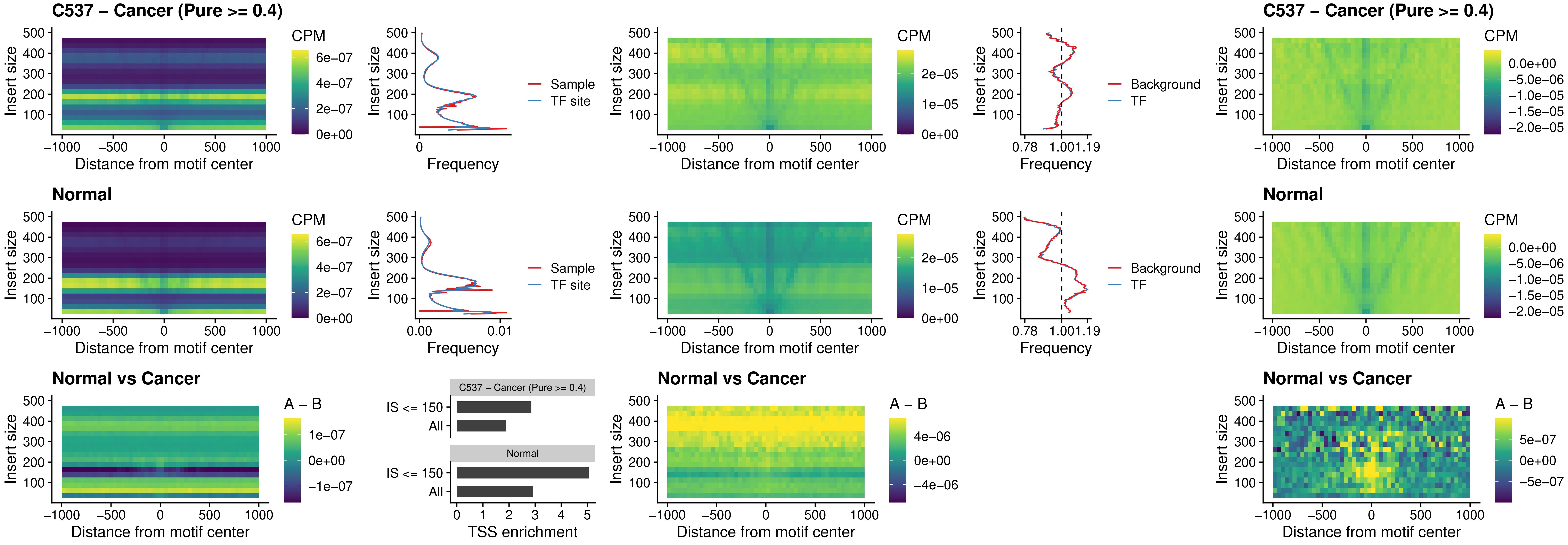

#### C537 – Cancer (Pure >= 0.4) – DMRT2_uc_dTSS_nPEAK

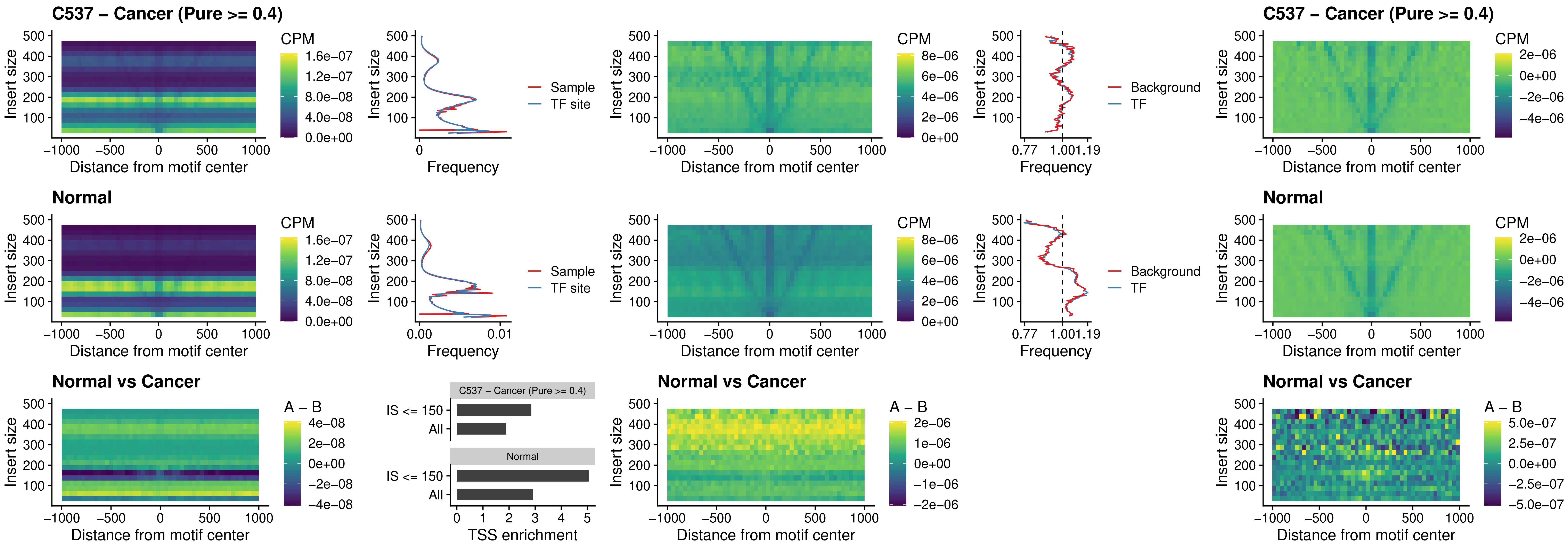

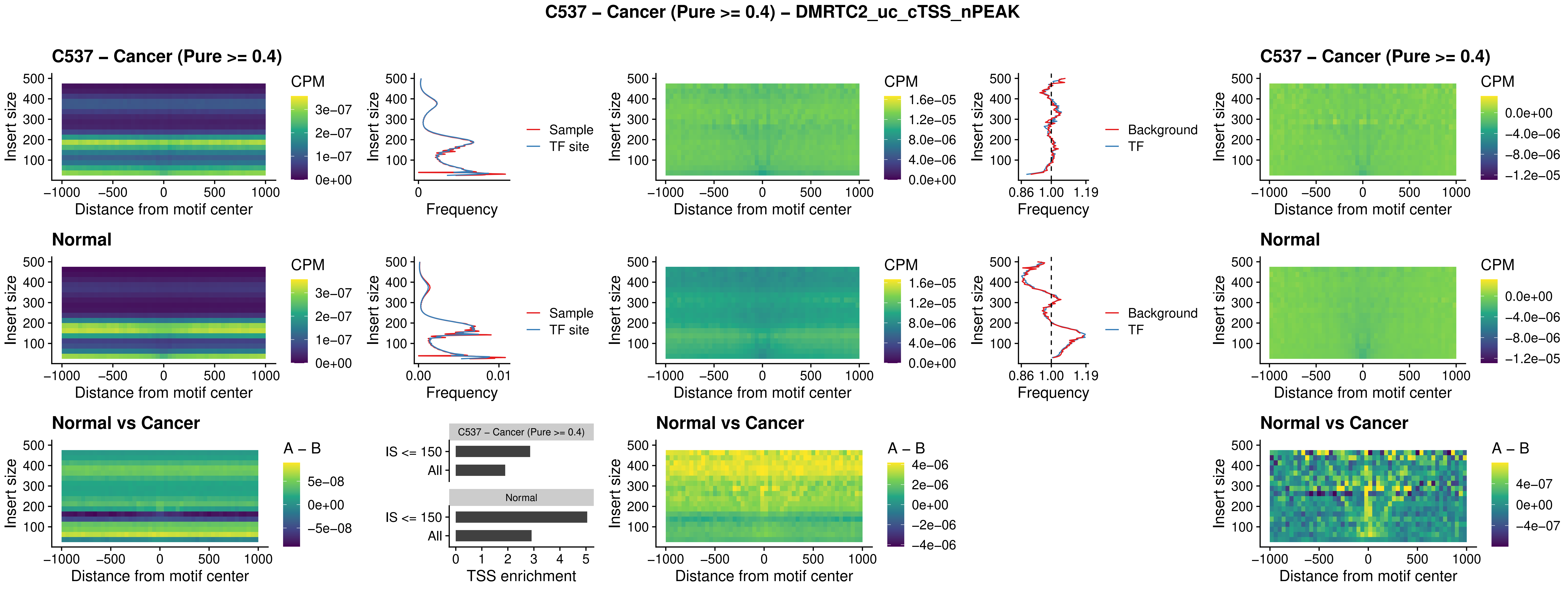
C537 – Cancer (Pure >= 0.4) – DMRTC2_uc_cTSS_nPEAK
C537 – Cancer (Pure >= 0.4)
Normal
Normal vs Cancer
Insert size
Distance from motif center
Frequency
CPM
A – B
Sample
TF site
Background
TF
C537 – Cancer (Pure >= 0.4)
Normal
IS <= 150
All
TSS enrichment

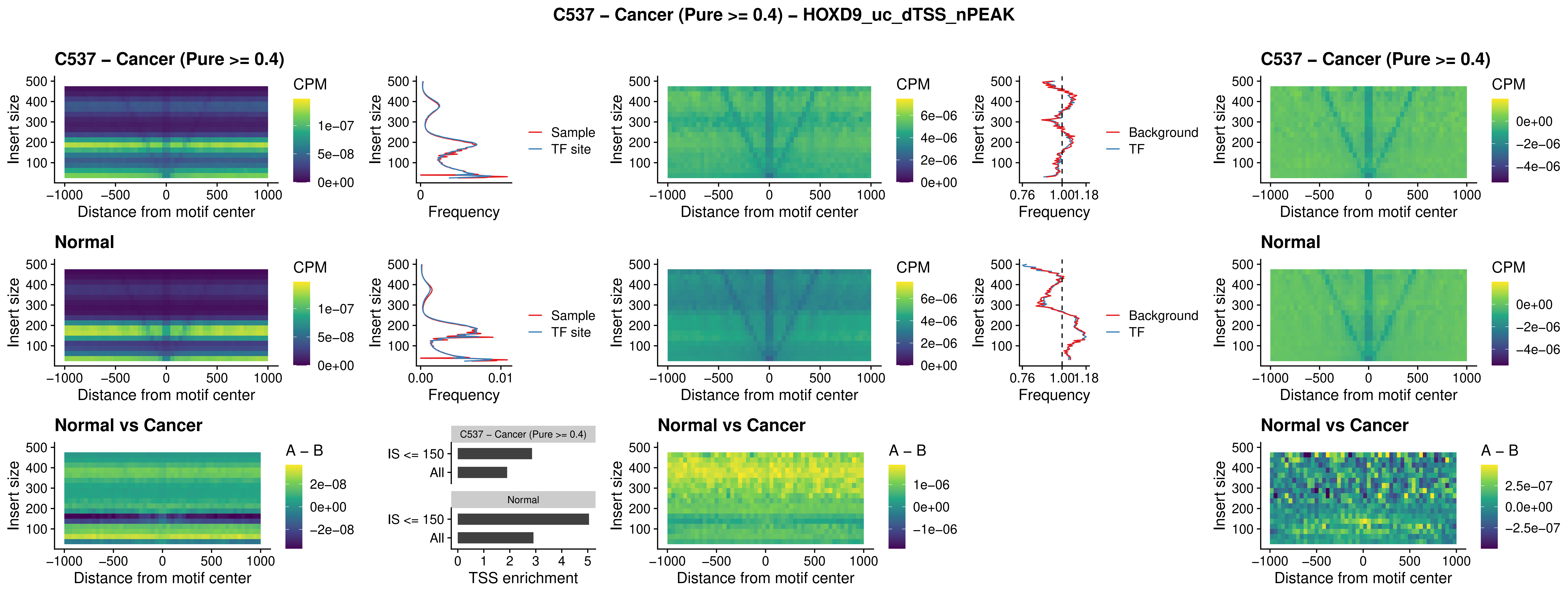

C537 – Cancer (Pure >= 0.4) – HOXD9_uc_dTSS_nPEAK
C537 – Cancer (Pure >= 0.4)
Normal
Normal vs Cancer
Insert size
Distance from motif center
Frequency
CPM
A – B
Sample
TF site
Background
TF
C537 – Cancer (Pure >= 0.4)
Normal
IS <= 150
All
TSS enrichment

#### C537 – Cancer (Pure >= 0.4) – HOXD4_uc_dTSS_nPEAK

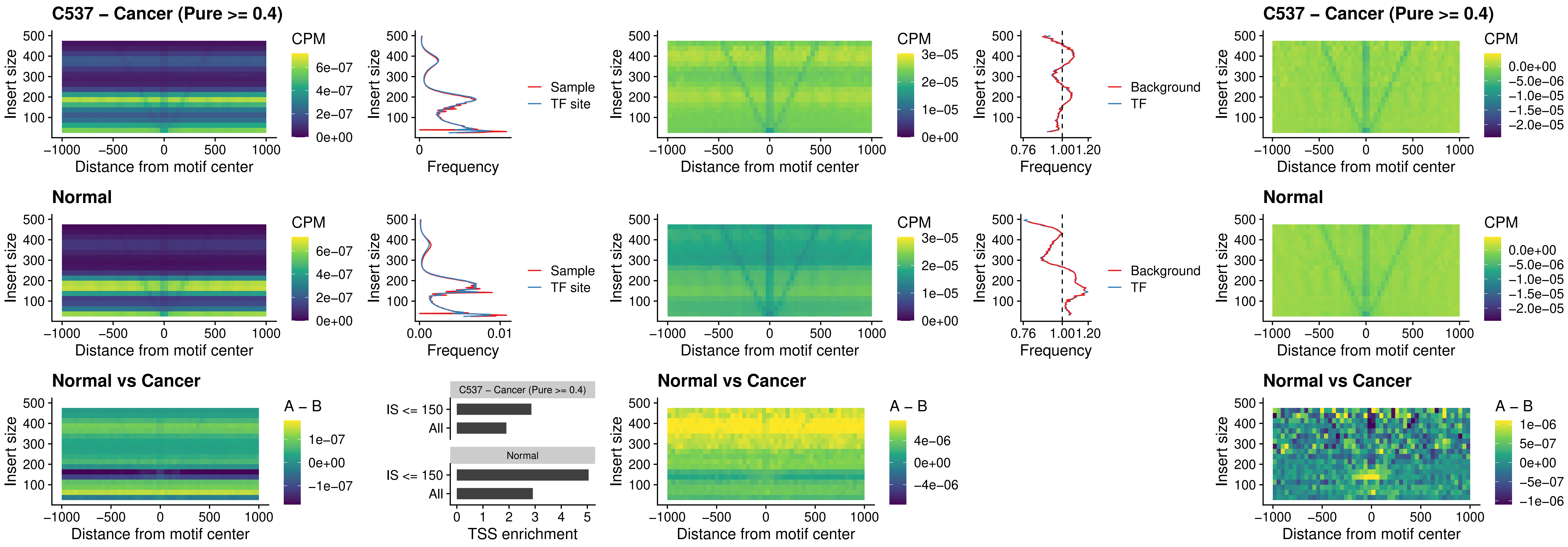

#### C537 – Cancer (Pure >= 0.4) – FOXP3_uc_cTSS_nPEAK

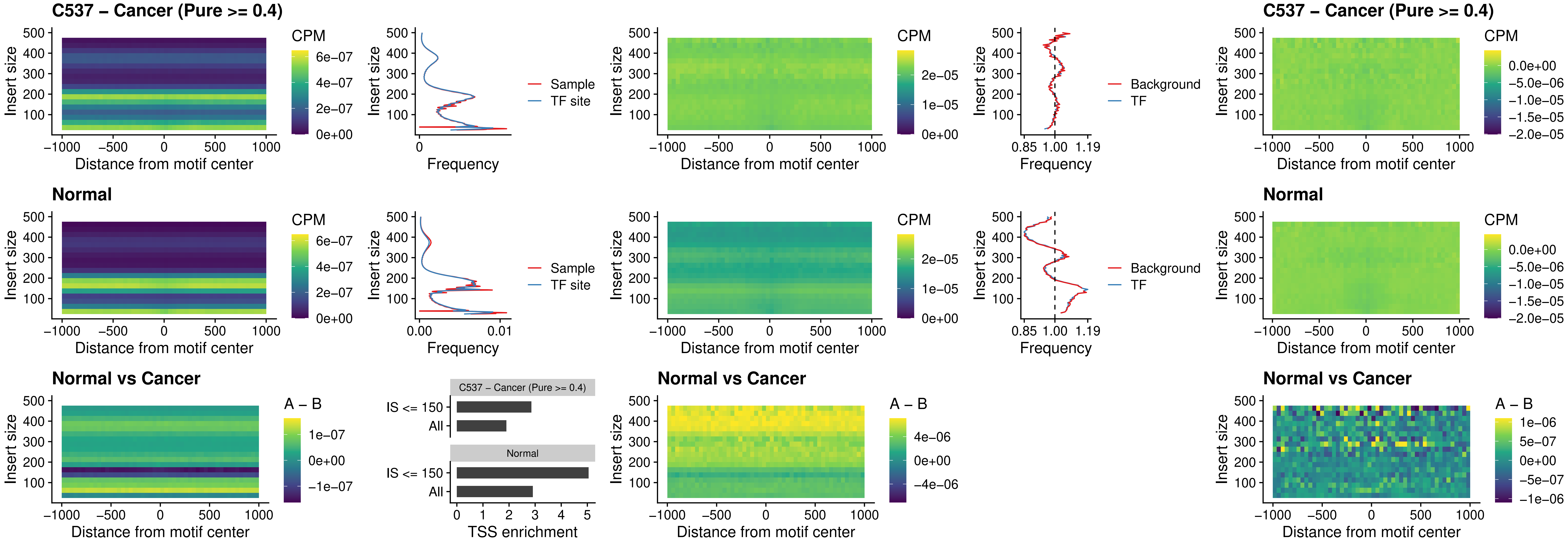

#### C537 – Cancer (Pure >= 0.4) – FOXD4L6_uc_cTSS_nPEAK

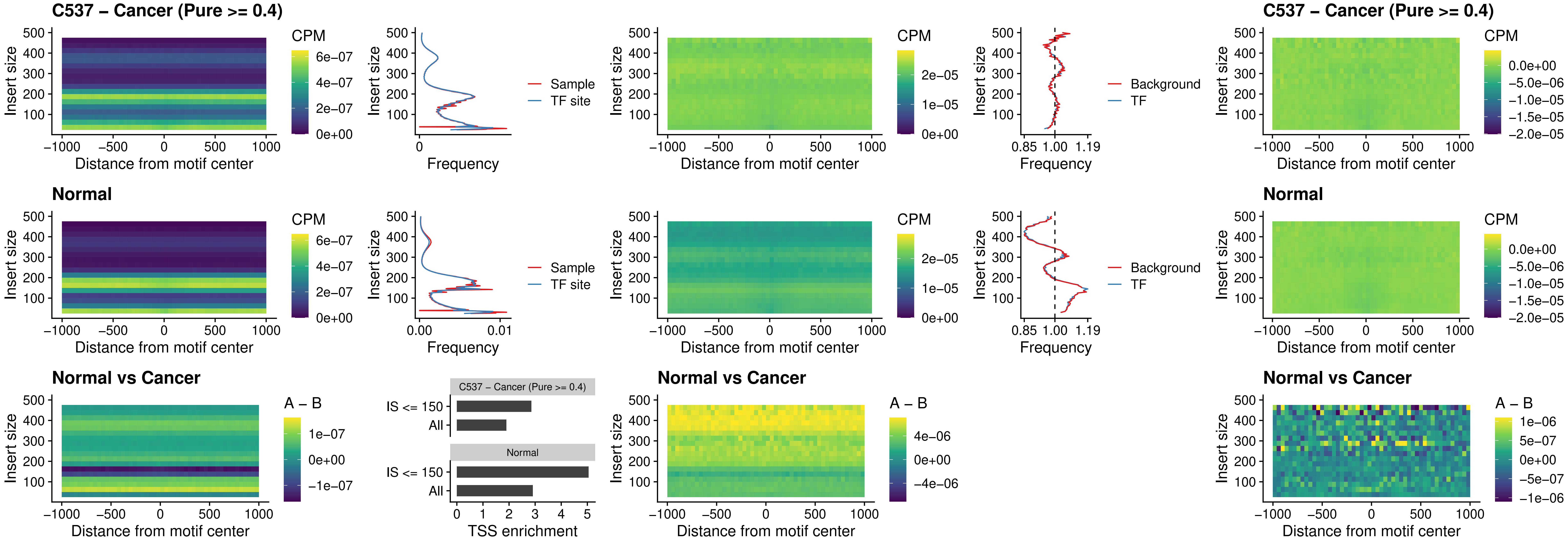

#### C537 – Cancer (Pure >= 0.4) – FOXD4L5_uc_cTSS_nPEAK

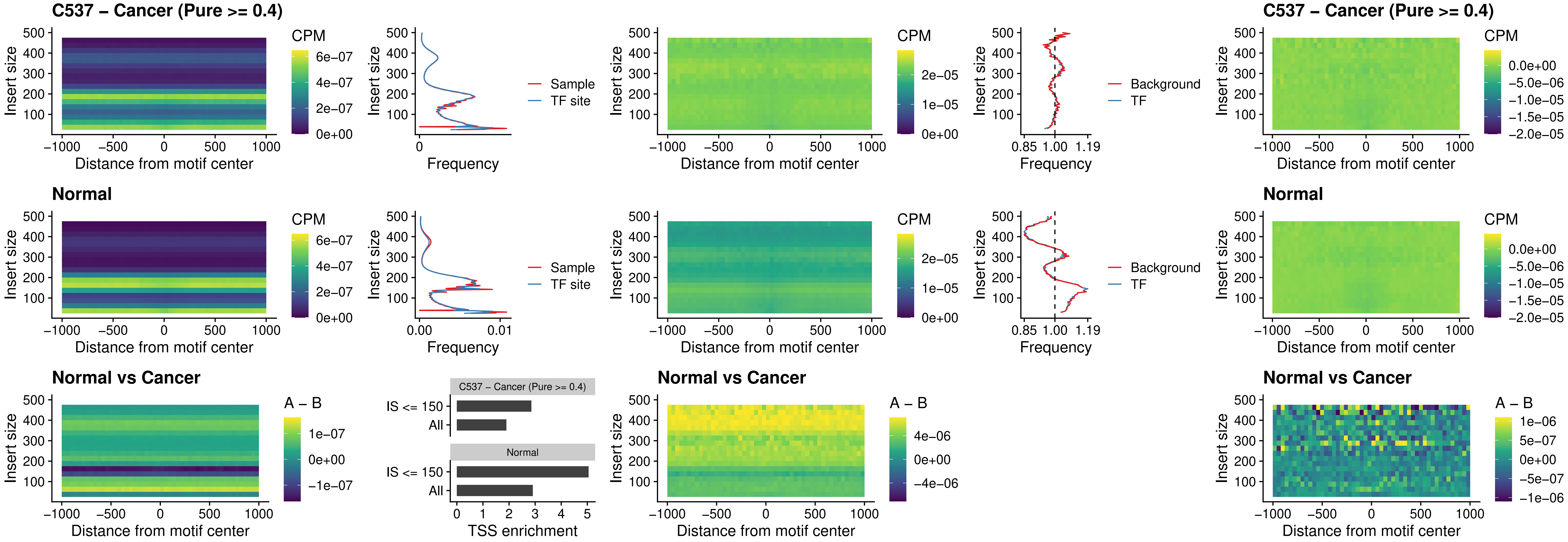

#### C537 – Cancer (Pure >= 0.4) – FOXD4L4_uc_cTSS_nPEAK

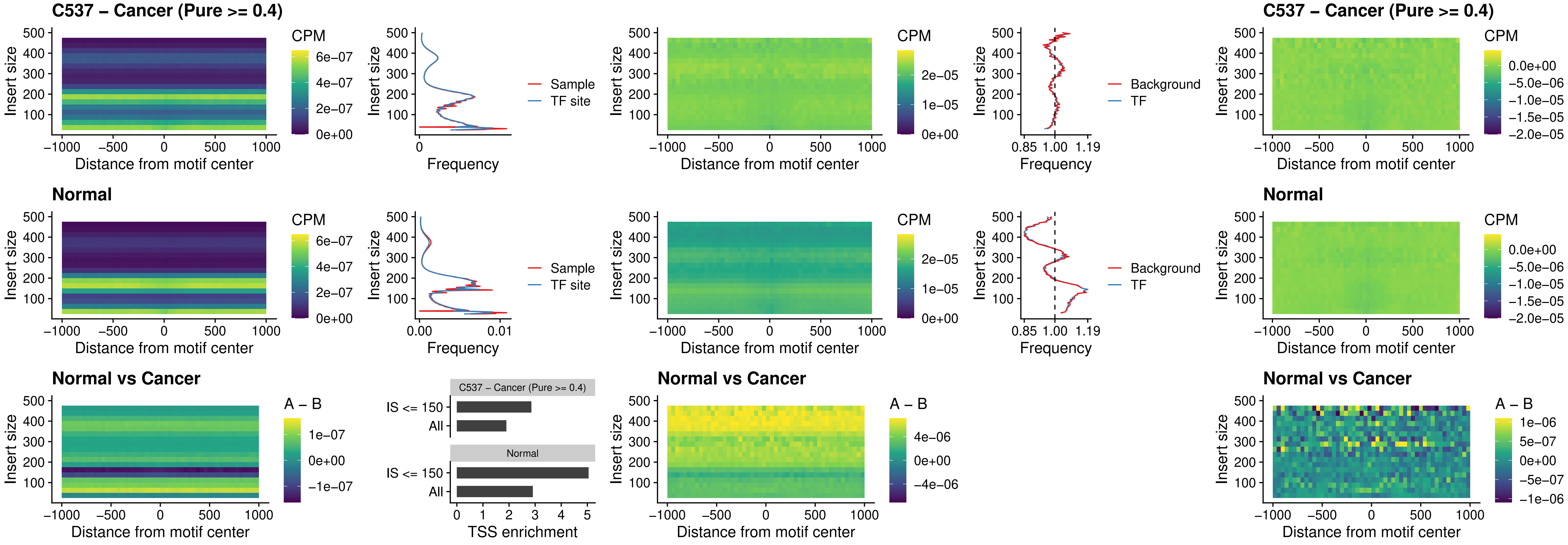

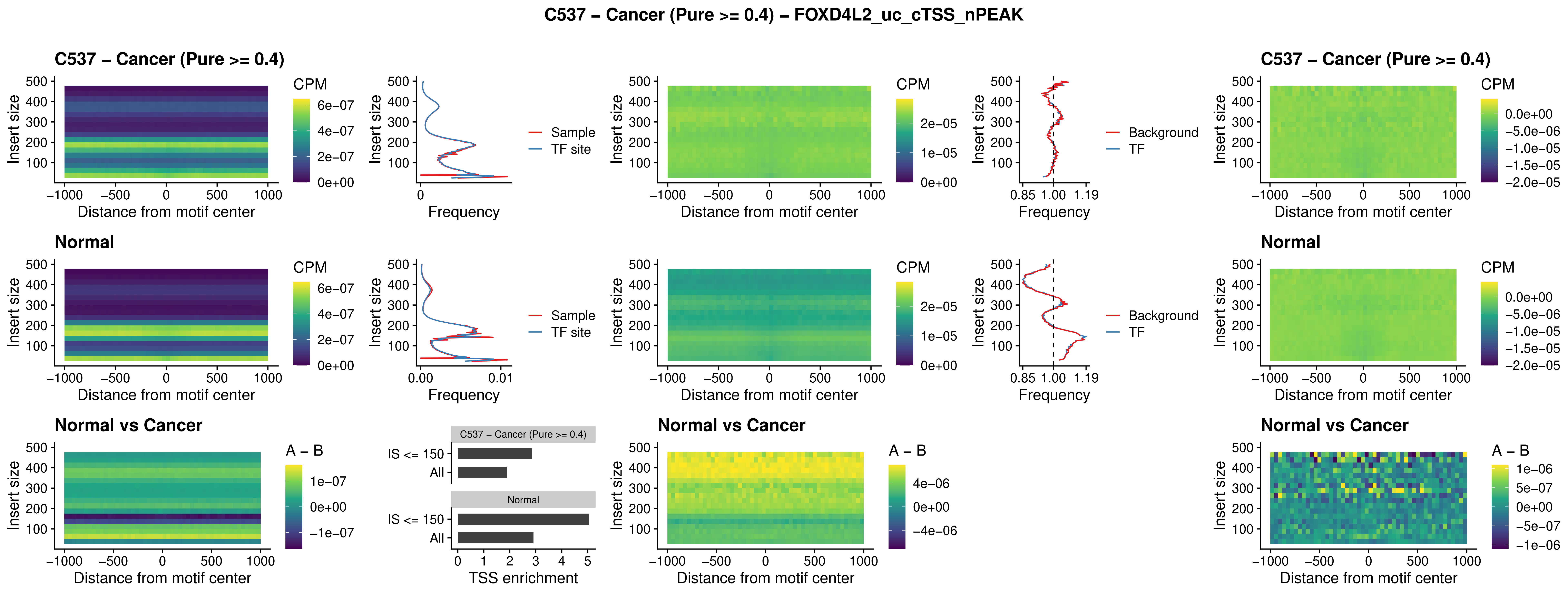
C537 – Cancer (Pure >= 0.4) – FOXD4L2_uc_cTSS_nPEAK
C537 – Cancer (Pure >= 0.4)
Normal
Normal vs Cancer
Insert size
Distance from motif center
Frequency
CPM
A – B
Sample
TF site
Background
TF
C537 – Cancer (Pure >= 0.4)
Normal
IS <= 150
All
TSS enrichment

#### C537 – Cancer (Pure >= 0.4) – FOXD4L3_uc_cTSS_nPEAK

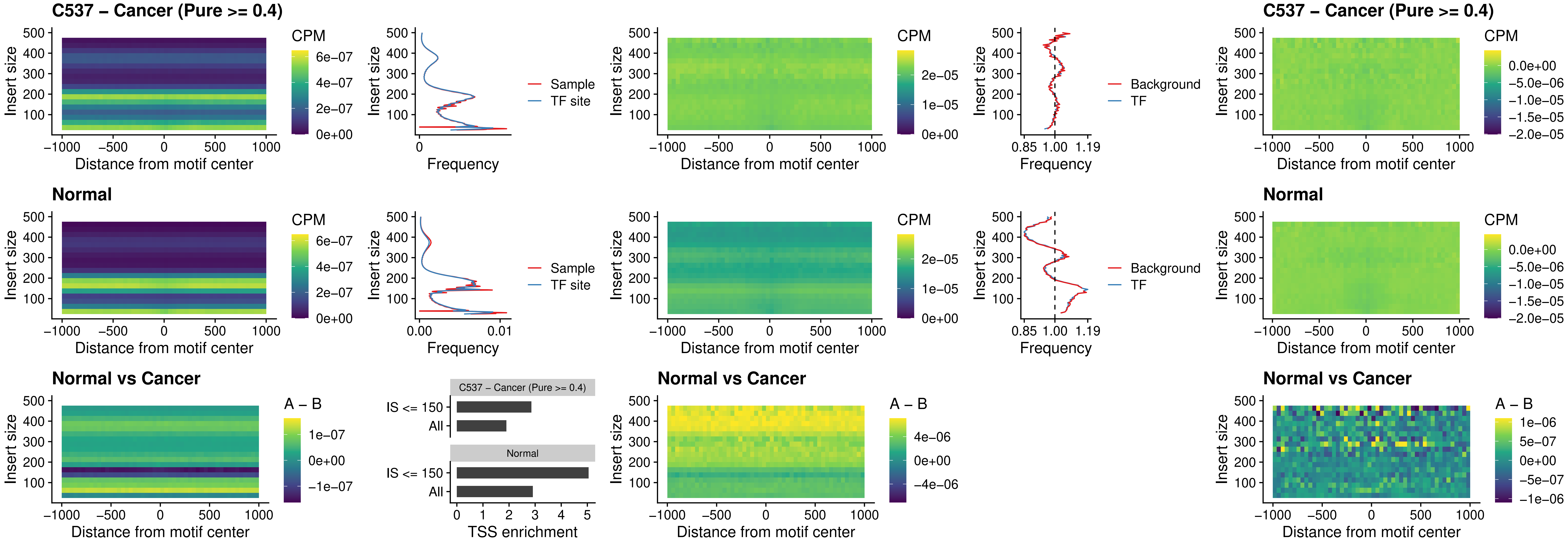

#### C537 – Cancer (Pure >= 0.4) – HOXD1_uc_dTSS_oPEAK

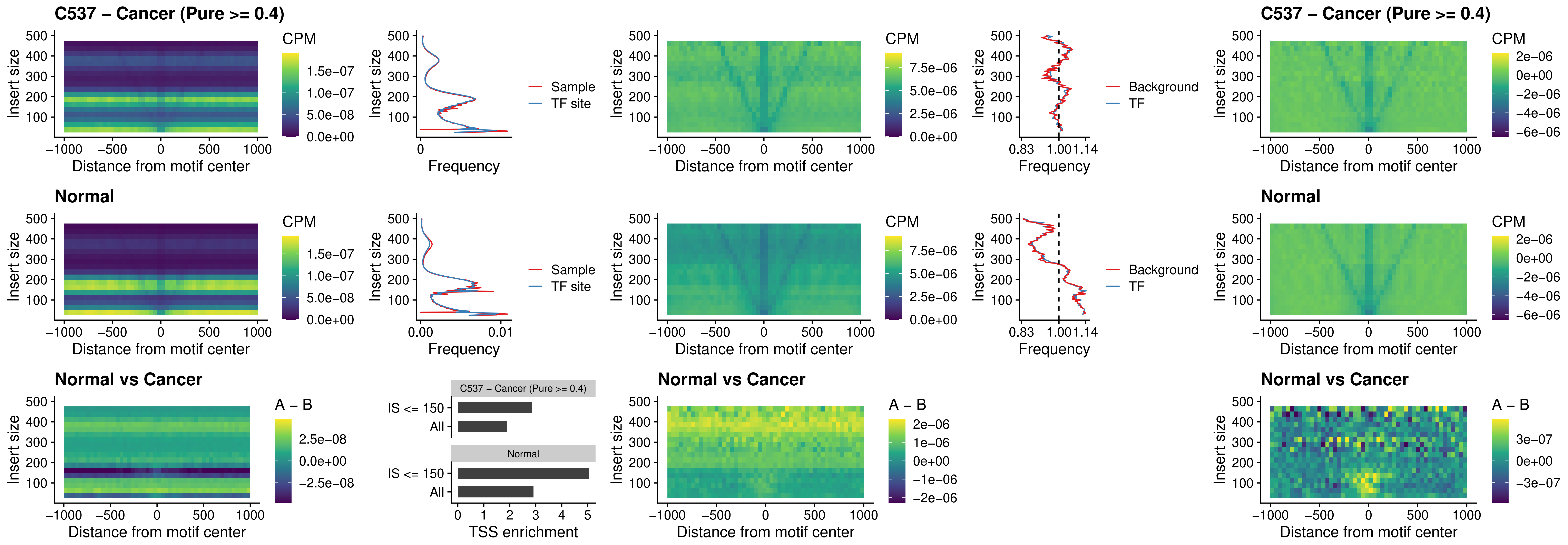

#### C537 – Cancer (Pure >= 0.4) – VAX1_uc_pTSS_nPEAK

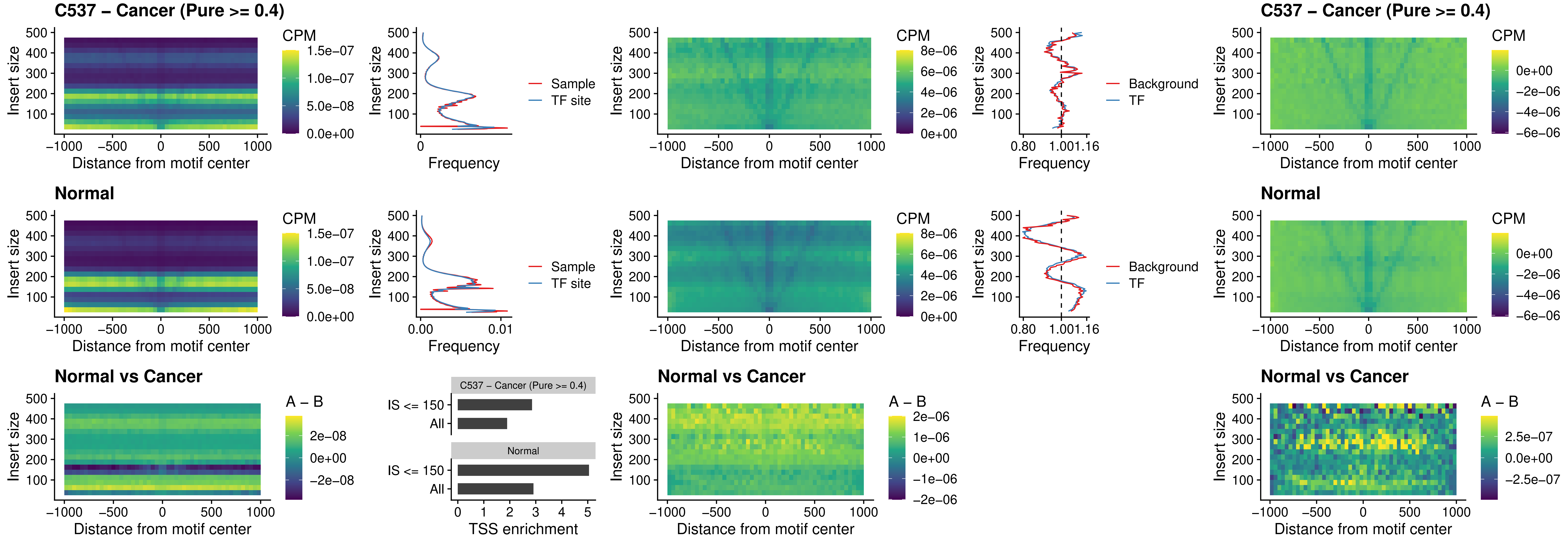

#### C537 – Cancer (Pure >= 0.4) – DLX3_uc_pTSS_nPEAK

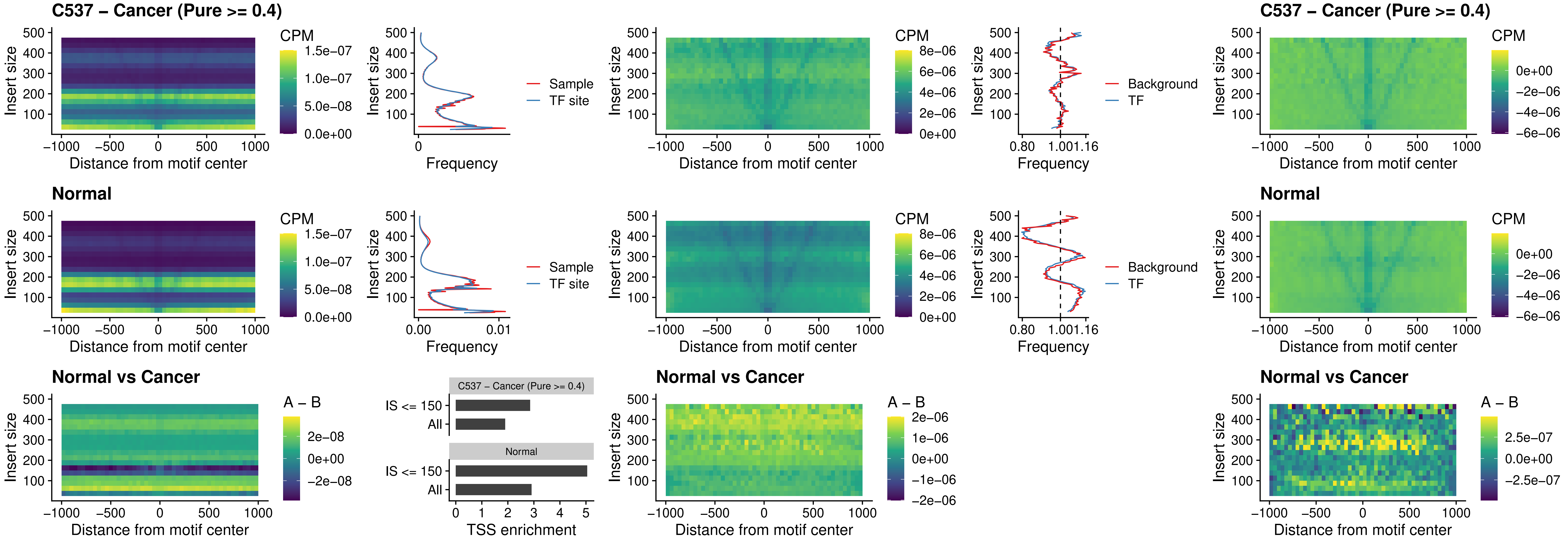

#### C537 – Cancer (Pure >= 0.4) – HOXD3_uc_pTSS_nPEAK

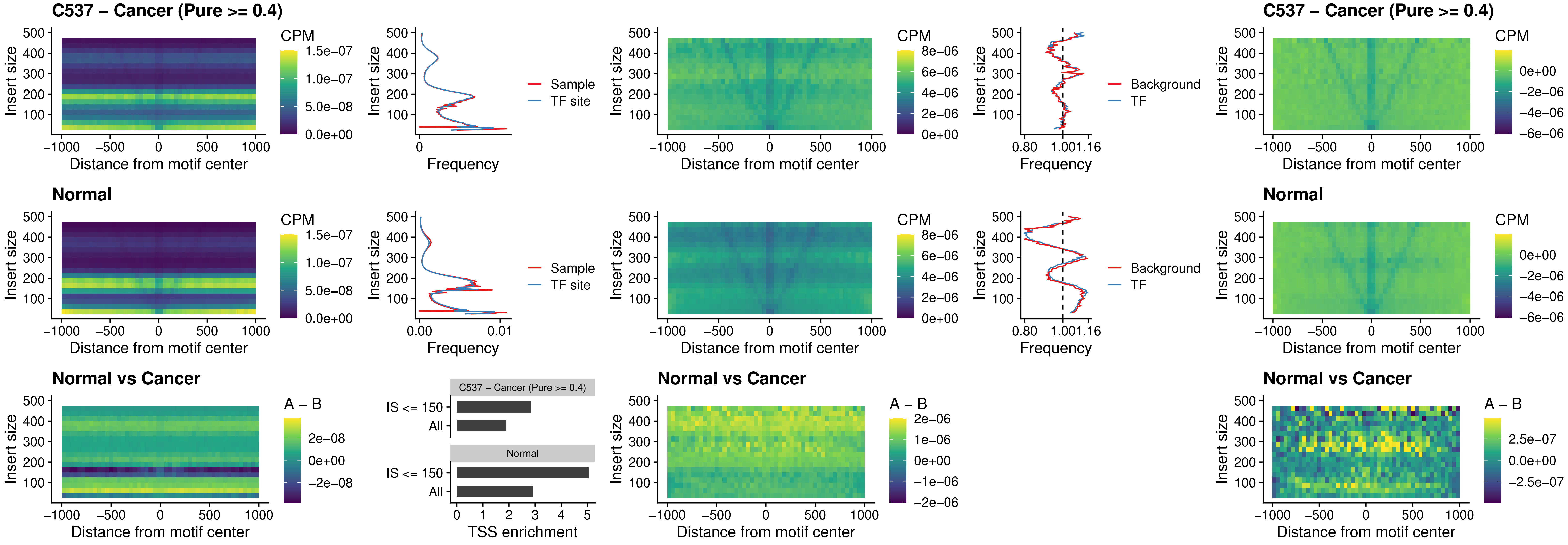

#### C537 – Cancer (Pure >= 0.4) – PDX1_uc_dTSS_nPEAK

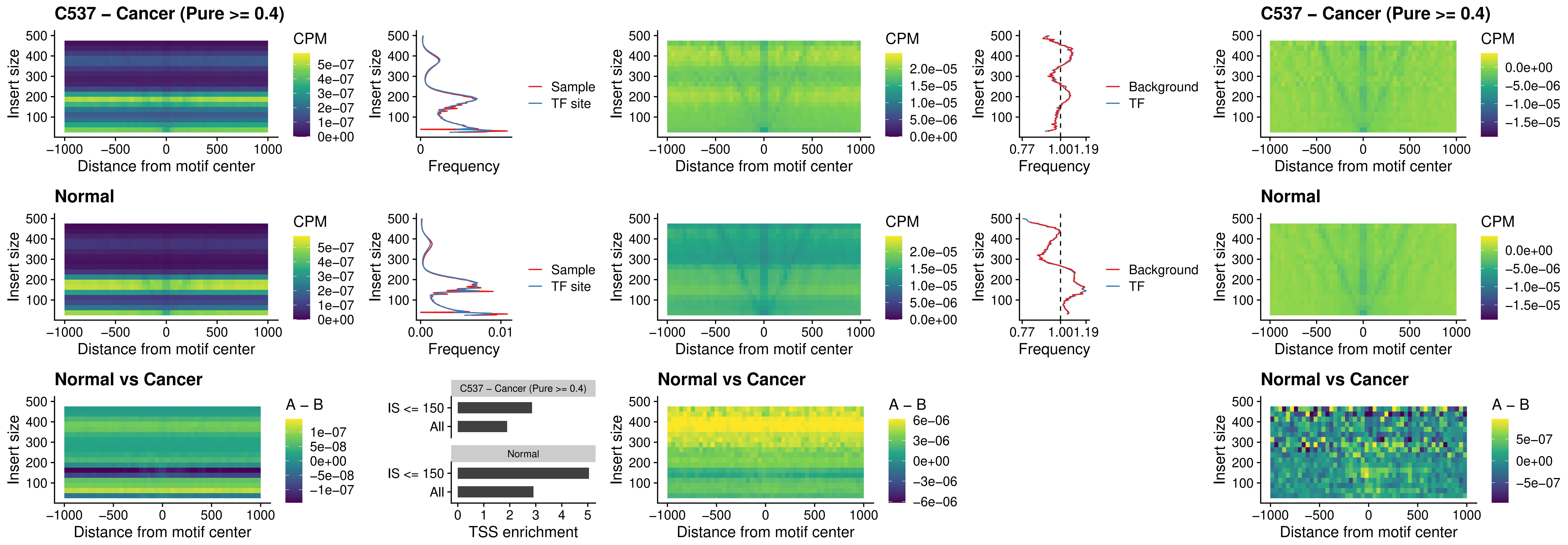

#### C537 – Cancer (Pure >= 0.4) – VAX2_uc_dTSS_nPEAK

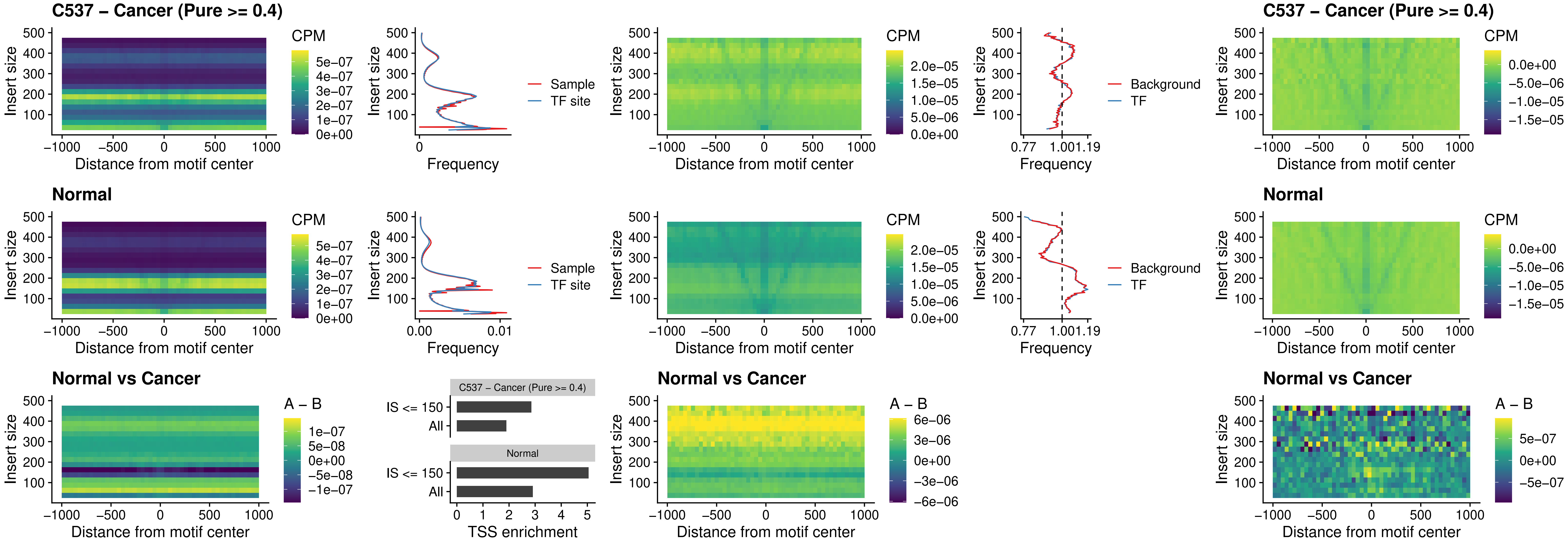

#### C537 – Cancer (Pure >= 0.4) – SOX5_uc_dTSS_nPEAK

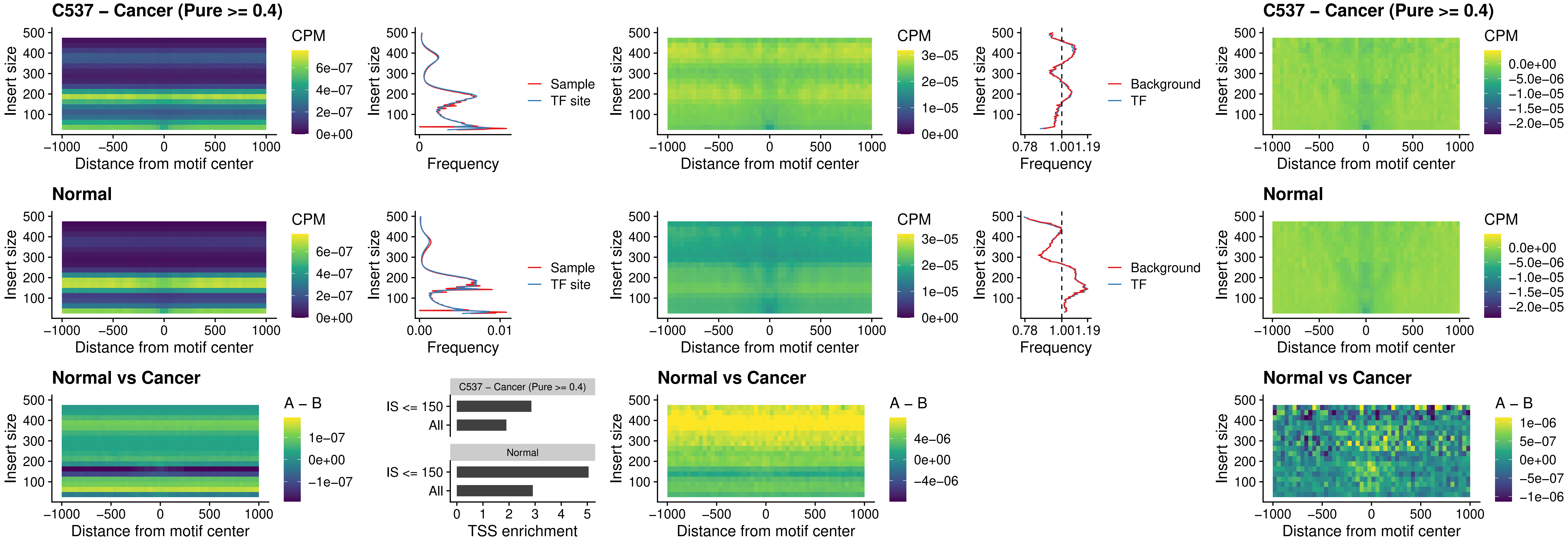

#### C537 – Cancer (Pure >= 0.4) – SOX6_uc_dTSS_nPEAK

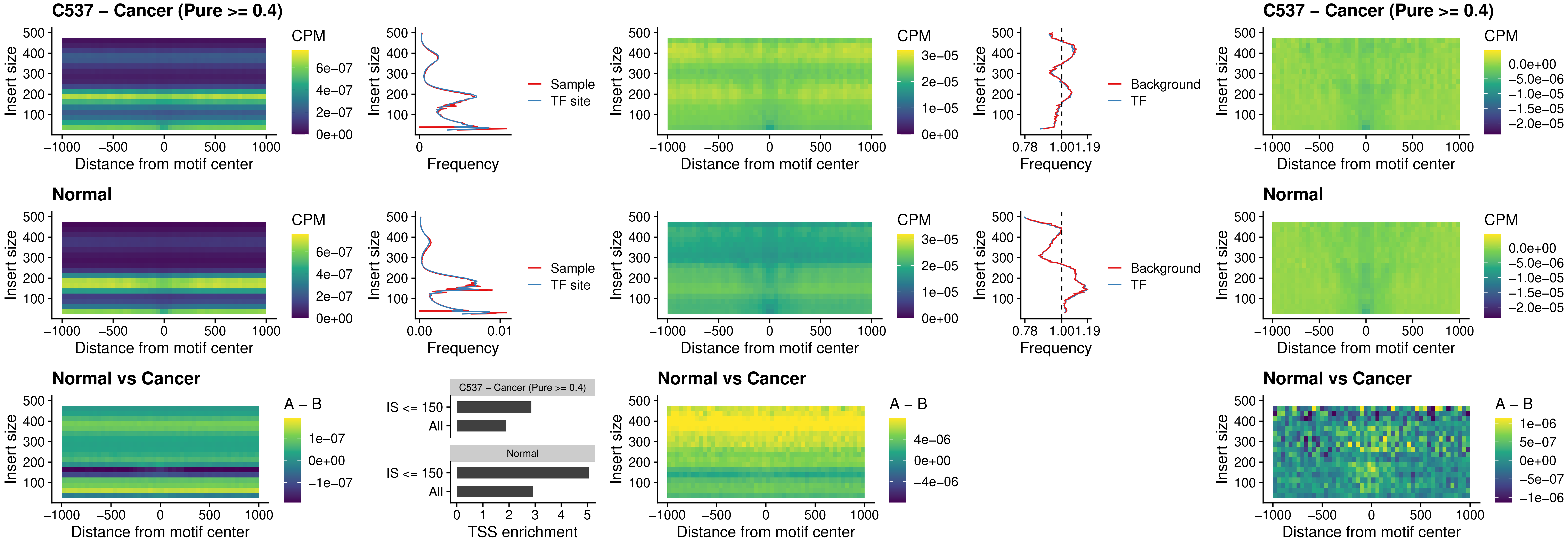

#### C537 – Cancer (Pure >= 0.4) – JDP2_uc_cTSS_nPEAK

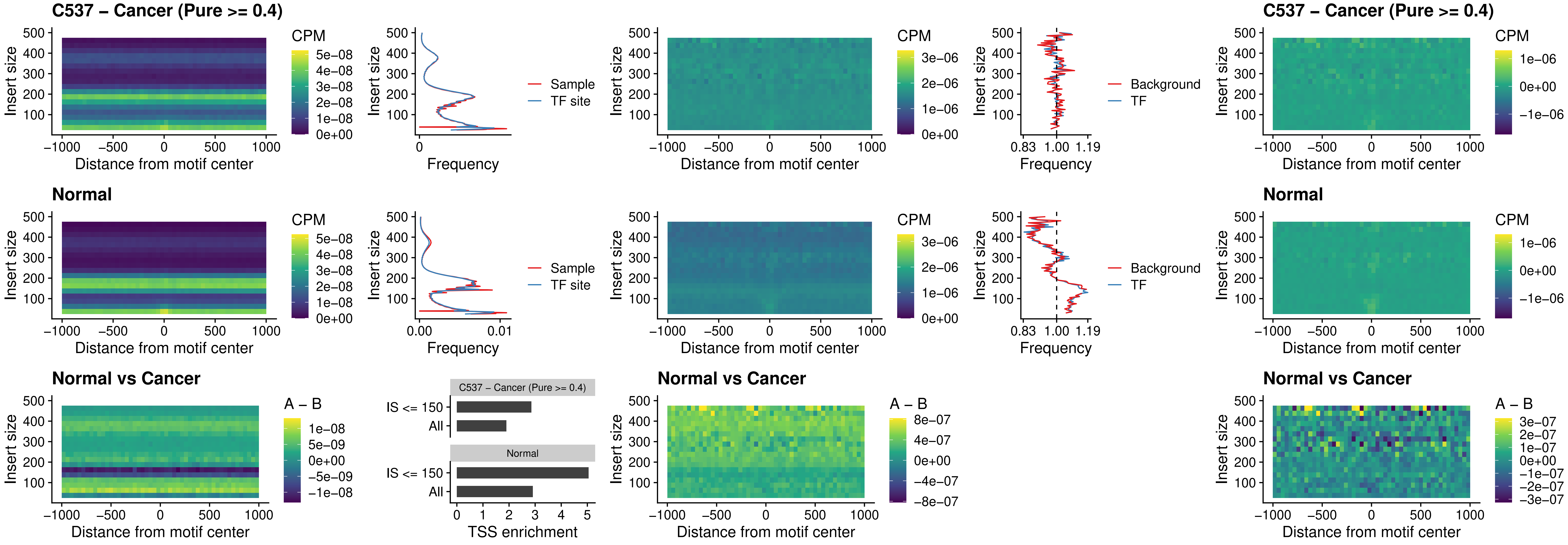

#### C537 – Cancer (Pure >= 0.4) – PURA_uc_dTSS_oPEAK

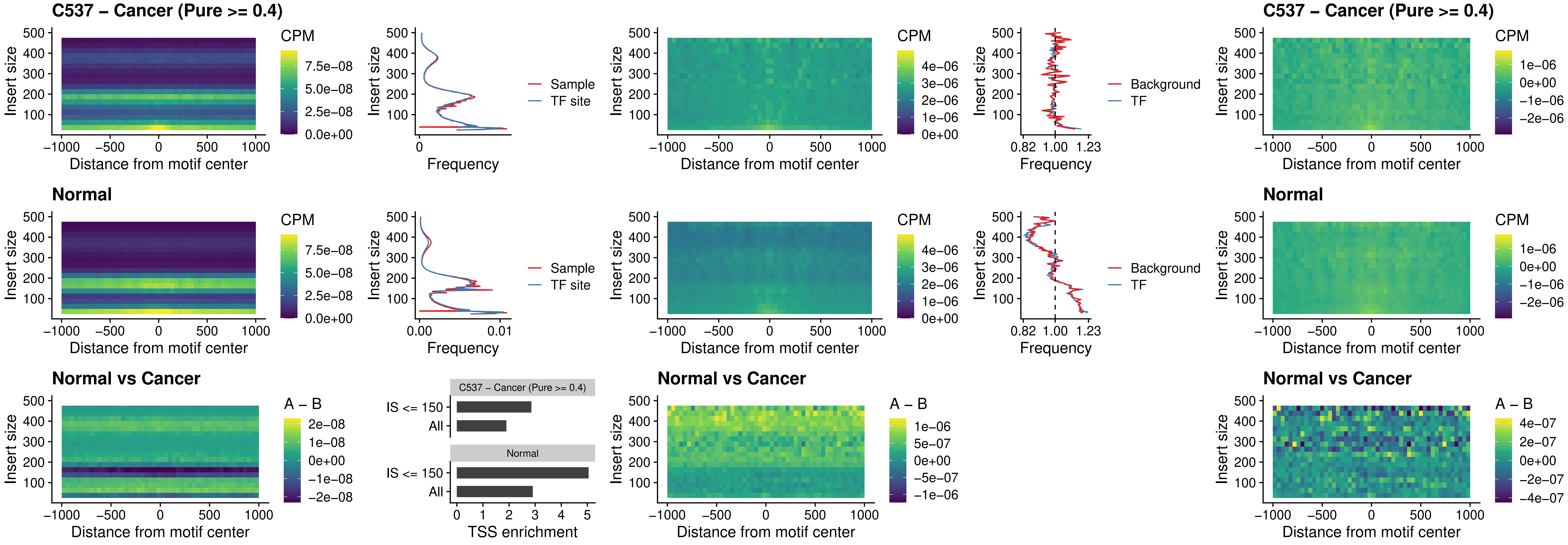

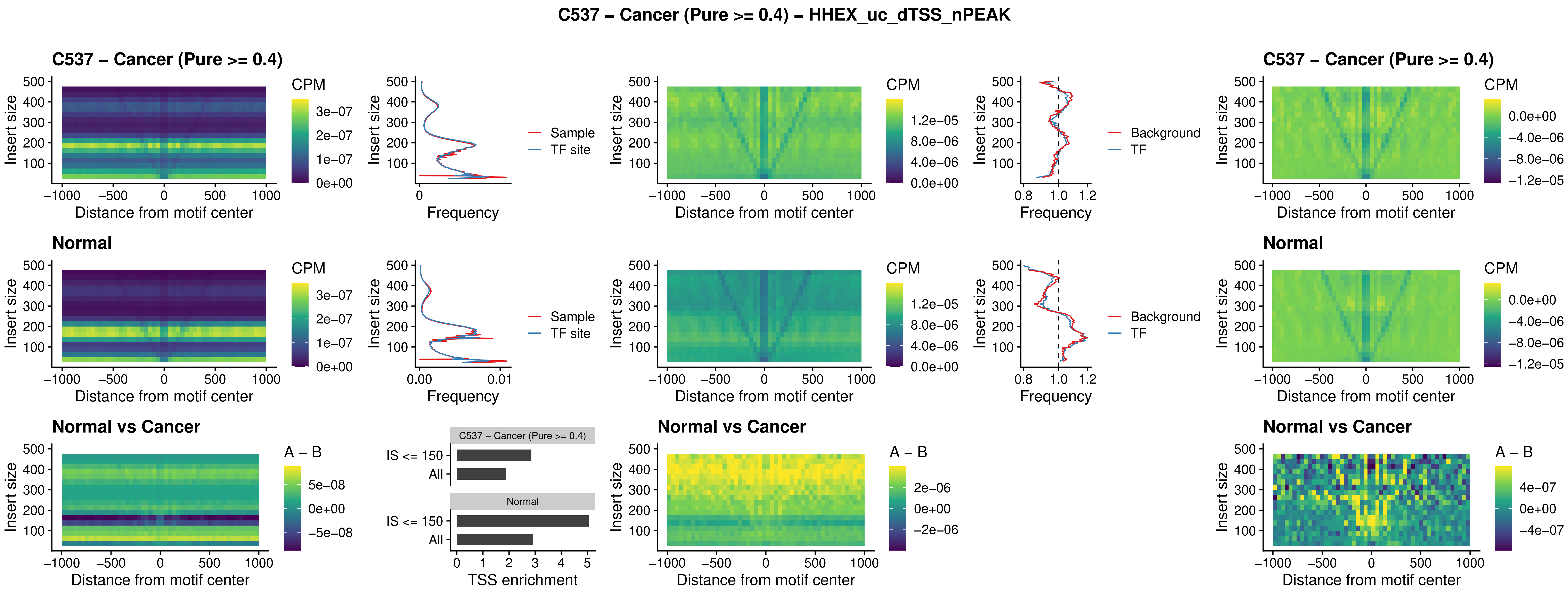
C537 – Cancer (Pure >= 0.4) – HHEX_uc_dTSS_nPEAK
C537 – Cancer (Pure >= 0.4)
Normal
Normal vs Cancer
Insert size
Distance from motif center
Frequency
CPM
A – B
Sample
TF site
Background
TF
C537 – Cancer (Pure >= 0.4)
Normal
IS <= 150
All
TSS enrichment

#### C537 – Cancer (Pure >= 0.4) – HMG20B_uc_dTSS_nPEAK

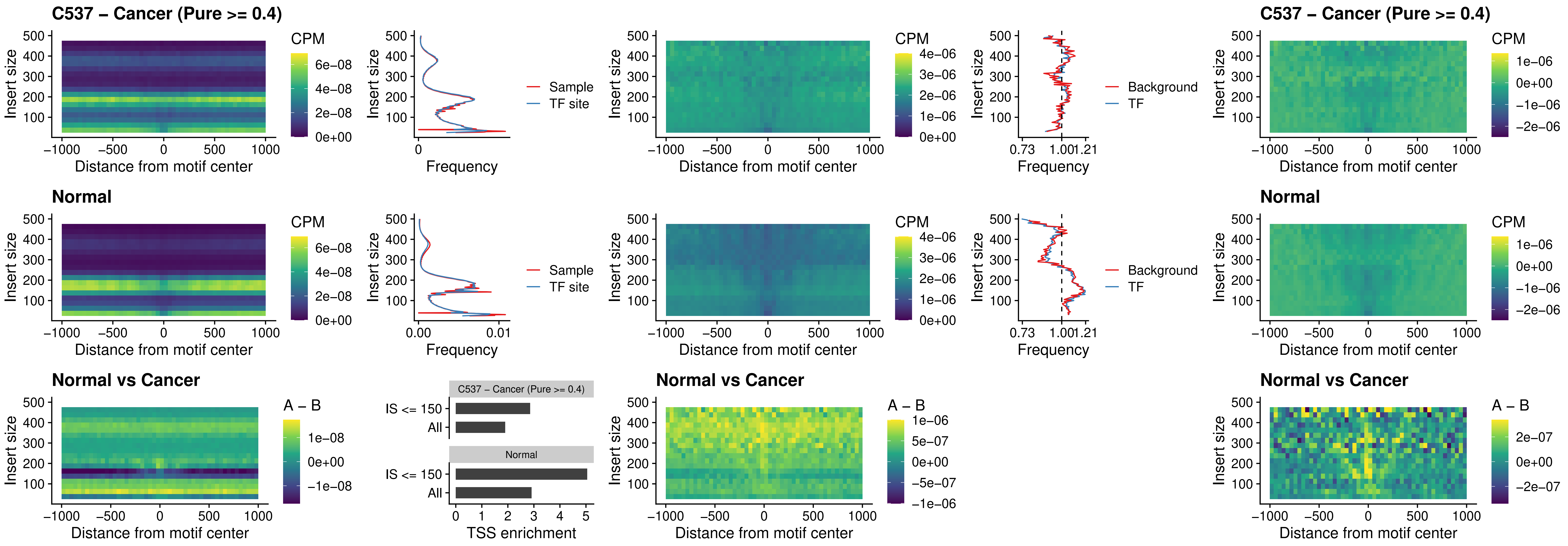

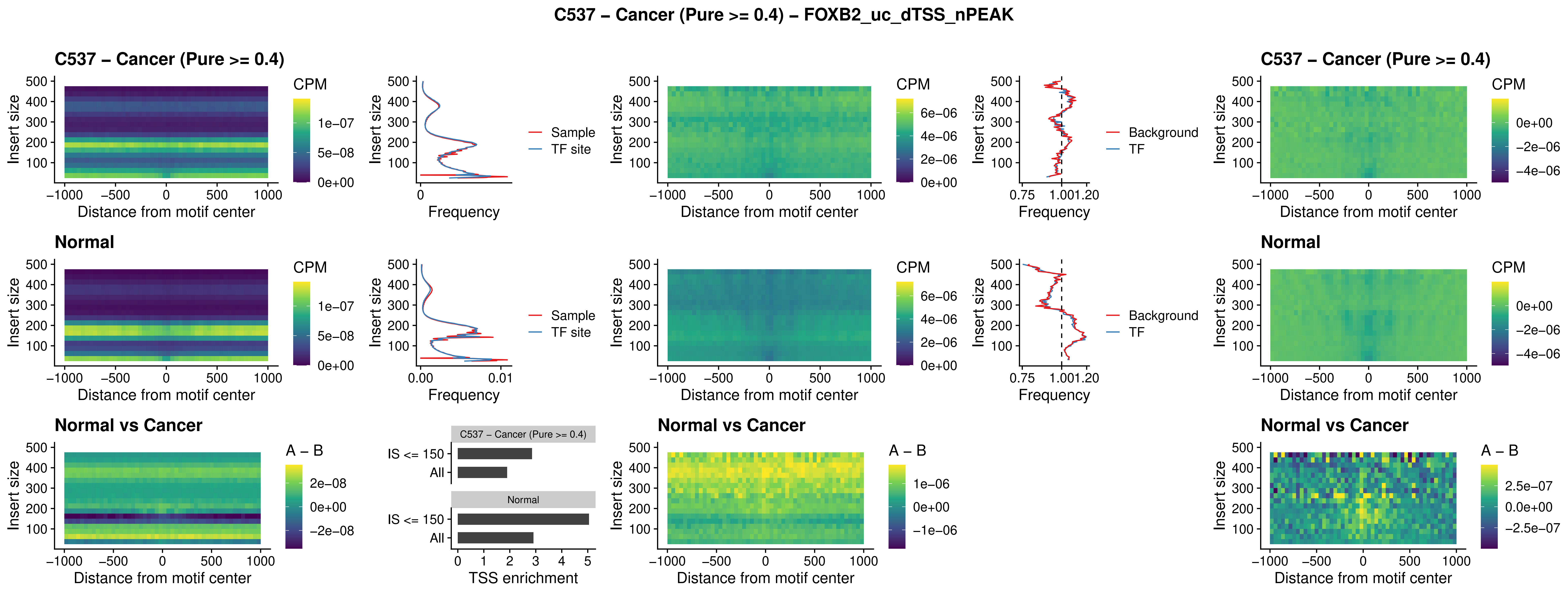
C537 – Cancer (Pure >= 0.4) – FOXB2_uc_dTSS_nPEAK
C537 – Cancer (Pure >= 0.4)
Insert size
Distance from motif center
CPM
Frequency
Sample
TF site
Background
TF
Normal
Normal vs Cancer
A – B
C537 – Cancer (Pure >= 0.4)
IS <= 150
All
TSS enrichment

### C537 – Cancer (Pure >= 0.4) – FOXL1_uc_cTSS_nPEAK

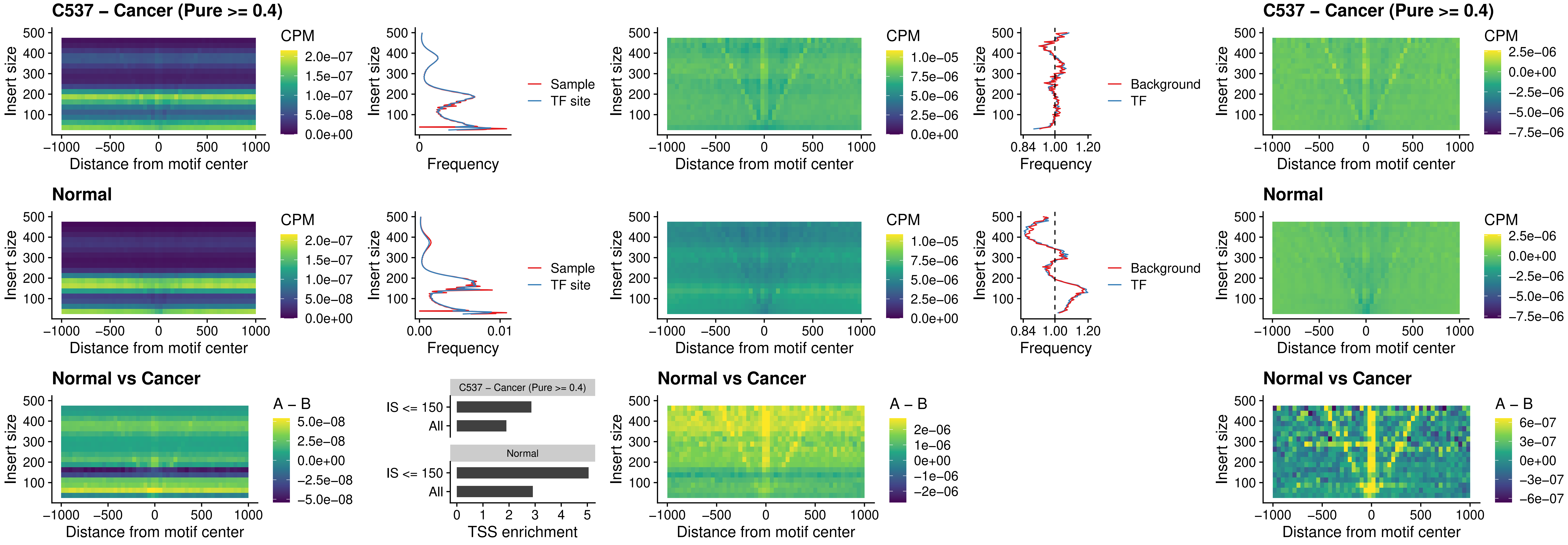

#### C537 – Cancer (Pure >= 0.4) – PBX3_uc_pTSS_oPEAK

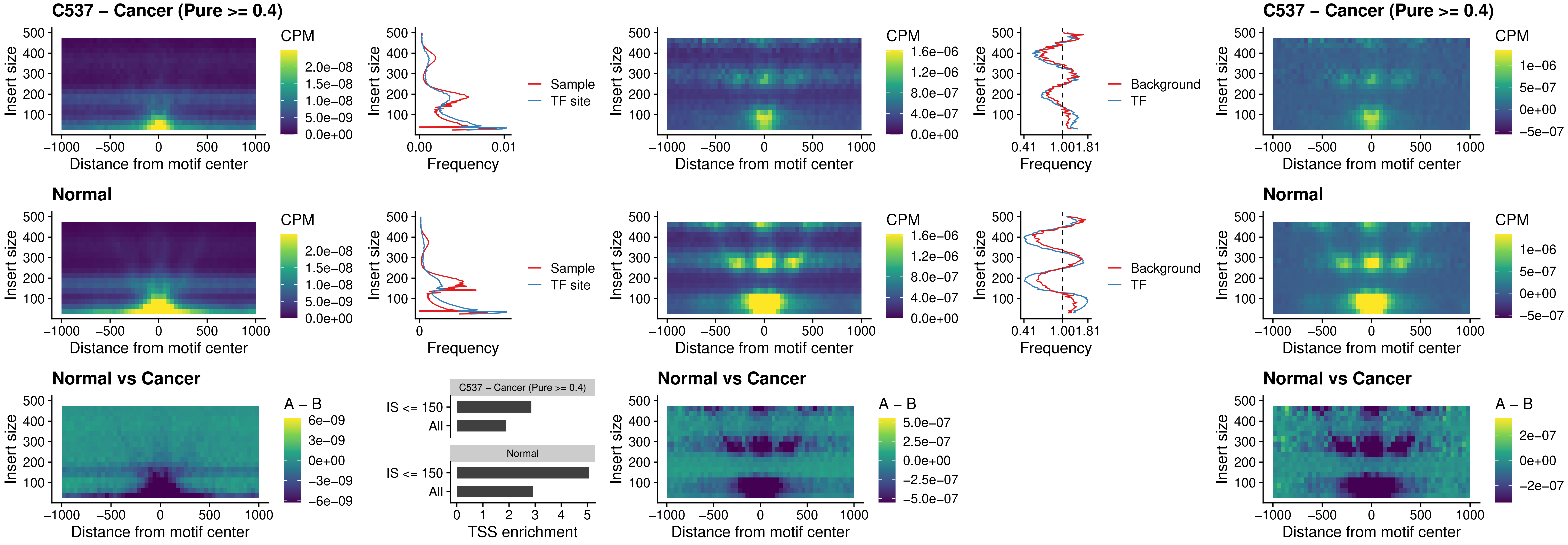

#### C537 – Cancer (Pure >= 0.4) – SNAI2_uc_cTSS_nPEAK

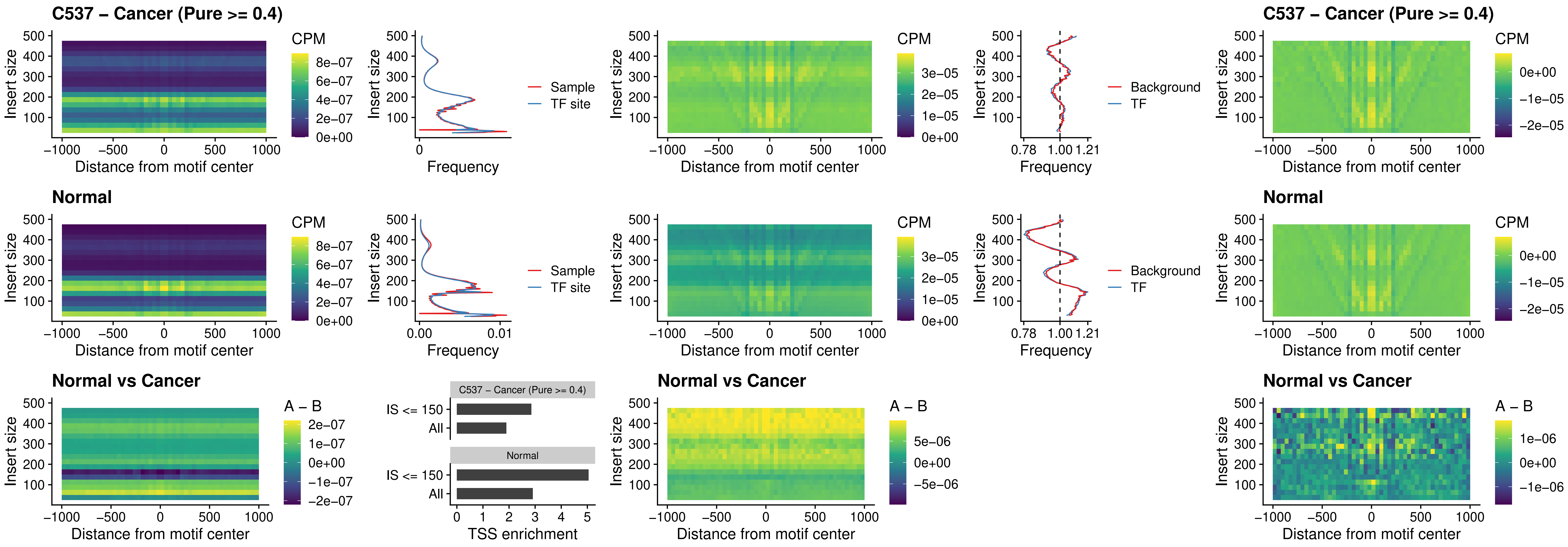

#### C537 – Cancer (Pure >= 0.4) – SNAI2_uc_dTSS_oPEAK

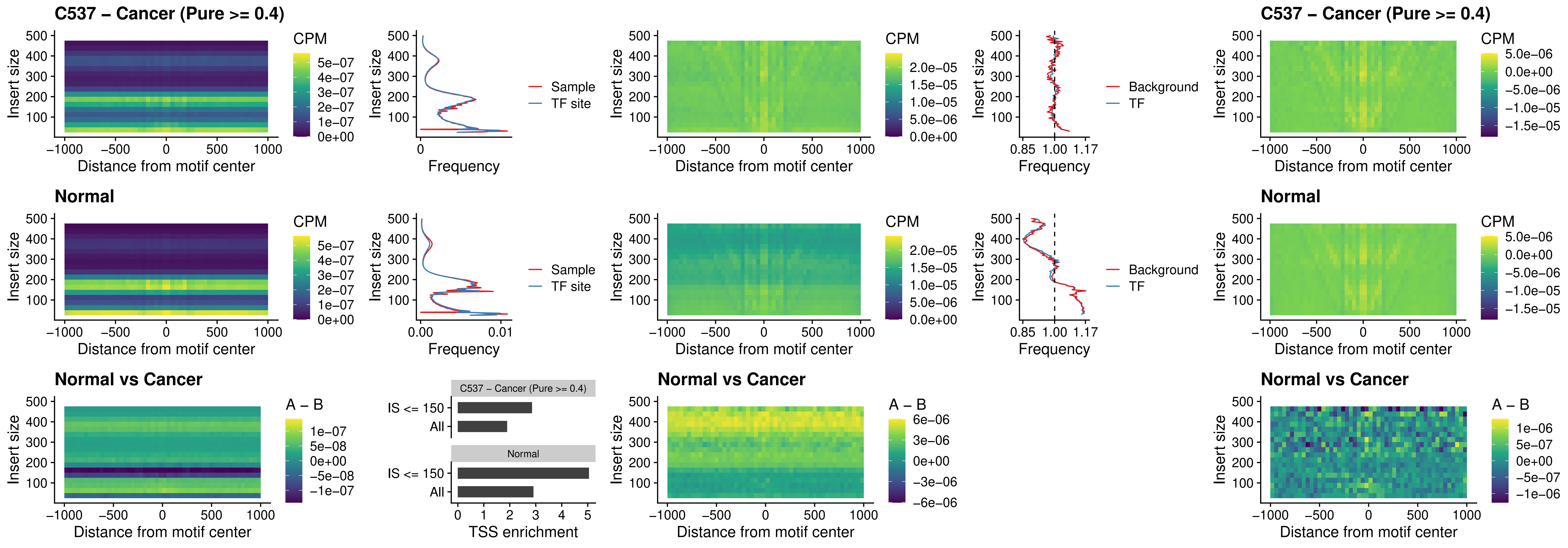

#### C537 – Cancer (Pure >= 0.4) – SPI1_uc_pTSS_nPEAK

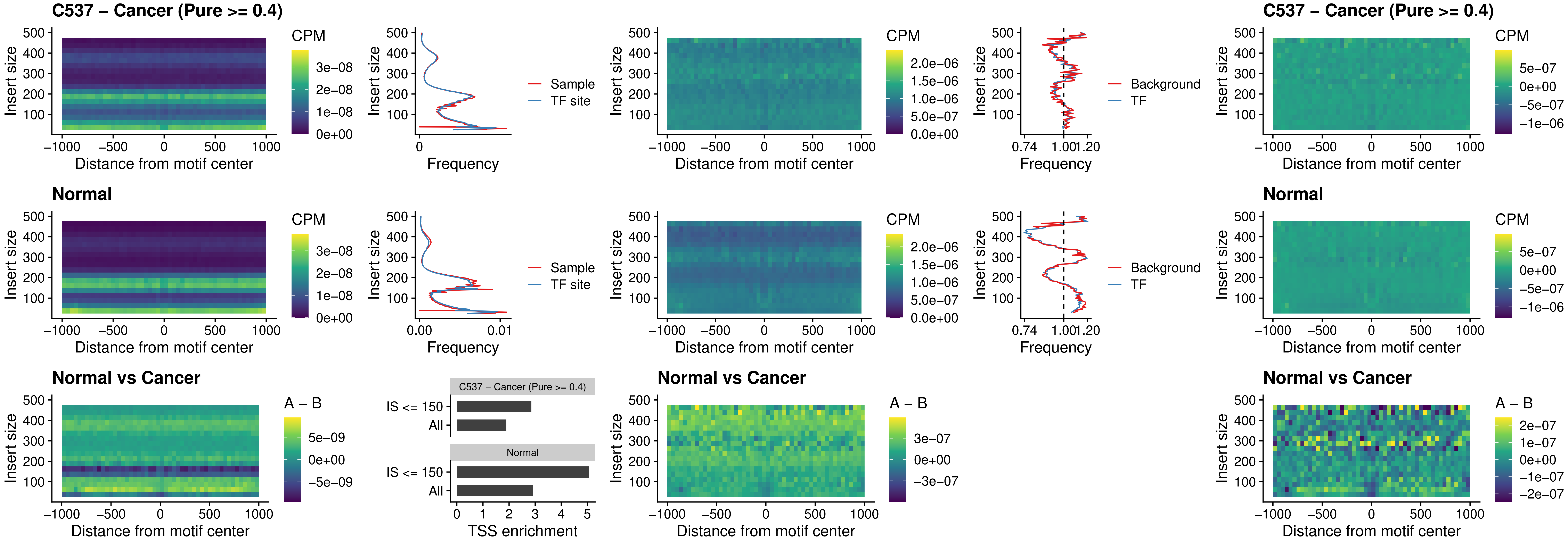

#### C537 – Cancer (Pure >= 0.4) – KLF16_uc_pTSS_nPEAK

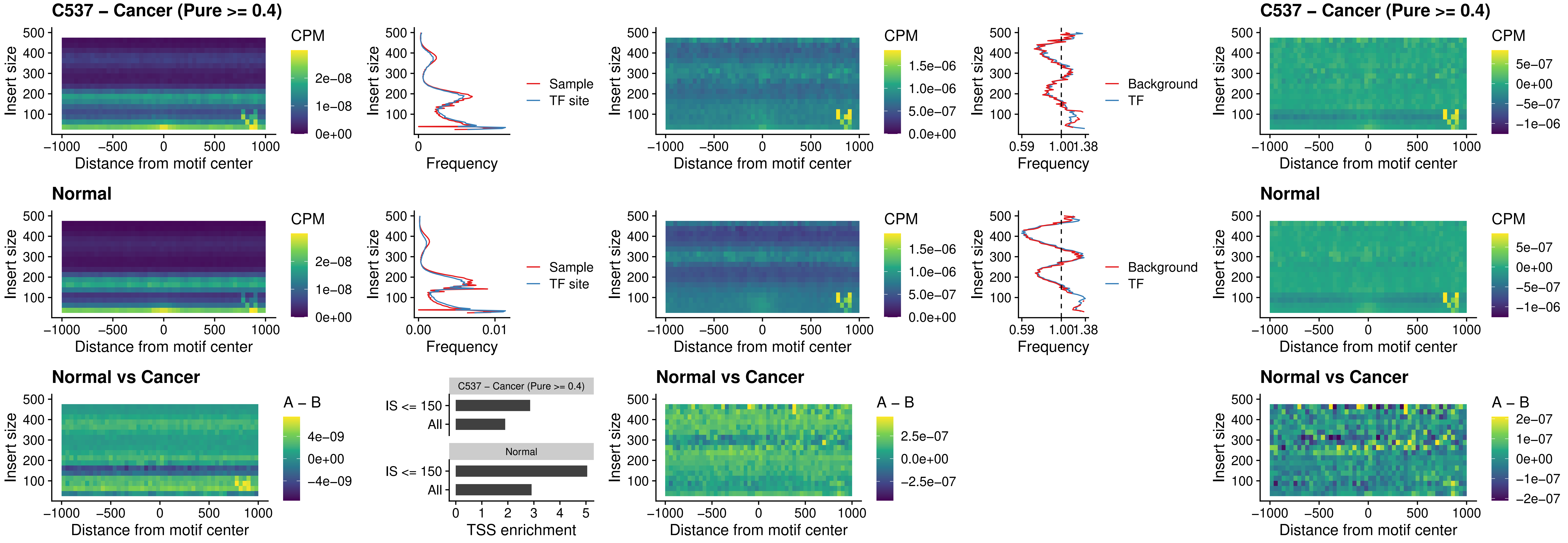

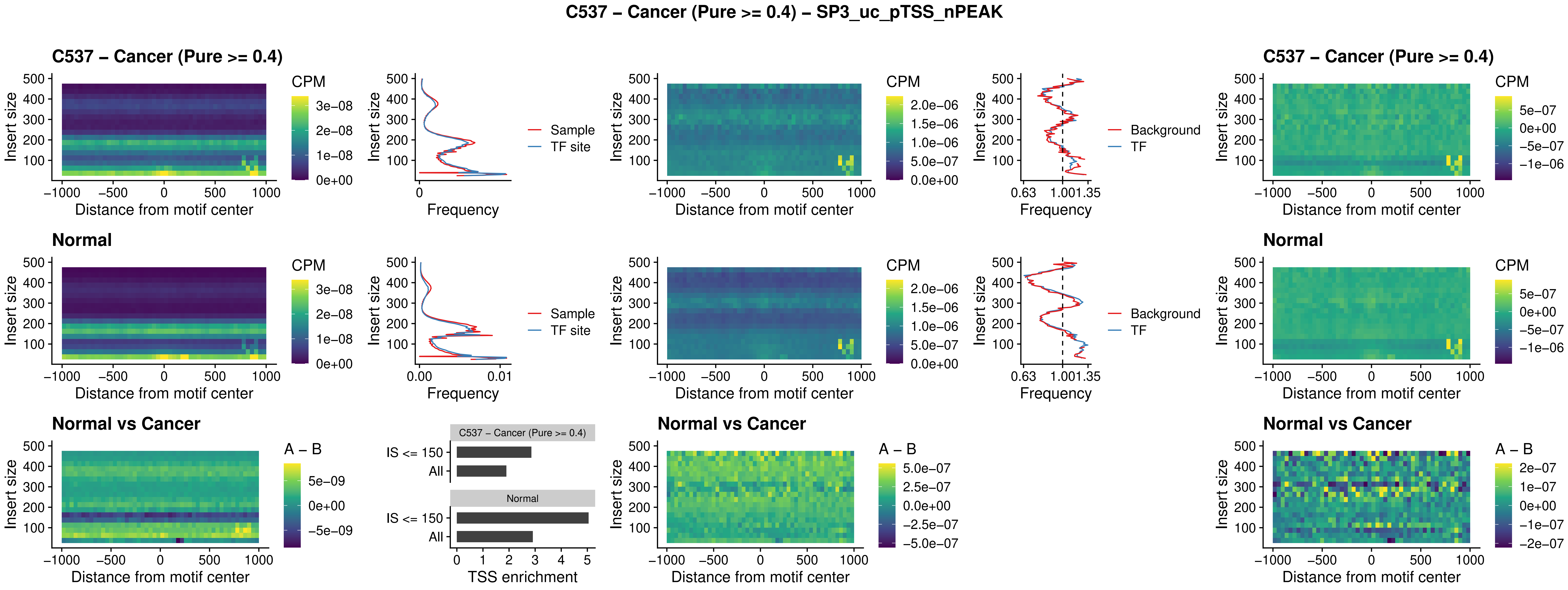
C537 – Cancer (Pure >= 0.4) – SP3_uc_pTSS_nPEAK
C537 – Cancer (Pure >= 0.4)
Normal
Normal vs Cancer
Insert size
Distance from motif center
Frequency
CPM
A – B
Sample
TF site
Background
TF
C537 – Cancer (Pure >= 0.4)
Normal
IS <= 150
All
TSS enrichment

#### C537 – Cancer (Pure >= 0.4) – NEUROD1_uc_dTSS_oPEAK

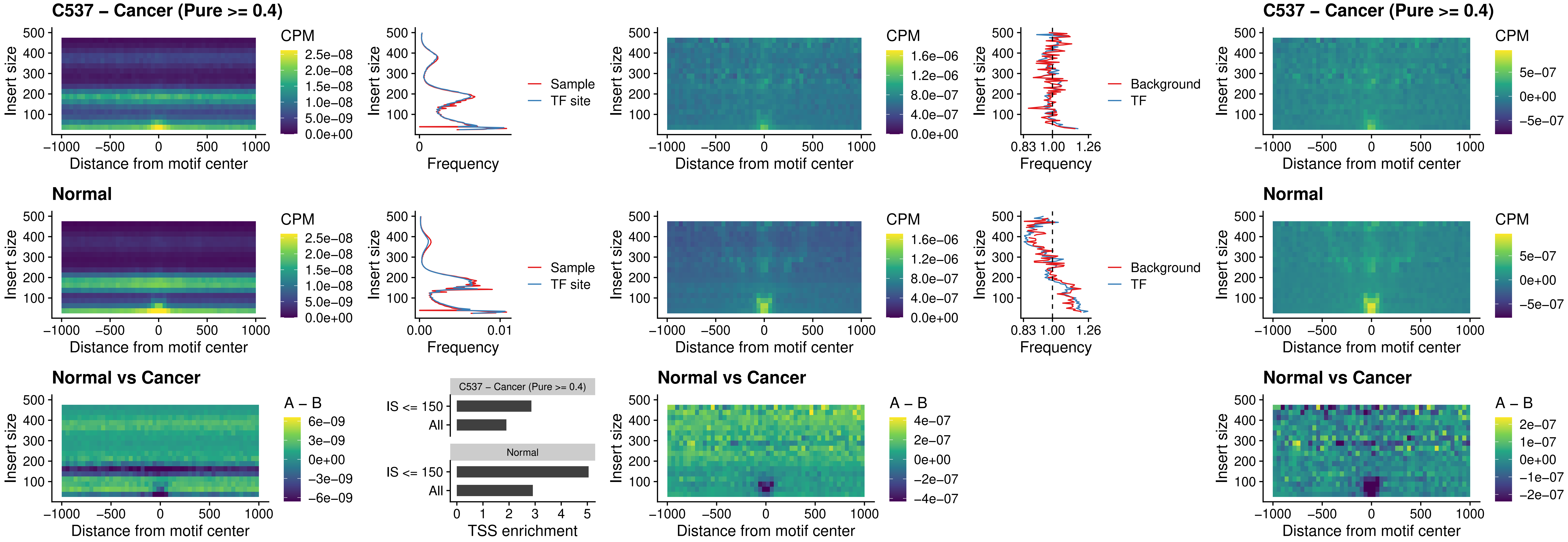

#### C537 – Cancer (Pure >= 0.4) – CTCF_uc_dTSS_oPEAK

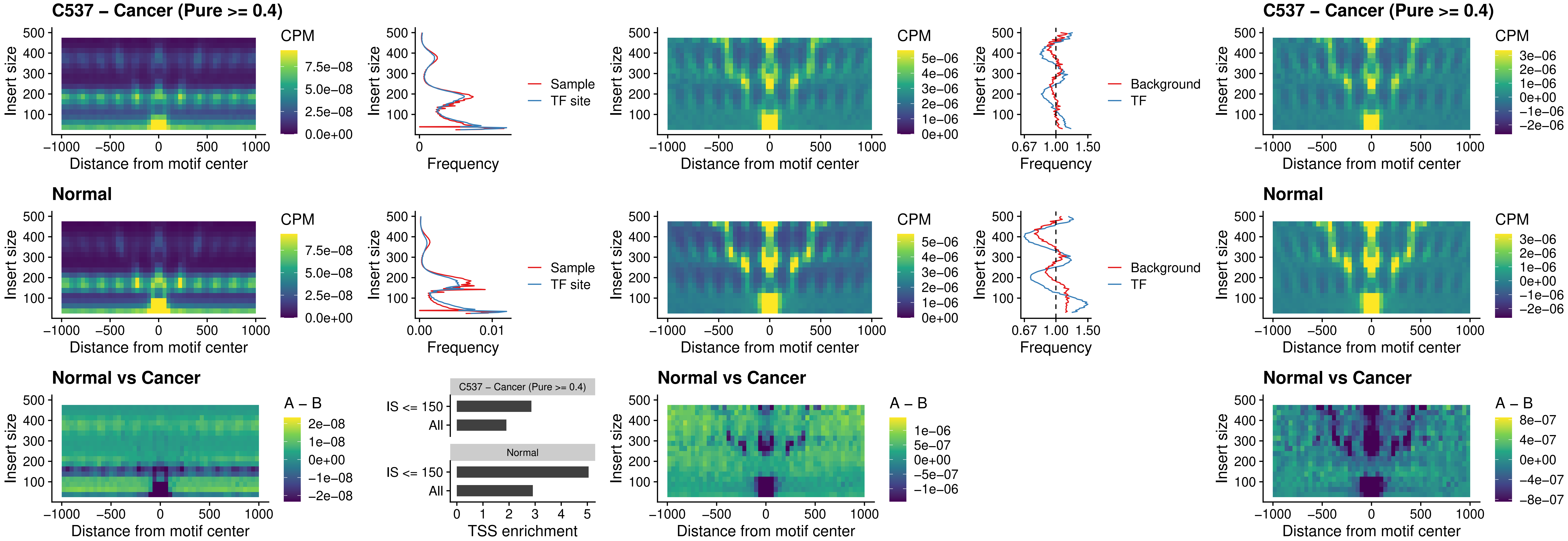

#### C537 – Cancer (Pure >= 0.4) – CTCF_uc_cTSS_oPEAK

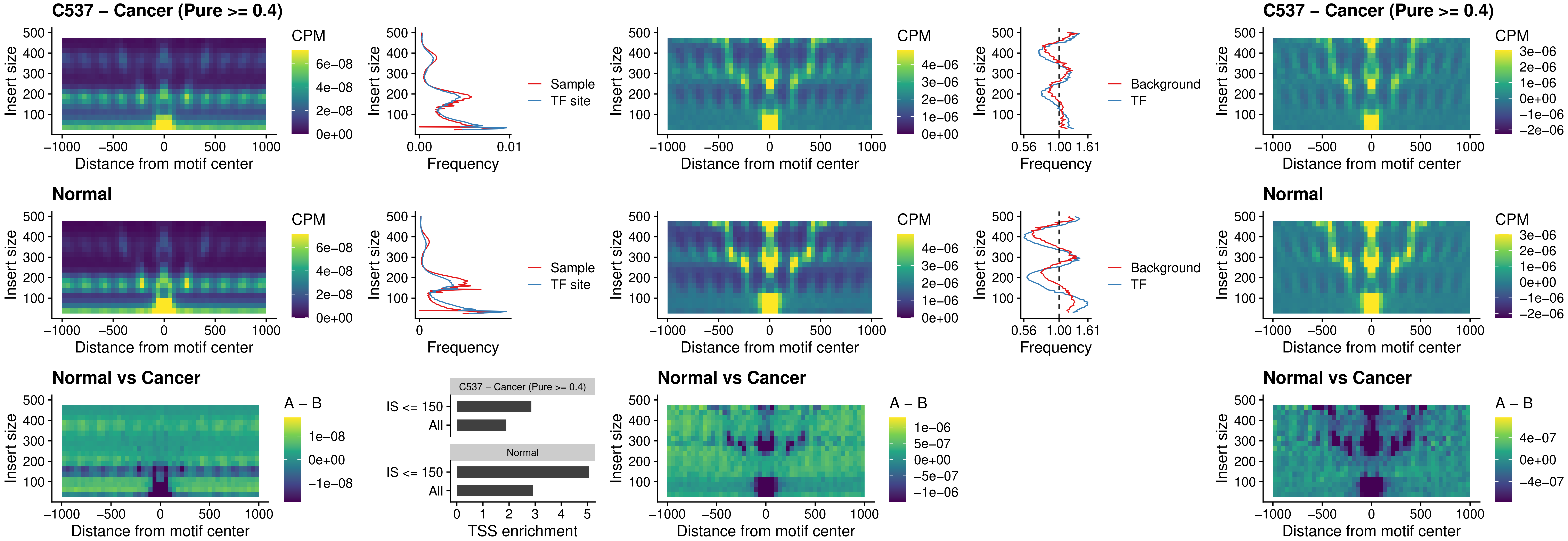

#### C537 – Cancer (Pure >= 0.4) – CTCFL_uc_dTSS_oPEAK

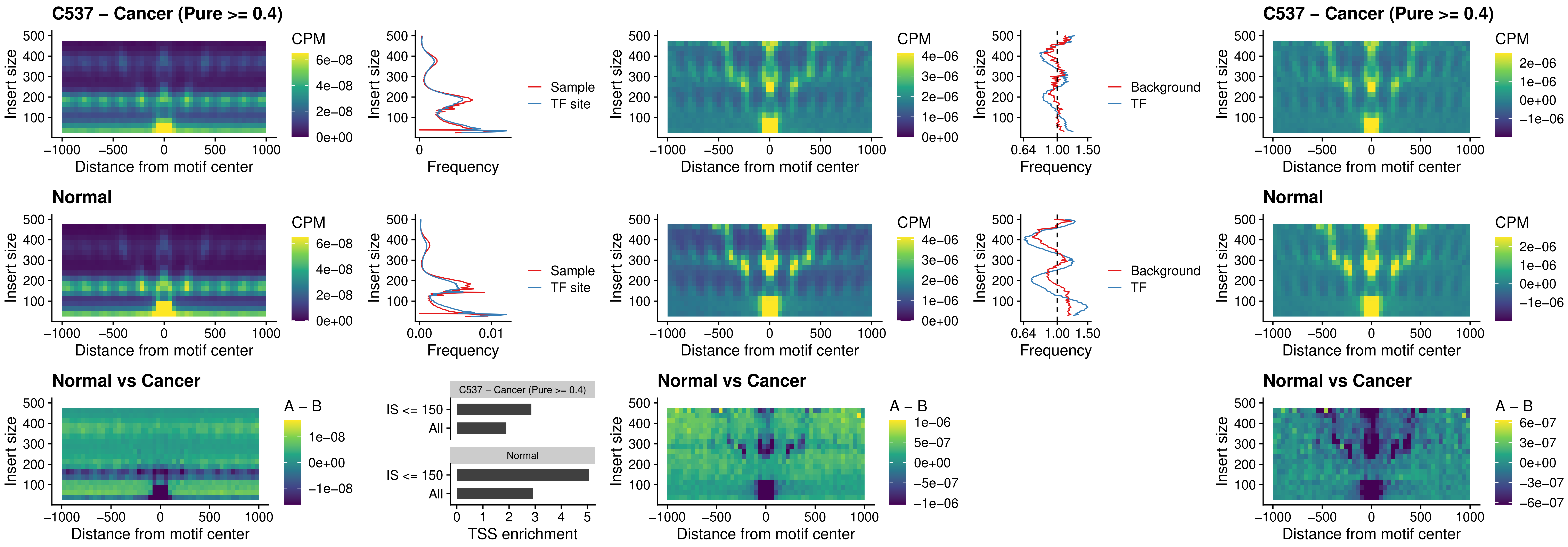

#### C537 – Cancer (Pure >= 0.4) – ENCODE_CTCF_sites_COLON

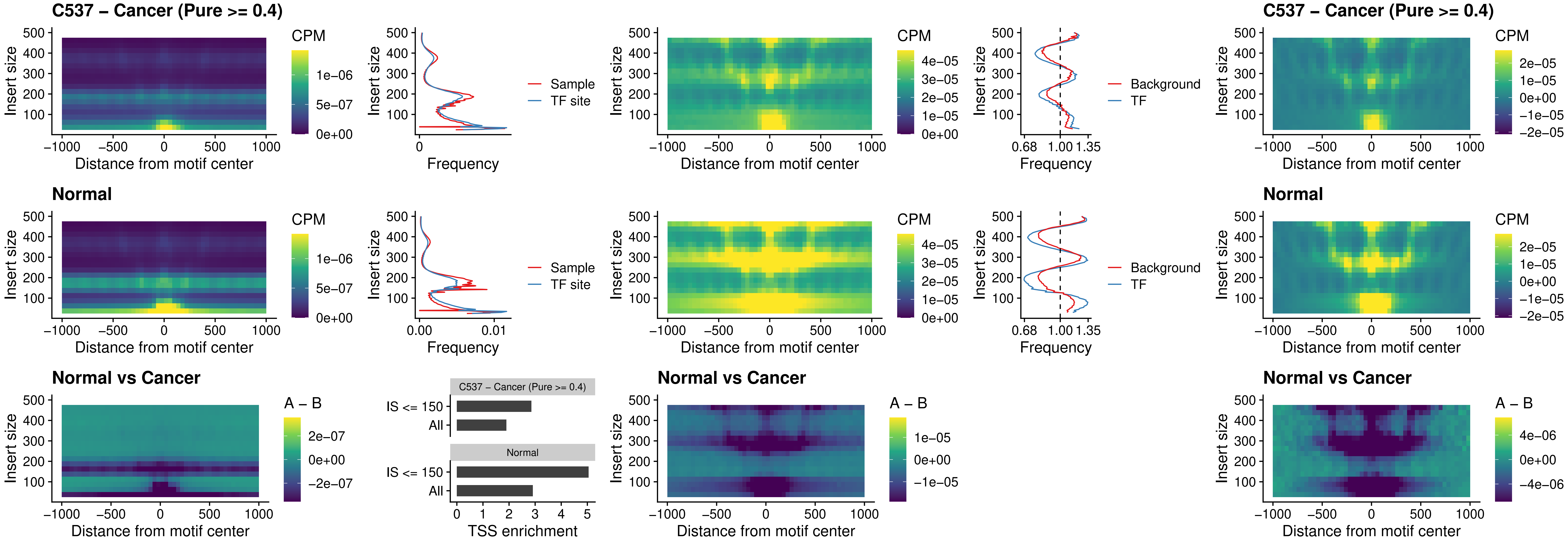

#### C537 – Cancer (Pure >= 0.4) – ENCODE_CTCF_sites_COLON_loop_ctcf

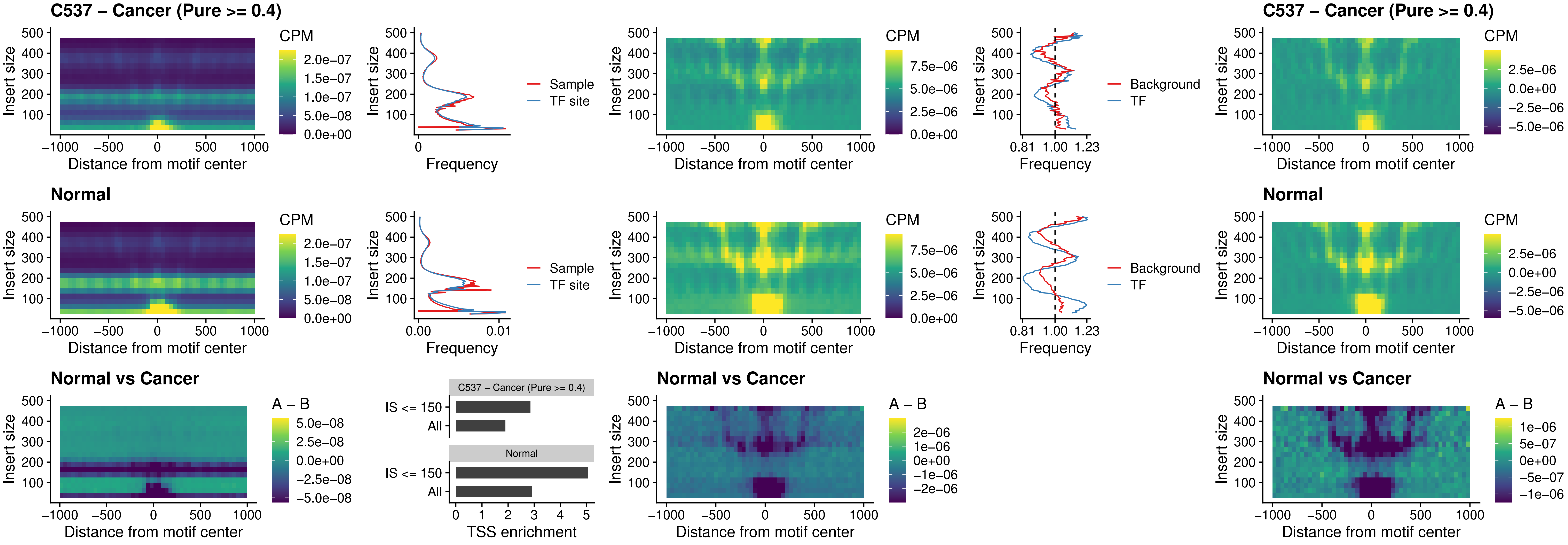

#### C537 – Cancer (Pure >= 0.4) – ENCODE_CTCF_sites_COLON_intergenic_ctcf

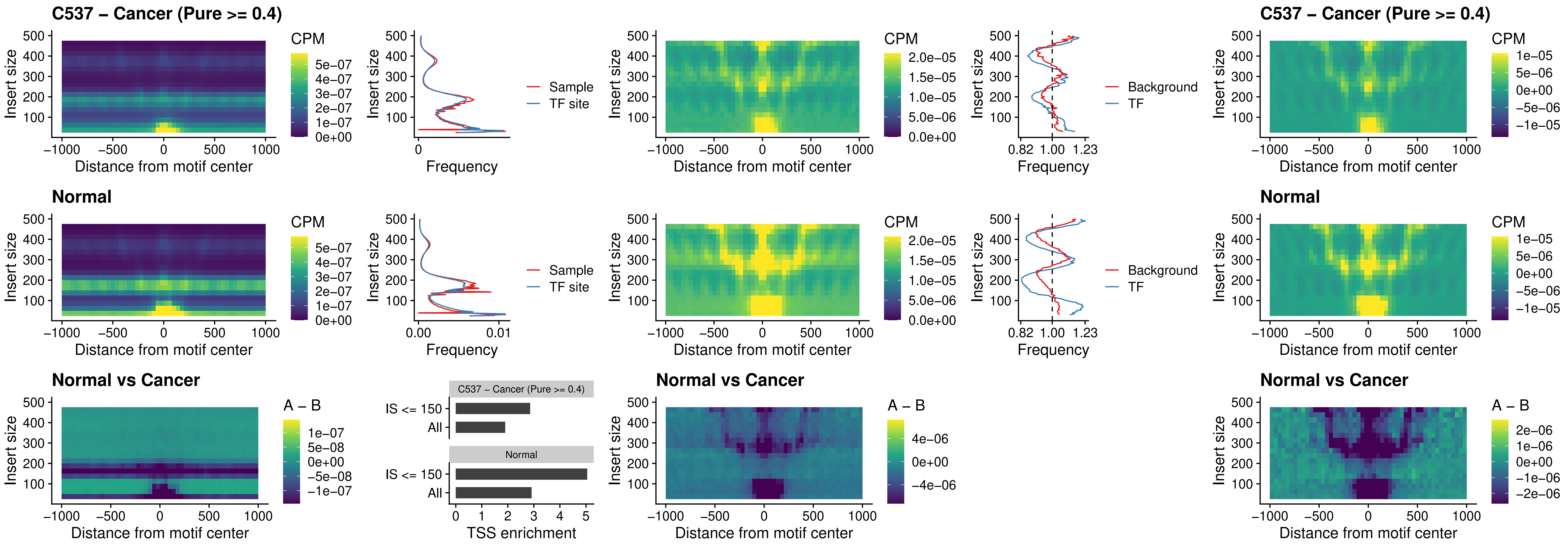

### C537 – Cancer (Pure >= 0.4) – ENCODE_CTCF_sites_COLON_nonpromotor

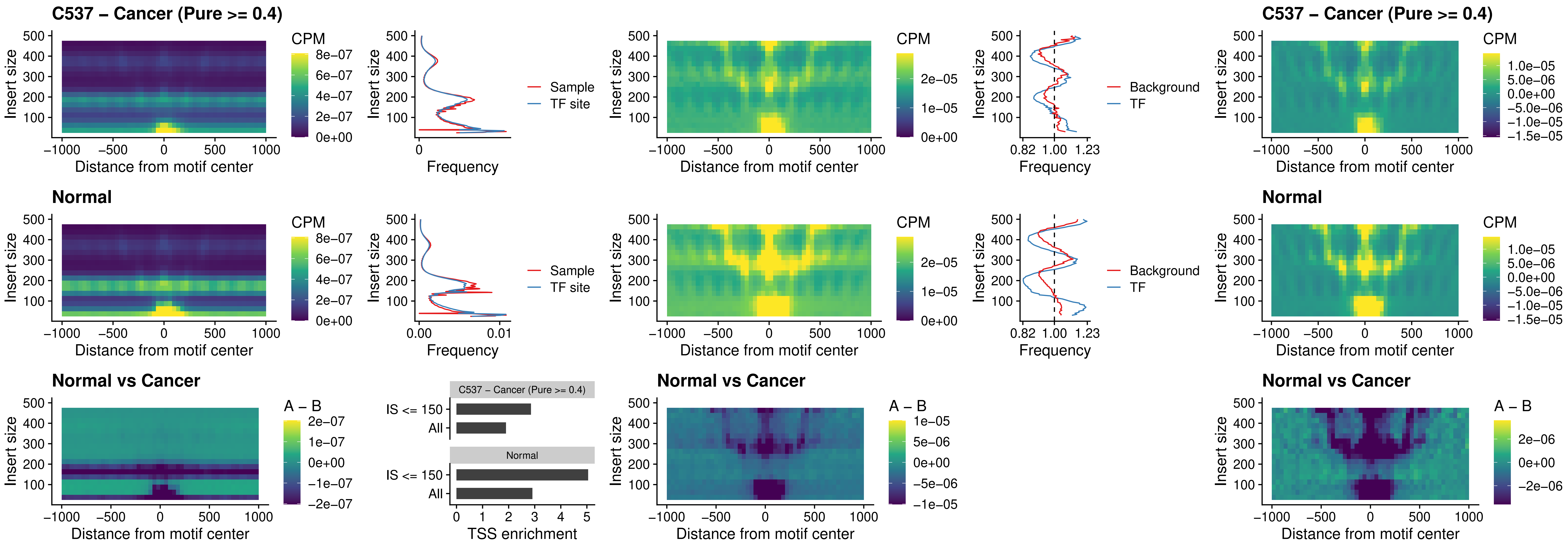

#### C537 – Cancer (Pure >= 0.4) – CTCFL_uc_cTSS_oPEAK

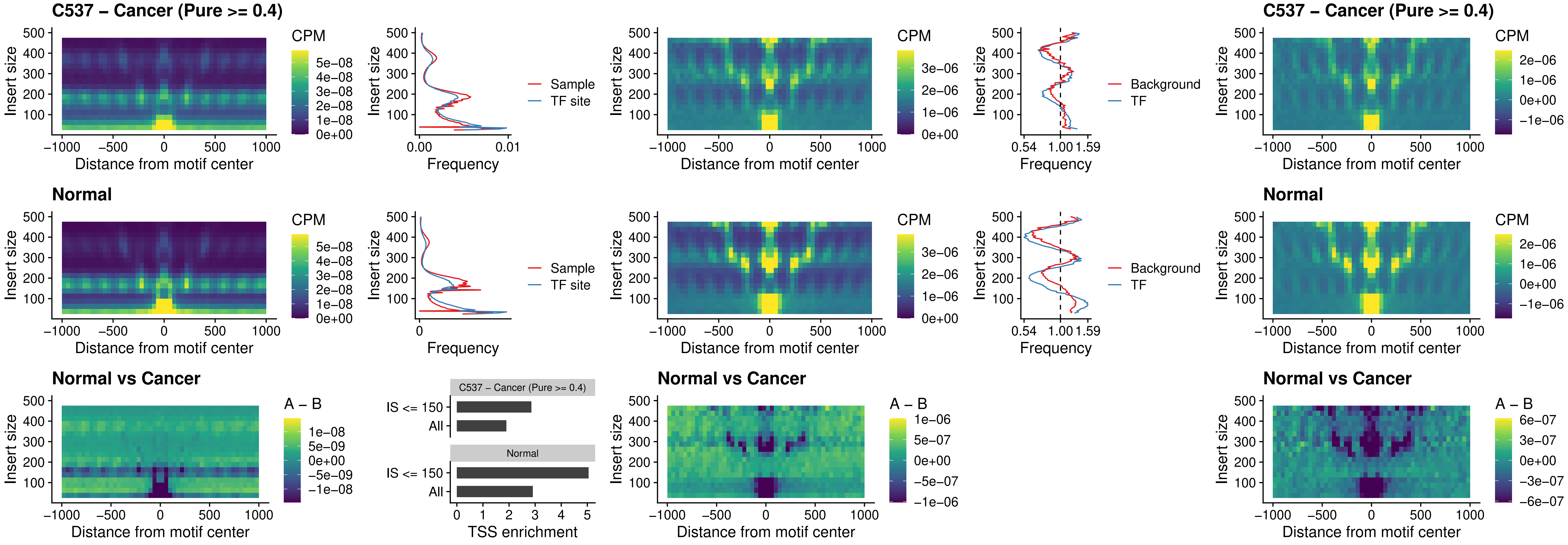

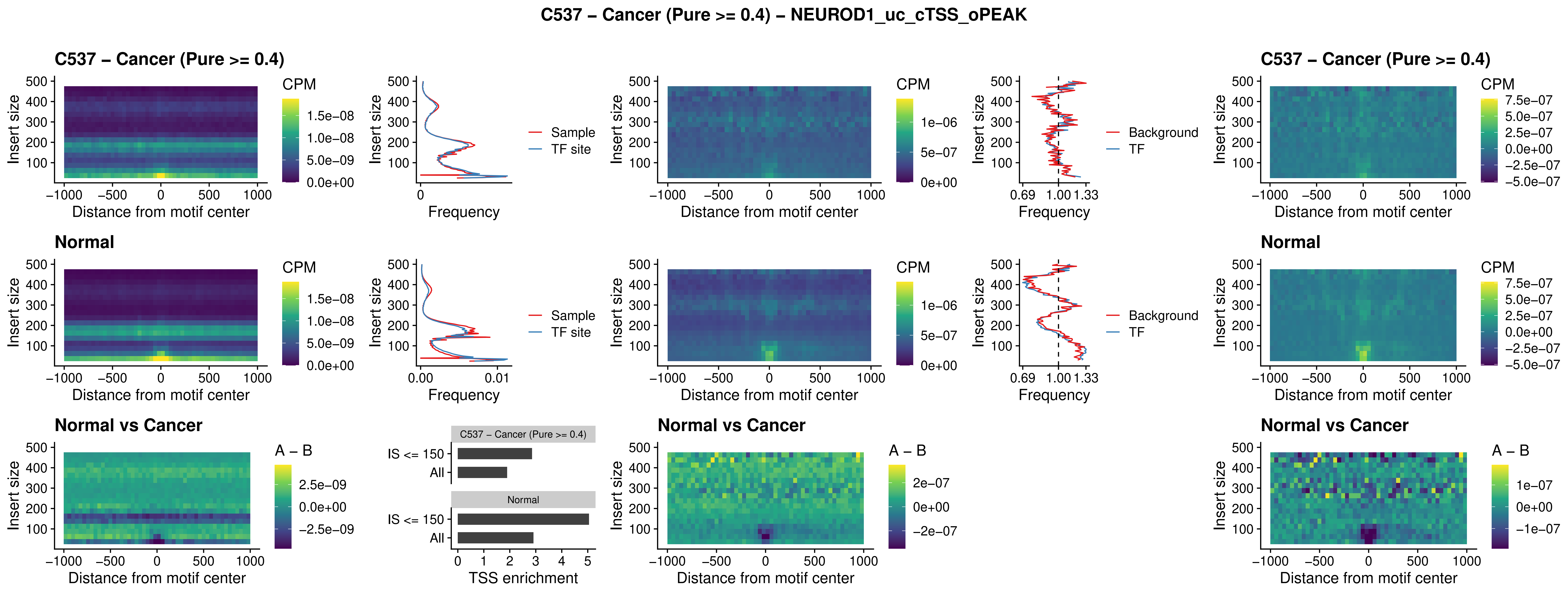
C537 – Cancer (Pure >= 0.4) – NEURODS1_uc_cTSS_oPEAK
C537 – Cancer (Pure >= 0.4)
Normal
Normal vs Cancer
CPM
A – B
Sample
TF site
Background
TF
Insert size
Frequency
Distance from motif center
TSS enrichment
IS <= 150
All
C537 – Cancer (Pure >= 0.4)
Normal

#### C537 – Cancer (Pure >= 0.4) – ENCODE_YY1_sites_COLON_nonpromotor

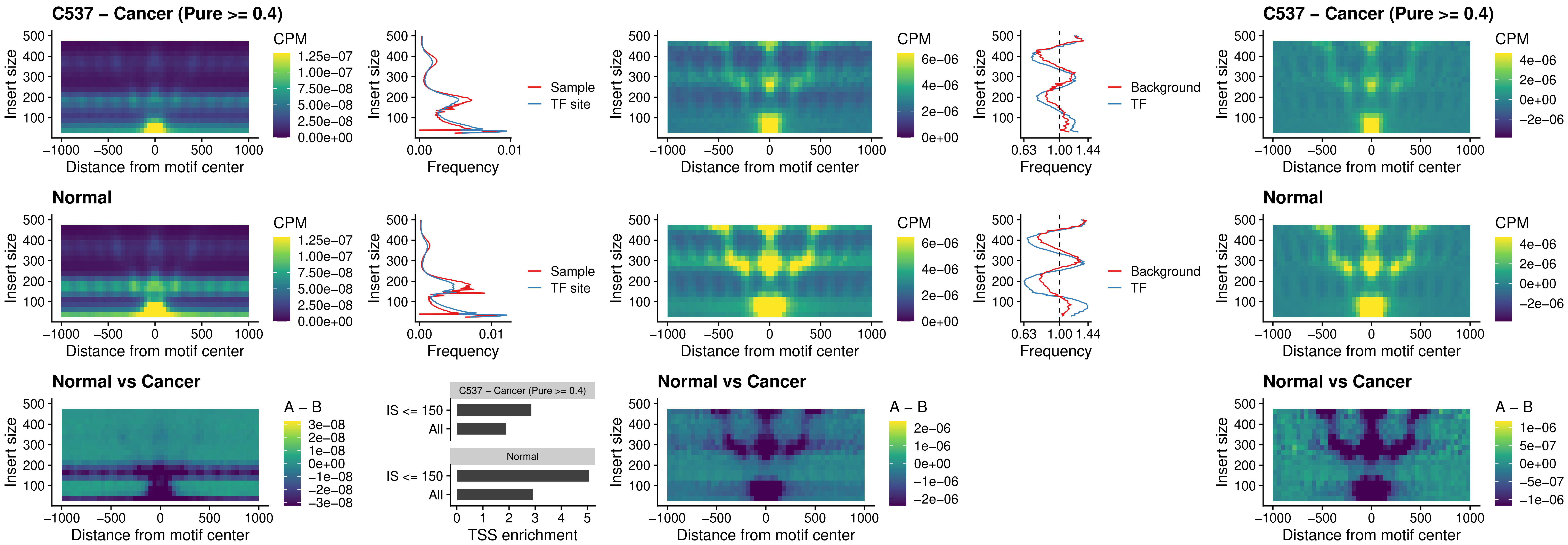

#### C537 – Cancer (Pure >= 0.4) – CTCF_uc_pTSS_oPEAK

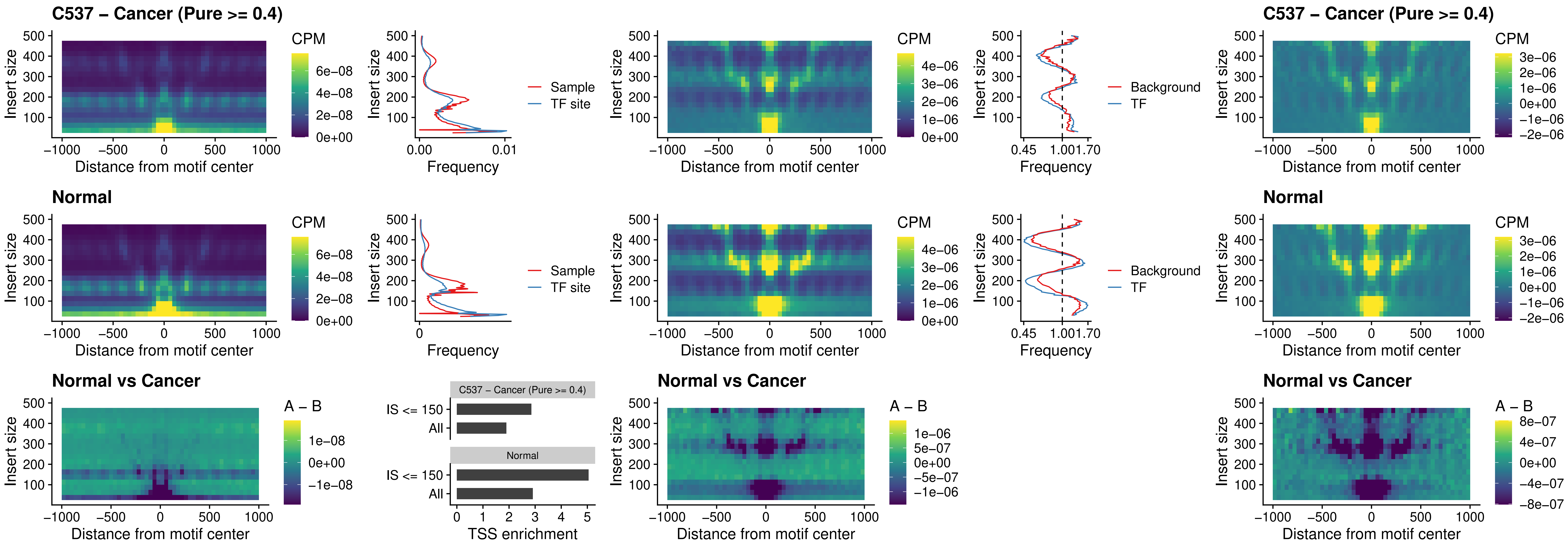

#### C537 – Cancer (Pure >= 0.4) – ENCODE_CTCF_sites_COLON_promotor

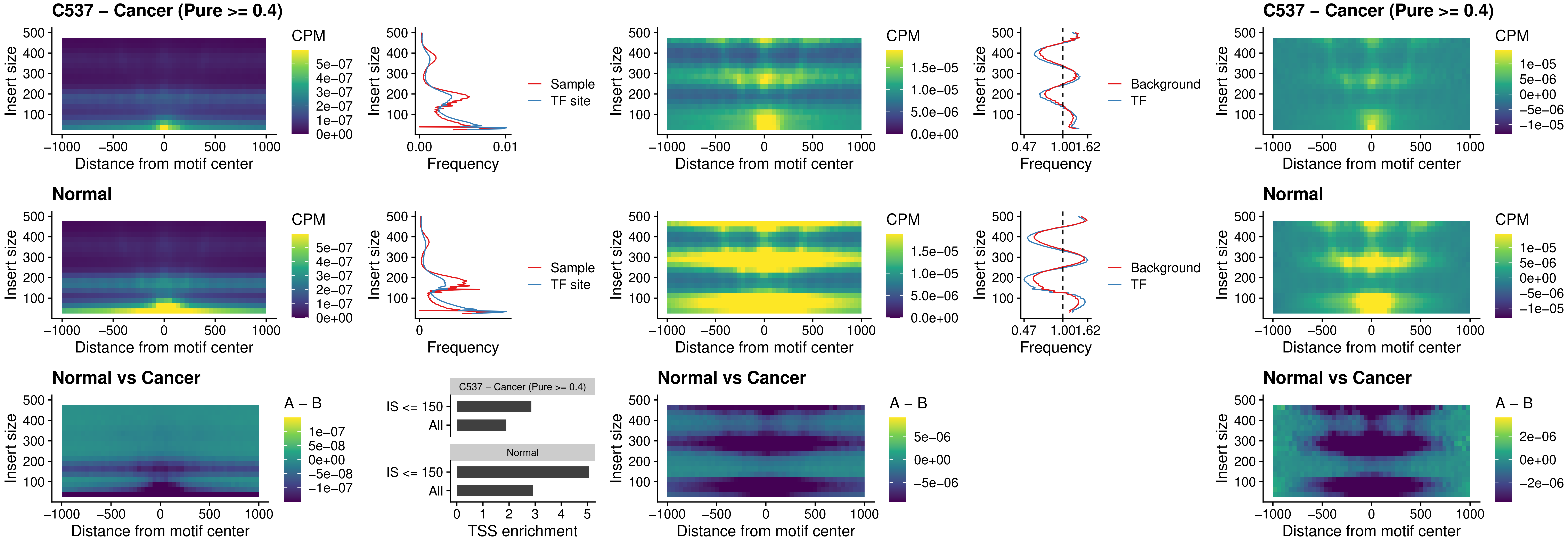

#### C537 – Cancer (Pure >= 0.4) – ENCODE_CTCF_sites_COLON_promotor_ctcf

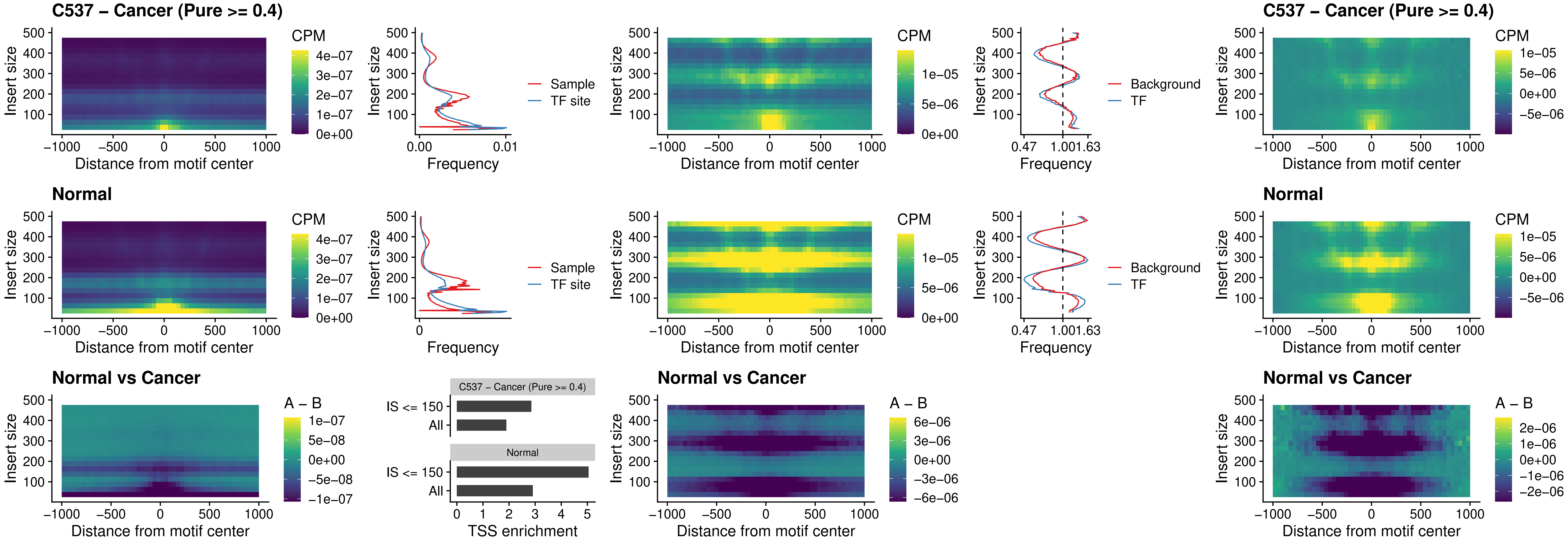

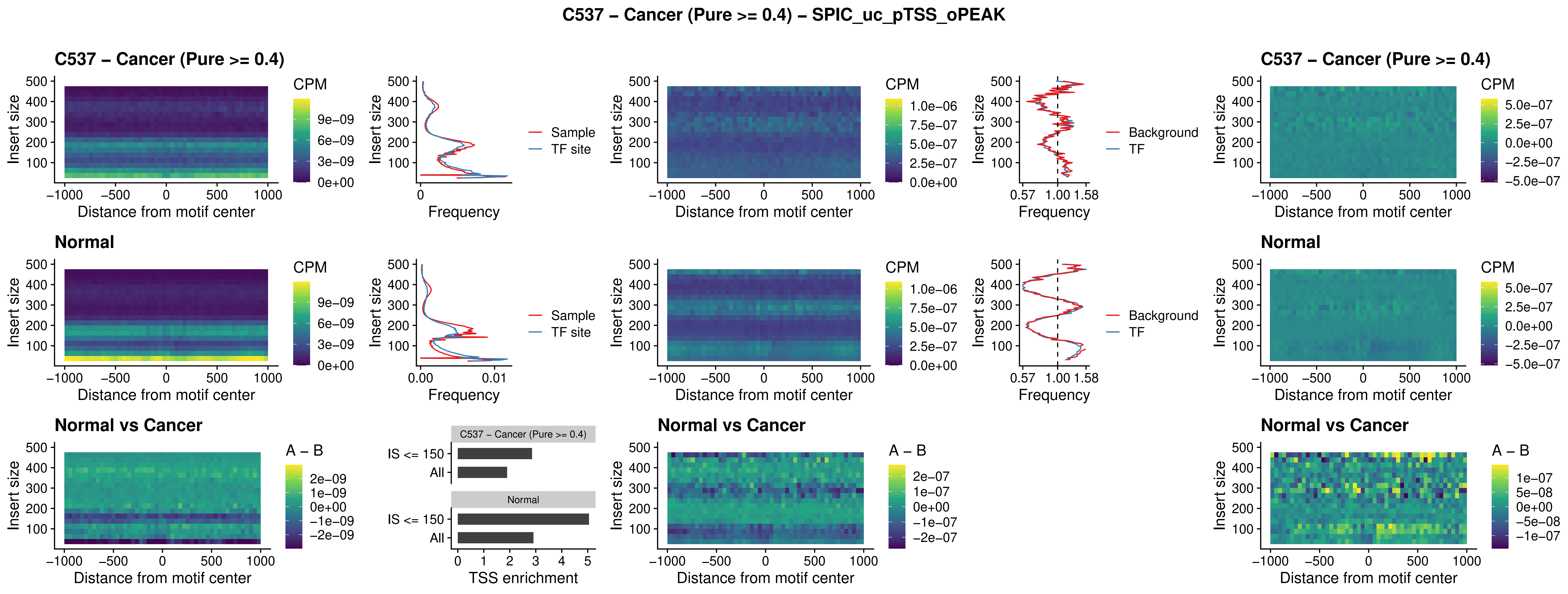
C537 – Cancer (Pure >= 0.4) – SPIC_uc_pTSS_oPEAK
C537 – Cancer (Pure >= 0.4)
Normal
Normal vs Cancer
Insert size
Distance from motif center
Frequency
CPM
A – B
Sample
TF site
Background
TF
C537 – Cancer (Pure >= 0.4)
Normal
IS <= 150
All
TSS enrichment

### C537 – Cancer (Pure >= 0.4) – SPI1_uc_dTSS_nPEAK

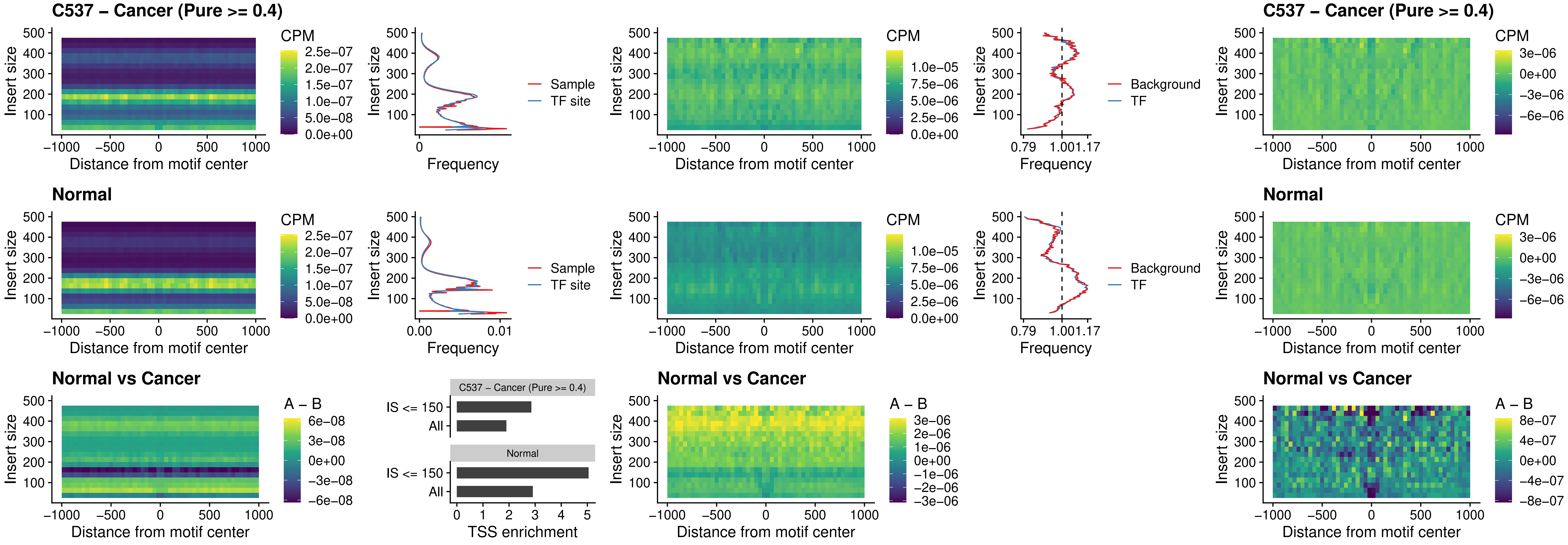

#### C537 – Cancer (Pure >= 0.4) – SPI1_uc_cTSS_nPEAK

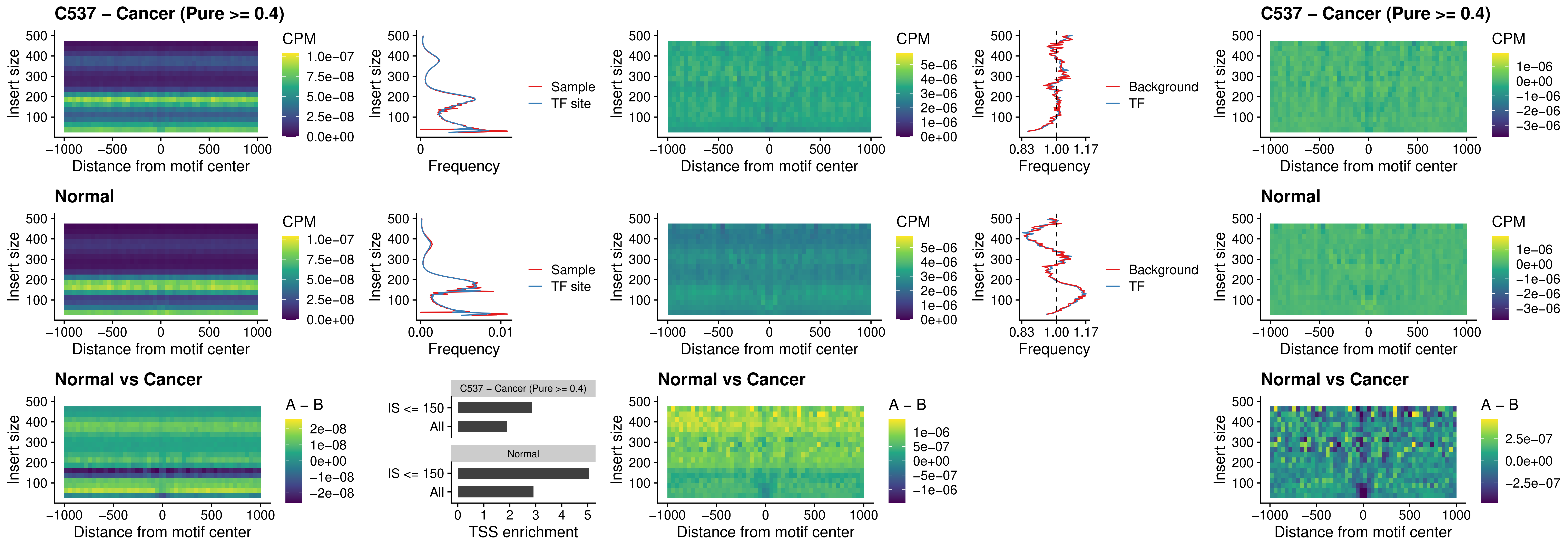

### C537 – Cancer (Pure >= 0.4) – BCL11A_uc_pTSS_nPEAK

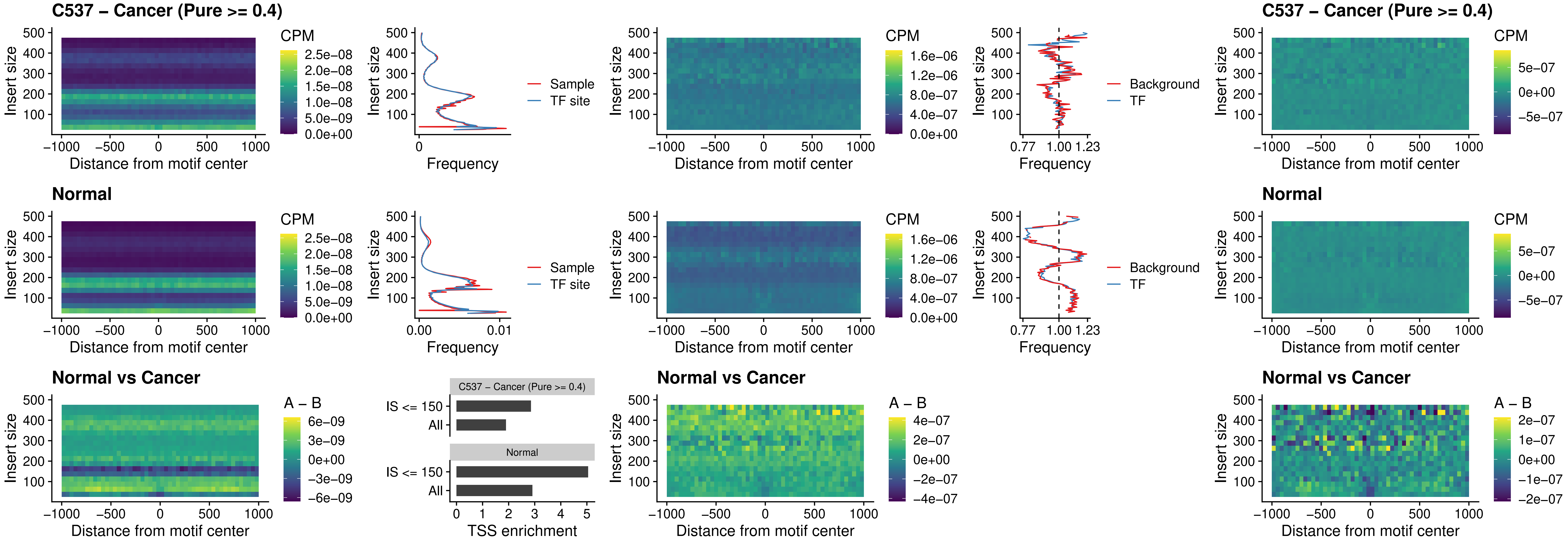

#### C537 – Cancer (Pure >= 0.4) – BCL11B_uc_pTSS_nPEAK

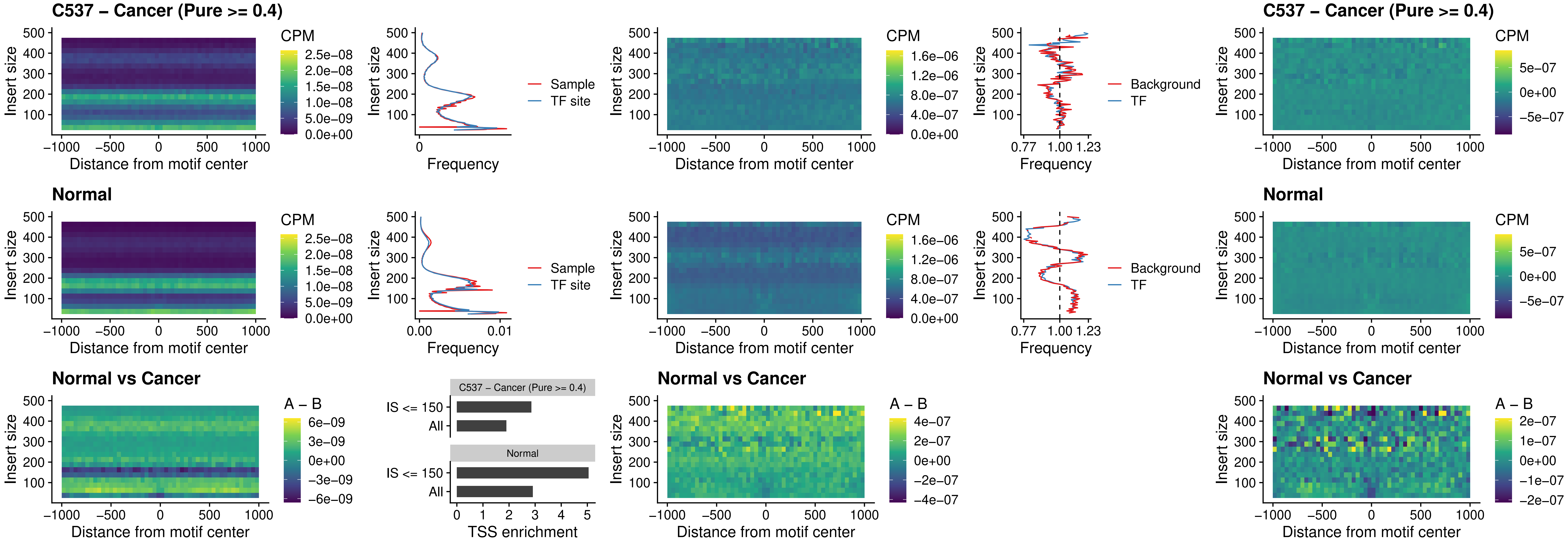

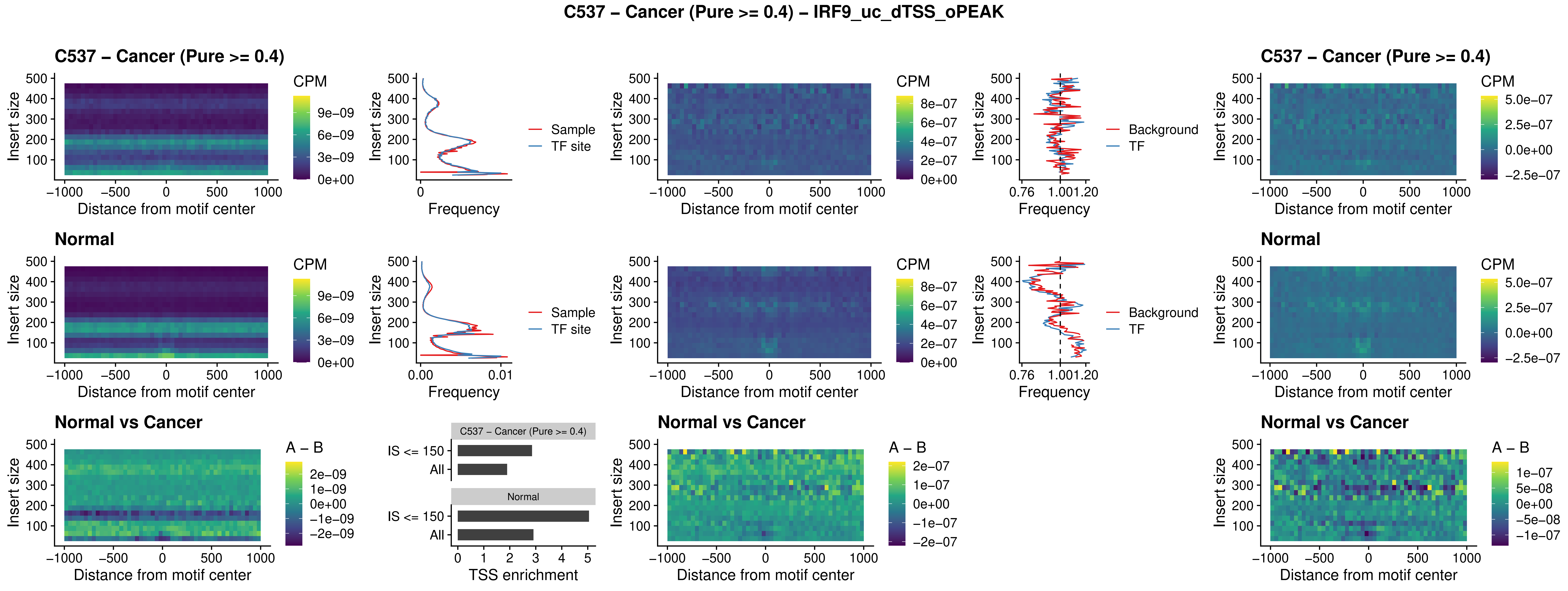
C537 – Cancer (Pure >= 0.4) – IRF9_uc_dTSS_oPEAK
C537 – Cancer (Pure >= 0.4)
Normal
Normal vs Cancer
Insert size
Distance from motif center
Frequency
CPM
A – B
Sample
TF site
Background
TF
C537 – Cancer (Pure >= 0.4)
Normal
IS <= 150
All
TSS enrichment

#### C537 – Cancer (Pure >= 0.4) – IRF9_uc_cTSS_oPEAK

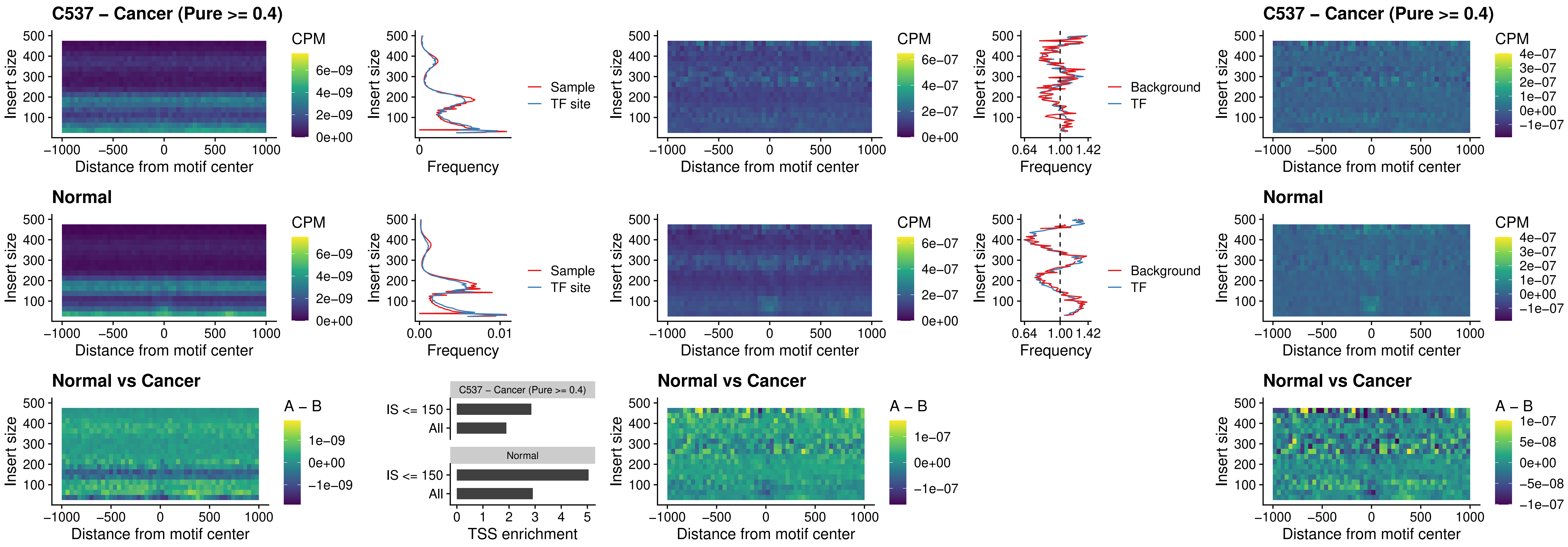

#### C537 – Cancer (Pure >= 0.4) – BCL11A_uc_cTSS_oPEAK

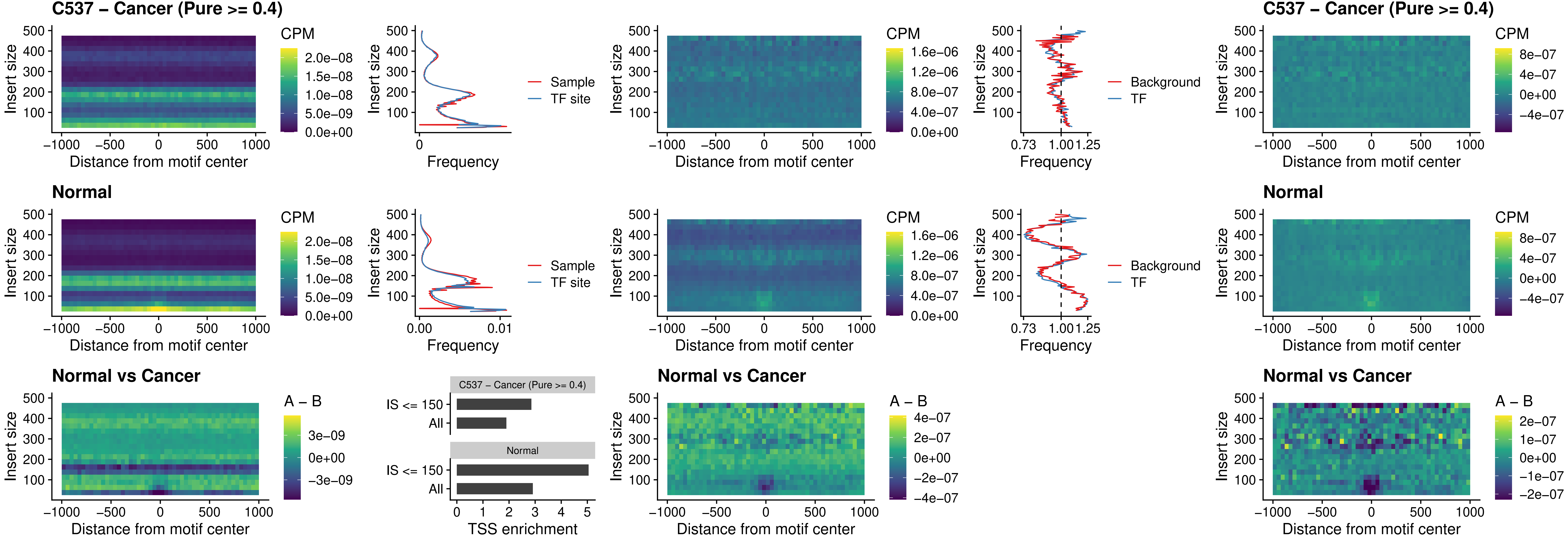

#### C537 – Cancer (Pure >= 0.4) – BCL11B_uc_cTSS_oPEAK

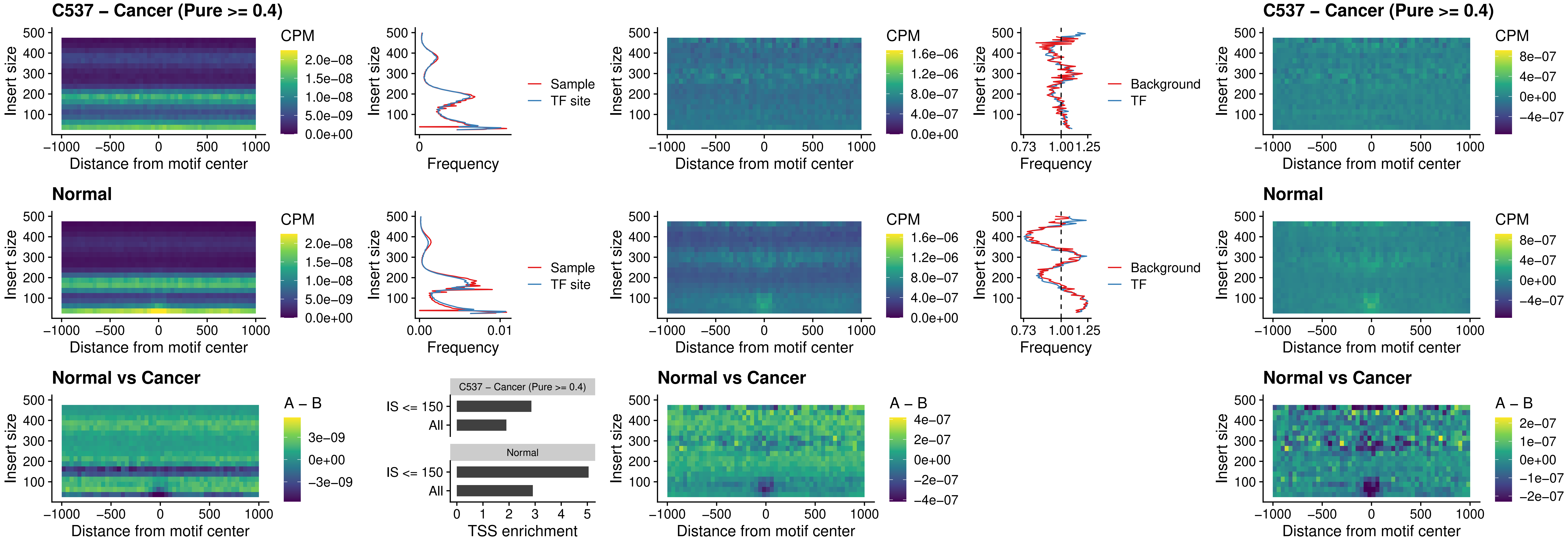

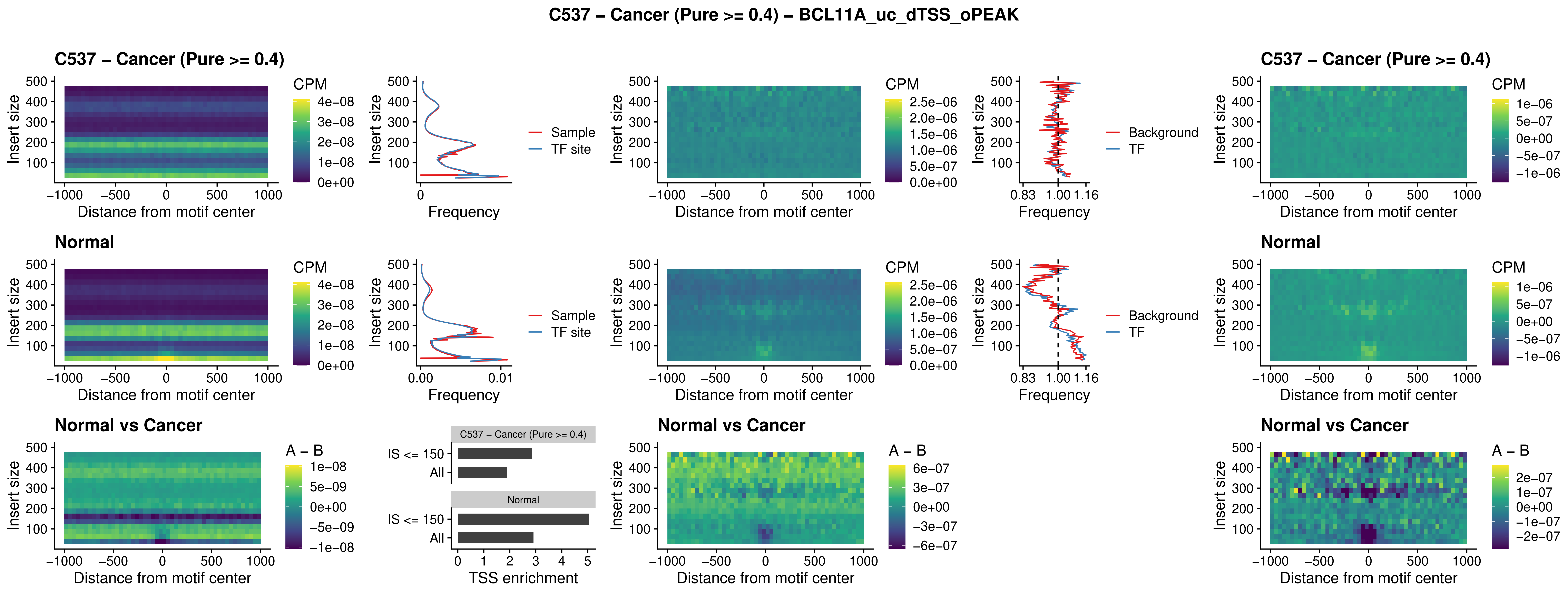
C537 – Cancer (Pure >= 0.4) – BCL11A_uc_dTSS_oPEAK
C537 – Cancer (Pure >= 0.4)
Normal
Normal vs Cancer
Insert size
Distance from motif center
Frequency
CPM
A – B
Sample
TF site
Background
TF
C537 – Cancer (Pure >= 0.4)
Normal
IS <= 150
All
TSS enrichment

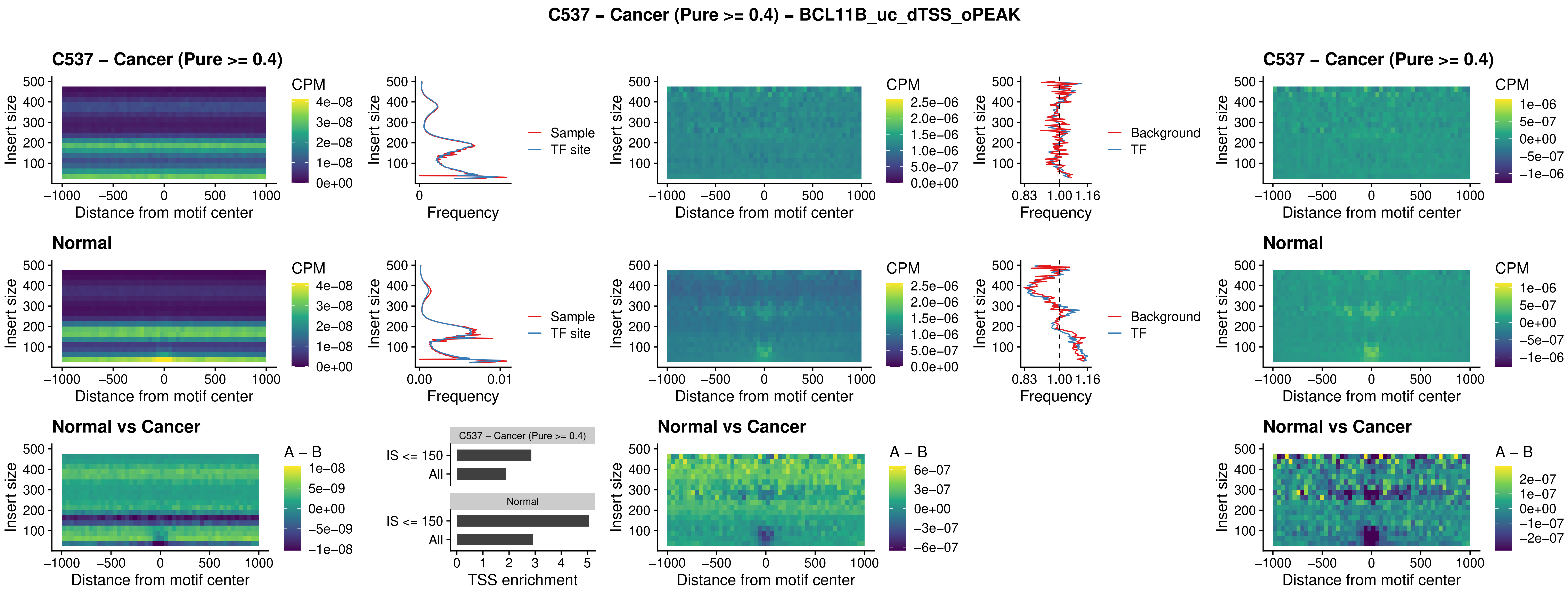
C537 – Cancer (Pure >= 0.4) – BCL11B_uc_dTSS_oPEAK
C537 – Cancer (Pure >= 0.4)
Normal
Normal vs Cancer
Insert size
Distance from motif center
Frequency
CPM
A – B
Sample
TF site
Background
TF
C537 – Cancer (Pure >= 0.4)
Normal
IS <= 150
All
TSS enrichment

#### C537 – Cancer (Pure >= 0.4) – SPIB_uc_cTSS_nPEAK

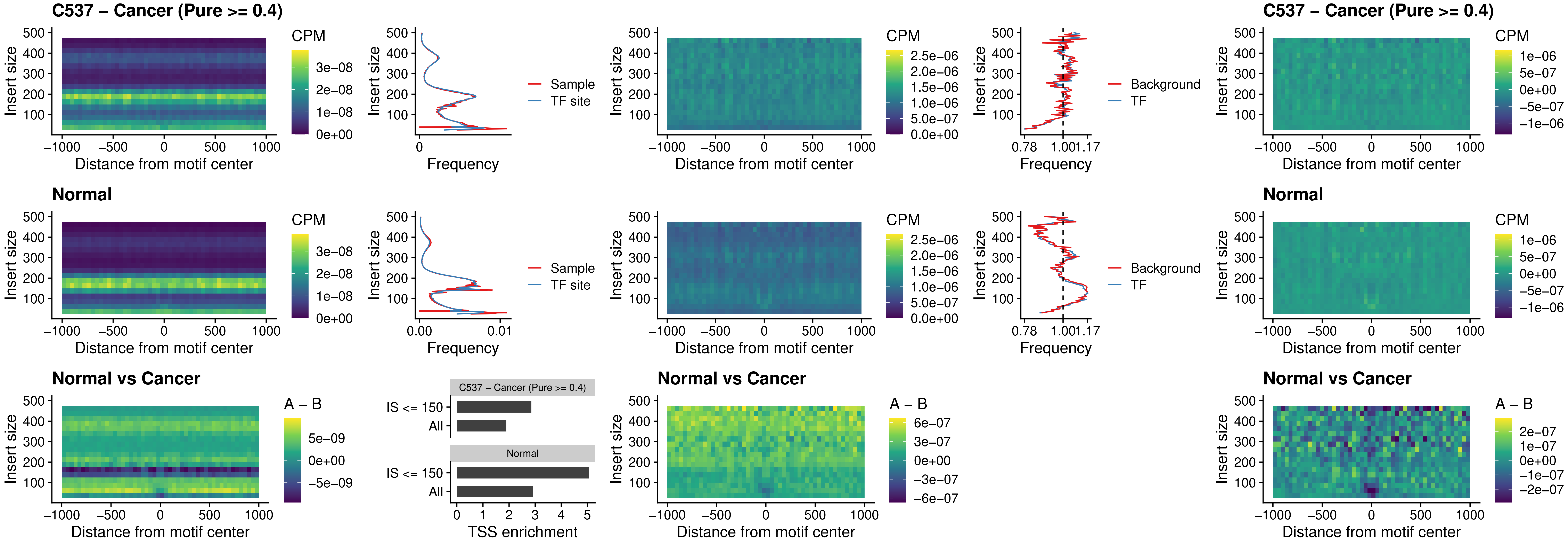

#### C537 – Cancer (Pure >= 0.4) – SPIC_uc_cTSS_nPEAK

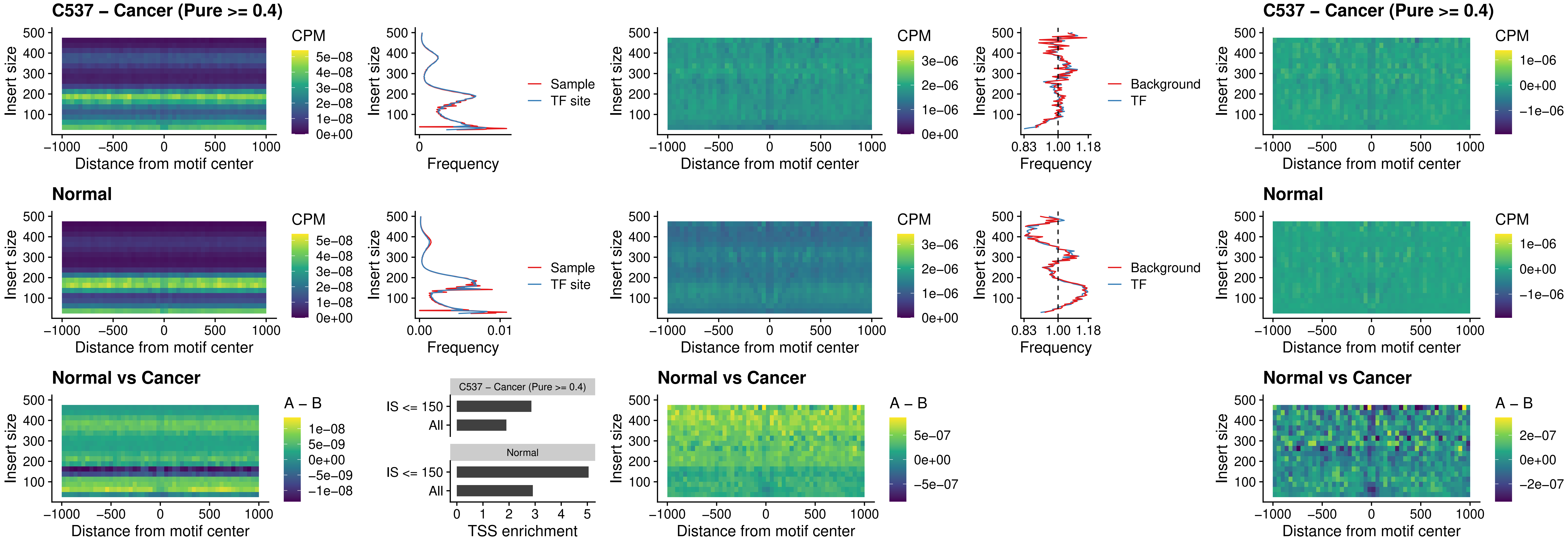

#### C537 – Cancer (Pure >= 0.4) – SPI1_uc_pTSS_oPEAK

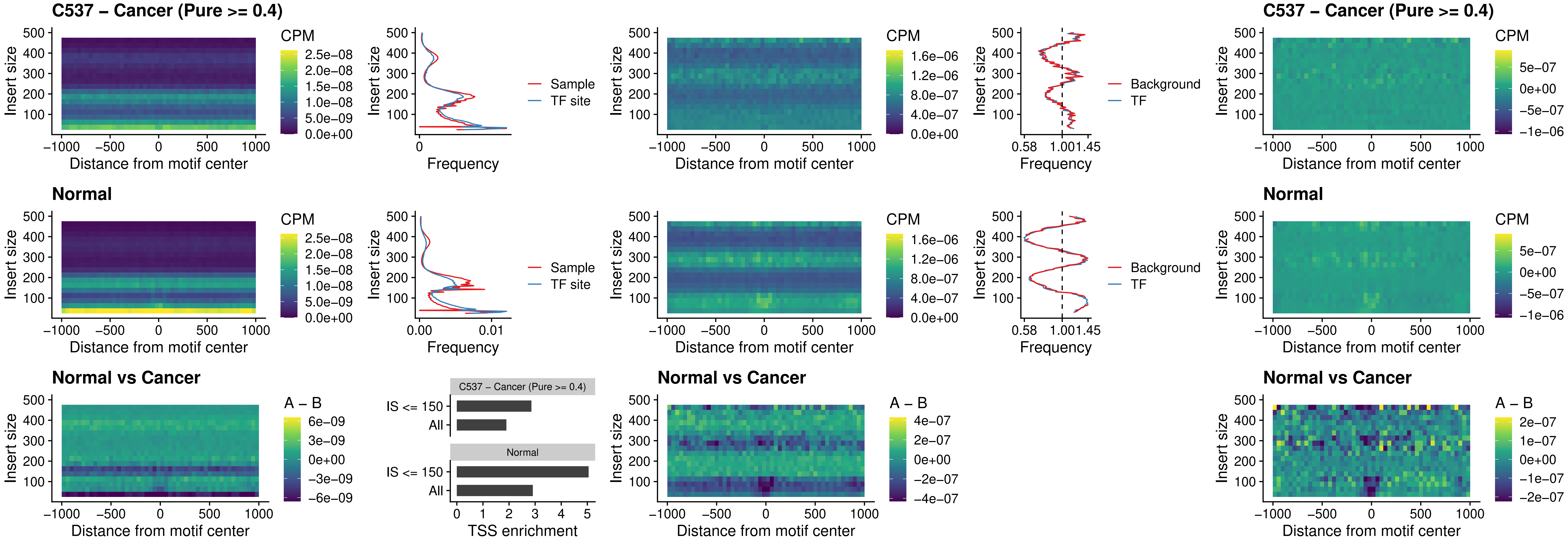

#### C537 – Cancer (Pure >= 0.4) – BCL11A_uc_pTSS_oPEAK

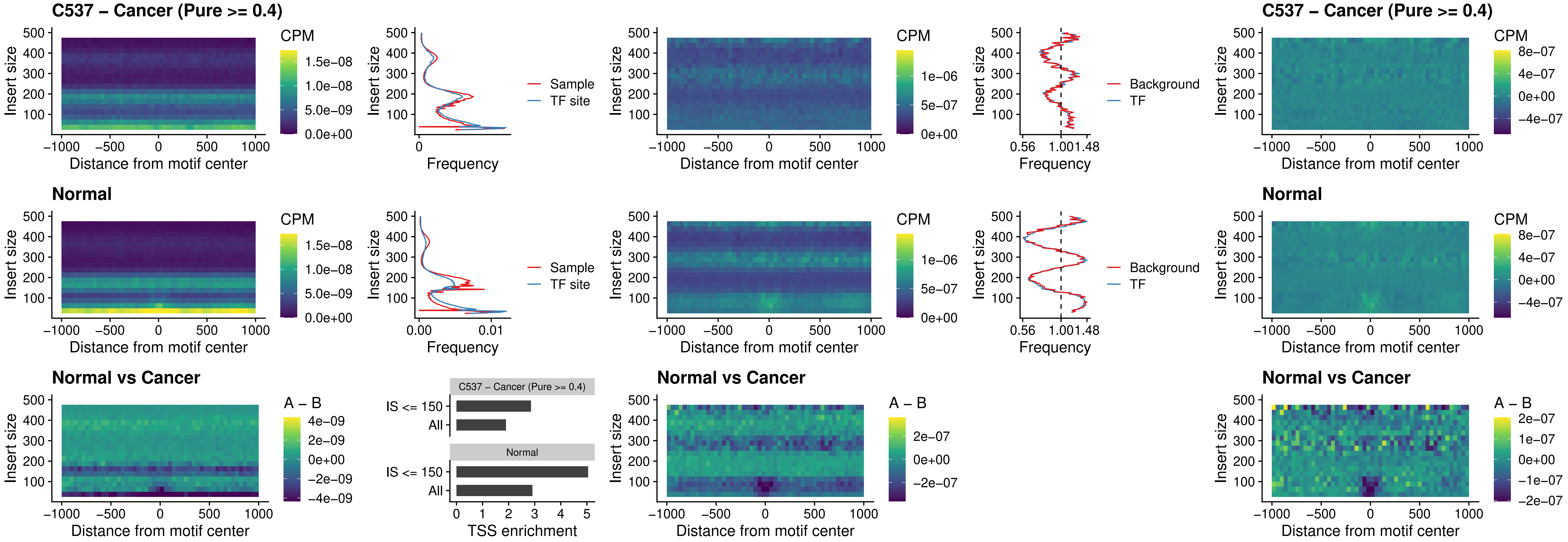

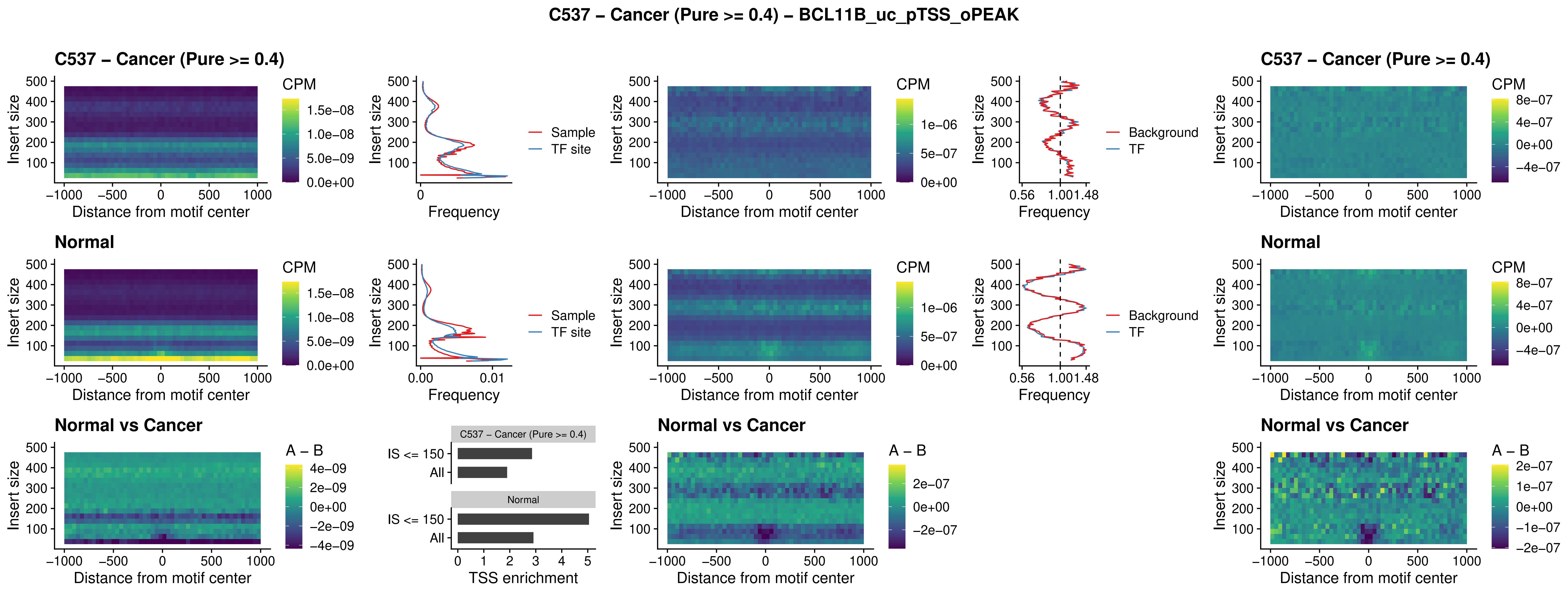
C537 – Cancer (Pure >= 0.4) – BCL11B_uc_pTSS_oPEAK
C537 – Cancer (Pure >= 0.4)
Normal
Normal vs Cancer
Insert size
Distance from motif center
CPM
A – B
Frequency
Sample
TF site
Background
TF
TSS enrichment
IS <= 150
All
Normal

#### C537 – Cancer (Pure >= 0.4) – SPI1_uc_cTSS_oPEAK

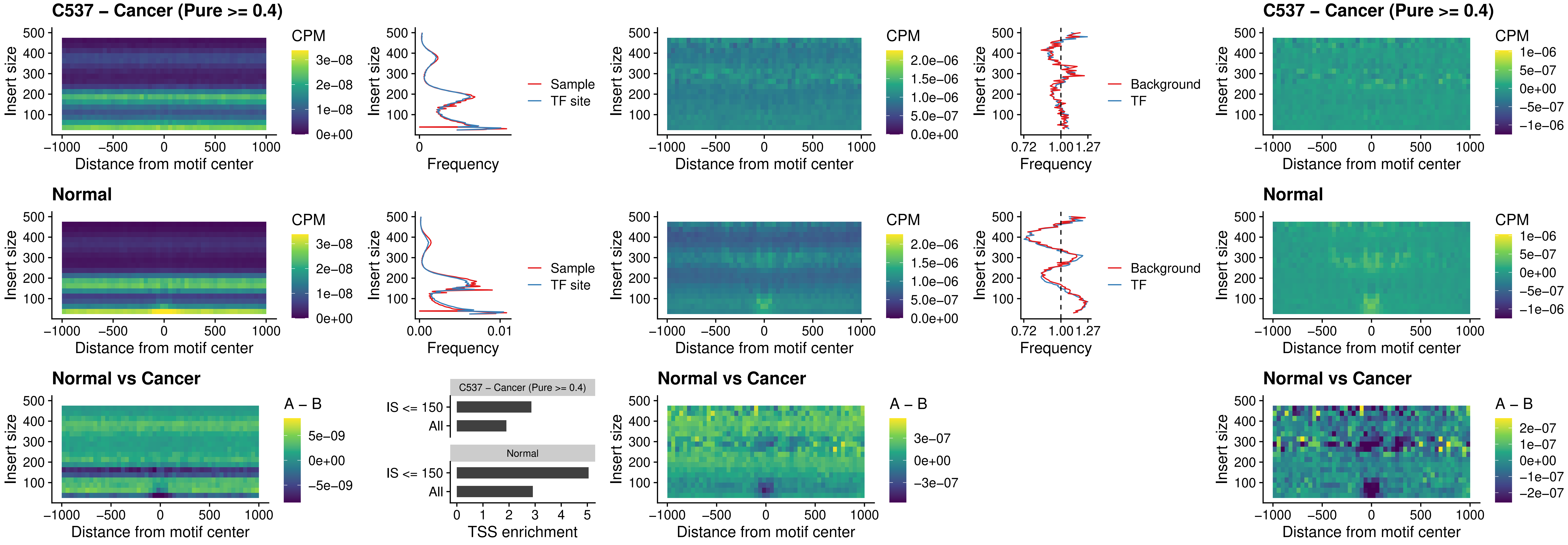

#### C537 – Cancer (Pure >= 0.4) – SPIB_uc_cTSS_oPEAK

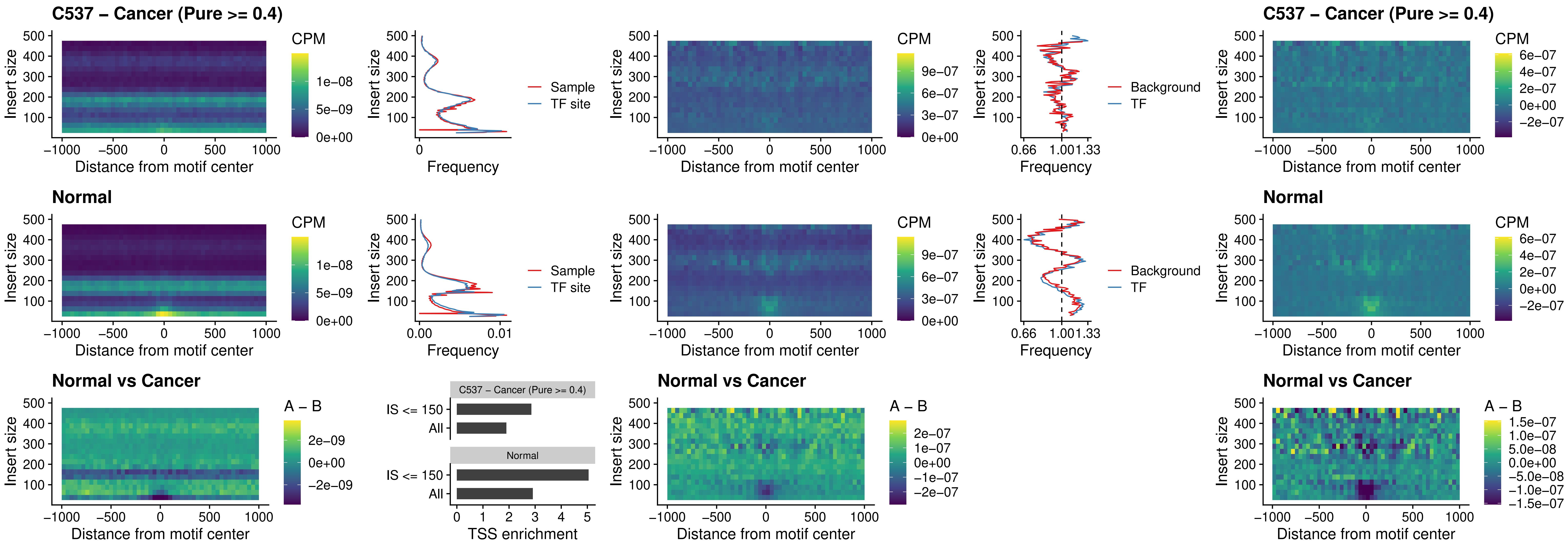

### C537 – Cancer (Pure >= 0.4) – SPIB_uc_pTSS_oPEAK

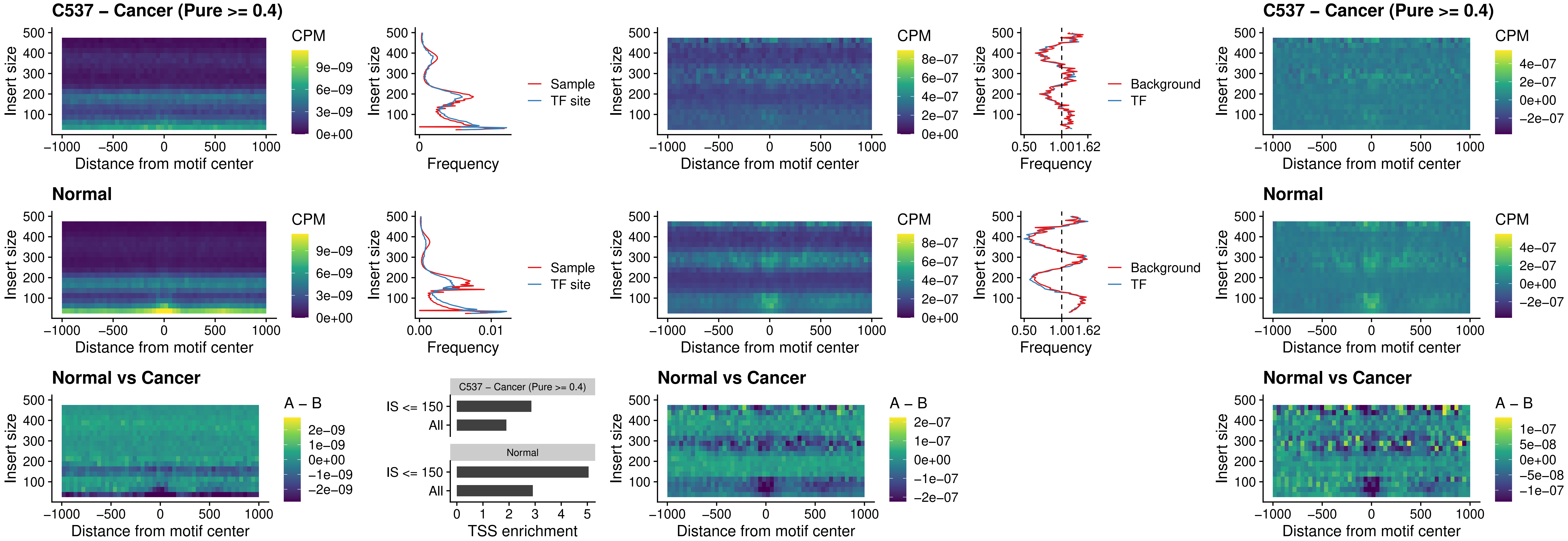

### C537 – Cancer (Pure >= 0.4) – SPI1_uc_dTSS_oPEAK

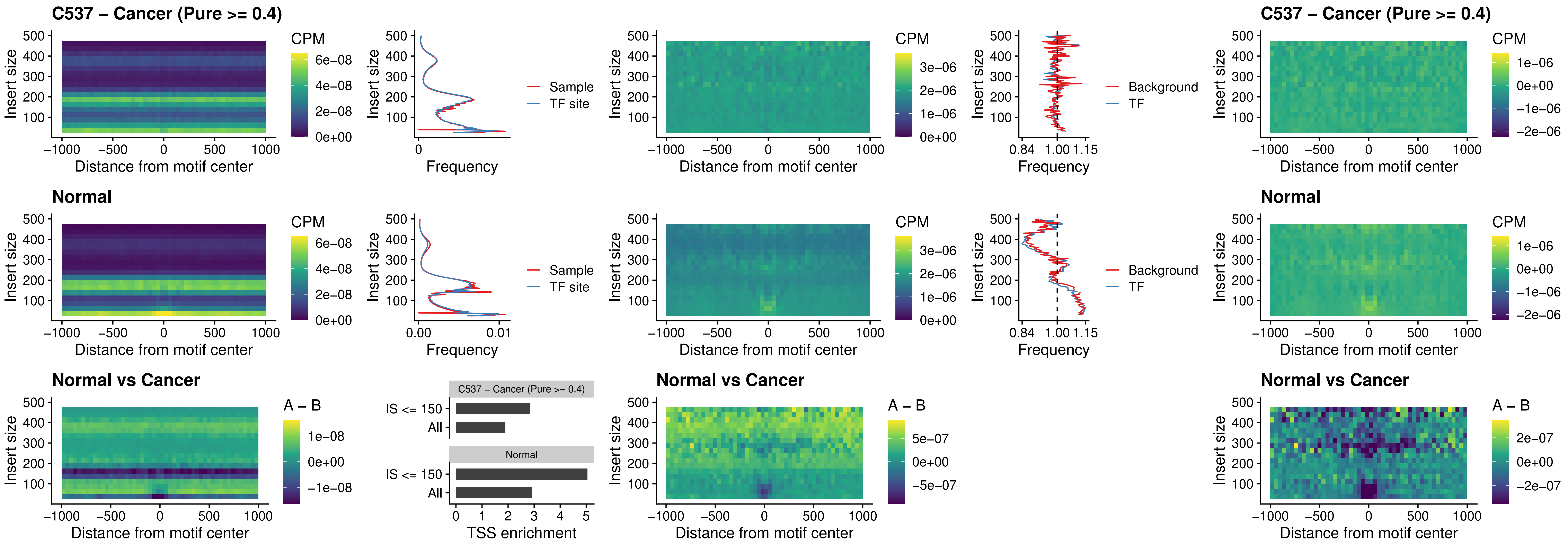

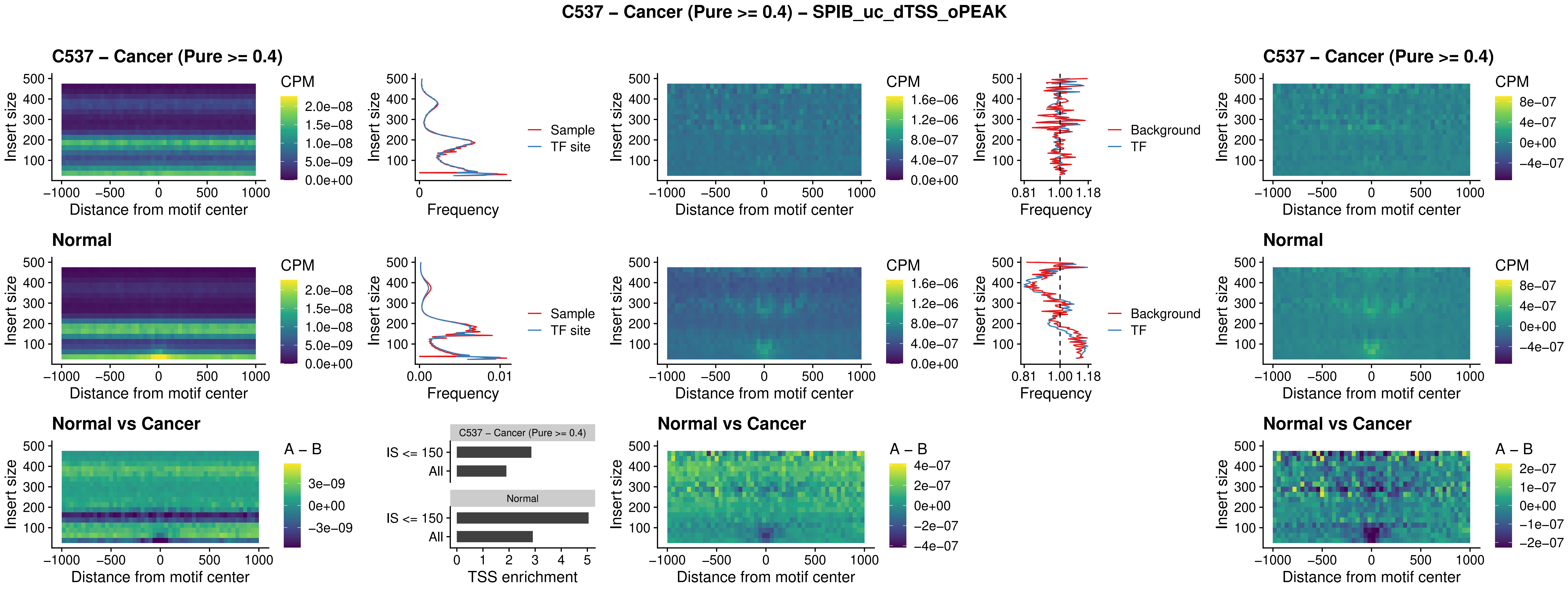
C537 – Cancer (Pure >= 0.4) – SPIB_uc_dTSS_oPEAK
C537 – Cancer (Pure >= 0.4)
Normal
Normal vs Cancer
Insert size
Distance from motif center
Frequency
CPM
A – B
Sample
TF site
Background
TF
C537 – Cancer (Pure >= 0.4)
Normal
IS <= 150
All
TSS enrichment

### C537 – Cancer (Pure >= 0.4) – IRF3_uc_pTSS_oPEAK

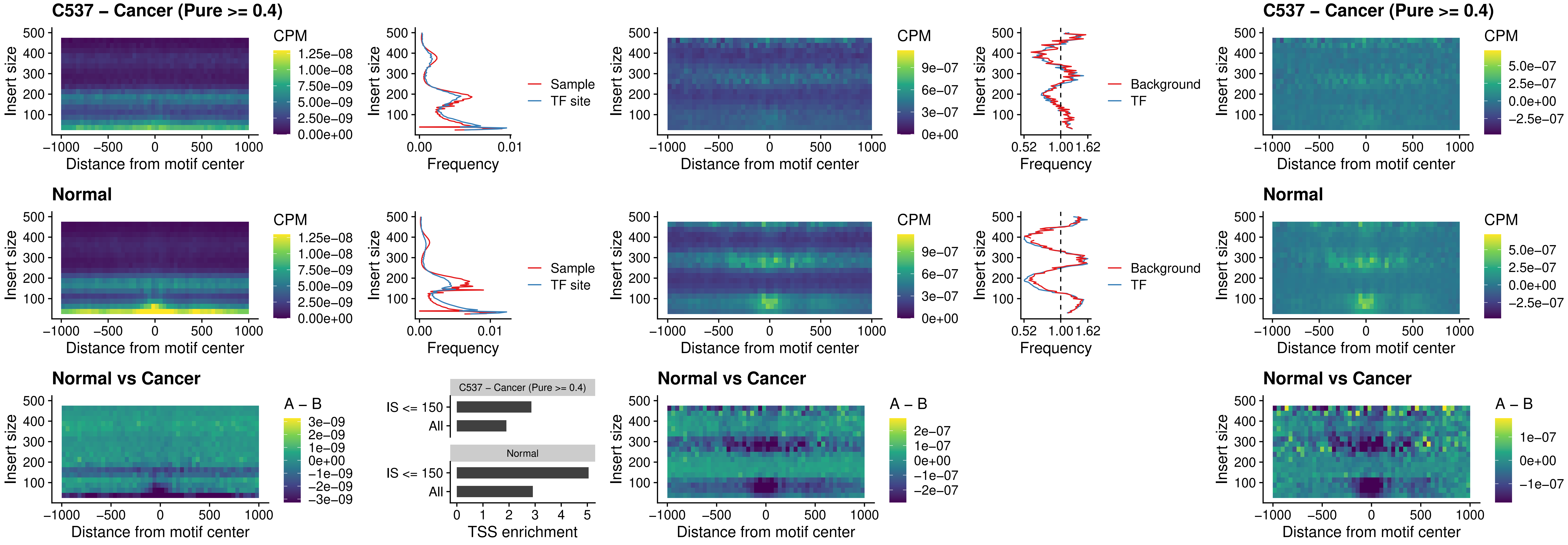

#### C537 – Cancer (Pure >= 0.4) – IRF2_uc_dTSS_oPEAK

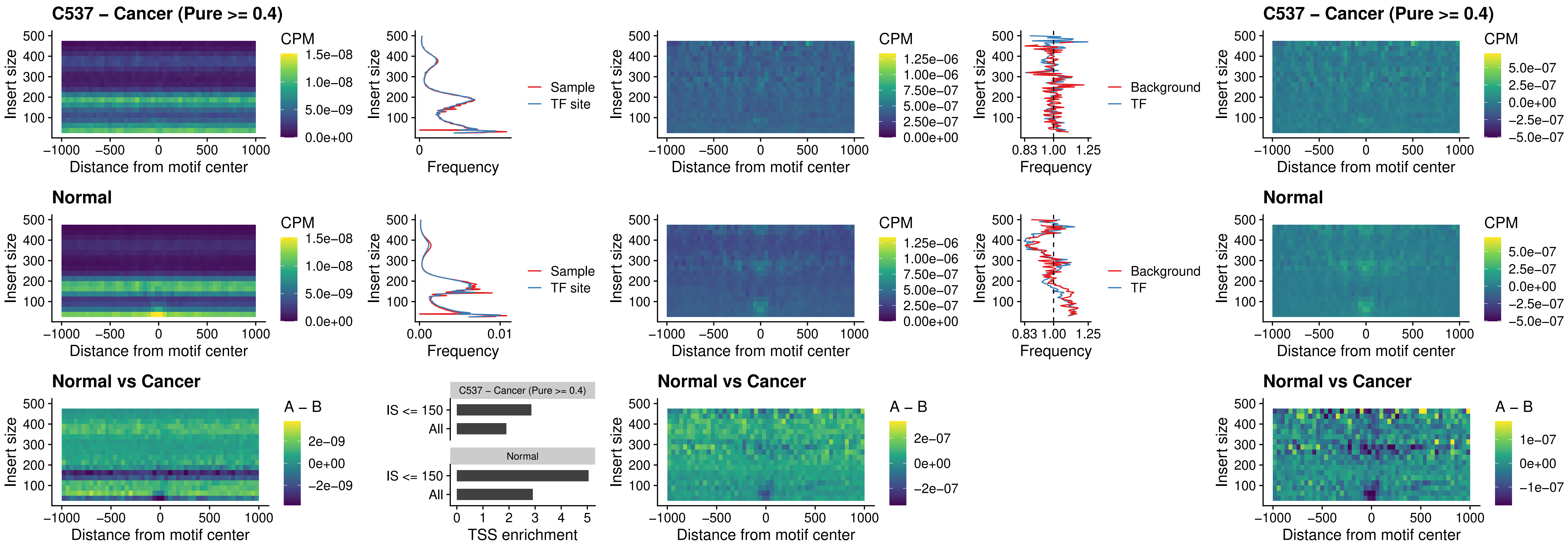

### C537 – Cancer (Pure >= 0.4) – IRF2_uc_pTSS_oPEAK

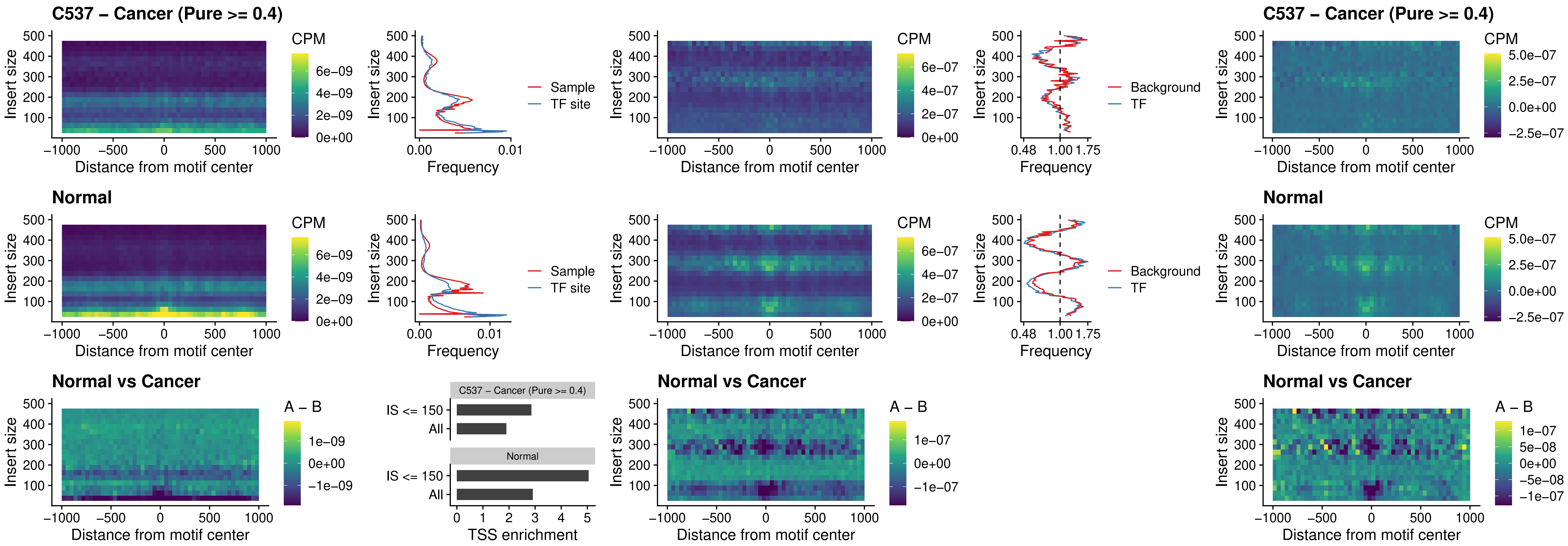

#### C537 – Cancer (Pure >= 0.4) – STAT2_uc_pTSS_oPEAK

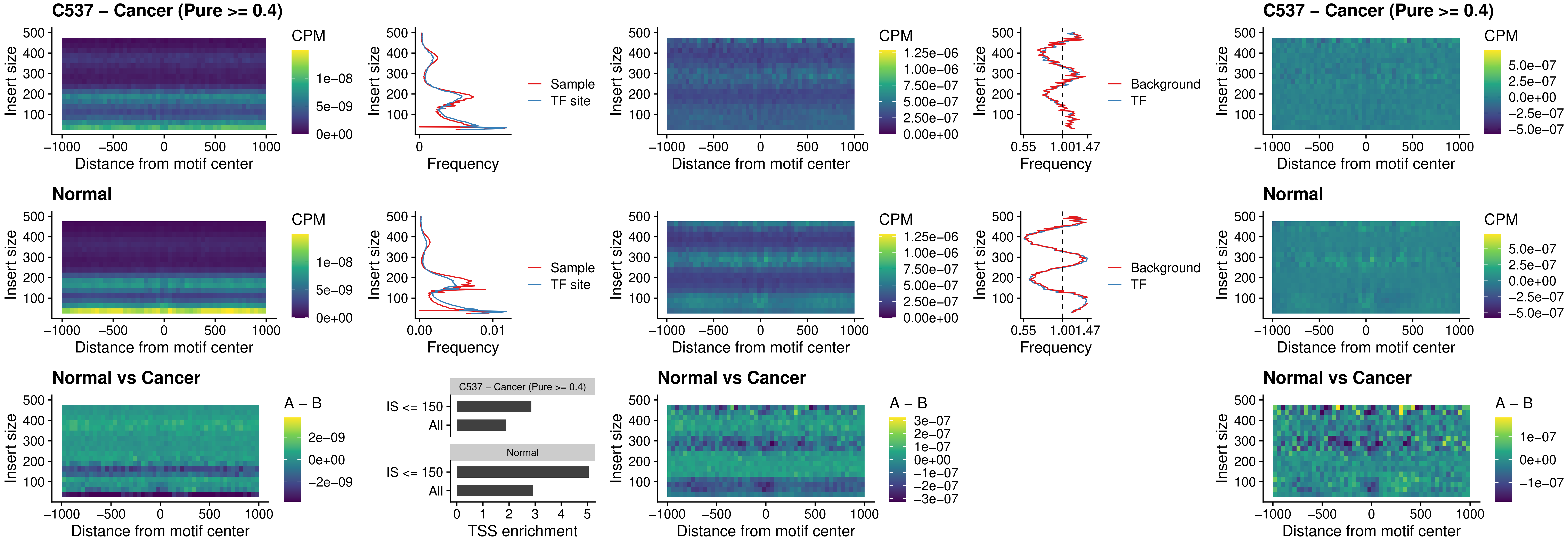

### C537 – Cancer (Pure >= 0.4) – SPIB_uc_dTSS_nPEAK

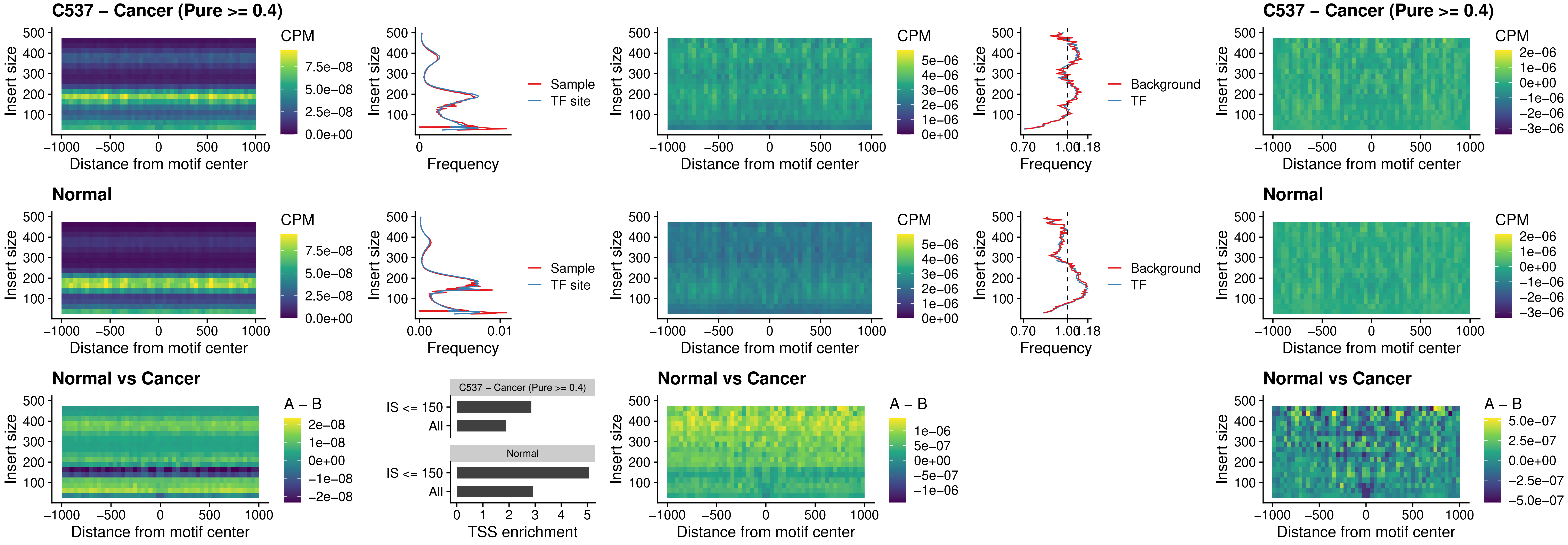

### C537 – Cancer (Pure >= 0.4) – SPIC_uc_cTSS_oPEAK

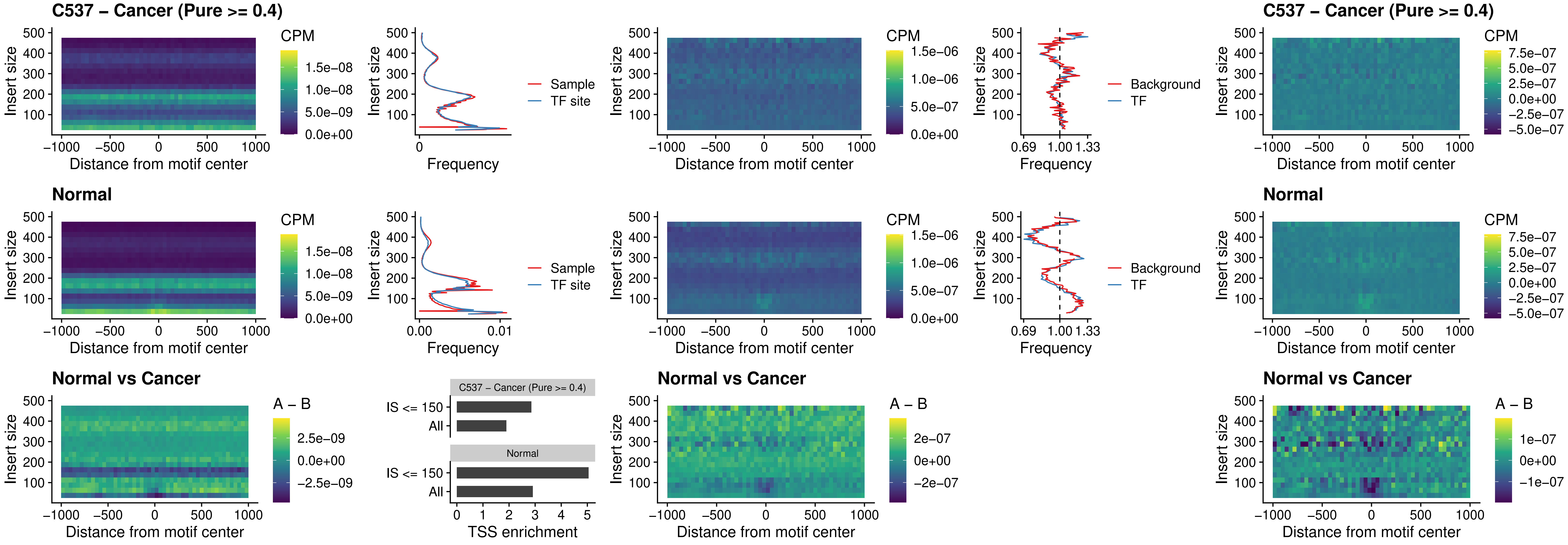

### C537 – Cancer (Pure >= 0.4) – PRDM1_uc_cTSS_oPEAK

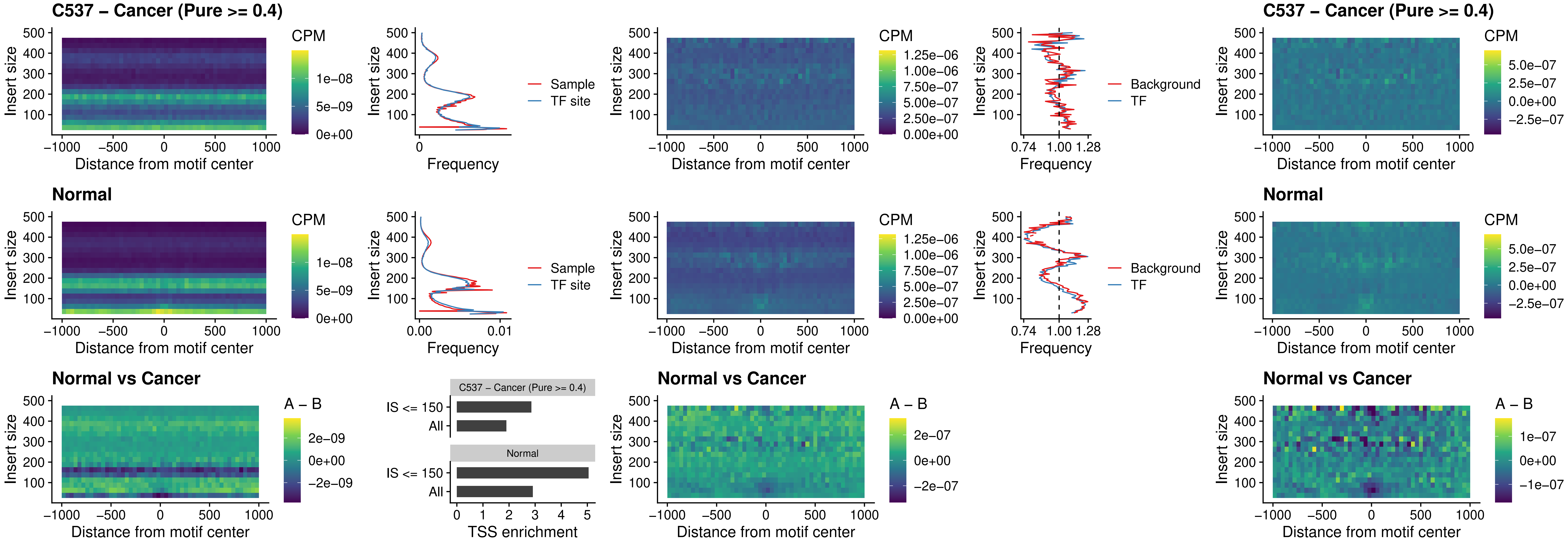

#### C537 – Cancer (Pure >= 0.4) – IRF3_uc_cTSS_oPEAK

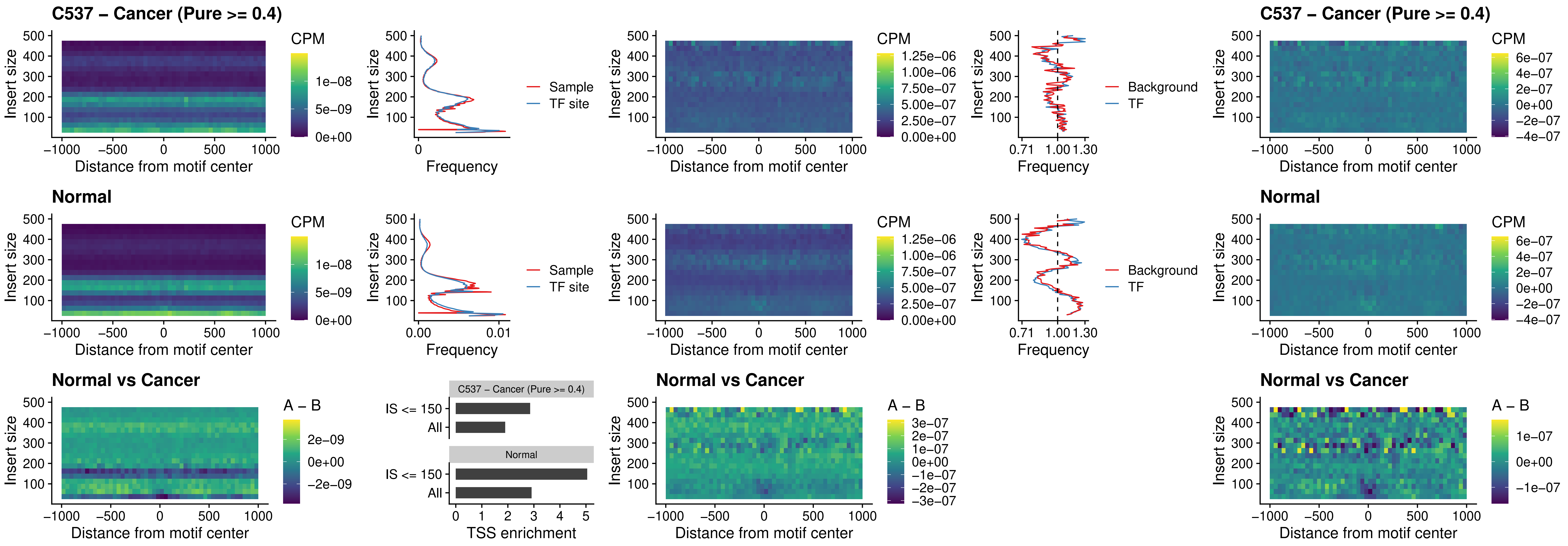

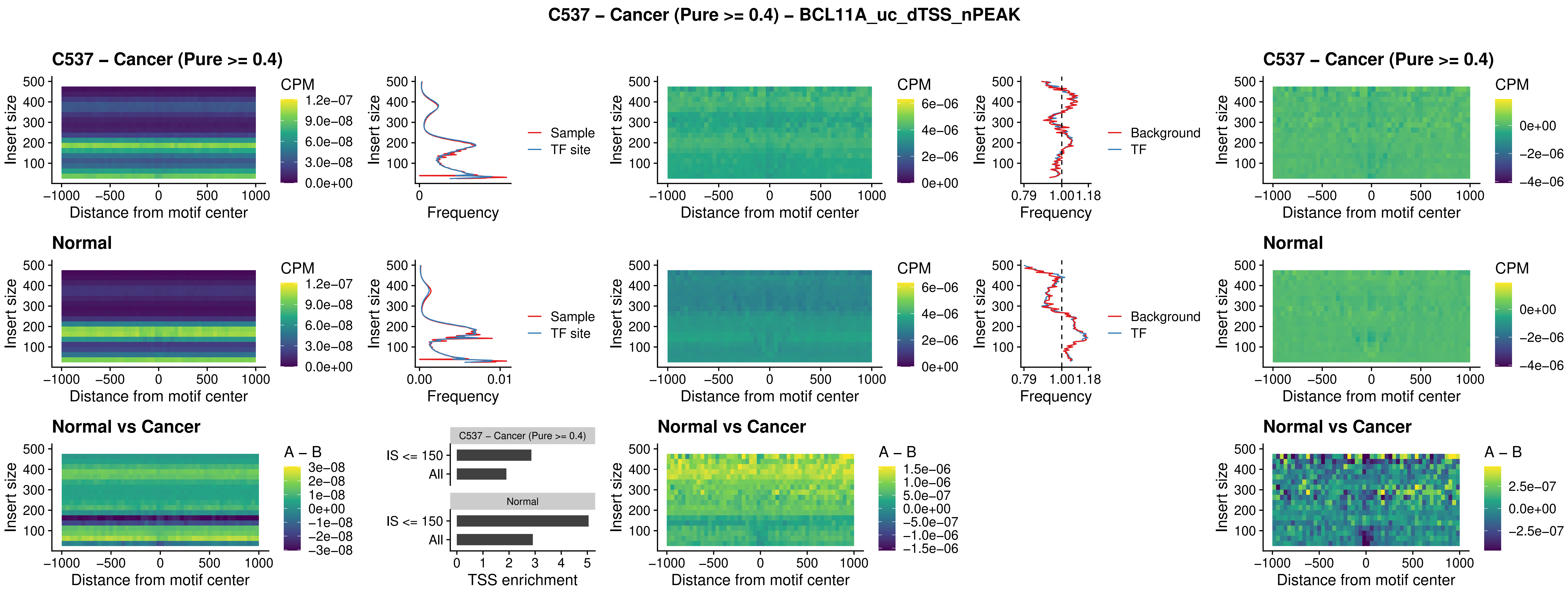
C537 – Cancer (Pure >= 0.4) – BCL11A_uc_dTSS_nPEAK
C537 – Cancer (Pure >= 0.4)
Normal
Normal vs Cancer
Insert size
Distance from motif center
CPM
A – B
Frequency
Sample
TF site
Background
TF
TSS enrichment
C537 – Cancer (Pure >= 0.4)
Normal
IS <= 150
All

#### C537 – Cancer (Pure >= 0.4) – BCL11B_uc_dTSS_nPEAK

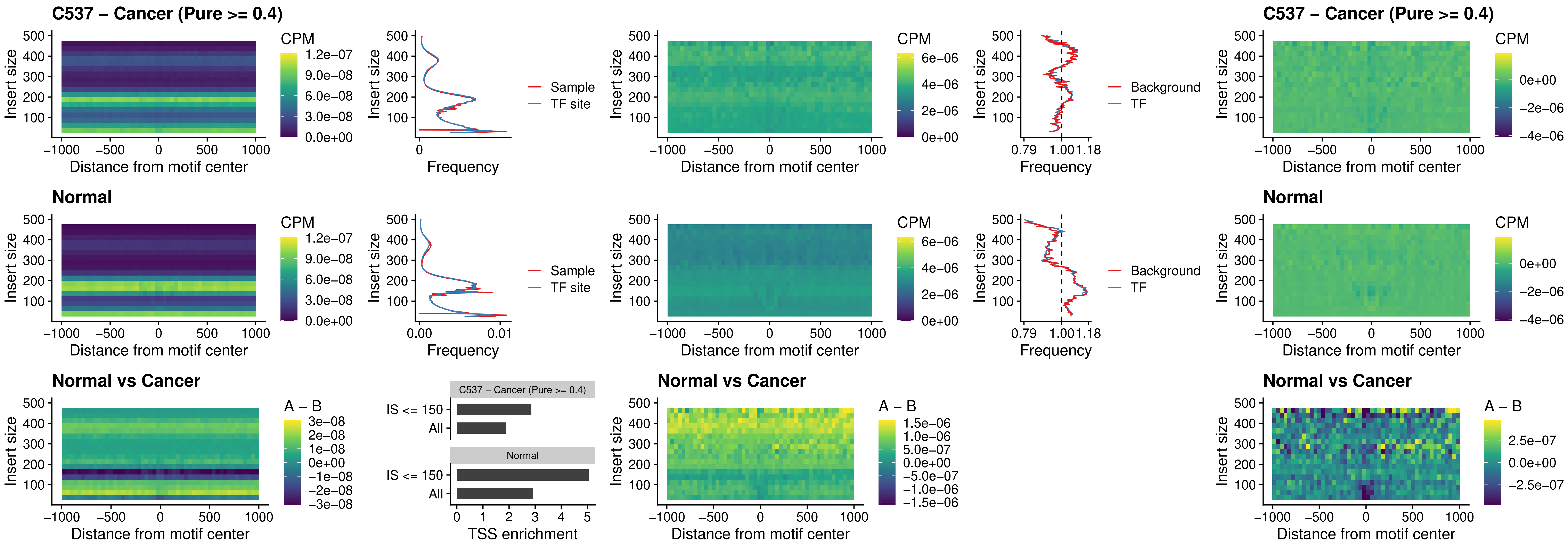

#### C537 – Cancer (Pure >= 0.4) – IRF1_uc_cTSS_oPEAK

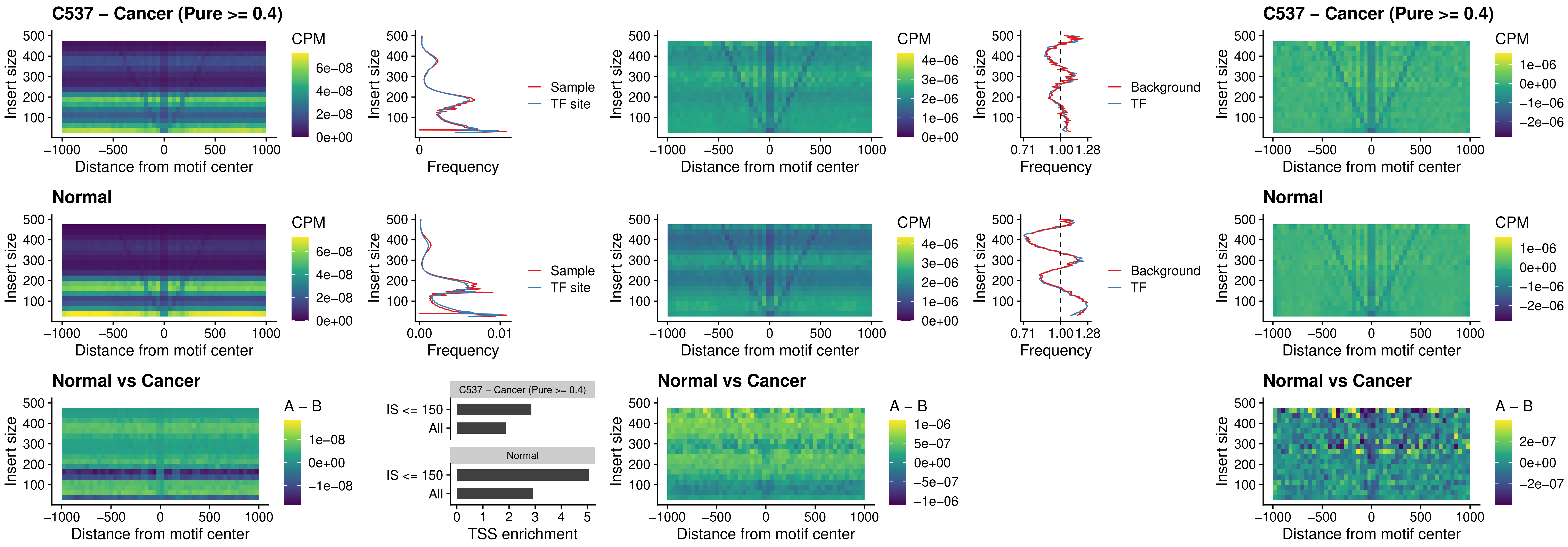

#### C537 – Cancer (Pure >= 0.4) – IRF1_uc_dTSS_oPEAK

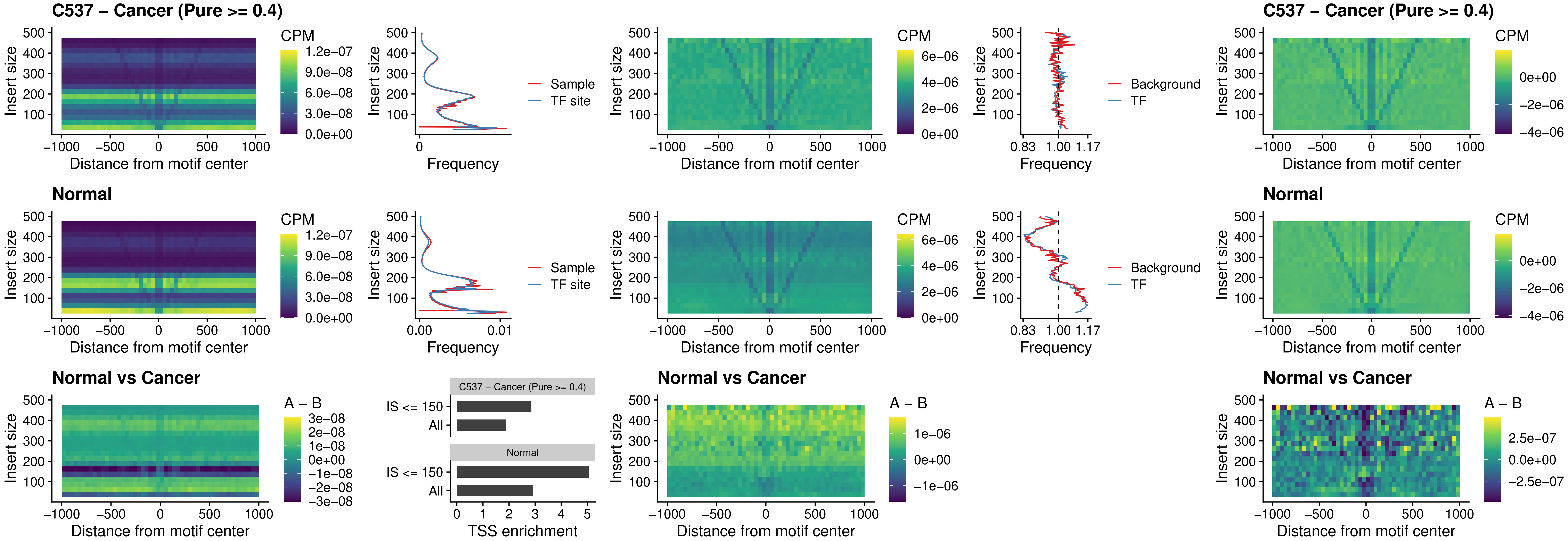

#### C537 – Cancer (Pure >= 0.4) – IRF1_uc_pTSS_oPEAK

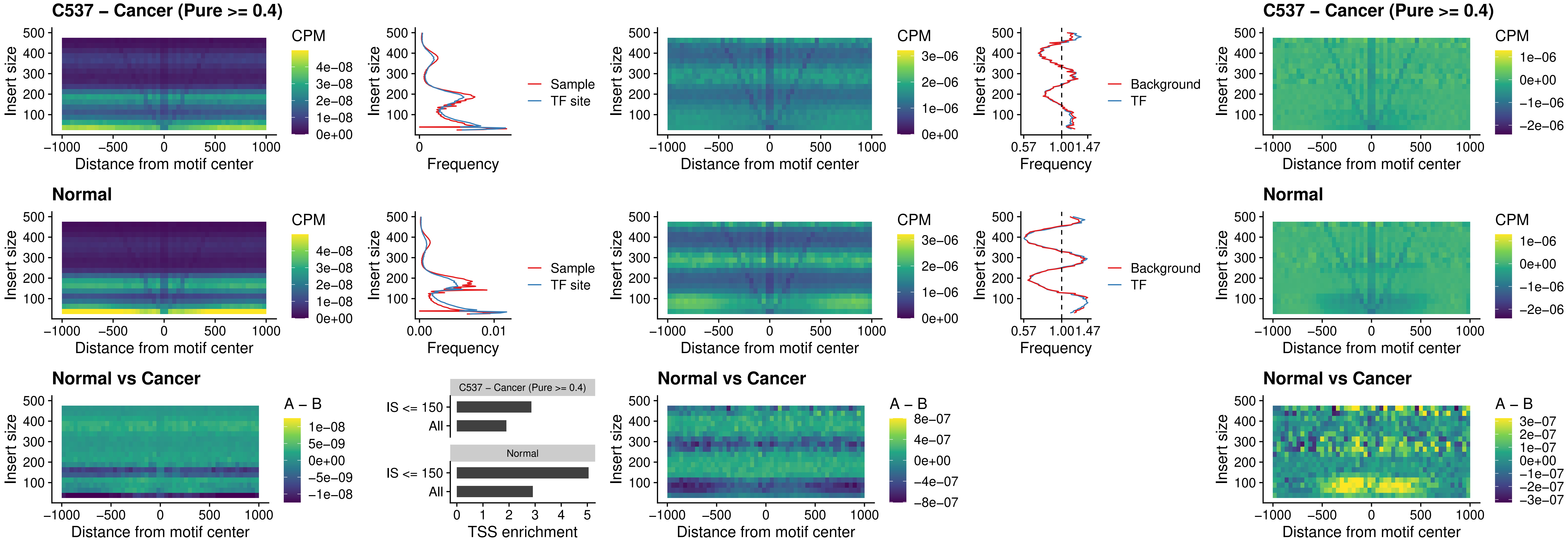

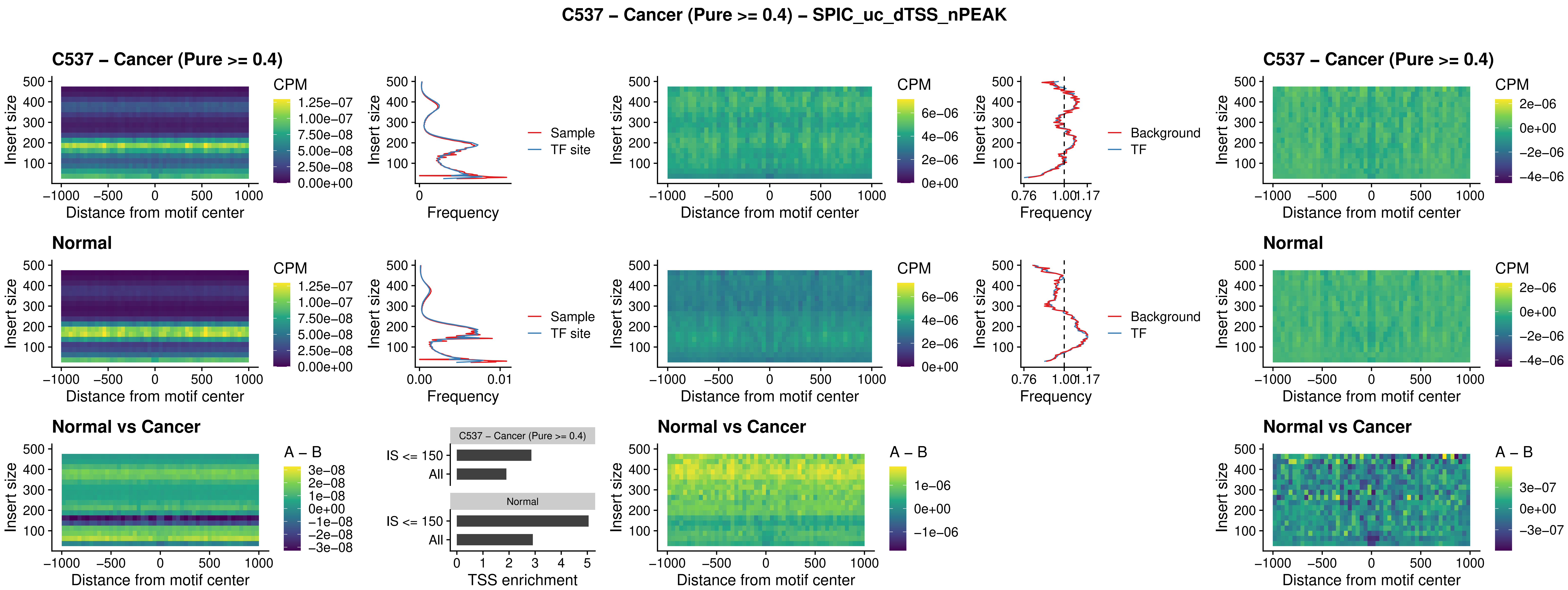
C537 – Cancer (Pure >= 0.4) – SPIC_uc_dTSS_nPEAK
C537 – Cancer (Pure >= 0.4)
Normal
Normal vs Cancer
Insert size
Distance from motif center
Frequency
CPM
A – B
Sample
TF site
Background
TF
C537 – Cancer (Pure >= 0.4)
IS <= 150
All
TSS enrichment

#### C537 – Cancer (Pure >= 0.4) – SPIC_uc_dTSS_oPEAK

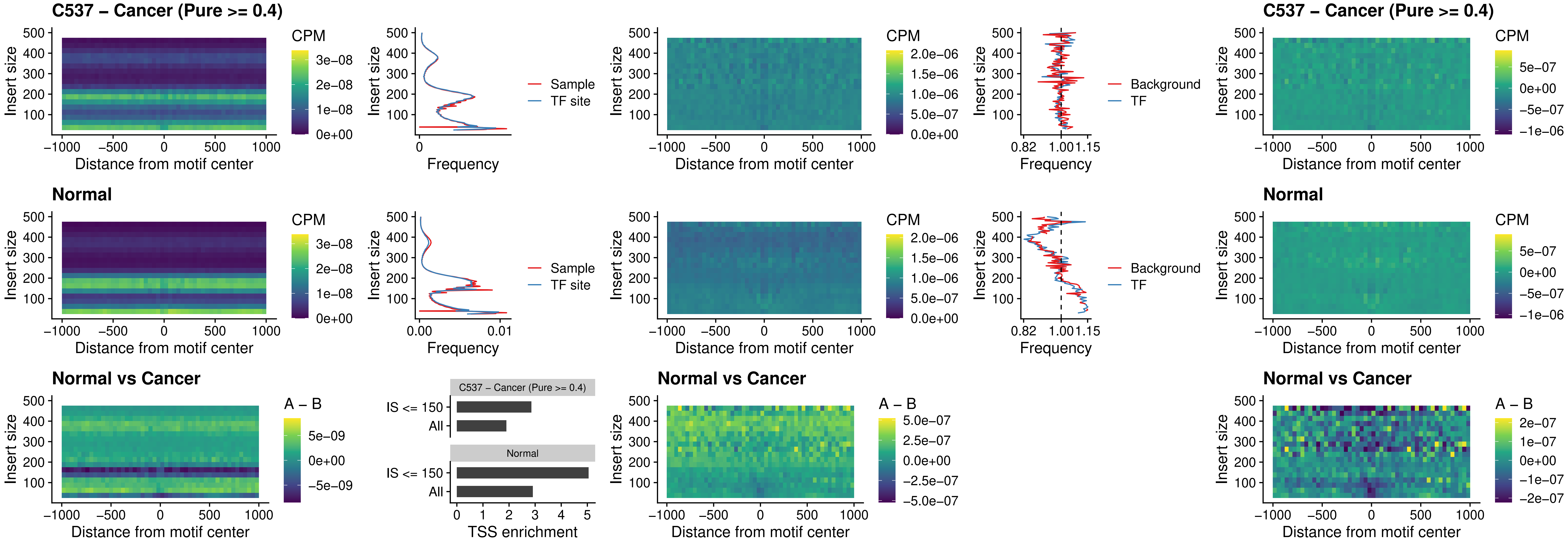

#### C537 – Cancer (Pure >= 0.4) – SPIB_uc_pTSS_nPEAK

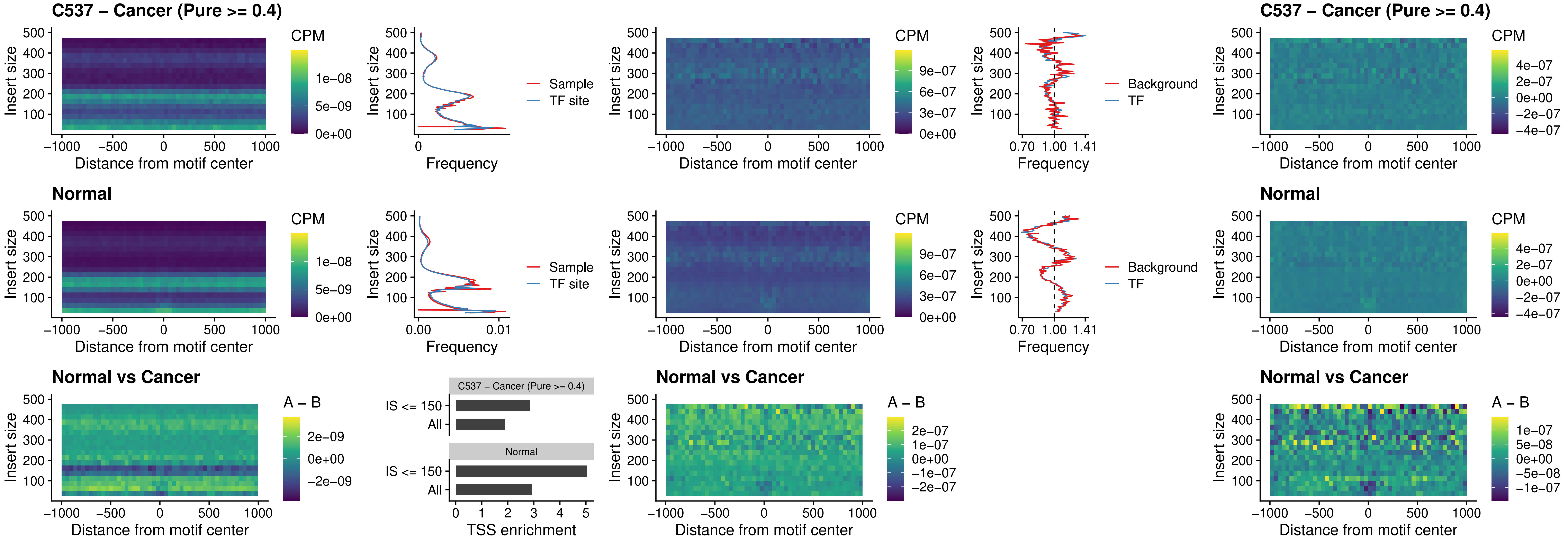

#### C537 – Cancer (Pure >= 0.4) – STAT2_uc_cTSS_oPEAK

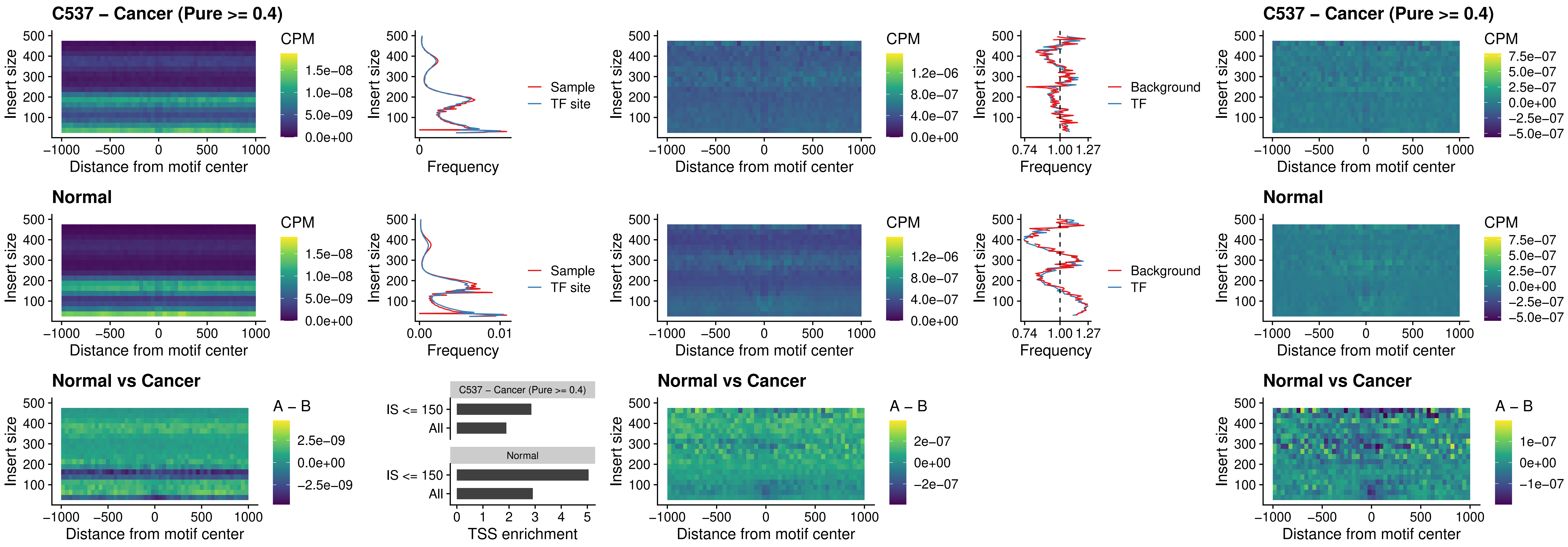

#### C537 – Cancer (Pure >= 0.4) – IRF8_uc_cTSS_oPEAK

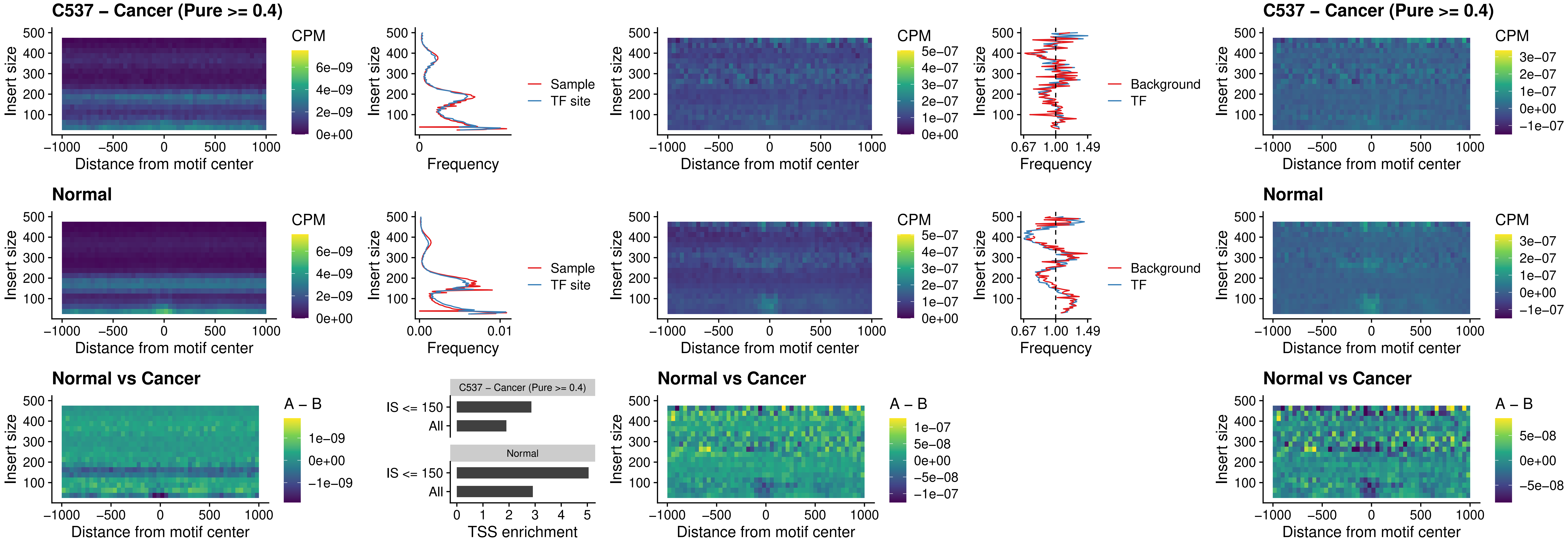

#### C537 – Cancer (Pure >= 0.4) – IRF3_uc_dTSS_oPEAK

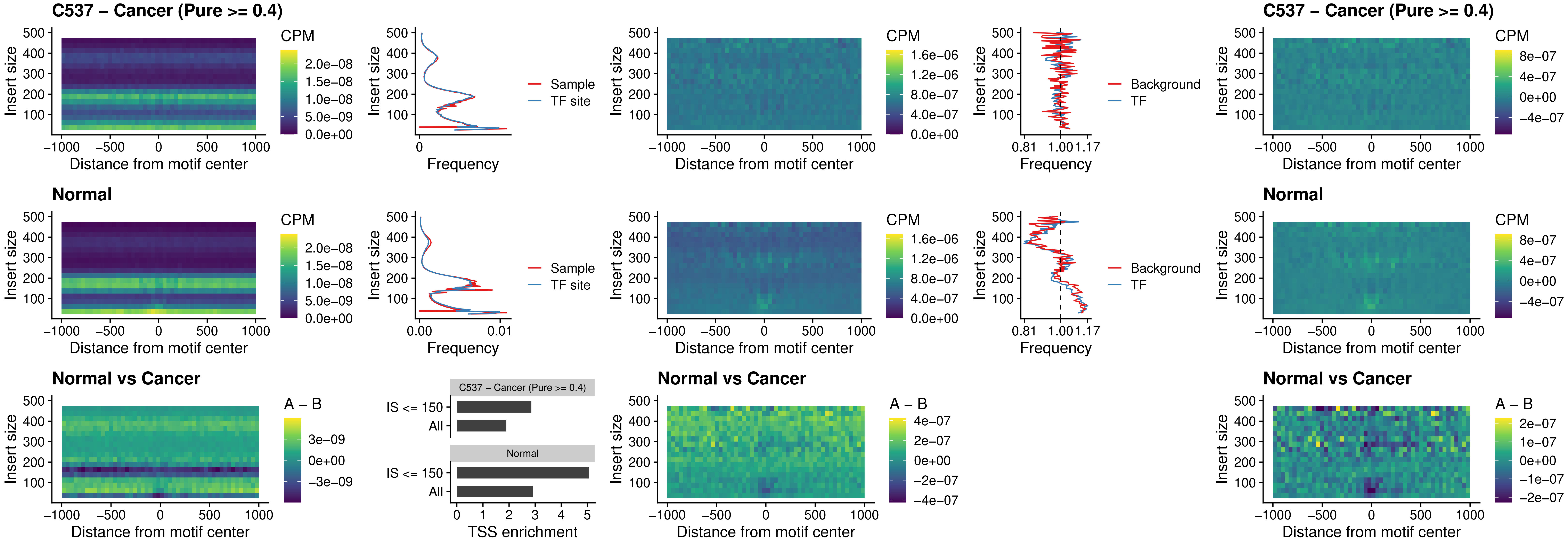

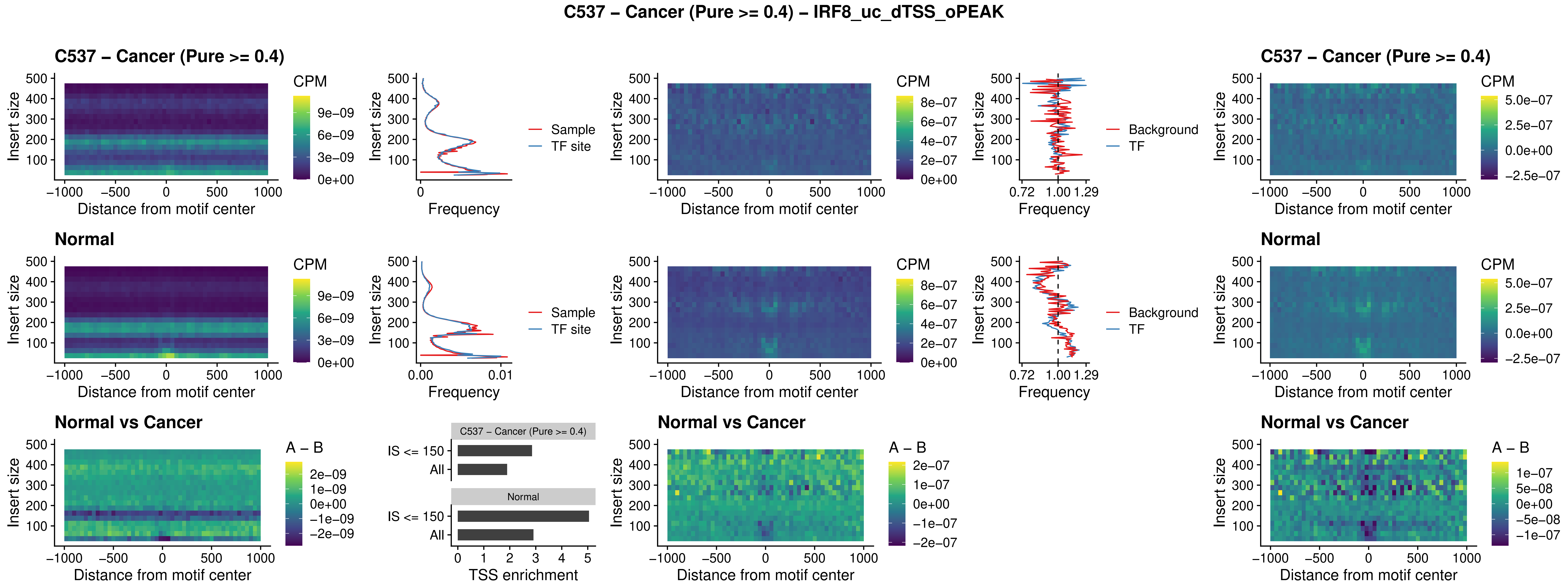
C537 – Cancer (Pure >= 0.4) – IRF8_uc_dTSS_oPEAK
C537 – Cancer (Pure >= 0.4)
Normal
Normal vs Cancer
Insert size
Distance from motif center
Frequency
CPM
A – B
Sample
TF site
Background
TF
C537 – Cancer (Pure >= 0.4)
Normal
IS <= 150
All
TSS enrichment

#### C537 – Cancer (Pure >= 0.4) – ETS2_uc_dTSS_oPEAK

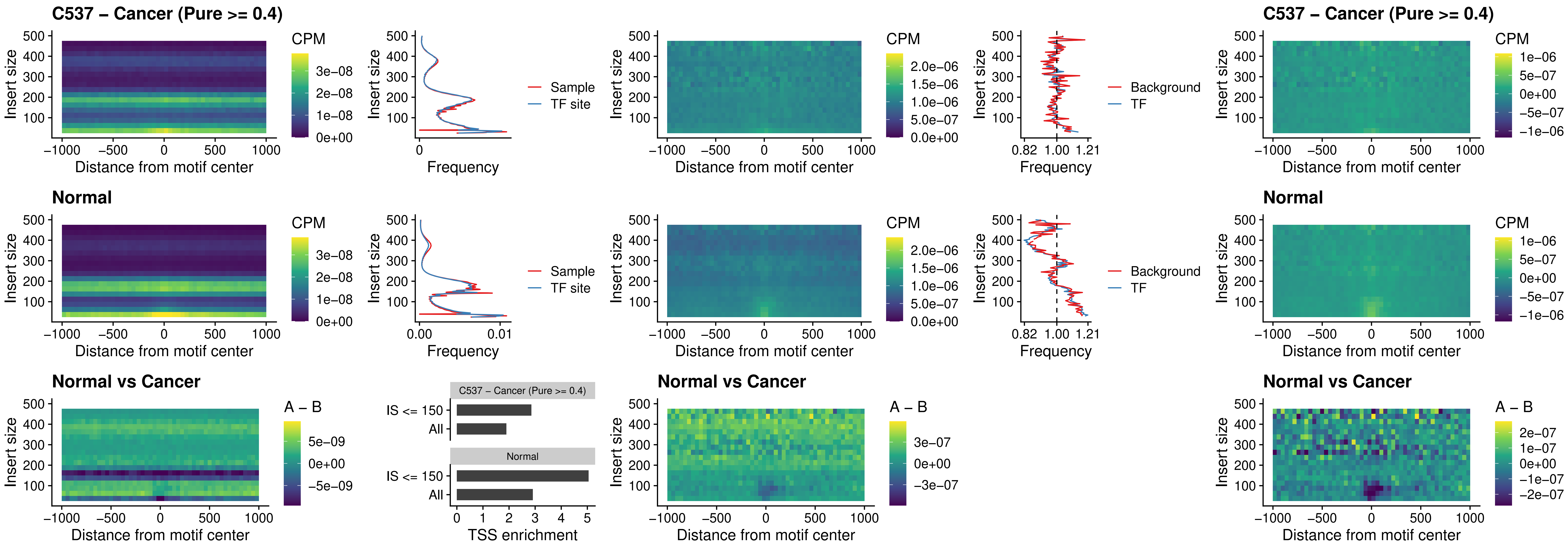

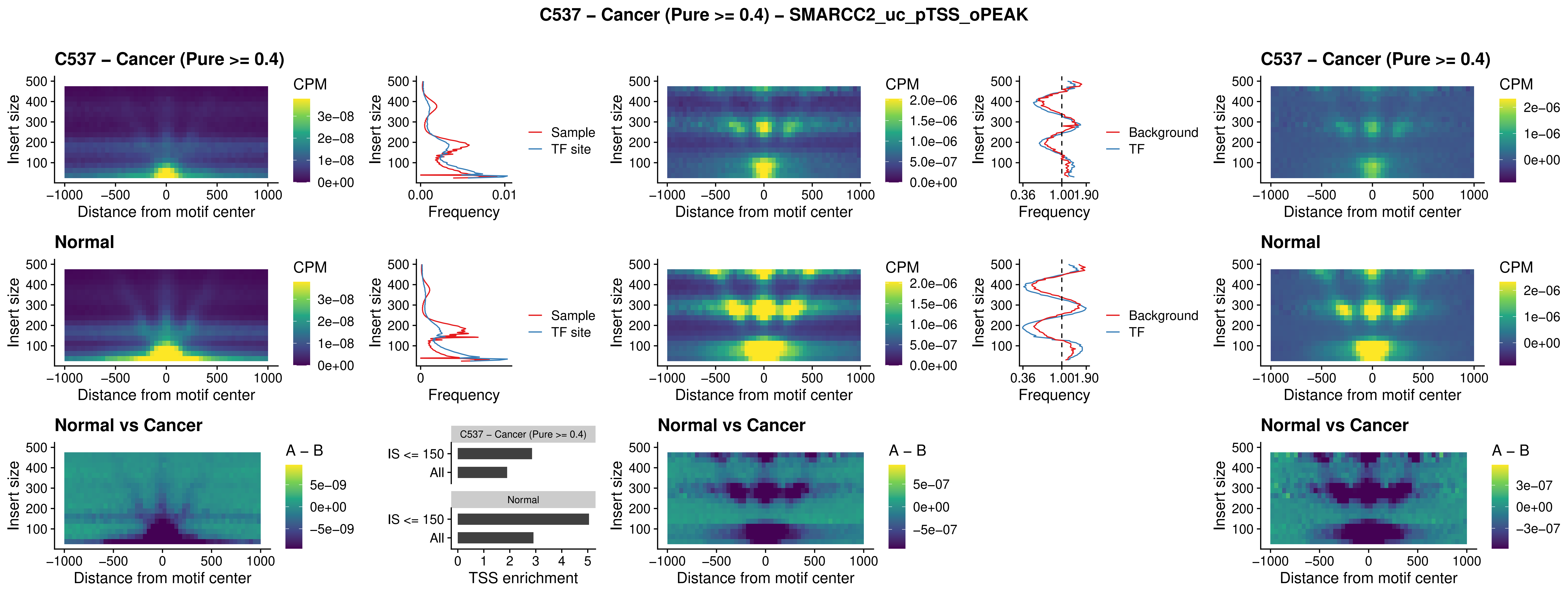

C537 – Cancer (Pure >= 0.4) – SMARCC2_uc_pTSS_oPEAK
C537 – Cancer (Pure >= 0.4)
Normal
Normal vs Cancer
Insert size
Distance from motif center
Frequency
CPM
A – B
Sample
TF site
Background
TF
TSS enrichment
IS <= 150
All
C537 – Cancer (Pure >= 0.4)
Normal

#### C537 – Cancer (Pure >= 0.4) – BCL11A_uc_cTSS_nPEAK

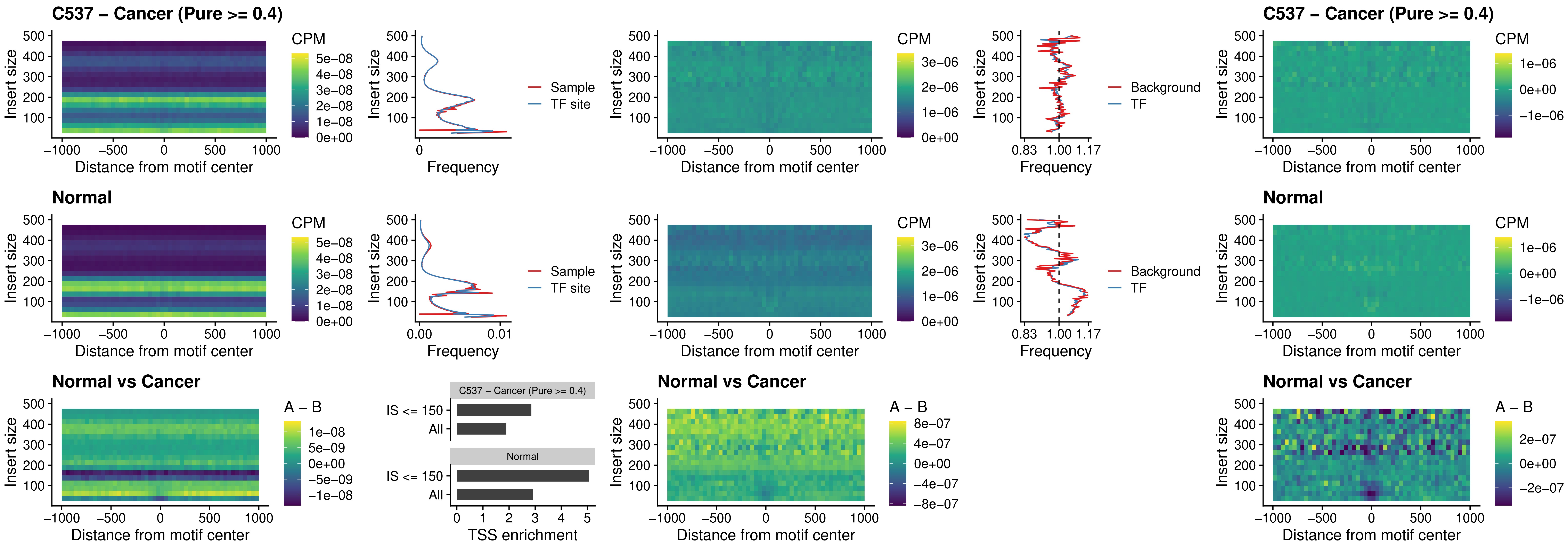

#### C537 – Cancer (Pure >= 0.4) – BCL11B_uc_cTSS_nPEAK

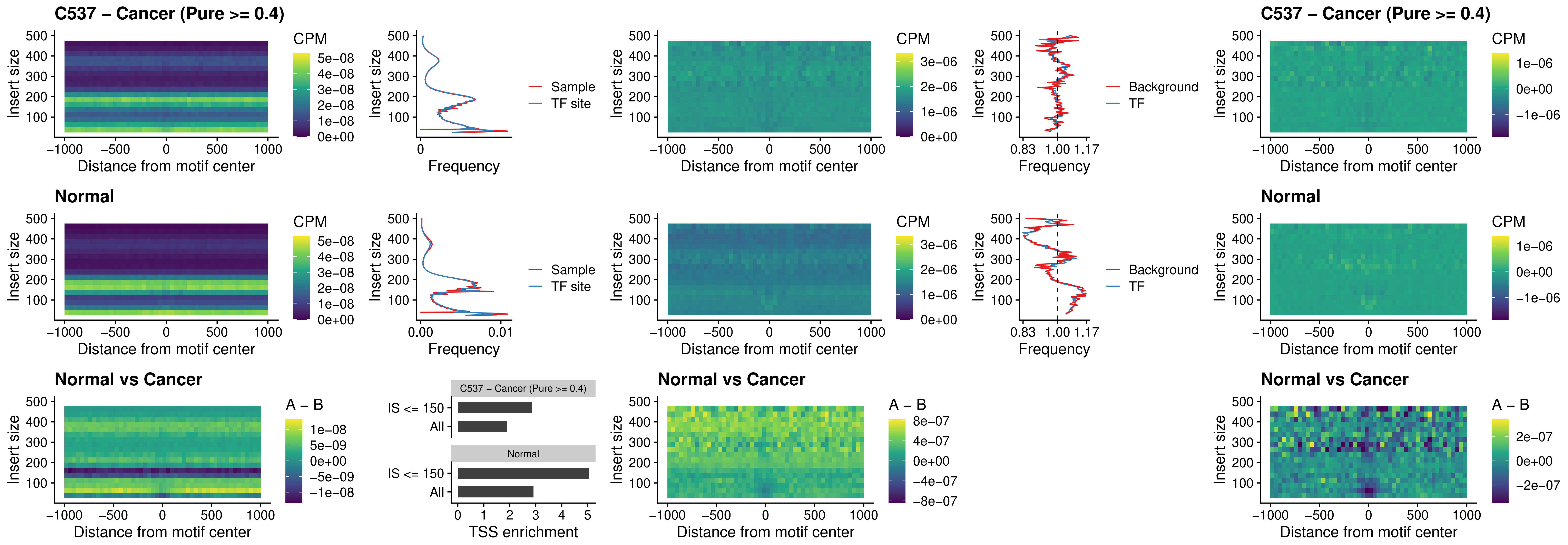

#### C537 – Cancer (Pure >= 0.4) – STAT2_uc_dTSS_oPEAK

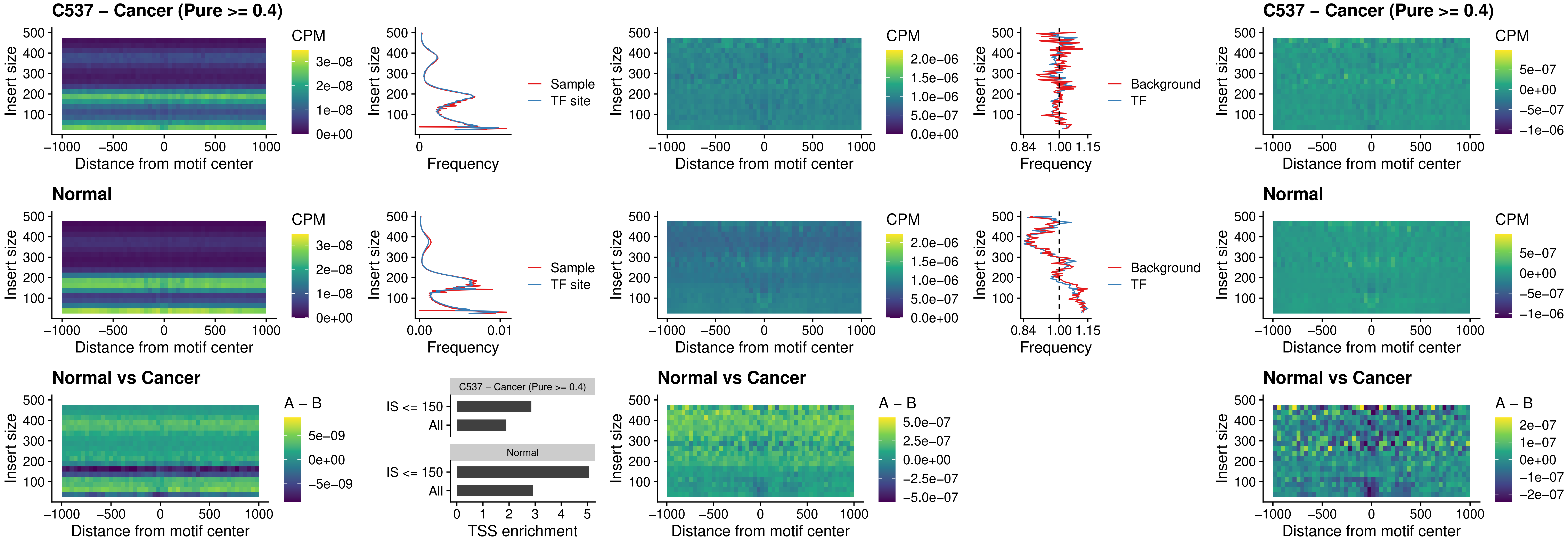

#### C537 – Cancer (Pure >= 0.4) – ZNF683_uc_cTSS_oPEAK

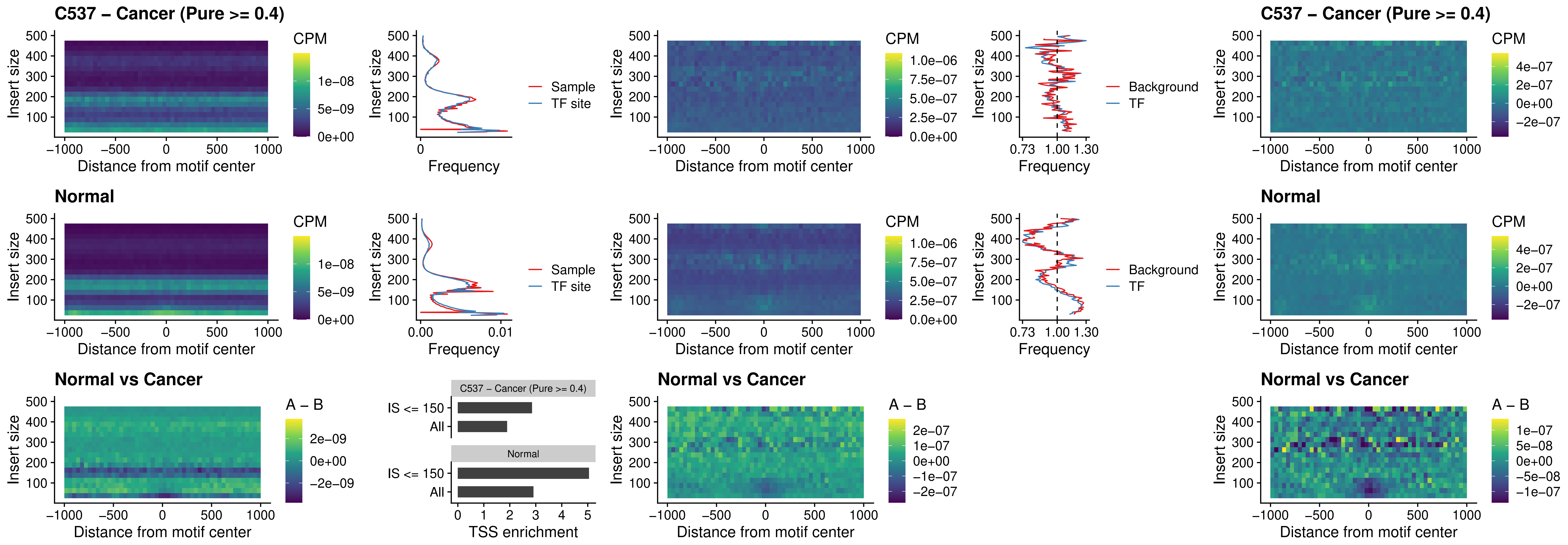

#### C537 – Cancer (Pure >= 0.4) – IRF8_uc_dTSS_nPEAK

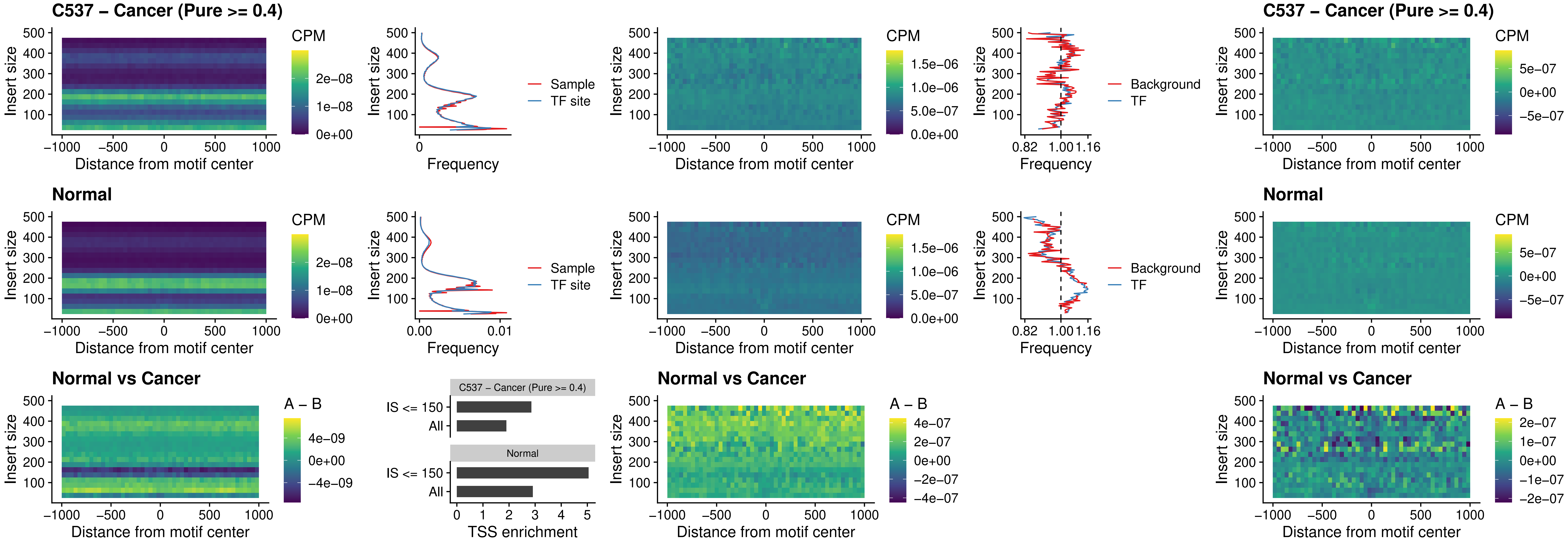

#### C537 – Cancer (Pure >= 0.4) – ETV7_uc_dTSS_oPEAK

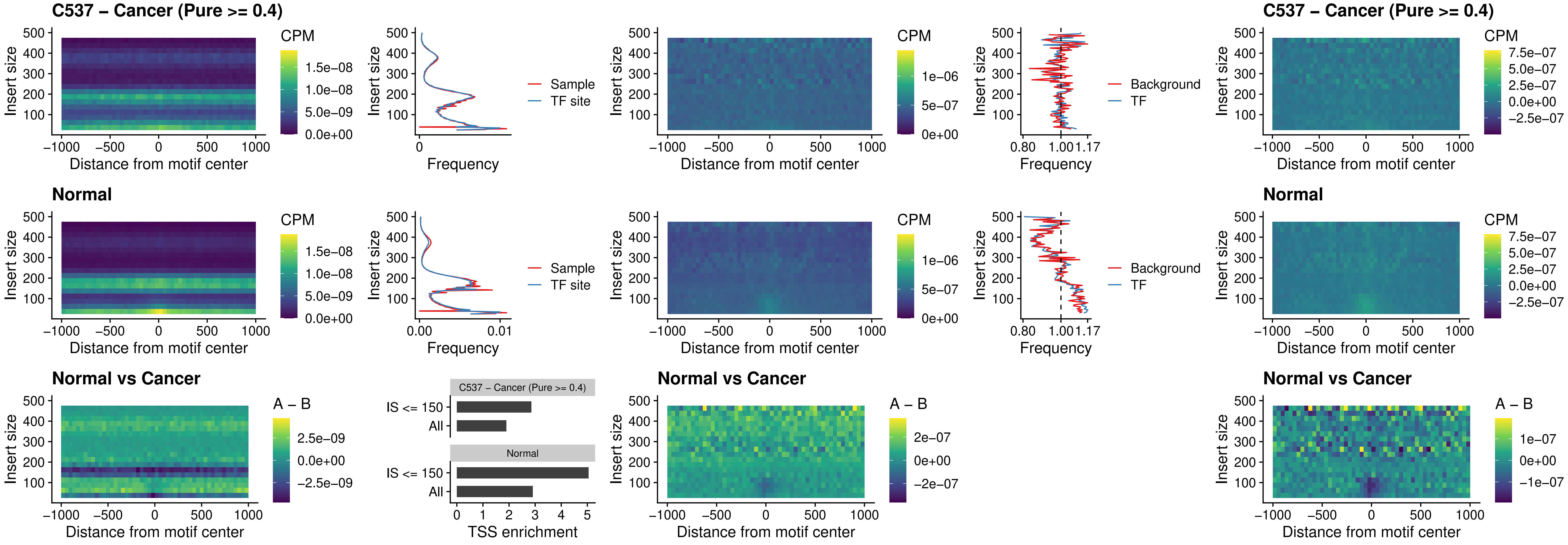

#### C538 – Cancer (Pure >= 0.4) – FOXL1_uc_pTSS_oPEAK

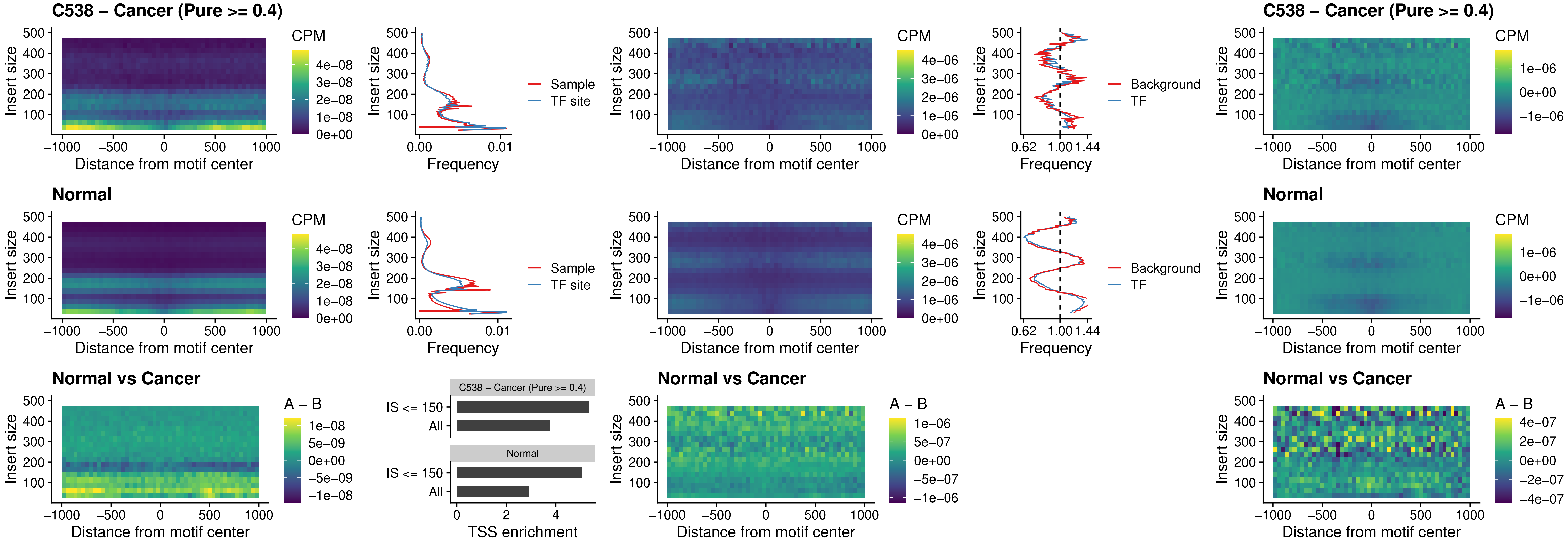

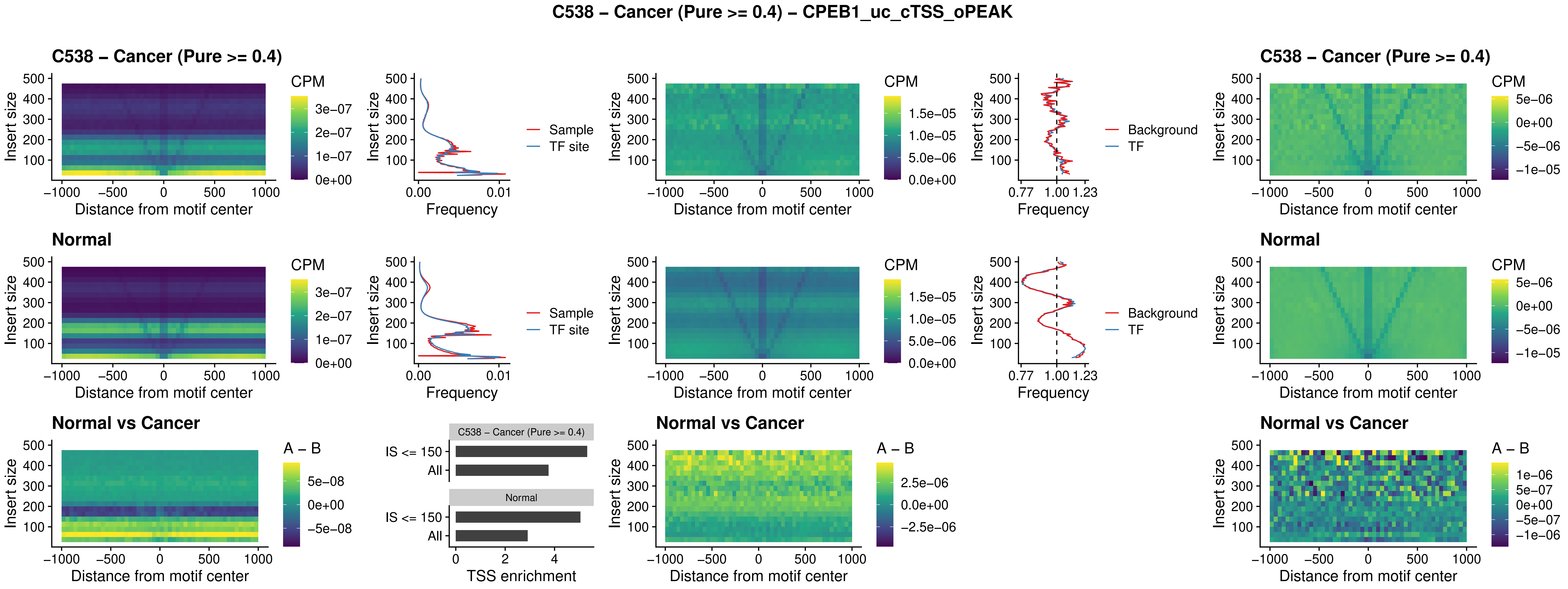
C538 – Cancer (Pure >= 0.4) – CPEB1_uc_cTSS_oPEAK
C538 – Cancer (Pure >= 0.4)
Normal
Normal vs Cancer
Insert size
Distance from motif center
CPM
A – B
Frequency
Sample
TF site
Background
TF
C538 – Cancer (Pure >= 0.4)
Normal
IS <= 150
All
TSS enrichment

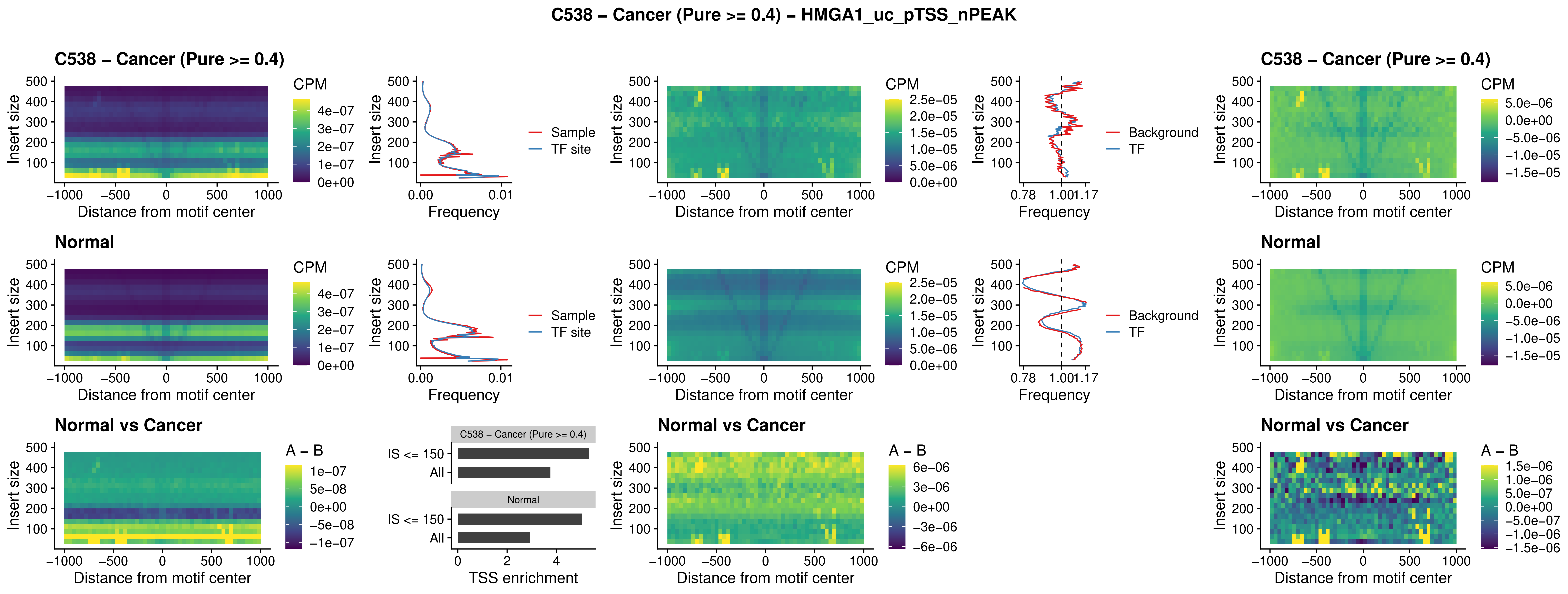
C538 – Cancer (Pure >= 0.4) – HMGA1_uc_pTSS_nPEAK
C538 – Cancer (Pure >= 0.4)
Normal
Normal vs Cancer
Insert size
Distance from motif center
Frequency
CPM
A – B
Sample
TF site
Background
TF
C538 – Cancer (Pure >= 0.4)
Normal
IS <= 150
All
TSS enrichment

#### C538 – Cancer (Pure >= 0.4) – SOX13_uc_cTSS_nPEAK

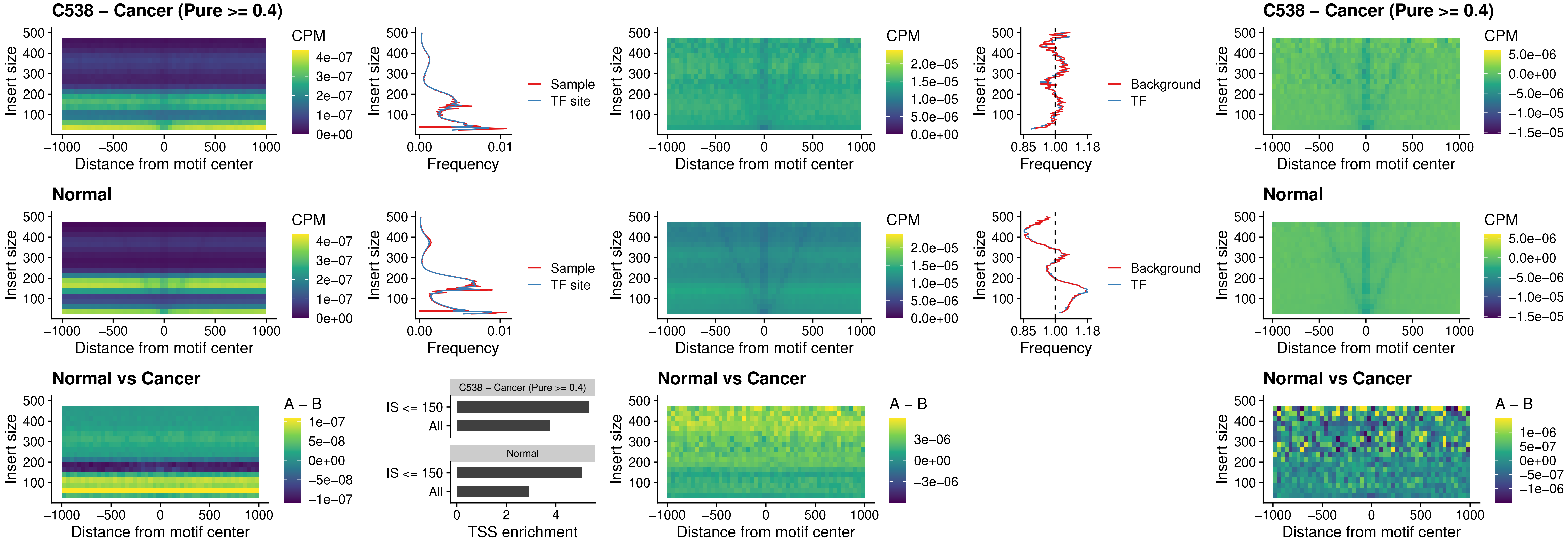

#### C538 – Cancer (Pure >= 0.4) – LHX4_uc_cTSS_nPEAK

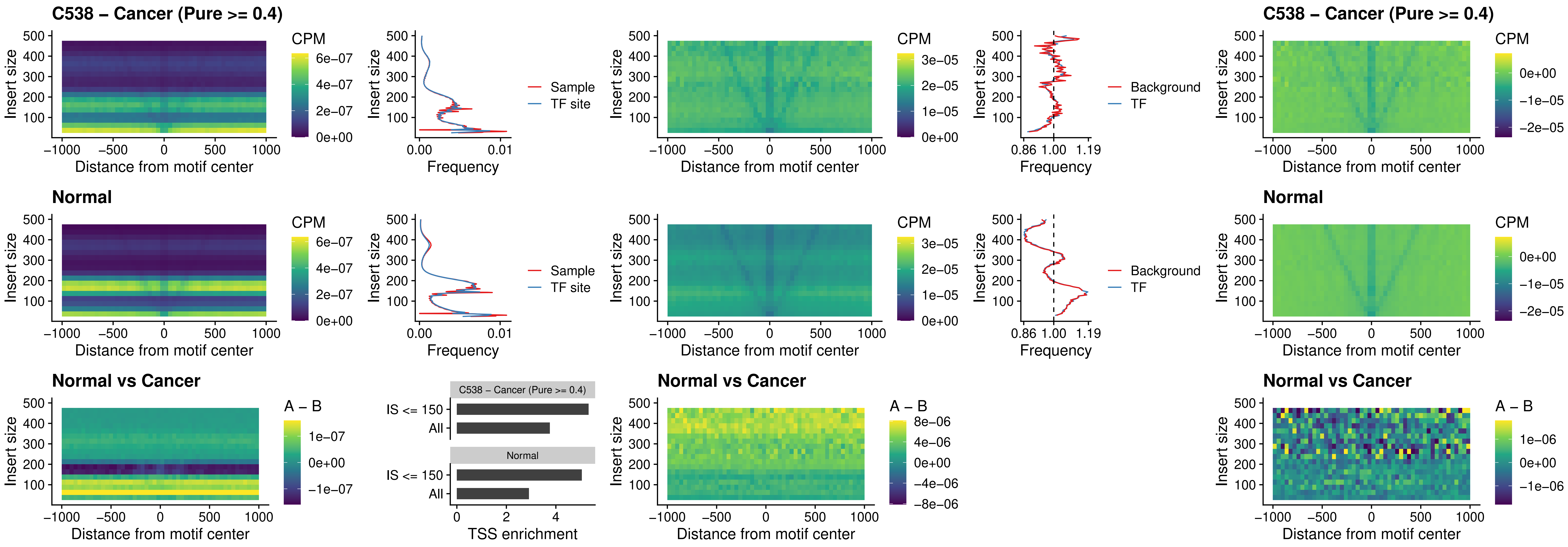

#### C538 – Cancer (Pure >= 0.4) – HOXA4_uc_cTSS_nPEAK

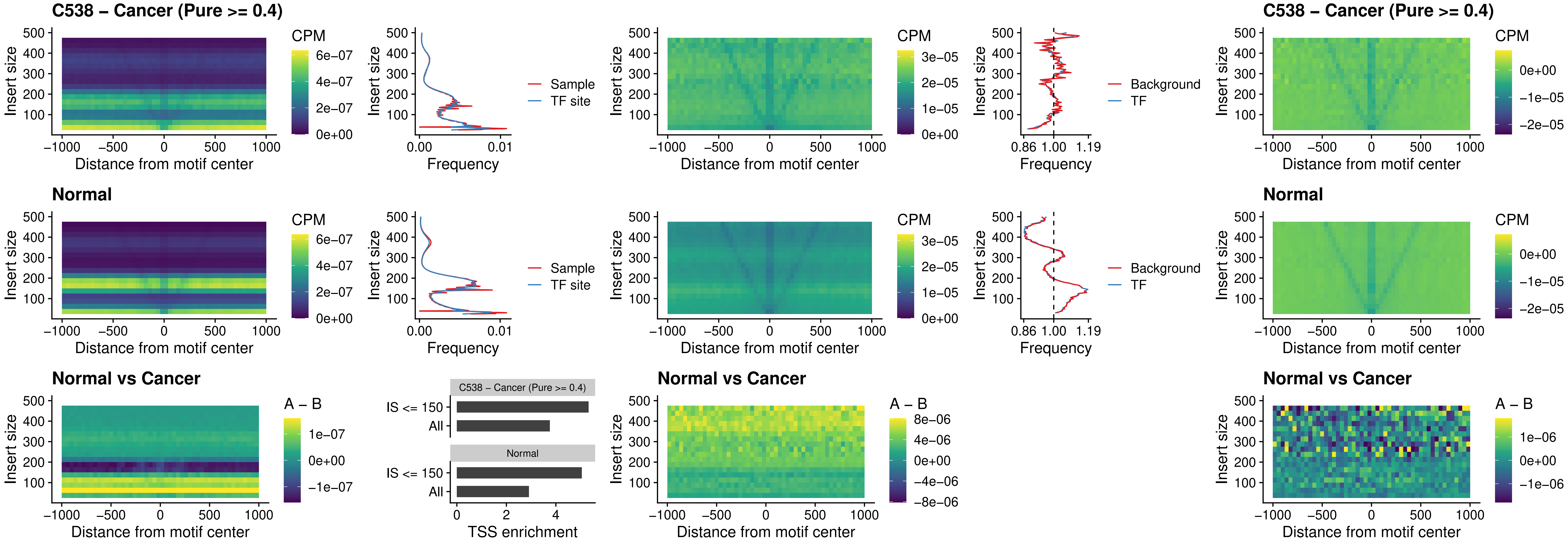

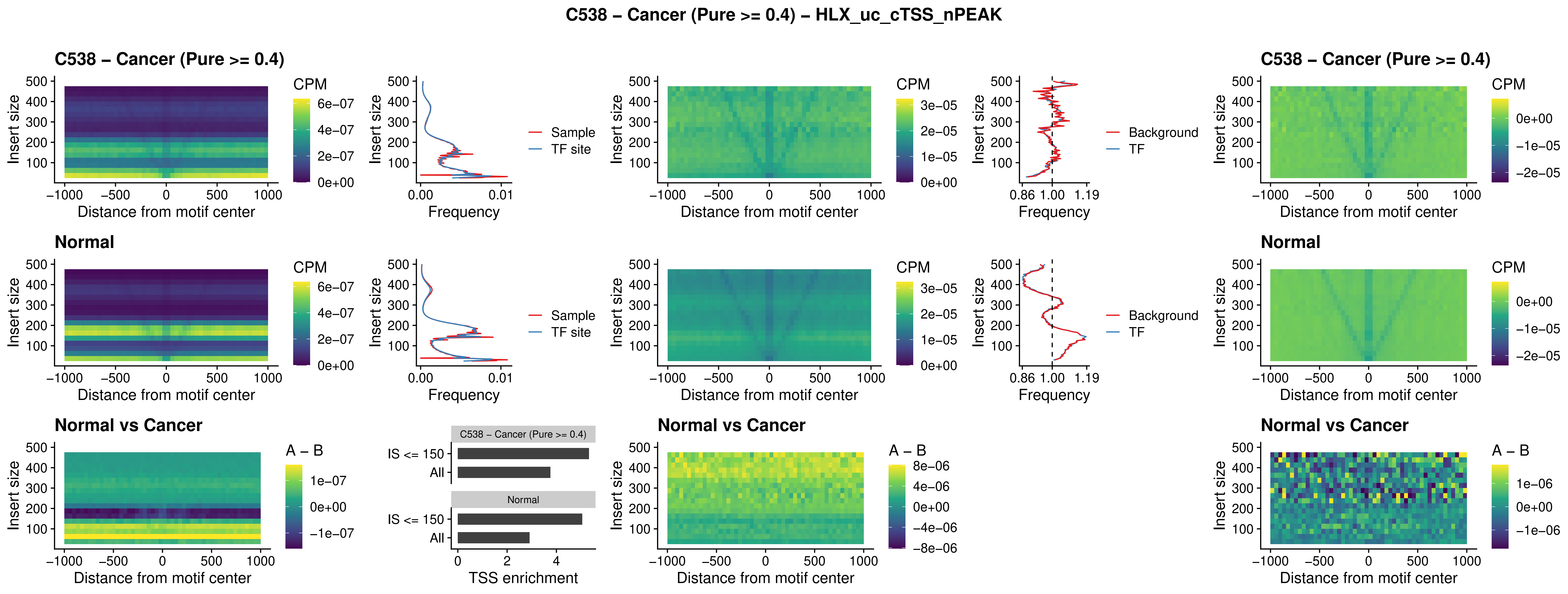
C538 – Cancer (Pure >= 0.4) – HLX_uc_cTSS_nPEAK
C538 – Cancer (Pure >= 0.4)
Normal
Normal vs Cancer
Insert size
Distance from motif center
Frequency
CPM
A – B
Sample
TF site
Background
TF
C538 – Cancer (Pure >= 0.4)
Normal
IS <= 150
All
TSS enrichment

#### C538 – Cancer (Pure >= 0.4) – AC0125311_uc_cTSS_nPEAK

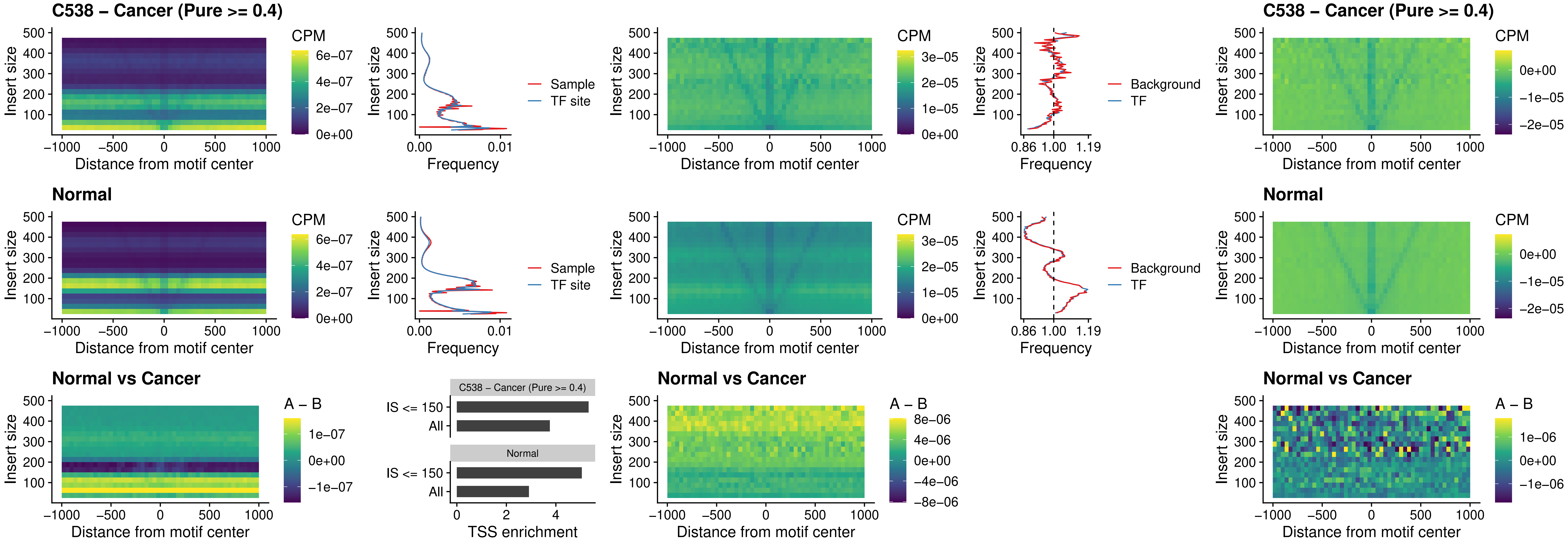

#### C538 – Cancer (Pure >= 0.4) – DBX2_uc_cTSS_nPEAK

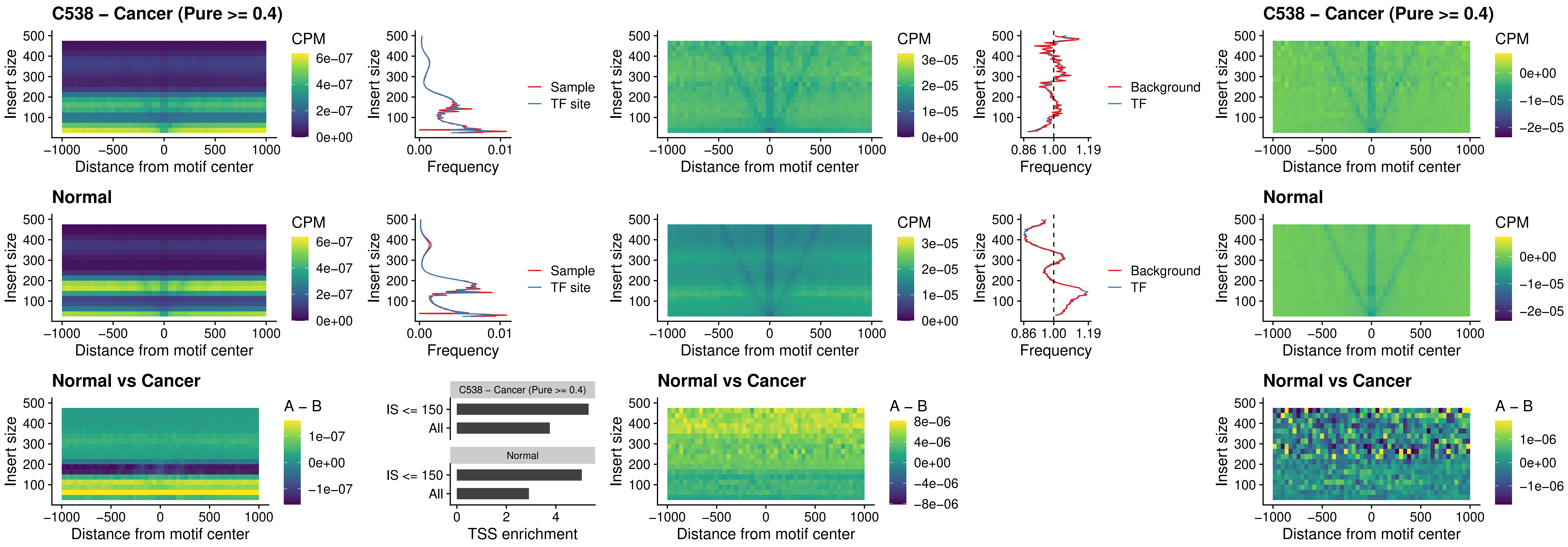

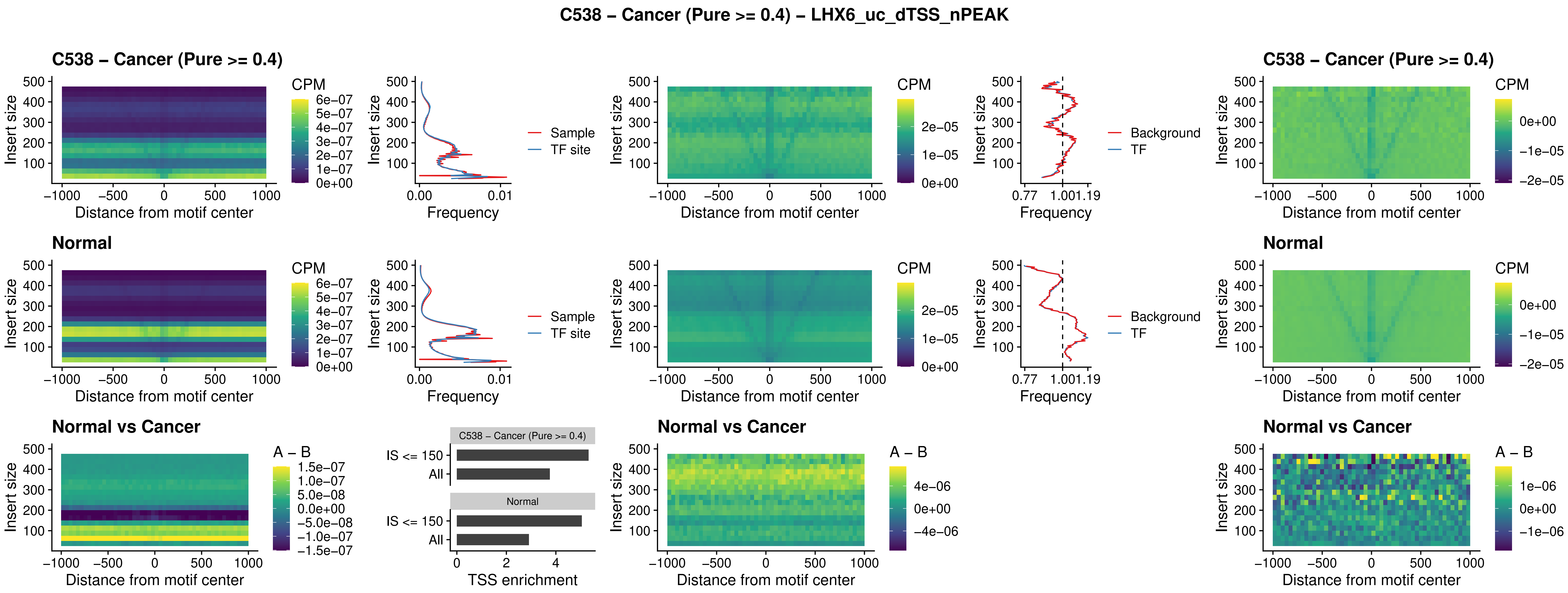
C538 – Cancer (Pure >= 0.4) – LHX6_uc_dTSS_nPEAK
C538 – Cancer (Pure >= 0.4)
Normal
Normal vs Cancer
Insert size
Distance from motif center
Frequency
CPM
A – B
Sample
TF site
Background
TF
C538 – Cancer (Pure >= 0.4)
Normal
IS <= 150
All
TSS enrichment

#### C538 – Cancer (Pure >= 0.4) – SOX1_uc_dTSS_nPEAK

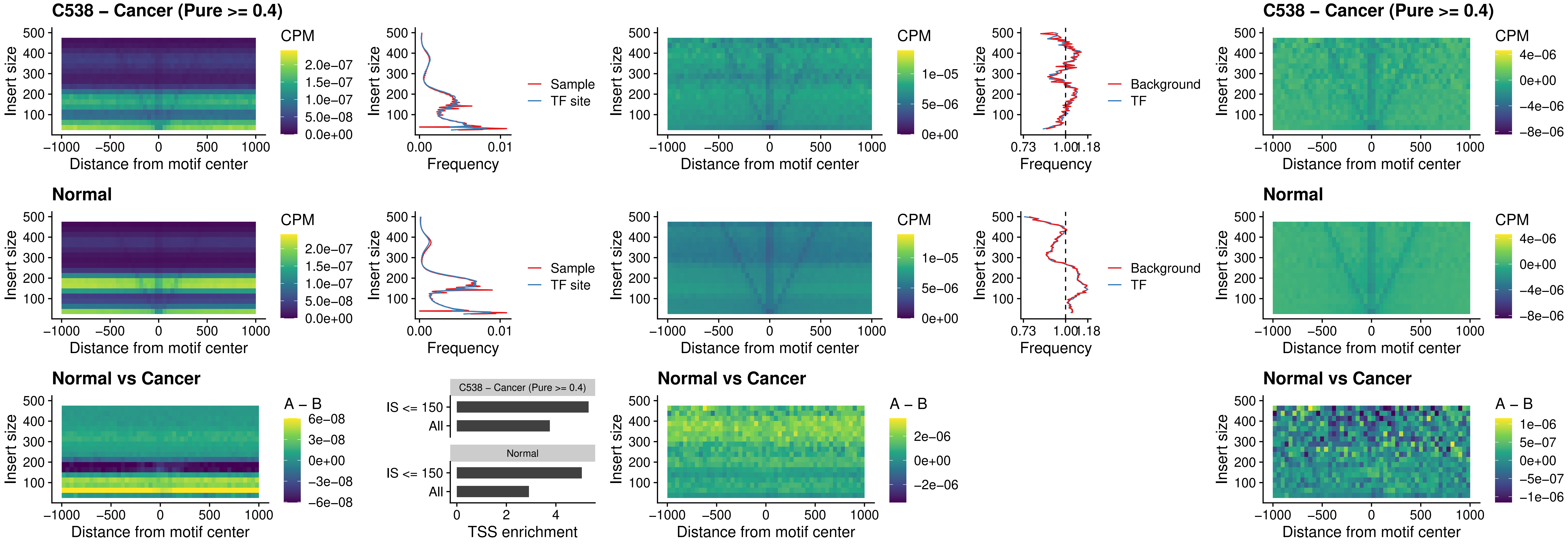

#### C538 – Cancer (Pure >= 0.4) – NKX24_uc_dTSS_nPEAK

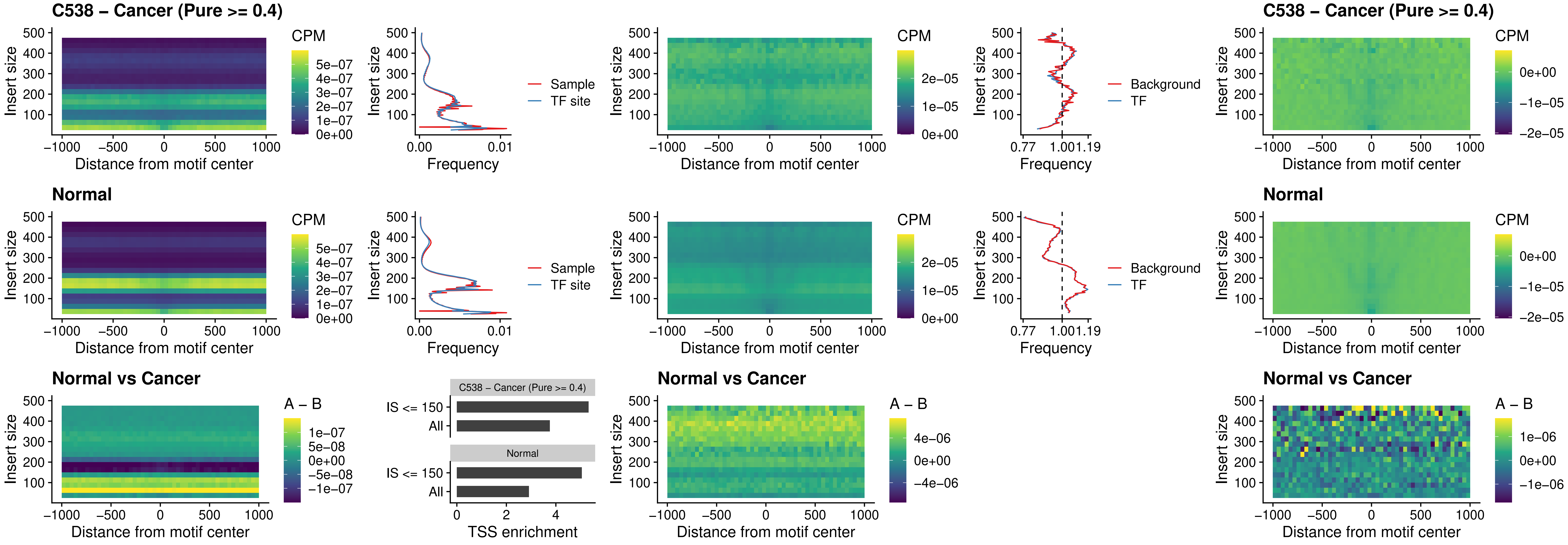

#### C538 – Cancer (Pure >= 0.4) – DLX3_uc_cTSS_nPEAK

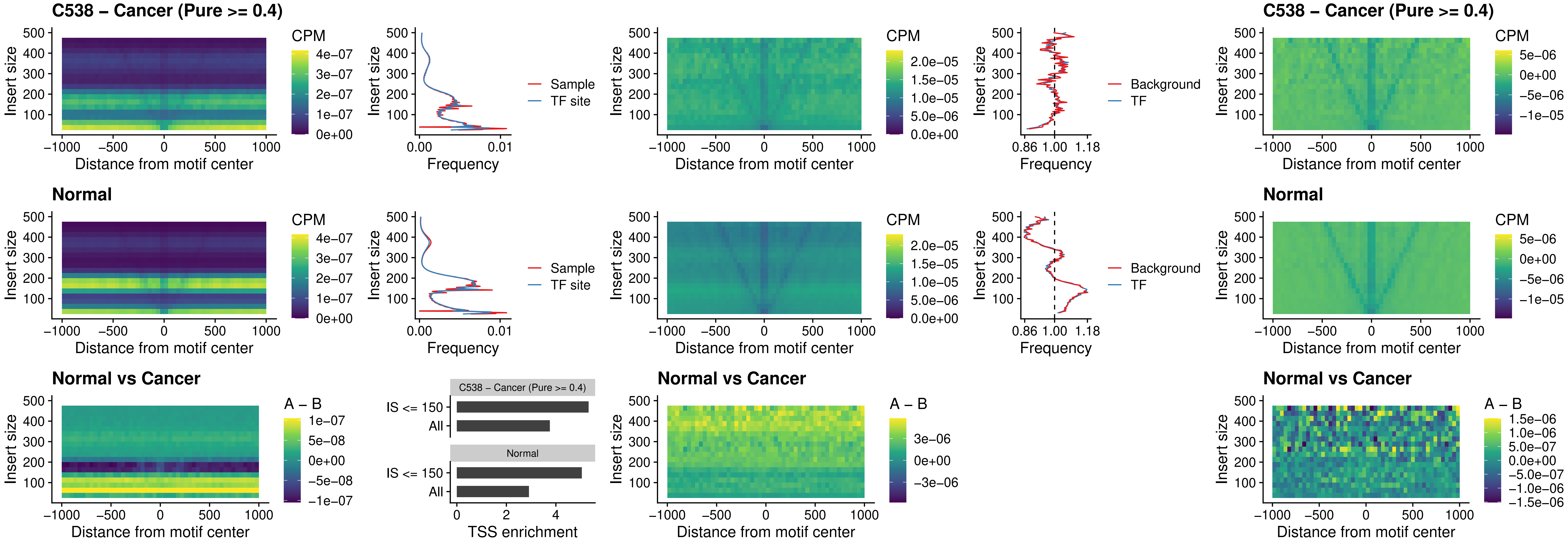

#### C538 – Cancer (Pure >= 0.4) – HOXD3_uc_cTSS_nPEAK

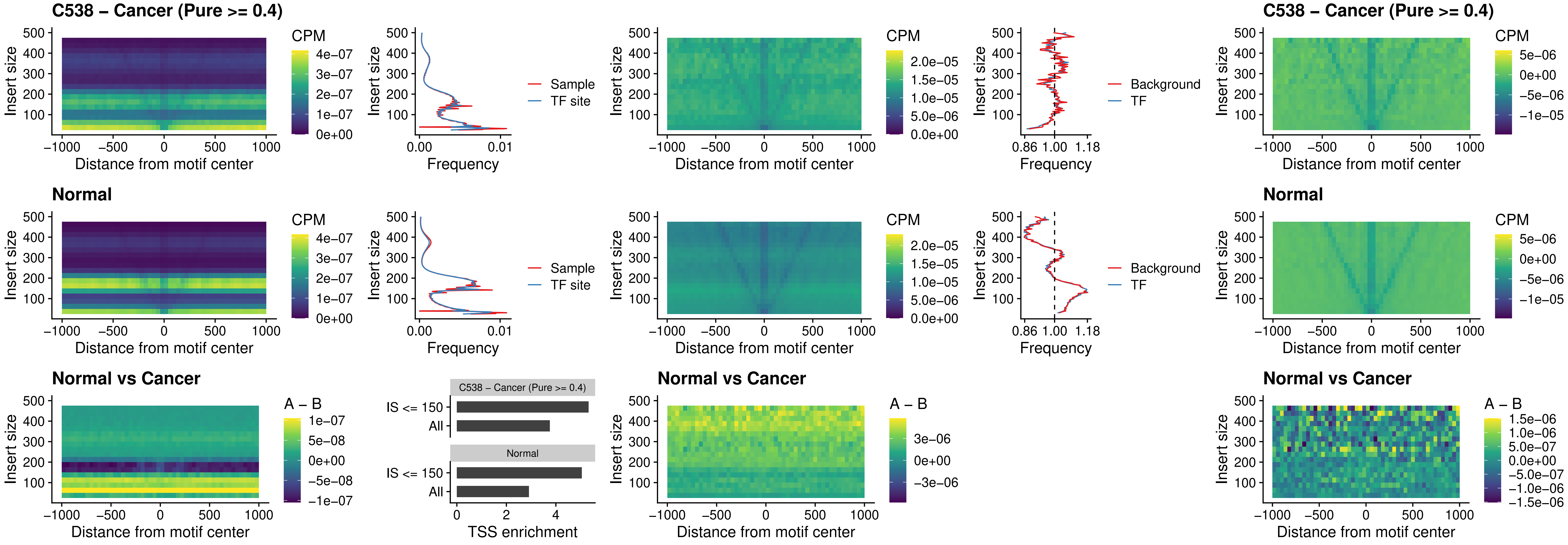

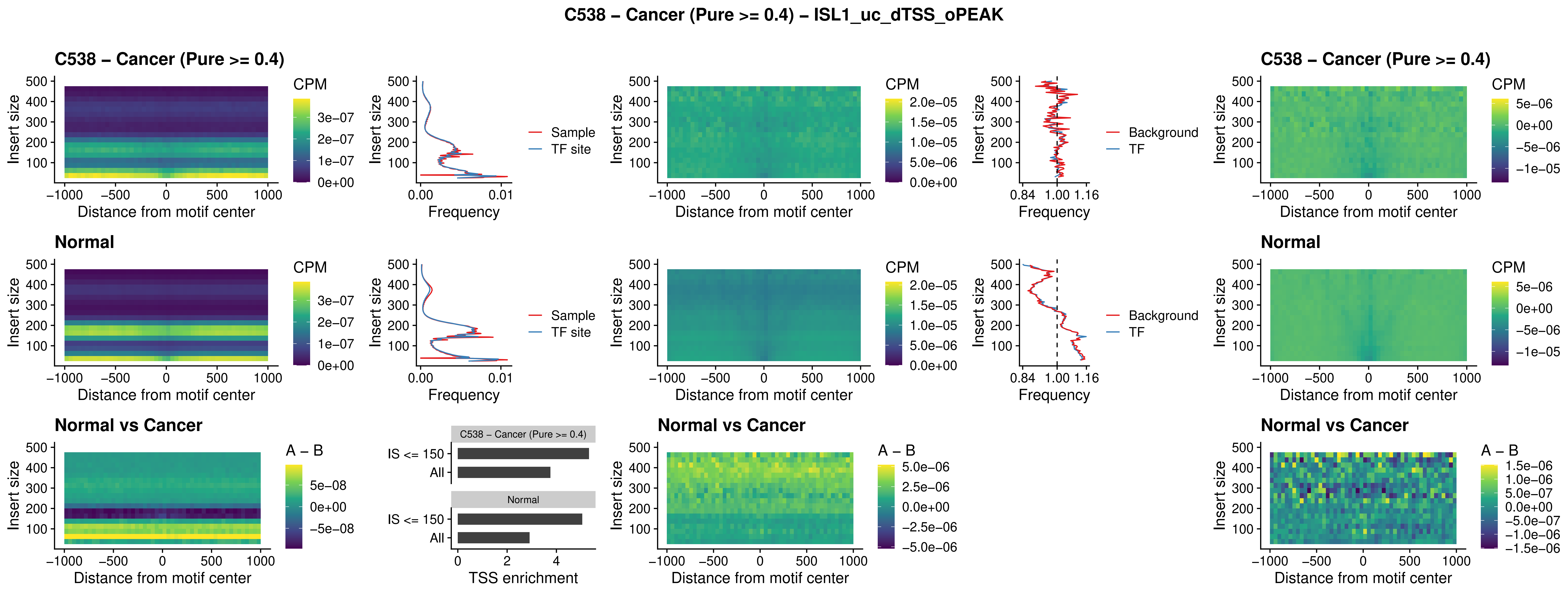

C538 – Cancer (Pure >= 0.4) – ISL1_uc_dTSS_oPEAK
C538 – Cancer (Pure >= 0.4)
Normal
Normal vs Cancer
Insert size
Distance from motif center
Frequency
CPM
A – B
Sample
TF site
Background
TF
C538 – Cancer (Pure >= 0.4)
Normal
IS <= 150
All
TSS enrichment

#### C538 – Cancer (Pure >= 0.4) – HOXB4_uc_dTSS_oPEAK

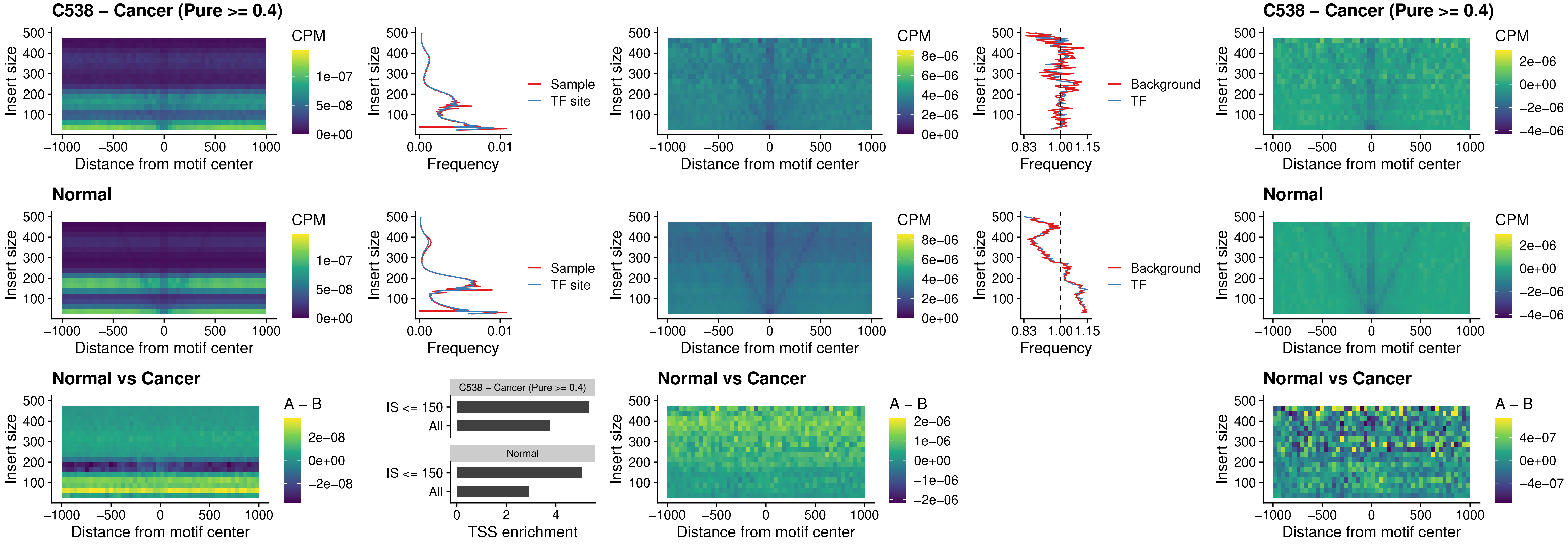

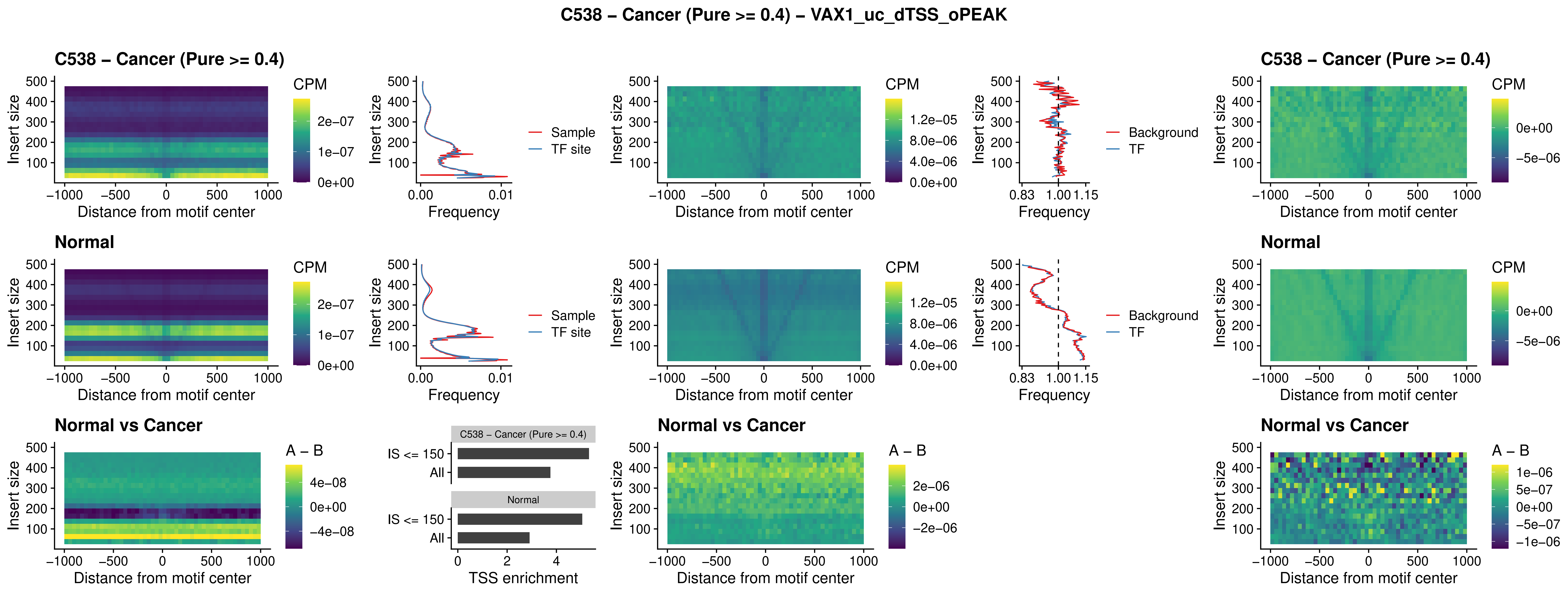
C538 – Cancer (Pure >= 0.4) – VAX1_uc_dTSS_oPEAK
C538 – Cancer (Pure >= 0.4)
Normal
Normal vs Cancer
Insert size
Distance from motif center
Frequency
CPM
A – B
Sample
TF site
Background
TF
C538 – Cancer (Pure >= 0.4)
Normal
IS <= 150
All
TSS enrichment

#### C538 – Cancer (Pure >= 0.4) – DLX3_uc_dTSS_oPEAK

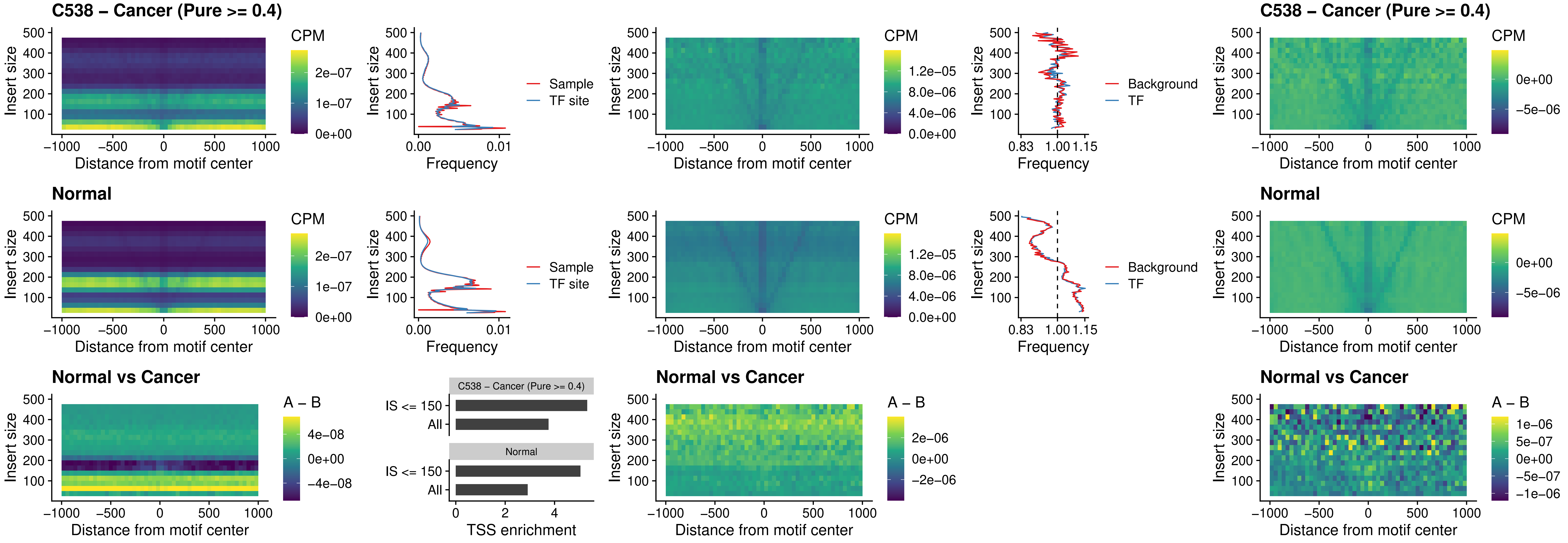

#### C538 – Cancer (Pure >= 0.4) – HOXD3_uc_dTSS_oPEAK

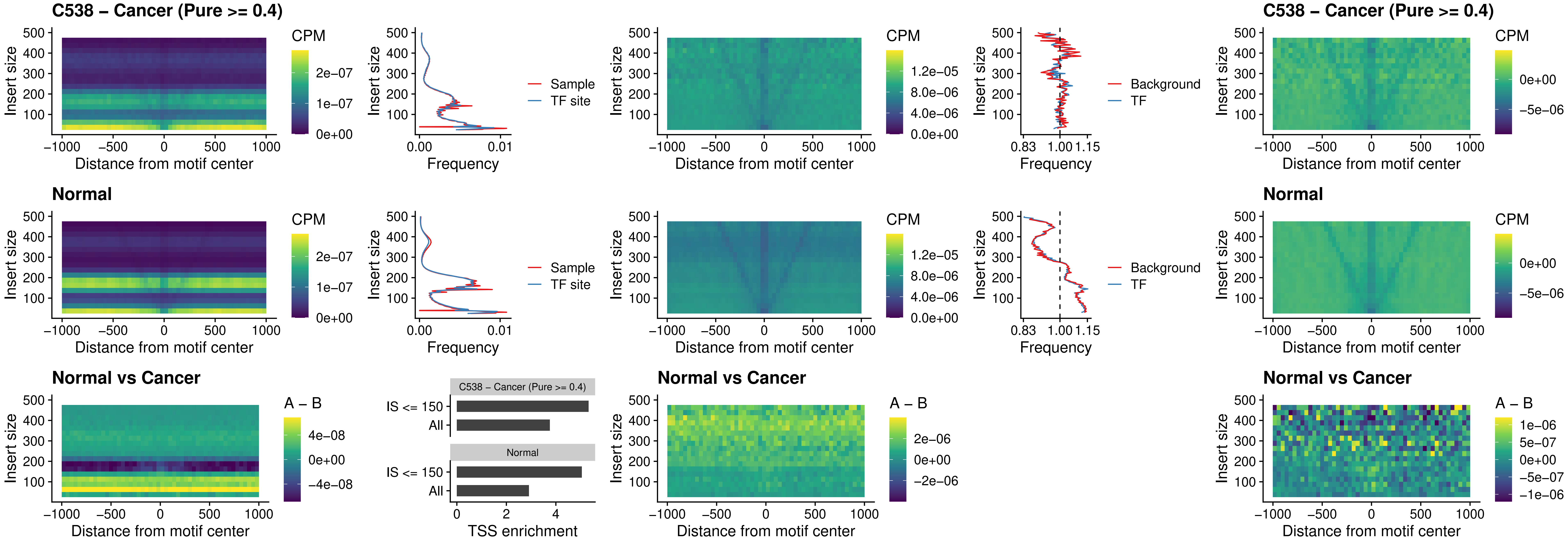

#### C538 – Cancer (Pure >= 0.4) – BBX_uc_dTSS_nPEAK

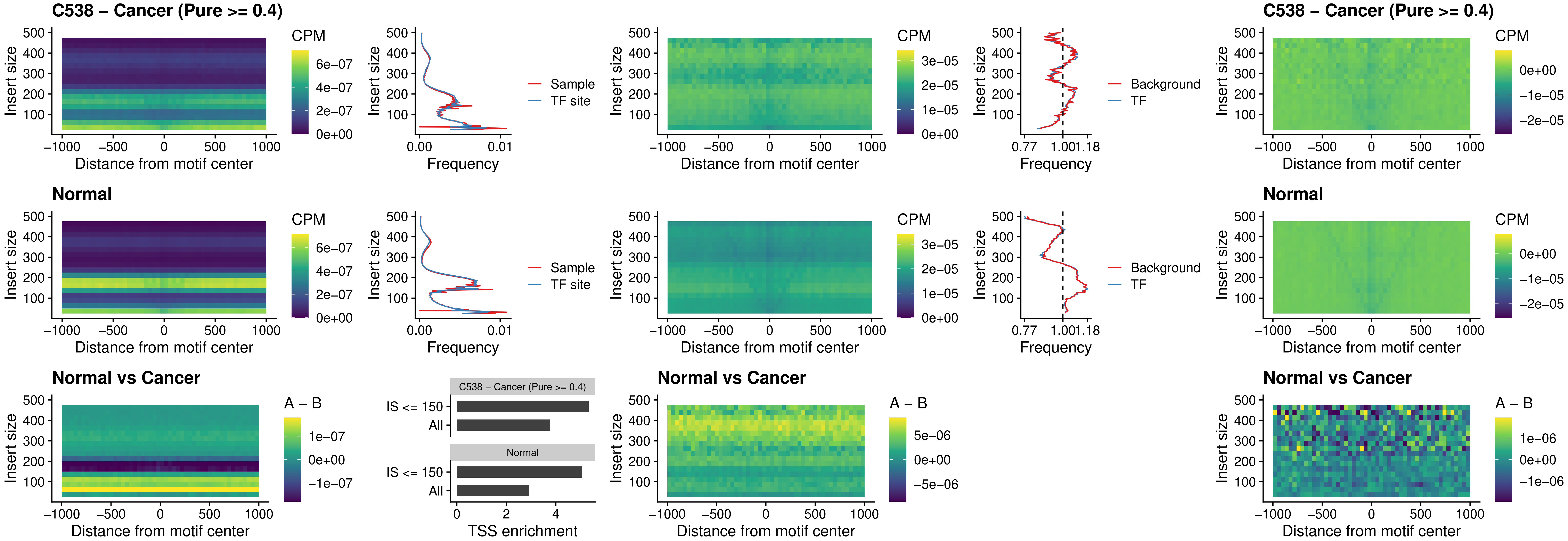

#### C538 – Cancer (Pure >= 0.4) – SOX3_uc_cTSS_nPEAK

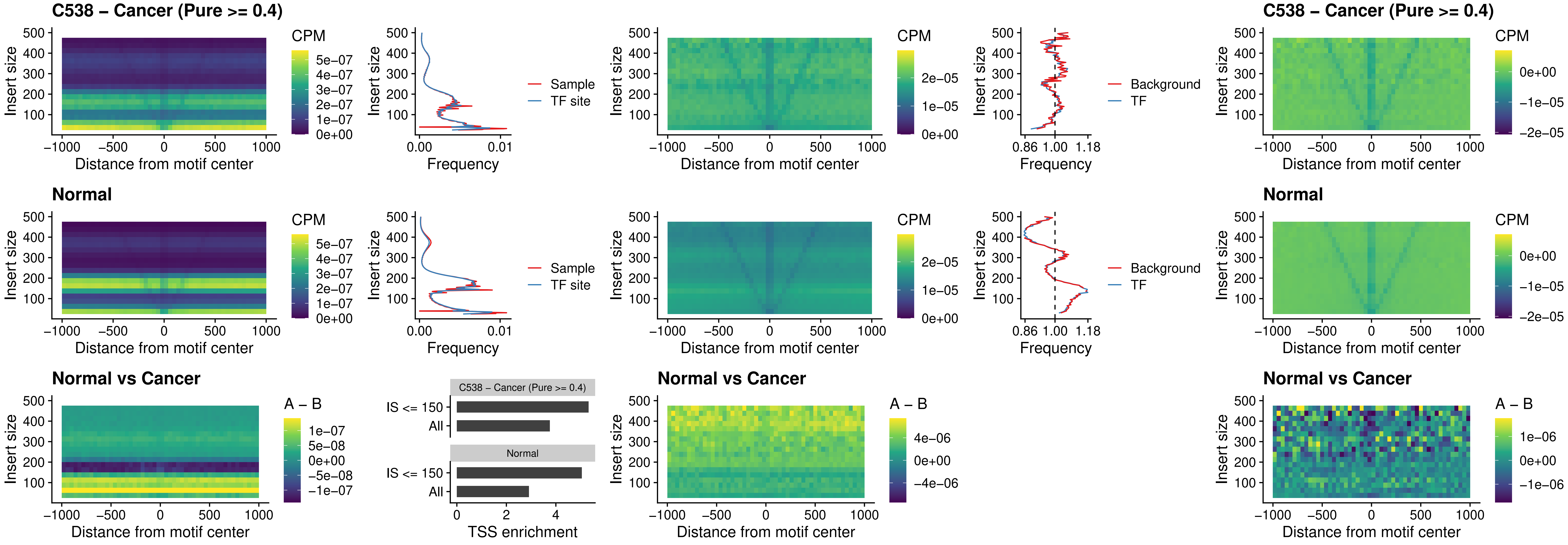

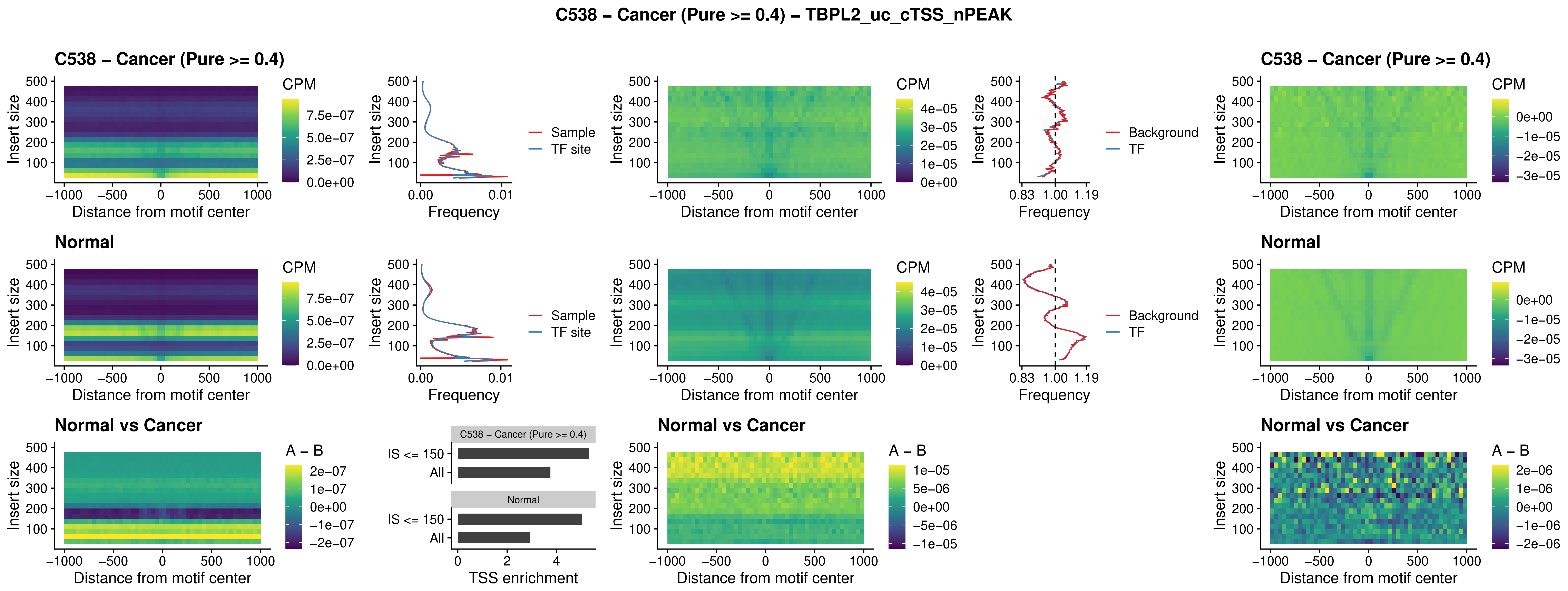
C538 – Cancer (Pure >= 0.4) – TBPL2_uc_cTSS_nPEAK
C538 – Cancer (Pure >= 0.4)
Normal
Normal vs Cancer
Insert size
Distance from motif center
CPM
A – B
Frequency
Sample
TF site
Background
TF
C538 – Cancer (Pure >= 0.4)
IS <= 150
All
TSS enrichment

### C538 – Cancer (Pure >= 0.4) – HOXA4_uc_cTSS_oPEAK

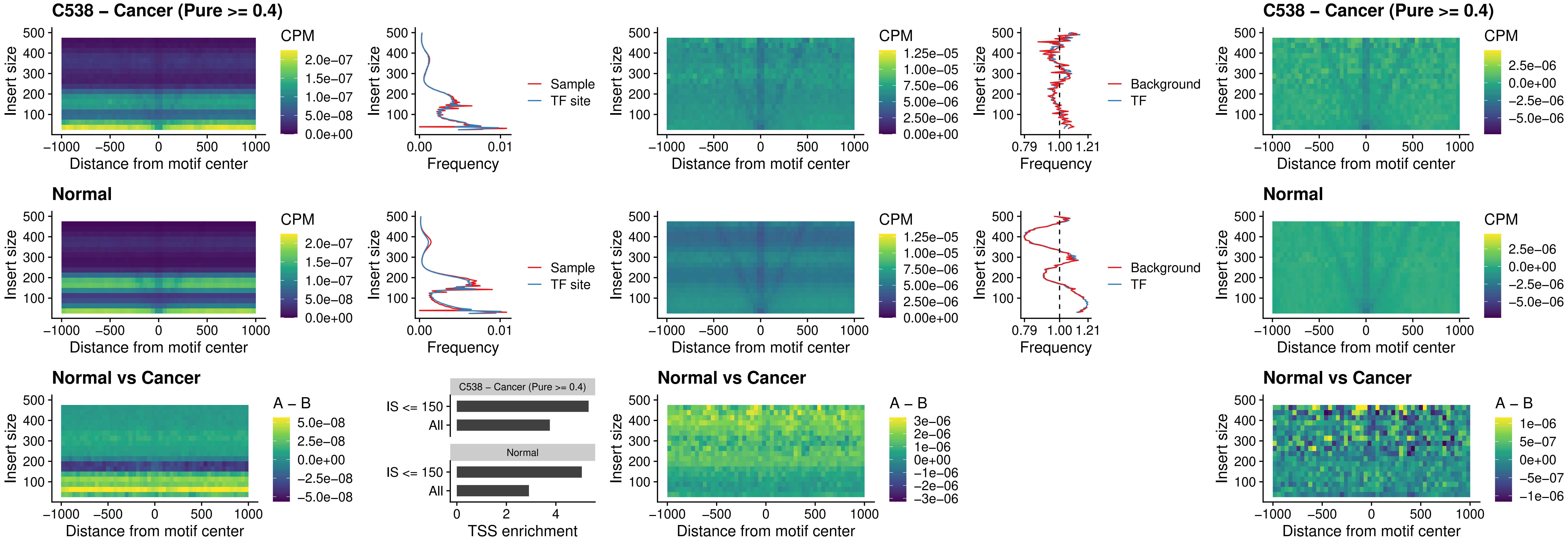

#### C538 – Cancer (Pure >= 0.4) – PDX1_uc_dTSS_oPEAK

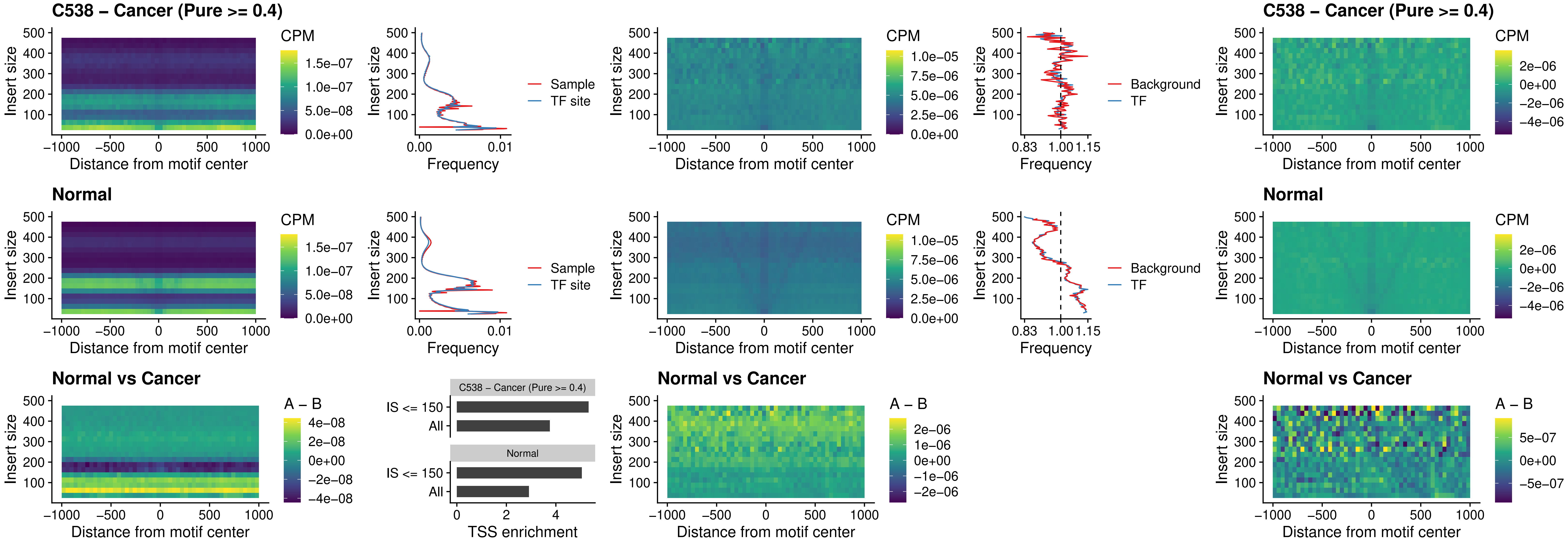

#### C538 – Cancer (Pure >= 0.4) – LHX4_uc_dTSS_oPEAK

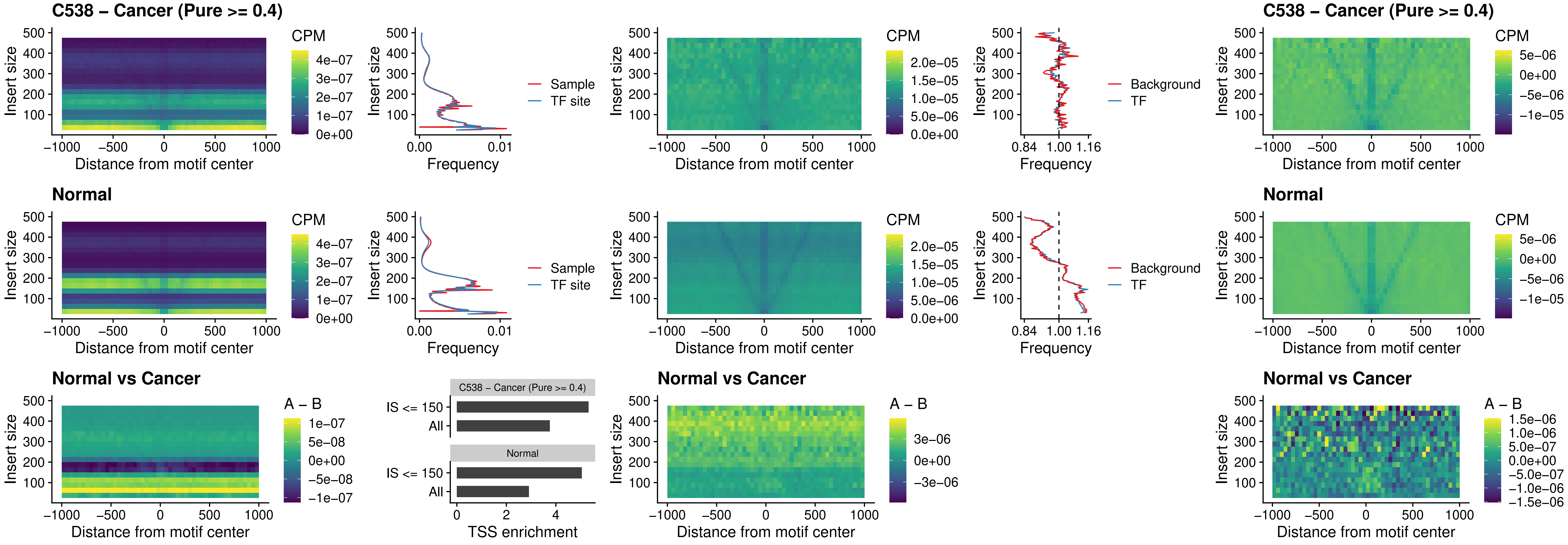

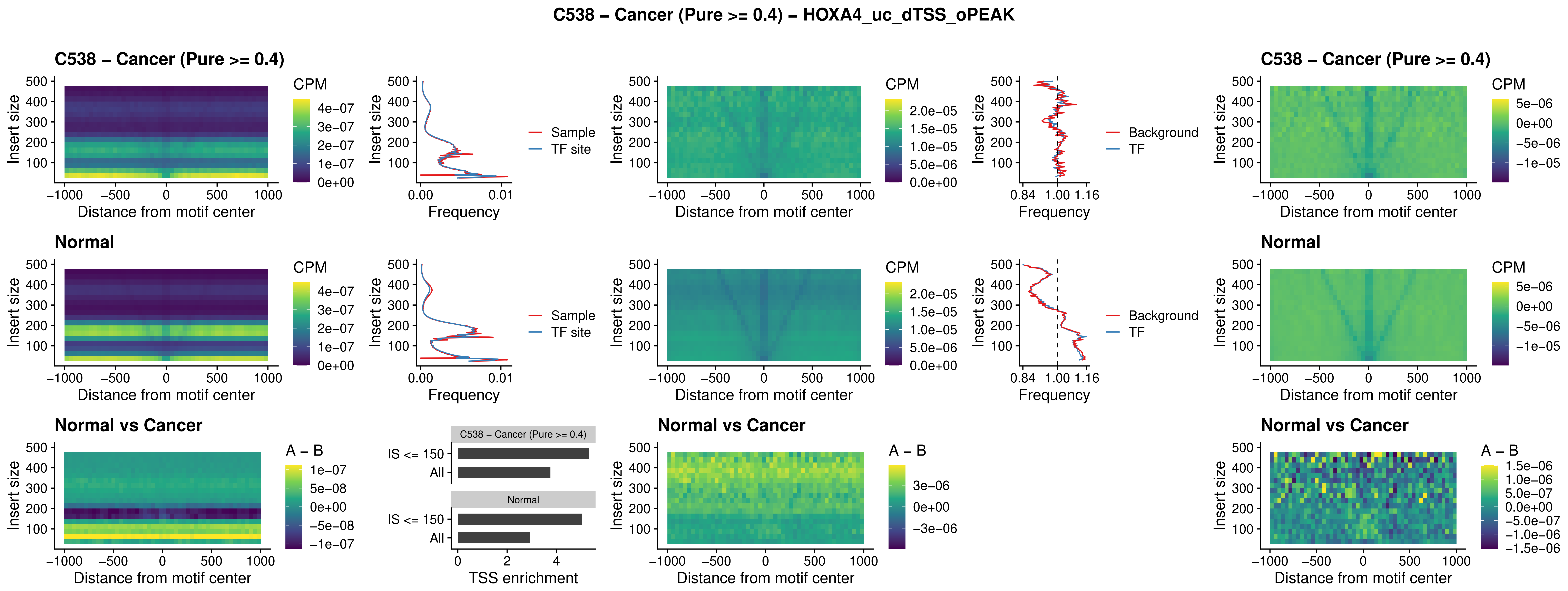
C538 – Cancer (Pure >= 0.4) – HOXA4_uc_dTSS_oPEAK
C538 – Cancer (Pure >= 0.4)
Normal
Normal vs Cancer
Insert size
Distance from motif center
Frequency
CPM
A – B
Sample
TF site
Background
TF
C538 – Cancer (Pure >= 0.4)
Normal
IS <= 150
All
TSS enrichment

#### C538 – Cancer (Pure >= 0.4) – HLX_uc_dTSS_oPEAK

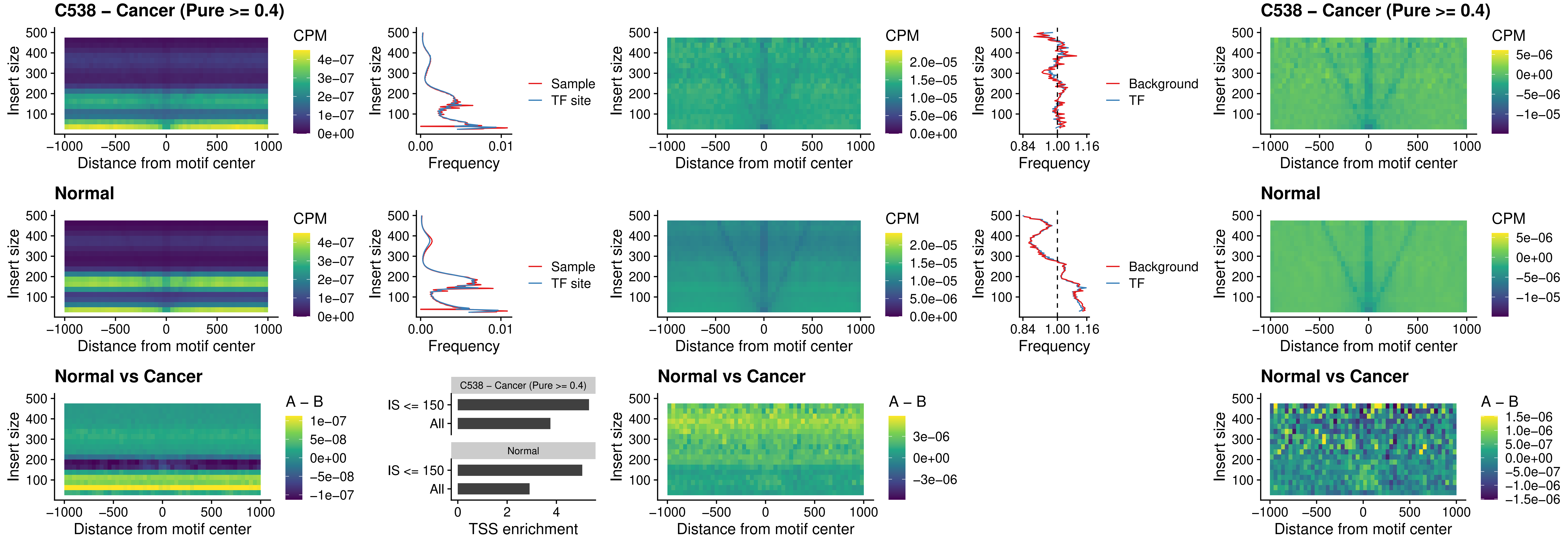

#### C538 – Cancer (Pure >= 0.4) – AC0125311_uc_dTSS_oPEAK

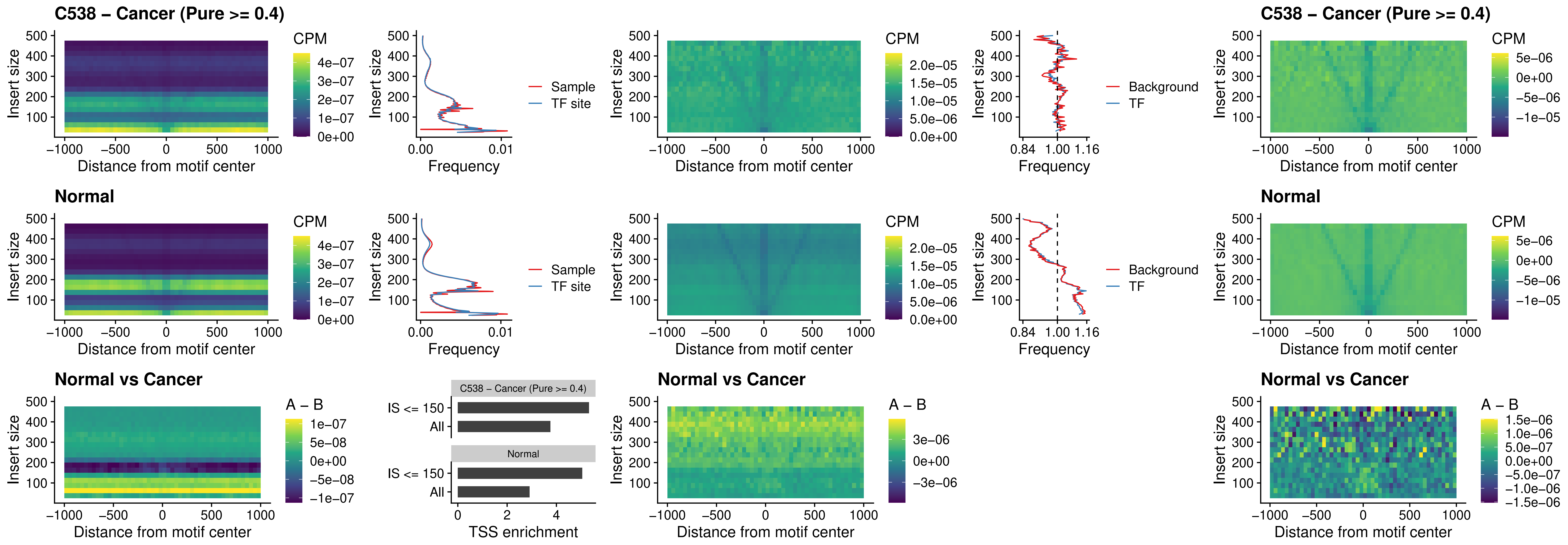

#### C538 – Cancer (Pure >= 0.4) – DBX2_uc_dTSS_oPEAK

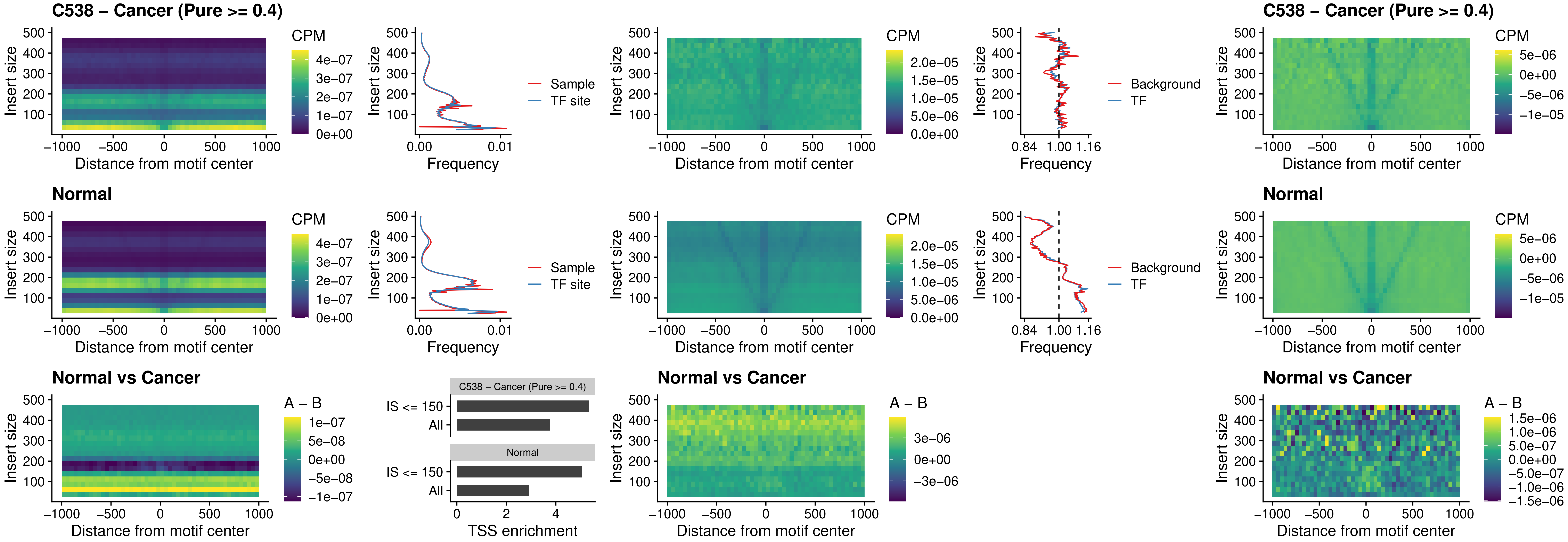

#### C538 – Cancer (Pure >= 0.4) – HOXA2_uc_dTSS_nPEAK

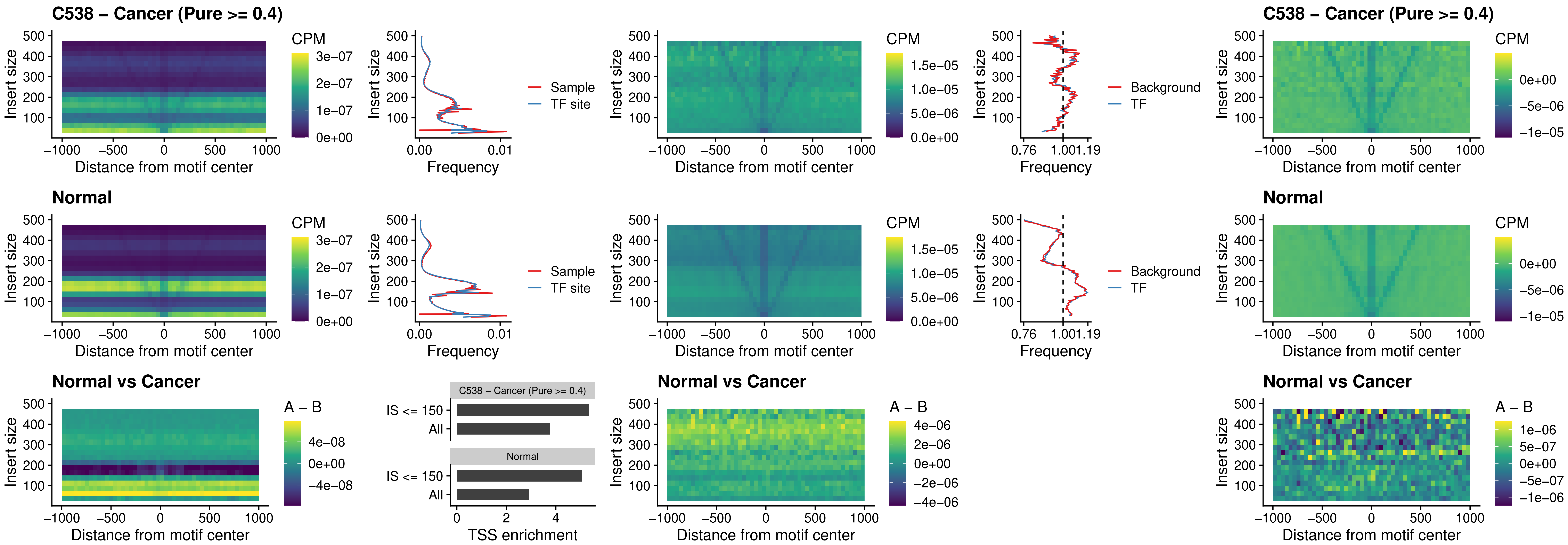

#### C538 – Cancer (Pure >= 0.4) – SHOX_uc_dTSS_nPEAK

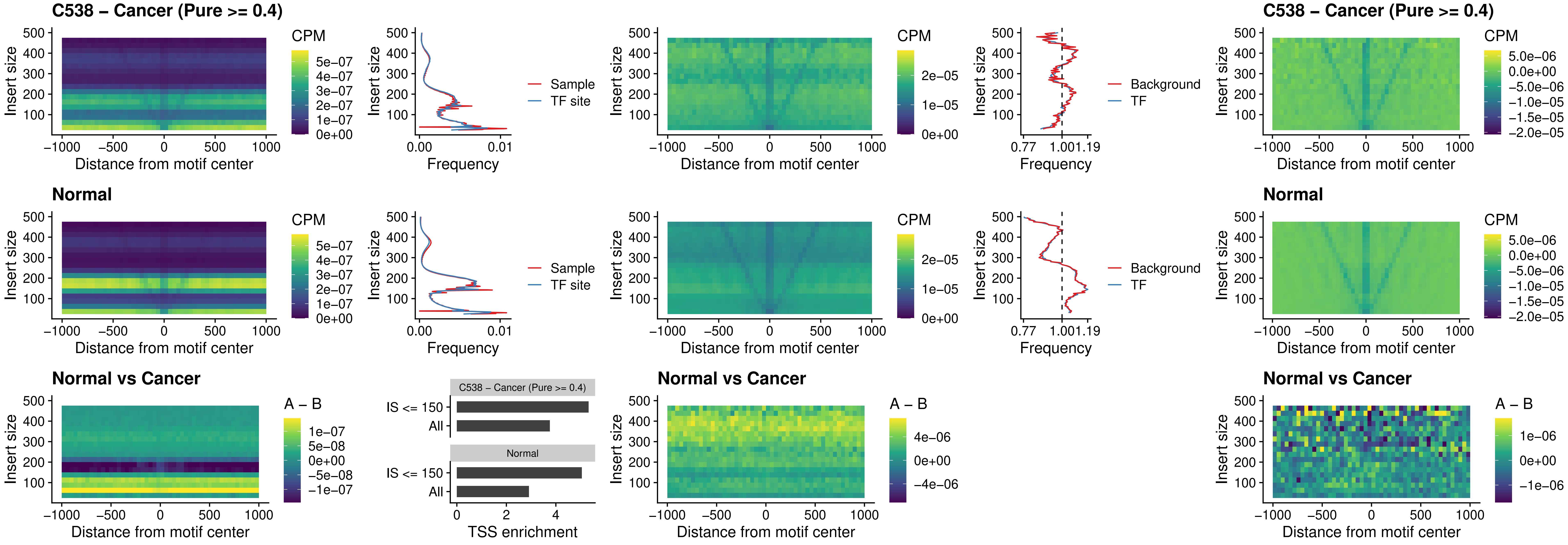

#### C538 – Cancer (Pure >= 0.4) – PRRX1_uc_dTSS_nPEAK

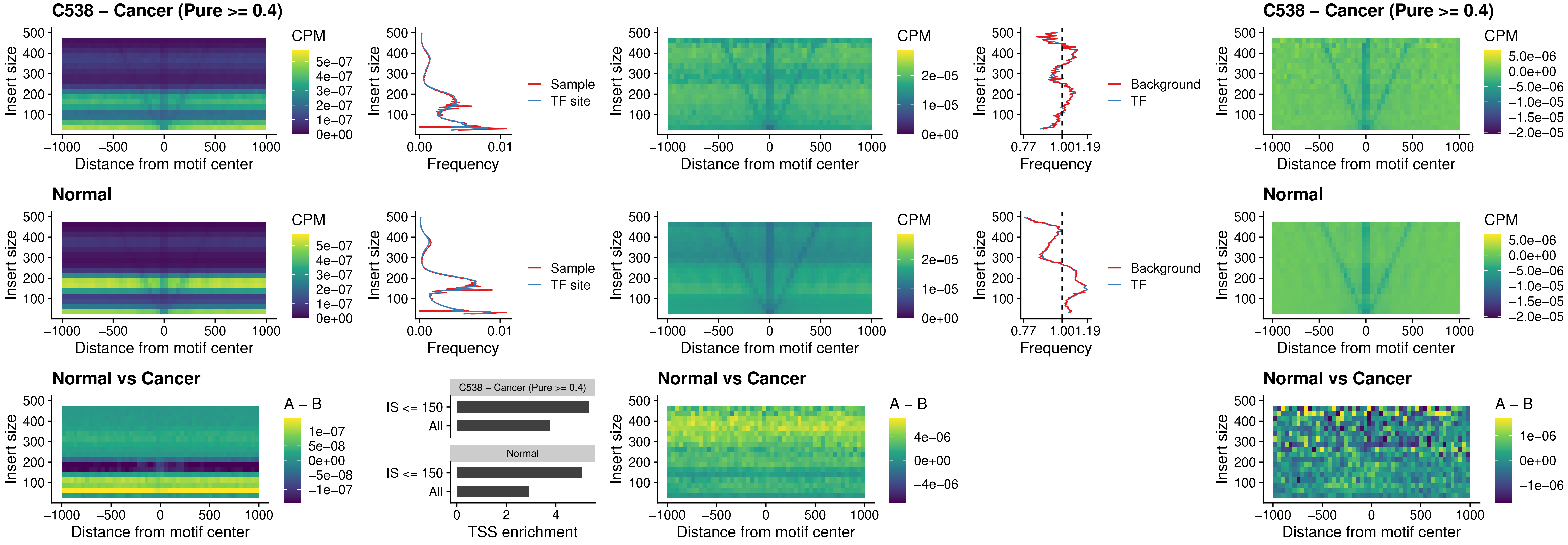

#### C538 – Cancer (Pure >= 0.4) – LMX1A_uc_dTSS_nPEAK

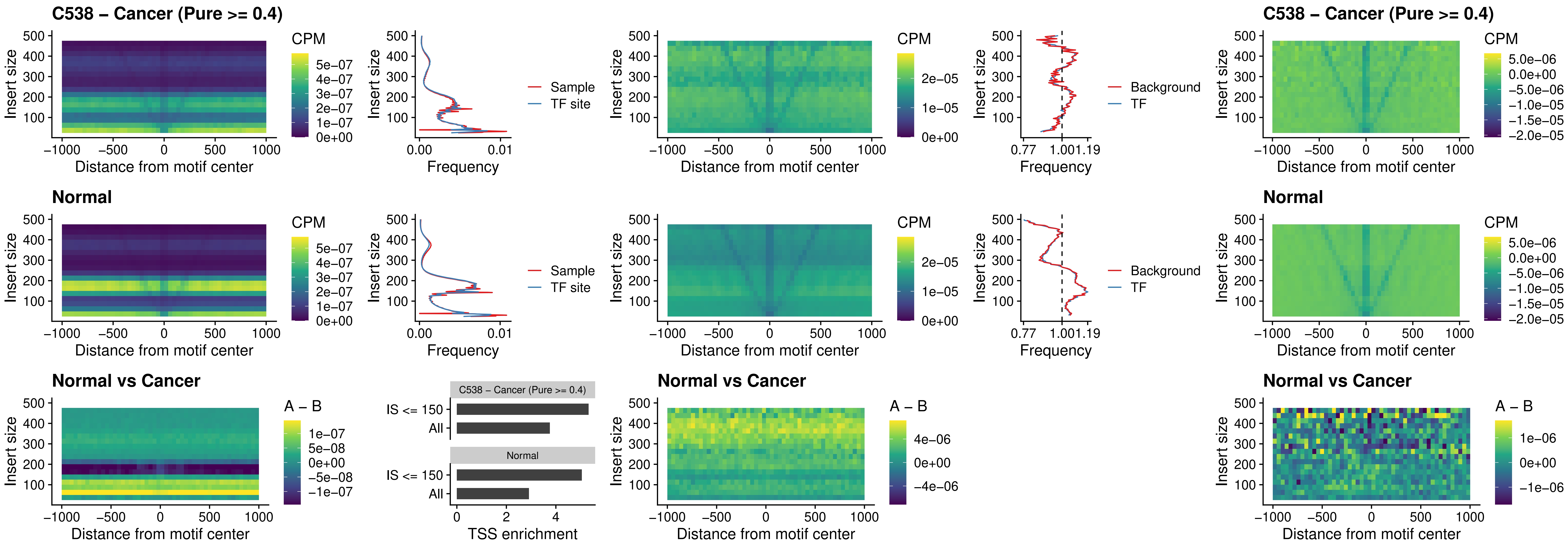

#### C538 – Cancer (Pure >= 0.4) – LMX1B_uc_dTSS_nPEAK

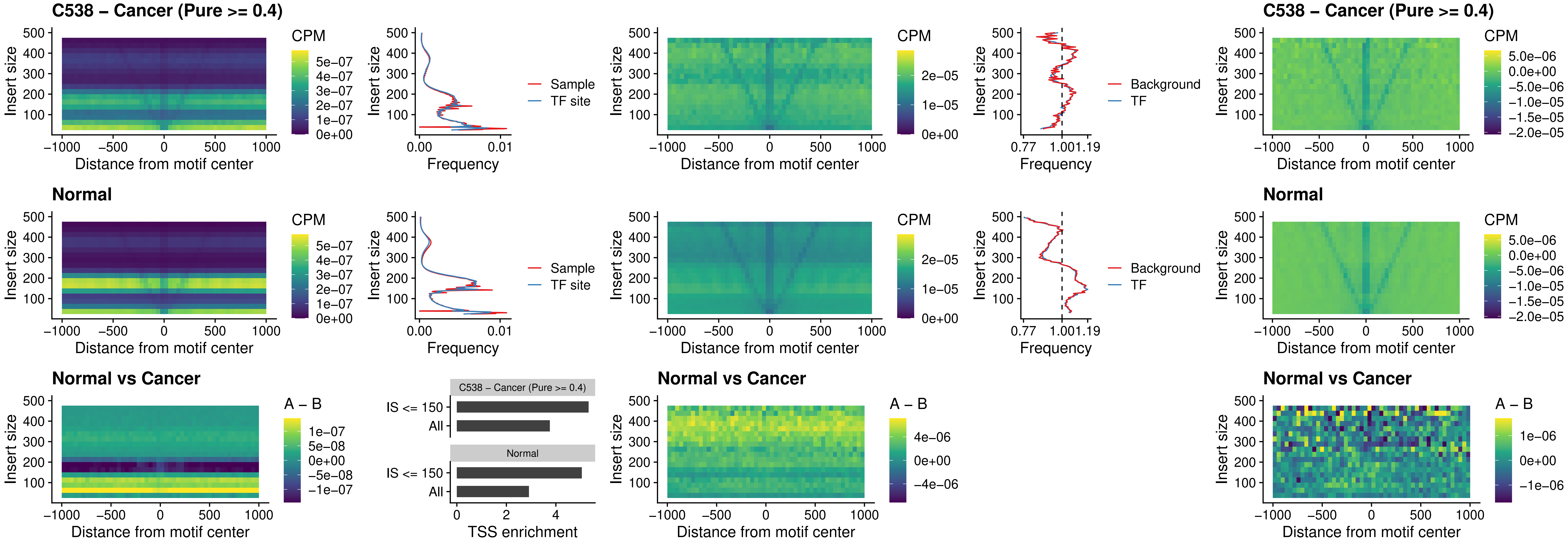

### C538 – Cancer (Pure >= 0.4) – DLX5_uc_dTSS_oPEAK

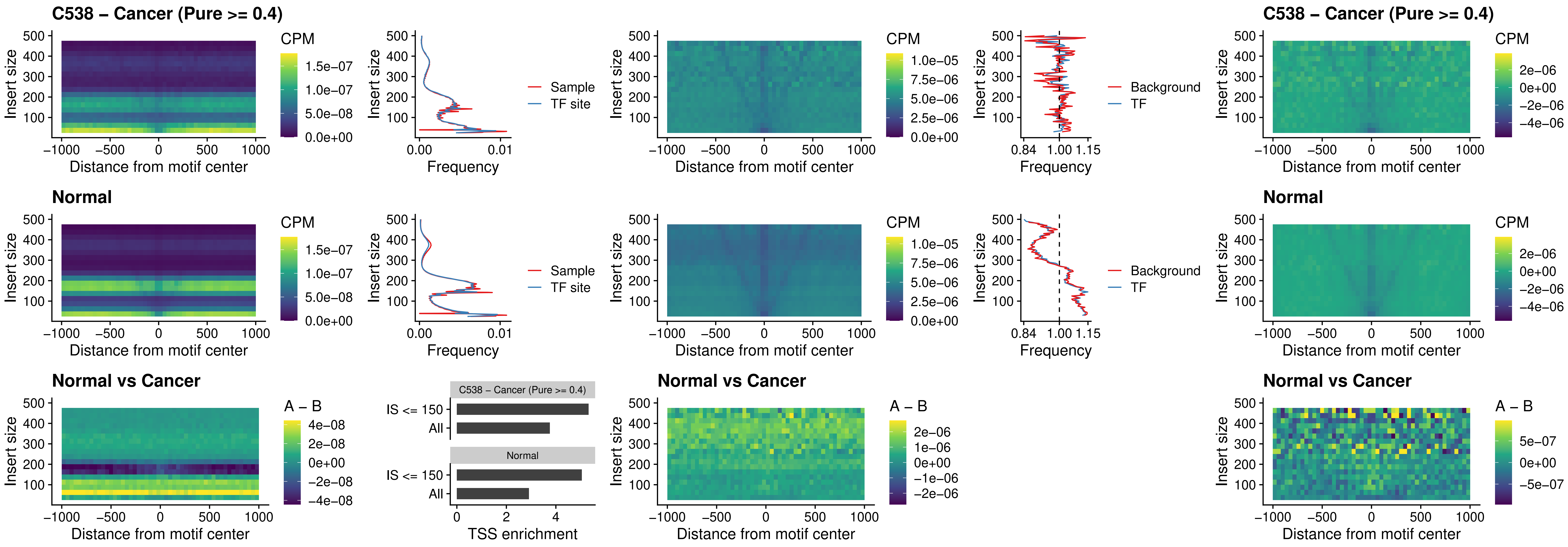

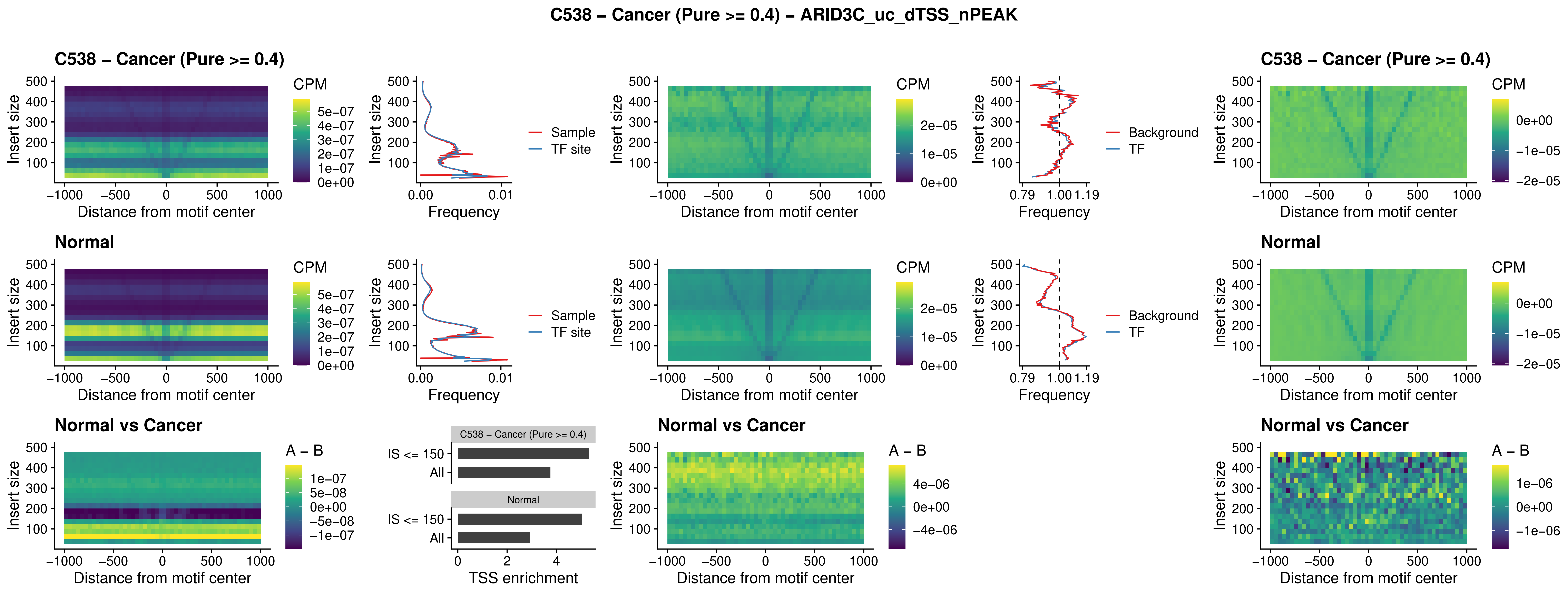
C538 – Cancer (Pure >= 0.4) – ARID3C_uc_dTSS_nPEAK
C538 – Cancer (Pure >= 0.4)
Normal
Normal vs Cancer
Insert size
Distance from motif center
Frequency
CPM
A – B
Sample
TF site
Background
TF
C538 – Cancer (Pure >= 0.4)
Normal
IS <= 150
All
TSS enrichment

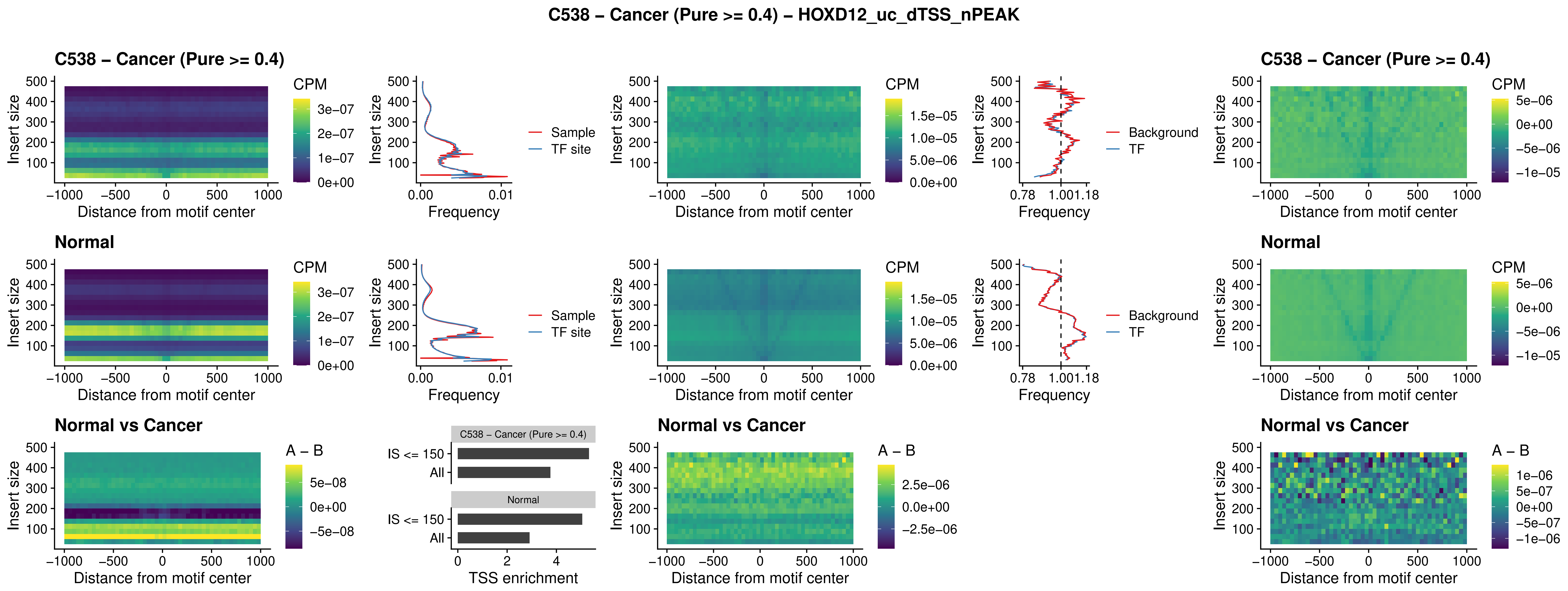
C538 – Cancer (Pure >= 0.4) – HOXD12_uc_dTSS_nPEAK
C538 – Cancer (Pure >= 0.4)
Normal
Normal vs Cancer
Insert size
Distance from motif center
Frequency
CPM
A – B
Sample
TF site
Background
TF
C538 – Cancer (Pure >= 0.4)
Normal
IS <= 150
All
TSS enrichment

#### C538 – Cancer (Pure >= 0.4) – CDX2_uc_dTSS_nPEAK

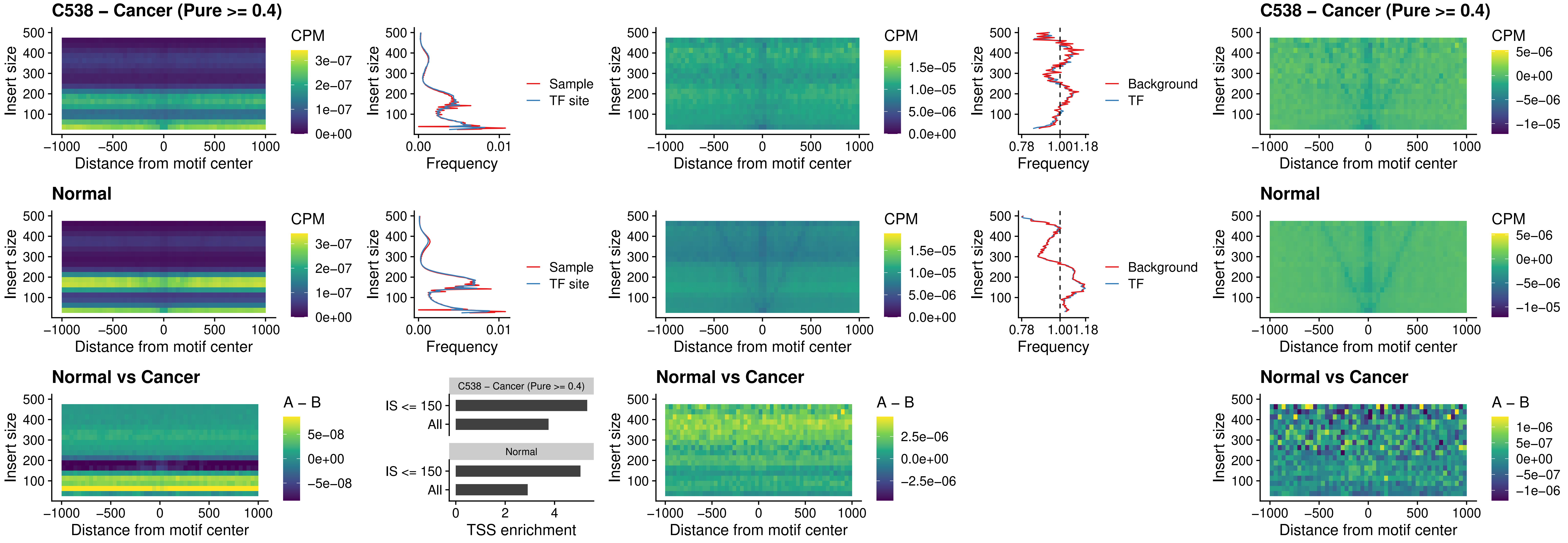

#### C538 – Cancer (Pure >= 0.4) – HOXC12_uc_dTSS_nPEAK

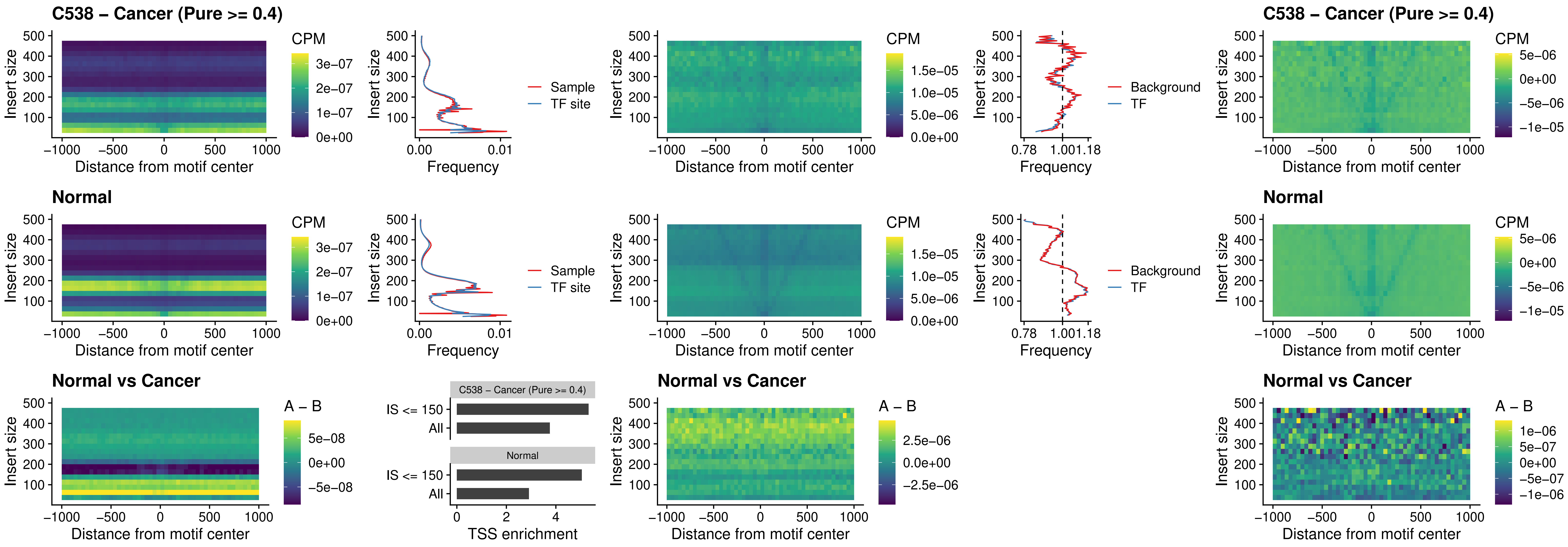

#### C538 – Cancer (Pure >= 0.4) – HHEX_uc_dTSS_oPEAK

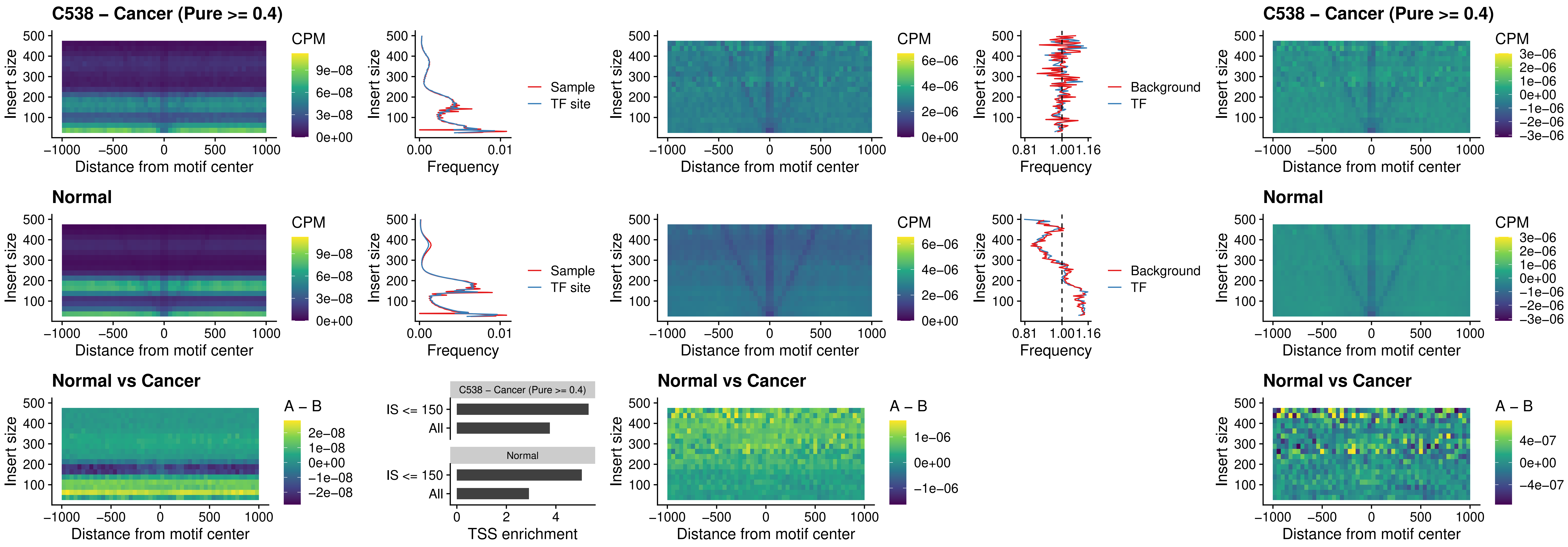

#### C538 – Cancer (Pure >= 0.4) – HOXA11_uc_dTSS_nPEAK

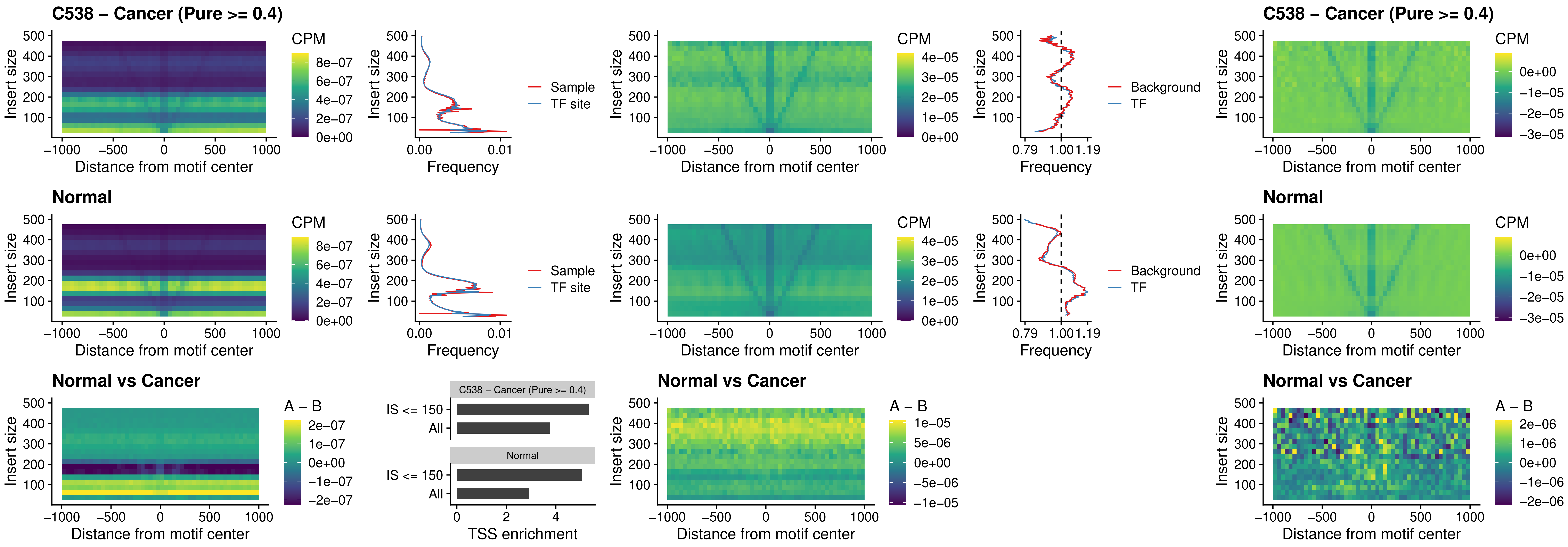

#### C538 – Cancer (Pure >= 0.4) – HOXA11_uc_cTSS_nPEAK

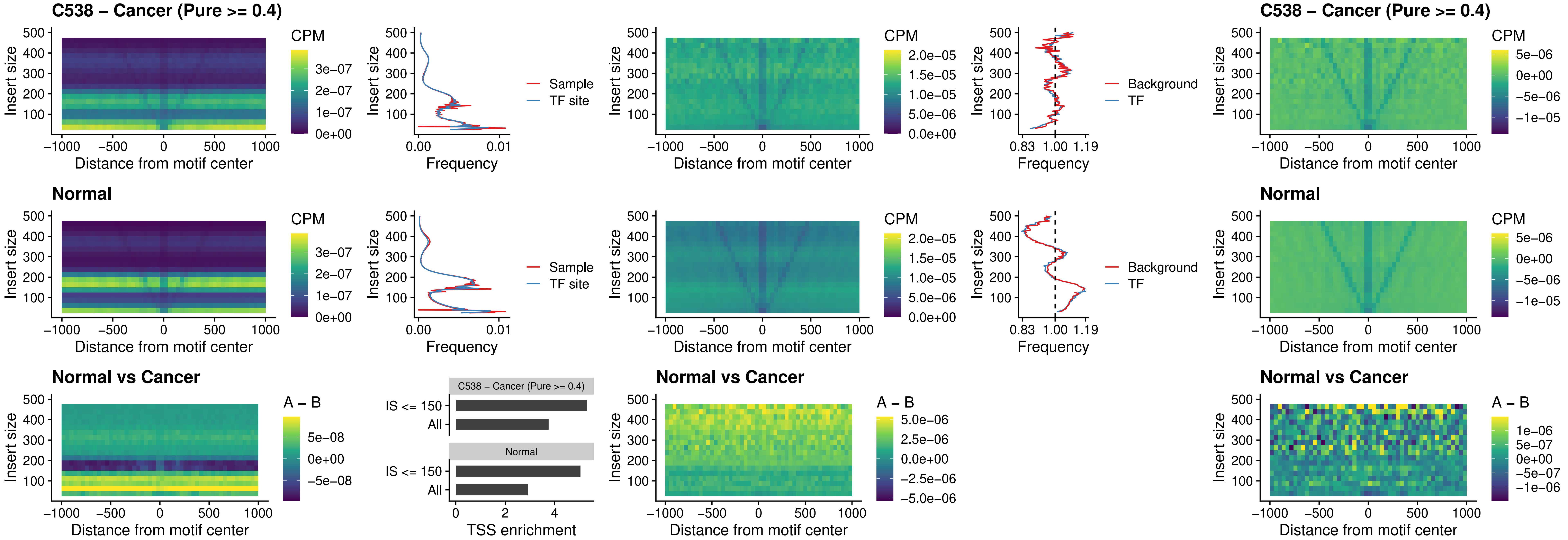

### C538 – Cancer (Pure >= 0.4) – ZFHX3_uc_dTSS_nPEAK

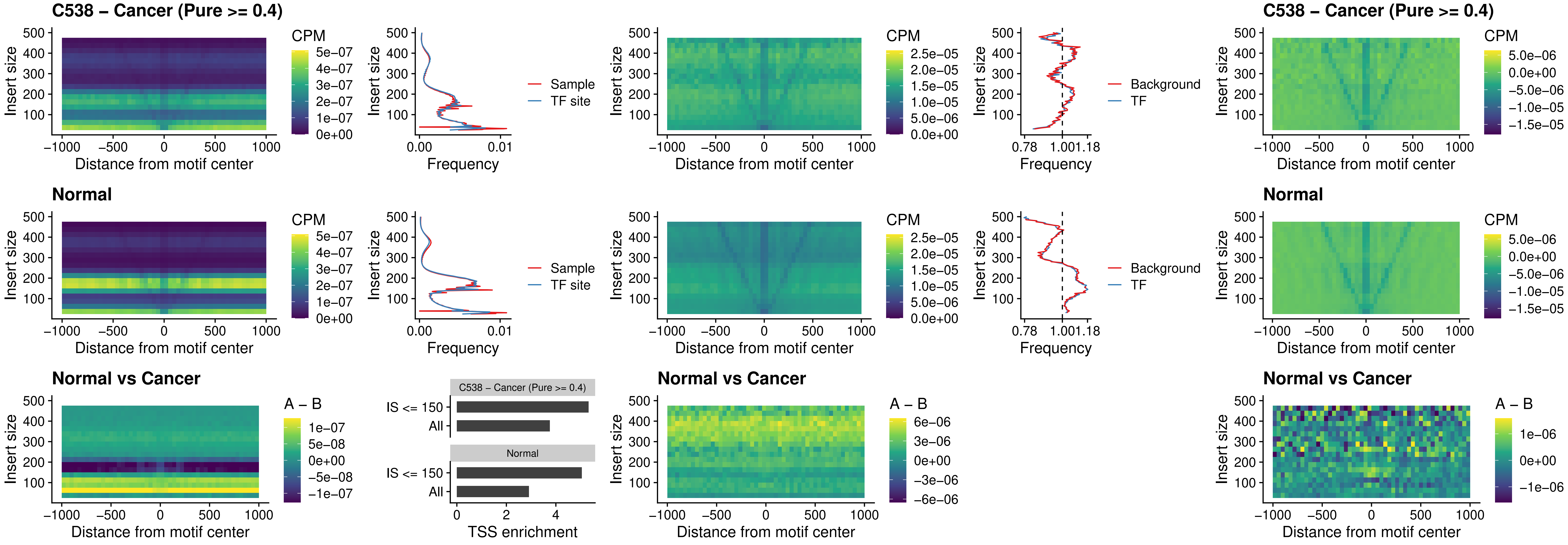

#### C538 – Cancer (Pure >= 0.4) – HOXB4_uc_dTSS_nPEAK

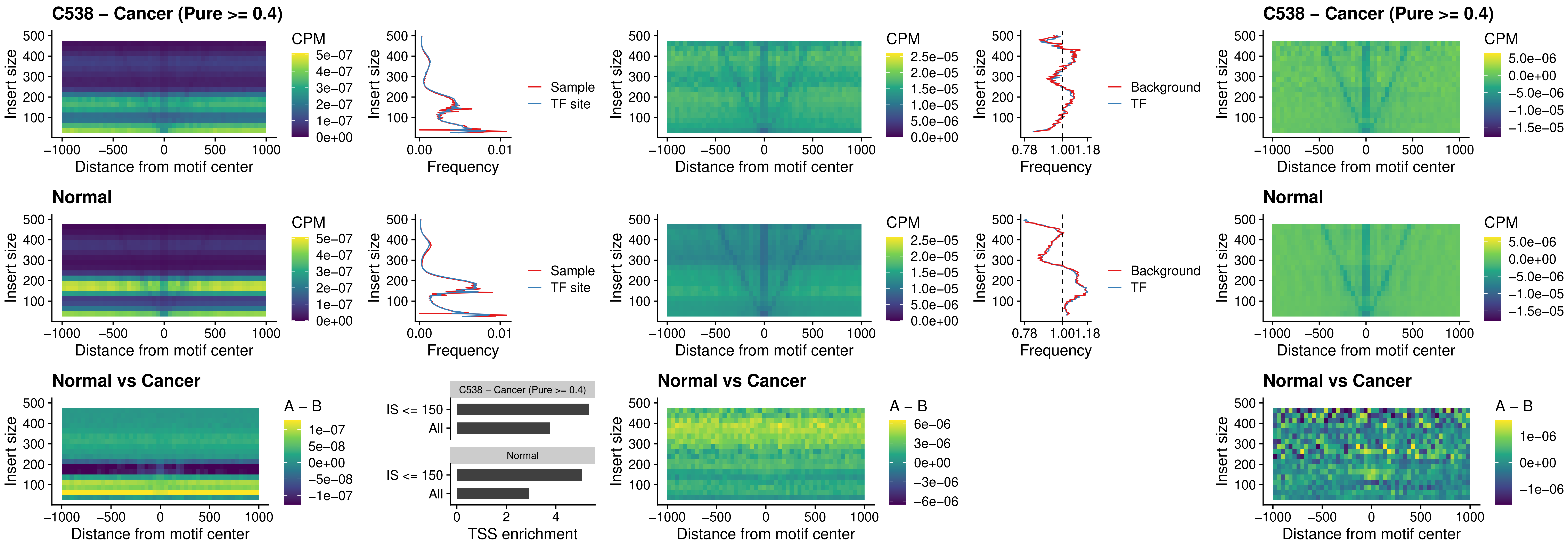

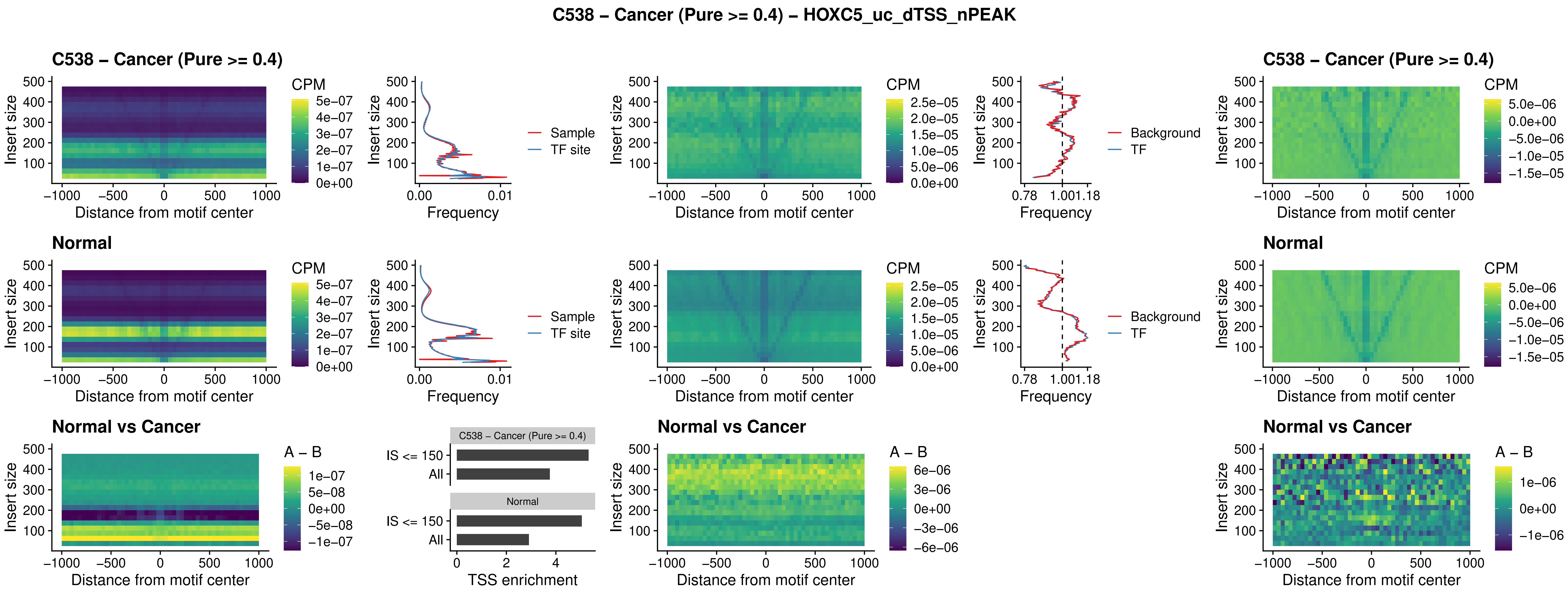
C538 – Cancer (Pure >= 0.4) – HOXC5_uc_dTSS_nPEAK
C538 – Cancer (Pure >= 0.4)
Normal
Normal vs Cancer
Insert size
Distance from motif center
Frequency
CPM
A – B
Sample
TF site
Background
TF
C538 – Cancer (Pure >= 0.4)
Normal
IS <= 150
All
TSS enrichment

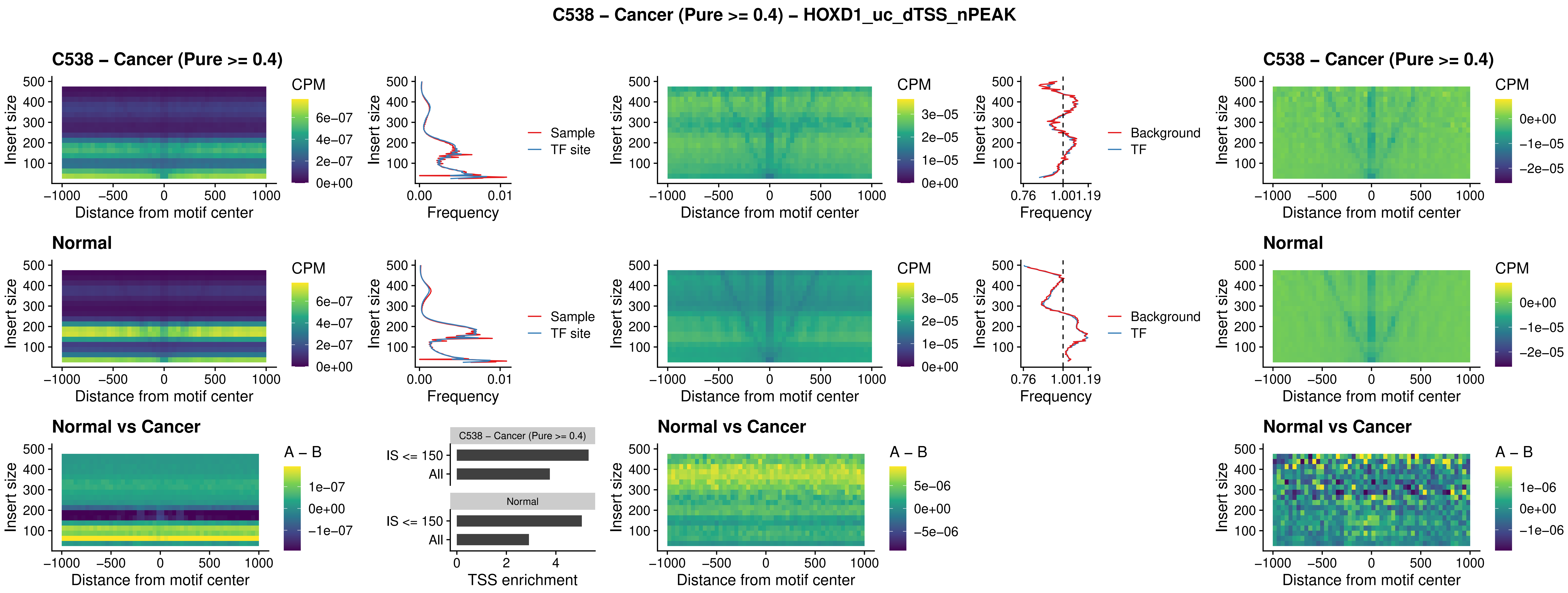
C538 – Cancer (Pure >= 0.4) – HOXD1_uc_dTSS_nPEAK
C538 – Cancer (Pure >= 0.4)
Normal
Normal vs Cancer
Insert size
Distance from motif center
Frequency
CPM
A – B
Sample
TF site
Background
TF
C538 – Cancer (Pure >= 0.4)
Normal
IS <= 150
All
TSS enrichment

#### C538 – Cancer (Pure >= 0.4) – OTP_uc_dTSS_nPEAK

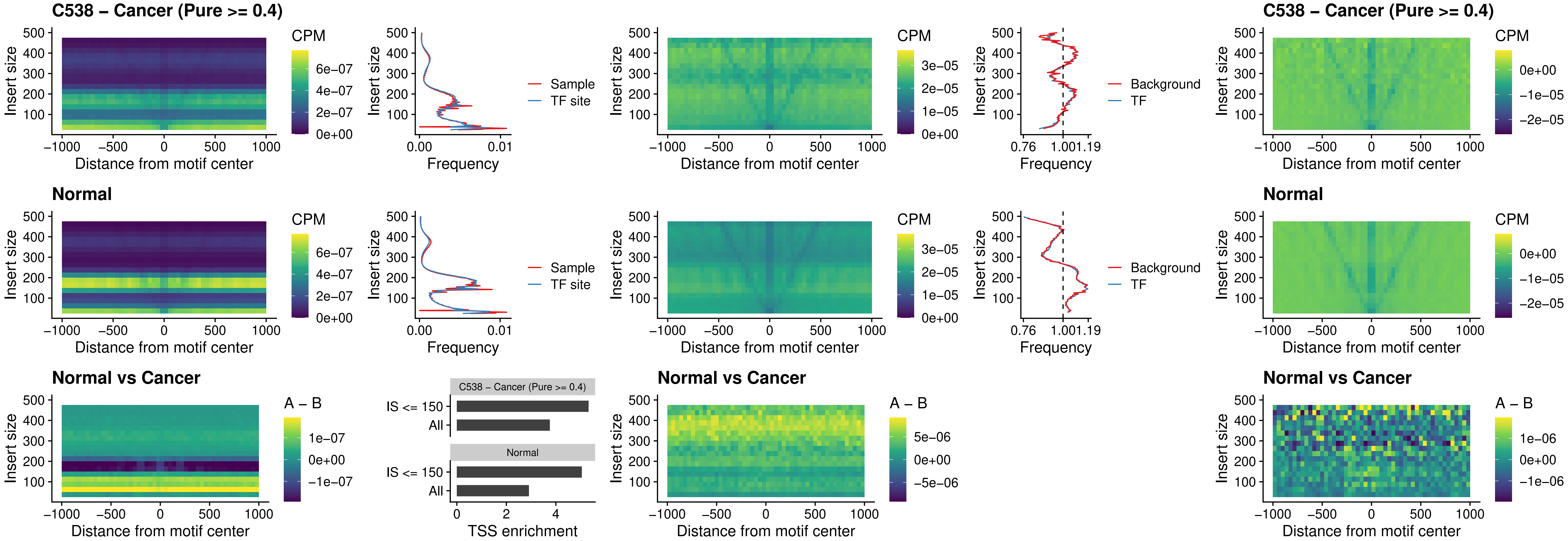

#### C538 – Cancer (Pure >= 0.4) – FOXL1_uc_dTSS_oPEAK

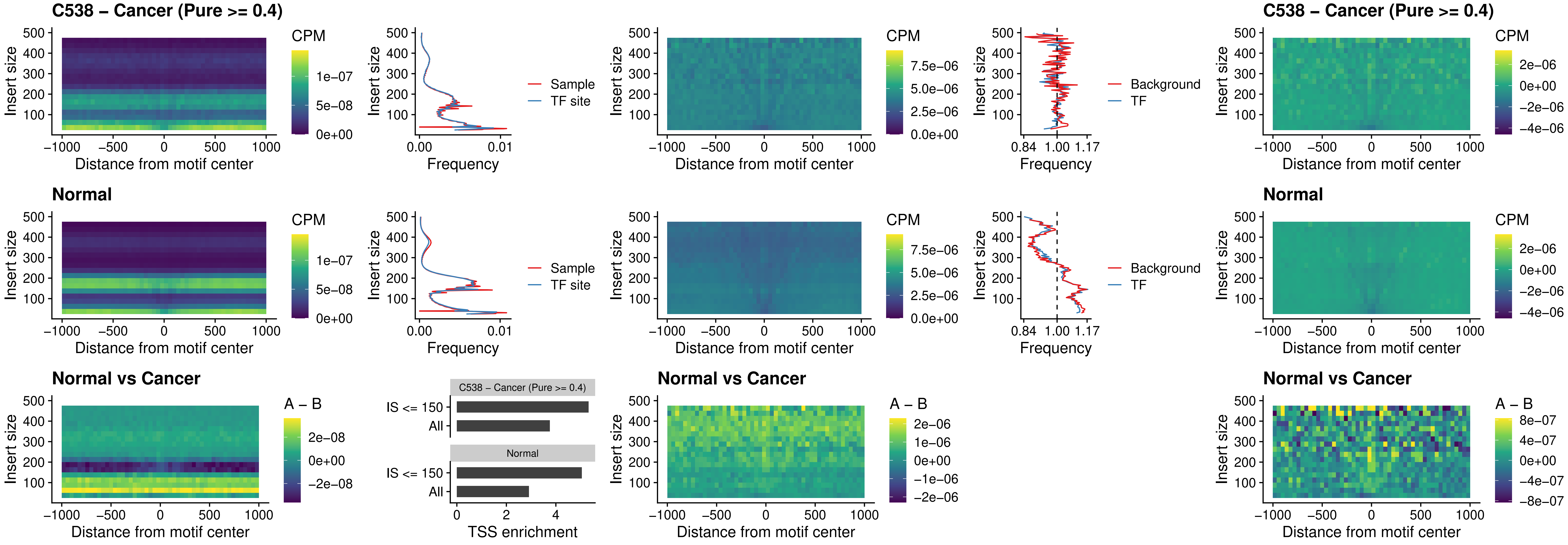

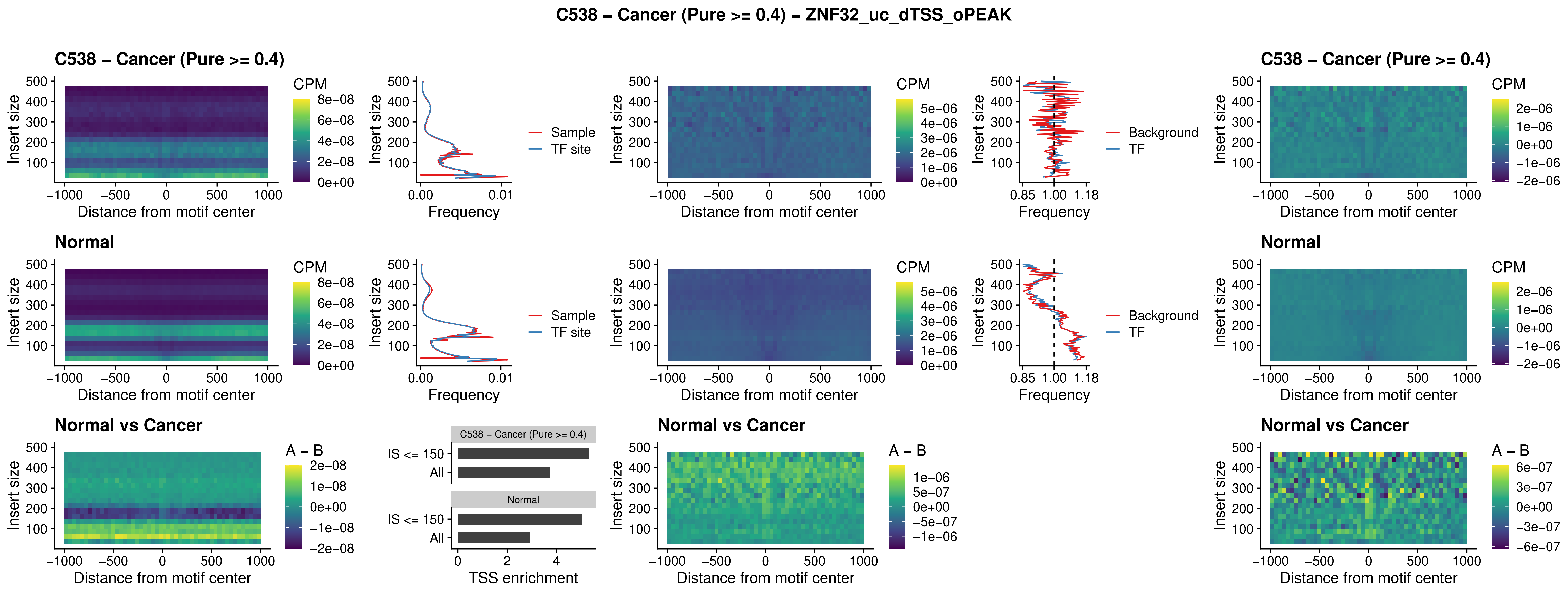
C538 – Cancer (Pure >= 0.4) – ZNF32_uc_dTSS_oPEAK
C538 – Cancer (Pure >= 0.4)
Normal
Normal vs Cancer
Insert size
Distance from motif center
Frequency
CPM
A – B
Sample
TF site
Background
TF
C538 – Cancer (Pure >= 0.4)
Normal
IS <= 150
All
TSS enrichment

#### C538 – Cancer (Pure >= 0.4) – FOXL1_uc_dTSS_nPEAK

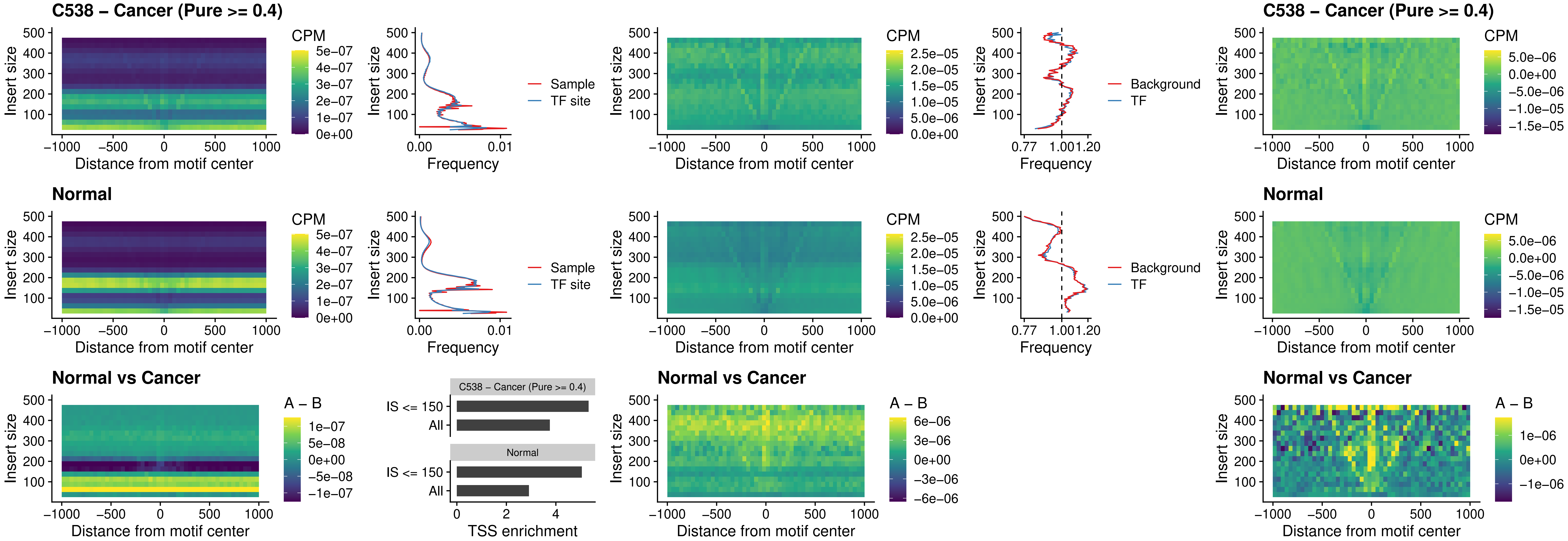

#### C538 – Cancer (Pure >= 0.4) – ZNF32_uc_dTSS_nPEAK

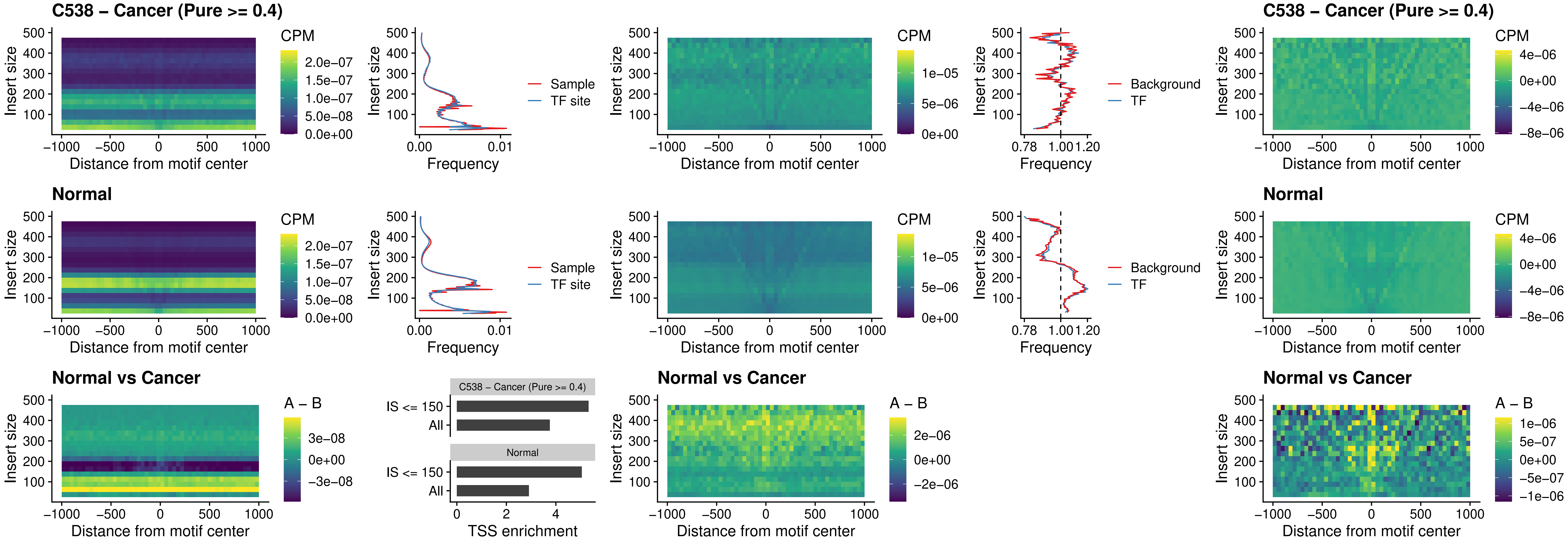

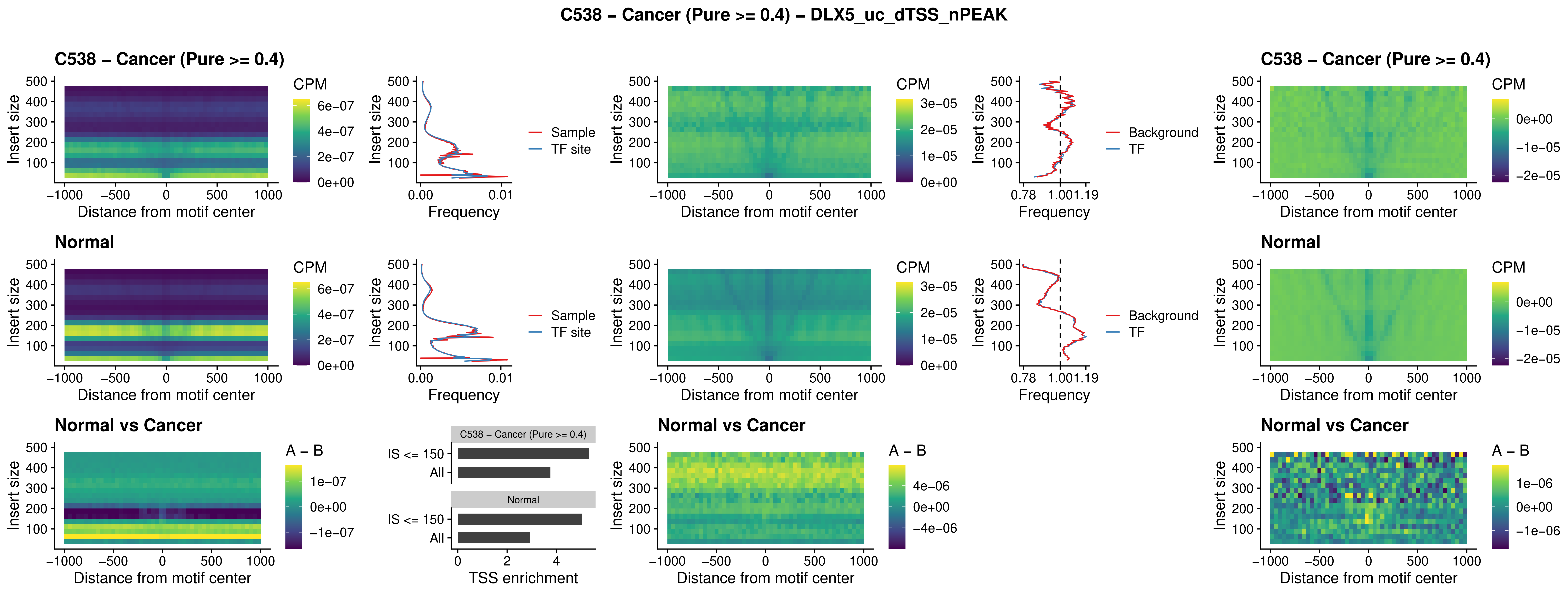
C538 – Cancer (Pure >= 0.4) – DLX5_uc_dTSS_nPEAK
C538 – Cancer (Pure >= 0.4)
Normal
Normal vs Cancer
Insert size
Distance from motif center
Frequency
CPM
A – B
Sample
TF site
Background
TF
C538 – Cancer (Pure >= 0.4)
Normal
IS <= 150
All
TSS enrichment

#### C538 – Cancer (Pure >= 0.4) – DMRT2_uc_dTSS_nPEAK

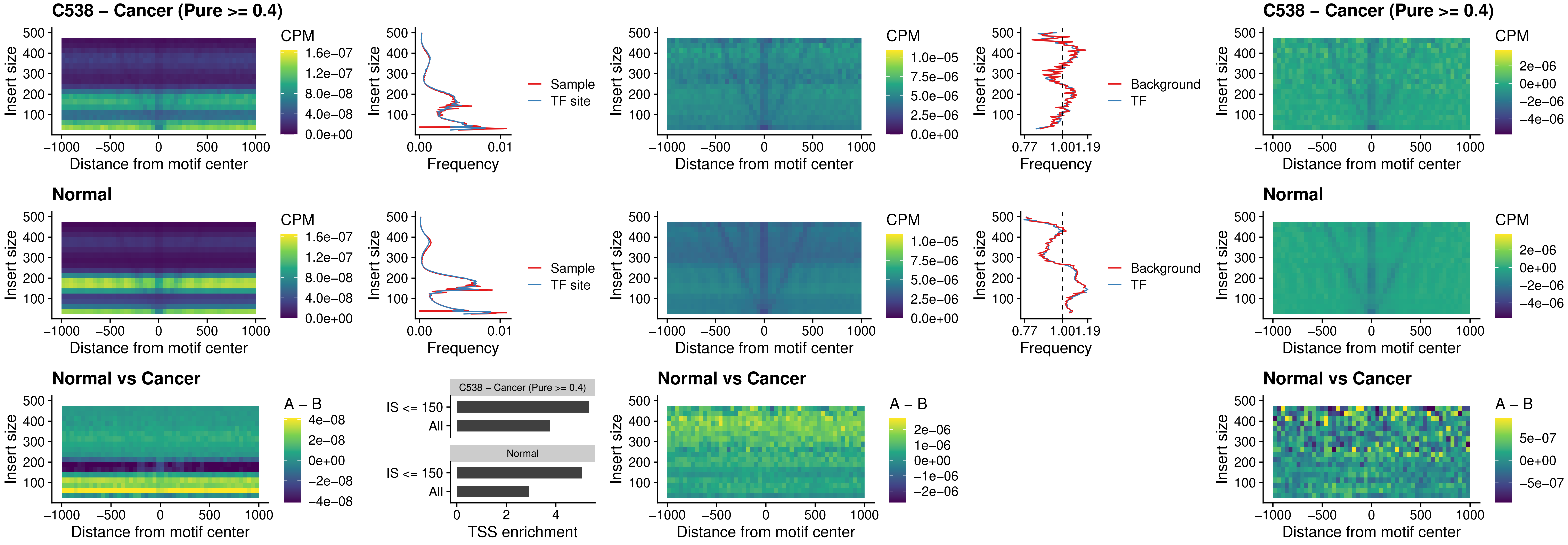

#### C538 – Cancer (Pure >= 0.4) – DMRTC2_uc_cTSS_nPEAK

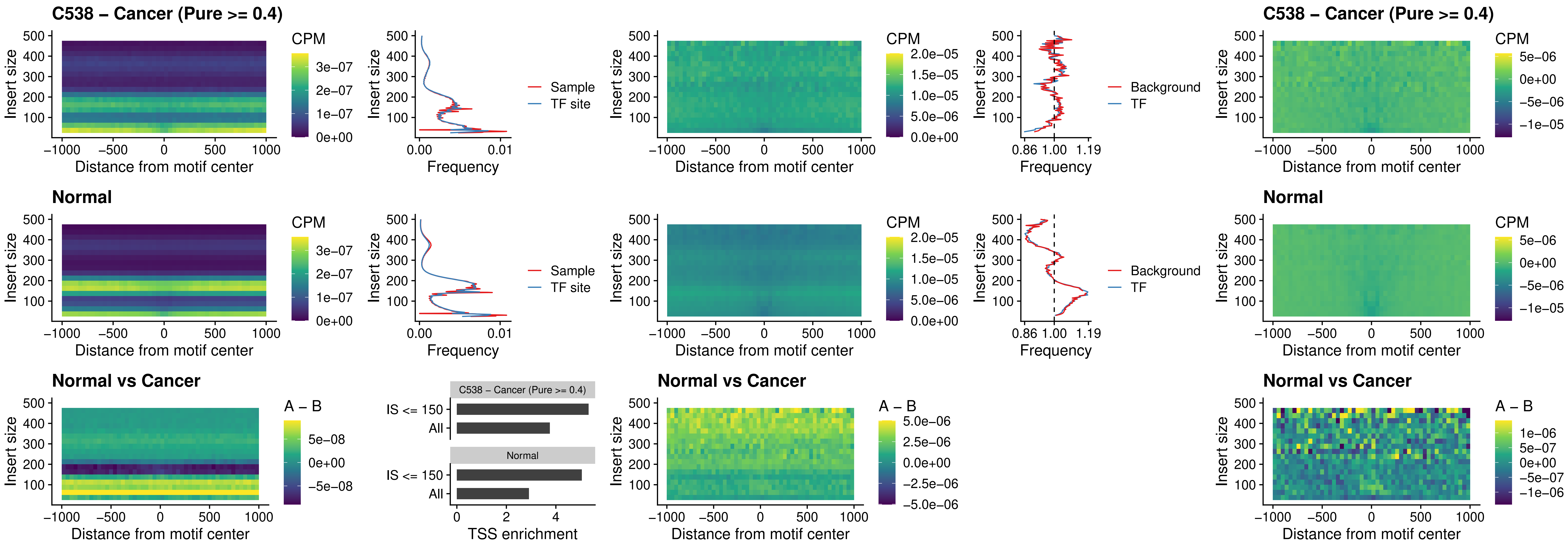

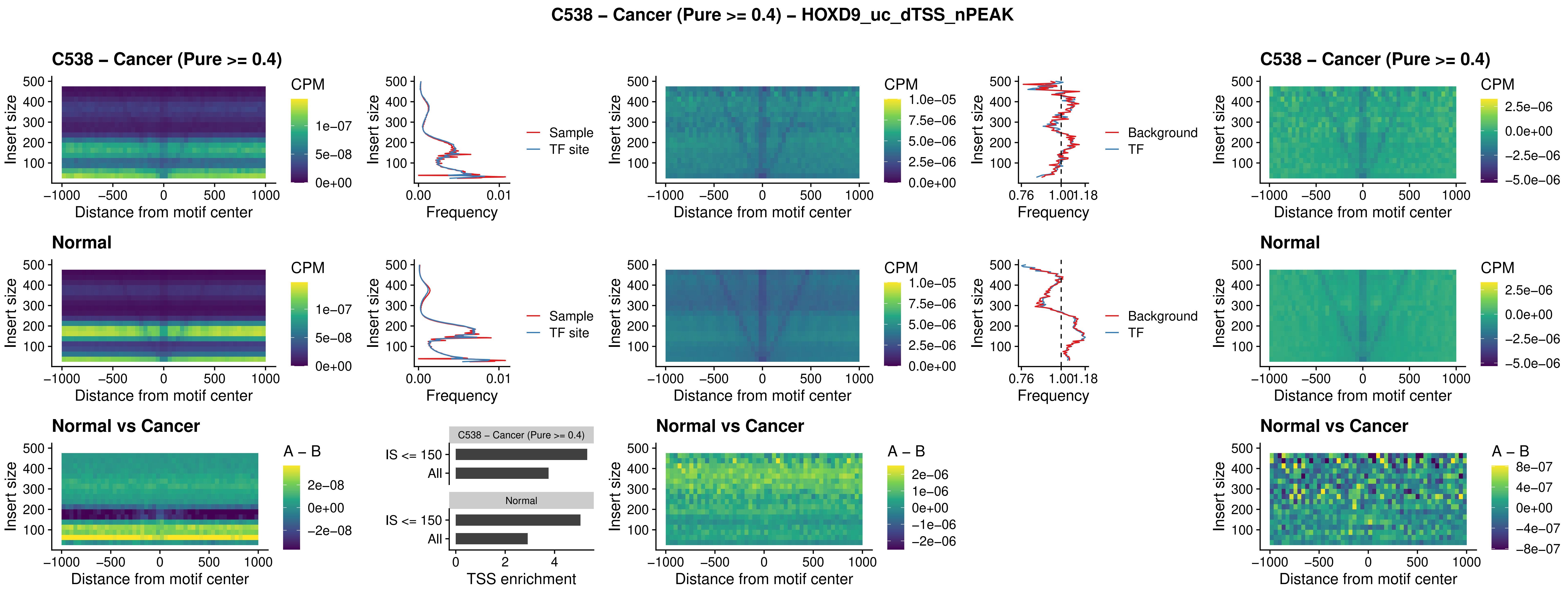
C538 – Cancer (Pure >= 0.4) – HOXD9_uc_dTSS_nPEAK
C538 – Cancer (Pure >= 0.4)
Normal
Normal vs Cancer
Insert size
Distance from motif center
Frequency
CPM
A – B
Sample
TF site
Background
TF
C538 – Cancer (Pure >= 0.4)
Normal
IS <= 150
All
TSS enrichment

#### C538 – Cancer (Pure >= 0.4) – HOXD4_uc_dTSS_nPEAK

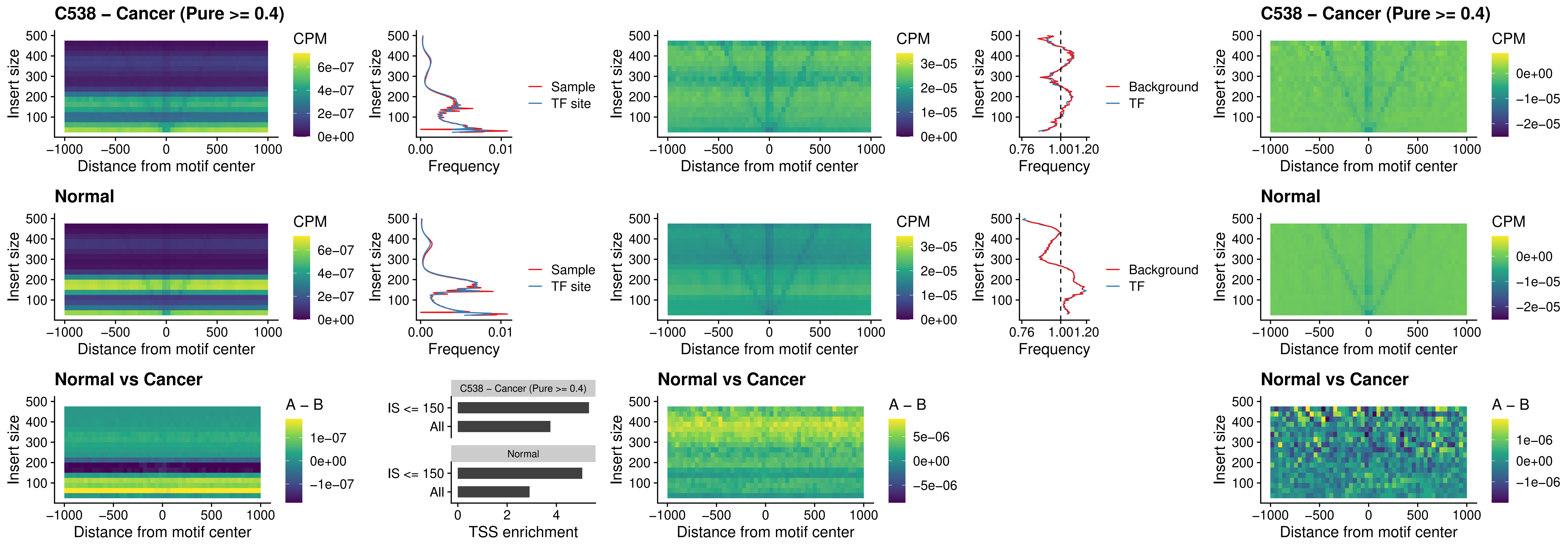

### C538 – Cancer (Pure >= 0.4) – FOXP3_uc_cTSS_nPEAK

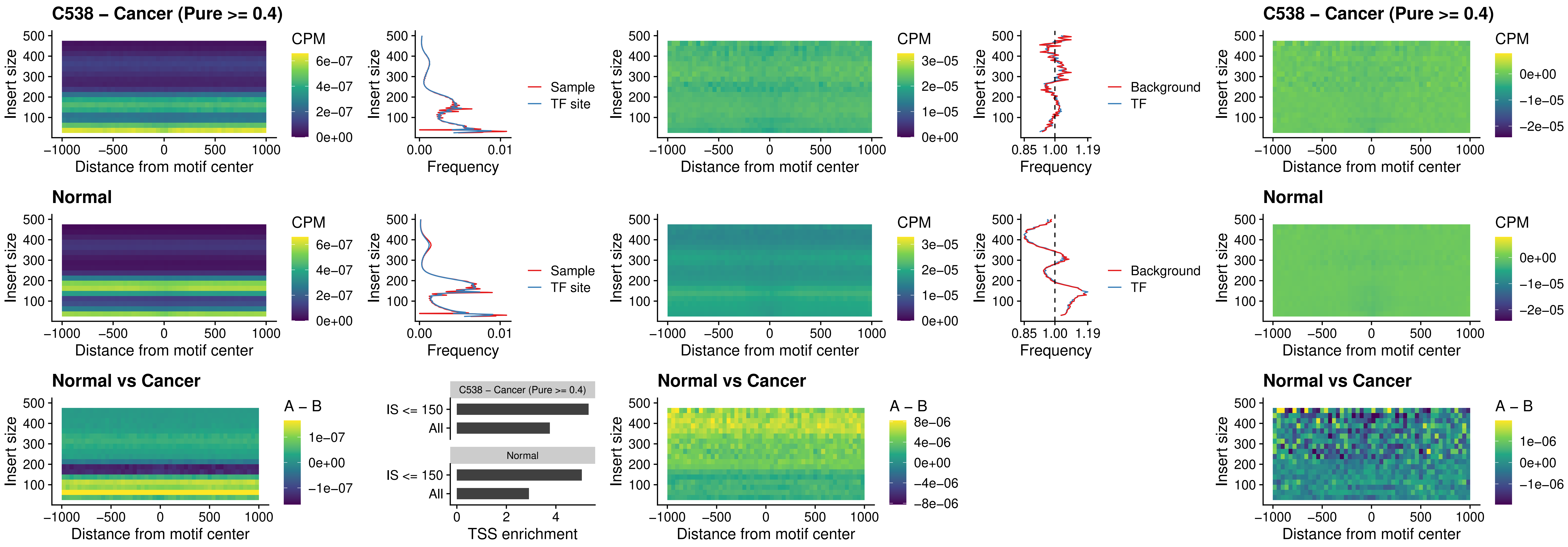

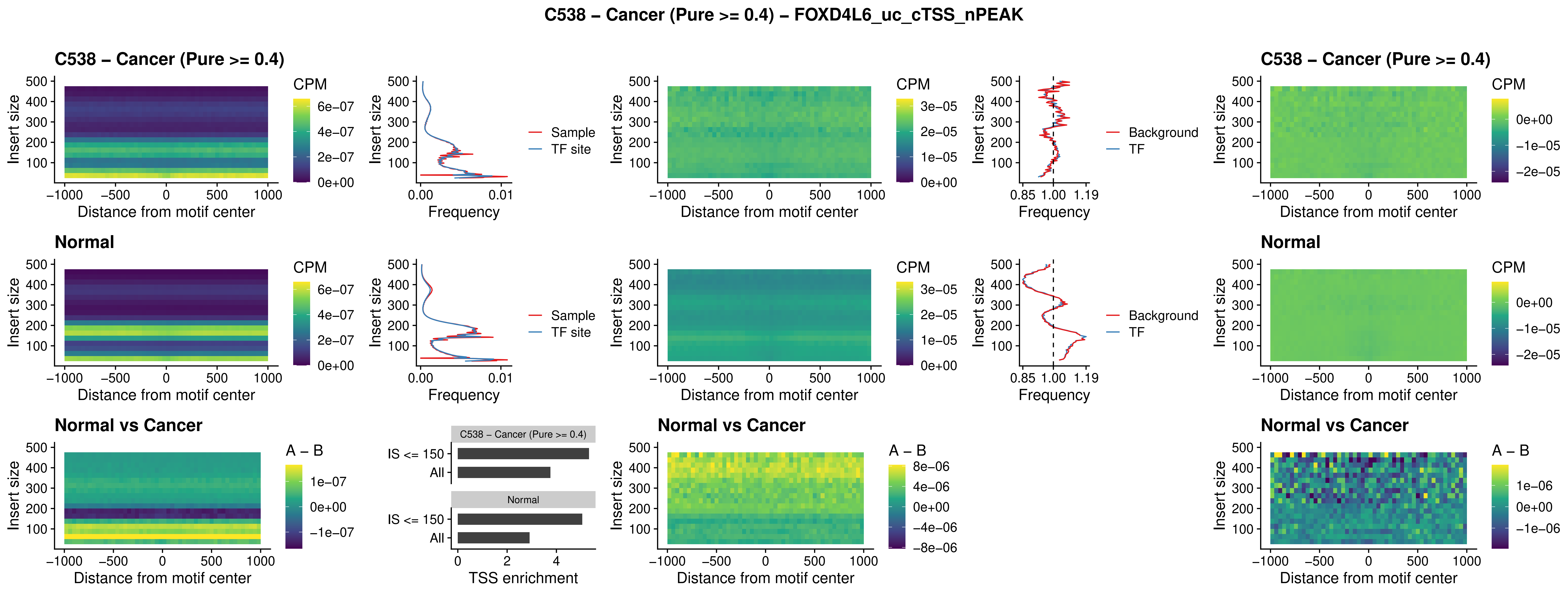
C538 – Cancer (Pure >= 0.4) – FOXD4L6_uc_cTSS_nPEAK
C538 – Cancer (Pure >= 0.4)
Normal
Normal vs Cancer
Insert size
Distance from motif center
Frequency
CPM
A – B
Sample
TF site
Background
TF
C538 – Cancer (Pure >= 0.4)
Normal
IS <= 150
All
TSS enrichment

#### C538 – Cancer (Pure >= 0.4) – FOXD4L5_uc_cTSS_nPEAK

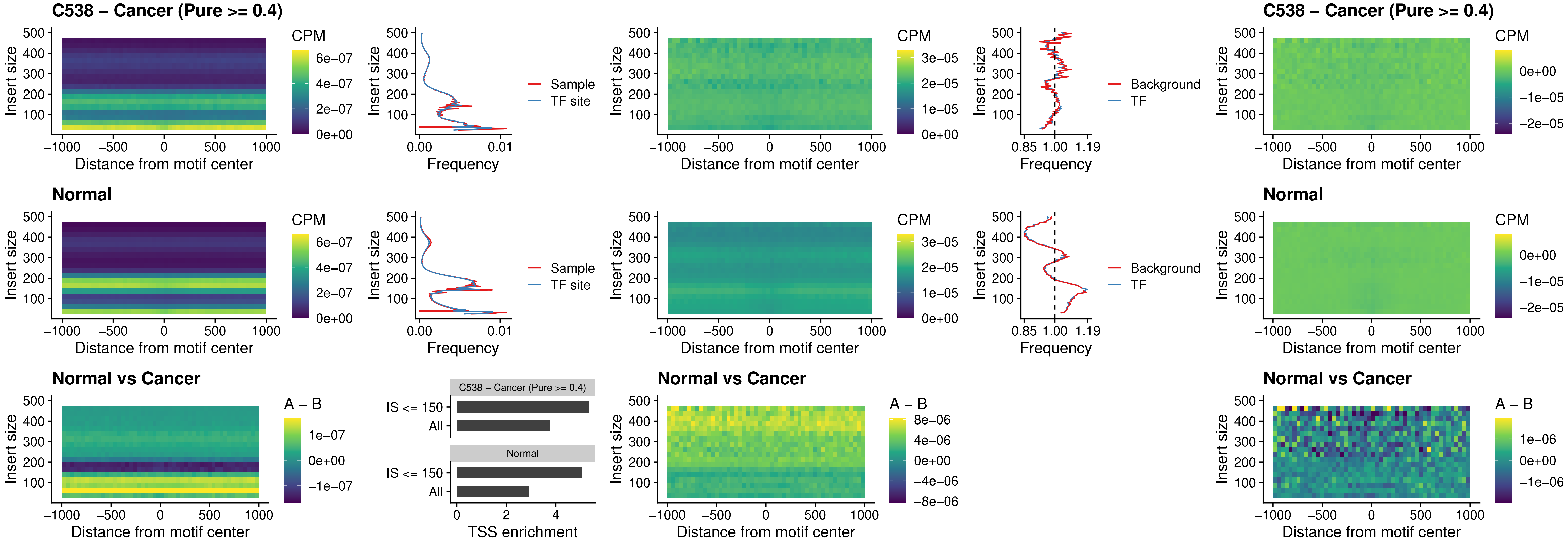

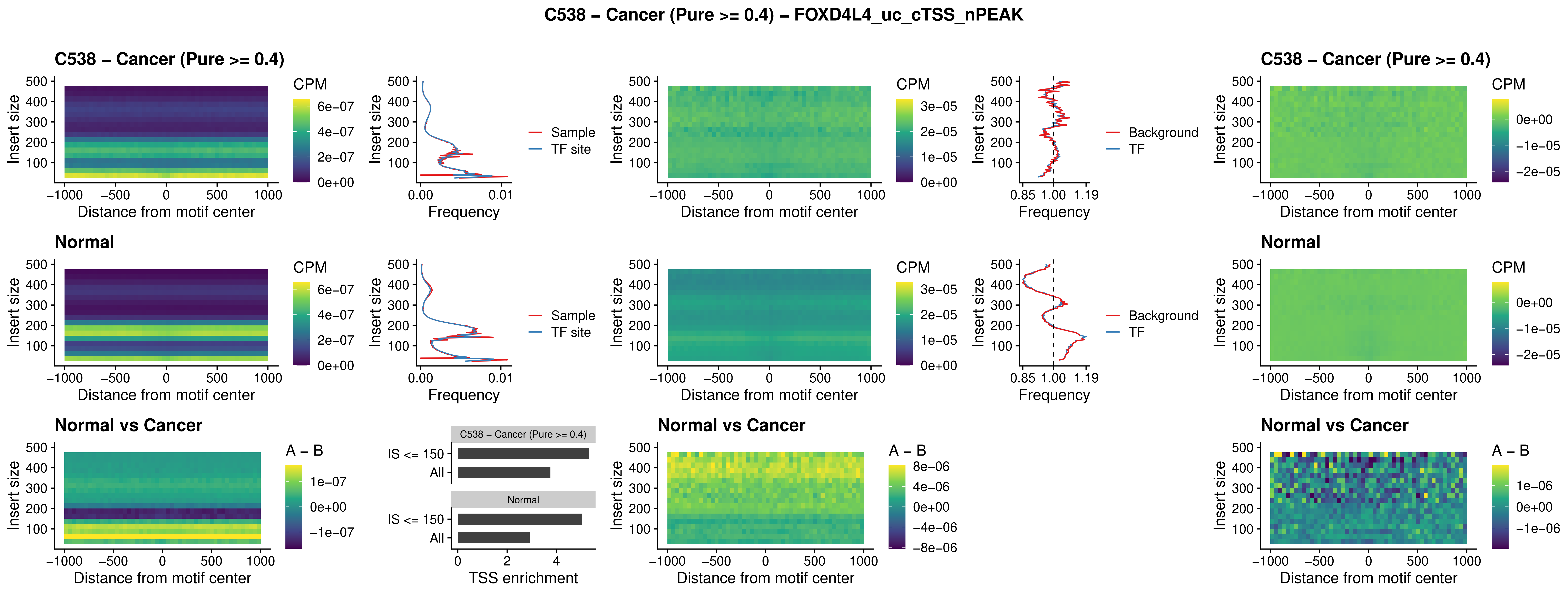

C538 – Cancer (Pure >= 0.4) – FOXD4L4_uc_cTSS_nPEAK
C538 – Cancer (Pure >= 0.4)
Normal
Normal vs Cancer
Insert size
Distance from motif center
Frequency
CPM
A – B
Sample
TF site
Background
TF
C538 – Cancer (Pure >= 0.4)
Normal
IS <= 150
All
TSS enrichment

#### C538 – Cancer (Pure >= 0.4) – FOXD4L2_uc_cTSS_nPEAK

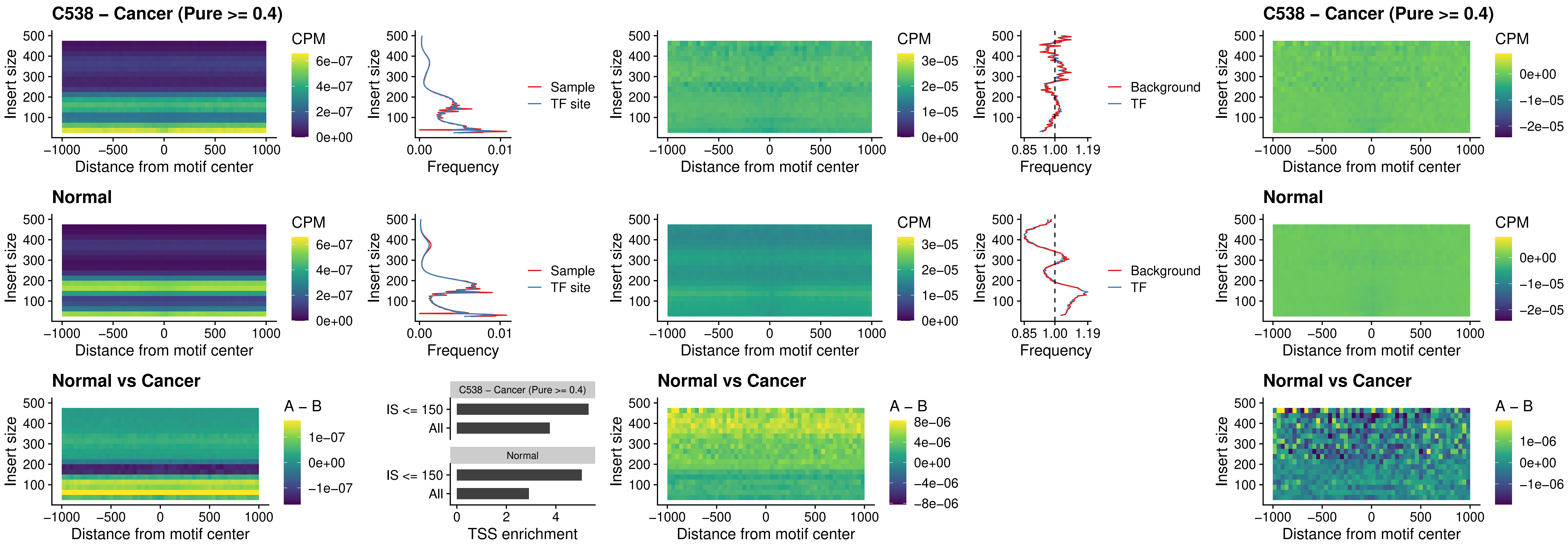

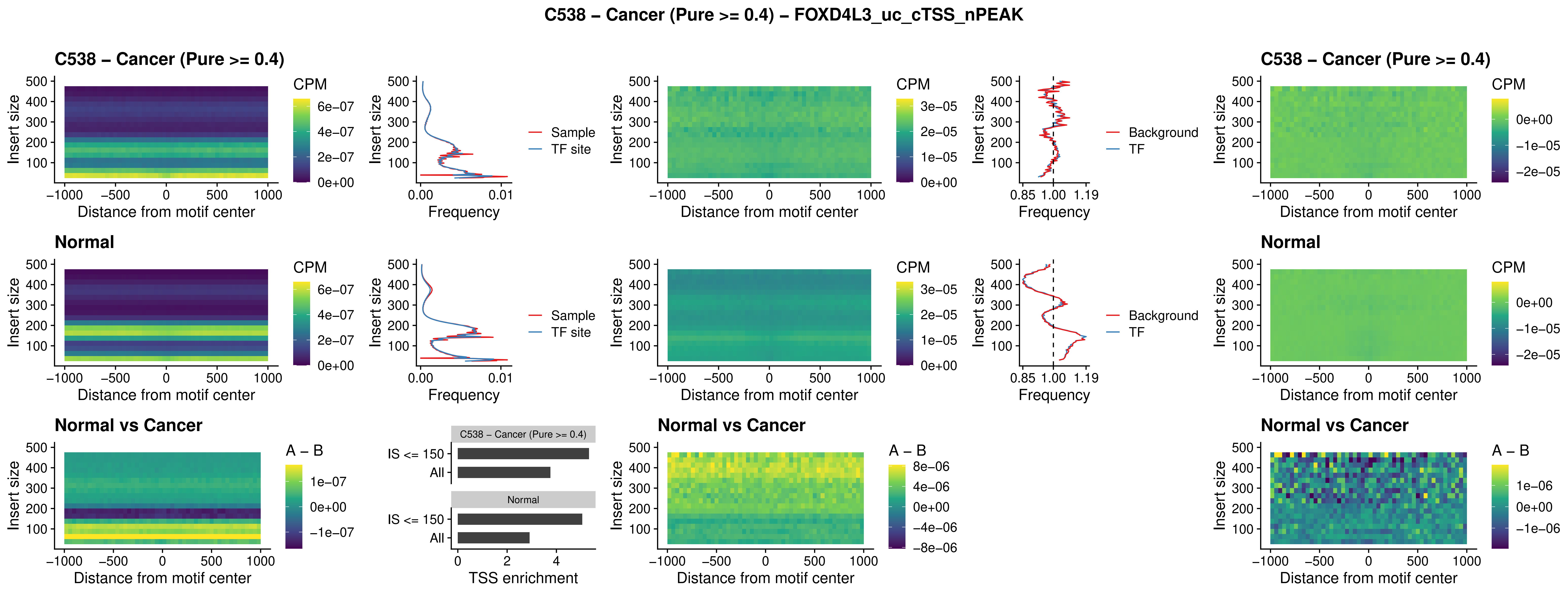
C538 – Cancer (Pure >= 0.4) – FOXD4L3_uc_cTSS_nPEAK
C538 – Cancer (Pure >= 0.4)
Normal
Normal vs Cancer
Insert size
Distance from motif center
Frequency
CPM
A – B
Sample
TF site
Background
TF
C538 – Cancer (Pure >= 0.4)
Normal
IS <= 150
All
TSS enrichment

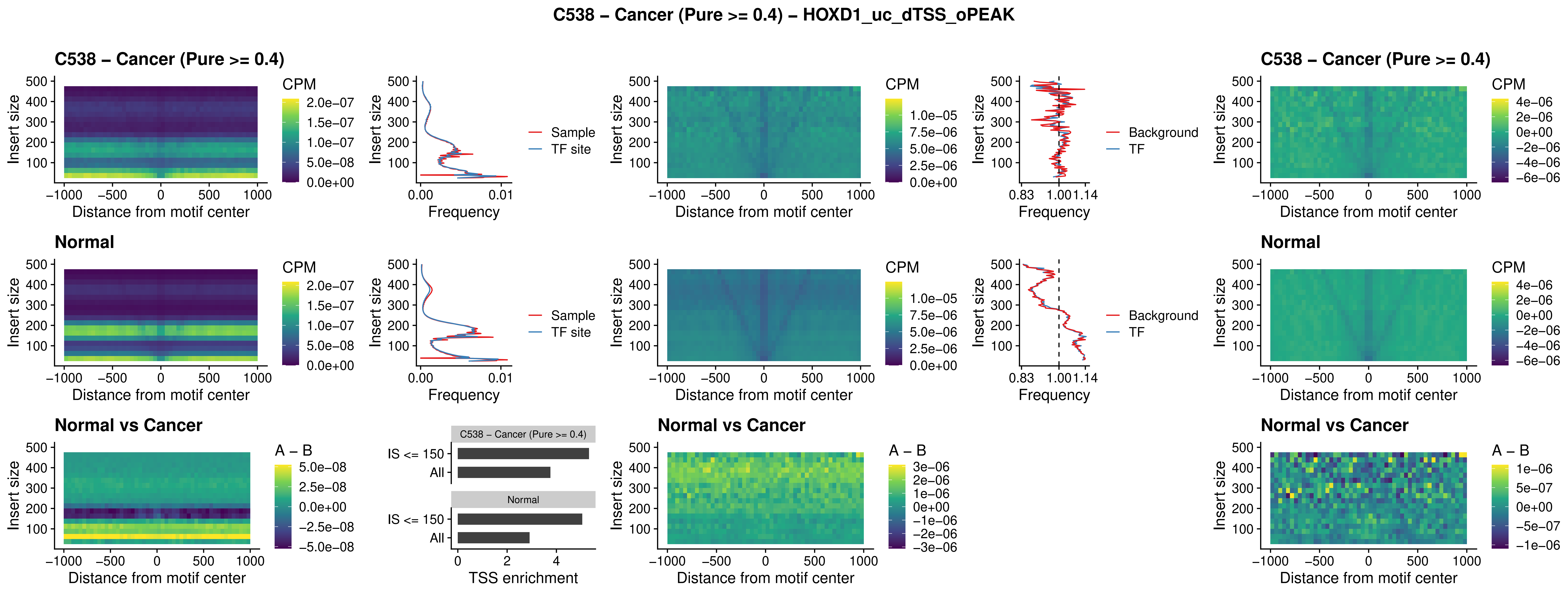
C538 – Cancer (Pure >= 0.4) – HOXD1_uc_dTSS_oPEAK
C538 – Cancer (Pure >= 0.4)
Normal
Normal vs Cancer
Insert size
Distance from motif center
Frequency
CPM
A – B
Sample
TF site
Background
TF
C538 – Cancer (Pure >= 0.4)
Normal
IS <= 150
All
TSS enrichment

#### C538 – Cancer (Pure >= 0.4) – VAX1_uc_pTSS_nPEAK

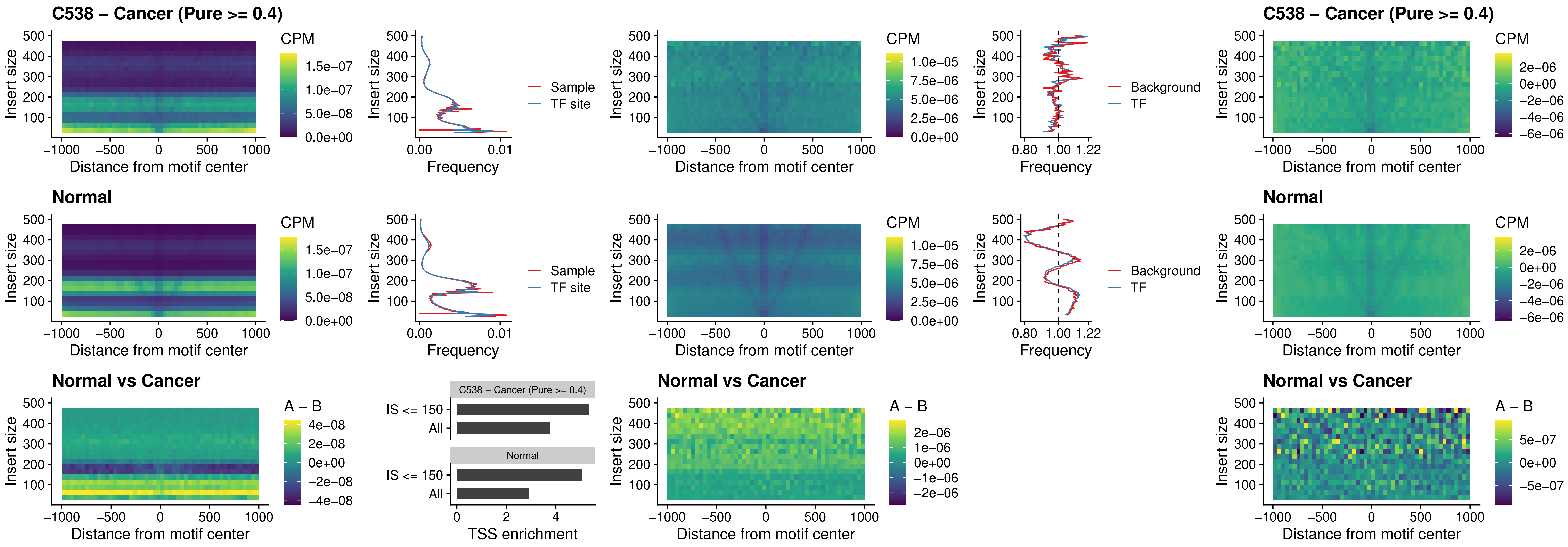

#### C538 – Cancer (Pure >= 0.4) – DLX3_uc_pTSS_nPEAK

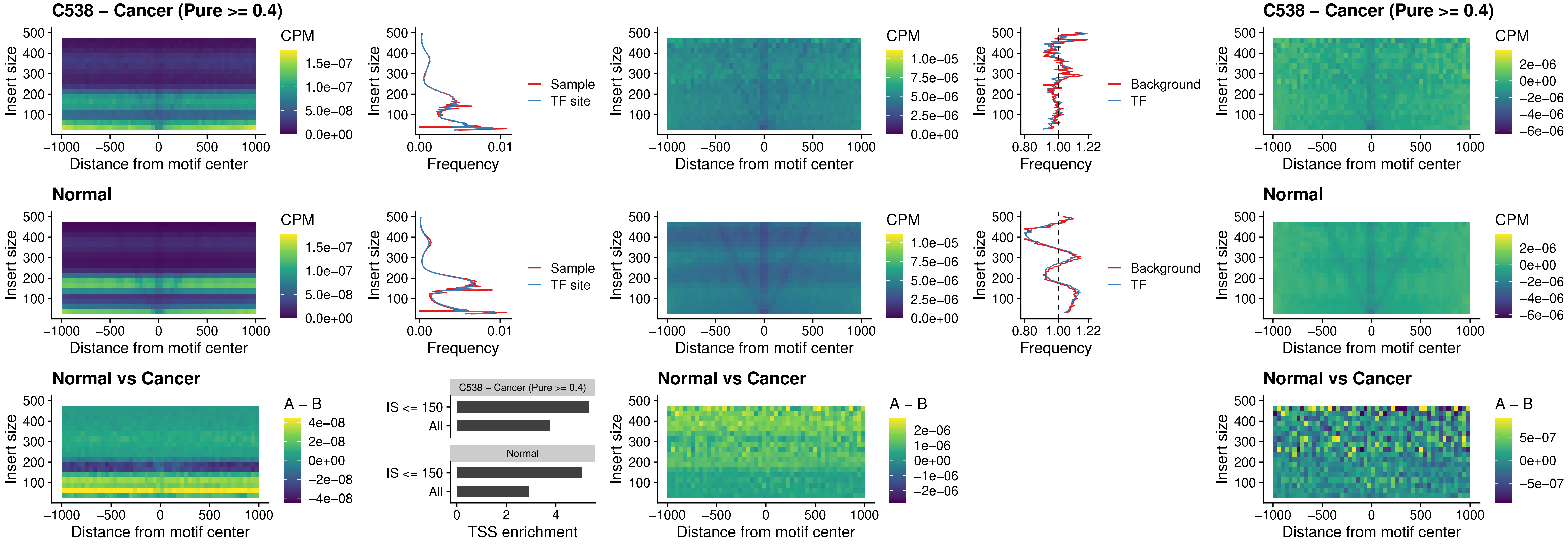

#### C538 – Cancer (Pure >= 0.4) – HOXD3_uc_pTSS_nPEAK

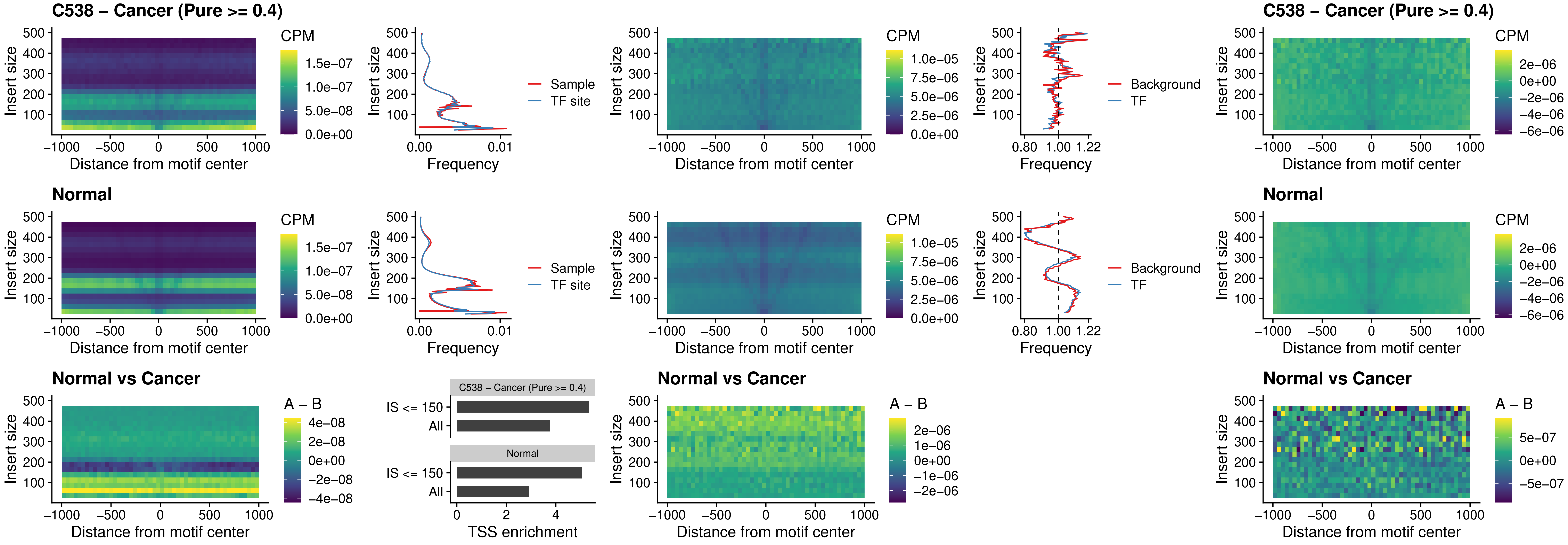

#### C538 – Cancer (Pure >= 0.4) – PDX1_uc_dTSS_nPEAK

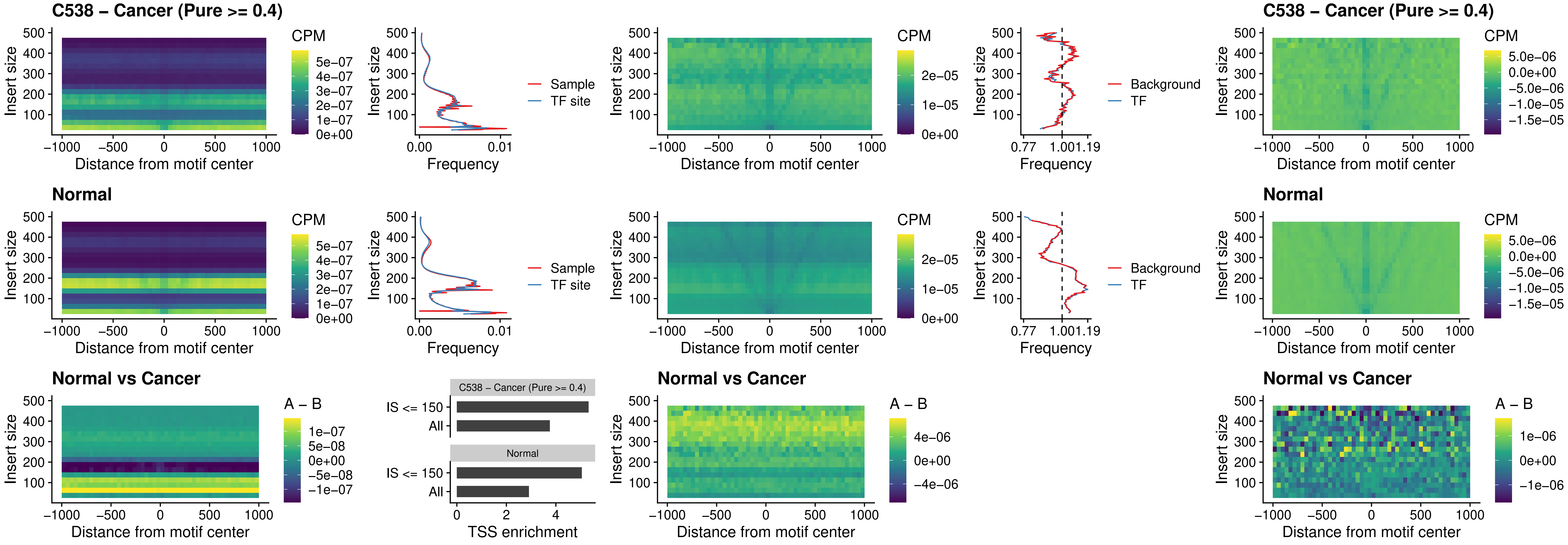

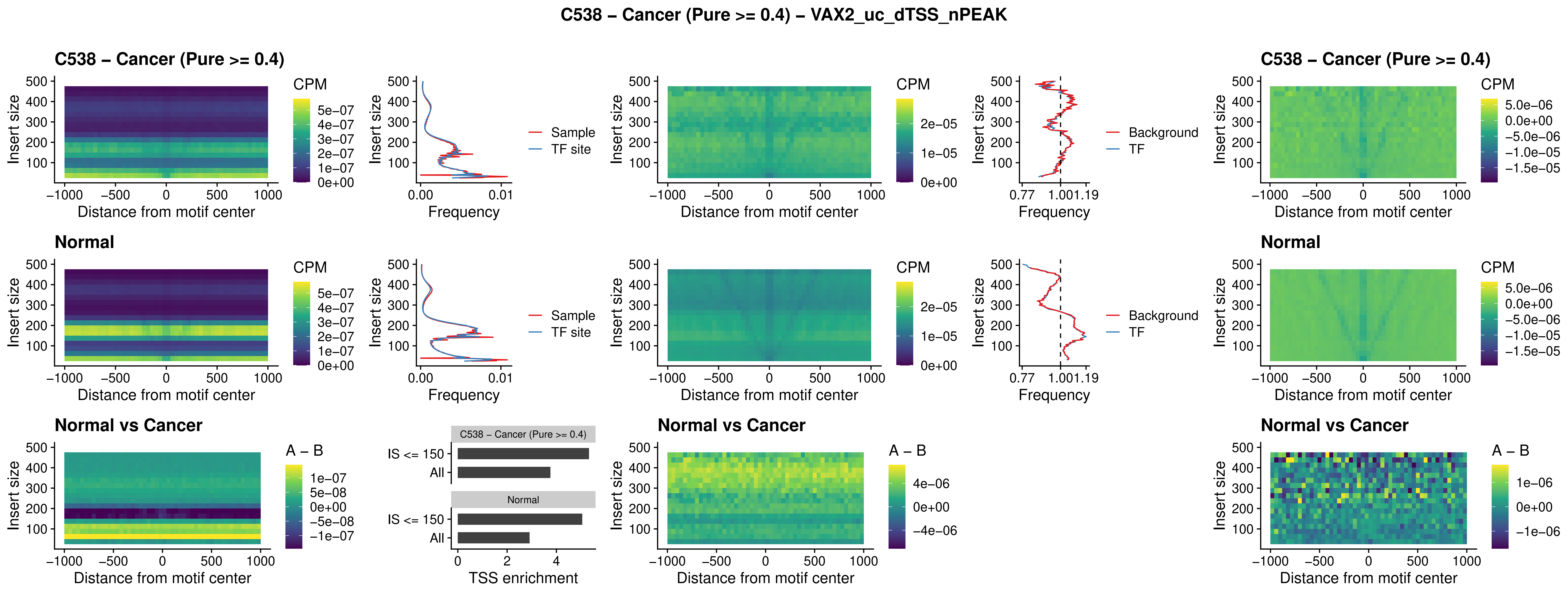

C538 – Cancer (Pure >= 0.4) – VAX2_uc_dTSS_nPEAK
C538 – Cancer (Pure >= 0.4)
Normal
Normal vs Cancer
Insert size
Distance from motif center
Frequency
CPM
A – B
Sample
TF site
Background
TF
C538 – Cancer (Pure >= 0.4)
Normal
IS <= 150
All
TSS enrichment

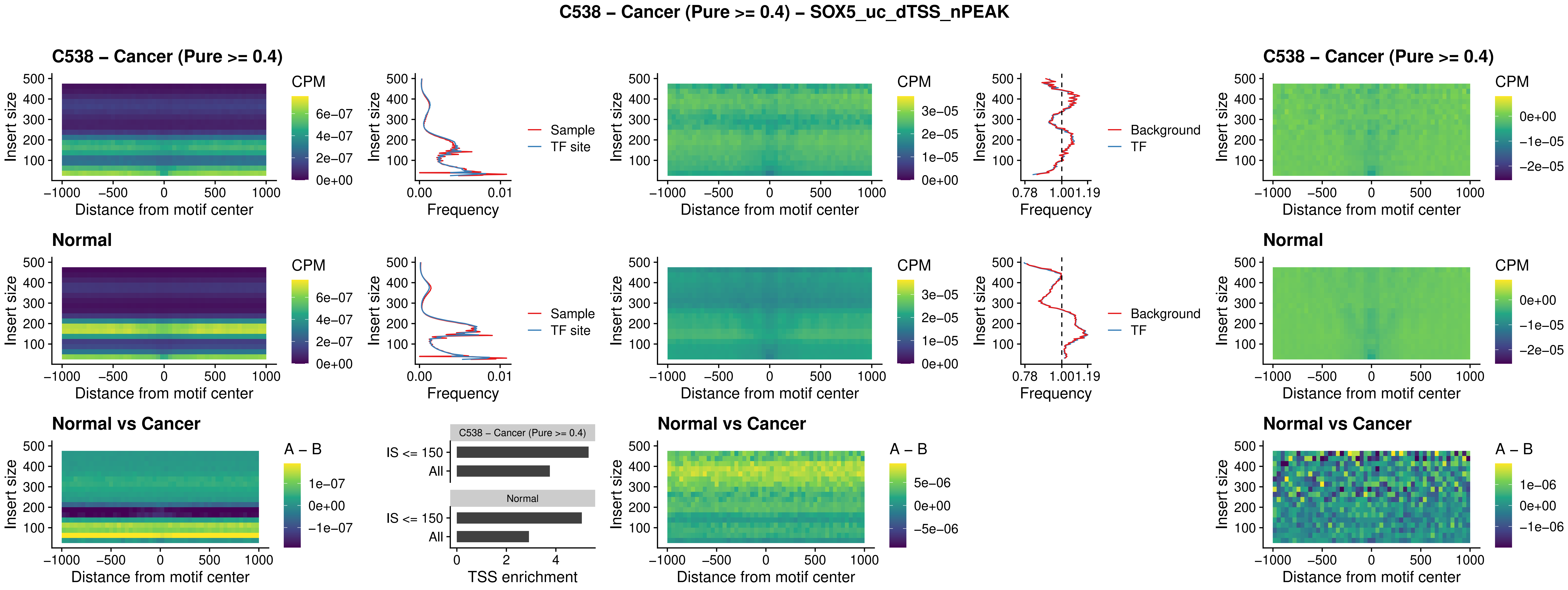
C538 – Cancer (Pure >= 0.4) – SOX5_uc_dTSS_nPEAK
C538 – Cancer (Pure >= 0.4)
Normal
Normal vs Cancer
Insert size
Distance from motif center
Frequency
CPM
A – B
Sample
TF site
Background
TF
C538 – Cancer (Pure >= 0.4)
Normal
IS <= 150
All
TSS enrichment

#### C538 – Cancer (Pure >= 0.4) – SOX6_uc_dTSS_nPEAK

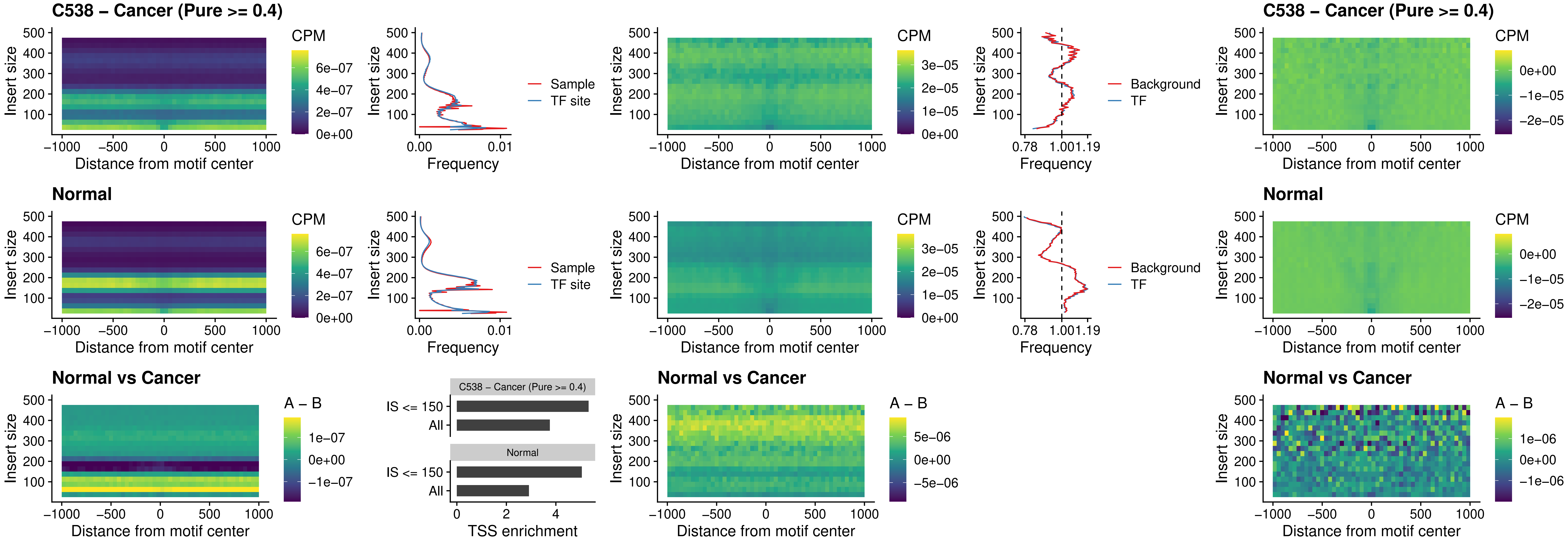

#### C538 – Cancer (Pure >= 0.4) – JDP2_uc_cTSS_nPEAK

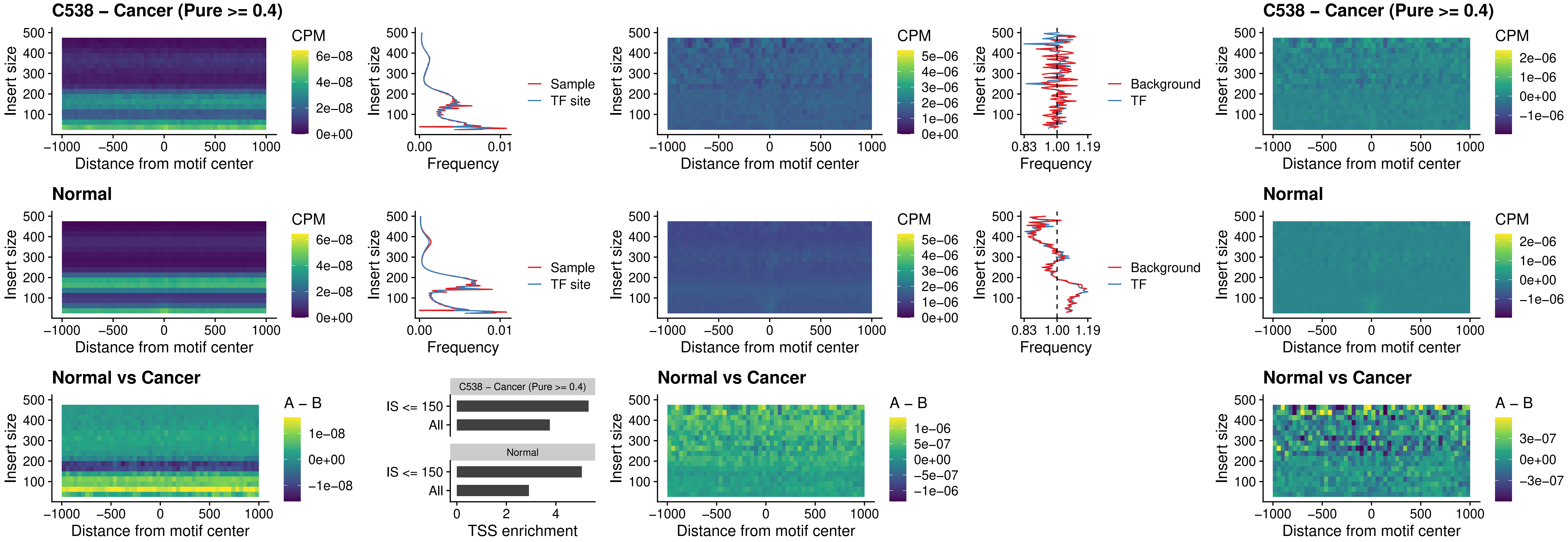

#### C538 – Cancer (Pure >= 0.4) – PURA_uc_dTSS_oPEAK

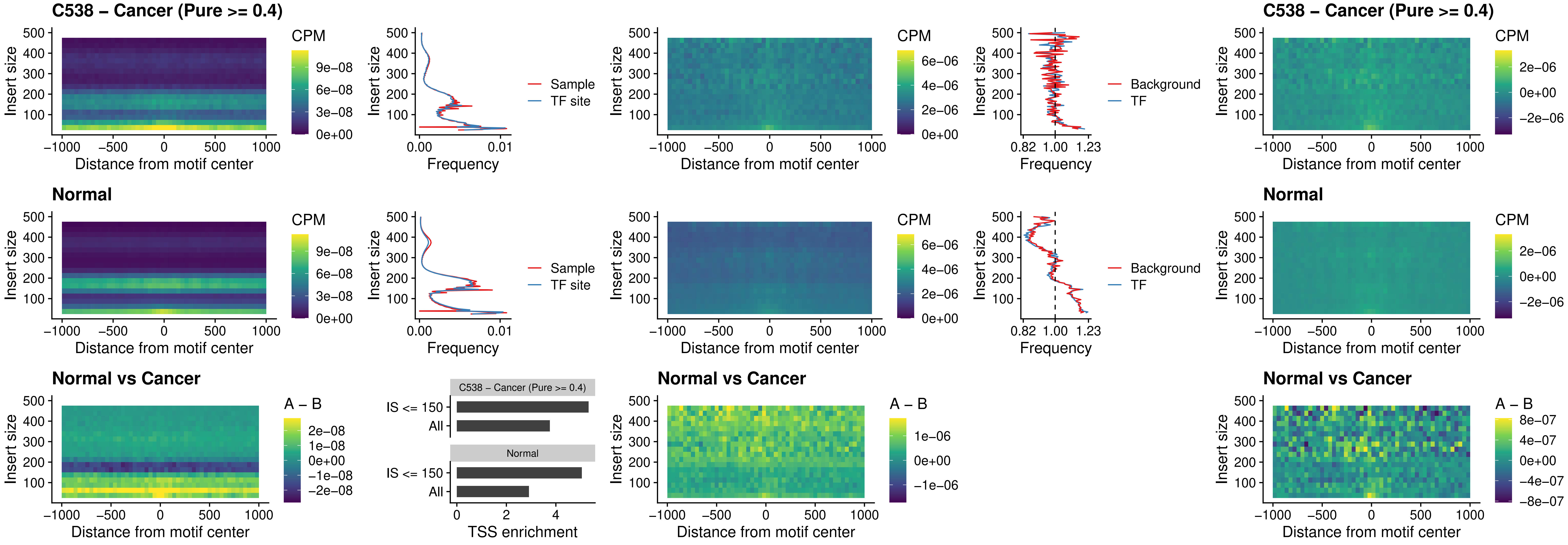

### C538 – Cancer (Pure >= 0.4) – HHEX_uc_dTSS_nPEAK

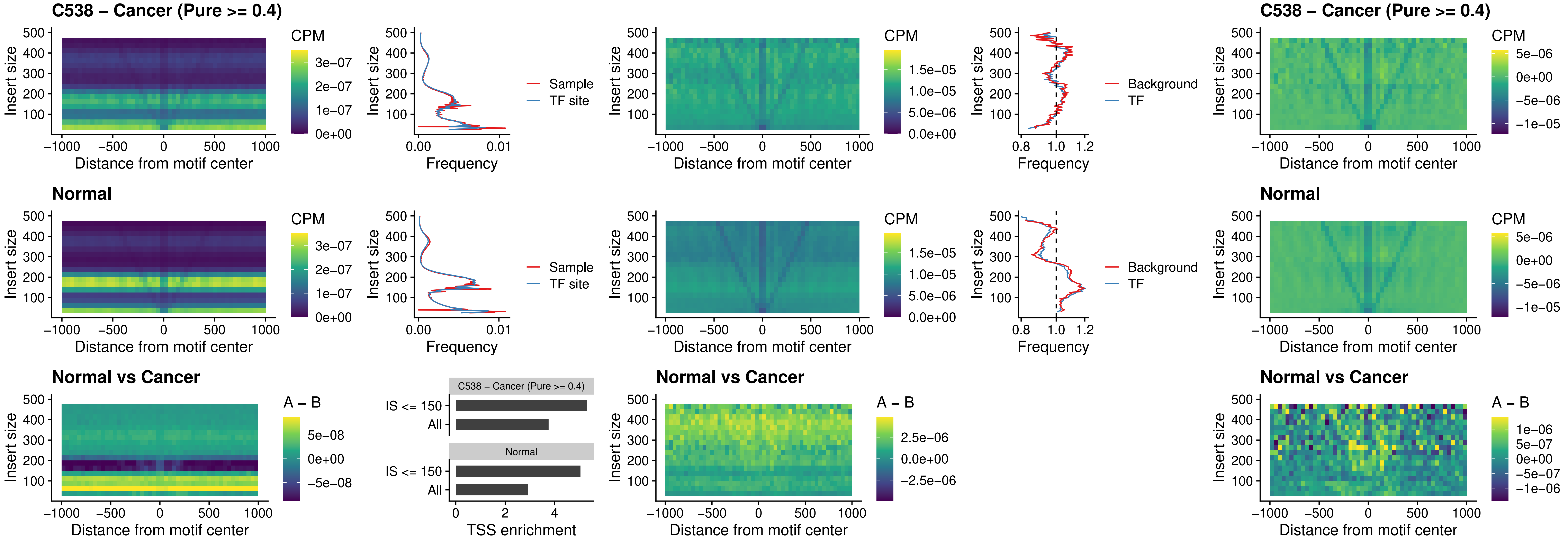

#### C538 – Cancer (Pure >= 0.4) – HMG20B_uc_dTSS_nPEAK

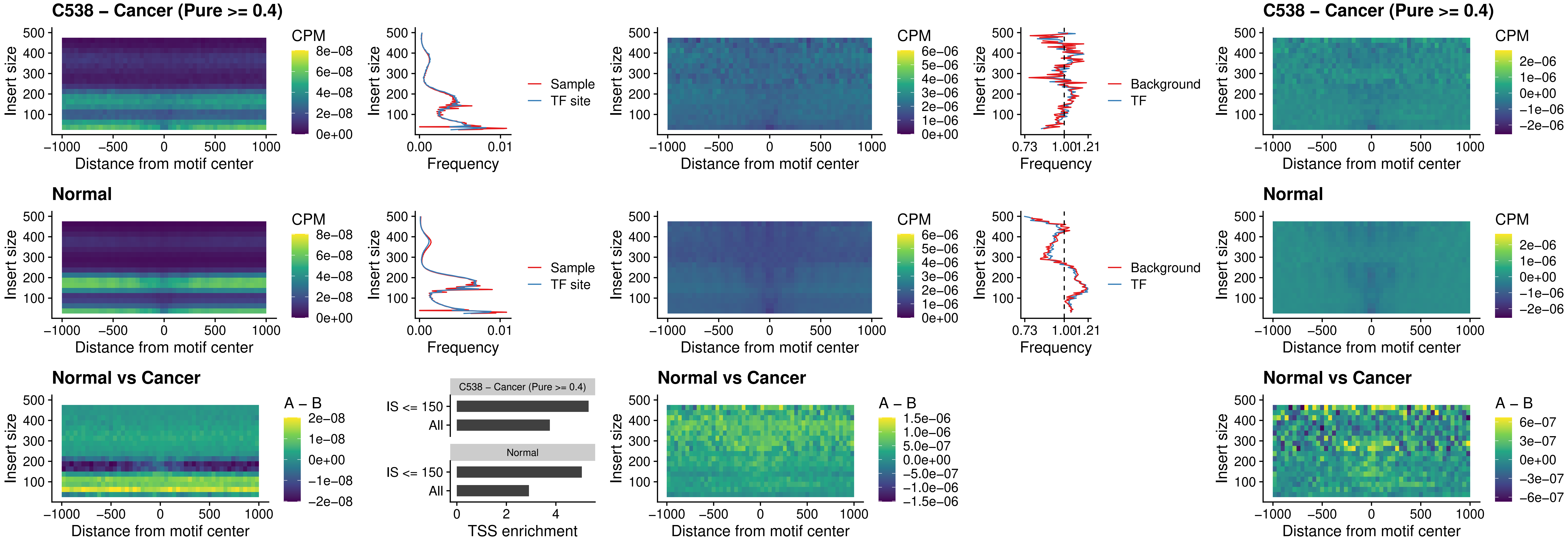

#### C538 – Cancer (Pure >= 0.4) – FOXB2_uc_dTSS_nPEAK

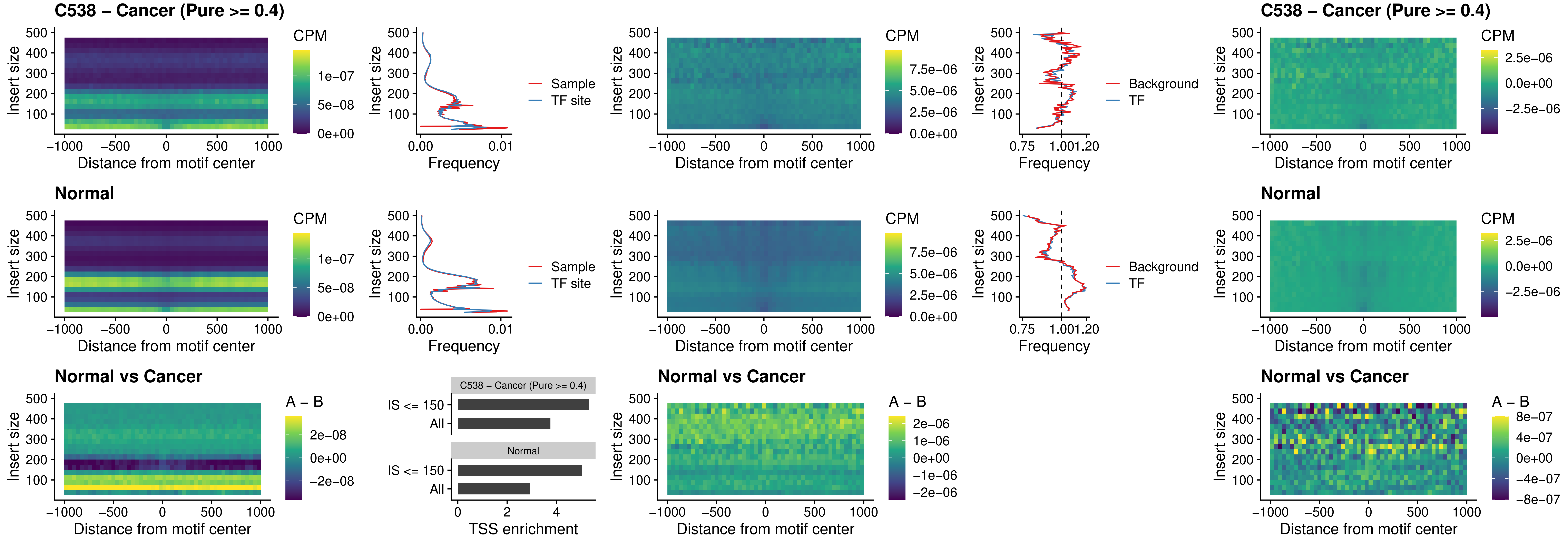

#### C538 – Cancer (Pure >= 0.4) – FOXL1_uc_cTSS_nPEAK

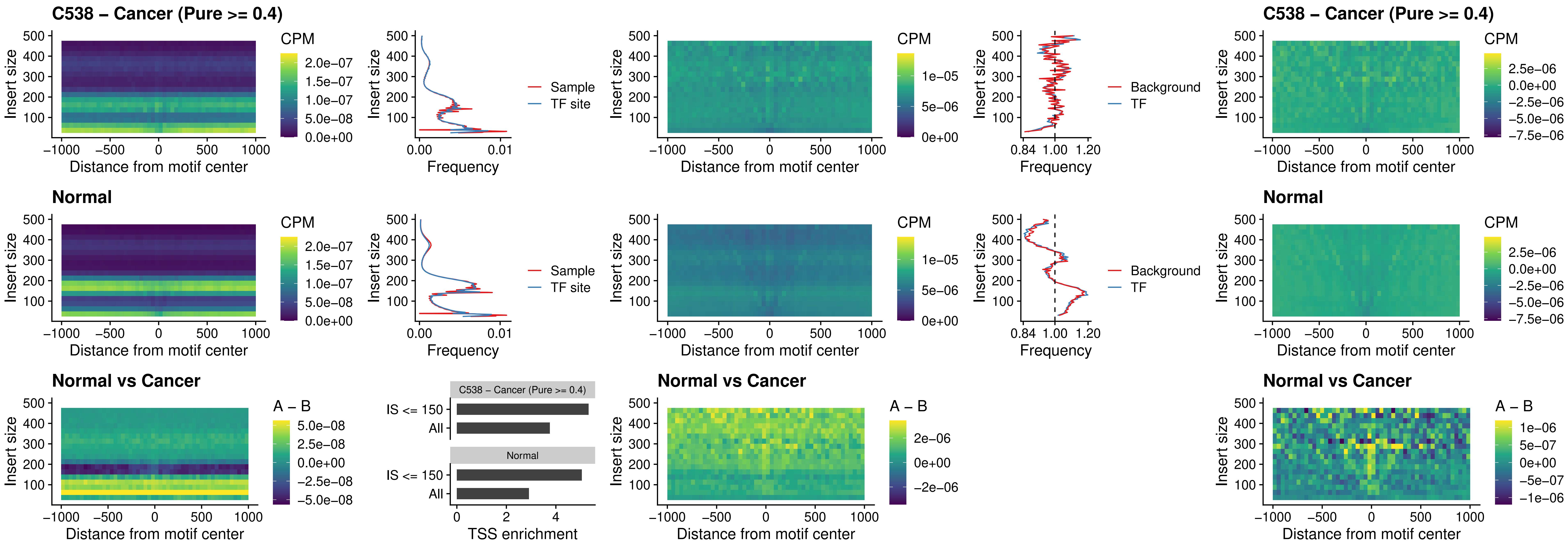

#### C538 – Cancer (Pure >= 0.4) – PBX3_uc_pTSS_oPEAK

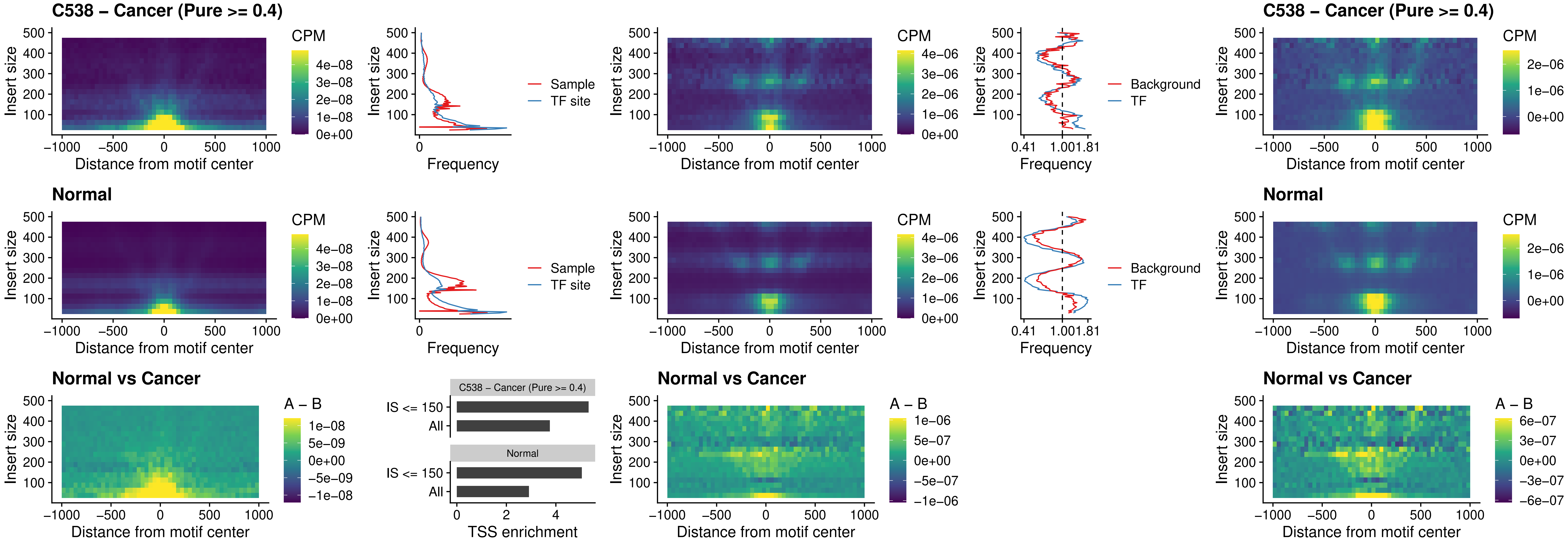

#### C538 – Cancer (Pure >= 0.4) – SNAI2_uc_cTSS_nPEAK

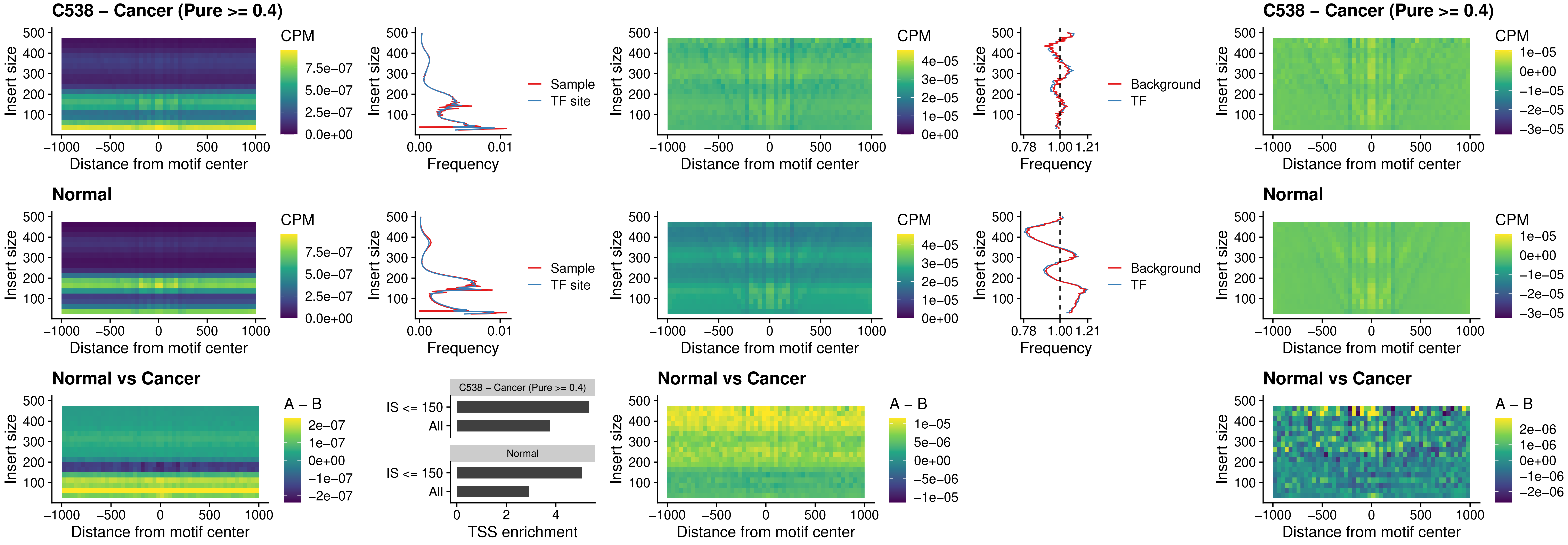

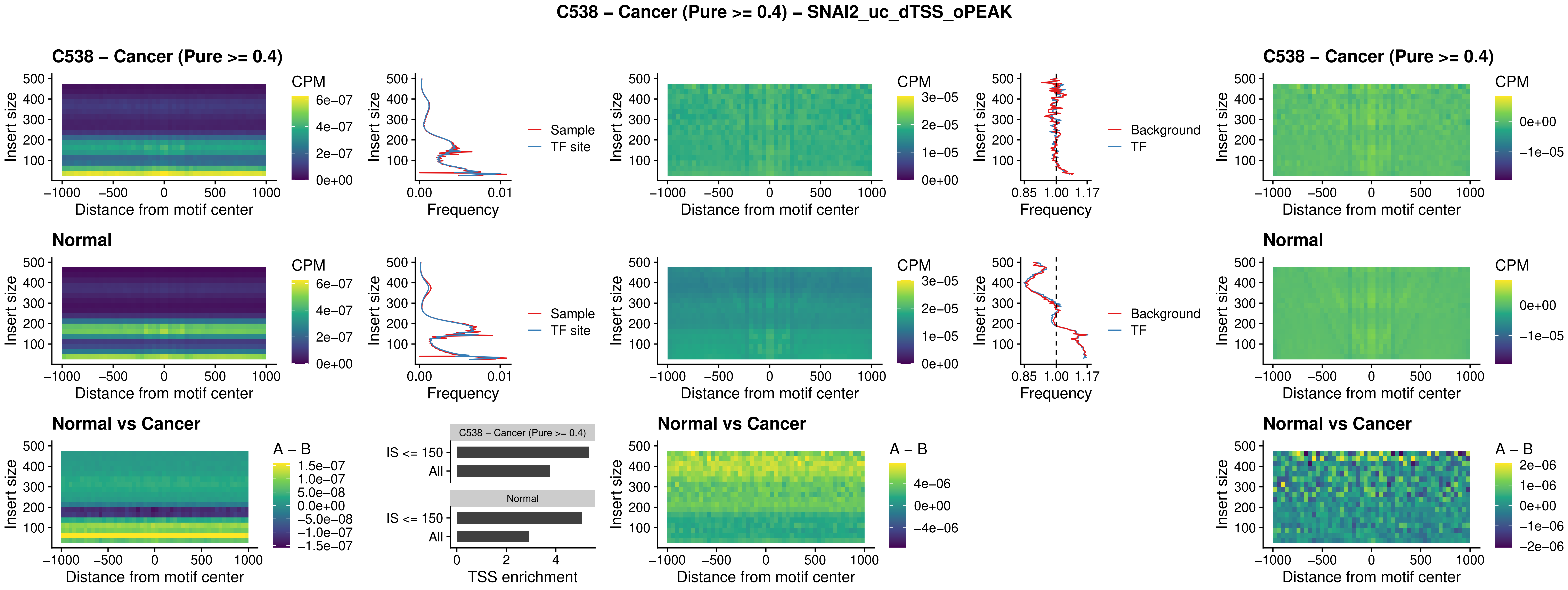
C538 – Cancer (Pure >= 0.4) – SNAI2_uc_dTSS_oPEAK
C538 – Cancer (Pure >= 0.4)
Normal
Normal vs Cancer
Insert size
Distance from motif center
Frequency
CPM
A – B
Sample
TF site
Background
TF
C538 – Cancer (Pure >= 0.4)
Normal
IS <= 150
All
TSS enrichment

#### C538 – Cancer (Pure >= 0.4) – SPI1_uc_pTSS_nPEAK

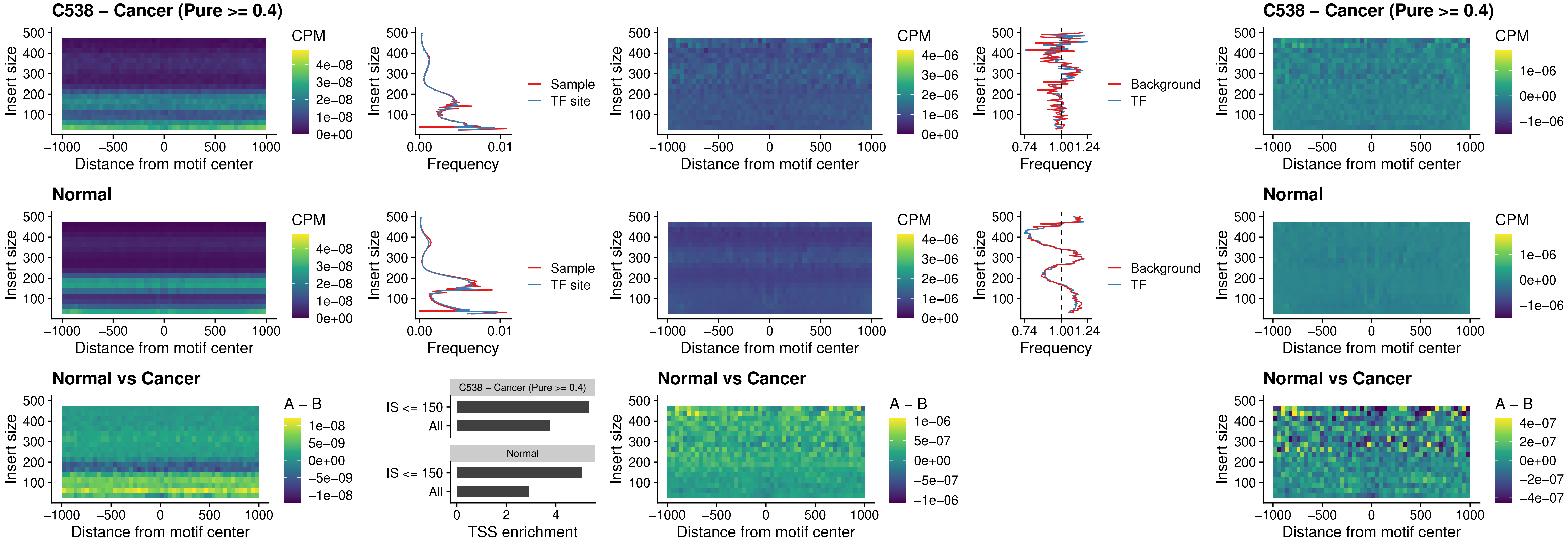

#### C538 – Cancer (Pure >= 0.4) – KLF16_uc_pTSS_nPEAK

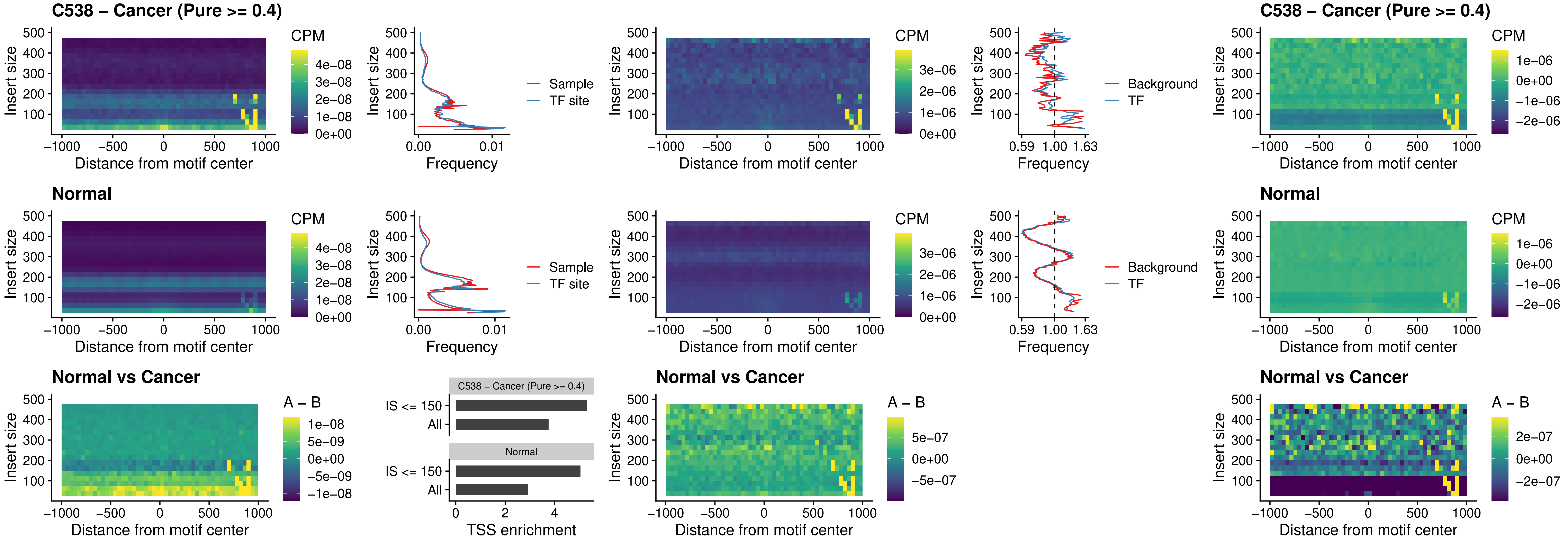

#### C538 – Cancer (Pure >= 0.4) – SP3_uc_pTSS_nPEAK

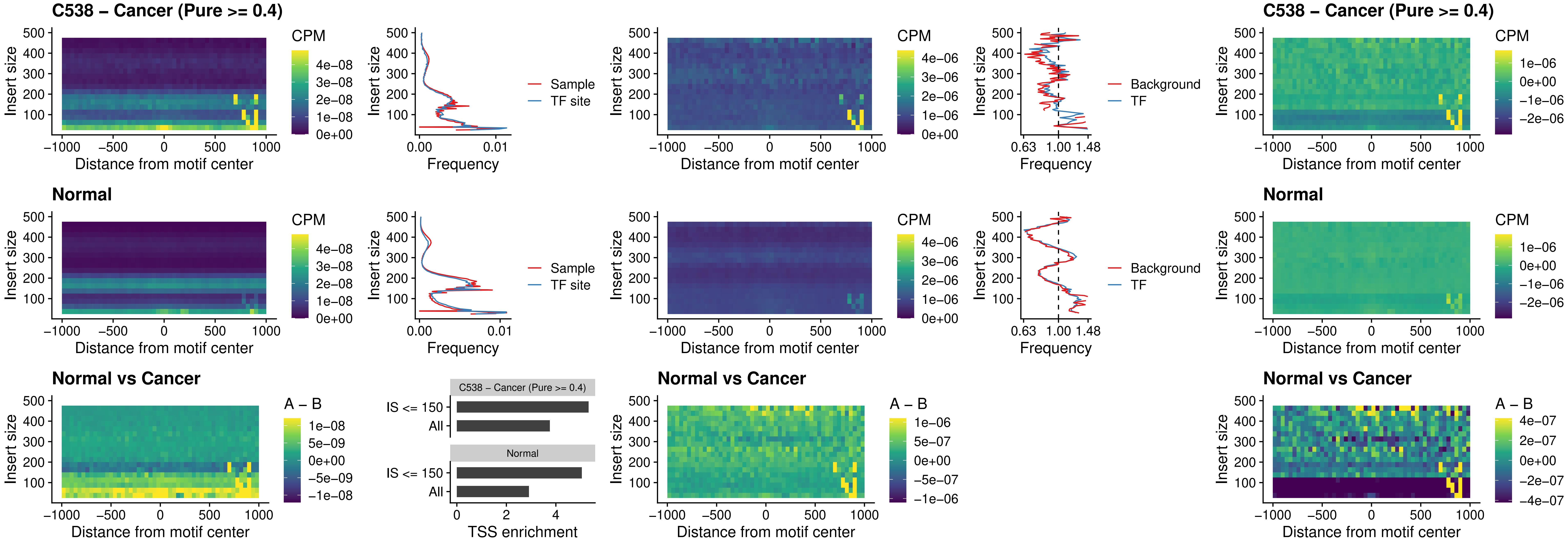

#### C538 – Cancer (Pure >= 0.4) – NEUROD1_uc_dTSS_oPEAK

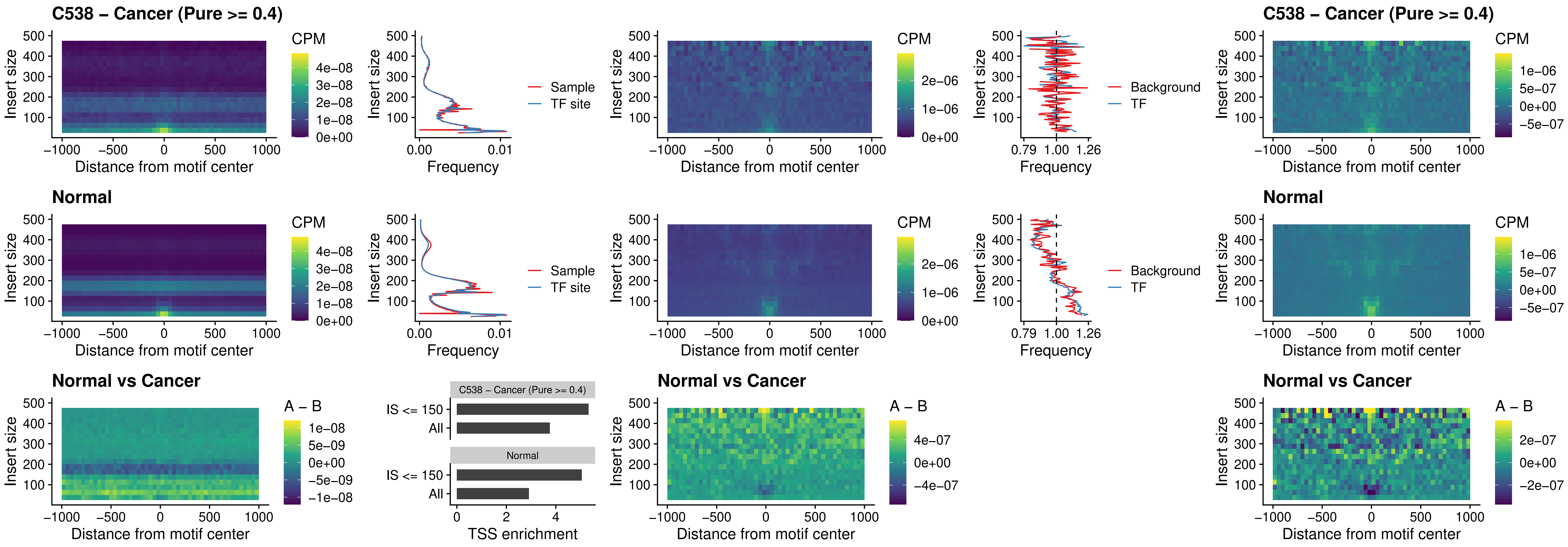

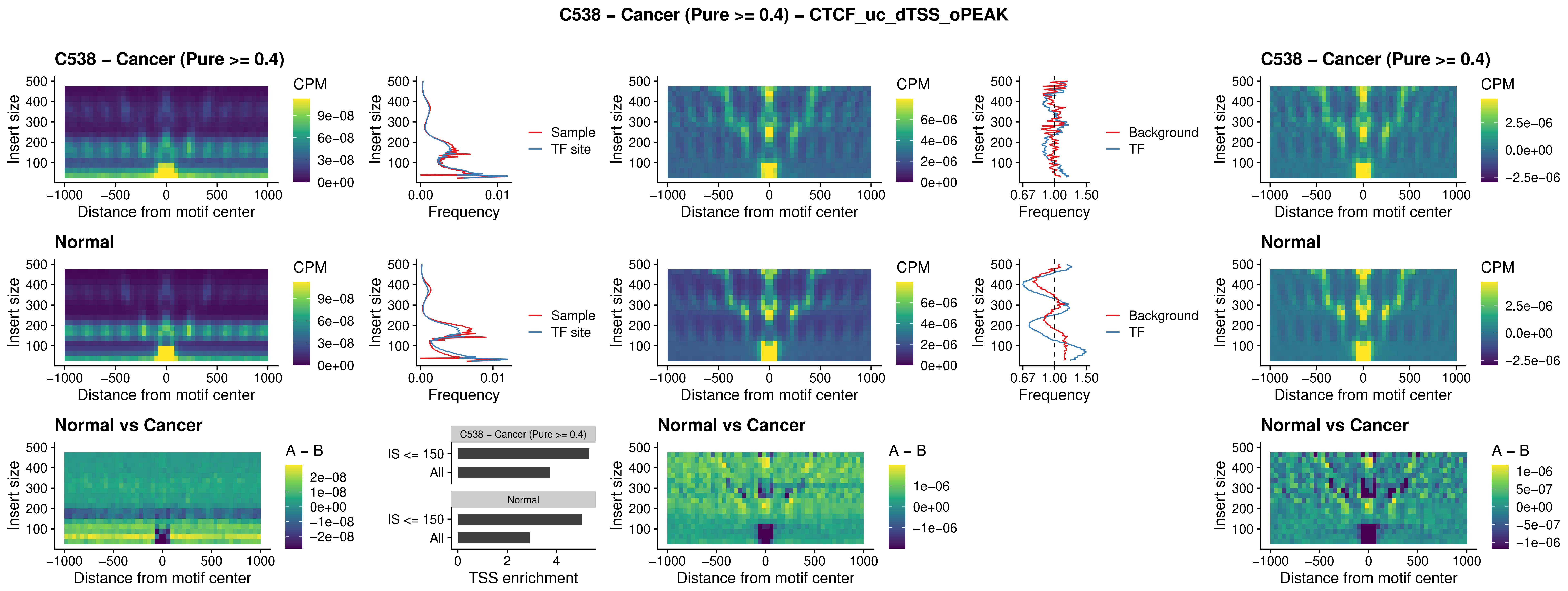

C538 – Cancer (Pure >= 0.4) – CTCF_uc_dTSS_oPEAK
C538 – Cancer (Pure >= 0.4)
Normal
Normal vs Cancer
Insert size
Distance from motif center
Frequency
CPM
A – B
Sample
TF site
Background
TF
C538 – Cancer (Pure >= 0.4)
IS <= 150
All
TSS enrichment

#### C538 – Cancer (Pure >= 0.4) – CTCF_uc_cTSS_oPEAK

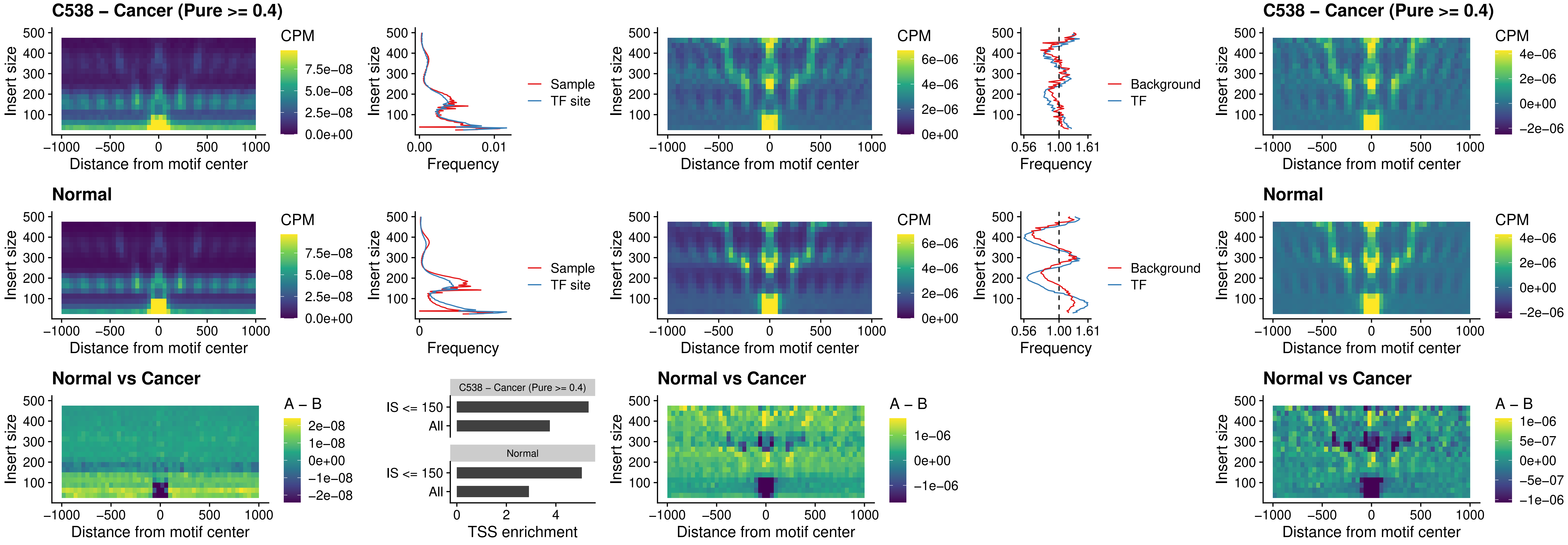

#### C538 – Cancer (Pure >= 0.4) – CTCFL_uc_dTSS_oPEAK

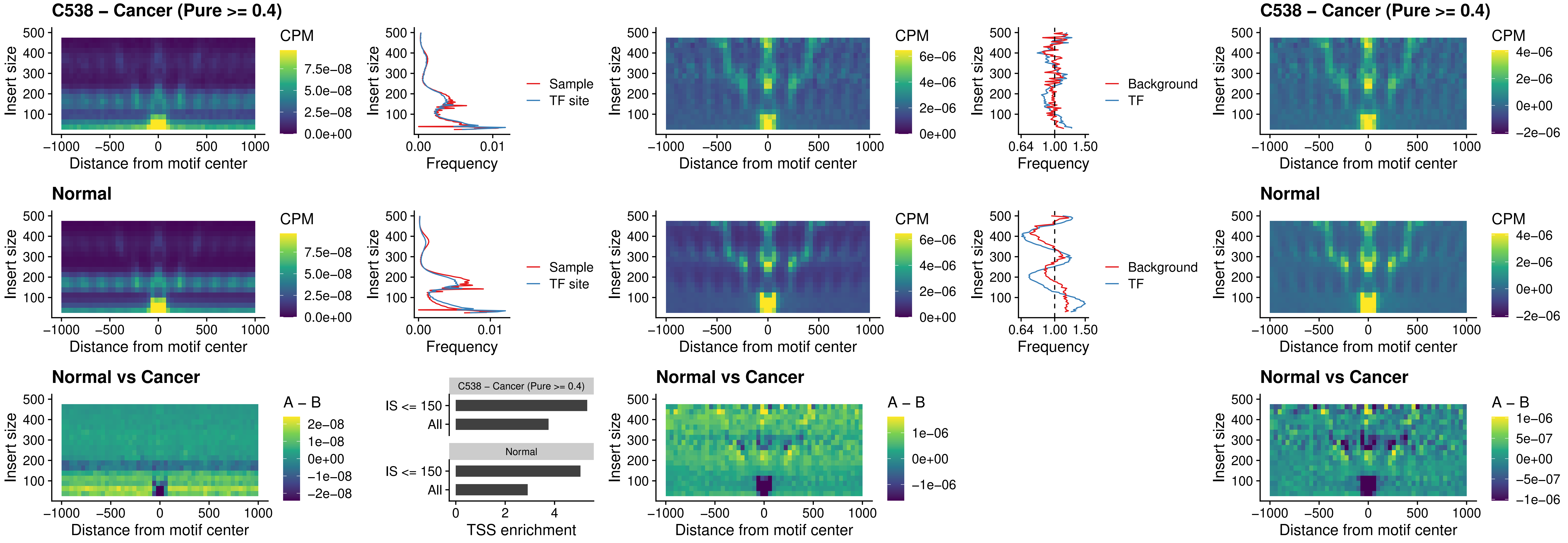

#### C538 – Cancer (Pure >= 0.4) – ENCODE_CTCF_sites_COLON

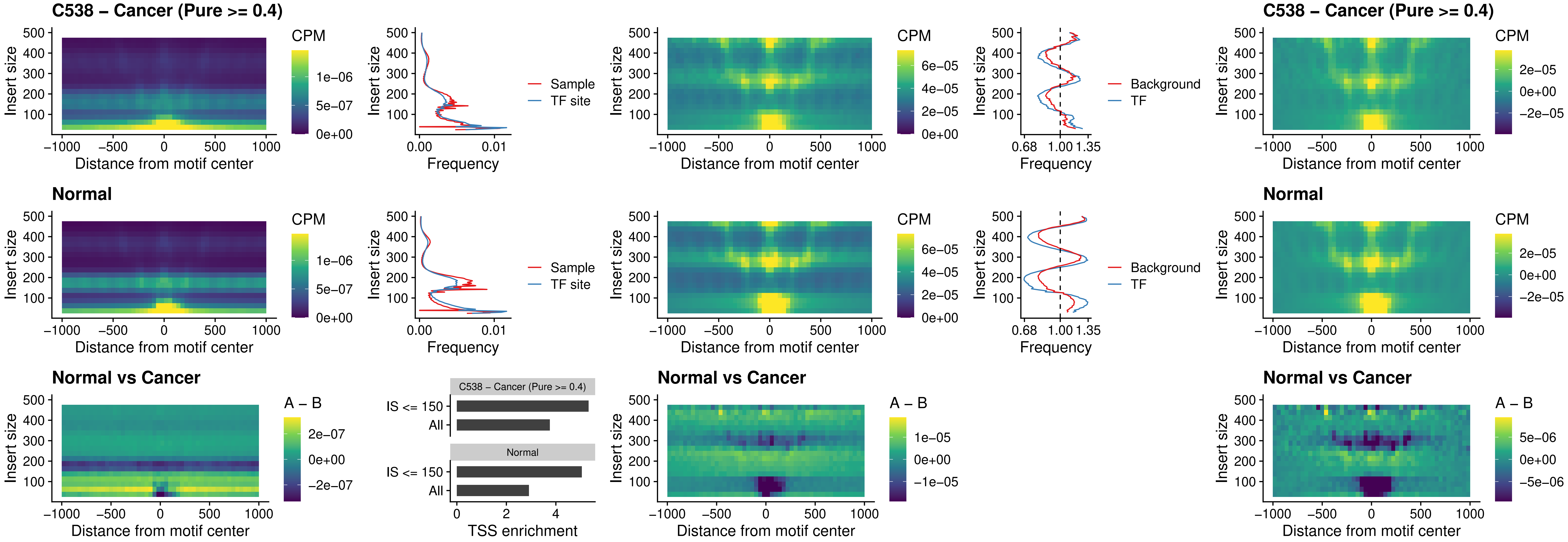

#### C538 – Cancer (Pure >= 0.4) – ENCODE_CTCF_sites_COLON_loop_ctcf

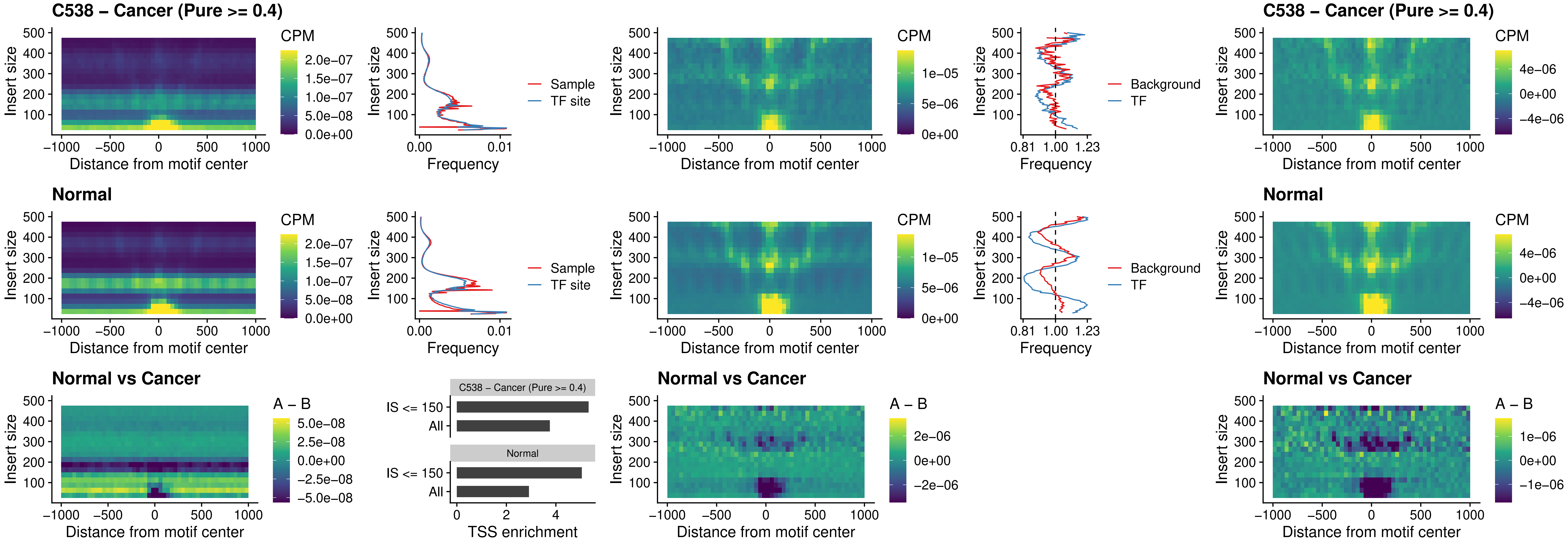

#### C538 – Cancer (Pure >= 0.4) – ENCODE_CTCF_sites_COLON_intergenic_ctcf

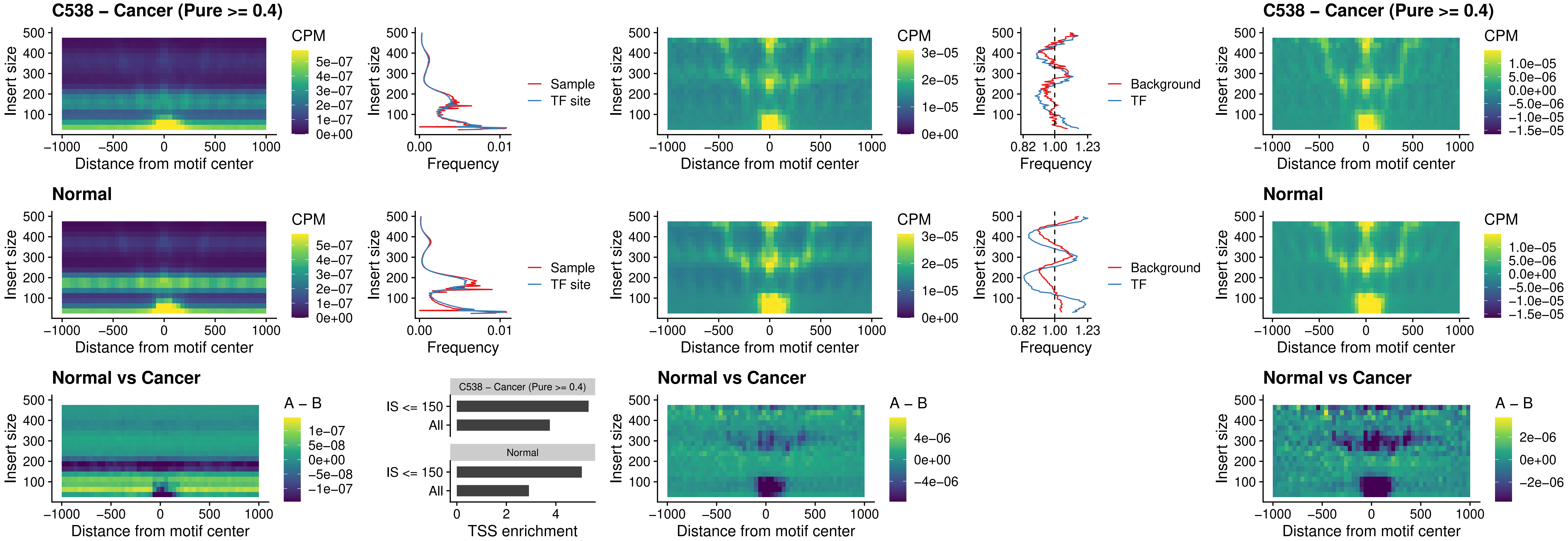

#### C538 – Cancer (Pure >= 0.4) – ENCODE_CTCF_sites_COLON_nonpromotor

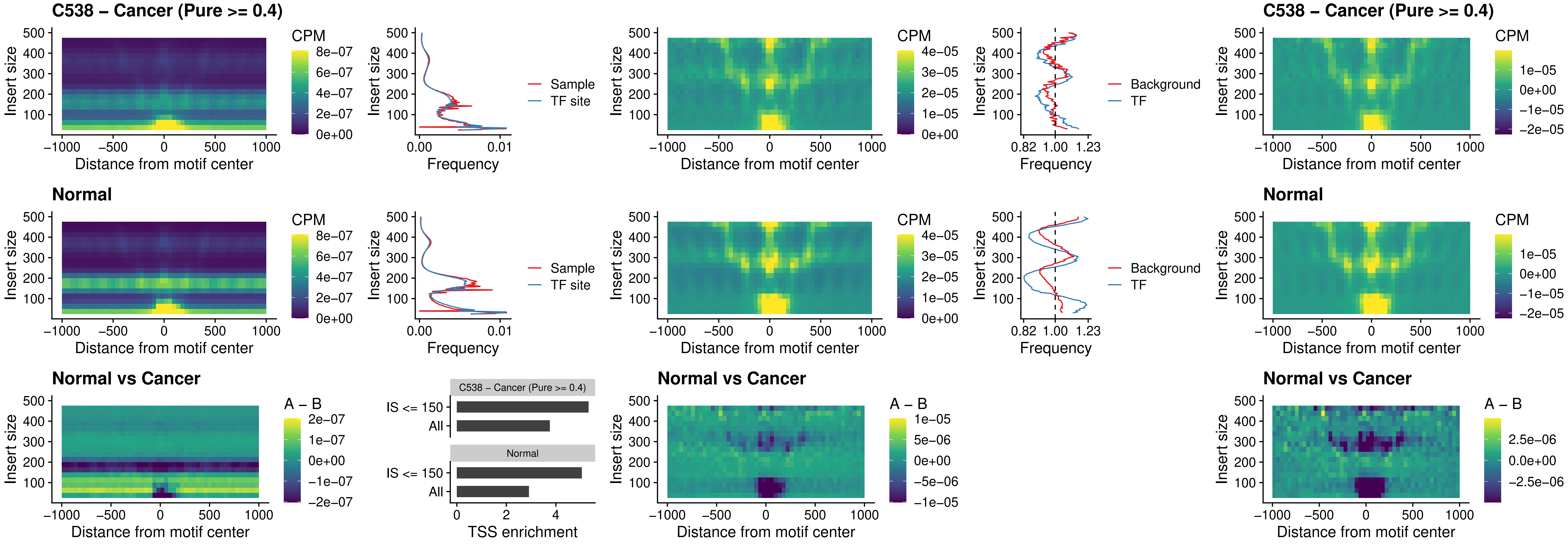

#### C538 – Cancer (Pure >= 0.4) – CTCFL_uc_cTSS_oPEAK

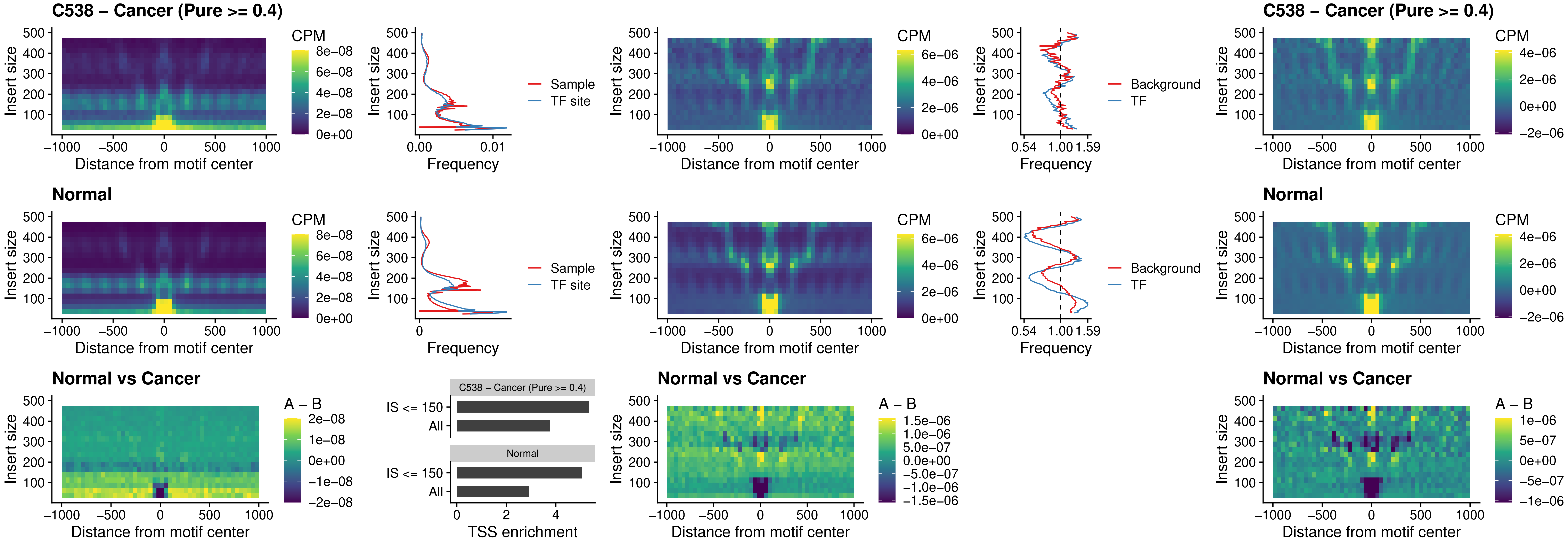

#### C538 – Cancer (Pure >= 0.4) – NEUROD1_uc_cTSS_oPEAK

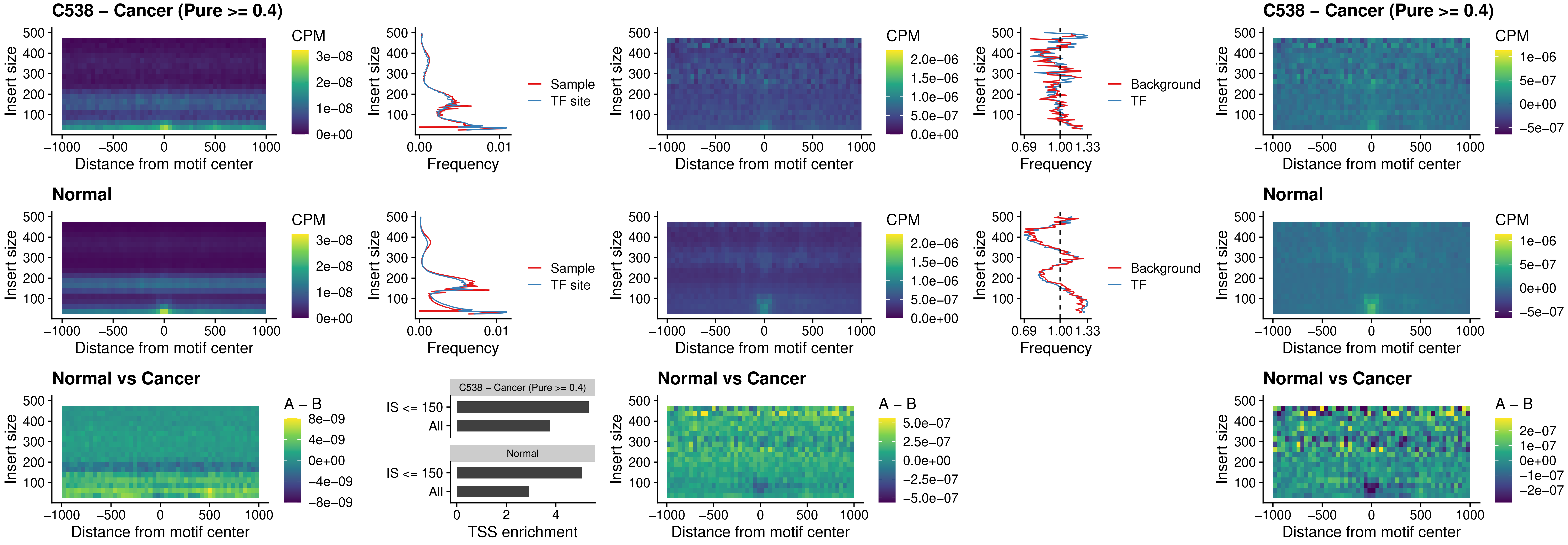

#### C538 – Cancer (Pure >= 0.4) – ENCODE_YY1_sites_COLON_nonpromotor

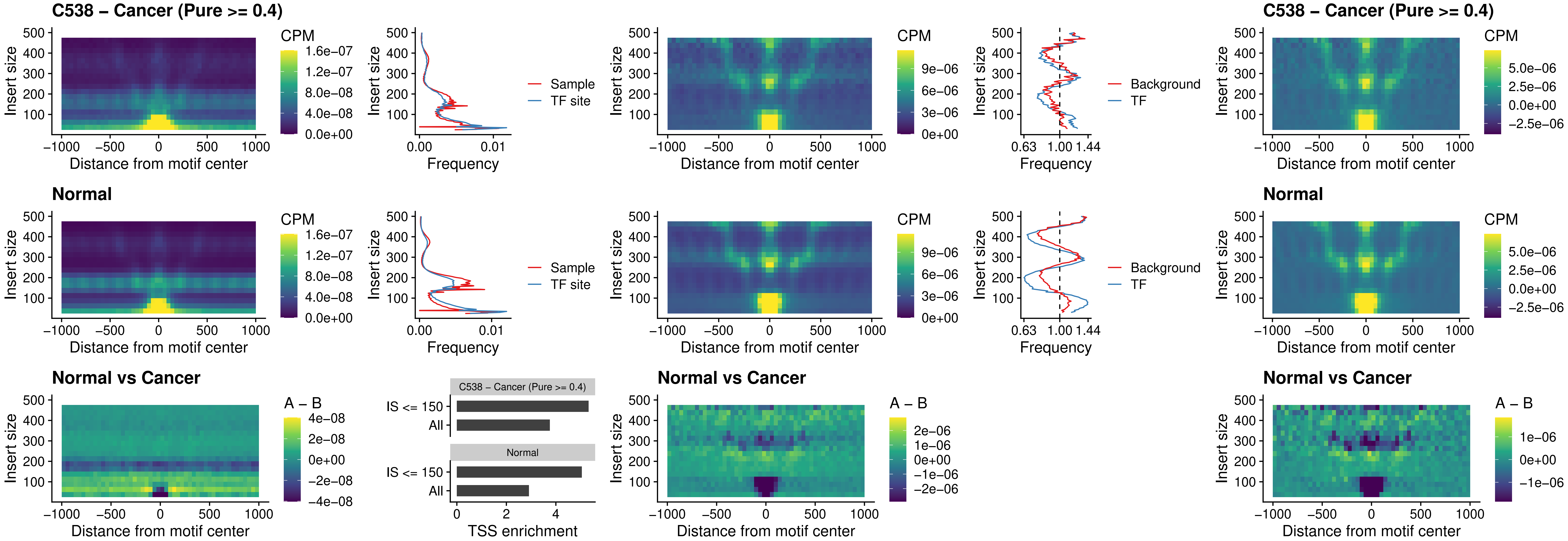

#### C538 – Cancer (Pure >= 0.4) – CTCF_uc_pTSS_oPEAK

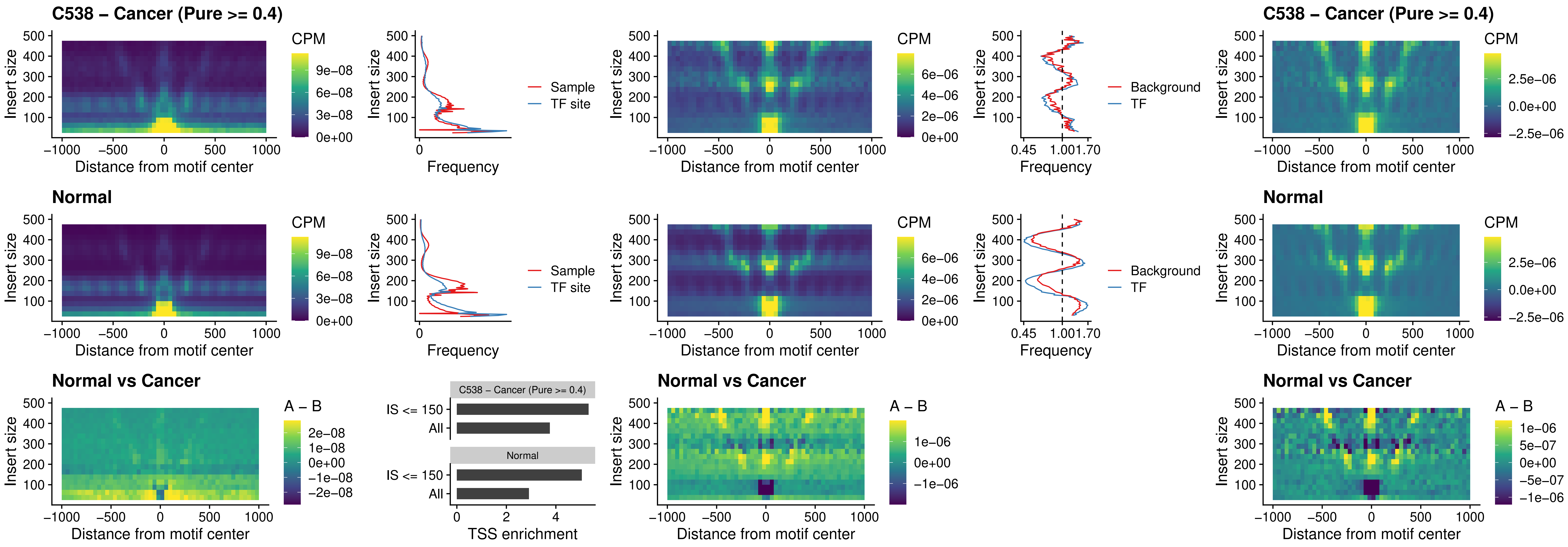

#### C538 – Cancer (Pure >= 0.4) – ENCODE_CTCF_sites_COLON_promotor

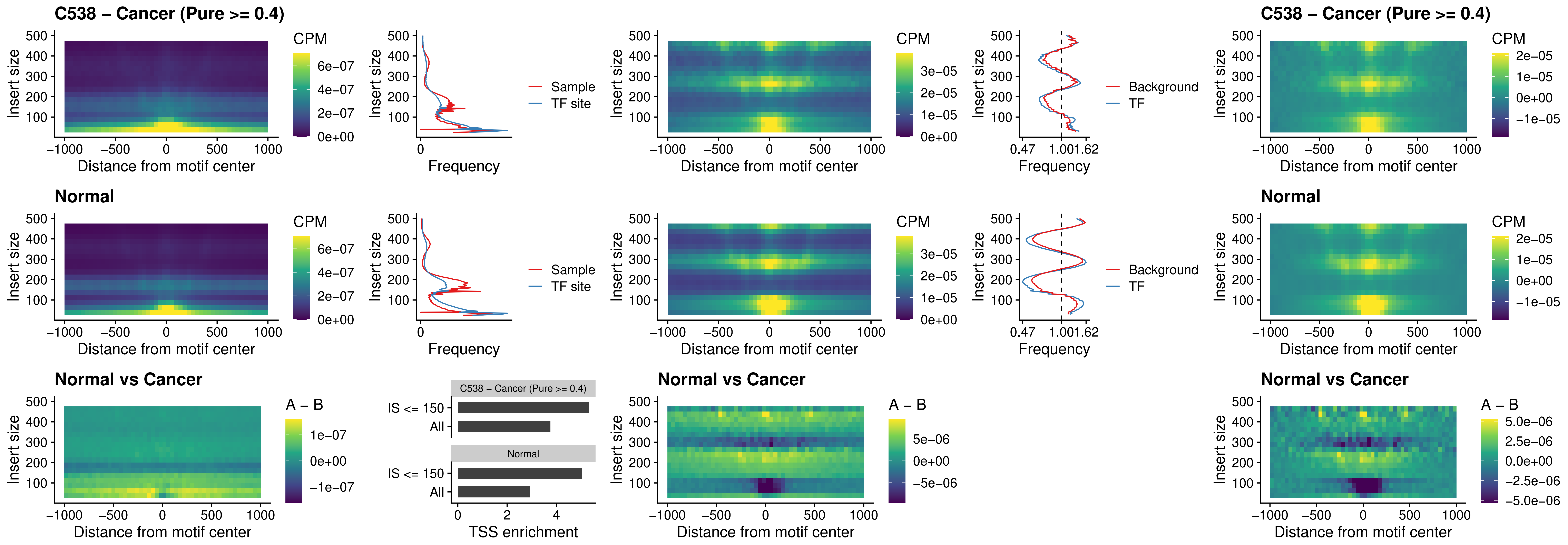

#### C538 – Cancer (Pure >= 0.4) – ENCODE_CTCF_sites_COLON_promotor_ctcf

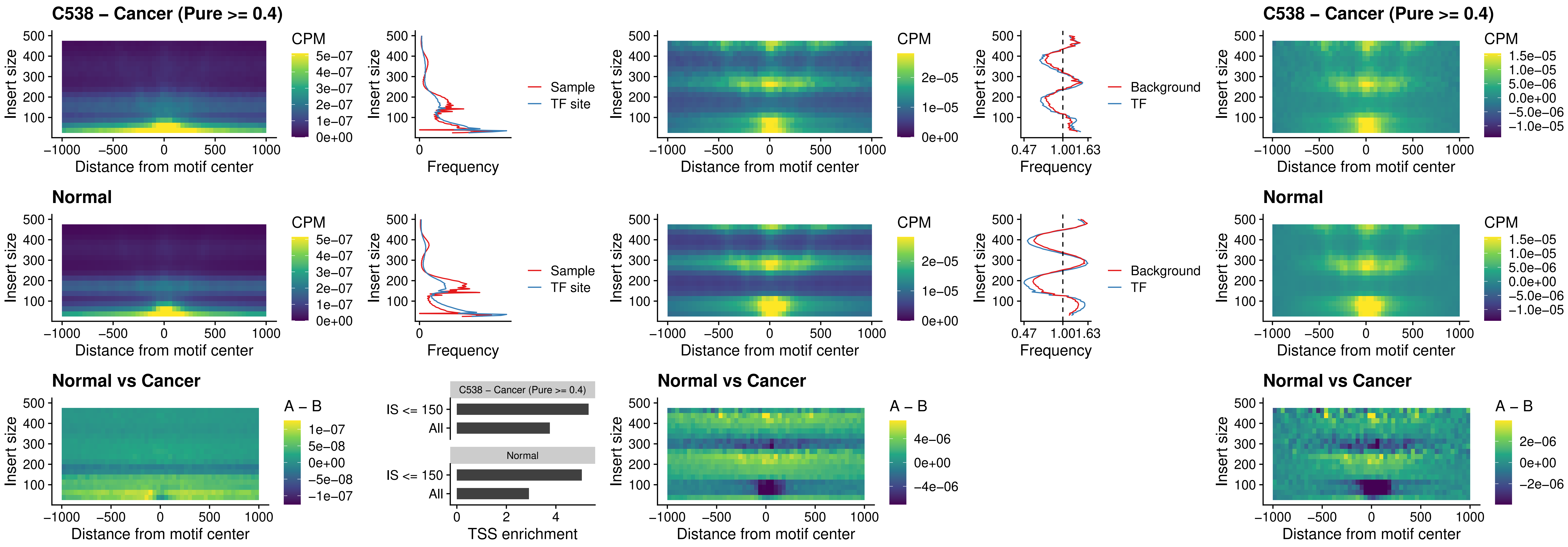

#### C538 – Cancer (Pure >= 0.4) – SPIC_uc_pTSS_oPEAK

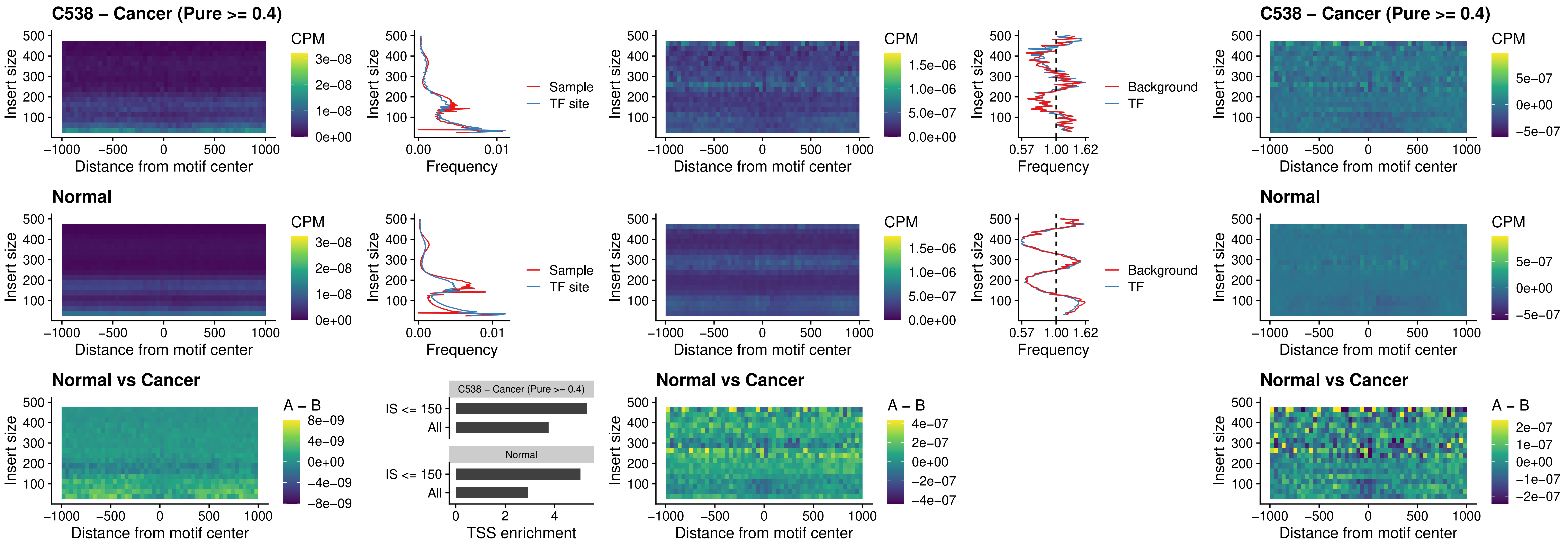

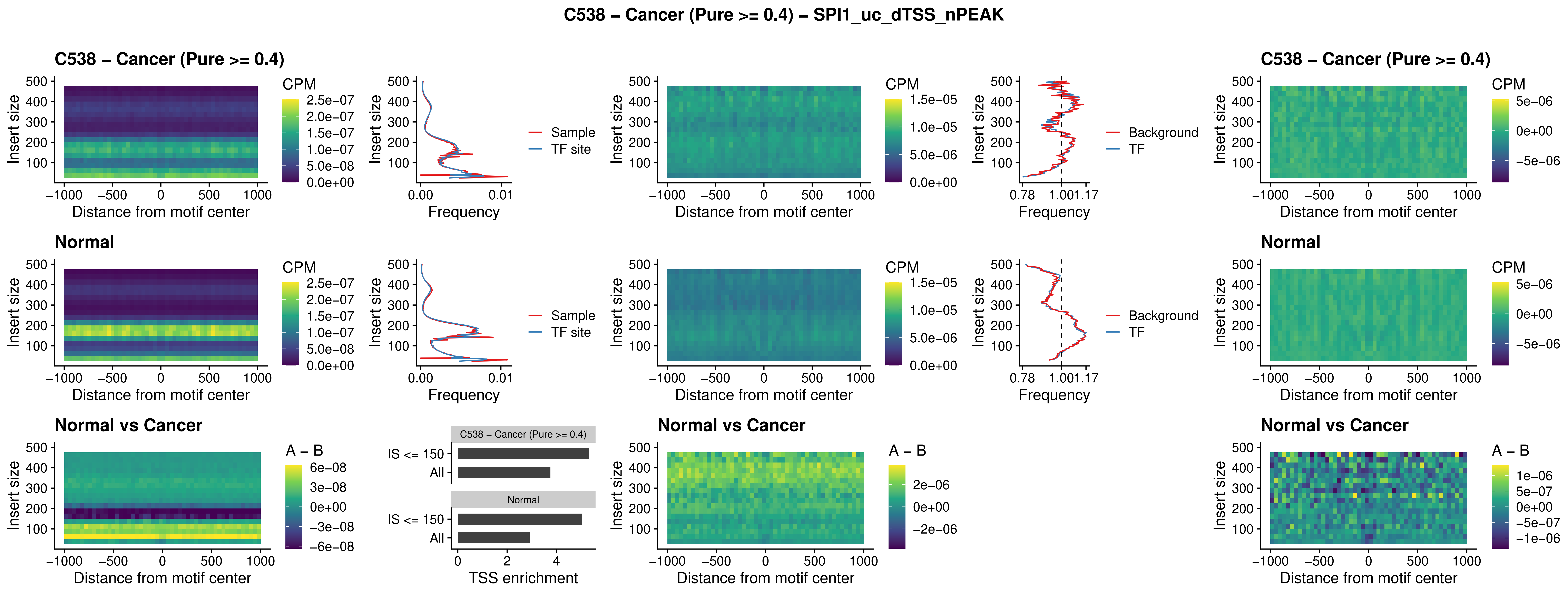
C538 – Cancer (Pure >= 0.4) – SPI1_uc_dTSS_nPEAK
C538 – Cancer (Pure >= 0.4)
Normal
Normal vs Cancer
Insert size
Distance from motif center
Frequency
CPM
A – B
Sample
TF site
Background
TF
C538 – Cancer (Pure >= 0.4)
Normal
IS <= 150
All
TSS enrichment

#### C538 – Cancer (Pure >= 0.4) – SPI1_uc_cTSS_nPEAK

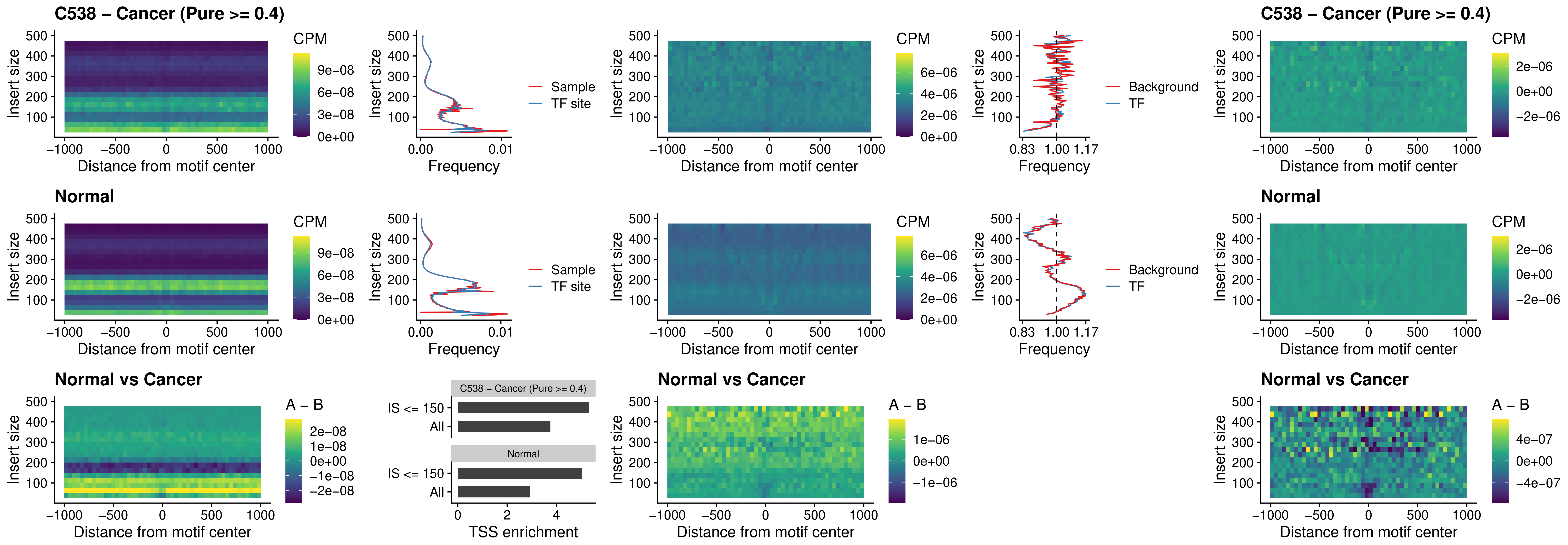

#### C538 – Cancer (Pure >= 0.4) – BCL11A_uc_pTSS_nPEAK

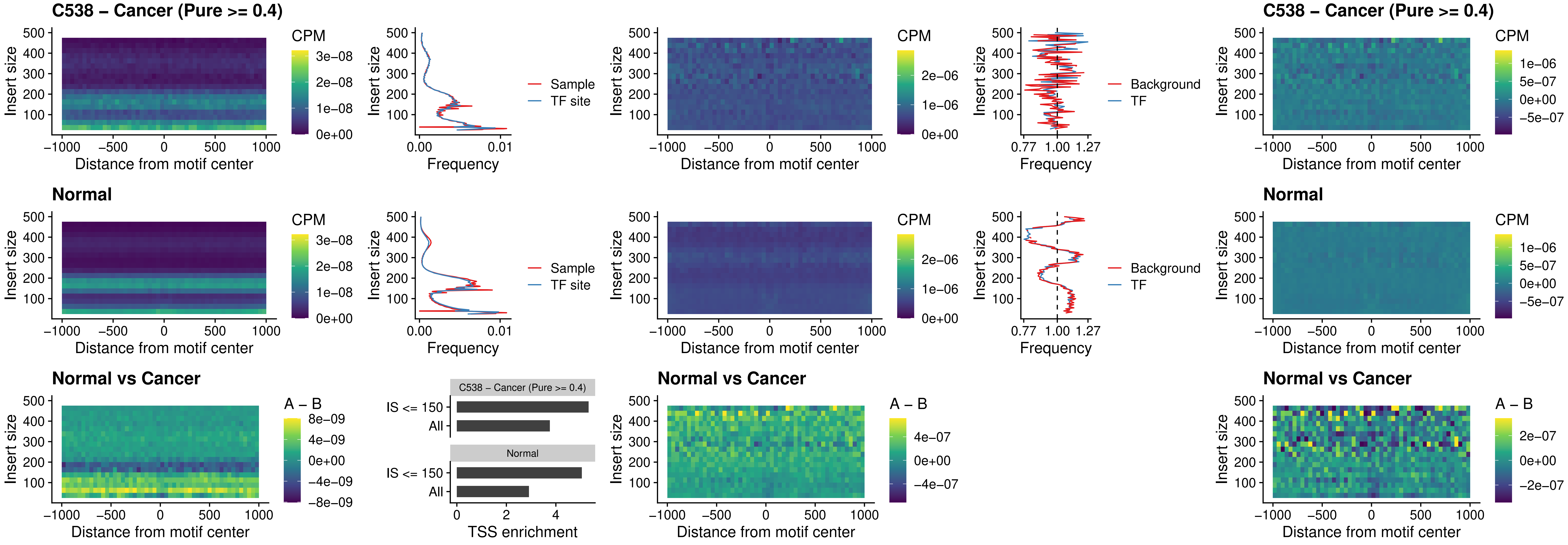

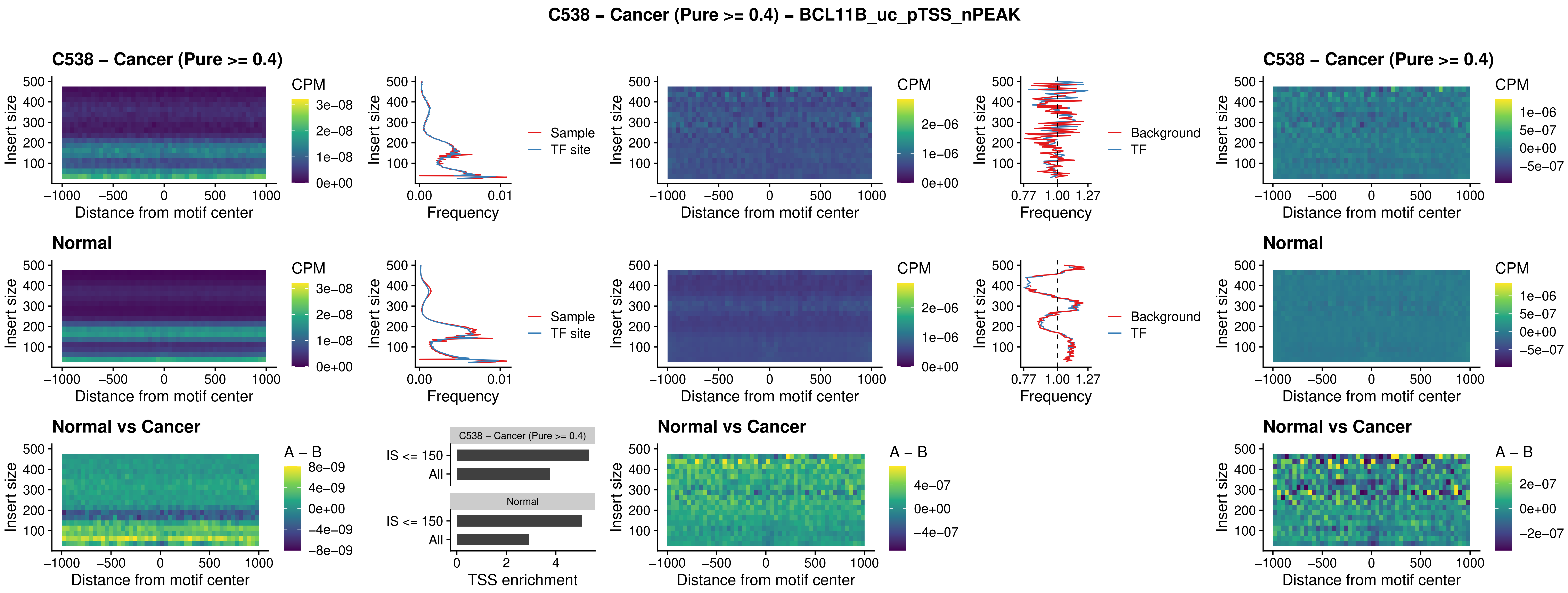

C538 – Cancer (Pure >= 0.4) – BCL11B_uc_pTSS_nPEAK
C538 – Cancer (Pure >= 0.4)
Normal
Normal vs Cancer
Insert size
Distance from motif center
CPM
A – B
Frequency
Sample
TF site
Background
TF
C538 – Cancer (Pure >= 0.4)
Normal
IS <= 150
All
TSS enrichment

#### C538 – Cancer (Pure >= 0.4) – IRF9_uc_dTSS_oPEAK

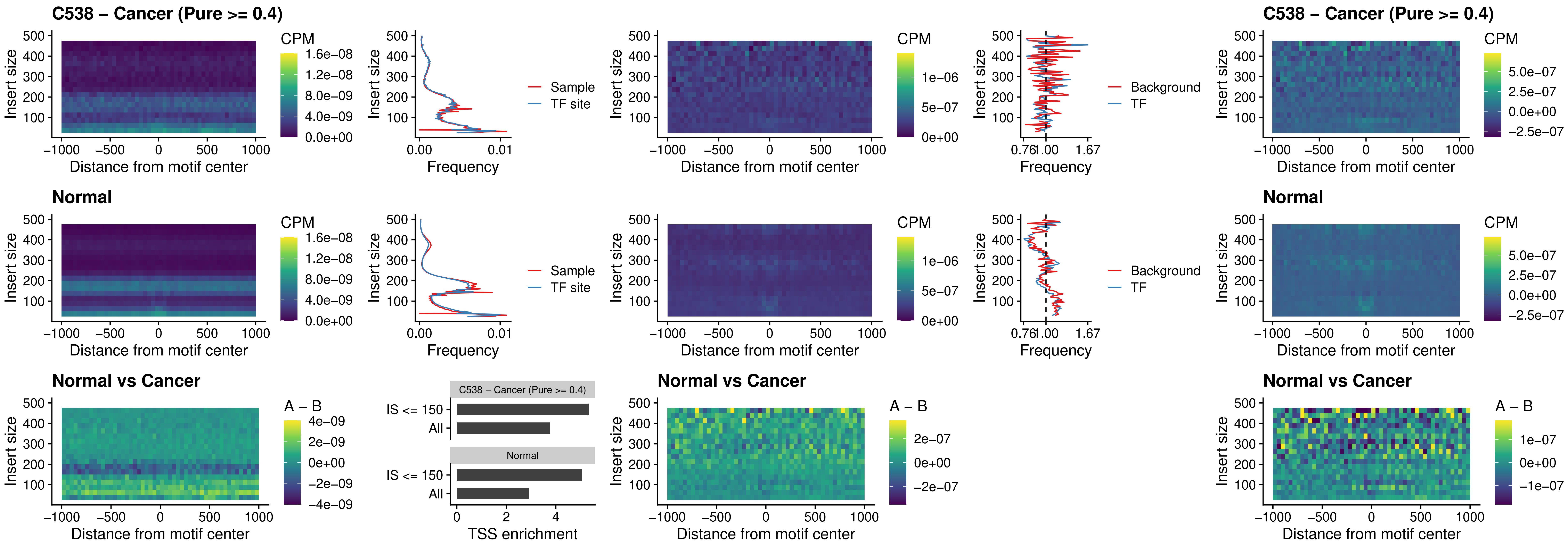

#### C538 – Cancer (Pure >= 0.4) – IRF9_uc_cTSS_oPEAK

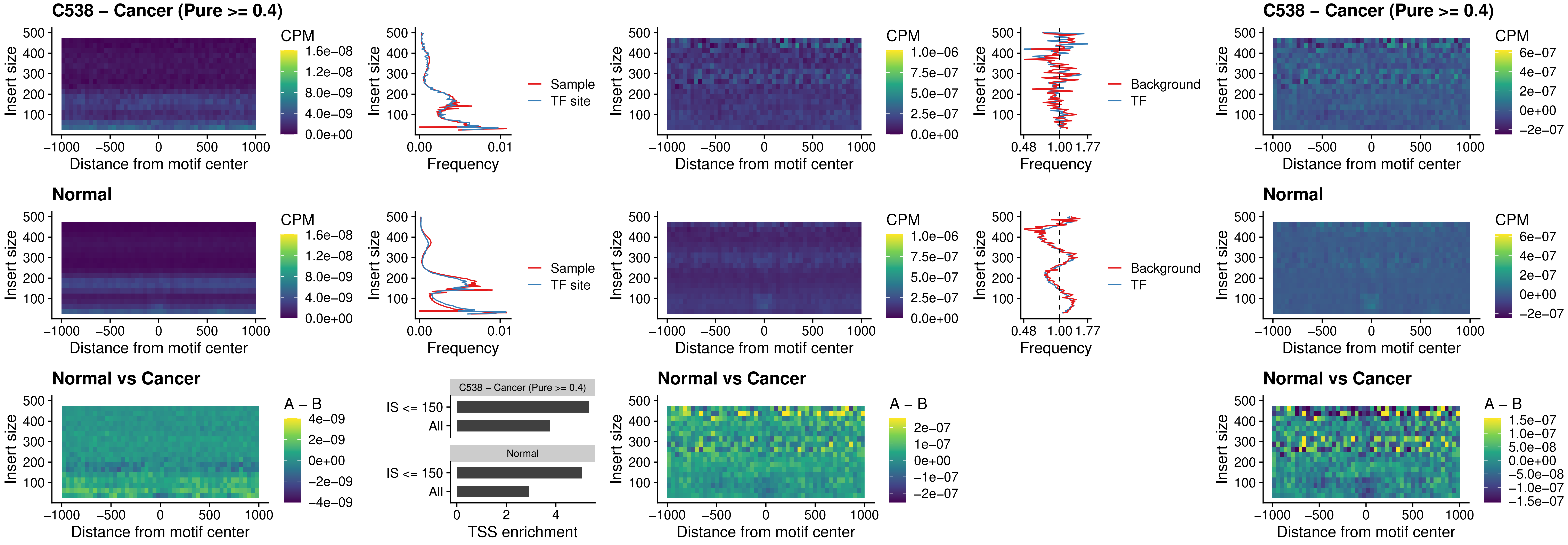

#### C538 – Cancer (Pure >= 0.4) – BCL11A_uc_cTSS_oPEAK

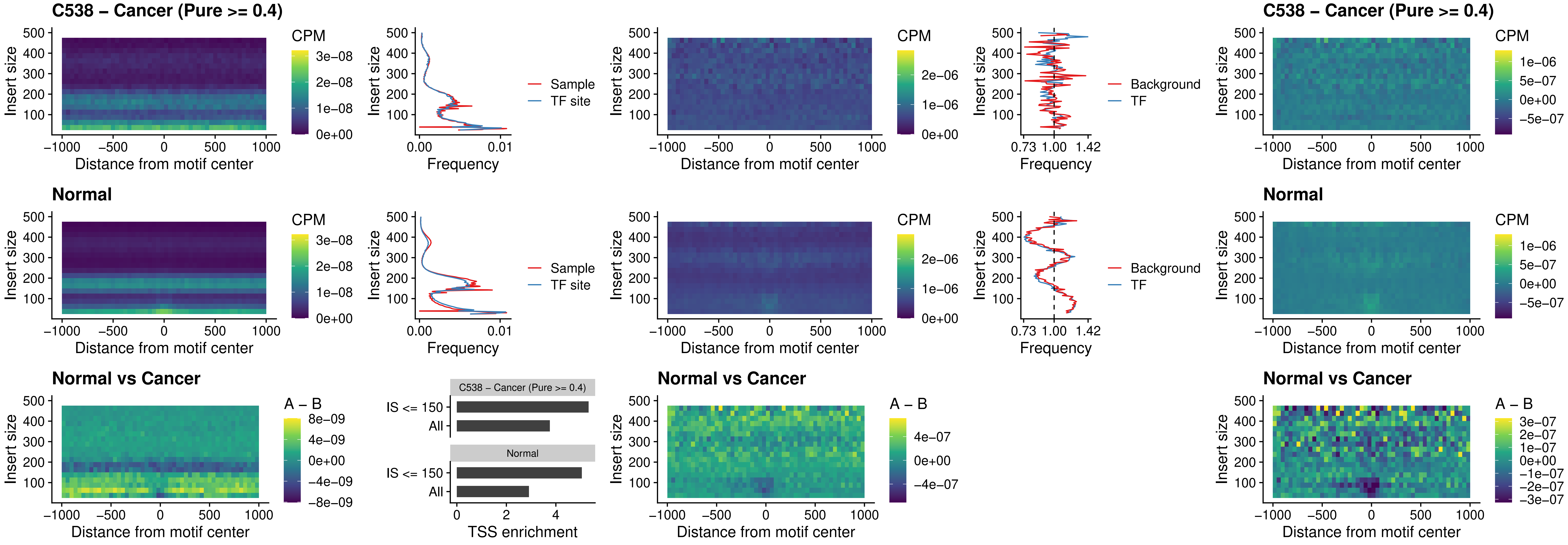

**C538 – Cancer (Pure >= 0.4) – BCL11B_uc_cTSS_oPEAK**

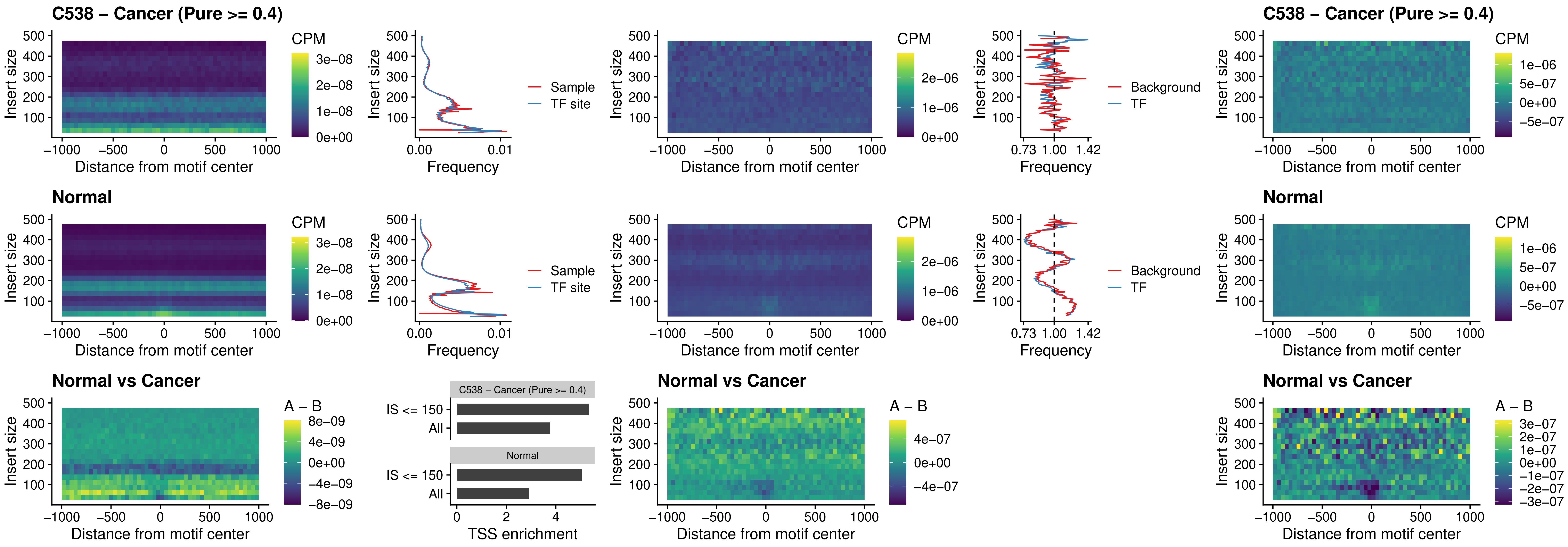

#### C538 – Cancer (Pure >= 0.4) – BCL11A_uc_dTSS_oPEAK

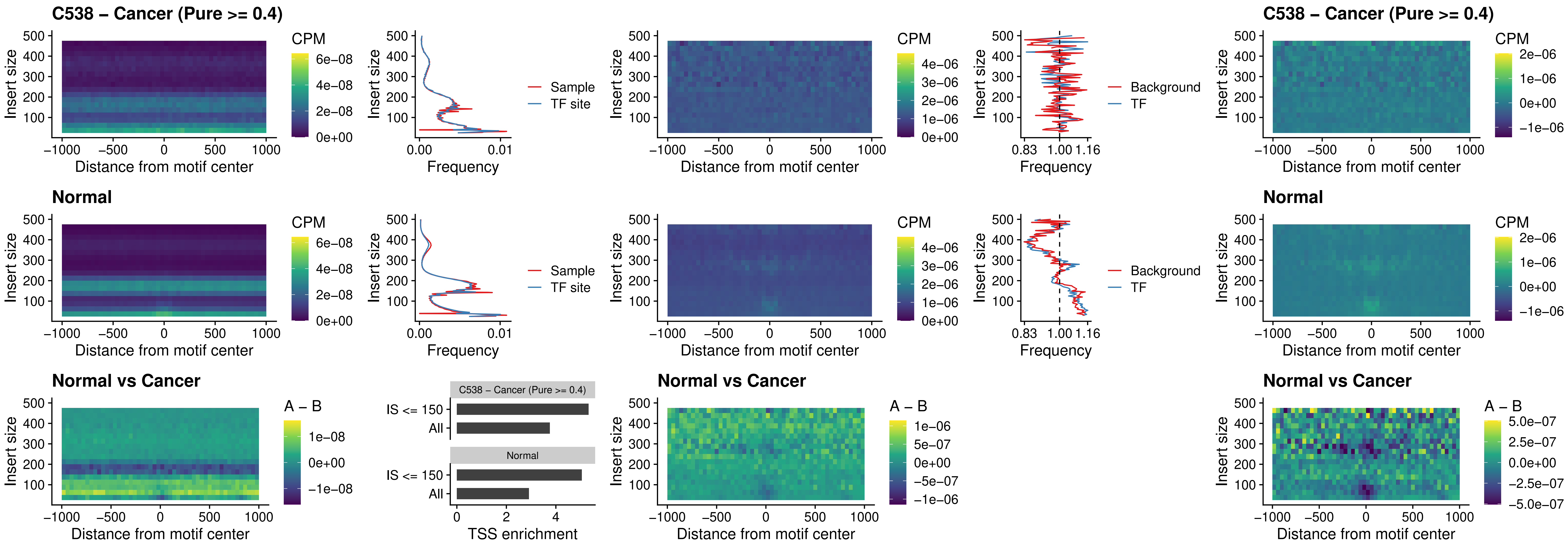

#### C538 – Cancer (Pure >= 0.4) – BCL11B_uc_dTSS_oPEAK

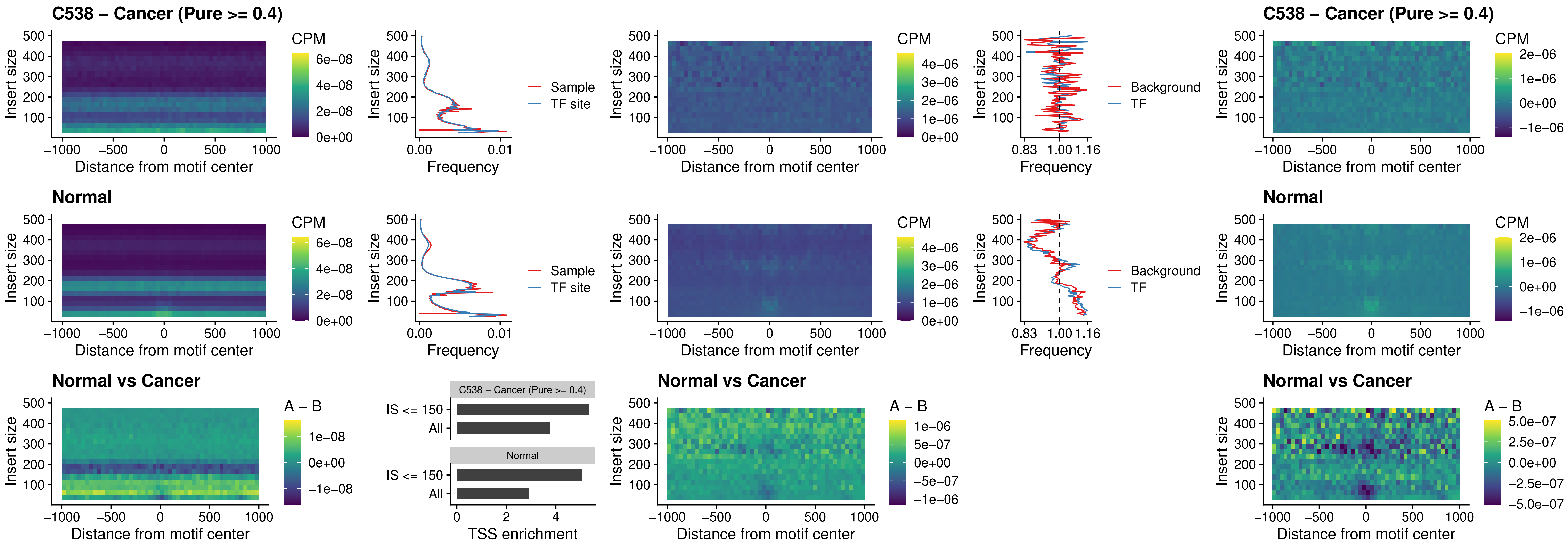

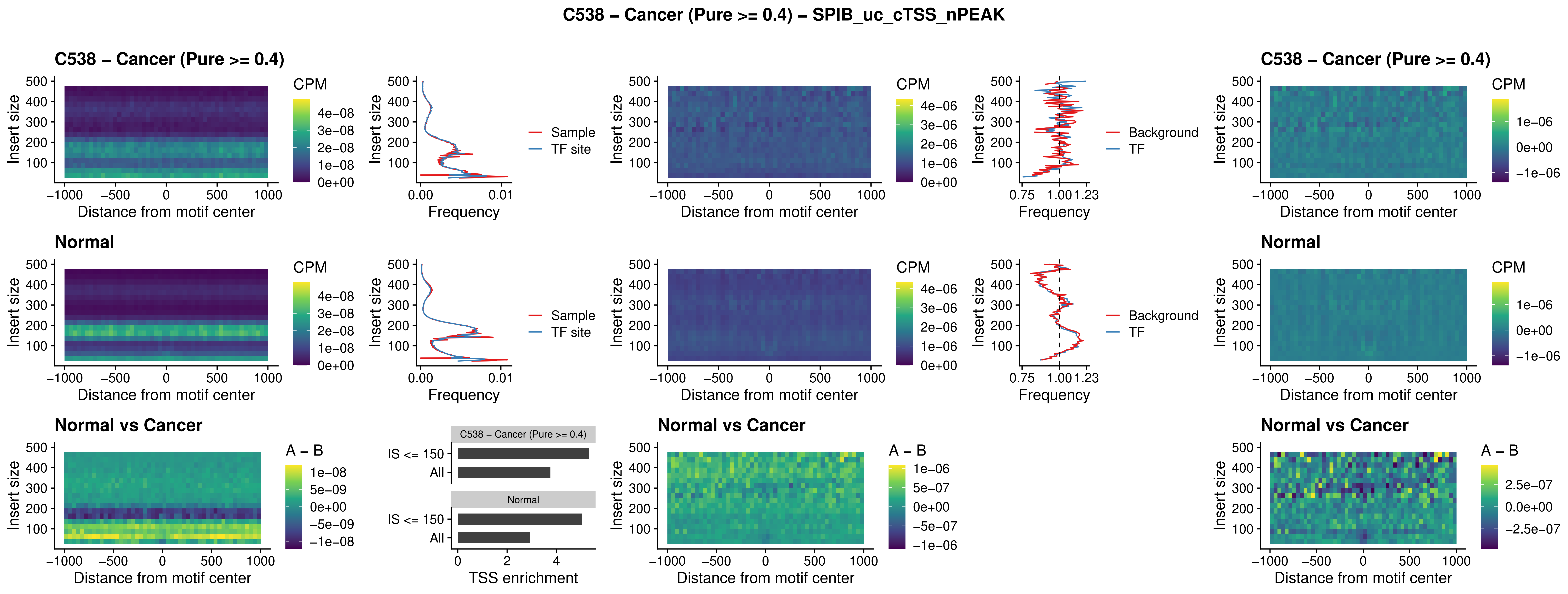
C538 – Cancer (Pure >= 0.4) – SPIB_uc_cTSS_nPEAK
C538 – Cancer (Pure >= 0.4)
Normal
Normal vs Cancer
Insert size
Distance from motif center
CPM
A – B
Frequency
Sample
TF site
Background
TF
C538 – Cancer (Pure >= 0.4)
IS <= 150
All
TSS enrichment

#### C538 – Cancer (Pure >= 0.4) – SPIC_uc_cTSS_nPEAK

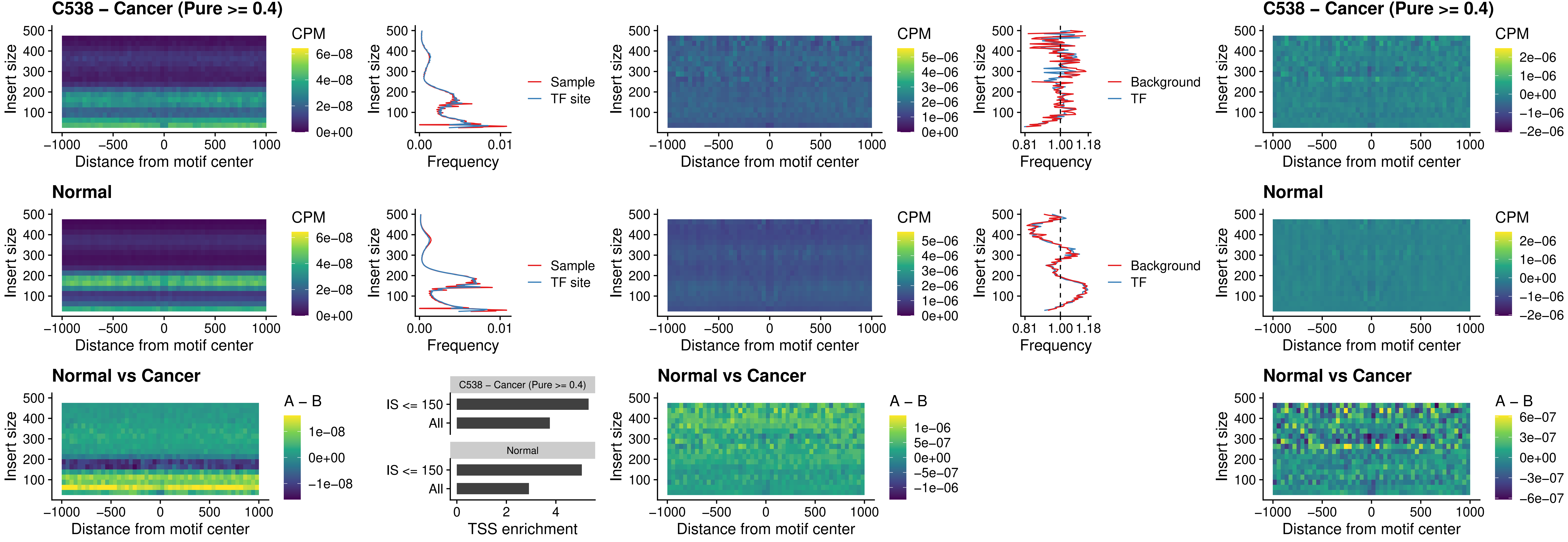

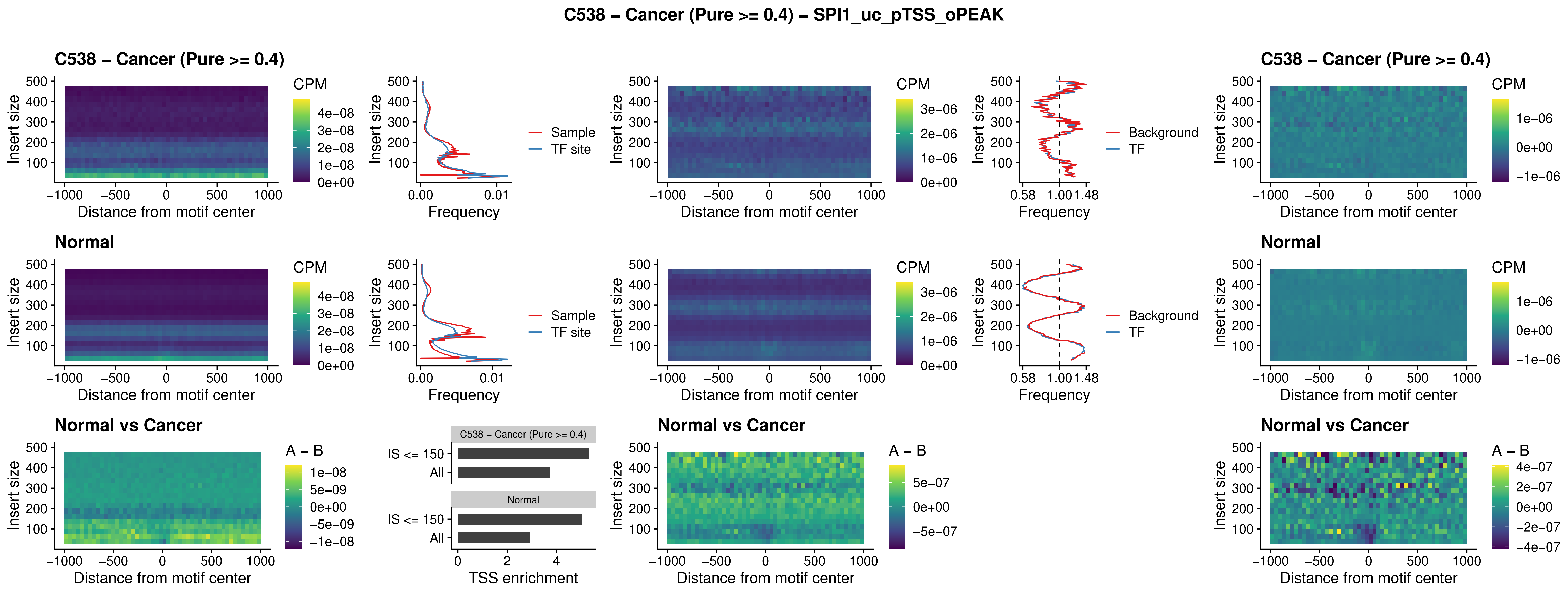
C538 – Cancer (Pure >= 0.4) – SPI1_uc_pTSS_oPEAK
C538 – Cancer (Pure >= 0.4)
Normal
Normal vs Cancer
Insert size
Distance from motif center
Frequency
CPM
A – B
Sample
TF site
Background
TF
C538 – Cancer (Pure >= 0.4)
Normal
IS <= 150
All
TSS enrichment

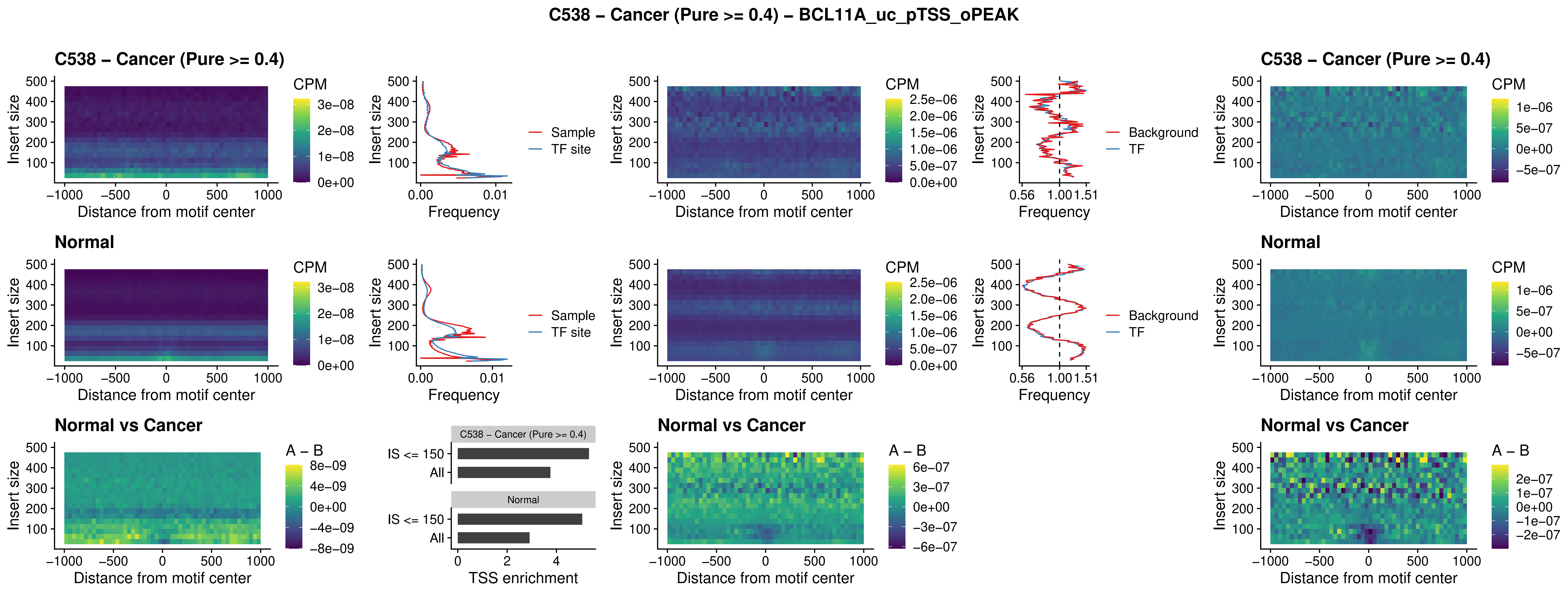
C538 – Cancer (Pure >= 0.4) – BCL11A_uc_pTSS_oPEAK
C538 – Cancer (Pure >= 0.4)
Normal
Normal vs Cancer
Insert size
Distance from motif center
Frequency
CPM
A – B
Sample
TF site
Background
TF
C538 – Cancer (Pure >= 0.4)
Normal
IS <= 150
All
TSS enrichment

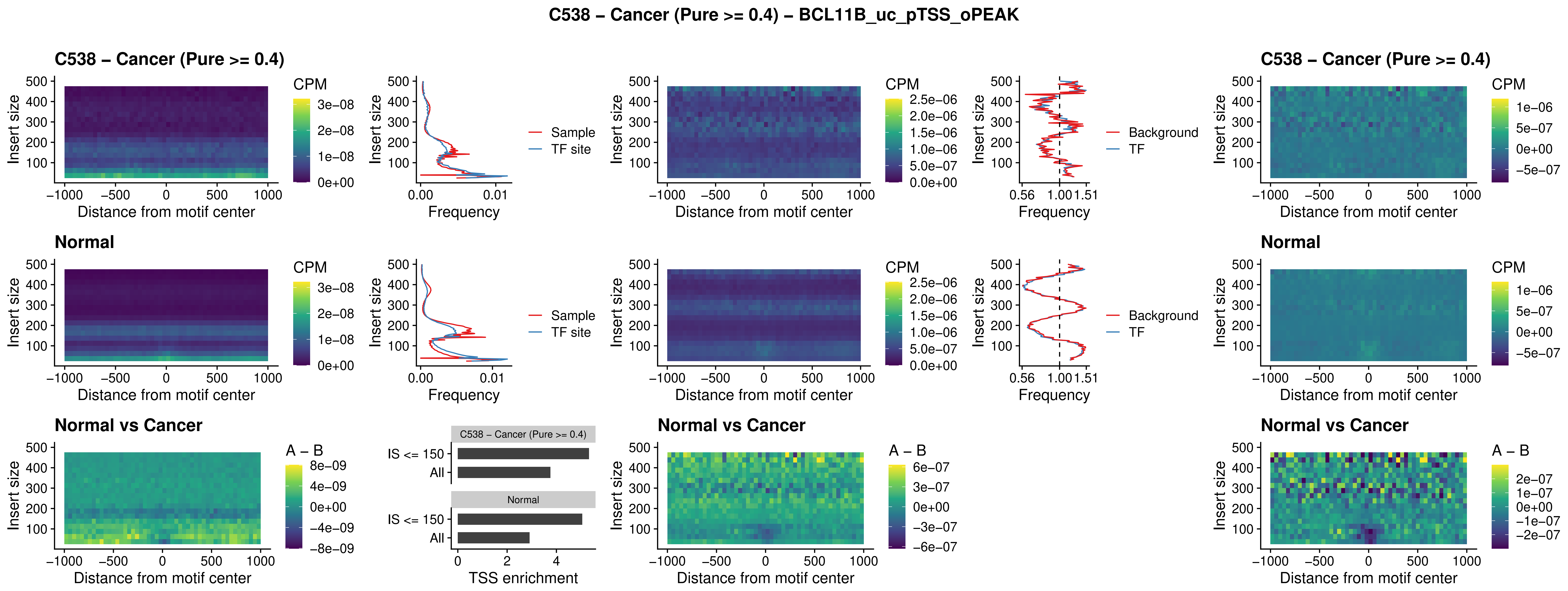
C538 – Cancer (Pure >= 0.4) – BCL11B_uc_pTSS_oPEAK
C538 – Cancer (Pure >= 0.4)
Normal
Normal vs Cancer
Insert size
Distance from motif center
Frequency
CPM
A – B
Sample
TF site
Background
TF
C538 – Cancer (Pure >= 0.4)
Normal
IS <= 150
All
TSS enrichment

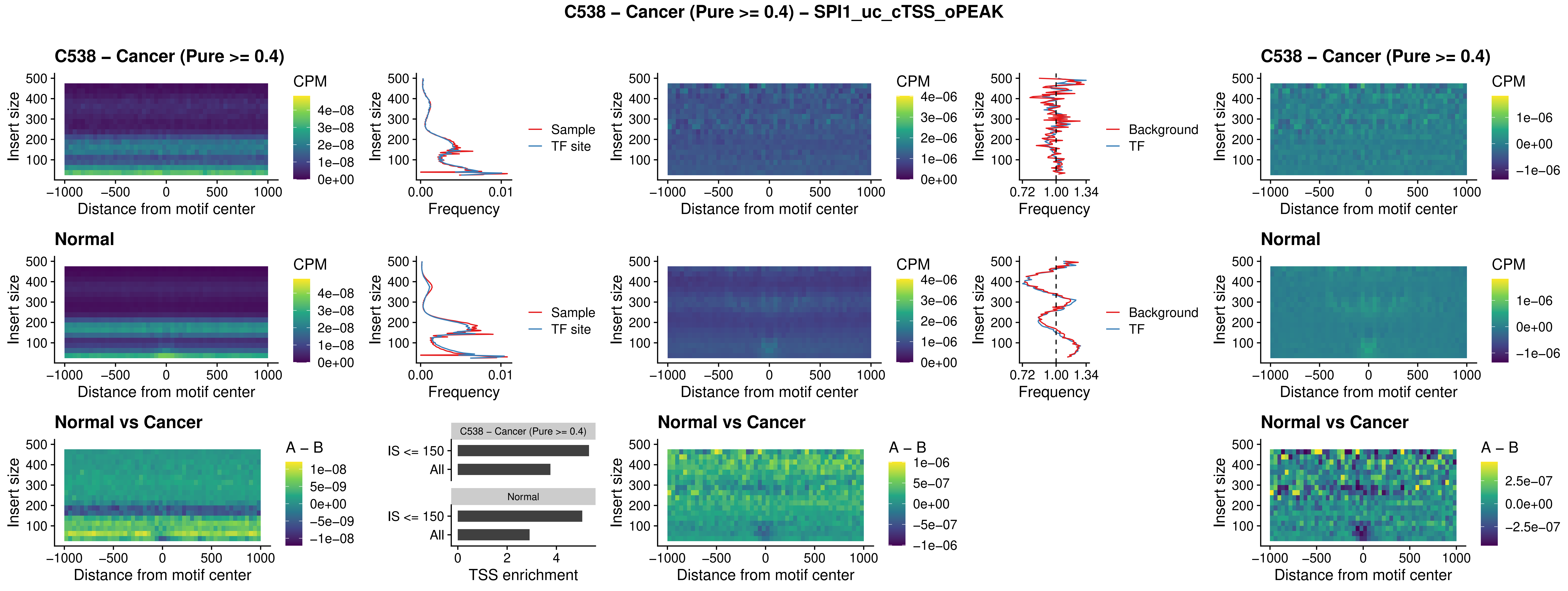
C538 – Cancer (Pure >= 0.4) – SPI1_uc_cTSS_oPEAK
C538 – Cancer (Pure >= 0.4)
Normal
Normal vs Cancer
Insert size
Distance from motif center
Frequency
CPM
A – B
Sample
TF site
Background
TF
C538 – Cancer (Pure >= 0.4)
Normal
IS <= 150
All
TSS enrichment

#### C538 – Cancer (Pure >= 0.4) – SPIB_uc_cTSS_oPEAK

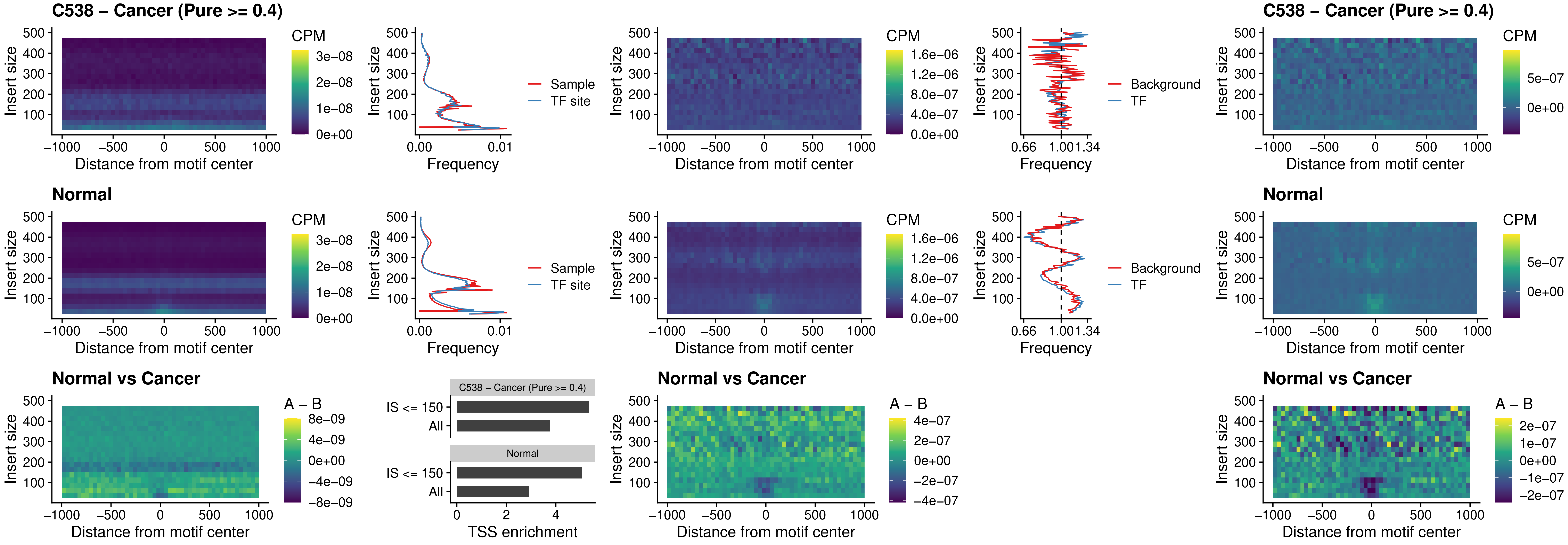

### C538 – Cancer (Pure >= 0.4) – SPIB_uc_pTSS_oPEAK

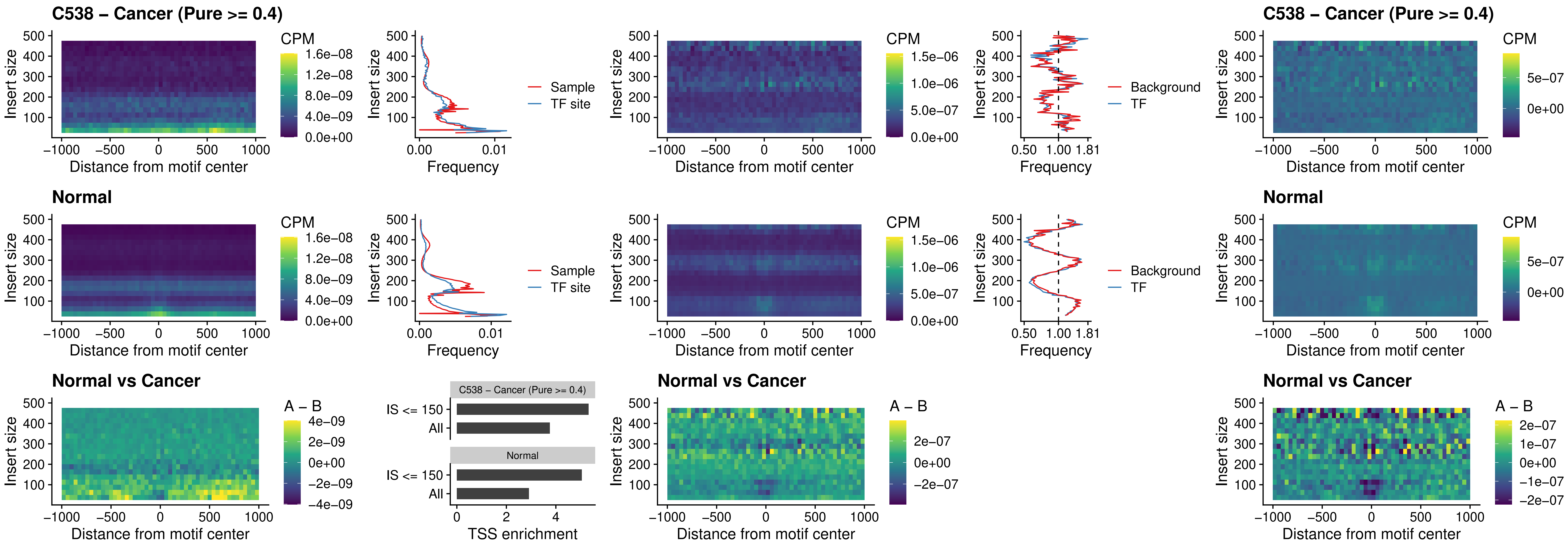

### C538 – Cancer (Pure >= 0.4) – SPI1_uc_dTSS_oPEAK

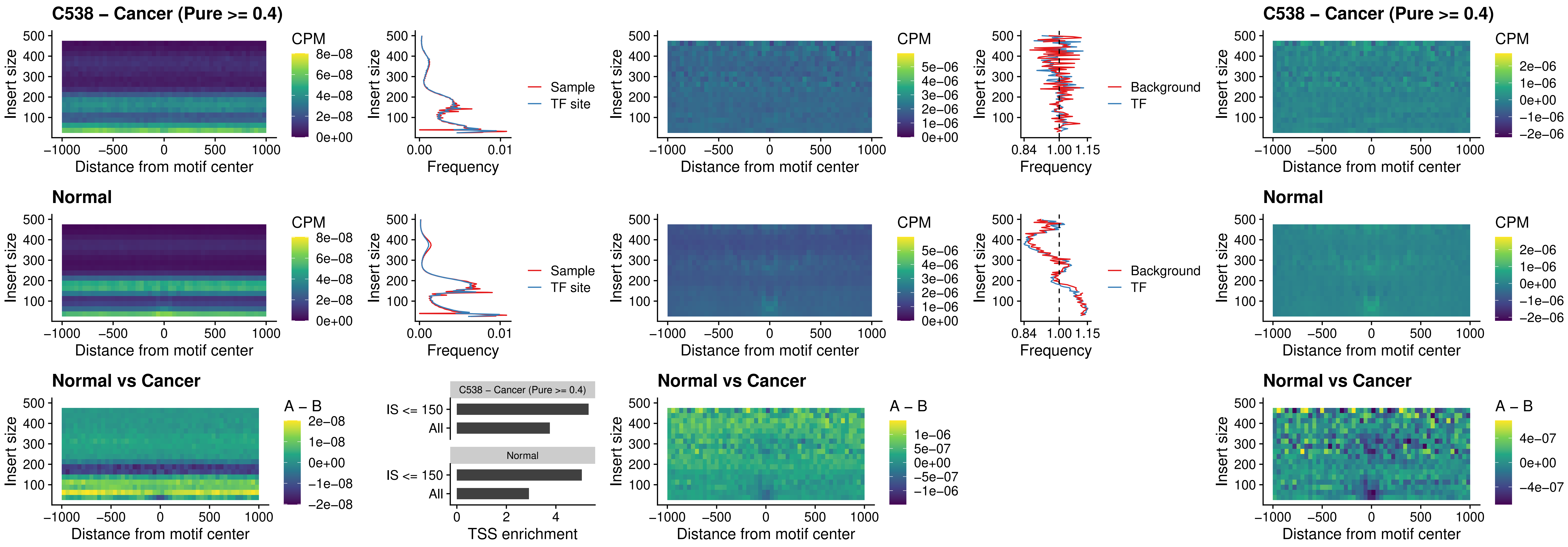

#### C538 – Cancer (Pure >= 0.4) – SPIB_uc_dTSS_oPEAK

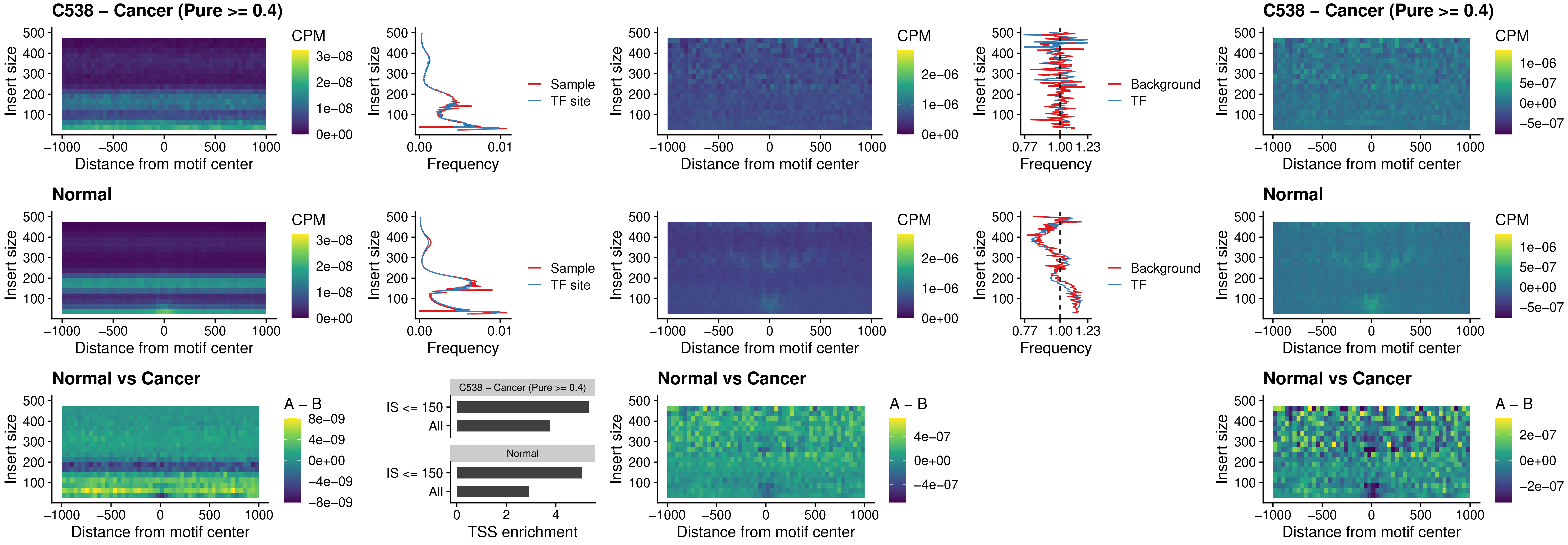

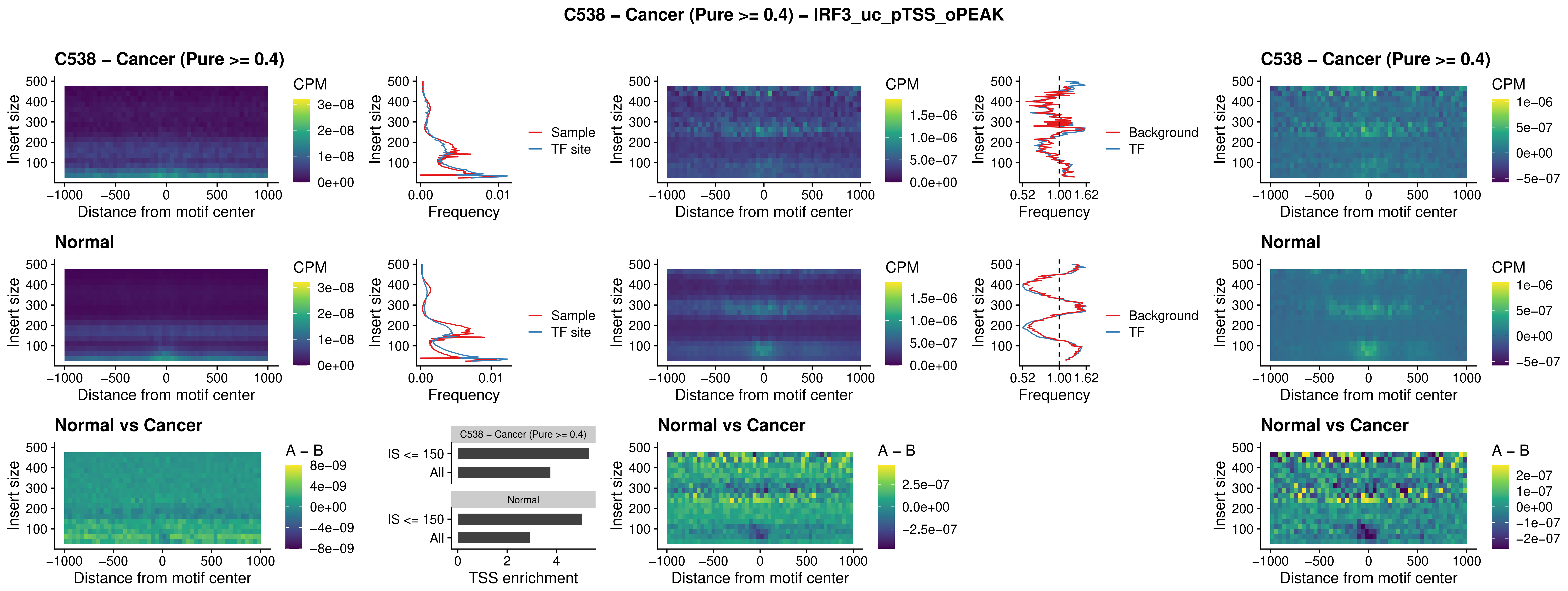

C538 – Cancer (Pure >= 0.4) – IRF3_uc_pTSS_oPEAK
C538 – Cancer (Pure >= 0.4)
Normal
Normal vs Cancer
Insert size
Distance from motif center
Frequency
CPM
A – B
Sample
TF site
Background
TF
C538 – Cancer (Pure >= 0.4)
Normal
IS <= 150
All
TSS enrichment

### C538 – Cancer (Pure >= 0.4) – IRF2_uc_dTSS_oPEAK

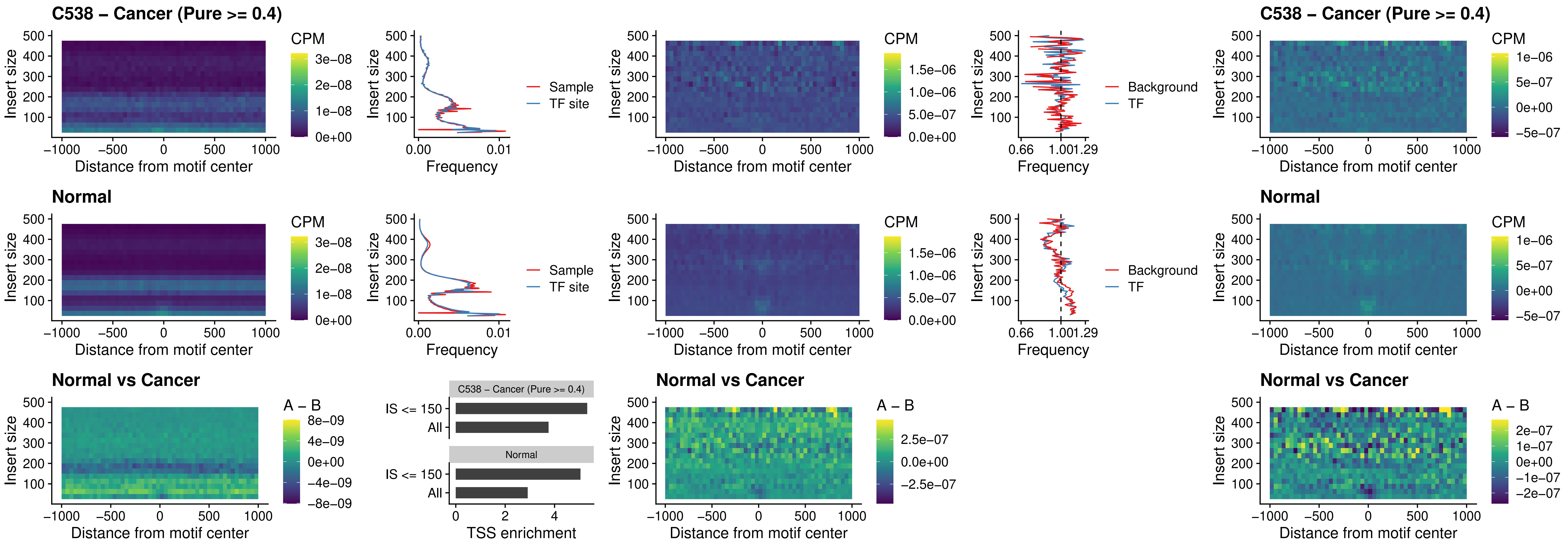

### C538 – Cancer (Pure >= 0.4) – IRF2_uc_pTSS_oPEAK

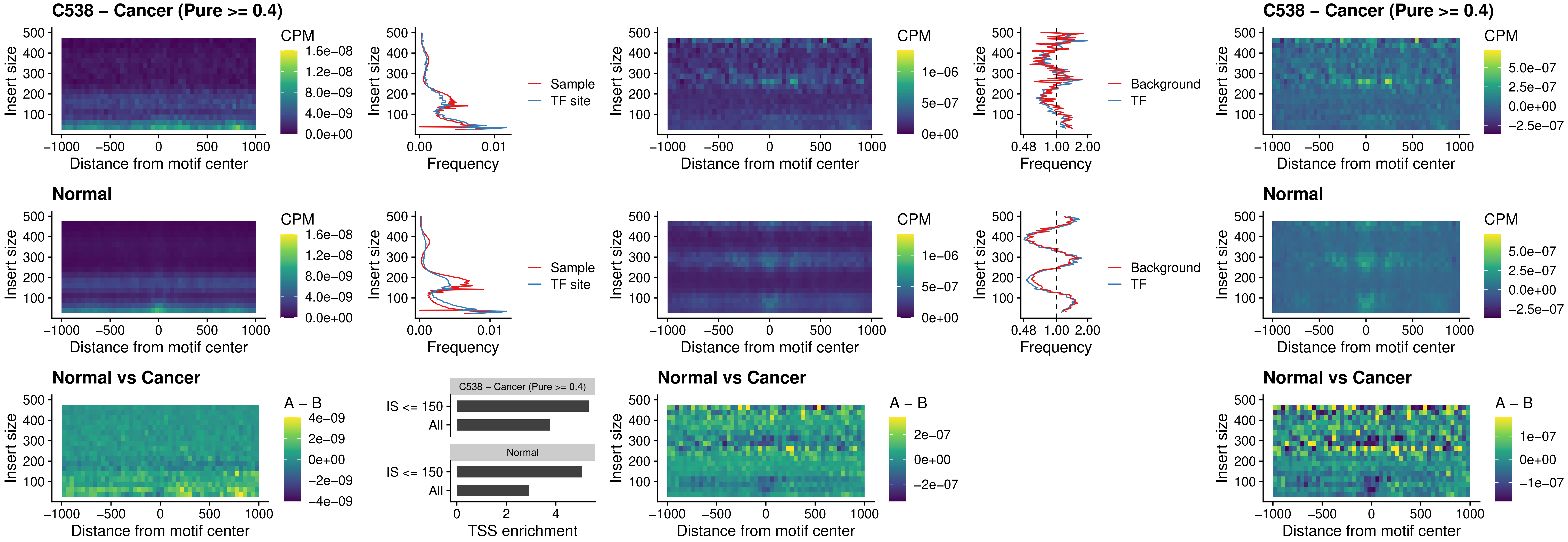

#### C538 – Cancer (Pure >= 0.4) – STAT2_uc_pTSS_oPEAK

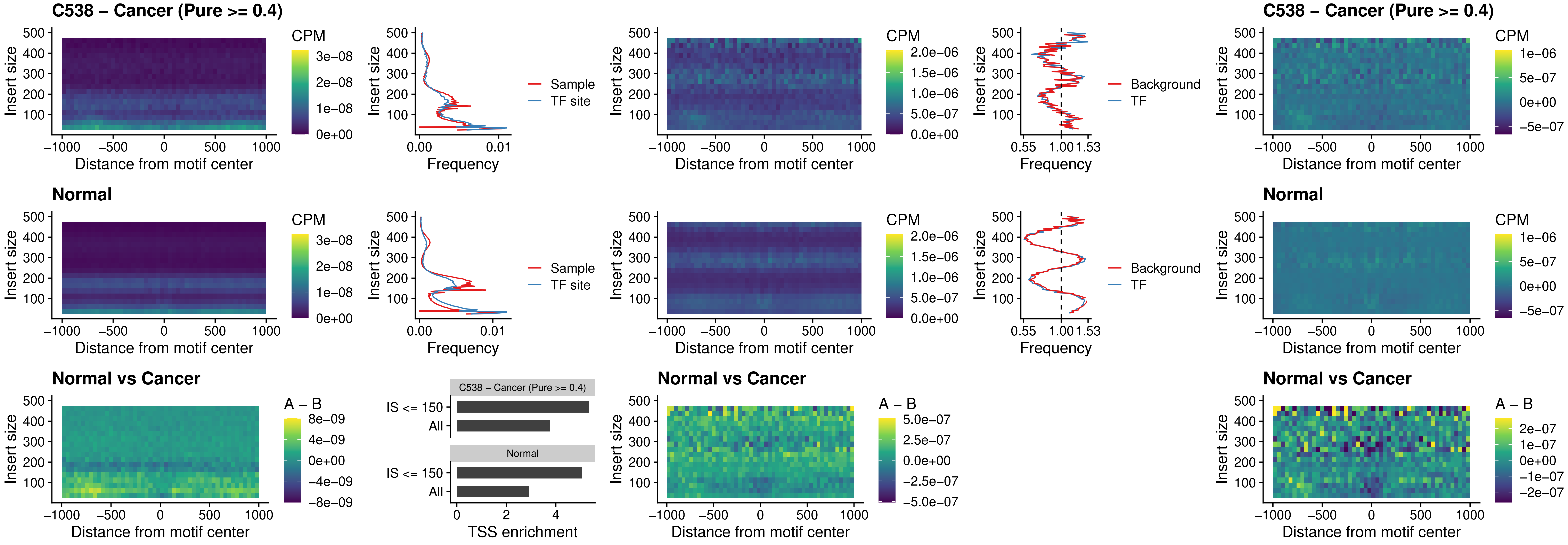

#### C538 – Cancer (Pure >= 0.4) – SPIB_uc_dTSS_nPEAK

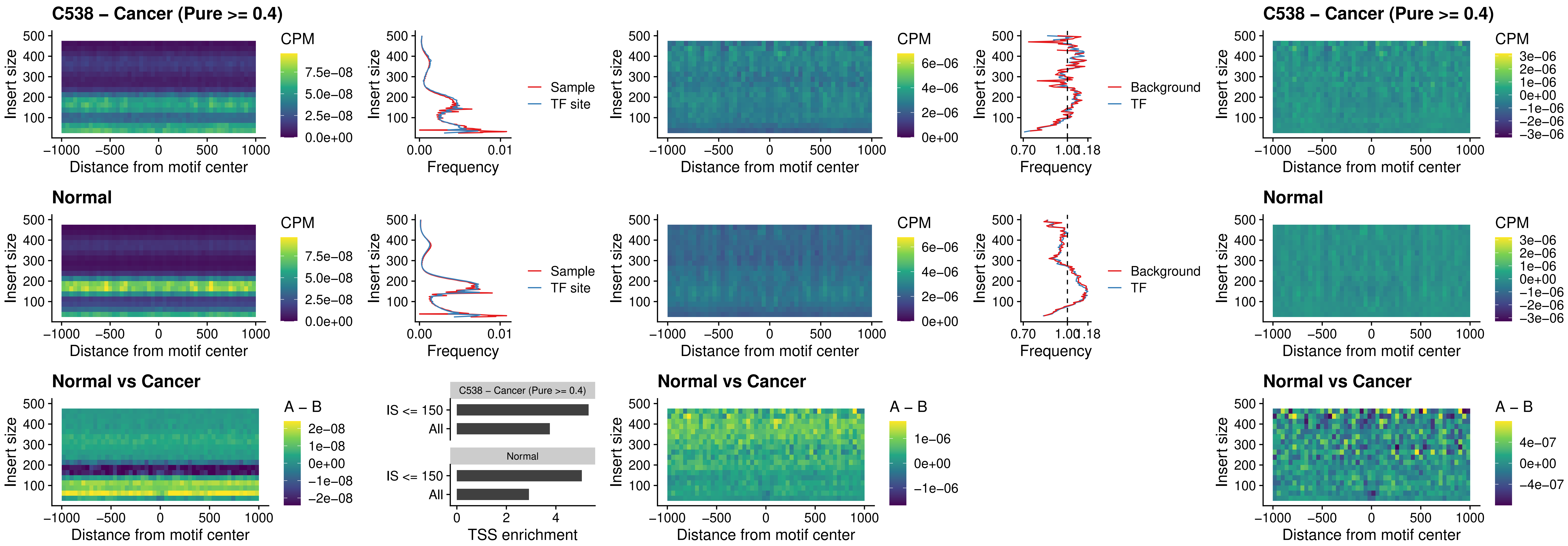

#### C538 – Cancer (Pure >= 0.4) – SPIC_uc_cTSS_oPEAK

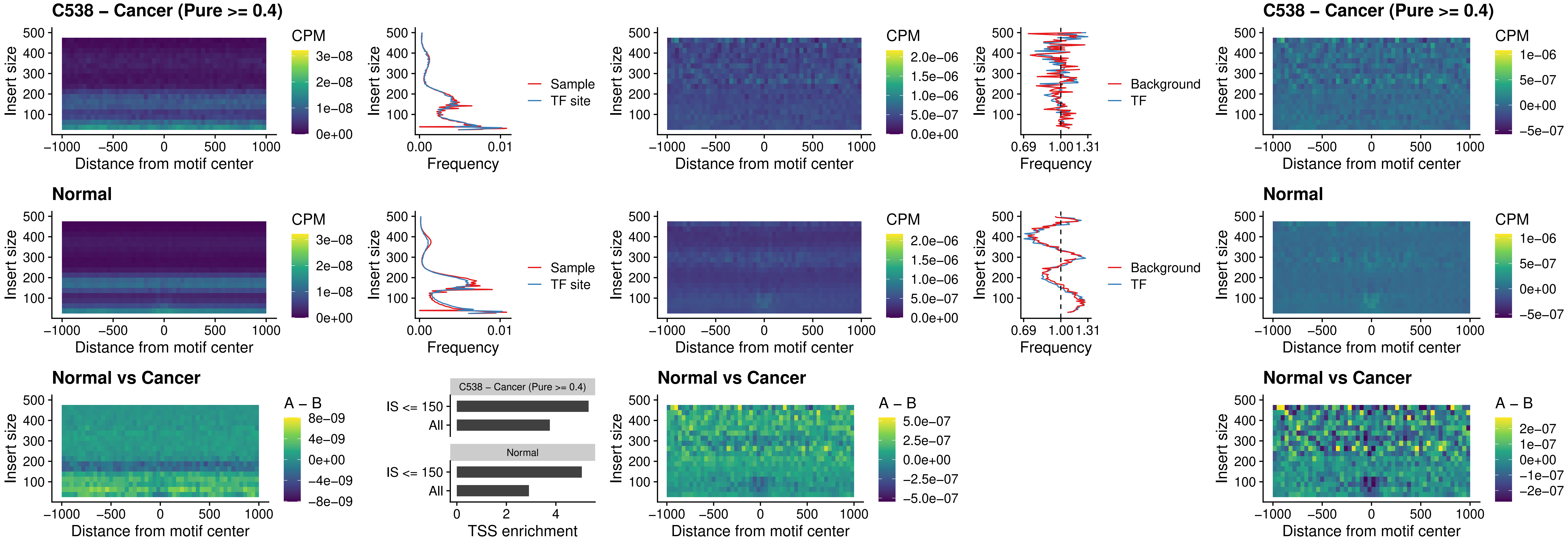

#### C538 – Cancer (Pure >= 0.4) – PRDM1_uc_cTSS_oPEAK

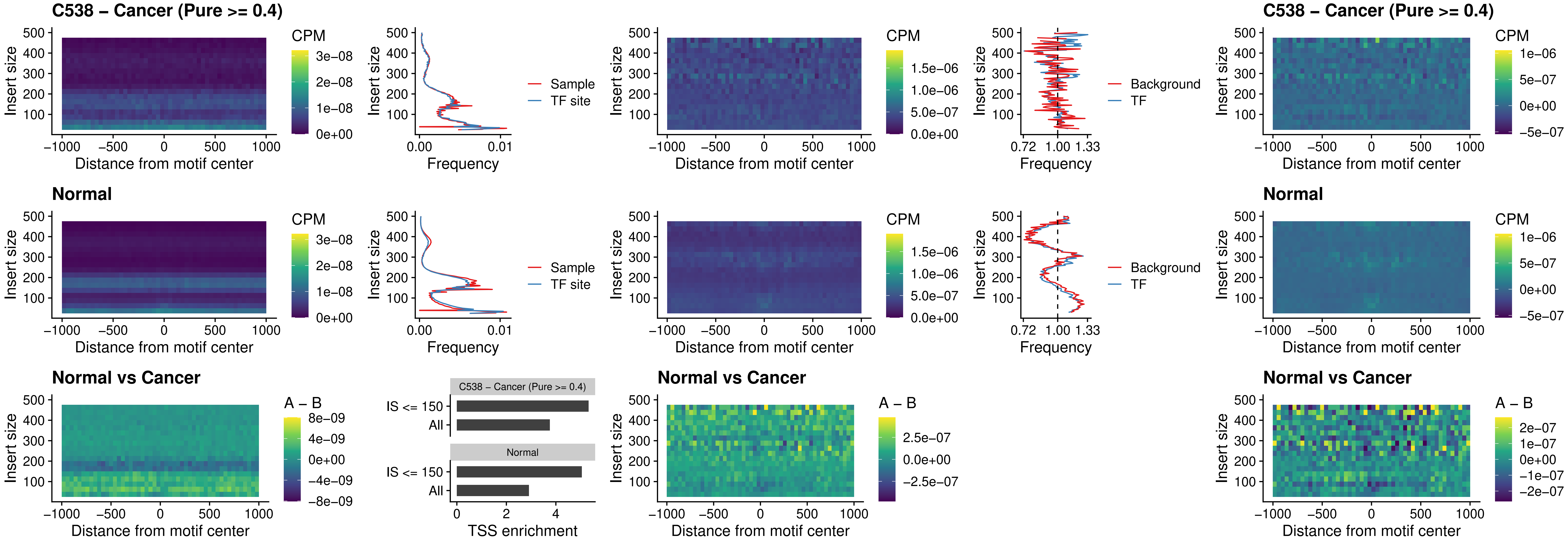

#### C538 – Cancer (Pure >= 0.4) – IRF3_uc_cTSS_oPEAK

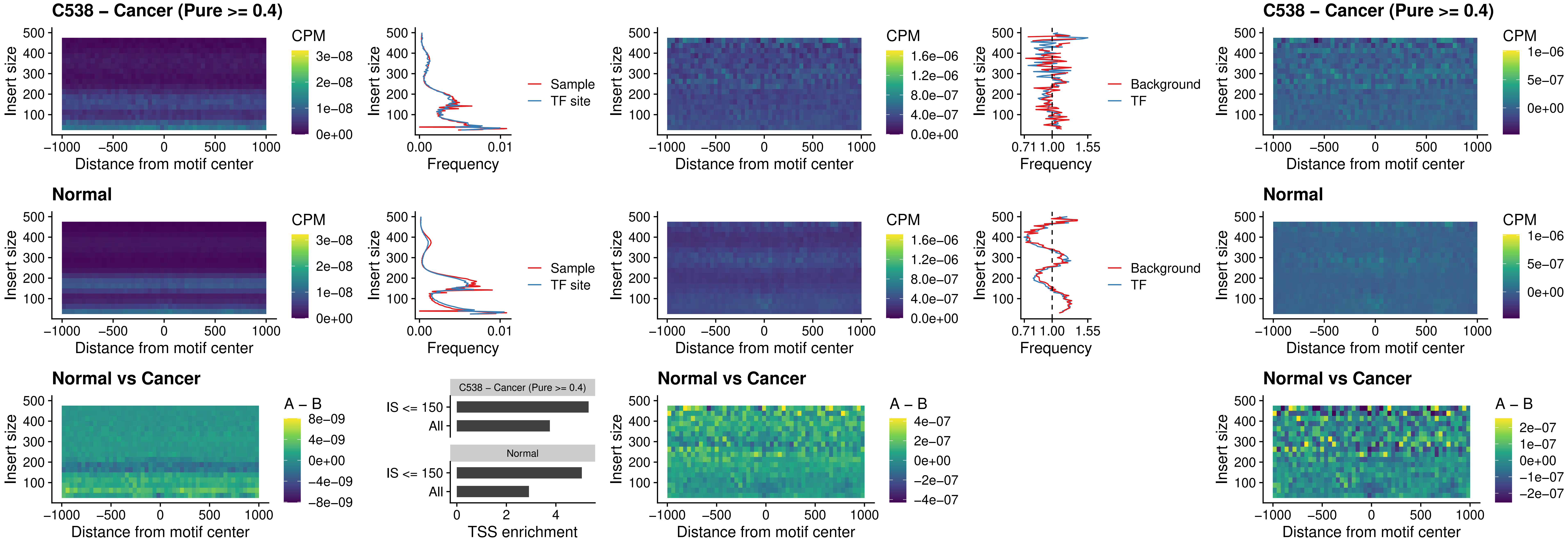

#### C538 – Cancer (Pure >= 0.4) – BCL11A_uc_dTSS_nPEAK

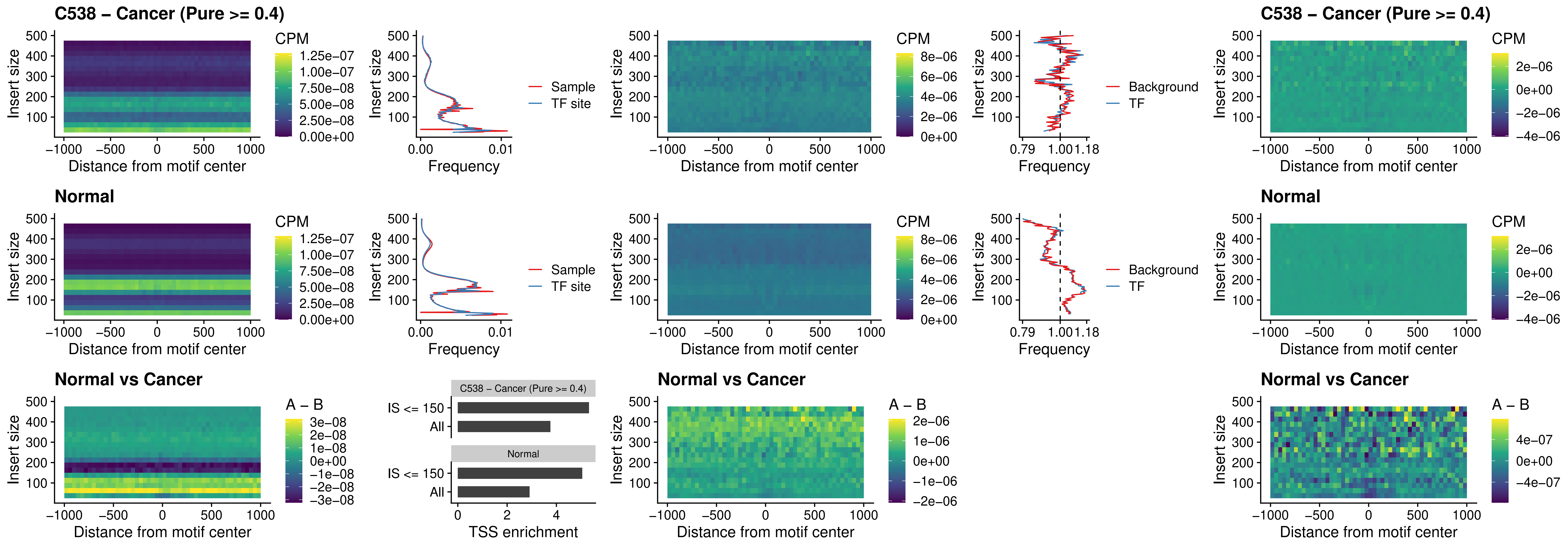

#### C538 – Cancer (Pure >= 0.4) – BCL11B_uc_dTSS_nPEAK

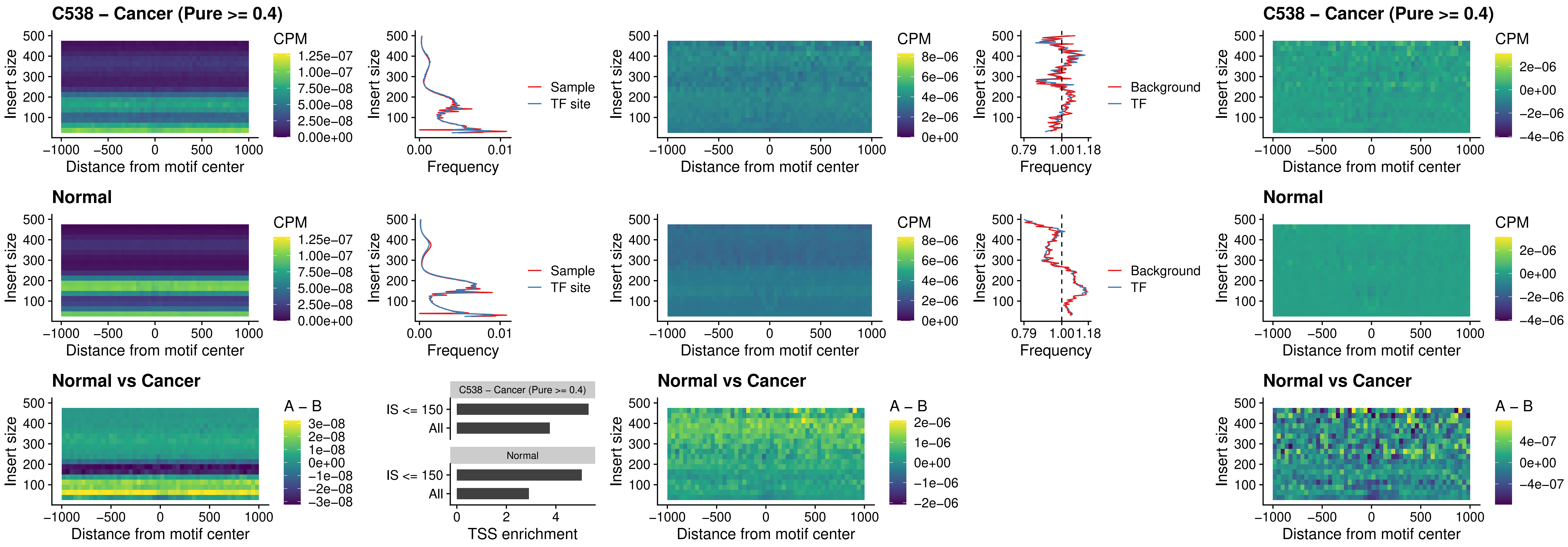

#### C538 – Cancer (Pure >= 0.4) – IRF1_uc_cTSS_oPEAK

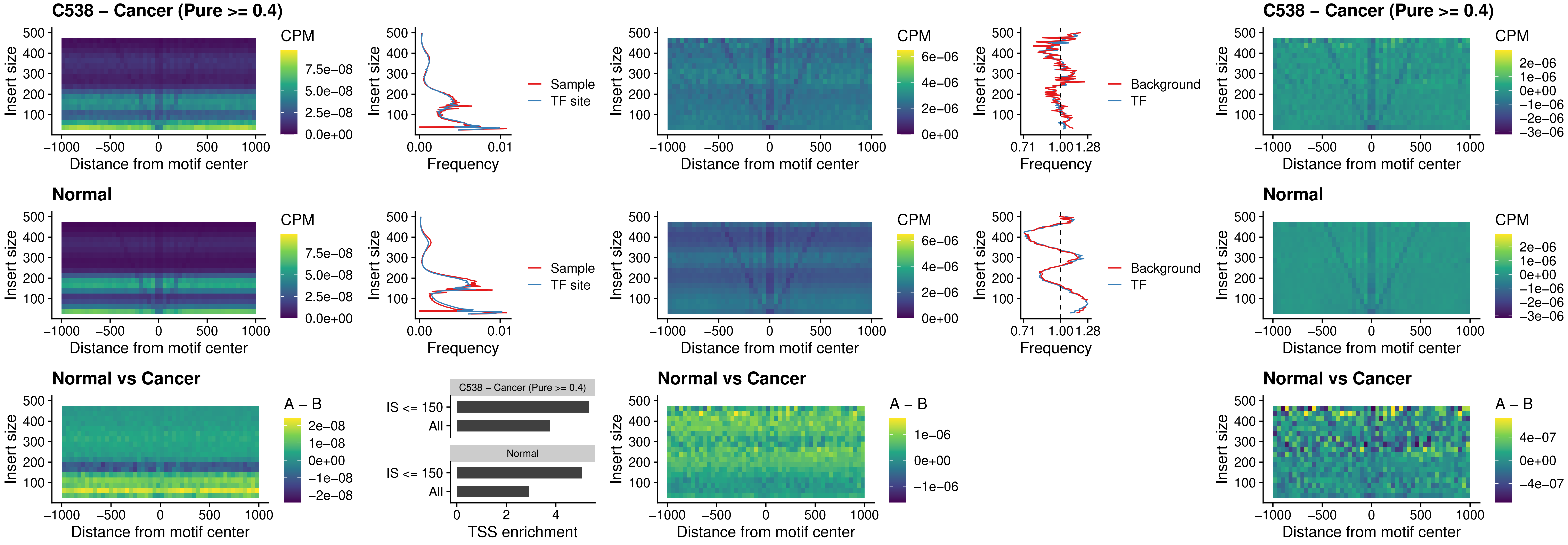

#### C538 – Cancer (Pure >= 0.4) – IRF1_uc_dTSS_oPEAK

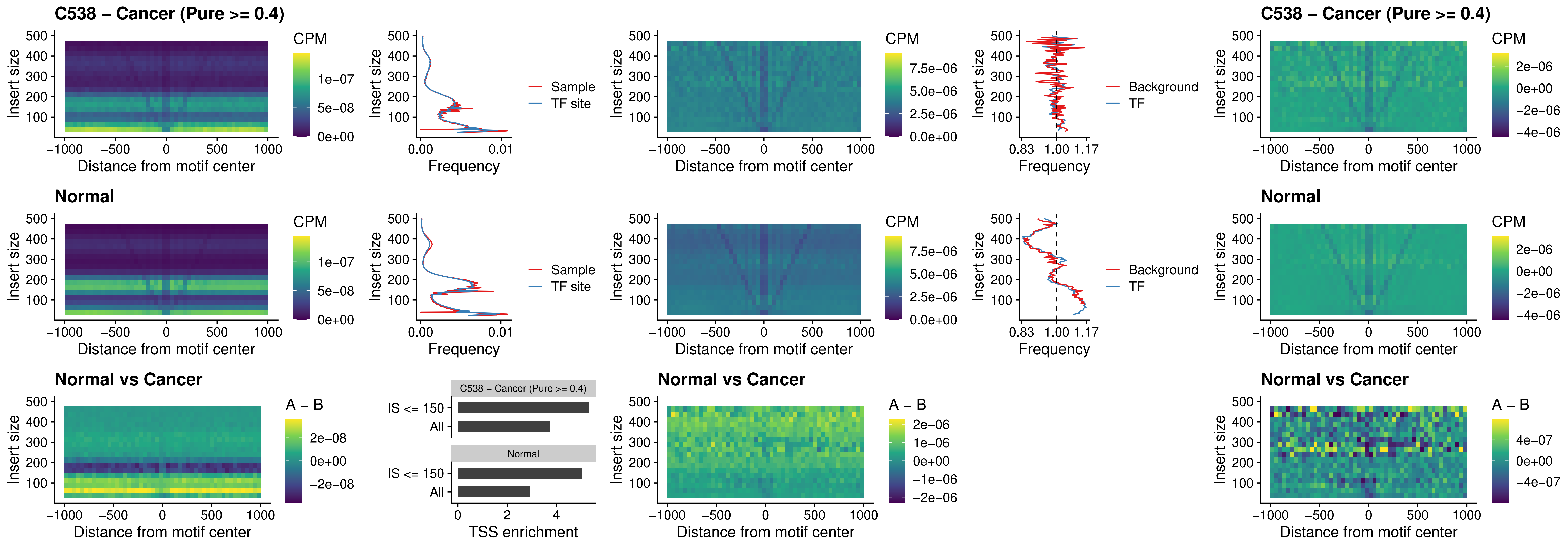

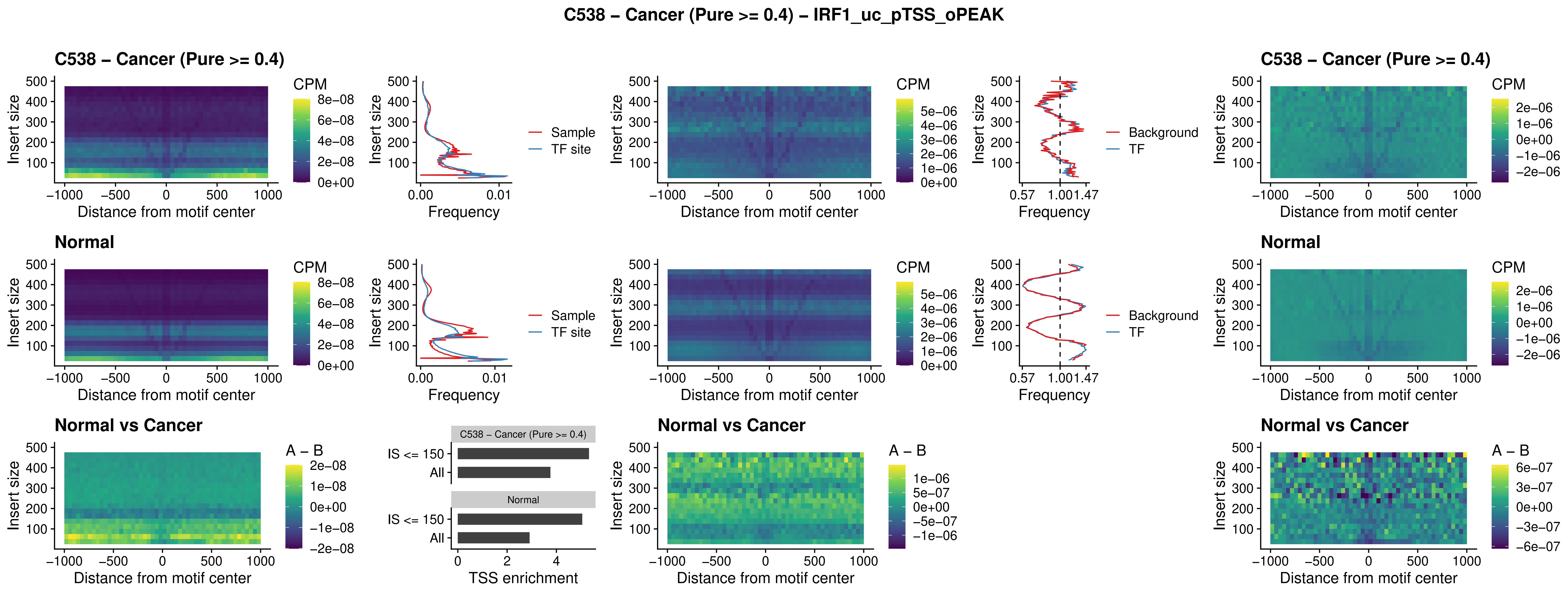

C538 – Cancer (Pure >= 0.4) – IRF1_uc_pTSS_oPEAK
C538 – Cancer (Pure >= 0.4)
Normal
Normal vs Cancer
Insert size
Distance from motif center
Frequency
CPM
A – B
Sample
TF site
Background
TF
C538 – Cancer (Pure >= 0.4)
Normal
IS <= 150
All
TSS enrichment

#### C538 – Cancer (Pure >= 0.4) – SPIC_uc_dTSS_nPEAK

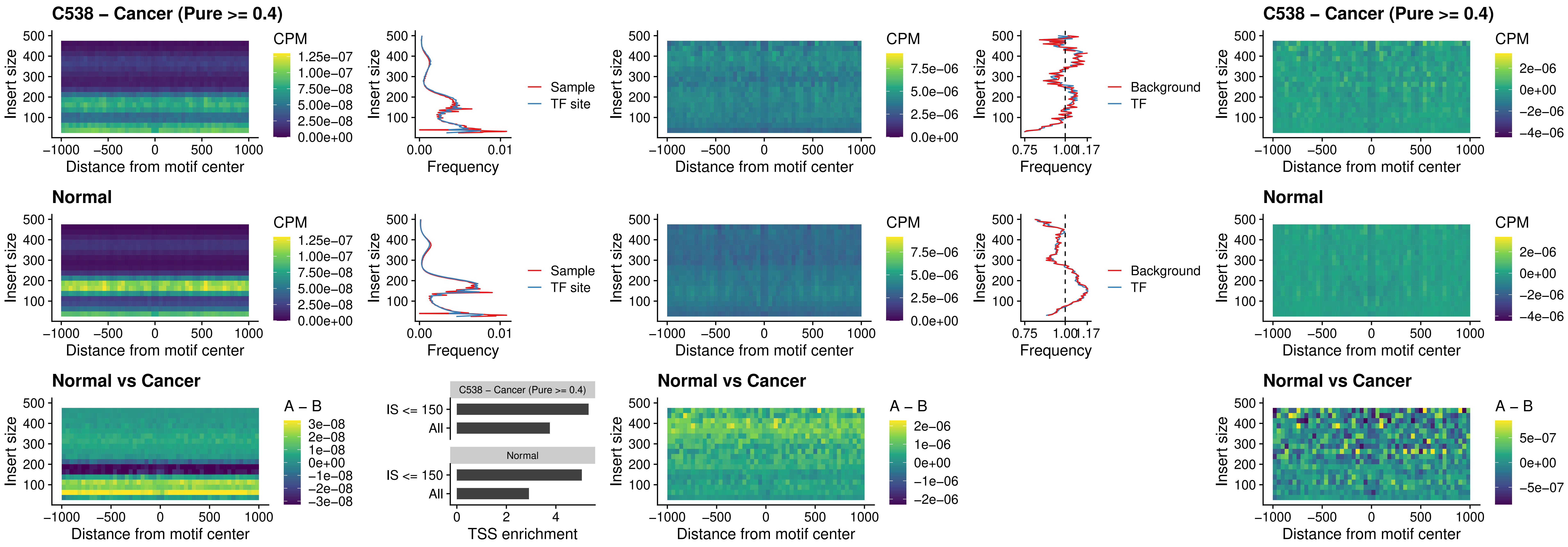

### C538 – Cancer (Pure >= 0.4) – SPIC_uc_dTSS_oPEAK

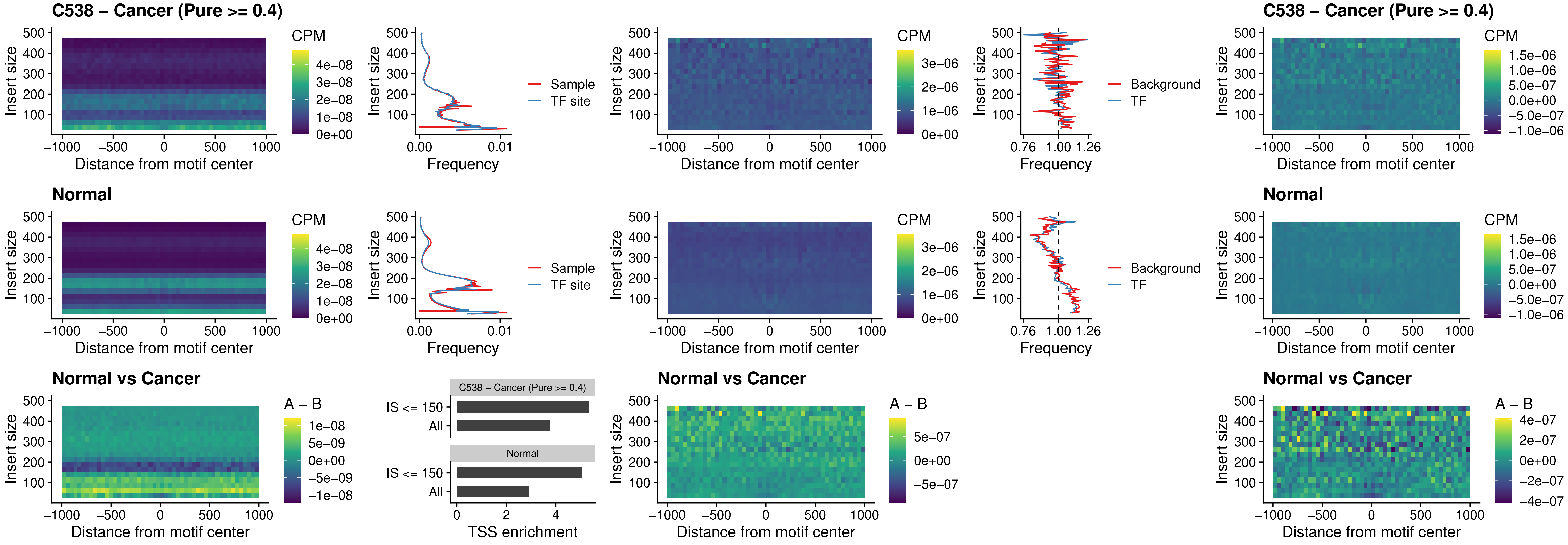

#### C538 – Cancer (Pure >= 0.4) – SPIB_uc_pTSS_nPEAK

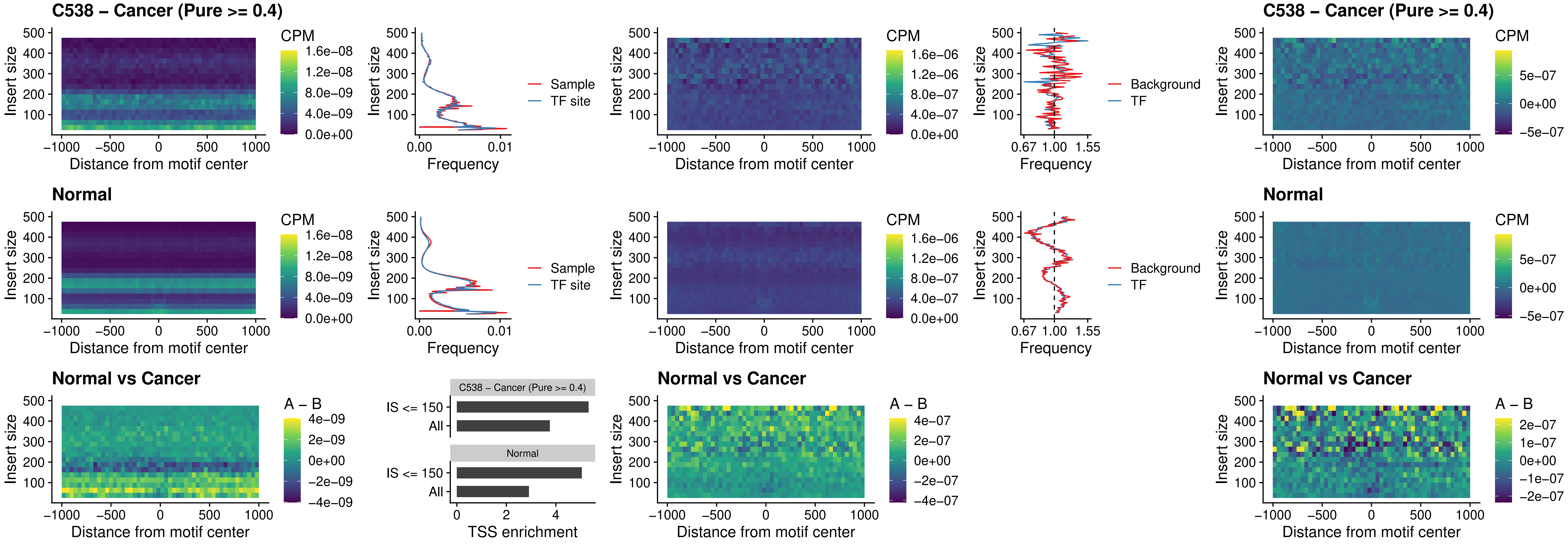

#### C538 – Cancer (Pure >= 0.4) – STAT2_uc_cTSS_oPEAK

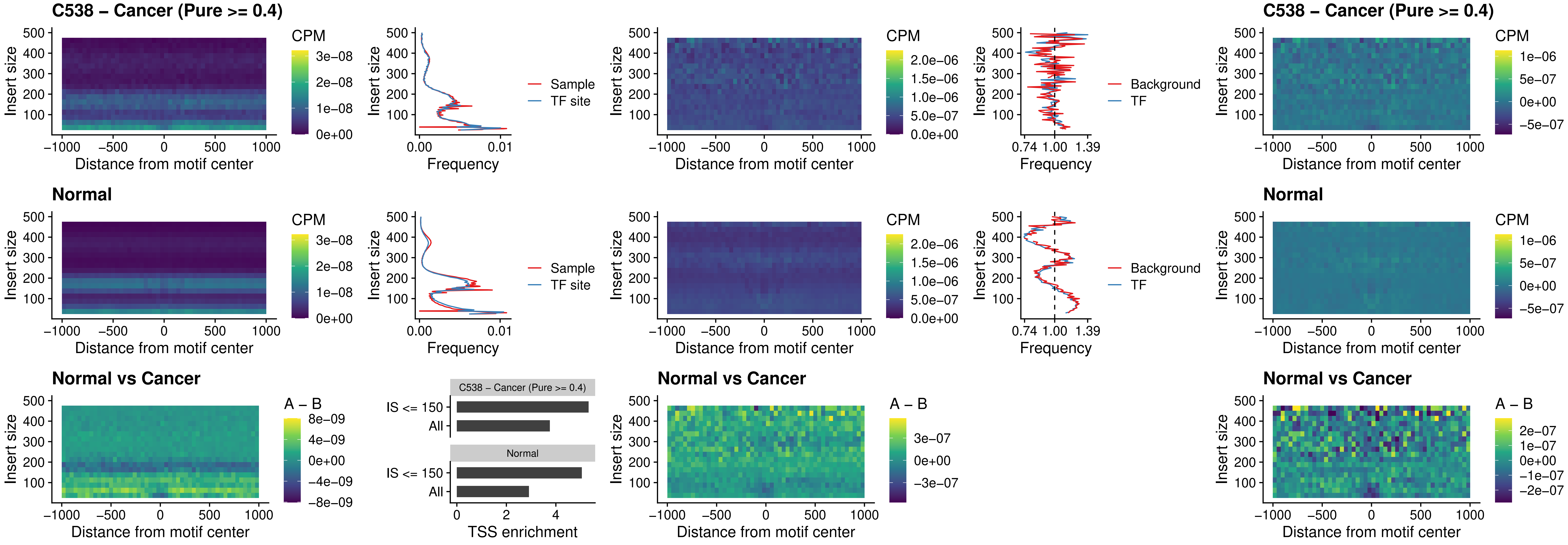

#### C538 – Cancer (Pure >= 0.4) – IRF8_uc_cTSS_oPEAK

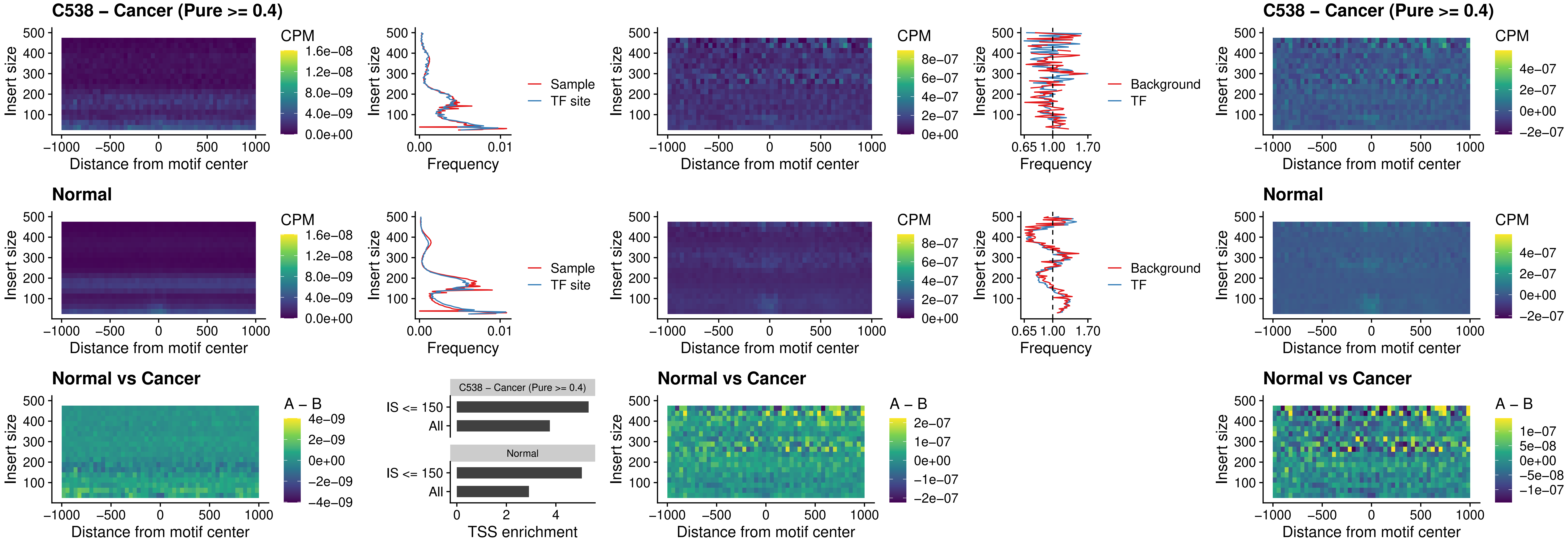

#### C538 – Cancer (Pure >= 0.4) – IRF3_uc_dTSS_oPEAK

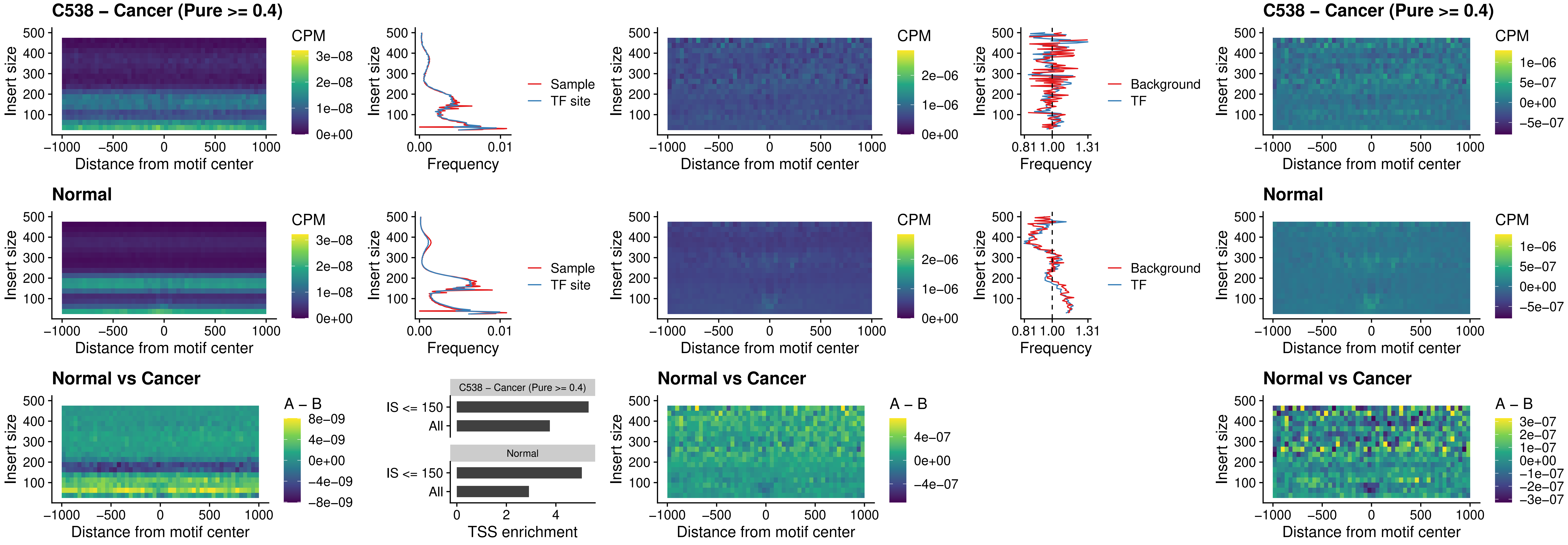

### C538 – Cancer (Pure >= 0.4) – IRF8_uc_dTSS_oPEAK

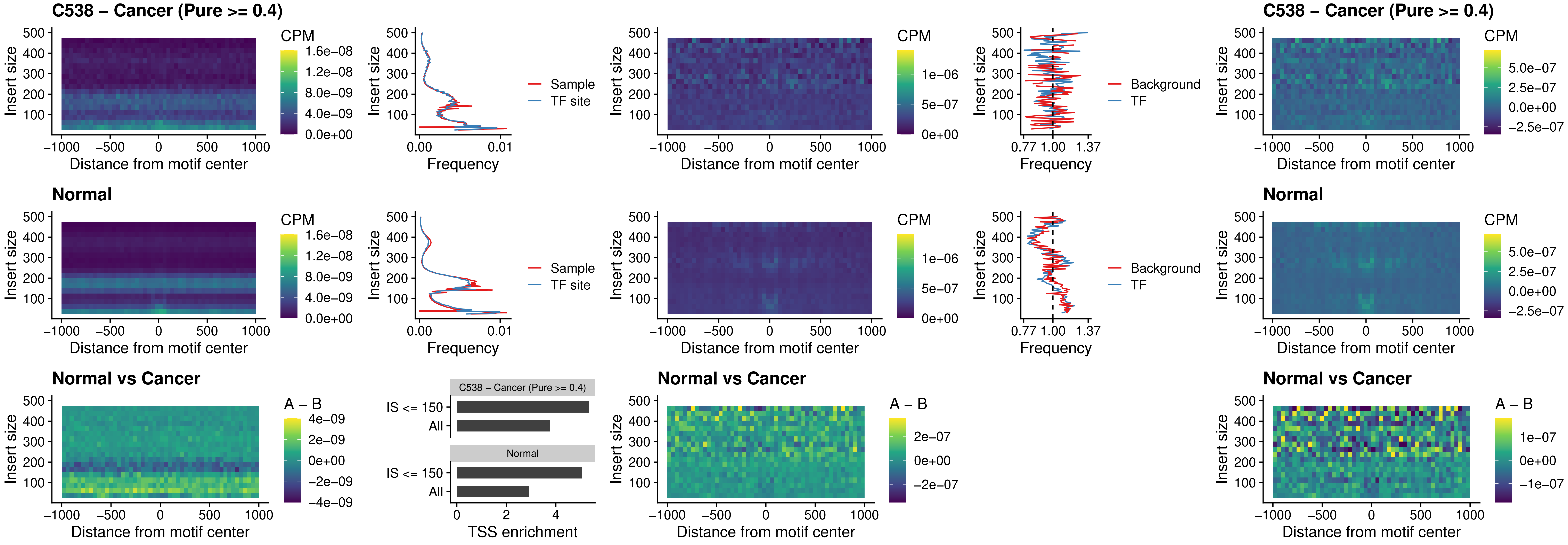

### C538 – Cancer (Pure >= 0.4) – ETS2_uc_dTSS_oPEAK

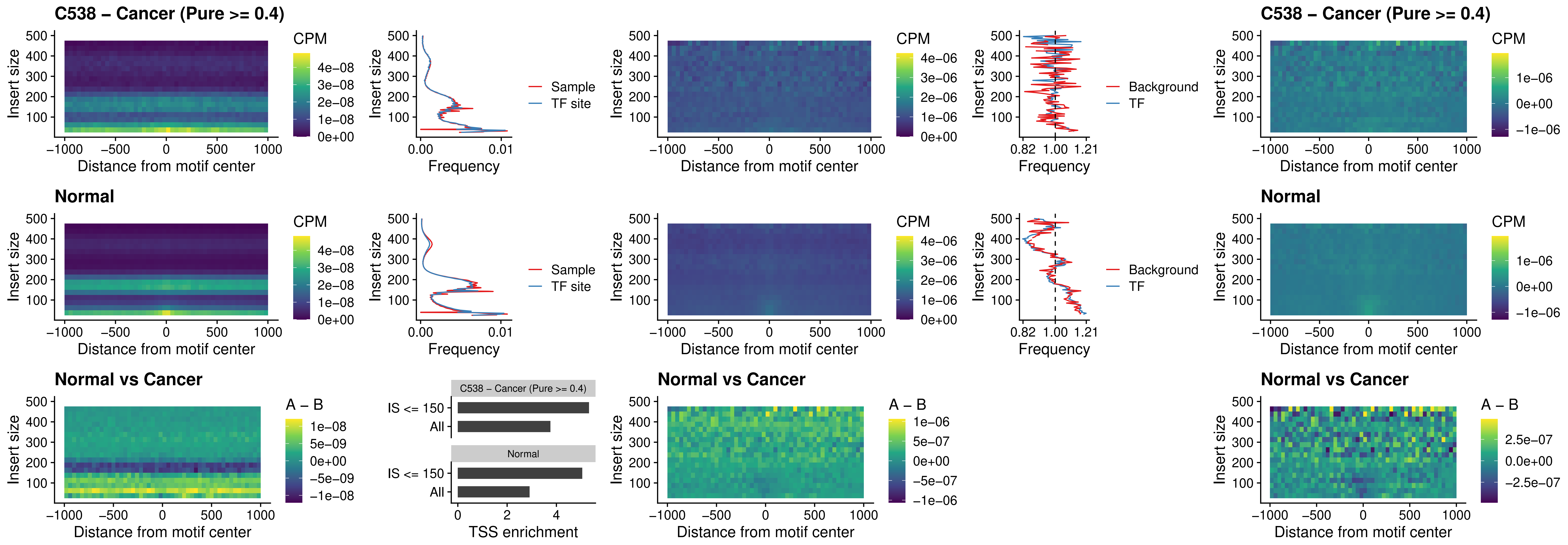

#### C538 – Cancer (Pure >= 0.4) – SMARCC2_uc_pTSS_oPEAK

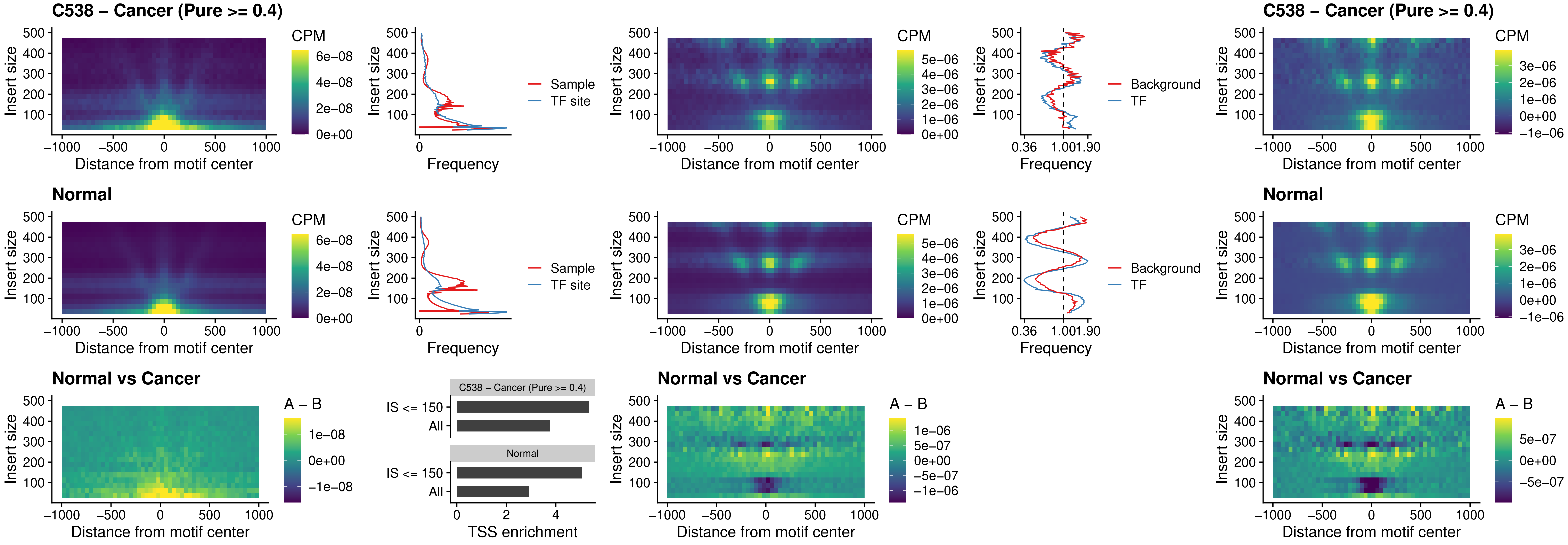

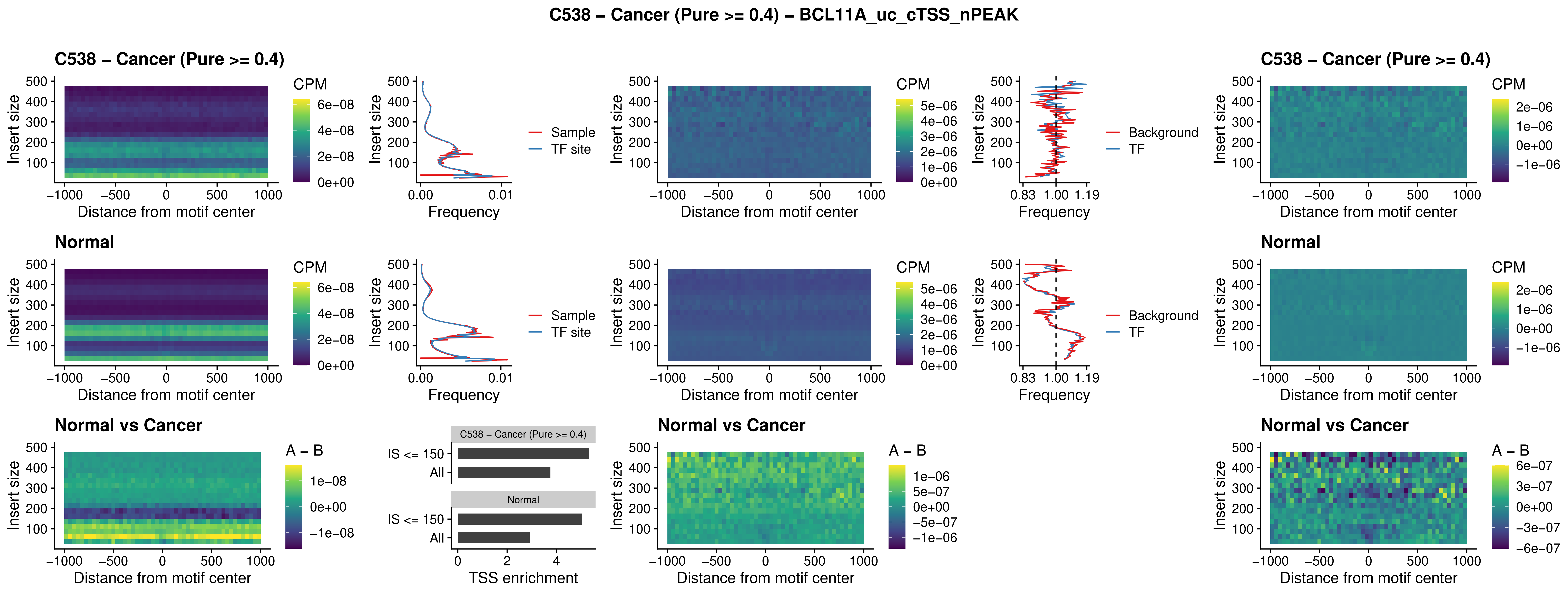
C538 – Cancer (Pure >= 0.4) – BCL11A_uc_cTSS_nPEAK
C538 – Cancer (Pure >= 0.4)
Normal
Normal vs Cancer
Insert size
Distance from motif center
CPM
A – B
Frequency
Sample
TF site
Background
TF
C538 – Cancer (Pure >= 0.4)
Normal
IS <= 150
All
TSS enrichment

#### C538 – Cancer (Pure >= 0.4) – BCL11B_uc_cTSS_nPEAK

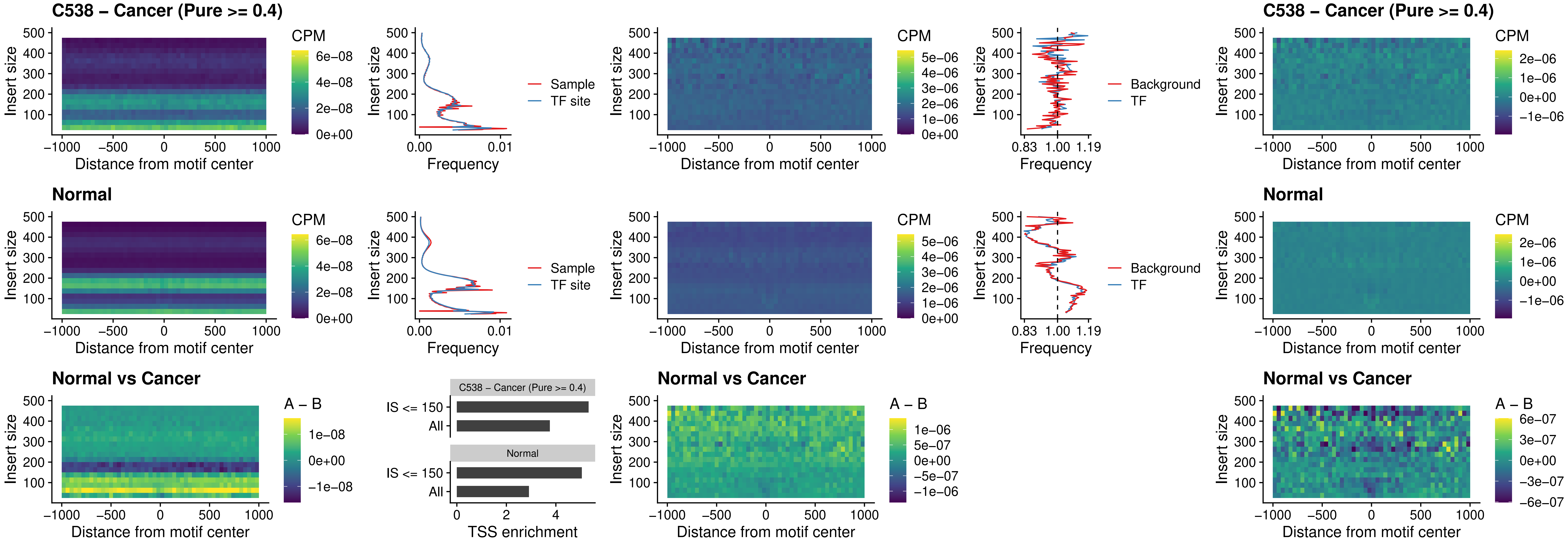

#### C538 – Cancer (Pure >= 0.4) – STAT2_uc_dTSS_oPEAK

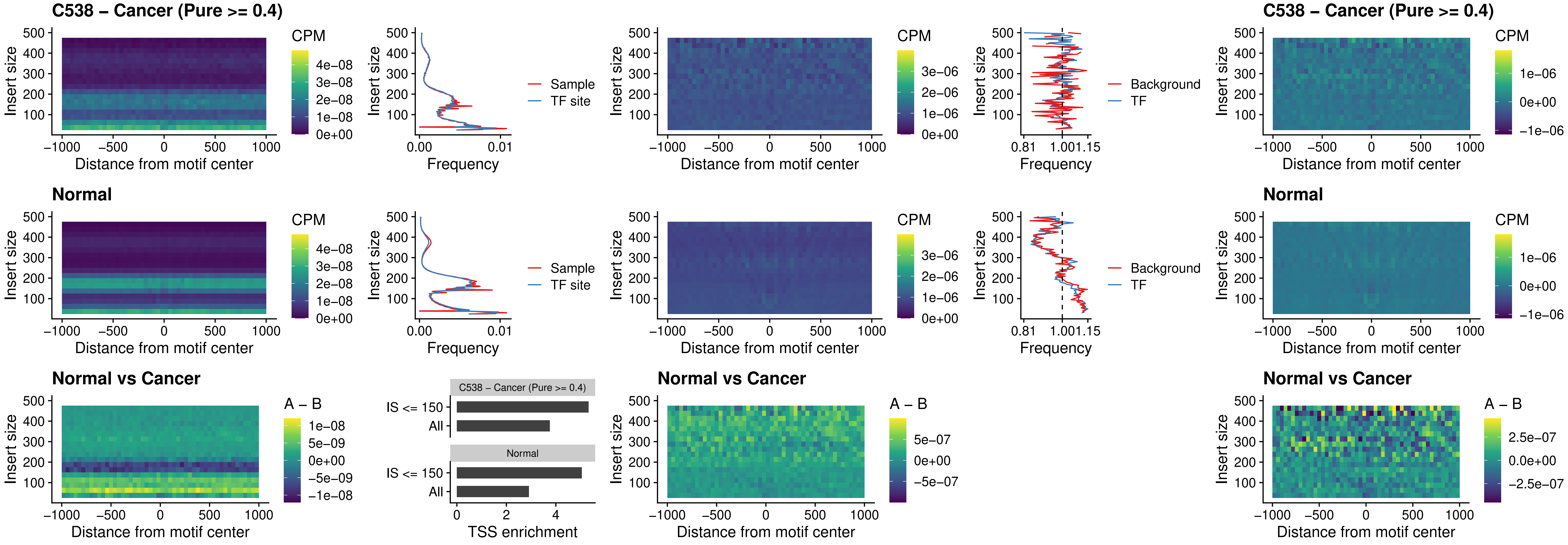

#### C538 – Cancer (Pure >= 0.4) – ZNF683_uc_cTSS_oPEAK

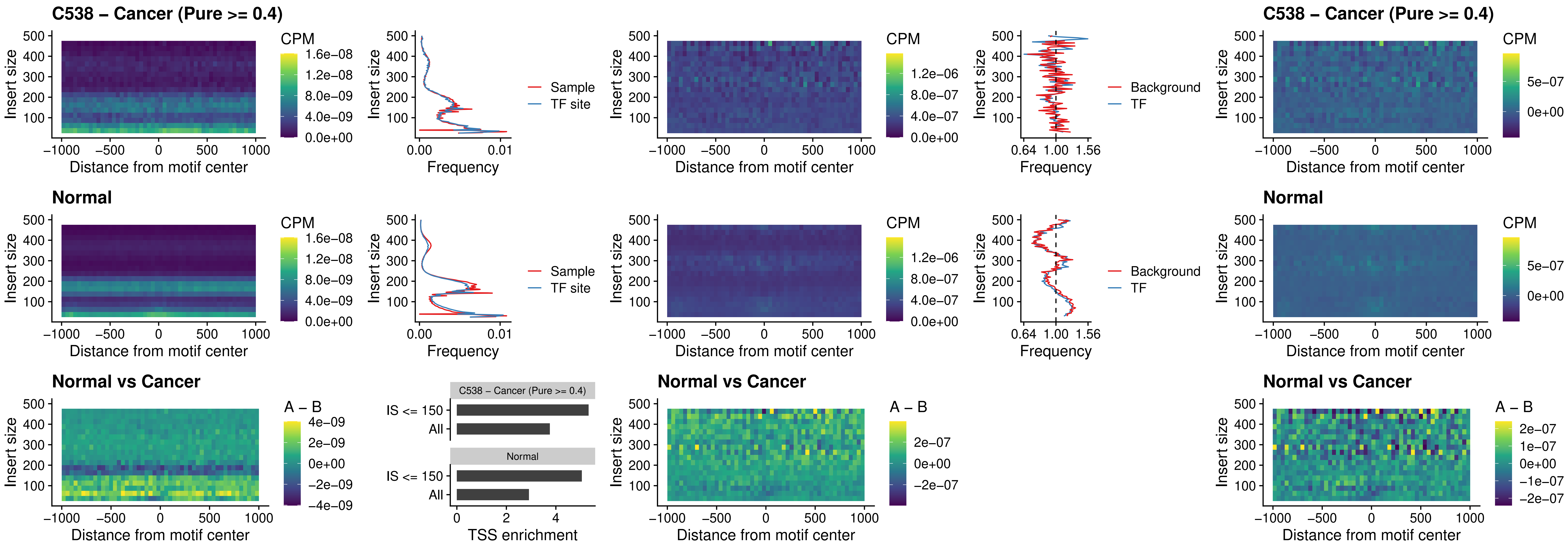

### C538 – Cancer (Pure >= 0.4) – IRF8_uc_dTSS_nPEAK

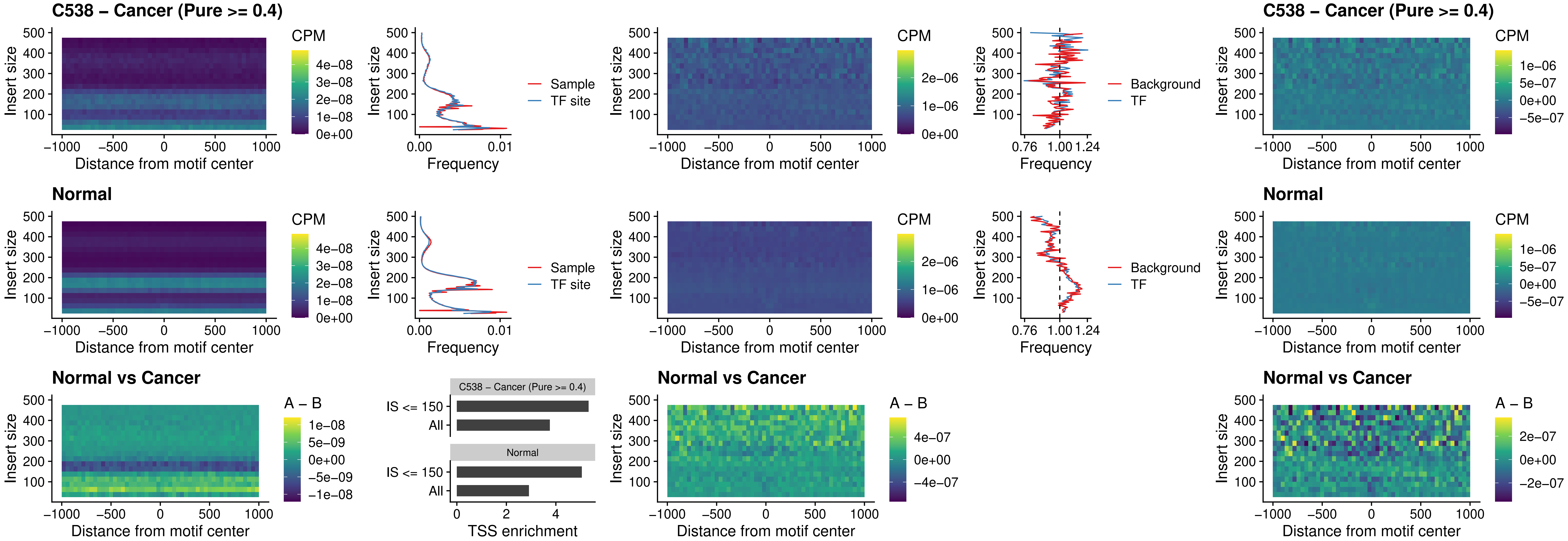

### C538 – Cancer (Pure >= 0.4) – ETV7_uc_dTSS_oPEAK

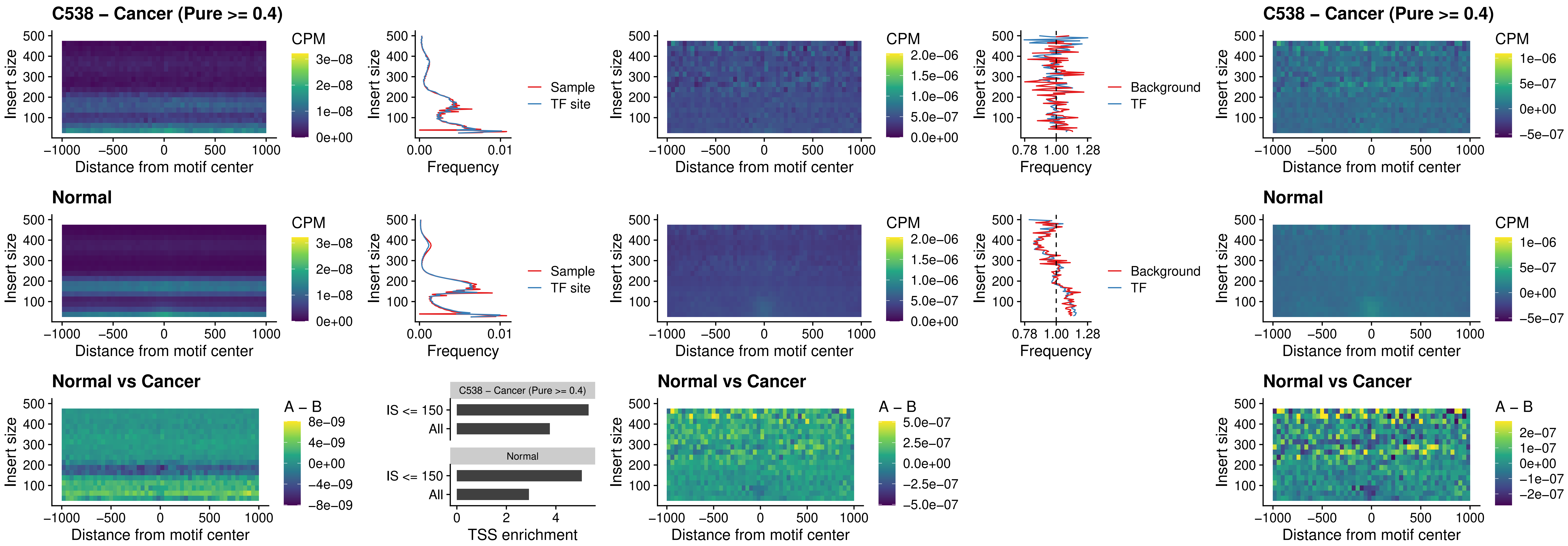

#### C539 – Cancer (Pure >= 0.4) – FOXL1_uc_pTSS_oPEAK

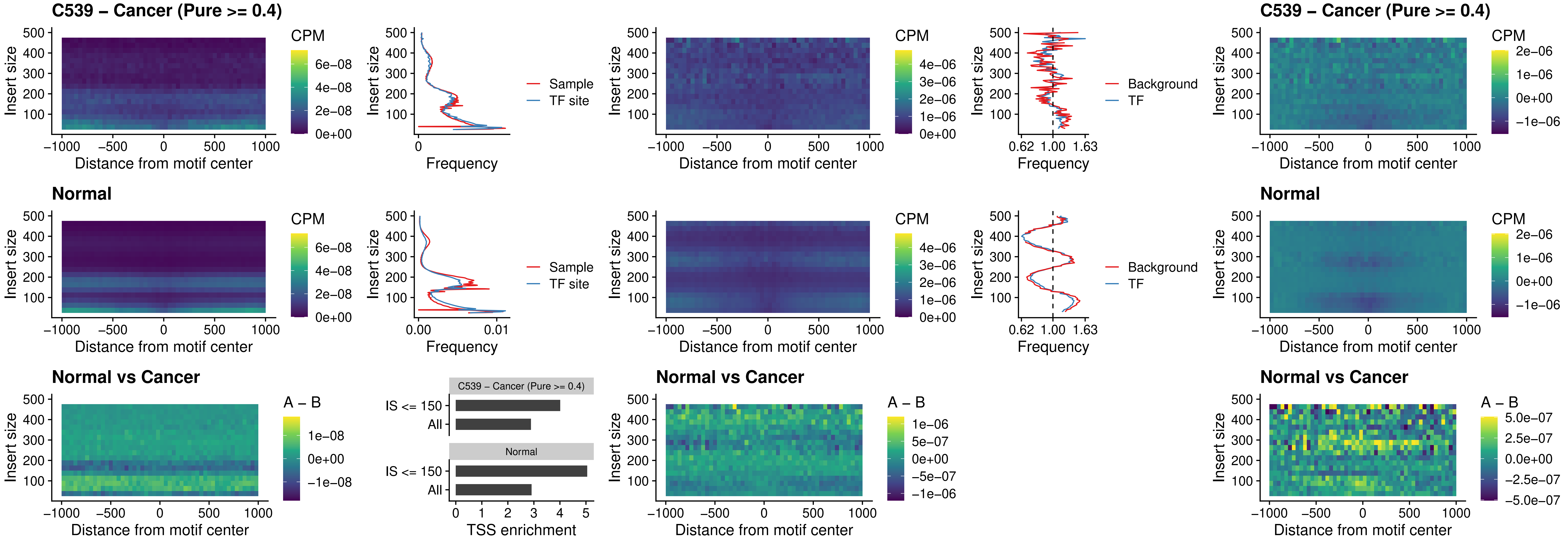

#### C539 – Cancer (Pure >= 0.4) – CPEB1_uc_cTSS_oPEAK

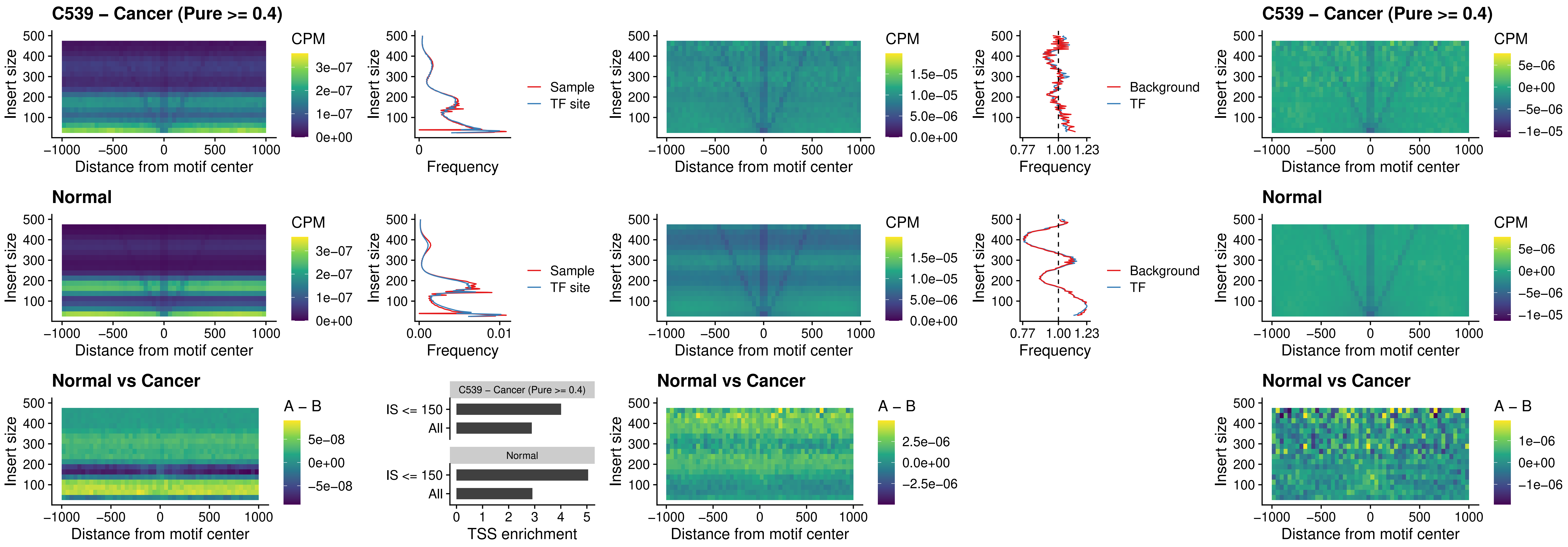

#### C539 – Cancer (Pure >= 0.4) – HMGA1_uc_pTSS_nPEAK

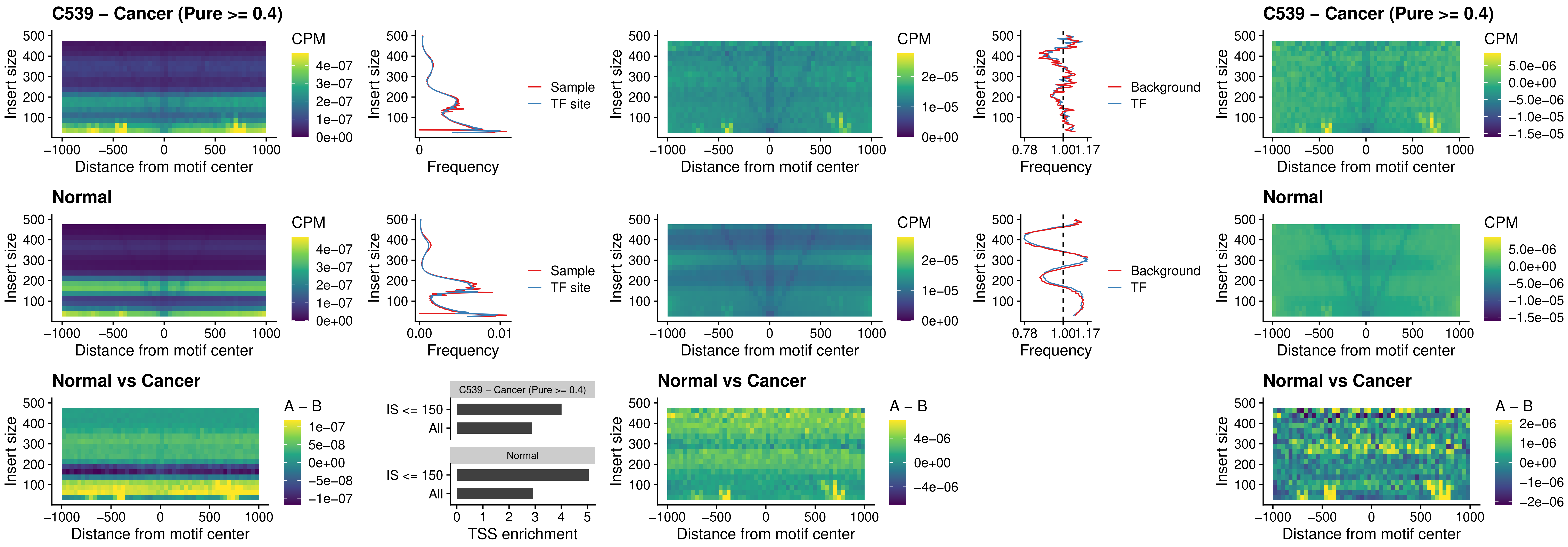

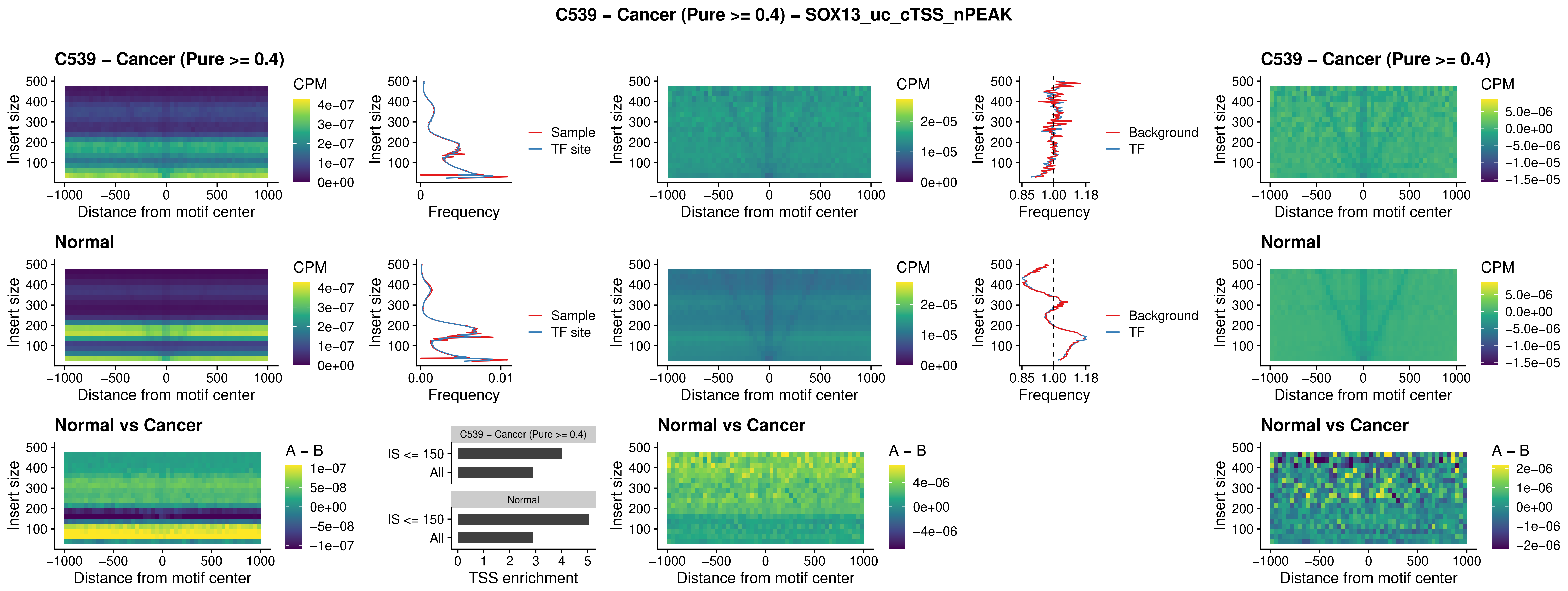
C539 – Cancer (Pure >= 0.4) – SOX13_uc_cTSS_nPEAK
C539 – Cancer (Pure >= 0.4)
Normal
Normal vs Cancer
Insert size
Distance from motif center
Frequency
CPM
A – B
Sample
TF site
Background
TF
C539 – Cancer (Pure >= 0.4)
Normal
IS <= 150
All
TSS enrichment

### C539 – Cancer (Pure >= 0.4) – LHX4_uc_cTSS_nPEAK

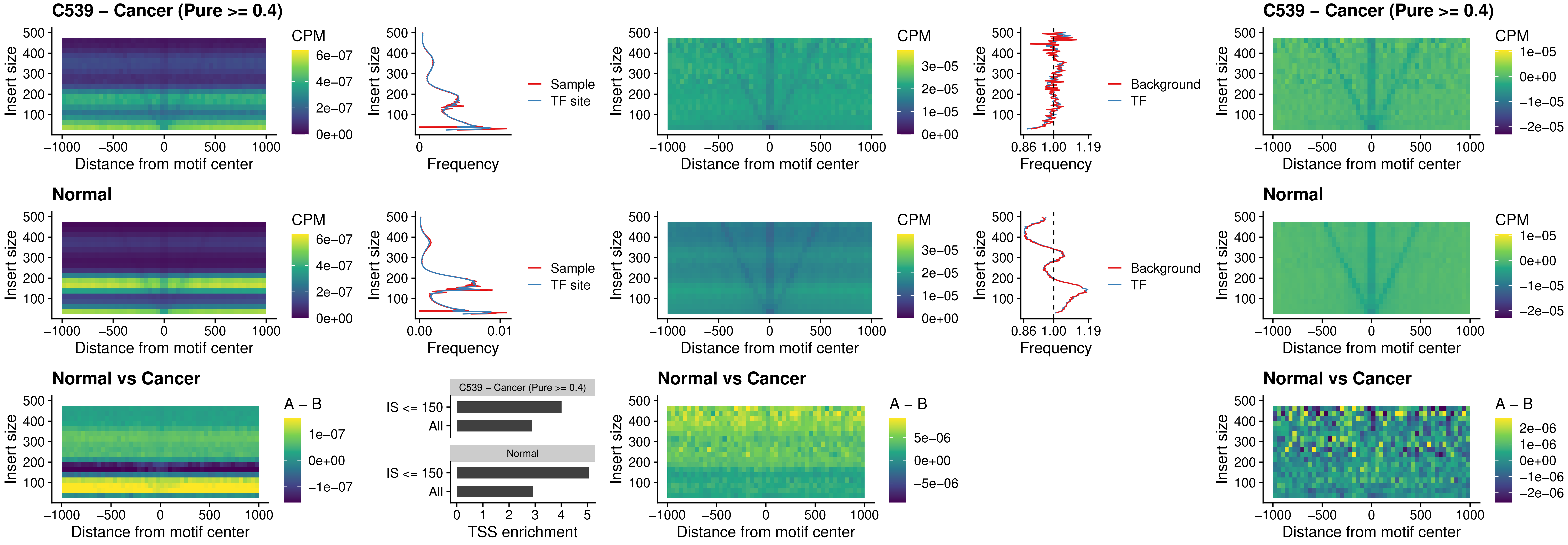

#### C539 – Cancer (Pure >= 0.4) – HOXA4_uc_cTSS_nPEAK

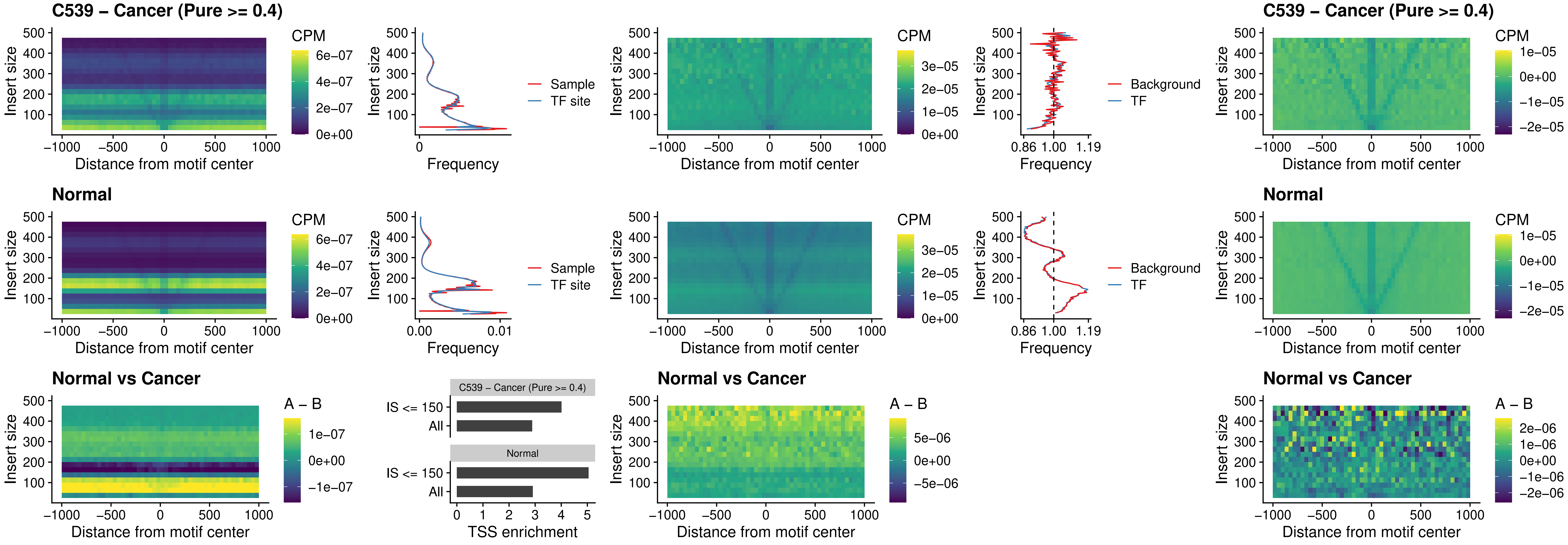

#### C539 – Cancer (Pure >= 0.4) – HLX_uc_cTSS_nPEAK

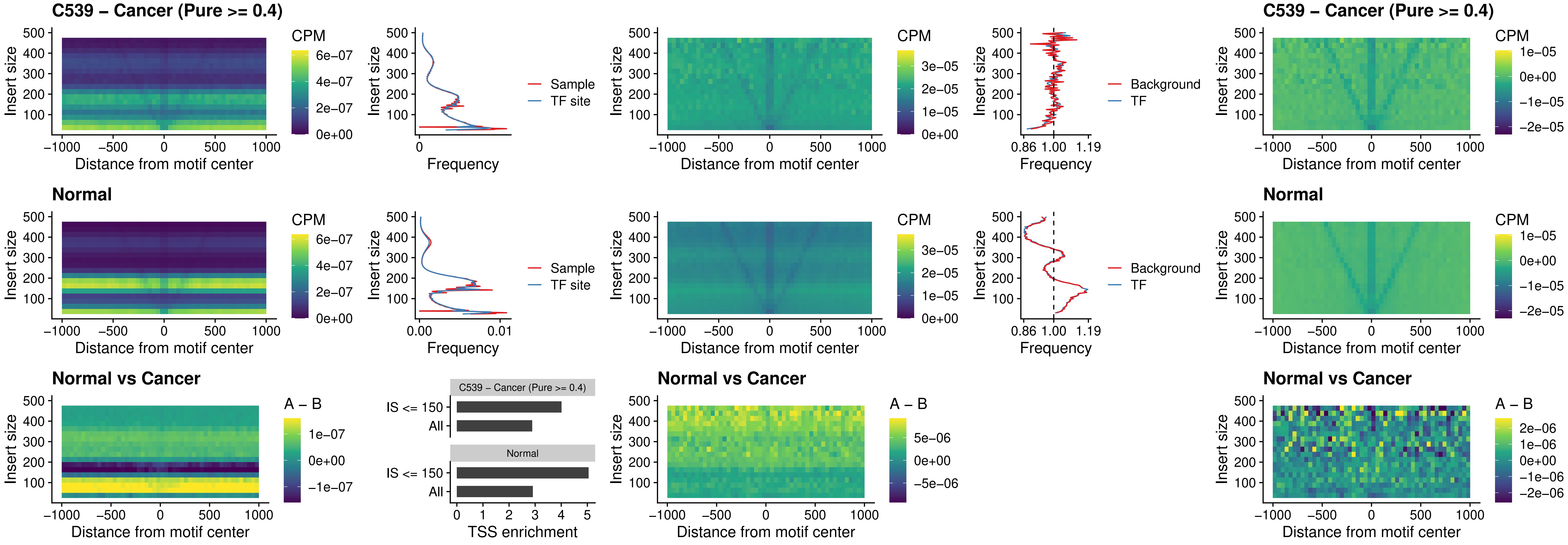

#### C539 – Cancer (Pure >= 0.4) – AC0125311_uc_cTSS_nPEAK

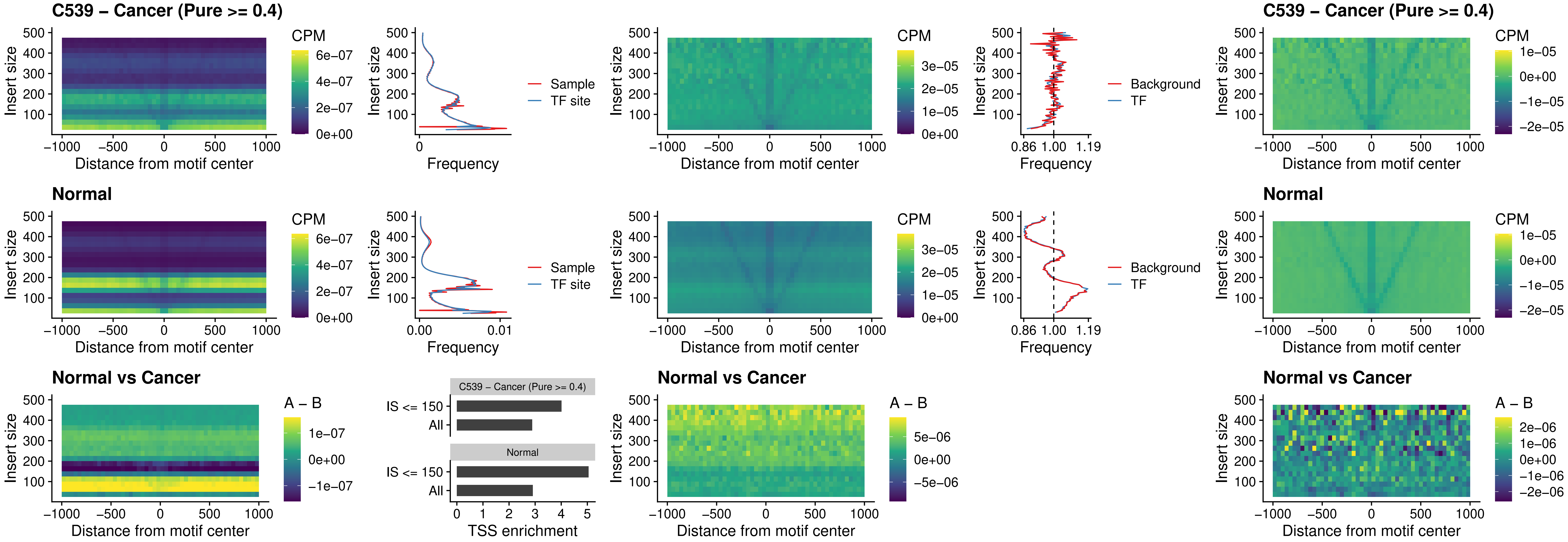

#### C539 – Cancer (Pure >= 0.4) – DBX2_uc_cTSS_nPEAK

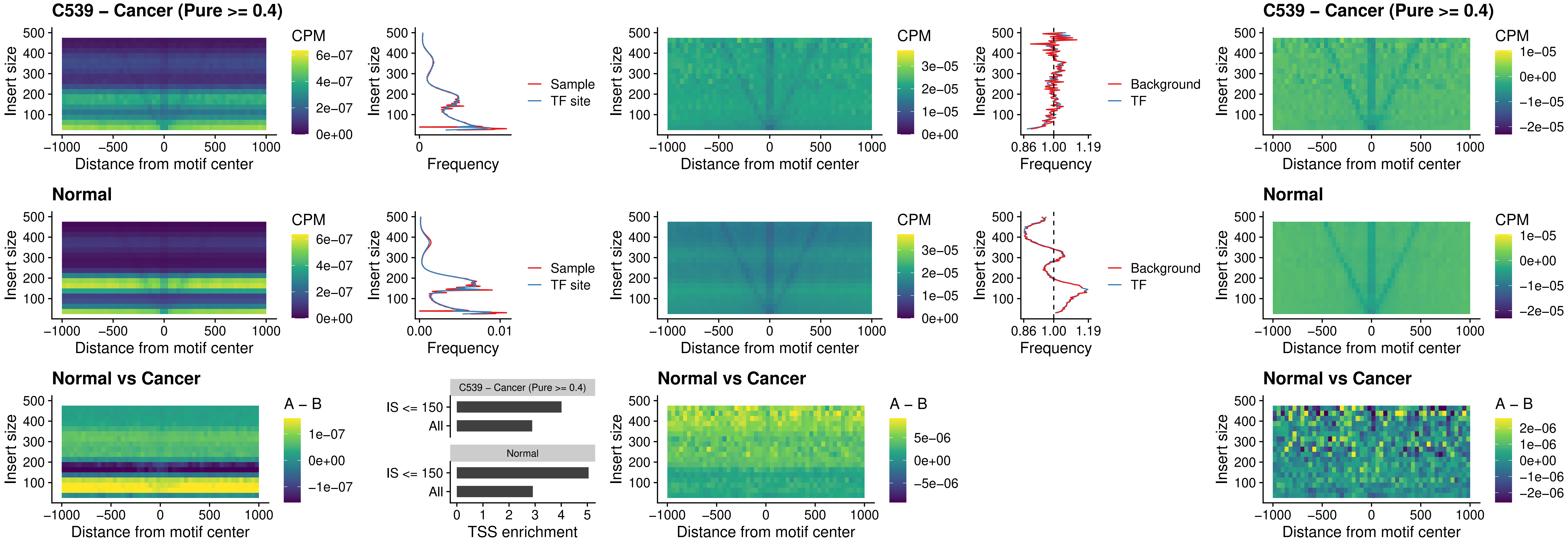

#### C539 – Cancer (Pure >= 0.4) – LHX6_uc_dTSS_nPEAK

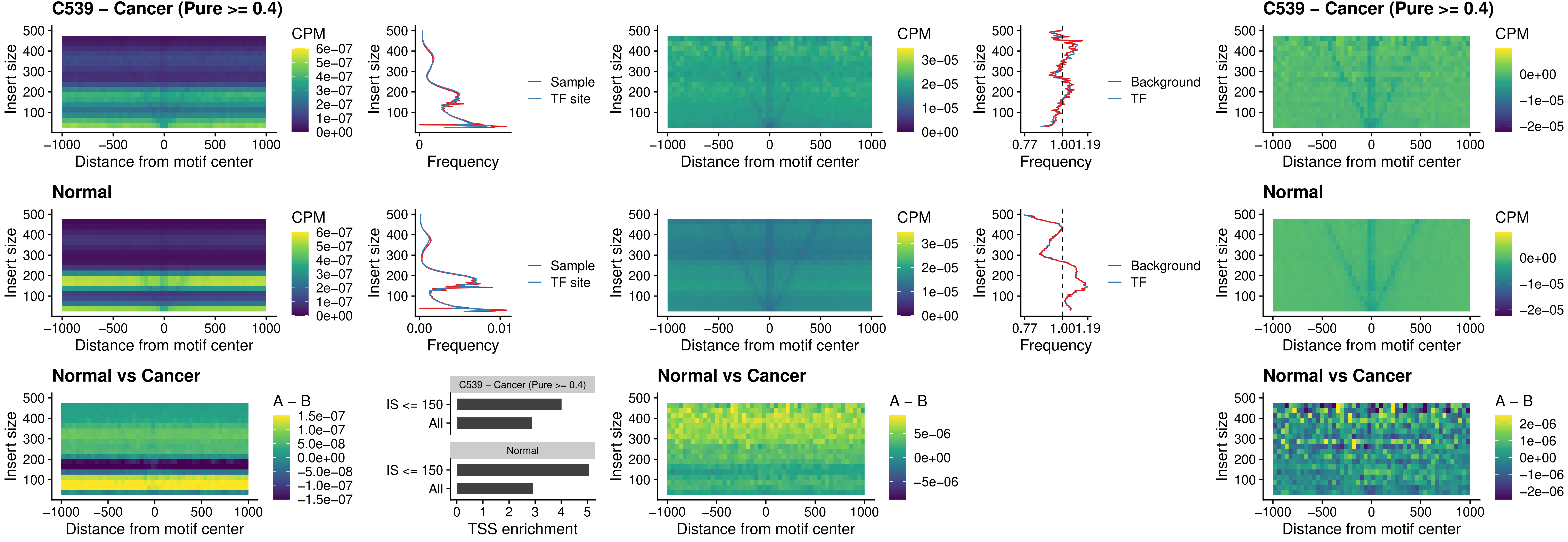

#### C539 – Cancer (Pure >= 0.4) – SOX1_uc_dTSS_nPEAK

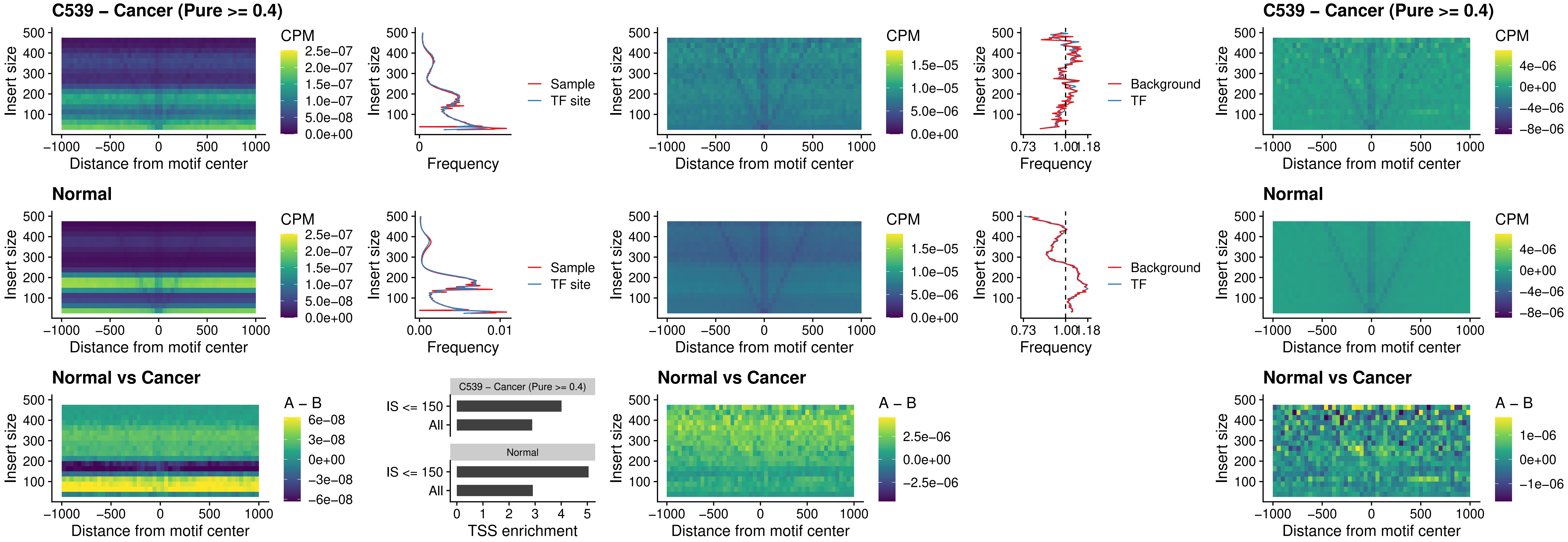

#### C539 – Cancer (Pure >= 0.4) – NKX24_uc_dTSS_nPEAK

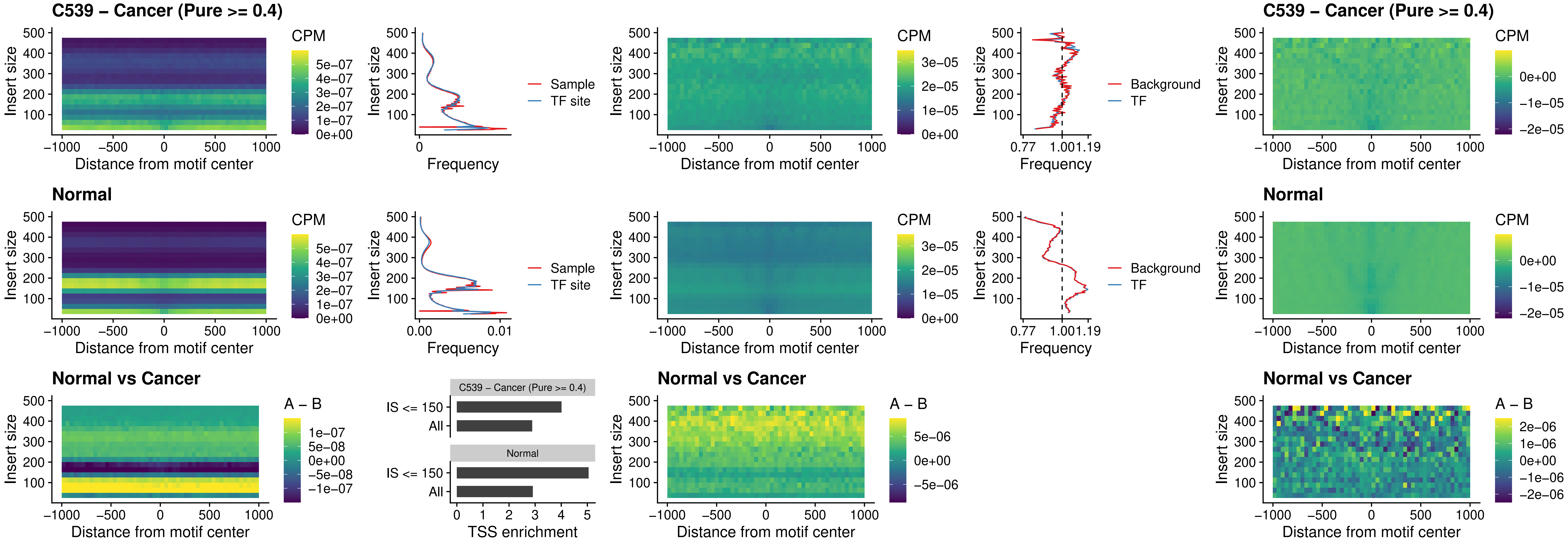

#### C539 – Cancer (Pure >= 0.4) – DLX3_uc_cTSS_nPEAK

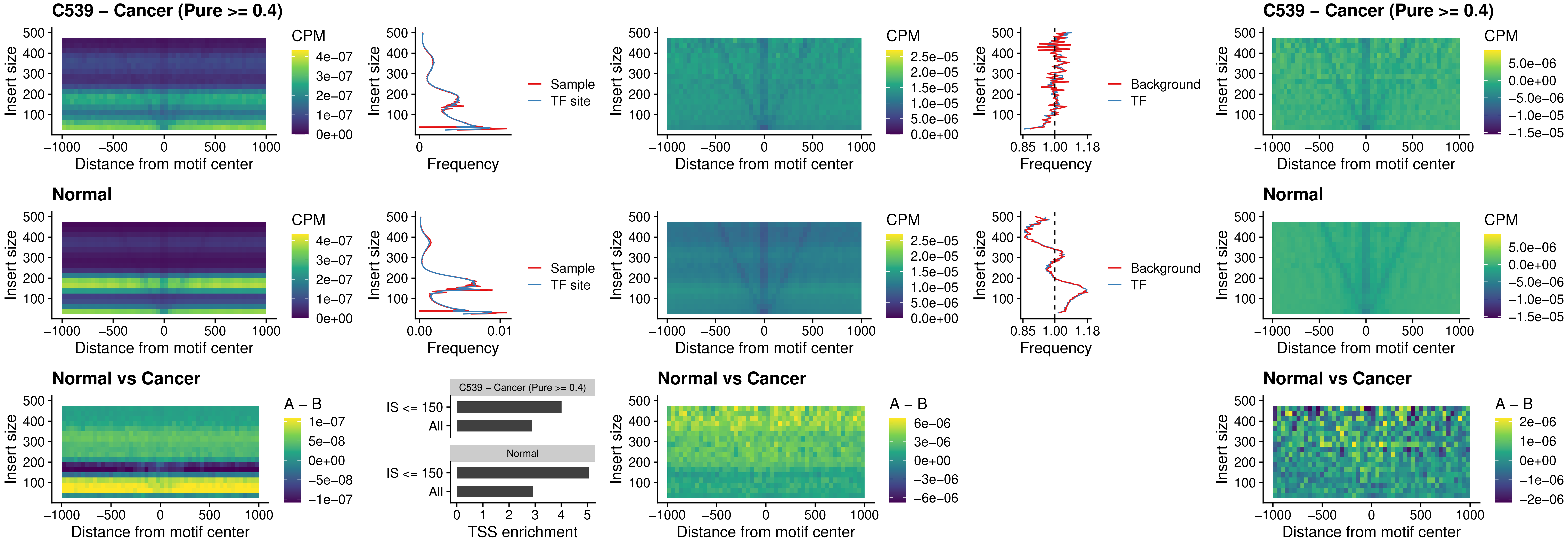

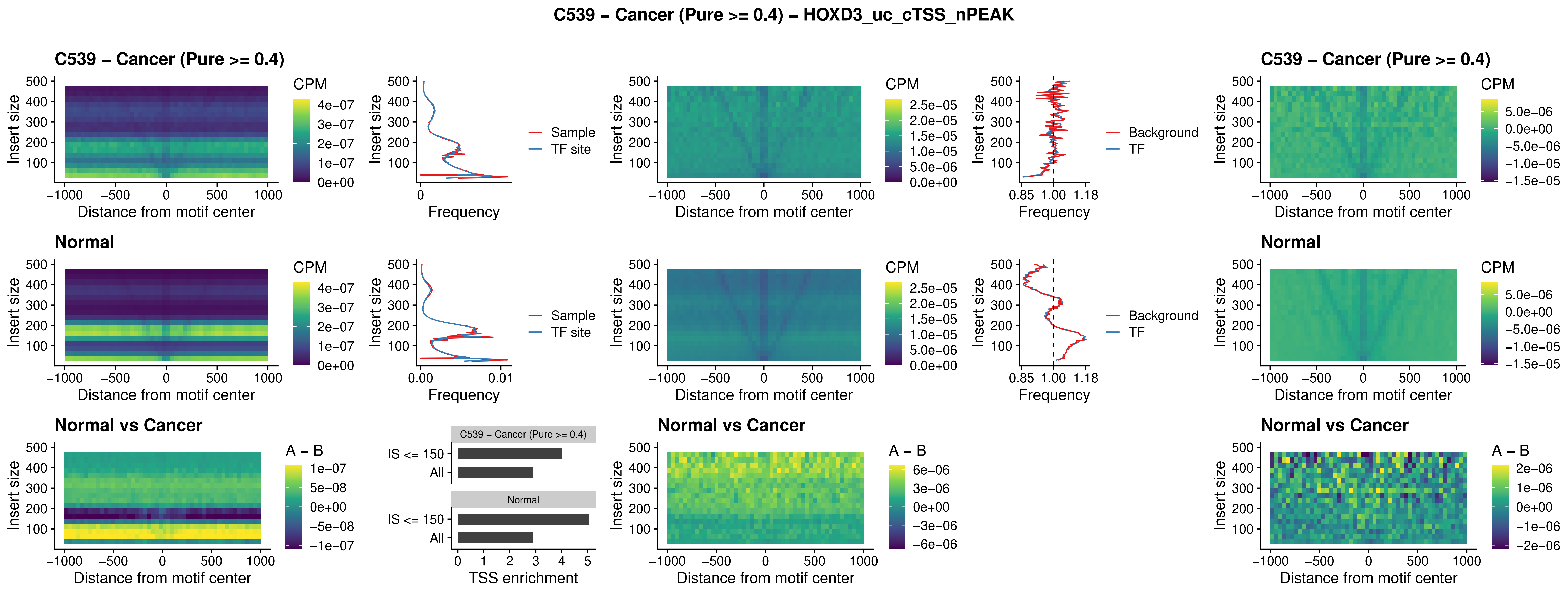

C539 – Cancer (Pure >= 0.4) – HOXD3_uc_cTSS_nPEAK
C539 – Cancer (Pure >= 0.4)
Normal
Normal vs Cancer
Insert size
Distance from motif center
Frequency
CPM
A – B
Sample
TF site
Background
TF
C539 – Cancer (Pure >= 0.4)
Normal
IS <= 150
All
TSS enrichment

#### C539 – Cancer (Pure >= 0.4) – ISL1_uc_dTSS_oPEAK

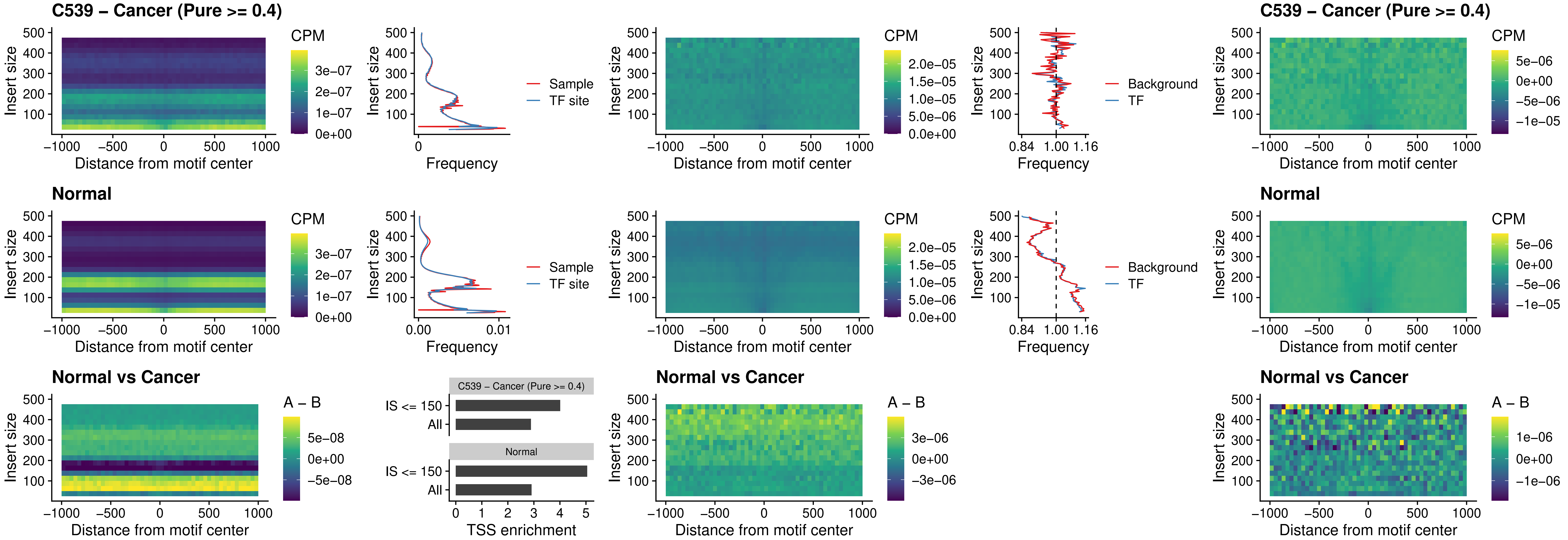

#### C539 – Cancer (Pure >= 0.4) – HOXB4_uc_dTSS_oPEAK

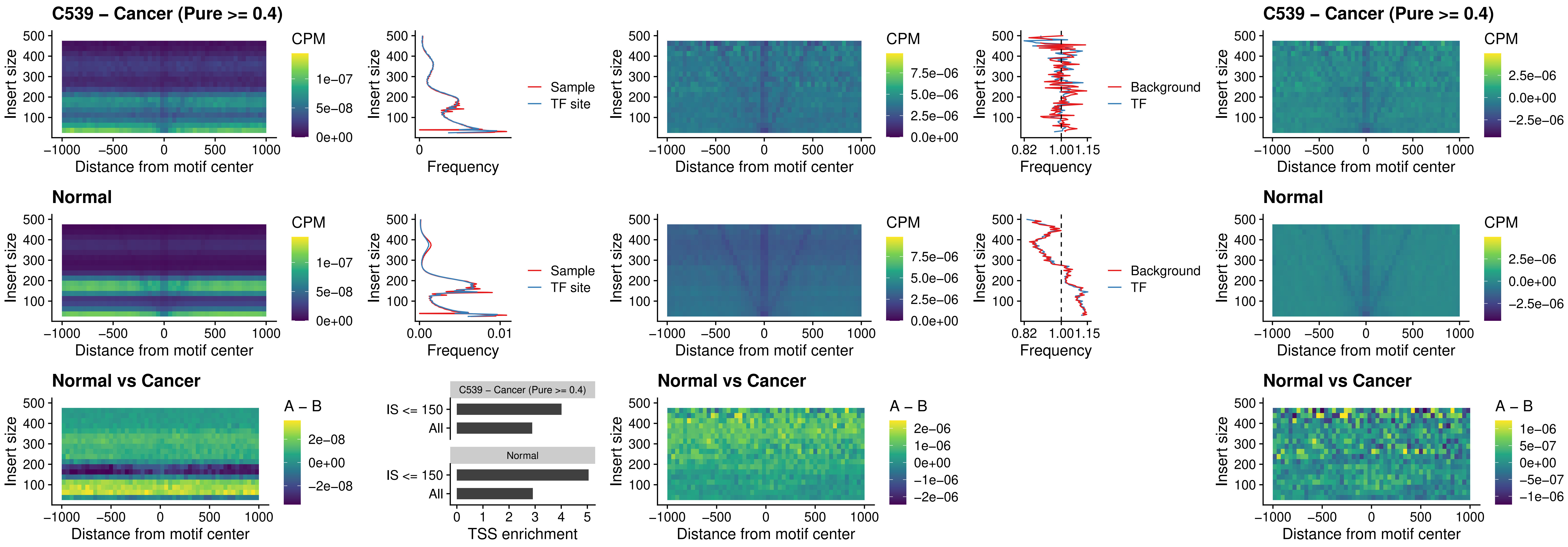

#### C539 – Cancer (Pure >= 0.4) – VAX1_uc_dTSS_oPEAK

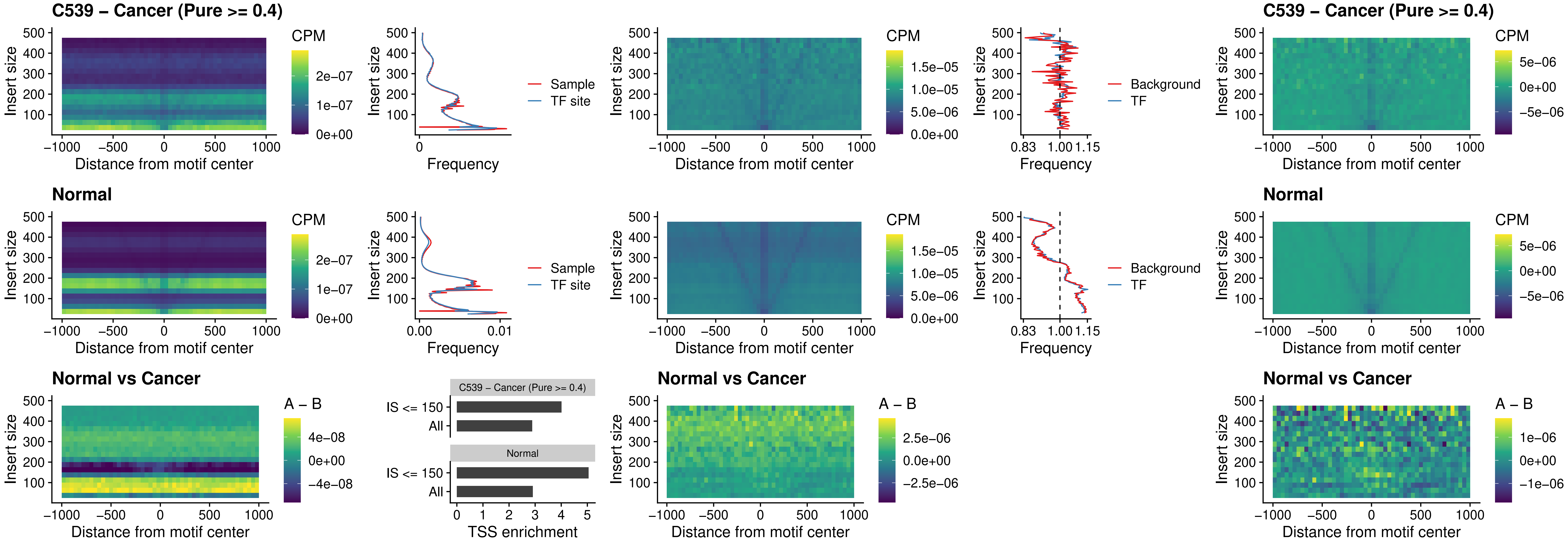

#### C539 – Cancer (Pure >= 0.4) – DLX3_uc_dTSS_oPEAK

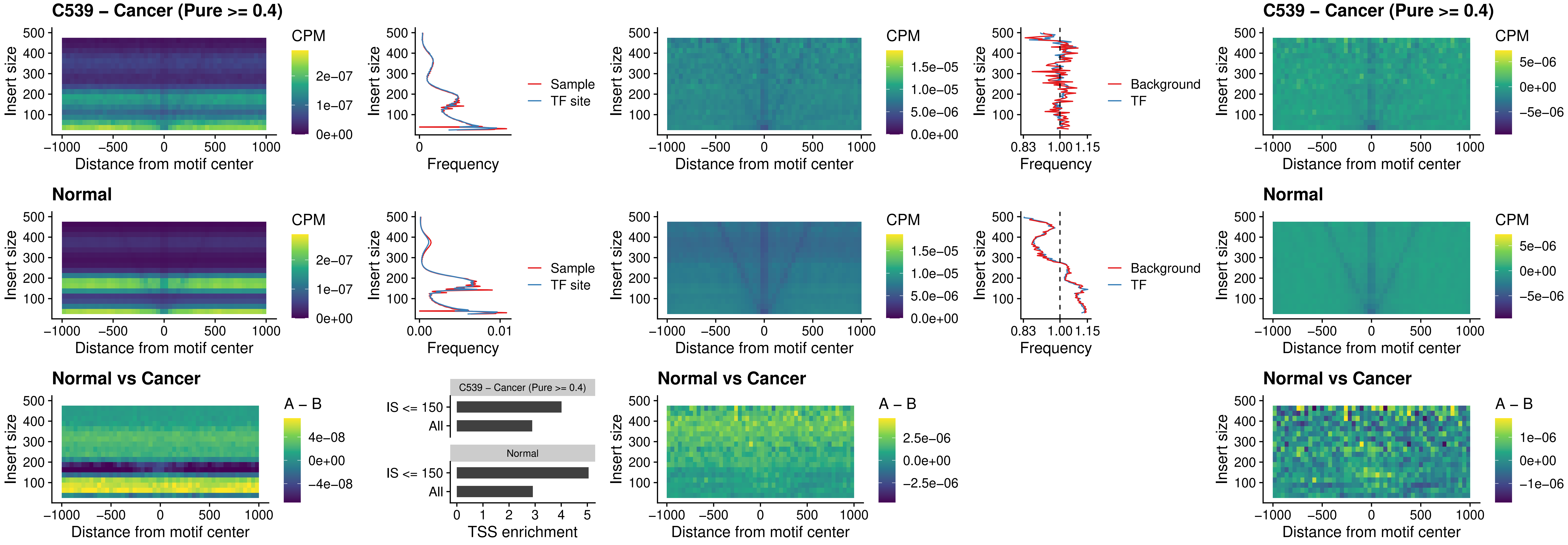

#### C539 – Cancer (Pure >= 0.4) – HOXD3_uc_dTSS_oPEAK

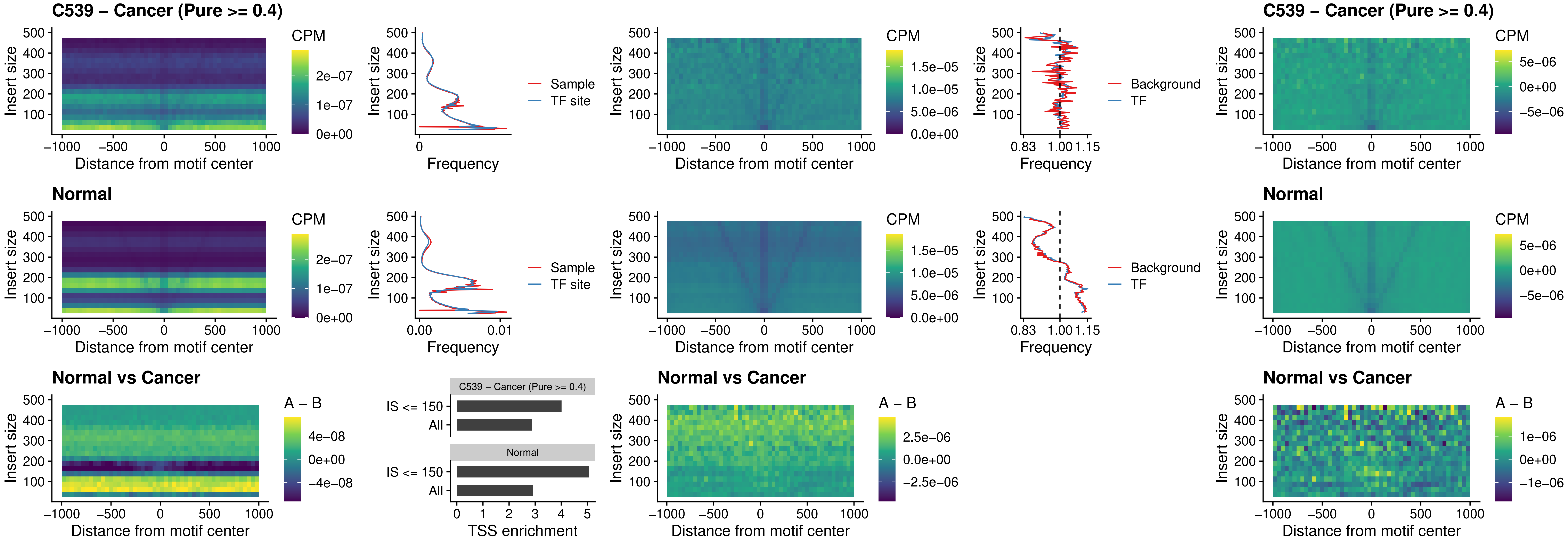

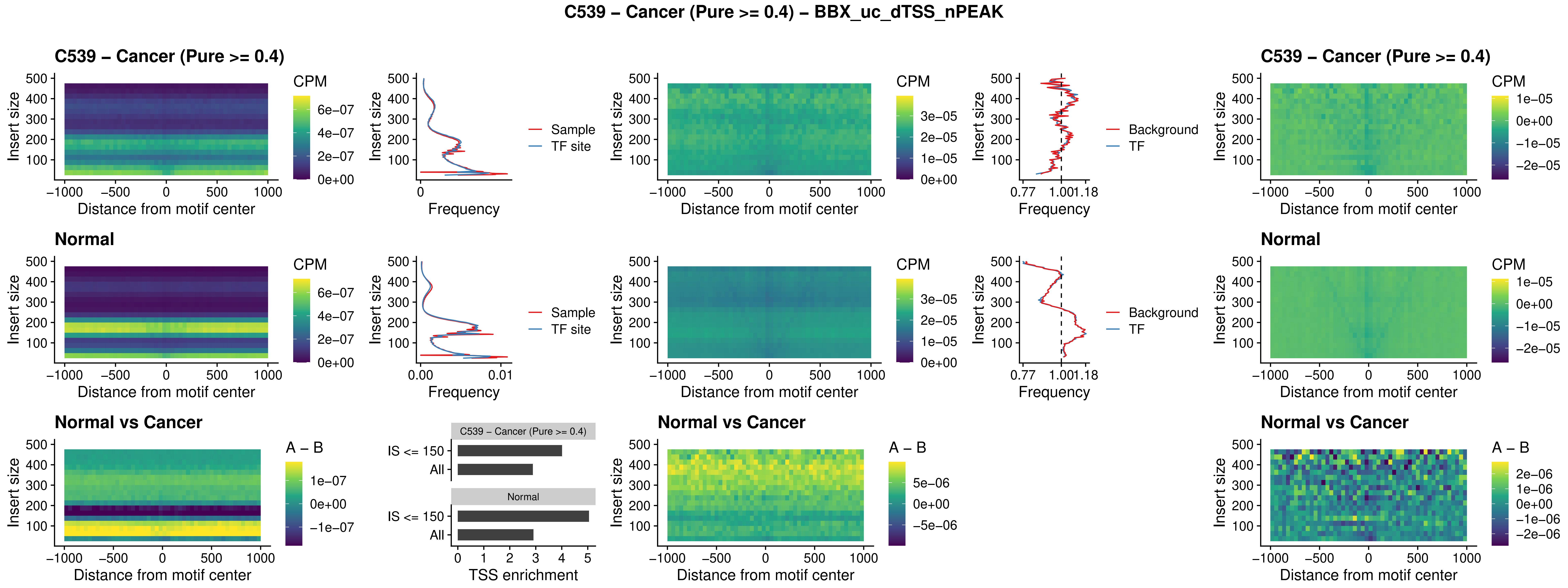
C539 – Cancer (Pure >= 0.4) – BBX_uc_dTSS_nPEAK
C539 – Cancer (Pure >= 0.4)
Normal
Normal vs Cancer
Insert size
Distance from motif center
Frequency
CPM
A – B
Sample
TF site
Background
TF
C539 – Cancer (Pure >= 0.4)
Normal
IS <= 150
All
TSS enrichment

#### C539 – Cancer (Pure >= 0.4) – SOX3_uc_cTSS_nPEAK

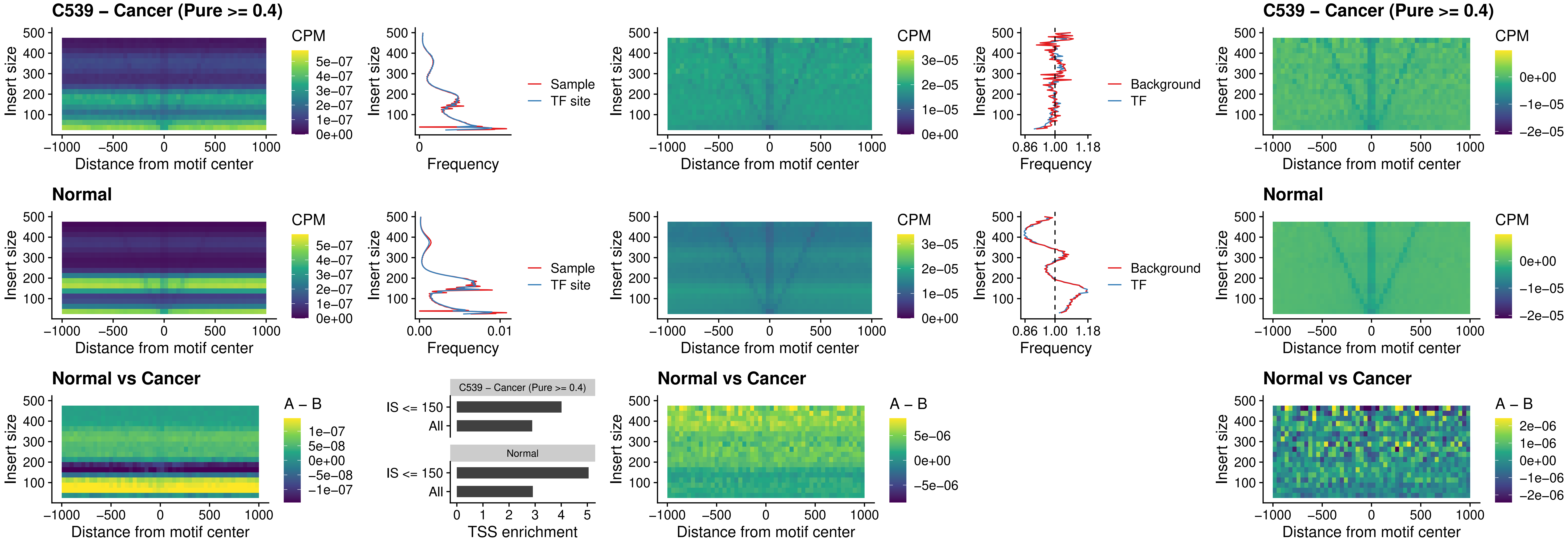

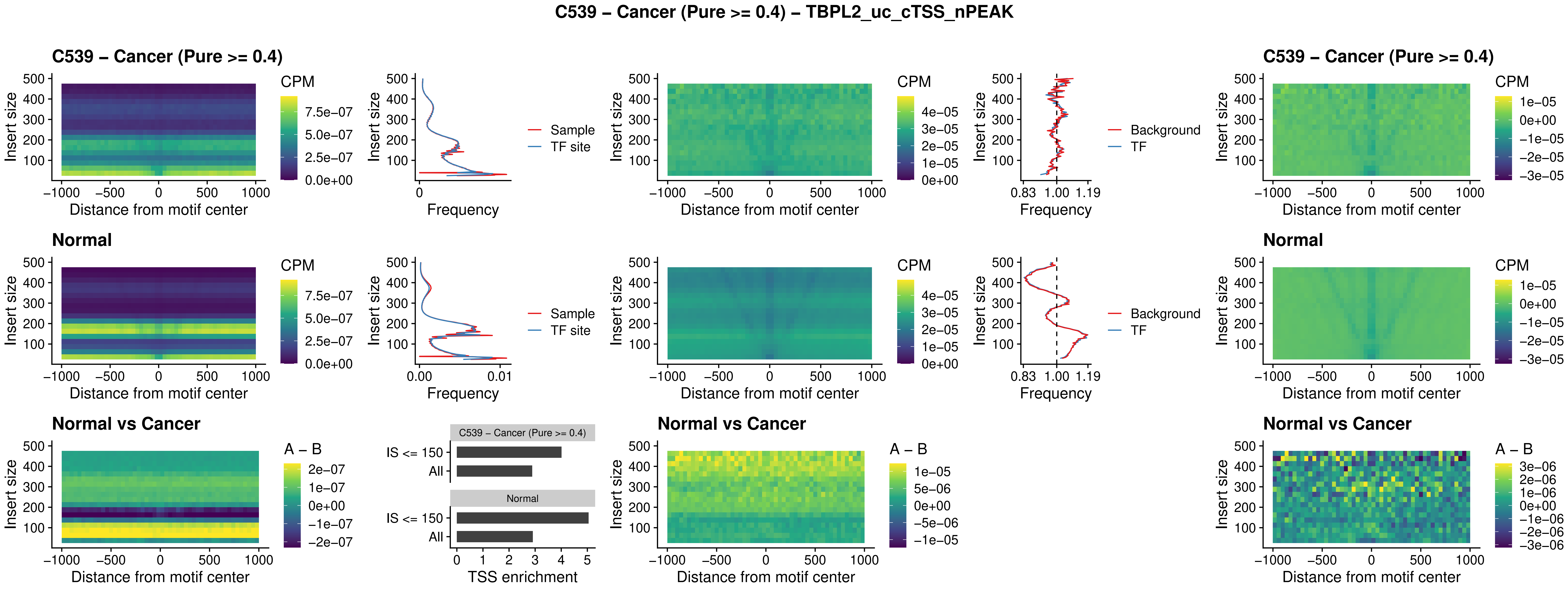
C539 – Cancer (Pure >= 0.4) – TBPL2_uc_cTSS_nPEAK
C539 – Cancer (Pure >= 0.4)
Normal
Normal vs Cancer
Insert size
Distance from motif center
Frequency
CPM
A – B
Sample
TF site
Background
TF
C539 – Cancer (Pure >= 0.4)
Normal
IS <= 150
All
TSS enrichment

#### C539 – Cancer (Pure >= 0.4) – HOXA4_uc_cTSS_oPEAK

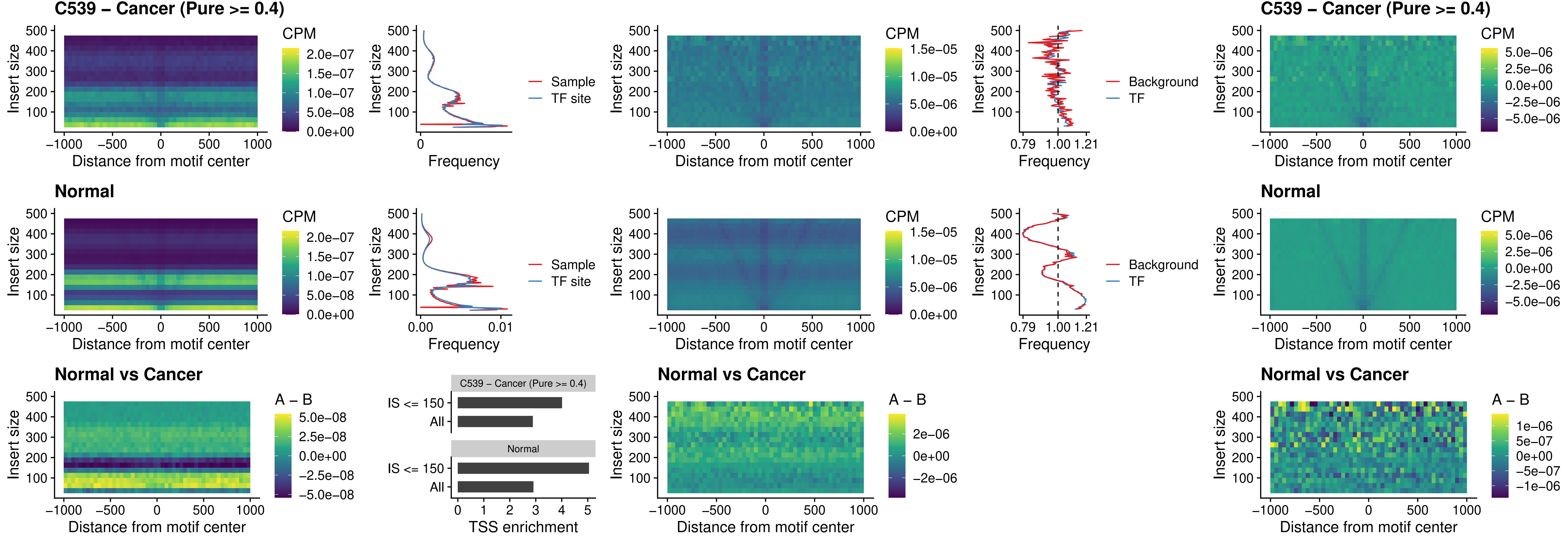

### C539 – Cancer (Pure >= 0.4) – PDX1_uc_dTSS_oPEAK

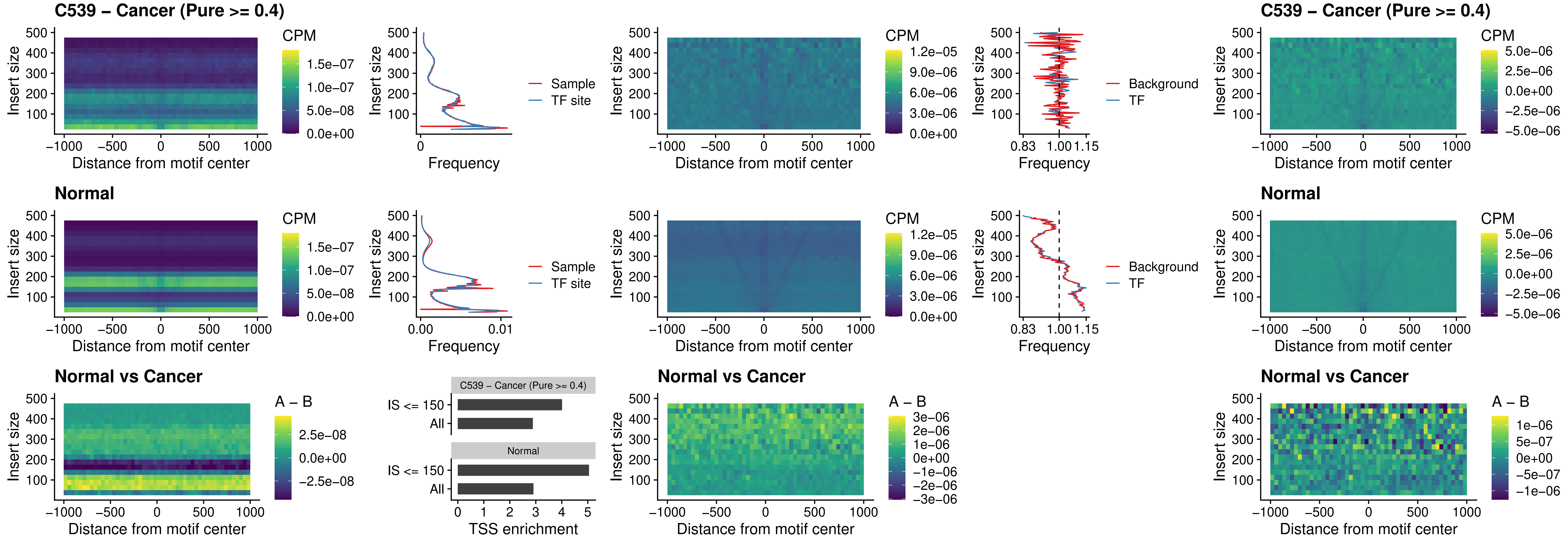

#### C539 – Cancer (Pure >= 0.4) – LHX4_uc_dTSS_oPEAK

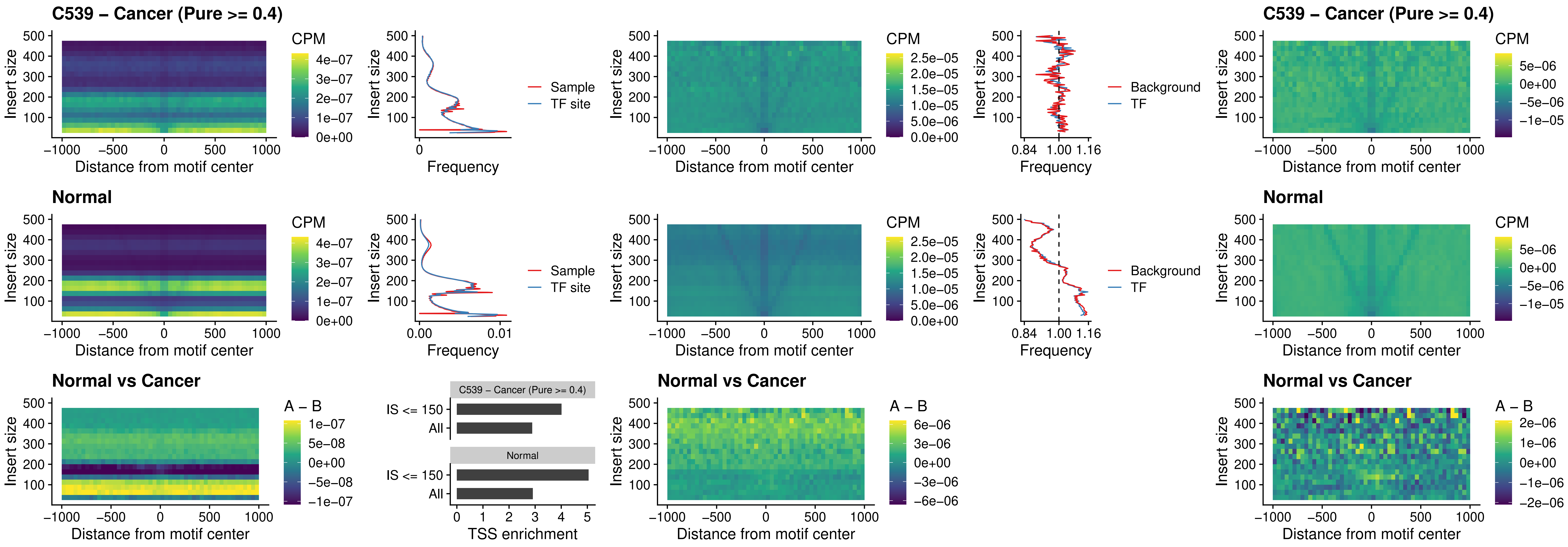

#### C539 – Cancer (Pure >= 0.4) – HOXA4_uc_dTSS_oPEAK

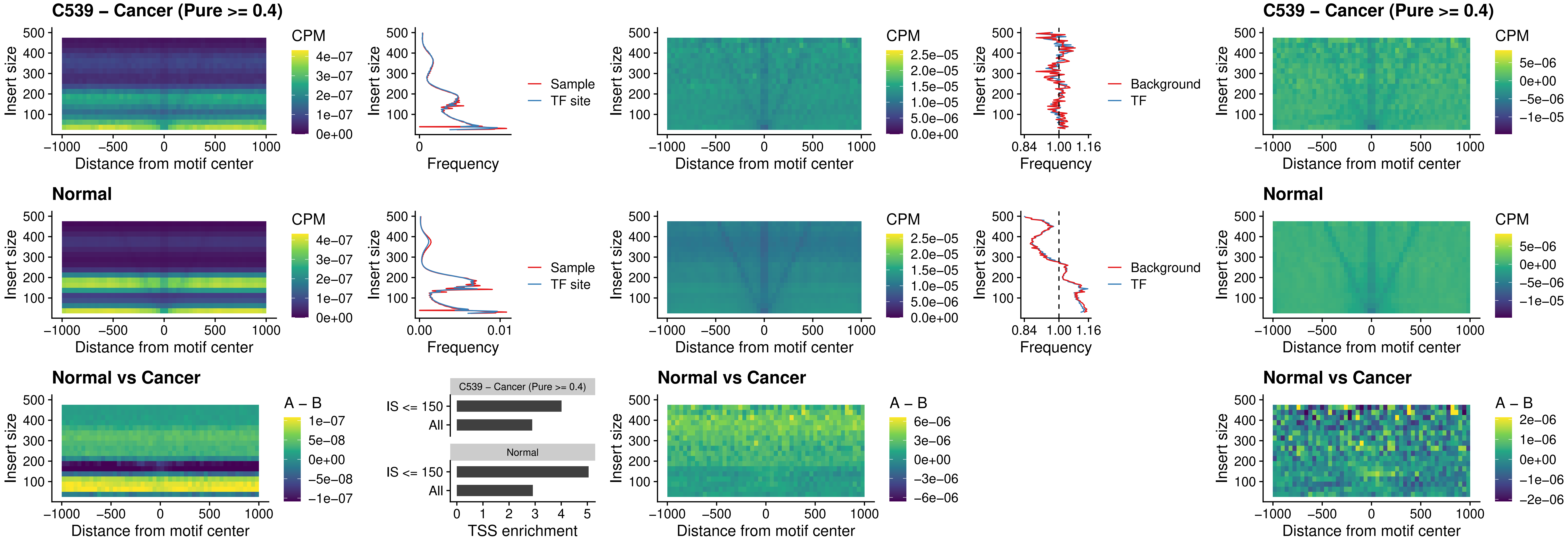

#### C539 – Cancer (Pure >= 0.4) – HLX_uc_dTSS_oPEAK

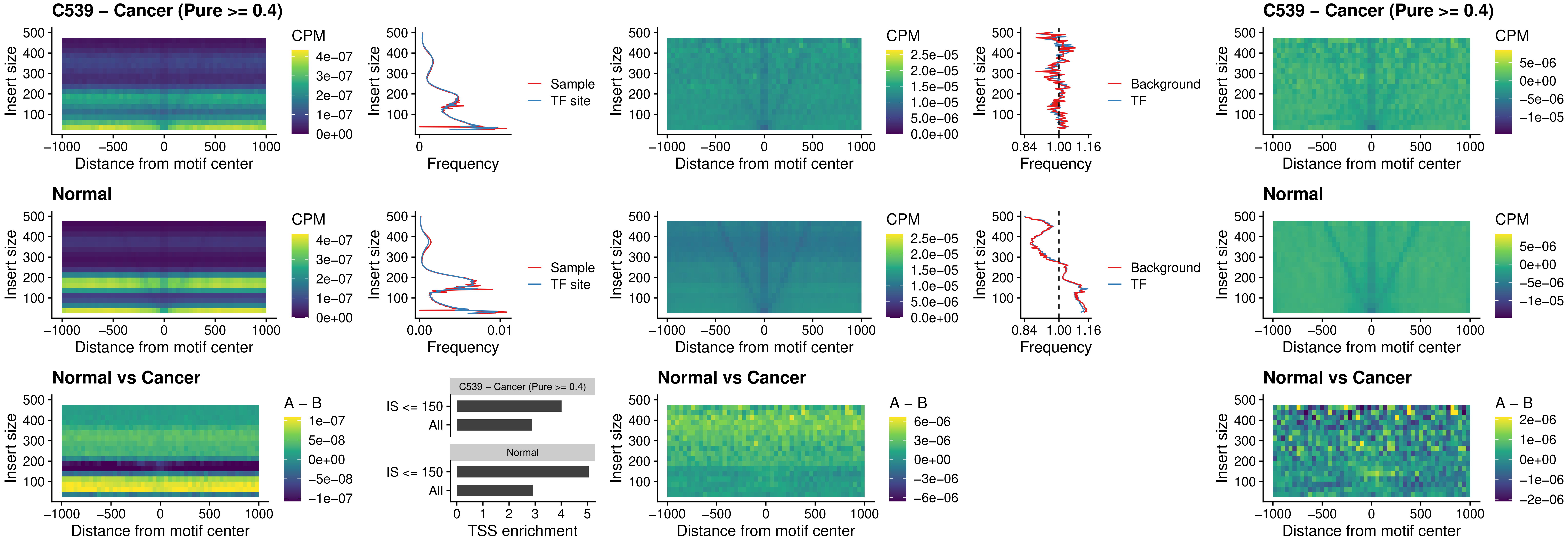

#### C539 – Cancer (Pure >= 0.4) – AC0125311_uc_dTSS_oPEAK

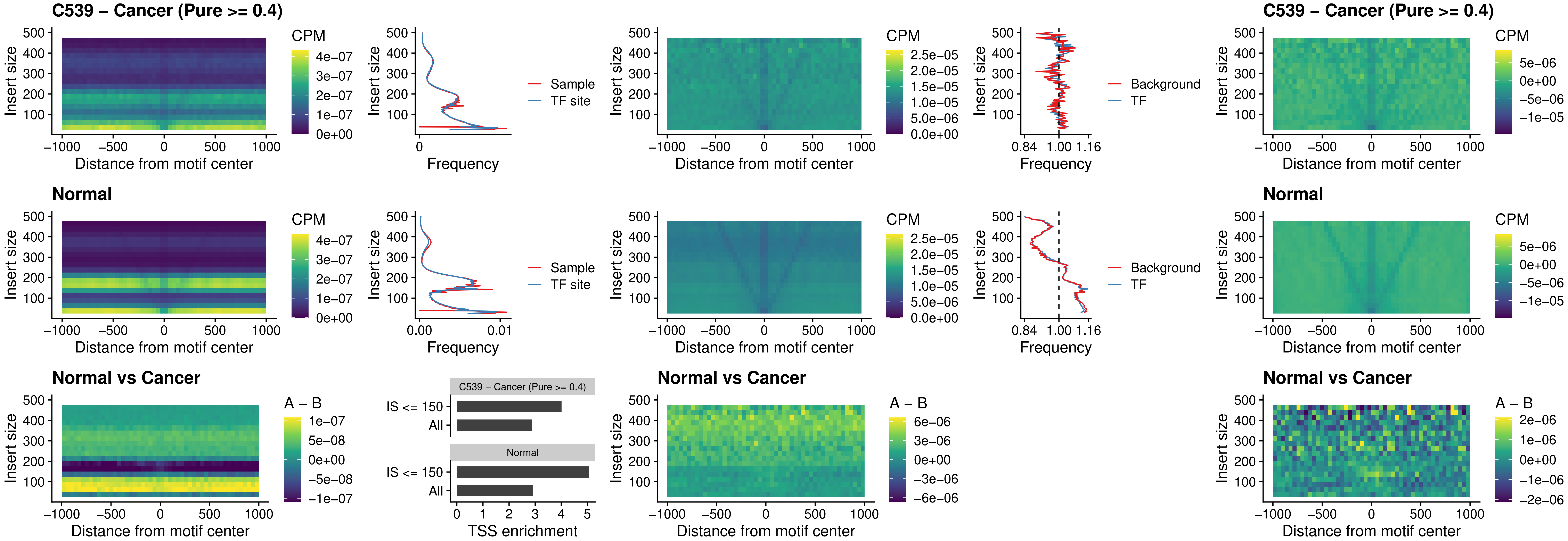

### C539 – Cancer (Pure >= 0.4) – DBX2_uc_dTSS_oPEAK

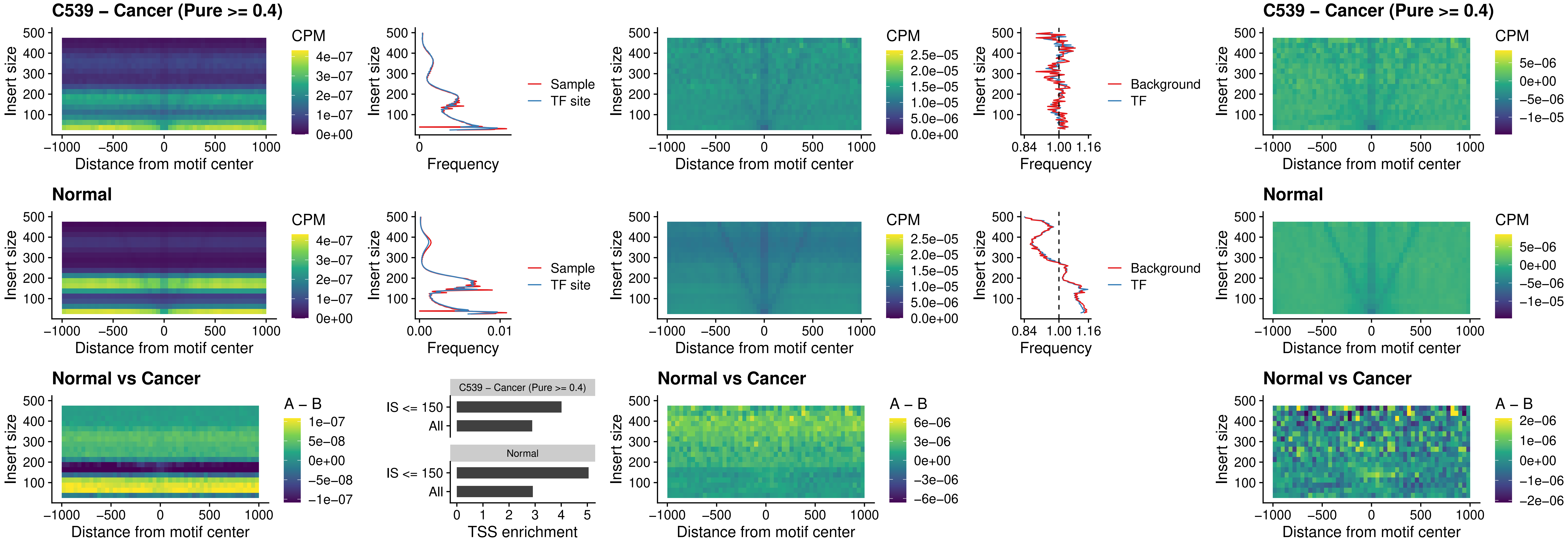

#### C539 – Cancer (Pure >= 0.4) – HOXA2_uc_dTSS_nPEAK

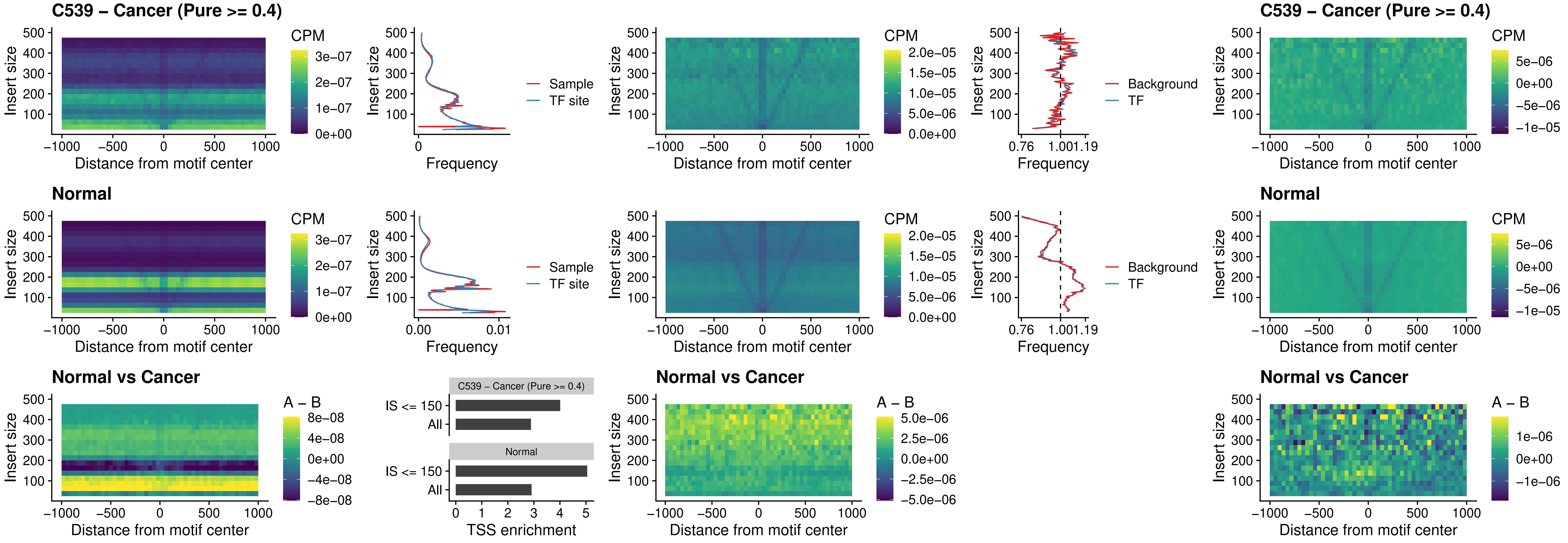

#### C539 – Cancer (Pure >= 0.4) – SHOX_uc_dTSS_nPEAK

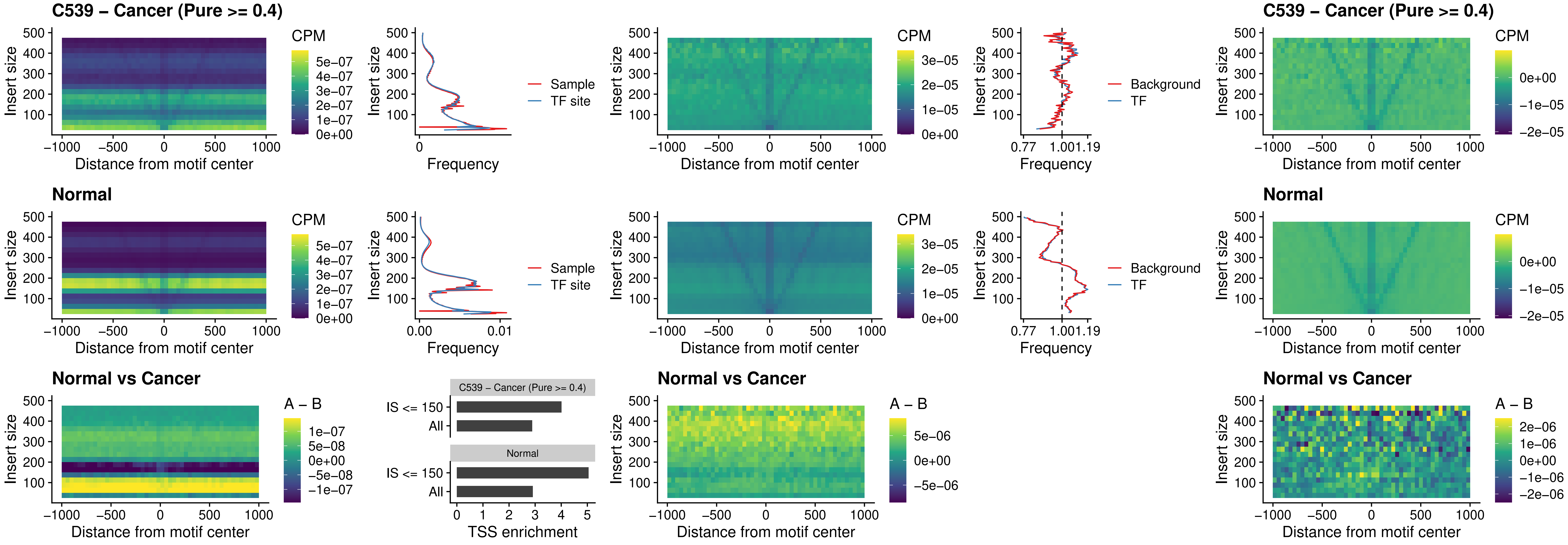

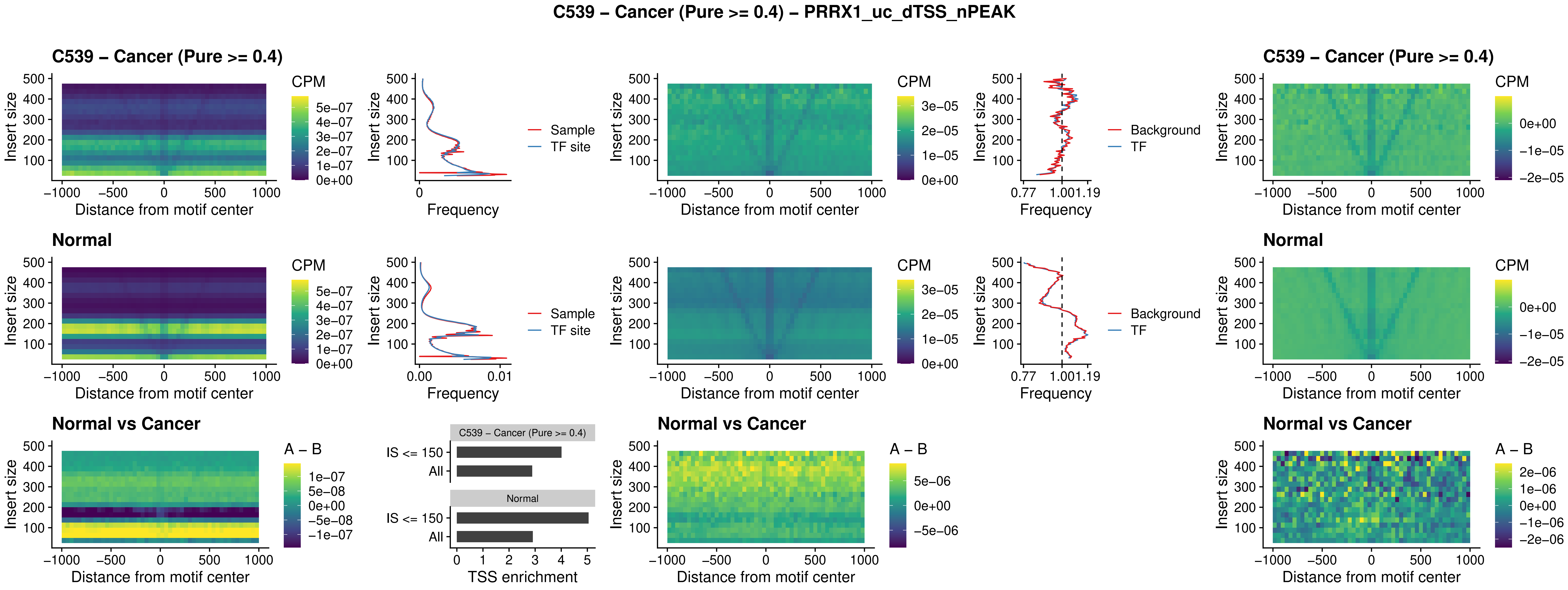
C539 – Cancer (Pure >= 0.4) – PRRX1_uc_dTSS_nPEAK
C539 – Cancer (Pure >= 0.4)
Normal
Normal vs Cancer
Insert size
Distance from motif center
Frequency
CPM
A – B
Sample
TF site
Background
TF
C539 – Cancer (Pure >= 0.4)
Normal
IS <= 150
All
TSS enrichment

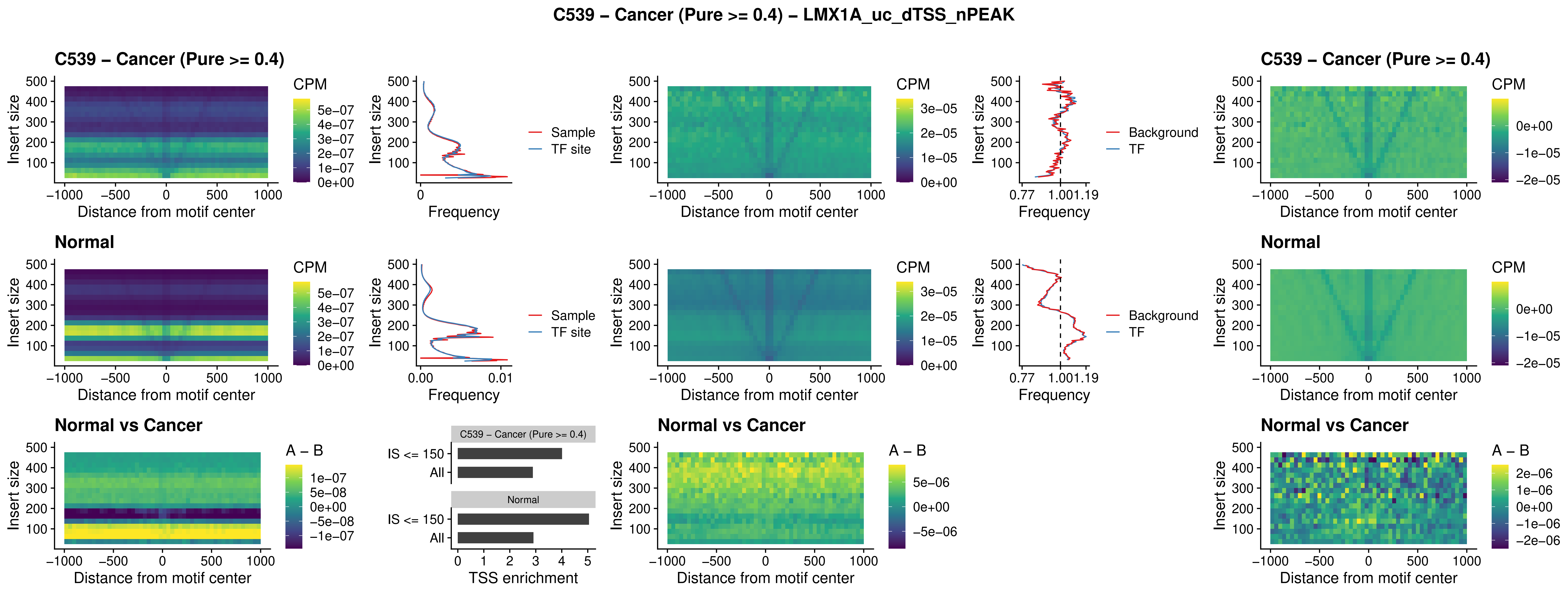

C539 – Cancer (Pure >= 0.4) – LMX1A_uc_dTSS_nPEAK
C539 – Cancer (Pure >= 0.4)
Normal
Normal vs Cancer
Insert size
Distance from motif center
CPM
A – B
Frequency
Sample
TF site
Background
TF
C539 – Cancer (Pure >= 0.4)
Normal
IS <= 150
All
TSS enrichment

#### C539 – Cancer (Pure >= 0.4) – LMX1B_uc_dTSS_nPEAK

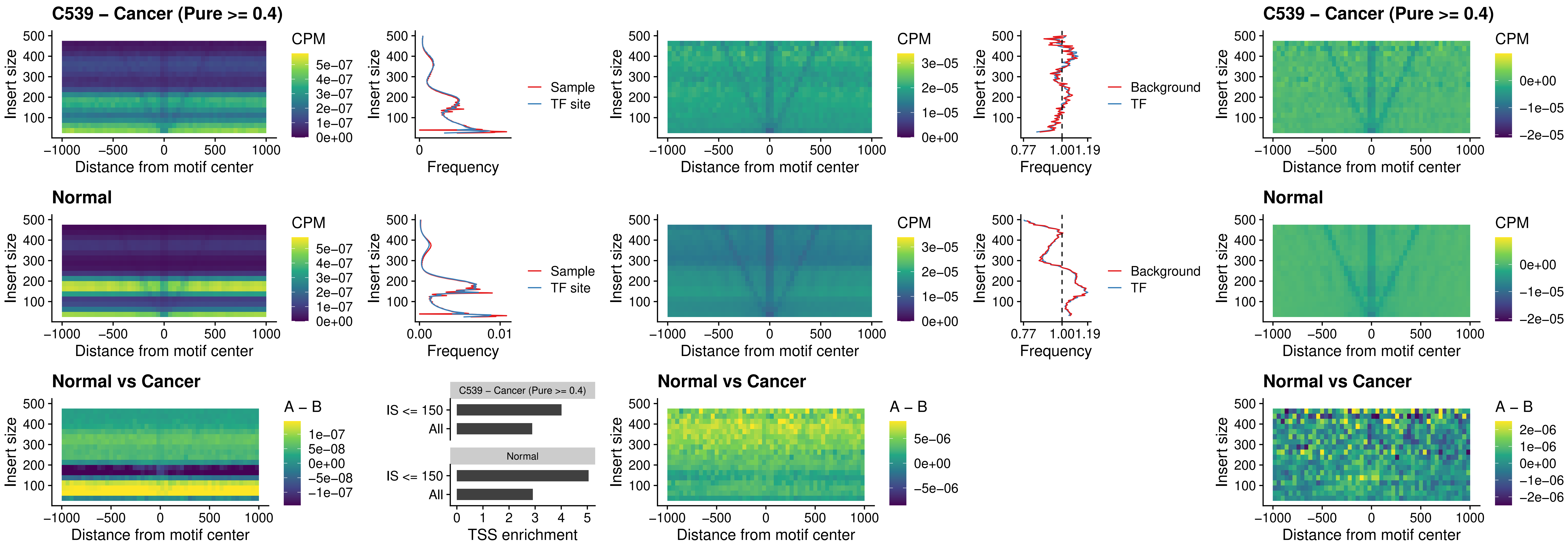

#### C539 – Cancer (Pure >= 0.4) – DLX5_uc_dTSS_oPEAK

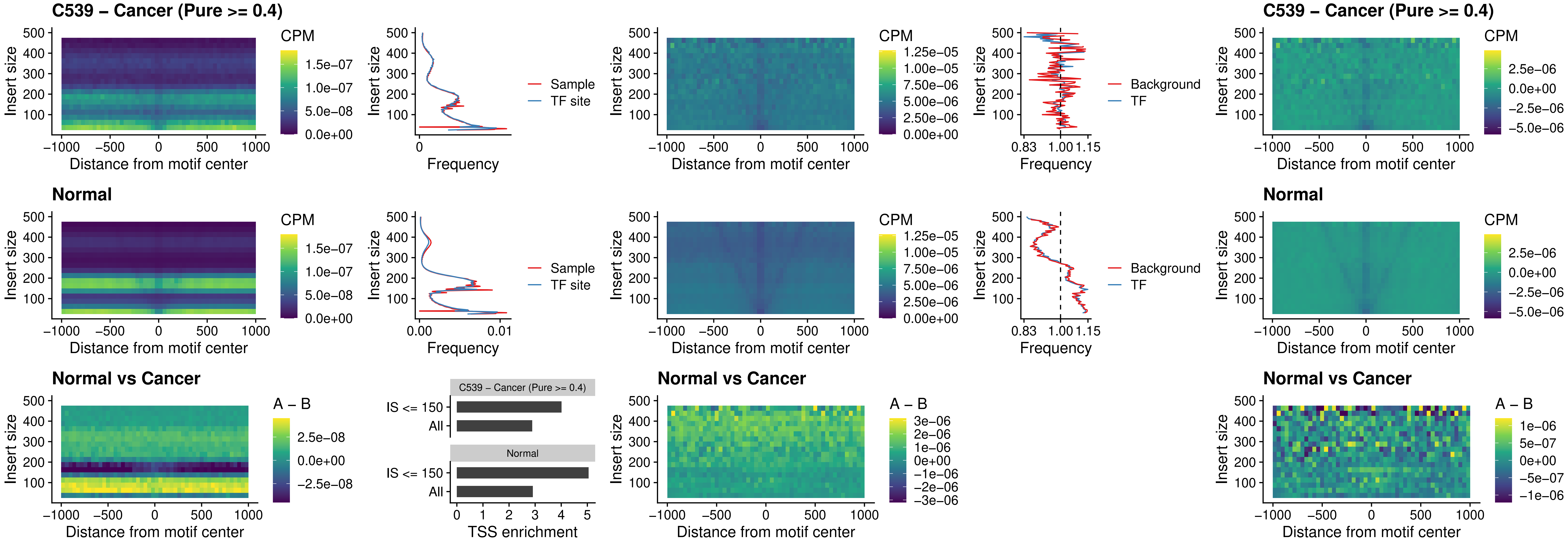

#### C539 – Cancer (Pure >= 0.4) – ARID3C_uc_dTSS_nPEAK

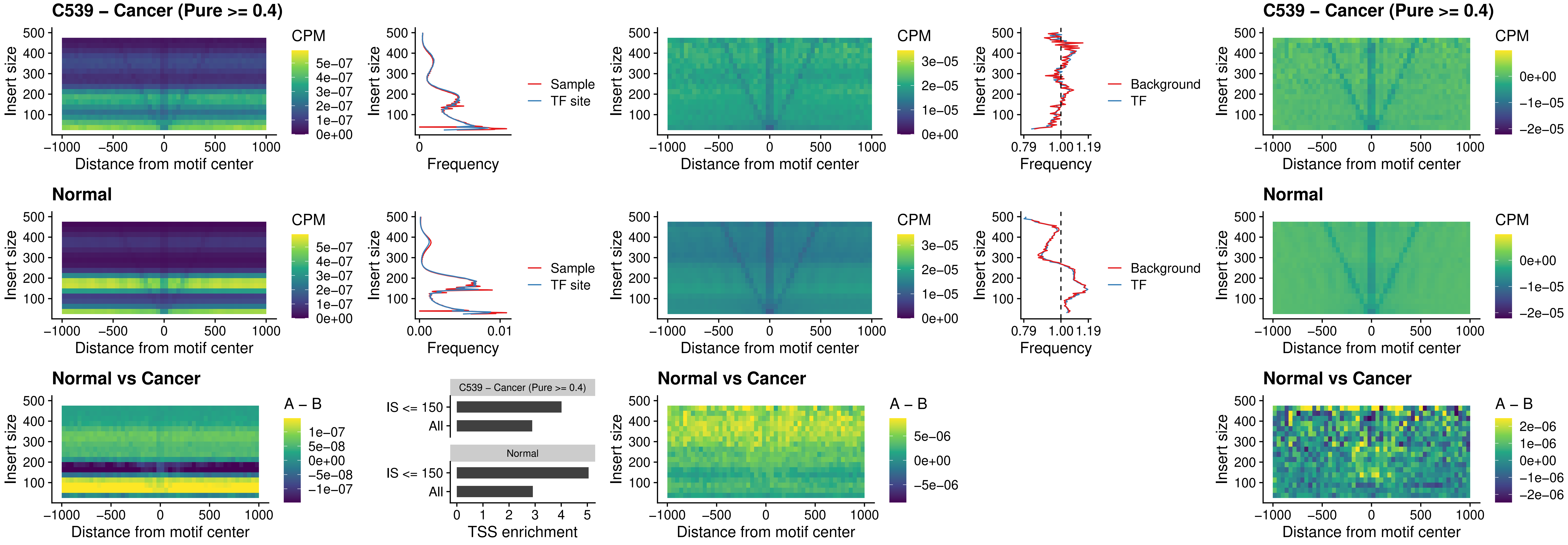

#### C539 – Cancer (Pure >= 0.4) – HOXD12_uc_dTSS_nPEAK

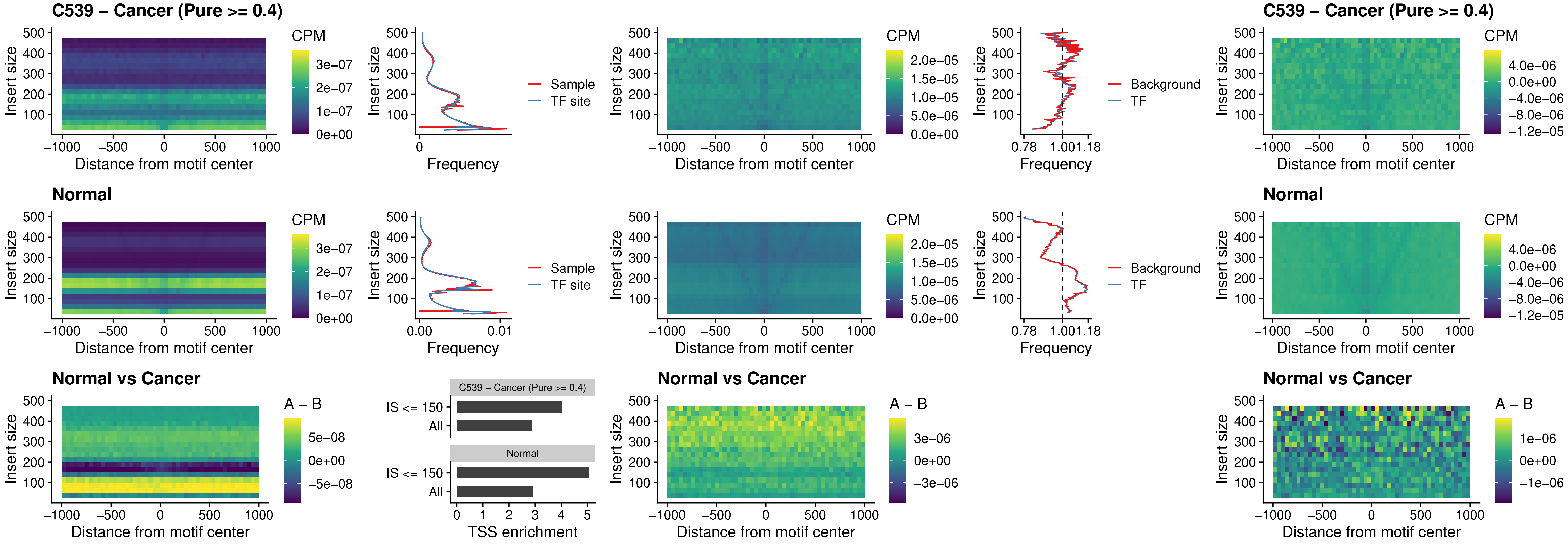

#### C539 – Cancer (Pure >= 0.4) – CDX2_uc_dTSS_nPEAK

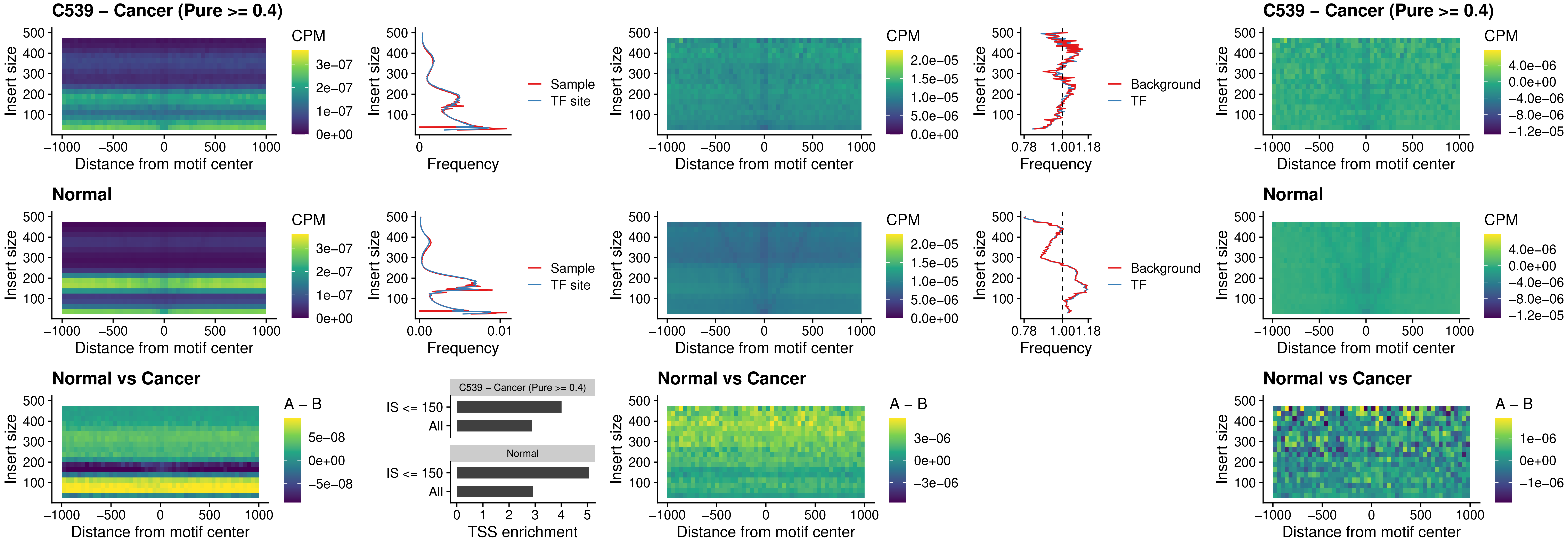

#### C539 – Cancer (Pure >= 0.4) – HOXC12_uc_dTSS_nPEAK

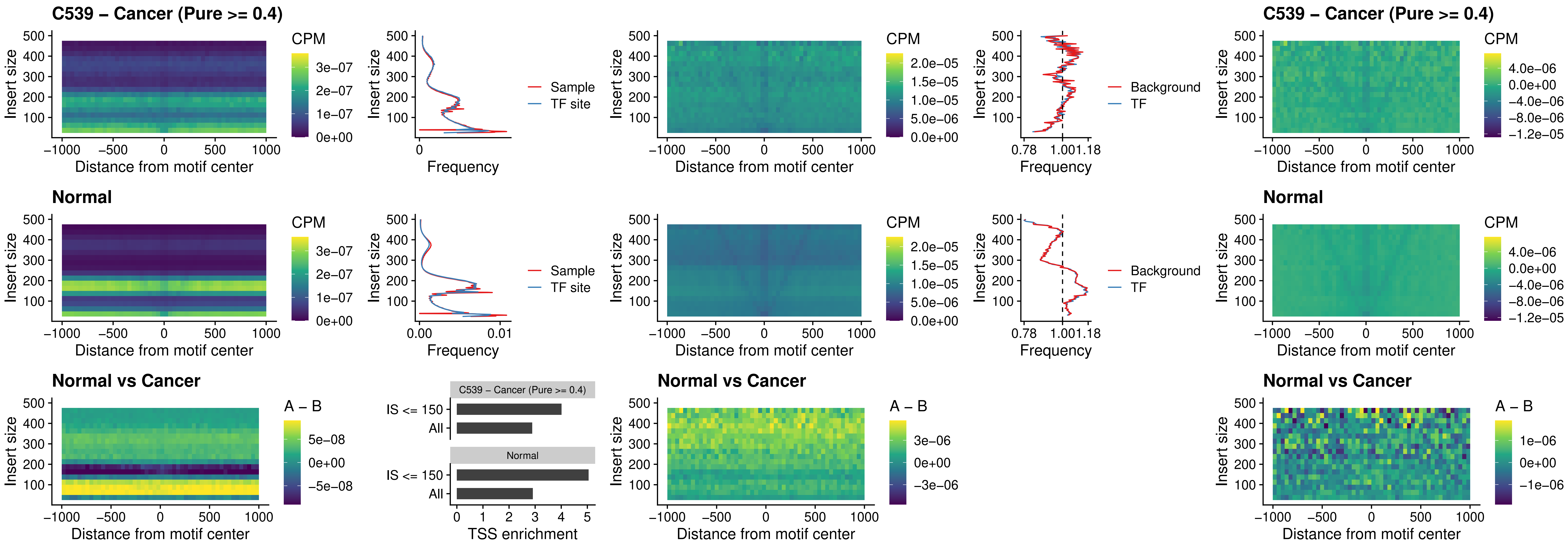

#### C539 – Cancer (Pure >= 0.4) – HHEX_uc_dTSS_oPEAK

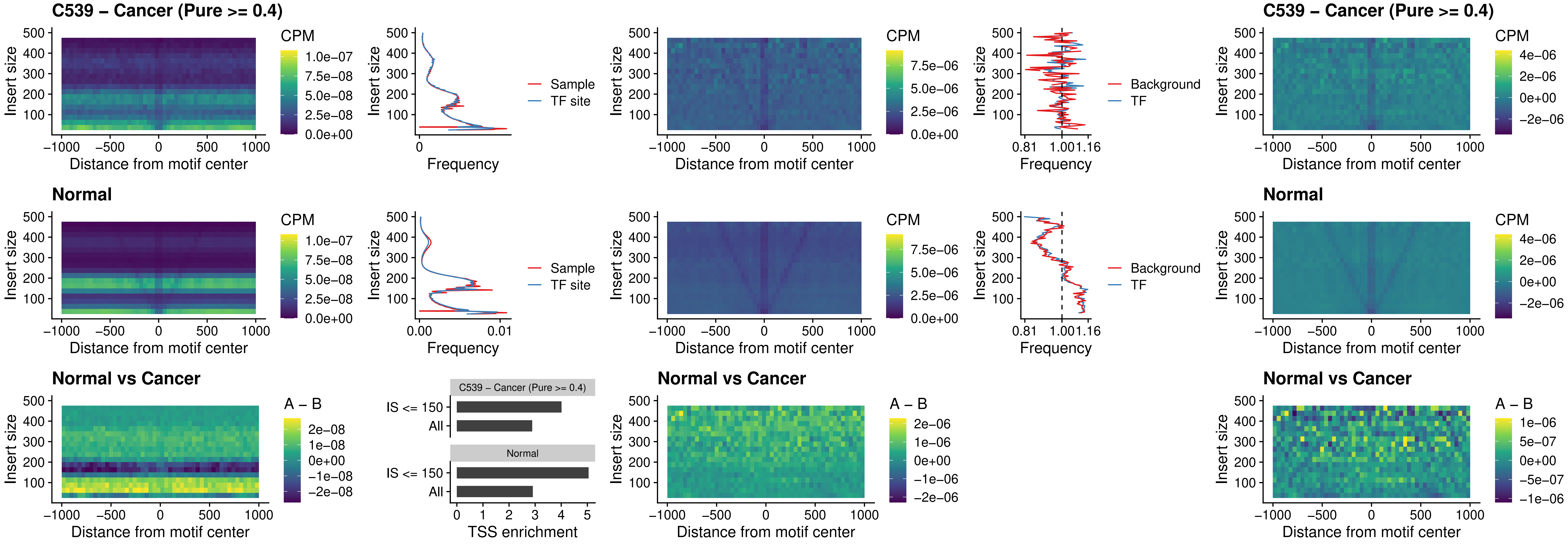

#### C539 – Cancer (Pure >= 0.4) – HOXA11_uc_dTSS_nPEAK

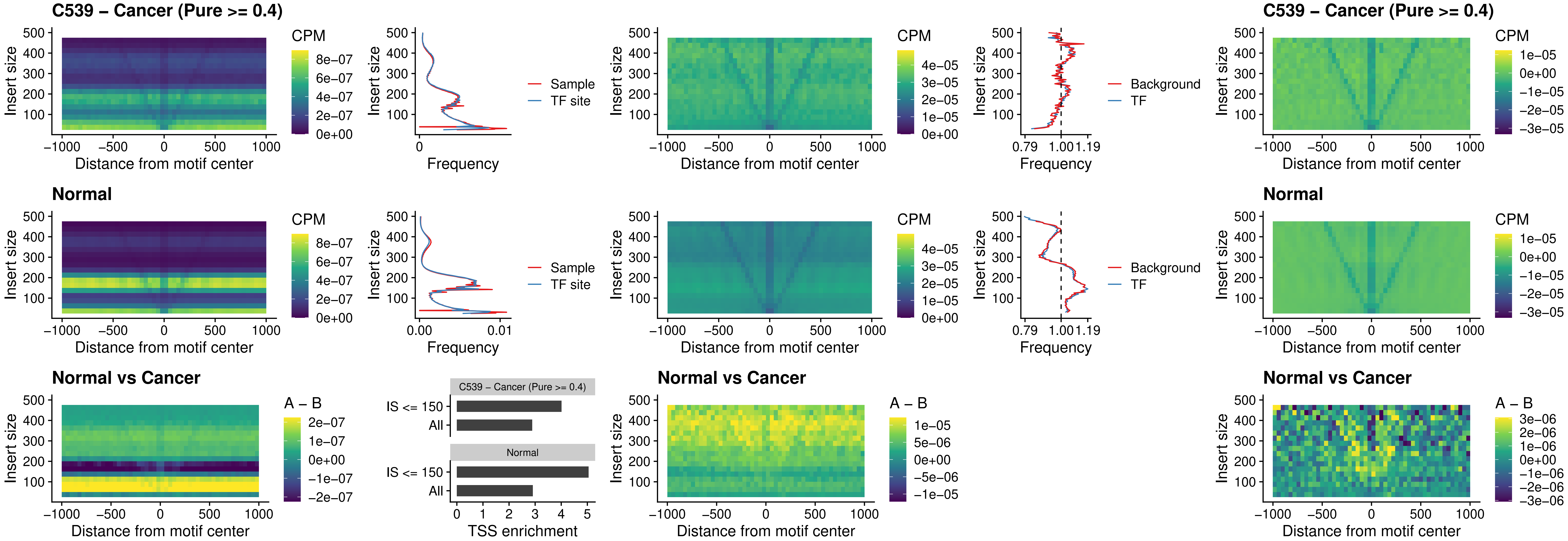

#### C539 – Cancer (Pure >= 0.4) – HOXA11_uc_cTSS_nPEAK

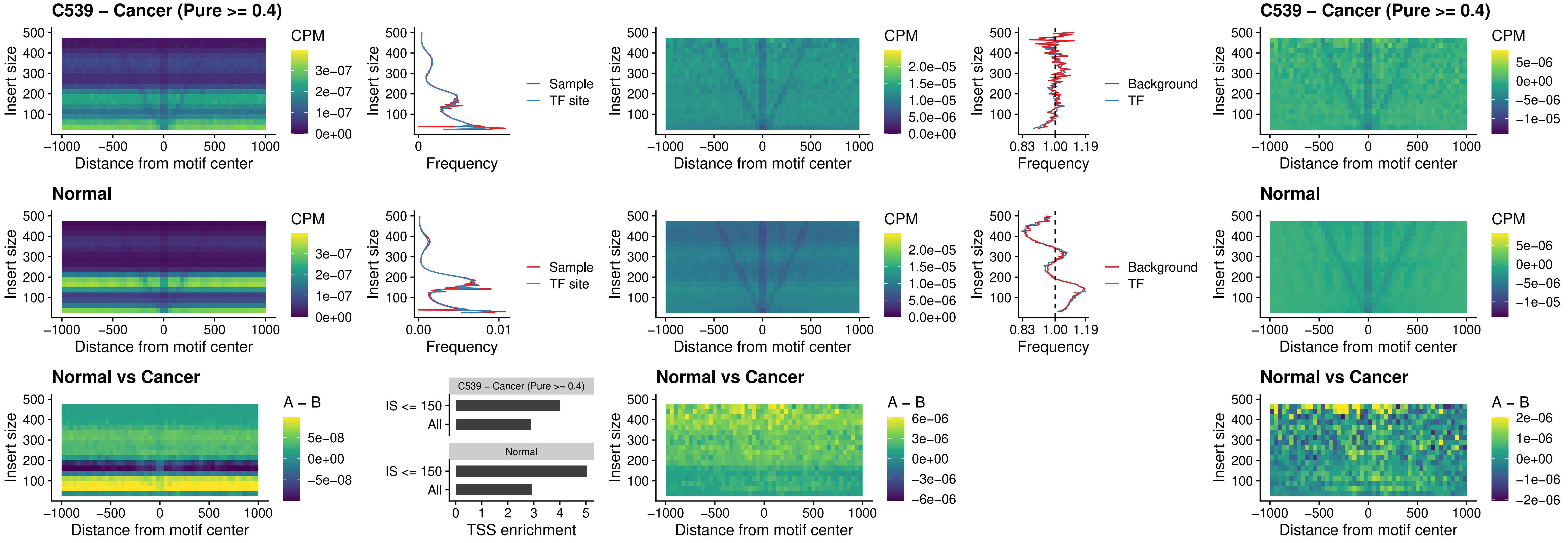

### C539 – Cancer (Pure >= 0.4) – ZFHX3_uc_dTSS_nPEAK

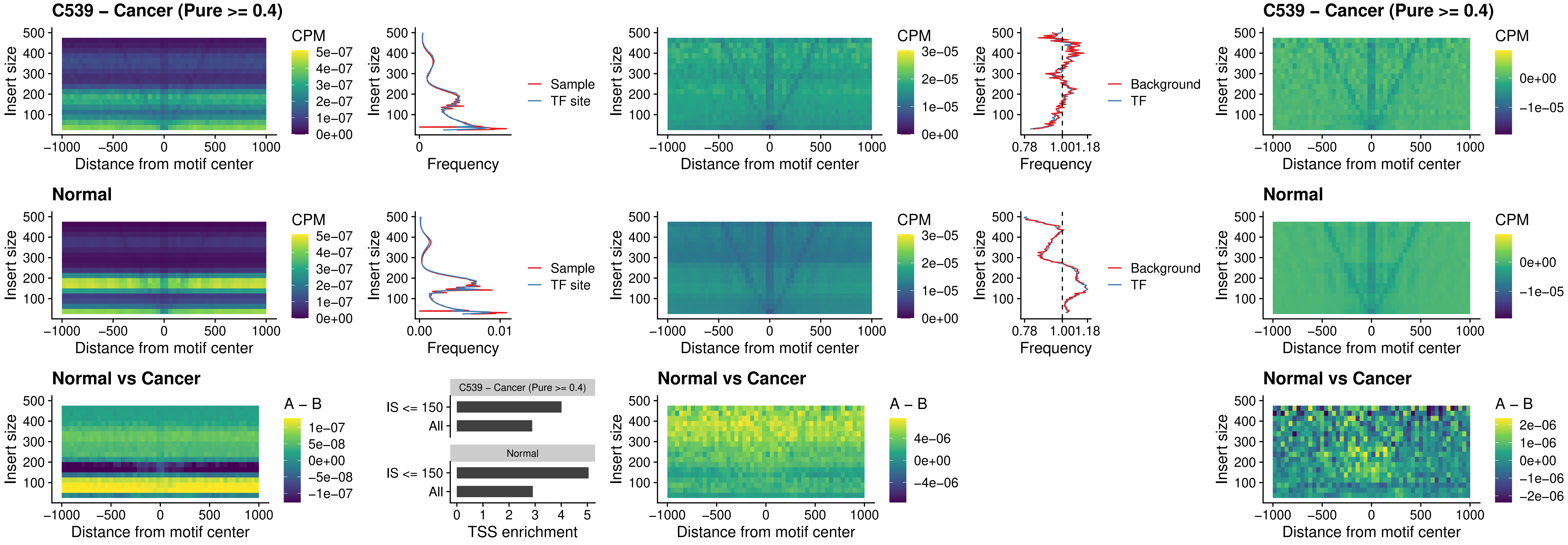

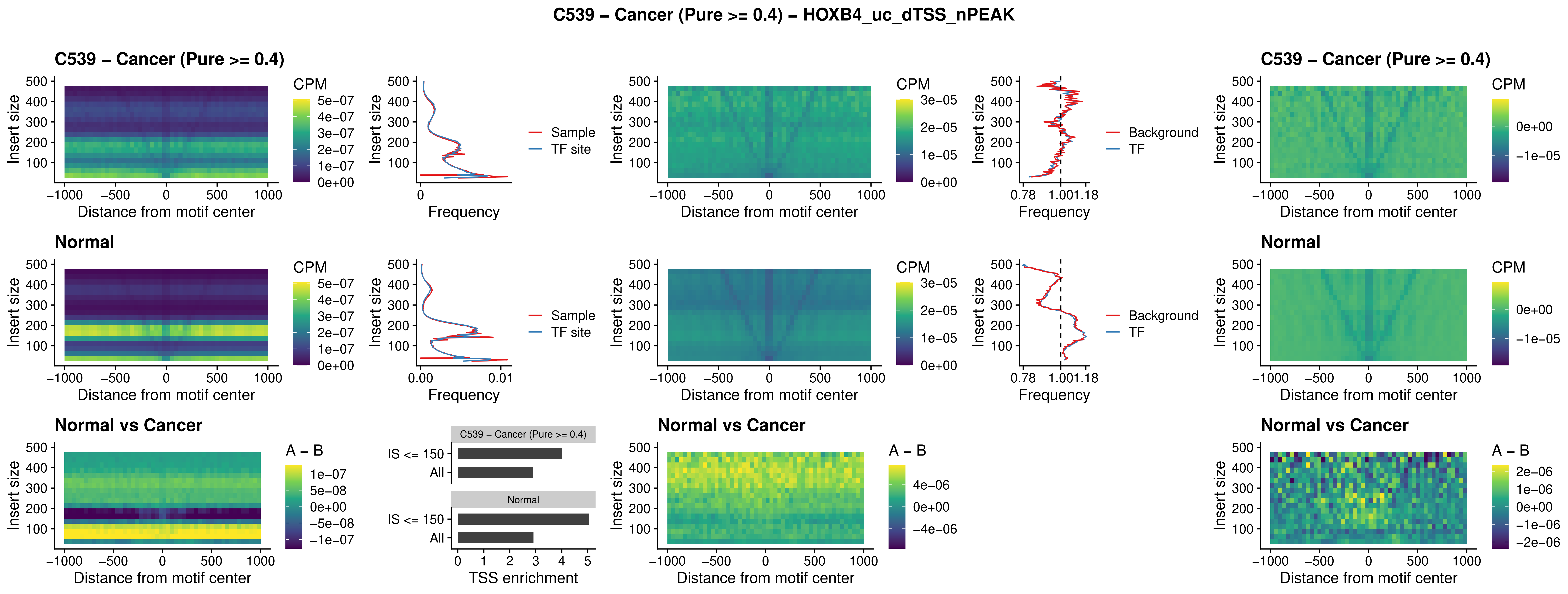

C539 – Cancer (Pure >= 0.4) – HOXB4_uc_dTSS_nPEAK
C539 – Cancer (Pure >= 0.4)
Normal
Normal vs Cancer
Insert size
Distance from motif center
Frequency
CPM
A – B
Sample
TF site
Background
TF
TSS enrichment
IS <= 150
All
C539 – Cancer (Pure >= 0.4)
Normal

#### C539 – Cancer (Pure >= 0.4) – HOXC5_uc_dTSS_nPEAK

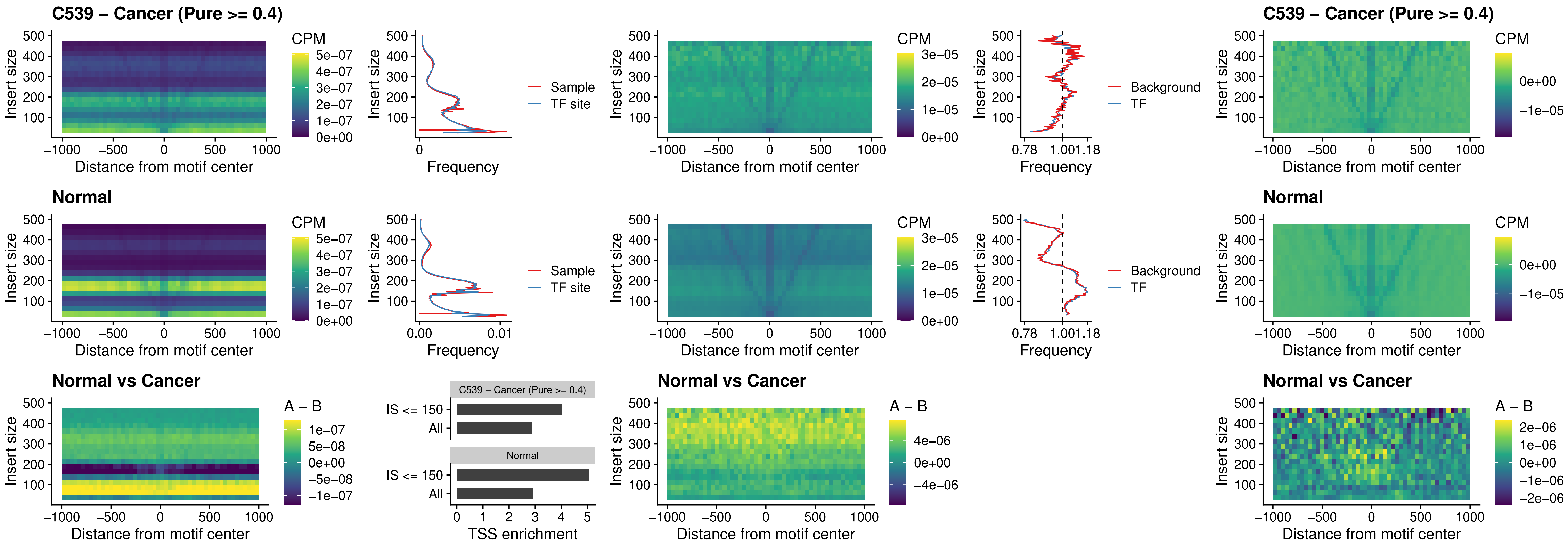

#### C539 – Cancer (Pure >= 0.4) – HOXD1_uc_dTSS_nPEAK

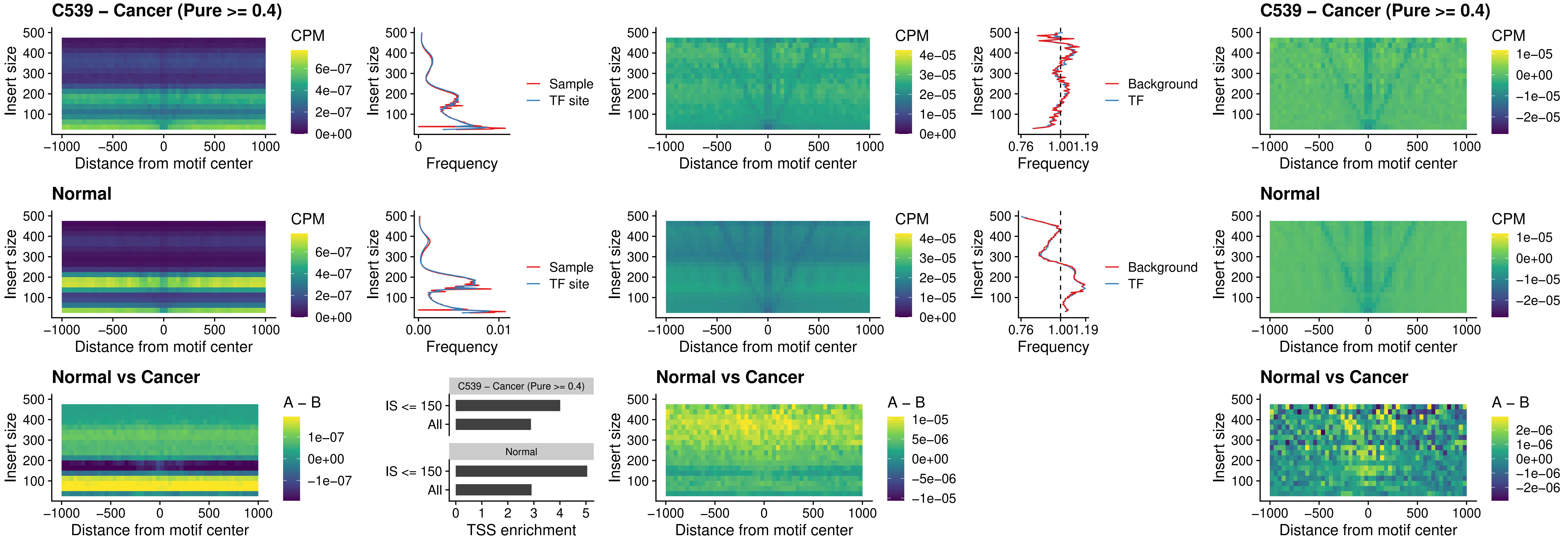

#### C539 – Cancer (Pure >= 0.4) – OTP_uc_dTSS_nPEAK

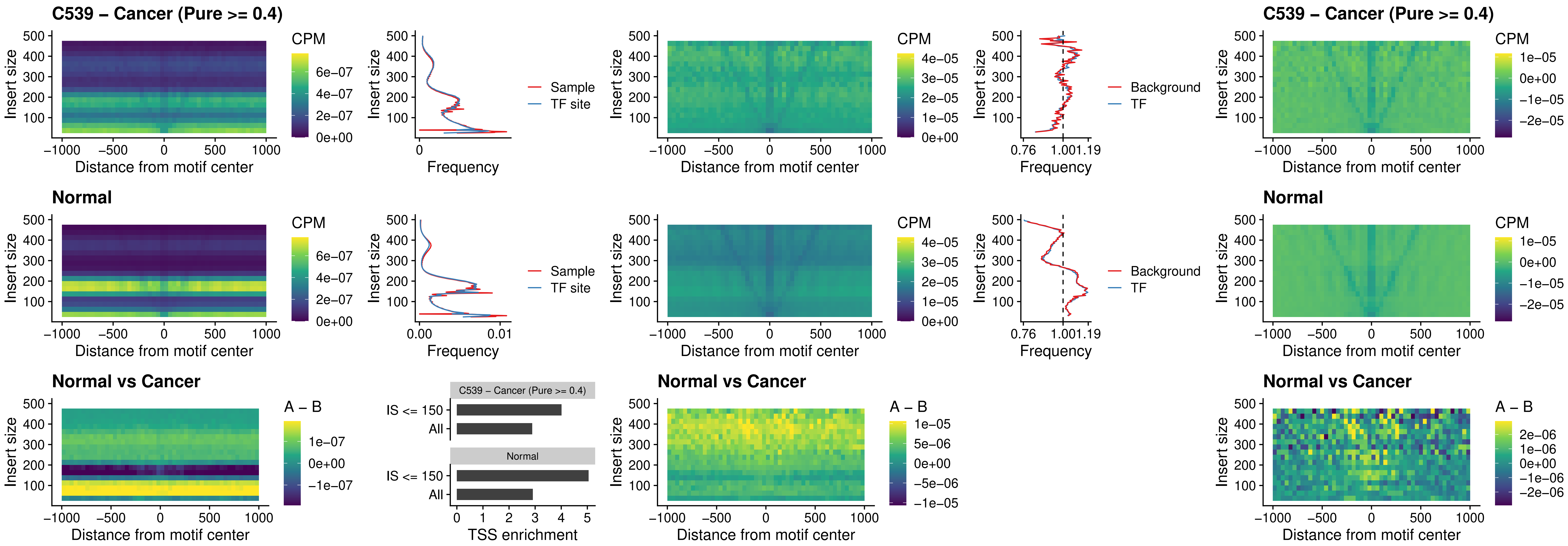

### C539 – Cancer (Pure >= 0.4) – FOXL1_uc_dTSS_oPEAK

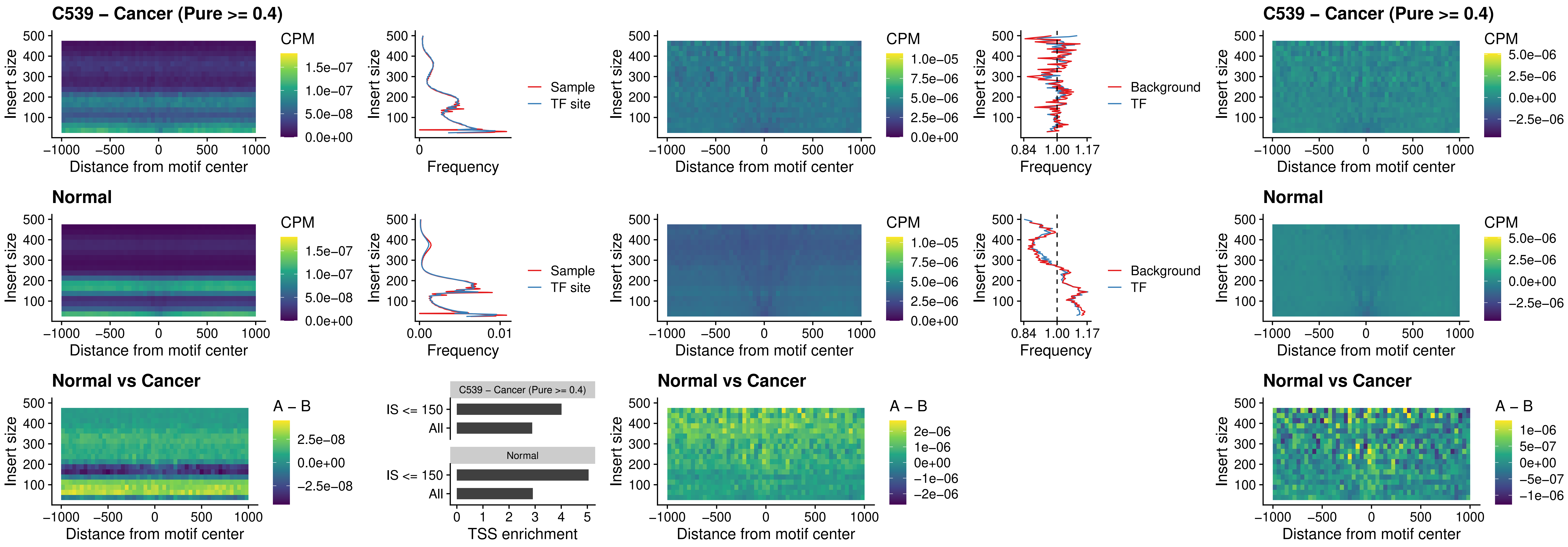

#### C539 – Cancer (Pure >= 0.4) – ZNF32_uc_dTSS_oPEAK

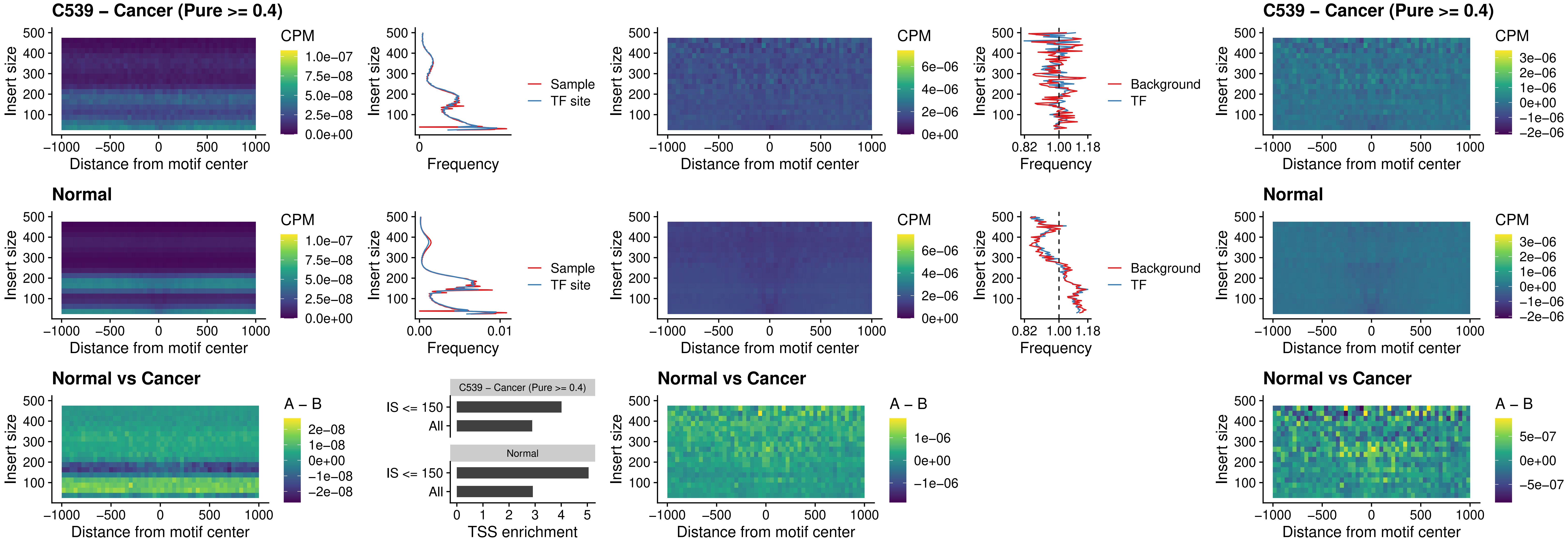

#### C539 – Cancer (Pure >= 0.4) – FOXL1_uc_dTSS_nPEAK

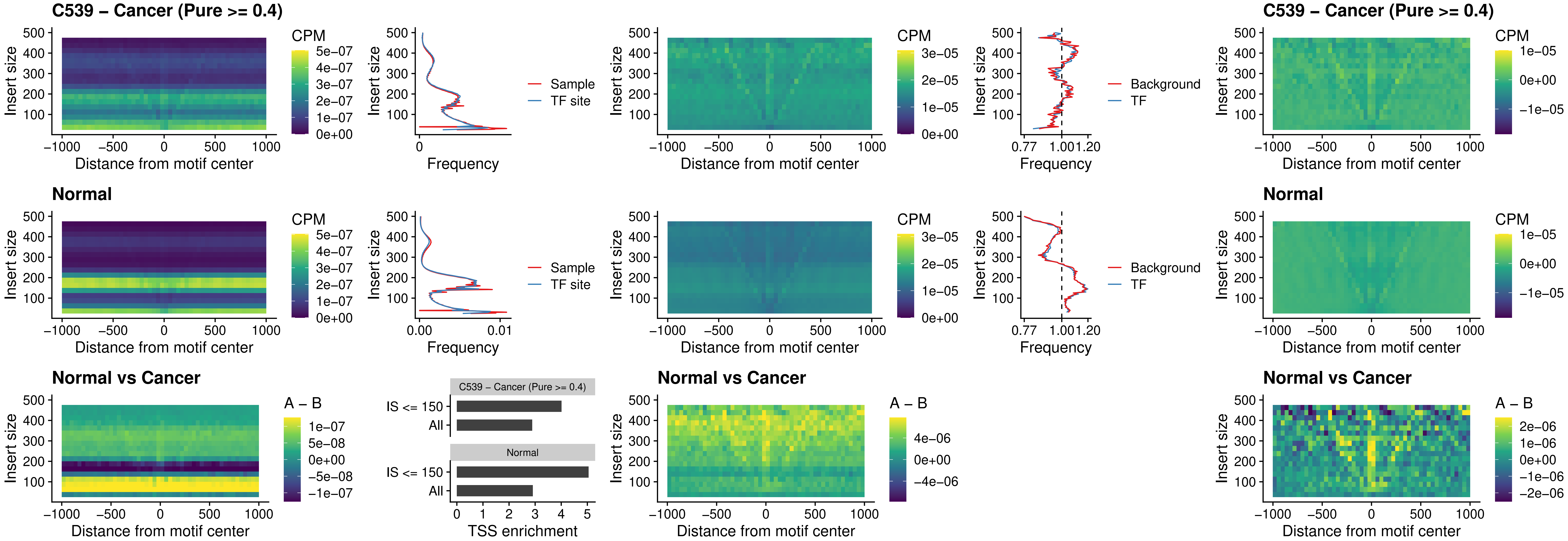

#### C539 – Cancer (Pure >= 0.4) – ZNF32_uc_dTSS_nPEAK

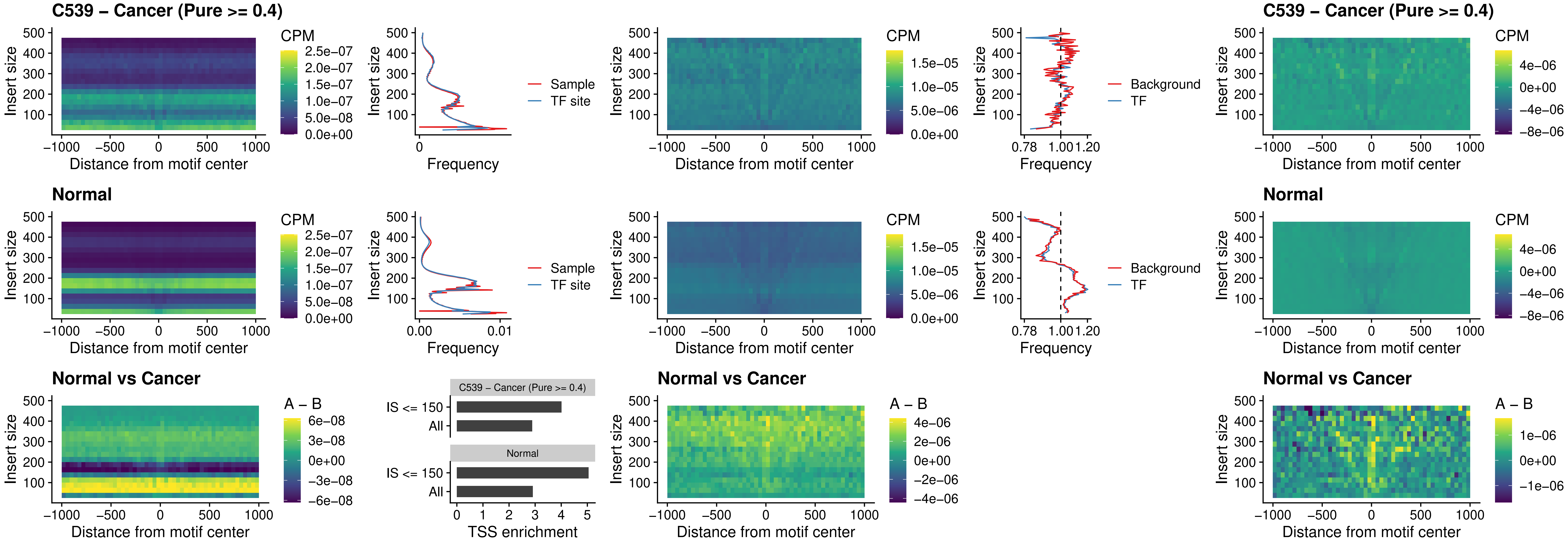

### C539 – Cancer (Pure >= 0.4) – DLX5_uc_dTSS_nPEAK

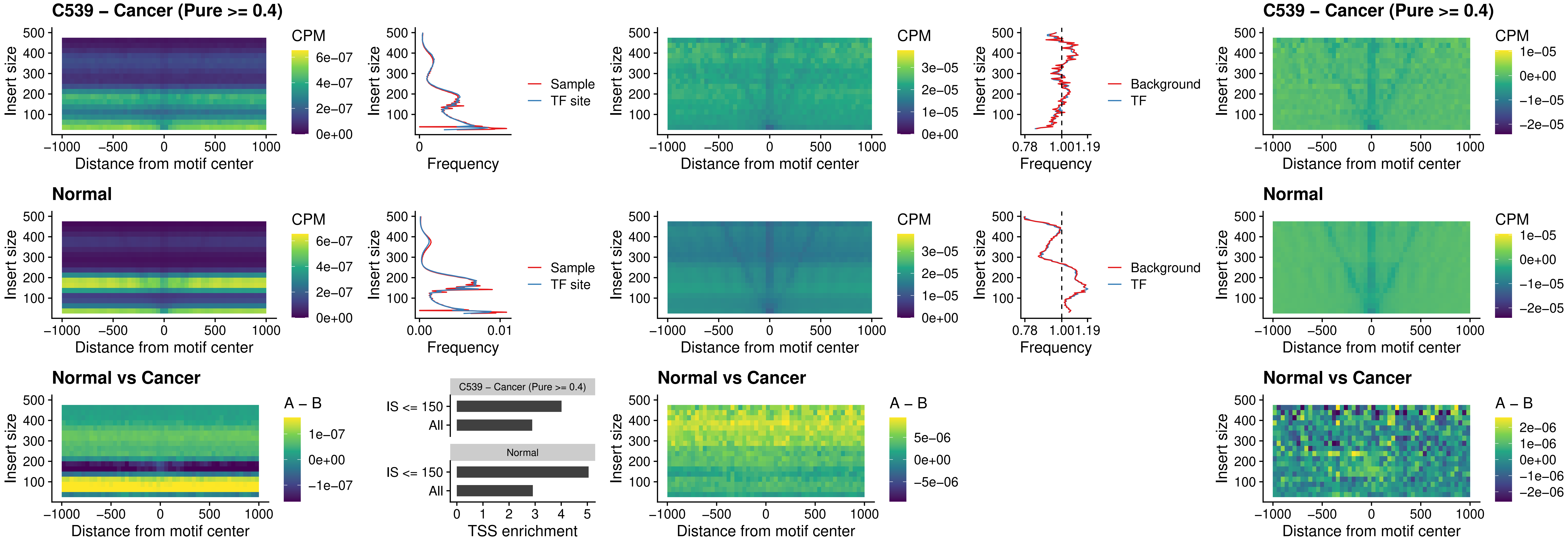

#### C539 – Cancer (Pure >= 0.4) – DMRT2_uc_dTSS_nPEAK

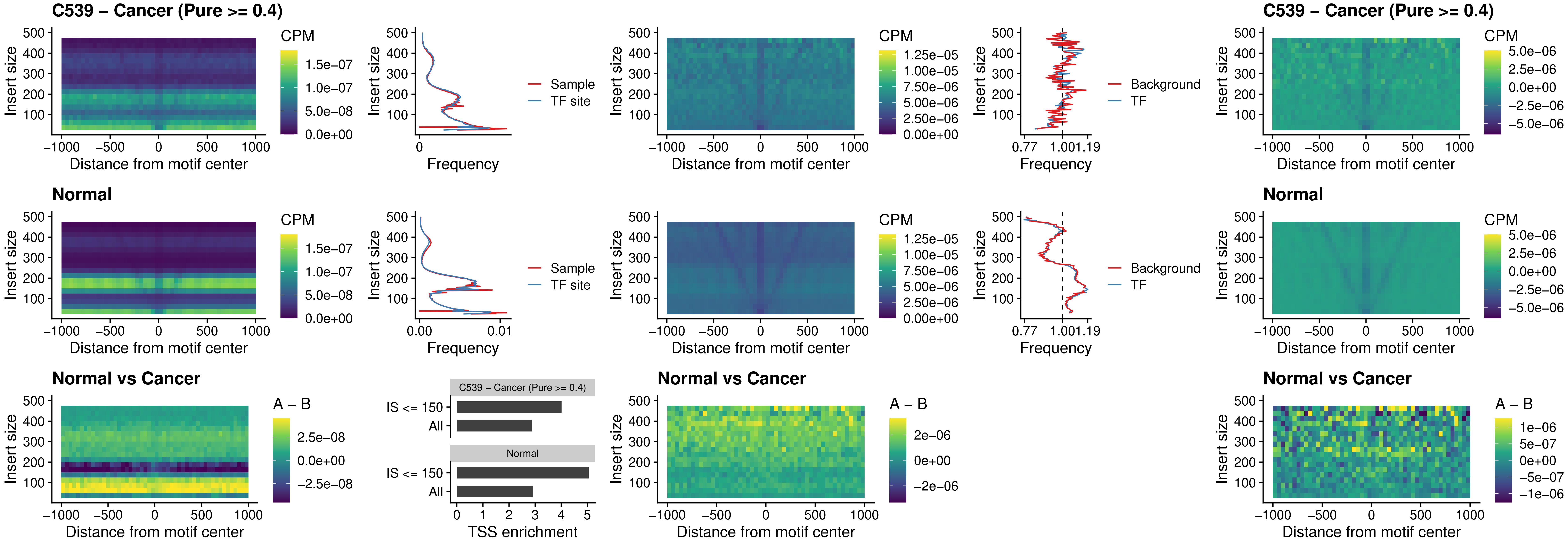

#### C539 – Cancer (Pure >= 0.4) – DMRTC2_uc_cTSS_nPEAK

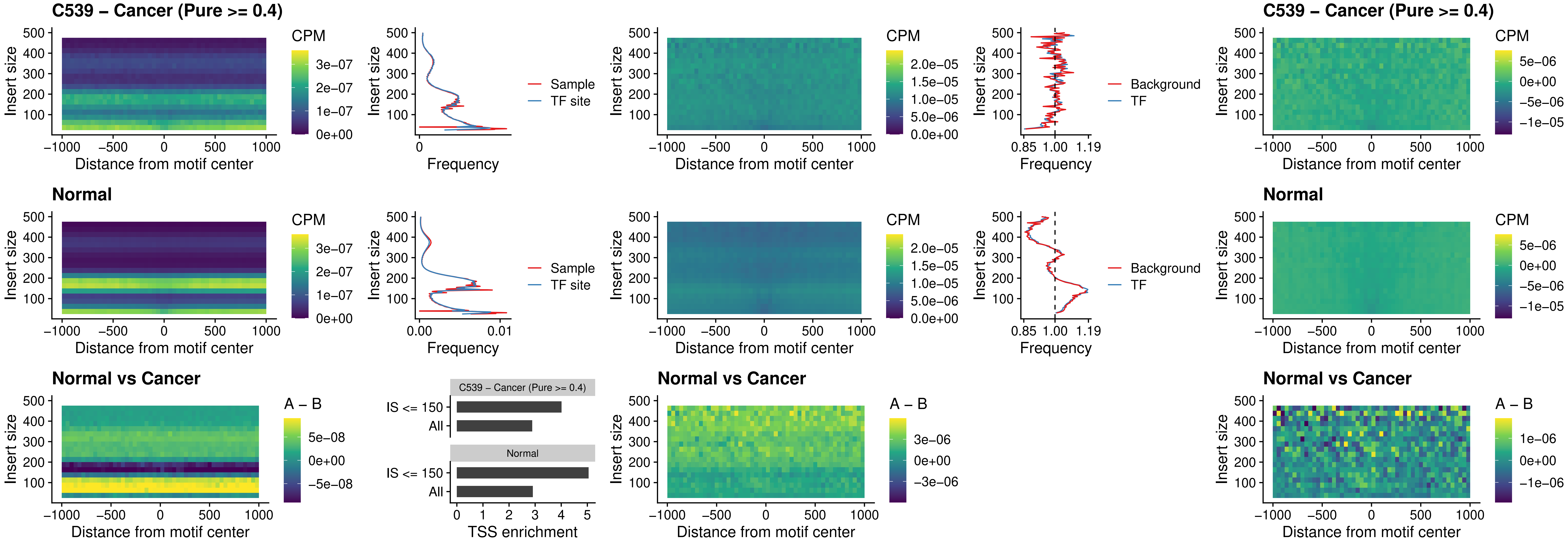

#### C539 – Cancer (Pure >= 0.4) – HOXD9_uc_dTSS_nPEAK

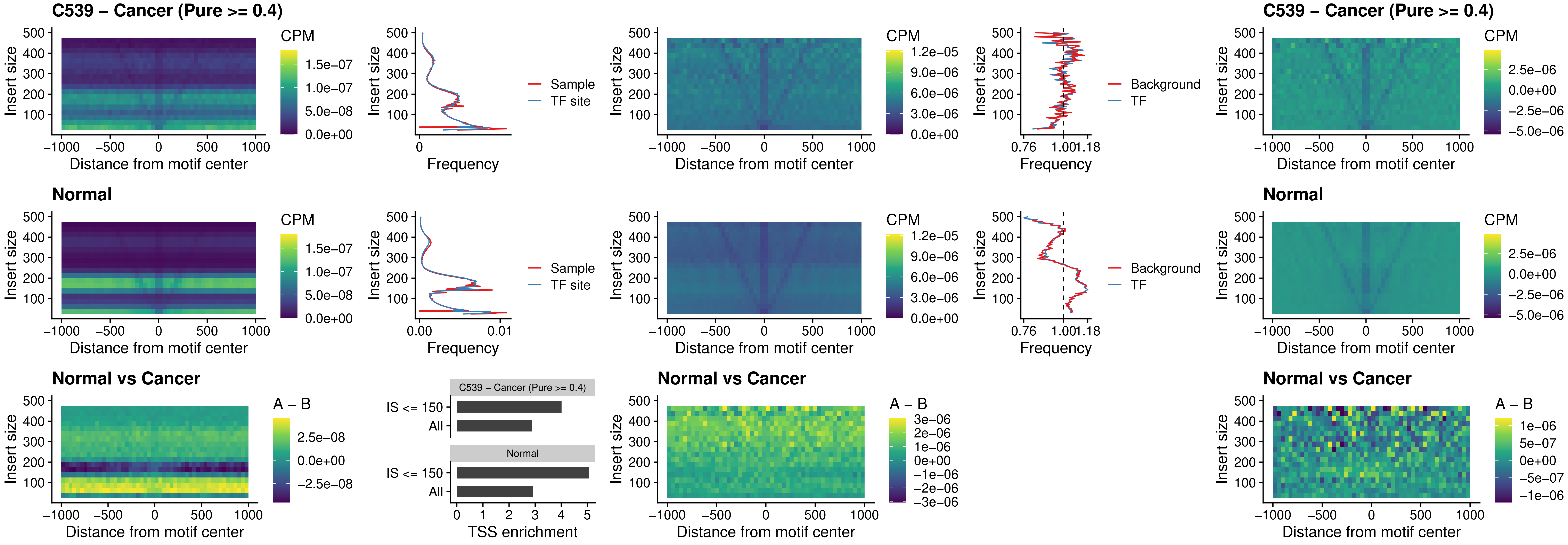

#### C539 – Cancer (Pure >= 0.4) – HOXD4_uc_dTSS_nPEAK

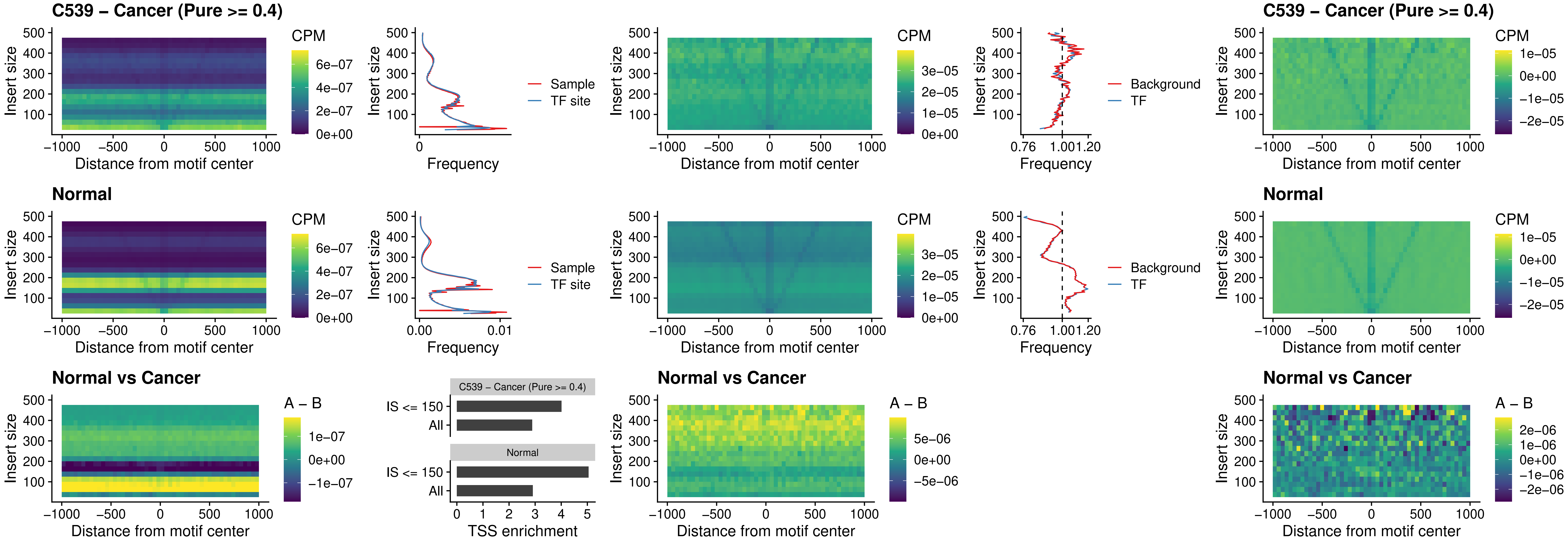

#### C539 – Cancer (Pure >= 0.4) – FOXP3_uc_cTSS_nPEAK

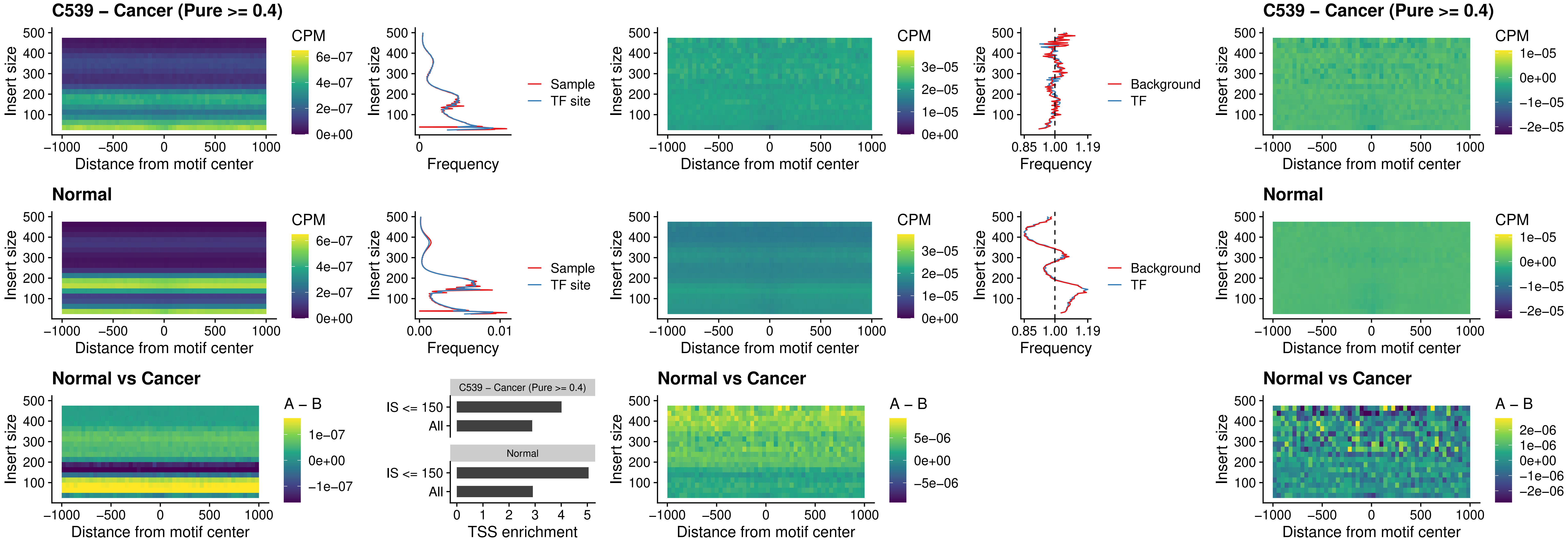

### C539 – Cancer (Pure >= 0.4) – FOXD4L6_uc_cTSS_nPEAK

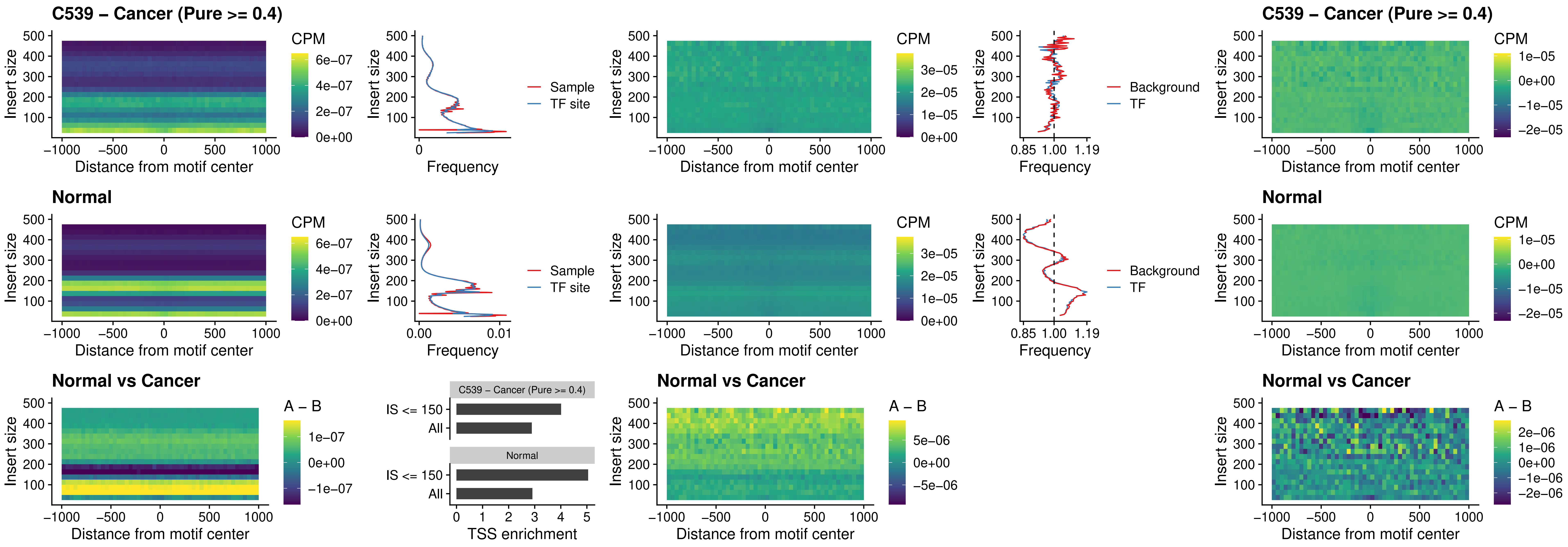

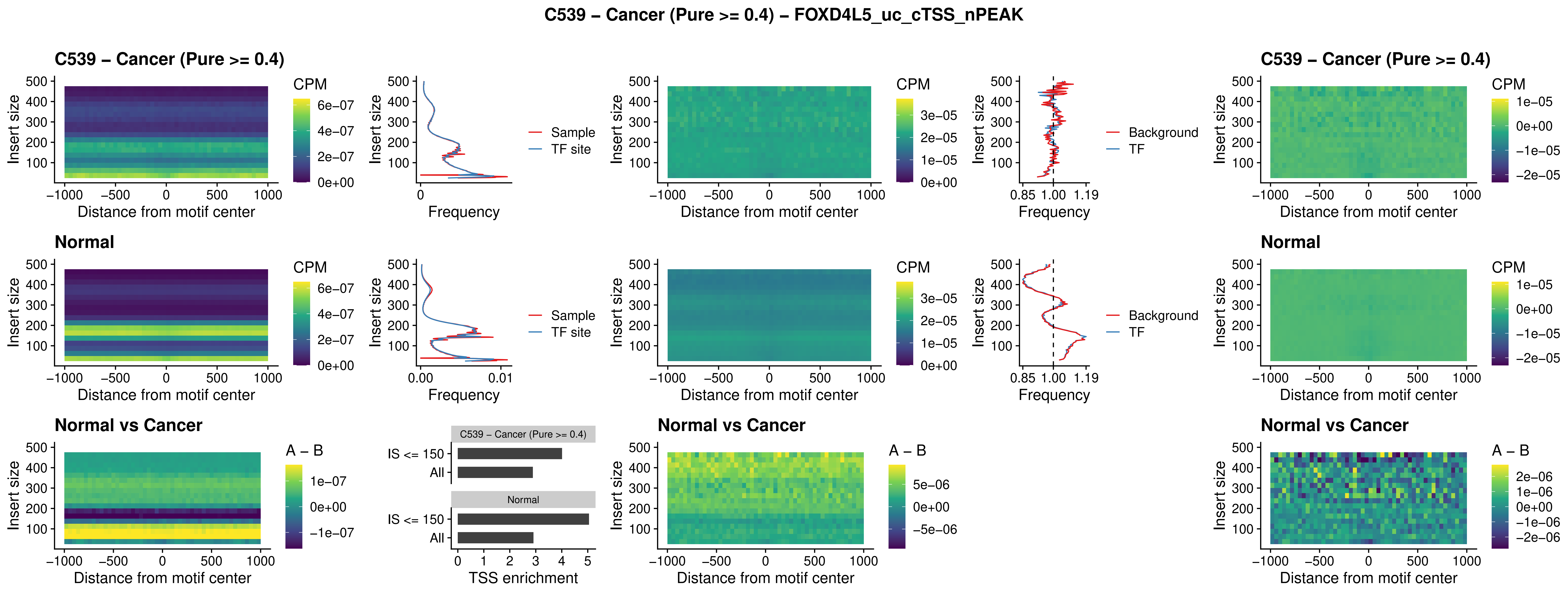
C539 – Cancer (Pure >= 0.4) – FOXD4L5_uc_cTSS_nPEAK
C539 – Cancer (Pure >= 0.4)
Normal
Normal vs Cancer
Insert size
Distance from motif center
Frequency
CPM
A – B
Sample
TF site
Background
TF
C539 – Cancer (Pure >= 0.4)
Normal
IS <= 150
All
TSS enrichment

#### C539 – Cancer (Pure >= 0.4) – FOXD4L4_uc_cTSS_nPEAK

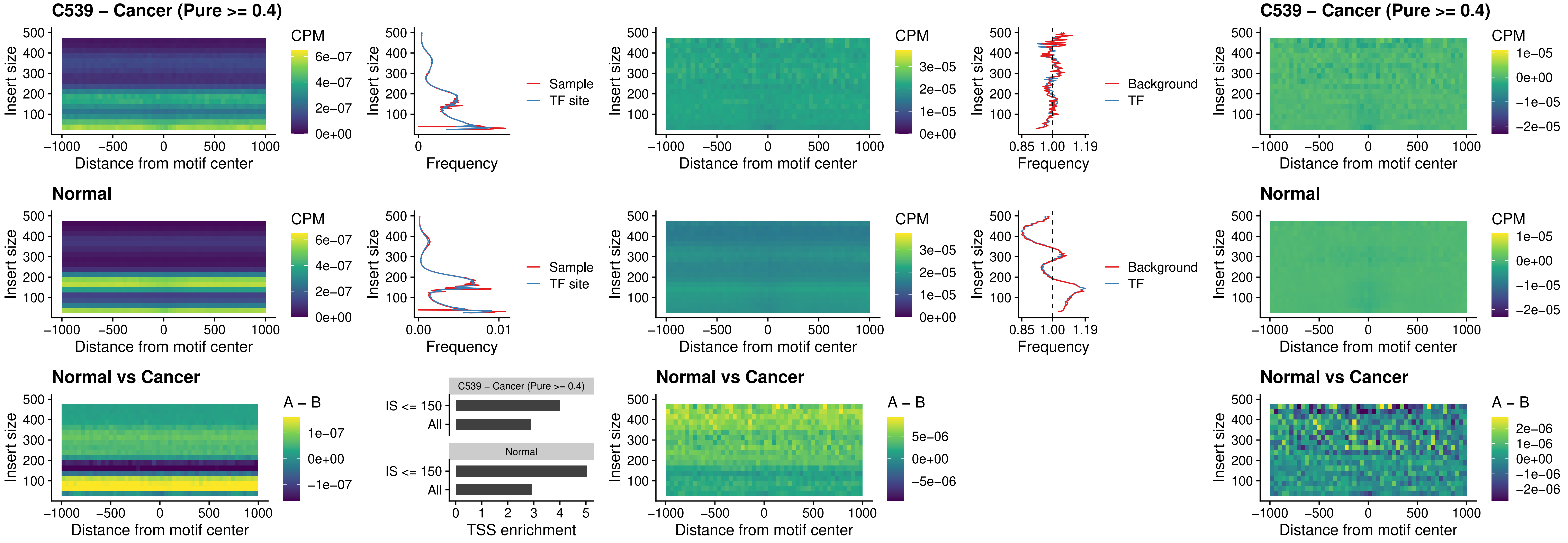

#### C539 – Cancer (Pure >= 0.4) – FOXD4L2_uc_cTSS_nPEAK

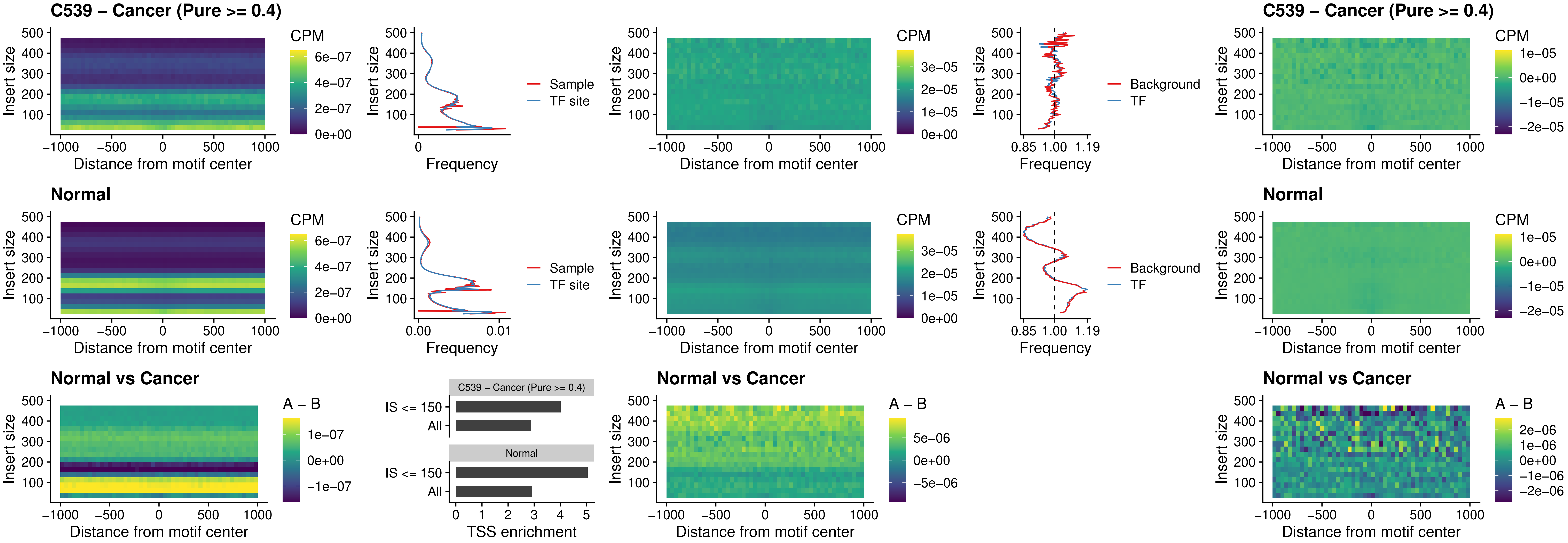

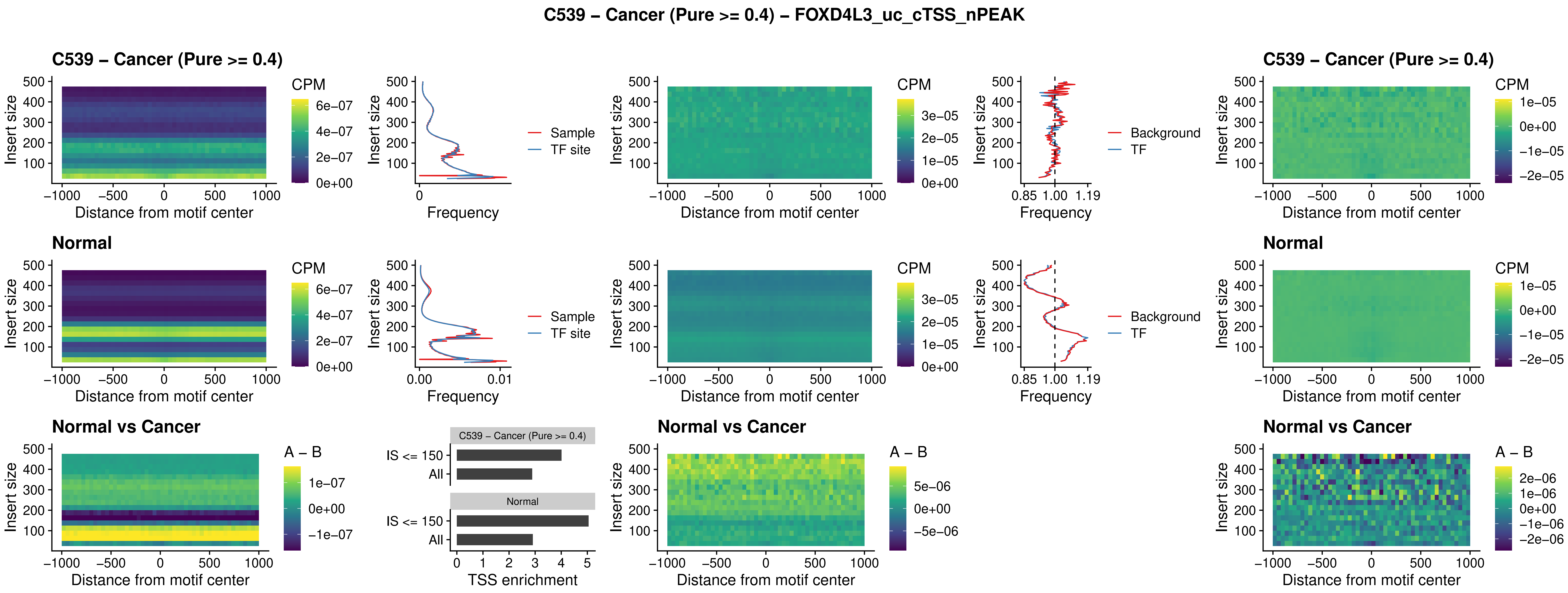

C539 – Cancer (Pure >= 0.4) – FOXD4L3_uc_cTSS_nPEAK
C539 – Cancer (Pure >= 0.4)
Normal
Normal vs Cancer
Insert size
Distance from motif center
Frequency
CPM
A – B
Sample
TF site
Background
TF
C539 – Cancer (Pure >= 0.4)
Normal
IS <= 150
All
TSS enrichment

#### C539 – Cancer (Pure >= 0.4) – HOXD1_uc_dTSS_oPEAK

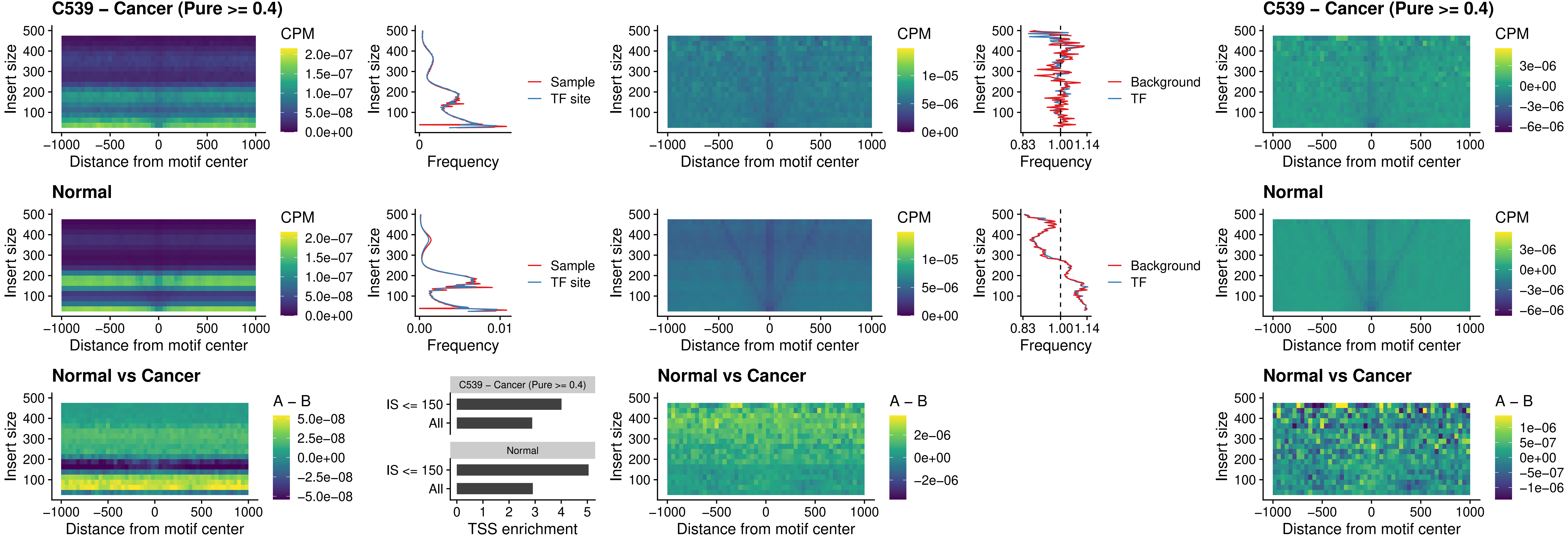

#### C539 – Cancer (Pure >= 0.4) – VAX1_uc_pTSS_nPEAK

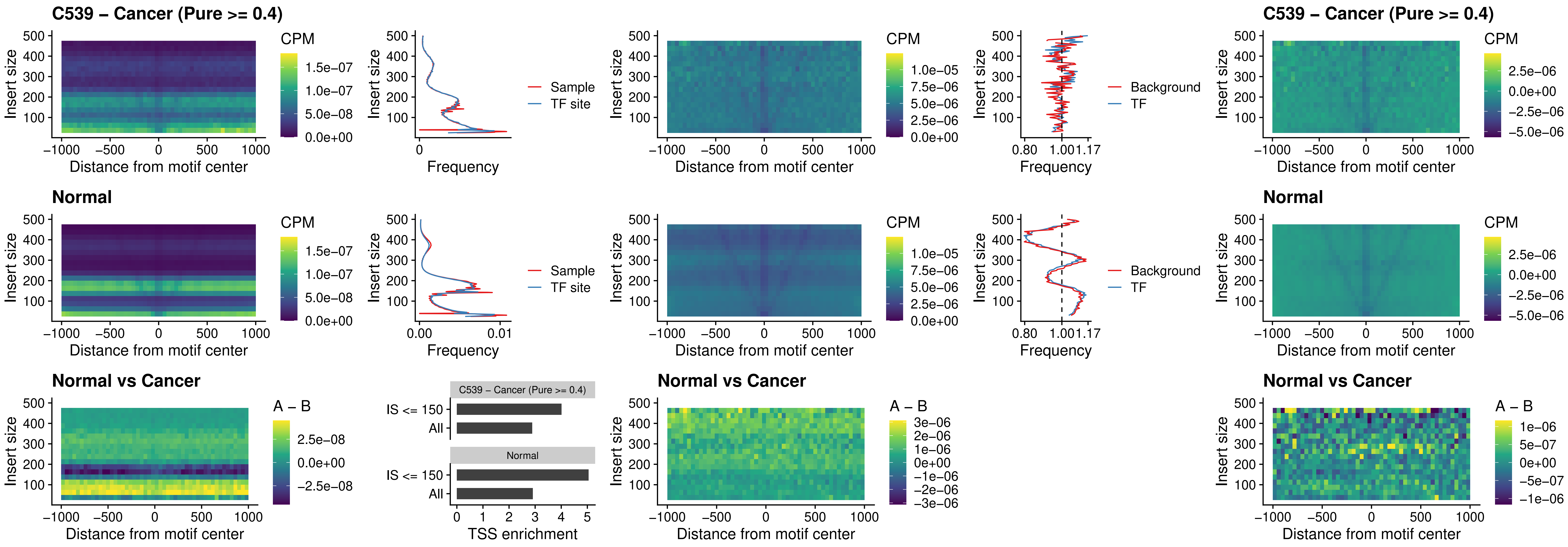

#### C539 – Cancer (Pure >= 0.4) – DLX3_uc_pTSS_nPEAK

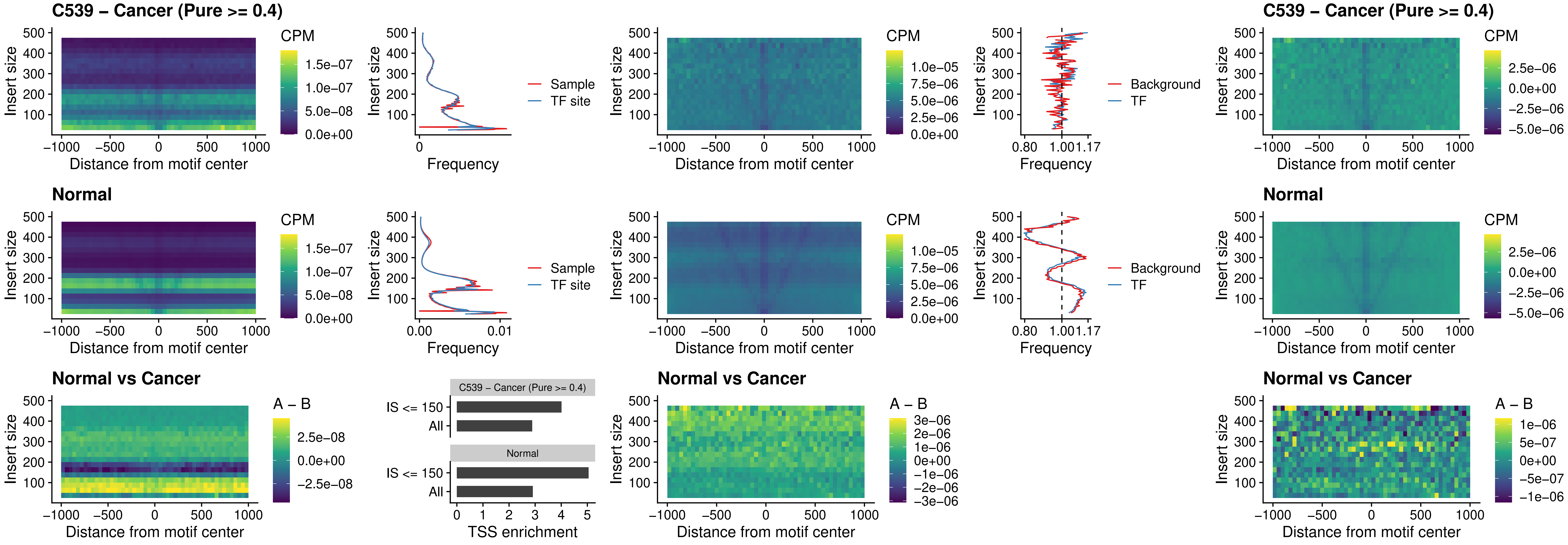

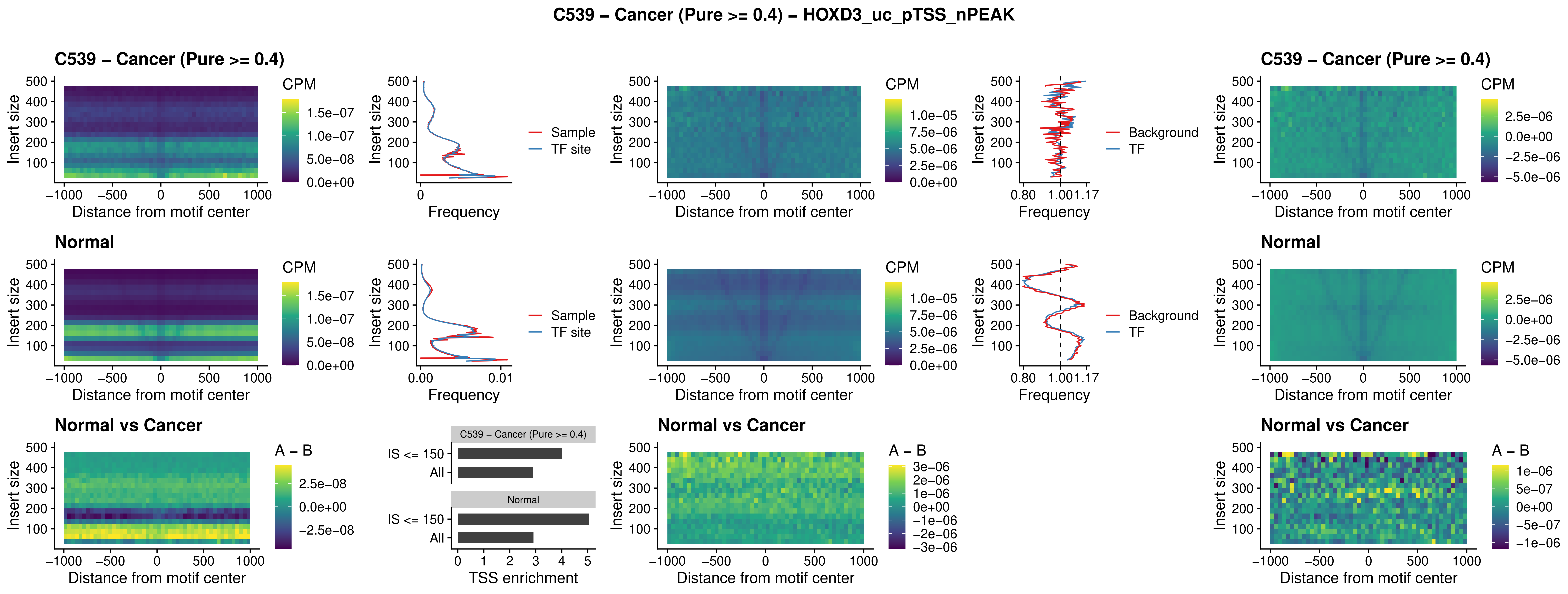

C539 – Cancer (Pure >= 0.4) – HOXD3_uc_pTSS_nPEAK
C539 – Cancer (Pure >= 0.4)
Normal
Normal vs Cancer
Insert size
Distance from motif center
Frequency
CPM
A – B
Sample
TF site
Background
TF
C539 – Cancer (Pure >= 0.4)
Normal
IS <= 150
All
TSS enrichment

#### C539 – Cancer (Pure >= 0.4) – PDX1_uc_dTSS_nPEAK

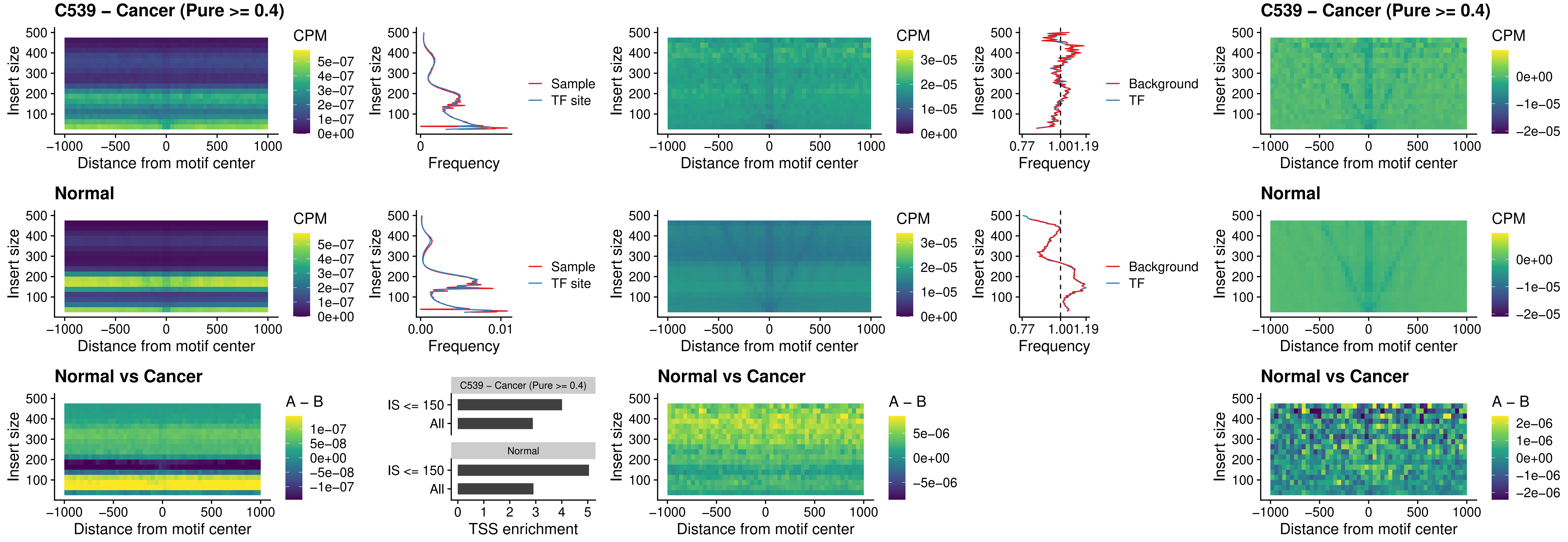

#### C539 – Cancer (Pure >= 0.4) – VAX2_uc_dTSS_nPEAK

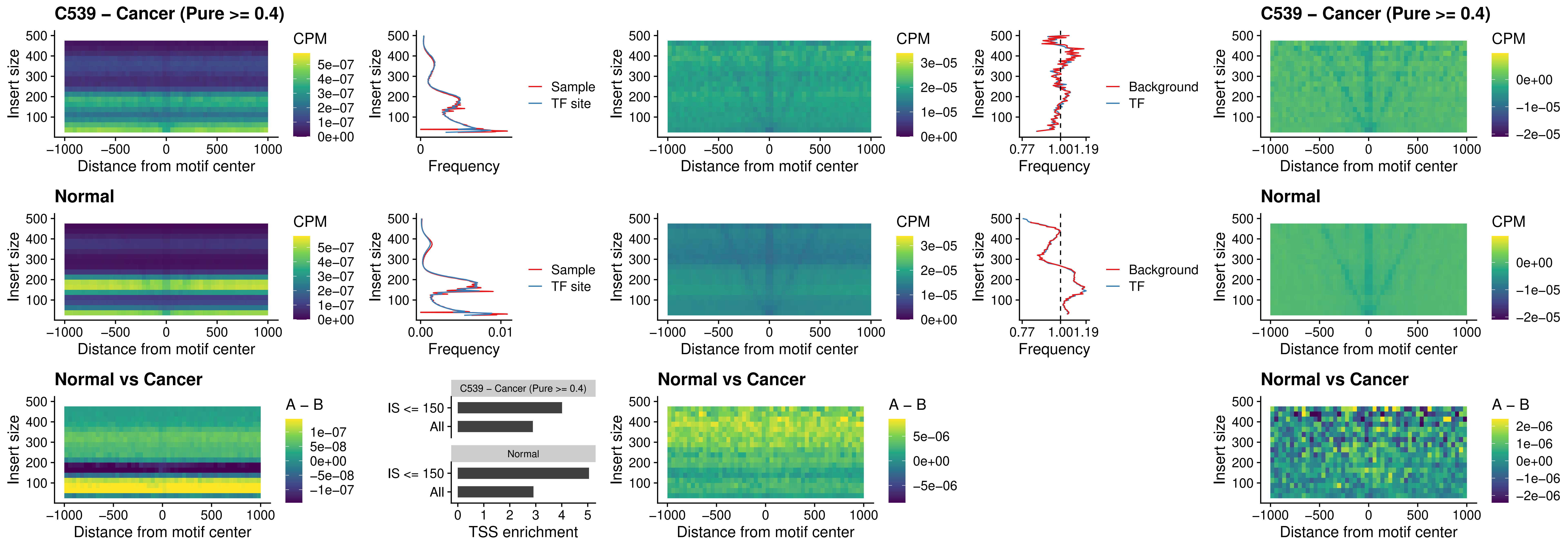

#### C539 – Cancer (Pure >= 0.4) – SOX5_uc_dTSS_nPEAK

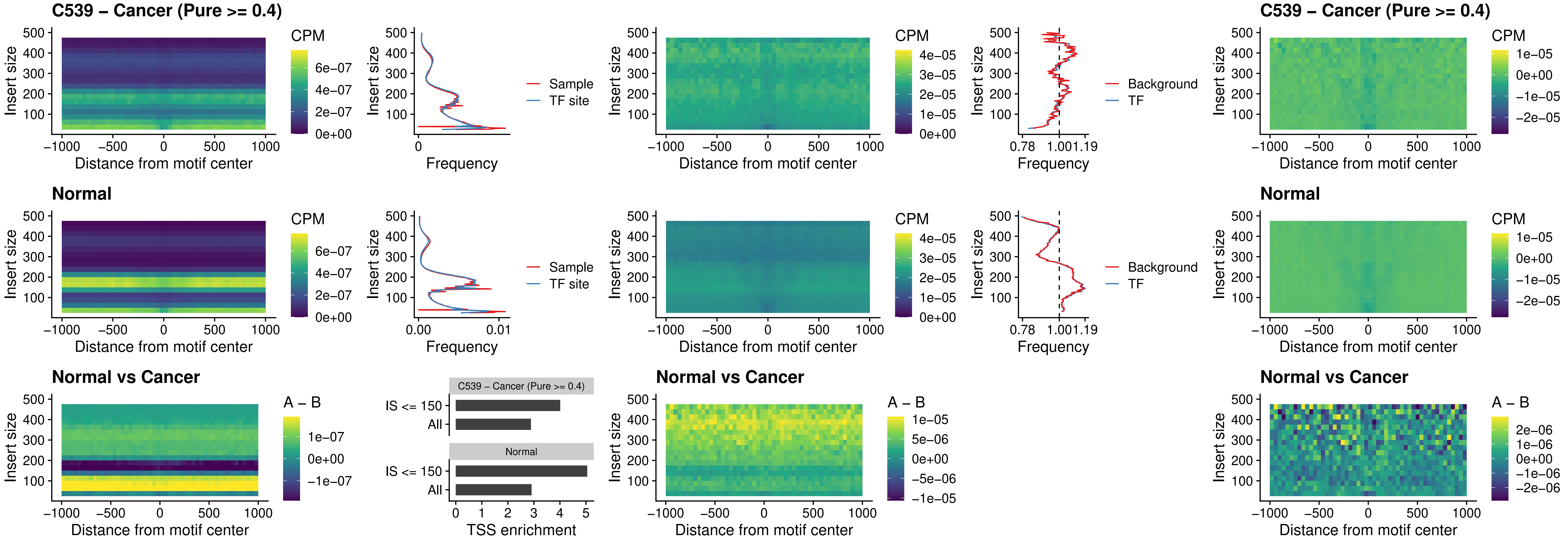

#### C539 – Cancer (Pure >= 0.4) – SOX6_uc_dTSS_nPEAK

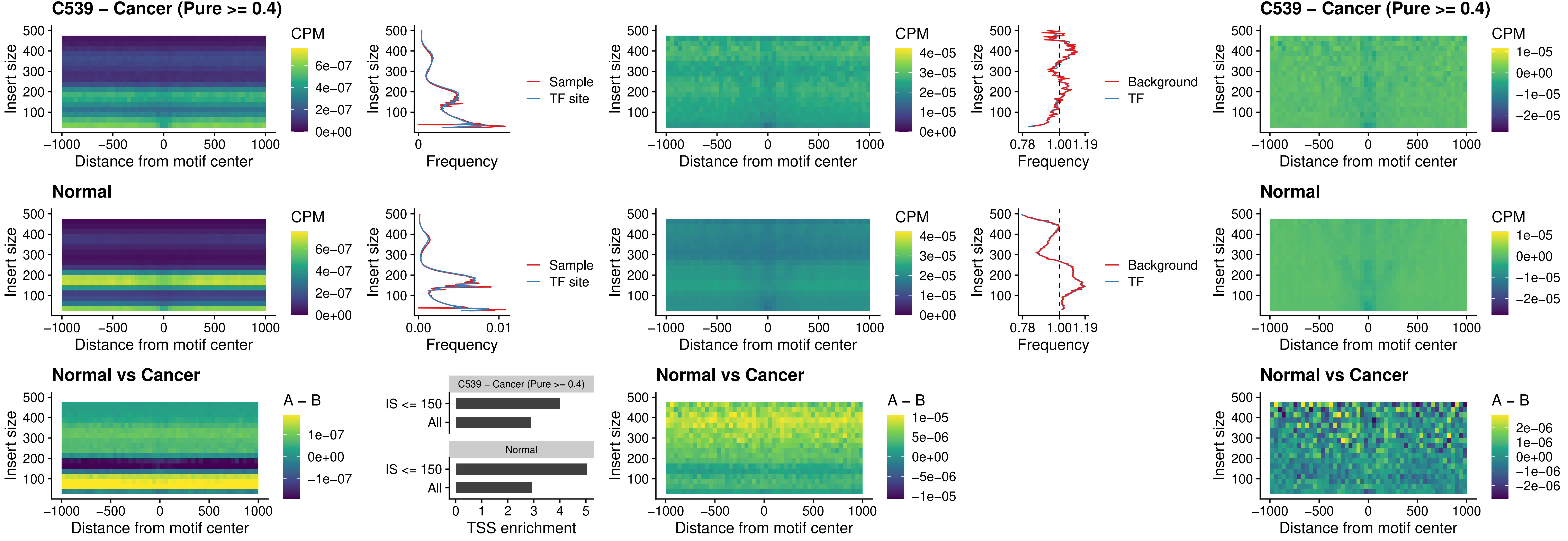

**C539 – Cancer (Pure >= 0.4) – JDP2_uc_cTSS_nPEAK**

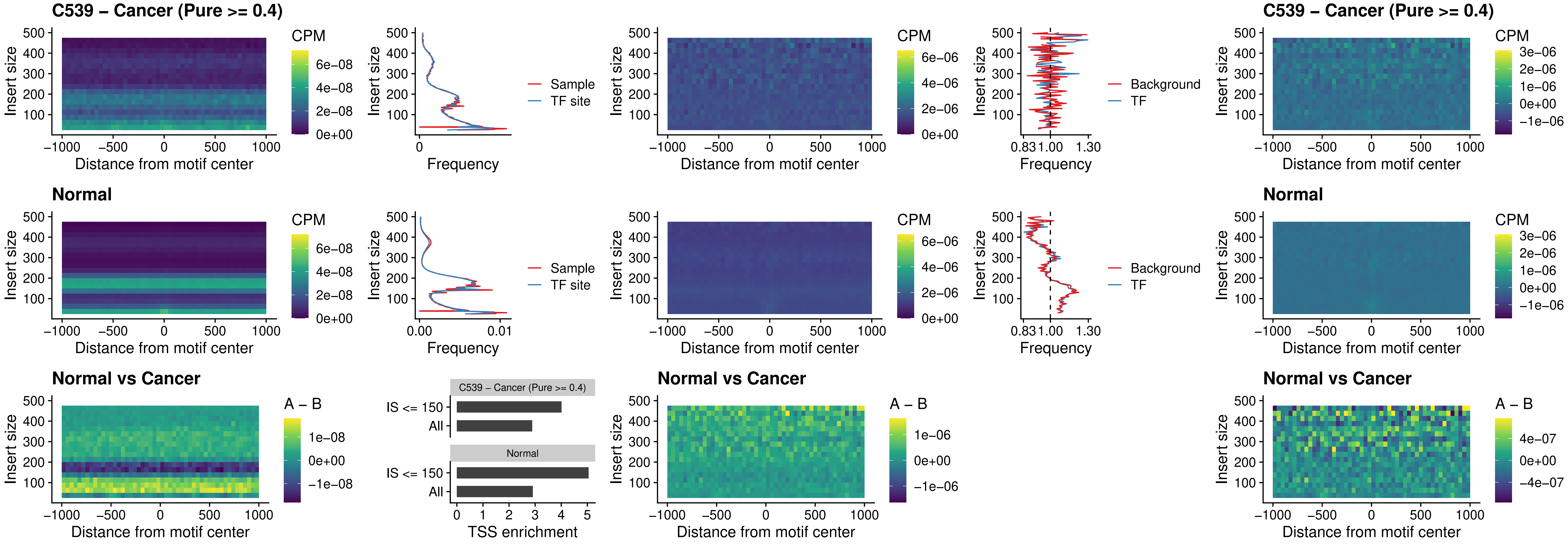

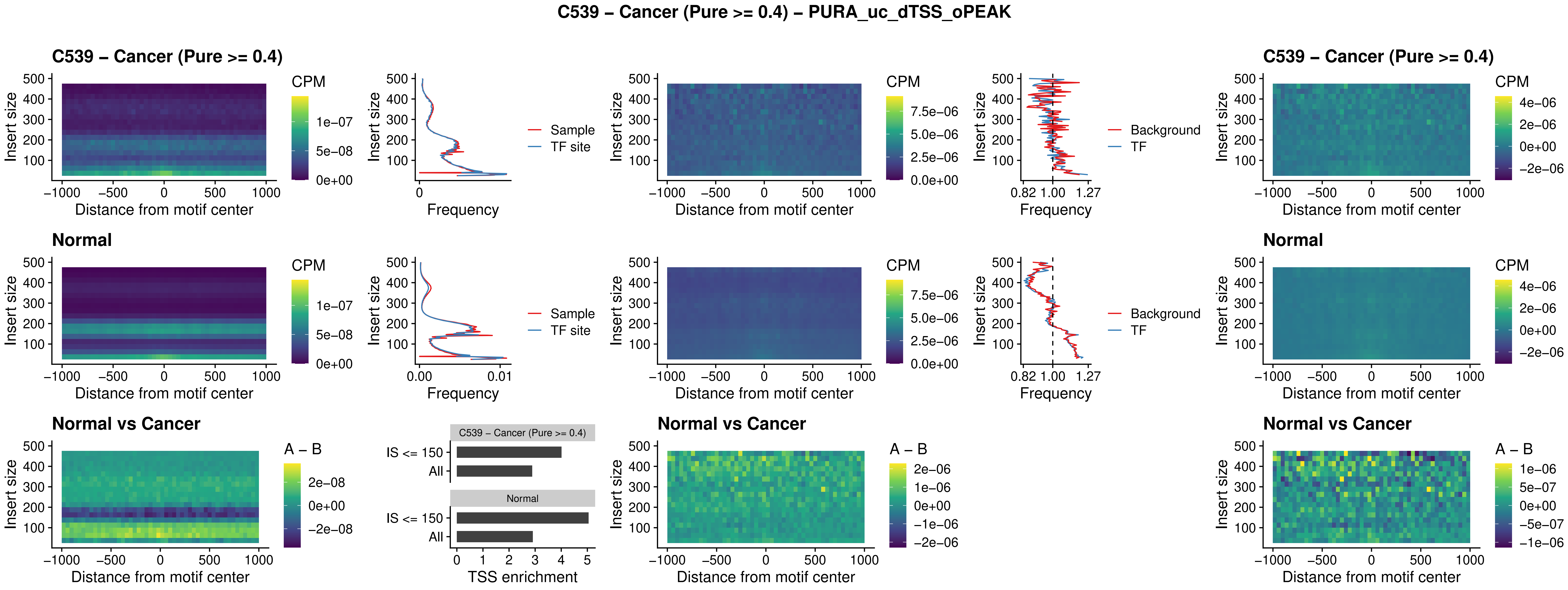
C539 – Cancer (Pure >= 0.4) – PURA_uc_dTSS_oPEAK
C539 – Cancer (Pure >= 0.4)
Normal
Normal vs Cancer
Insert size
Distance from motif center
CPM
A – B
Frequency
Sample
TF site
Background
TF
C539 – Cancer (Pure >= 0.4)
Normal
IS <= 150
All
TSS enrichment

#### C539 – Cancer (Pure >= 0.4) – HHEX_uc_dTSS_nPEAK

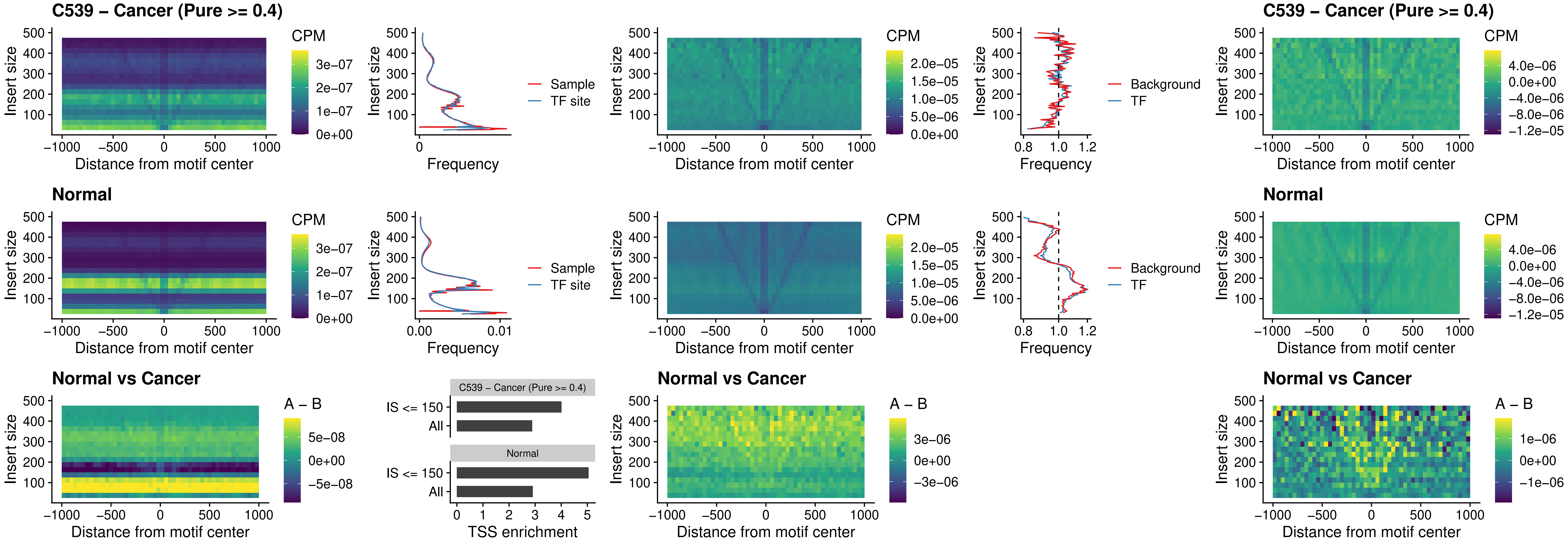

#### C539 – Cancer (Pure >= 0.4) – HMG20B_uc_dTSS_nPEAK

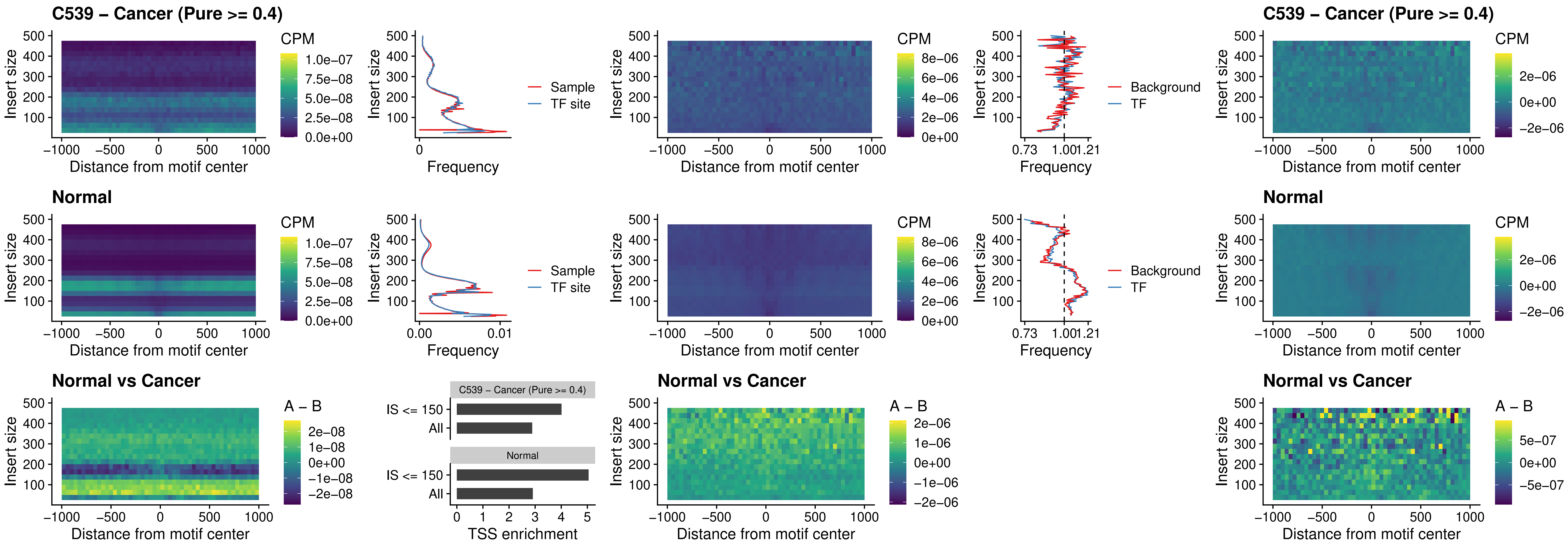

#### C539 – Cancer (Pure >= 0.4) – FOXB2_uc_dTSS_nPEAK

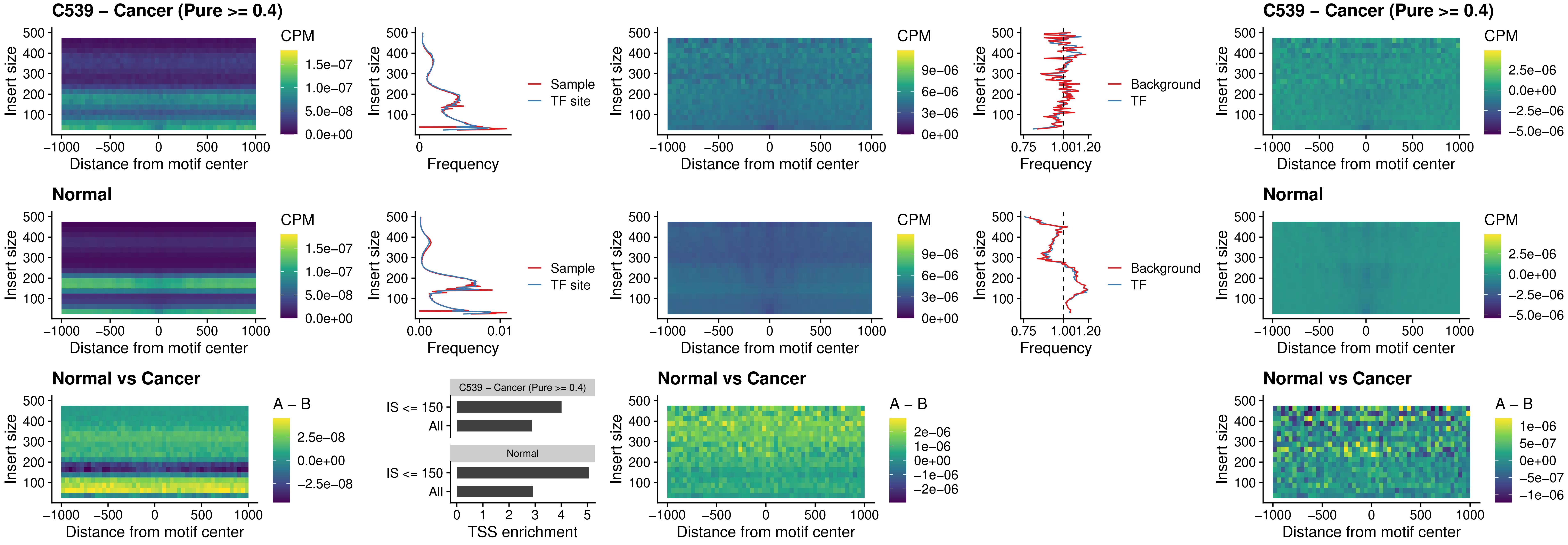

#### C539 – Cancer (Pure >= 0.4) – FOXL1_uc_cTSS_nPEAK

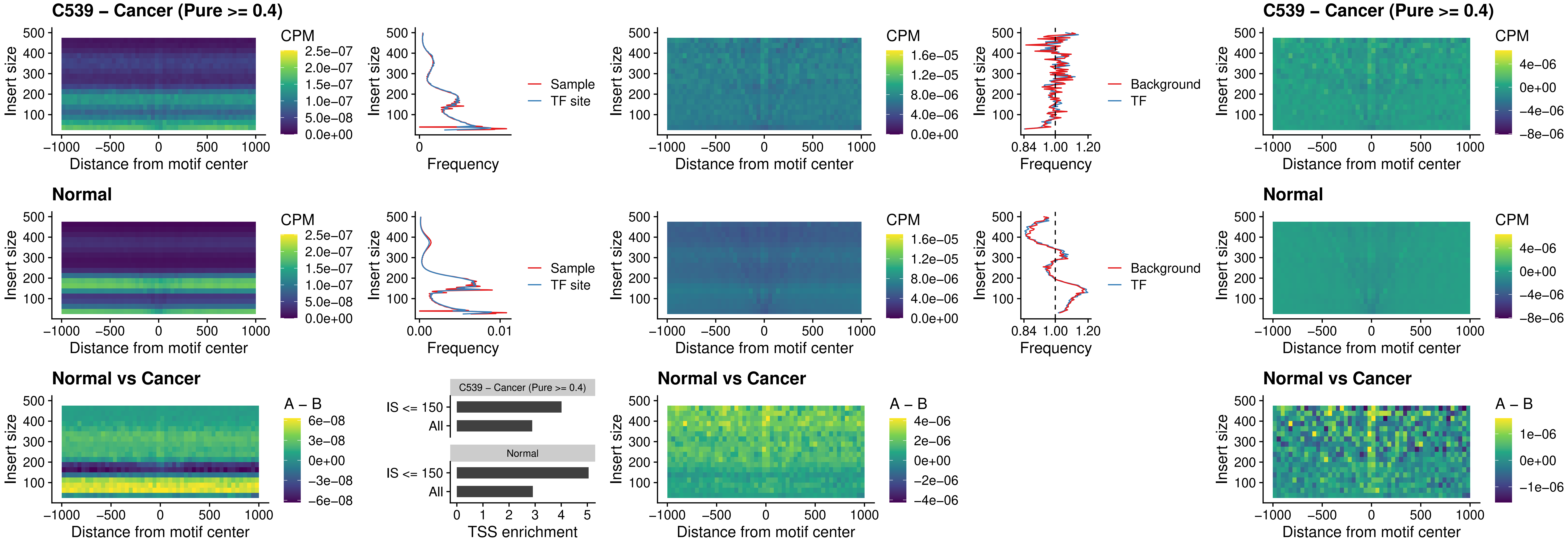

#### C539 – Cancer (Pure >= 0.4) – PBX3_uc_pTSS_oPEAK

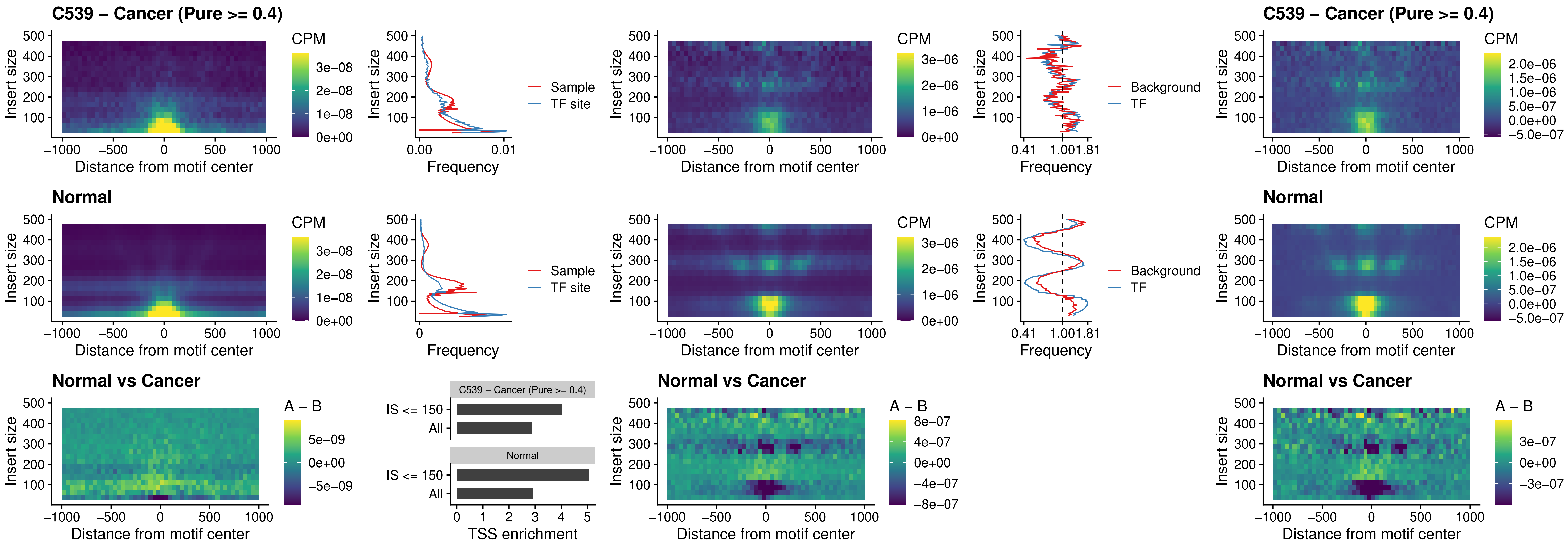

#### C539 – Cancer (Pure >= 0.4) – SNAI2_uc_cTSS_nPEAK

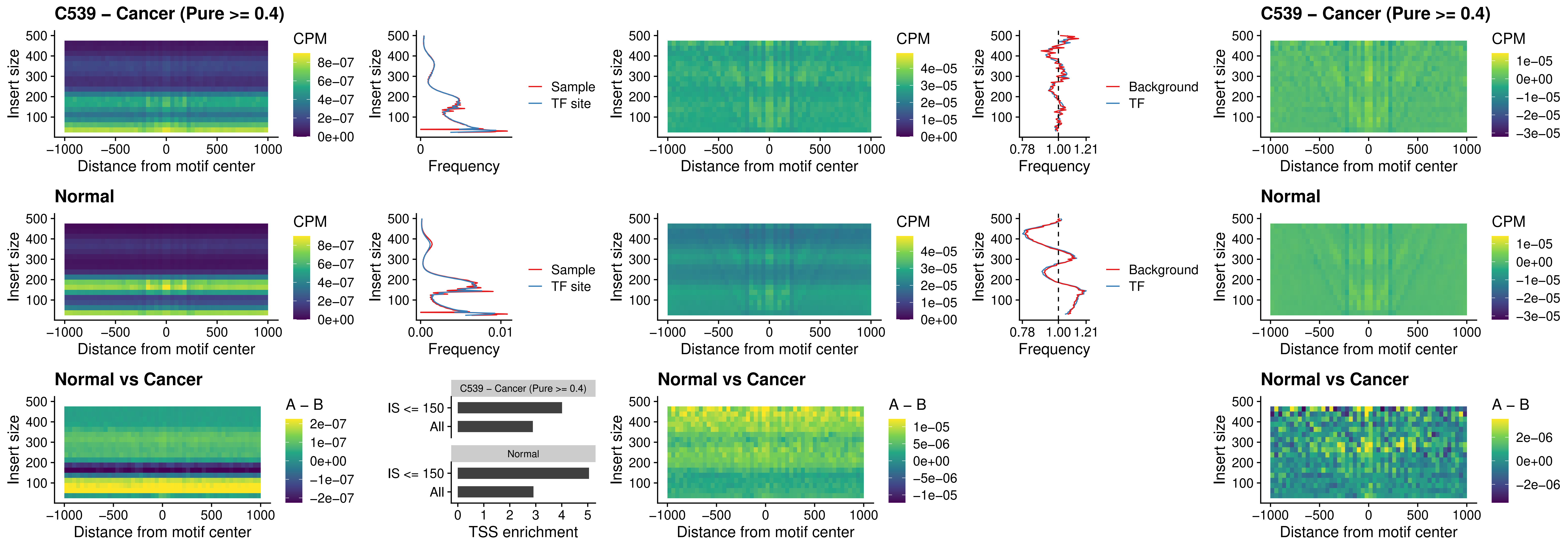

#### C539 – Cancer (Pure >= 0.4) – SNAI2_uc_dTSS_oPEAK

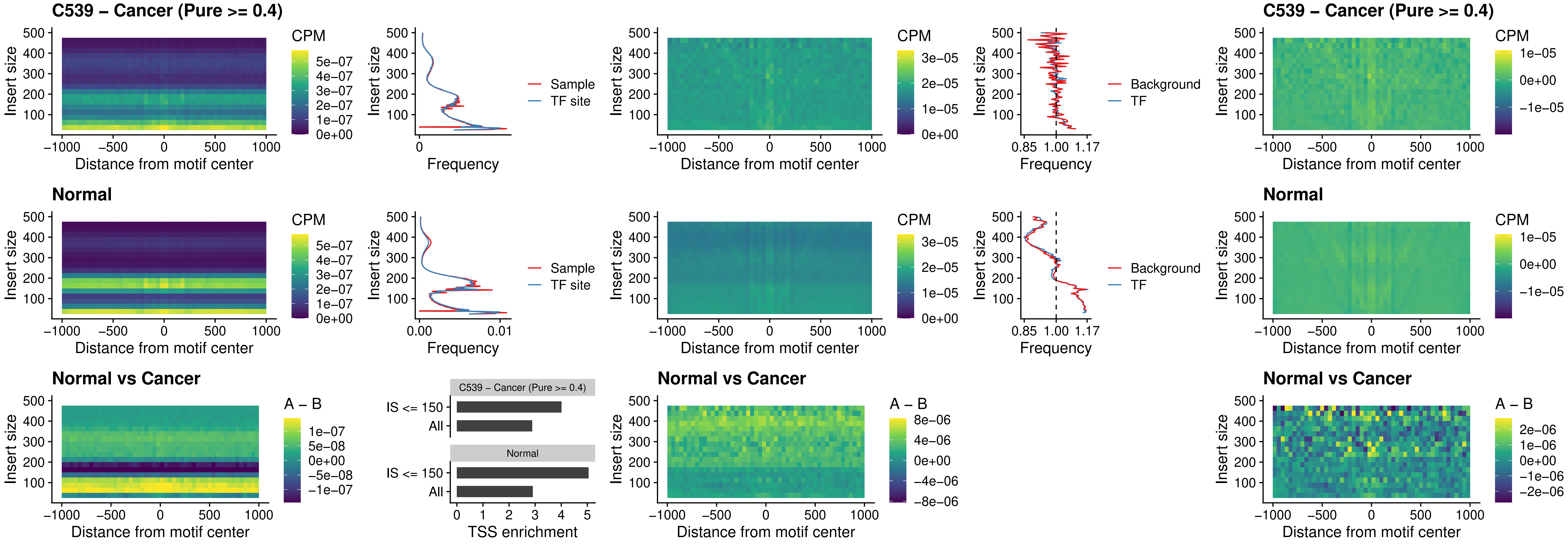

#### C539 – Cancer (Pure >= 0.4) – SPI1_uc_pTSS_nPEAK

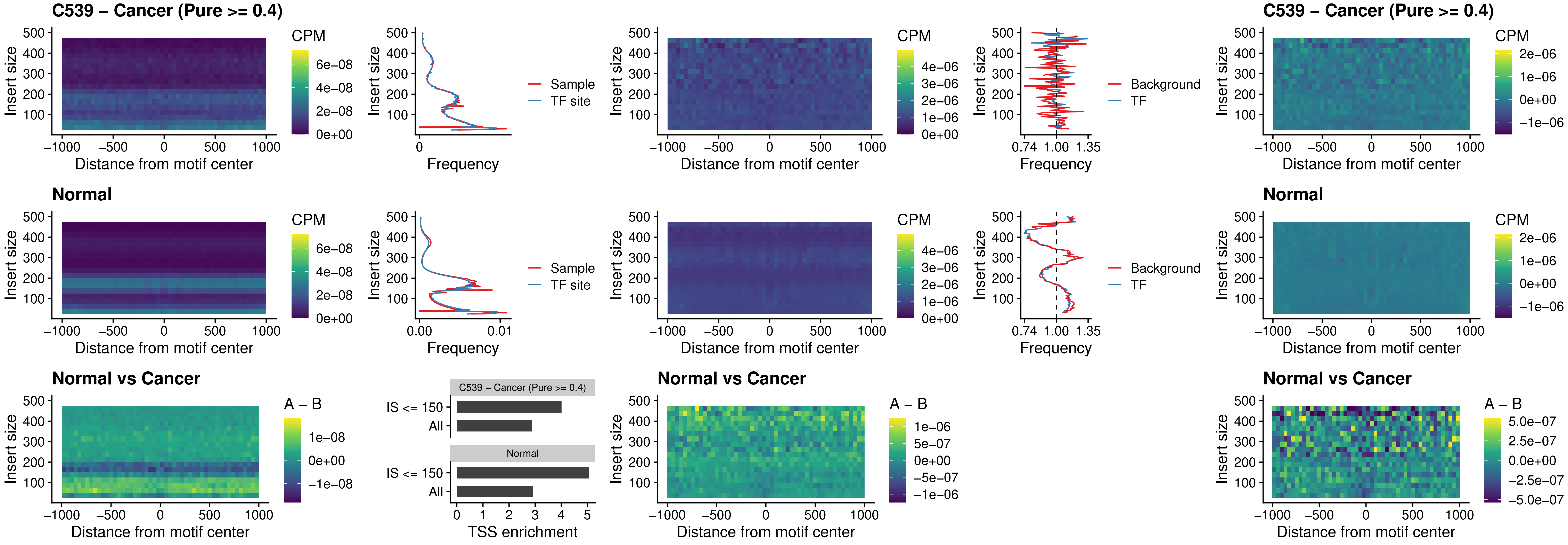

#### C539 – Cancer (Pure >= 0.4) – KLF16_uc_pTSS_nPEAK

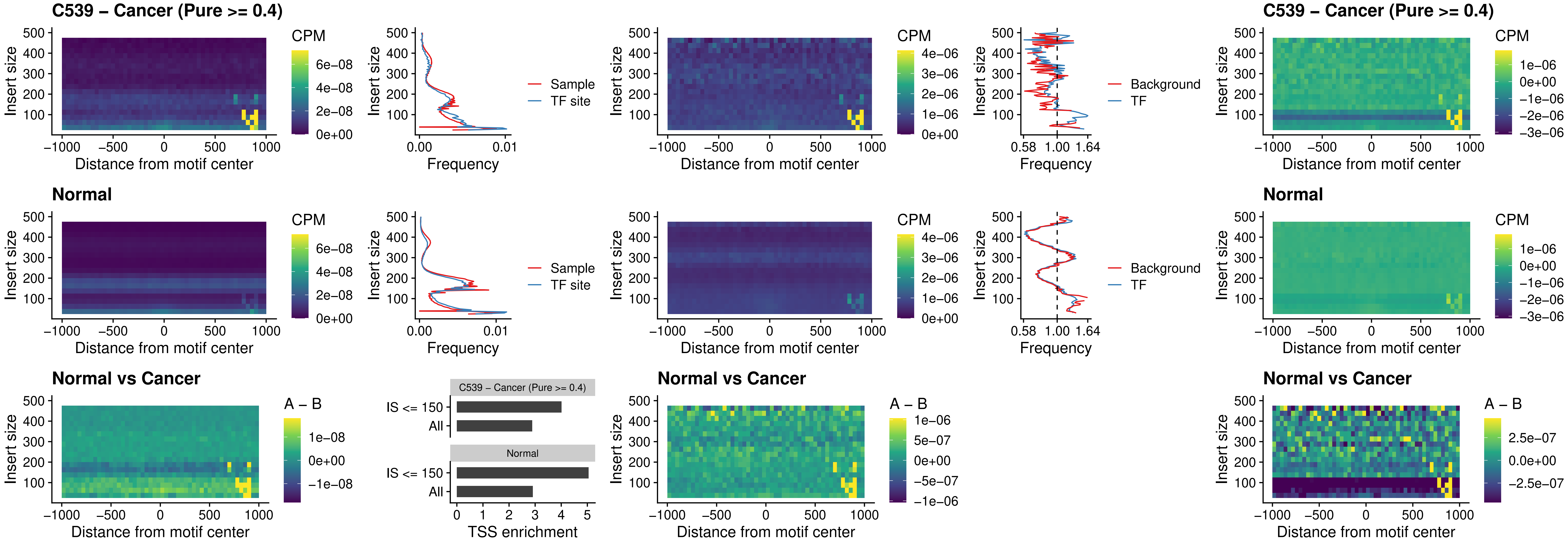

#### C539 – Cancer (Pure >= 0.4) – SP3_uc_pTSS_nPEAK

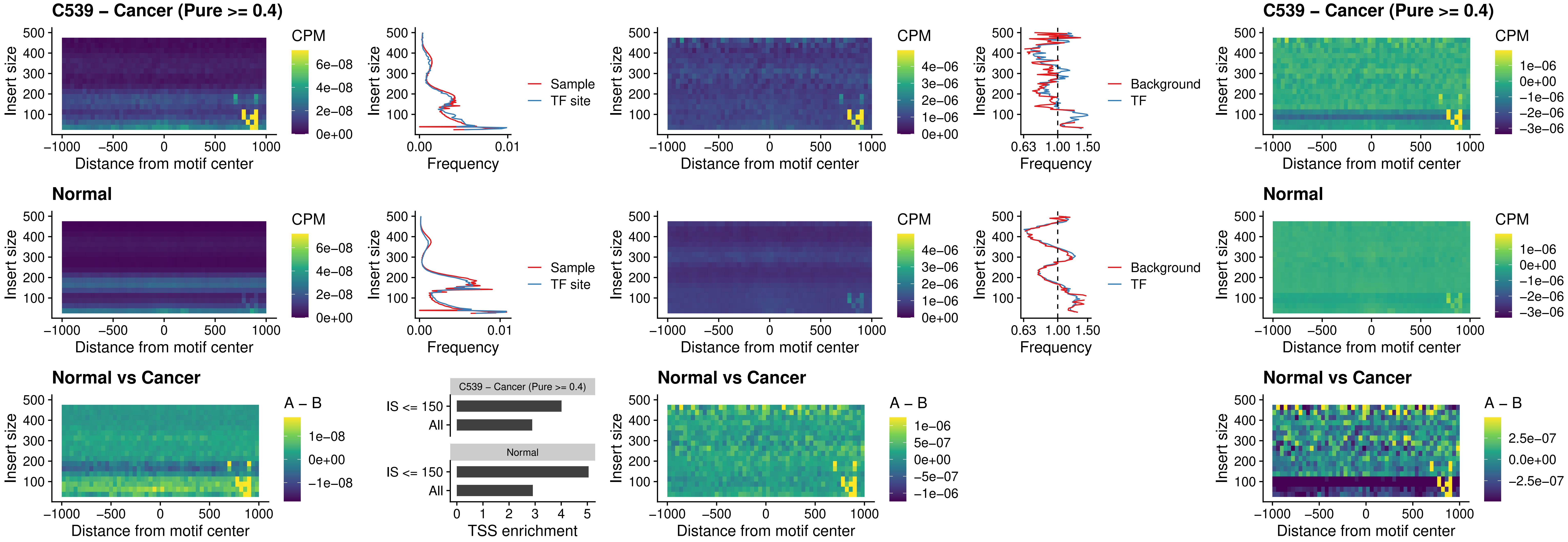

#### C539 – Cancer (Pure >= 0.4) – NEUROD1_uc_dTSS_oPEAK

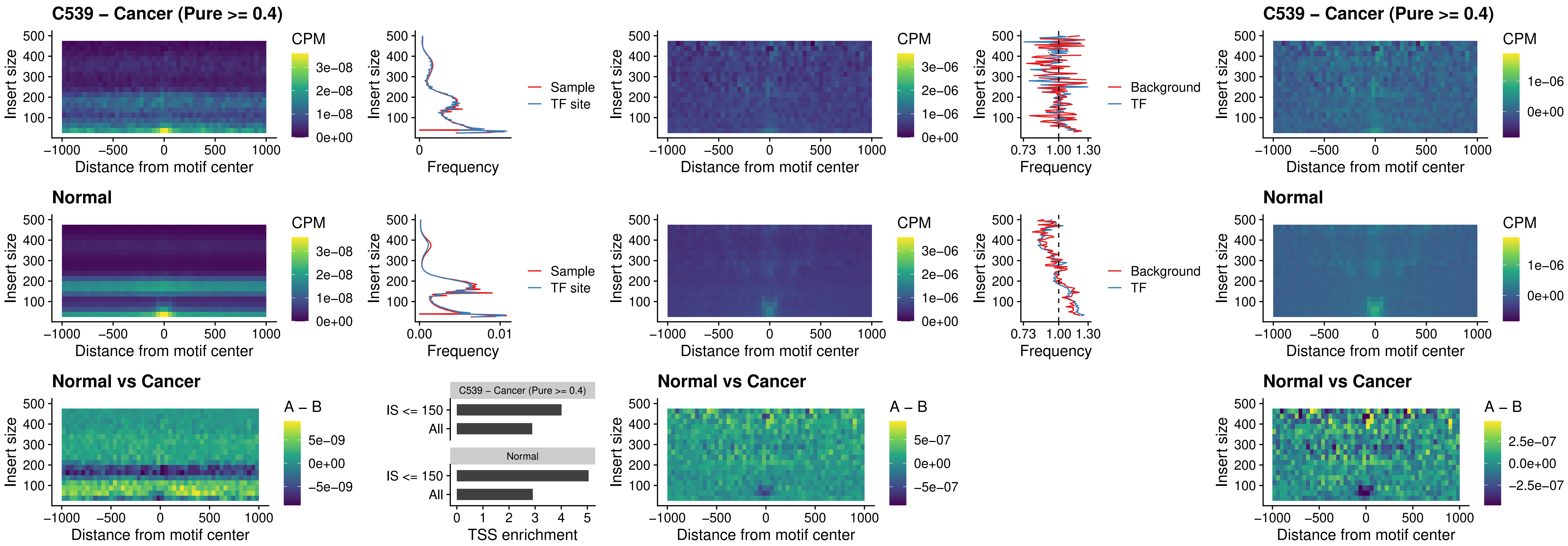

#### C539 – Cancer (Pure >= 0.4) – CTCF_uc_dTSS_oPEAK

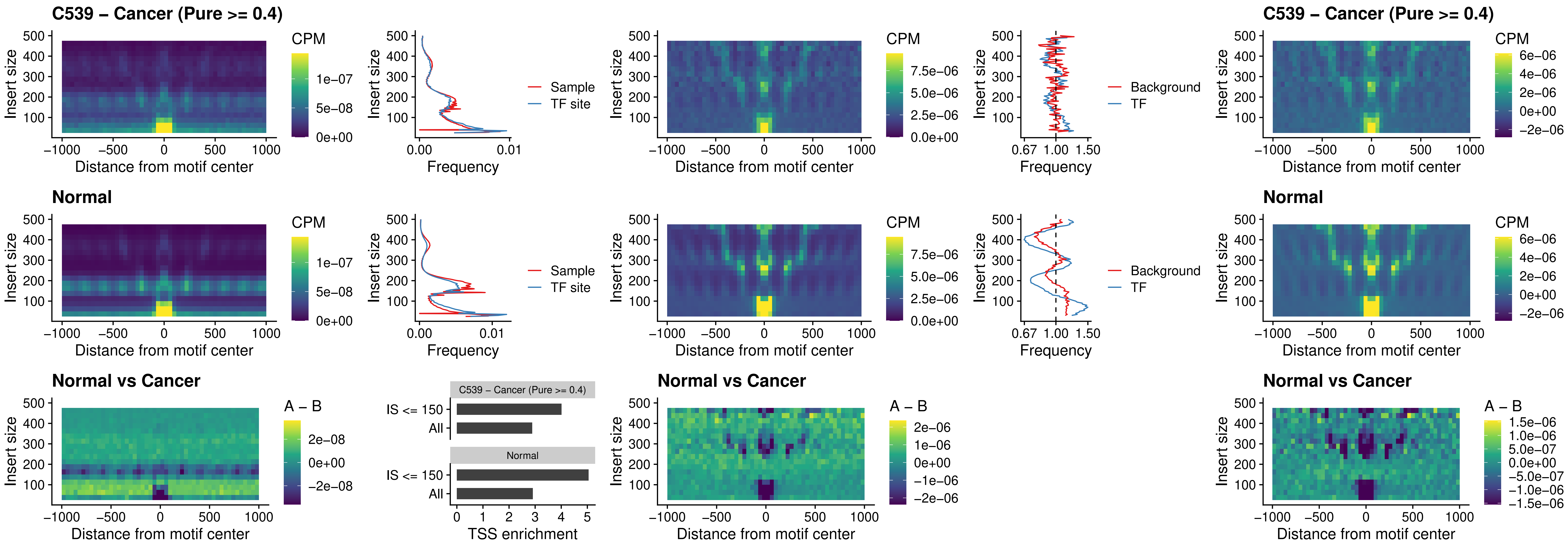

#### C539 – Cancer (Pure >= 0.4) – CTCF_uc_cTSS_oPEAK

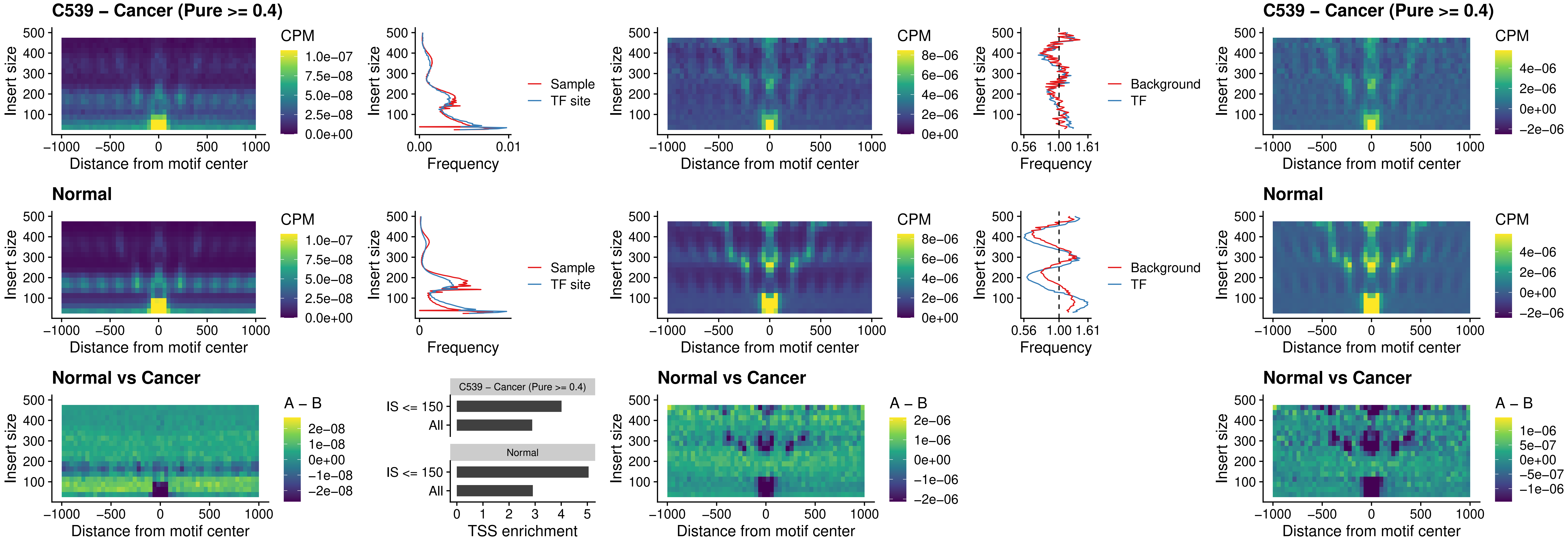

#### C539 – Cancer (Pure >= 0.4) – CTCFL_uc_dTSS_oPEAK

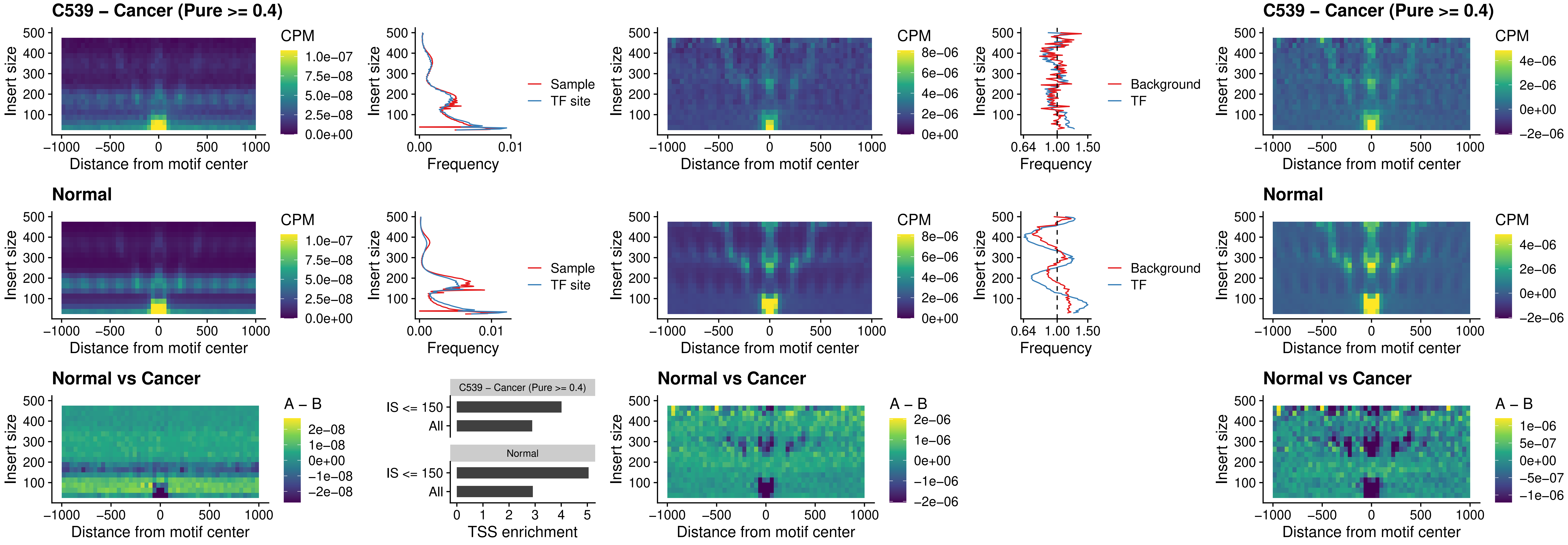

#### C539 – Cancer (Pure >= 0.4) – ENCODE_CTCF_sites_COLON

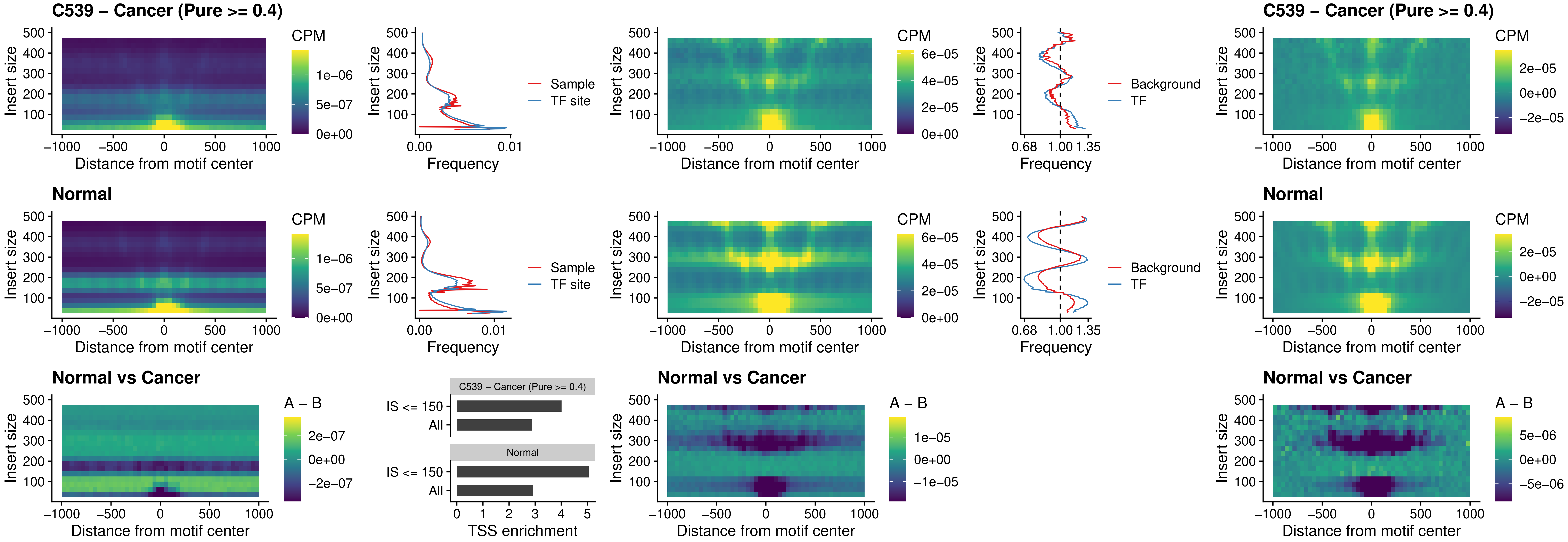

#### C539 – Cancer (Pure >= 0.4) – ENCODE_CTCF_sites_COLON_loop_ctcf

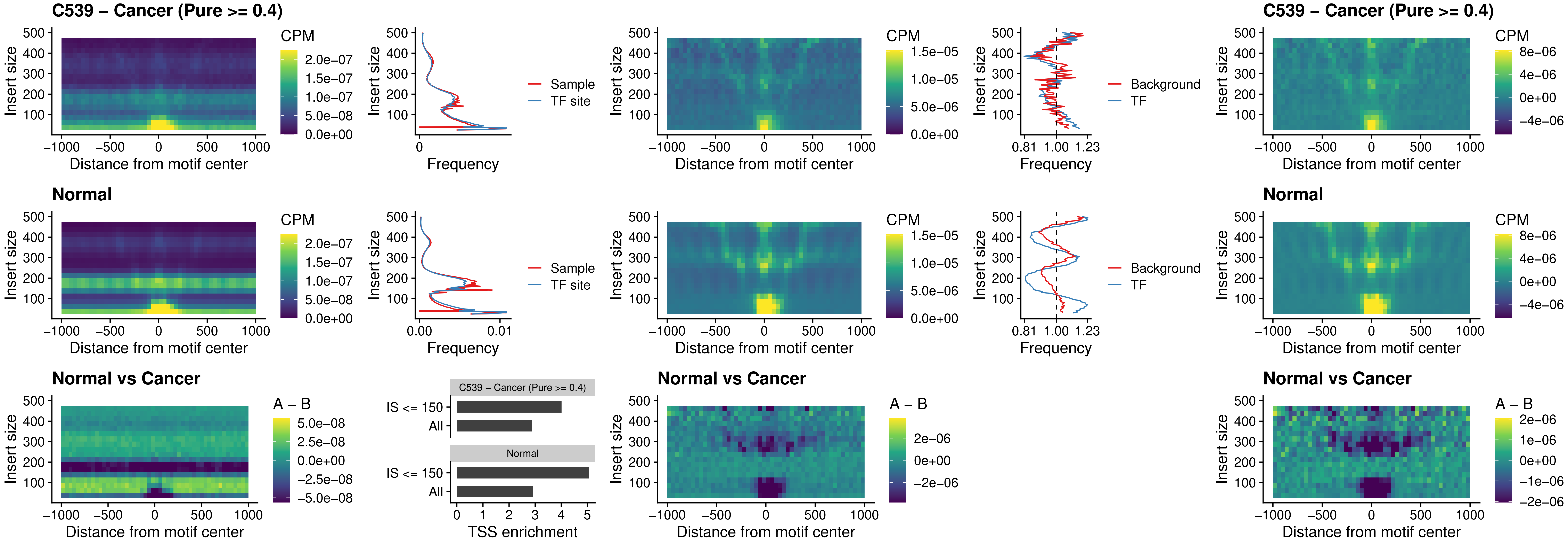

#### C539 – Cancer (Pure >= 0.4) – ENCODE_CTCF_sites_COLON_intergenic_ctcf

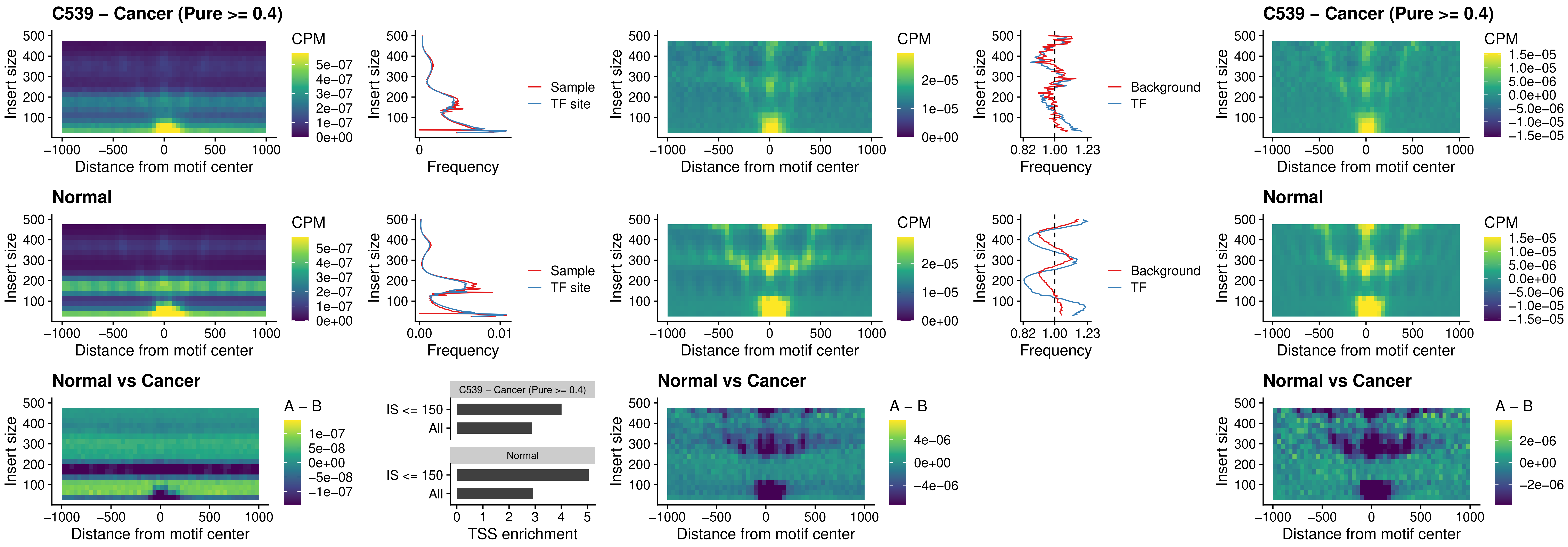

#### C539 – Cancer (Pure >= 0.4) – ENCODE_CTCF_sites_COLON_nonpromotor

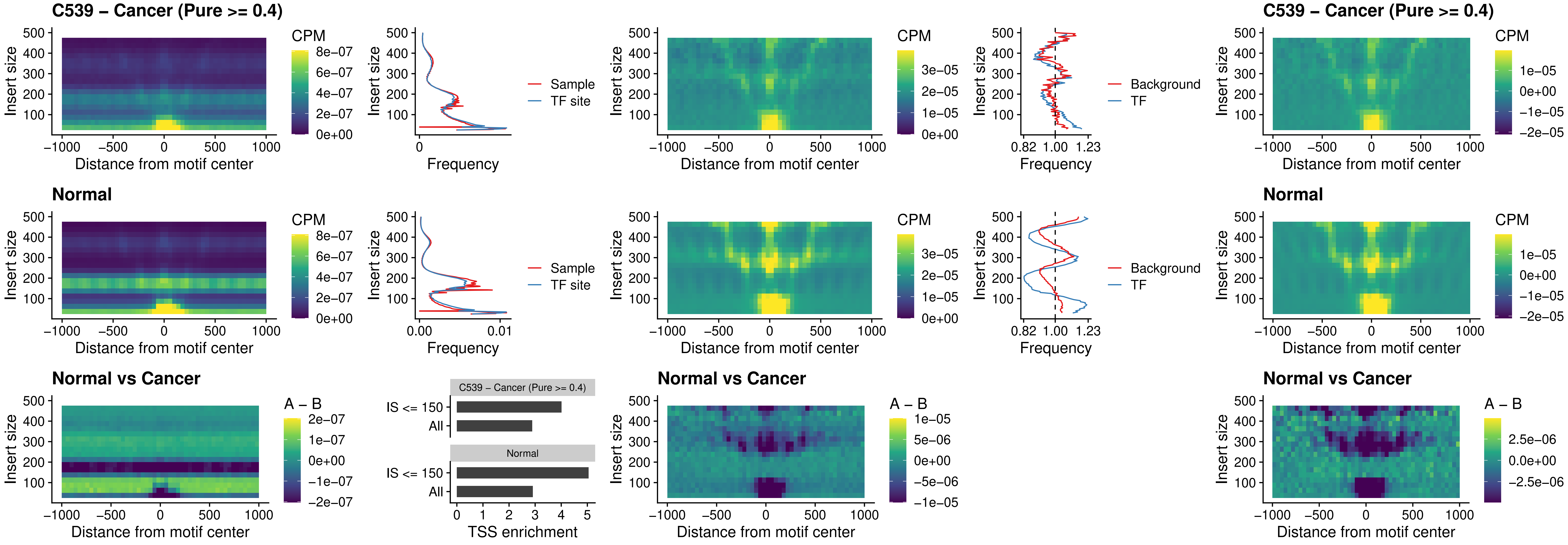

### C539 – Cancer (Pure >= 0.4) – CTCFL_uc_cTSS_oPEAK

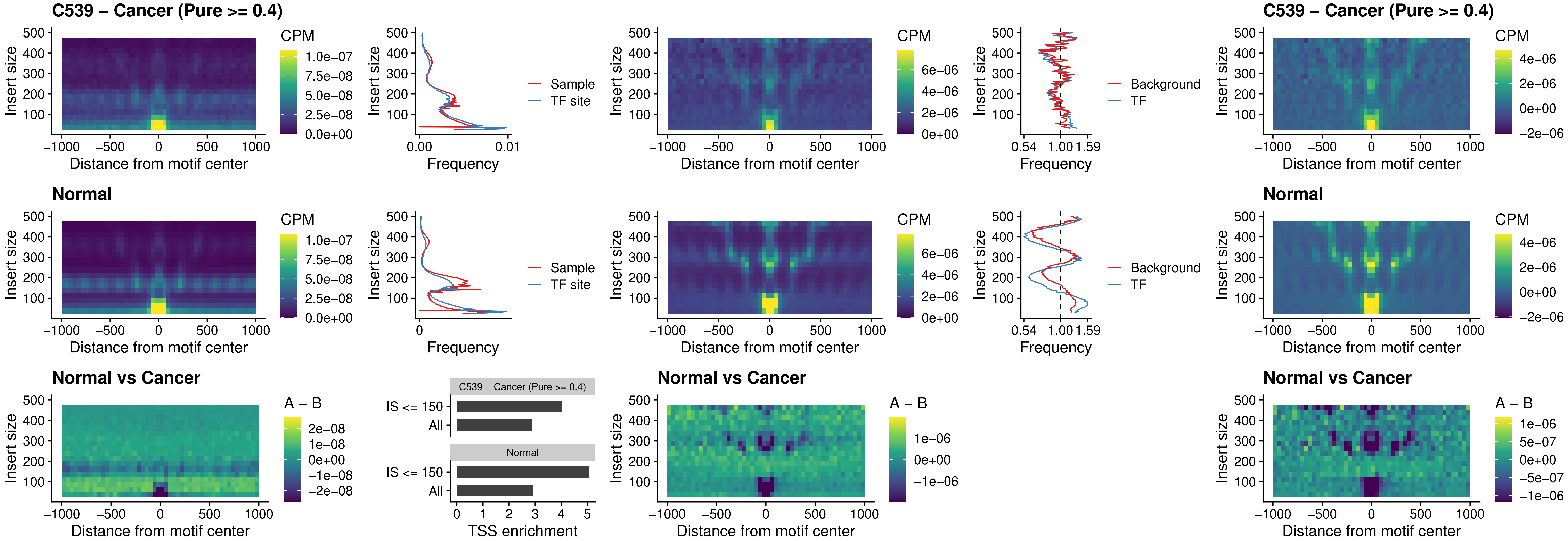

#### C539 – Cancer (Pure >= 0.4) – NEUROD1_uc_cTSS_oPEAK

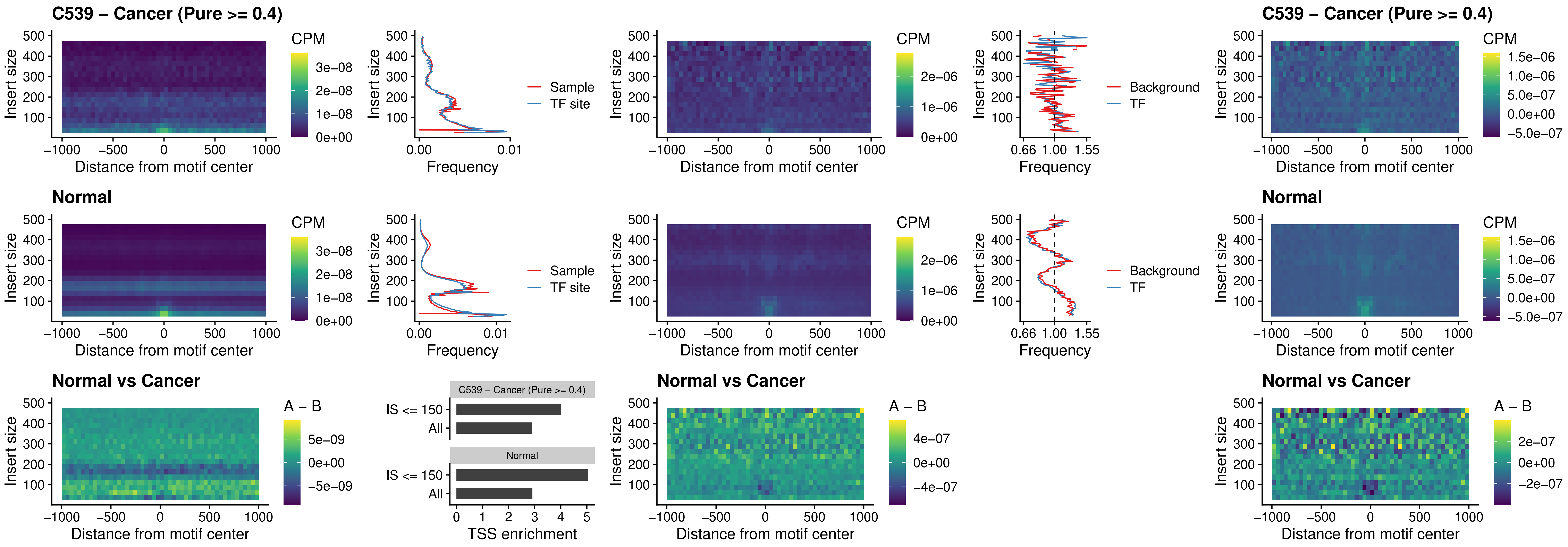

#### C539 – Cancer (Pure >= 0.4) – ENCODE_YY1_sites_COLON_nonpromotor

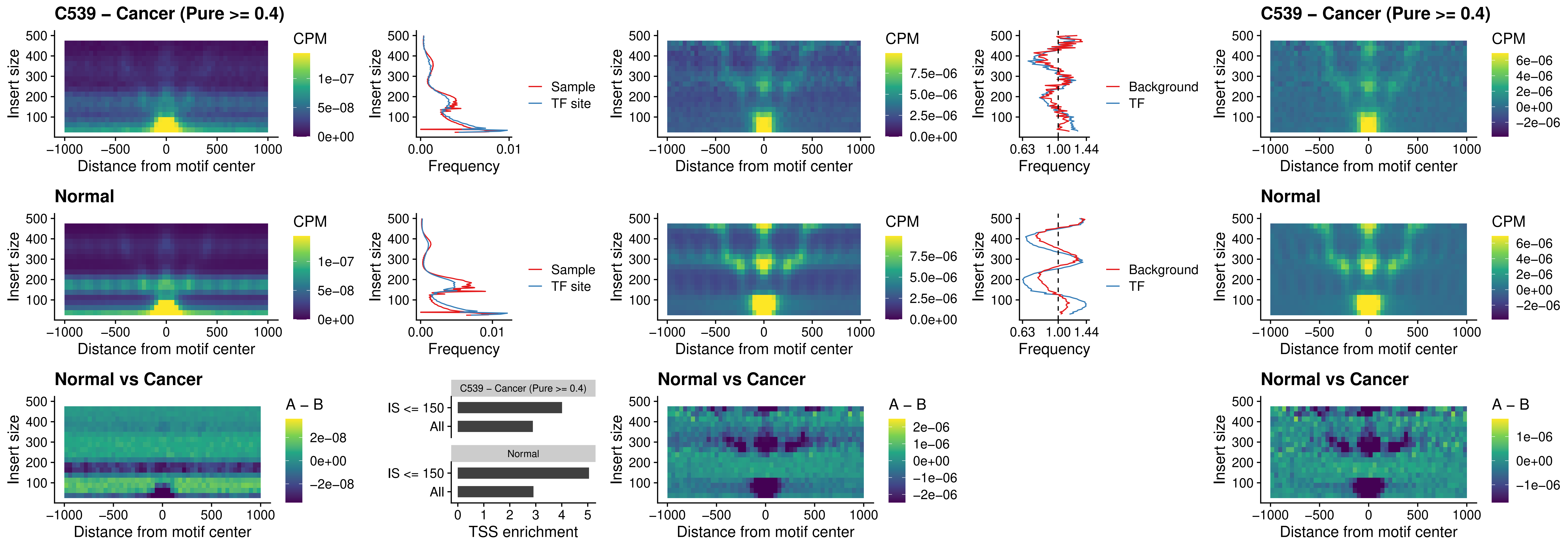

#### C539 – Cancer (Pure >= 0.4) – CTCF_uc_pTSS_oPEAK

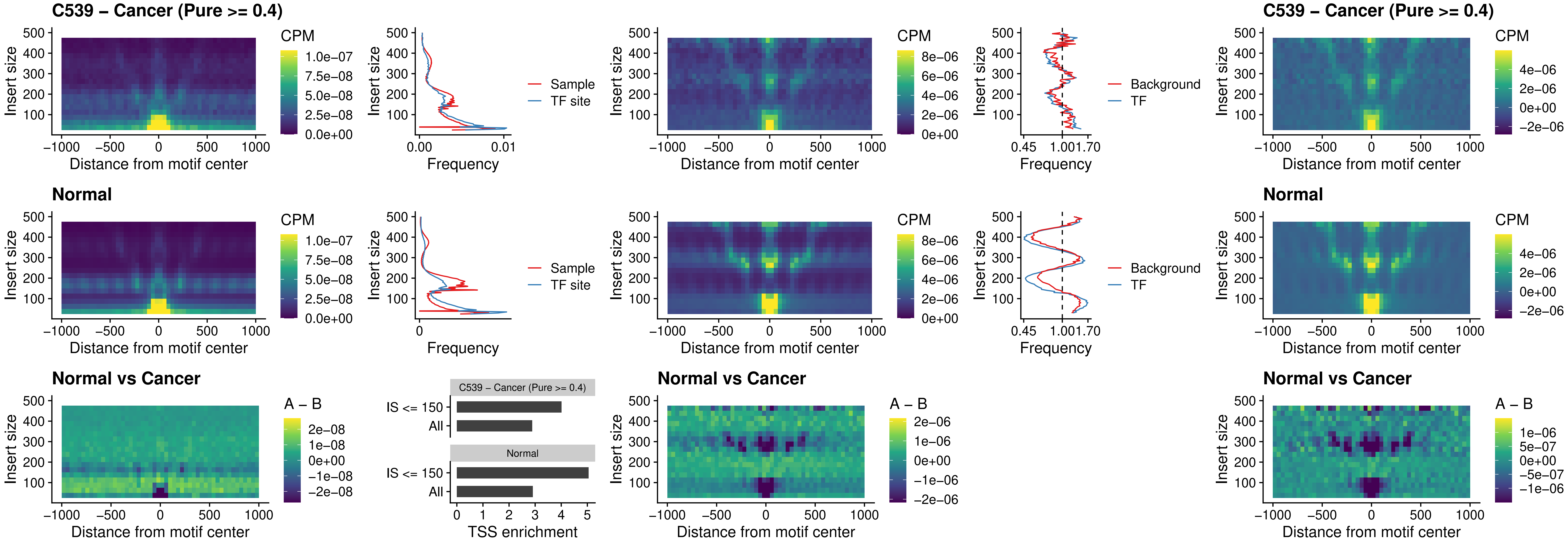

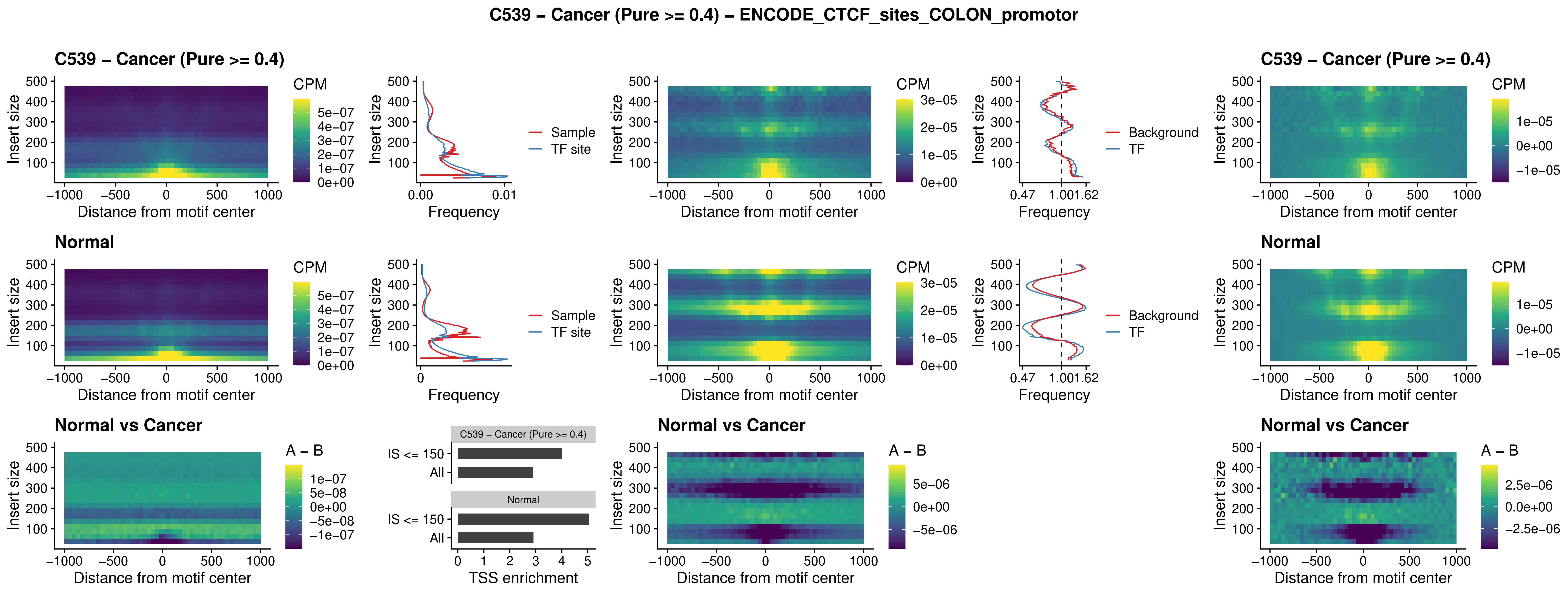
C539 – Cancer (Pure >= 0.4) – ENCODE_CTCF_sites_COLON_promotor
C539 – Cancer (Pure >= 0.4)
Normal
Normal vs Cancer
Insert size
Distance from motif center
CPM
A – B
Frequency
Sample
TF site
Background
TF
C539 – Cancer (Pure >= 0.4)
Normal
IS <= 150
All
TSS enrichment

#### C539 – Cancer (Pure >= 0.4) – ENCODE_CTCF_sites_COLON_promotor_ctcf

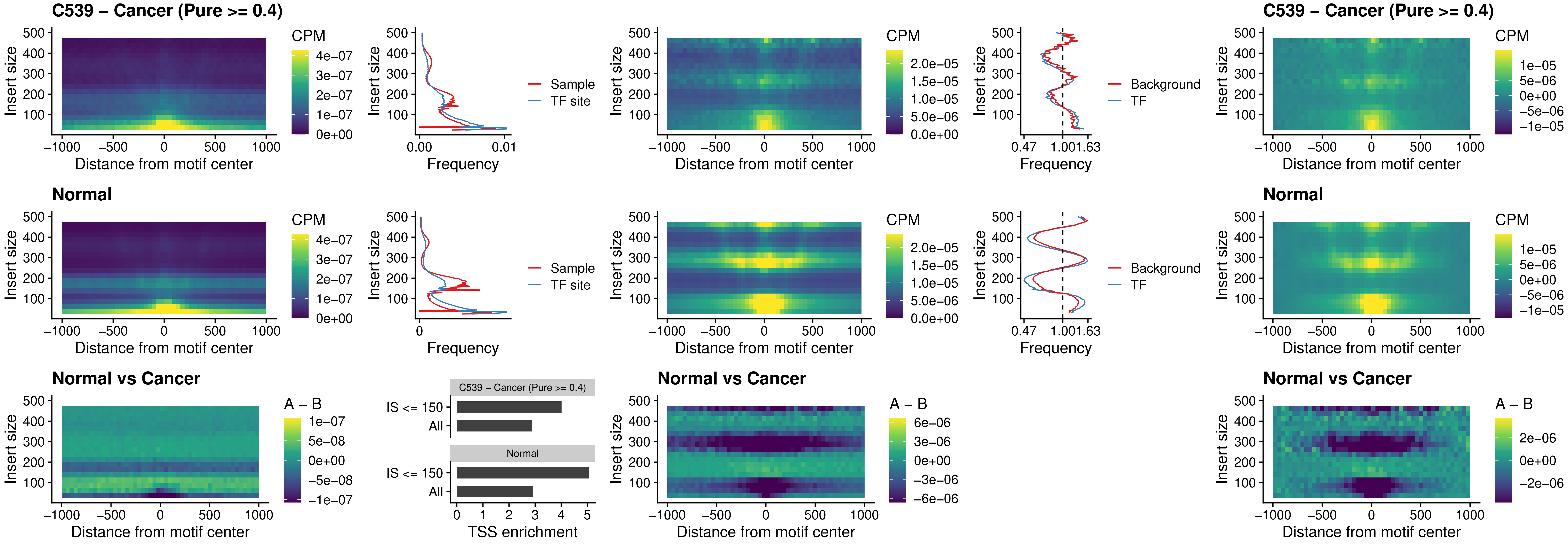

#### C539 – Cancer (Pure >= 0.4) – SPIC_uc_pTSS_oPEAK

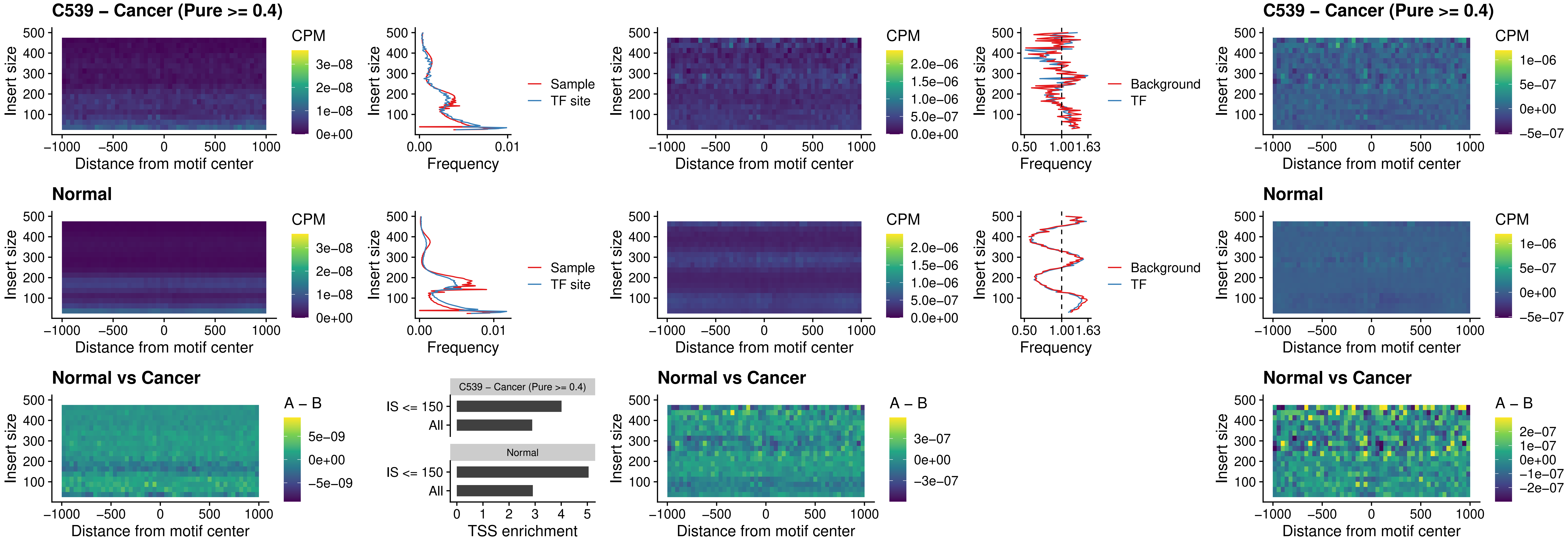

#### C539 – Cancer (Pure >= 0.4) – SPI1_uc_dTSS_nPEAK

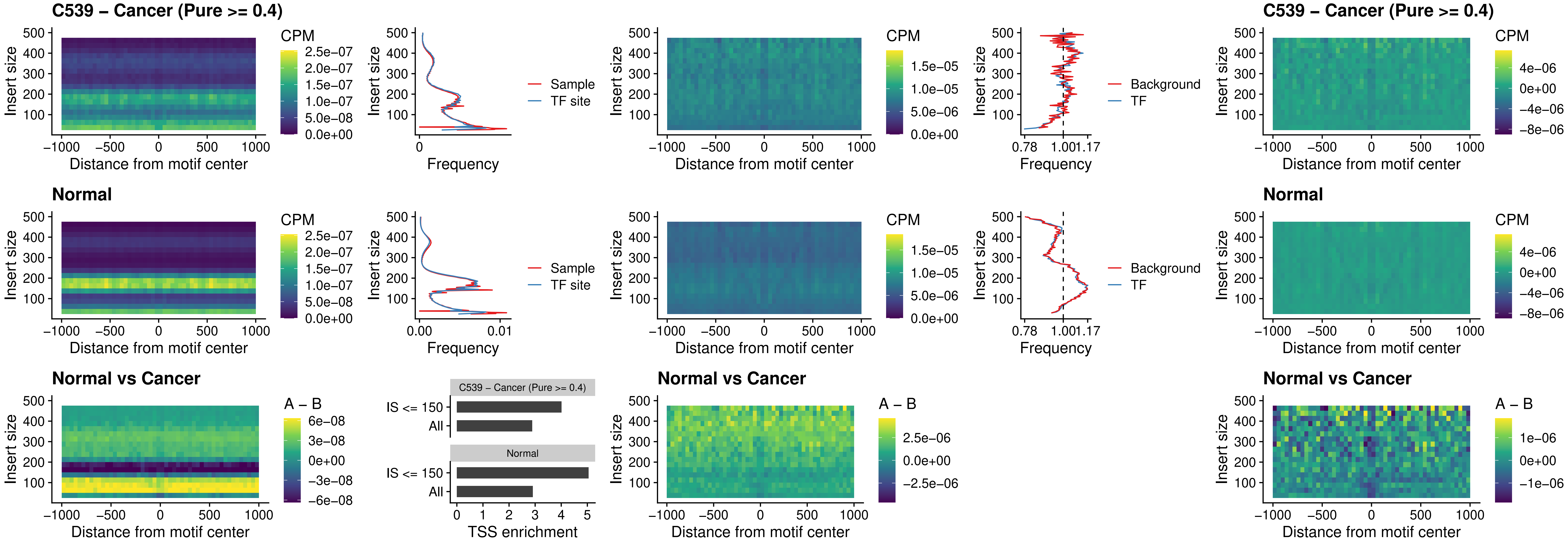

#### C539 – Cancer (Pure >= 0.4) – SPI1_uc_cTSS_nPEAK

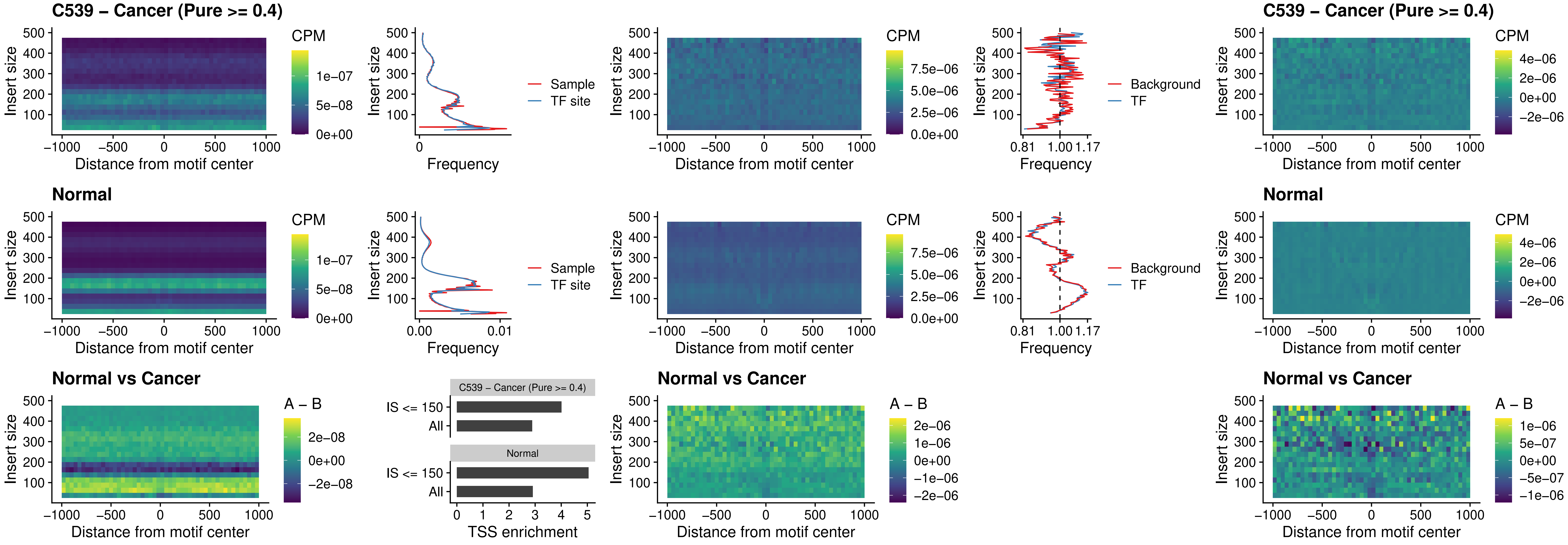

#### C539 – Cancer (Pure >= 0.4) – BCL11A_uc_pTSS_nPEAK

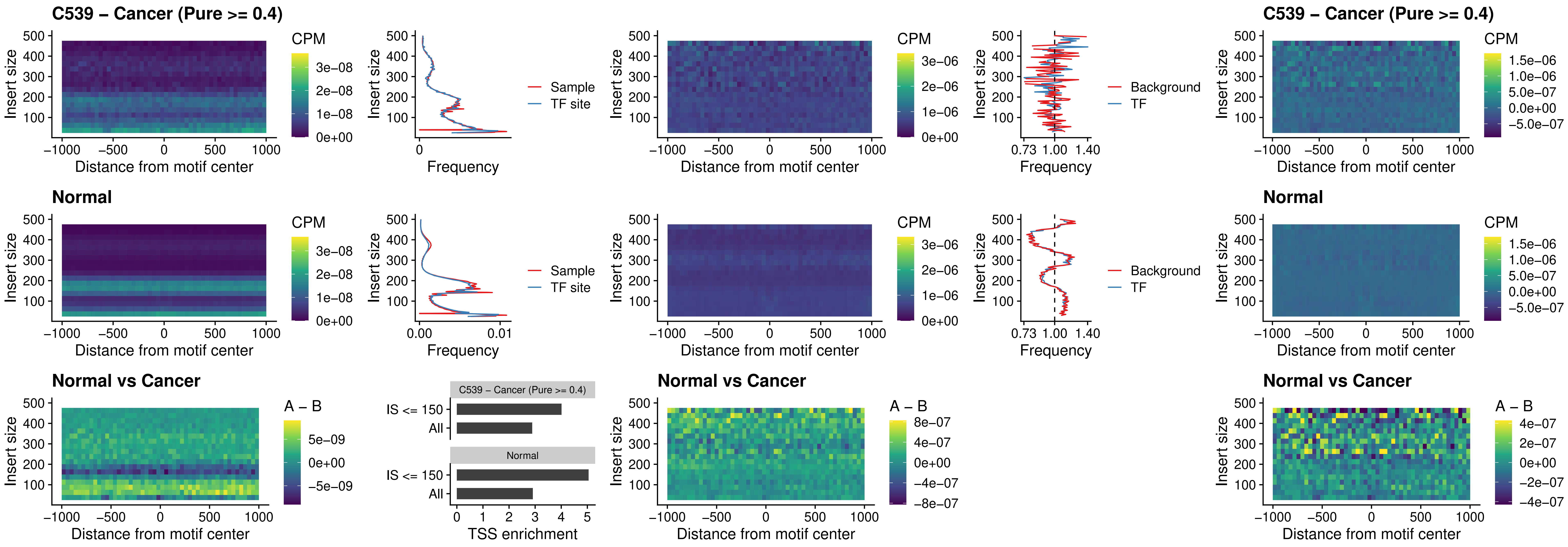

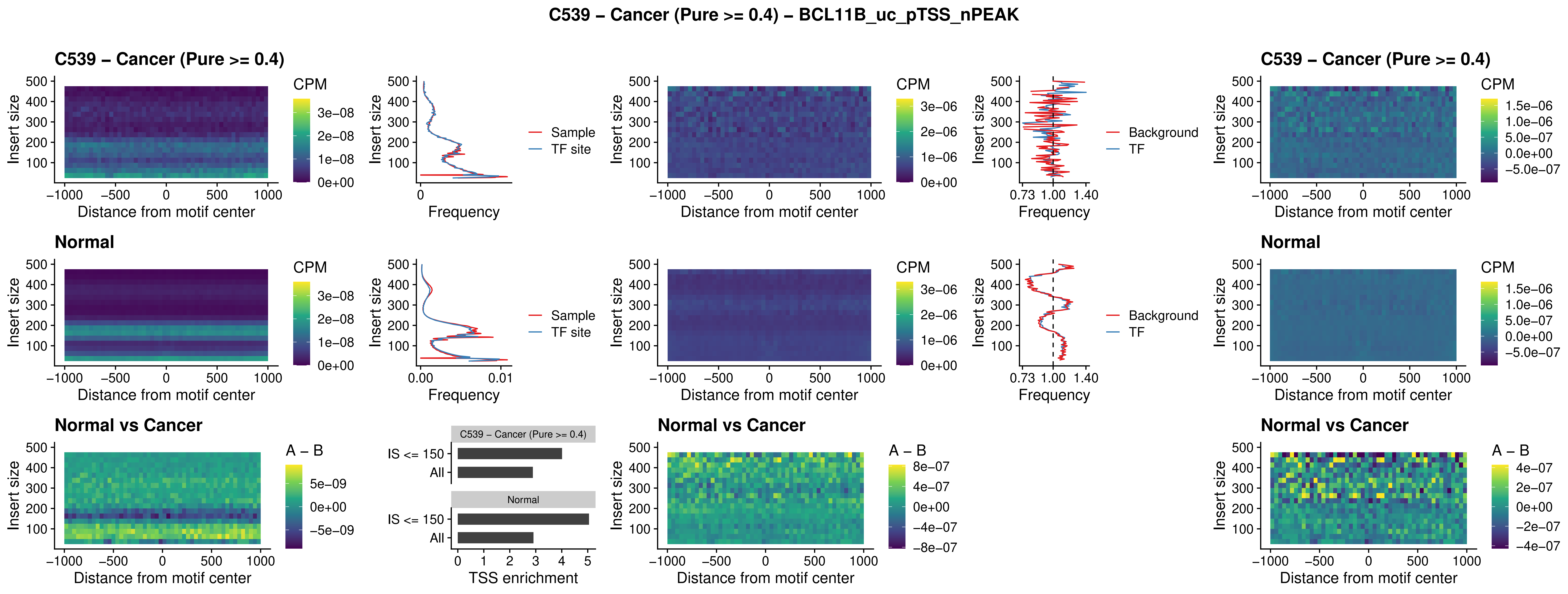
C539 – Cancer (Pure >= 0.4) – BCL11B_uc_pTSS_nPEAK
C539 – Cancer (Pure >= 0.4)
Normal
Normal vs Cancer
Insert size
Distance from motif center
Frequency
CPM
A – B
Sample
TF site
Background
TF
C539 – Cancer (Pure >= 0.4)
Normal
IS <= 150
All
TSS enrichment

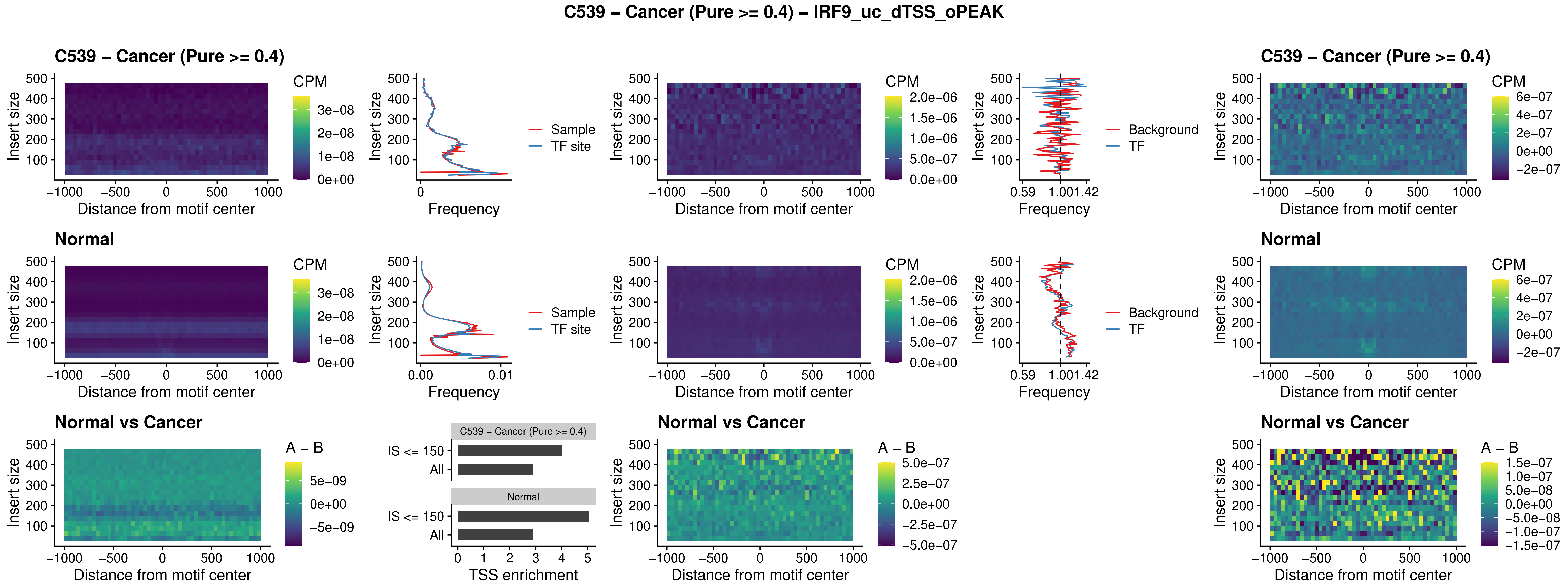
C539 – Cancer (Pure >= 0.4) – IRF9_uc_dTSS_oPEAK
C539 – Cancer (Pure >= 0.4)
Normal
Normal vs Cancer
Insert size
Distance from motif center
Frequency
CPM
A – B
Sample
TF site
Background
TF
C539 – Cancer (Pure >= 0.4)
Normal
IS <= 150
All
TSS enrichment

#### C539 – Cancer (Pure >= 0.4) – IRF9_uc_cTSS_oPEAK

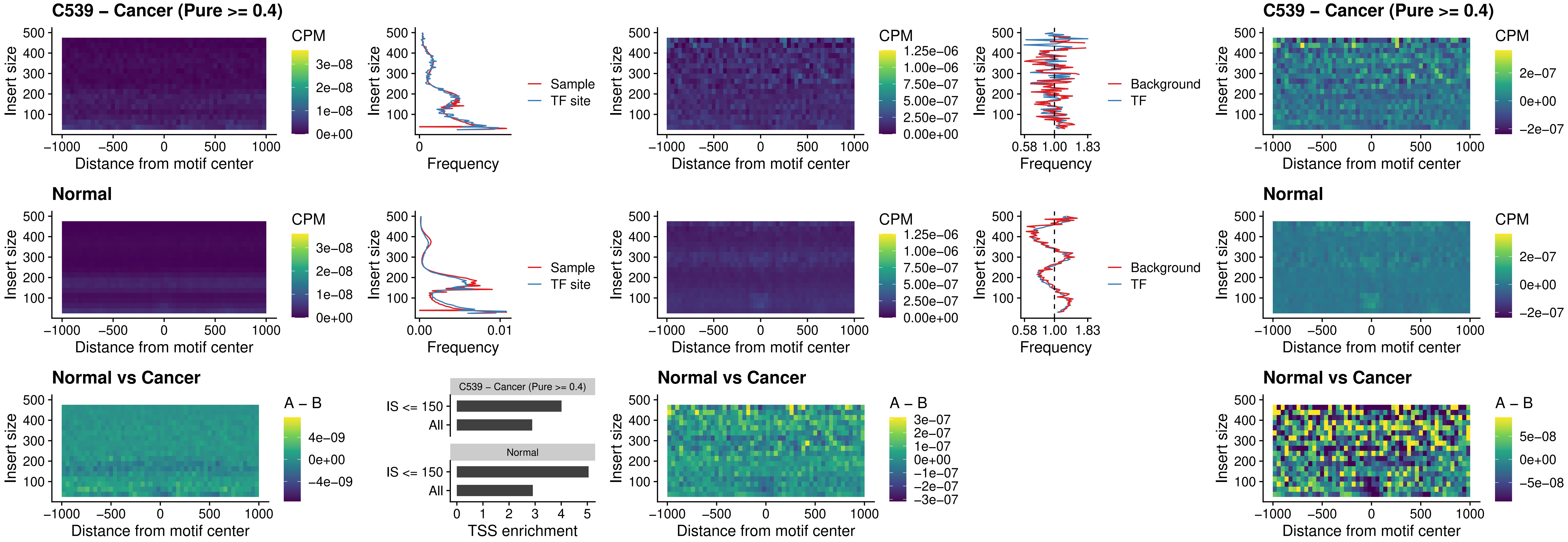

#### C539 – Cancer (Pure >= 0.4) – BCL11A_uc_cTSS_oPEAK

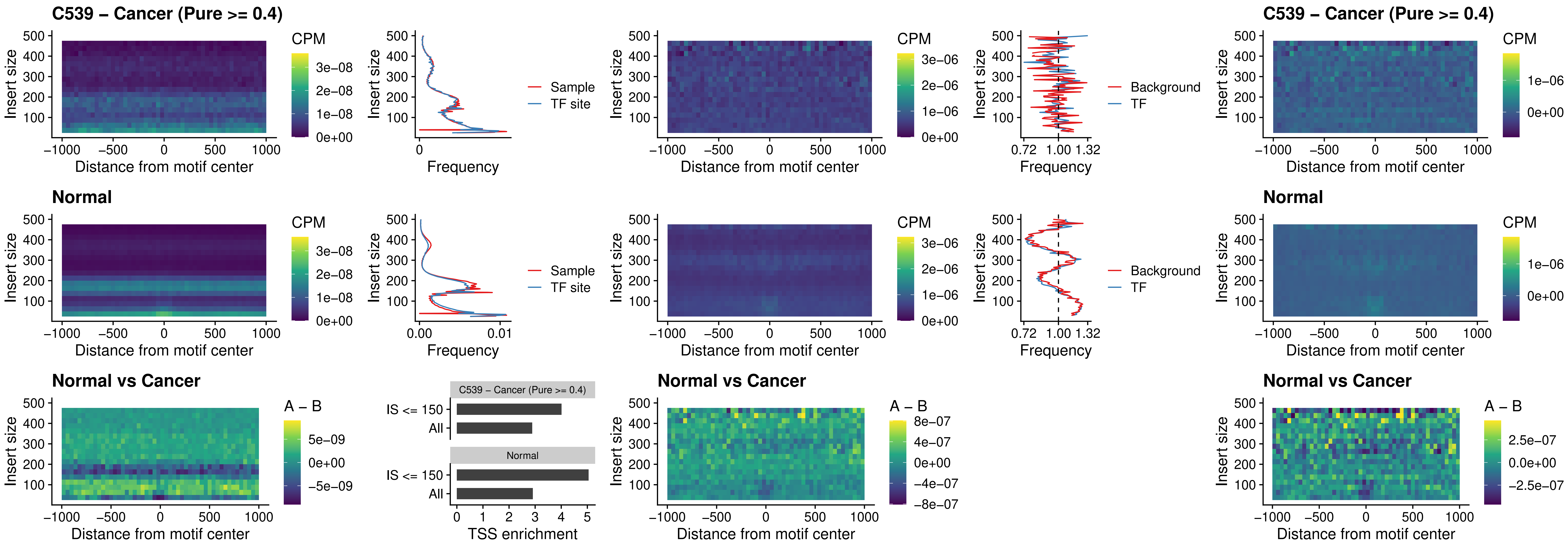

#### C539 – Cancer (Pure >= 0.4) – BCL11B_uc_cTSS_oPEAK

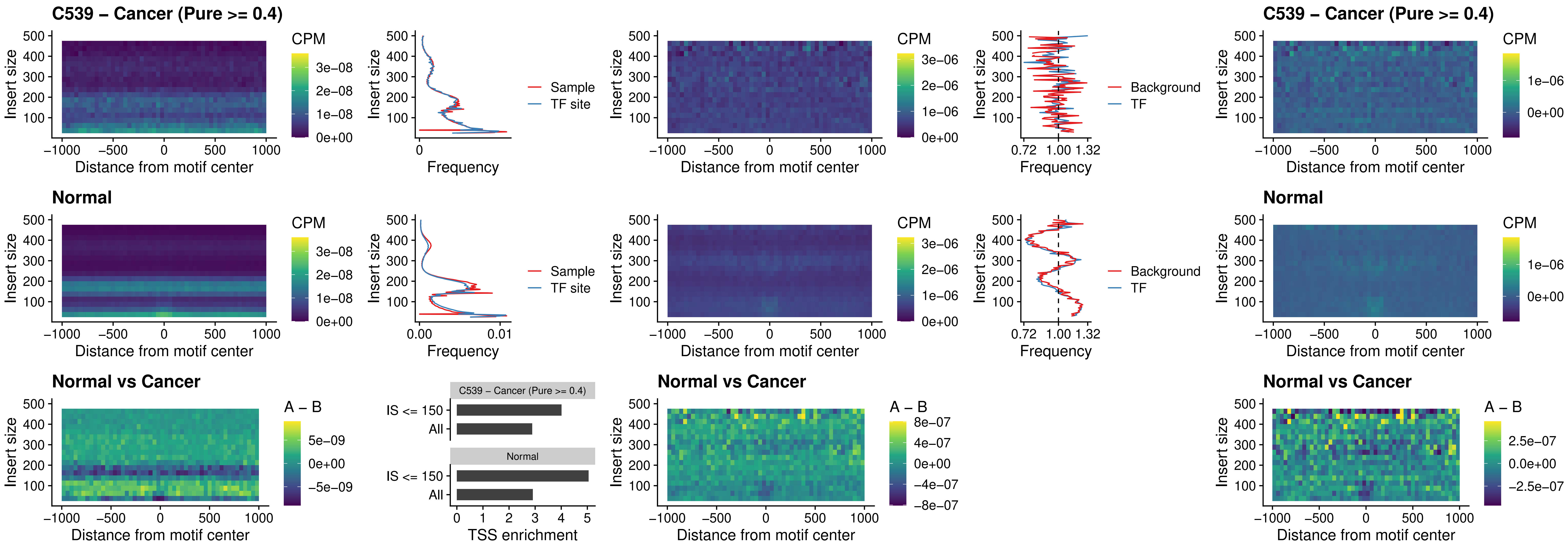

#### C539 – Cancer (Pure >= 0.4) – BCL11A_uc_dTSS_oPEAK

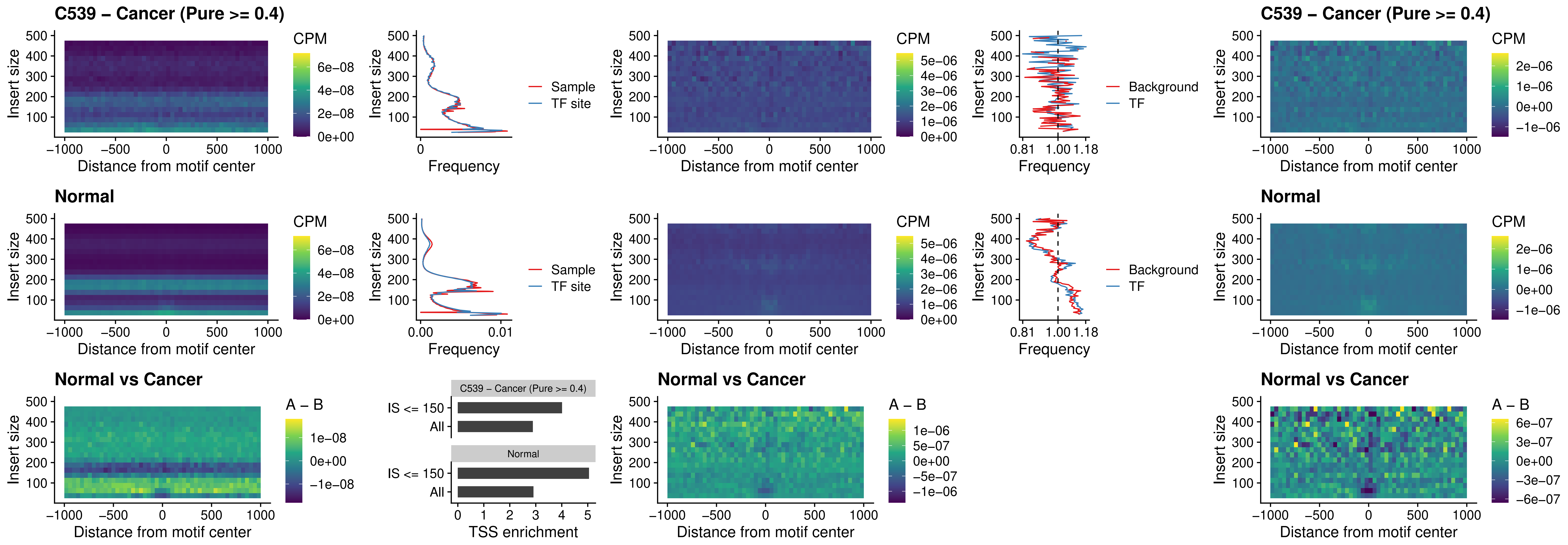

#### C539 – Cancer (Pure >= 0.4) – BCL11B_uc_dTSS_oPEAK

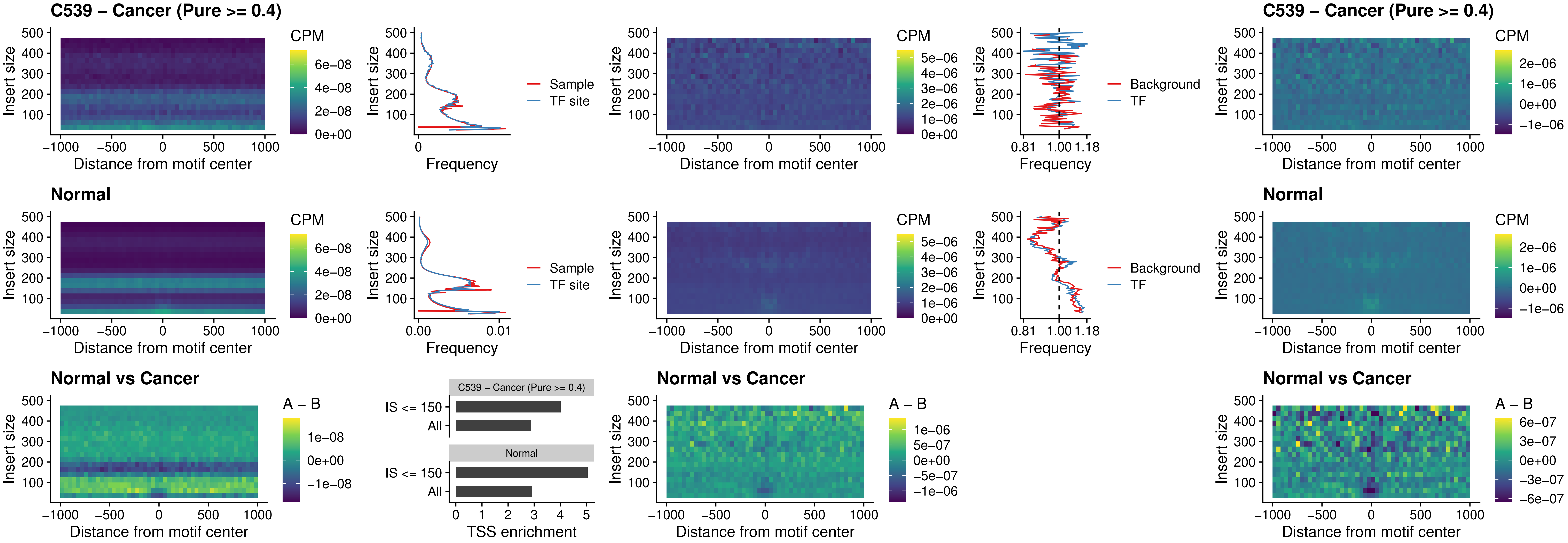

#### C539 – Cancer (Pure >= 0.4) – SPIB_uc_cTSS_nPEAK

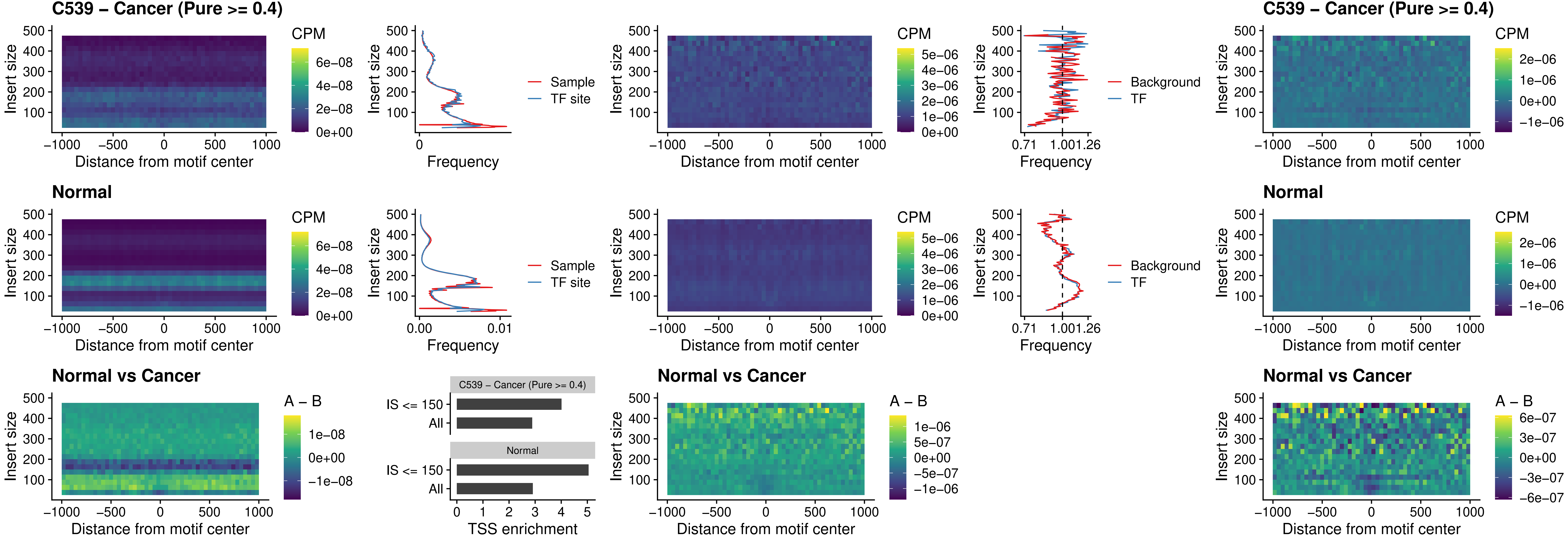

#### C539 – Cancer (Pure >= 0.4) – SPIC_uc_cTSS_nPEAK

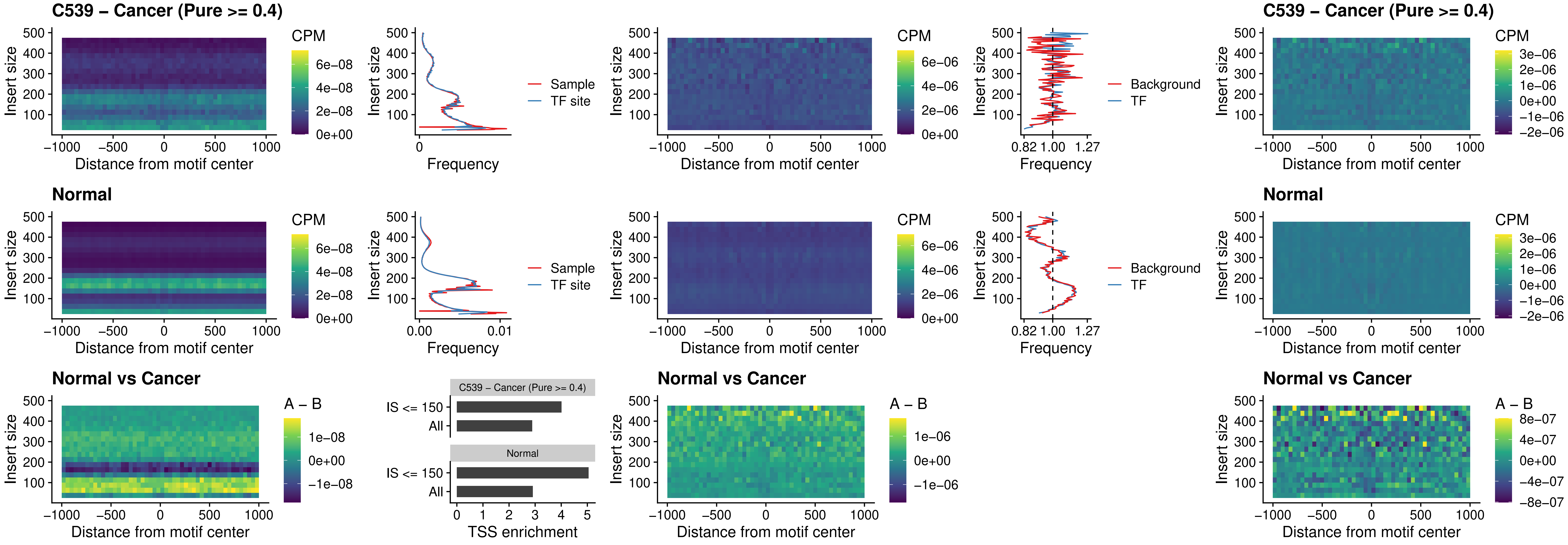

#### C539 – Cancer (Pure >= 0.4) – SPI1_uc_pTSS_oPEAK

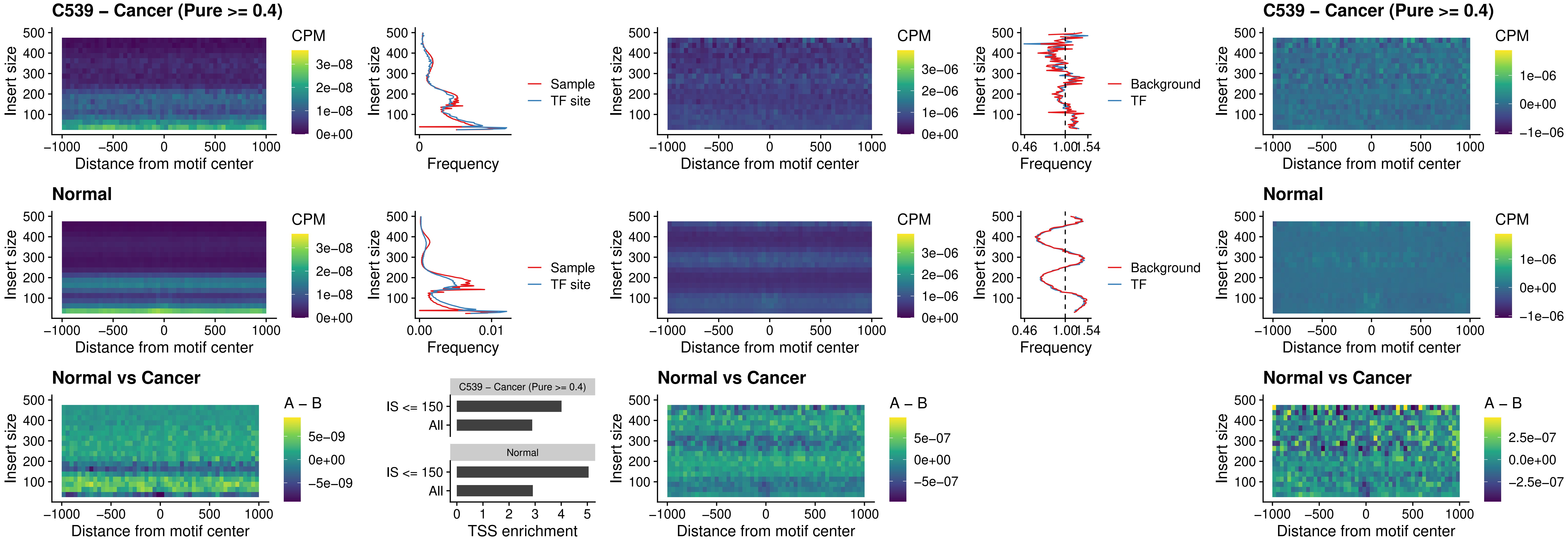

#### C539 – Cancer (Pure >= 0.4) – BCL11A_uc_pTSS_oPEAK

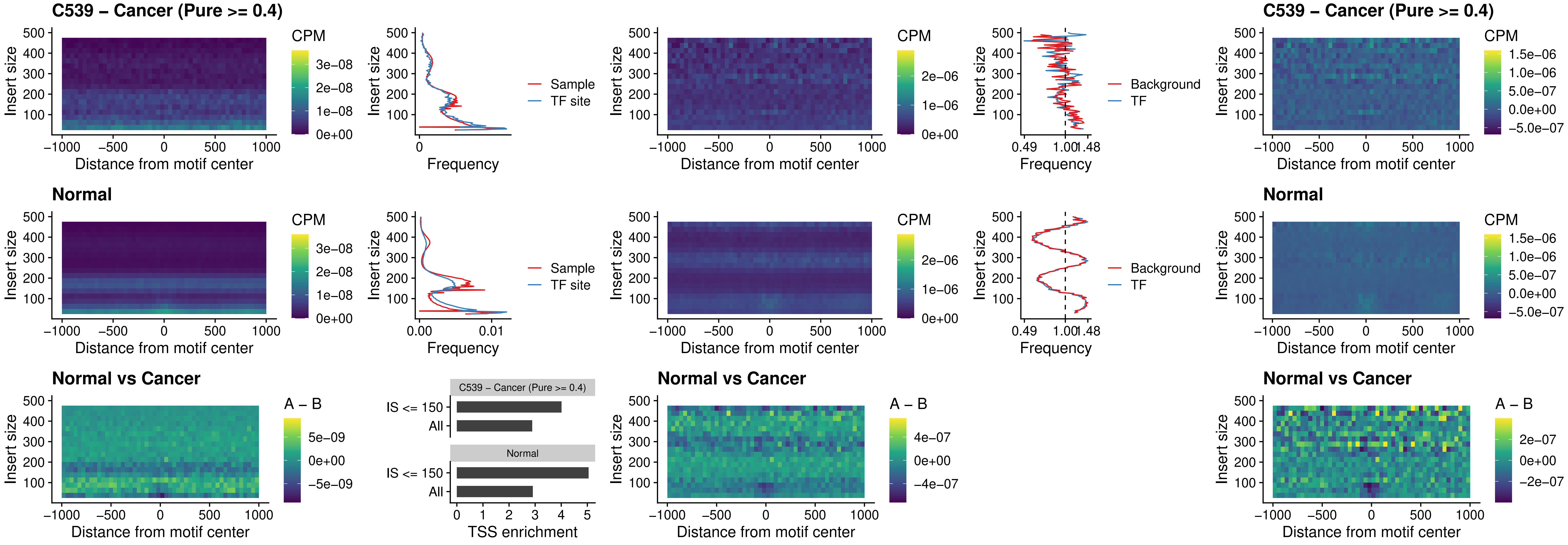

#### C539 – Cancer (Pure >= 0.4) – BCL11B_uc_pTSS_oPEAK

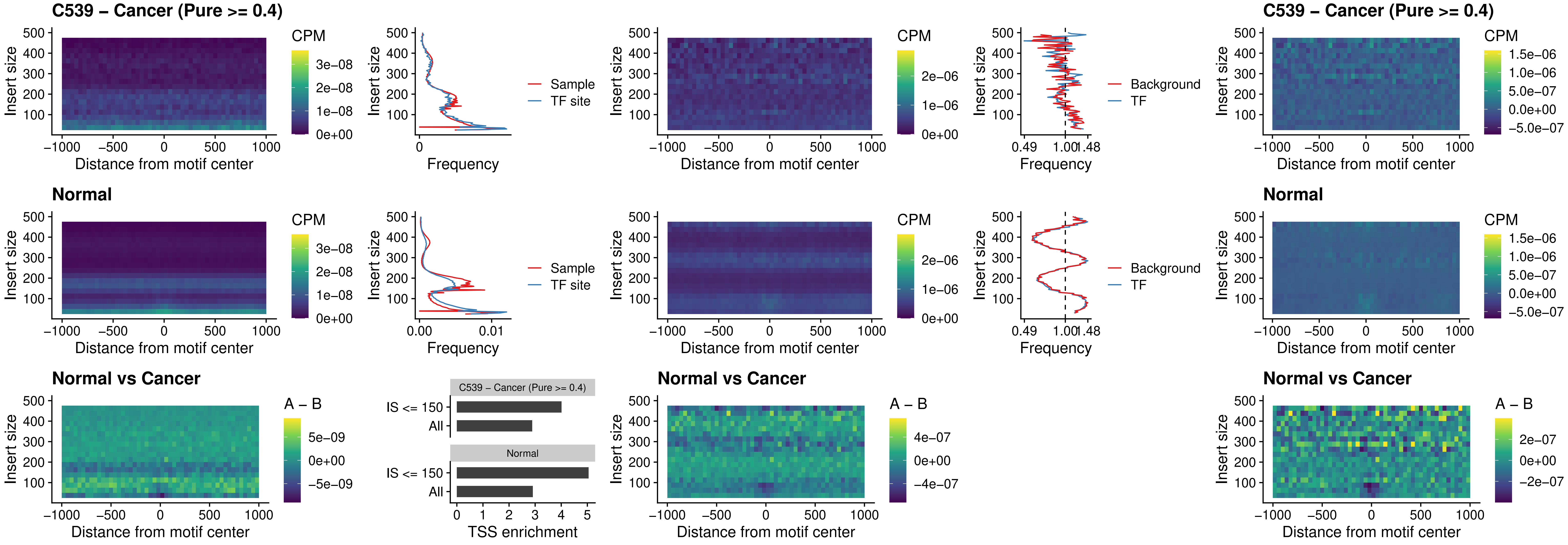

#### C539 – Cancer (Pure >= 0.4) – SPI1_uc_cTSS_oPEAK

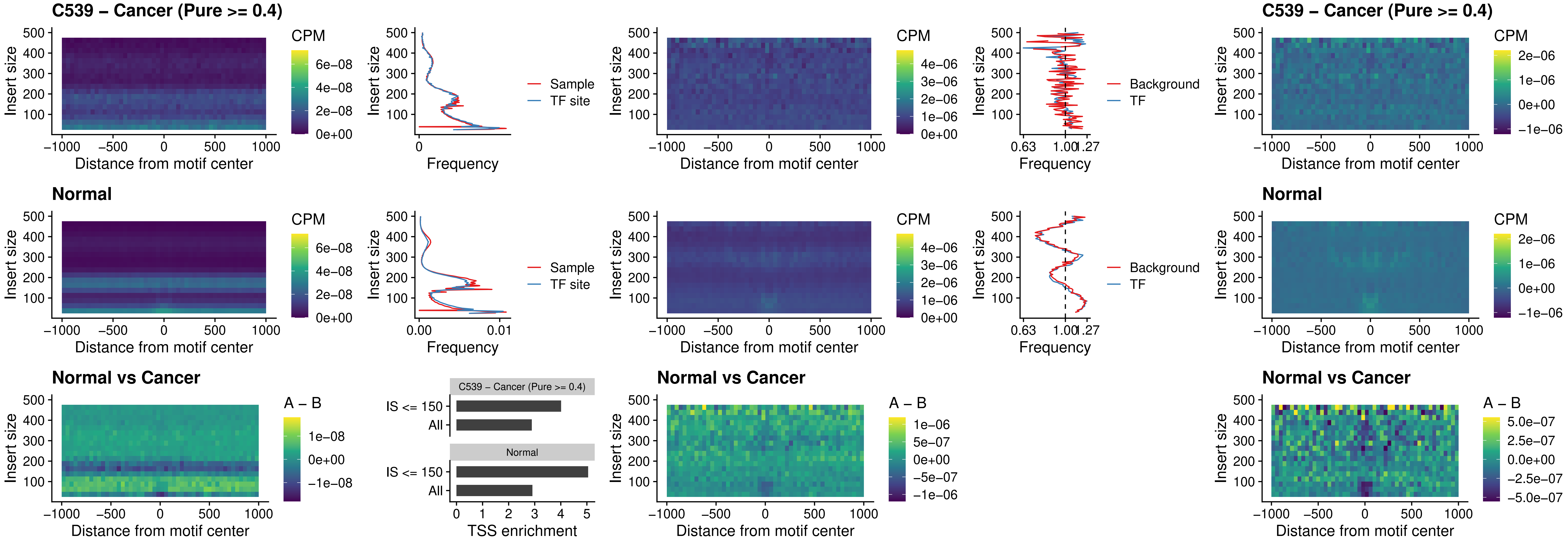

#### C539 – Cancer (Pure >= 0.4) – SPIB_uc_cTSS_oPEAK

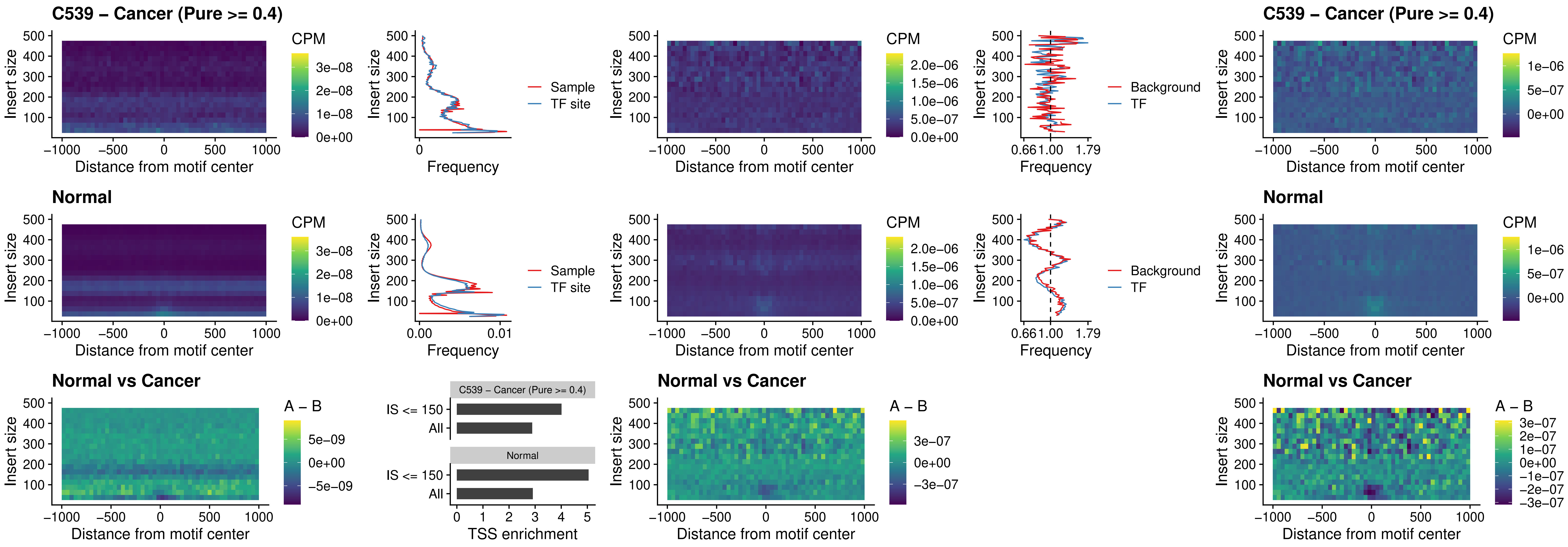

### C539 – Cancer (Pure >= 0.4) – SPIB_uc_pTSS_oPEAK

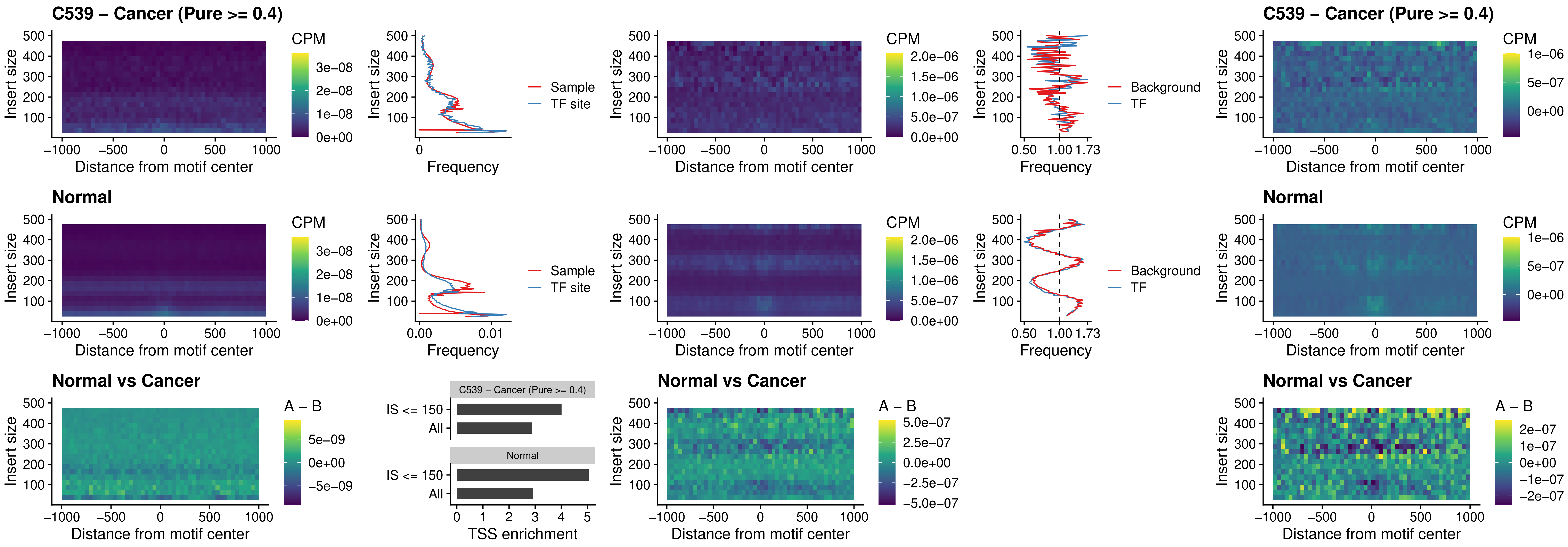

### C539 – Cancer (Pure >= 0.4) – SPI1_uc_dTSS_oPEAK

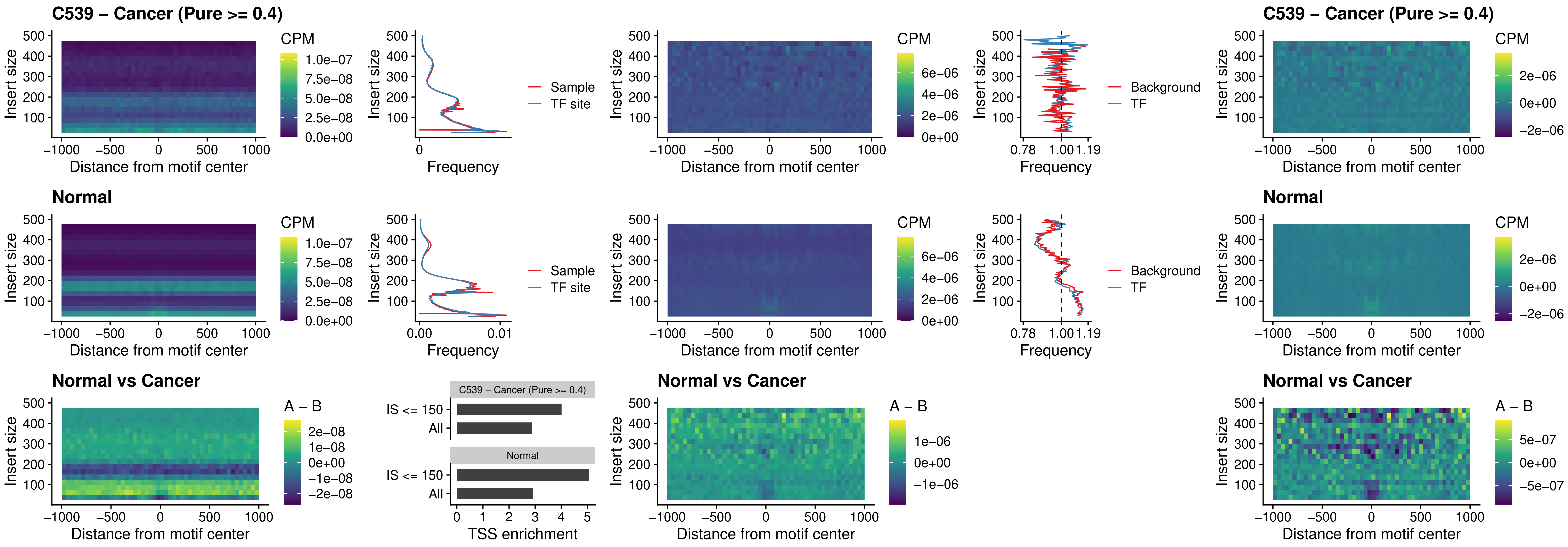

### C539 – Cancer (Pure >= 0.4) – SPIB_uc_dTSS_oPEAK

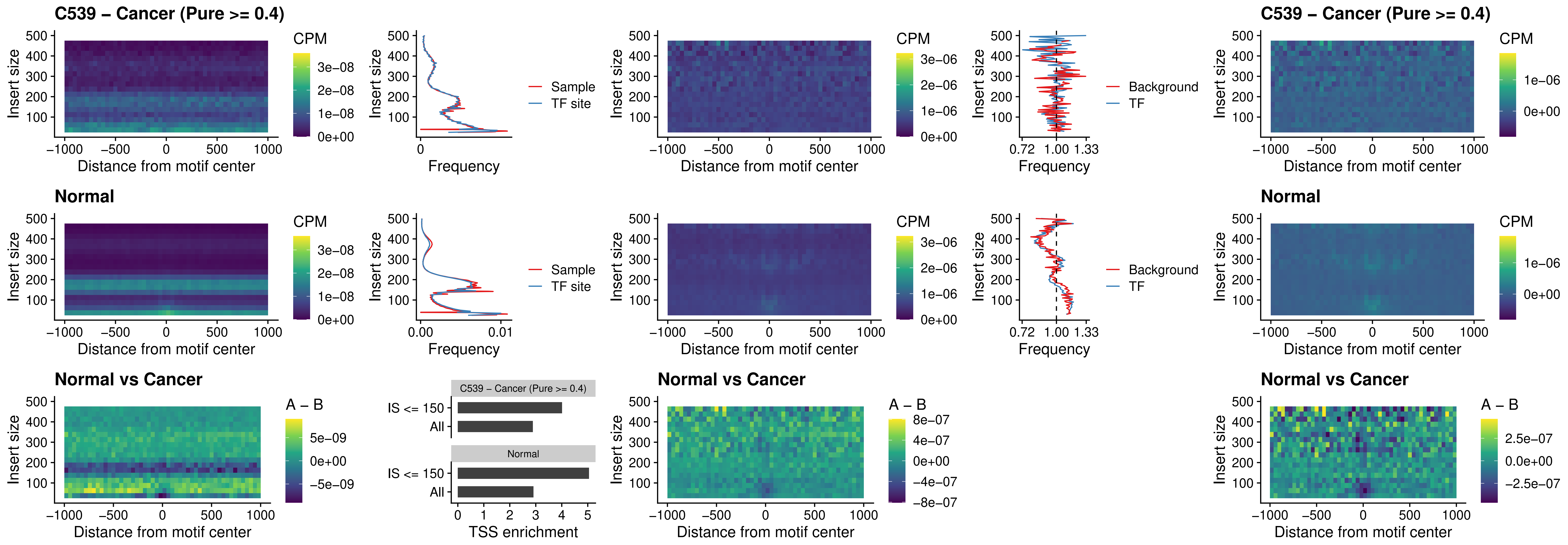

#### C539 – Cancer (Pure >= 0.4) – IRF3_uc_pTSS_oPEAK

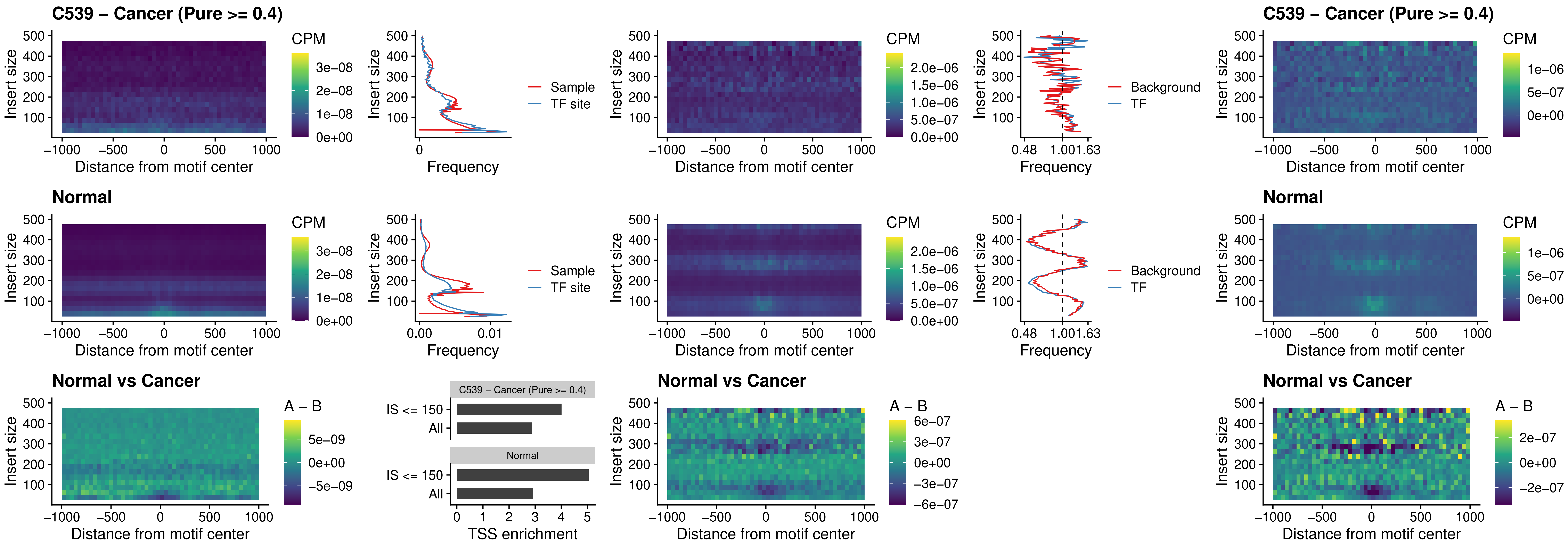

#### C539 – Cancer (Pure >= 0.4) – IRF2_uc_dTSS_oPEAK

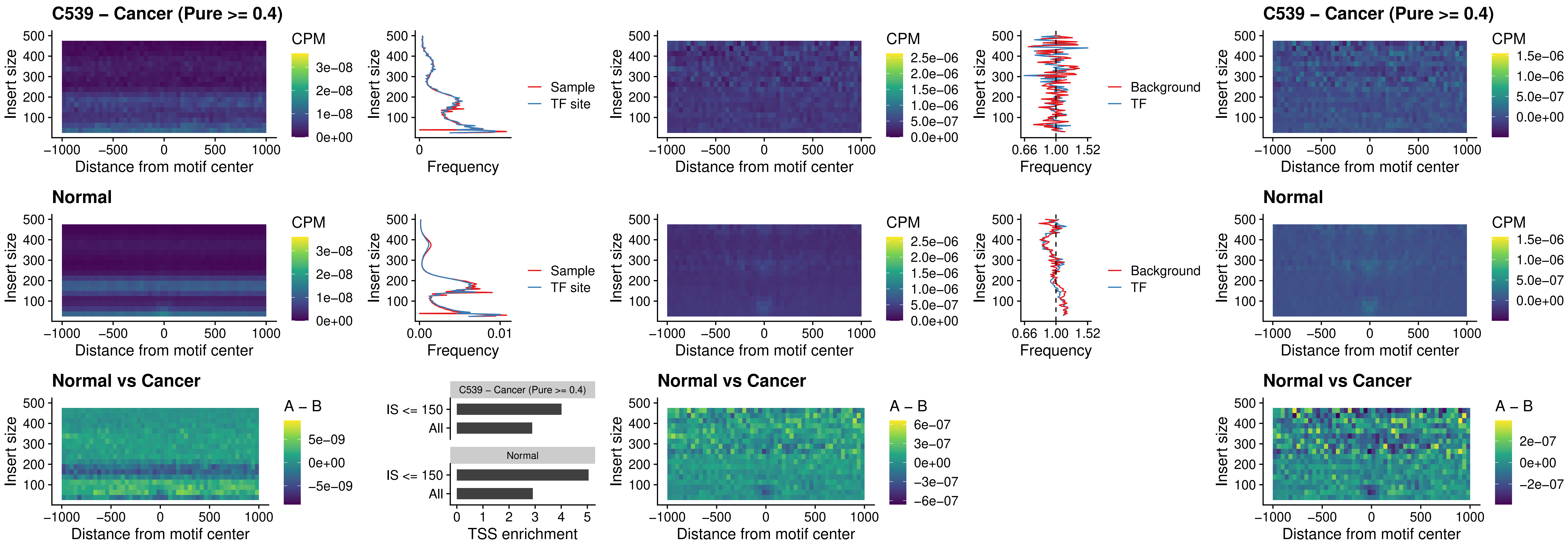

#### C539 – Cancer (Pure >= 0.4) – IRF2_uc_pTSS_oPEAK

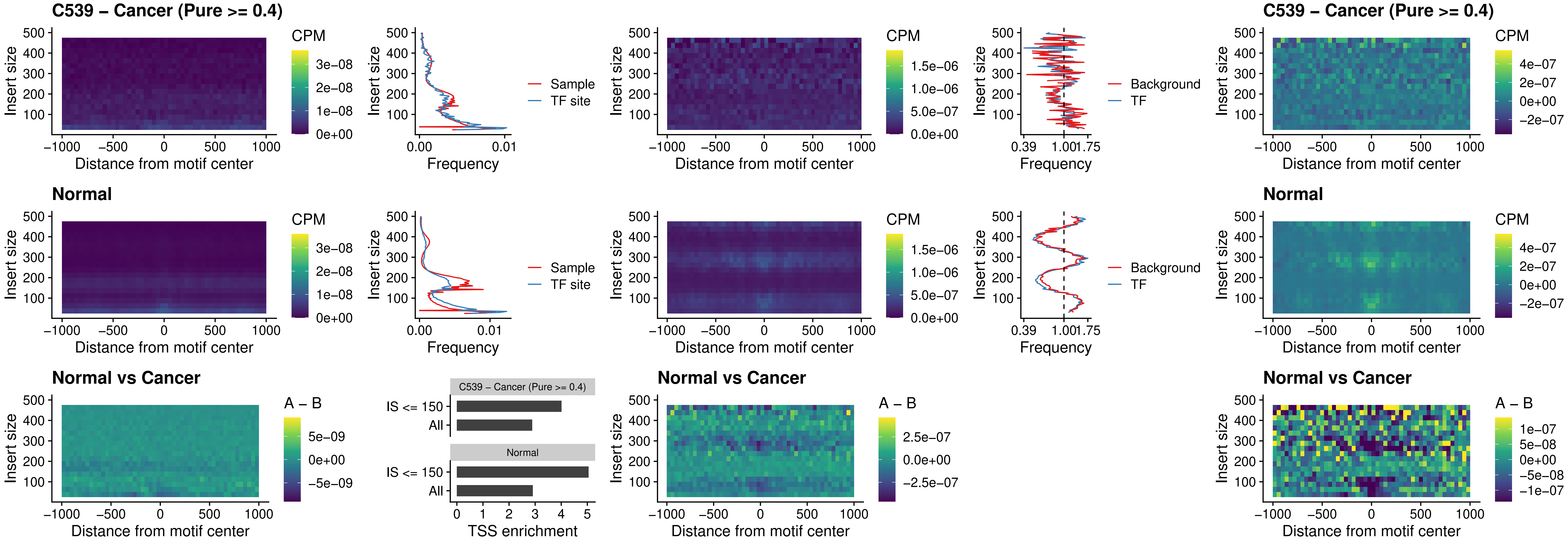

#### C539 – Cancer (Pure >= 0.4) – STAT2_uc_pTSS_oPEAK

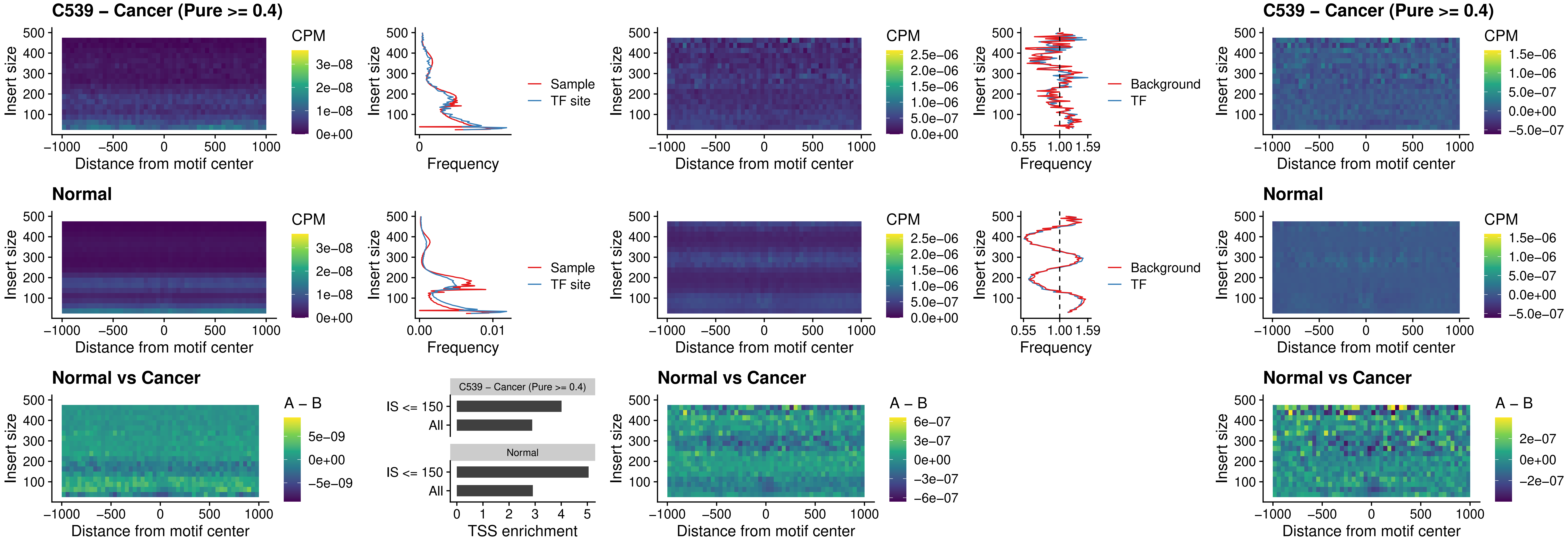

#### C539 – Cancer (Pure >= 0.4) – SPIB_uc_dTSS_nPEAK

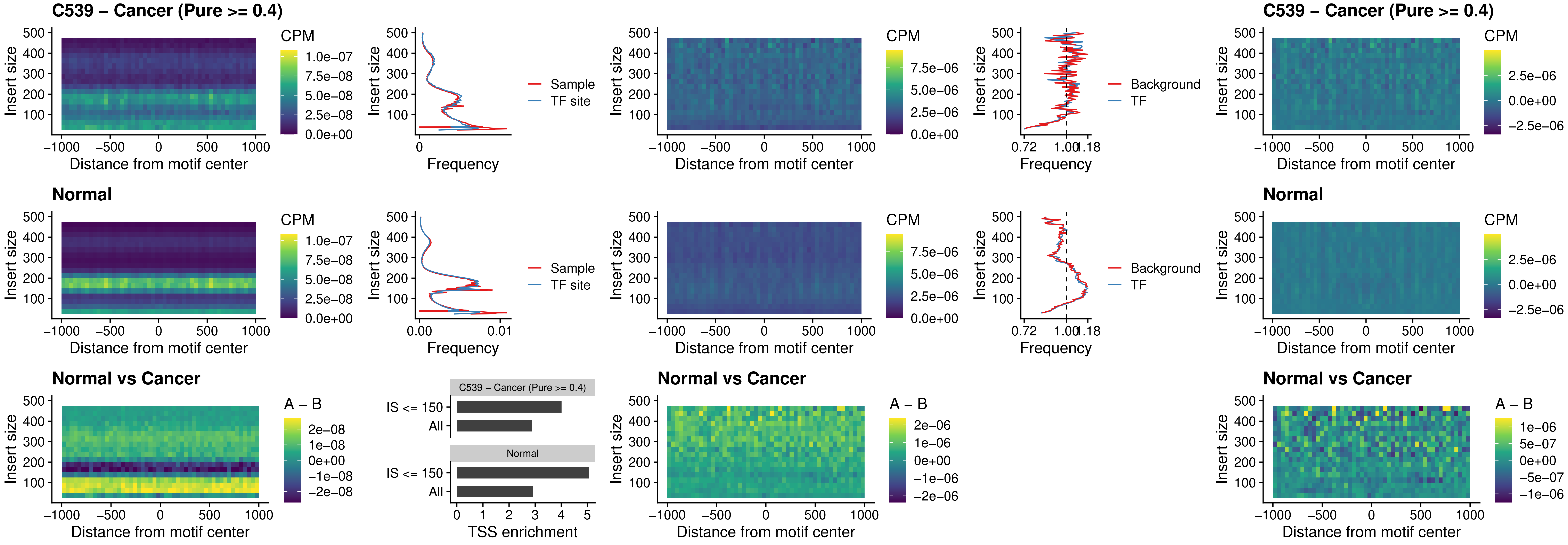

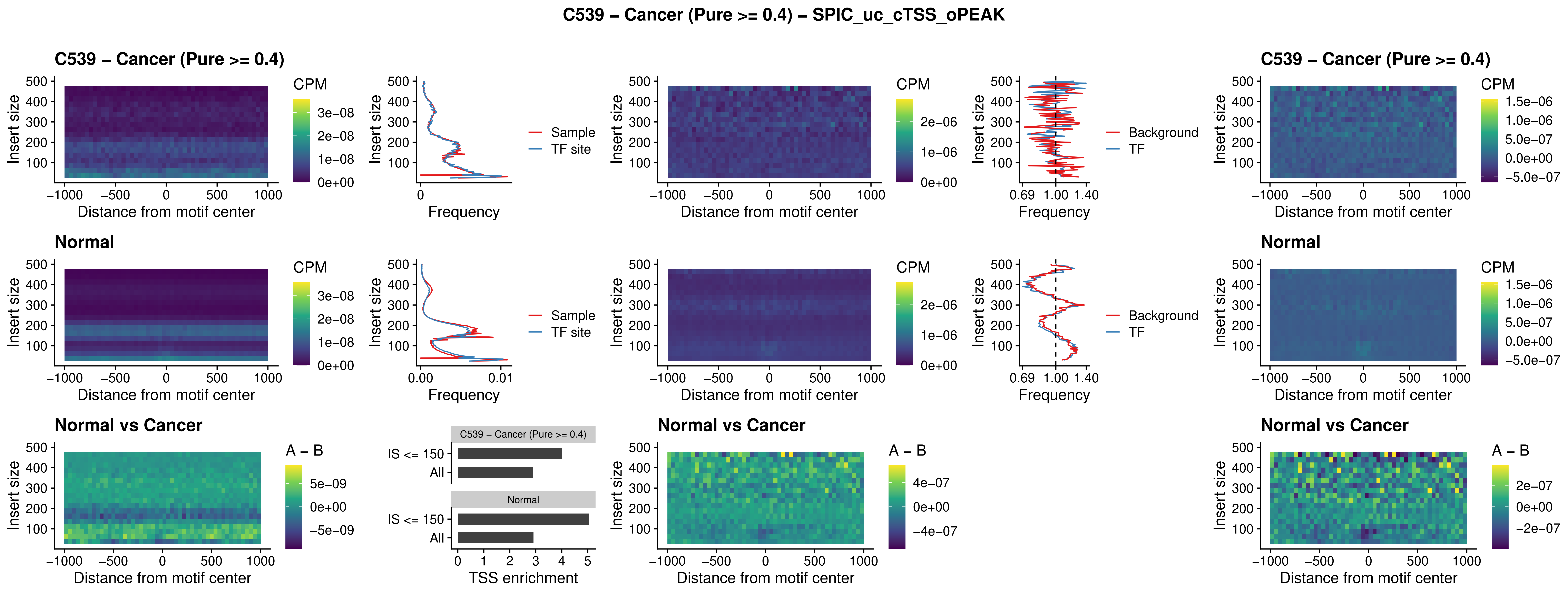
C539 – Cancer (Pure >= 0.4) – SPIC_uc_cTSS_oPEAK
C539 – Cancer (Pure >= 0.4)
Normal
Normal vs Cancer
Insert size
Distance from motif center
Frequency
CPM
A – B
Sample
TF site
Background
TF
C539 – Cancer (Pure >= 0.4)
Normal
IS <= 150
All
TSS enrichment

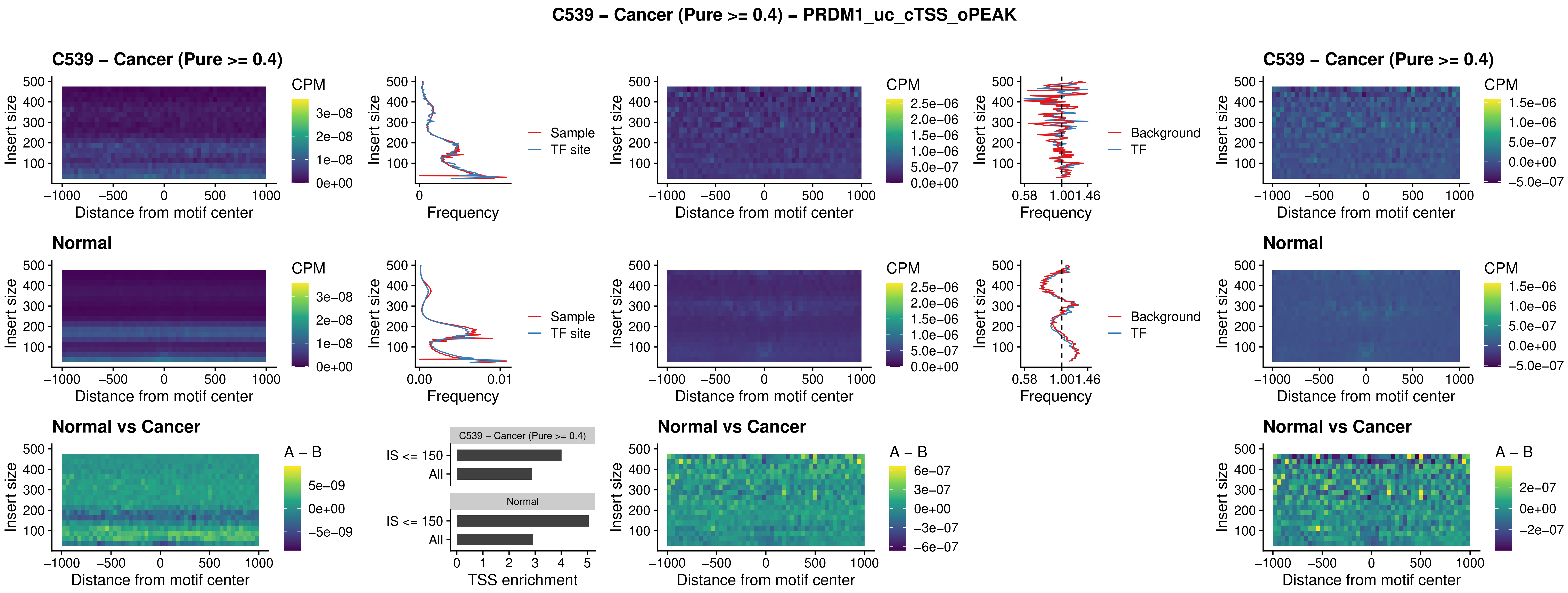
C539 – Cancer (Pure >= 0.4) – PRDM1_uc_cTSS_oPEAK
C539 – Cancer (Pure >= 0.4)
Normal
Normal vs Cancer
Insert size
Distance from motif center
Frequency
CPM
A – B
Sample
TF site
Background
TF
C539 – Cancer (Pure >= 0.4)
Normal
IS <= 150
All
TSS enrichment

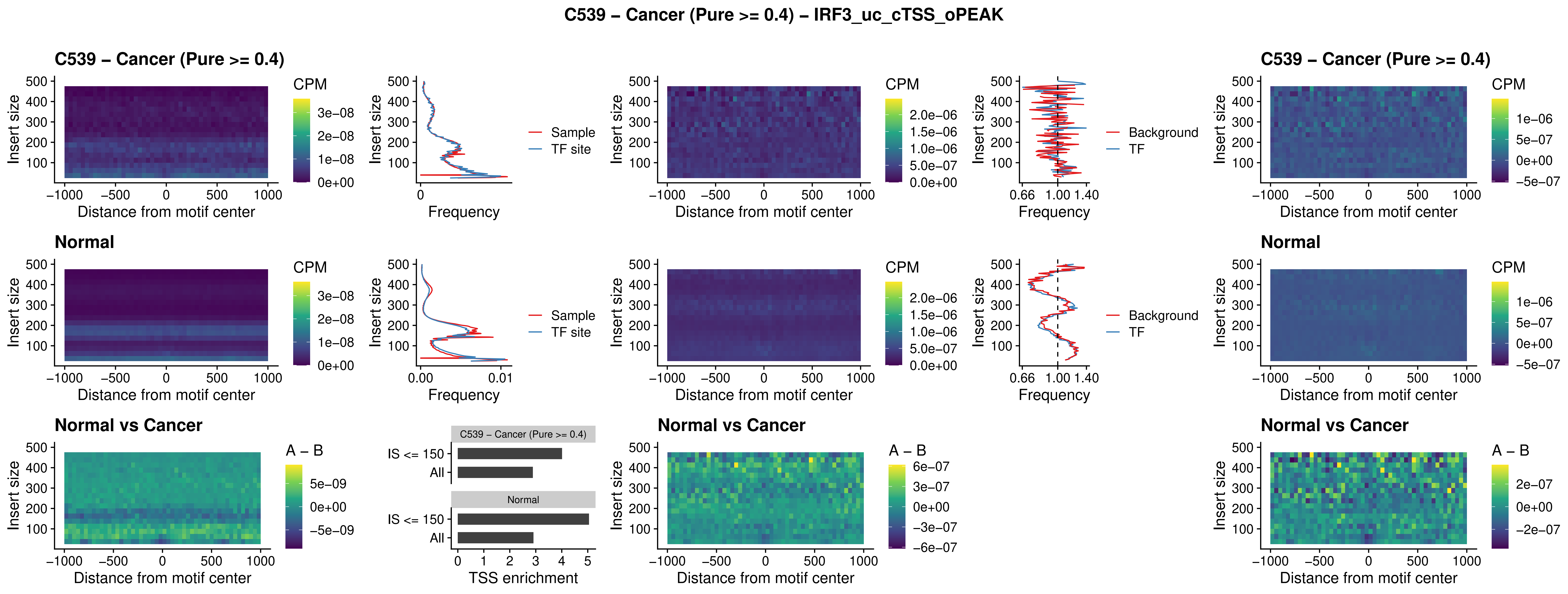
C539 – Cancer (Pure >= 0.4) – IRF3_uc_cTSS_oPEAK
C539 – Cancer (Pure >= 0.4)
Normal
Normal vs Cancer
Insert size
Distance from motif center
Frequency
CPM
A – B
Sample
TF site
Background
TF
C539 – Cancer (Pure >= 0.4)
IS <= 150
All
TSS enrichment

#### C539 – Cancer (Pure >= 0.4) – BCL11A_uc_dTSS_nPEAK

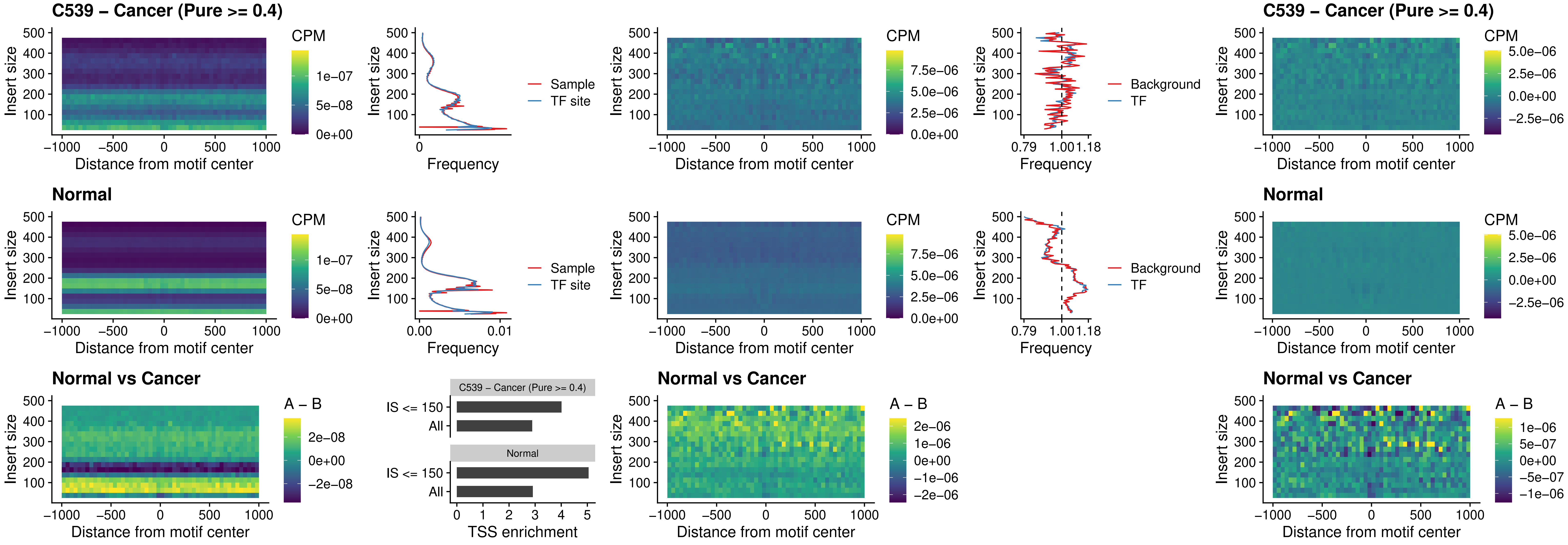

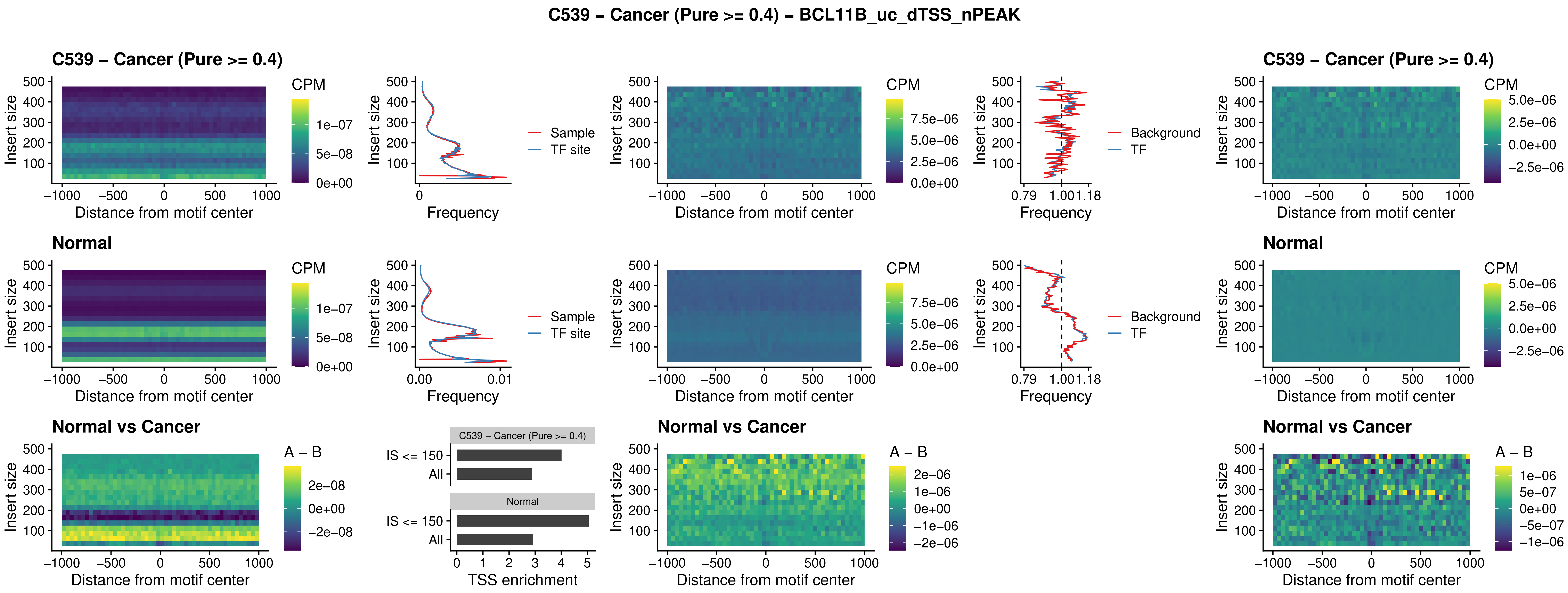
C539 – Cancer (Pure >= 0.4) – BCL11B_uc_dTSS_nPEAK
C539 – Cancer (Pure >= 0.4)
Normal
Normal vs Cancer
Insert size
Distance from motif center
Frequency
CPM
A – B
Sample
TF site
Background
TF
TSS enrichment
IS <= 150
All
C539 – Cancer (Pure >= 0.4)
Normal

#### C539 – Cancer (Pure >= 0.4) – IRF1_uc_cTSS_oPEAK

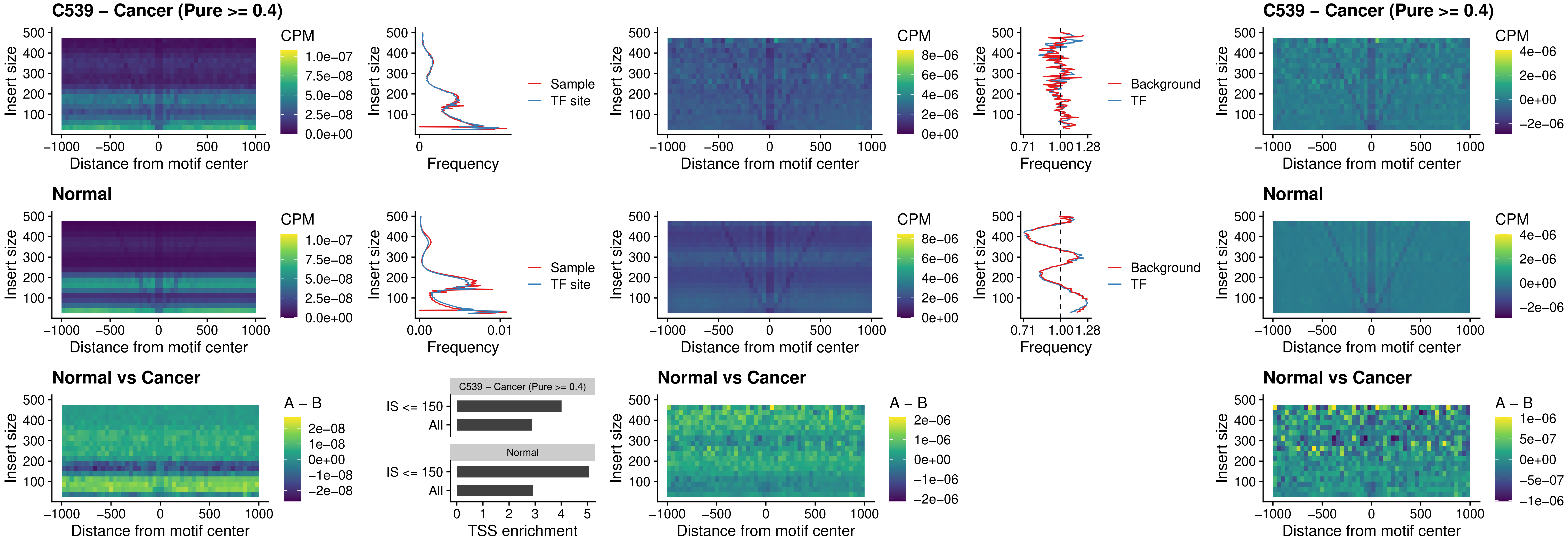

#### C539 – Cancer (Pure >= 0.4) – IRF1_uc_dTSS_oPEAK

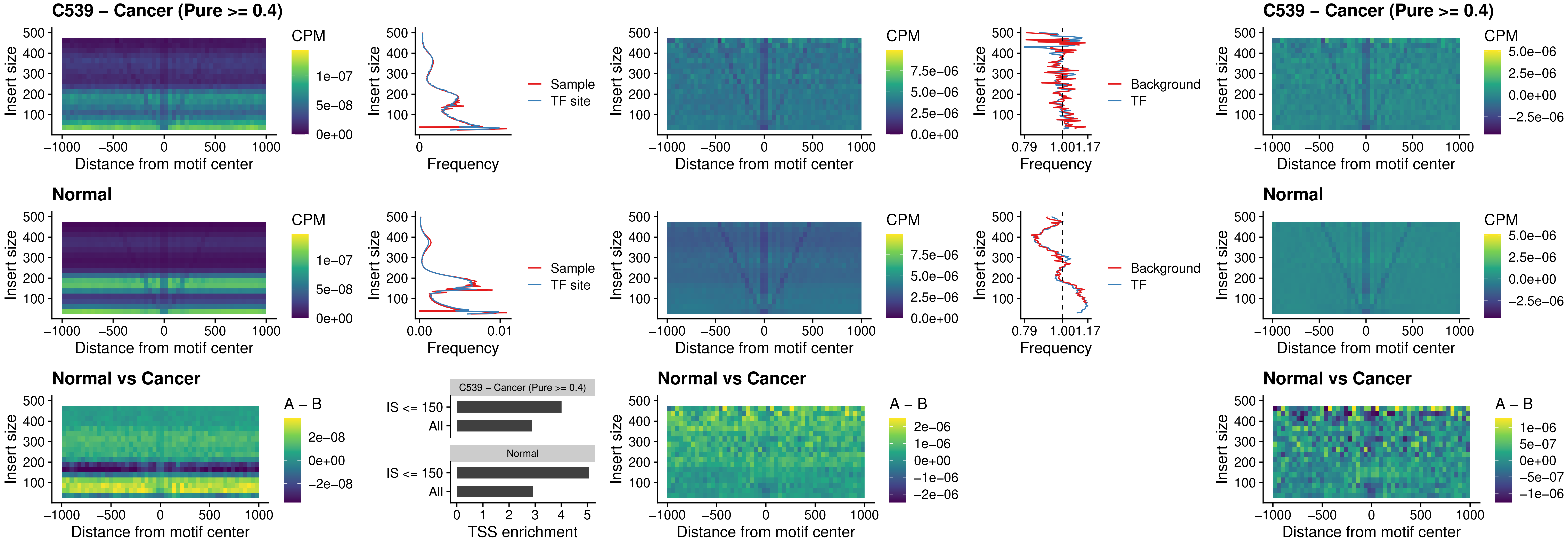

### C539 – Cancer (Pure >= 0.4) – IRF1_uc_pTSS_oPEAK

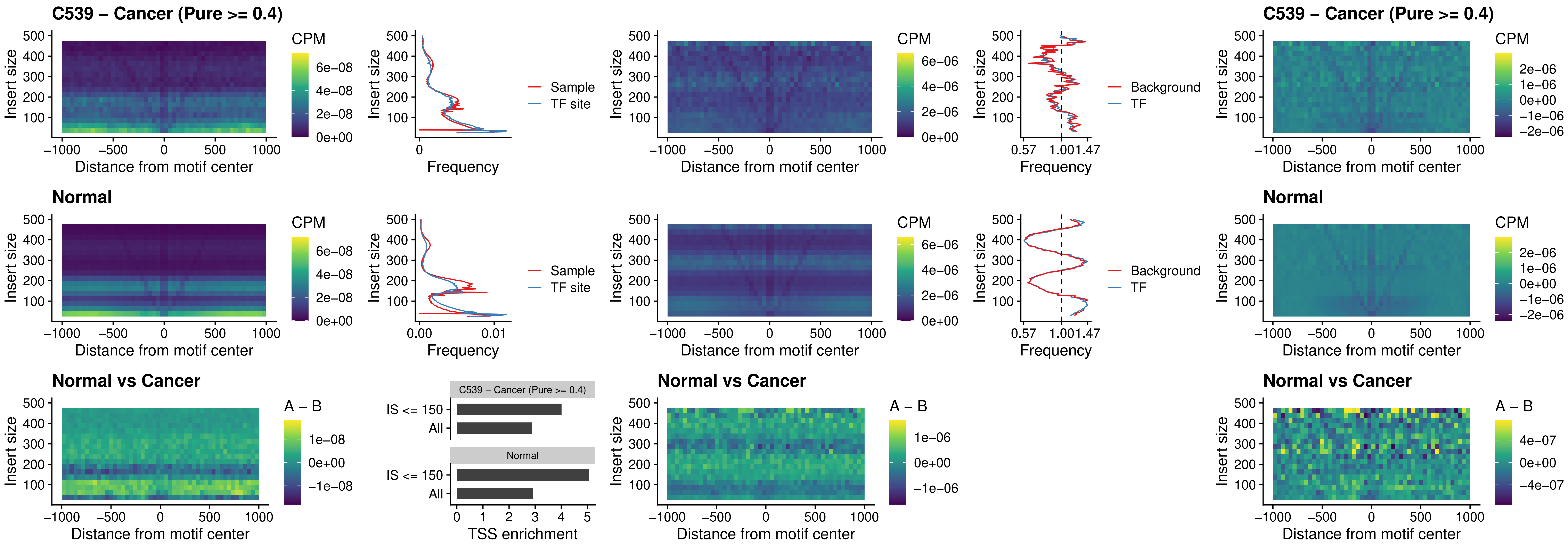

#### C539 – Cancer (Pure >= 0.4) – SPIC_uc_dTSS_nPEAK

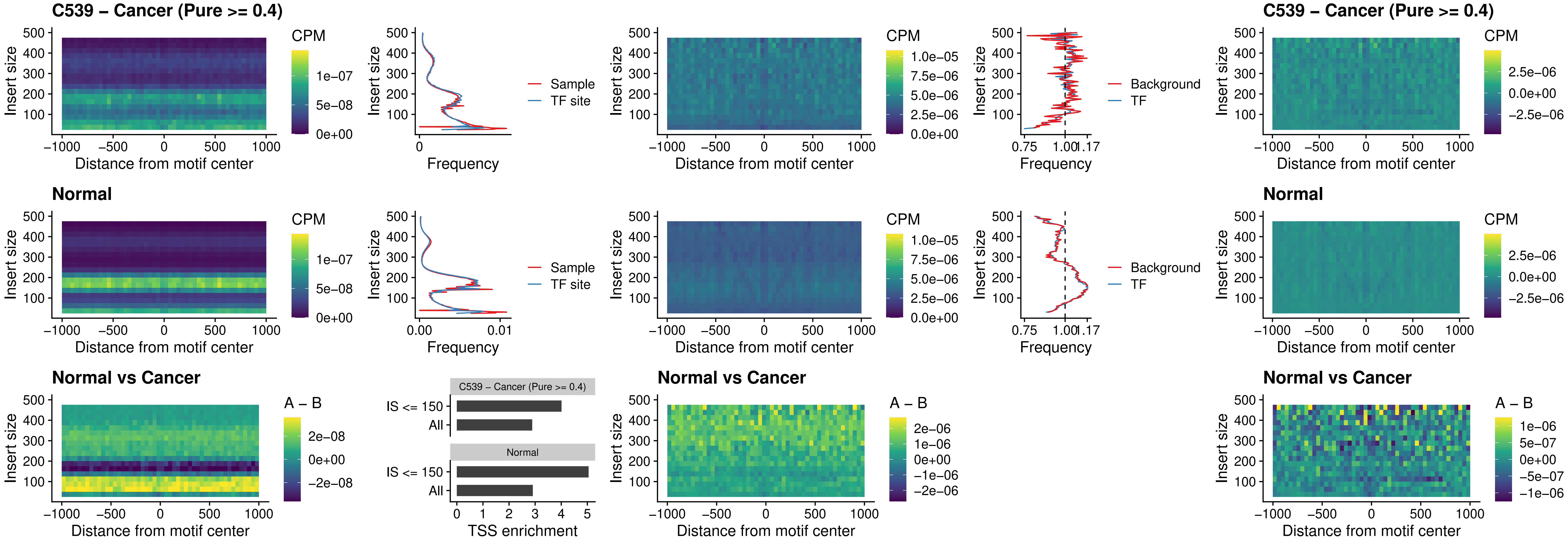

#### C539 – Cancer (Pure >= 0.4) – SPIC_uc_dTSS_oPEAK

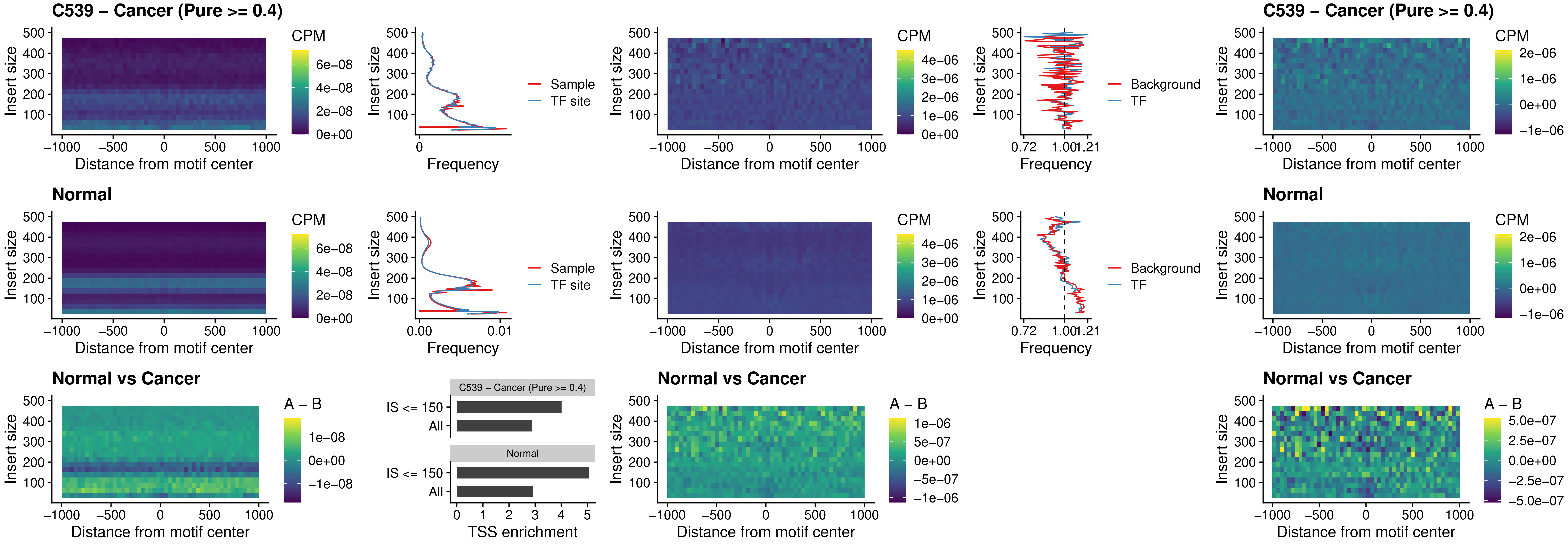

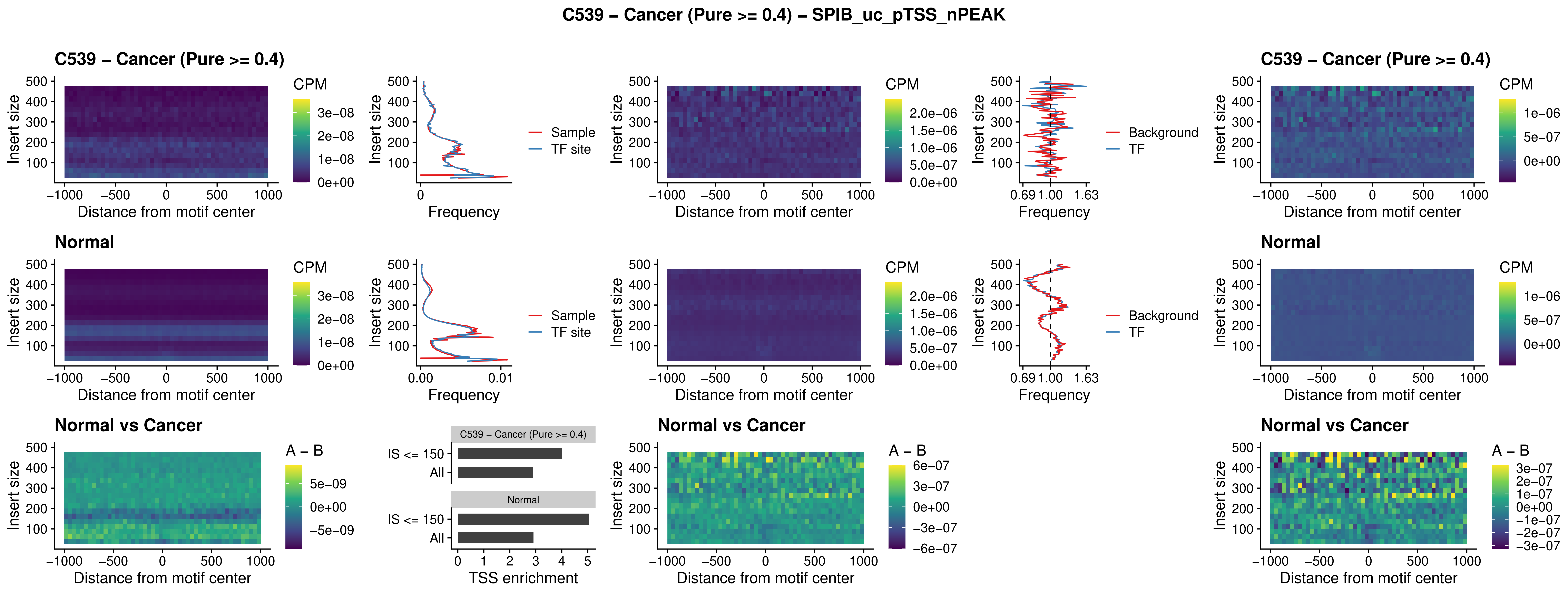
C539 – Cancer (Pure >= 0.4) – SPIB_uc_pTSS_nPEAK
C539 – Cancer (Pure >= 0.4)
Normal
Normal vs Cancer
Insert size
Distance from motif center
CPM
A – B
Frequency
Sample
TF site
Background
TF
C539 – Cancer (Pure >= 0.4)
Normal
IS <= 150
All
TSS enrichment

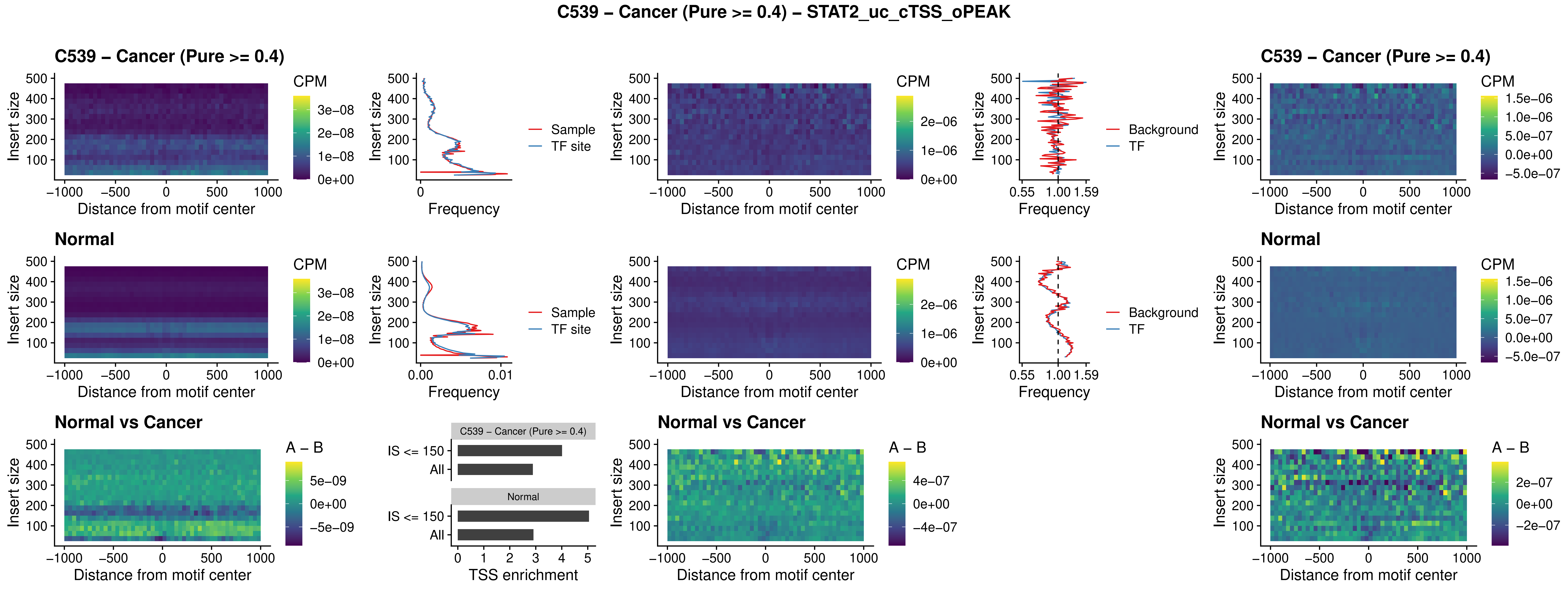
C539 – Cancer (Pure >= 0.4) – STAT2_uc_cTSS_oPEAK
C539 – Cancer (Pure >= 0.4)
Normal
Normal vs Cancer
Insert size
Distance from motif center
CPM
A – B
Frequency
Sample
TF site
Background
TF
C539 – Cancer (Pure >= 0.4)
Normal
IS <= 150
All
TSS enrichment

#### C539 – Cancer (Pure >= 0.4) – IRF8_uc_cTSS_oPEAK

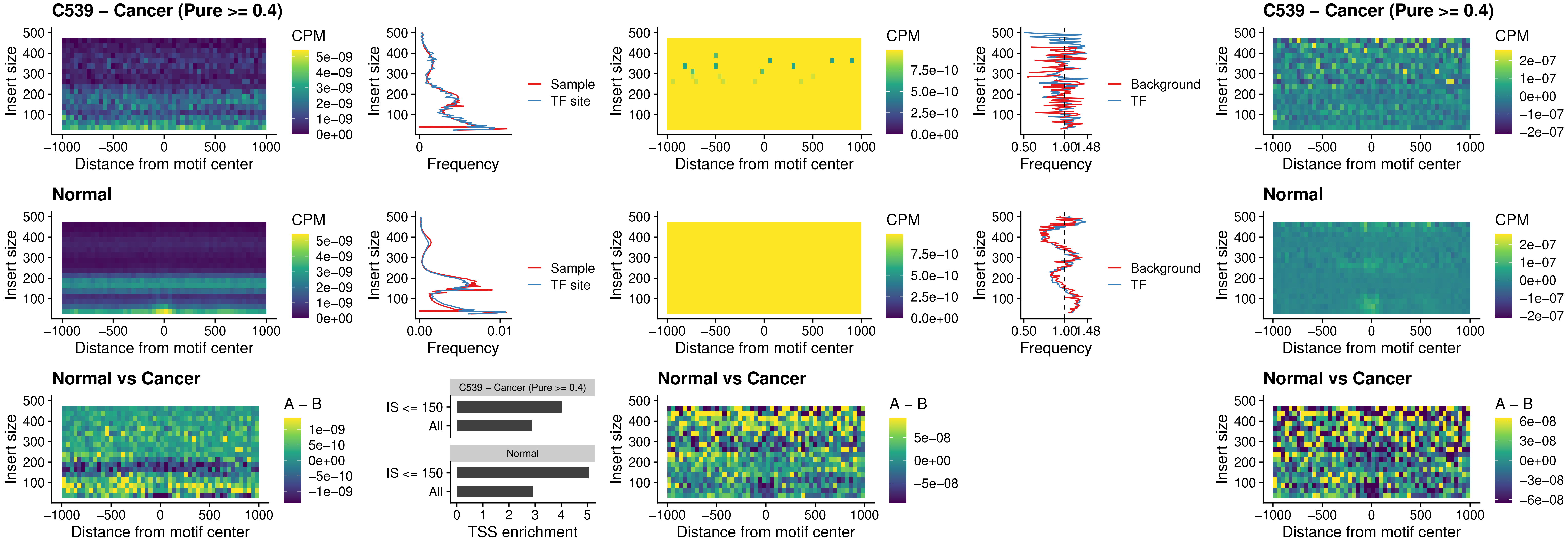

### C539 – Cancer (Pure >= 0.4) – IRF3_uc_dTSS_oPEAK

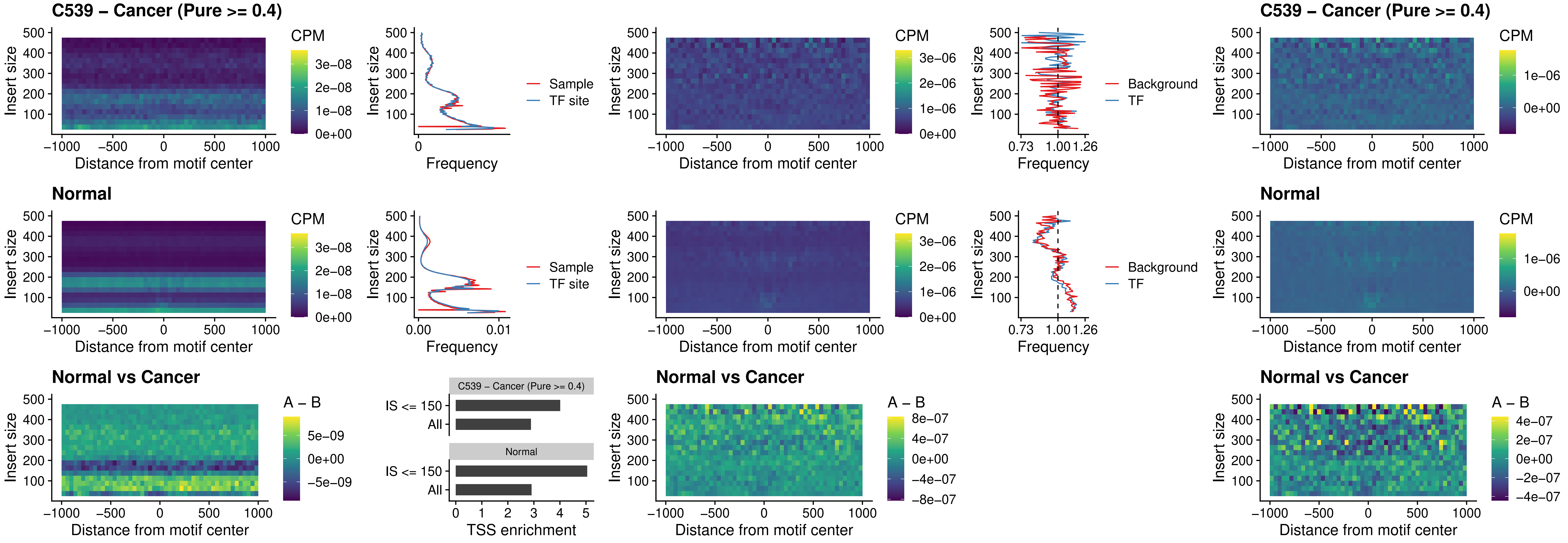

#### C539 – Cancer (Pure >= 0.4) – IRF8_uc_dTSS_oPEAK

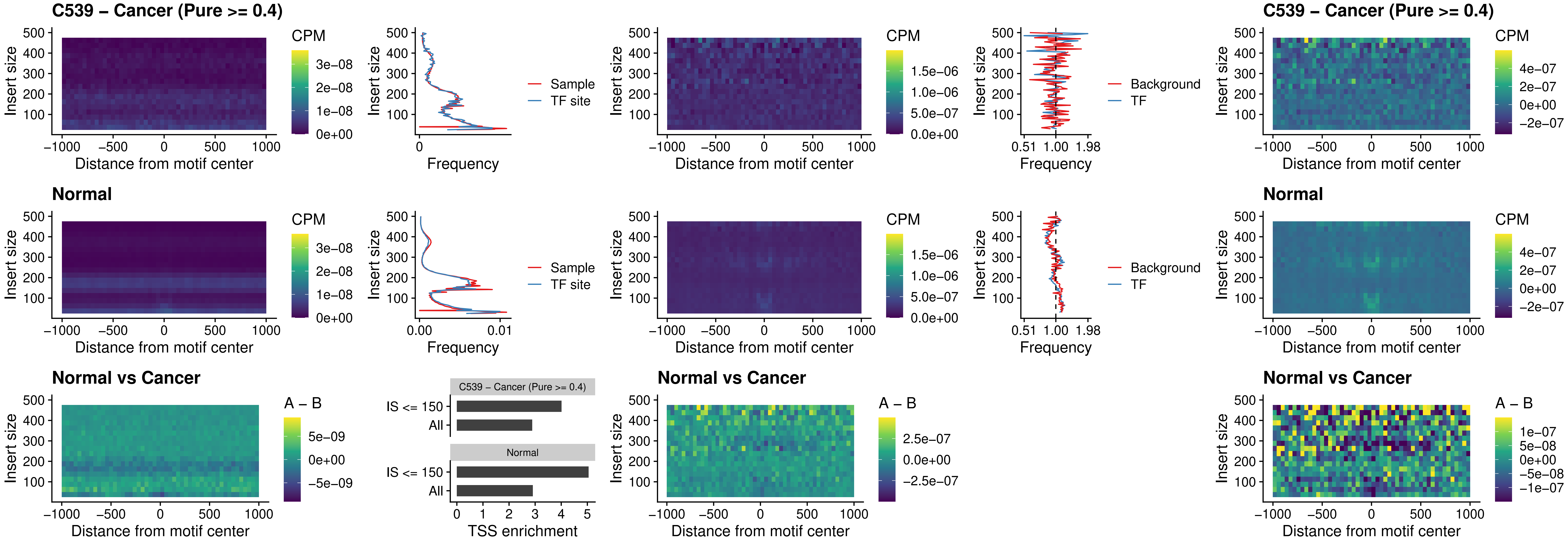

#### C539 – Cancer (Pure >= 0.4) – ETS2_uc_dTSS_oPEAK

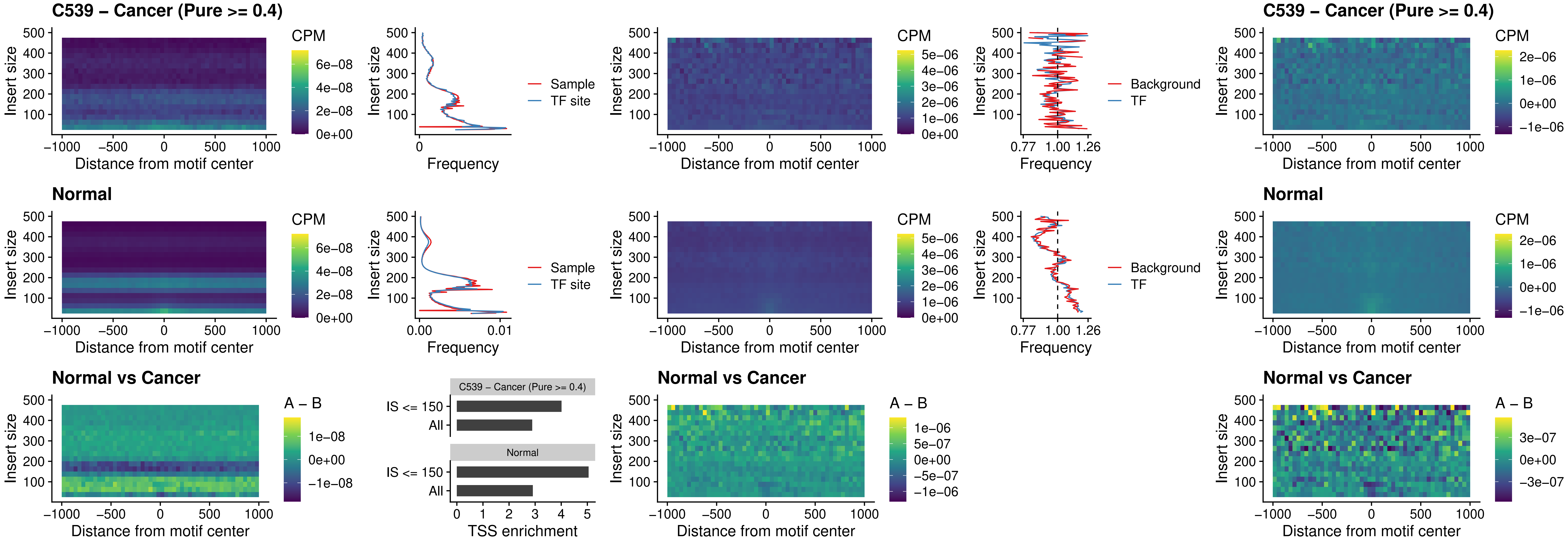

#### C539 – Cancer (Pure >= 0.4) – SMARCC2_uc_pTSS_oPEAK

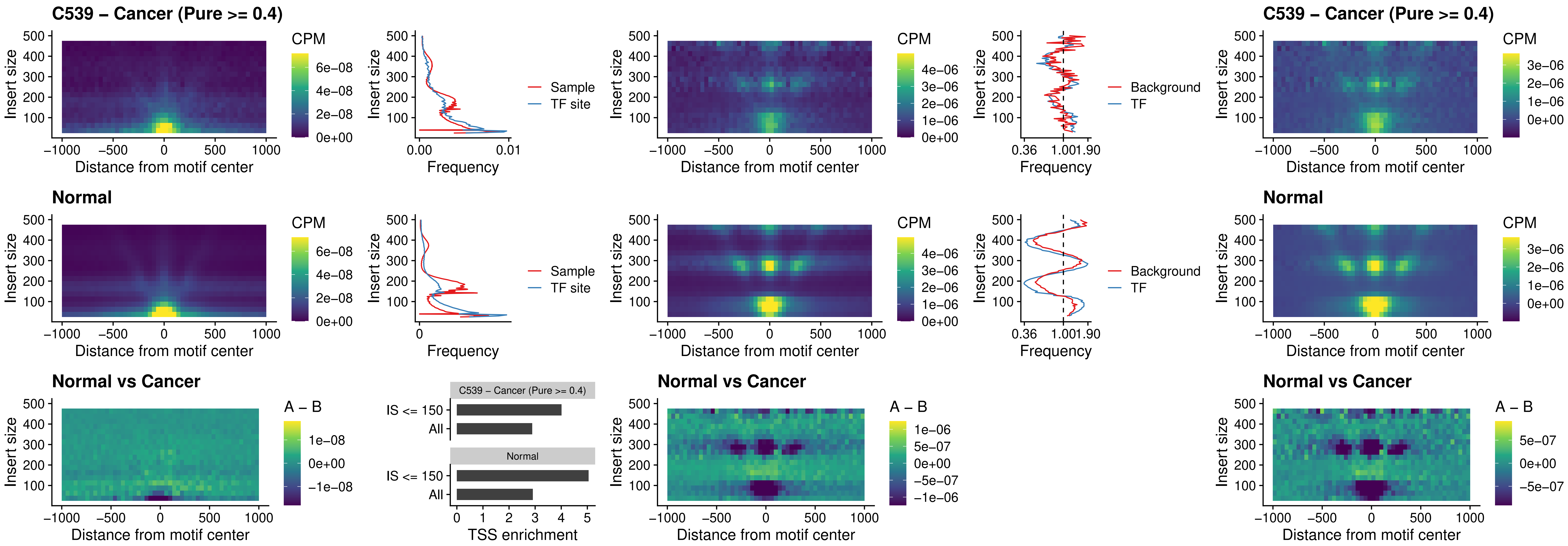

#### C539 – Cancer (Pure >= 0.4) – BCL11A_uc_cTSS_nPEAK

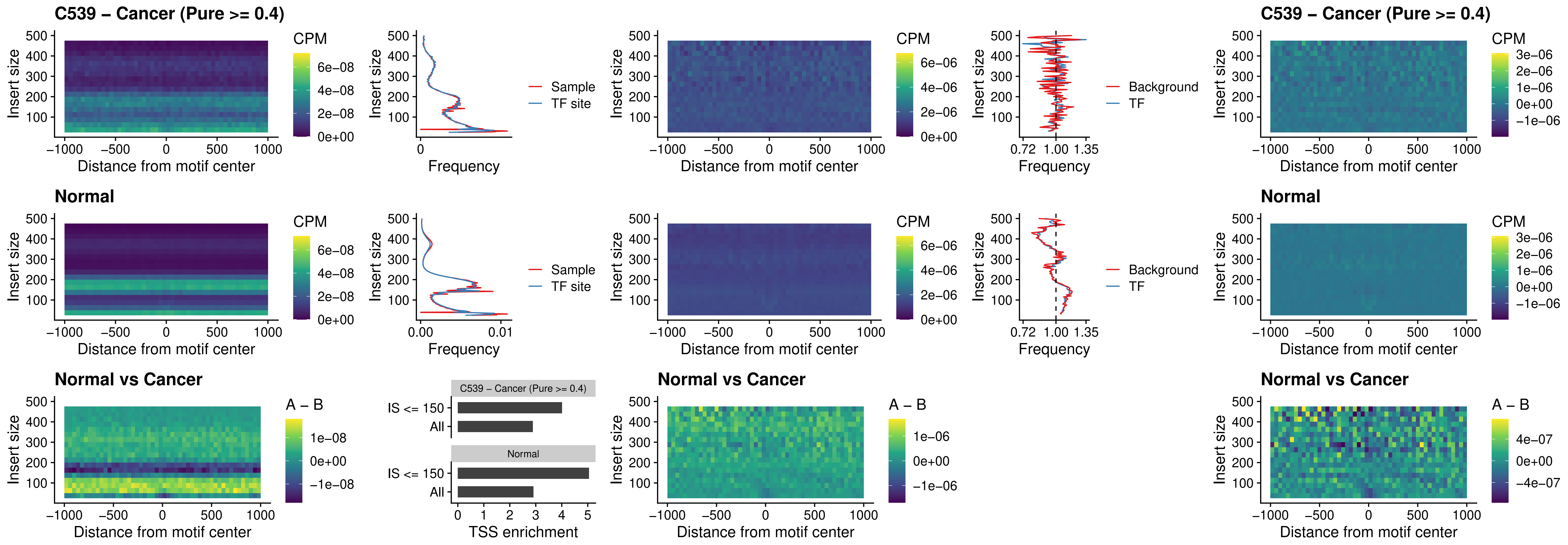

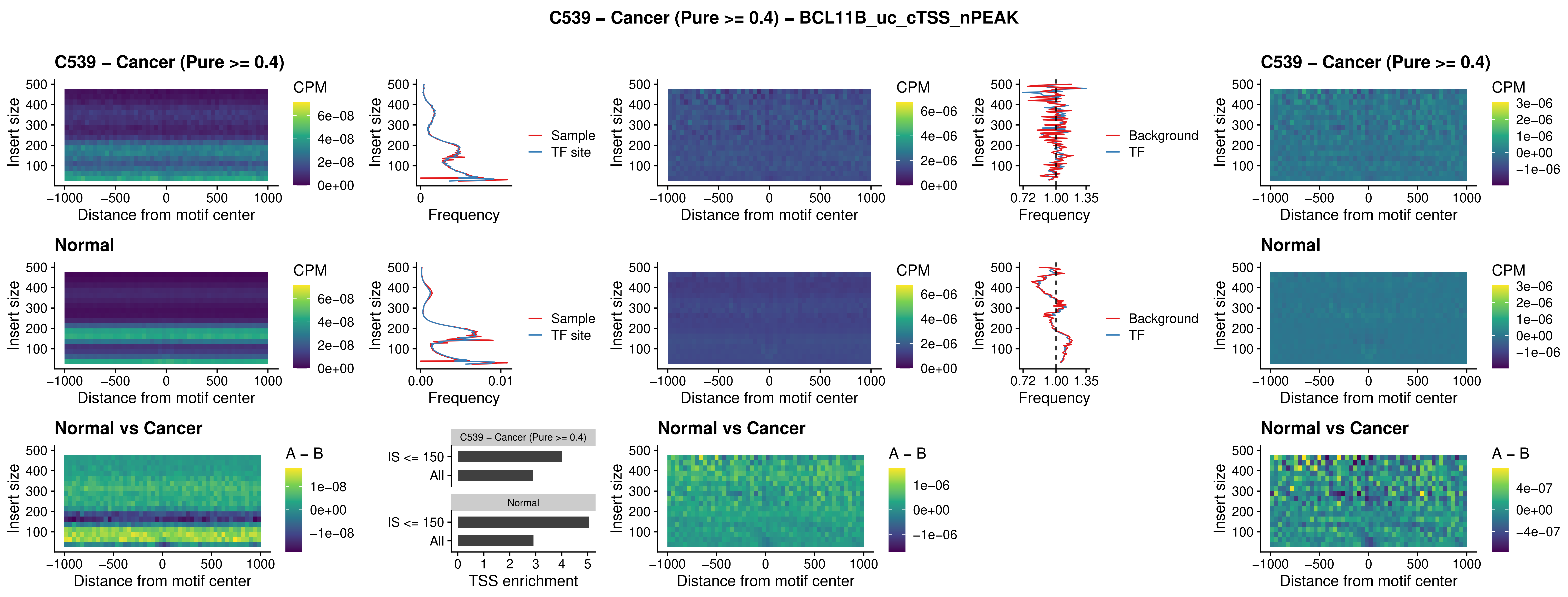
C539 – Cancer (Pure >= 0.4) – BCL11B_uc_cTSS_nPEAK
C539 – Cancer (Pure >= 0.4)
Normal
Normal vs Cancer
Insert size
Distance from motif center
Frequency
CPM
A – B
Sample
TF site
Background
TF
C539 – Cancer (Pure >= 0.4)
Normal
IS <= 150
All
TSS enrichment

**C539 – Cancer (Pure >= 0.4) – STAT2_uc_dTSS_oPEAK**

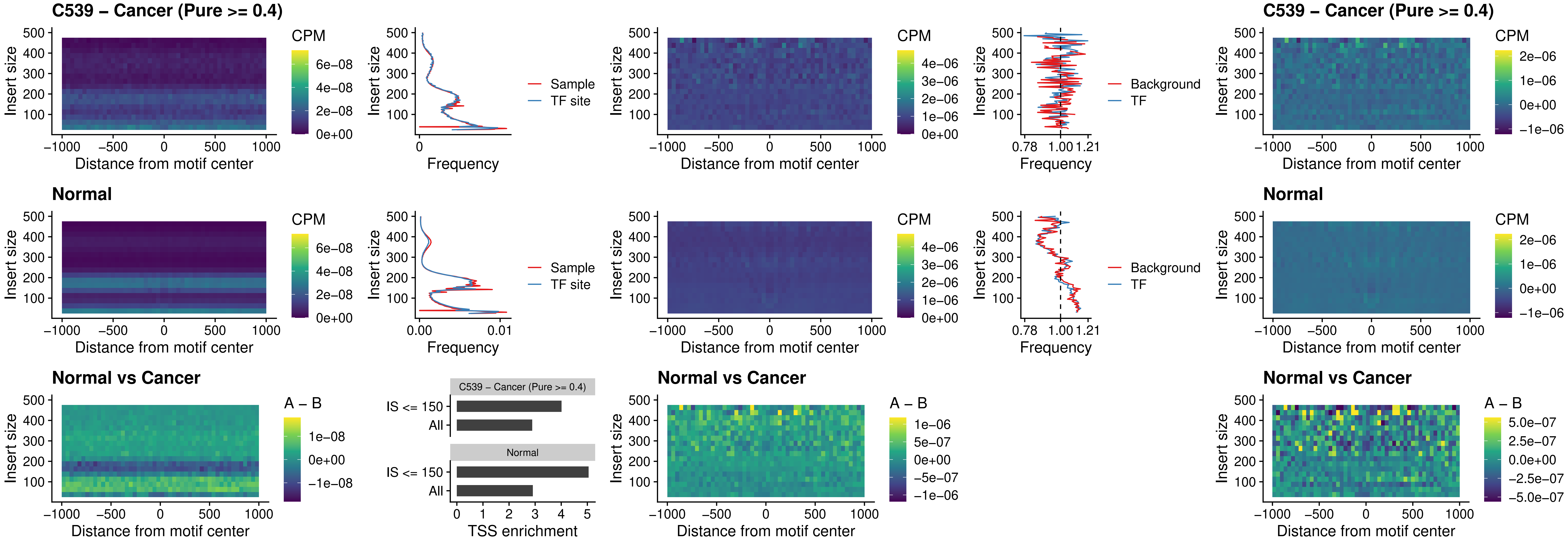

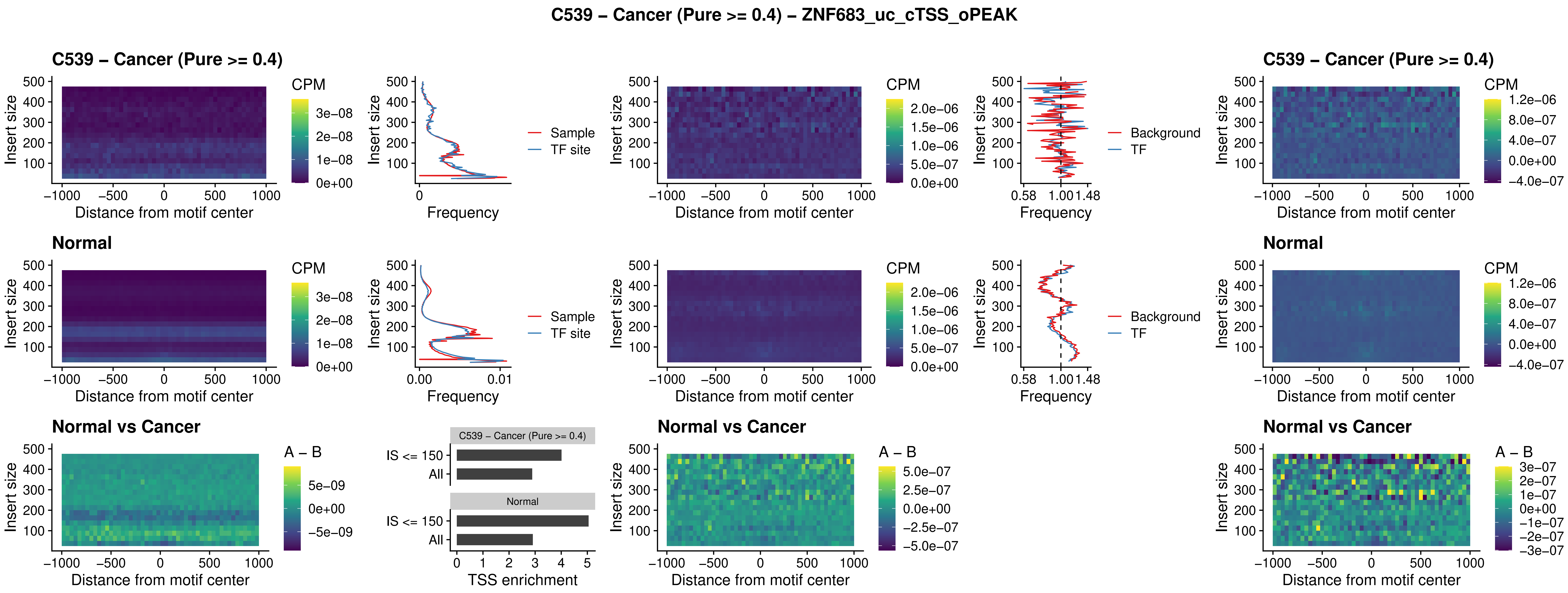
C539 – Cancer (Pure >= 0.4) – ZNF683_uc_cTSS_oPEAK
C539 – Cancer (Pure >= 0.4)
Normal
Normal vs Cancer
Insert size
Distance from motif center
Frequency
CPM
A – B
Sample
TF site
Background
TF
C539 – Cancer (Pure >= 0.4)
Normal
IS <= 150
All
TSS enrichment

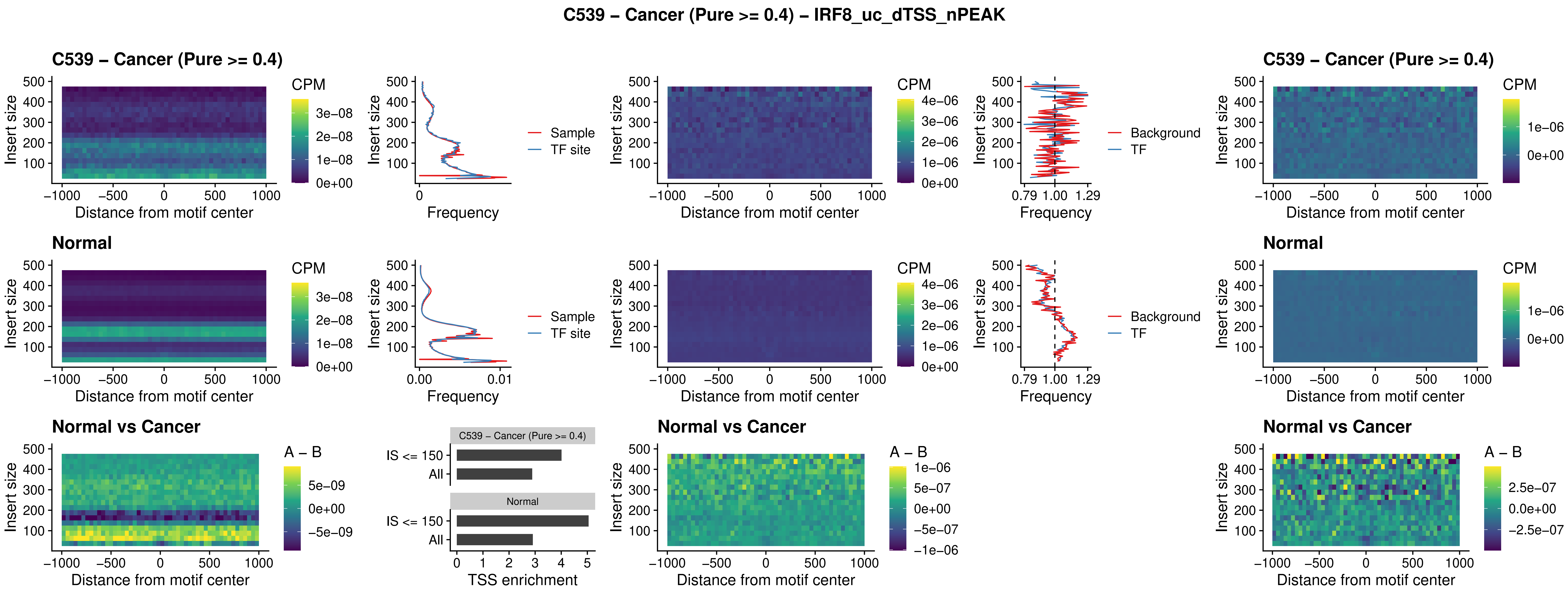
C539 – Cancer (Pure >= 0.4) – IRF8_uc_dTSS_nPEAK
C539 – Cancer (Pure >= 0.4)
Normal
Normal vs Cancer
Insert size
Distance from motif center
Frequency
CPM
Sample
TF site
Background
TF
A – B
C539 – Cancer (Pure >= 0.4)
Normal
IS <= 150
All
TSS enrichment

### C539 – Cancer (Pure >= 0.4) – ETV7_uc_dTSS_oPEAK

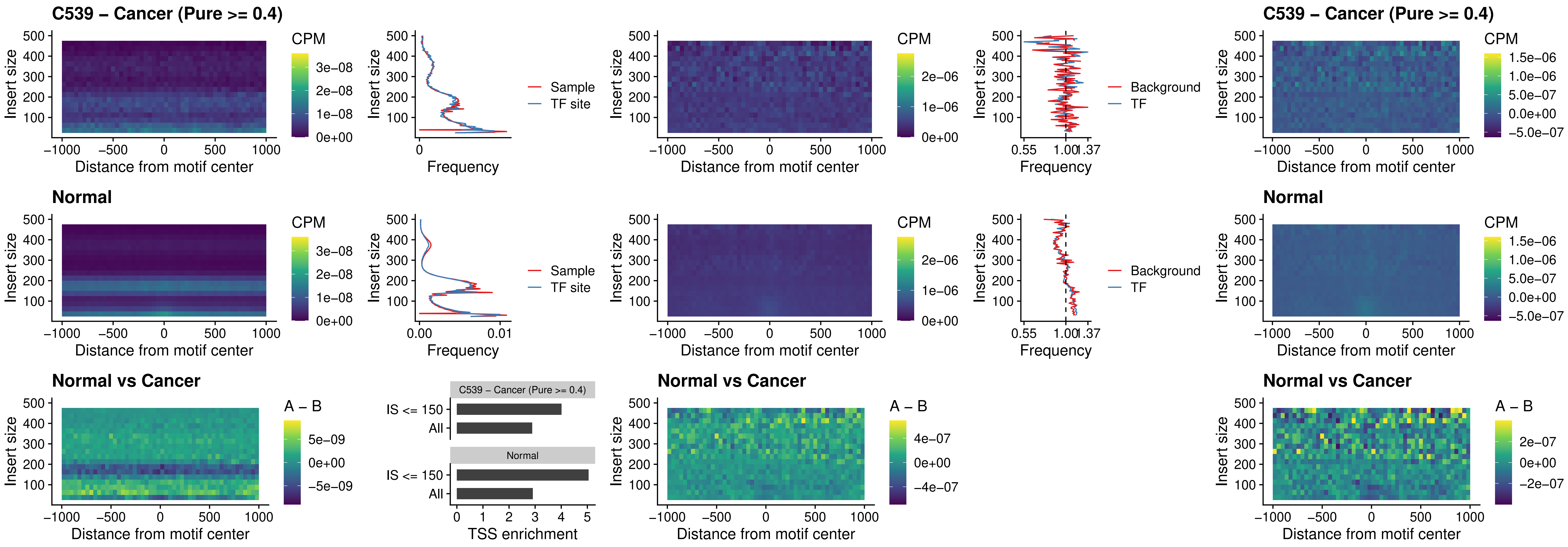

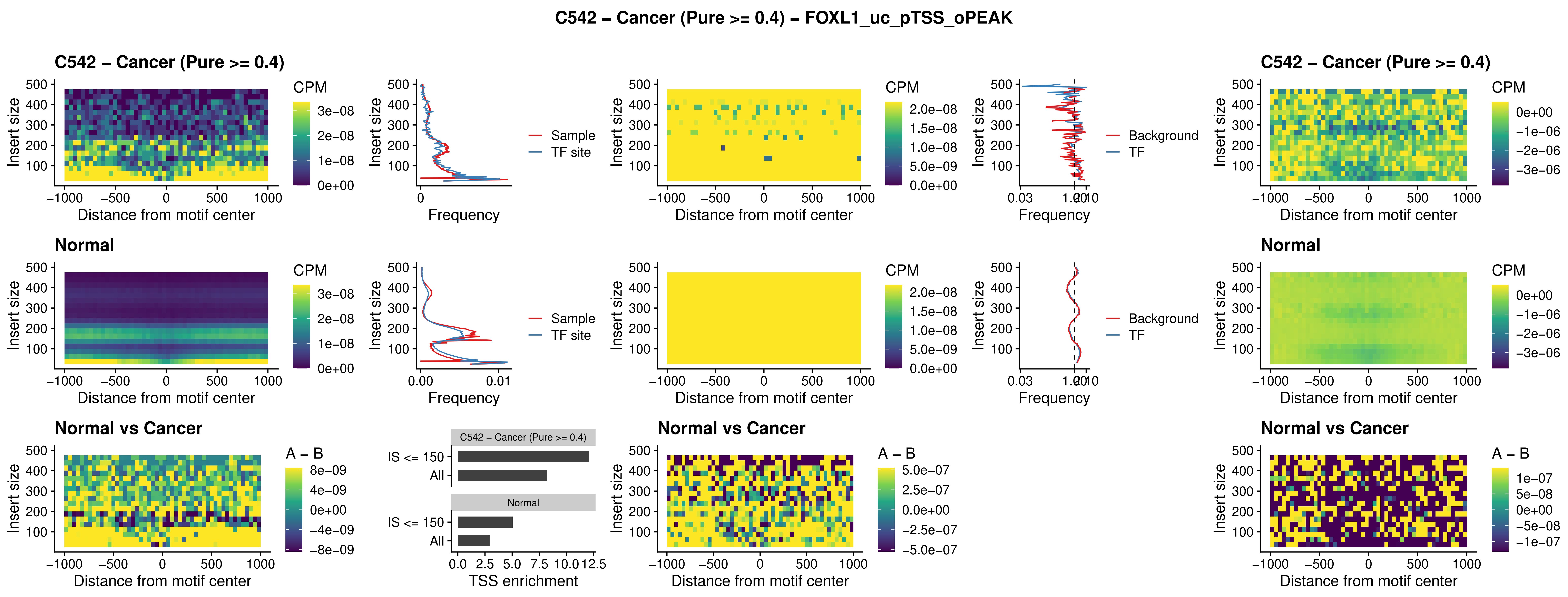
C542 – Cancer (Pure >= 0.4) – FOXL1_uc_pTSS_oPEAK
C542 – Cancer (Pure >= 0.4)
Normal
Normal vs Cancer
Insert size
Distance from motif center
CPM
A – B
Frequency
Sample
TF site
Background
TF
C542 – Cancer (Pure >= 0.4)
Normal
IS <= 150
All
TSS enrichment

#### C542 – Cancer (Pure >= 0.4) – CPEB1_uc_cTSS_oPEAK

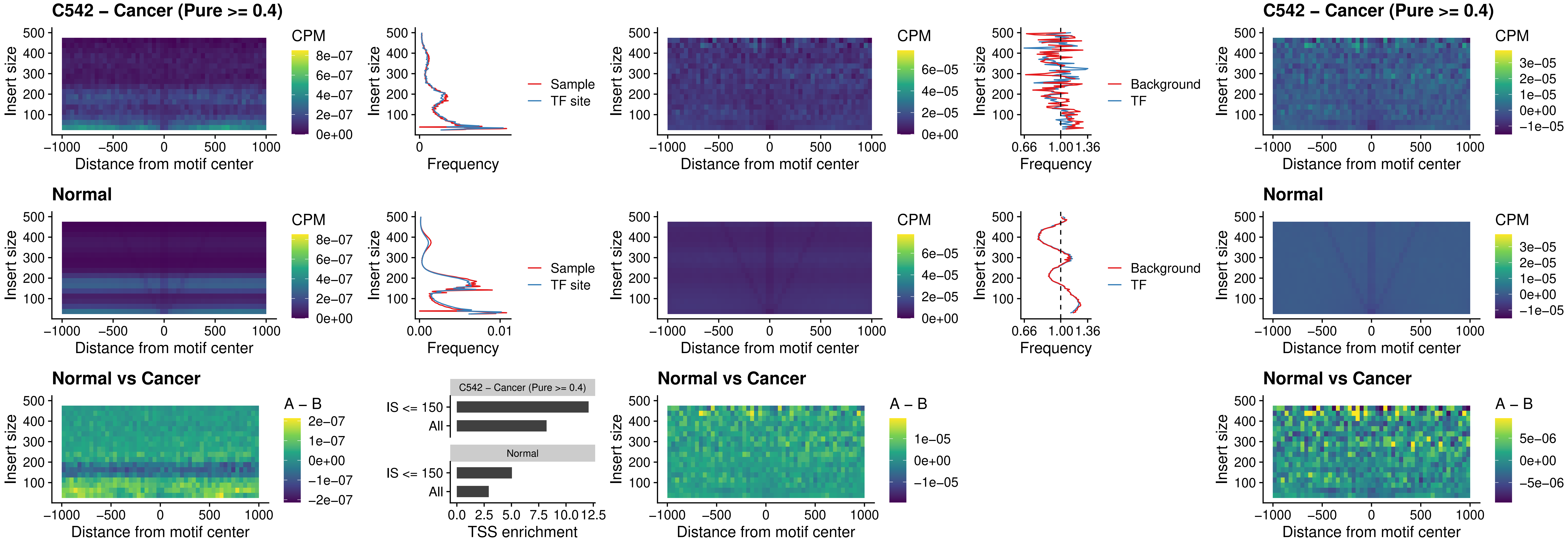

#### C542 – Cancer (Pure >= 0.4) – HMGA1_uc_pTSS_nPEAK

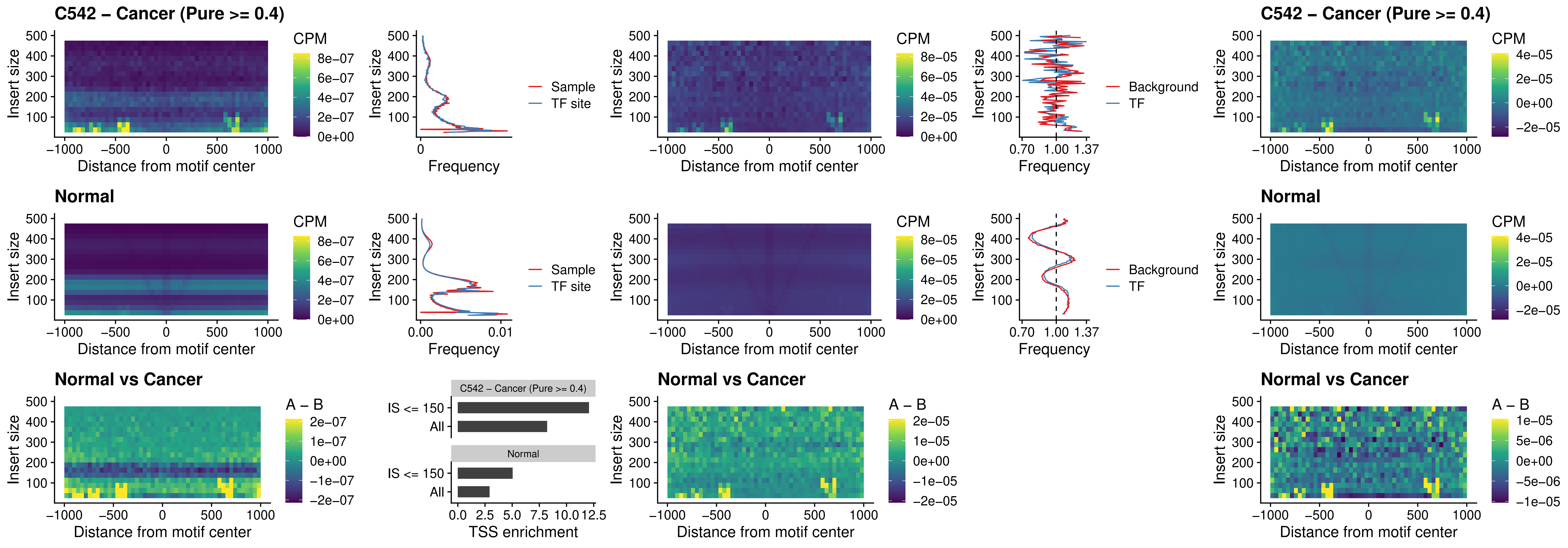

#### C542 – Cancer (Pure >= 0.4) – SOX13_uc_cTSS_nPEAK

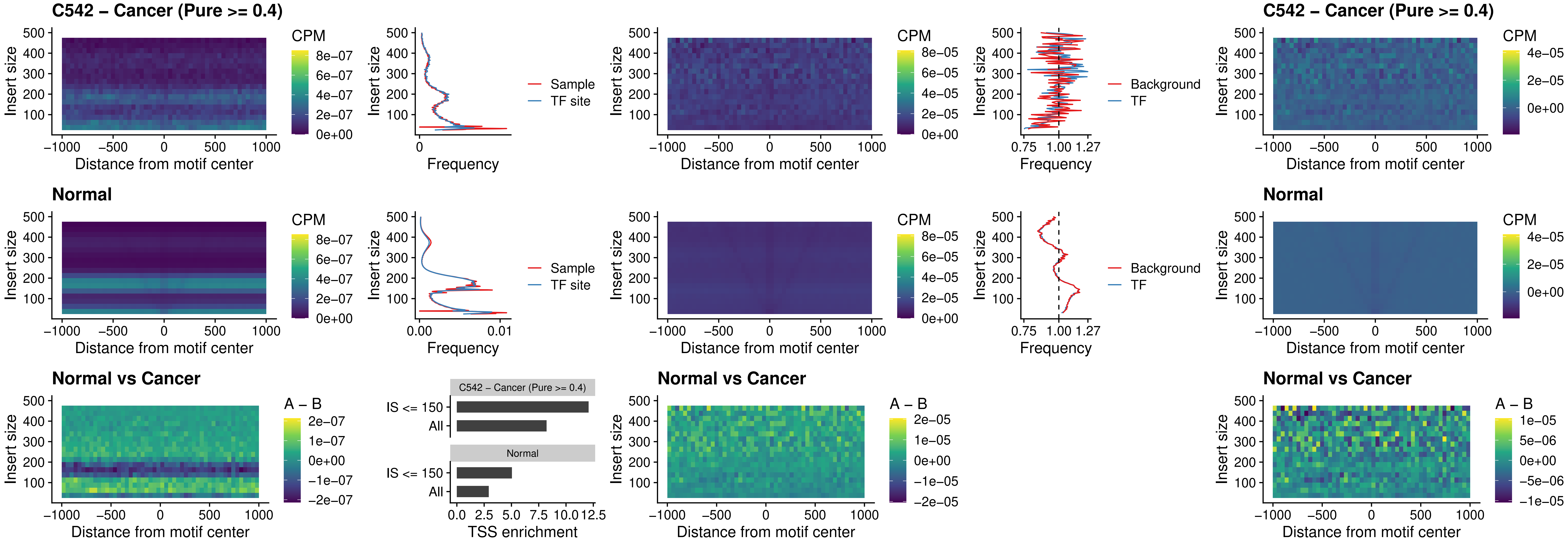

#### C542 – Cancer (Pure >= 0.4) – LHX4_uc_cTSS_nPEAK

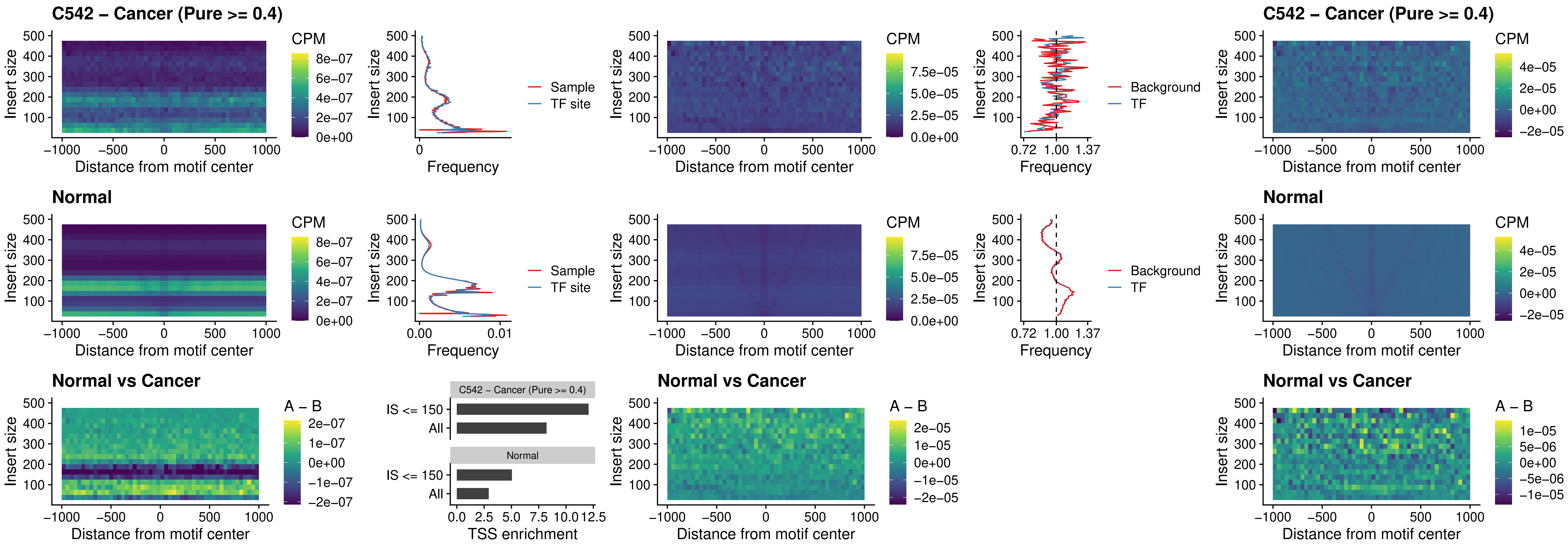

#### C542 – Cancer (Pure >= 0.4) – HOXA4_uc_cTSS_nPEAK

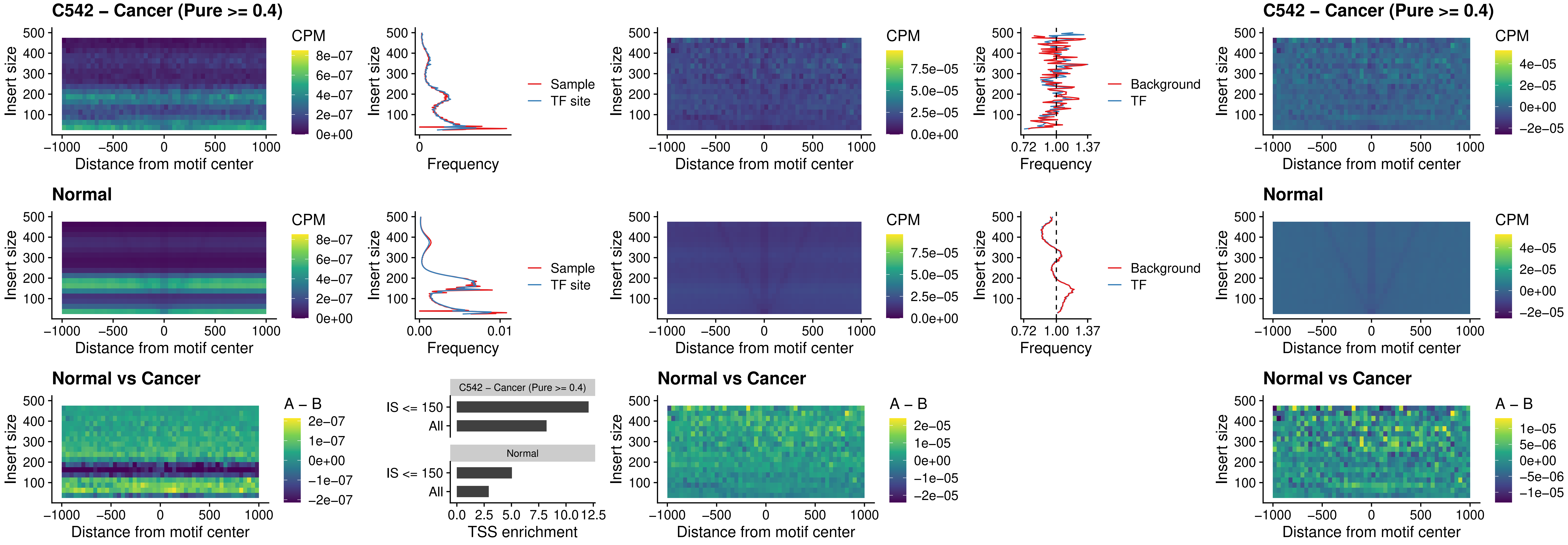

#### C542 – Cancer (Pure >= 0.4) – HLX_uc_cTSS_nPEAK

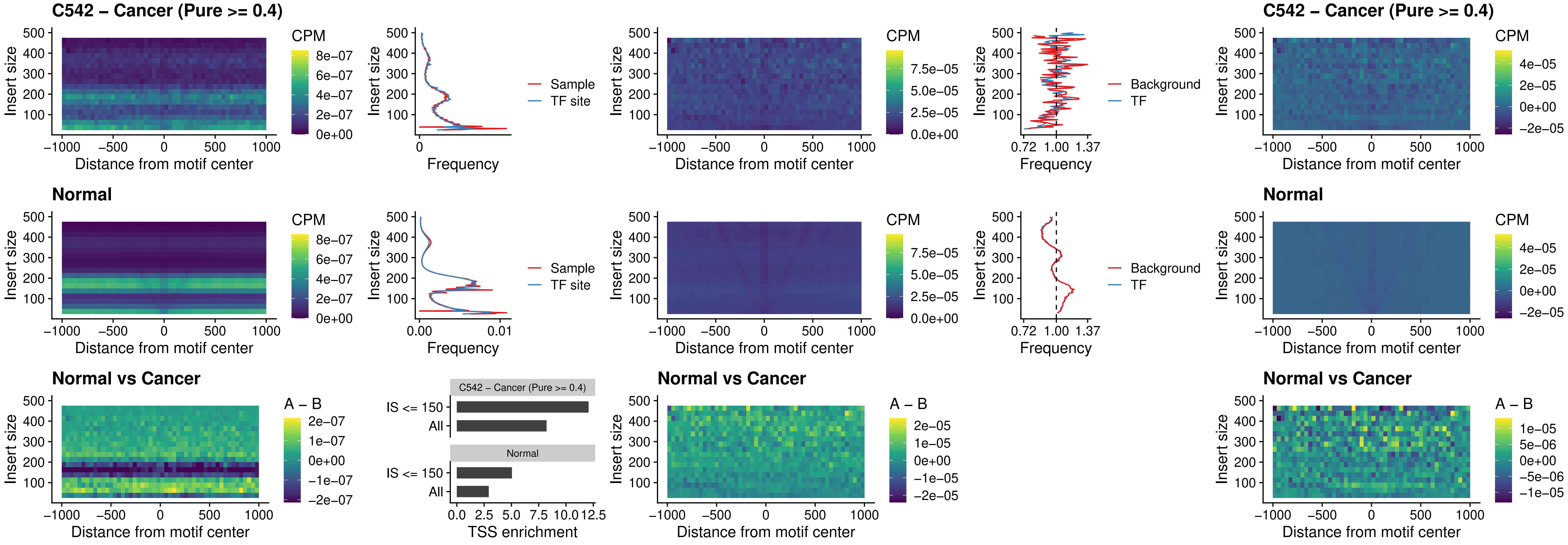

#### C542 – Cancer (Pure >= 0.4) – AC0125311_uc_cTSS_nPEAK

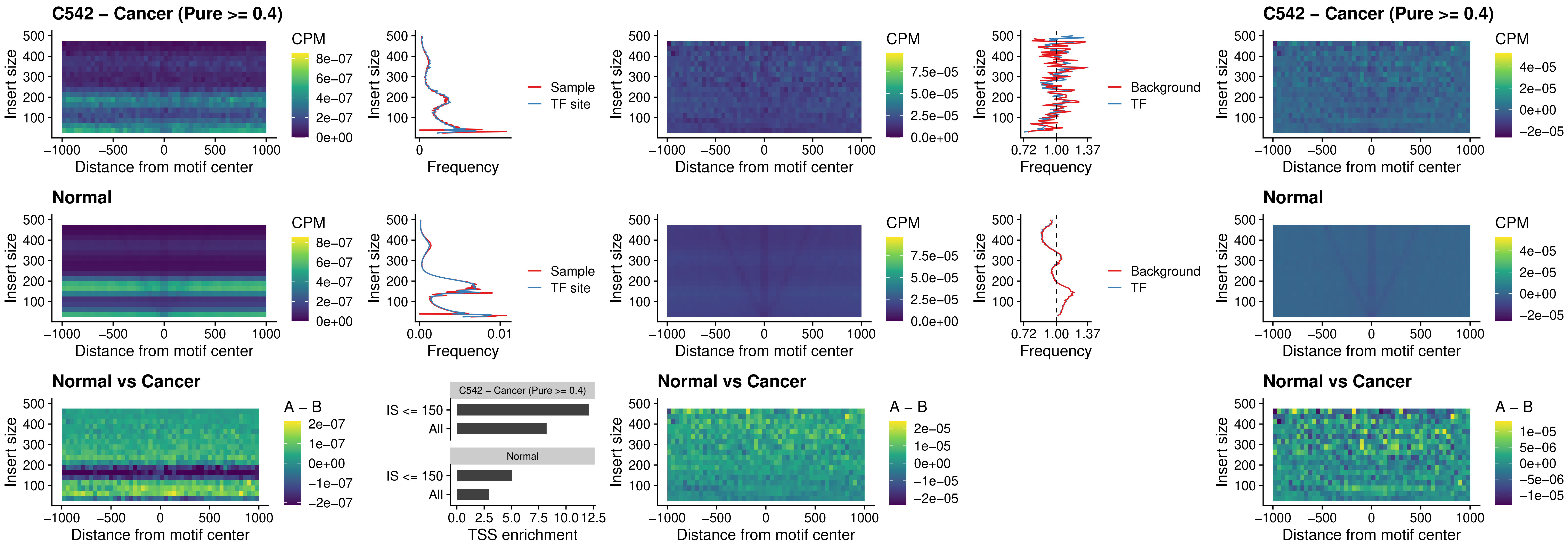

#### C542 – Cancer (Pure >= 0.4) – DBX2_uc_cTSS_nPEAK

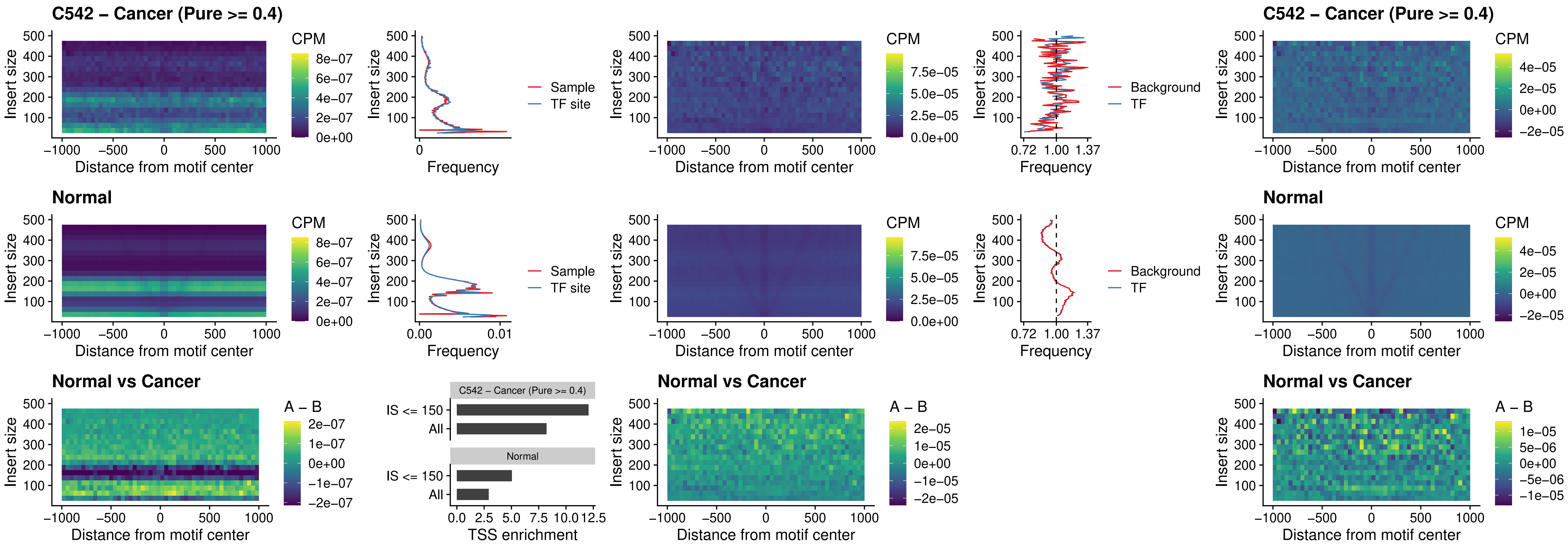

### C542 – Cancer (Pure >= 0.4) – LHX6_uc_dTSS_nPEAK

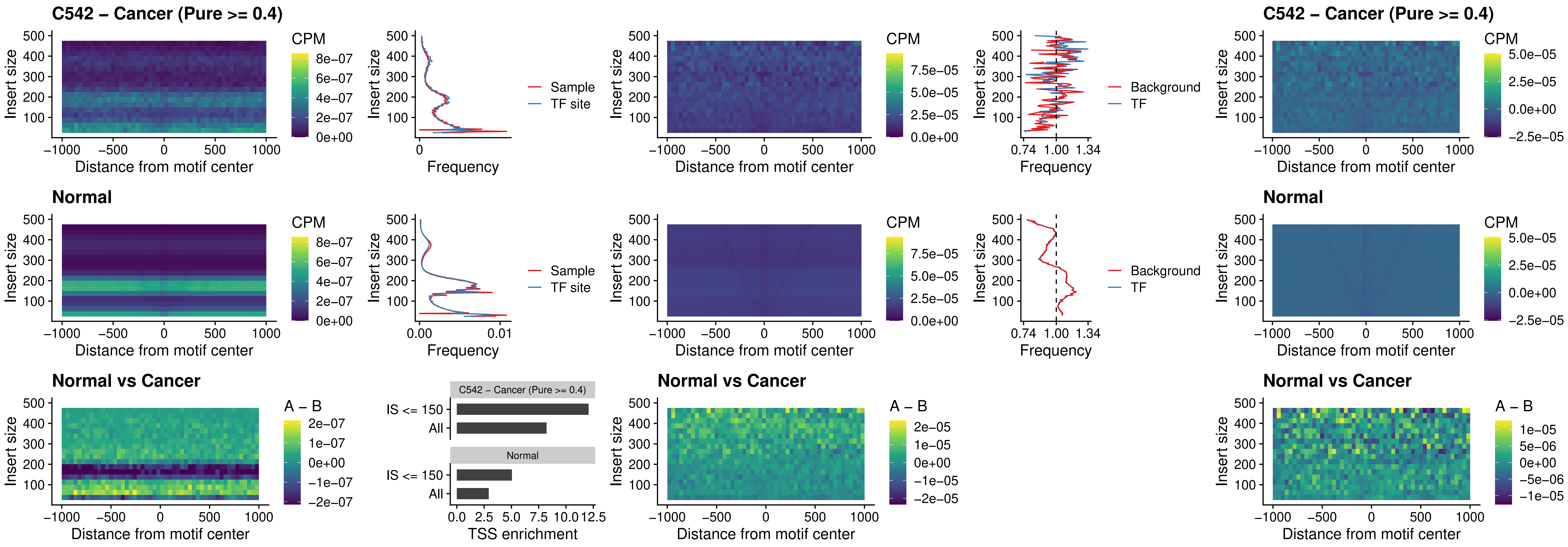

#### C542 – Cancer (Pure >= 0.4) – SOX1_uc_dTSS_nPEAK

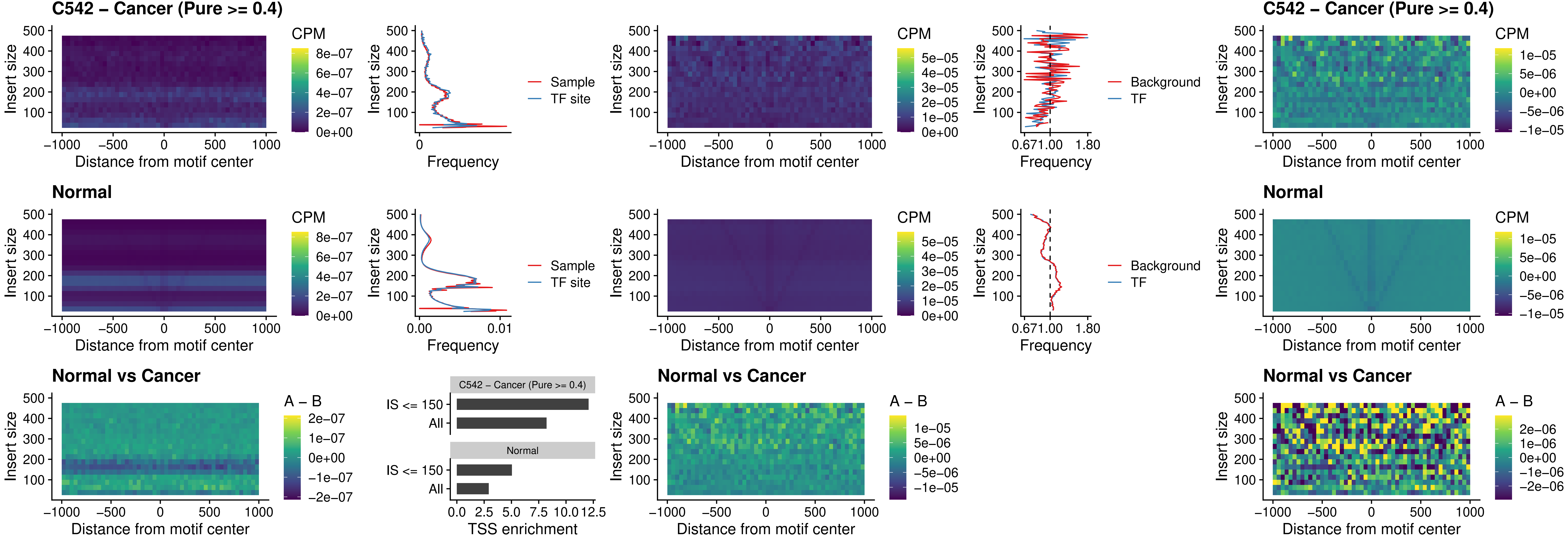

#### C542 – Cancer (Pure >= 0.4) – NKX24_uc_dTSS_nPEAK

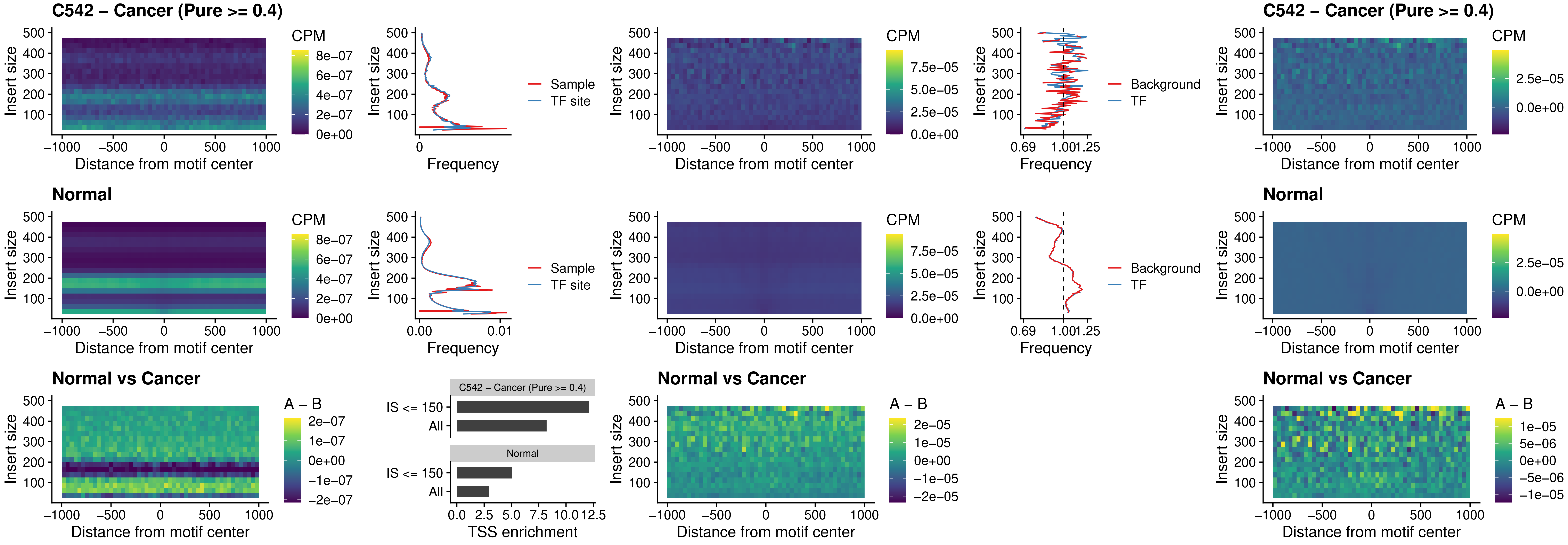

#### C542 – Cancer (Pure >= 0.4) – DLX3_uc_cTSS_nPEAK

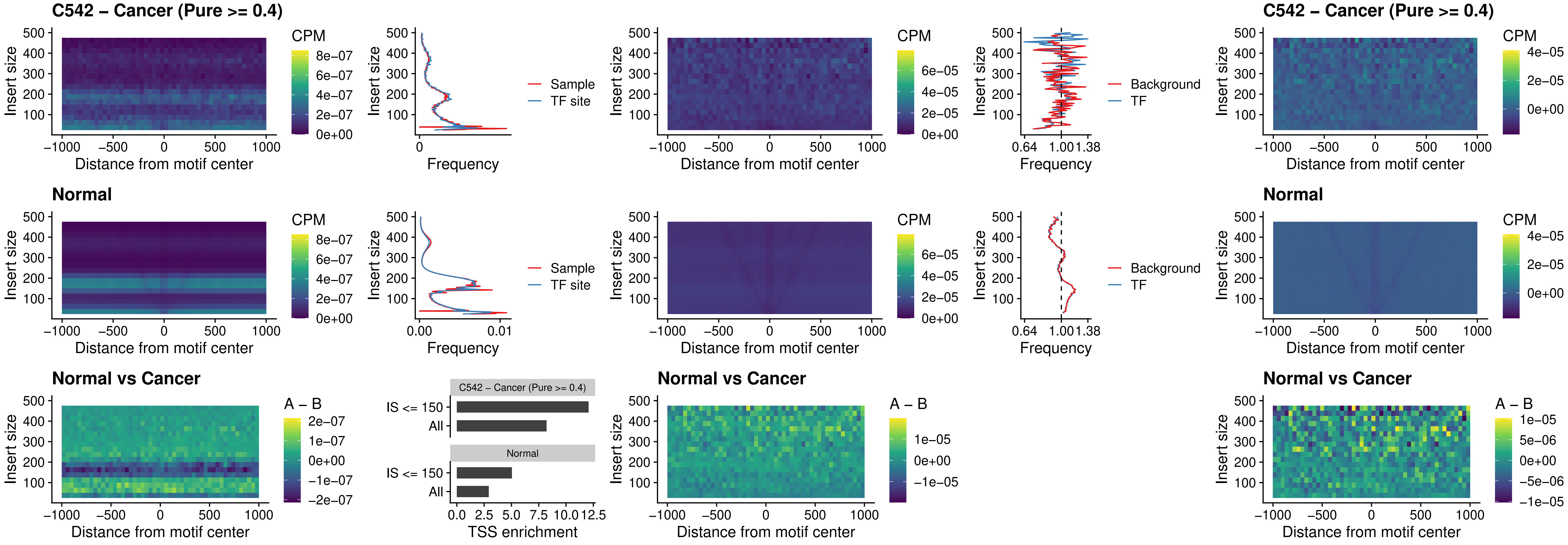

### C542 – Cancer (Pure >= 0.4) – HOXD3_uc_cTSS_nPEAK

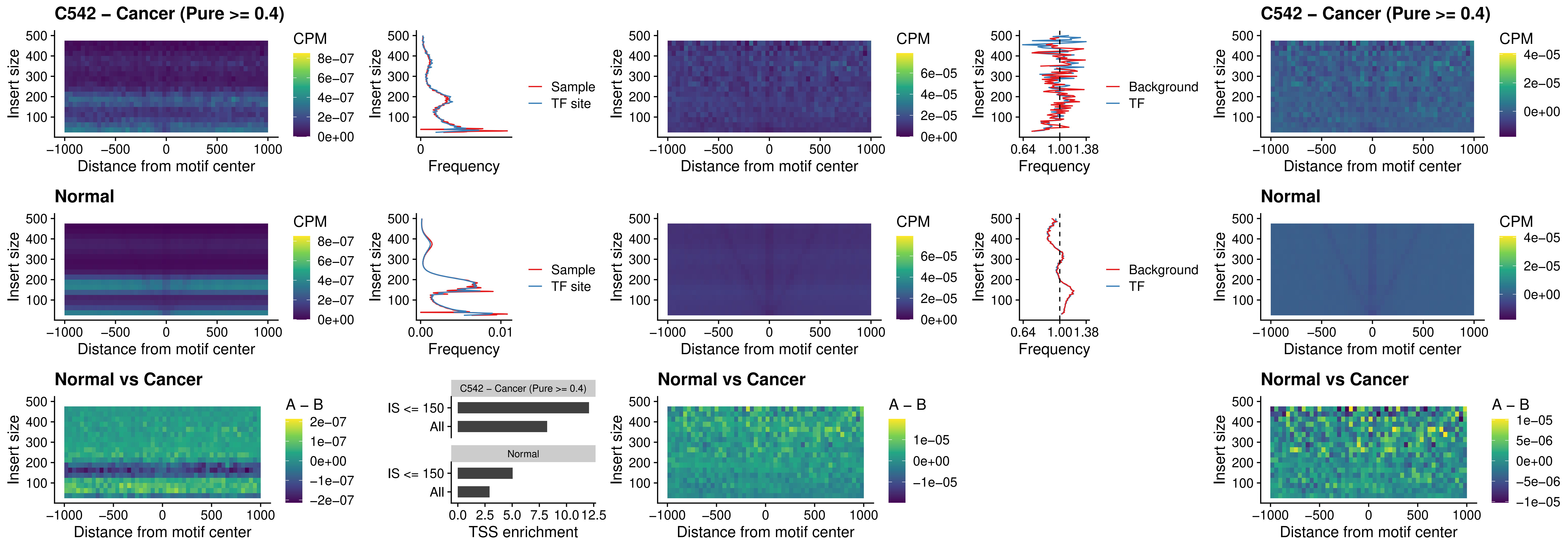

#### C542 – Cancer (Pure >= 0.4) – ISL1_uc_dTSS_oPEAK

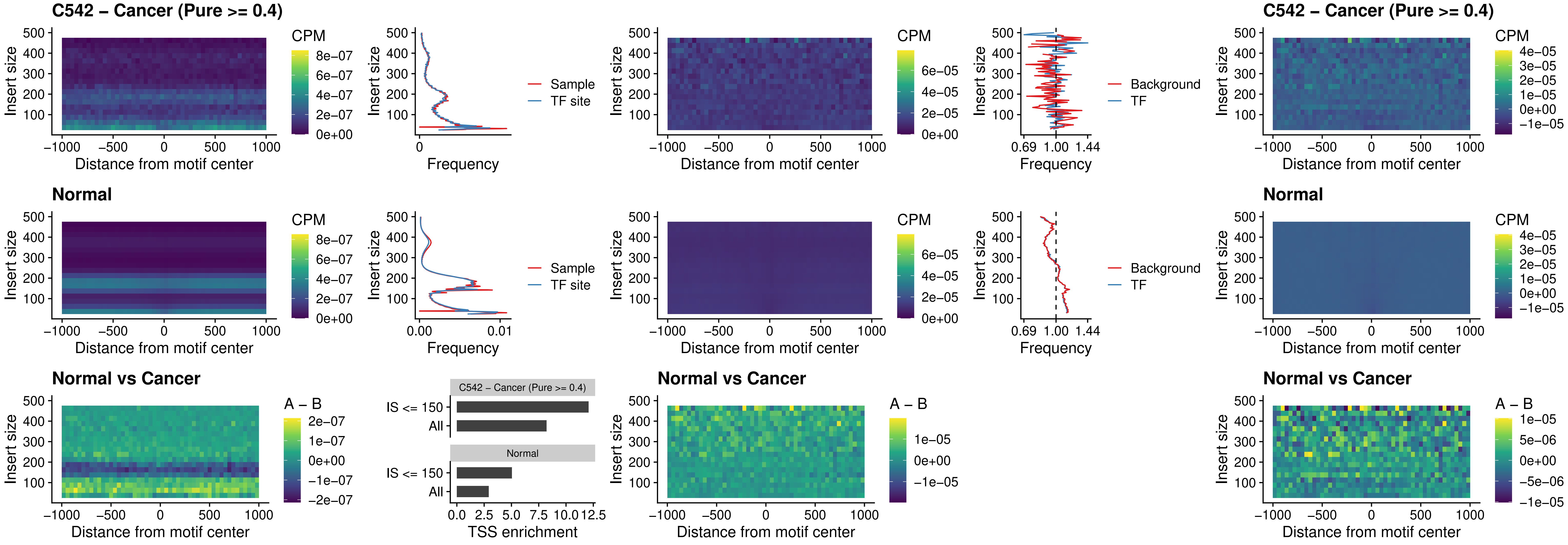

#### C542 – Cancer (Pure >= 0.4) – HOXB4_uc_dTSS_oPEAK

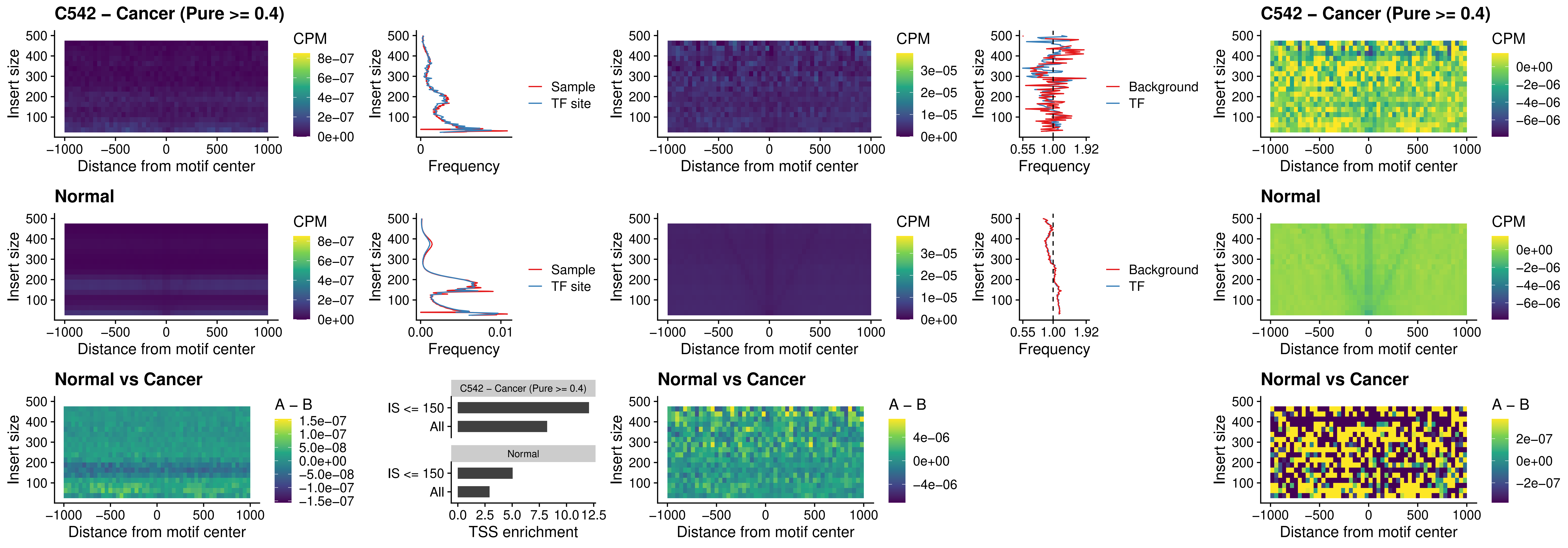

#### C542 – Cancer (Pure >= 0.4) – VAX1_uc_dTSS_oPEAK

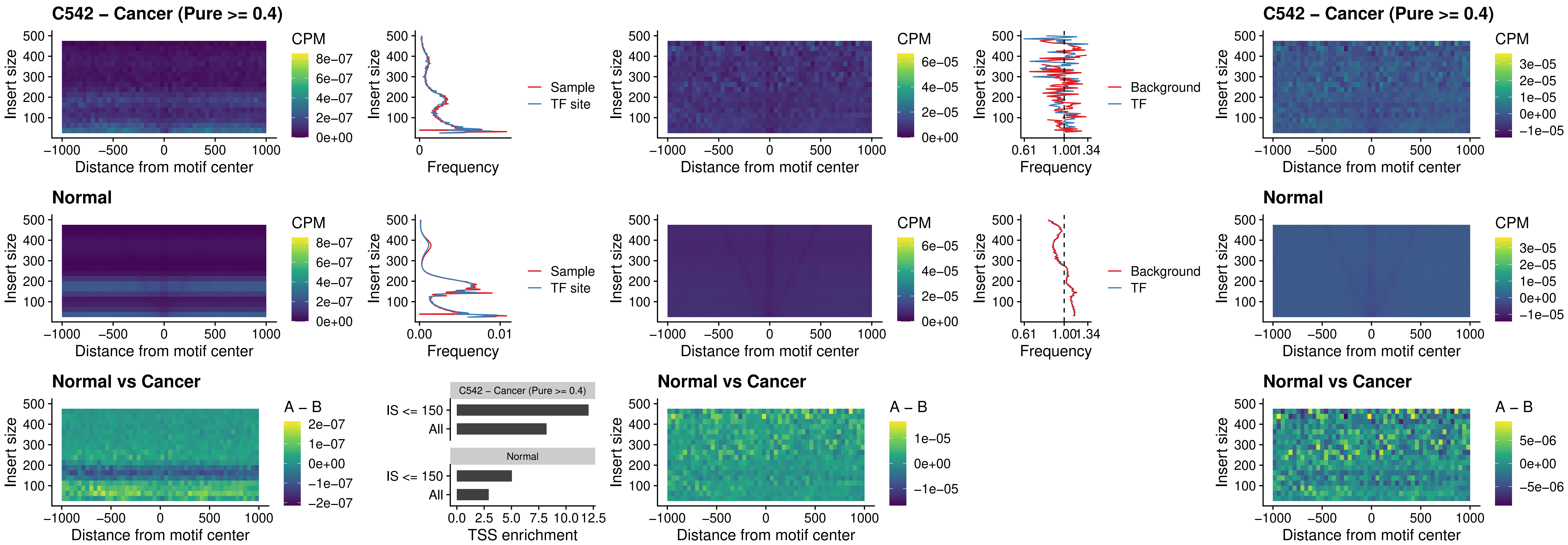

#### C542 – Cancer (Pure >= 0.4) – DLX3_uc_dTSS_oPEAK

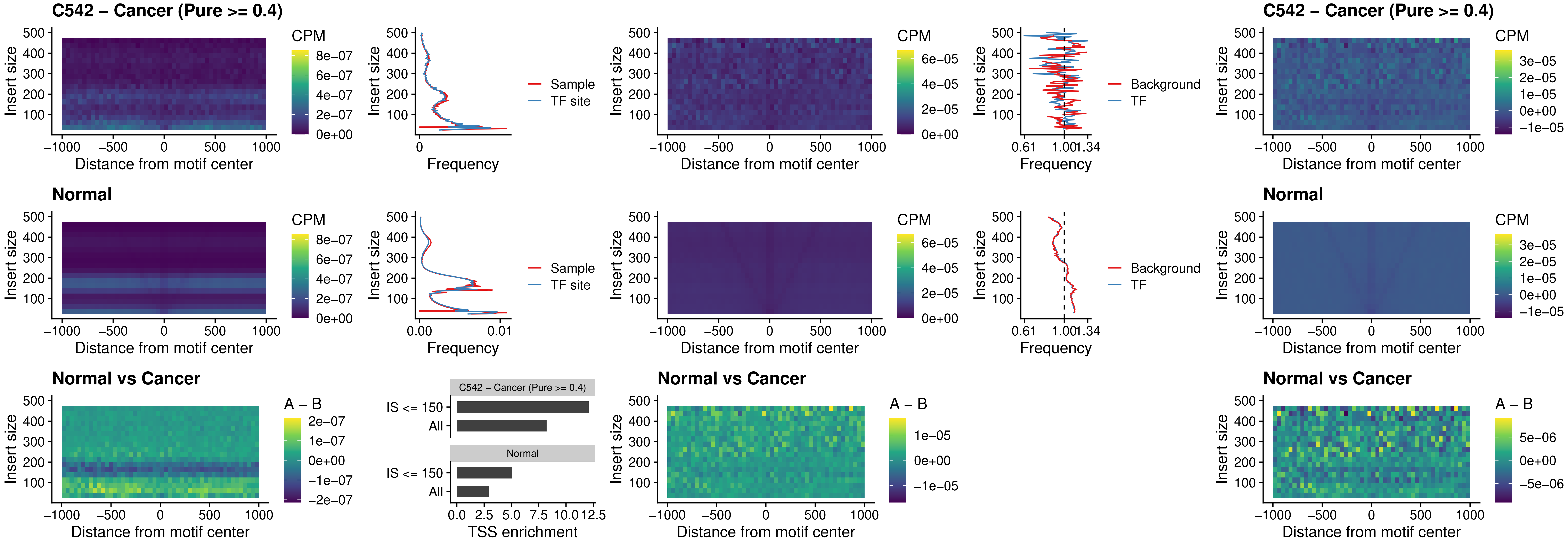

#### C542 – Cancer (Pure >= 0.4) – HOXD3_uc_dTSS_oPEAK

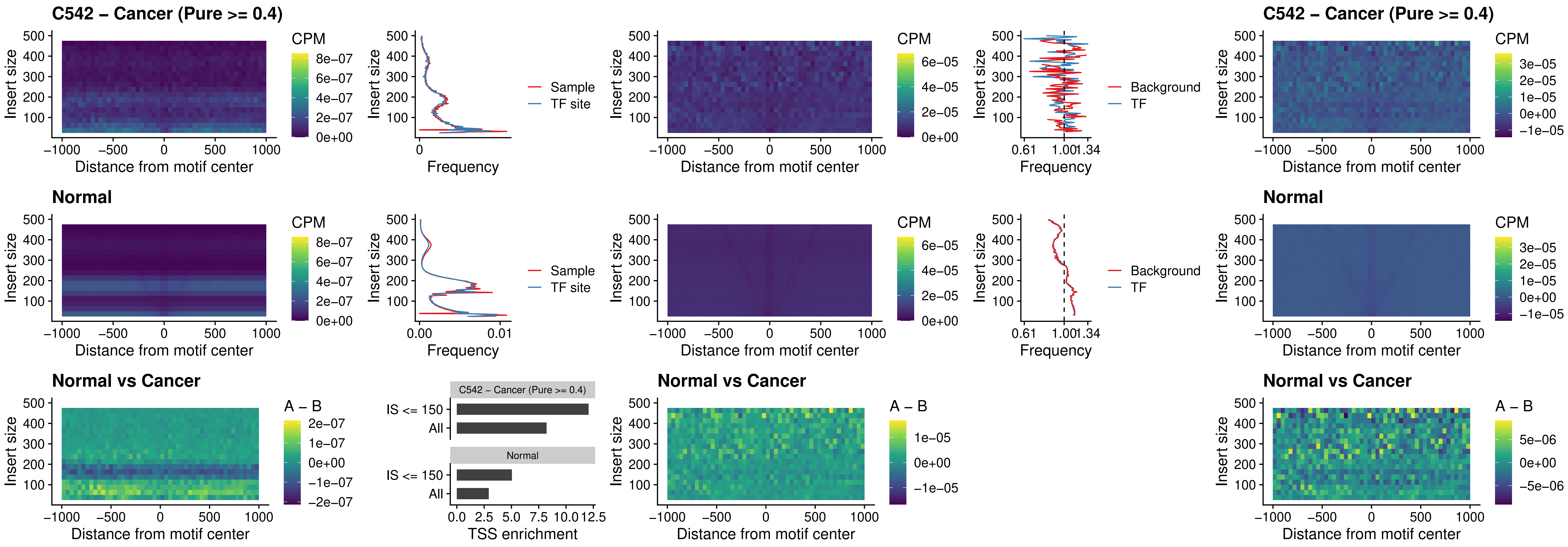

#### C542 – Cancer (Pure >= 0.4) – BBX_uc_dTSS_nPEAK

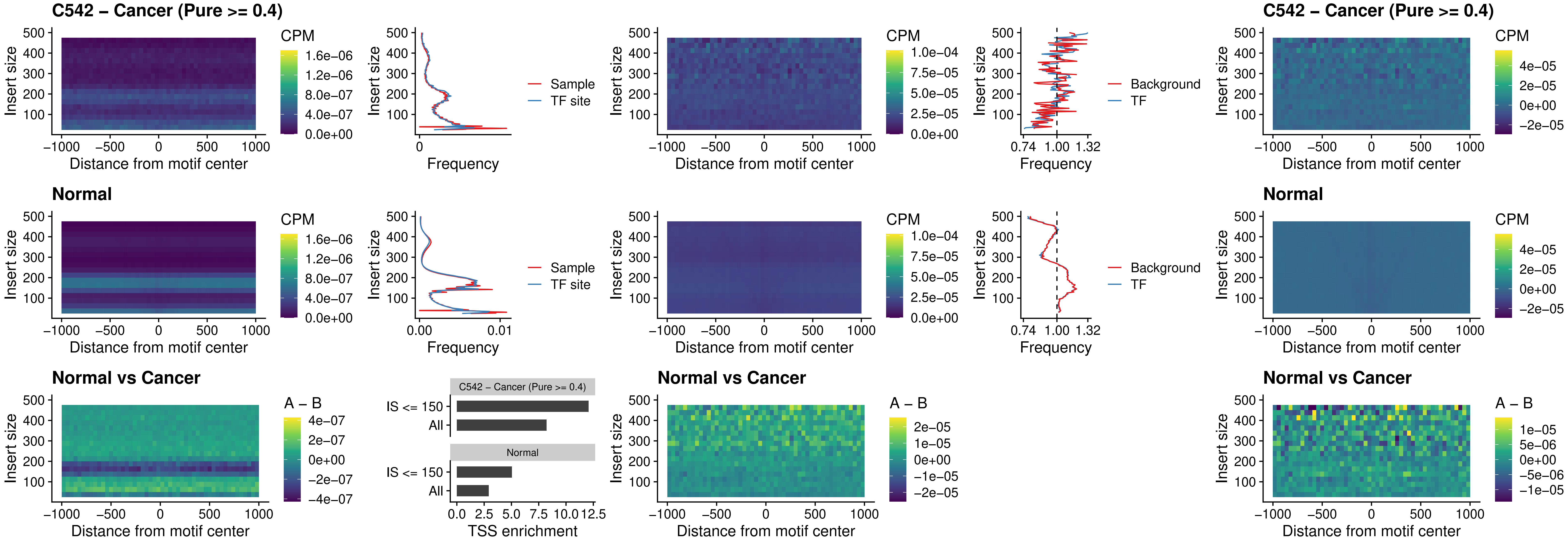

#### C542 – Cancer (Pure >= 0.4) – SOX3_uc_cTSS_nPEAK

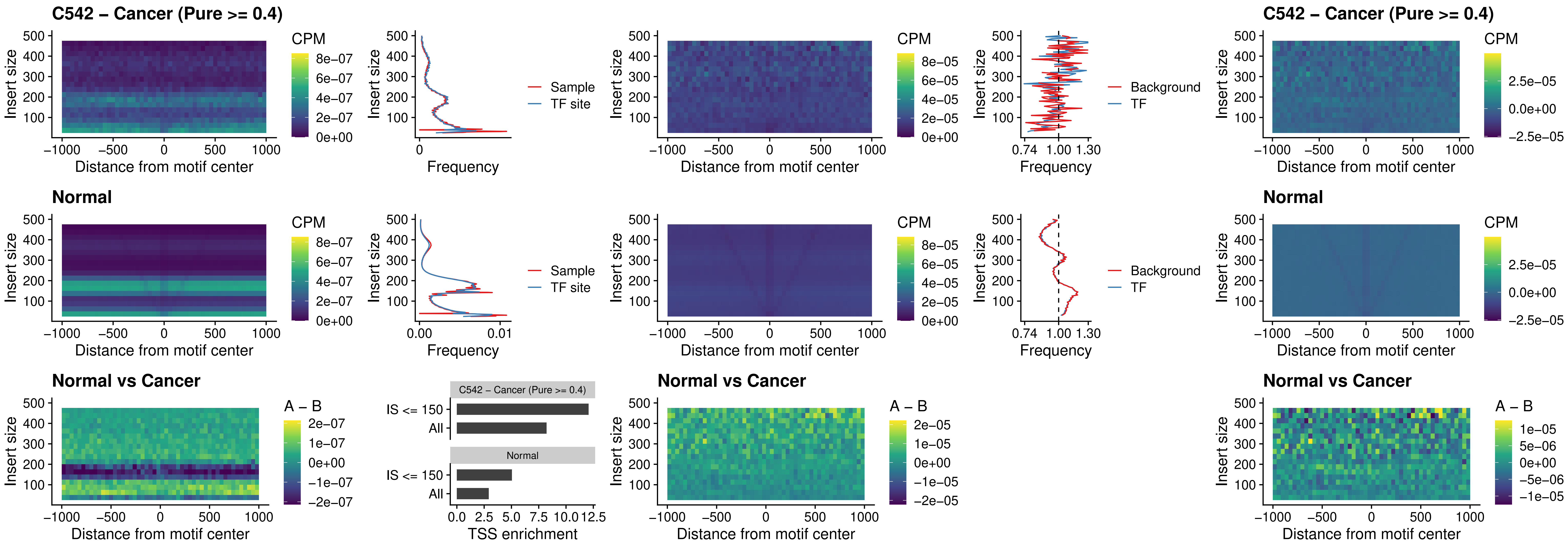

#### C542 – Cancer (Pure >= 0.4) – TBPL2_uc_cTSS_nPEAK

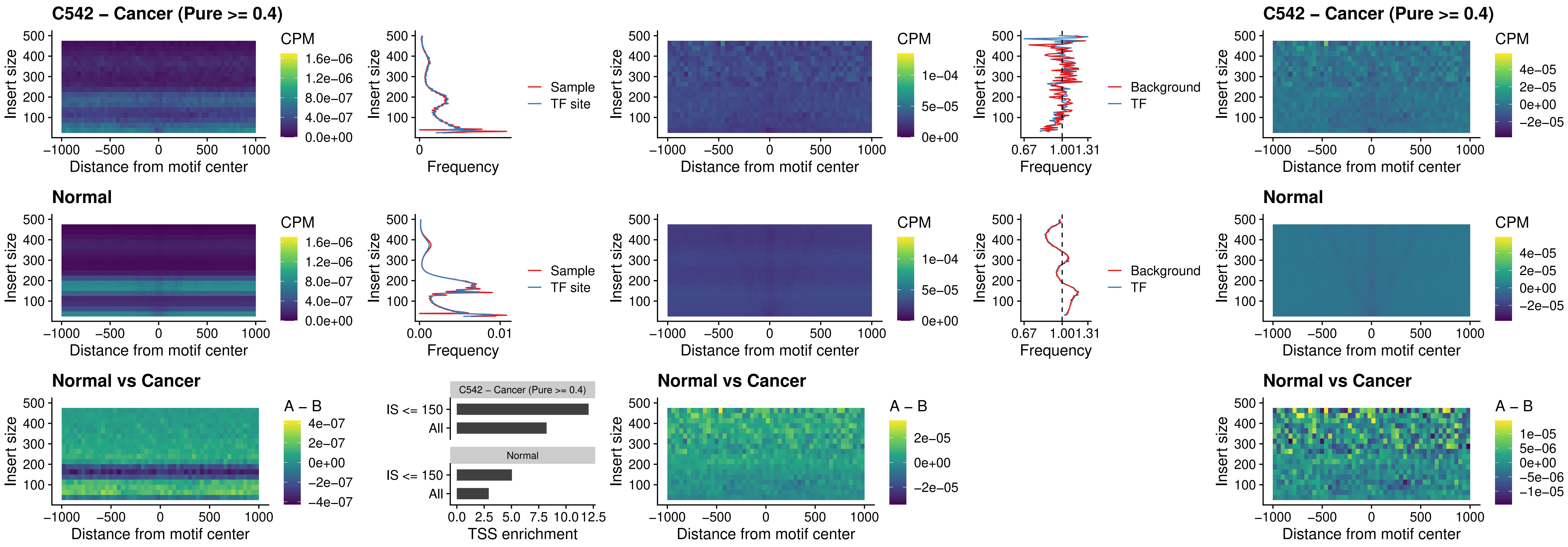

#### C542 – Cancer (Pure >= 0.4) – HOXA4_uc_cTSS_oPEAK

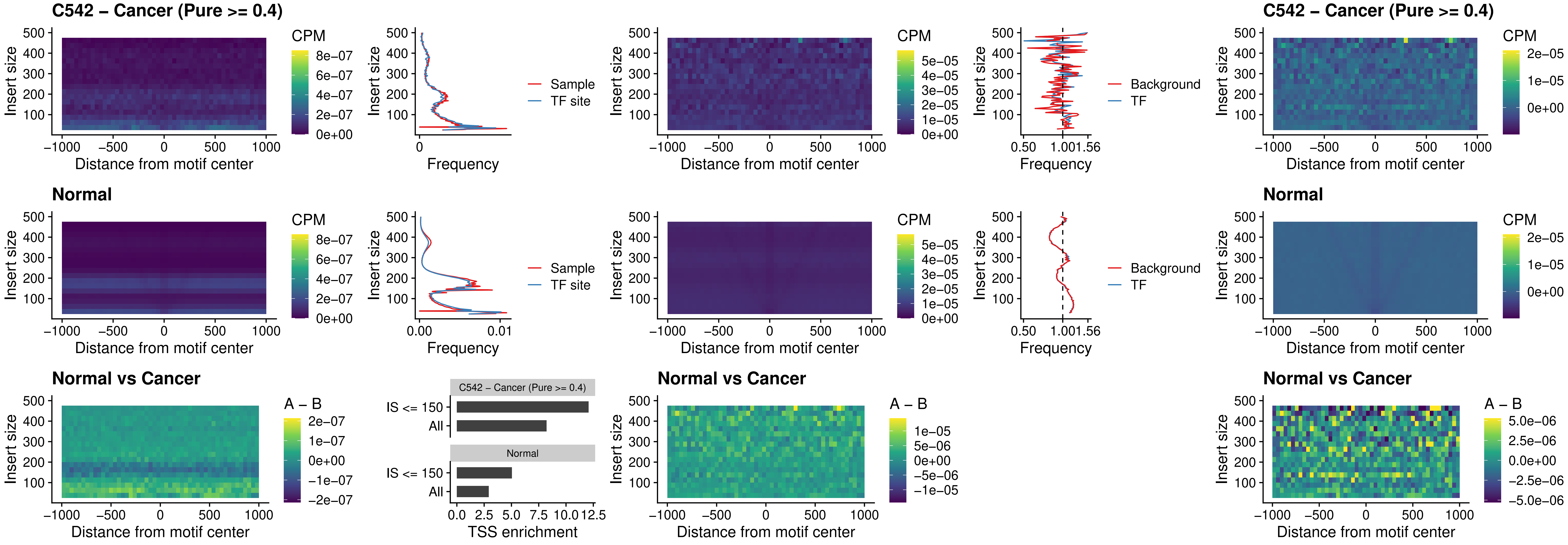

#### C542 – Cancer (Pure >= 0.4) – PDX1_uc_dTSS_oPEAK

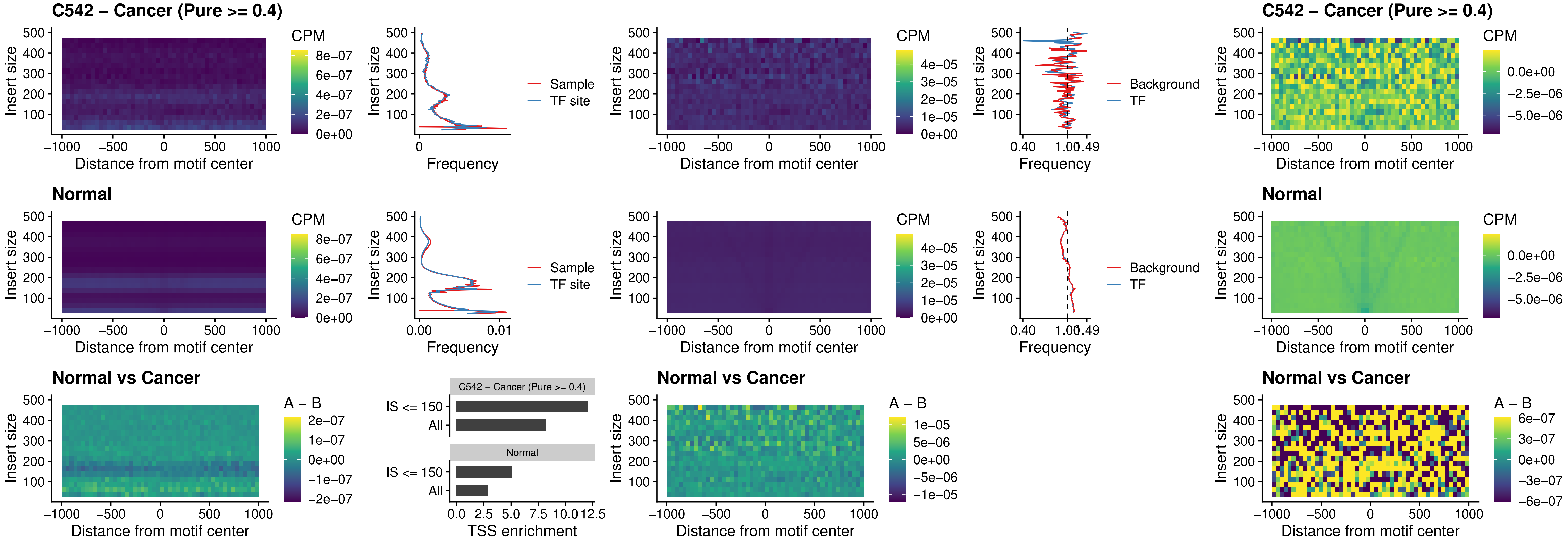

#### C542 – Cancer (Pure >= 0.4) – LHX4_uc_dTSS_oPEAK

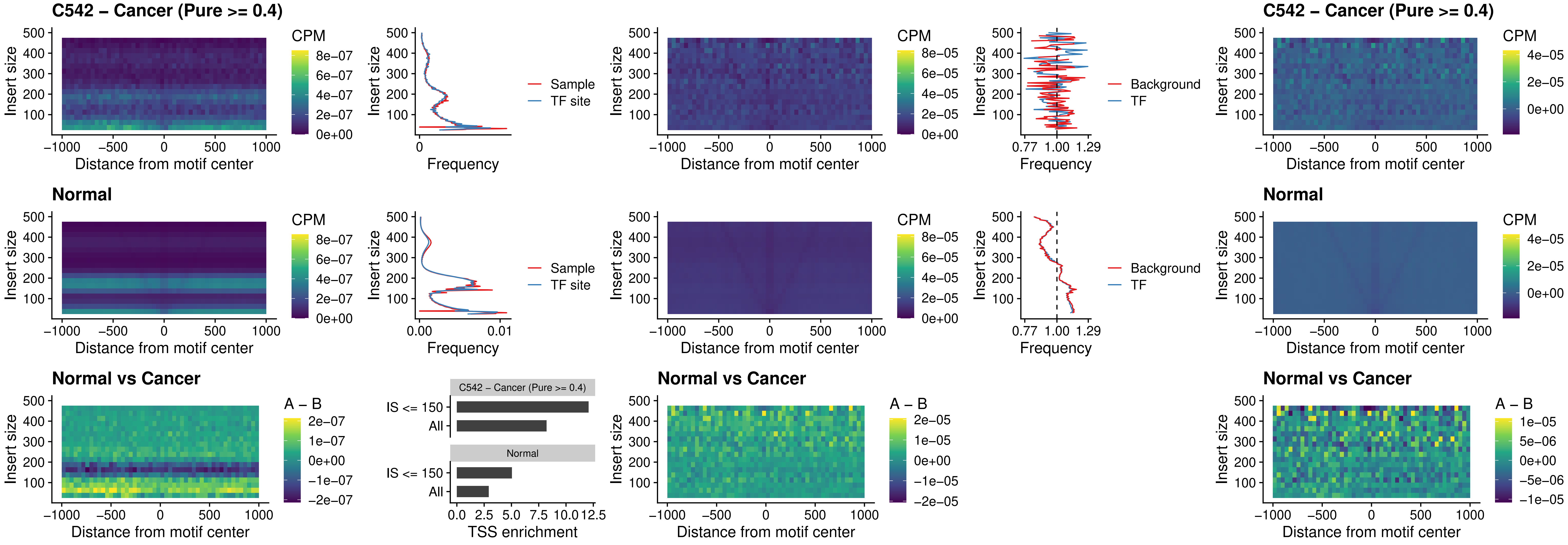

#### C542 – Cancer (Pure >= 0.4) – HOXA4_uc_dTSS_oPEAK

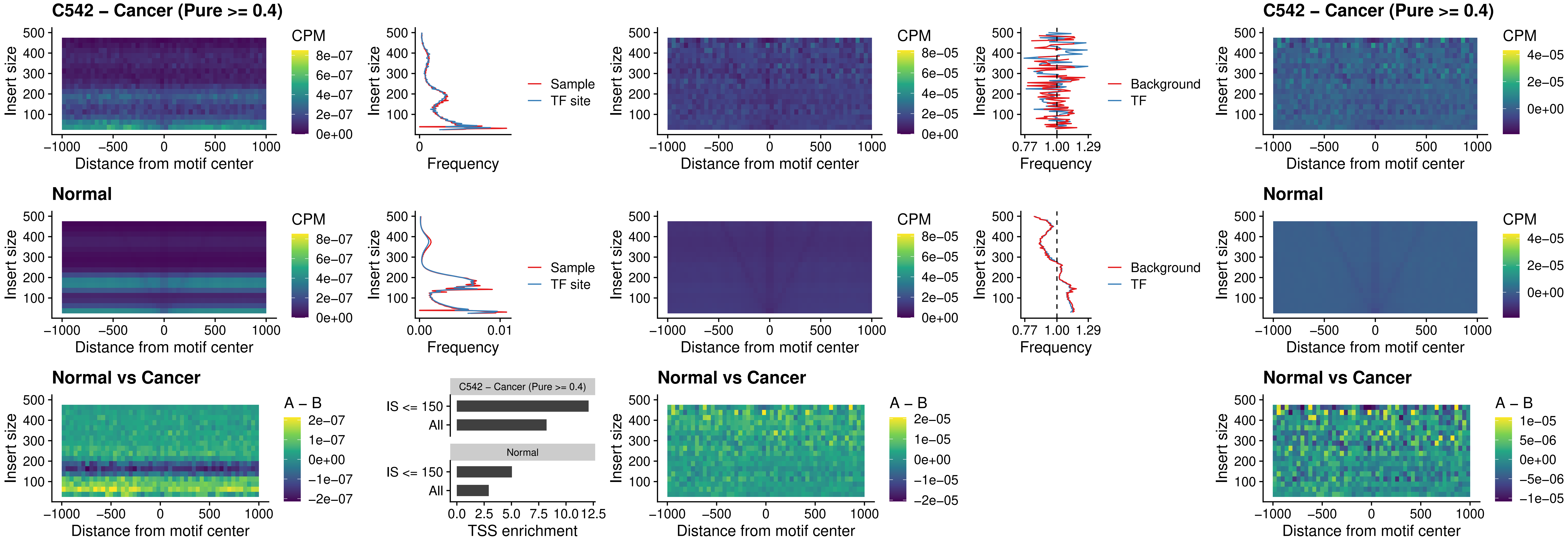

#### C542 – Cancer (Pure >= 0.4) – HLX_uc_dTSS_oPEAK

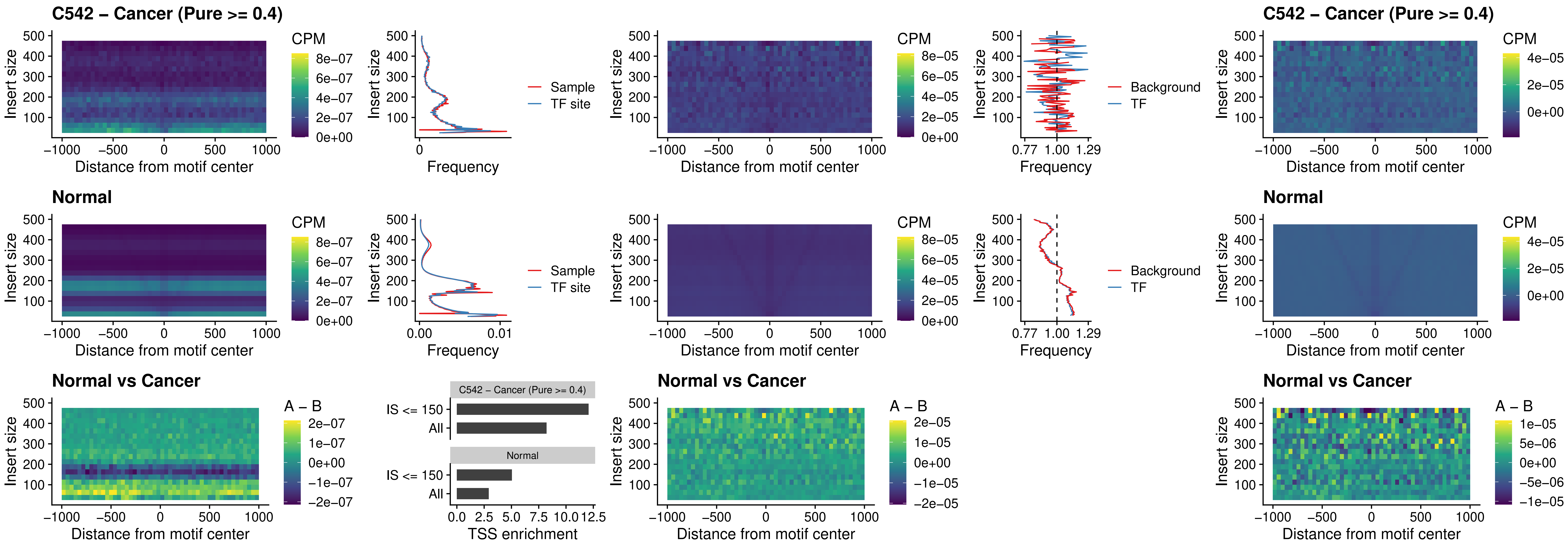

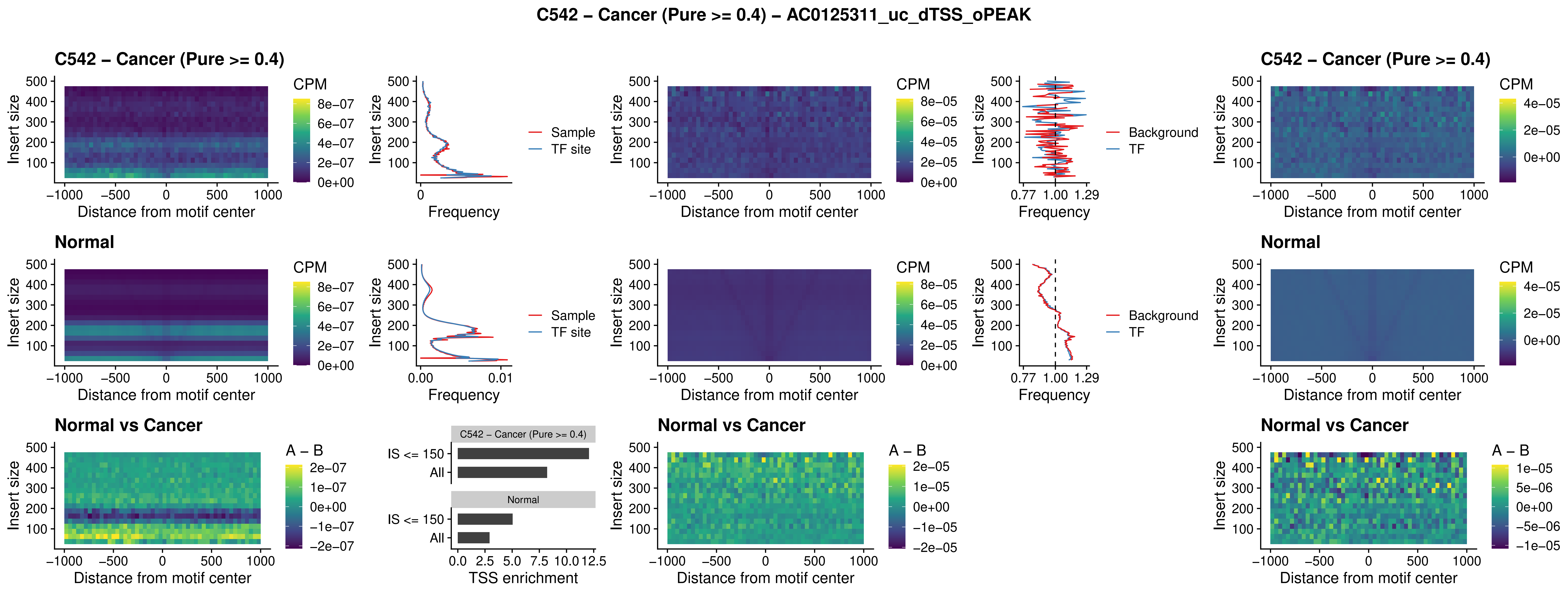
C542 – Cancer (Pure >= 0.4) – AC0125311_uc_dTSS_oPEAK
C542 – Cancer (Pure >= 0.4)
Normal
Normal vs Cancer
Insert size
Distance from motif center
Frequency
CPM
A – B
Sample
TF site
Background
TF
C542 – Cancer (Pure >= 0.4)
Normal
IS <= 150
All
TSS enrichment

### C542 – Cancer (Pure >= 0.4) – DBX2_uc_dTSS_oPEAK

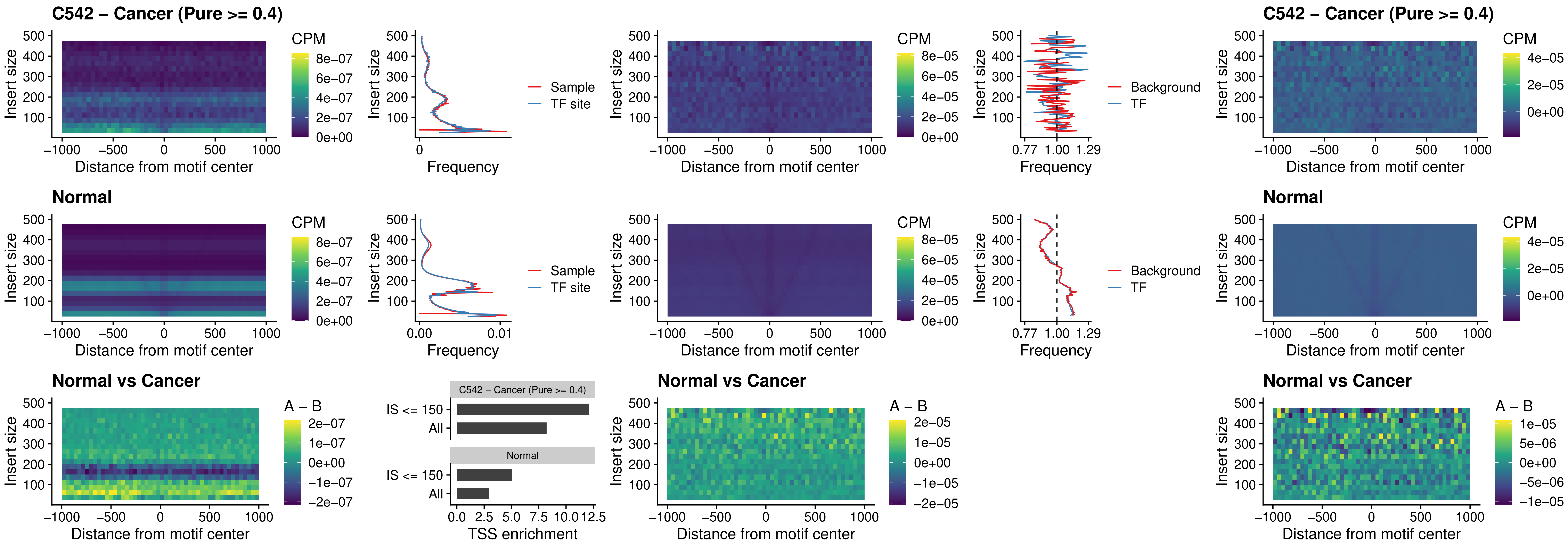

#### C542 – Cancer (Pure >= 0.4) – HOXA2_uc_dTSS_nPEAK

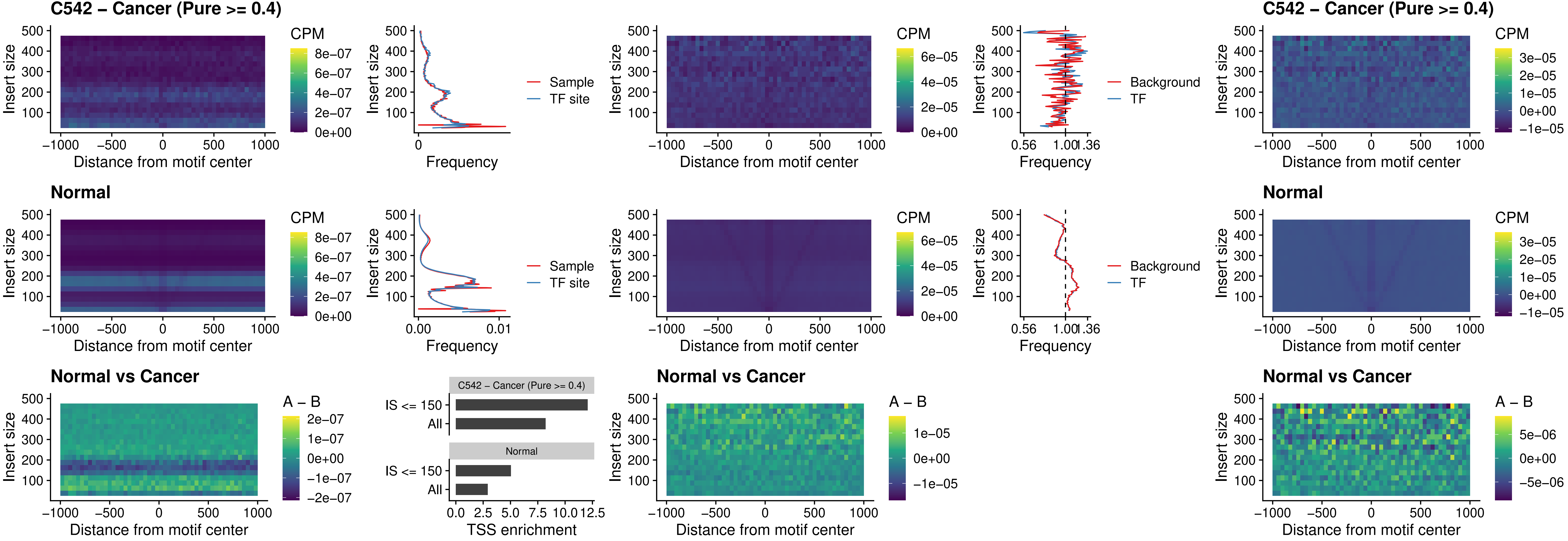

#### C542 – Cancer (Pure >= 0.4) – SHOX_uc_dTSS_nPEAK

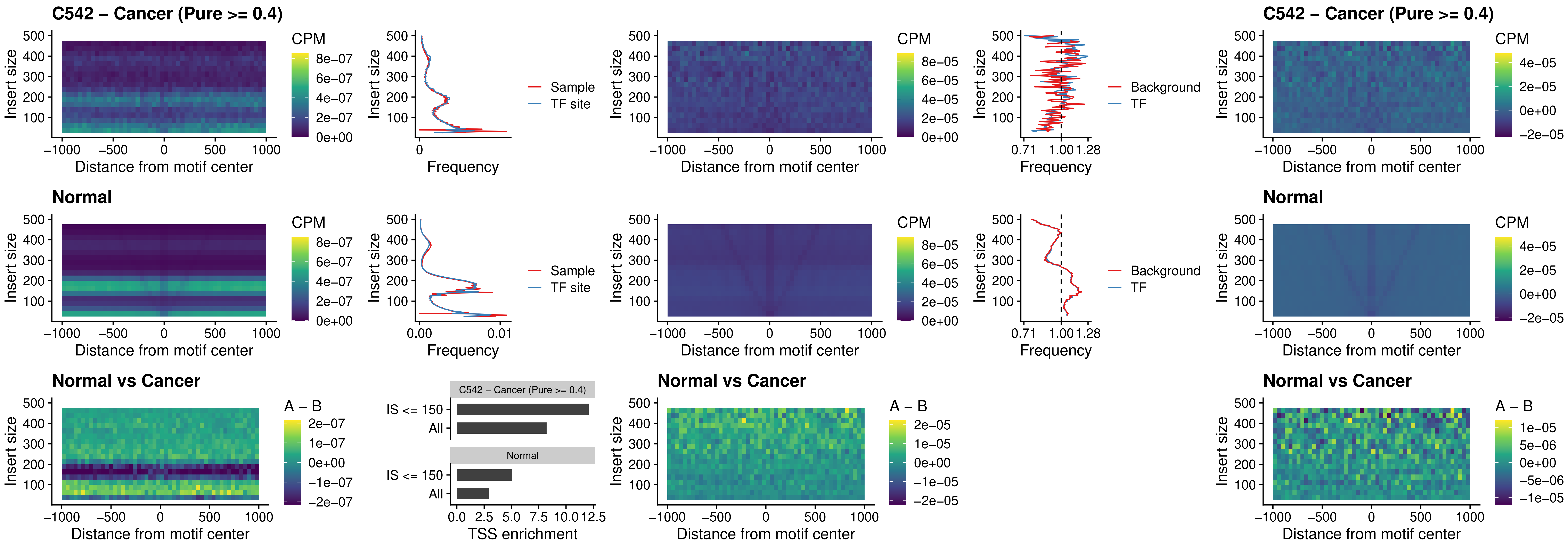

#### C542 – Cancer (Pure >= 0.4) – PRRX1_uc_dTSS_nPEAK

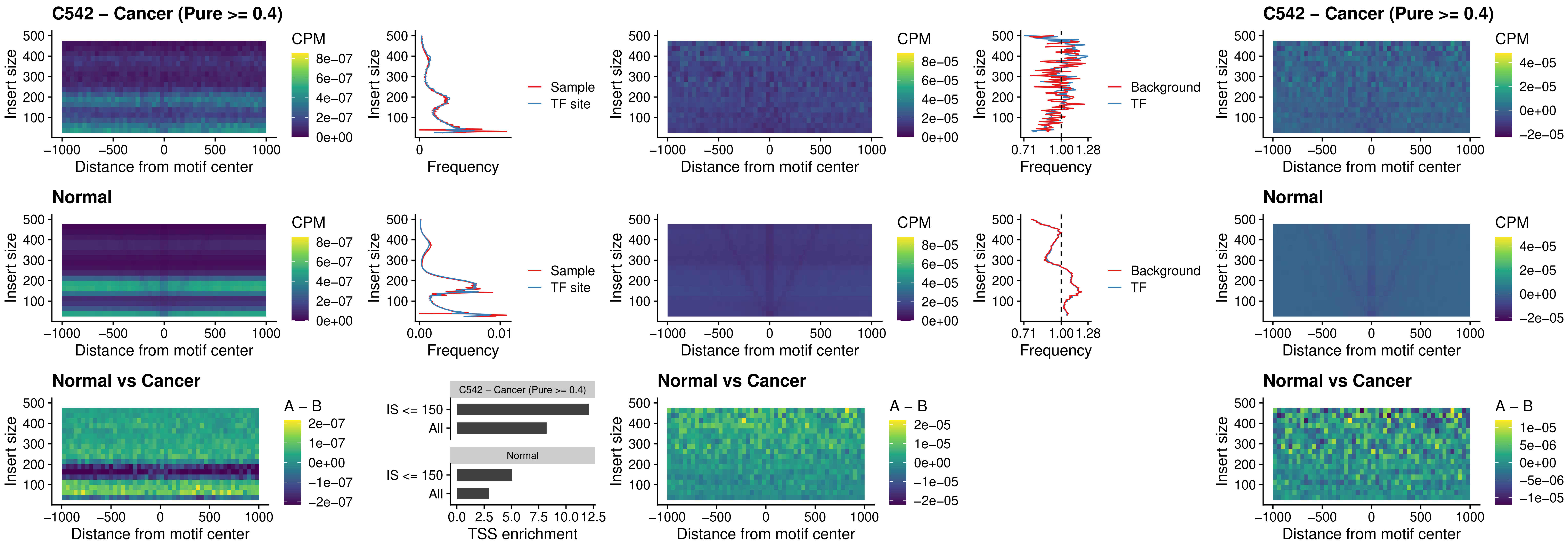

#### C542 – Cancer (Pure >= 0.4) – LMX1A_uc_dTSS_nPEAK

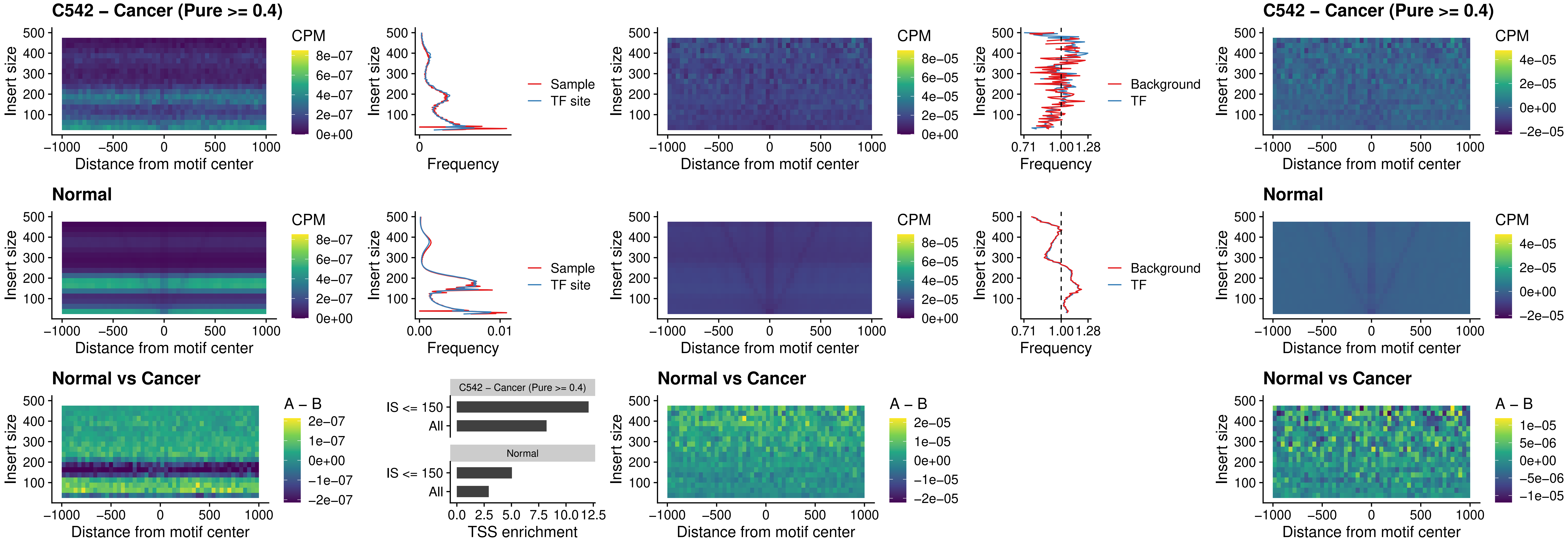

#### C542 – Cancer (Pure >= 0.4) – LMX1B_uc_dTSS_nPEAK

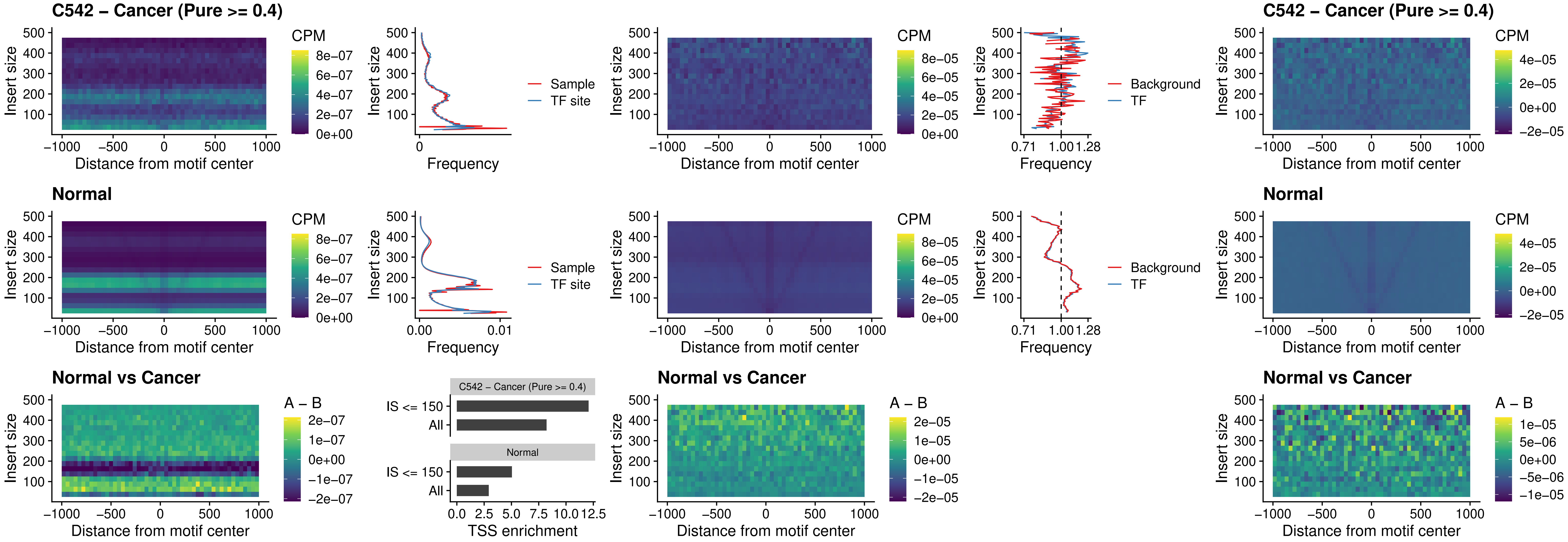

#### C542 – Cancer (Pure >= 0.4) – DLX5_uc_dTSS_oPEAK

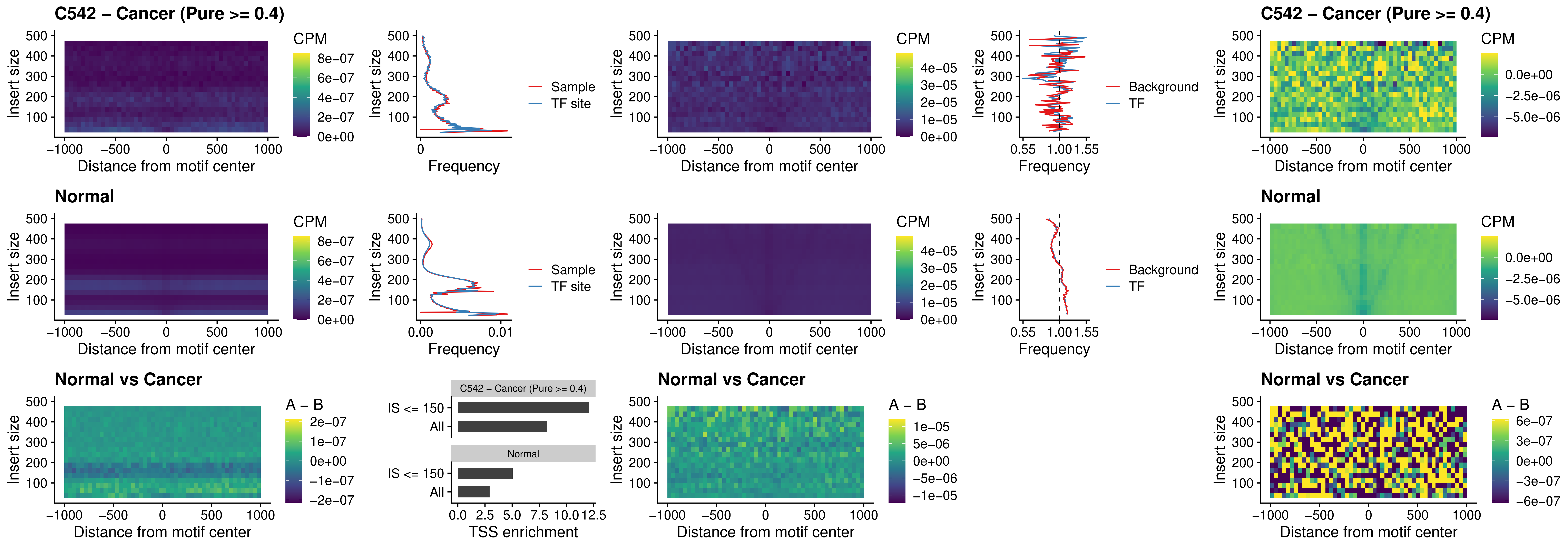

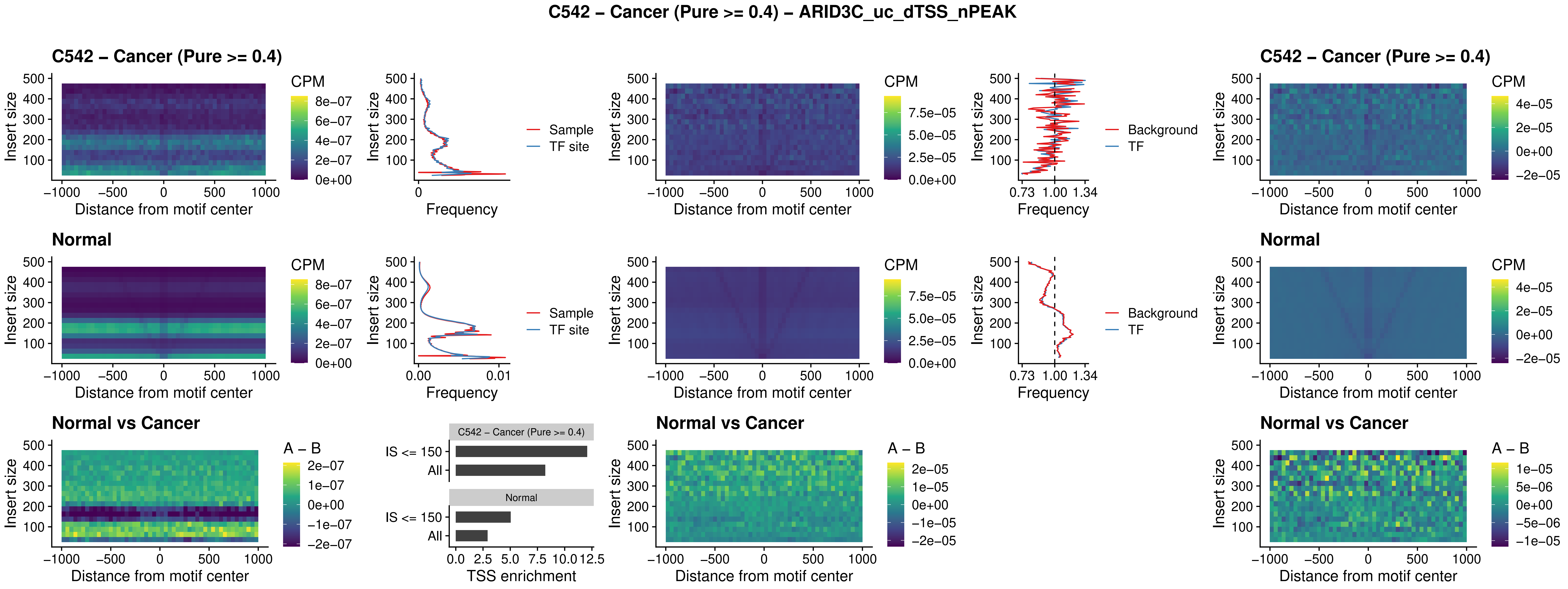
C542 – Cancer (Pure >= 0.4) – ARID3C_uc_dTSS_nPEAK
C542 – Cancer (Pure >= 0.4)
Normal
Normal vs Cancer
Insert size
Distance from motif center
Frequency
CPM
A – B
Sample
TF site
Background
TF
C542 – Cancer (Pure >= 0.4)
Normal
IS <= 150
All
TSS enrichment

### C542 – Cancer (Pure >= 0.4) – HOXD12_uc_dTSS_nPEAK

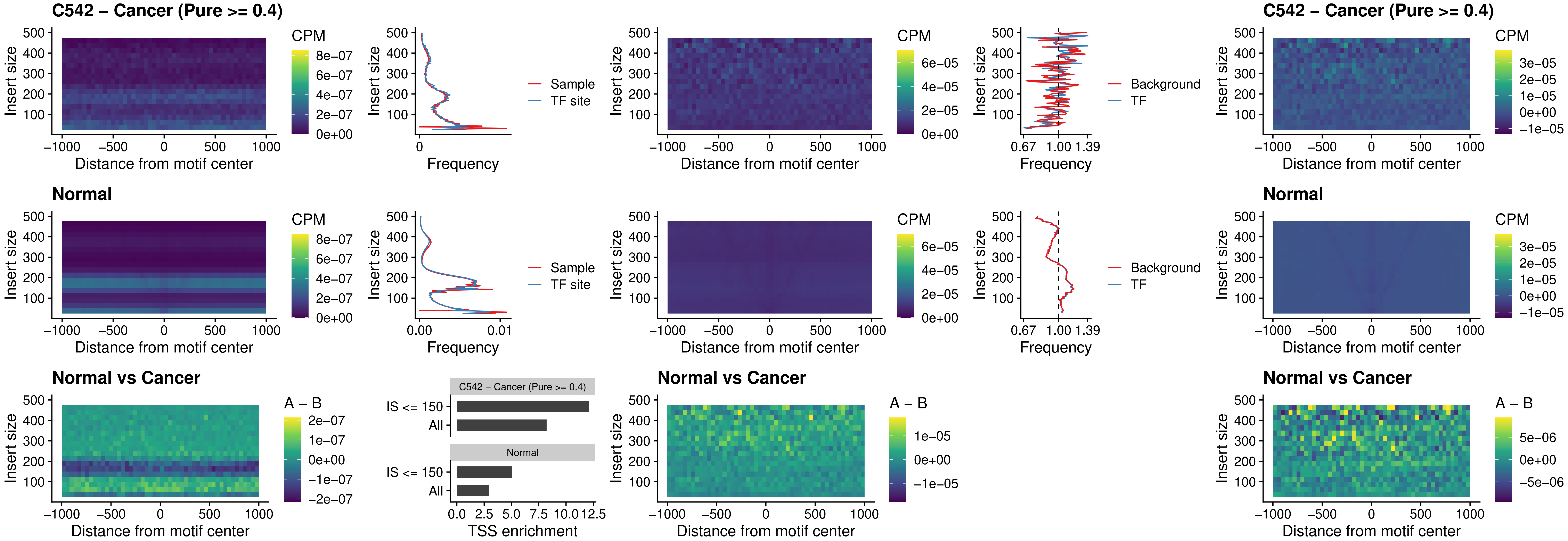

#### C542 – Cancer (Pure >= 0.4) – CDX2_uc_dTSS_nPEAK

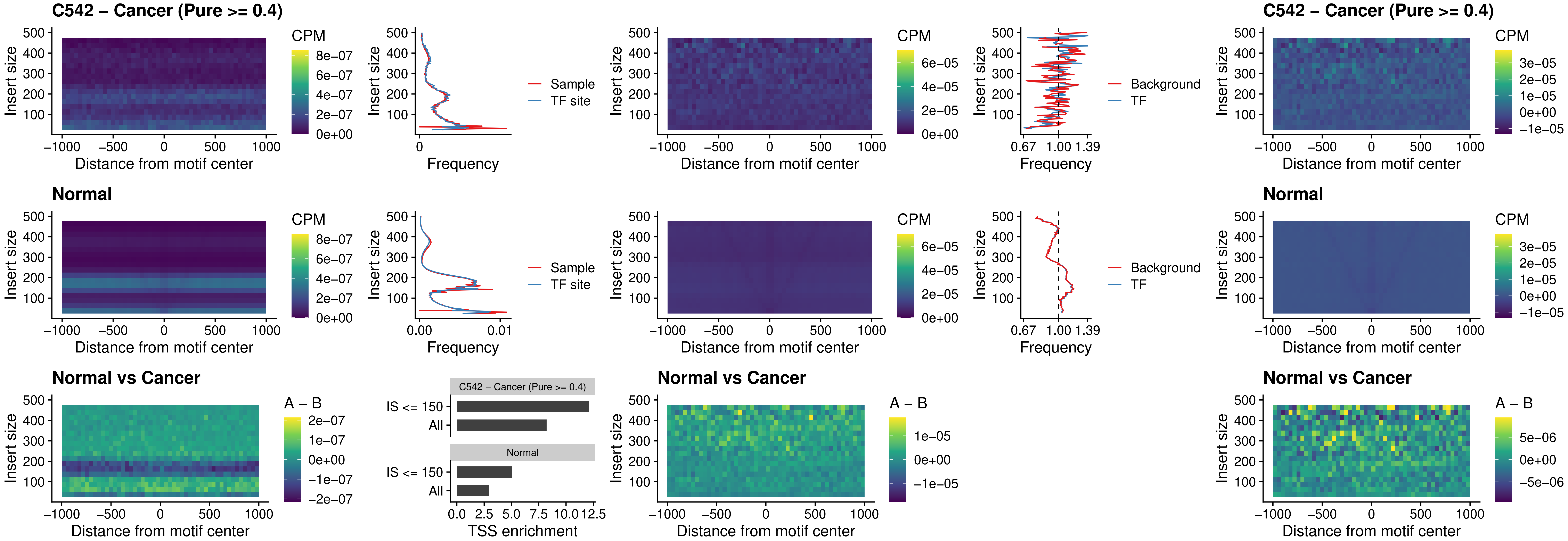

#### C542 – Cancer (Pure >= 0.4) – HOXC12_uc_dTSS_nPEAK

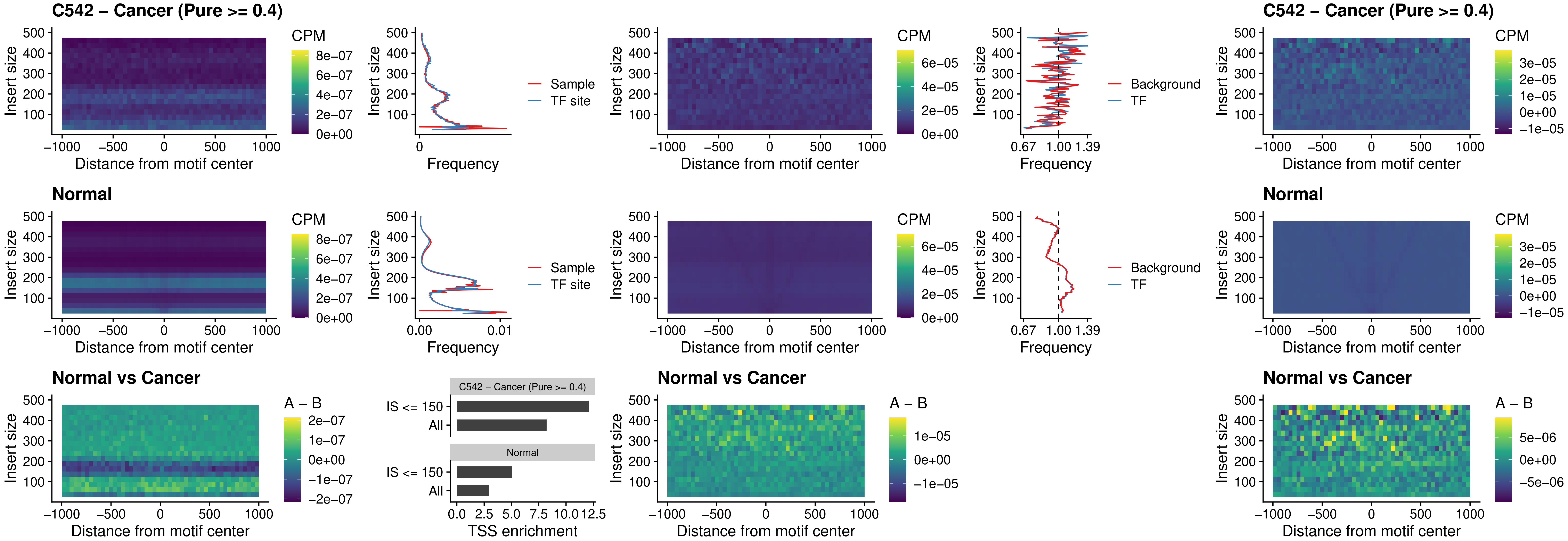

#### C542 – Cancer (Pure >= 0.4) – HHEX_uc_dTSS_oPEAK

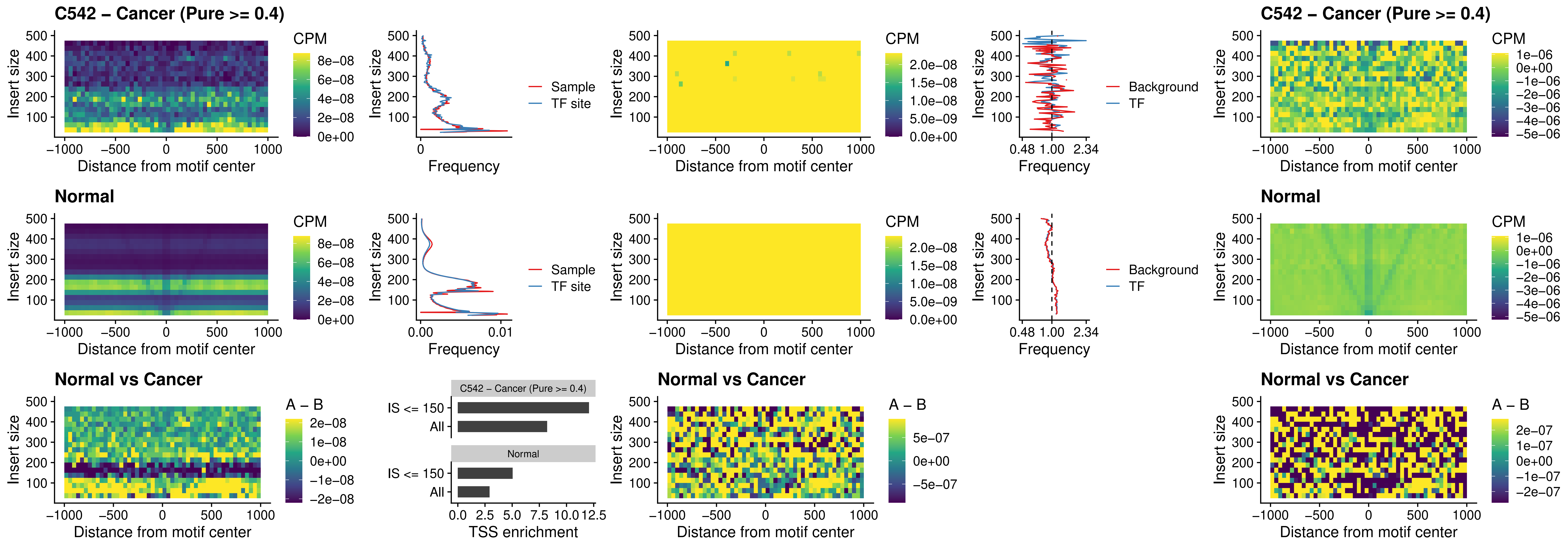

#### C542 – Cancer (Pure >= 0.4) – HOXA11_uc_dTSS_nPEAK

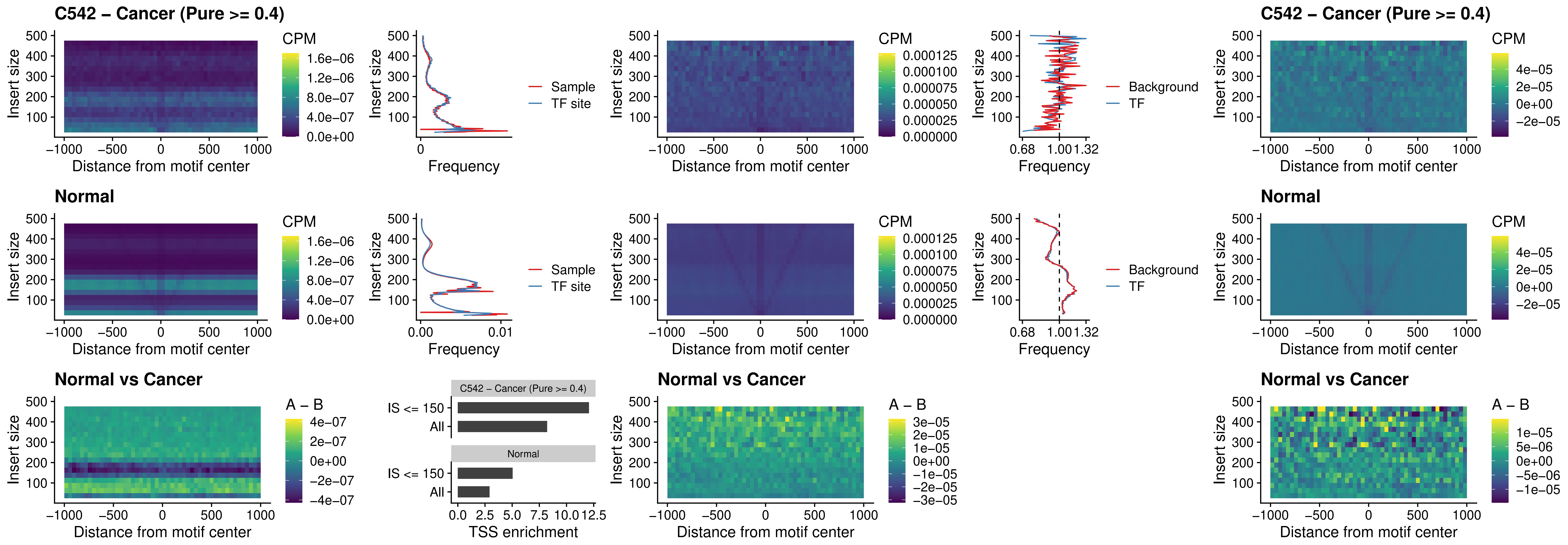

### C542 – Cancer (Pure >= 0.4) – HOXA11_uc_cTSS_nPEAK

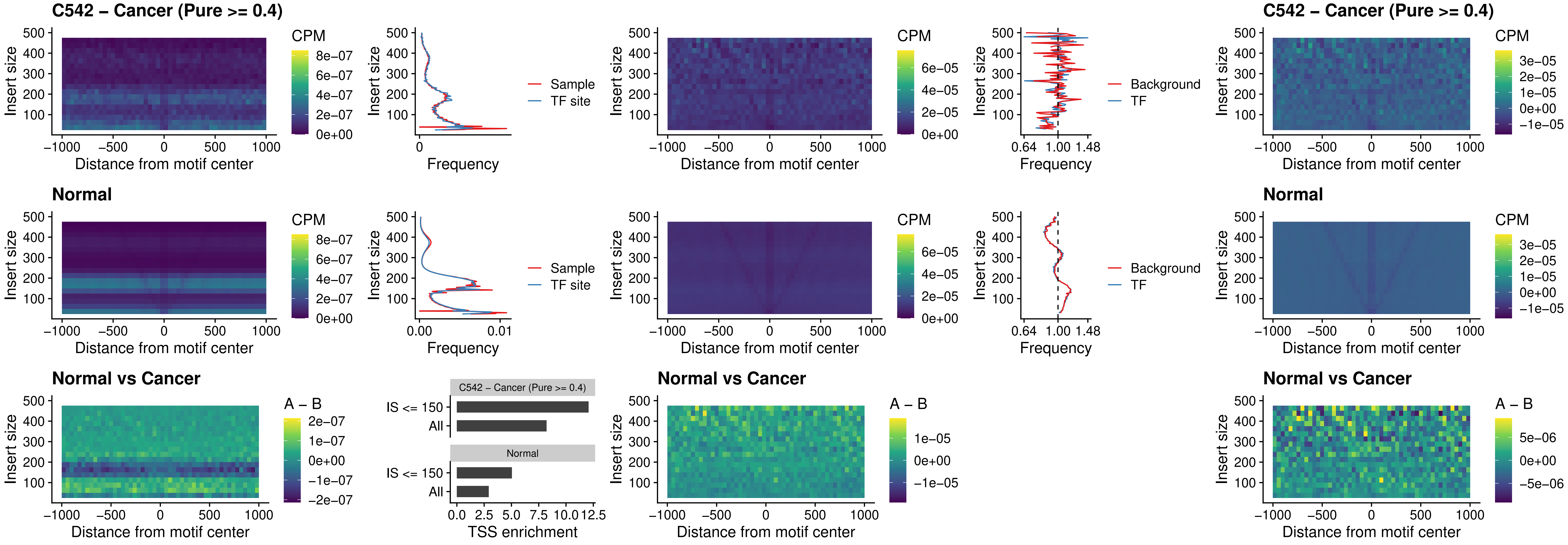

#### C542 – Cancer (Pure >= 0.4) – ZFHX3_uc_dTSS_nPEAK

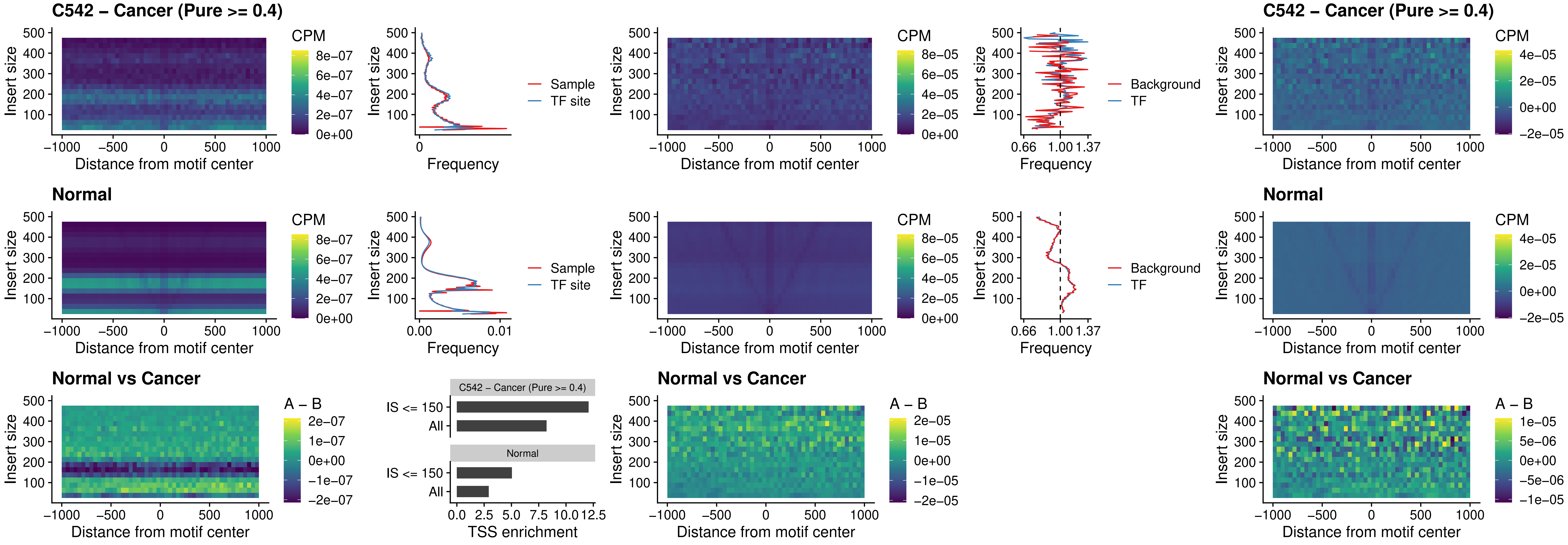

#### C542 – Cancer (Pure >= 0.4) – HOXB4_uc_dTSS_nPEAK

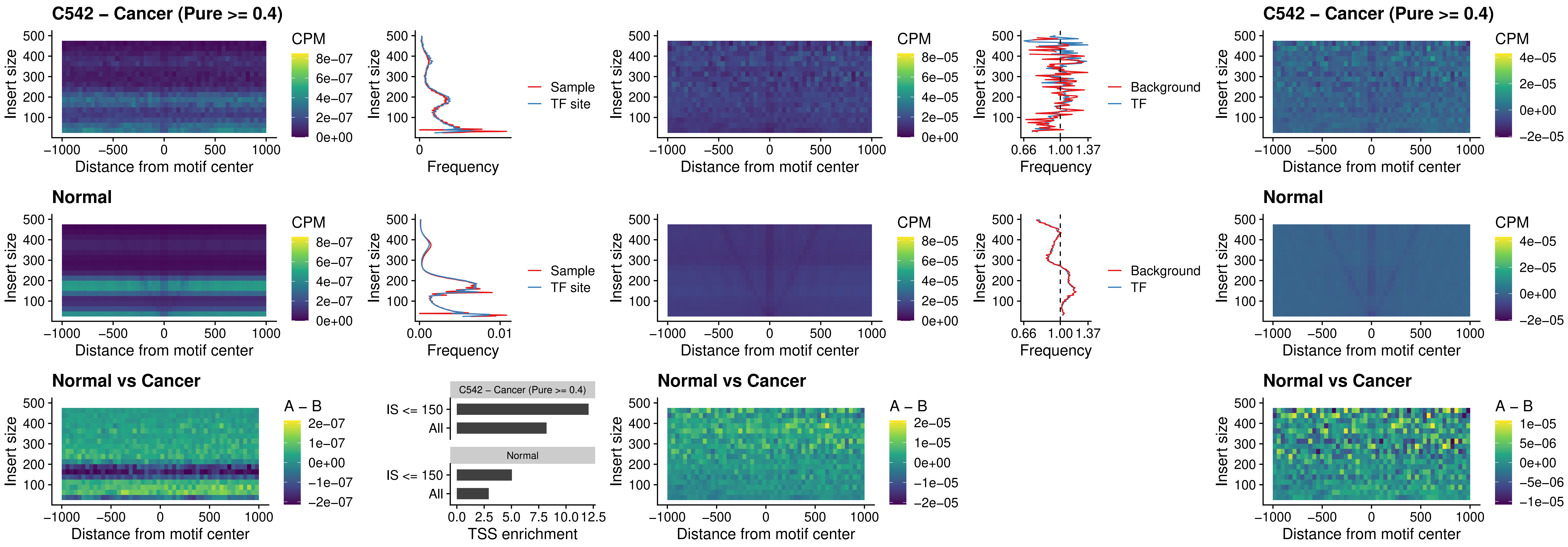

#### C542 – Cancer (Pure >= 0.4) – HOXC5_uc_dTSS_nPEAK

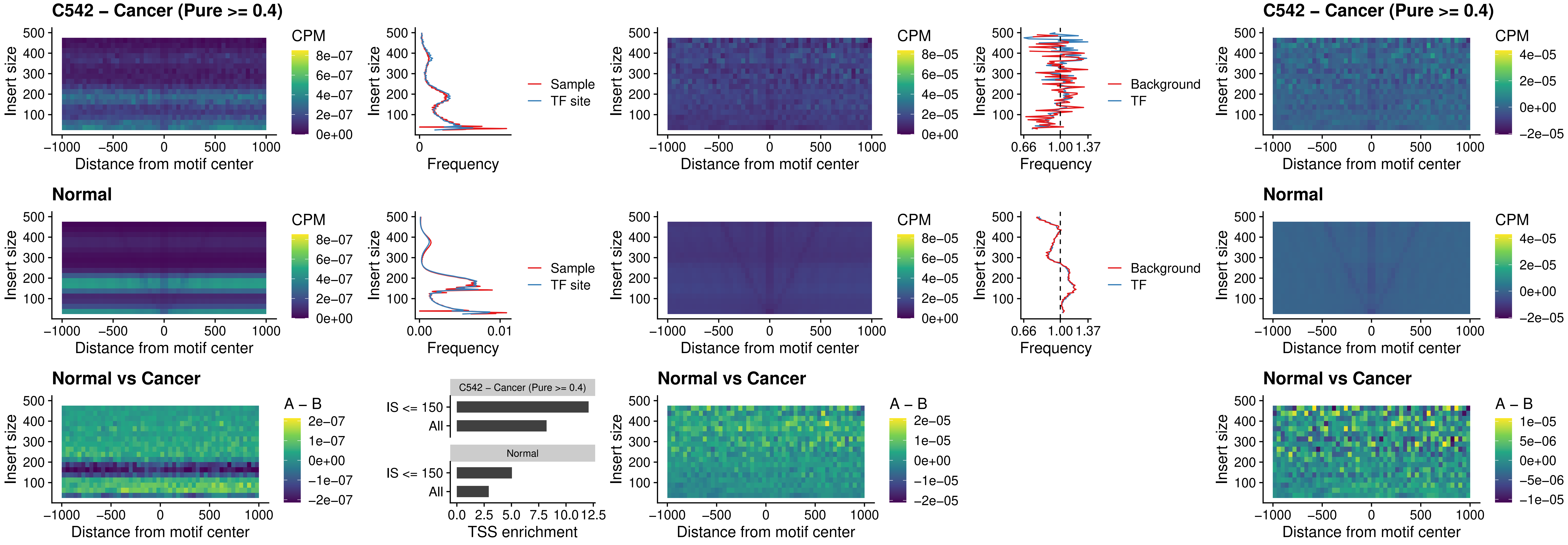

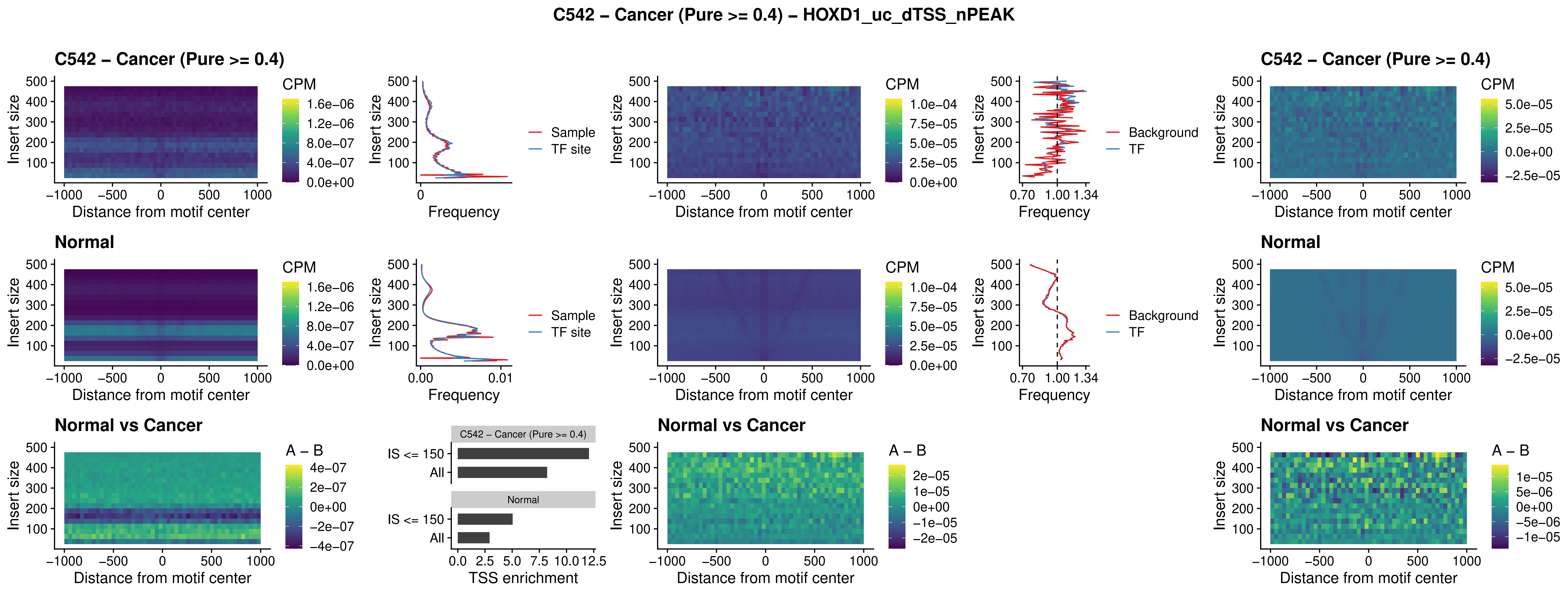
C542 – Cancer (Pure >= 0.4) – HOXD1_uc_dTSS_nPEAK
C542 – Cancer (Pure >= 0.4)
Normal
Normal vs Cancer
Insert size
Distance from motif center
Frequency
CPM
A – B
Sample
TF site
Background
TF
C542 – Cancer (Pure >= 0.4)
Normal
IS <= 150
All
TSS enrichment

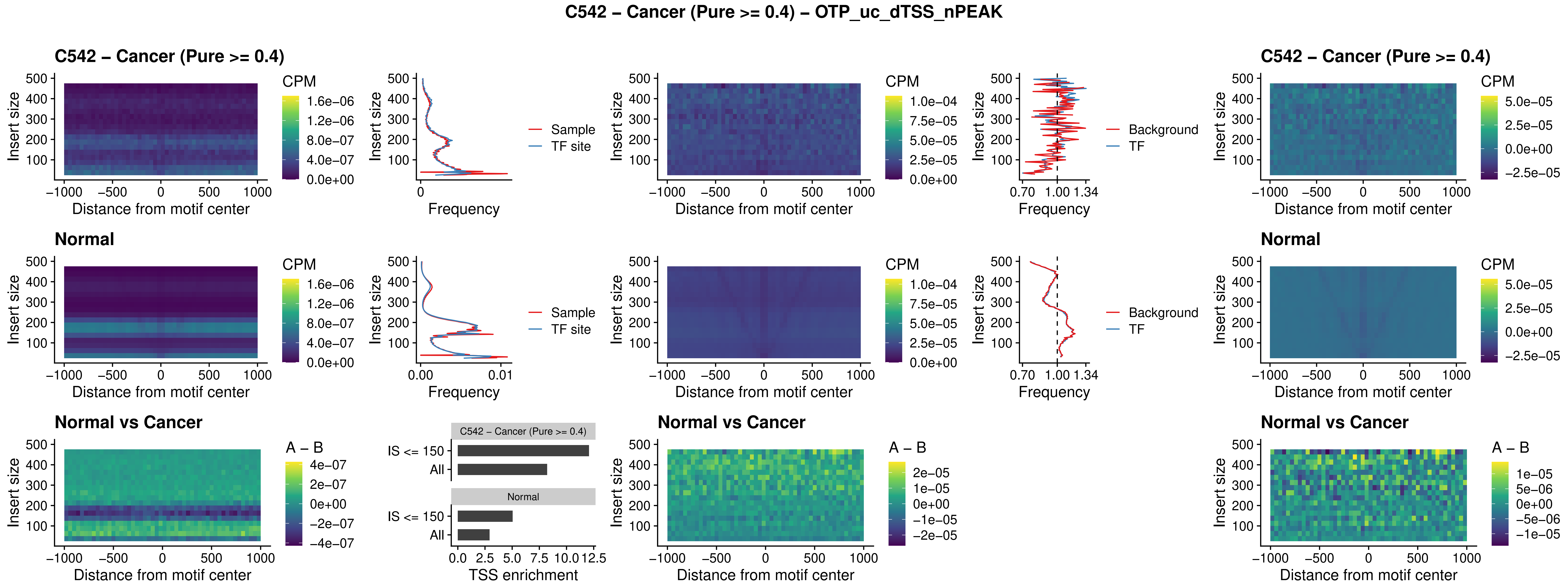
C542 – Cancer (Pure >= 0.4) – OTP_uc_dTSS_nPEAK
C542 – Cancer (Pure >= 0.4)
Normal
Normal vs Cancer
Insert size
Distance from motif center
Frequency
CPM
A – B
Sample
TF site
Background
TF
C542 – Cancer (Pure >= 0.4)
Normal
IS <= 150
All
TSS enrichment

#### C542 – Cancer (Pure >= 0.4) – FOXL1_uc_dTSS_oPEAK

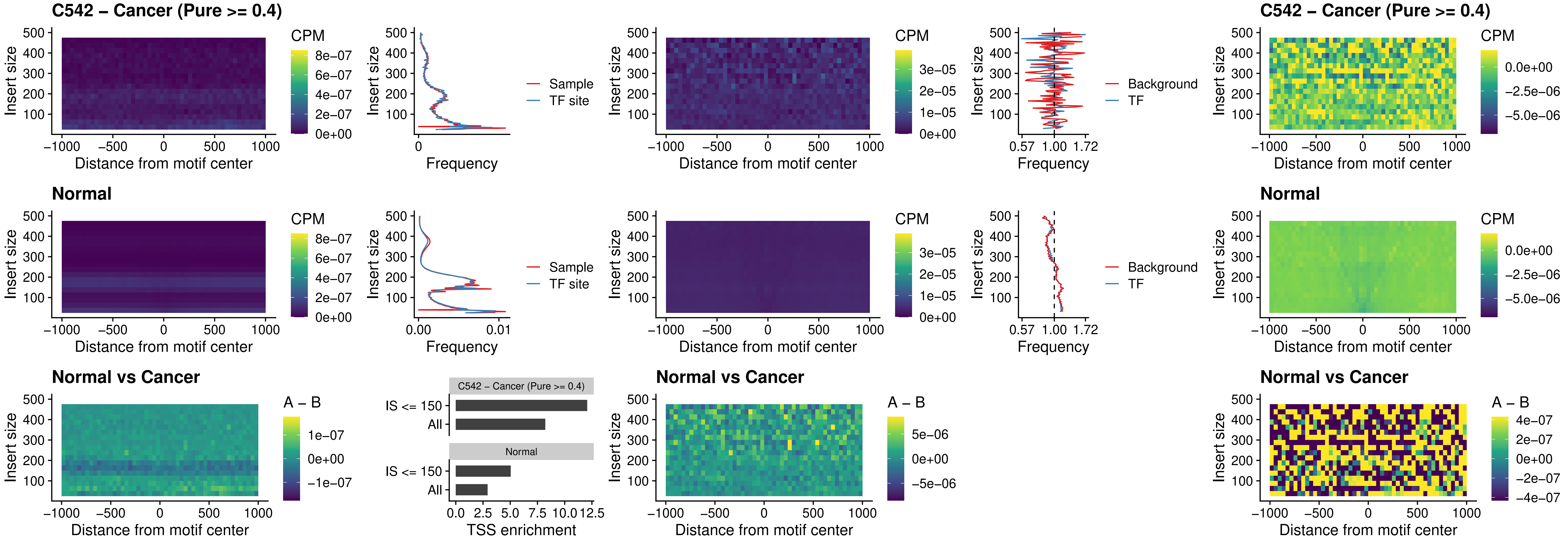

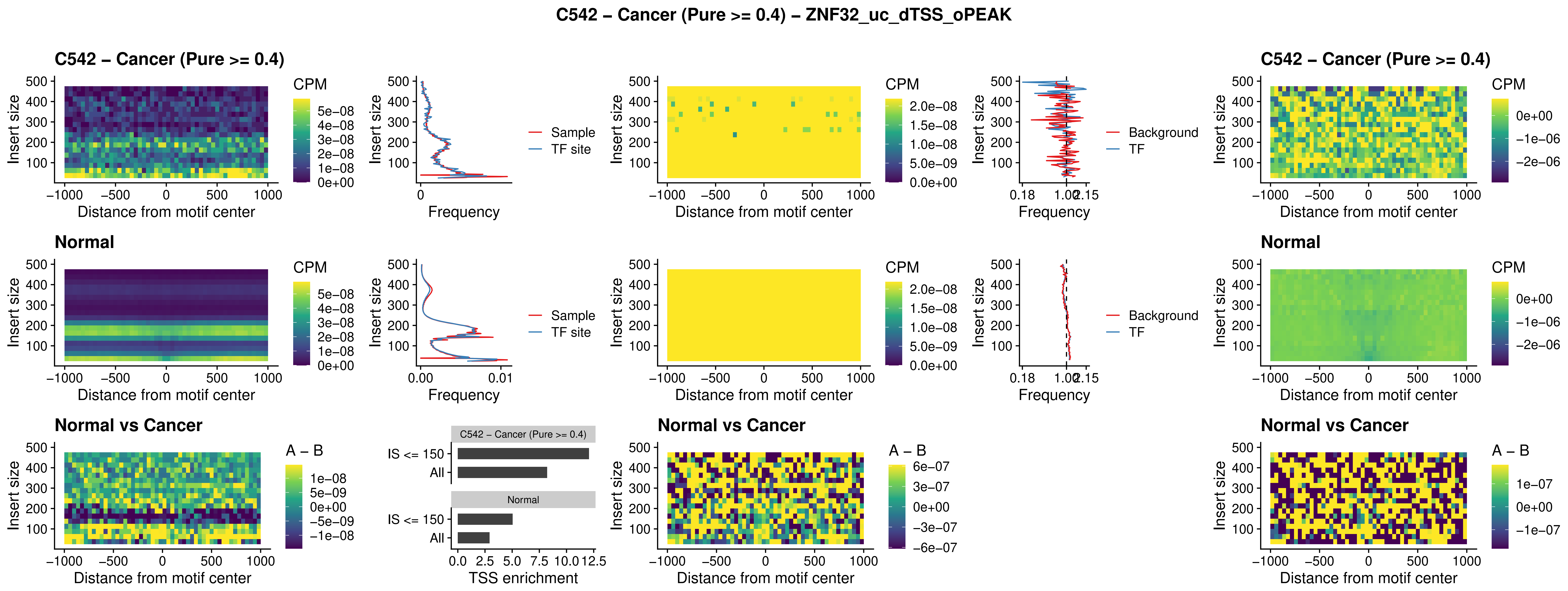

C542 – Cancer (Pure >= 0.4) – ZNF32_uc_dTSS_oPEAK
C542 – Cancer (Pure >= 0.4)
Normal
Normal vs Cancer
Insert size
Distance from motif center
Frequency
CPM
A – B
Sample
TF site
Background
TF
C542 – Cancer (Pure >= 0.4)
IS <= 150
All
TSS enrichment

#### C542 – Cancer (Pure >= 0.4) – FOXL1_uc_dTSS_nPEAK

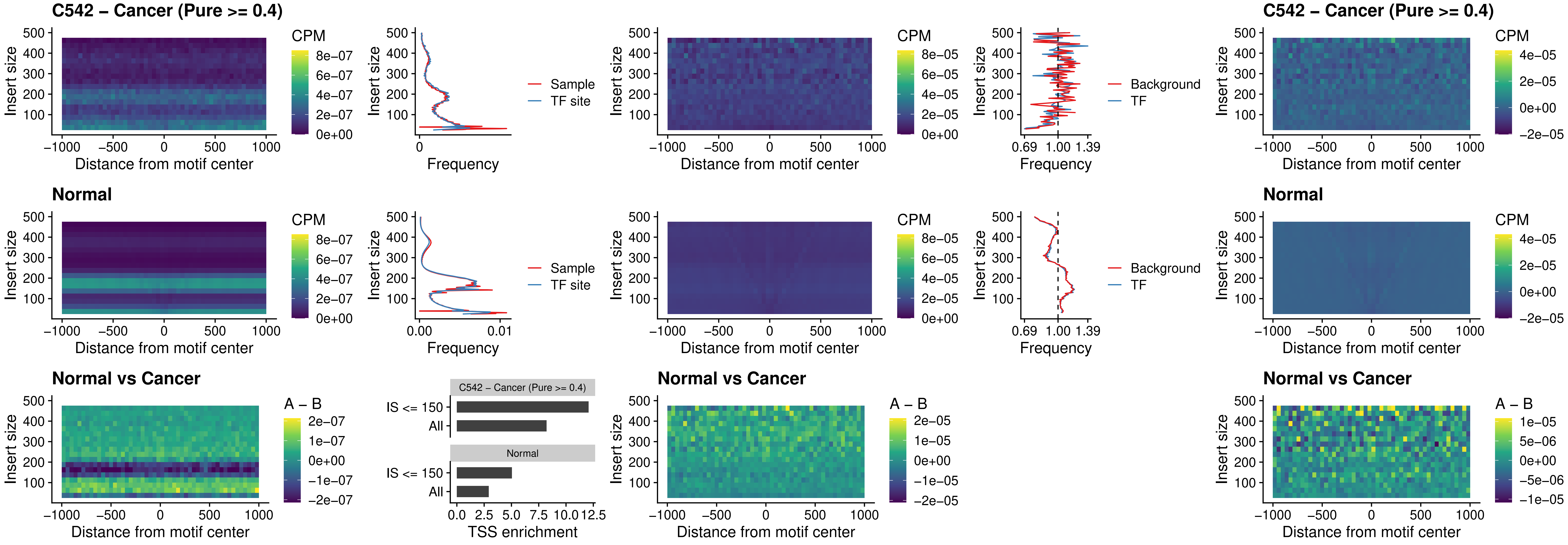

#### C542 – Cancer (Pure >= 0.4) – ZNF32_uc_dTSS_nPEAK

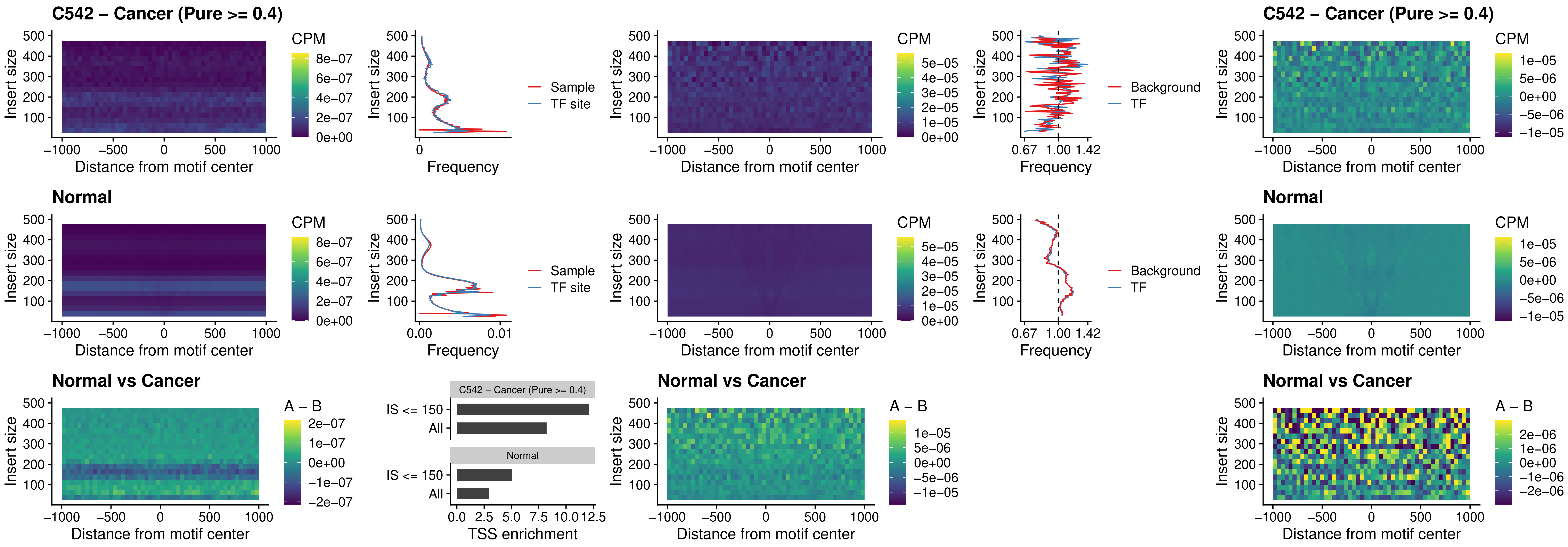

### C542 – Cancer (Pure >= 0.4) – DLX5_uc_dTSS_nPEAK

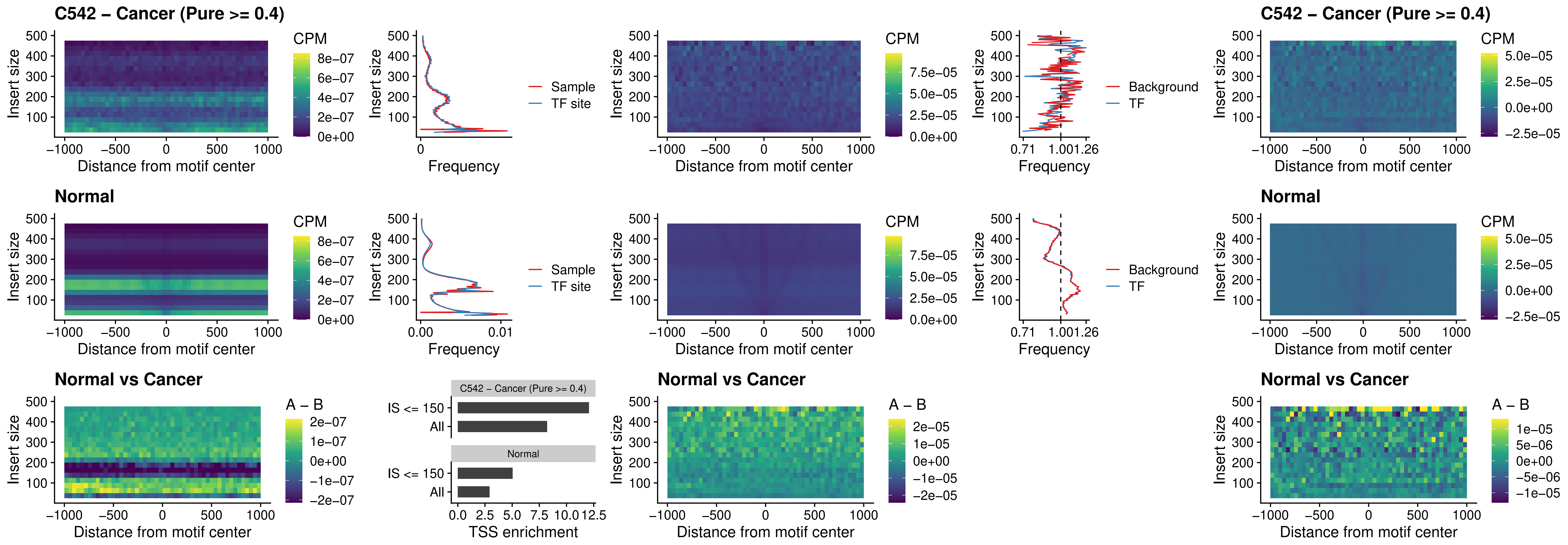

#### C542 – Cancer (Pure >= 0.4) – DMRT2_uc_dTSS_nPEAK

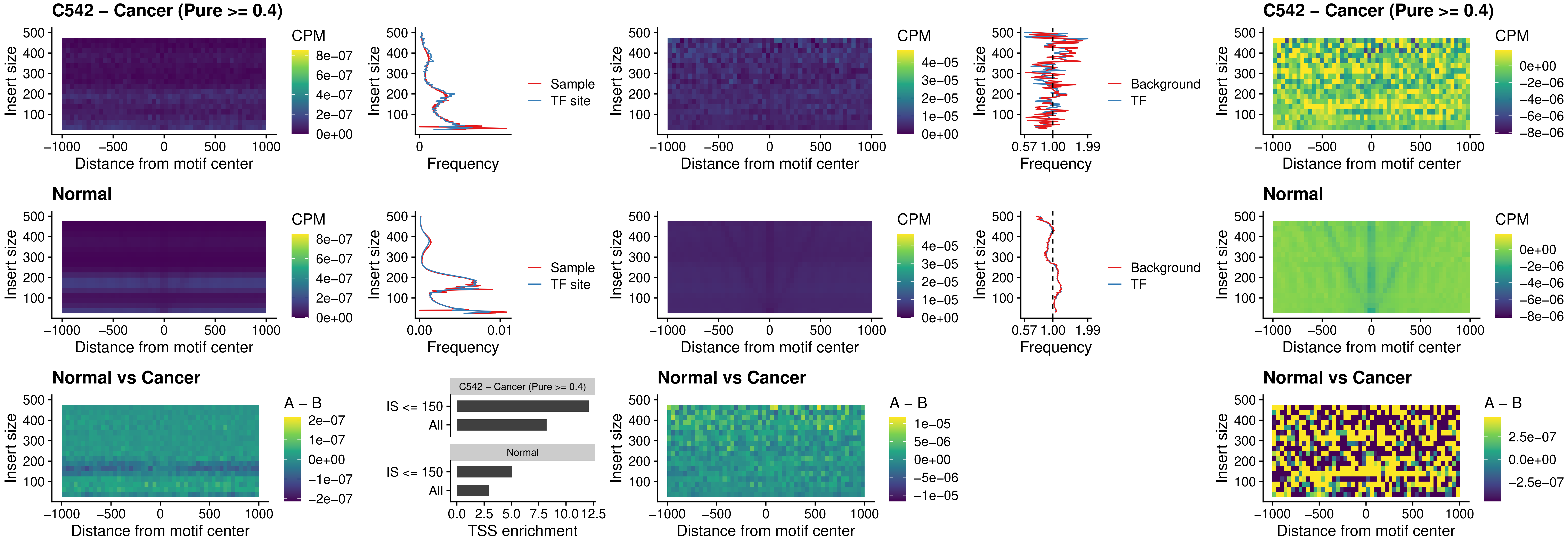

#### C542 – Cancer (Pure >= 0.4) – DMRTC2_uc_cTSS_nPEAK

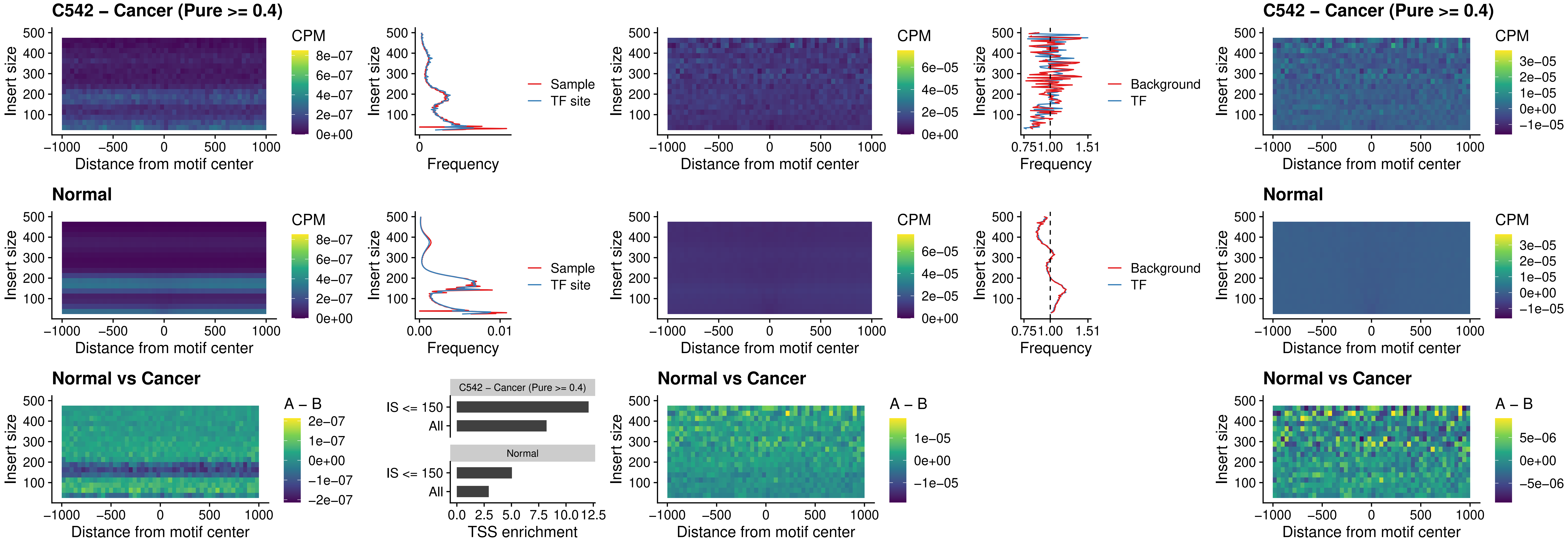

#### C542 – Cancer (Pure >= 0.4) – HOXD9_uc_dTSS_nPEAK

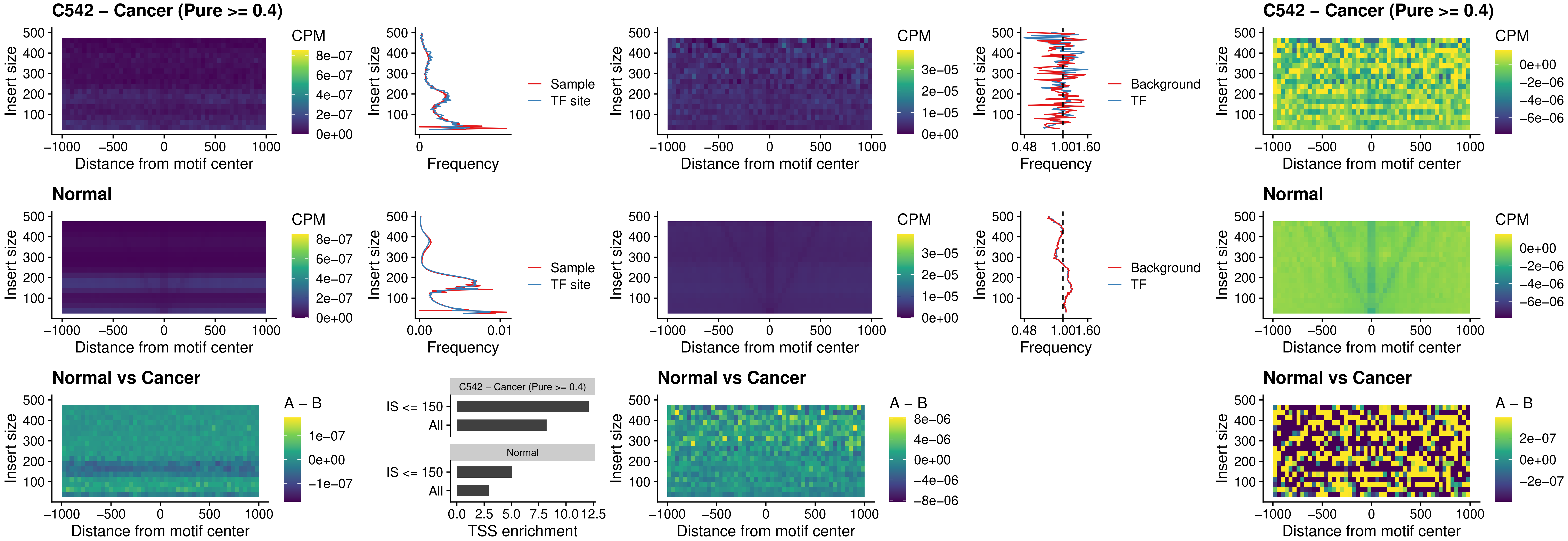

#### C542 – Cancer (Pure >= 0.4) – HOXD4_uc_dTSS_nPEAK

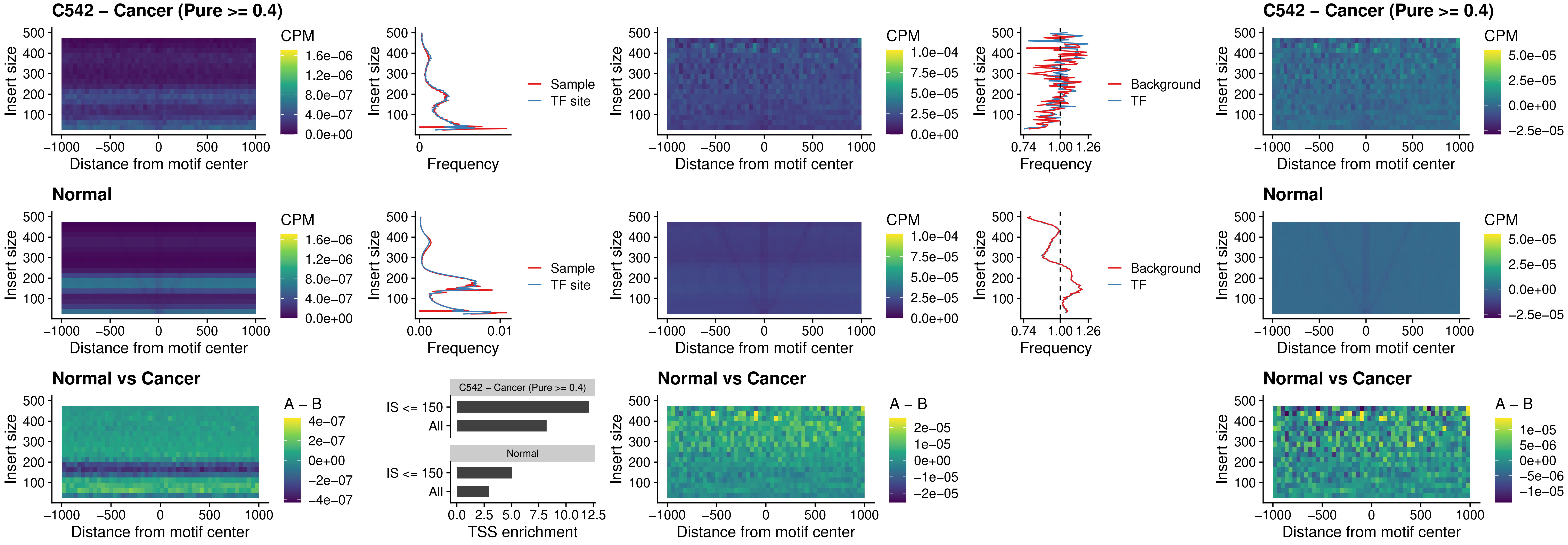

### C542 – Cancer (Pure >= 0.4) – FOXP3_uc_cTSS_nPEAK

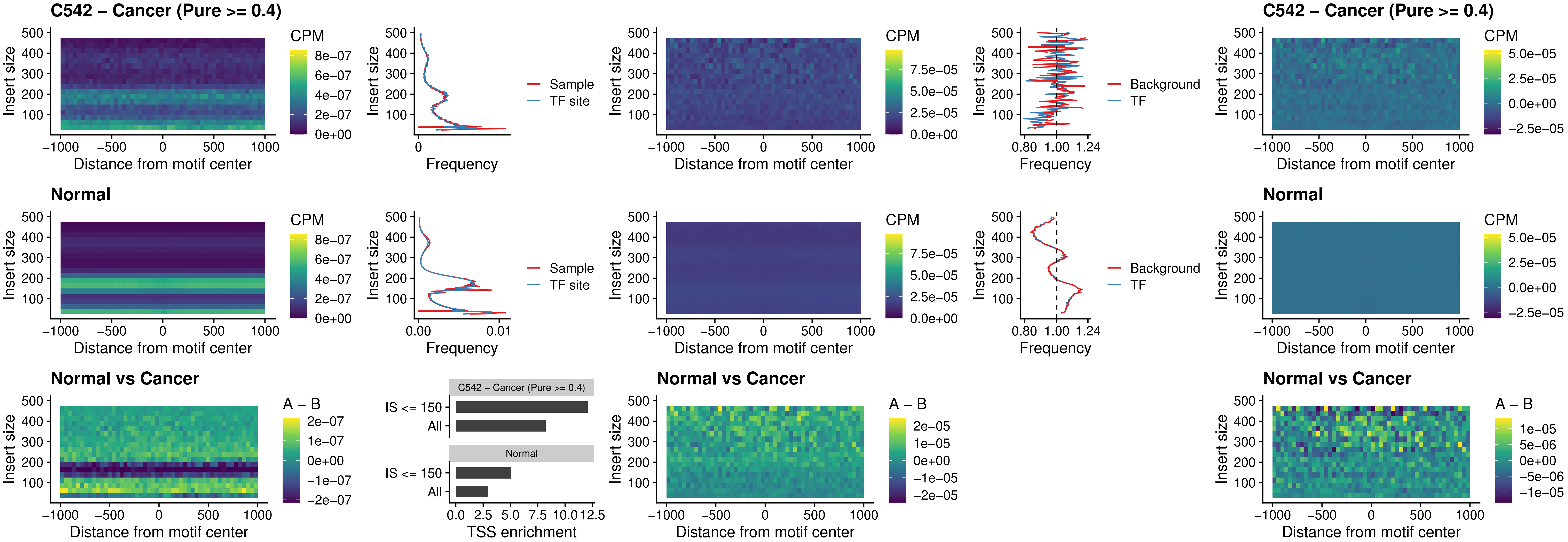

#### C542 – Cancer (Pure >= 0.4) – FOXD4L6_uc_cTSS_nPEAK

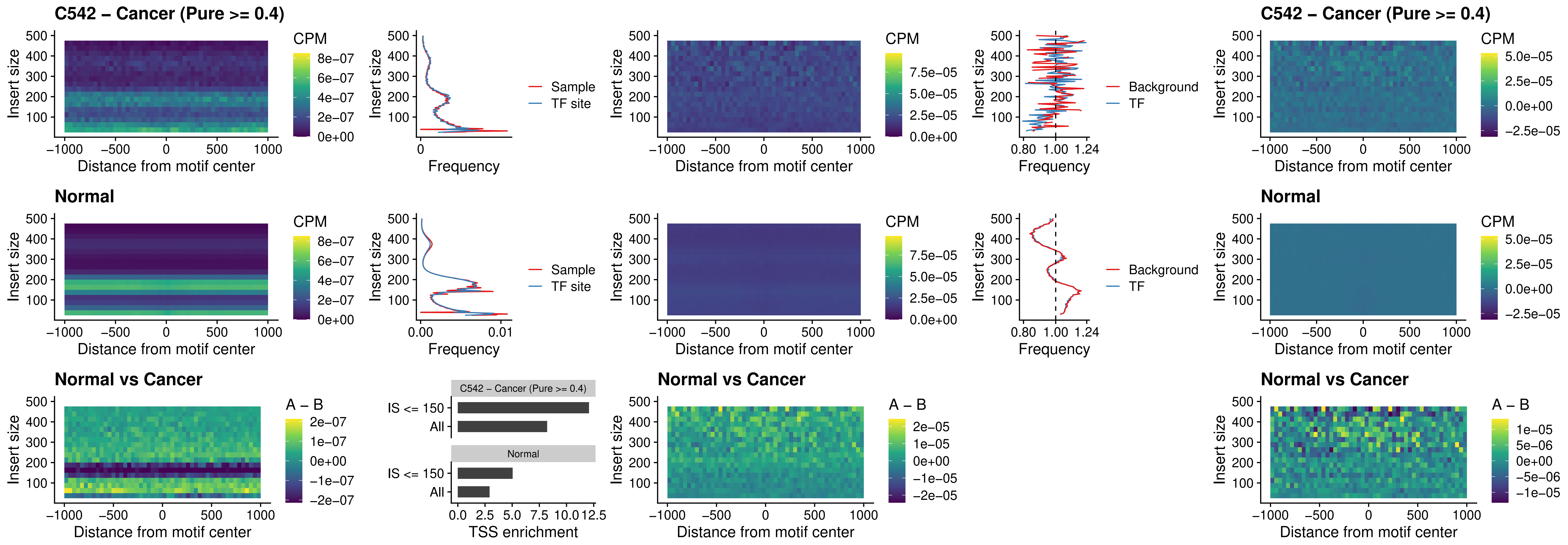

#### C542 – Cancer (Pure >= 0.4) – FOXD4L5_uc_cTSS_nPEAK

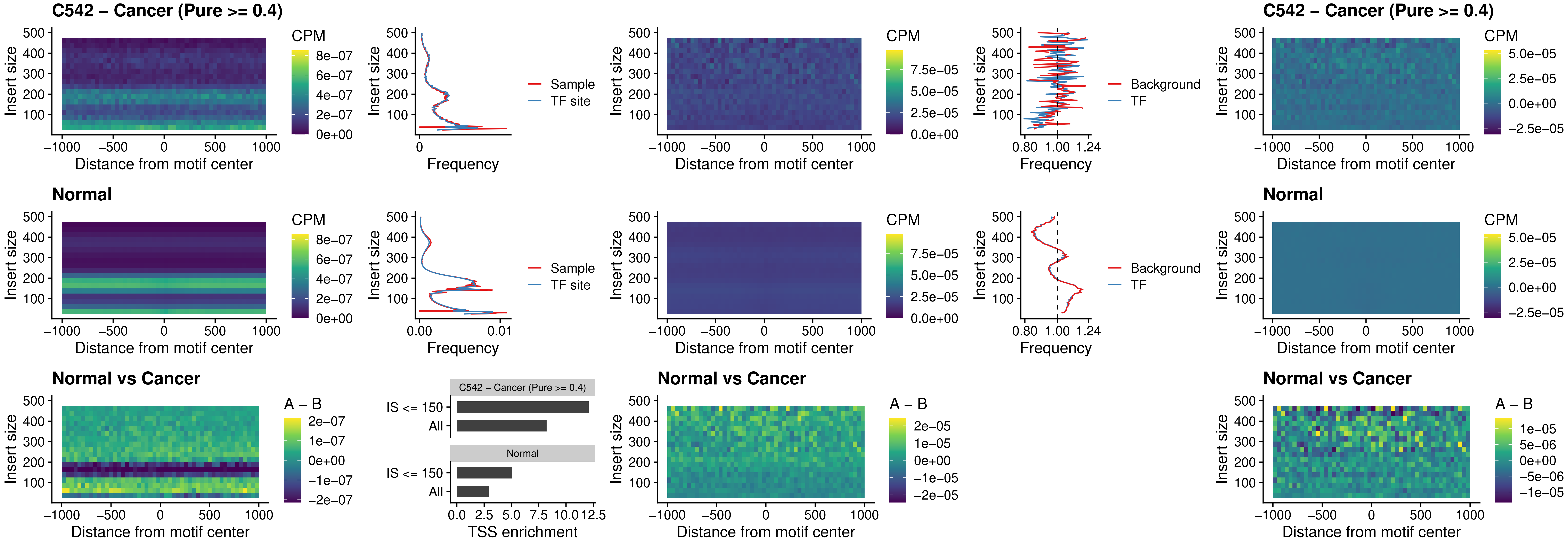

#### C542 – Cancer (Pure >= 0.4) – FOXD4L4_uc_cTSS_nPEAK

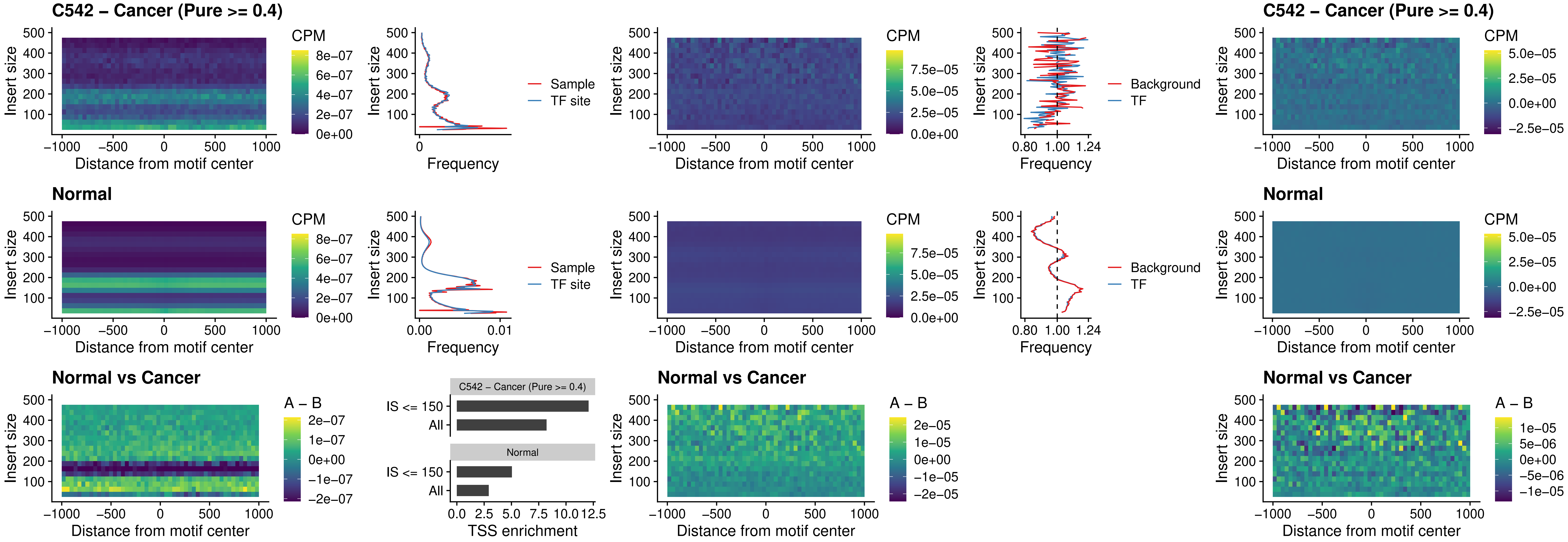

#### C542 – Cancer (Pure >= 0.4) – FOXD4L2_uc_cTSS_nPEAK

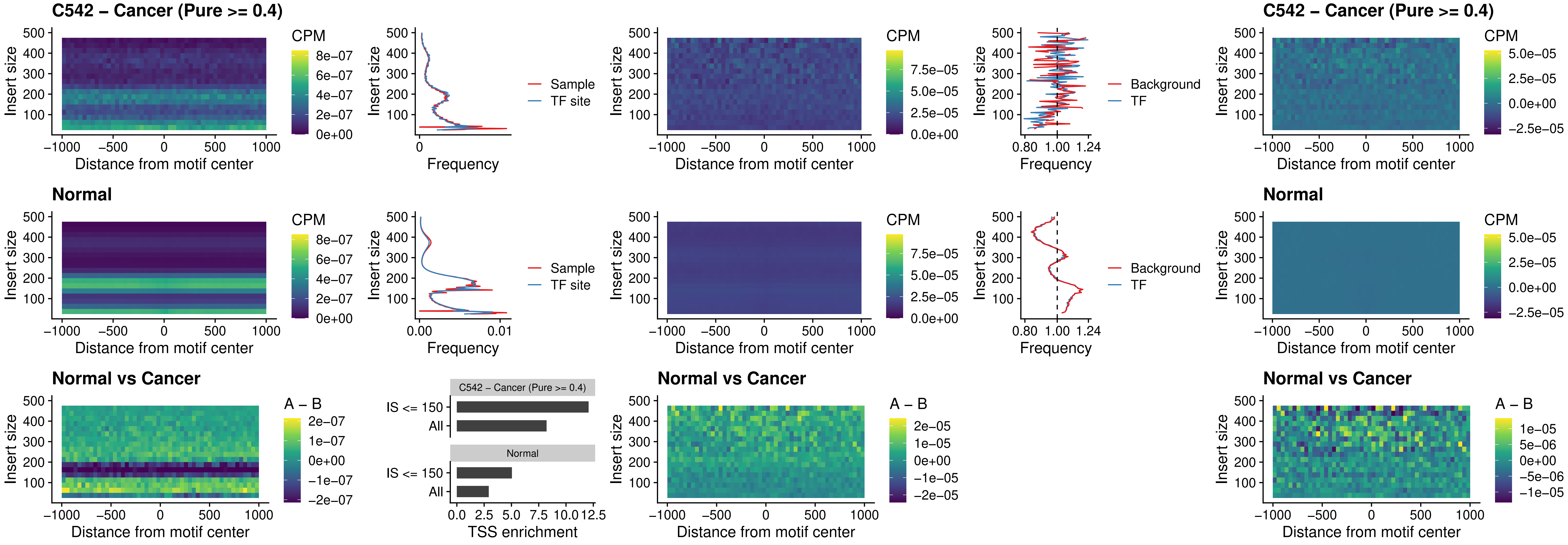

#### C542 – Cancer (Pure >= 0.4) – FOXD4L3_uc_cTSS_nPEAK

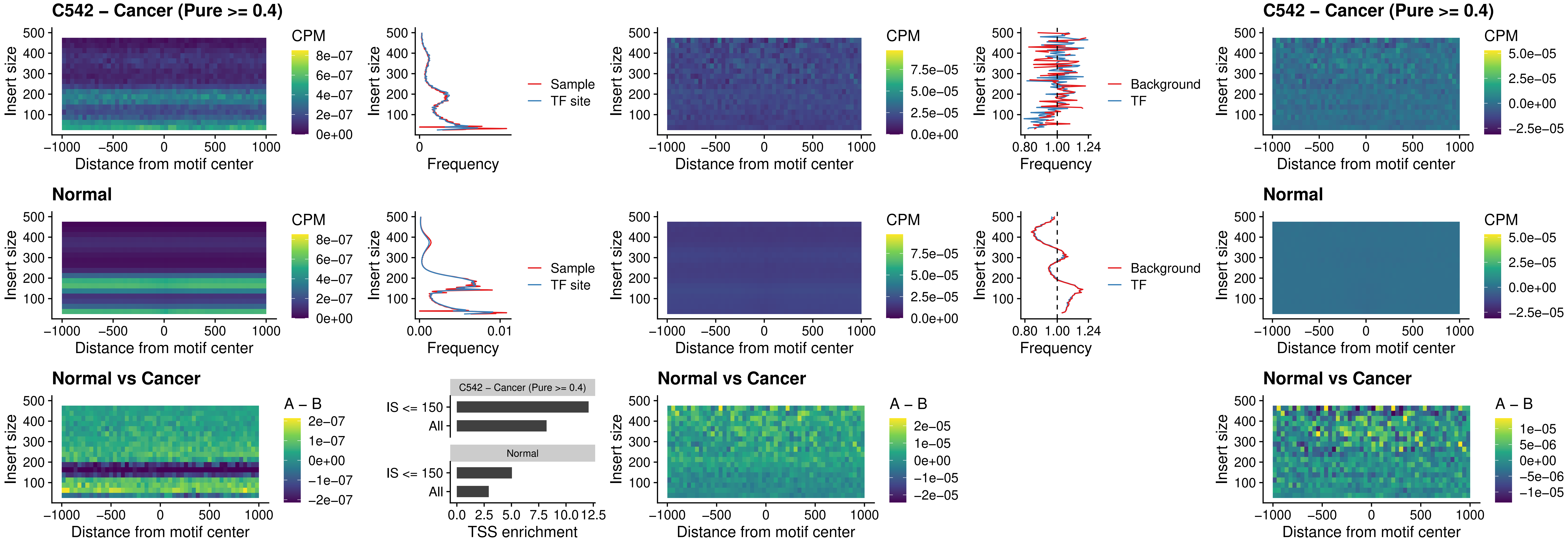

#### C542 – Cancer (Pure >= 0.4) – HOXD1_uc_dTSS_oPEAK

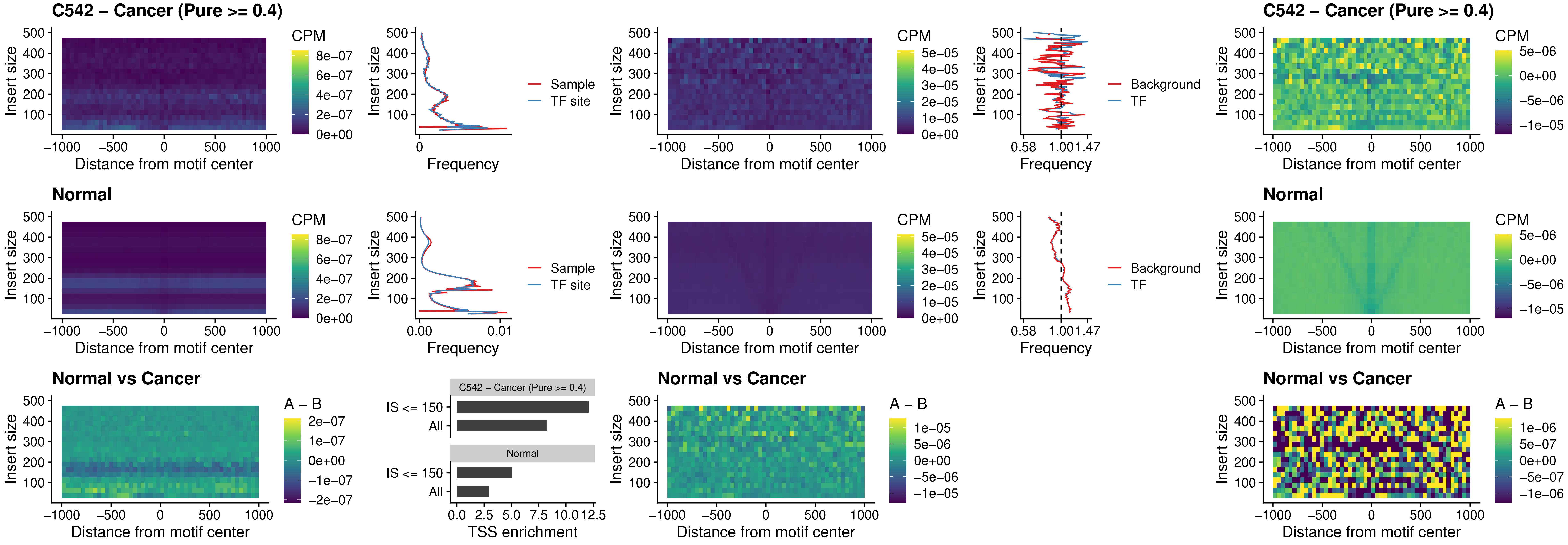

### C542 – Cancer (Pure >= 0.4) – VAX1_uc_pTSS_nPEAK

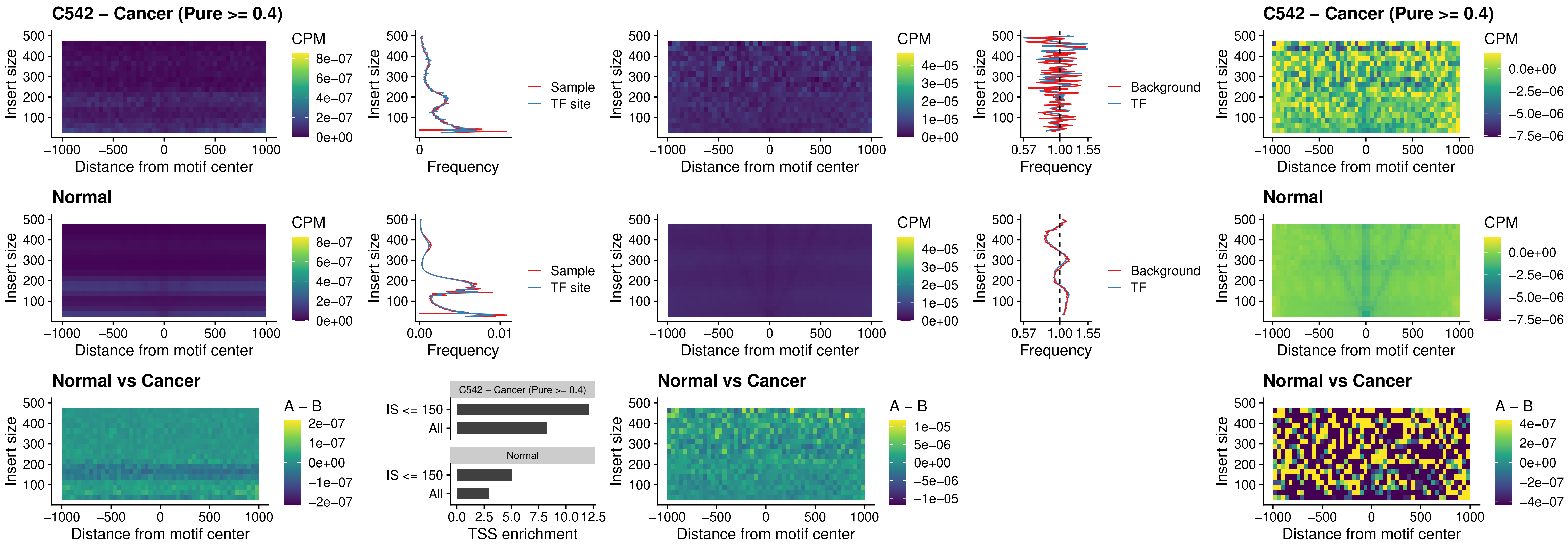

#### C542 – Cancer (Pure >= 0.4) – DLX3_uc_pTSS_nPEAK

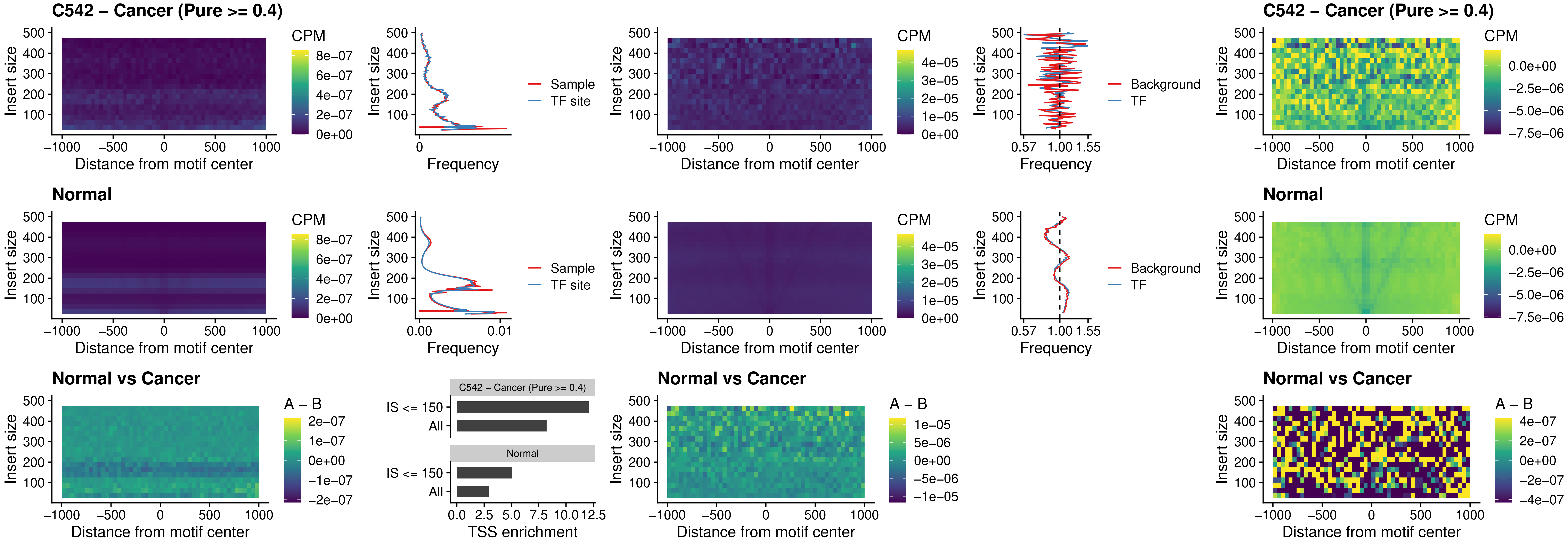

#### C542 – Cancer (Pure >= 0.4) – HOXD3_uc_pTSS_nPEAK

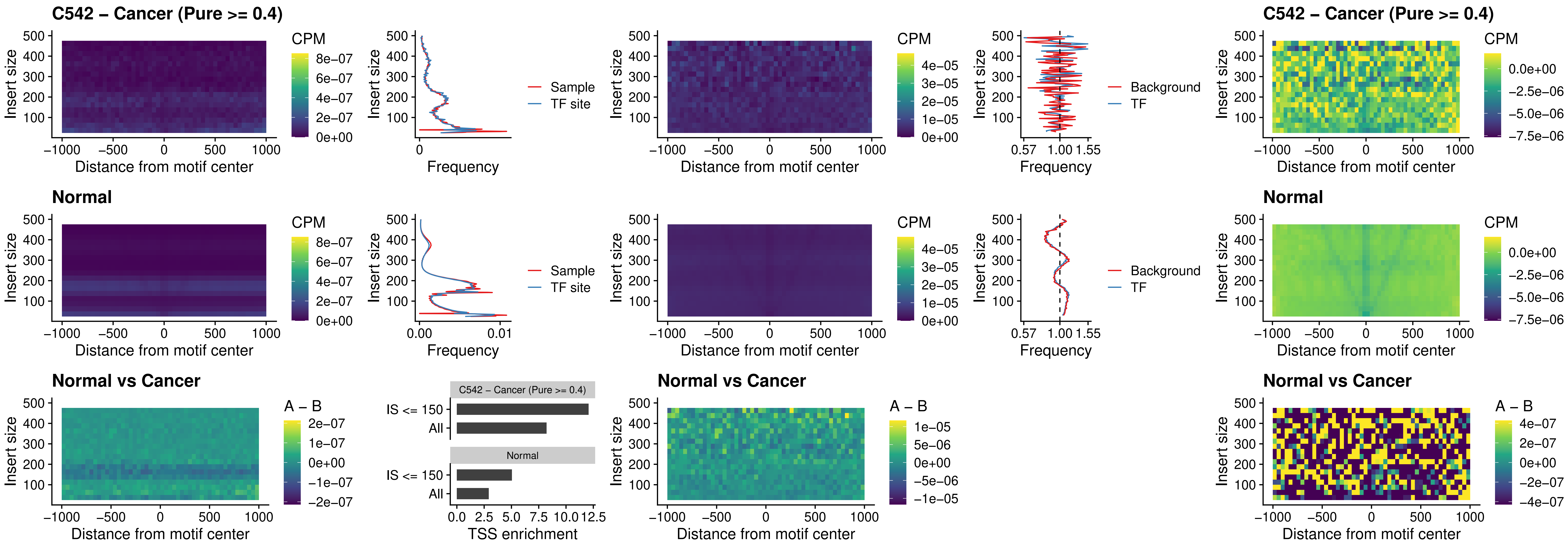

#### C542 – Cancer (Pure >= 0.4) – PDX1_uc_dTSS_nPEAK

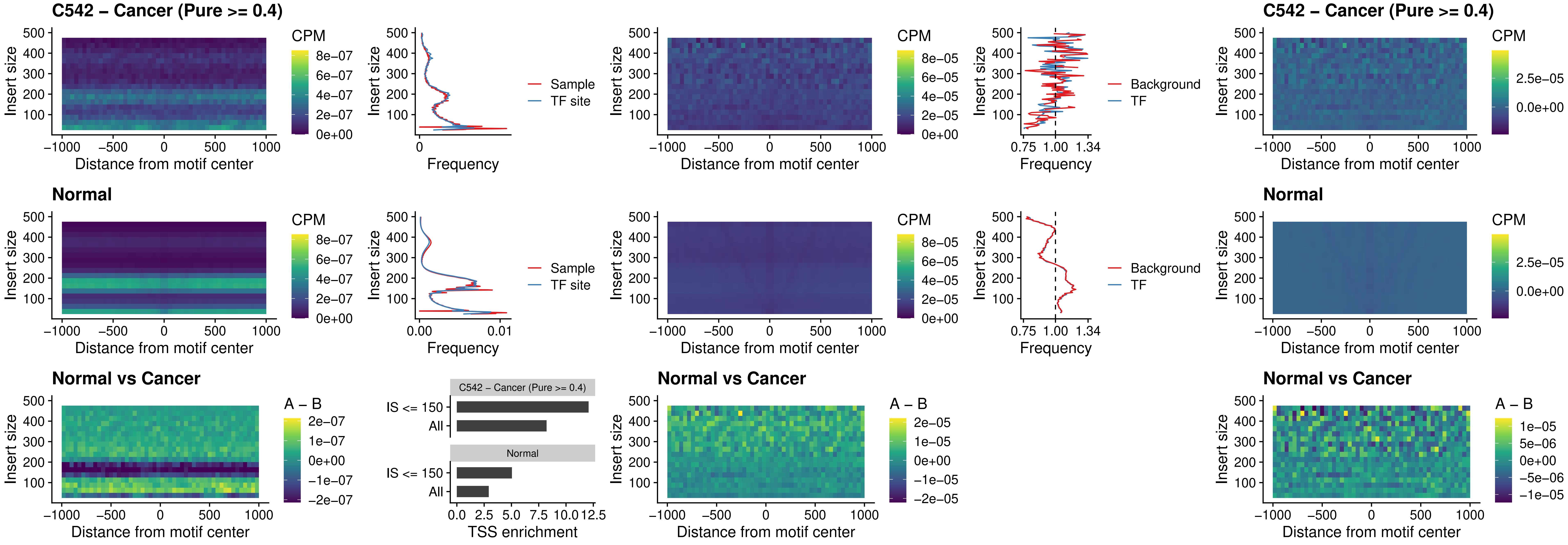

#### C542 – Cancer (Pure >= 0.4) – VAX2_uc_dTSS_nPEAK

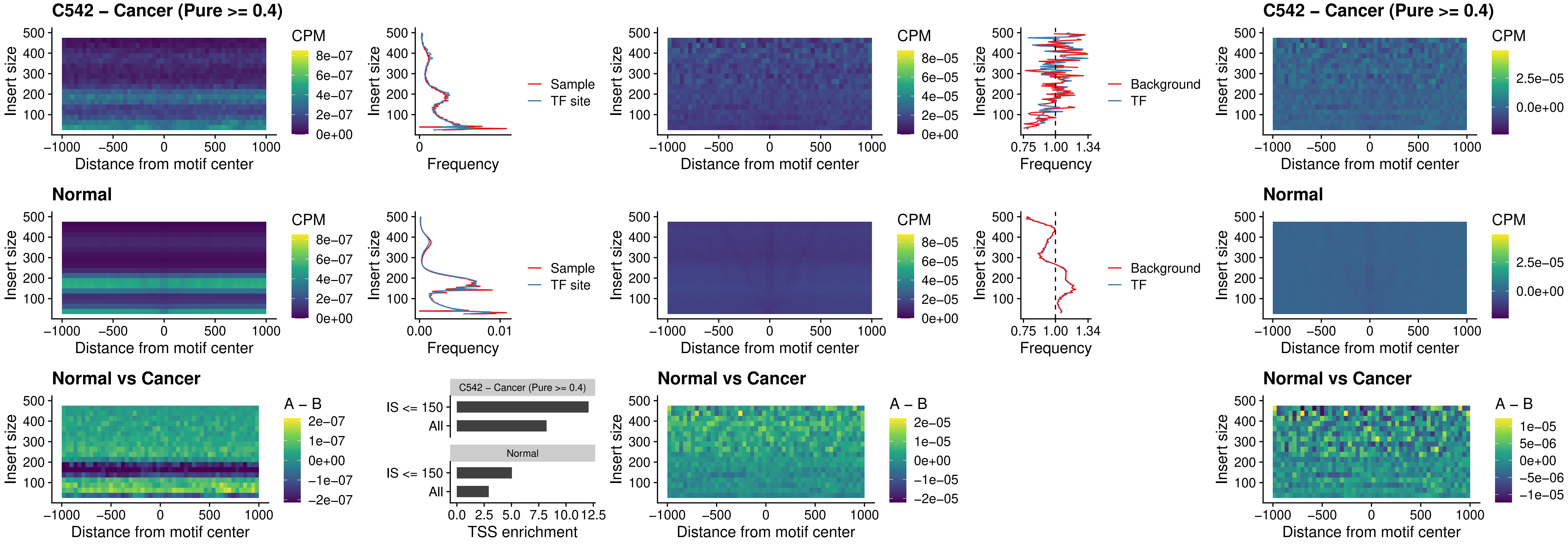

#### C542 – Cancer (Pure >= 0.4) – SOX5_uc_dTSS_nPEAK

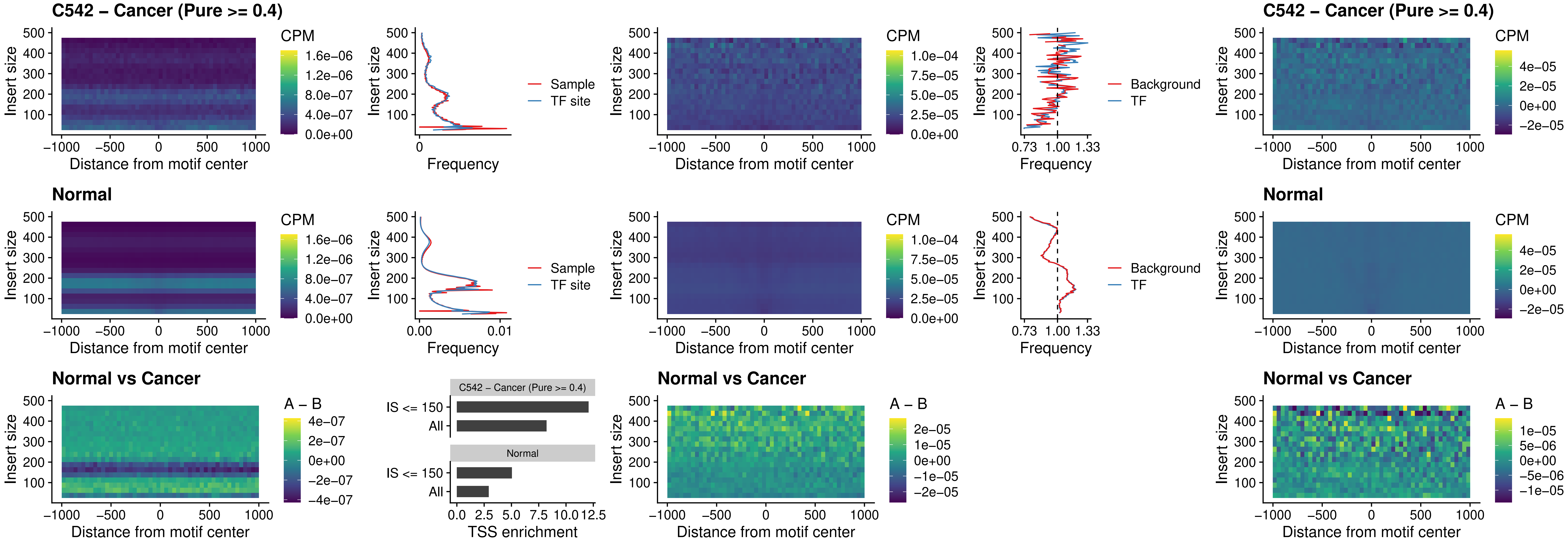

#### C542 – Cancer (Pure >= 0.4) – SOX6_uc_dTSS_nPEAK

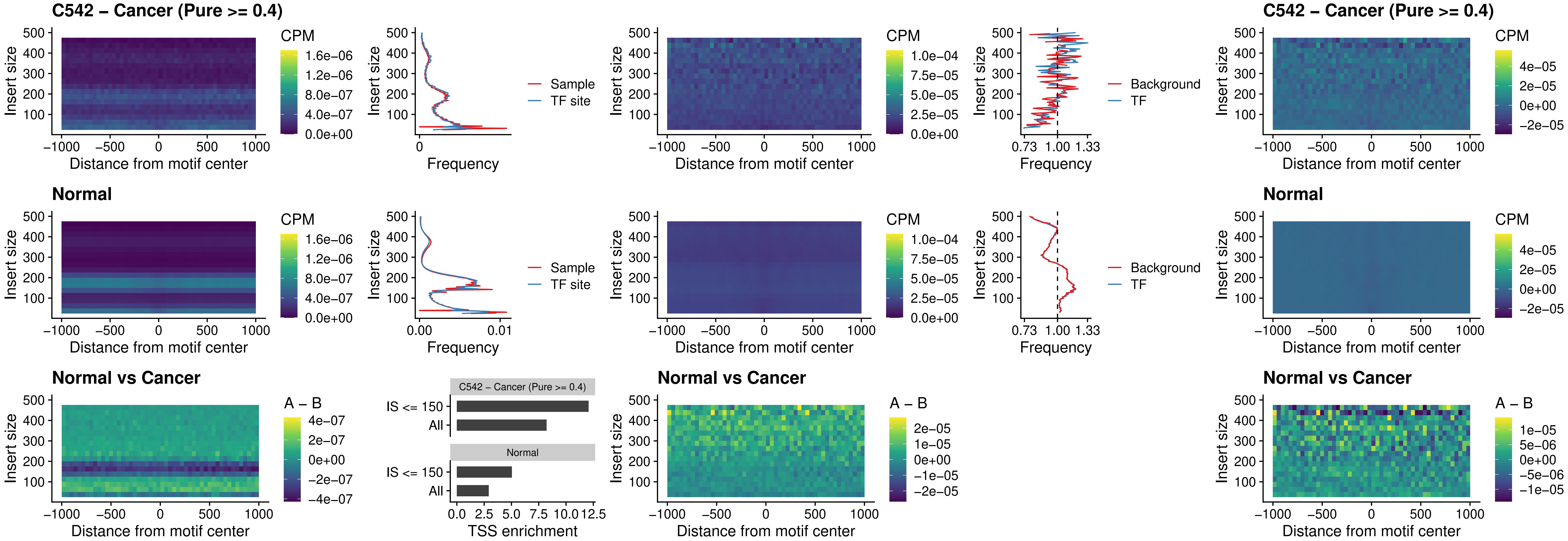

#### C542 – Cancer (Pure >= 0.4) – JDP2_uc_cTSS_nPEAK

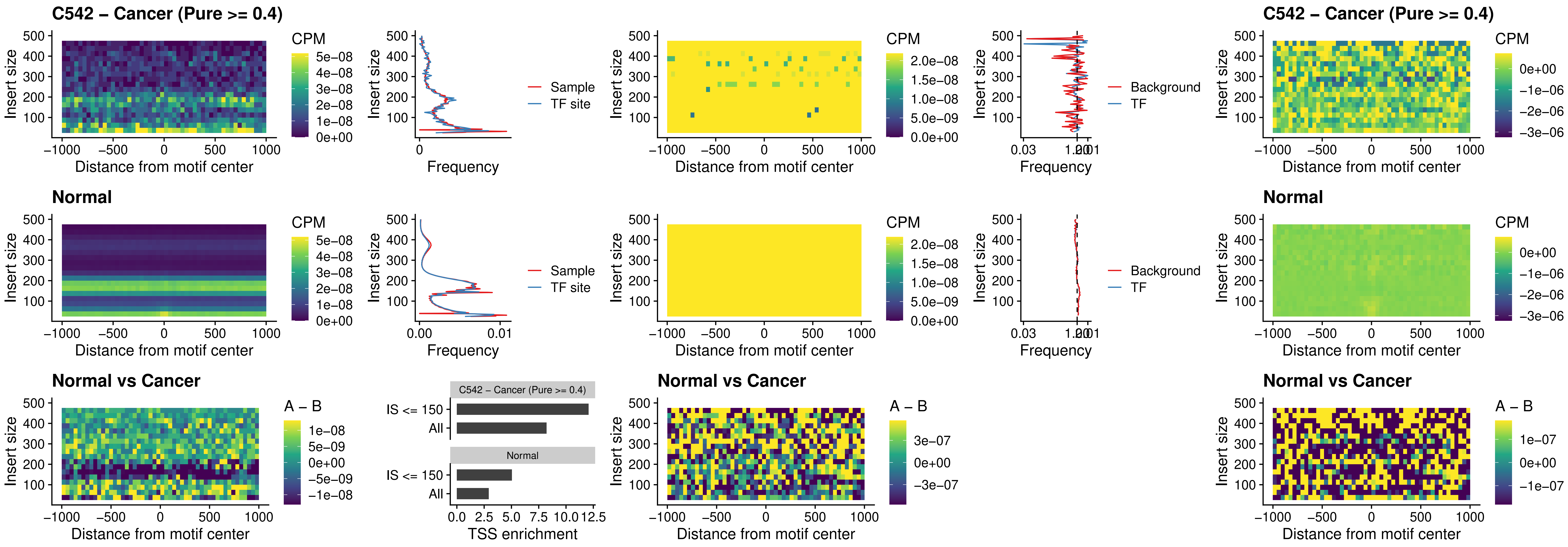

#### C542 – Cancer (Pure >= 0.4) – PURA_uc_dTSS_oPEAK

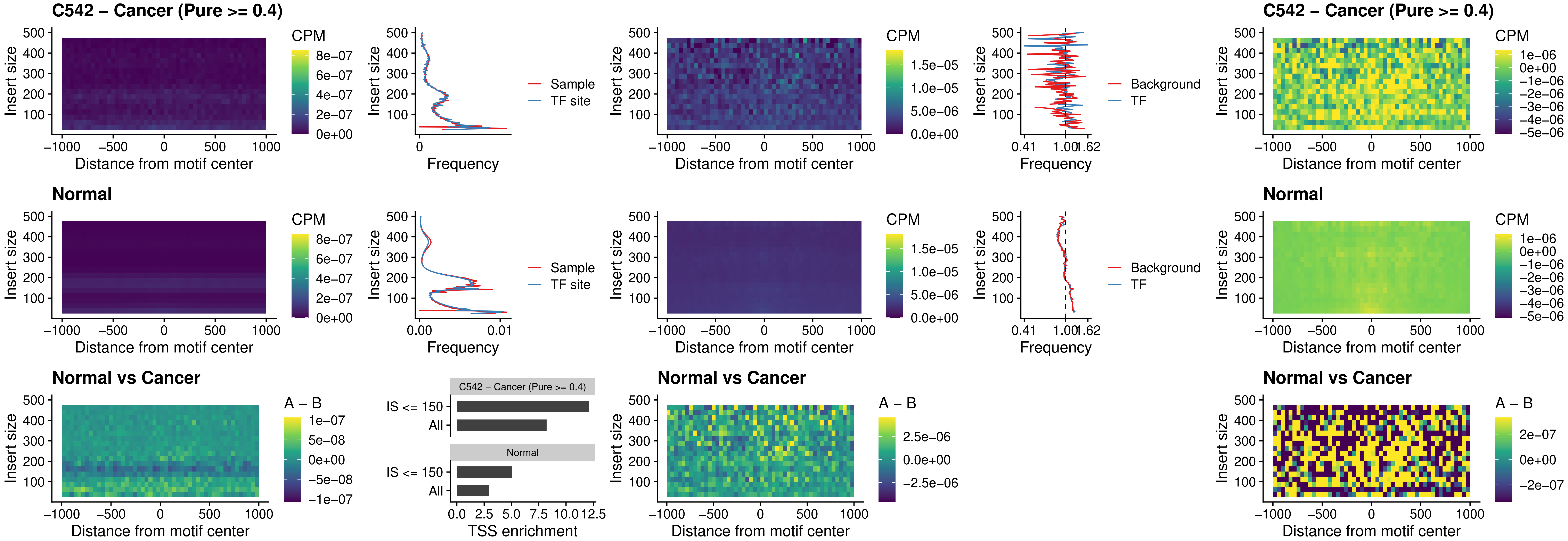

### C542 – Cancer (Pure >= 0.4) – HHEX_uc_dTSS_nPEAK

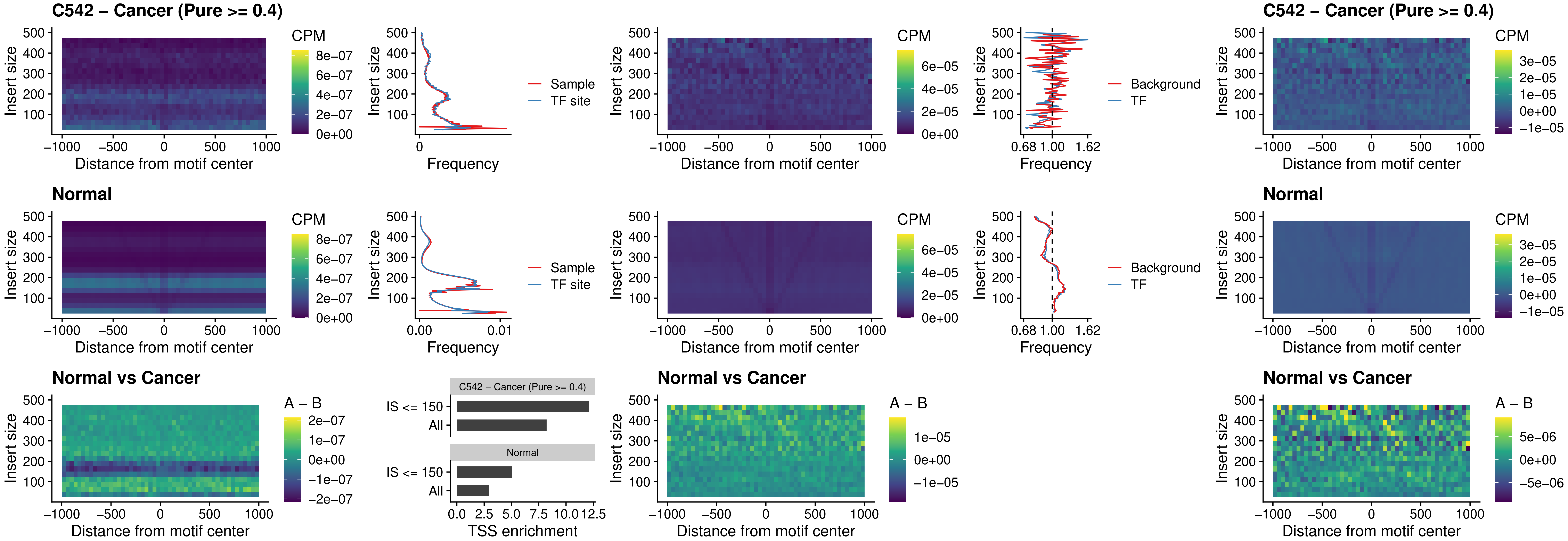

#### C542 – Cancer (Pure >= 0.4) – HMG20B_uc_dTSS_nPEAK

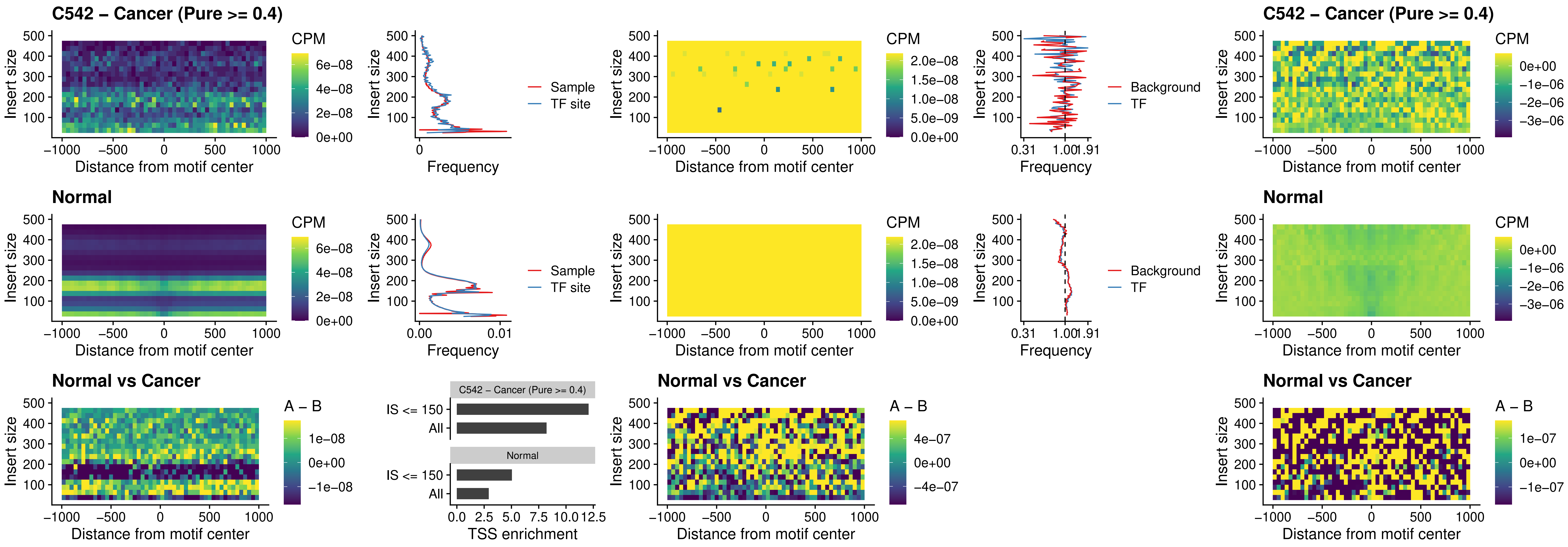

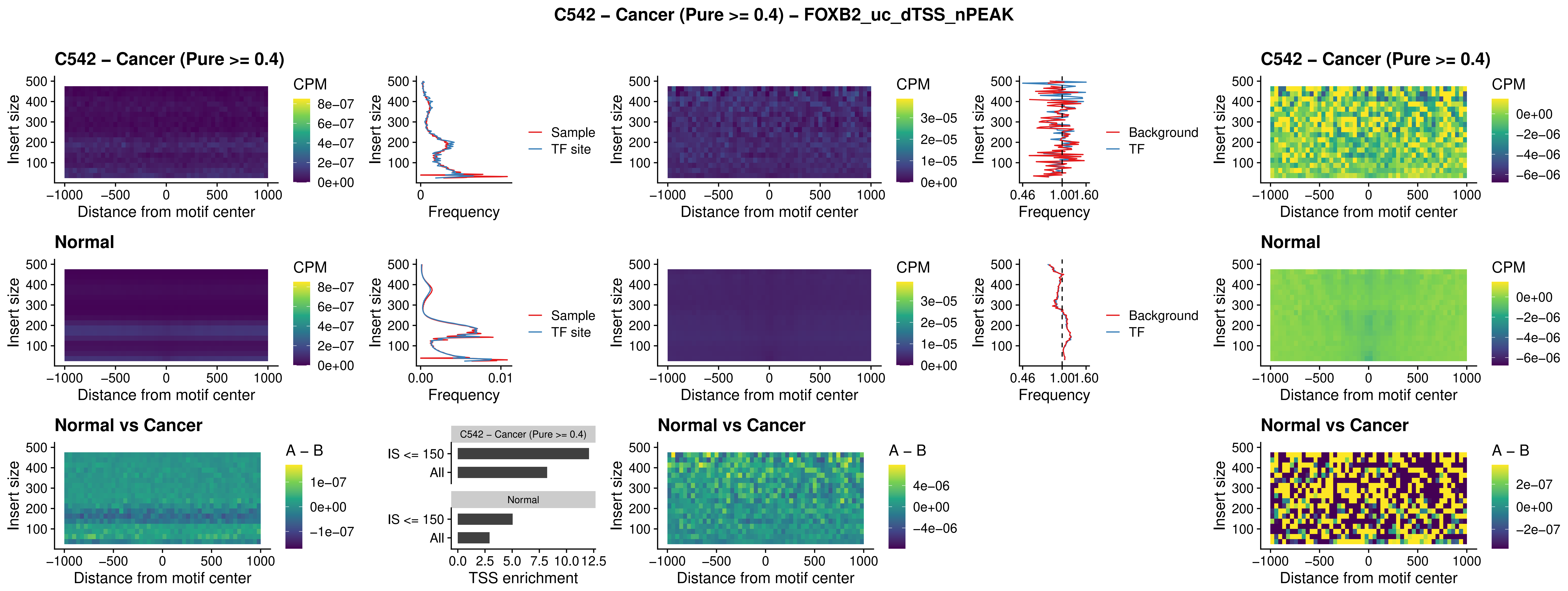
C542 – Cancer (Pure >= 0.4) – FOXB2_uc_dTSS_nPEAK
C542 – Cancer (Pure >= 0.4)
Normal
Normal vs Cancer
Insert size
Distance from motif center
Frequency
CPM
A – B
Sample
TF site
Background
TF
C542 – Cancer (Pure >= 0.4)
Normal
IS <= 150
All
TSS enrichment

#### C542 – Cancer (Pure >= 0.4) – FOXL1_uc_cTSS_nPEAK

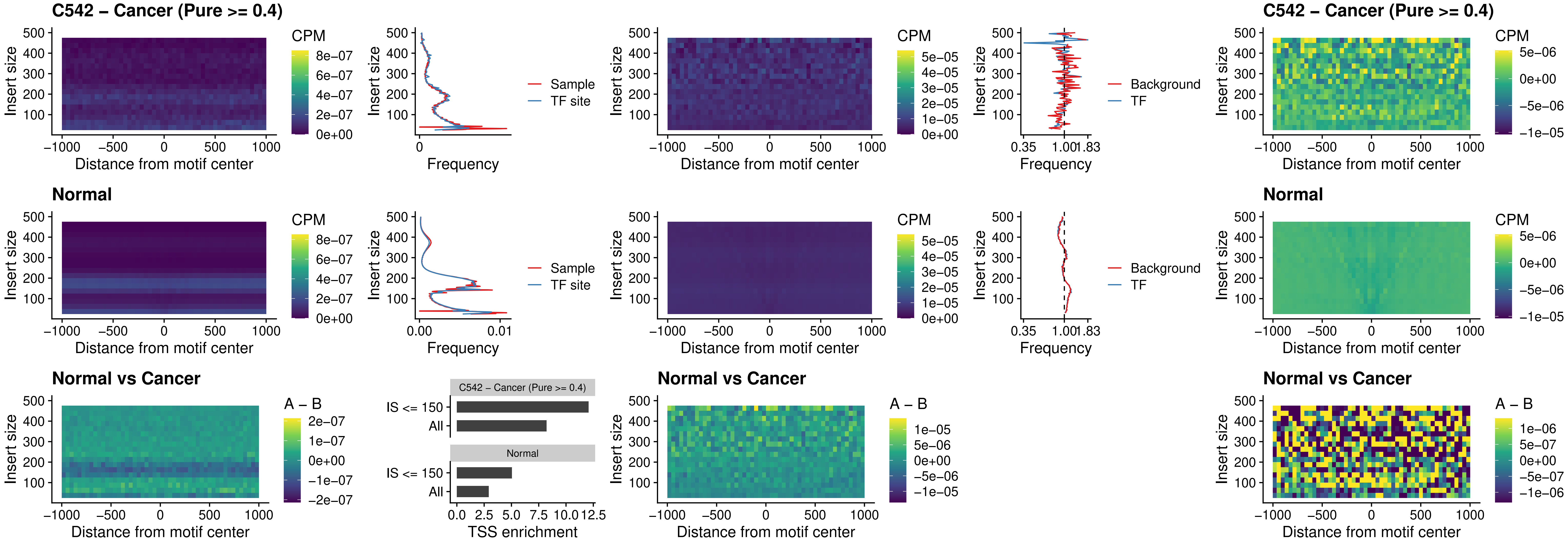

### C542 – Cancer (Pure >= 0.4) – PBX3_uc_pTSS_oPEAK

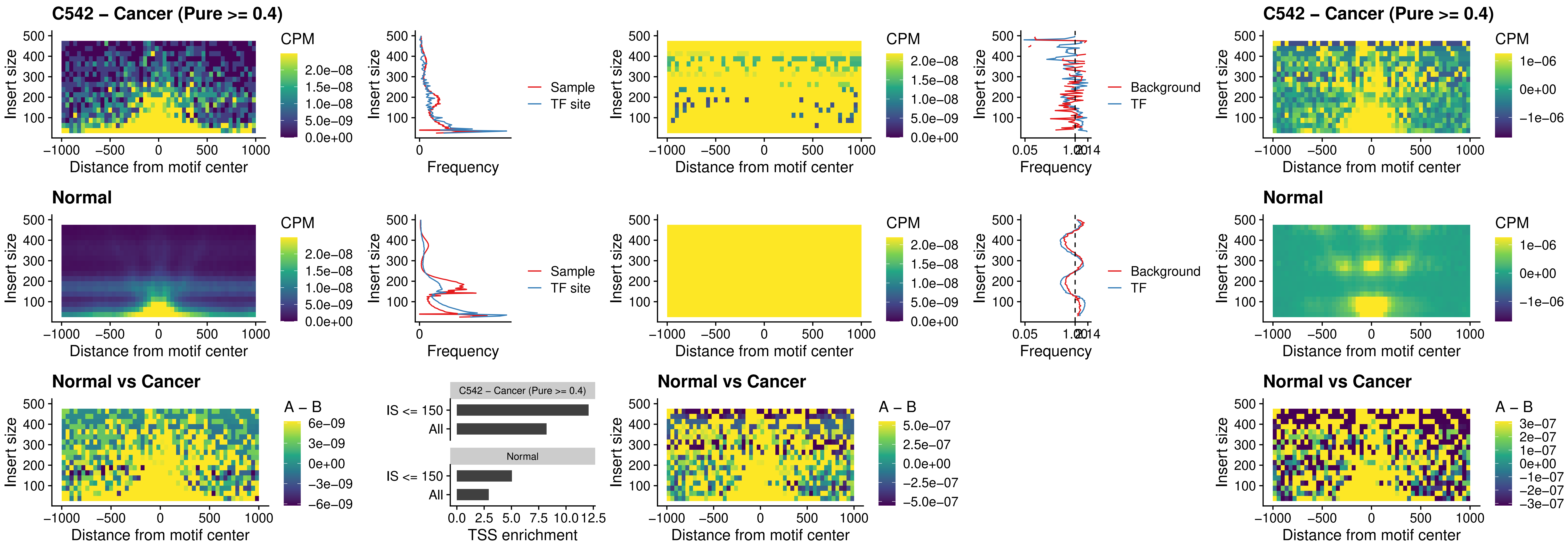

#### C542 – Cancer (Pure >= 0.4) – SNAI2_uc_cTSS_nPEAK

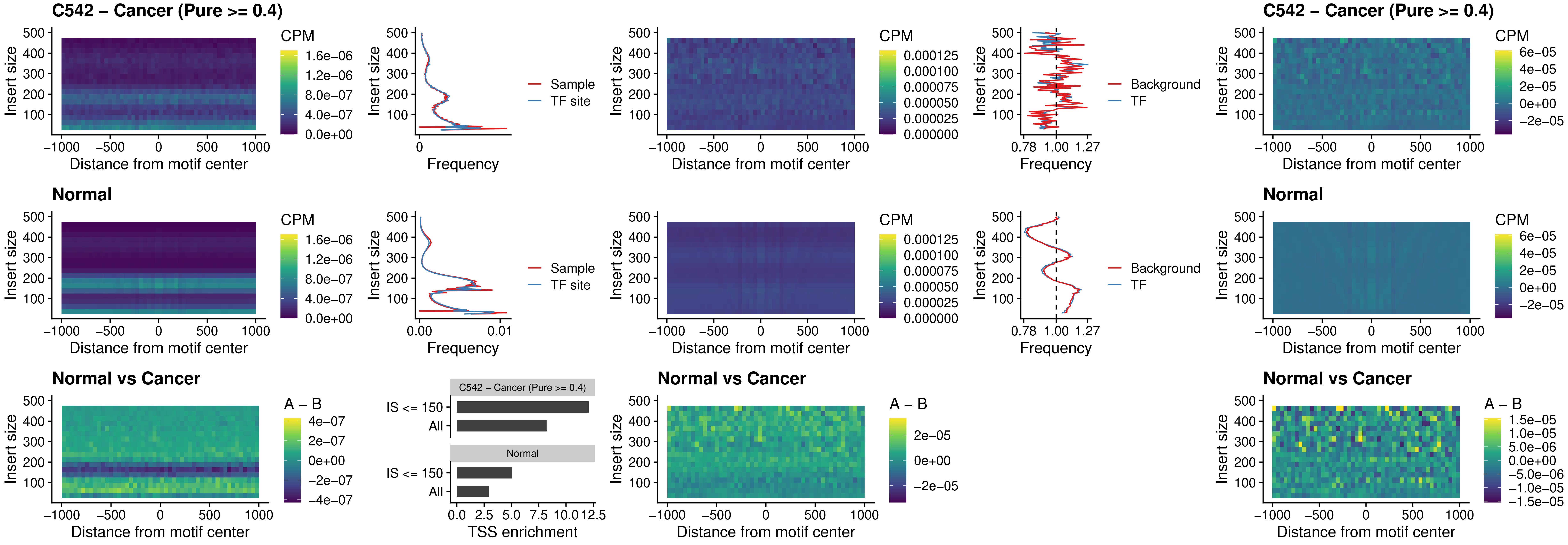

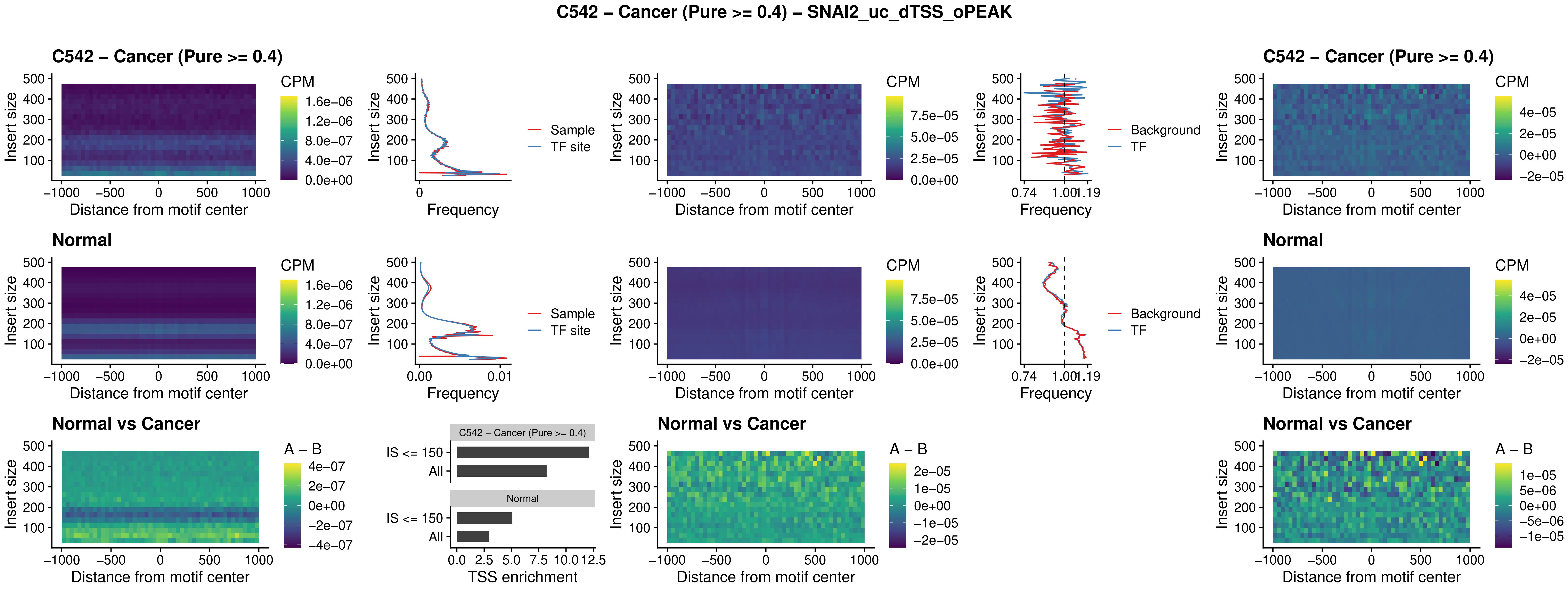
C542 – Cancer (Pure >= 0.4) – SNAI2_uc_dTSS_oPEAK
C542 – Cancer (Pure >= 0.4)
Normal
Normal vs Cancer
Insert size
Distance from motif center
Frequency
CPM
A – B
Sample
TF site
Background
TF
C542 – Cancer (Pure >= 0.4)
Normal
IS <= 150
All
TSS enrichment

#### C542 – Cancer (Pure >= 0.4) – SPI1_uc_pTSS_nPEAK

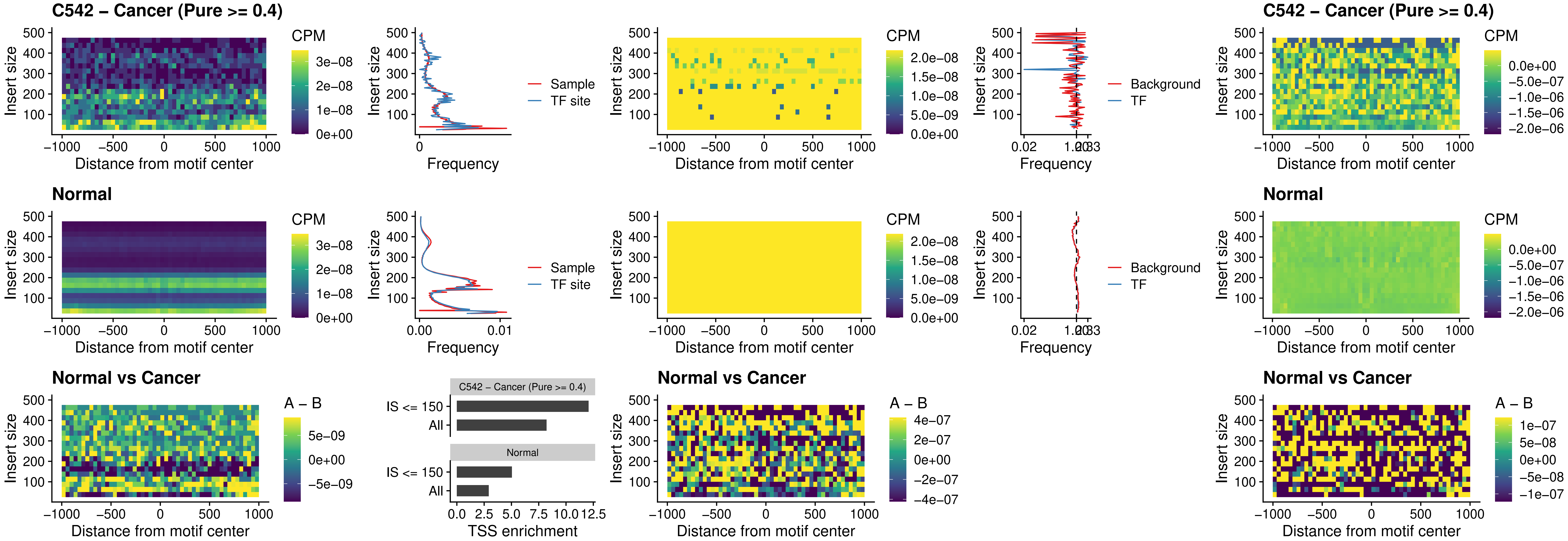

#### C542 – Cancer (Pure >= 0.4) – KLF16_uc_pTSS_nPEAK

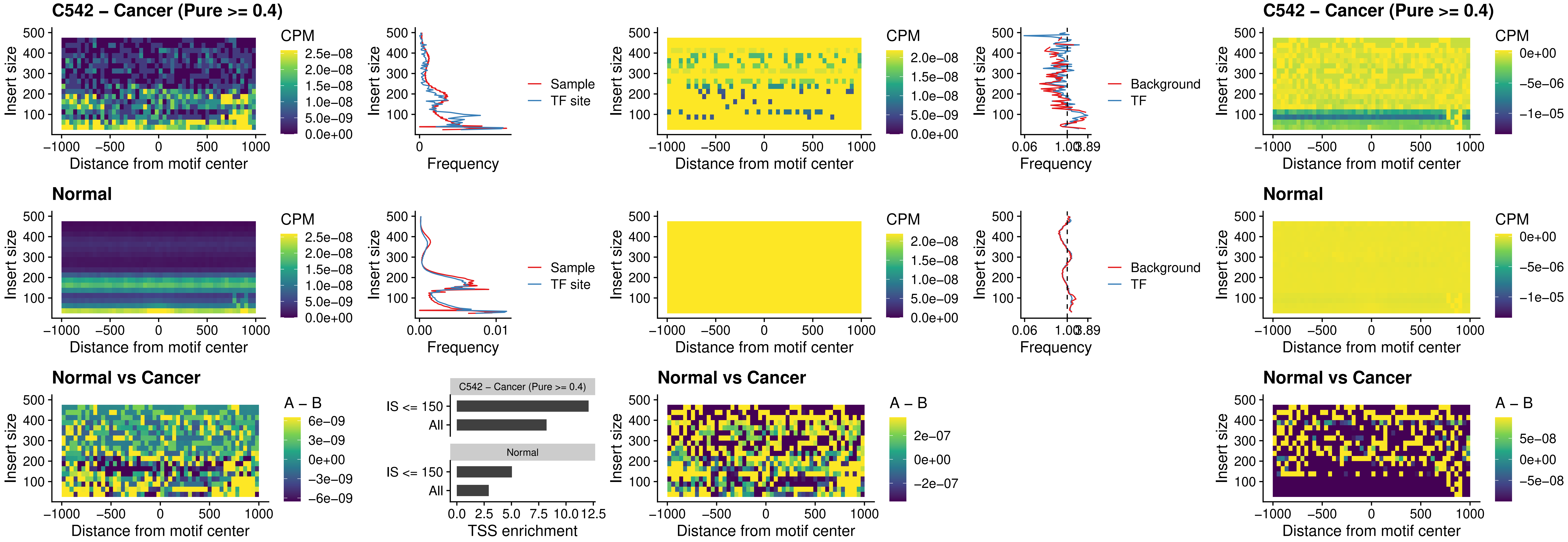

#### C542 – Cancer (Pure >= 0.4) – SP3_uc_pTSS_nPEAK

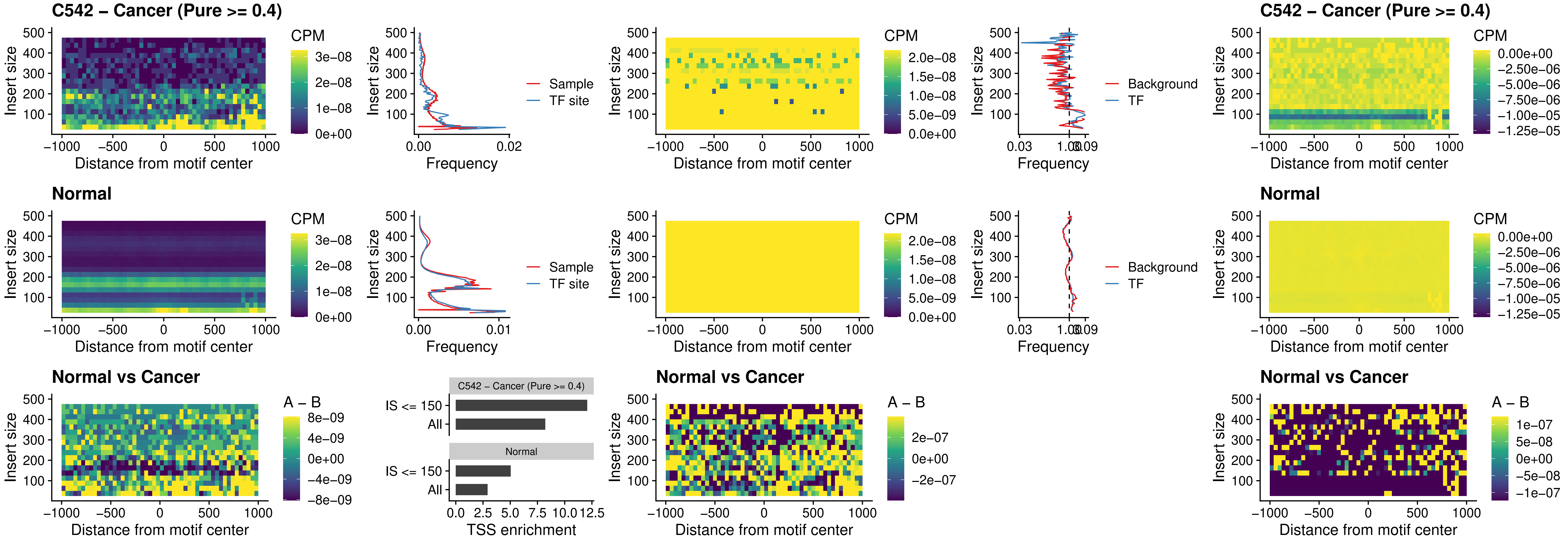

### C542 – Cancer (Pure >= 0.4) – NEURODI_uc_dTSS_oPEAK

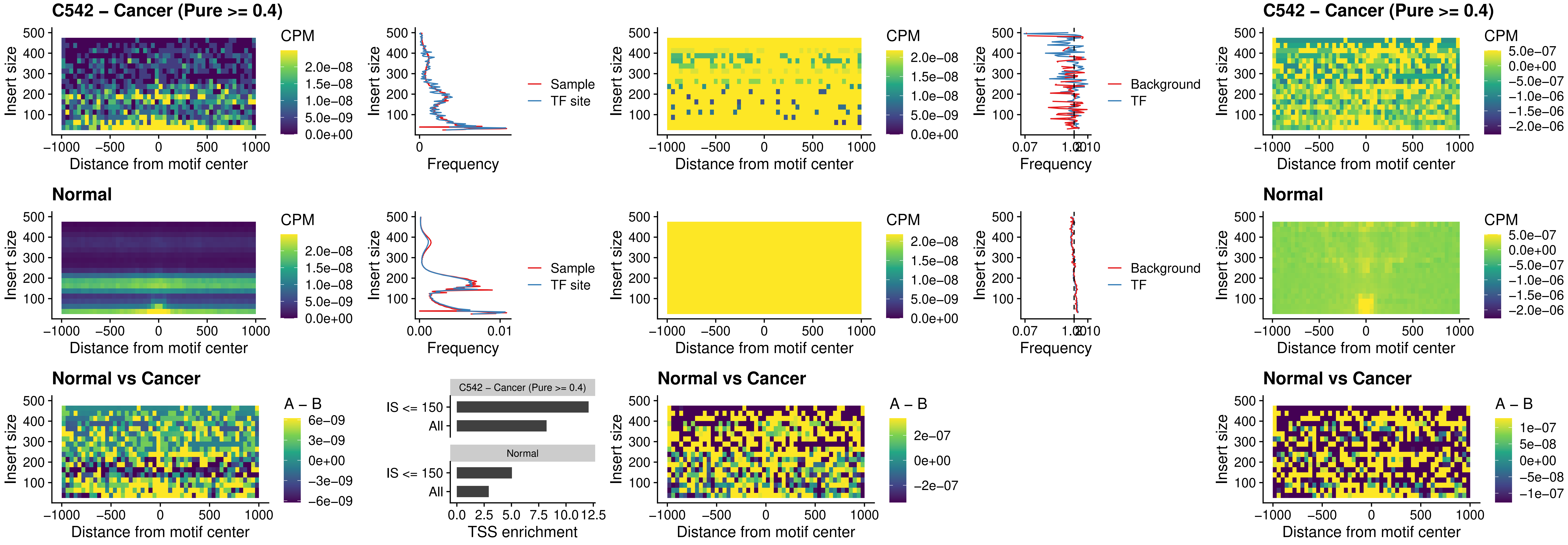

#### C542 – Cancer (Pure >= 0.4) – CTCF_uc_dTSS_oPEAK

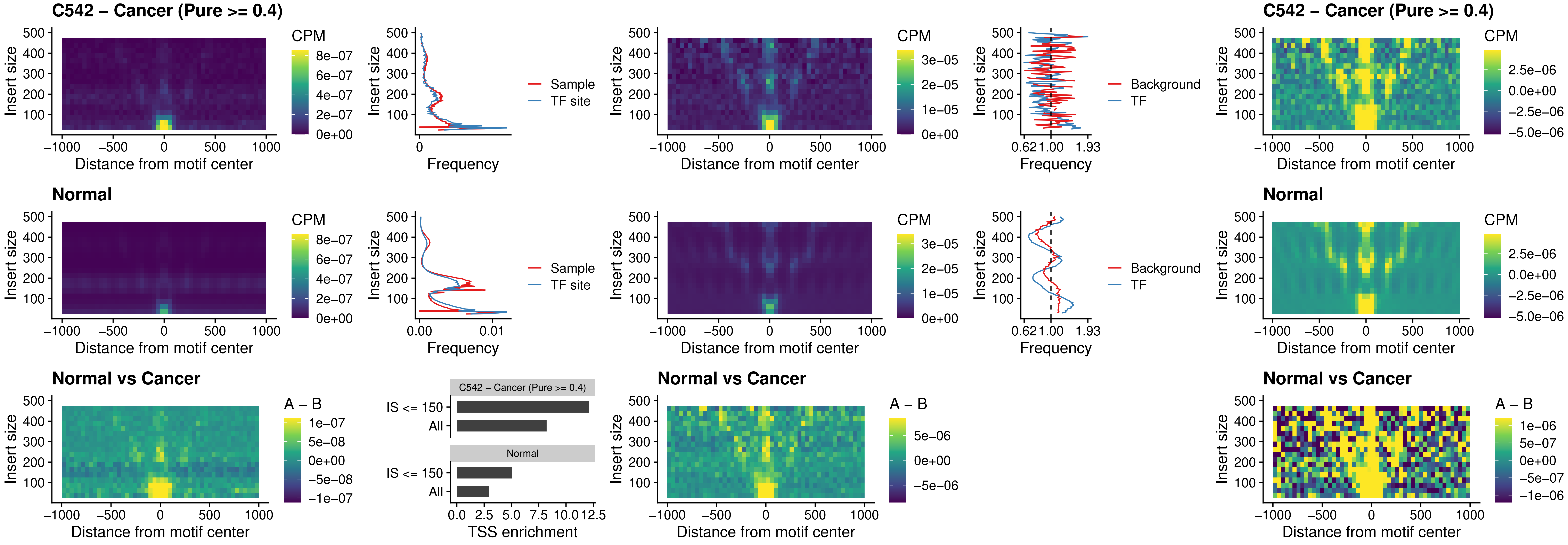

#### C542 – Cancer (Pure >= 0.4) – CTCF_uc_cTSS_oPEAK

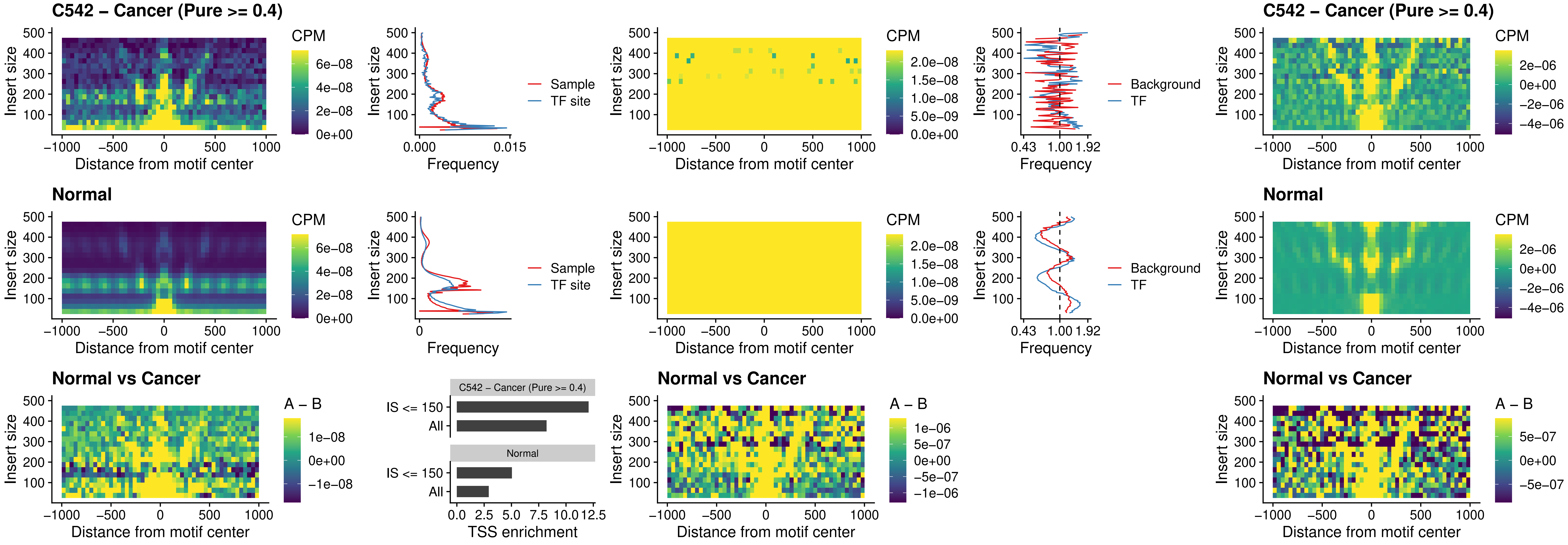

### C542 – Cancer (Pure >= 0.4) – CTCFL_uc_dTSS_oPEAK

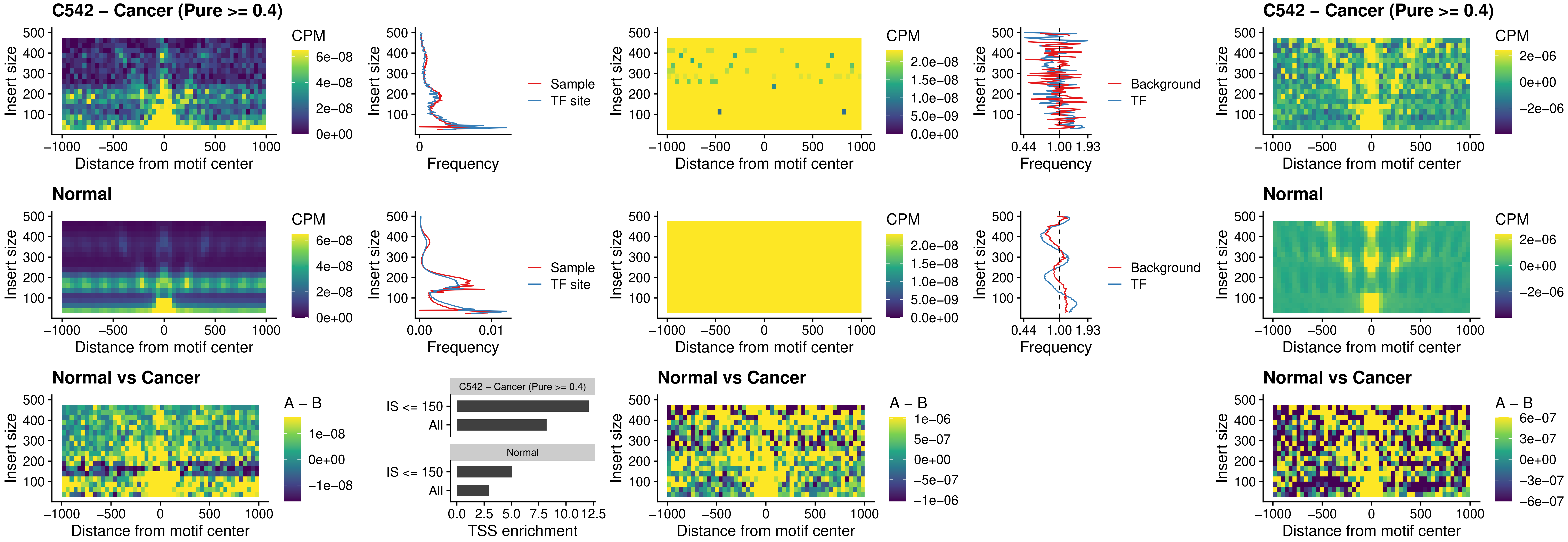

#### C542 – Cancer (Pure >= 0.4) – ENCODE_CTCF_sites_COLON

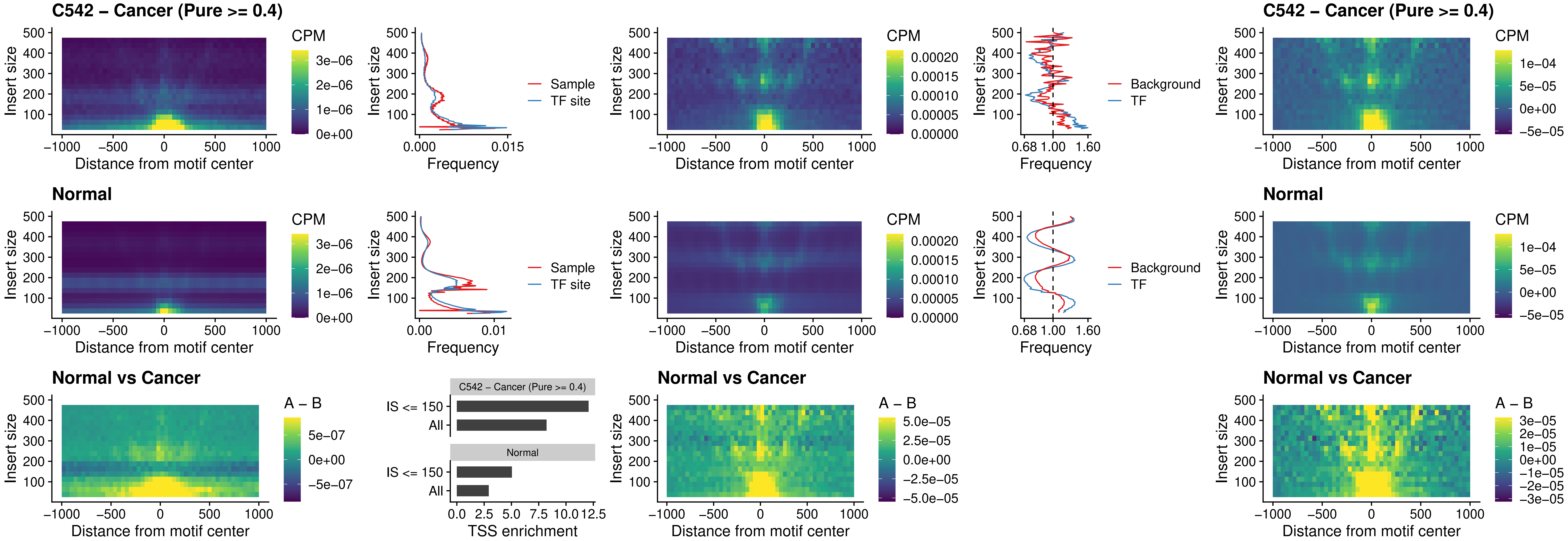

#### C542 – Cancer (Pure >= 0.4) – ENCODE_CTCF_sites_COLON_loop_ctcf

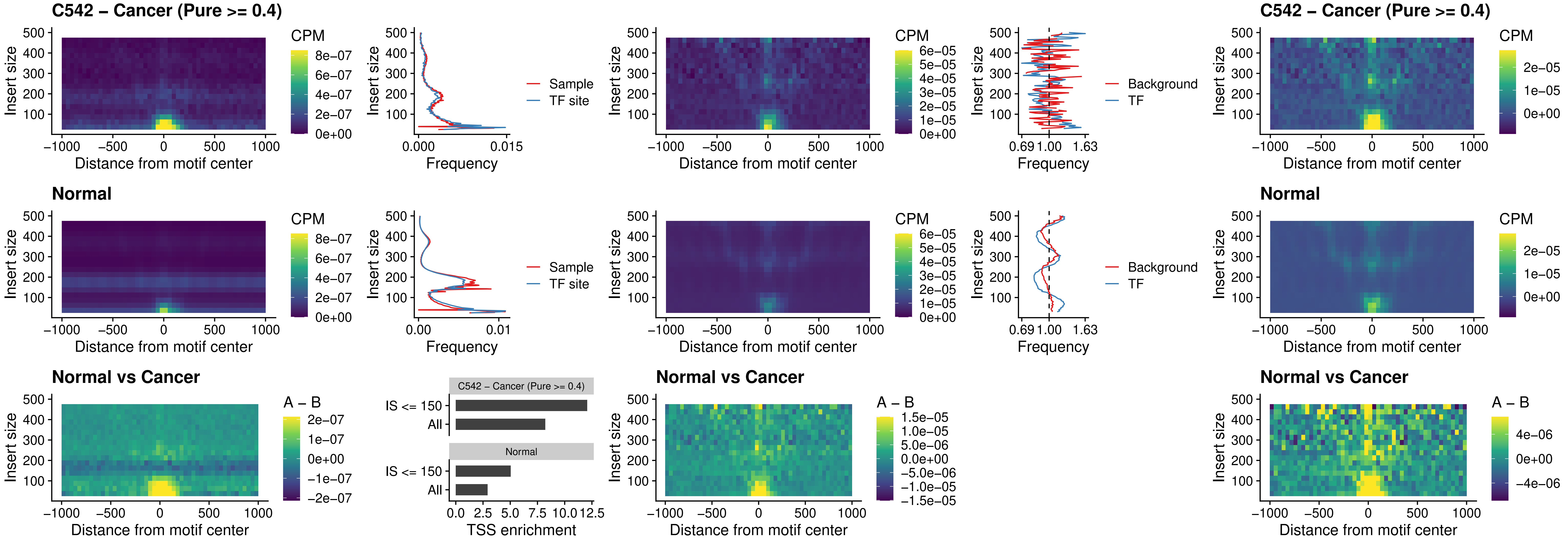

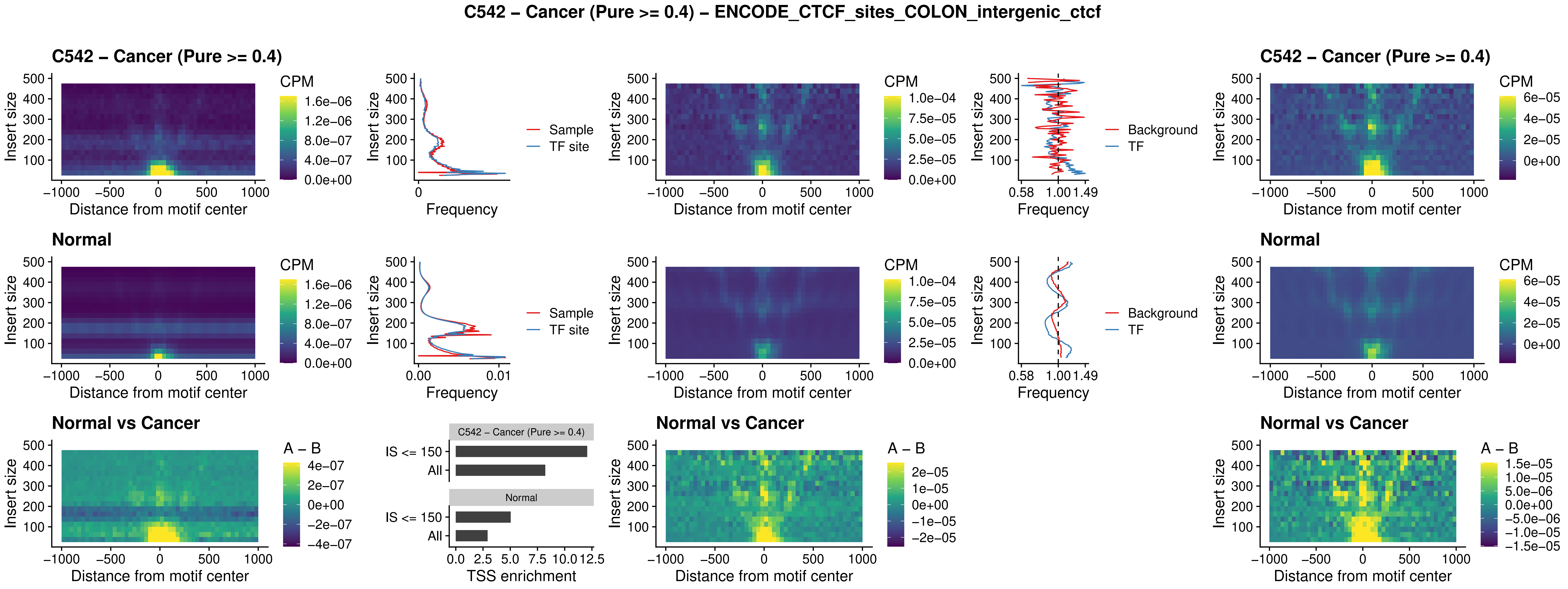

C542 – Cancer (Pure >= 0.4) – ENCODE_CTCF_sites_COLON_intergenic_ctcf
C542 – Cancer (Pure >= 0.4)
Normal
Normal vs Cancer
Insert size
Distance from motif center
Frequency
CPM
A – B
Sample
TF site
Background
TF
C542 – Cancer (Pure >= 0.4)
Normal
IS <= 150
All
TSS enrichment

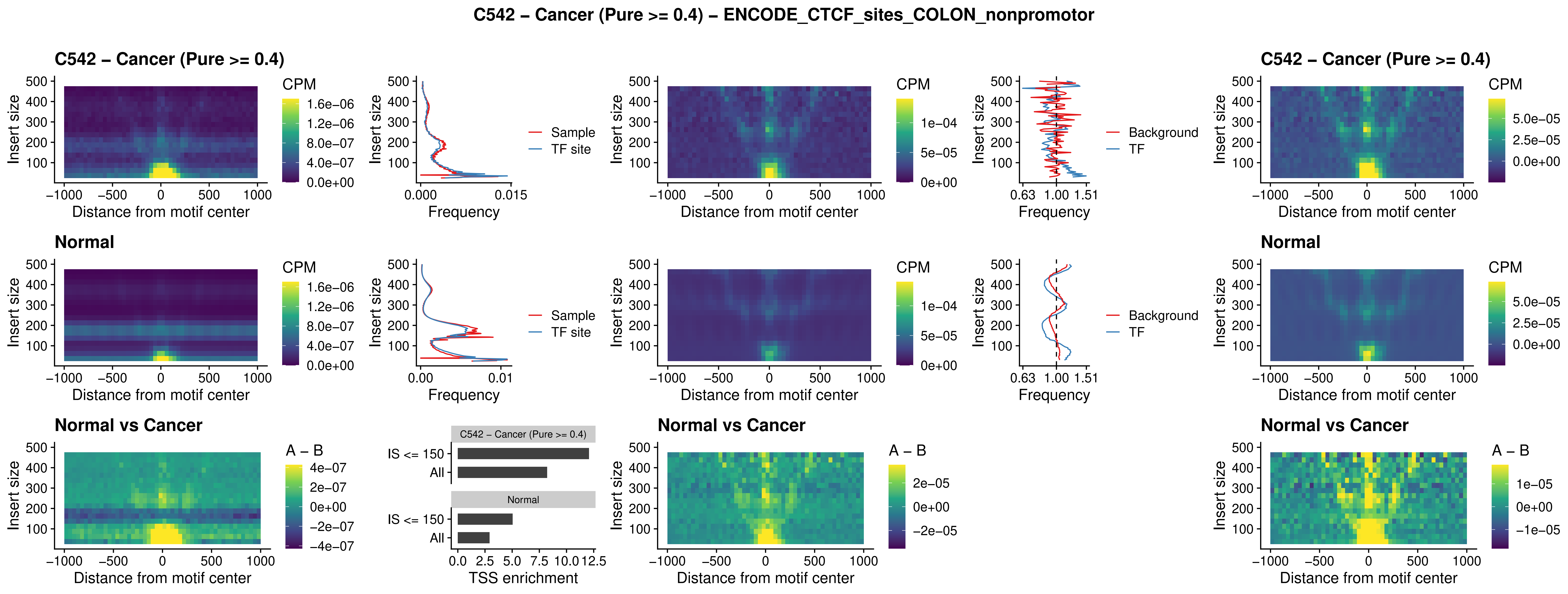
C542 – Cancer (Pure >= 0.4) – ENCODE_CTCF_sites_COLON_nonpromotor
C542 – Cancer (Pure >= 0.4)
Normal
Normal vs Cancer
Insert size
Distance from motif center
Frequency
CPM
A – B
Sample
TF site
Background
TF
C542 – Cancer (Pure >= 0.4)
Normal
IS <= 150
All
TSS enrichment

#### C542 – Cancer (Pure >= 0.4) – CTCFL_uc_cTSS_oPEAK

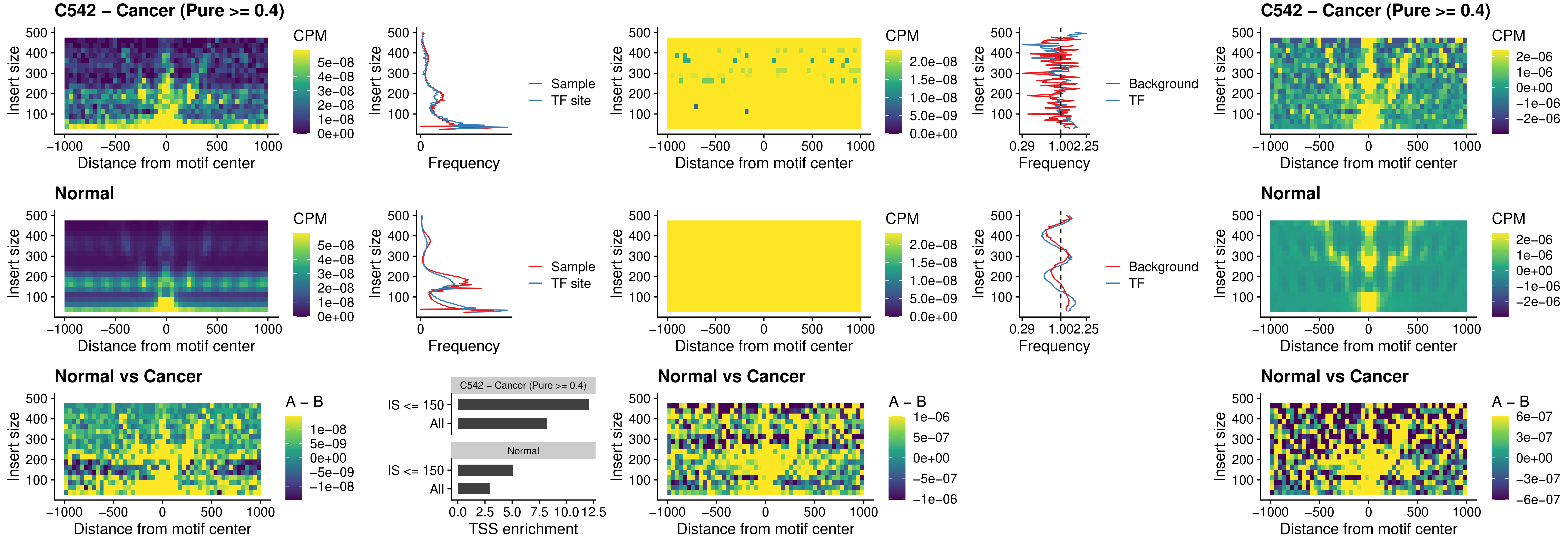

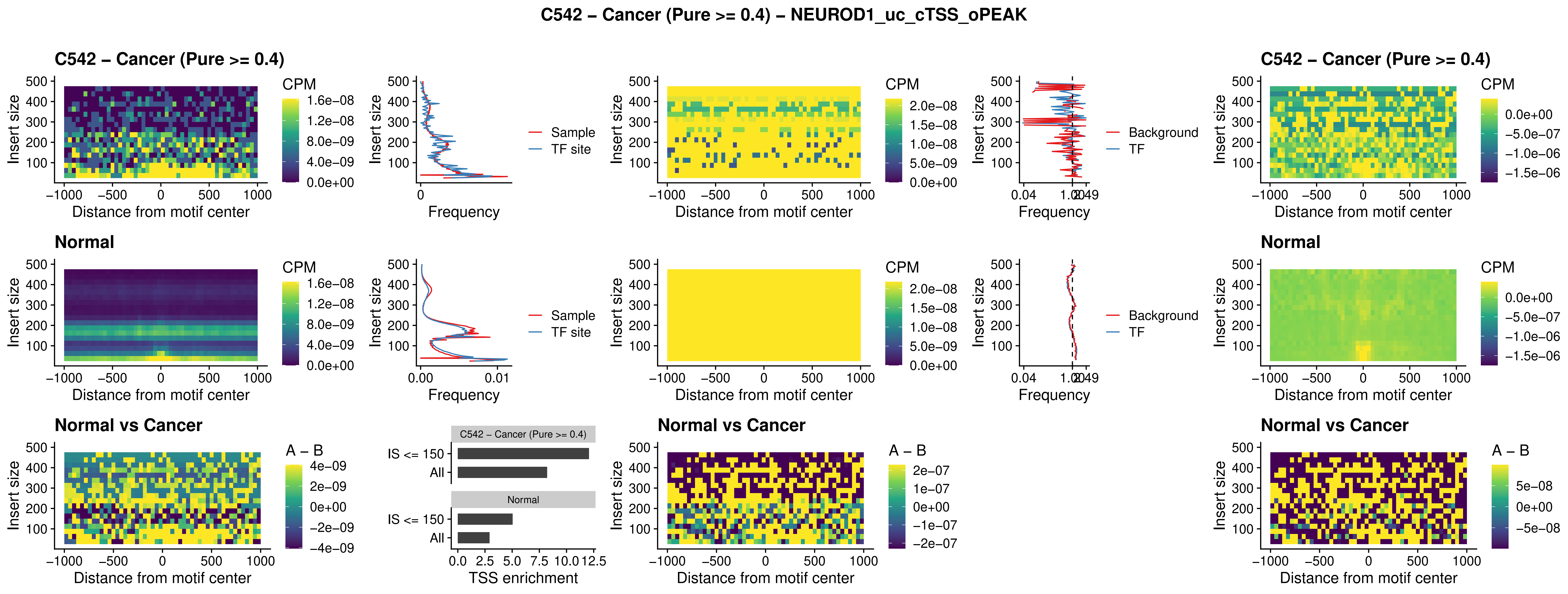
C542 – Cancer (Pure >= 0.4) – NEUROD1_uc_cTSS_oPEAK
C542 – Cancer (Pure >= 0.4)
Normal
Normal vs Cancer
Insert size
Distance from motif center
Frequency
CPM
A – B
Sample
TF site
Background
TF
C542 – Cancer (Pure >= 0.4)
Normal
IS <= 150
All
TSS enrichment

#### C542 – Cancer (Pure >= 0.4) – ENCODE_YY1_sites_COLON_nonpromotor

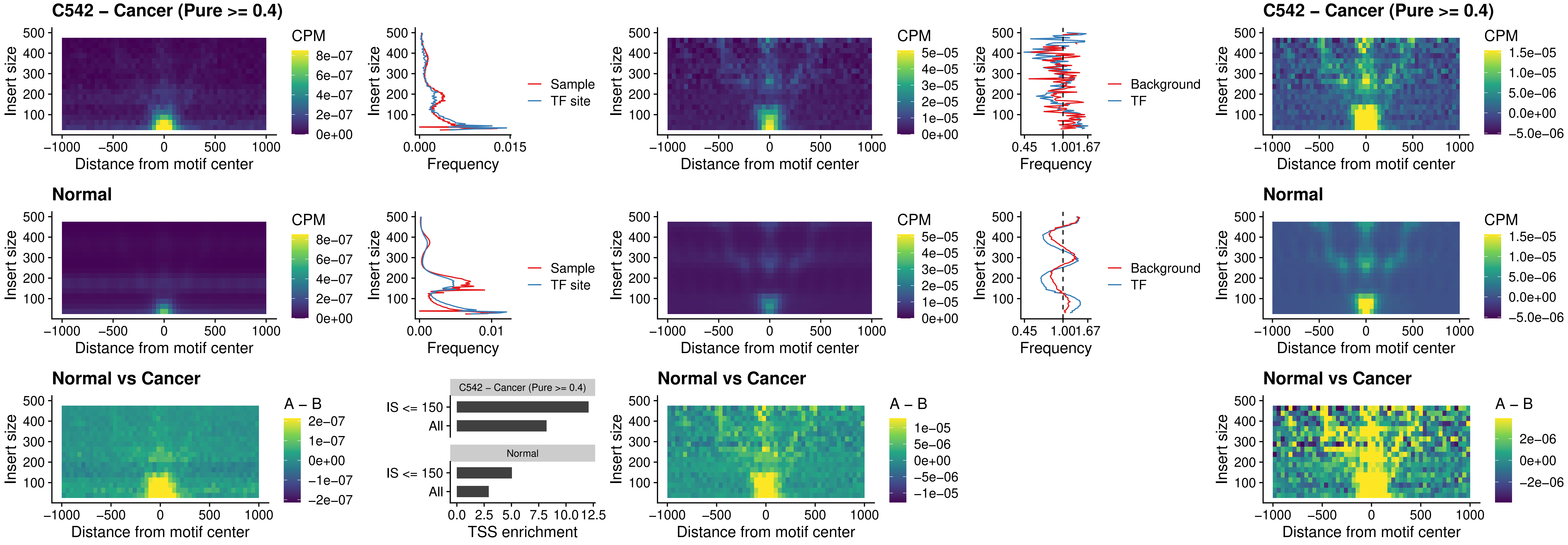

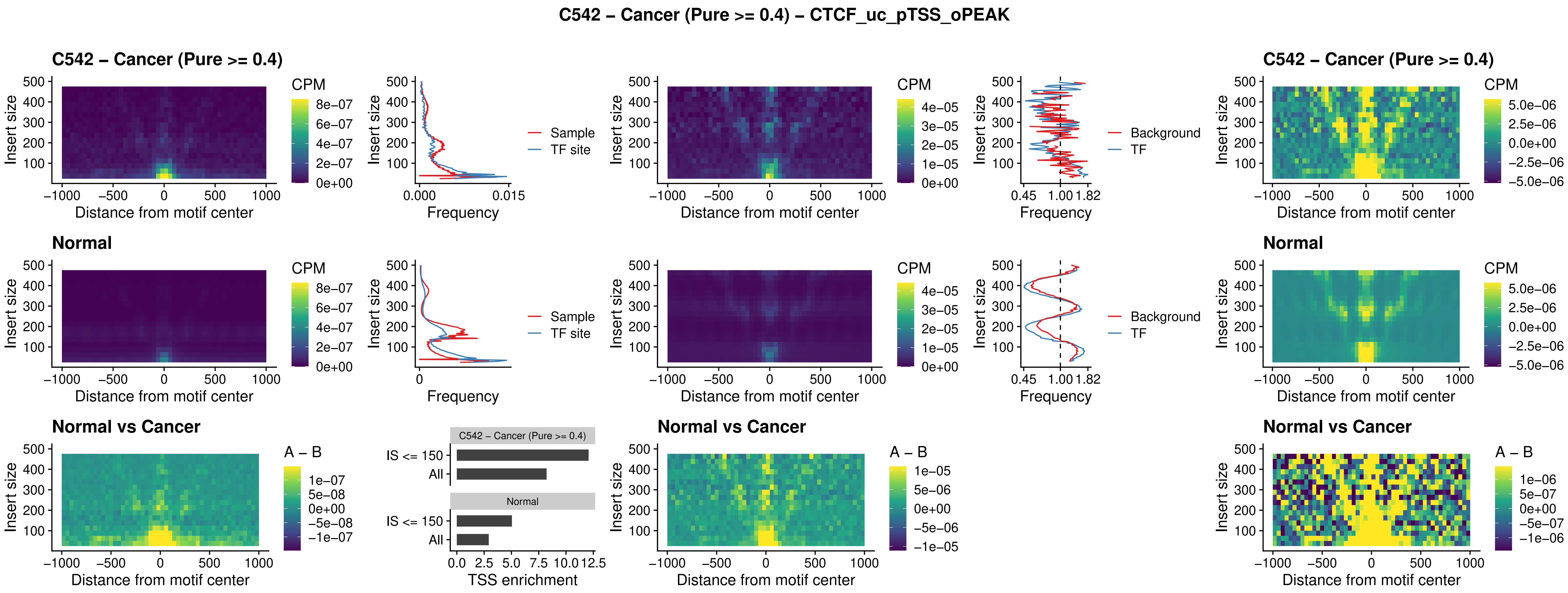

C542 – Cancer (Pure >= 0.4) – CTCF_uc_pTSS_oPEAK
C542 – Cancer (Pure >= 0.4)
Normal
Normal vs Cancer
Insert size
Distance from motif center
Frequency
CPM
A – B
Sample
TF site
Background
TF
C542 – Cancer (Pure >= 0.4)
Normal
IS <= 150
All
TSS enrichment

#### C542 – Cancer (Pure >= 0.4) – ENCODE_CTCF_sites_COLON_promotor

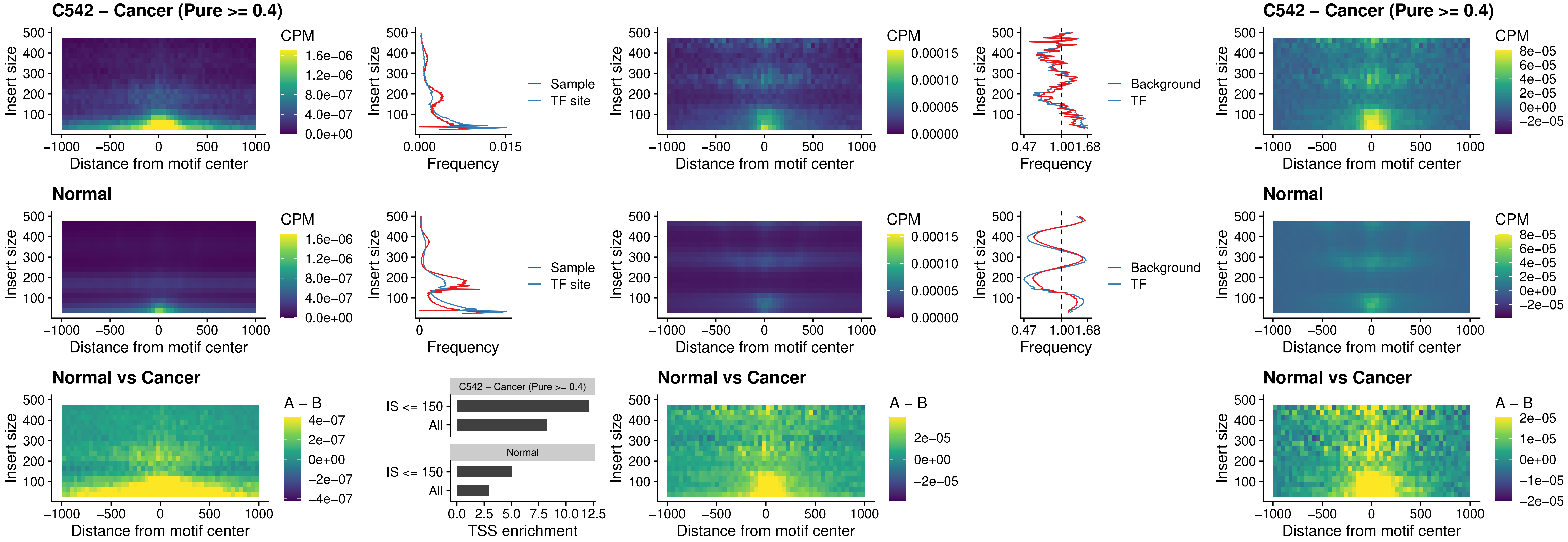

#### C542 – Cancer (Pure >= 0.4) – ENCODE_CTCF_sites_COLON_promotor_ctcf

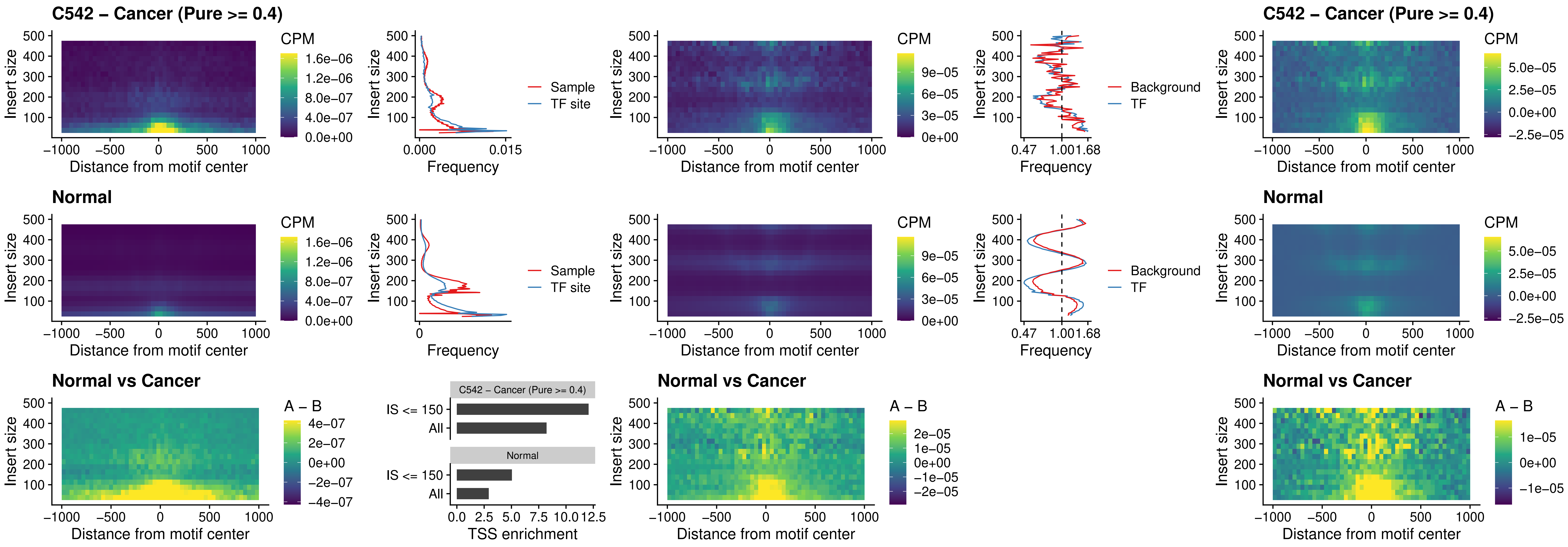

### C542 – Cancer (Pure >= 0.4) – SPIC_uc_pTSS_oPEAK

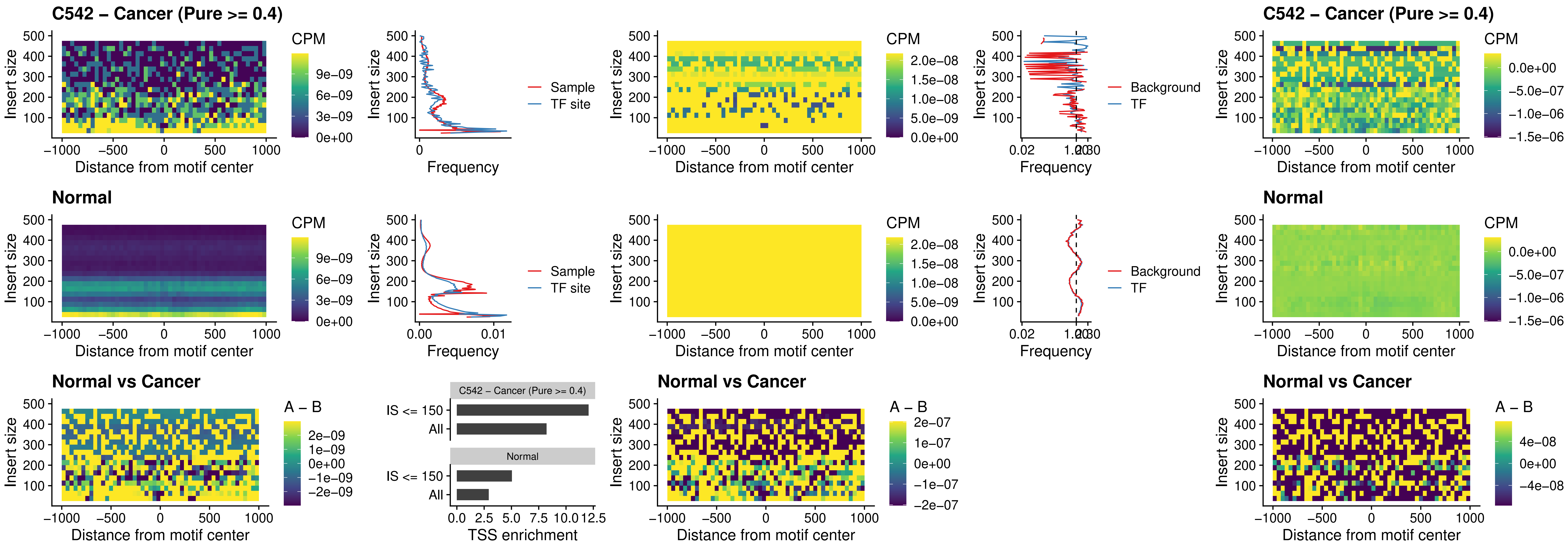

#### C542 – Cancer (Pure >= 0.4) – SPI1_uc_dTSS_nPEAK

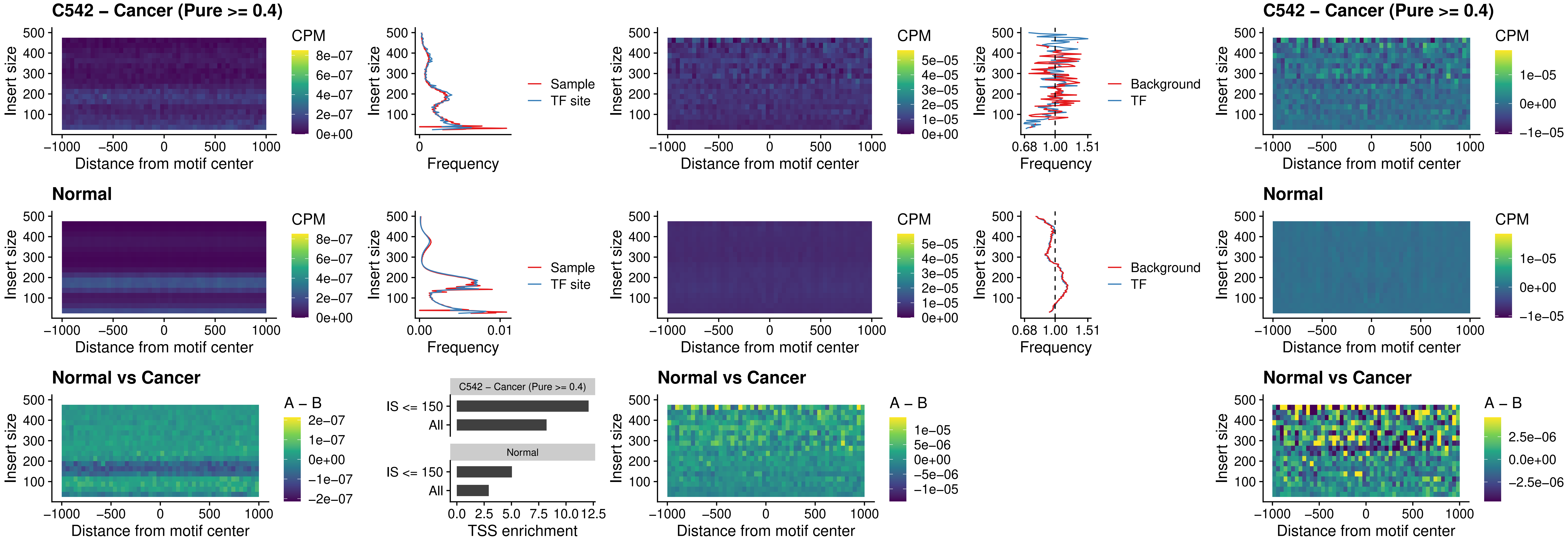

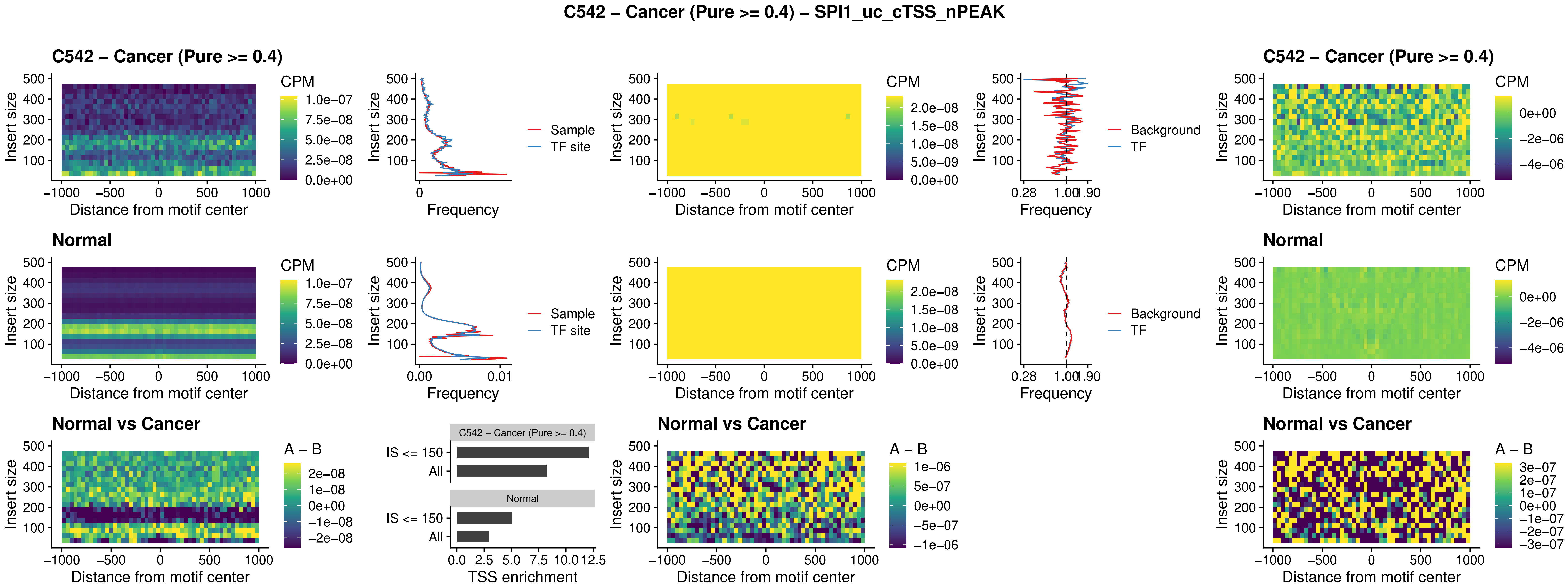
C542 – Cancer (Pure >= 0.4) – SPI1_uc_cTSS_nPEAK
C542 – Cancer (Pure >= 0.4)
Normal
Normal vs Cancer
Insert size
Distance from motif center
Frequency
CPM
A – B
Sample
TF site
Background
TF
C542 – Cancer (Pure >= 0.4)
IS <= 150
All
TSS enrichment

#### C542 – Cancer (Pure >= 0.4) – BCL11A_uc_pTSS_nPEAK

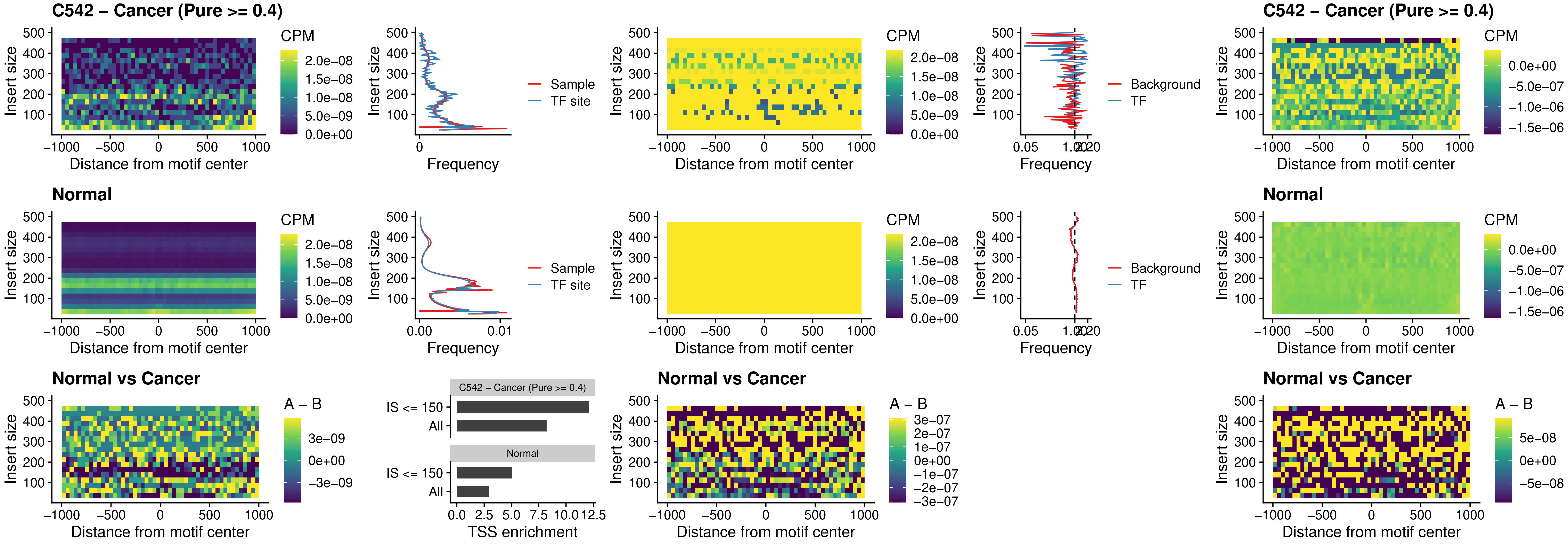

#### C542 – Cancer (Pure >= 0.4) – BCL11B_uc_pTSS_nPEAK

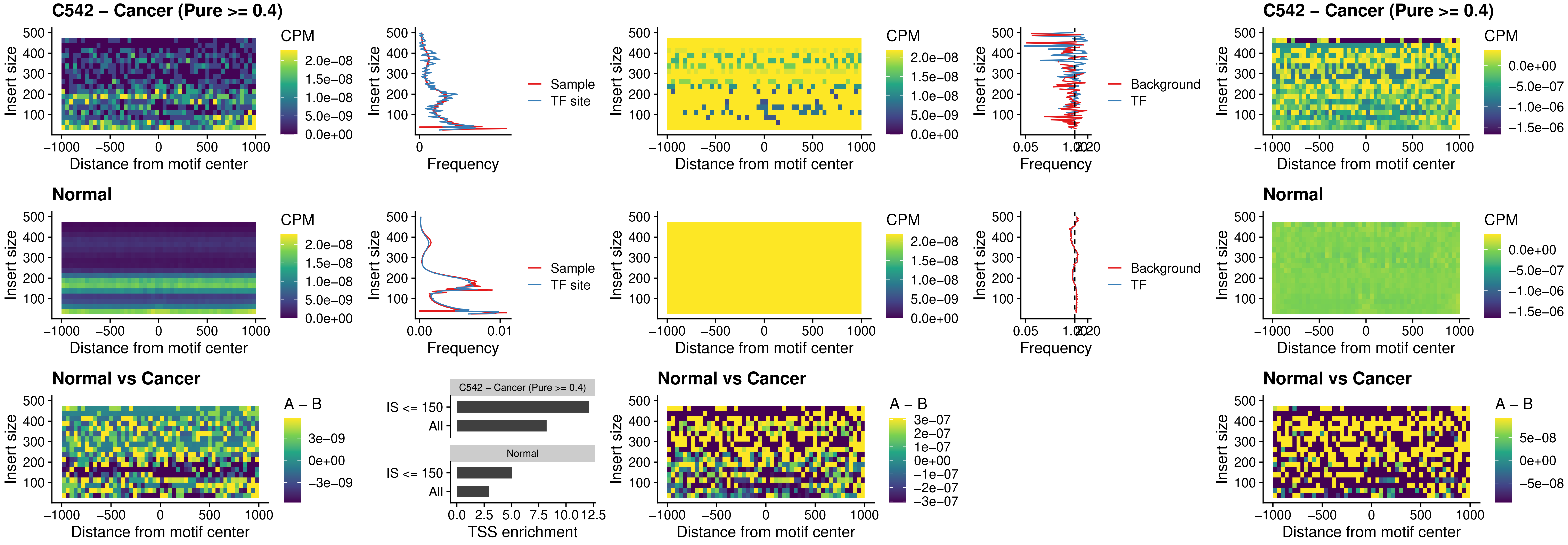

#### C542 – Cancer (Pure >= 0.4) – IRF9_uc_dTSS_oPEAK

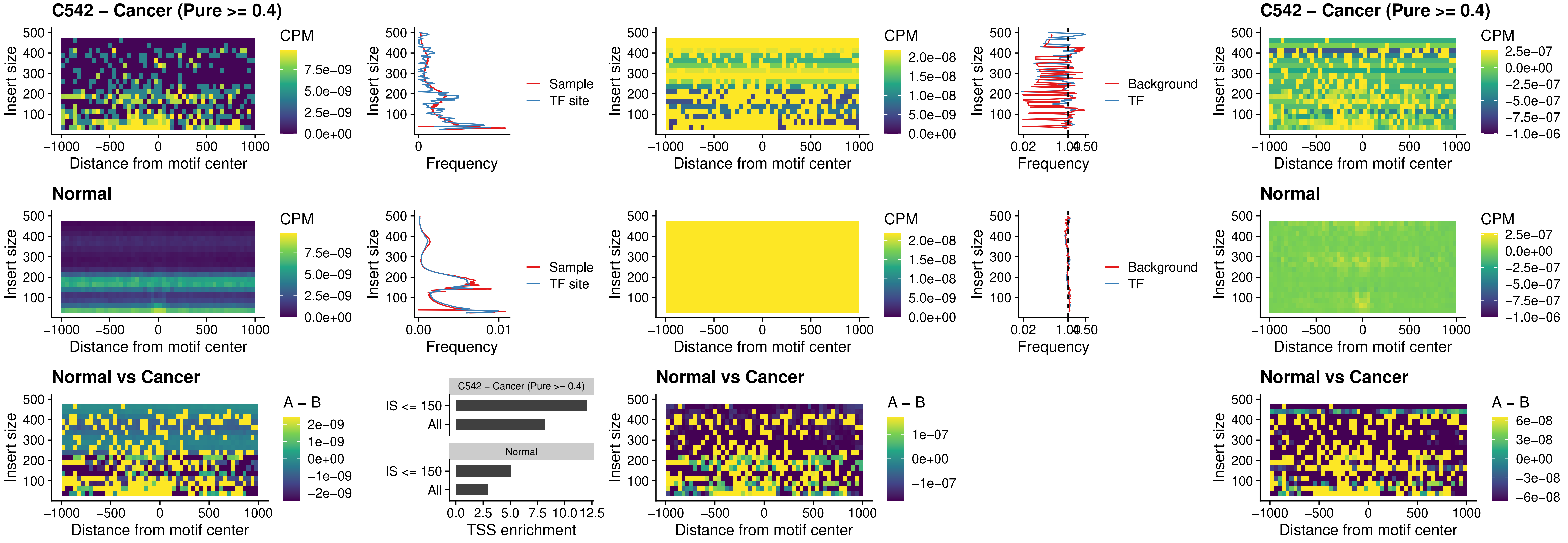

#### C542 – Cancer (Pure >= 0.4) – IRF9_uc_cTSS_oPEAK

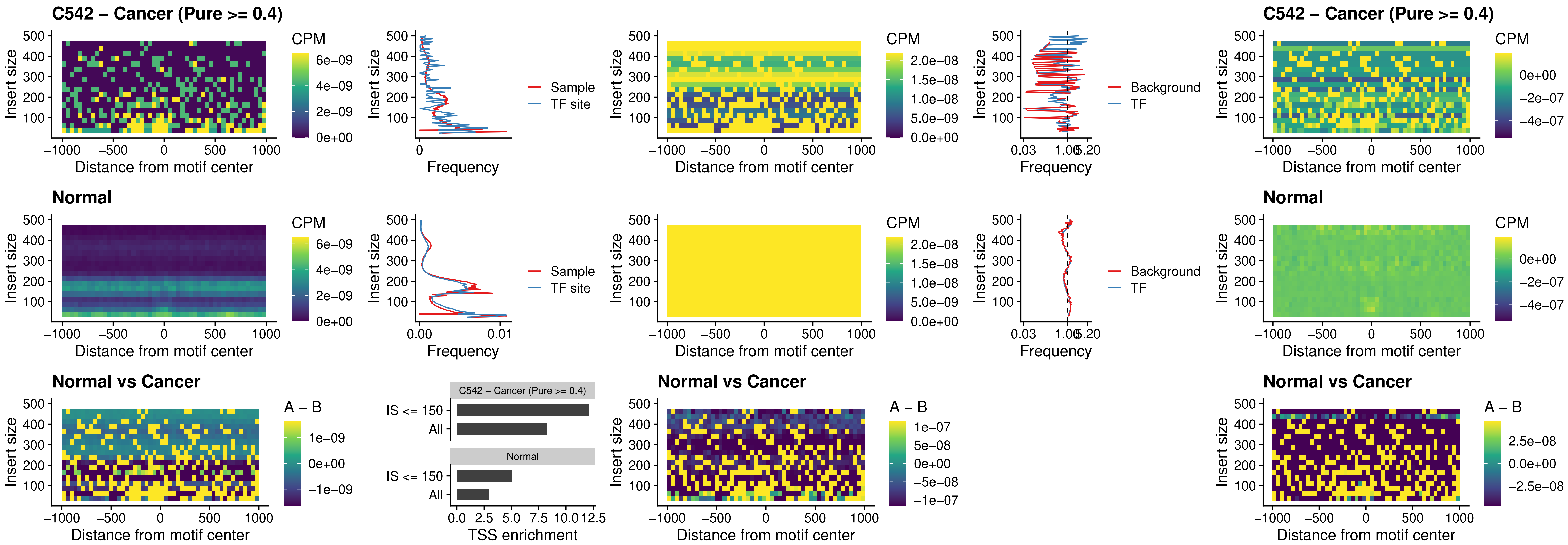

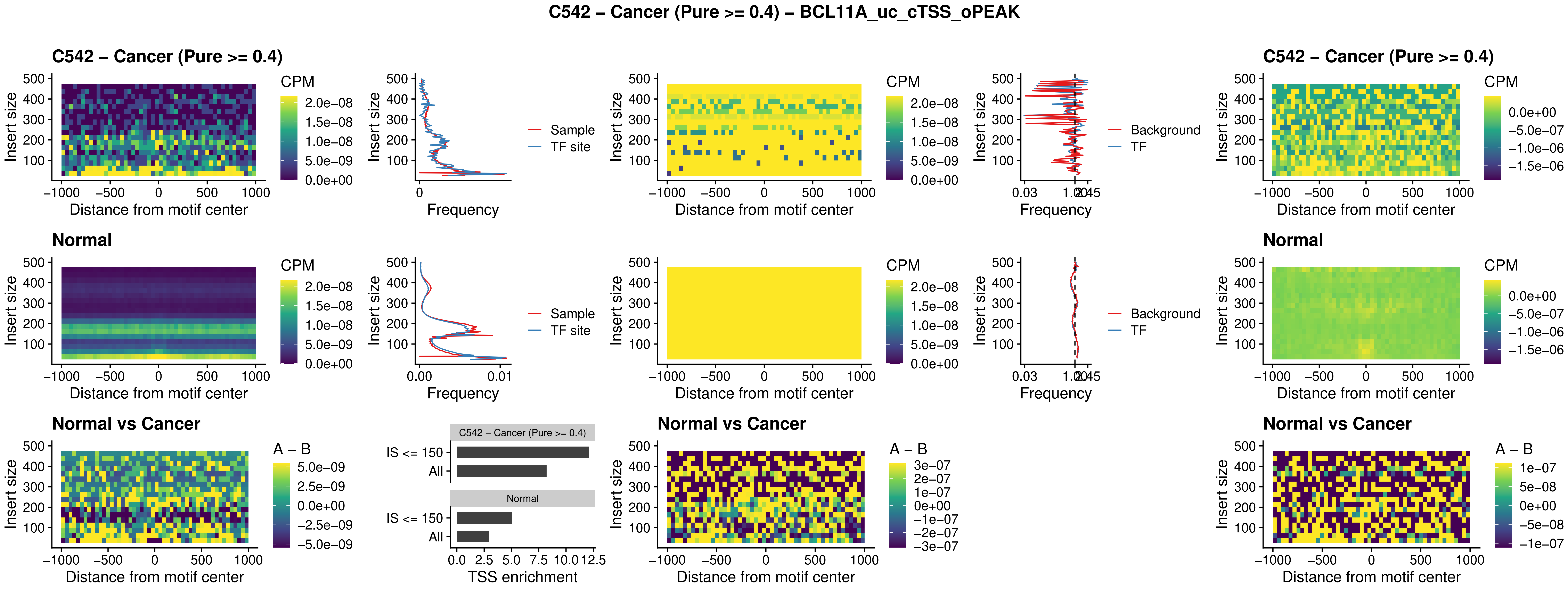
C542 – Cancer (Pure >= 0.4) – BCL11A_uc_cTSS_oPEAK
C542 – Cancer (Pure >= 0.4)
Normal
Normal vs Cancer
Insert size
Distance from motif center
Frequency
CPM
A – B
Sample
TF site
Background
TF
C542 – Cancer (Pure >= 0.4)
Normal
IS <= 150
All
TSS enrichment

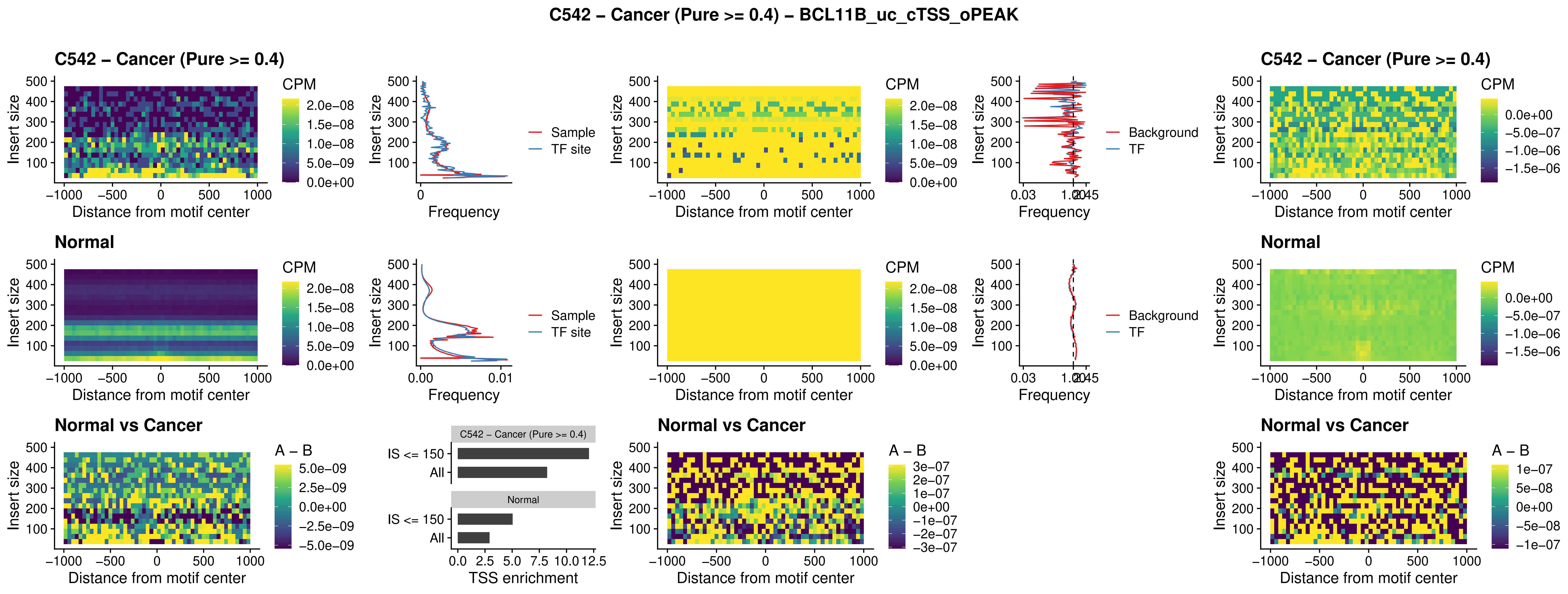
C542 – Cancer (Pure >= 0.4) – BCL11B_uc_cTSS_oPEAK
C542 – Cancer (Pure >= 0.4)
Normal
Normal vs Cancer
Insert size
Distance from motif center
Frequency
CPM
A – B
Sample
TF site
Background
TF
C542 – Cancer (Pure >= 0.4)
Normal
IS <= 150
All
TSS enrichment

### C542 – Cancer (Pure >= 0.4) – BCL11A_uc_dTSS_oPEAK

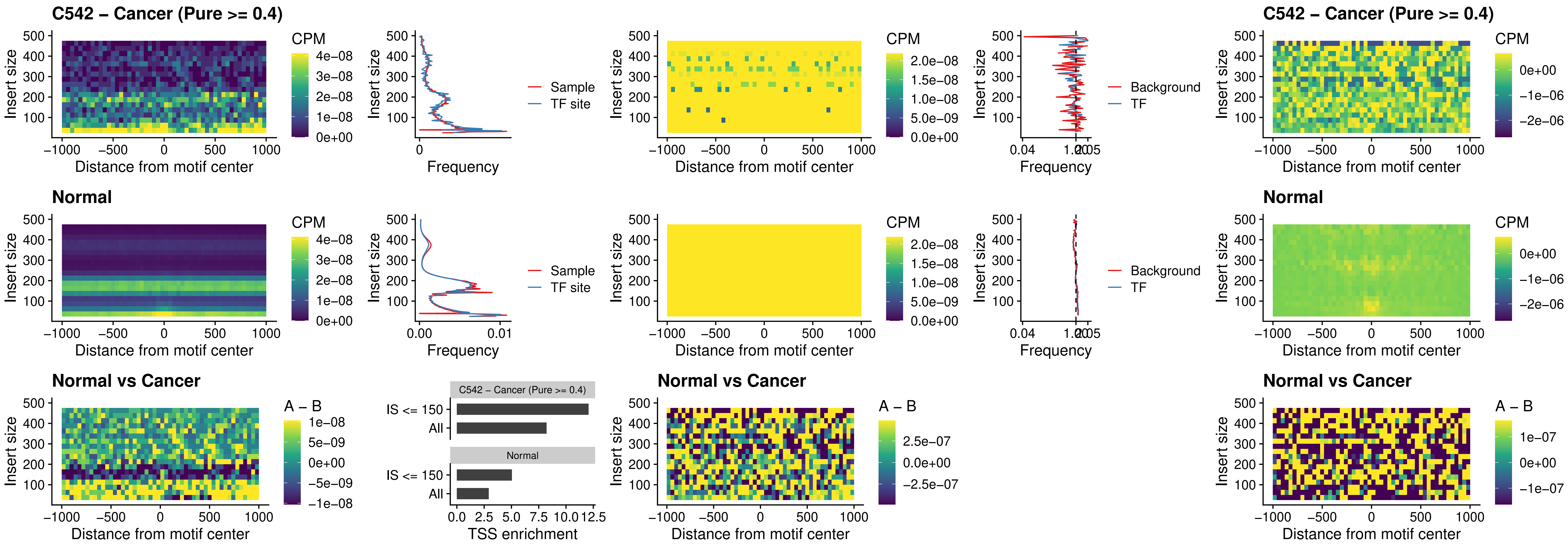

### C542 – Cancer (Pure >= 0.4) – BCL11B_uc_dTSS_oPEAK

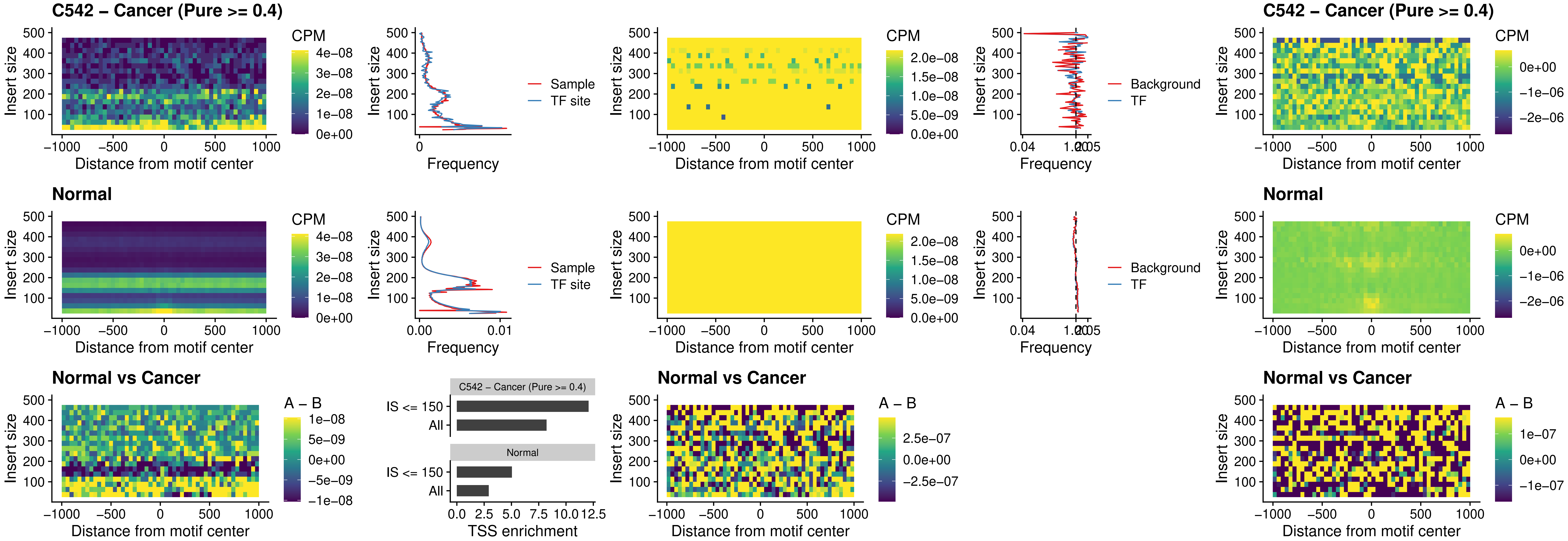

#### C542 – Cancer (Pure >= 0.4) – SPIB_uc_cTSS_nPEAK

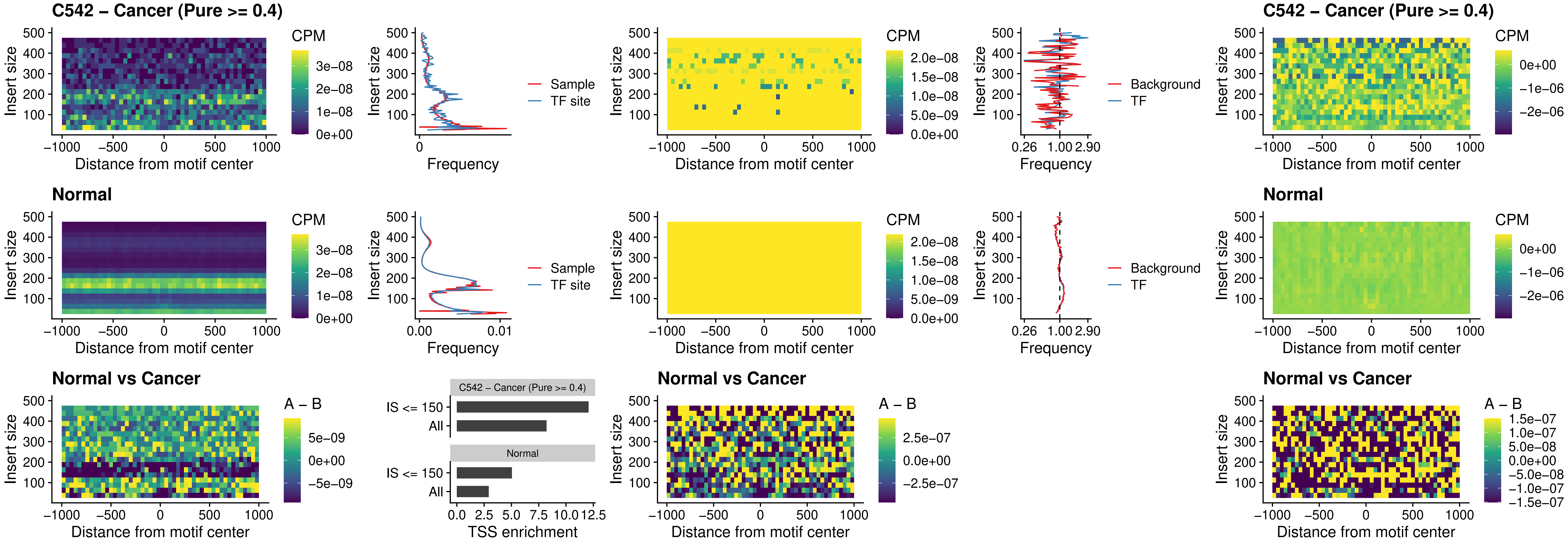

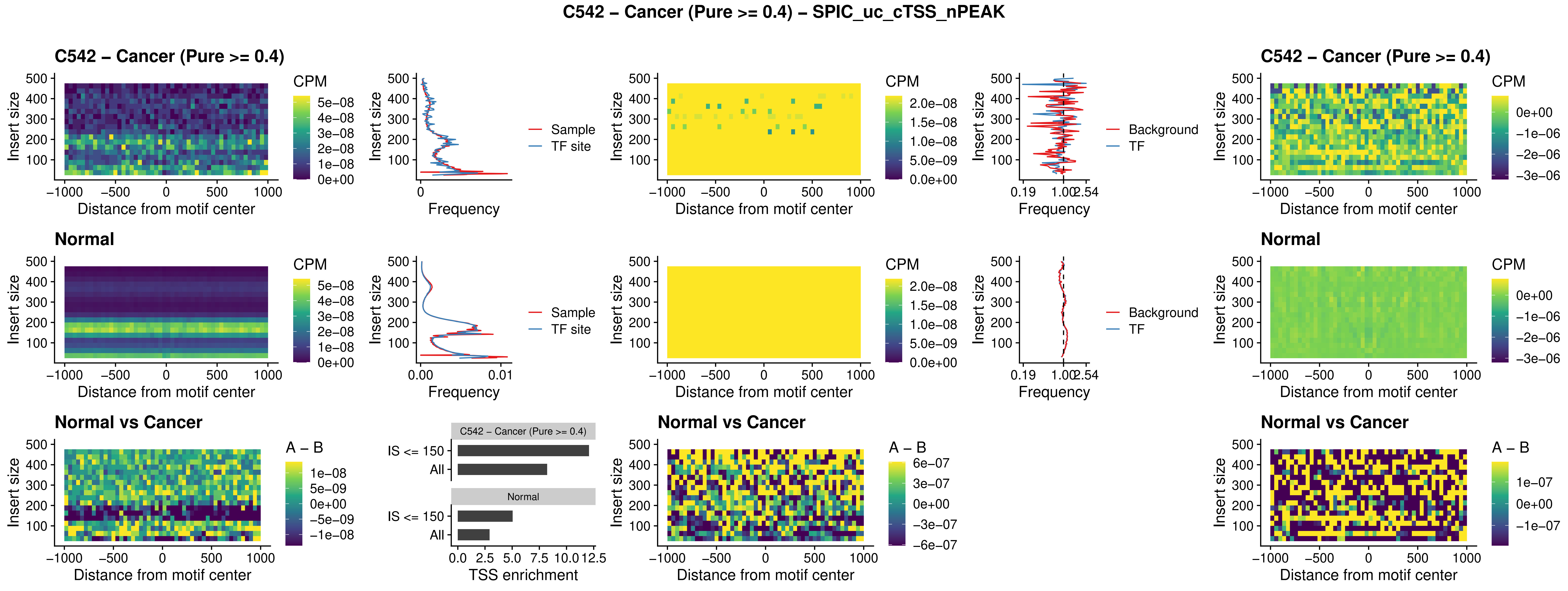
C542 – Cancer (Pure >= 0.4) – SPIC_uc_cTSS_nPEAK
C542 – Cancer (Pure >= 0.4)
Normal
Normal vs Cancer
Insert size
Distance from motif center
Frequency
CPM
A – B
Sample
TF site
Background
TF
C542 – Cancer (Pure >= 0.4)
Normal
IS <= 150
All
TSS enrichment

#### C542 – Cancer (Pure >= 0.4) – SPI1_uc_pTSS_oPEAK

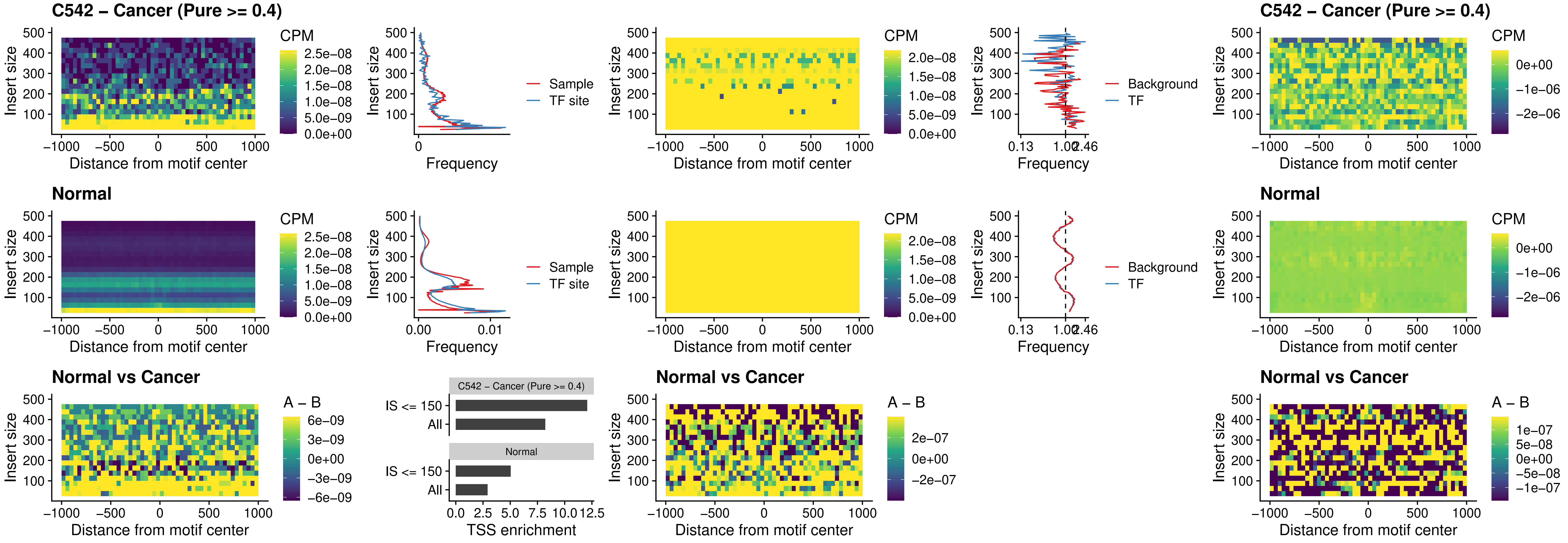

#### C542 – Cancer (Pure >= 0.4) – BCL11A_uc_pTSS_oPEAK

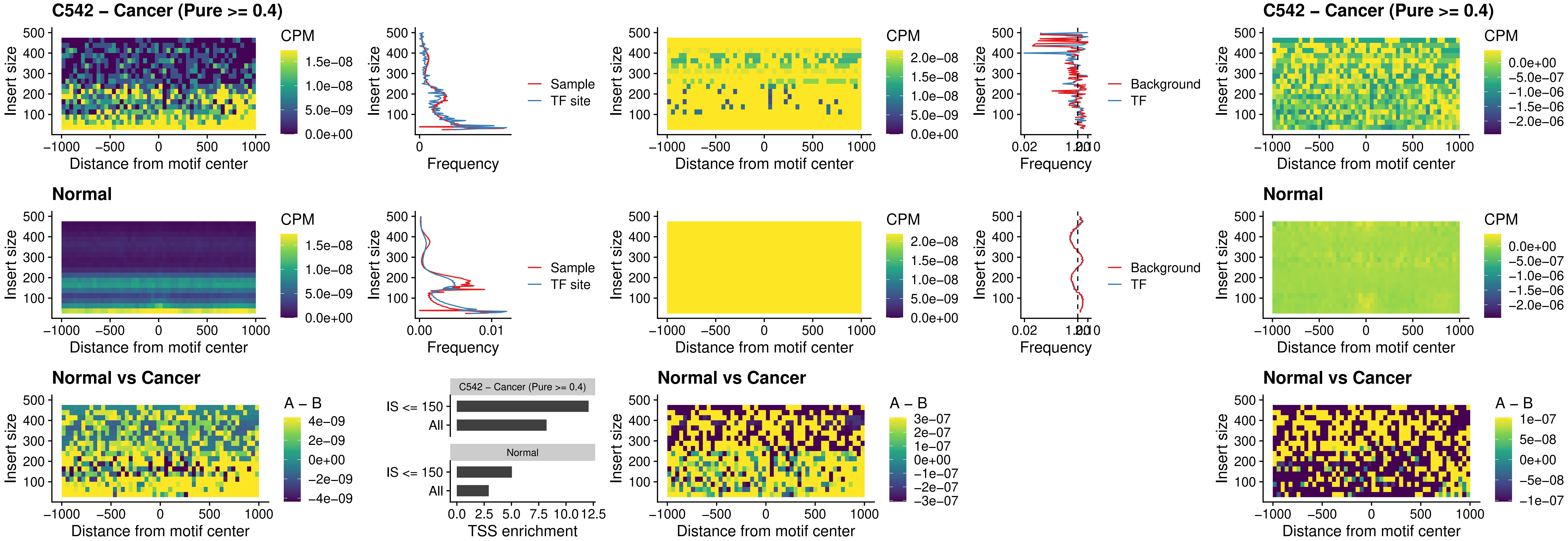

#### C542 – Cancer (Pure >= 0.4) – BCL11B_uc_pTSS_oPEAK

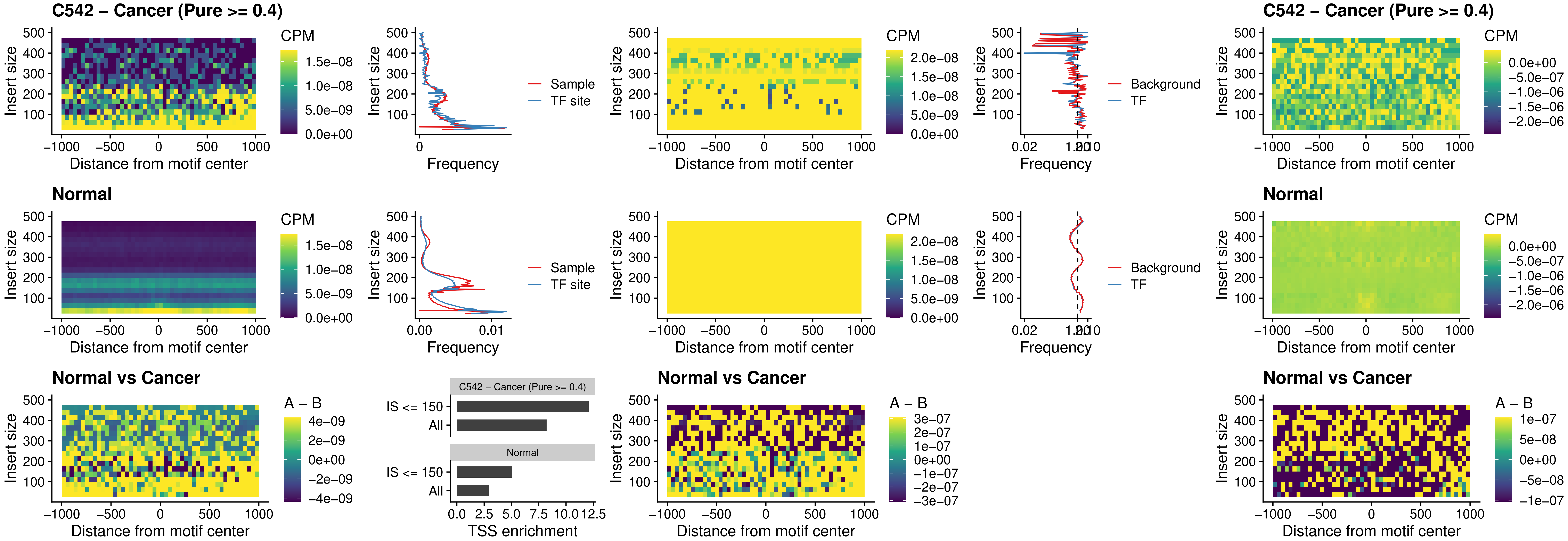

#### C542 – Cancer (Pure >= 0.4) – SPI1_uc_cTSS_oPEAK

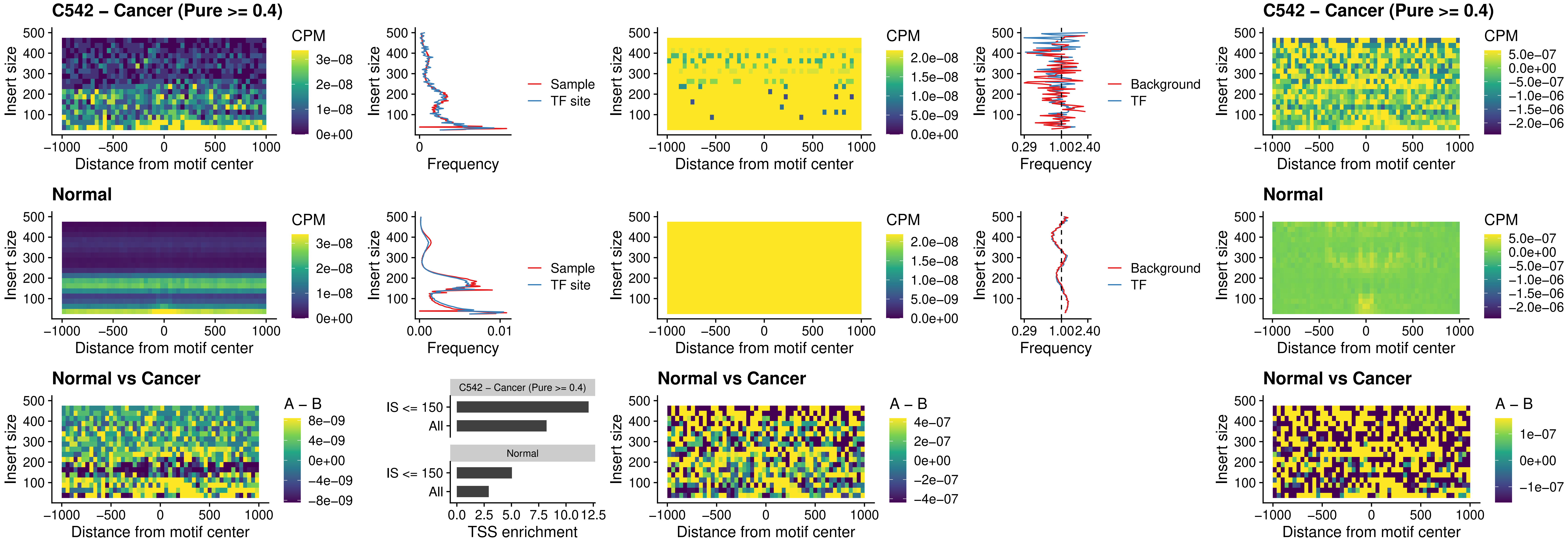

### C542 – Cancer (Pure >= 0.4) – SPIB_uc_cTSS_oPEAK

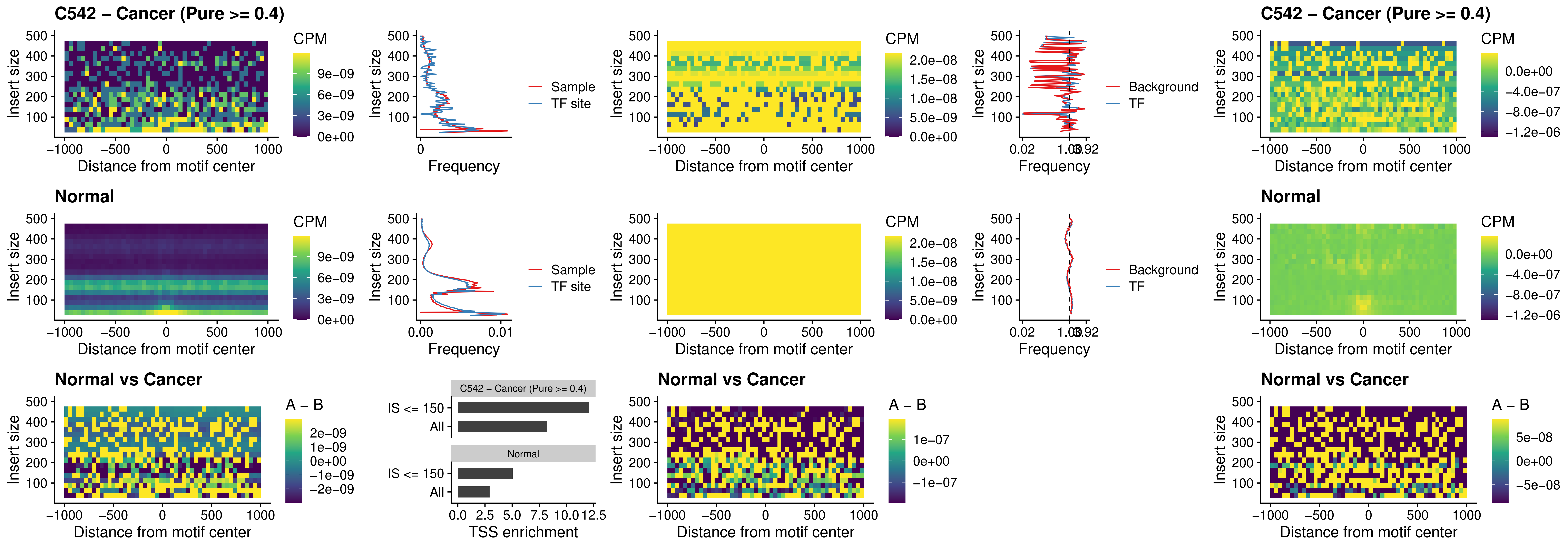

#### C542 – Cancer (Pure >= 0.4) – SPIB_uc_pTSS_oPEAK

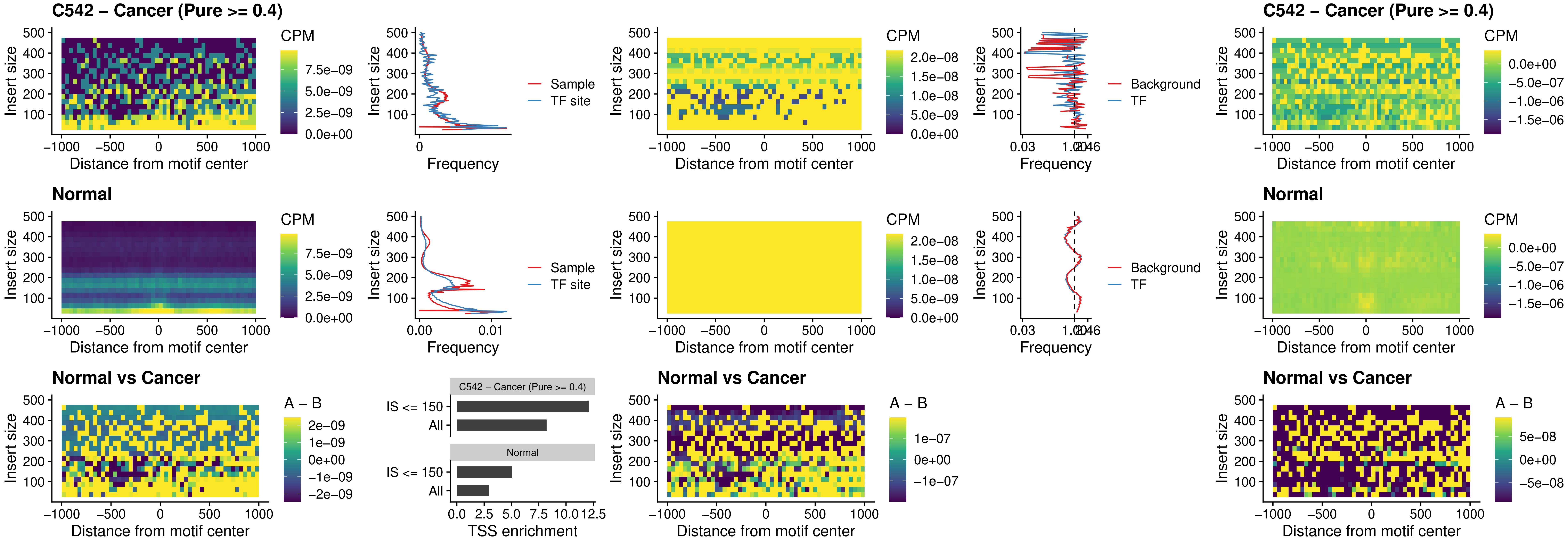

#### C542 – Cancer (Pure >= 0.4) – SPI1_uc_dTSS_oPEAK

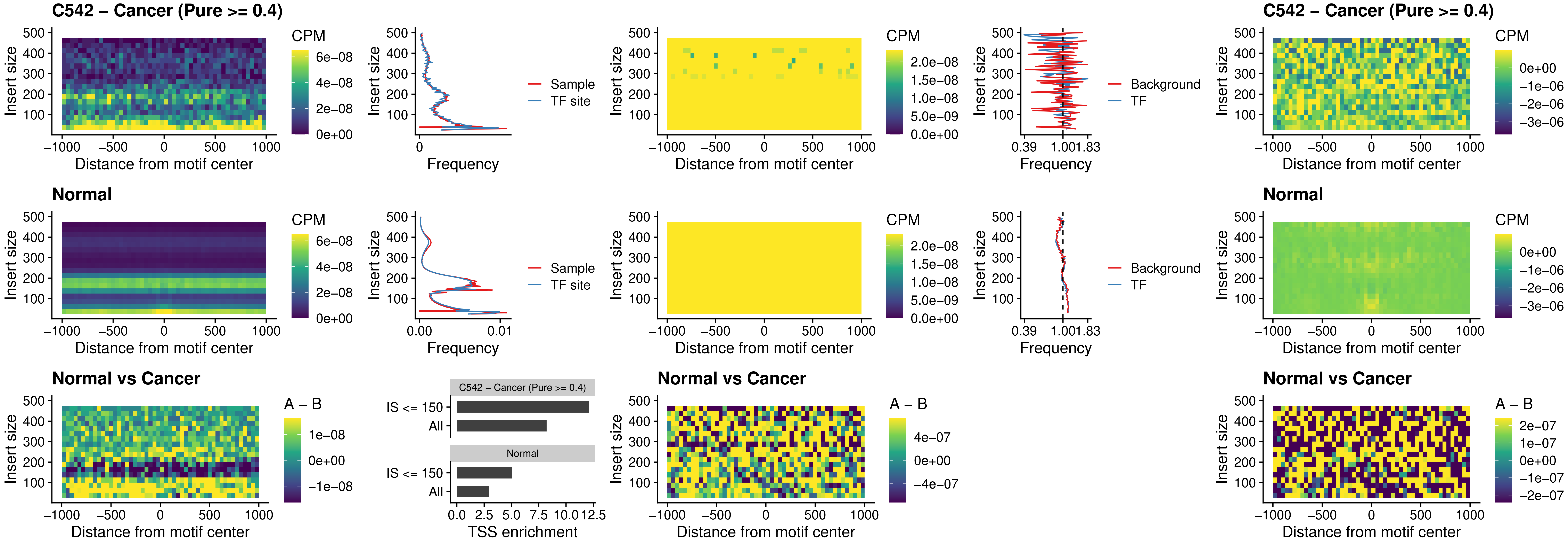

### C542 – Cancer (Pure >= 0.4) – SPIB_uc_dTSS_oPEAK

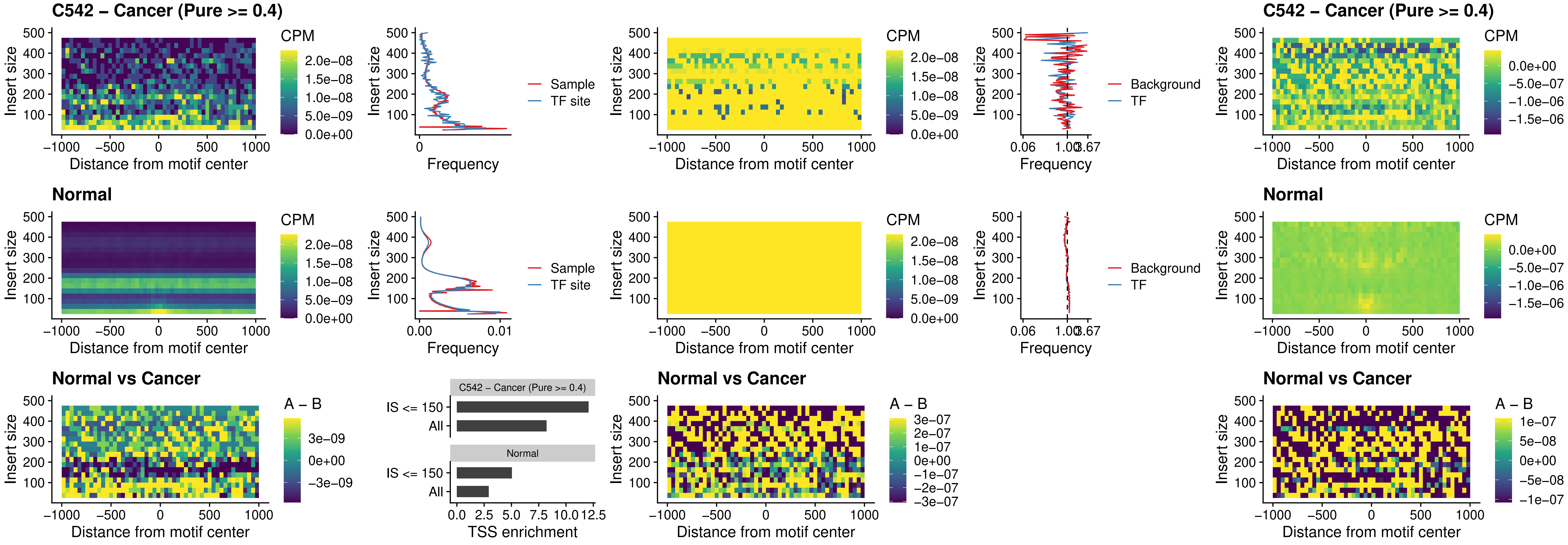

#### C542 – Cancer (Pure >= 0.4) – IRF3_uc_pTSS_oPEAK

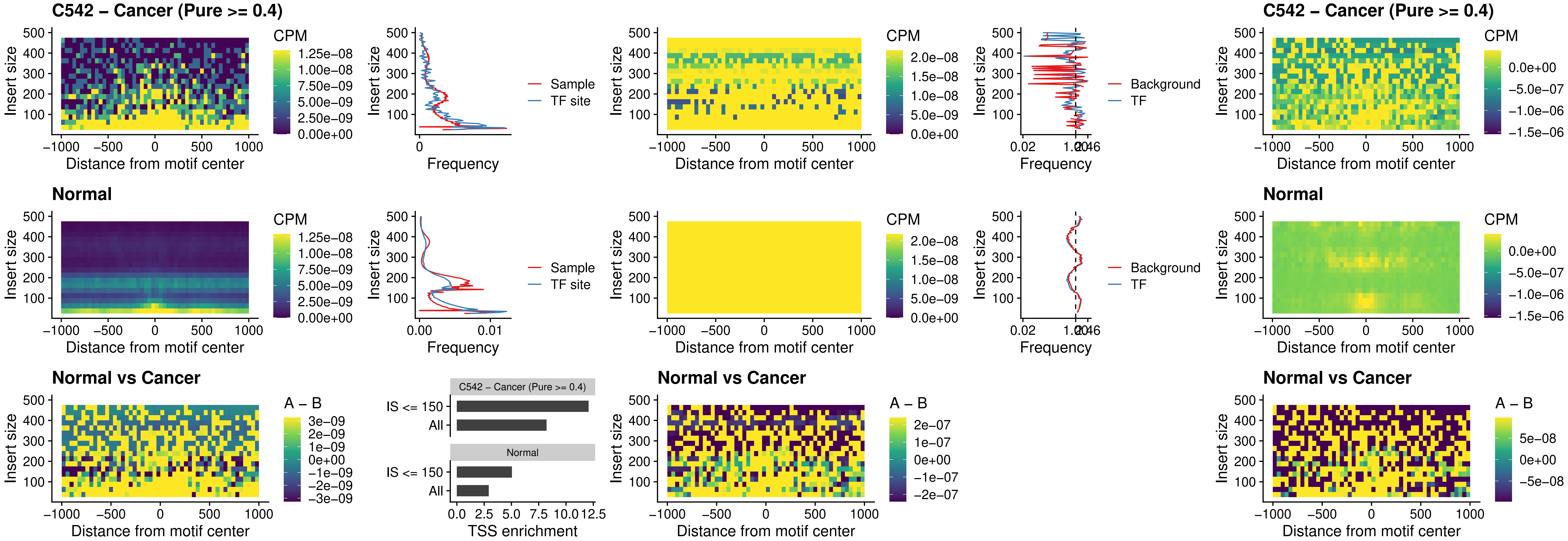

### C542 – Cancer (Pure >= 0.4) – IRF2_uc_dTSS_oPEAK

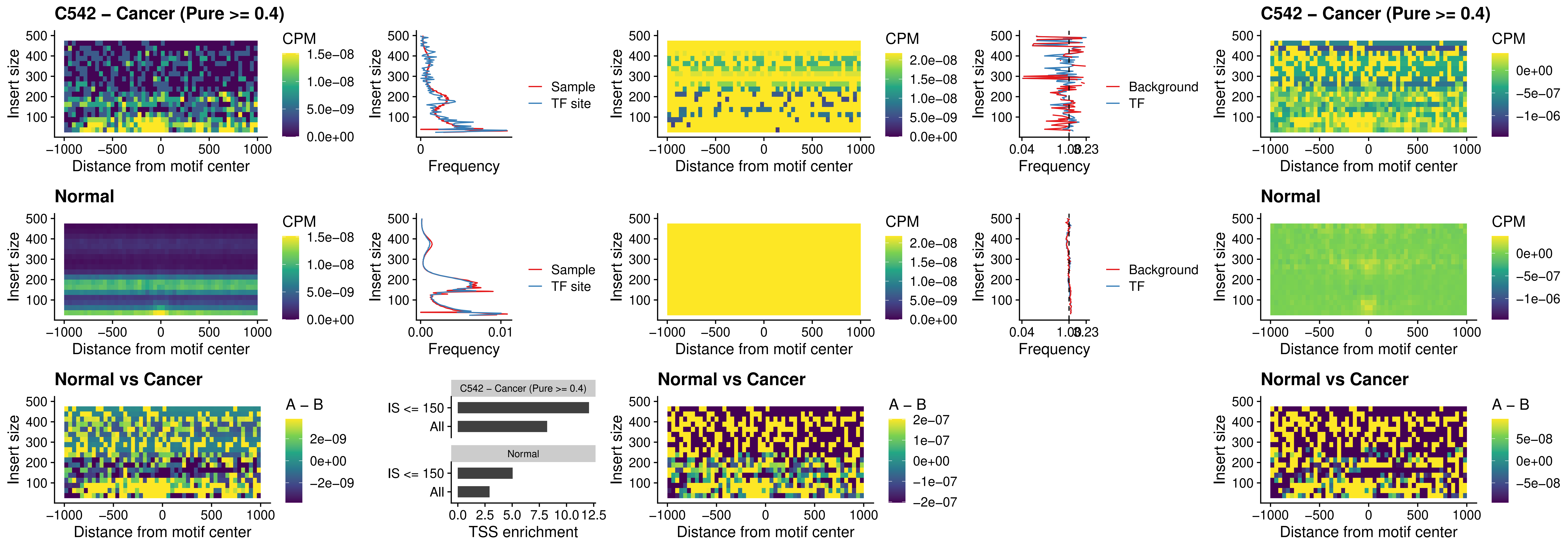

#### C542 – Cancer (Pure >= 0.4) – IRF2_uc_pTSS_oPEAK

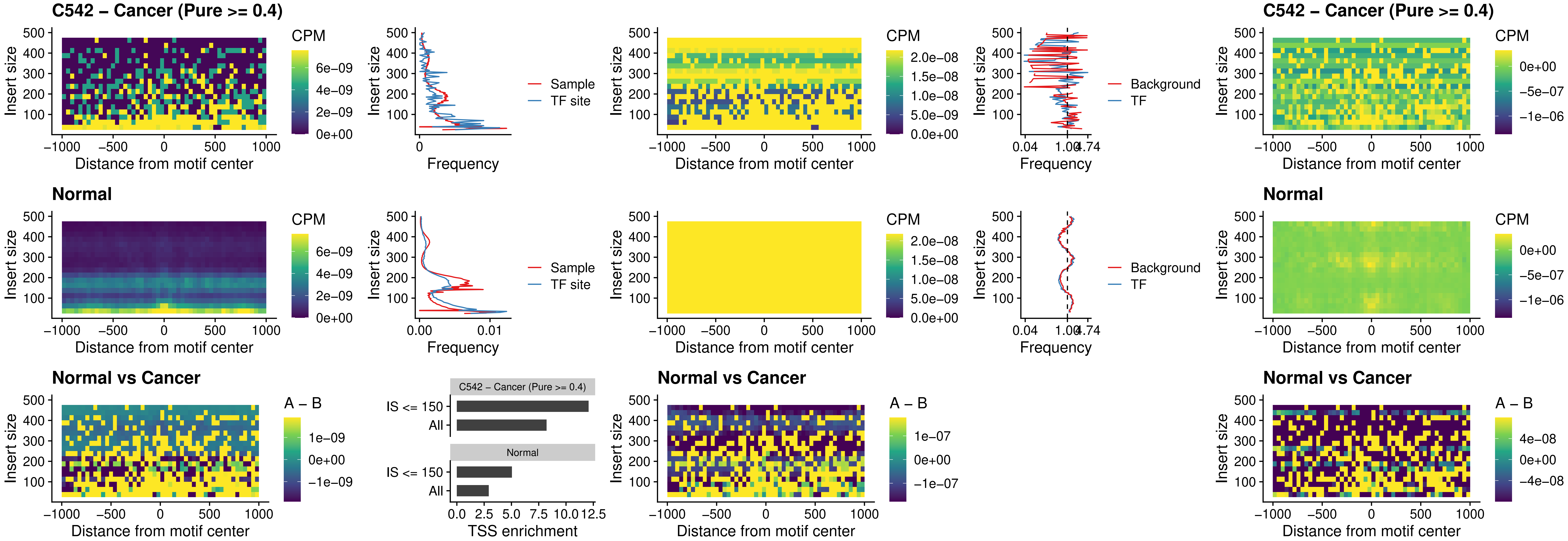

#### C542 – Cancer (Pure >= 0.4) – STAT2_uc_pTSS_oPEAK

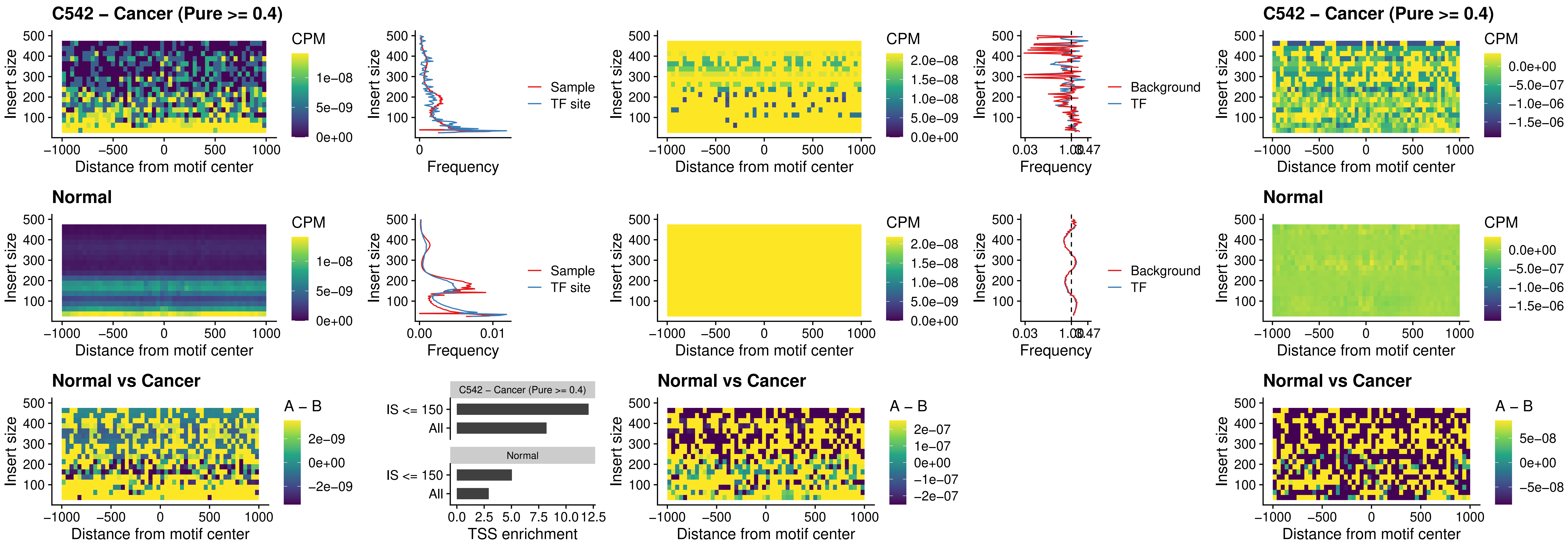

#### C542 – Cancer (Pure >= 0.4) – SPIB_uc_dTSS_nPEAK

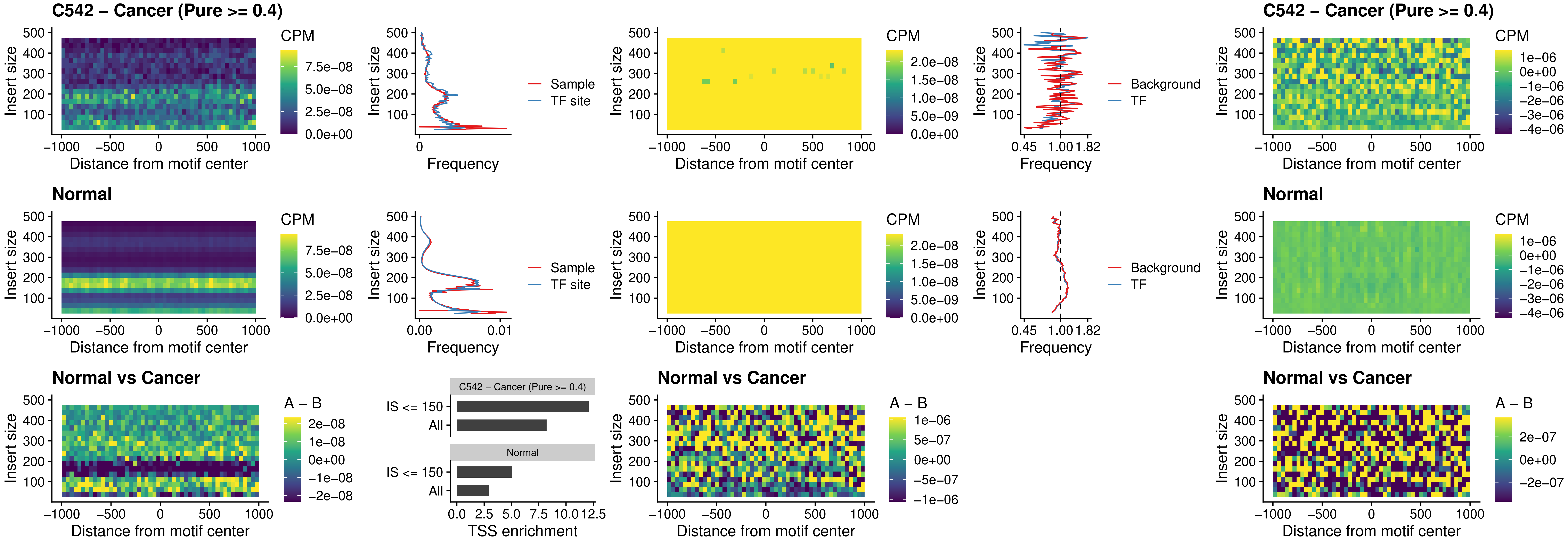

#### C542 – Cancer (Pure >= 0.4) – SPIC_uc_cTSS_oPEAK

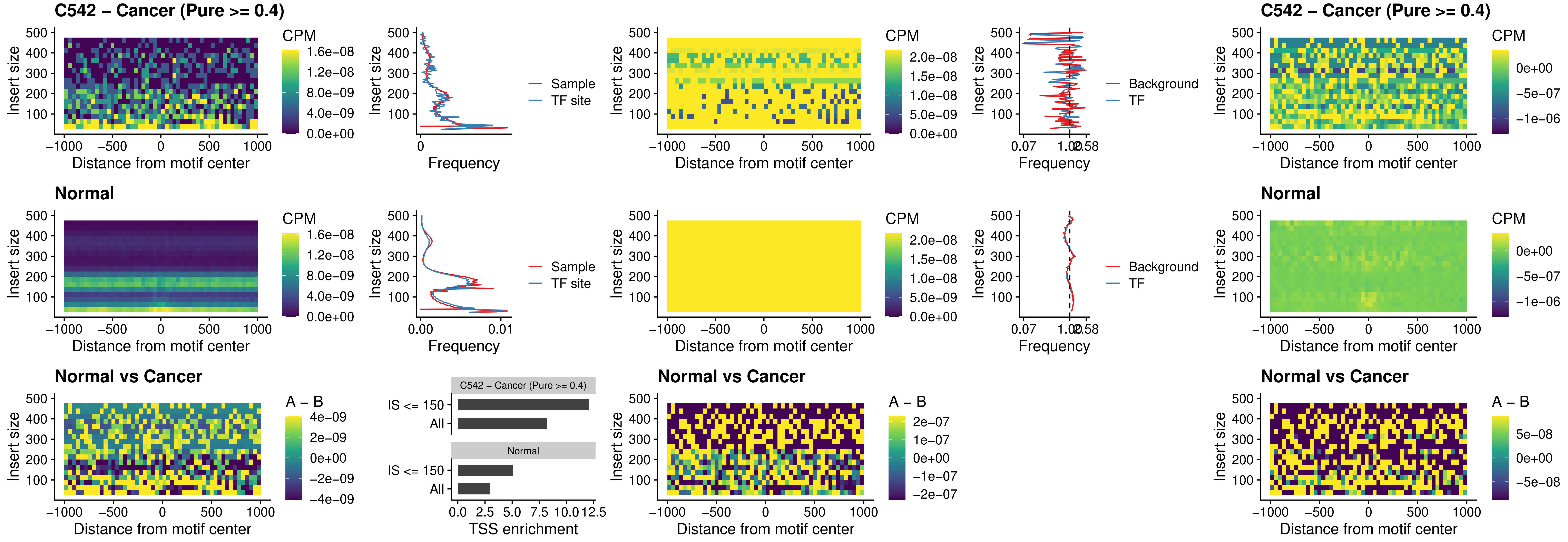

#### C542 – Cancer (Pure >= 0.4) – PRDM1_uc_cTSS_oPEAK

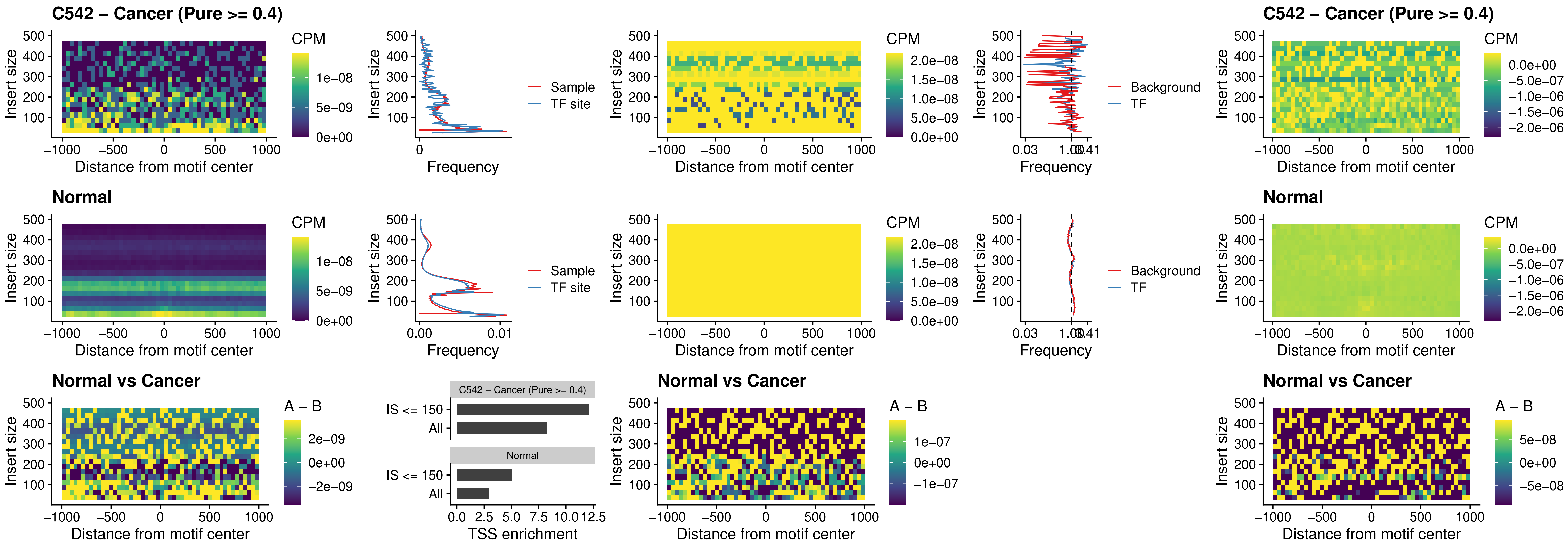

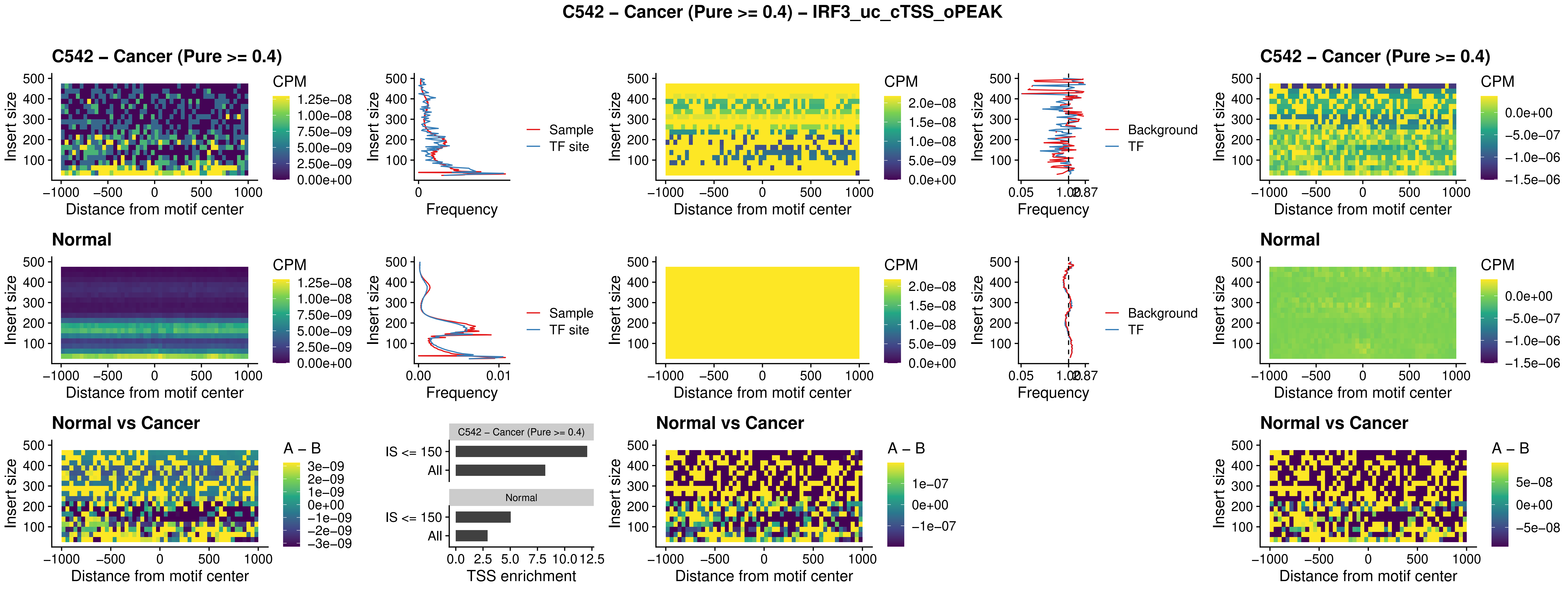
C542 – Cancer (Pure >= 0.4) – IRF3_uc_cTSS_oPEAK
C542 – Cancer (Pure >= 0.4)
Normal
Normal vs Cancer
Insert size
Distance from motif center
Frequency
CPM
Sample
TF site
Background
TF
A – B
C542 – Cancer (Pure >= 0.4)
Normal
IS <= 150
All
TSS enrichment

#### C542 – Cancer (Pure >= 0.4) – BCL11A_uc_dTSS_nPEAK

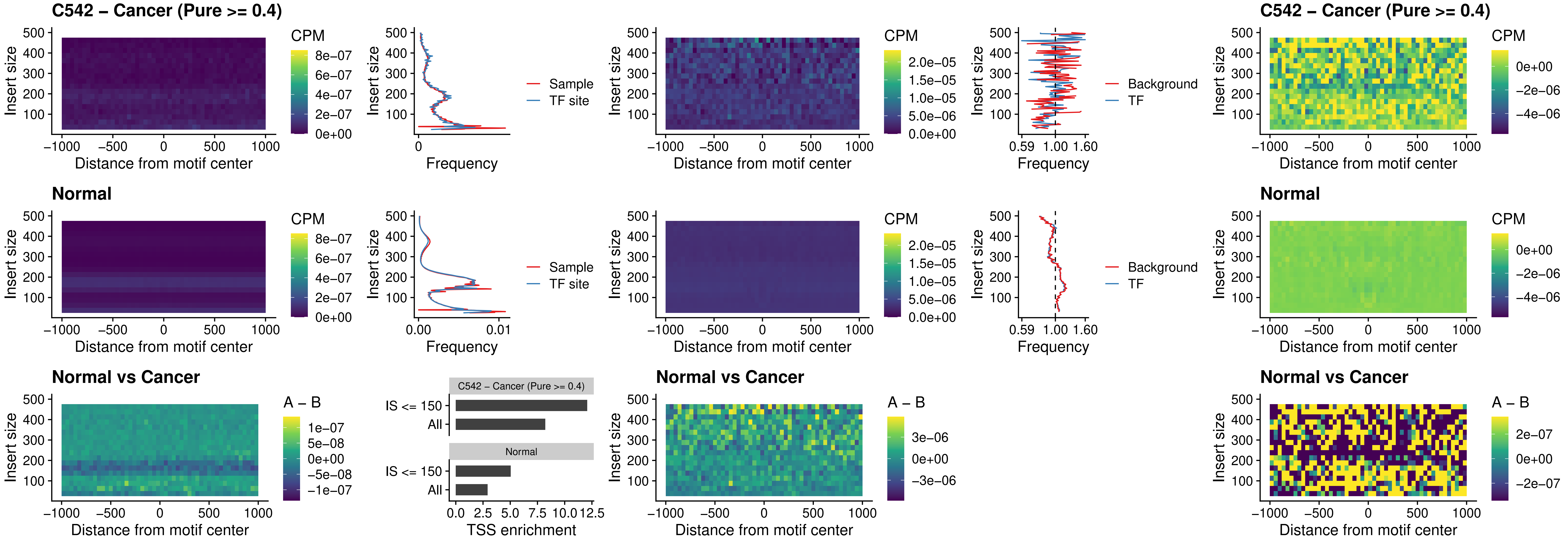

#### C542 – Cancer (Pure >= 0.4) – BCL11B_uc_dTSS_nPEAK

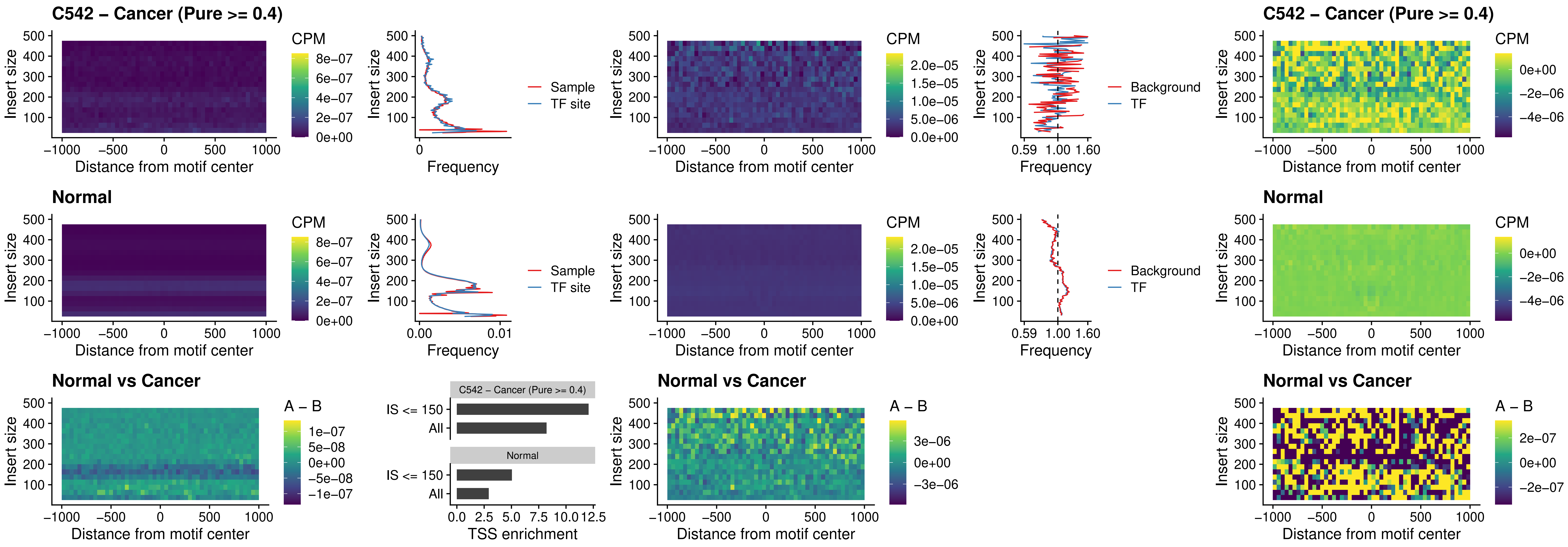

#### C542 – Cancer (Pure >= 0.4) – IRF1_uc_cTSS_oPEAK

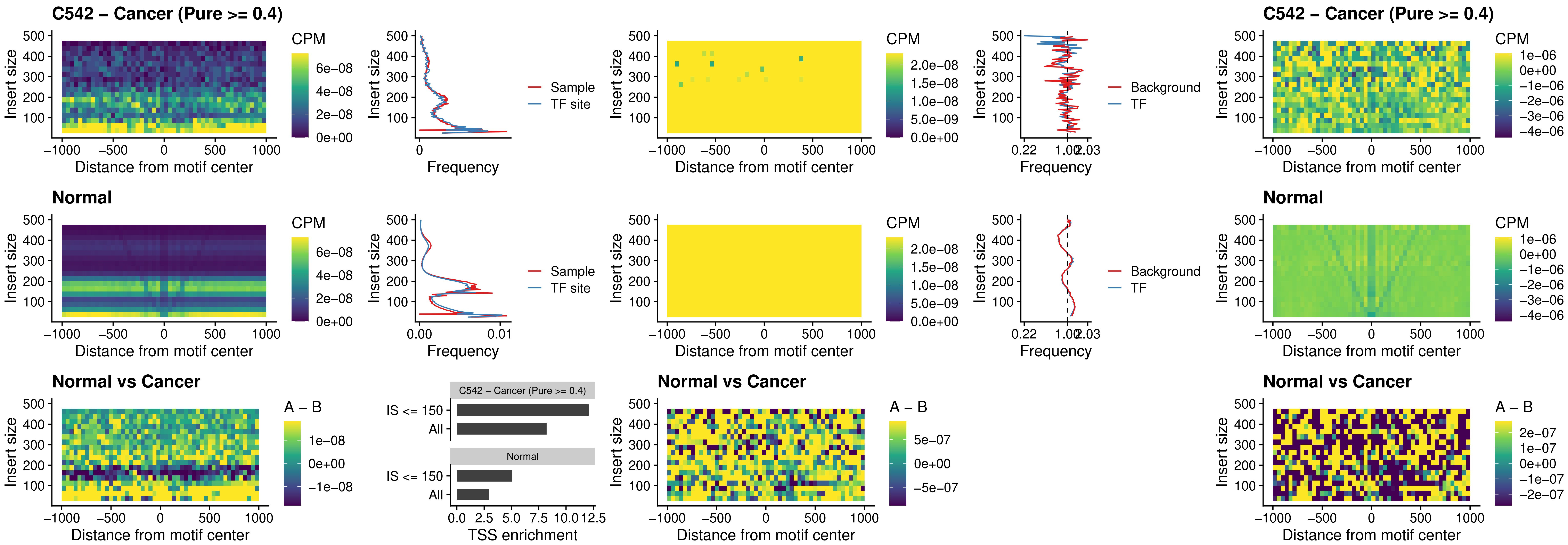

#### C542 – Cancer (Pure >= 0.4) – IRF1_uc_dTSS_oPEAK

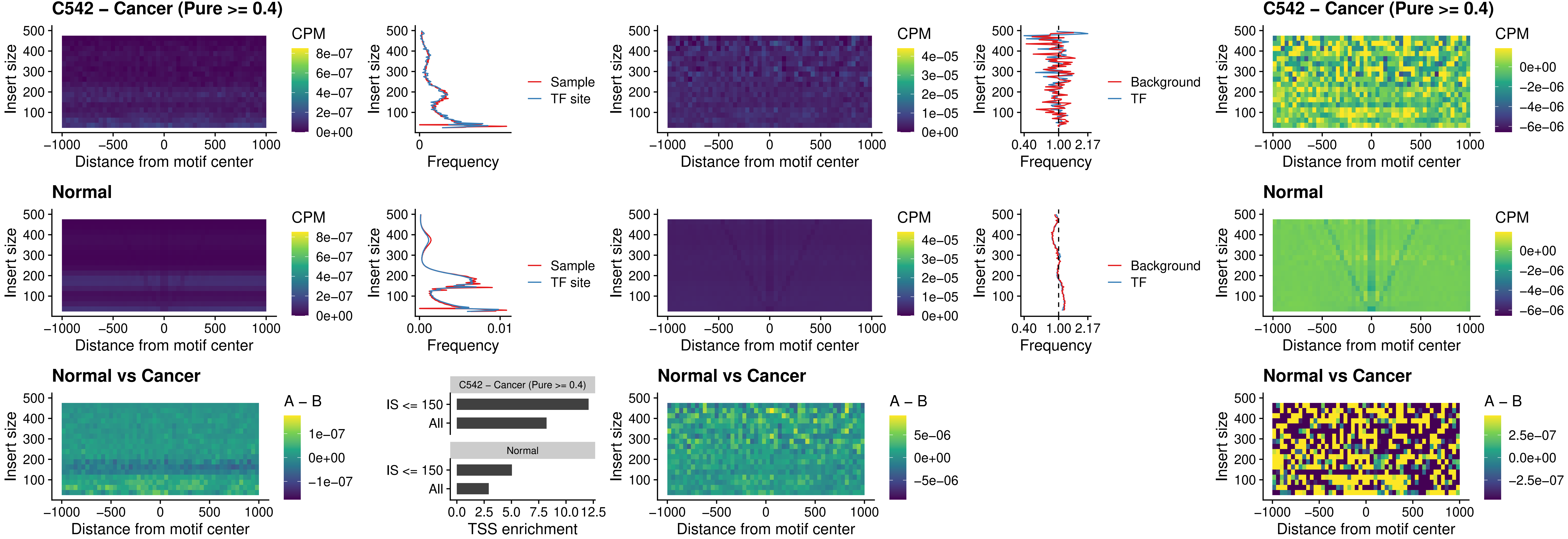

#### C542 – Cancer (Pure >= 0.4) – IRF1_uc_pTSS_oPEAK

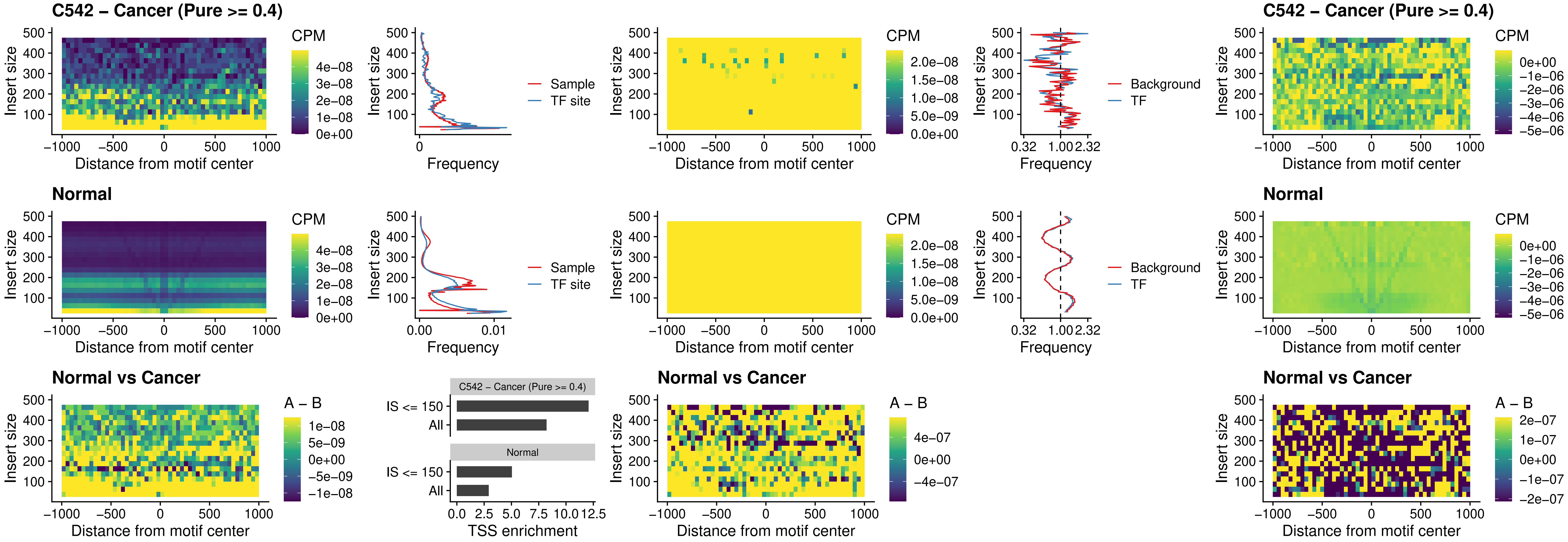

#### C542 – Cancer (Pure >= 0.4) – SPIC_uc_dTSS_nPEAK

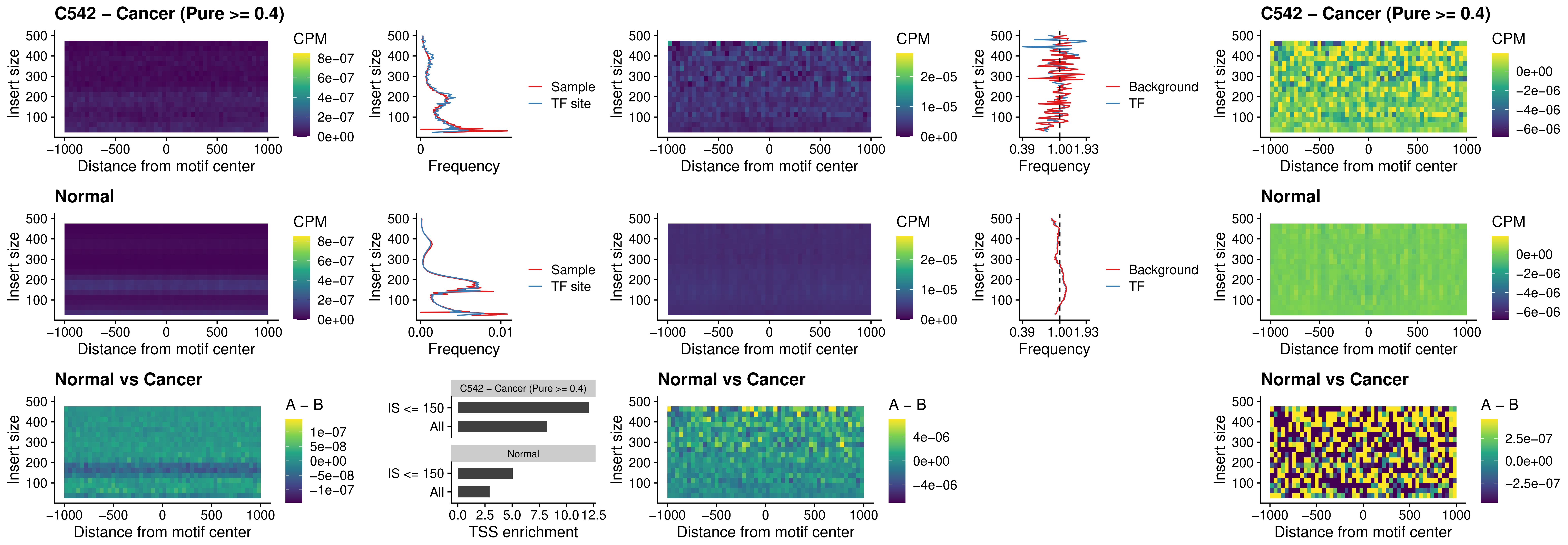

#### C542 – Cancer (Pure >= 0.4) – SPIC_uc_dTSS_oPEAK

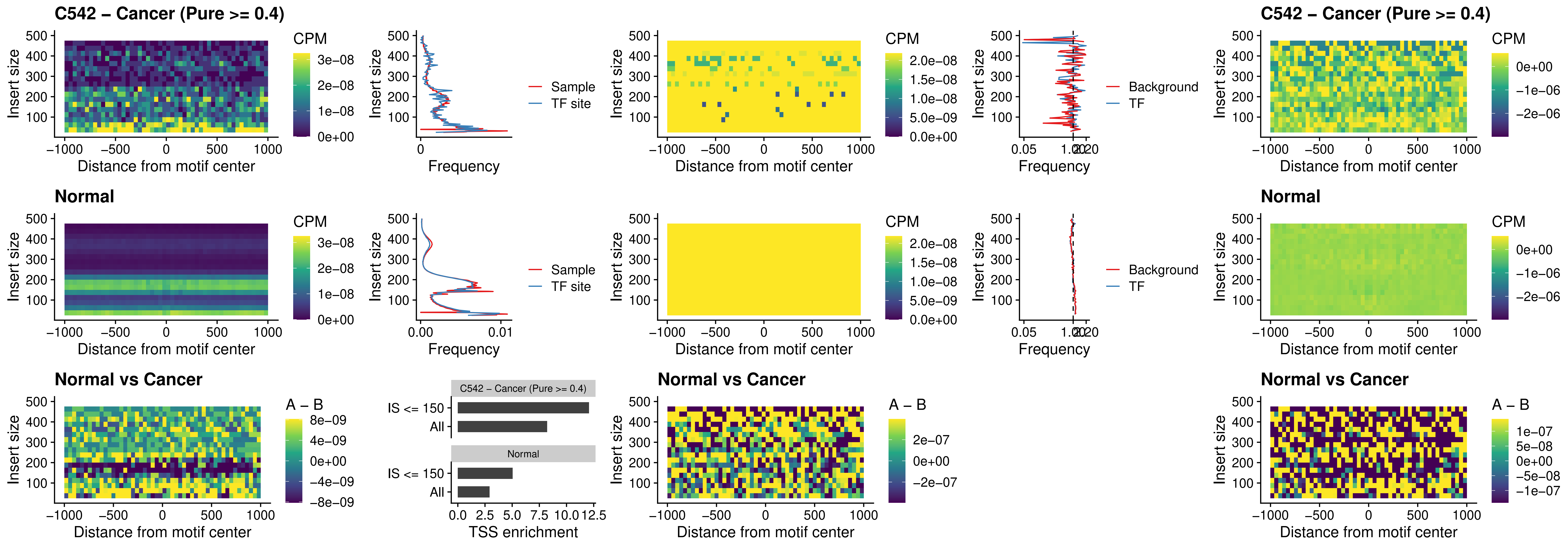

#### C542 – Cancer (Pure >= 0.4) – SPIB_uc_pTSS_nPEAK

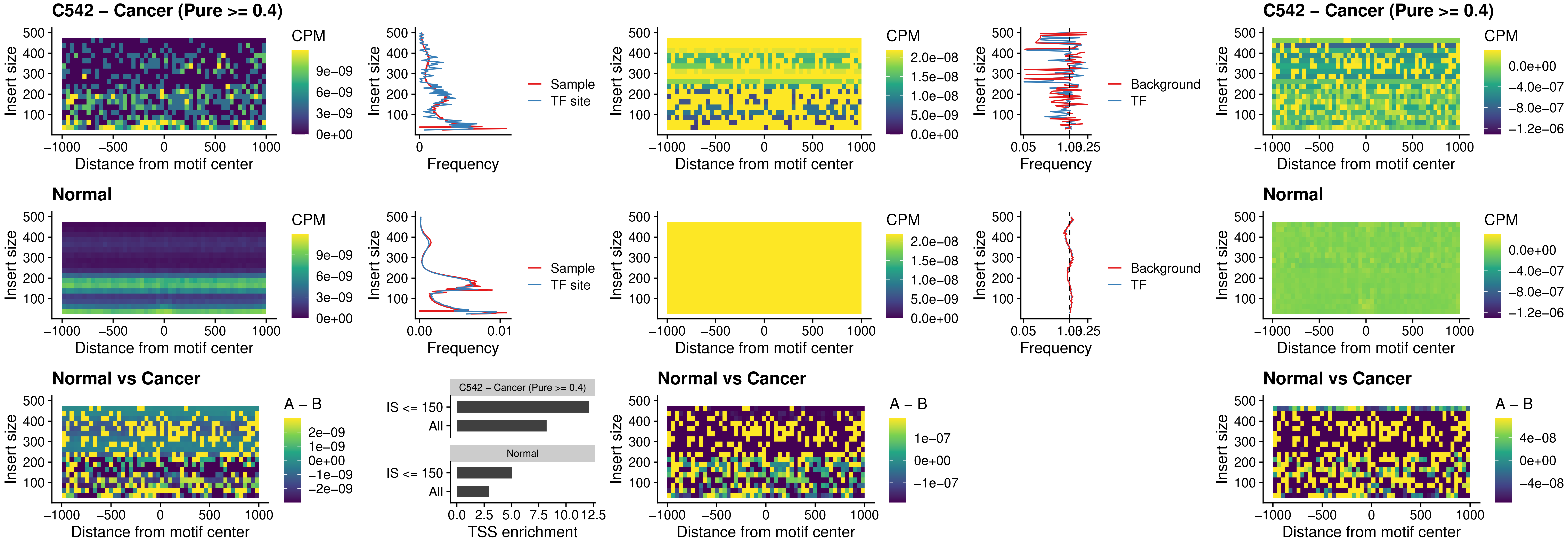

### C542 – Cancer (Pure >= 0.4) – STAT2_uc_cTSS_oPEAK

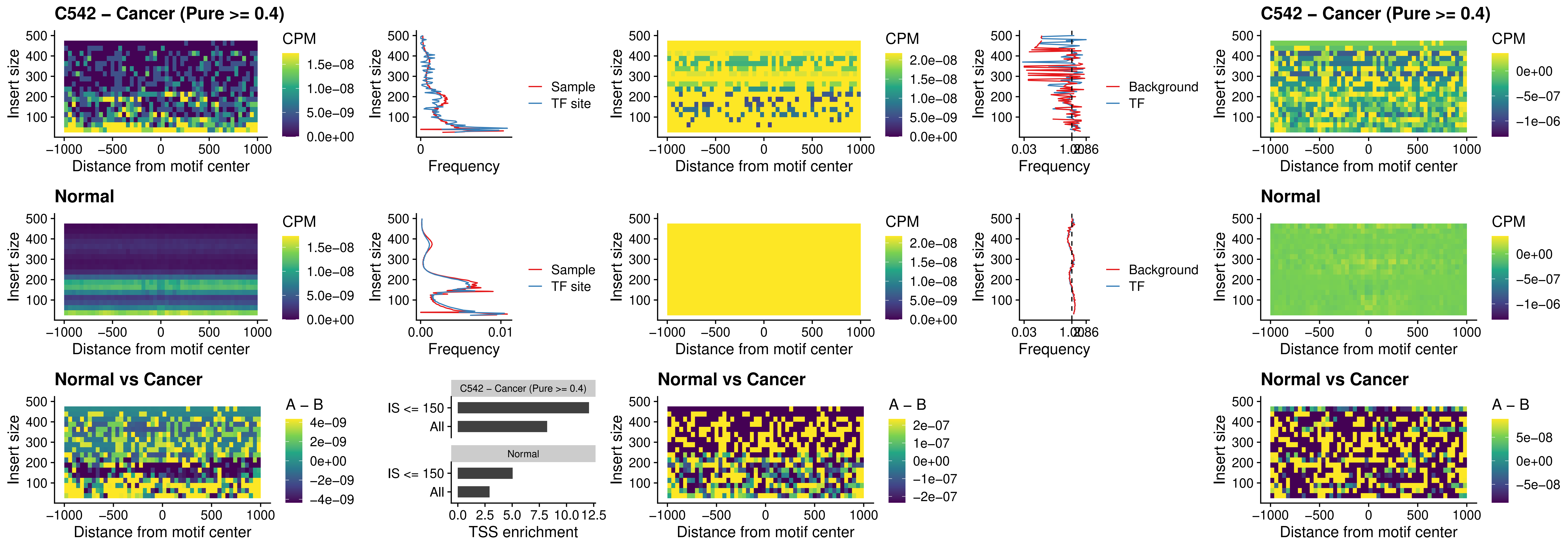

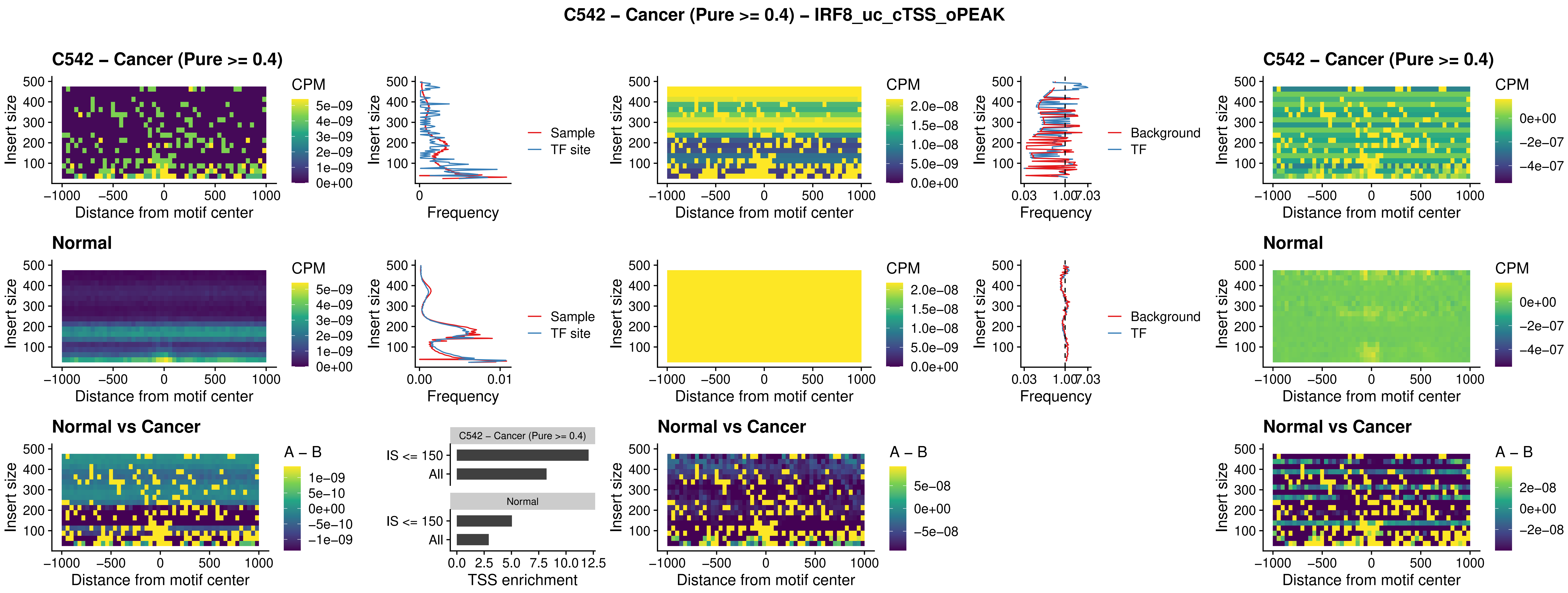

C542 – Cancer (Pure >= 0.4) – IRF8_uc_cTSS_oPEAK
C542 – Cancer (Pure >= 0.4)
Normal
Normal vs Cancer
Insert size
Distance from motif center
Frequency
CPM
A – B
Sample
TF site
Background
TF
C542 – Cancer (Pure >= 0.4)
Normal
IS <= 150
All
TSS enrichment

#### C542 – Cancer (Pure >= 0.4) – IRF3_uc_dTSS_oPEAK

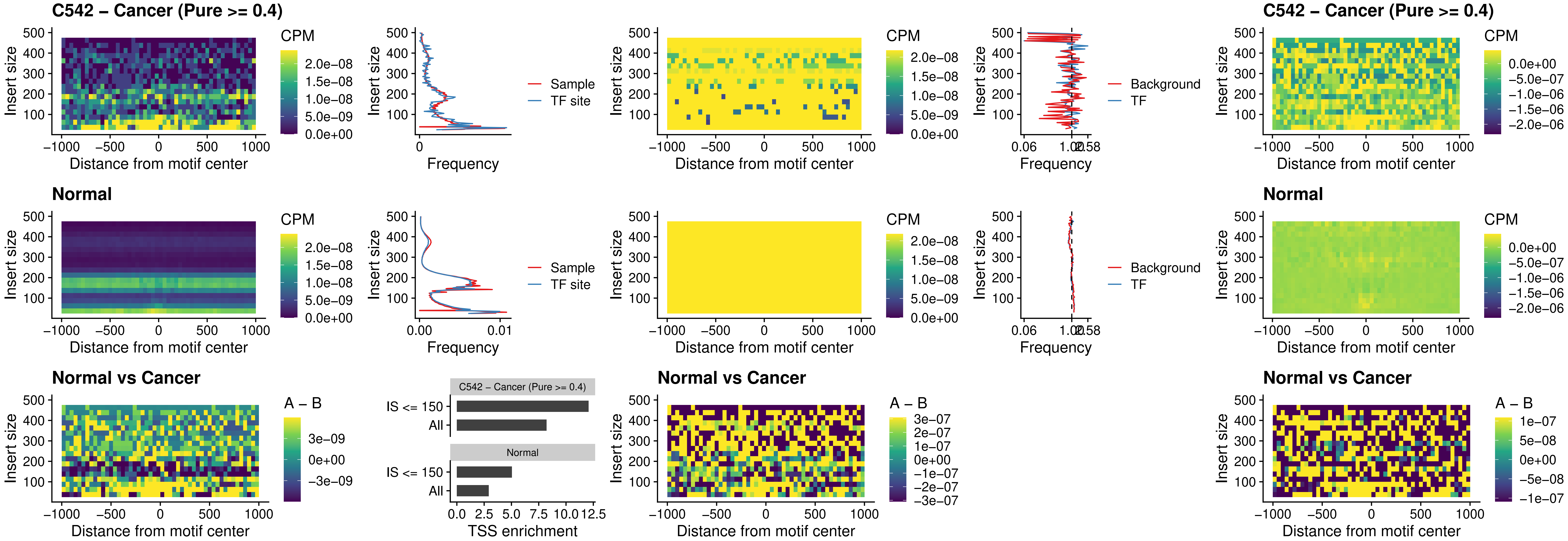

### C542 – Cancer (Pure >= 0.4) – IRF8_uc_dTSS_oPEAK

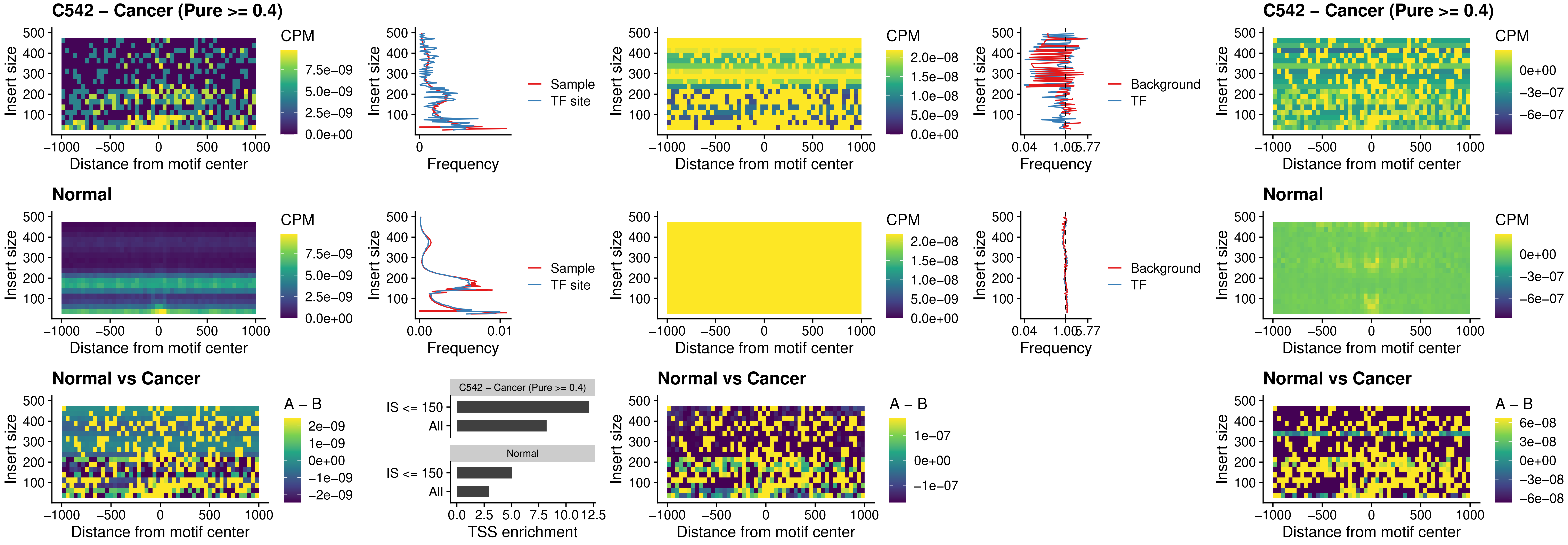

### C542 – Cancer (Pure >= 0.4) – ETS2_uc_dTSS_oPEAK

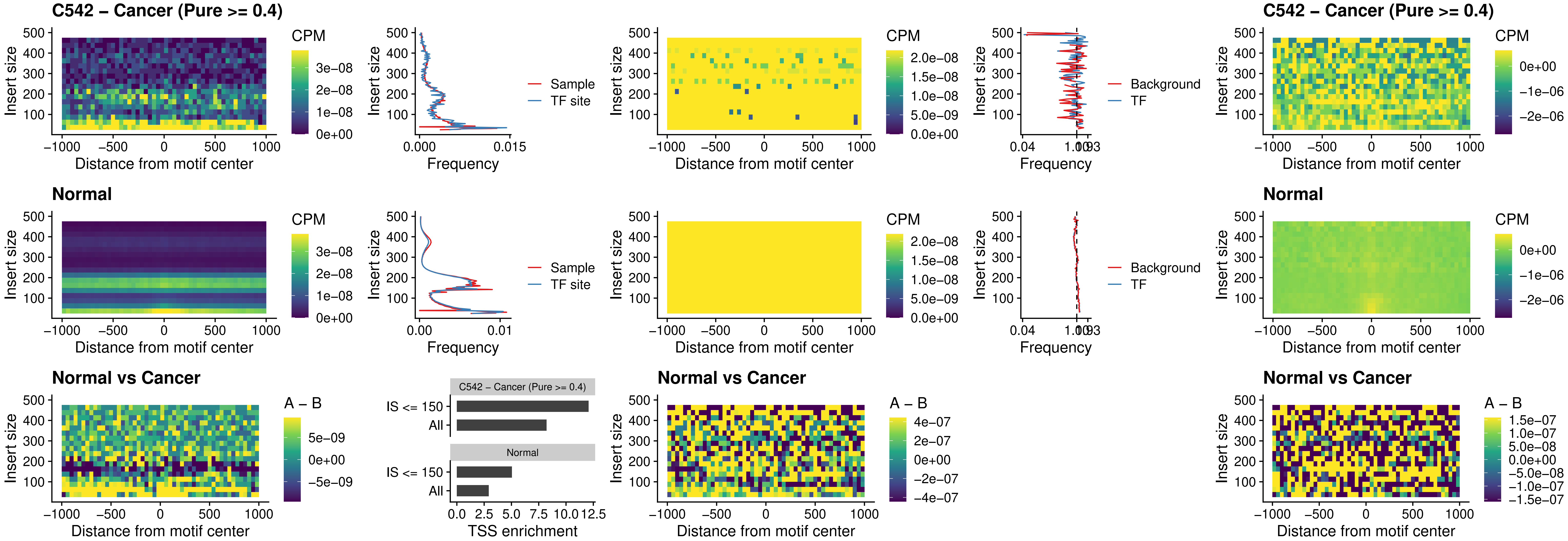

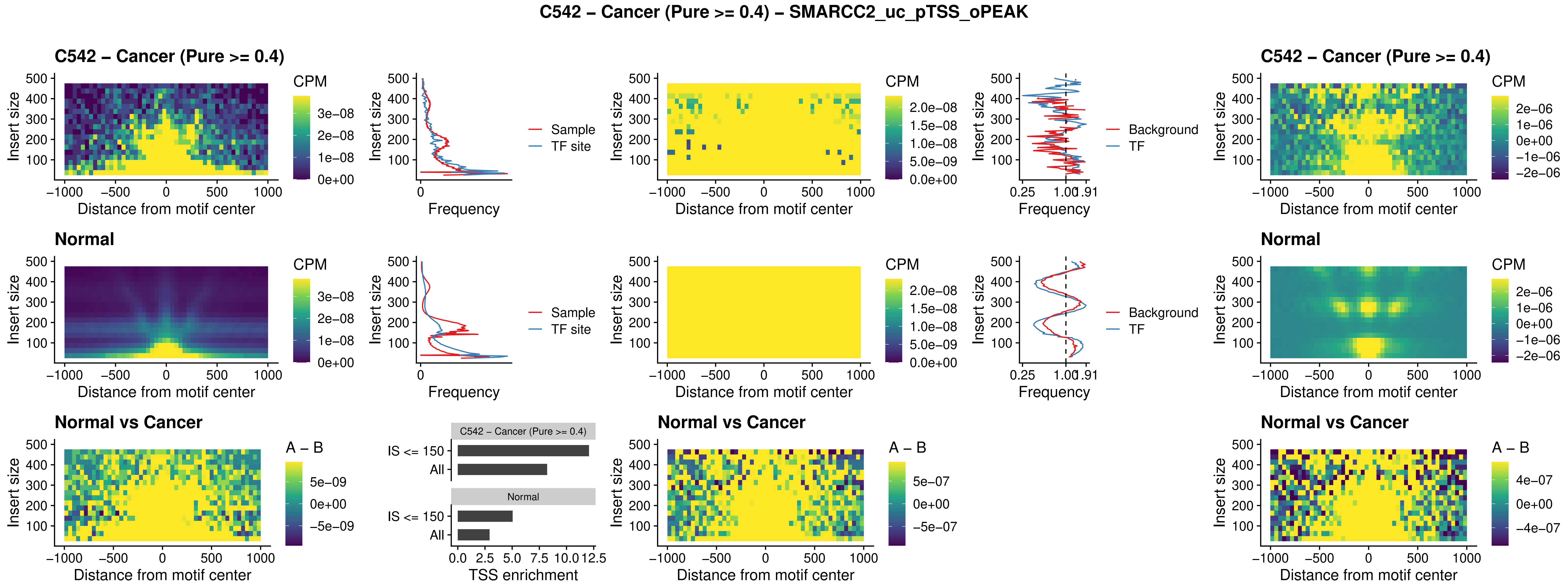
C542 – Cancer (Pure >= 0.4) – SMARCC2_uc_pTSS_oPEAK
C542 – Cancer (Pure >= 0.4)
Normal
Normal vs Cancer
Insert size
Distance from motif center
Frequency
CPM
A – B
Sample
TF site
Background
TF
C542 – Cancer (Pure >= 0.4)
Normal
IS <= 150
All
TSS enrichment

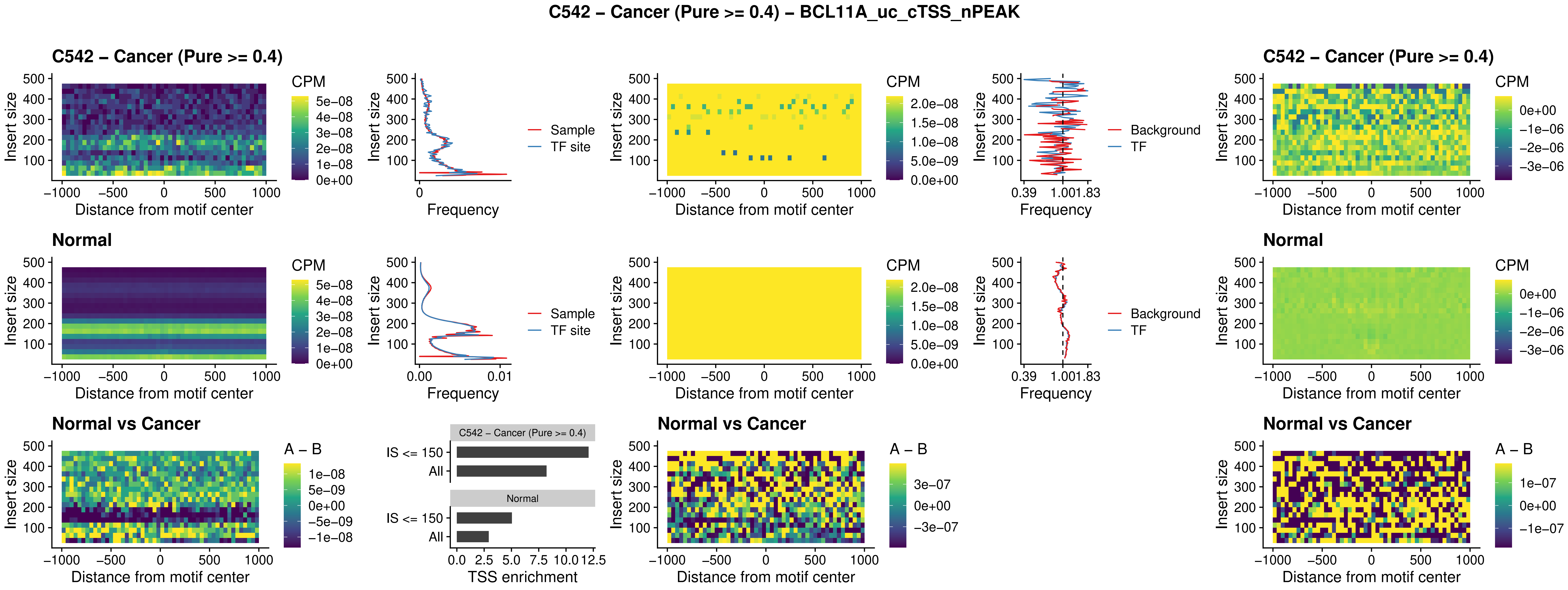
C542 – Cancer (Pure >= 0.4) – BCL11A_uc_cTSS_nPEAK
C542 – Cancer (Pure >= 0.4)
Normal
Normal vs Cancer
Insert size
Distance from motif center
CPM
A – B
Frequency
Sample
TF site
Background
TF
C542 – Cancer (Pure >= 0.4)
Normal
IS <= 150
All
TSS enrichment

#### C542 – Cancer (Pure >= 0.4) – BCL11B_uc_cTSS_nPEAK

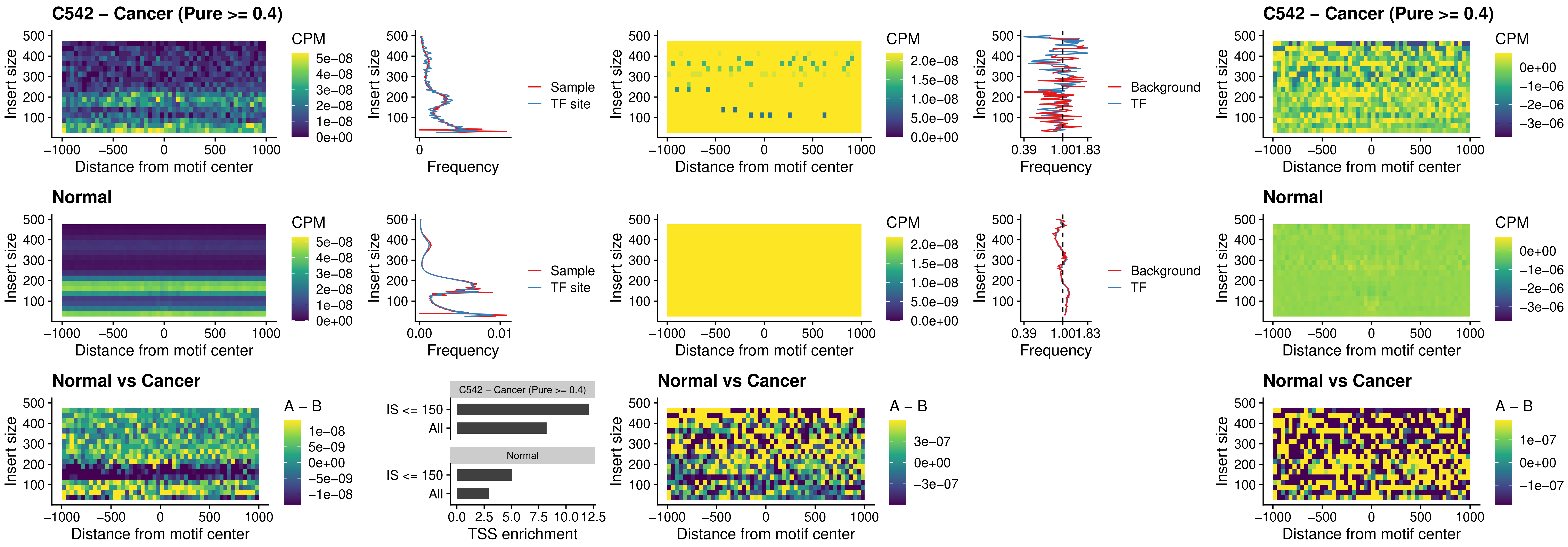

#### C542 – Cancer (Pure >= 0.4) – STAT2_uc_dTSS_oPEAK

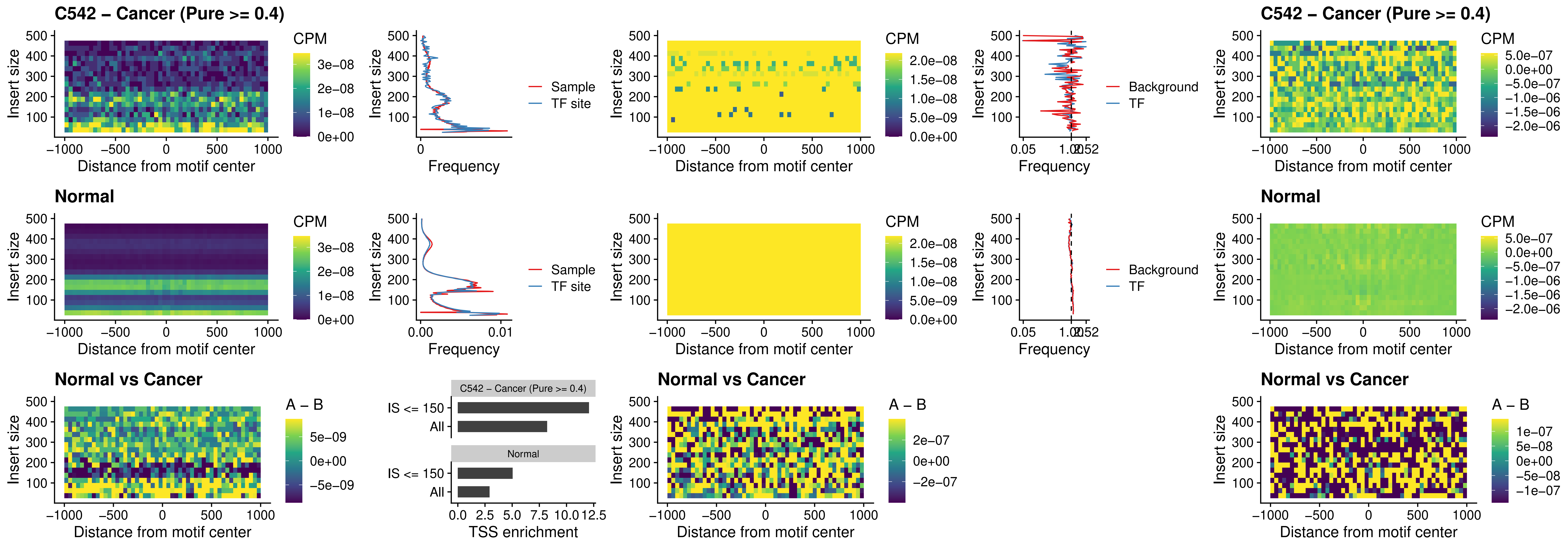

#### C542 – Cancer (Pure >= 0.4) – ZNF683_uc_cTSS_oPEAK

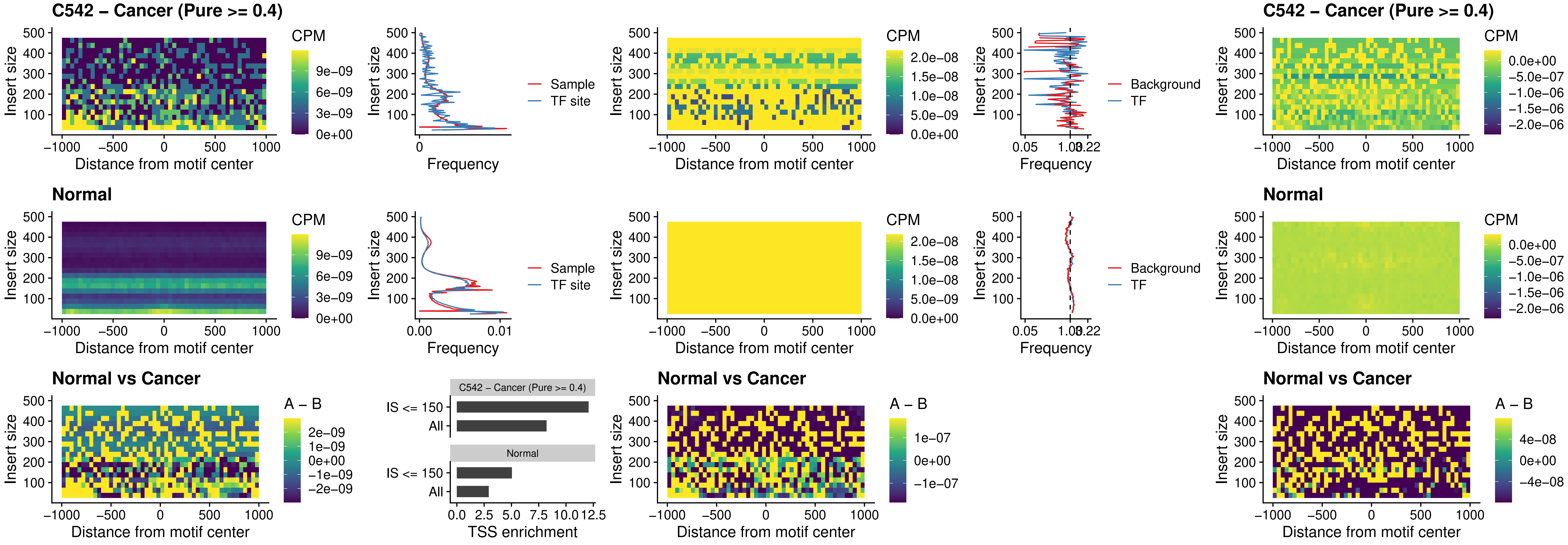

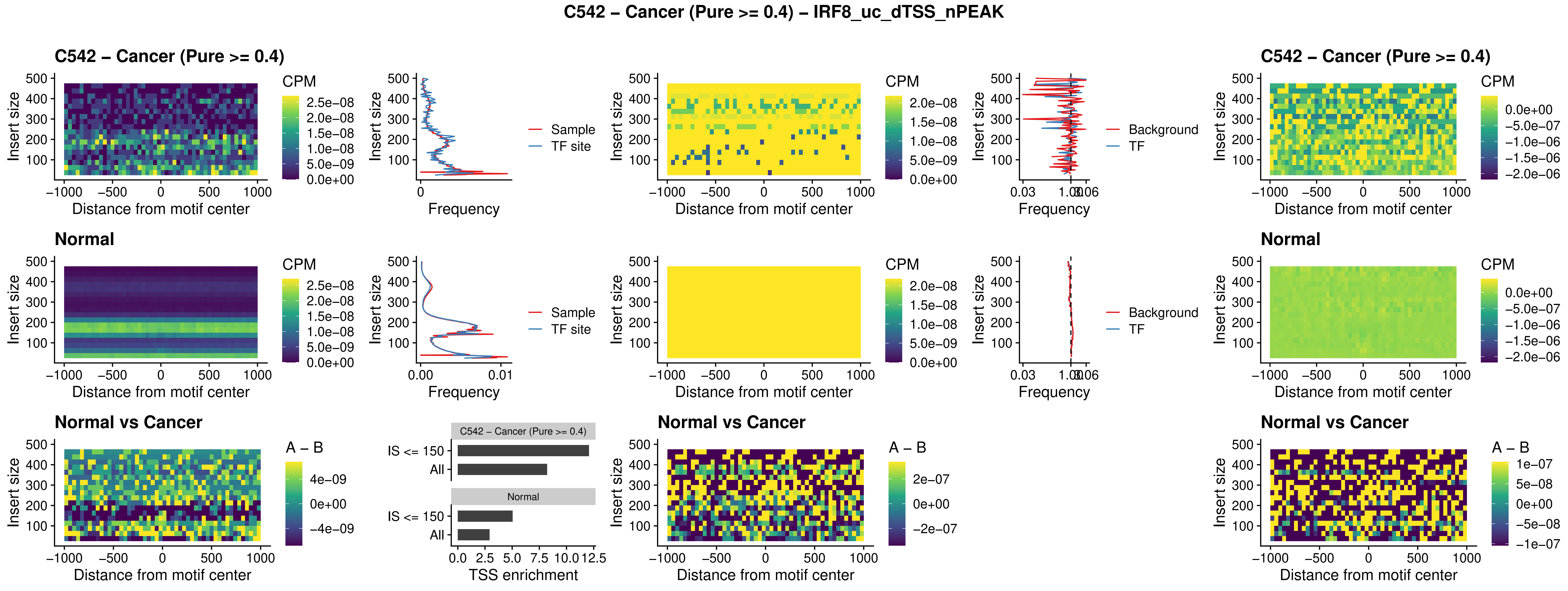
C542 – Cancer (Pure >= 0.4) – IRF8_uc_dTSS_nPEAK
C542 – Cancer (Pure >= 0.4)
Normal
Normal vs Cancer
Insert size
Distance from motif center
Frequency
CPM
A – B
Sample
TF site
Background
TF
C542 – Cancer (Pure >= 0.4)
Normal
IS <= 150
All
TSS enrichment

### C542 – Cancer (Pure >= 0.4) – ETV7_uc_dTSS_oPEAK

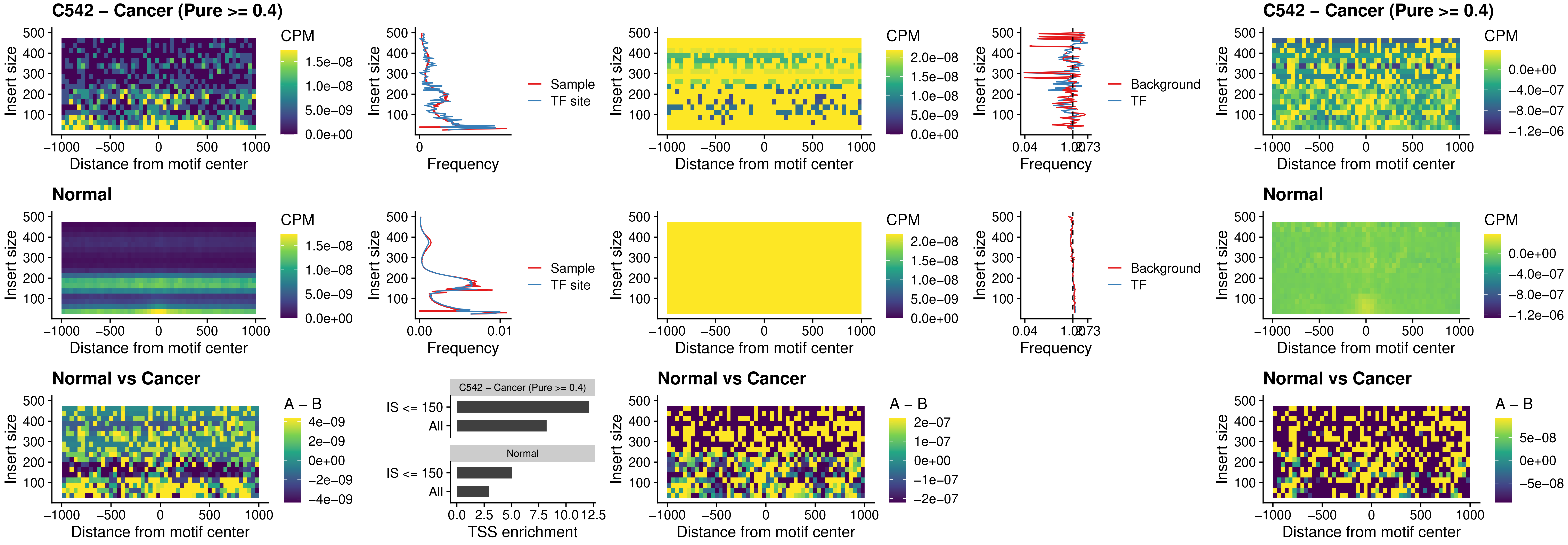

### C543 – Cancer (Pure >= 0.4) – FOXL1_uc_pTSS_oPEAK

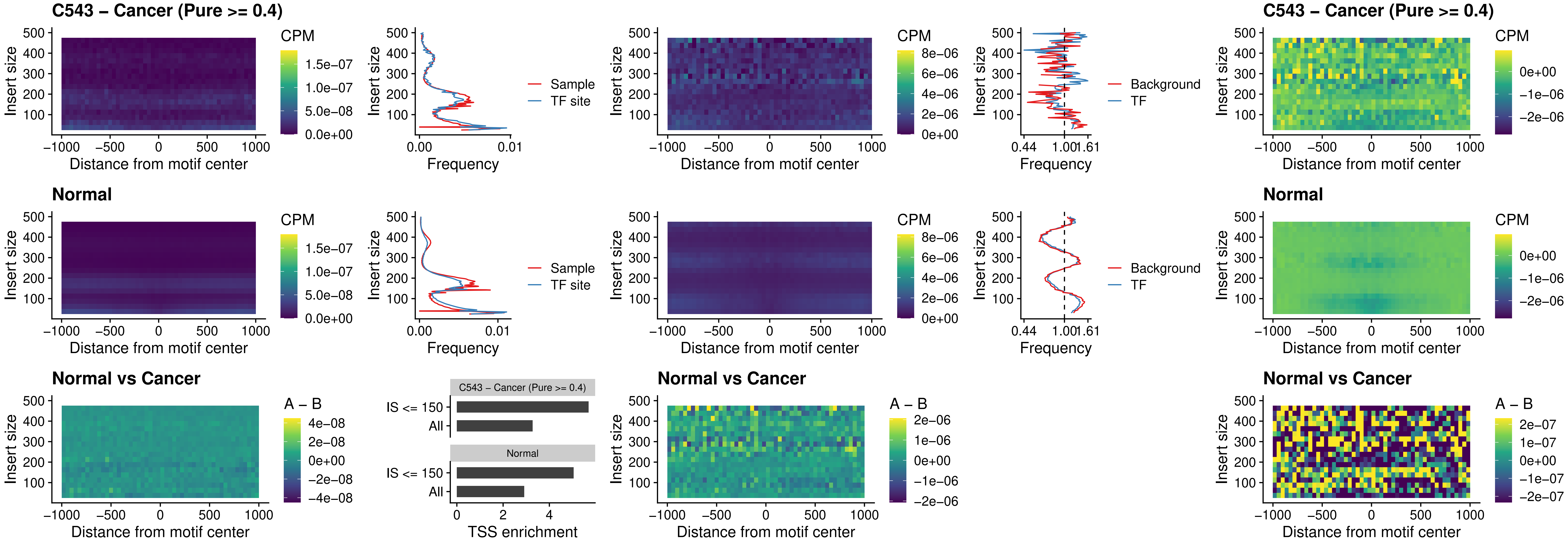

### C543 – Cancer (Pure >= 0.4) – CPEB1_uc_cTSS_oPEAK

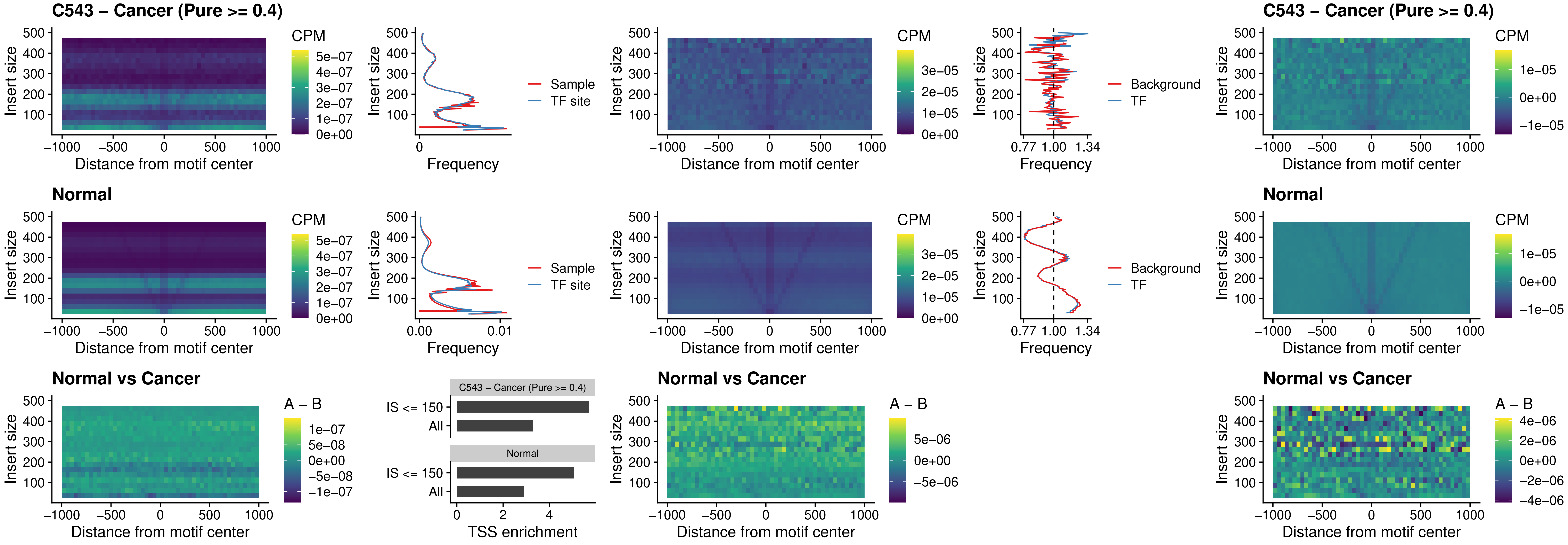

**C543 – Cancer (Pure >= 0.4) – HMGA1_uc_pTSS_nPEAK**

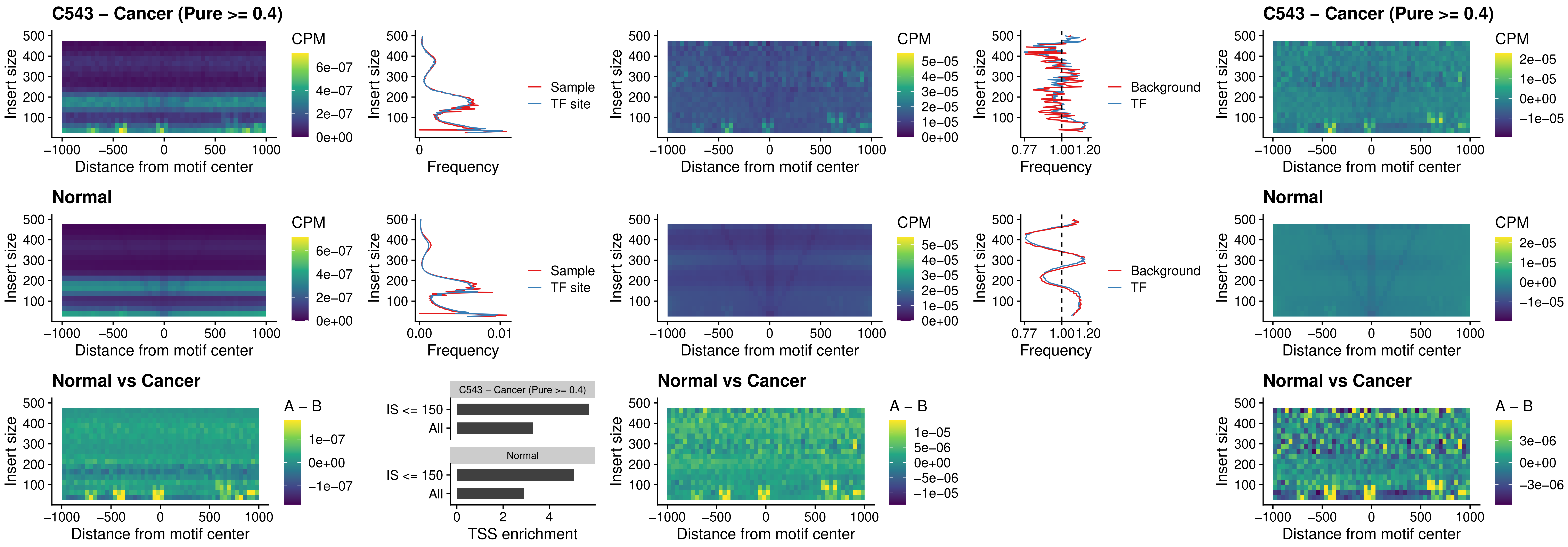

### C543 – Cancer (Pure >= 0.4) – SOX13_uc_cTSS_nPEAK

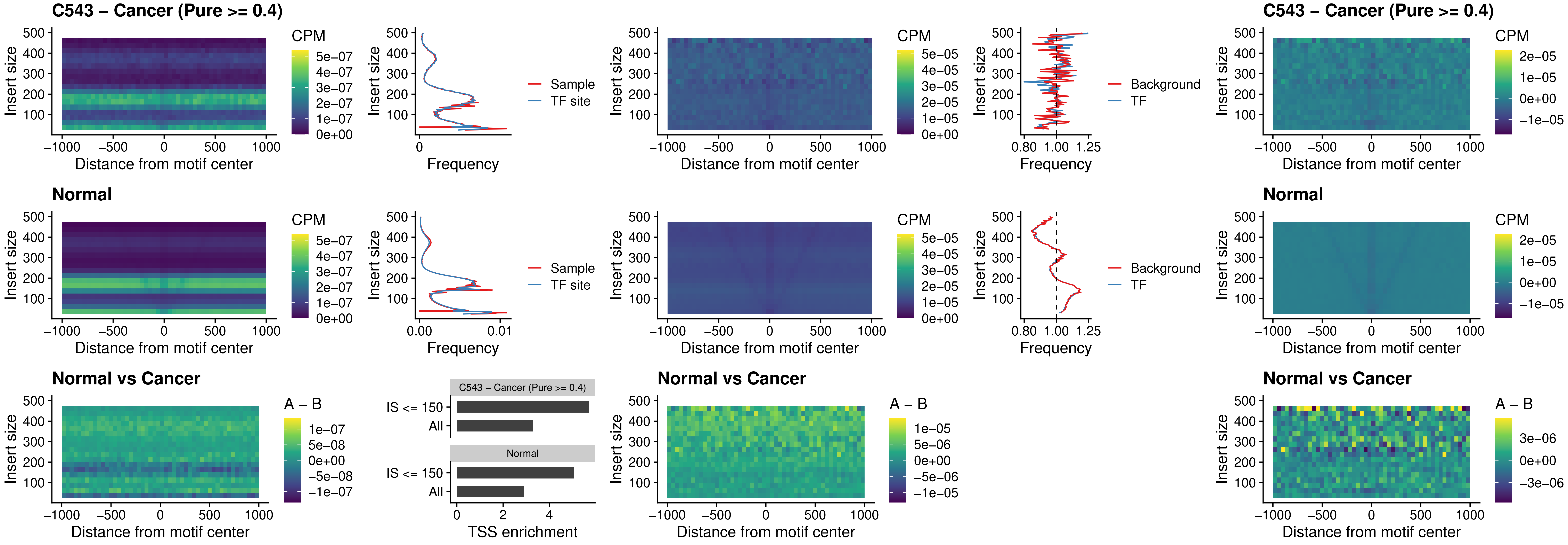

#### C543 – Cancer (Pure >= 0.4) – LHX4_uc_cTSS_nPEAK

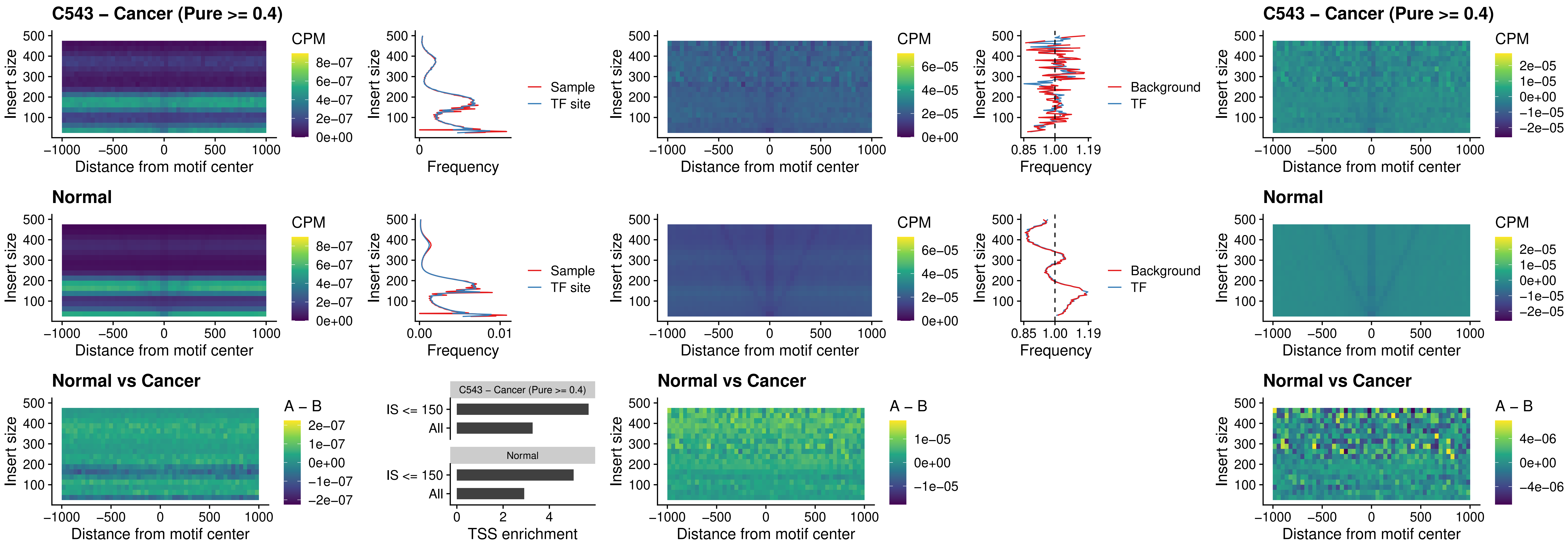

#### C543 – Cancer (Pure >= 0.4) – HOXA4_uc_cTSS_nPEAK

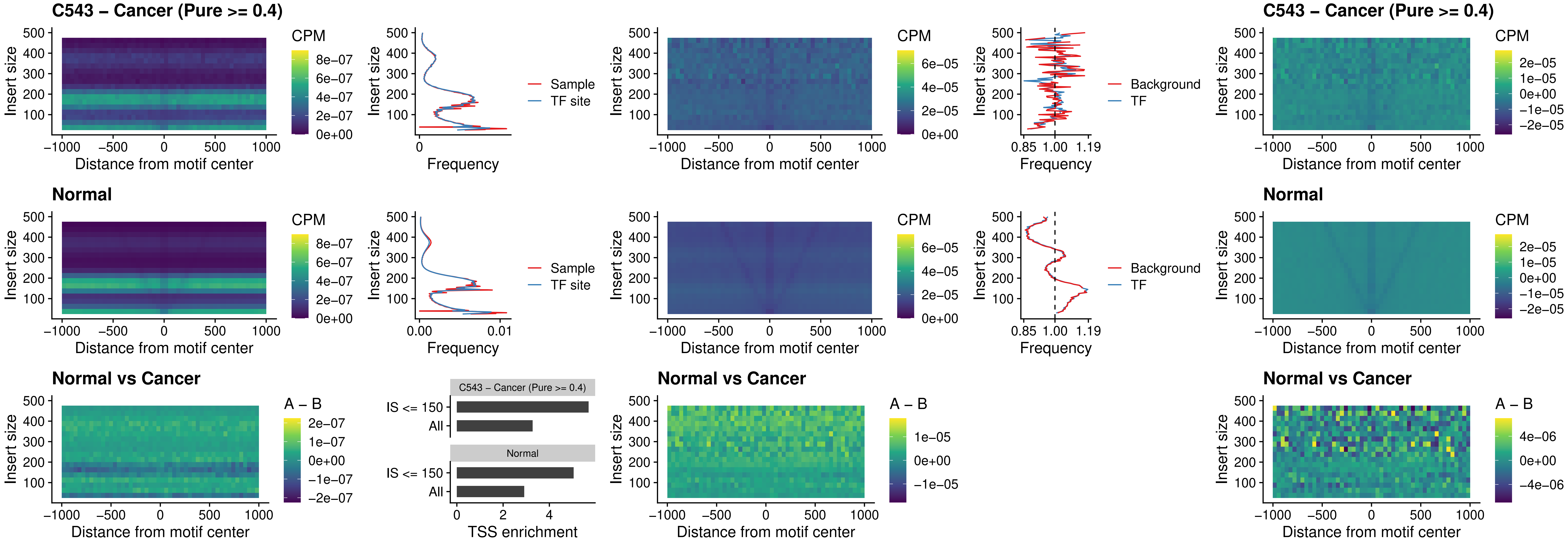

#### C543 – Cancer (Pure >= 0.4) – HLX_uc_cTSS_nPEAK

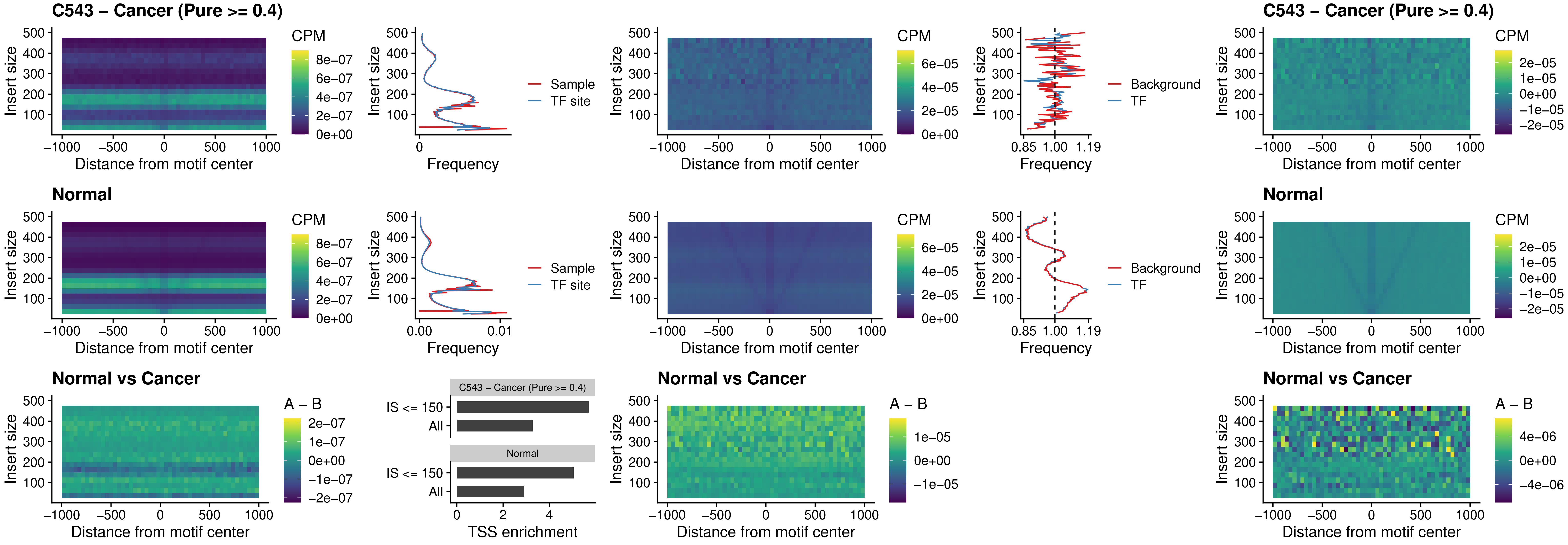

#### C543 – Cancer (Pure >= 0.4) – AC0125311_uc_cTSS_nPEAK

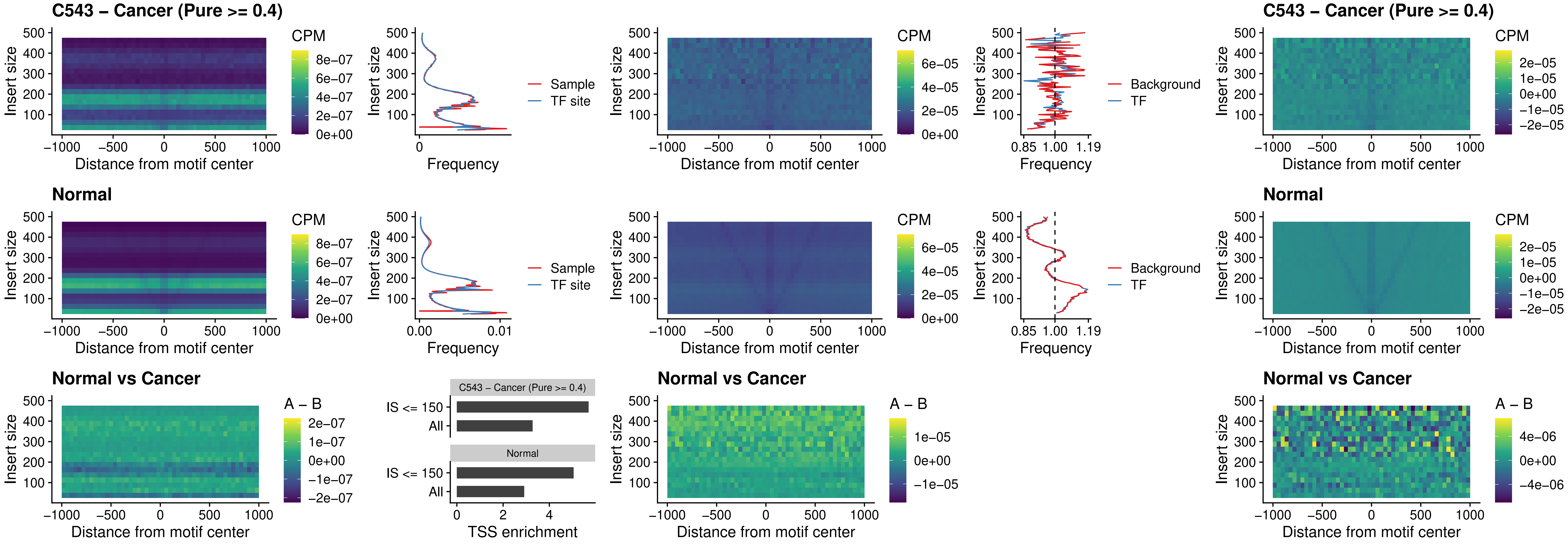

#### C543 – Cancer (Pure >= 0.4) – DBX2_uc_cTSS_nPEAK

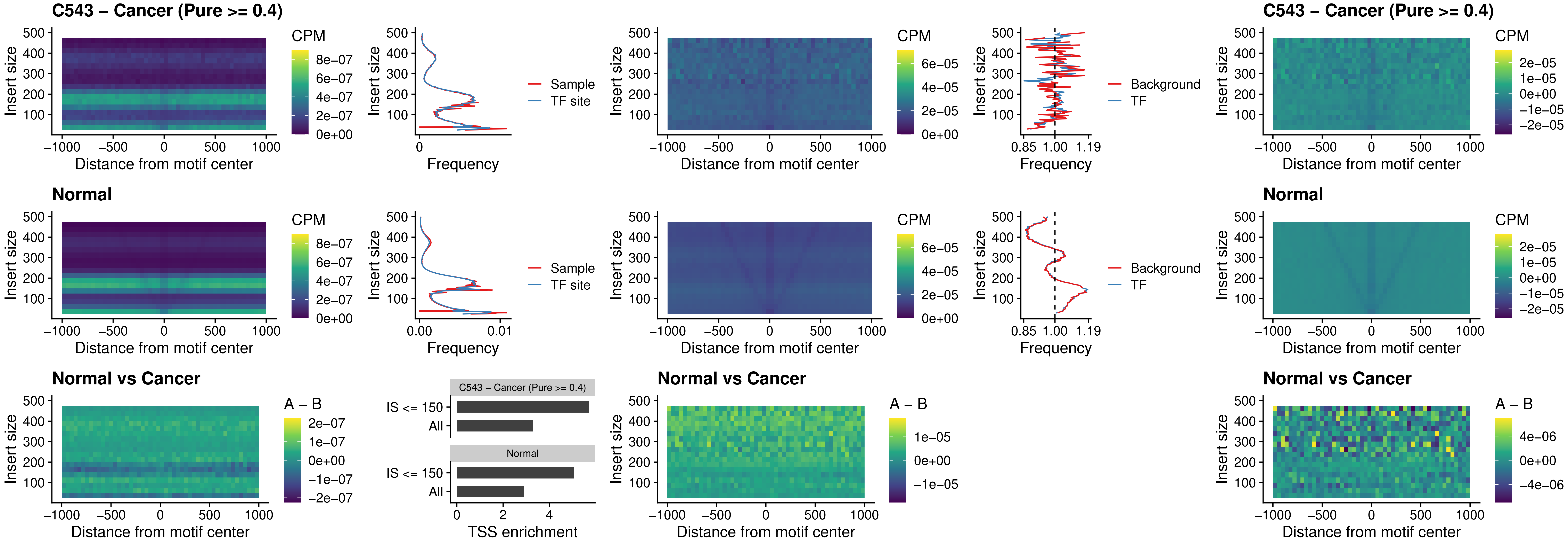

#### C543 – Cancer (Pure >= 0.4) – LHX6_uc_dTSS_nPEAK

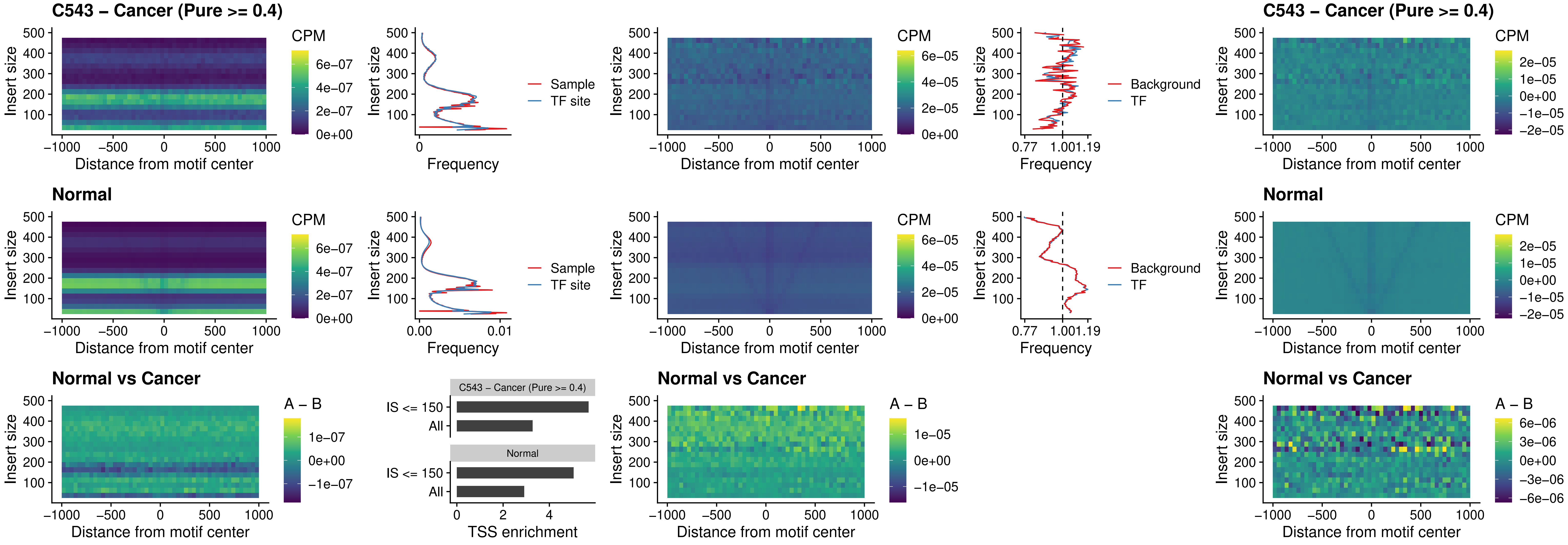

#### C543 – Cancer (Pure >= 0.4) – SOX1_uc_dTSS_nPEAK

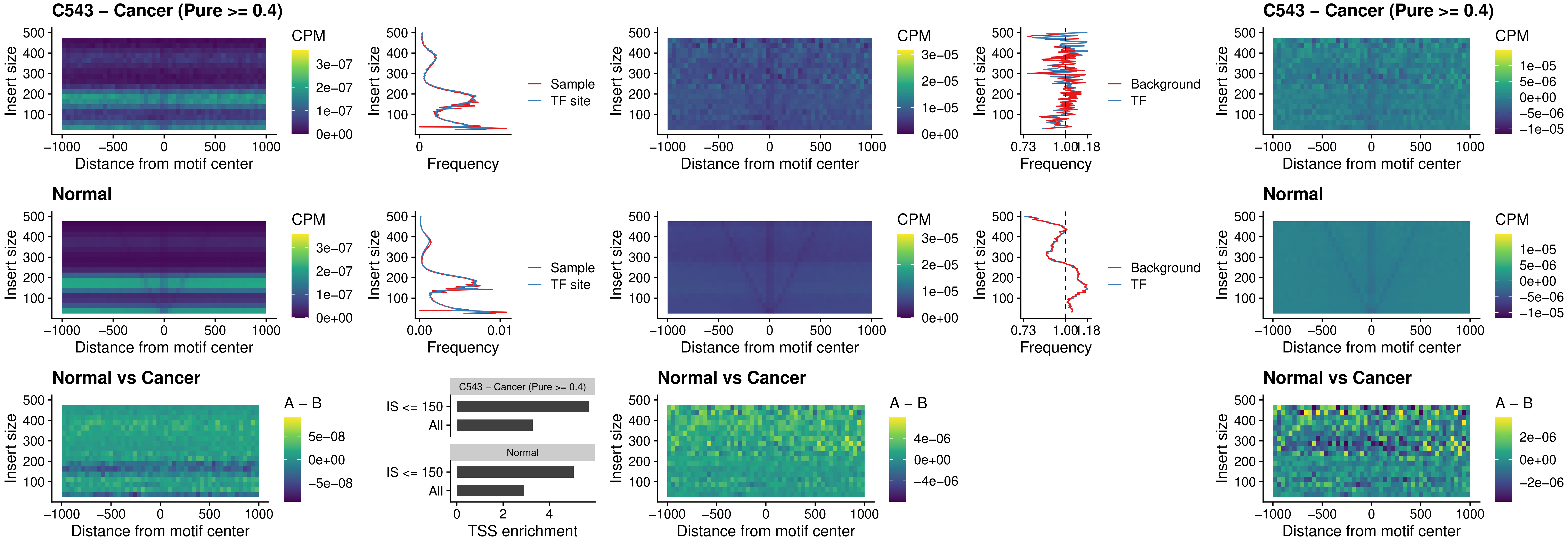

#### C543 – Cancer (Pure >= 0.4) – NKX24_uc_dTSS_nPEAK

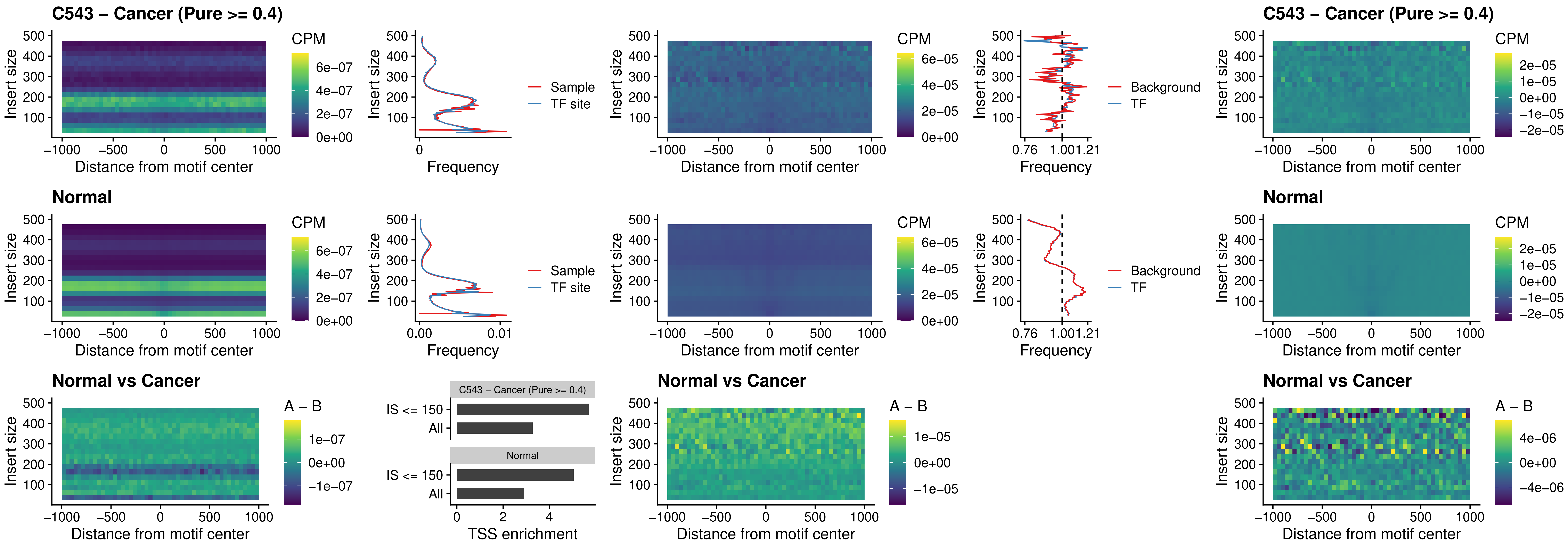

### C543 – Cancer (Pure >= 0.4) – DLX3_uc_cTSS_nPEAK

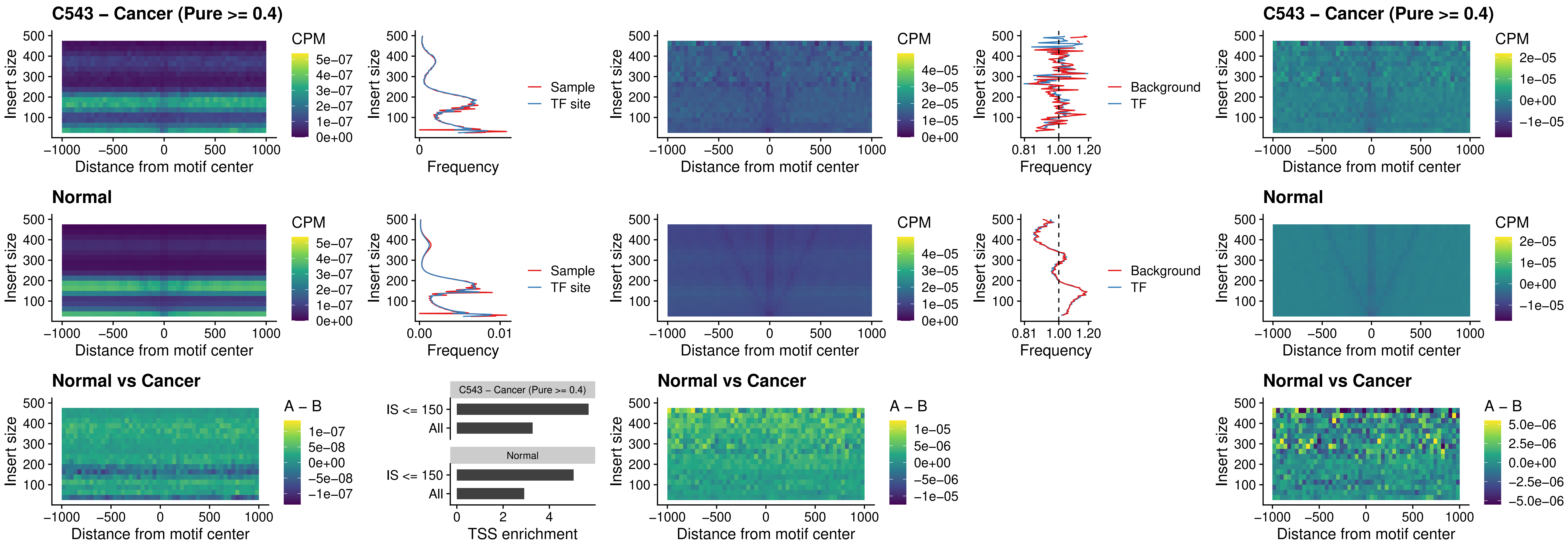

#### C543 – Cancer (Pure >= 0.4) – HOXD3_uc_cTSS_nPEAK

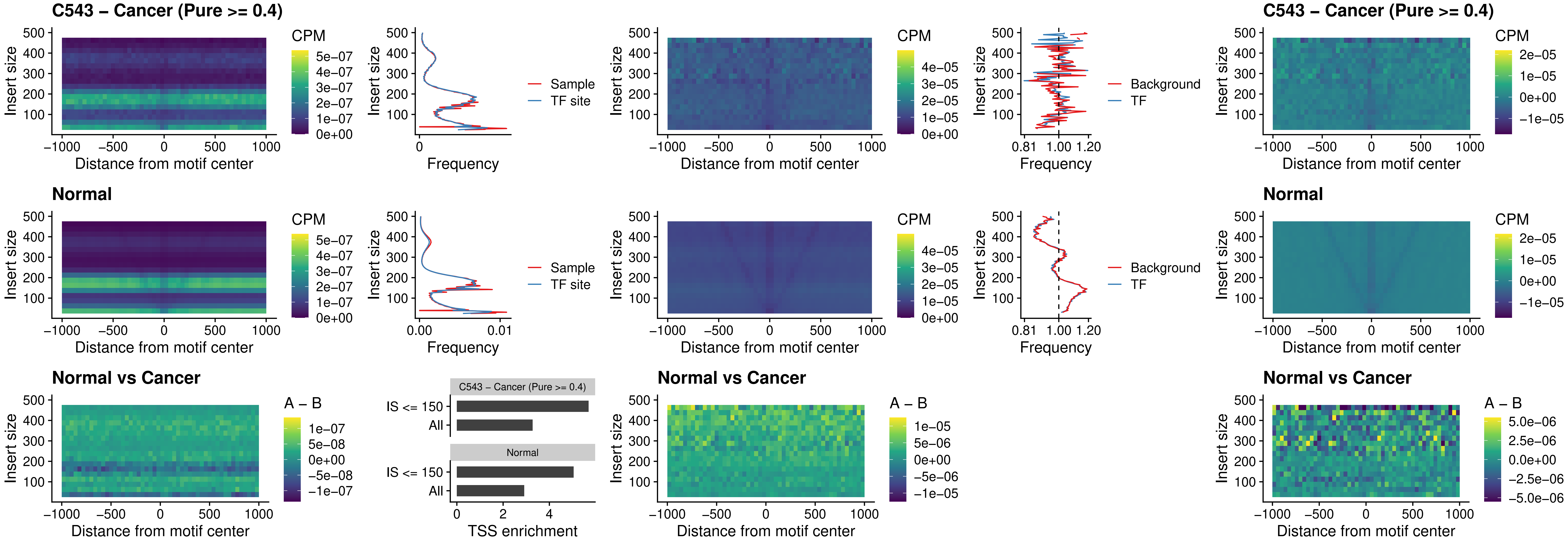

#### C543 – Cancer (Pure >= 0.4) – ISL1_uc_dTSS_oPEAK

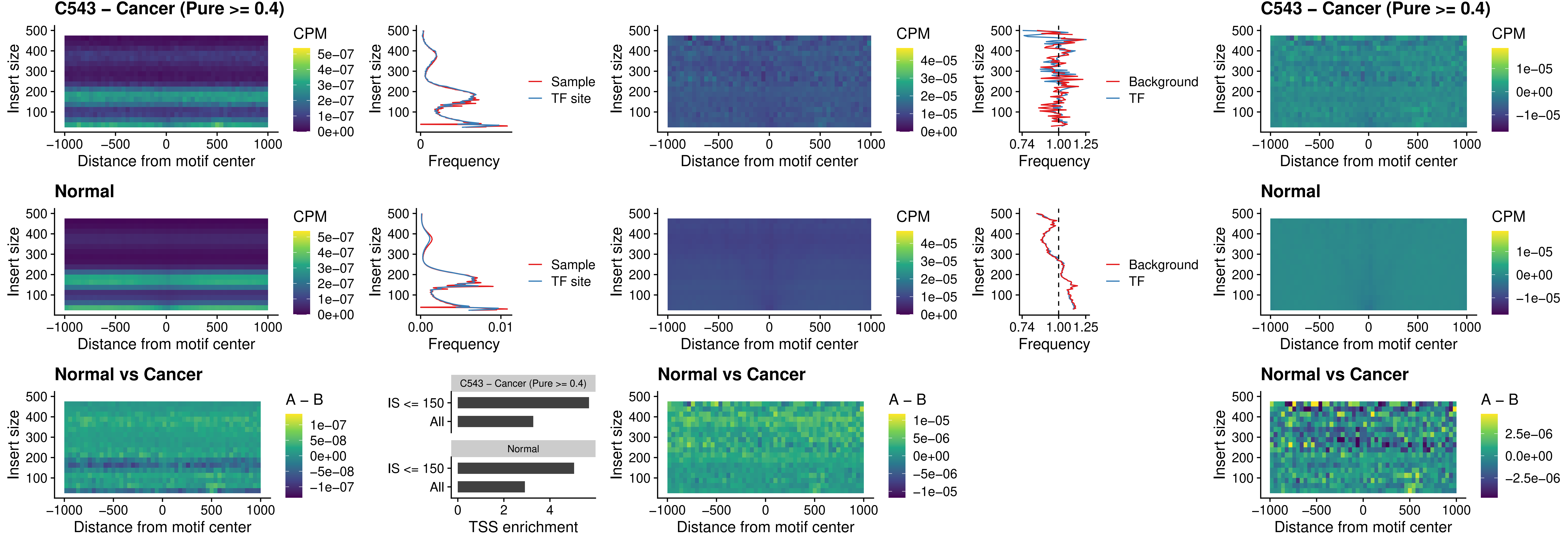

#### C543 – Cancer (Pure >= 0.4) – HOXB4_uc_dTSS_oPEAK

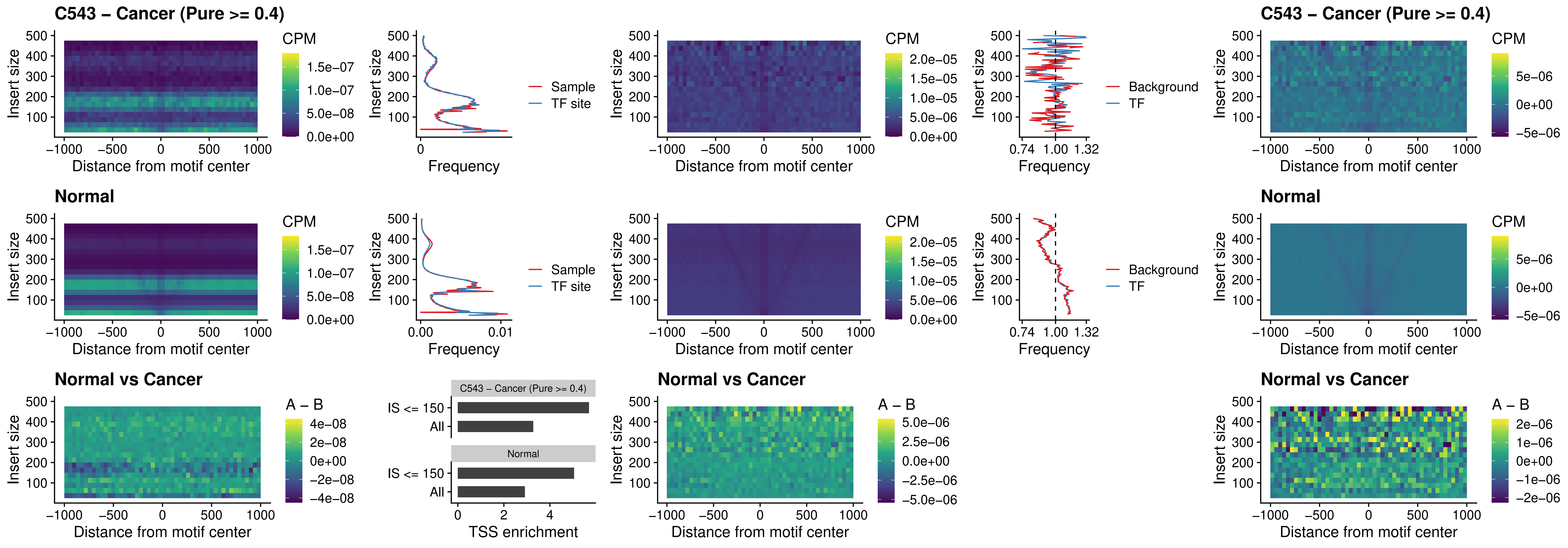

#### C543 – Cancer (Pure >= 0.4) – VAX1_uc_dTSS_oPEAK

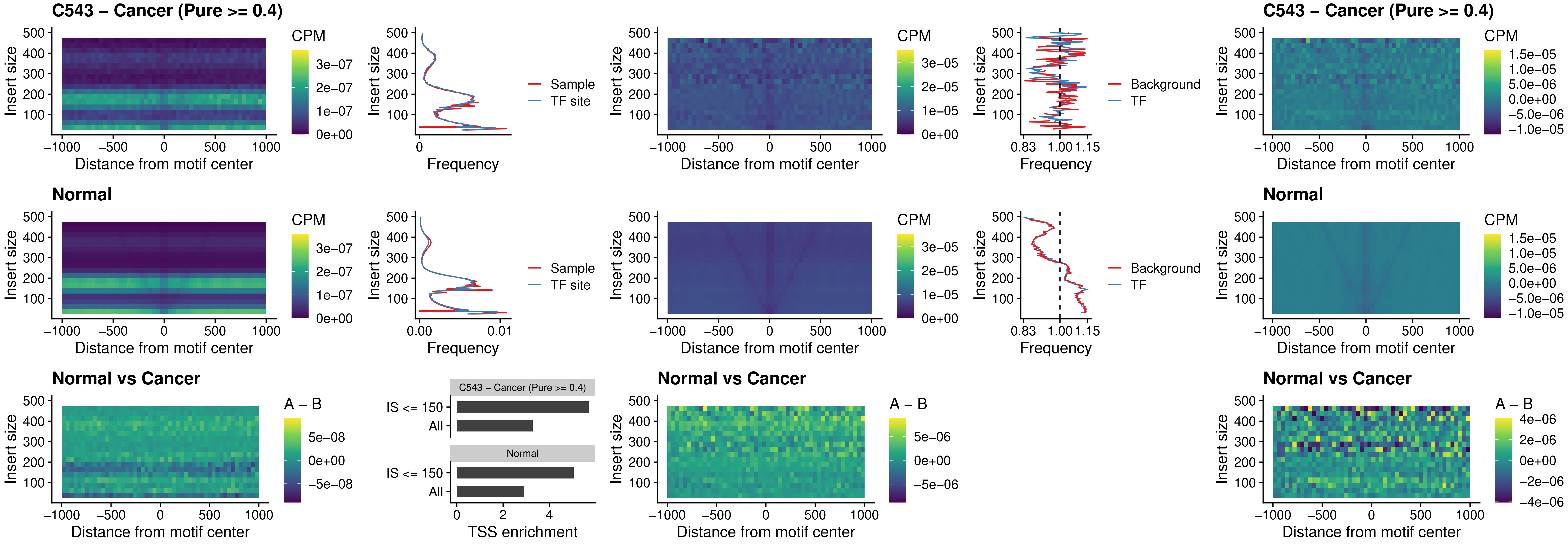

#### C543 – Cancer (Pure >= 0.4) – DLX3_uc_dTSS_oPEAK

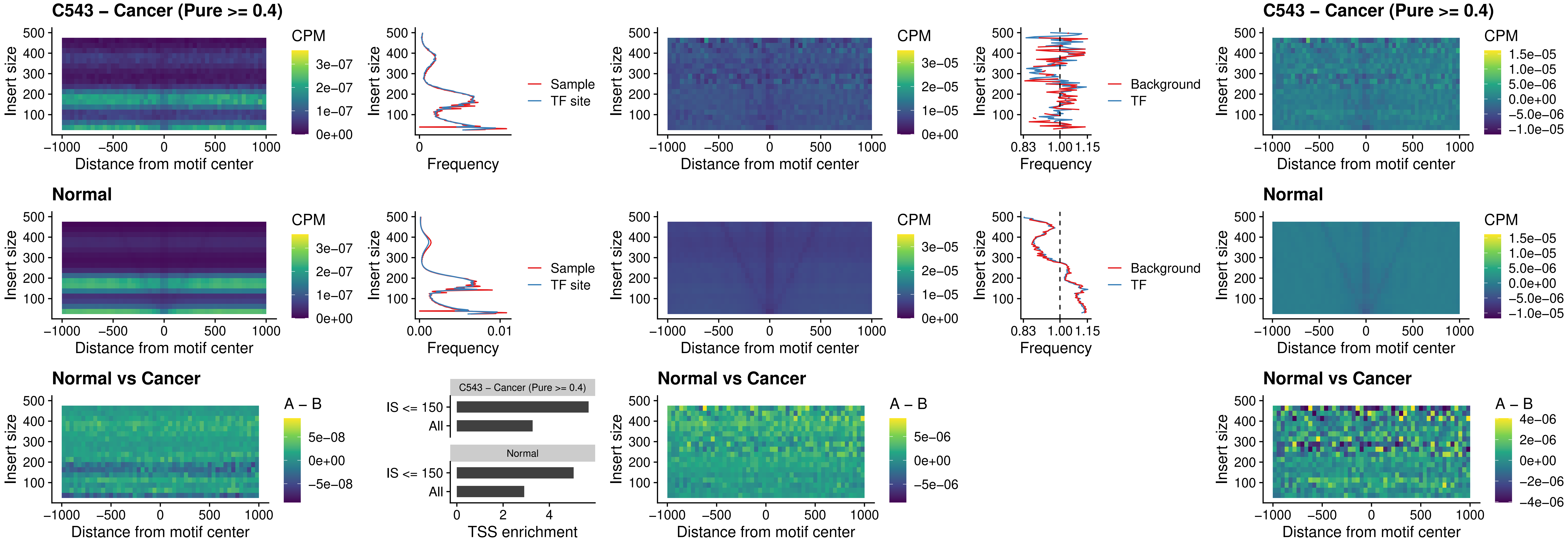

#### C543 – Cancer (Pure >= 0.4) – HOXD3_uc_dTSS_oPEAK

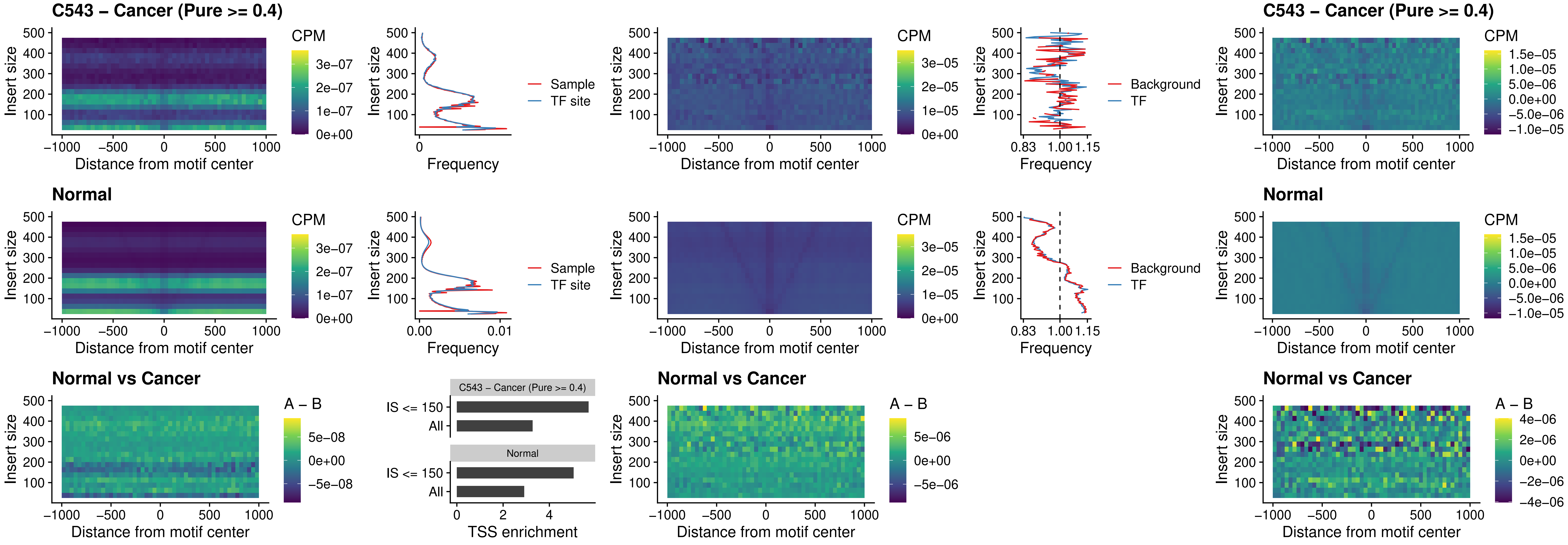

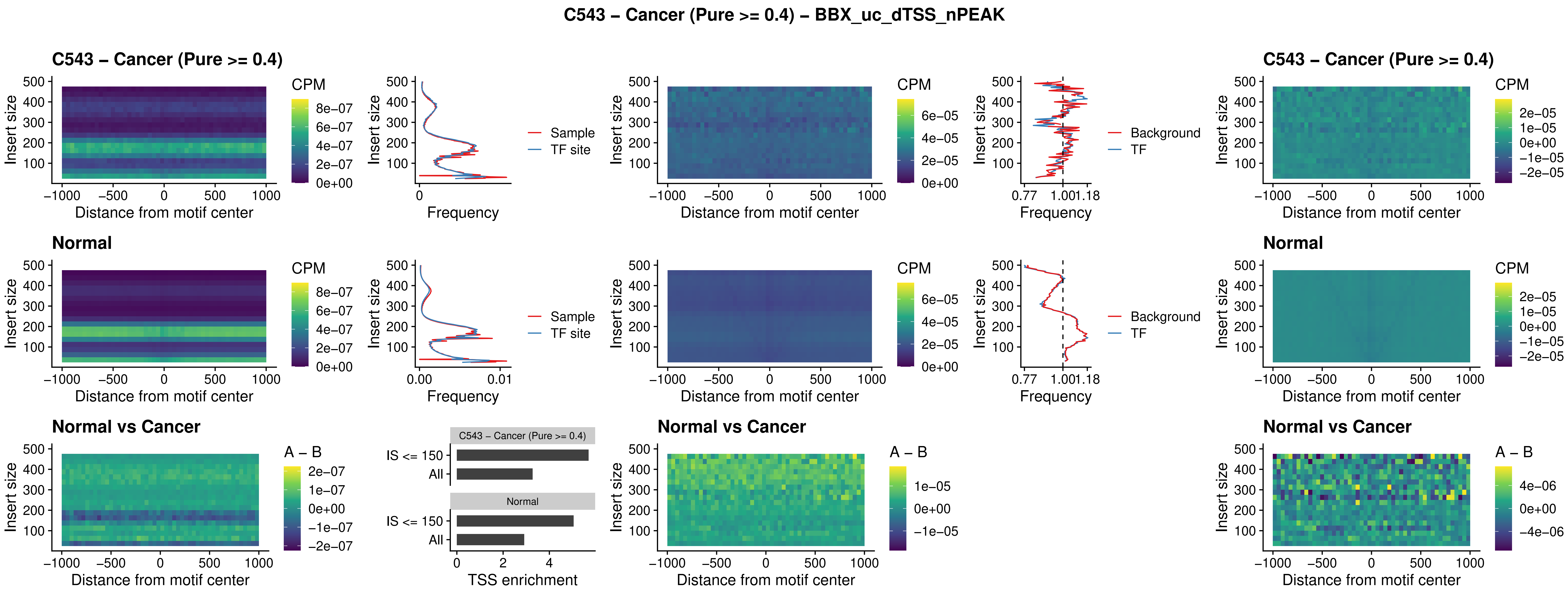
C543 – Cancer (Pure >= 0.4) – BBX_uc_dTSS_nPEAK
C543 – Cancer (Pure >= 0.4)
Normal
Normal vs Cancer
Insert size
Distance from motif center
Frequency
CPM
A – B
Sample
TF site
Background
TF
C543 – Cancer (Pure >= 0.4)
Normal
IS <= 150
All
TSS enrichment

#### C543 – Cancer (Pure >= 0.4) – SOX3_uc_cTSS_nPEAK

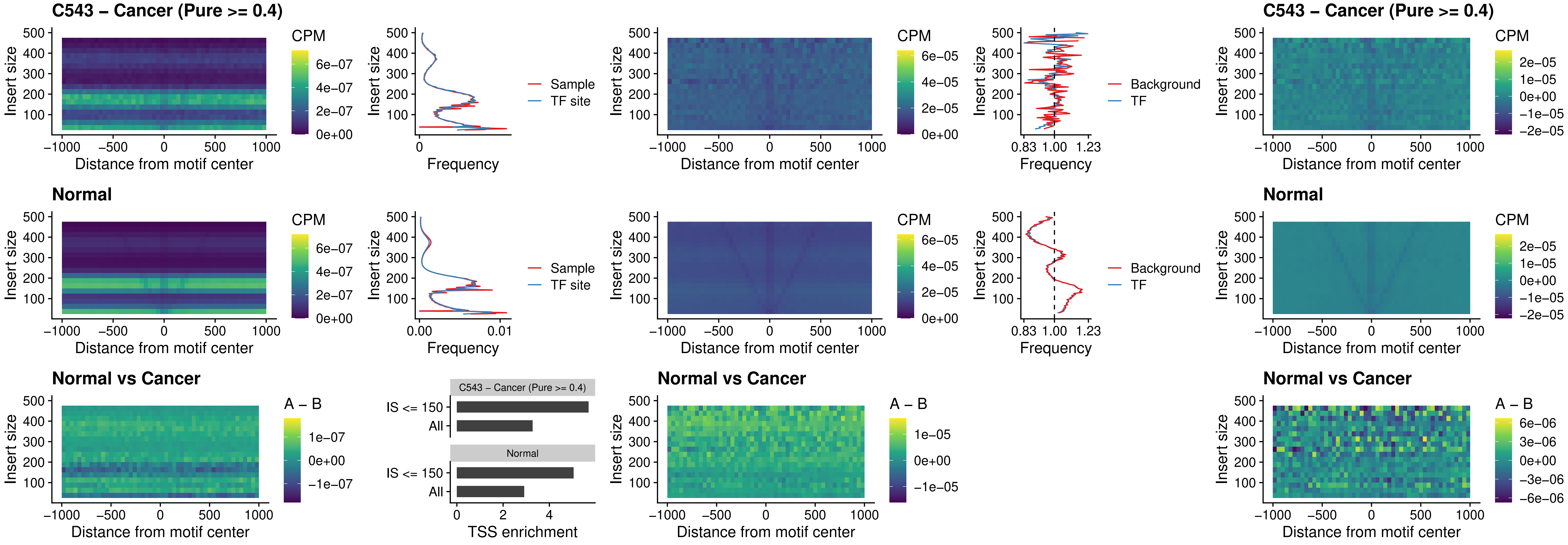

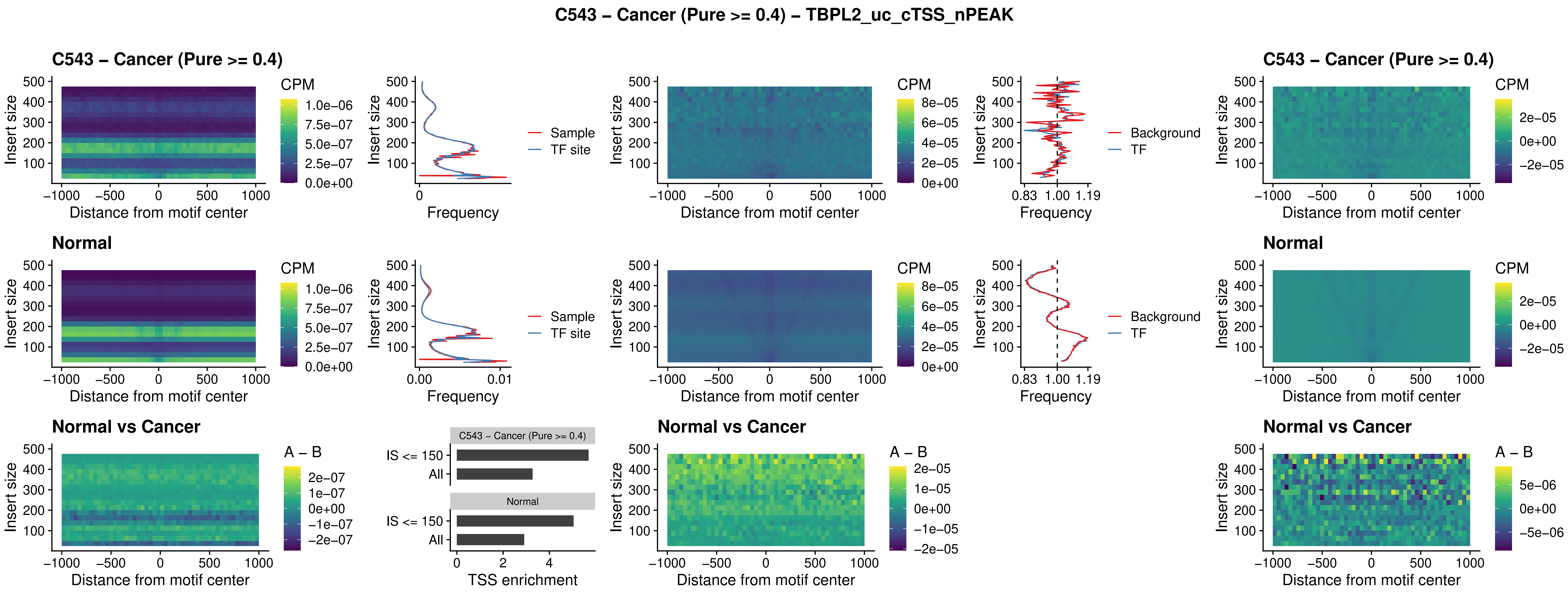
C543 – Cancer (Pure >= 0.4) – TBPL2_uc_cTSS_nPEAK
C543 – Cancer (Pure >= 0.4)
Normal
Normal vs Cancer
Insert size
Distance from motif center
Frequency
CPM
A – B
Sample
TF site
Background
TF
C543 – Cancer (Pure >= 0.4)
IS <= 150
All
TSS enrichment

#### C543 – Cancer (Pure >= 0.4) – HOXA4_uc_cTSS_oPEAK

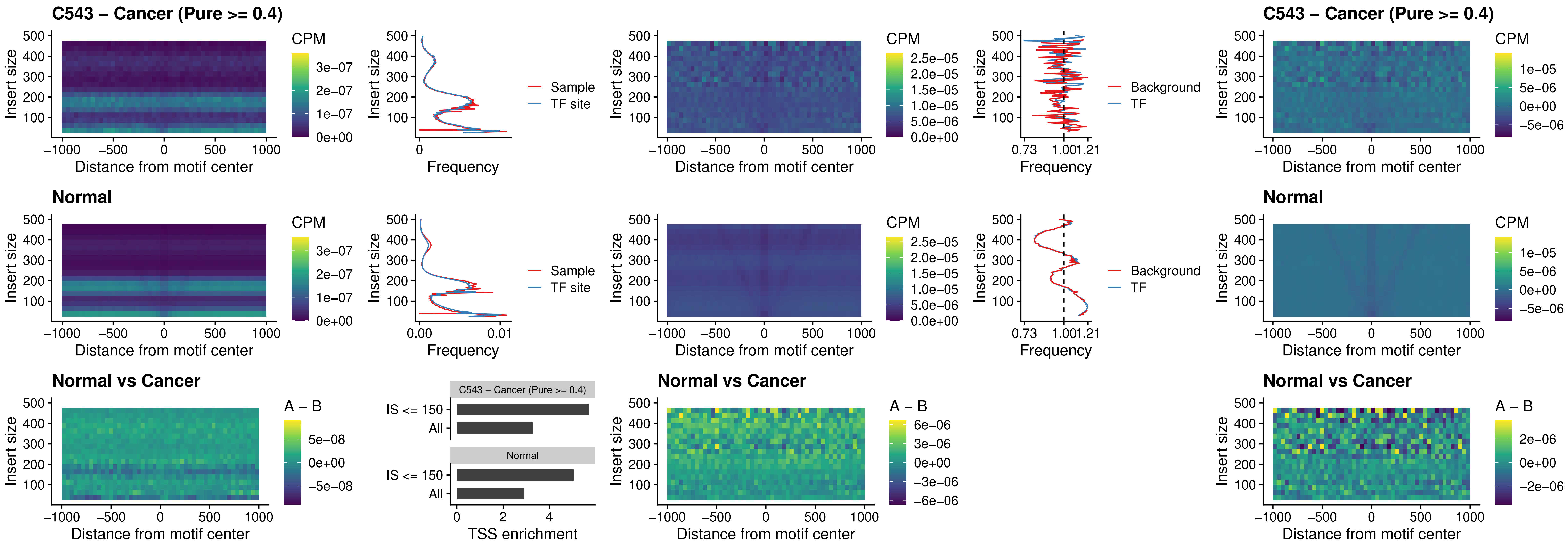

### C543 – Cancer (Pure >= 0.4) – PDX1_uc_dTSS_oPEAK

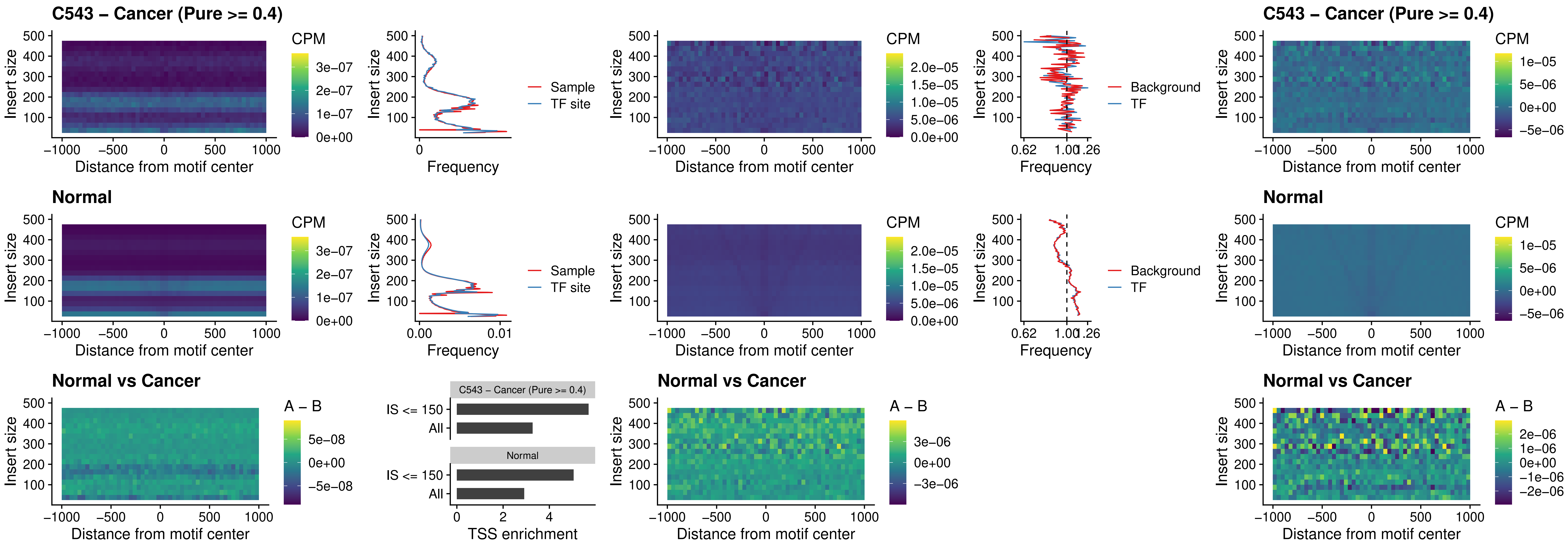

#### C543 – Cancer (Pure >= 0.4) – LHX4_uc_dTSS_oPEAK

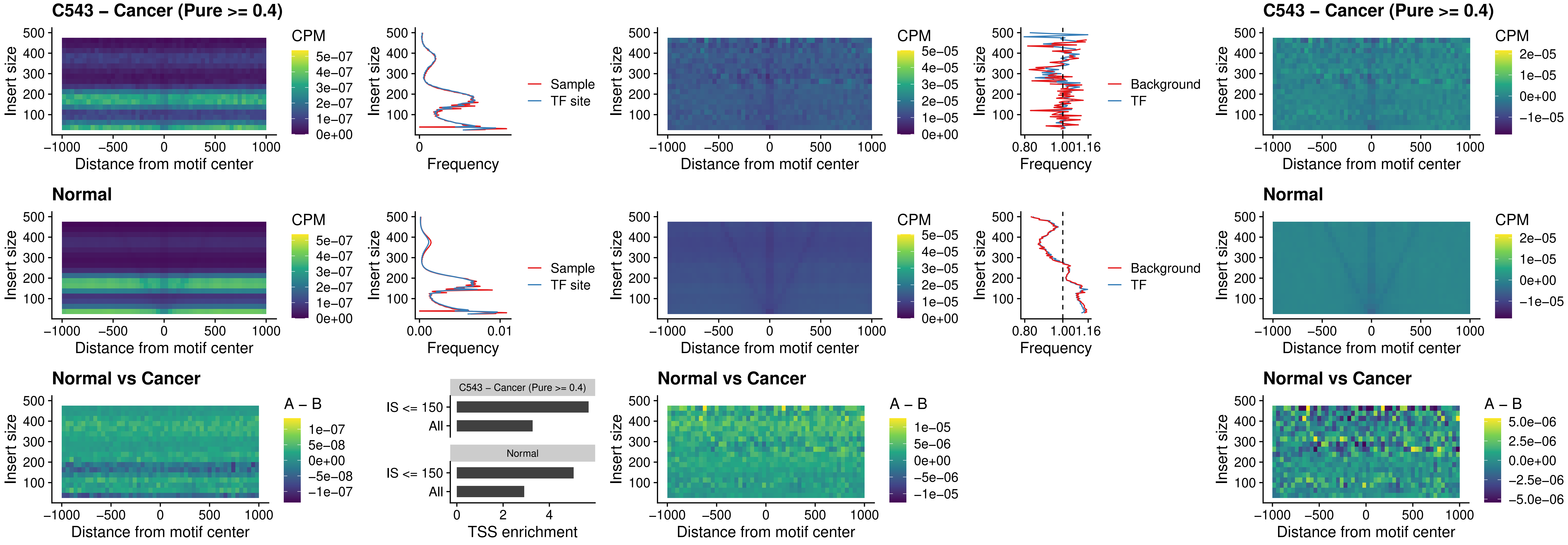

#### C543 – Cancer (Pure >= 0.4) – HOXA4_uc_dTSS_oPEAK

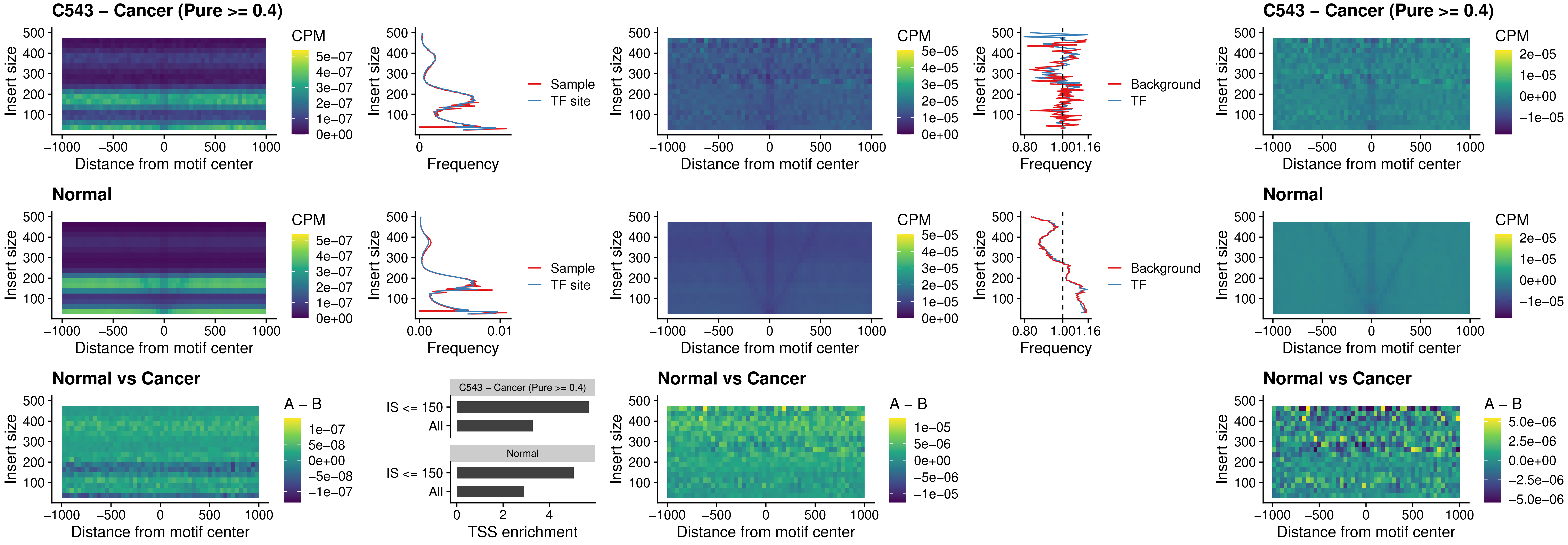

#### C543 – Cancer (Pure >= 0.4) – HLX_uc_dTSS_oPEAK

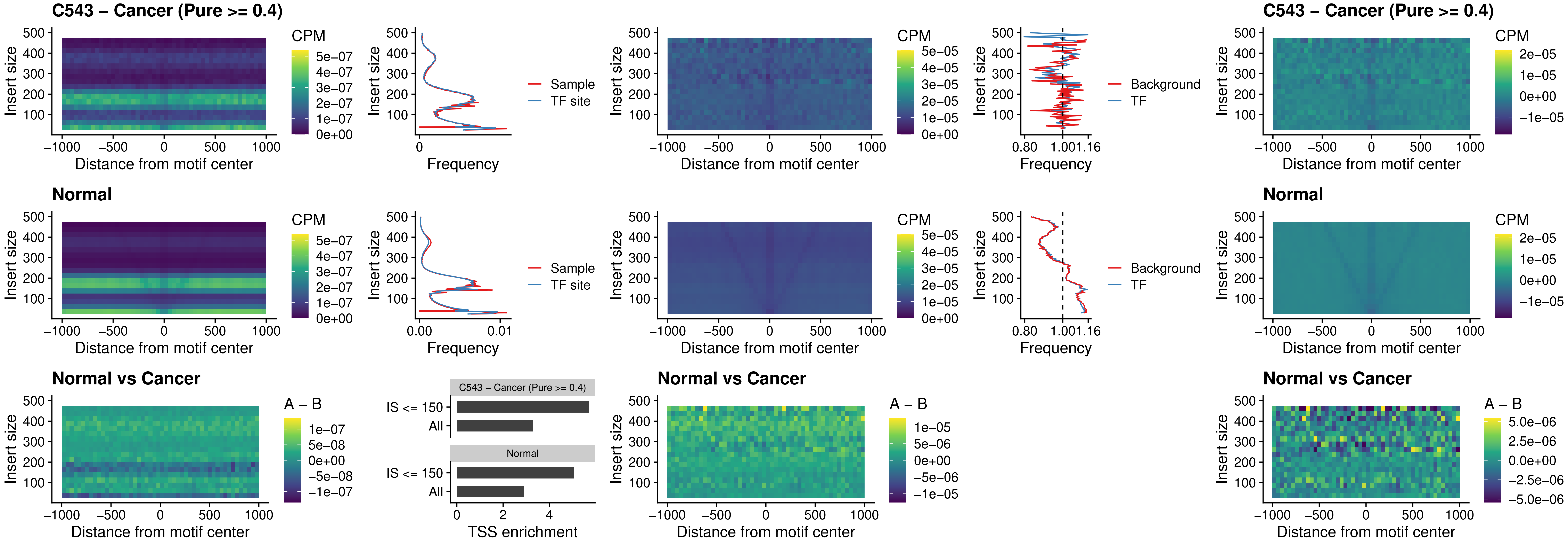

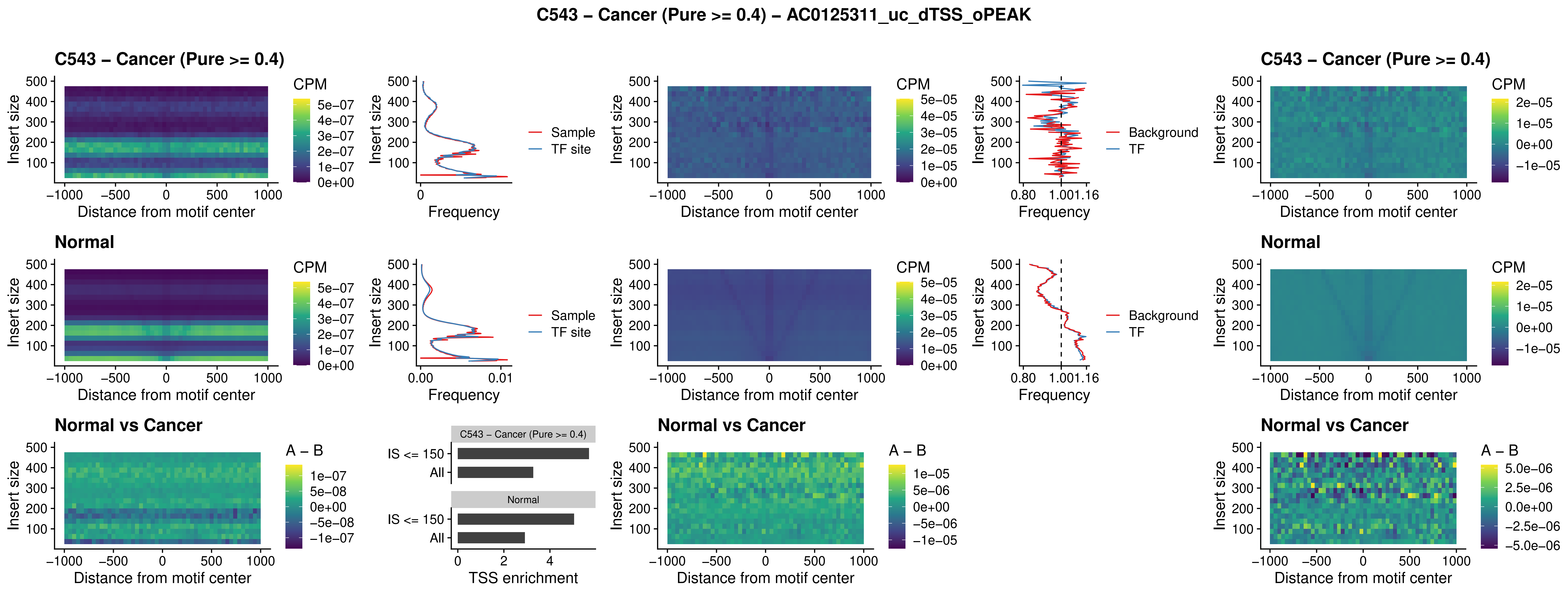
C543 – Cancer (Pure >= 0.4) – AC0125311_uc_dTSS_oPEAK
C543 – Cancer (Pure >= 0.4)
Normal
Normal vs Cancer
Insert size
Distance from motif center
CPM
A – B
Frequency
Sample
TF site
Background
TF
C543 – Cancer (Pure >= 0.4)
Normal
IS <= 150
All
TSS enrichment

#### C543 – Cancer (Pure >= 0.4) – DBX2_uc_dTSS_oPEAK

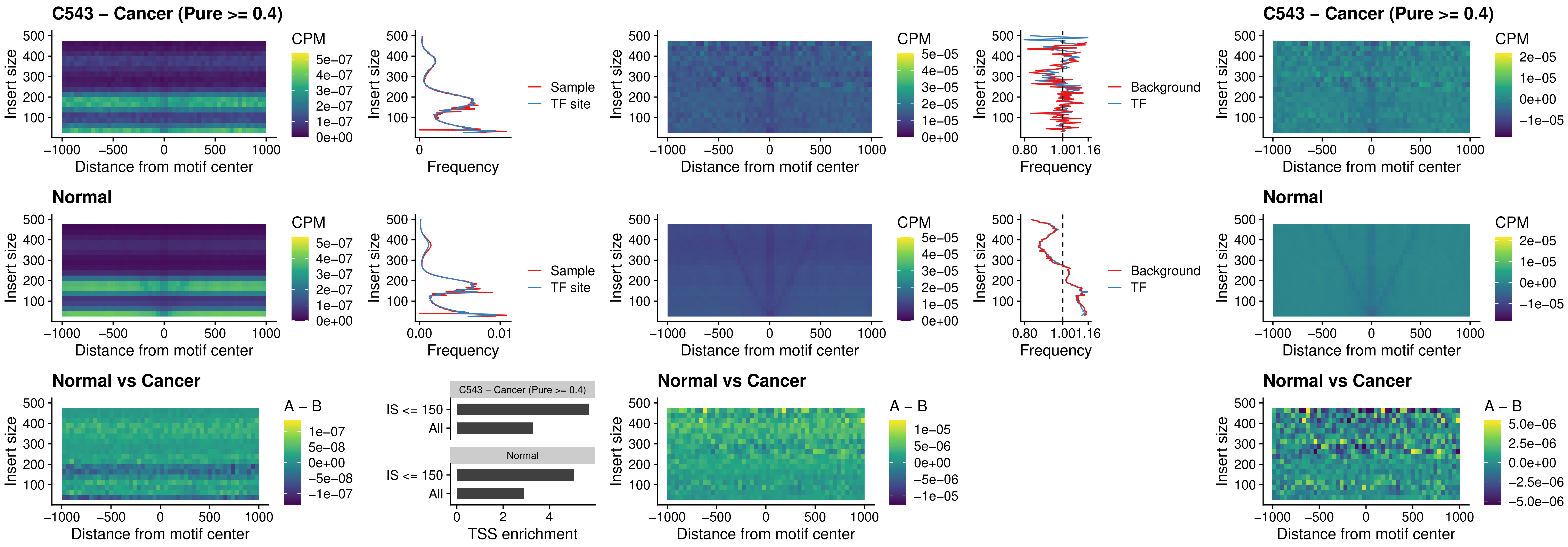

#### C543 – Cancer (Pure >= 0.4) – HOXA2_uc_dTSS_nPEAK

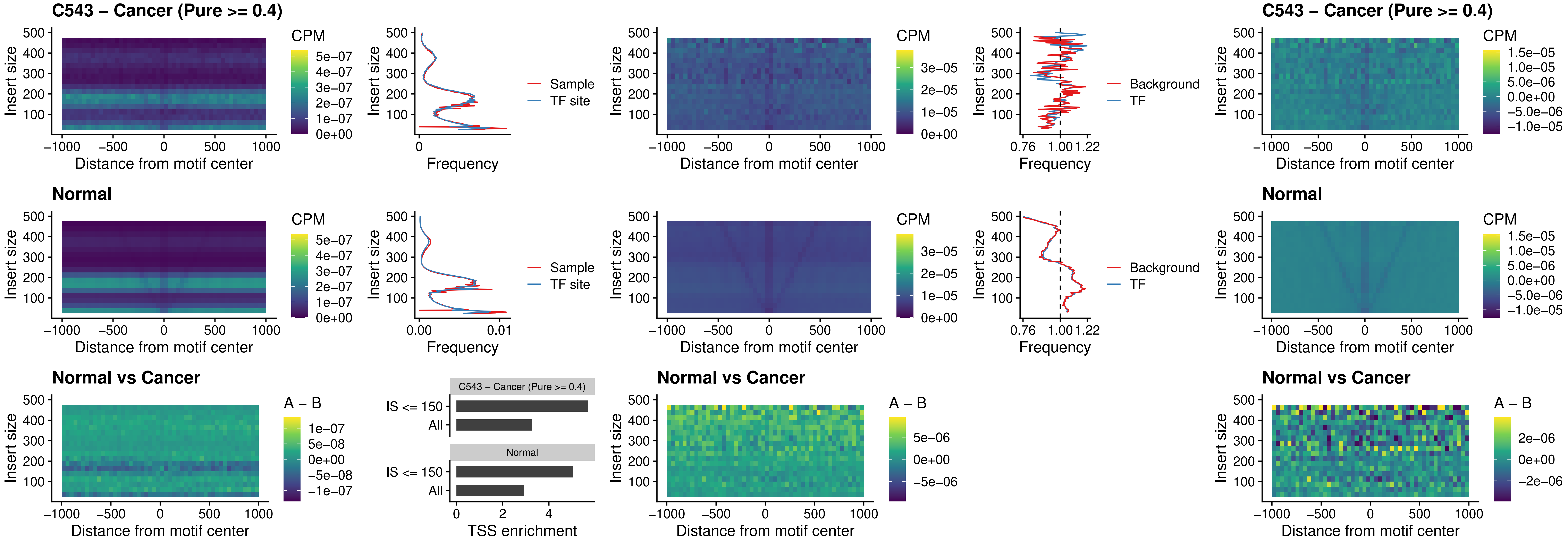

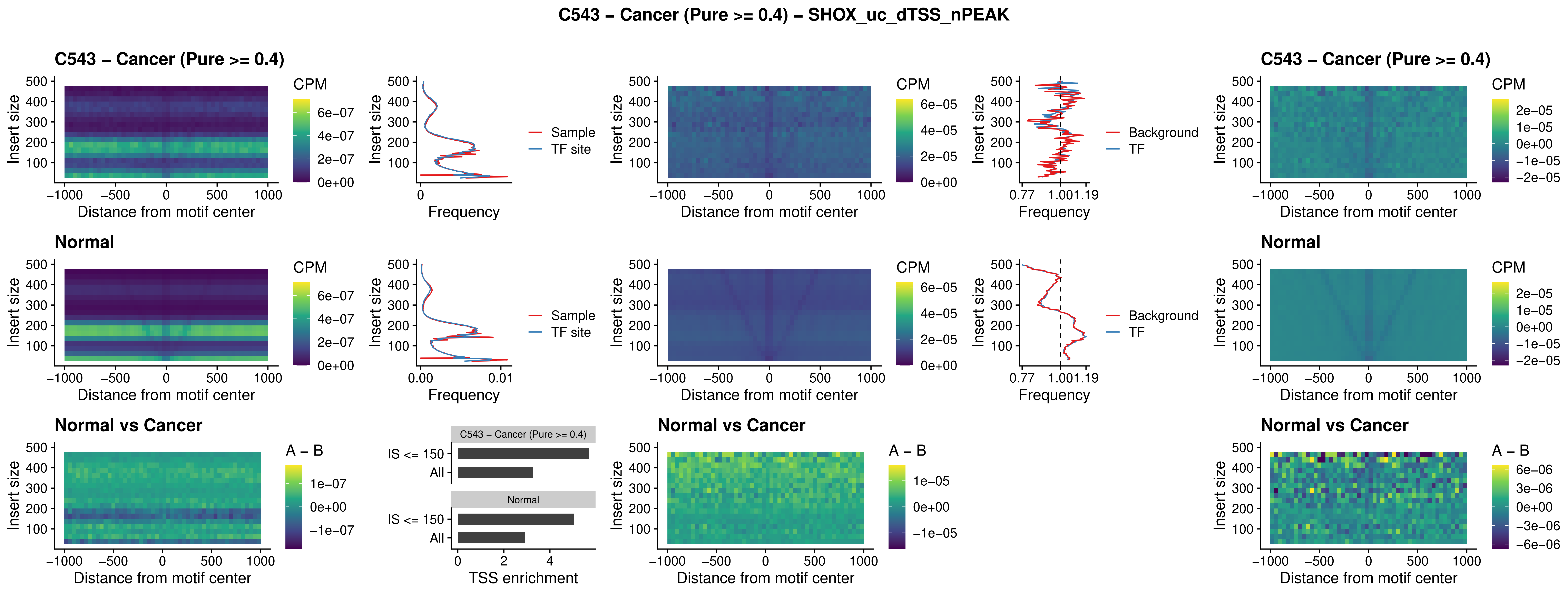
C543 – Cancer (Pure >= 0.4) – SHOX_uc_dTSS_nPEAK
C543 – Cancer (Pure >= 0.4)
Normal
Normal vs Cancer
Insert size
Distance from motif center
Frequency
CPM
A – B
Sample
TF site
Background
TF
C543 – Cancer (Pure >= 0.4)
Normal
IS <= 150
All
TSS enrichment

#### C543 – Cancer (Pure >= 0.4) – PRRX1_uc_dTSS_nPEAK

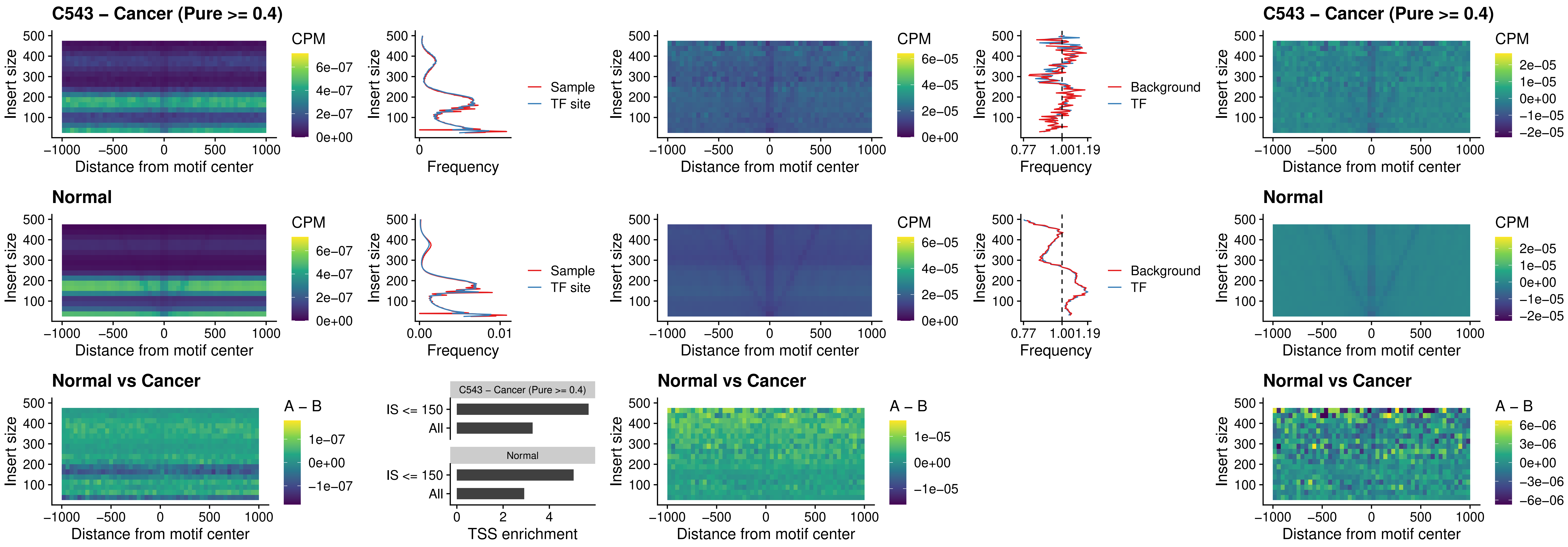

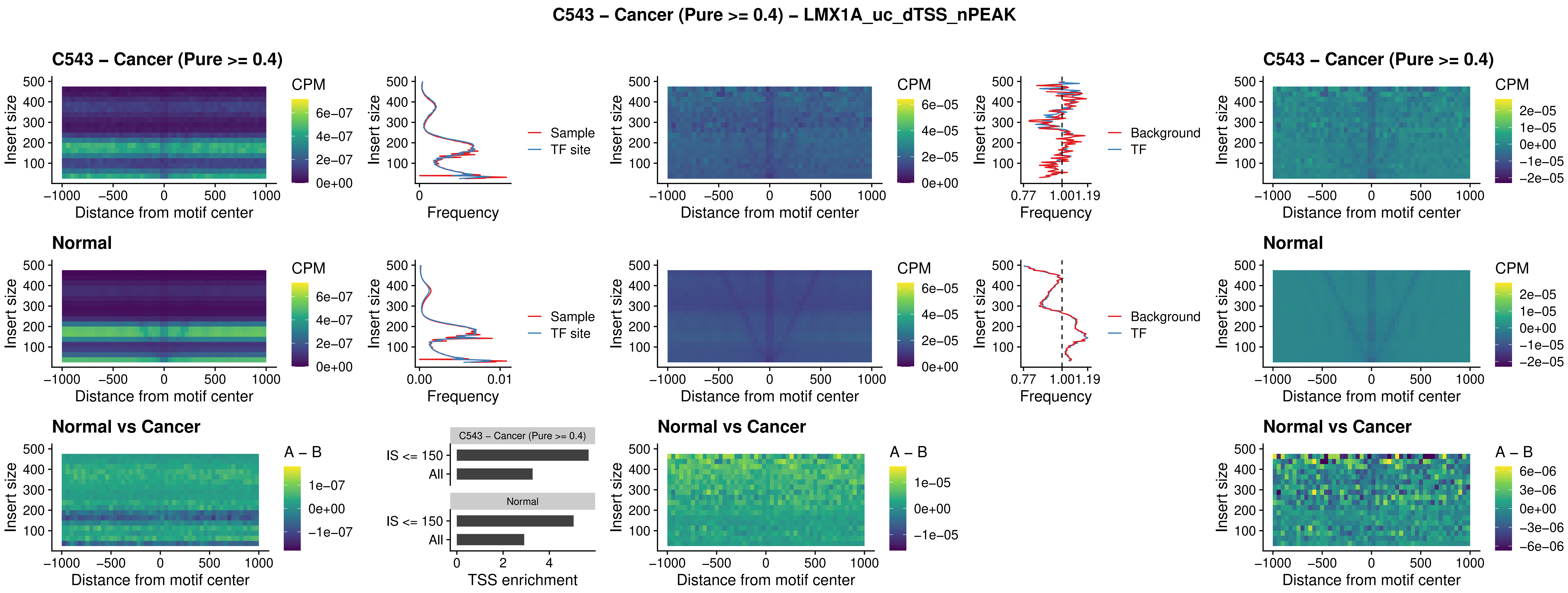
C543 – Cancer (Pure >= 0.4) – LMX1A_uc_dTSS_nPEAK
C543 – Cancer (Pure >= 0.4)
Normal
Normal vs Cancer
Insert size
Distance from motif center
CPM
A – B
Frequency
Sample
TF site
Background
TF
C543 – Cancer (Pure >= 0.4)
Normal
IS <= 150
All
TSS enrichment

#### C543 – Cancer (Pure >= 0.4) – LMX1B_uc_dTSS_nPEAK

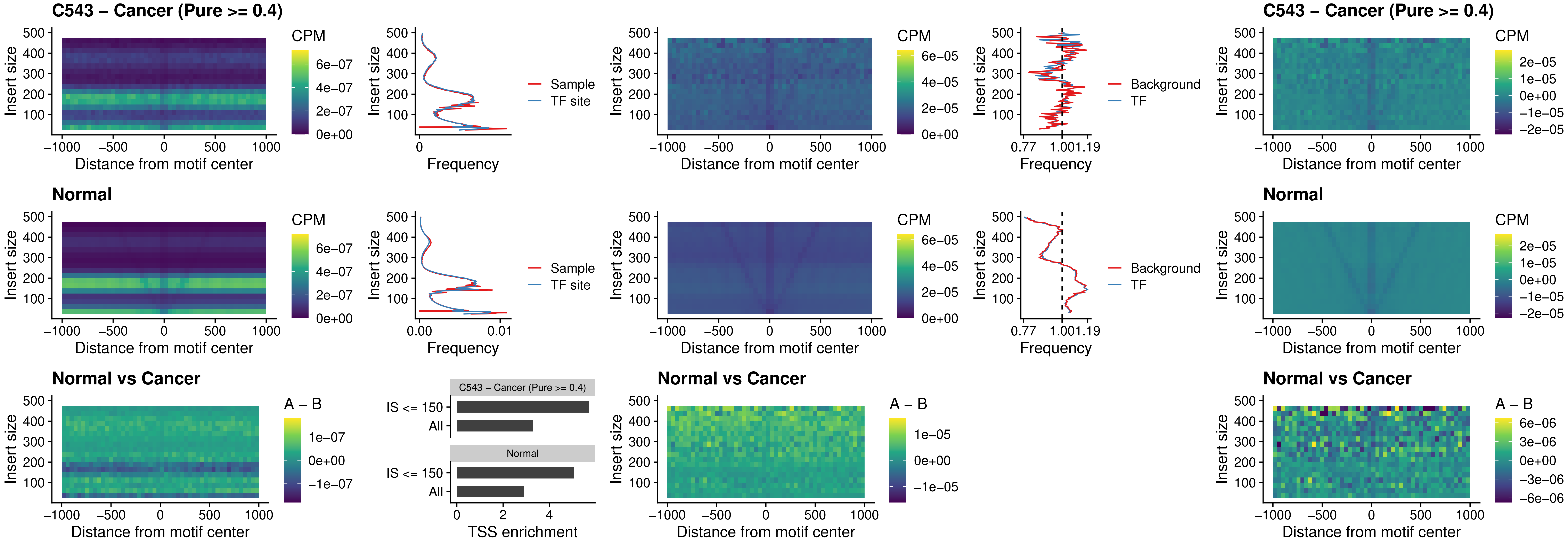

#### C543 – Cancer (Pure >= 0.4) – DLX5_uc_dTSS_oPEAK

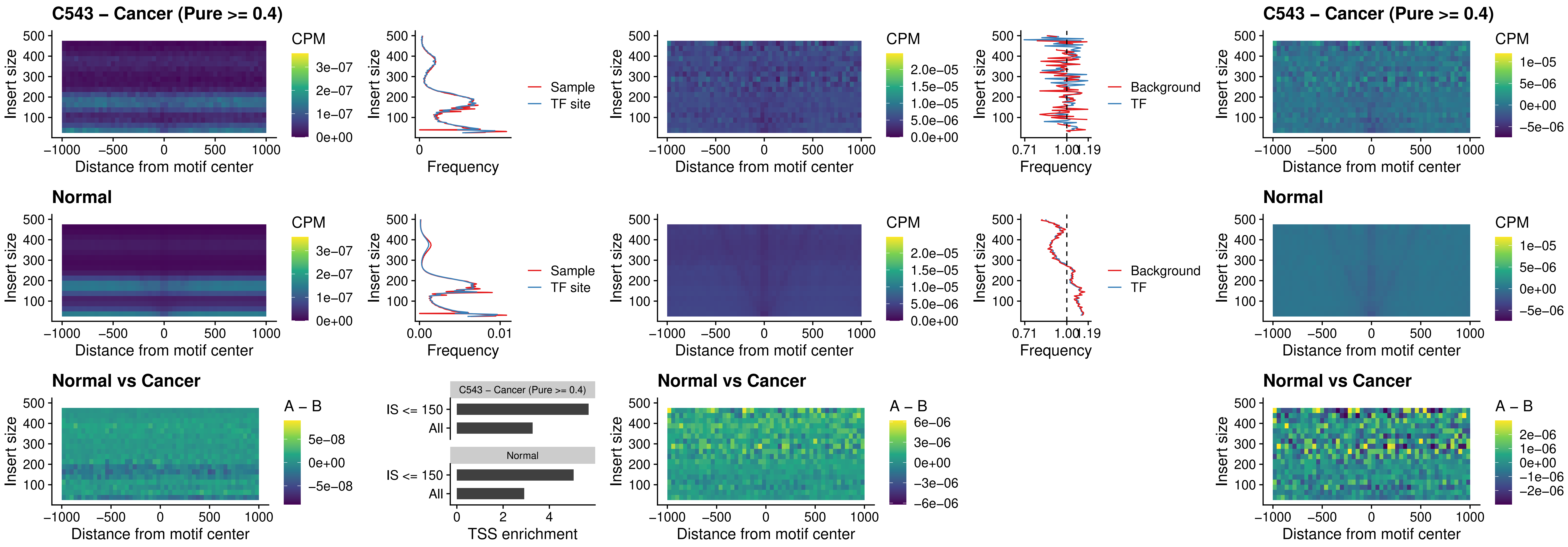

#### C543 – Cancer (Pure >= 0.4) – ARID3C_uc_dTSS_nPEAK

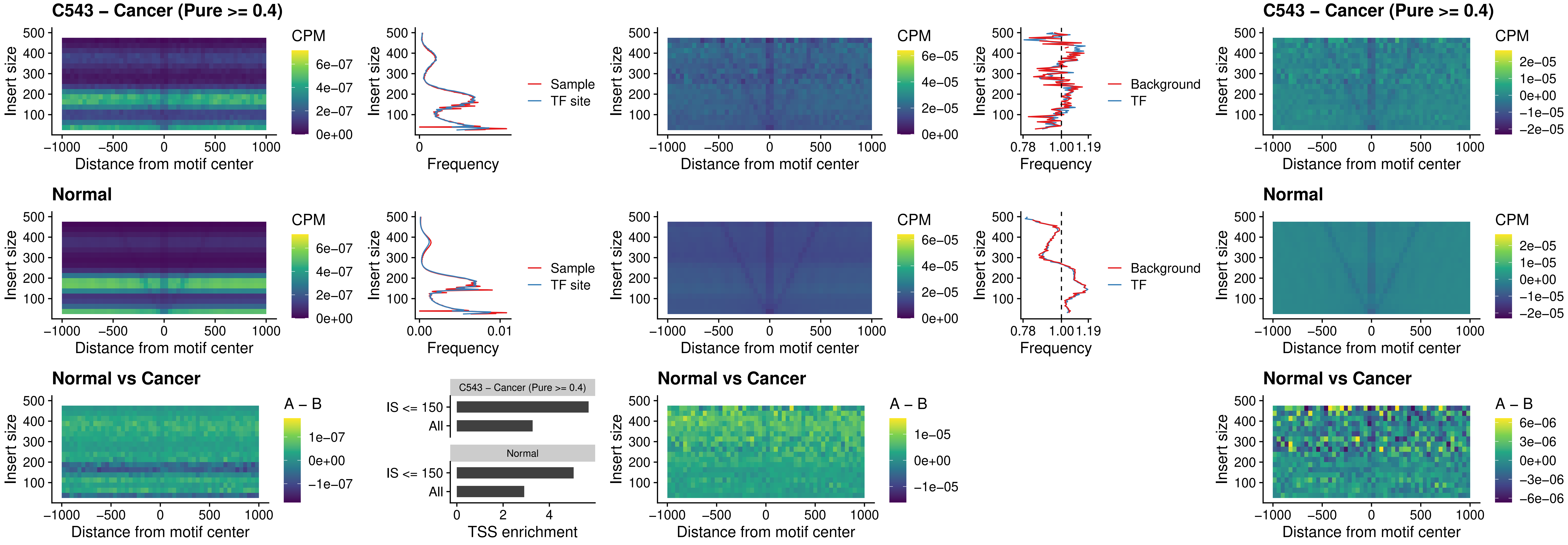

#### C543 – Cancer (Pure >= 0.4) – HOXD12_uc_dTSS_nPEAK

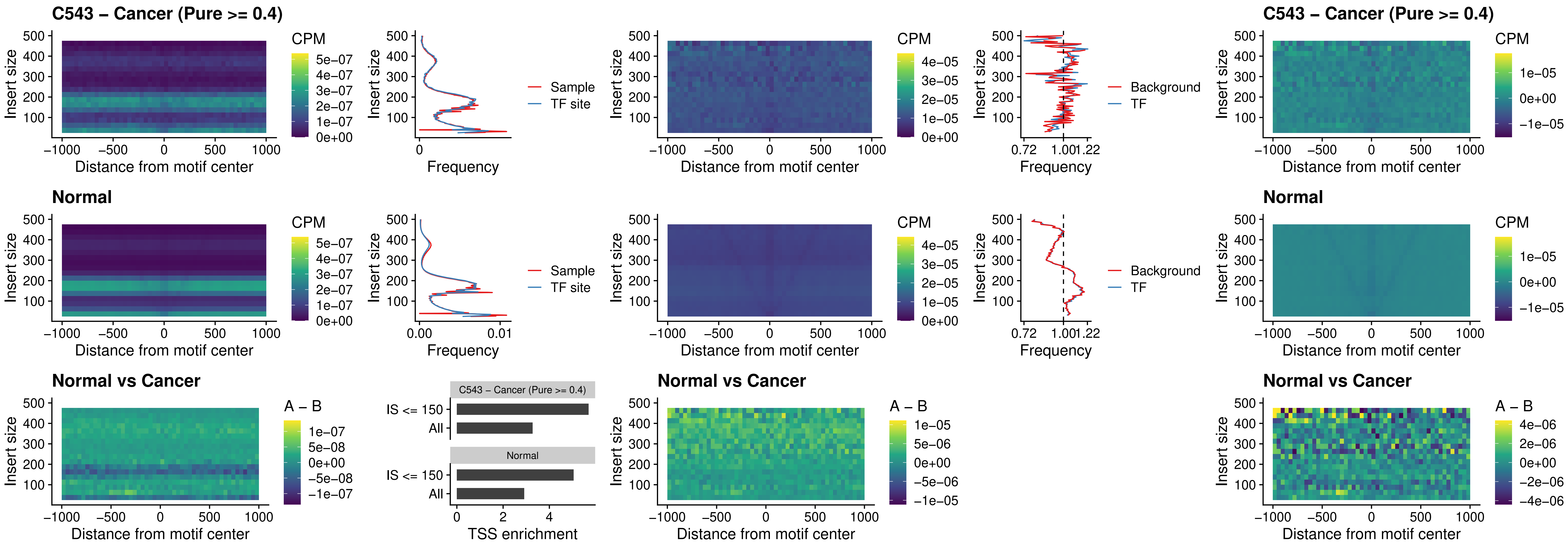

#### C543 – Cancer (Pure >= 0.4) – CDX2_uc_dTSS_nPEAK

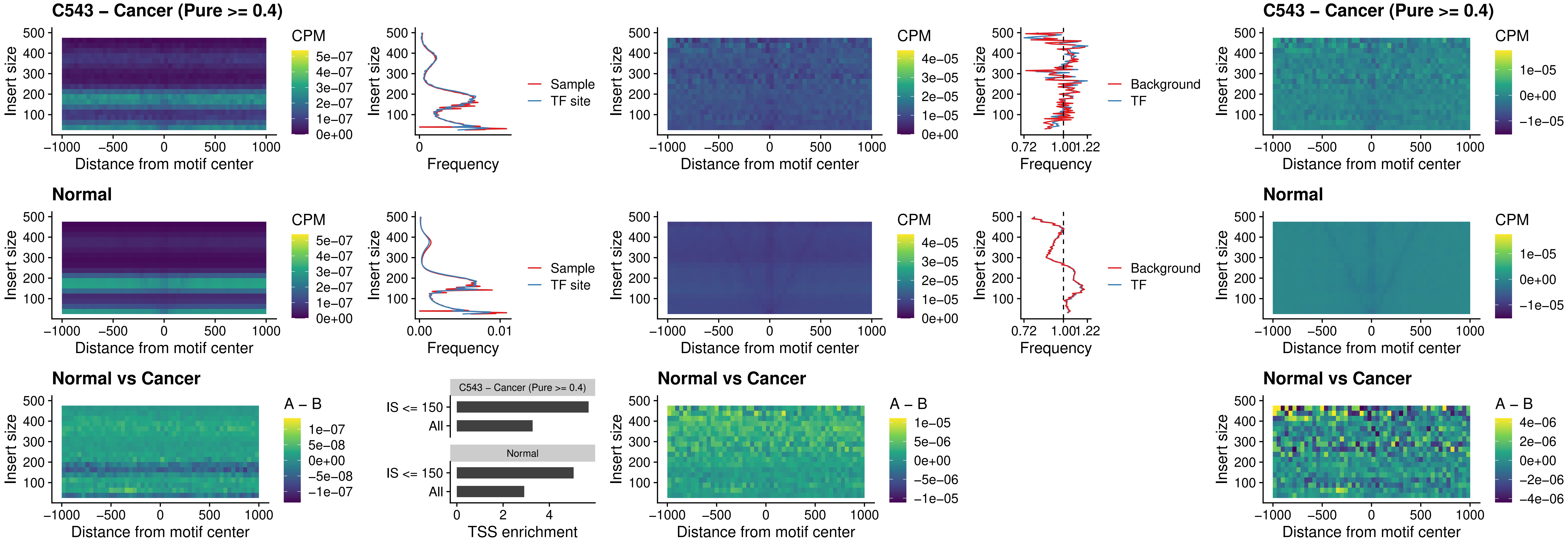

#### C543 – Cancer (Pure >= 0.4) – HOXC12_uc_dTSS_nPEAK

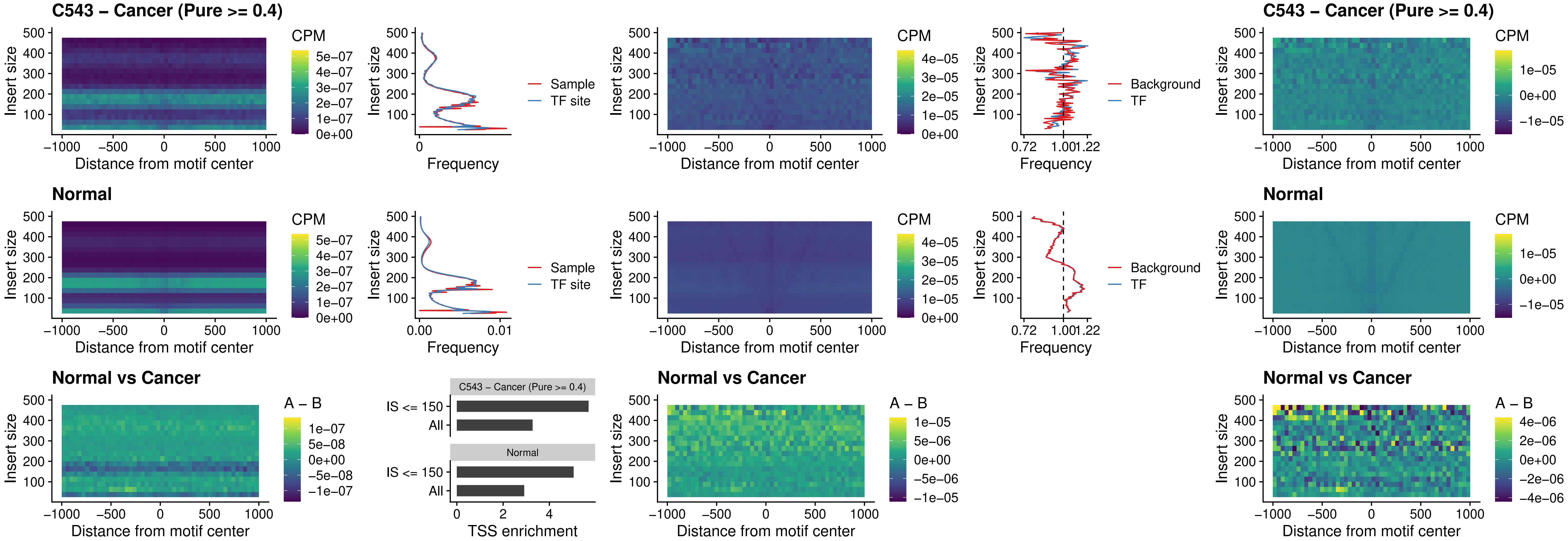

### C543 – Cancer (Pure >= 0.4) – HHEX_uc_dTSS_oPEAK

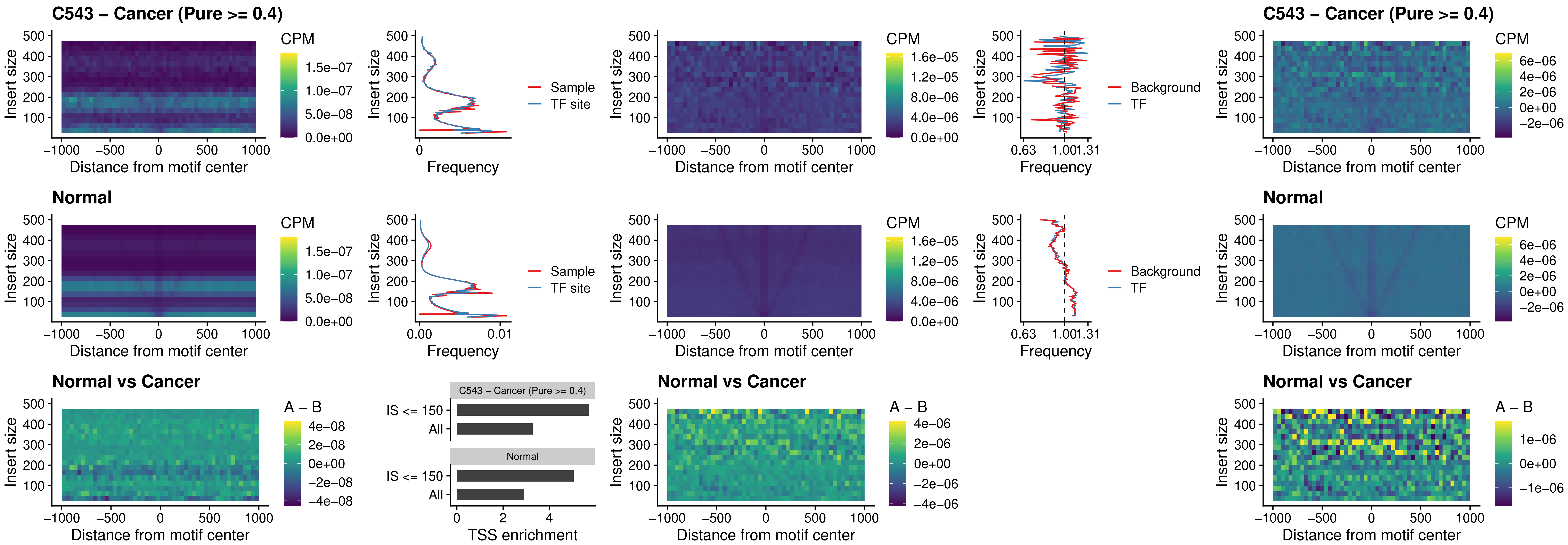

#### C543 – Cancer (Pure >= 0.4) – HOXA11_uc_dTSS_nPEAK

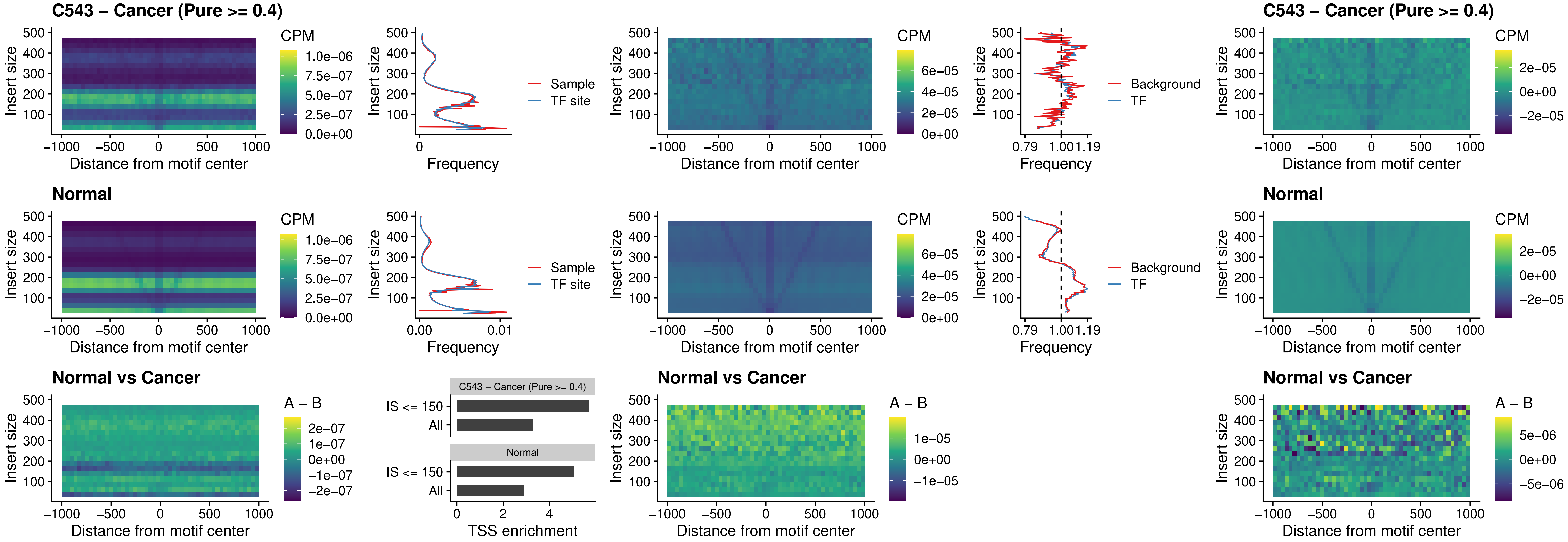

#### C543 – Cancer (Pure >= 0.4) – HOXA11_uc_cTSS_nPEAK

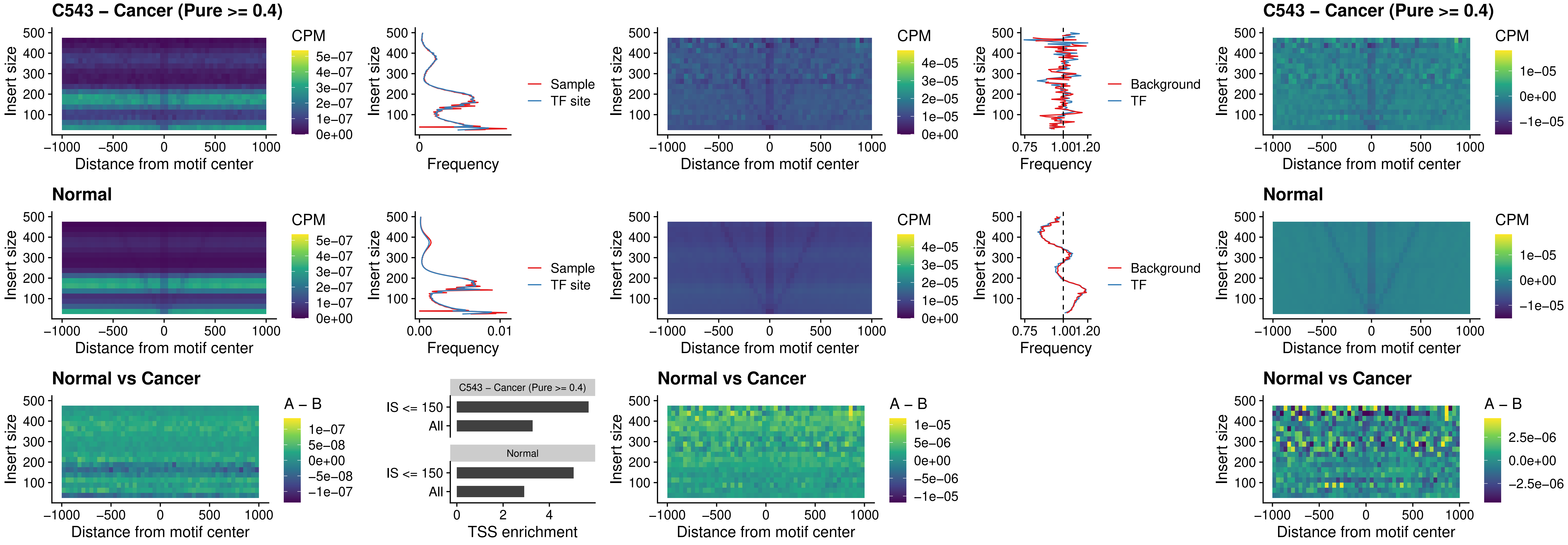

#### C543 – Cancer (Pure >= 0.4) – ZFHX3_uc_dTSS_nPEAK

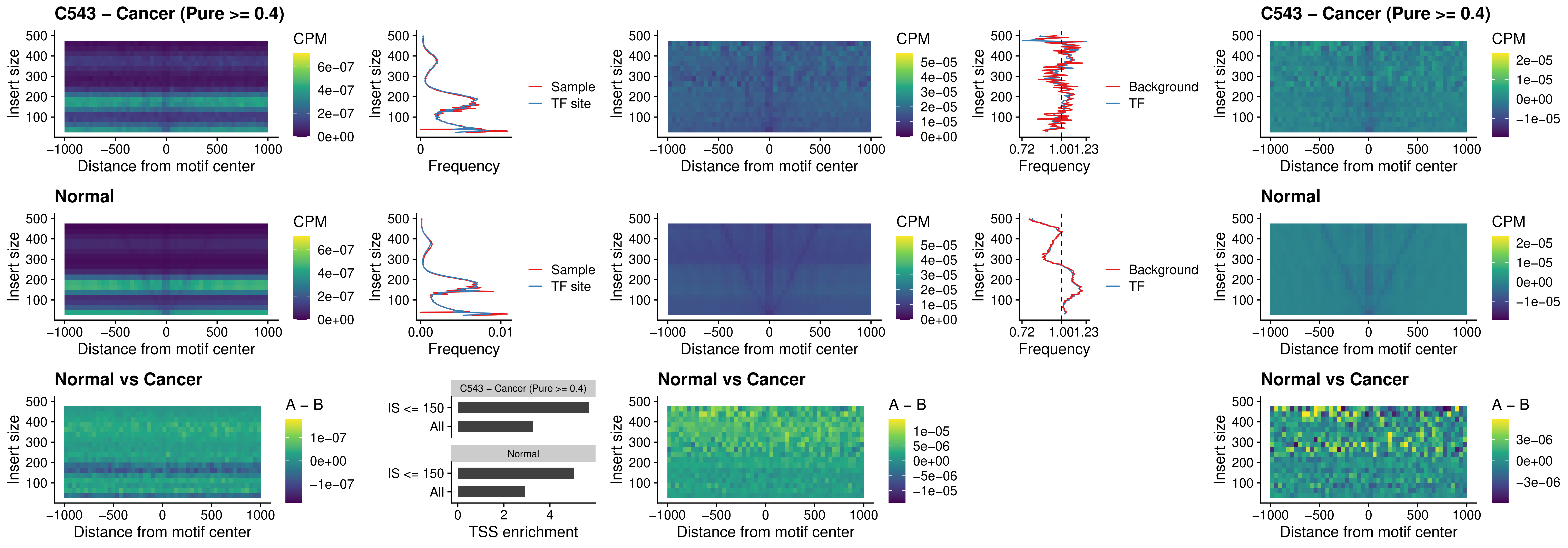

#### C543 – Cancer (Pure >= 0.4) – HOXB4_uc_dTSS_nPEAK

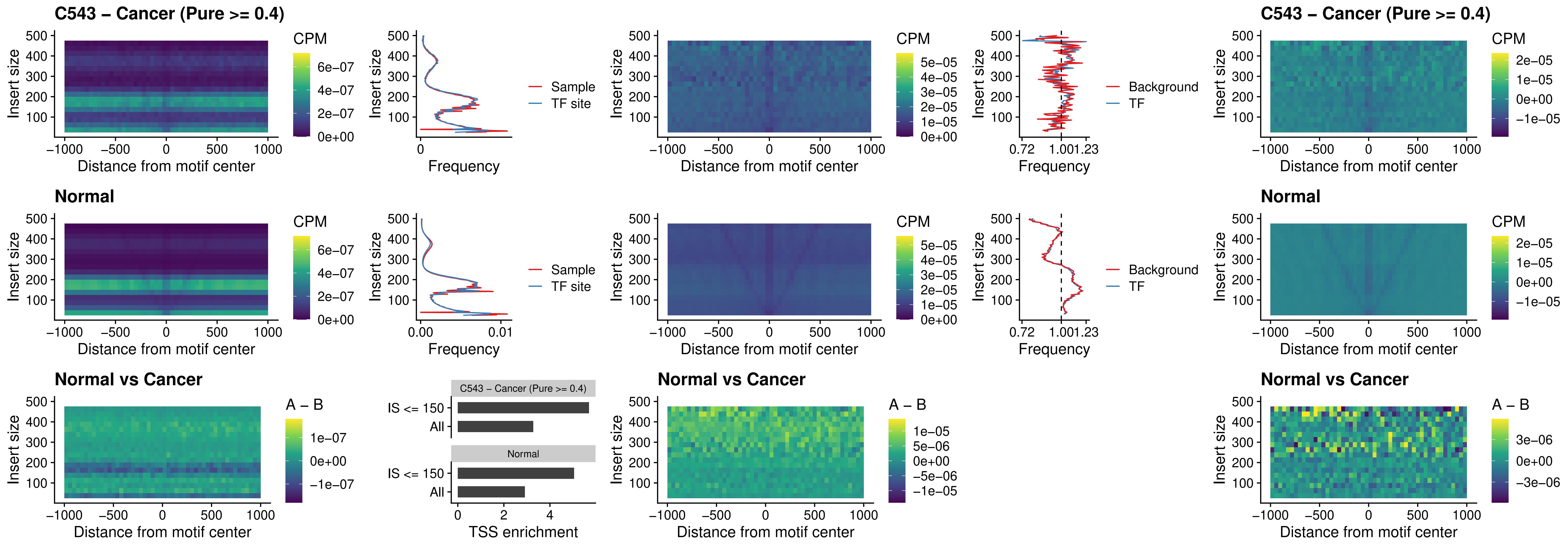

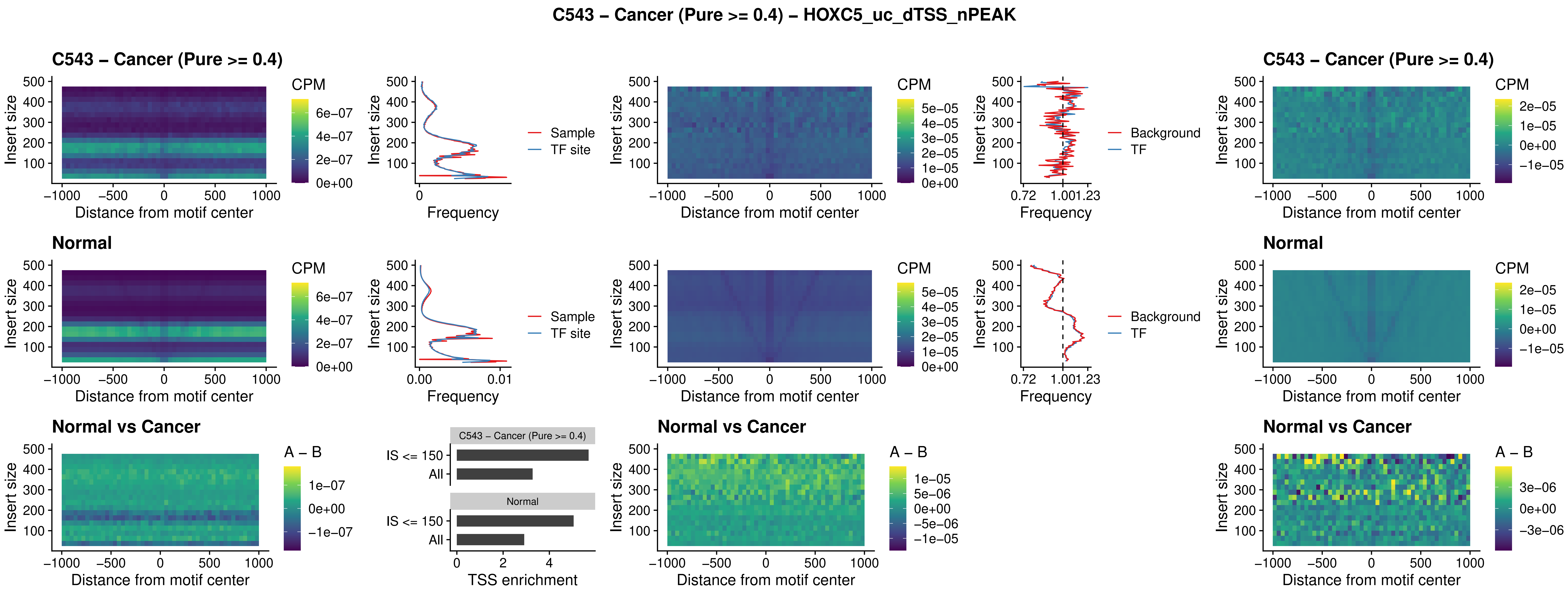
C543 – Cancer (Pure >= 0.4) – HOXC5_uc_dTSS_nPEAK
C543 – Cancer (Pure >= 0.4)
Normal
Normal vs Cancer
Insert size
Distance from motif center
Frequency
CPM
A – B
Sample
TF site
Background
TF
C543 – Cancer (Pure >= 0.4)
Normal
IS <= 150
All
TSS enrichment

#### C543 – Cancer (Pure >= 0.4) – HOXD1_uc_dTSS_nPEAK

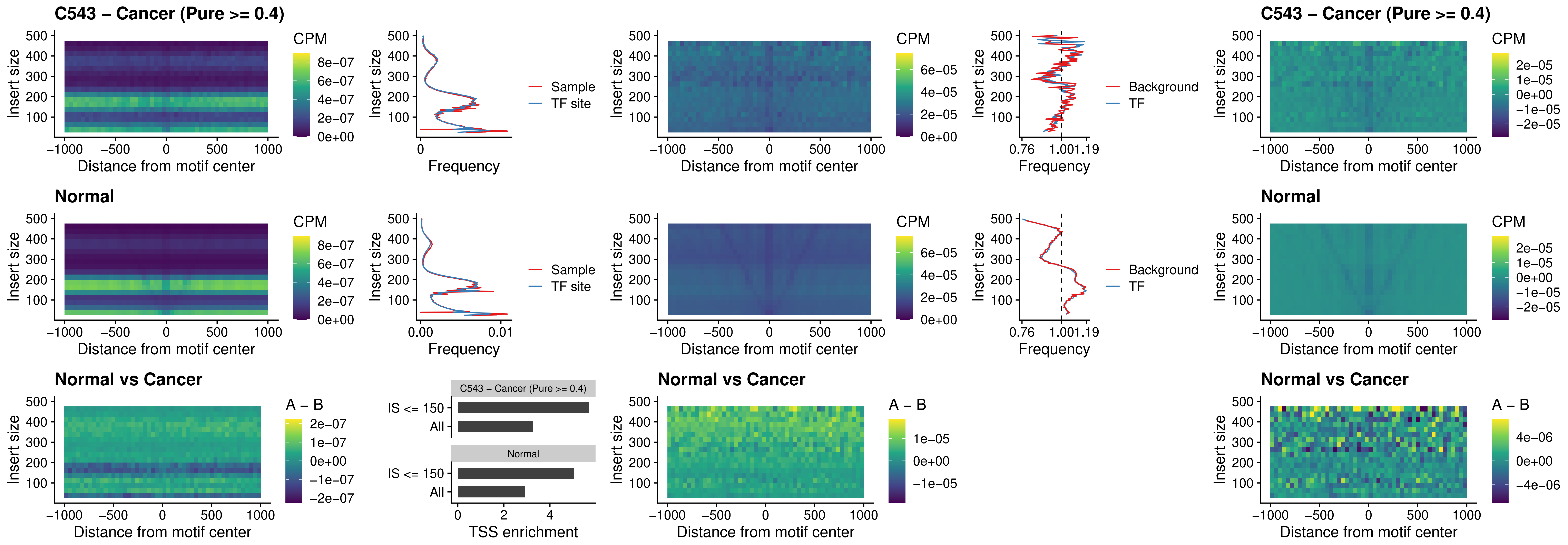

#### C543 – Cancer (Pure >= 0.4) – OTP_uc_dTSS_nPEAK

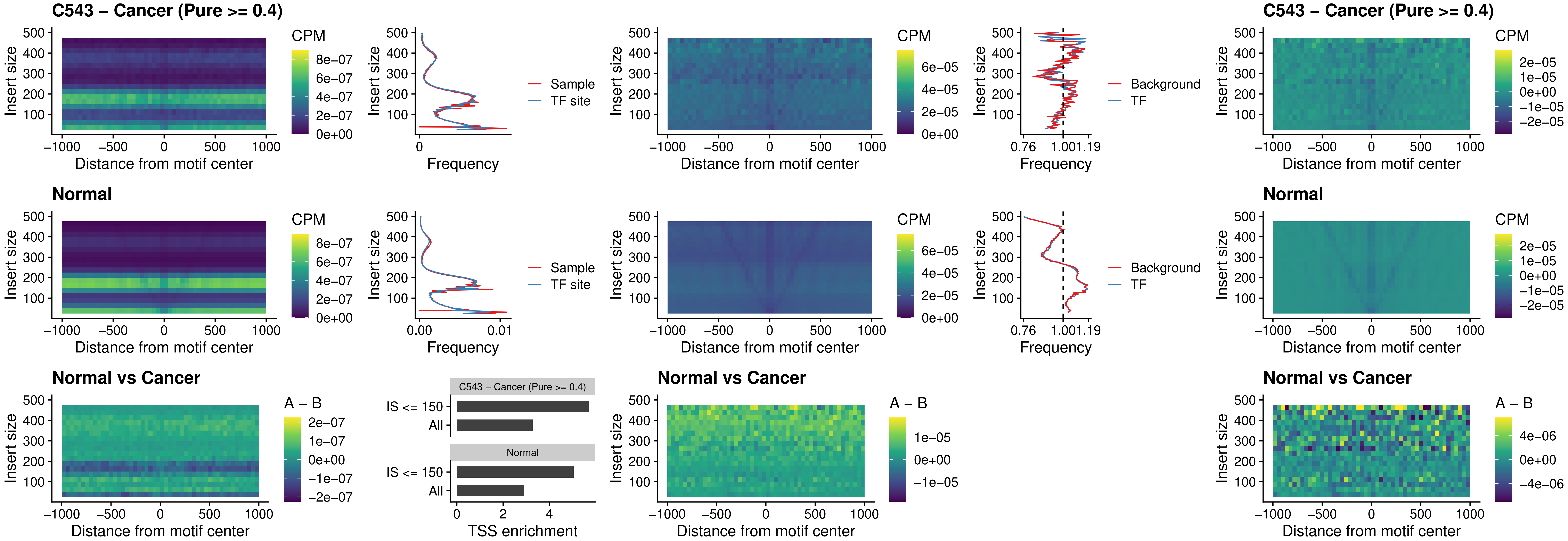

#### C543 – Cancer (Pure >= 0.4) – FOXL1_uc_dTSS_oPEAK

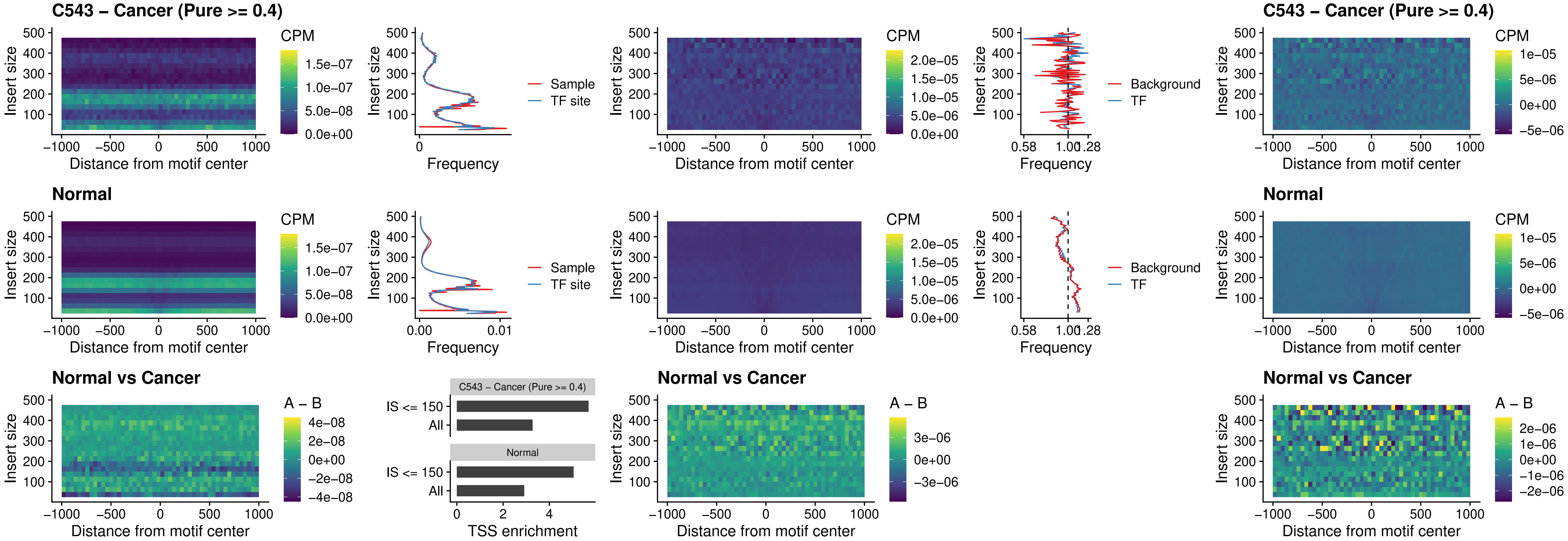

### C543 – Cancer (Pure >= 0.4) – ZNF32_uc_dTSS_oPEAK

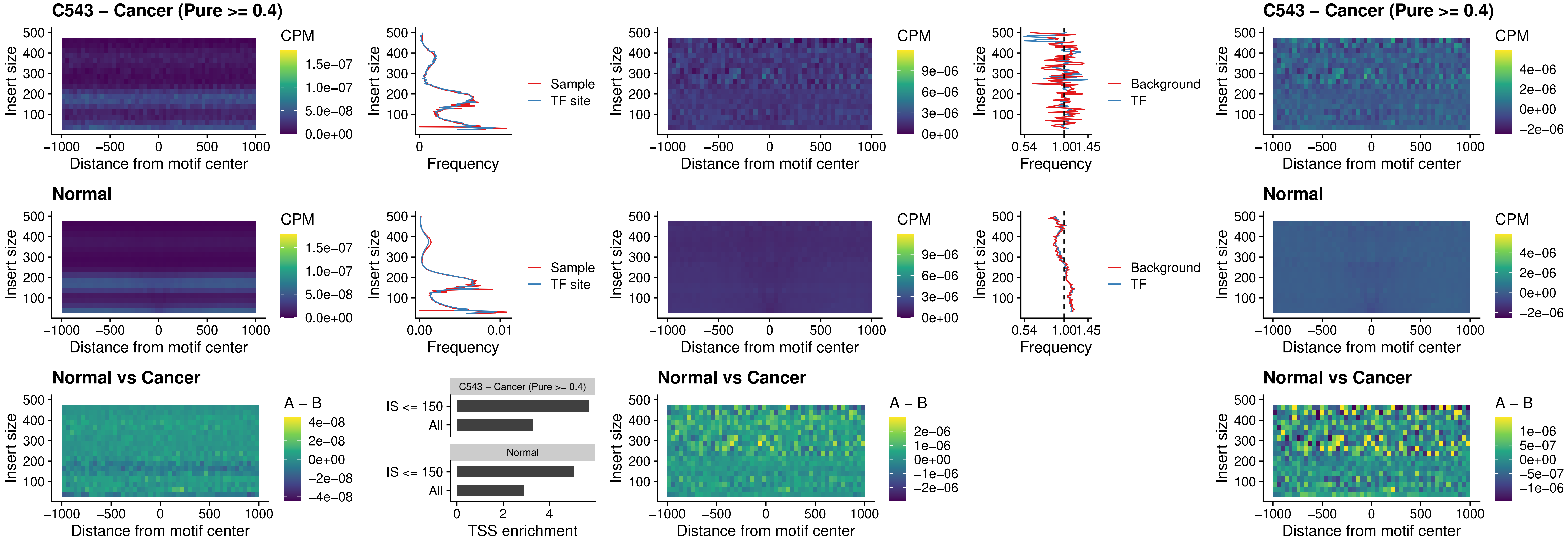

#### C543 – Cancer (Pure >= 0.4) – FOXL1_uc_dTSS_nPEAK

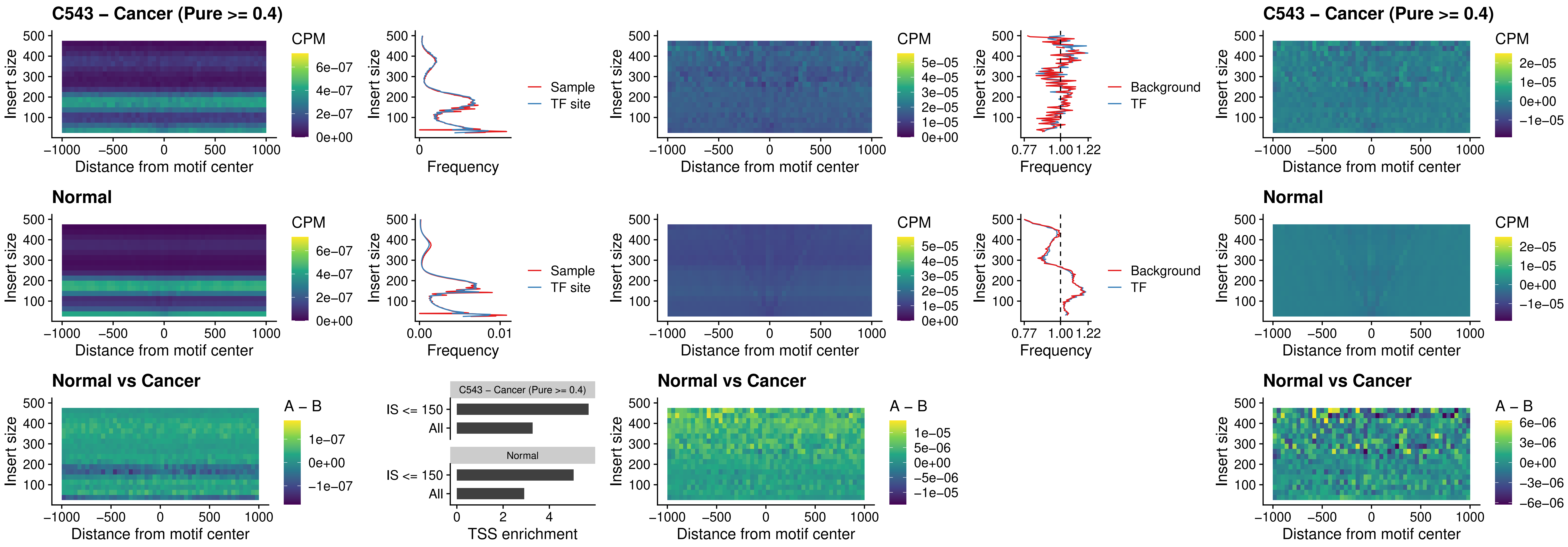

### C543 – Cancer (Pure >= 0.4) – ZNF32_uc_dTSS_nPEAK

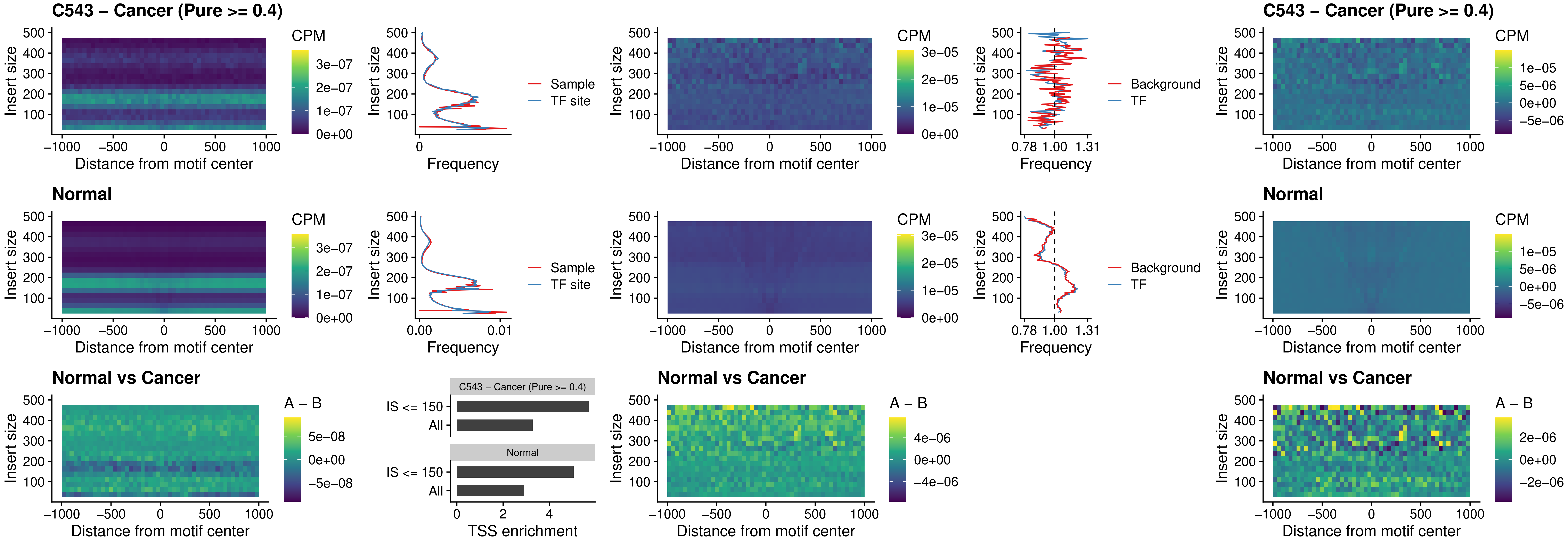

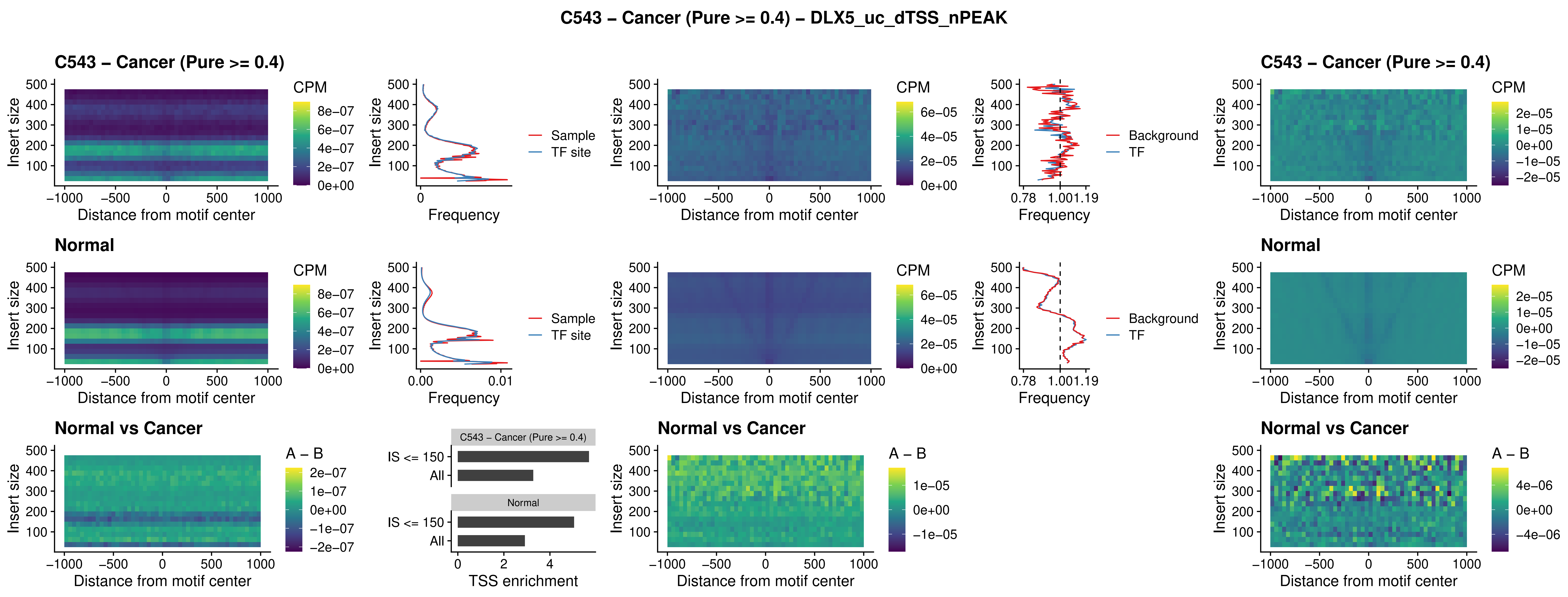
C543 – Cancer (Pure >= 0.4) – DLX5_uc_dTSS_nPEAK
C543 – Cancer (Pure >= 0.4)
Normal
Normal vs Cancer
Insert size
Distance from motif center
CPM
A – B
Frequency
Sample
TF site
Background
TF
C543 – Cancer (Pure >= 0.4)
Normal
IS <= 150
All
TSS enrichment

#### C543 – Cancer (Pure >= 0.4) – DMRT2_uc_dTSS_nPEAK

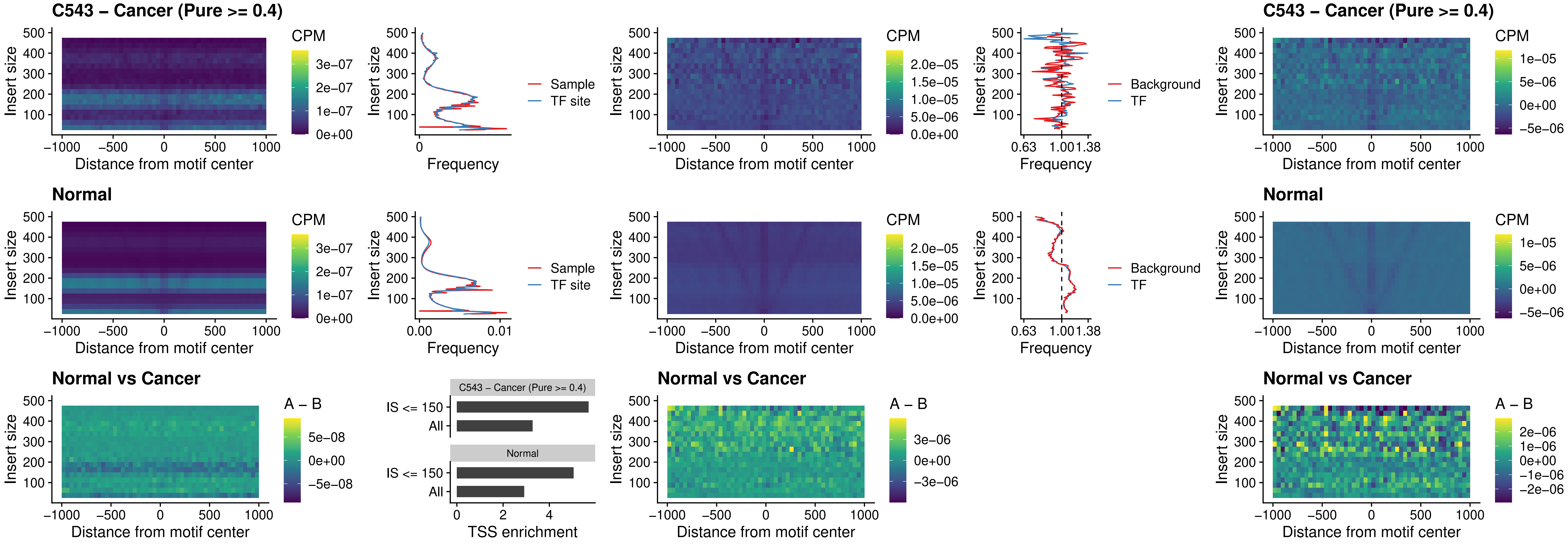

#### C543 – Cancer (Pure >= 0.4) – DMRTC2_uc_cTSS_nPEAK

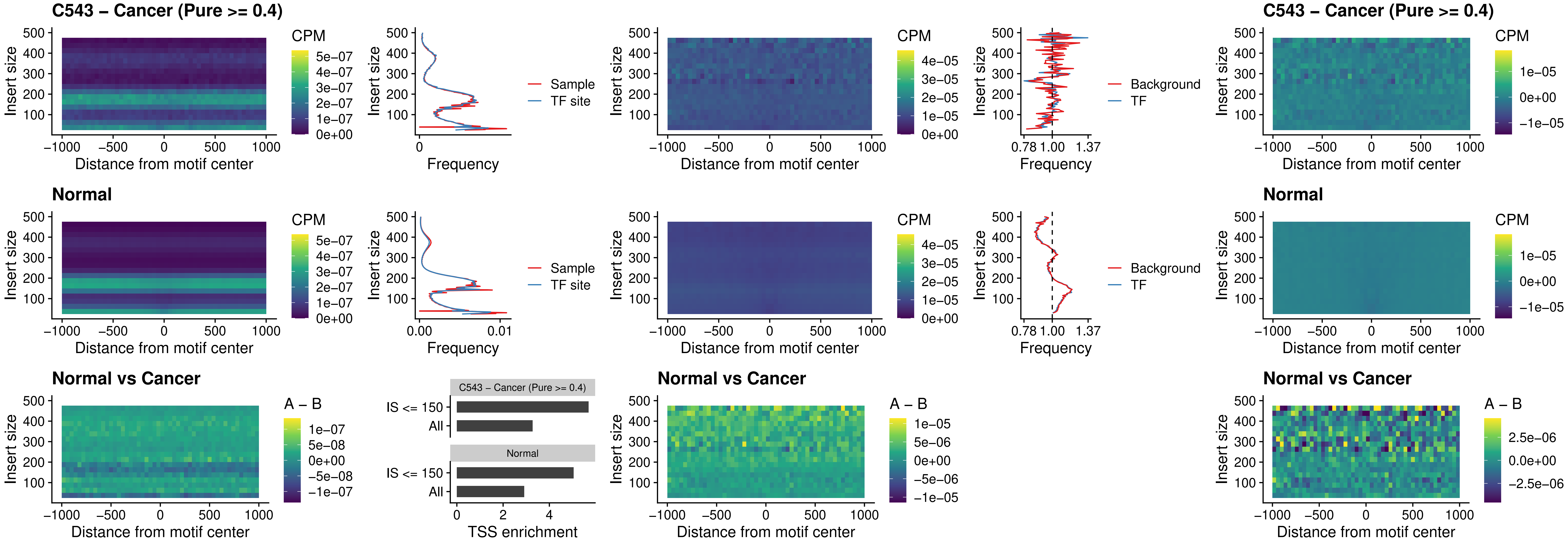

#### C543 – Cancer (Pure >= 0.4) – HOXD9_uc_dTSS_nPEAK

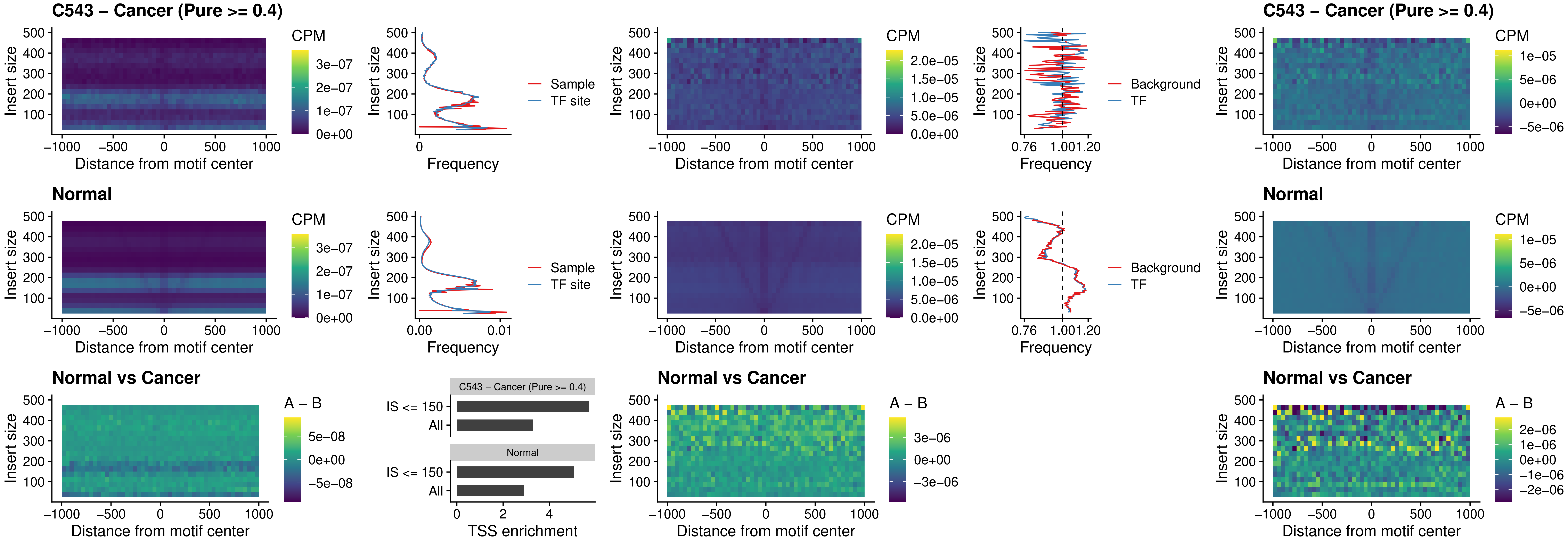

#### C543 – Cancer (Pure >= 0.4) – HOXD4_uc_dTSS_nPEAK

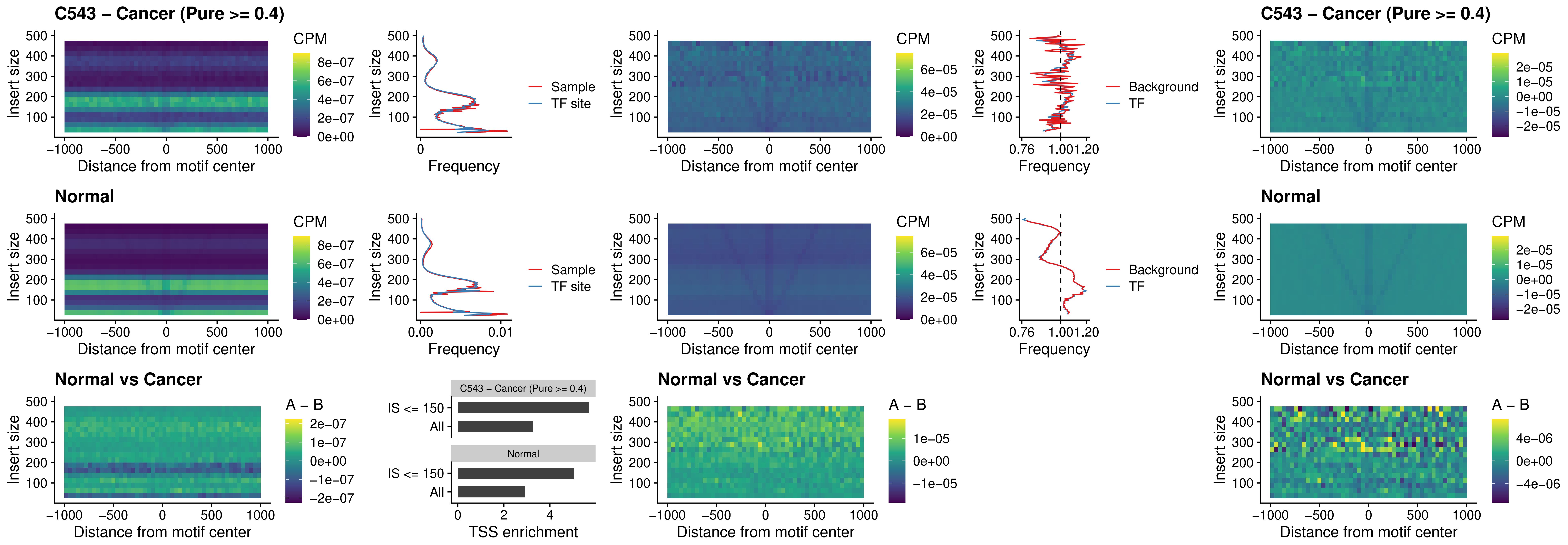

#### C543 – Cancer (Pure >= 0.4) – FOXP3_uc_cTSS_nPEAK

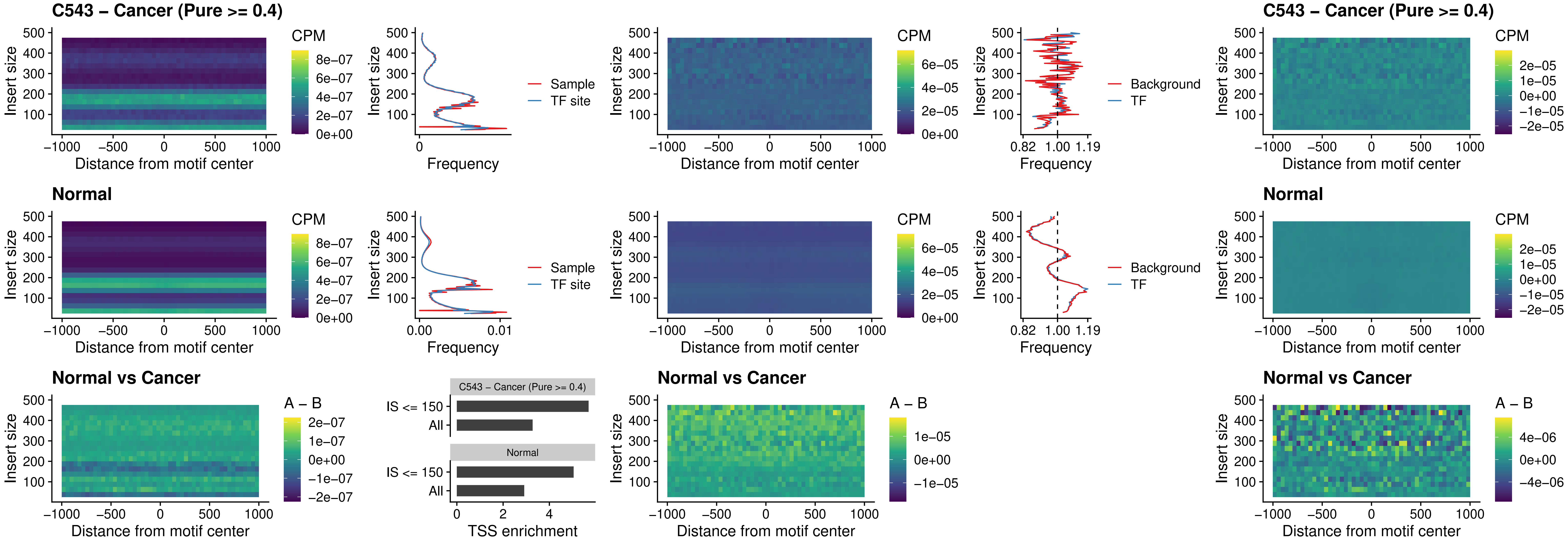

#### C543 – Cancer (Pure >= 0.4) – FOXD4L6_uc_cTSS_nPEAK

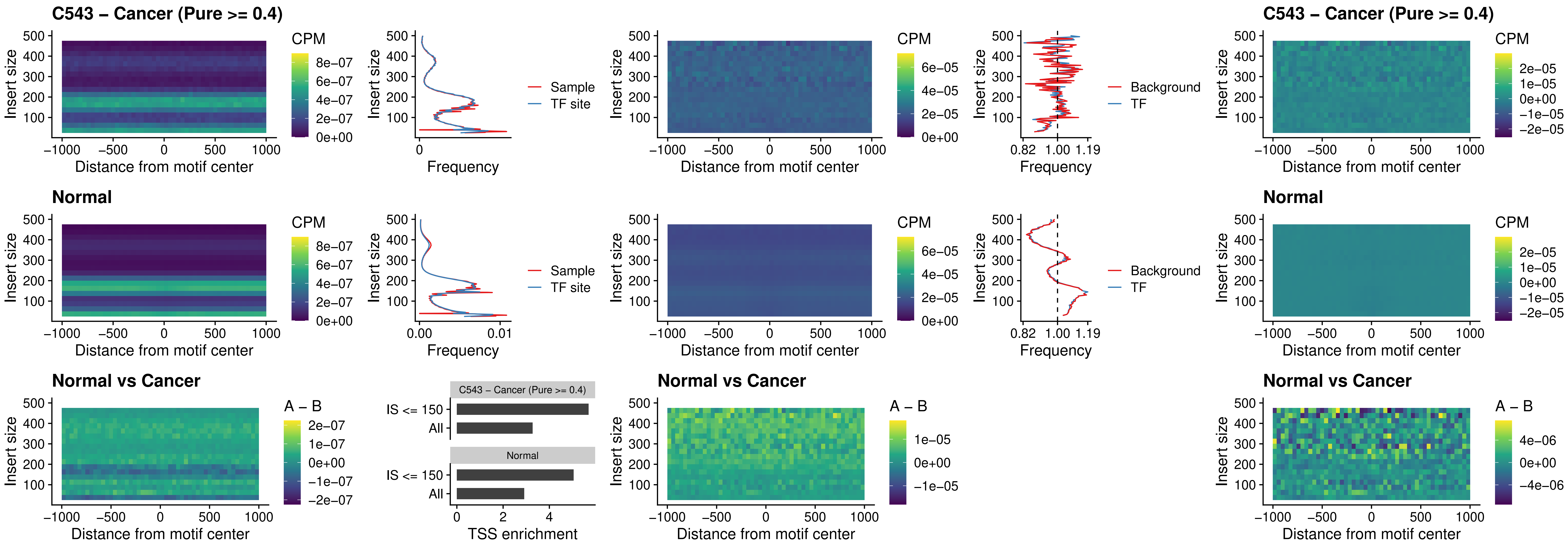

### C543 – Cancer (Pure >= 0.4) – FOXD4L5_uc_cTSS_nPEAK

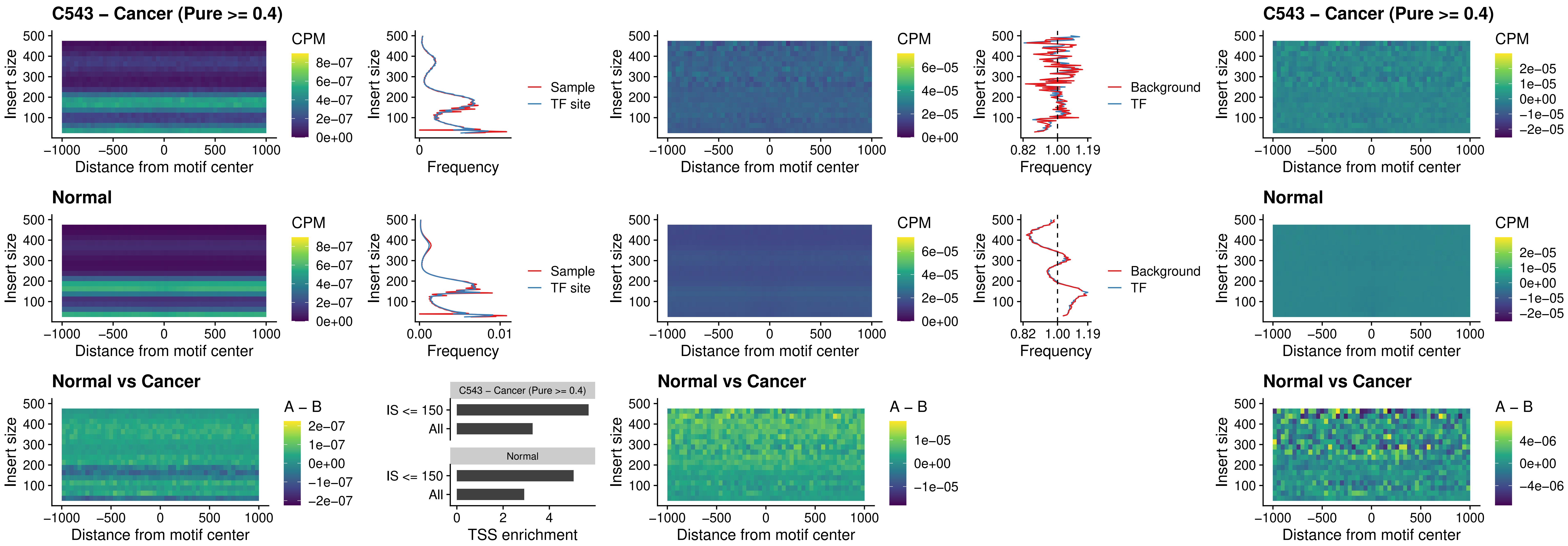

#### C543 – Cancer (Pure >= 0.4) – FOXD4L4_uc_cTSS_nPEAK

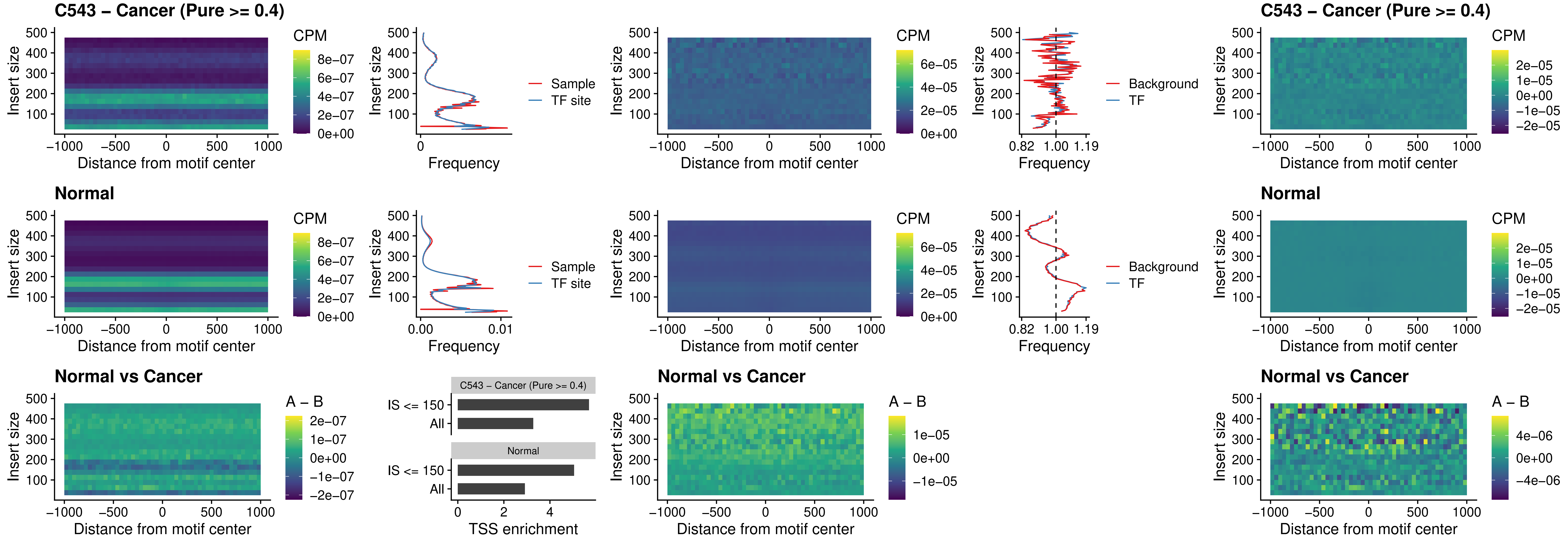

### C543 – Cancer (Pure >= 0.4) – FOXD4L2_uc_cTSS_nPEAK

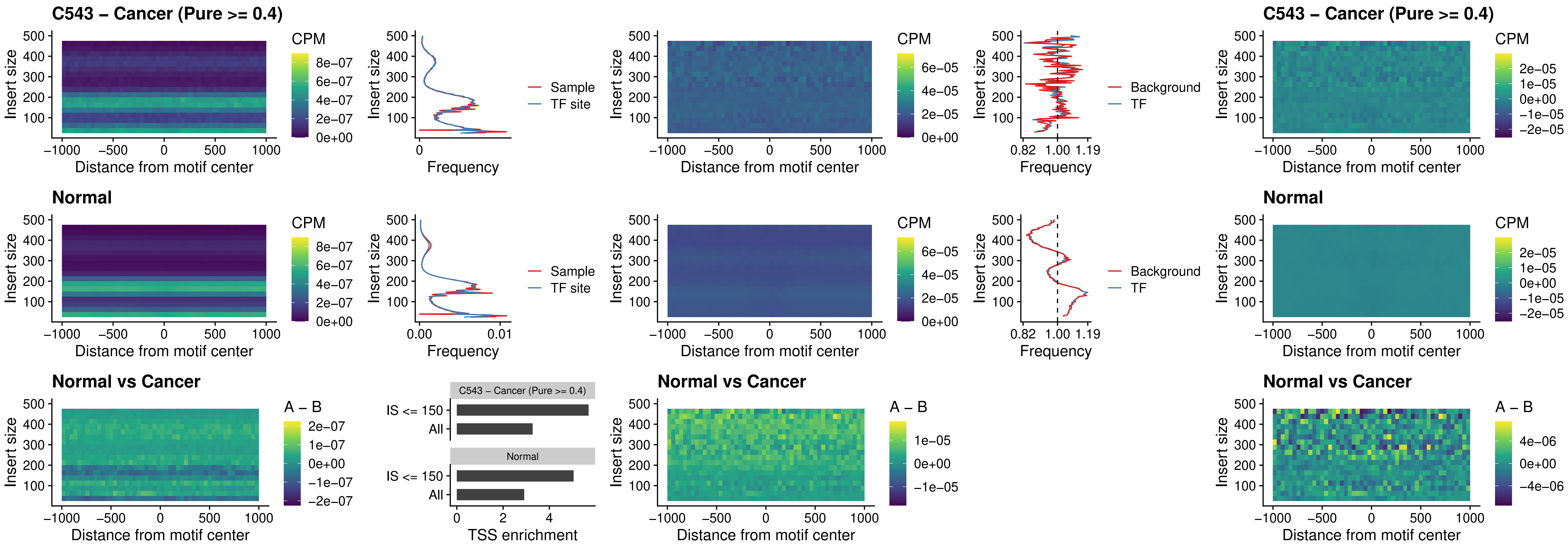

#### C543 – Cancer (Pure >= 0.4) – FOXD4L3_uc_cTSS_nPEAK

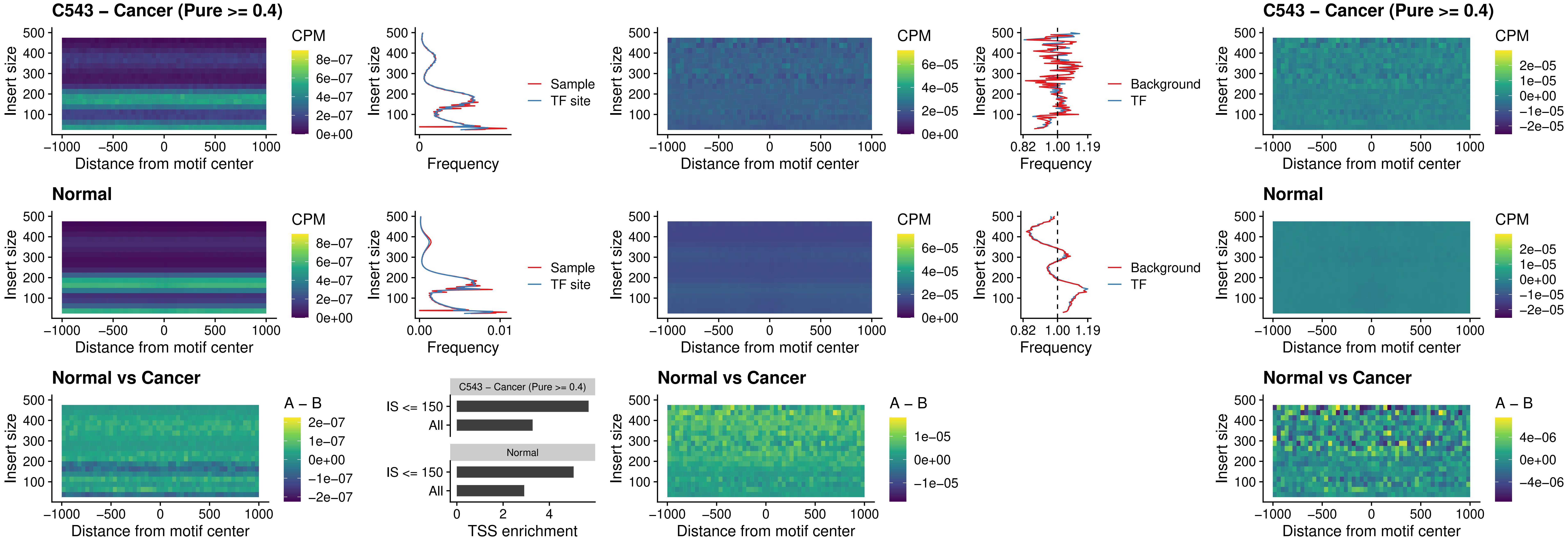

#### C543 – Cancer (Pure >= 0.4) – HOXD1_uc_dTSS_oPEAK

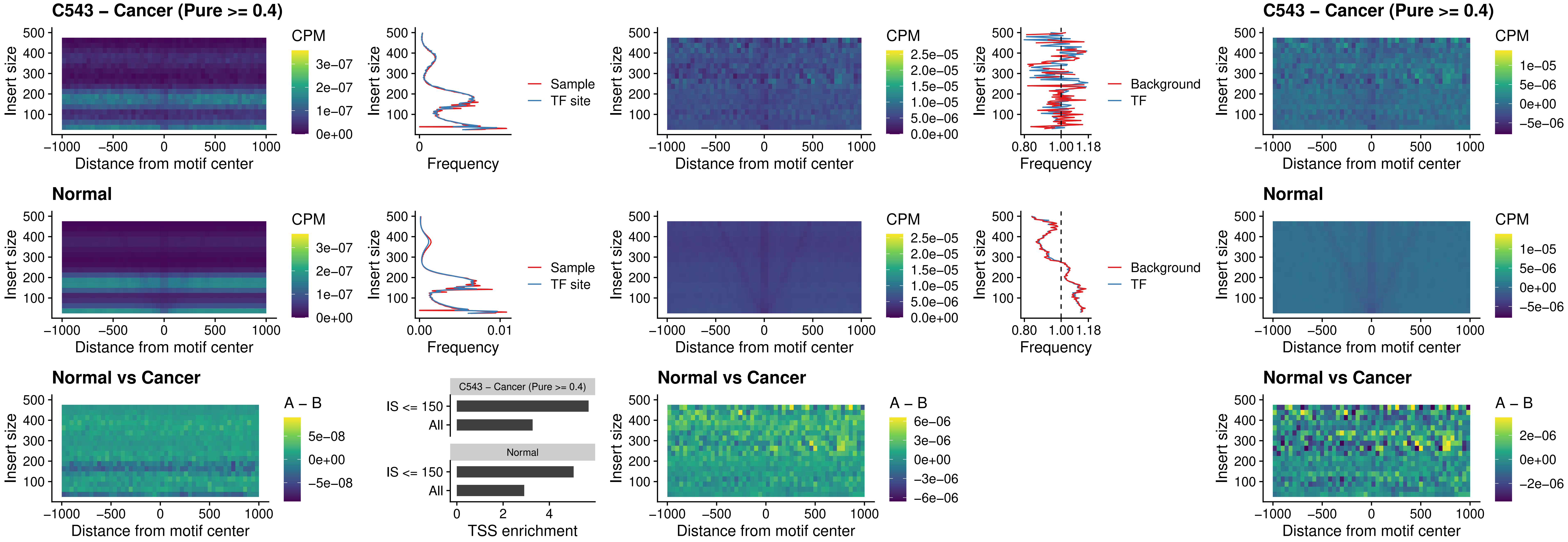

#### C543 – Cancer (Pure >= 0.4) – VAX1_uc_pTSS_nPEAK

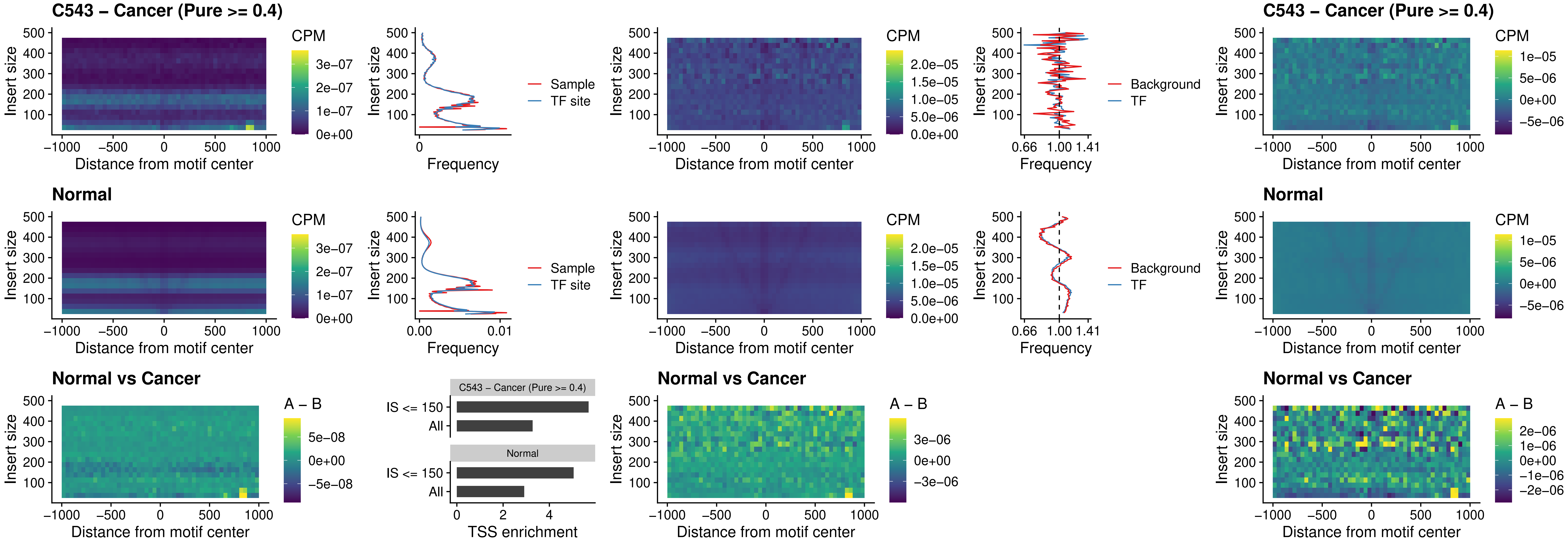

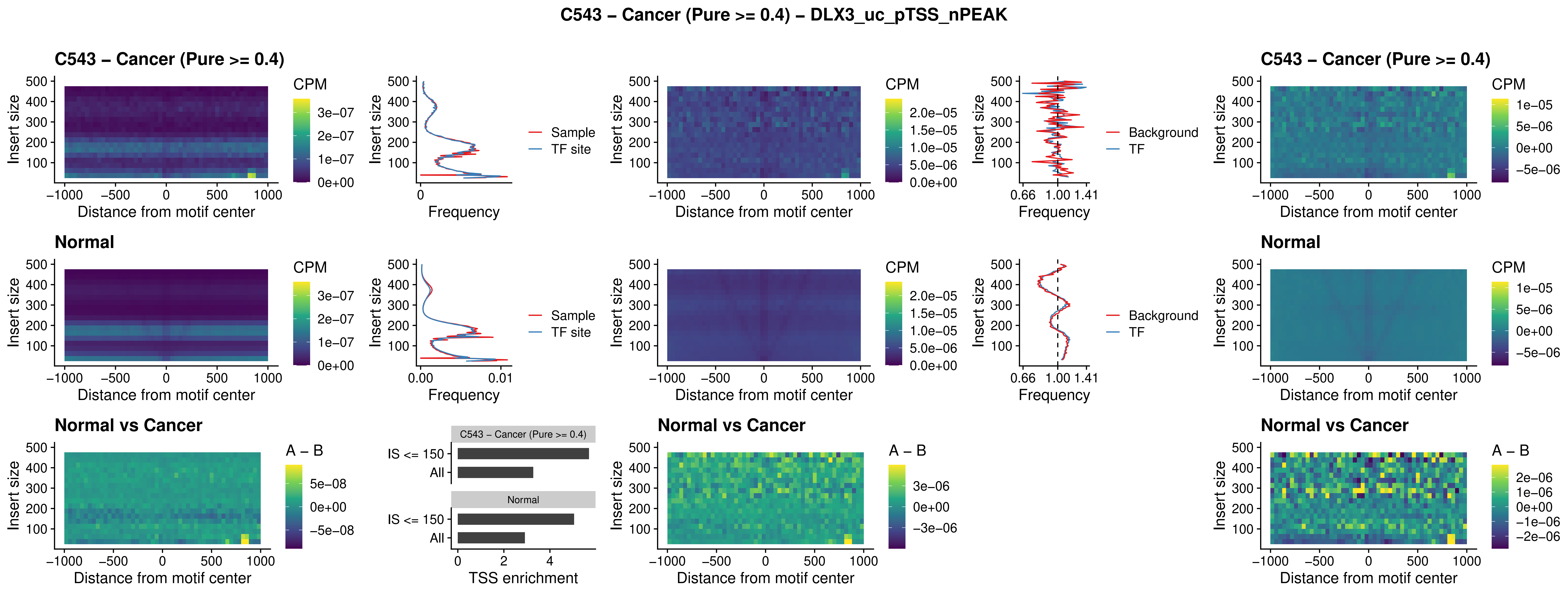
C543 – Cancer (Pure >= 0.4) – DLX3_uc_pTSS_nPEAK
C543 – Cancer (Pure >= 0.4)
Normal
Normal vs Cancer
Insert size
Distance from motif center
CPM
A – B
Frequency
Sample
TF site
Background
TF
C543 – Cancer (Pure >= 0.4)
Normal
IS <= 150
All
TSS enrichment

#### C543 – Cancer (Pure >= 0.4) – HOXD3_uc_pTSS_nPEAK

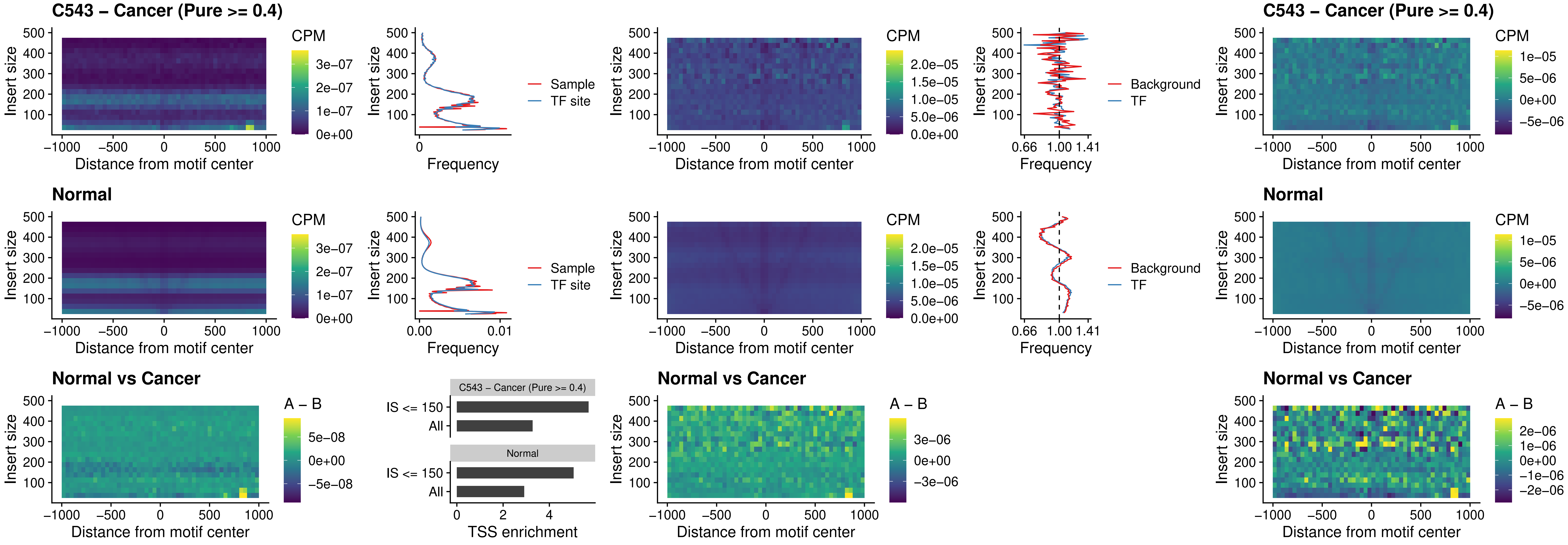

#### C543 – Cancer (Pure >= 0.4) – PDX1_uc_dTSS_nPEAK

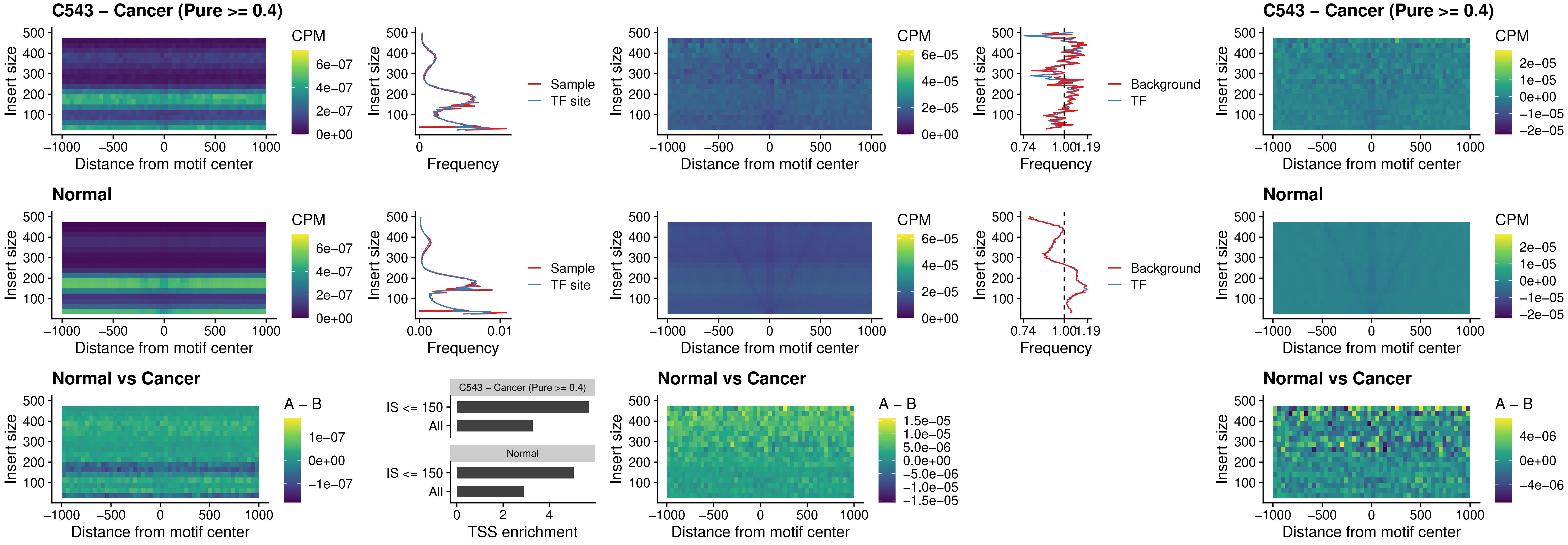

### C543 – Cancer (Pure >= 0.4) – VAX2_uc_dTSS_nPEAK

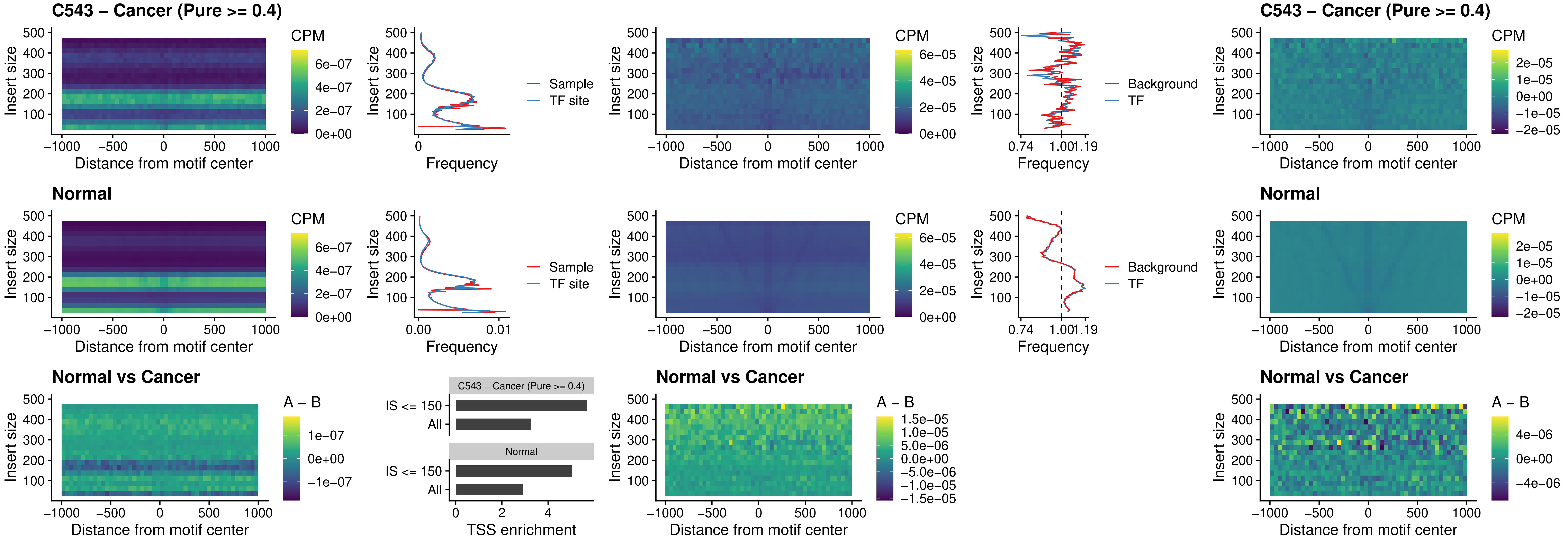

#### C543 – Cancer (Pure >= 0.4) – SOX5_uc_dTSS_nPEAK

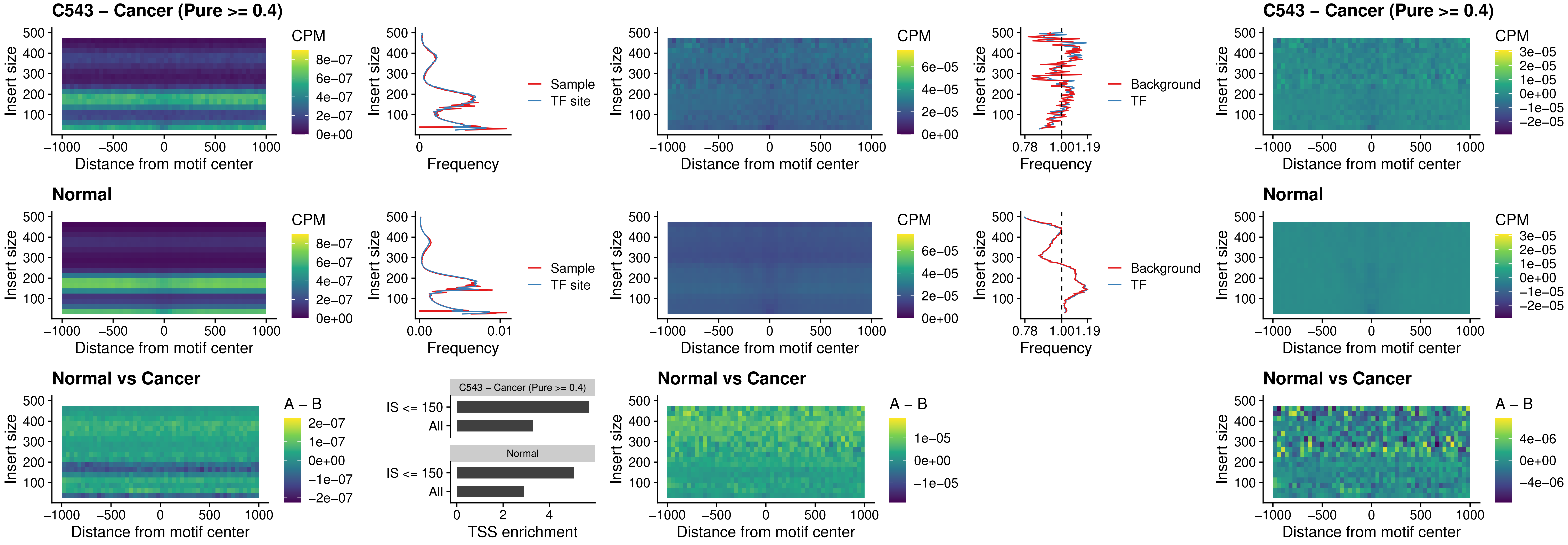

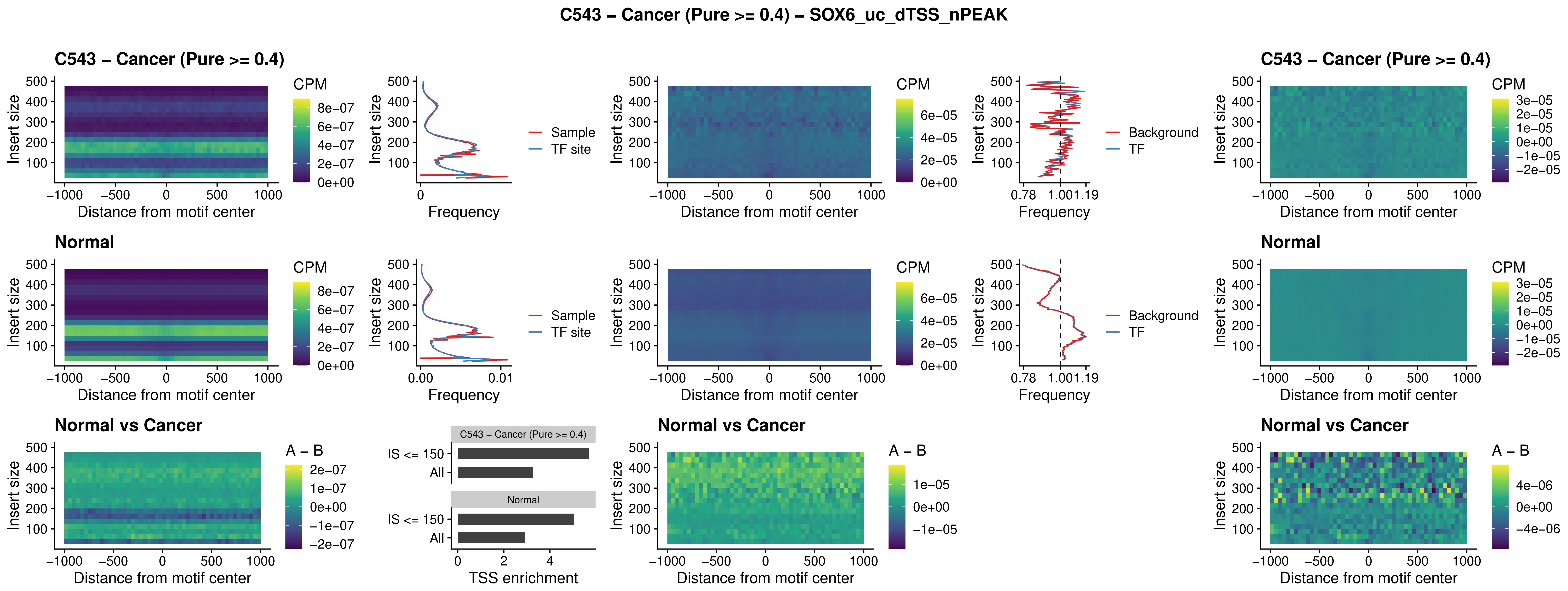
C543 – Cancer (Pure >= 0.4) – SOX6_uc_dTSS_nPEAK
C543 – Cancer (Pure >= 0.4)
Normal
Normal vs Cancer
Insert size
Distance from motif center
CPM
A – B
Frequency
Sample
TF site
Background
TF
C543 – Cancer (Pure >= 0.4)
IS <= 150
All
TSS enrichment

#### C543 – Cancer (Pure >= 0.4) – JDP2_uc_cTSS_nPEAK

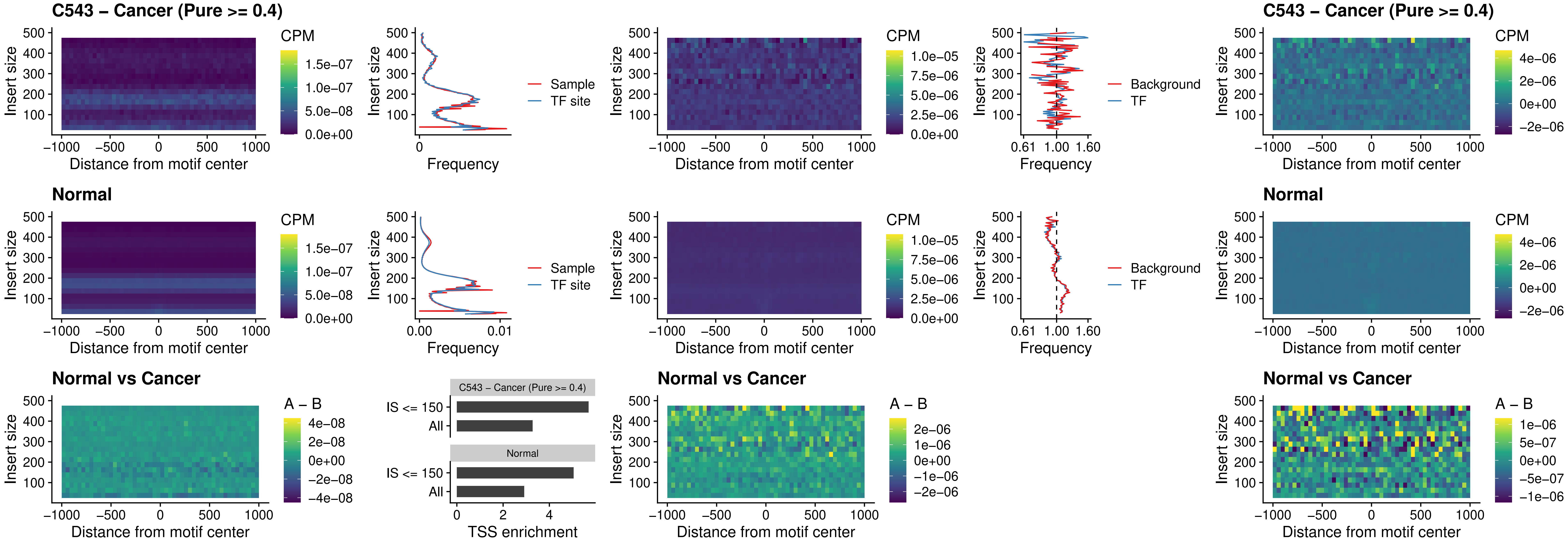

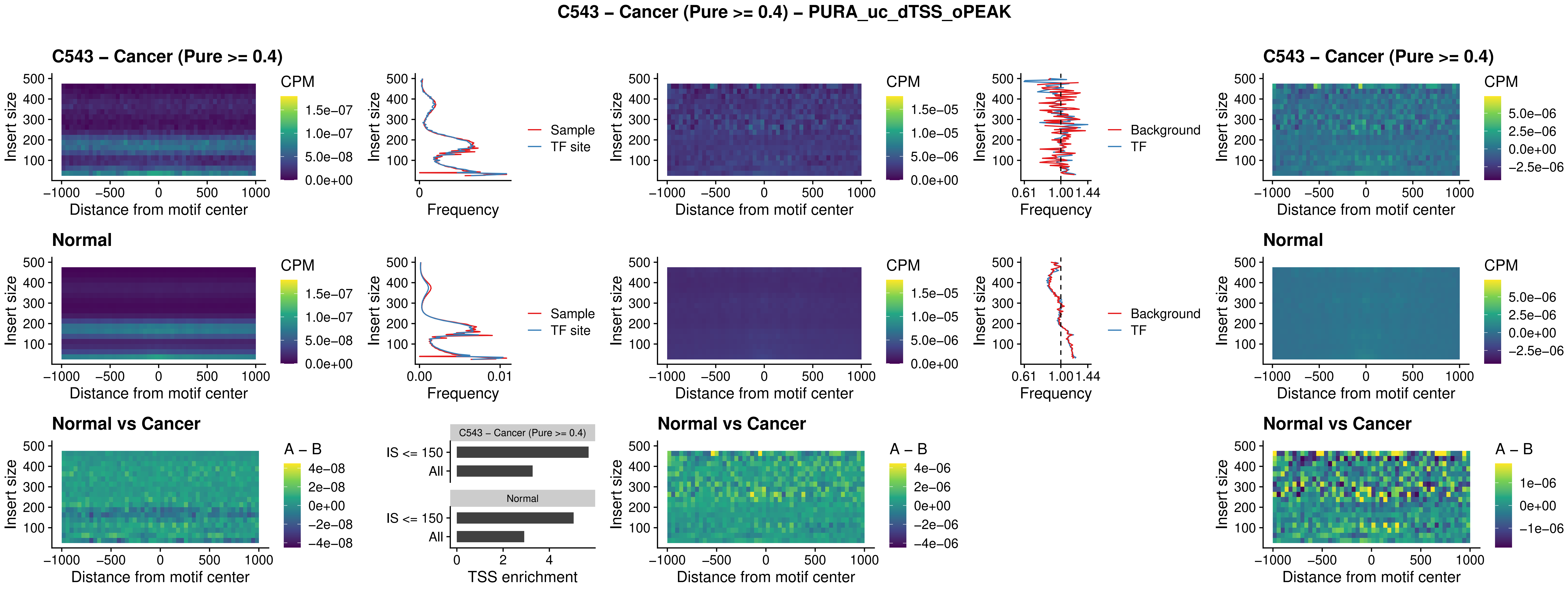
C543 – Cancer (Pure >= 0.4) – PURA_uc_dTSS_oPEAK
C543 – Cancer (Pure >= 0.4)
Normal
Normal vs Cancer
Insert size
Distance from motif center
CPM
A – B
Frequency
Sample
TF site
Background
TF
TSS enrichment
IS <= 150
All
C543 – Cancer (Pure >= 0.4)
Normal

### C543 – Cancer (Pure >= 0.4) – HHEX_uc_dTSS_nPEAK

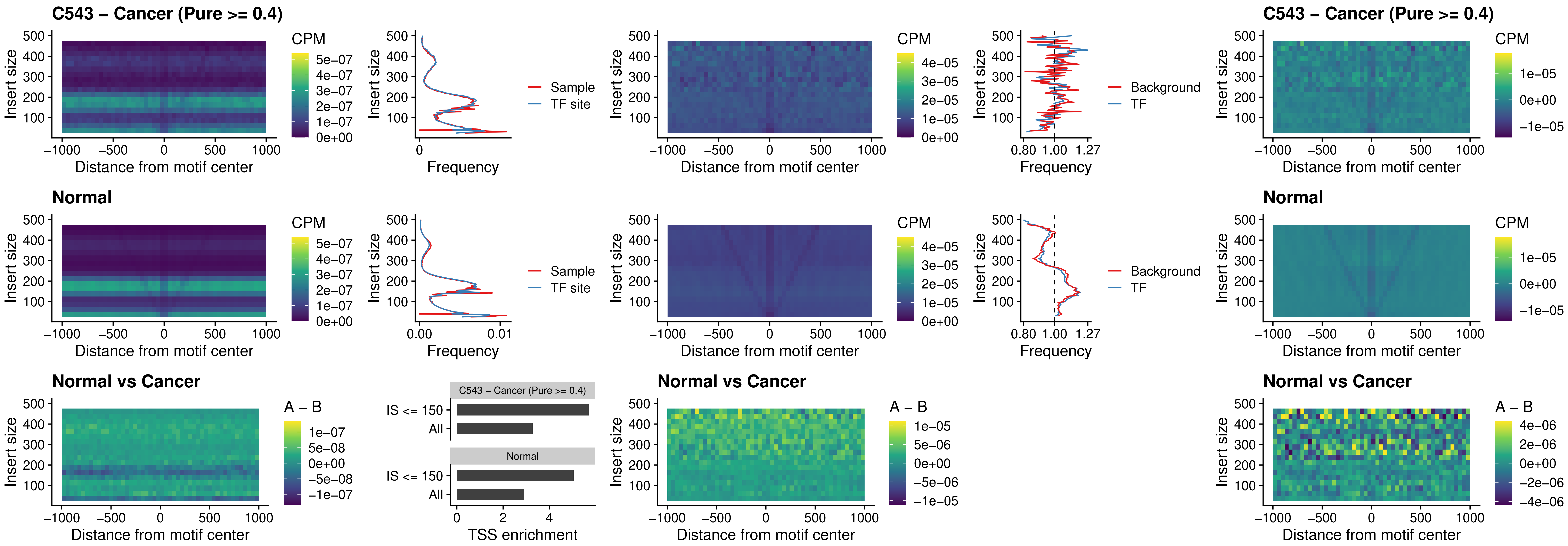

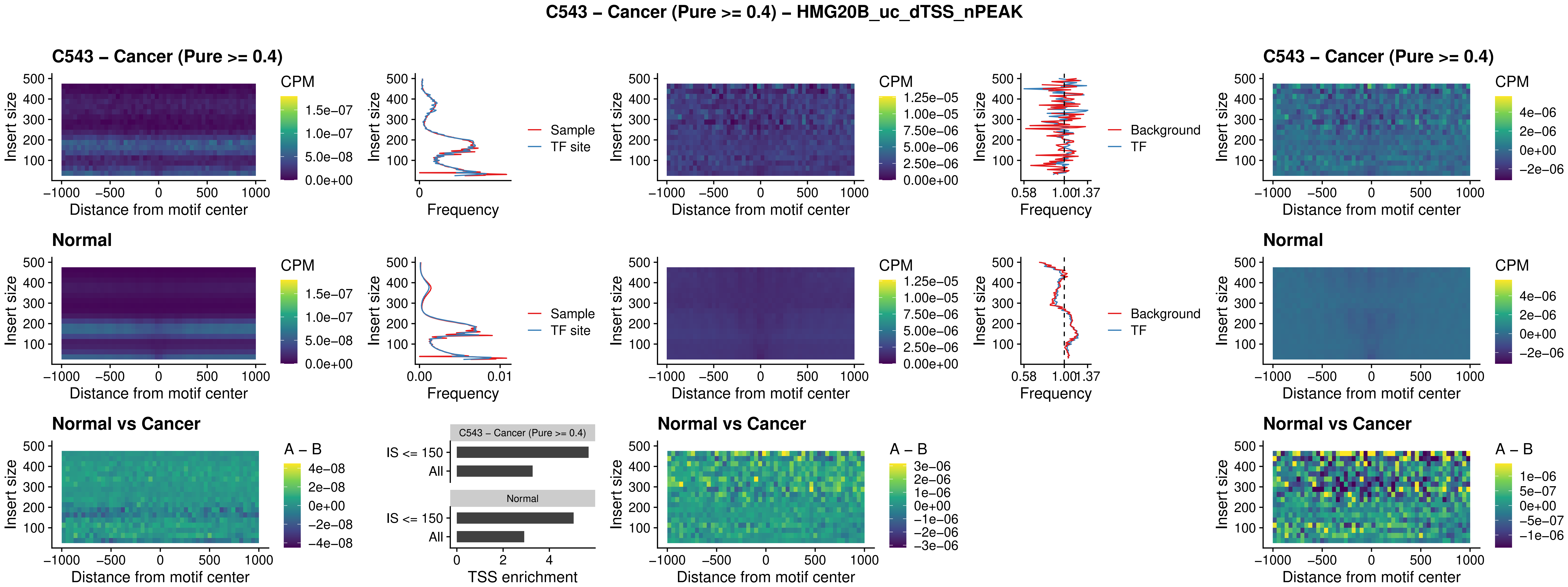

C543 – Cancer (Pure >= 0.4) – HMG20B_uc_dTSS_nPEAK
C543 – Cancer (Pure >= 0.4)
Normal
Normal vs Cancer
Insert size
Distance from motif center
Frequency
CPM
A – B
Sample
TF site
Background
TF
C543 – Cancer (Pure >= 0.4)
IS <= 150
All
TSS enrichment

#### C543 – Cancer (Pure >= 0.4) – FOXB2_uc_dTSS_nPEAK

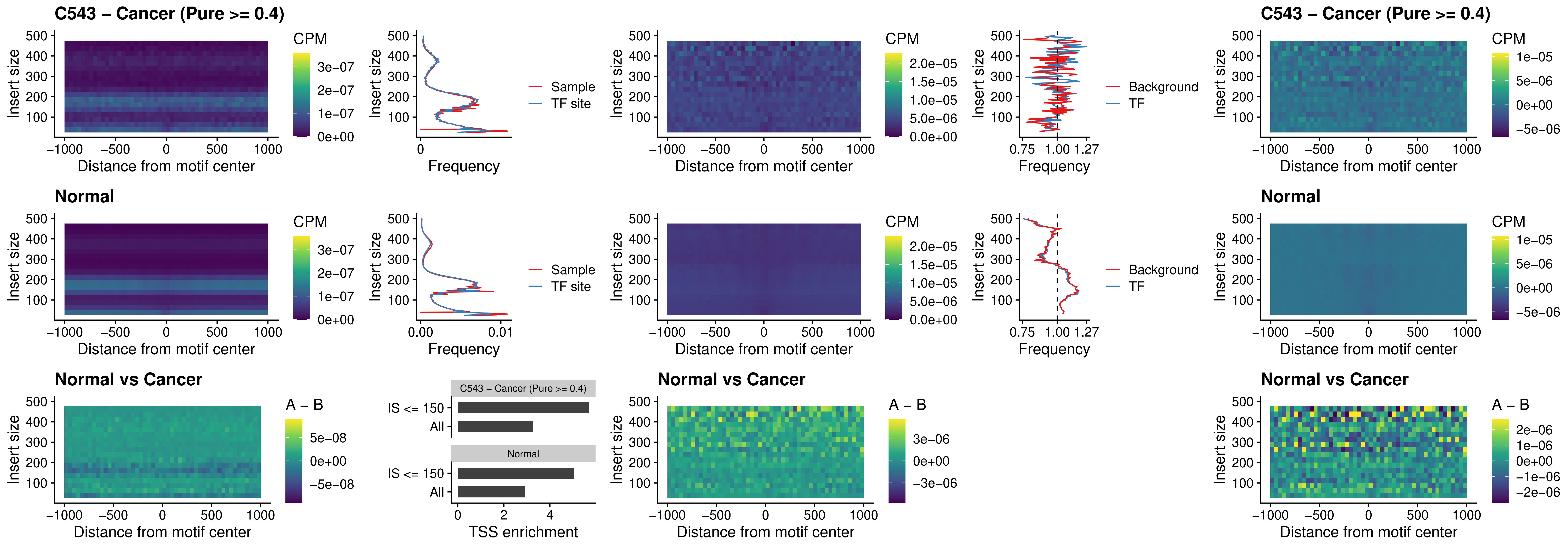

### C543 – Cancer (Pure >= 0.4) – FOXL1_uc_cTSS_nPEAK

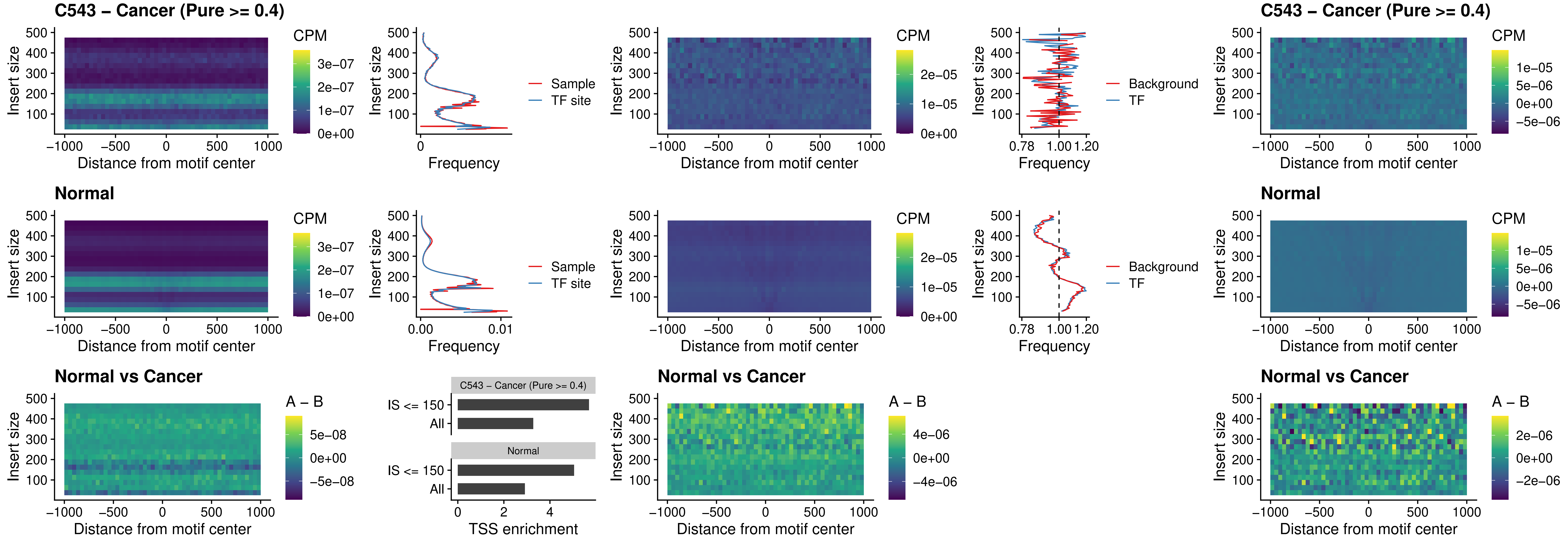

#### C543 – Cancer (Pure >= 0.4) – PBX3_uc_pTSS_oPEAK

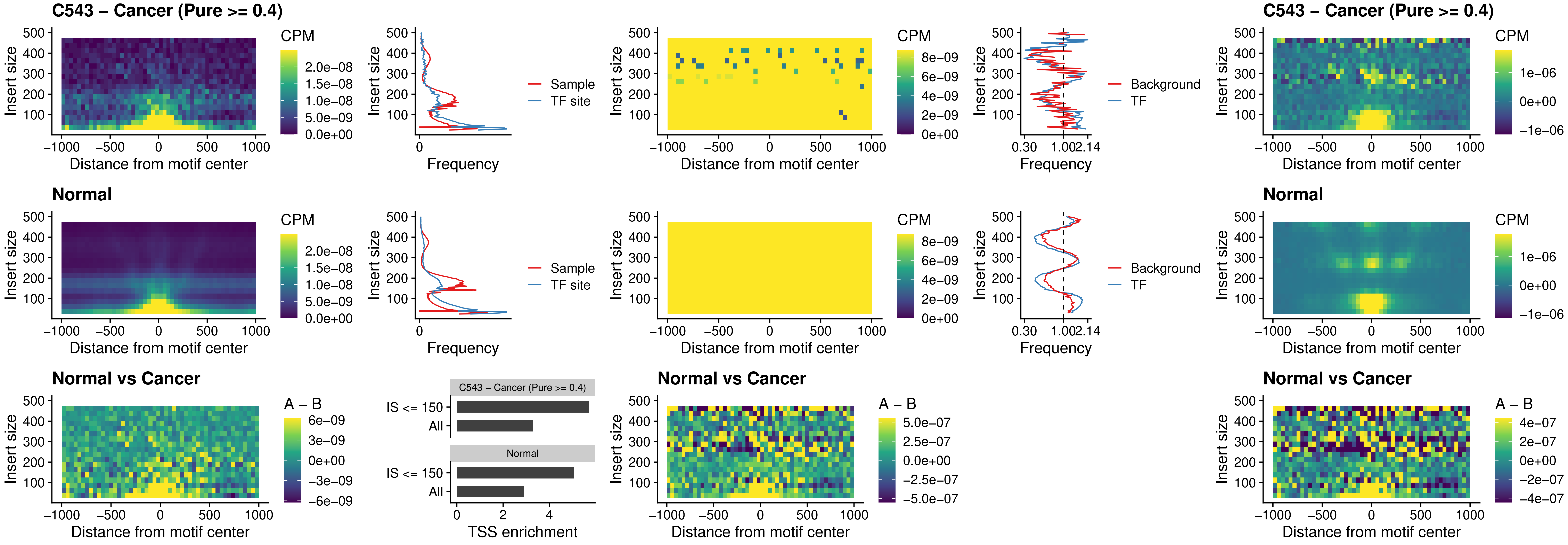

#### C543 – Cancer (Pure >= 0.4) – SNAI2_uc_cTSS_nPEAK

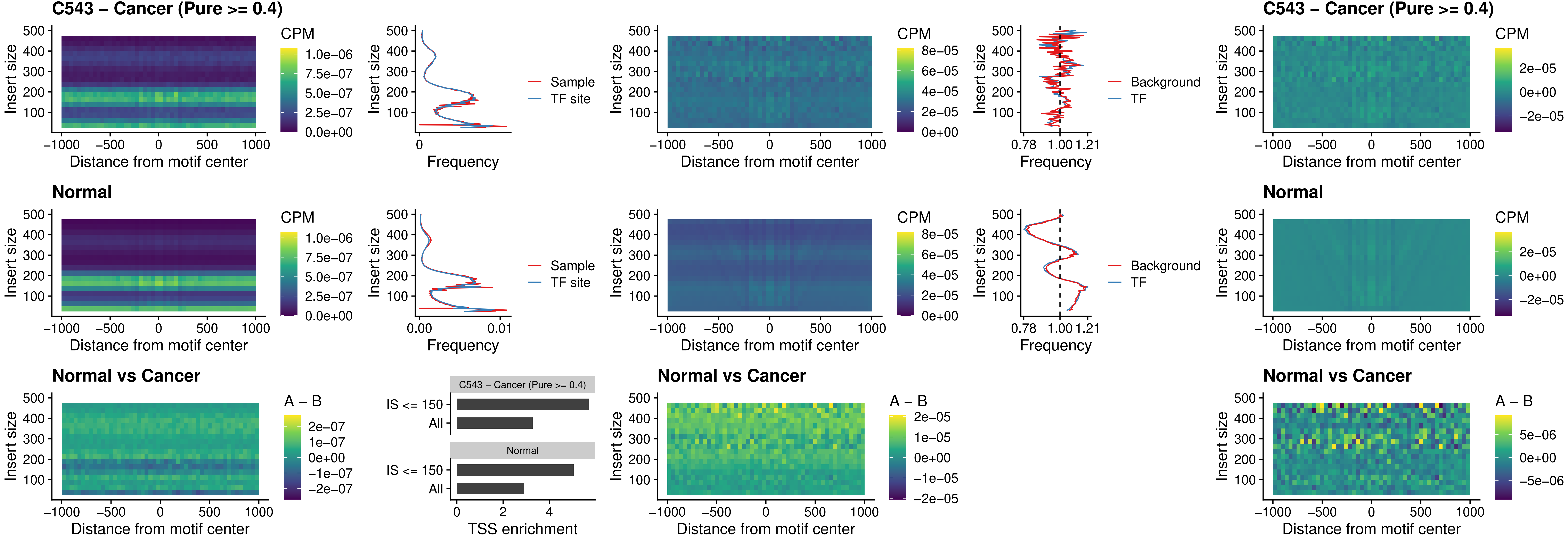

#### C543 – Cancer (Pure >= 0.4) – SNAI2_uc_dTSS_oPEAK

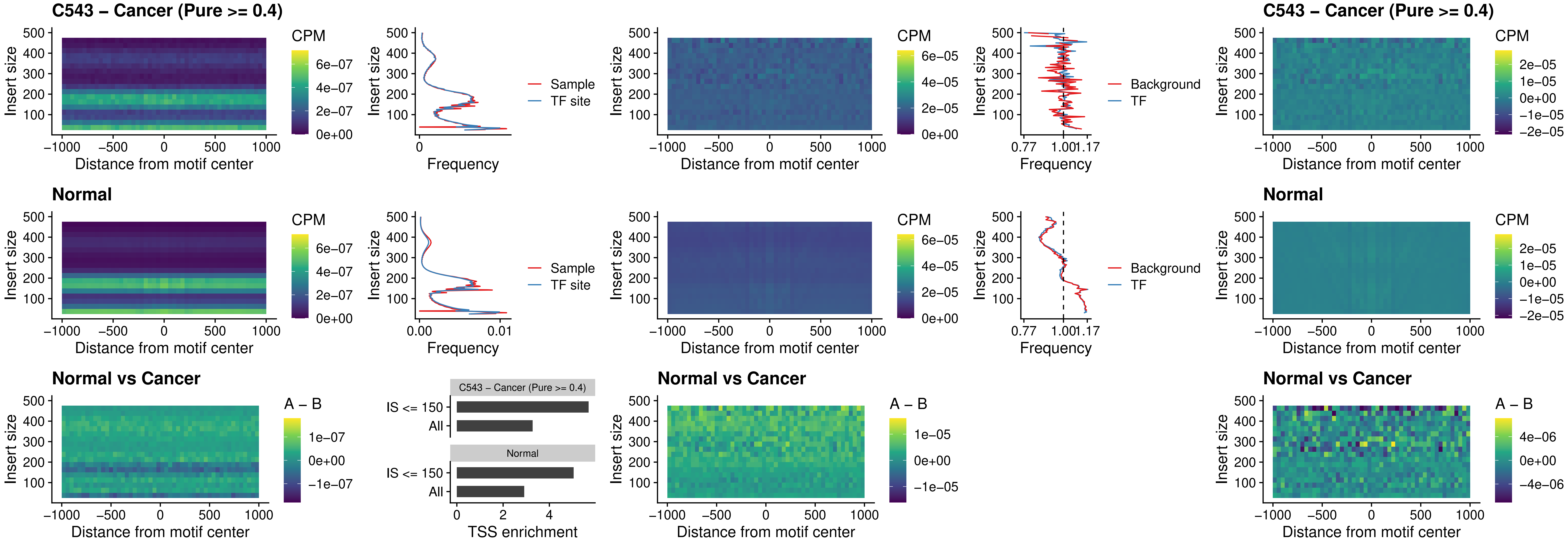

#### C543 – Cancer (Pure >= 0.4) – SPI1_uc_pTSS_nPEAK

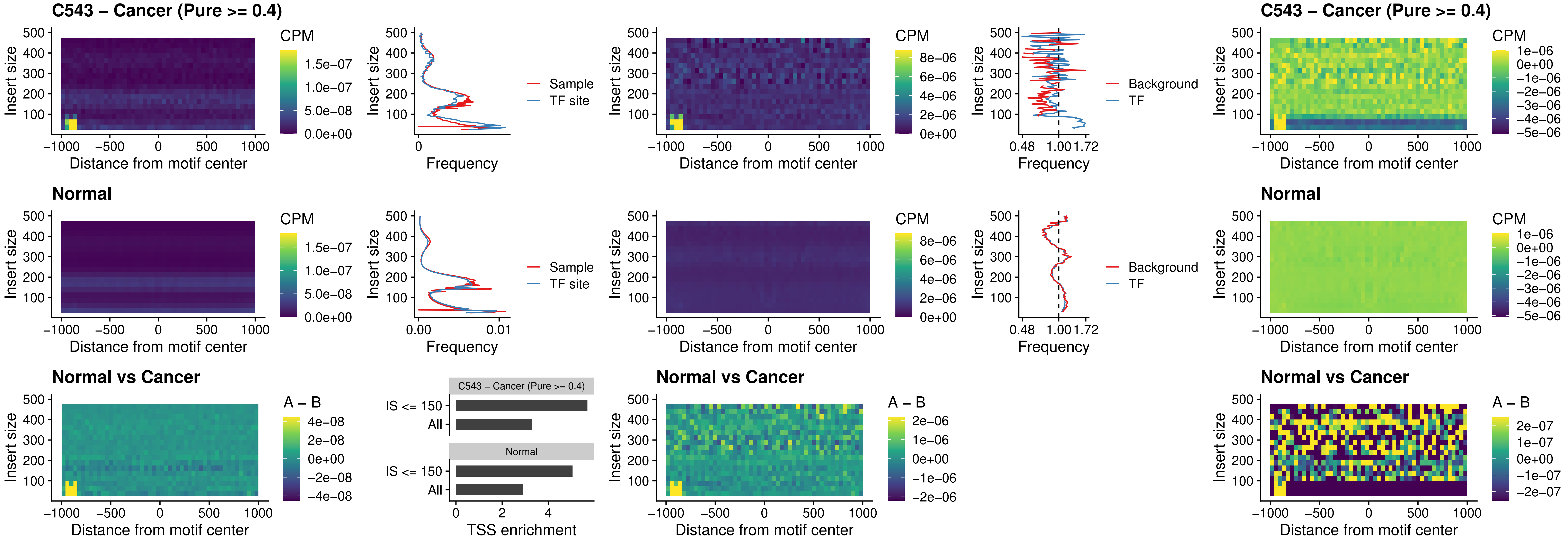

#### C543 – Cancer (Pure >= 0.4) – KLF16_uc_pTSS_nPEAK

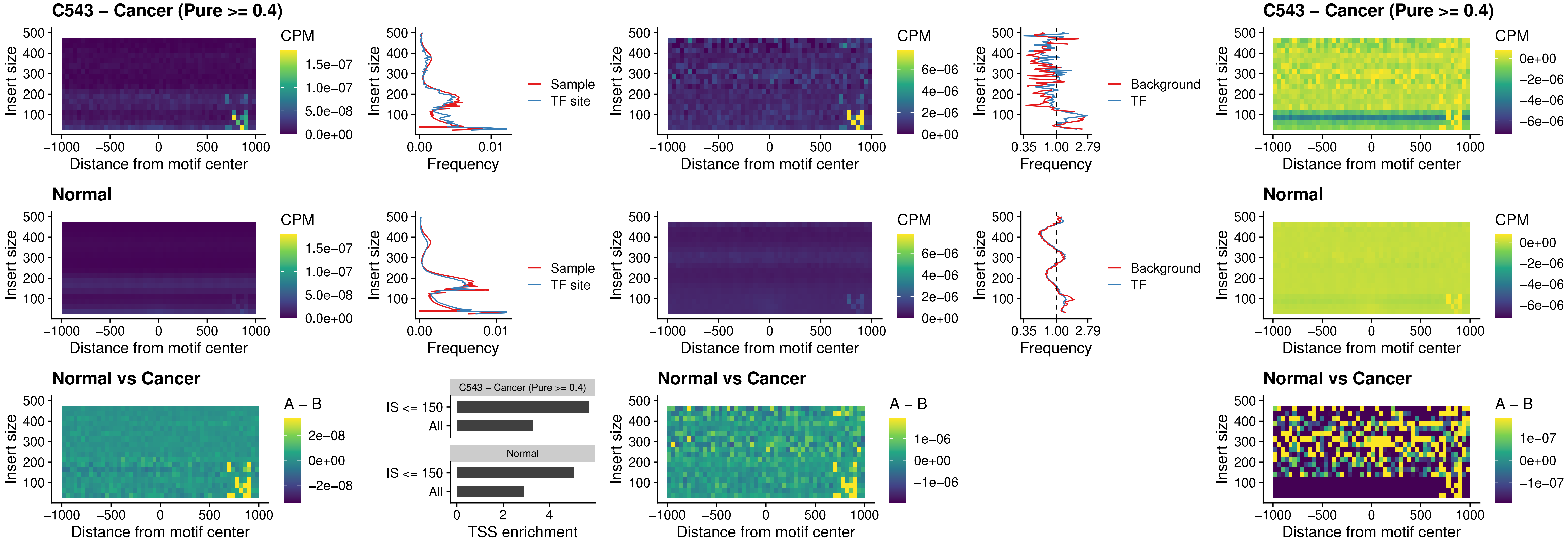

#### C543 – Cancer (Pure >= 0.4) – SP3_uc_pTSS_nPEAK

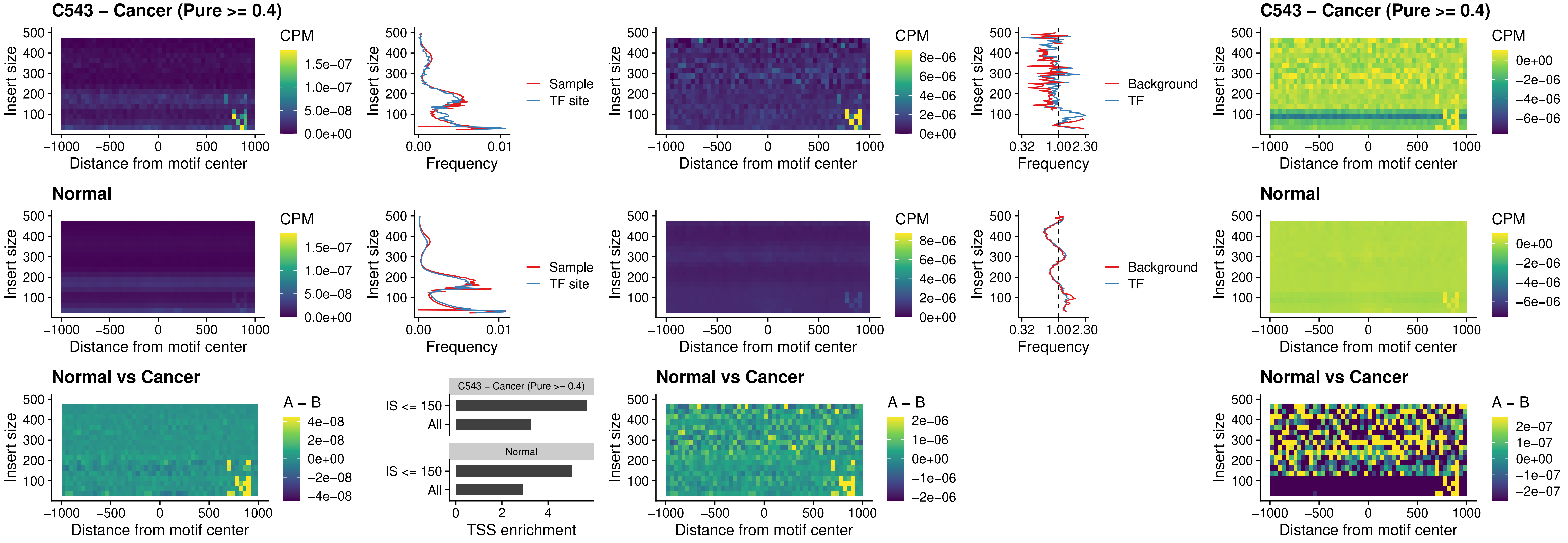

### C543 – Cancer (Pure >= 0.4) – NEUROD1_uc_dTSS_oPEAK

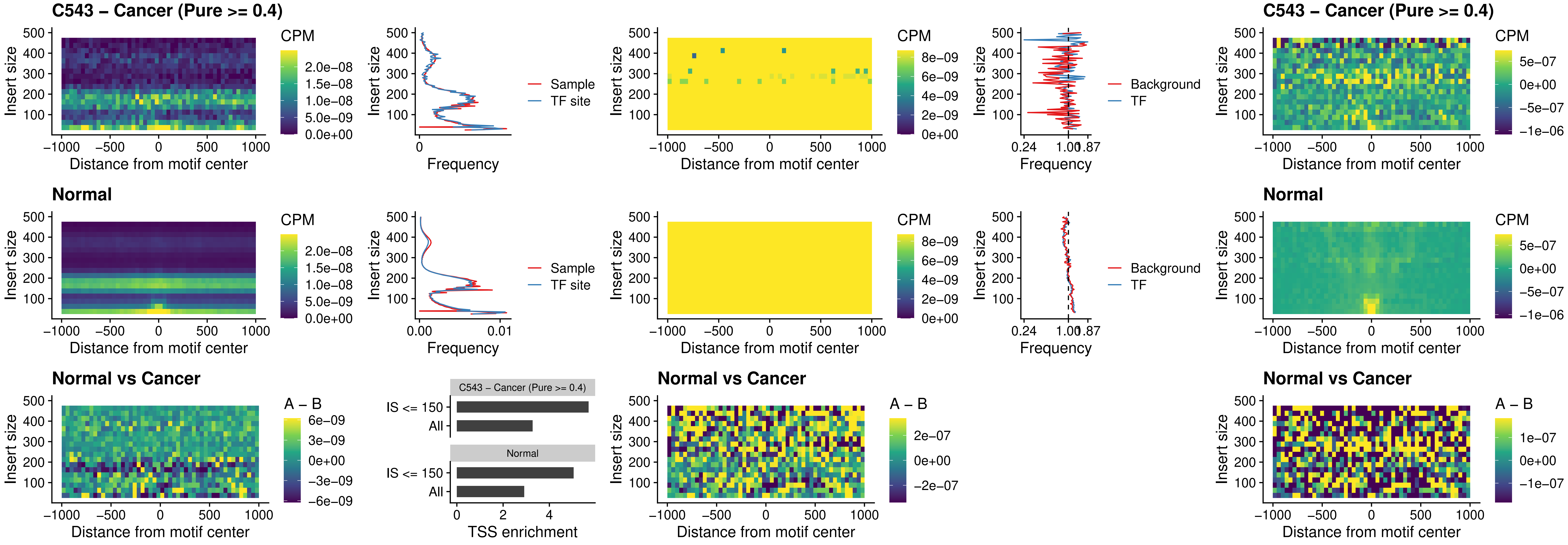

#### C543 – Cancer (Pure >= 0.4) – CTCF_uc_dTSS_oPEAK

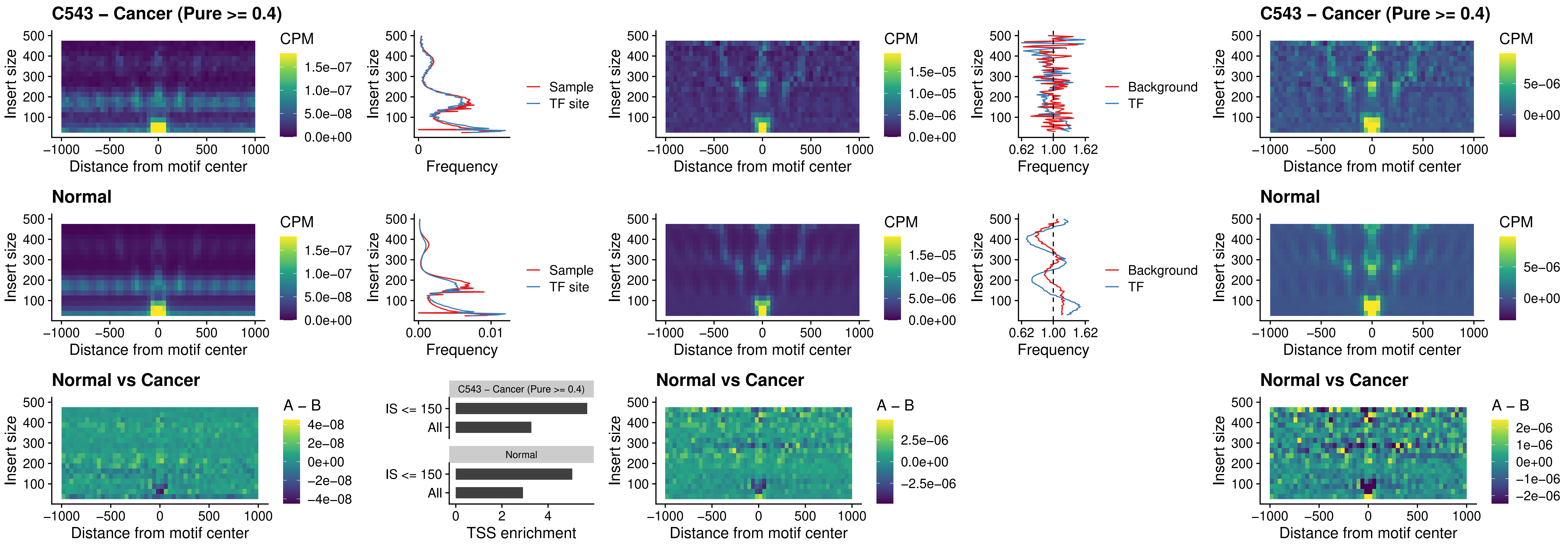

#### C543 – Cancer (Pure >= 0.4) – CTCF_uc_cTSS_oPEAK

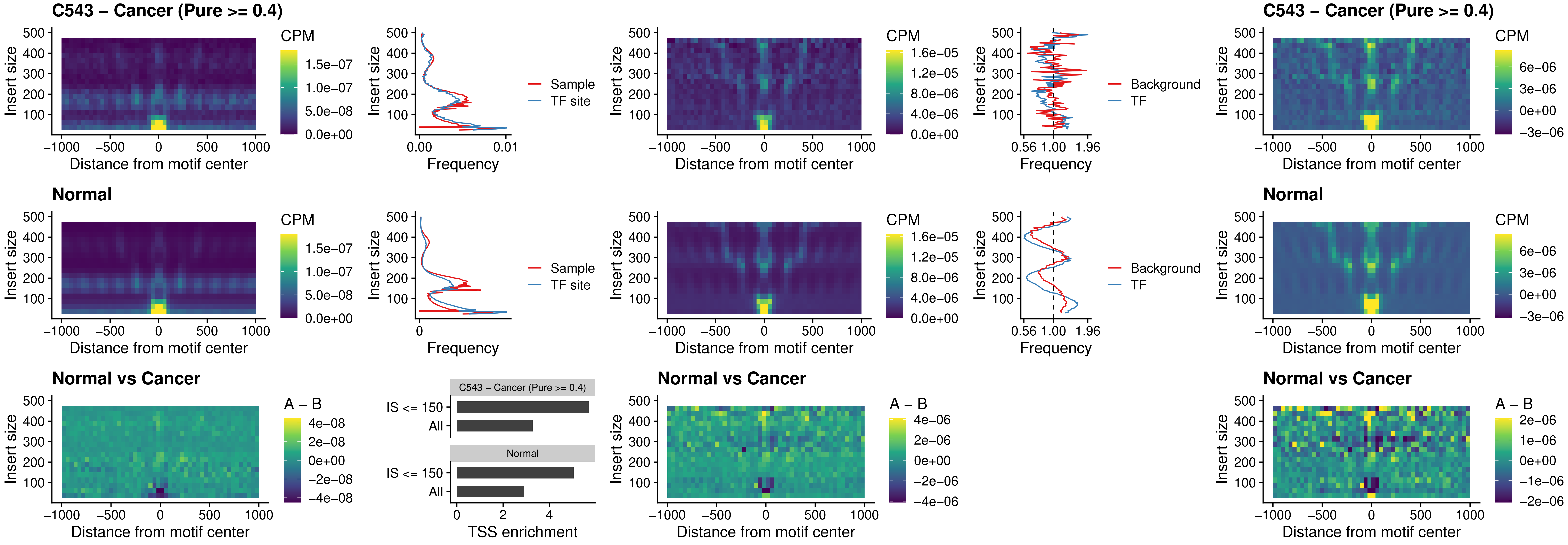

#### C543 – Cancer (Pure >= 0.4) – CTCFL_uc_dTSS_oPEAK

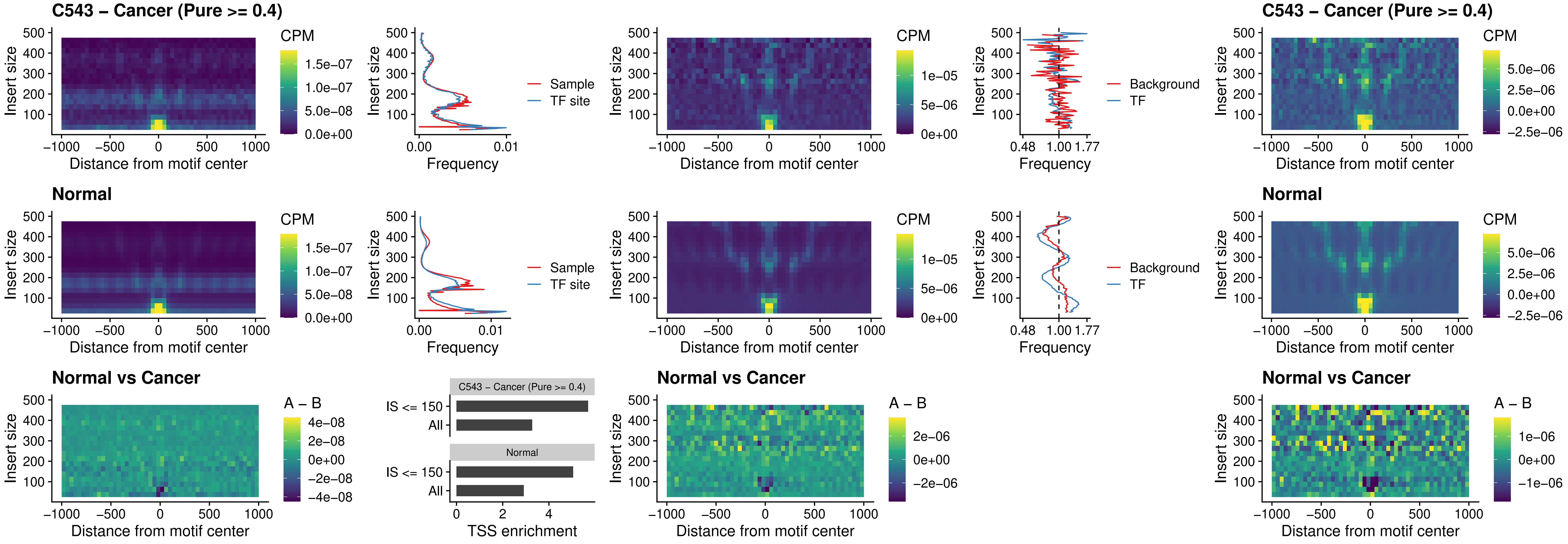

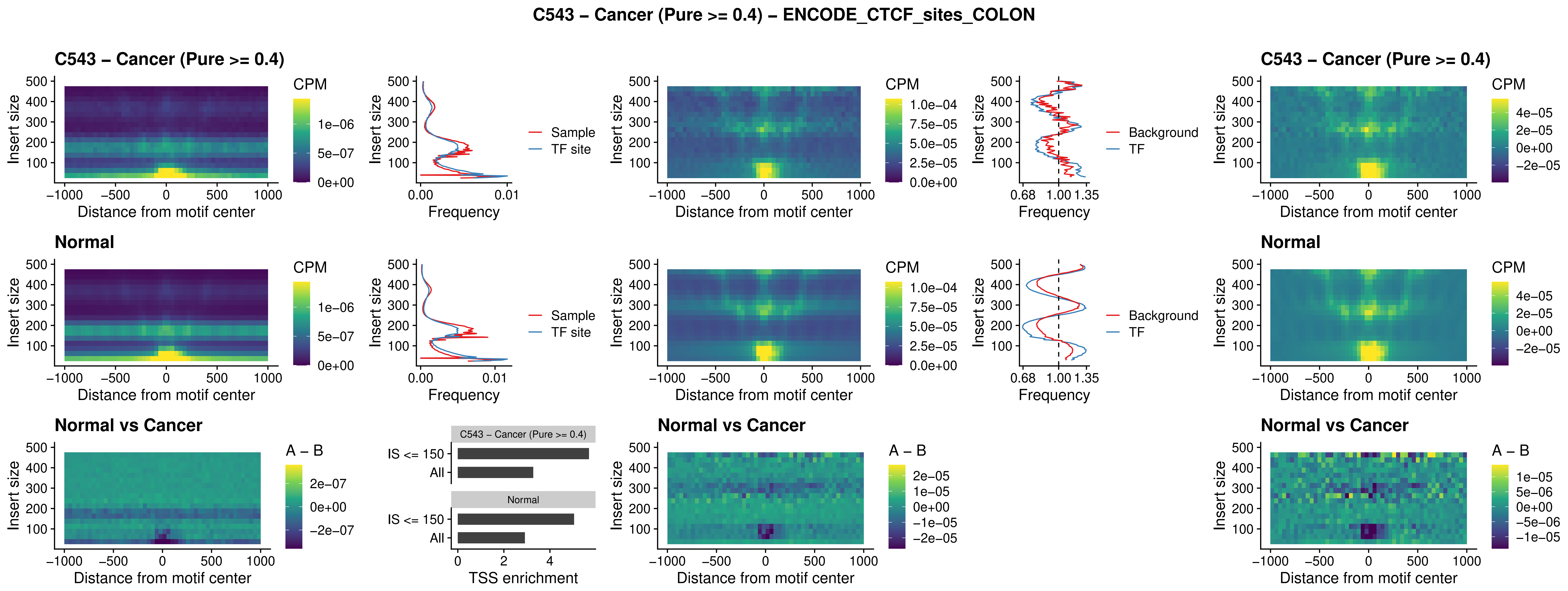
C543 – Cancer (Pure >= 0.4) – ENCODE_CTCF_sites_COLON
C543 – Cancer (Pure >= 0.4)
Normal
Normal vs Cancer
Insert size
Distance from motif center
Frequency
CPM
A – B
Sample
TF site
Background
TF
C543 – Cancer (Pure >= 0.4)
Normal
IS <= 150
All
TSS enrichment

**C543 – Cancer (Pure >= 0.4) – ENCODE_CTCF_sites_COLON_loop_ctcf**

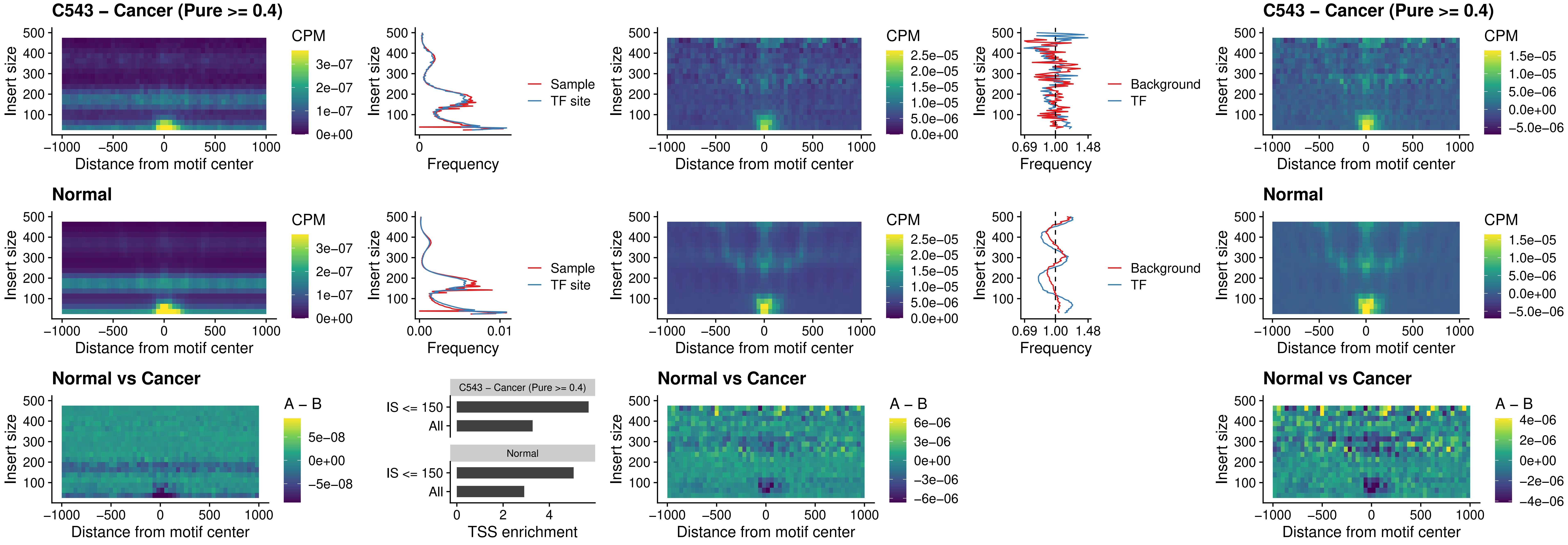

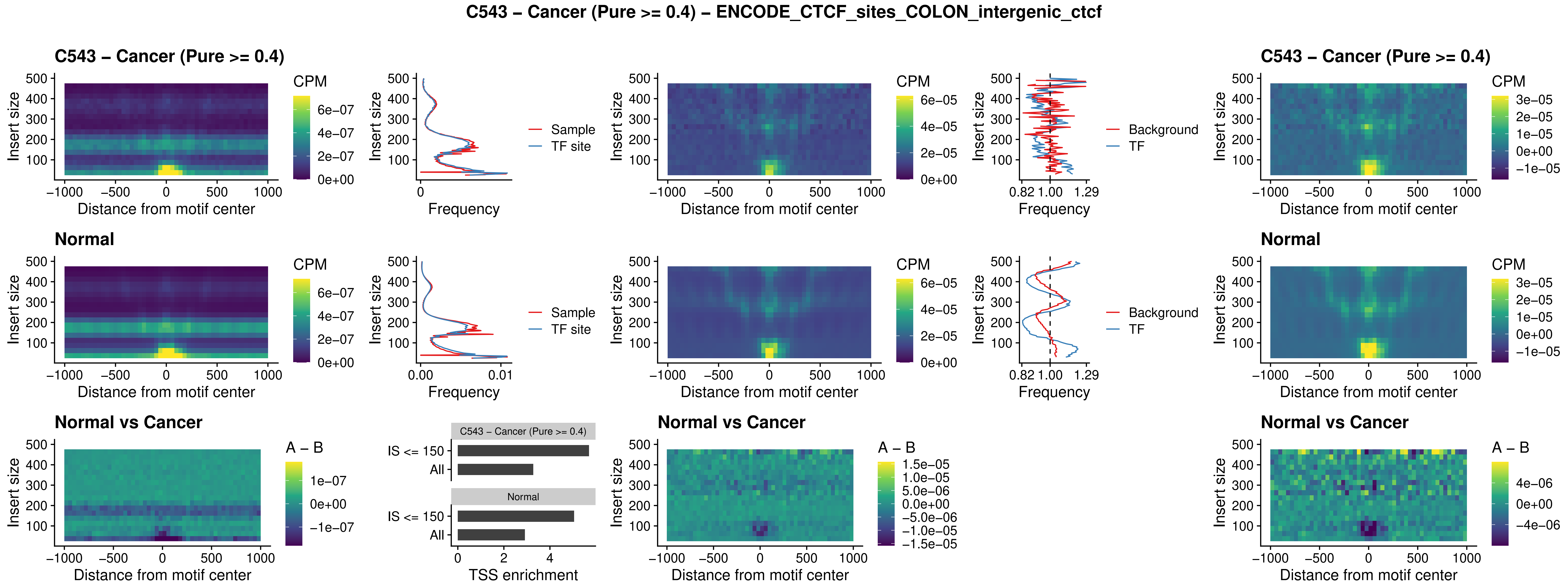
C543 – Cancer (Pure >= 0.4) – ENCODE_CTCF_sites_COLON_intergenic_ctcf
C543 – Cancer (Pure >= 0.4)
Normal
Normal vs Cancer
Insert size
Distance from motif center
Frequency
CPM
A – B
Sample
TF site
Background
TF
C543 – Cancer (Pure >= 0.4)
Normal
IS <= 150
All
TSS enrichment

#### C543 – Cancer (Pure >= 0.4) – ENCODE_CTCF_sites_COLON_nonpromotor

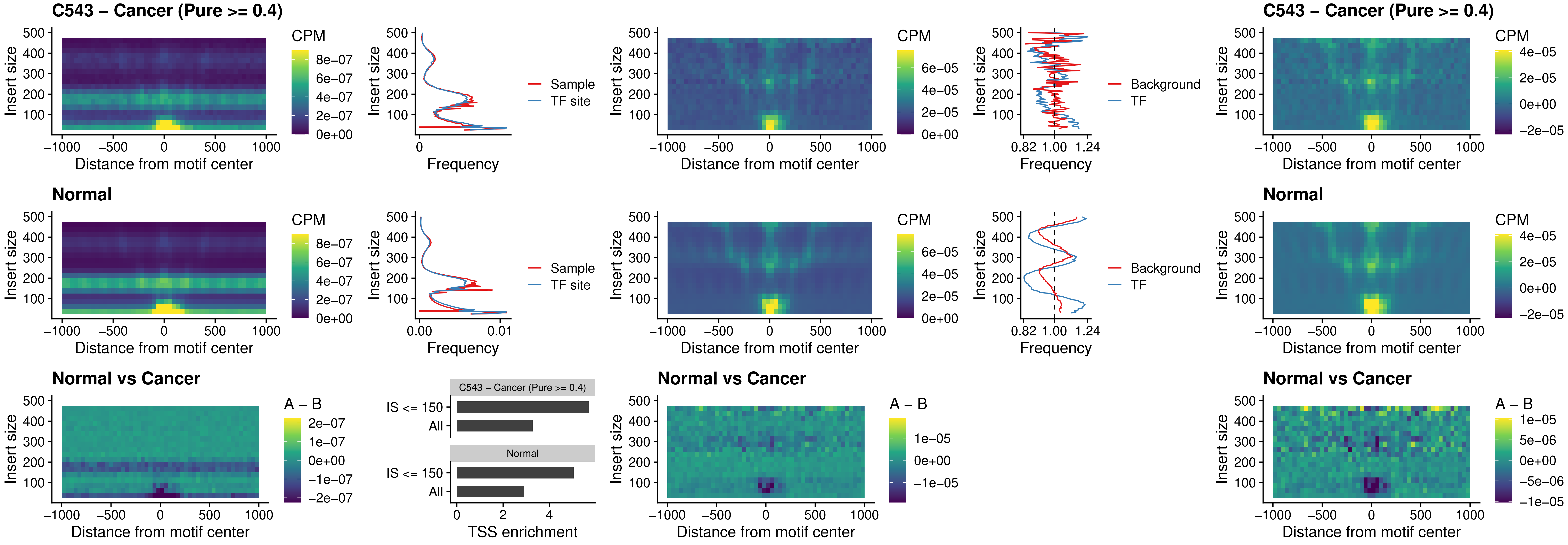

#### C543 – Cancer (Pure >= 0.4) – CTCFL_uc_cTSS_oPEAK

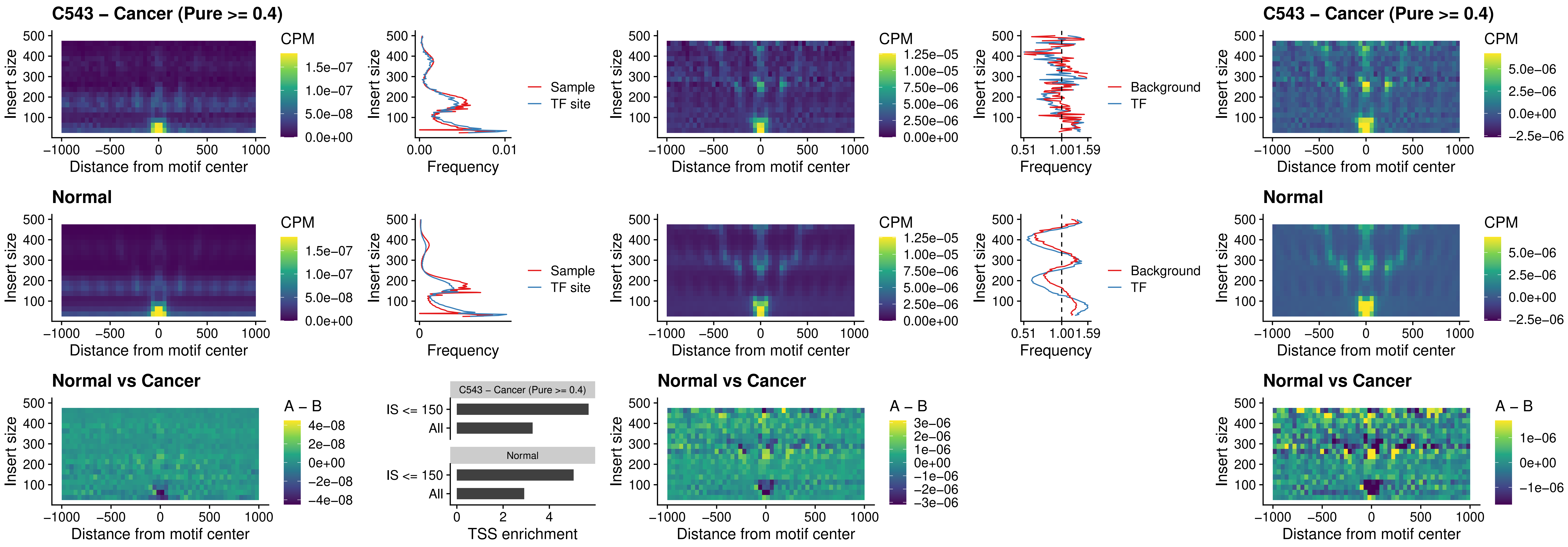

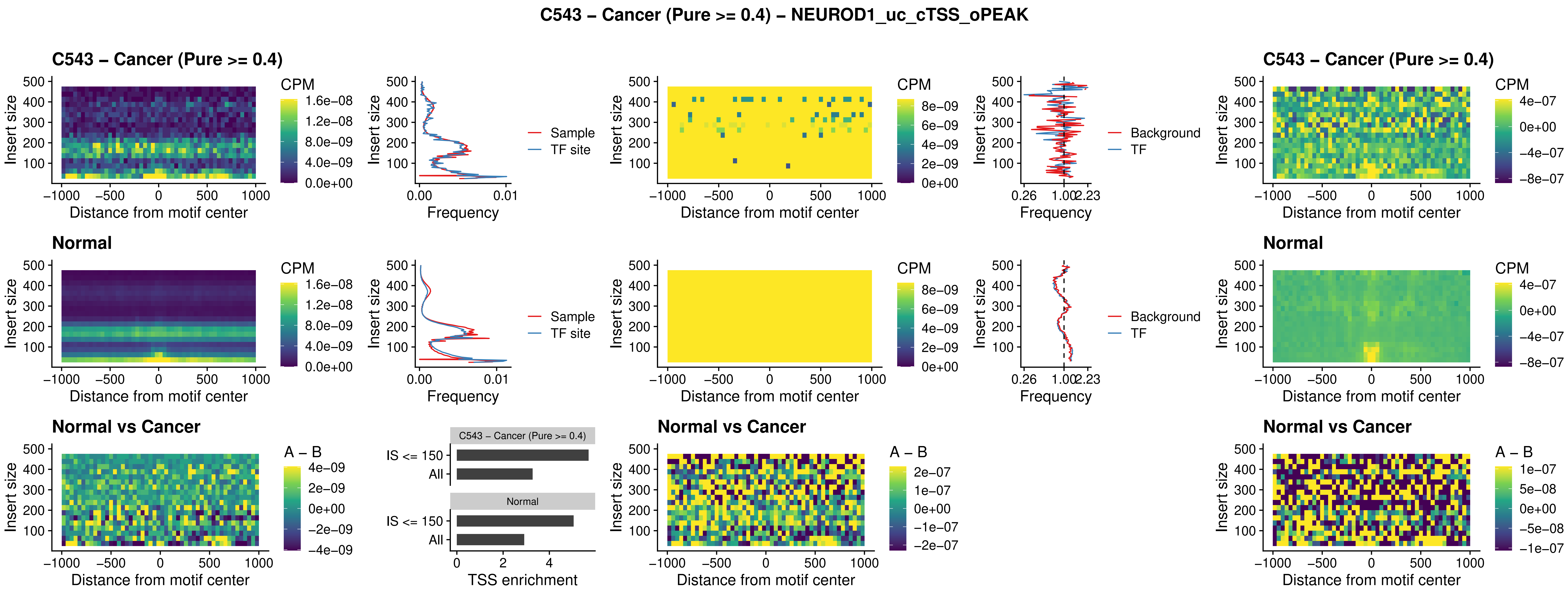

C543 – Cancer (Pure >= 0.4) – NEUROD1_uc_cTSS_oPEAK
C543 – Cancer (Pure >= 0.4)
Normal
Normal vs Cancer
Insert size
Distance from motif center
CPM
A – B
Frequency
Sample
TF site
Background
TF
C543 – Cancer (Pure >= 0.4)
Normal
IS <= 150
All
TSS enrichment

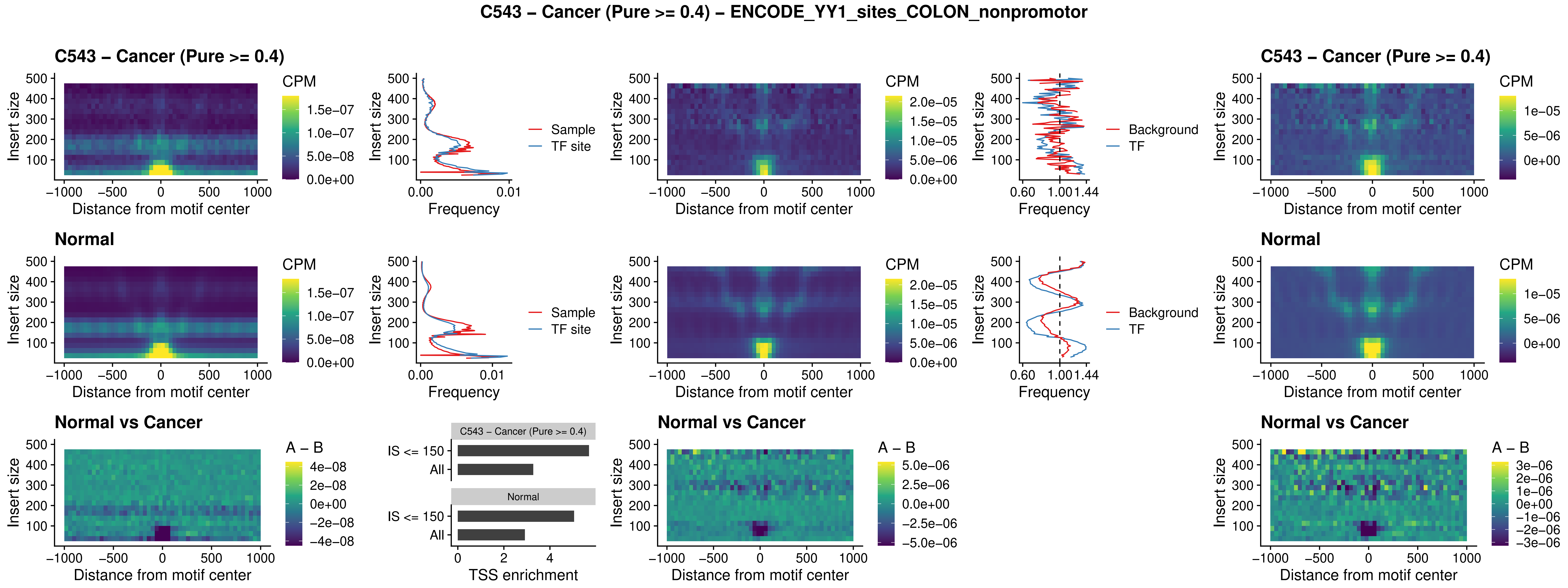
C543 – Cancer (Pure >= 0.4) – ENCODE_YY1_sites_COLON_nonpromotor
C543 – Cancer (Pure >= 0.4)
Normal
Normal vs Cancer
Insert size
Distance from motif center
CPM
A – B
Frequency
Sample
TF site
Background
TF
C543 – Cancer (Pure >= 0.4)
Normal
IS <= 150
All
TSS enrichment

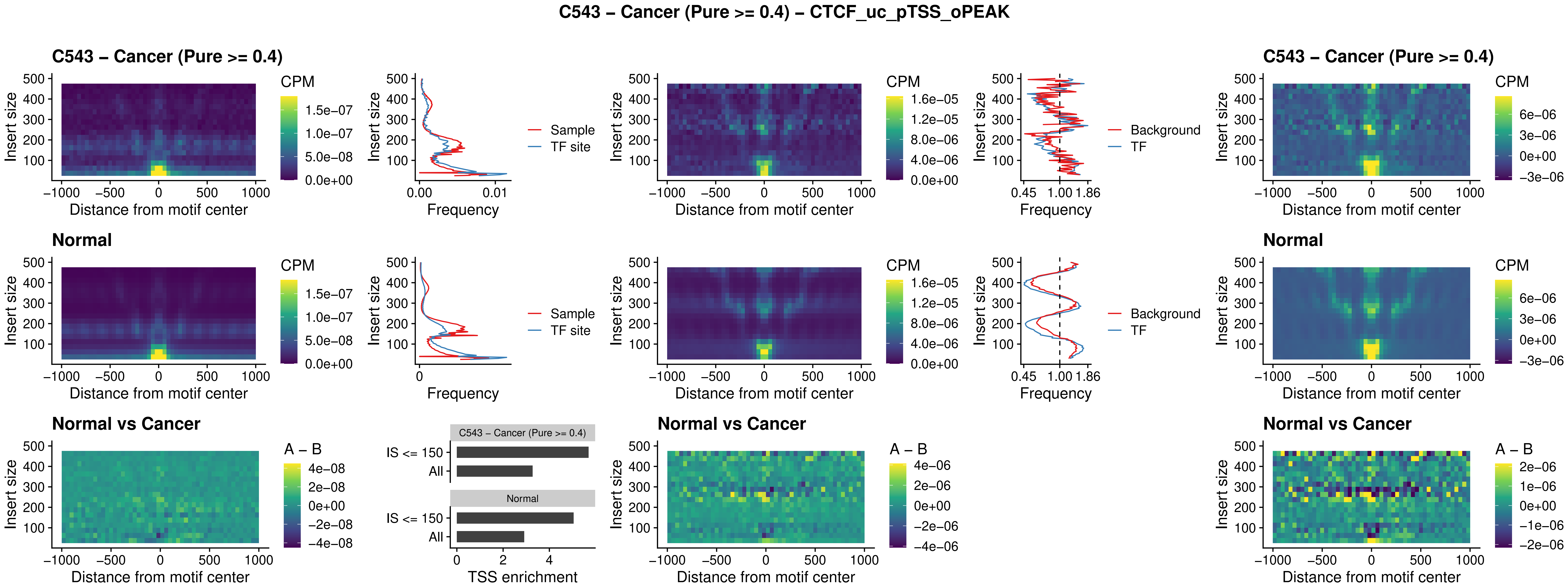
C543 – Cancer (Pure >= 0.4) – CTCF_uc_pTSS_oPEAK
C543 – Cancer (Pure >= 0.4)
Normal
Normal vs Cancer
Insert size
Distance from motif center
Frequency
CPM
A – B
Sample
TF site
Background
TF
C543 – Cancer (Pure >= 0.4)
Normal
IS <= 150
All
TSS enrichment

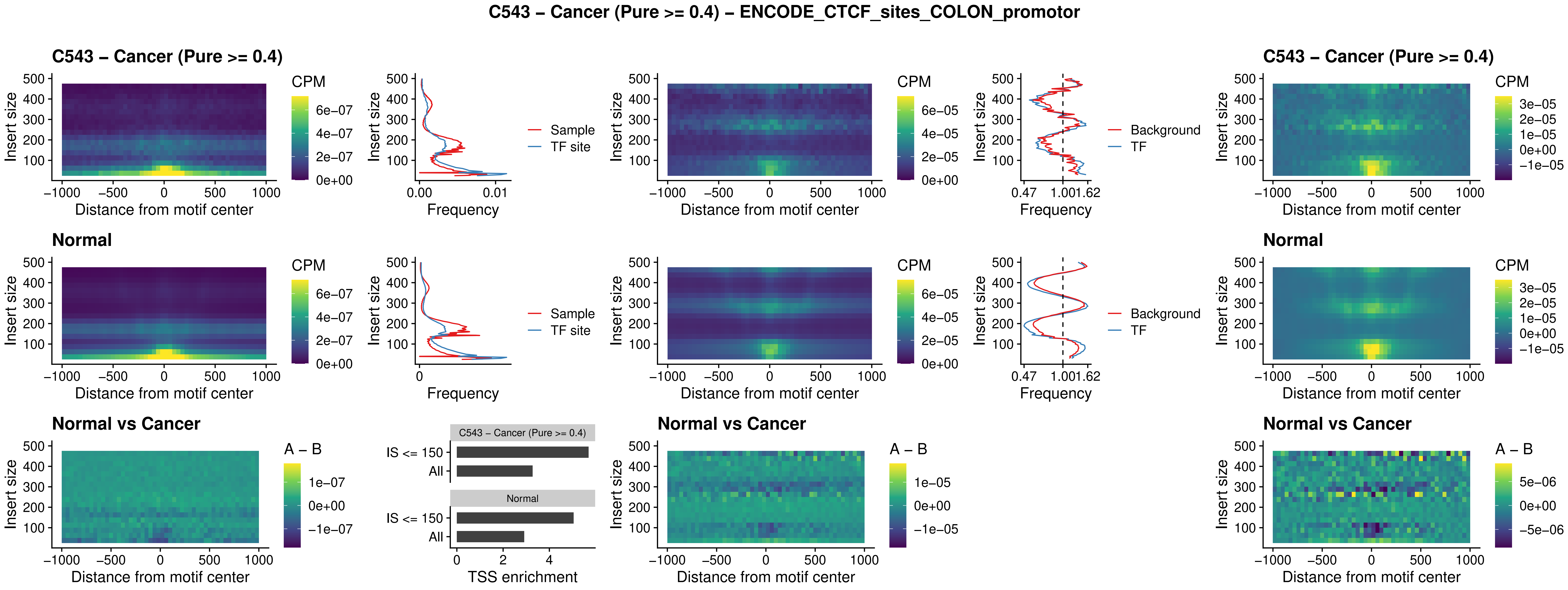
C543 – Cancer (Pure >= 0.4) – ENCODE_CTCF_sites_COLON_promotor
C543 – Cancer (Pure >= 0.4)
Normal
Normal vs Cancer
Insert size
Distance from motif center
Frequency
CPM
A – B
Sample
TF site
Background
TF
C543 – Cancer (Pure >= 0.4)
Normal
IS <= 150
All
TSS enrichment

#### C543 – Cancer (Pure >= 0.4) – ENCODE_CTCF_sites_COLON_promotor_ctcf

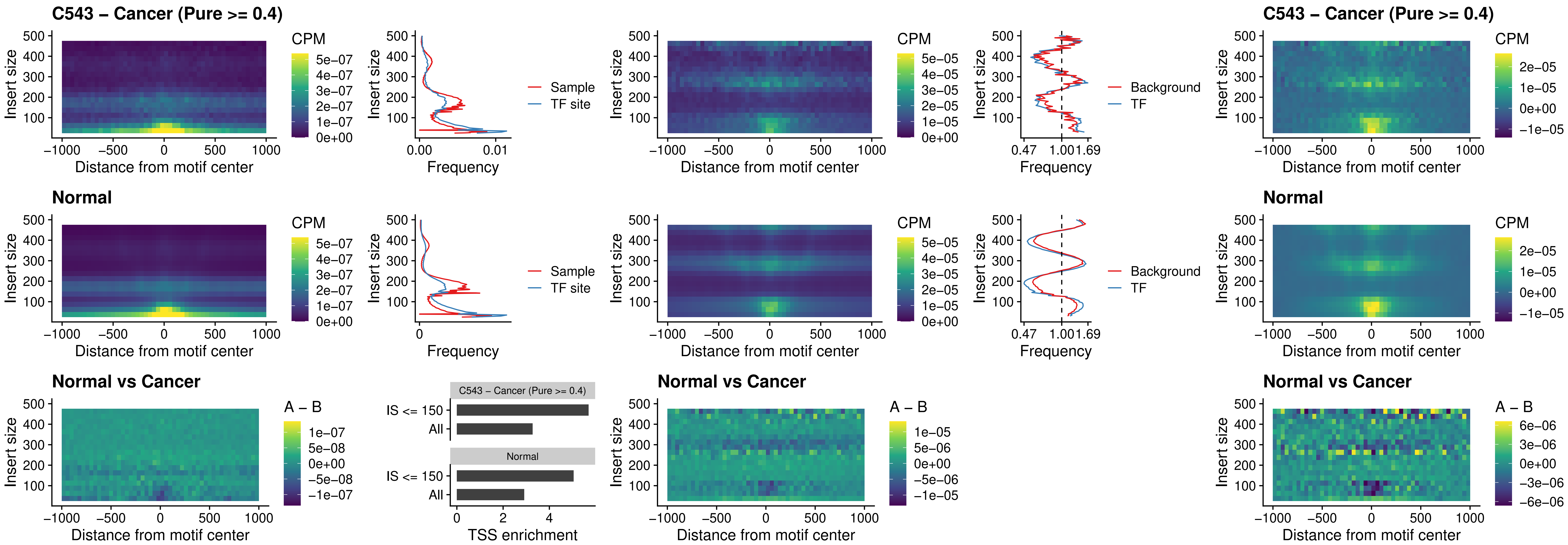

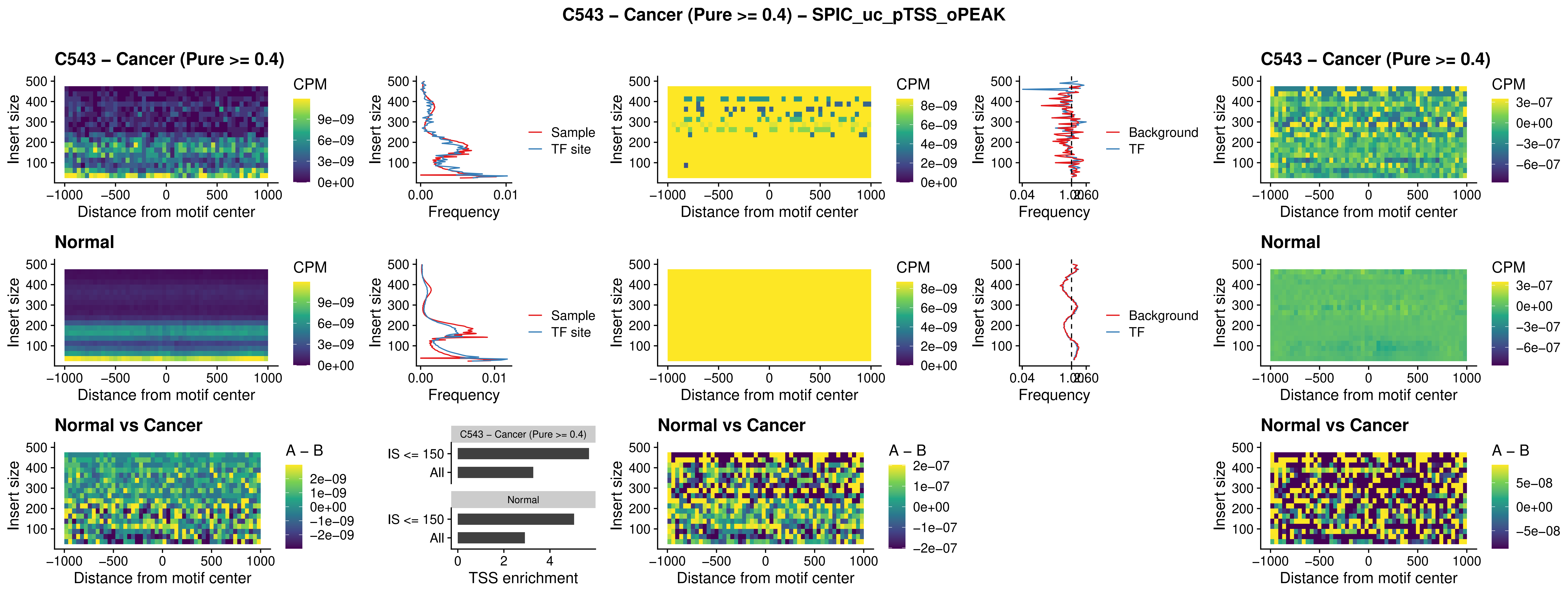
C543 – Cancer (Pure >= 0.4) – SPIC_uc_pTSS_oPEAK
C543 – Cancer (Pure >= 0.4)
Normal
Normal vs Cancer
Insert size
Distance from motif center
CPM
A – B
Frequency
Sample
TF site
Background
TF
C543 – Cancer (Pure >= 0.4)
Normal
IS <= 150
All
TSS enrichment

### C543 – Cancer (Pure >= 0.4) – SPI1_uc_dTSS_nPEAK

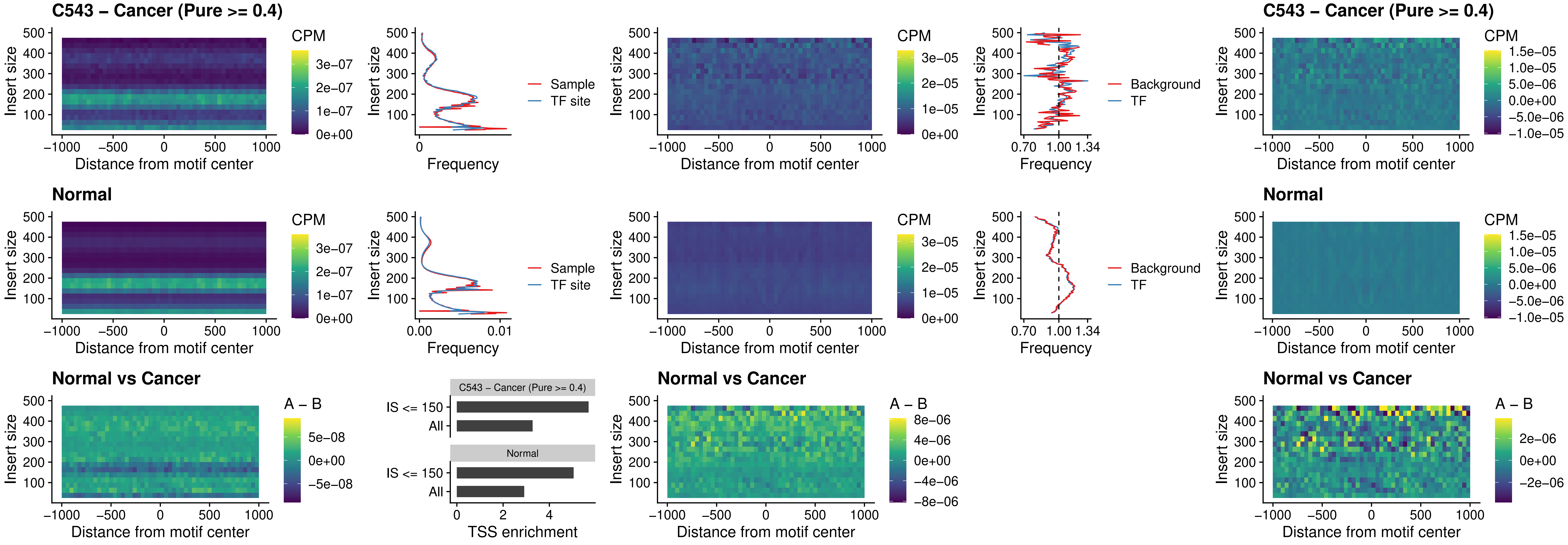

#### C543 – Cancer (Pure >= 0.4) – SPI1_uc_cTSS_nPEAK

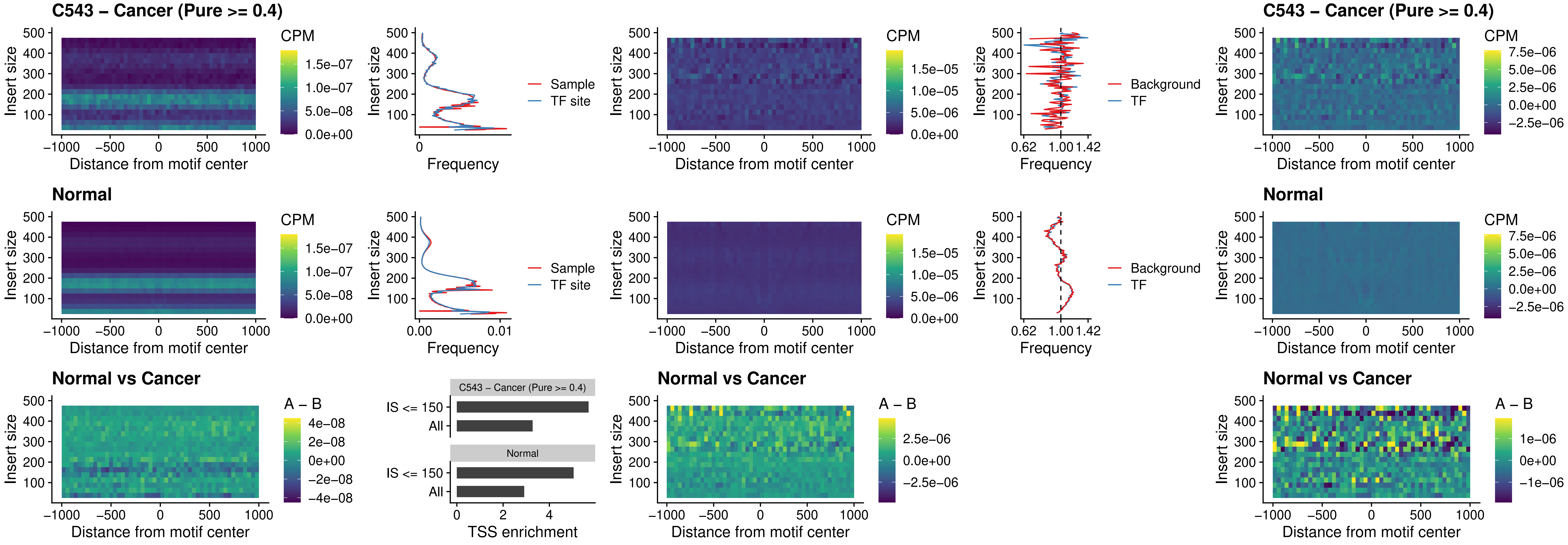

#### C543 – Cancer (Pure >= 0.4) – BCL11A_uc_pTSS_nPEAK

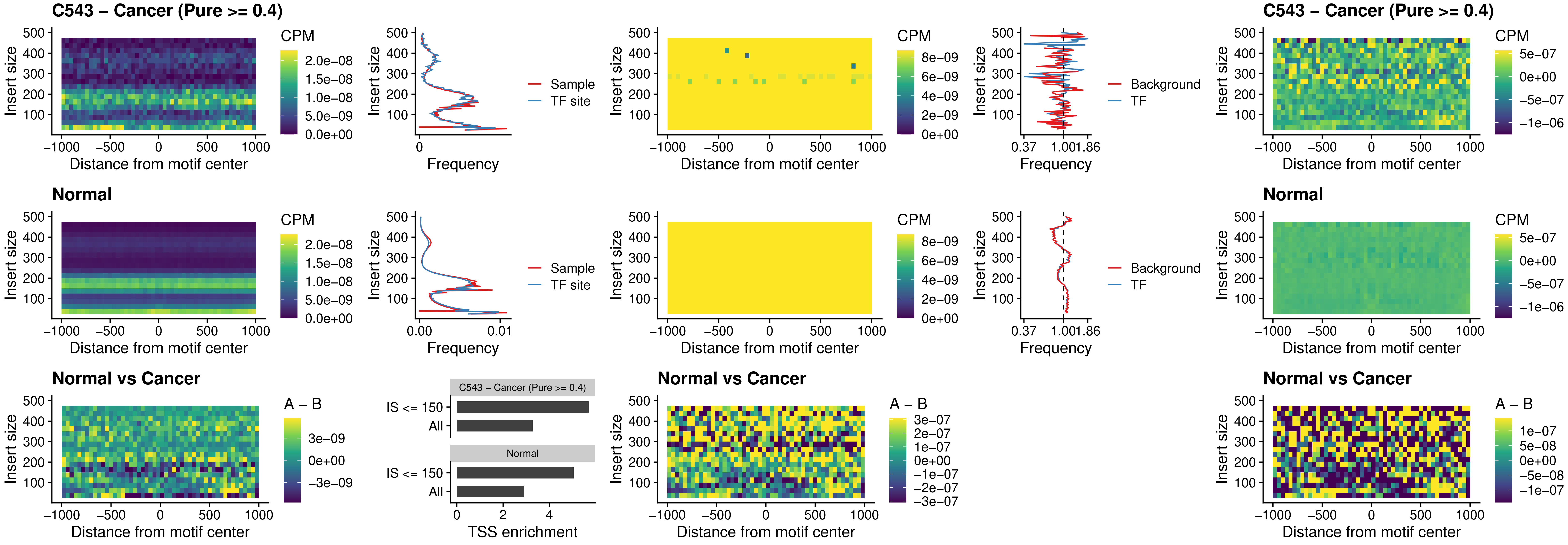

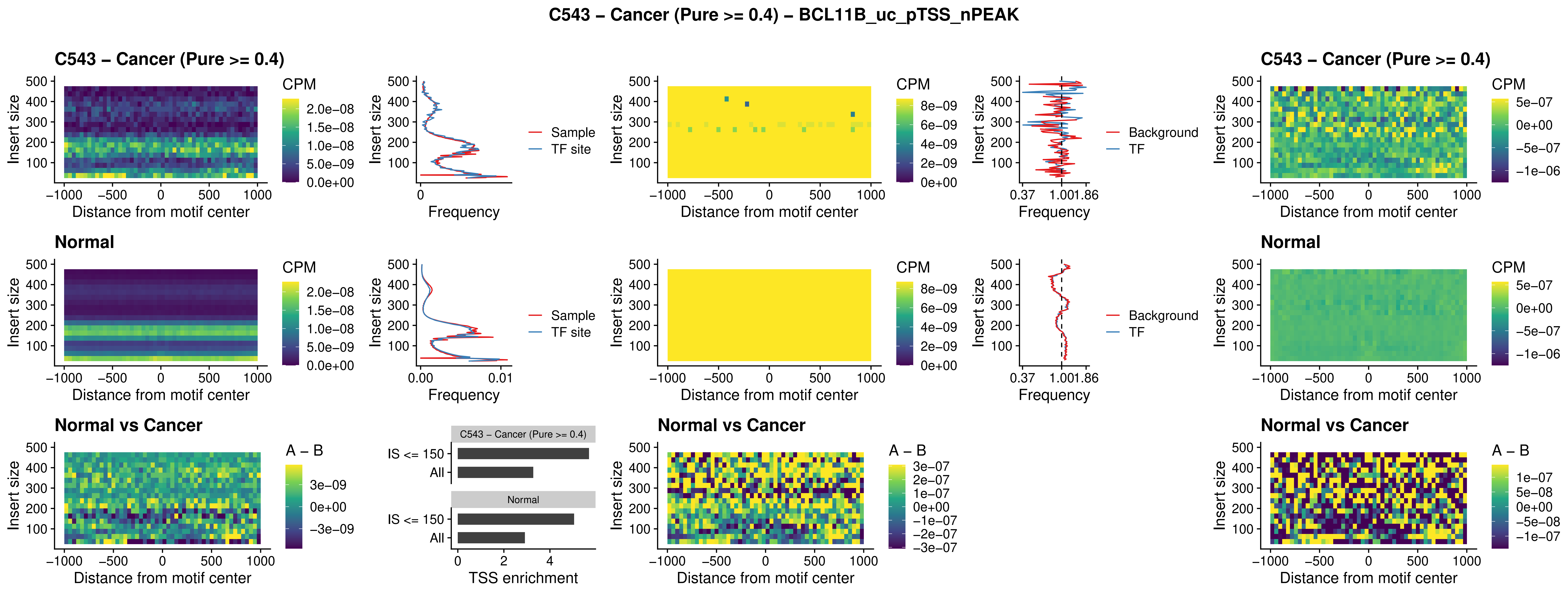
C543 – Cancer (Pure >= 0.4) – BCL11B_uc_pTSS_nPEAK
C543 – Cancer (Pure >= 0.4)
Normal
Normal vs Cancer
Insert size
Distance from motif center
CPM
A – B
Frequency
Sample
TF site
Background
TF
TSS enrichment
C543 – Cancer (Pure >= 0.4)
Normal
IS <= 150
All

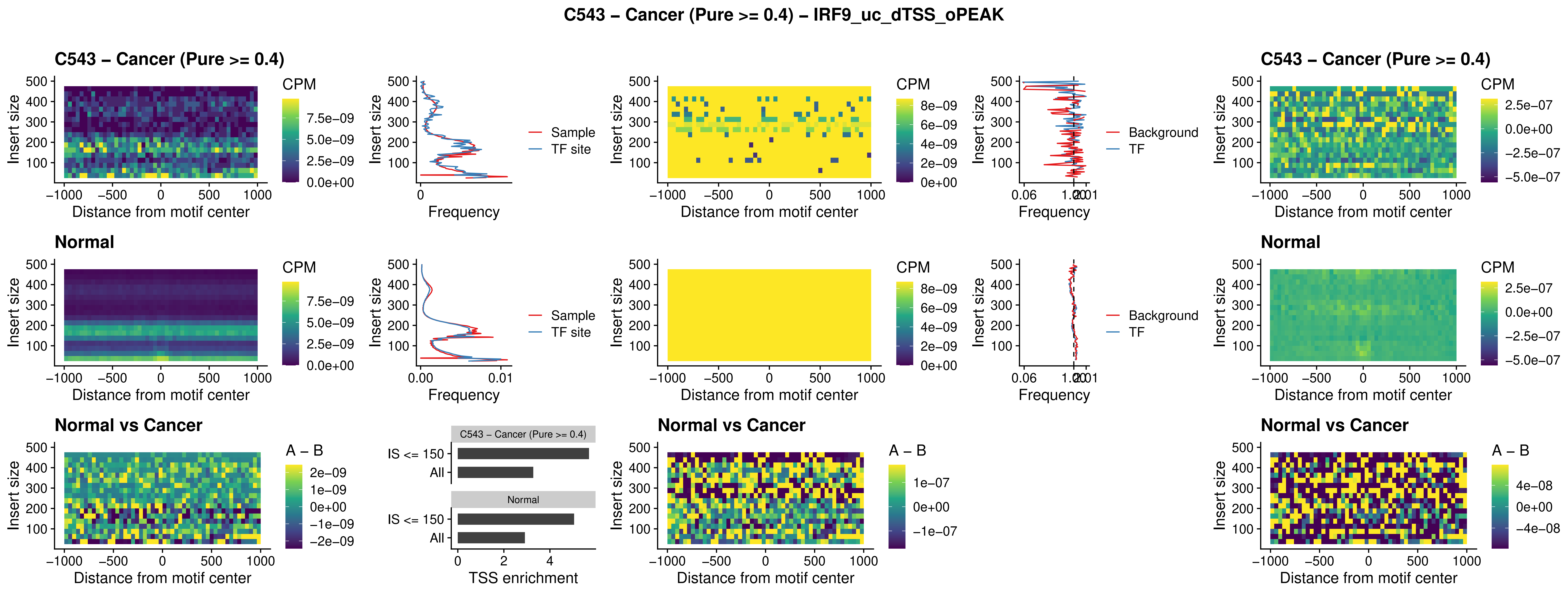

C543 – Cancer (Pure >= 0.4) – IRF9_uc_dTSS_oPEAK
C543 – Cancer (Pure >= 0.4)
Normal
Normal vs Cancer
Insert size
Distance from motif center
CPM
A – B
Frequency
Sample
TF site
Background
TF
TSS enrichment
IS <= 150
All

#### C543 – Cancer (Pure >= 0.4) – IRF9_uc_cTSS_oPEAK

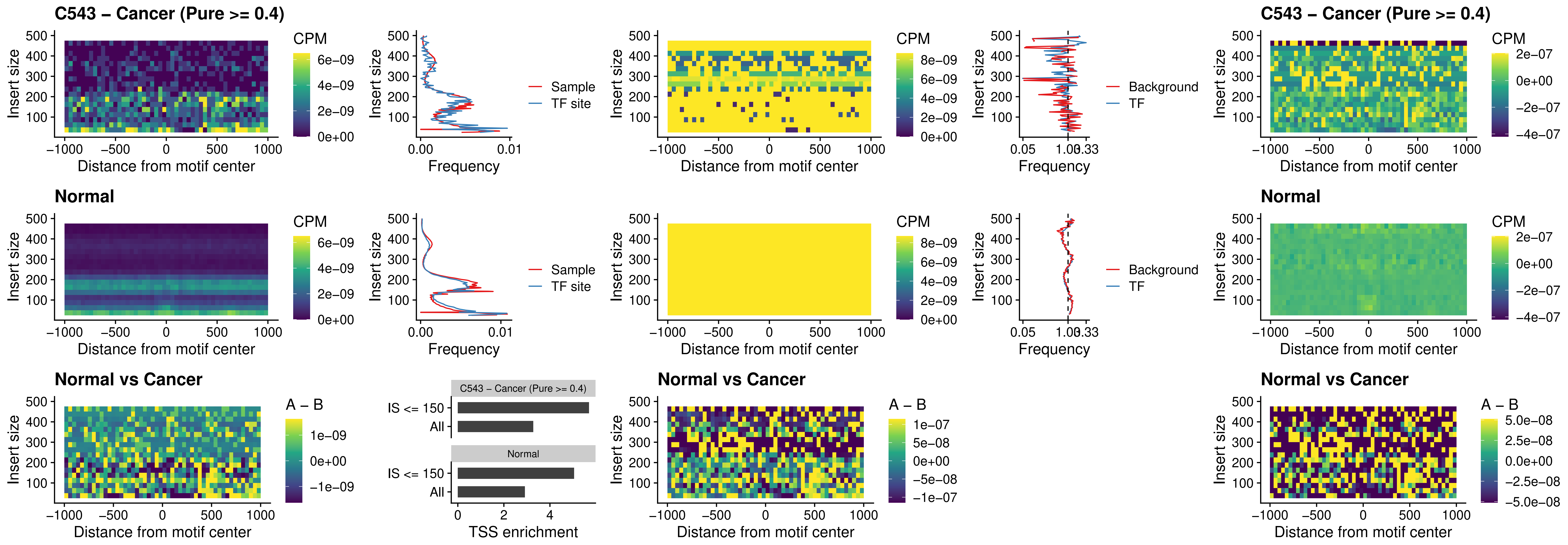

#### C543 – Cancer (Pure >= 0.4) – BCL11A_uc_cTSS_oPEAK

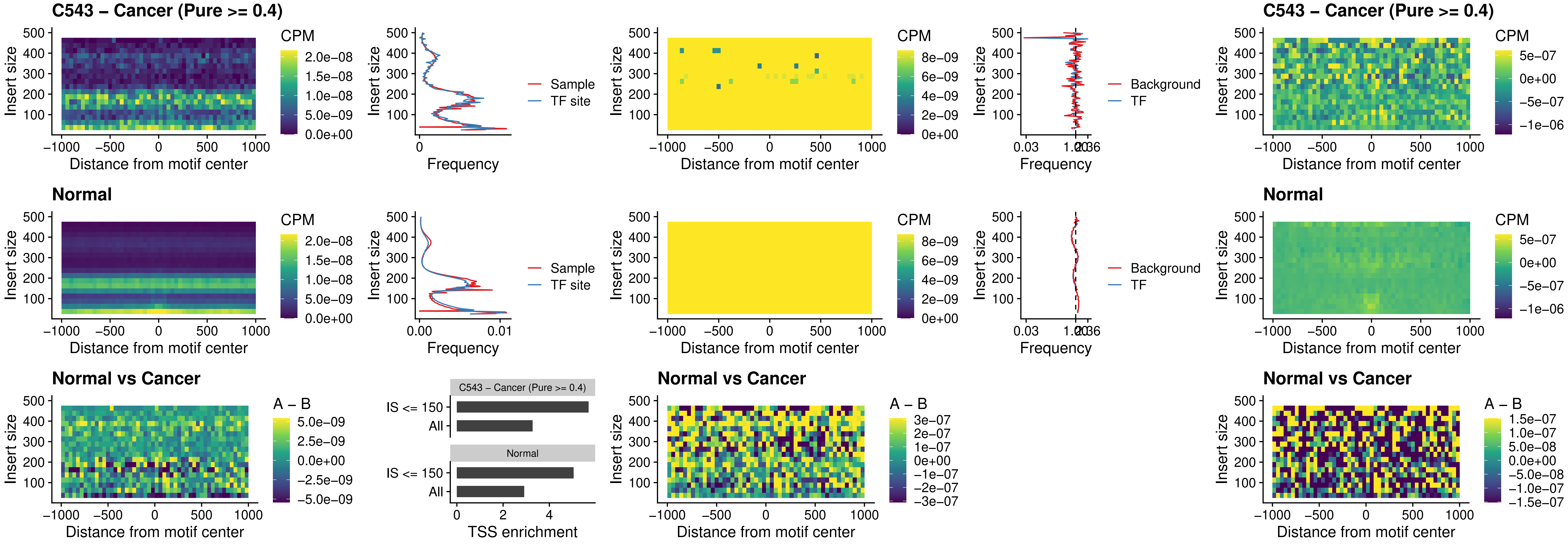

#### C543 – Cancer (Pure >= 0.4) – BCL11B_uc_cTSS_oPEAK

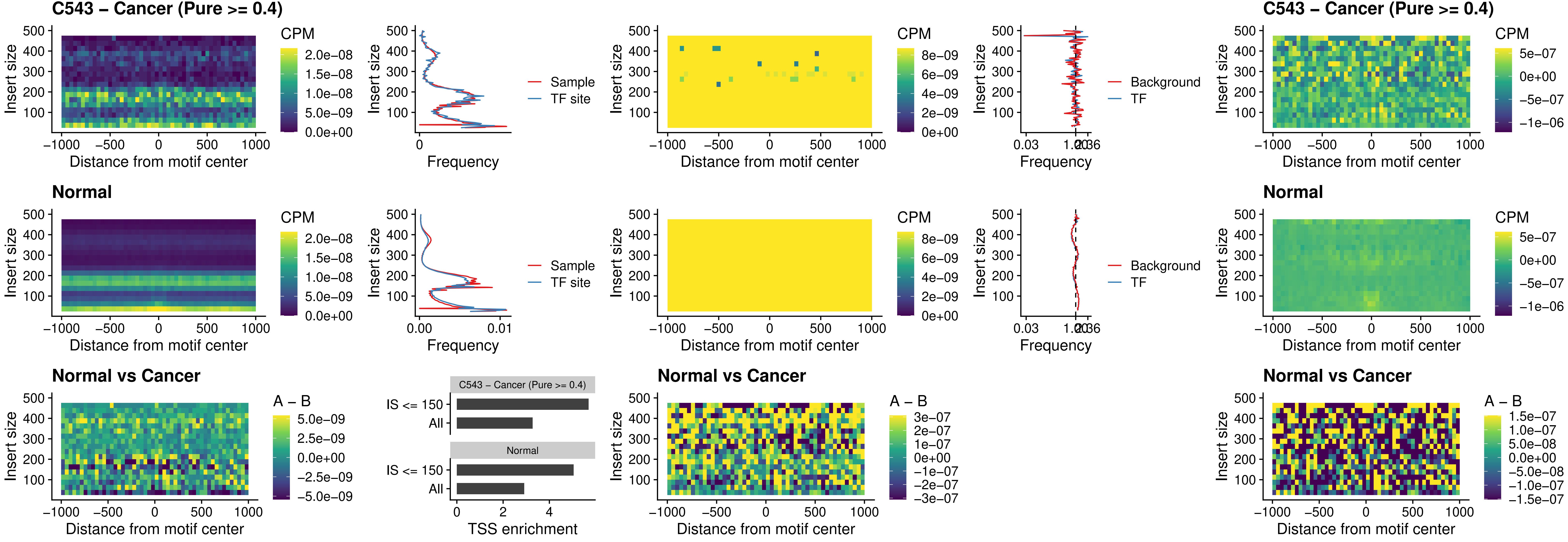

#### C543 – Cancer (Pure >= 0.4) – BCL11A_uc_dTSS_oPEAK

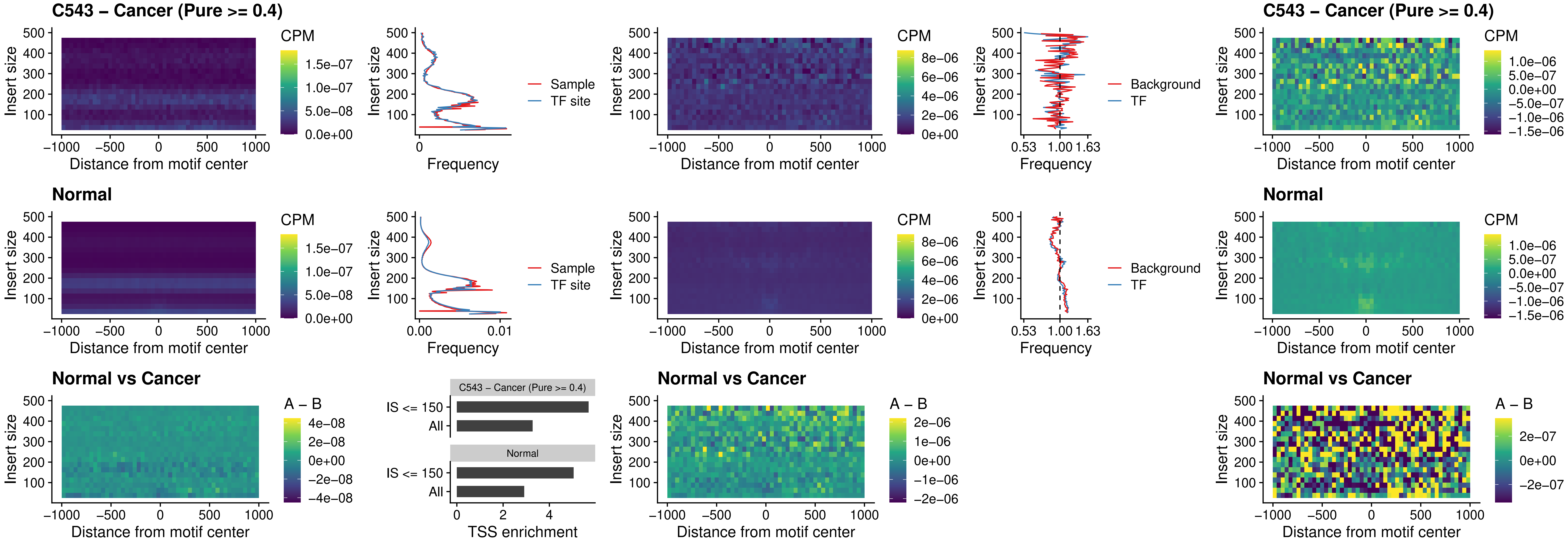

#### C543 – Cancer (Pure >= 0.4) – BCL11B_uc_dTSS_oPEAK

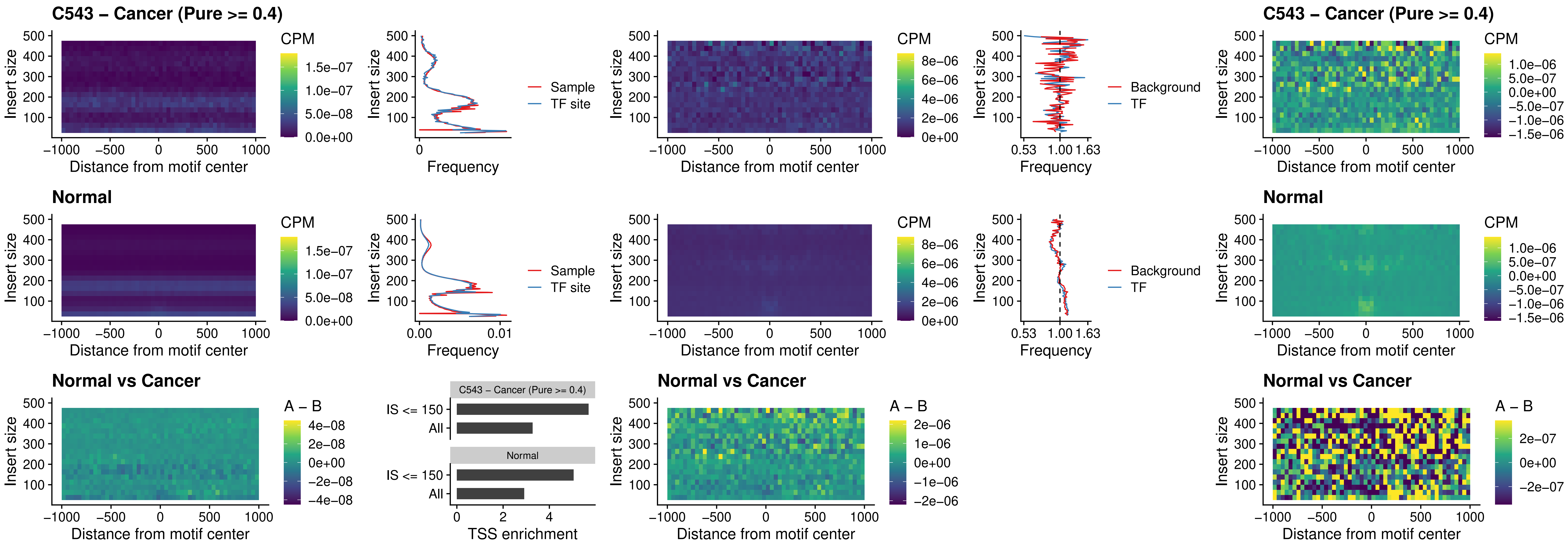

#### C543 – Cancer (Pure >= 0.4) – SPIB_uc_cTSS_nPEAK

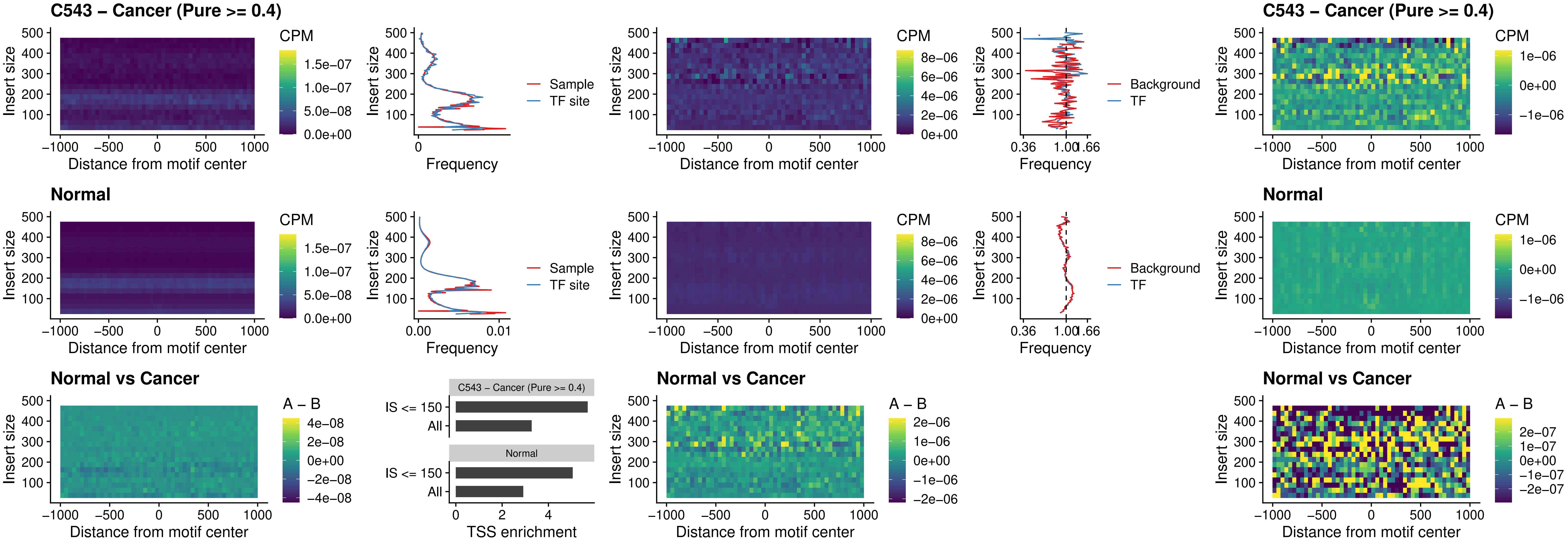

#### C543 – Cancer (Pure >= 0.4) – SPIC_uc_cTSS_nPEAK

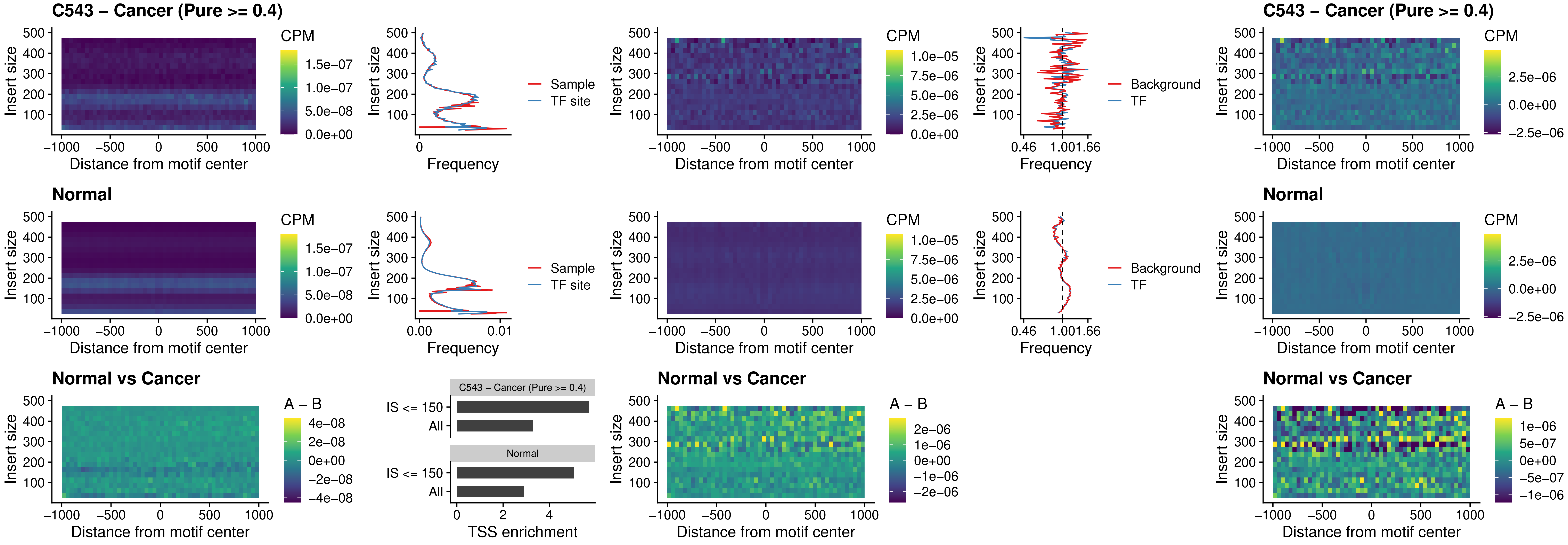

#### C543 – Cancer (Pure >= 0.4) – SPI1_uc_pTSS_oPEAK

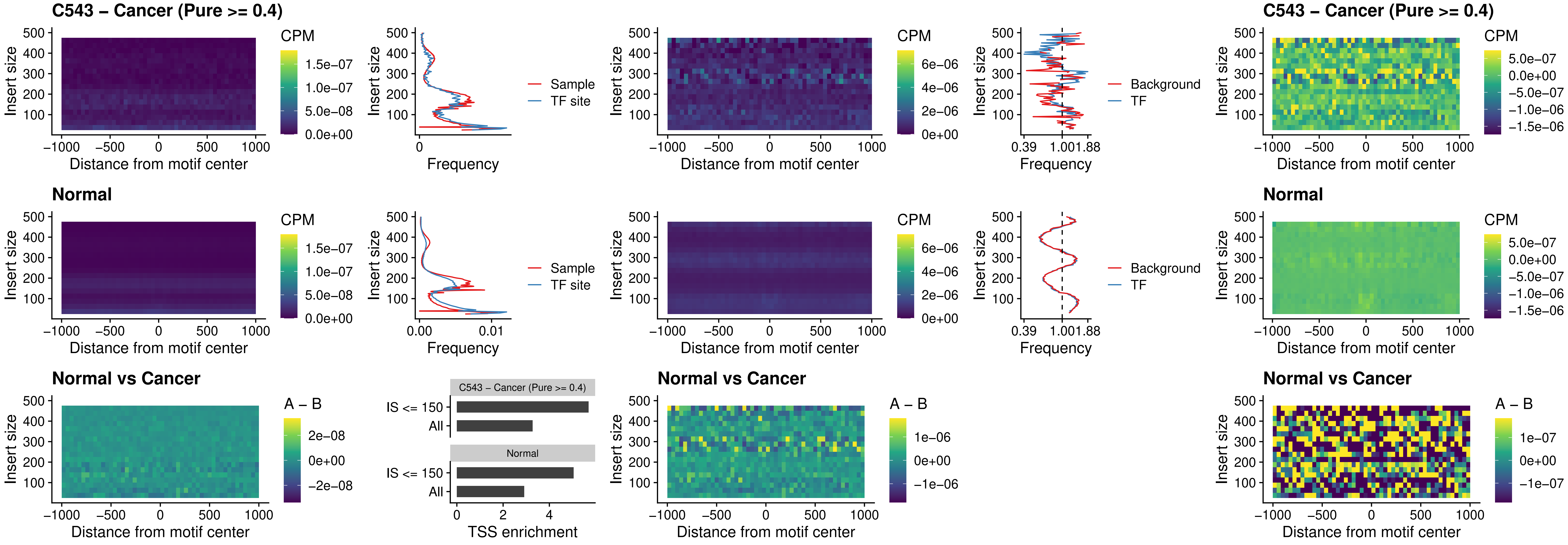

#### C543 – Cancer (Pure >= 0.4) – BCL11A_uc_pTSS_oPEAK

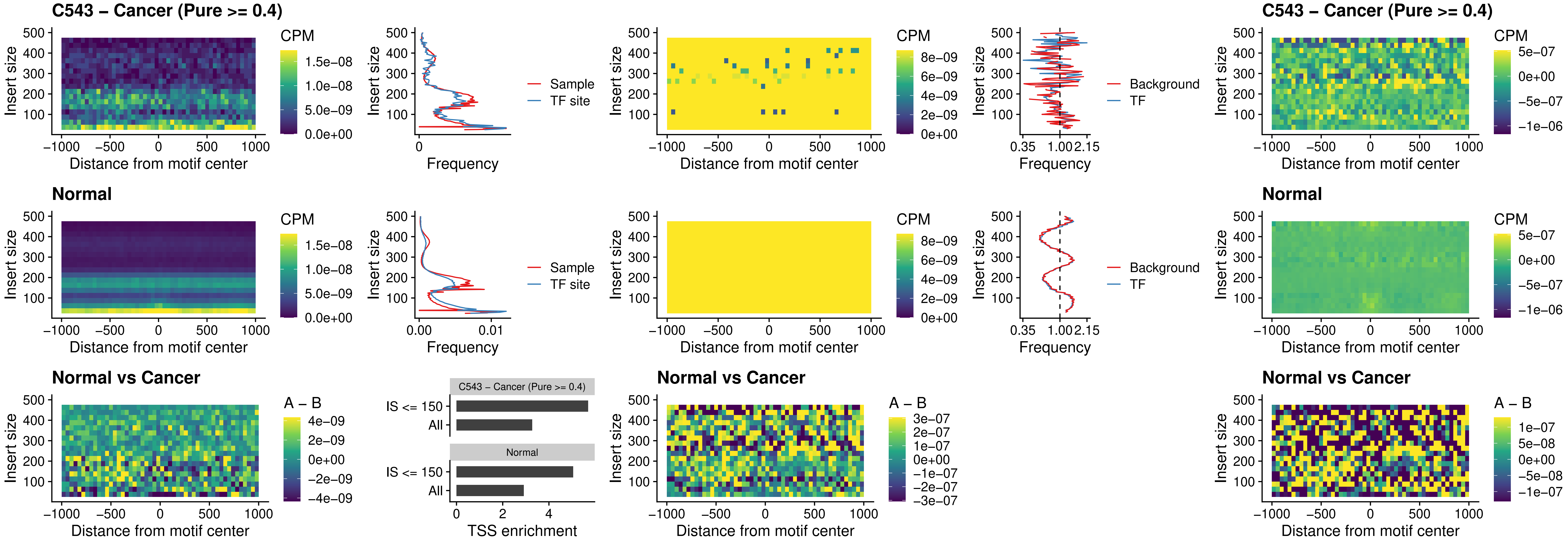

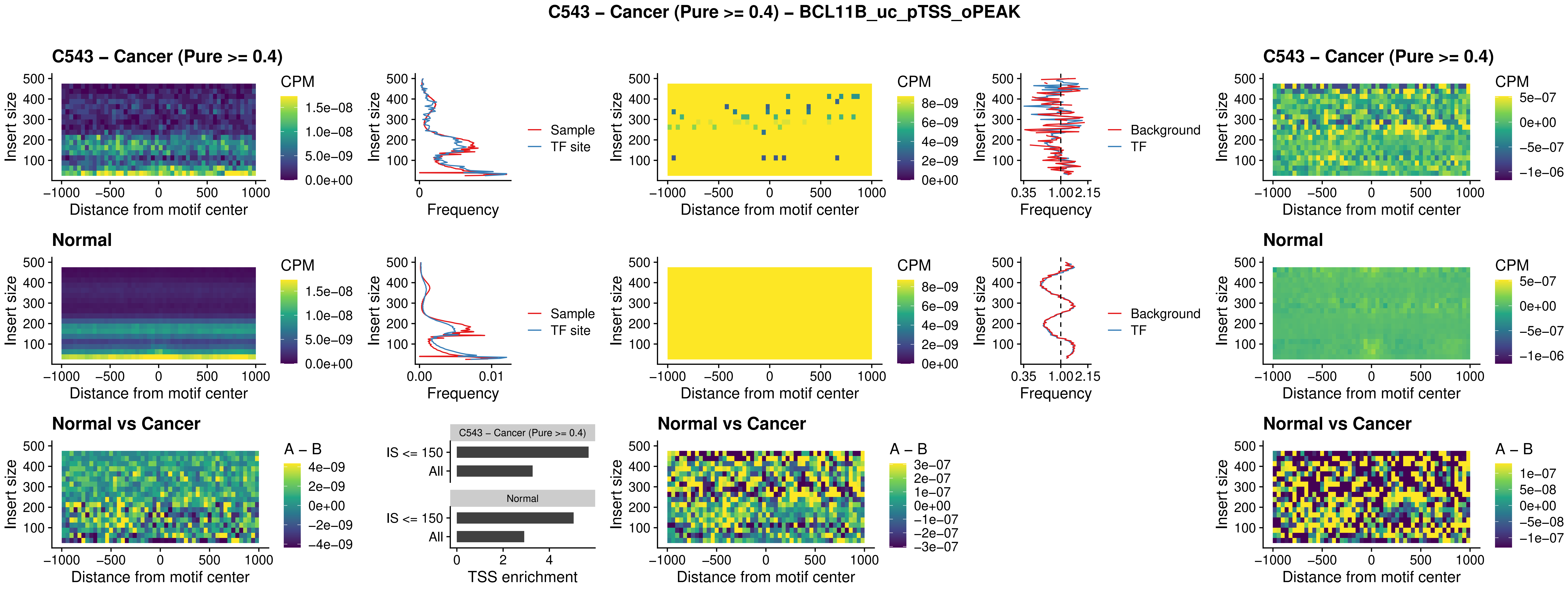
C543 – Cancer (Pure >= 0.4) – BCL11B_uc_pTSS_oPEAK
C543 – Cancer (Pure >= 0.4)
Normal
Normal vs Cancer
Insert size
Distance from motif center
Frequency
CPM
A – B
Sample
TF site
Background
TF
C543 – Cancer (Pure >= 0.4)
Normal
IS <= 150
All
TSS enrichment

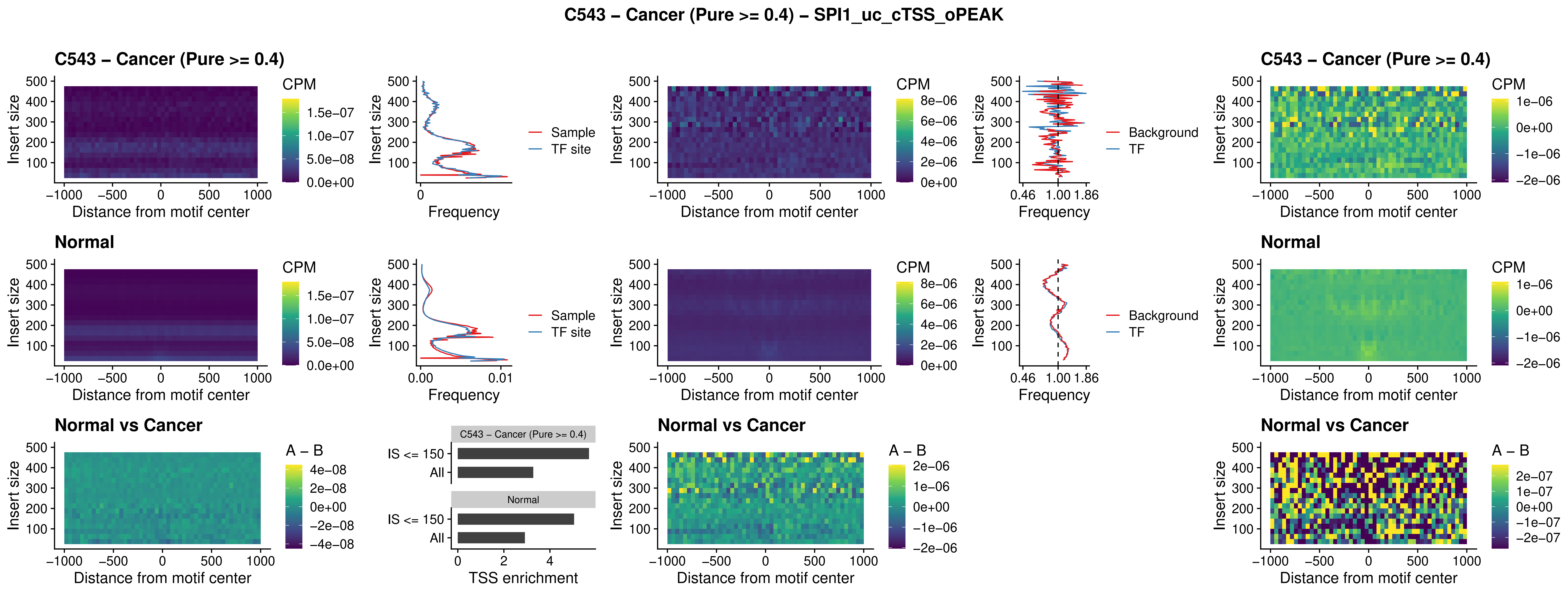
C543 – Cancer (Pure >= 0.4) – SPI1_uc_cTSS_oPEAK
C543 – Cancer (Pure >= 0.4)
Normal
Normal vs Cancer
Insert size
Distance from motif center
Frequency
CPM
A – B
Sample
TF site
Background
TF
C543 – Cancer (Pure >= 0.4)
Normal
IS <= 150
All
TSS enrichment

#### C543 – Cancer (Pure >= 0.4) – SPIB_uc_cTSS_oPEAK

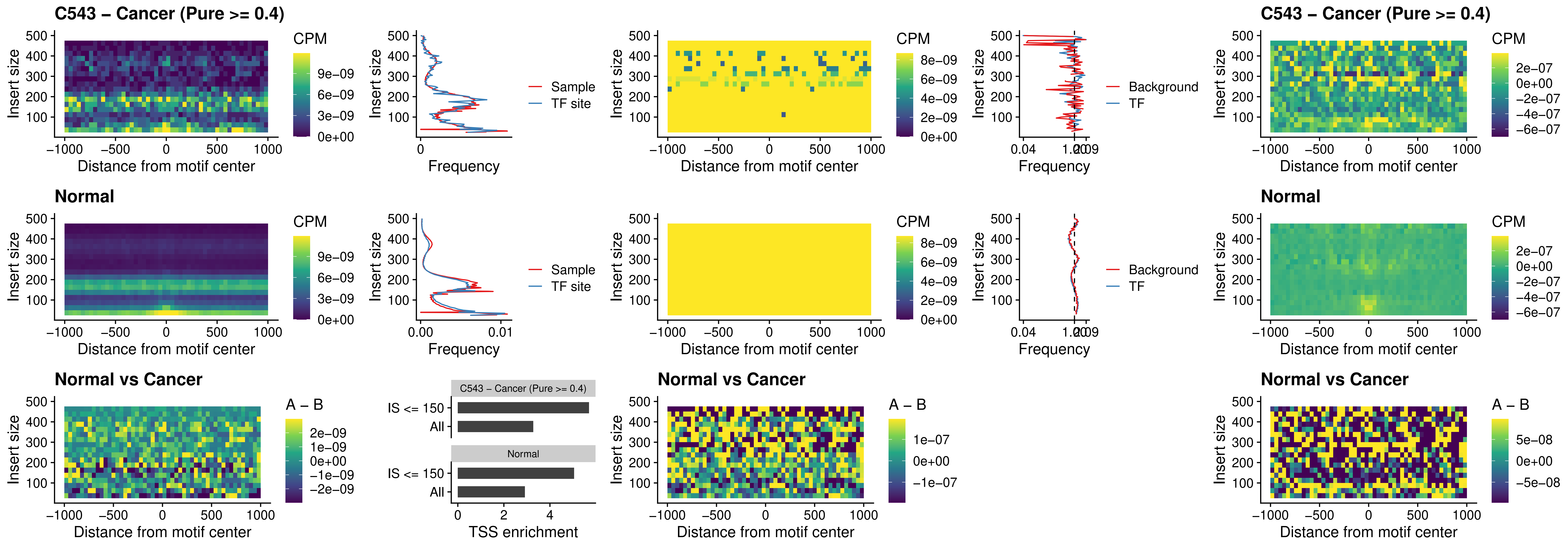

#### C543 – Cancer (Pure >= 0.4) – SPIB_uc_pTSS_oPEAK

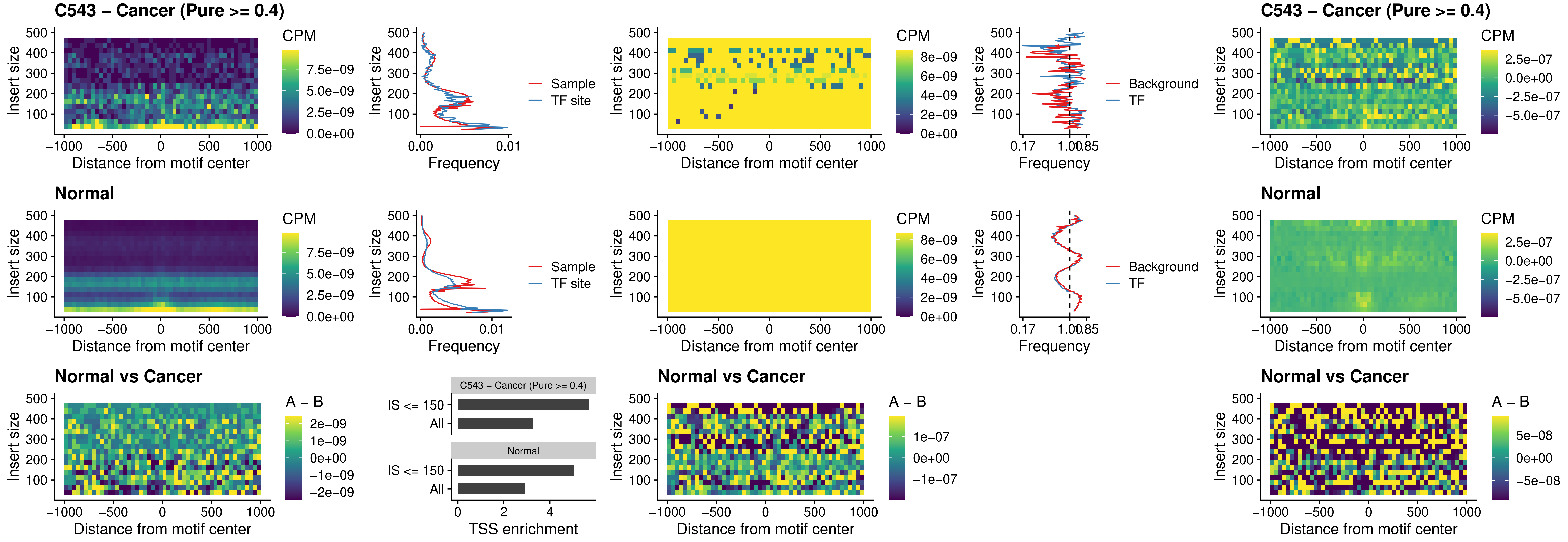

#### C543 – Cancer (Pure >= 0.4) – SPI1_uc_dTSS_oPEAK

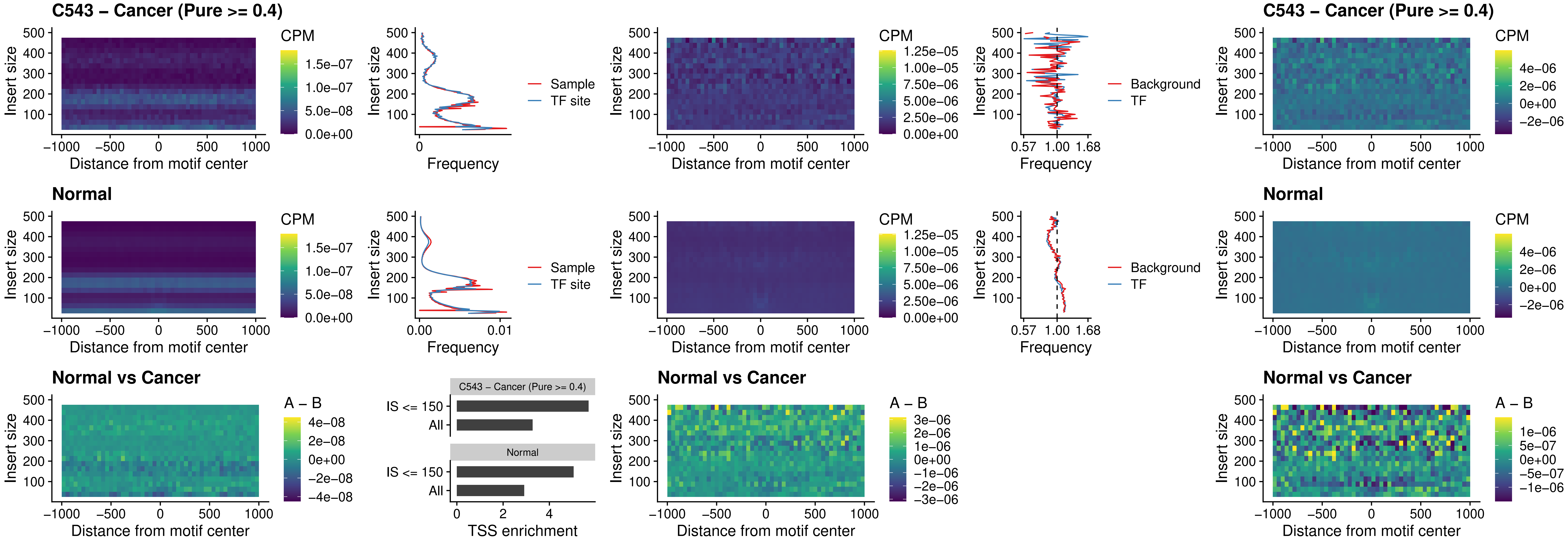

#### C543 – Cancer (Pure >= 0.4) – SPIB_uc_dTSS_oPEAK

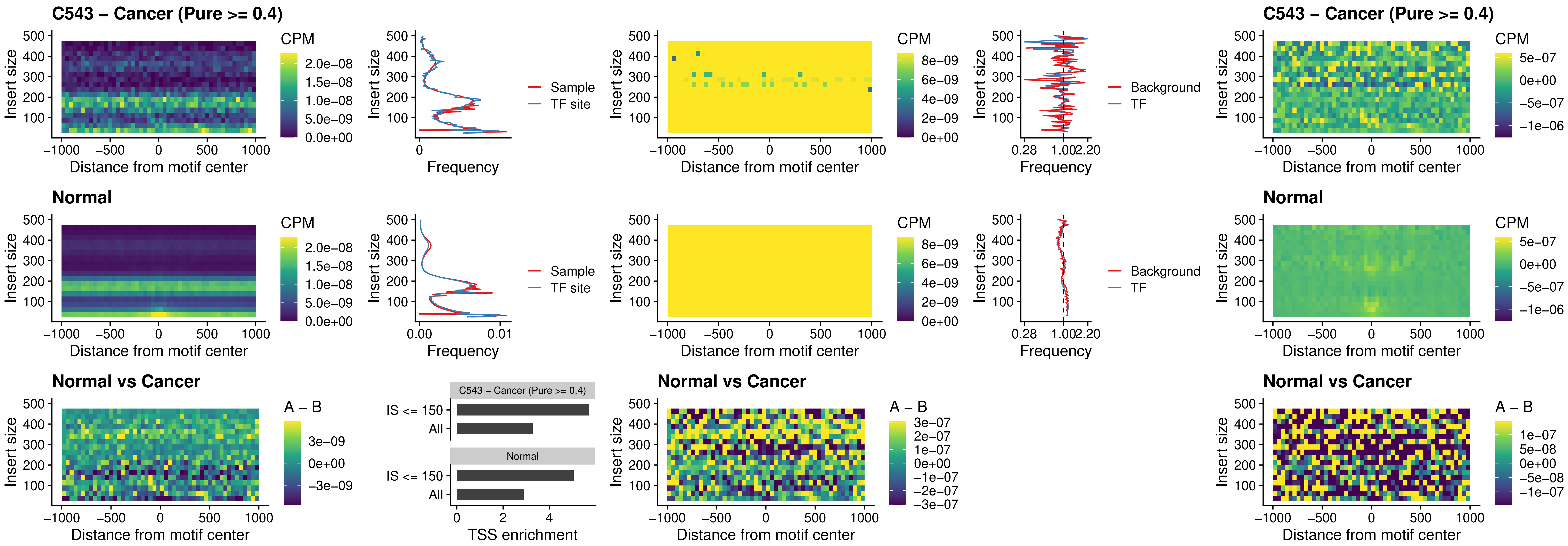

### C543 – Cancer (Pure >= 0.4) – IRF3_uc_pTSS_oPEAK

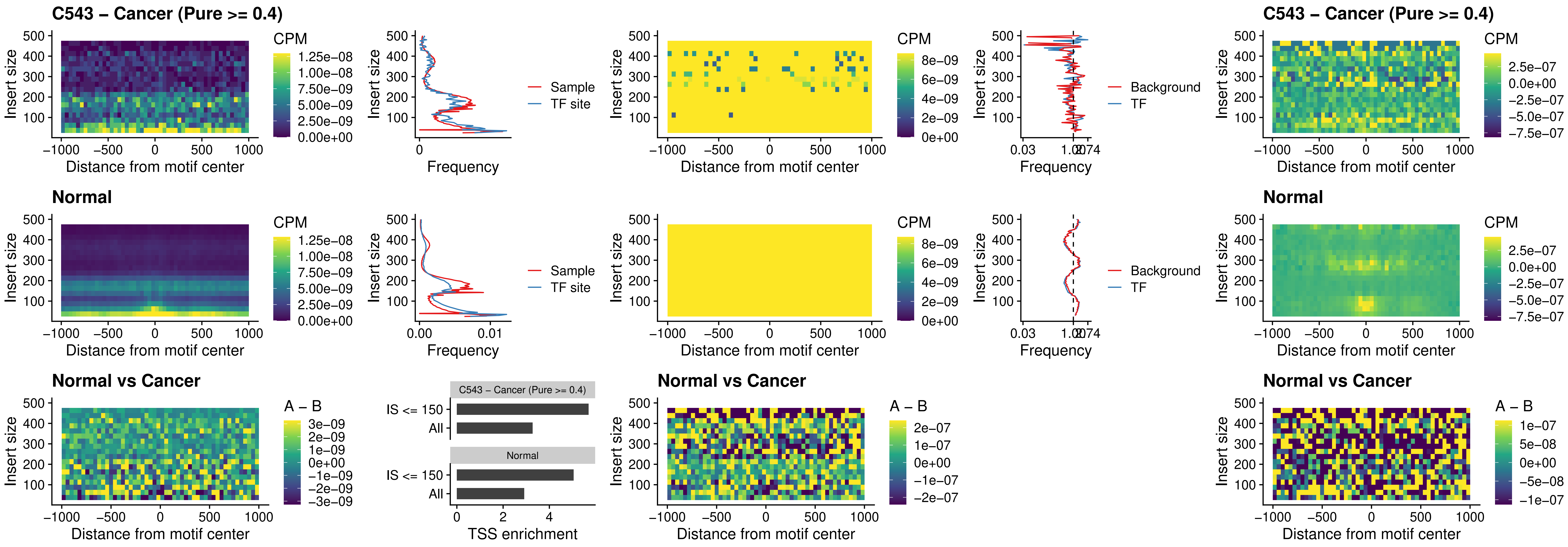

#### C543 – Cancer (Pure >= 0.4) – IRF2_uc_dTSS_oPEAK

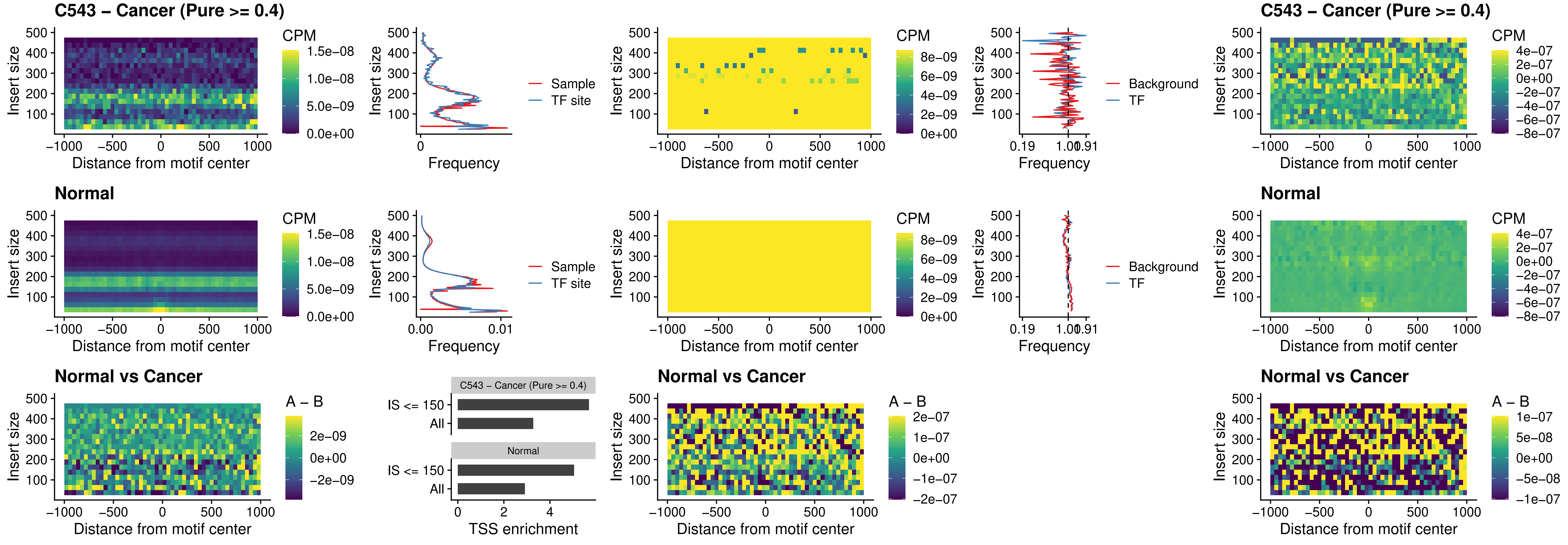

#### C543 – Cancer (Pure >= 0.4) – IRF2_uc_pTSS_oPEAK

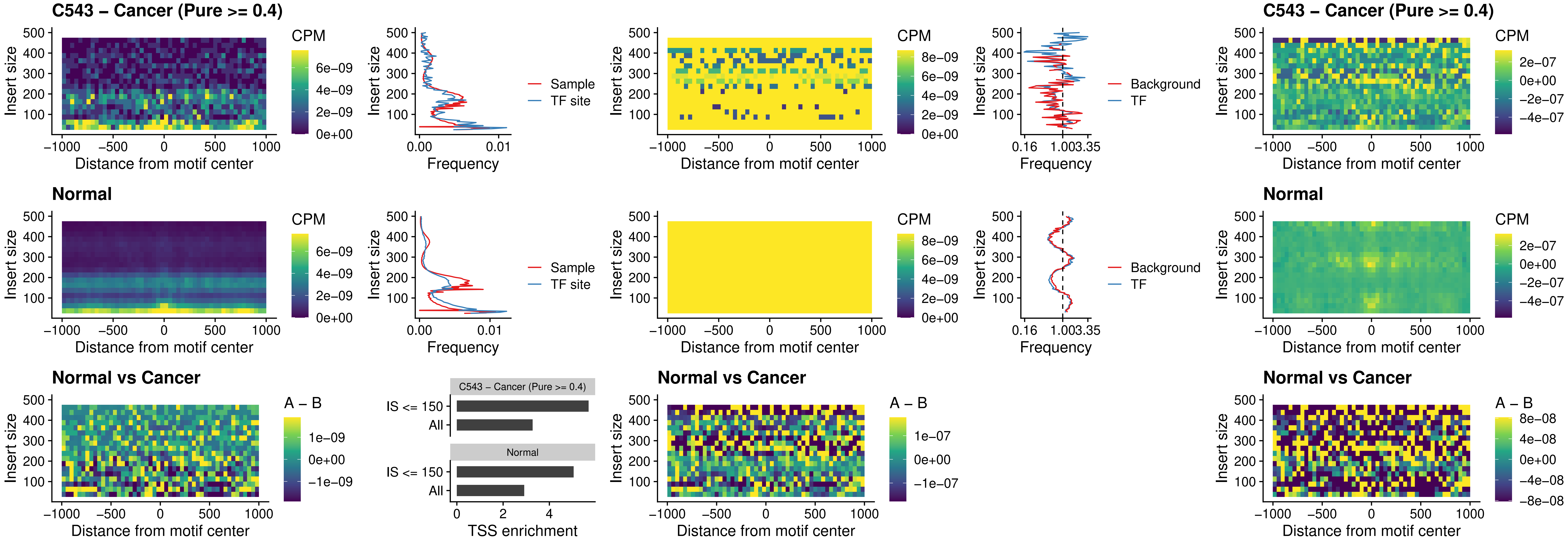

#### C543 – Cancer (Pure >= 0.4) – STAT2_uc_pTSS_oPEAK

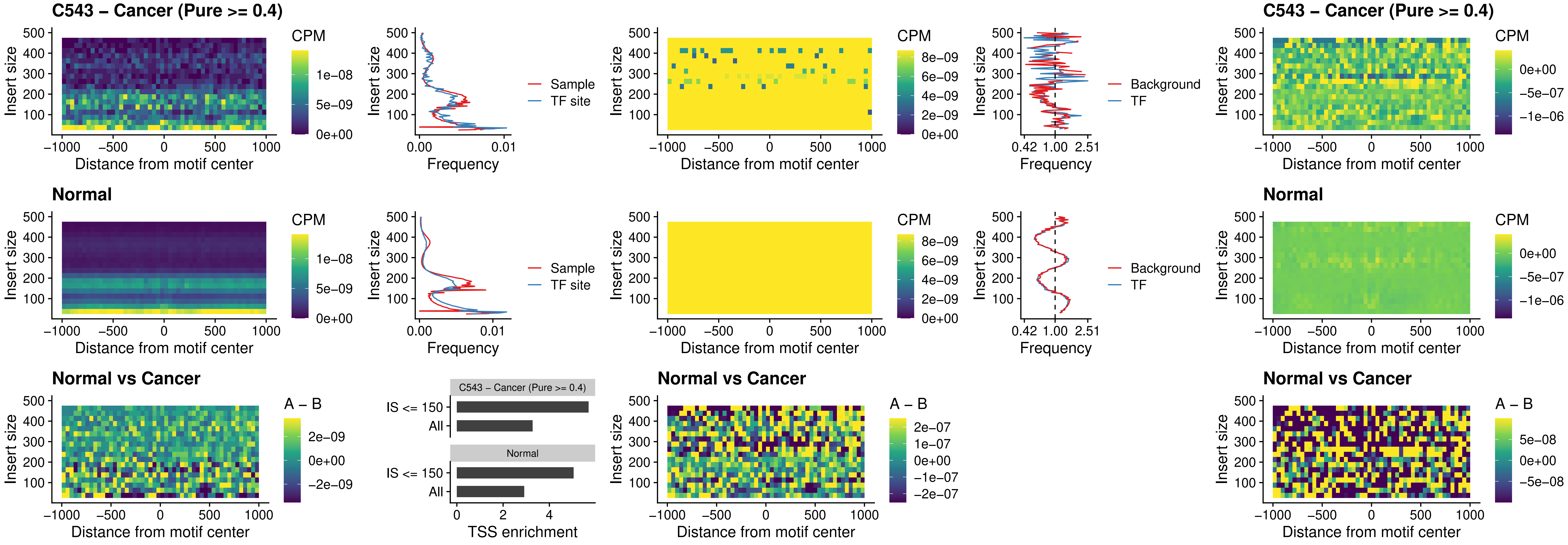

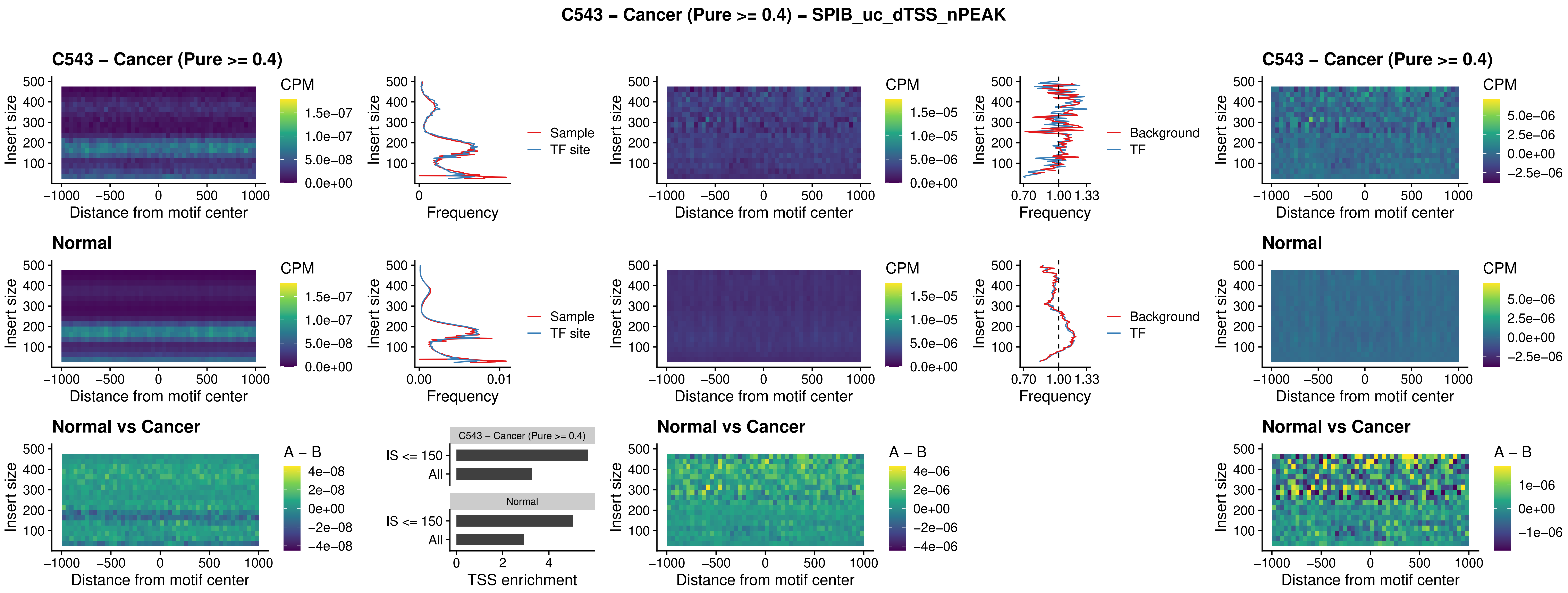
C543 – Cancer (Pure >= 0.4) – SPIB_uc_dTSS_nPEAK
C543 – Cancer (Pure >= 0.4)
Normal
Normal vs Cancer
Insert size
Distance from motif center
Frequency
CPM
A – B
Sample
TF site
Background
TF
TSS enrichment
IS <= 150
All
C543 – Cancer (Pure >= 0.4)
Normal

#### C543 – Cancer (Pure >= 0.4) – SPIC_uc_cTSS_oPEAK

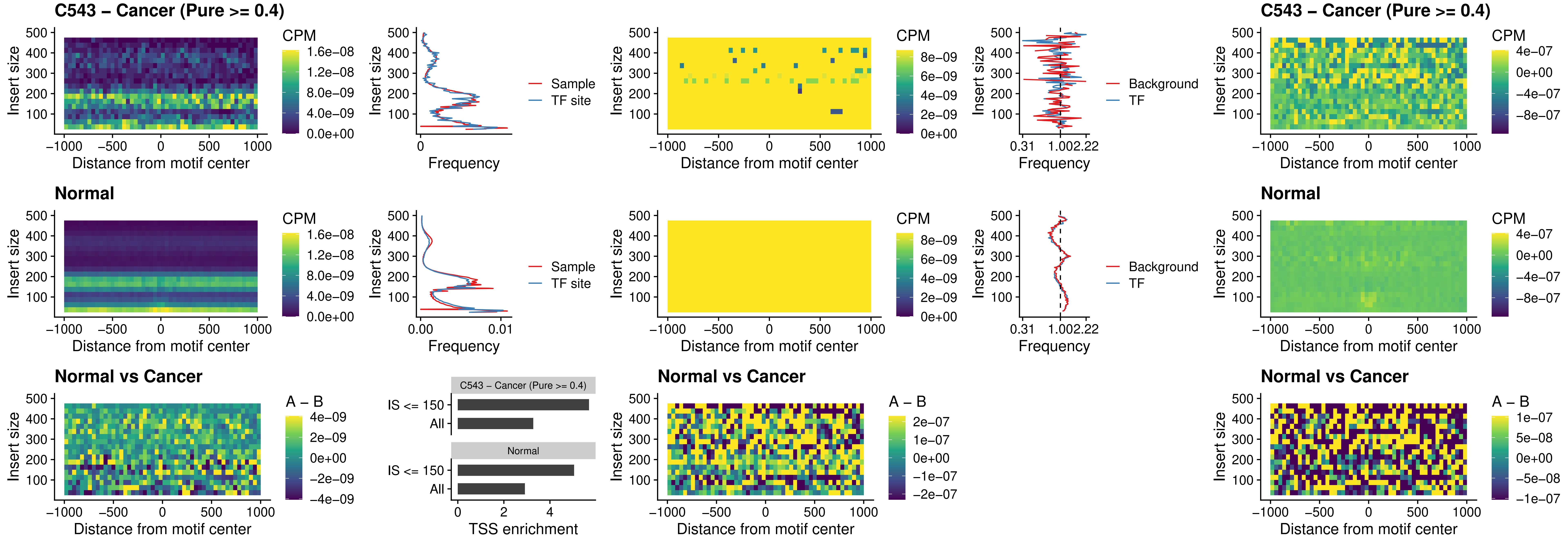

#### C543 – Cancer (Pure >= 0.4) – PRDM1_uc_cTSS_oPEAK

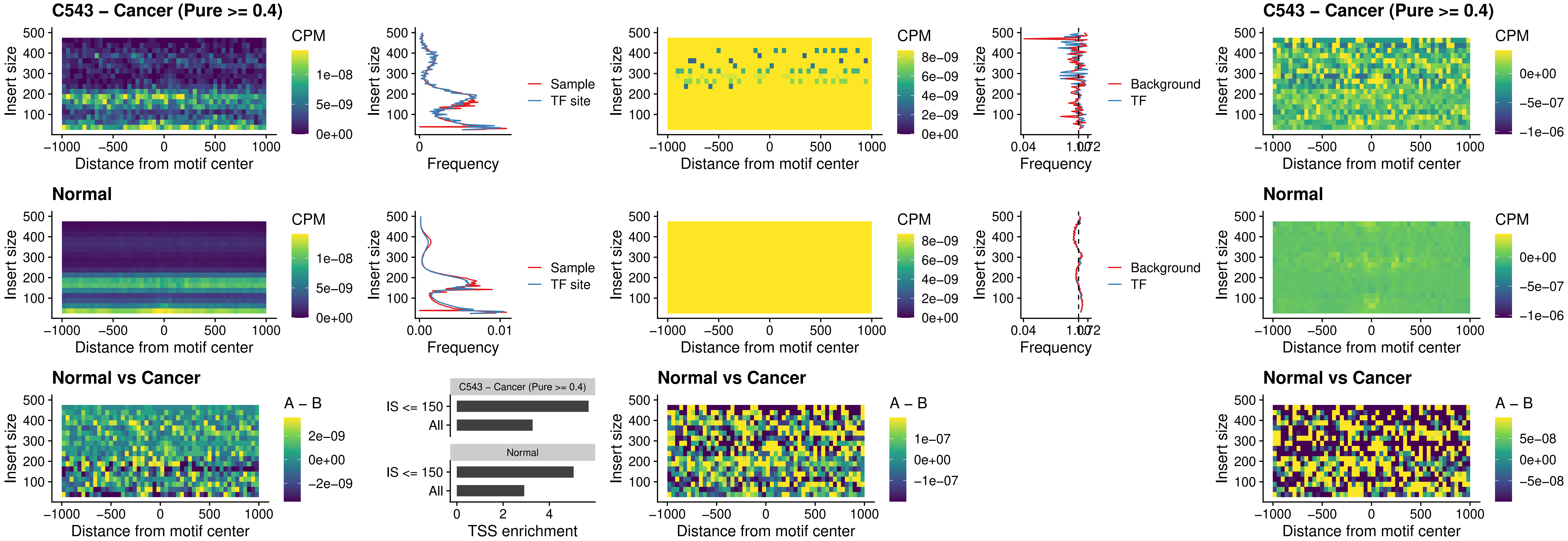

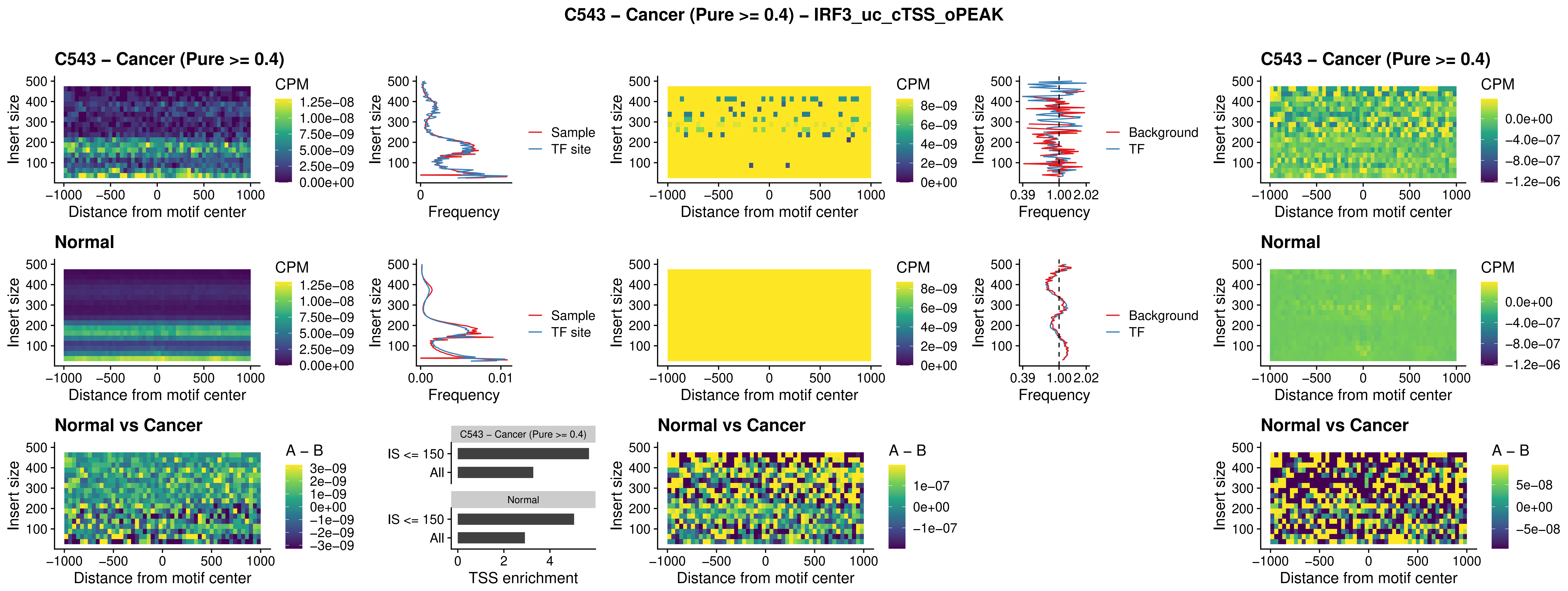
C543 – Cancer (Pure >= 0.4) – IRF3_uc_cTSS_oPEAK
C543 – Cancer (Pure >= 0.4)
Normal
Normal vs Cancer
Insert size
Distance from motif center
Frequency
CPM
A – B
Sample
TF site
Background
TF
TSS enrichment
IS <= 150
All
C543 – Cancer (Pure >= 0.4)
Normal

#### C543 – Cancer (Pure >= 0.4) – BCL11A_uc_dTSS_nPEAK

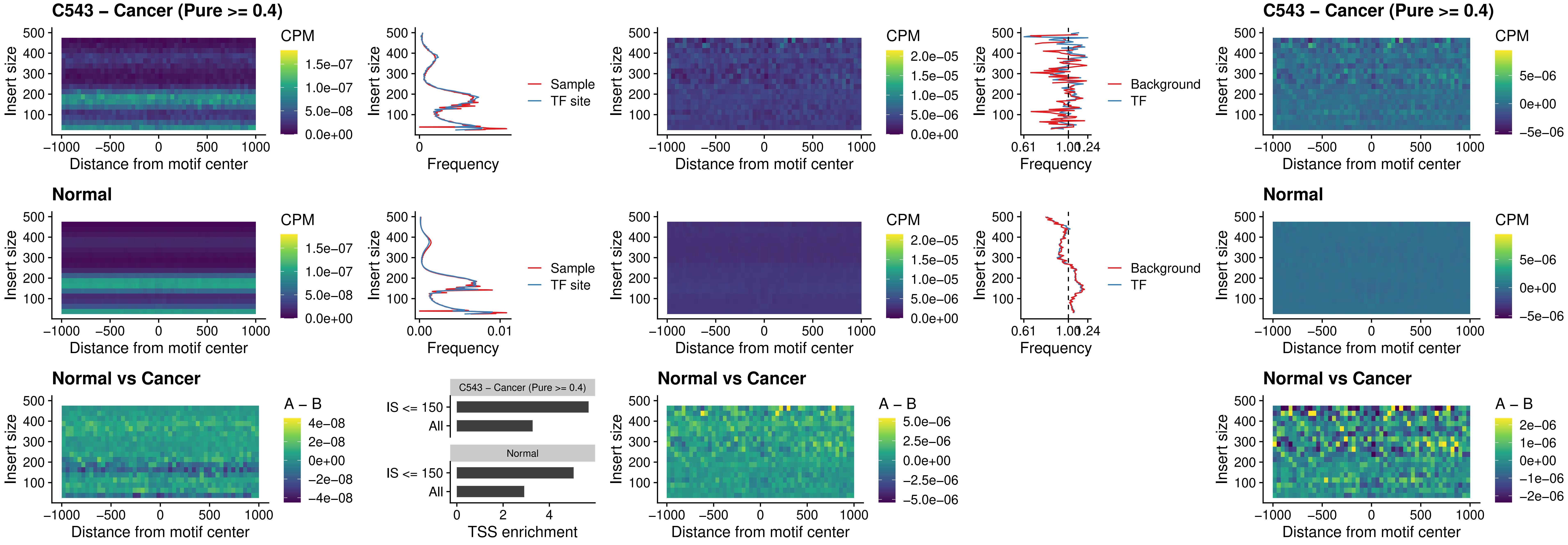

#### C543 – Cancer (Pure >= 0.4) – BCL11B_uc_dTSS_nPEAK

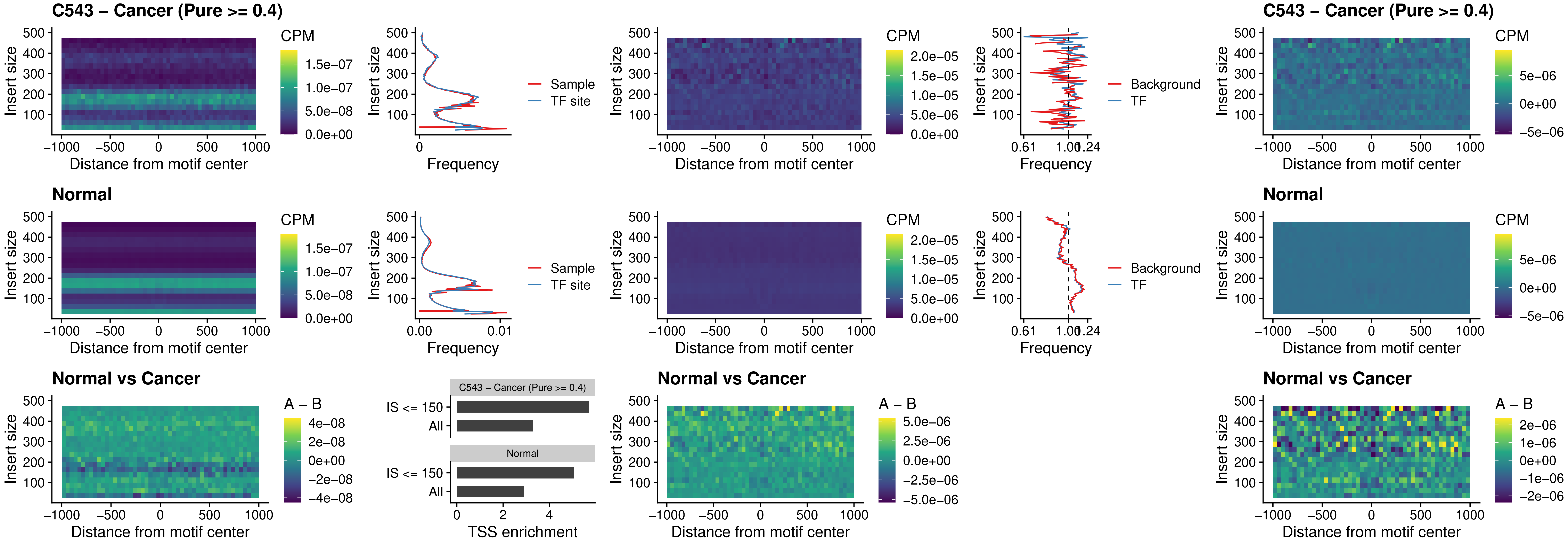

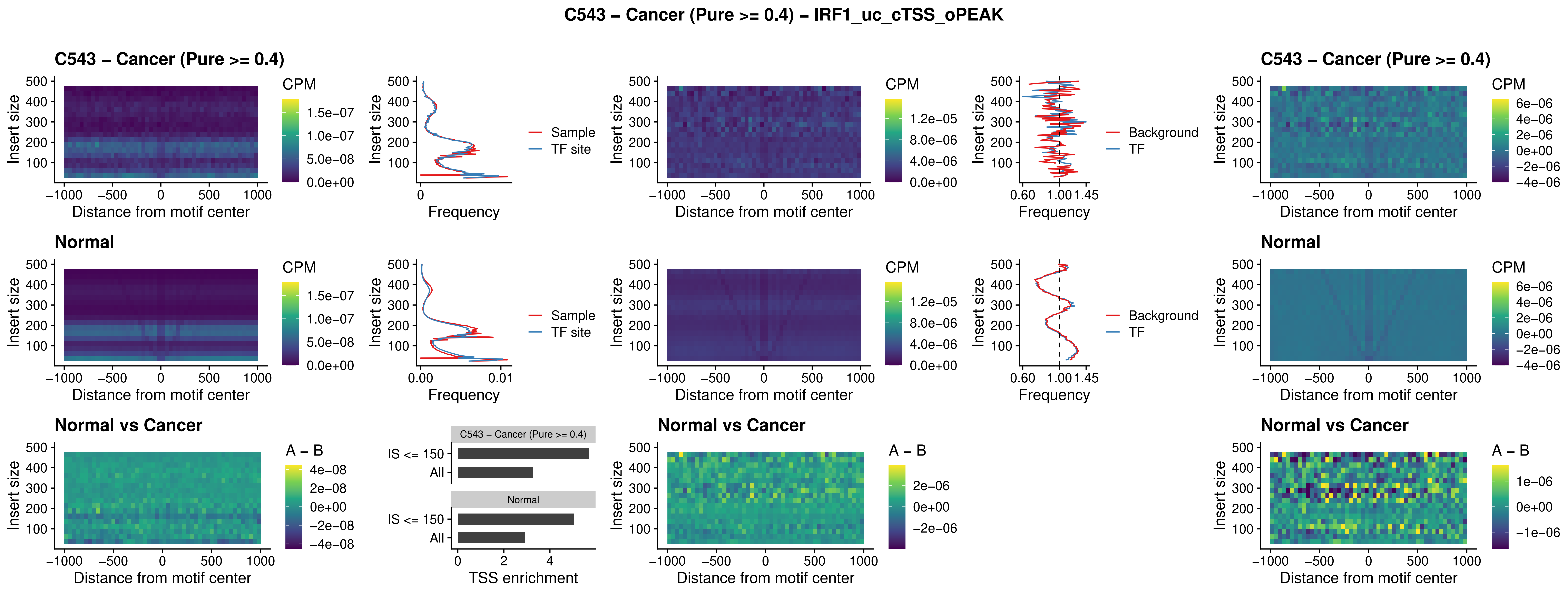
C543 – Cancer (Pure >= 0.4) – IRF1_uc_cTSS_oPEAK
C543 – Cancer (Pure >= 0.4)
Normal
Normal vs Cancer
Insert size
Distance from motif center
CPM
A – B
Frequency
Sample
TF site
Background
TF
C543 – Cancer (Pure >= 0.4)
Normal
IS <= 150
All
TSS enrichment

### C543 – Cancer (Pure >= 0.4) – IRF1_uc_dTSS_oPEAK

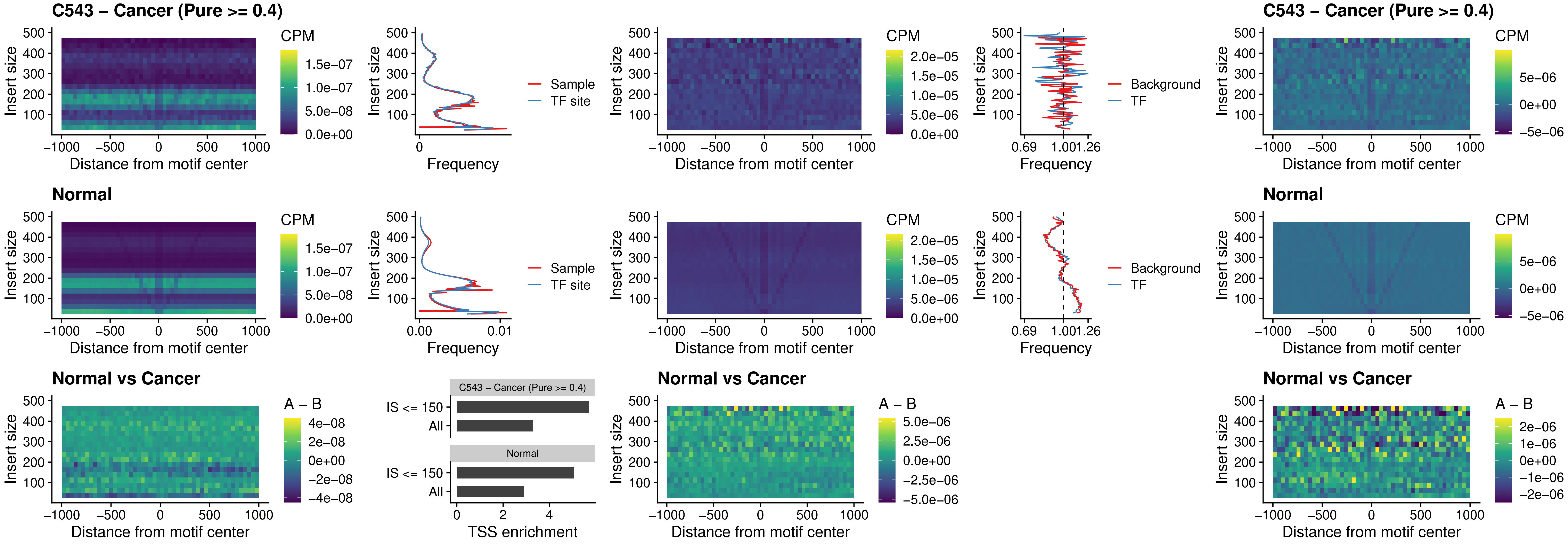

#### C543 – Cancer (Pure >= 0.4) – IRF1_uc_pTSS_oPEAK

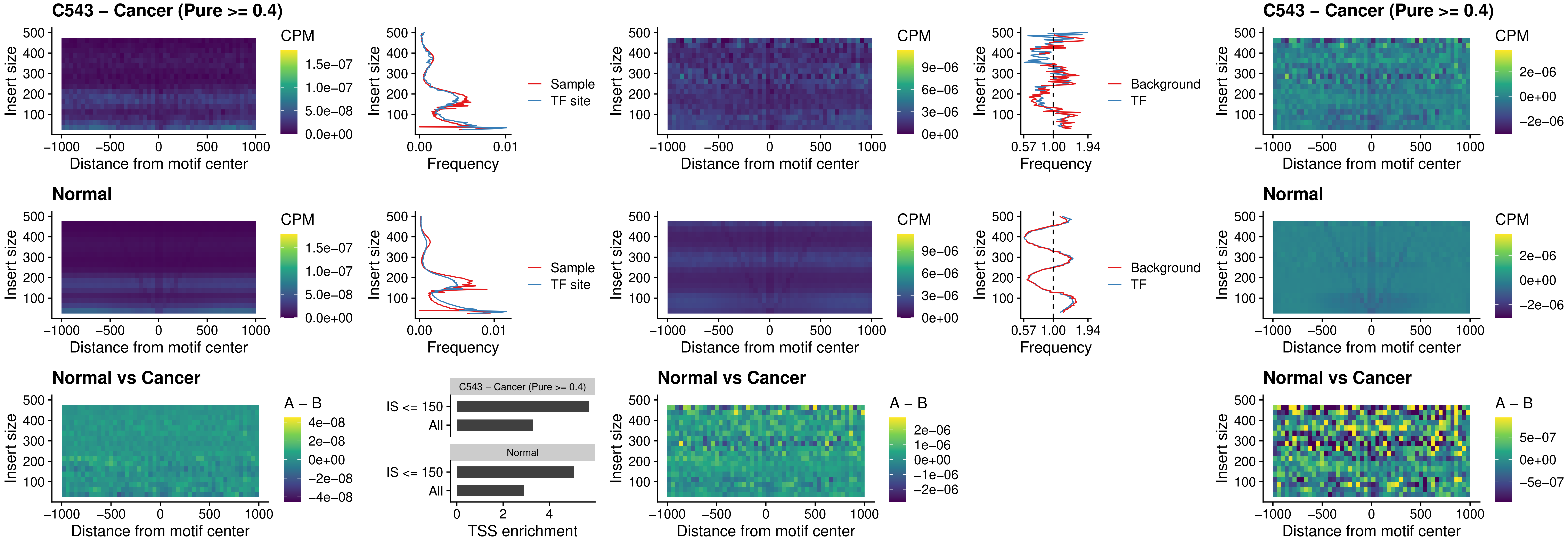

#### C543 – Cancer (Pure >= 0.4) – SPIC_uc_dTSS_nPEAK

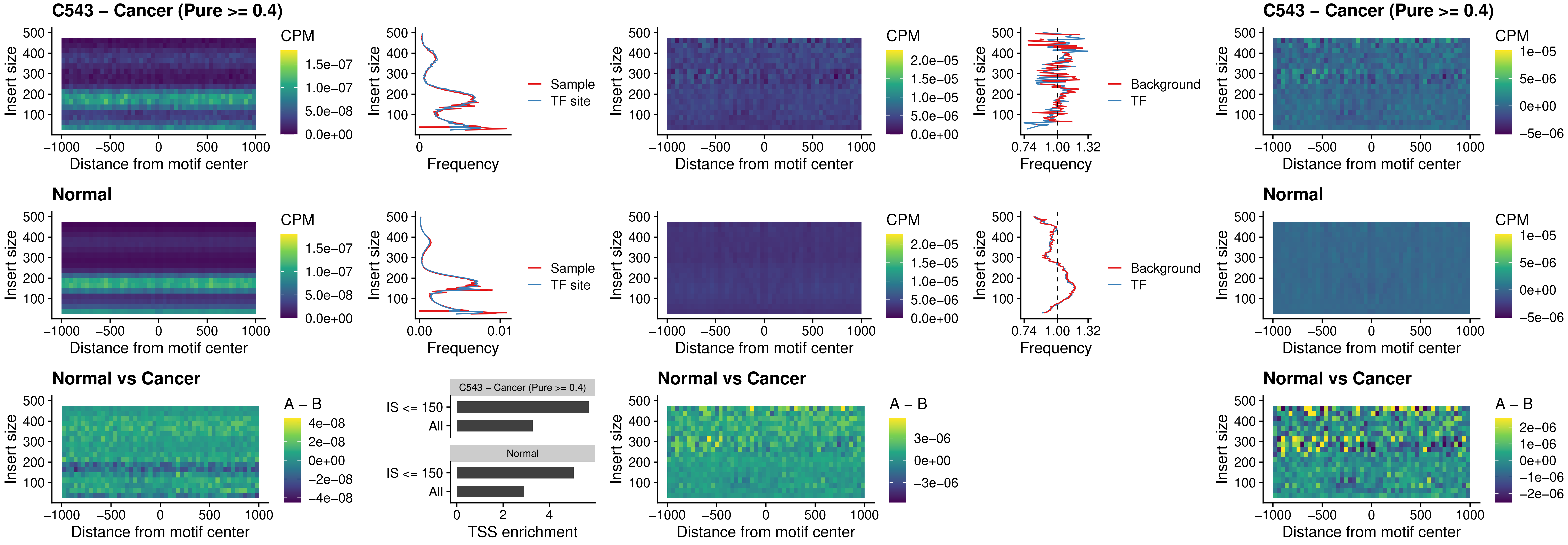

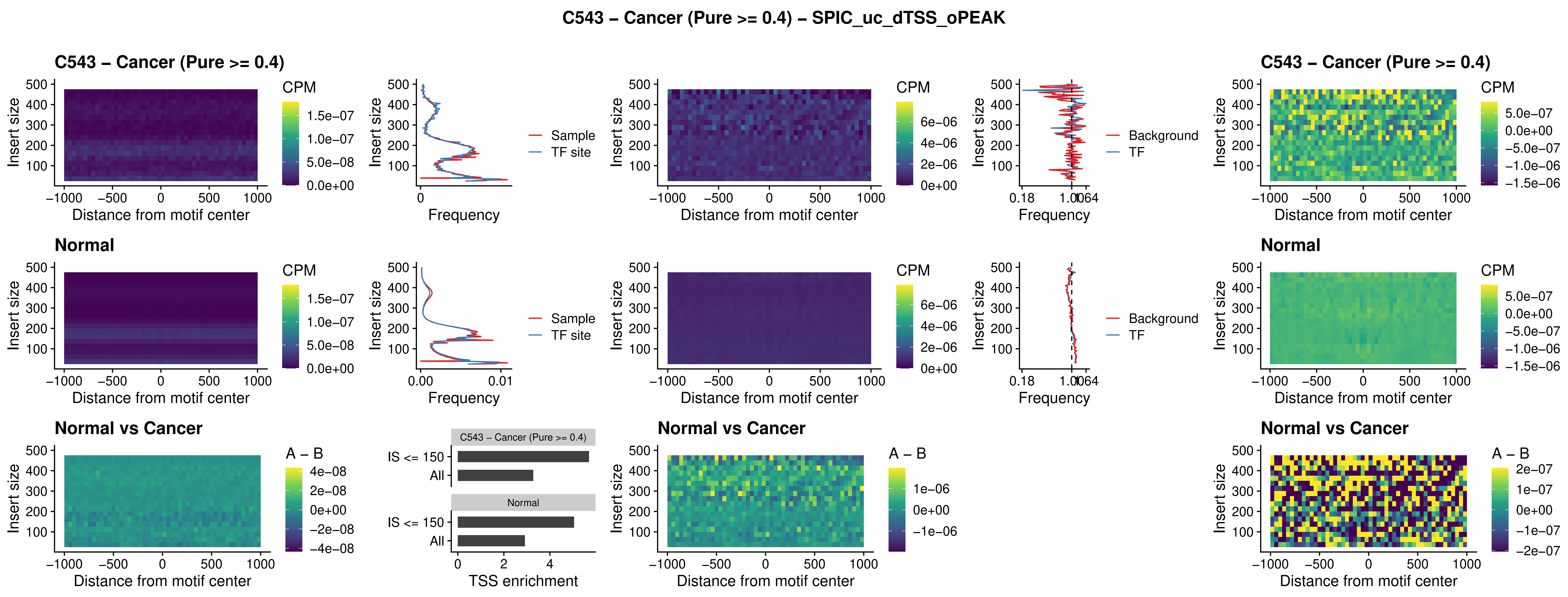
C543 – Cancer (Pure >= 0.4) – SPIC_uc_dTSS_oPEAK
C543 – Cancer (Pure >= 0.4)
Normal
Normal vs Cancer
Insert size
Distance from motif center
Frequency
CPM
A – B
Sample
TF site
Background
TF
C543 – Cancer (Pure >= 0.4)
Normal
IS <= 150
All
TSS enrichment

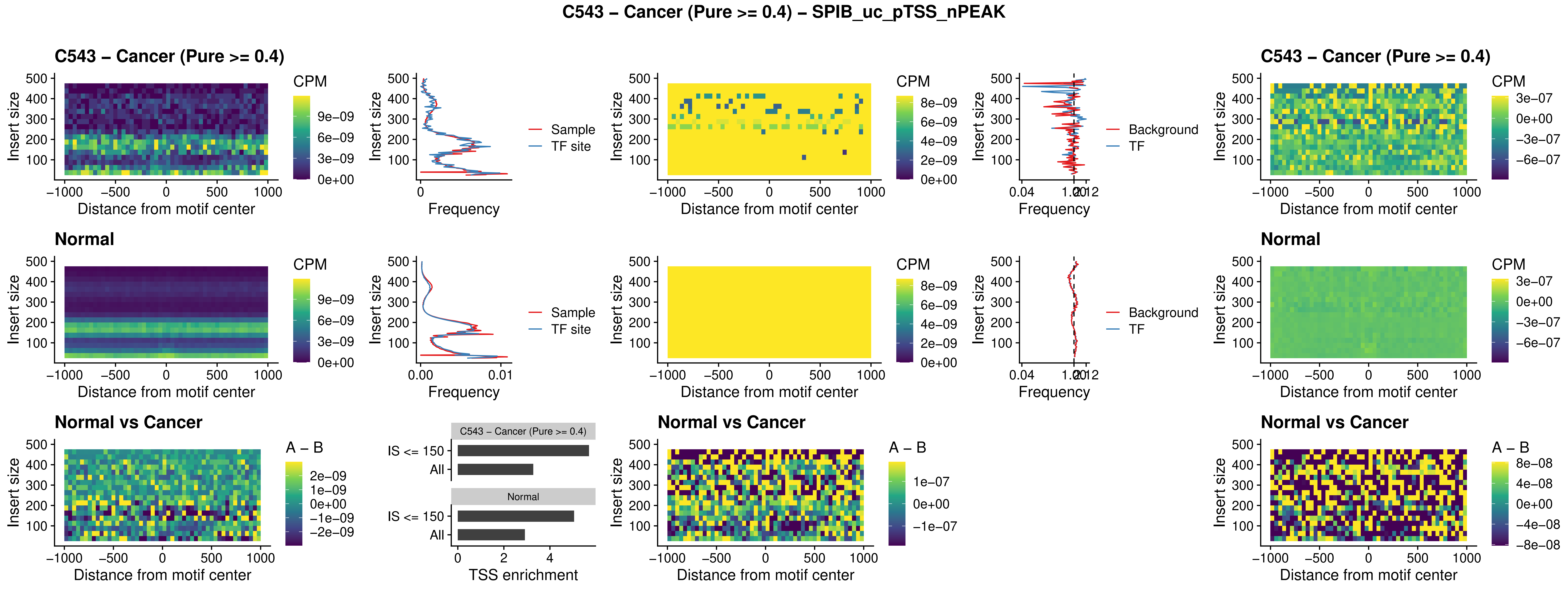
C543 – Cancer (Pure >= 0.4) – SPIB_uc_pTSS_nPEAK
C543 – Cancer (Pure >= 0.4)
Normal
Normal vs Cancer
Insert size
Distance from motif center
Frequency
CPM
A – B
Sample
TF site
Background
TF
C543 – Cancer (Pure >= 0.4)
Normal
IS <= 150
All
TSS enrichment

#### C543 – Cancer (Pure >= 0.4) – STAT2_uc_cTSS_oPEAK

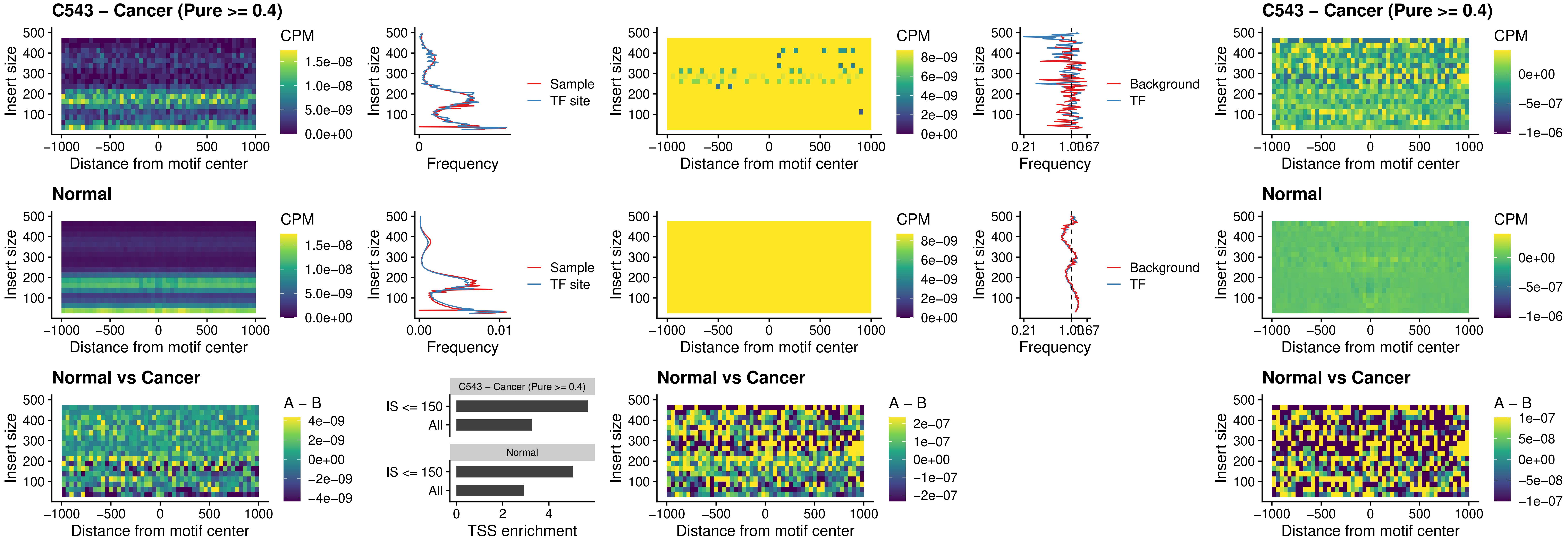

#### C543 – Cancer (Pure >= 0.4) – IRF8_uc_cTSS_oPEAK

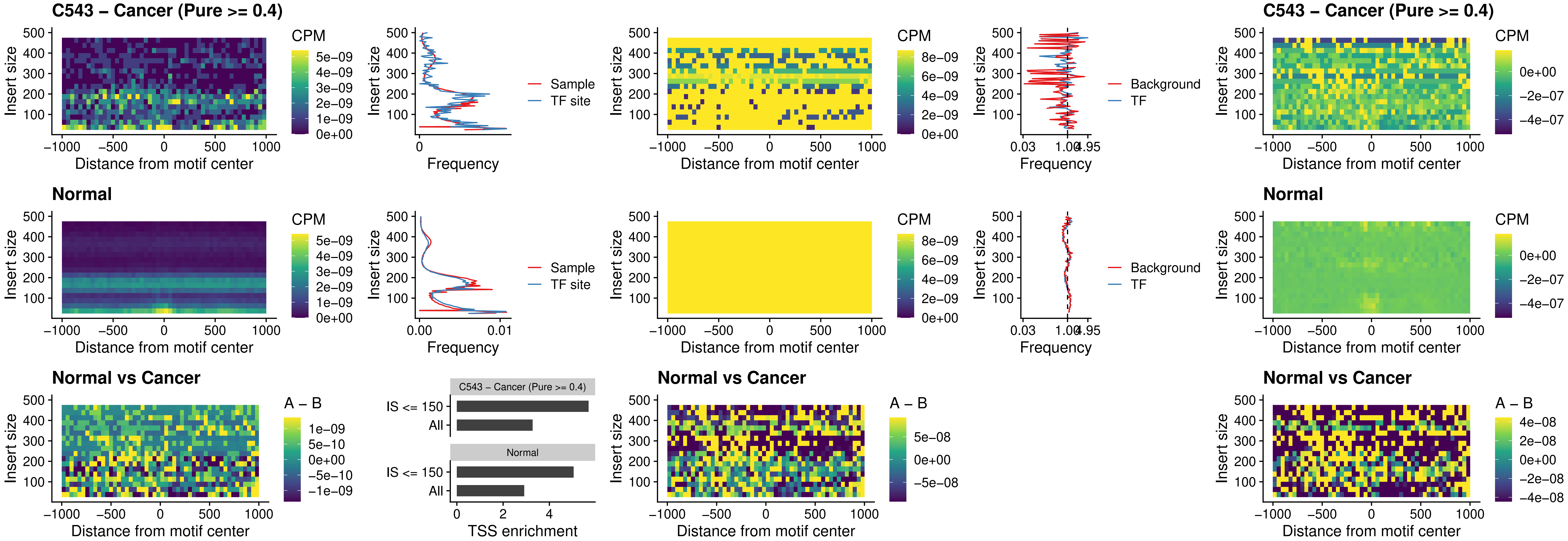

#### C543 – Cancer (Pure >= 0.4) – IRF3_uc_dTSS_oPEAK

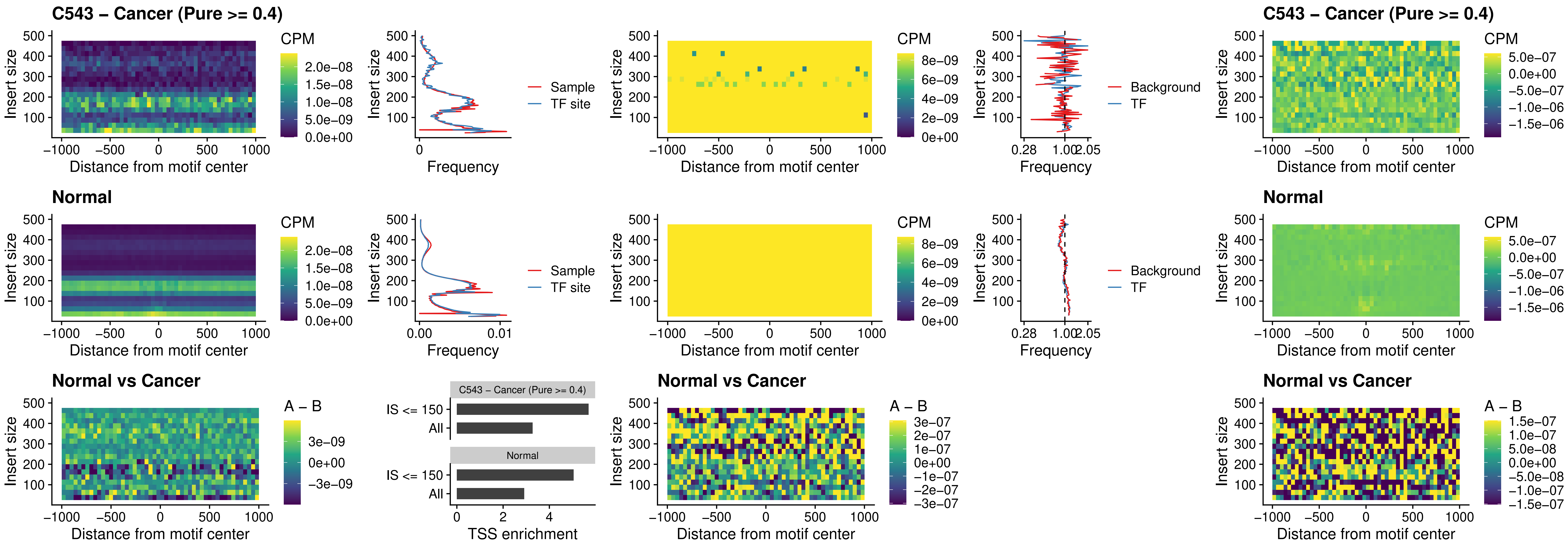

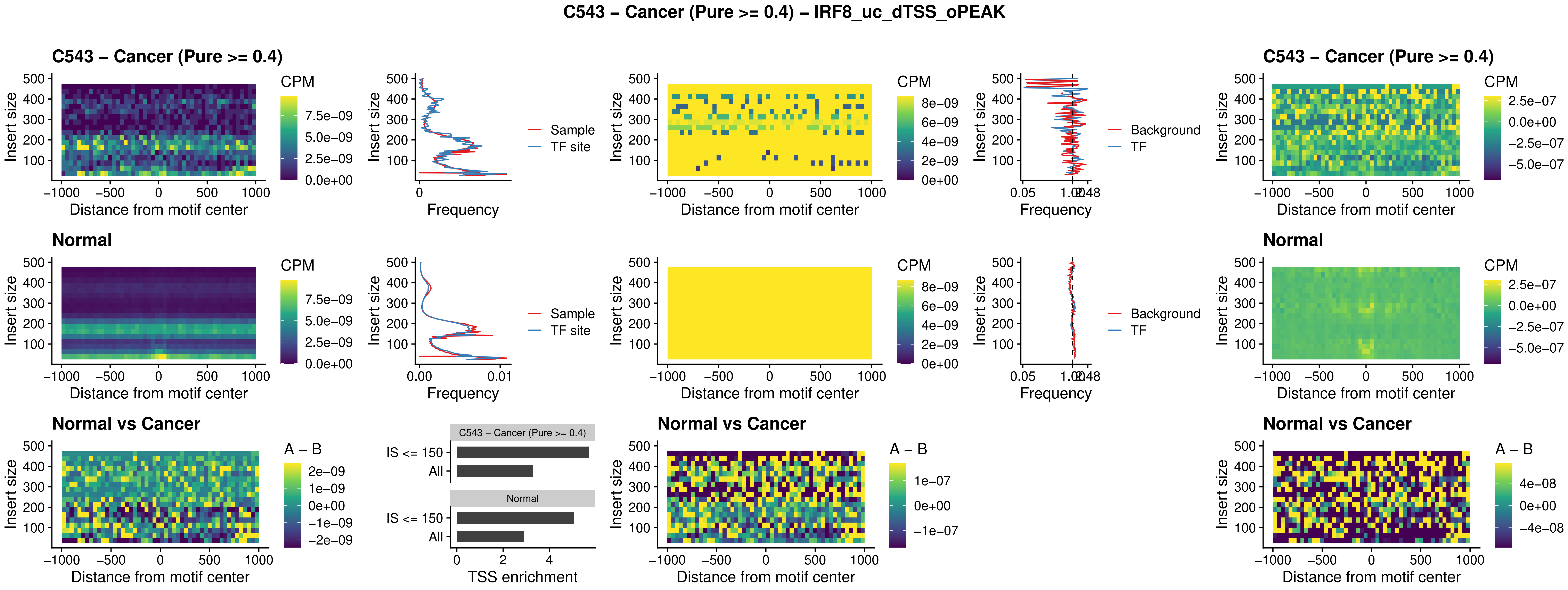

C543 – Cancer (Pure >= 0.4) – IRF8_uc_dTSS_oPEAK
C543 – Cancer (Pure >= 0.4)
Normal
Normal vs Cancer
Insert size
Distance from motif center
Frequency
CPM
A – B
Sample
TF site
Background
TF
C543 – Cancer (Pure >= 0.4)
Normal
IS <= 150
All
TSS enrichment

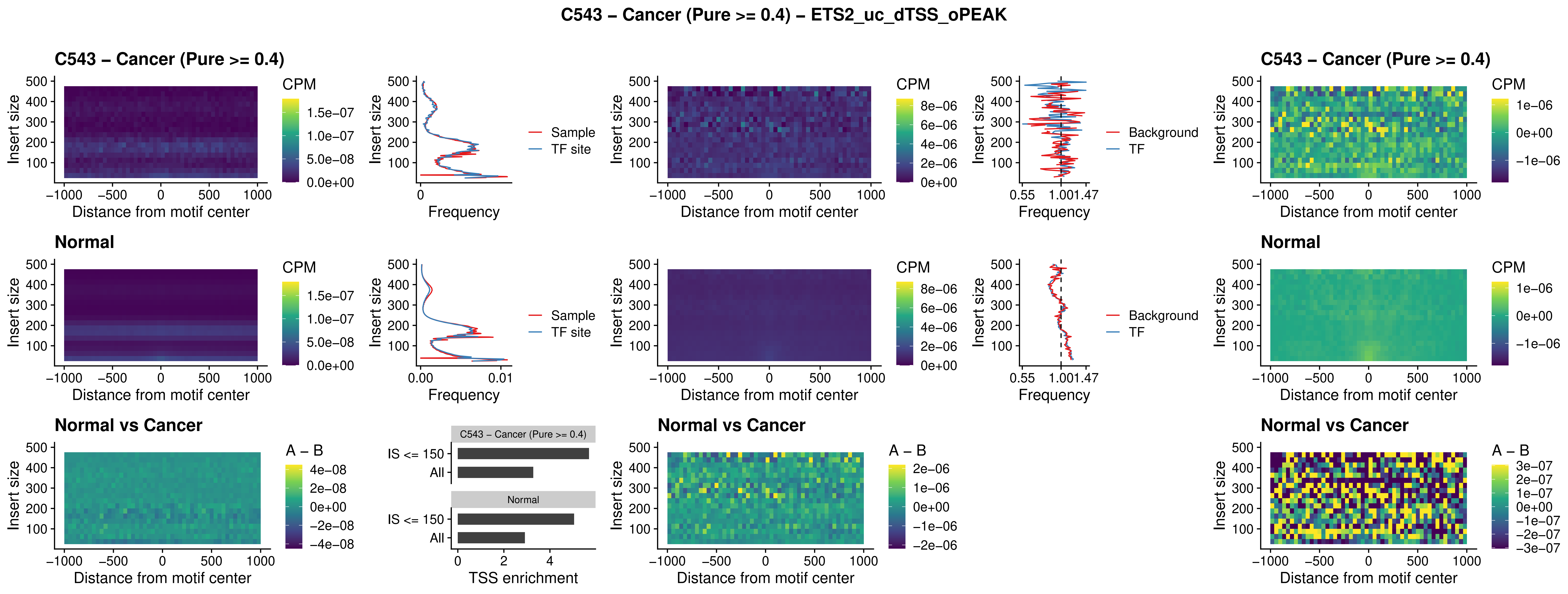
C543 – Cancer (Pure >= 0.4) – ETS2_uc_dTSS_oPEAK
C543 – Cancer (Pure >= 0.4)
Normal
Normal vs Cancer
Insert size
Distance from motif center
Frequency
CPM
A – B
Sample
TF site
Background
TF
C543 – Cancer (Pure >= 0.4)
Normal
IS <= 150
All
TSS enrichment

#### C543 – Cancer (Pure >= 0.4) – SMARCC2_uc_pTSS_oPEAK

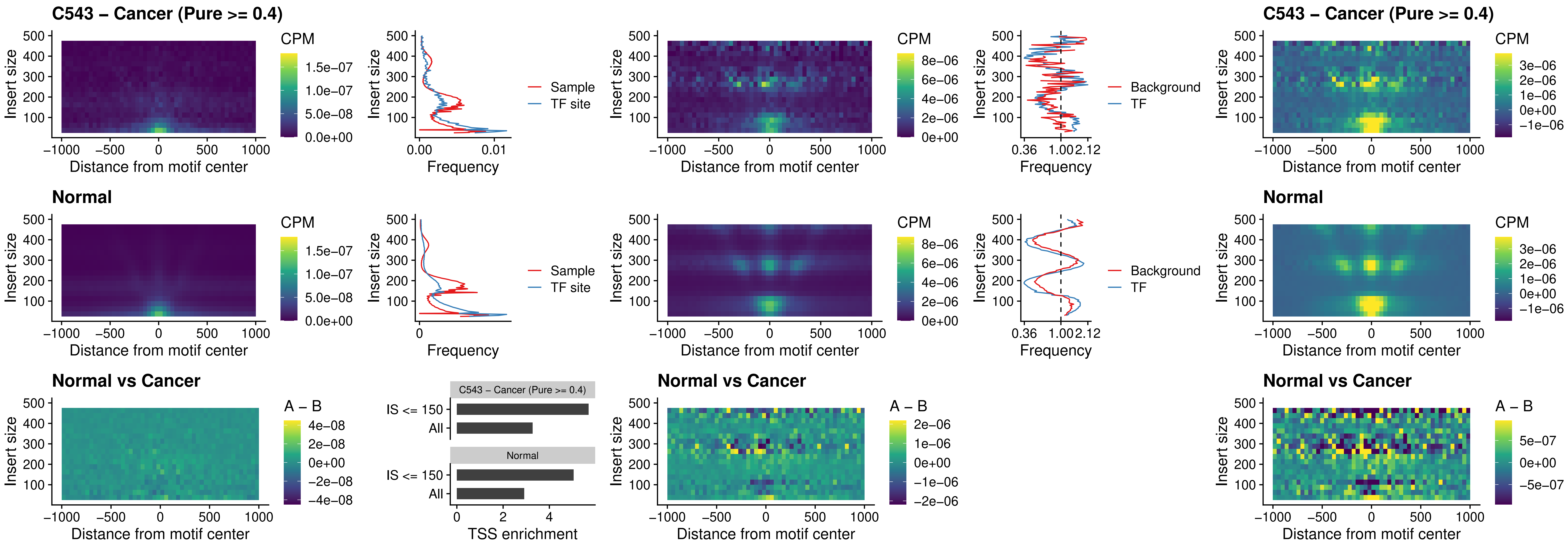

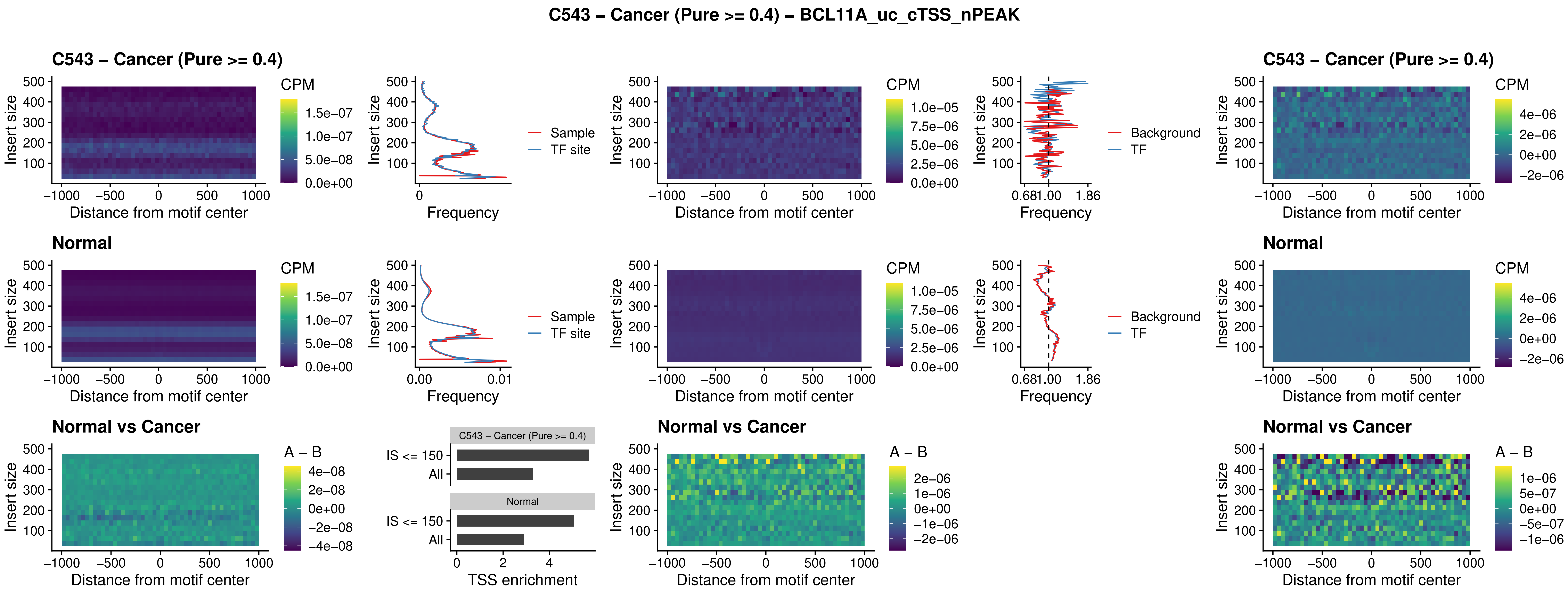
C543 – Cancer (Pure >= 0.4) – BCL11A_uc_cTSS_nPEAK
C543 – Cancer (Pure >= 0.4)
Normal
Normal vs Cancer
Insert size
Distance from motif center
Frequency
CPM
A – B
Sample
TF site
Background
TF
TSS enrichment
IS <= 150
All
C543 – Cancer (Pure >= 0.4)
Normal

### C543 – Cancer (Pure >= 0.4) – BCL11B_uc_cTSS_nPEAK

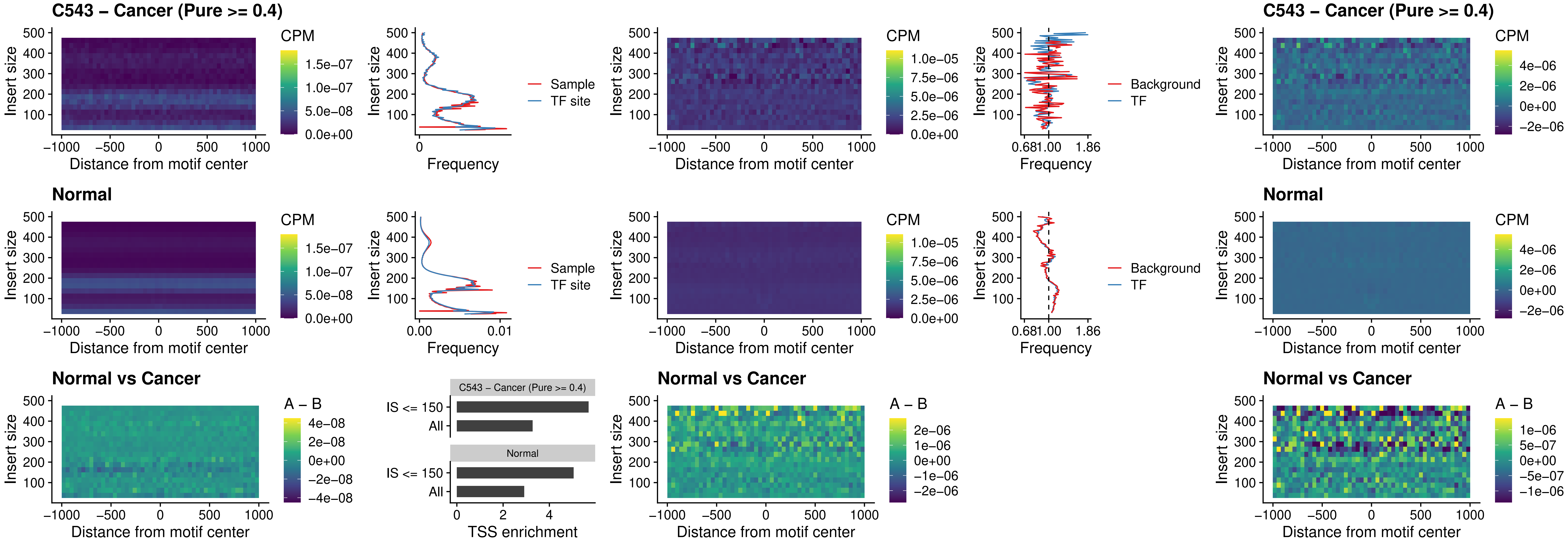

#### C543 – Cancer (Pure >= 0.4) – STAT2_uc_dTSS_oPEAK

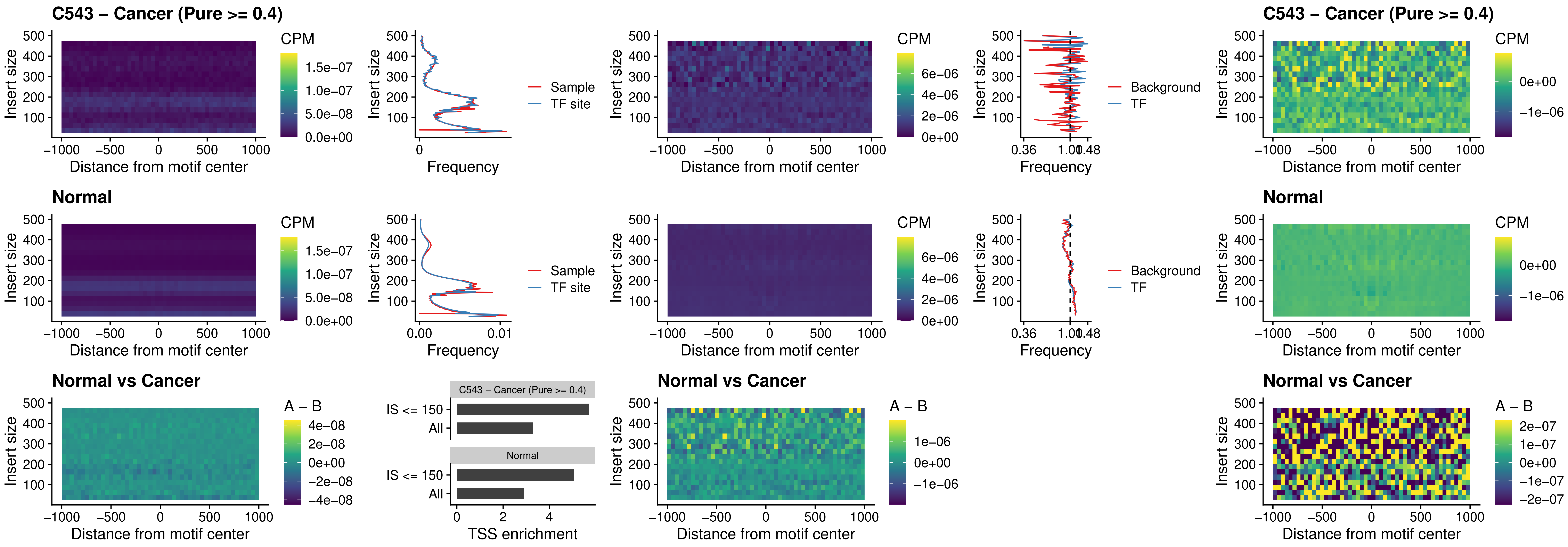

#### C543 – Cancer (Pure >= 0.4) – ZNF683_uc_cTSS_oPEAK

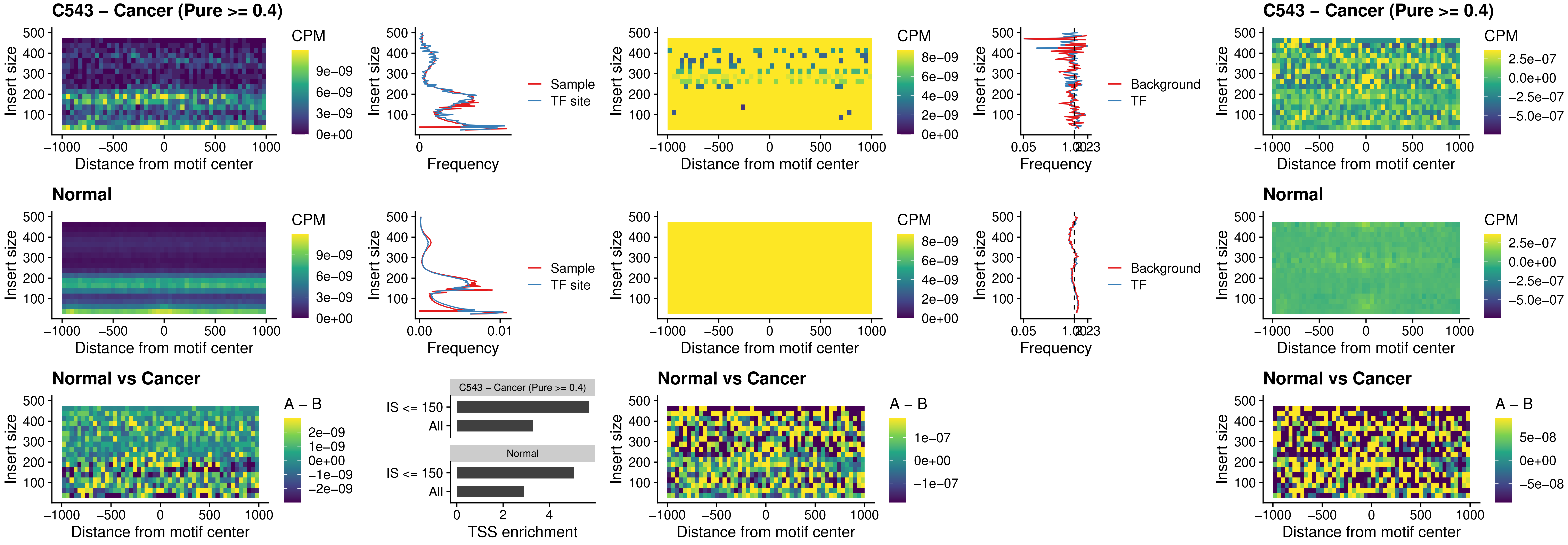

#### C543 – Cancer (Pure >= 0.4) – IRF8_uc_dTSS_nPEAK

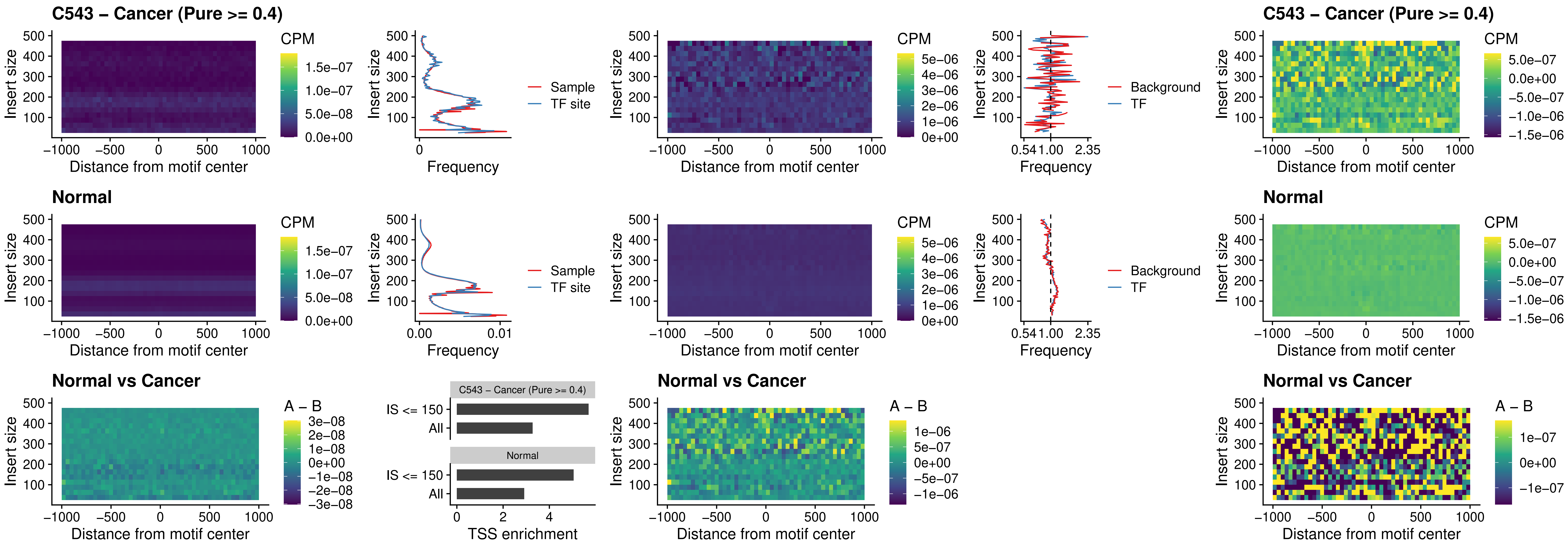

#### C543 – Cancer (Pure >= 0.4) – ETV7_uc_dTSS_oPEAK

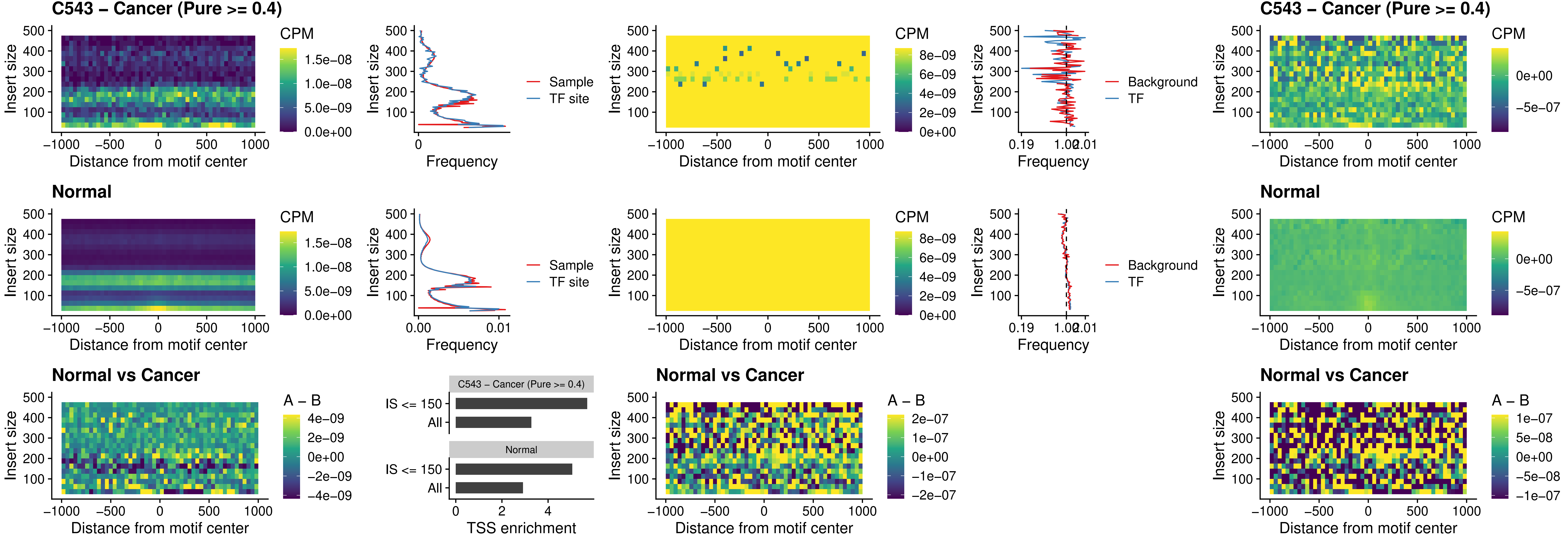

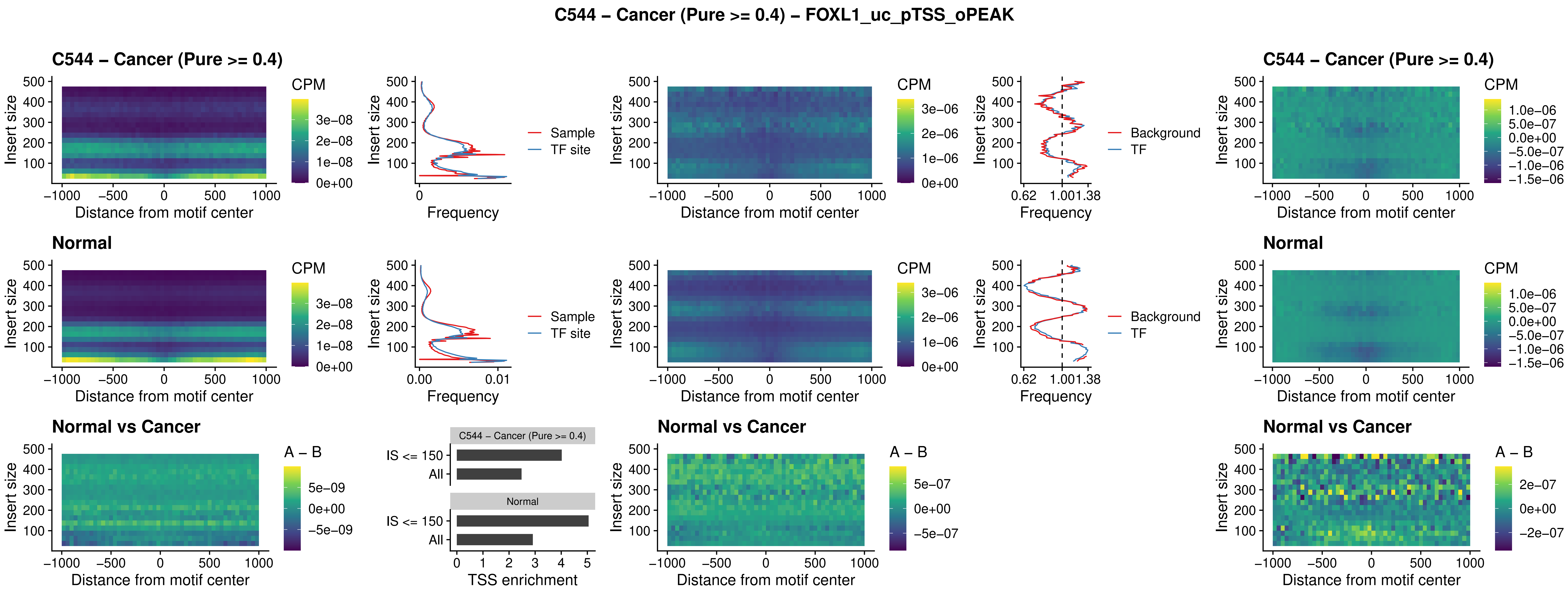
C544 – Cancer (Pure >= 0.4) – FOXL1_uc_pTSS_oPEAK
C544 – Cancer (Pure >= 0.4)
Normal
Normal vs Cancer
Insert size
Distance from motif center
Frequency
CPM
A – B
Sample
TF site
Background
TF
C544 – Cancer (Pure >= 0.4)
Normal
IS <= 150
All
TSS enrichment

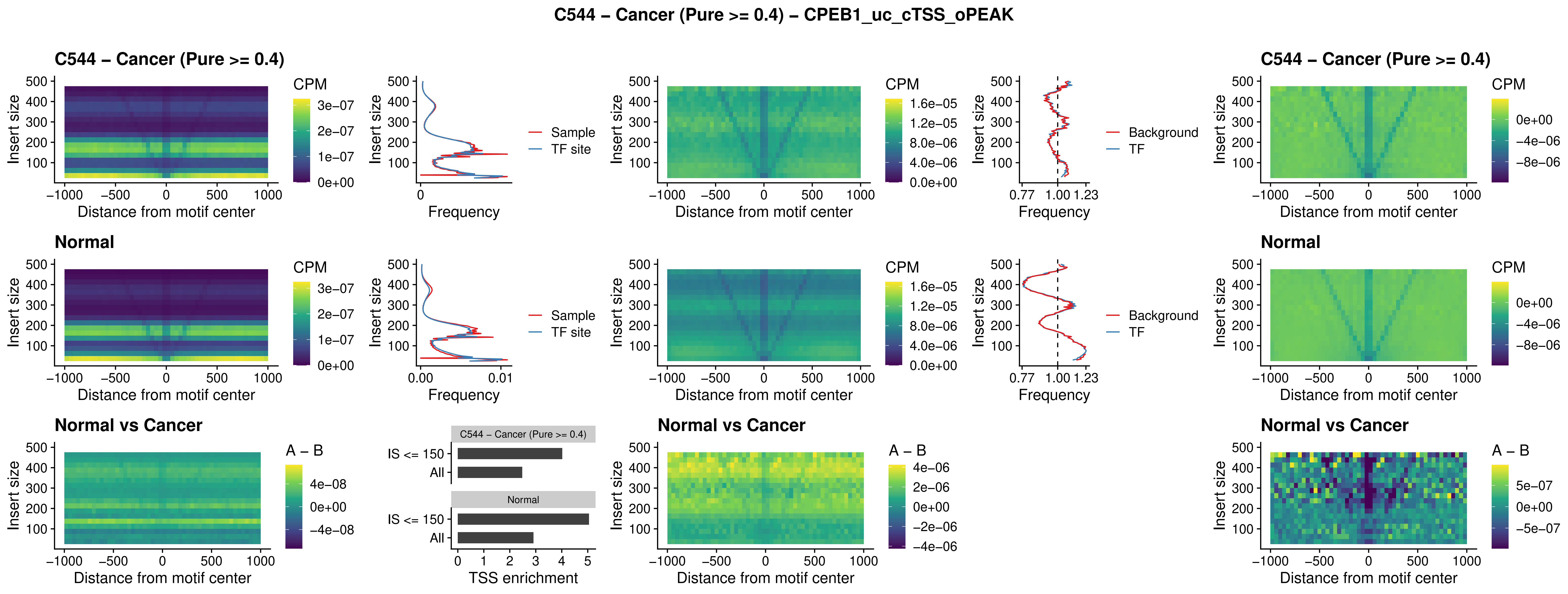
C544 – Cancer (Pure >= 0.4) – CPEB1_uc_cTSS_oPEAK
C544 – Cancer (Pure >= 0.4)
Normal
Normal vs Cancer
Insert size
Distance from motif center
Frequency
CPM
A – B
Sample
TF site
Background
TF
C544 – Cancer (Pure >= 0.4)
Normal
IS <= 150
All
TSS enrichment

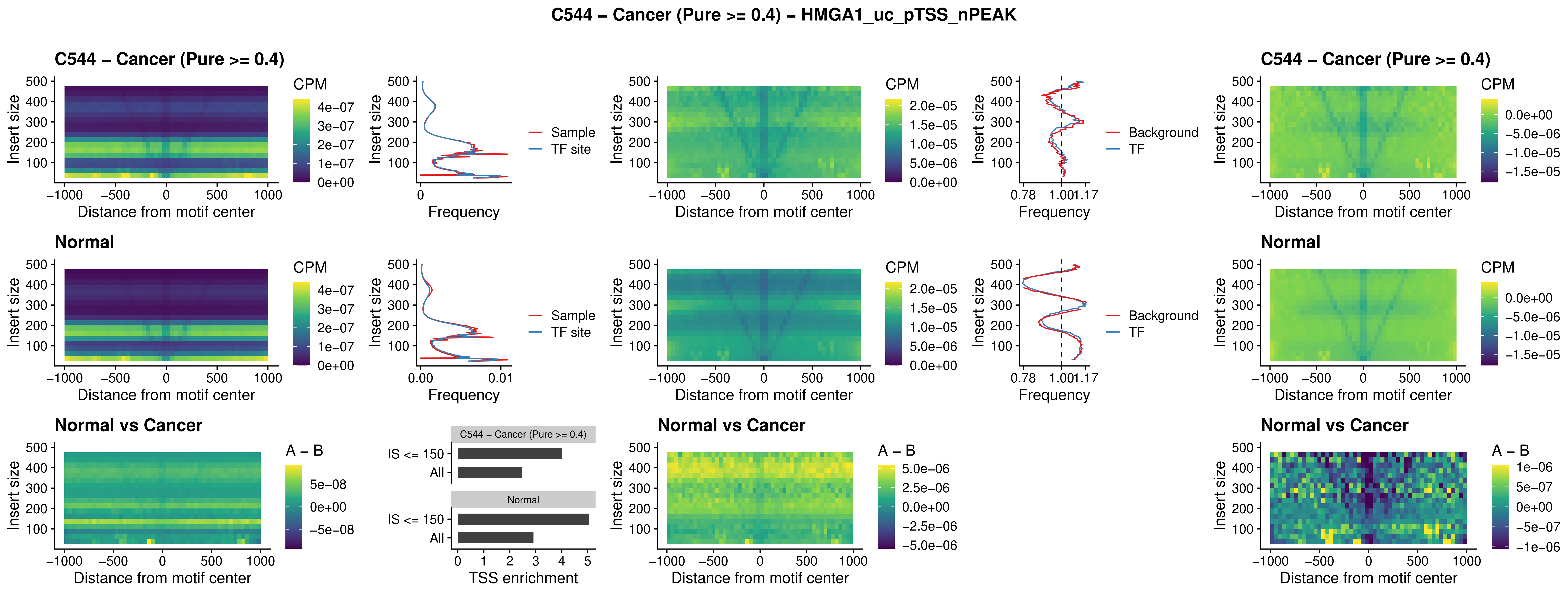
C544 – Cancer (Pure >= 0.4) – HMGA1_uc_pTSS_nPEAK
C544 – Cancer (Pure >= 0.4)
Normal
Normal vs Cancer
Insert size
Distance from motif center
Frequency
CPM
A – B
Sample
TF site
Background
TF
C544 – Cancer (Pure >= 0.4)
Normal
IS <= 150
All
TSS enrichment

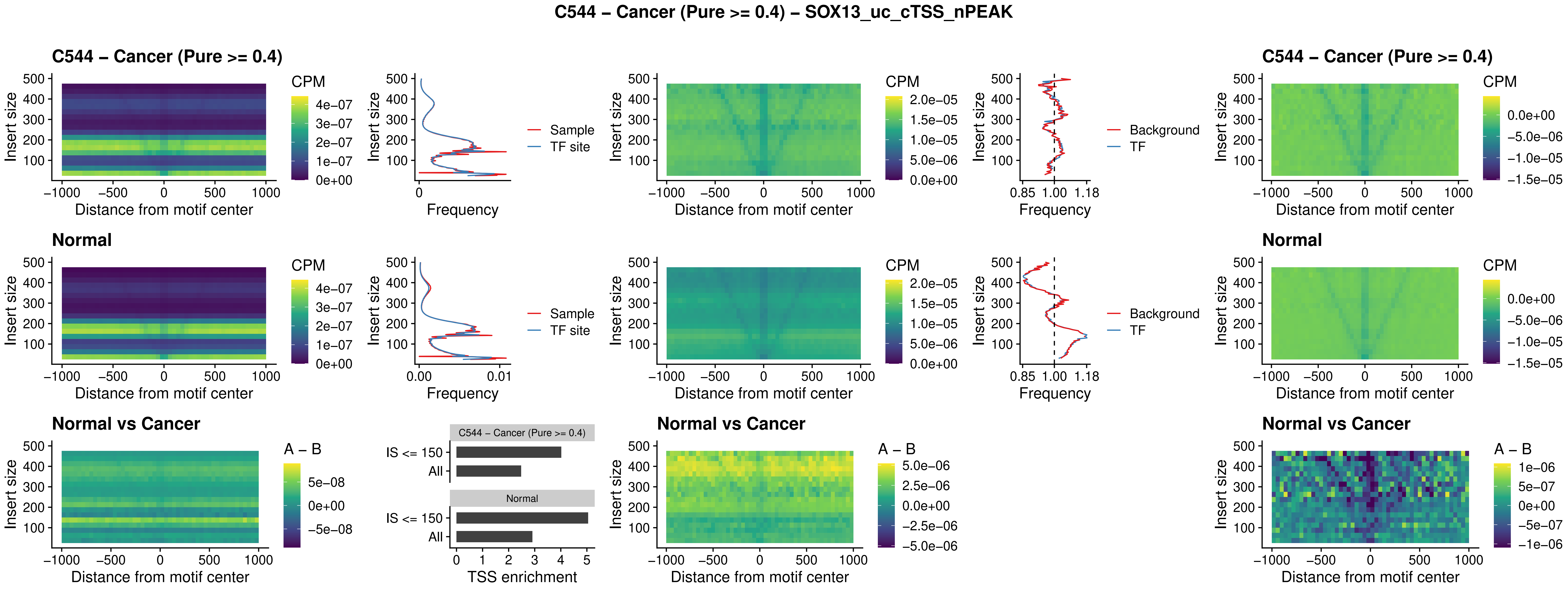
C544 – Cancer (Pure >= 0.4) – SOX13_uc_cTSS_nPEAK
C544 – Cancer (Pure >= 0.4)
Normal
Normal vs Cancer
Insert size
Distance from motif center
Frequency
CPM
A – B
Sample
TF site
Background
TF
C544 – Cancer (Pure >= 0.4)
Normal
IS <= 150
All
TSS enrichment

### C544 – Cancer (Pure >= 0.4) – LHX4_uc_cTSS_nPEAK

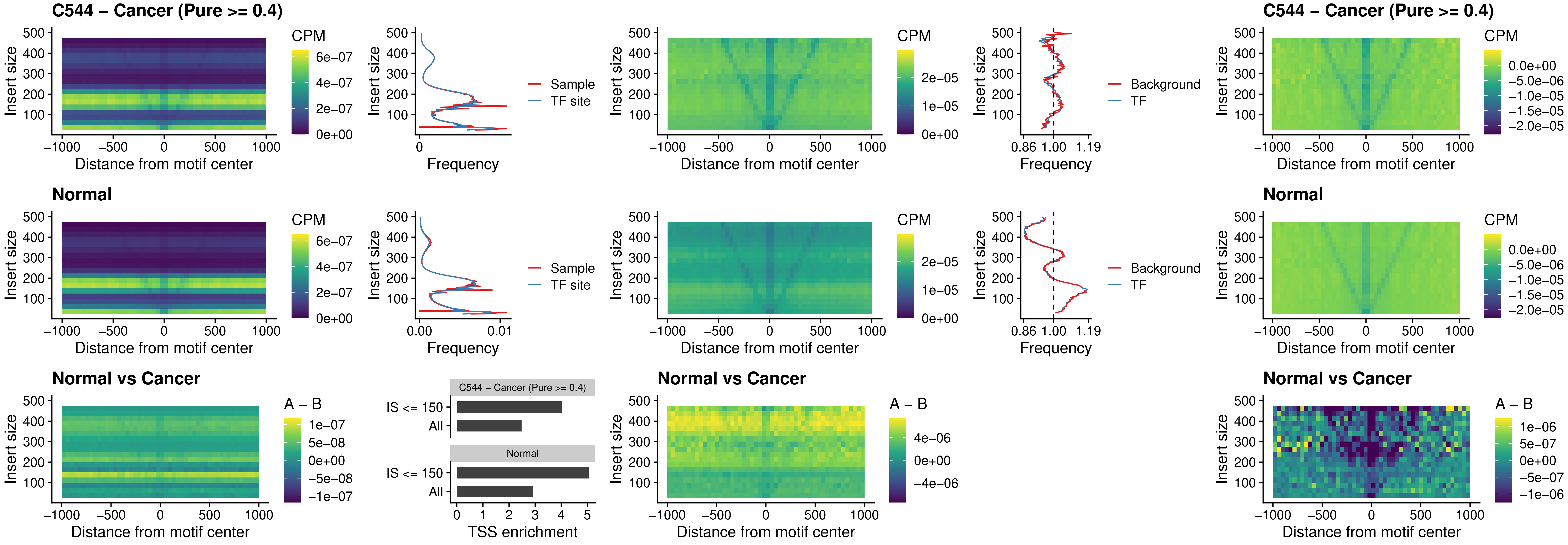

#### C544 – Cancer (Pure >= 0.4) – HOXA4_uc_cTSS_nPEAK

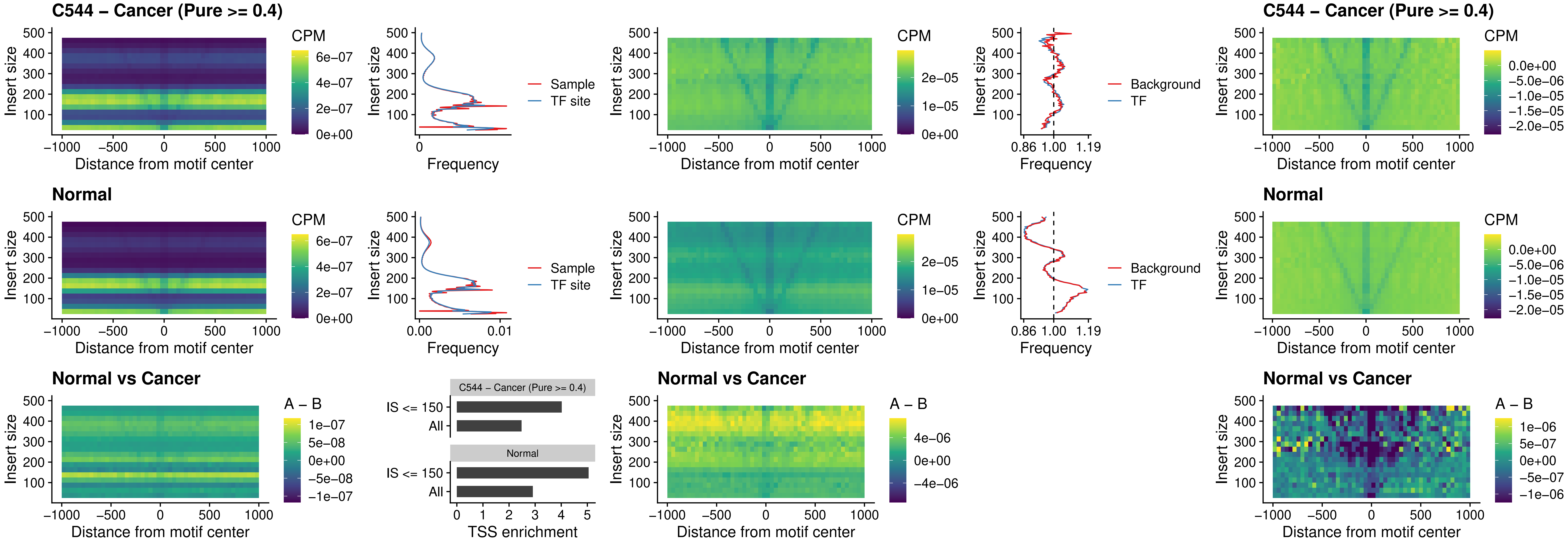

#### C544 – Cancer (Pure >= 0.4) – HLX_uc_cTSS_nPEAK

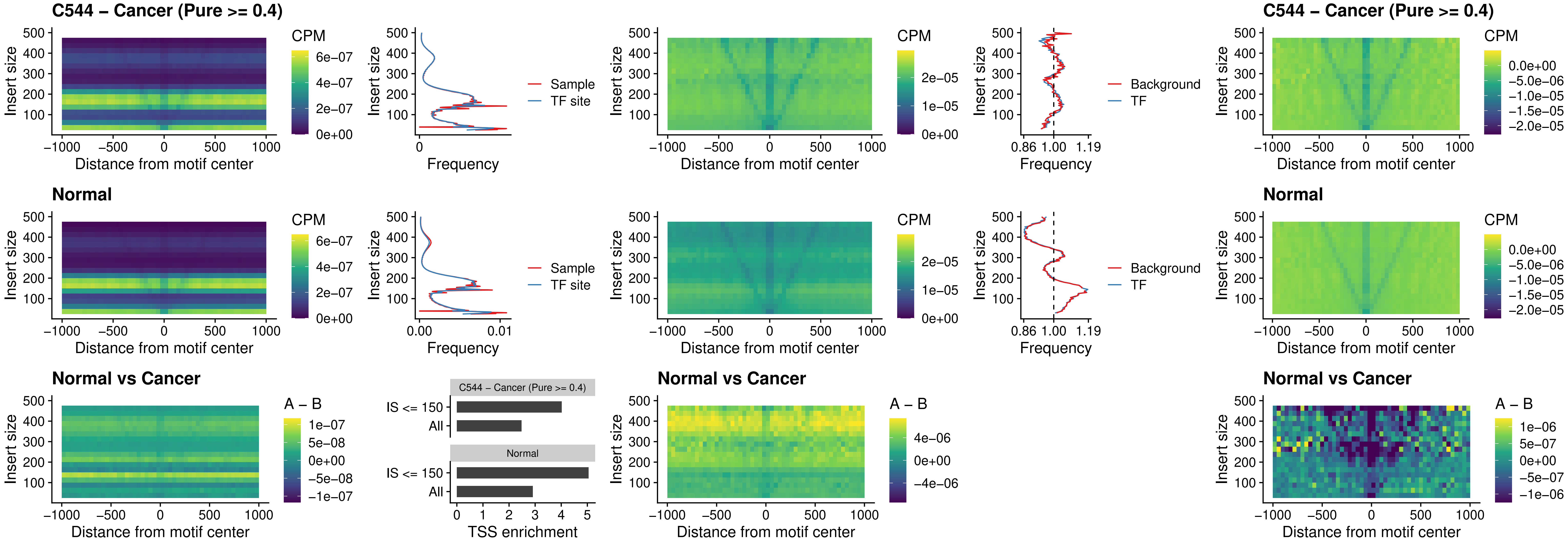

#### C544 – Cancer (Pure >= 0.4) – AC0125311_uc_cTSS_nPEAK

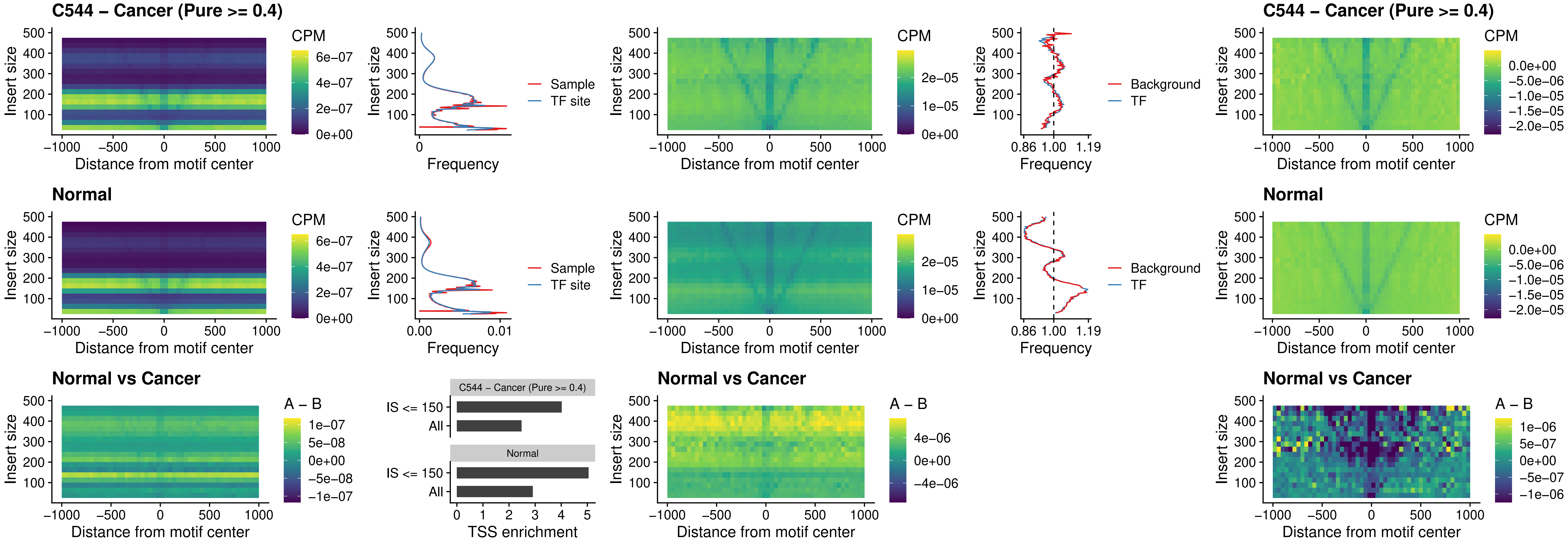

#### C544 – Cancer (Pure >= 0.4) – DBX2_uc_cTSS_nPEAK

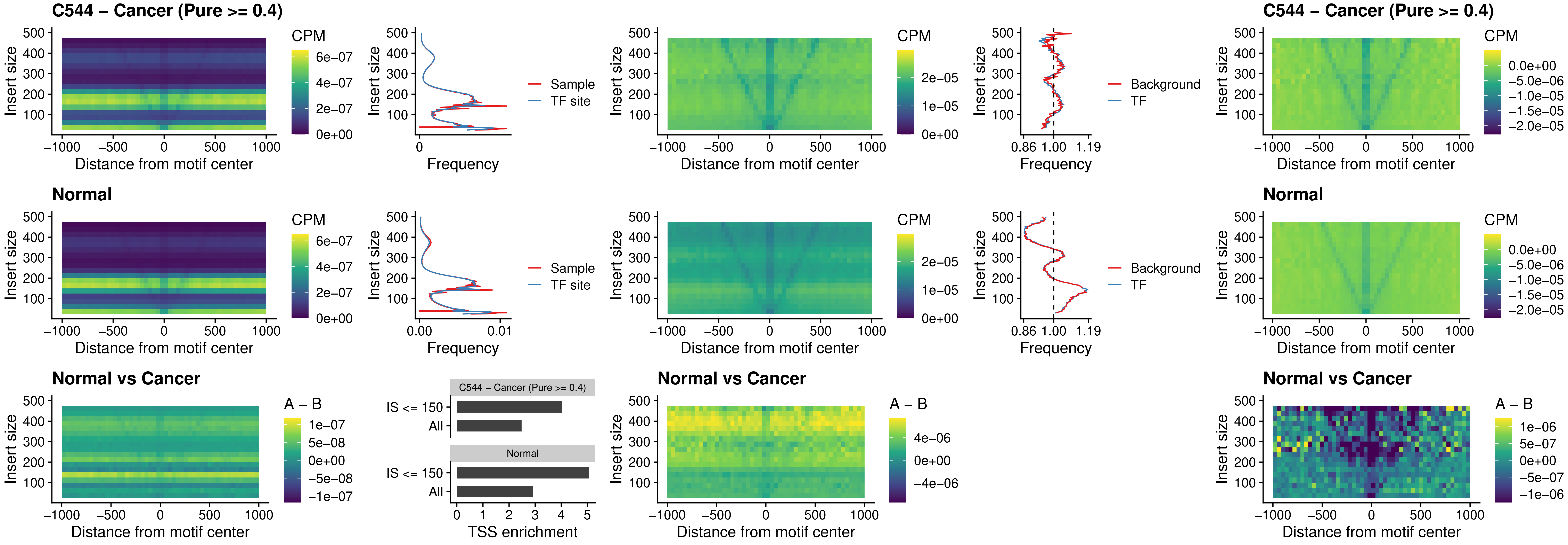

#### C544 – Cancer (Pure >= 0.4) – LHX6_uc_dTSS_nPEAK

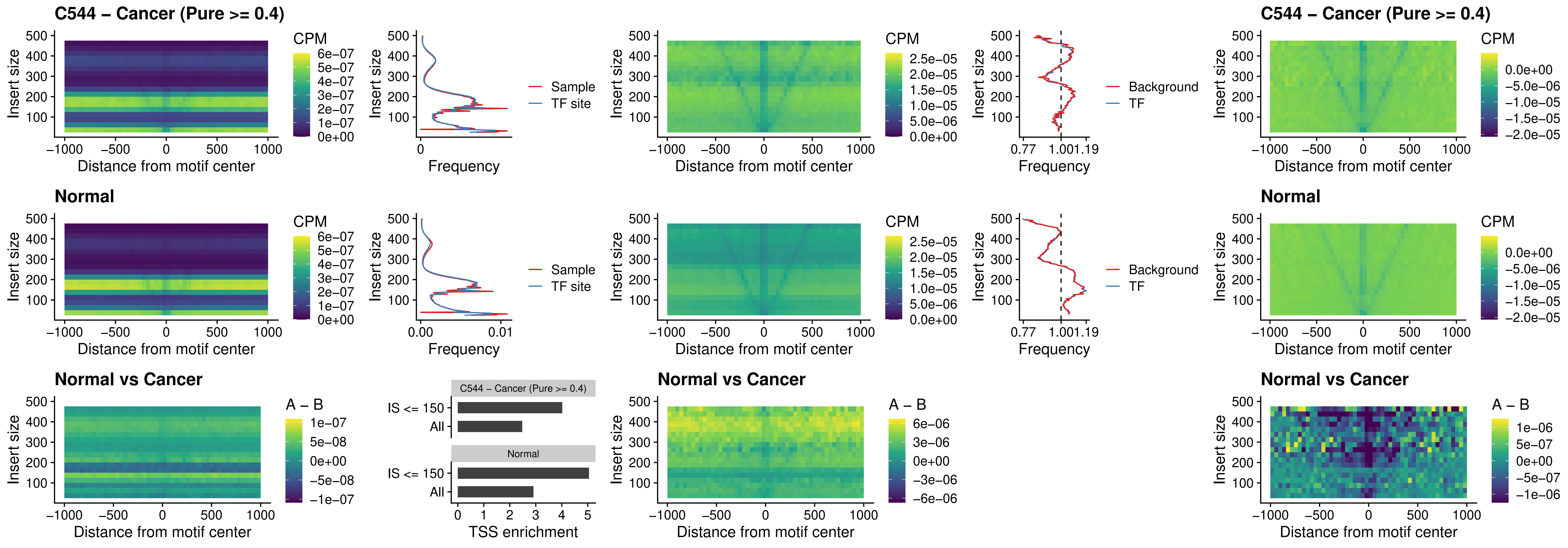

#### C544 – Cancer (Pure >= 0.4) – SOX1_uc_dTSS_nPEAK

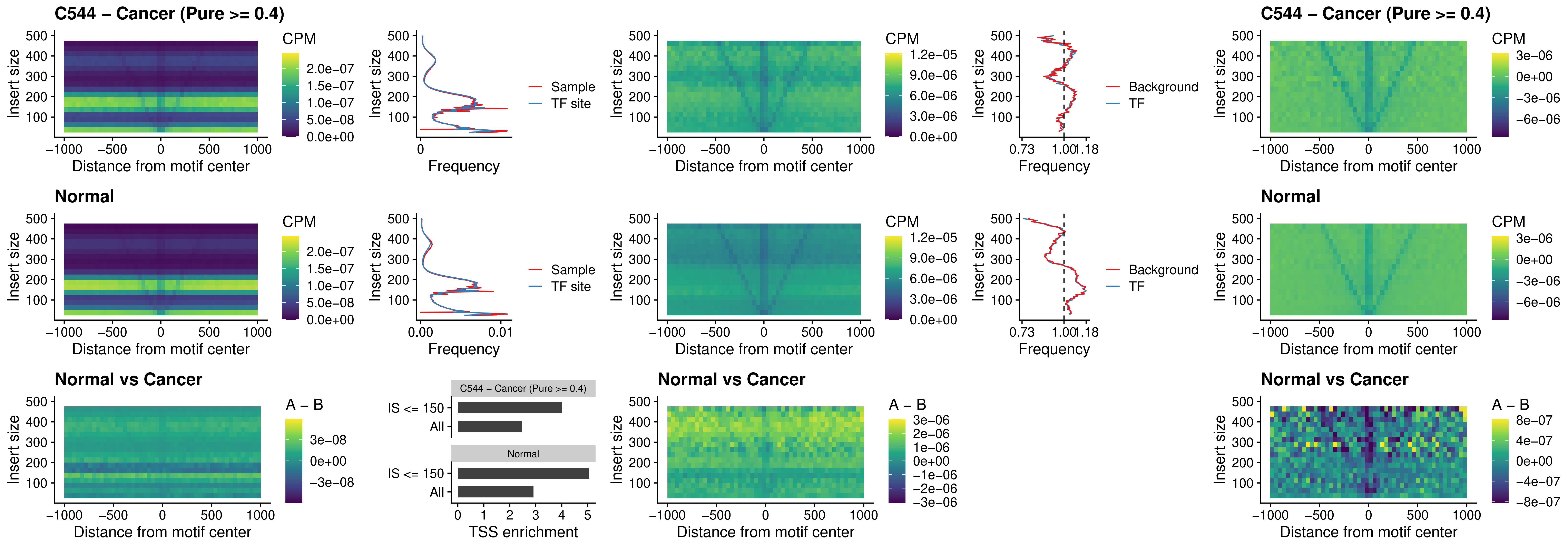

#### C544 – Cancer (Pure >= 0.4) – NKX24_uc_dTSS_nPEAK

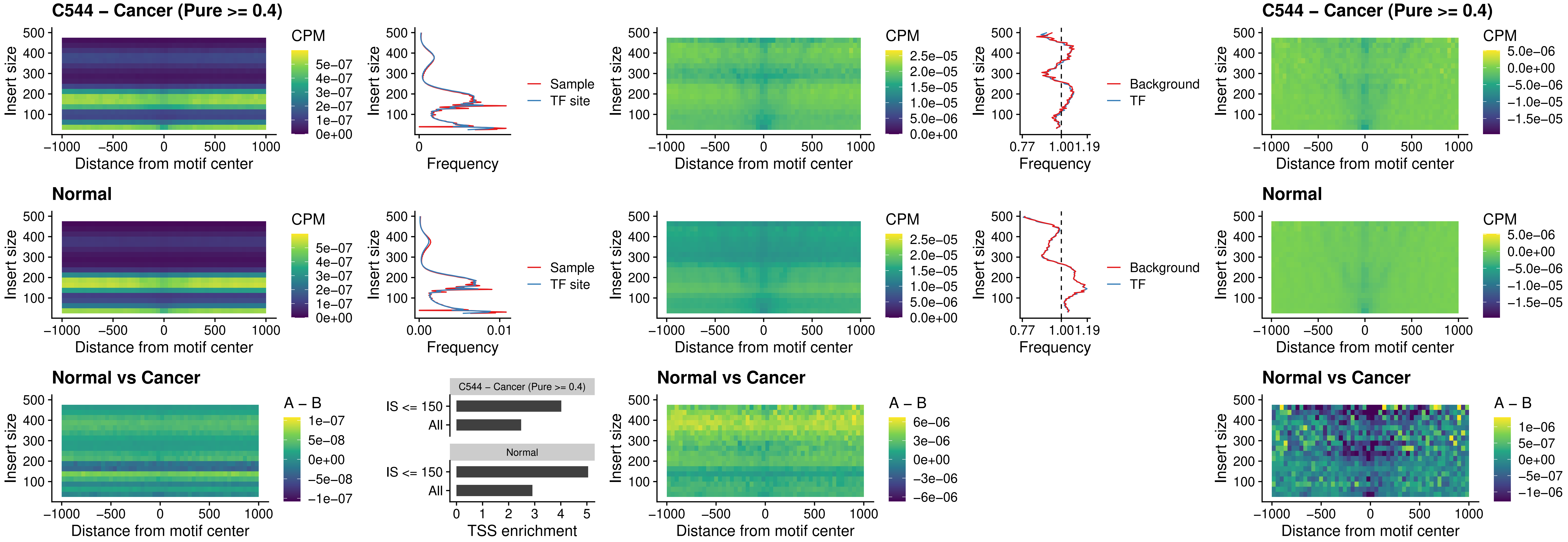

#### C544 – Cancer (Pure >= 0.4) – DLX3_uc_cTSS_nPEAK

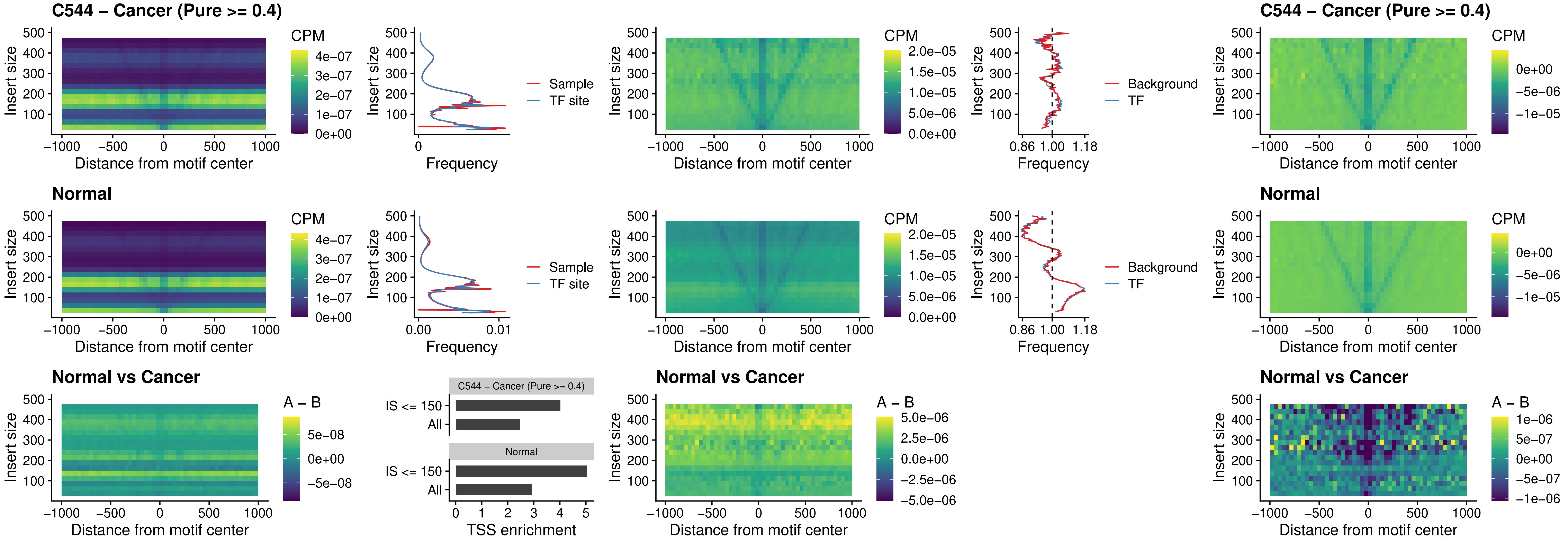

#### C544 – Cancer (Pure >= 0.4) – HOXD3_uc_cTSS_nPEAK

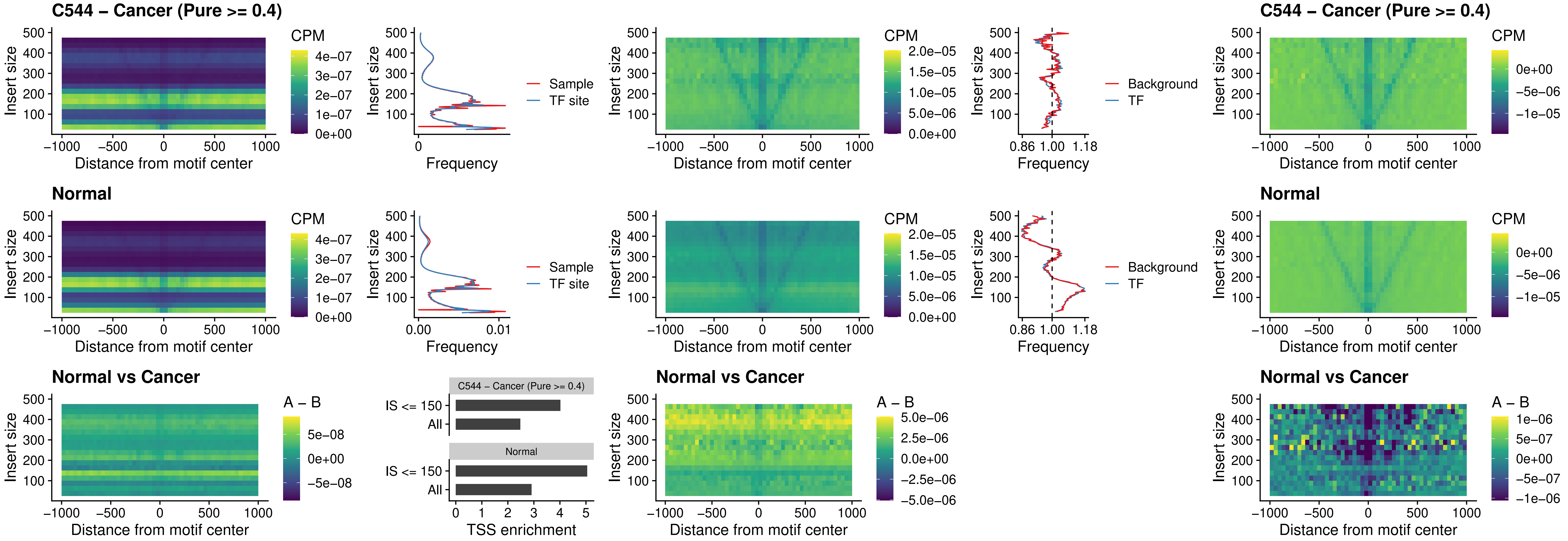

### C544 – Cancer (Pure >= 0.4) – ISL1_uc_dTSS_oPEAK

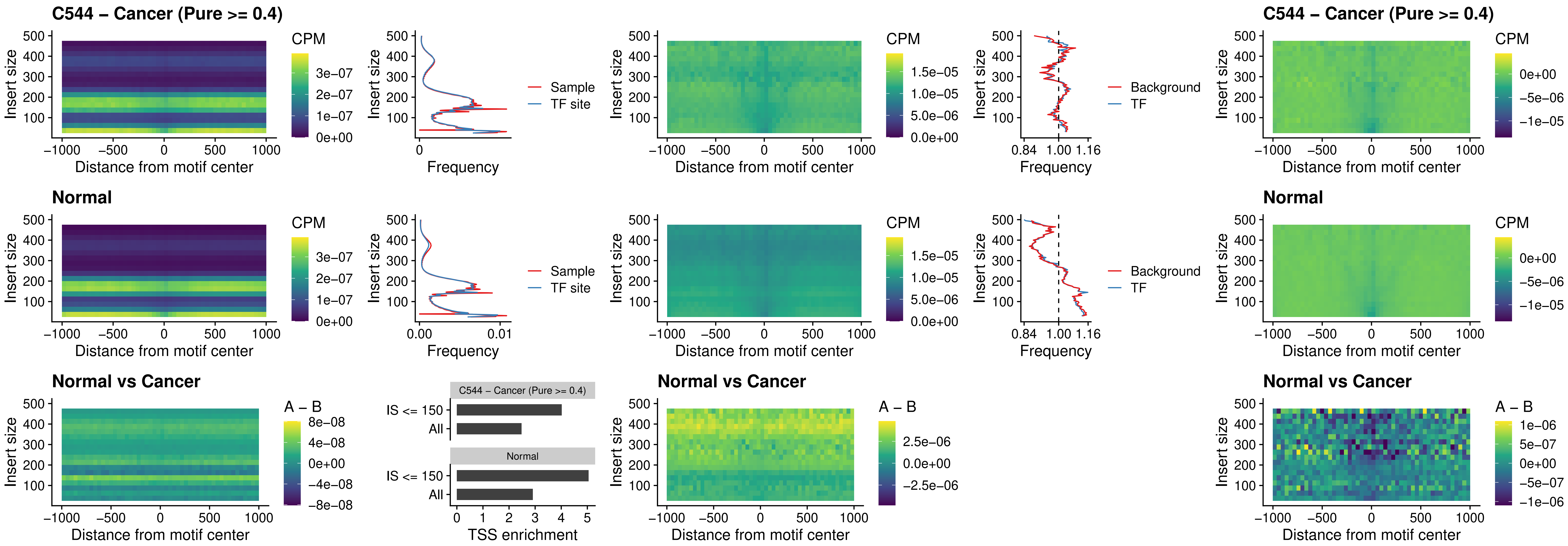

#### C544 – Cancer (Pure >= 0.4) – HOXB4_uc_dTSS_oPEAK

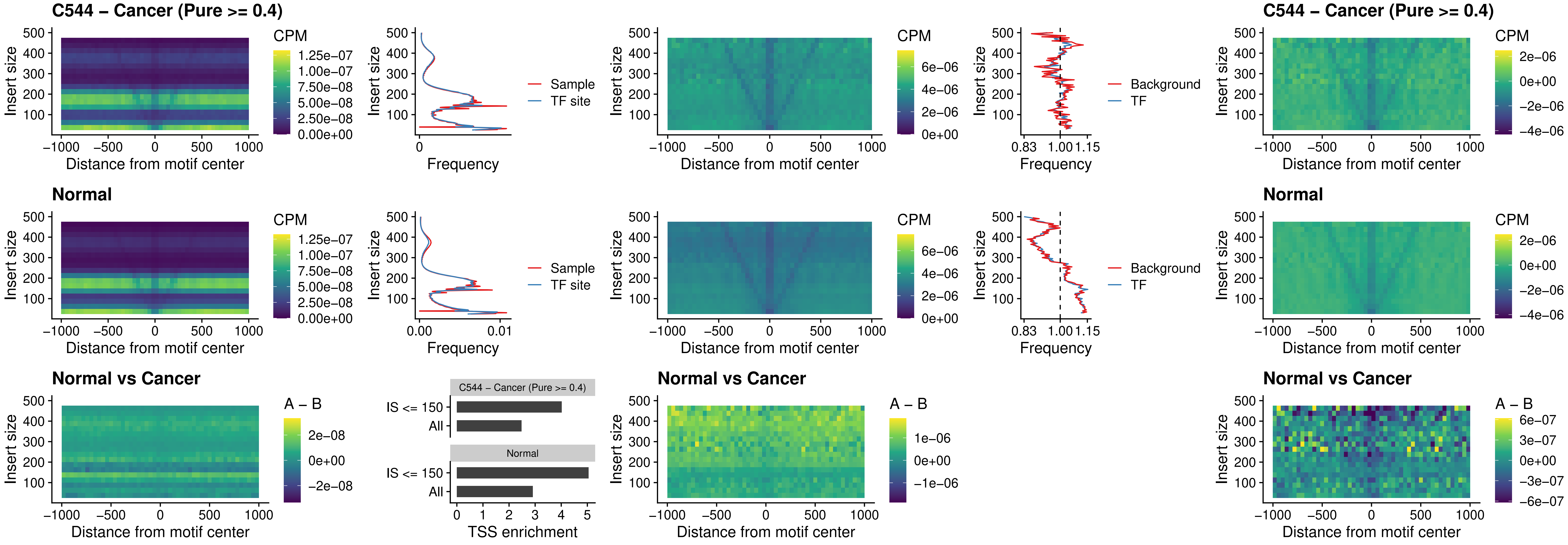

#### C544 – Cancer (Pure >= 0.4) – VAX1_uc_dTSS_oPEAK

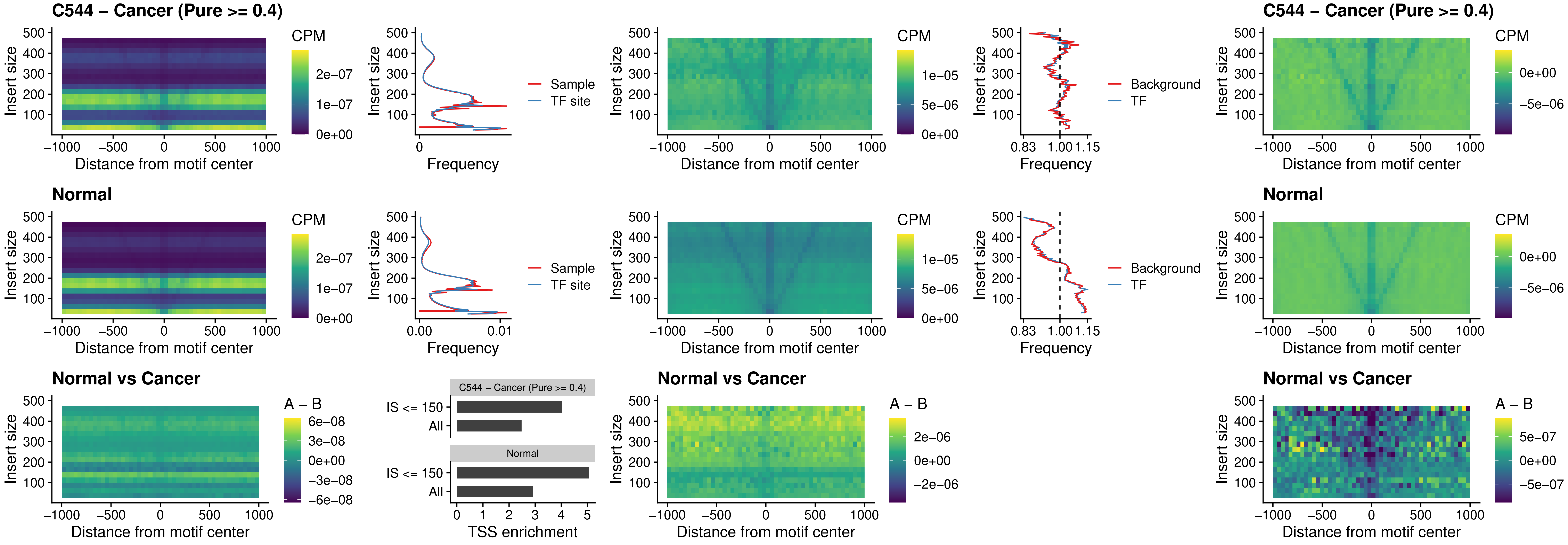

#### C544 – Cancer (Pure >= 0.4) – DLX3_uc_dTSS_oPEAK

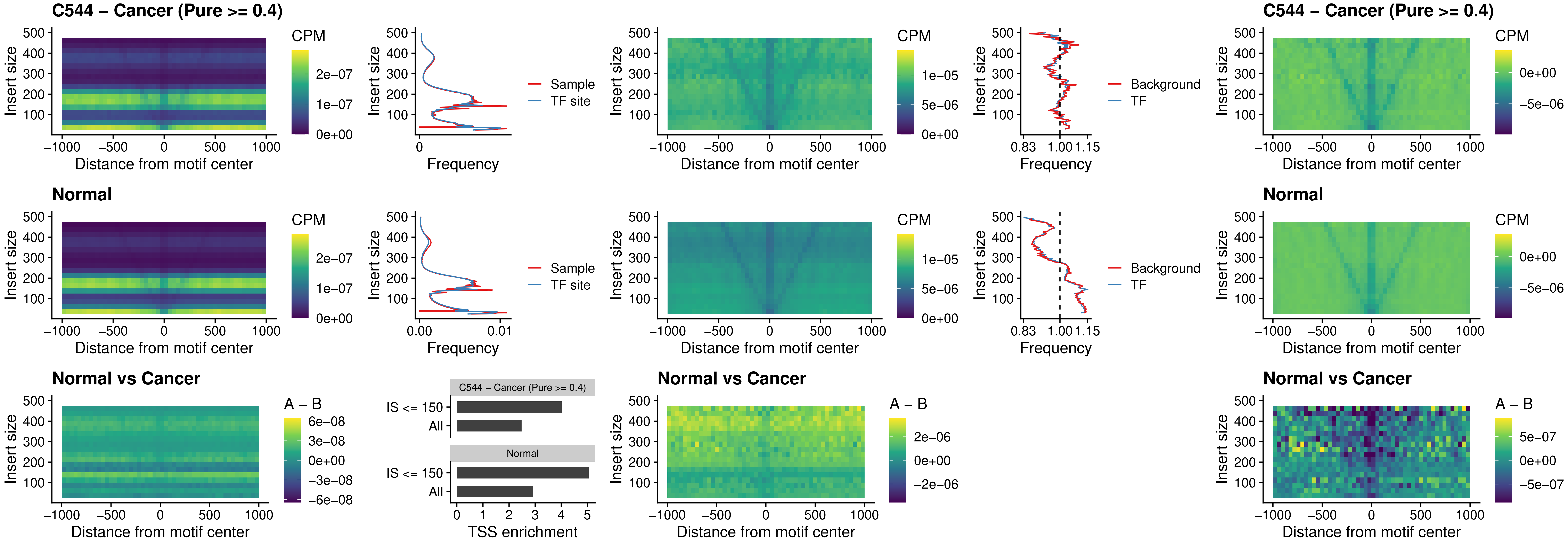

### C544 – Cancer (Pure >= 0.4) – HOXD3_uc_dTSS_oPEAK

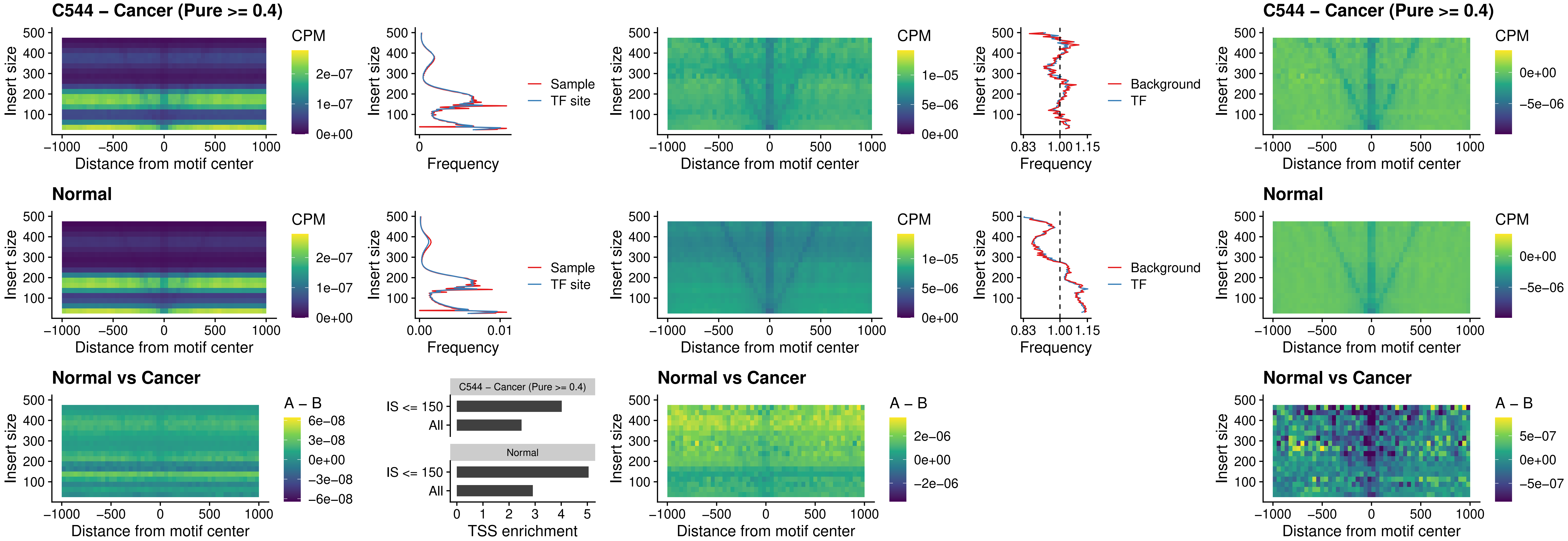

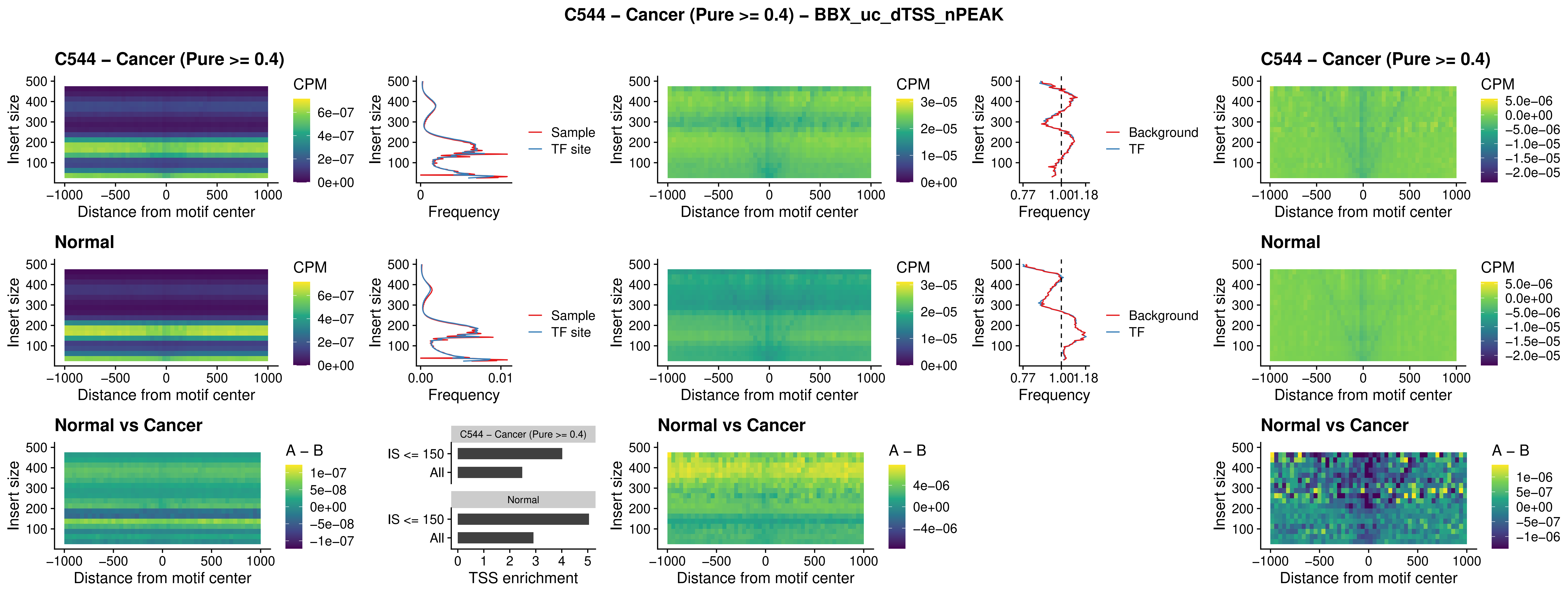
C544 – Cancer (Pure >= 0.4) – BBX_uc_dTSS_nPEAK
C544 – Cancer (Pure >= 0.4)
Normal
Normal vs Cancer
Insert size
Distance from motif center
Frequency
CPM
A – B
Sample
TF site
Background
TF
C544 – Cancer (Pure >= 0.4)
Normal
IS <= 150
All
TSS enrichment

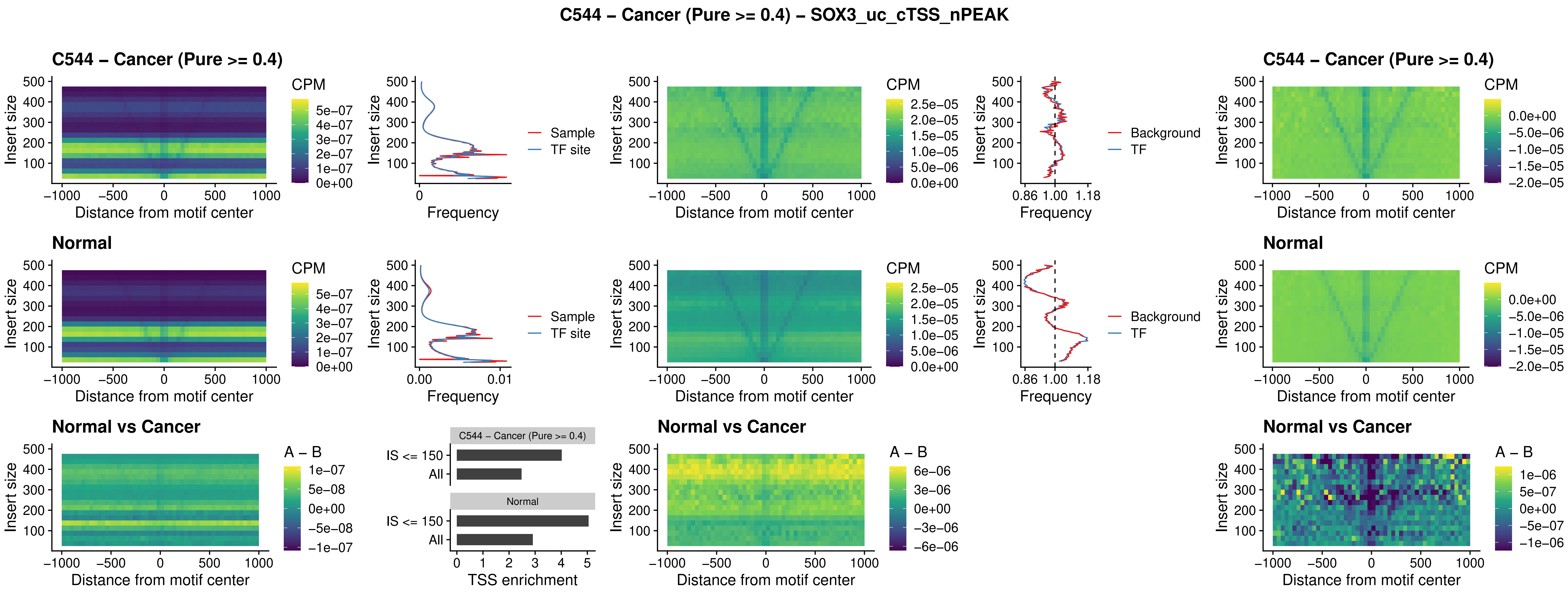
C544 – Cancer (Pure >= 0.4) – SOX3_uc_cTSS_nPEAK
C544 – Cancer (Pure >= 0.4)
Normal
Normal vs Cancer
Insert size
Distance from motif center
Frequency
CPM
A – B
Sample
TF site
Background
TF
C544 – Cancer (Pure >= 0.4)
Normal
IS <= 150
All
TSS enrichment

#### C544 – Cancer (Pure >= 0.4) – TBPL2_uc_cTSS_nPEAK

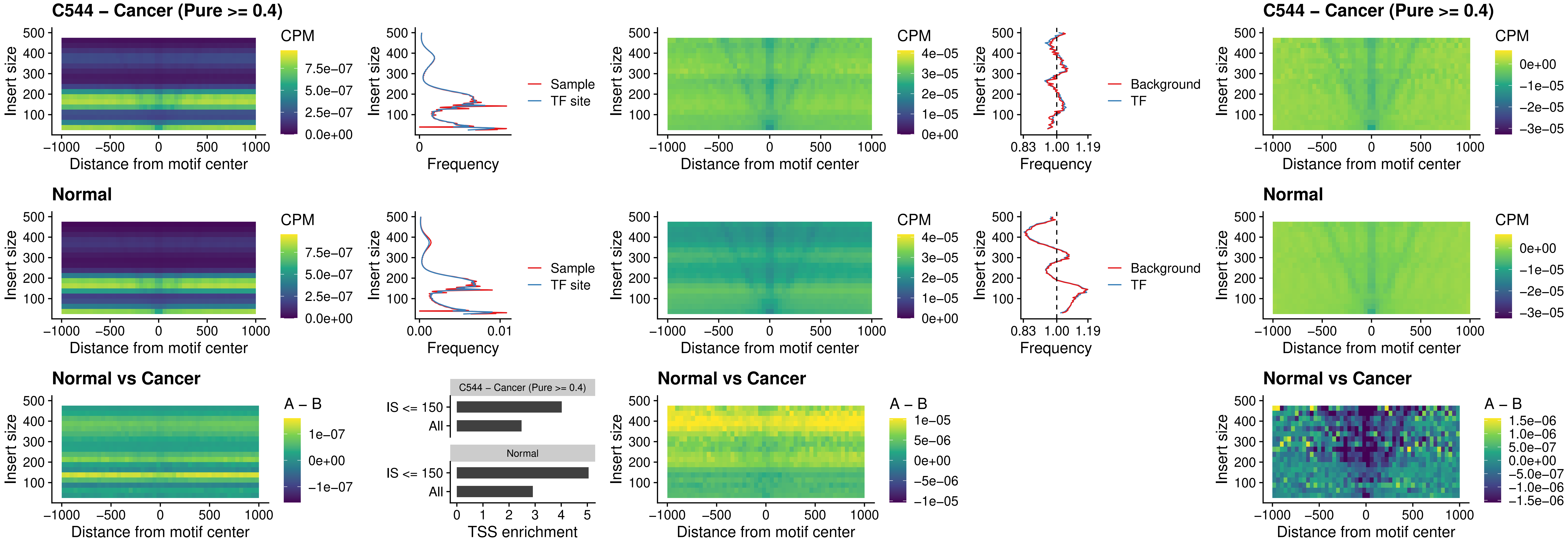

#### C544 – Cancer (Pure >= 0.4) – HOXA4_uc_cTSS_oPEAK

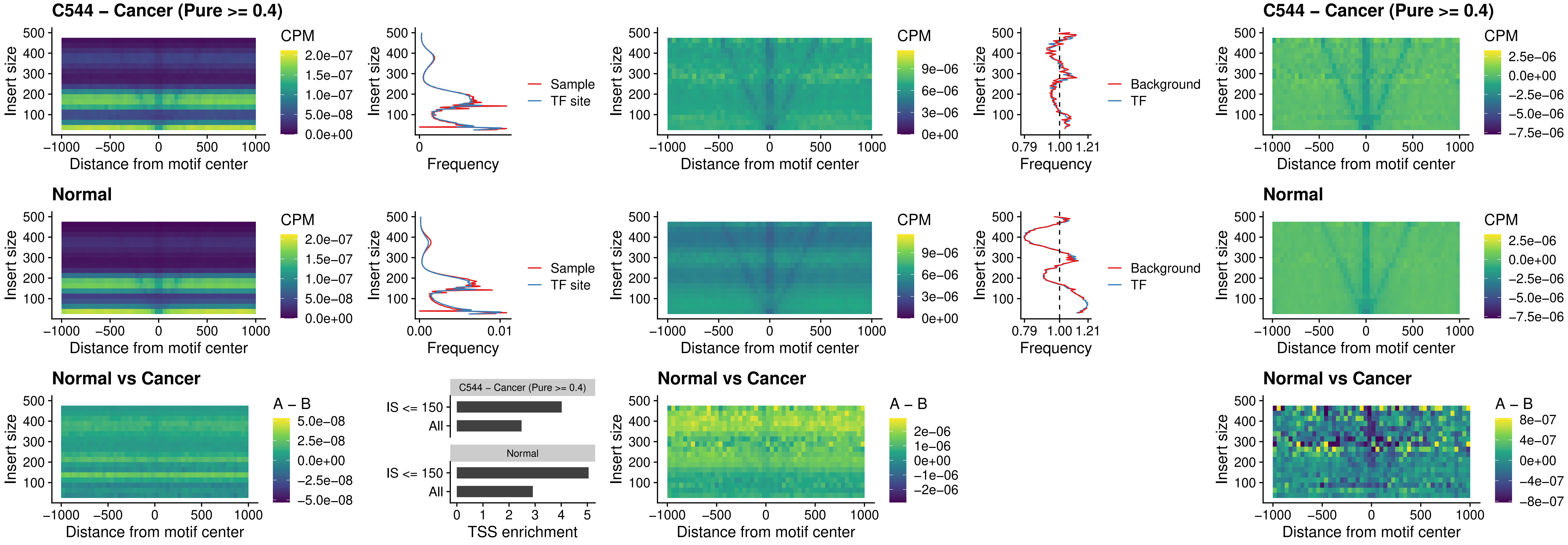

#### C544 – Cancer (Pure >= 0.4) – PDX1_uc_dTSS_oPEAK

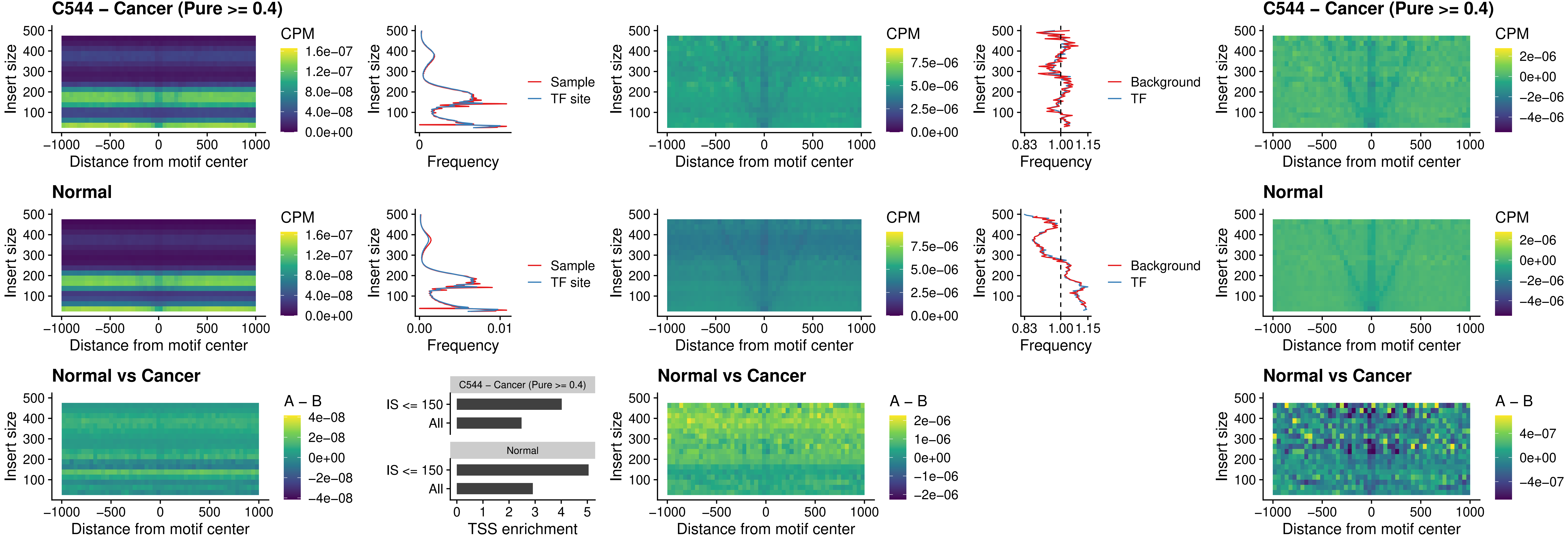

#### C544 – Cancer (Pure >= 0.4) – LHX4_uc_dTSS_oPEAK

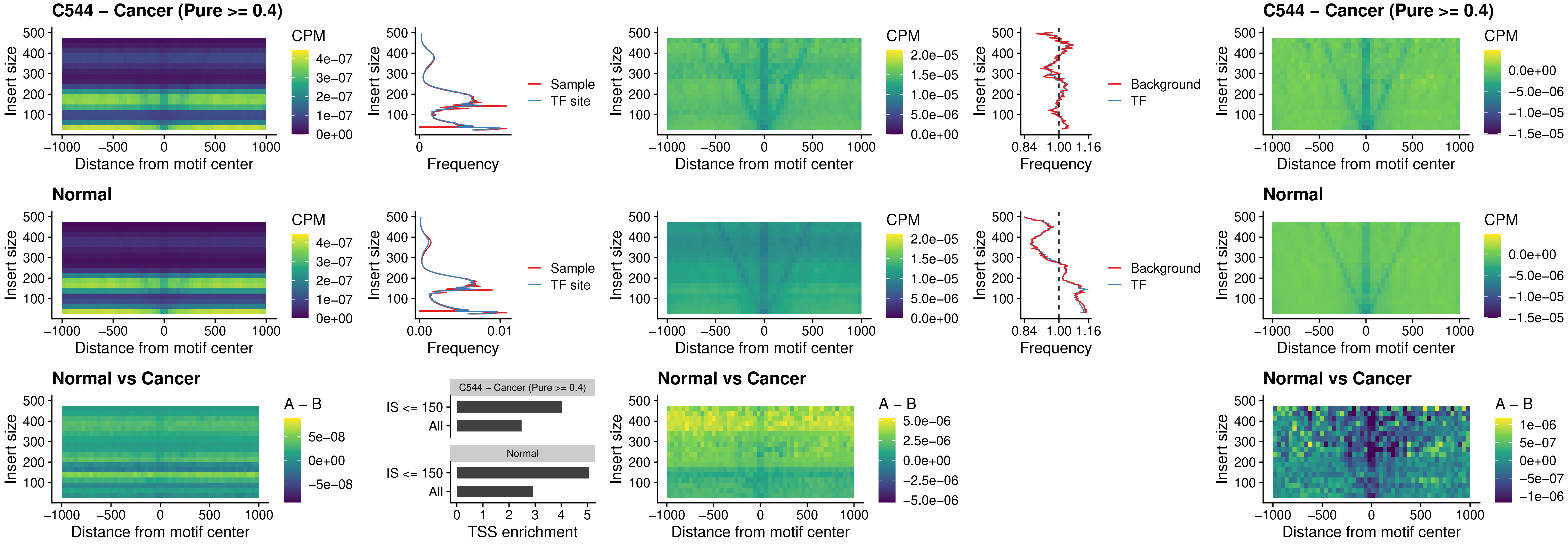

#### C544 – Cancer (Pure >= 0.4) – HOXA4_uc_dTSS_oPEAK

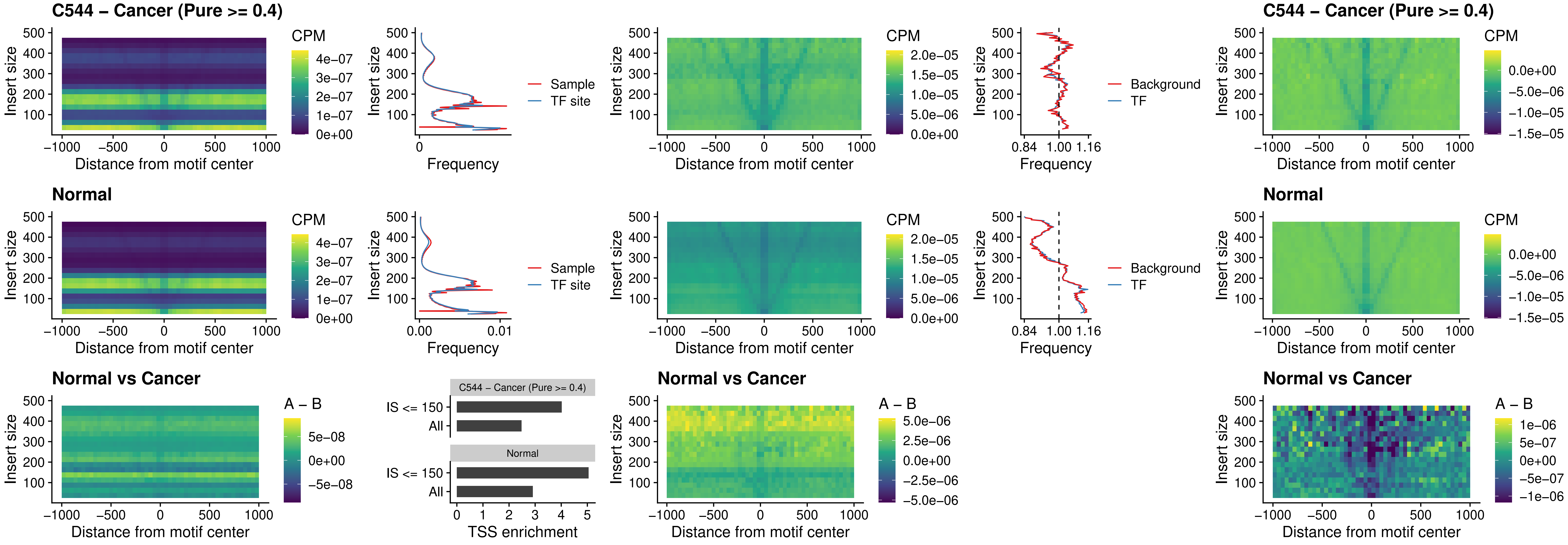

#### C544 – Cancer (Pure >= 0.4) – HLX_uc_dTSS_oPEAK

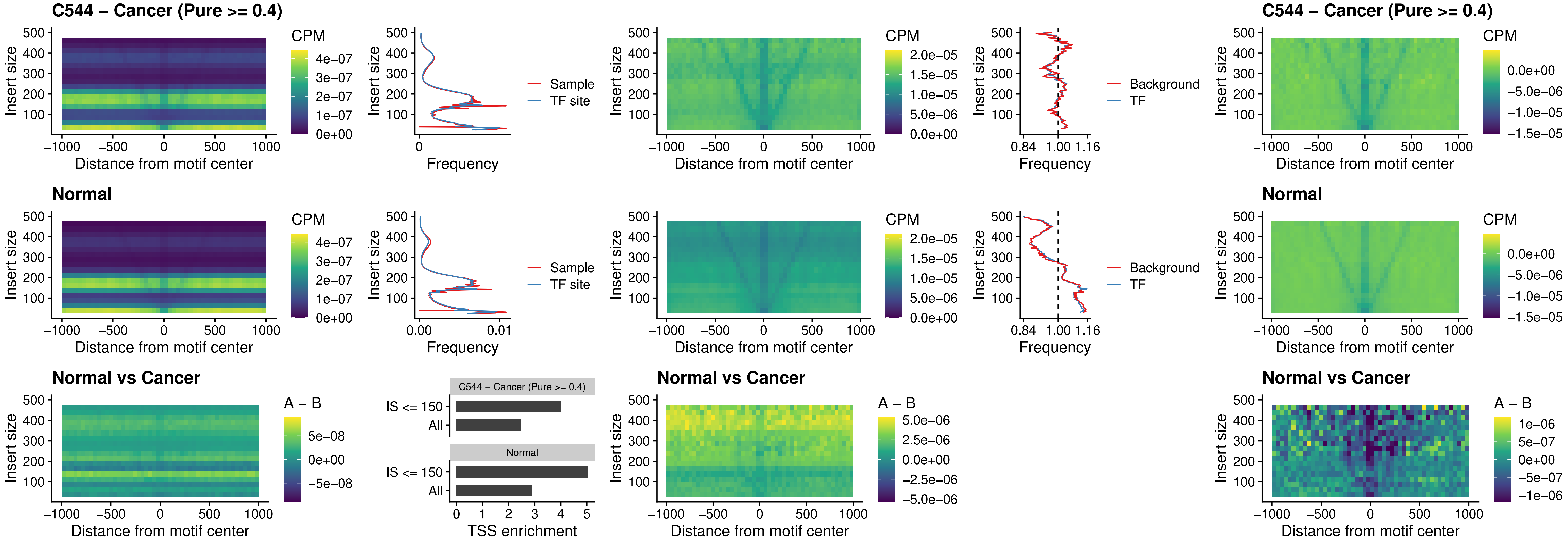

#### C544 – Cancer (Pure >= 0.4) – AC0125311_uc_dTSS_oPEAK

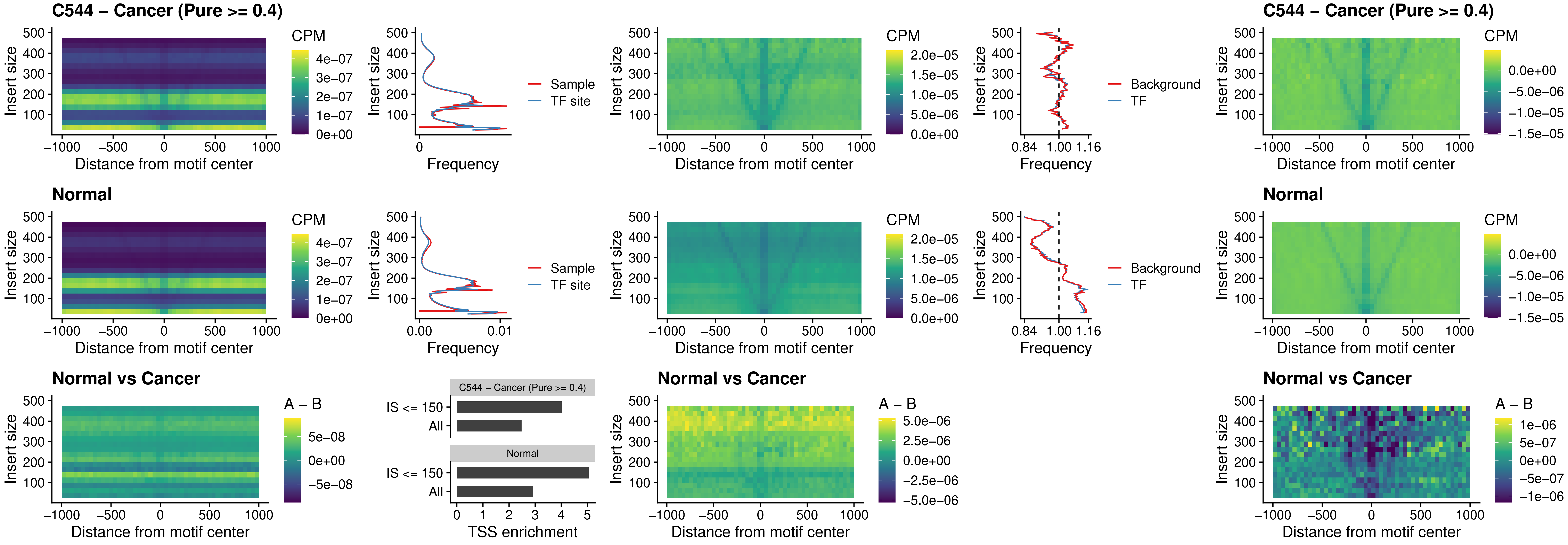

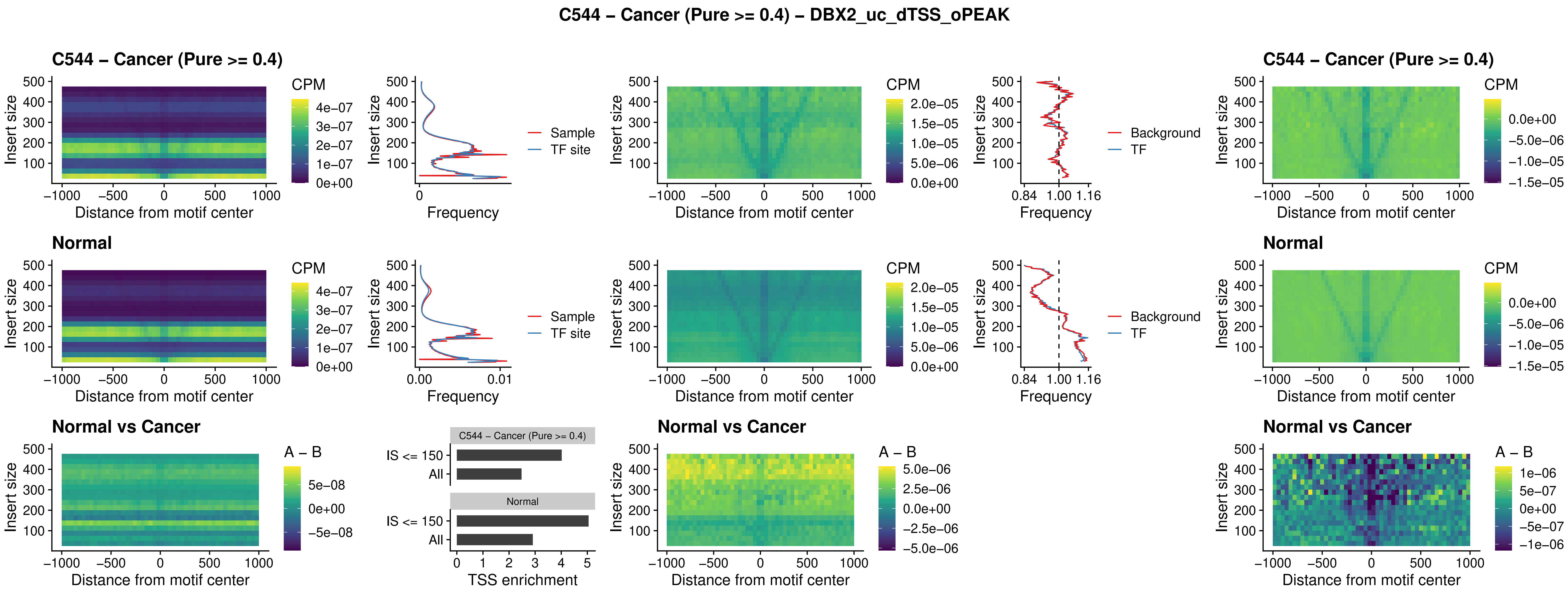
C544 – Cancer (Pure >= 0.4) – DBX2_uc_dTSS_oPEAK
C544 – Cancer (Pure >= 0.4)
Normal
Normal vs Cancer
Insert size
Distance from motif center
CPM
A – B
Frequency
Sample
TF site
Background
TF
C544 – Cancer (Pure >= 0.4)
IS <= 150
All
TSS enrichment

#### C544 – Cancer (Pure >= 0.4) – HOXA2_uc_dTSS_nPEAK

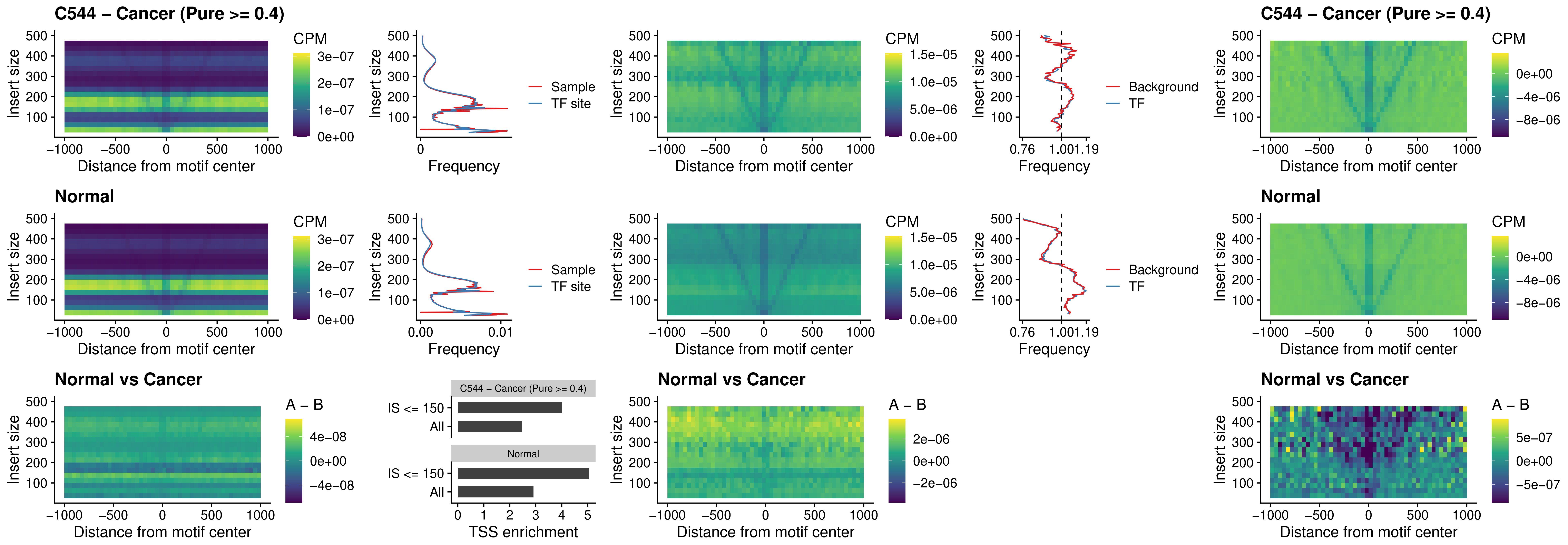

### C544 – Cancer (Pure >= 0.4) – SHOX_uc_dTSS_nPEAK

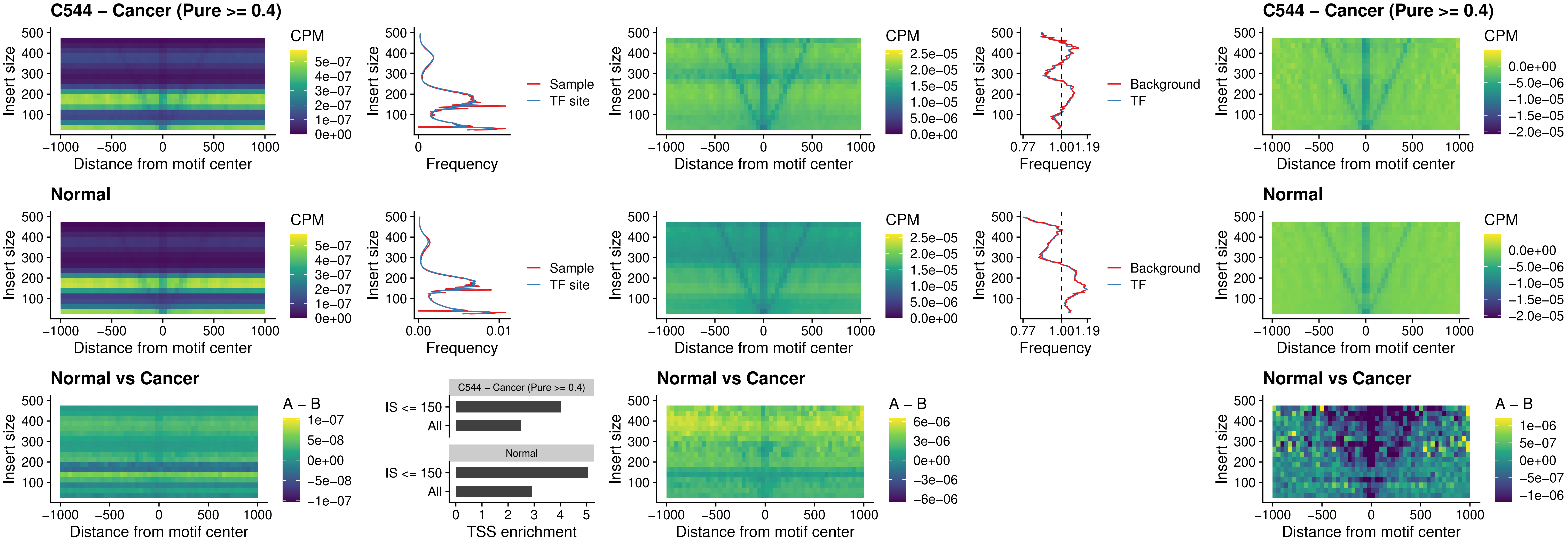

#### C544 – Cancer (Pure >= 0.4) – PRRX1_uc_dTSS_nPEAK

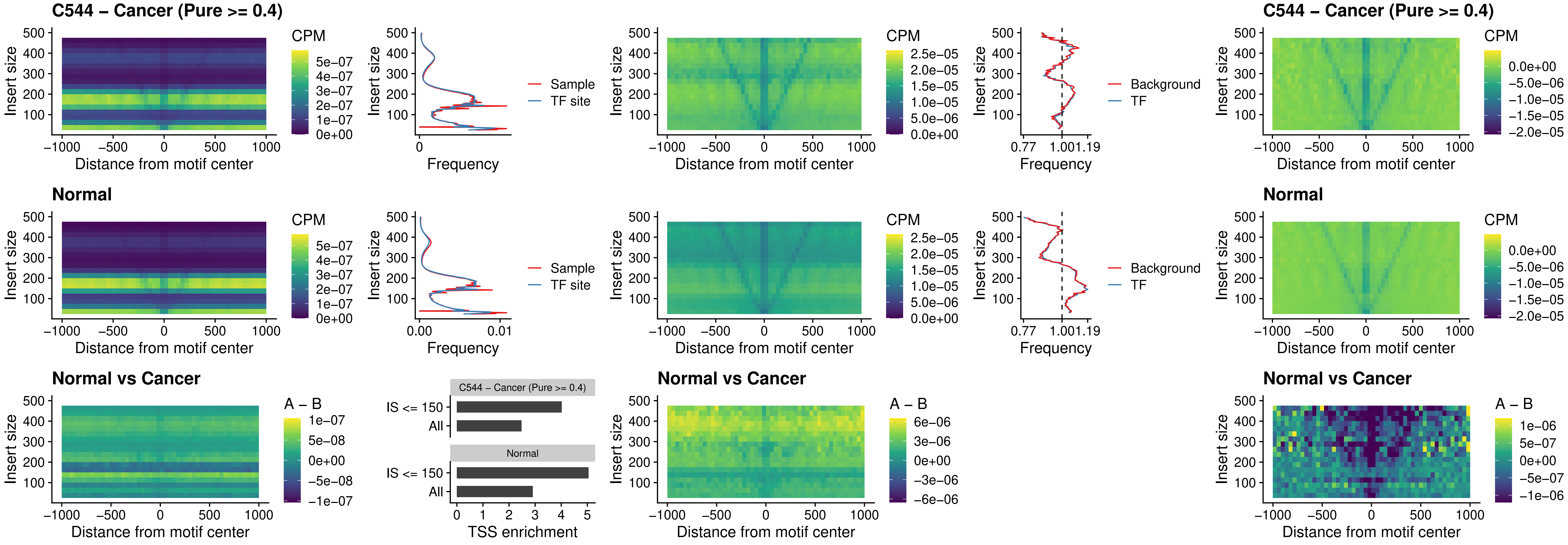

#### C544 – Cancer (Pure >= 0.4) – LMX1A_uc_dTSS_nPEAK

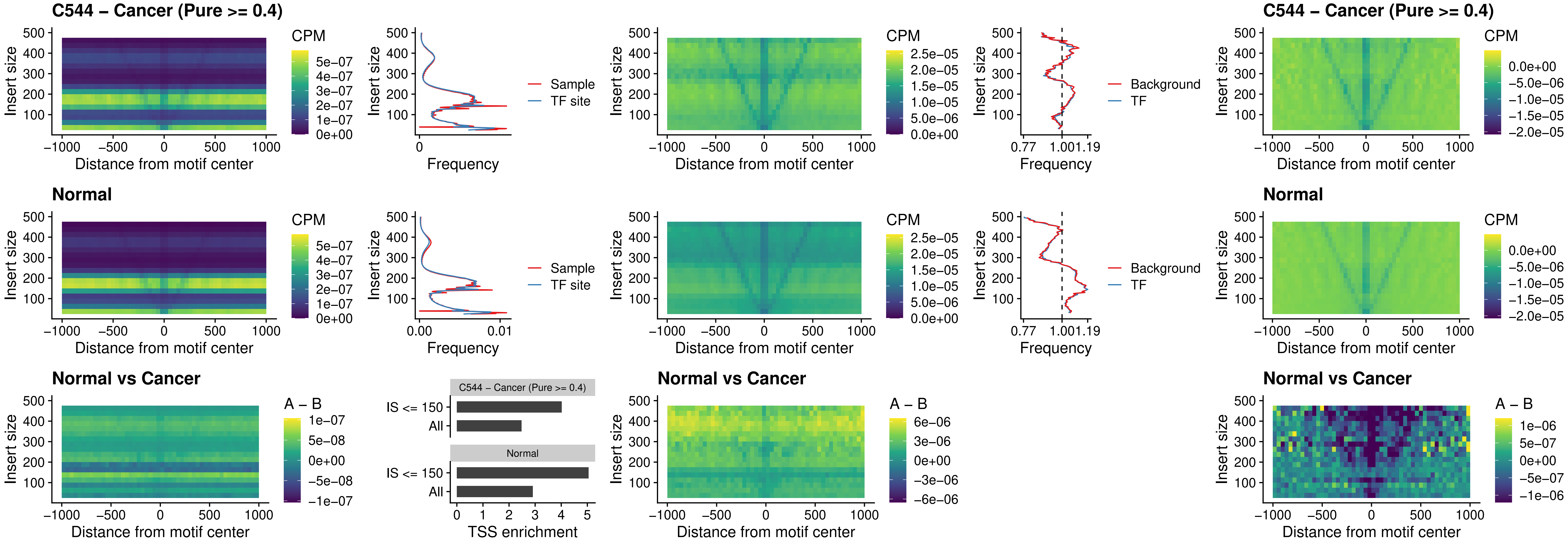

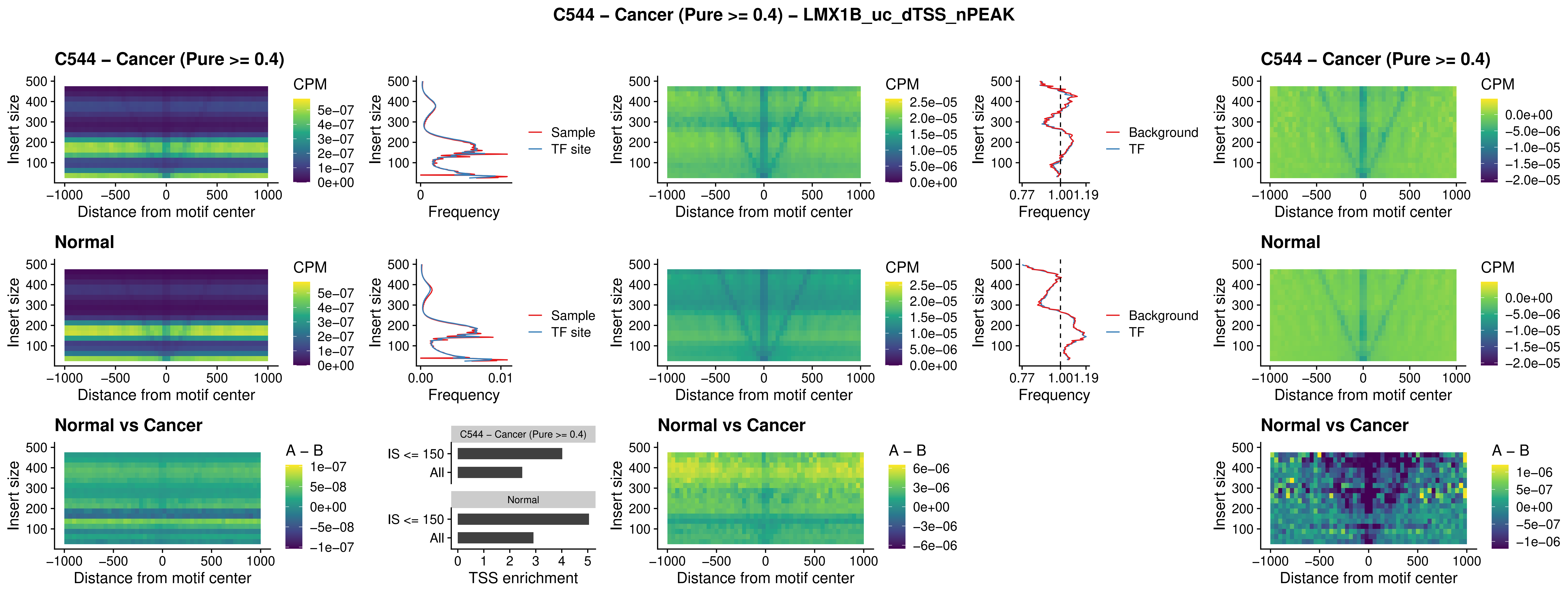

C544 – Cancer (Pure >= 0.4) – LMX1B_uc_dTSS_nPEAK
C544 – Cancer (Pure >= 0.4)
Normal
Normal vs Cancer
Insert size
Distance from motif center
Frequency
CPM
A – B
Sample
TF site
Background
TF
C544 – Cancer (Pure >= 0.4)
Normal
IS <= 150
All
TSS enrichment

### C544 – Cancer (Pure >= 0.4) – DLX5_uc_dTSS_oPEAK

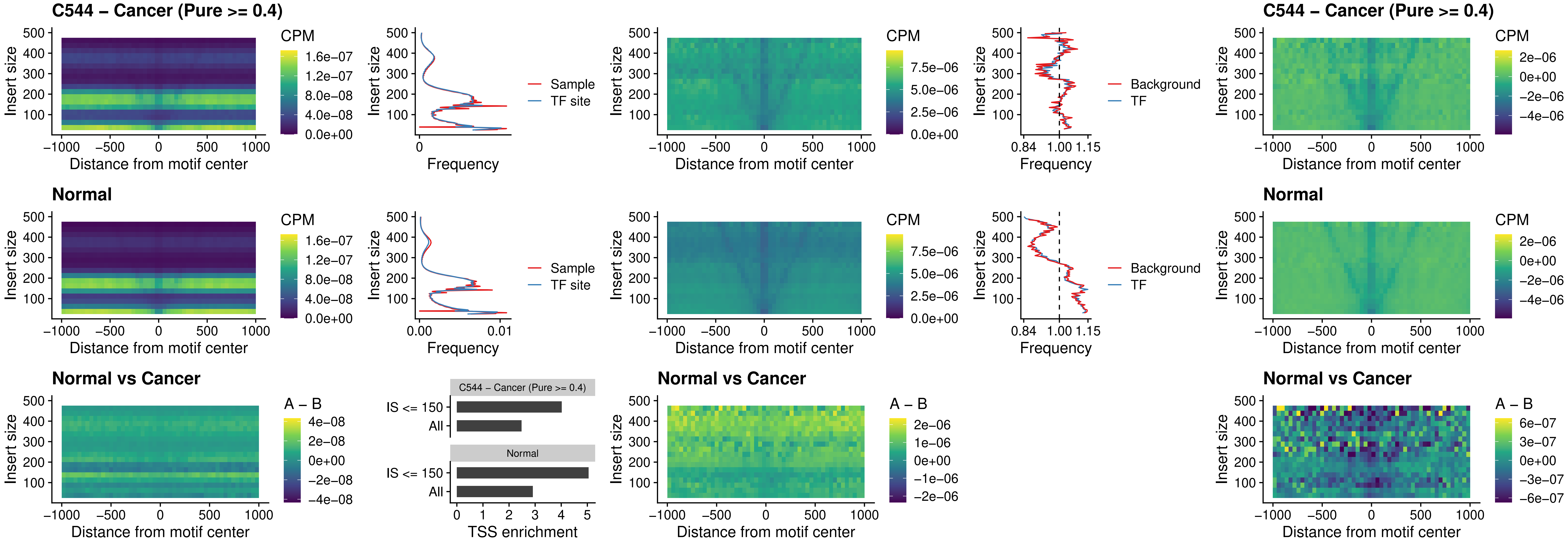

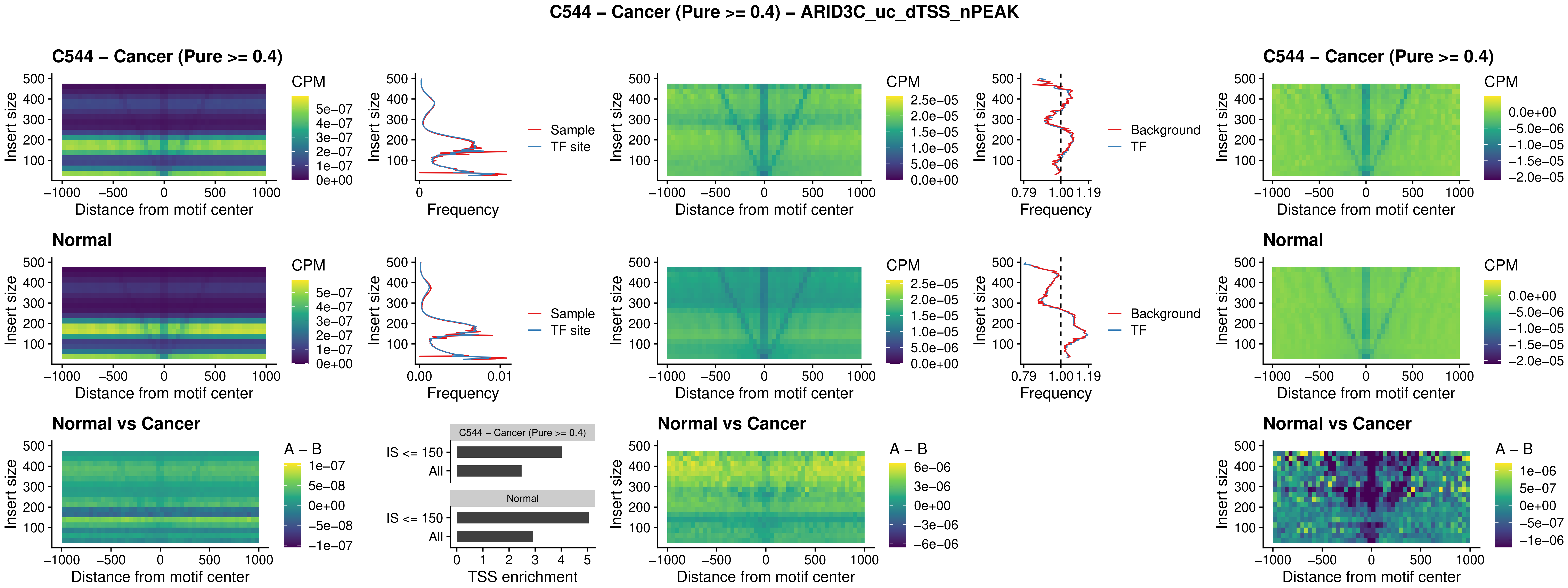
C544 – Cancer (Pure >= 0.4) – ARID3C_uc_dTSS_nPEAK
C544 – Cancer (Pure >= 0.4)
Normal
Normal vs Cancer
Insert size
Distance from motif center
Frequency
CPM
A – B
Sample
TF site
Background
TF
C544 – Cancer (Pure >= 0.4)
Normal
IS <= 150
All
TSS enrichment

#### C544 – Cancer (Pure >= 0.4) – HOXD12_uc_dTSS_nPEAK

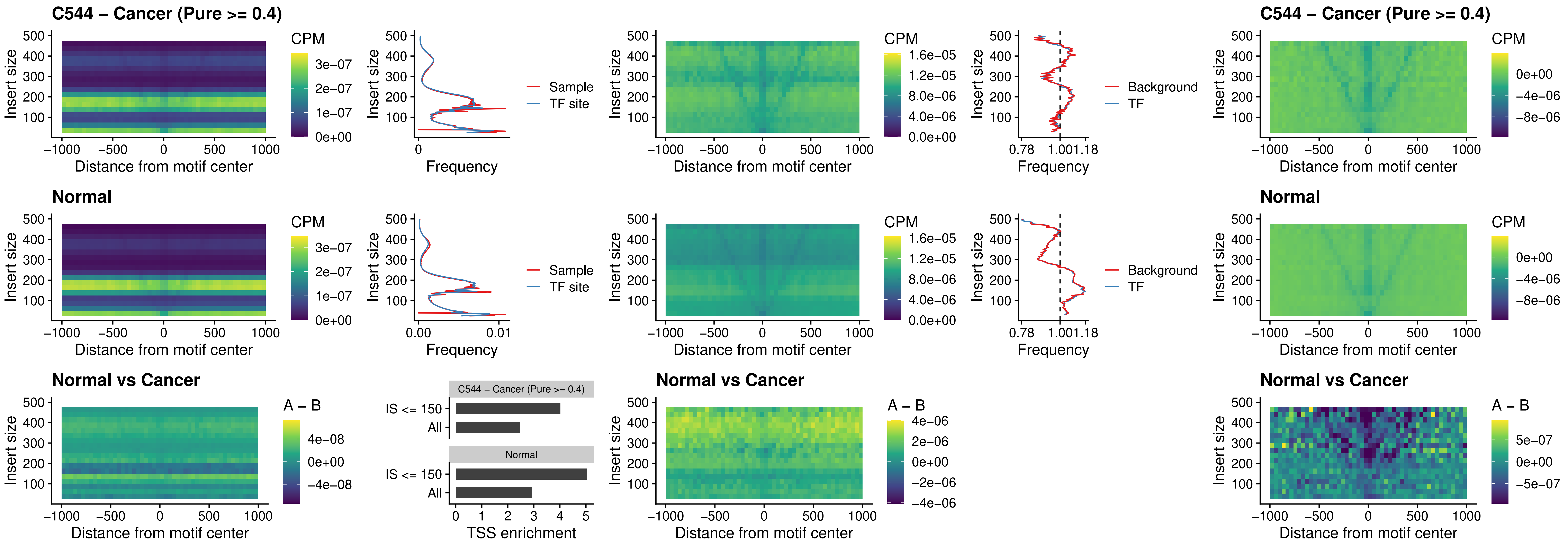

#### C544 – Cancer (Pure >= 0.4) – CDX2_uc_dTSS_nPEAK

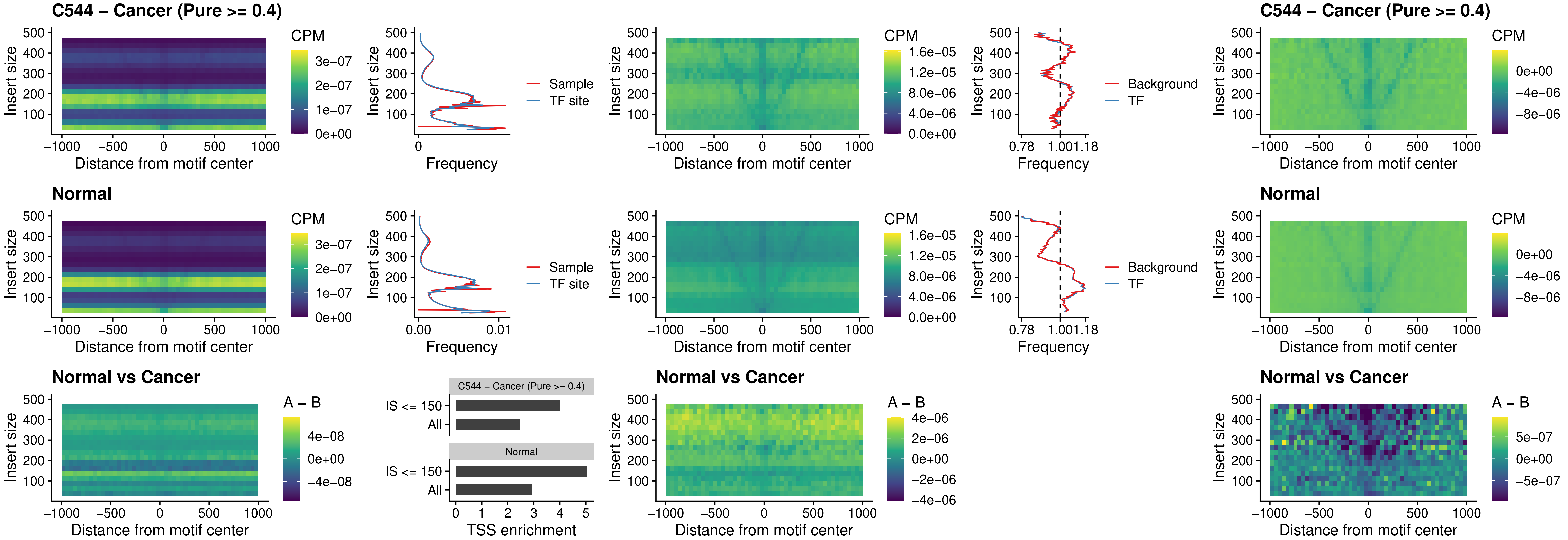

#### C544 – Cancer (Pure >= 0.4) – HOXC12_uc_dTSS_nPEAK

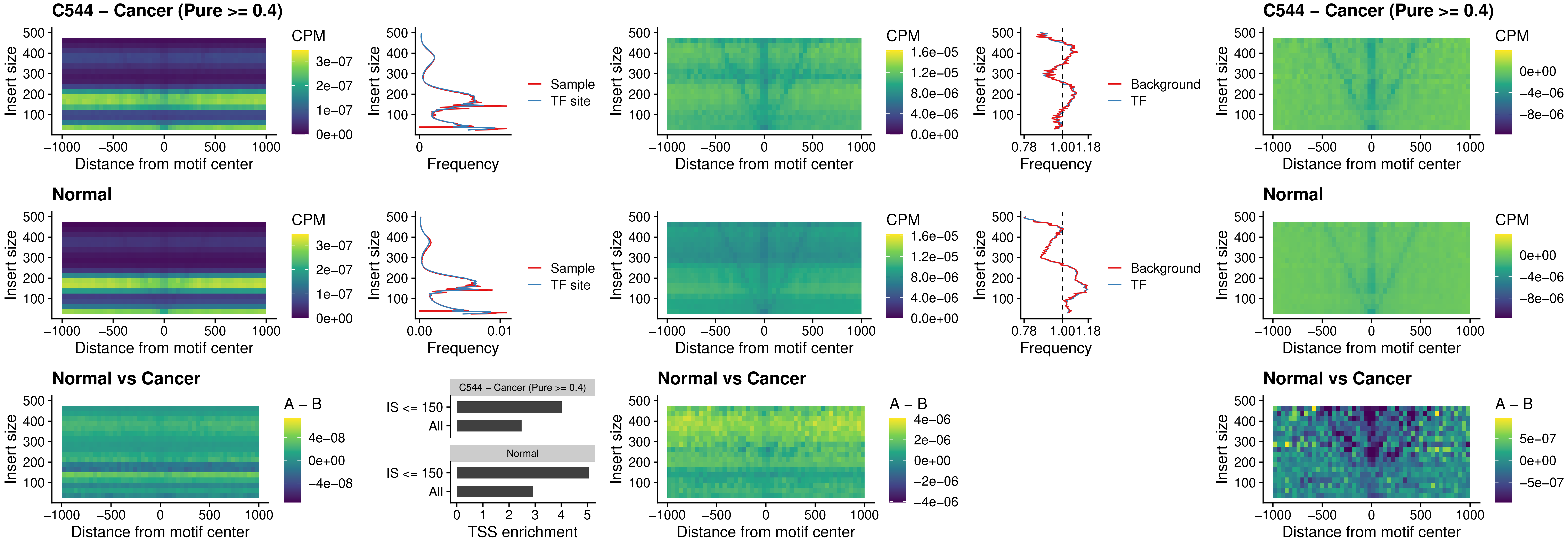

#### C544 – Cancer (Pure >= 0.4) – HHEX_uc_dTSS_oPEAK

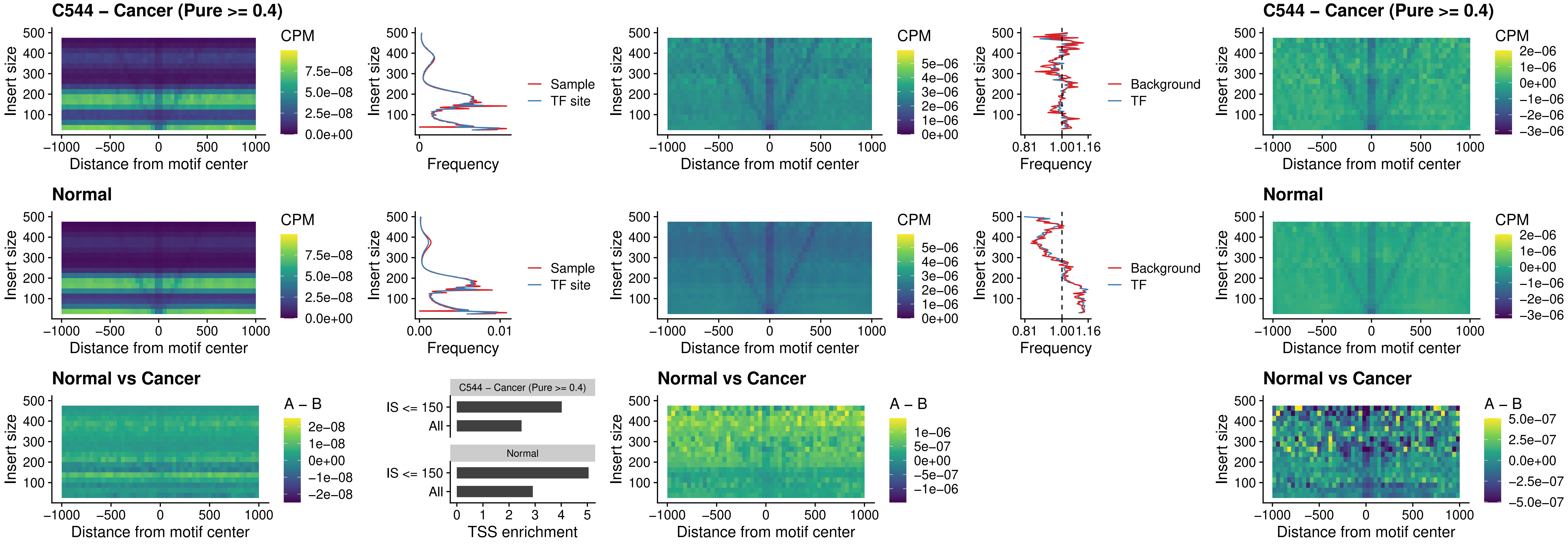

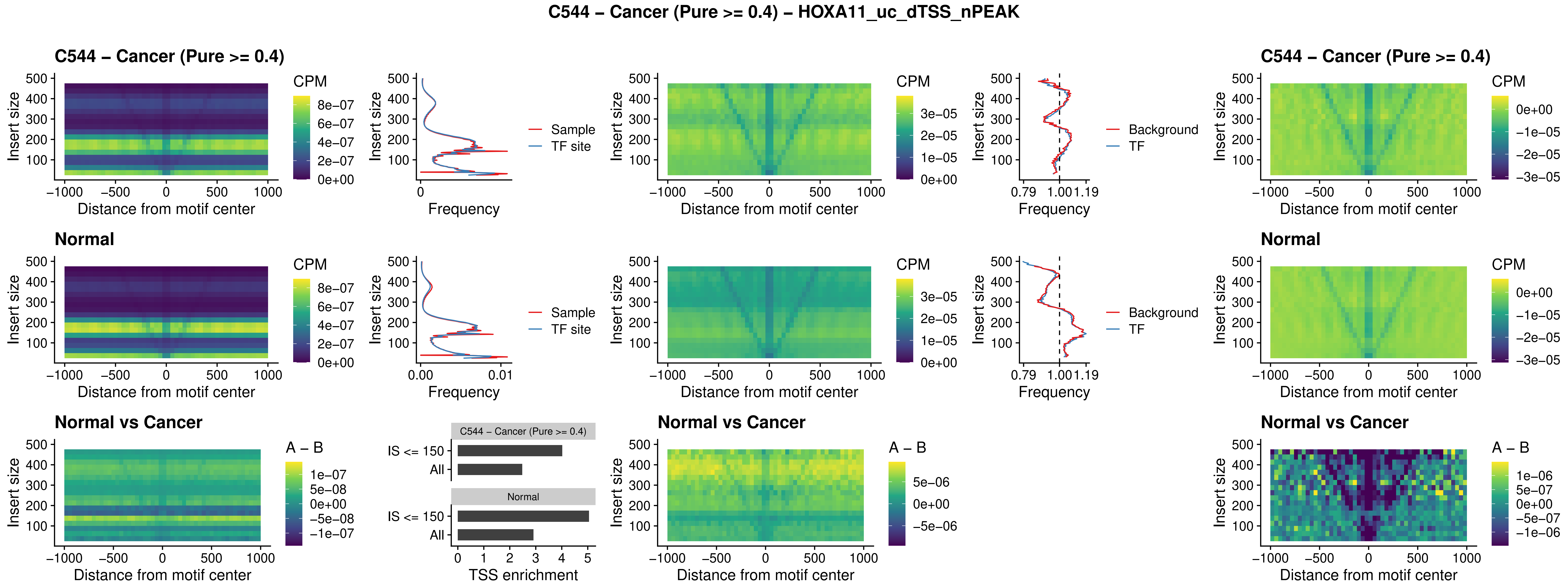

C544 – Cancer (Pure >= 0.4) – HOXA11_uc_dTSS_nPEAK
C544 – Cancer (Pure >= 0.4)
Normal
Normal vs Cancer
Insert size
Distance from motif center
Frequency
CPM
A – B
Sample
TF site
Background
TF
C544 – Cancer (Pure >= 0.4)
Normal
IS <= 150
All
TSS enrichment

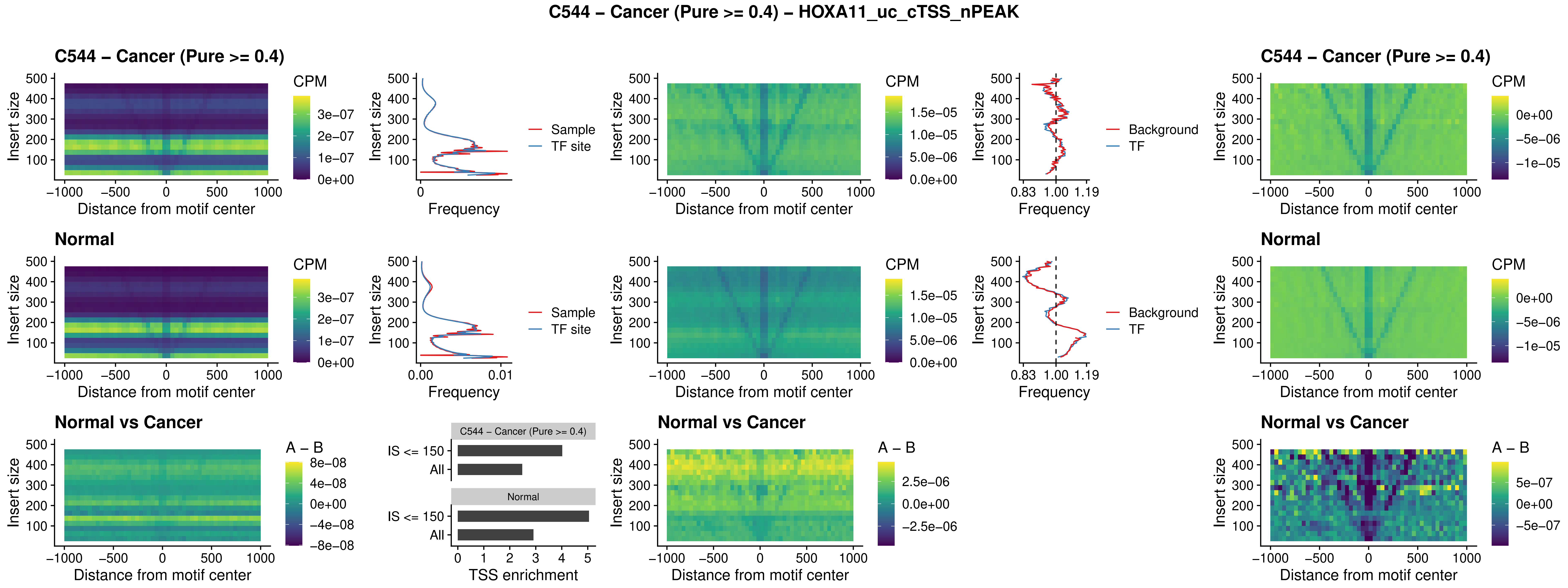
C544 – Cancer (Pure >= 0.4) – HOXA11_uc_cTSS_nPEAK
C544 – Cancer (Pure >= 0.4)
Normal
Normal vs Cancer
Insert size
Distance from motif center
Frequency
CPM
A – B
Sample
TF site
Background
TF
C544 – Cancer (Pure >= 0.4)
Normal
IS <= 150
All
TSS enrichment

#### C544 – Cancer (Pure >= 0.4) – ZFHX3_uc_dTSS_nPEAK

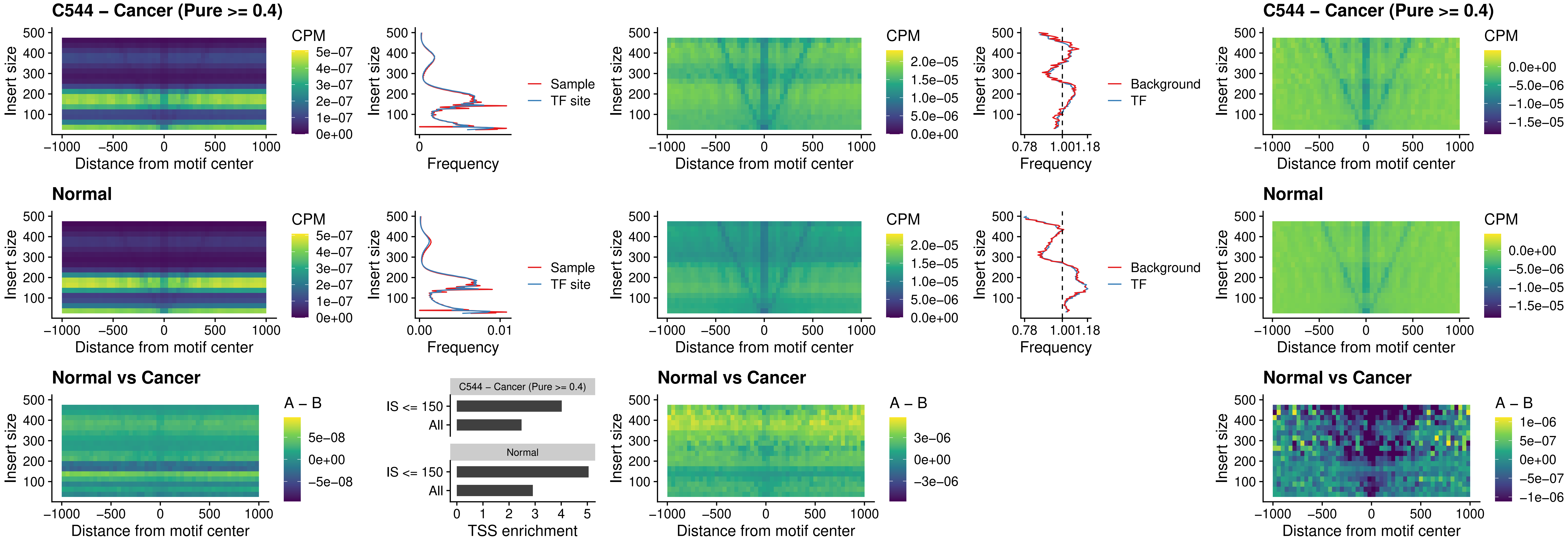

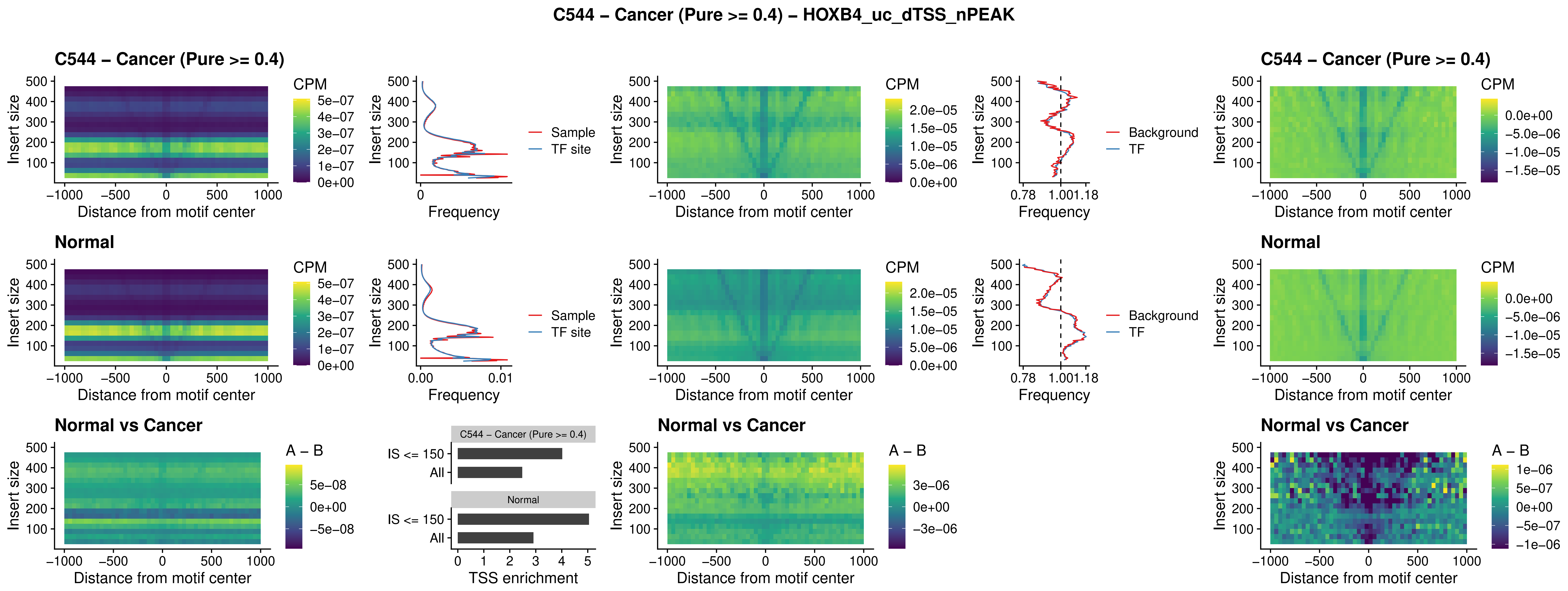
C544 – Cancer (Pure >= 0.4) – HOXB4_uc_dTSS_nPEAK
C544 – Cancer (Pure >= 0.4)
Normal
Normal vs Cancer
Insert size
Distance from motif center
Frequency
CPM
A – B
Sample
TF site
Background
TF
C544 – Cancer (Pure >= 0.4)
Normal
IS <= 150
All
TSS enrichment

#### C544 – Cancer (Pure >= 0.4) – HOXC5_uc_dTSS_nPEAK

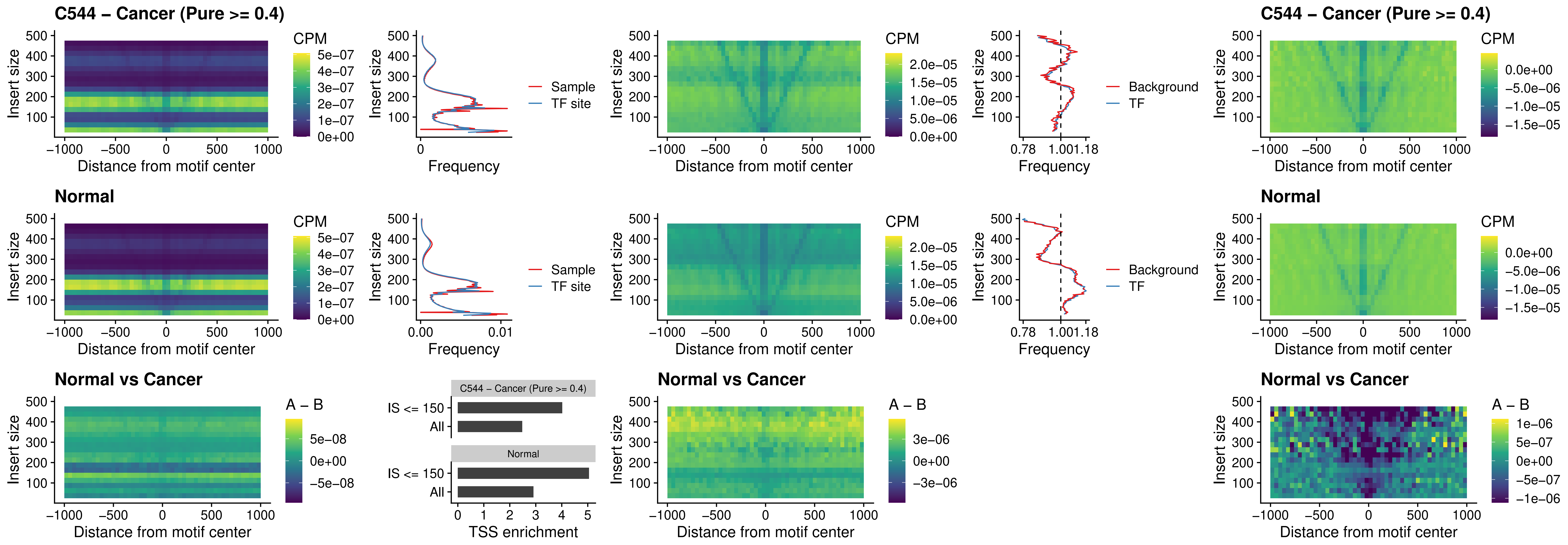

#### C544 – Cancer (Pure >= 0.4) – HOXD1_uc_dTSS_nPEAK

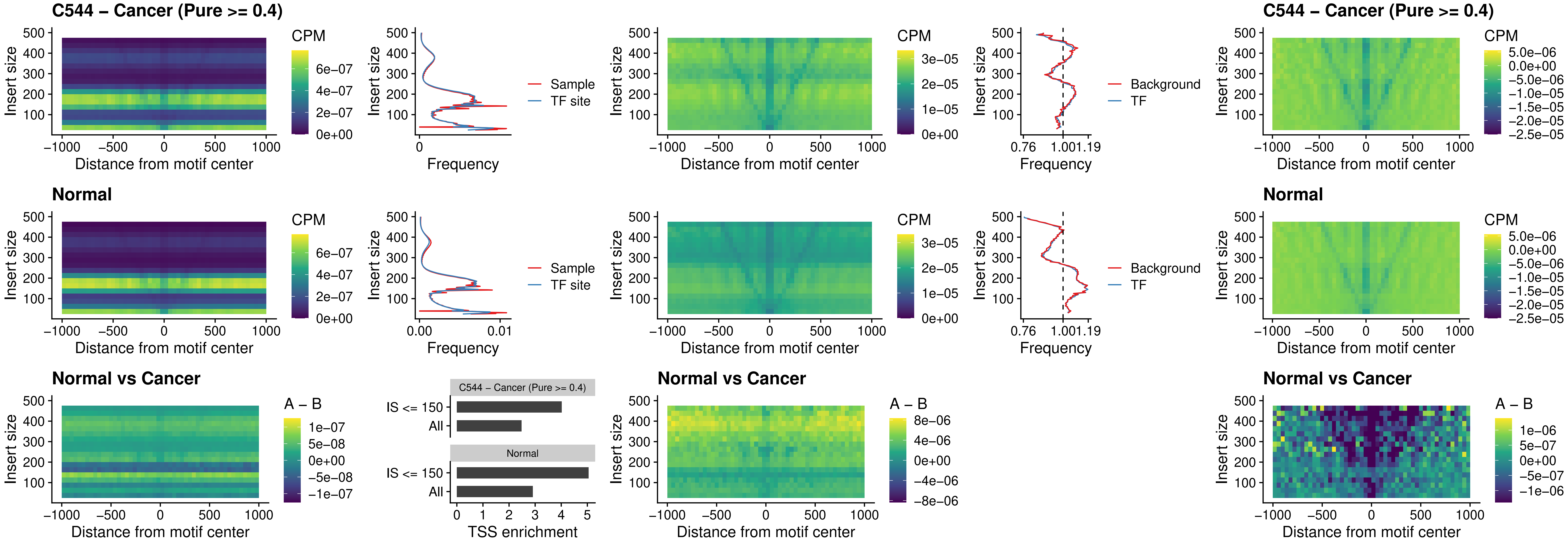

#### C544 – Cancer (Pure >= 0.4) – OTP_uc_dTSS_nPEAK

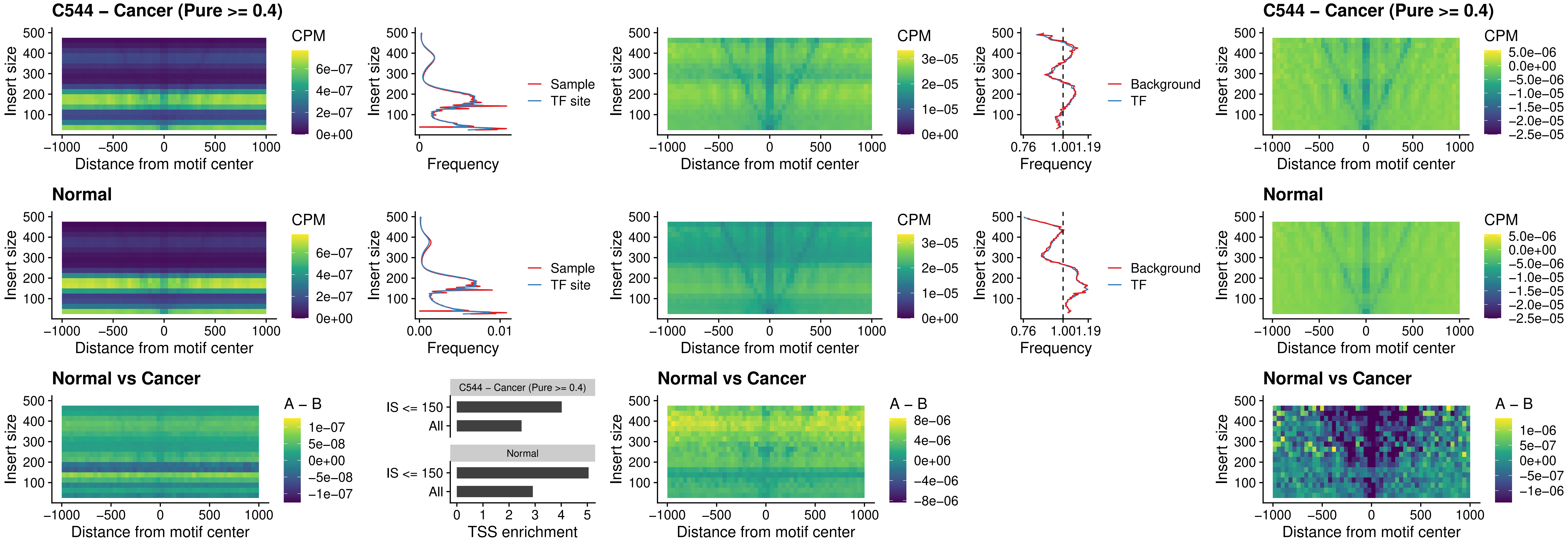

#### C544 – Cancer (Pure >= 0.4) – FOXL1_uc_dTSS_oPEAK

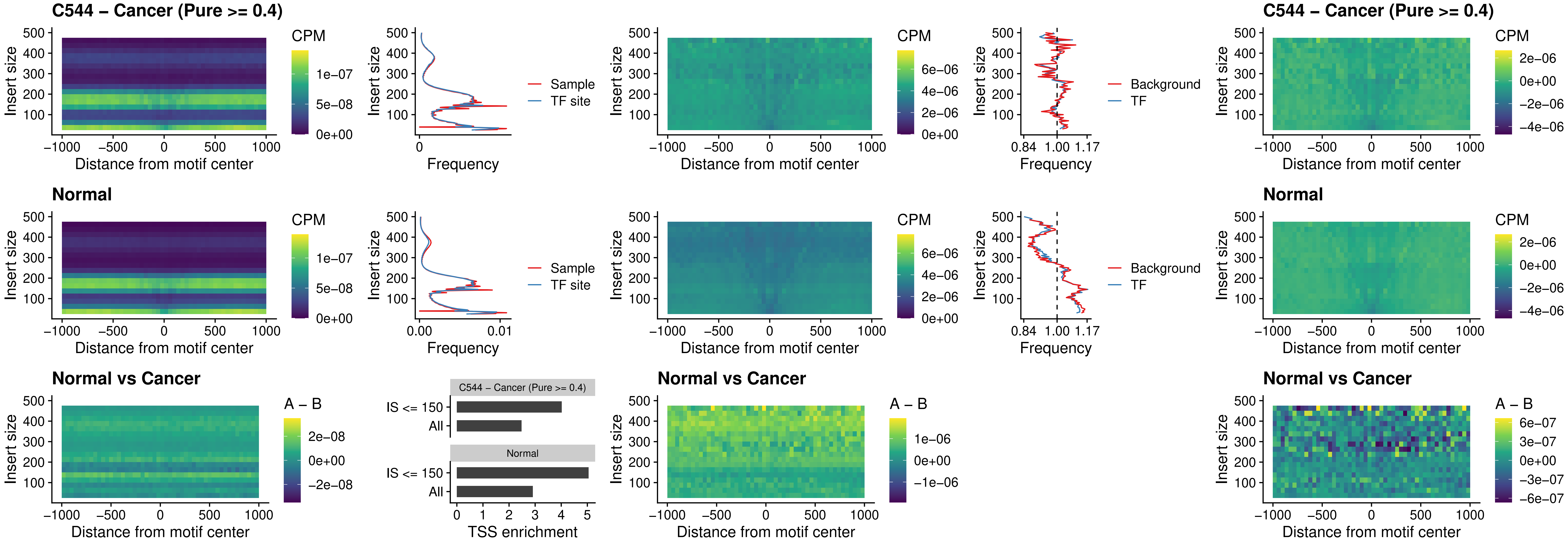

#### C544 – Cancer (Pure >= 0.4) – ZNF32_uc_dTSS_oPEAK

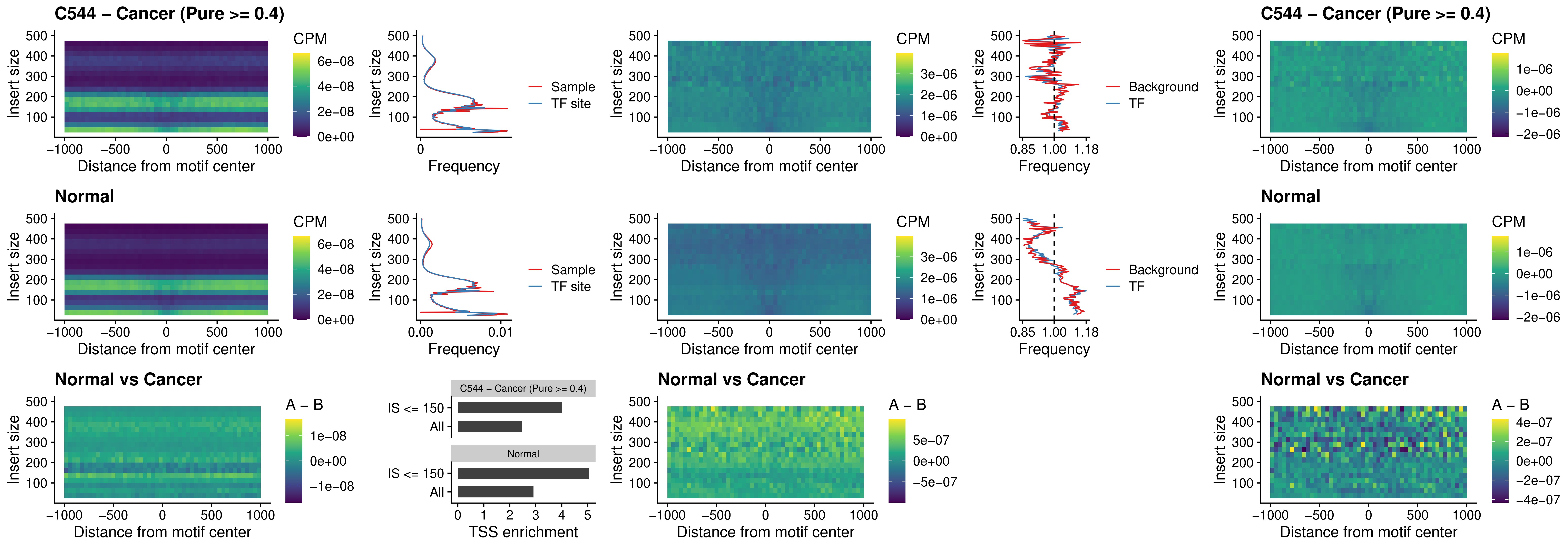

#### C544 – Cancer (Pure >= 0.4) – FOXL1_uc_dTSS_nPEAK

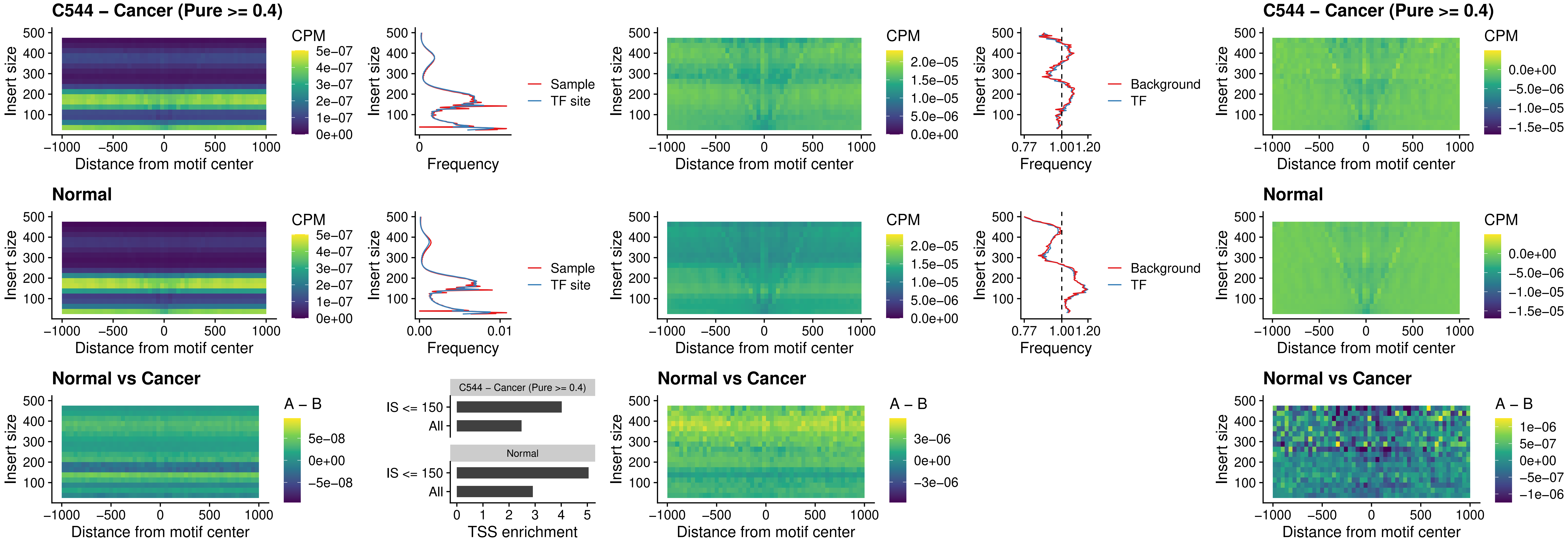

### C544 – Cancer (Pure >= 0.4) – ZNF32_uc_dTSS_nPEAK

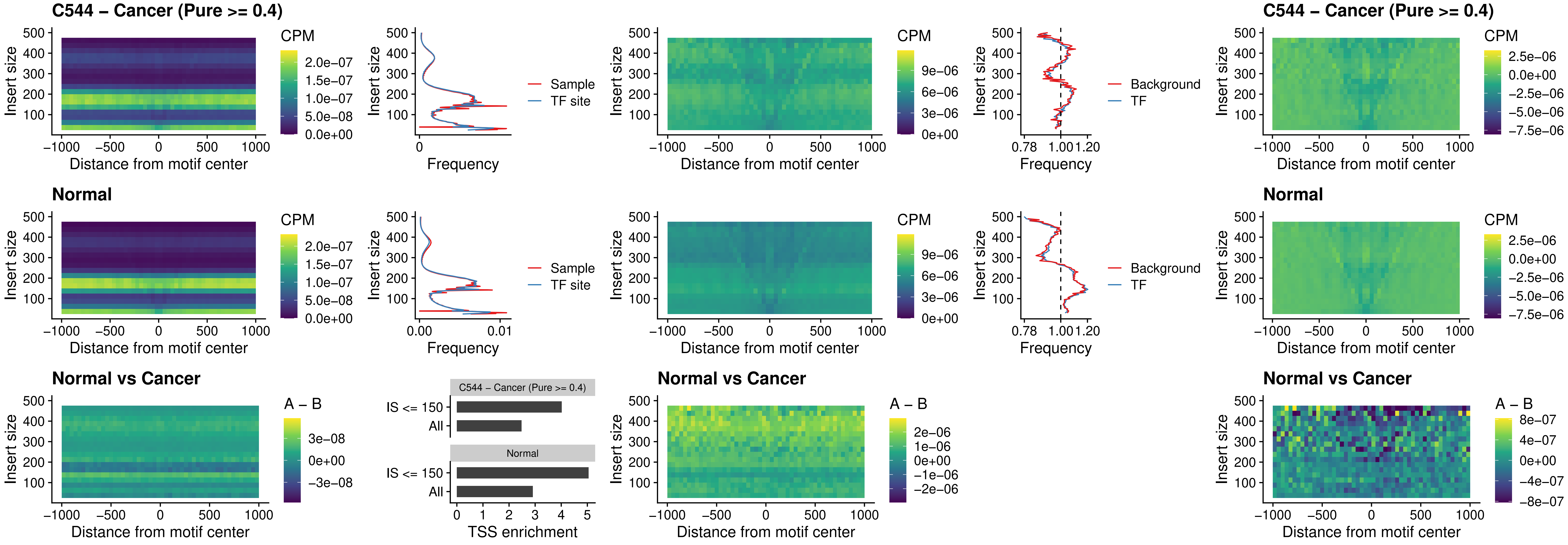

#### C544 – Cancer (Pure >= 0.4) – DLX5_uc_dTSS_nPEAK

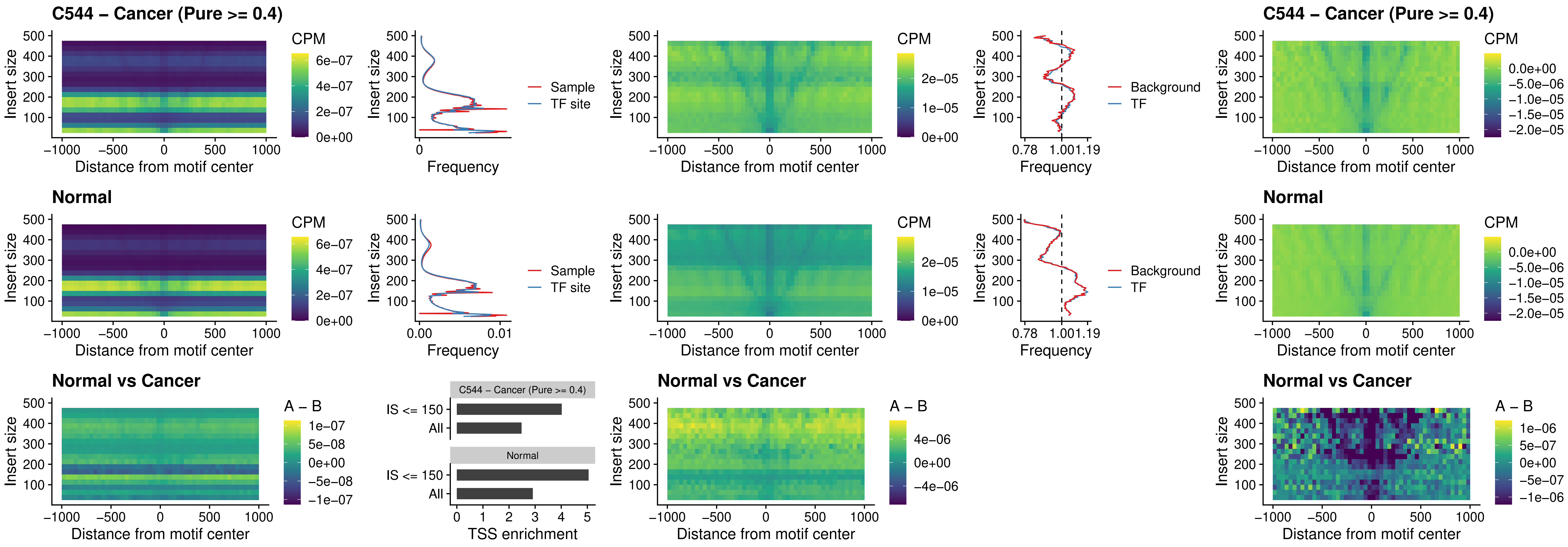

#### C544 – Cancer (Pure >= 0.4) – DMRT2_uc_dTSS_nPEAK

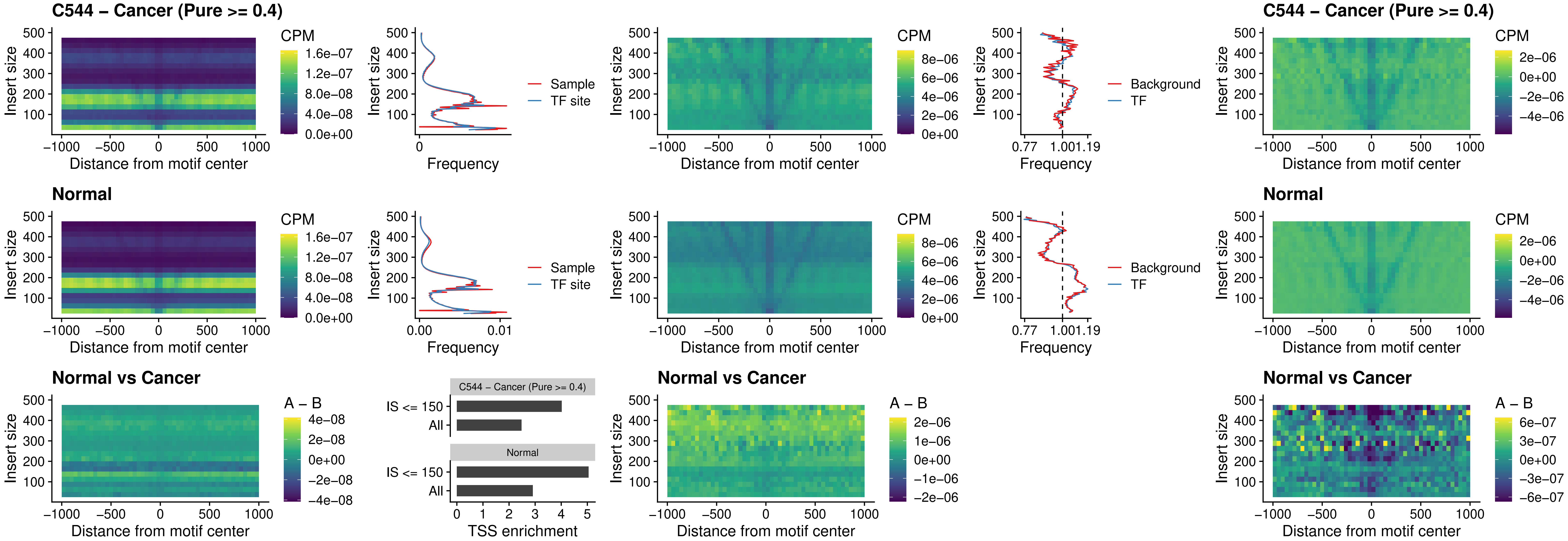

### C544 – Cancer (Pure >= 0.4) – DMRTC2_uc_cTSS_nPEAK

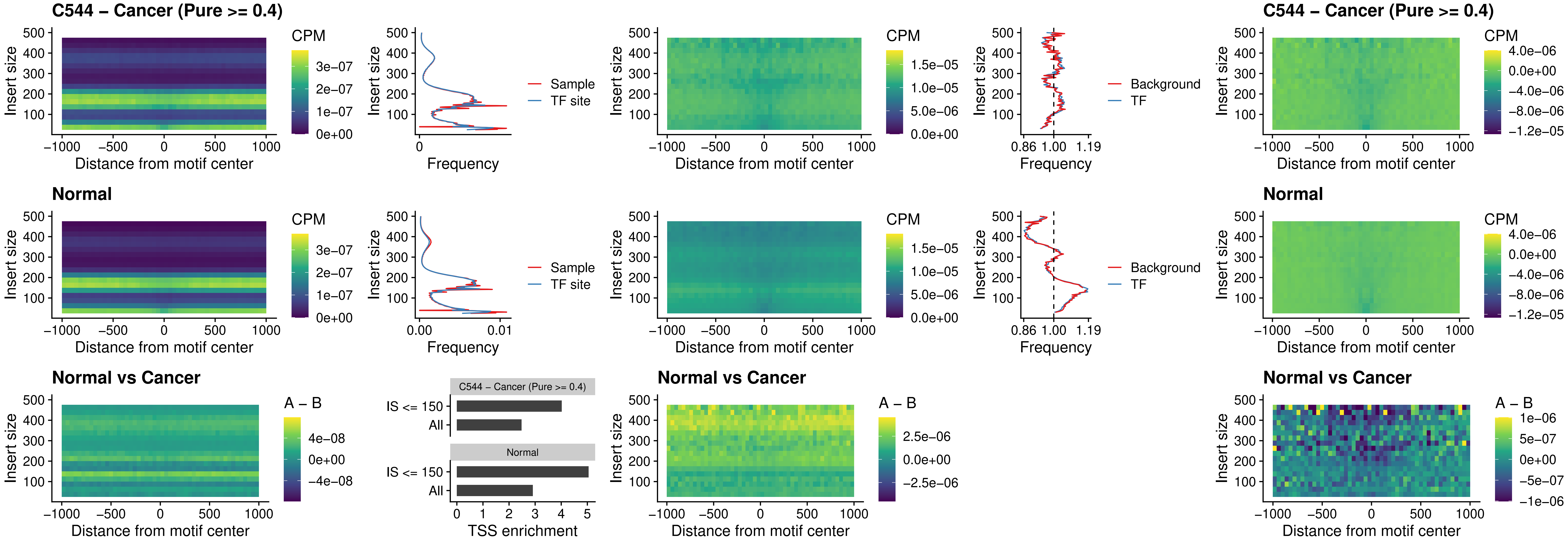

#### C544 – Cancer (Pure >= 0.4) – HOXD9_uc_dTSS_nPEAK

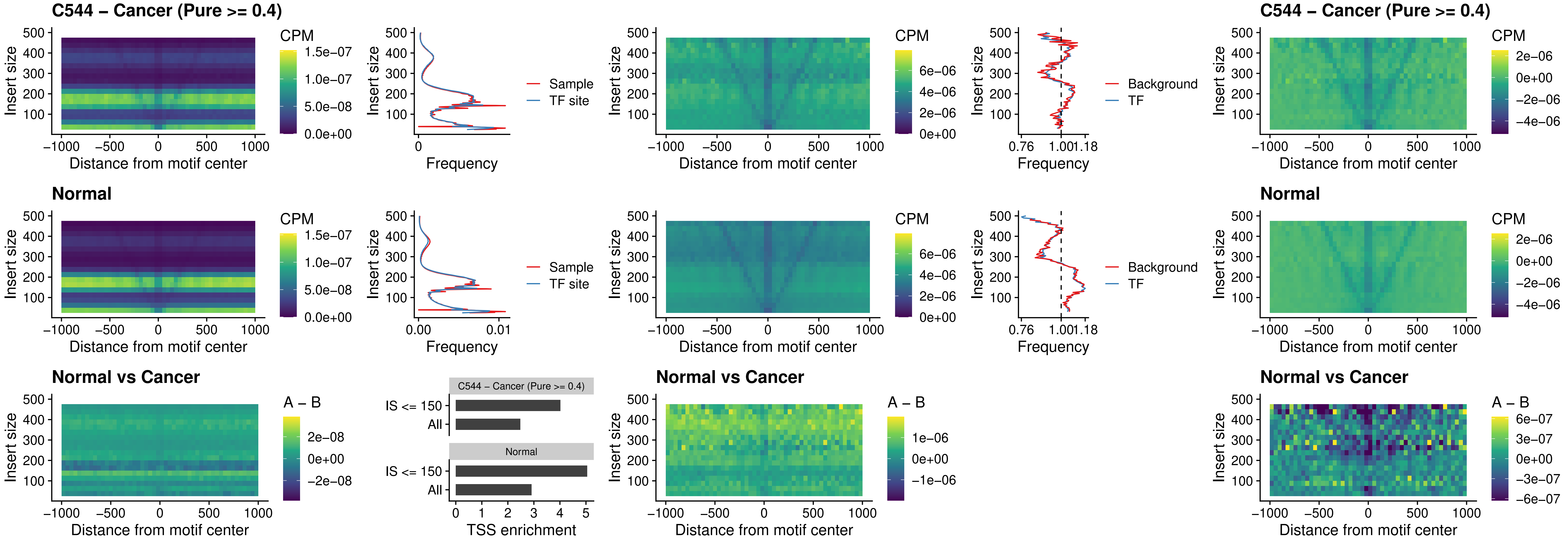

#### C544 – Cancer (Pure >= 0.4) – HOXD4_uc_dTSS_nPEAK

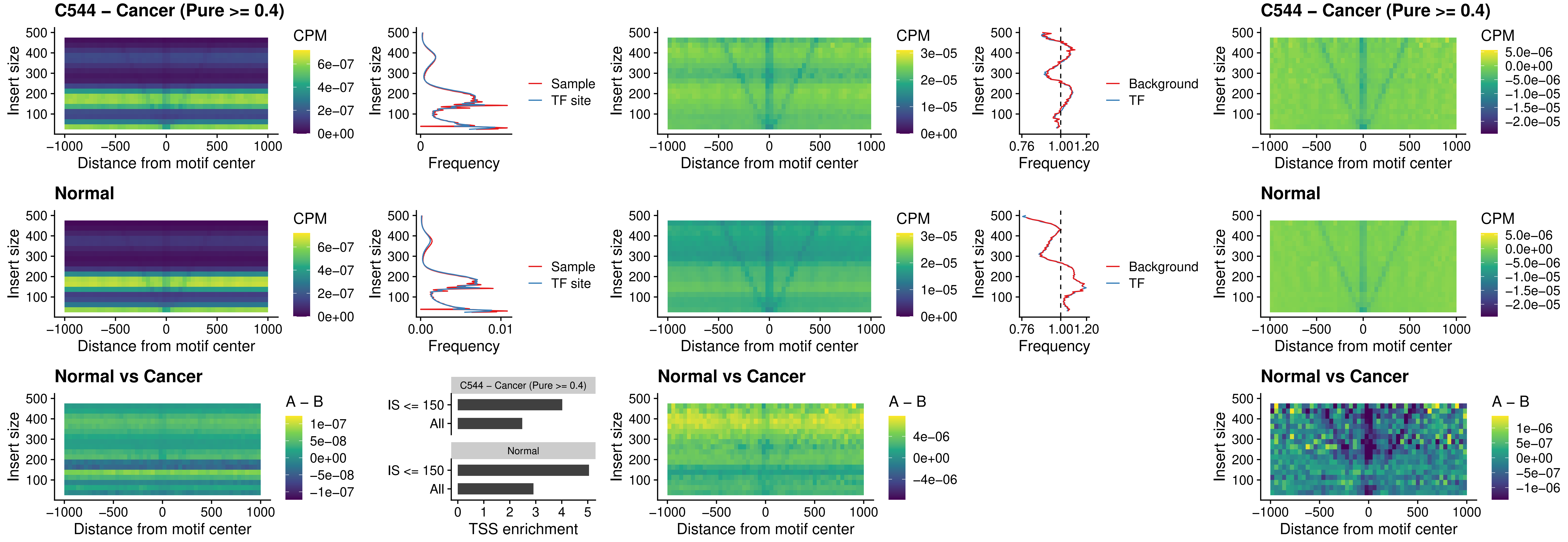

#### C544 – Cancer (Pure >= 0.4) – FOXP3_uc_cTSS_nPEAK

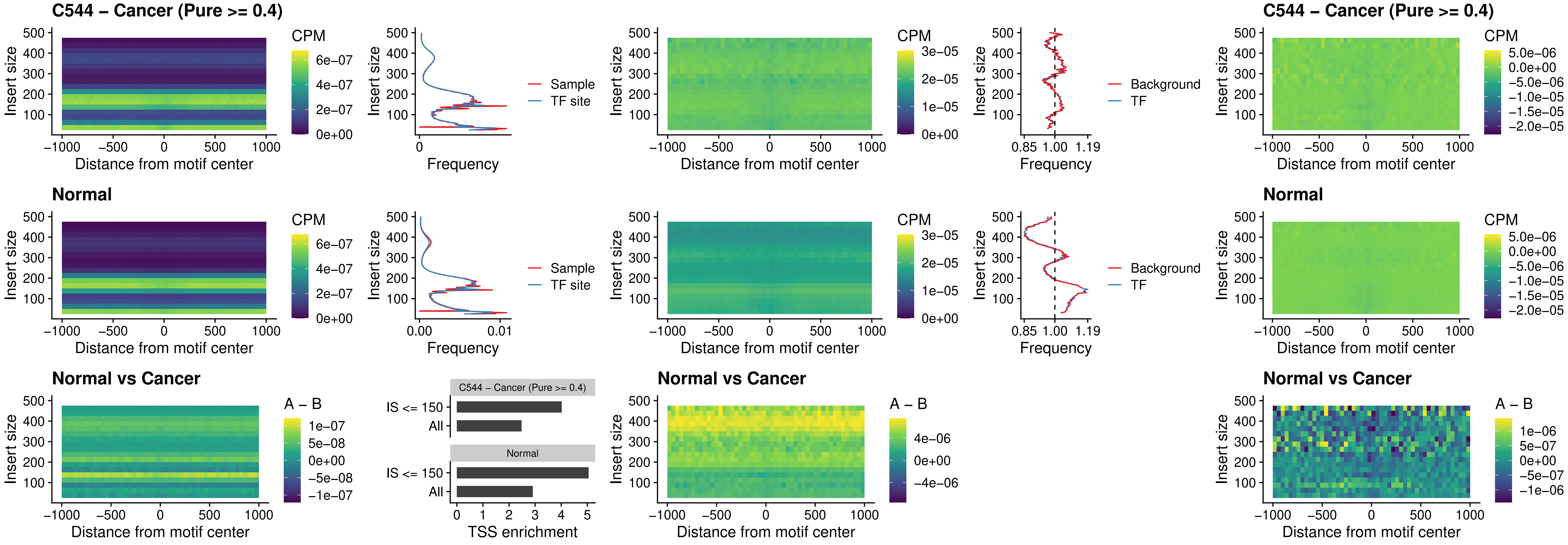

### C544 – Cancer (Pure >= 0.4) – FOXD4L6_uc_cTSS_nPEAK

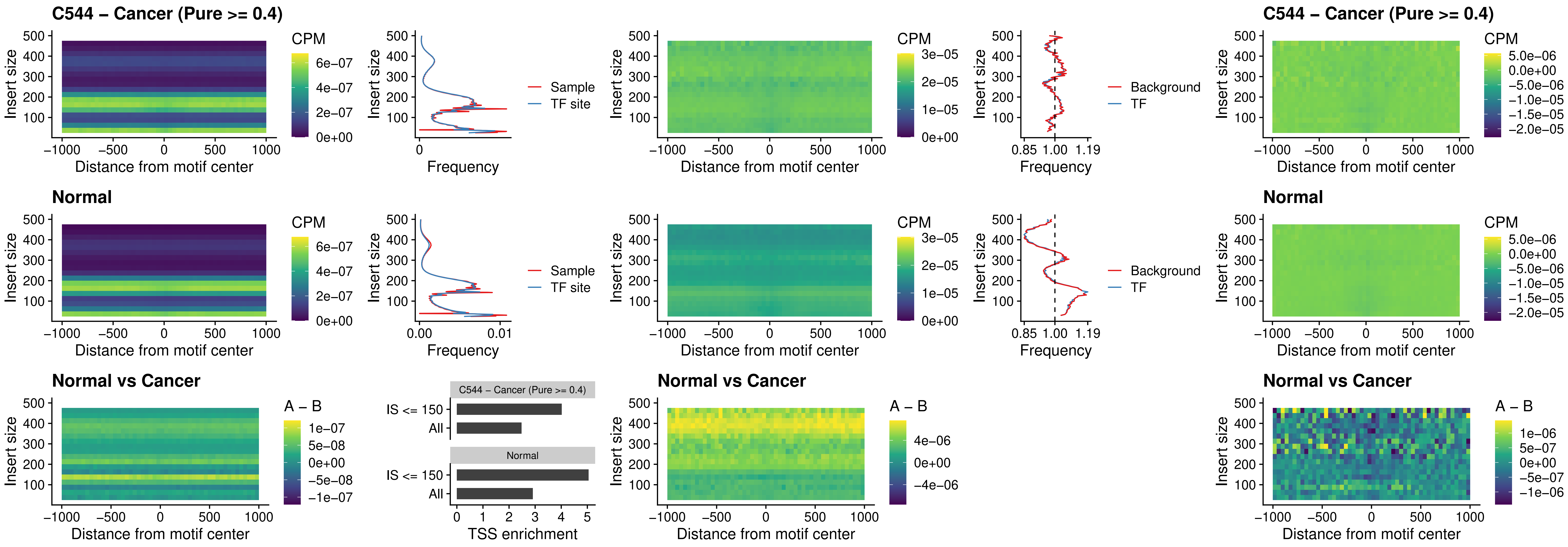

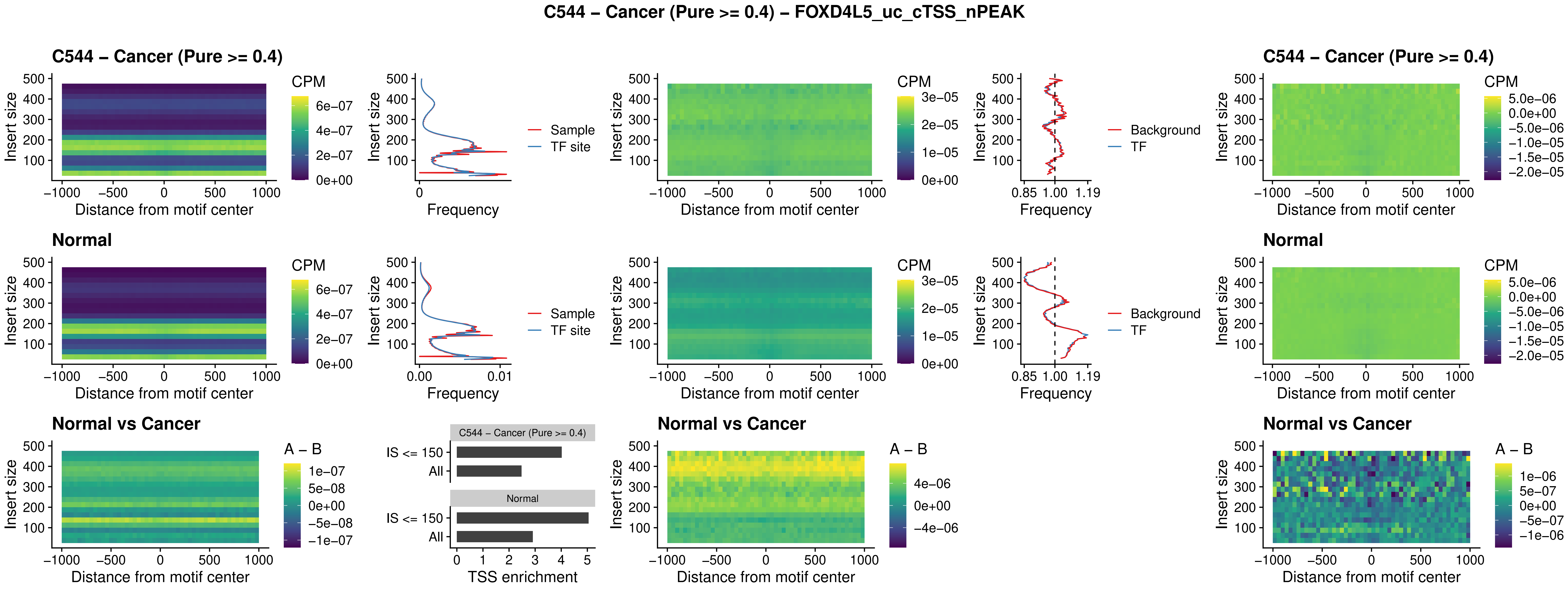
C544 – Cancer (Pure >= 0.4) – FOXD4L5_uc_cTSS_nPEAK
C544 – Cancer (Pure >= 0.4)
Normal
Normal vs Cancer
Insert size
Distance from motif center
CPM
A – B
Frequency
Sample
TF site
Background
TF
C544 – Cancer (Pure >= 0.4)
Normal
IS <= 150
All
TSS enrichment

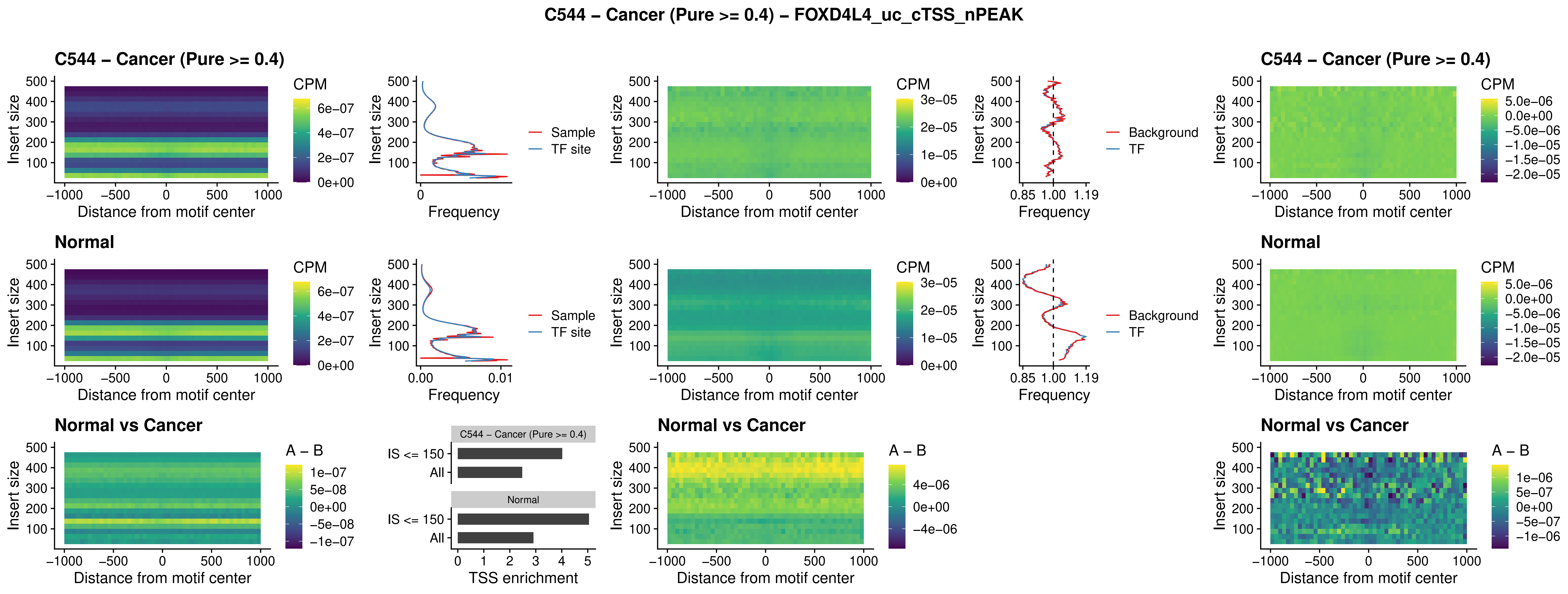
C544 – Cancer (Pure >= 0.4) – FOXD4L4_uc_cTSS_nPEAK
C544 – Cancer (Pure >= 0.4)
Normal
Normal vs Cancer
Insert size
Distance from motif center
Frequency
CPM
A – B
Sample
TF site
Background
TF
TSS enrichment
IS <= 150
All
C544 – Cancer (Pure >= 0.4)
Normal

#### C544 – Cancer (Pure >= 0.4) – FOXD4L2_uc_cTSS_nPEAK

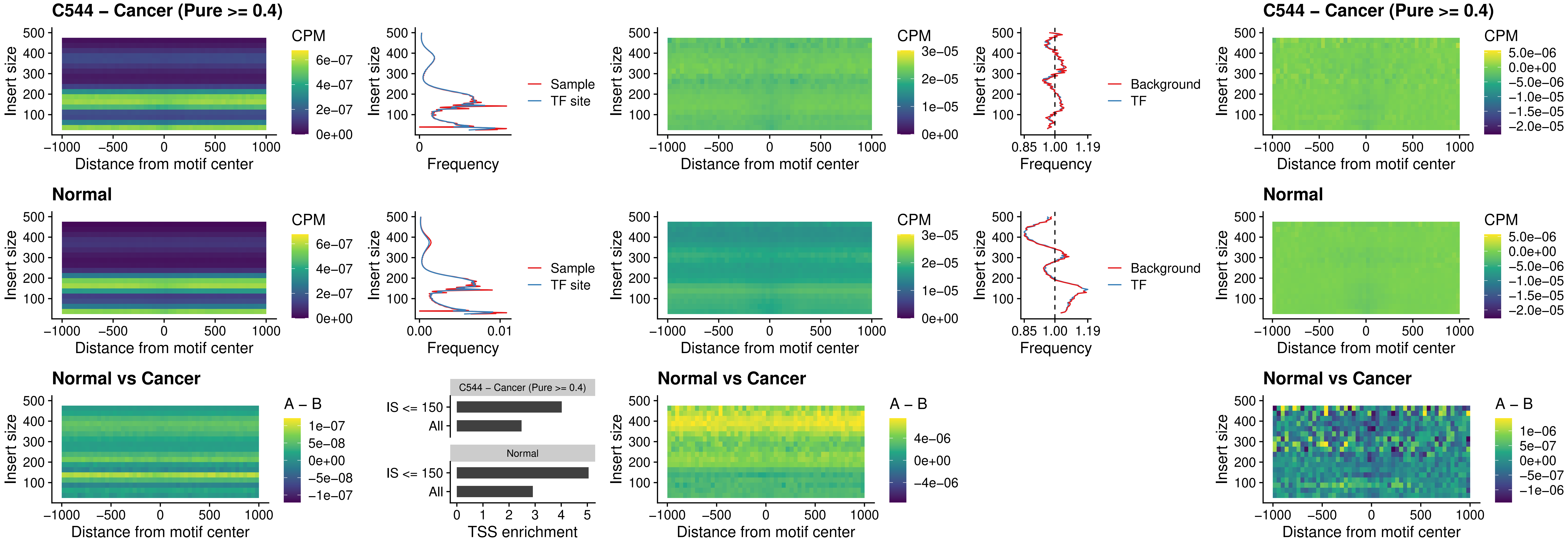

#### C544 – Cancer (Pure >= 0.4) – FOXD4L3_uc_cTSS_nPEAK

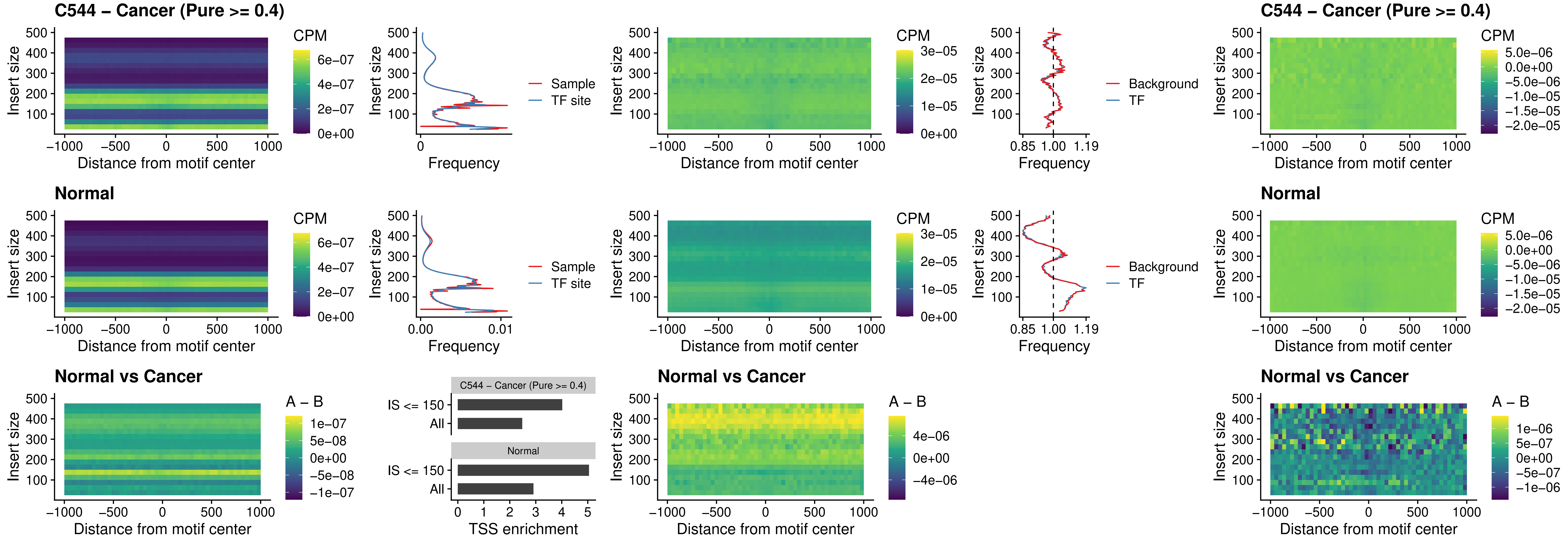

### C544 – Cancer (Pure >= 0.4) – HOXD1_uc_dTSS_oPEAK

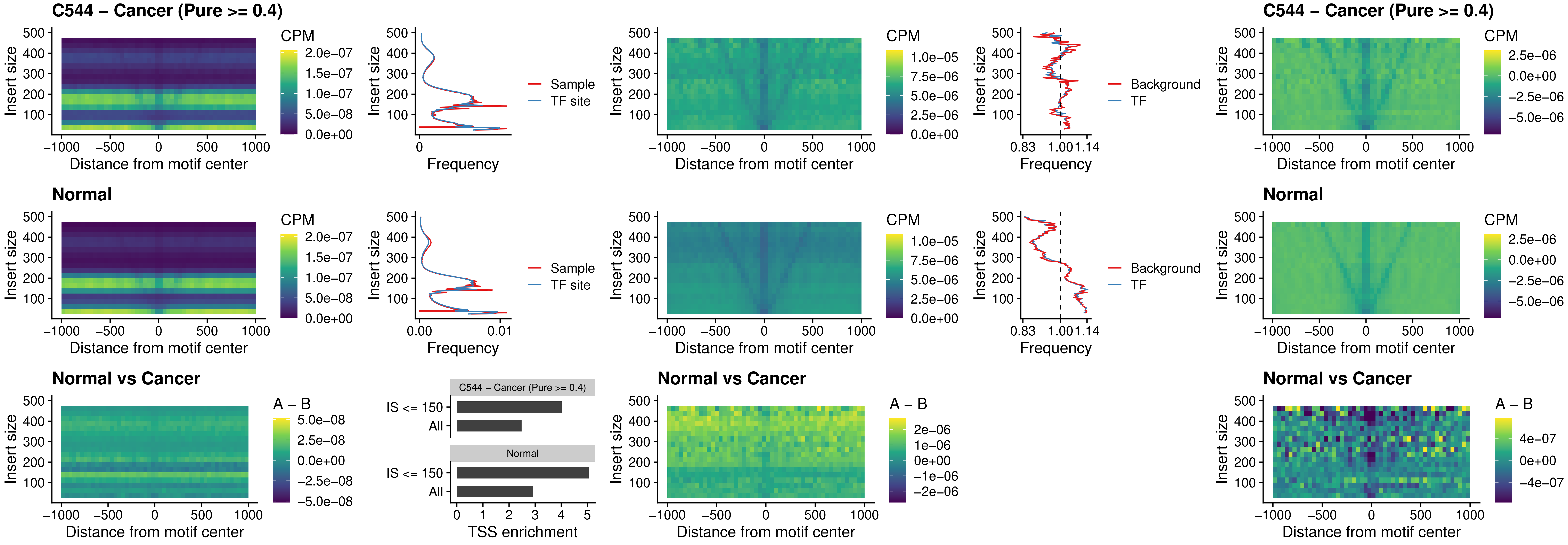

#### C544 – Cancer (Pure >= 0.4) – VAX1_uc_pTSS_nPEAK

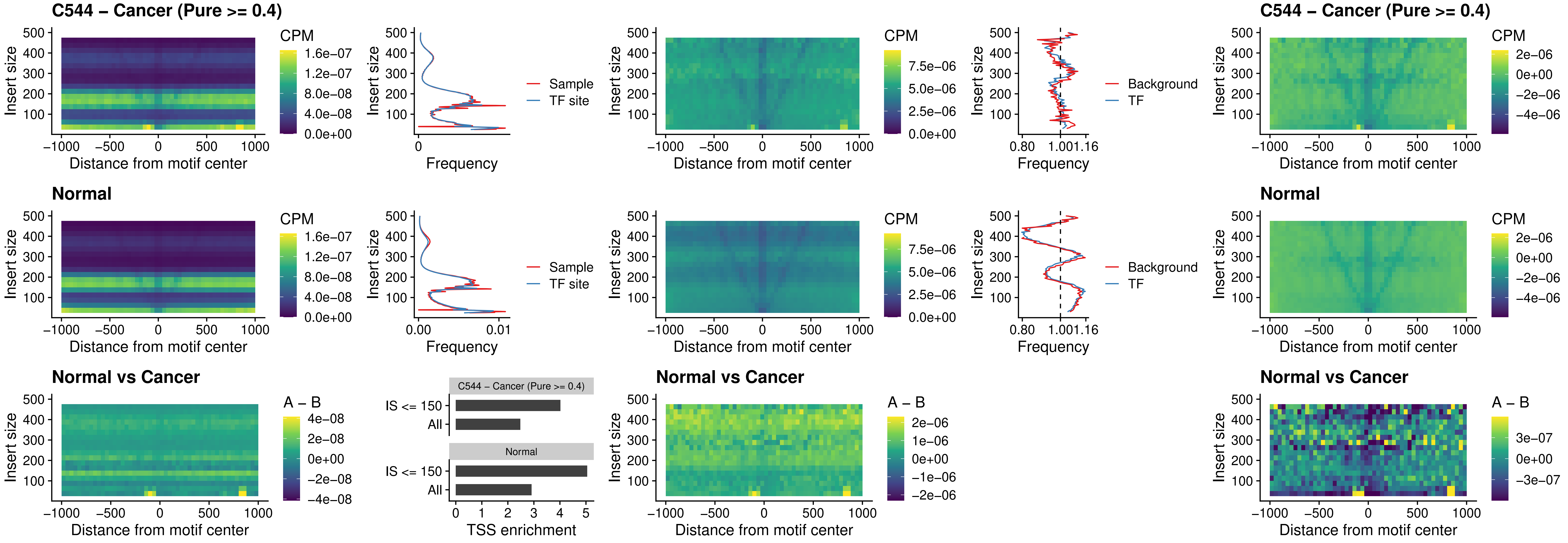

#### C544 – Cancer (Pure >= 0.4) – DLX3_uc_pTSS_nPEAK

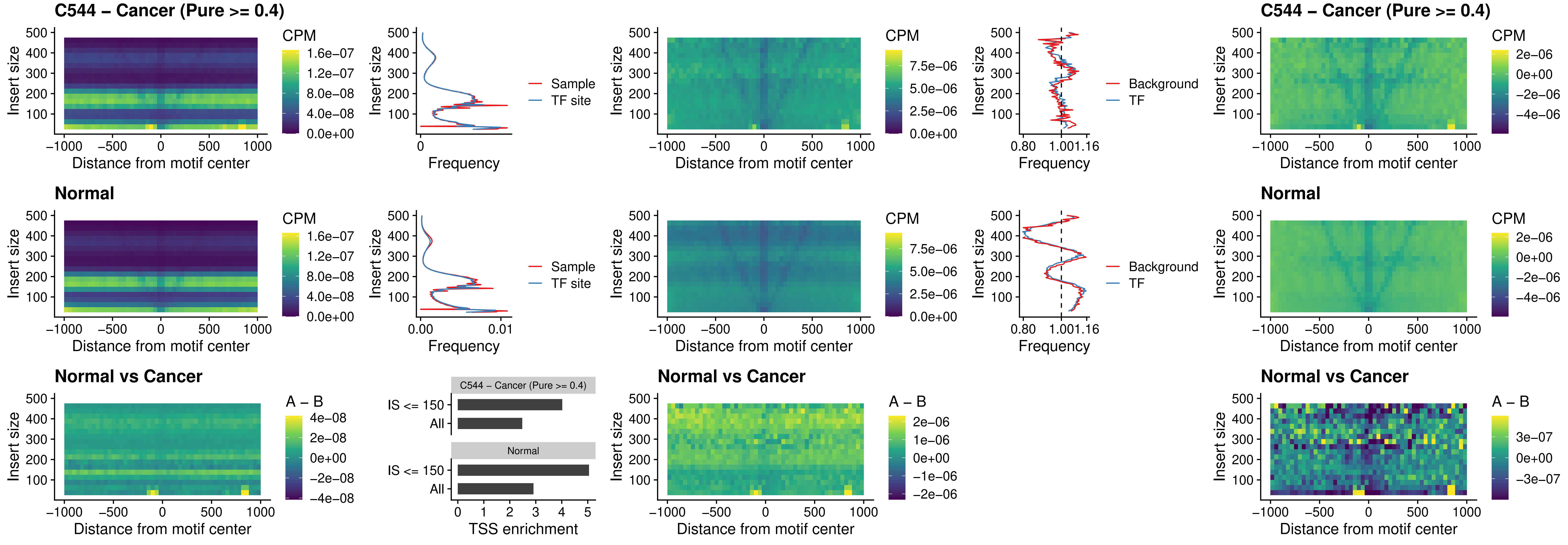

#### C544 – Cancer (Pure >= 0.4) – HOXD3_uc_pTSS_nPEAK

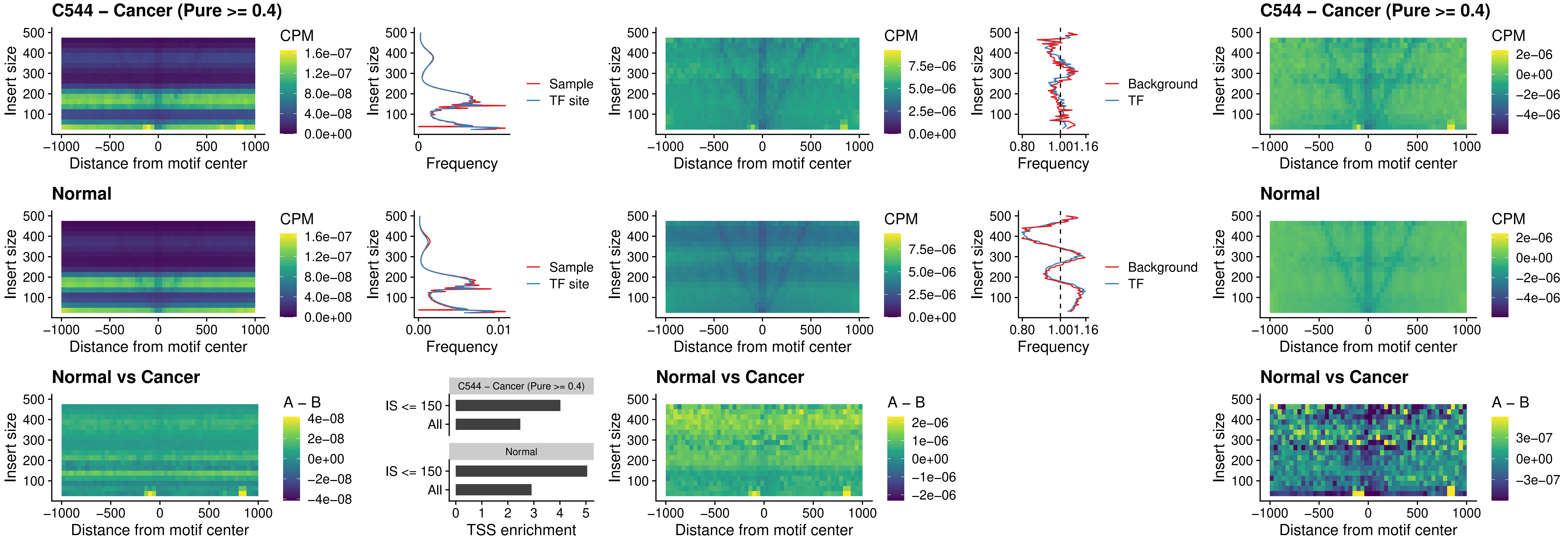

#### C544 – Cancer (Pure >= 0.4) – PDX1_uc_dTSS_nPEAK

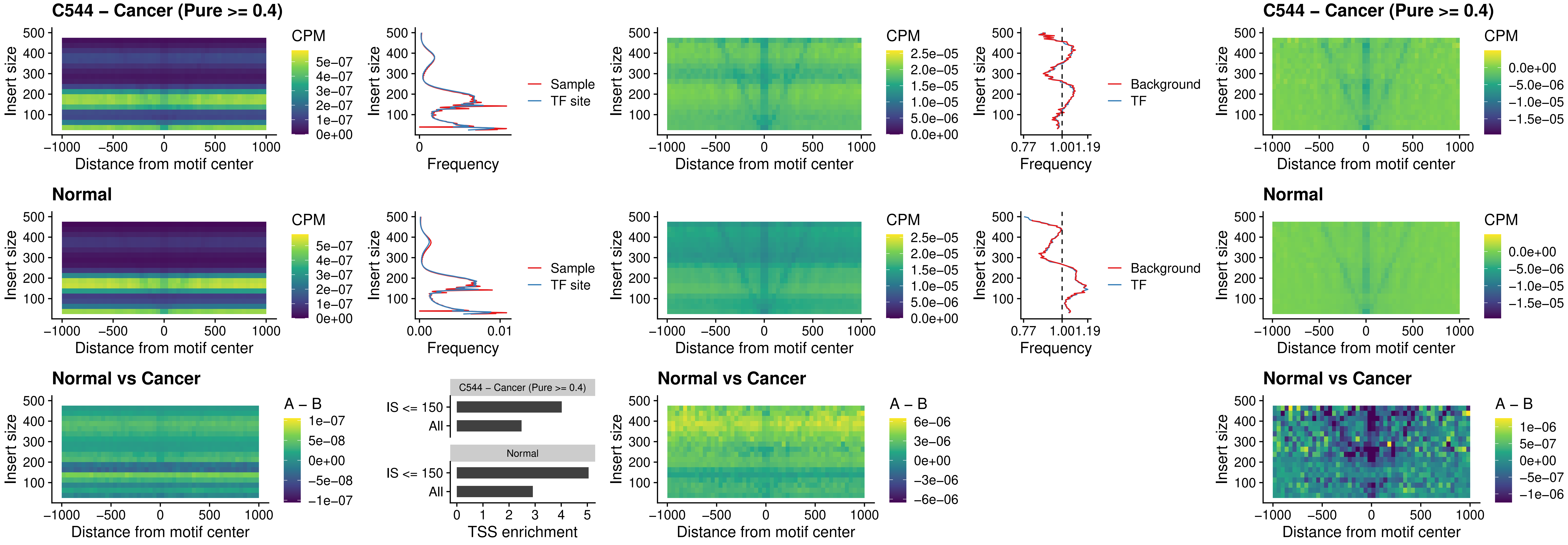

#### C544 – Cancer (Pure >= 0.4) – VAX2_uc_dTSS_nPEAK

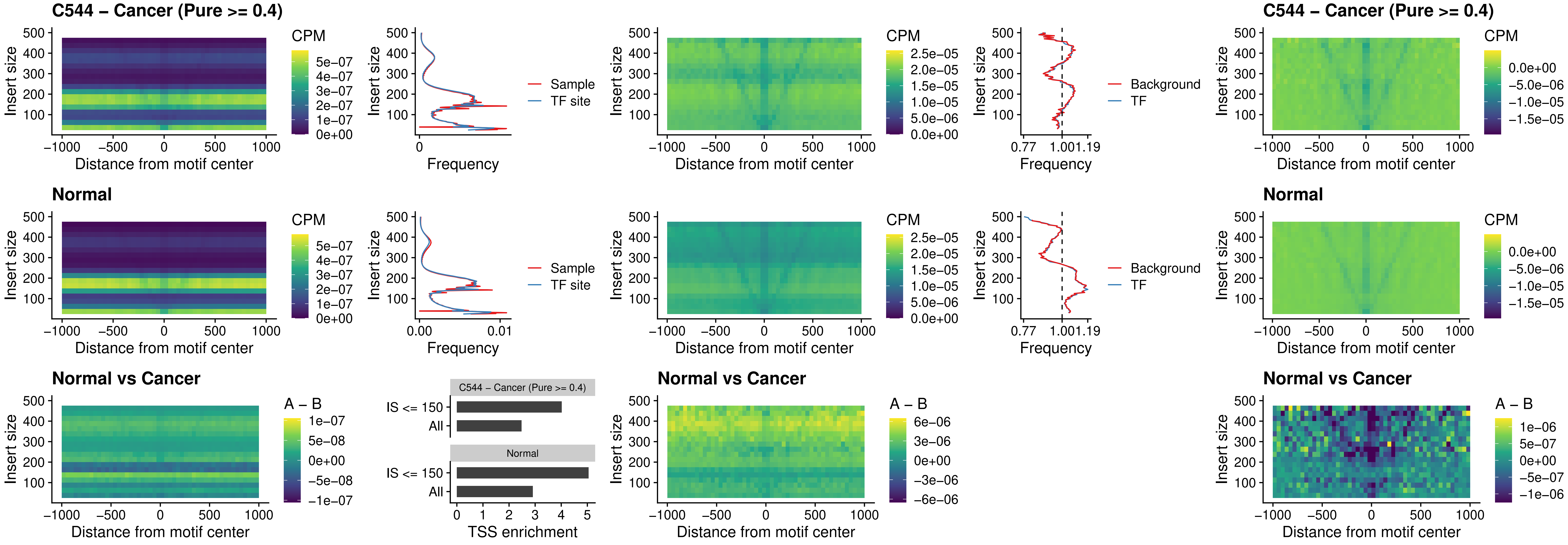

### C544 – Cancer (Pure >= 0.4) – SOX5_uc_dTSS_nPEAK

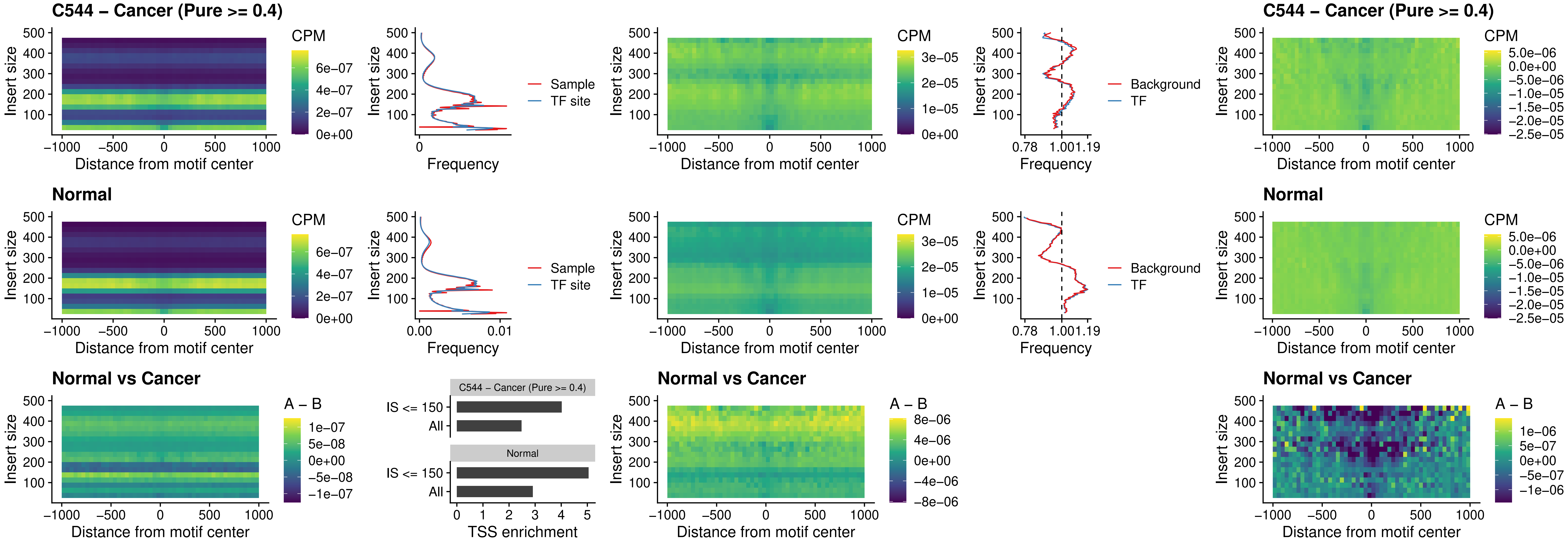

### C544 – Cancer (Pure >= 0.4) – SOX6_uc_dTSS_nPEAK

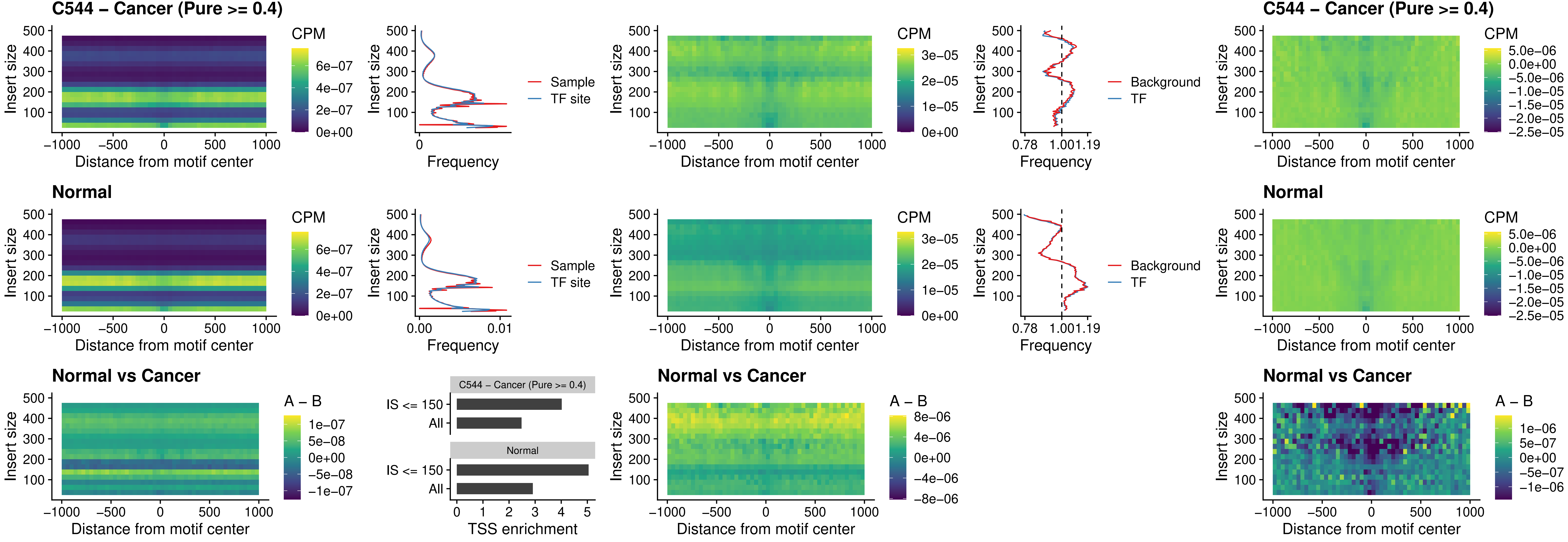

### C544 – Cancer (Pure >= 0.4) – JDP2_uc_cTSS_nPEAK

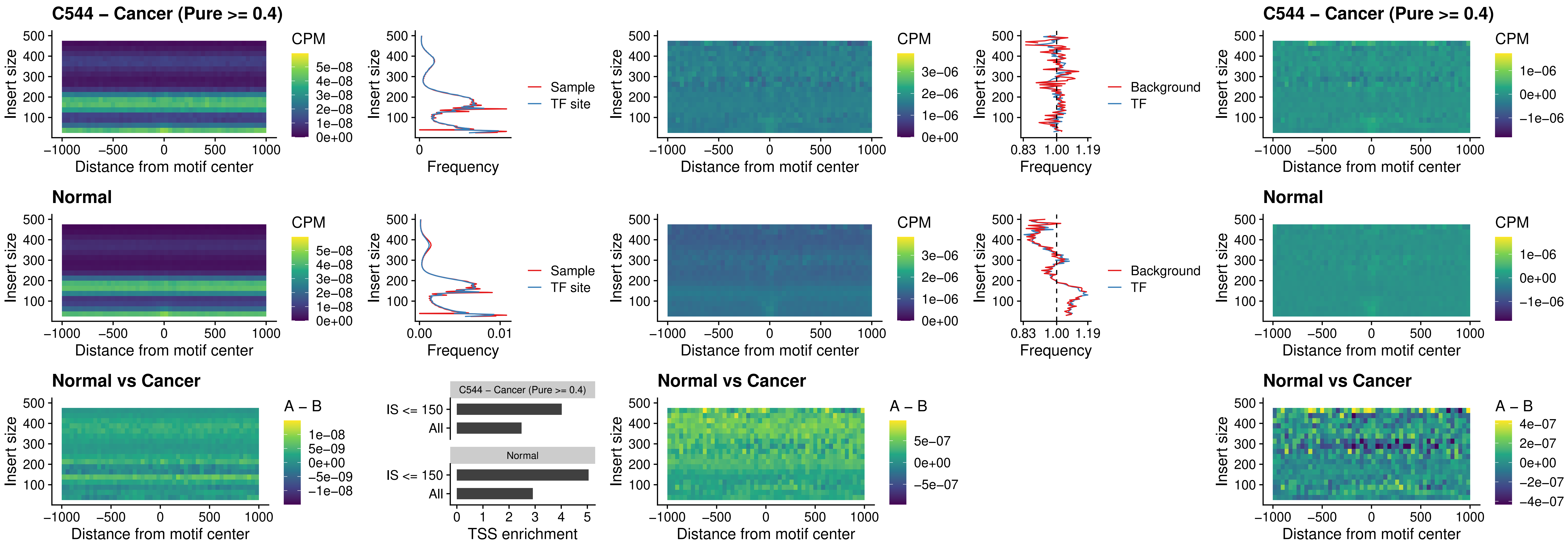

#### C544 – Cancer (Pure >= 0.4) – PURA_uc_dTSS_oPEAK

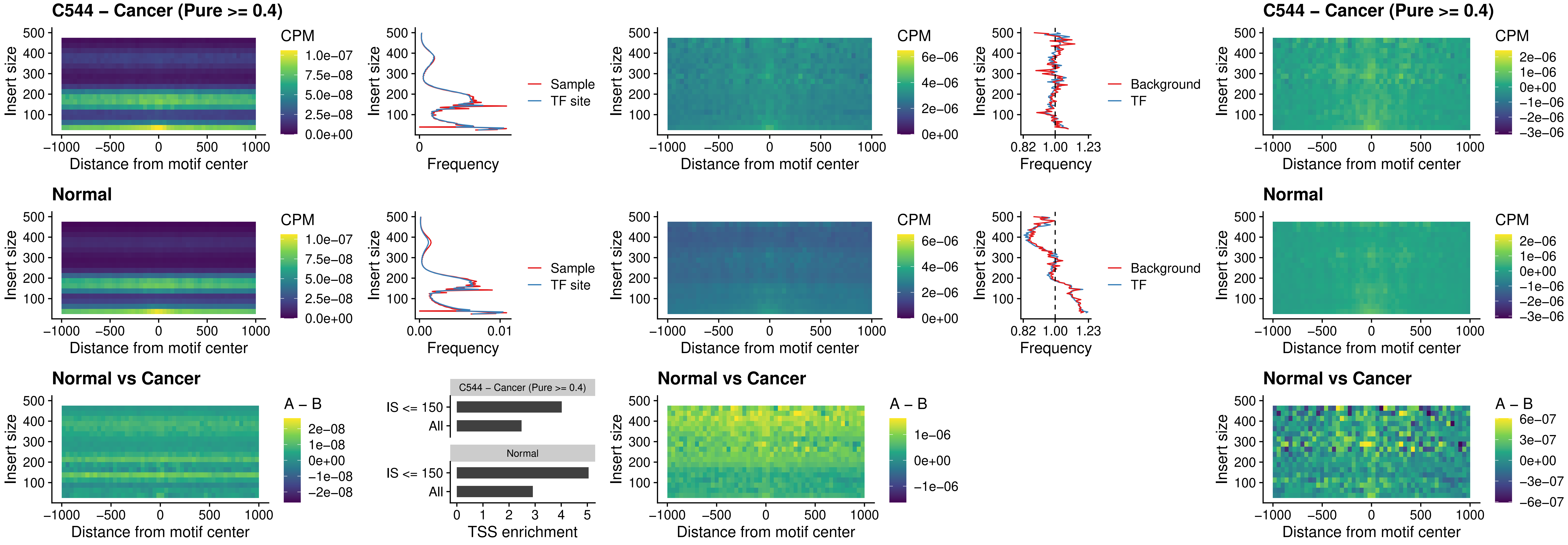

#### C544 – Cancer (Pure >= 0.4) – HHEX_uc_dTSS_nPEAK

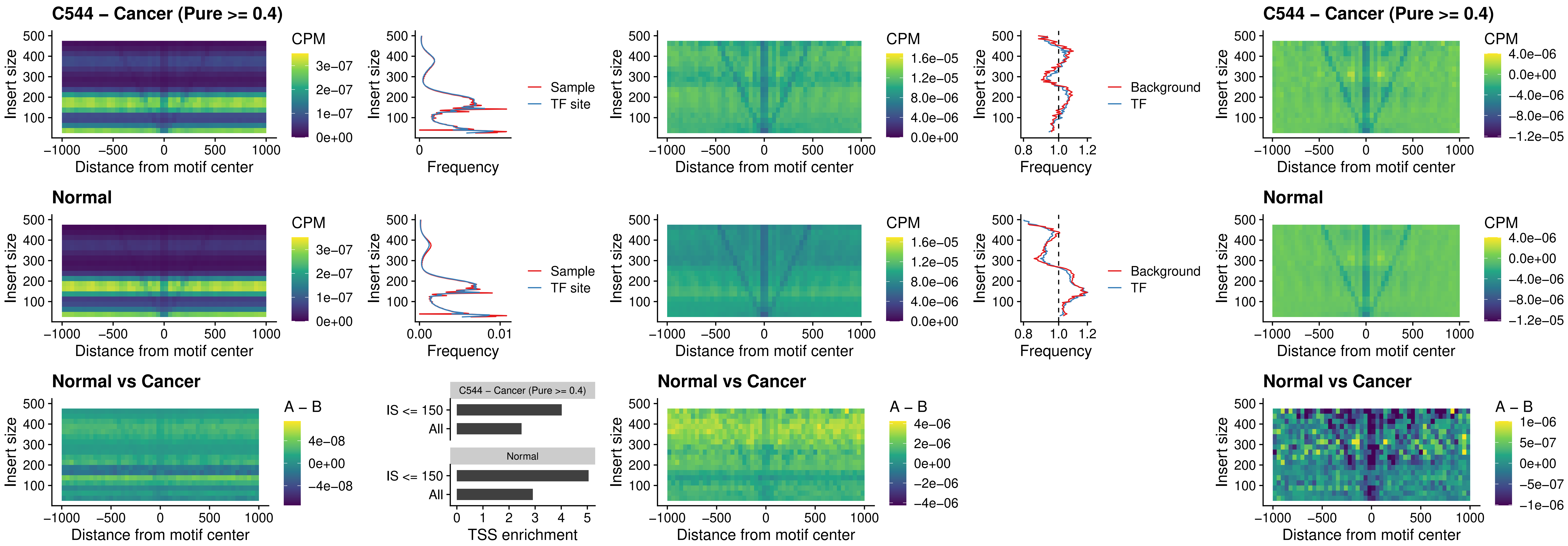

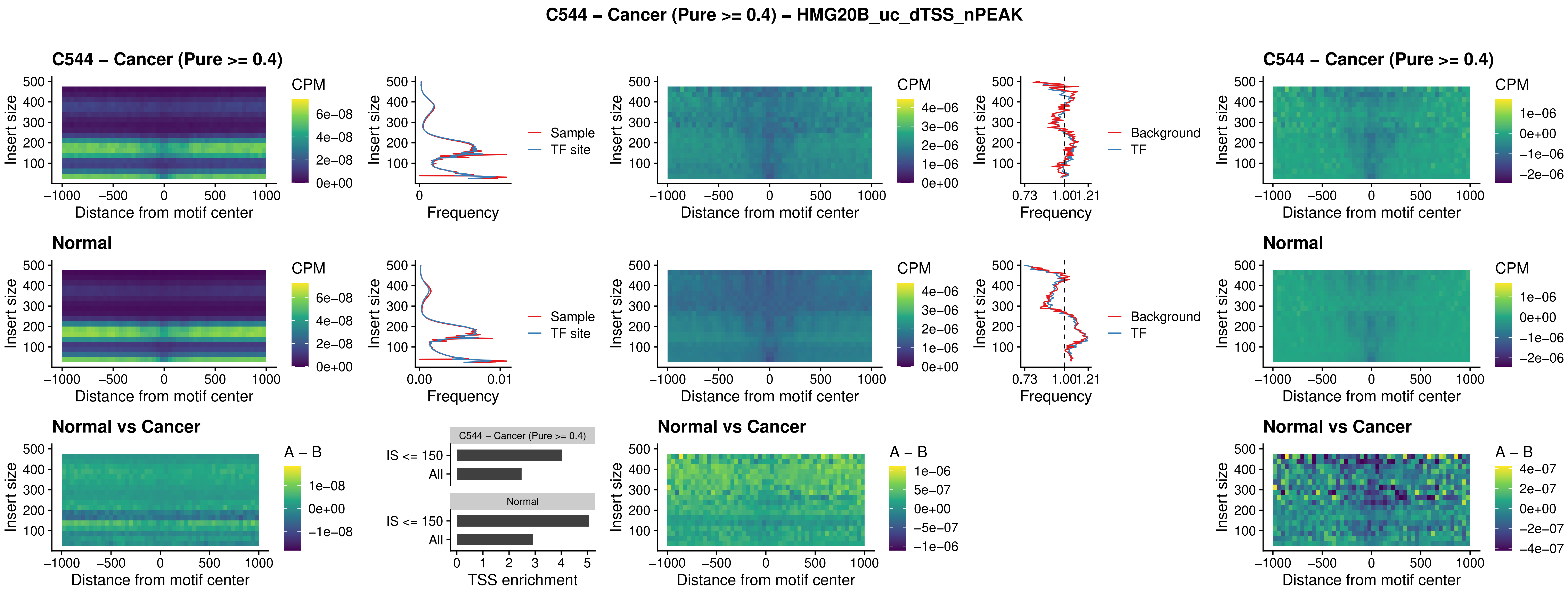
C544 – Cancer (Pure >= 0.4) – HMG20B_uc_dTSS_nPEAK
C544 – Cancer (Pure >= 0.4)
Normal
Normal vs Cancer
Insert size
Distance from motif center
Frequency
CPM
A – B
Sample
TF site
Background
TF
C544 – Cancer (Pure >= 0.4)
Normal
IS <= 150
All
TSS enrichment

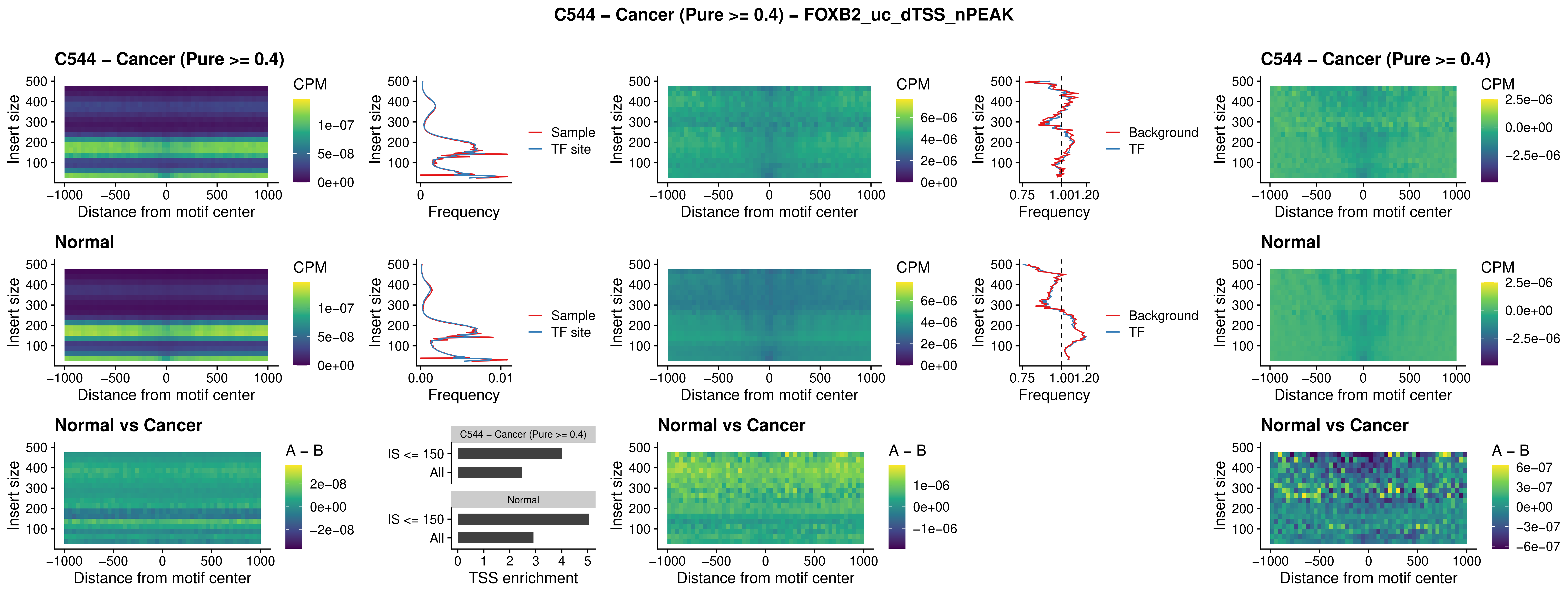

C544 – Cancer (Pure >= 0.4) – FOXB2_uc_dTSS_nPEAK
C544 – Cancer (Pure >= 0.4)
Normal
Normal vs Cancer
Insert size
Distance from motif center
Frequency
CPM
A – B
Sample
TF site
Background
TF
C544 – Cancer (Pure >= 0.4)
Normal
IS <= 150
All
TSS enrichment

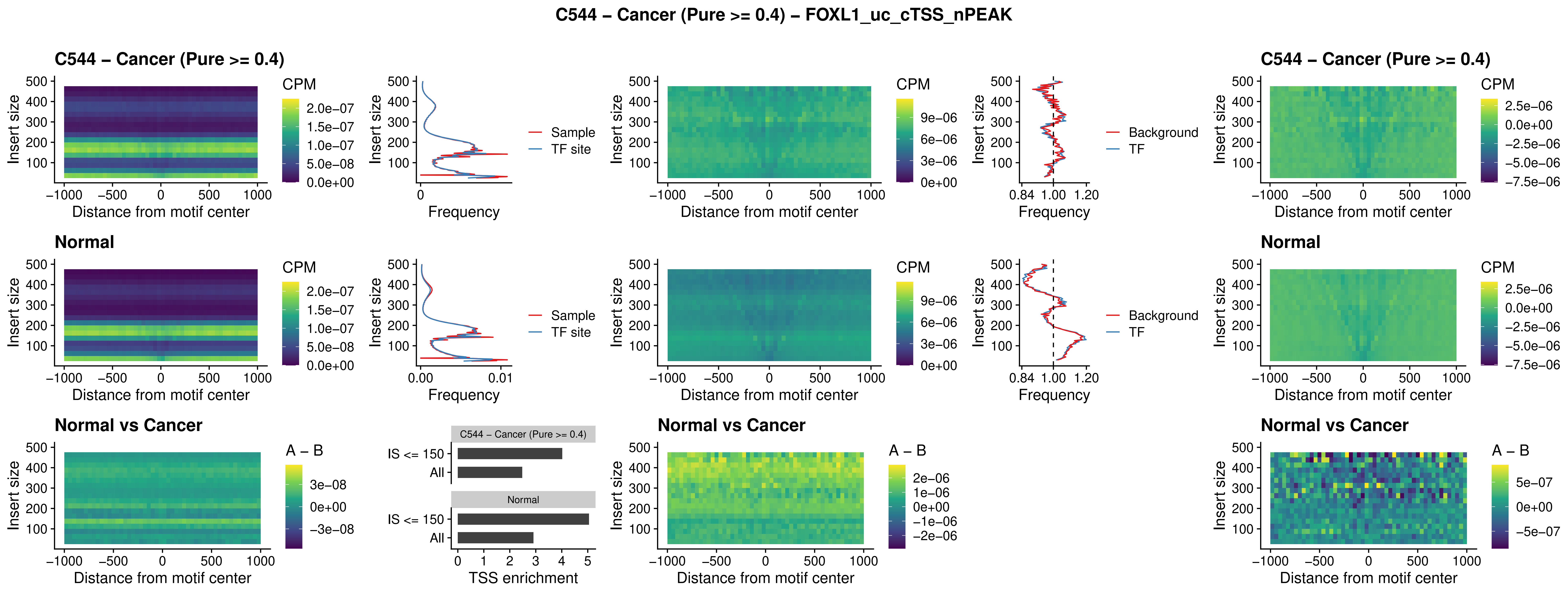

C544 – Cancer (Pure >= 0.4) – FOXL1_uc_cTSS_nPEAK
C544 – Cancer (Pure >= 0.4)
Normal
Normal vs Cancer
Insert size
Distance from motif center
Frequency
CPM
A – B
Sample
TF site
Background
TF
C544 – Cancer (Pure >= 0.4)
Normal
IS <= 150
All
TSS enrichment

#### C544 – Cancer (Pure >= 0.4) – PBX3_uc_pTSS_oPEAK

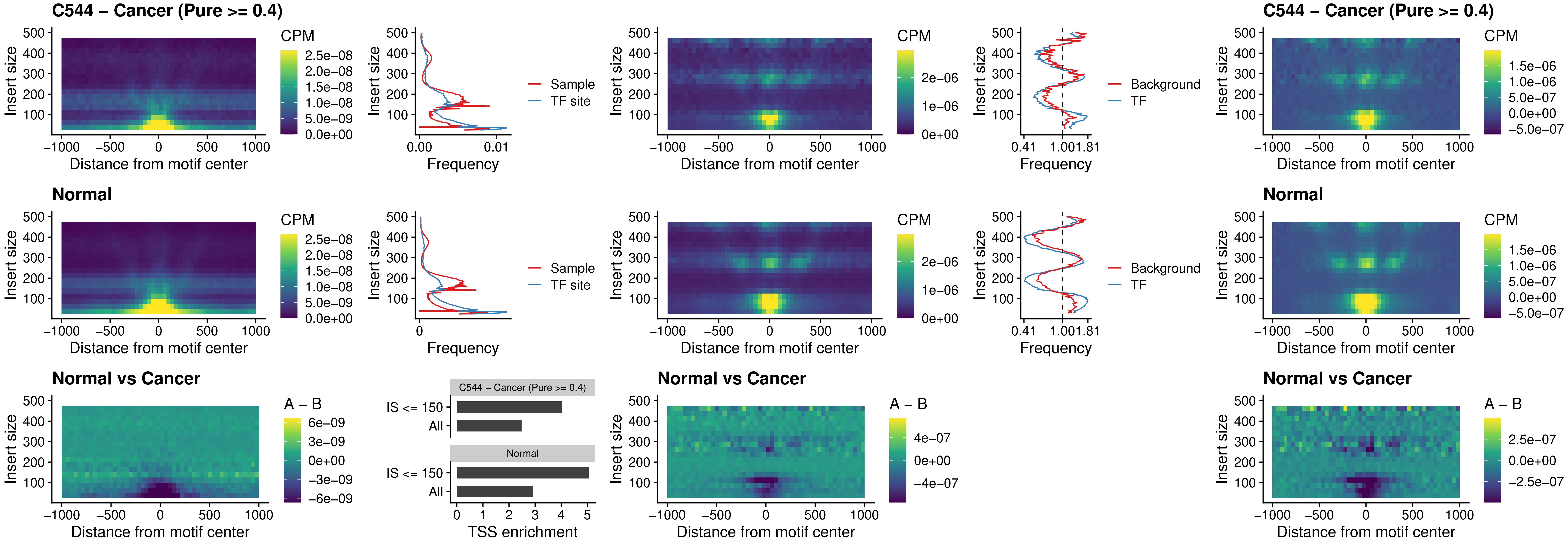

#### C544 – Cancer (Pure >= 0.4) – SNAI2_uc_cTSS_nPEAK

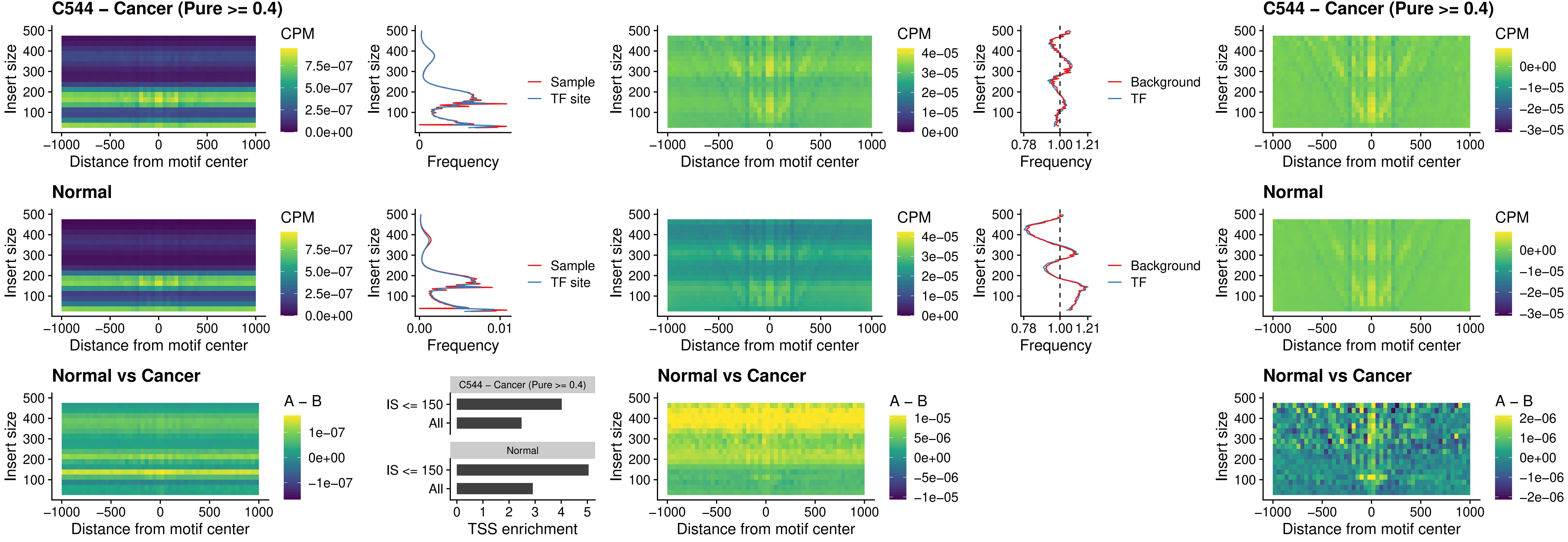

### C544 – Cancer (Pure >= 0.4) – SNAI2_uc_dTSS_oPEAK

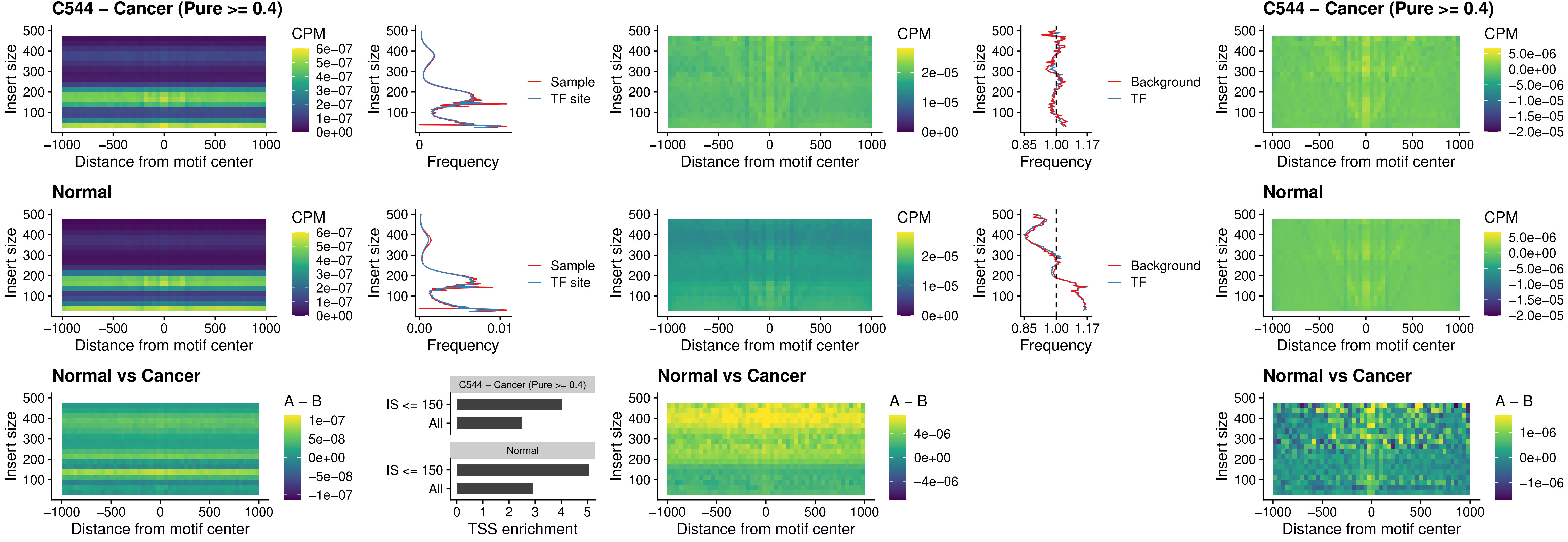

#### C544 – Cancer (Pure >= 0.4) – SPI1_uc_pTSS_nPEAK

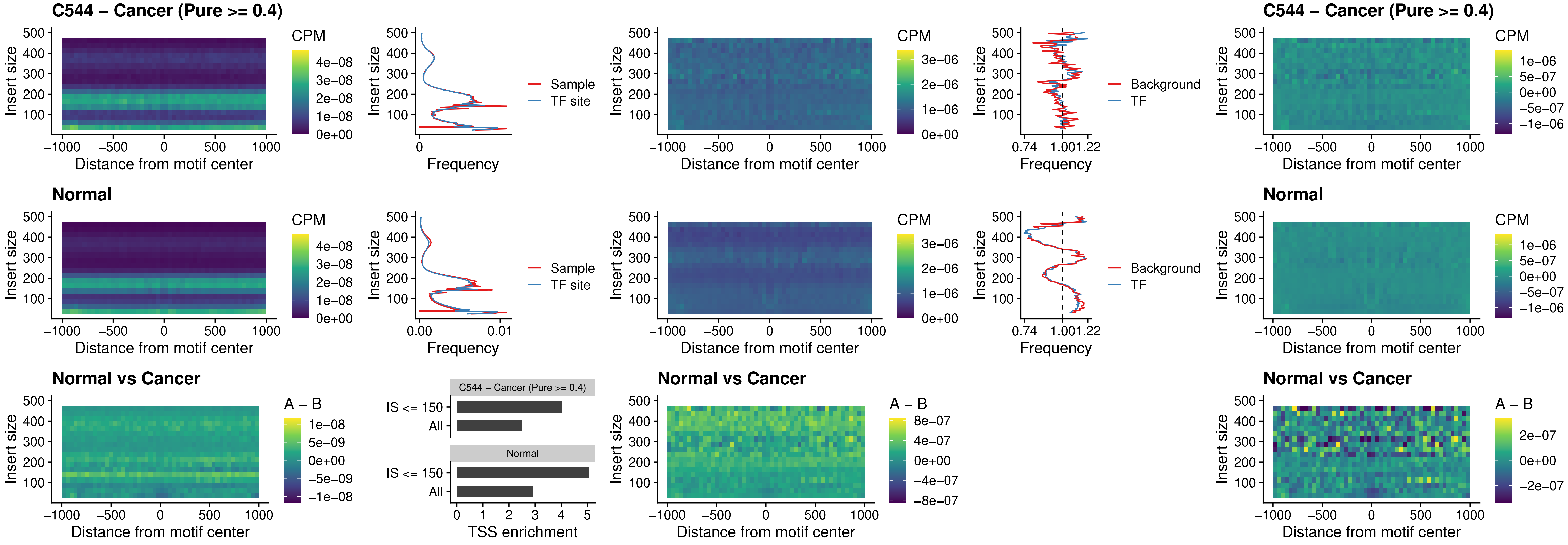

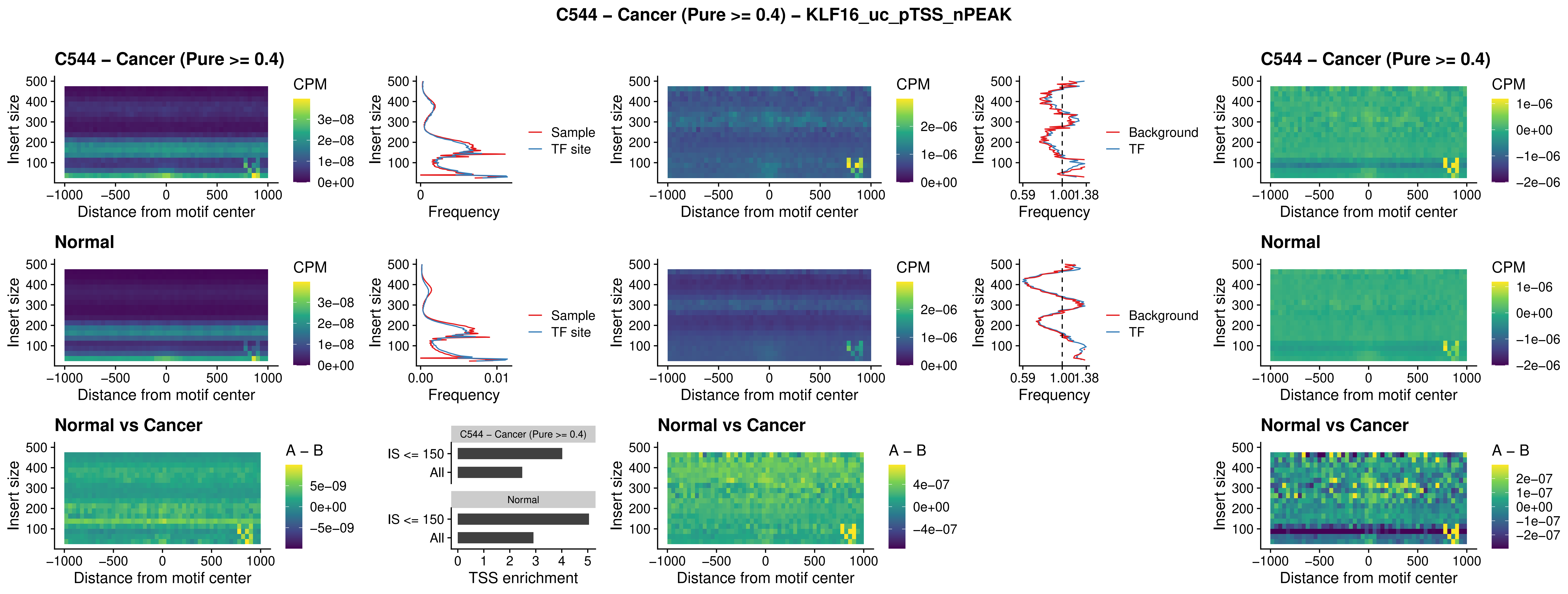
C544 – Cancer (Pure >= 0.4) – KLF16_uc_pTSS_nPEAK
C544 – Cancer (Pure >= 0.4)
Normal
Normal vs Cancer
Insert size
Distance from motif center
Frequency
CPM
A – B
Sample
TF site
Background
TF
C544 – Cancer (Pure >= 0.4)
IS <= 150
All
TSS enrichment

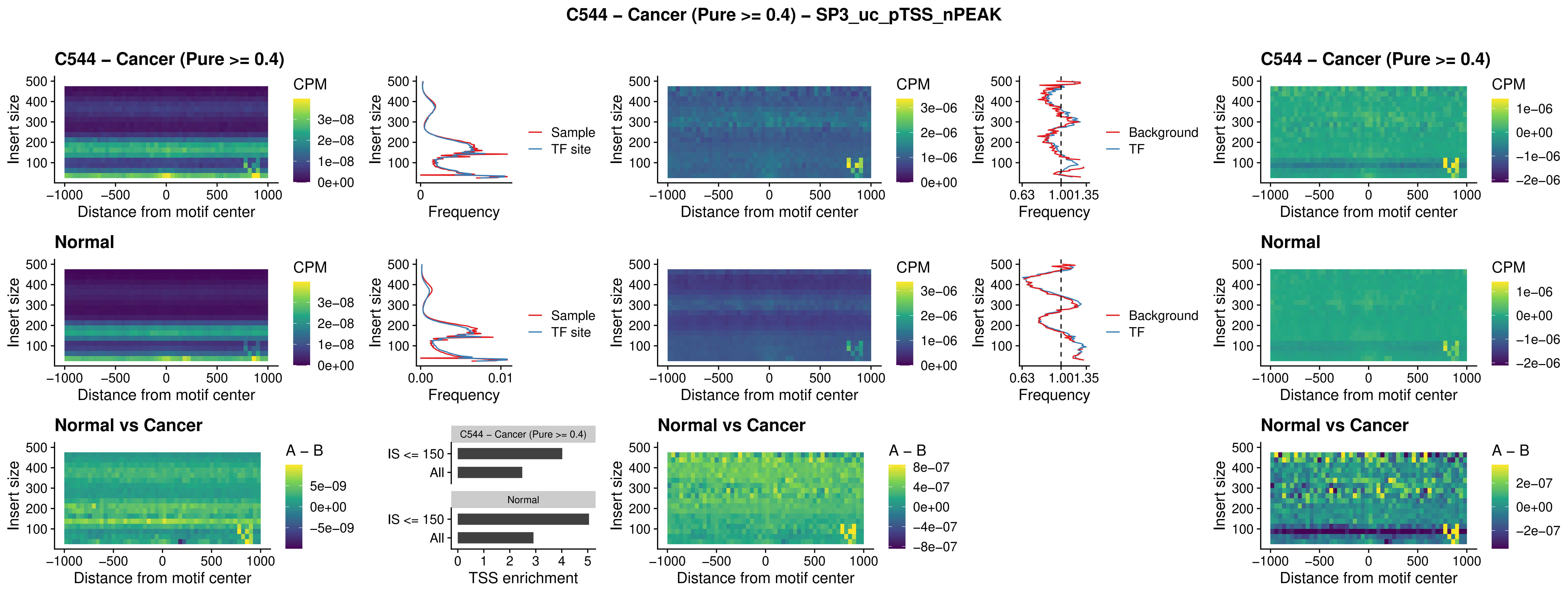
C544 – Cancer (Pure >= 0.4) – SP3_uc_pTSS_nPEAK
C544 – Cancer (Pure >= 0.4)
Normal
Normal vs Cancer
Insert size
Distance from motif center
CPM
A – B
Frequency
Sample
TF site
Background
TF
C544 – Cancer (Pure >= 0.4)
Normal
IS <= 150
All
TSS enrichment

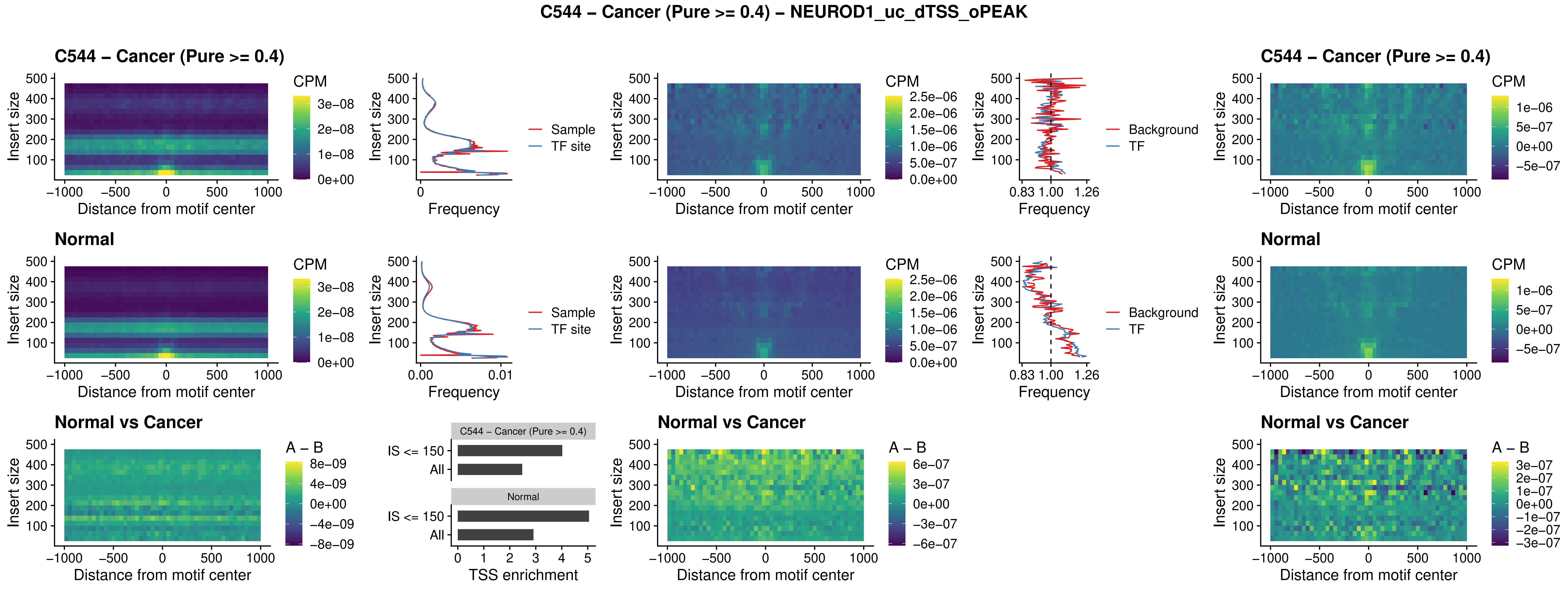
C544 – Cancer (Pure >= 0.4) – NEURОD1_uc_dTSS_oPEAK
C544 – Cancer (Pure >= 0.4)
Normal
Normal vs Cancer
Insert size
Distance from motif center
Frequency
CPM
A – B
Sample
TF site
Background
TF
C544 – Cancer (Pure >= 0.4)
Normal
IS <= 150
All
TSS enrichment

#### C544 – Cancer (Pure >= 0.4) – CTCF_uc_dTSS_oPEAK

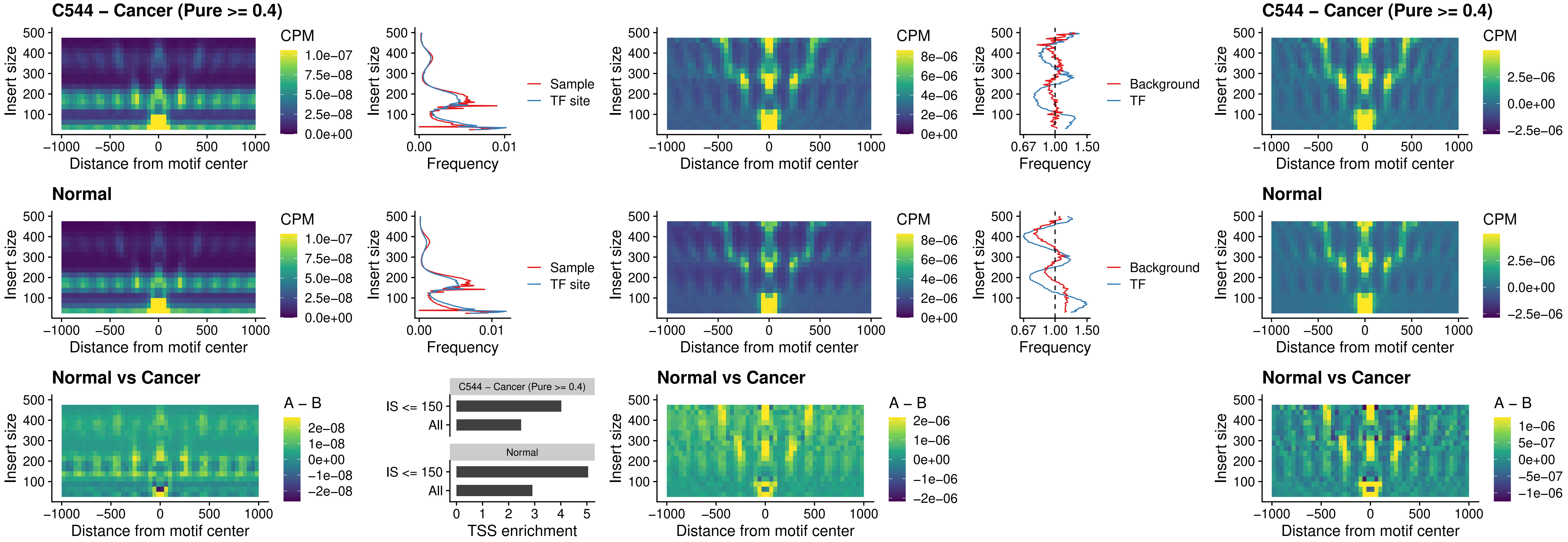

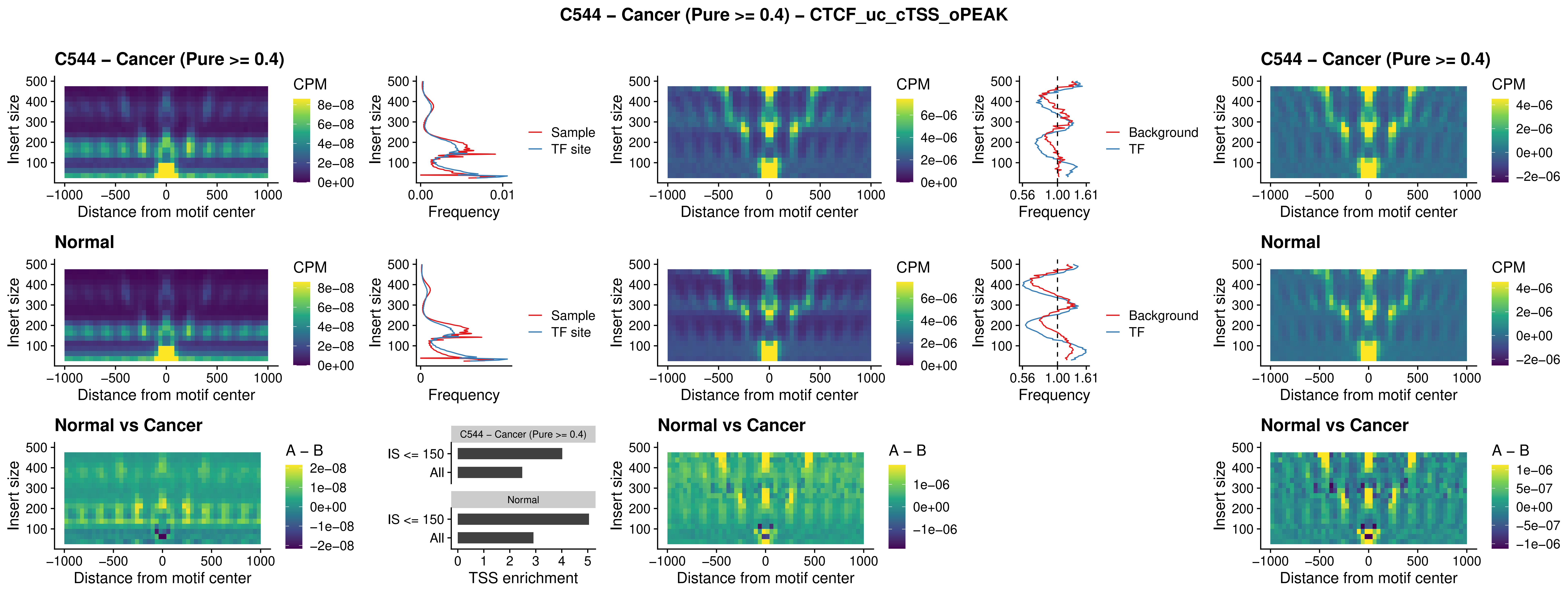

C544 – Cancer (Pure >= 0.4) – CTCF_uc_cTSS_oPEAK
C544 – Cancer (Pure >= 0.4)
Normal
Normal vs Cancer
Insert size
Distance from motif center
CPM
A – B
Frequency
Sample
TF site
Background
TF
C544 – Cancer (Pure >= 0.4)
Normal
IS <= 150
All
TSS enrichment

### C544 – Cancer (Pure >= 0.4) – CTCFL_uc_dTSS_oPEAK

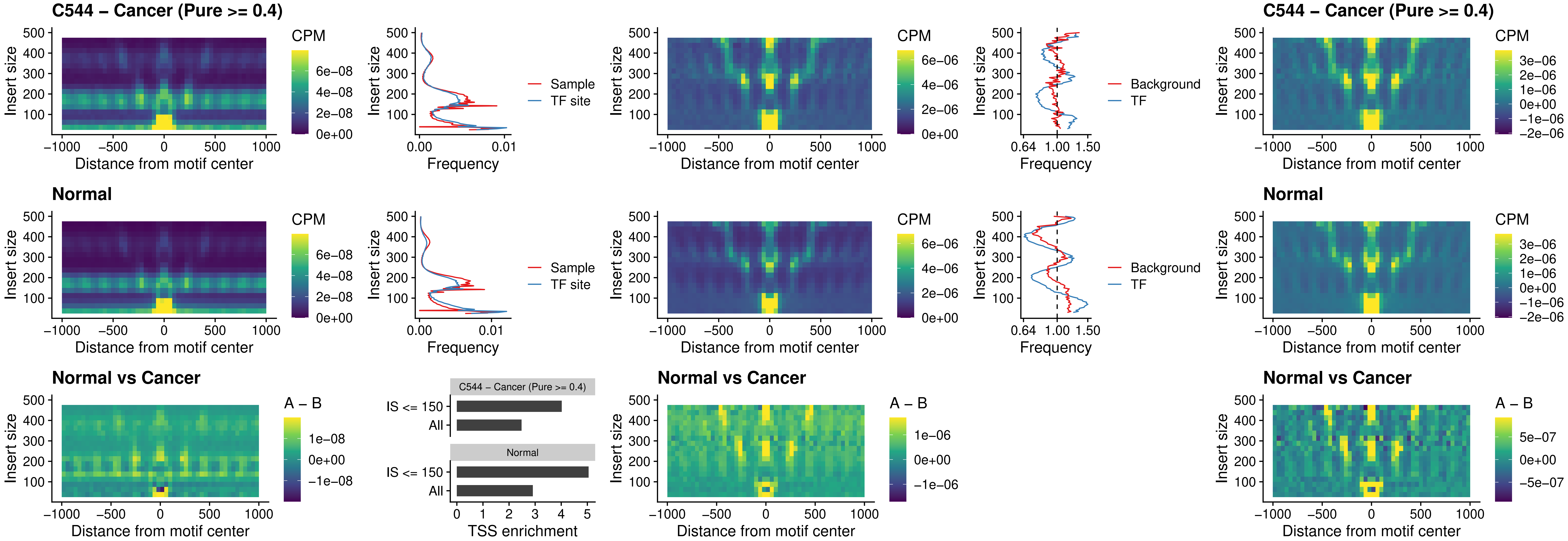

#### C544 – Cancer (Pure >= 0.4) – ENCODE_CTCF_sites_COLON

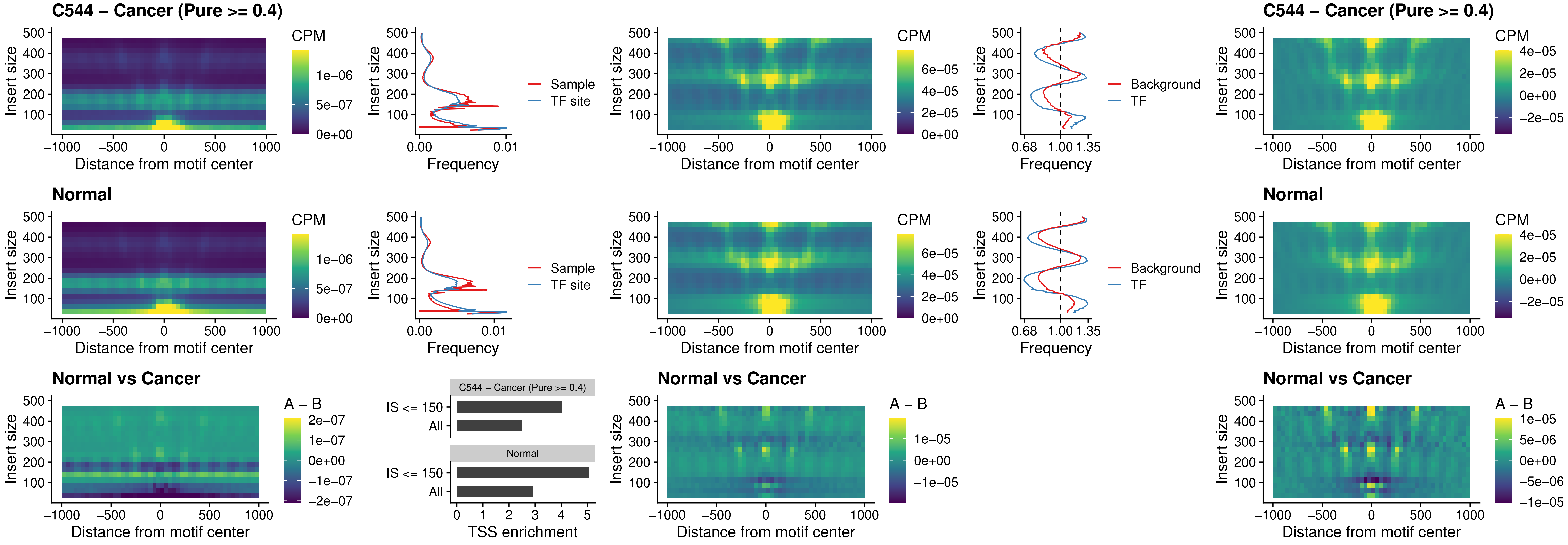

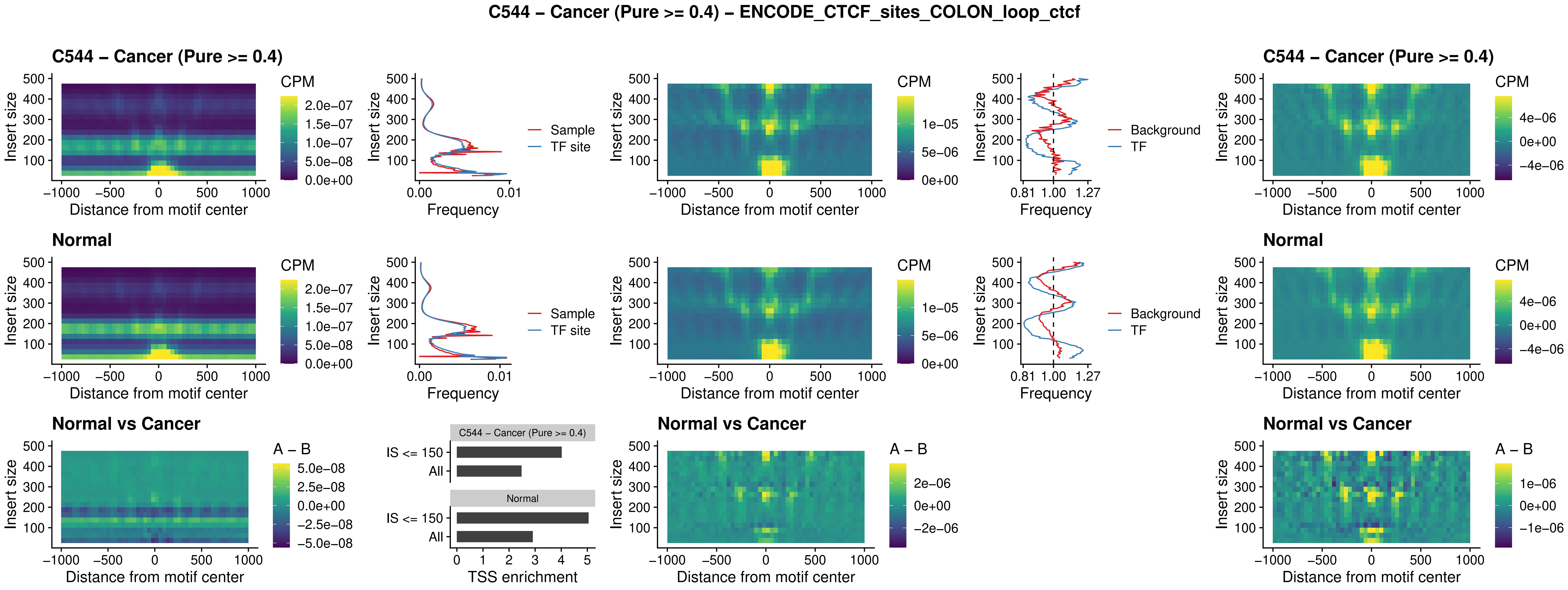
C544 – Cancer (Pure >= 0.4) – ENCODE_CTCF_sites_COLON_loop_ctcf
C544 – Cancer (Pure >= 0.4)
Normal
Normal vs Cancer
Insert size
Distance from motif center
Frequency
CPM
A – B
Sample
TF site
Background
TF
IS <= 150
All
Normal
TSS enrichment

#### C544 – Cancer (Pure >= 0.4) – ENCODE_CTCF_sites_COLON_intergenic_ctcf

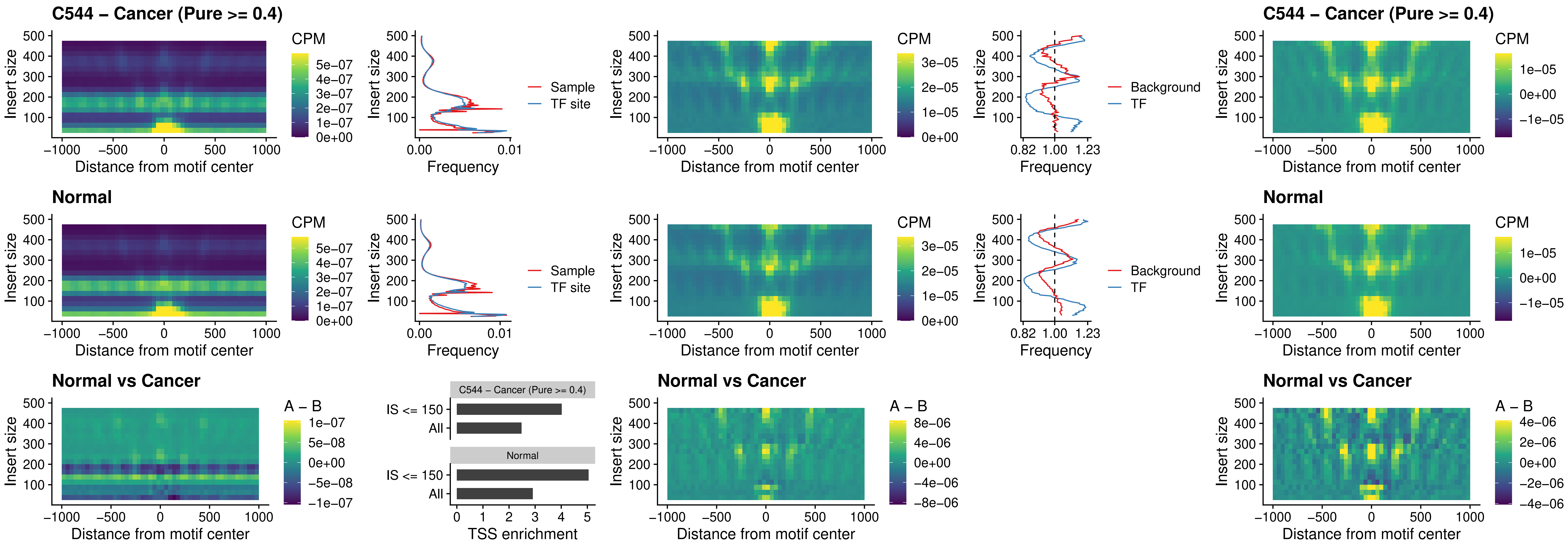

#### C544 – Cancer (Pure >= 0.4) – ENCODE_CTCF_sites_COLON_nonpromotor

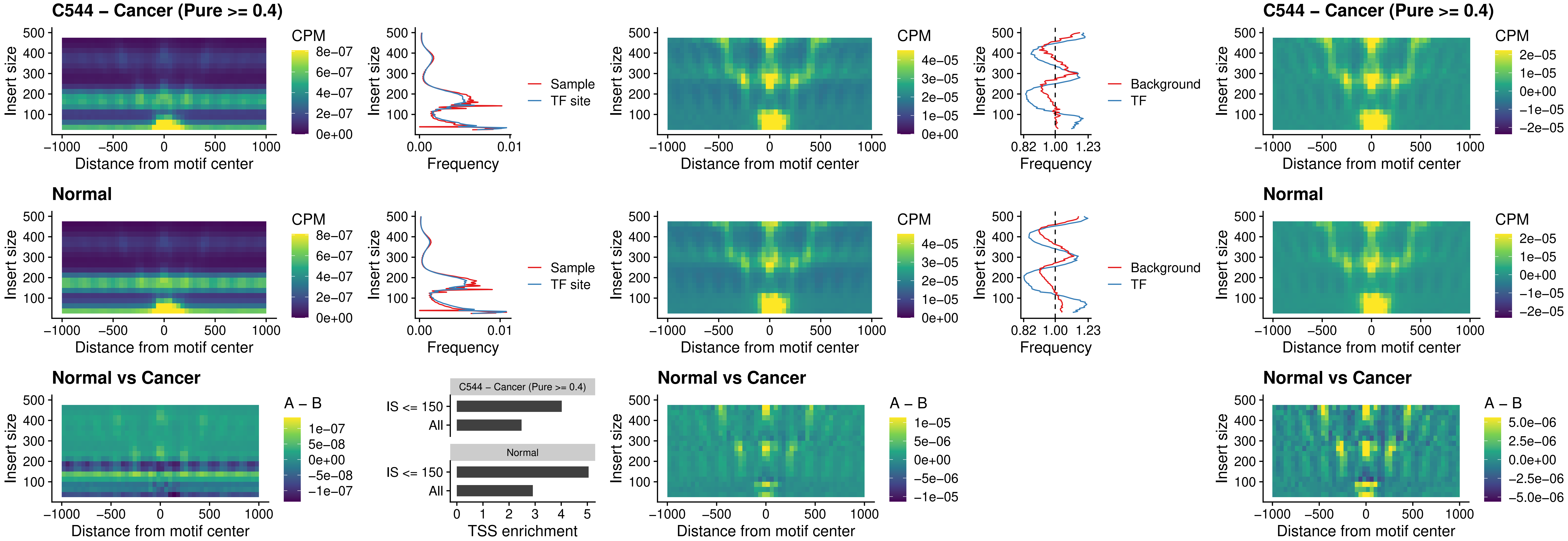

#### C544 – Cancer (Pure >= 0.4) – CTCFL_uc_cTSS_oPEAK

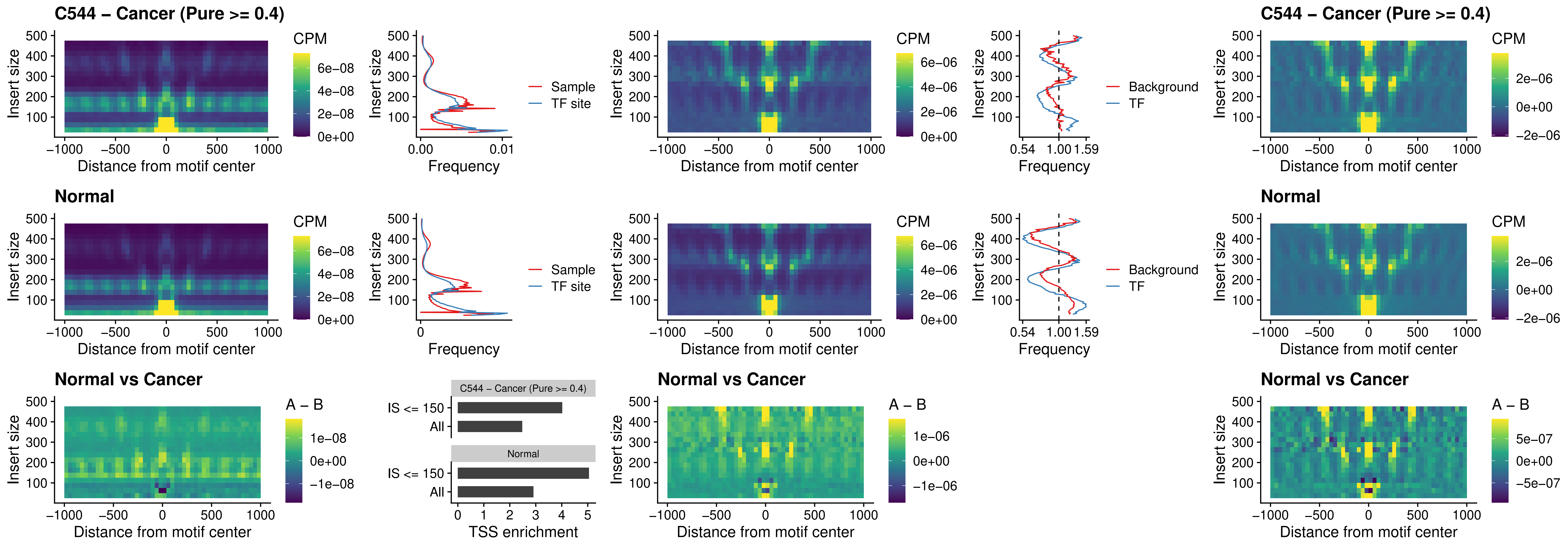

#### C544 – Cancer (Pure >= 0.4) – NEUROD1_uc_cTSS_oPEAK

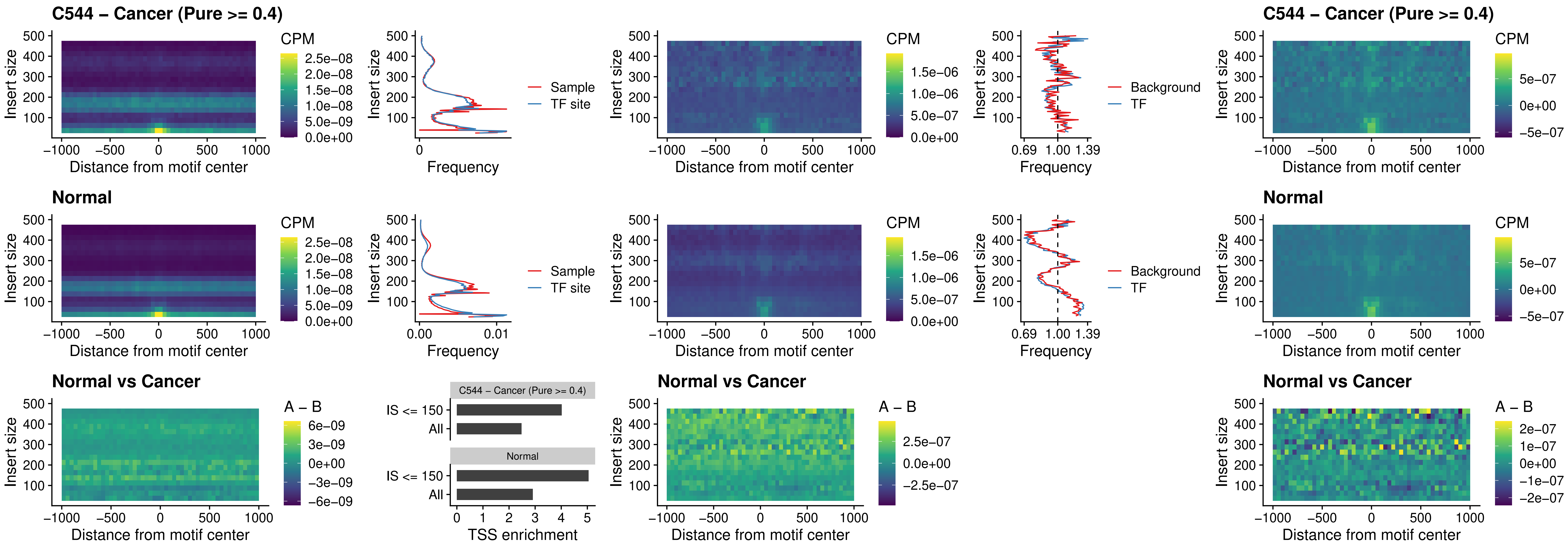

#### C544 – Cancer (Pure >= 0.4) – ENCODE_YY1_sites_COLON_nonpromotor

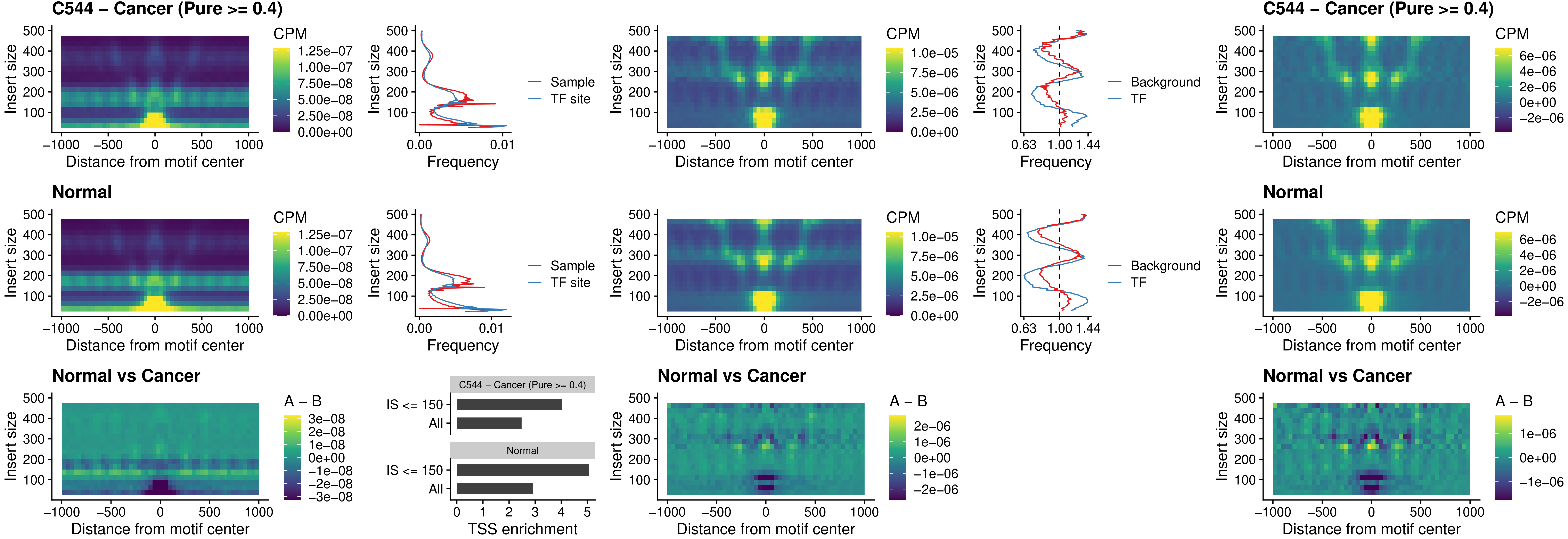

#### C544 – Cancer (Pure >= 0.4) – CTCF_uc_pTSS_oPEAK

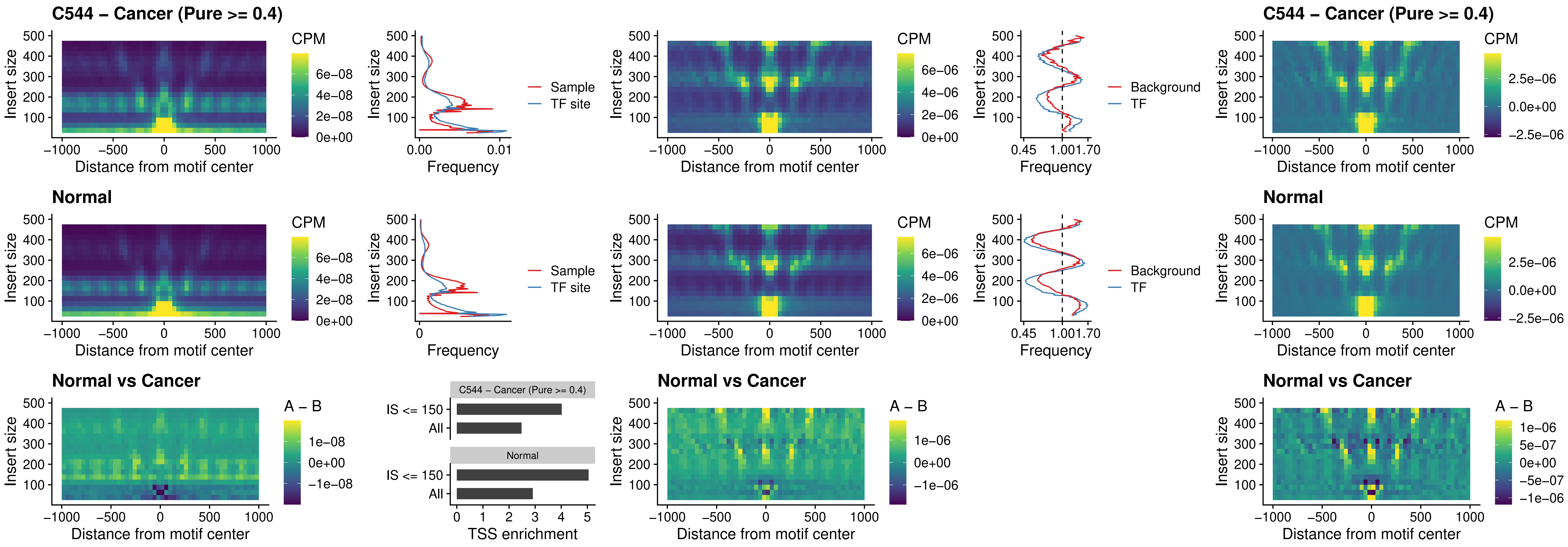

### C544 – Cancer (Pure >= 0.4) – ENCODE_CTCF_sites_COLON_promotor

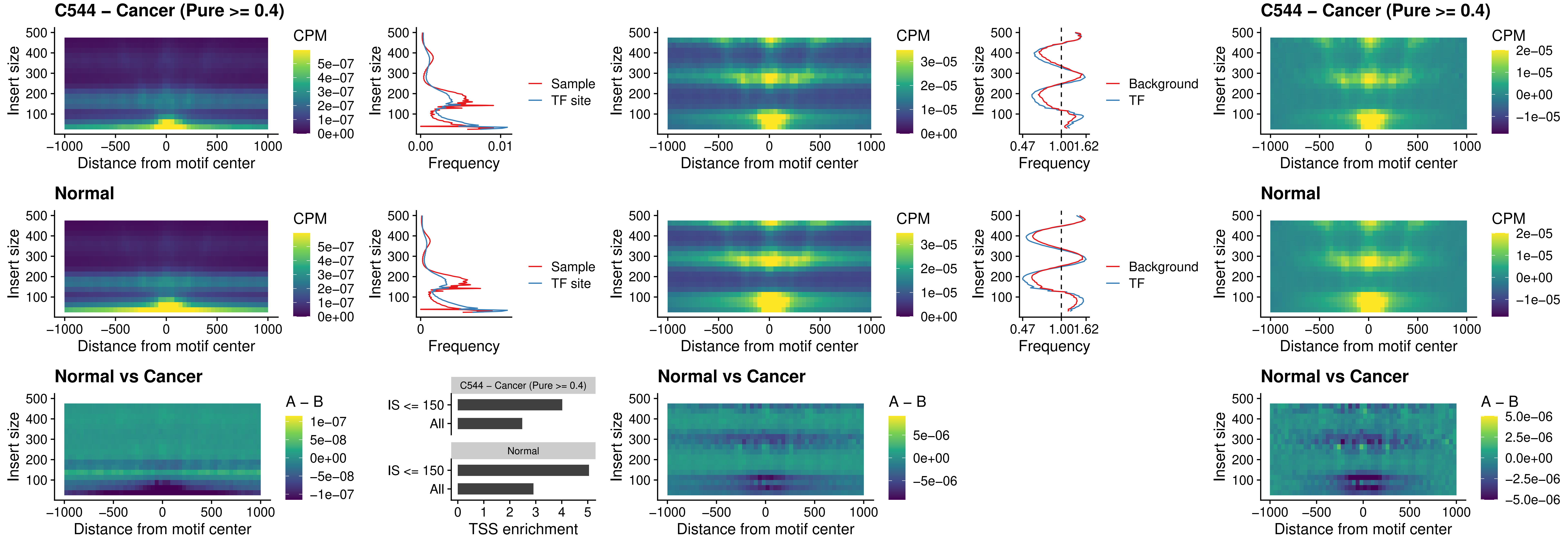

#### C544 – Cancer (Pure >= 0.4) – ENCODE_CTCF_sites_COLON_promotor_ctcf

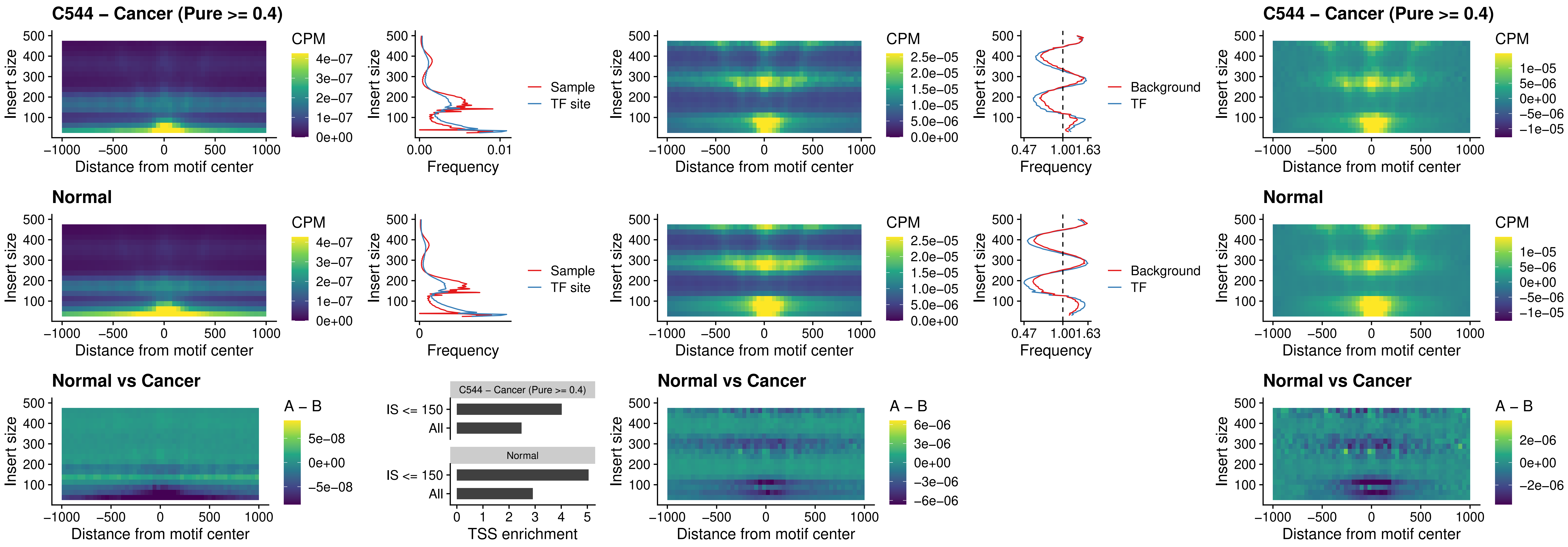

#### C544 – Cancer (Pure >= 0.4) – SPIC_uc_pTSS_oPEAK

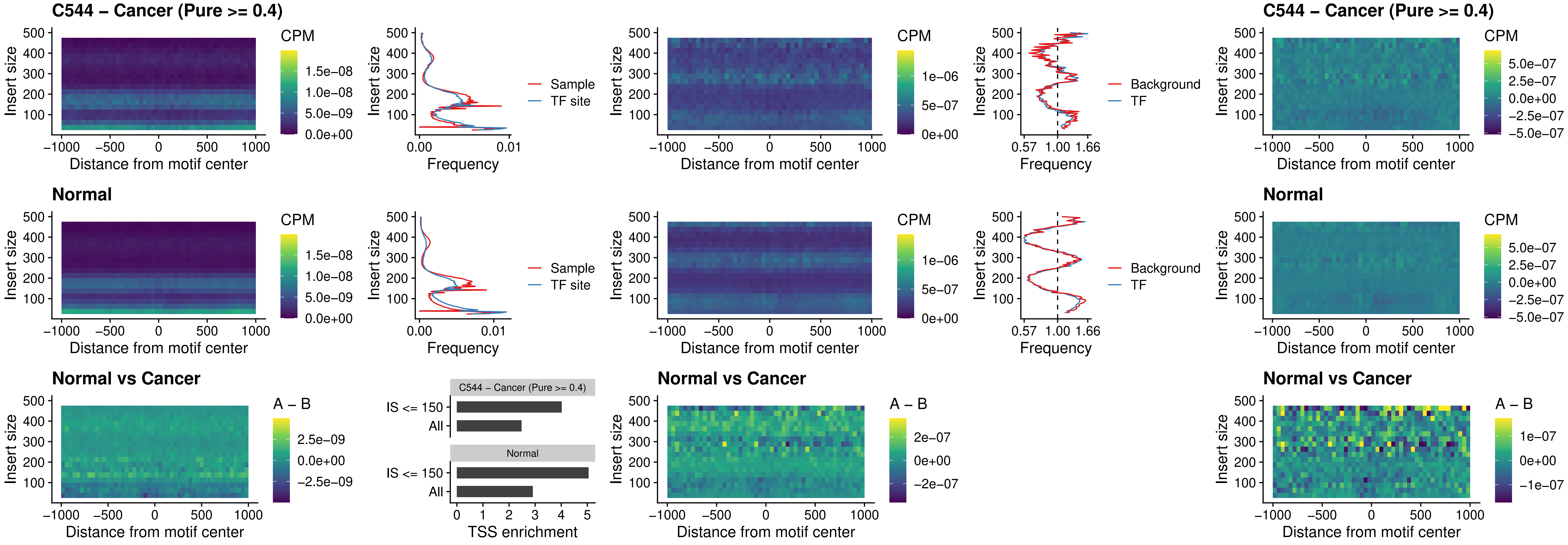

#### C544 – Cancer (Pure >= 0.4) – SPI1_uc_dTSS_nPEAK

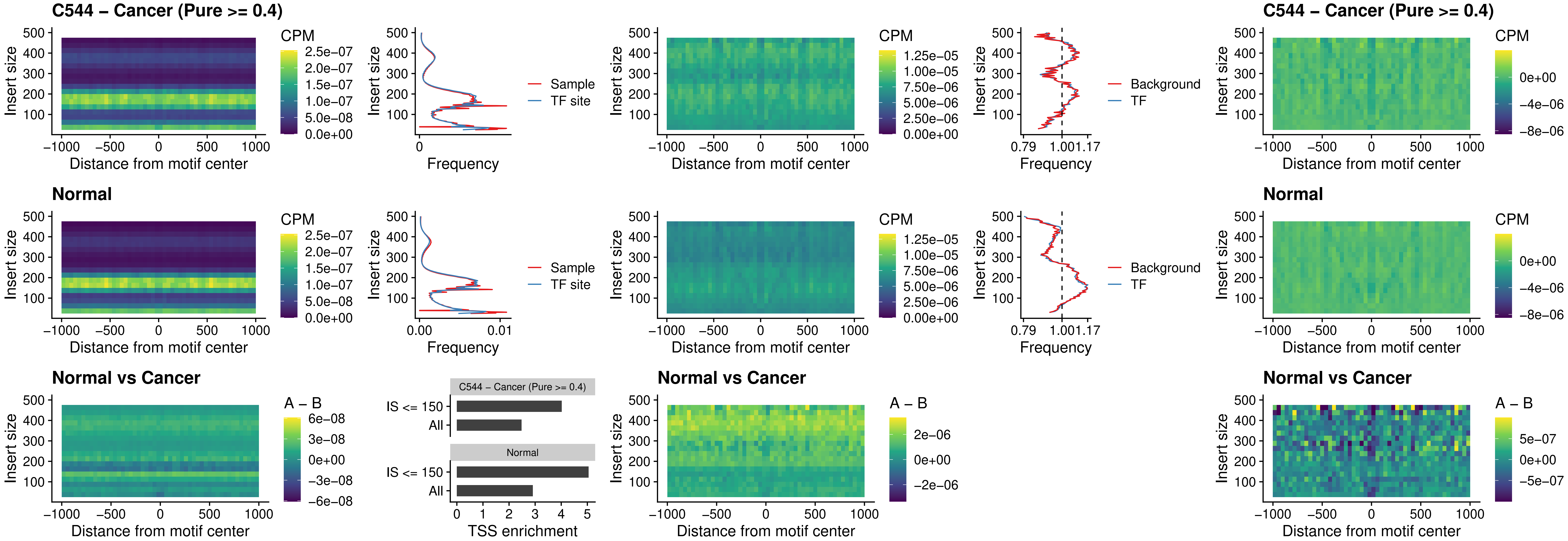

#### C544 – Cancer (Pure >= 0.4) – SPI1_uc_cTSS_nPEAK

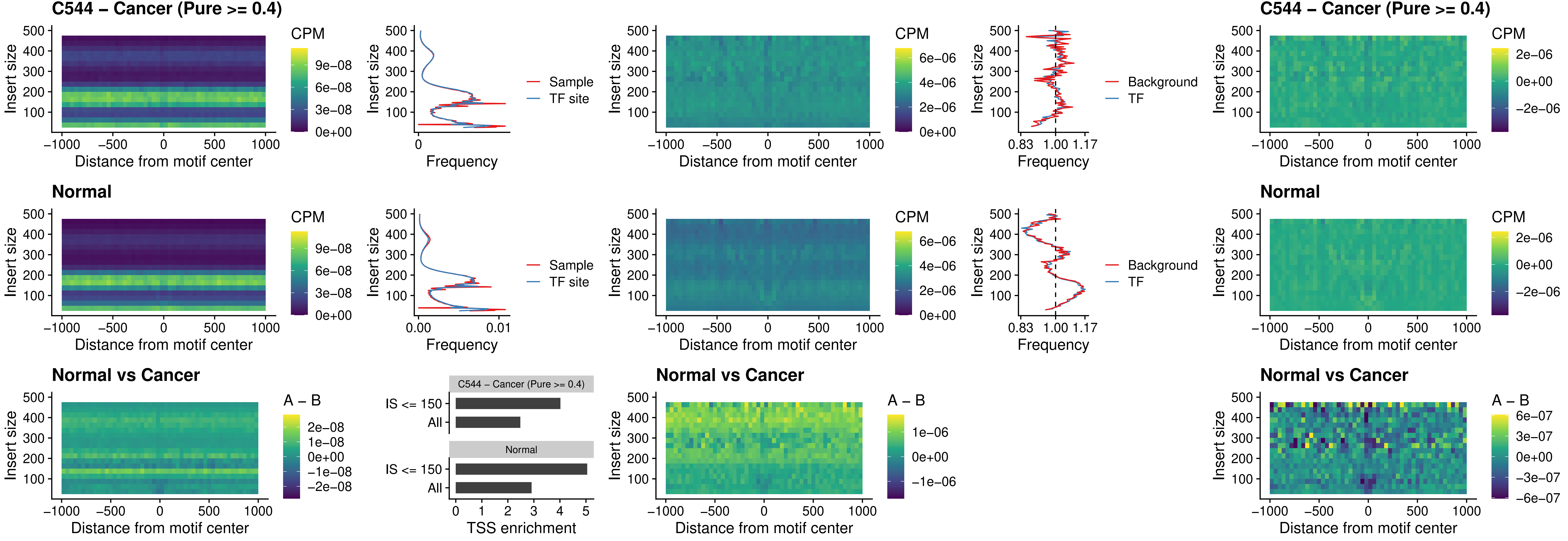

#### C544 – Cancer (Pure >= 0.4) – BCL11A_uc_pTSS_nPEAK

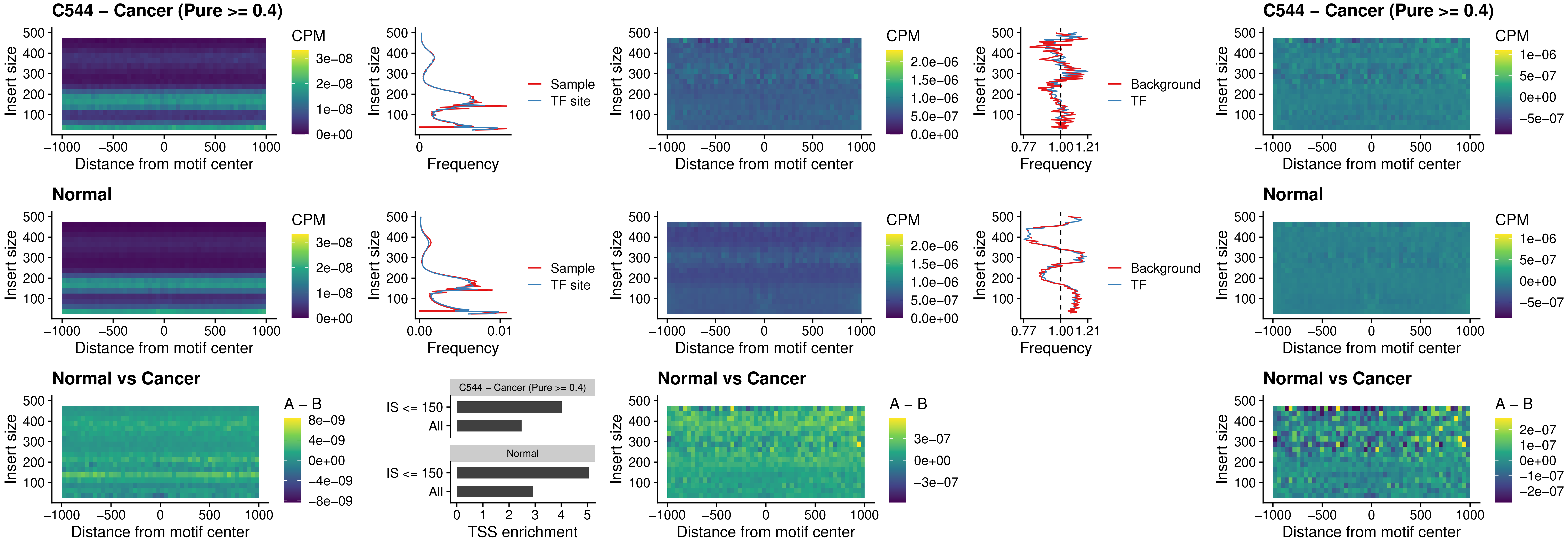

#### C544 – Cancer (Pure >= 0.4) – BCL11B_uc_pTSS_nPEAK

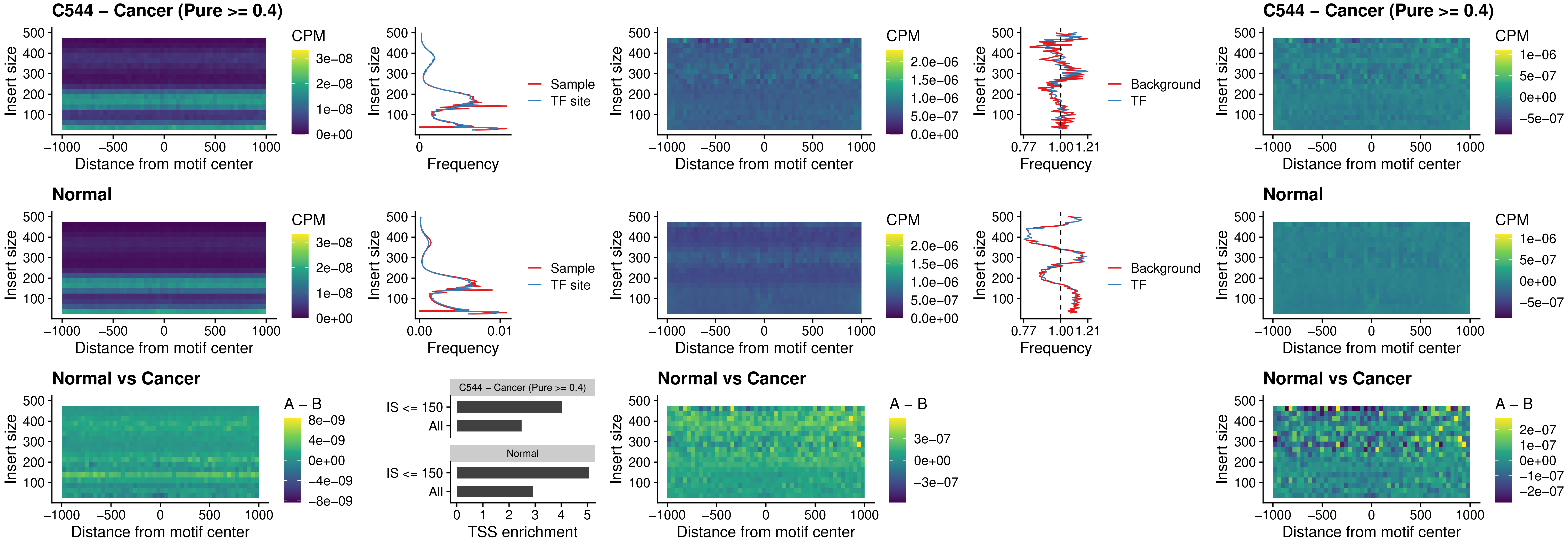

#### C544 – Cancer (Pure >= 0.4) – IRF9_uc_dTSS_oPEAK

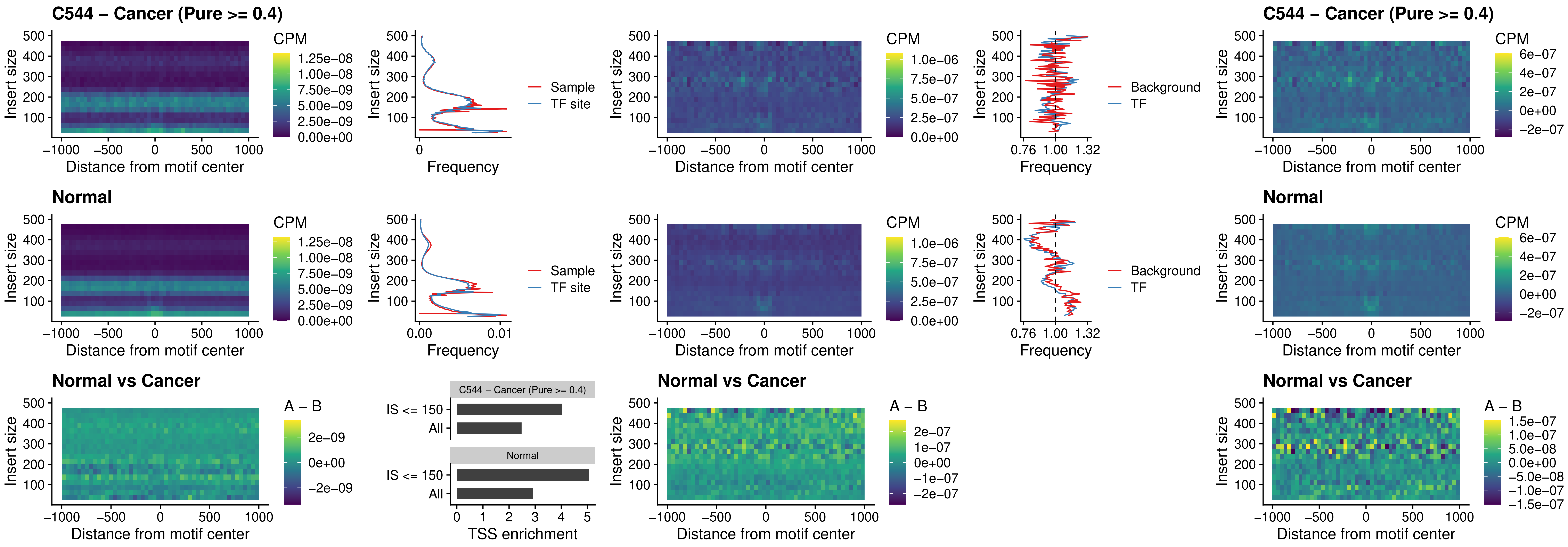

#### C544 – Cancer (Pure >= 0.4) – IRF9_uc_cTSS_oPEAK

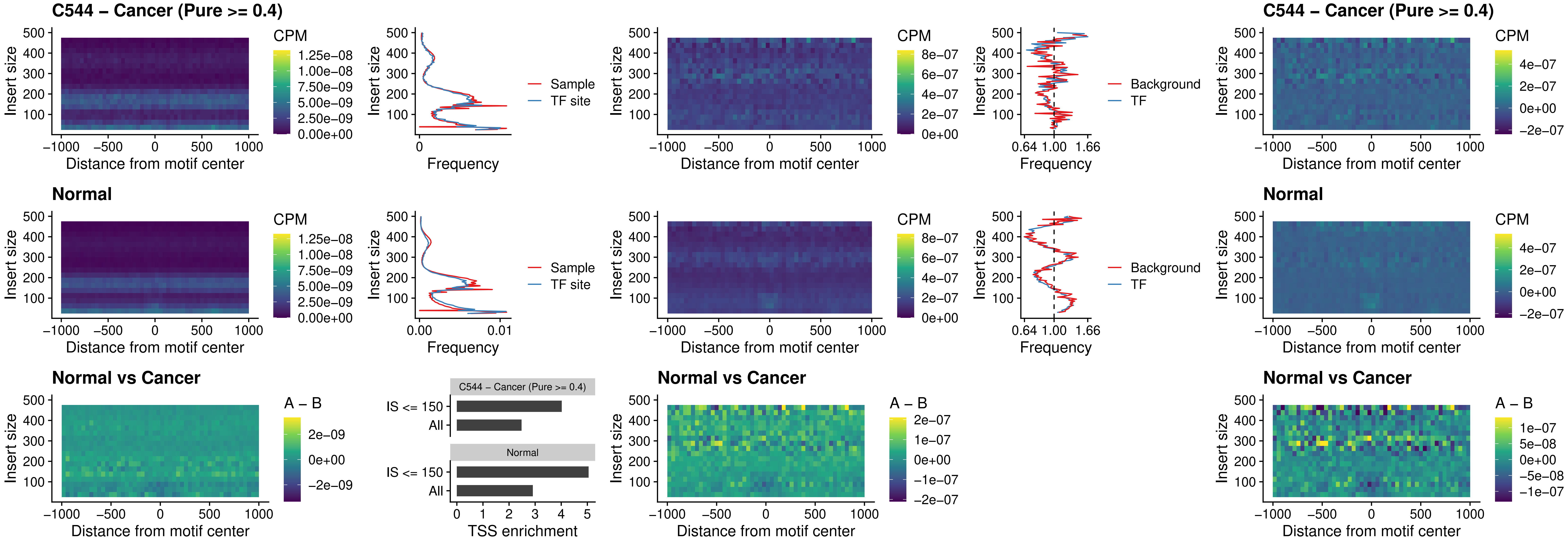

#### C544 – Cancer (Pure >= 0.4) – BCL11A_uc_cTSS_oPEAK

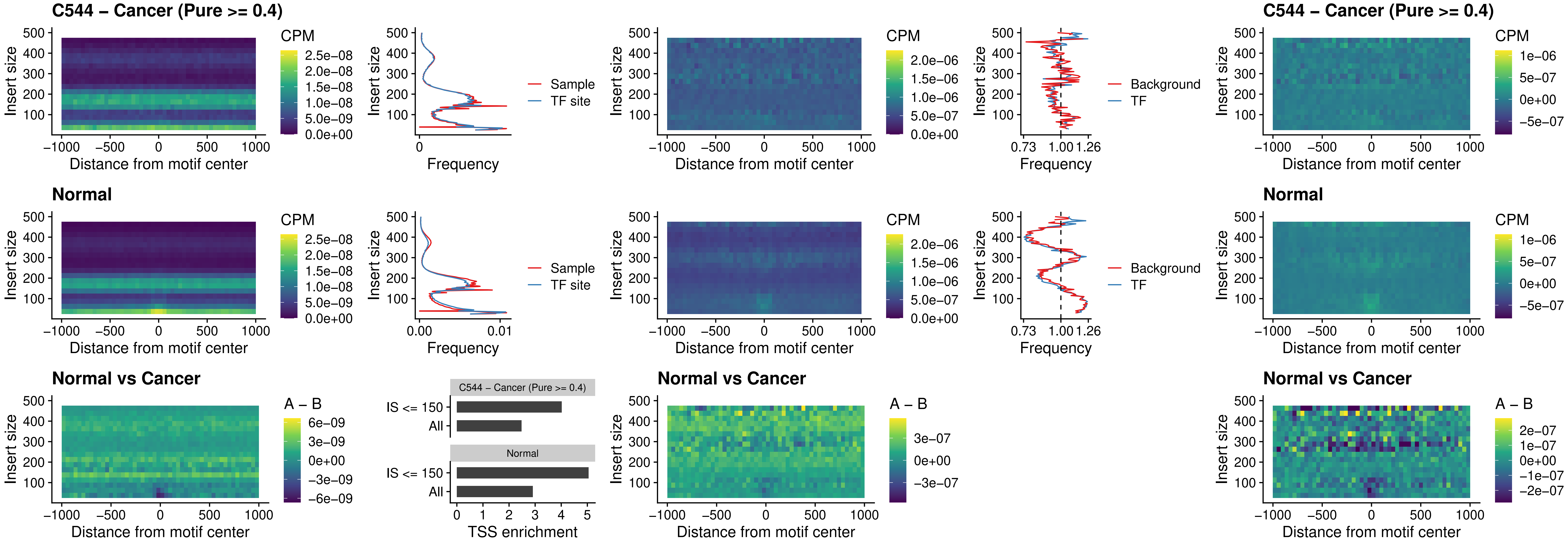

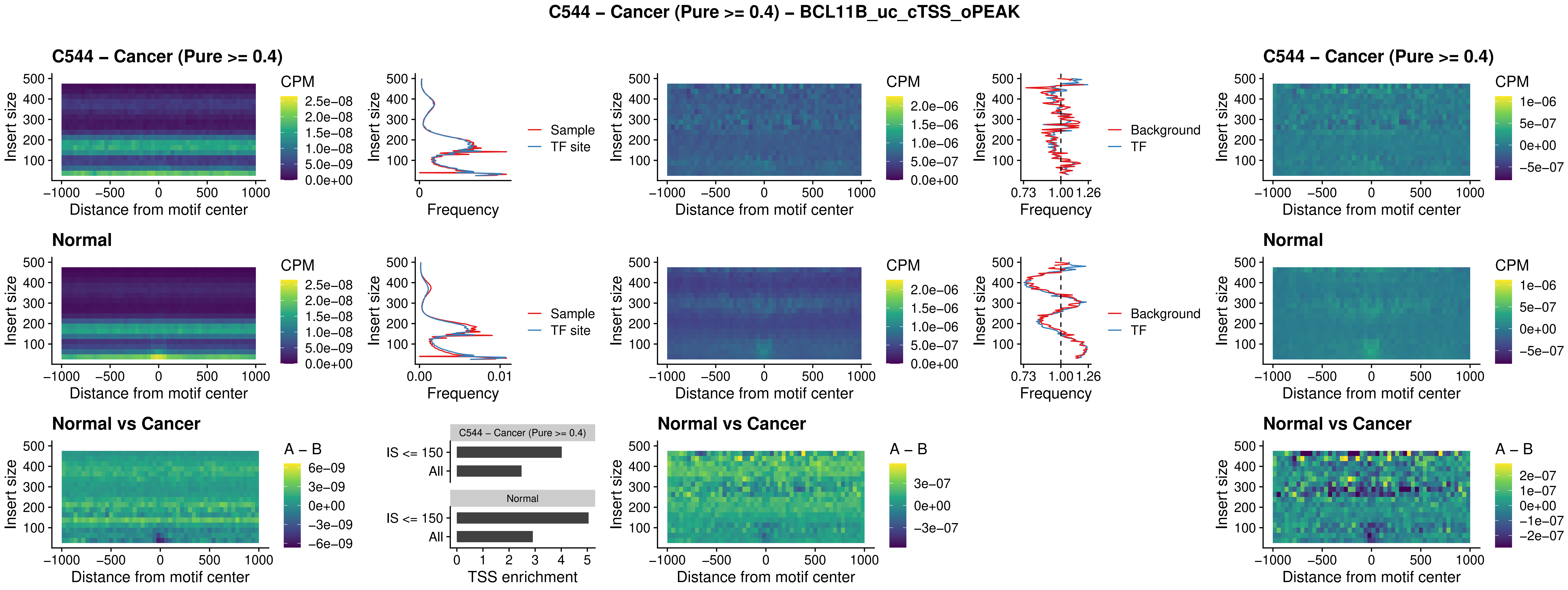
C544 – Cancer (Pure >= 0.4) – BCL11B_uc_cTSS_oPEAK
C544 – Cancer (Pure >= 0.4)
Normal
Normal vs Cancer
Insert size
Distance from motif center
Frequency
CPM
A – B
Sample
TF site
Background
TF
C544 – Cancer (Pure >= 0.4)
Normal
IS <= 150
All
TSS enrichment

### C544 – Cancer (Pure >= 0.4) – BCL11A_uc_dTSS_oPEAK

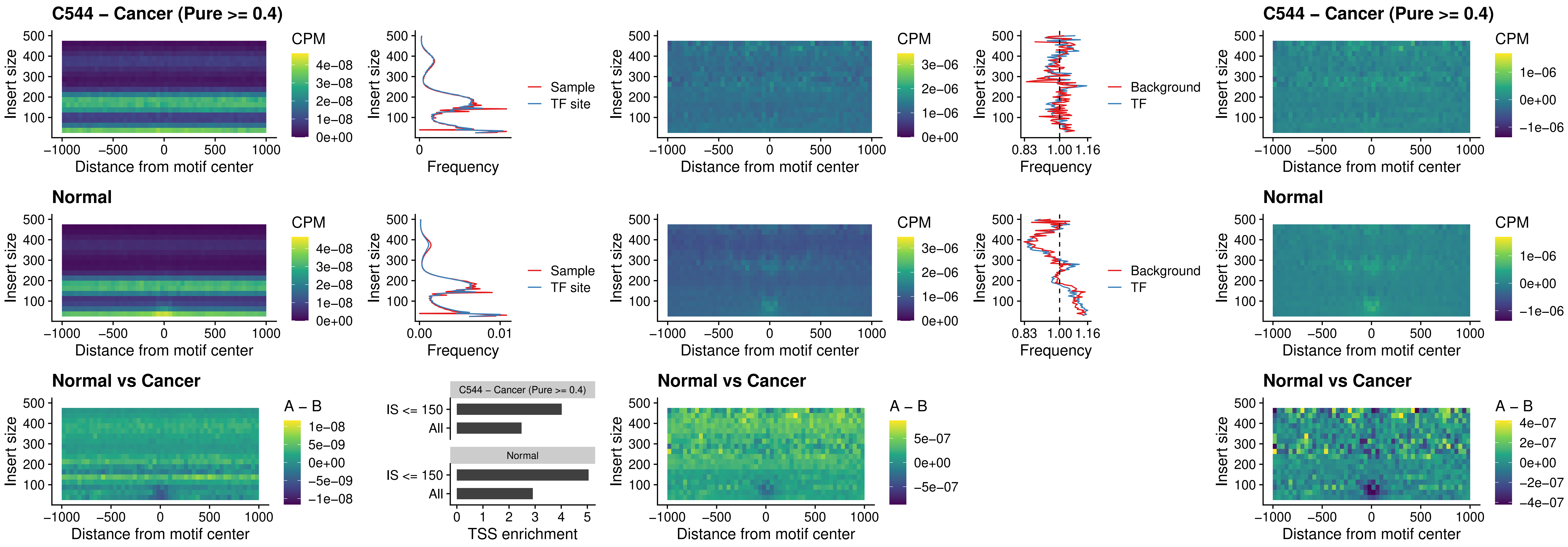

#### C544 – Cancer (Pure >= 0.4) – BCL11B_uc_dTSS_oPEAK

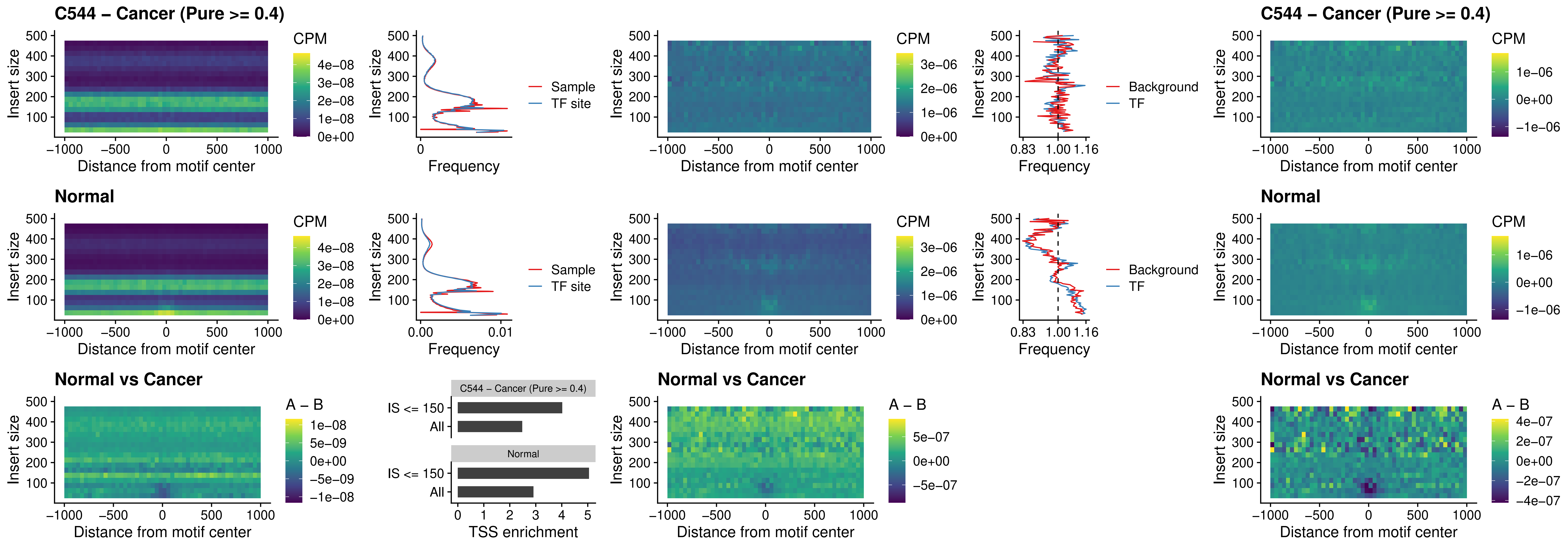

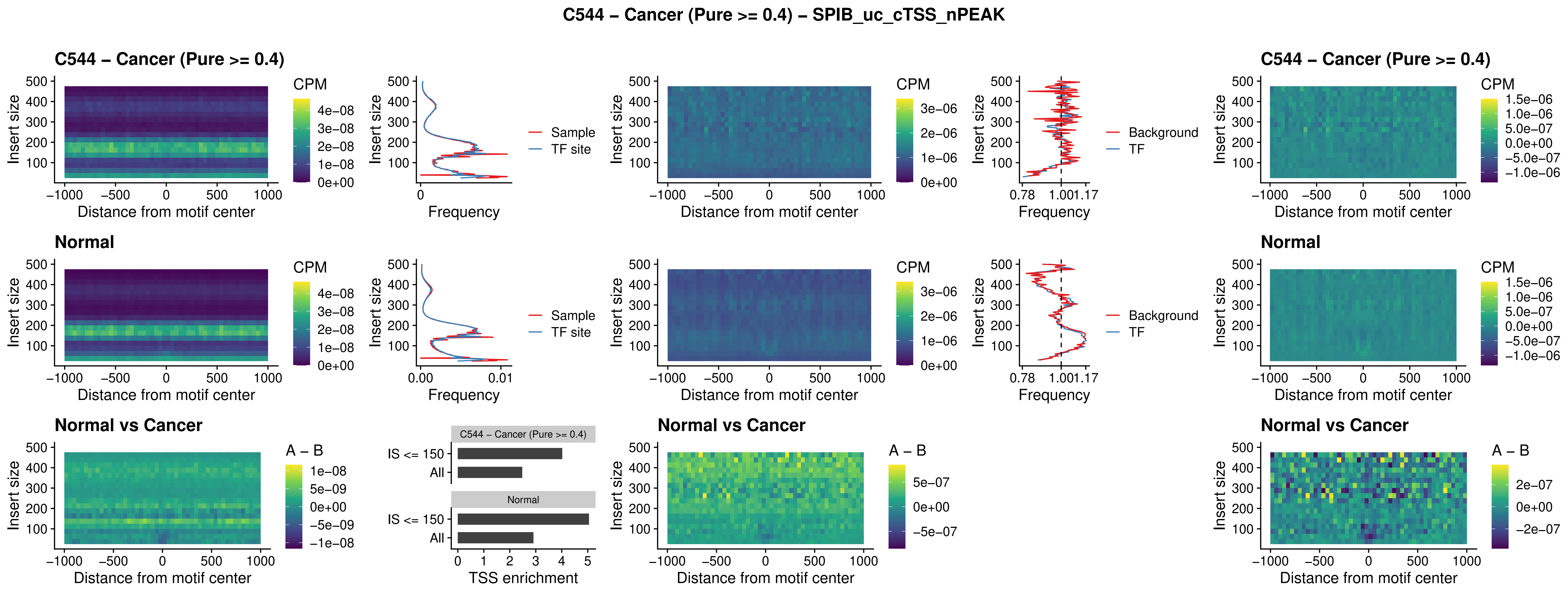
C544 – Cancer (Pure >= 0.4) – SPIB_uc_cTSS_nPEAK
C544 – Cancer (Pure >= 0.4)
Normal
Normal vs Cancer
Insert size
Distance from motif center
Frequency
CPM
A – B
Sample
TF site
Background
TF
TSS enrichment
IS <= 150
All

#### C544 – Cancer (Pure >= 0.4) – SPIC_uc_cTSS_nPEAK

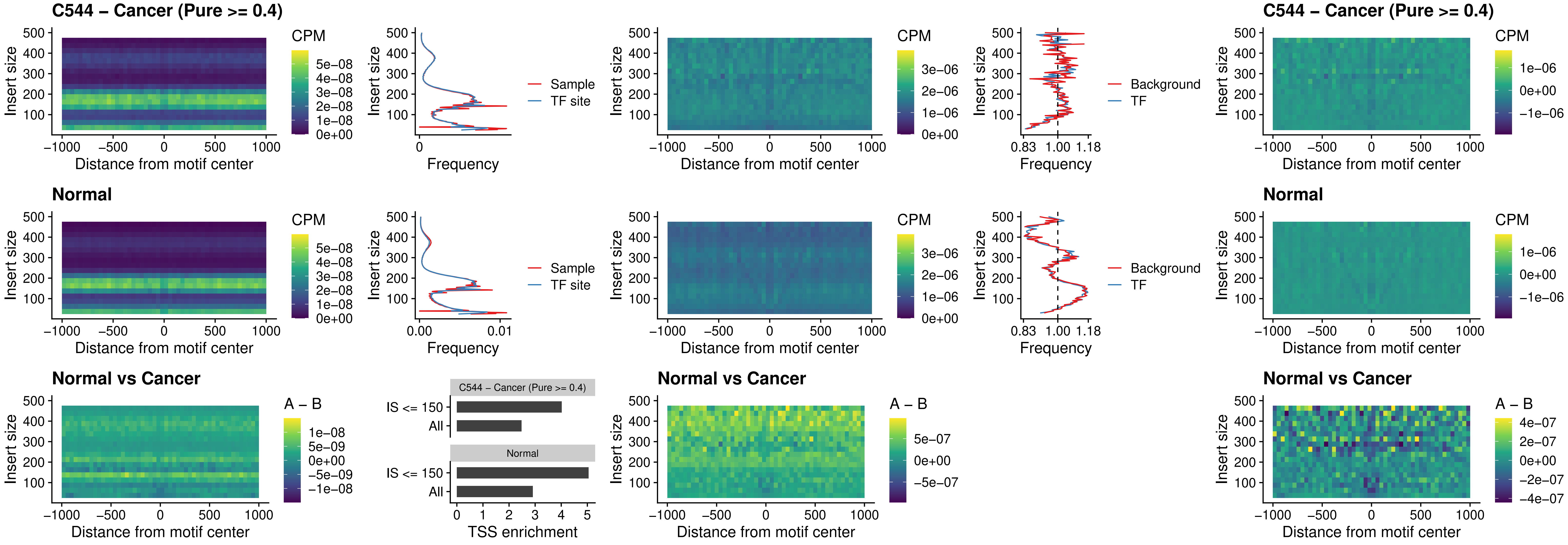

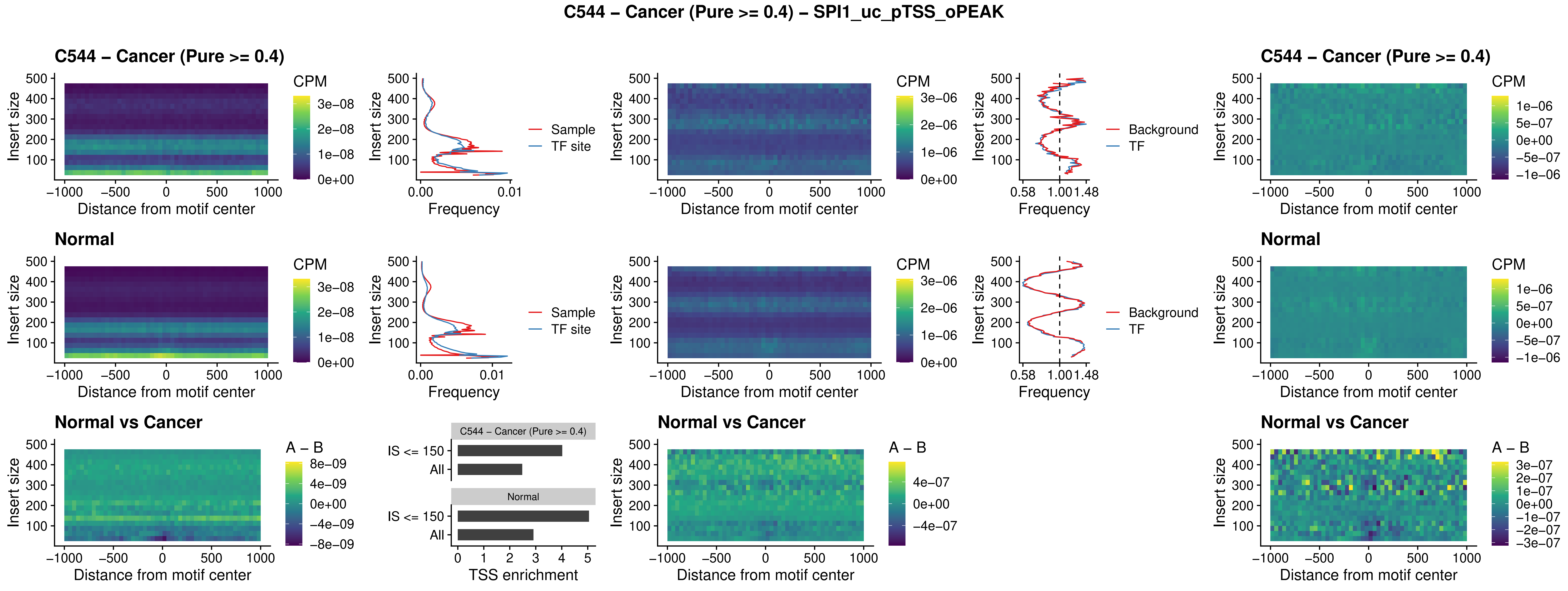
C544 – Cancer (Pure >= 0.4) – SPI1_uc_pTSS_oPEAK
C544 – Cancer (Pure >= 0.4)
Normal
Normal vs Cancer
Insert size
Distance from motif center
Frequency
CPM
A – B
Sample
TF site
Background
TF
C544 – Cancer (Pure >= 0.4)
Normal
IS <= 150
All
TSS enrichment

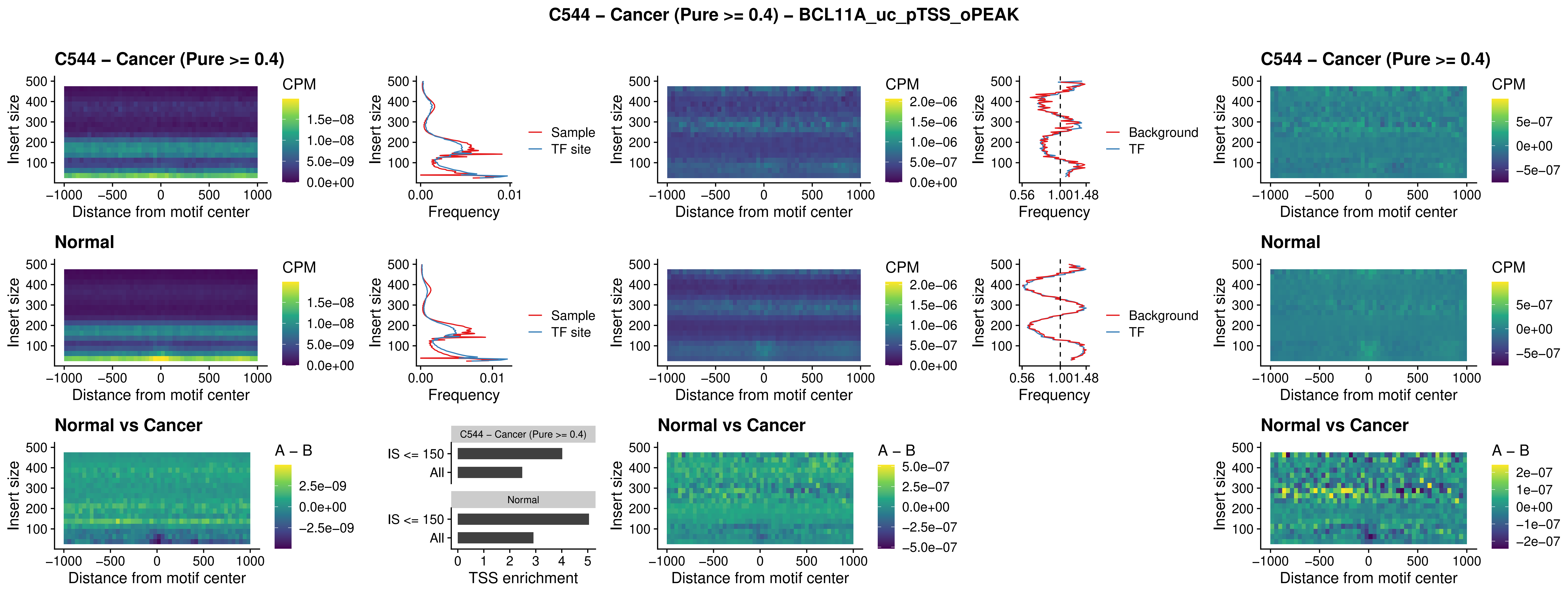
C544 – Cancer (Pure >= 0.4) – BCL11A_uc_pTSS_oPEAK
C544 – Cancer (Pure >= 0.4)
Normal
Normal vs Cancer
Insert size
Distance from motif center
Frequency
CPM
A – B
Sample
TF site
Background
TF
C544 – Cancer (Pure >= 0.4)
Normal
IS <= 150
All
TSS enrichment

#### C544 – Cancer (Pure >= 0.4) – BCL11B_uc_pTSS_oPEAK

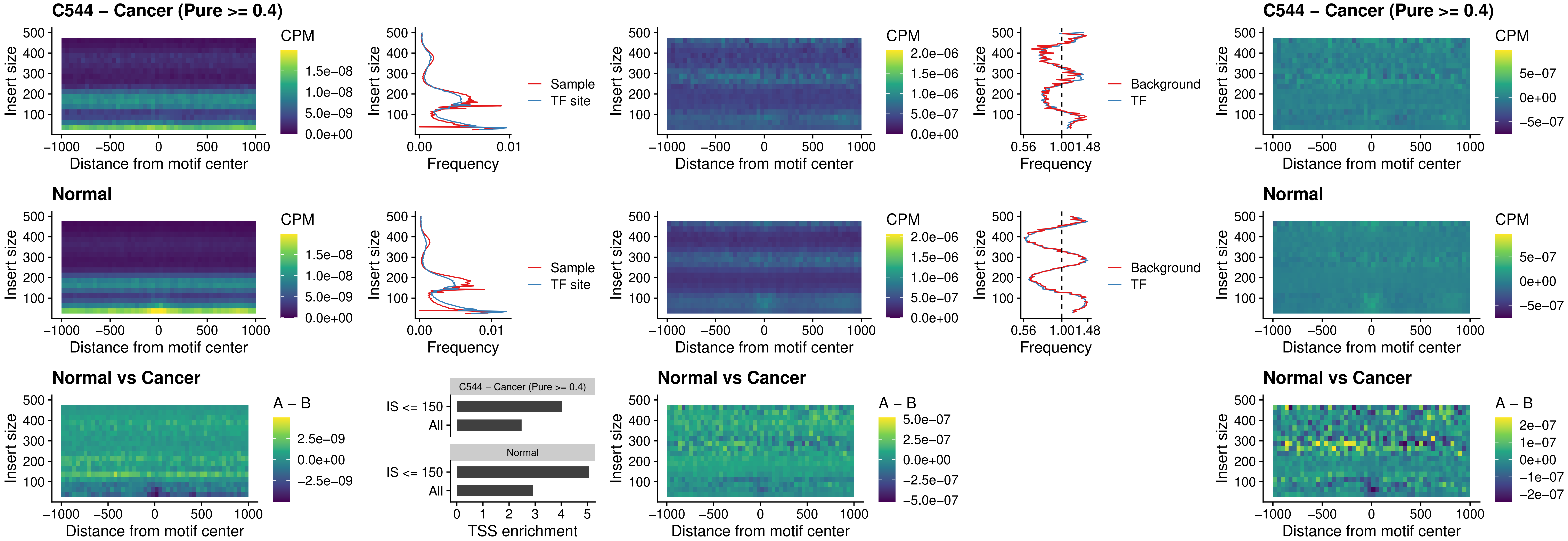

#### C544 – Cancer (Pure >= 0.4) – SPI1_uc_cTSS_oPEAK

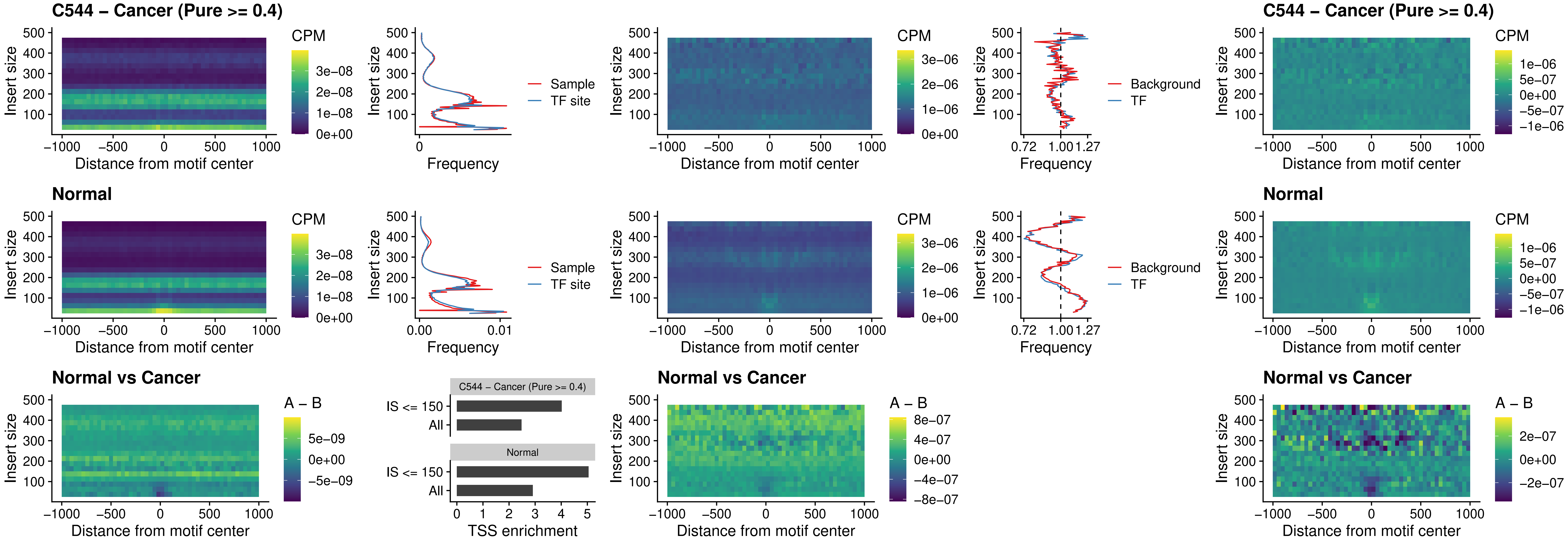

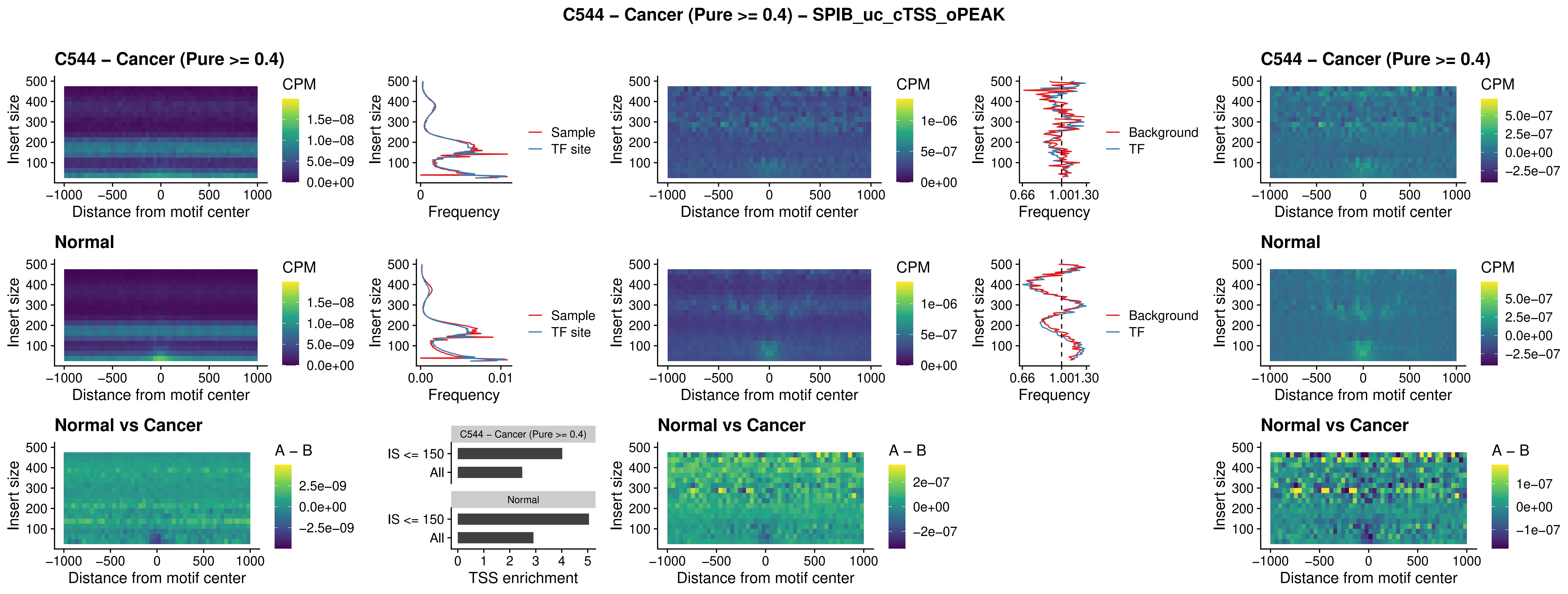
C544 – Cancer (Pure >= 0.4) – SPIB_uc_cTSS_oPEAK
C544 – Cancer (Pure >= 0.4)
Normal
Normal vs Cancer
Insert size
Distance from motif center
Frequency
CPM
A – B
Sample
TF site
Background
TF
C544 – Cancer (Pure >= 0.4)
Normal
IS <= 150
All
TSS enrichment

#### C544 – Cancer (Pure >= 0.4) – SPIB_uc_pTSS_oPEAK

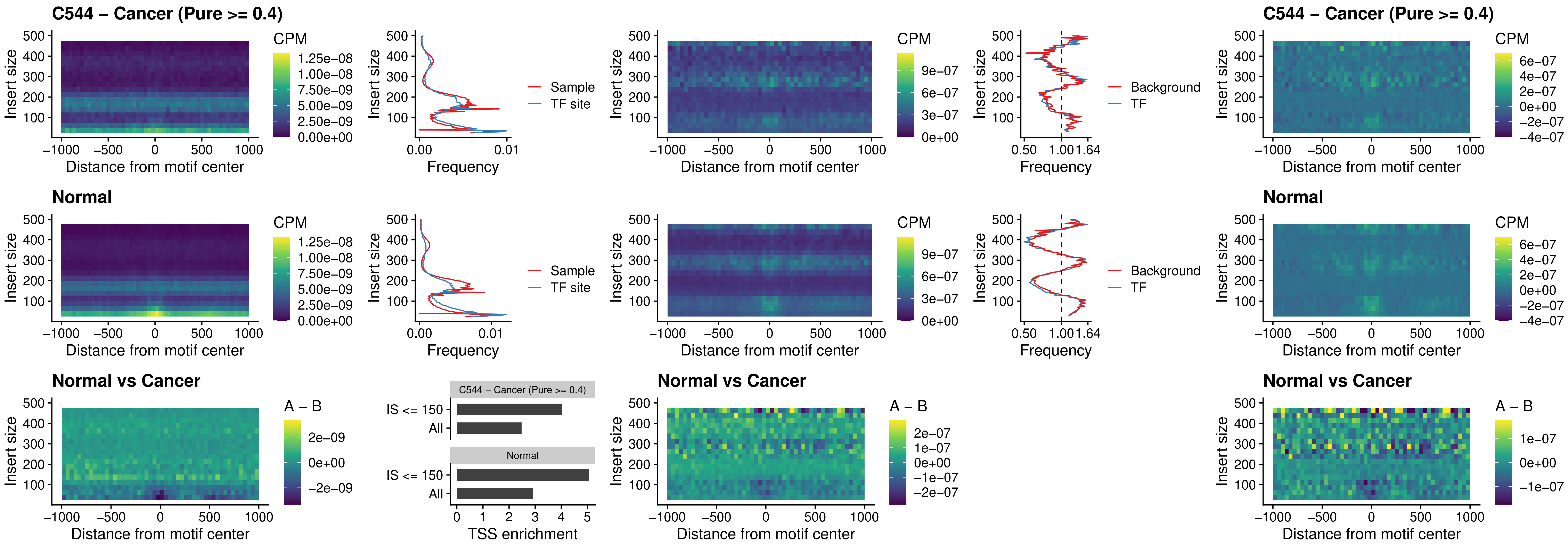

#### C544 – Cancer (Pure >= 0.4) – SPI1_uc_dTSS_oPEAK

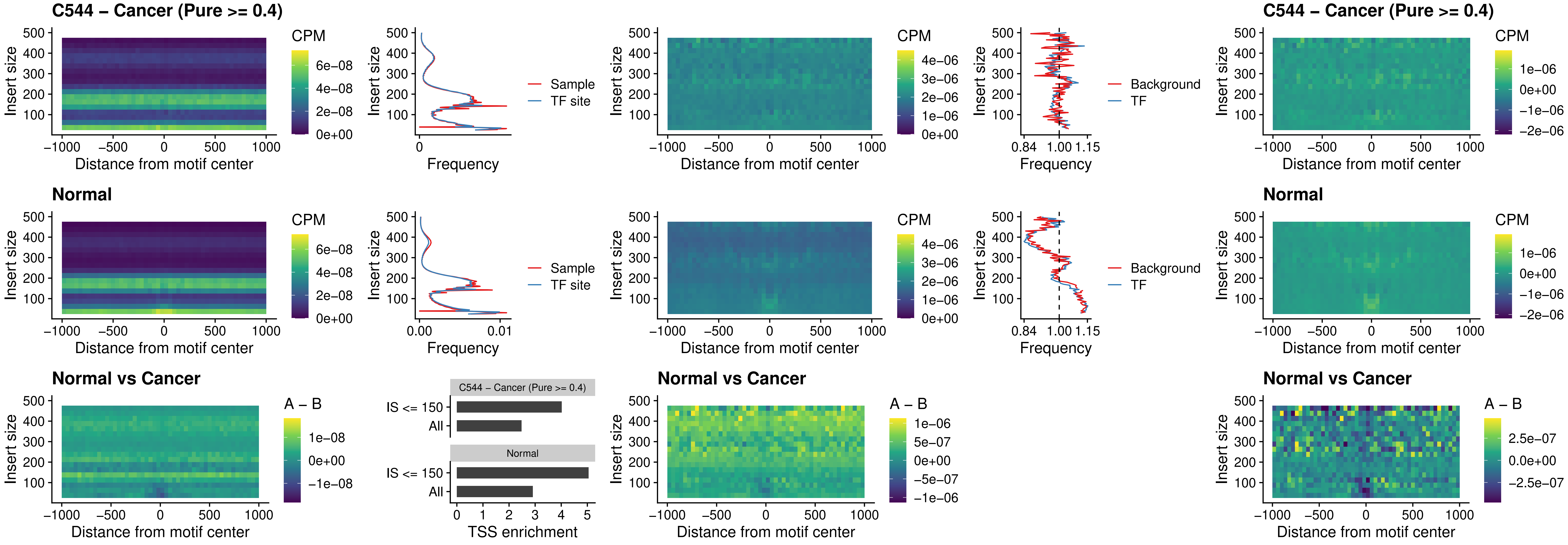

#### C544 – Cancer (Pure >= 0.4) – SPIB_uc_dTSS_oPEAK

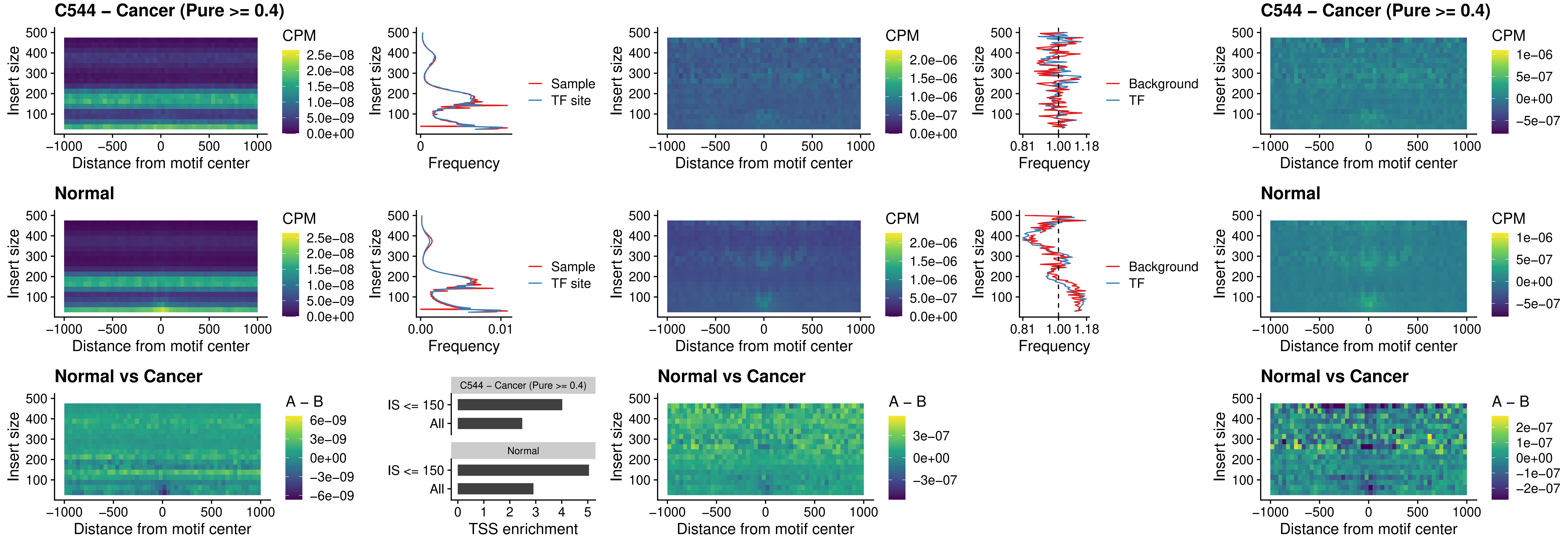

#### C544 – Cancer (Pure >= 0.4) – IRF3_uc_pTSS_oPEAK

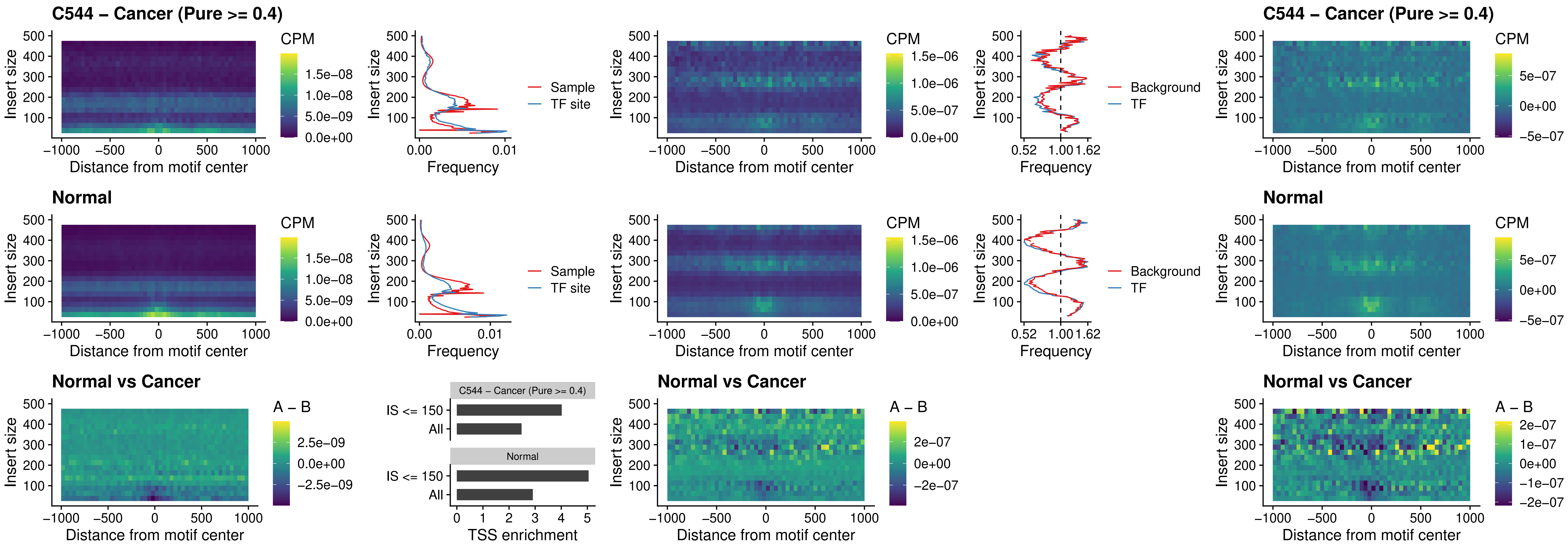

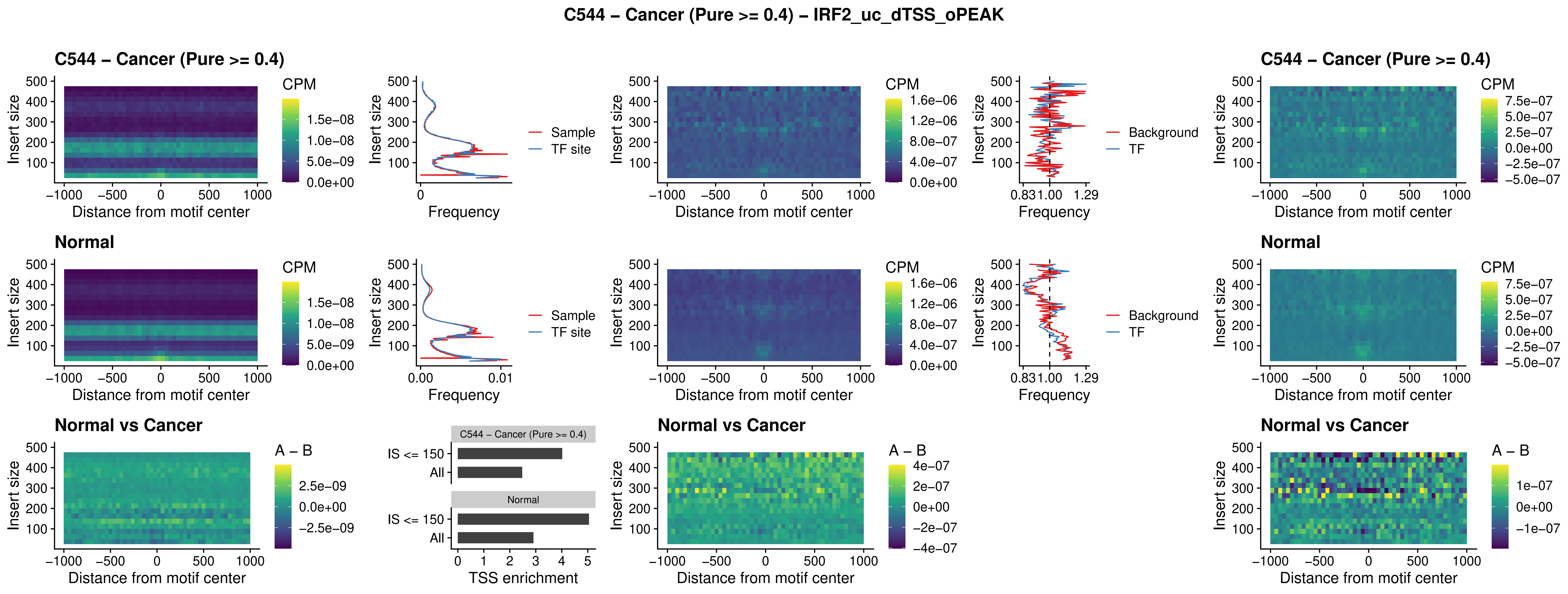
C544 – Cancer (Pure >= 0.4) – IRF2_uc_dTSS_oPEAK
C544 – Cancer (Pure >= 0.4)
Normal
Normal vs Cancer
Insert size
Distance from motif center
Frequency
CPM
A – B
Sample
TF site
Background
TF
TSS enrichment
IS <= 150
All
C544 – Cancer (Pure >= 0.4)
Normal

#### C544 – Cancer (Pure >= 0.4) – IRF2_uc_pTSS_oPEAK

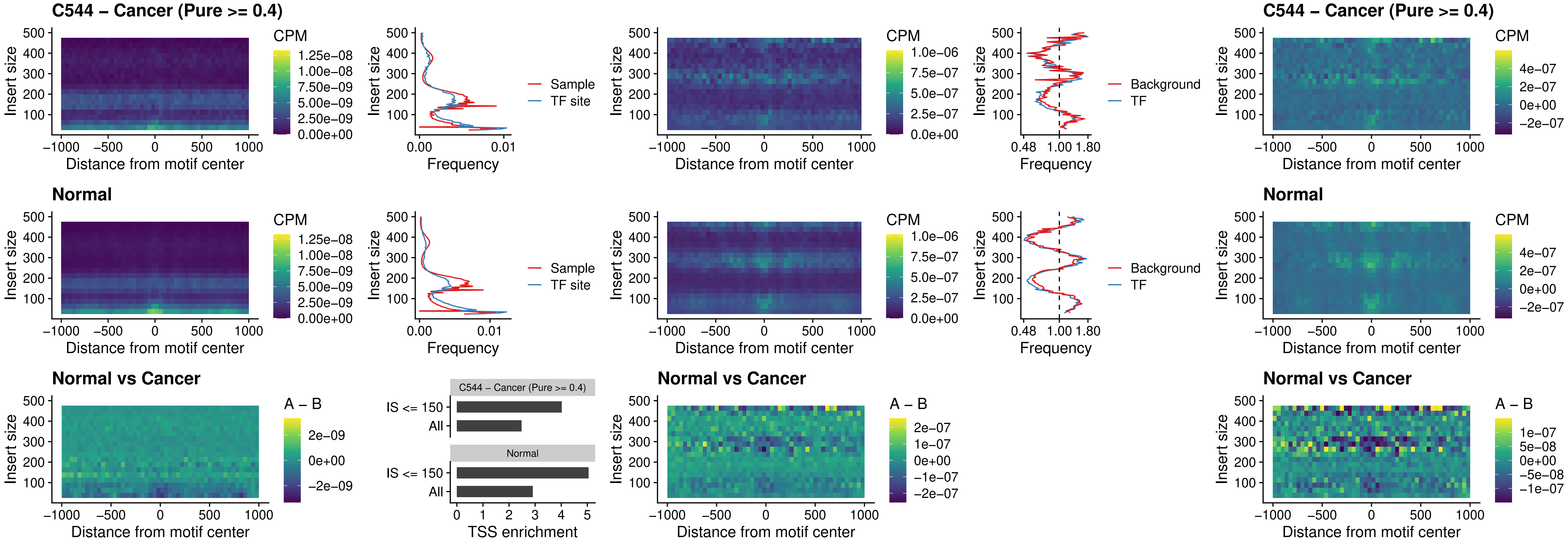

#### C544 – Cancer (Pure >= 0.4) – STAT2_uc_pTSS_oPEAK

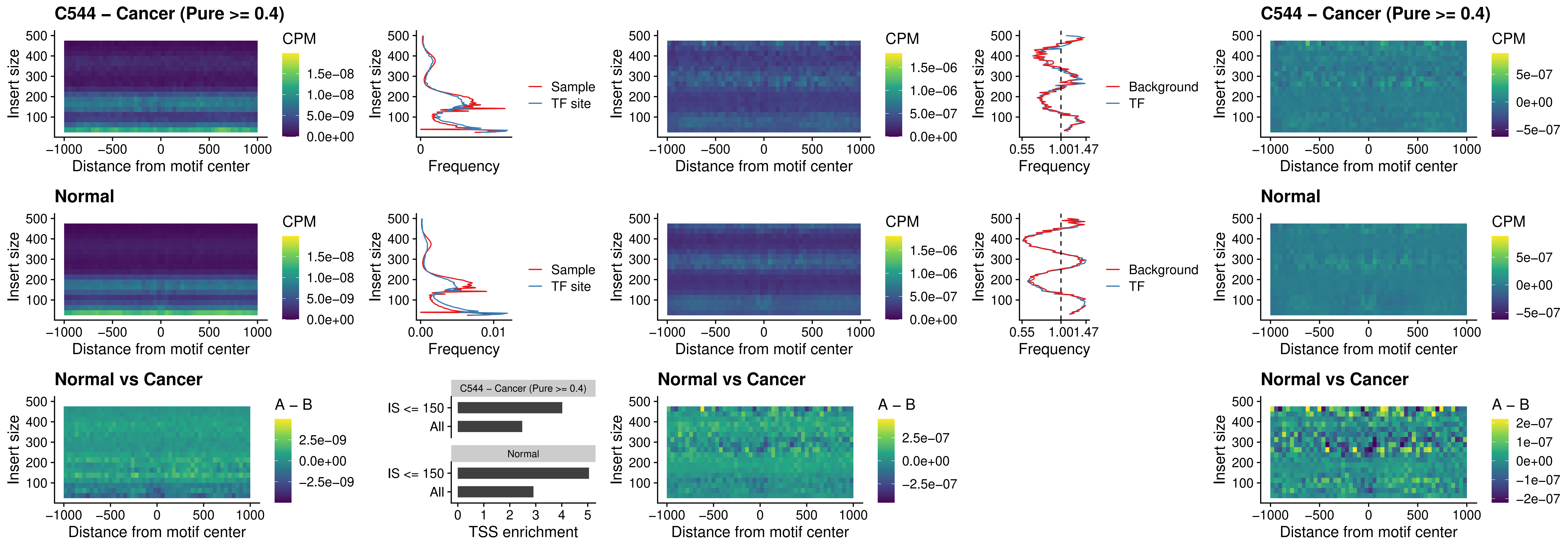

#### C544 – Cancer (Pure >= 0.4) – SPIB_uc_dTSS_nPEAK

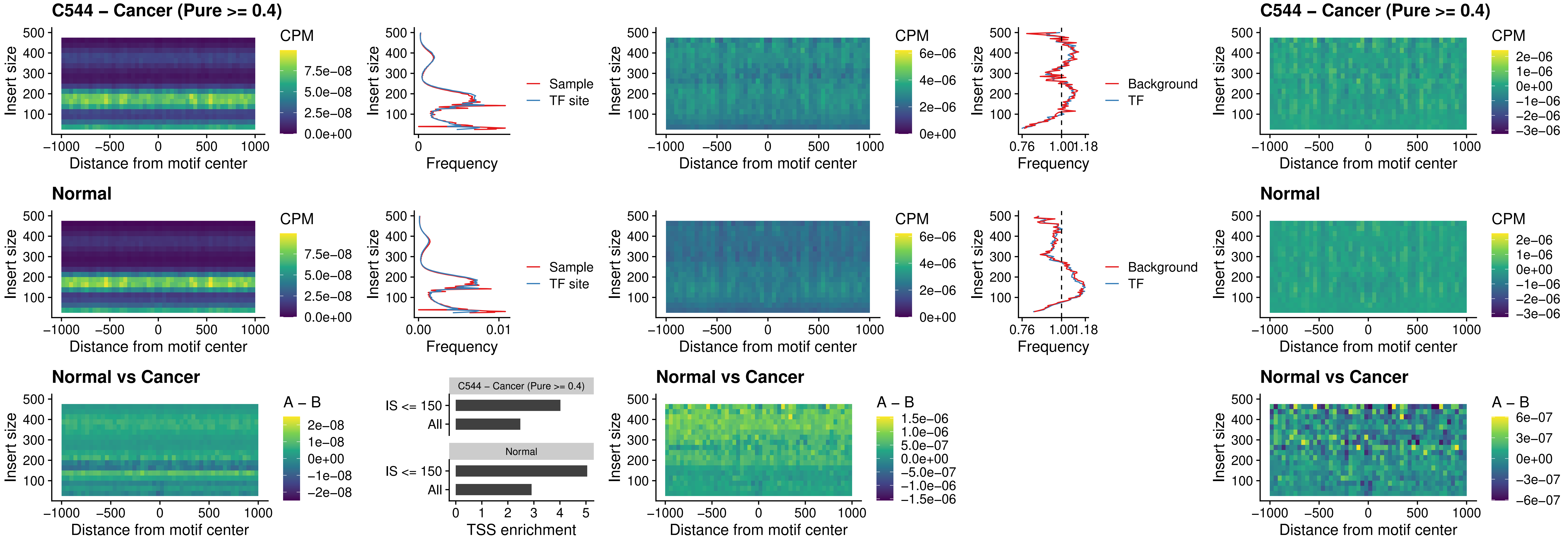

#### C544 – Cancer (Pure >= 0.4) – SPIC_uc_cTSS_oPEAK

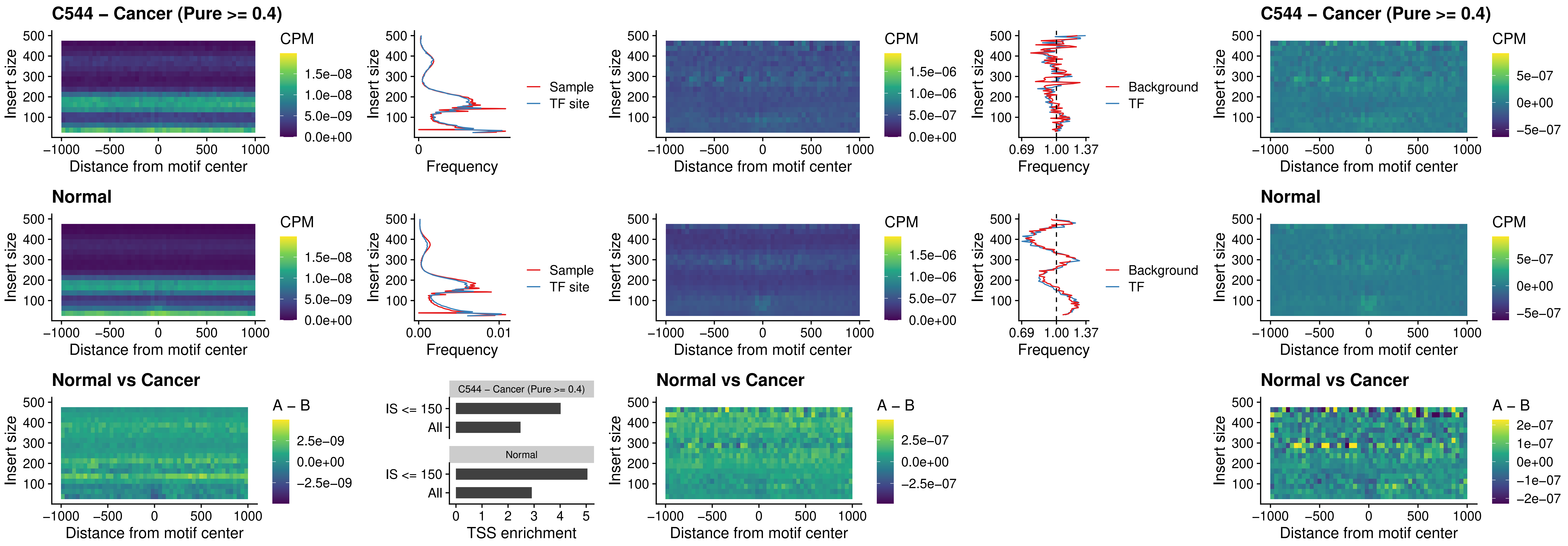

#### C544 – Cancer (Pure >= 0.4) – PRDM1_uc_cTSS_oPEAK

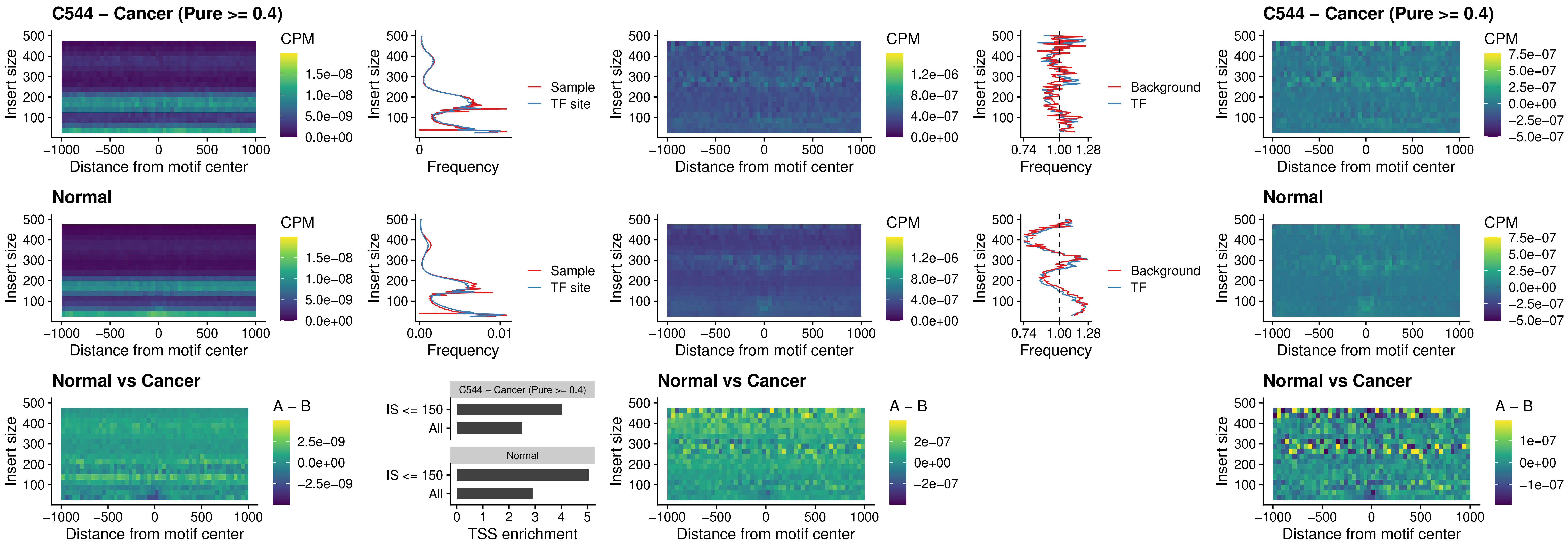

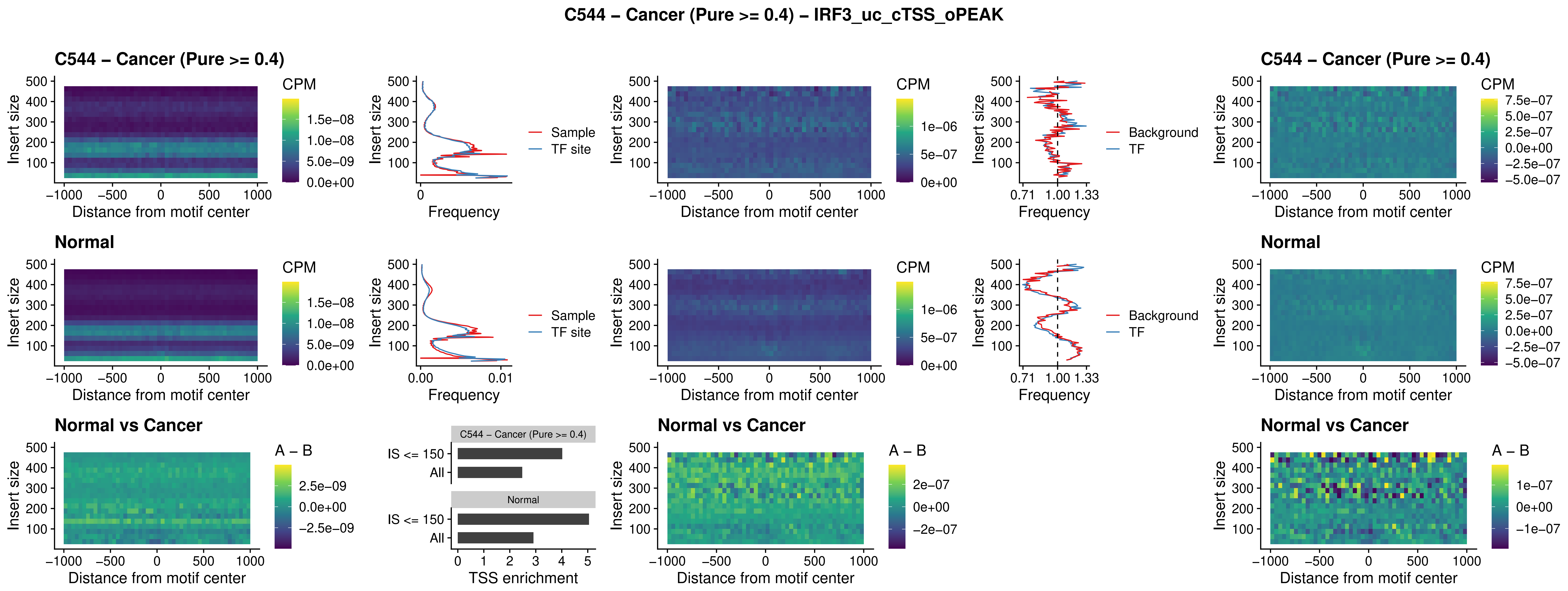
C544 – Cancer (Pure >= 0.4) – IRF3_uc_cTSS_oPEAK
C544 – Cancer (Pure >= 0.4)
Normal
Normal vs Cancer
Insert size
Distance from motif center
CPM
A – B
Frequency
Sample
TF site
Background
TF
C544 – Cancer (Pure >= 0.4)
Normal
IS <= 150
All
TSS enrichment

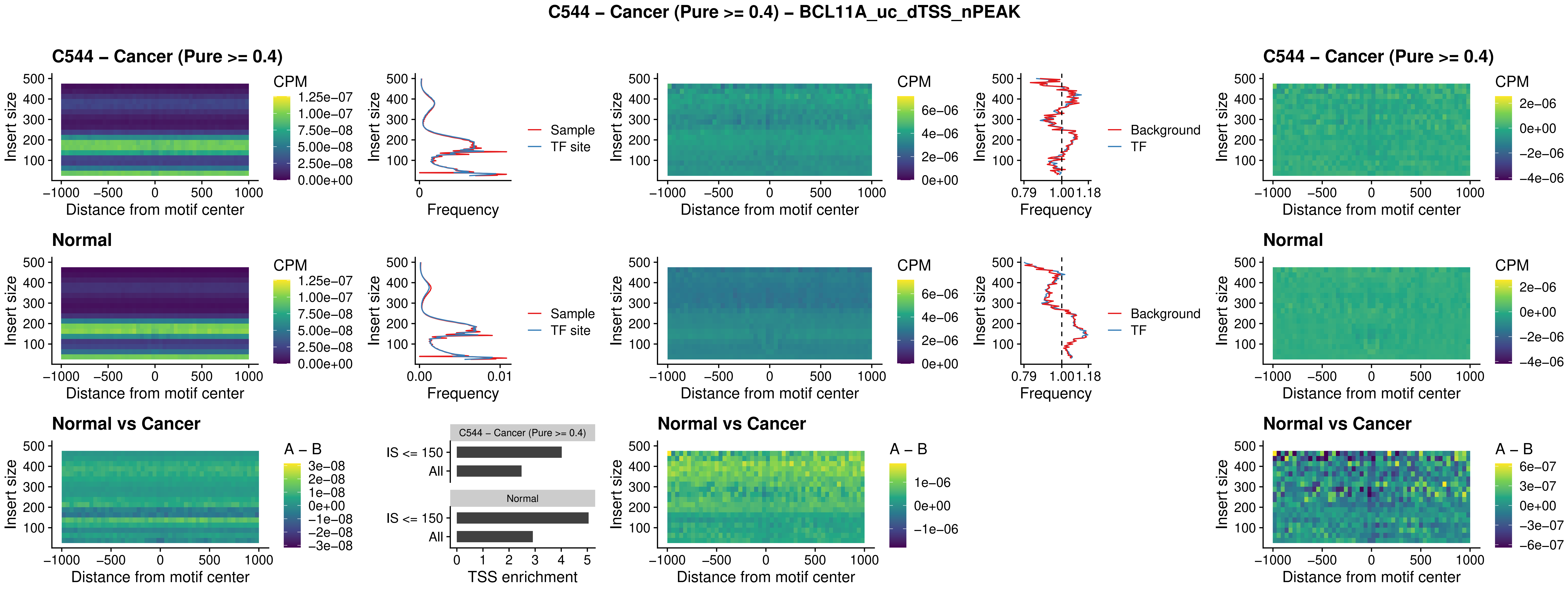
C544 – Cancer (Pure >= 0.4) – BCL11A_uc_dTSS_nPEAK
C544 – Cancer (Pure >= 0.4)
Normal
Normal vs Cancer
Insert size
Distance from motif center
Frequency
CPM
A – B
Sample
TF site
Background
TF
C544 – Cancer (Pure >= 0.4)
Normal
IS <= 150
All
TSS enrichment
0.79
1.00
1.18
1.25e–07
1.00e–07
7.50e–08
5.00e–08
2.50e–08
0.00e+00
6e–06
4e–06
2e–06
0e+00
2e–06
0e+00
–2e–06
–4e–06
3e–08
2e–08
1e–08
0e+00
–1e–08
–2e–08
–3e–08
1e–06
0e+00
–1e–06
6e–07
3e–07
0e+00
–3e–07
–6e–07
0.00
0.01

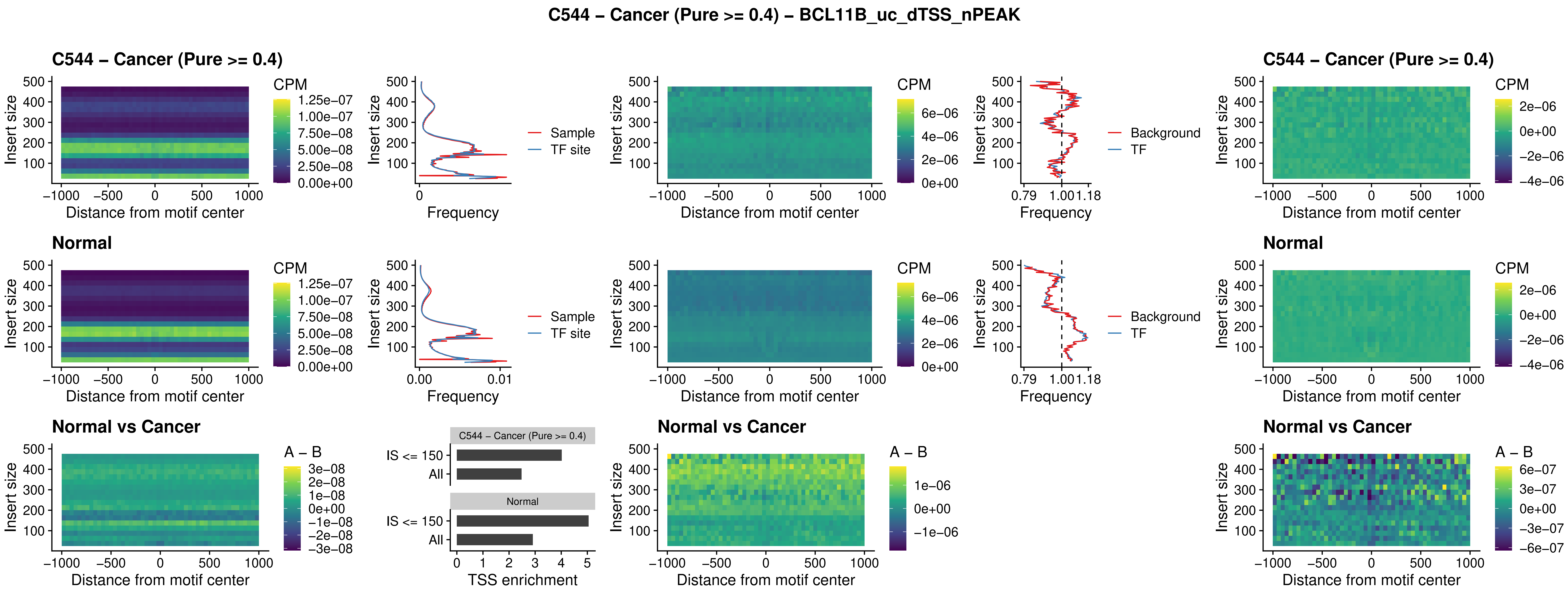
C544 – Cancer (Pure >= 0.4) – BCL11B_uc_dTSS_nPEAK
C544 – Cancer (Pure >= 0.4)
Normal
Normal vs Cancer
Insert size
Distance from motif center
Frequency
CPM
A – B
Sample
TF site
Background
TF
TSS enrichment
IS <= 150
All
C544 – Cancer (Pure >= 0.4)
Normal

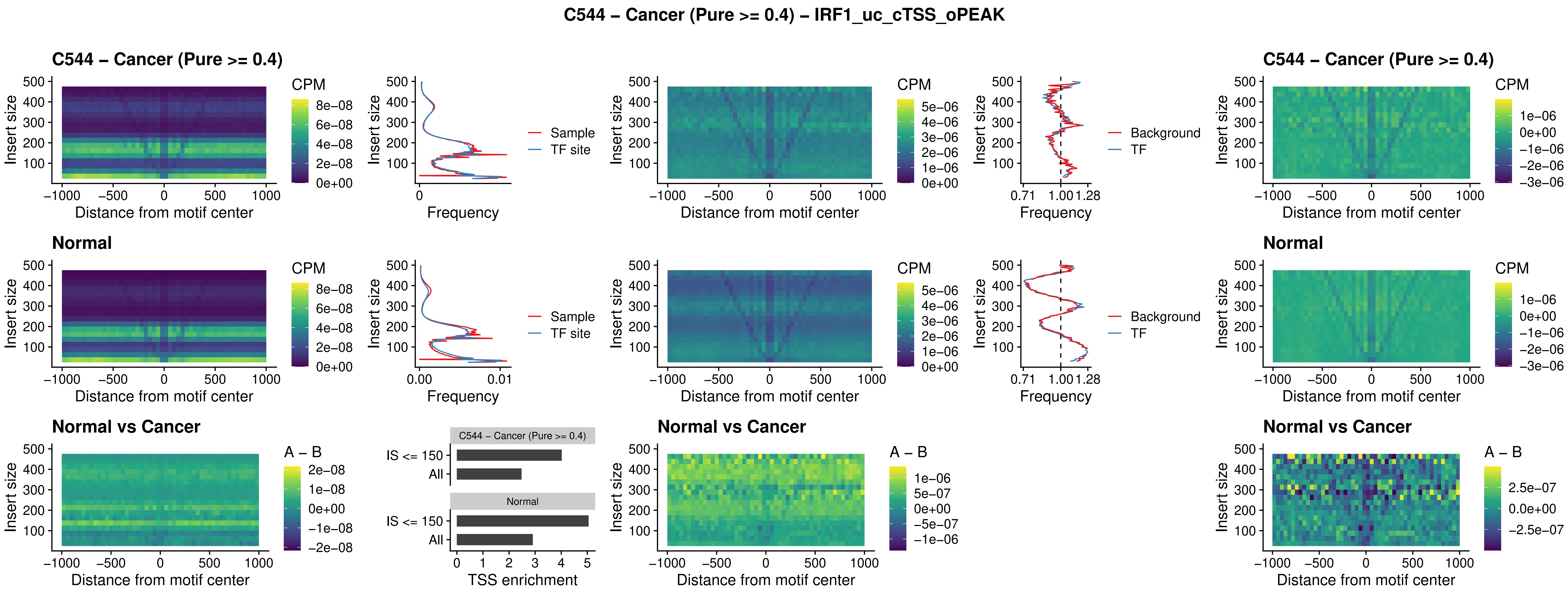
C544 – Cancer (Pure >= 0.4) – IRF1_uc_cTSS_oPEAK
C544 – Cancer (Pure >= 0.4)
Normal
Normal vs Cancer
Insert size
Distance from motif center
Frequency
CPM
A – B
Sample
TF site
Background
TF
C544 – Cancer (Pure >= 0.4)
Normal
IS <= 150
All
TSS enrichment

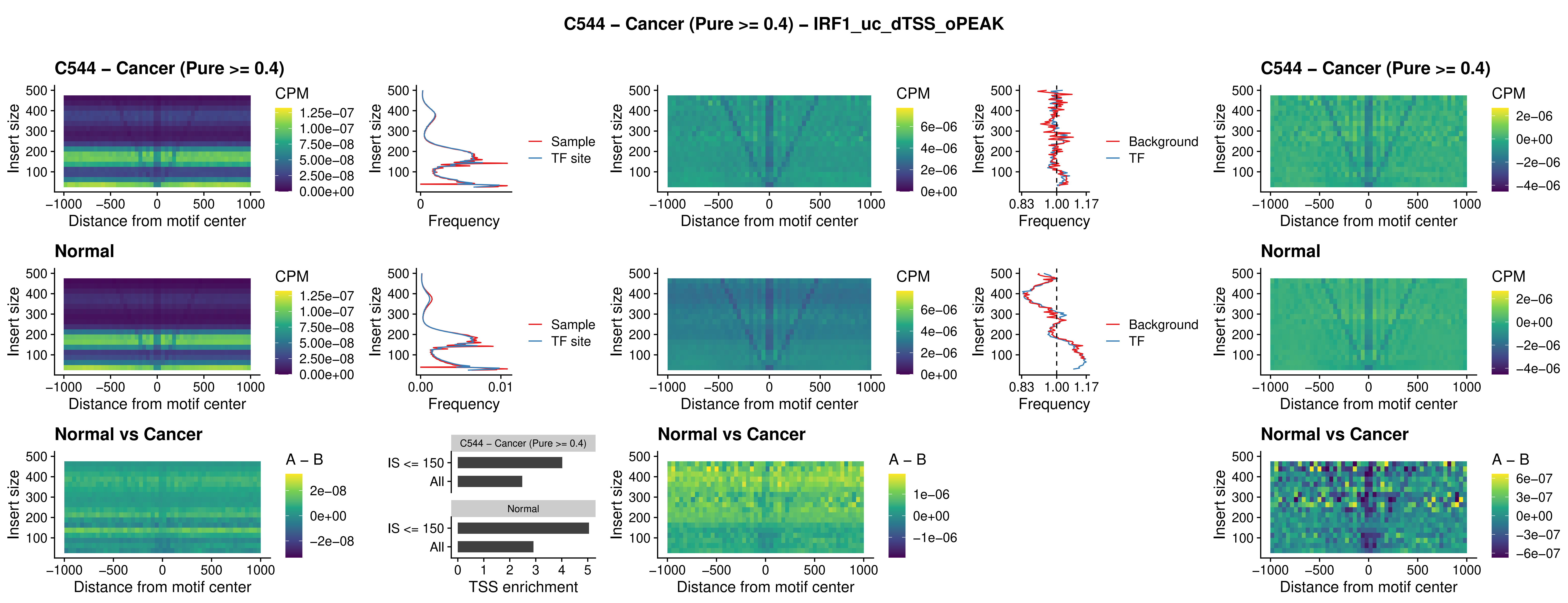
C544 – Cancer (Pure >= 0.4) – IRF1_uc_dTSS_oPEAK
C544 – Cancer (Pure >= 0.4)
Normal
Normal vs Cancer
Insert size
Distance from motif center
Frequency
CPM
A – B
Sample
TF site
Background
TF
C544 – Cancer (Pure >= 0.4)
Normal
IS <= 150
All
TSS enrichment

#### C544 – Cancer (Pure >= 0.4) – IRF1_uc_pTSS_oPEAK

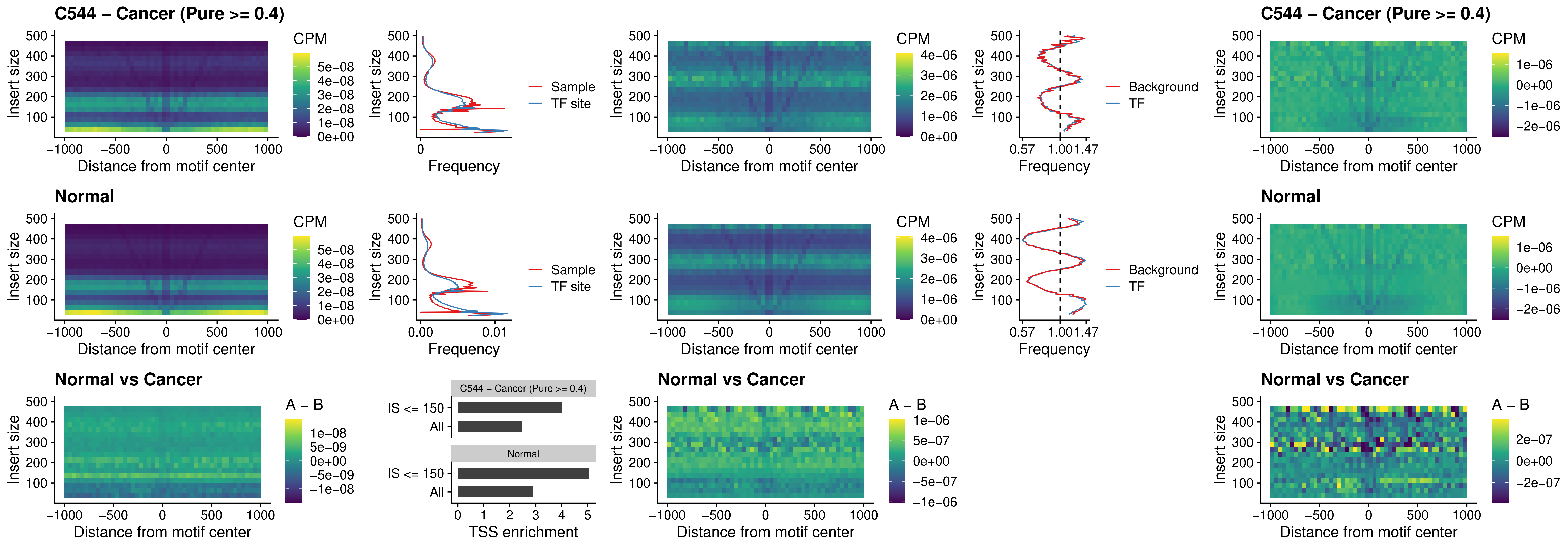

#### C544 – Cancer (Pure >= 0.4) – SPIC_uc_dTSS_nPEAK

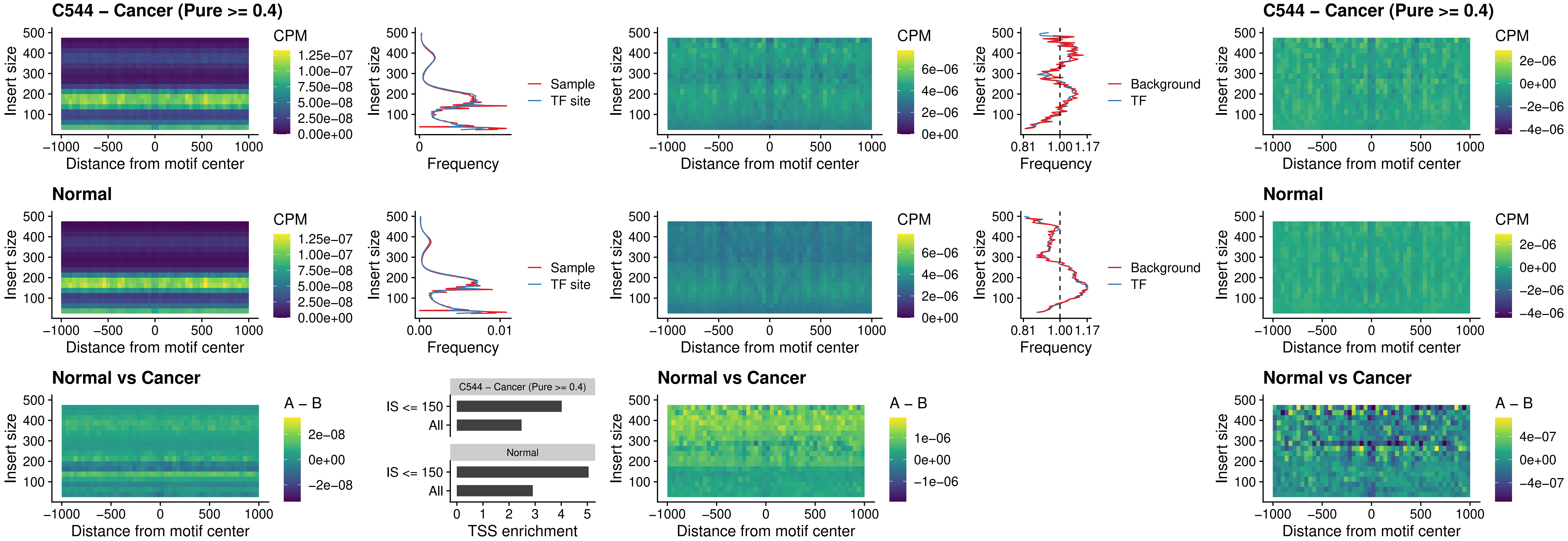

### C544 – Cancer (Pure >= 0.4) – SPIC_uc_dTSS_oPEAK

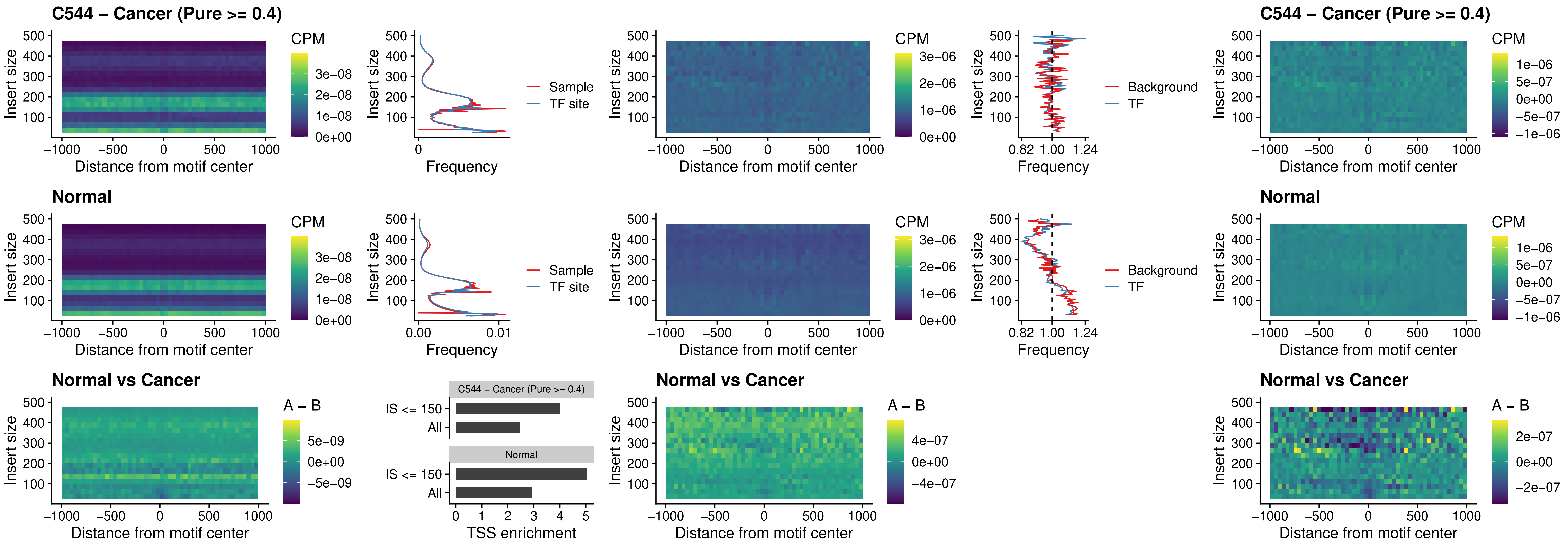

#### C544 – Cancer (Pure >= 0.4) – SPIB_uc_pTSS_nPEAK

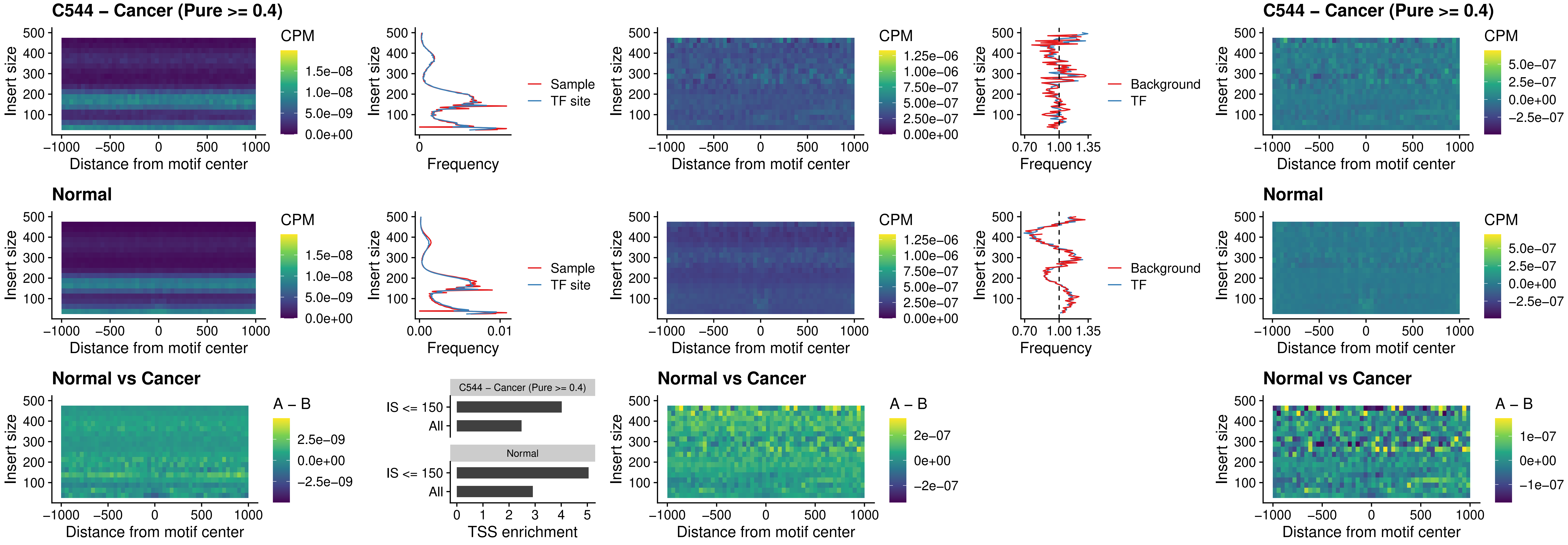

#### C544 – Cancer (Pure >= 0.4) – STAT2_uc_cTSS_oPEAK

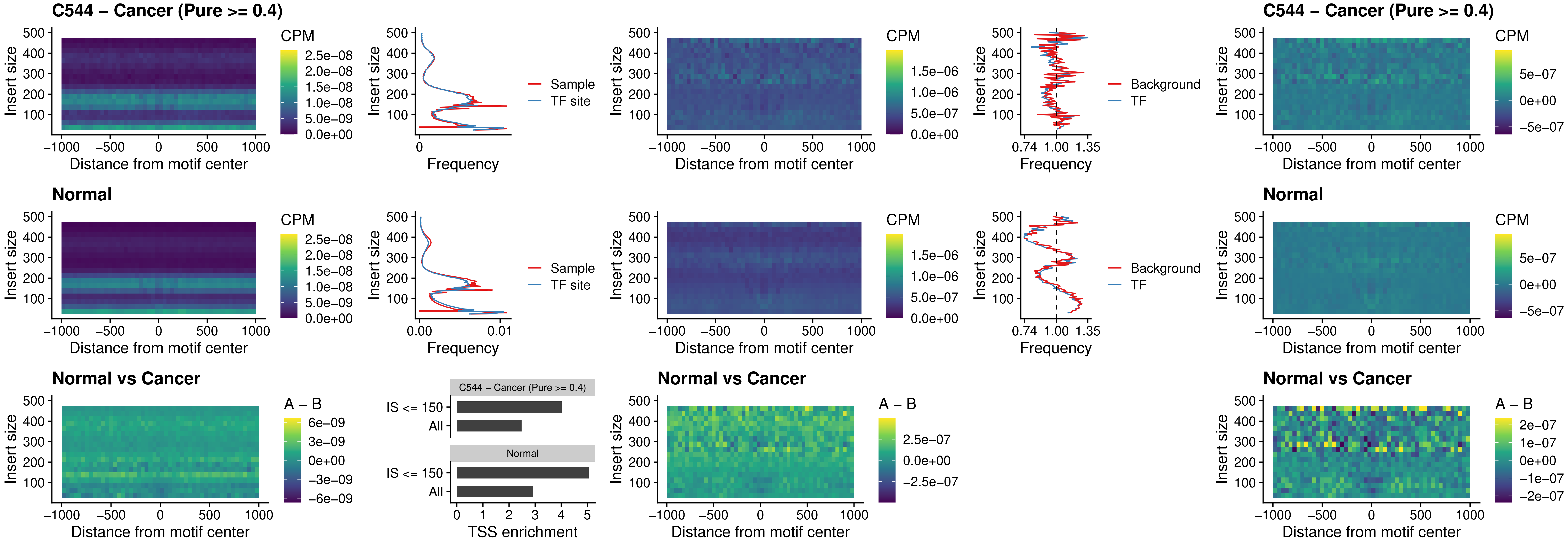

#### C544 – Cancer (Pure >= 0.4) – IRF8_uc_cTSS_oPEAK

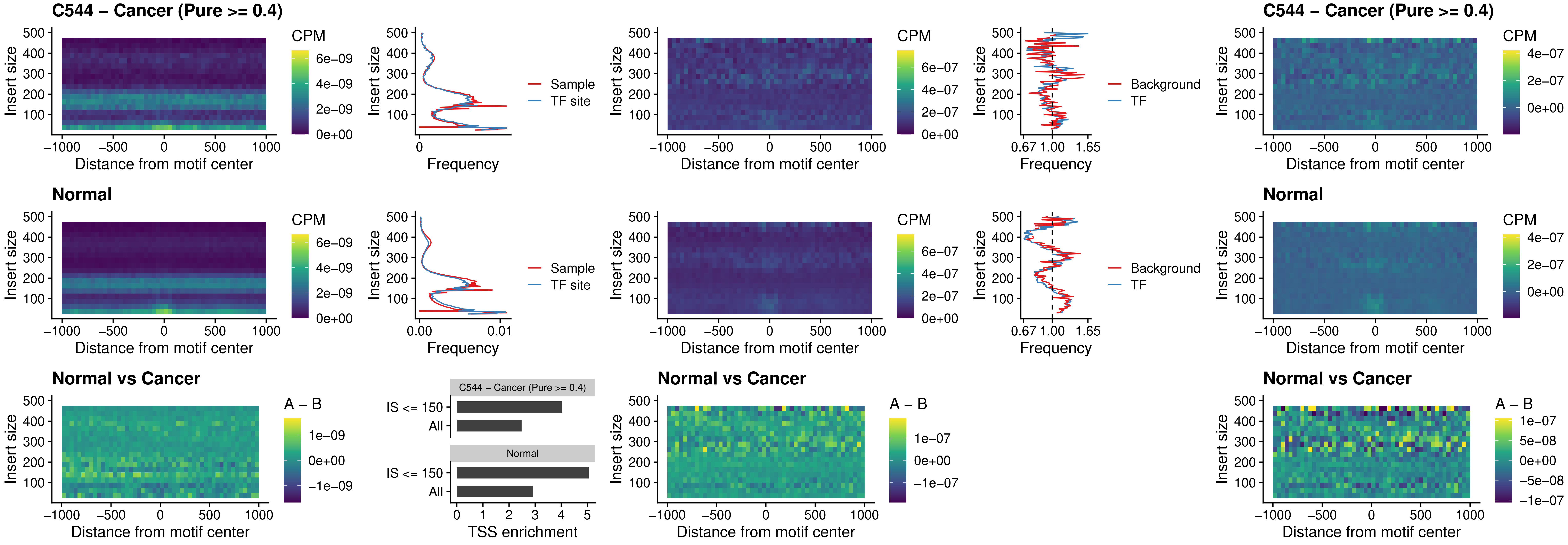

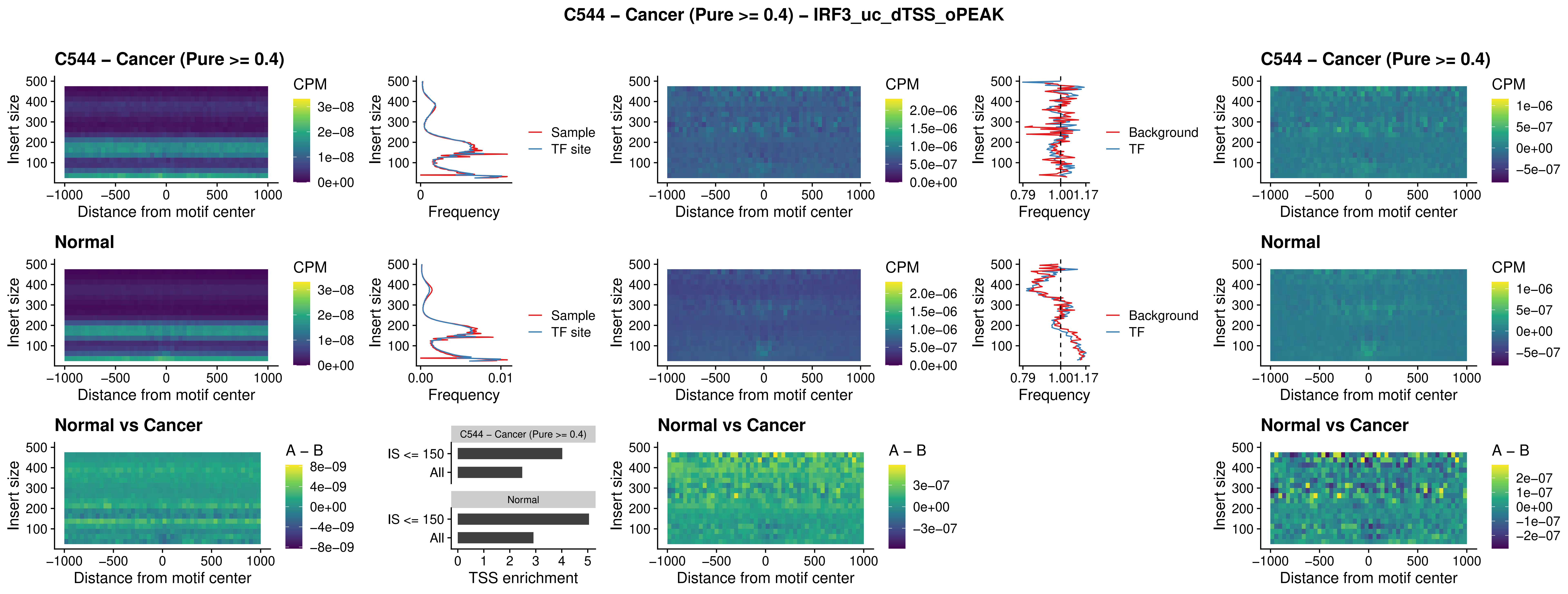
C544 – Cancer (Pure >= 0.4) – IRF3_uc_dTSS_oPEAK
C544 – Cancer (Pure >= 0.4)
Normal
Normal vs Cancer
Insert size
Distance from motif center
CPM
A – B
Frequency
Sample
TF site
Background
TF
C544 – Cancer (Pure >= 0.4)
Normal
IS <= 150
All
TSS enrichment

#### C544 – Cancer (Pure >= 0.4) – IRF8_uc_dTSS_oPEAK

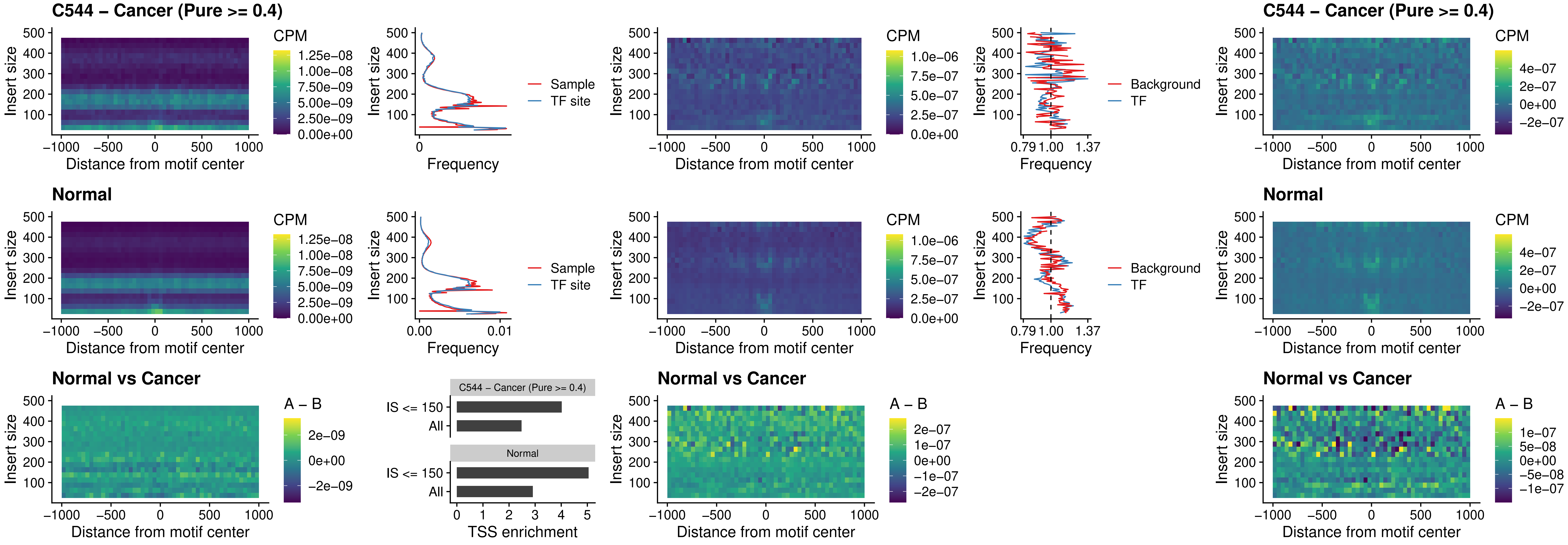

#### C544 – Cancer (Pure >= 0.4) – ETS2_uc_dTSS_oPEAK

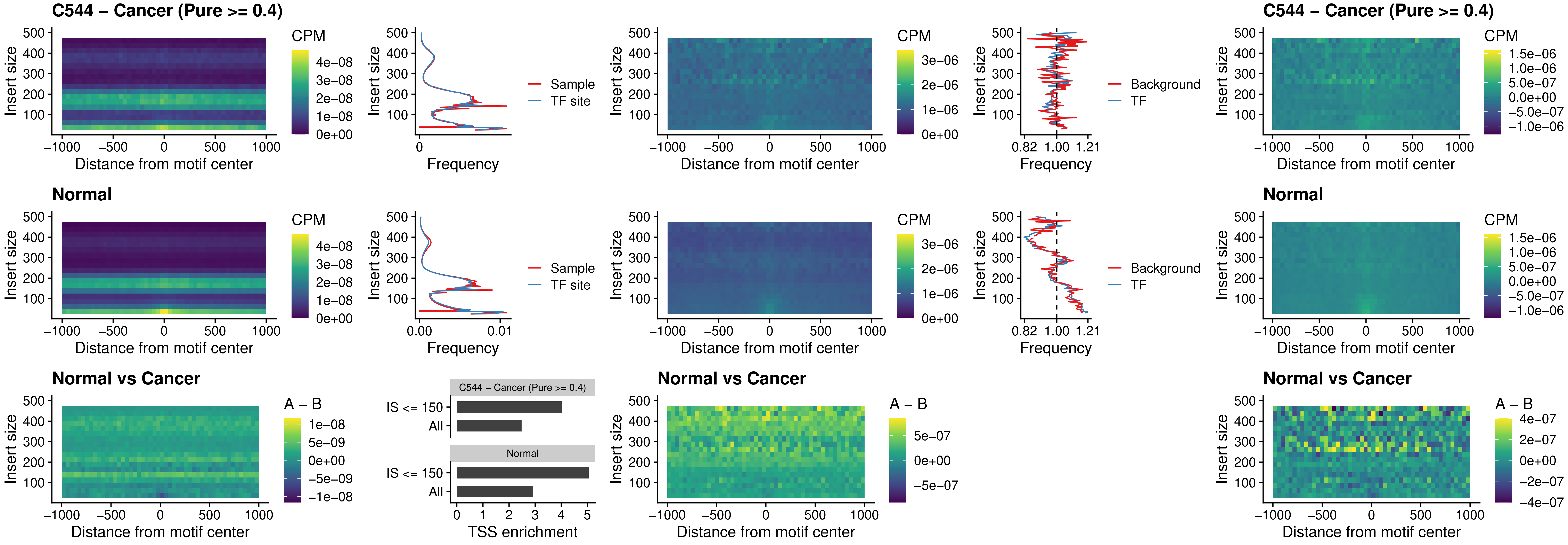

#### C544 – Cancer (Pure >= 0.4) – SMARCC2_uc_pTSS_oPEAK

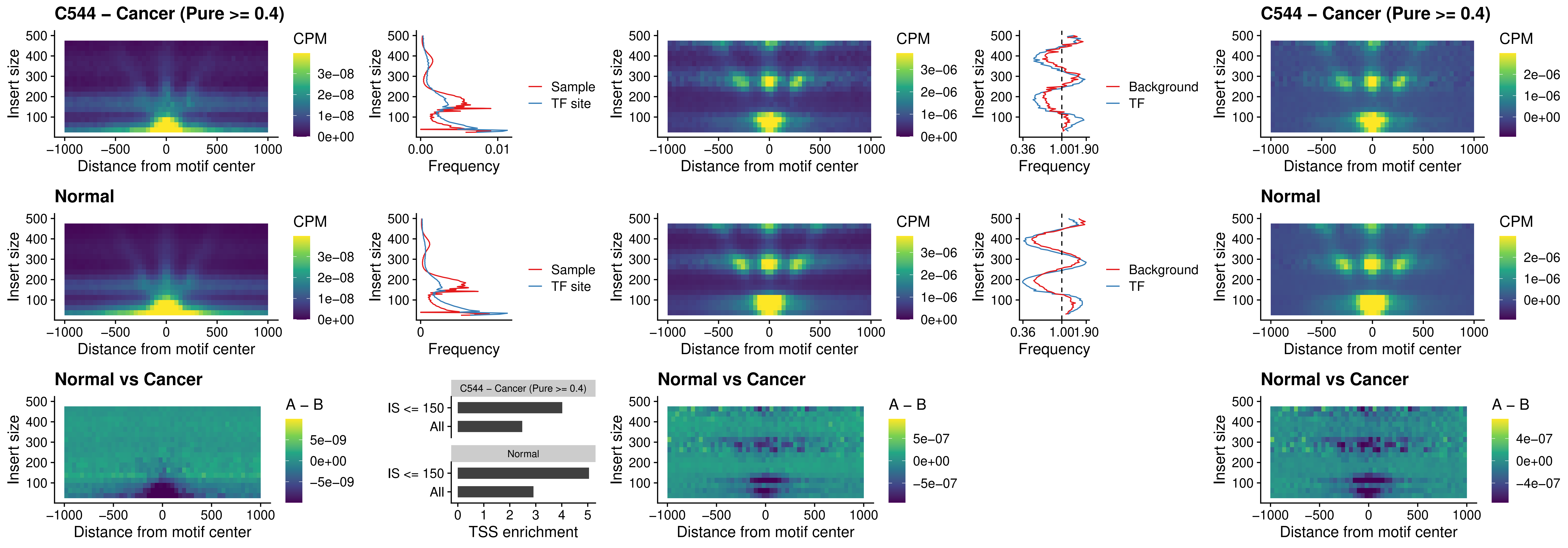

#### C544 – Cancer (Pure >= 0.4) – BCL11A_uc_cTSS_nPEAK

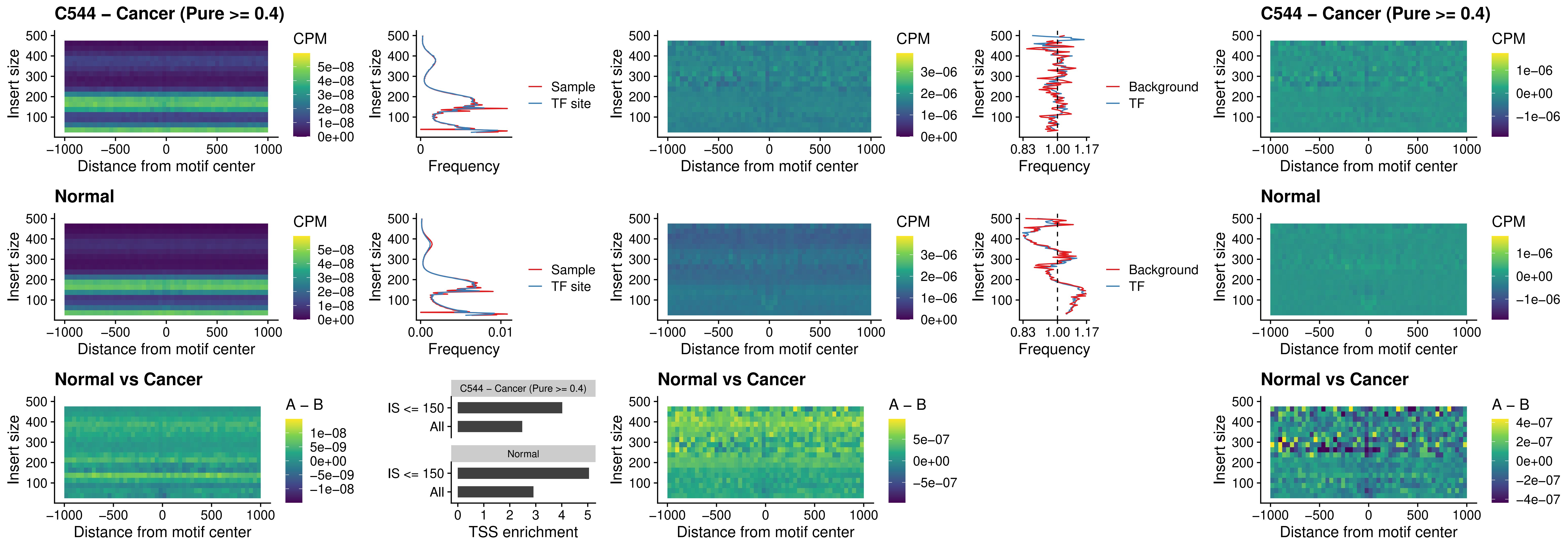

#### C544 – Cancer (Pure >= 0.4) – BCL11B_uc_cTSS_nPEAK

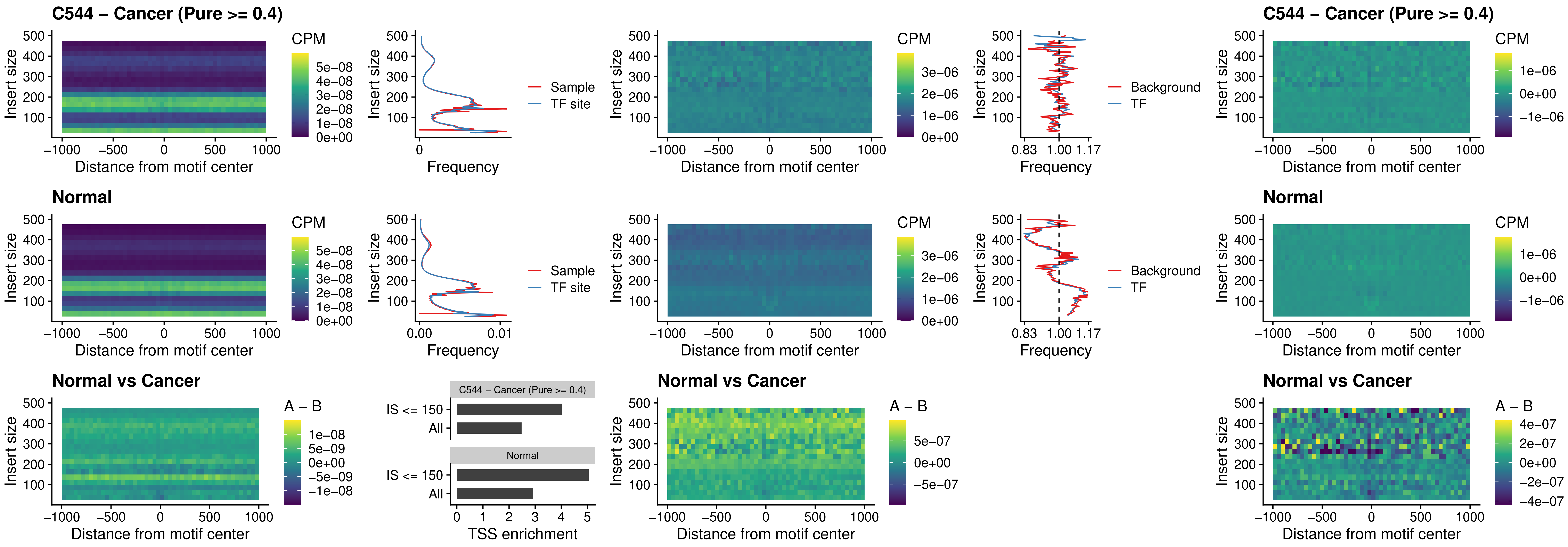

#### C544 – Cancer (Pure >= 0.4) – STAT2_uc_dTSS_oPEAK

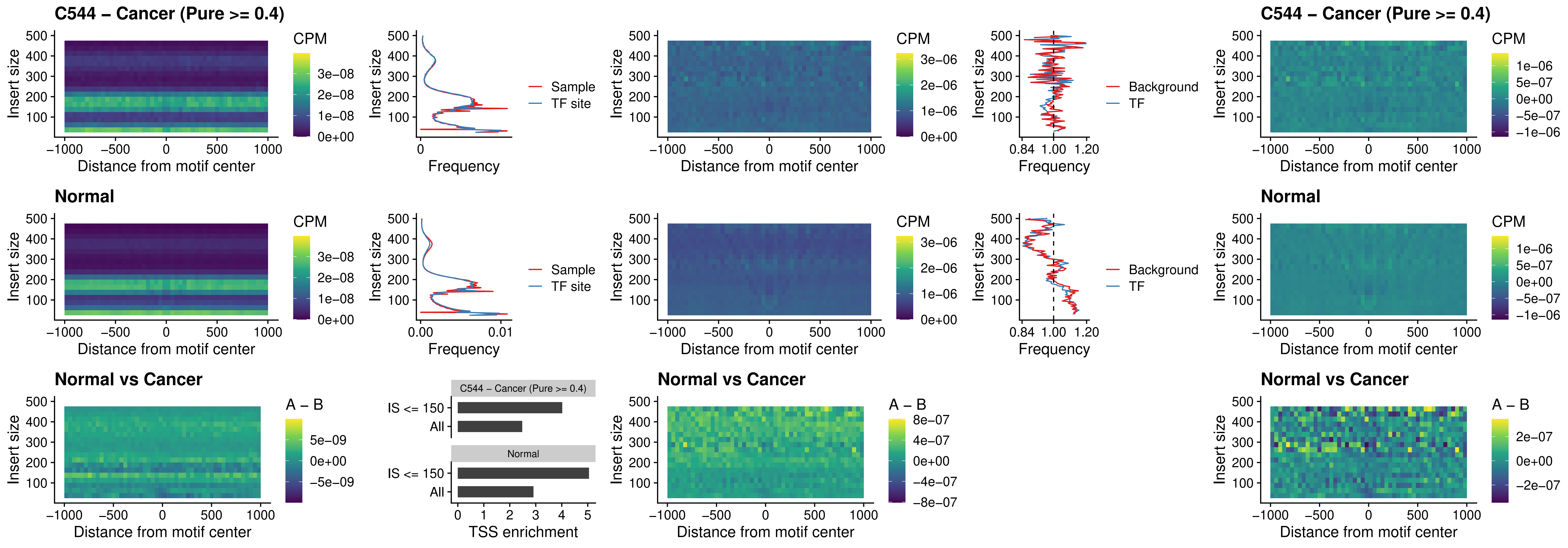

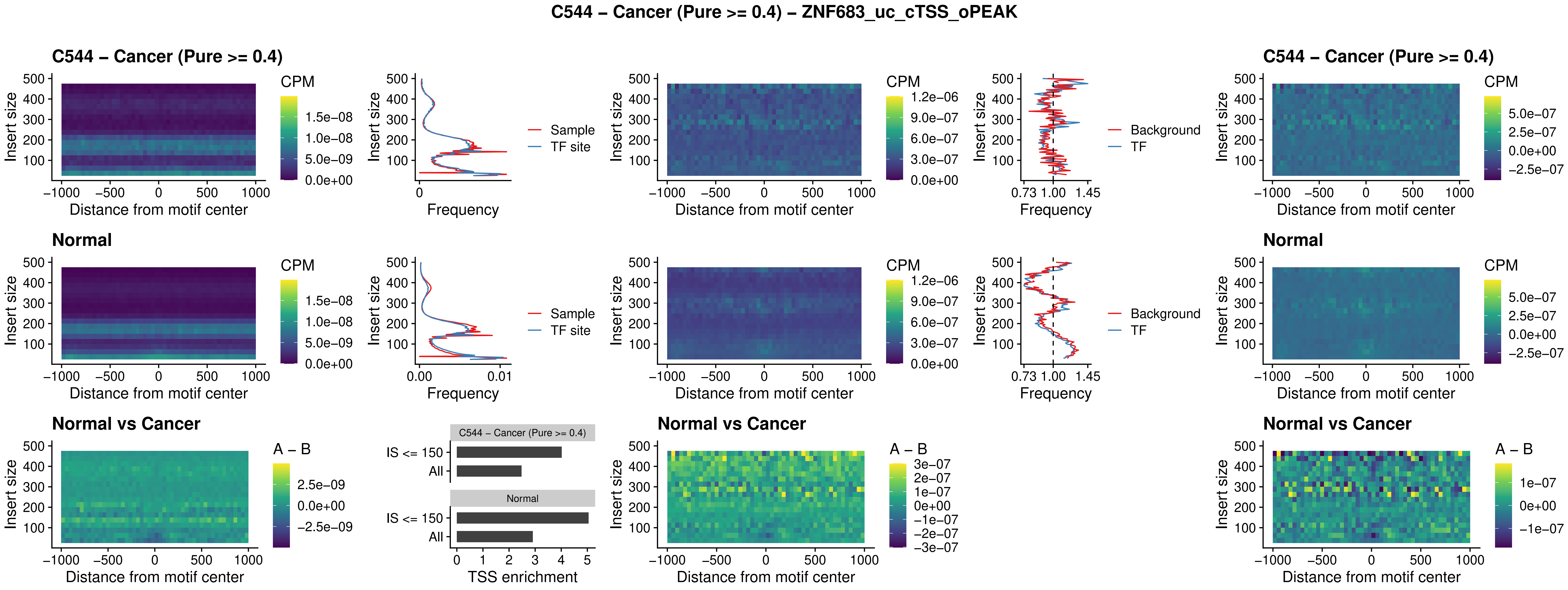
C544 – Cancer (Pure >= 0.4) – ZNF683_uc_cTSS_oPEAK
C544 – Cancer (Pure >= 0.4)
Normal
Normal vs Cancer
Insert size
Distance from motif center
Frequency
CPM
A – B
Sample
TF site
Background
TF
C544 – Cancer (Pure >= 0.4)
Normal
IS <= 150
All
TSS enrichment

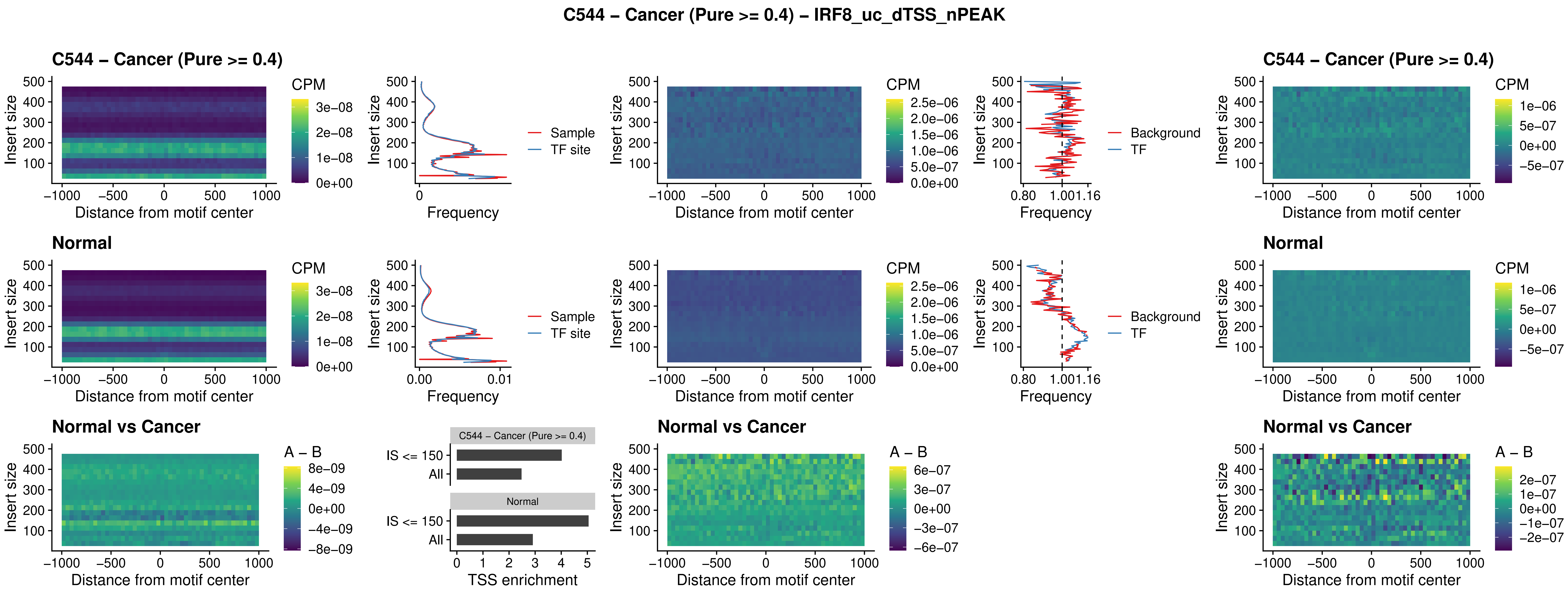
C544 – Cancer (Pure >= 0.4) – IRF8_uc_dTSS_nPEAK
C544 – Cancer (Pure >= 0.4)
Normal
Normal vs Cancer
Insert size
Distance from motif center
Frequency
CPM
A – B
Sample
TF site
Background
TF
C544 – Cancer (Pure >= 0.4)
Normal
IS <= 150
All
TSS enrichment

#### C544 – Cancer (Pure >= 0.4) – ETV7_uc_dTSS_oPEAK

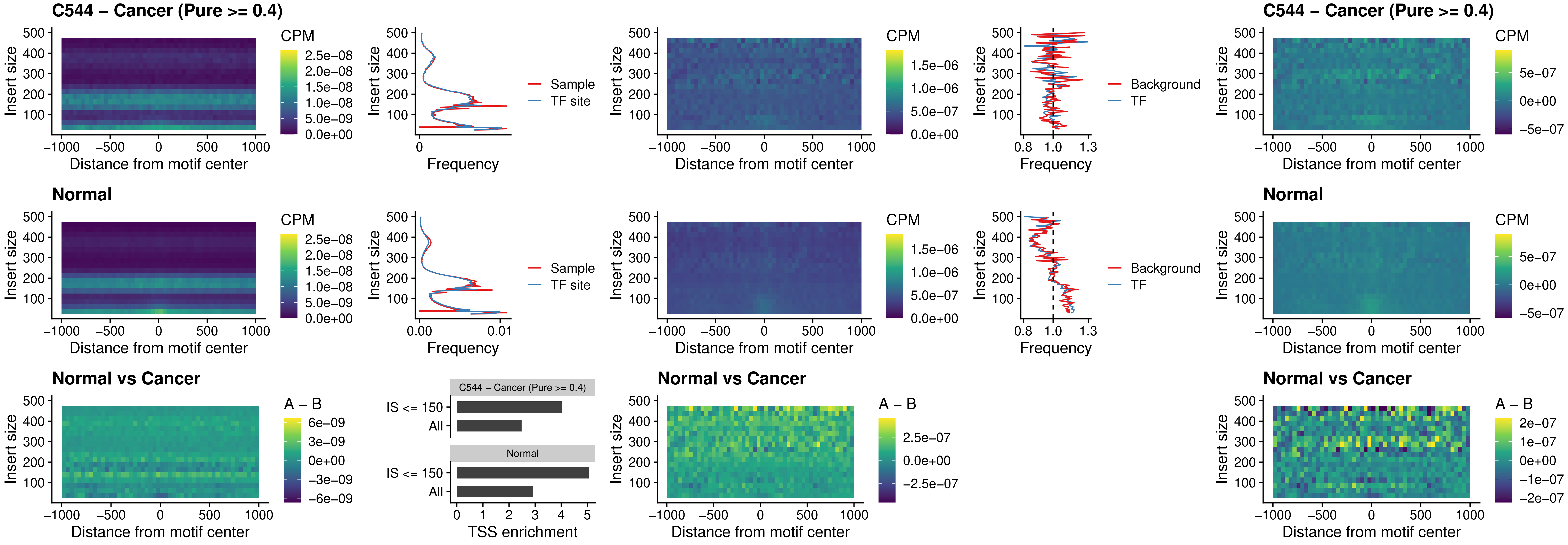

#### C548 – Cancer (Pure >= 0.4) – FOXL1_uc_pTSS_oPEAK

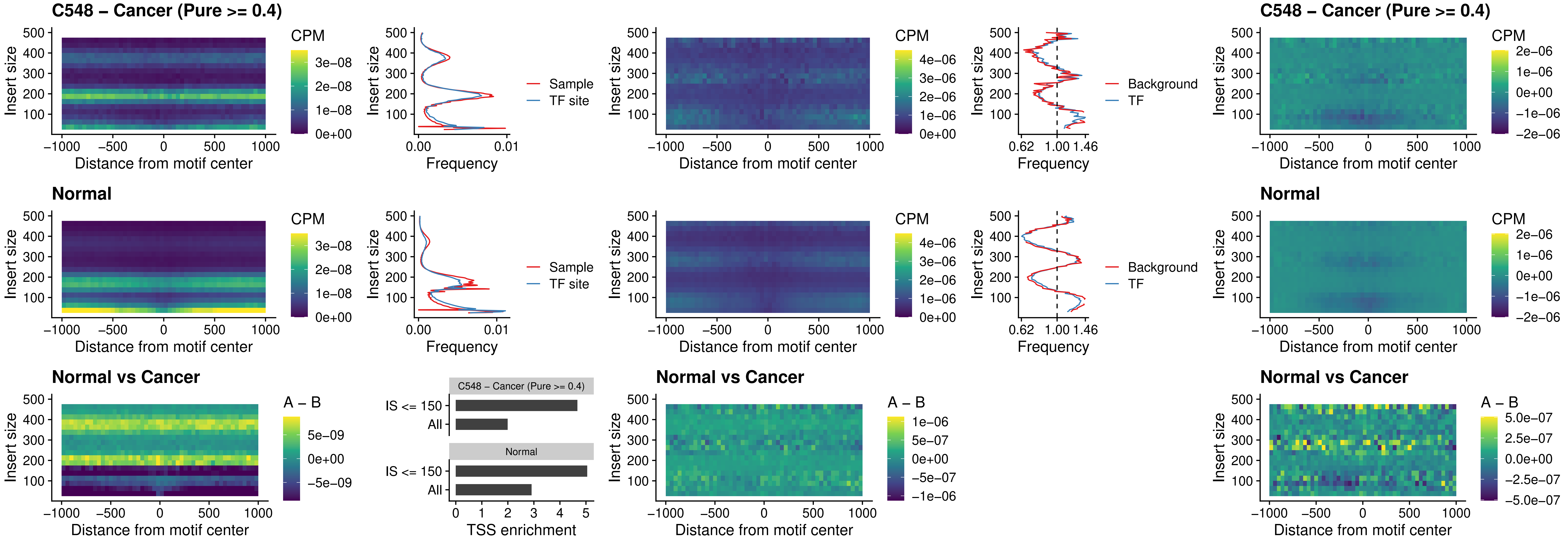

### C548 – Cancer (Pure >= 0.4) – CPEB1_uc_cTSS_oPEAK

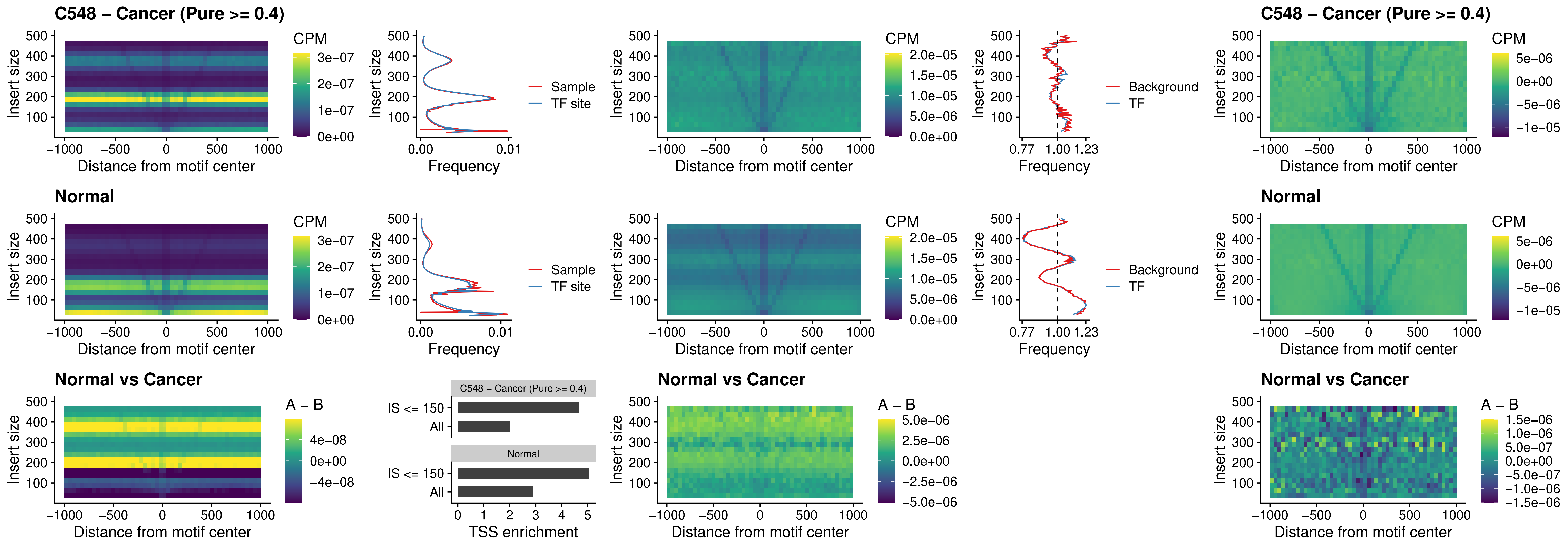

#### C548 – Cancer (Pure >= 0.4) – HMGA1_uc_pTSS_nPEAK

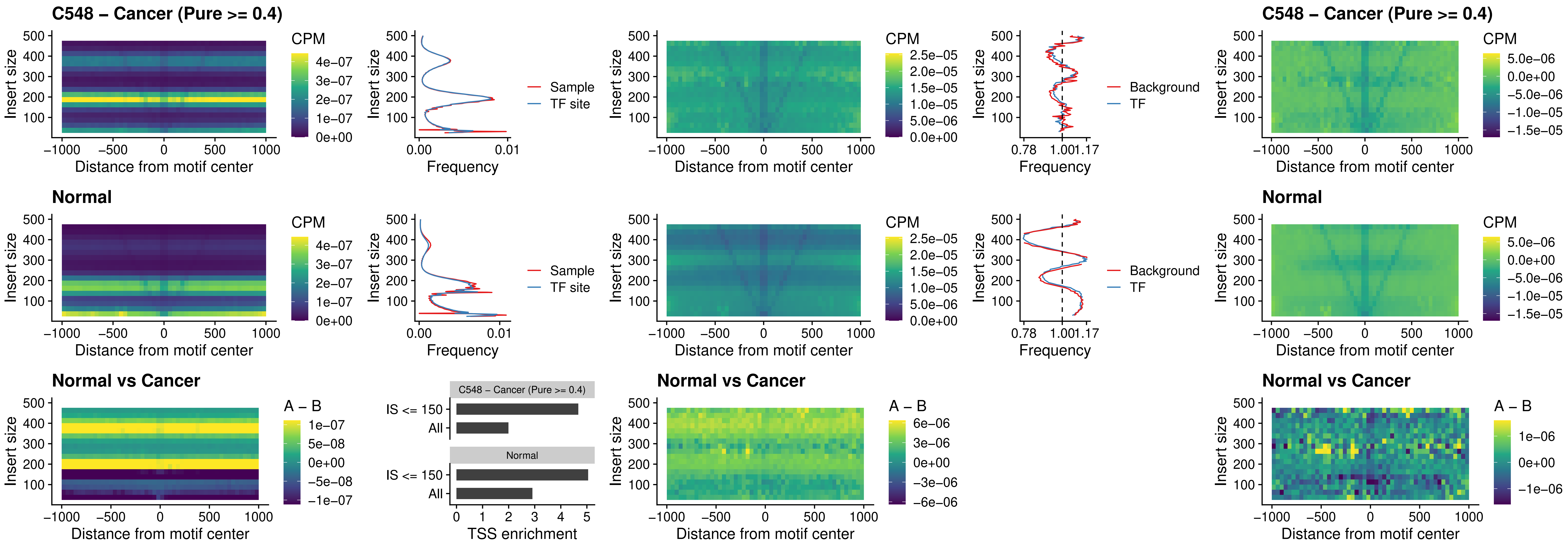

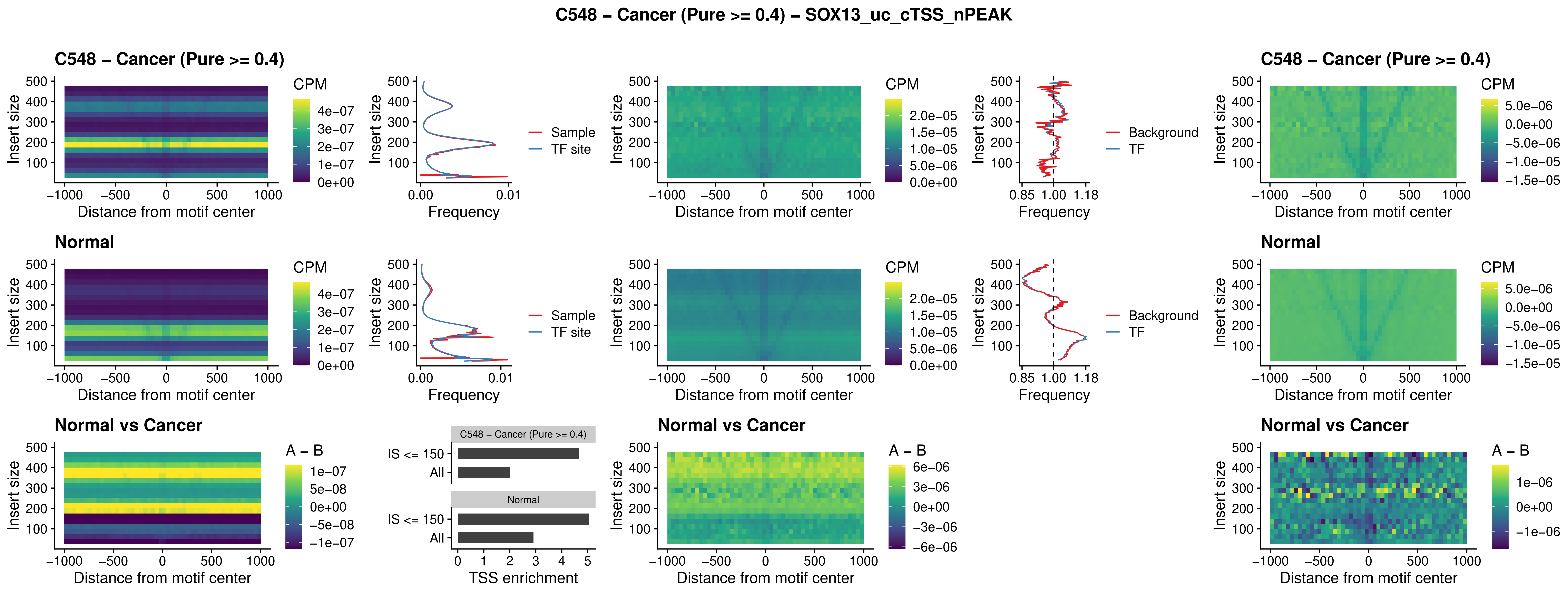

C548 – Cancer (Pure >= 0.4) – SOX13_uc_cTSS_nPEAK
C548 – Cancer (Pure >= 0.4)
Normal
Normal vs Cancer
Insert size
Distance from motif center
CPM
A – B
Frequency
Sample
TF site
Background
TF
C548 – Cancer (Pure >= 0.4)
Normal
IS <= 150
All
TSS enrichment

### C548 – Cancer (Pure >= 0.4) – LHX4_uc_cTSS_nPEAK

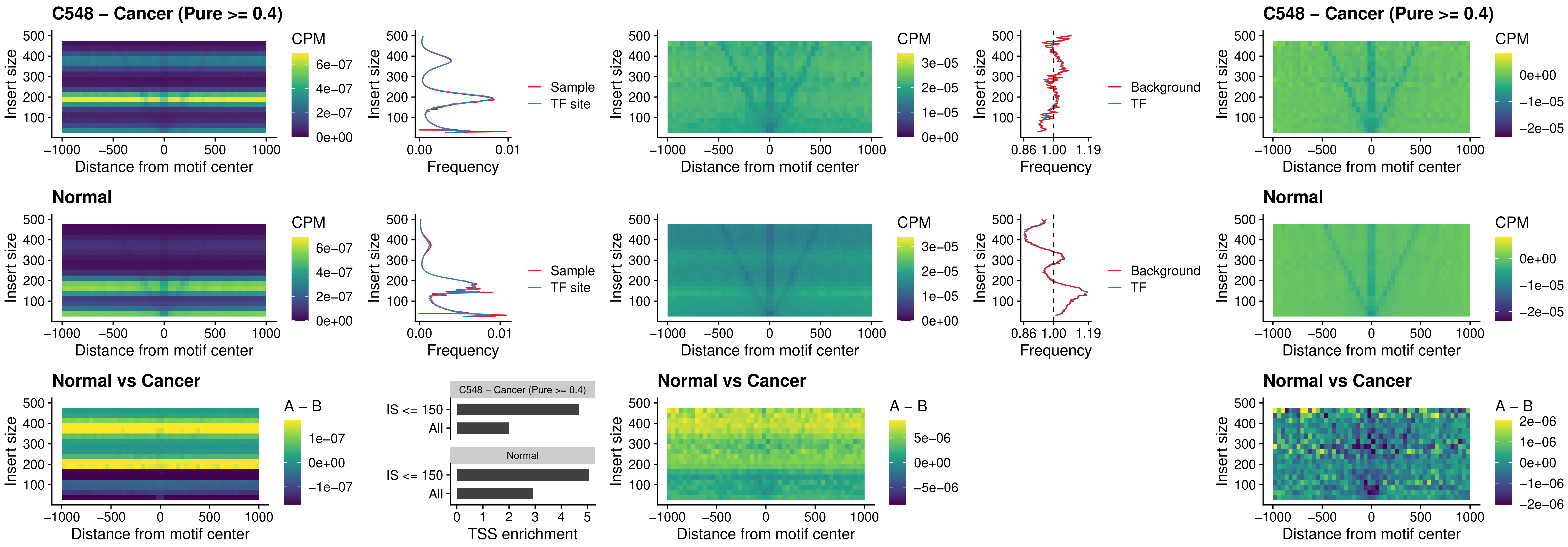

#### C548 – Cancer (Pure >= 0.4) – HOXA4_uc_cTSS_nPEAK

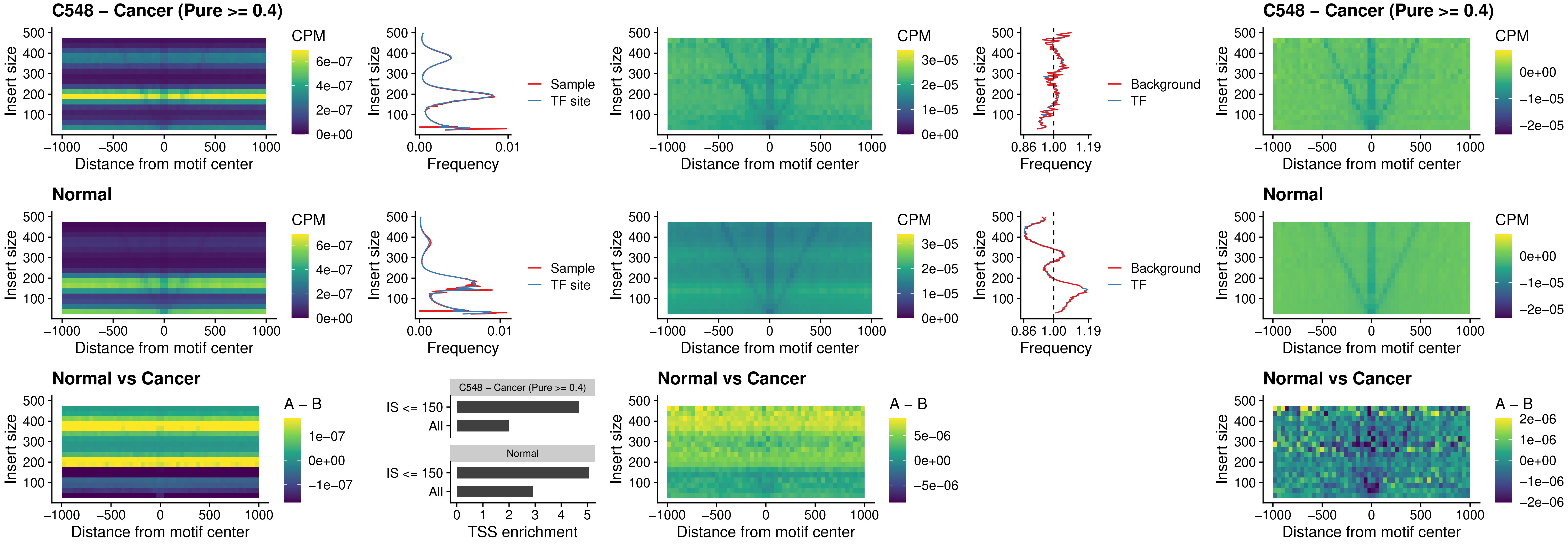

#### C548 – Cancer (Pure >= 0.4) – HLX_uc_cTSS_nPEAK

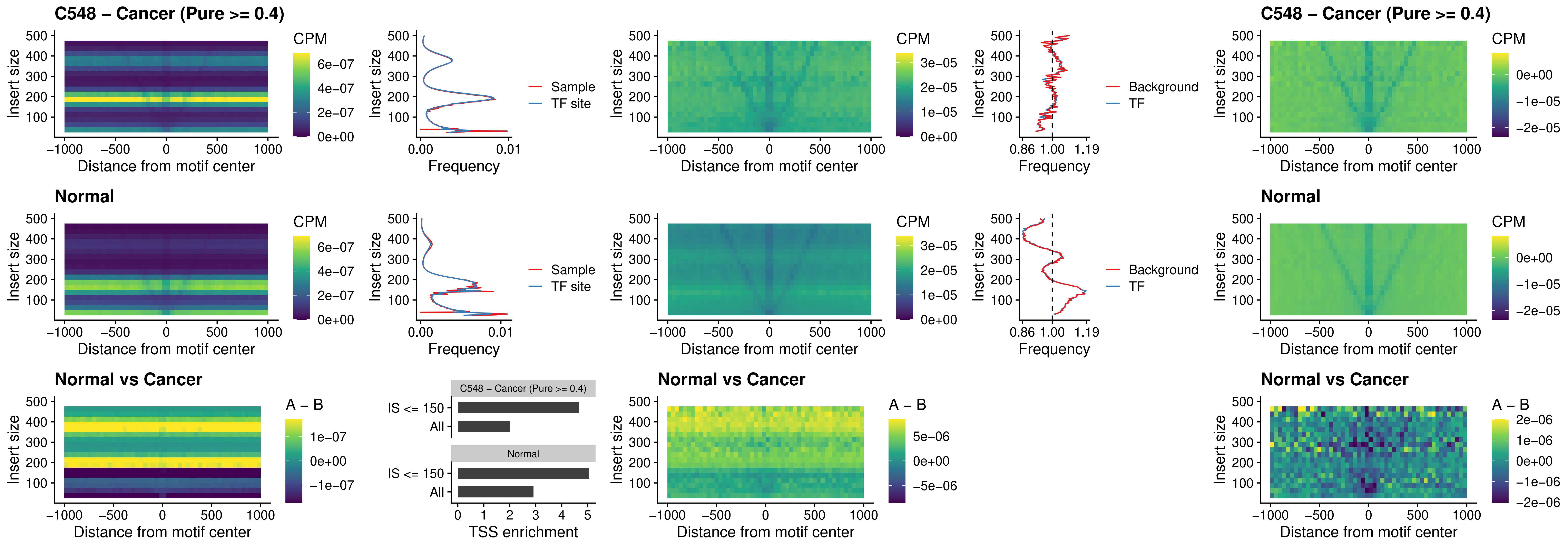

#### C548 – Cancer (Pure >= 0.4) – AC0125311_uc_cTSS_nPEAK

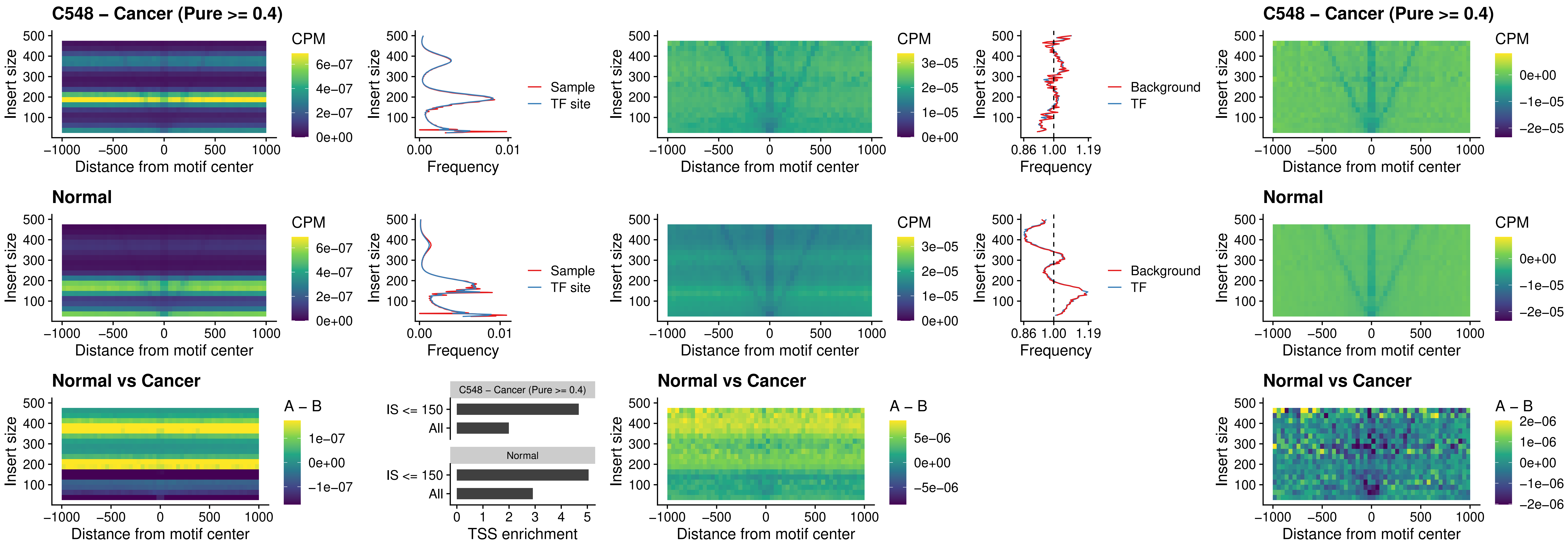

#### C548 – Cancer (Pure >= 0.4) – DBX2_uc_cTSS_nPEAK

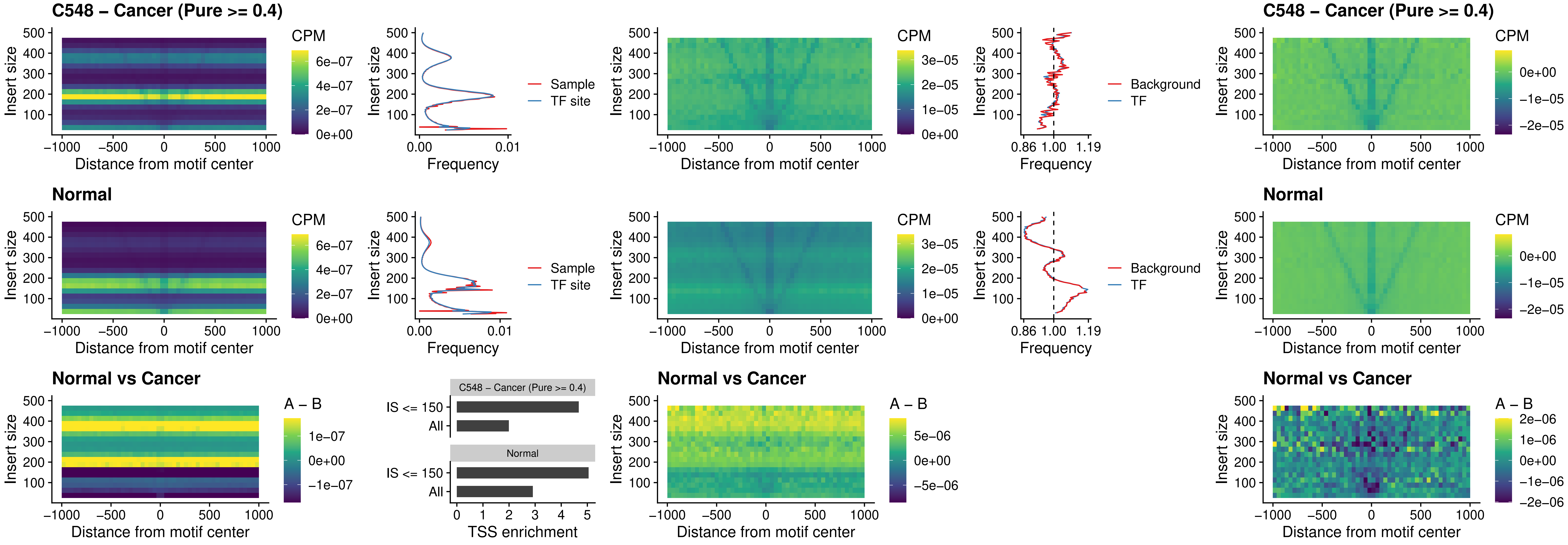

#### C548 – Cancer (Pure >= 0.4) – LHX6_uc_dTSS_nPEAK

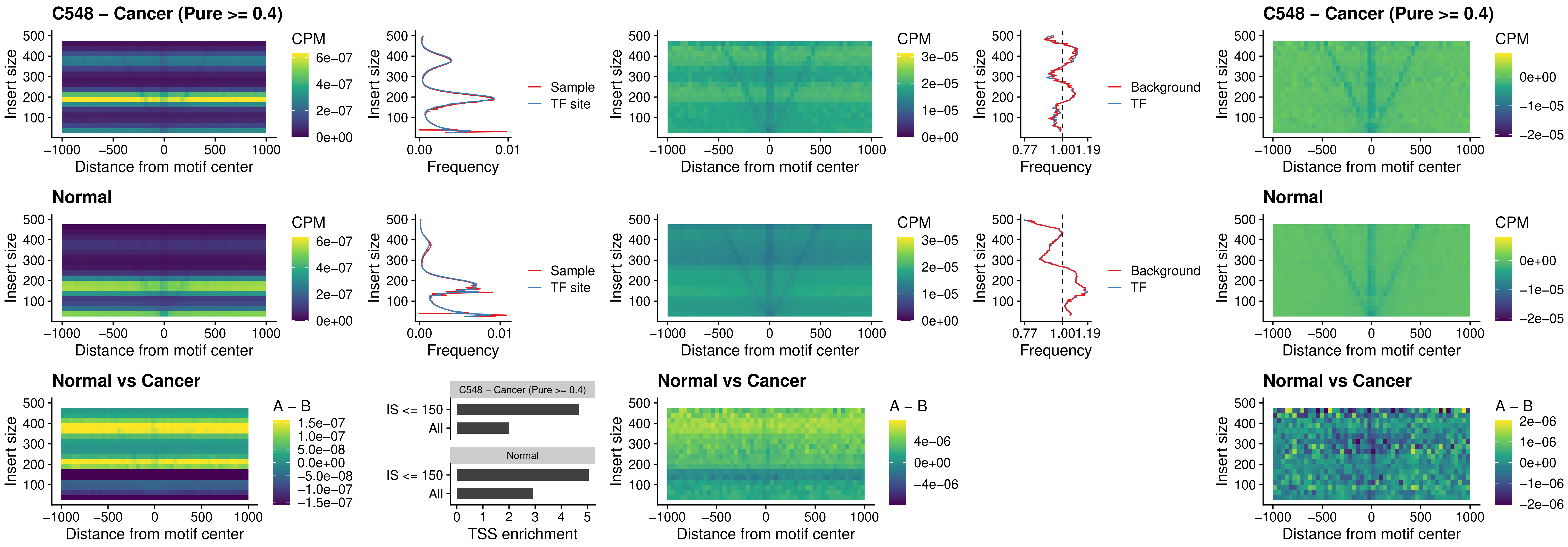

#### C548 – Cancer (Pure >= 0.4) – SOX1_uc_dTSS_nPEAK

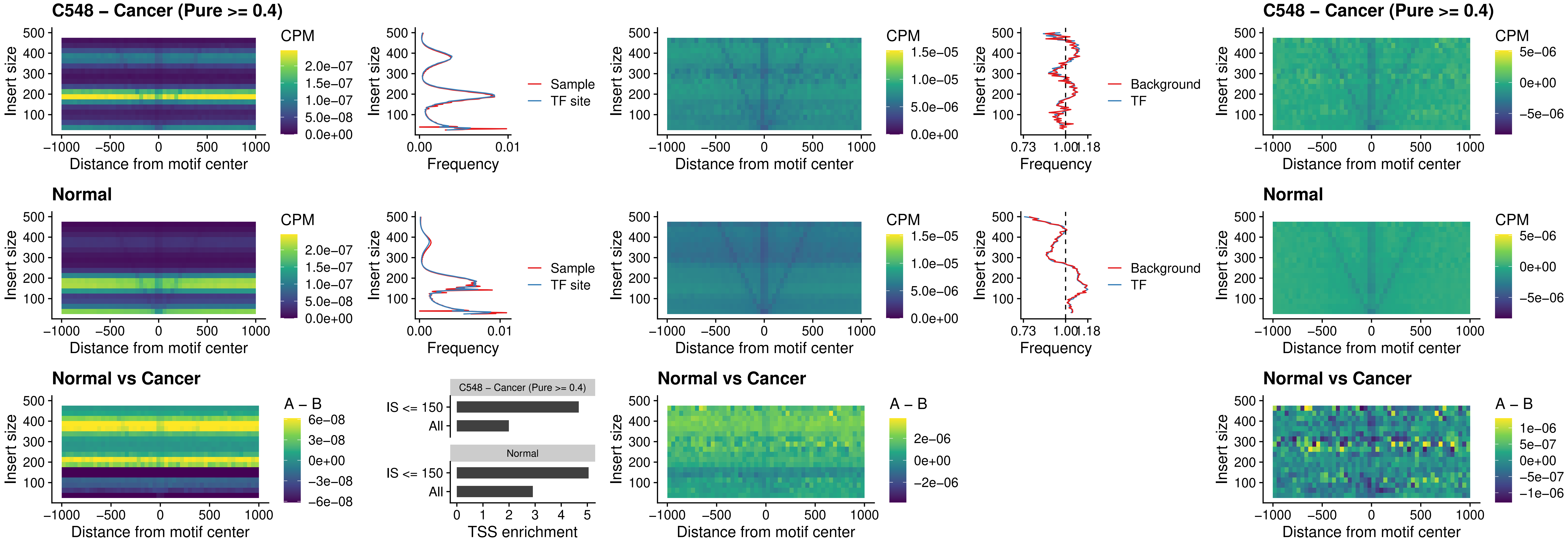

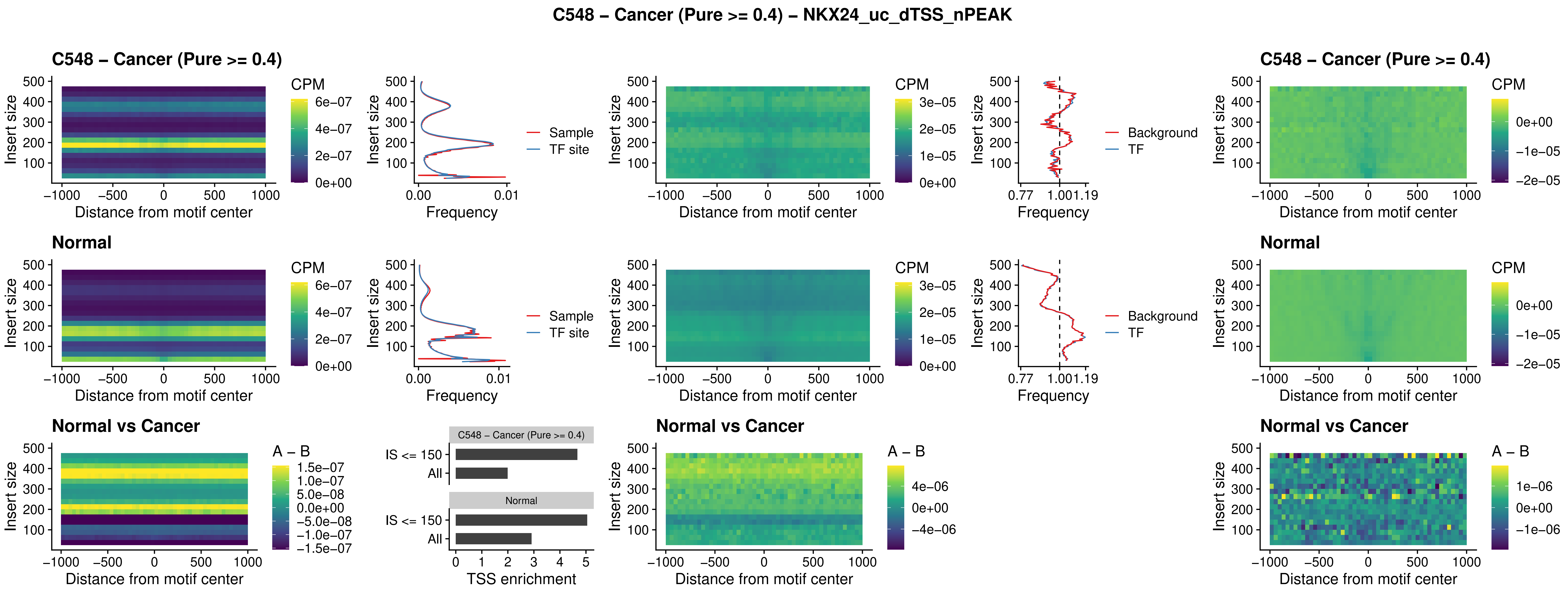
C548 – Cancer (Pure >= 0.4) – NKX24_uc_dTSS_nPEAK
C548 – Cancer (Pure >= 0.4)
Normal
Normal vs Cancer
Insert size
Distance from motif center
Frequency
CPM
A – B
Sample
TF site
Background
TF
C548 – Cancer (Pure >= 0.4)
Normal
IS <= 150
All
TSS enrichment

#### C548 – Cancer (Pure >= 0.4) – DLX3_uc_cTSS_nPEAK

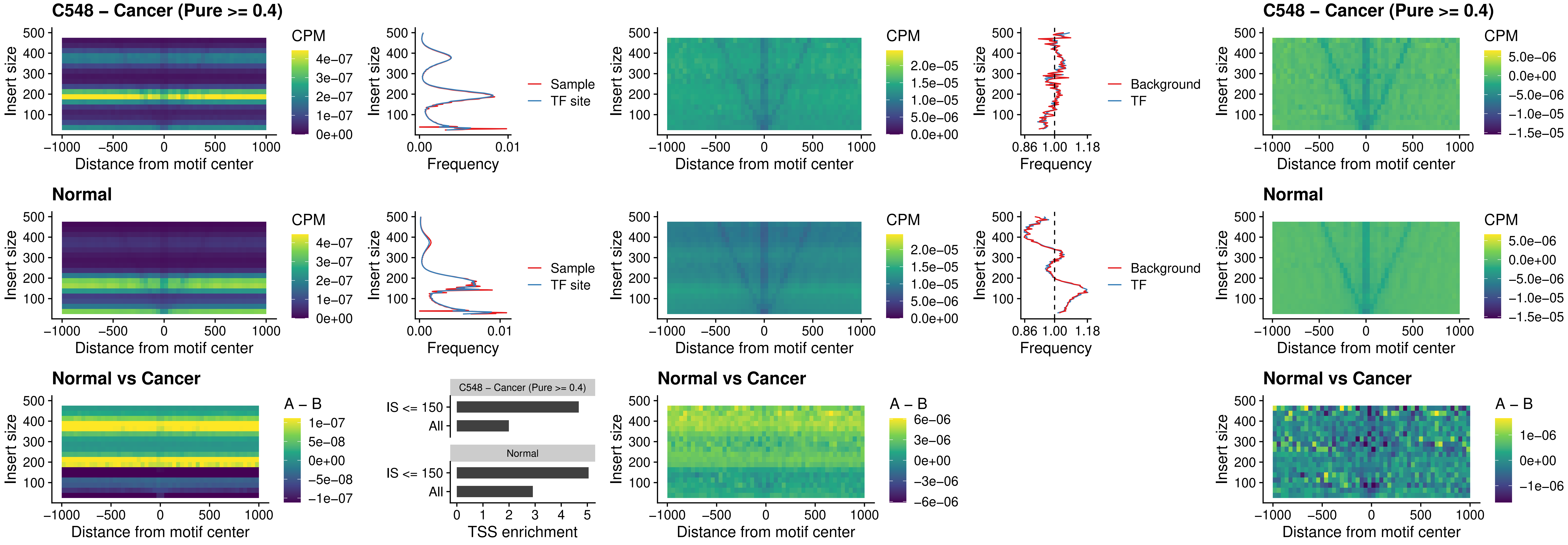

#### C548 – Cancer (Pure >= 0.4) – HOXD3_uc_cTSS_nPEAK

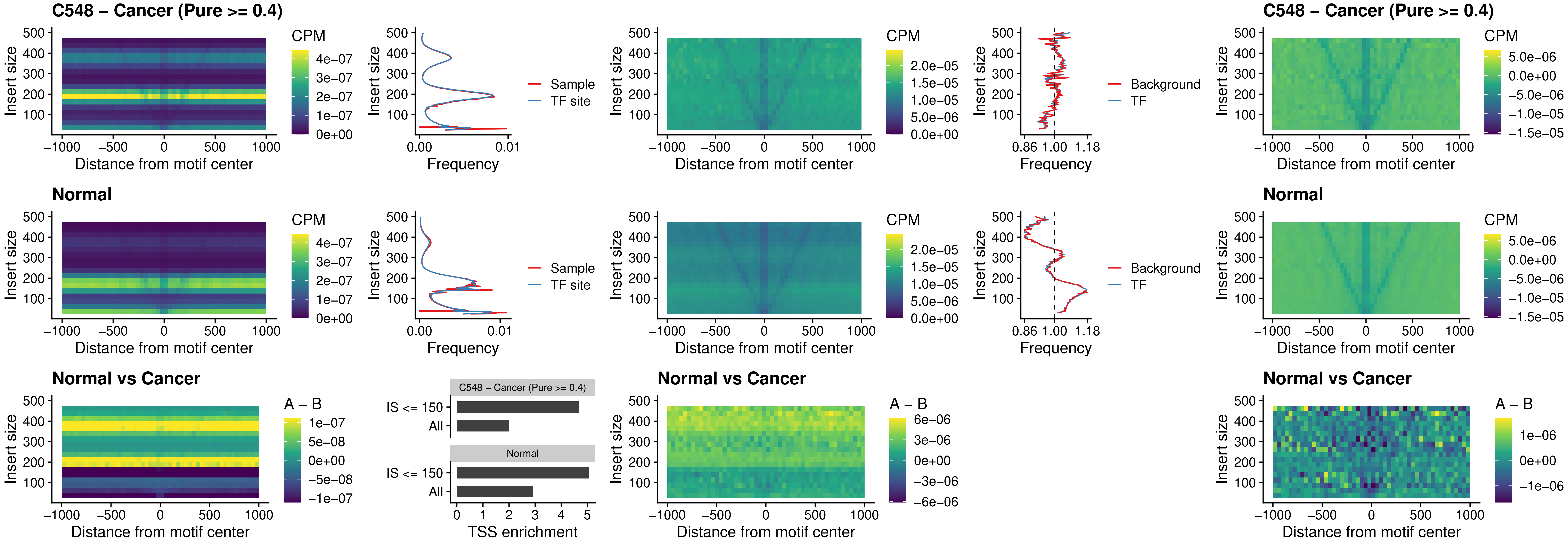

#### C548 – Cancer (Pure >= 0.4) – ISL1_uc_dTSS_oPEAK

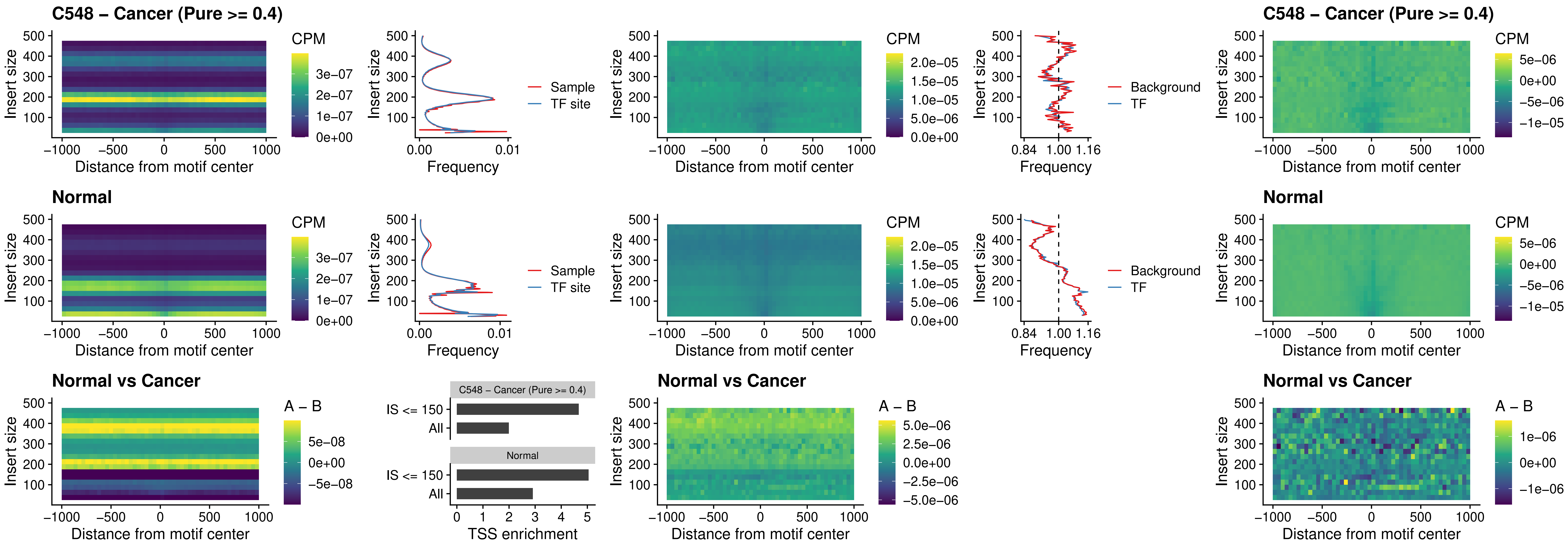

### C548 – Cancer (Pure >= 0.4) – HOXB4_uc_dTSS_oPEAK

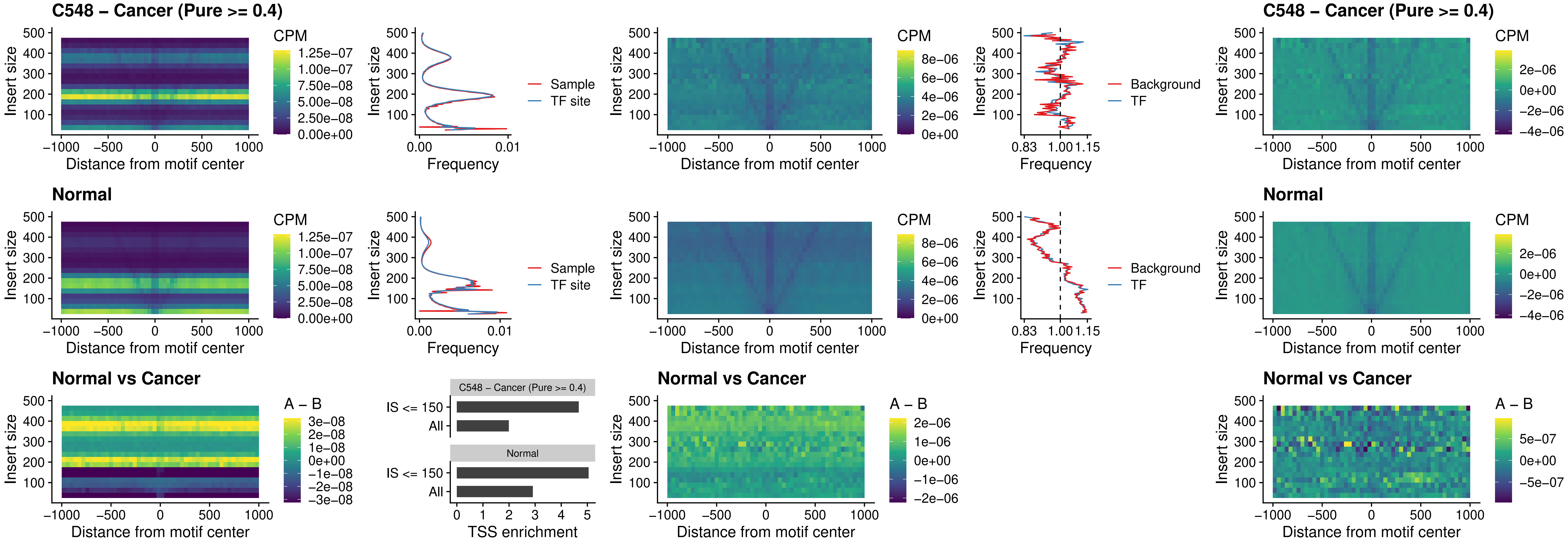

**C548 – Cancer (Pure >= 0.4) – VAX1_uc_dTSS_oPEAK**

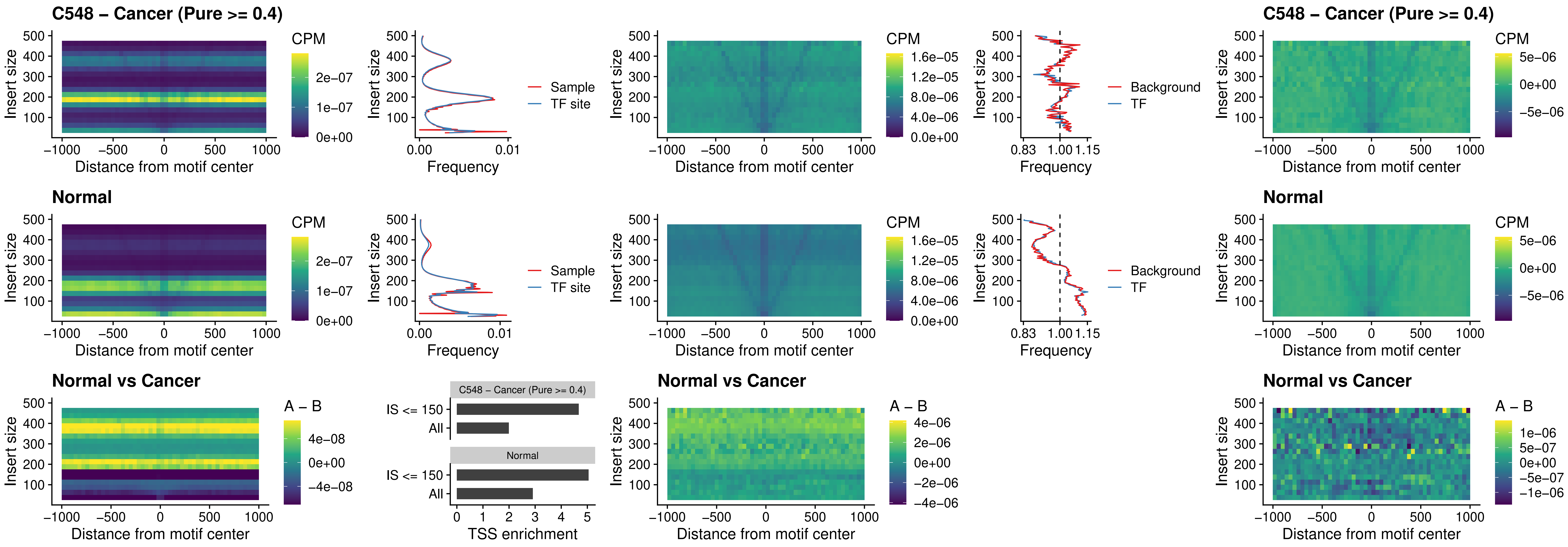

#### C548 – Cancer (Pure >= 0.4) – DLX3_uc_dTSS_oPEAK

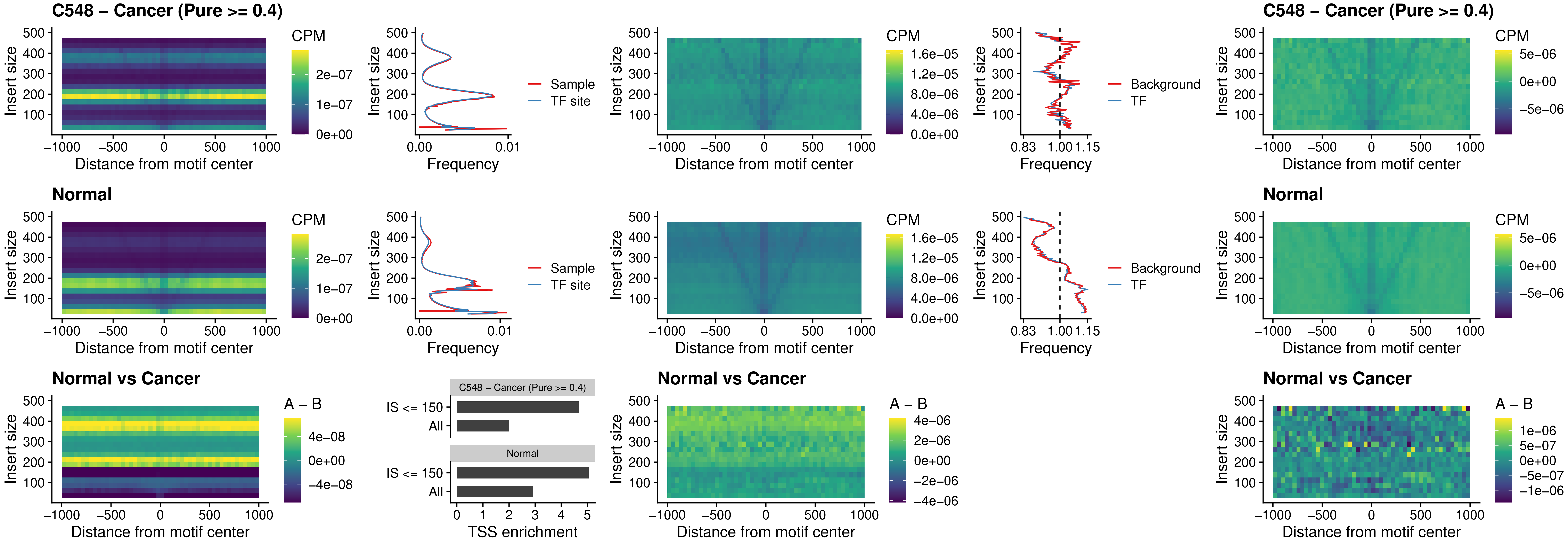

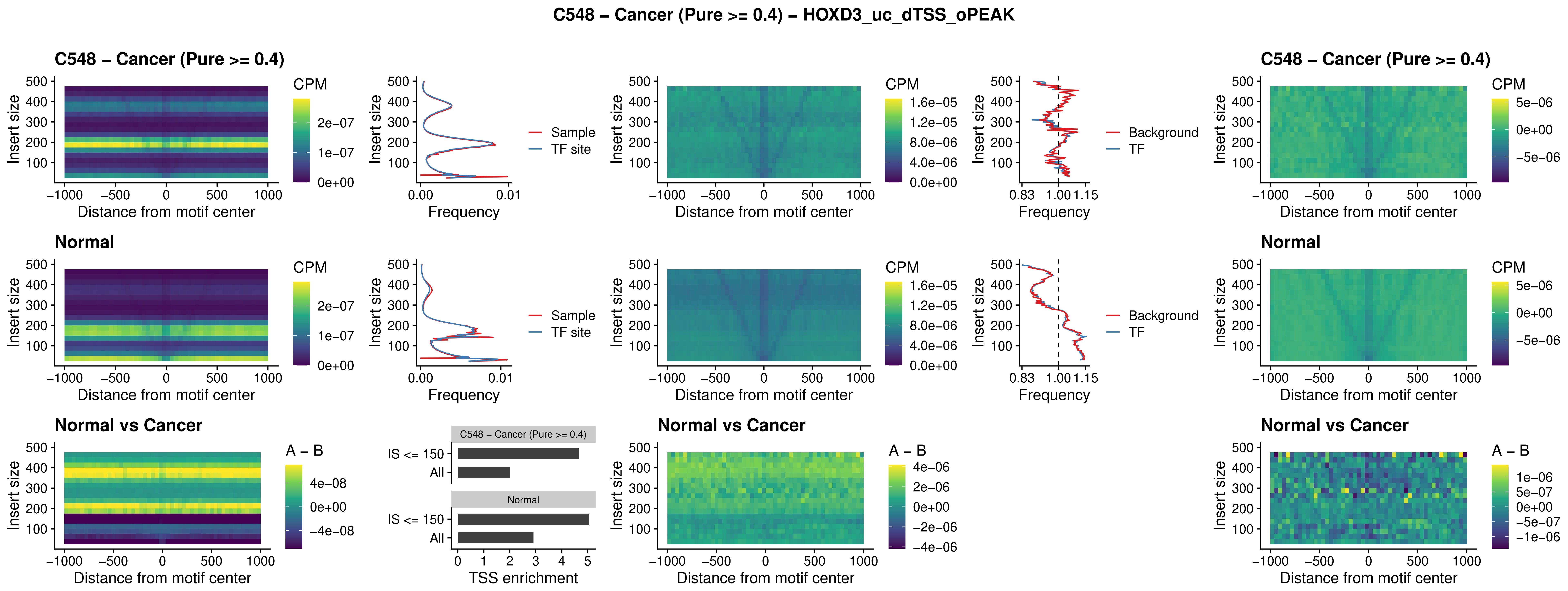

C548 – Cancer (Pure >= 0.4) – HOXD3_uc_dTSS_oPEAK
C548 – Cancer (Pure >= 0.4)
Normal
Normal vs Cancer
Insert size
Distance from motif center
Frequency
CPM
A – B
Sample
TF site
Background
TF
C548 – Cancer (Pure >= 0.4)
Normal
IS <= 150
All
TSS enrichment

#### C548 – Cancer (Pure >= 0.4) – BBX_uc_dTSS_nPEAK

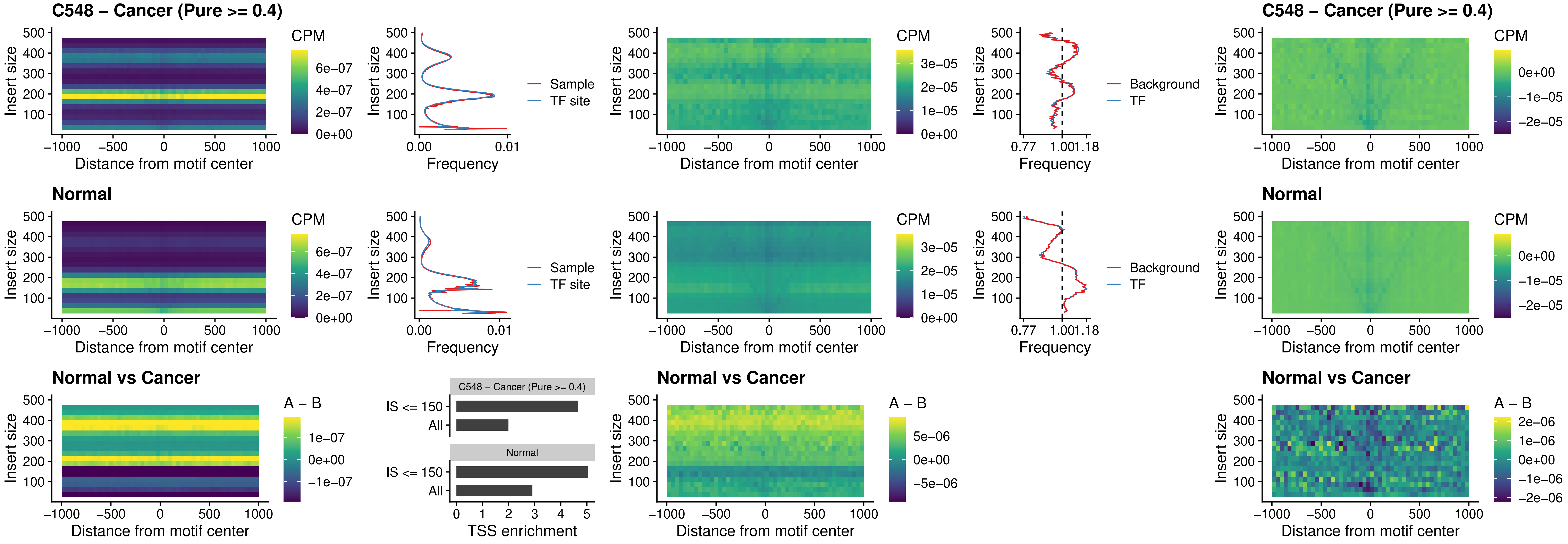

### C548 – Cancer (Pure >= 0.4) – SOX3_uc_cTSS_nPEAK

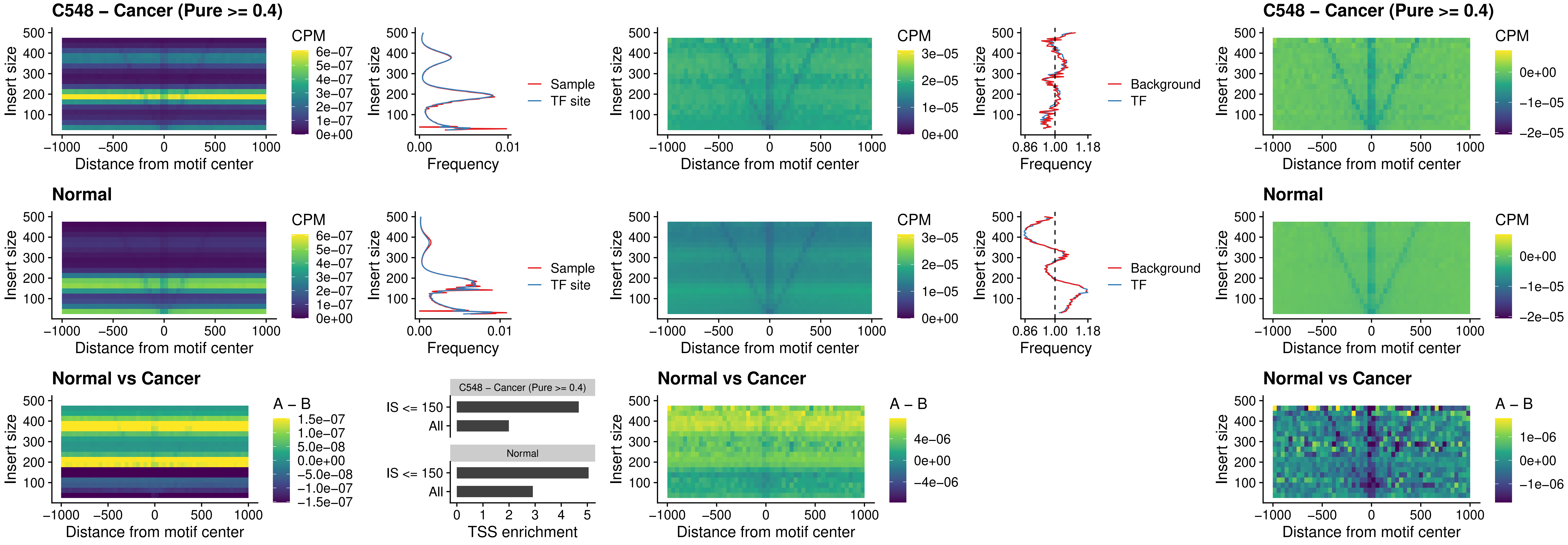

#### C548 – Cancer (Pure >= 0.4) – TBPL2_uc_cTSS_nPEAK

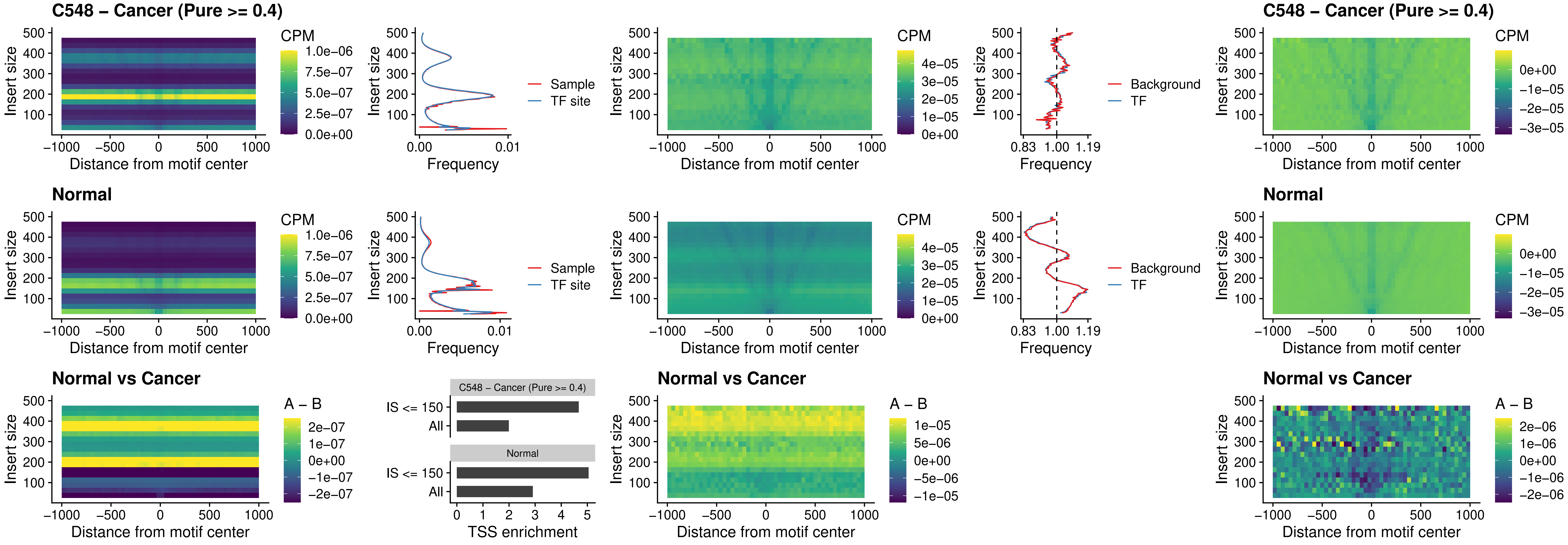

#### C548 – Cancer (Pure >= 0.4) – HOXA4_uc_cTSS_oPEAK

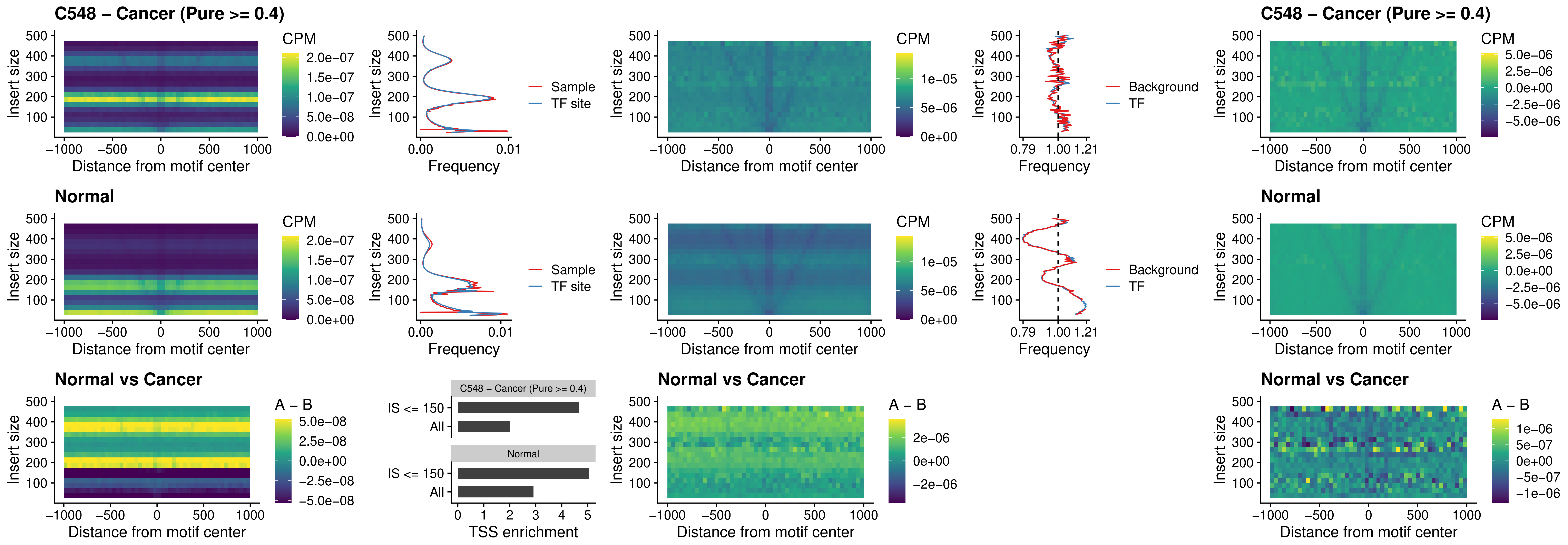

#### C548 – Cancer (Pure >= 0.4) – PDX1_uc_dTSS_oPEAK

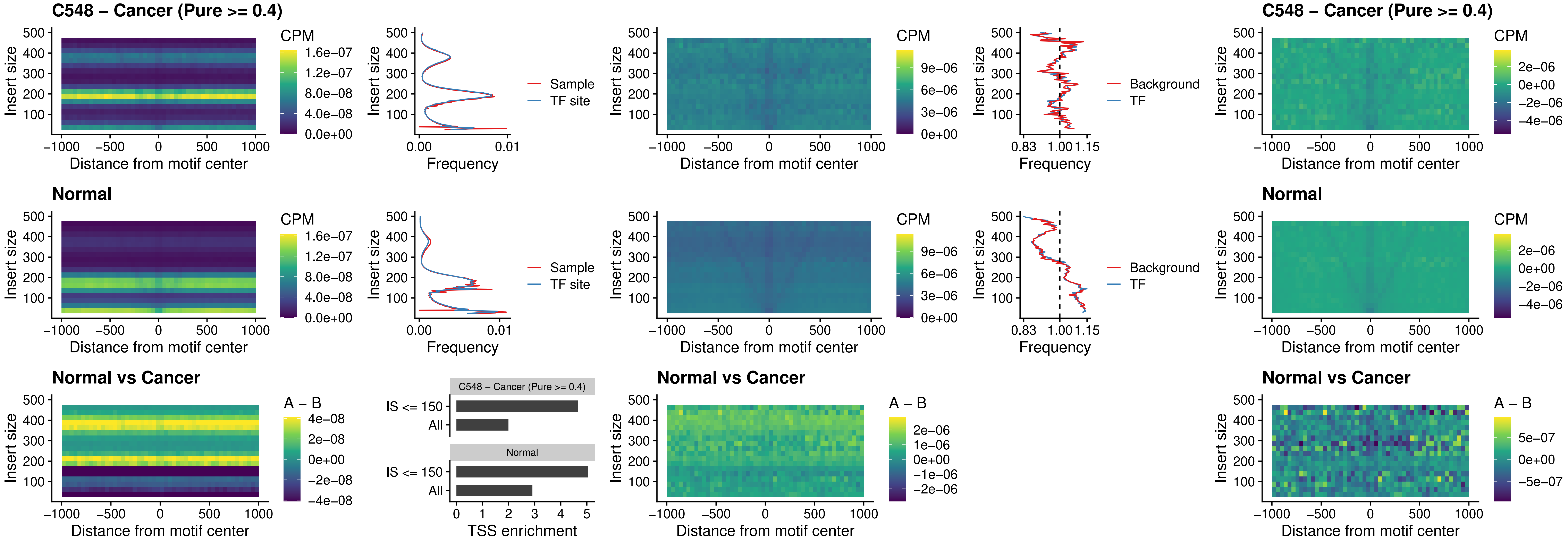

#### C548 – Cancer (Pure >= 0.4) – LHX4_uc_dTSS_oPEAK

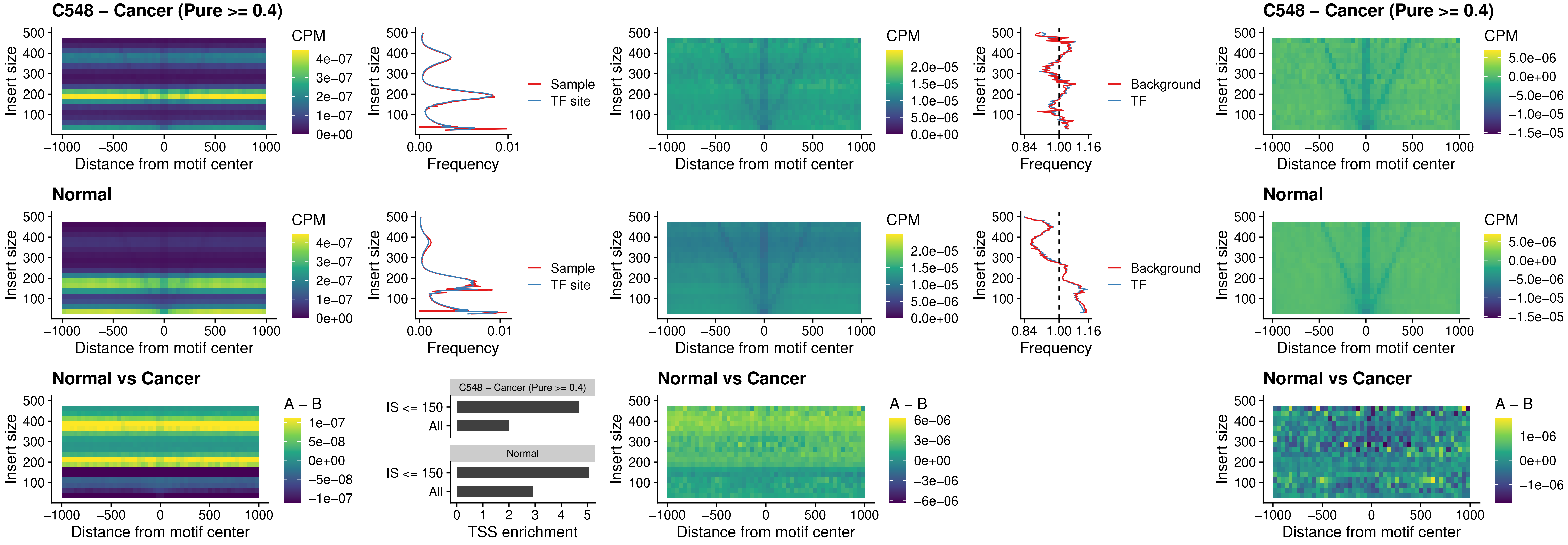

#### C548 – Cancer (Pure >= 0.4) – HOXA4_uc_dTSS_oPEAK

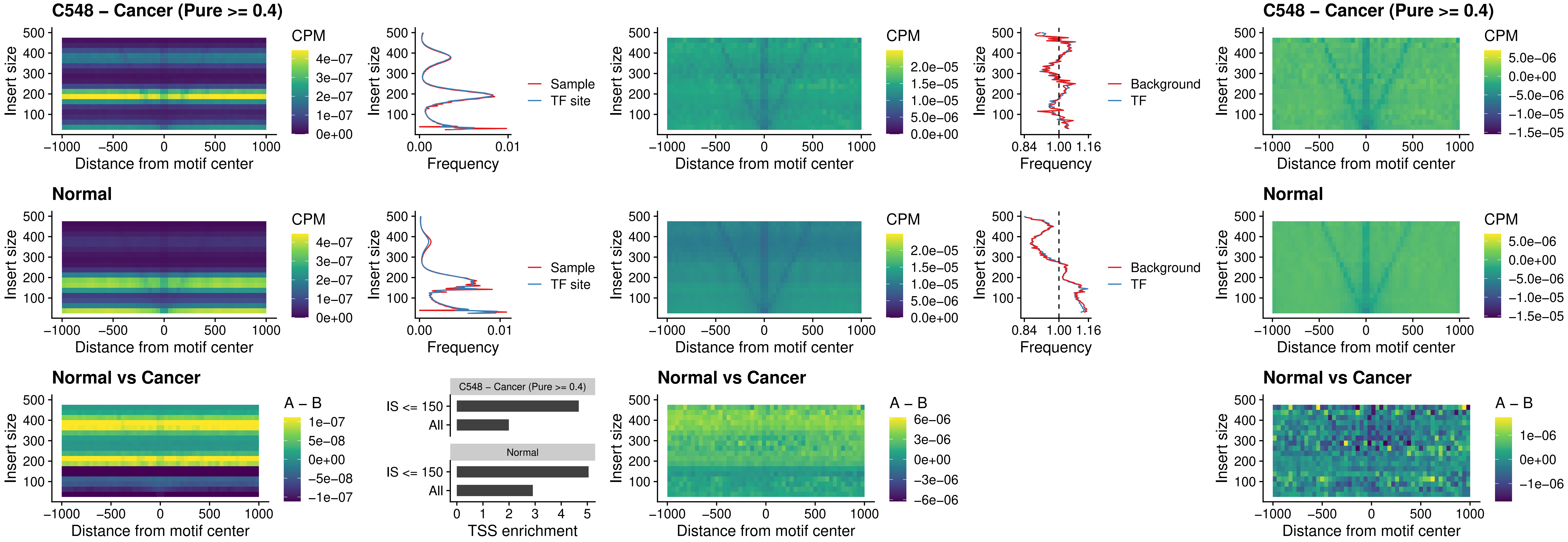

#### C548 – Cancer (Pure >= 0.4) – HLX_uc_dTSS_oPEAK

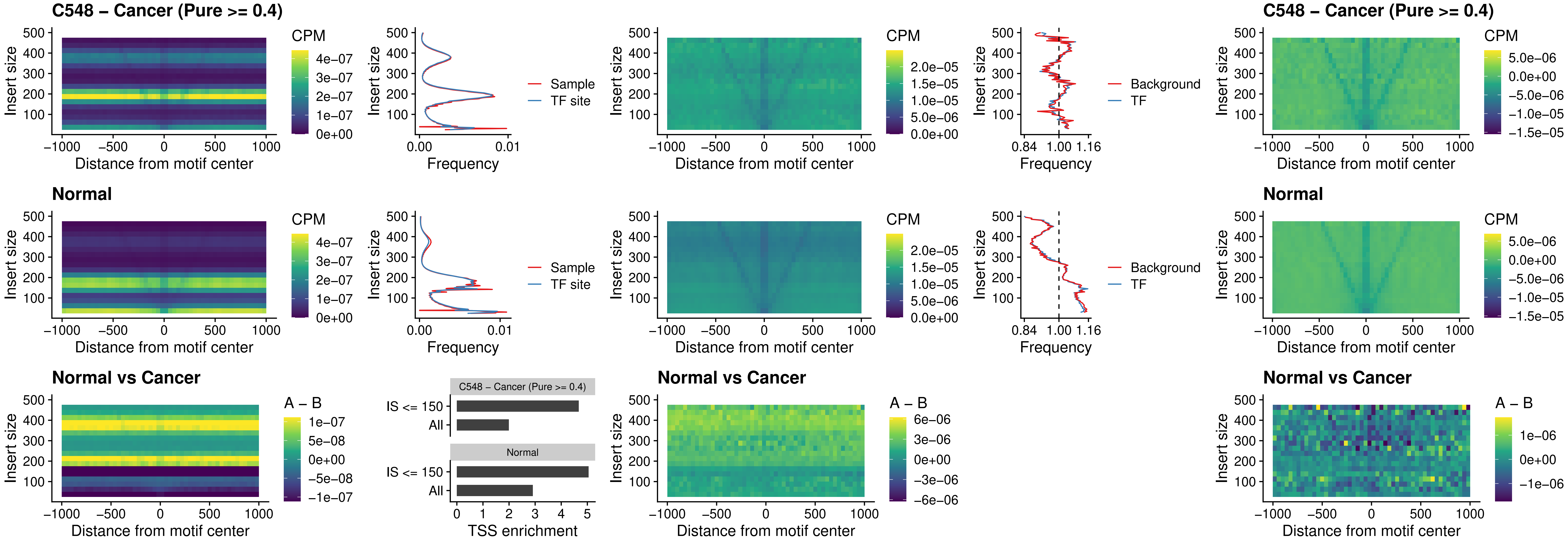

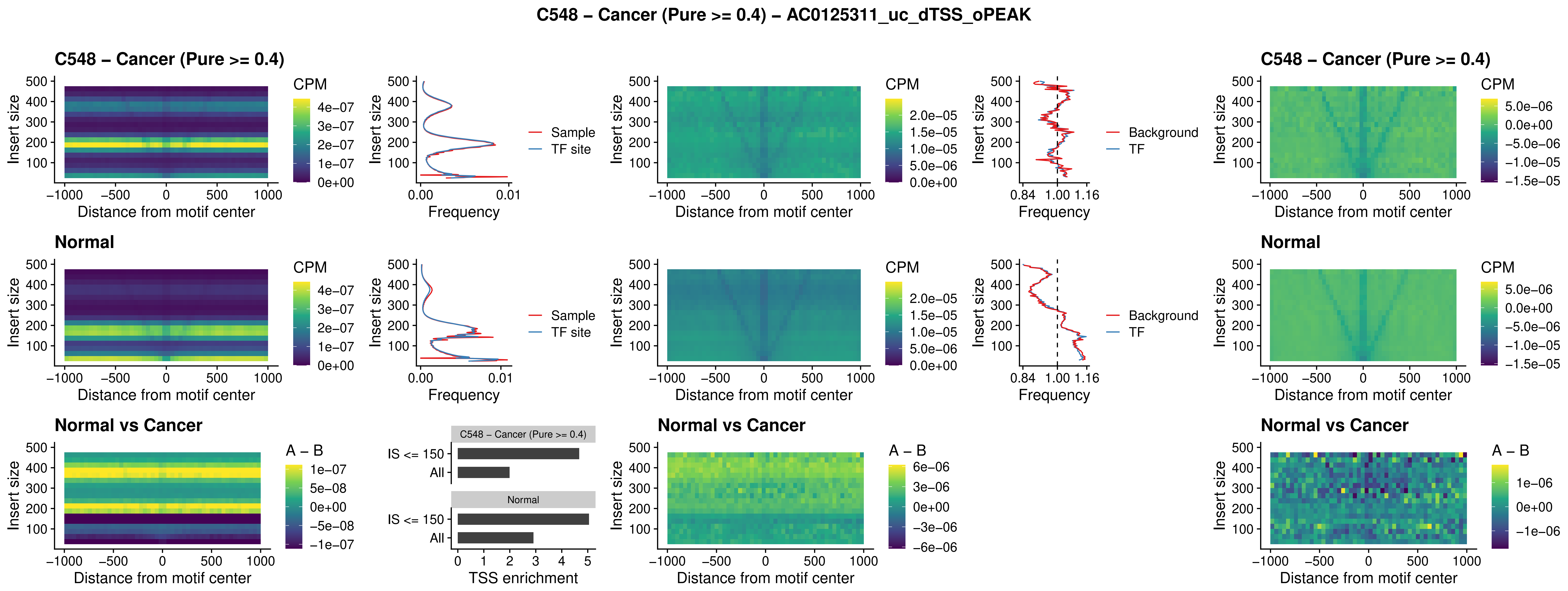
C548 – Cancer (Pure >= 0.4) – AC0125311_uc_dTSS_oPEAK
C548 – Cancer (Pure >= 0.4)
Normal
Normal vs Cancer
Insert size
Distance from motif center
Frequency
CPM
A – B
Sample
TF site
Background
TF
C548 – Cancer (Pure >= 0.4)
Normal
IS <= 150
All
TSS enrichment

#### C548 – Cancer (Pure >= 0.4) – DBX2_uc_dTSS_oPEAK

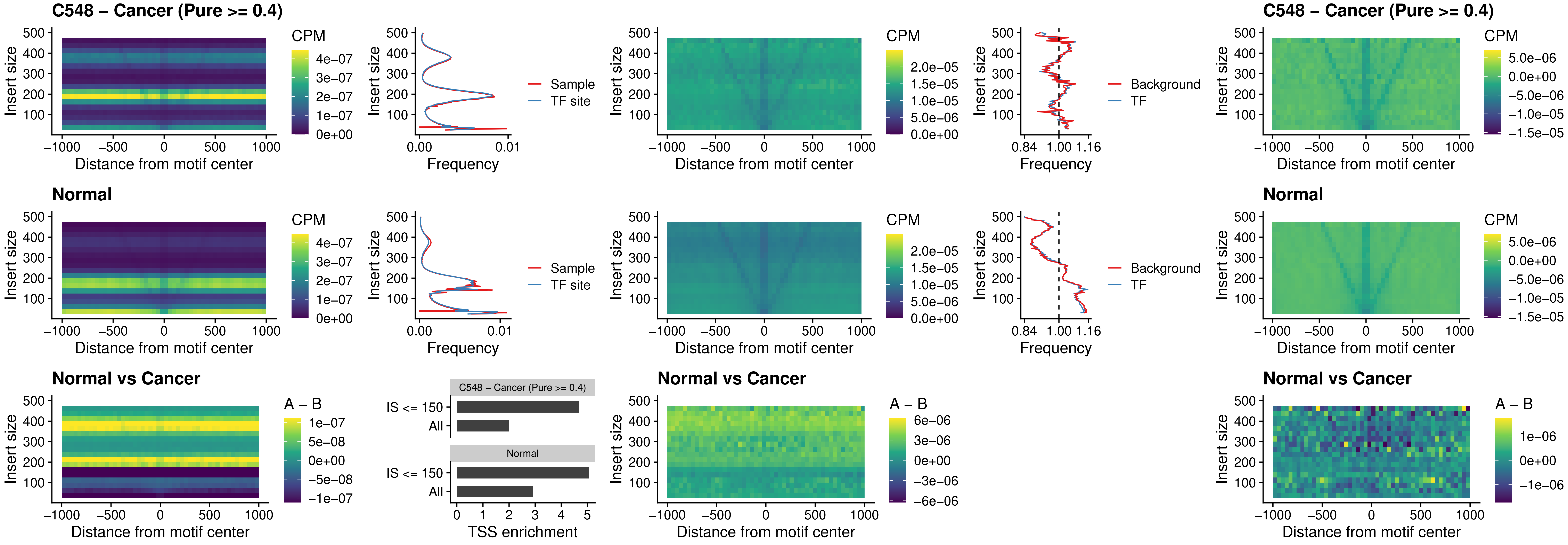

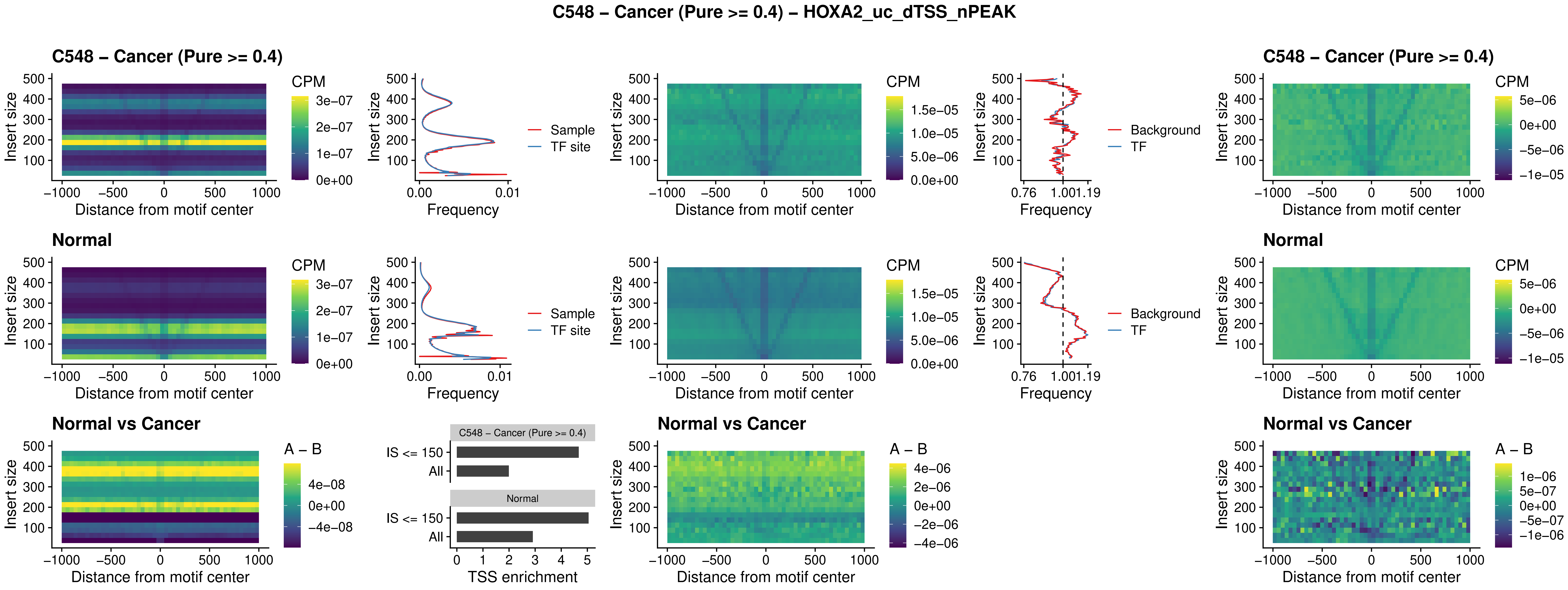
C548 – Cancer (Pure >= 0.4) – HOXA2_uc_dTSS_nPEAK
C548 – Cancer (Pure >= 0.4)
Normal
Normal vs Cancer
Insert size
Distance from motif center
Frequency
CPM
A – B
Sample
TF site
Background
TF
C548 – Cancer (Pure >= 0.4)
Normal
IS <= 150
All
TSS enrichment

#### C548 – Cancer (Pure >= 0.4) – SHOX_uc_dTSS_nPEAK

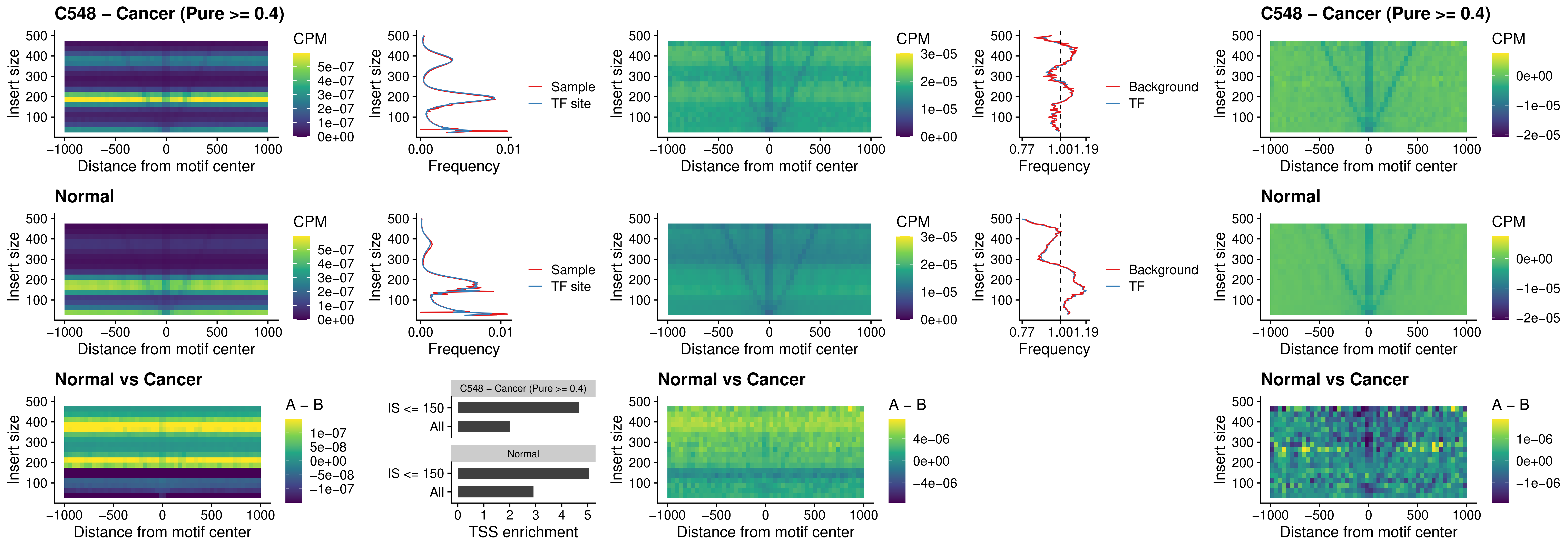

#### C548 – Cancer (Pure >= 0.4) – PRRX1_uc_dTSS_nPEAK

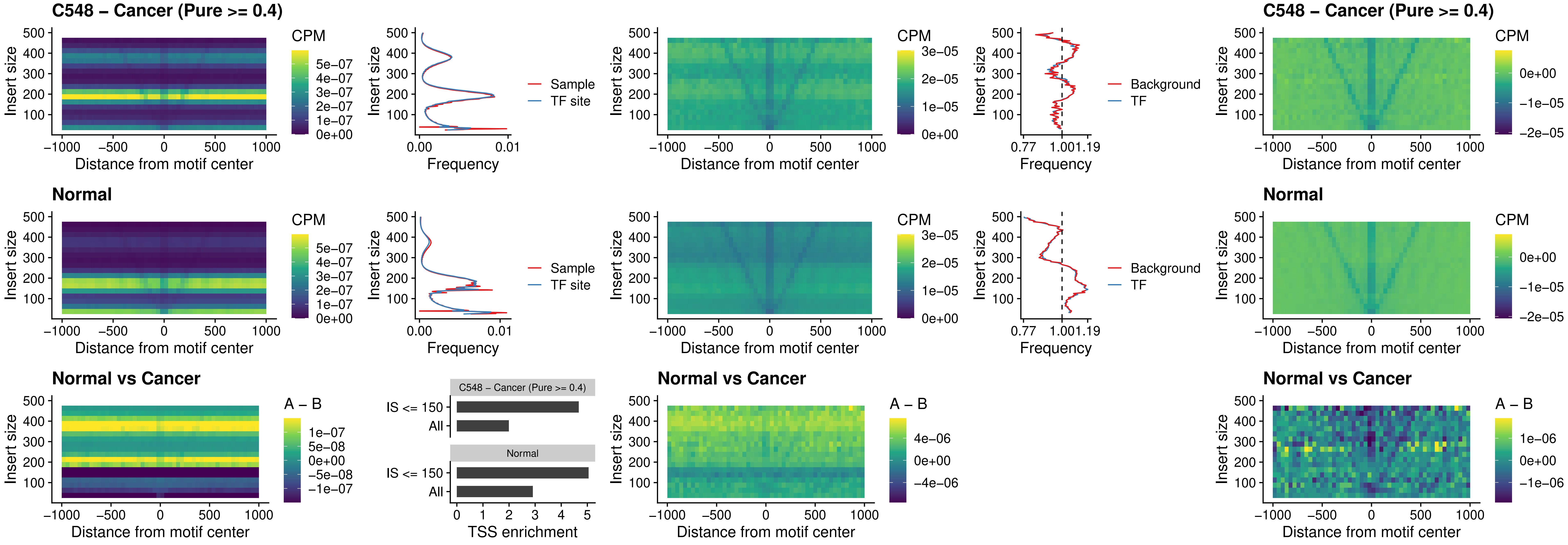

#### C548 – Cancer (Pure >= 0.4) – LMX1A_uc_dTSS_nPEAK

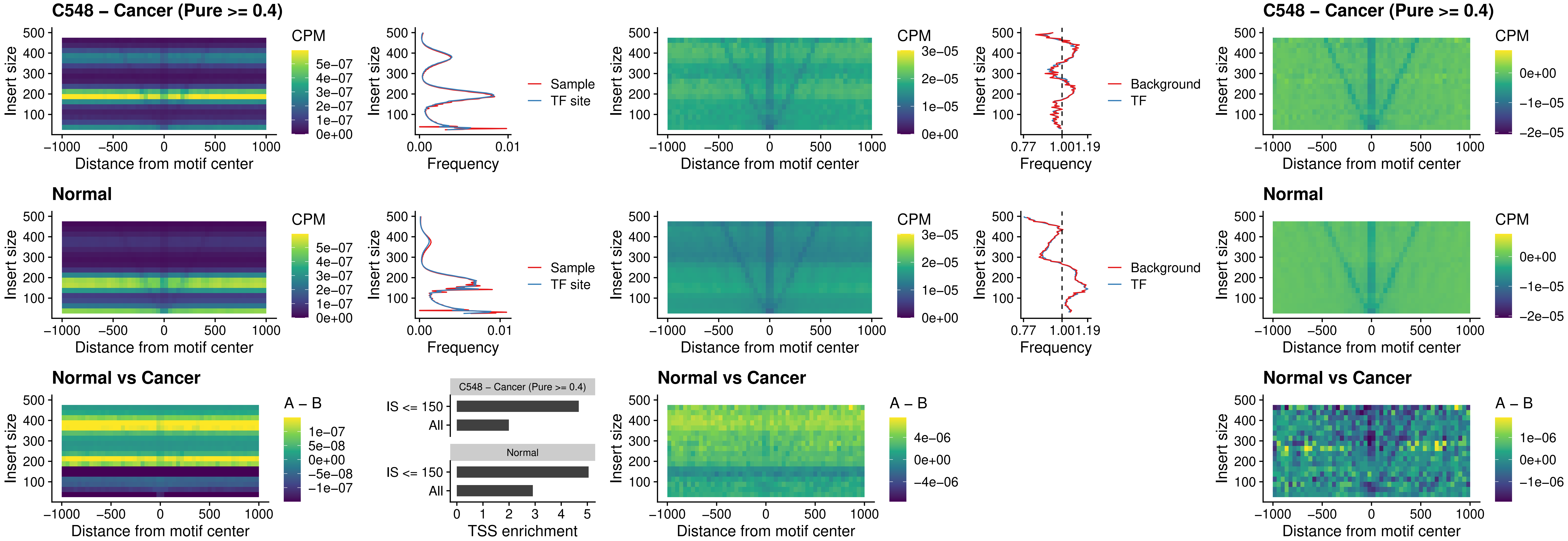

#### C548 – Cancer (Pure >= 0.4) – LMX1B_uc_dTSS_nPEAK

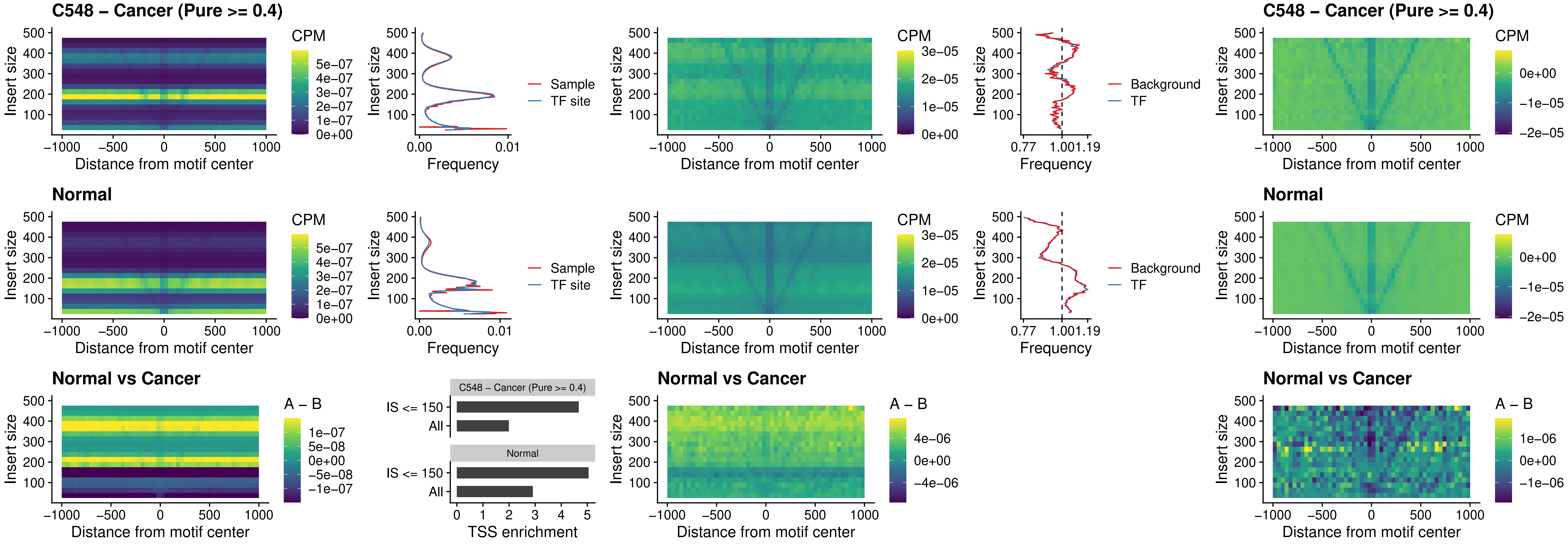

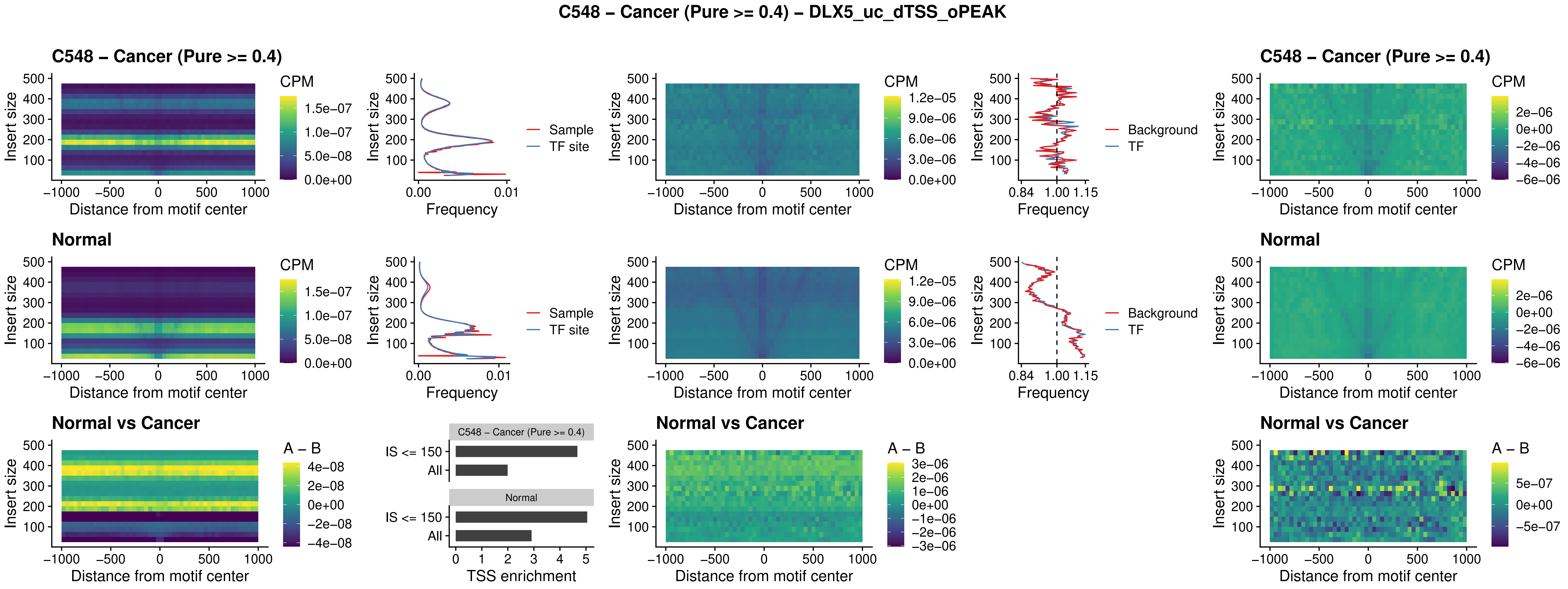

C548 – Cancer (Pure >= 0.4) – DLX5_uc_dTSS_oPEAK
C548 – Cancer (Pure >= 0.4)
Normal
Normal vs Cancer
Insert size
Distance from motif center
Frequency
CPM
A – B
Sample
TF site
Background
TF
C548 – Cancer (Pure >= 0.4)
Normal
IS <= 150
All
TSS enrichment

#### C548 – Cancer (Pure >= 0.4) – ARID3C_uc_dTSS_nPEAK

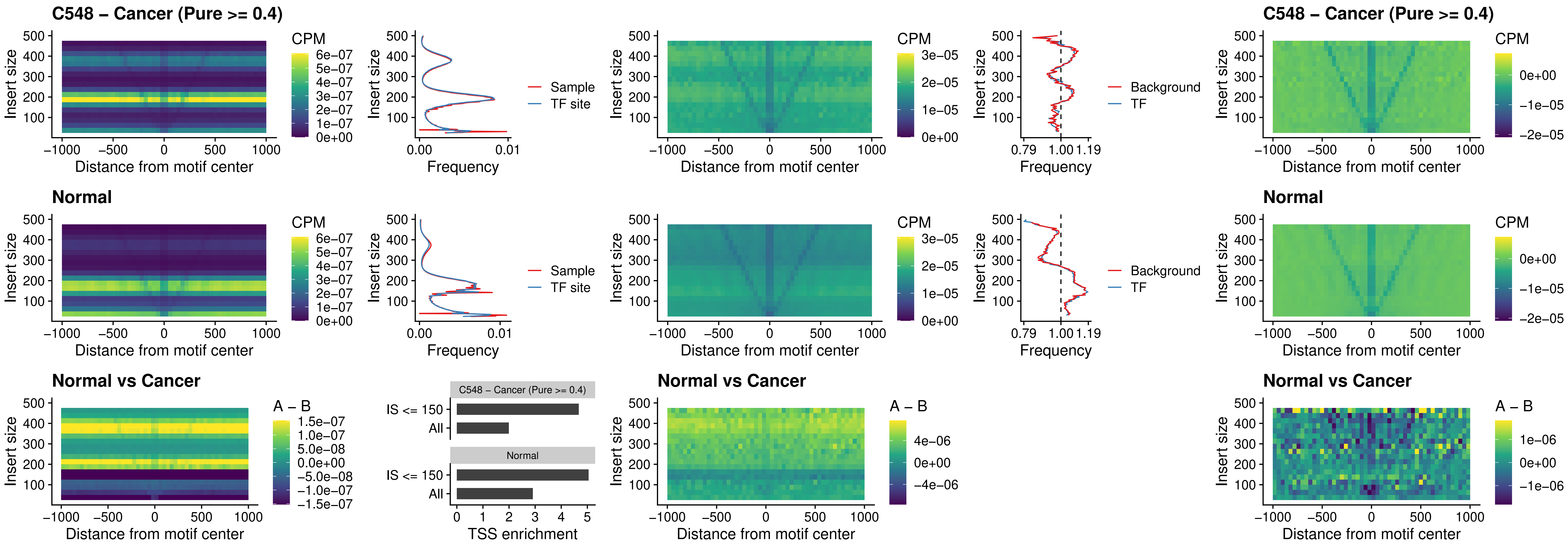

#### C548 – Cancer (Pure >= 0.4) – HOXD12_uc_dTSS_nPEAK

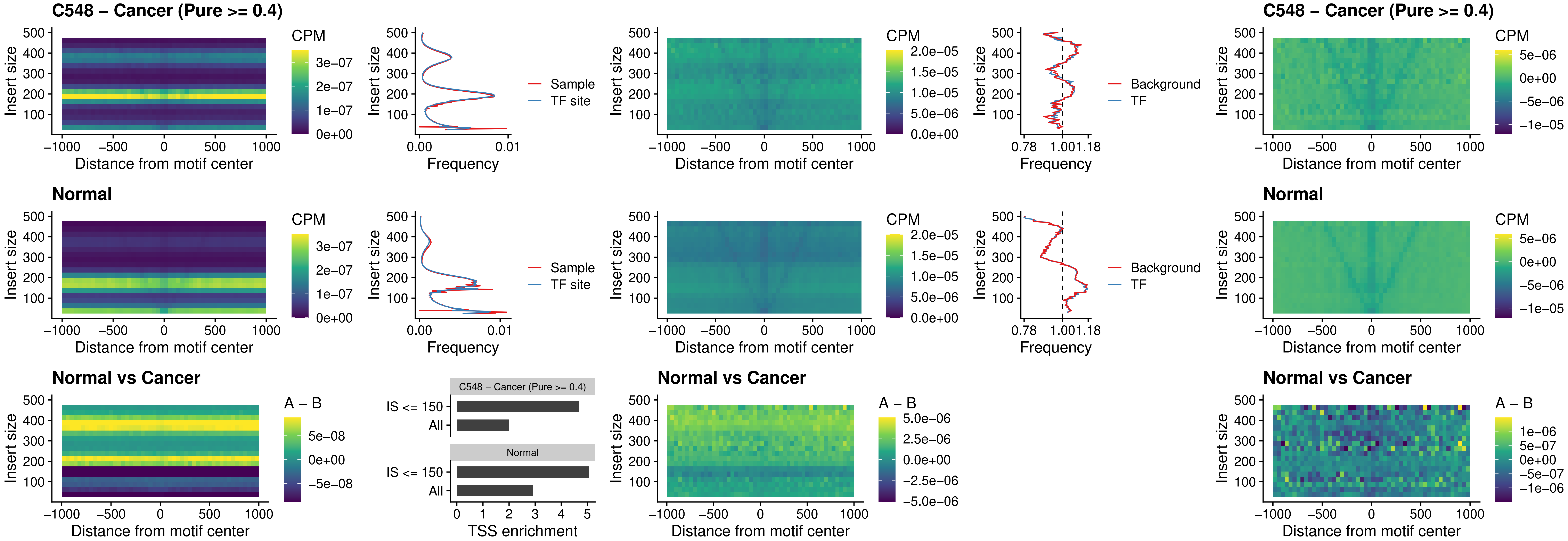

#### C548 – Cancer (Pure >= 0.4) – CDX2_uc_dTSS_nPEAK

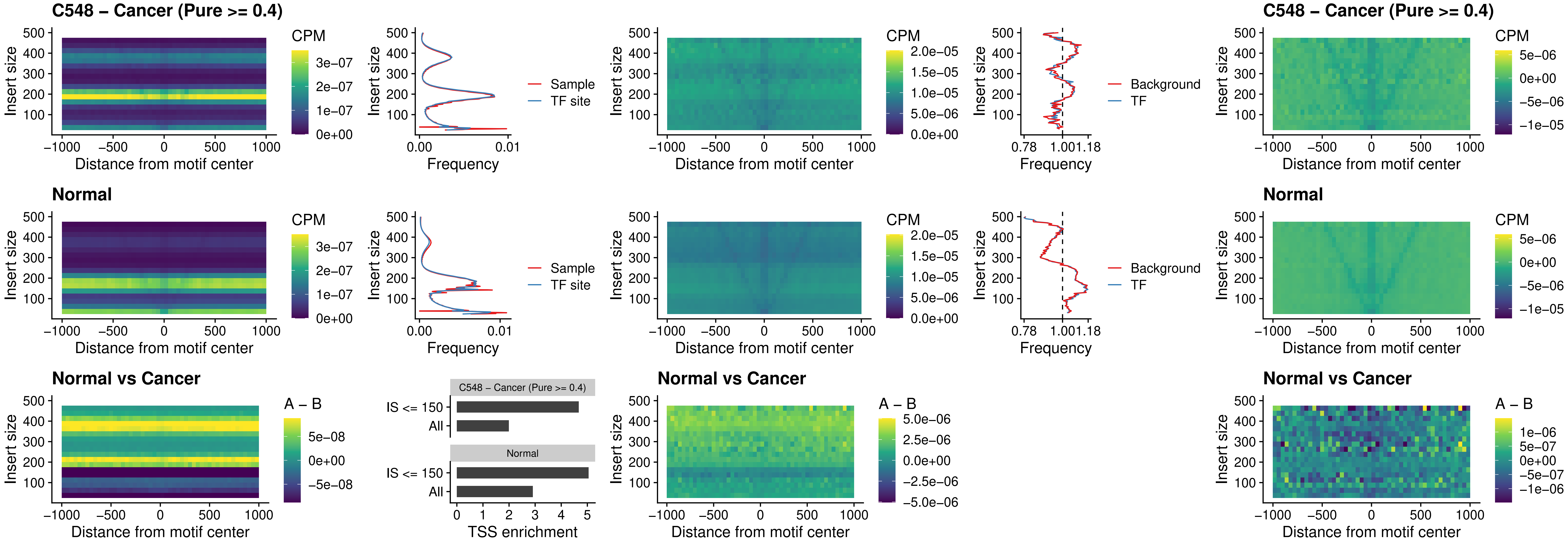

### C548 – Cancer (Pure >= 0.4) – HOXC12_uc_dTSS_nPEAK

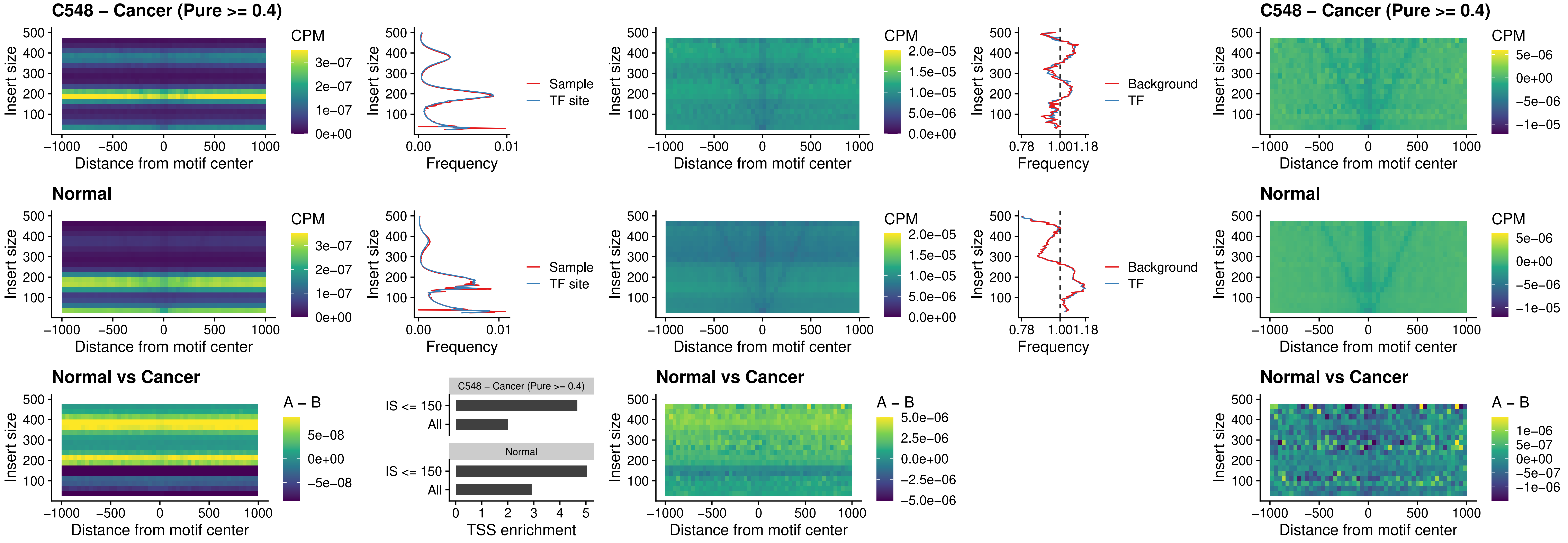

#### C548 – Cancer (Pure >= 0.4) – HHEX_uc_dTSS_oPEAK

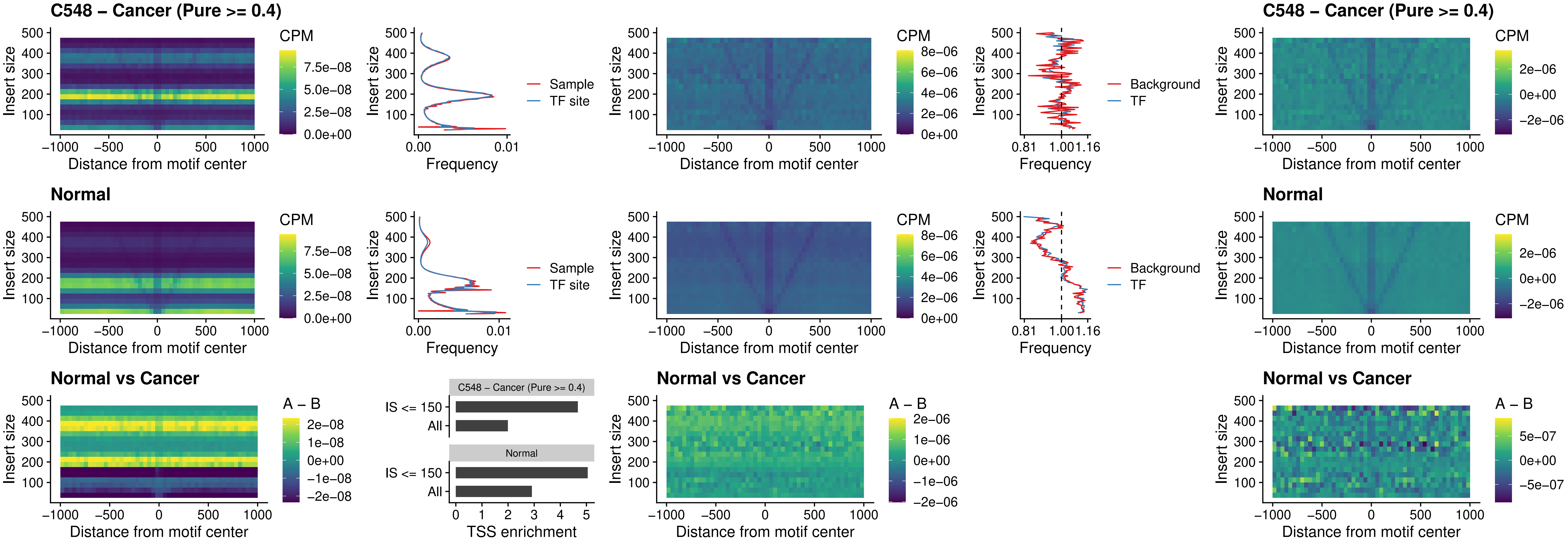

#### C548 – Cancer (Pure >= 0.4) – HOXA11_uc_dTSS_nPEAK

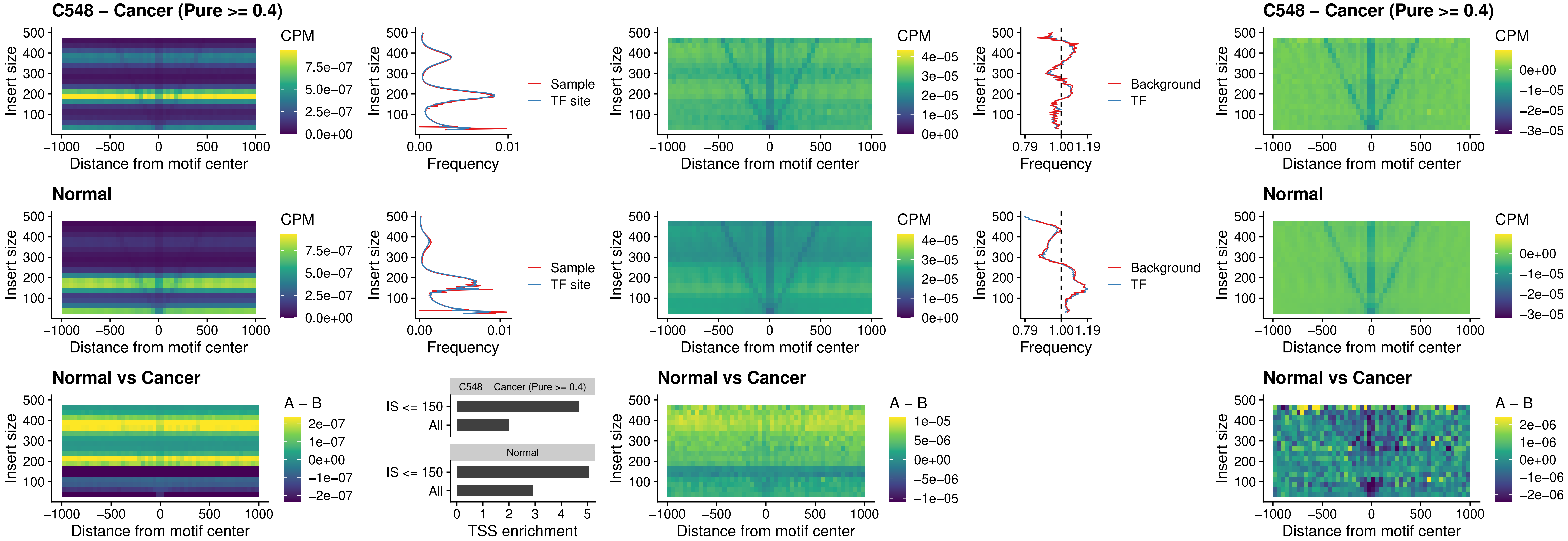

#### C548 – Cancer (Pure >= 0.4) – HOXA11_uc_cTSS_nPEAK

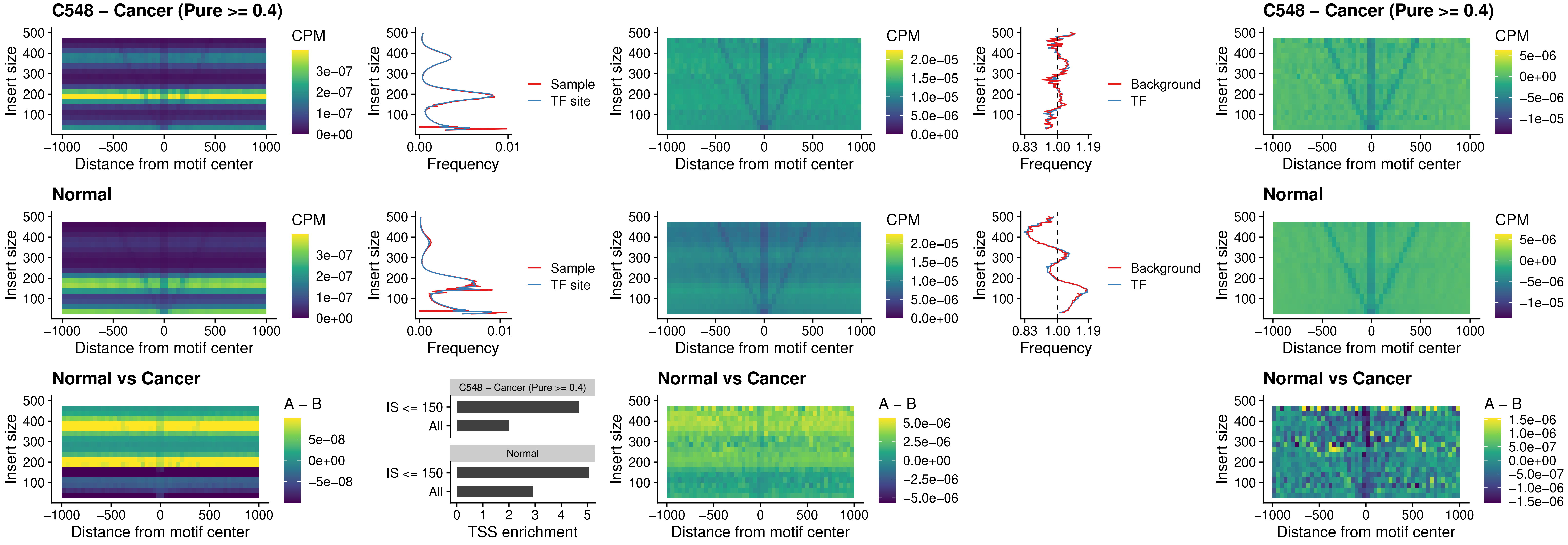

#### C548 – Cancer (Pure >= 0.4) – ZFHX3_uc_dTSS_nPEAK

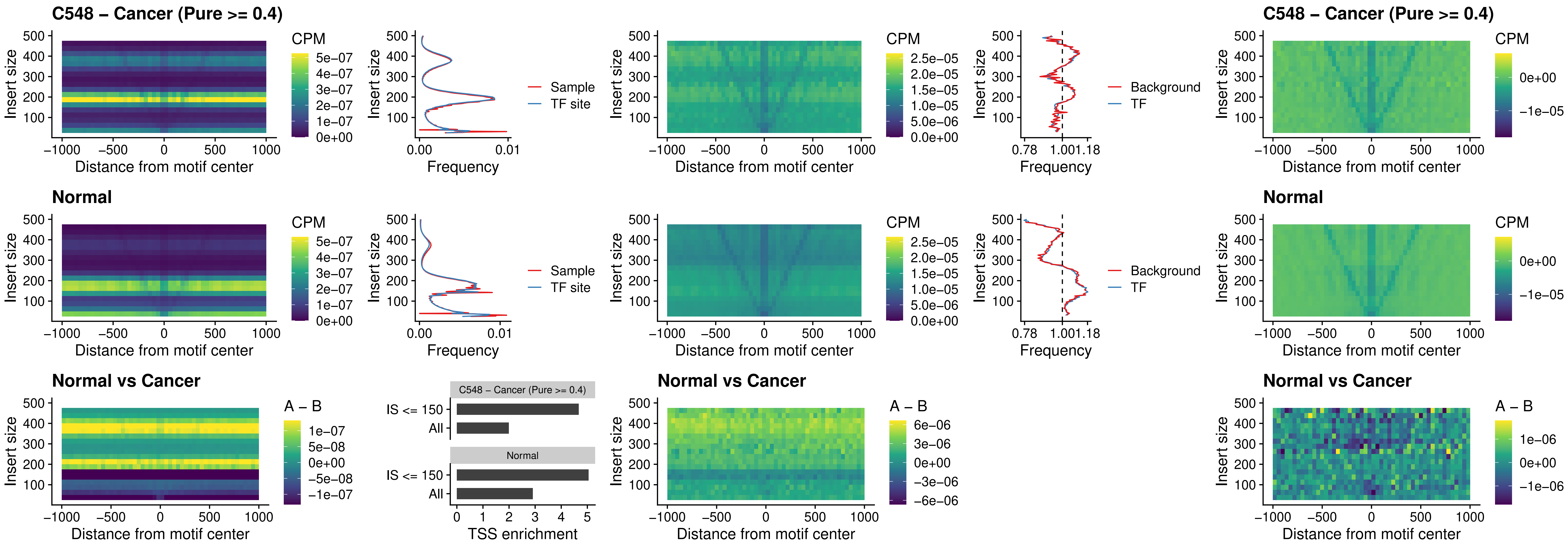

#### C548 – Cancer (Pure >= 0.4) – HOXB4_uc_dTSS_nPEAK

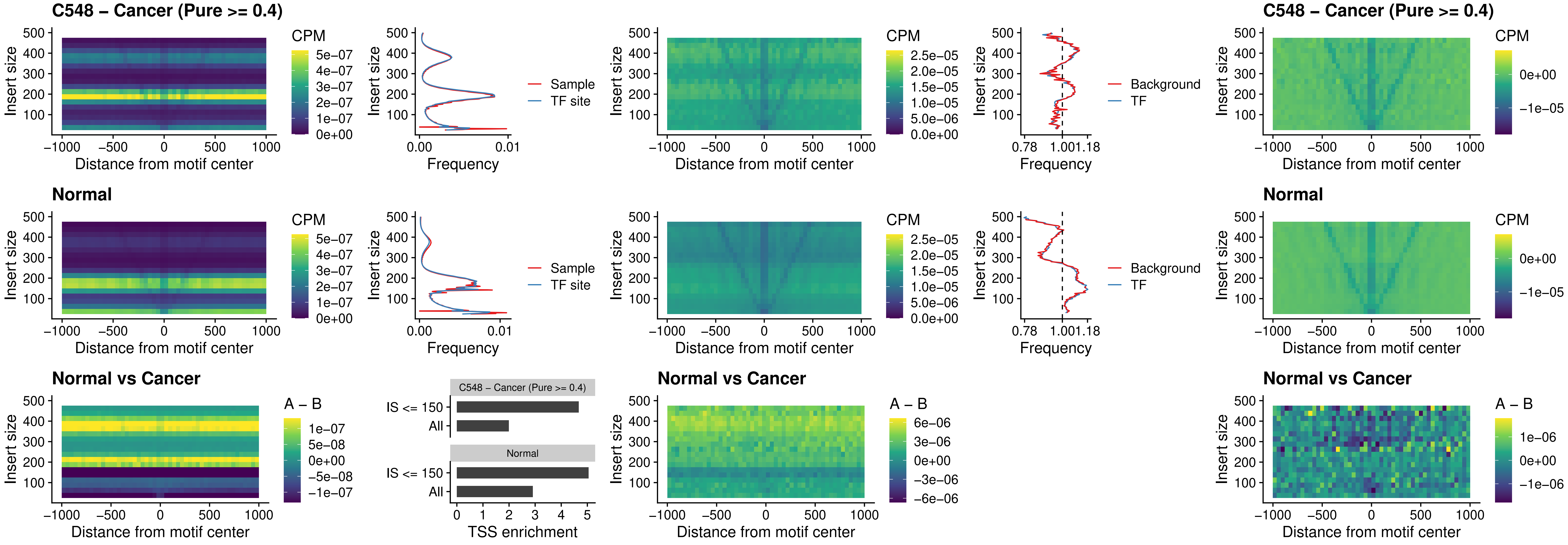

### C548 – Cancer (Pure >= 0.4) – HOXC5_uc_dTSS_nPEAK

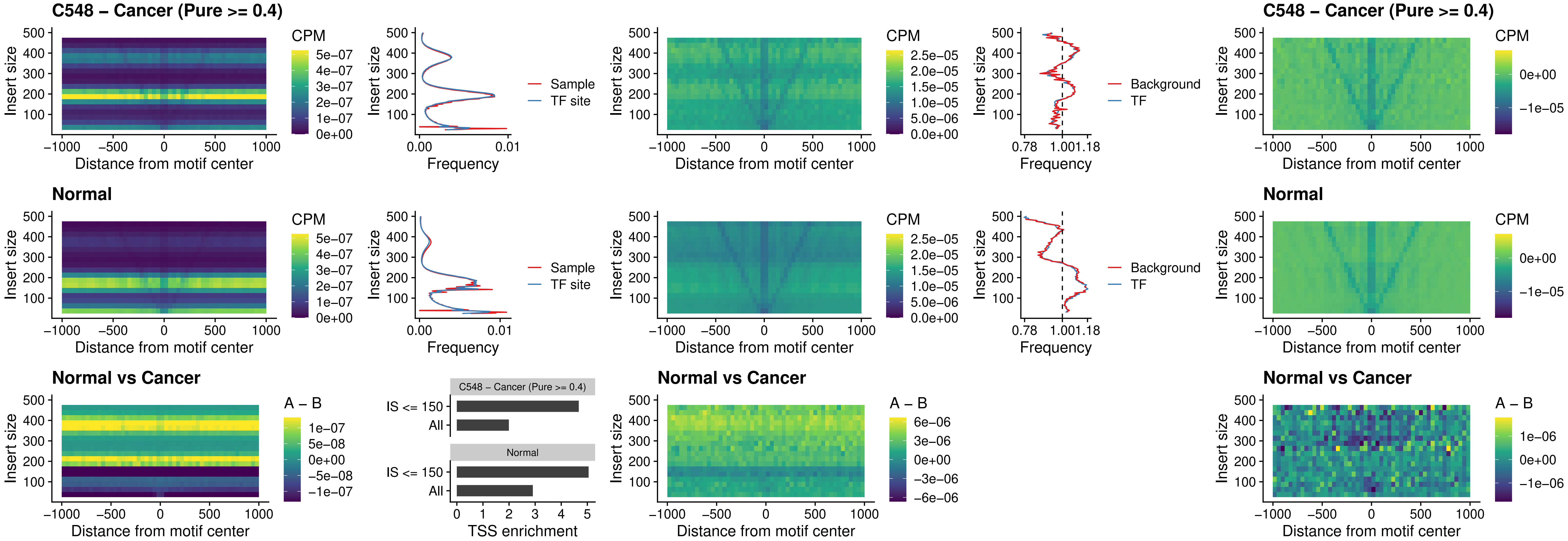

#### C548 – Cancer (Pure >= 0.4) – HOXD1_uc_dTSS_nPEAK

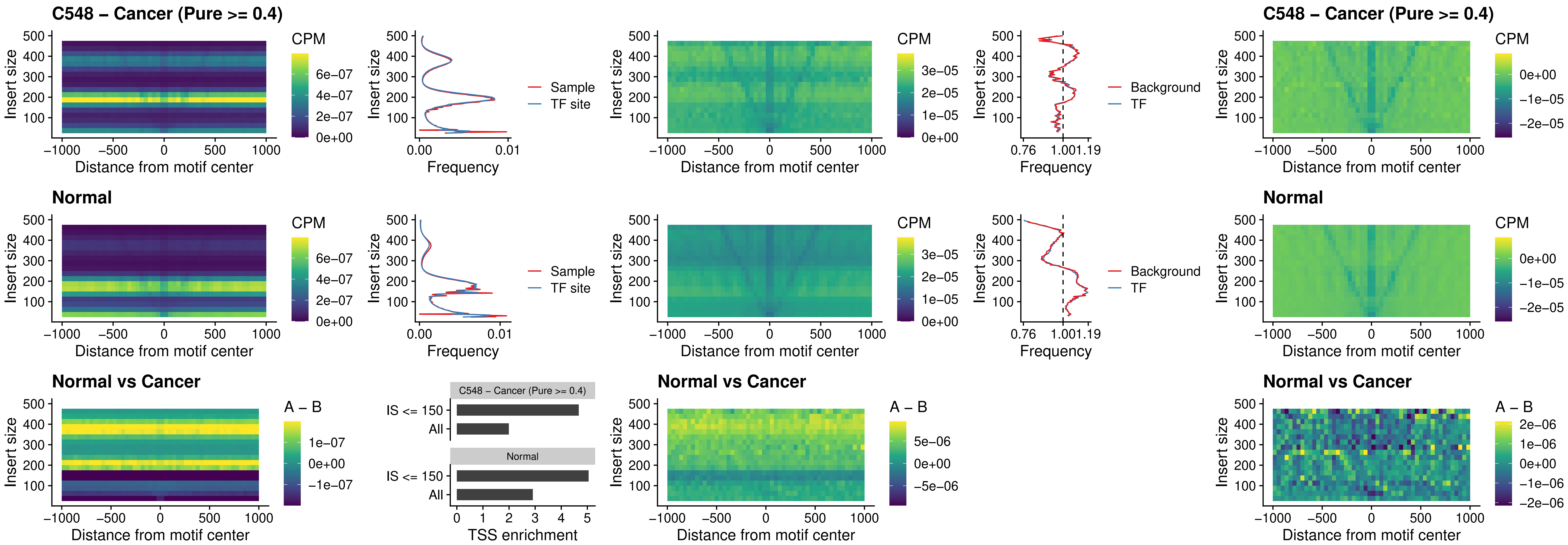

#### C548 – Cancer (Pure >= 0.4) – OTP_uc_dTSS_nPEAK

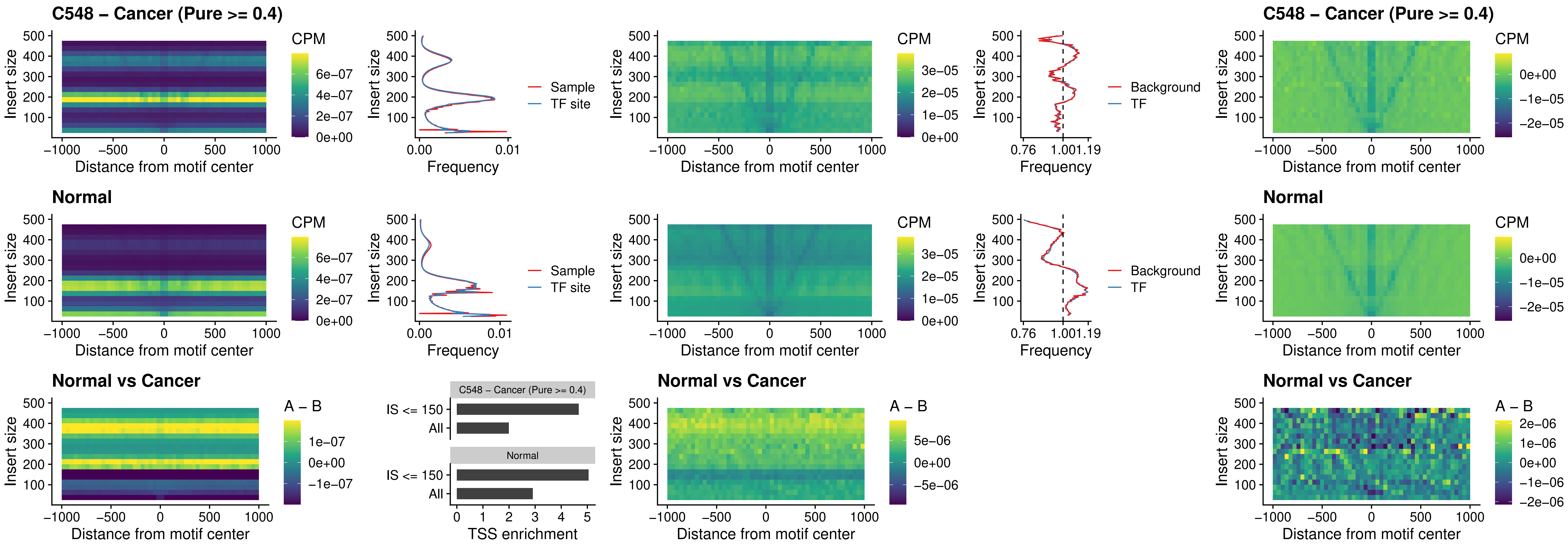

#### C548 – Cancer (Pure >= 0.4) – FOXL1_uc_dTSS_oPEAK

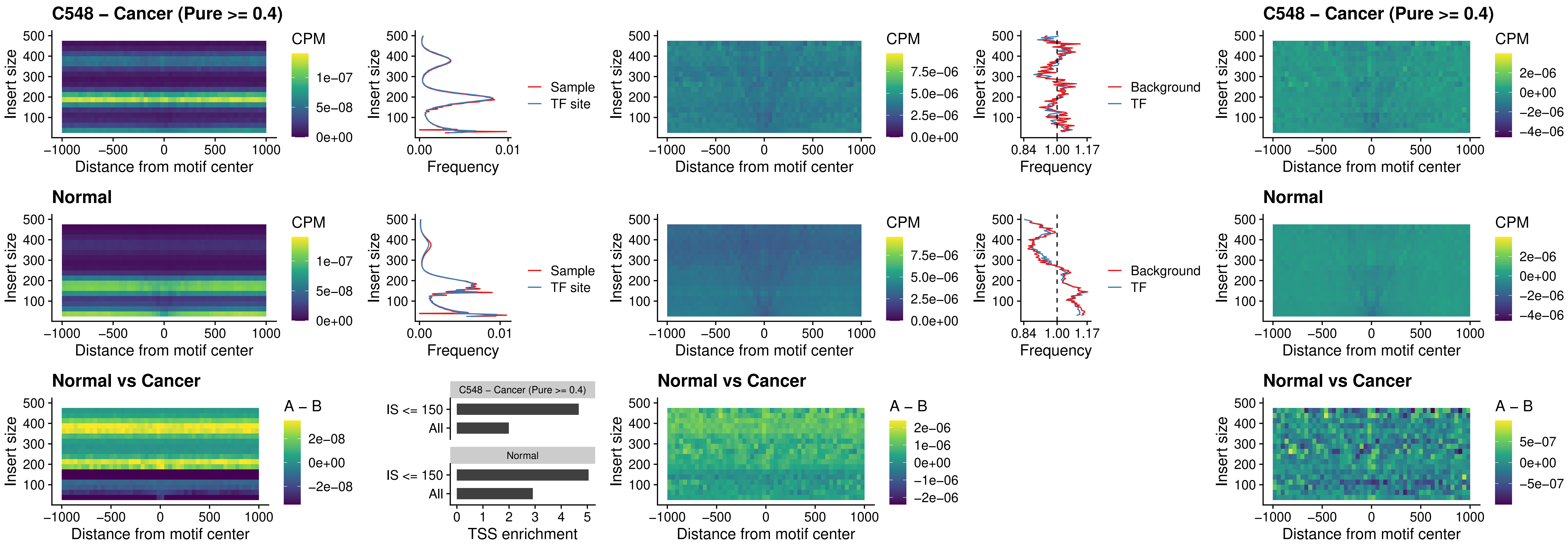

### C548 – Cancer (Pure >= 0.4) – ZNF32_uc_dTSS_oPEAK

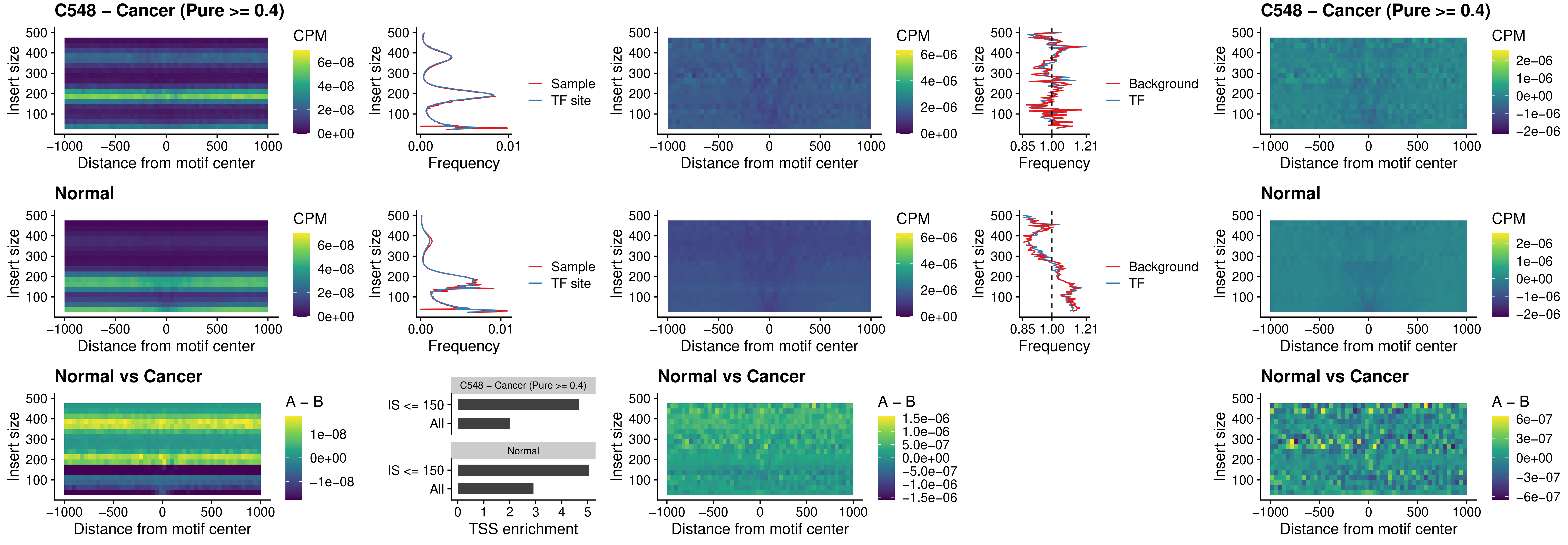

#### C548 – Cancer (Pure >= 0.4) – FOXL1_uc_dTSS_nPEAK

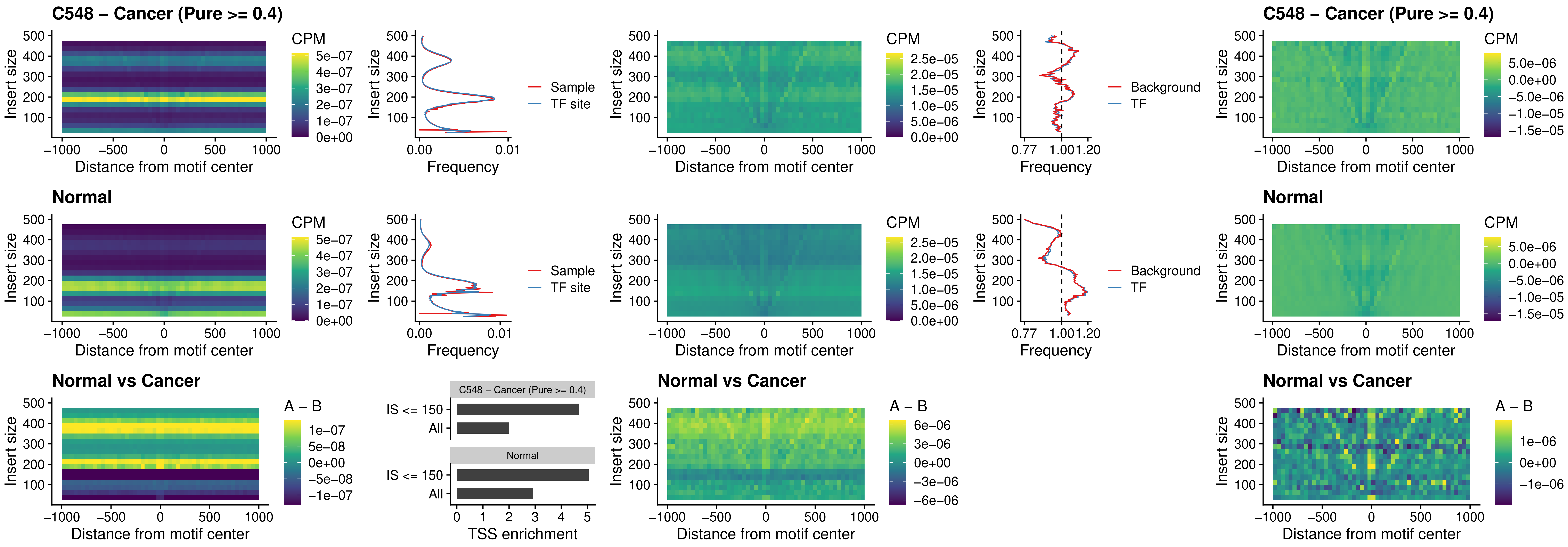

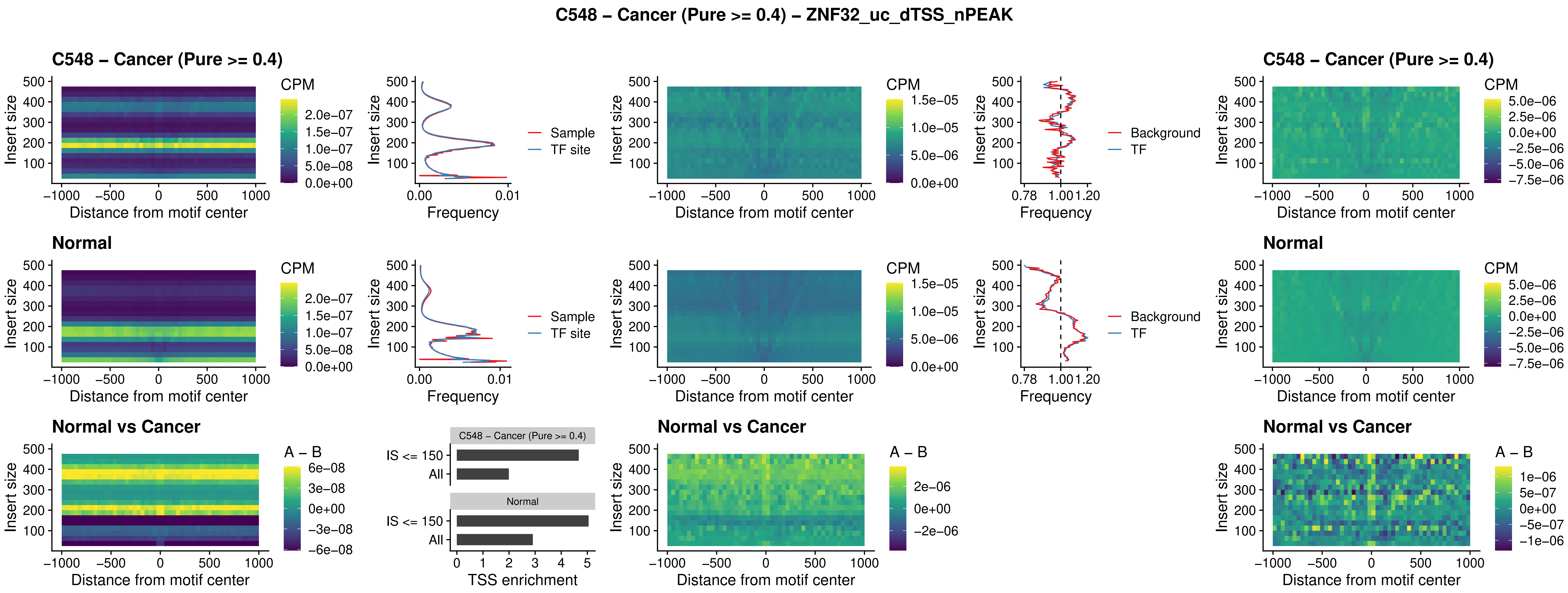
C548 – Cancer (Pure >= 0.4) – ZNF32_uc_dTSS_nPEAK
C548 – Cancer (Pure >= 0.4)
Normal
Normal vs Cancer
Insert size
Distance from motif center
Frequency
CPM
A – B
Sample
TF site
Background
TF
C548 – Cancer (Pure >= 0.4)
Normal
IS <= 150
All
TSS enrichment

### C548 – Cancer (Pure >= 0.4) – DLX5_uc_dTSS_nPEAK

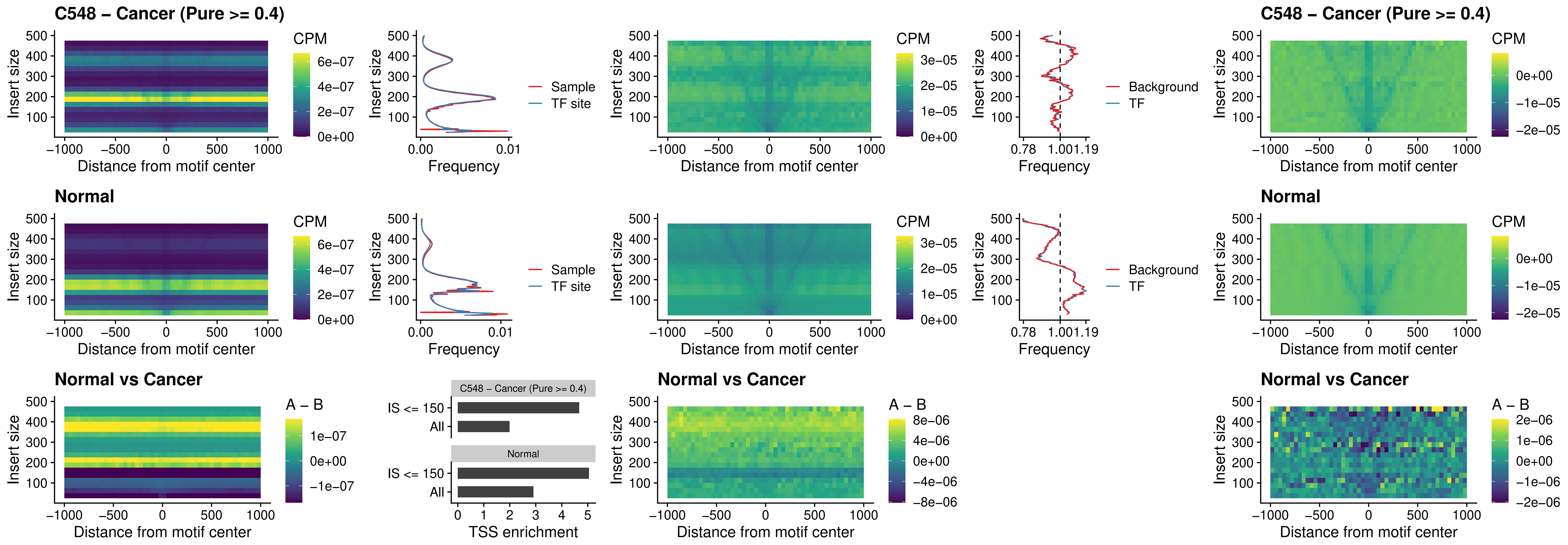

### C548 – Cancer (Pure >= 0.4) – DMRT2_uc_dTSS_nPEAK

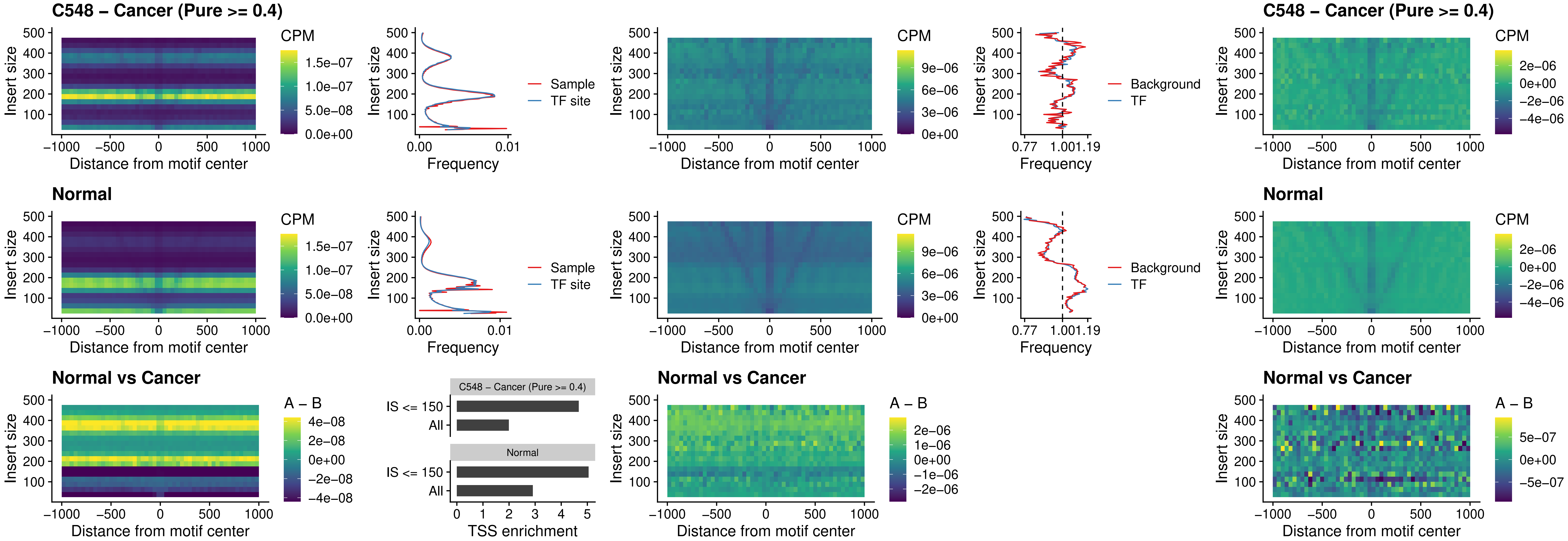

#### C548 – Cancer (Pure >= 0.4) – DMRTC2_uc_cTSS_nPEAK

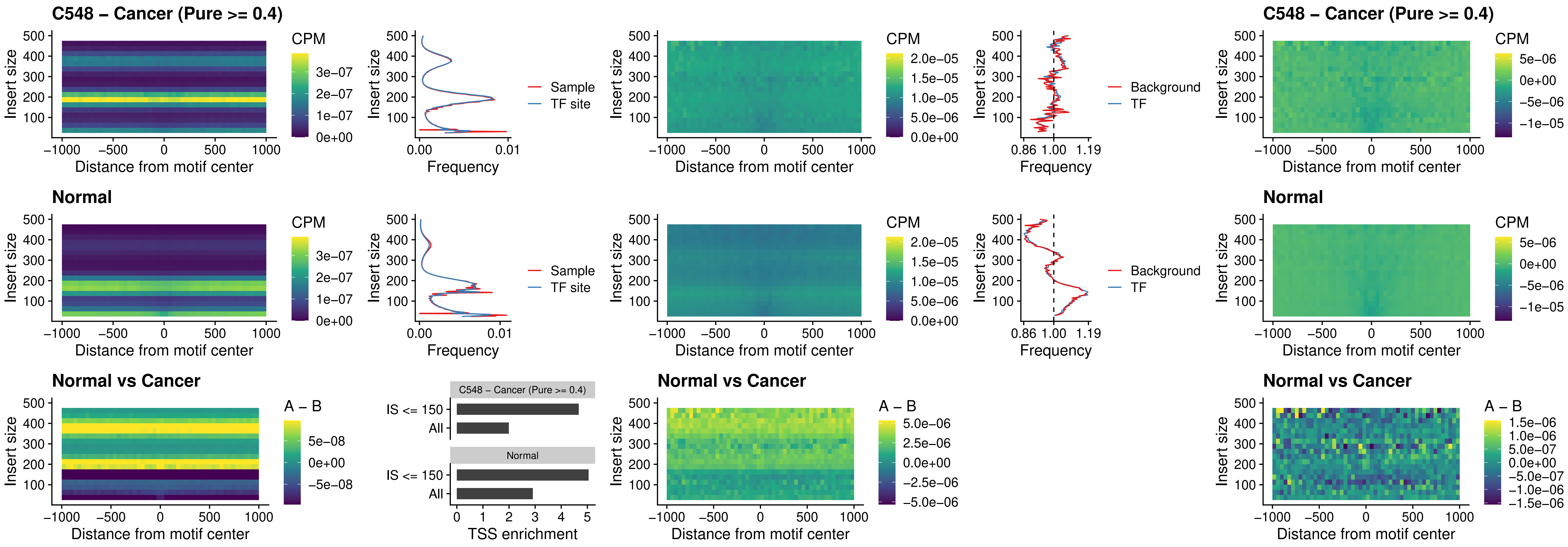

#### C548 – Cancer (Pure >= 0.4) – HOXD9_uc_dTSS_nPEAK

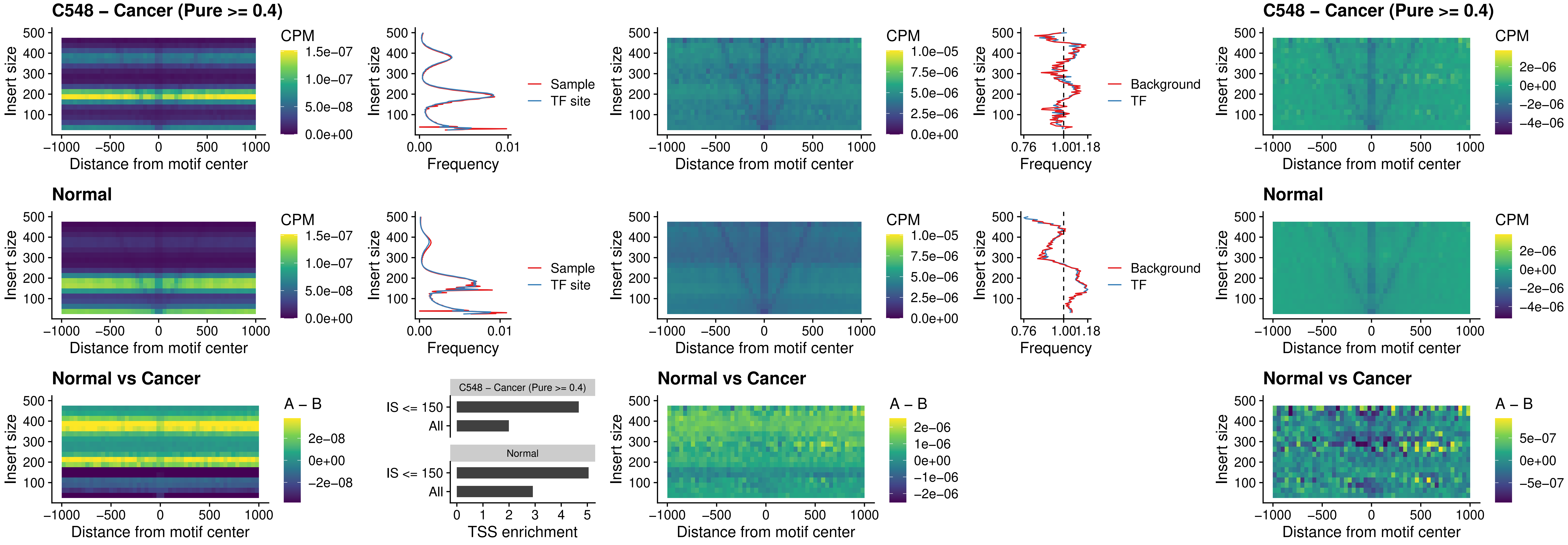

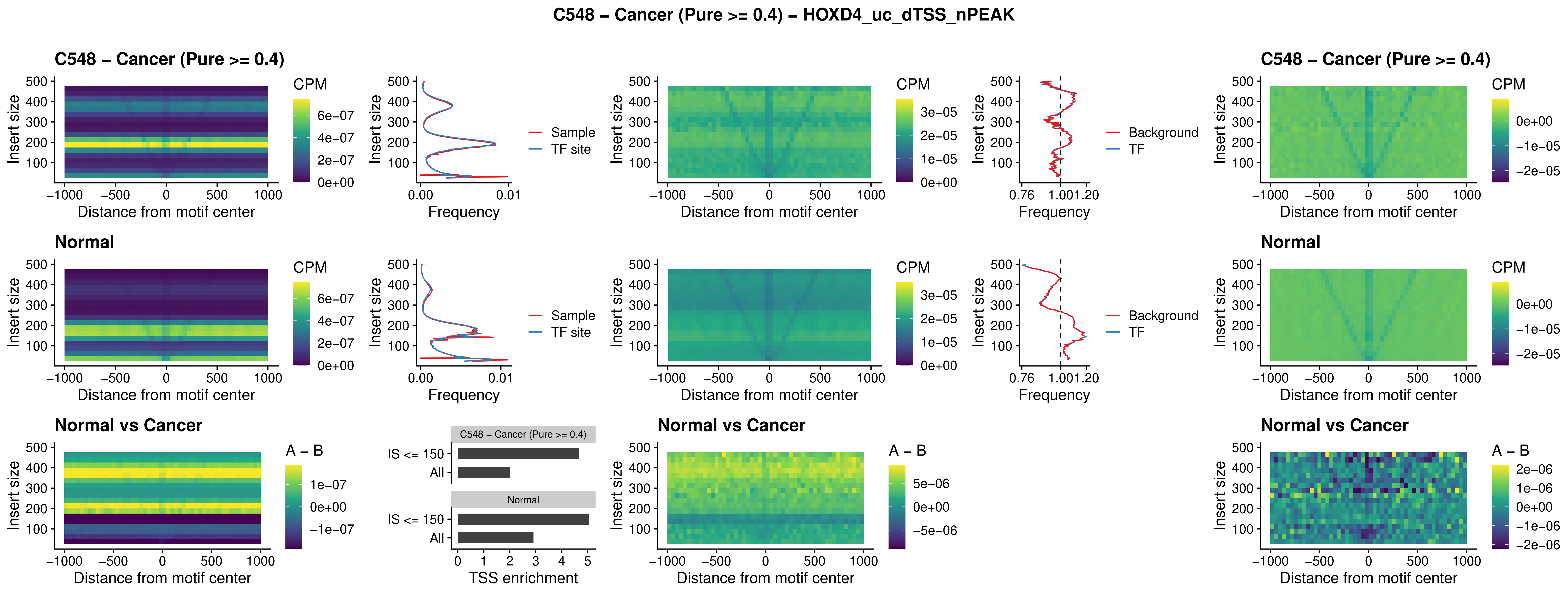
C548 – Cancer (Pure >= 0.4) – HOXD4_uc_dTSS_nPEAK
C548 – Cancer (Pure >= 0.4)
Normal
Normal vs Cancer
Insert size
Distance from motif center
Frequency
CPM
A – B
Sample
TF site
Background
TF
C548 – Cancer (Pure >= 0.4)
Normal
IS <= 150
All
TSS enrichment

#### C548 – Cancer (Pure >= 0.4) – FOXP3_uc_cTSS_nPEAK

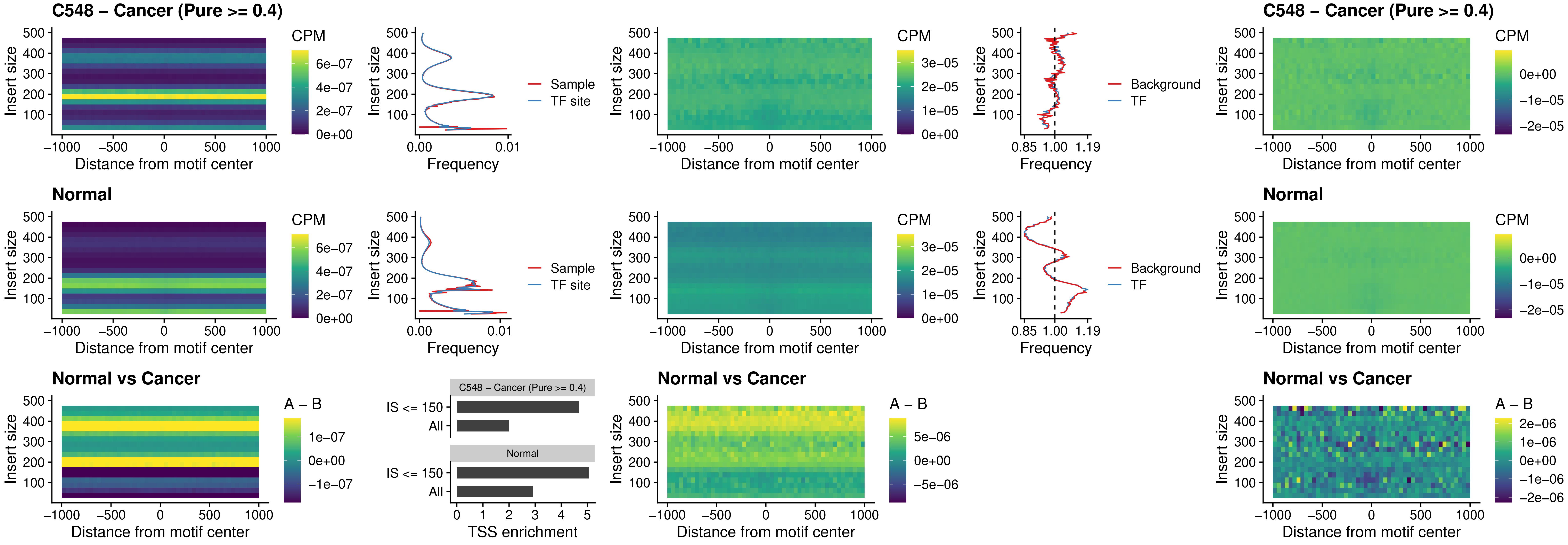

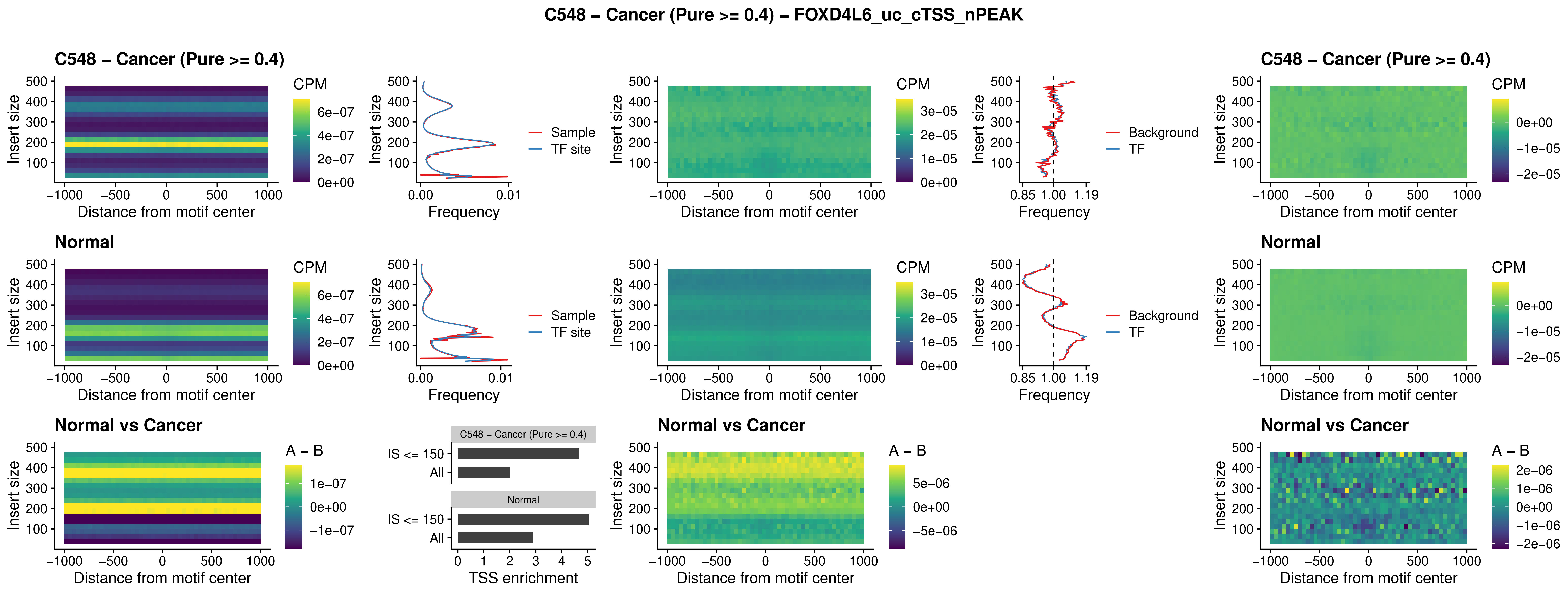
C548 – Cancer (Pure >= 0.4) – FOXD4L6_uc_cTSS_nPEAK
C548 – Cancer (Pure >= 0.4)
Normal
Normal vs Cancer
Insert size
Distance from motif center
Frequency
CPM
A – B
Sample
TF site
Background
TF
C548 – Cancer (Pure >= 0.4)
Normal
IS <= 150
All
TSS enrichment

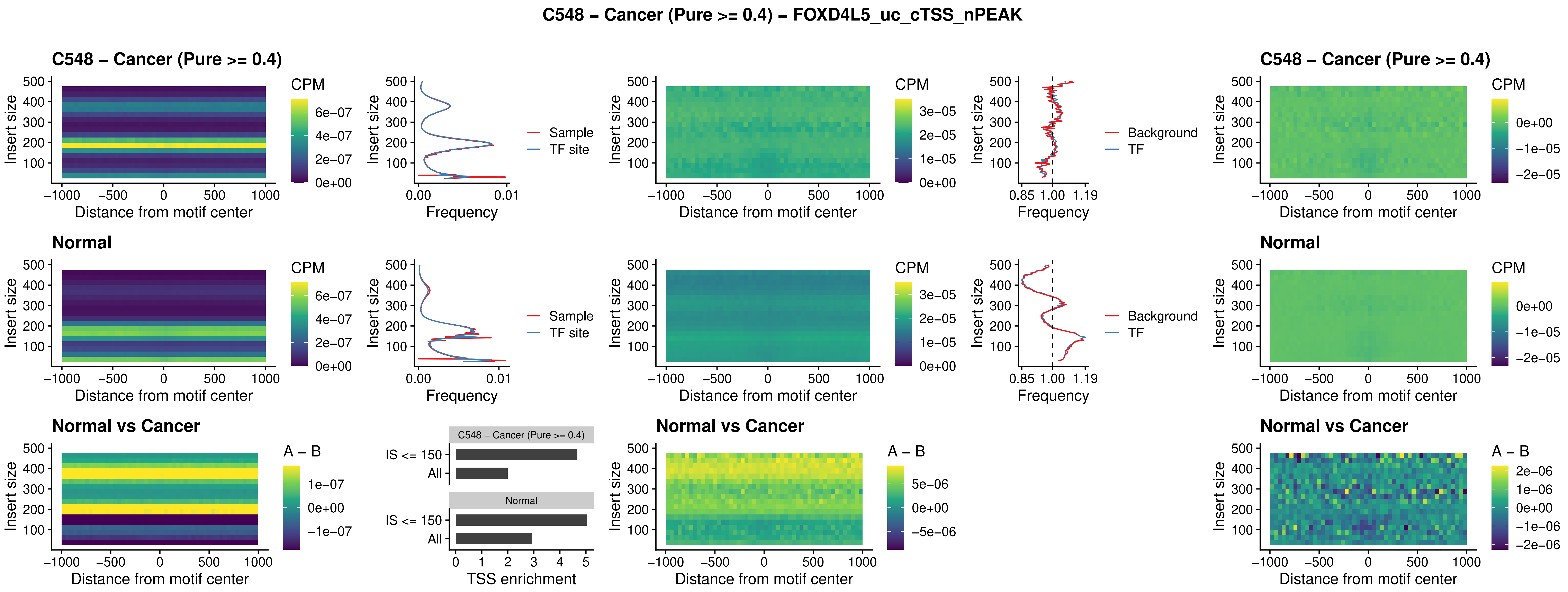
C548 – Cancer (Pure >= 0.4) – FOXD4L5_uc_cTSS_nPEAK
C548 – Cancer (Pure >= 0.4)
Normal
Normal vs Cancer
Insert size
Distance from motif center
Frequency
CPM
A – B
Sample
TF site
Background
TF
C548 – Cancer (Pure >= 0.4)
Normal
IS <= 150
All
TSS enrichment

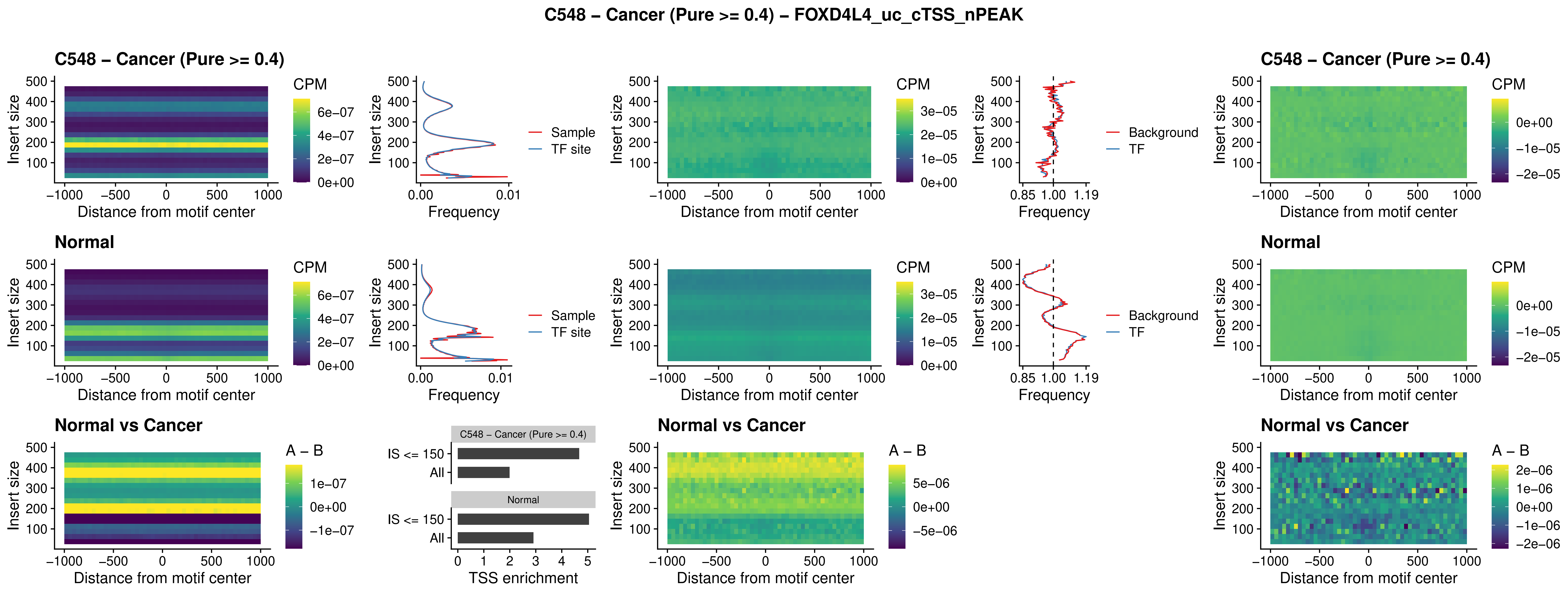

C548 – Cancer (Pure >= 0.4) – FOXD4L4_uc_cTSS_nPEAK
C548 – Cancer (Pure >= 0.4)
Normal
Normal vs Cancer
Insert size
Distance from motif center
Frequency
CPM
A – B
Sample
TF site
Background
TF
C548 – Cancer (Pure >= 0.4)
Normal
IS <= 150
All
TSS enrichment

#### C548 – Cancer (Pure >= 0.4) – FOXD4L2_uc_cTSS_nPEAK

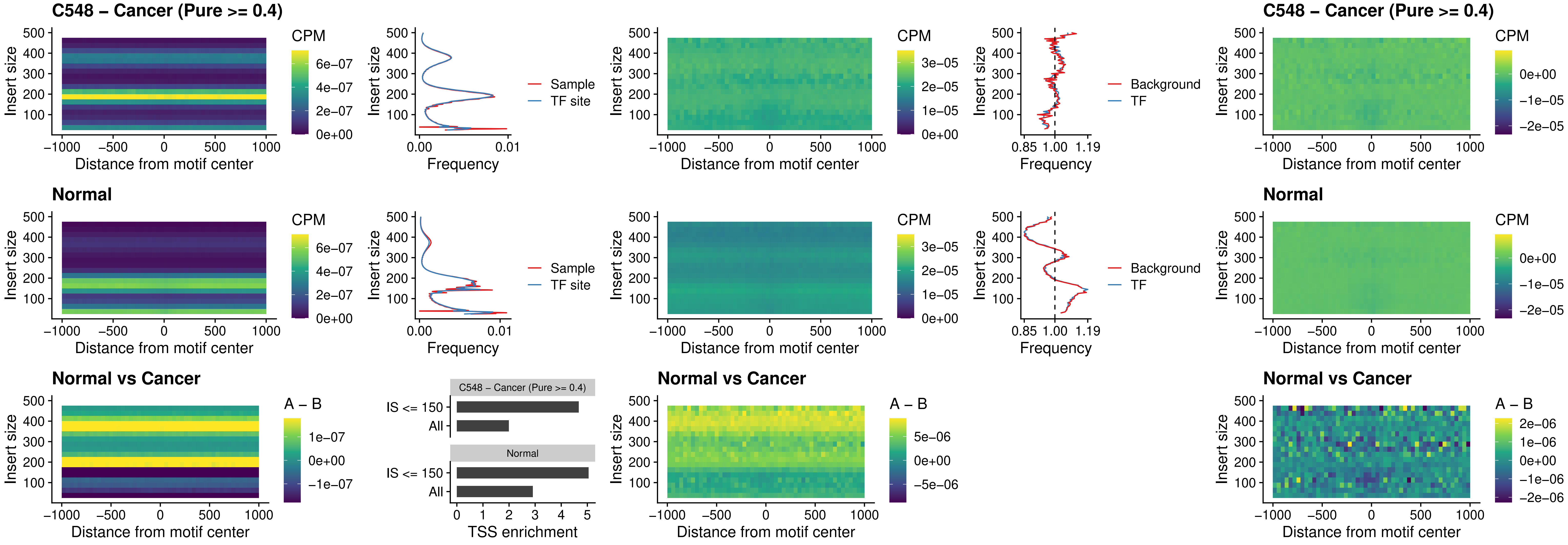

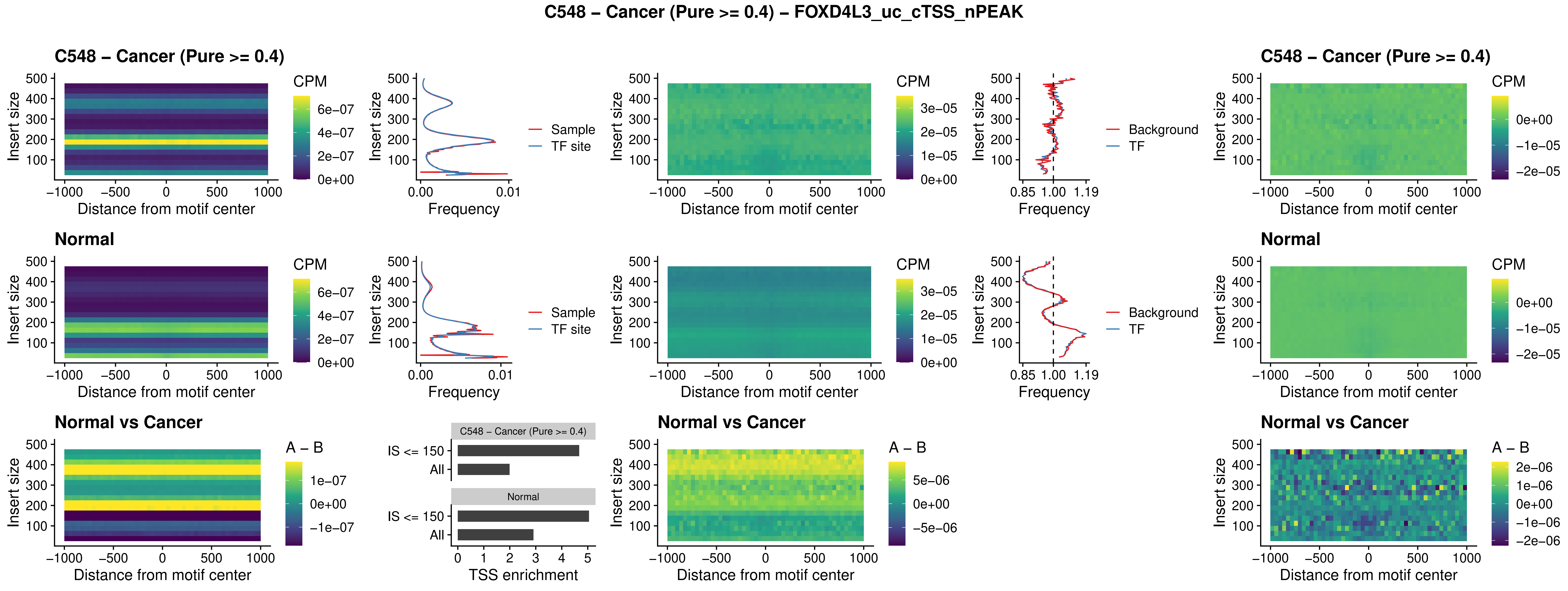

C548 – Cancer (Pure >= 0.4) – FOXD4L3_uc_cTSS_nPEAK
C548 – Cancer (Pure >= 0.4)
Normal
Normal vs Cancer
Insert size
Distance from motif center
Frequency
CPM
A – B
Sample
TF site
Background
TF
C548 – Cancer (Pure >= 0.4)
Normal
IS <= 150
All
TSS enrichment

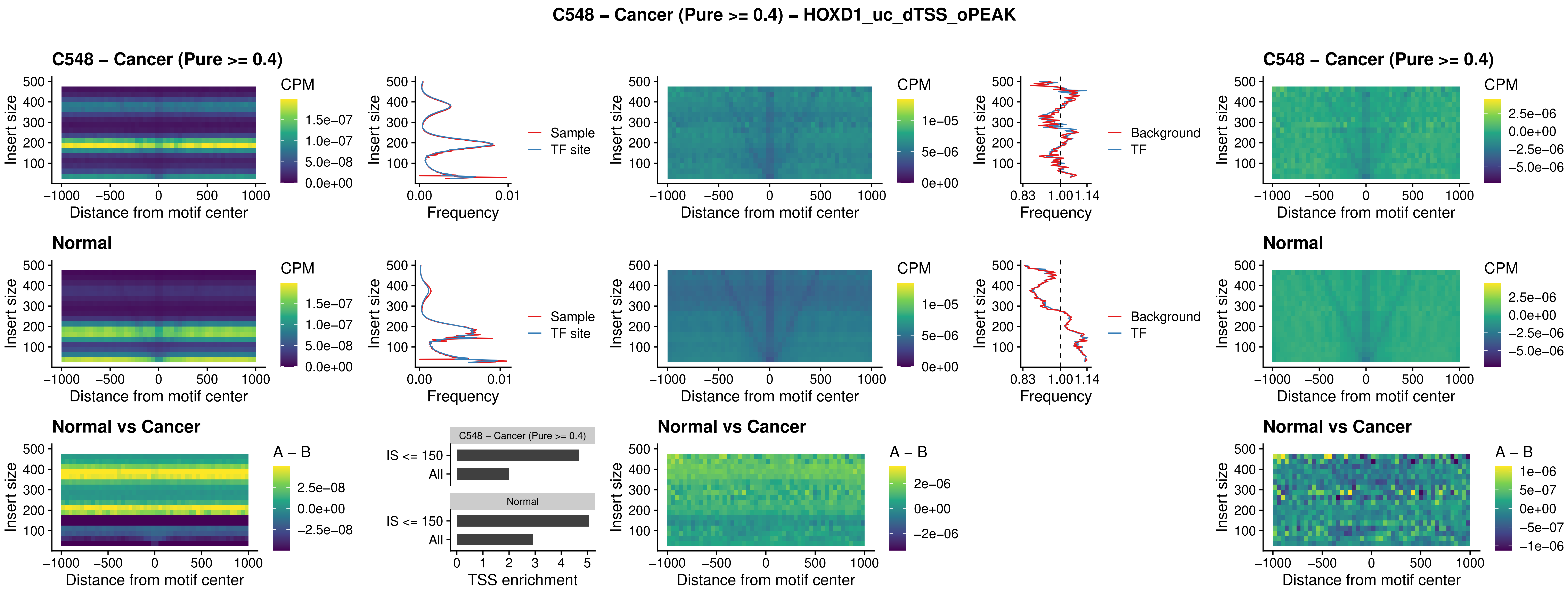
C548 – Cancer (Pure >= 0.4) – HOXD1_uc_dTSS_oPEAK
C548 – Cancer (Pure >= 0.4)
Normal
Normal vs Cancer
Insert size
Distance from motif center
Frequency
CPM
A – B
Sample
TF site
Background
TF
C548 – Cancer (Pure >= 0.4)
IS <= 150
All
TSS enrichment

#### C548 – Cancer (Pure >= 0.4) – VAX1_uc_pTSS_nPEAK

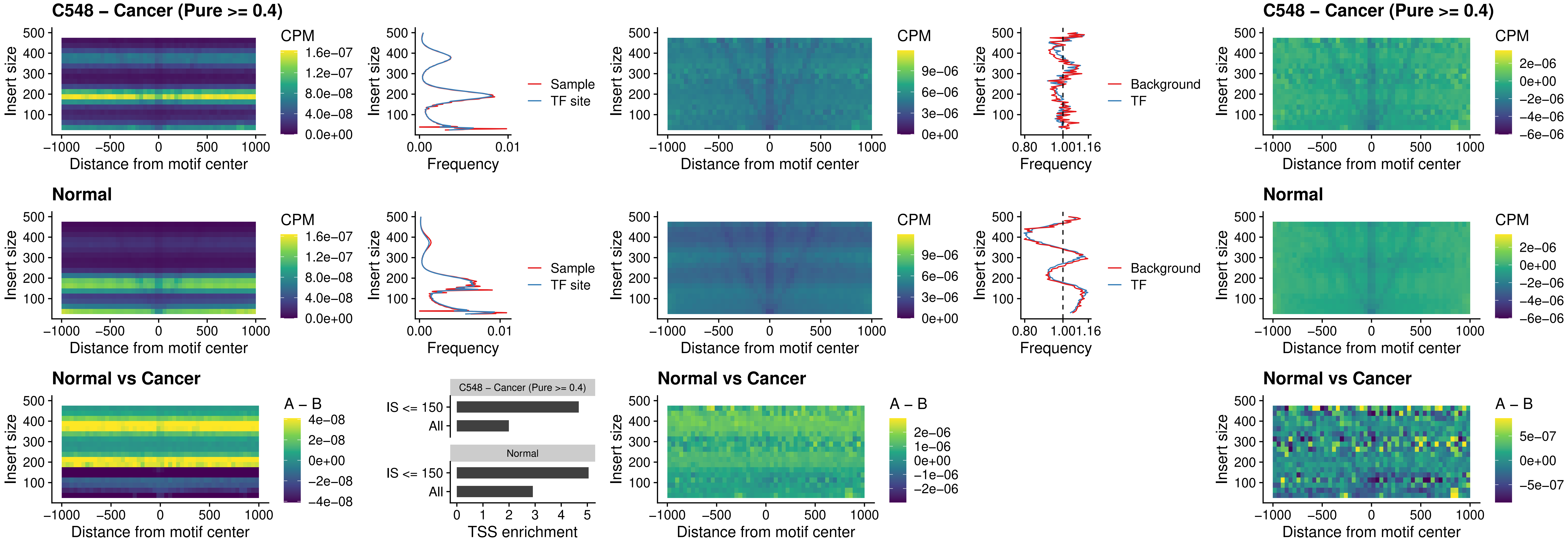

#### C548 – Cancer (Pure >= 0.4) – DLX3_uc_pTSS_nPEAK

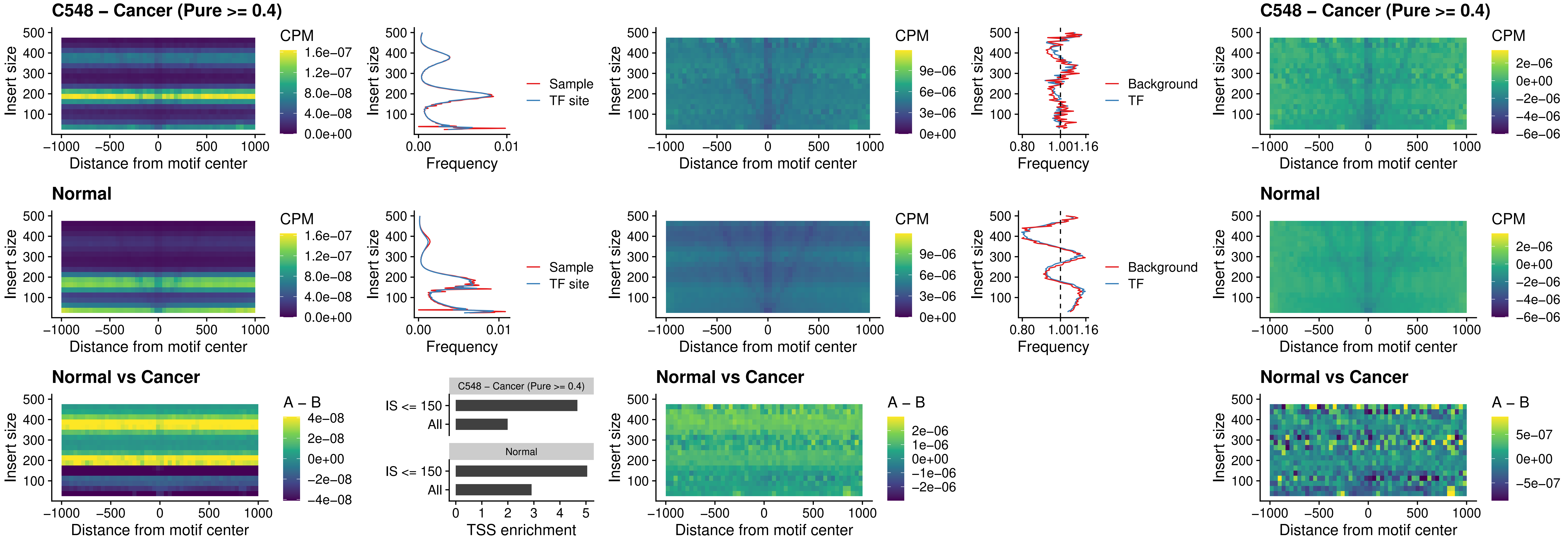

#### C548 – Cancer (Pure >= 0.4) – HOXD3_uc_pTSS_nPEAK

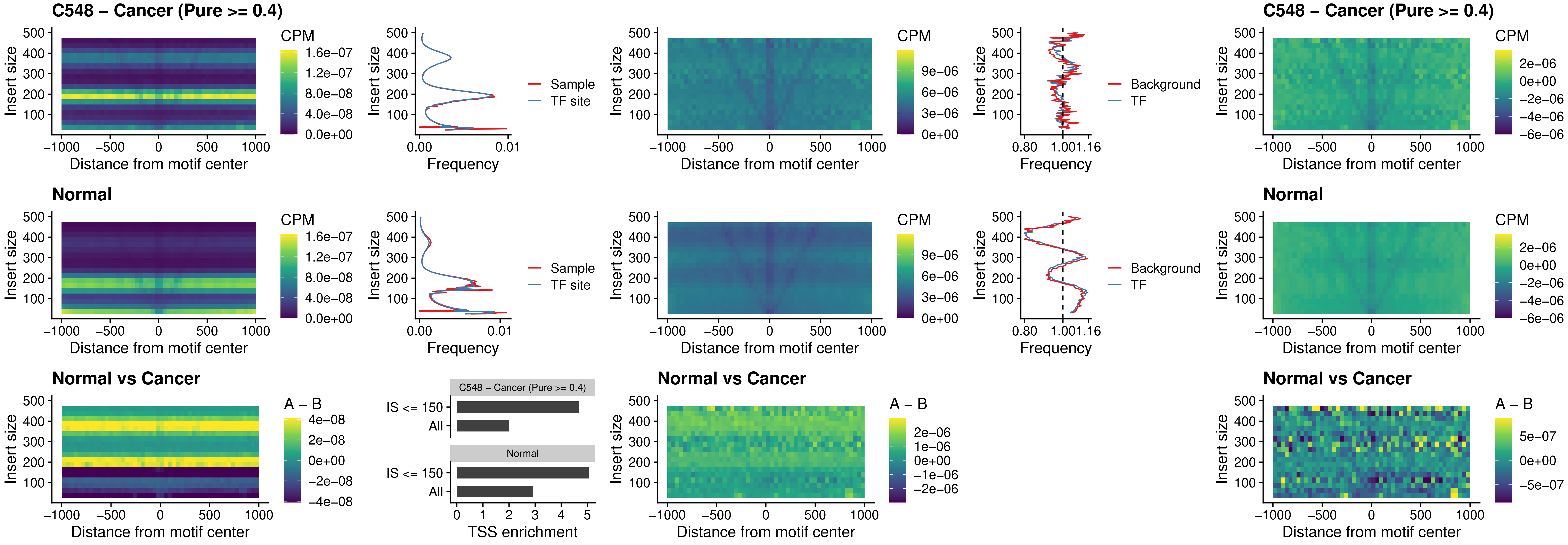

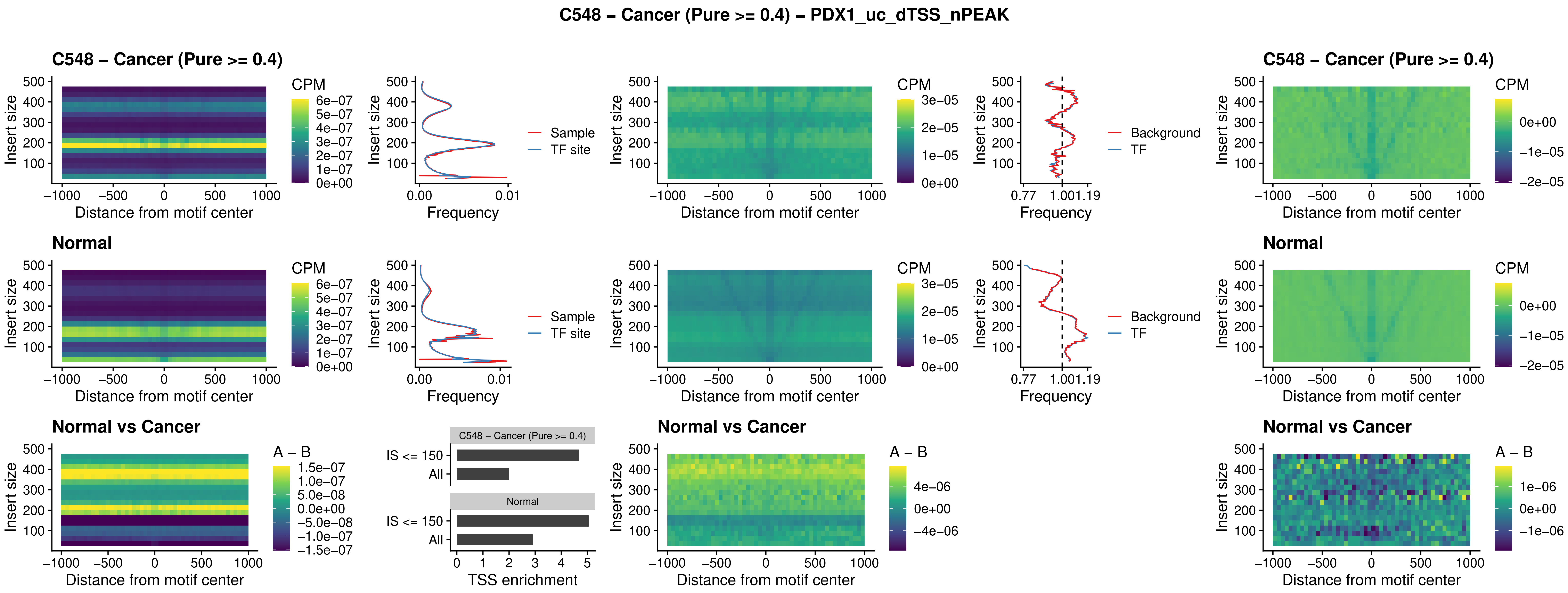
C548 – Cancer (Pure >= 0.4) – PDX1_uc_dTSS_nPEAK
C548 – Cancer (Pure >= 0.4)
Normal
Normal vs Cancer
Insert size
Distance from motif center
Frequency
CPM
A – B
Sample
TF site
Background
TF
C548 – Cancer (Pure >= 0.4)
Normal
IS <= 150
All
TSS enrichment

#### C548 – Cancer (Pure >= 0.4) – VAX2_uc_dTSS_nPEAK

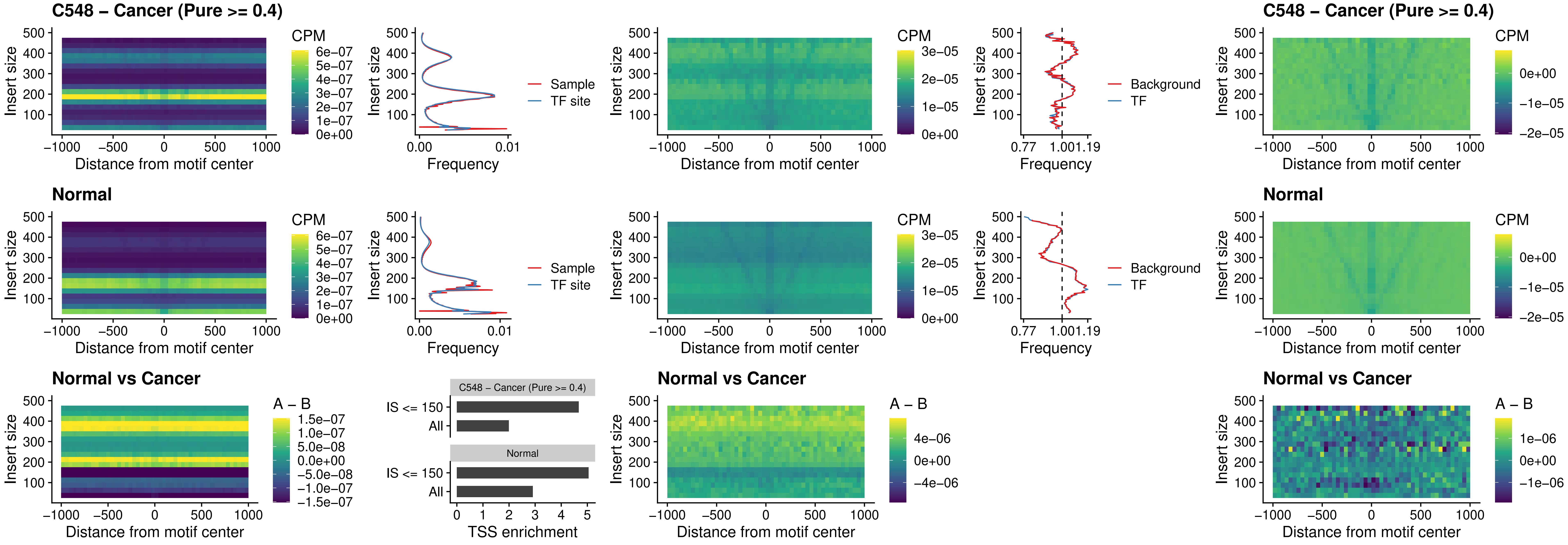

#### C548 – Cancer (Pure >= 0.4) – SOX5_uc_dTSS_nPEAK

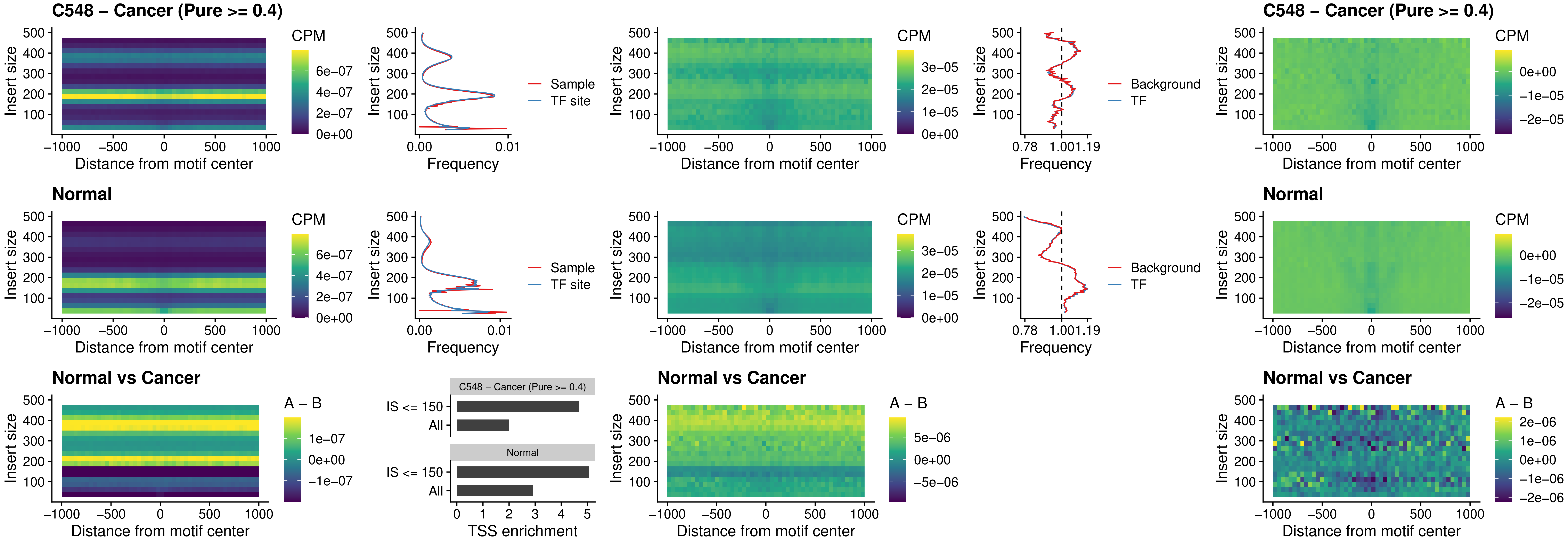

### C548 – Cancer (Pure >= 0.4) – SOX6_uc_dTSS_nPEAK

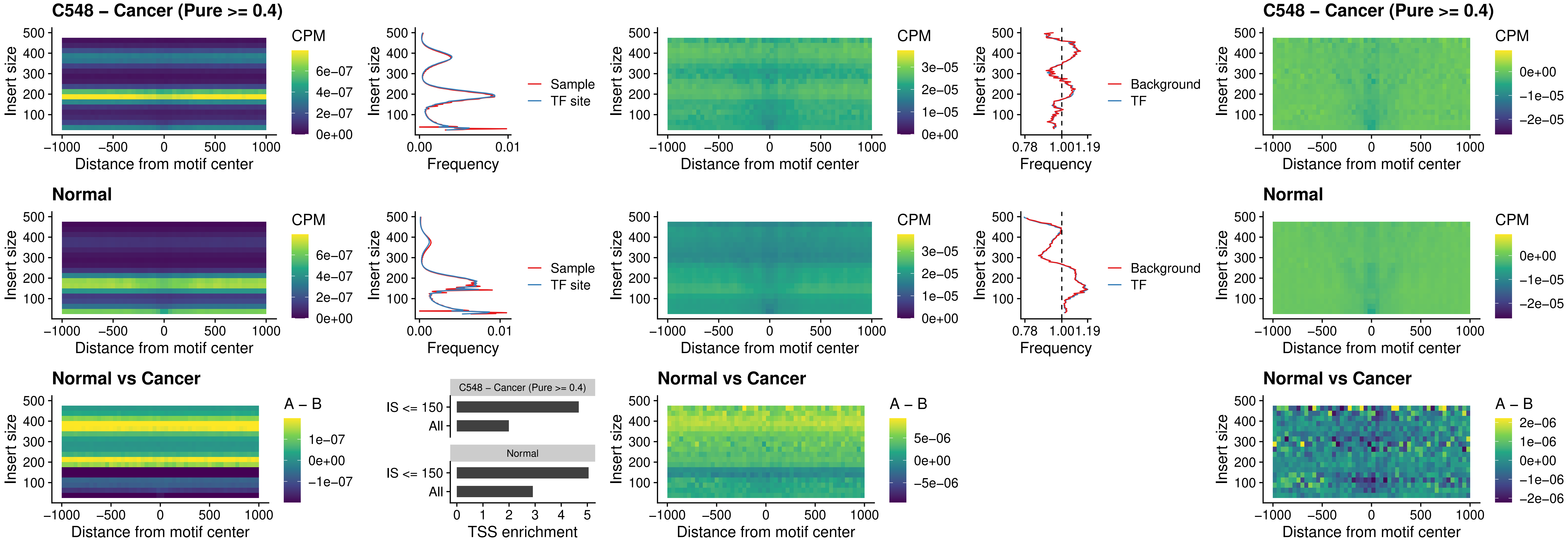

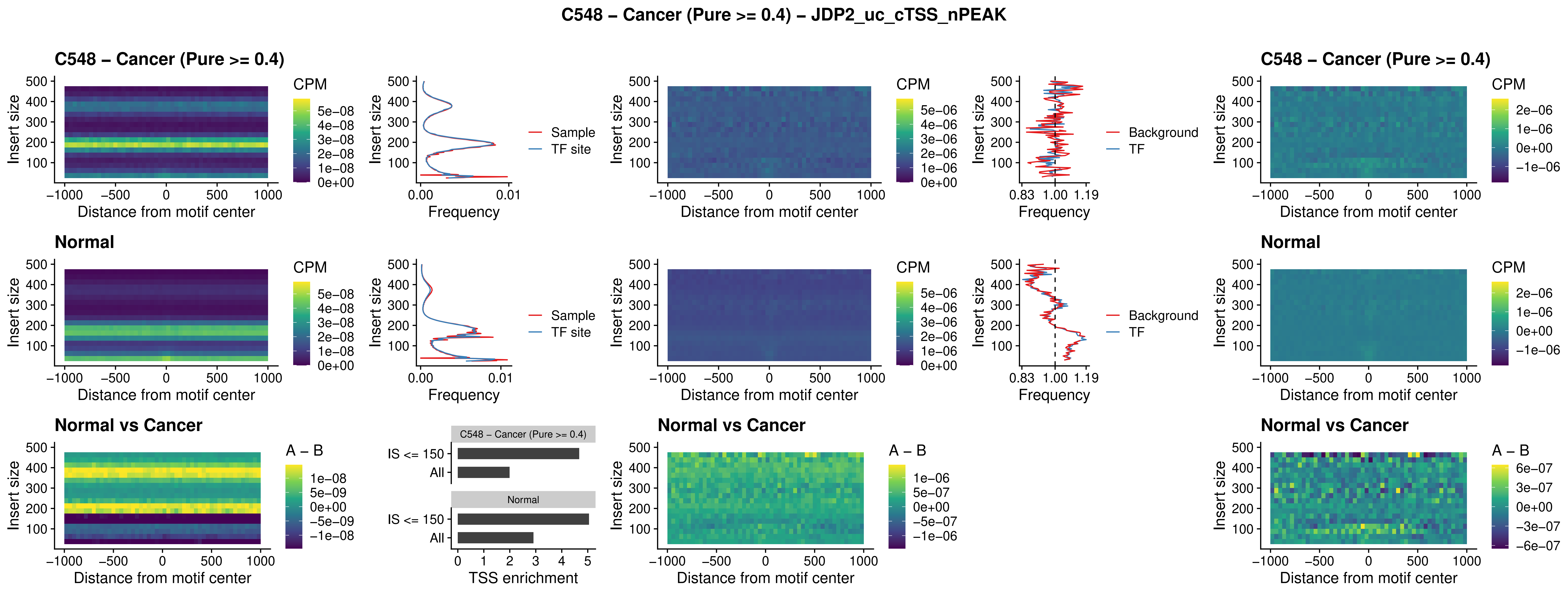

C548 – Cancer (Pure >= 0.4) – JDP2_uc_cTSS_nPEAK
C548 – Cancer (Pure >= 0.4)
Normal
Normal vs Cancer
Insert size
Distance from motif center
Frequency
CPM
A – B
Sample
TF site
Background
TF
C548 – Cancer (Pure >= 0.4)
Normal
IS <= 150
All
TSS enrichment

#### C548 – Cancer (Pure >= 0.4) – PURA_uc_dTSS_oPEAK

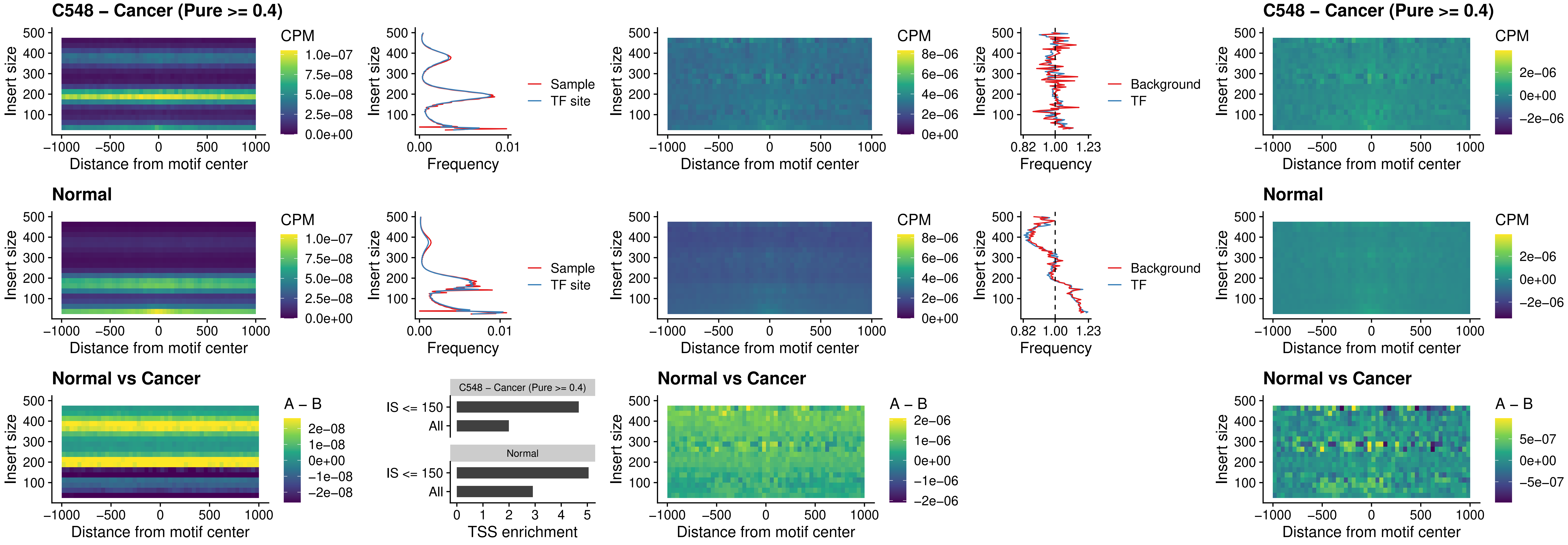

#### C548 – Cancer (Pure >= 0.4) – HHEX_uc_dTSS_nPEAK

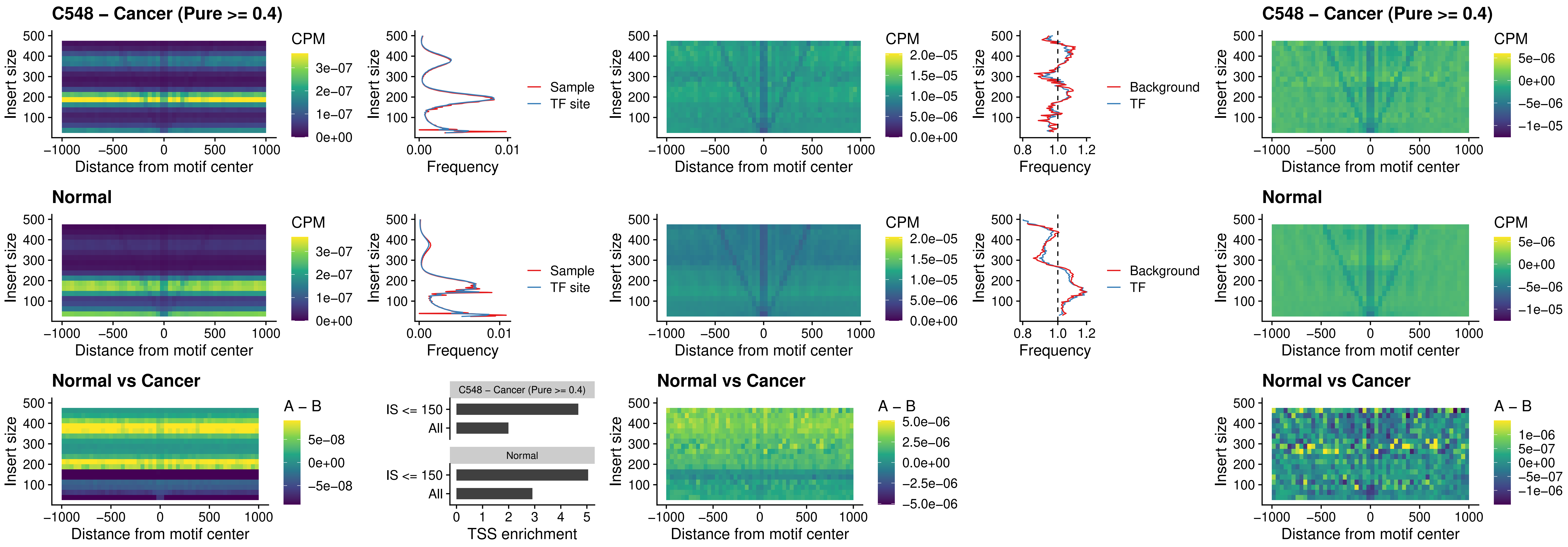

#### C548 – Cancer (Pure >= 0.4) – HMG20B_uc_dTSS_nPEAK

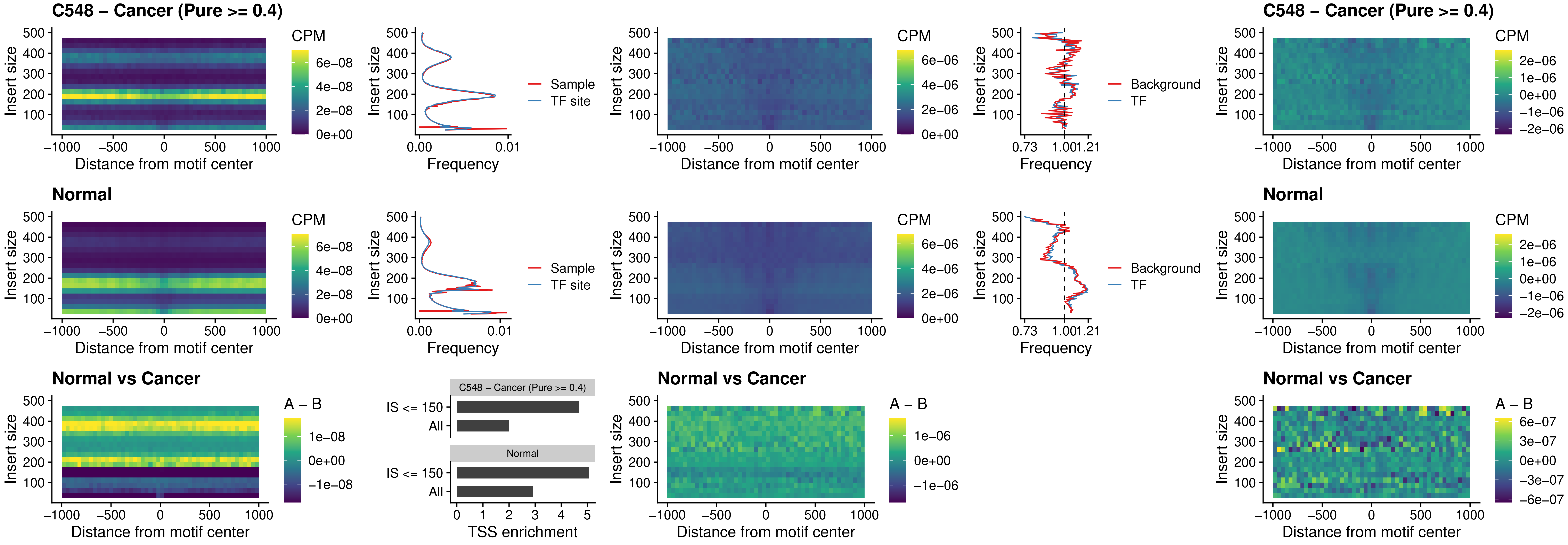

#### C548 – Cancer (Pure >= 0.4) – FOXB2_uc_dTSS_nPEAK

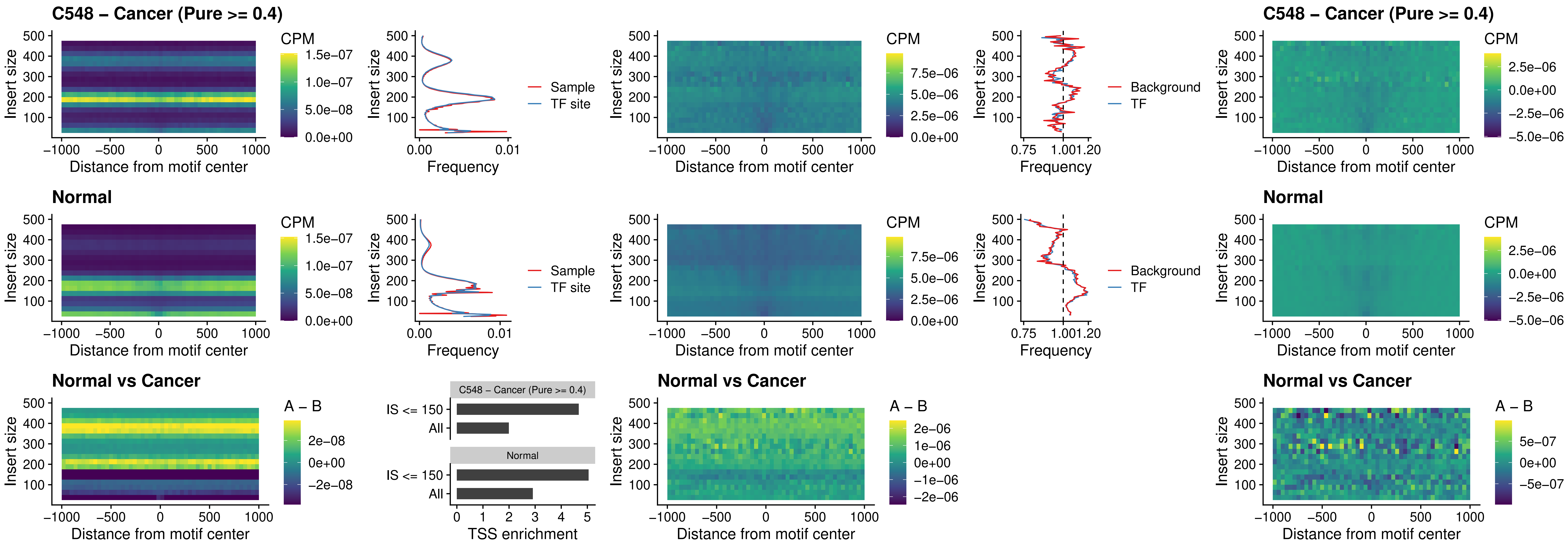

#### C548 – Cancer (Pure >= 0.4) – FOXL1_uc_cTSS_nPEAK

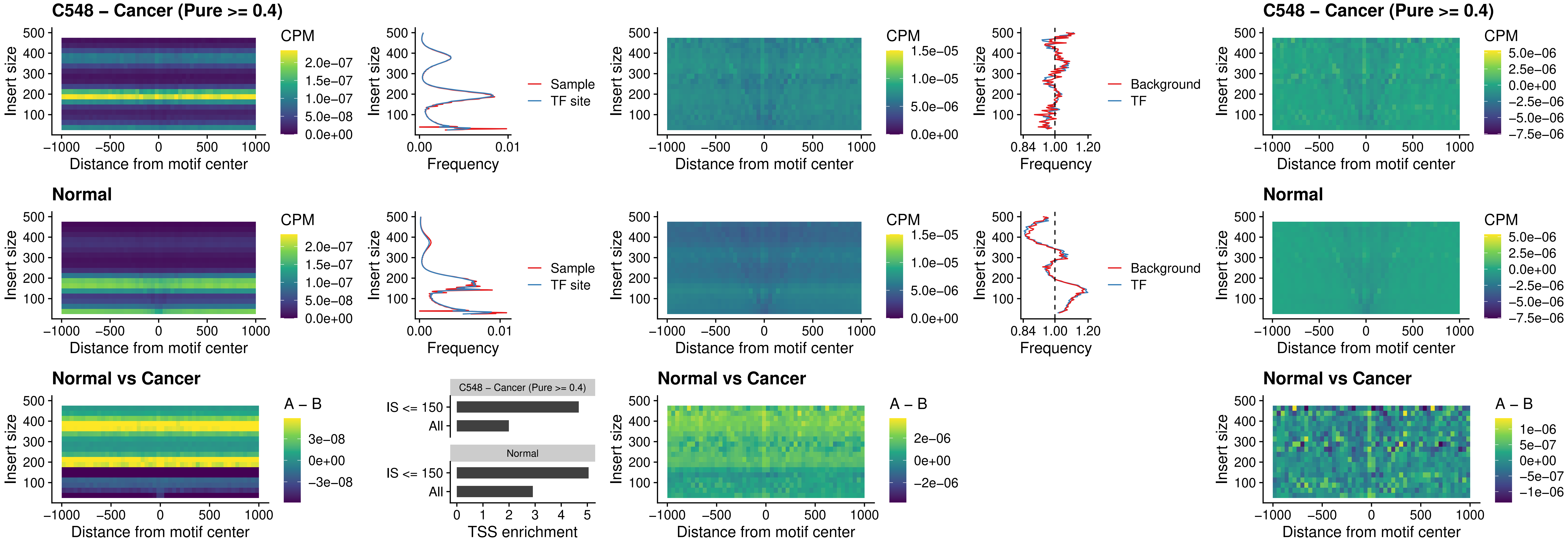

#### C548 – Cancer (Pure >= 0.4) – PBX3_uc_pTSS_oPEAK

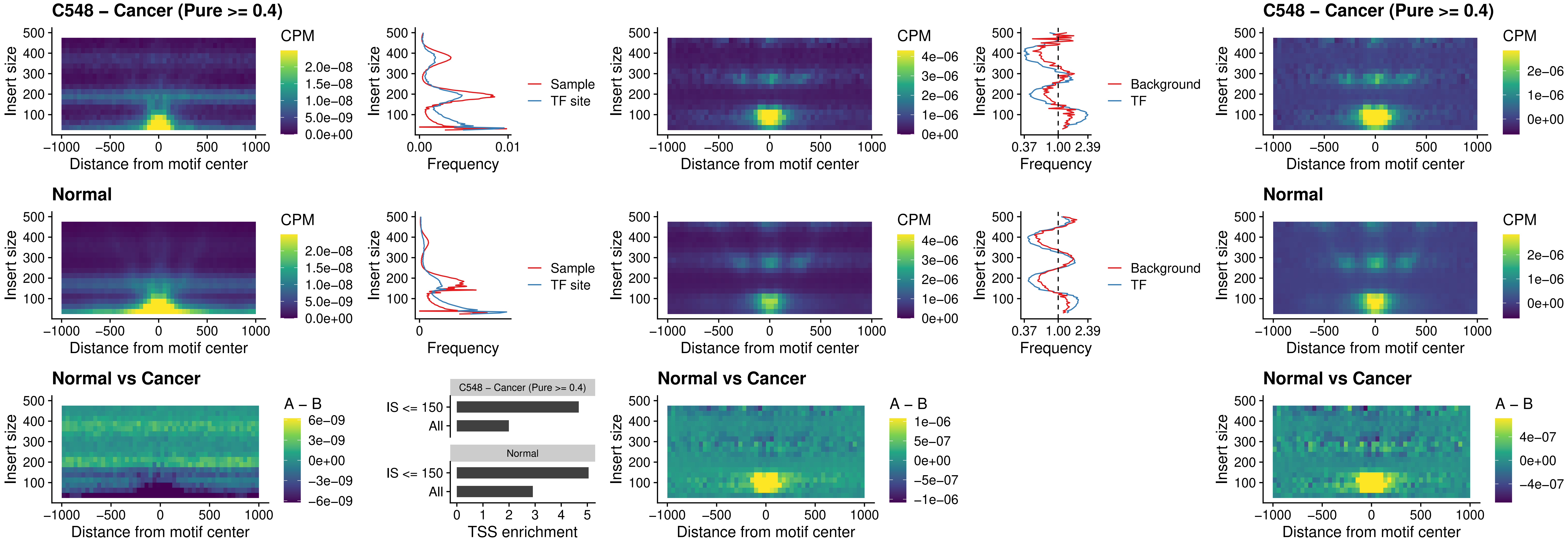

#### C548 – Cancer (Pure >= 0.4) – SNAI2_uc_cTSS_nPEAK

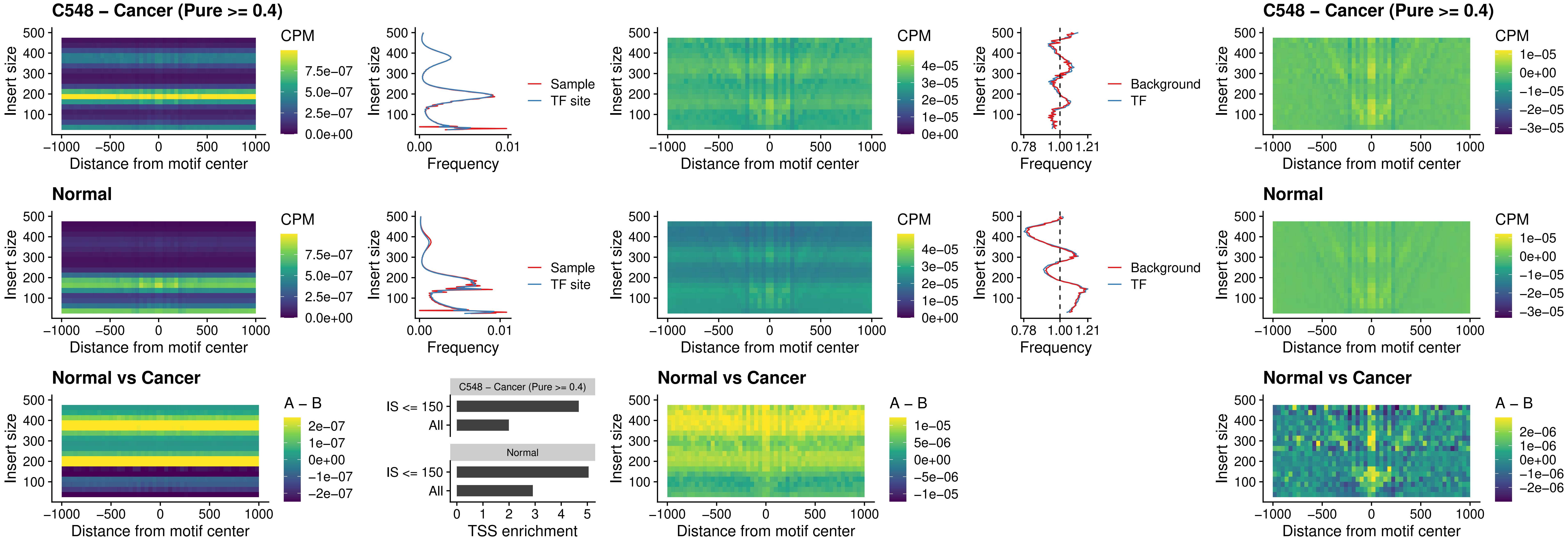

#### C548 – Cancer (Pure >= 0.4) – SNAI2_uc_dTSS_oPEAK

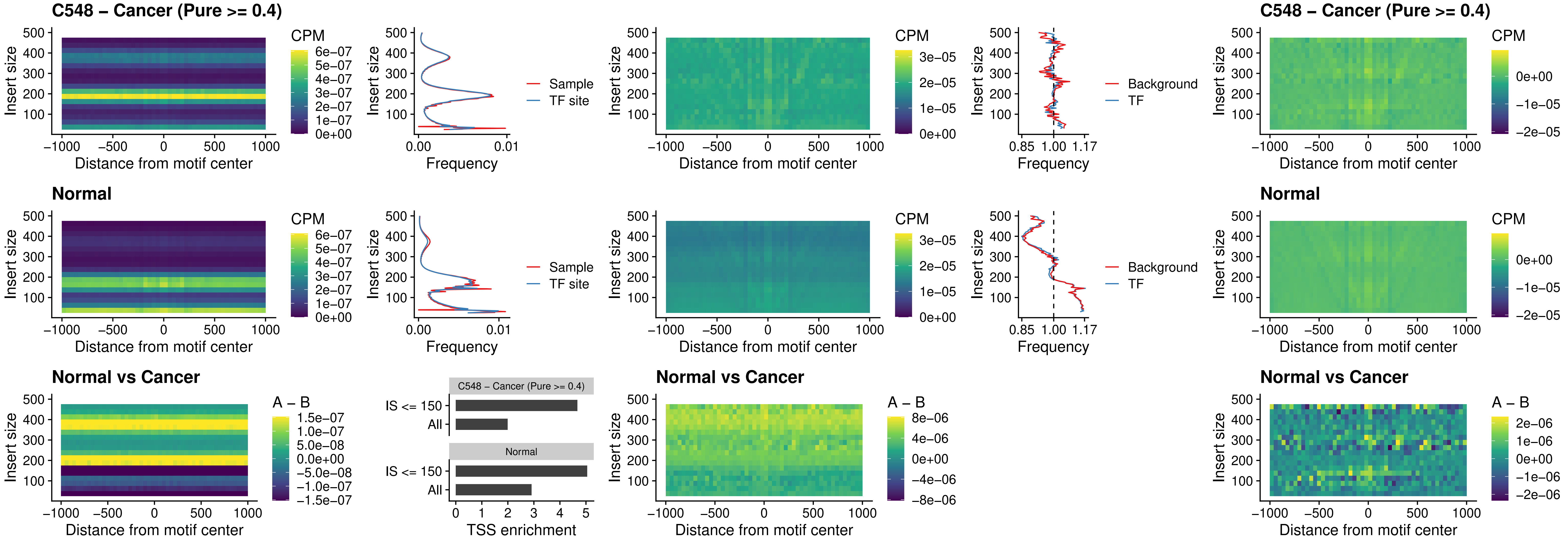

### C548 – Cancer (Pure >= 0.4) – SPI1_uc_pTSS_nPEAK

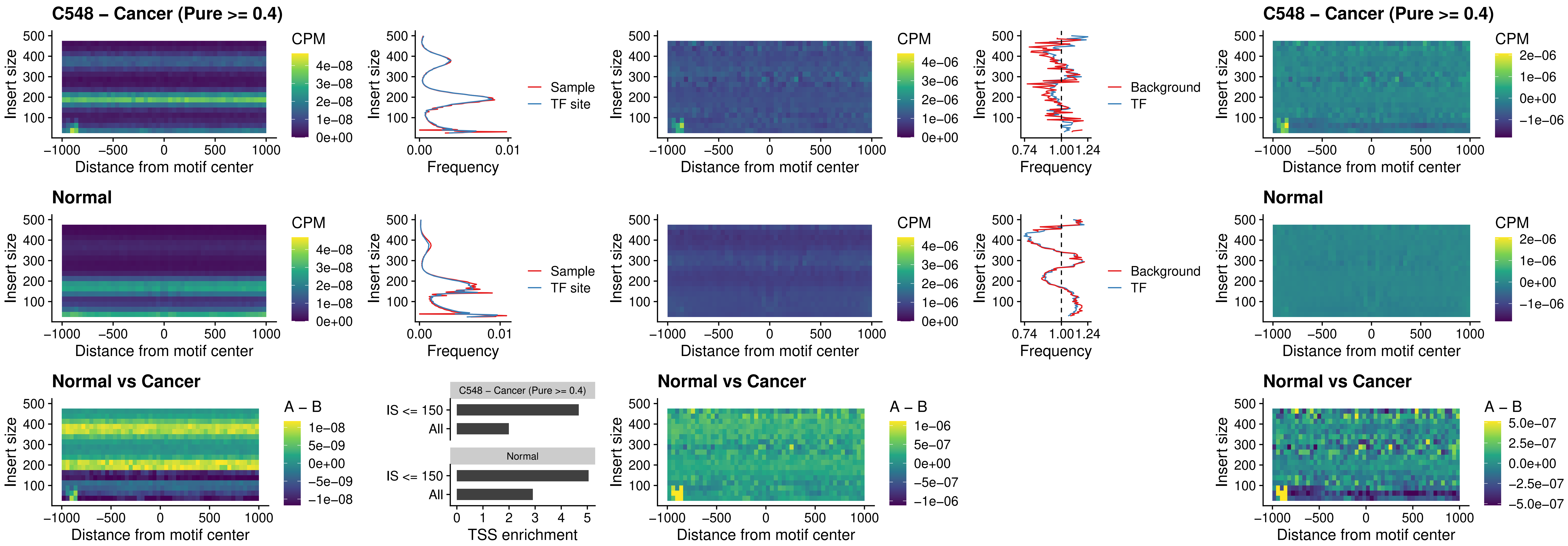

#### C548 – Cancer (Pure >= 0.4) – KLF16_uc_pTSS_nPEAK

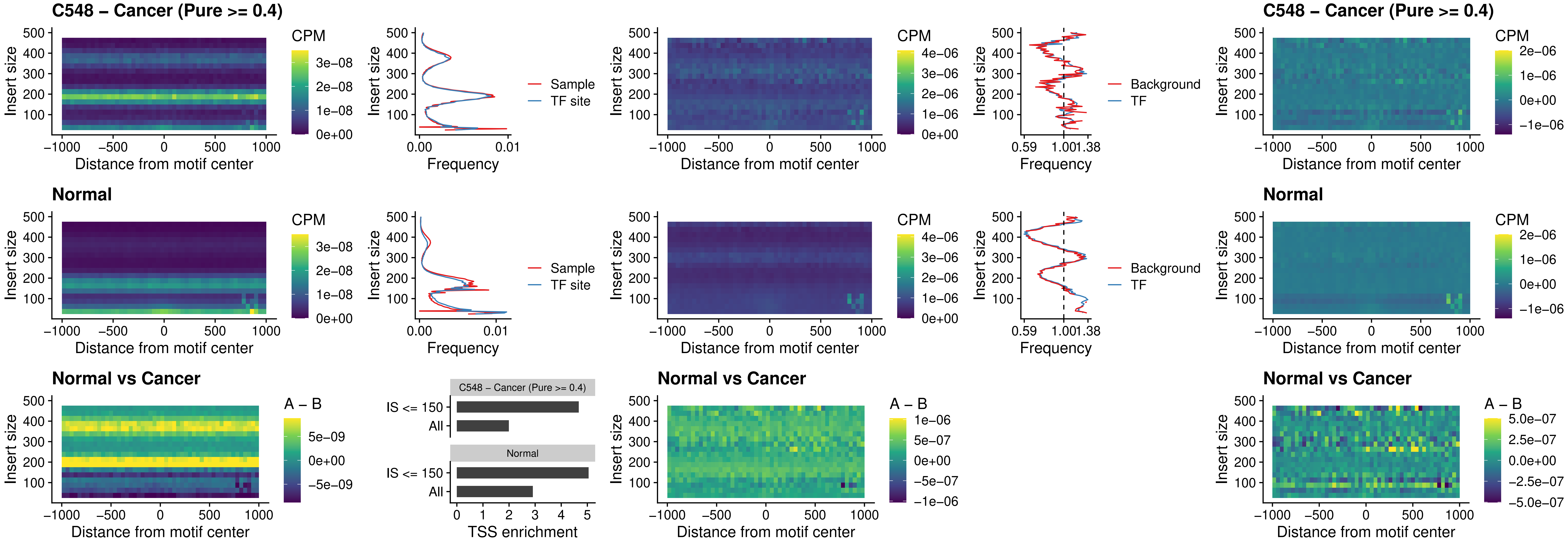

#### C548 – Cancer (Pure >= 0.4) – SP3_uc_pTSS_nPEAK

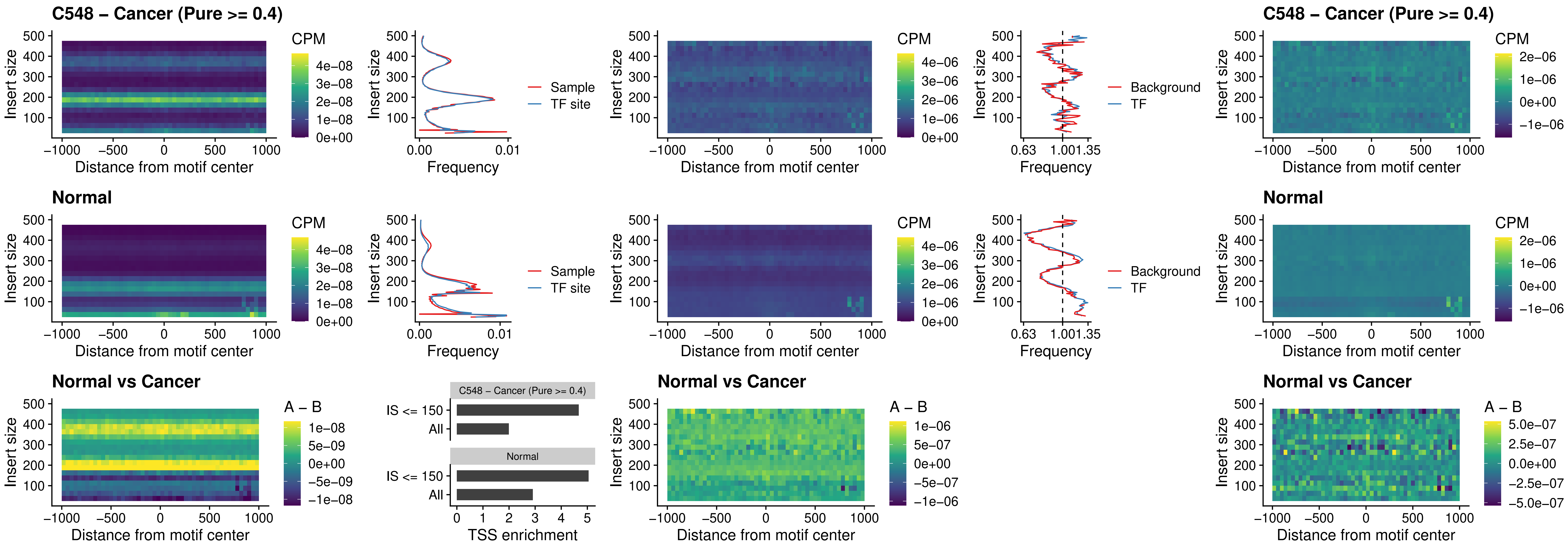

#### C548 – Cancer (Pure >= 0.4) – NEUROD1_uc_dTSS_oPEAK

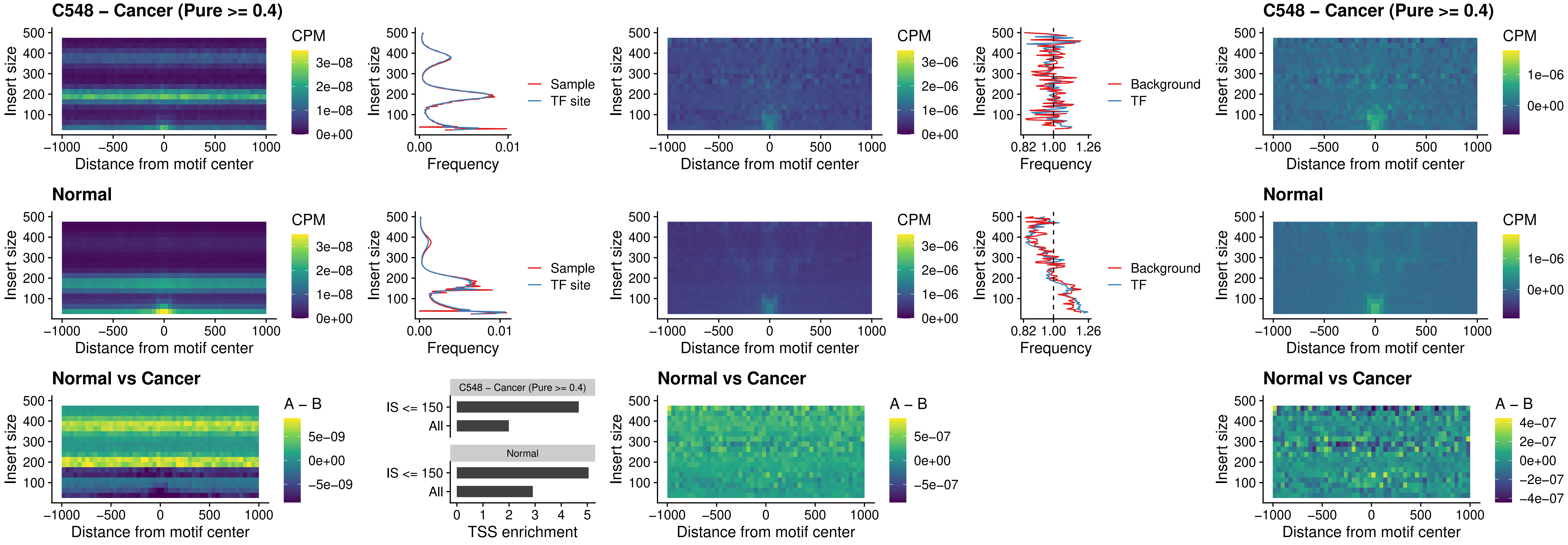

### C548 – Cancer (Pure >= 0.4) – CTCF_uc_dTSS_oPEAK

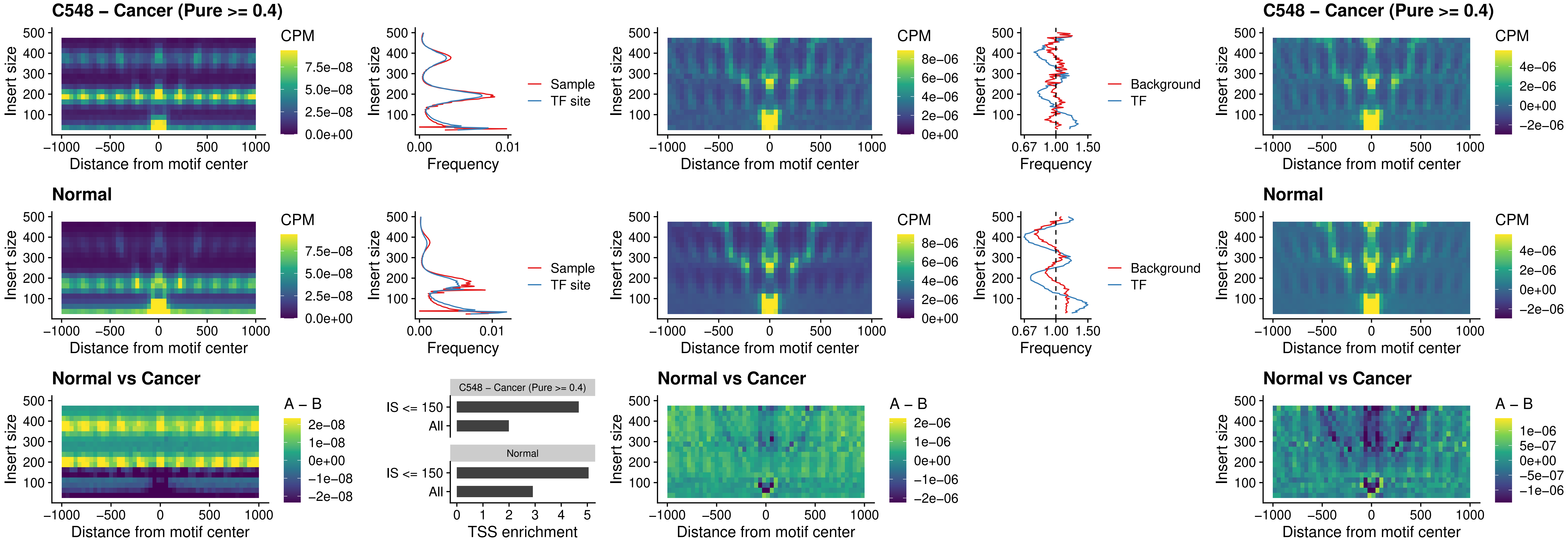

#### C548 – Cancer (Pure >= 0.4) – CTCF_uc_cTSS_oPEAK

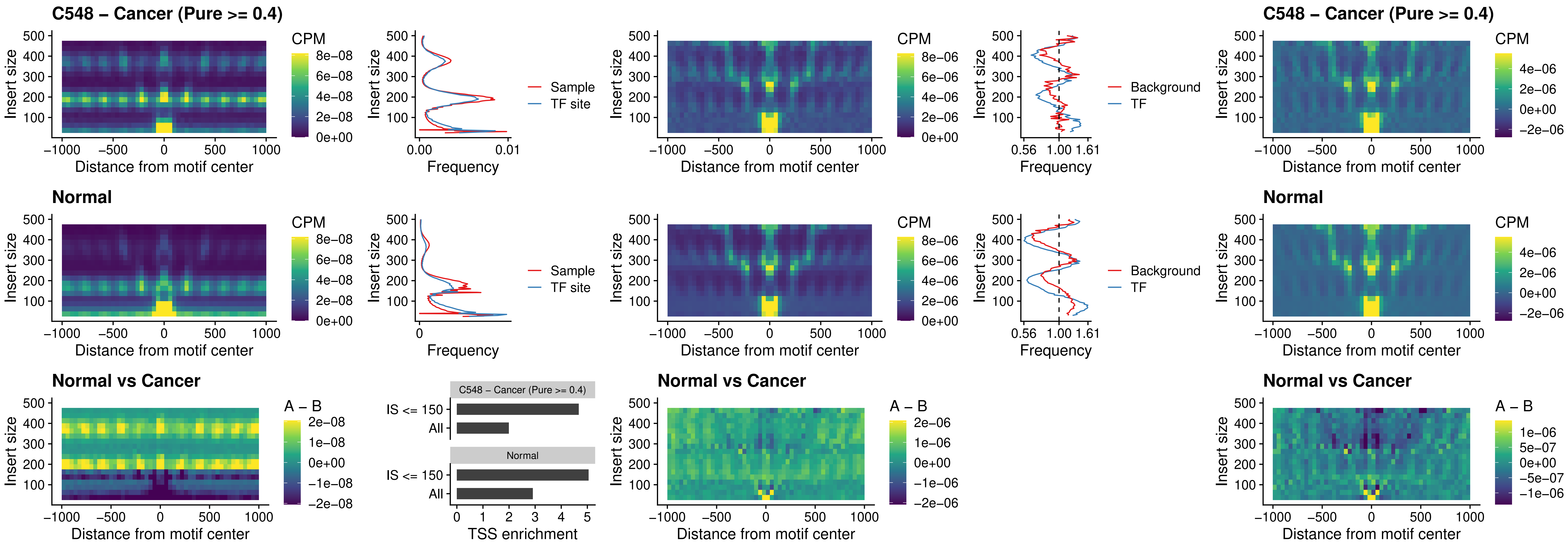

### C548 – Cancer (Pure >= 0.4) – CTCFL_uc_dTSS_oPEAK

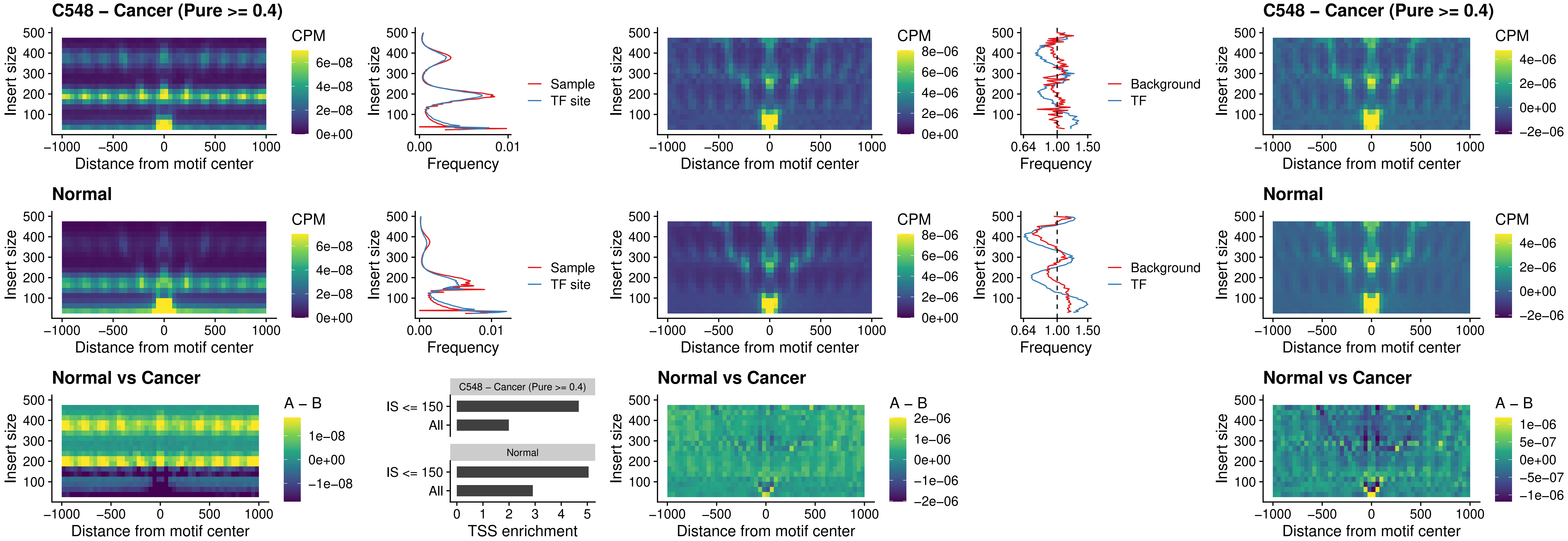

#### C548 – Cancer (Pure >= 0.4) – ENCODE_CTCF_sites_COLON

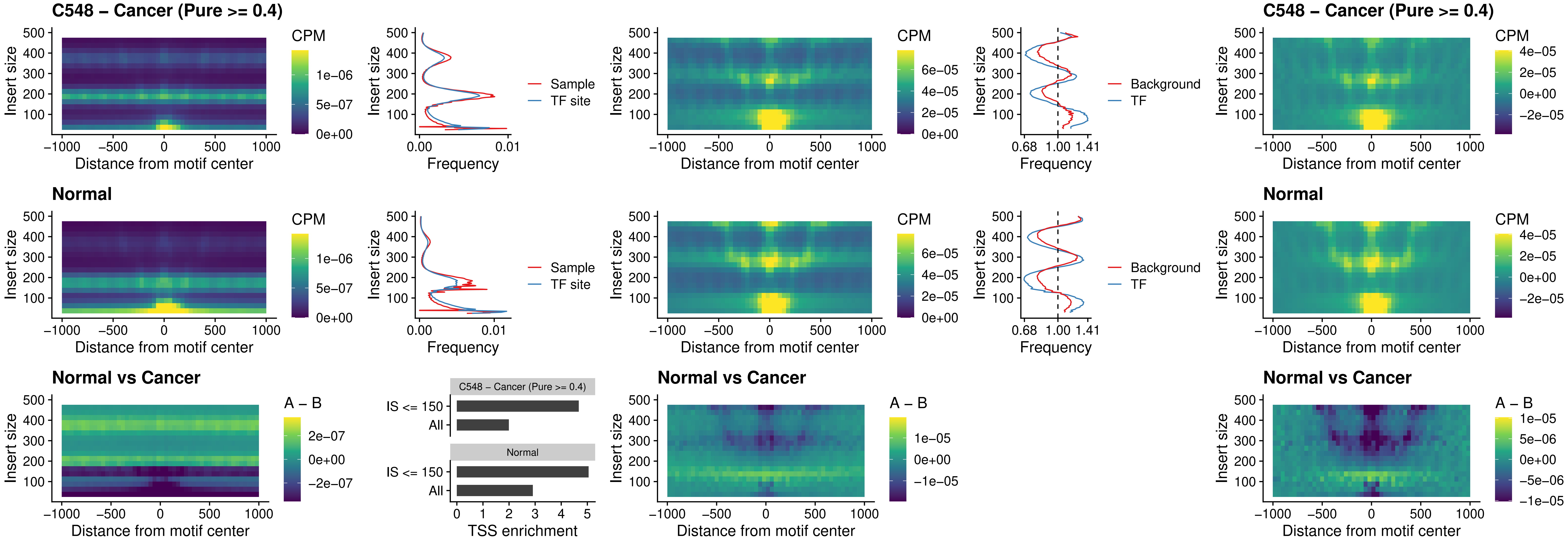

#### C548 – Cancer (Pure >= 0.4) – ENCODE_CTCF_sites_COLON_loop_ctcf

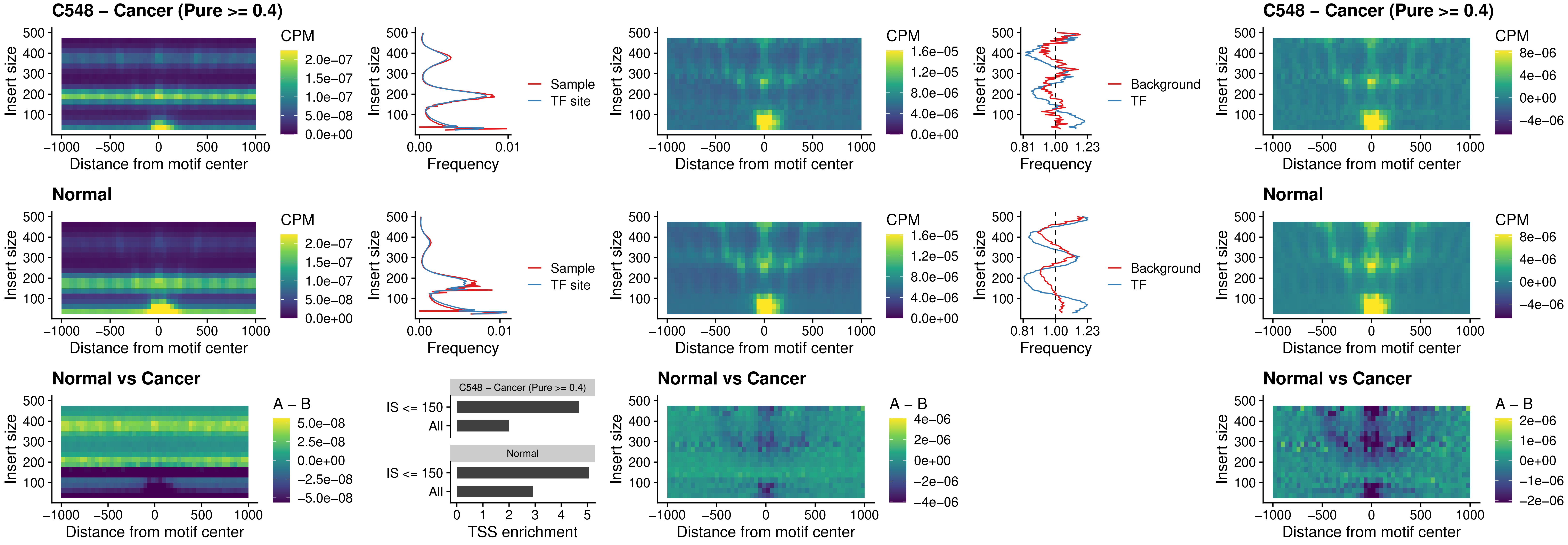

#### C548 – Cancer (Pure >= 0.4) – ENCODE_CTCF_sites_COLON_intergenic_ctcf

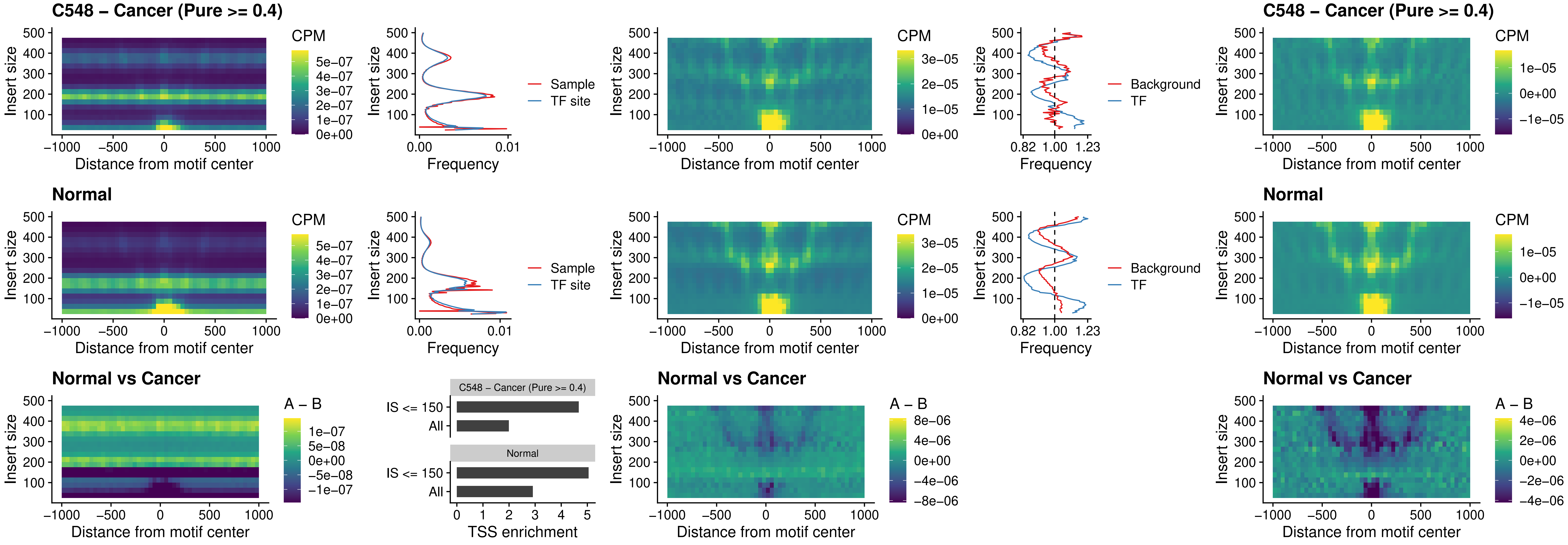

#### C548 – Cancer (Pure >= 0.4) – ENCODE_CTCF_sites_COLON_nonpromotor

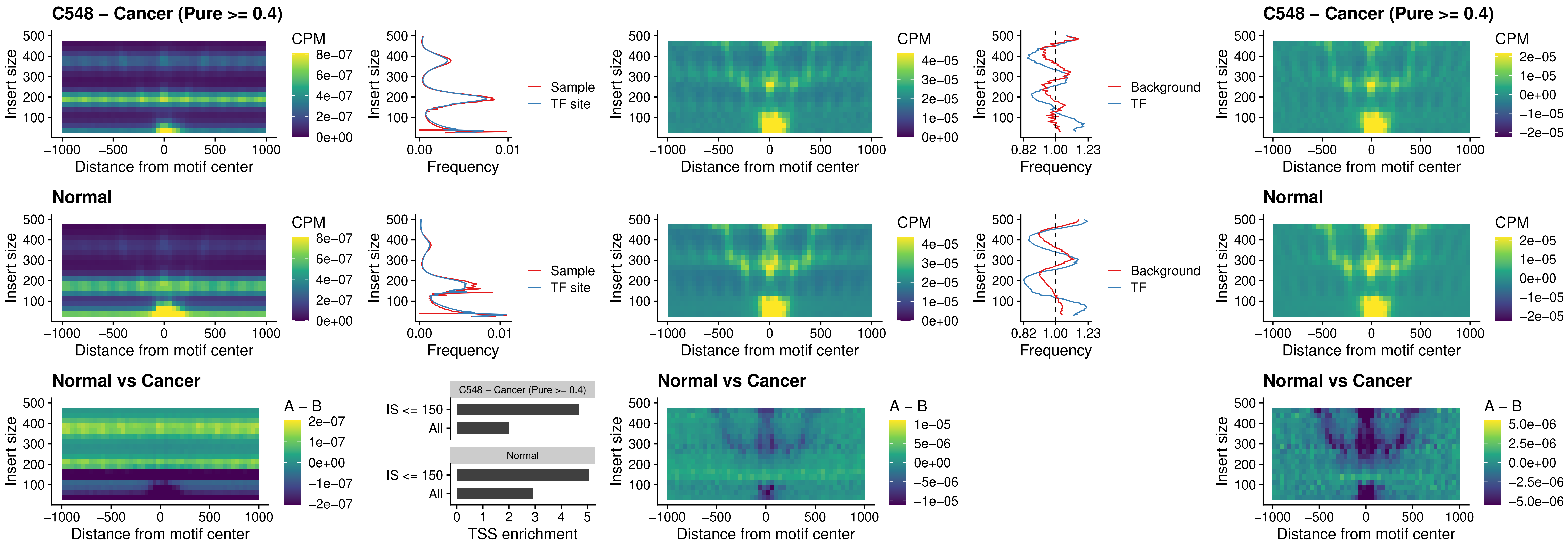

#### C548 – Cancer (Pure >= 0.4) – CTCFL_uc_cTSS_oPEAK

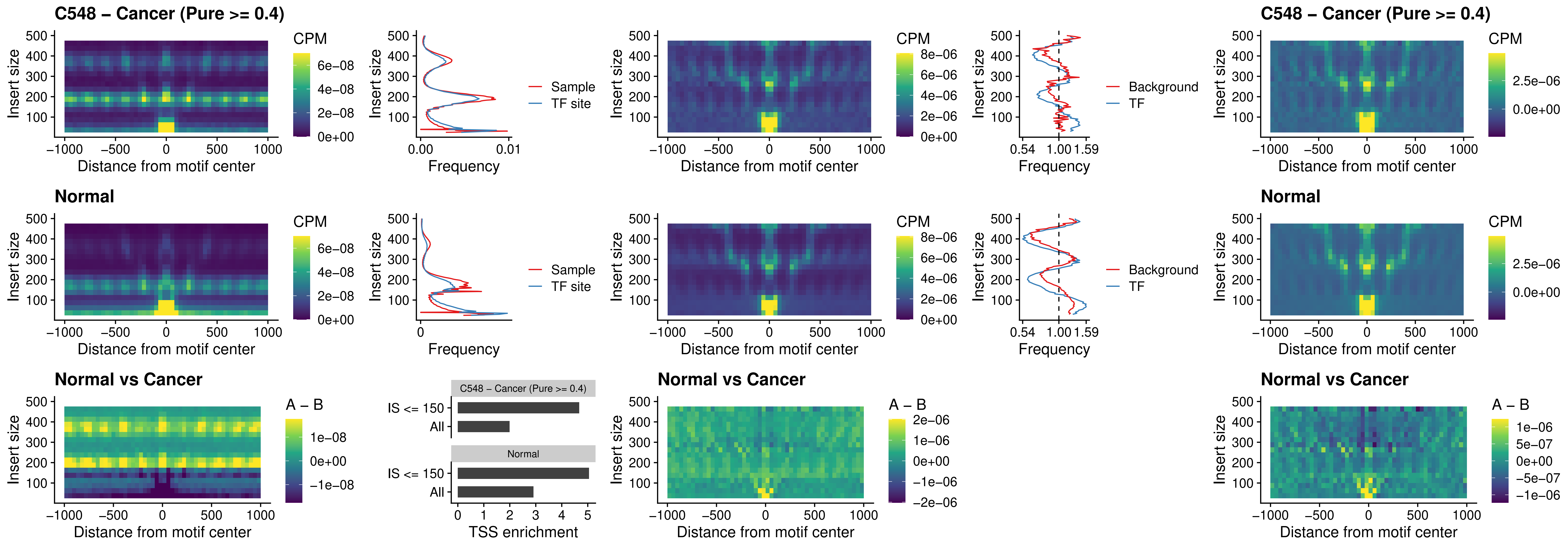

#### C548 – Cancer (Pure >= 0.4) – NEURODL1_uc_cTSS_oPEAK

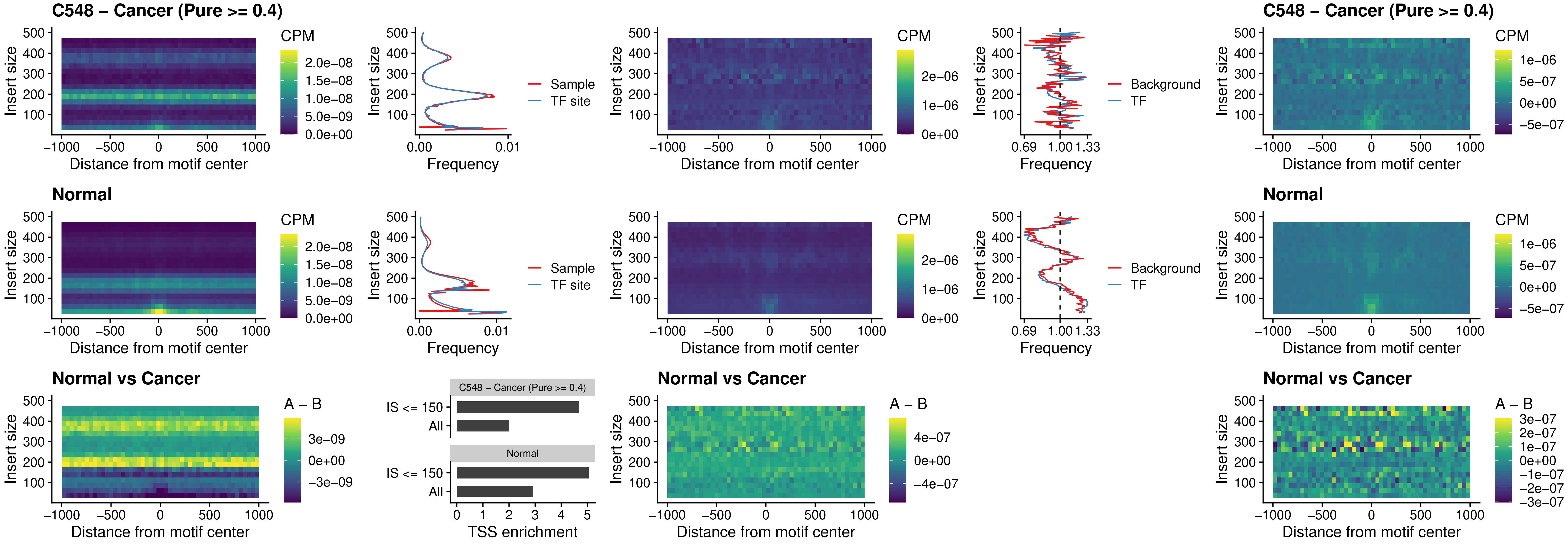

#### C548 – Cancer (Pure >= 0.4) – ENCODE_YY1_sites_COLON_nonpromotor

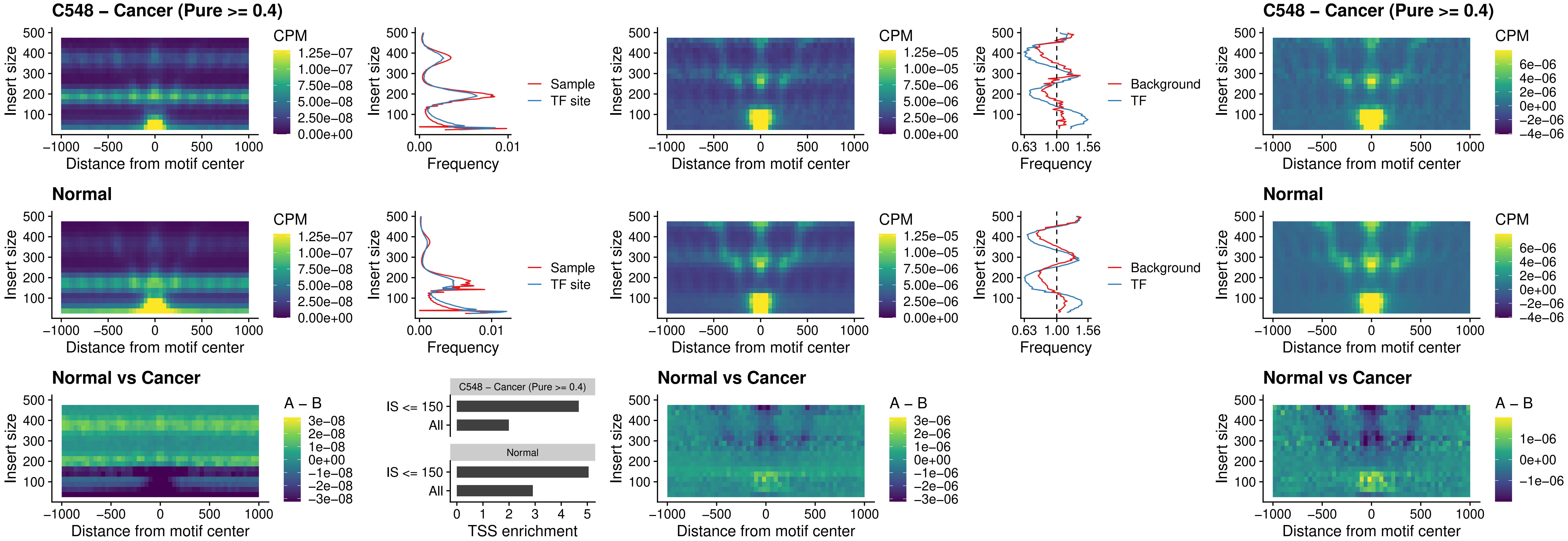

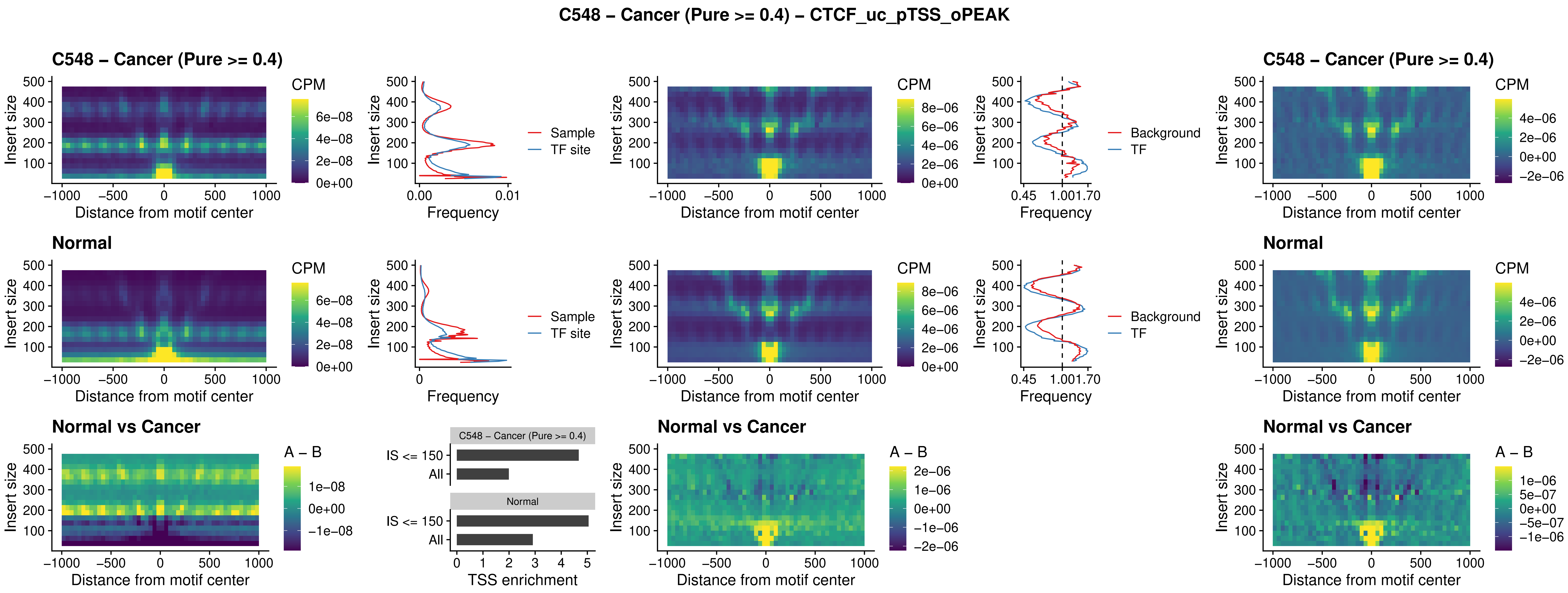
C548 – Cancer (Pure >= 0.4) – CTCF_uc_pTSS_oPEAK
C548 – Cancer (Pure >= 0.4)
Normal
Normal vs Cancer
Insert size
Distance from motif center
CPM
A – B
Frequency
Sample
TF site
Background
TF
C548 – Cancer (Pure >= 0.4)
Normal
IS <= 150
All
TSS enrichment

#### C548 – Cancer (Pure >= 0.4) – ENCODE_CTCF_sites_COLON_promotor

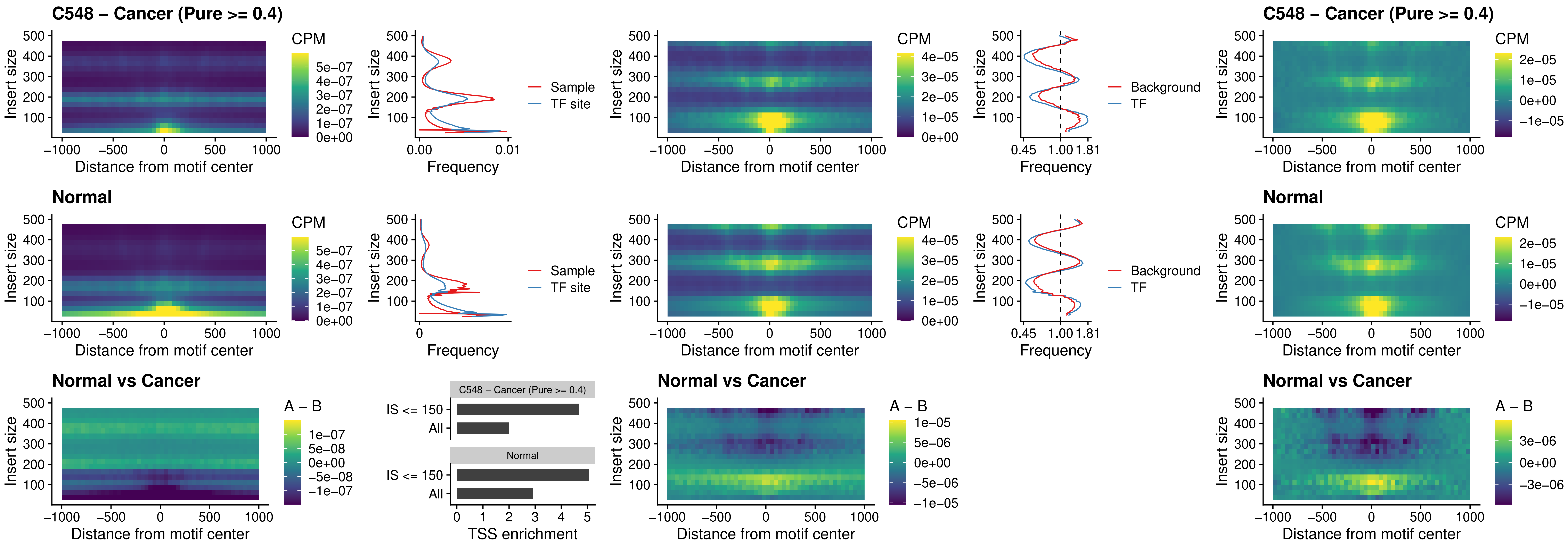

### C548 – Cancer (Pure >= 0.4) – ENCODE_CTCF_sites_COLON_promotor_ctcf

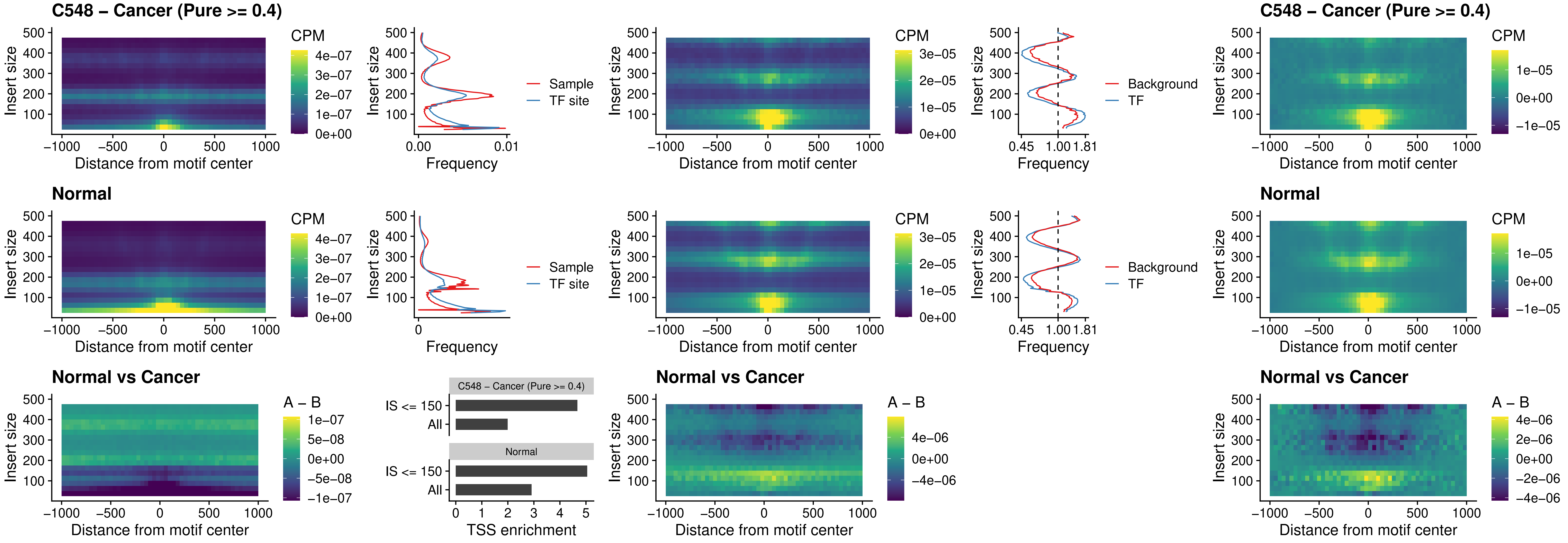

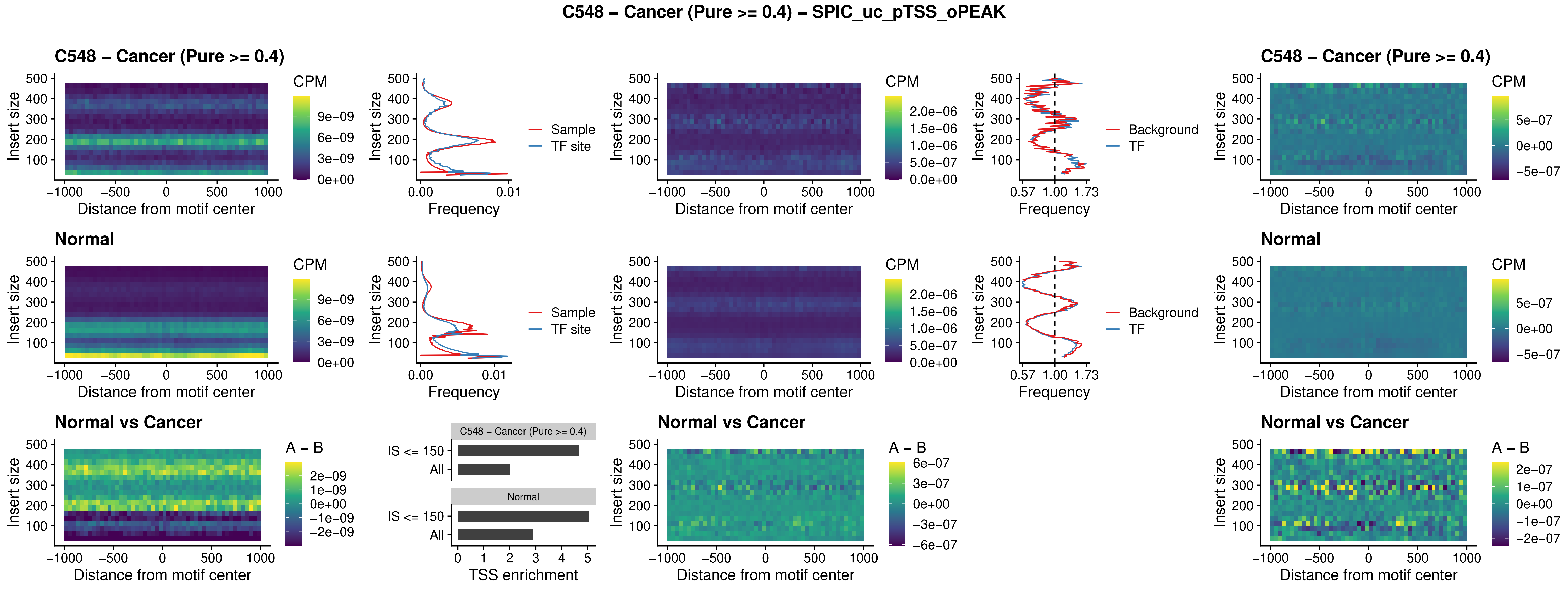
C548 – Cancer (Pure >= 0.4) – SPIC_uc_pTSS_oPEAK
C548 – Cancer (Pure >= 0.4)
Normal
Normal vs Cancer
Insert size
Distance from motif center
Frequency
CPM
A – B
Sample
TF site
Background
TF
C548 – Cancer (Pure >= 0.4)
Normal
IS <= 150
All
TSS enrichment

#### C548 – Cancer (Pure >= 0.4) – SPI1_uc_dTSS_nPEAK

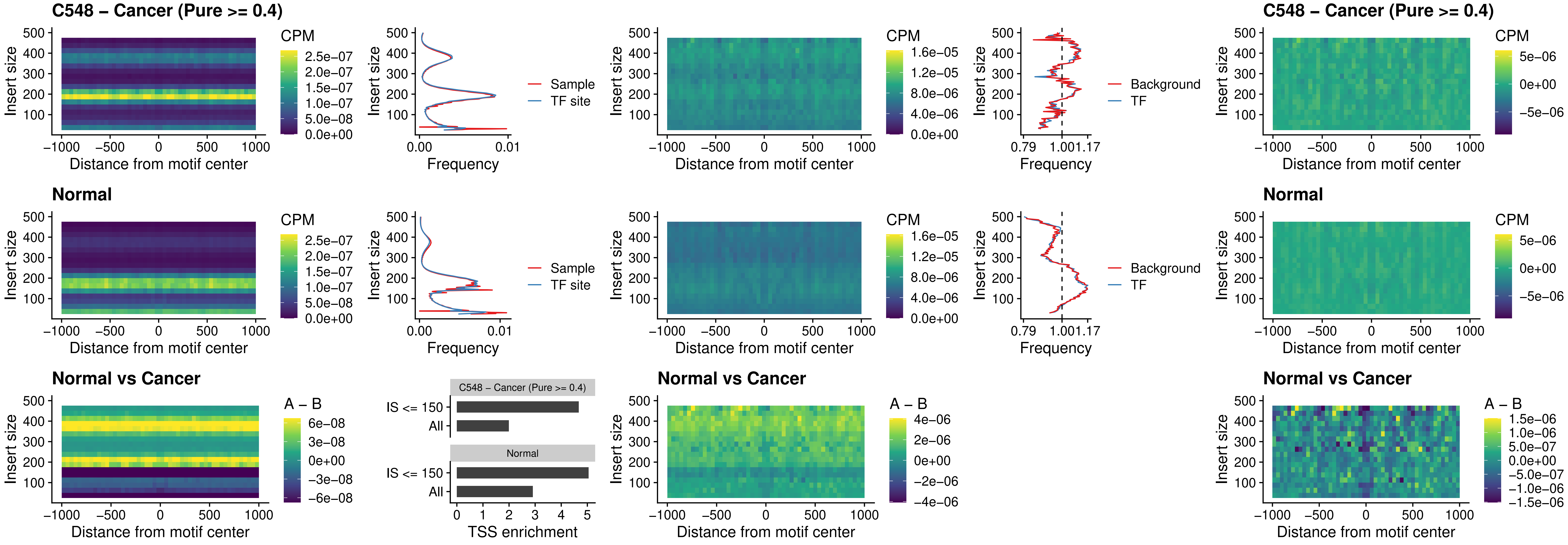

#### C548 – Cancer (Pure >= 0.4) – SPI1_uc_cTSS_nPEAK

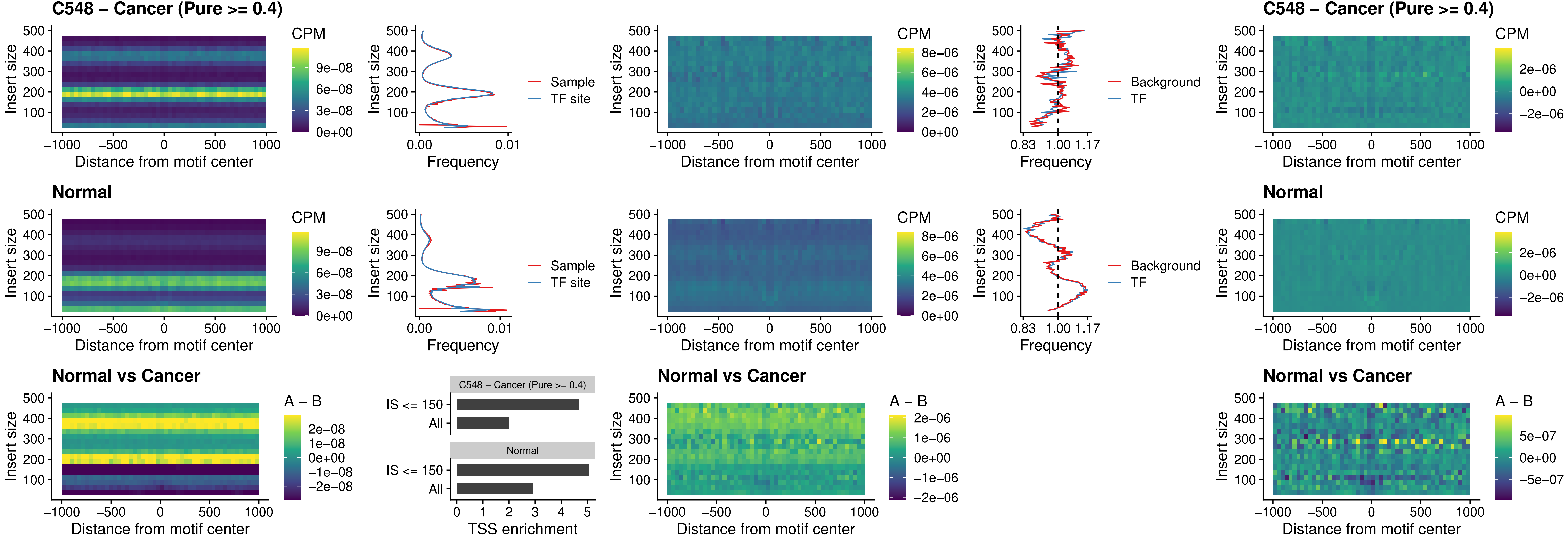

#### C548 – Cancer (Pure >= 0.4) – BCL11A_uc_pTSS_nPEAK

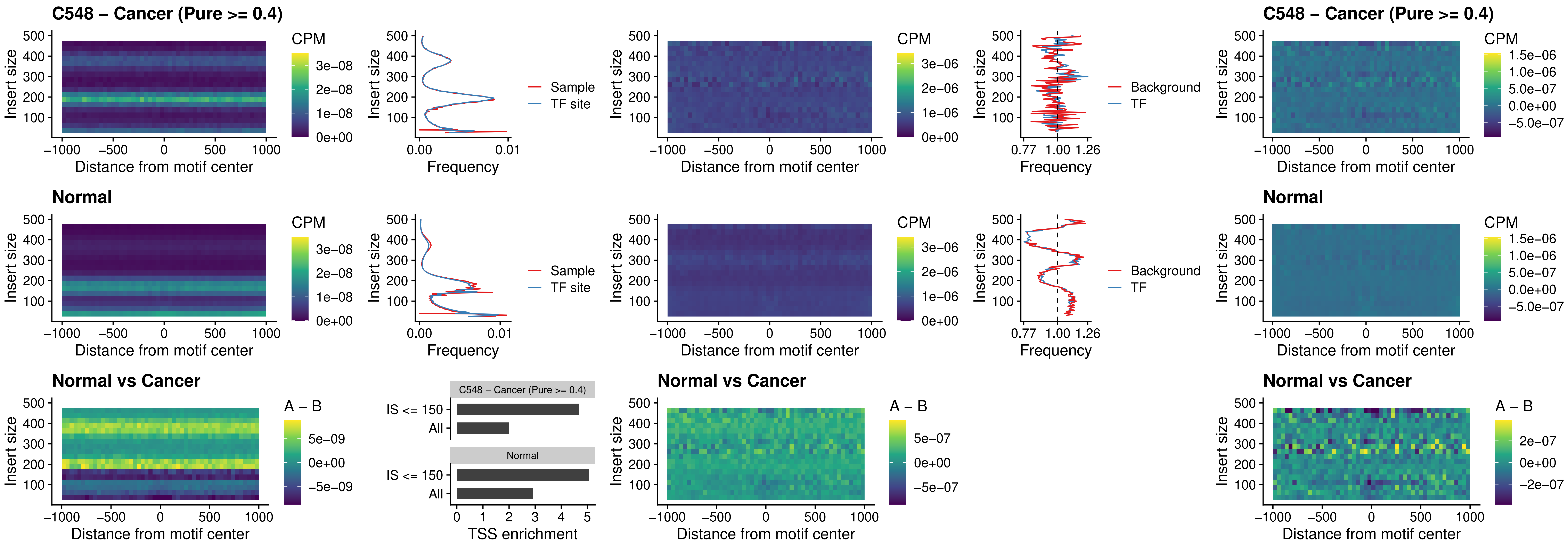

#### C548 – Cancer (Pure >= 0.4) – BCL11B_uc_pTSS_nPEAK

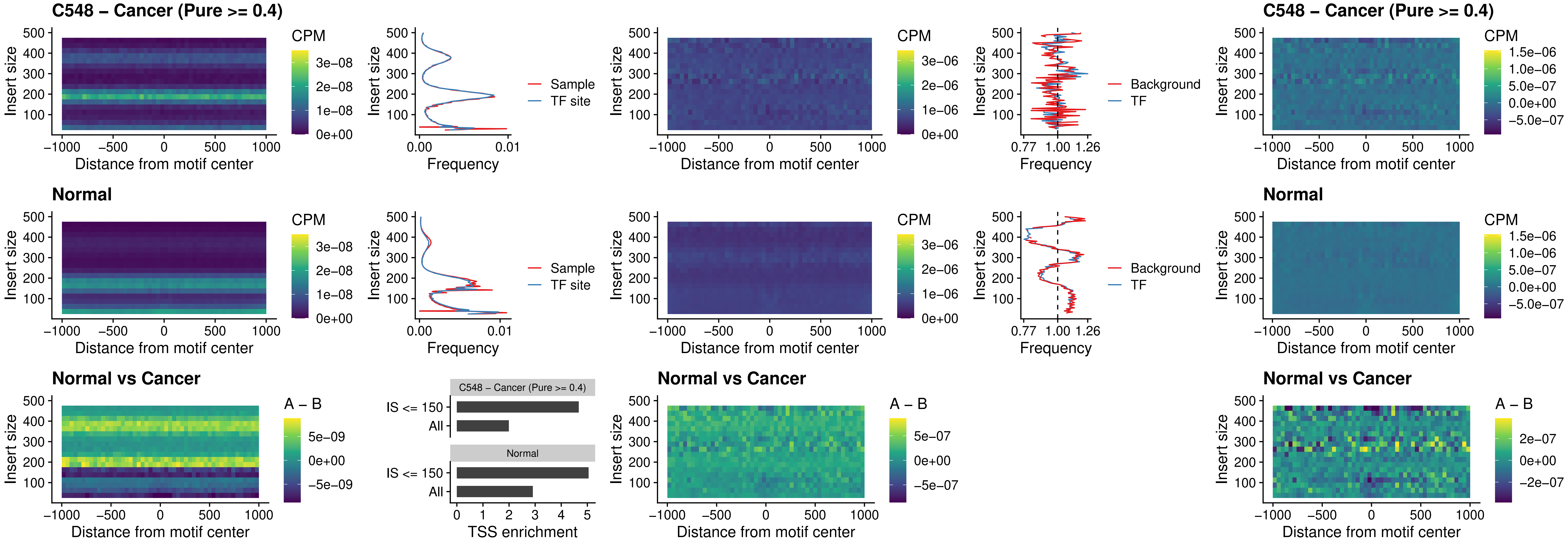

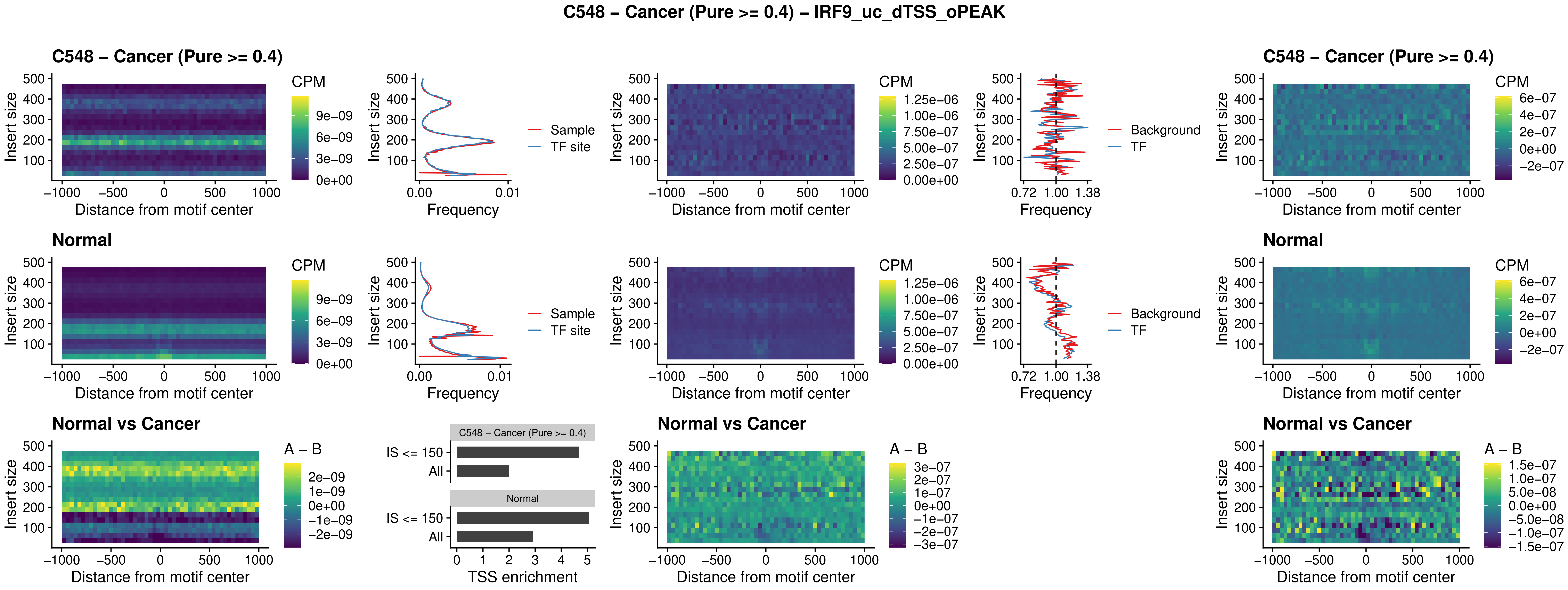
C548 – Cancer (Pure >= 0.4) – IRF9_uc_dTSS_oPEAK
C548 – Cancer (Pure >= 0.4)
Normal
Normal vs Cancer
Insert size
Distance from motif center
CPM
A – B
Frequency
Sample
TF site
Background
TF
TSS enrichment
IS <= 150
All
C548 – Cancer (Pure >= 0.4)
Normal

#### C548 – Cancer (Pure >= 0.4) – IRF9_uc_cTSS_oPEAK

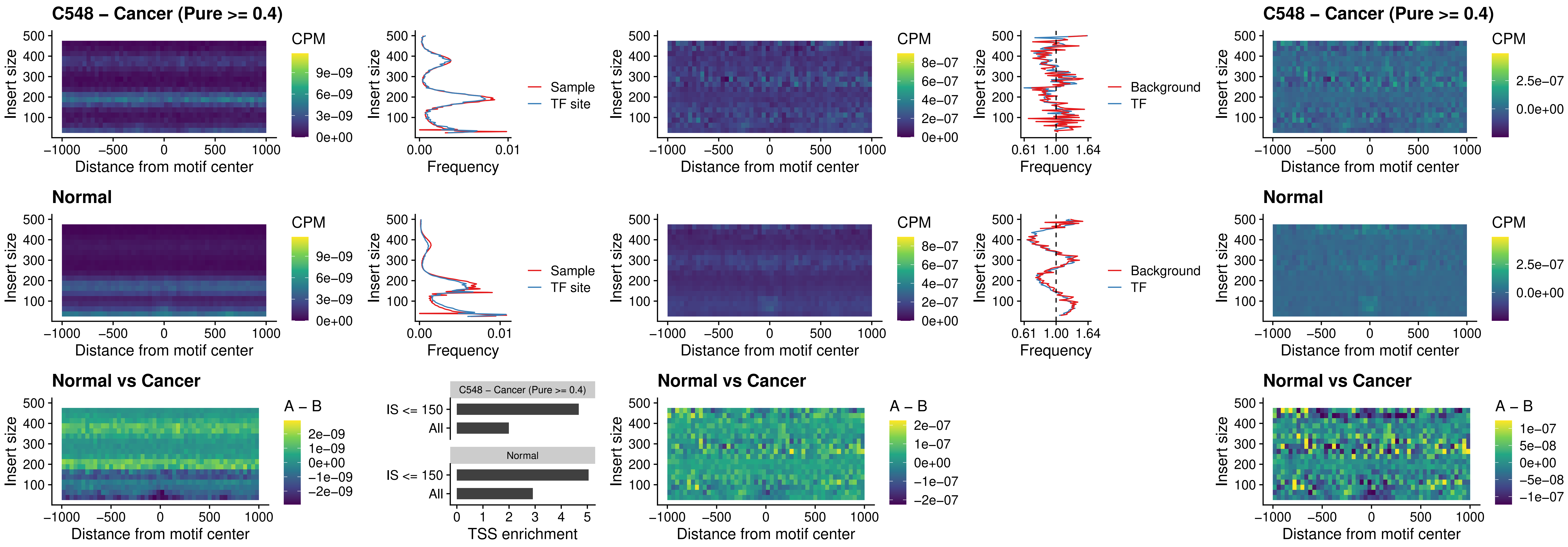

#### C548 – Cancer (Pure >= 0.4) – BCL11A_uc_cTSS_oPEAK

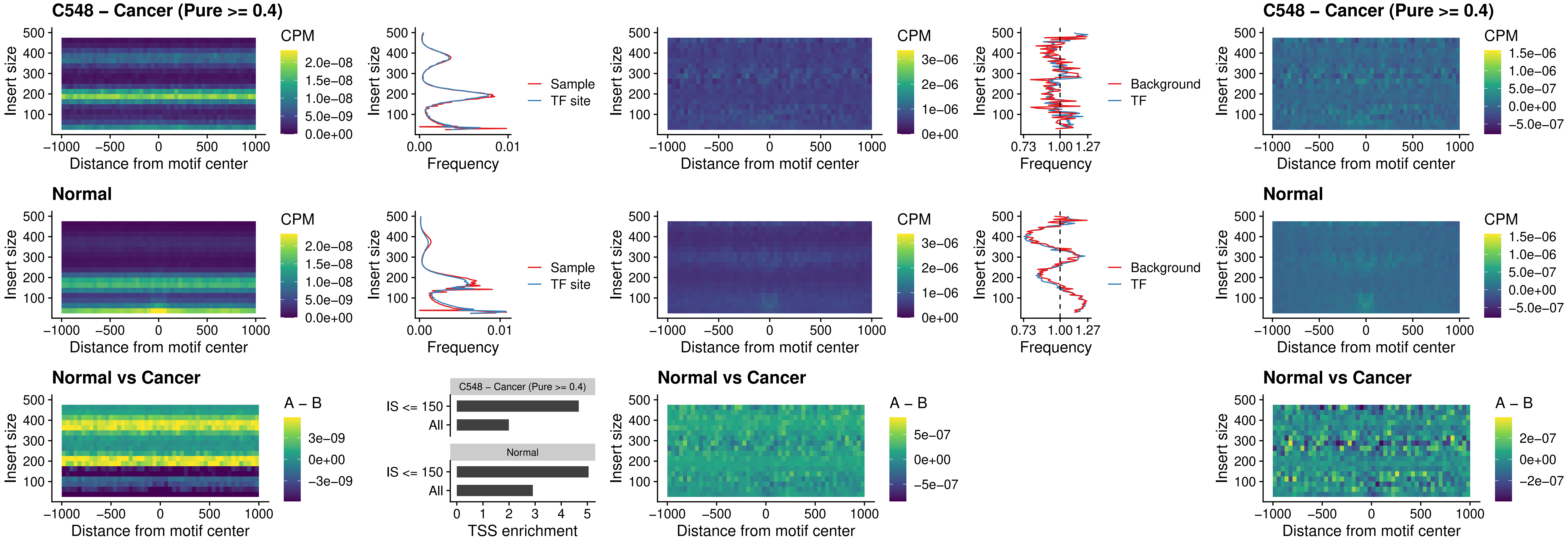

#### C548 – Cancer (Pure >= 0.4) – BCL11B_uc_cTSS_oPEAK

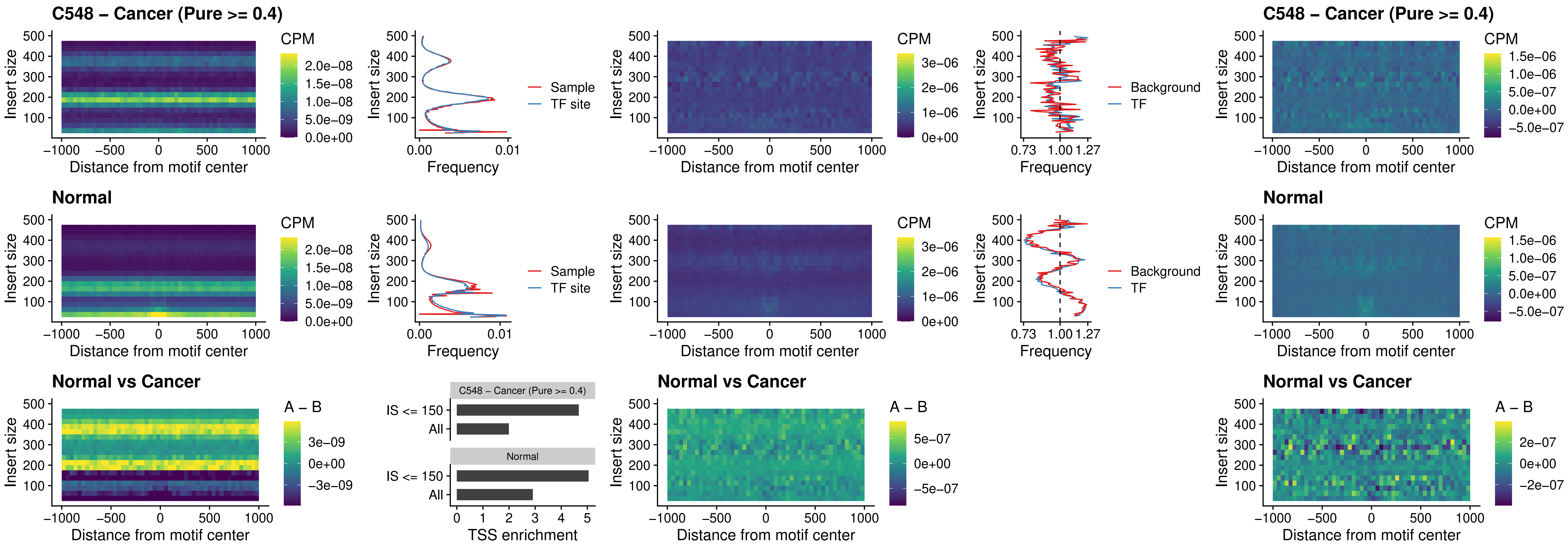

#### C548 – Cancer (Pure >= 0.4) – BCL11A_uc_dTSS_oPEAK

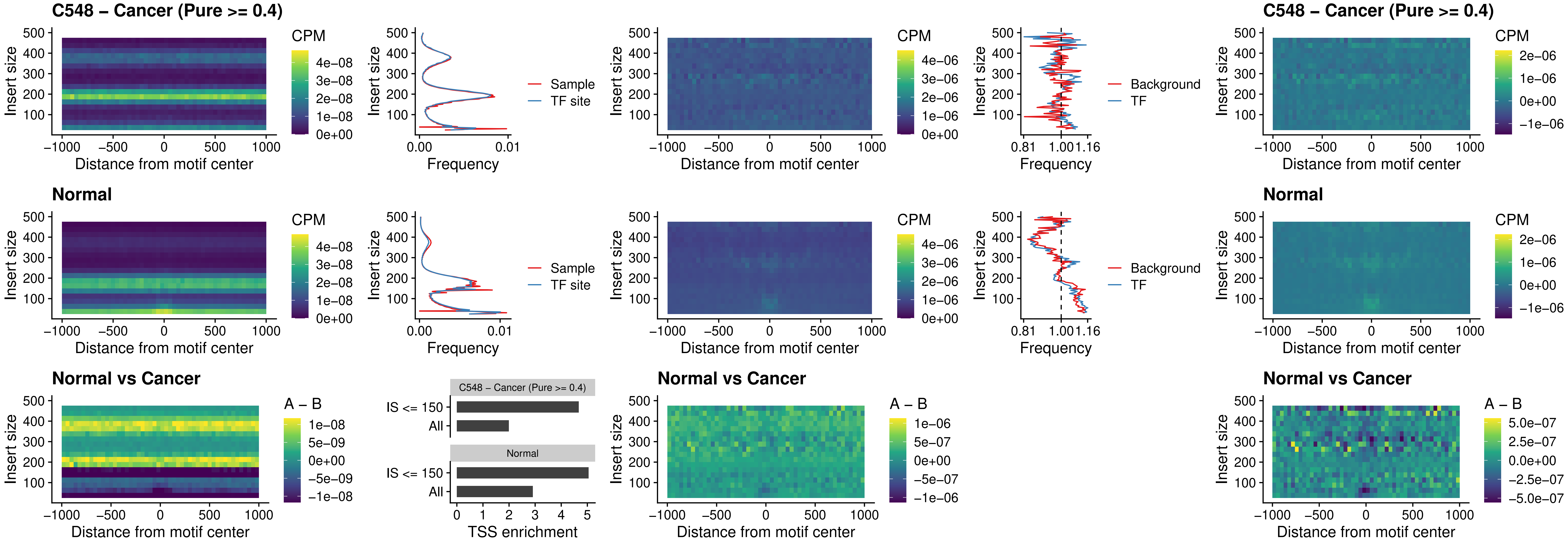

#### C548 – Cancer (Pure >= 0.4) – BCL11B_uc_dTSS_oPEAK

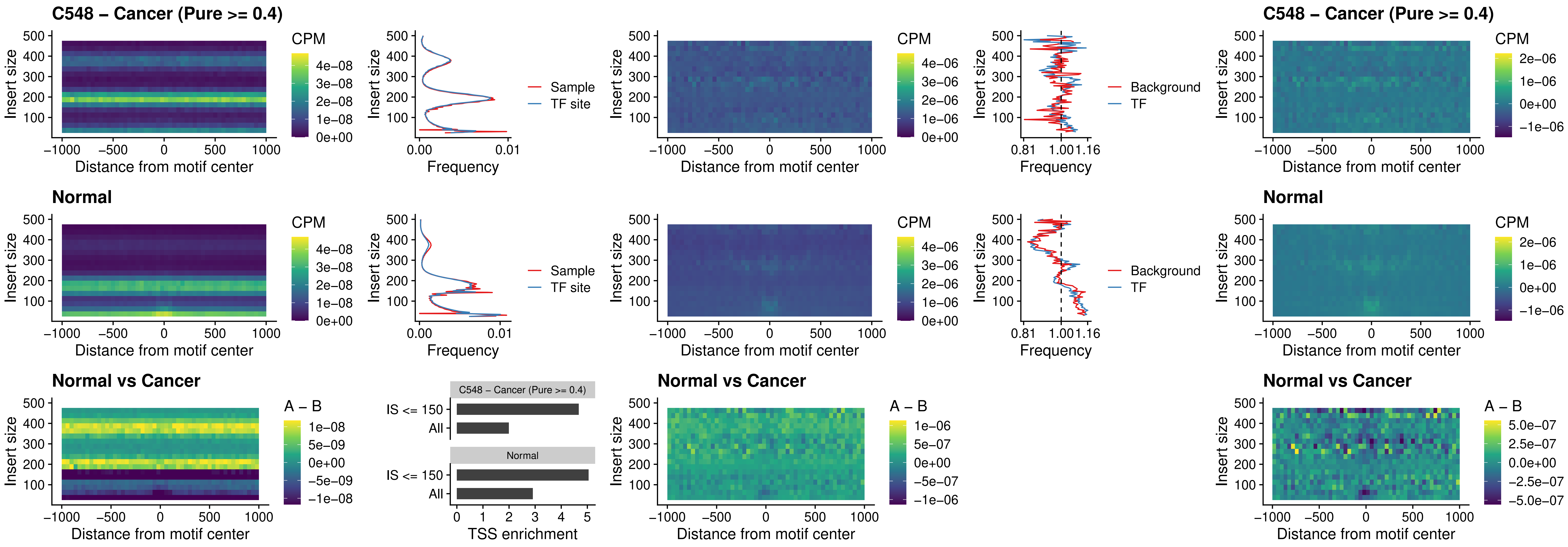

#### C548 – Cancer (Pure >= 0.4) – SPIB_uc_cTSS_nPEAK

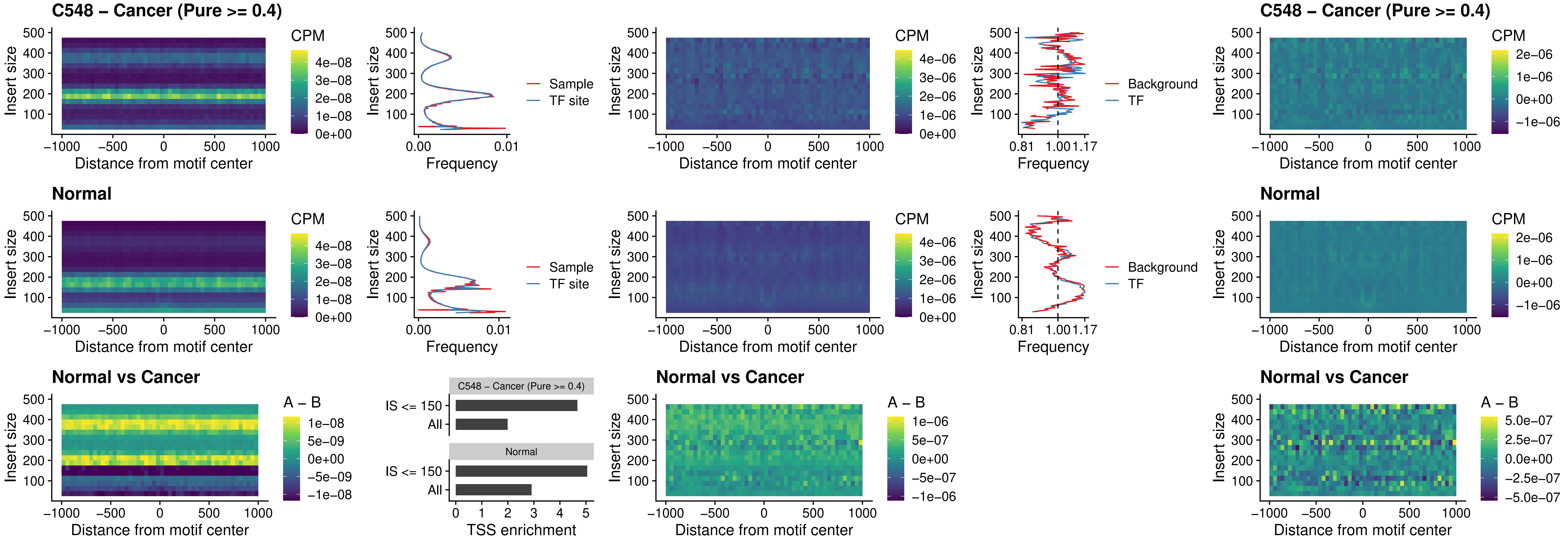

#### C548 – Cancer (Pure >= 0.4) – SPIC_uc_cTSS_nPEAK

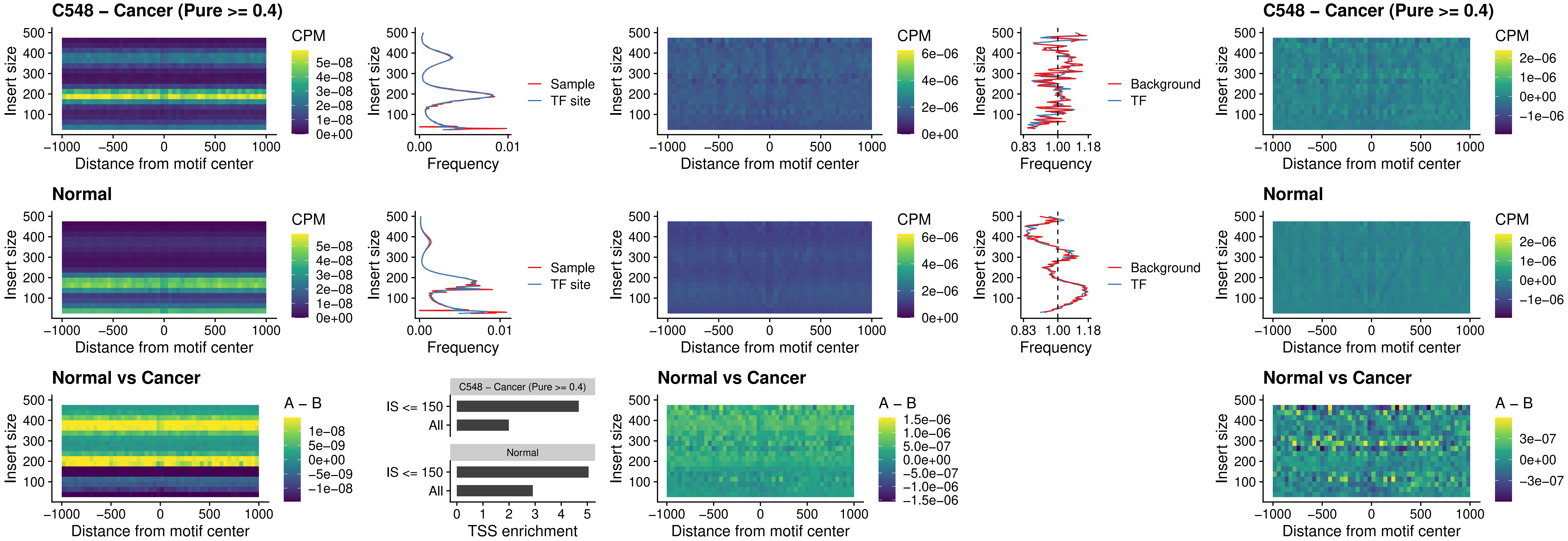

#### C548 – Cancer (Pure >= 0.4) – SPI1_uc_pTSS_oPEAK

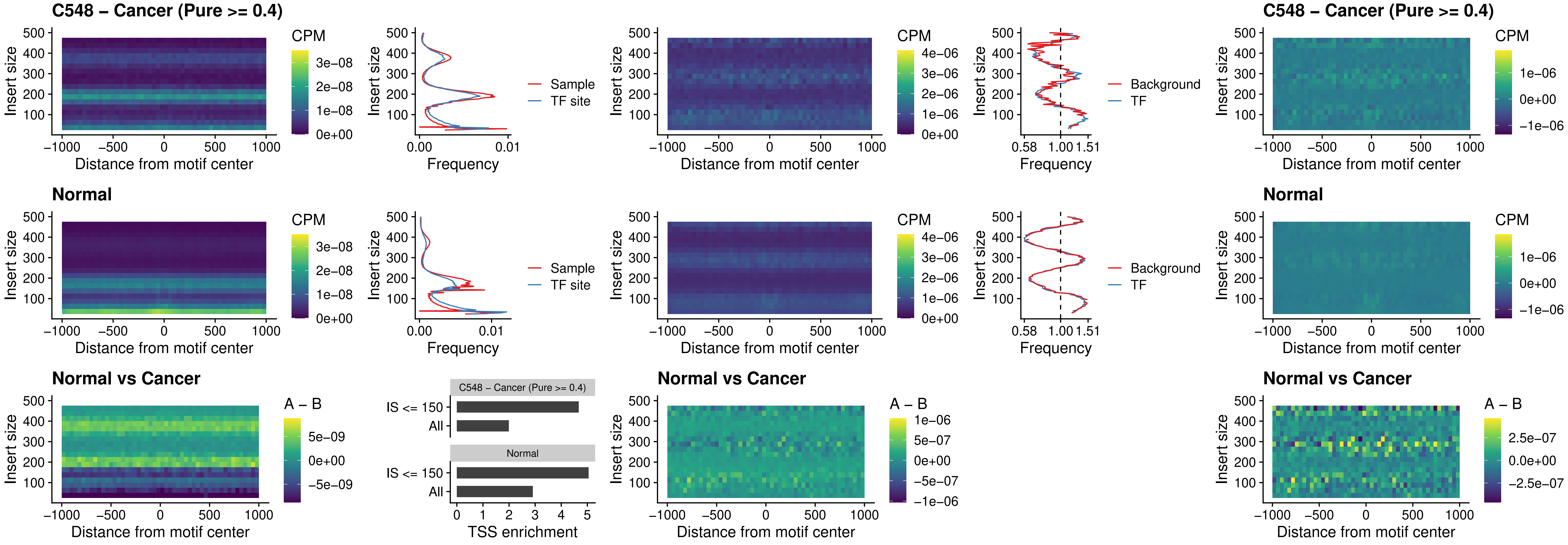

#### C548 – Cancer (Pure >= 0.4) – BCL11A_uc_pTSS_oPEAK

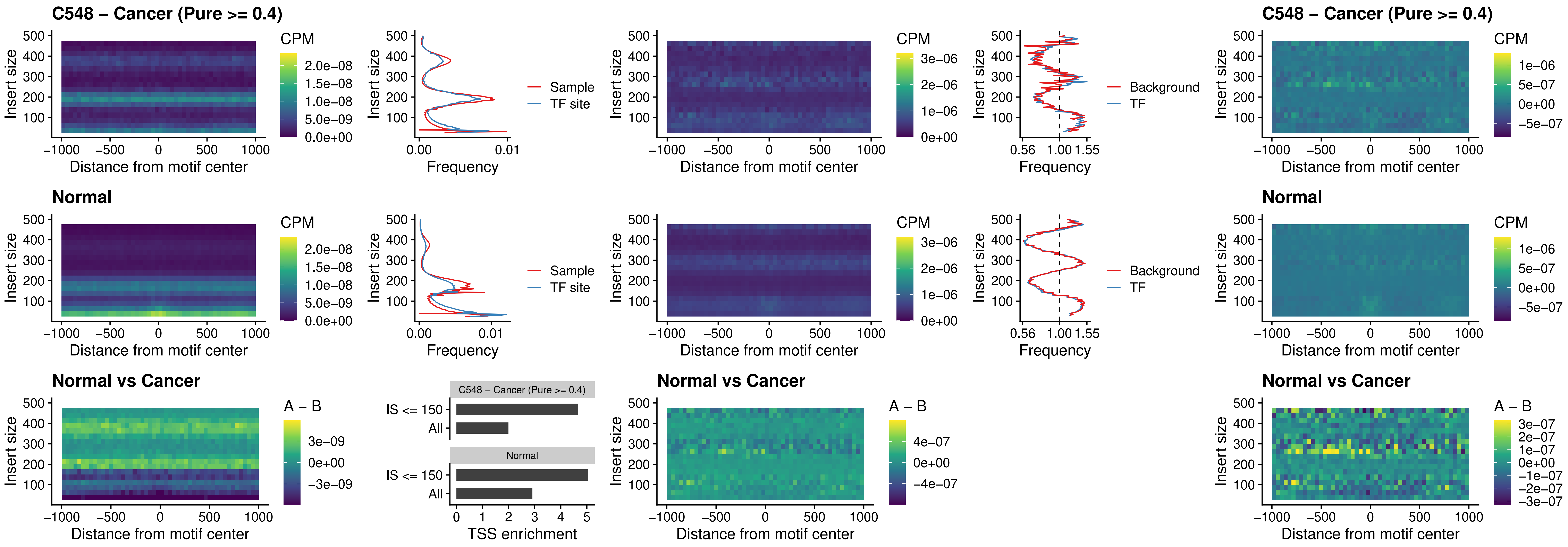

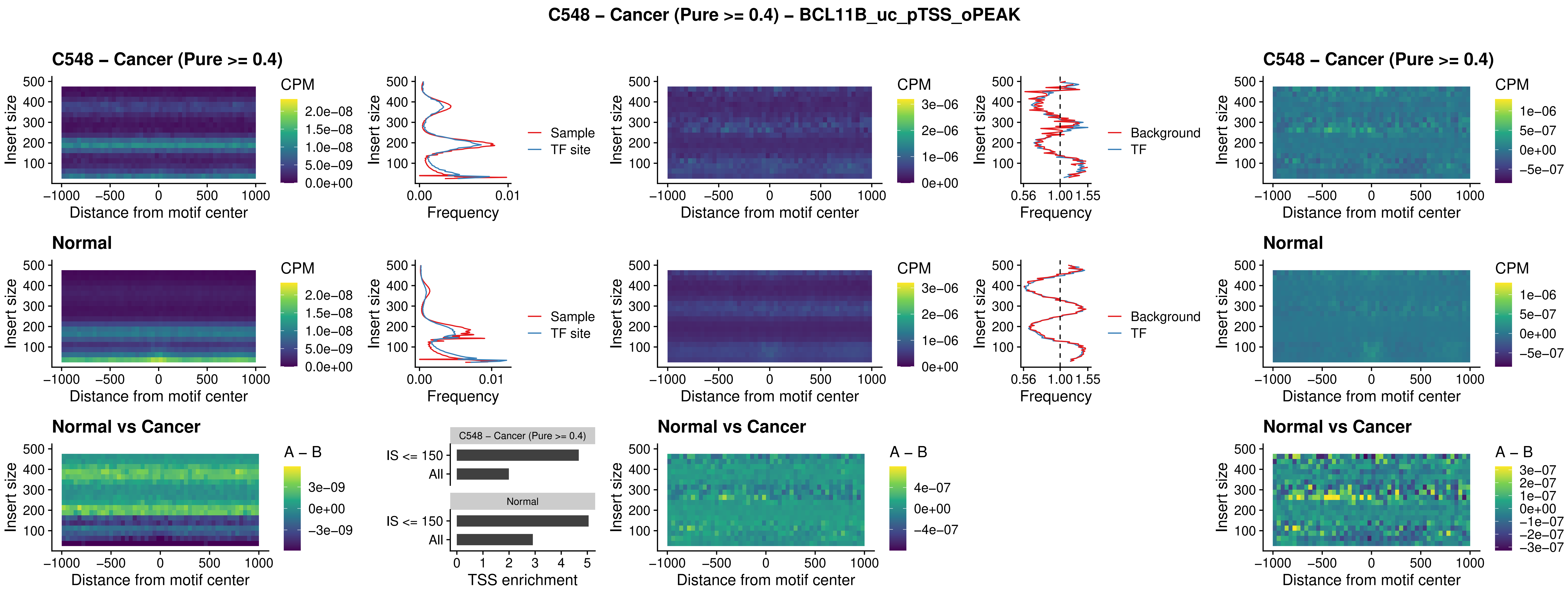
C548 – Cancer (Pure >= 0.4) – BCL11B_uc_pTSS_oPEAK
C548 – Cancer (Pure >= 0.4)
Normal
Normal vs Cancer
Insert size
Distance from motif center
Frequency
CPM
A – B
Sample
TF site
Background
TF
C548 – Cancer (Pure >= 0.4)
Normal
IS <= 150
All
TSS enrichment

#### C548 – Cancer (Pure >= 0.4) – SPI1_uc_cTSS_oPEAK

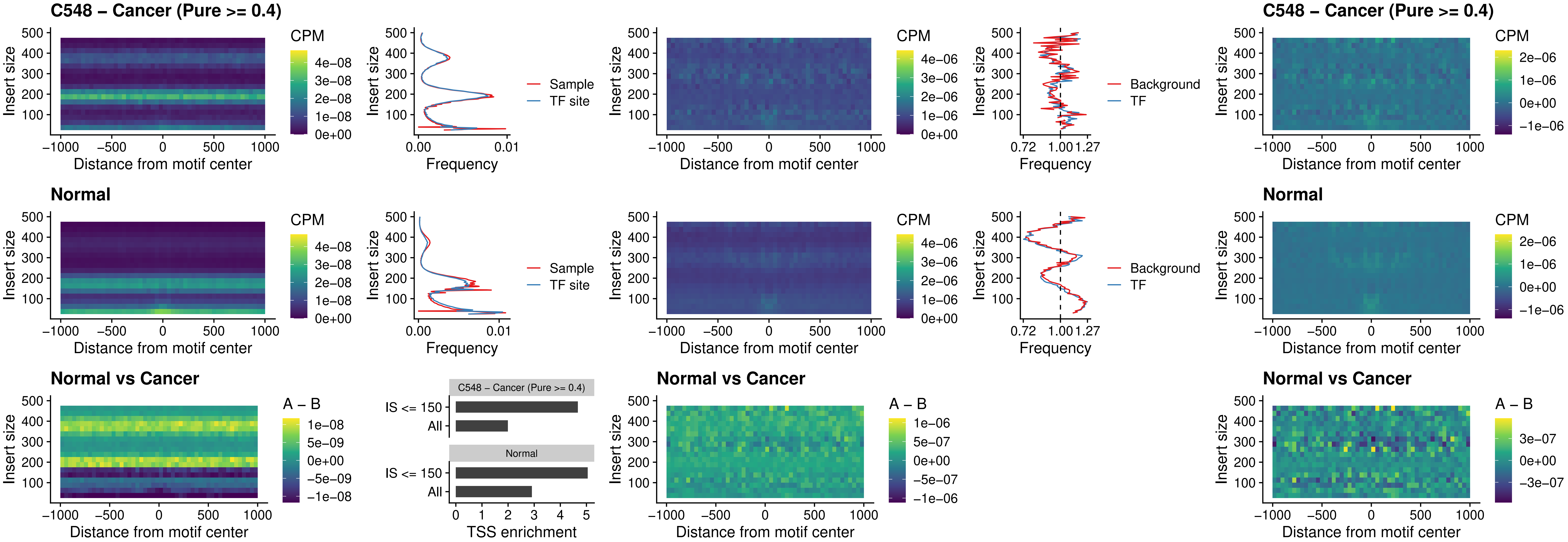

#### C548 – Cancer (Pure >= 0.4) – SPIB_uc_cTSS_oPEAK

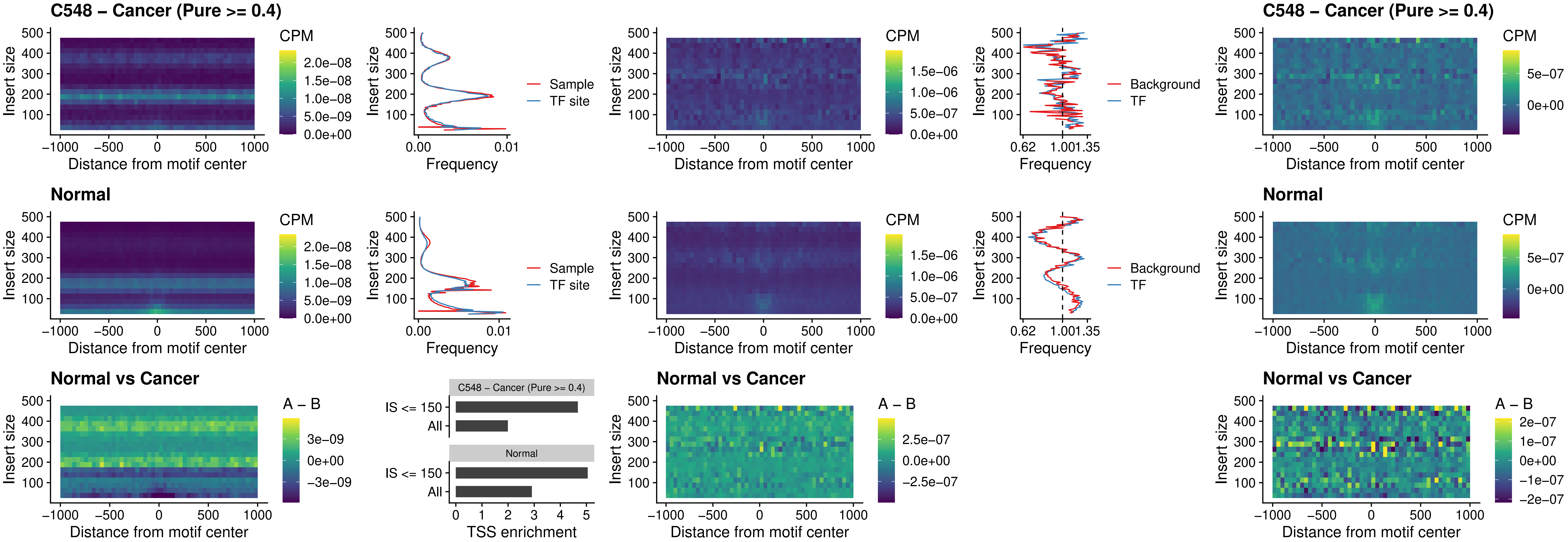

#### C548 – Cancer (Pure >= 0.4) – SPIB_uc_pTSS_oPEAK

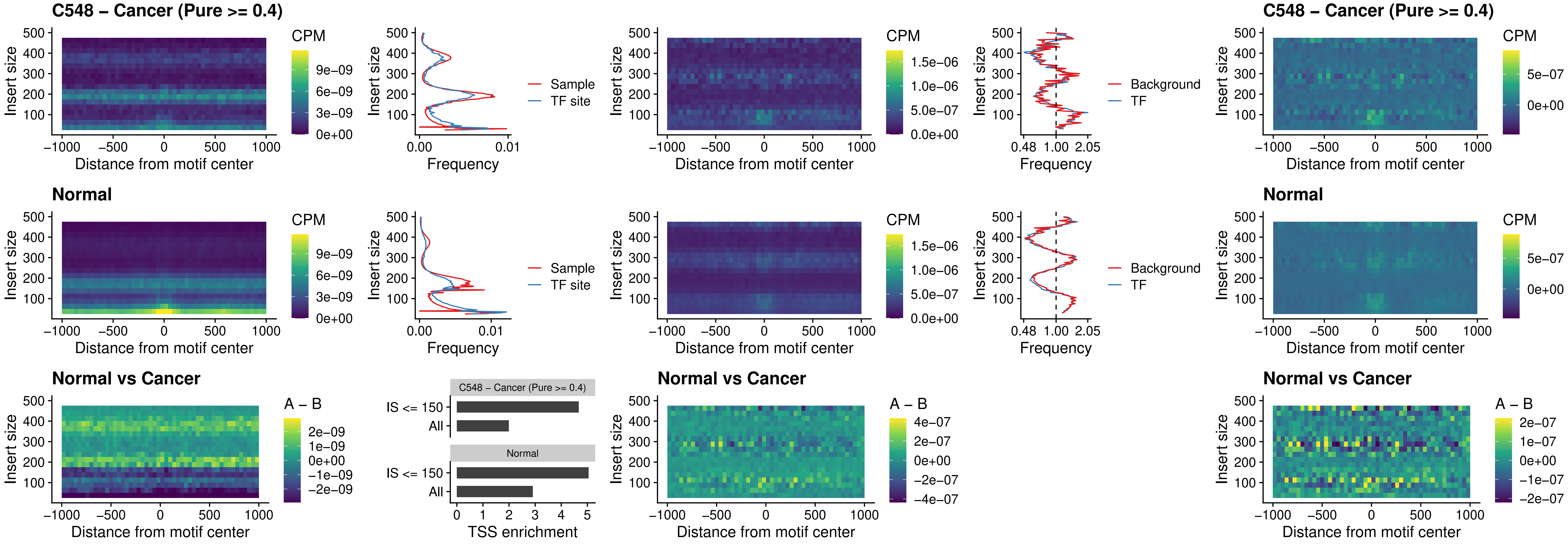

#### C548 – Cancer (Pure >= 0.4) – SPI1_uc_dTSS_oPEAK

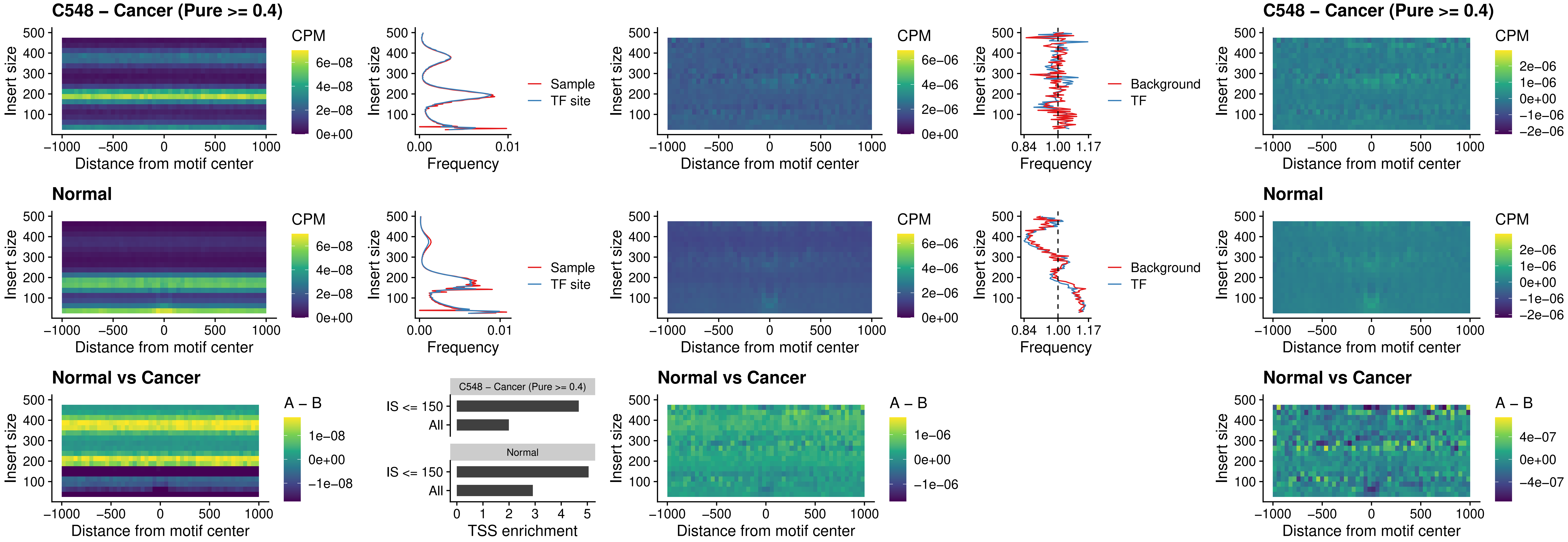

#### C548 – Cancer (Pure >= 0.4) – SPIB_uc_dTSS_oPEAK

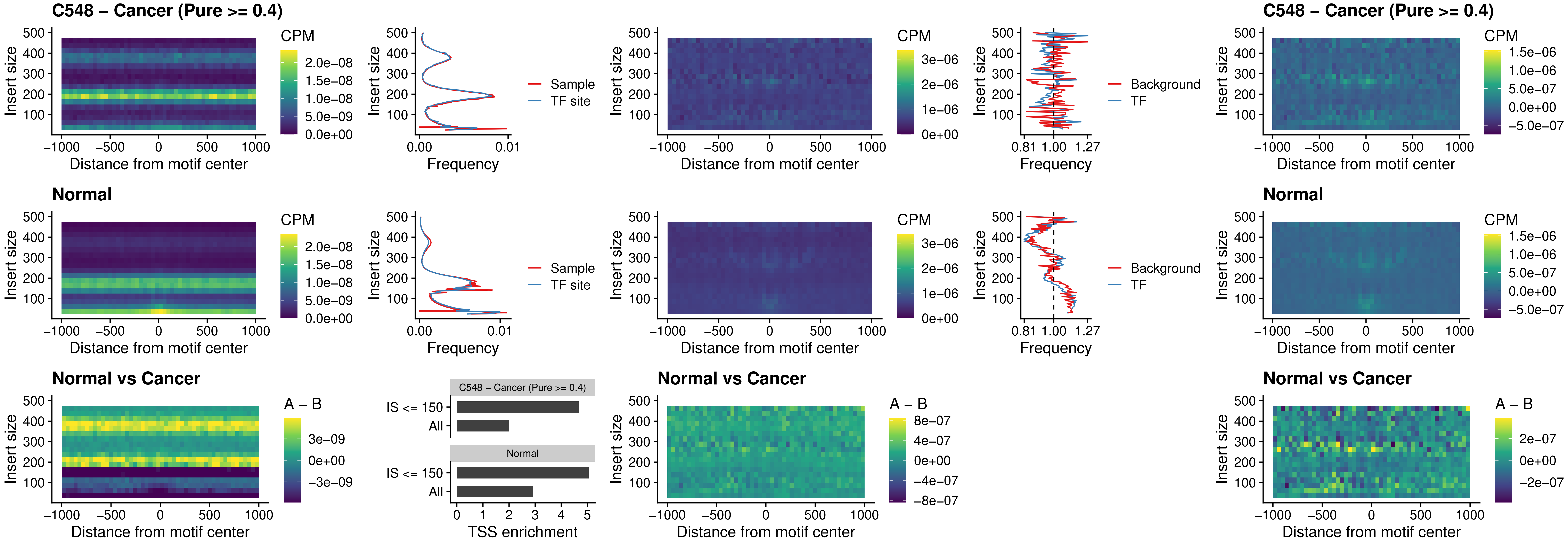

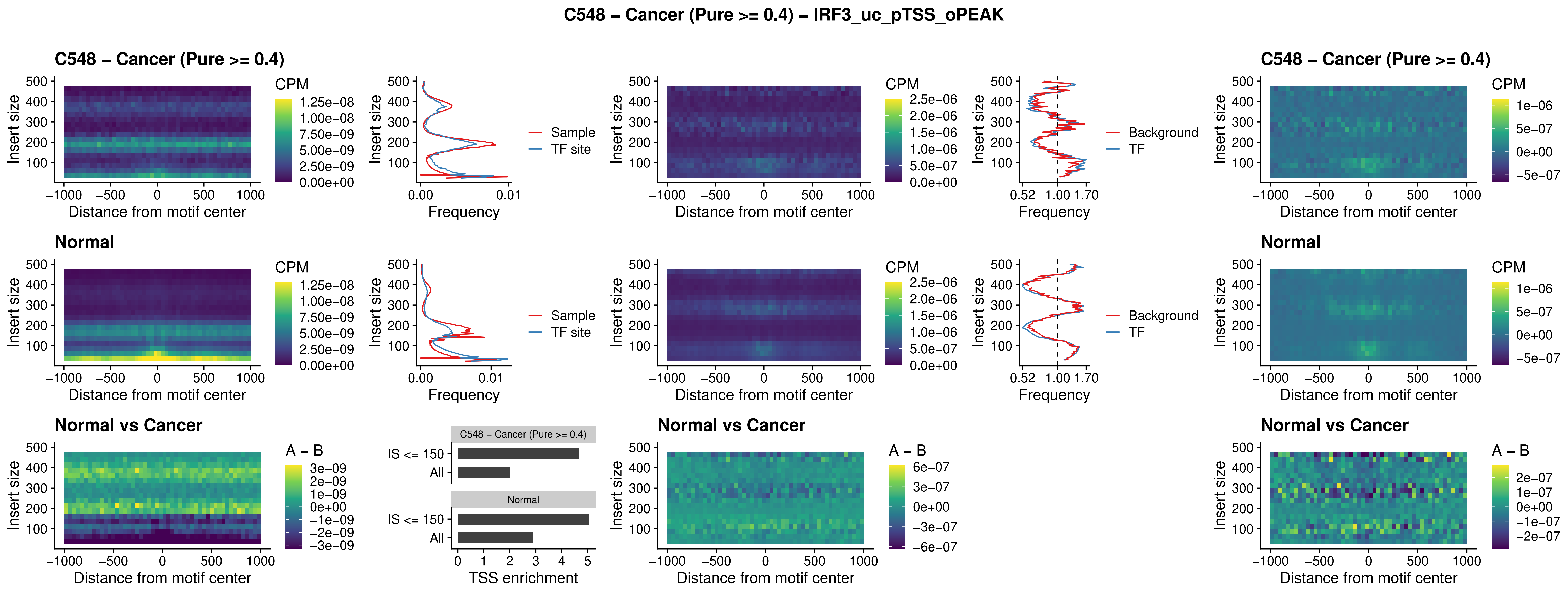
C548 – Cancer (Pure >= 0.4) – IRF3_uc_pTSS_oPEAK
C548 – Cancer (Pure >= 0.4)
Normal
Normal vs Cancer
Insert size
Distance from motif center
CPM
A – B
Frequency
Sample
TF site
Background
TF
C548 – Cancer (Pure >= 0.4)
Normal
IS <= 150
All
TSS enrichment

#### C548 – Cancer (Pure >= 0.4) – IRF2_uc_dTSS_oPEAK

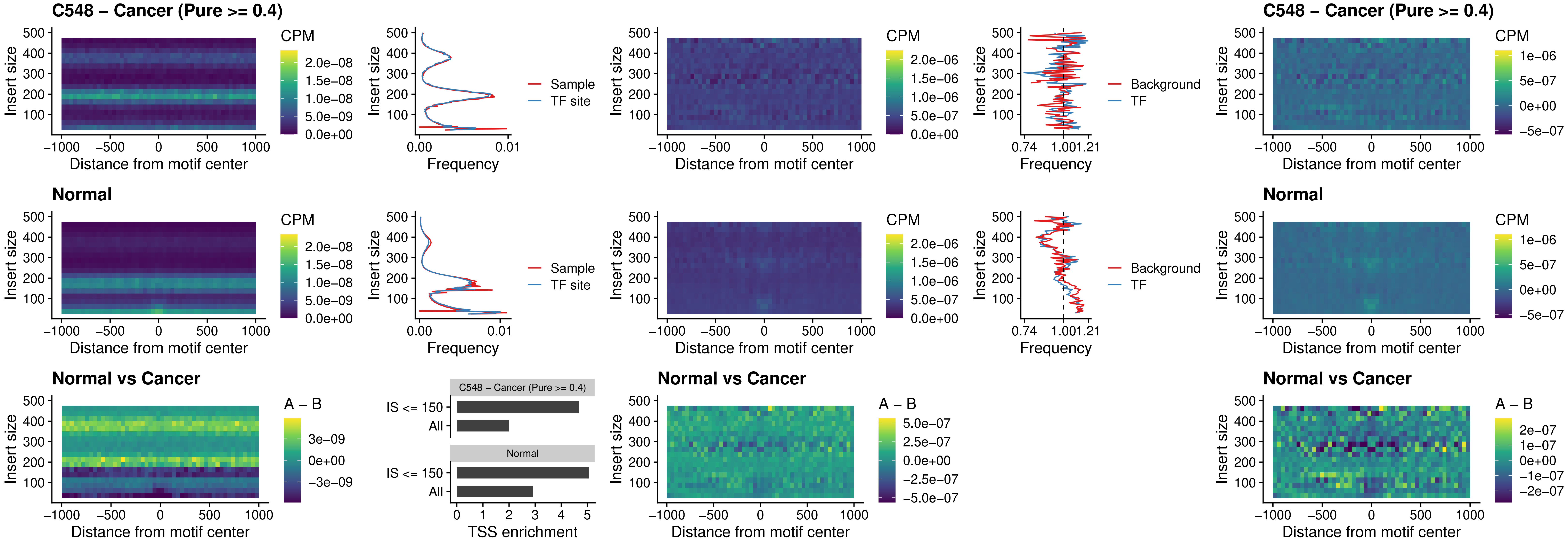

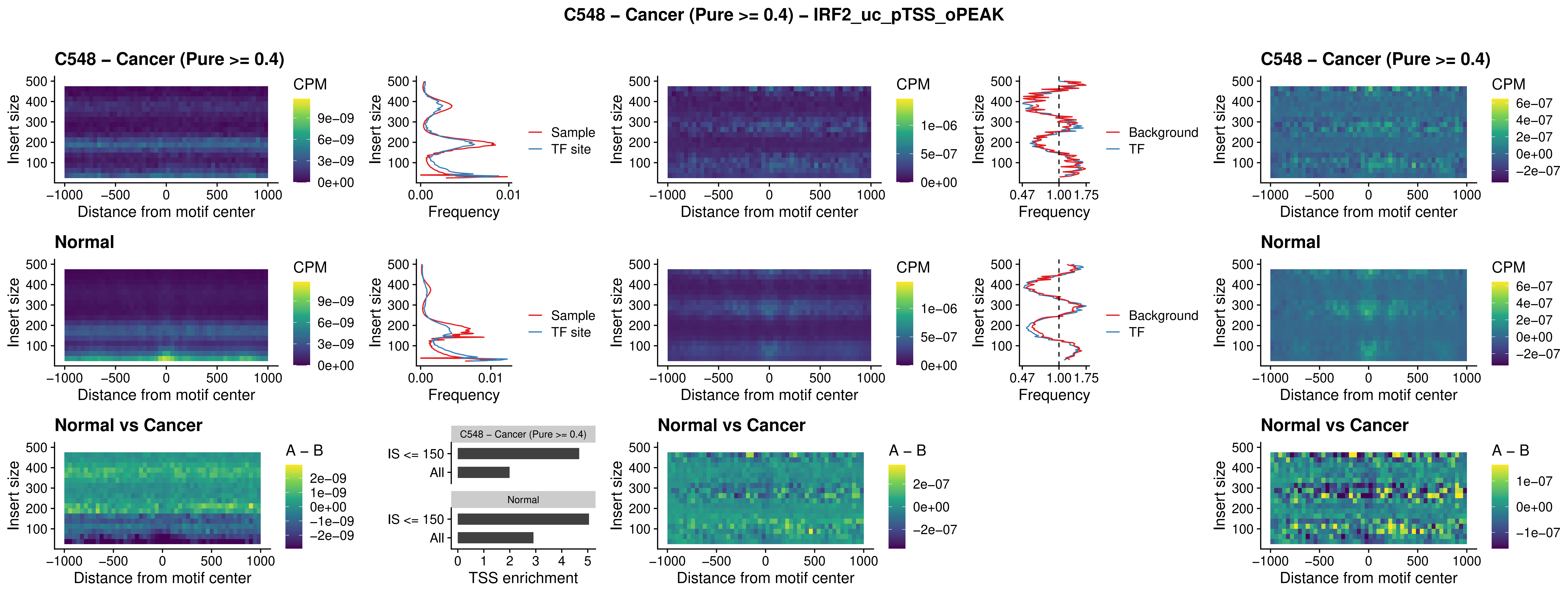
C548 – Cancer (Pure >= 0.4) – IRF2_uc_pTSS_oPEAK
C548 – Cancer (Pure >= 0.4)
Normal
Normal vs Cancer
Insert size
Distance from motif center
CPM
A – B
Frequency
Sample
TF site
Background
TF
C548 – Cancer (Pure >= 0.4)
Normal
IS <= 150
All
TSS enrichment

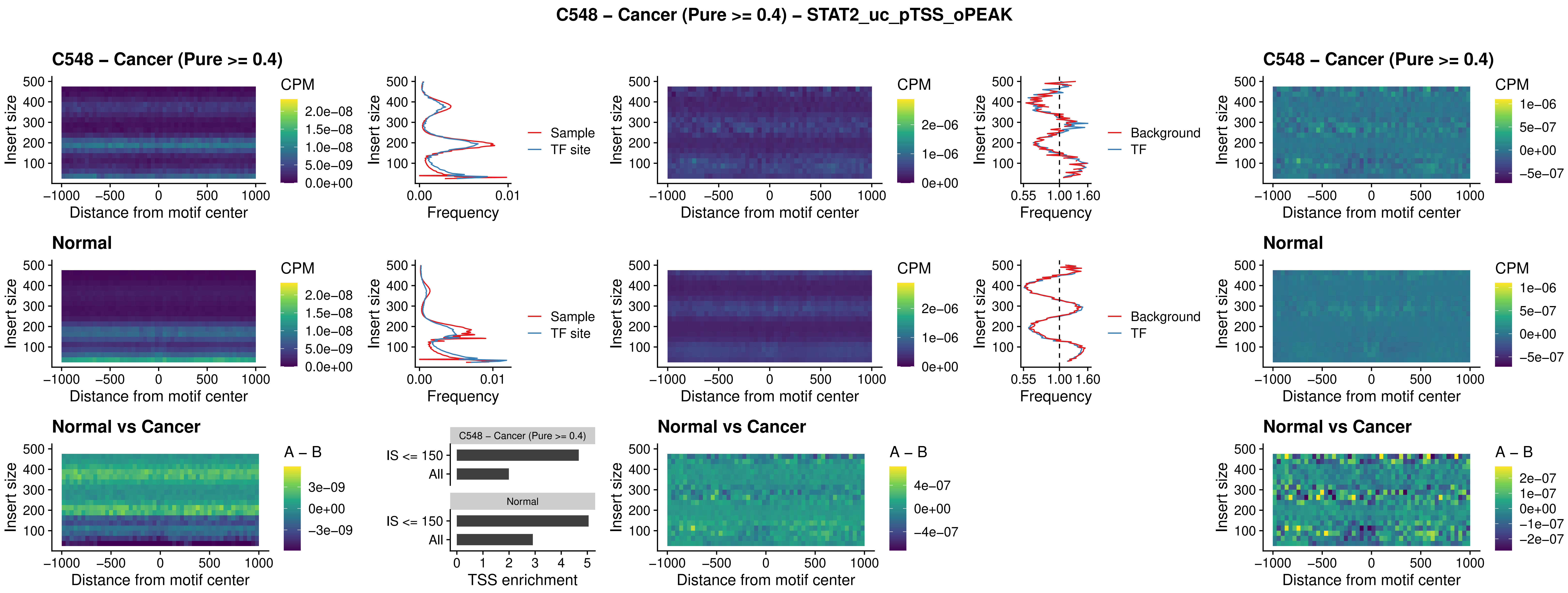
C548 – Cancer (Pure >= 0.4) – STAT2_uc_pTSS_oPEAK
C548 – Cancer (Pure >= 0.4)
Normal
Normal vs Cancer
Insert size
Distance from motif center
CPM
A – B
Frequency
Sample
TF site
Background
TF
IS <= 150
All
TSS enrichment

#### C548 – Cancer (Pure >= 0.4) – SPIB_uc_dTSS_nPEAK

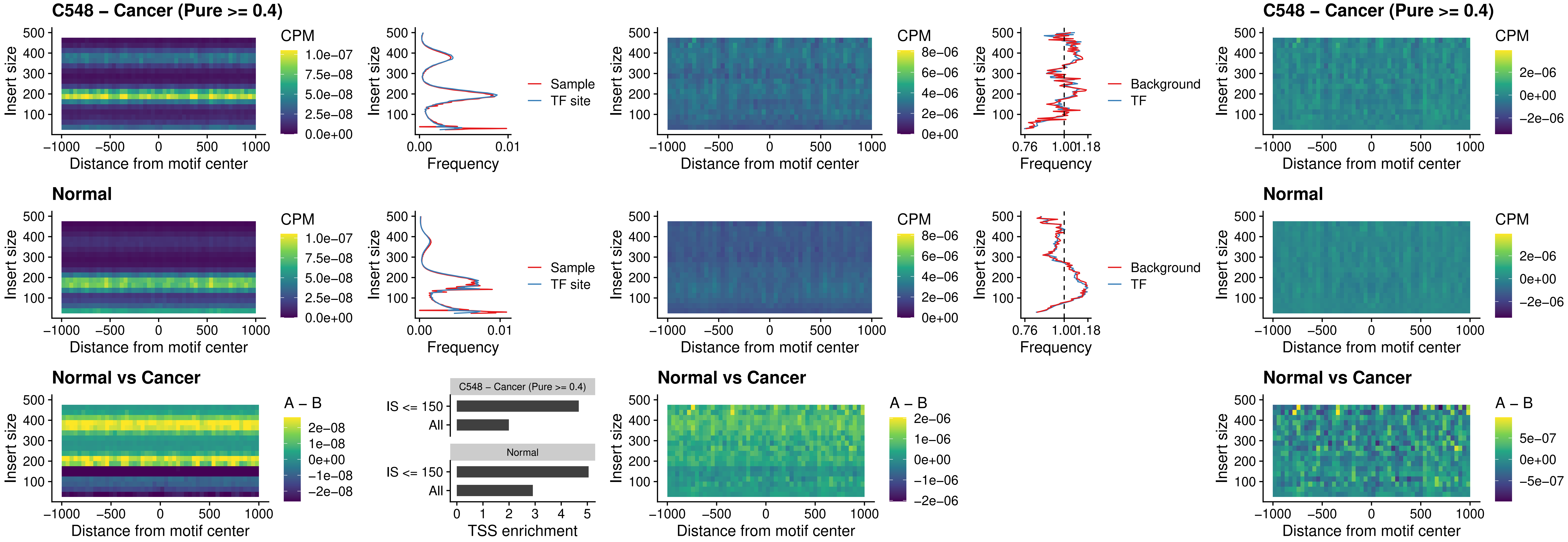

#### C548 – Cancer (Pure >= 0.4) – SPIC_uc_cTSS_oPEAK

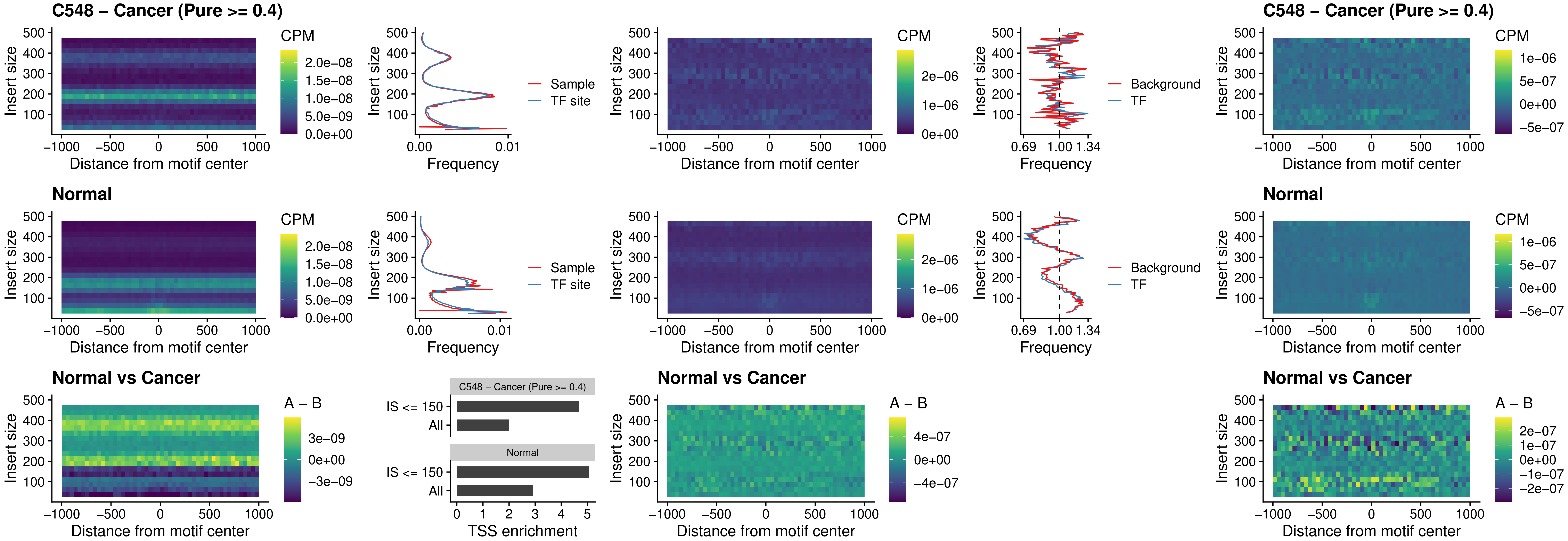

#### C548 – Cancer (Pure >= 0.4) – PRDM1_uc_cTSS_oPEAK

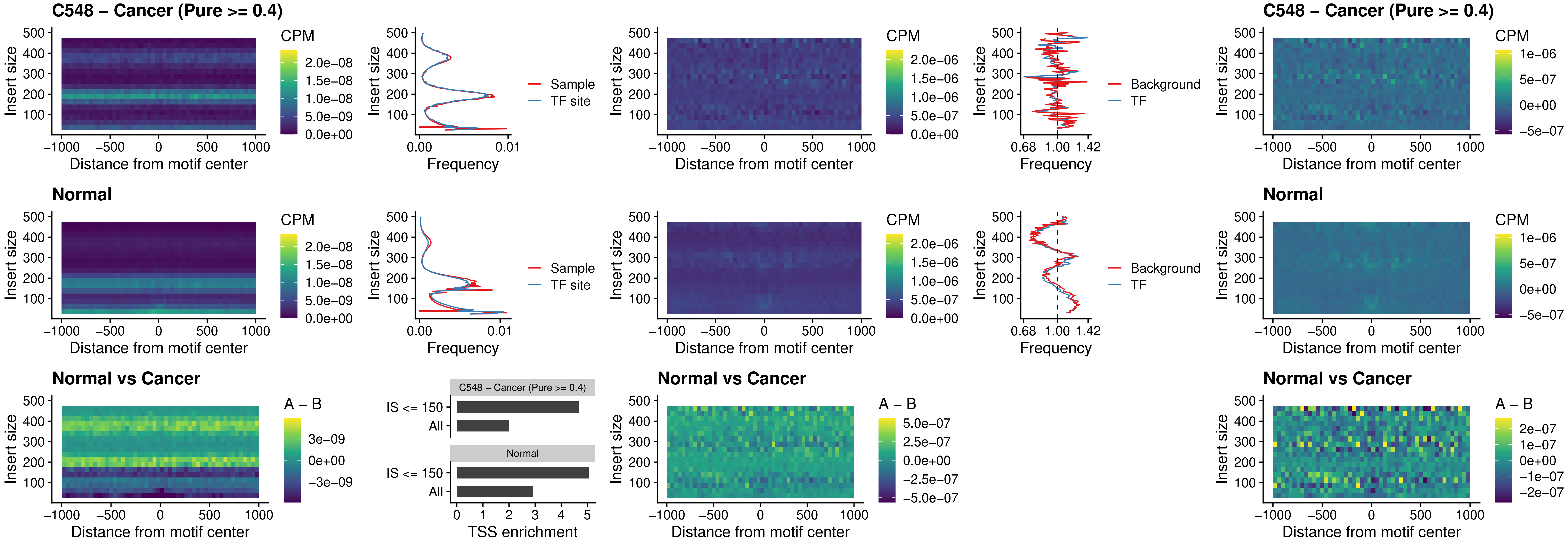

#### C548 – Cancer (Pure >= 0.4) – IRF3_uc_cTSS_oPEAK

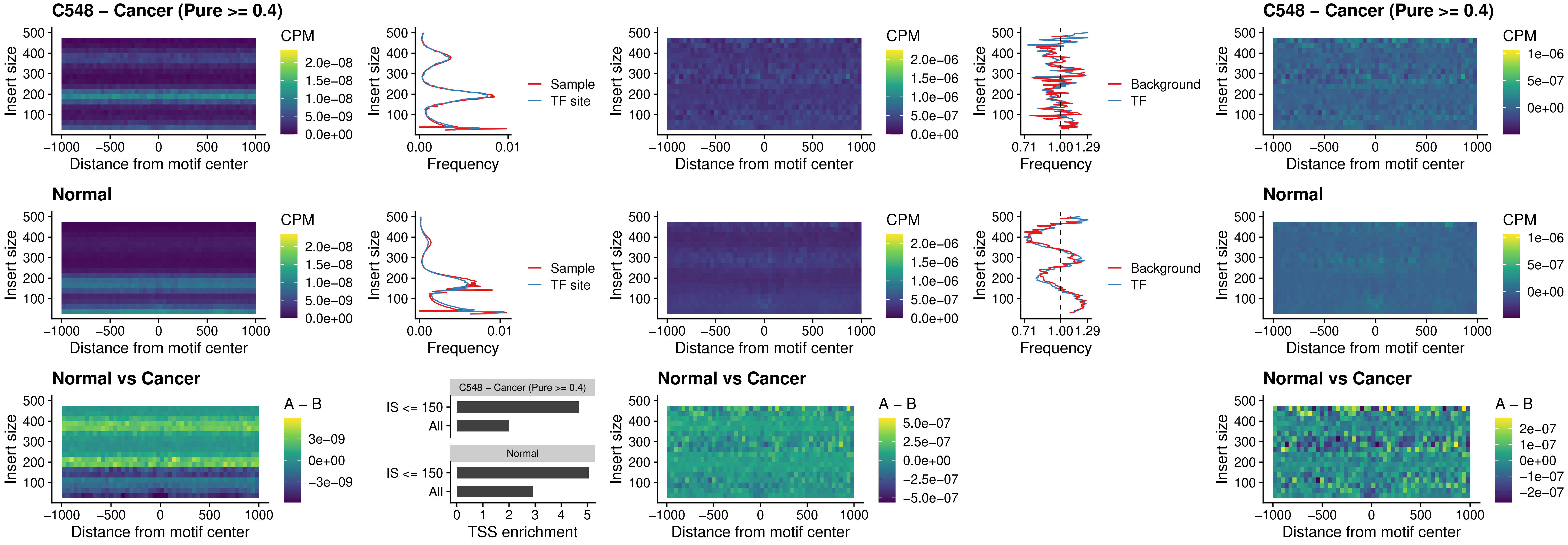

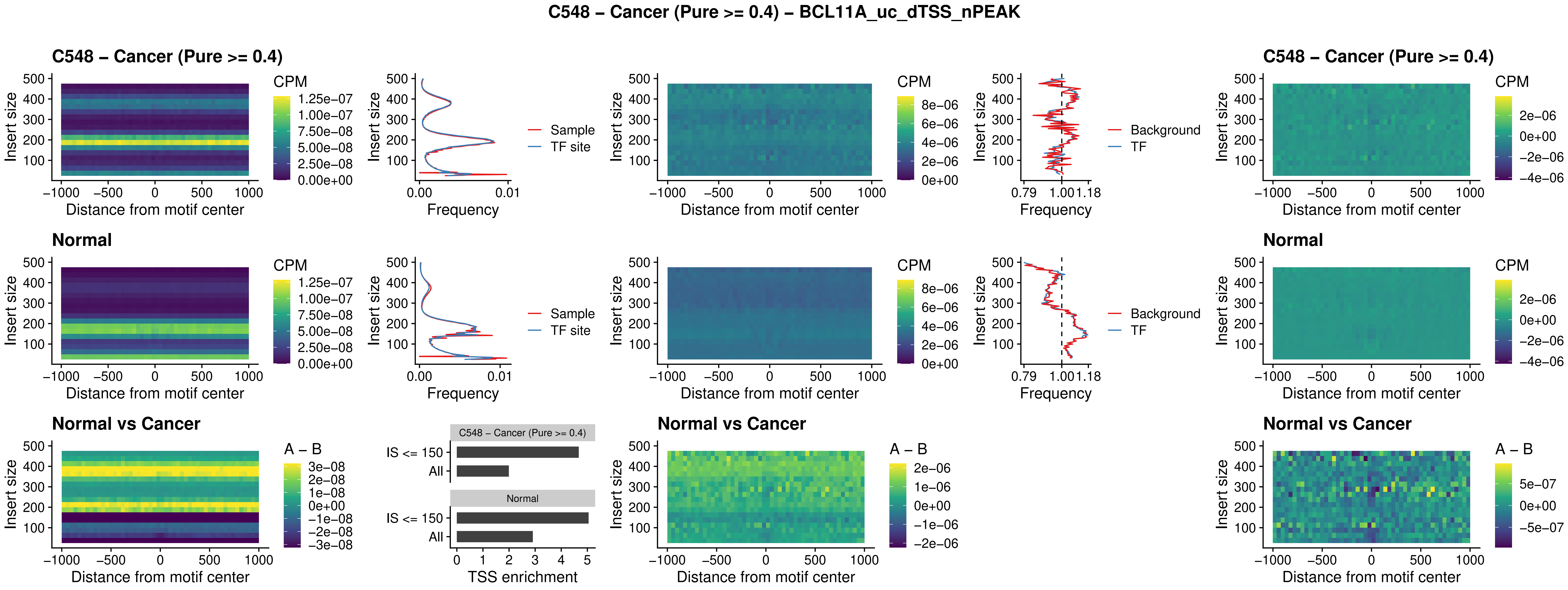
C548 – Cancer (Pure >= 0.4) – BCL11A_uc_dTSS_nPEAK
C548 – Cancer (Pure >= 0.4)
Normal
Normal vs Cancer
Insert size
Distance from motif center
Frequency
CPM
A – B
Sample
TF site
Background
TF
TSS enrichment
IS <= 150
All

#### C548 – Cancer (Pure >= 0.4) – BCL11B_uc_dTSS_nPEAK

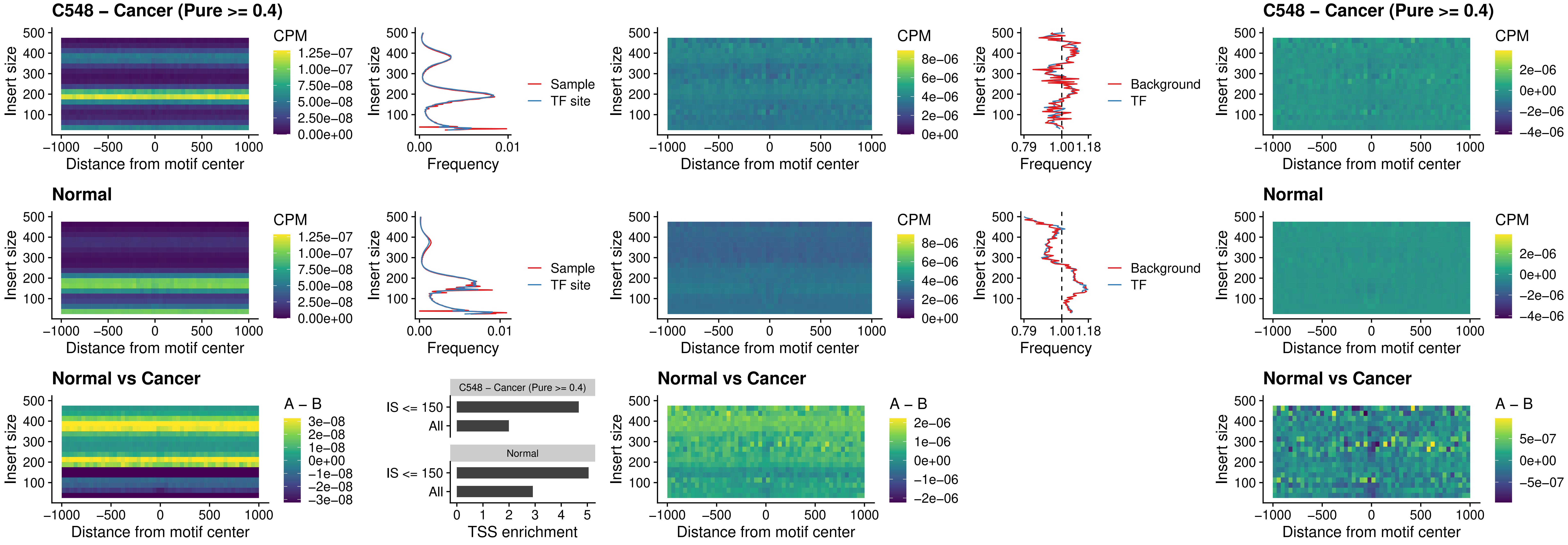

#### C548 – Cancer (Pure >= 0.4) – IRF1_uc_cTSS_oPEAK

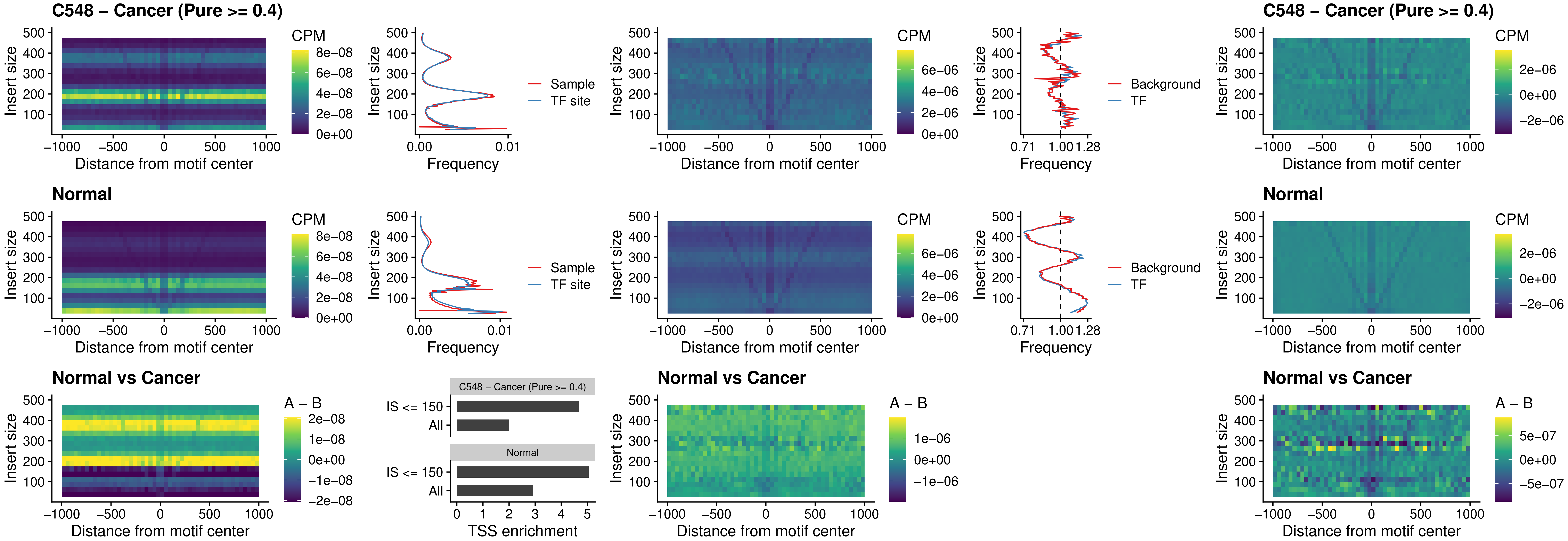

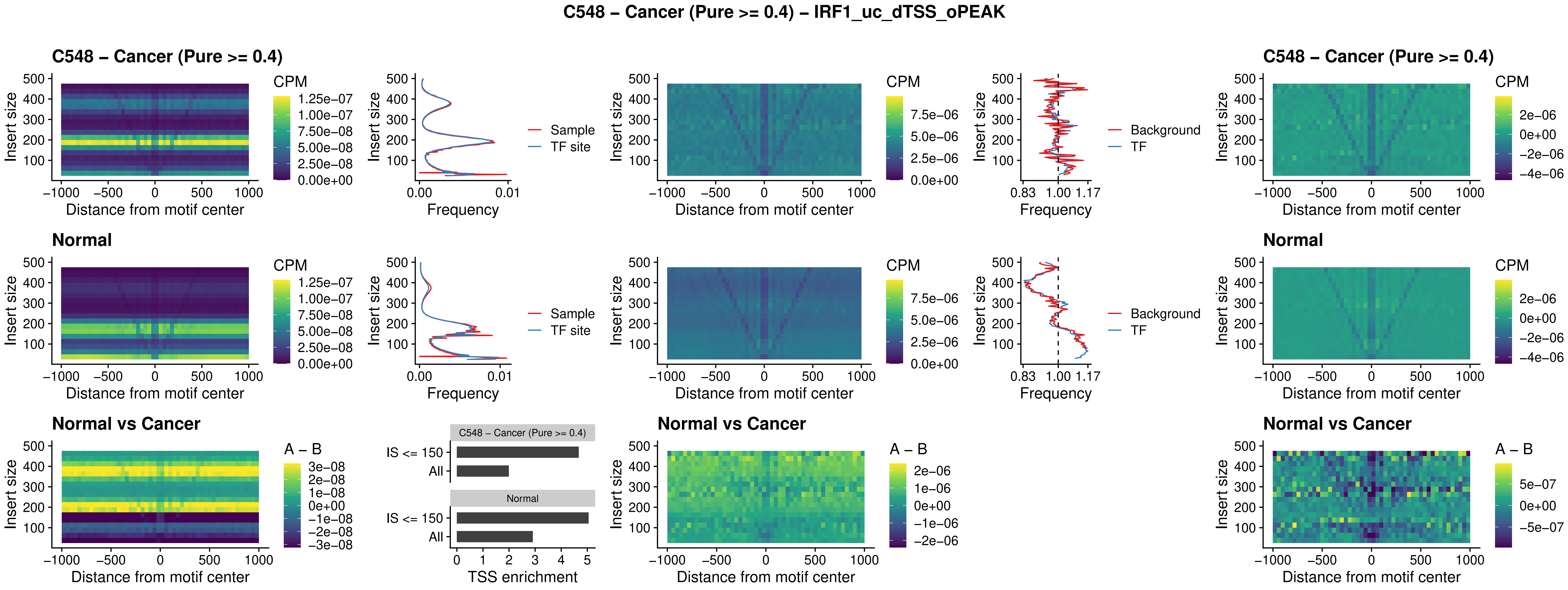
C548 – Cancer (Pure >= 0.4) – IRF1_uc_dTSS_oPEAK
C548 – Cancer (Pure >= 0.4)
Normal
Normal vs Cancer
Insert size
Distance from motif center
Frequency
CPM
A – B
Sample
TF site
Background
TF
C548 – Cancer (Pure >= 0.4)
Normal
IS <= 150
All
TSS enrichment

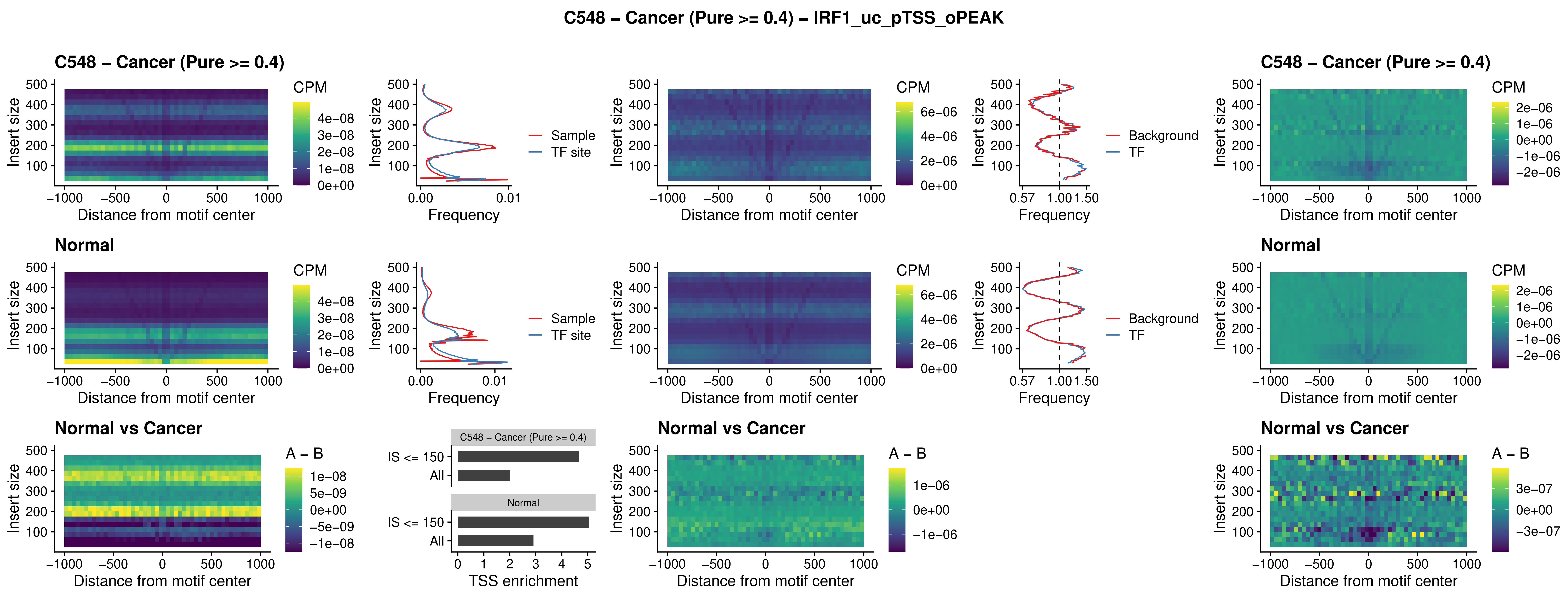
C548 – Cancer (Pure >= 0.4) – IRF1_uc_pTSS_oPEAK
C548 – Cancer (Pure >= 0.4)
Normal
Normal vs Cancer
Insert size
Distance from motif center
Frequency
CPM
A – B
Sample
TF site
Background
TF
C548 – Cancer (Pure >= 0.4)
Normal
IS <= 150
All
TSS enrichment

#### C548 – Cancer (Pure >= 0.4) – SPIC_uc_dTSS_nPEAK

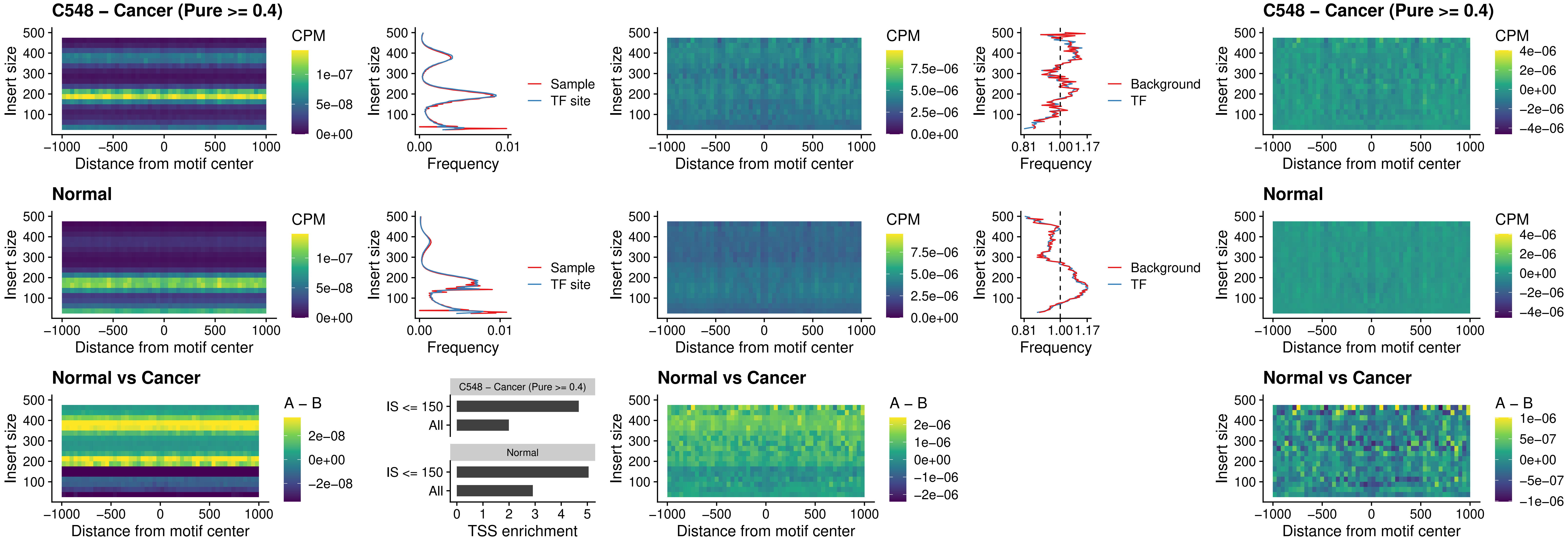

#### C548 – Cancer (Pure >= 0.4) – SPIC_uc_dTSS_oPEAK

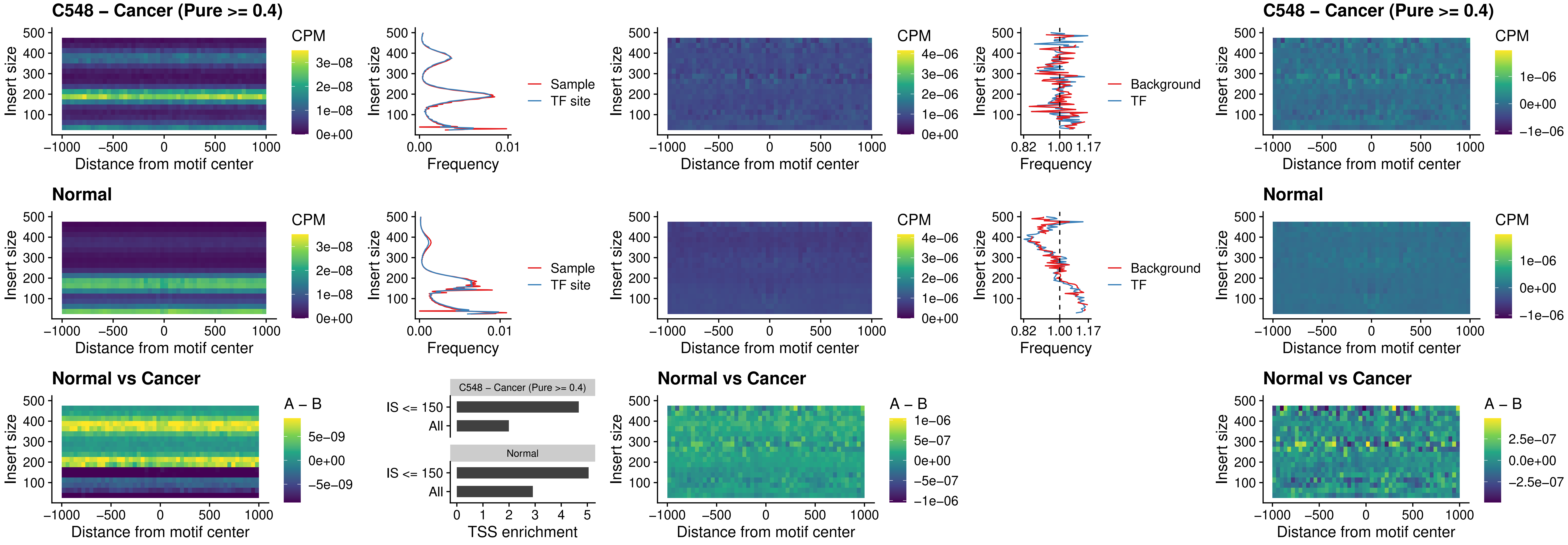

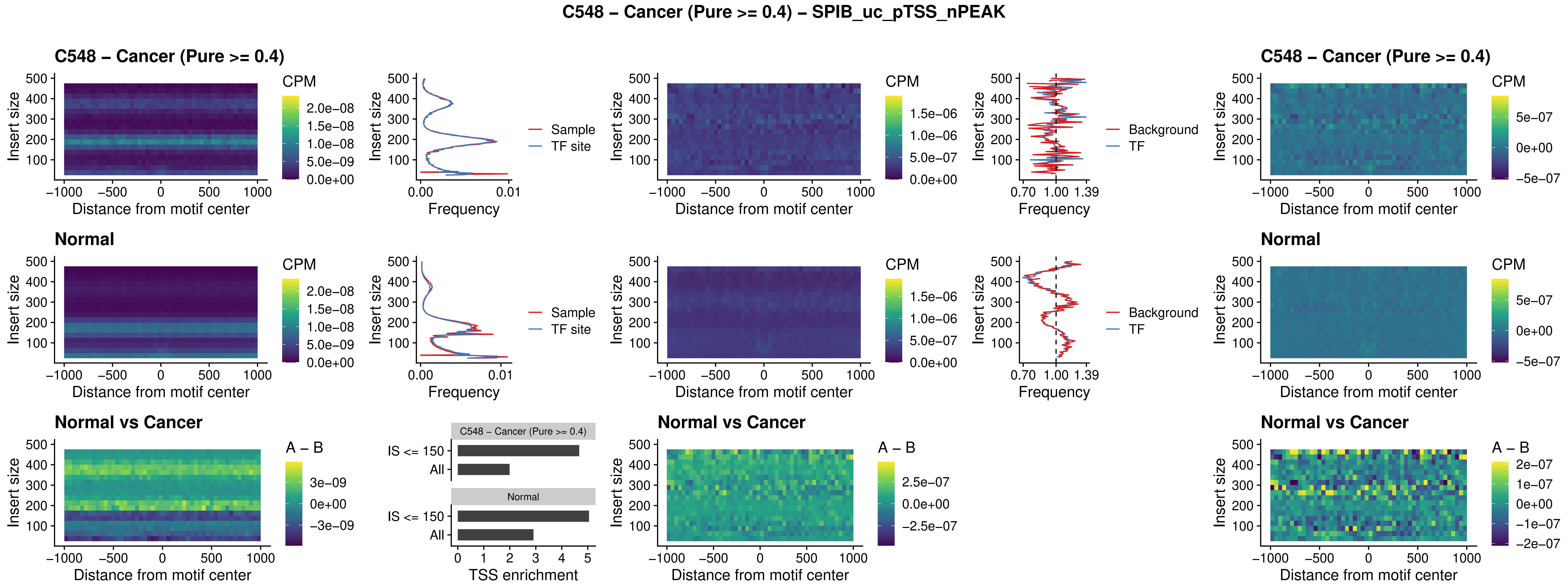
C548 – Cancer (Pure >= 0.4) – SPIB_uc_pTSS_nPEAK
C548 – Cancer (Pure >= 0.4)
Normal
Normal vs Cancer
Insert size
Distance from motif center
Frequency
CPM
A – B
Sample
TF site
Background
TF
C548 – Cancer (Pure >= 0.4)
Normal
IS <= 150
All
TSS enrichment

#### C548 – Cancer (Pure >= 0.4) – STAT2_uc_cTSS_oPEAK

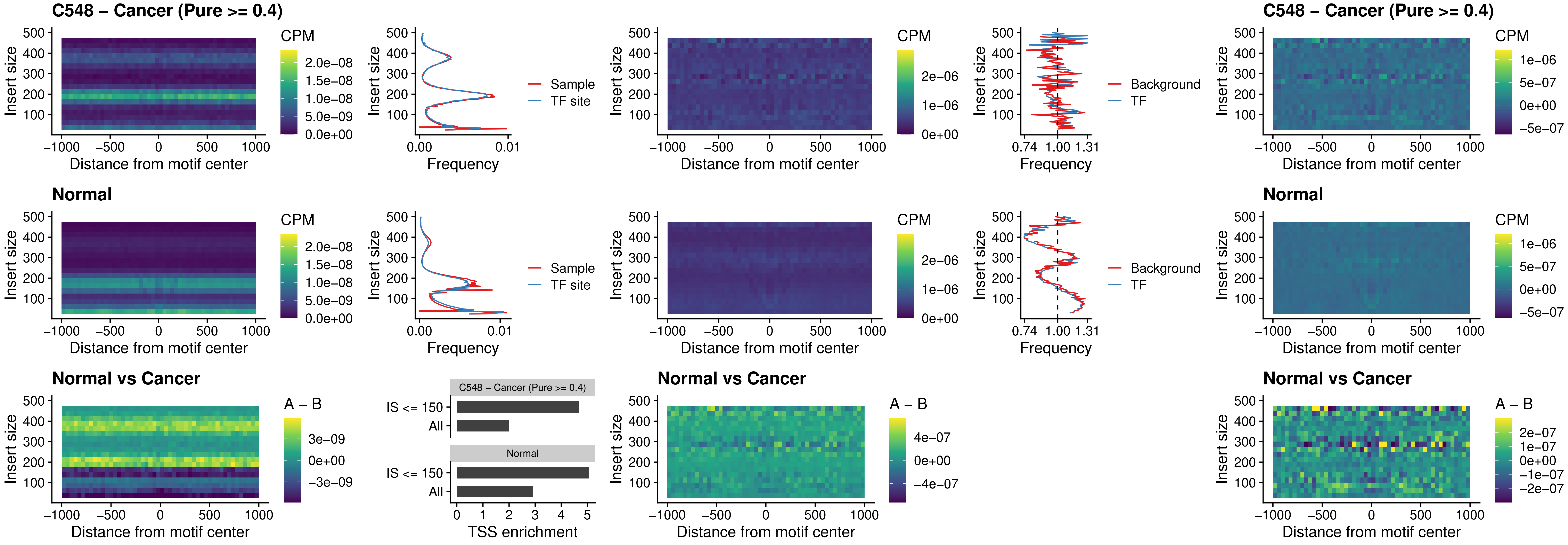

#### C548 – Cancer (Pure >= 0.4) – IRF8_uc_cTSS_oPEAK

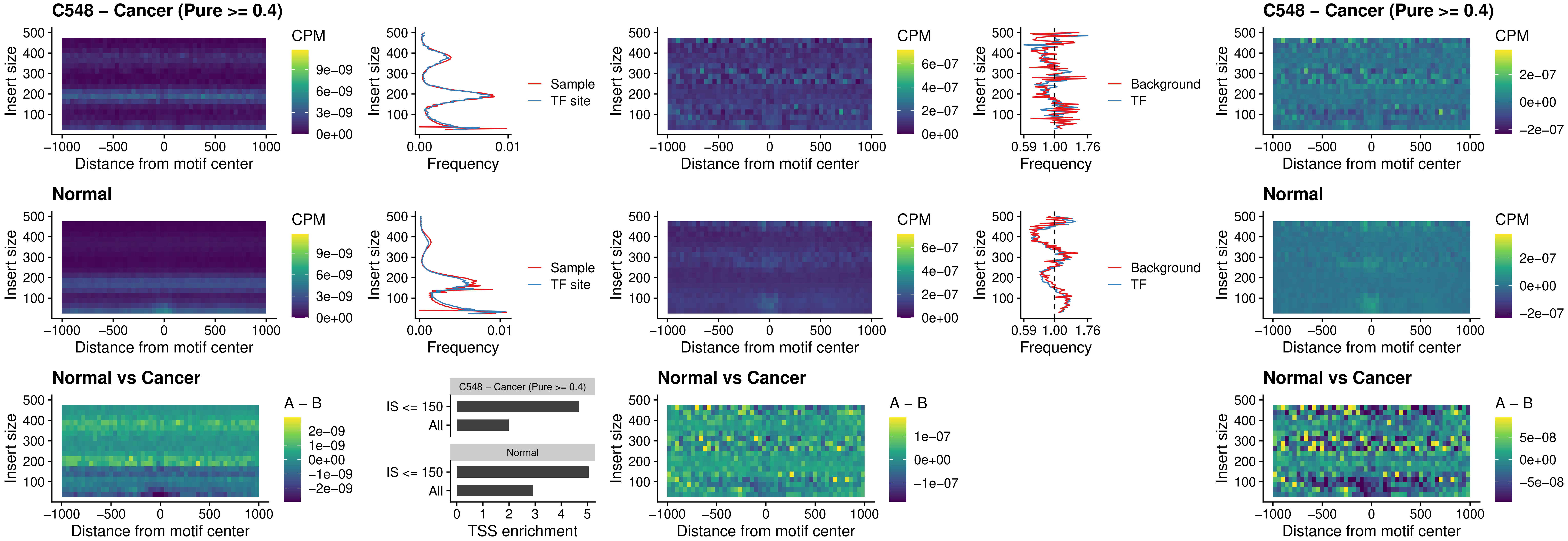

### C548 – Cancer (Pure >= 0.4) – IRF3_uc_dTSS_oPEAK

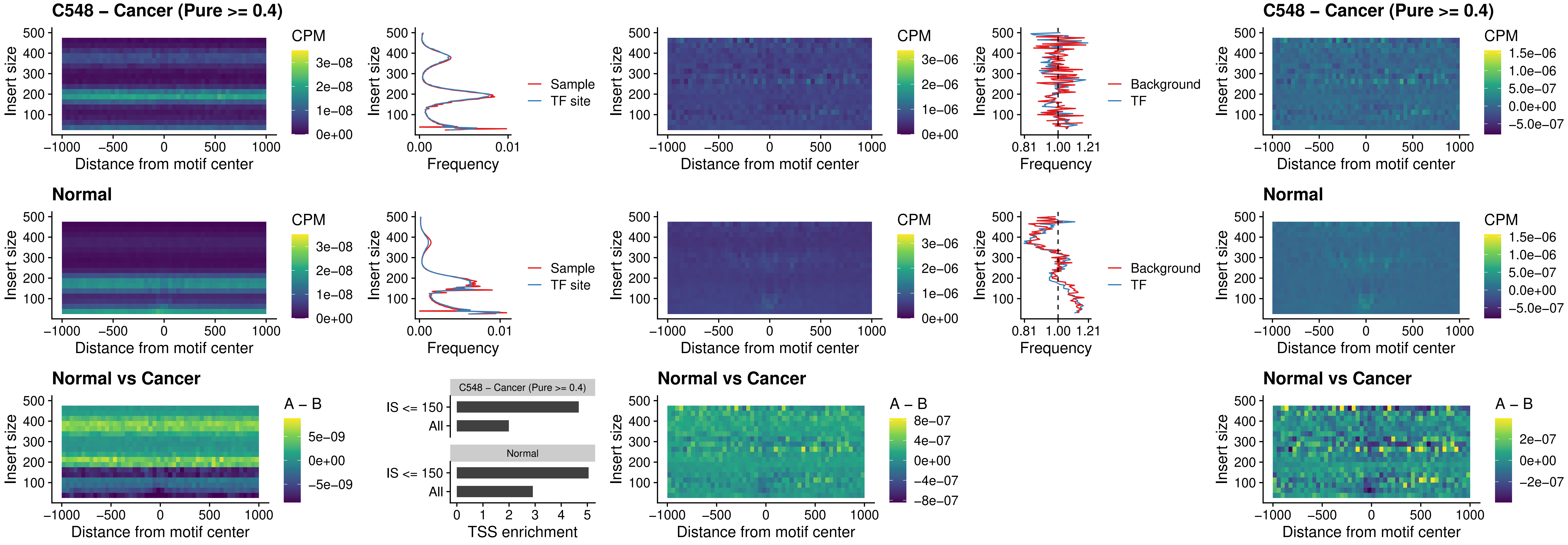

#### C548 – Cancer (Pure >= 0.4) – IRF8_uc_dTSS_oPEAK

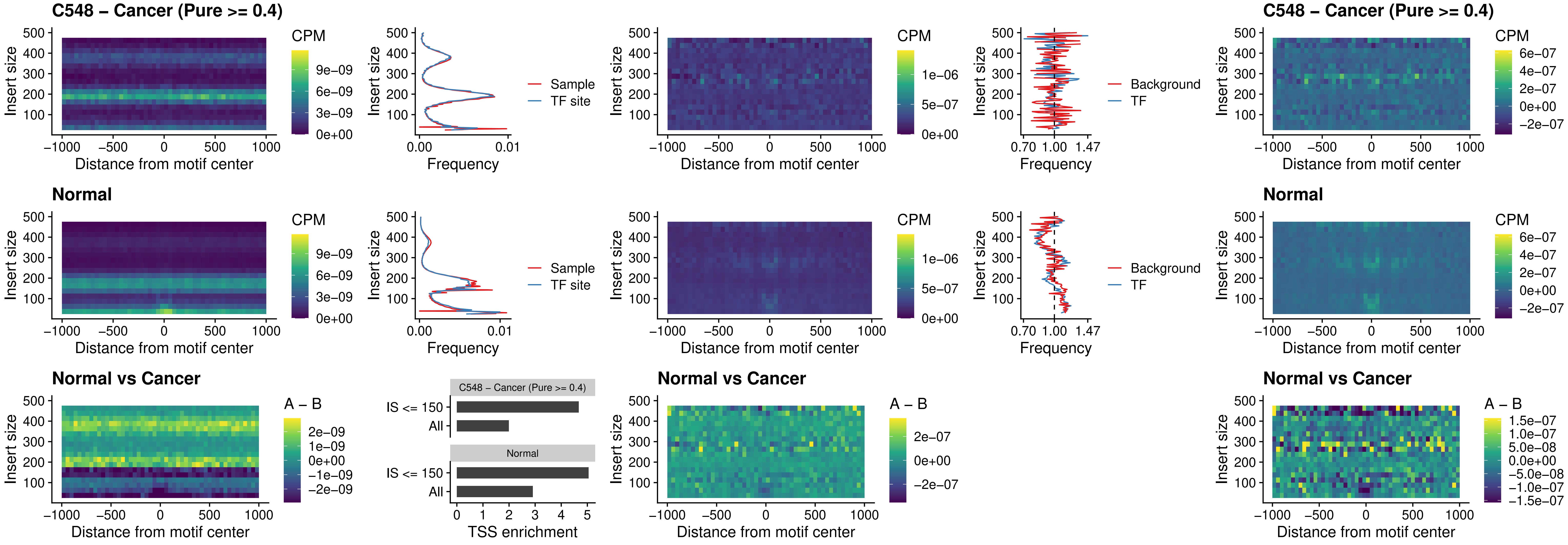

### C548 – Cancer (Pure >= 0.4) – ETS2_uc_dTSS_oPEAK

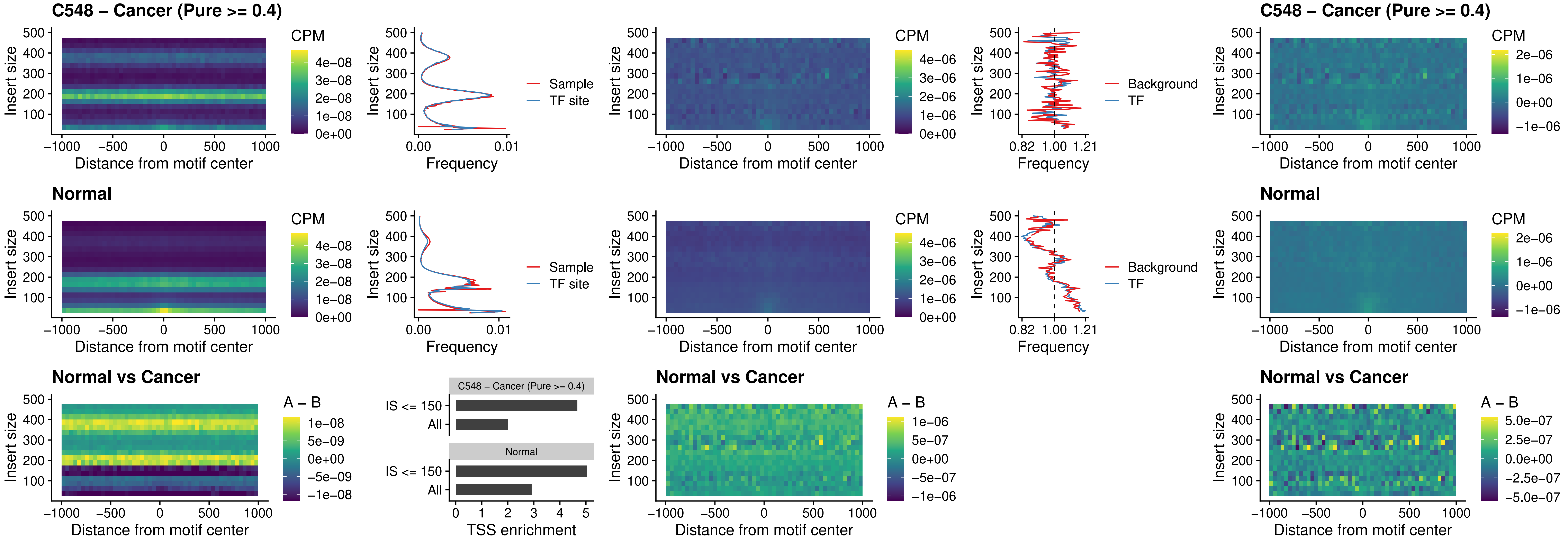

#### C548 – Cancer (Pure >= 0.4) – SMARCC2_uc_pTSS_oPEAK

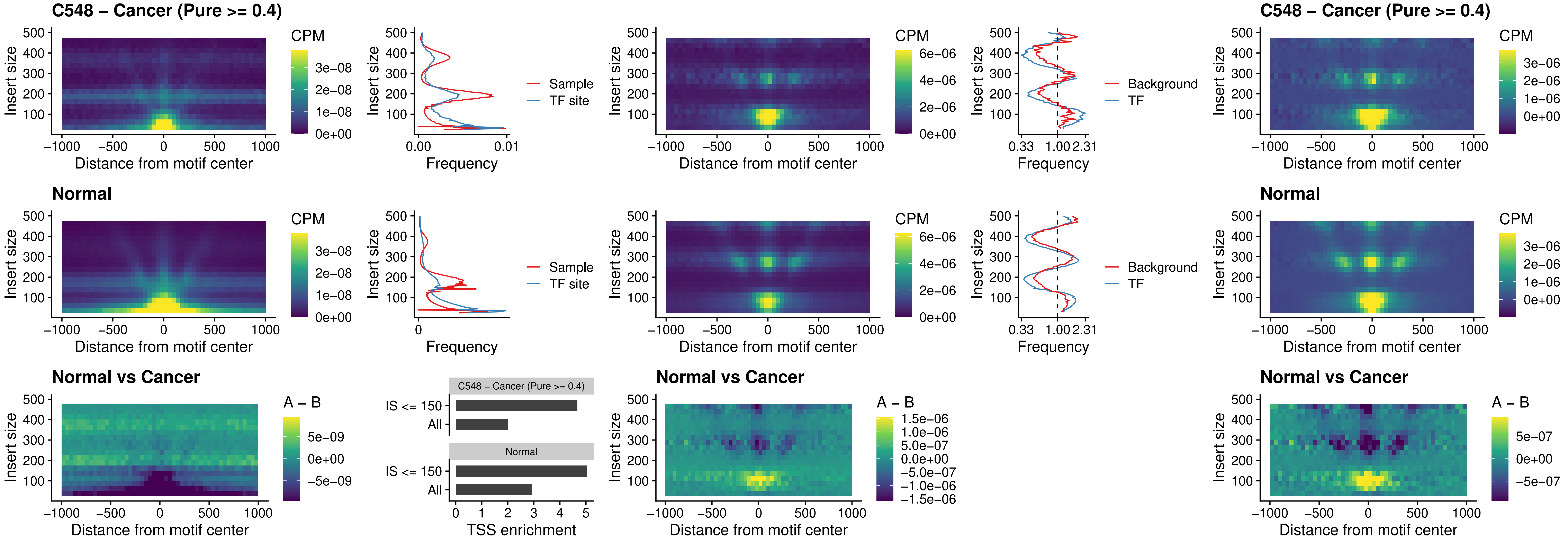

#### C548 – Cancer (Pure >= 0.4) – BCL11A_uc_cTSS_nPEAK

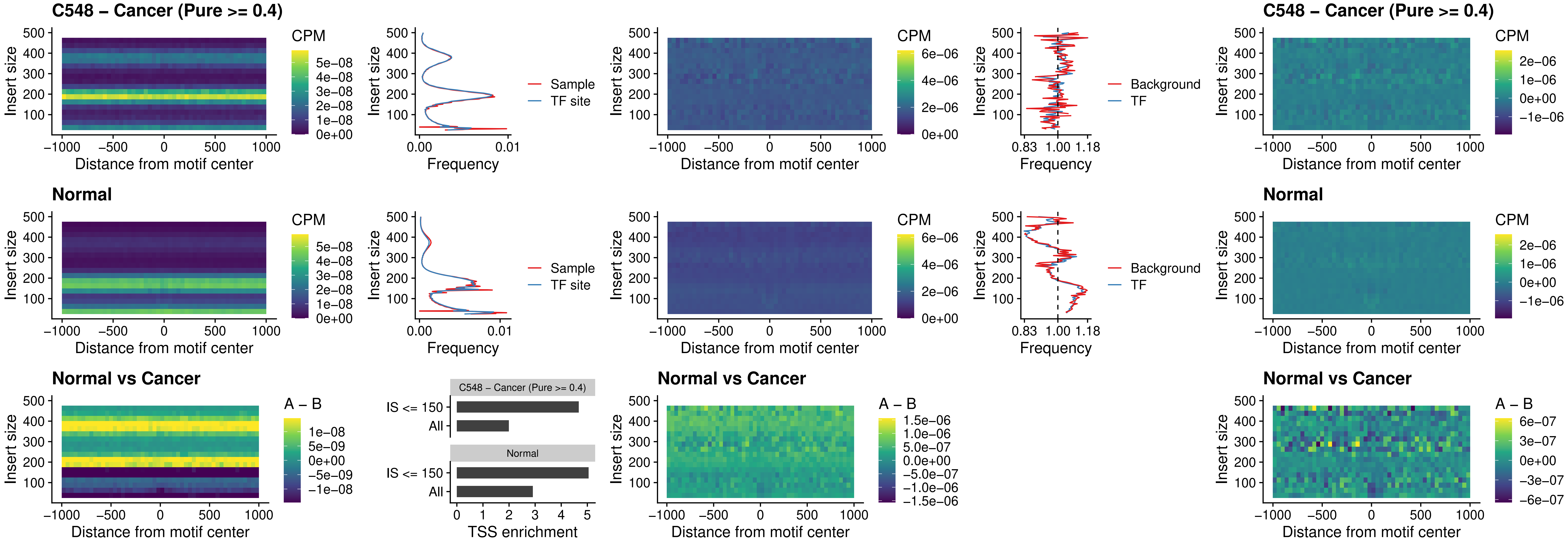

#### C548 – Cancer (Pure >= 0.4) – BCL11B_uc_cTSS_nPEAK

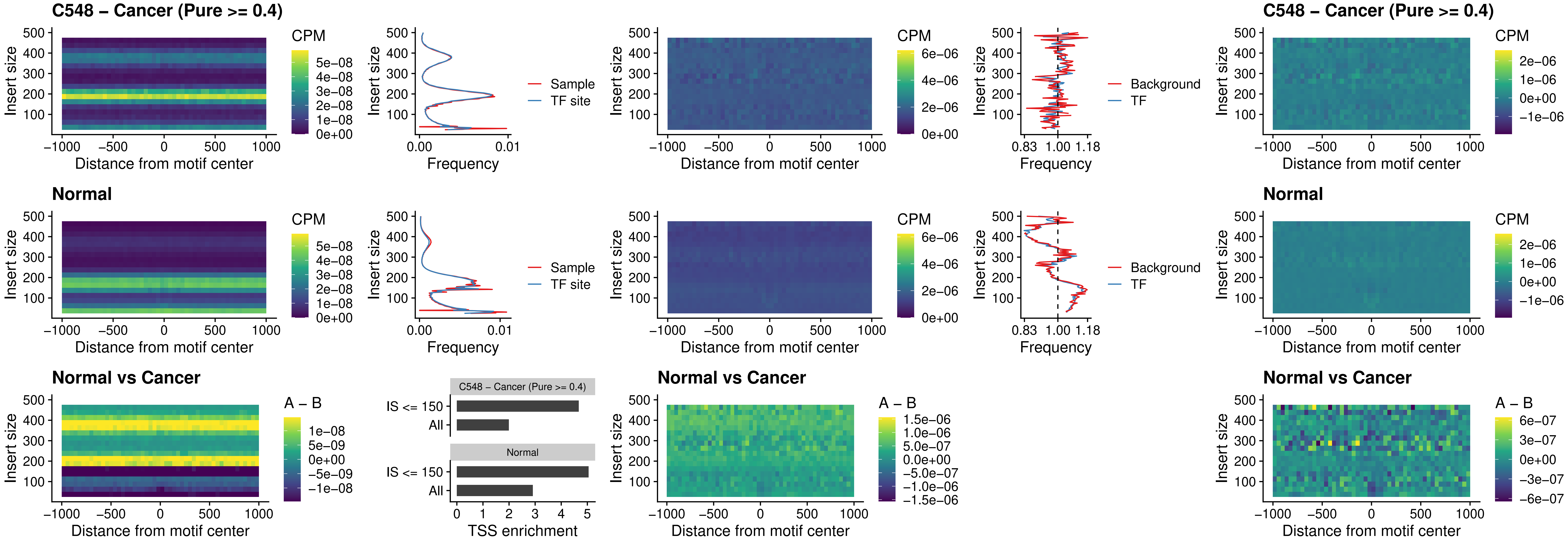

### C548 – Cancer (Pure >= 0.4) – STAT2_uc_dTSS_oPEAK

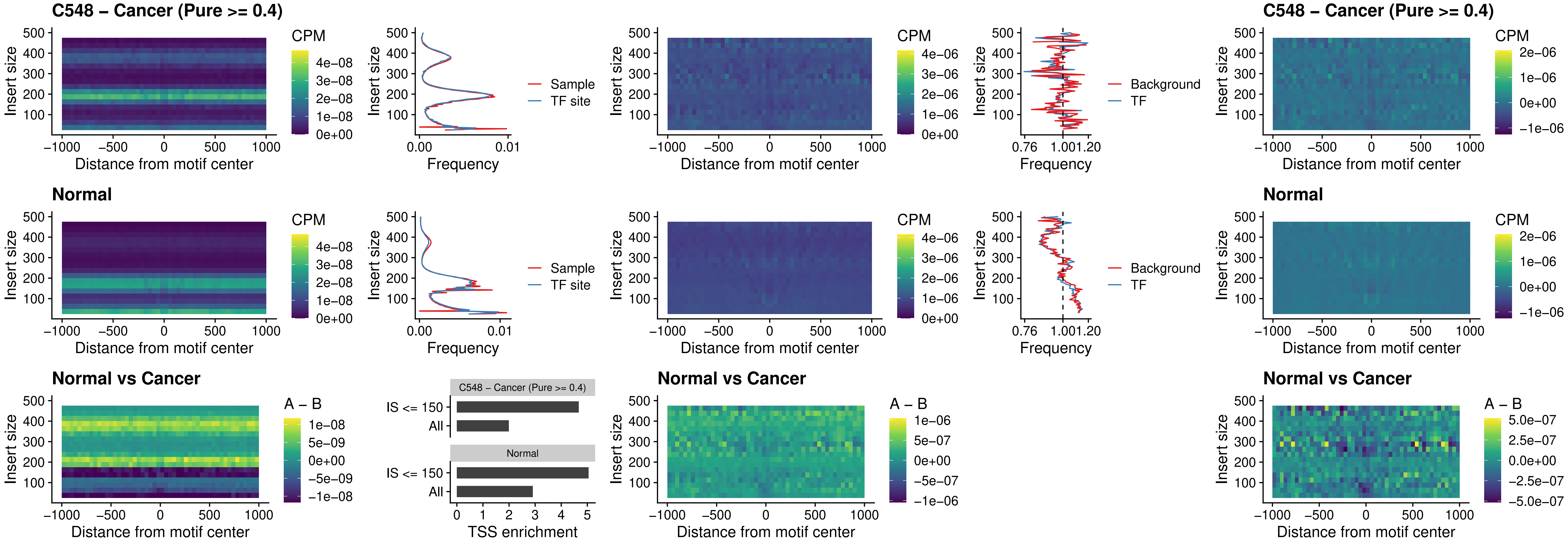

#### C548 – Cancer (Pure >= 0.4) – ZNF683_uc_cTSS_oPEAK

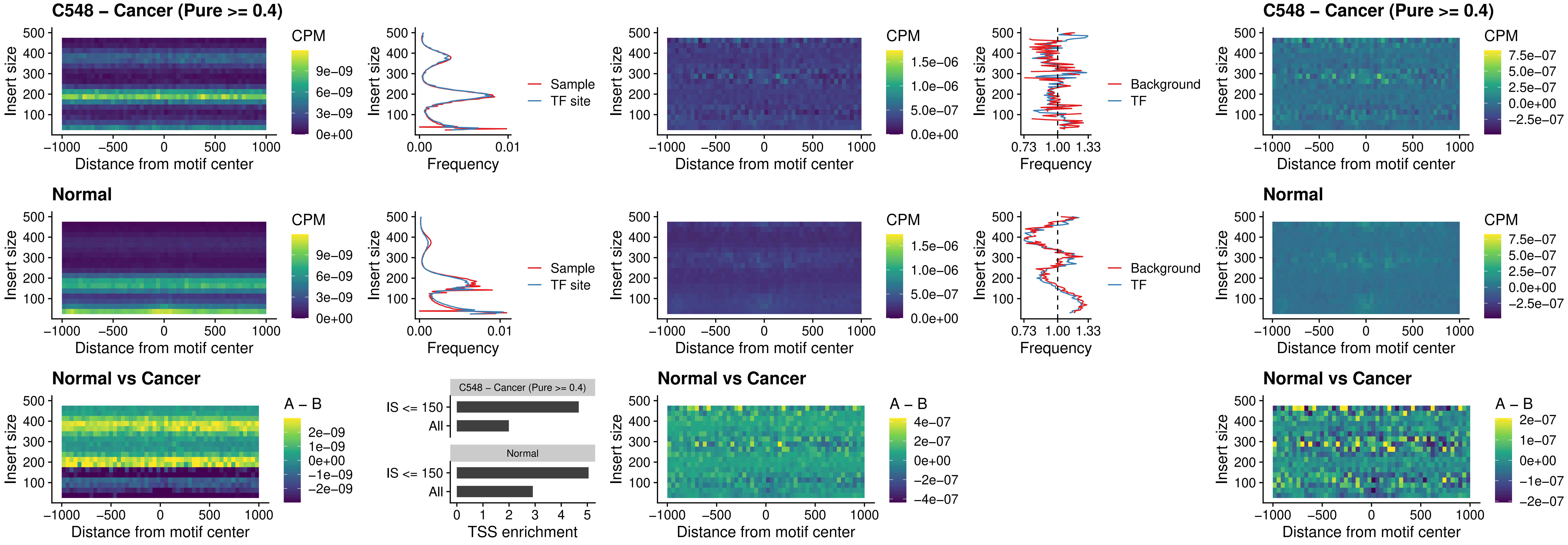

### C548 – Cancer (Pure >= 0.4) – IRF8_uc_dTSS_nPEAK

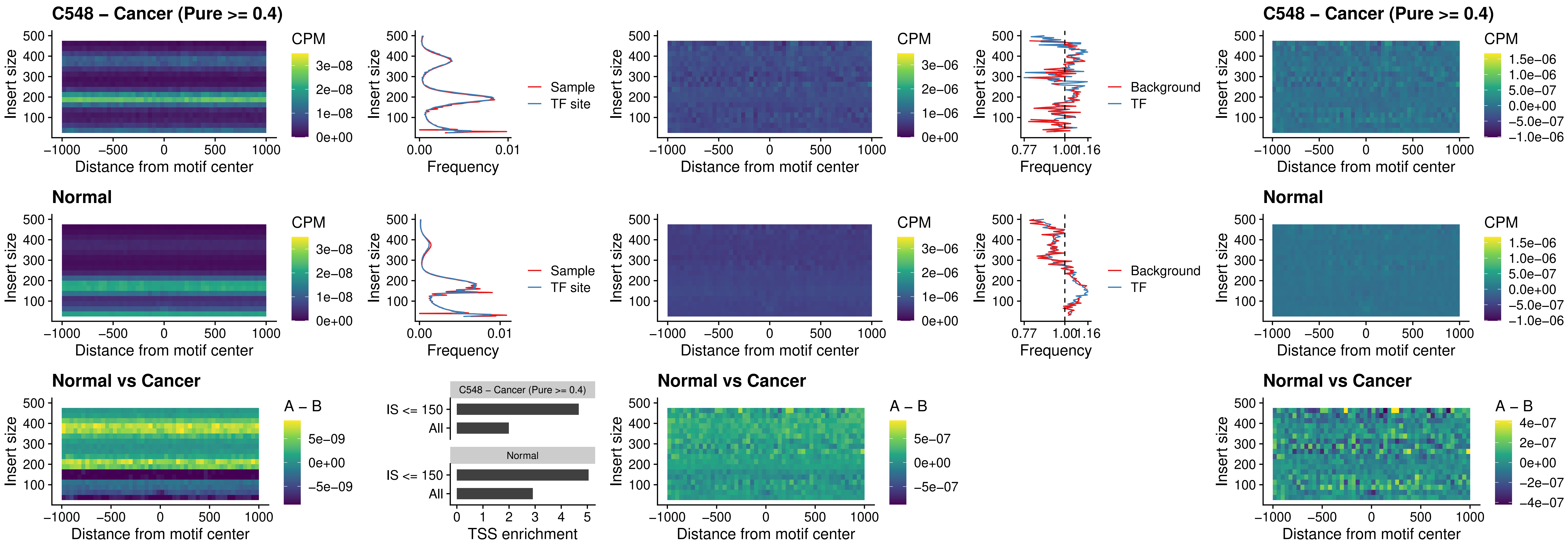

#### C548 – Cancer (Pure >= 0.4) – ETV7_uc_dTSS_oPEAK

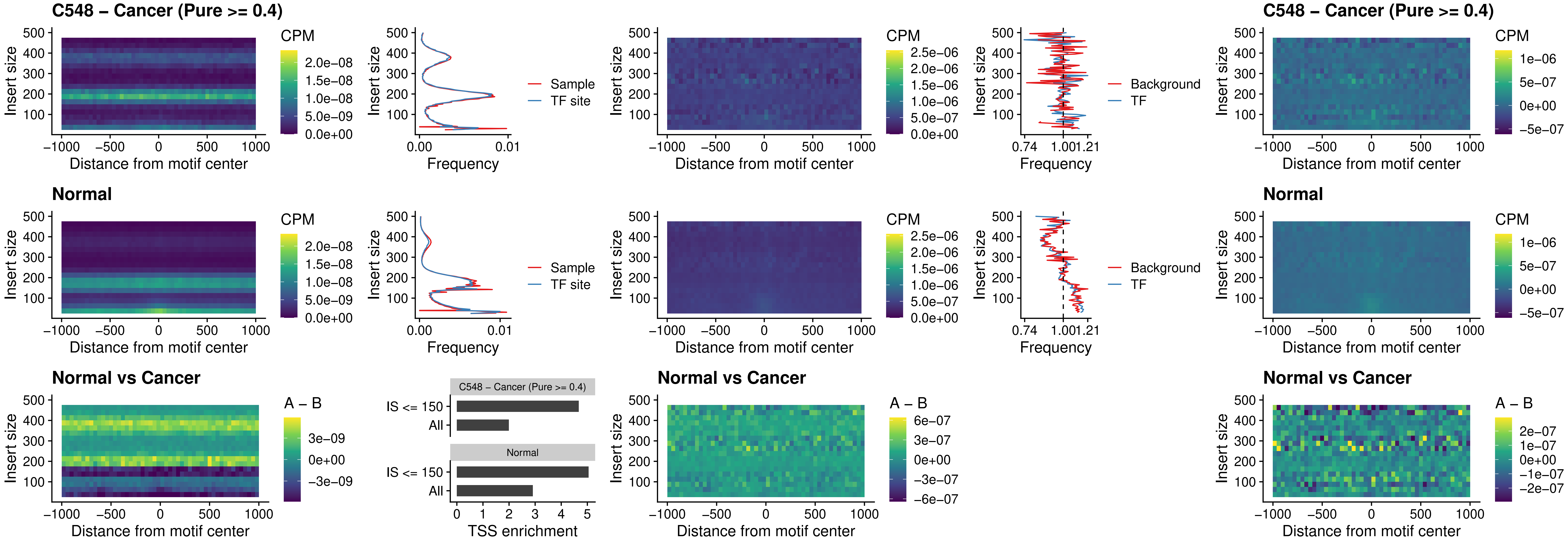

#### C550 – Cancer (Pure >= 0.4) – FOXL1_uc_pTSS_oPEAK

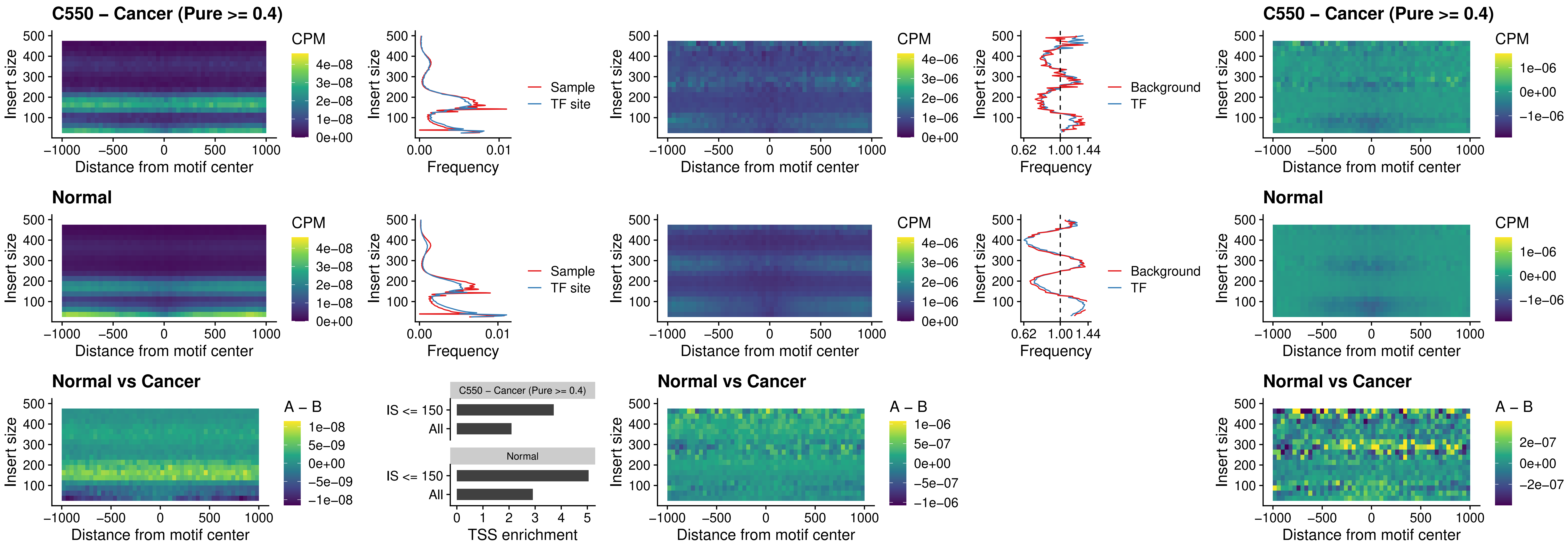

#### C550 – Cancer (Pure >= 0.4) – CPEB1_uc_cTSS_oPEAK

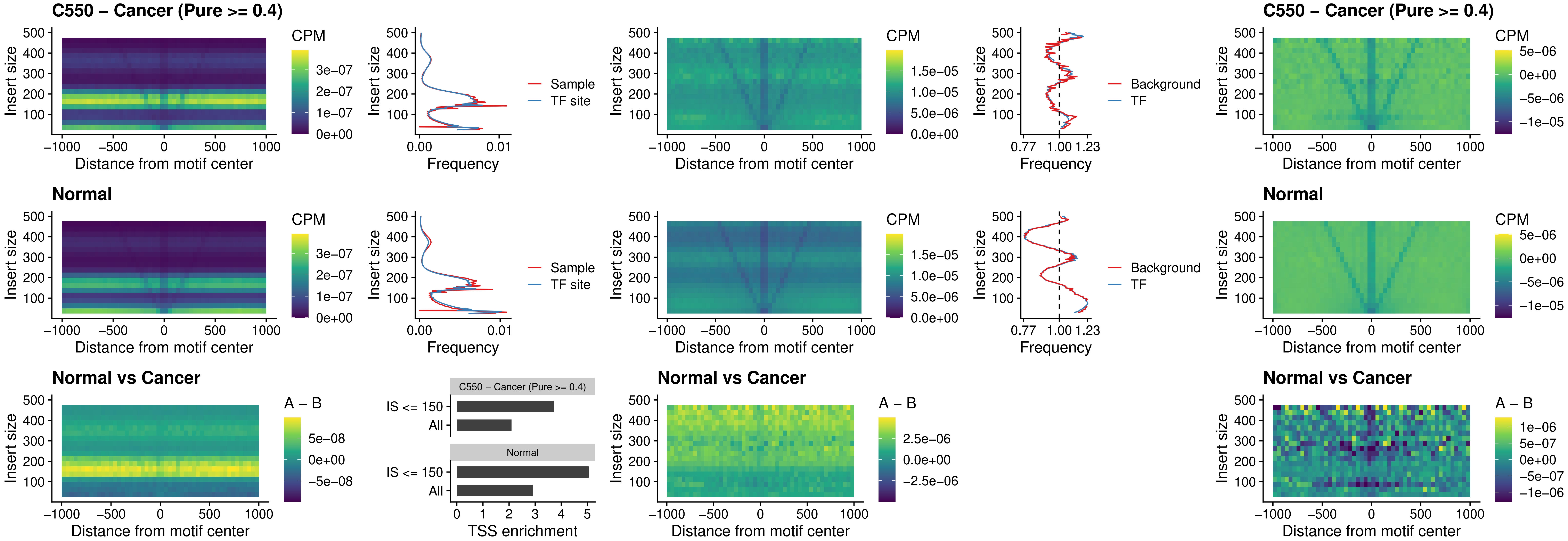

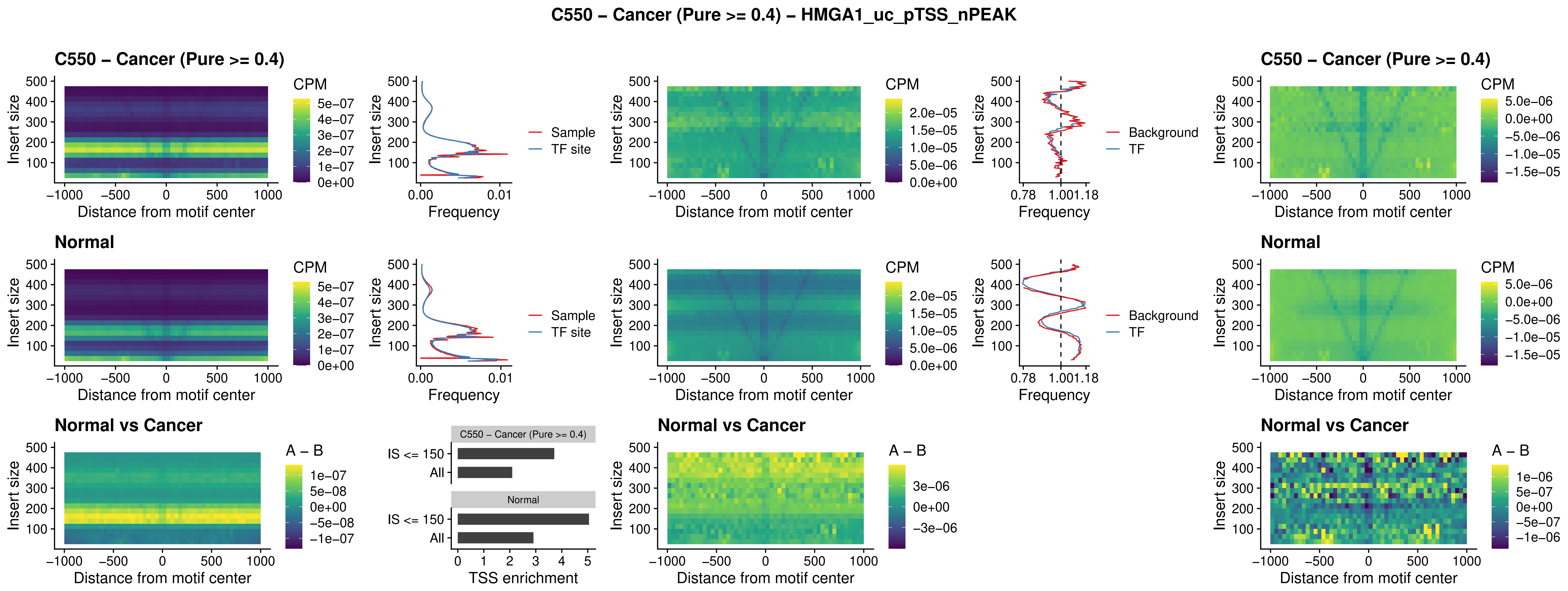
C550 – Cancer (Pure >= 0.4) – HMGA1_uc_pTSS_nPEAK
C550 – Cancer (Pure >= 0.4)
Normal
Normal vs Cancer
Insert size
Distance from motif center
CPM
A – B
Frequency
Sample
TF site
Background
TF
C550 – Cancer (Pure >= 0.4)
Normal
IS <= 150
All
TSS enrichment

### C550 – Cancer (Pure >= 0.4) – SOX13_uc_cTSS_nPEAK

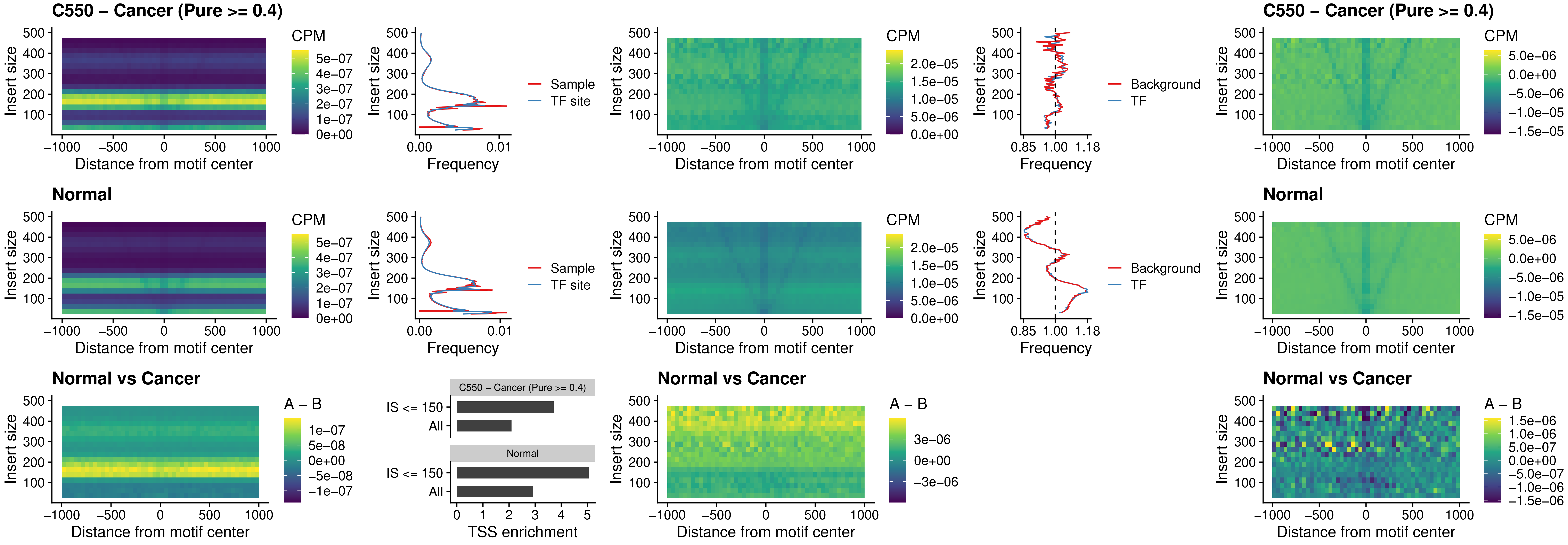

#### C550 – Cancer (Pure >= 0.4) – LHX4_uc_cTSS_nPEAK

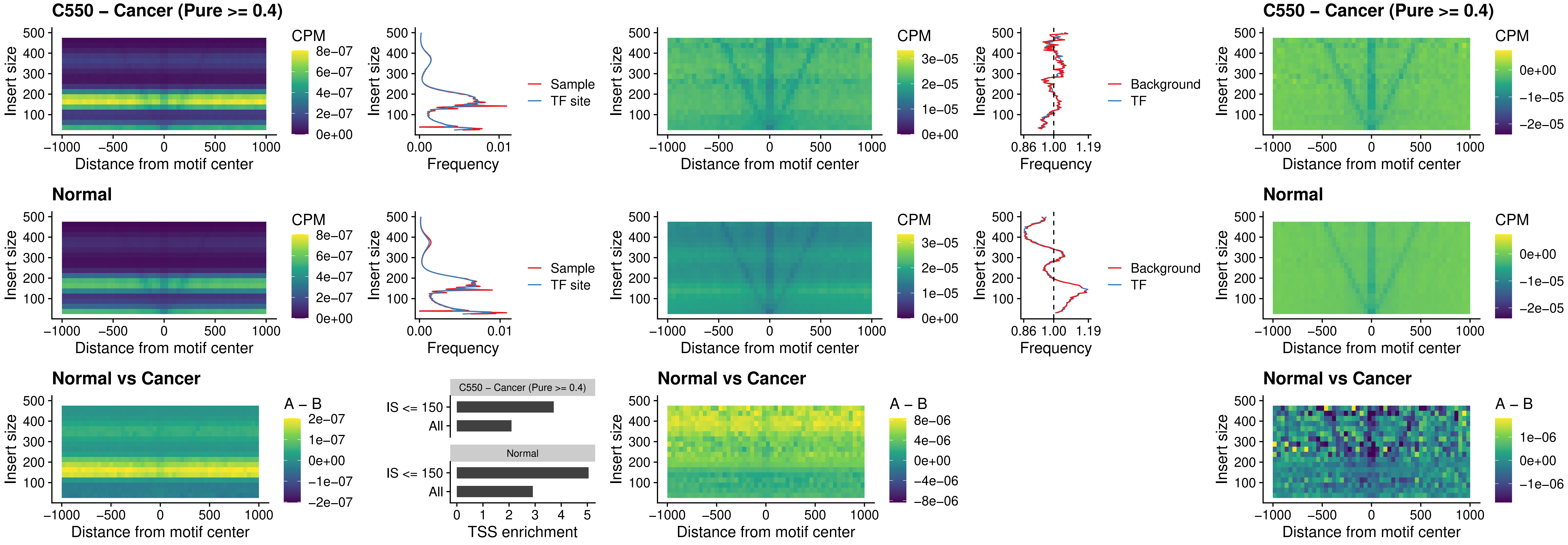

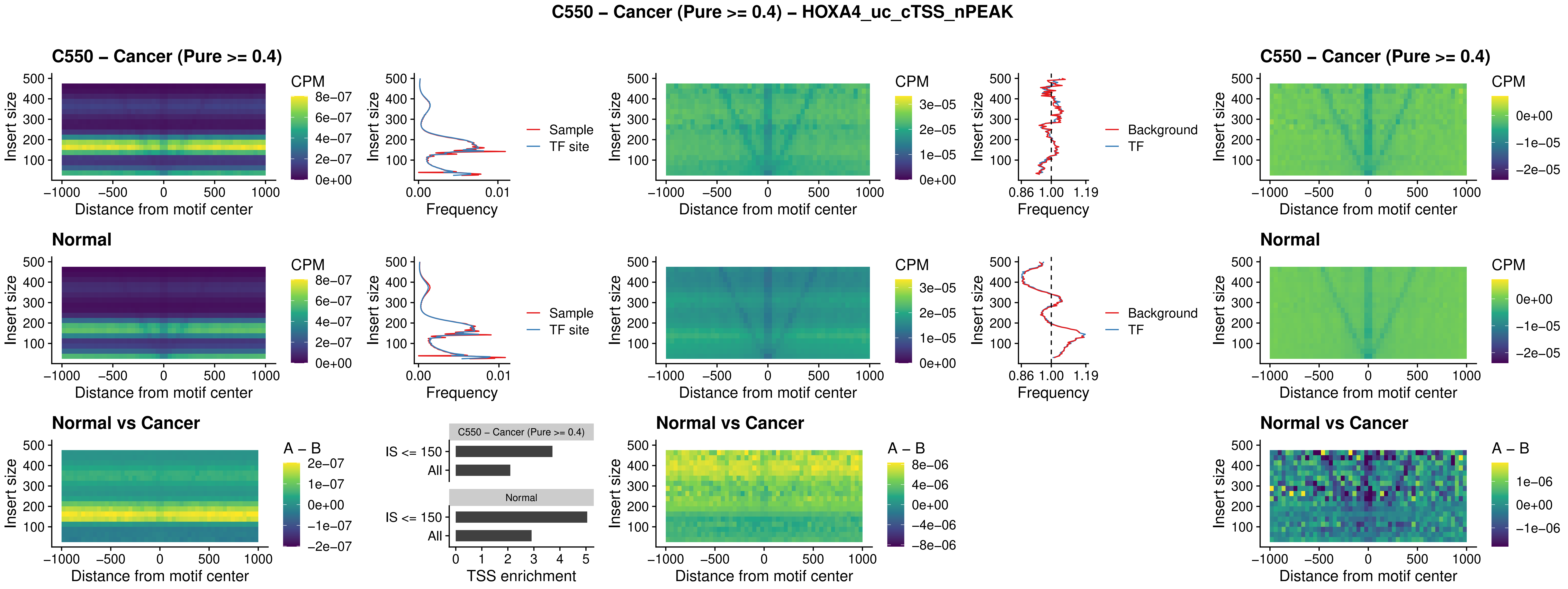
C550 – Cancer (Pure >= 0.4) – HOXA4_uc_cTSS_nPEAK
C550 – Cancer (Pure >= 0.4)
Normal
Normal vs Cancer
Insert size
Distance from motif center
Frequency
CPM
A – B
Sample
TF site
Background
TF
C550 – Cancer (Pure >= 0.4)
Normal
IS <= 150
All
TSS enrichment

#### C550 – Cancer (Pure >= 0.4) – HLX_uc_cTSS_nPEAK

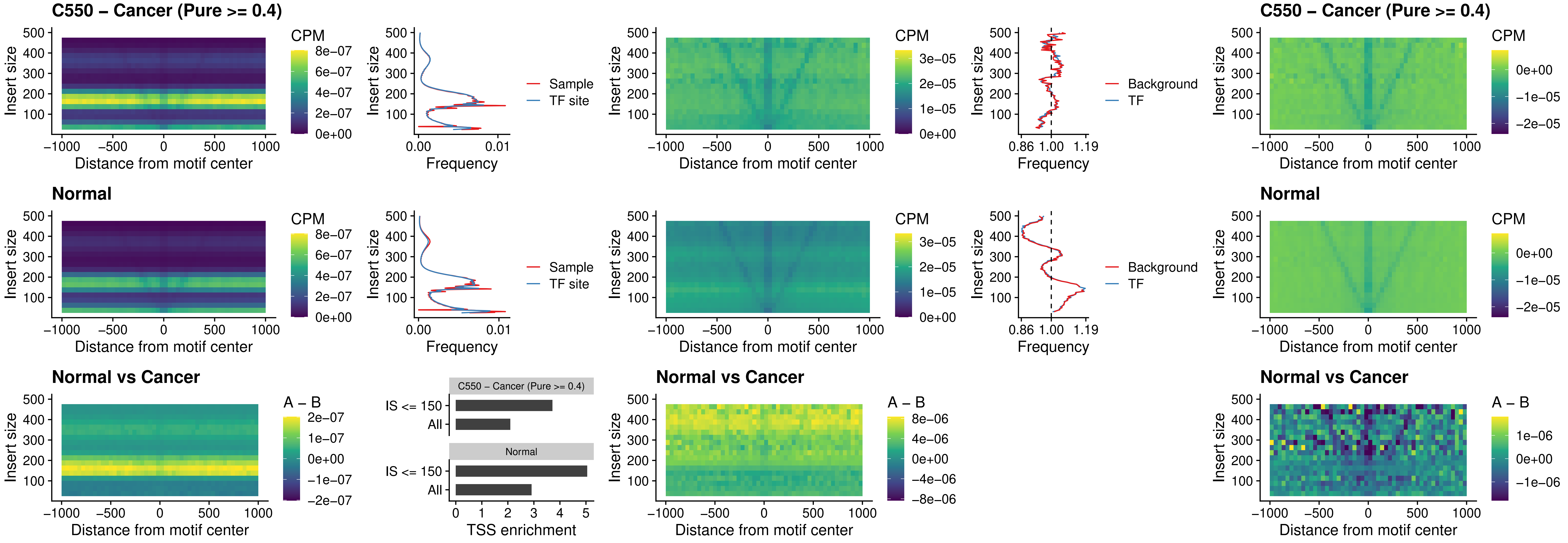

#### C550 – Cancer (Pure >= 0.4) – AC0125311_uc_cTSS_nPEAK

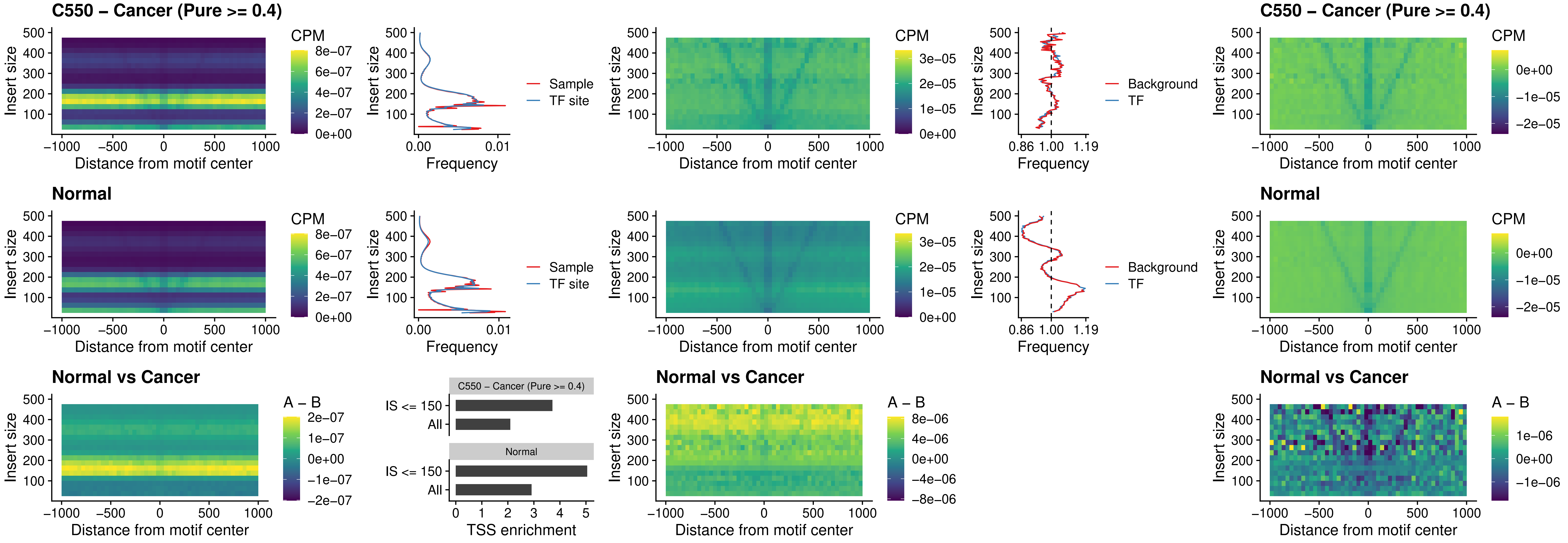

#### C550 – Cancer (Pure >= 0.4) – DBX2_uc_cTSS_nPEAK

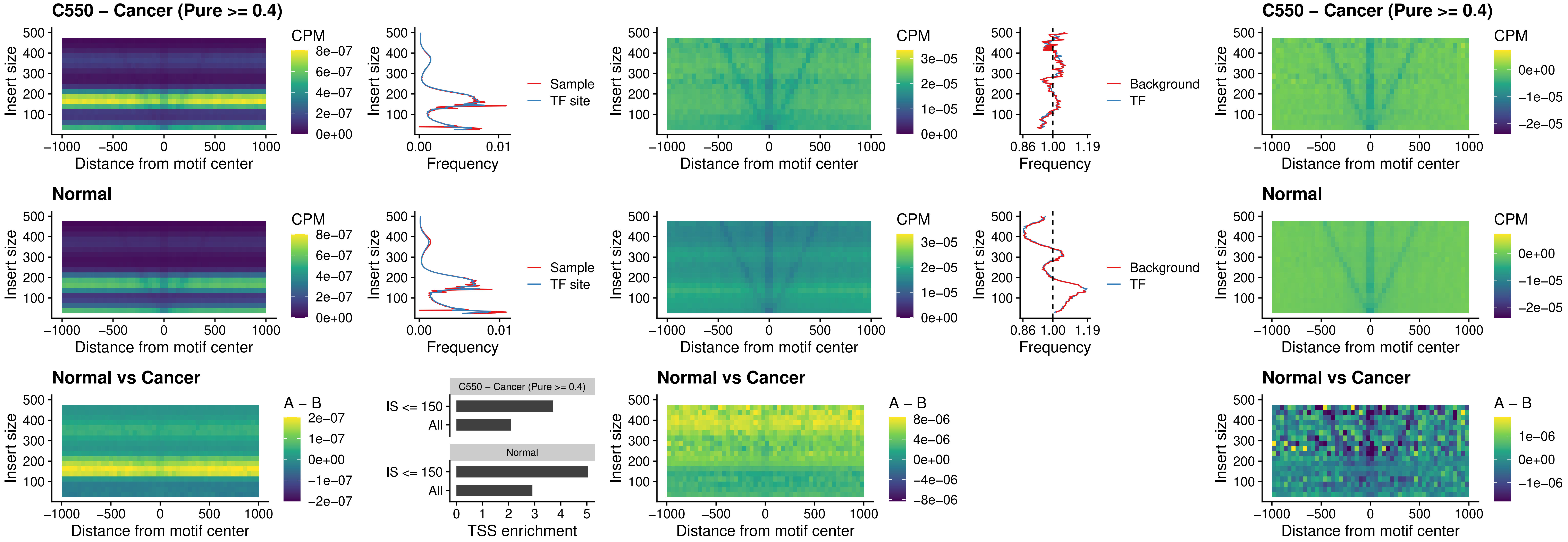

#### C550 – Cancer (Pure >= 0.4) – LHX6_uc_dTSS_nPEAK

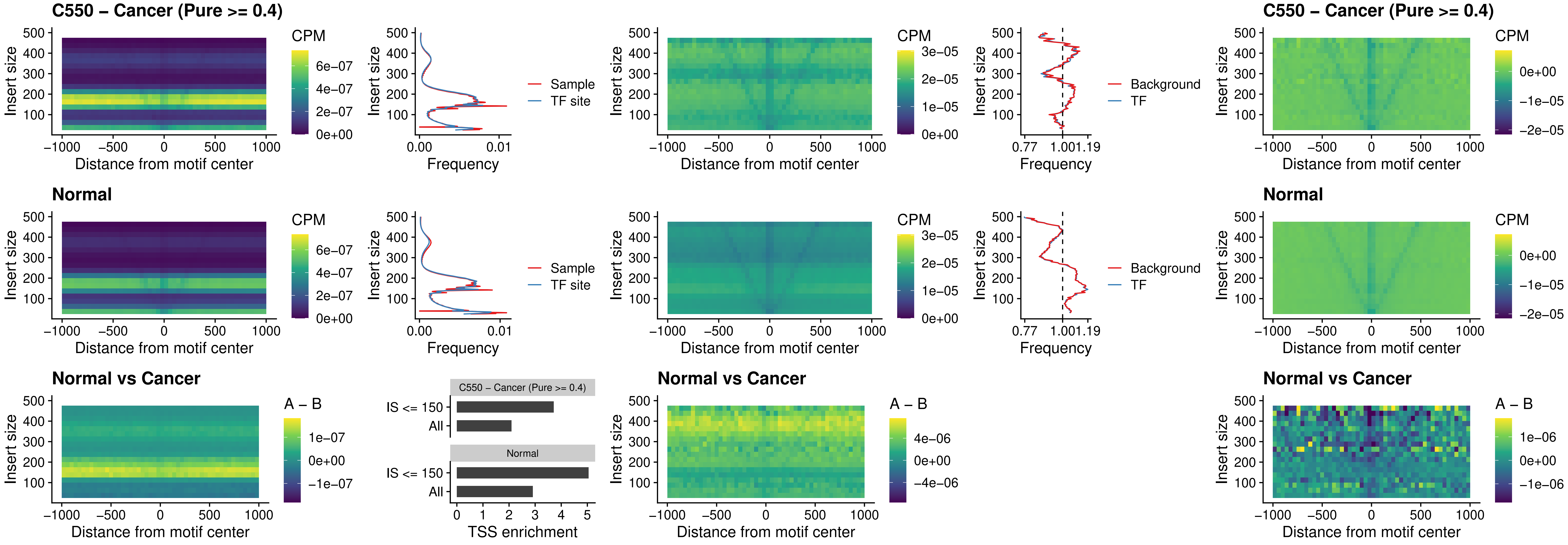

#### C550 – Cancer (Pure >= 0.4) – SOX1_uc_dTSS_nPEAK

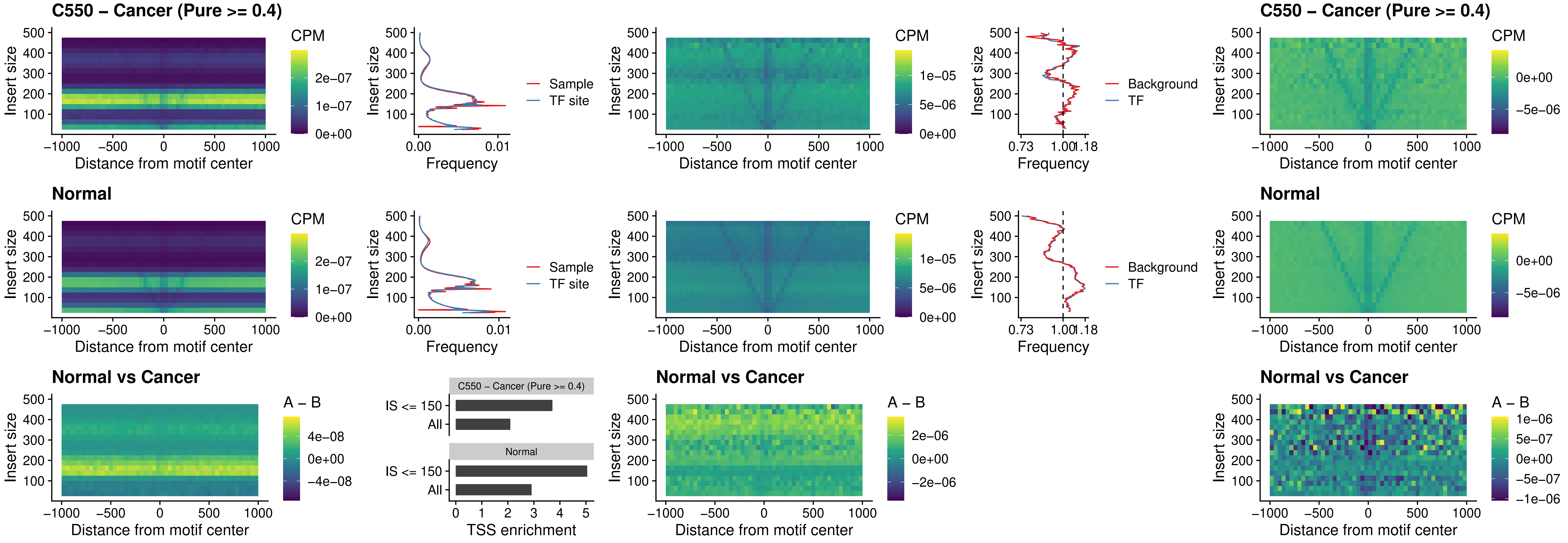

#### C550 – Cancer (Pure >= 0.4) – NKX24_uc_dTSS_nPEAK

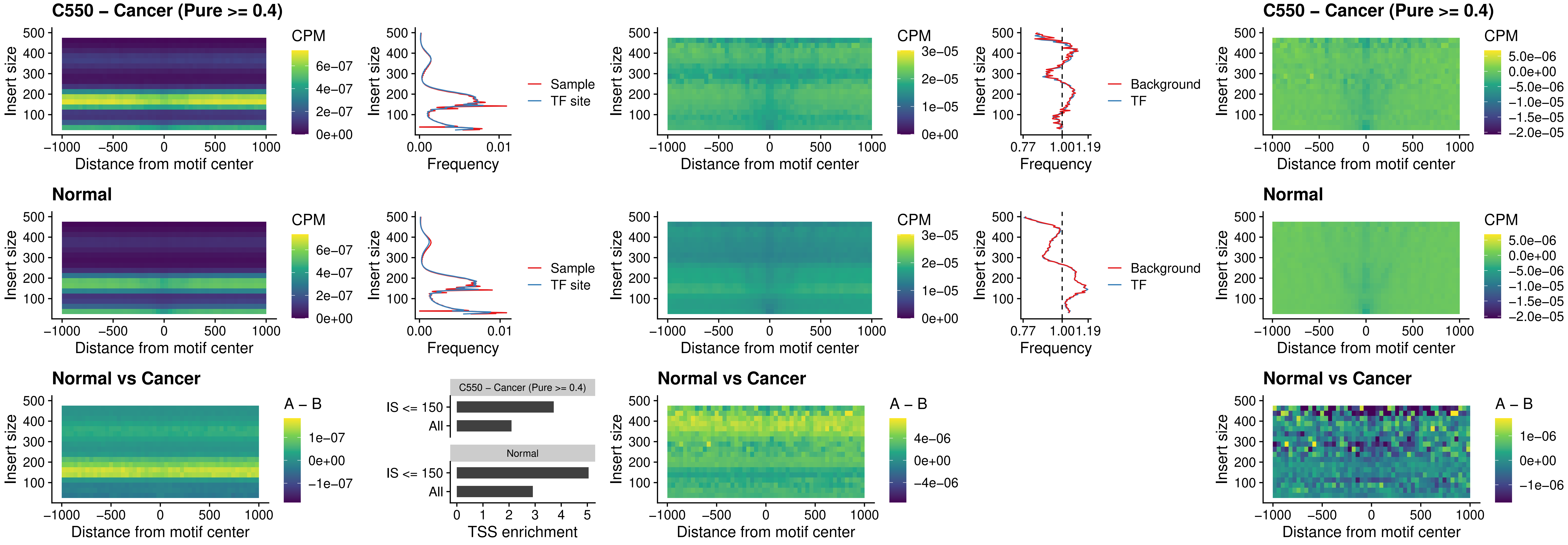

#### C550 – Cancer (Pure >= 0.4) – DLX3_uc_cTSS_nPEAK

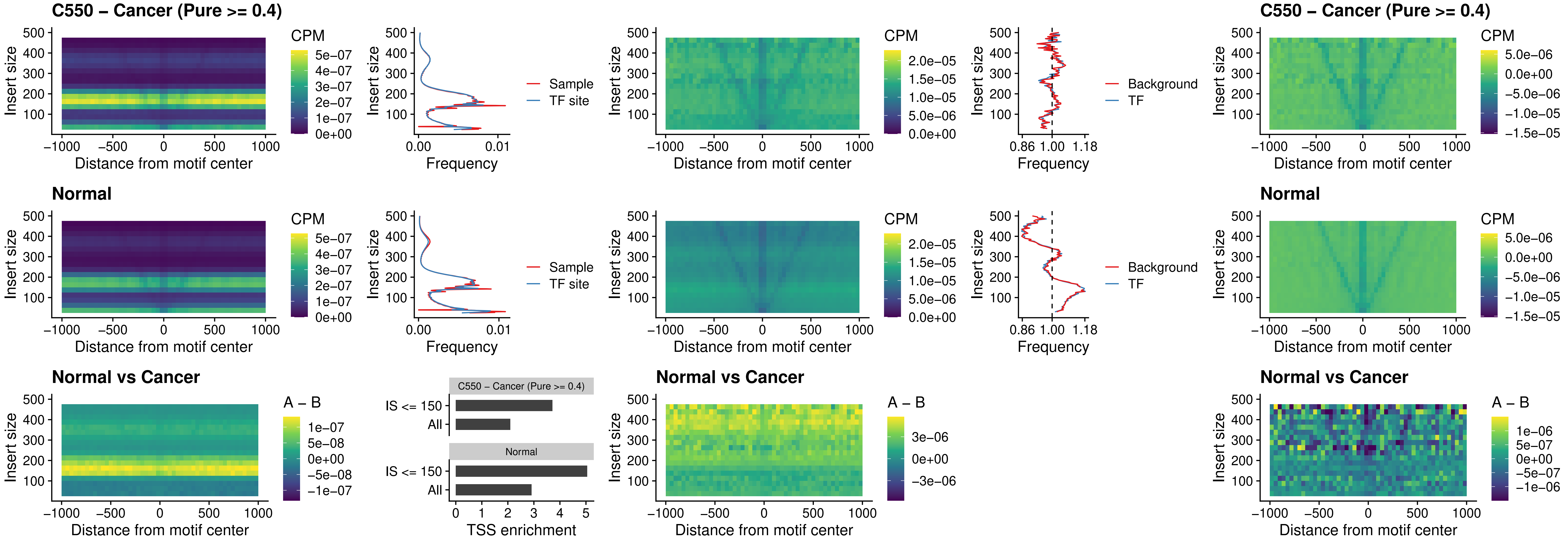

### C550 – Cancer (Pure >= 0.4) – HOXD3_uc_cTSS_nPEAK

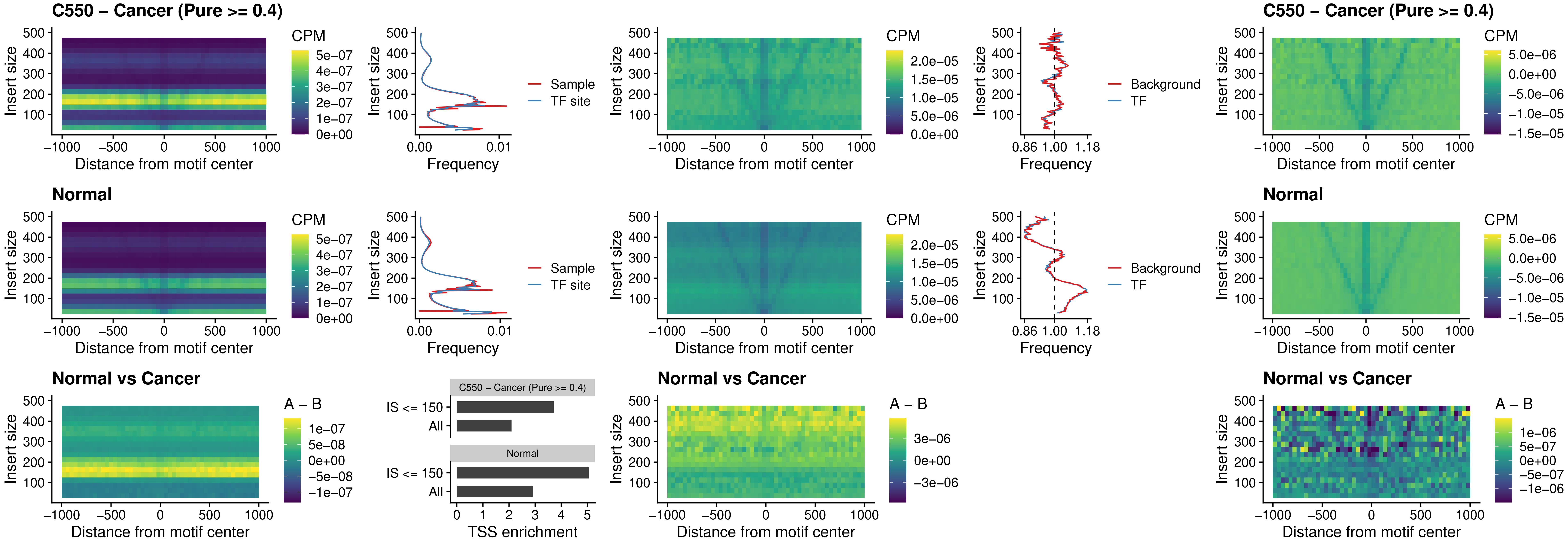

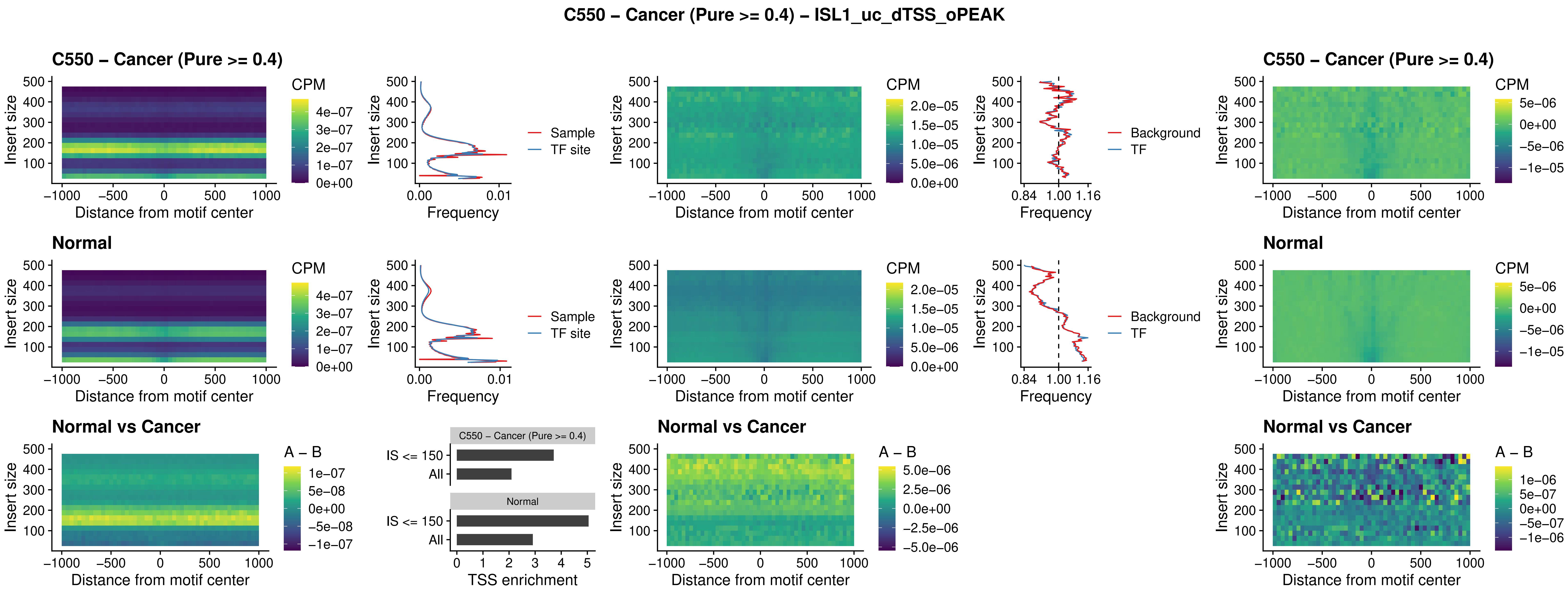
C550 – Cancer (Pure >= 0.4) – ISL1_uc_dTSS_oPEAK
C550 – Cancer (Pure >= 0.4)
Normal
Normal vs Cancer
Insert size
Distance from motif center
Frequency
CPM
A – B
Sample
TF site
Background
TF
C550 – Cancer (Pure >= 0.4)
Normal
IS <= 150
All
TSS enrichment

### C550 – Cancer (Pure >= 0.4) – HOXB4_uc_dTSS_oPEAK

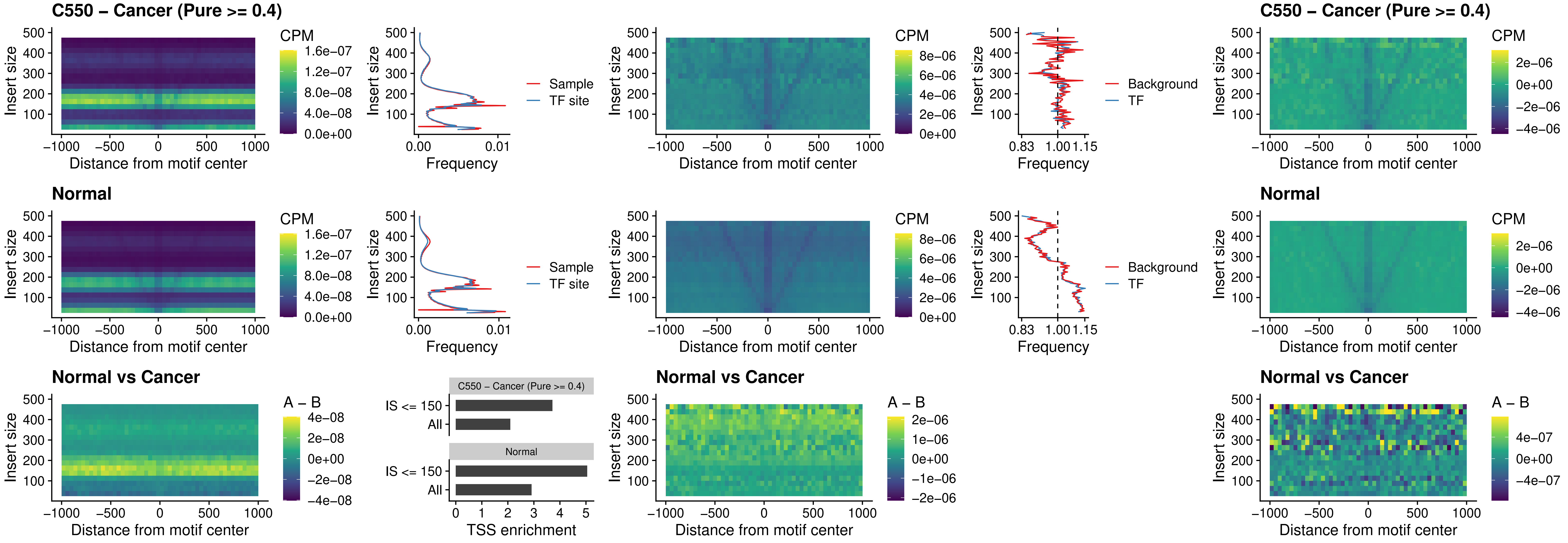

#### C550 – Cancer (Pure >= 0.4) – VAX1_uc_dTSS_oPEAK

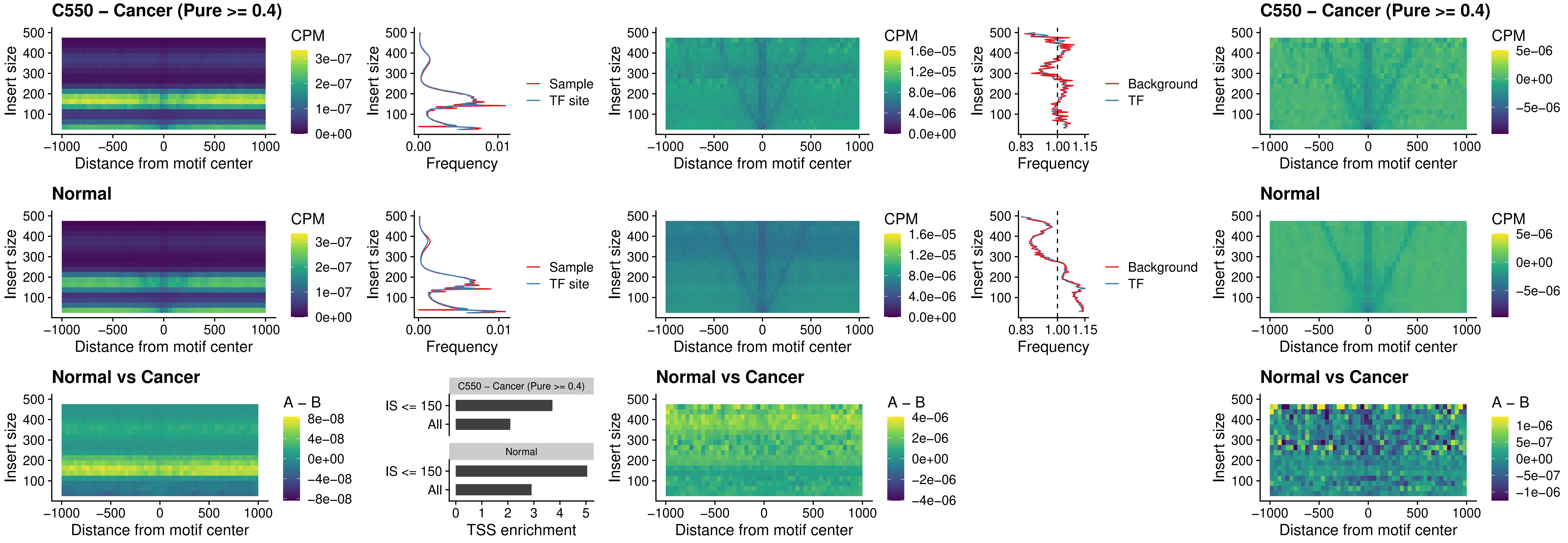

### C550 – Cancer (Pure >= 0.4) – DLX3_uc_dTSS_oPEAK

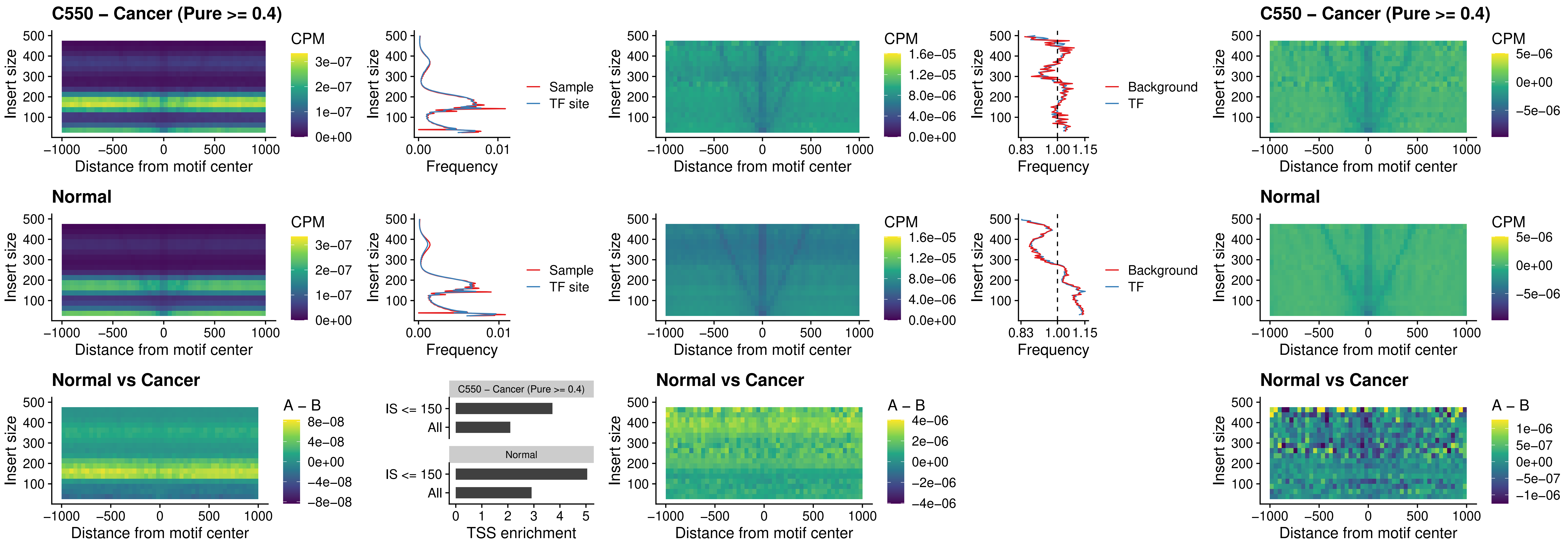

#### C550 – Cancer (Pure >= 0.4) – HOXD3_uc_dTSS_oPEAK

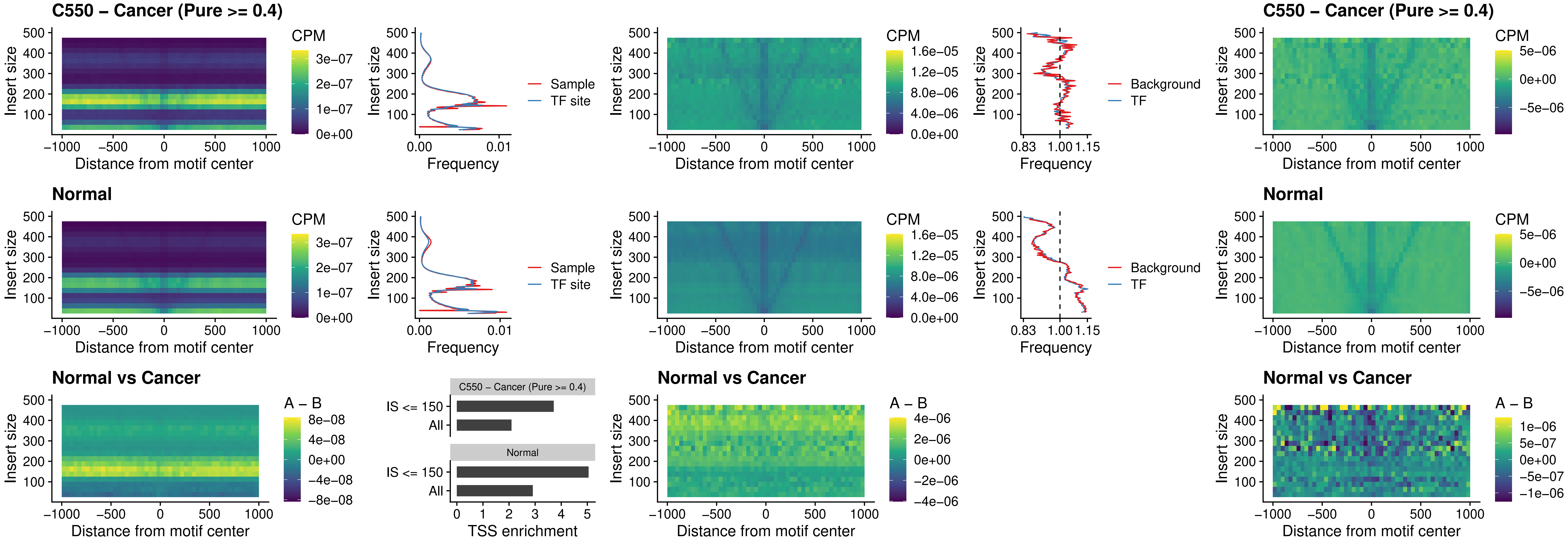

#### C550 – Cancer (Pure >= 0.4) – BBX_uc_dTSS_nPEAK

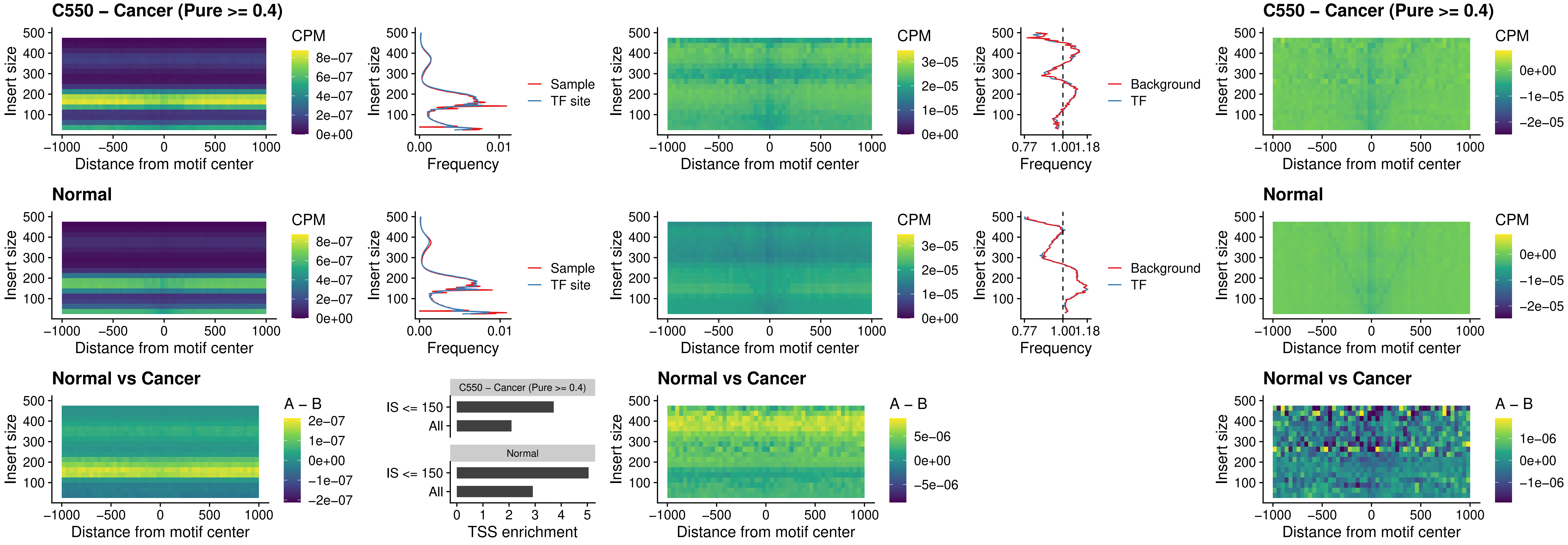

#### C550 – Cancer (Pure >= 0.4) – SOX3_uc_cTSS_nPEAK

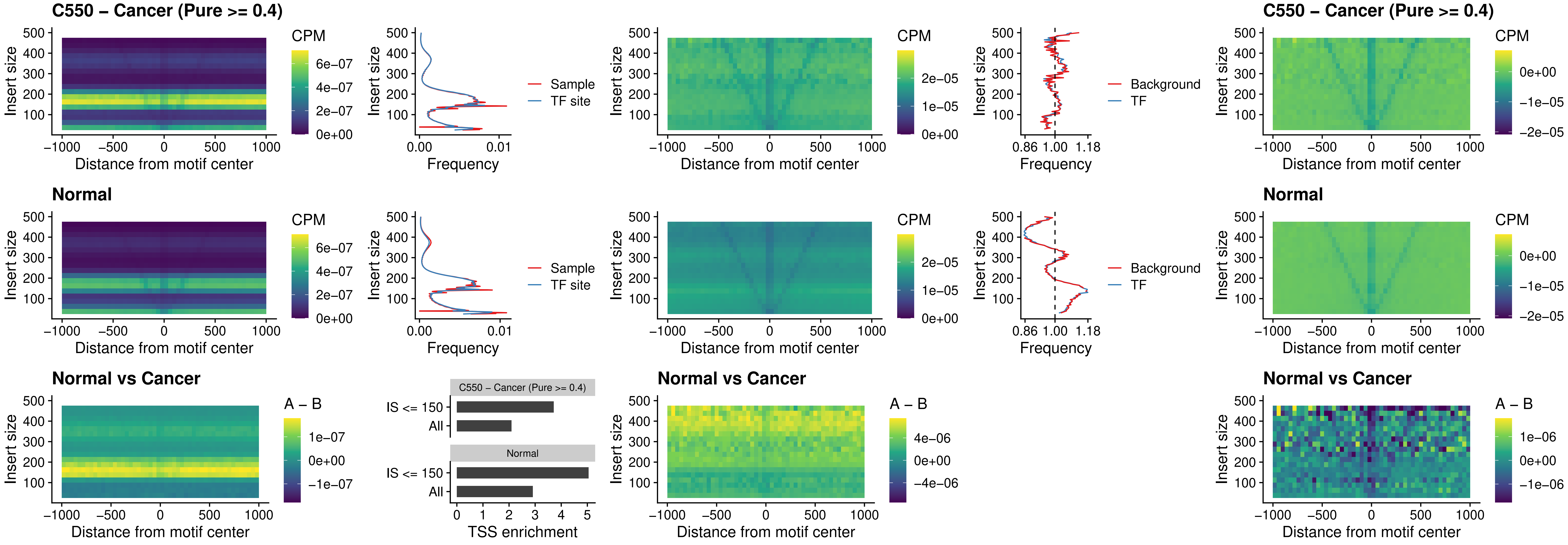

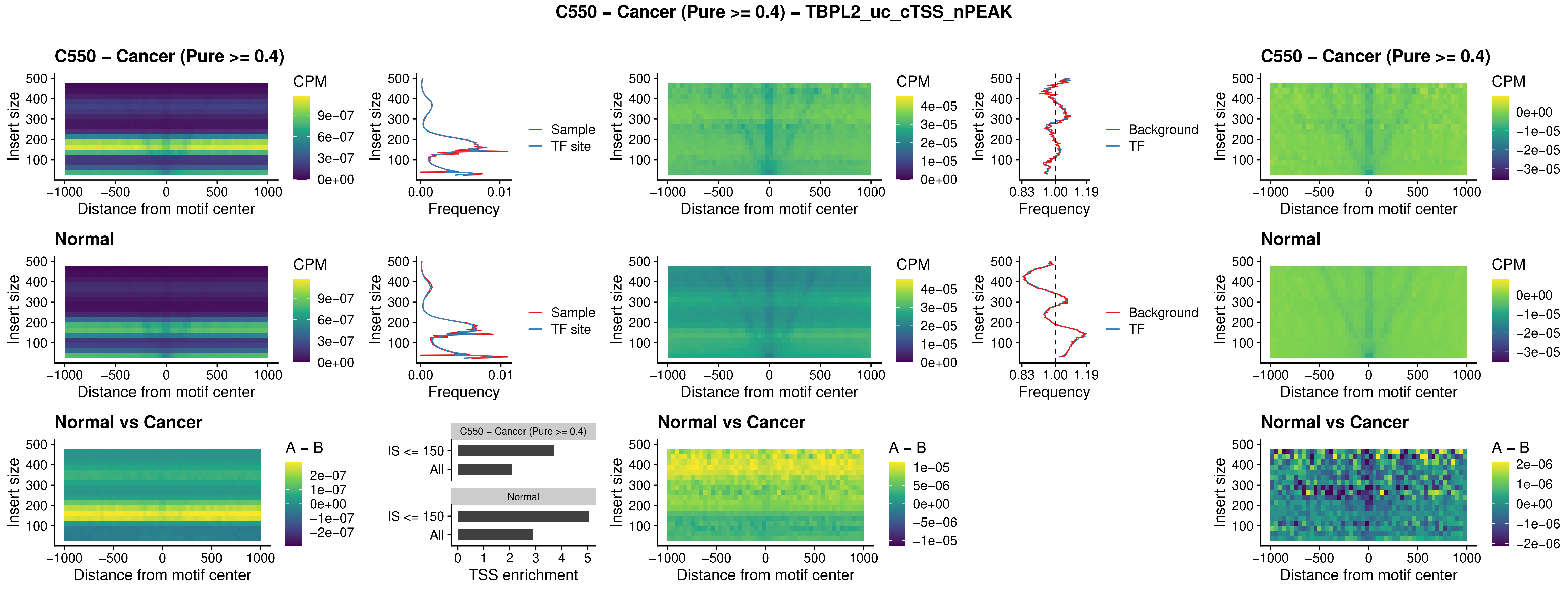

C550 – Cancer (Pure >= 0.4) – TBPL2_uc_cTSS_nPEAK
C550 – Cancer (Pure >= 0.4)
Normal
Normal vs Cancer
Insert size
Distance from motif center
CPM
A – B
Frequency
Sample
TF site
Background
TF
C550 – Cancer (Pure >= 0.4)
Normal
IS <= 150
All
TSS enrichment

### C550 – Cancer (Pure >= 0.4) – HOXA4_uc_cTSS_oPEAK

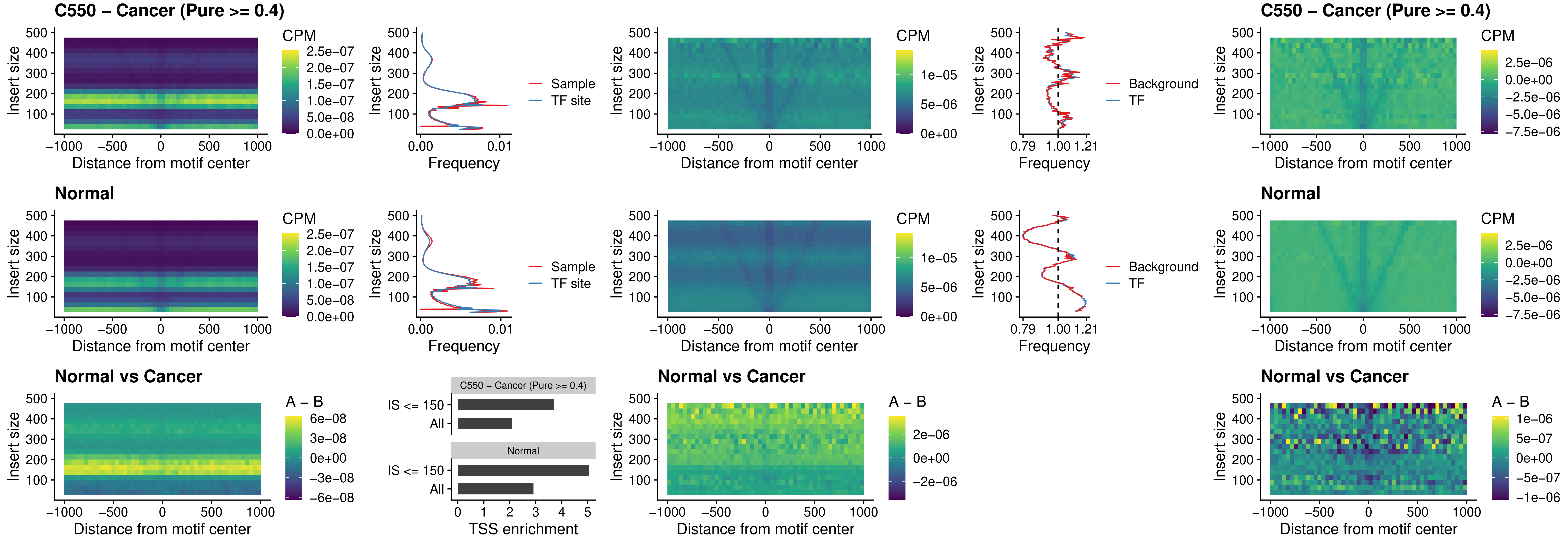

#### C550 – Cancer (Pure >= 0.4) – PDX1_uc_dTSS_oPEAK

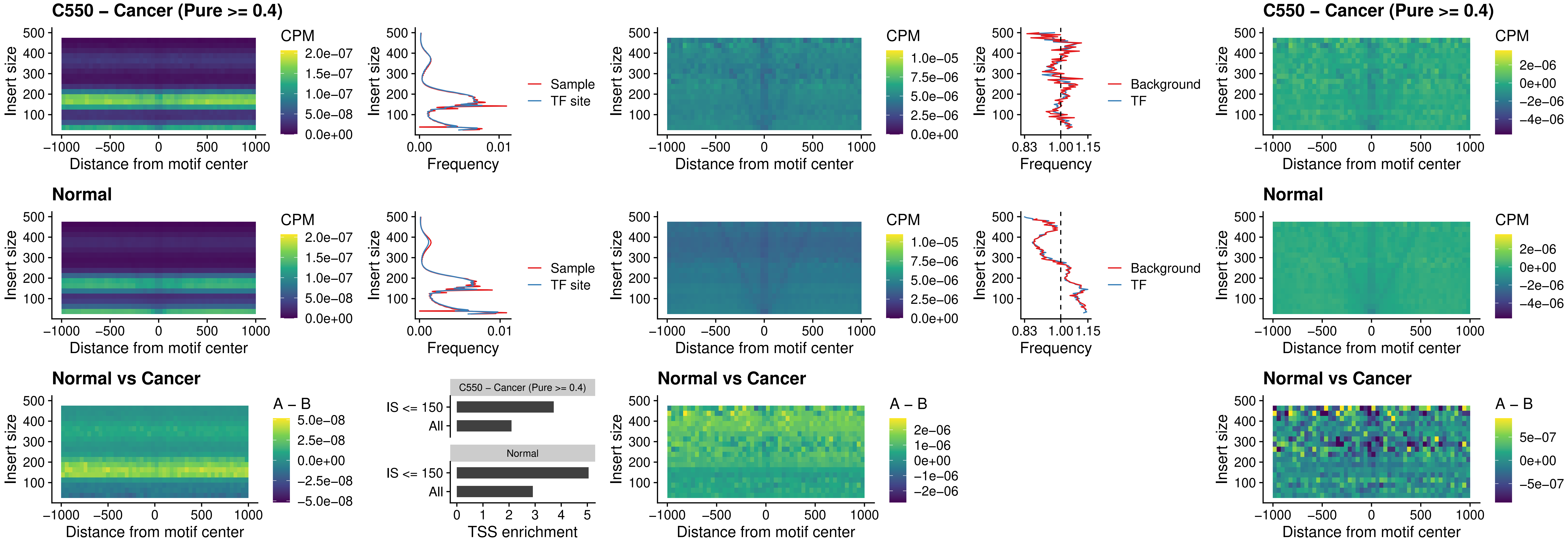

#### C550 – Cancer (Pure >= 0.4) – LHX4_uc_dTSS_oPEAK

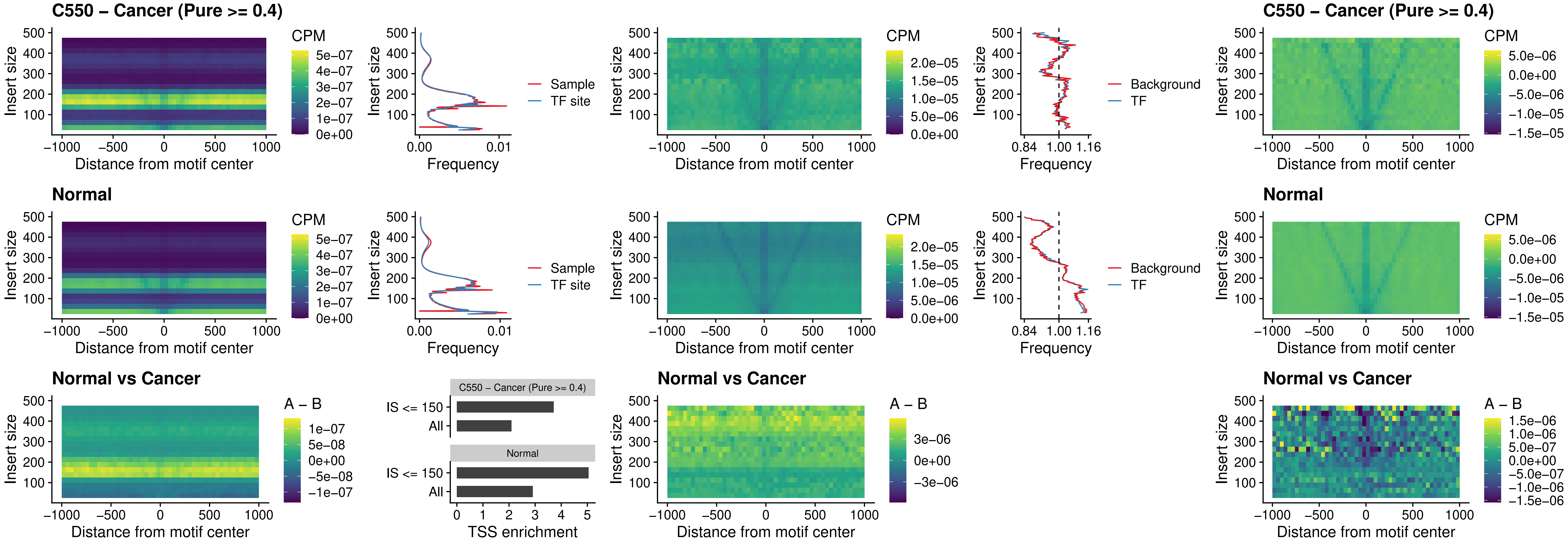

#### C550 – Cancer (Pure >= 0.4) – HOXA4_uc_dTSS_oPEAK

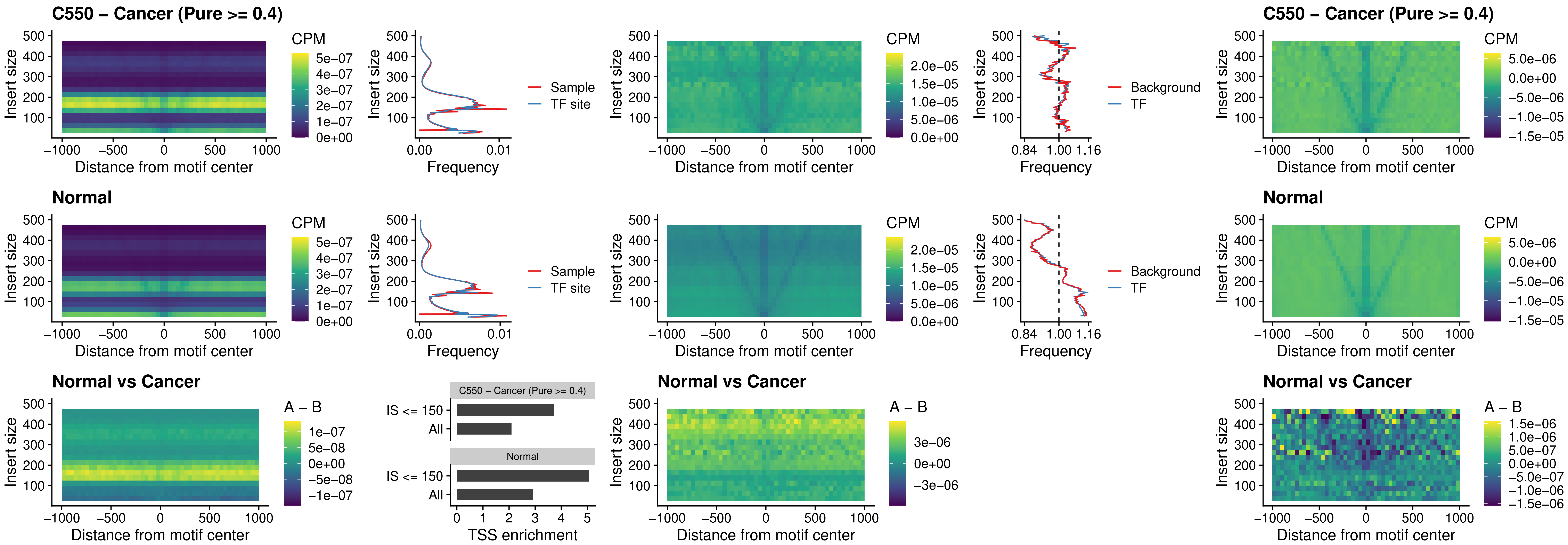

### C550 – Cancer (Pure >= 0.4) – HLX_uc_dTSS_oPEAK

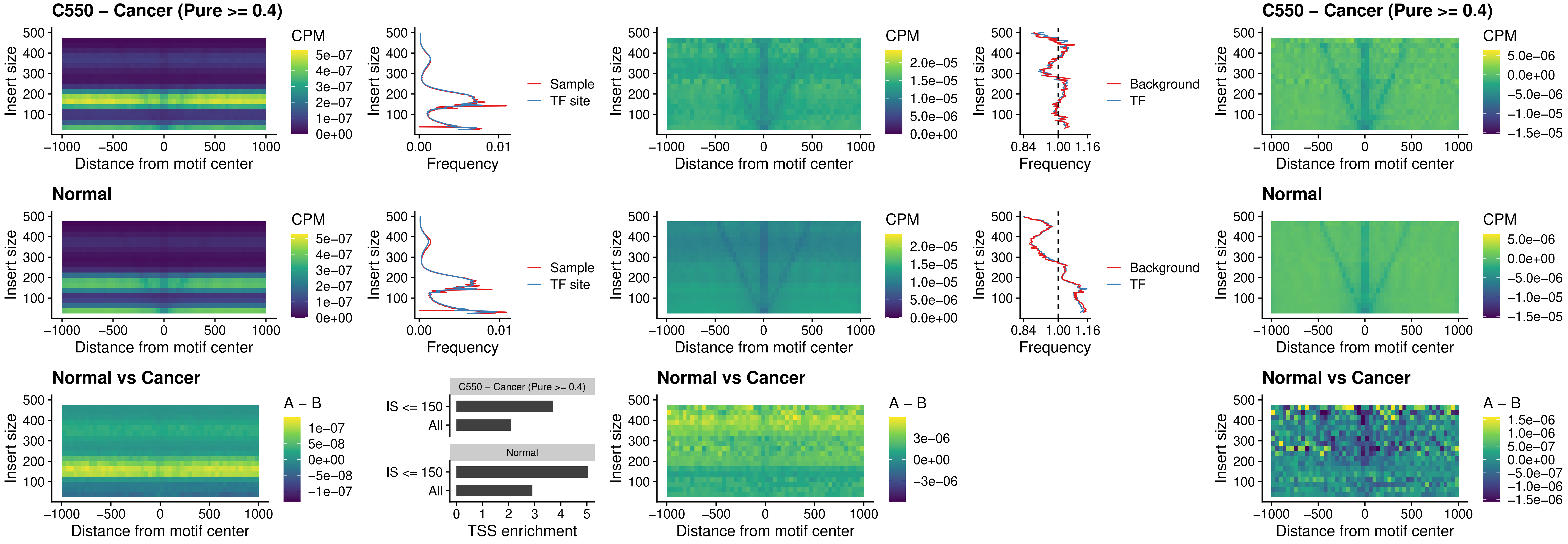

#### C550 – Cancer (Pure >= 0.4) – AC0125311_uc_dTSS_oPEAK

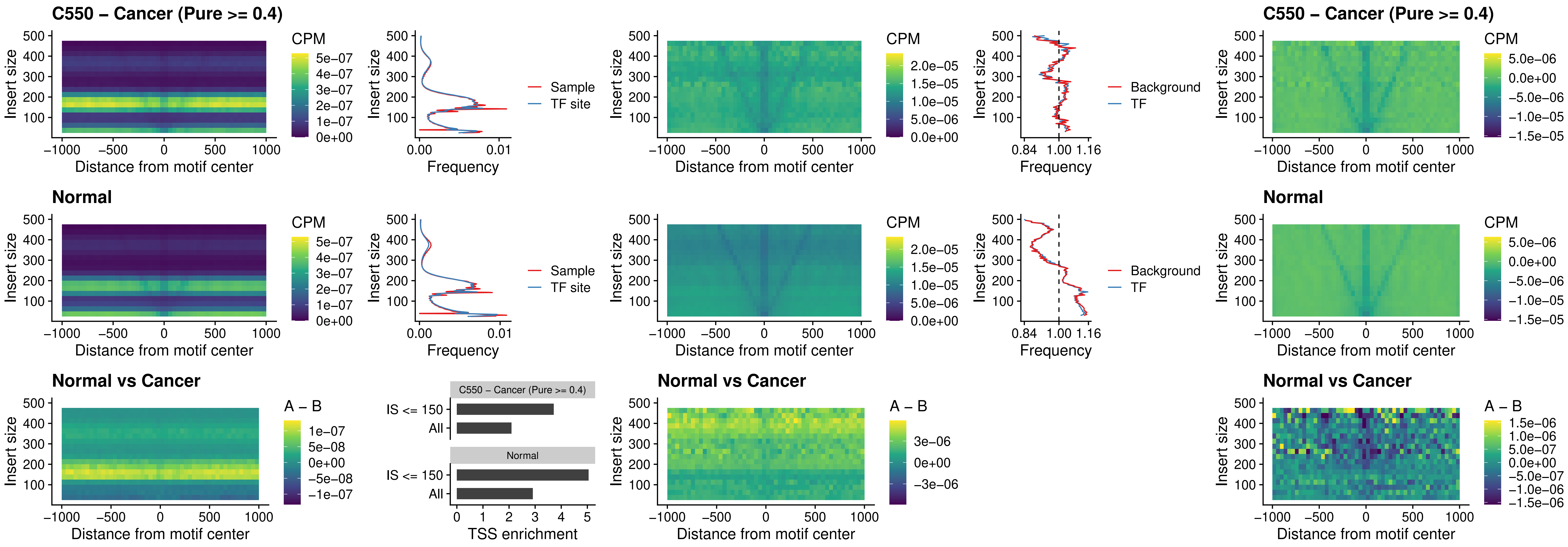

#### C550 – Cancer (Pure >= 0.4) – DBX2_uc_dTSS_oPEAK

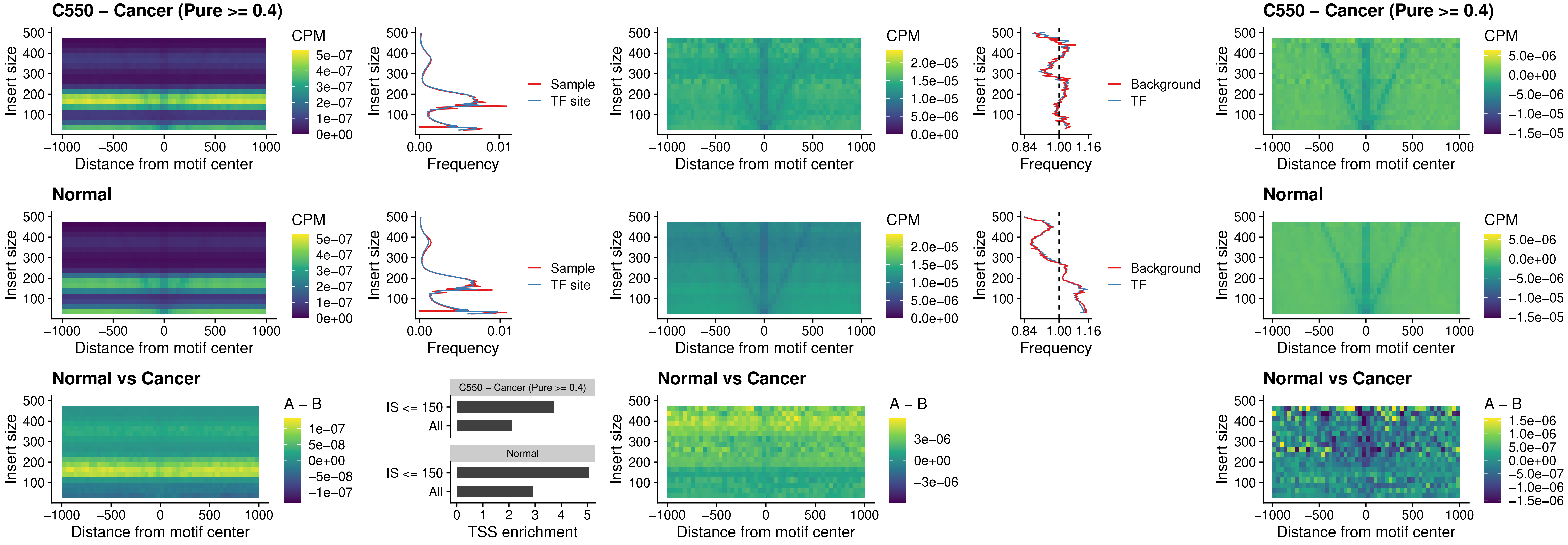

#### C550 – Cancer (Pure >= 0.4) – HOXA2_uc_dTSS_nPEAK

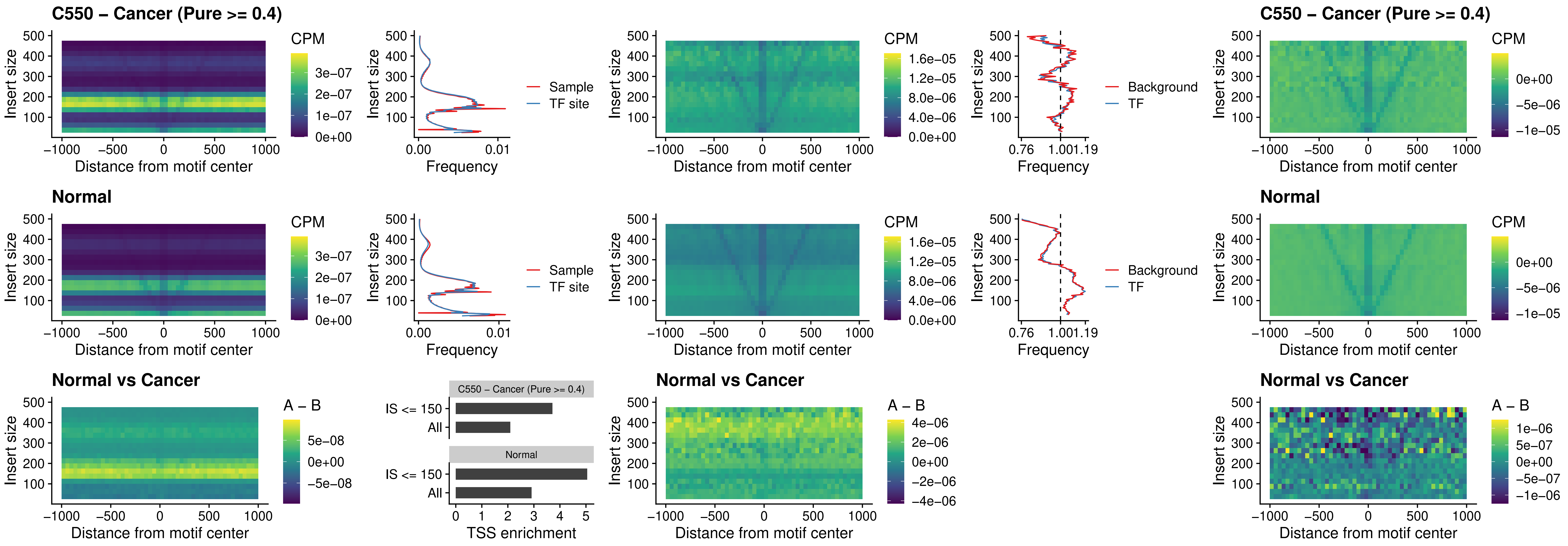

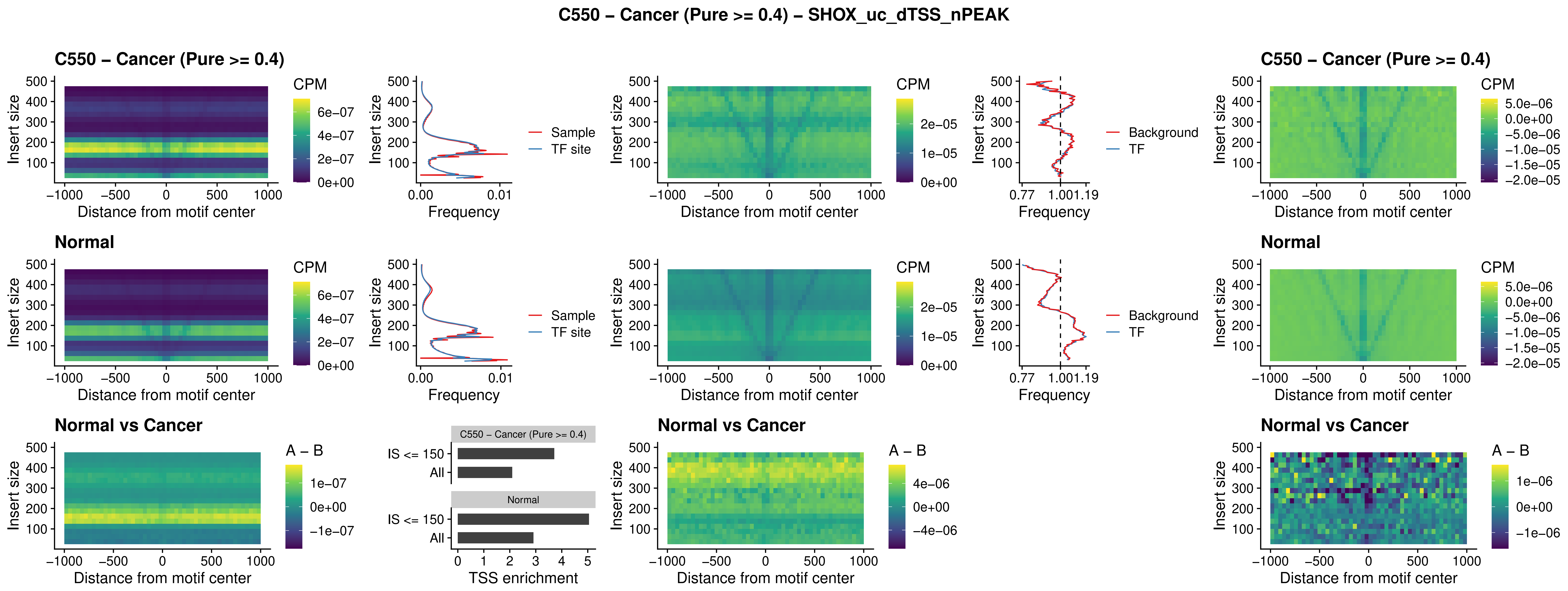
C550 – Cancer (Pure >= 0.4) – SHOX_uc_dTSS_nPEAK
C550 – Cancer (Pure >= 0.4)
Normal
Normal vs Cancer
Insert size
Distance from motif center
Frequency
CPM
A – B
Sample
TF site
Background
TF
C550 – Cancer (Pure >= 0.4)
Normal
IS <= 150
All
TSS enrichment

#### C550 – Cancer (Pure >= 0.4) – PRRX1_uc_dTSS_nPEAK

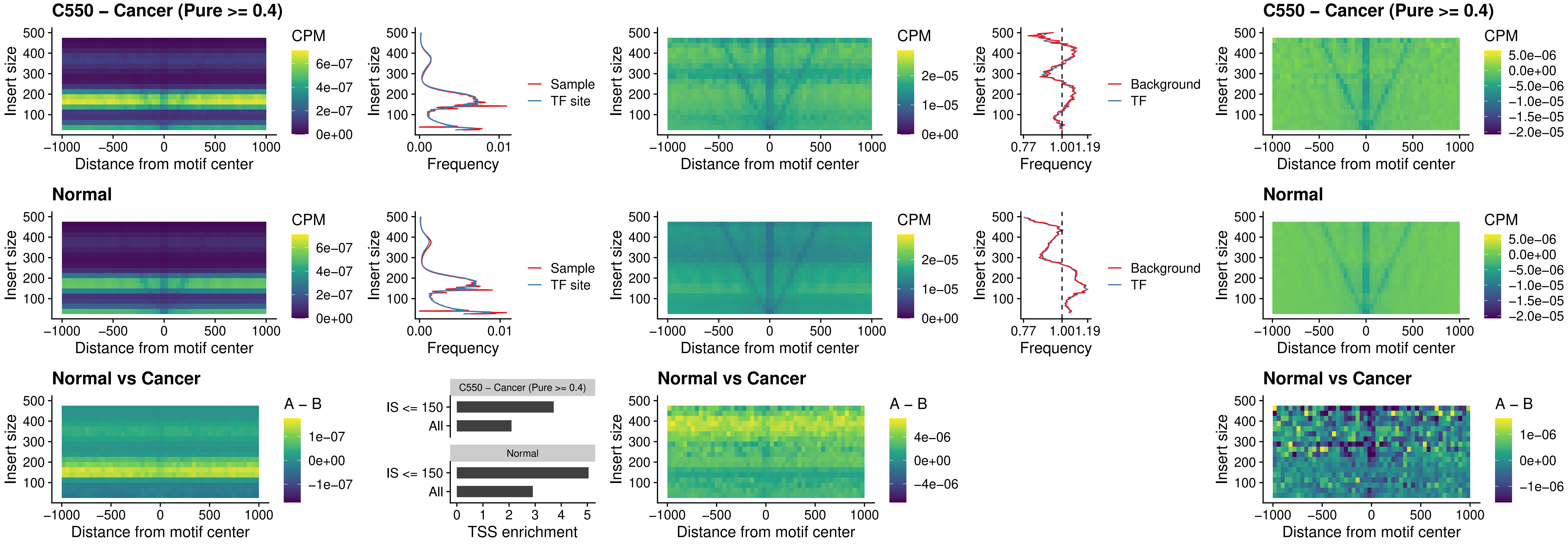

#### C550 – Cancer (Pure >= 0.4) – LMX1A_uc_dTSS_nPEAK

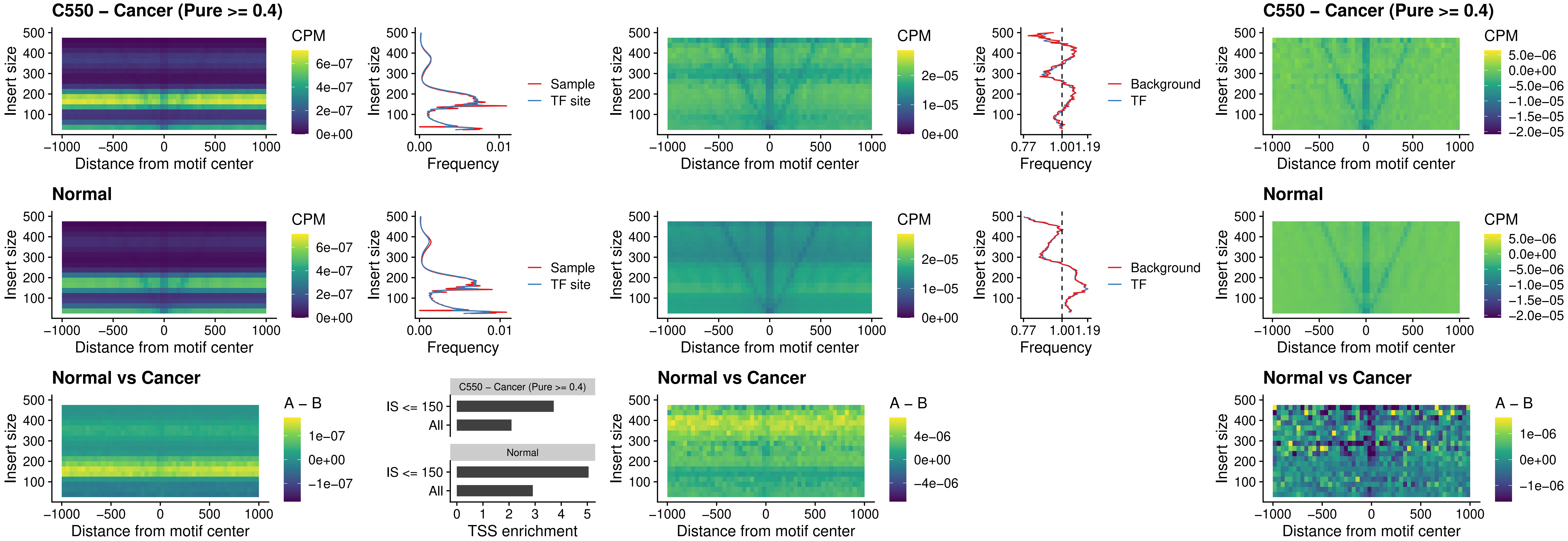

#### C550 – Cancer (Pure >= 0.4) – LMX1B_uc_dTSS_nPEAK

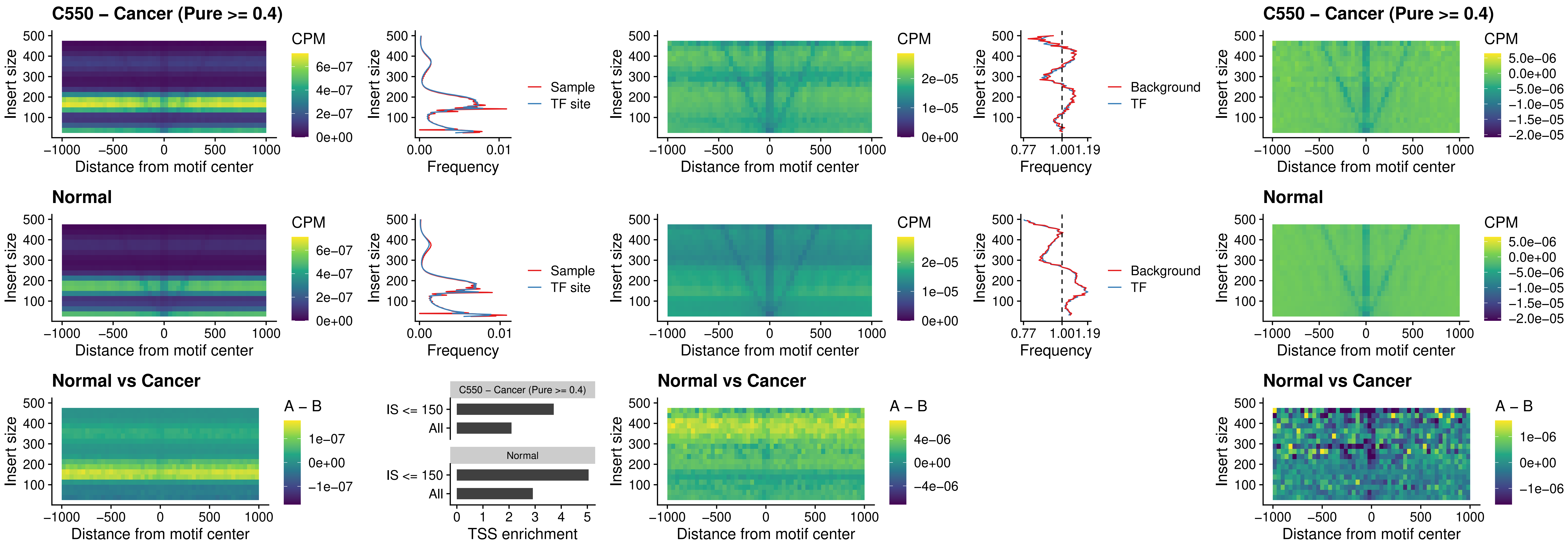

#### C550 – Cancer (Pure >= 0.4) – DLX5_uc_dTSS_oPEAK

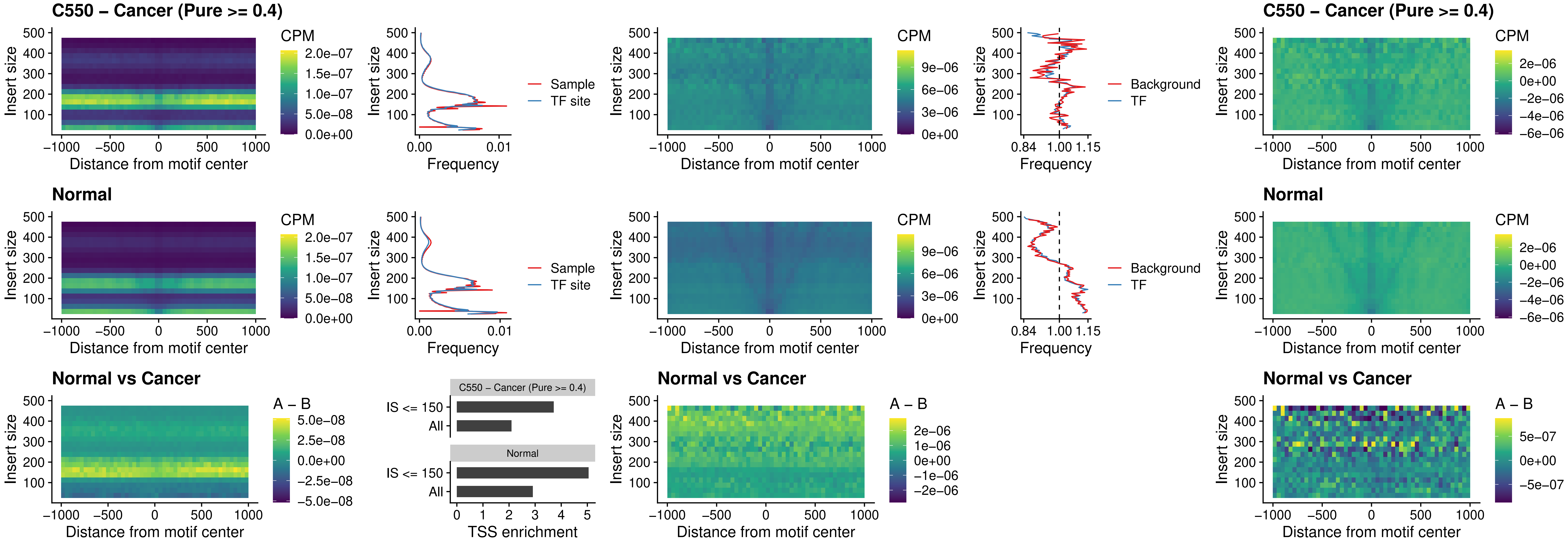

#### C550 – Cancer (Pure >= 0.4) – ARID3C_uc_dTSS_nPEAK

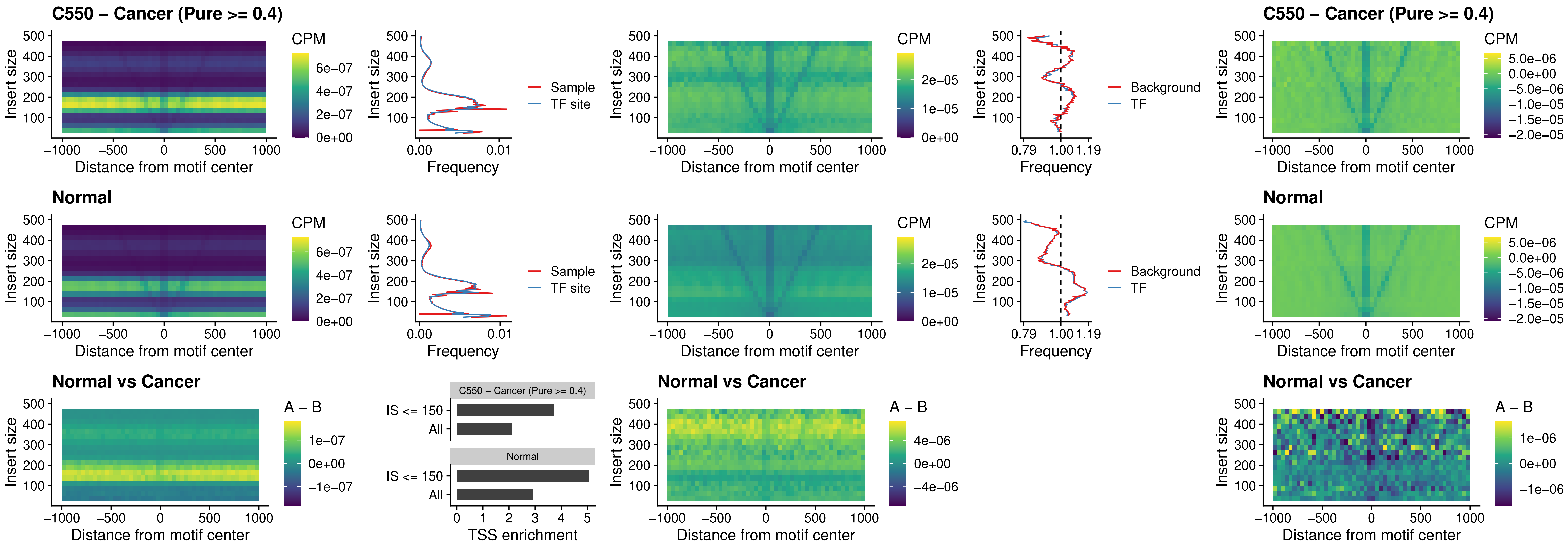

### C550 – Cancer (Pure >= 0.4) – HOXD12_uc_dTSS_nPEAK

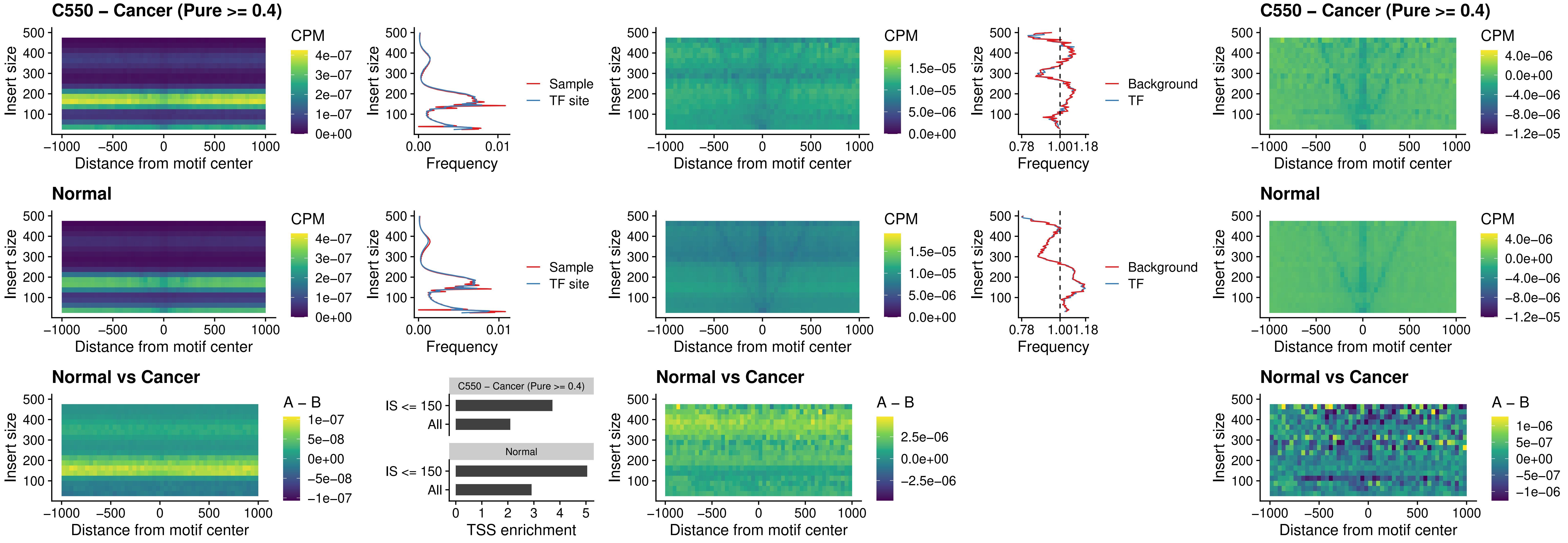

#### C550 – Cancer (Pure >= 0.4) – CDX2_uc_dTSS_nPEAK

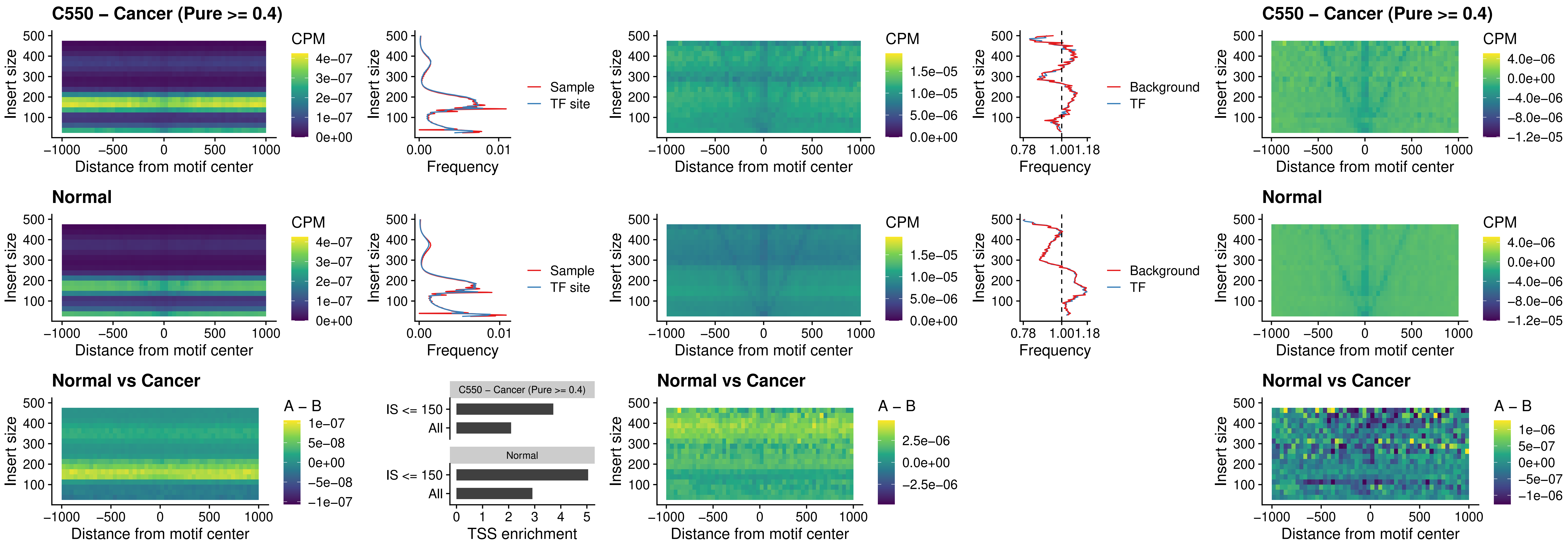

#### C550 – Cancer (Pure >= 0.4) – HOXC12_uc_dTSS_nPEAK

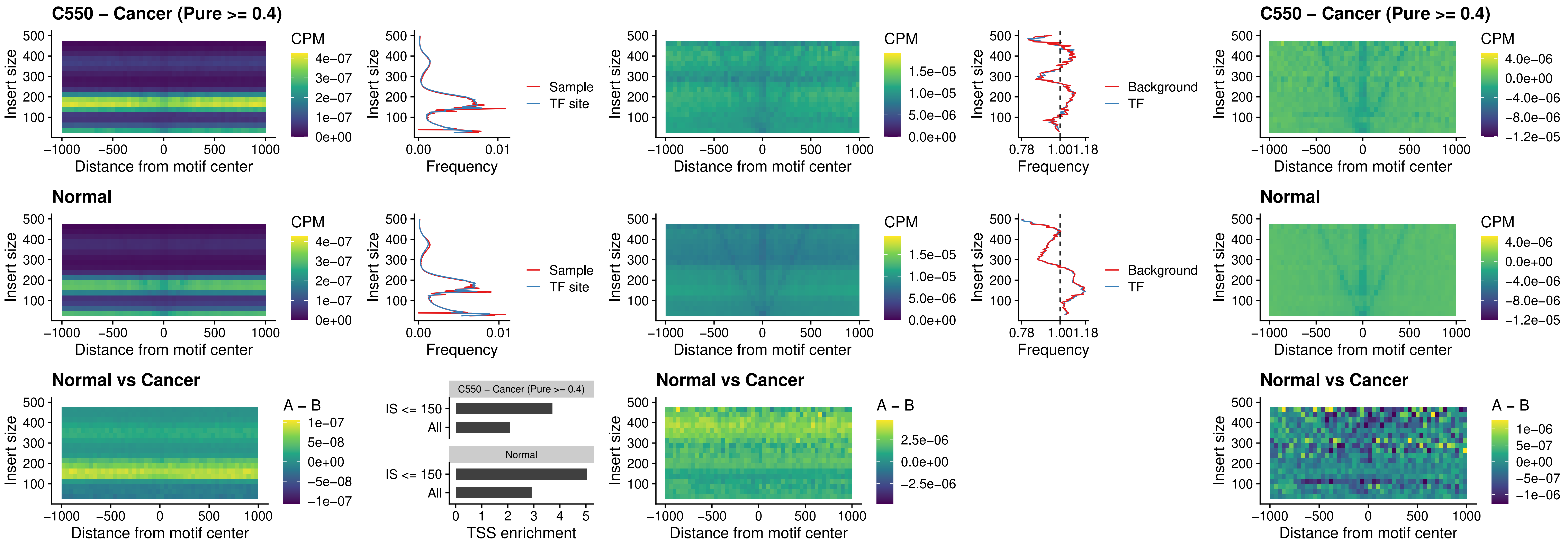

### C550 – Cancer (Pure >= 0.4) – HHEX_uc_dTSS_oPEAK

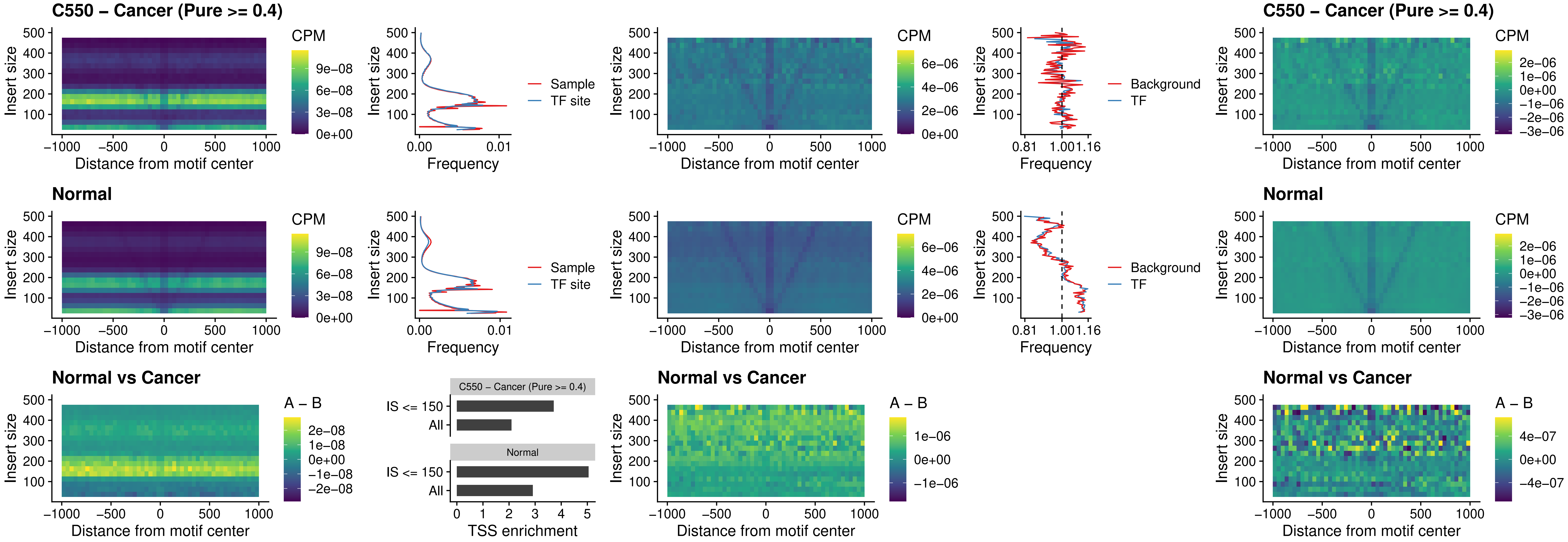

#### C550 – Cancer (Pure >= 0.4) – HOXA11_uc_dTSS_nPEAK

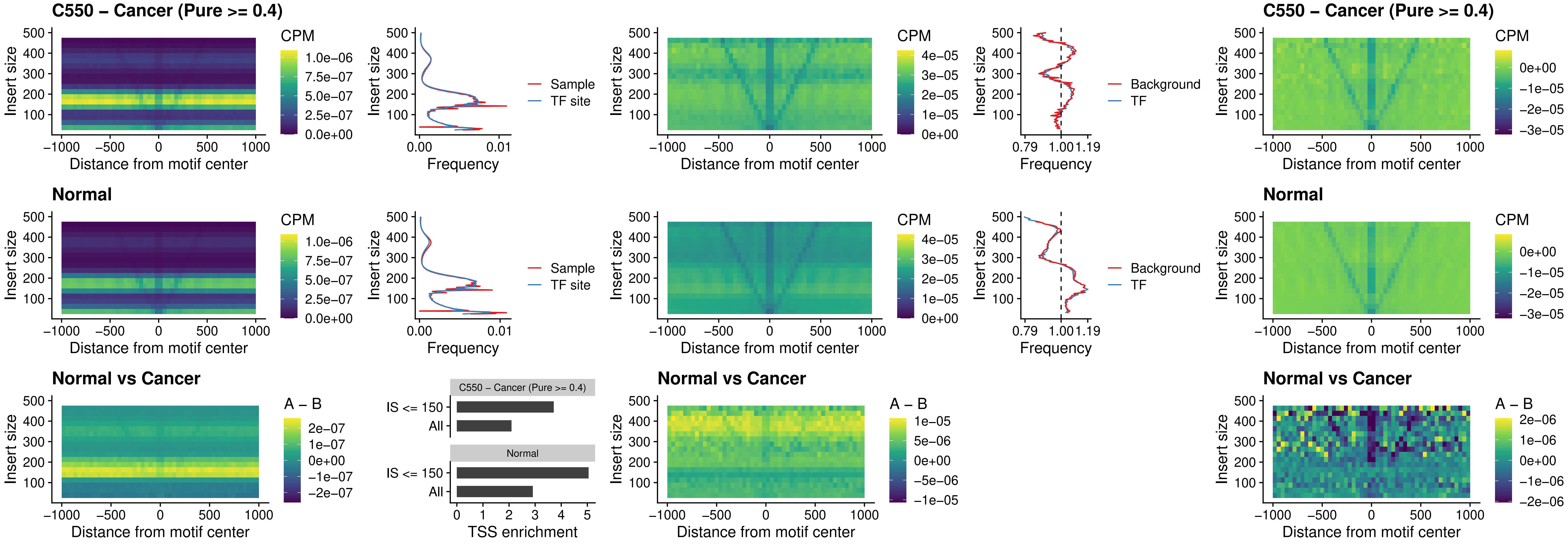

#### C550 – Cancer (Pure >= 0.4) – HOXA11_uc_cTSS_nPEAK

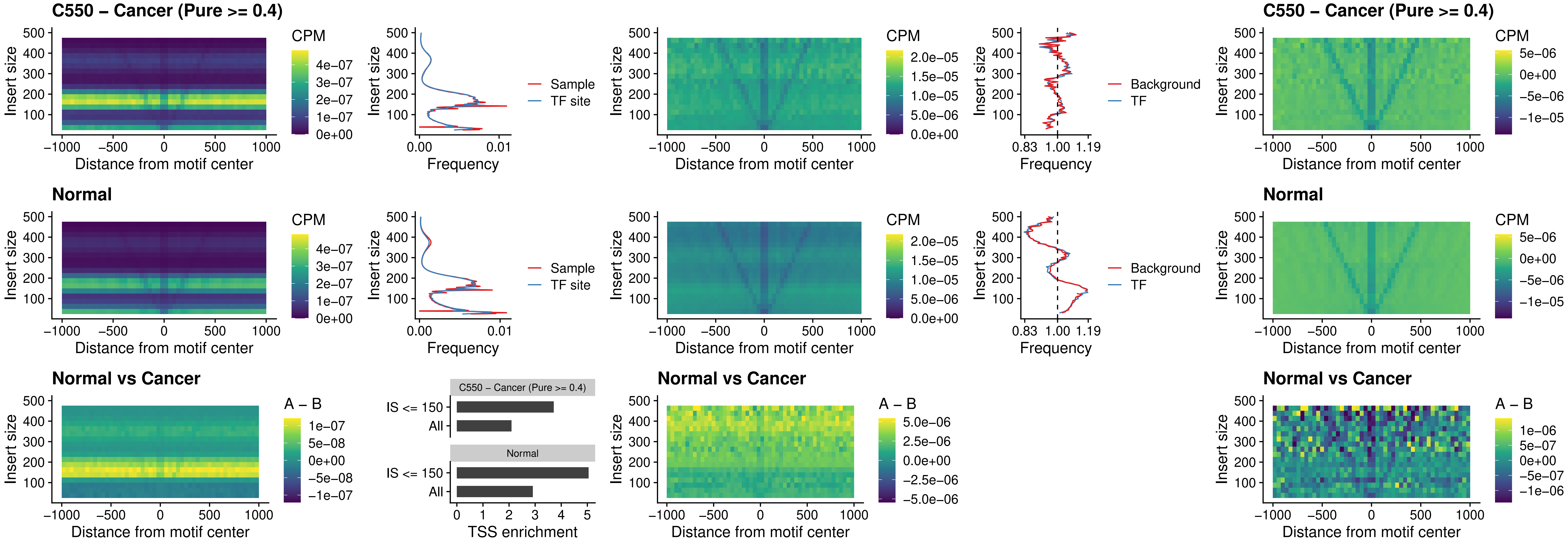

### C550 – Cancer (Pure >= 0.4) – ZFHX3_uc_dTSS_nPEAK

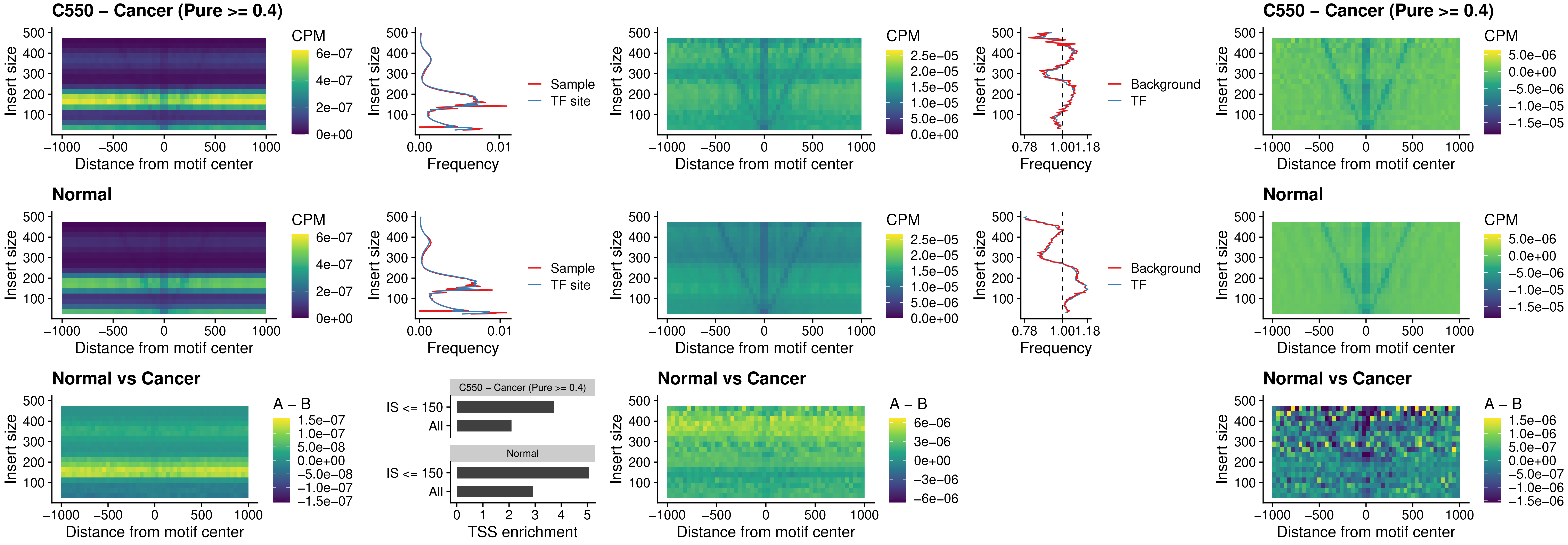

#### C550 – Cancer (Pure >= 0.4) – HOXB4_uc_dTSS_nPEAK

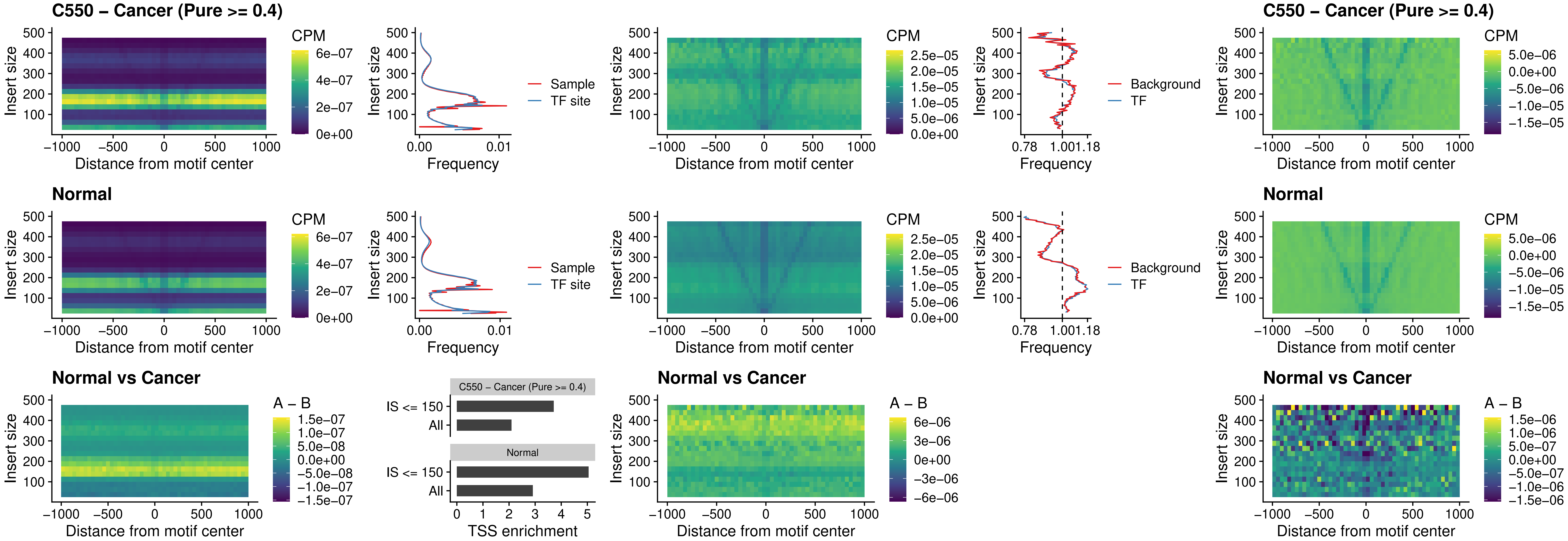

**C550 – Cancer (Pure >= 0.4) – HOXC5_uc_dTSS_nPEAK**

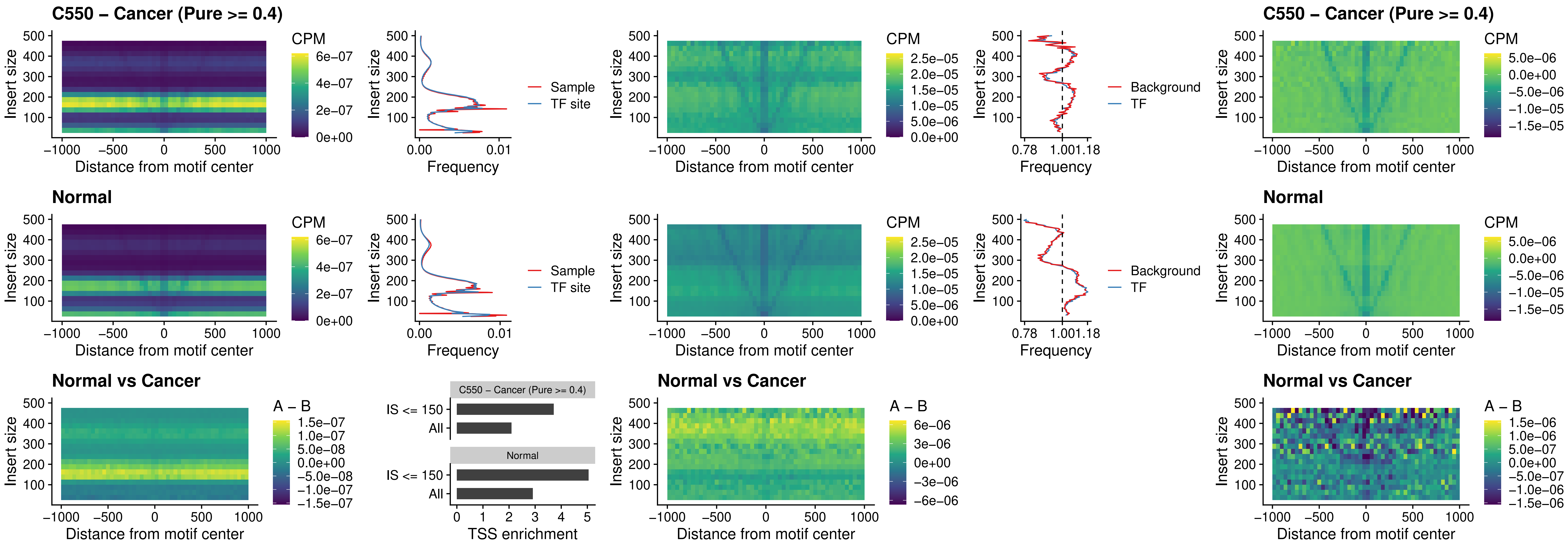

#### C550 – Cancer (Pure >= 0.4) – HOXD1_uc_dTSS_nPEAK

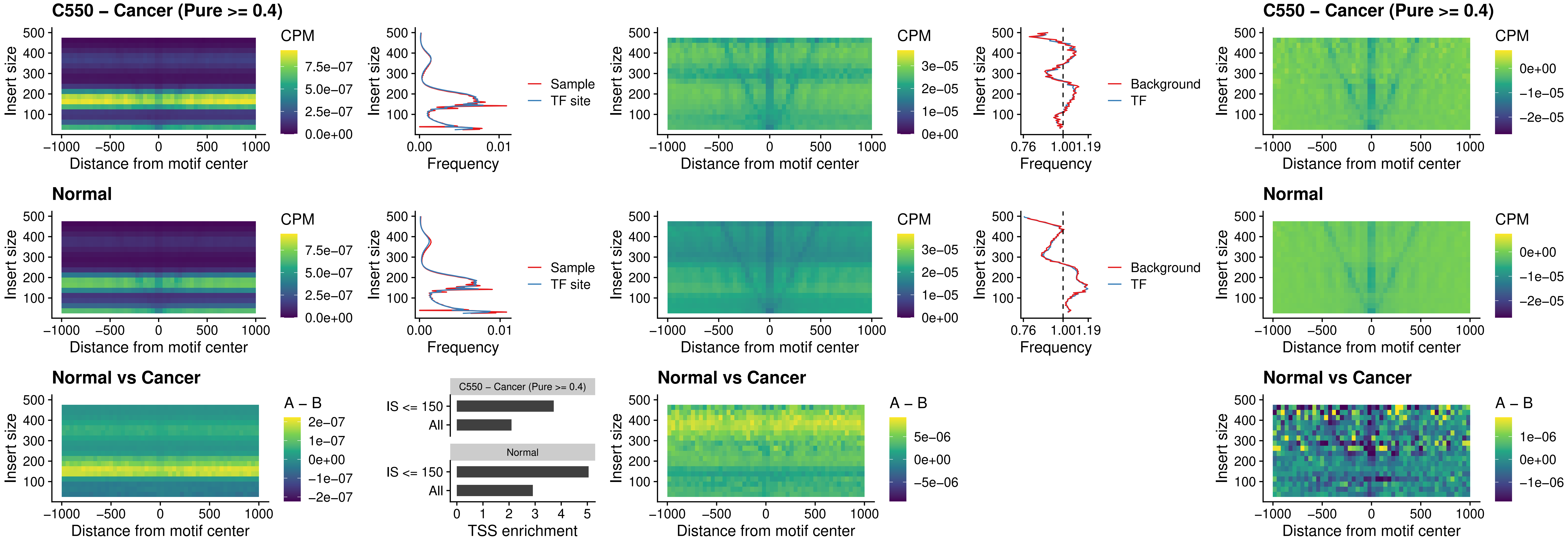

#### C550 – Cancer (Pure >= 0.4) – OTP_uc_dTSS_nPEAK

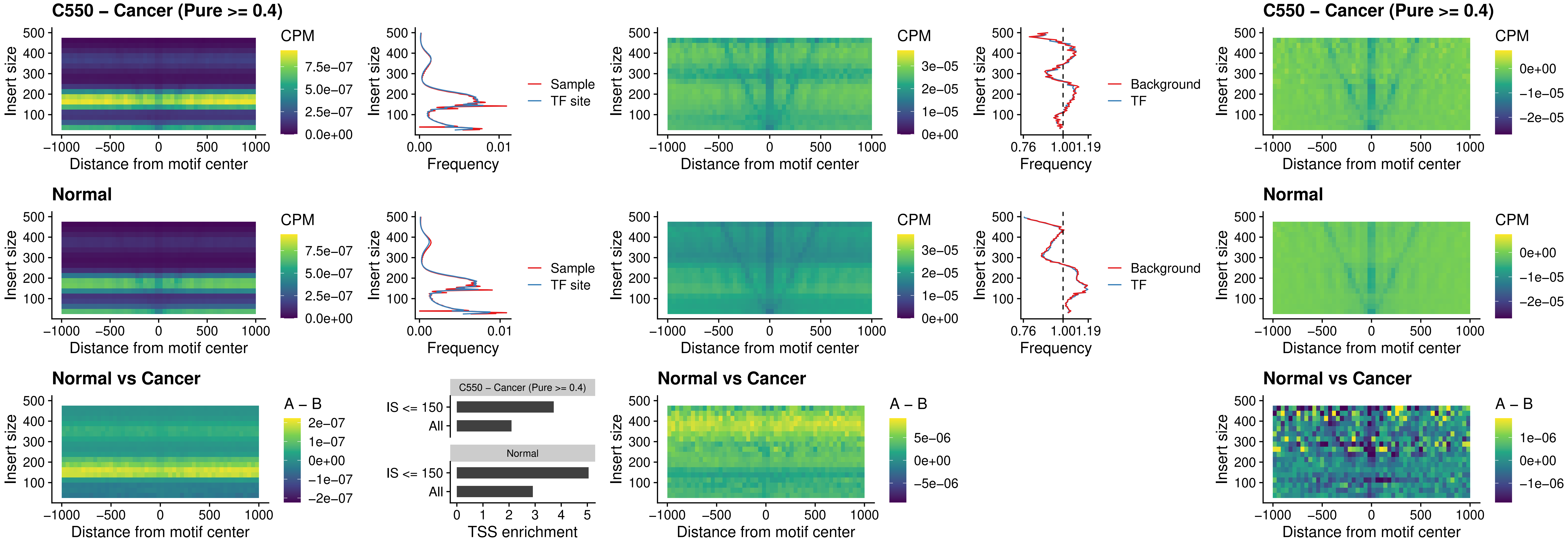

### C550 – Cancer (Pure >= 0.4) – FOXL1_uc_dTSS_oPEAK

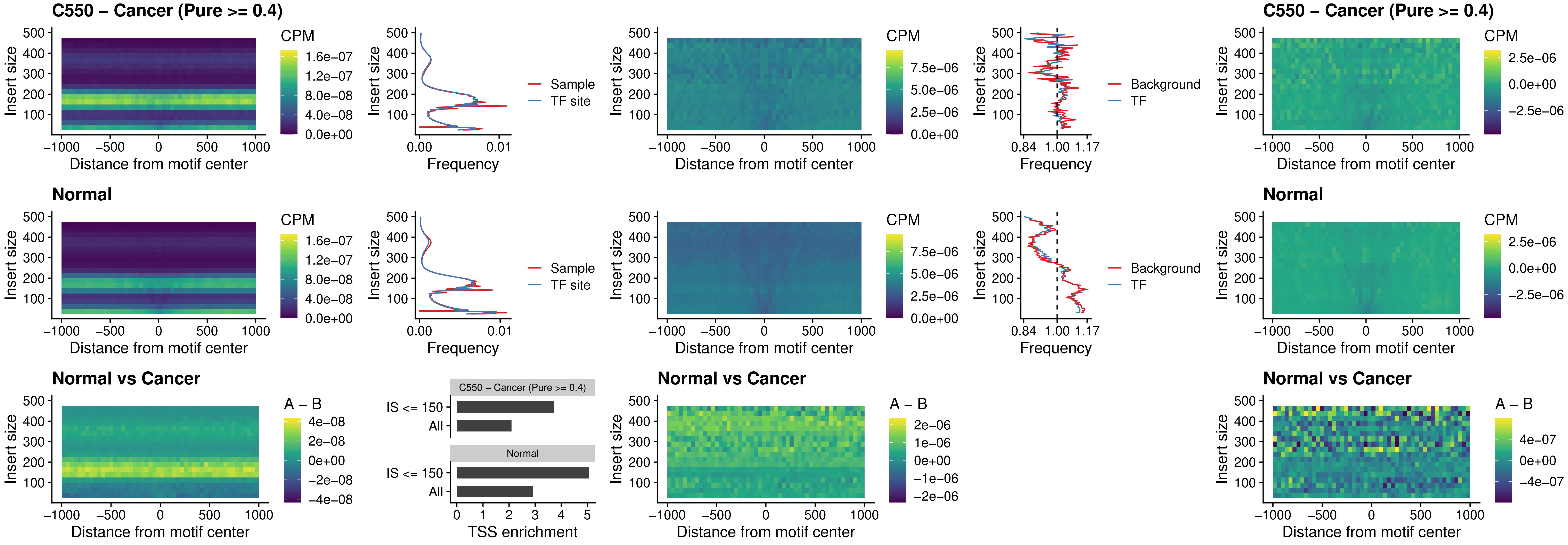

### C550 – Cancer (Pure >= 0.4) – ZNF32_uc_dTSS_oPEAK

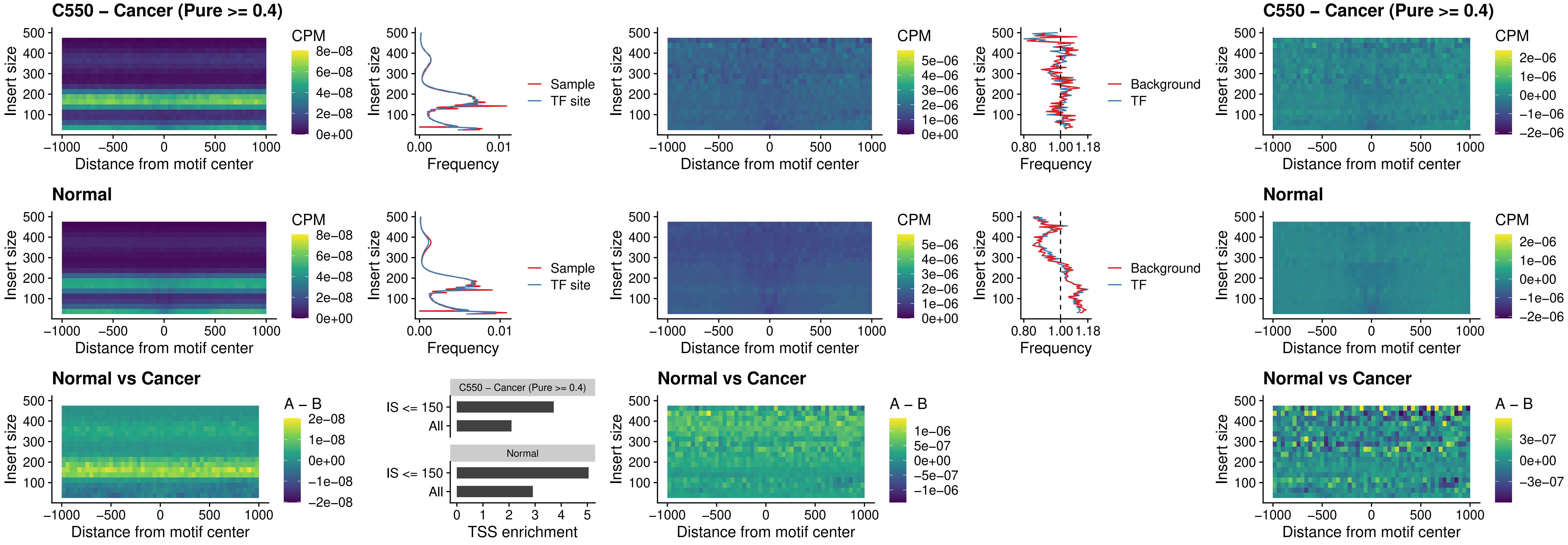

### C550 – Cancer (Pure >= 0.4) – FOXL1_uc_dTSS_nPEAK

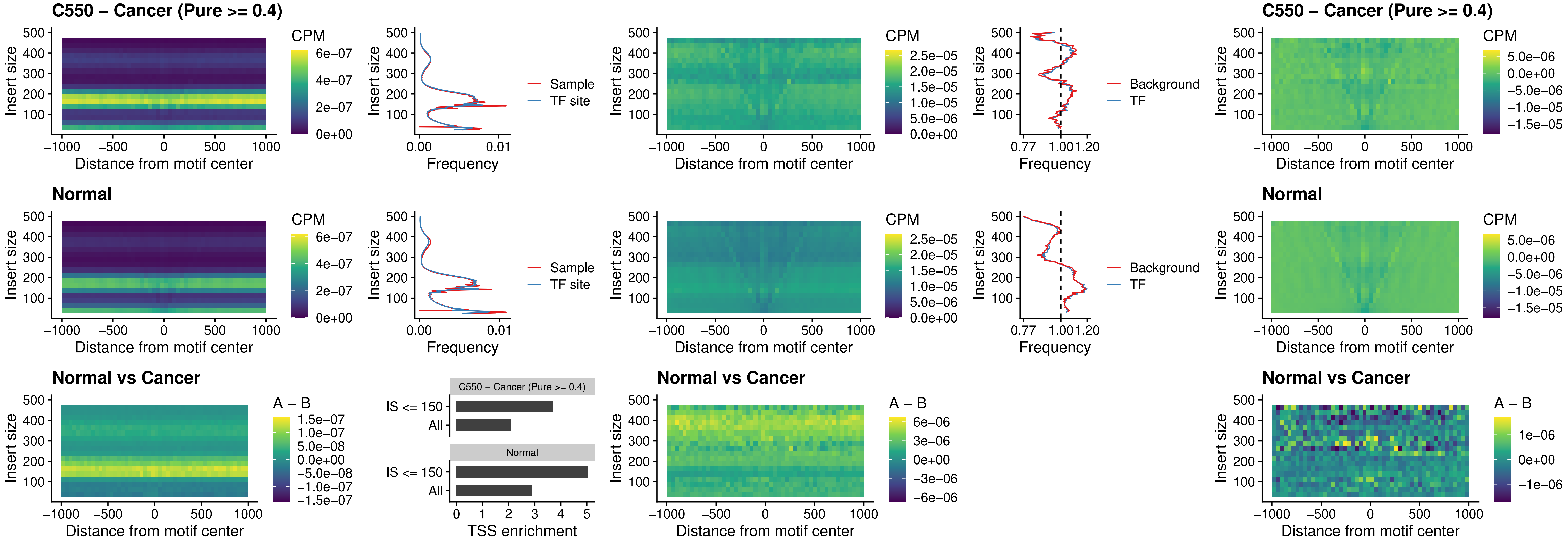

#### C550 – Cancer (Pure >= 0.4) – ZNF32_uc_dTSS_nPEAK

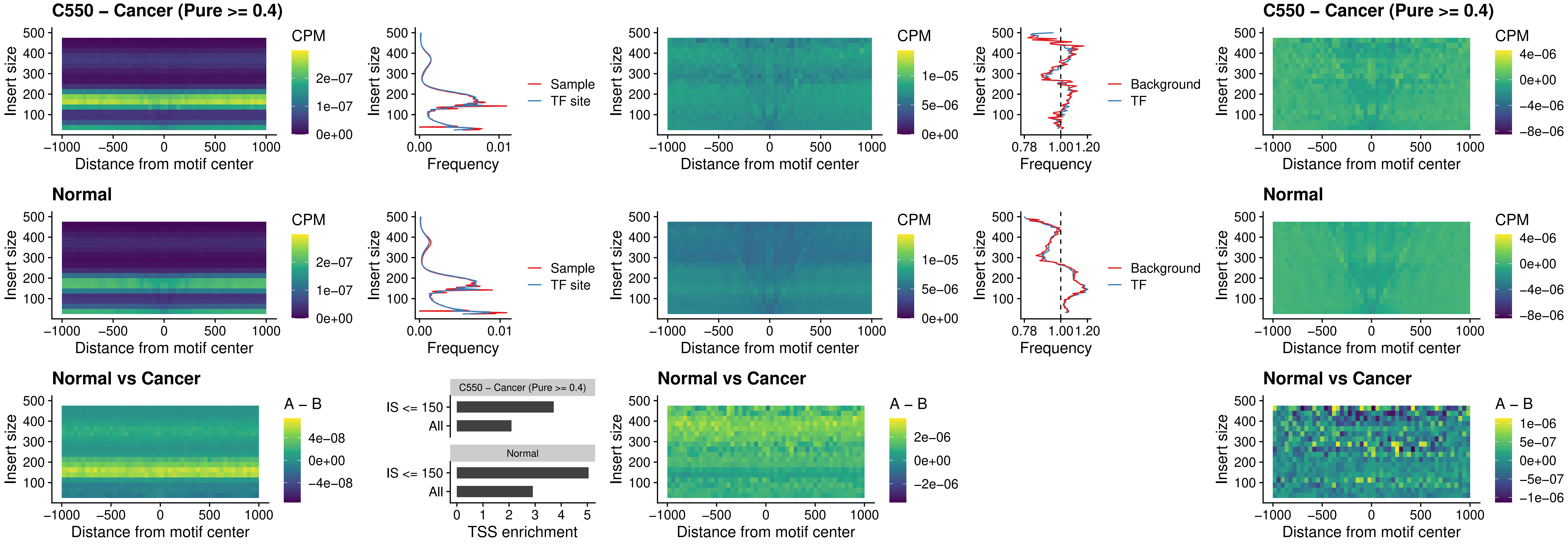

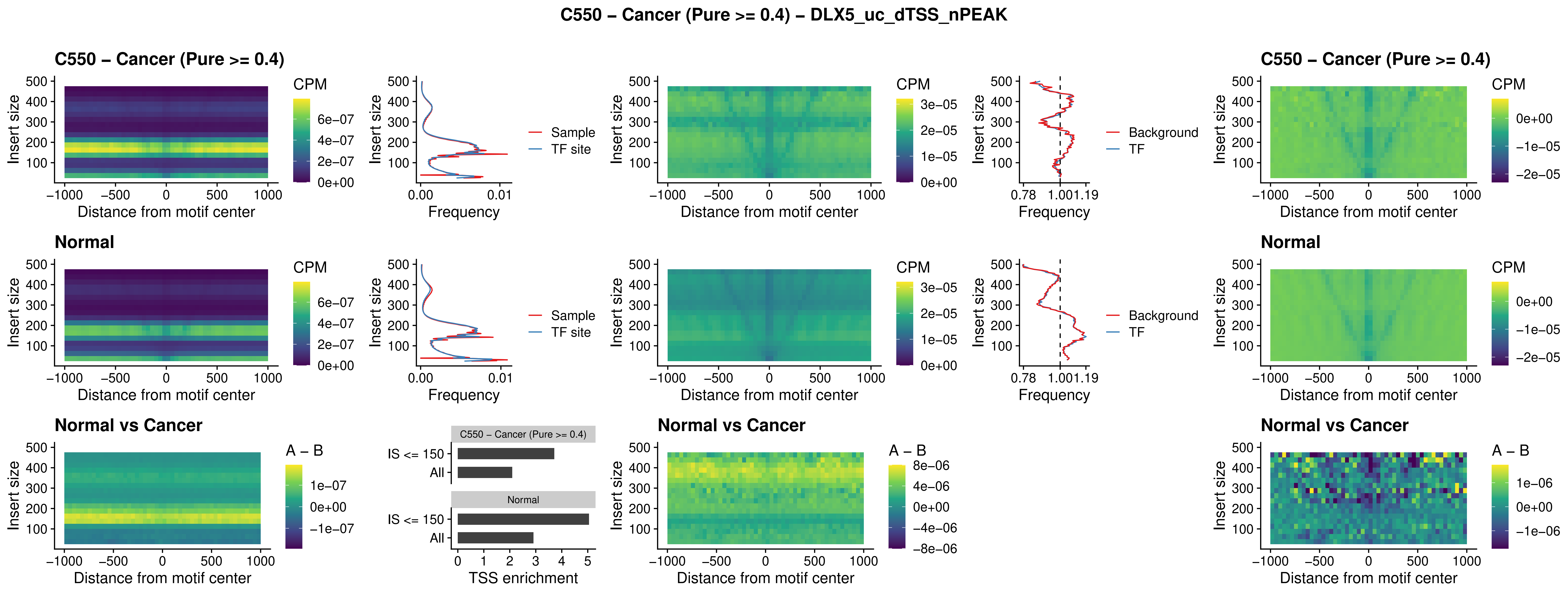

C550 – Cancer (Pure >= 0.4) – DLX5_uc_dTSS_nPEAK
C550 – Cancer (Pure >= 0.4)
Normal
Normal vs Cancer
Insert size
Distance from motif center
Frequency
CPM
A – B
Sample
TF site
Background
TF
C550 – Cancer (Pure >= 0.4)
IS <= 150
All
TSS enrichment

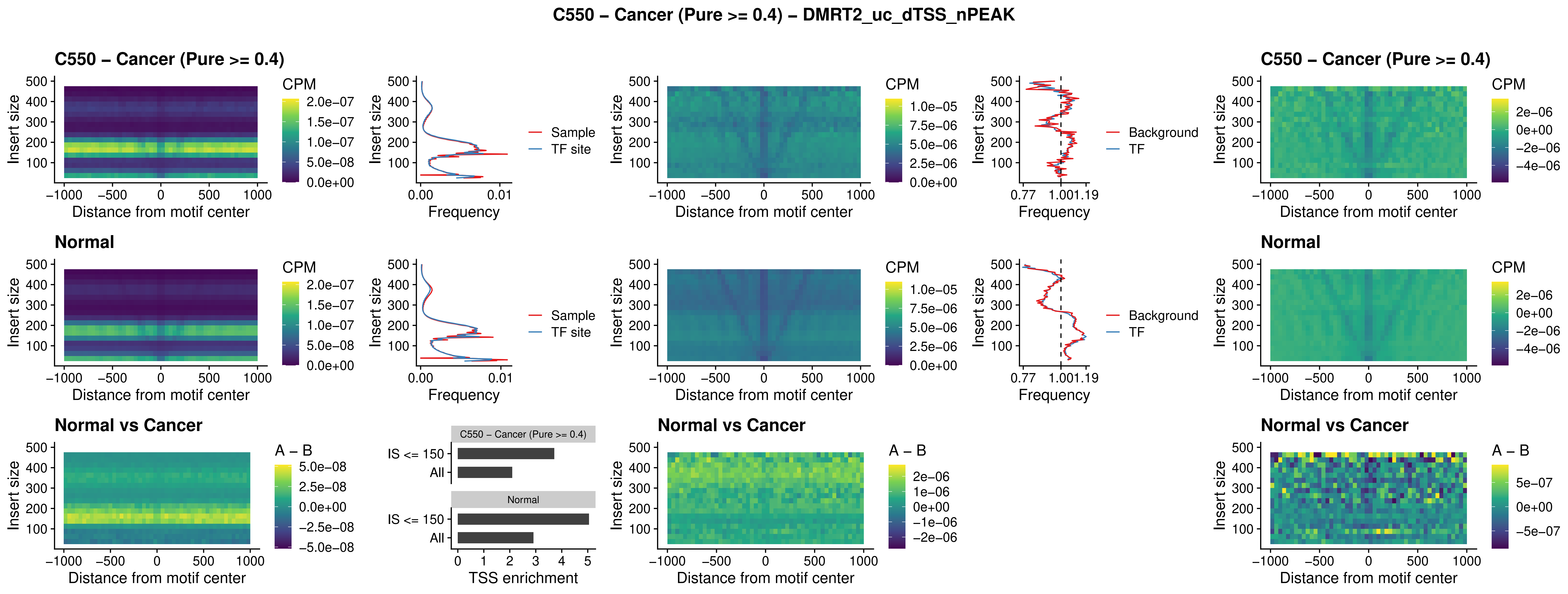

C550 – Cancer (Pure >= 0.4) – DMRT2_uc_dTSS_nPEAK
C550 – Cancer (Pure >= 0.4)
Normal
Normal vs Cancer
Insert size
Distance from motif center
Frequency
CPM
A – B
Sample
TF site
Background
TF
C550 – Cancer (Pure >= 0.4)
Normal
IS <= 150
All
TSS enrichment

#### C550 – Cancer (Pure >= 0.4) – DMRTC2_uc_cTSS_nPEAK

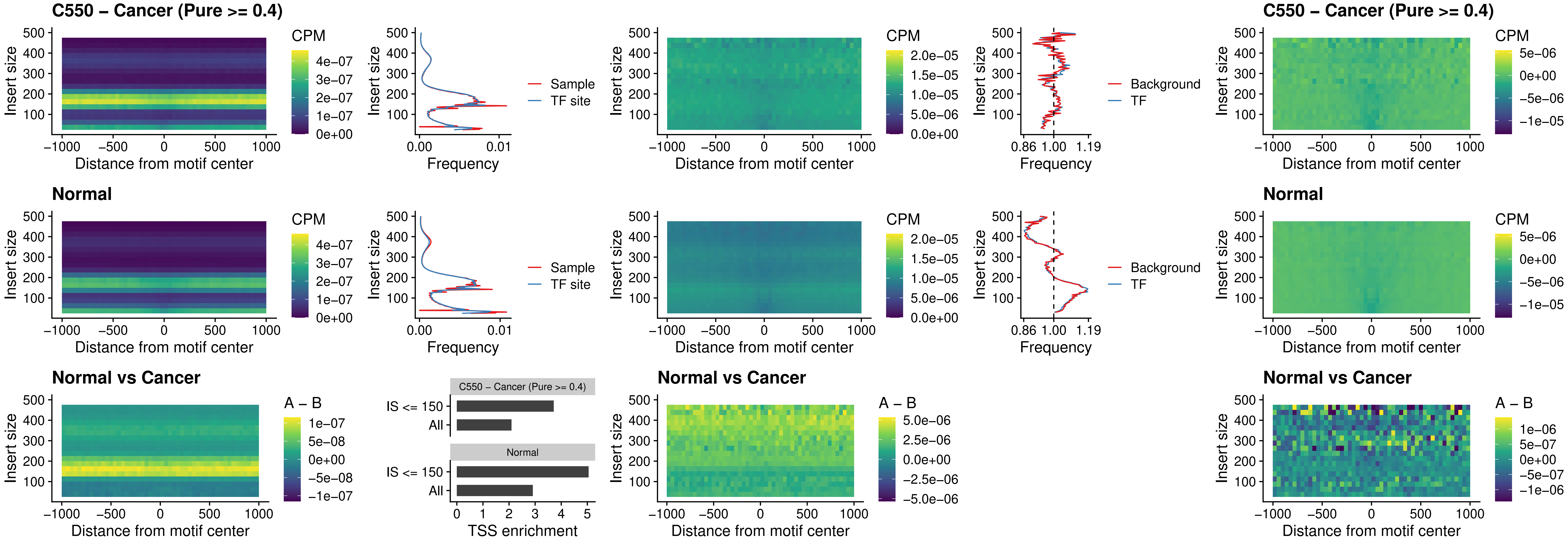

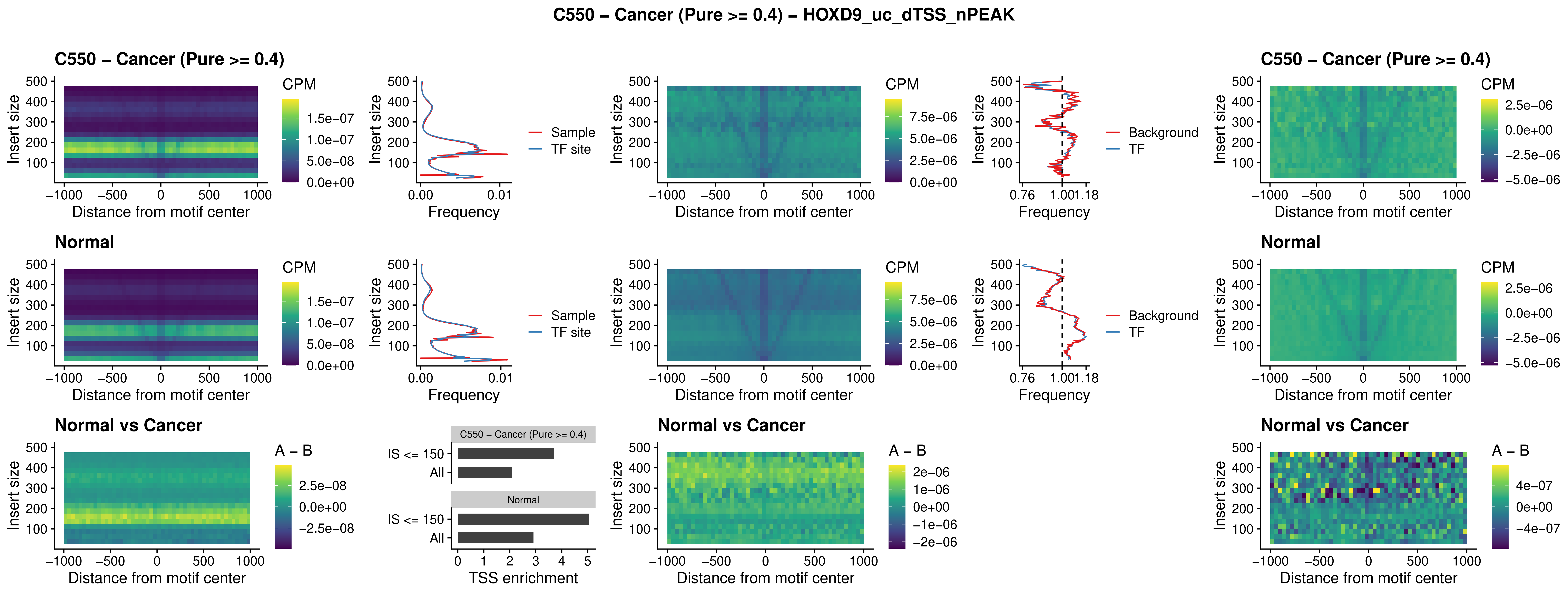
C550 – Cancer (Pure >= 0.4) – HOXD9_uc_dTSS_nPEAK
C550 – Cancer (Pure >= 0.4)
Normal
Normal vs Cancer
Insert size
Distance from motif center
CPM
A – B
Frequency
Sample
TF site
Background
TF
TSS enrichment
IS <= 150
All
C550 – Cancer (Pure >= 0.4)
Normal

#### C550 – Cancer (Pure >= 0.4) – HOXD4_uc_dTSS_nPEAK

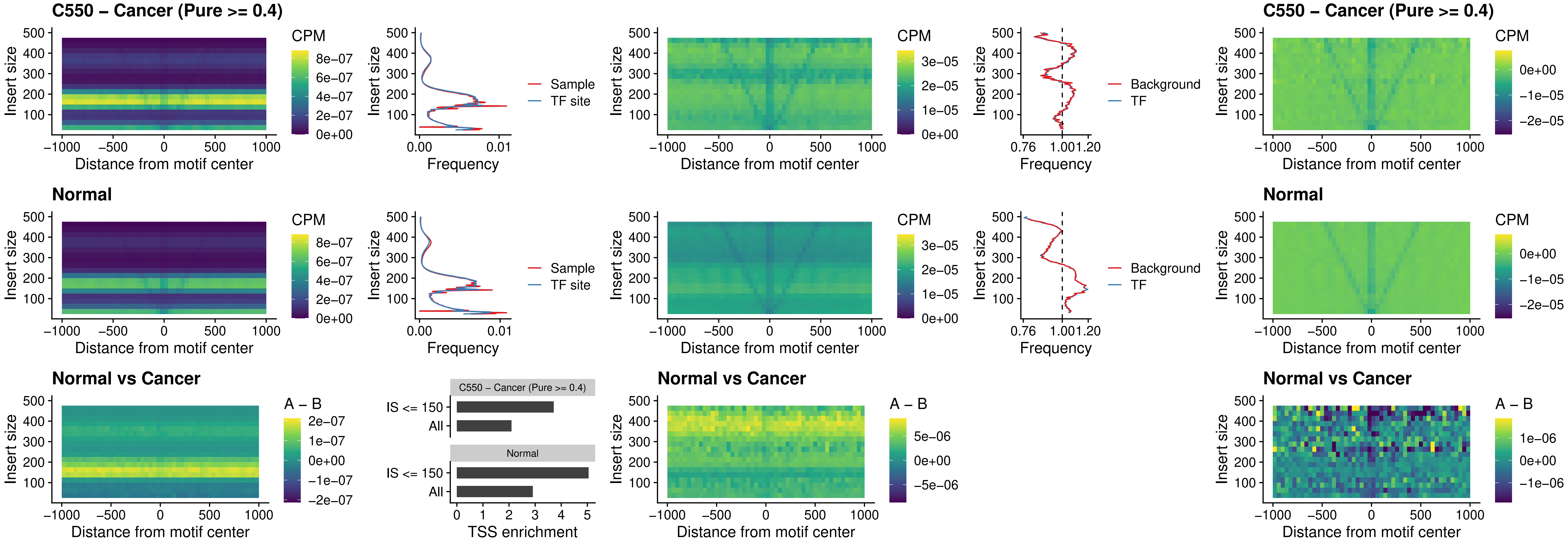

### C550 – Cancer (Pure >= 0.4) – FOXP3_uc_cTSS_nPEAK

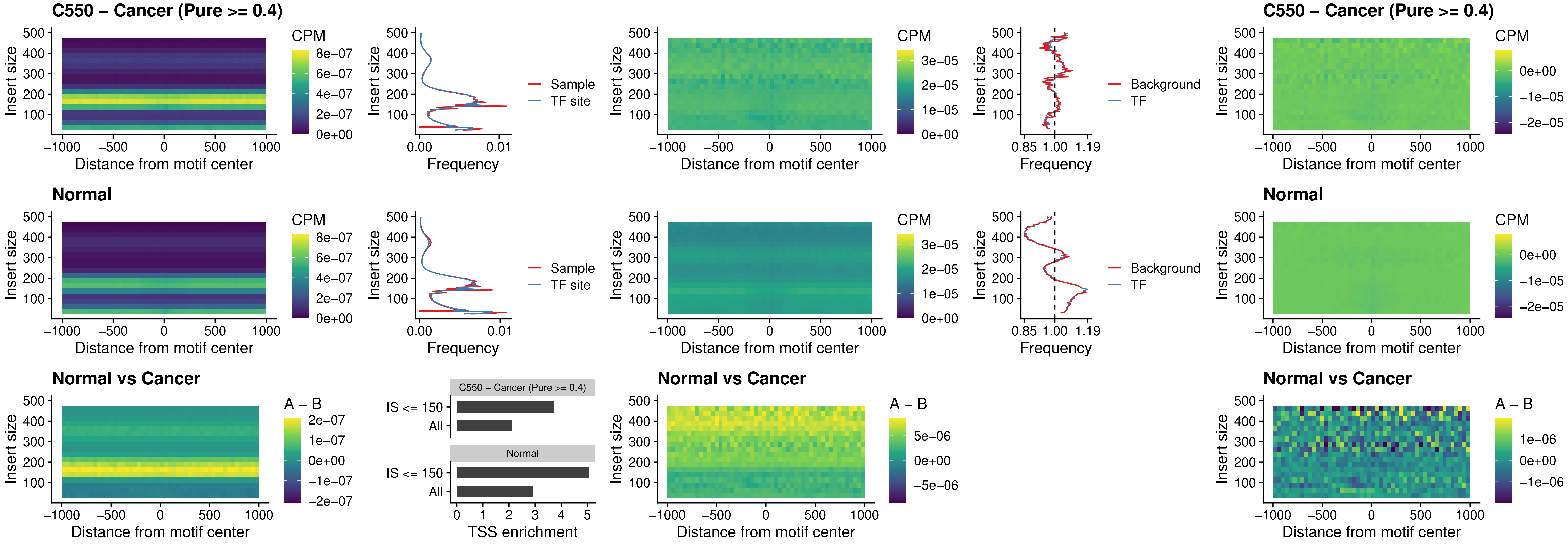

#### C550 – Cancer (Pure >= 0.4) – FOXD4L6_uc_cTSS_nPEAK

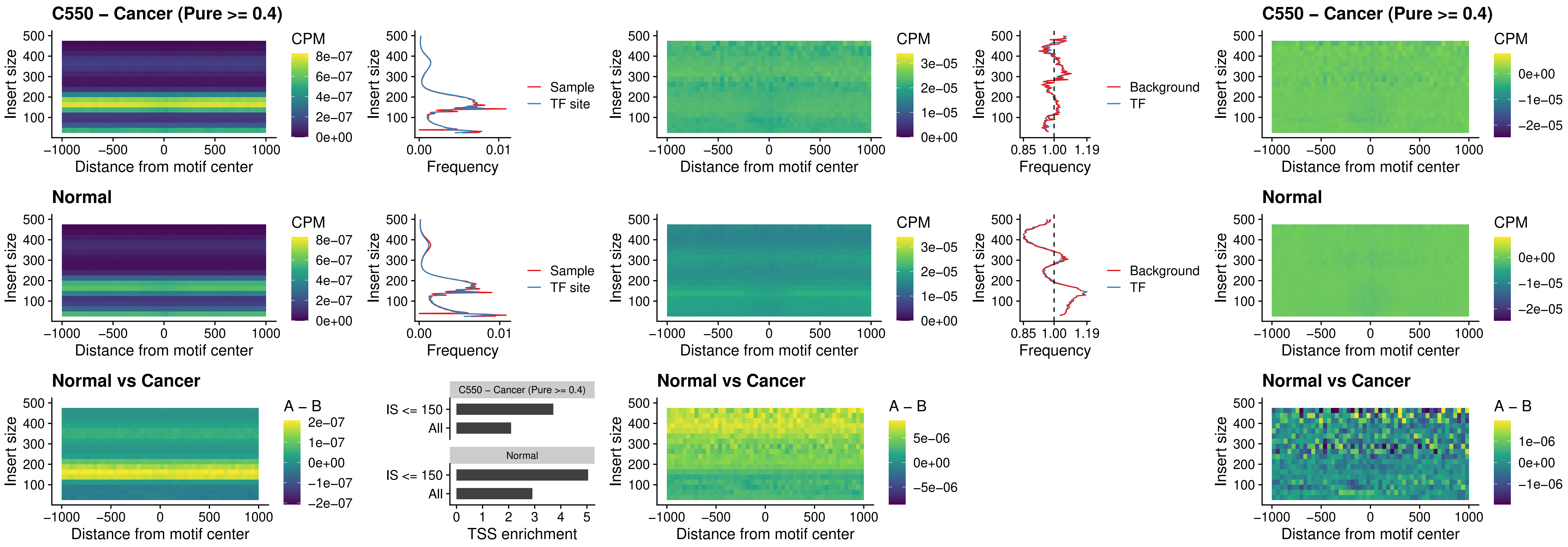

#### C550 – Cancer (Pure >= 0.4) – FOXD4L5_uc_cTSS_nPEAK

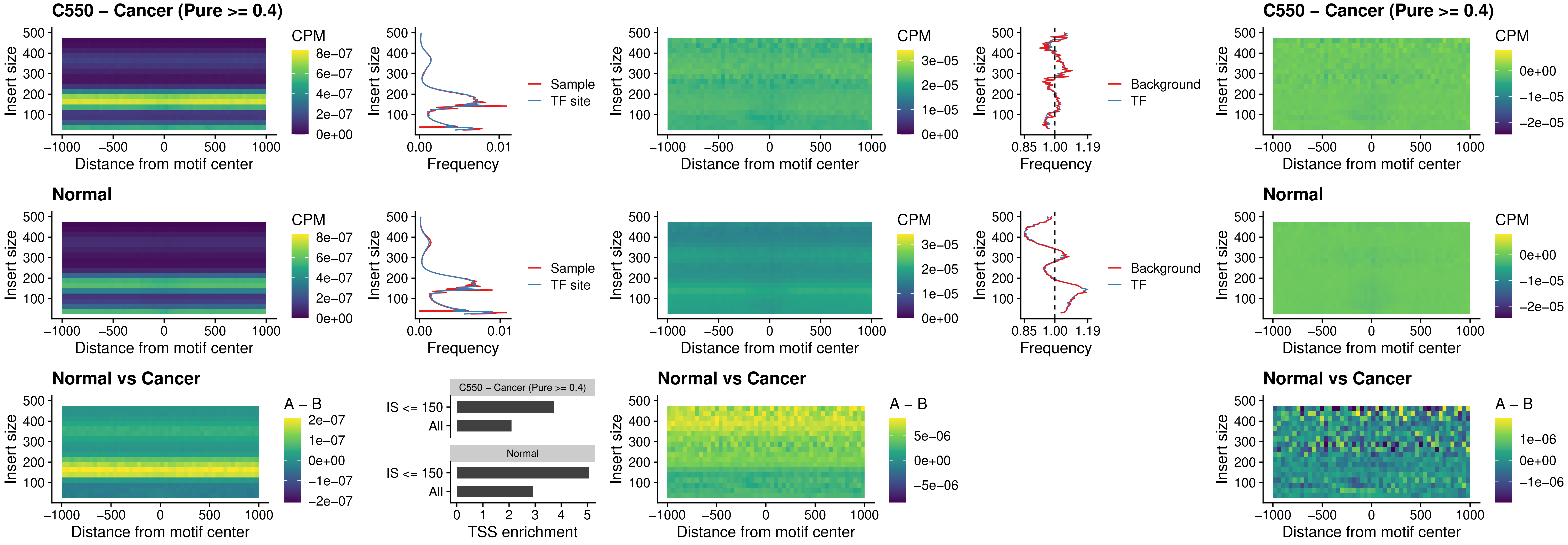

#### C550 – Cancer (Pure >= 0.4) – FOXD4L4_uc_cTSS_nPEAK

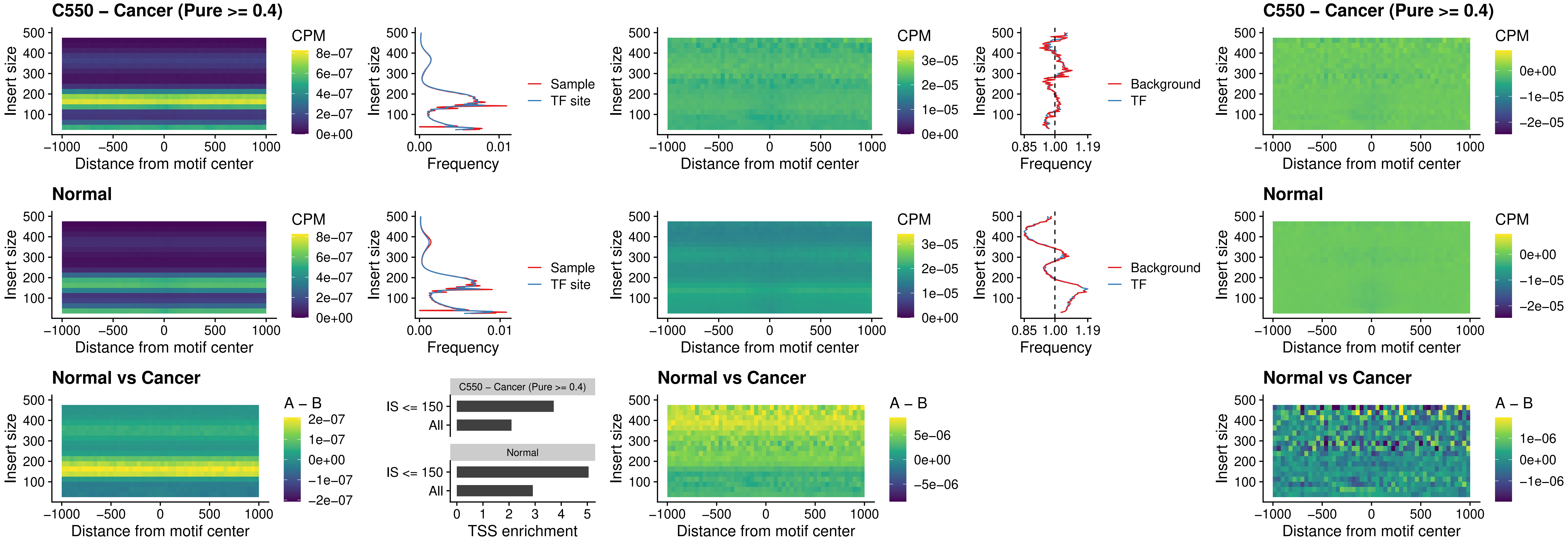

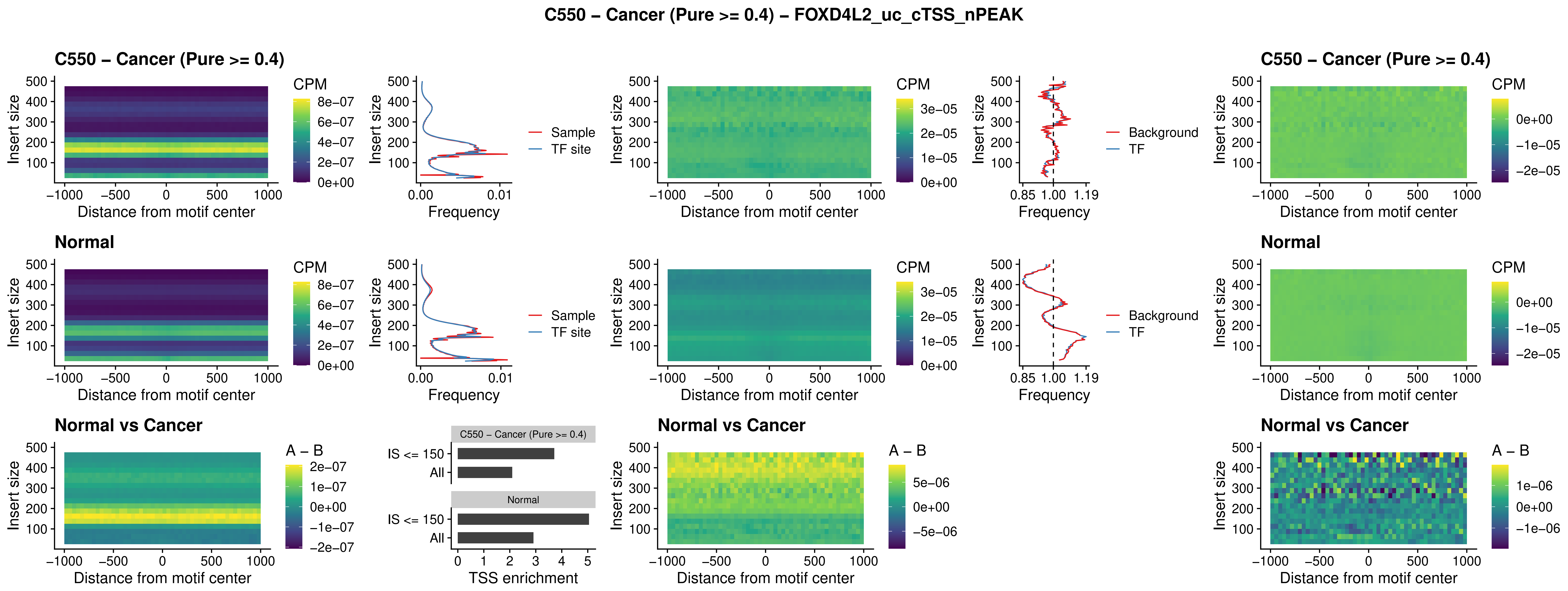
C550 – Cancer (Pure >= 0.4) – FOXD4L2_uc_cTSS_nPEAK
C550 – Cancer (Pure >= 0.4)
Normal
Normal vs Cancer
Insert size
Distance from motif center
Frequency
CPM
A – B
Sample
TF site
Background
TF
C550 – Cancer (Pure >= 0.4)
Normal
IS <= 150
All
TSS enrichment

#### C550 – Cancer (Pure >= 0.4) – FOXD4L3_uc_cTSS_nPEAK

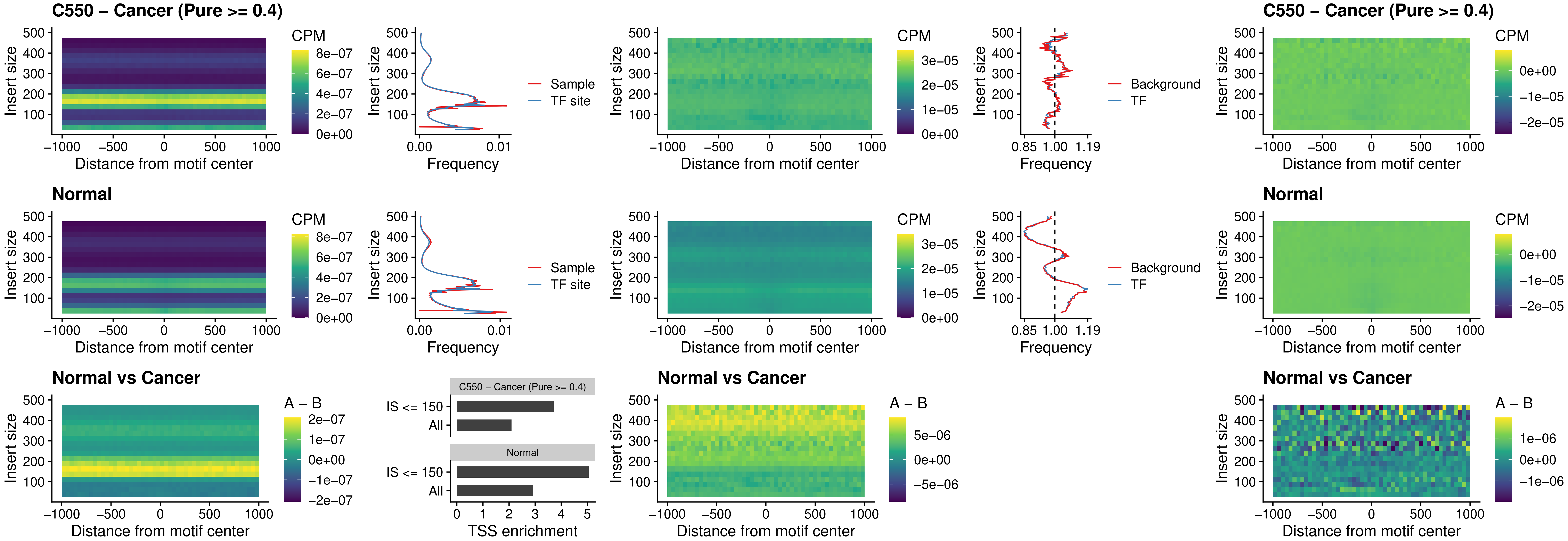

#### C550 – Cancer (Pure >= 0.4) – HOXD1_uc_dTSS_oPEAK

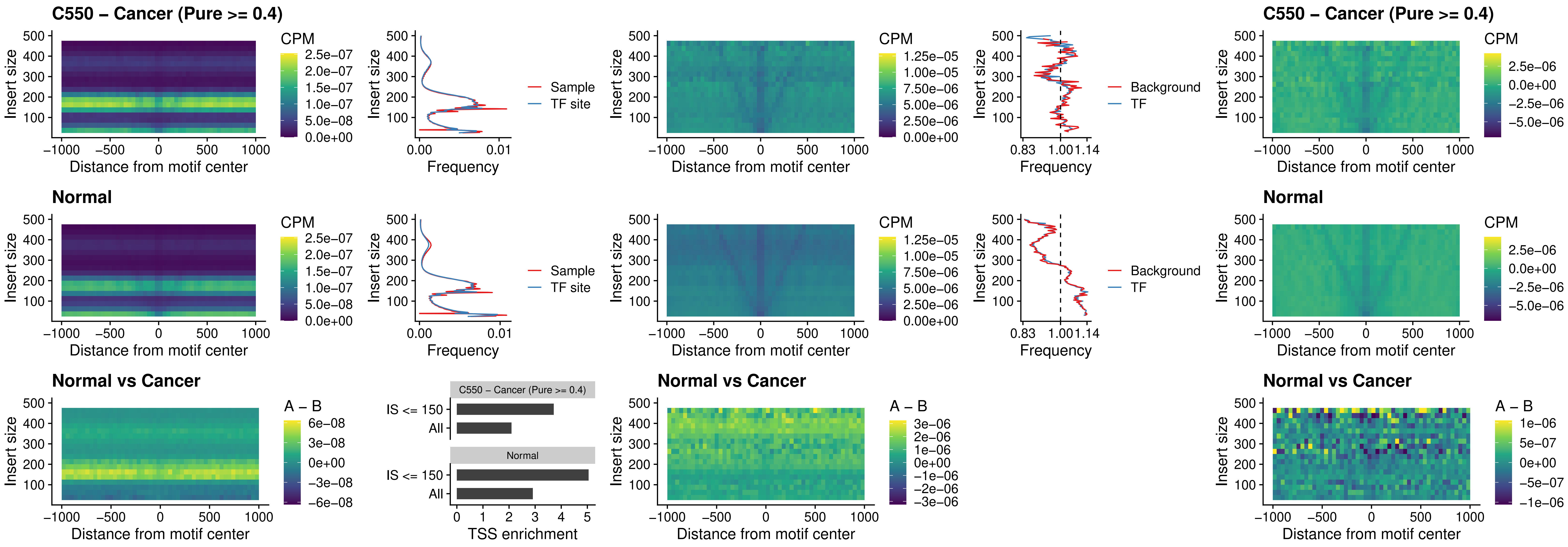

#### C550 – Cancer (Pure >= 0.4) – VAX1_uc_pTSS_nPEAK

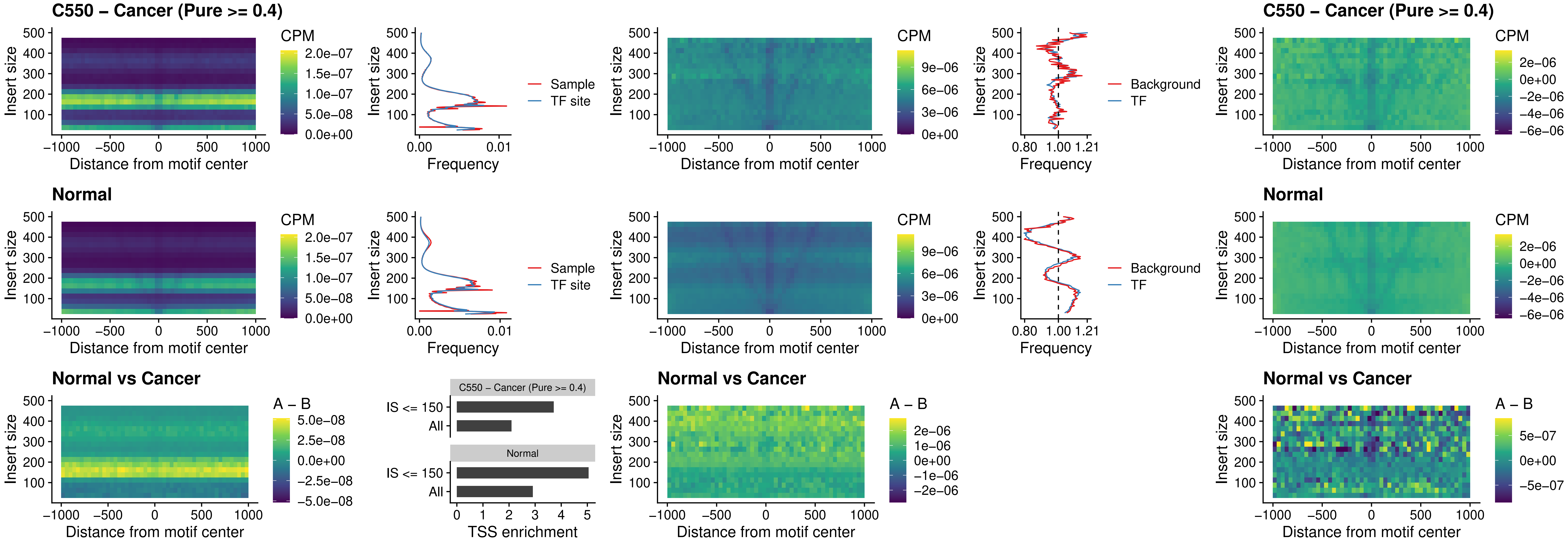

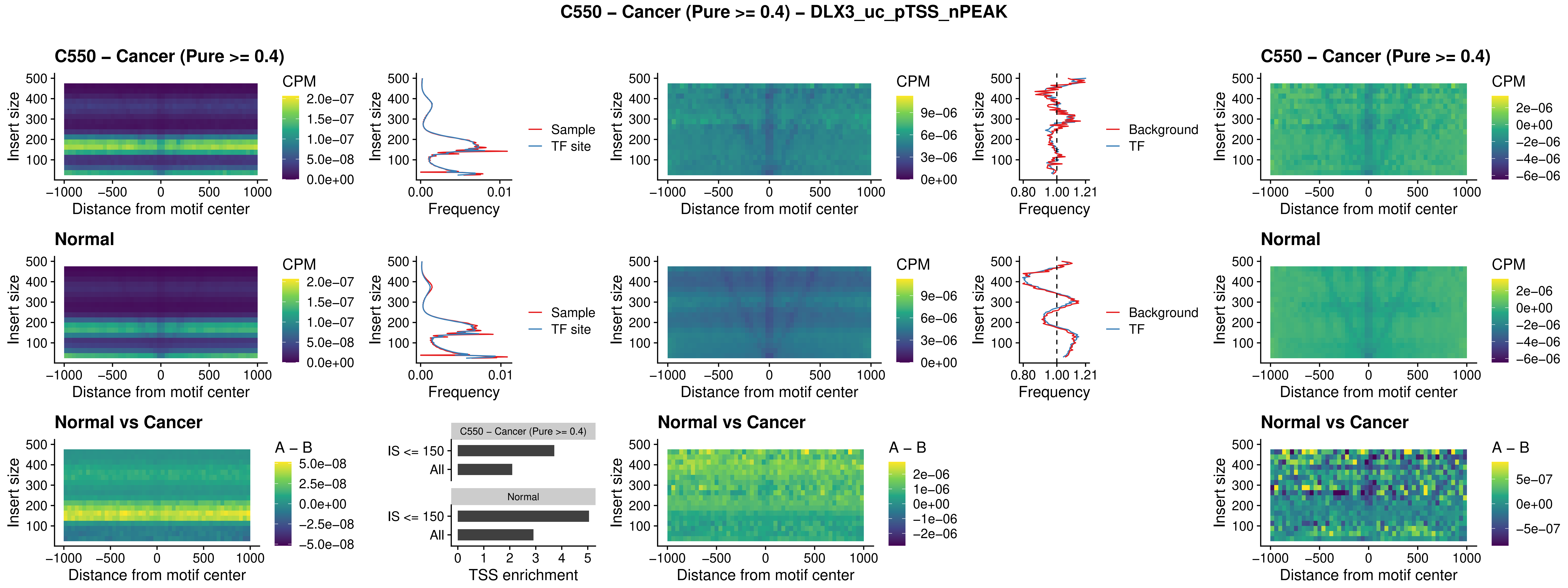

C550 – Cancer (Pure >= 0.4) – DLX3_uc_pTSS_nPEAK
C550 – Cancer (Pure >= 0.4)
Normal
Normal vs Cancer
Insert size
Distance from motif center
Frequency
CPM
A – B
Sample
TF site
Background
TF
C550 – Cancer (Pure >= 0.4)
Normal
IS <= 150
All
TSS enrichment

#### C550 – Cancer (Pure >= 0.4) – HOXD3_uc_pTSS_nPEAK

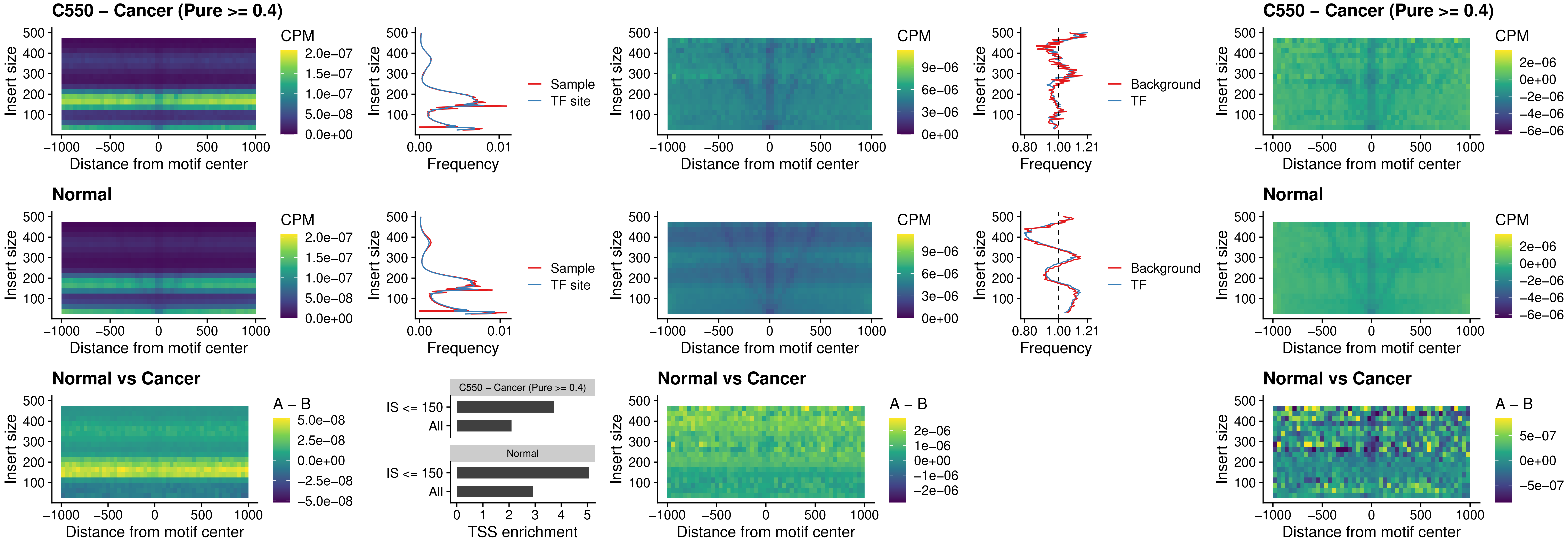

### C550 – Cancer (Pure >= 0.4) – PDX1_uc_dTSS_nPEAK

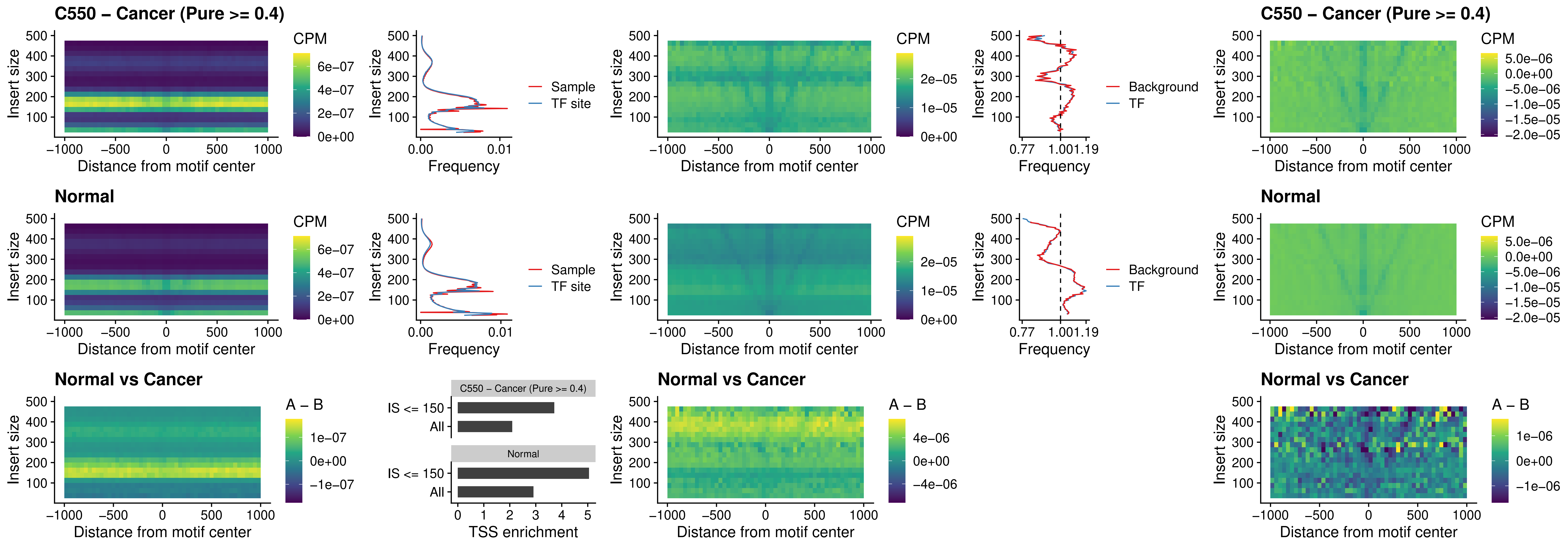

### C550 – Cancer (Pure >= 0.4) – VAX2_uc_dTSS_nPEAK

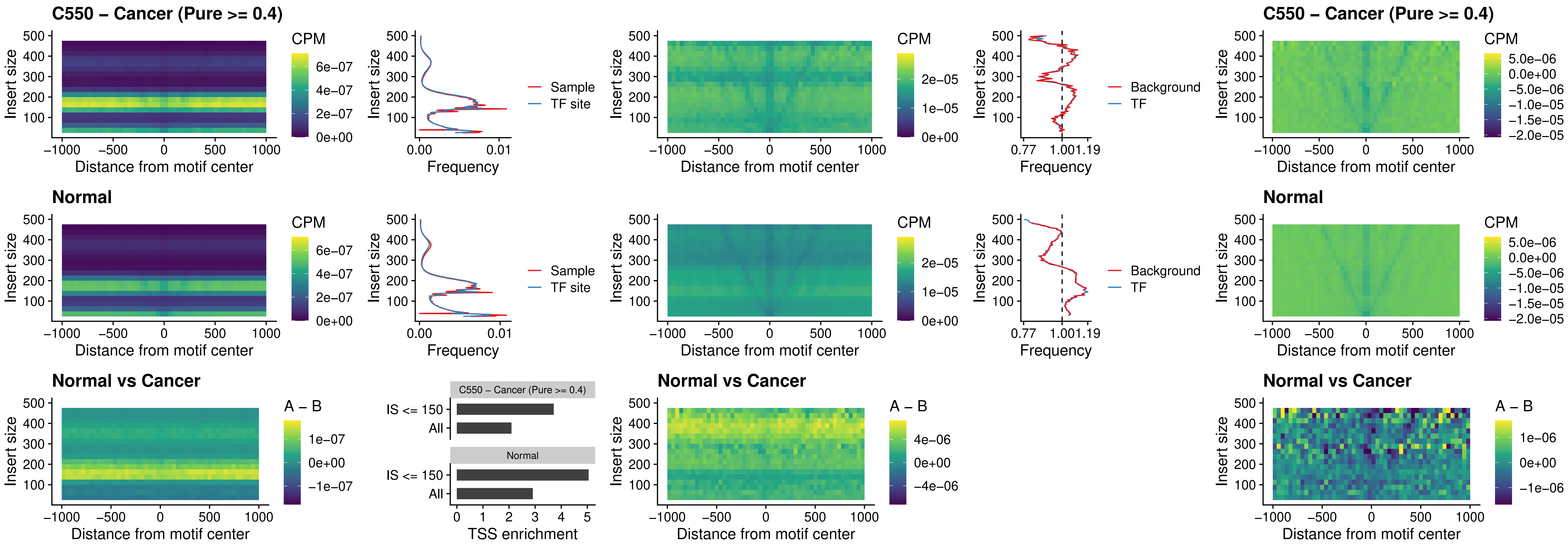

#### C550 – Cancer (Pure >= 0.4) – SOX5_uc_dTSS_nPEAK

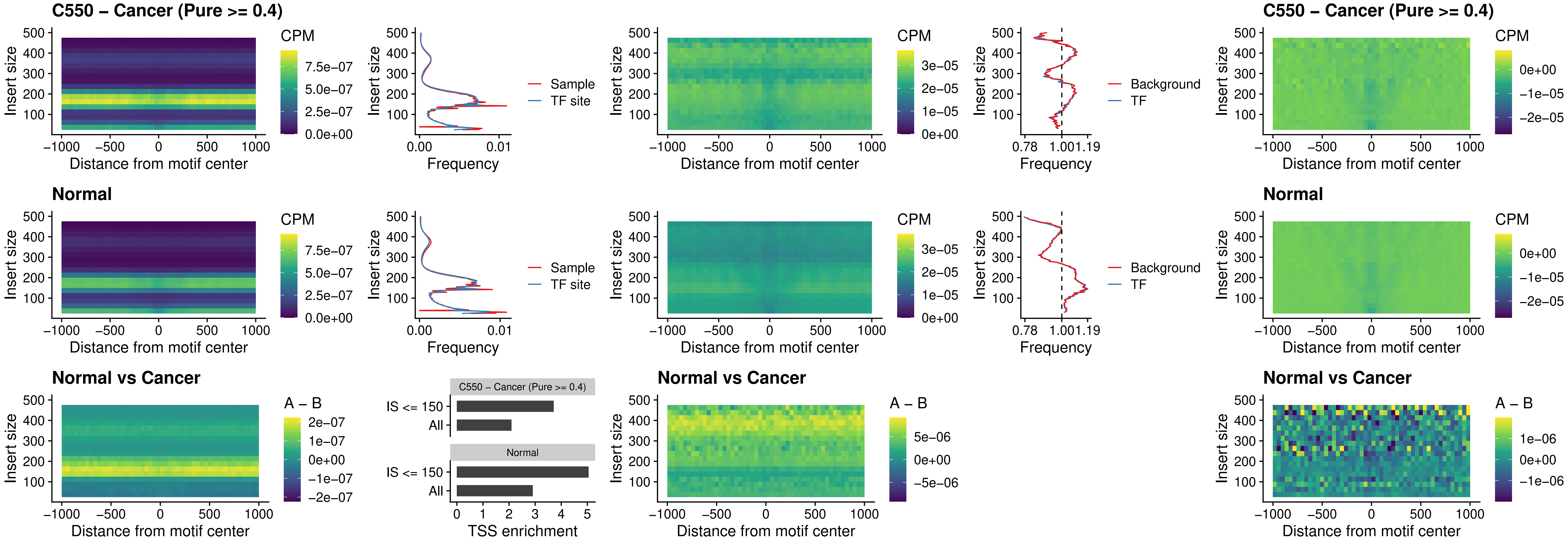

#### C550 – Cancer (Pure >= 0.4) – SOX6_uc_dTSS_nPEAK

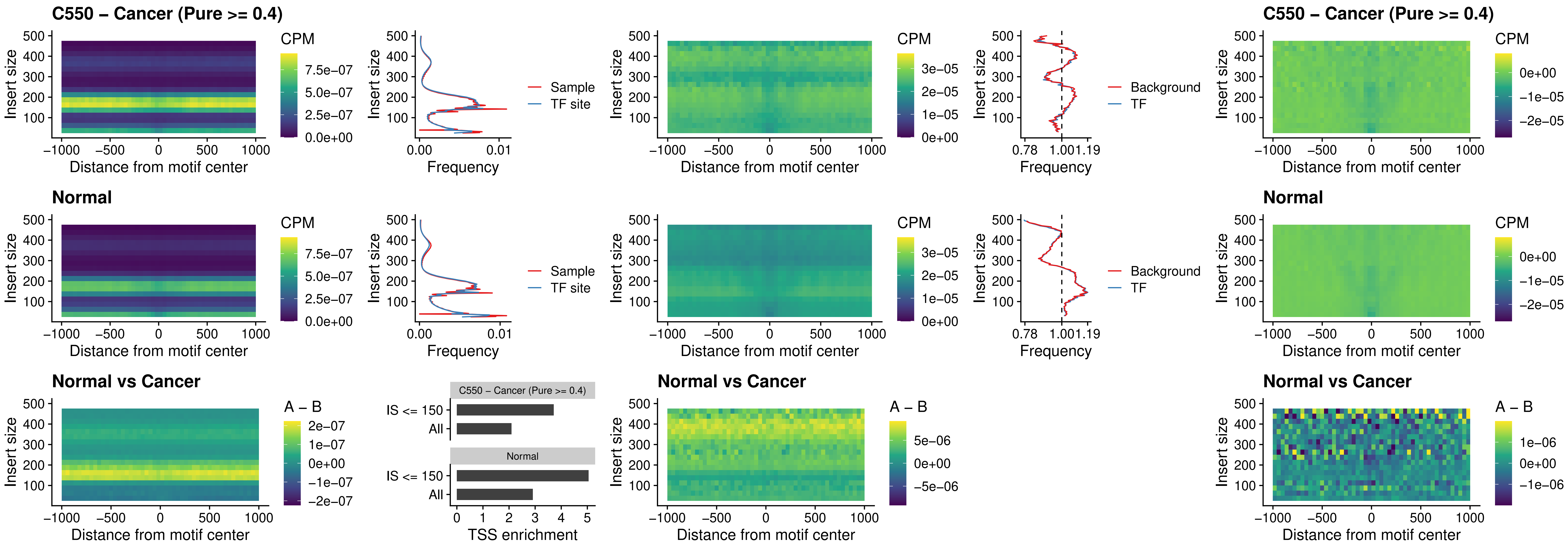

#### C550 – Cancer (Pure >= 0.4) – JDP2_uc_cTSS_nPEAK

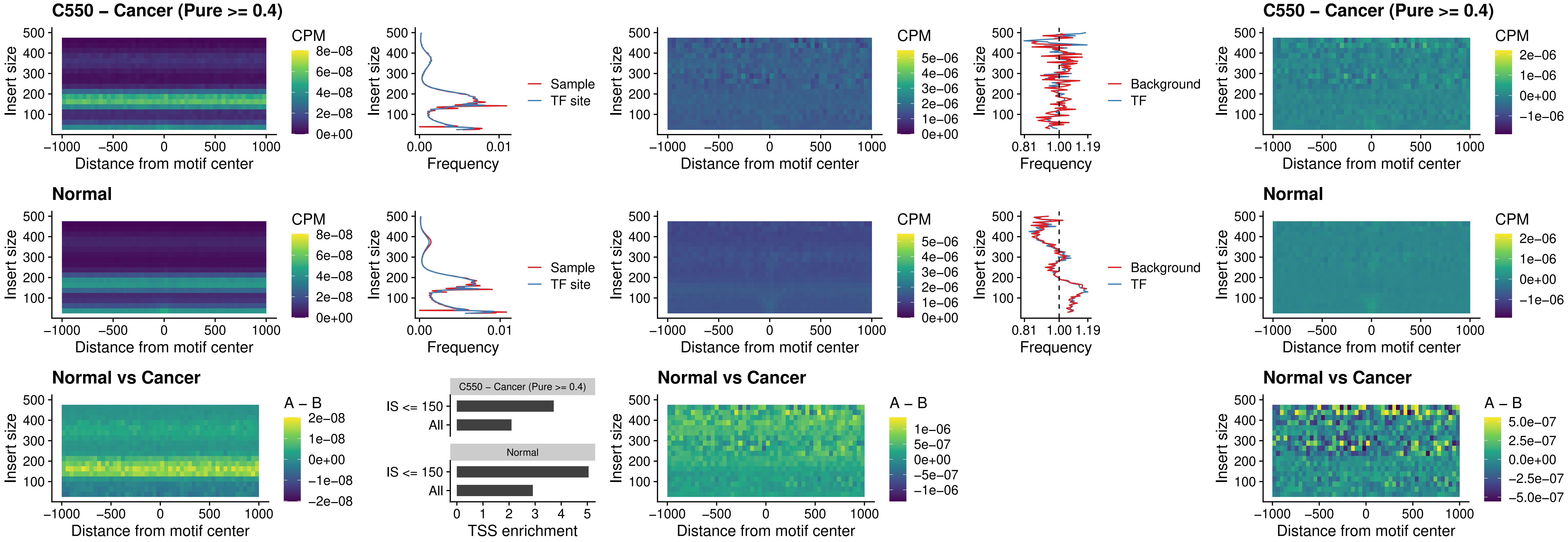

### C550 – Cancer (Pure >= 0.4) – PURA_uc_dTSS_oPEAK

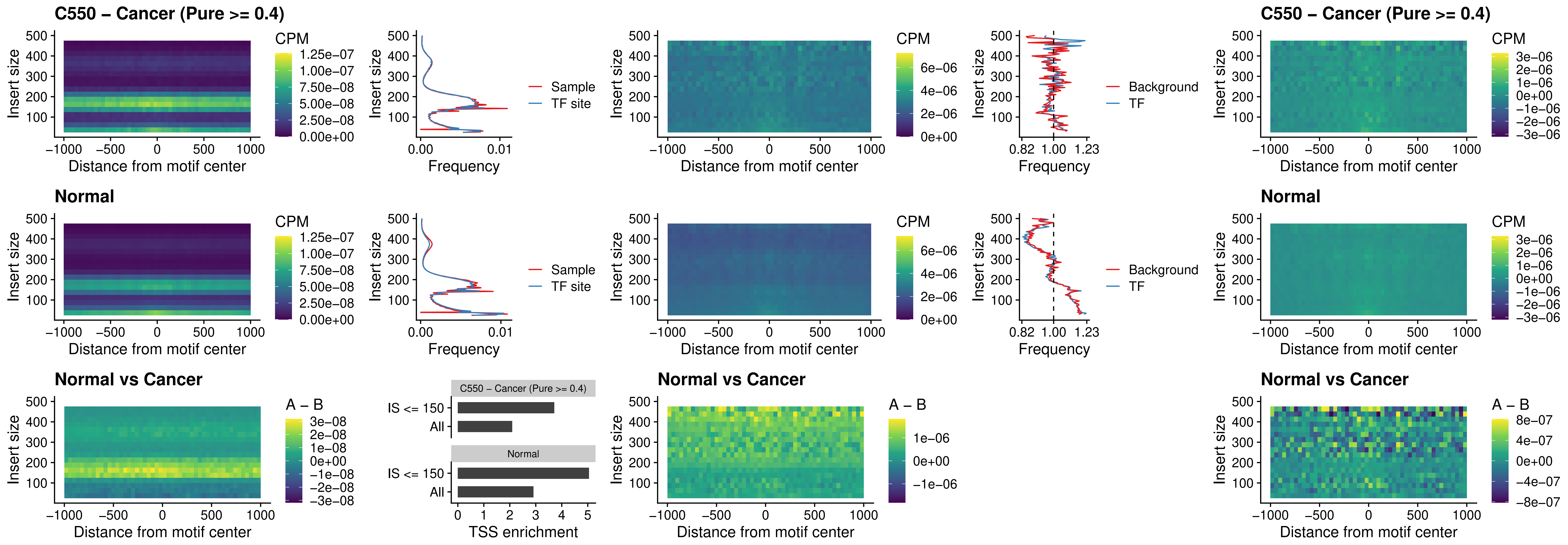

#### C550 – Cancer (Pure >= 0.4) – HHEX_uc_dTSS_nPEAK

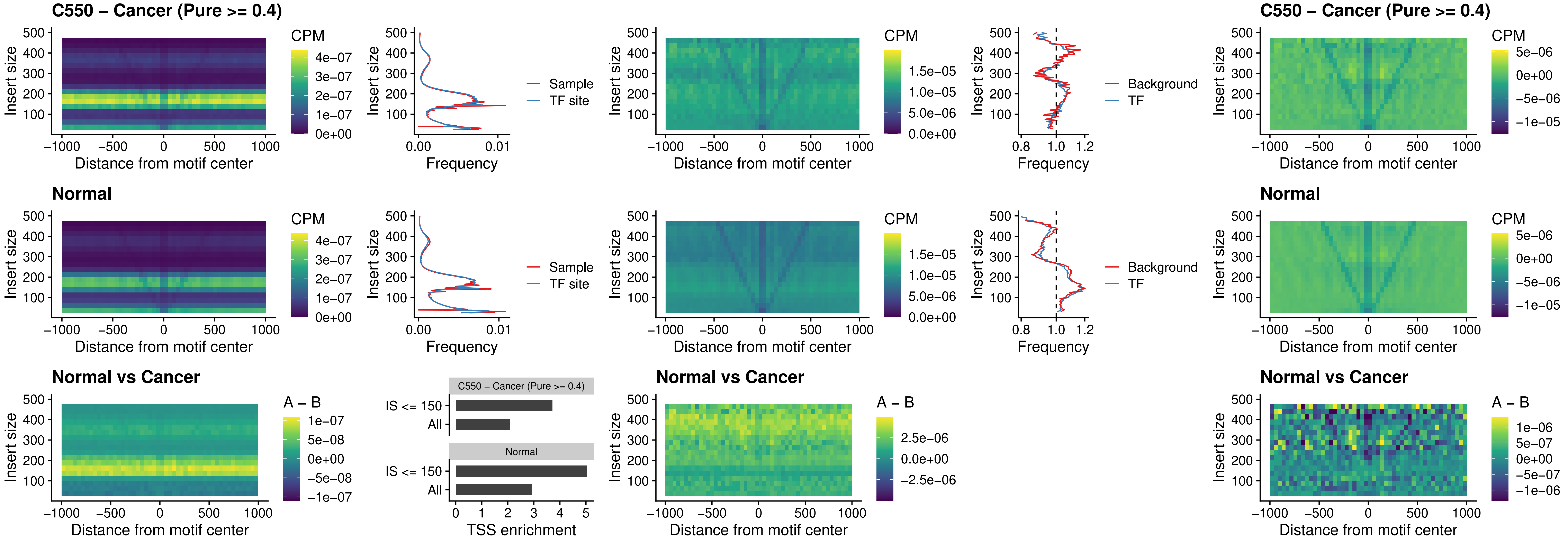

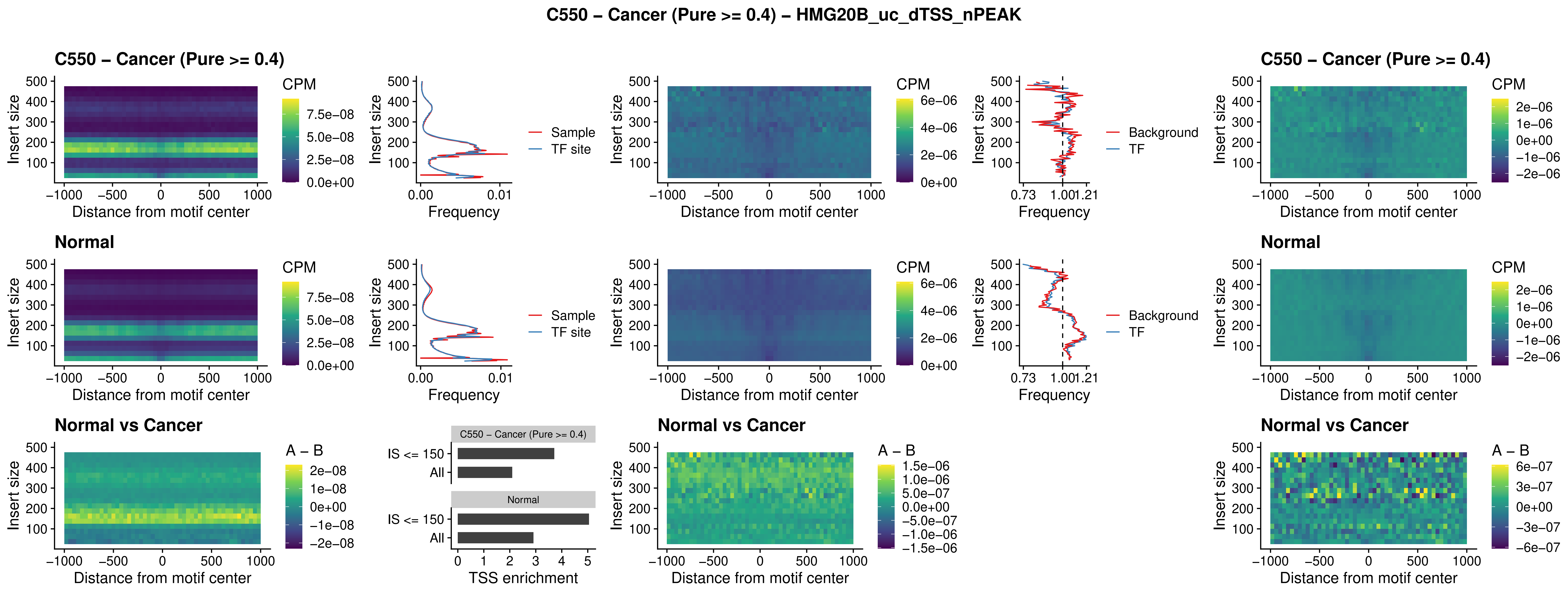

C550 – Cancer (Pure >= 0.4) – HMG20B_uc_dTSS_nPEAK
C550 – Cancer (Pure >= 0.4)
Normal
Normal vs Cancer
Insert size
Distance from motif center
CPM
A – B
Frequency
Sample
TF site
Background
TF
C550 – Cancer (Pure >= 0.4)
Normal
IS <= 150
All
TSS enrichment

#### C550 – Cancer (Pure >= 0.4) – FOXB2_uc_dTSS_nPEAK

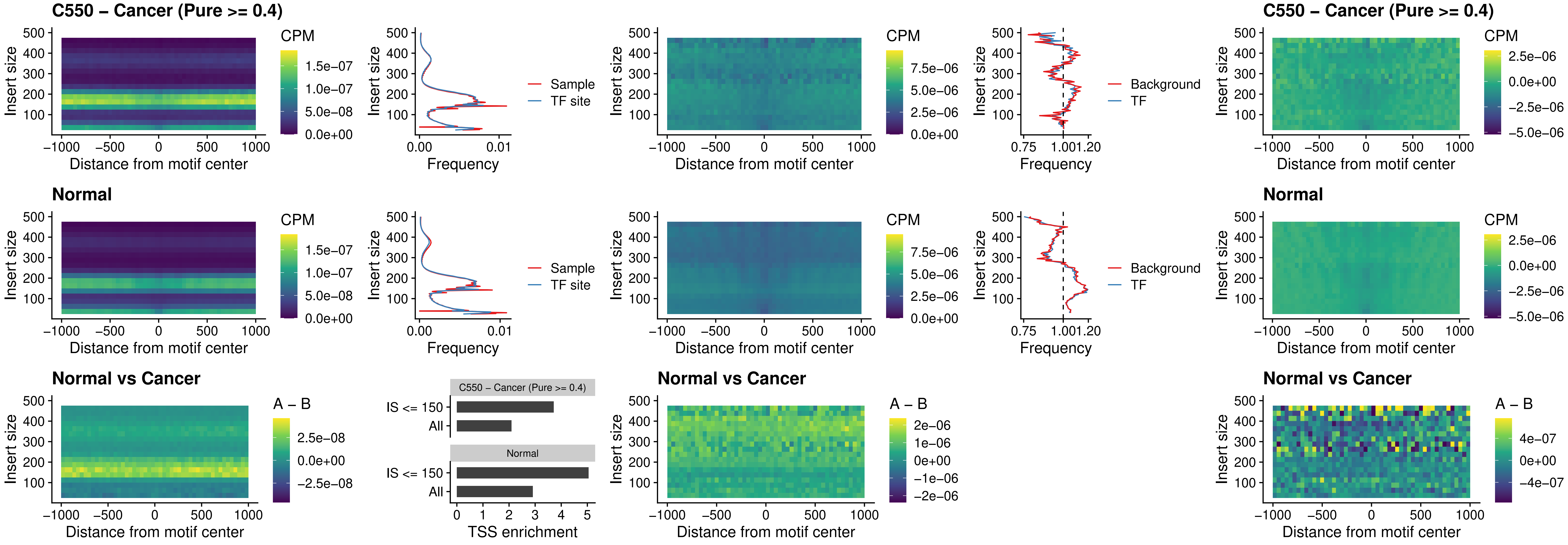

### C550 – Cancer (Pure >= 0.4) – FOXL1_uc_cTSS_nPEAK

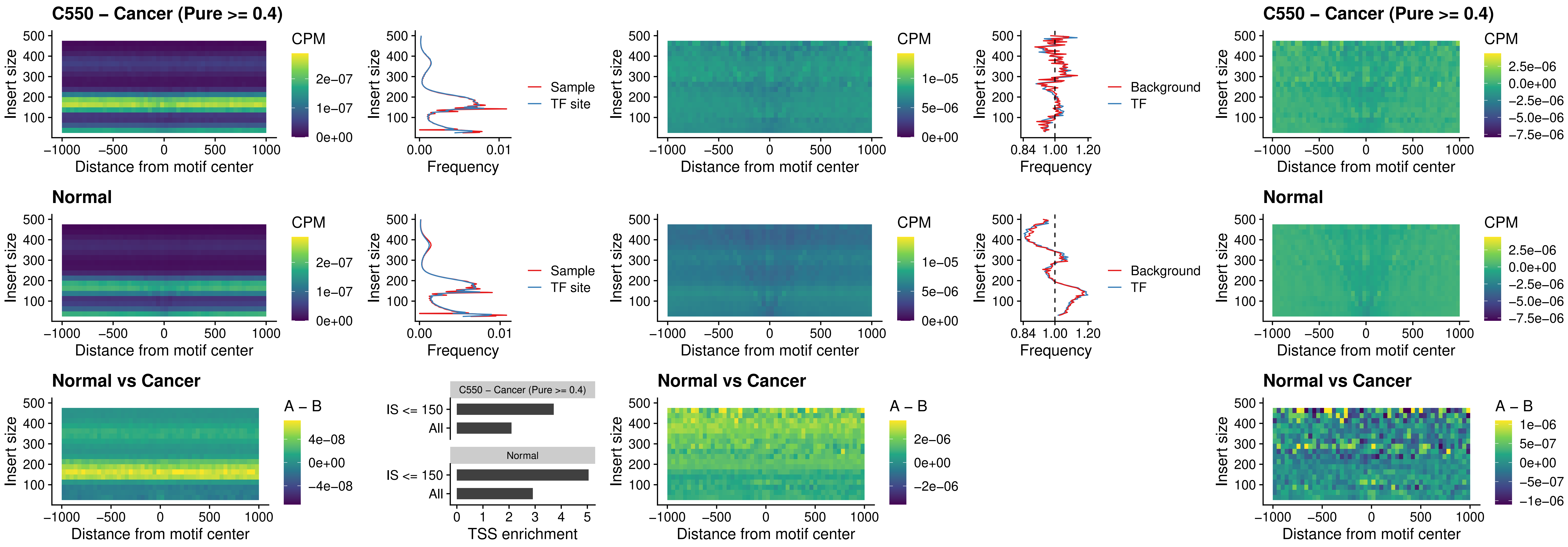

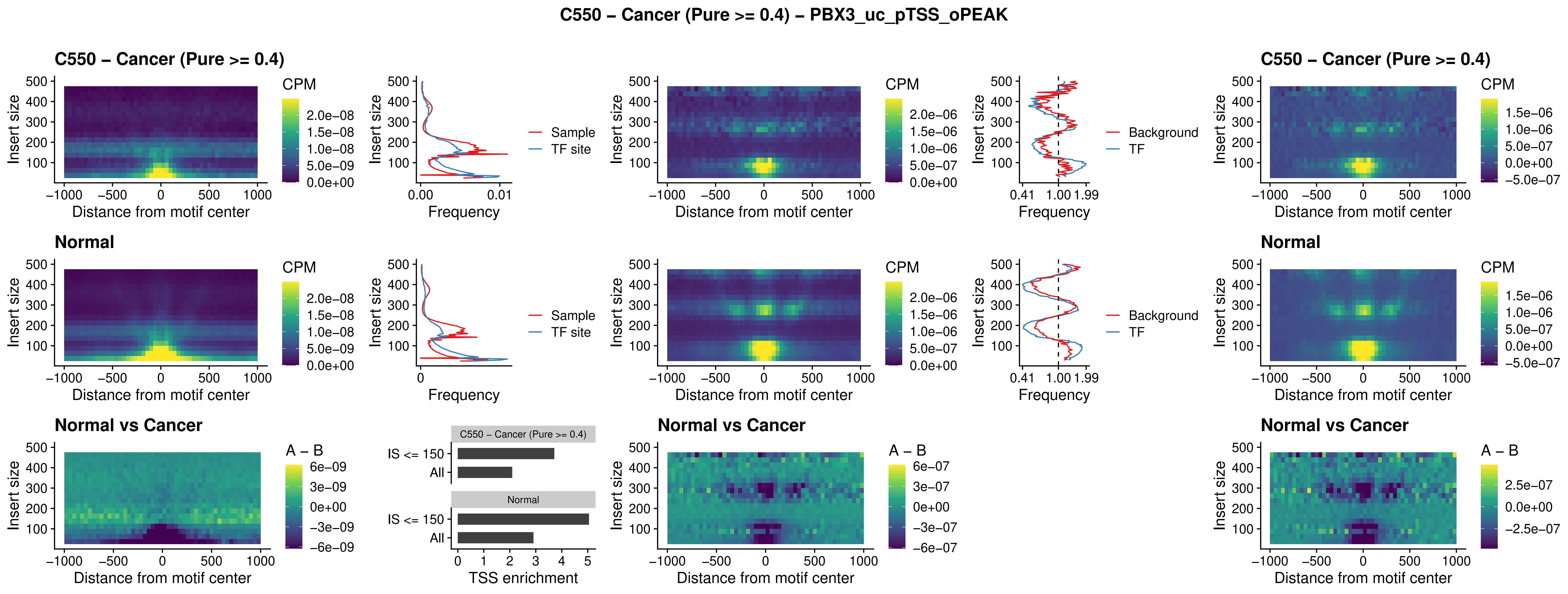
C550 – Cancer (Pure >= 0.4) – PBX3_uc_pTSS_oPEAK
C550 – Cancer (Pure >= 0.4)
Normal
Normal vs Cancer
Insert size
Distance from motif center
CPM
A – B
Frequency
Sample
TF site
Background
TF
C550 – Cancer (Pure >= 0.4)
Normal
IS <= 150
All
TSS enrichment

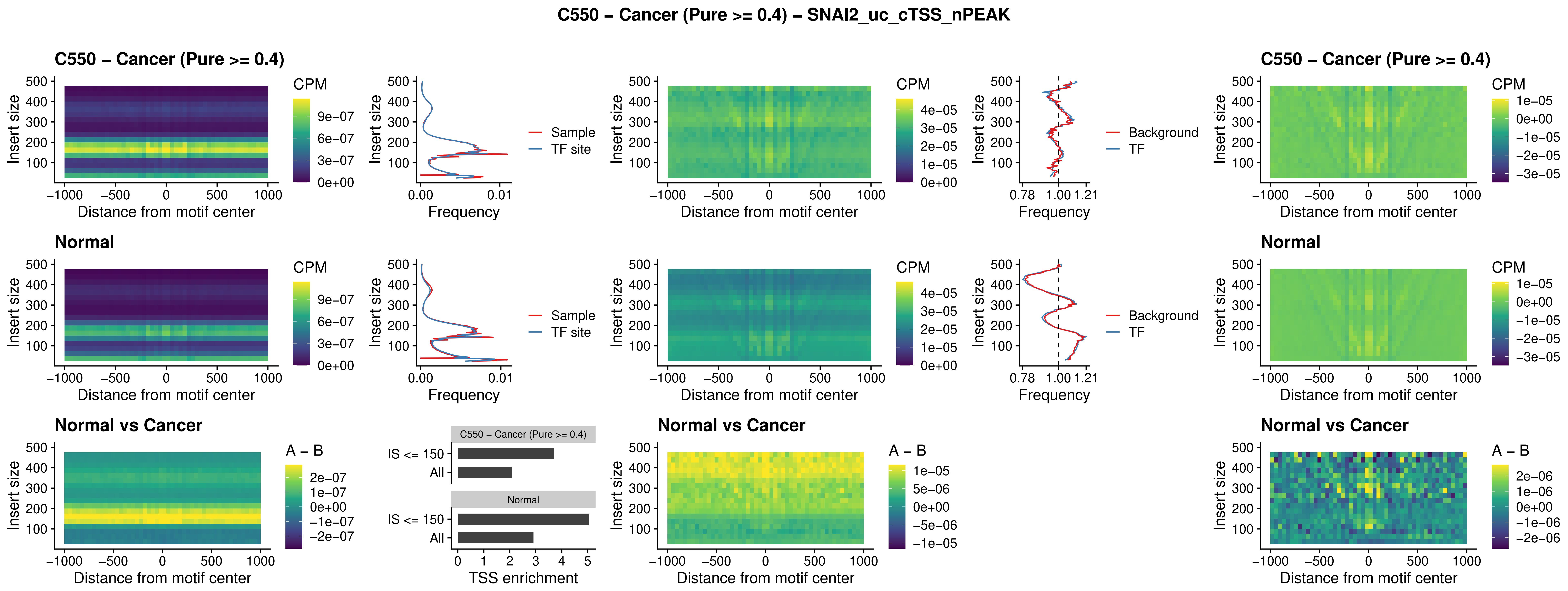
C550 – Cancer (Pure >= 0.4) – SNAI2_uc_cTSS_nPEAK
C550 – Cancer (Pure >= 0.4)
Normal
Normal vs Cancer
Insert size
Distance from motif center
Frequency
CPM
A – B
Sample
TF site
Background
TF
C550 – Cancer (Pure >= 0.4)
Normal
IS <= 150
All
TSS enrichment

#### C550 – Cancer (Pure >= 0.4) – SNAI2_uc_dTSS_oPEAK

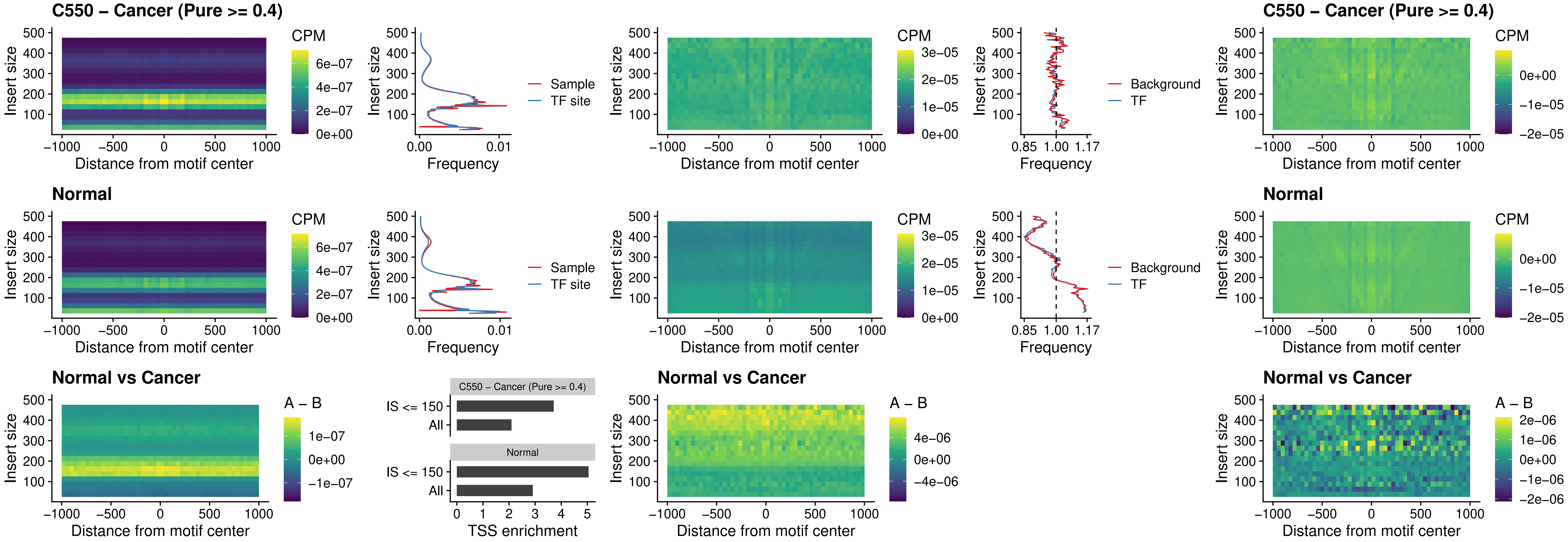

#### C550 – Cancer (Pure >= 0.4) – SPI1_uc_pTSS_nPEAK

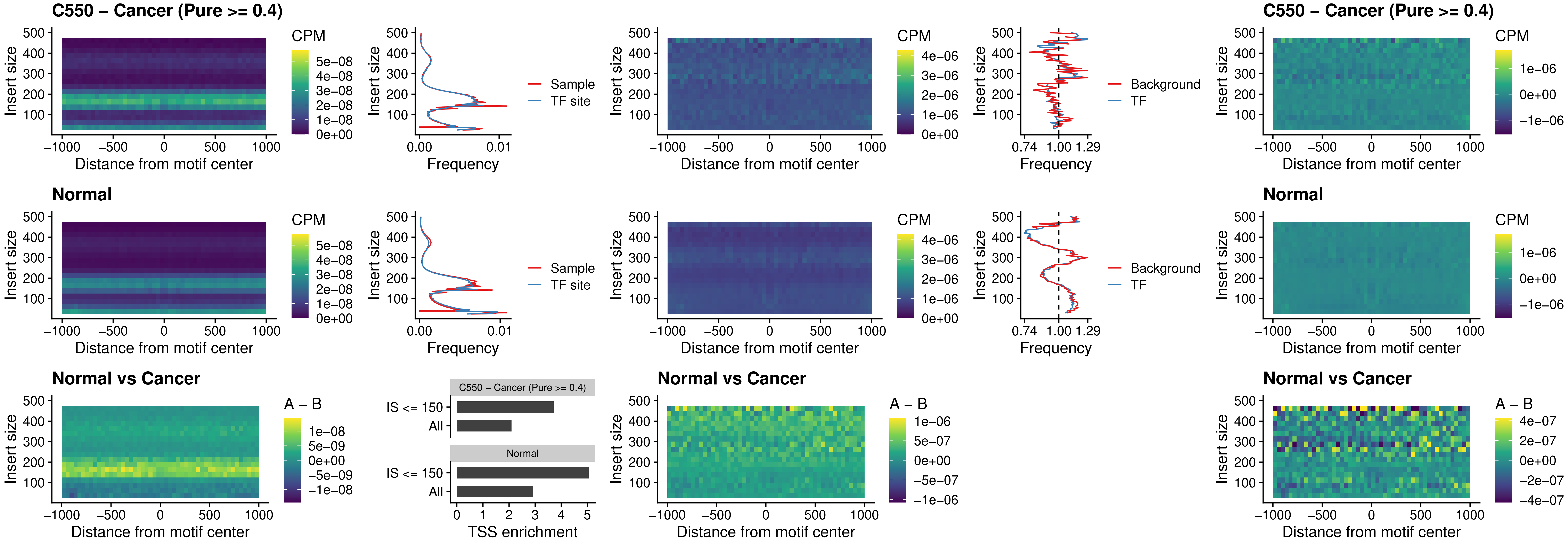

#### C550 – Cancer (Pure >= 0.4) – KLF16_uc_pTSS_nPEAK

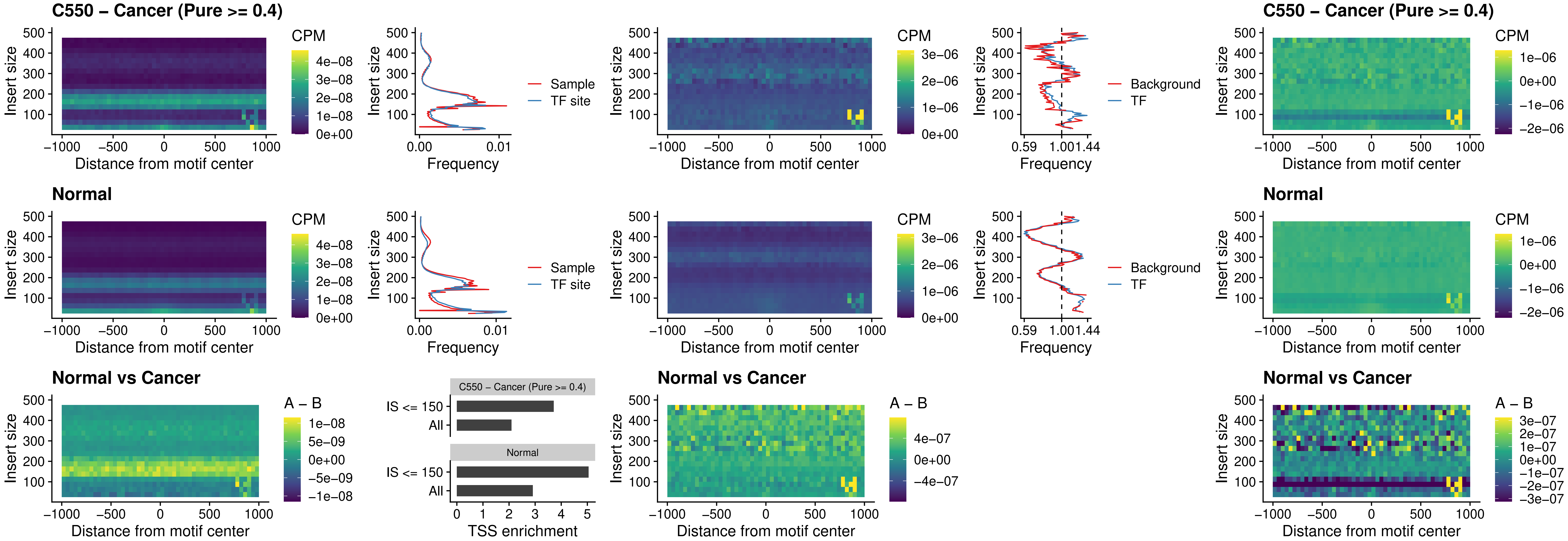

#### C550 – Cancer (Pure >= 0.4) – SP3_uc_pTSS_nPEAK

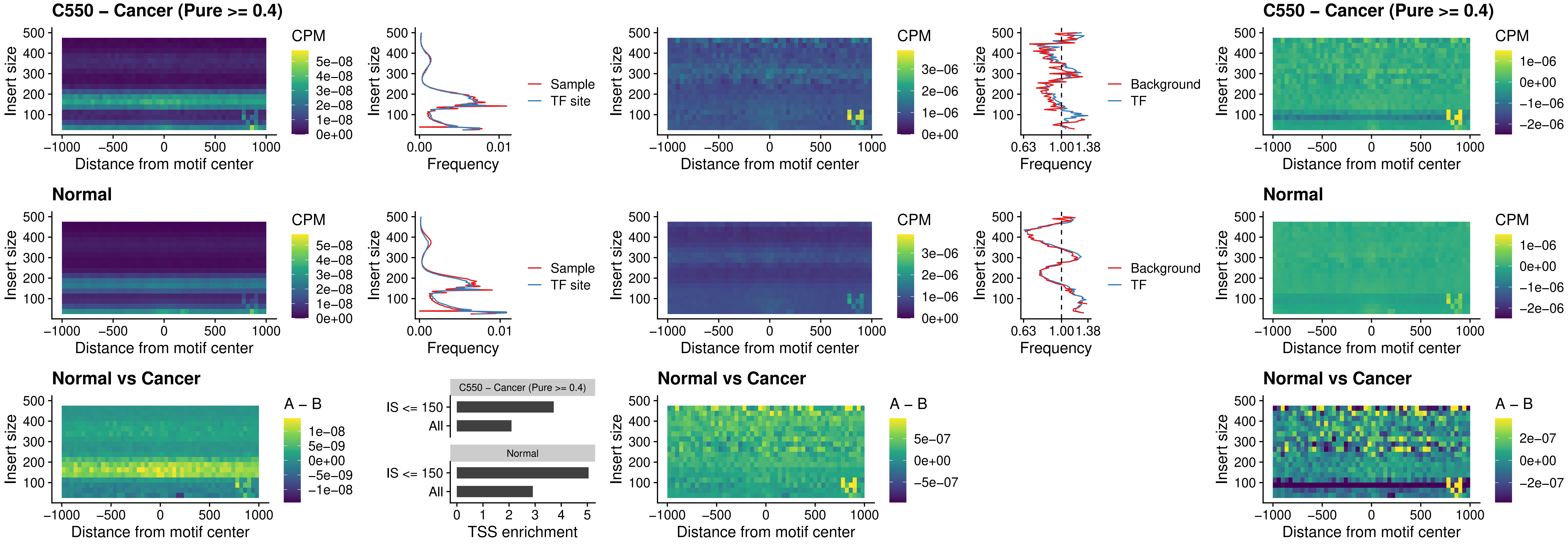

#### C550 – Cancer (Pure >= 0.4) – NEUROD1_uc_dTSS_oPEAK

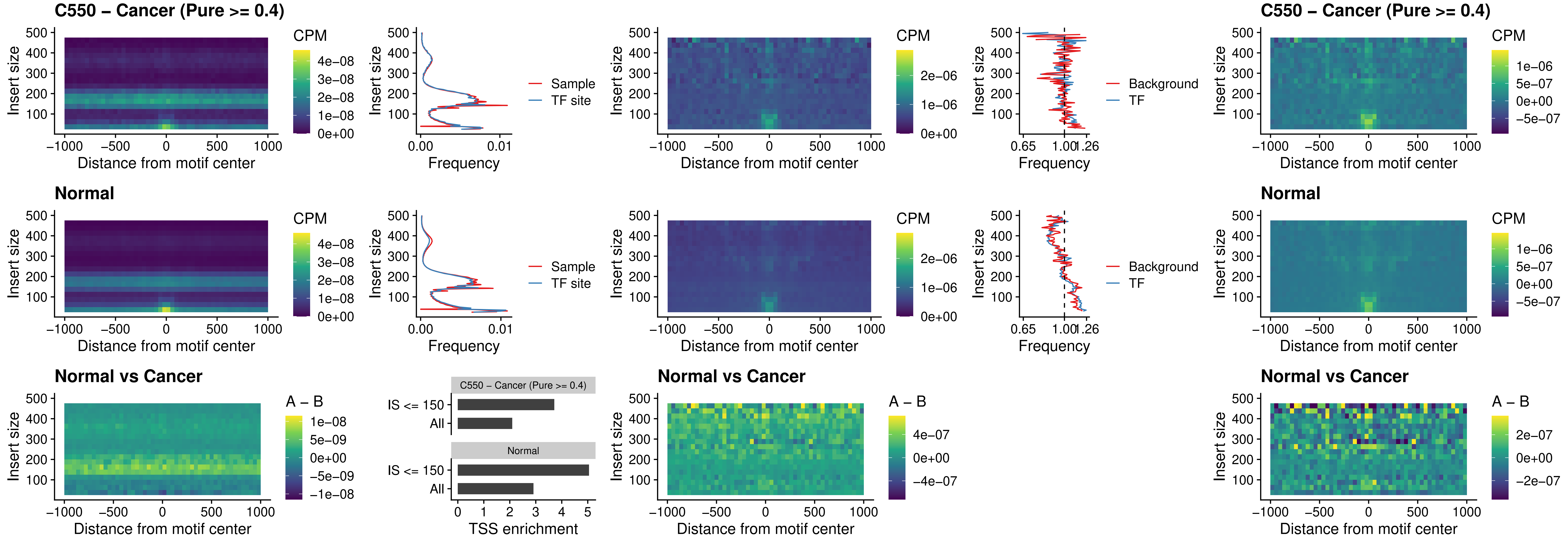

#### C550 – Cancer (Pure >= 0.4) – CTCF_uc_dTSS_oPEAK

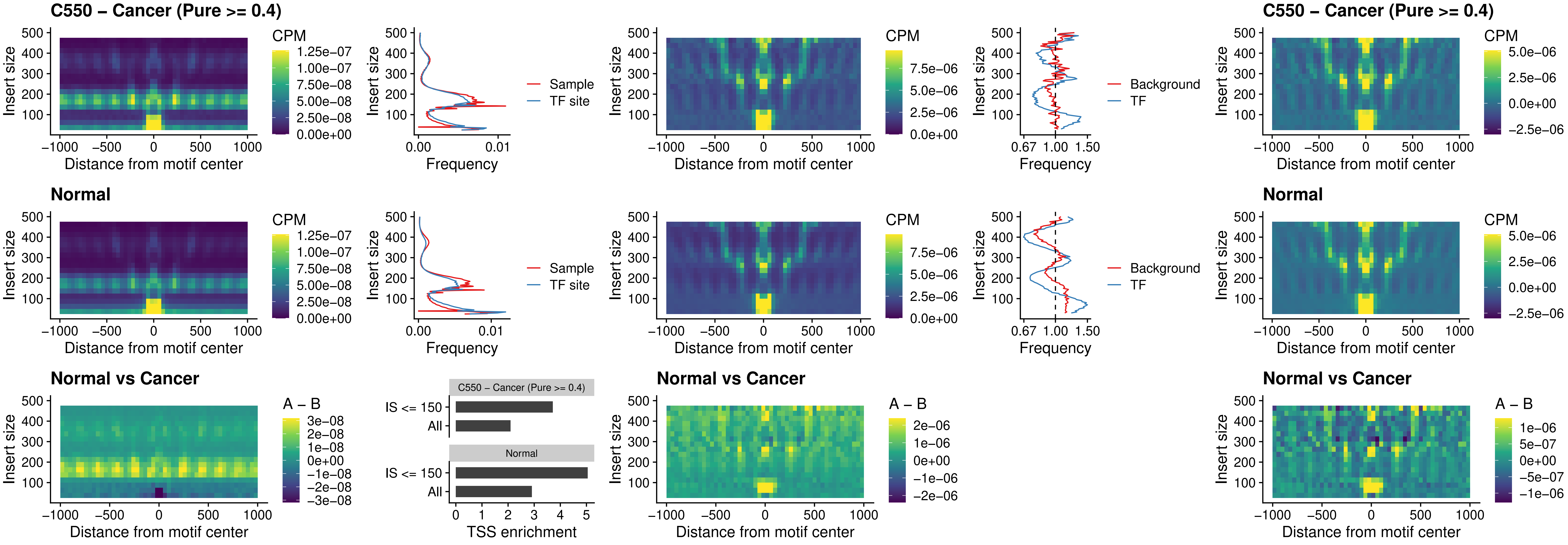

#### C550 – Cancer (Pure >= 0.4) – CTCF_uc_cTSS_oPEAK

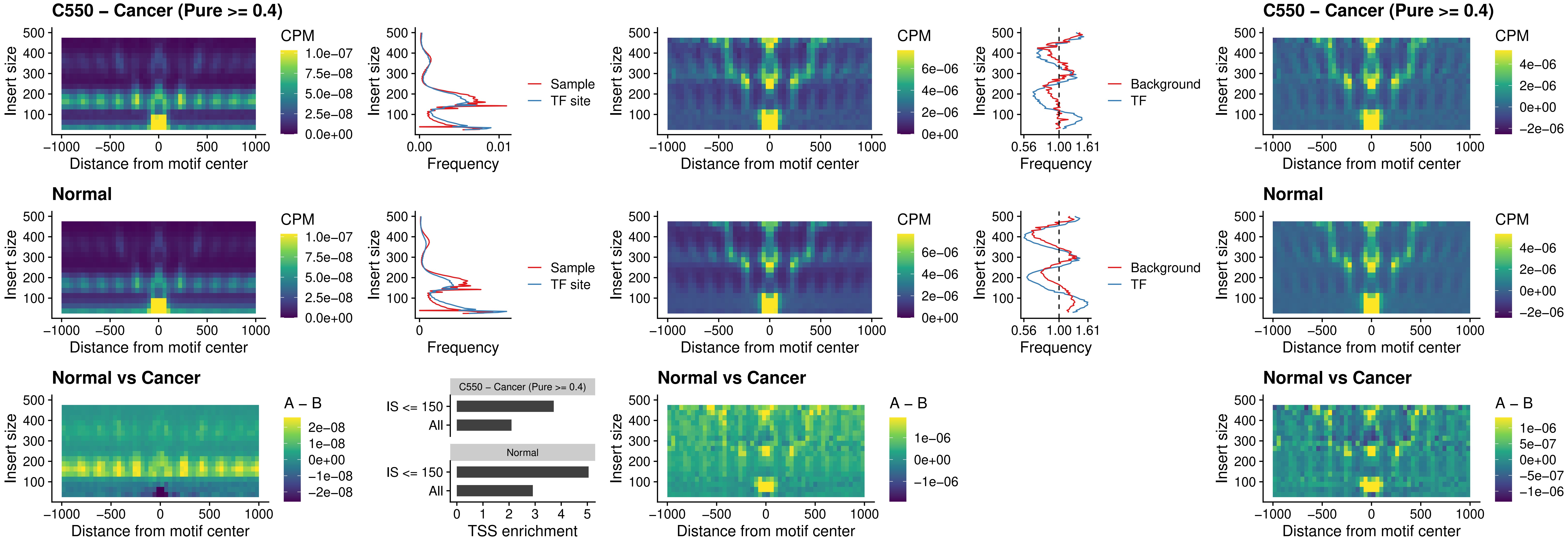

#### C550 – Cancer (Pure >= 0.4) – CTCFL_uc_dTSS_oPEAK

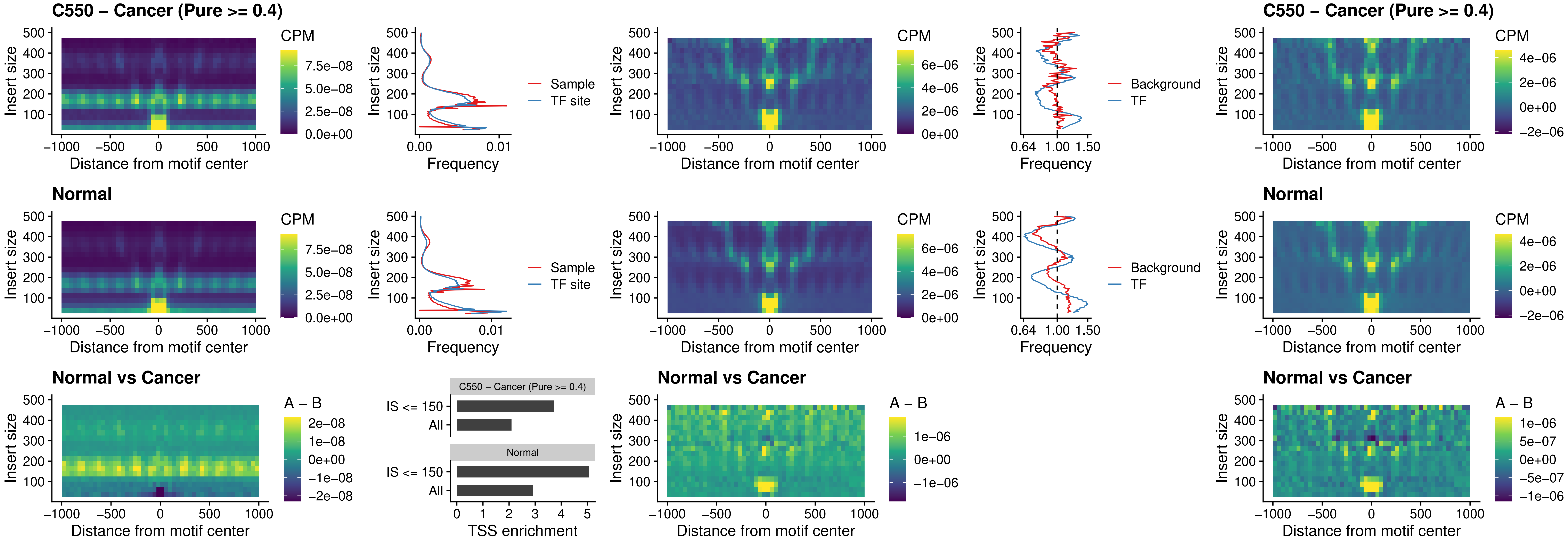

#### C550 – Cancer (Pure >= 0.4) – ENCODE_CTCF_sites_COLON

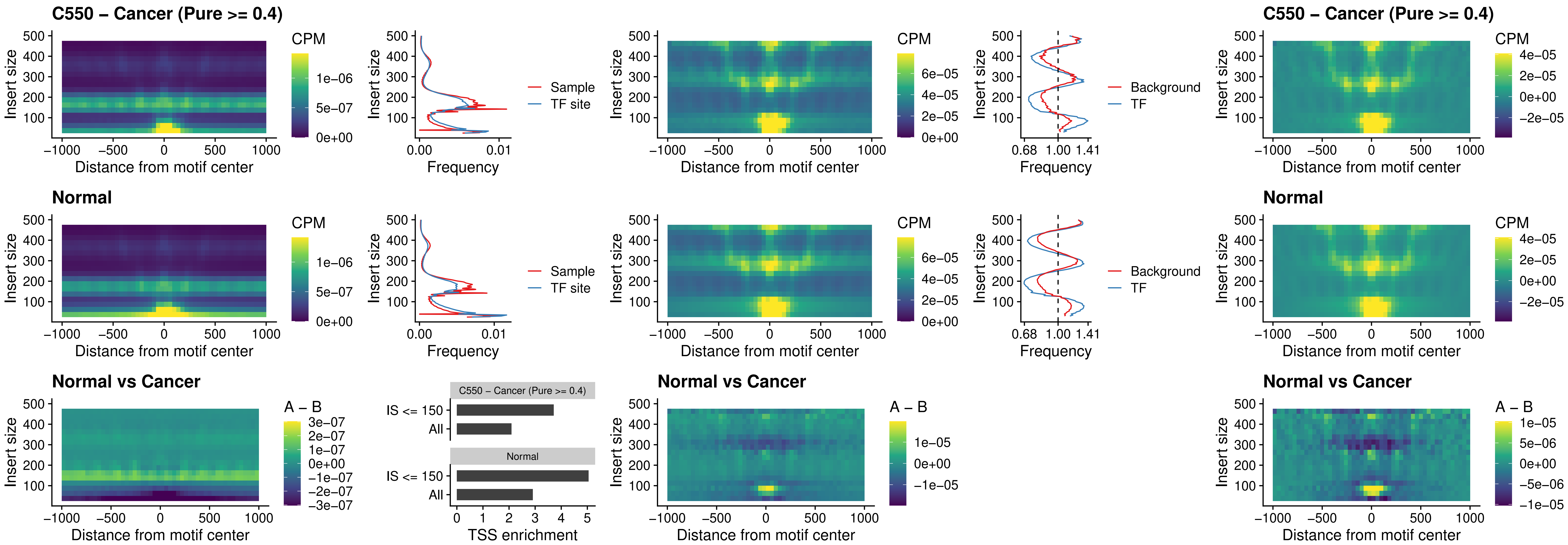

#### C550 – Cancer (Pure >= 0.4) – ENCODE_CTCF_sites_COLON_loop_ctcf

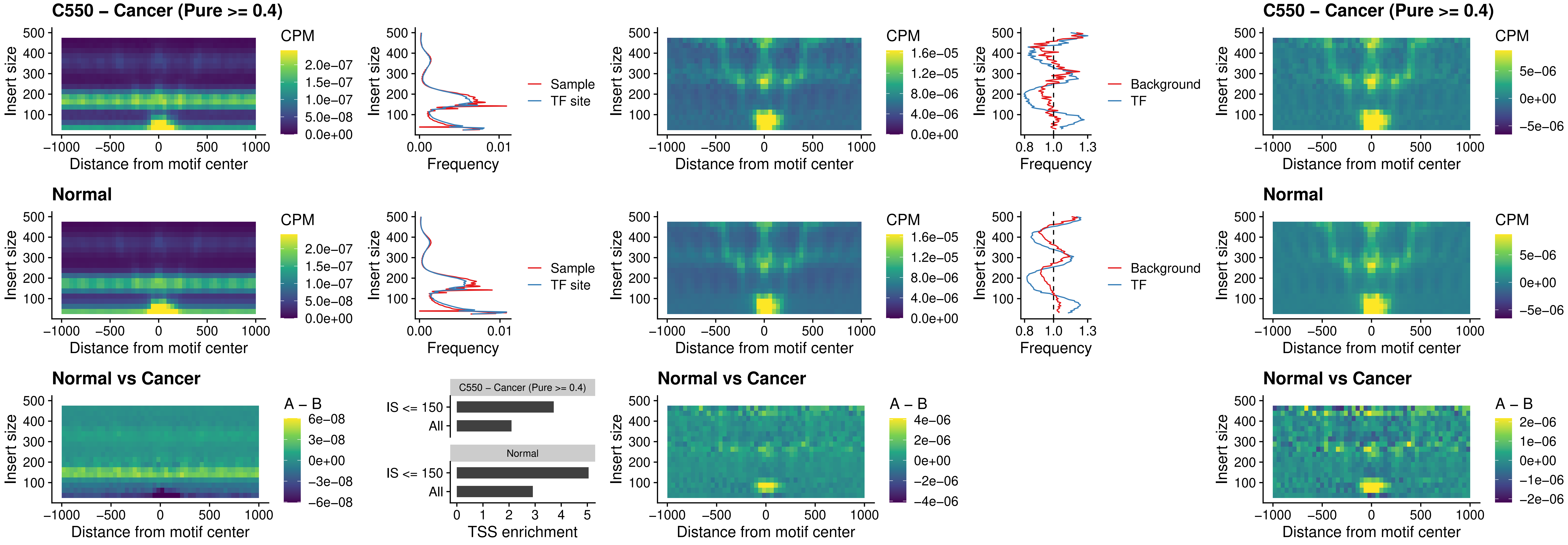

#### C550 – Cancer (Pure >= 0.4) – ENCODE_CTCF_sites_COLON_intergenic_ctcf

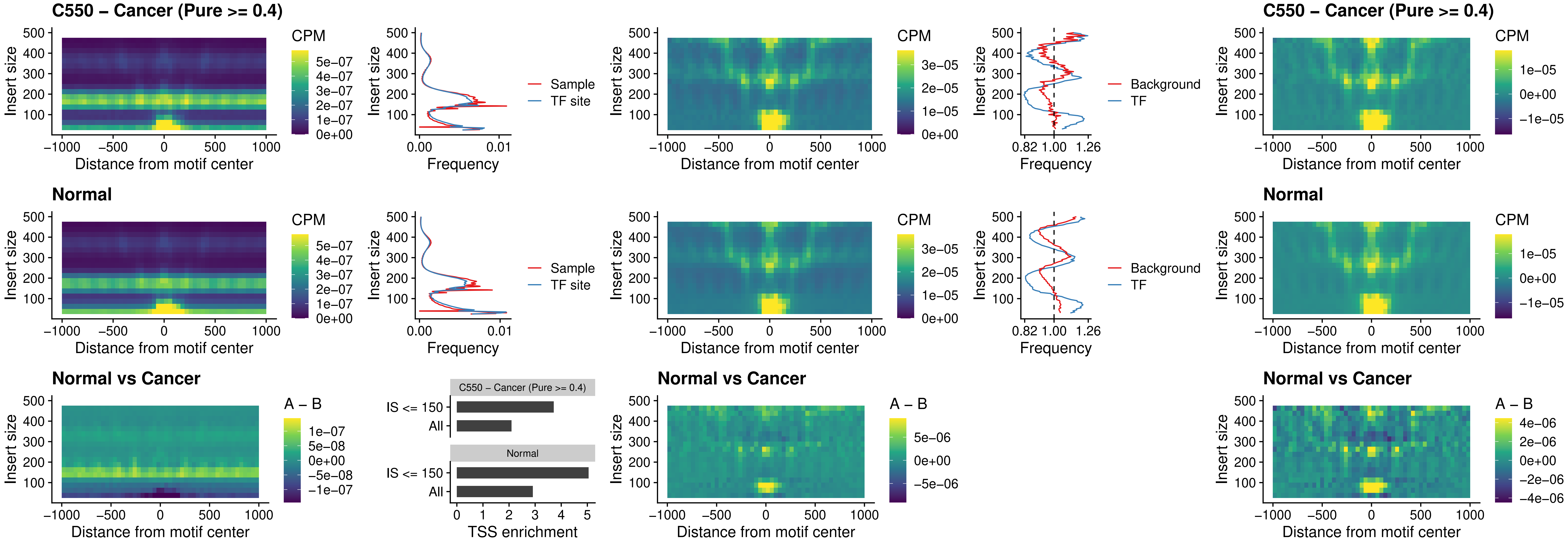

#### C550 – Cancer (Pure >= 0.4) – ENCODE_CTCF_sites_COLON_nonpromotor

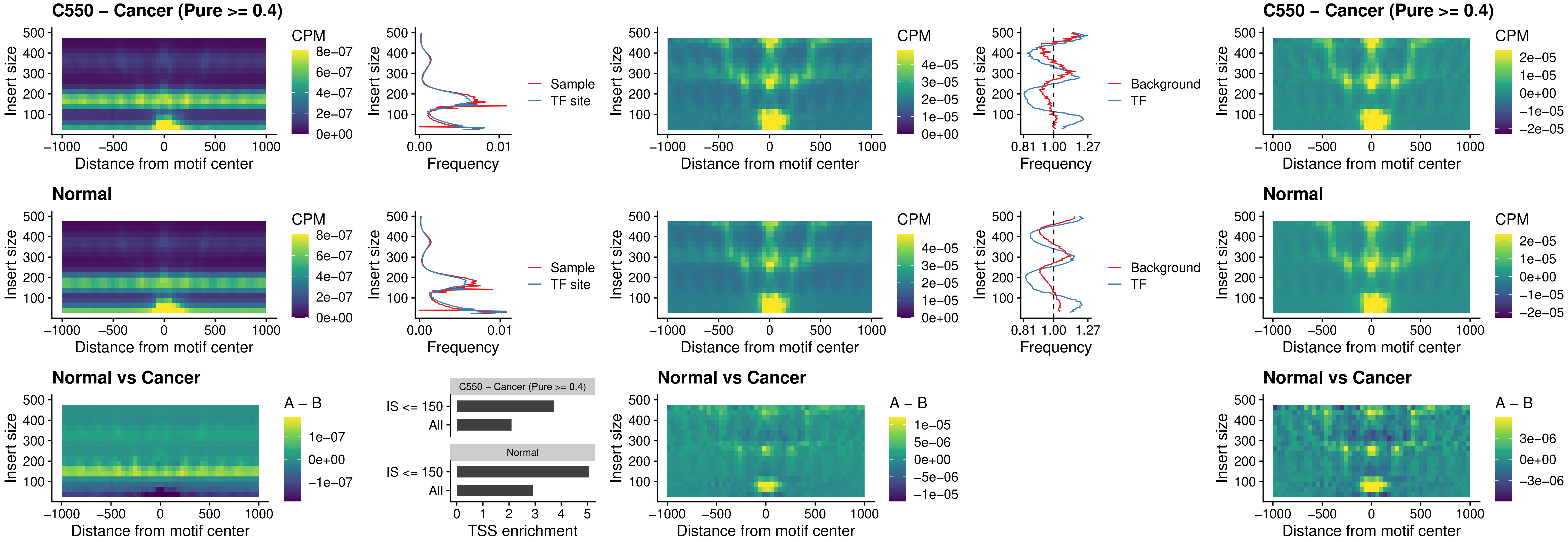

#### C550 – Cancer (Pure >= 0.4) – CTCFL_uc_cTSS_oPEAK

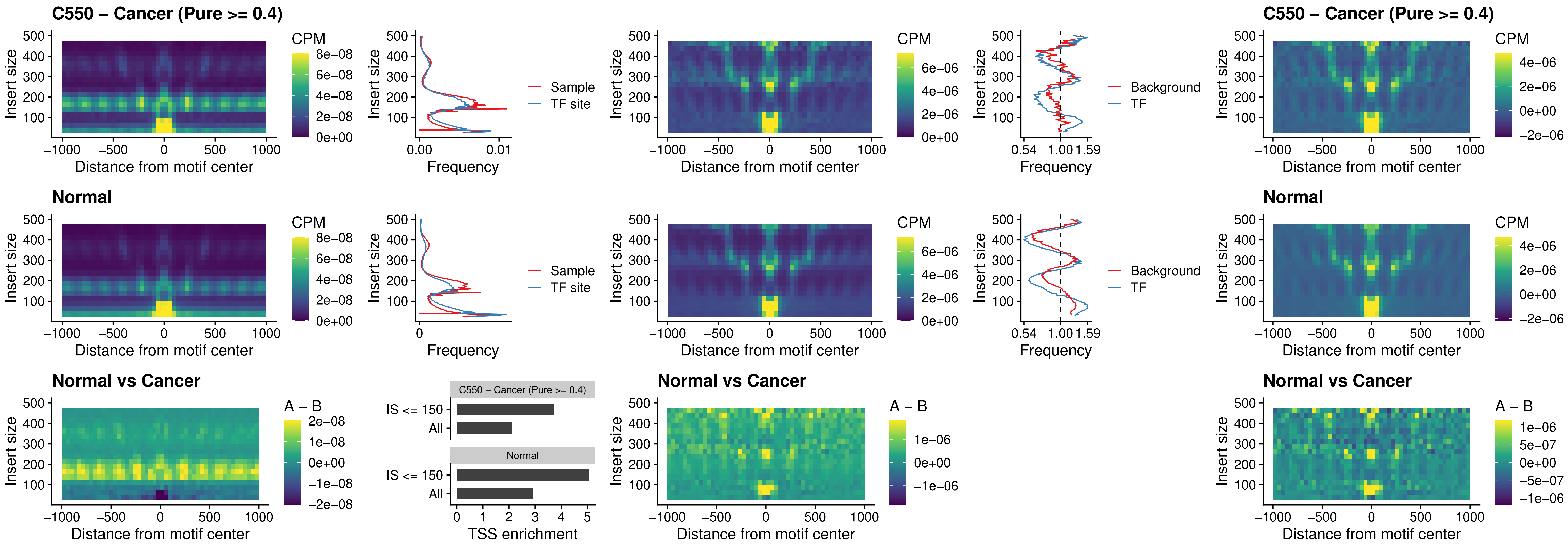

#### C550 – Cancer (Pure >= 0.4) – NEUROD1_uc_cTSS_oPEAK

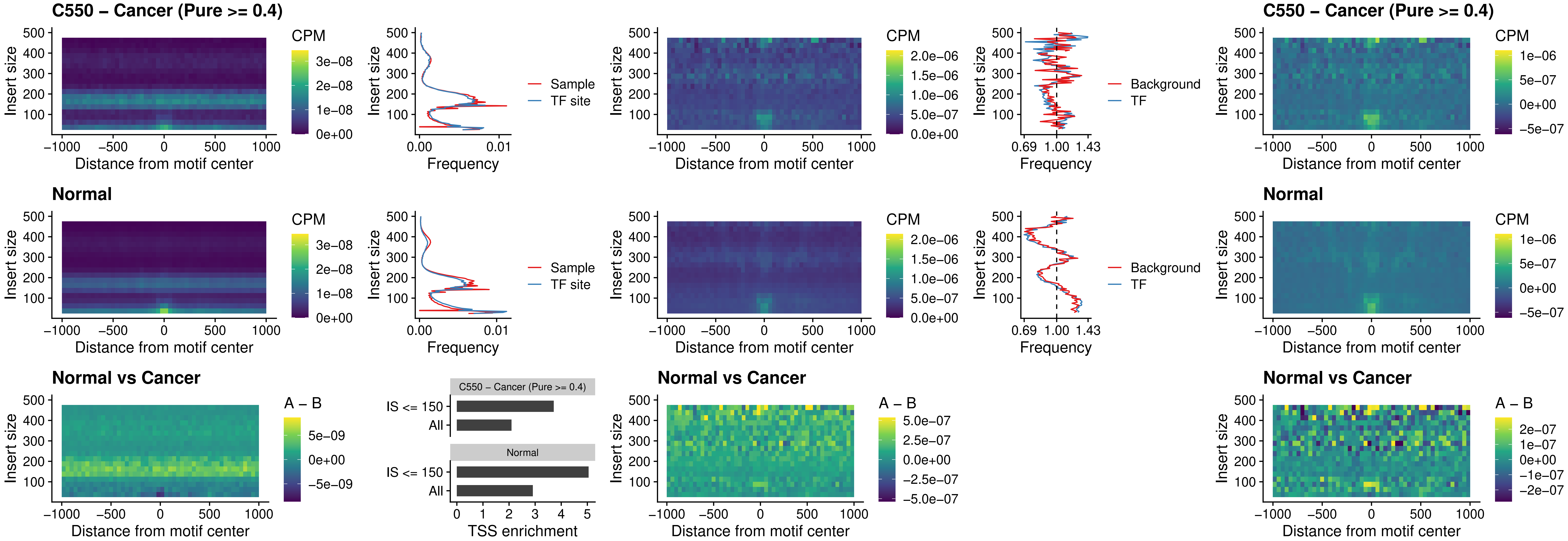

#### C550 – Cancer (Pure >= 0.4) – ENCODE_YY1_sites_COLON_nonpromotor

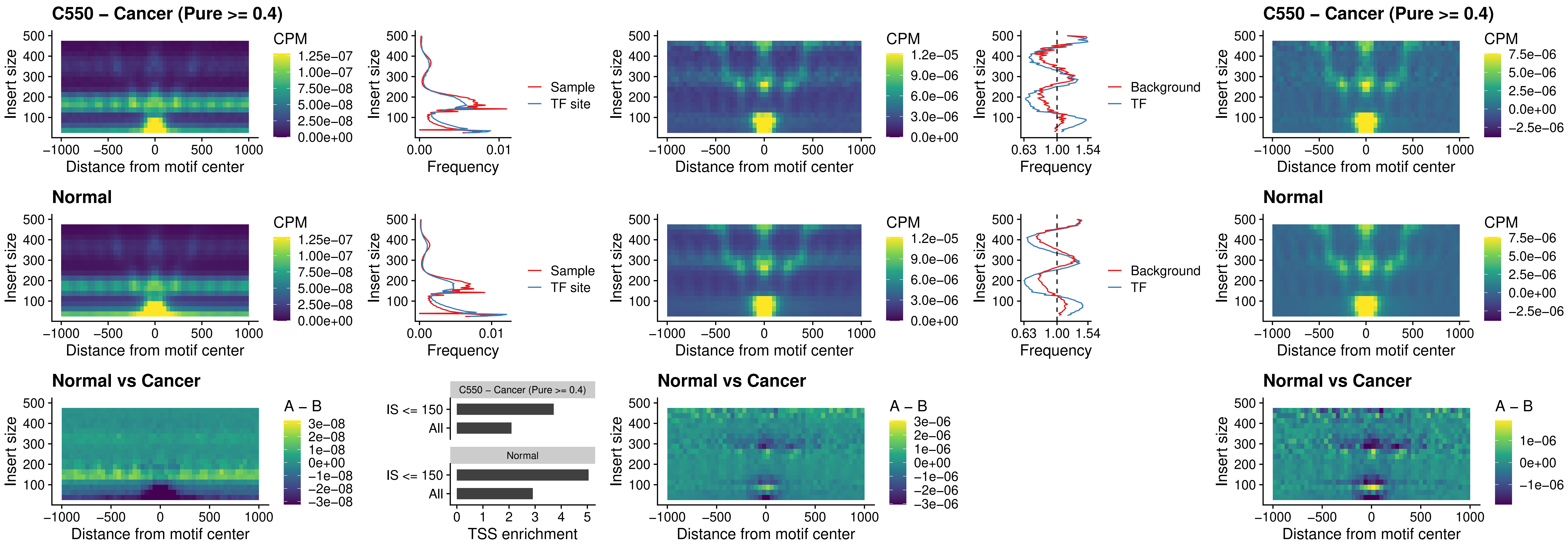

#### C550 – Cancer (Pure >= 0.4) – CTCF_uc_pTSS_oPEAK

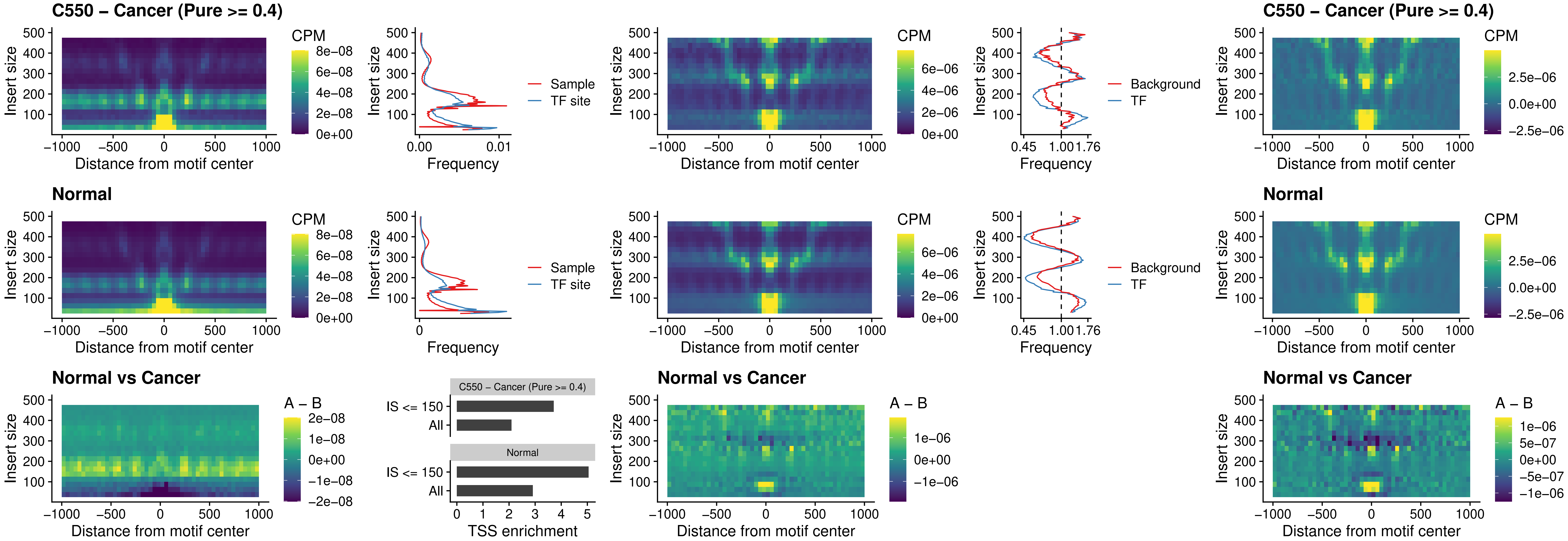

#### C550 – Cancer (Pure >= 0.4) – ENCODE_CTCF_sites_COLON_promotor

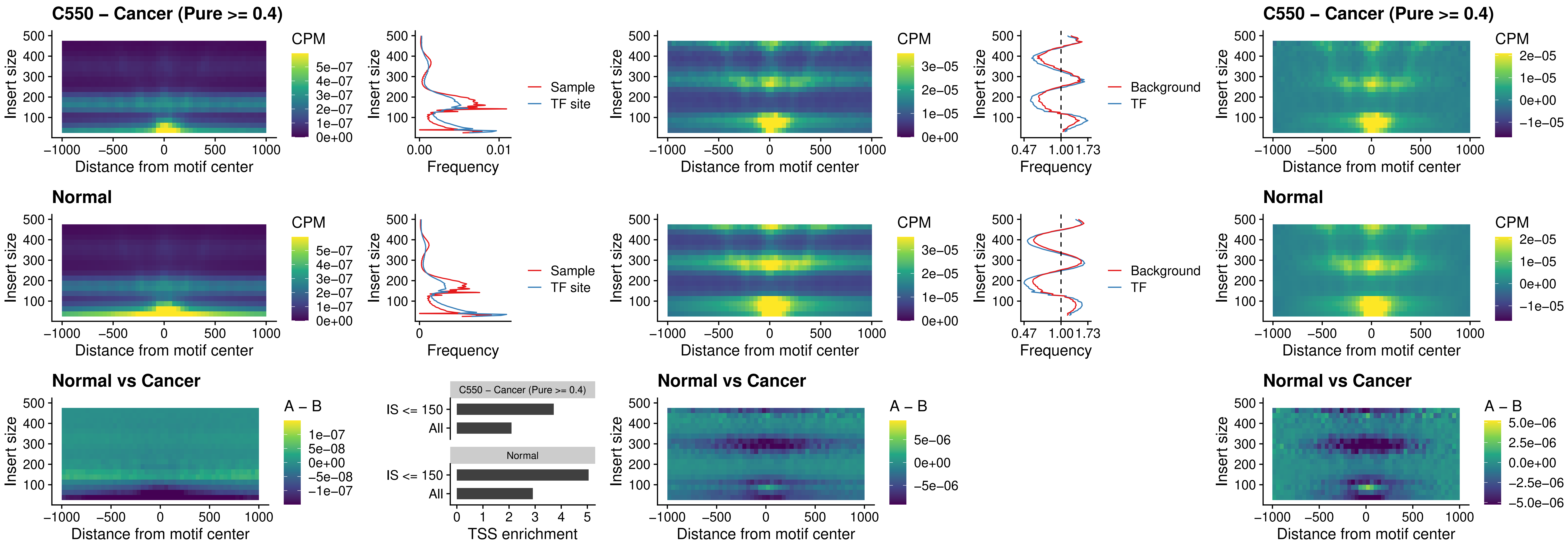

#### C550 – Cancer (Pure >= 0.4) – ENCODE_CTCF_sites_COLON_promotor_ctcf

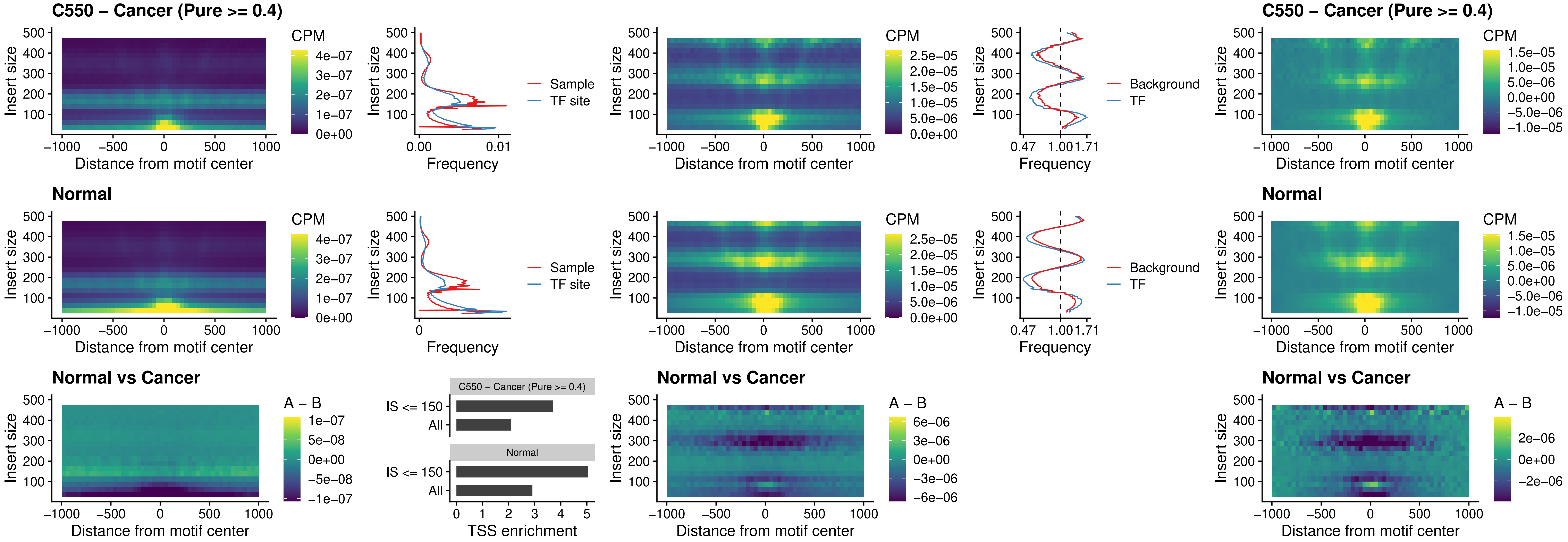

#### C550 – Cancer (Pure >= 0.4) – SPIC_uc_pTSS_oPEAK

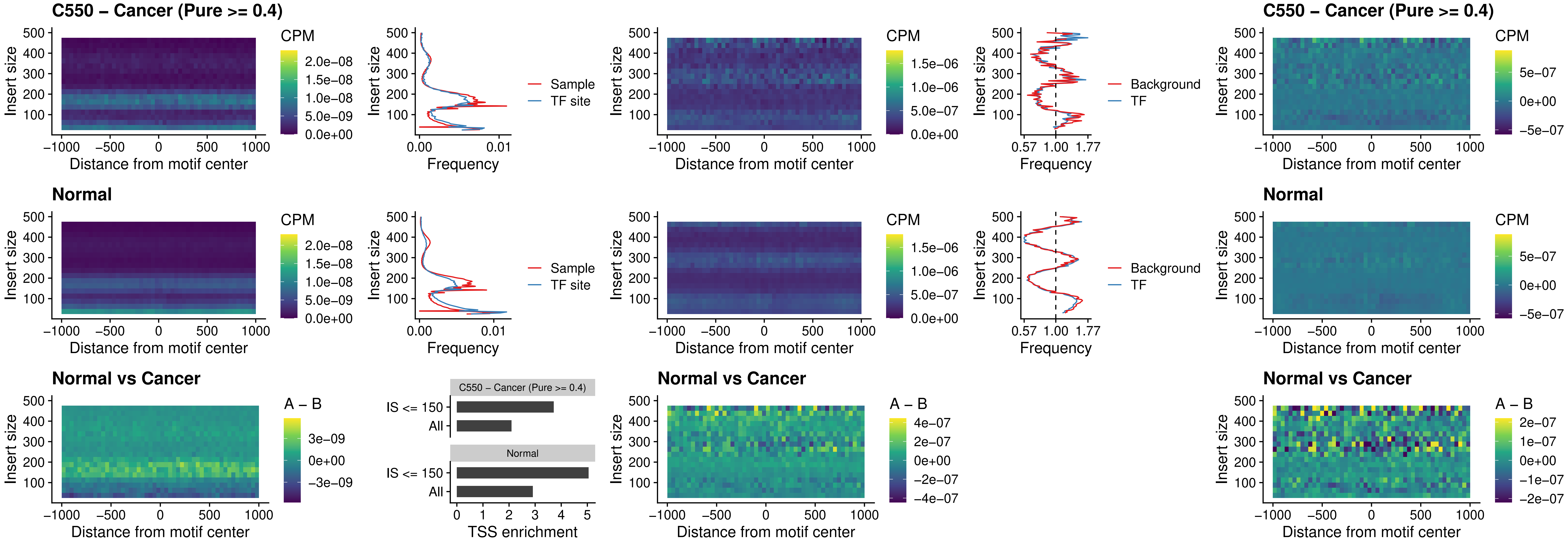

#### C550 – Cancer (Pure >= 0.4) – SPI1_uc_dTSS_nPEAK

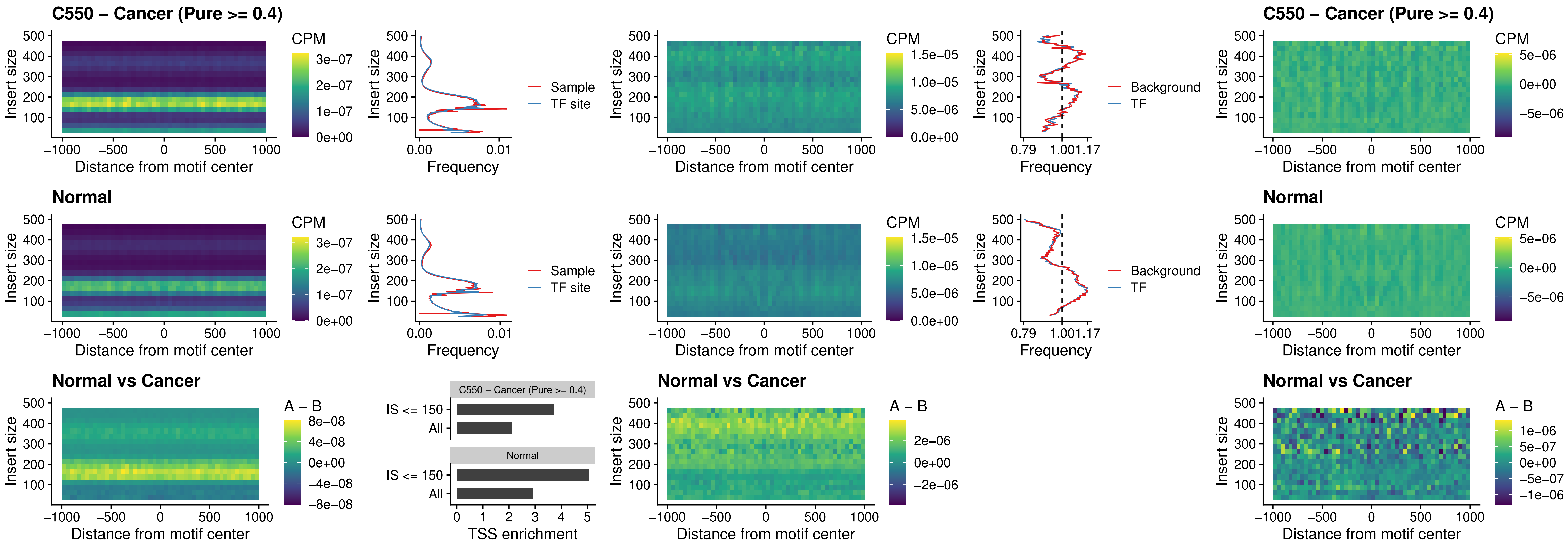

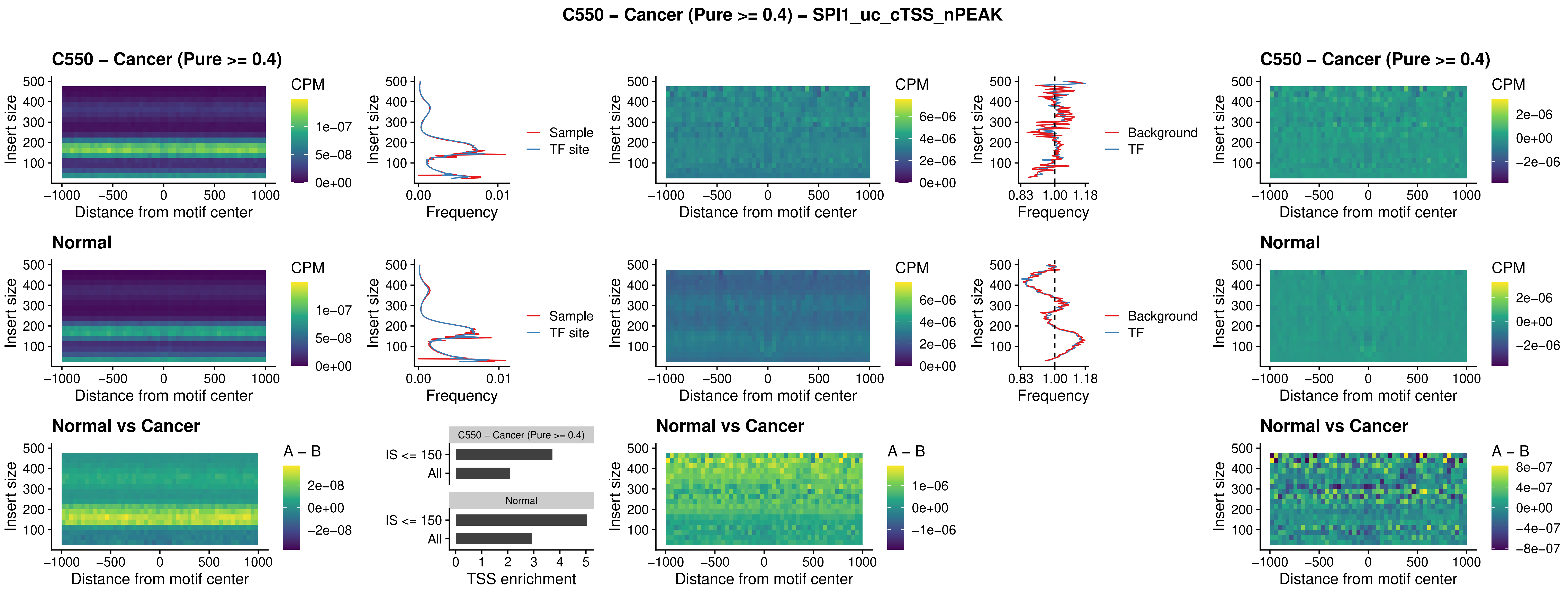
C550 – Cancer (Pure >= 0.4) – SPI1_uc_cTSS_nPEAK
C550 – Cancer (Pure >= 0.4)
Normal
Normal vs Cancer
Insert size
Distance from motif center
CPM
A – B
Frequency
Sample
TF site
Background
TF
C550 – Cancer (Pure >= 0.4)
Normal
IS <= 150
All
TSS enrichment

#### C550 – Cancer (Pure >= 0.4) – BCL11A_uc_pTSS_nPEAK

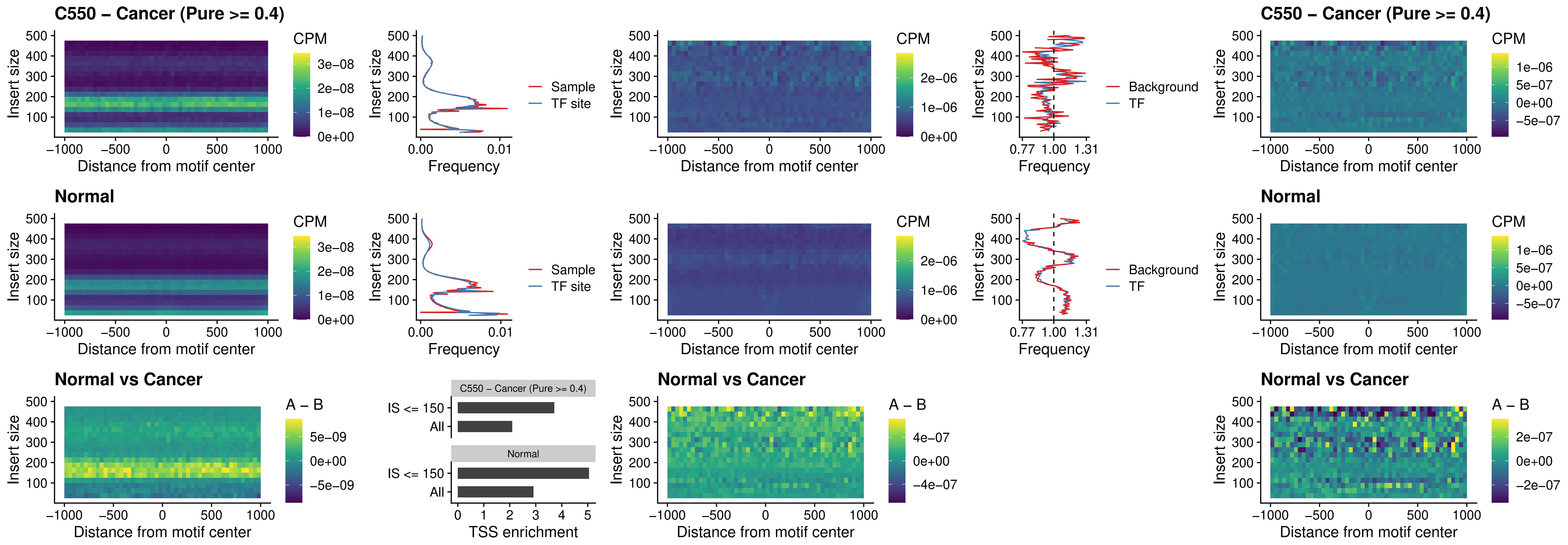

#### C550 – Cancer (Pure >= 0.4) – BCL11B_uc_pTSS_nPEAK

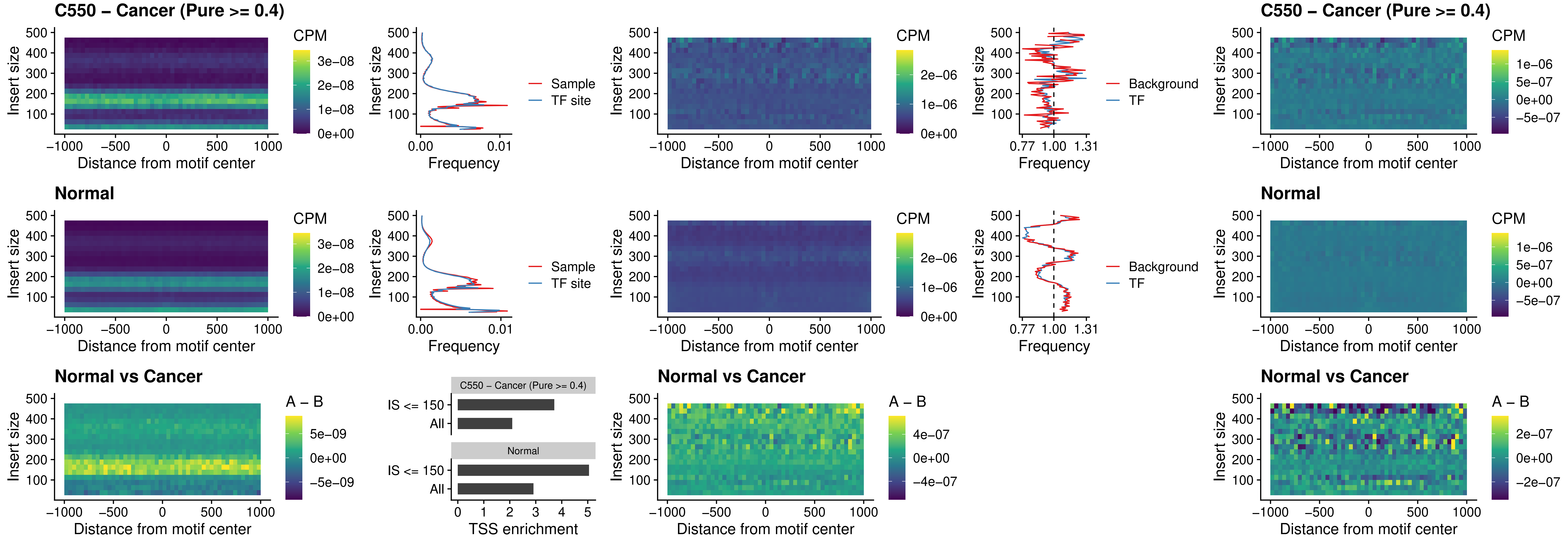

#### C550 – Cancer (Pure >= 0.4) – IRF9_uc_dTSS_oPEAK

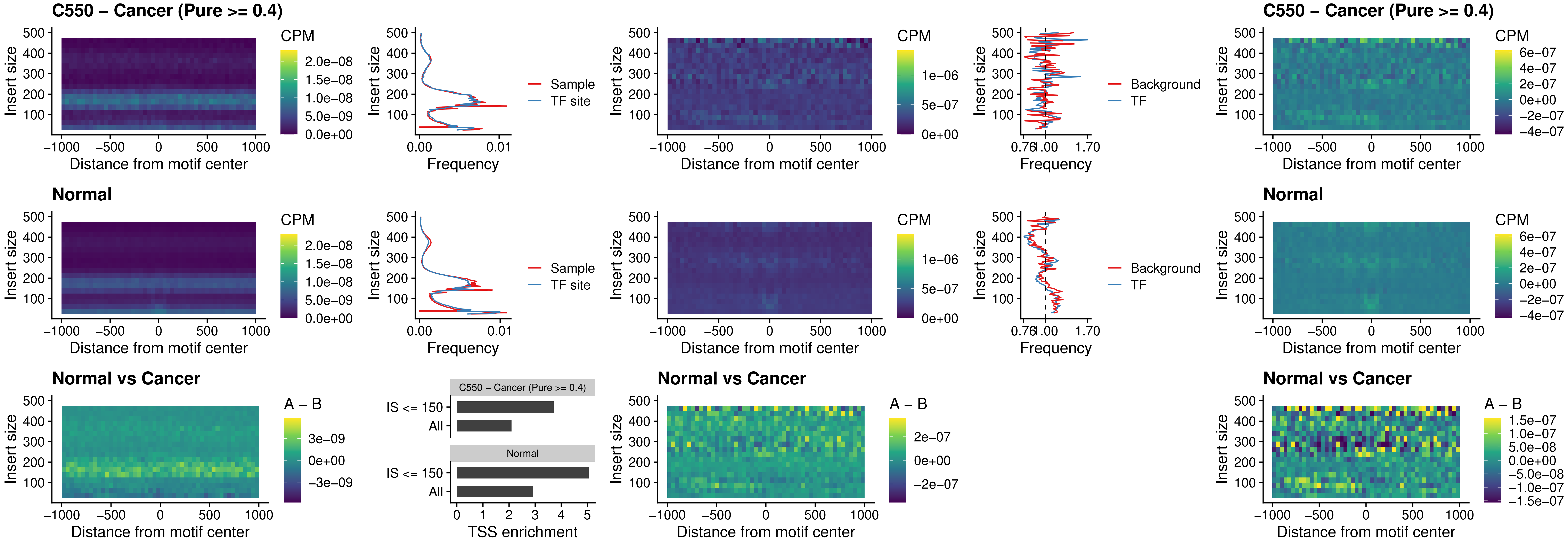

#### C550 – Cancer (Pure >= 0.4) – IRF9_uc_cTSS_oPEAK

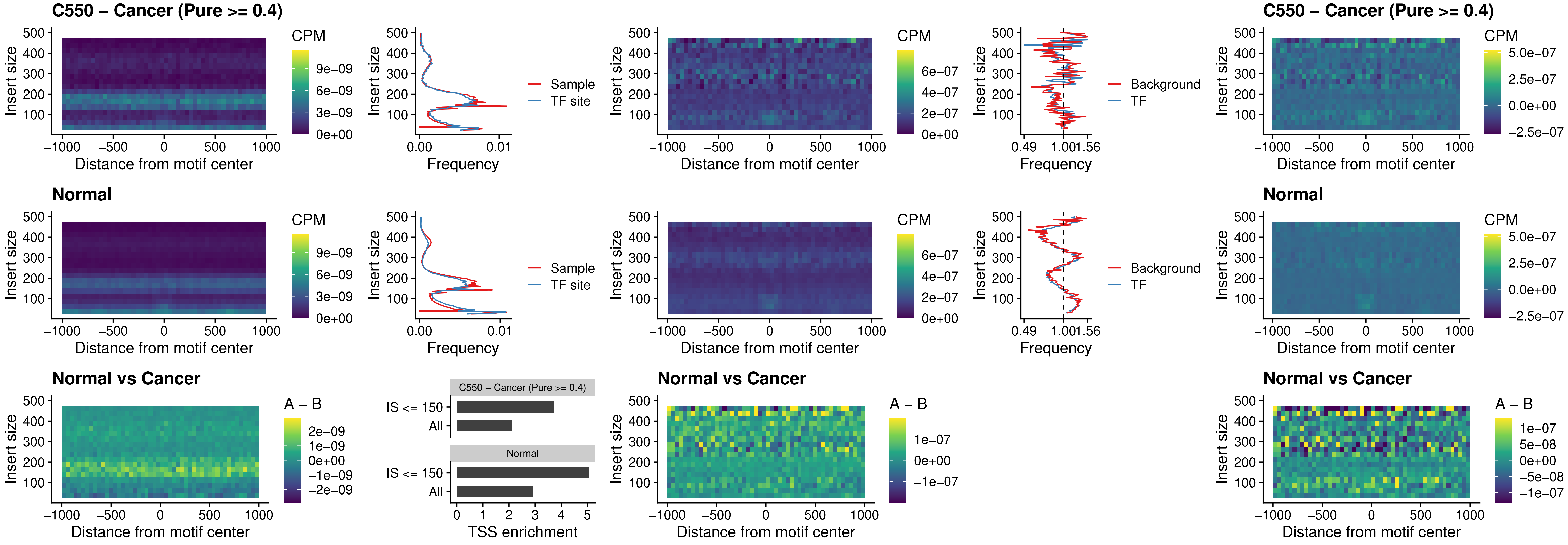

#### C550 – Cancer (Pure >= 0.4) – BCL11A_uc_cTSS_oPEAK

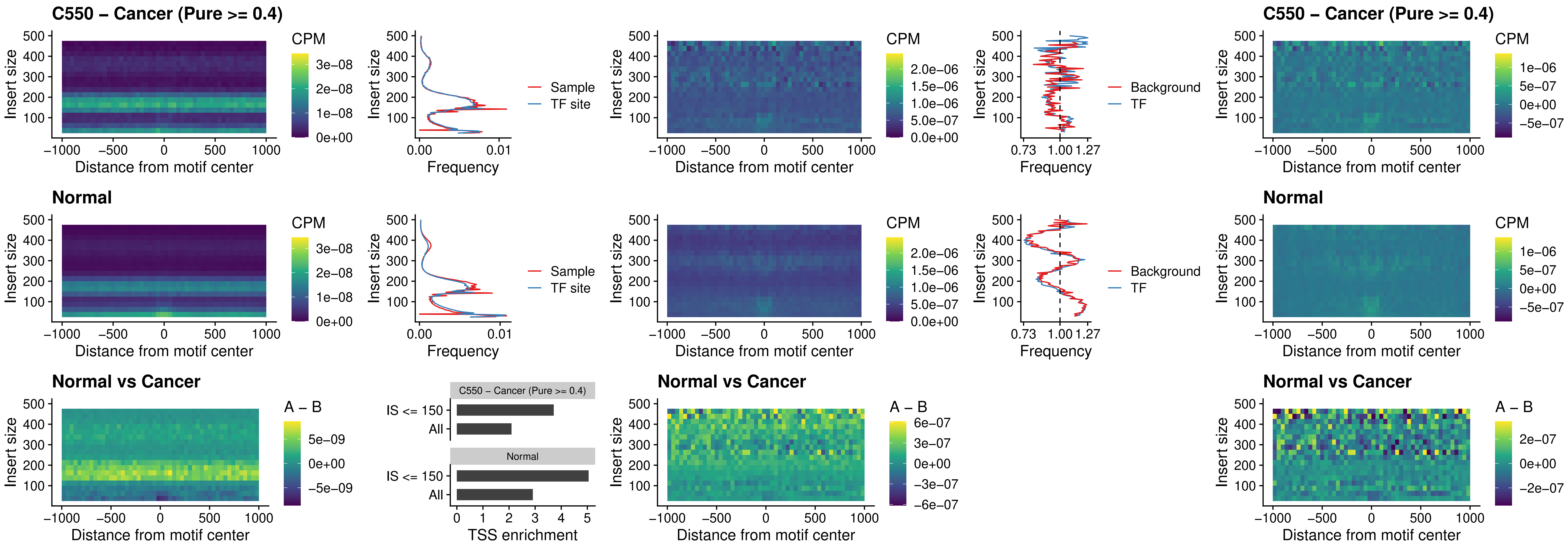

#### C550 – Cancer (Pure >= 0.4) – BCL11B_uc_cTSS_oPEAK

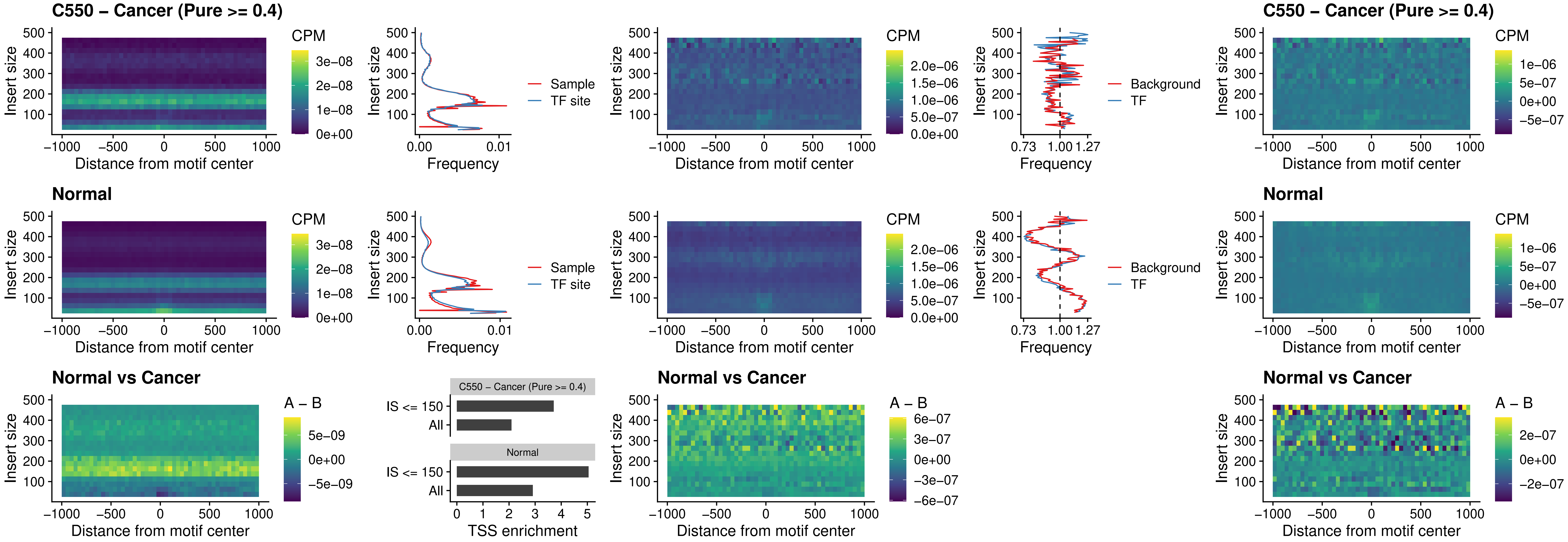

### C550 – Cancer (Pure >= 0.4) – BCL11A_uc_dTSS_oPEAK

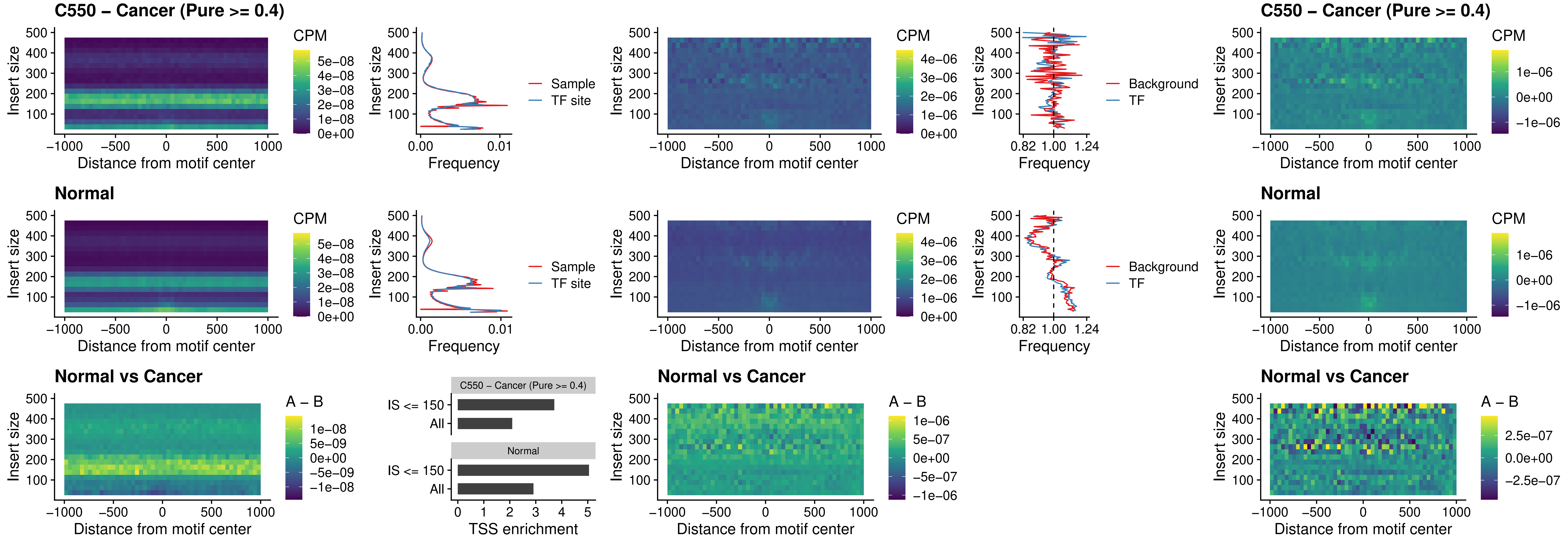

### C550 – Cancer (Pure >= 0.4) – BCL11B_uc_dTSS_oPEAK

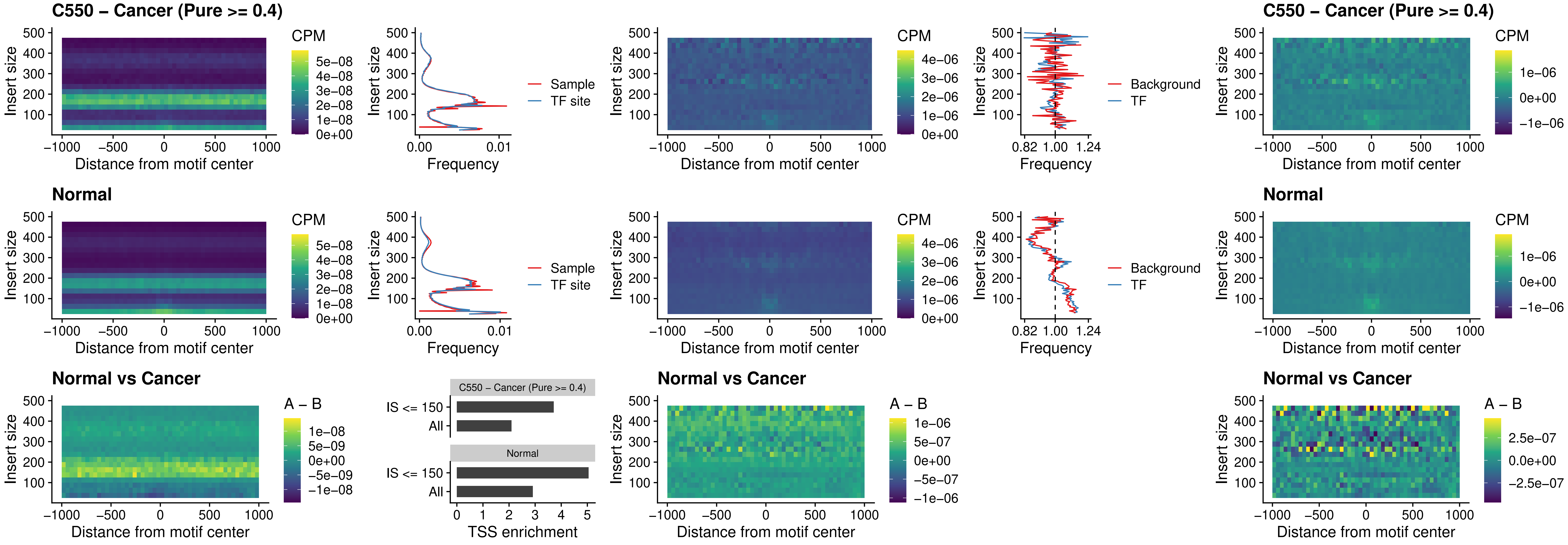

### C550 – Cancer (Pure >= 0.4) – SPIB_uc_cTSS_nPEAK

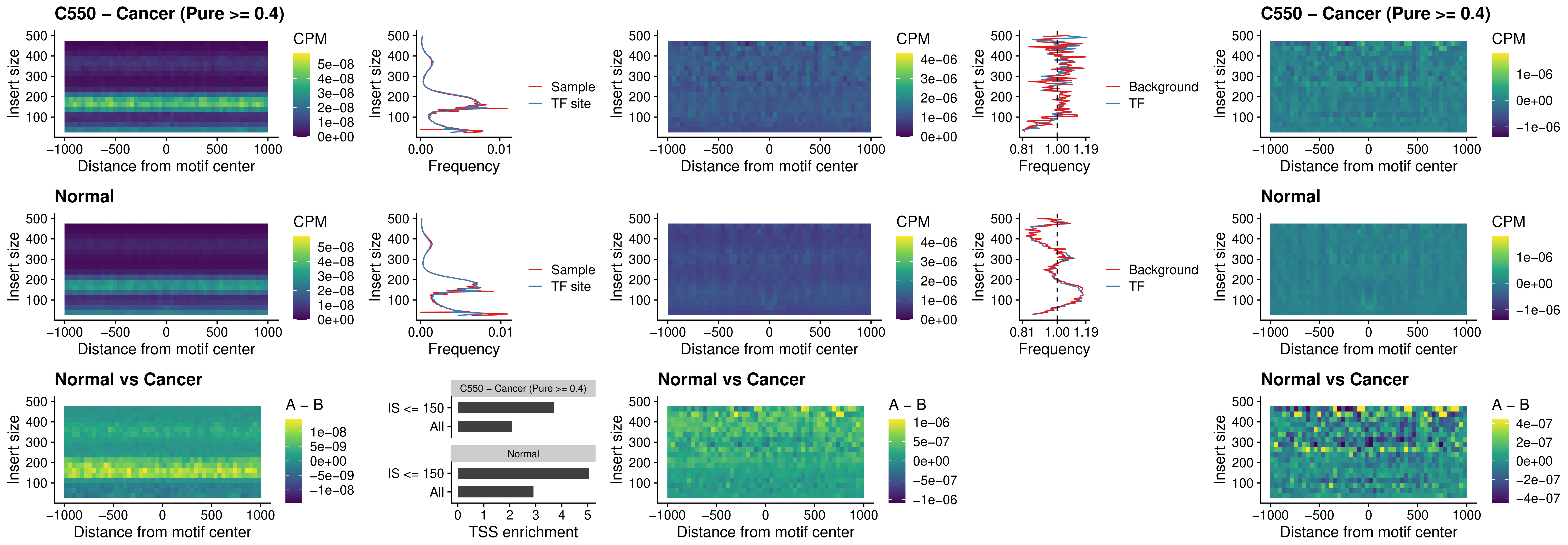

### C550 – Cancer (Pure >= 0.4) – SPIC_uc_cTSS_nPEAK

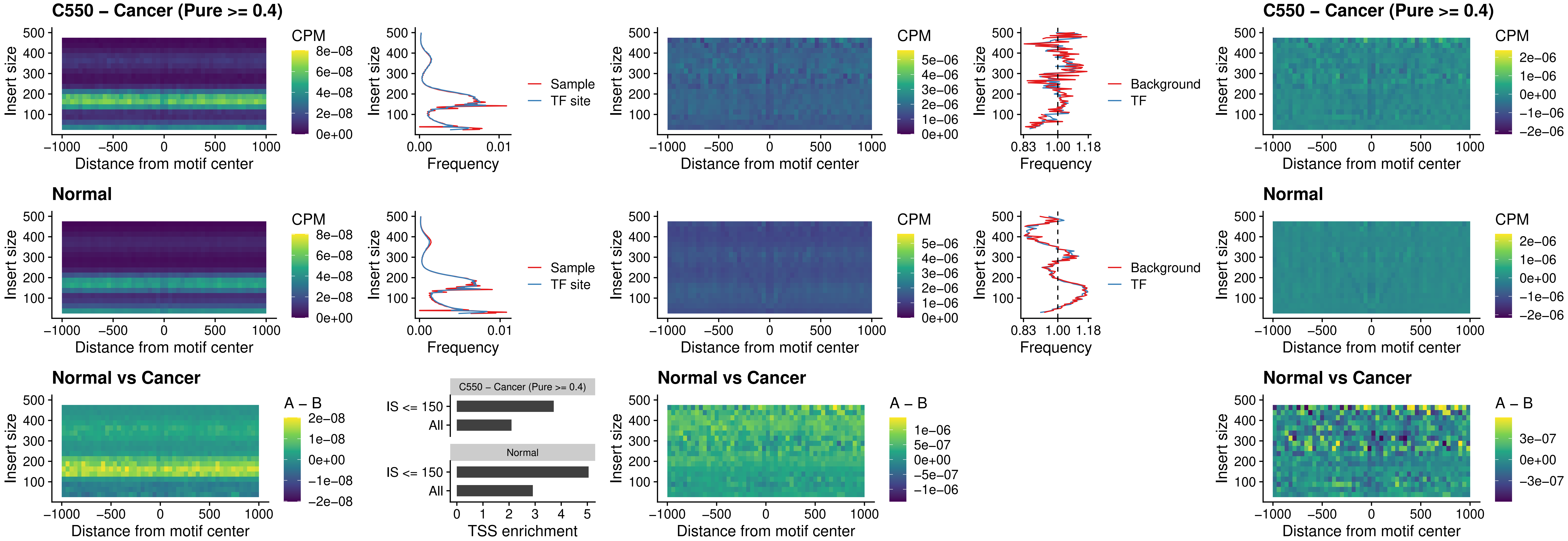

#### C550 – Cancer (Pure >= 0.4) – SPI1_uc_pTSS_oPEAK

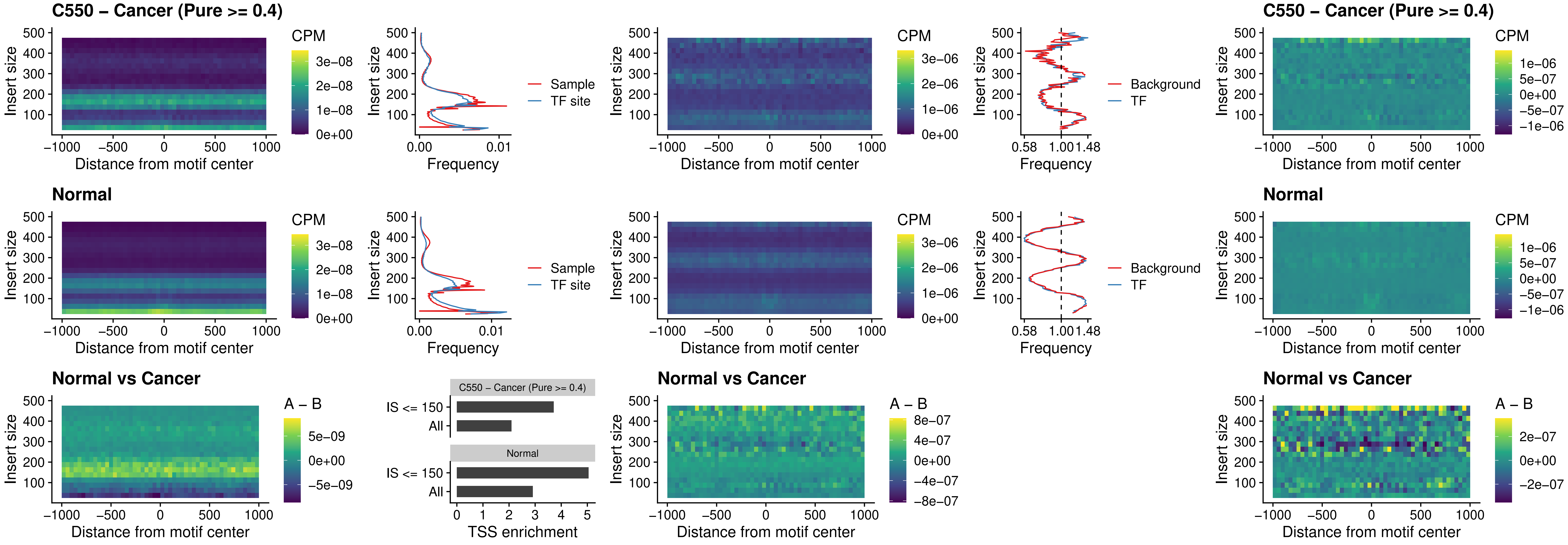

#### C550 – Cancer (Pure >= 0.4) – BCL11A_uc_pTSS_oPEAK

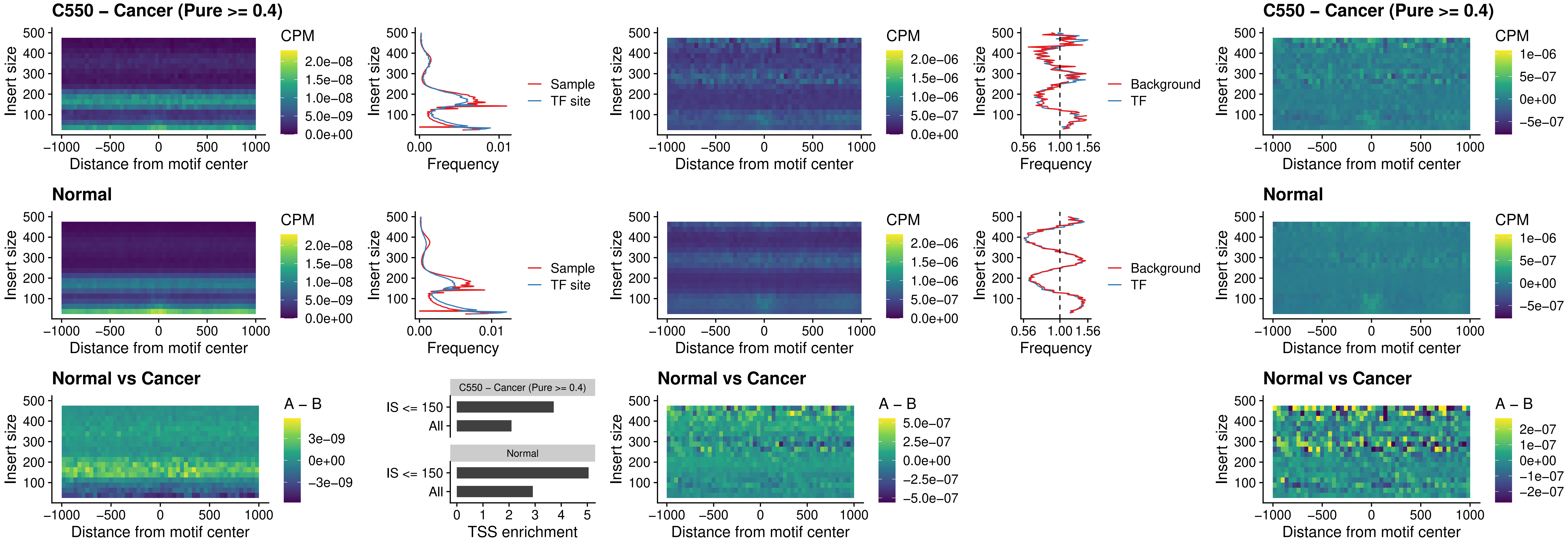

#### C550 – Cancer (Pure >= 0.4) – BCL11B_uc_pTSS_oPEAK

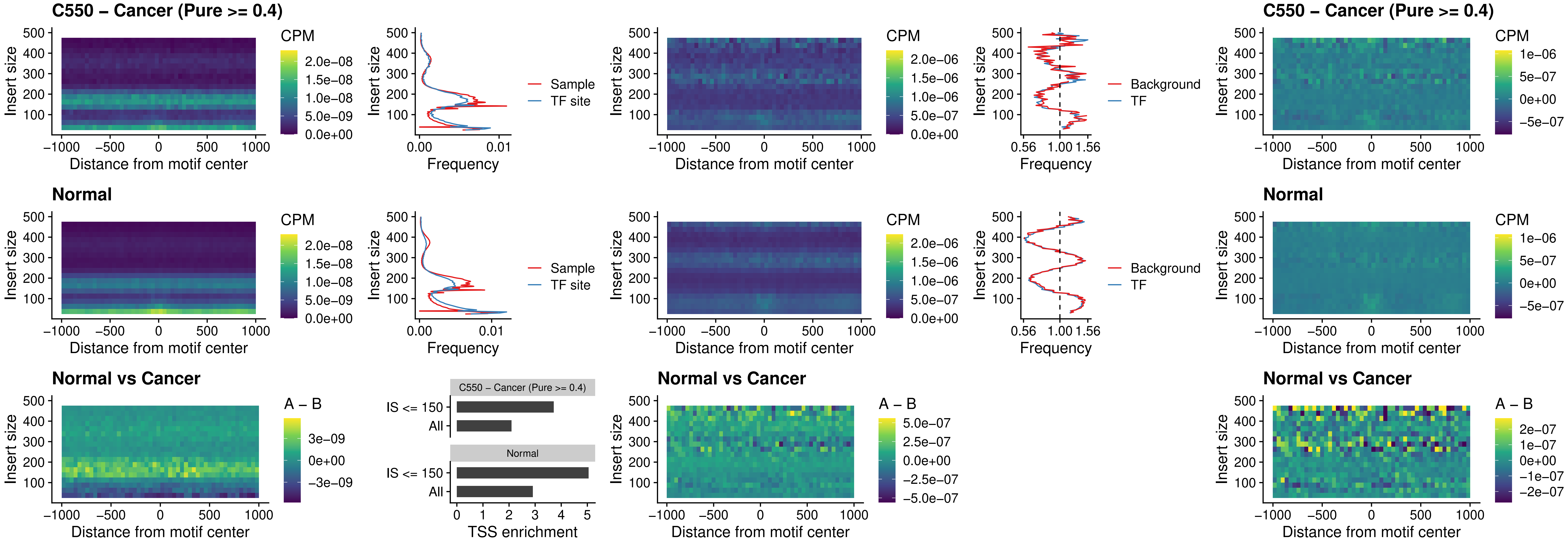

#### C550 – Cancer (Pure >= 0.4) – SPI1_uc_cTSS_oPEAK

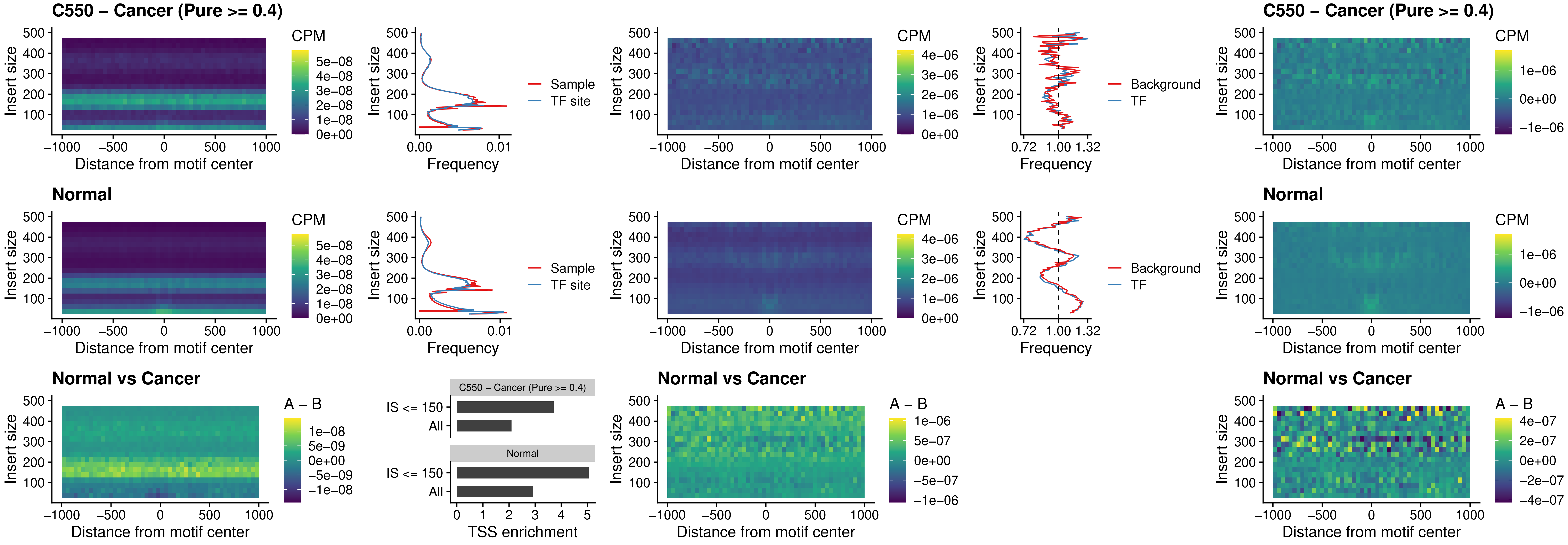

#### C550 – Cancer (Pure >= 0.4) – SPIB_uc_cTSS_oPEAK

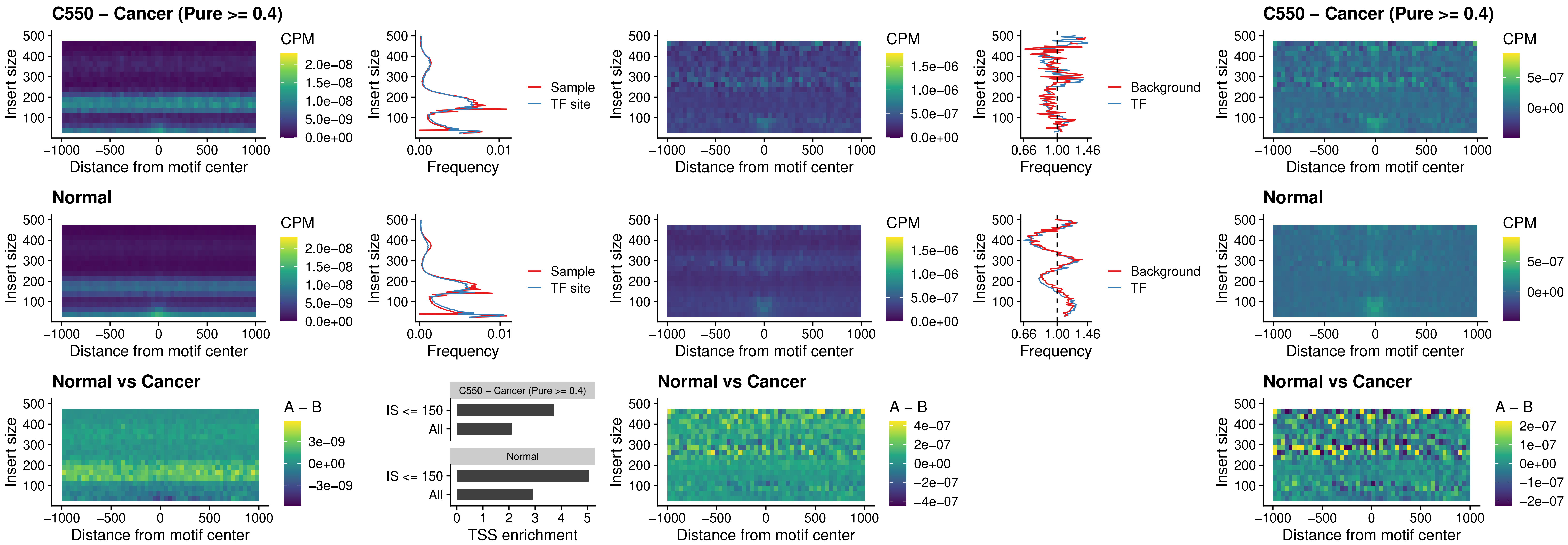

#### C550 – Cancer (Pure >= 0.4) – SPIB_uc_pTSS_oPEAK

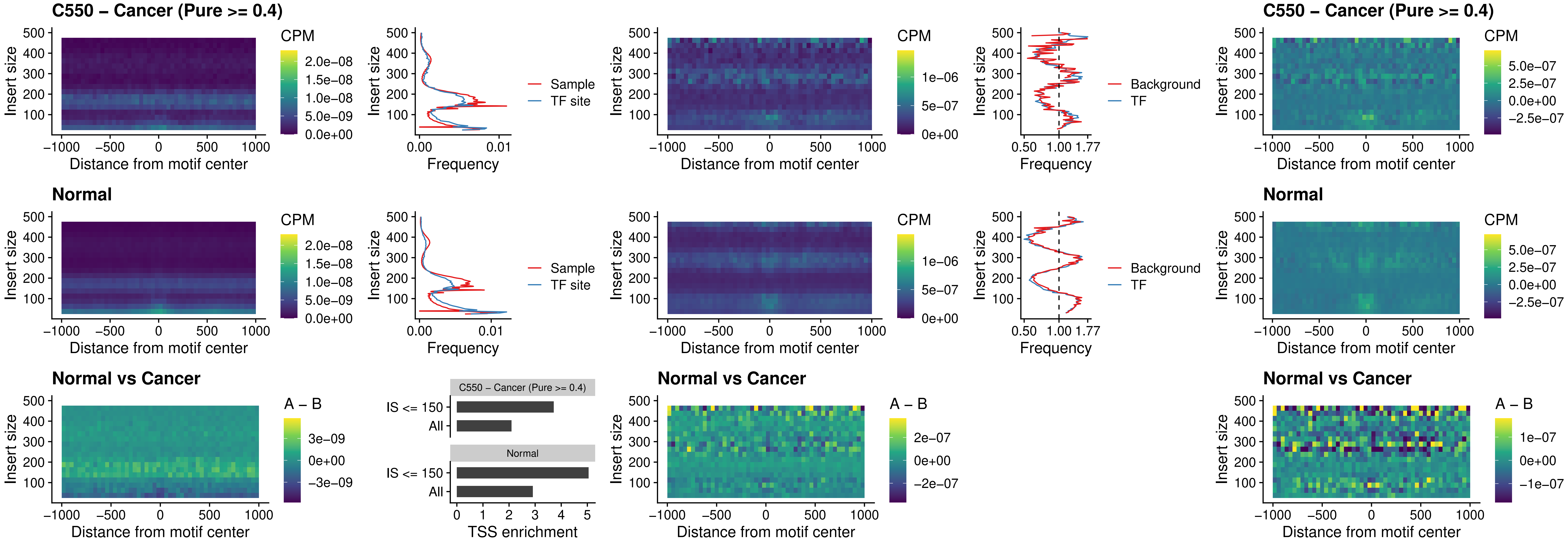

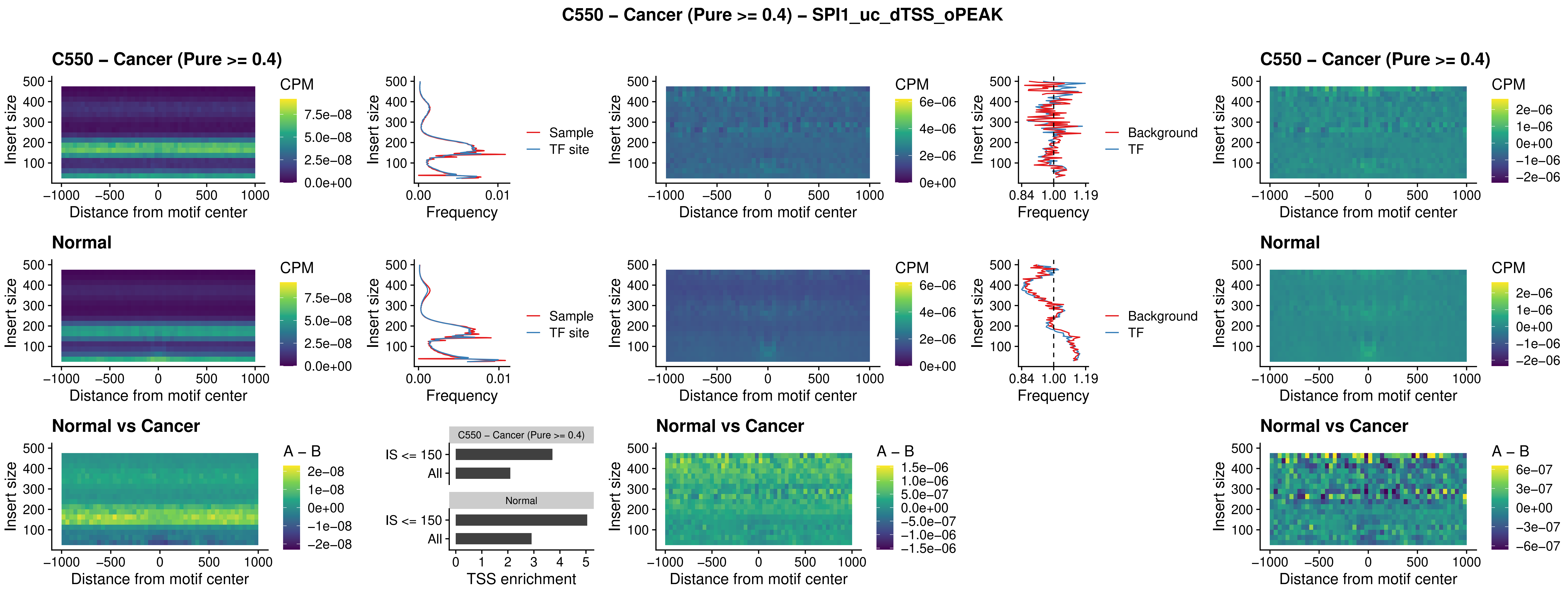
C550 – Cancer (Pure >= 0.4) – SPI1_uc_dTSS_oPEAK
C550 – Cancer (Pure >= 0.4)
Normal
Normal vs Cancer
Insert size
Distance from motif center
CPM
A – B
Frequency
Sample
TF site
Background
TF
C550 – Cancer (Pure >= 0.4)
Normal
IS <= 150
All
TSS enrichment

#### C550 – Cancer (Pure >= 0.4) – SPIB_uc_dTSS_oPEAK

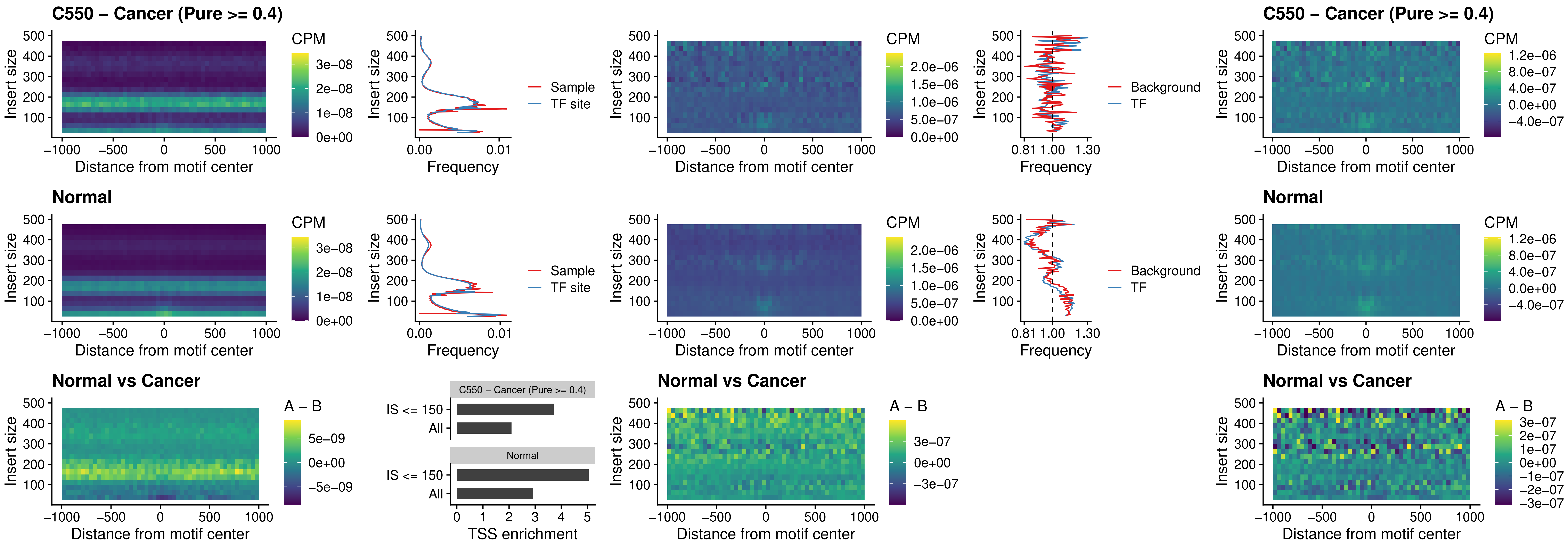

#### C550 – Cancer (Pure >= 0.4) – IRF3_uc_pTSS_oPEAK

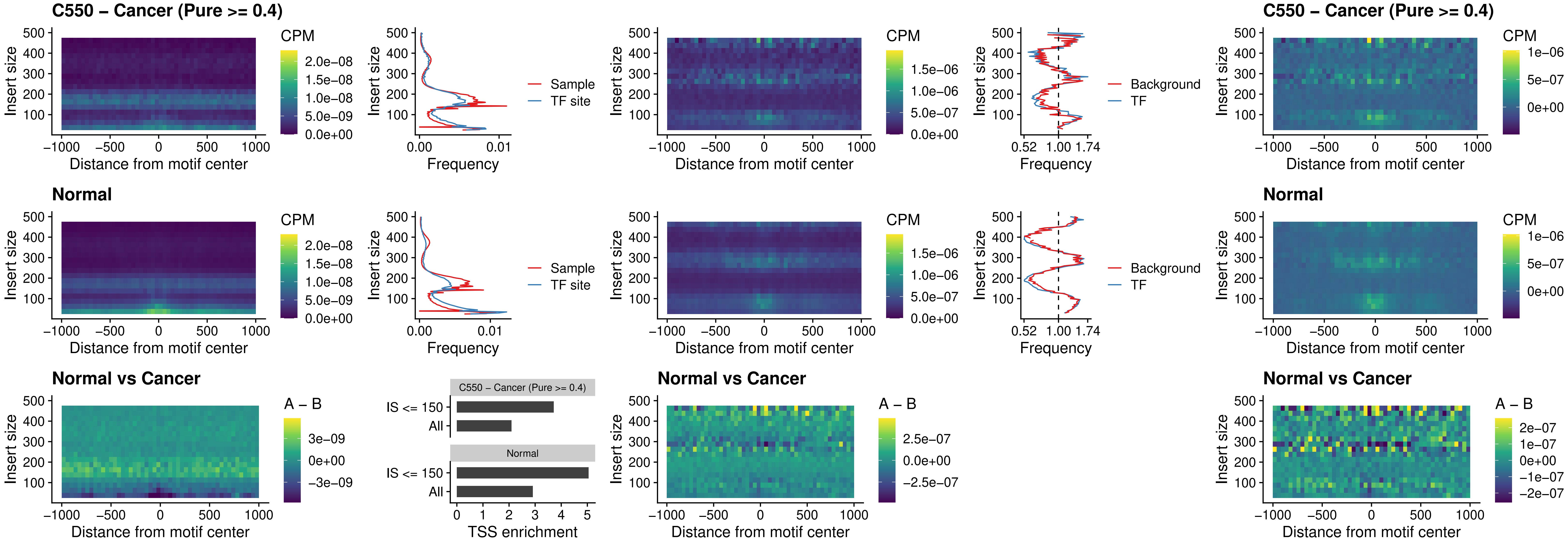

#### C550 – Cancer (Pure >= 0.4) – IRF2_uc_dTSS_oPEAK

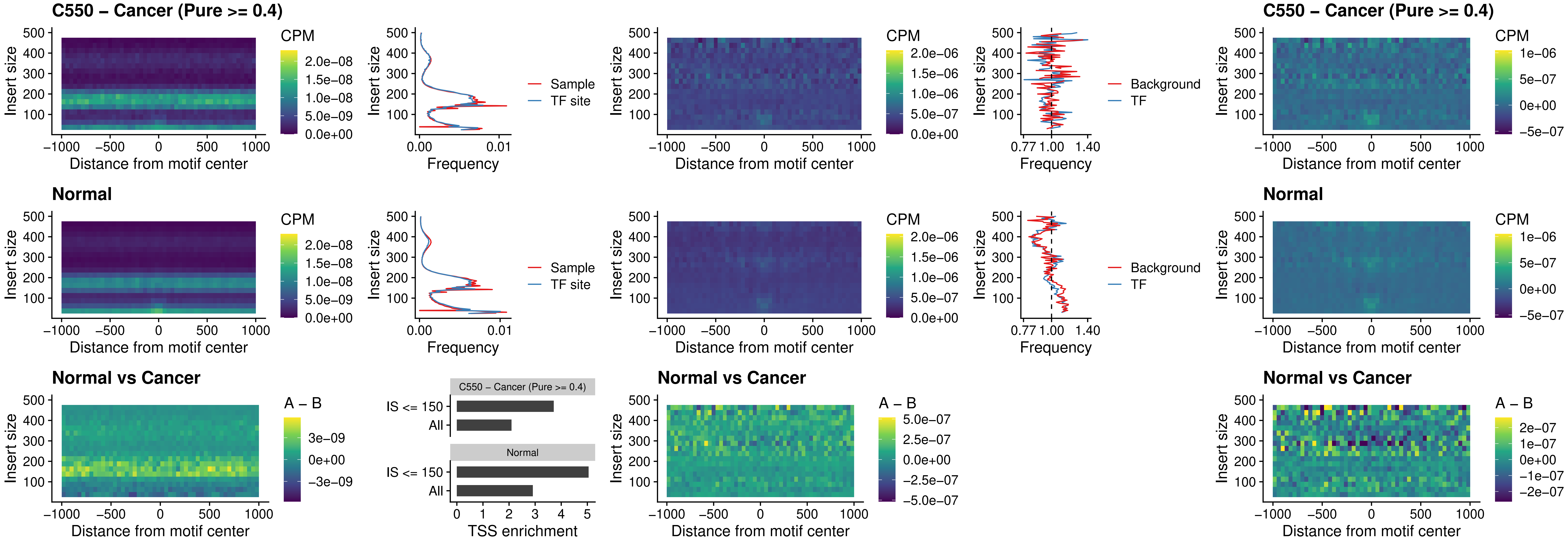

#### C550 – Cancer (Pure >= 0.4) – IRF2_uc_pTSS_oPEAK

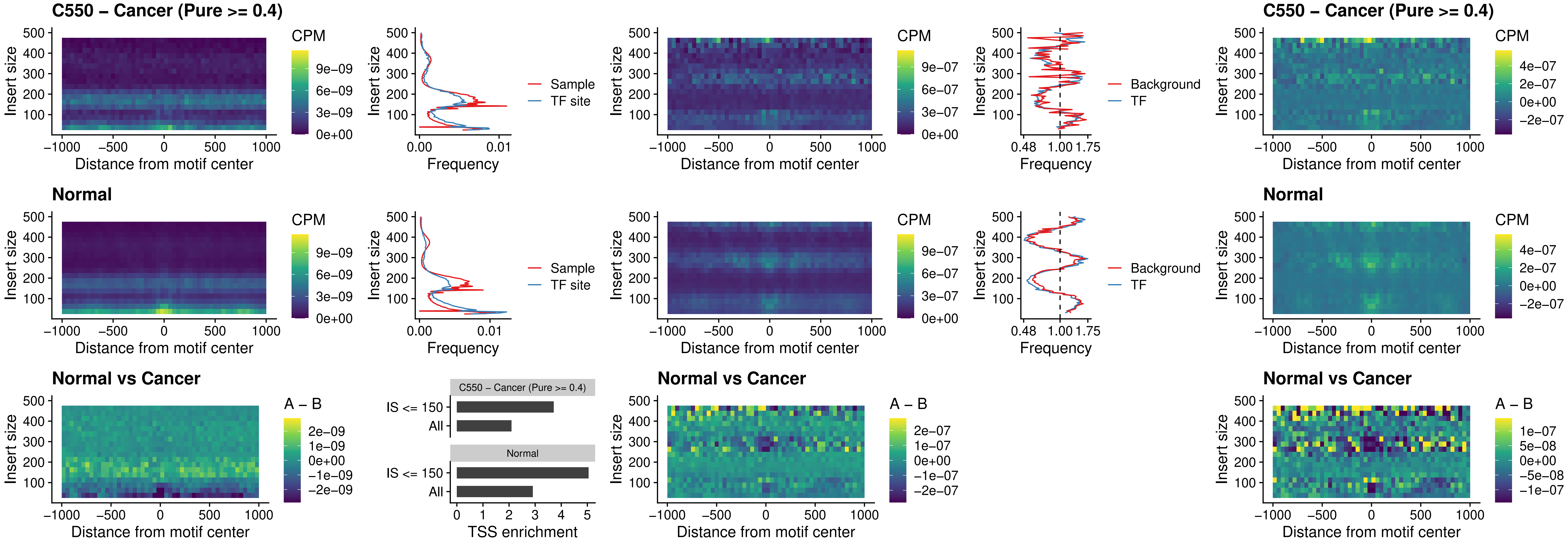

#### C550 – Cancer (Pure >= 0.4) – STAT2_uc_pTSS_oPEAK

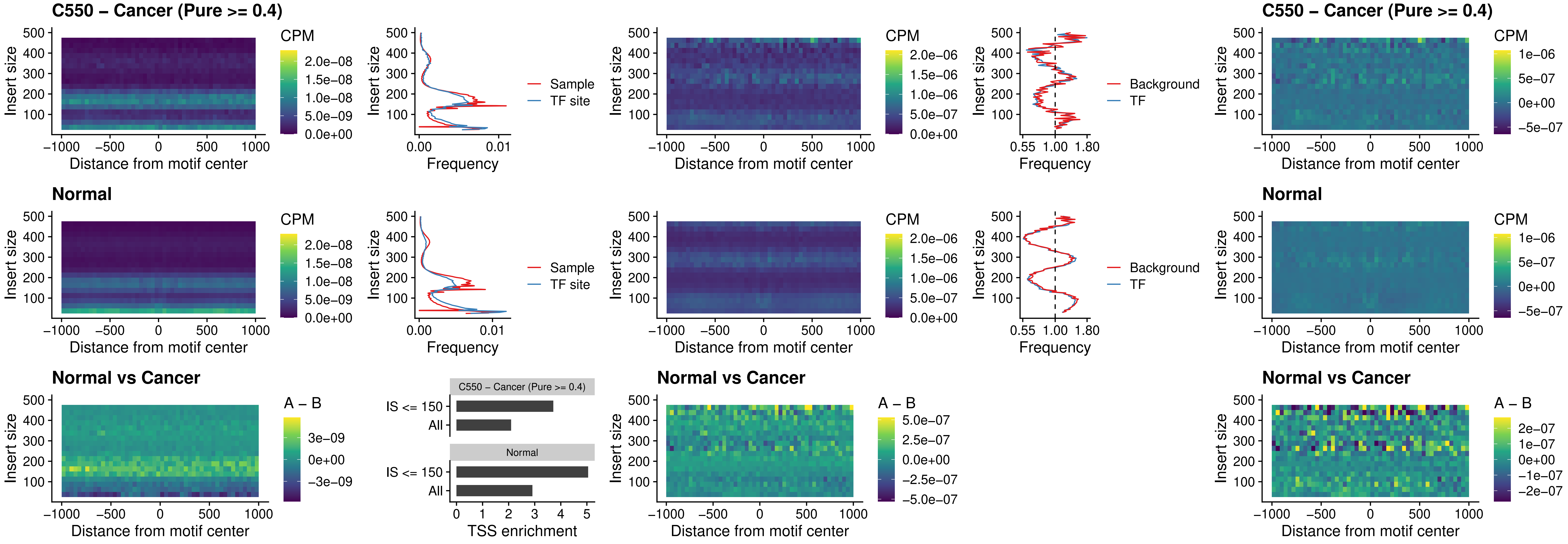

### C550 – Cancer (Pure >= 0.4) – SPIB_uc_dTSS_nPEAK

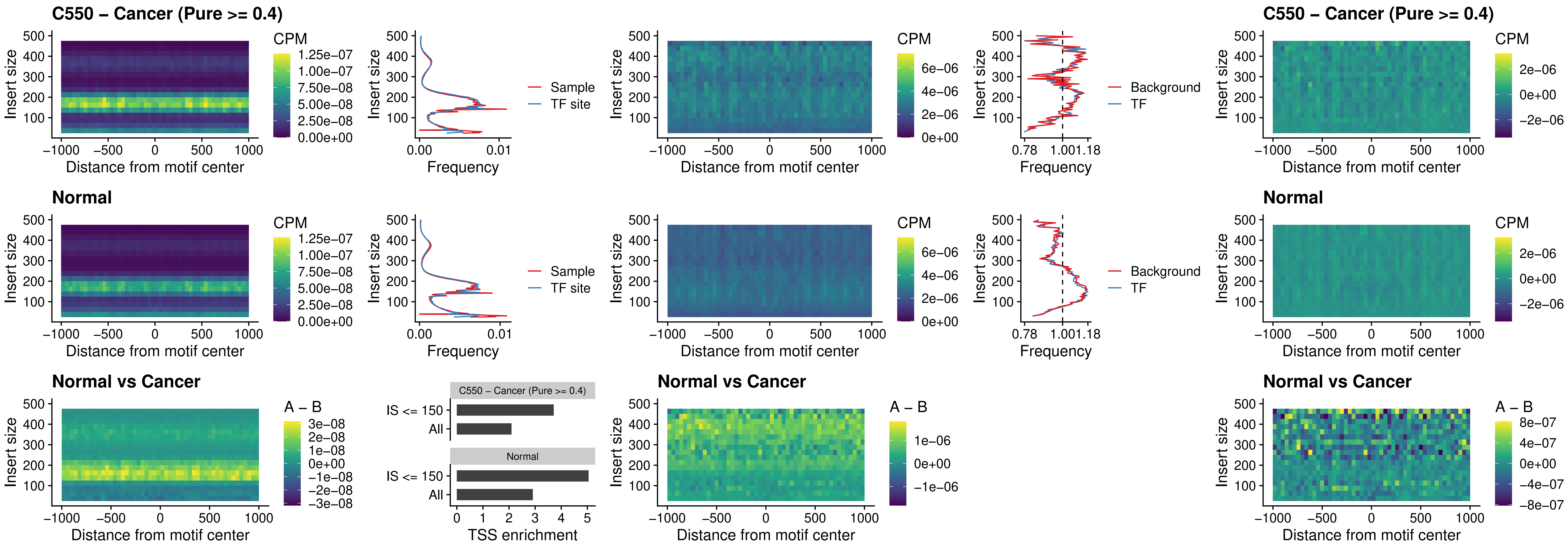

#### C550 – Cancer (Pure >= 0.4) – SPIC_uc_cTSS_oPEAK

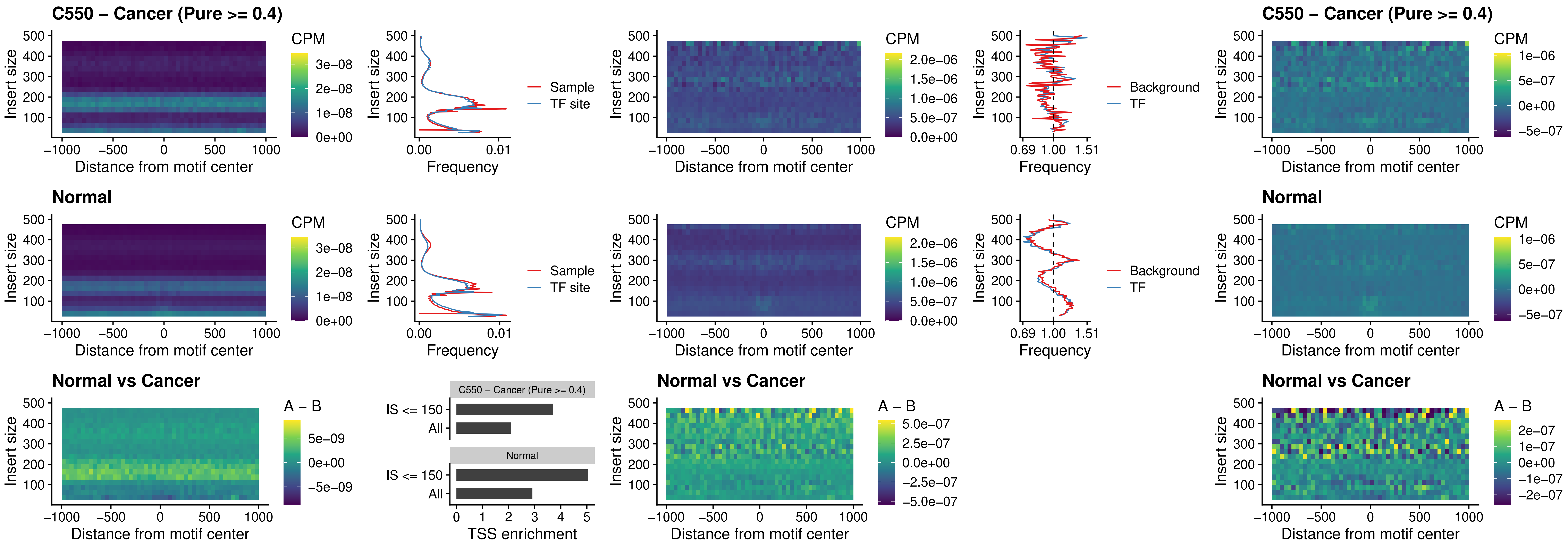

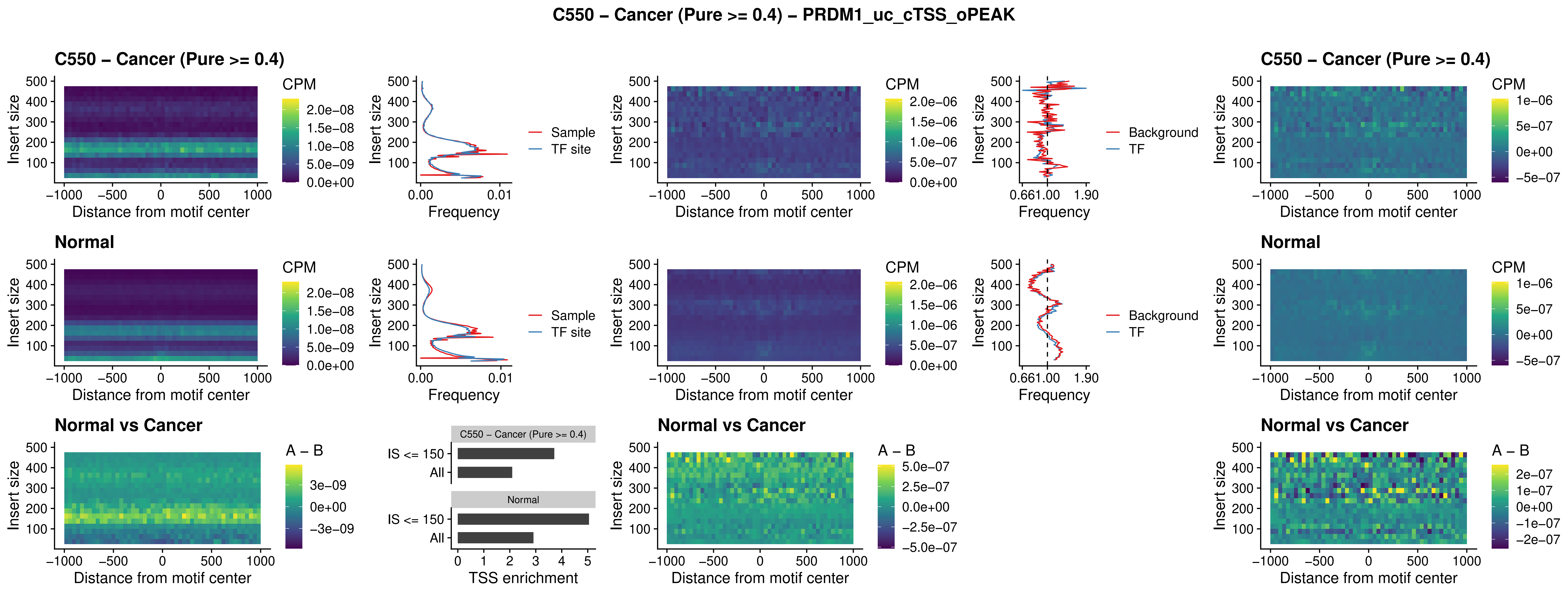
C550 – Cancer (Pure >= 0.4) – PRDM1_uc_cTSS_oPEAK
C550 – Cancer (Pure >= 0.4)
Normal
Normal vs Cancer
Insert size
Distance from motif center
Frequency
CPM
A – B
Sample
TF site
Background
TF
C550 – Cancer (Pure >= 0.4)
Normal
IS <= 150
All
TSS enrichment

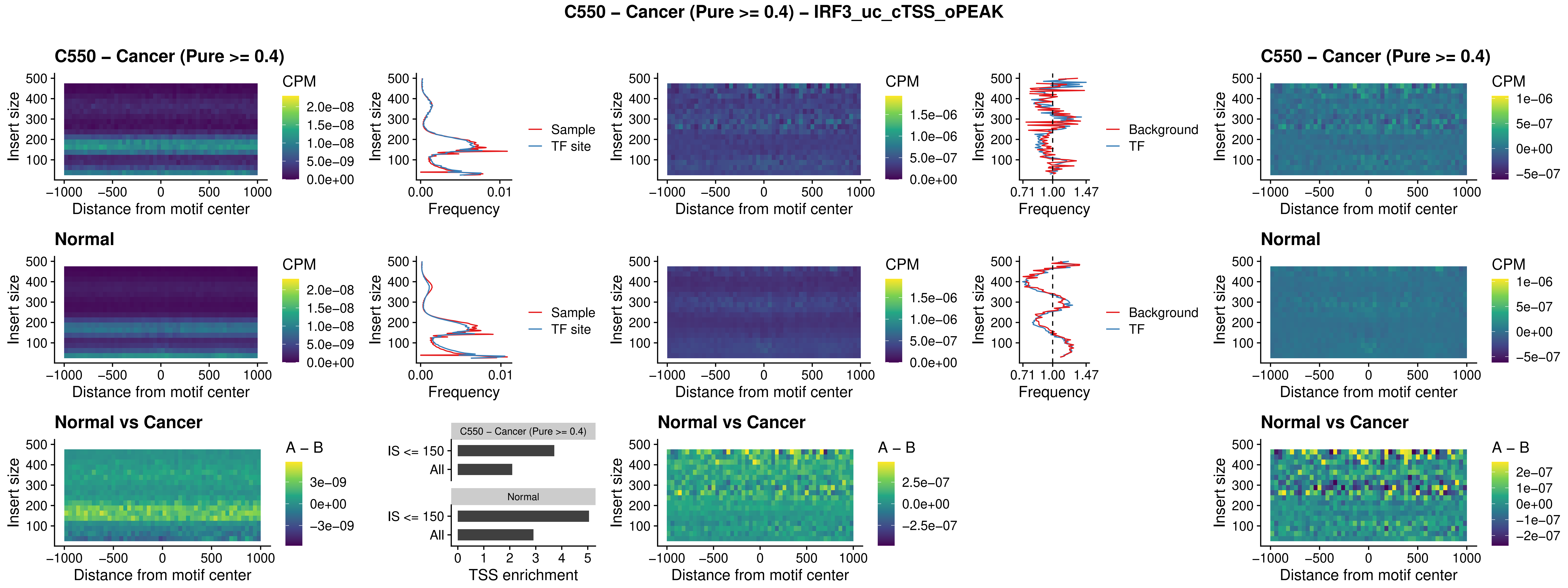
C550 – Cancer (Pure >= 0.4) – IRF3_uc_cTSS_oPEAK
C550 – Cancer (Pure >= 0.4)
Normal
Normal vs Cancer
Insert size
Distance from motif center
CPM
A – B
Frequency
Sample
TF site
Background
TF
C550 – Cancer (Pure >= 0.4)
Normal
IS <= 150
All
TSS enrichment

#### C550 – Cancer (Pure >= 0.4) – BCL11A_uc_dTSS_nPEAK

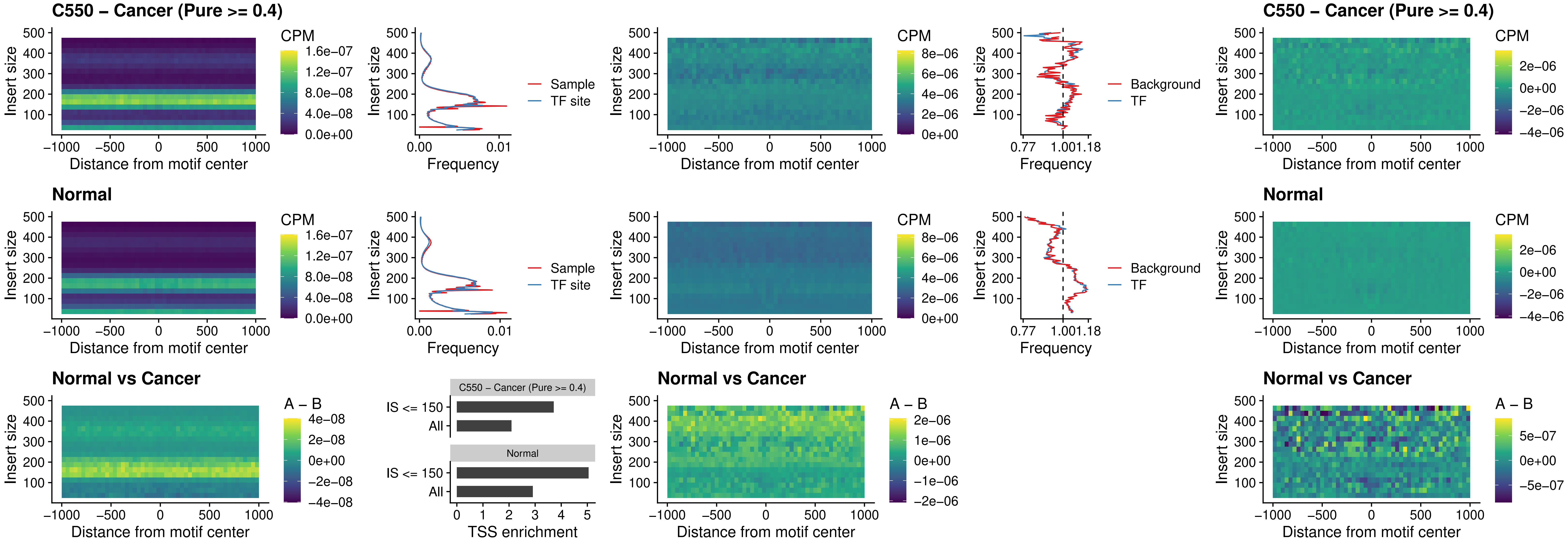

### C550 – Cancer (Pure >= 0.4) – BCL11B_uc_dTSS_nPEAK

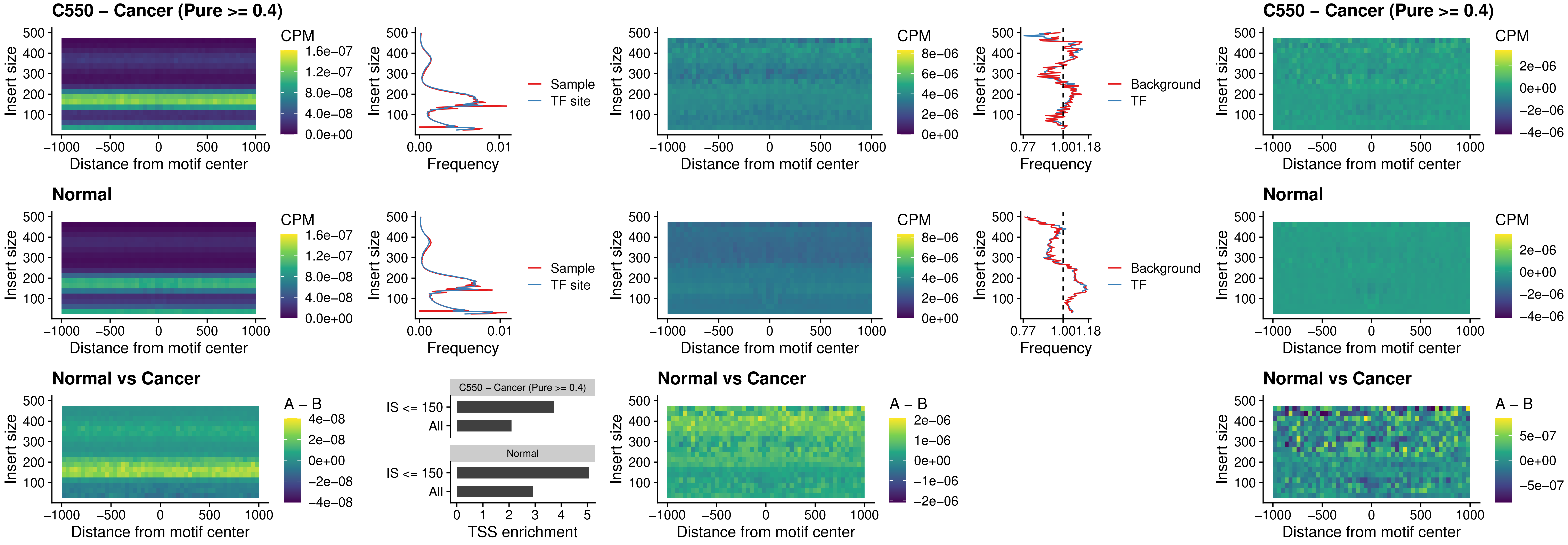

### C550 – Cancer (Pure >= 0.4) – IRF1_uc_cTSS_oPEAK

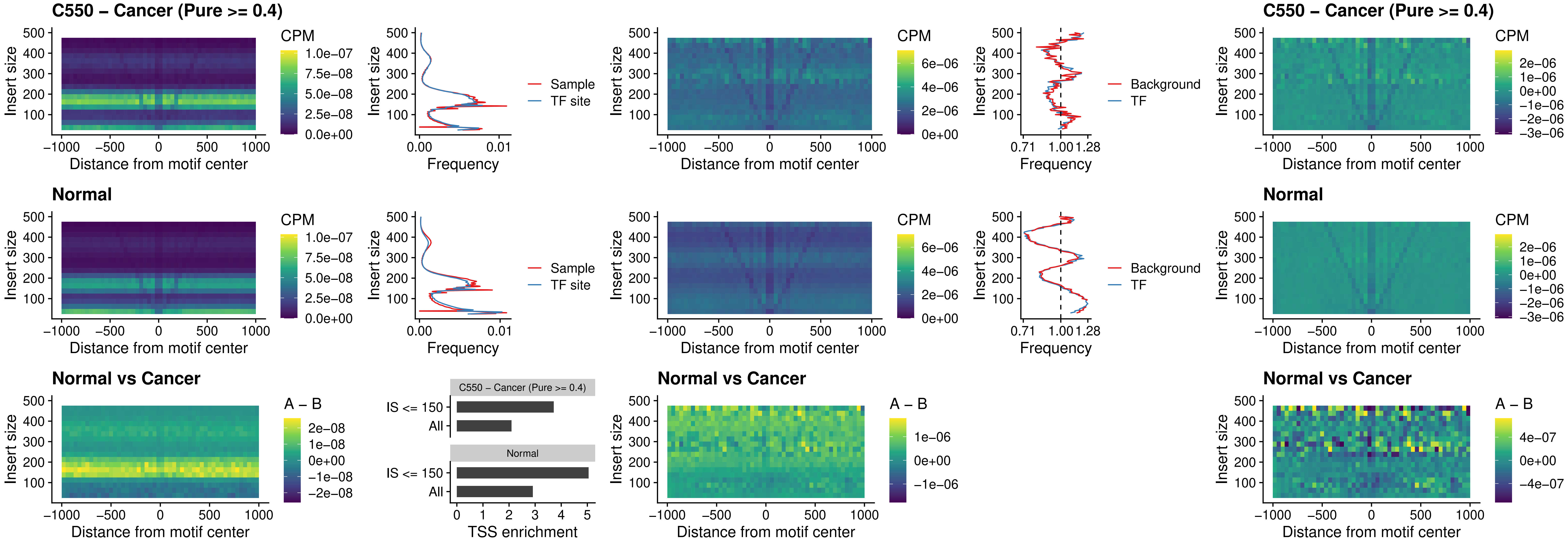

#### C550 – Cancer (Pure >= 0.4) – IRF1_uc_dTSS_oPEAK

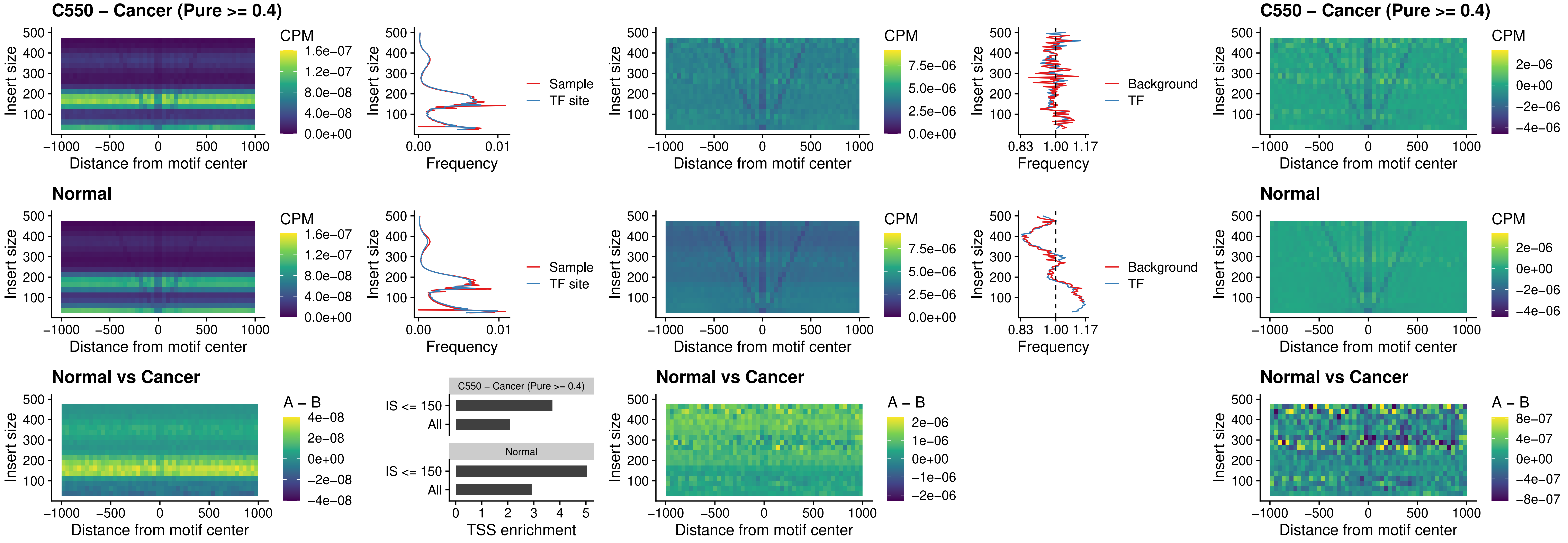

#### C550 – Cancer (Pure >= 0.4) – IRF1_uc_pTSS_oPEAK

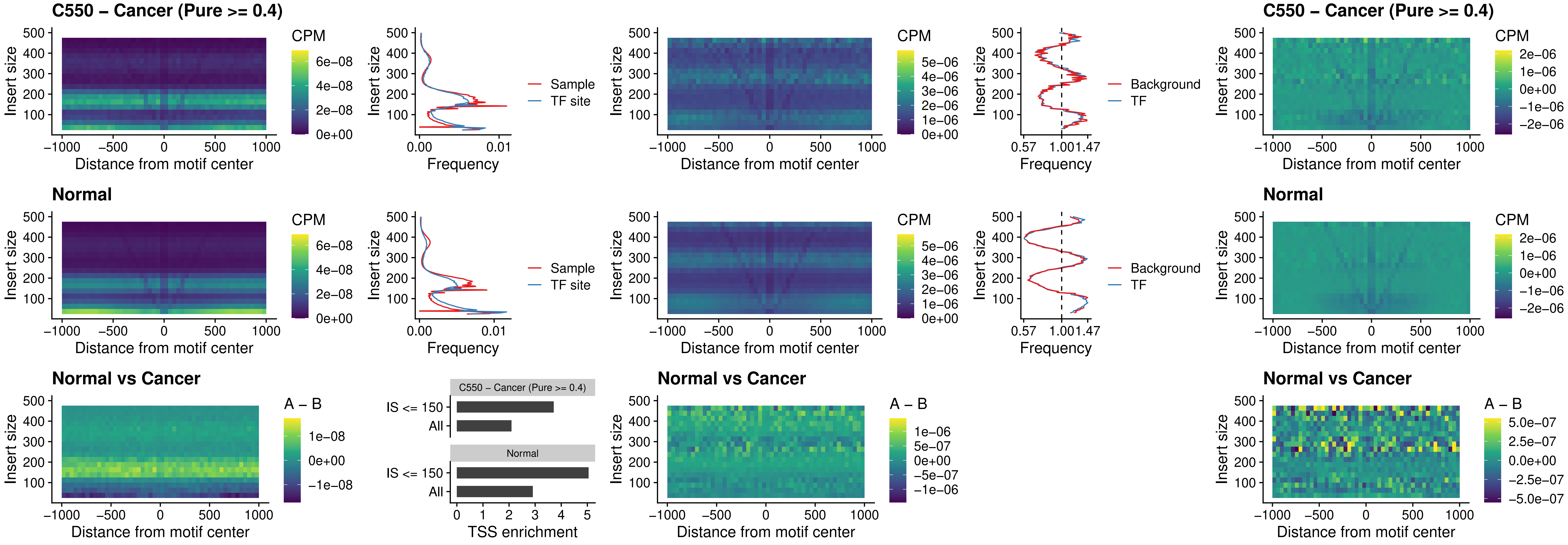

### C550 – Cancer (Pure >= 0.4) – SPIC_uc_dTSS_nPEAK

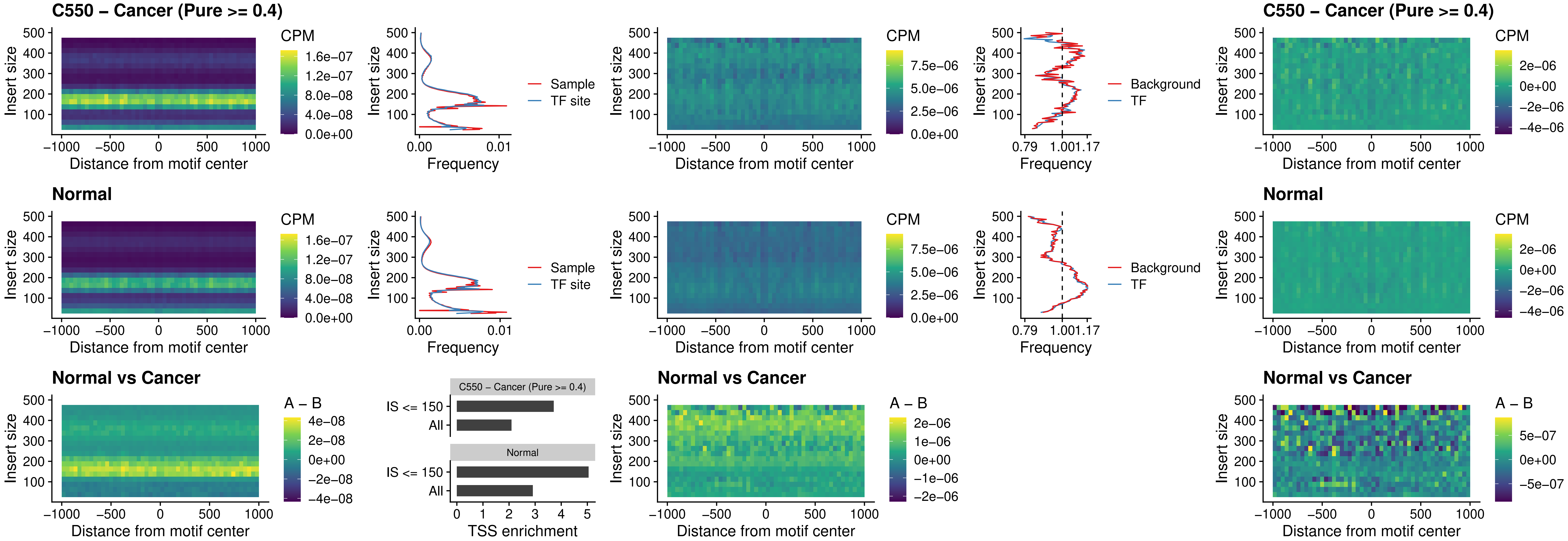

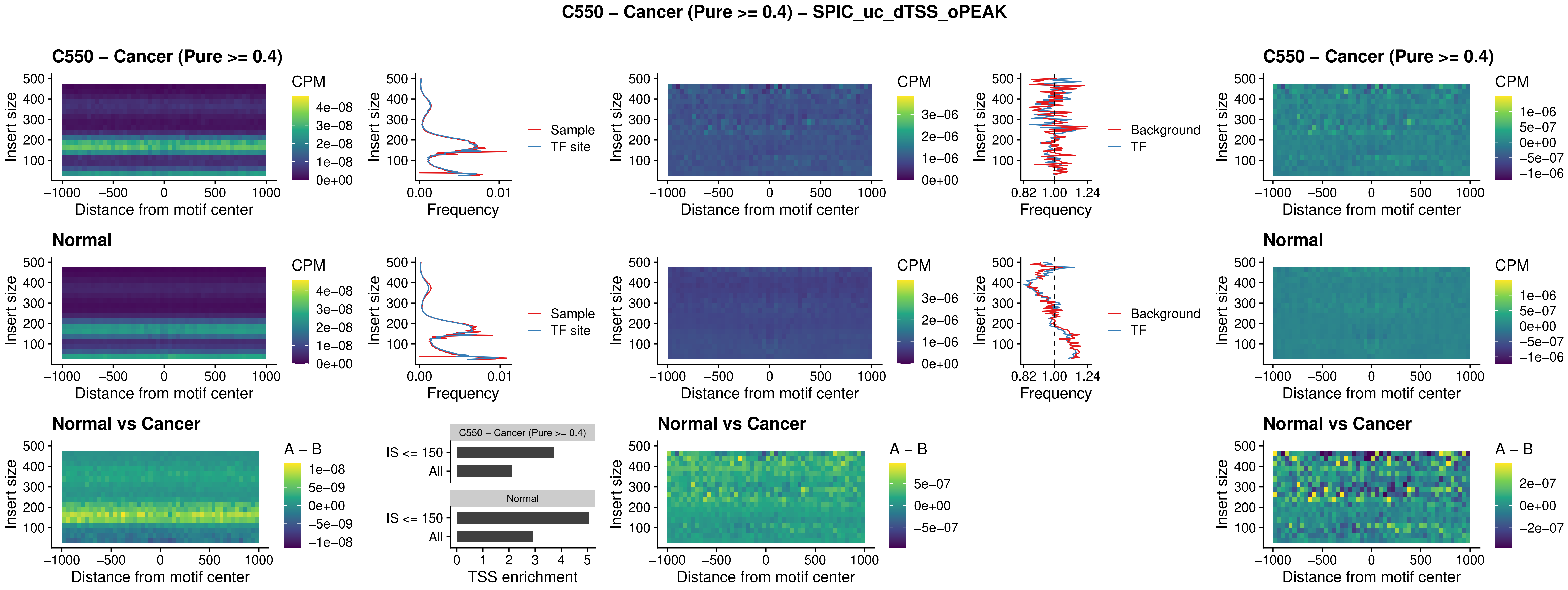
C550 – Cancer (Pure >= 0.4) – SPIC_uc_dTSS_oPEAK
C550 – Cancer (Pure >= 0.4)
Normal
Normal vs Cancer
Insert size
Distance from motif center
Frequency
CPM
A – B
Sample
TF site
Background
TF
C550 – Cancer (Pure >= 0.4)
Normal
IS <= 150
All
TSS enrichment

### C550 – Cancer (Pure >= 0.4) – SPIB_uc_pTSS_nPEAK

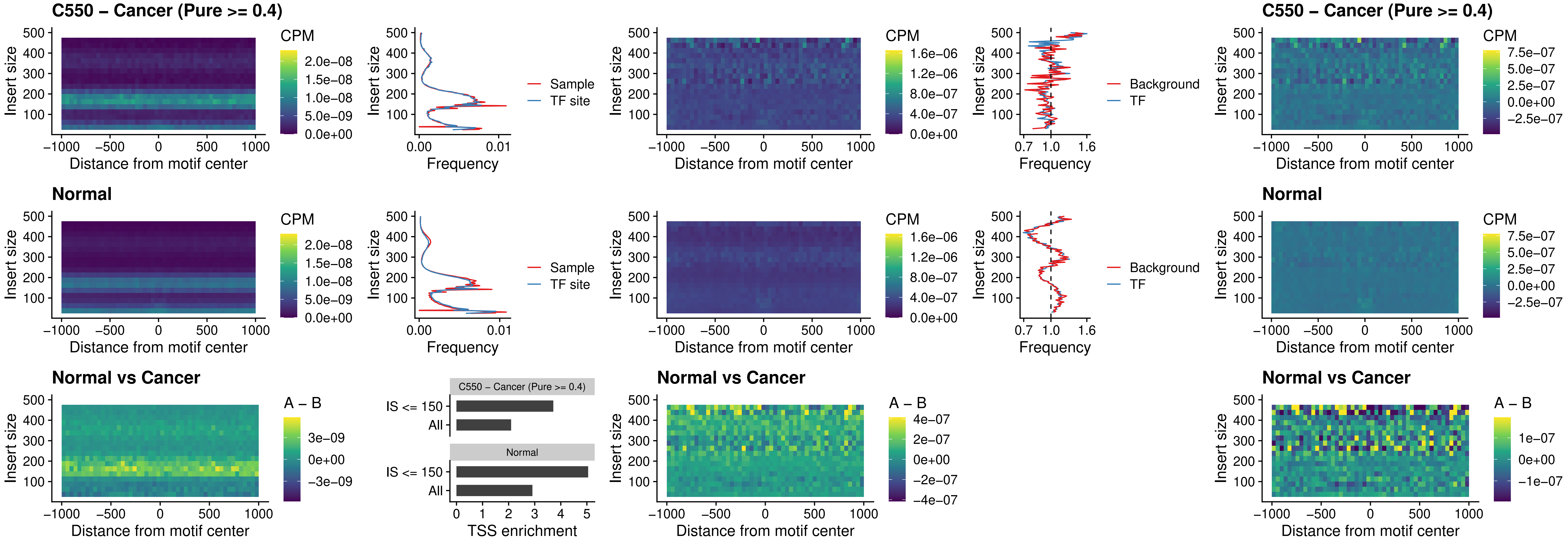

#### C550 – Cancer (Pure >= 0.4) – STAT2_uc_cTSS_oPEAK

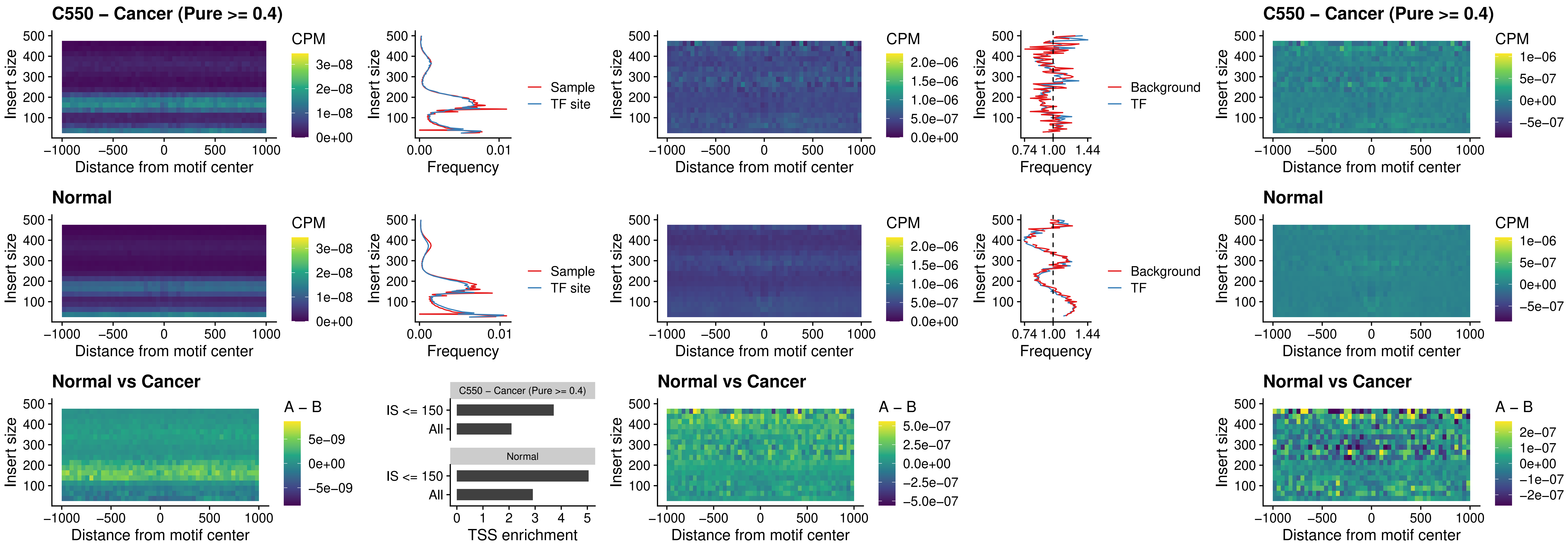

#### C550 – Cancer (Pure >= 0.4) – IRF8_uc_cTSS_oPEAK

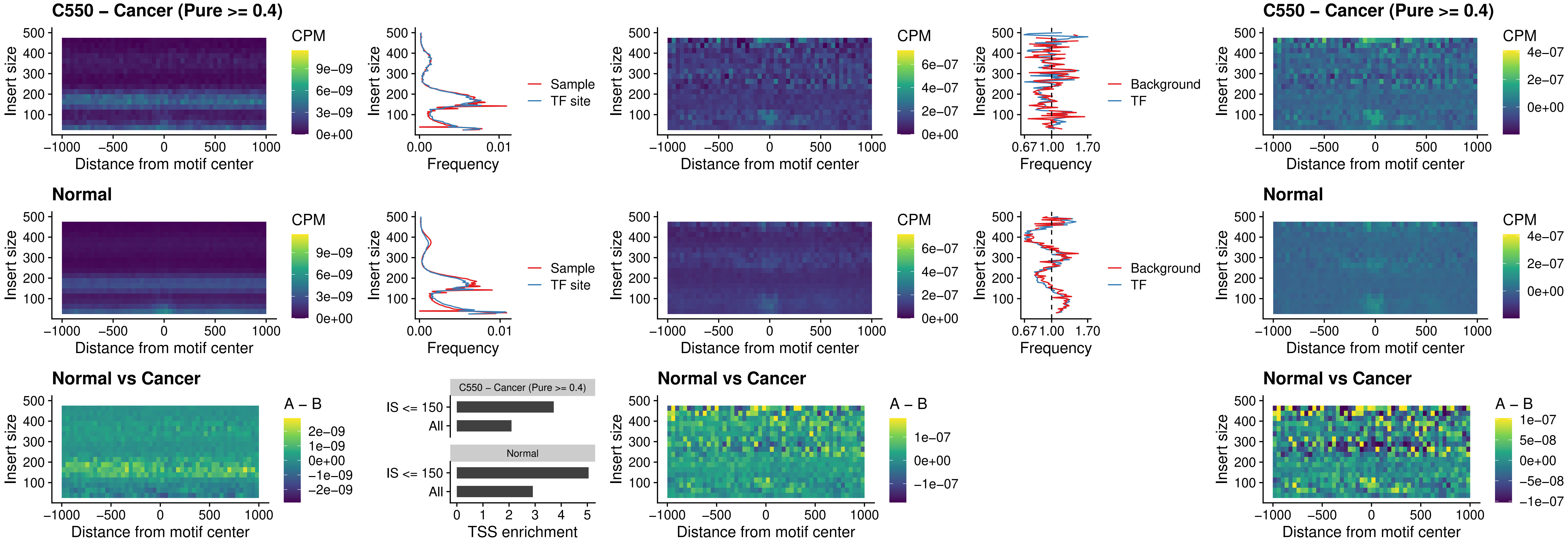

#### C550 – Cancer (Pure >= 0.4) – IRF3_uc_dTSS_oPEAK

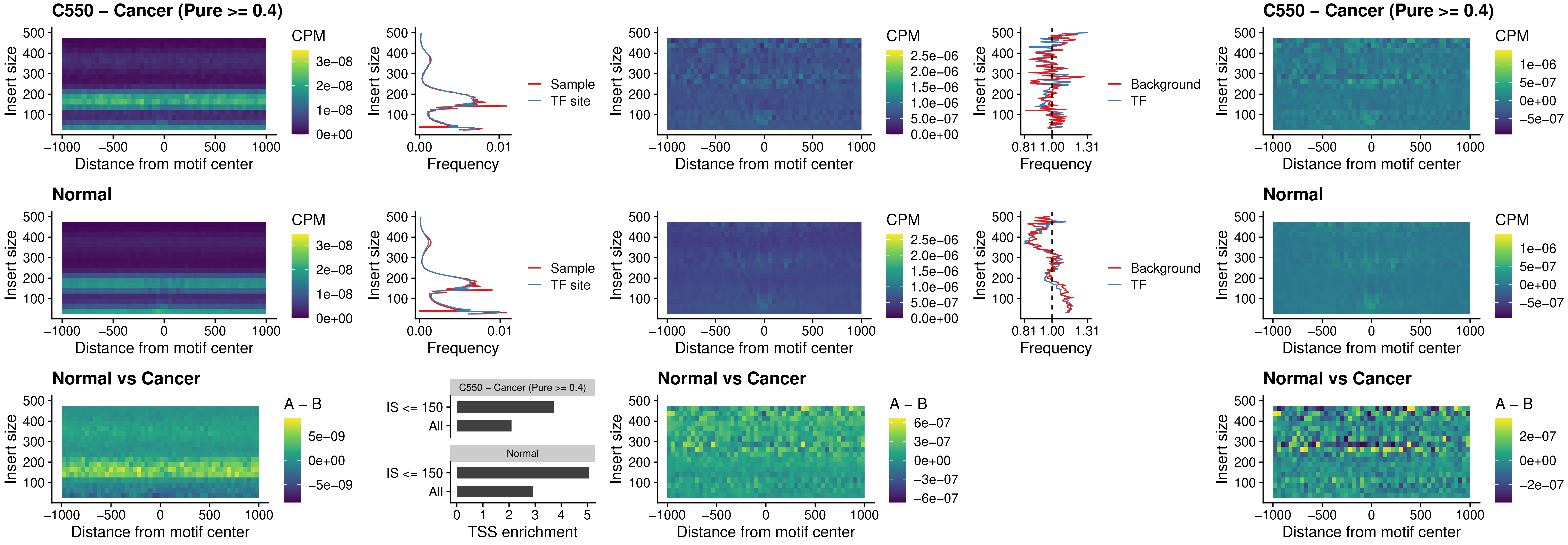

#### C550 – Cancer (Pure >= 0.4) – IRF8_uc_dTSS_oPEAK

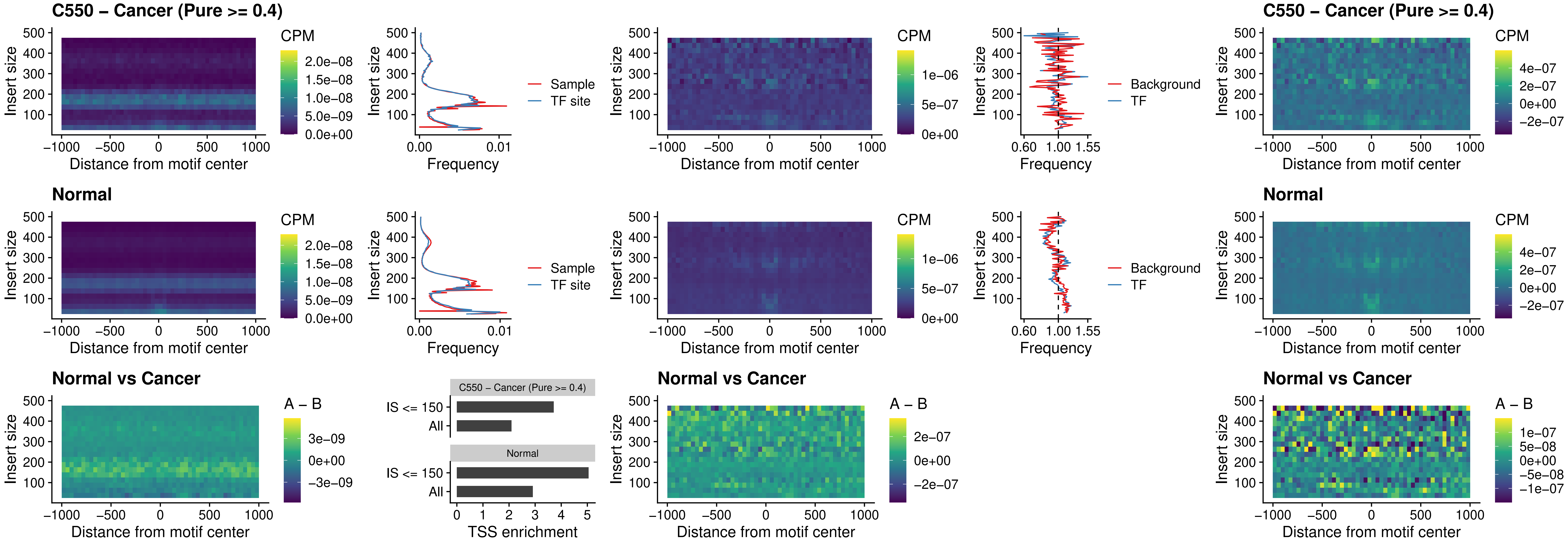

#### C550 – Cancer (Pure >= 0.4) – ETS2_uc_dTSS_oPEAK

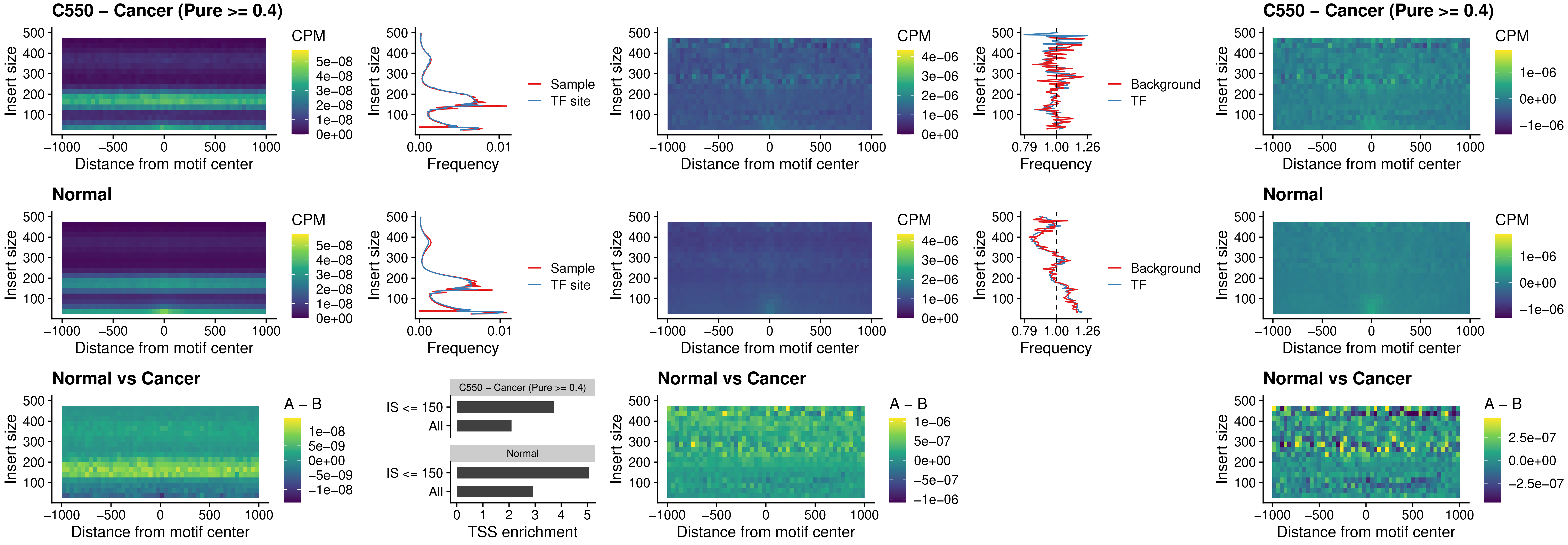

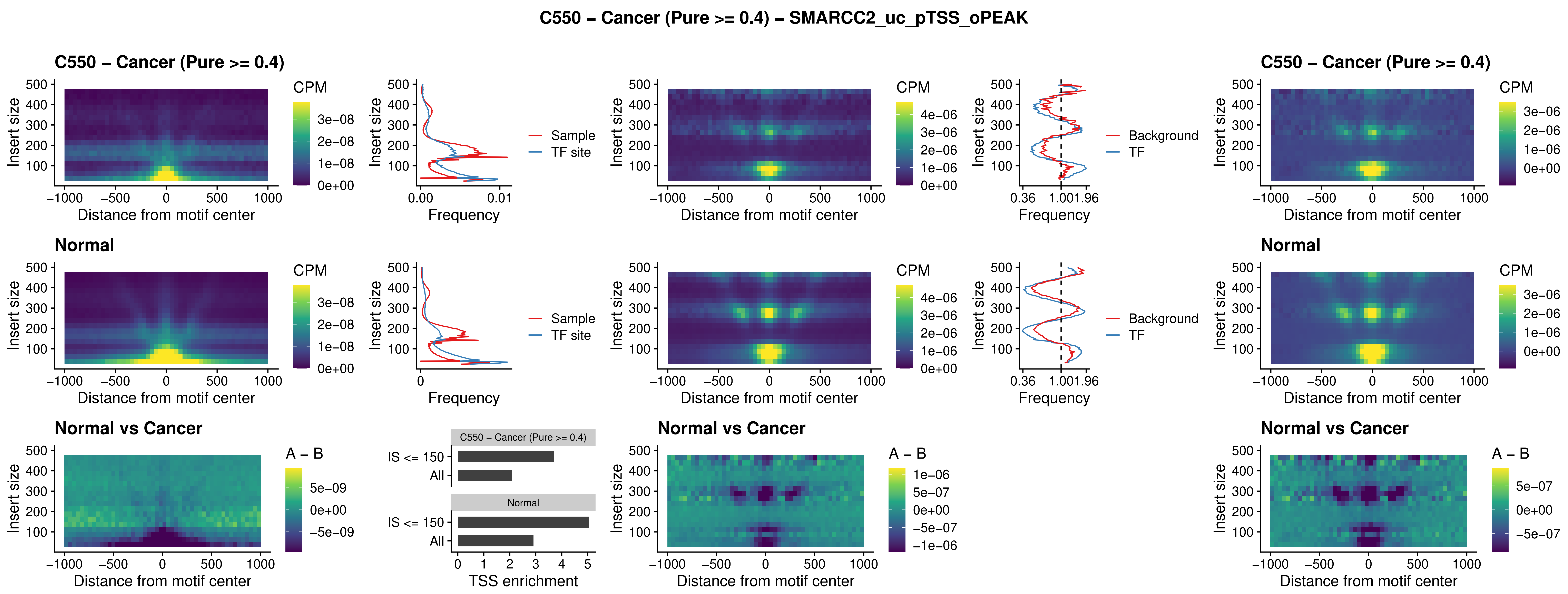
C550 – Cancer (Pure >= 0.4) – SMARCC2_uc_pTSS_oPEAK
C550 – Cancer (Pure >= 0.4)
Normal
Normal vs Cancer
Insert size
Distance from motif center
CPM
A – B
Frequency
Sample
TF site
Background
TF
TSS enrichment
IS <= 150
All
C550 – Cancer (Pure >= 0.4)
Normal

**C550 – Cancer (Pure >= 0.4) – BCL11A_uc_cTSS_nPEAK**

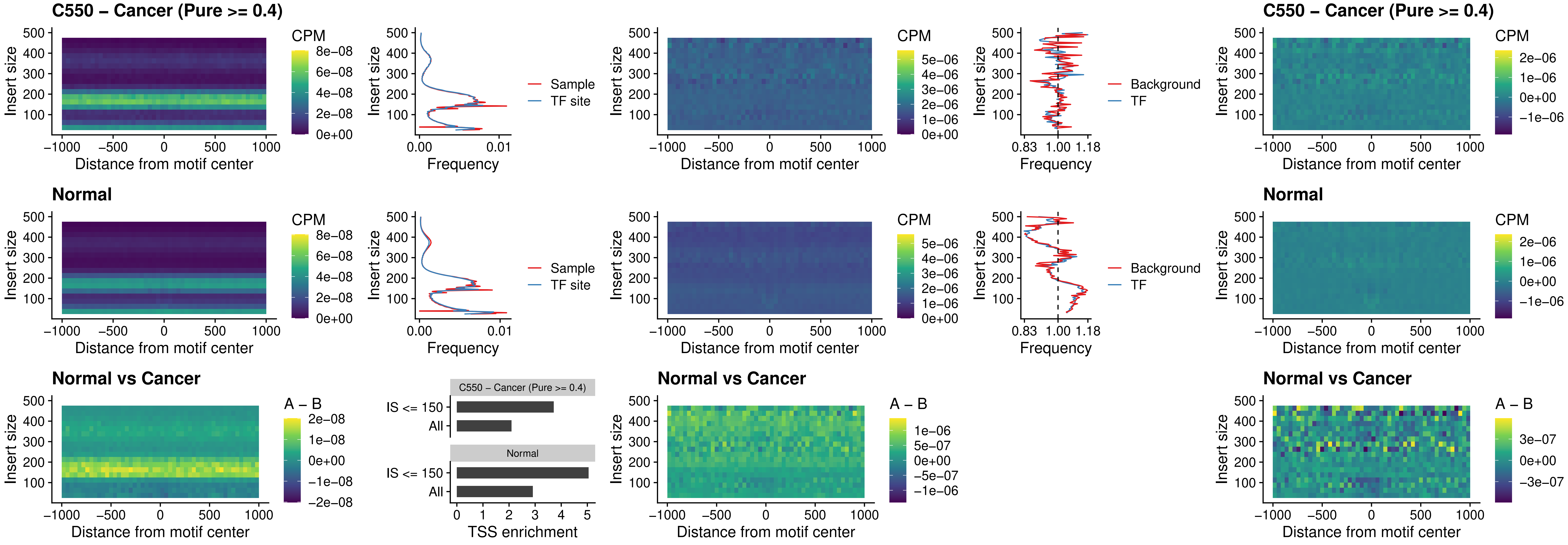

#### C550 – Cancer (Pure >= 0.4) – BCL11B_uc_cTSS_nPEAK

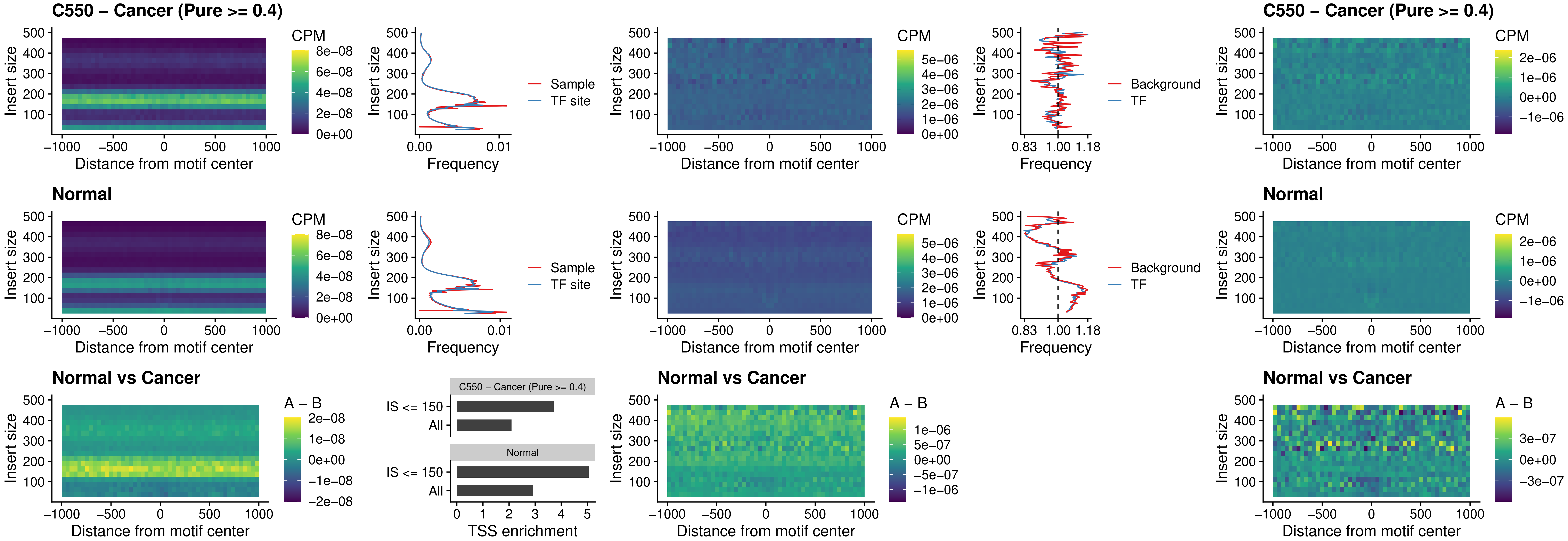

#### C550 – Cancer (Pure >= 0.4) – STAT2_uc_dTSS_oPEAK

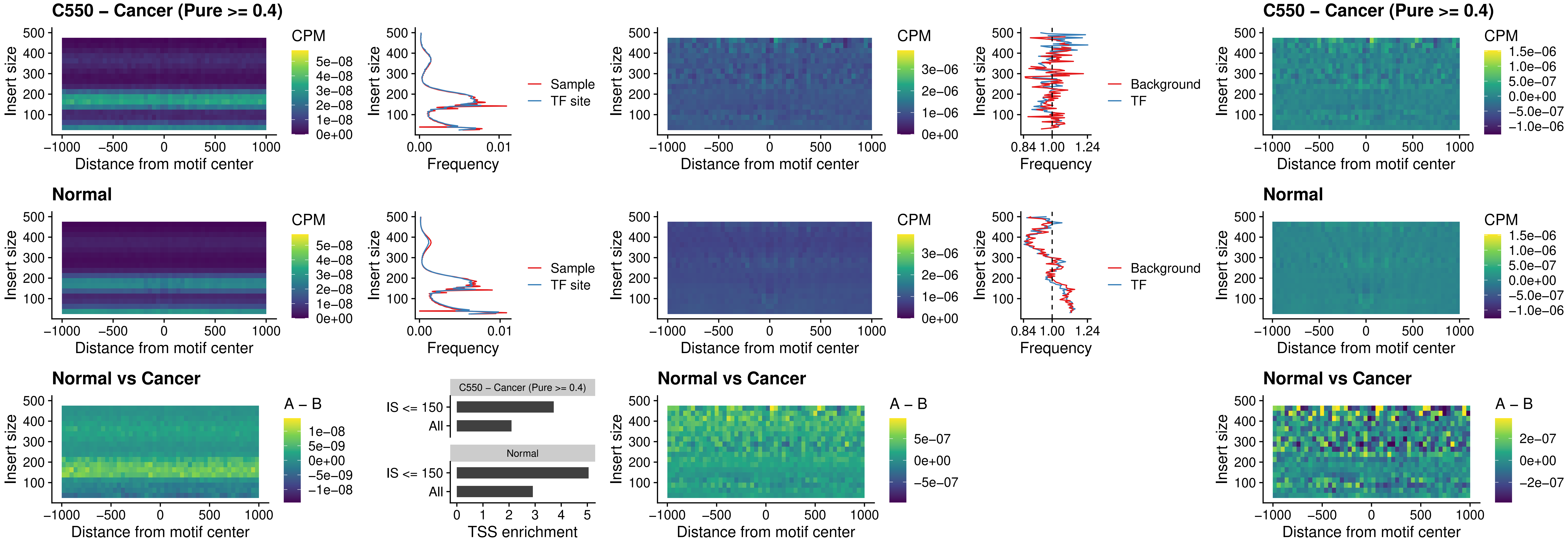

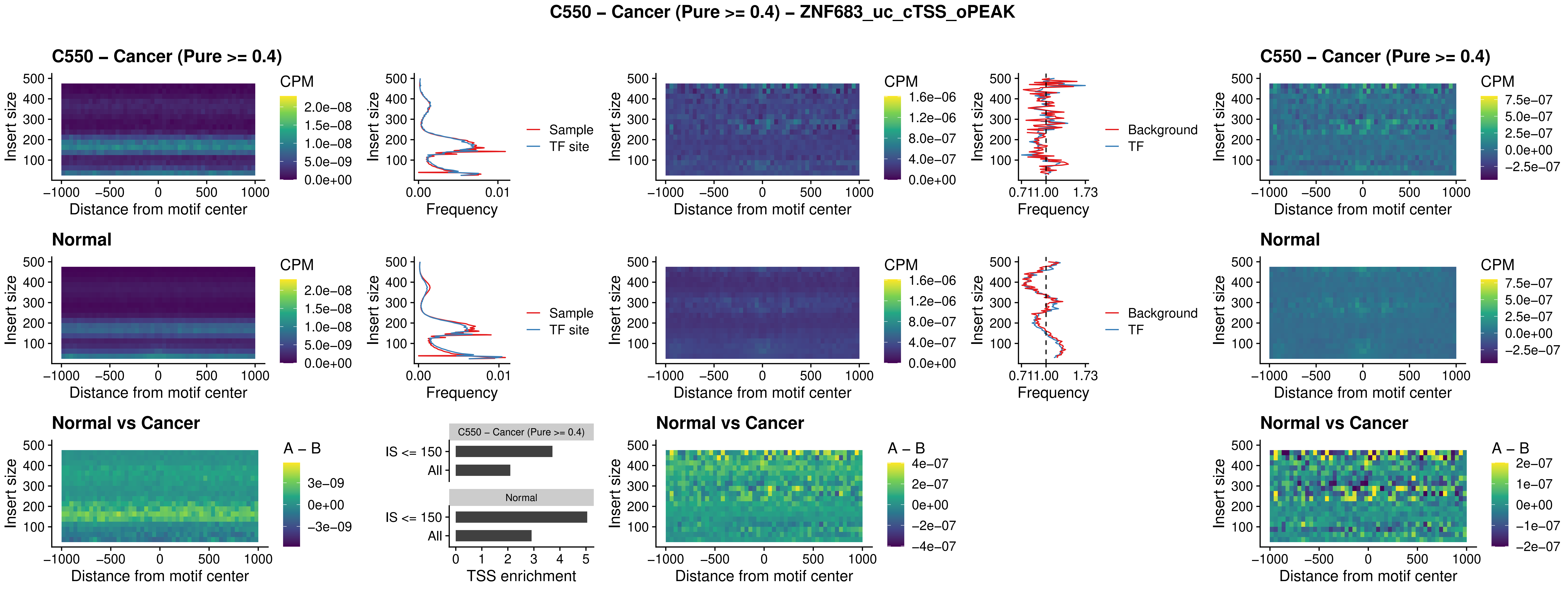
C550 – Cancer (Pure >= 0.4) – ZNF683_uc_cTSS_oPEAK
C550 – Cancer (Pure >= 0.4)
Normal
Normal vs Cancer
Insert size
Distance from motif center
Frequency
CPM
A – B
Sample
TF site
Background
TF
C550 – Cancer (Pure >= 0.4)
Normal
IS <= 150
All
TSS enrichment

#### C550 – Cancer (Pure >= 0.4) – IRF8_uc_dTSS_nPEAK

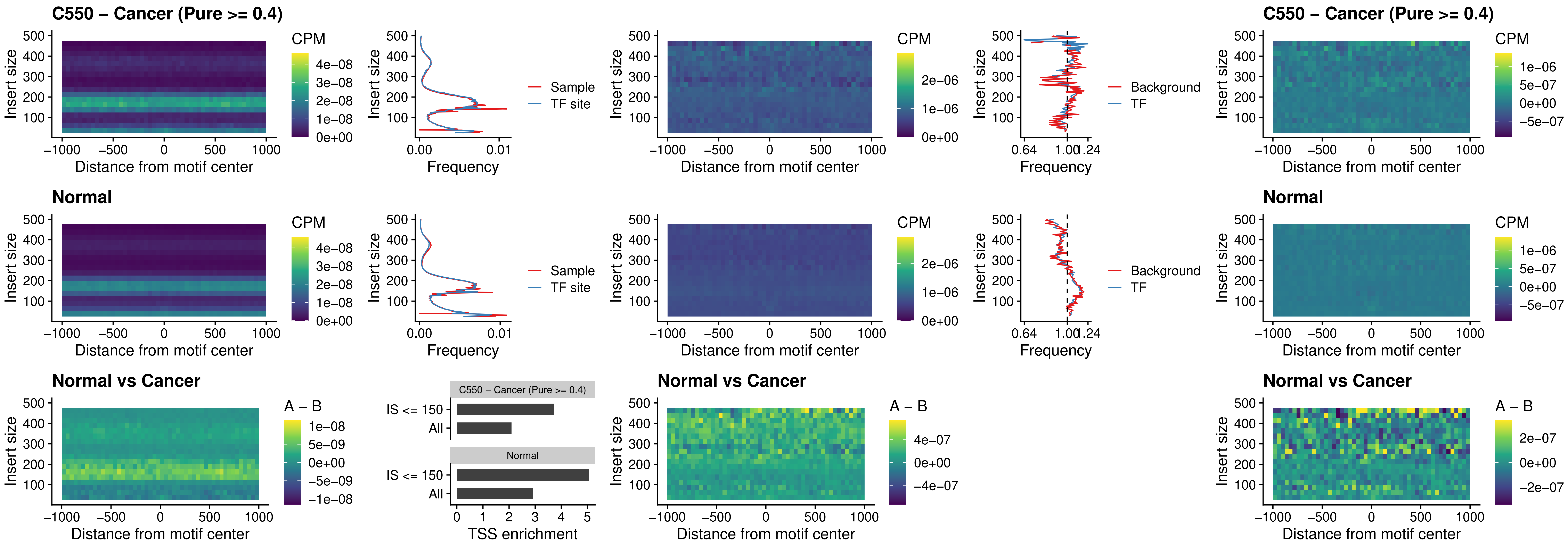

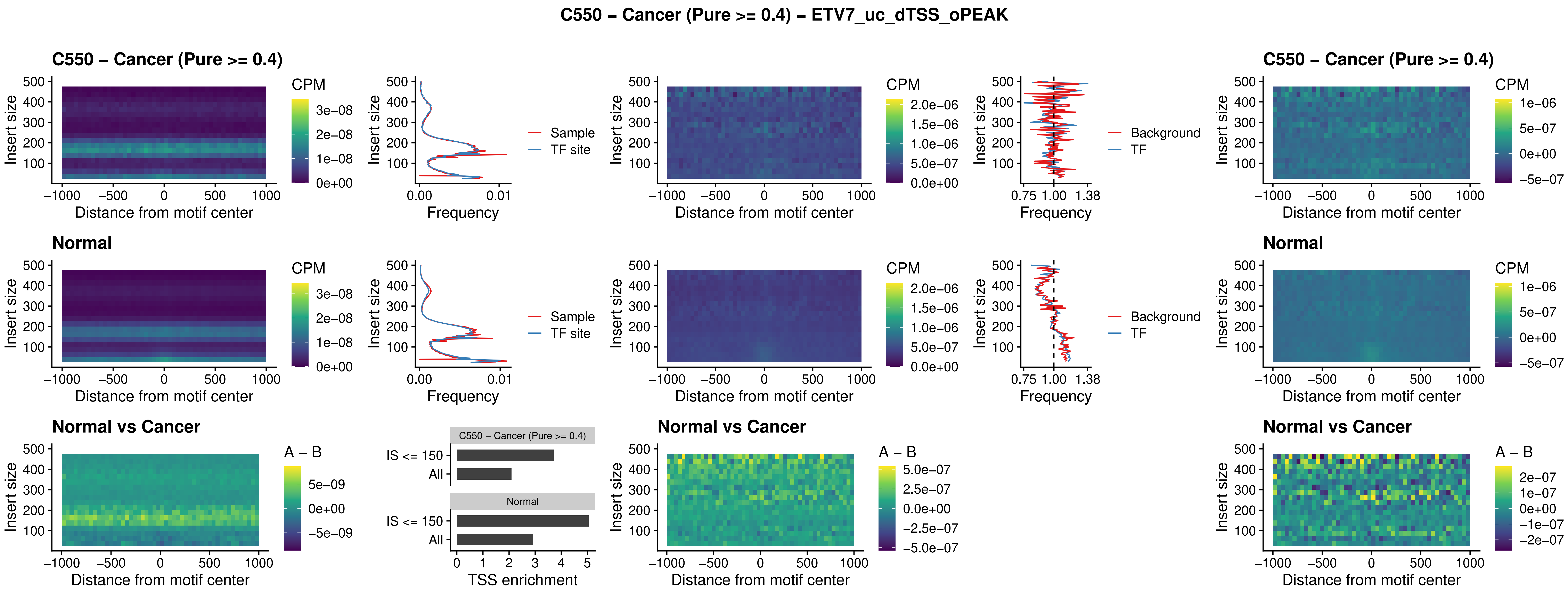
C550 – Cancer (Pure >= 0.4) – ETV7_uc_dTSS_oPEAK
C550 – Cancer (Pure >= 0.4)
Normal
Normal vs Cancer
Insert size
Distance from motif center
Frequency
CPM
A – B
Sample
TF site
Background
TF
C550 – Cancer (Pure >= 0.4)
Normal
IS <= 150
All
TSS enrichment

#### C551 – Cancer (Pure >= 0.4) – FOXL1_uc_pTSS_oPEAK

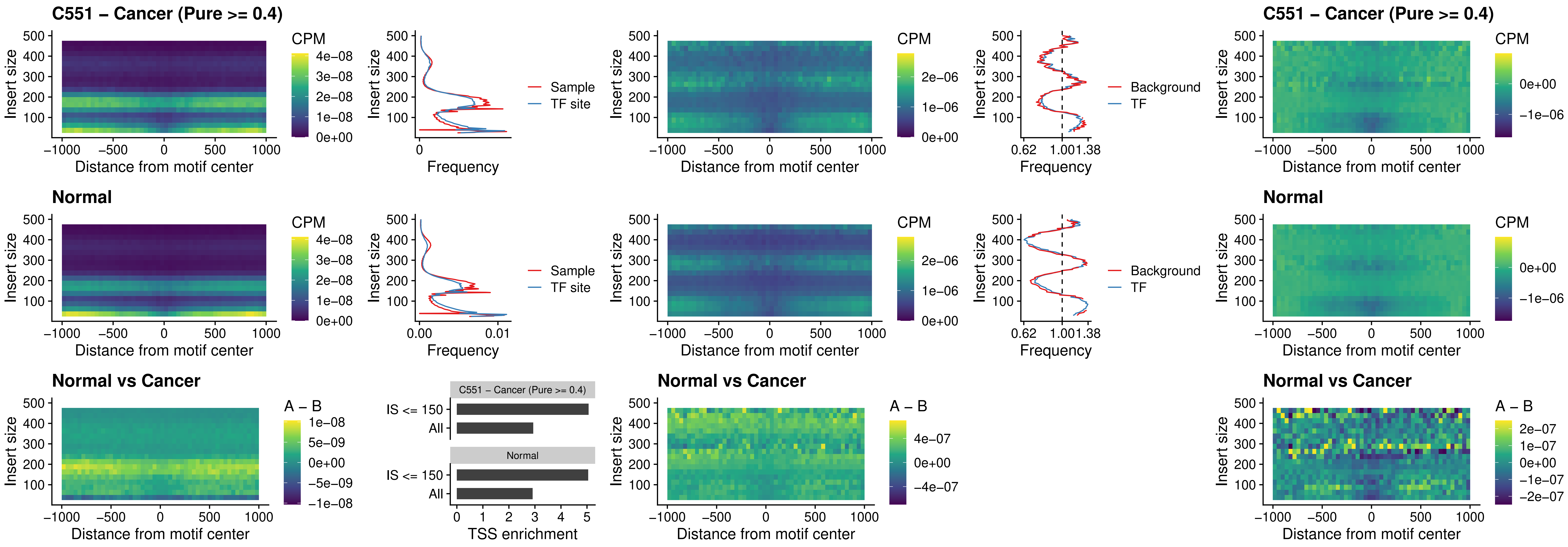

#### C551 – Cancer (Pure >= 0.4) – CPEB1_uc_cTSS_oPEAK

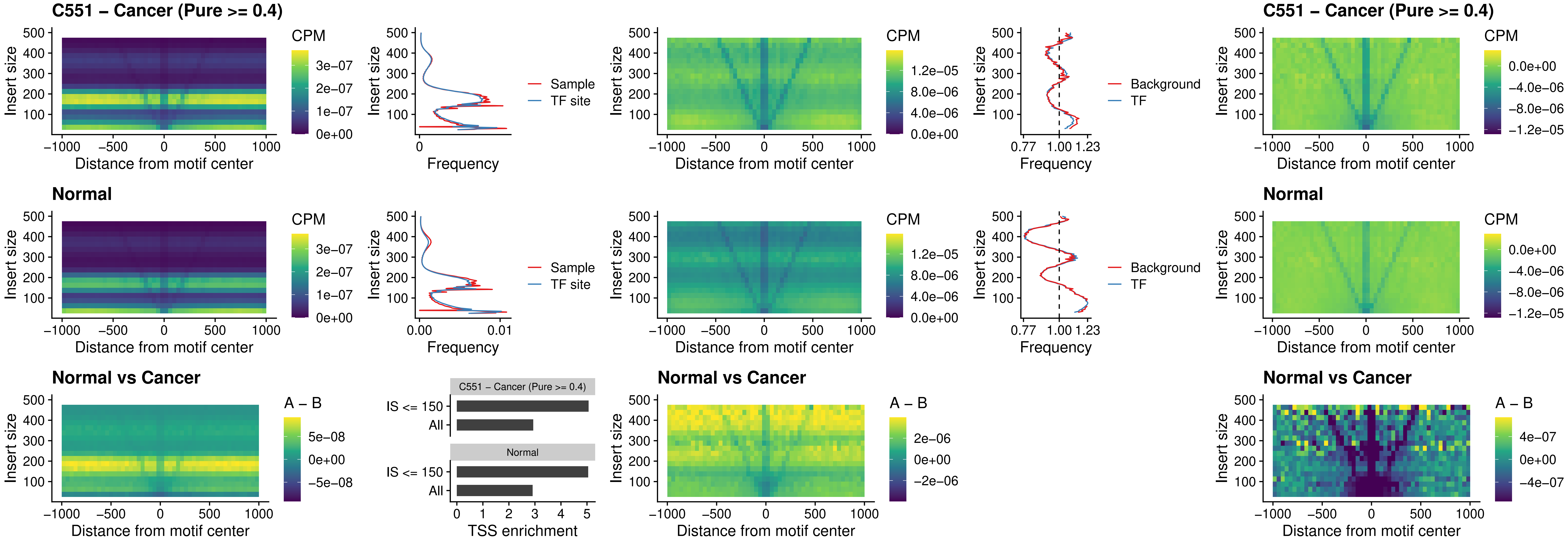

#### C551 – Cancer (Pure >= 0.4) – HMGA1_uc_pTSS_nPEAK

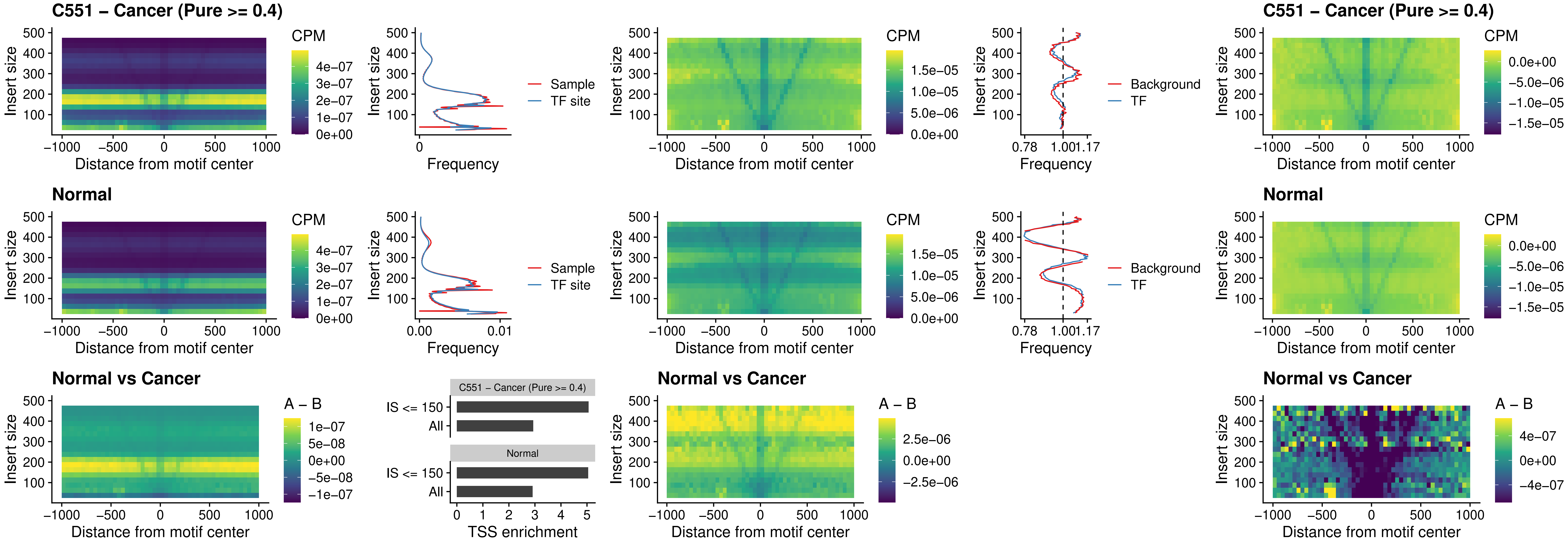

#### C551 – Cancer (Pure >= 0.4) – SOX13_uc_cTSS_nPEAK

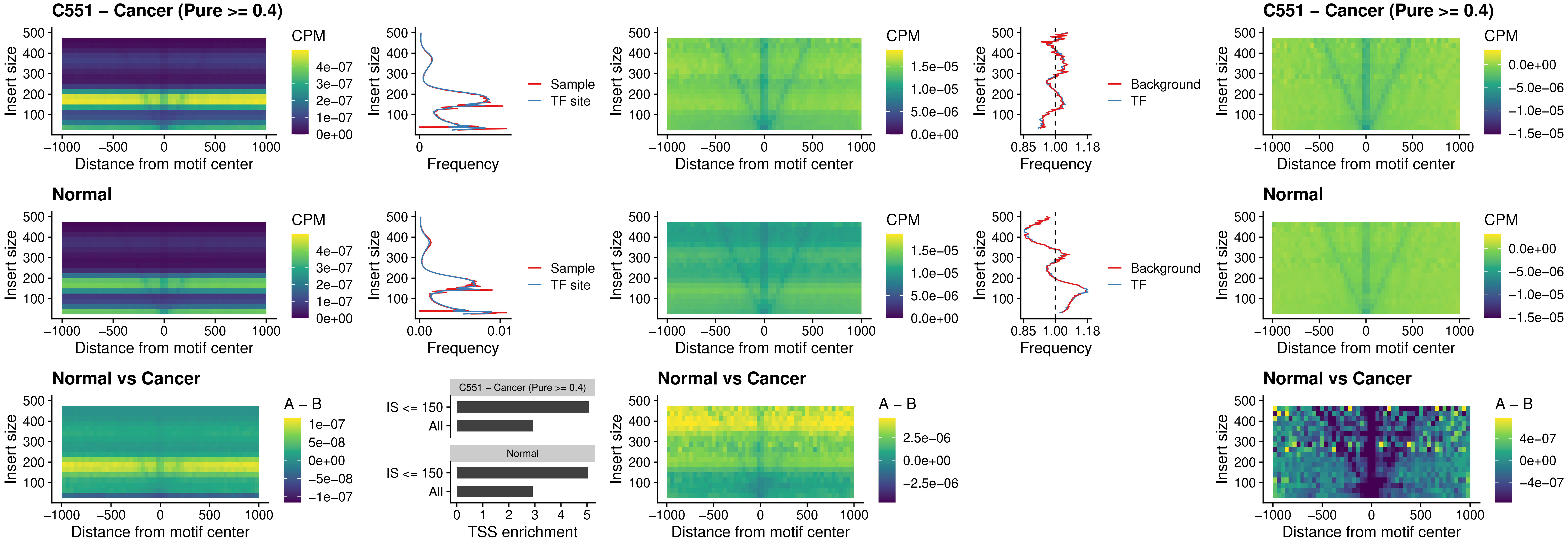

### C551 – Cancer (Pure >= 0.4) – LHX4_uc_cTSS_nPEAK

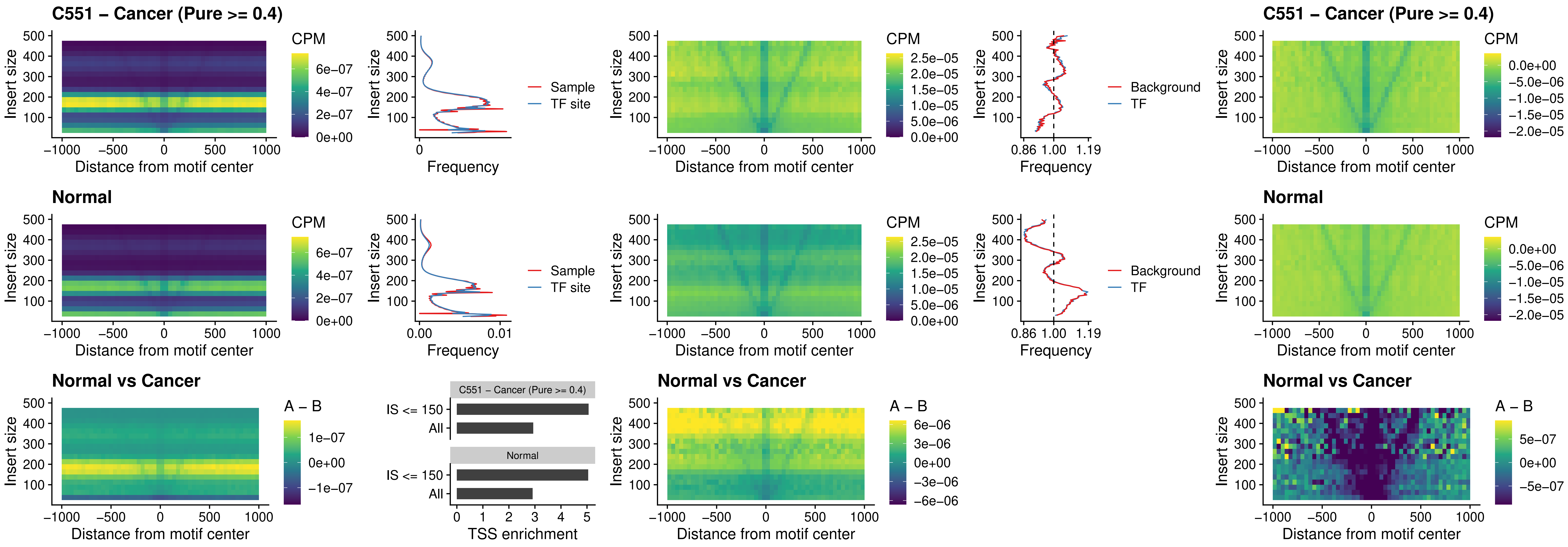

#### C551 – Cancer (Pure >= 0.4) – HOXA4_uc_cTSS_nPEAK

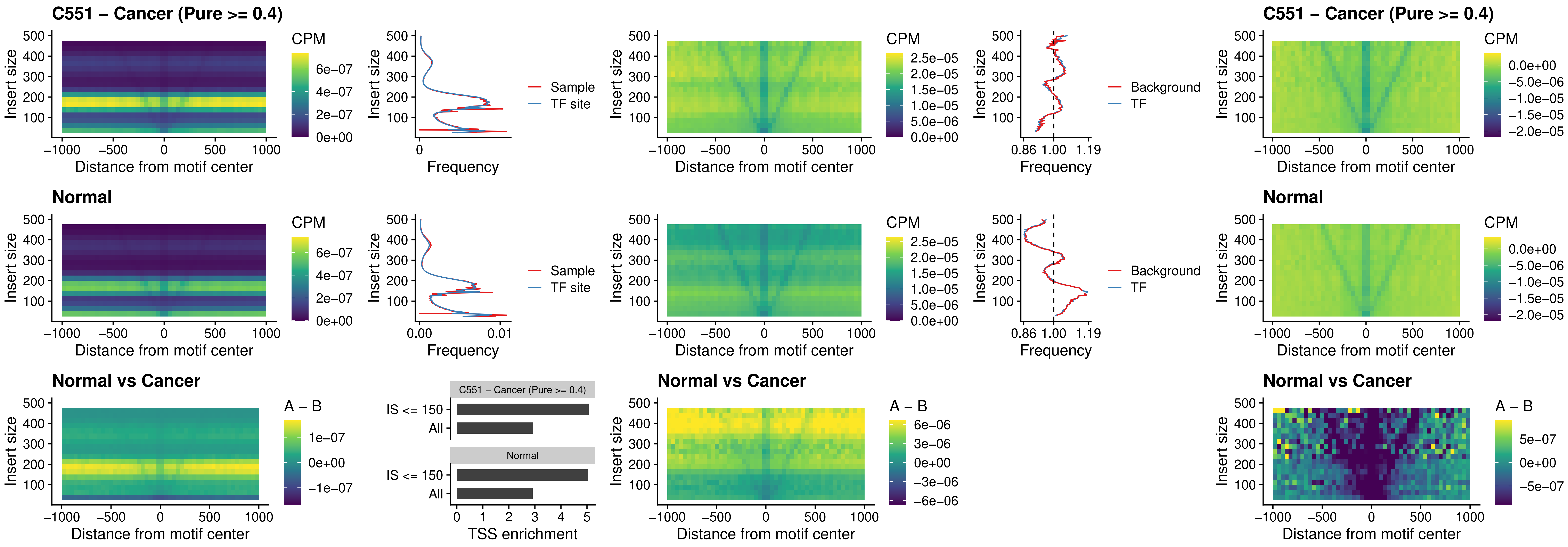

#### C551 – Cancer (Pure >= 0.4) – HLX_uc_cTSS_nPEAK

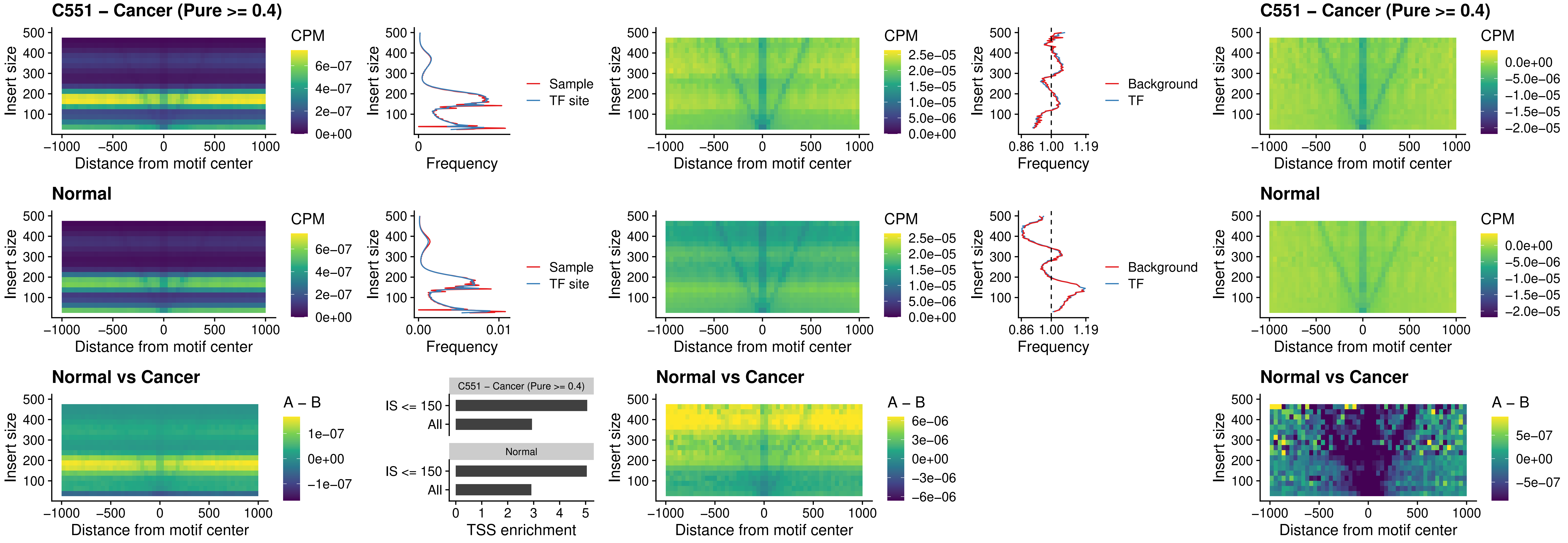

#### C551 – Cancer (Pure >= 0.4) – AC0125311_uc_cTSS_nPEAK

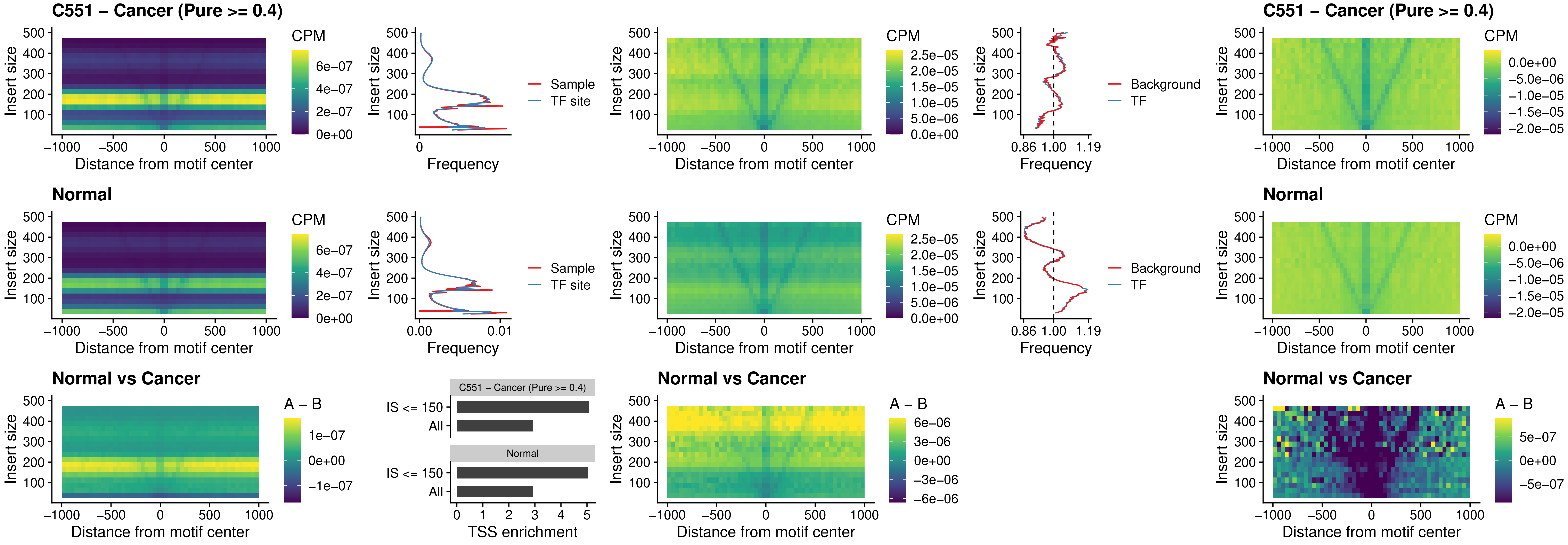

### C551 – Cancer (Pure >= 0.4) – DBX2_uc_cTSS_nPEAK

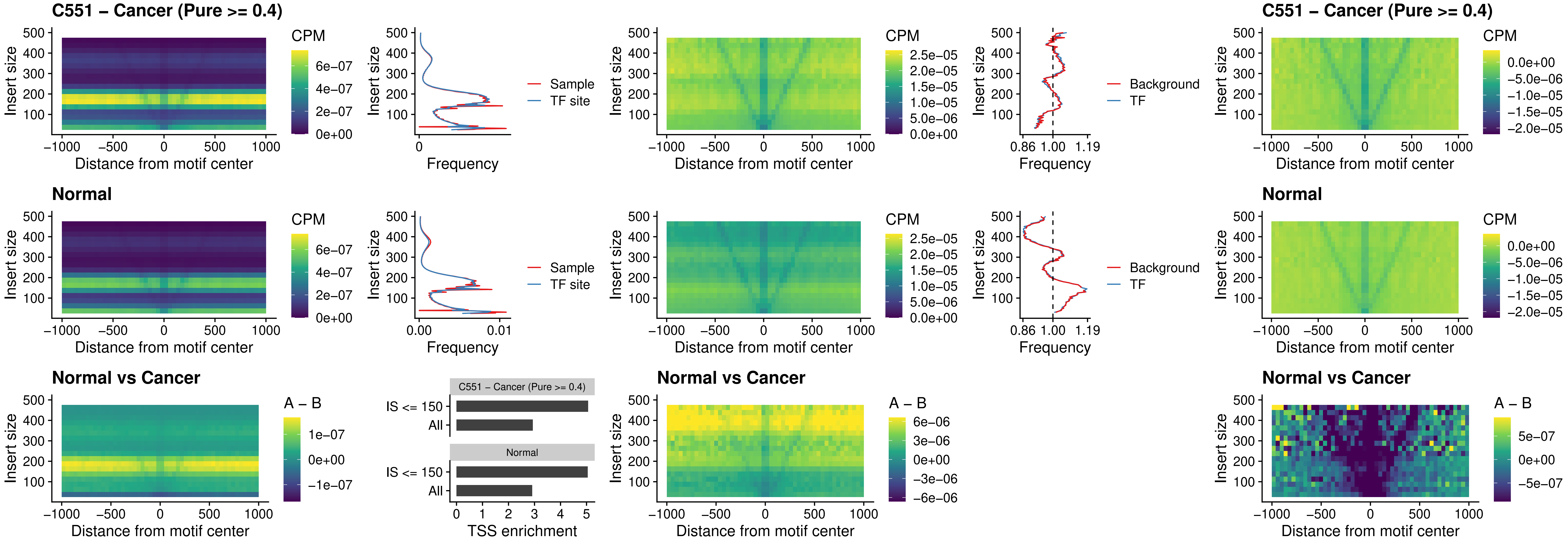

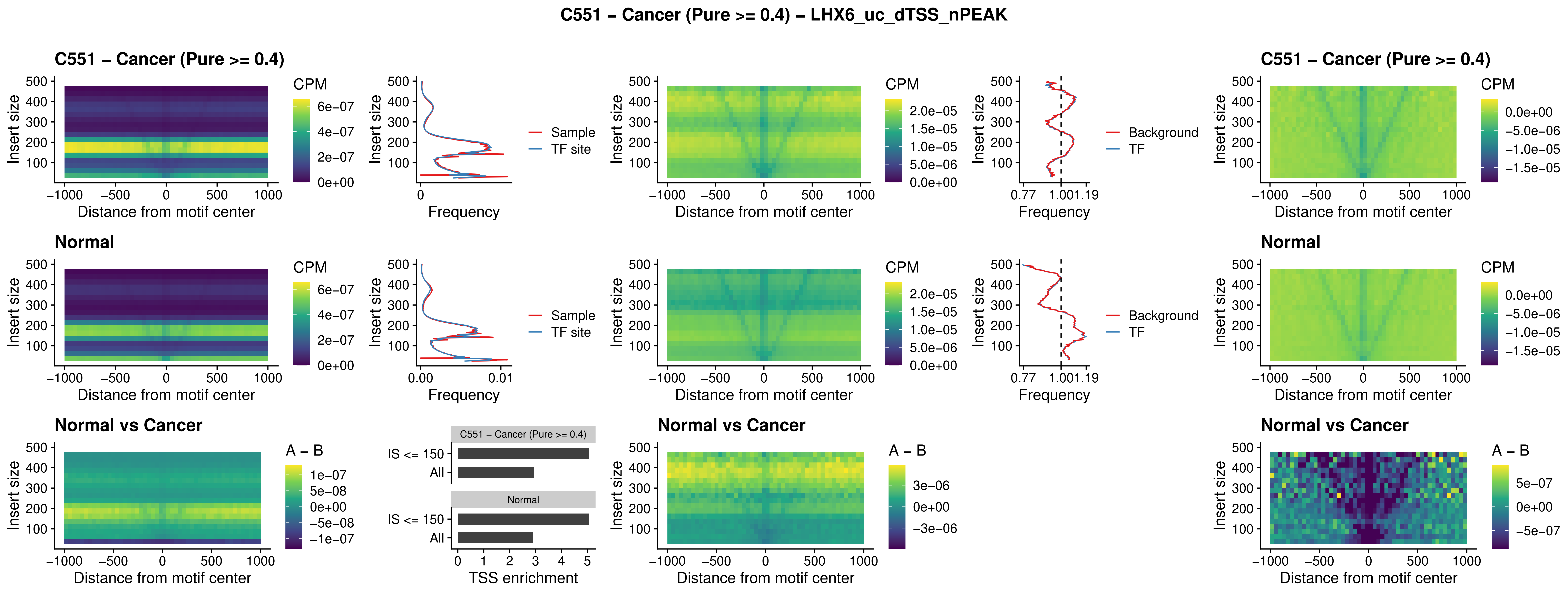
C551 – Cancer (Pure >= 0.4) – LHX6_uc_dTSS_nPEAK
C551 – Cancer (Pure >= 0.4)
Normal
Normal vs Cancer
Insert size
Distance from motif center
Frequency
CPM
A – B
Sample
TF site
Background
TF
C551 – Cancer (Pure >= 0.4)
Normal
IS <= 150
All
TSS enrichment

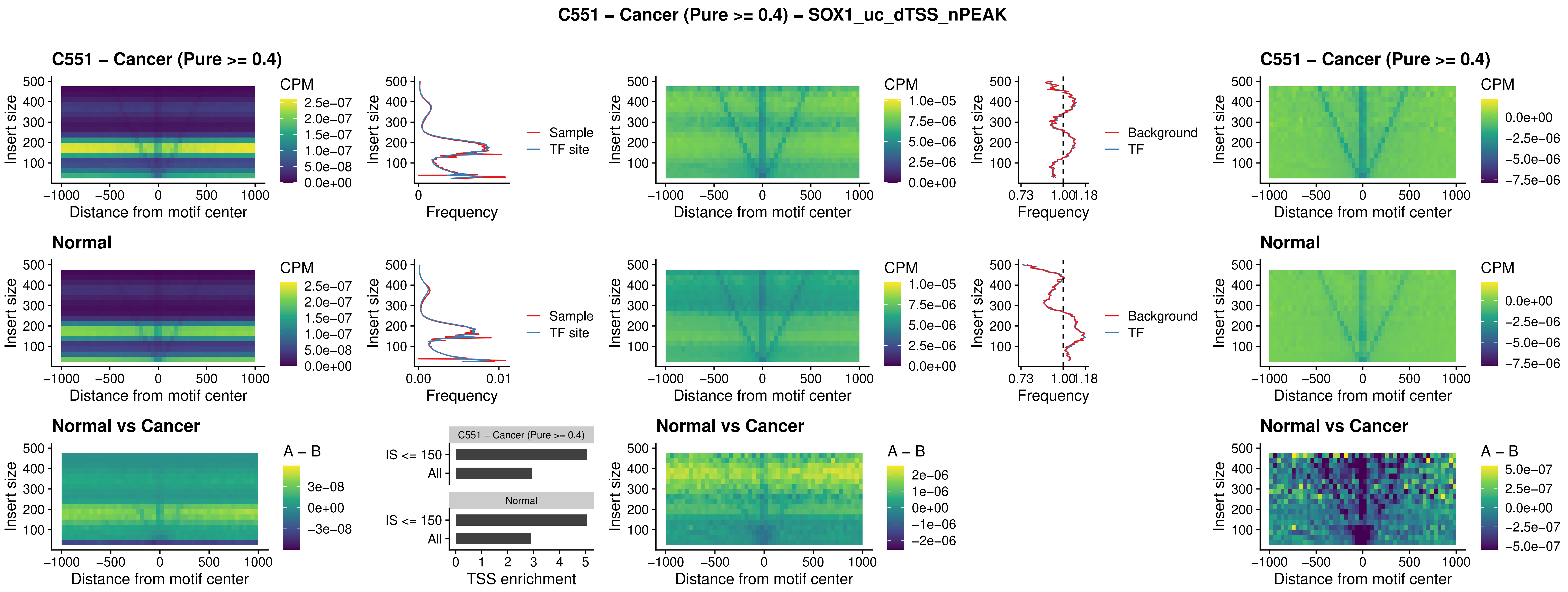
C551 – Cancer (Pure >= 0.4) – SOX1_uc_dTSS_nPEAK
C551 – Cancer (Pure >= 0.4)
Normal
Normal vs Cancer
Insert size
Distance from motif center
CPM
A – B
Frequency
Sample
TF site
Background
TF
C551 – Cancer (Pure >= 0.4)
Normal
IS <= 150
All
TSS enrichment

### C551 – Cancer (Pure >= 0.4) – NKX24_uc_dTSS_nPEAK

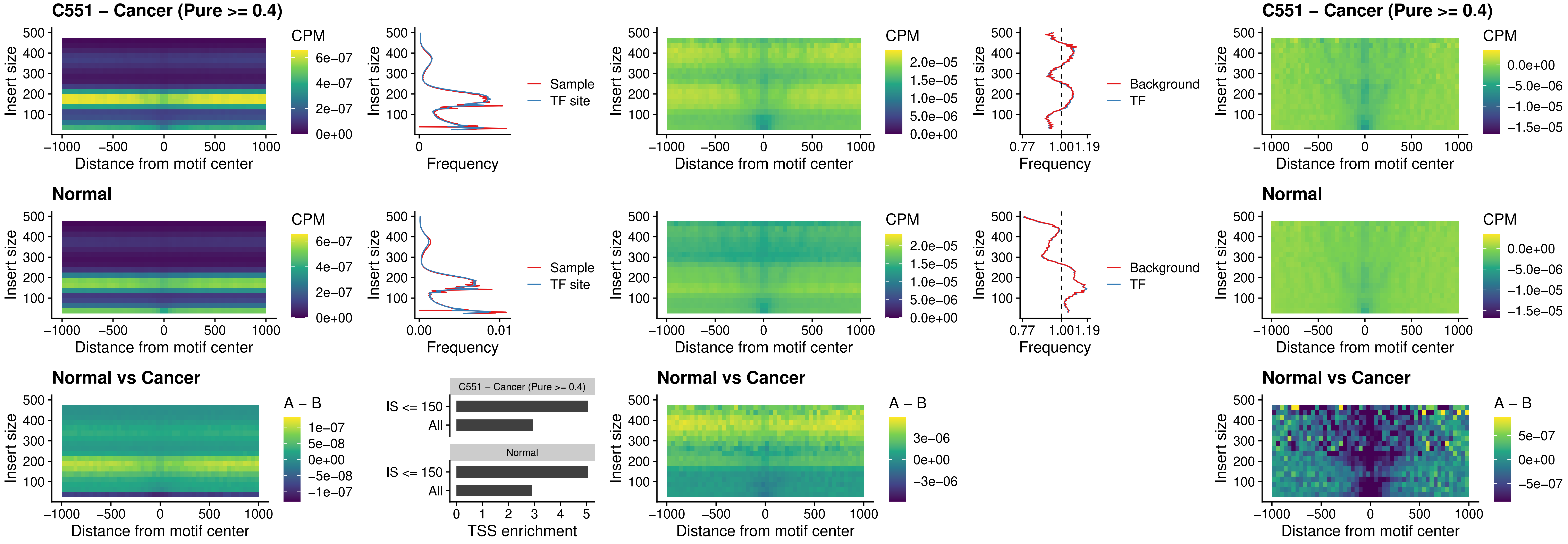

#### C551 – Cancer (Pure >= 0.4) – DLX3_uc_cTSS_nPEAK

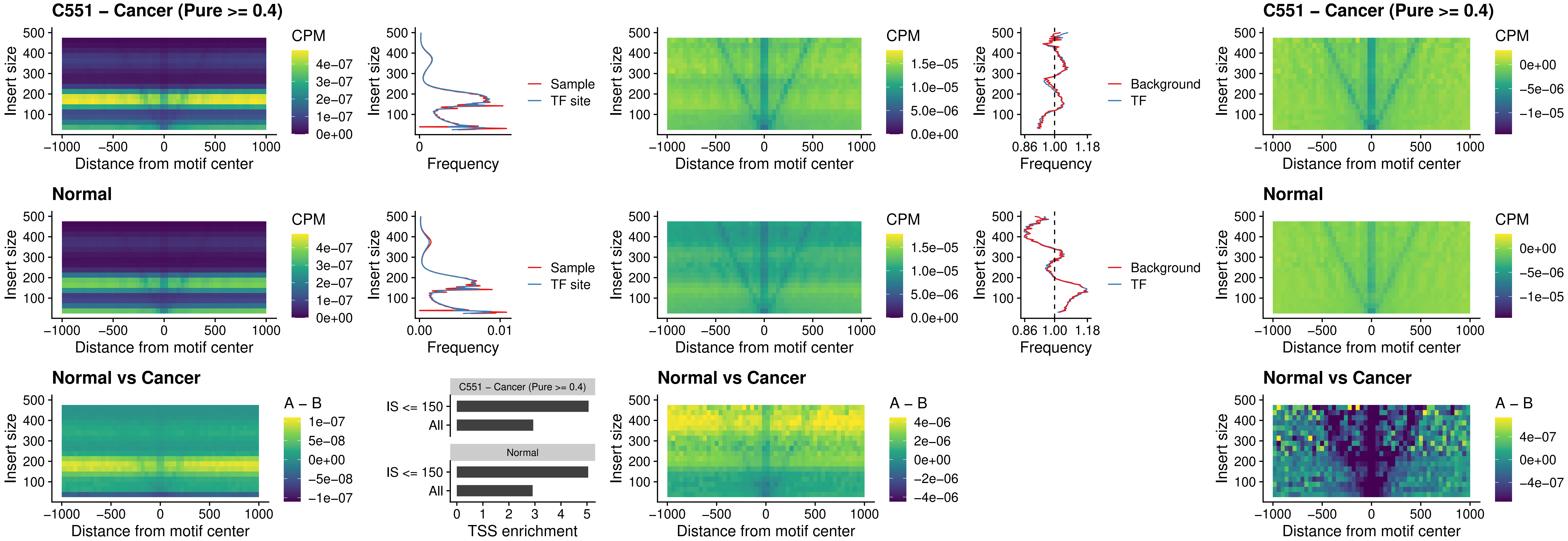

#### C551 – Cancer (Pure >= 0.4) – HOXD3_uc_cTSS_nPEAK

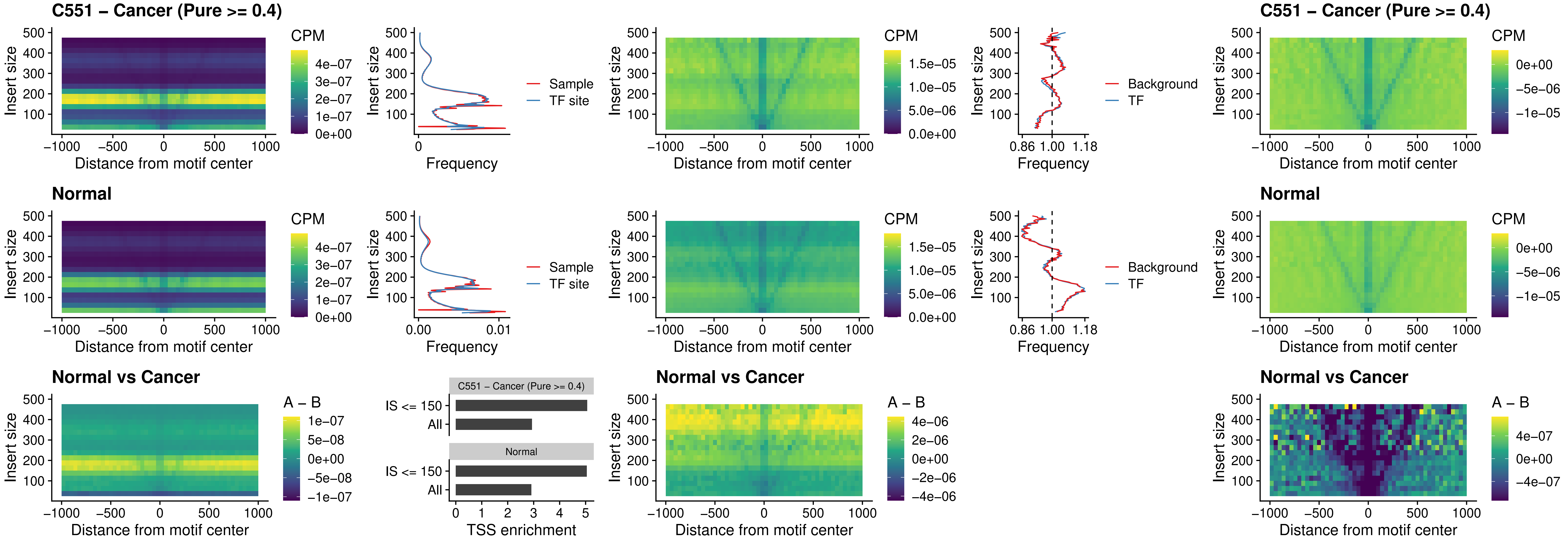

### C551 – Cancer (Pure >= 0.4) – ISL1_uc_dTSS_oPEAK

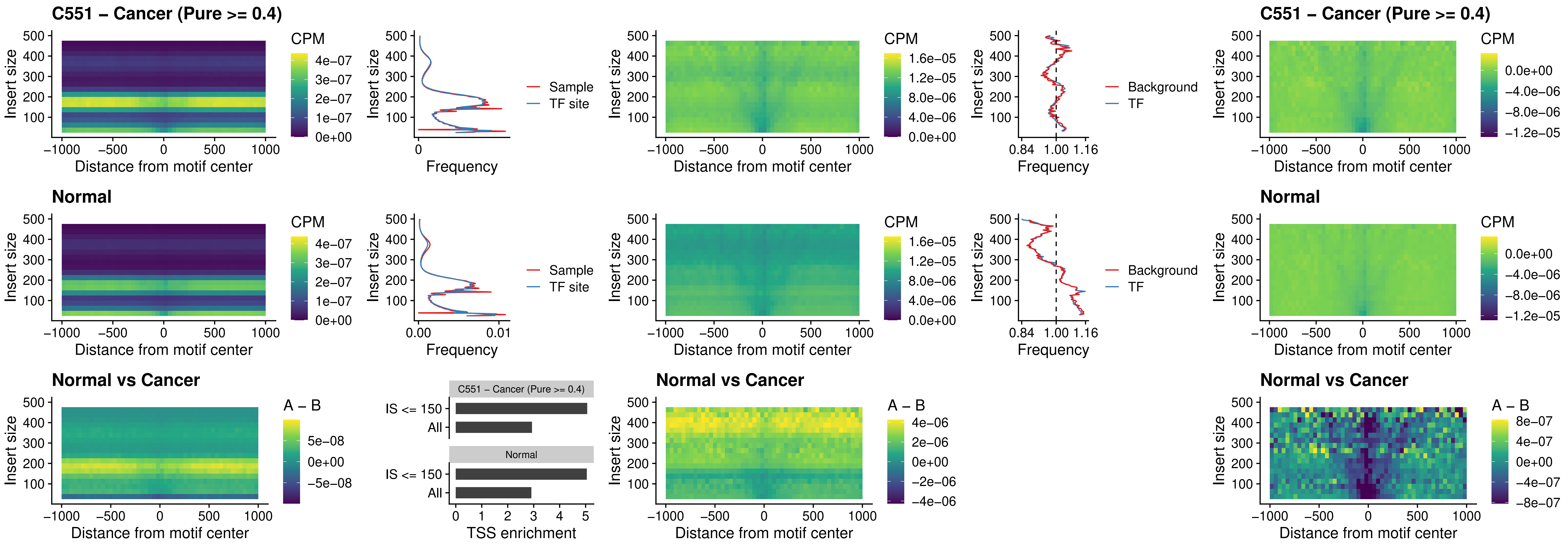

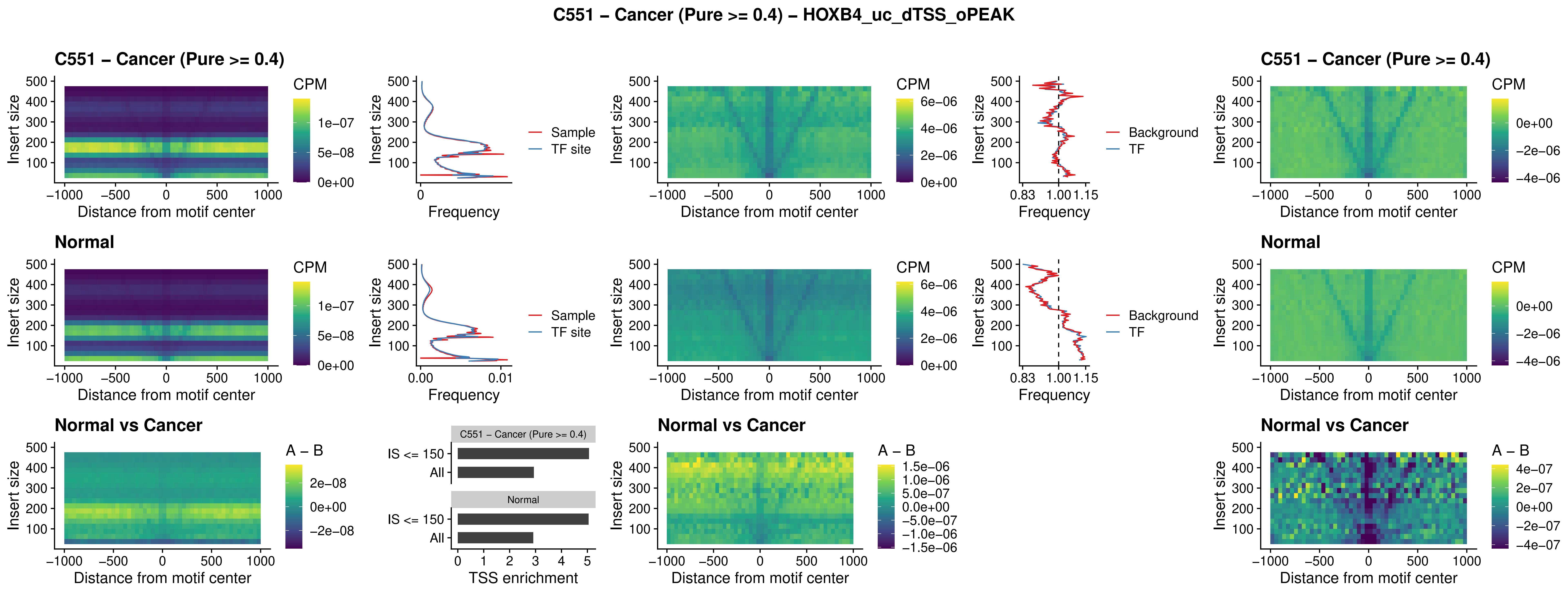

C551 – Cancer (Pure >= 0.4) – HOXB4_uc_dTSS_oPEAK
C551 – Cancer (Pure >= 0.4)
Normal
Normal vs Cancer
Insert size
Distance from motif center
Frequency
CPM
A – B
Sample
TF site
Background
TF
C551 – Cancer (Pure >= 0.4)
Normal
IS <= 150
All
TSS enrichment

#### C551 – Cancer (Pure >= 0.4) – VAX1_uc_dTSS_oPEAK

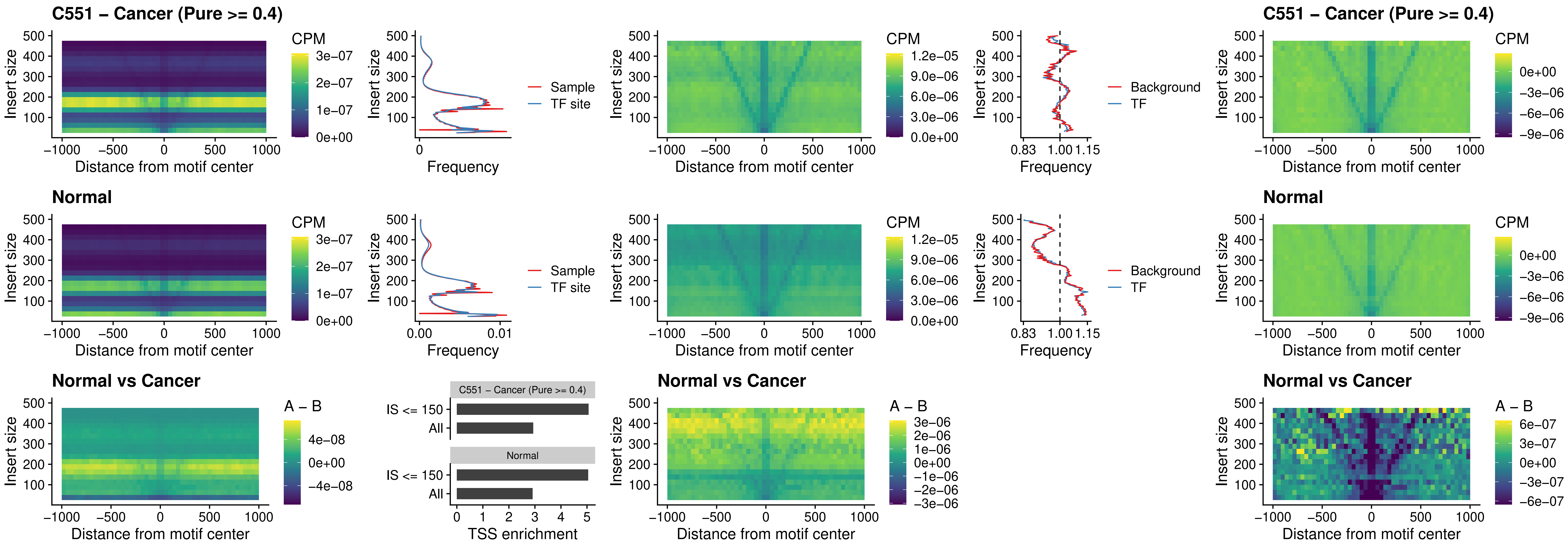

### C551 – Cancer (Pure >= 0.4) – DLX3_uc_dTSS_oPEAK

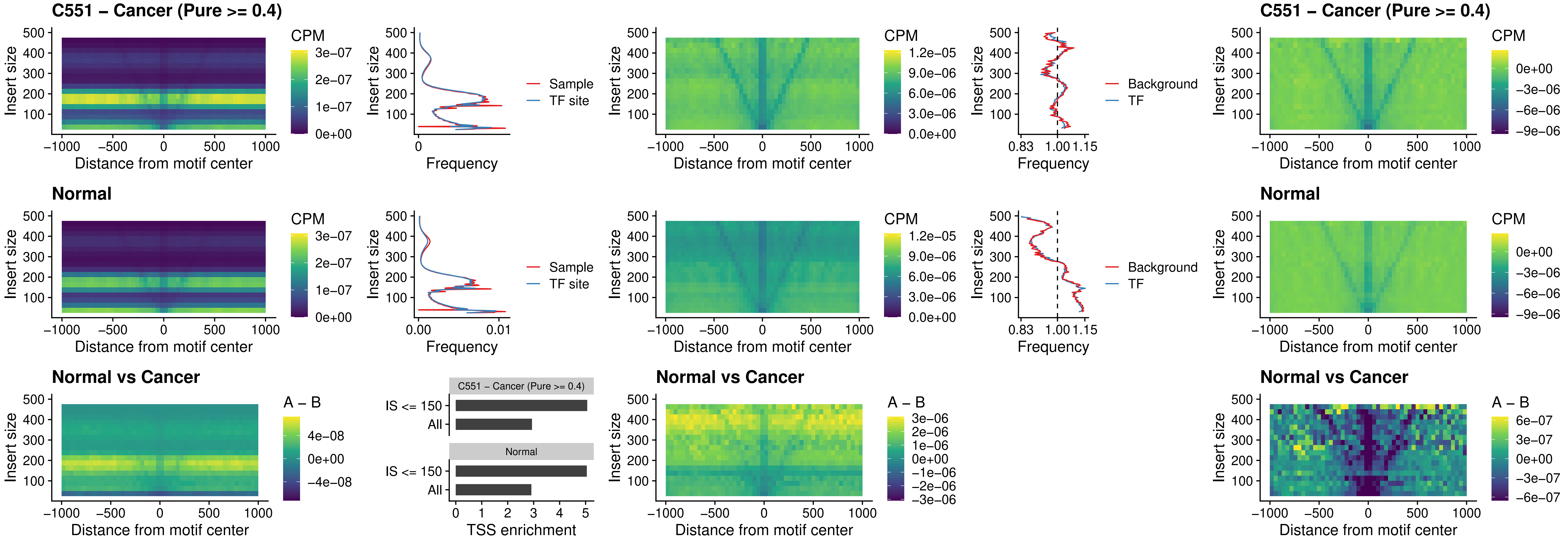

#### C551 – Cancer (Pure >= 0.4) – HOXD3_uc_dTSS_oPEAK

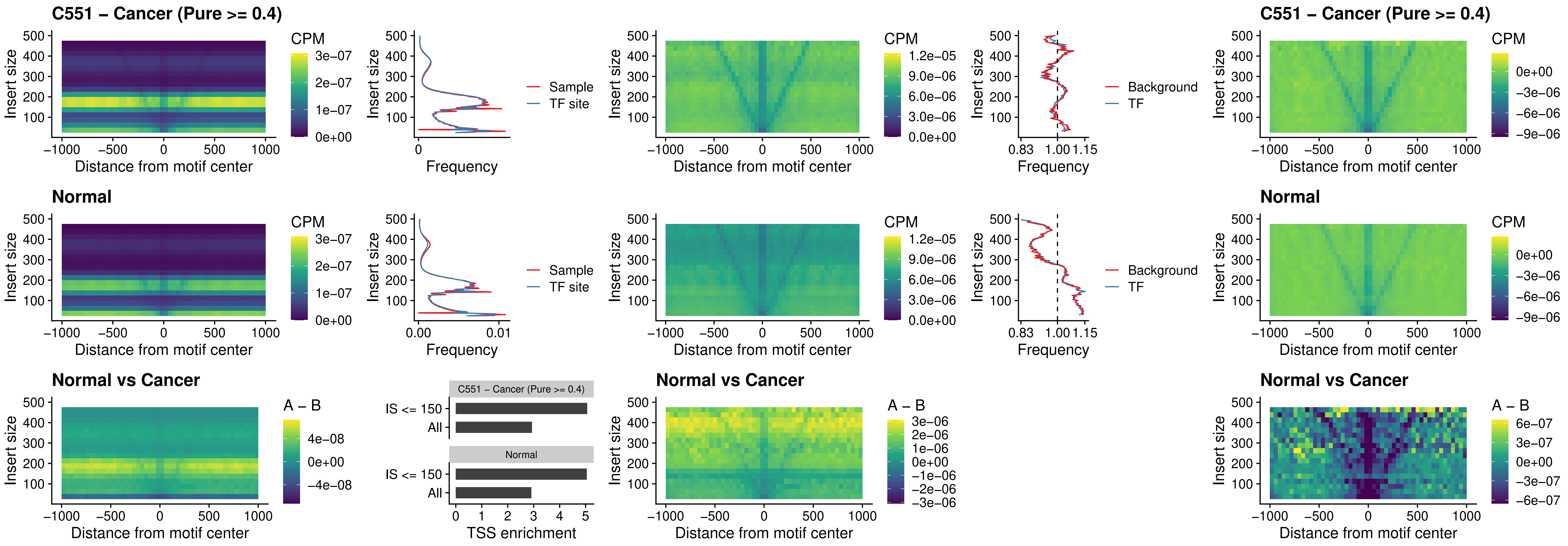

### C551 – Cancer (Pure >= 0.4) – BBX_uc_dTSS_nPEAK

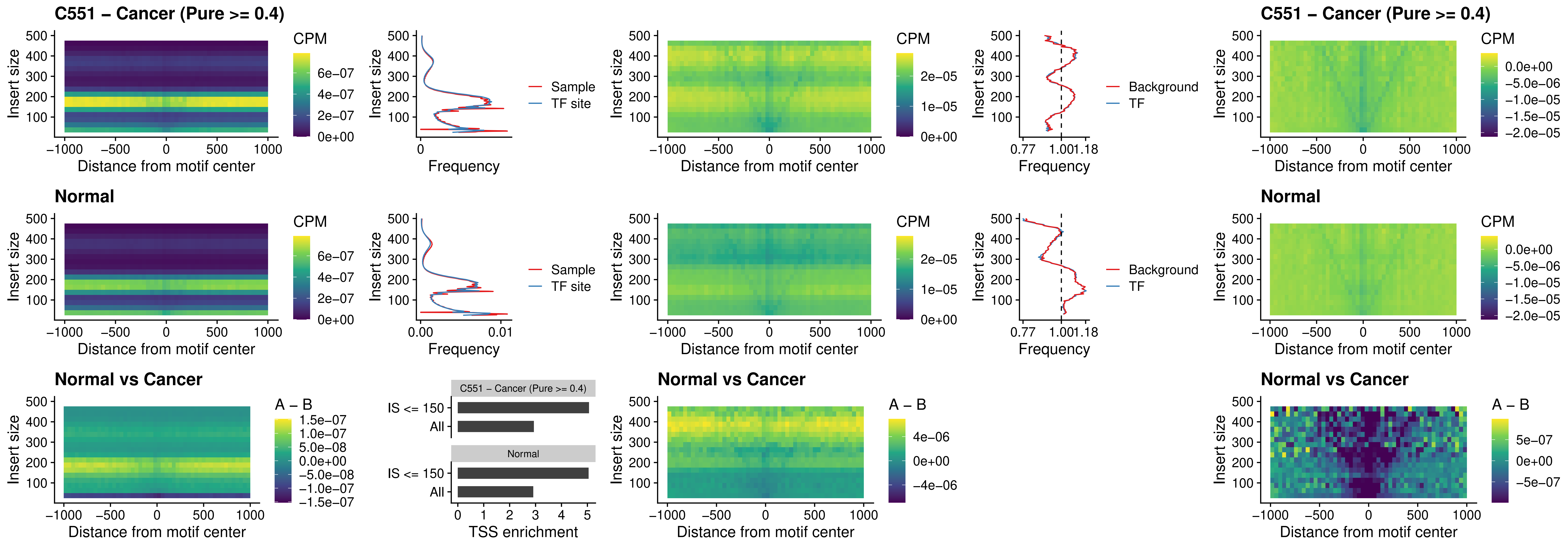

#### C551 – Cancer (Pure >= 0.4) – SOX3_uc_cTSS_nPEAK

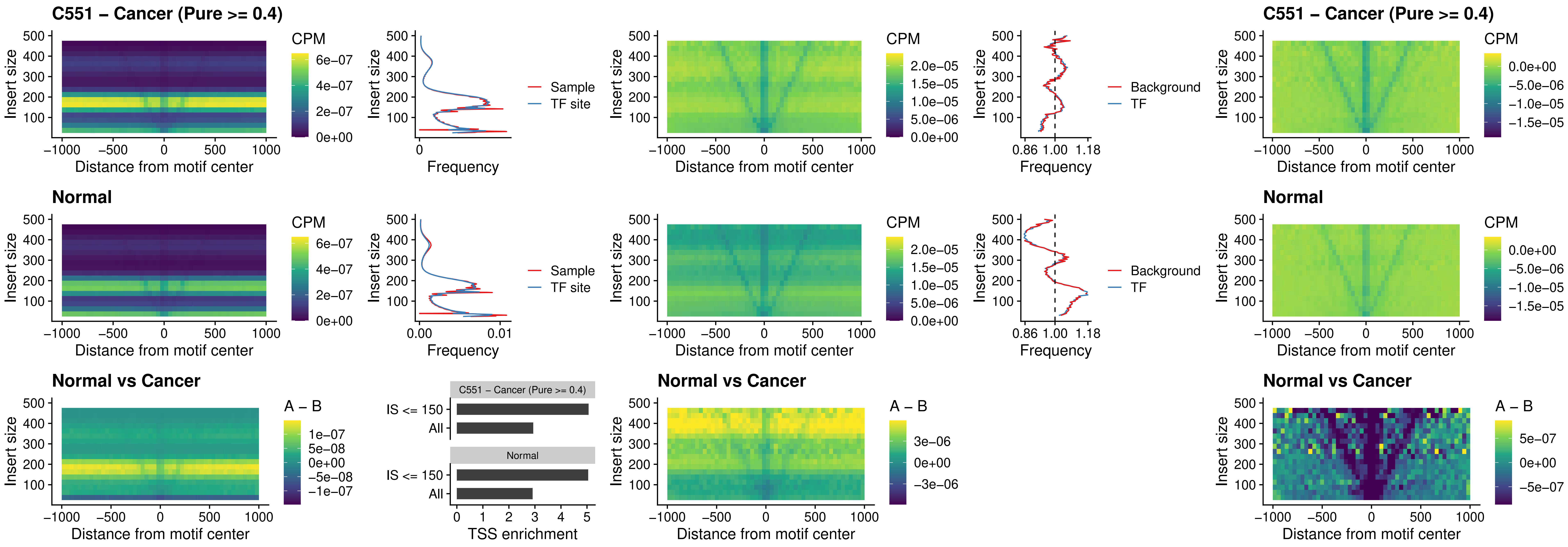

#### C551 – Cancer (Pure >= 0.4) – TBPL2_uc_cTSS_nPEAK

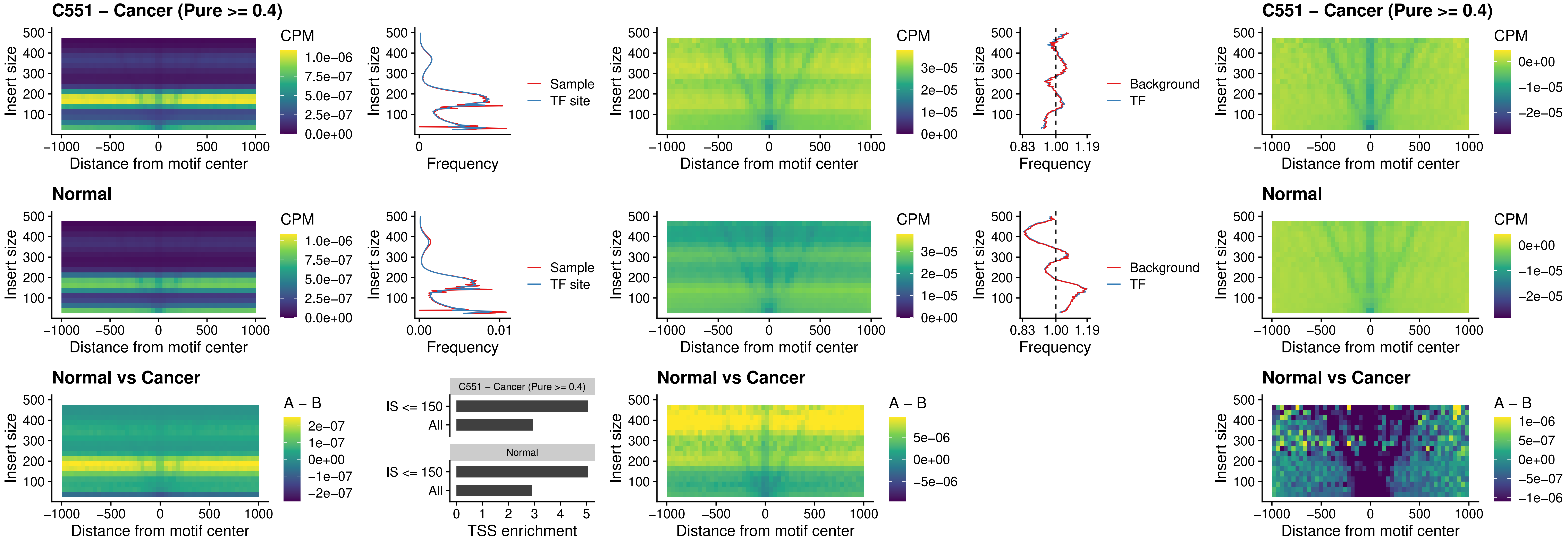

#### C551 – Cancer (Pure >= 0.4) – HOXA4_uc_cTSS_oPEAK

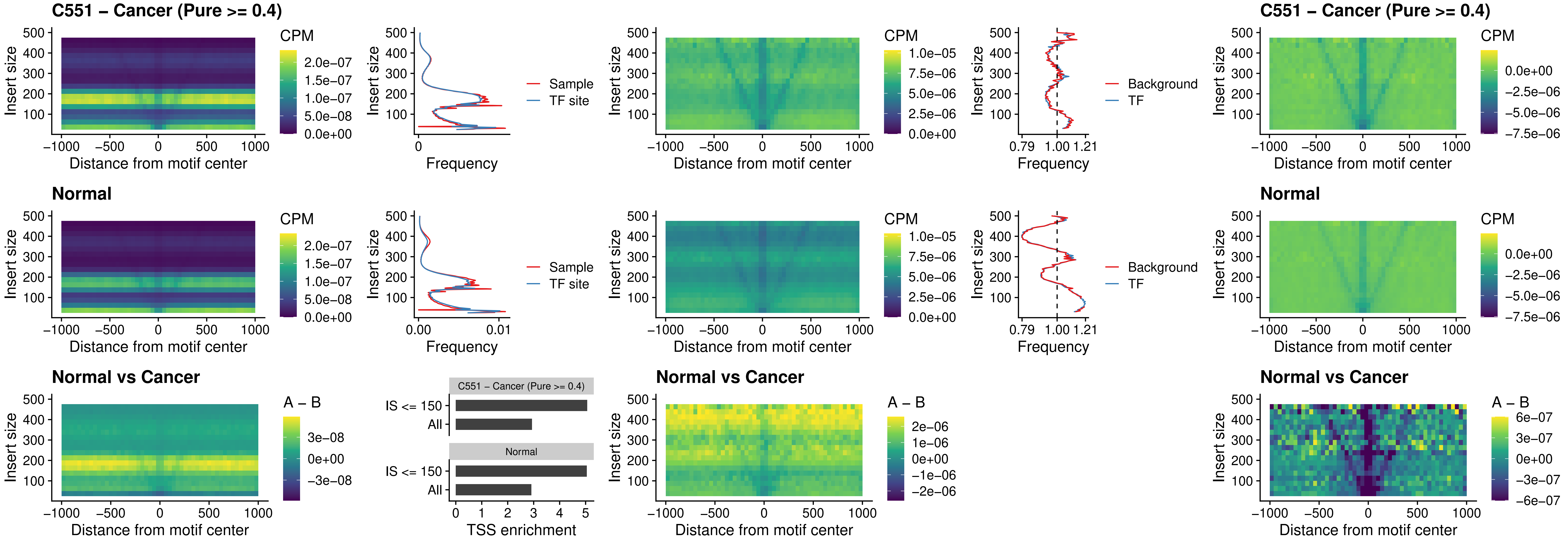

#### C551 – Cancer (Pure >= 0.4) – PDX1_uc_dTSS_oPEAK

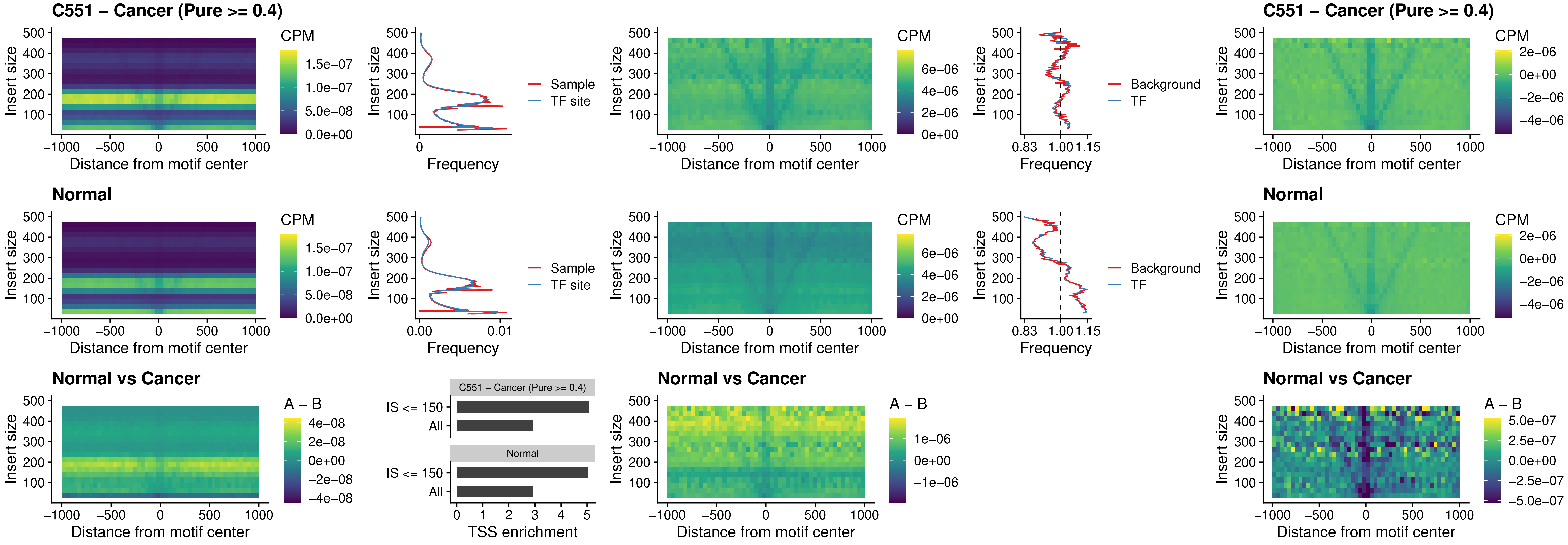

#### C551 – Cancer (Pure >= 0.4) – LHX4_uc_dTSS_oPEAK

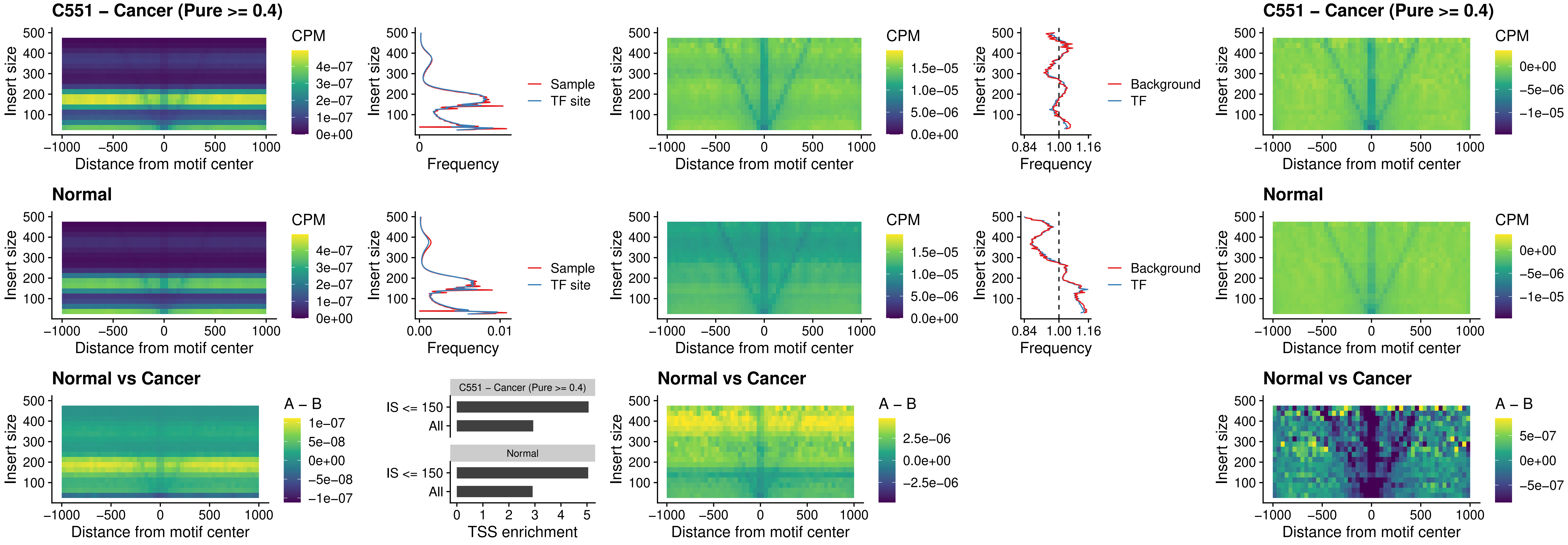

#### C551 – Cancer (Pure >= 0.4) – HOXA4_uc_dTSS_oPEAK

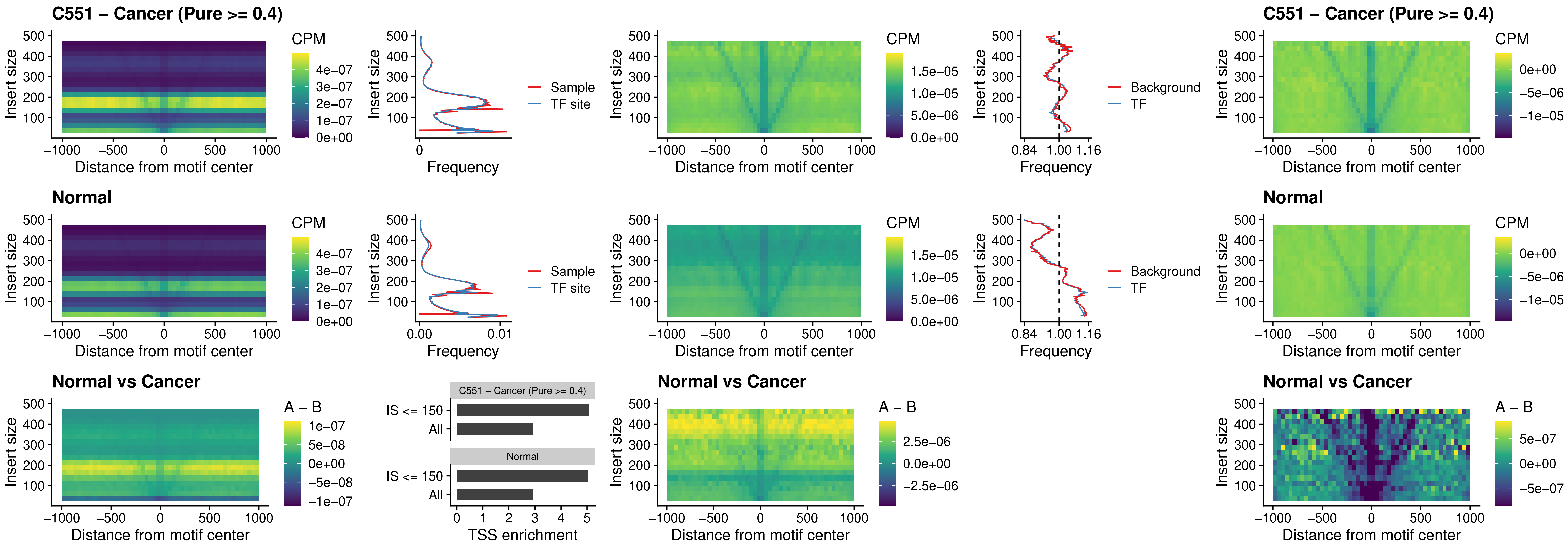

#### C551 – Cancer (Pure >= 0.4) – HLX_uc_dTSS_oPEAK

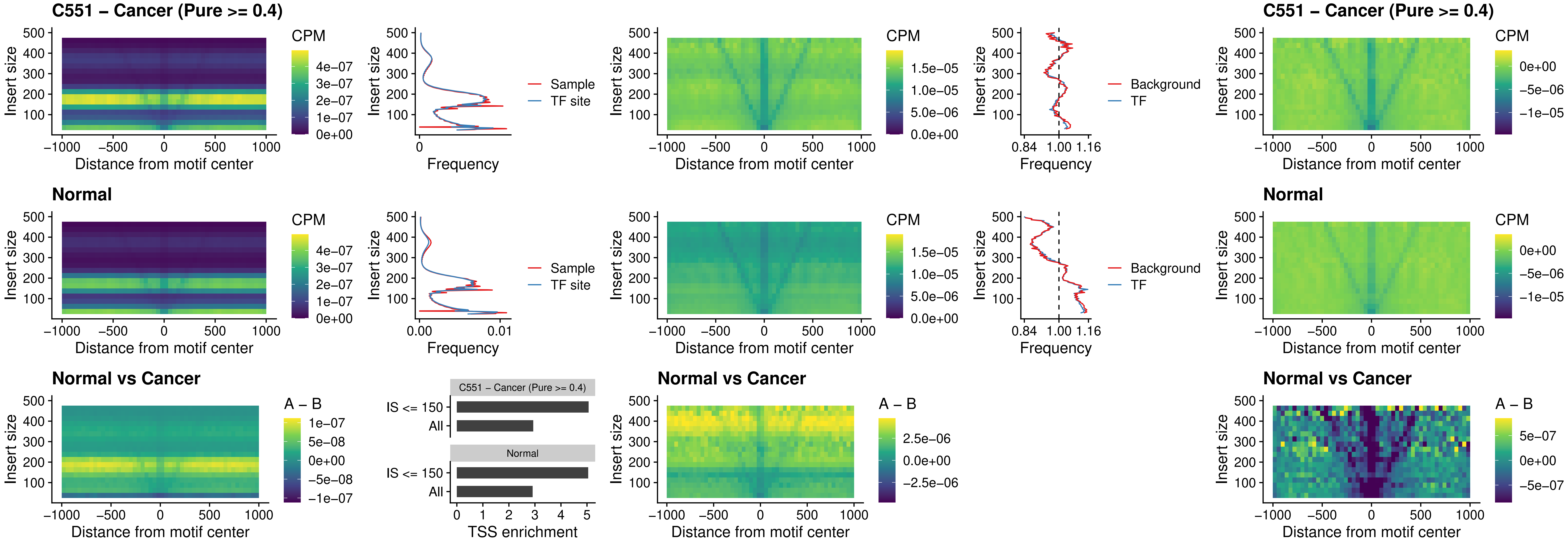

#### C551 – Cancer (Pure >= 0.4) – AC0125311_uc_dTSS_oPEAK

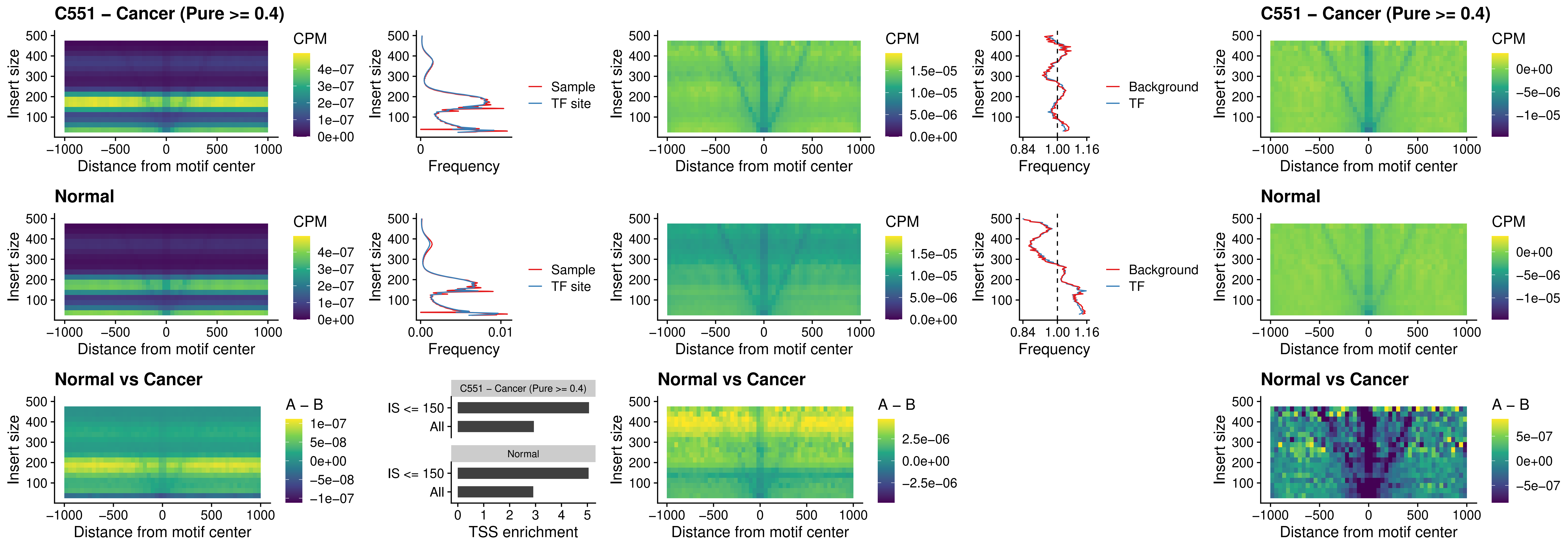

#### C551 – Cancer (Pure >= 0.4) – DBX2_uc_dTSS_oPEAK

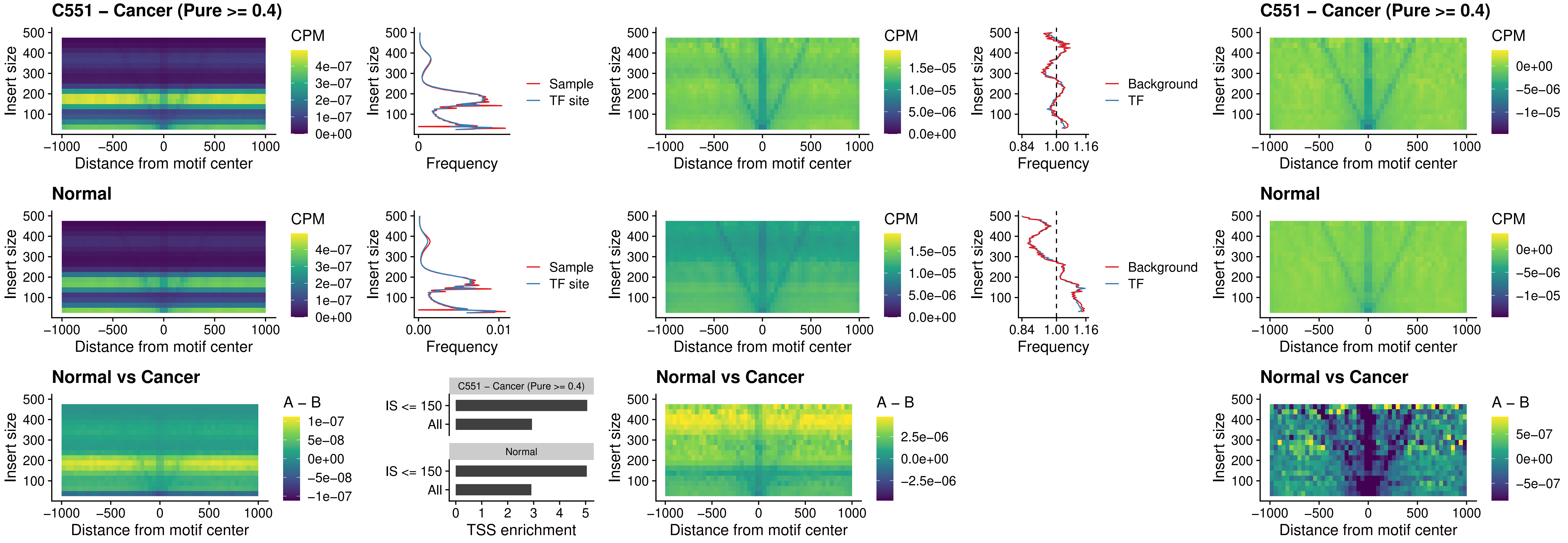

#### C551 – Cancer (Pure >= 0.4) – HOXA2_uc_dTSS_nPEAK

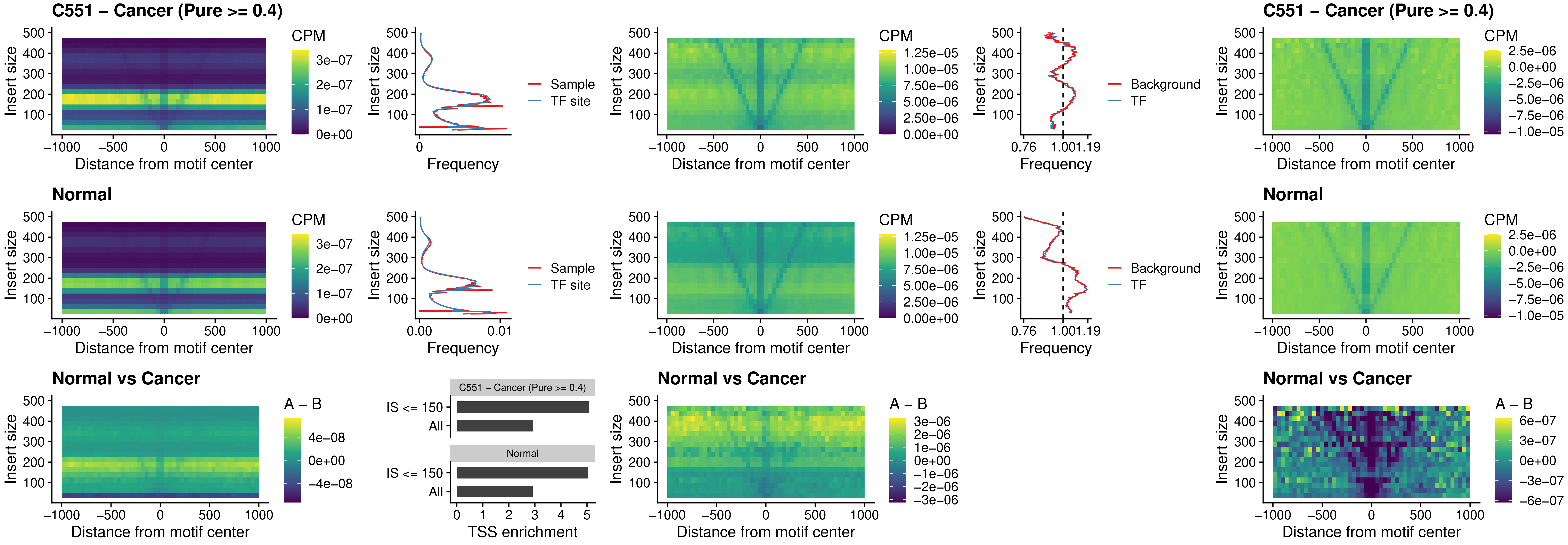

#### C551 – Cancer (Pure >= 0.4) – SHOX_uc_dTSS_nPEAK

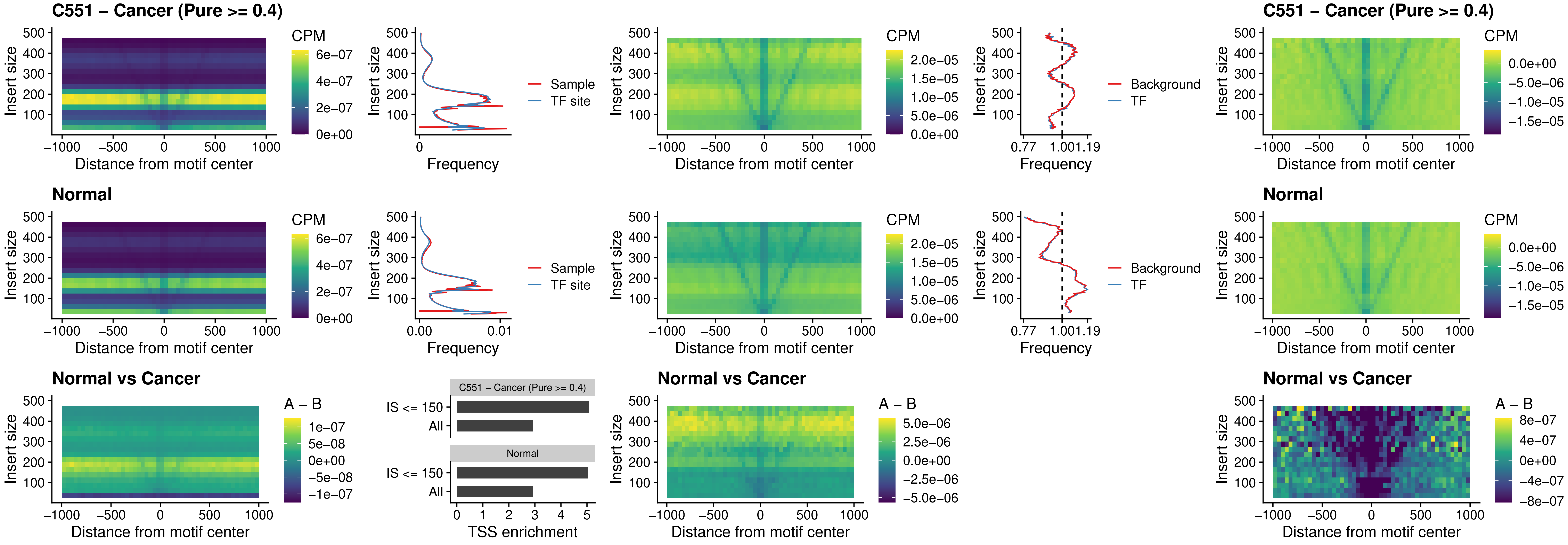

### C551 – Cancer (Pure >= 0.4) – PRRX1_uc_dTSS_nPEAK

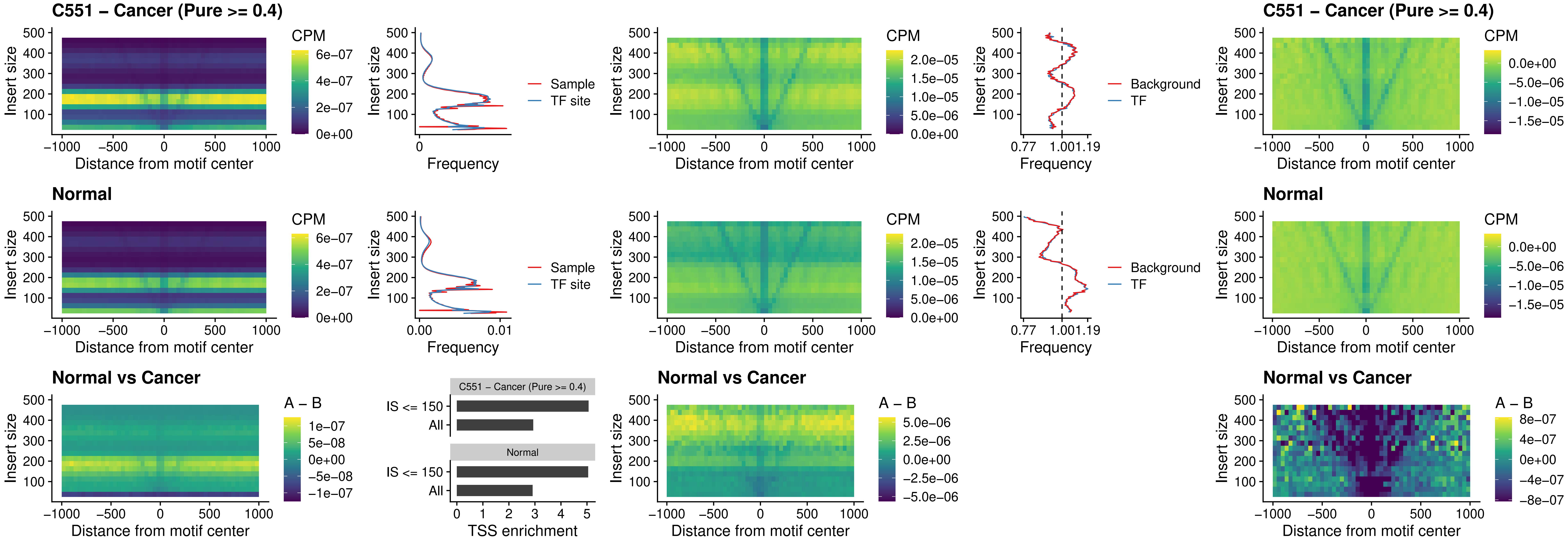

#### C551 – Cancer (Pure >= 0.4) – LMX1A_uc_dTSS_nPEAK

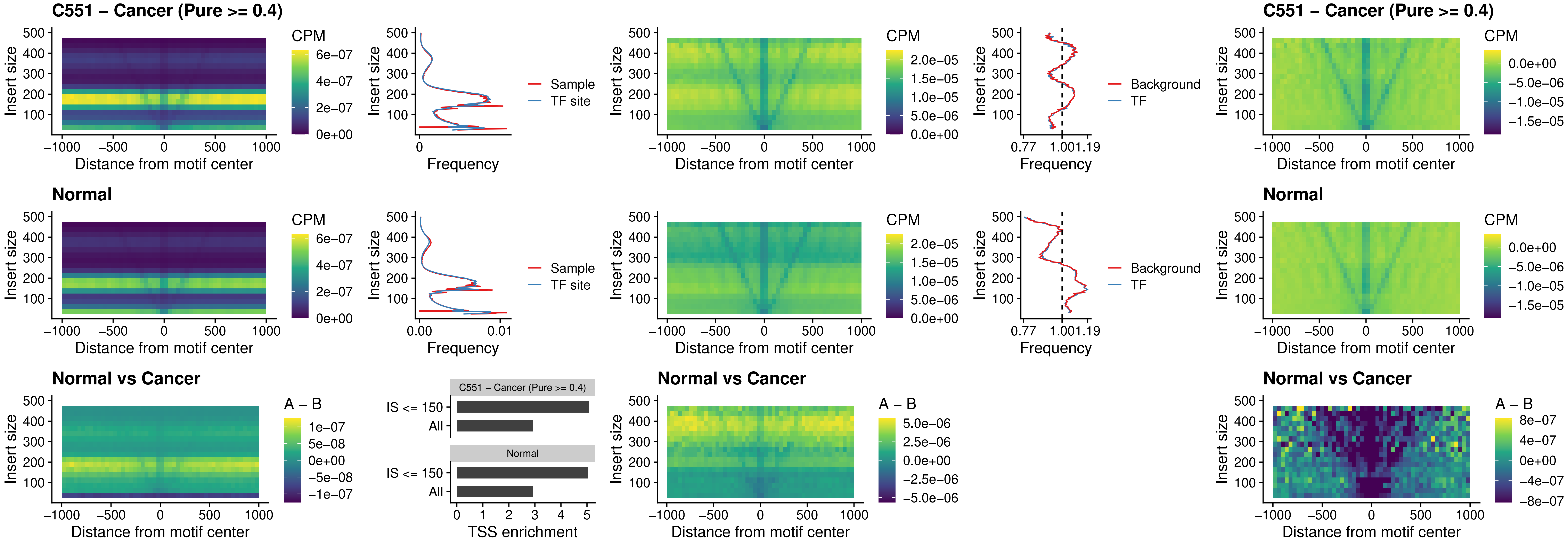

#### C551 – Cancer (Pure >= 0.4) – LMX1B_uc_dTSS_nPEAK

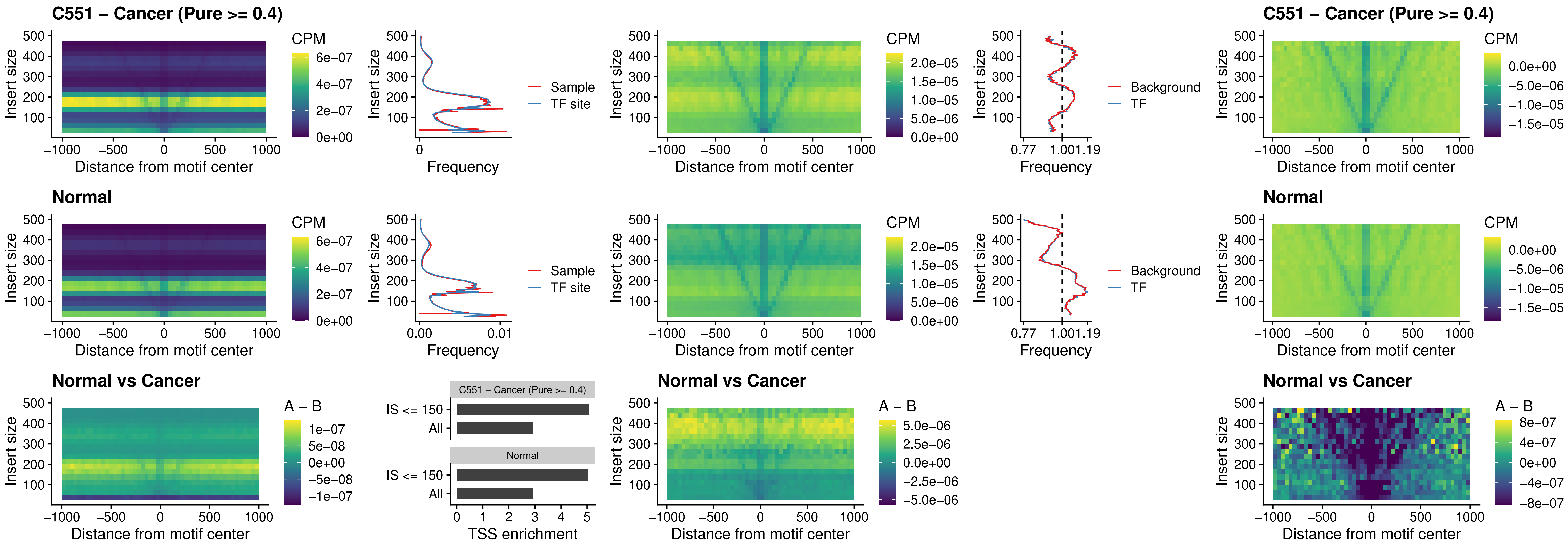

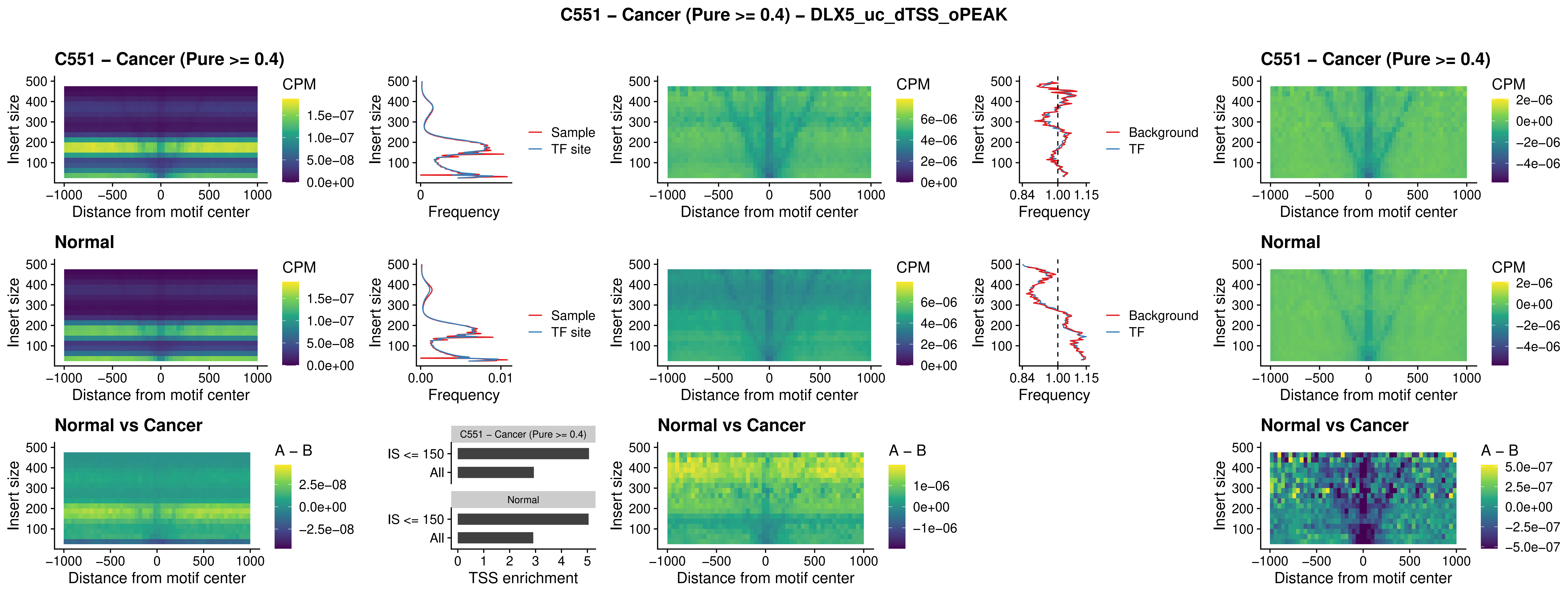

C551 – Cancer (Pure >= 0.4) – DLX5_uc_dTSS_oPEAK
C551 – Cancer (Pure >= 0.4)
Normal
Normal vs Cancer
Insert size
Distance from motif center
Frequency
CPM
A – B
Sample
TF site
Background
TF
C551 – Cancer (Pure >= 0.4)
Normal
IS <= 150
All
TSS enrichment

#### C551 – Cancer (Pure >= 0.4) – ARID3C_uc_dTSS_nPEAK

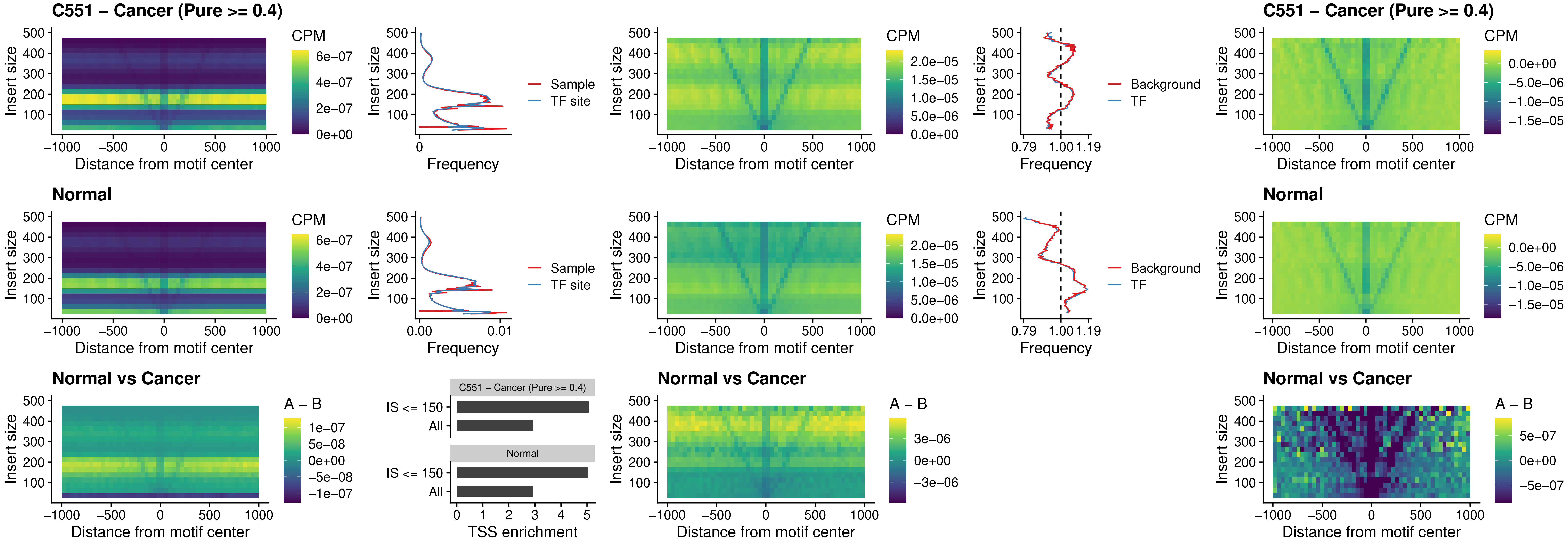

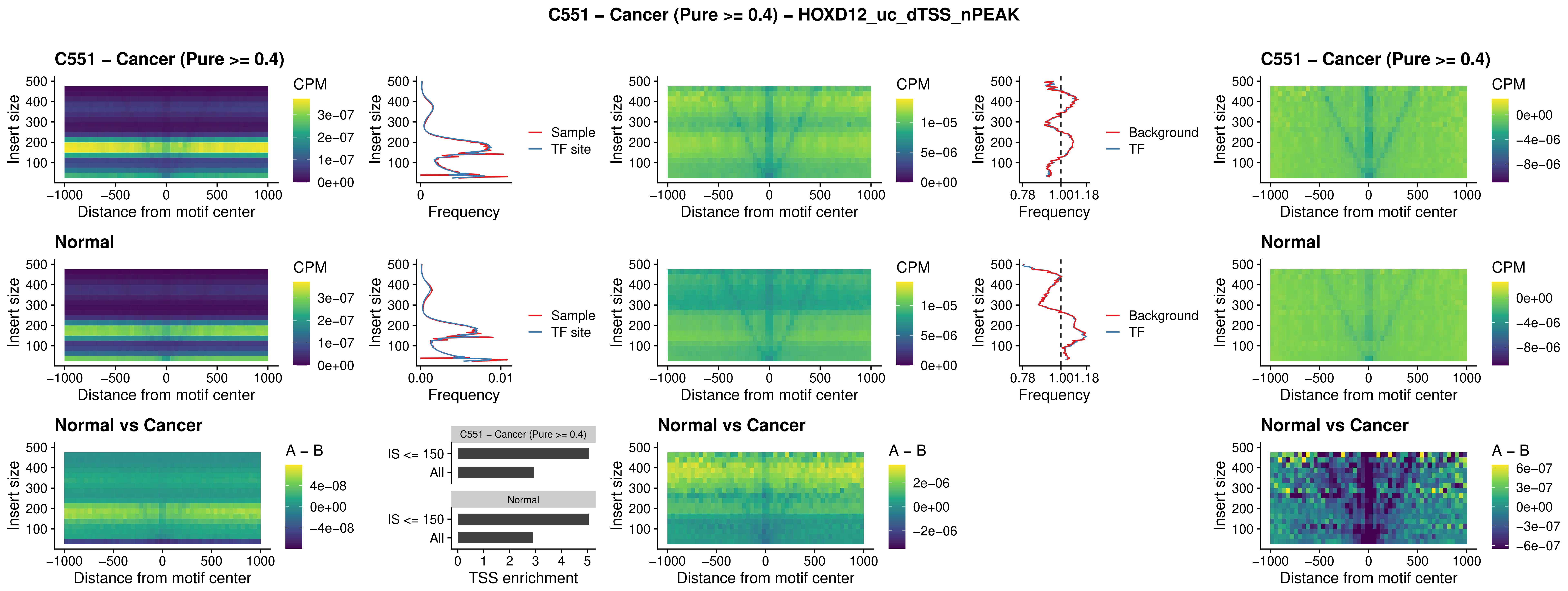
C551 – Cancer (Pure >= 0.4) – HOXD12_uc_dTSS_nPEAK
C551 – Cancer (Pure >= 0.4)
Normal
Normal vs Cancer
Insert size
Distance from motif center
Frequency
CPM
A – B
Sample
TF site
Background
TF
C551 – Cancer (Pure >= 0.4)
Normal
IS <= 150
All
TSS enrichment

#### C551 – Cancer (Pure >= 0.4) – CDX2_uc_dTSS_nPEAK

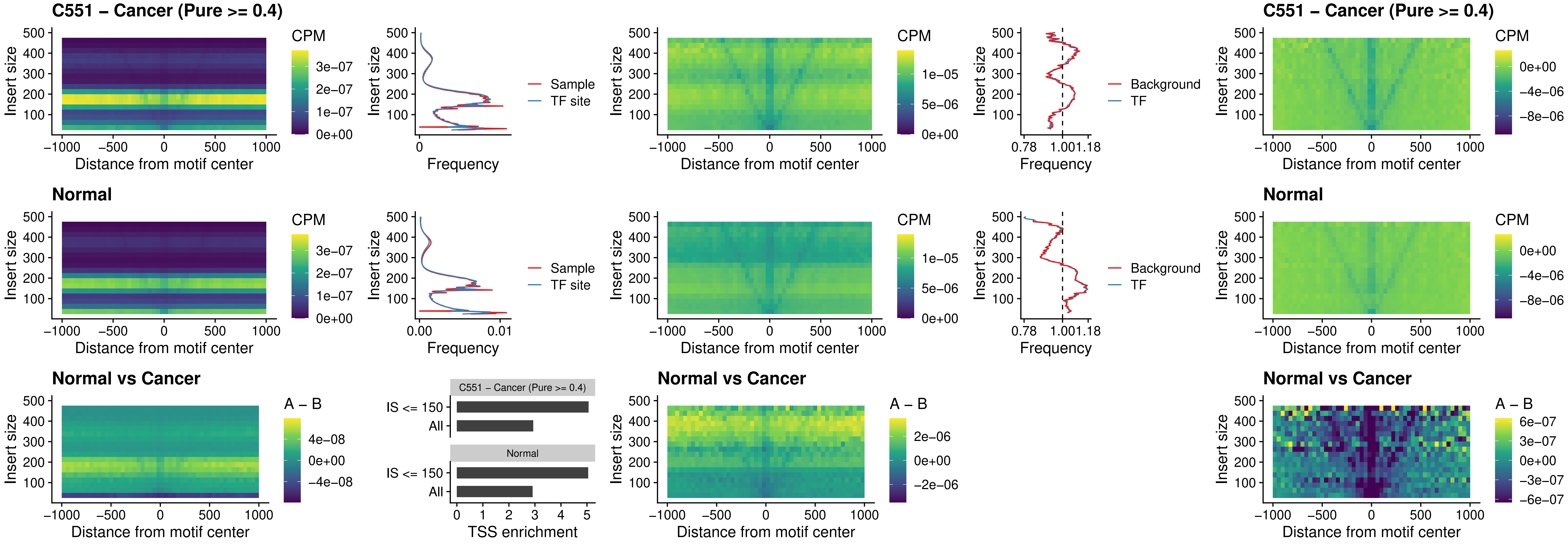

#### C551 – Cancer (Pure >= 0.4) – HOXC12_uc_dTSS_nPEAK

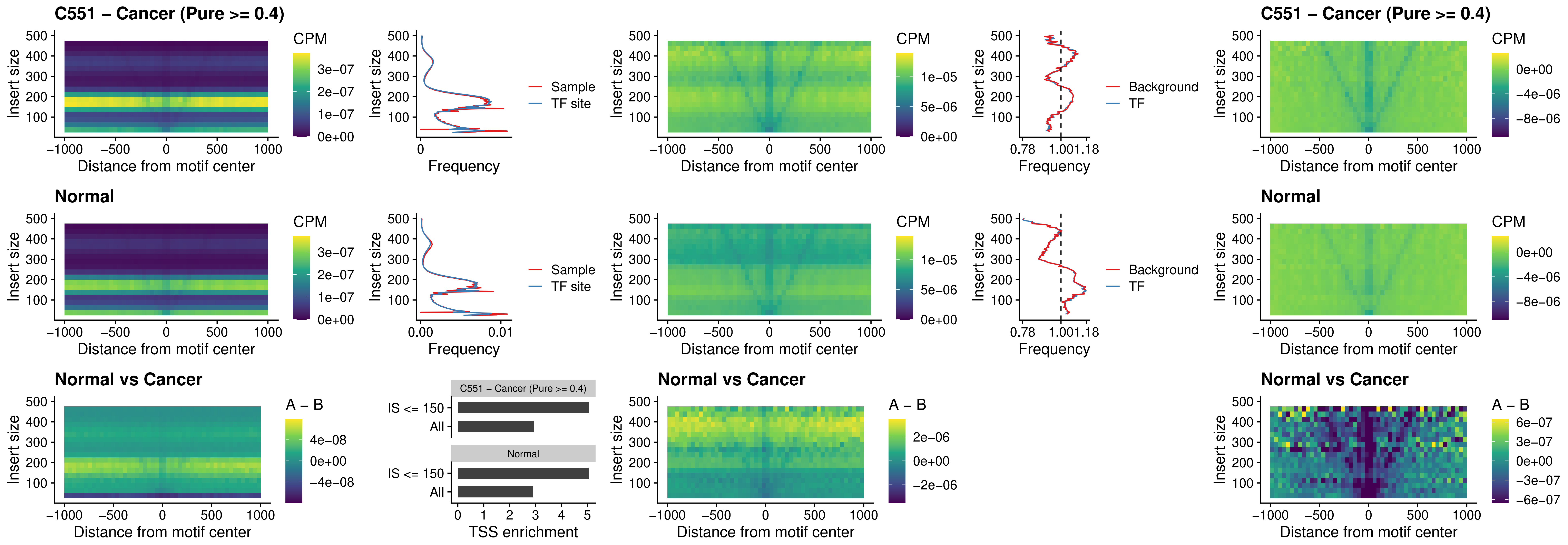

### C551 – Cancer (Pure >= 0.4) – HHEX_uc_dTSS_oPEAK

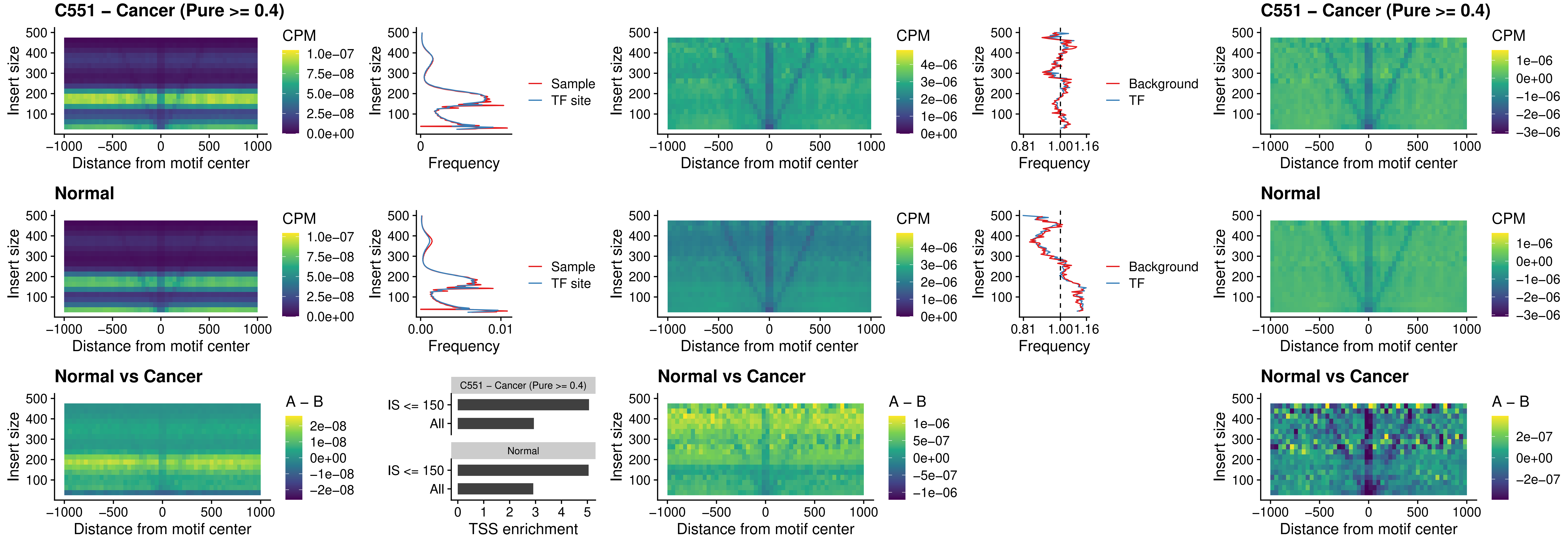

#### C551 – Cancer (Pure >= 0.4) – HOXA11_uc_dTSS_nPEAK

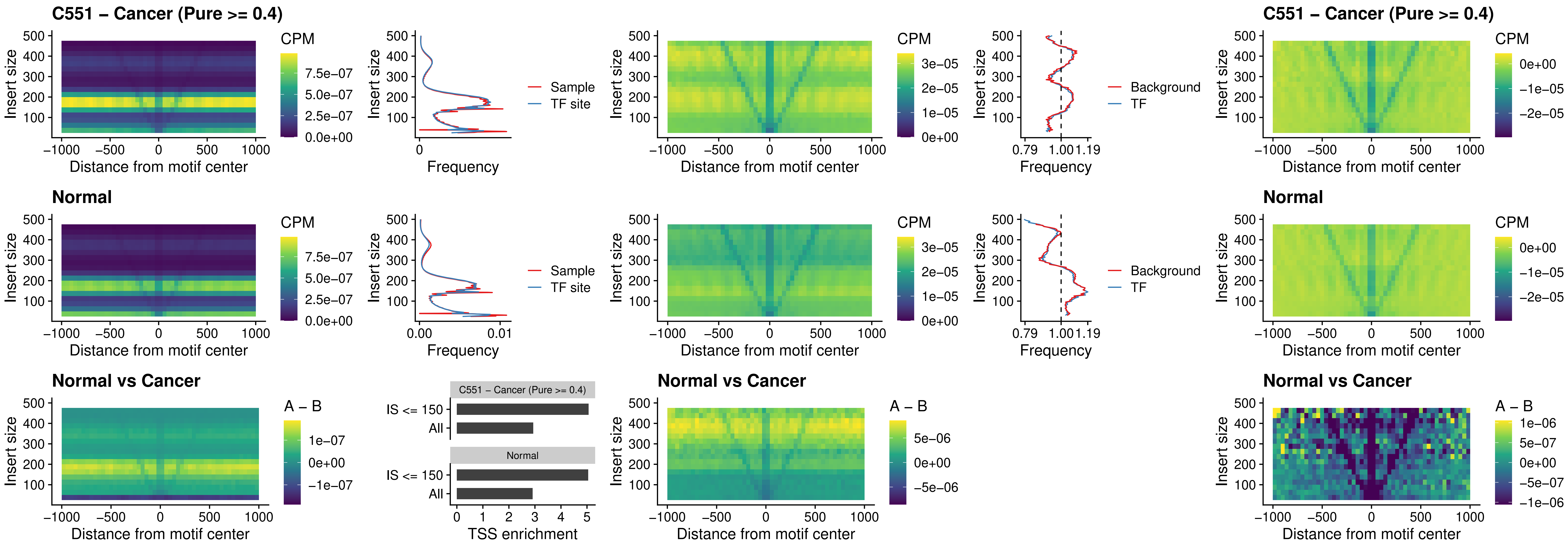

#### C551 – Cancer (Pure >= 0.4) – HOXA11_uc_cTSS_nPEAK

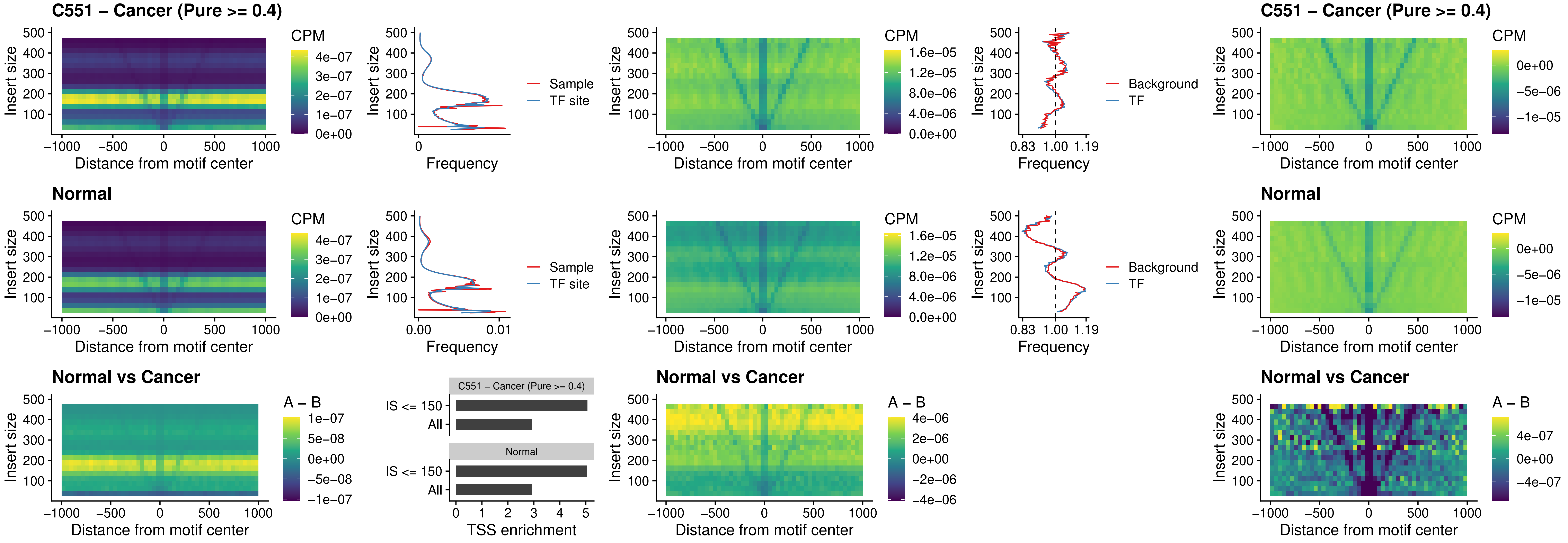

#### C551 – Cancer (Pure >= 0.4) – ZFHX3_uc_dTSS_nPEAK

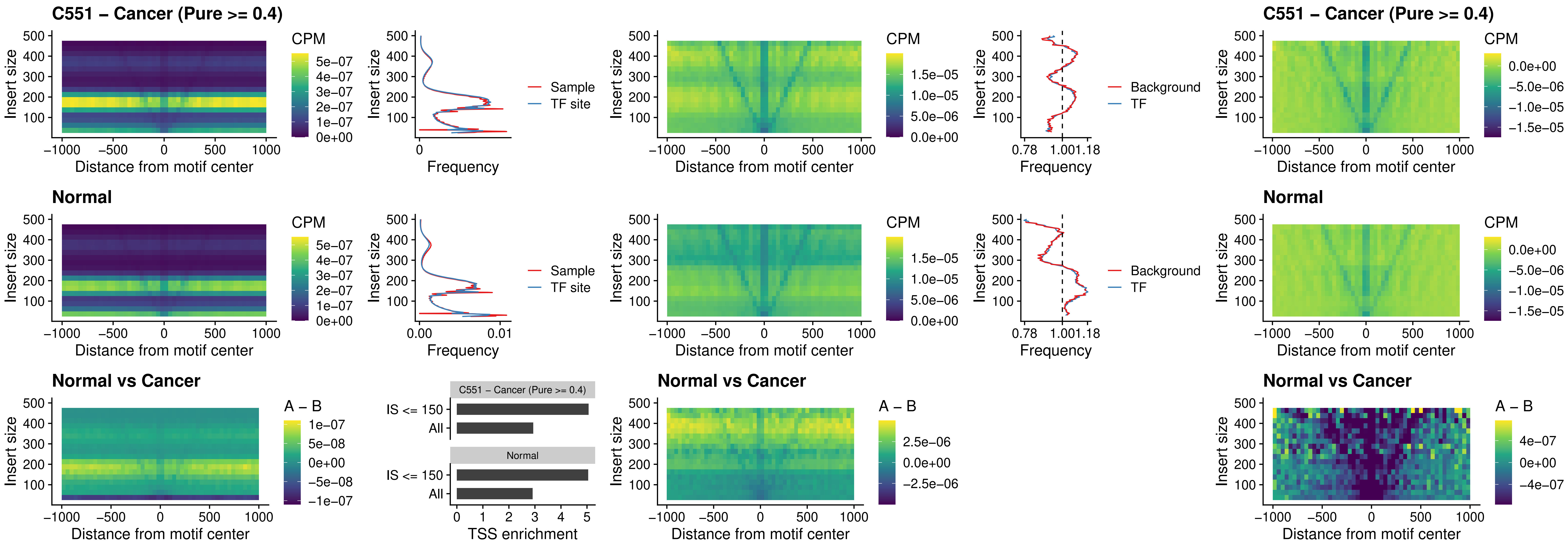

#### C551 – Cancer (Pure >= 0.4) – HOXB4_uc_dTSS_nPEAK

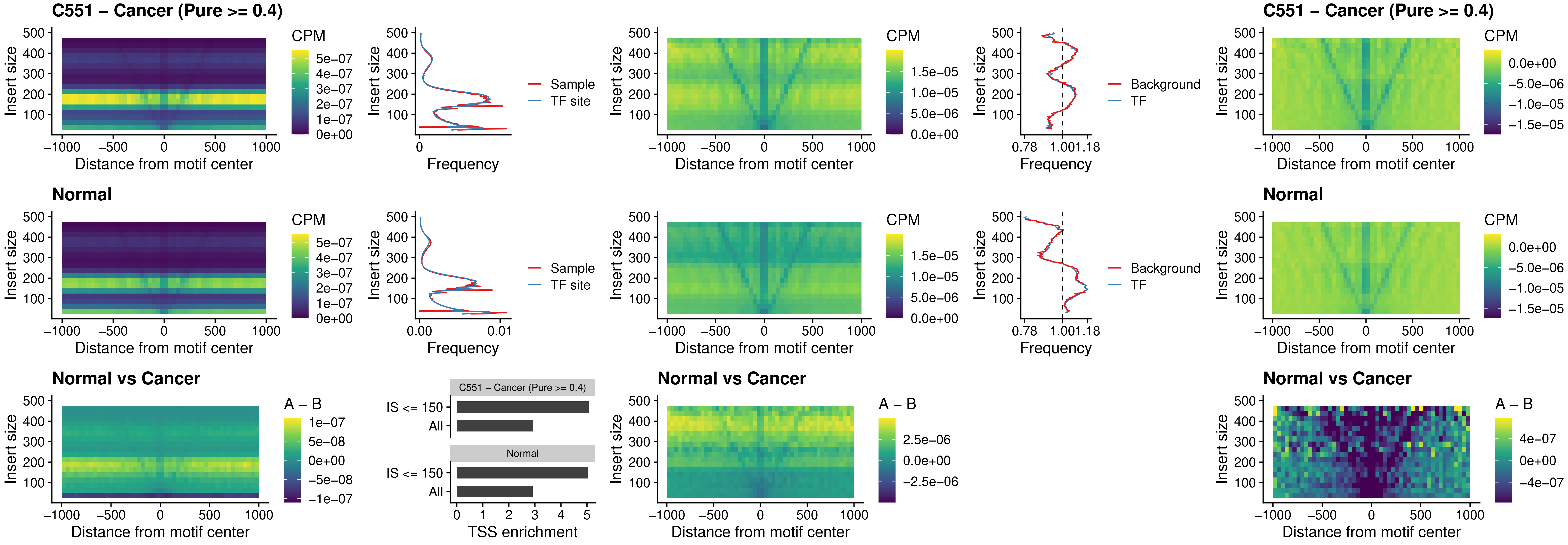

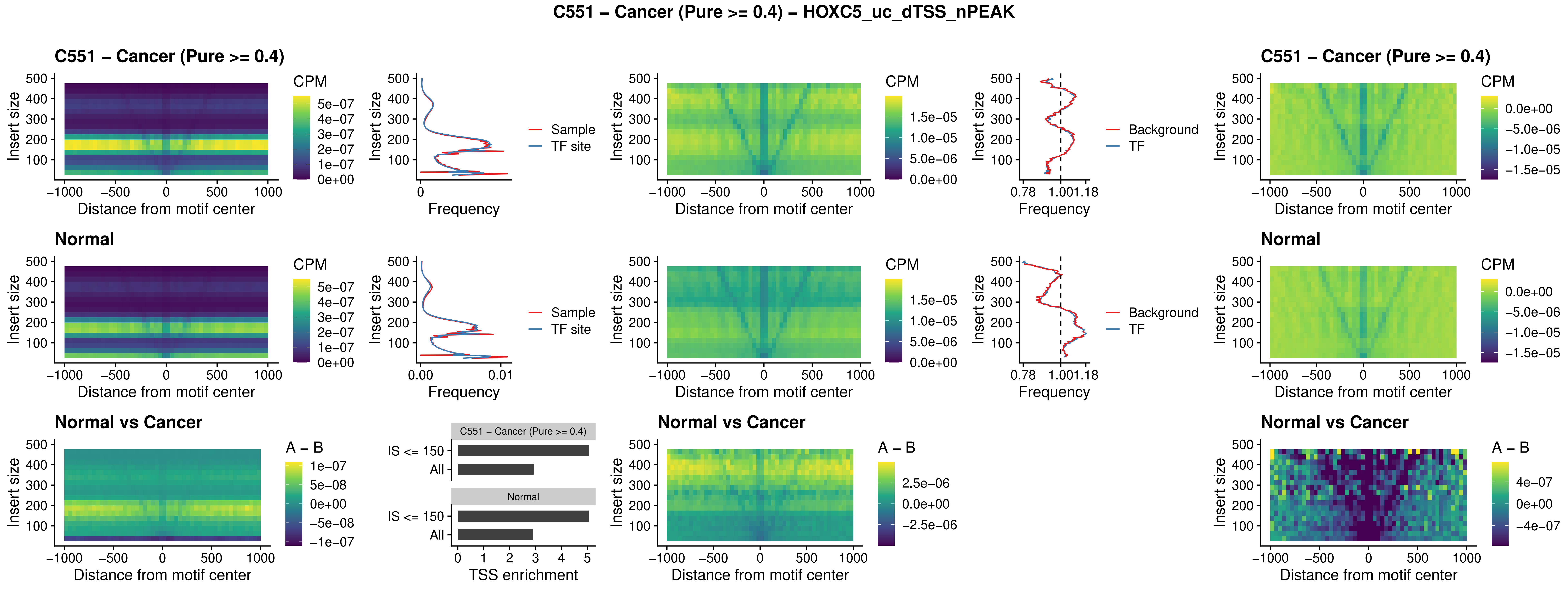

C551 – Cancer (Pure >= 0.4) – HOXC5_uc_dTSS_nPEAK
C551 – Cancer (Pure >= 0.4)
Normal
Normal vs Cancer
Insert size
Distance from motif center
Frequency
CPM
A – B
Sample
TF site
Background
TF
C551 – Cancer (Pure >= 0.4)
Normal
IS <= 150
All
TSS enrichment

#### C551 – Cancer (Pure >= 0.4) – HOXD1_uc_dTSS_nPEAK

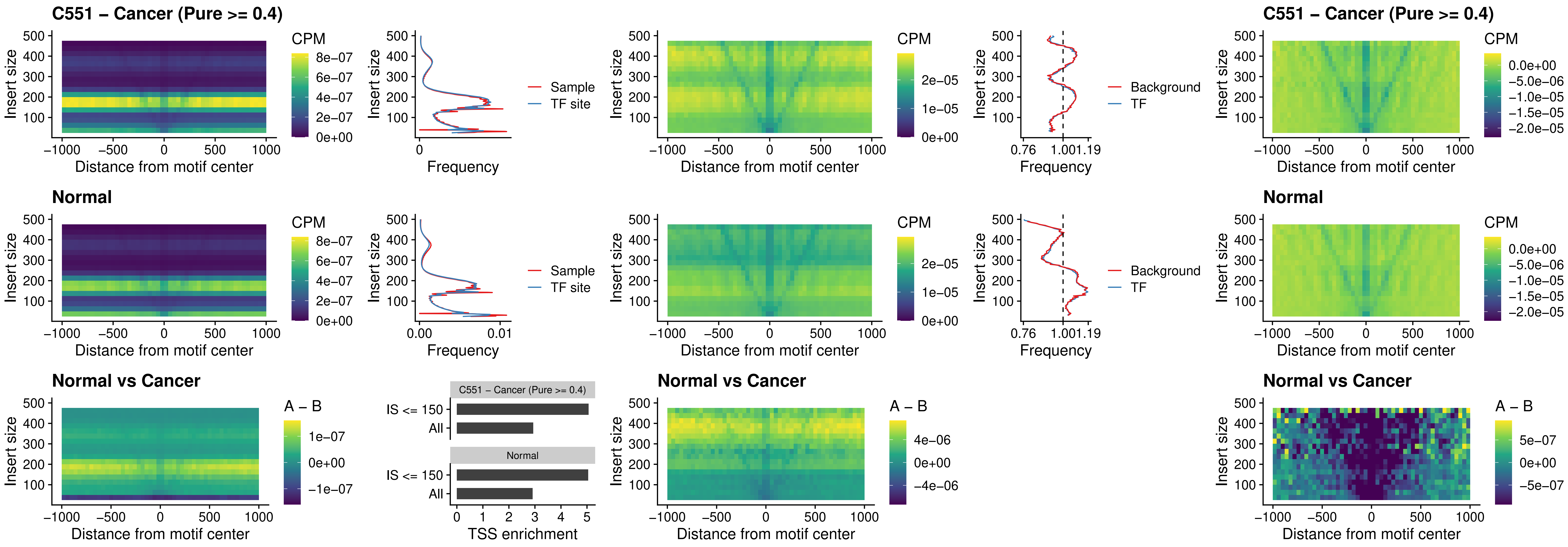

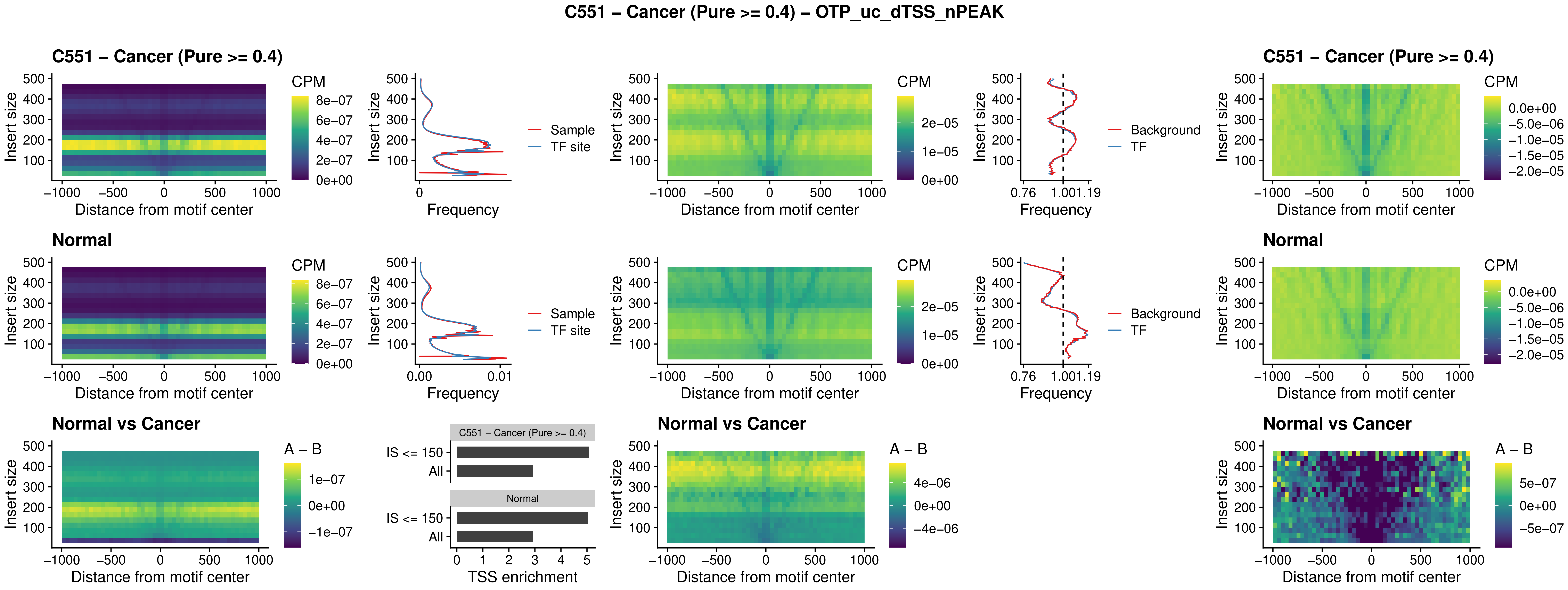
C551 – Cancer (Pure >= 0.4) – OTP_uc_dTSS_nPEAK
C551 – Cancer (Pure >= 0.4)
Normal
Normal vs Cancer
Insert size
Distance from motif center
CPM
A – B
Frequency
Sample
TF site
Background
TF
TSS enrichment
IS <= 150
All
C551 – Cancer (Pure >= 0.4)
Normal

#### C551 – Cancer (Pure >= 0.4) – FOXL1_uc_dTSS_oPEAK

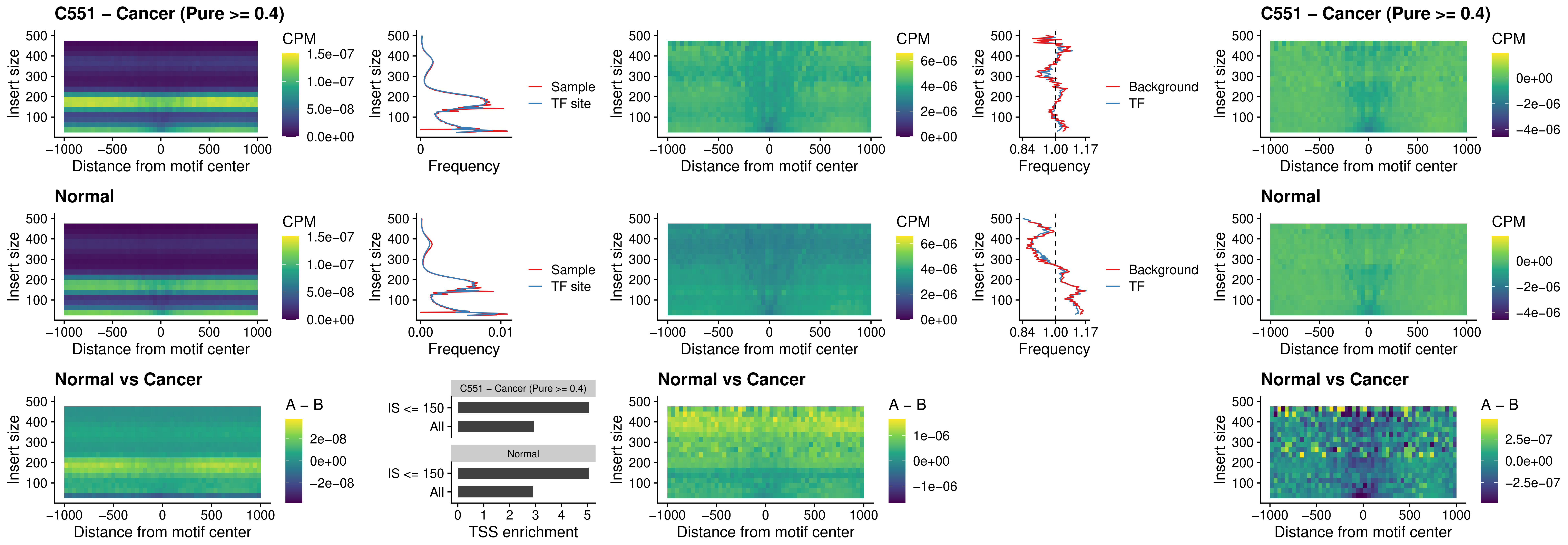

### C551 – Cancer (Pure >= 0.4) – ZNF32_uc_dTSS_oPEAK

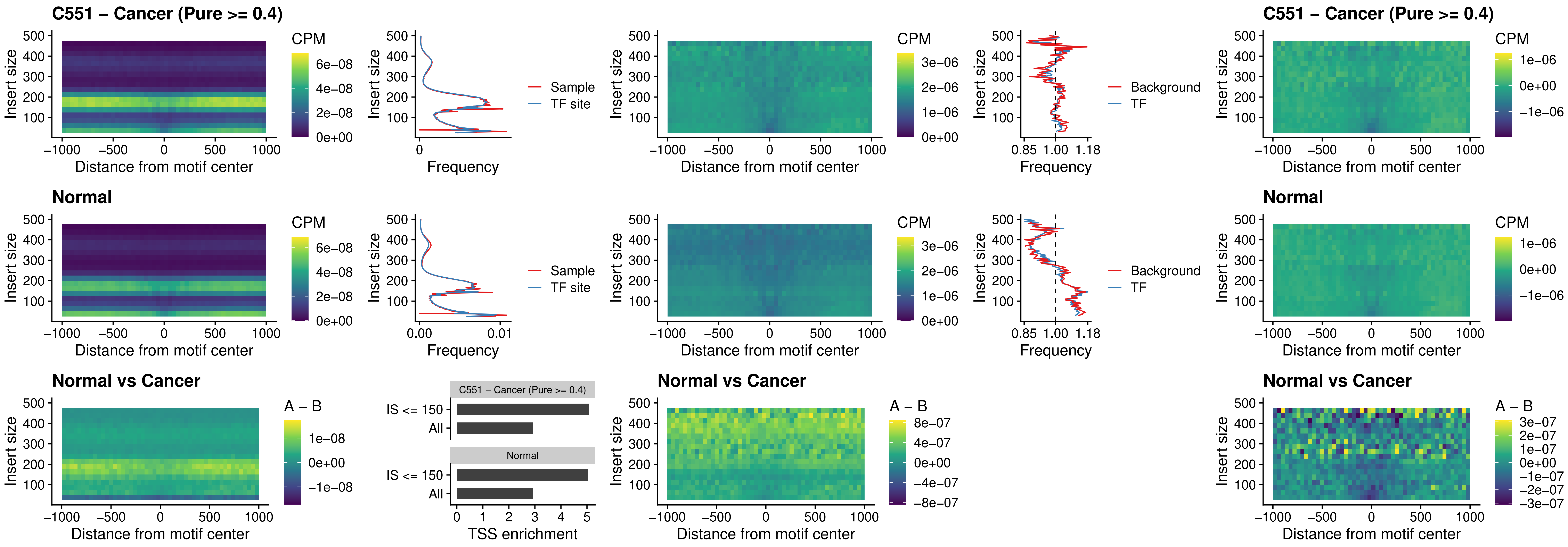

#### C551 – Cancer (Pure >= 0.4) – FOXL1_uc_dTSS_nPEAK

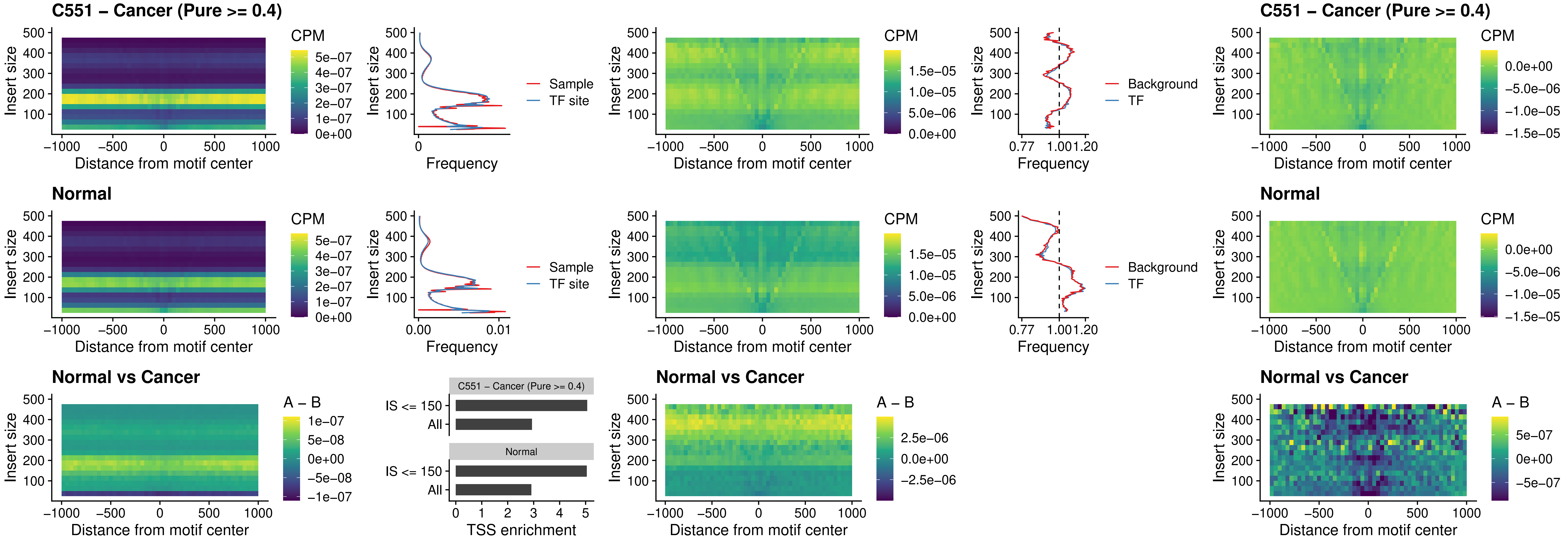

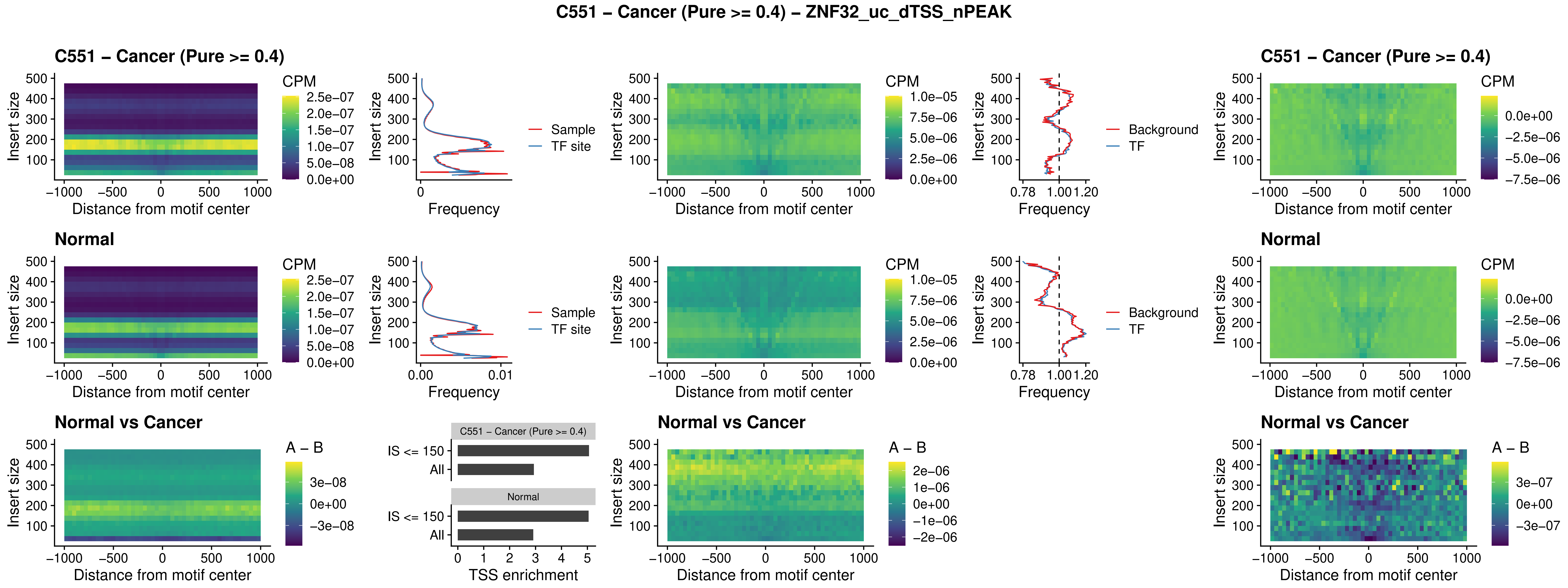
C551 – Cancer (Pure >= 0.4) – ZNF32_uc_dTSS_nPEAK
C551 – Cancer (Pure >= 0.4)
Normal
Normal vs Cancer
Insert size
Distance from motif center
Frequency
CPM
A – B
Sample
TF site
Background
TF
TSS enrichment
IS <= 150
All
C551 – Cancer (Pure >= 0.4)
Normal

#### C551 – Cancer (Pure >= 0.4) – DLX5_uc_dTSS_nPEAK

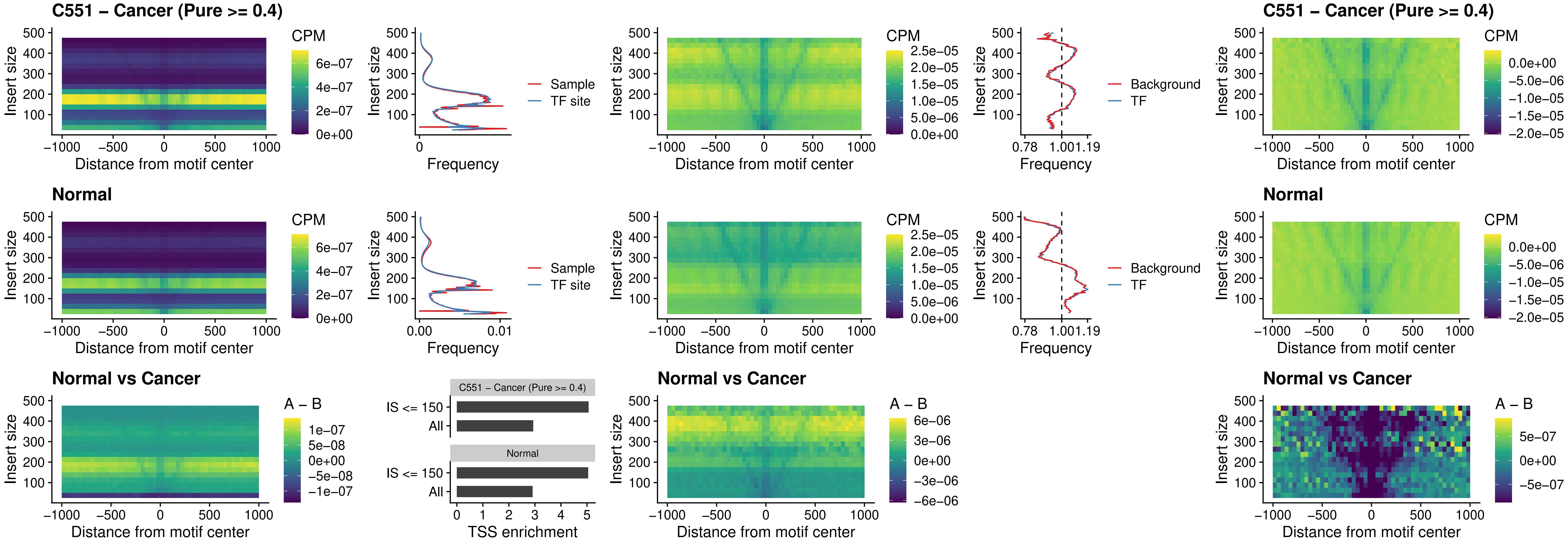

#### C551 – Cancer (Pure >= 0.4) – DMRT2_uc_dTSS_nPEAK

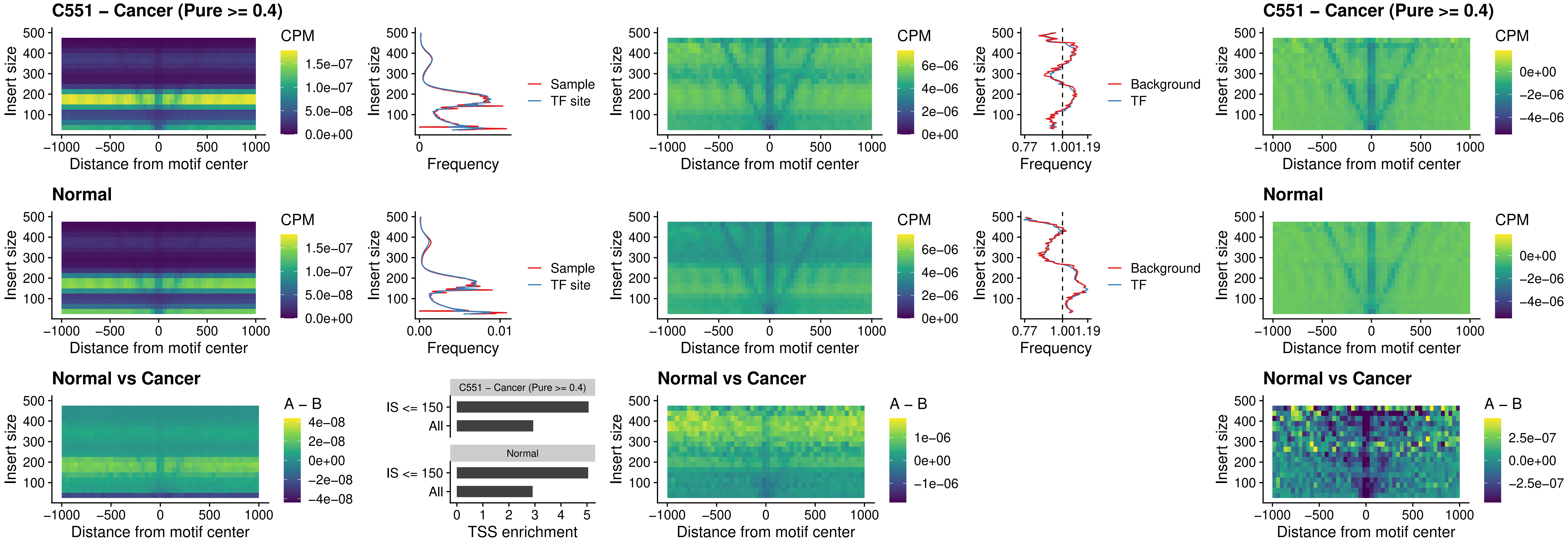

### C551 – Cancer (Pure >= 0.4) – DMRTC2_uc_cTSS_nPEAK

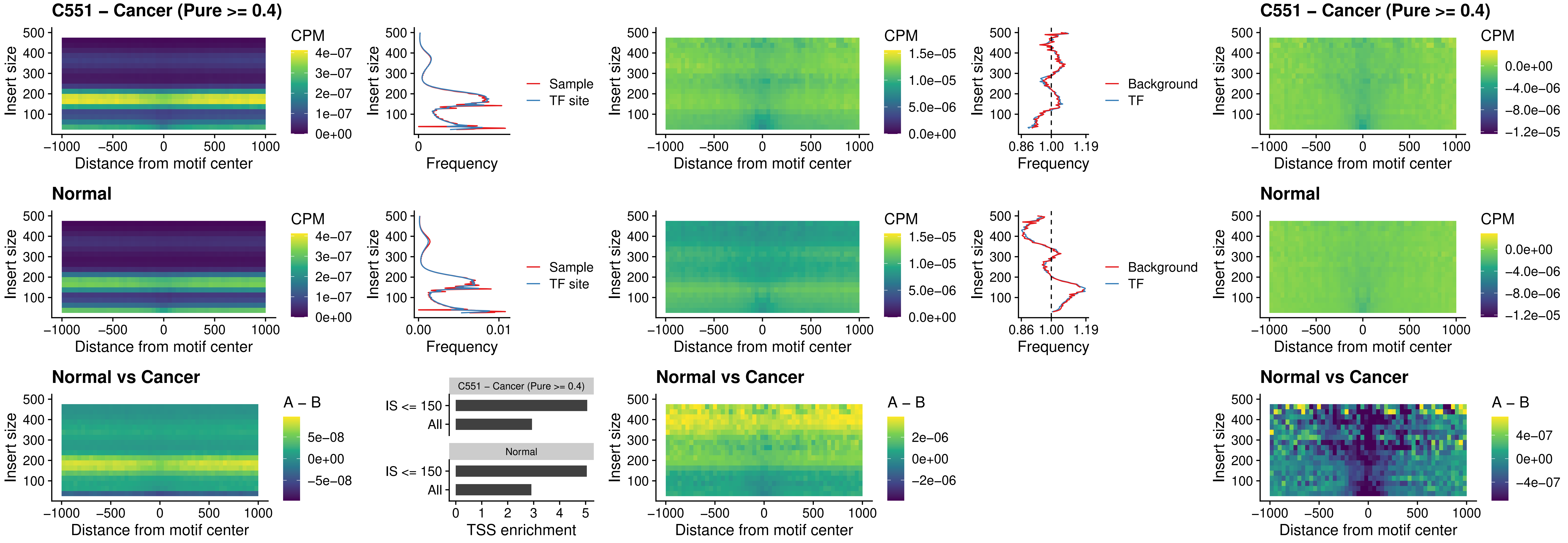

#### C551 – Cancer (Pure >= 0.4) – HOXD9_uc_dTSS_nPEAK

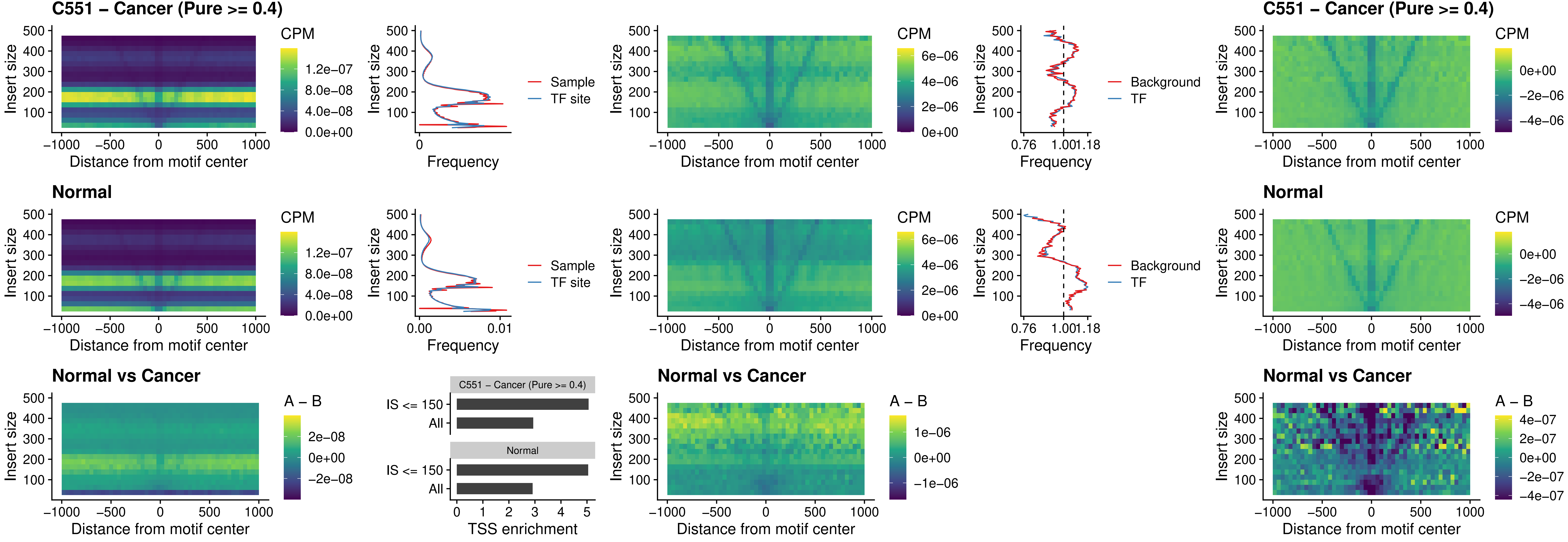

#### C551 – Cancer (Pure >= 0.4) – HOXD4_uc_dTSS_nPEAK

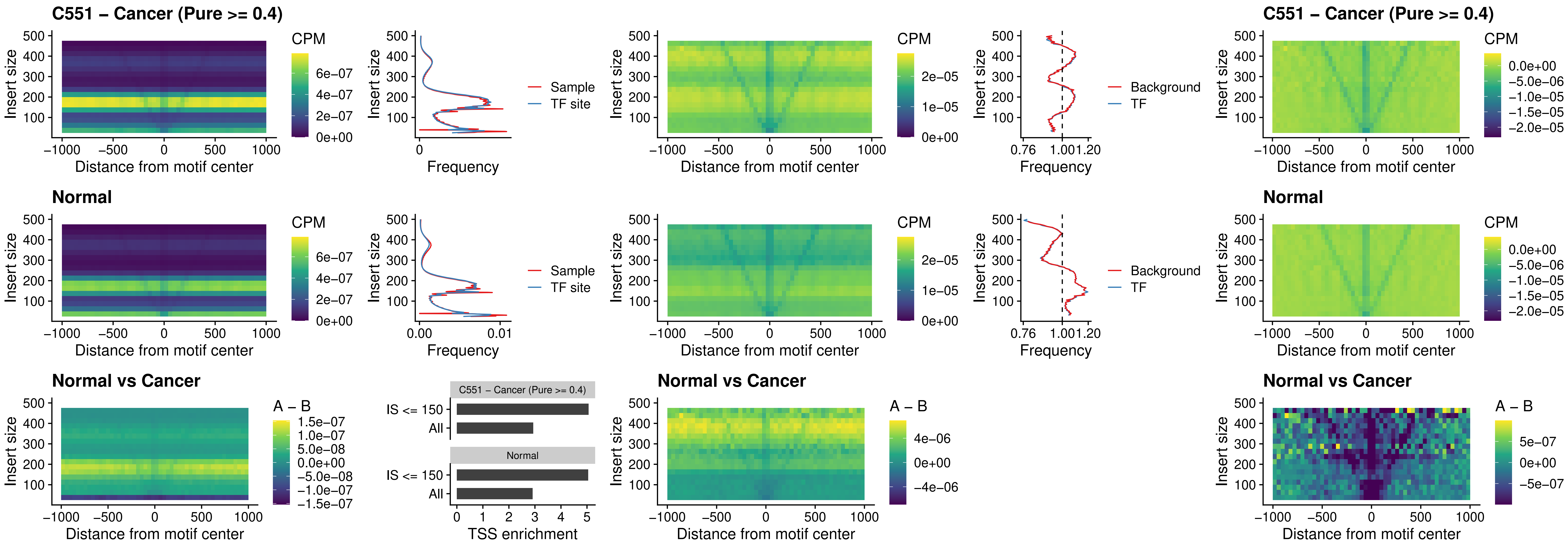

#### C551 – Cancer (Pure >= 0.4) – FOXP3_uc_cTSS_nPEAK

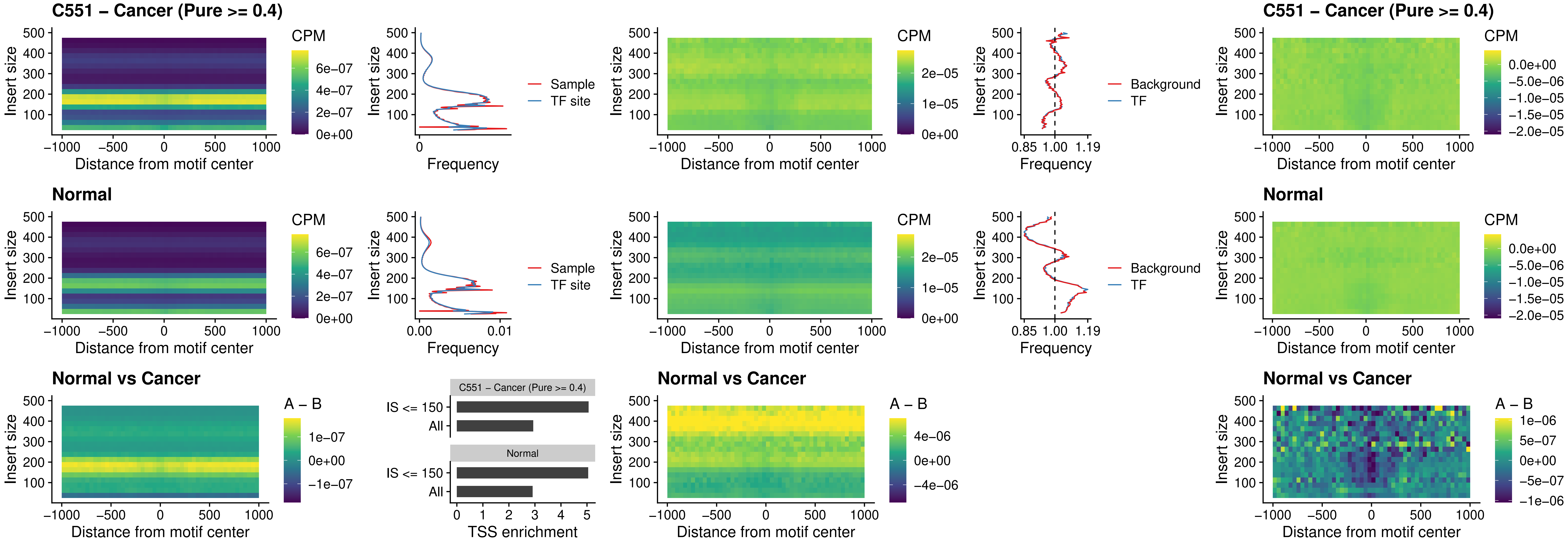

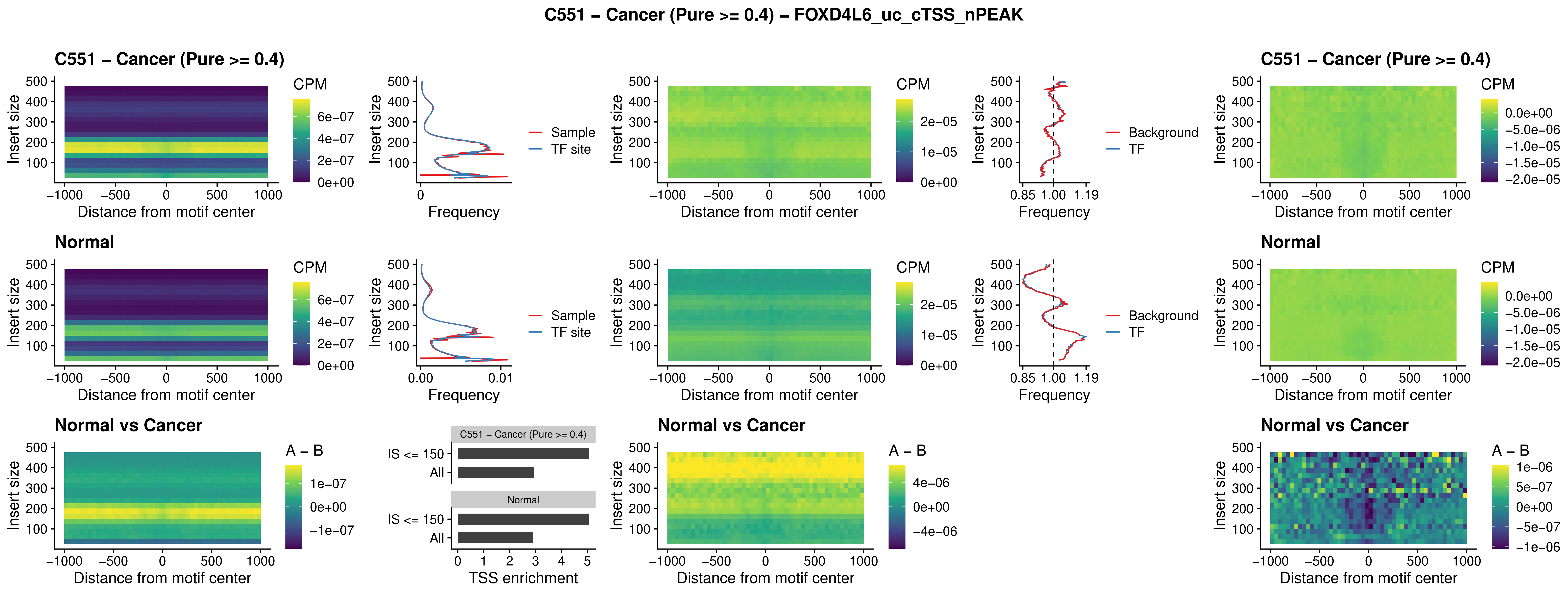
C551 – Cancer (Pure >= 0.4) – FOXD4L6_uc_cTSS_nPEAK
C551 – Cancer (Pure >= 0.4)
Normal
Normal vs Cancer
Insert size
Distance from motif center
Frequency
CPM
A – B
Sample
TF site
Background
TF
C551 – Cancer (Pure >= 0.4)
Normal
IS <= 150
All
TSS enrichment

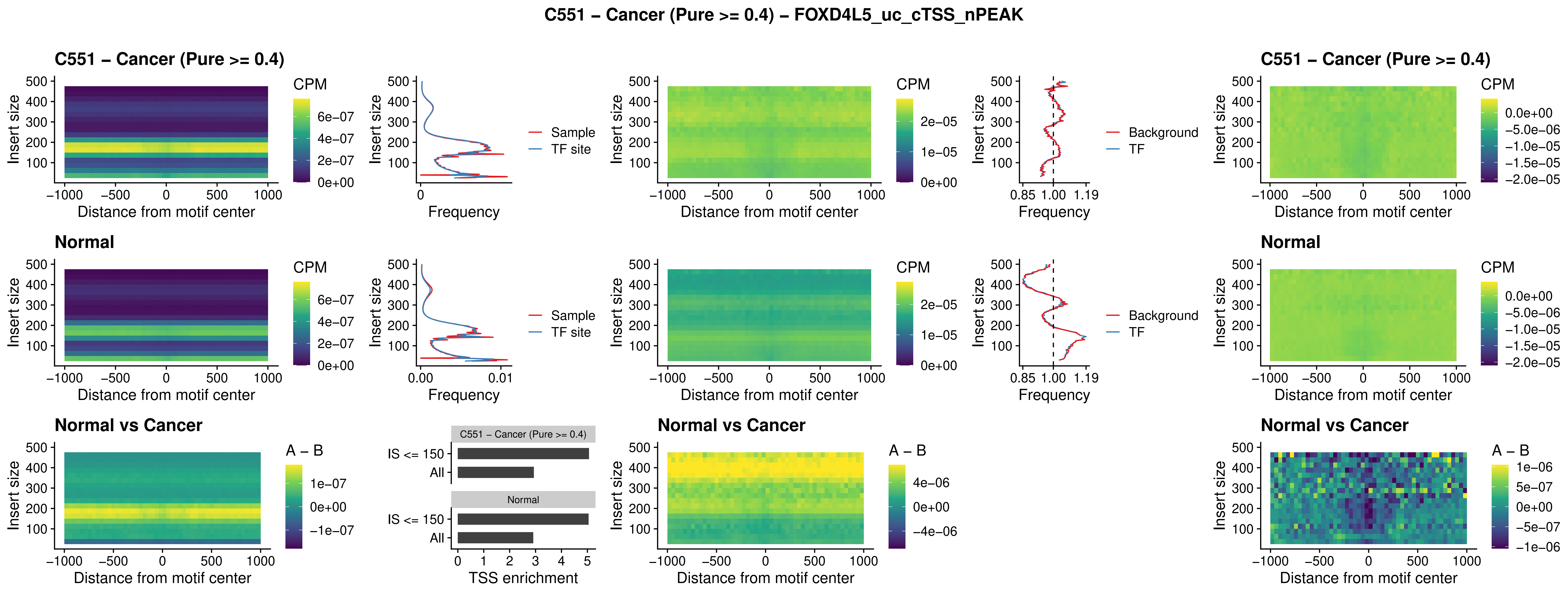
C551 – Cancer (Pure >= 0.4) – FOXD4L5_uc_cTSS_nPEAK
C551 – Cancer (Pure >= 0.4)
Normal
Normal vs Cancer
Insert size
Distance from motif center
CPM
A – B
Frequency
Sample
TF site
Background
TF
C551 – Cancer (Pure >= 0.4)
Normal
IS <= 150
All
TSS enrichment

#### C551 – Cancer (Pure >= 0.4) – FOXD4L4_uc_cTSS_nPEAK

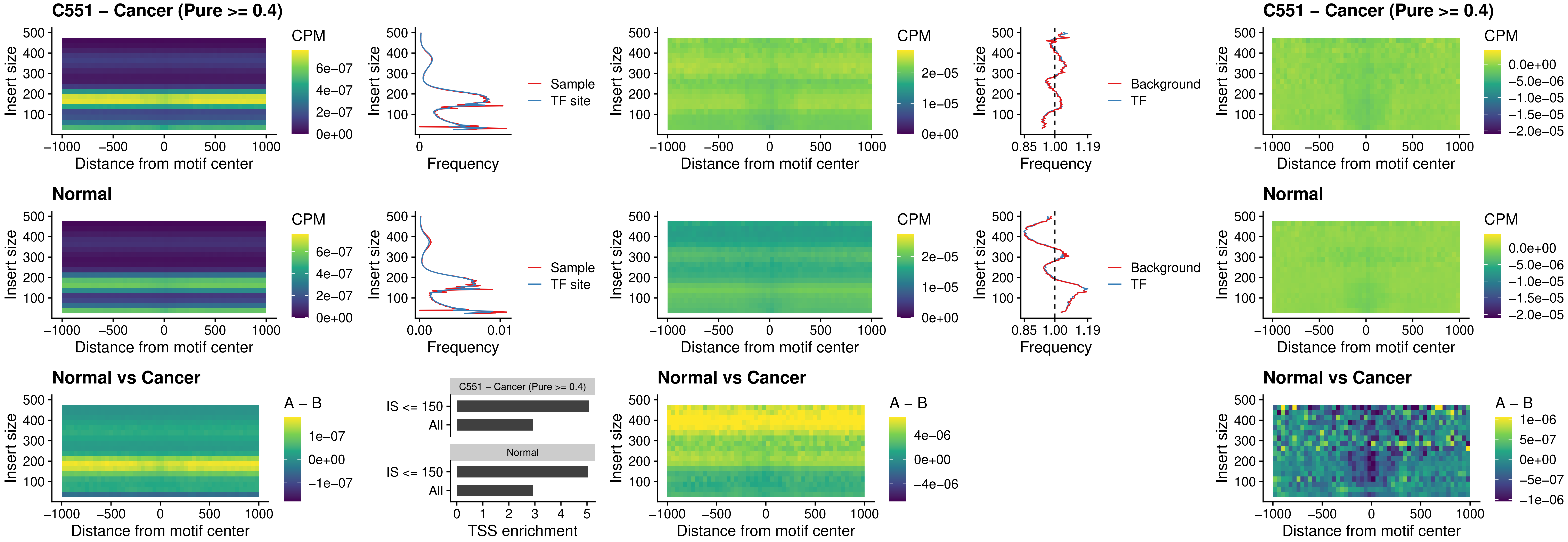

#### C551 – Cancer (Pure >= 0.4) – FOXD4L2_uc_cTSS_nPEAK

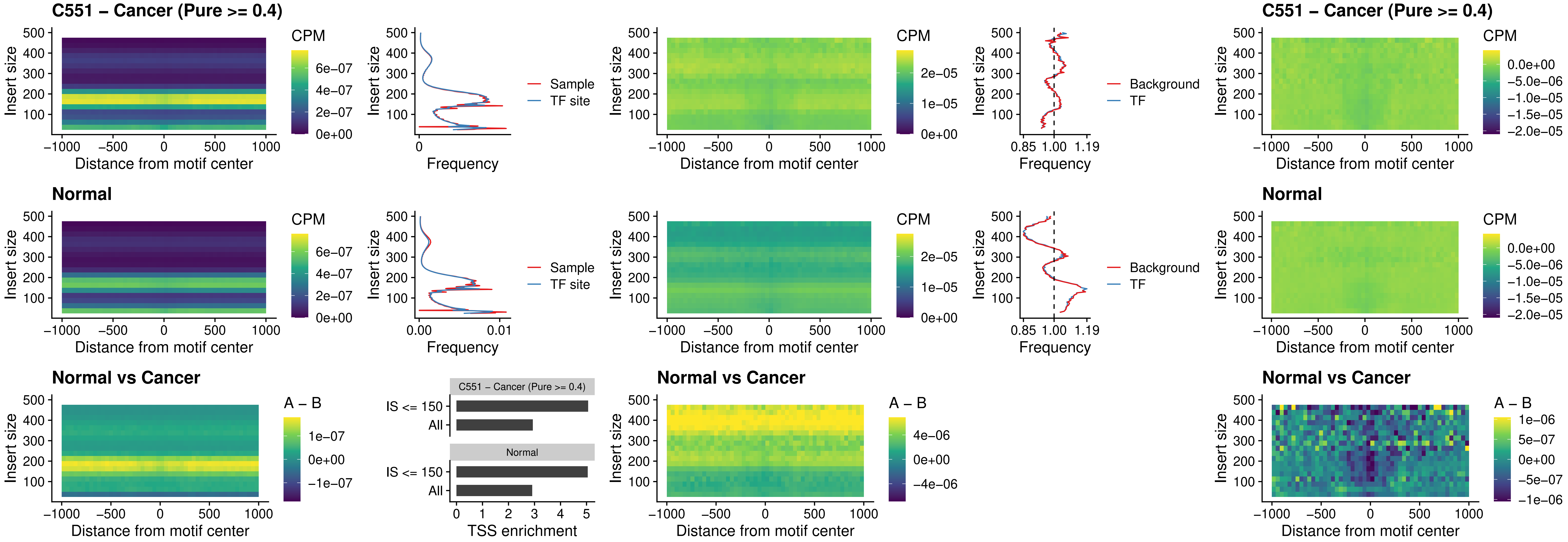

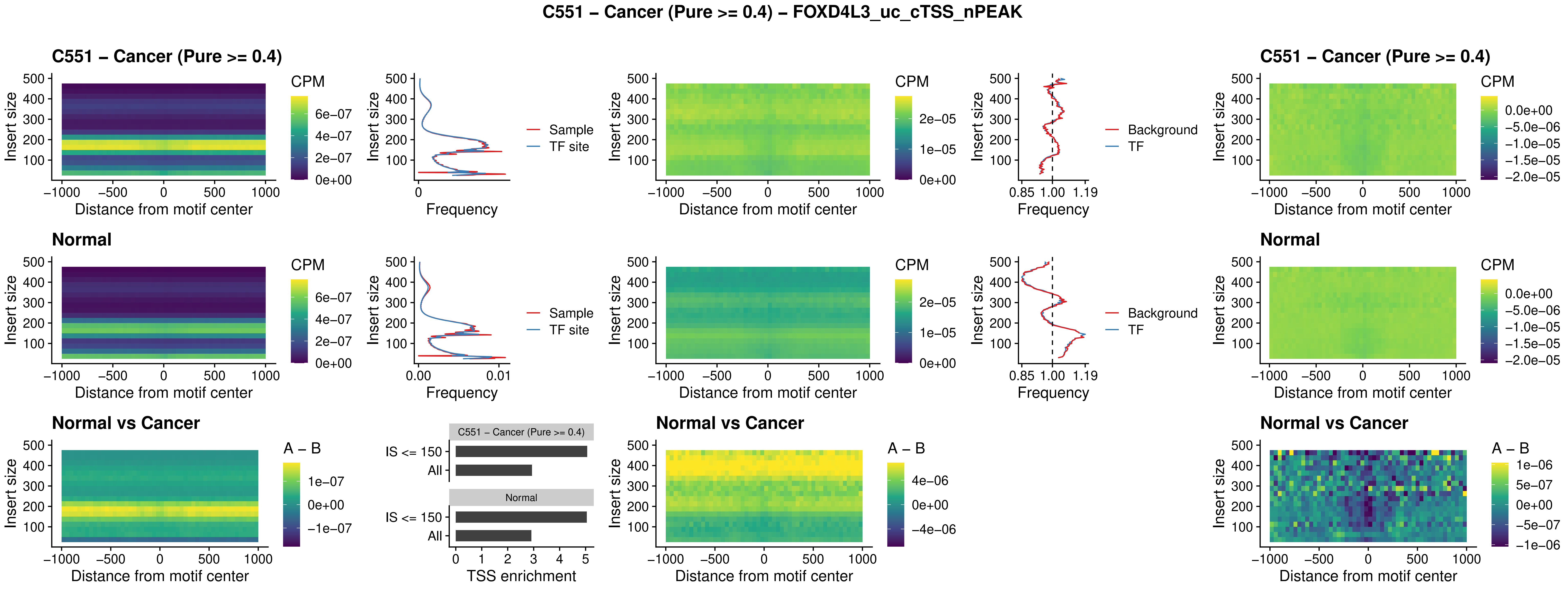
C551 – Cancer (Pure >= 0.4) – FOXD4L3_uc_cTSS_nPEAK
C551 – Cancer (Pure >= 0.4)
Normal
Normal vs Cancer
Insert size
Distance from motif center
Frequency
CPM
A – B
Sample
TF site
Background
TF
C551 – Cancer (Pure >= 0.4)
Normal
IS <= 150
All
TSS enrichment

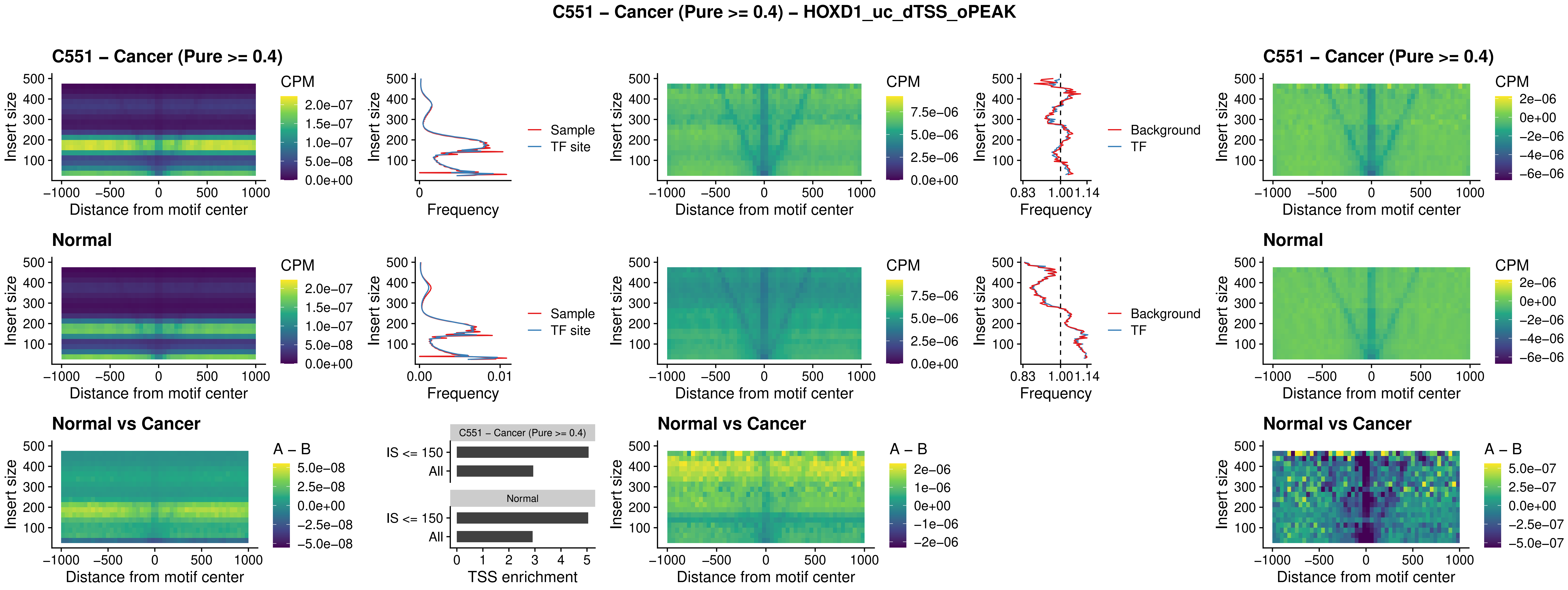
C551 – Cancer (Pure >= 0.4) – HOXD1_uc_dTSS_oPEAK
C551 – Cancer (Pure >= 0.4)
Normal
Normal vs Cancer
Insert size
Distance from motif center
CPM
A – B
Frequency
Sample
TF site
Background
TF
C551 – Cancer (Pure >= 0.4)
Normal
IS <= 150
All
TSS enrichment

#### C551 – Cancer (Pure >= 0.4) – VAX1_uc_pTSS_nPEAK

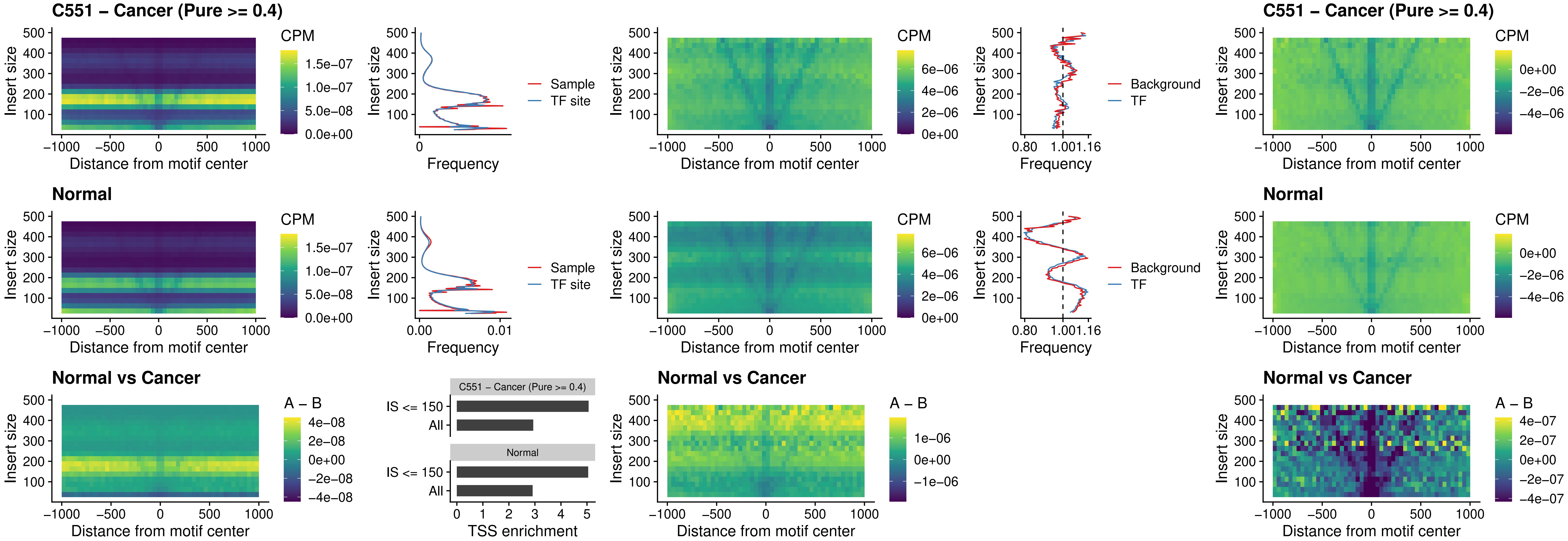

#### C551 – Cancer (Pure >= 0.4) – DLX3_uc_pTSS_nPEAK

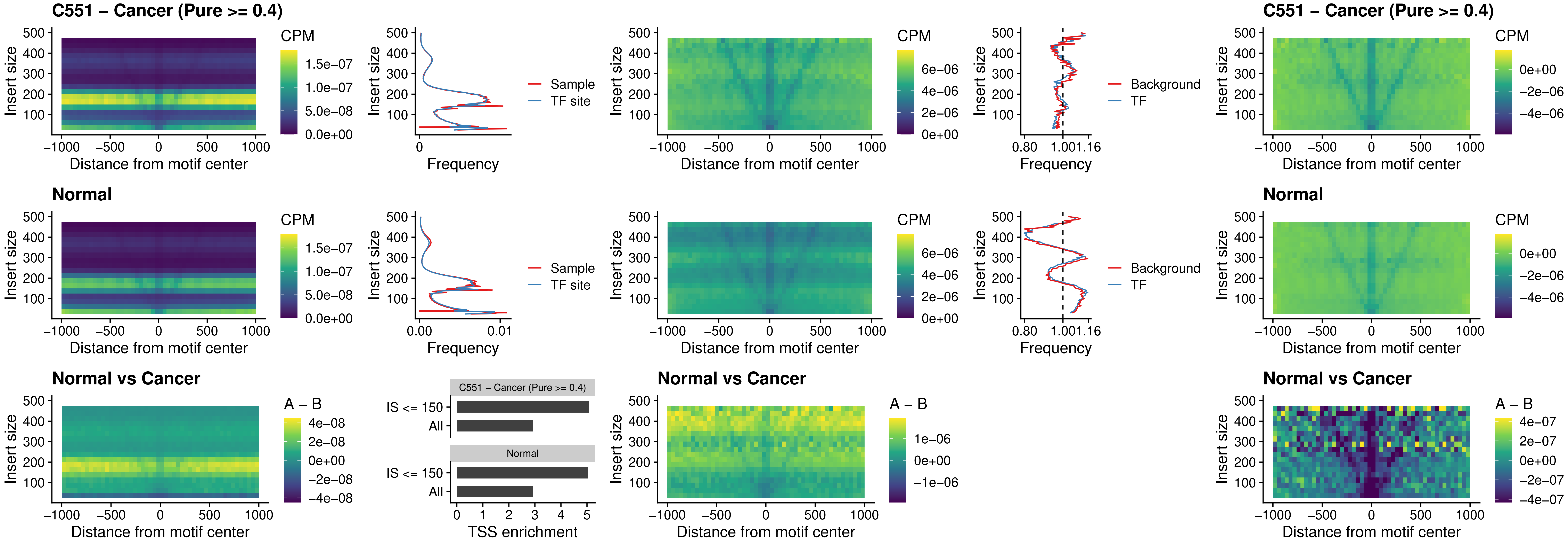

### C551 – Cancer (Pure >= 0.4) – HOXD3_uc_pTSS_nPEAK

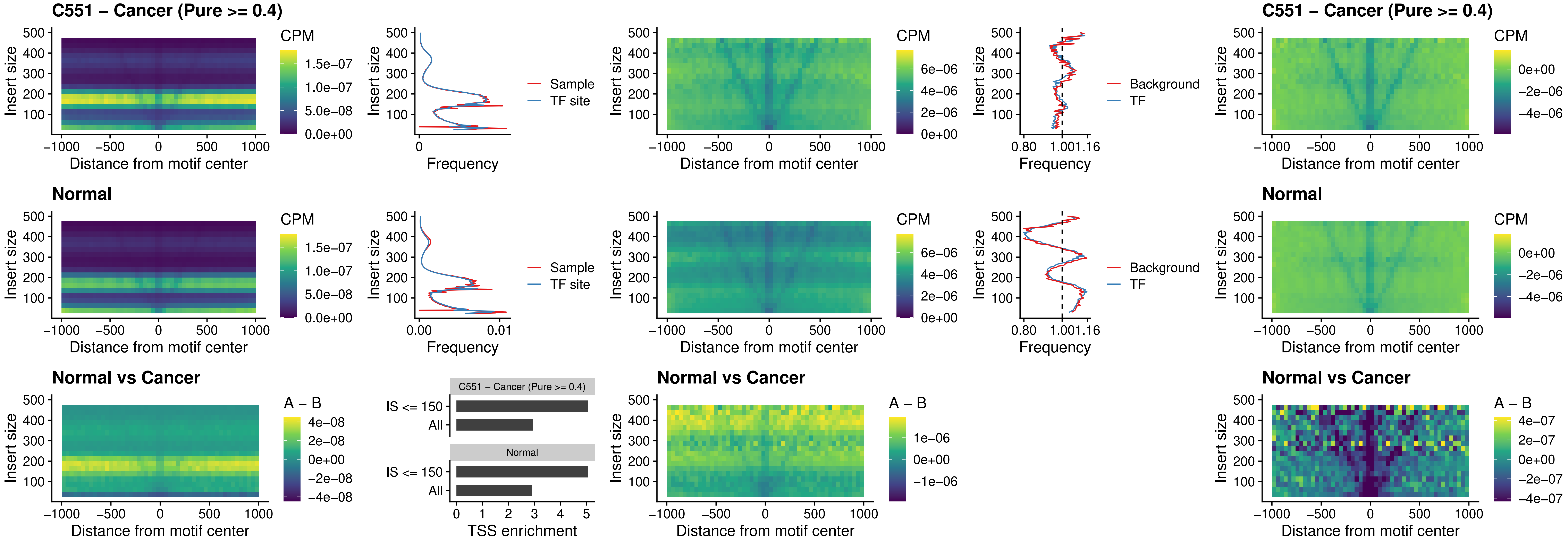

#### C551 – Cancer (Pure >= 0.4) – PDX1_uc_dTSS_nPEAK

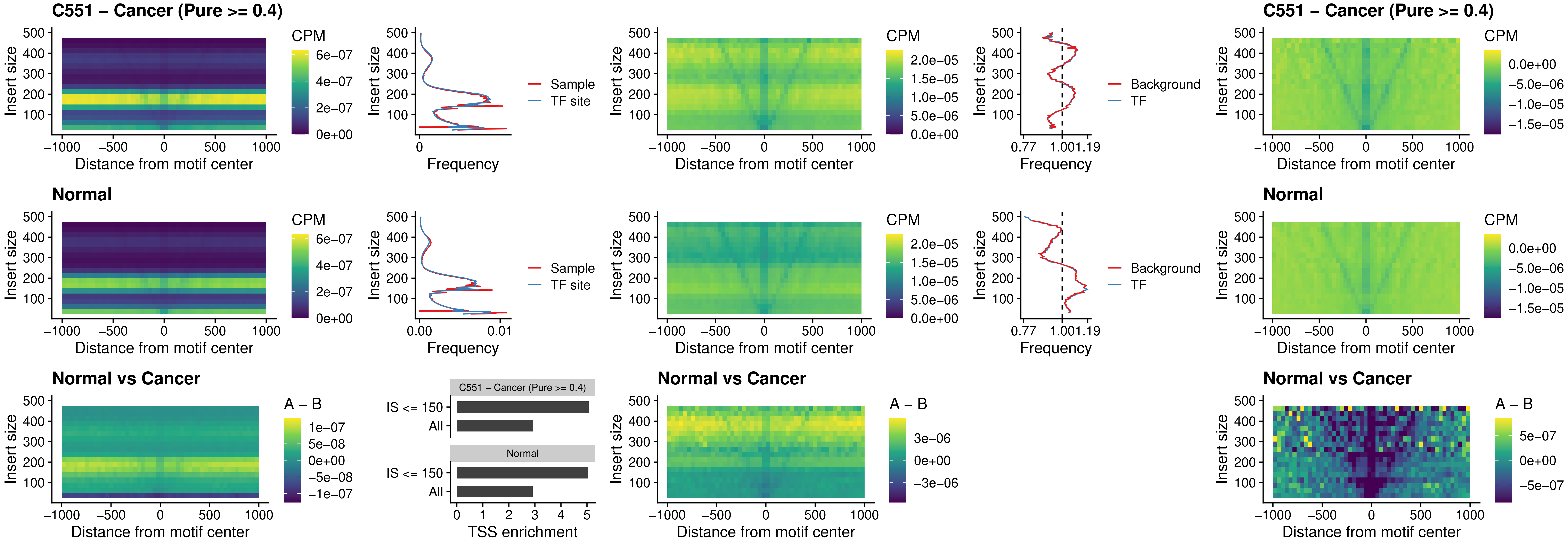

#### C551 – Cancer (Pure >= 0.4) – VAX2_uc_dTSS_nPEAK

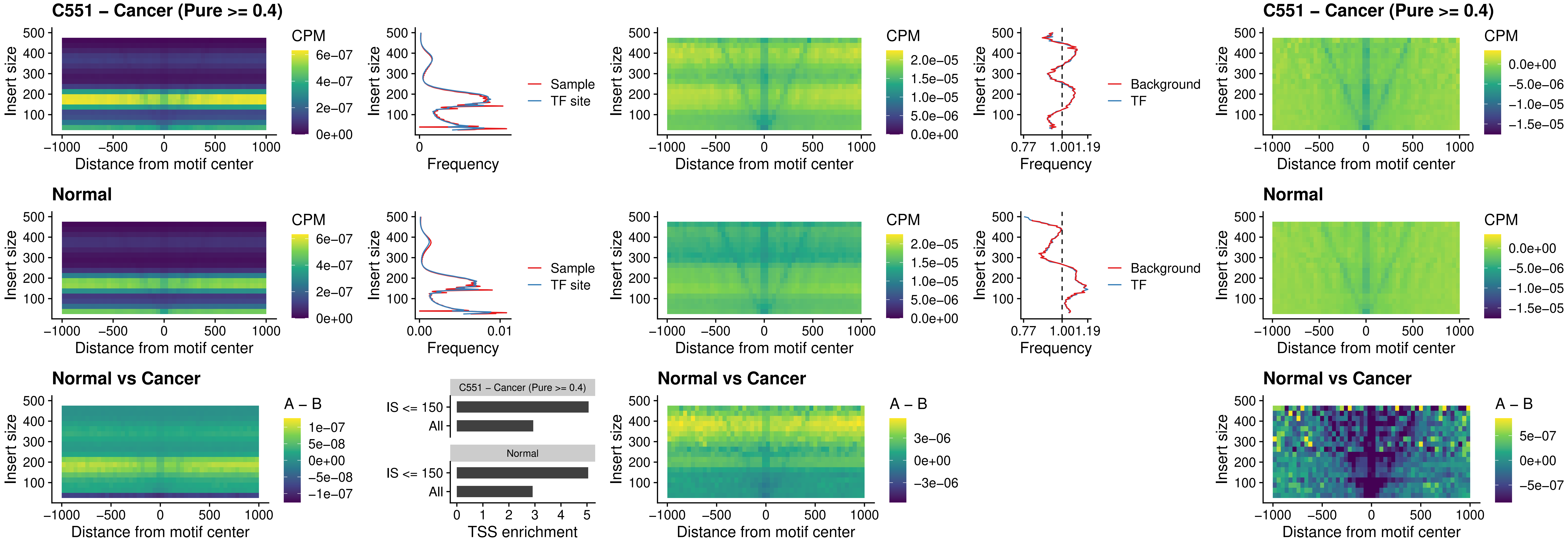

#### C551 – Cancer (Pure >= 0.4) – SOX5_uc_dTSS_nPEAK

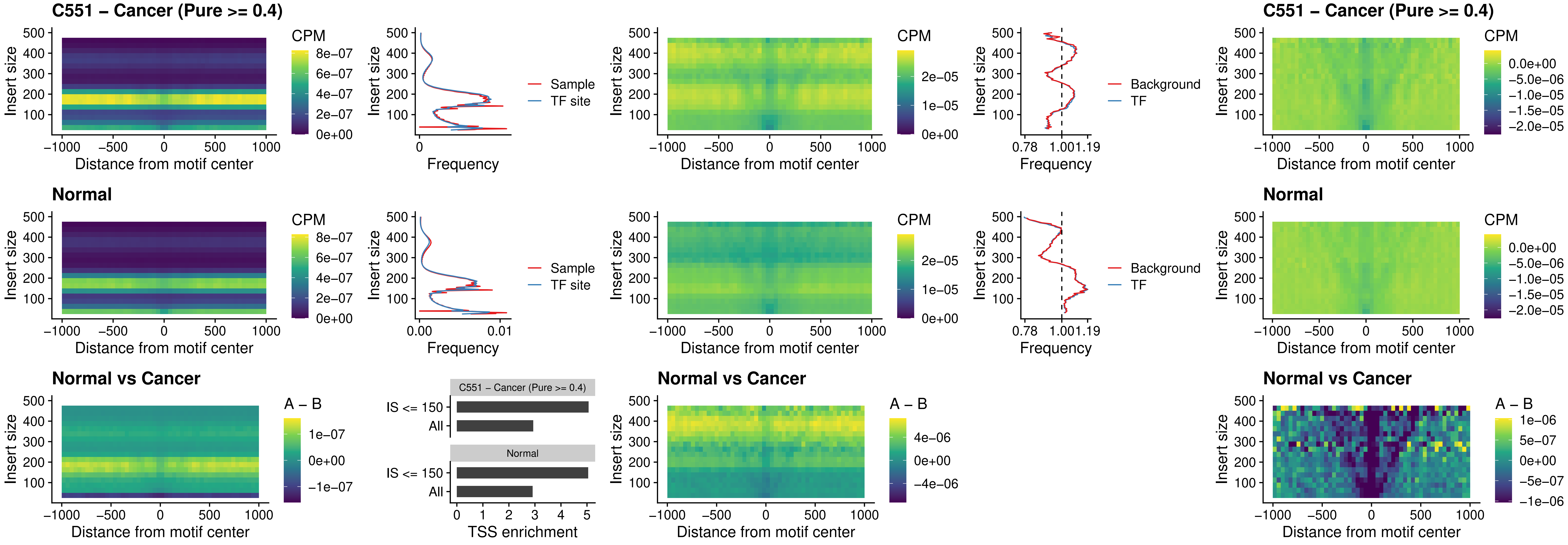

#### C551 – Cancer (Pure >= 0.4) – SOX6_uc_dTSS_nPEAK

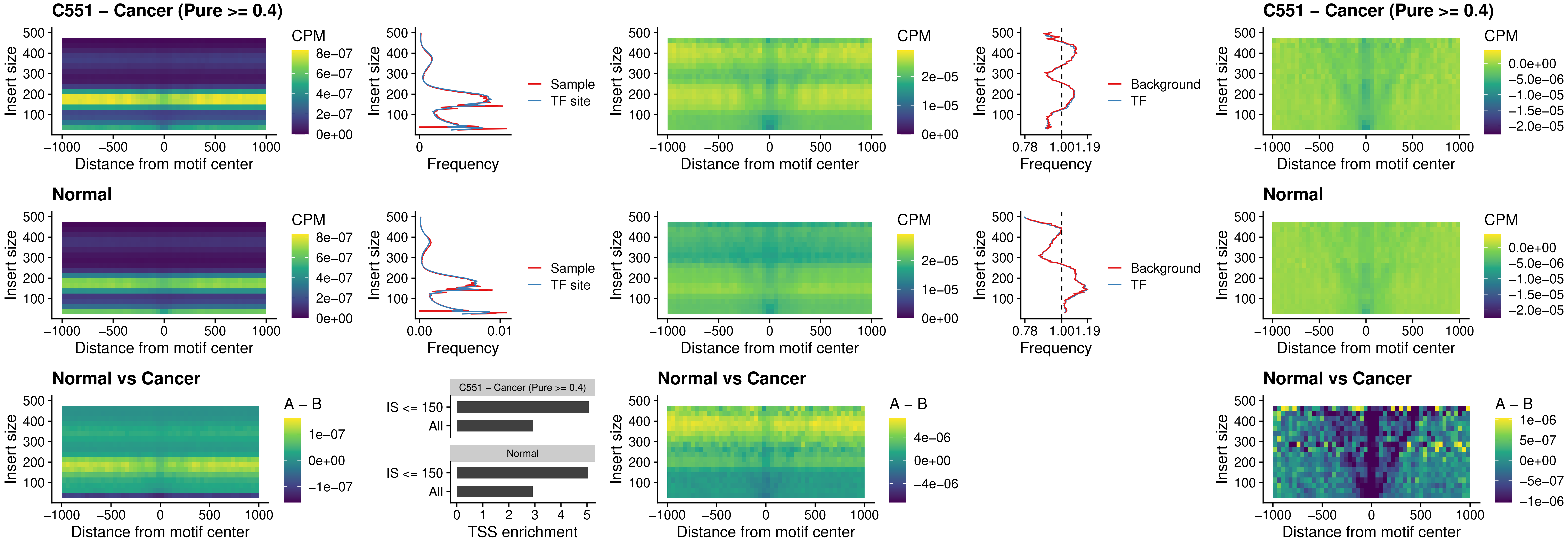

### C551 – Cancer (Pure >= 0.4) – JDP2_uc_cTSS_nPEAK

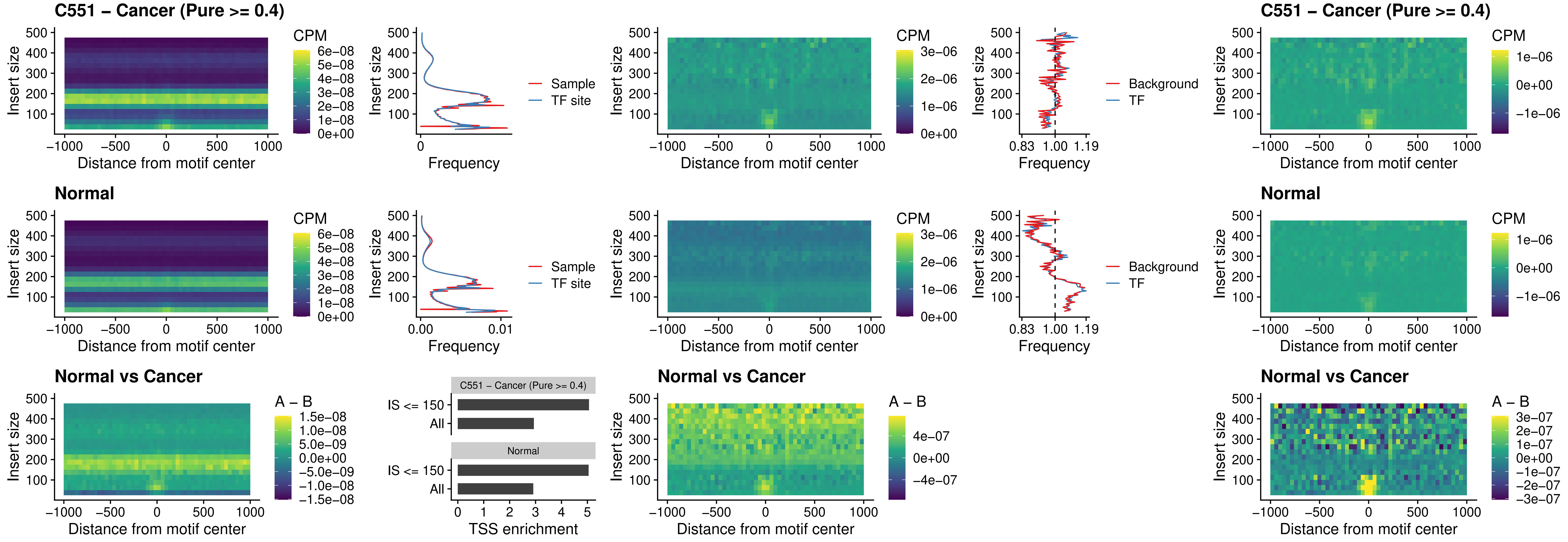

#### C551 – Cancer (Pure >= 0.4) – PURA_uc_dTSS_oPEAK

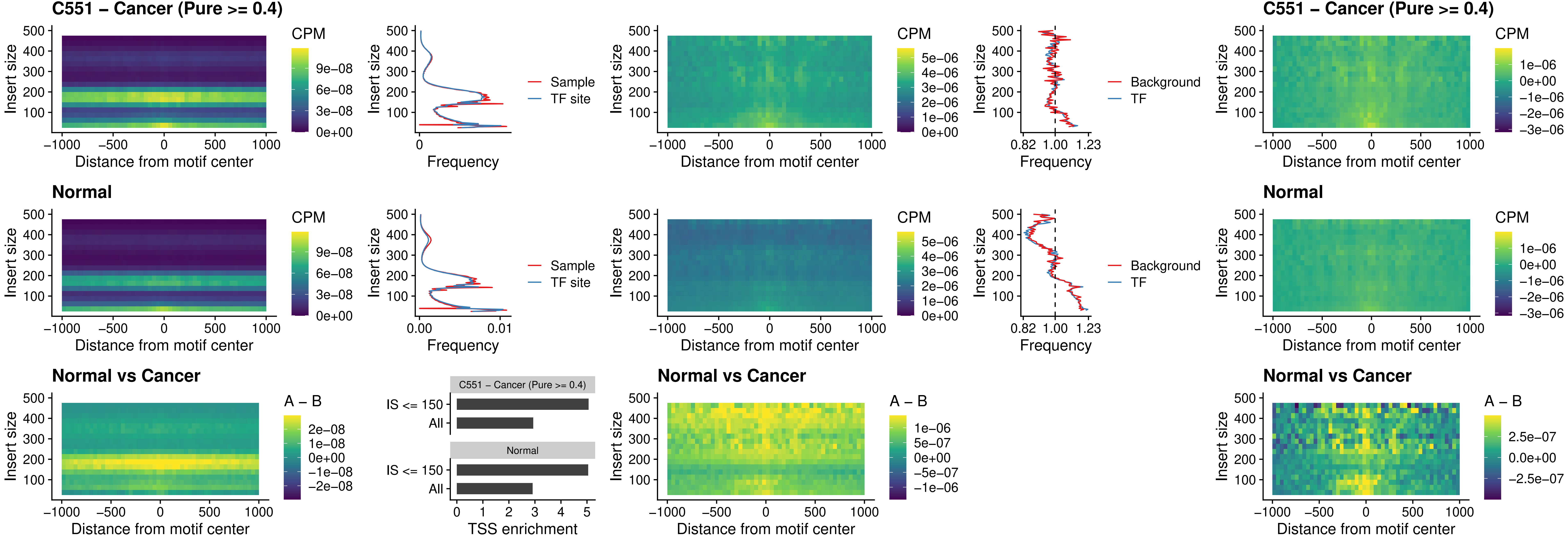

#### C551 – Cancer (Pure >= 0.4) – HHEX_uc_dTSS_nPEAK

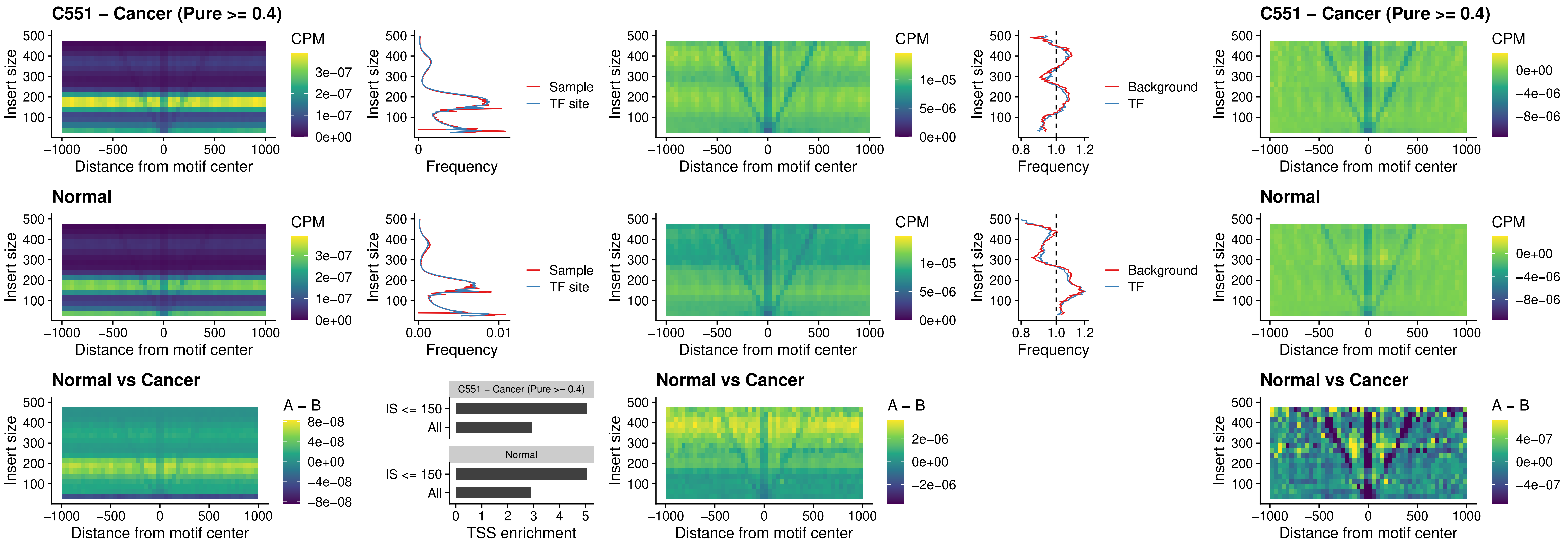

#### C551 – Cancer (Pure >= 0.4) – HMG20B_uc_dTSS_nPEAK

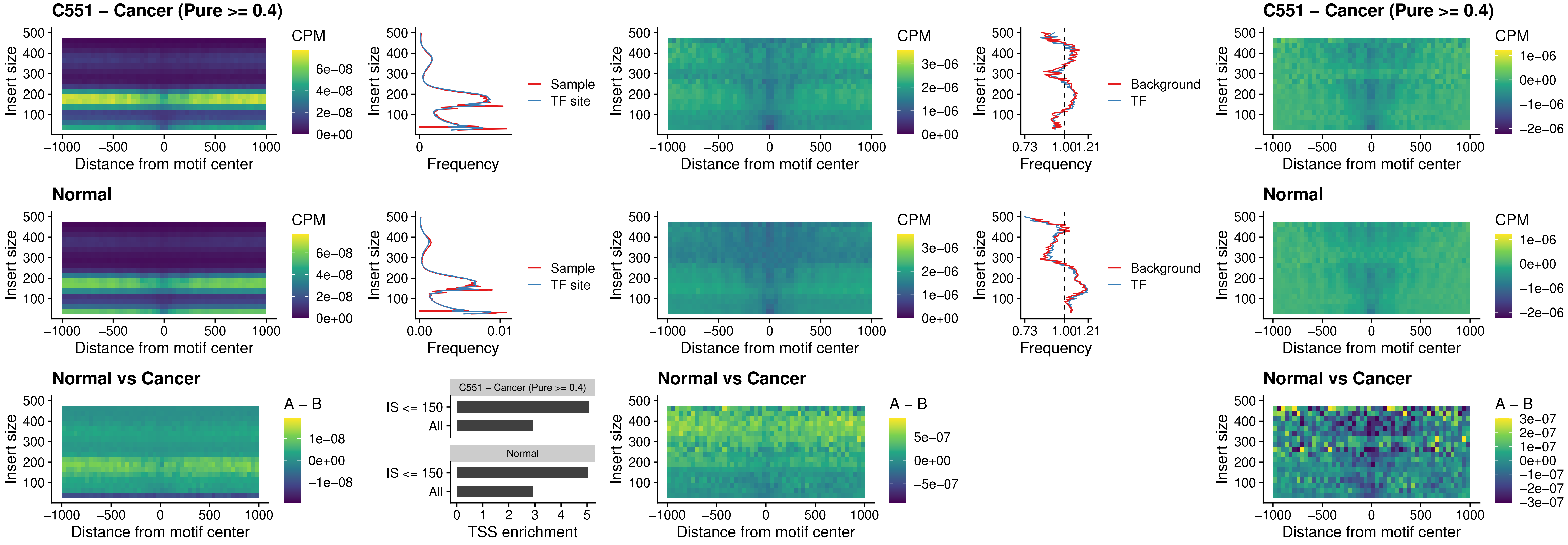

#### C551 – Cancer (Pure >= 0.4) – FOXB2_uc_dTSS_nPEAK

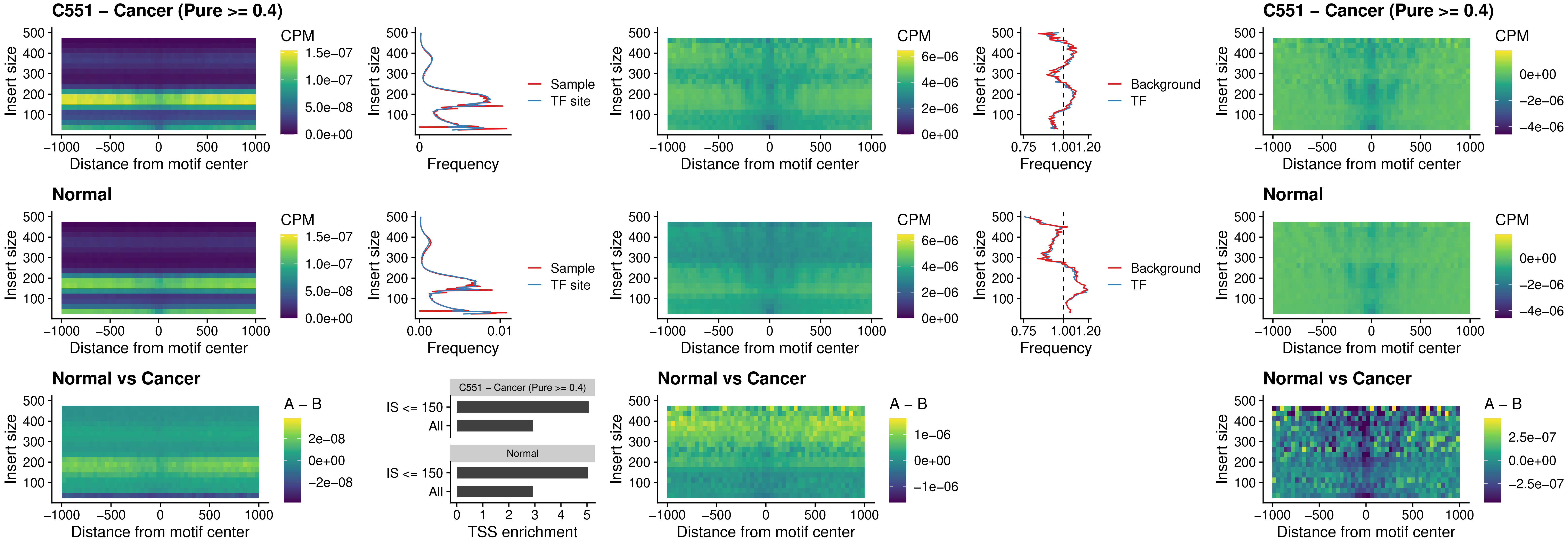

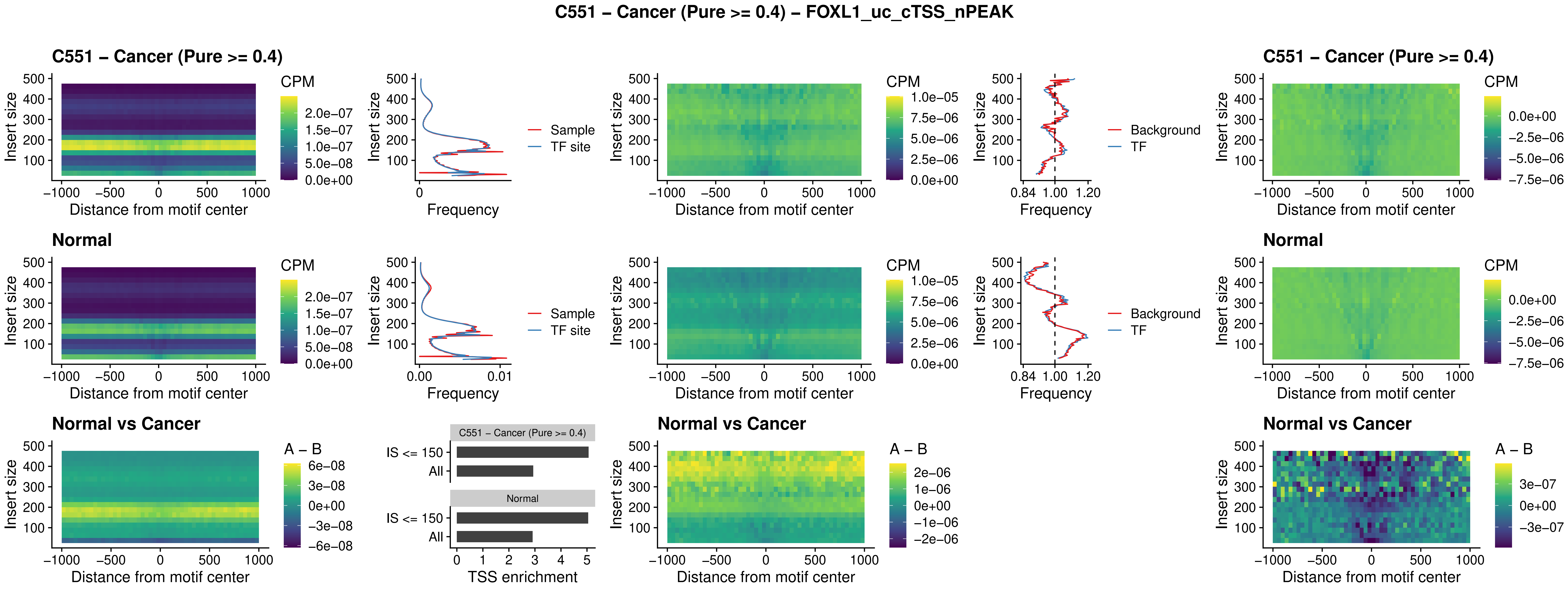
C551 – Cancer (Pure >= 0.4) – FOXL1_uc_cTSS_nPEAK
C551 – Cancer (Pure >= 0.4)
Normal
Normal vs Cancer
Insert size
Distance from motif center
Frequency
CPM
A – B
Sample
TF site
Background
TF
C551 – Cancer (Pure >= 0.4)
Normal
IS <= 150
All
TSS enrichment

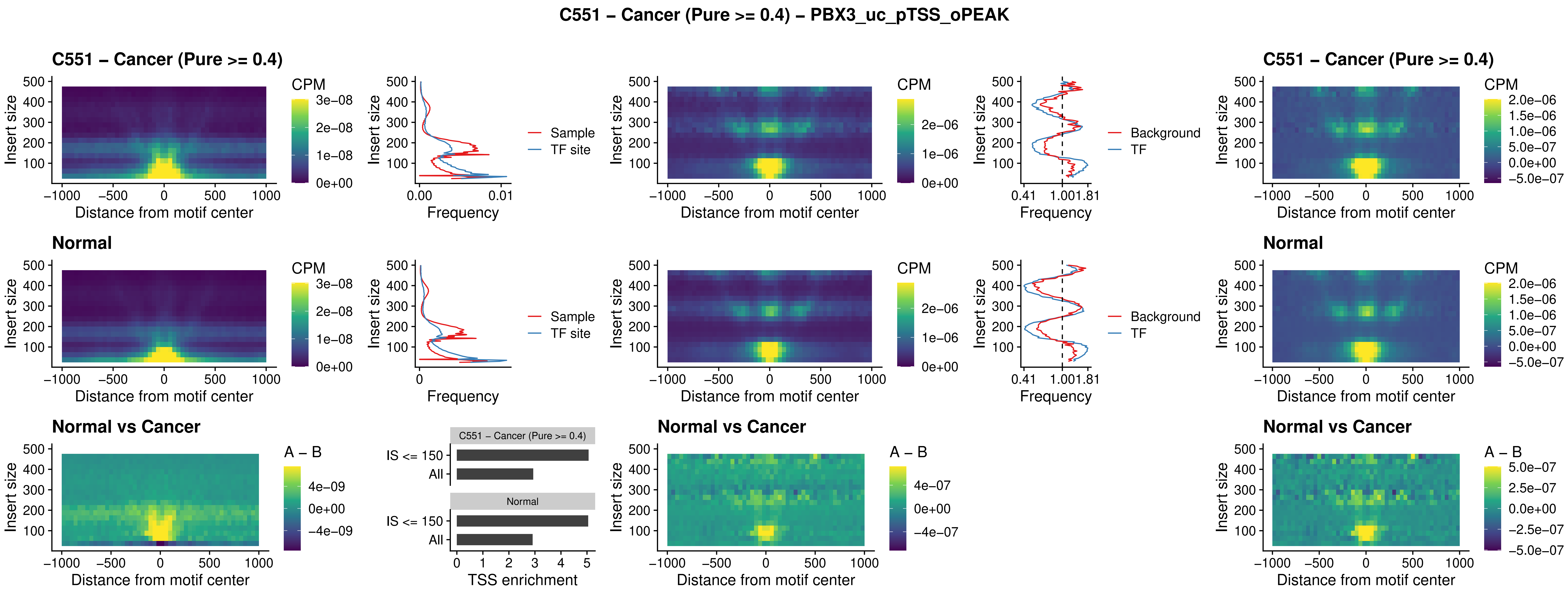
C551 – Cancer (Pure >= 0.4) – PBX3_uc_pTSS_oPEAK
C551 – Cancer (Pure >= 0.4)
Normal
Normal vs Cancer
Insert size
Distance from motif center
CPM
A – B
Frequency
Sample
TF site
Background
TF
C551 – Cancer (Pure >= 0.4)
IS <= 150
All
TSS enrichment

#### C551 – Cancer (Pure >= 0.4) – SNAI2_uc_cTSS_nPEAK

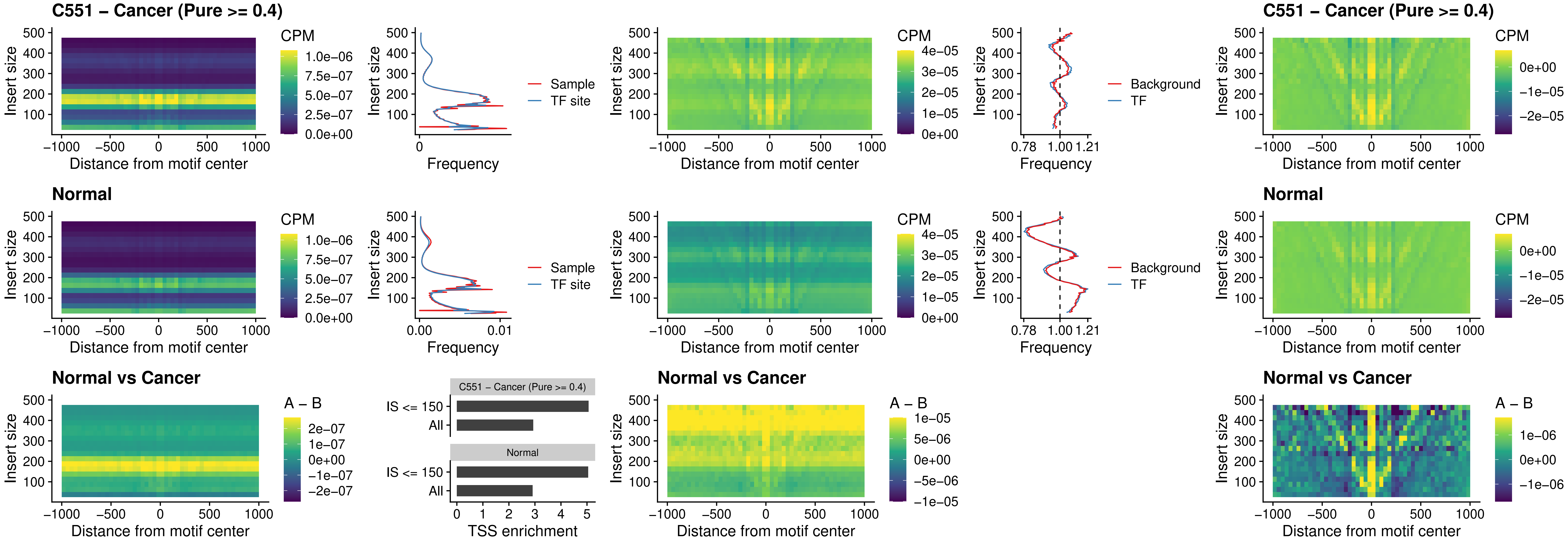

### C551 – Cancer (Pure >= 0.4) – SNAI2_uc_dTSS_oPEAK

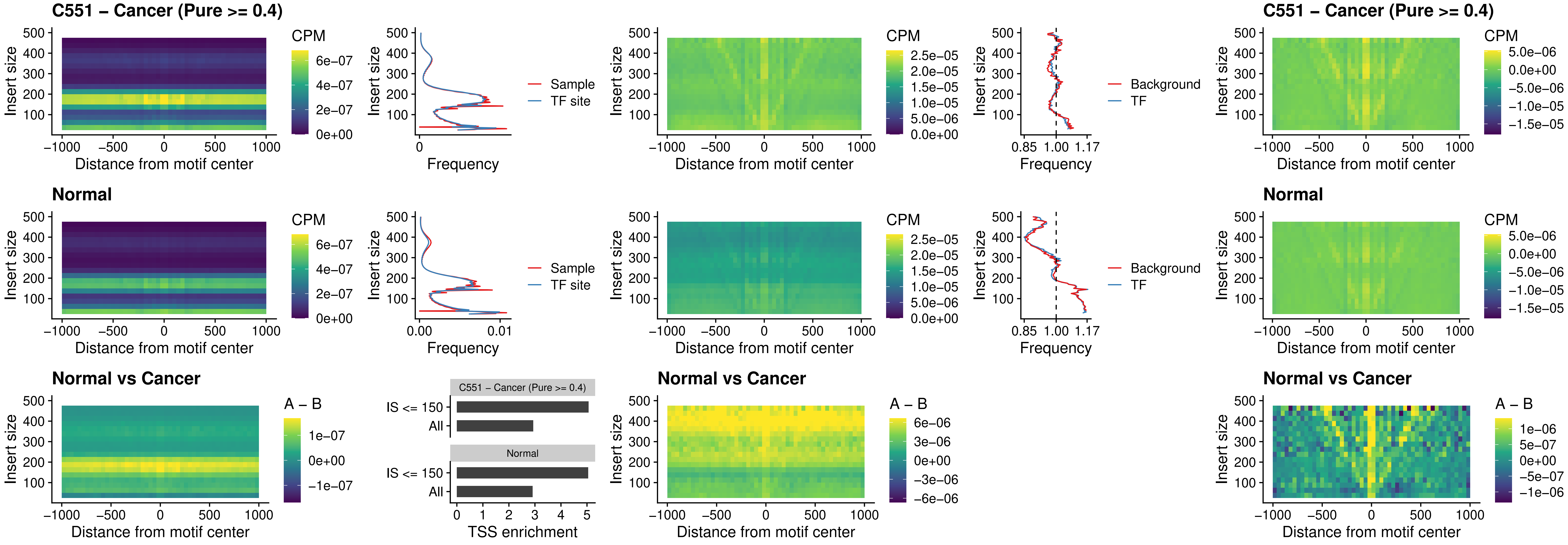

#### C551 – Cancer (Pure >= 0.4) – SPI1_uc_pTSS_nPEAK

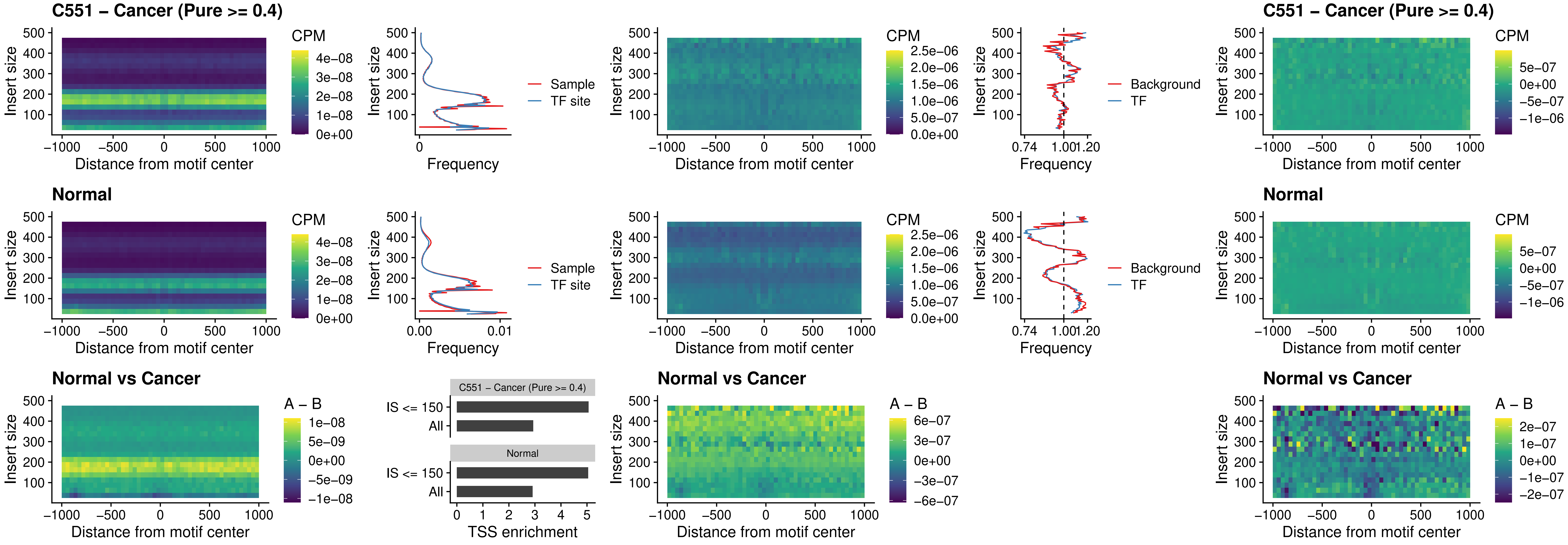

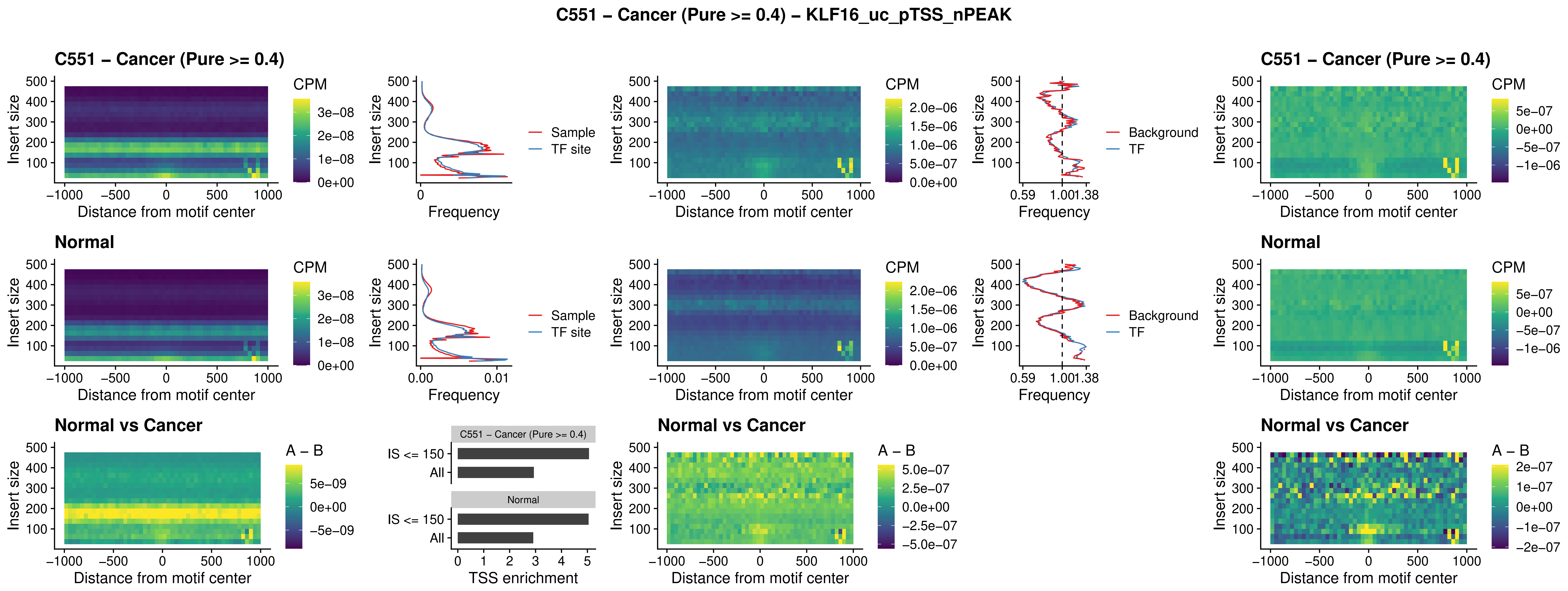
C551 – Cancer (Pure >= 0.4) – KLF16_uc_pTSS_nPEAK
C551 – Cancer (Pure >= 0.4)
Normal
Normal vs Cancer
Insert size
Distance from motif center
CPM
A – B
Frequency
Sample
TF site
Background
TF
TSS enrichment
IS <= 150
All
C551 – Cancer (Pure >= 0.4)
Normal

### C551 – Cancer (Pure >= 0.4) – SP3_uc_pTSS_nPEAK

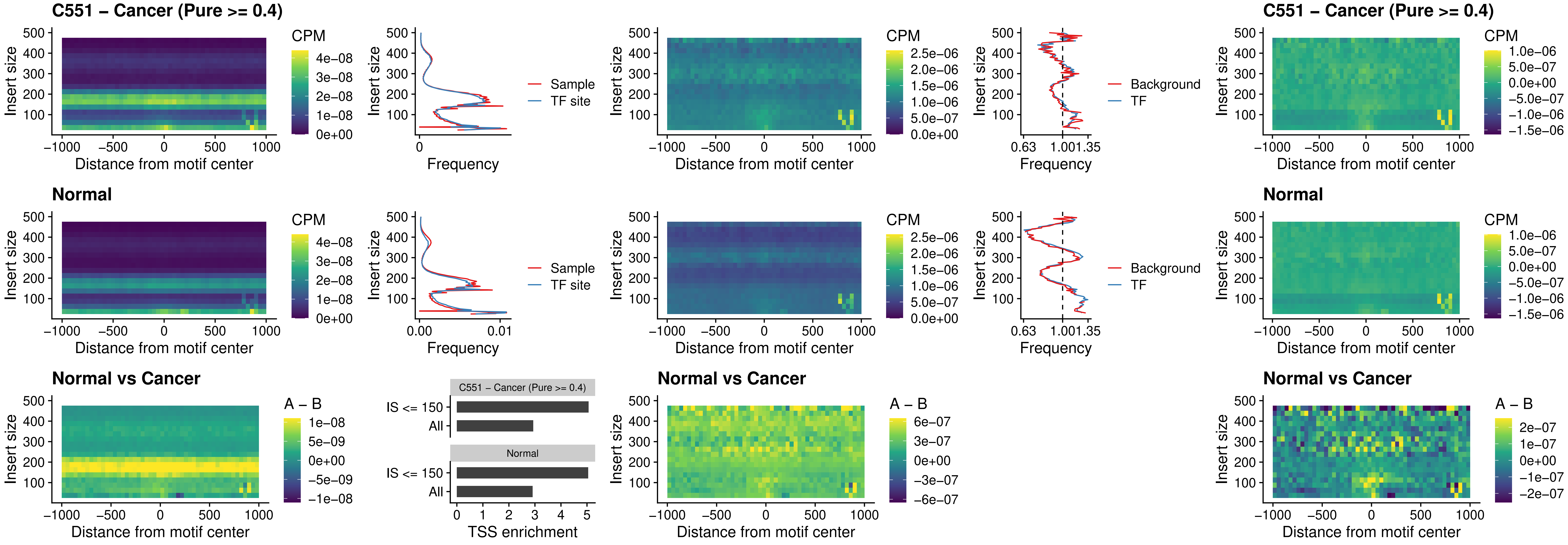

#### C551 – Cancer (Pure >= 0.4) – NEUROD1_uc_dTSS_oPEAK

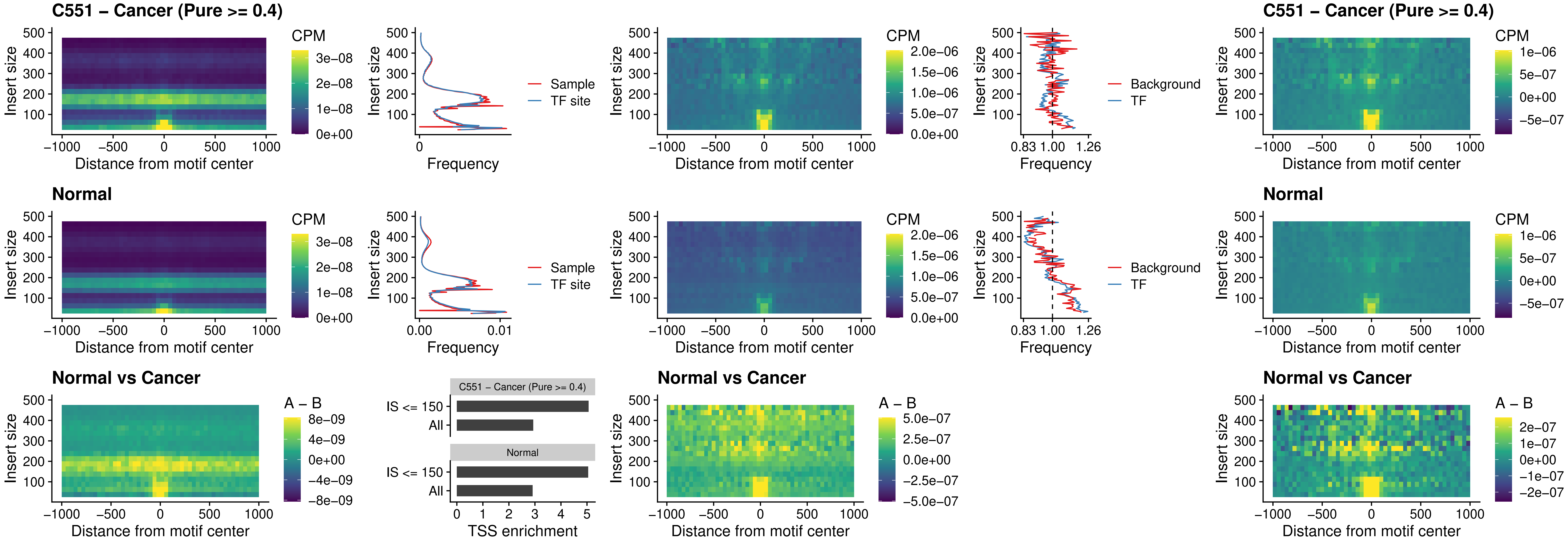

### C551 – Cancer (Pure >= 0.4) – CTCF_uc_dTSS_oPEAK

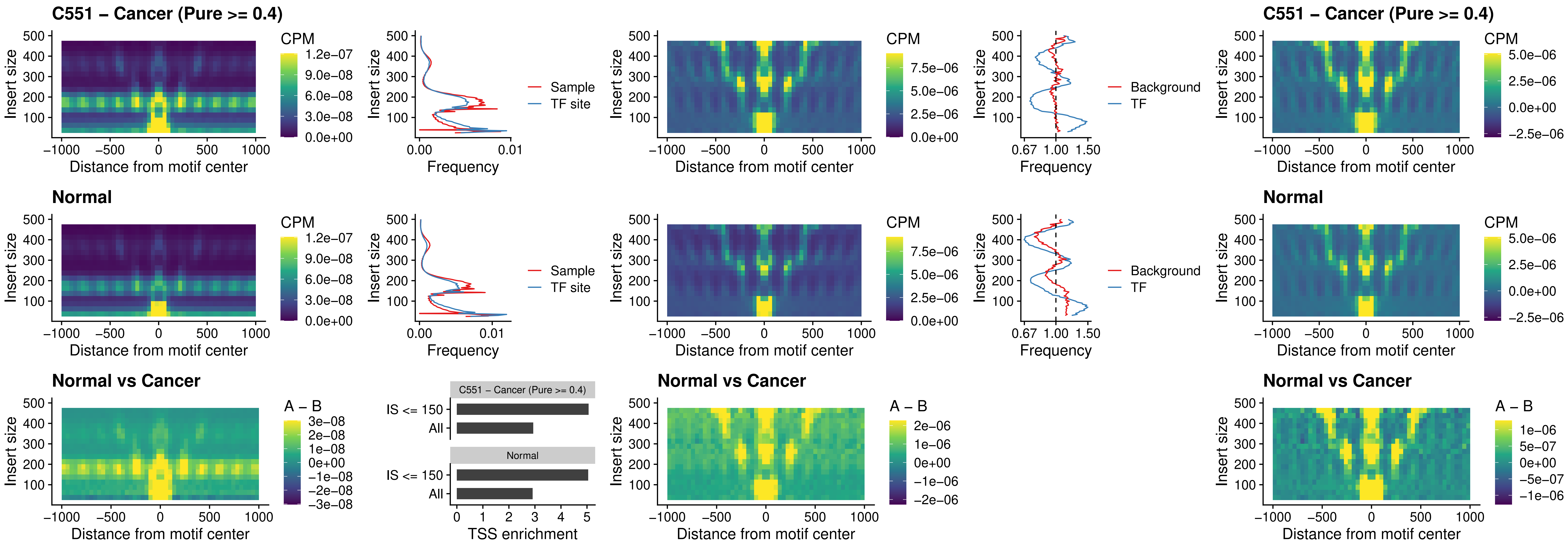

#### C551 – Cancer (Pure >= 0.4) – CTCF_uc_cTSS_oPEAK

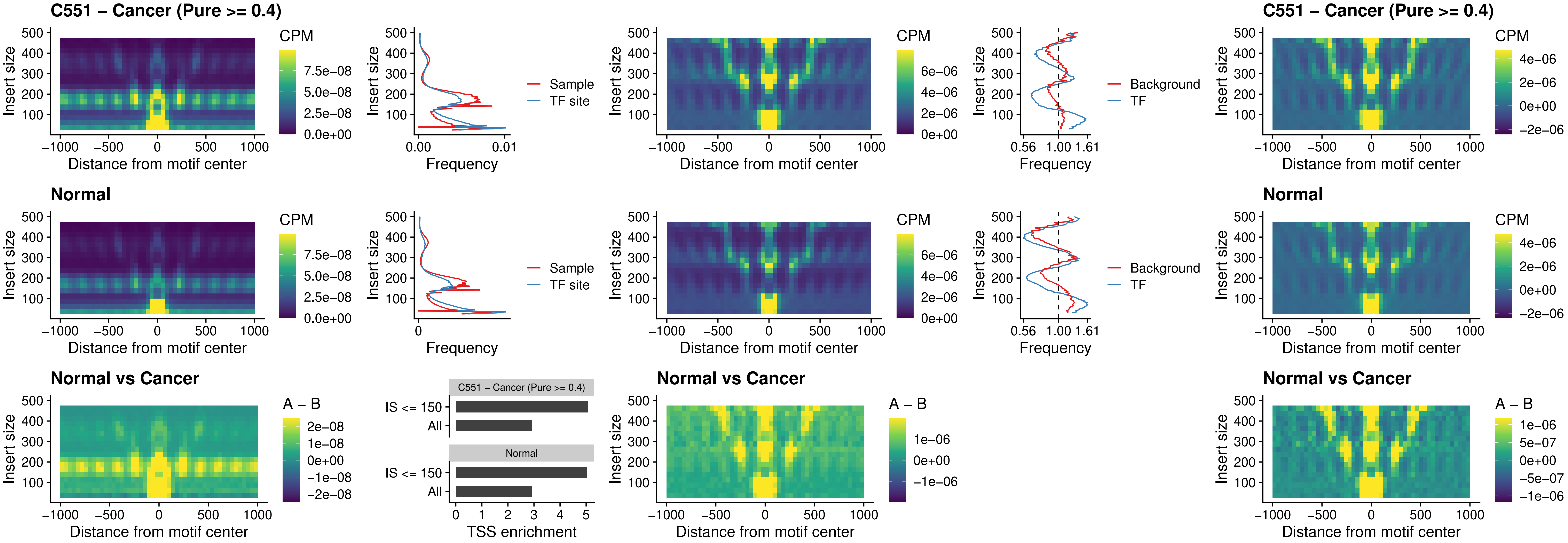

#### C551 – Cancer (Pure >= 0.4) – CTCFL_uc_dTSS_oPEAK

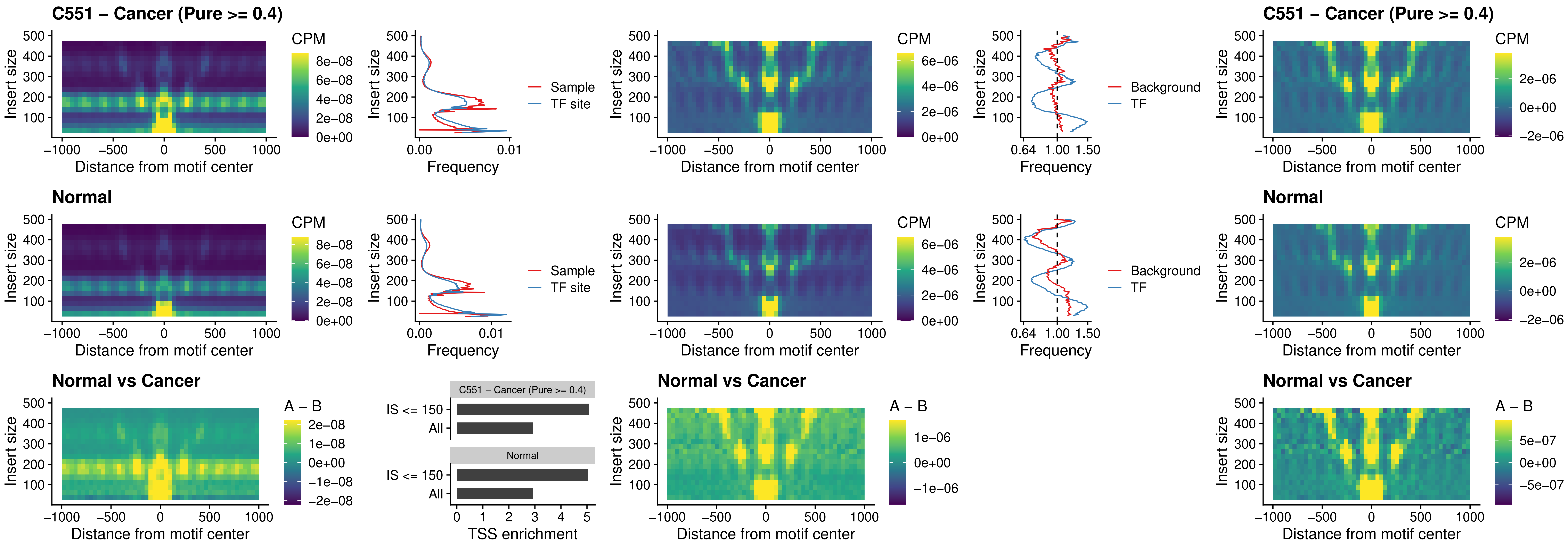

#### C551 – Cancer (Pure >= 0.4) – ENCODE_CTCF_sites_COLON

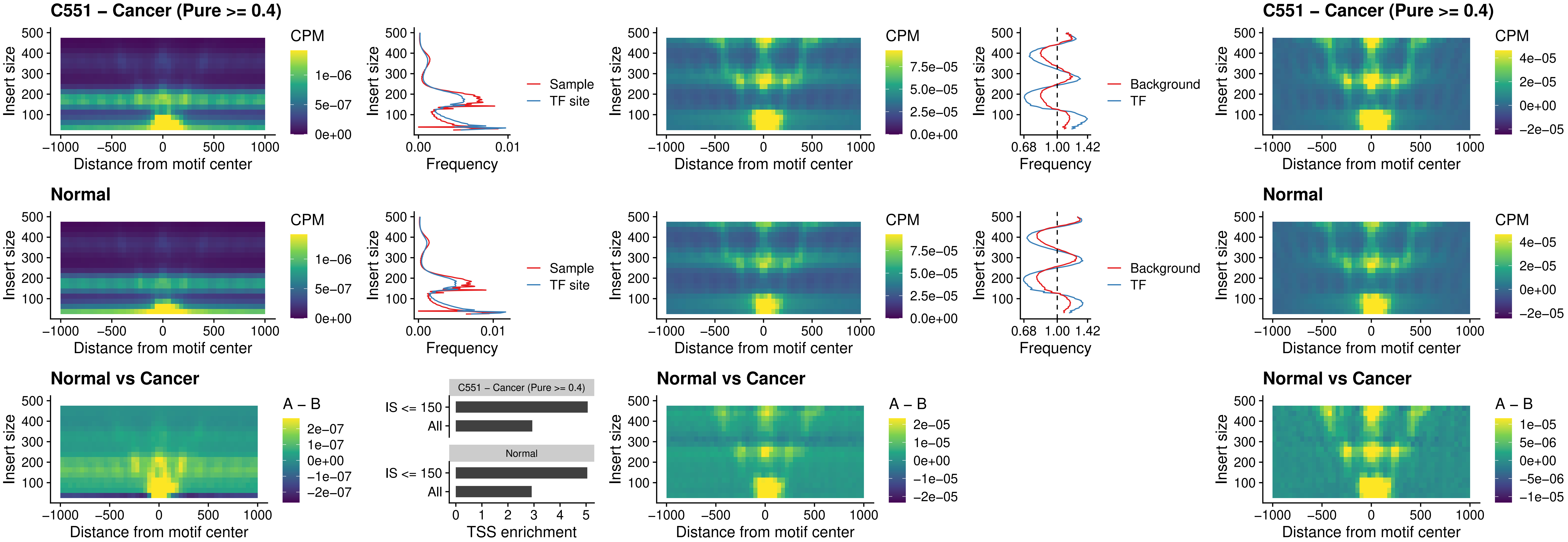

### C551 – Cancer (Pure >= 0.4) – ENCODE_CTCF_sites_COLON_loop_ctcf

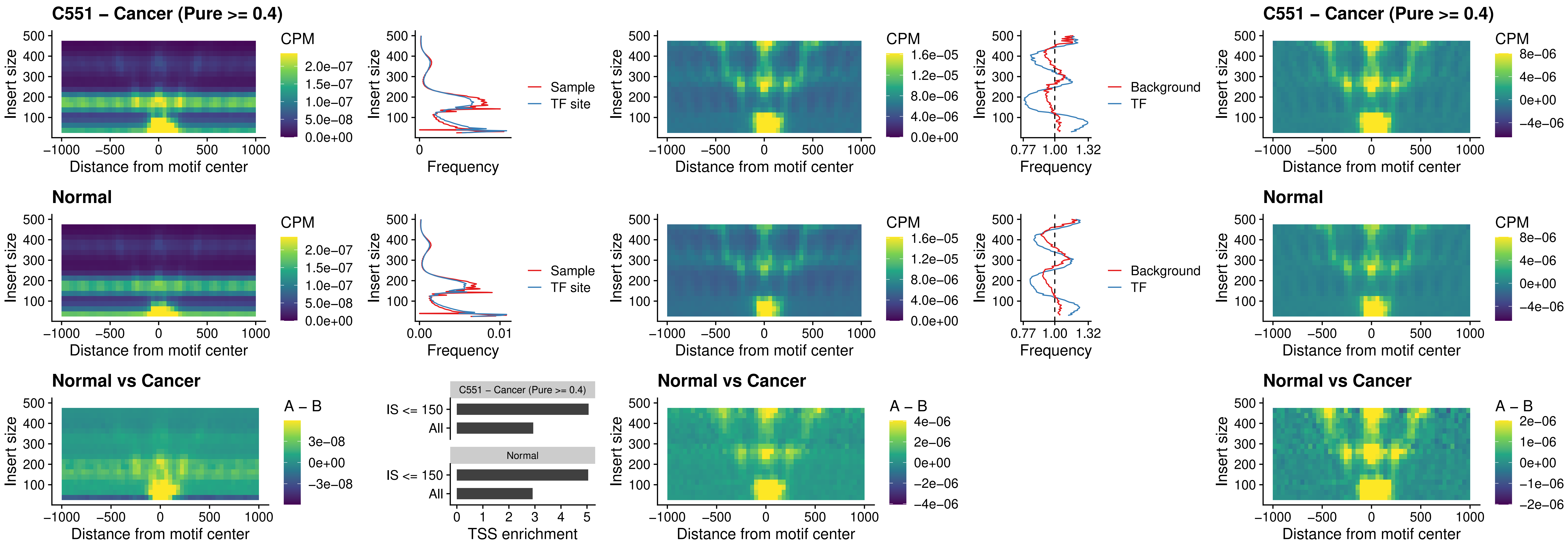

#### C551 – Cancer (Pure >= 0.4) – ENCODE_CTCF_sites_COLON_intergenic_ctcf

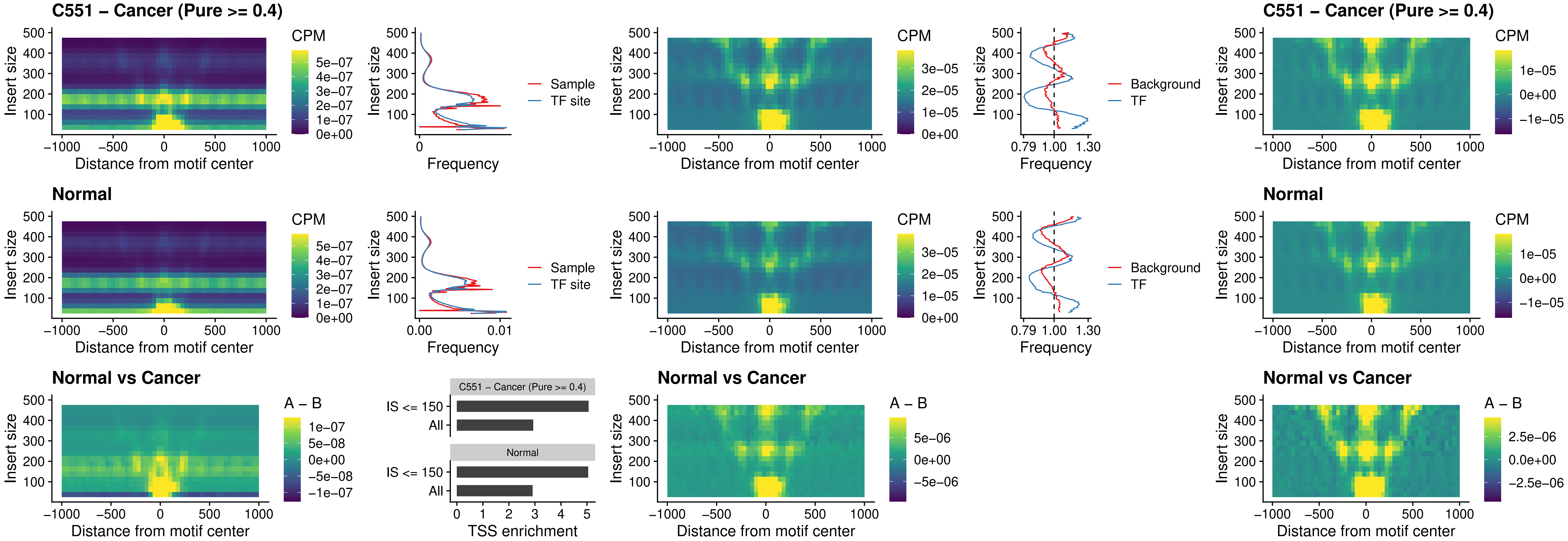

#### C551 – Cancer (Pure >= 0.4) – ENCODE_CTCF_sites_COLON_nonpromotor

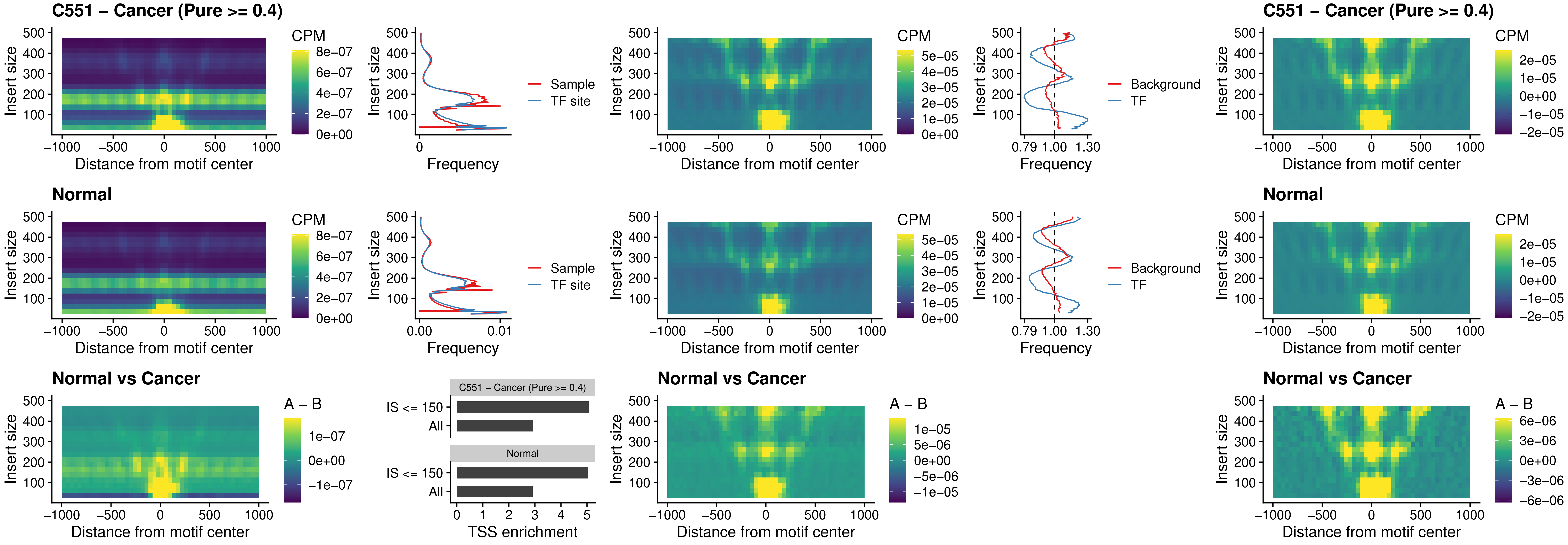

#### C551 – Cancer (Pure >= 0.4) – CTCFL_uc_cTSS_oPEAK

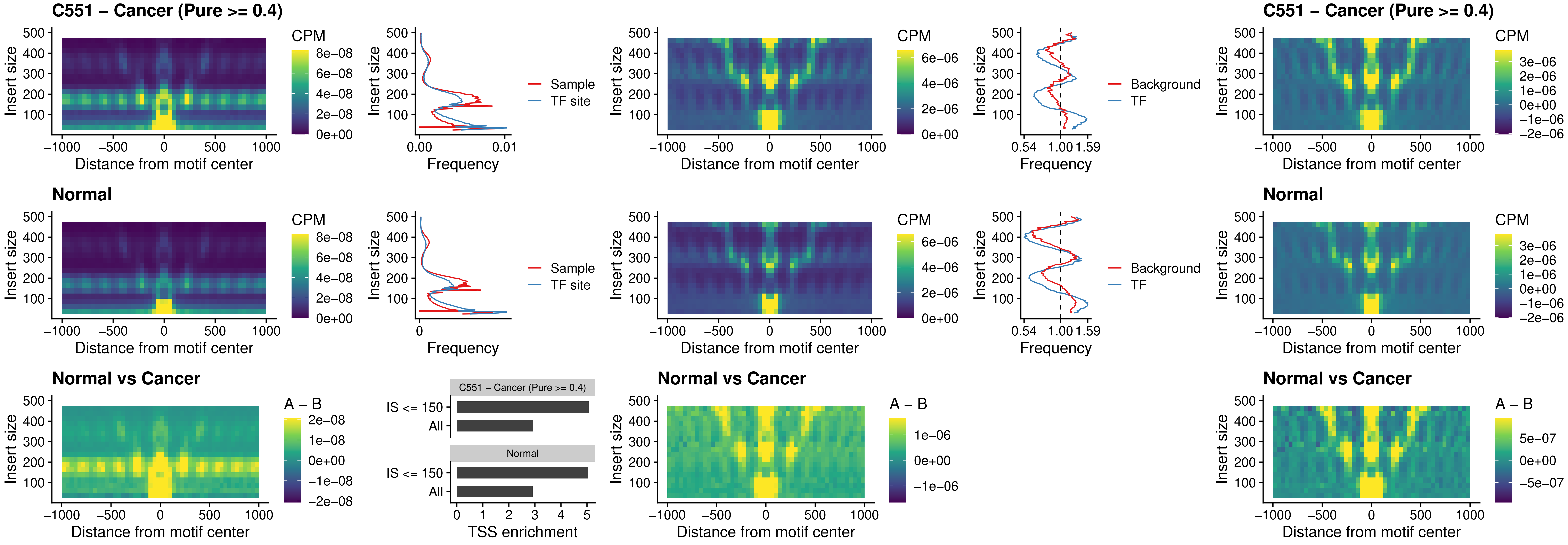

#### C551 – Cancer (Pure >= 0.4) – NEUROD1_uc_cTSS_oPEAK

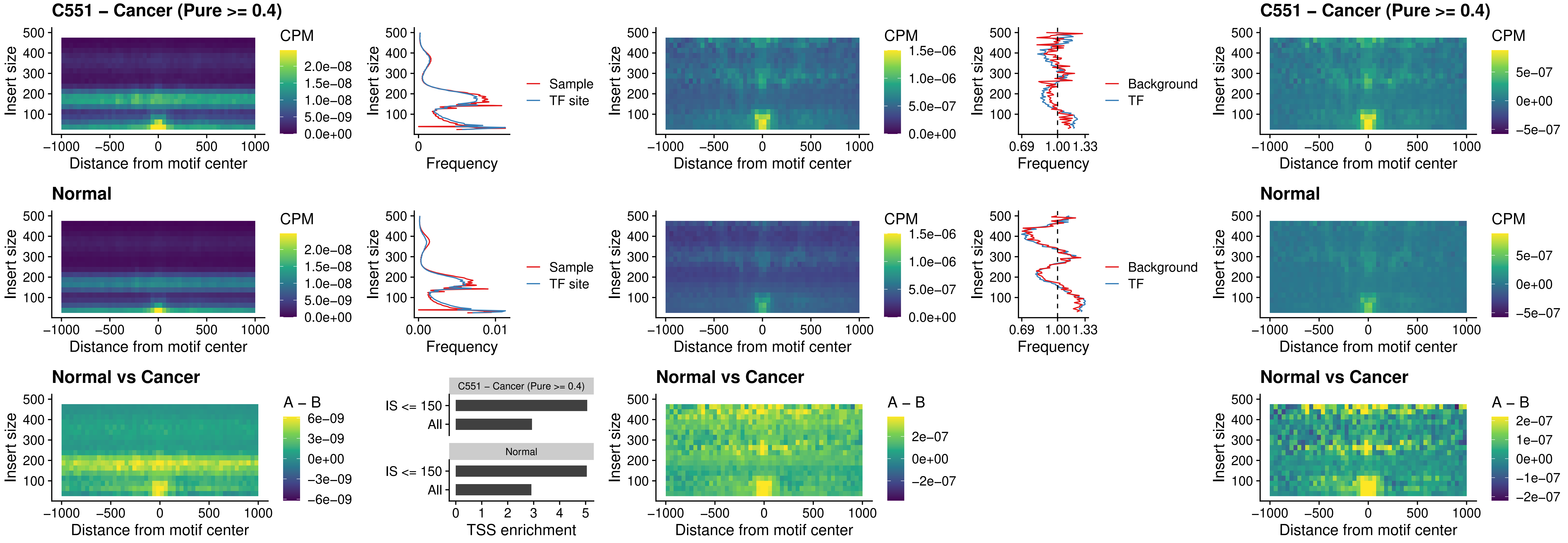

#### C551 – Cancer (Pure >= 0.4) – ENCODE_YY1_sites_COLON_nonpromotor

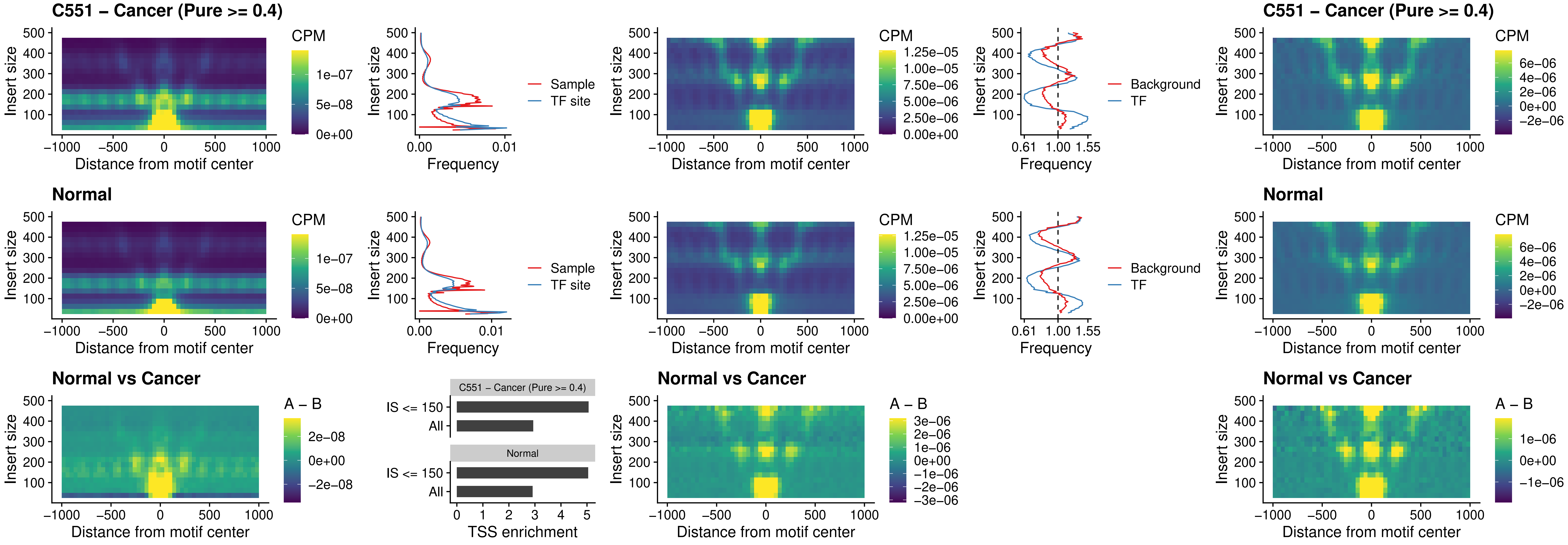

#### C551 – Cancer (Pure >= 0.4) – CTCF_uc_pTSS_oPEAK

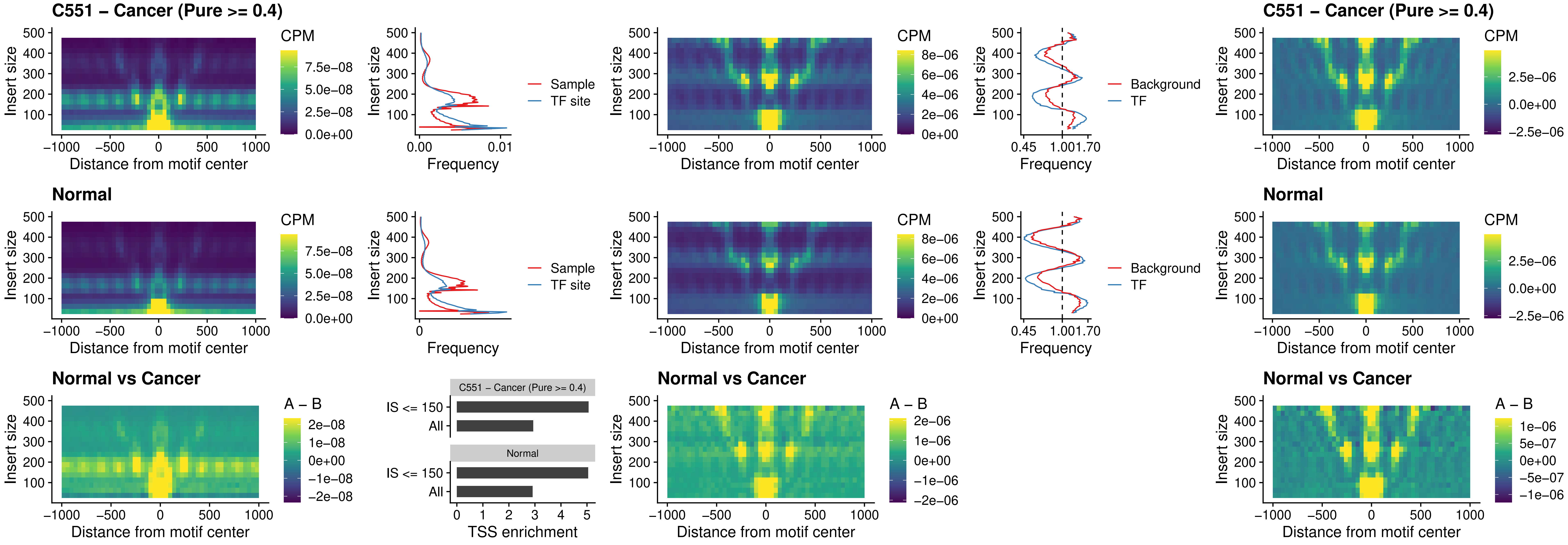

#### C551 – Cancer (Pure >= 0.4) – ENCODE_CTCF_sites_COLON_promotor

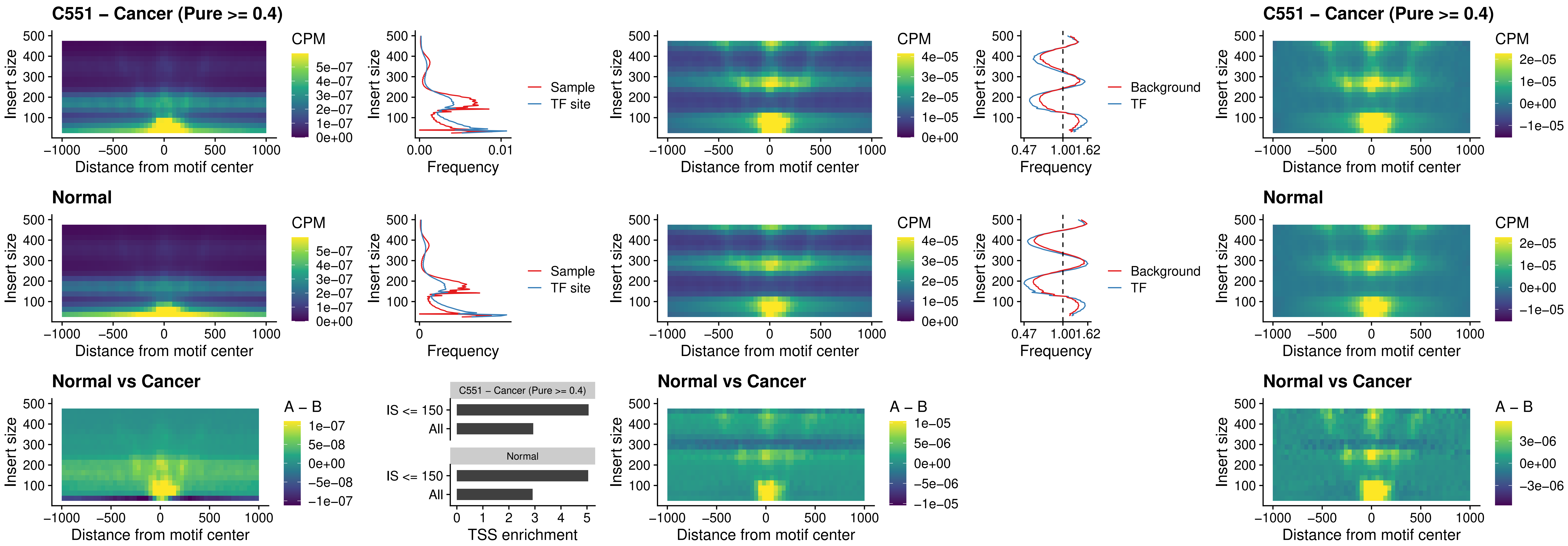

#### C551 – Cancer (Pure >= 0.4) – ENCODE_CTCF_sites_COLON_promotor_ctcf

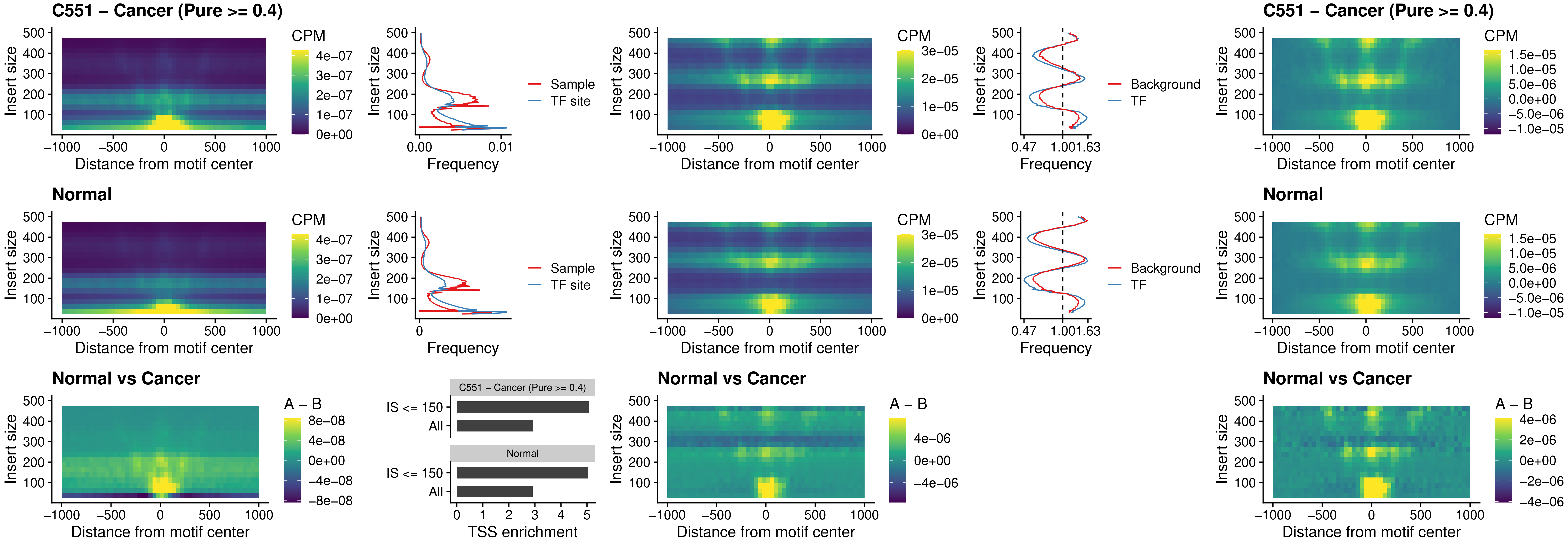

#### C551 – Cancer (Pure >= 0.4) – SPIC_uc_pTSS_oPEAK

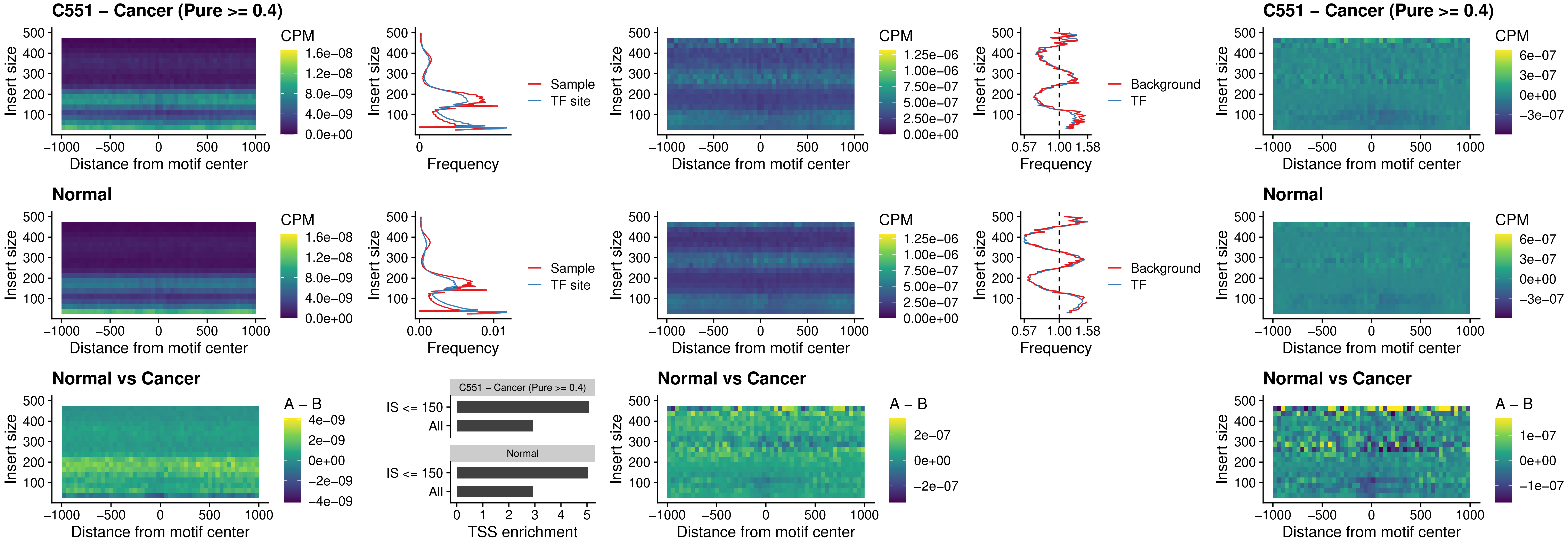

#### C551 – Cancer (Pure >= 0.4) – SPI1_uc_dTSS_nPEAK

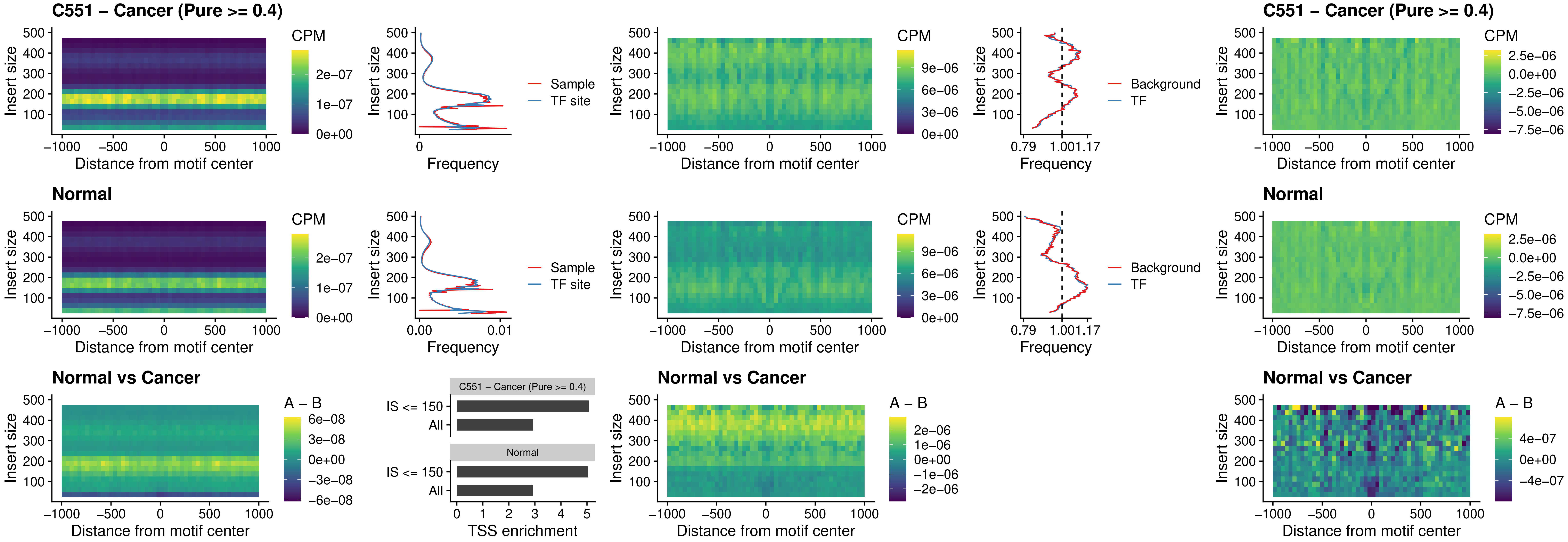

### C551 – Cancer (Pure >= 0.4) – SPI1_uc_cTSS_nPEAK

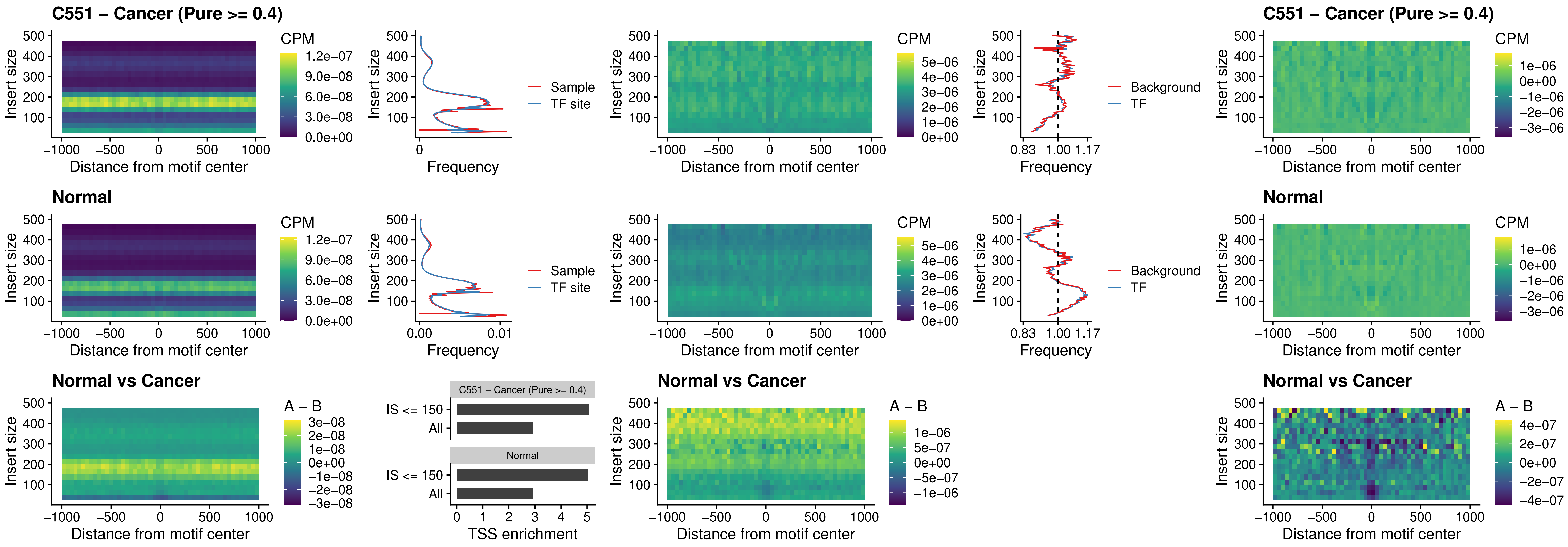

#### C551 – Cancer (Pure >= 0.4) – BCL11A_uc_pTSS_nPEAK

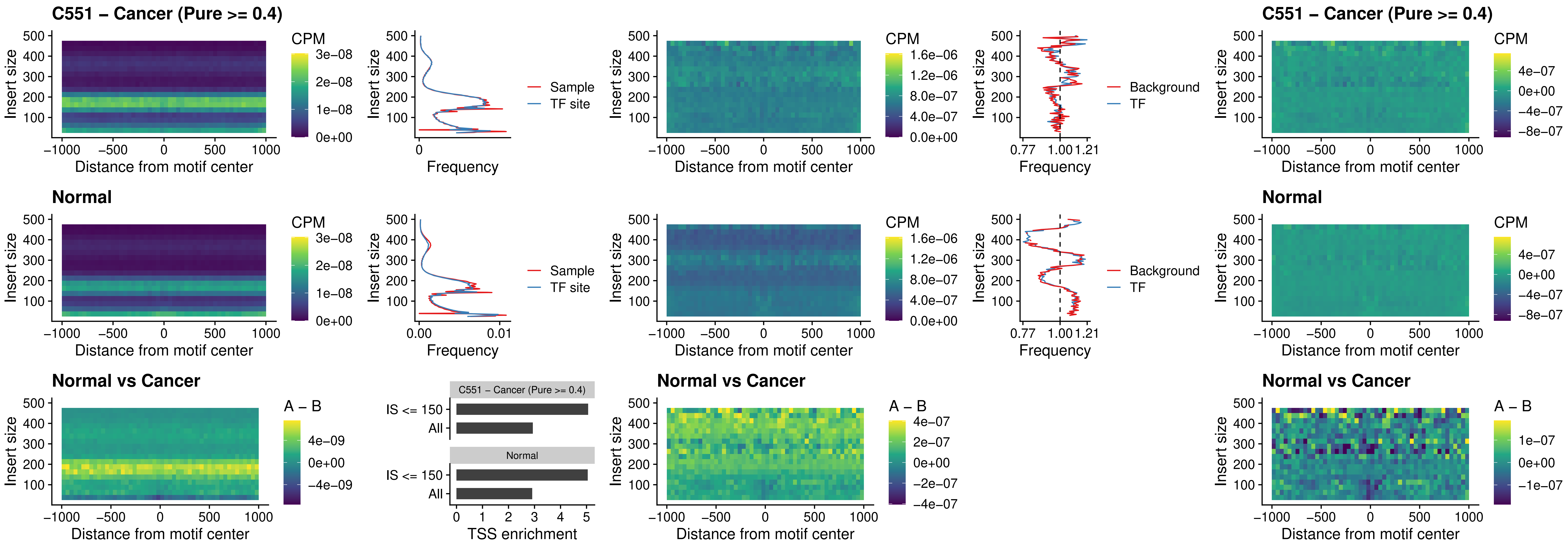

#### C551 – Cancer (Pure >= 0.4) – BCL11B_uc_pTSS_nPEAK

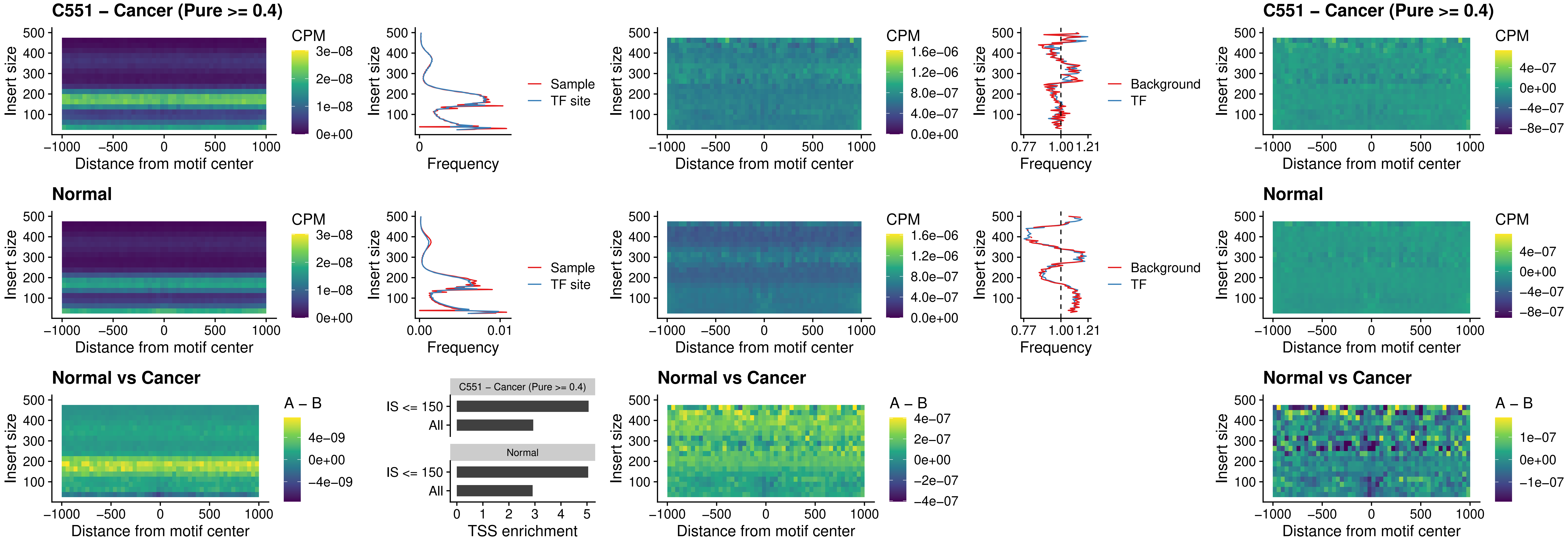

#### C551 – Cancer (Pure >= 0.4) – IRF9_uc_dTSS_oPEAK

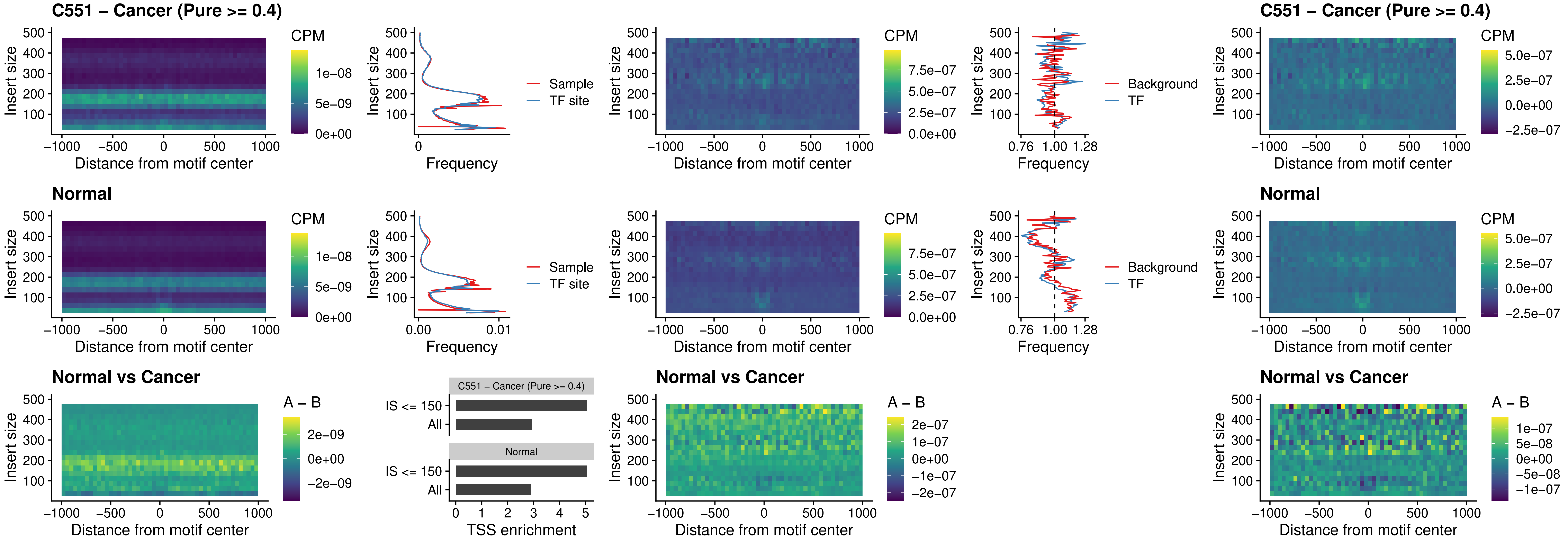

#### C551 – Cancer (Pure >= 0.4) – IRF9_uc_cTSS_oPEAK

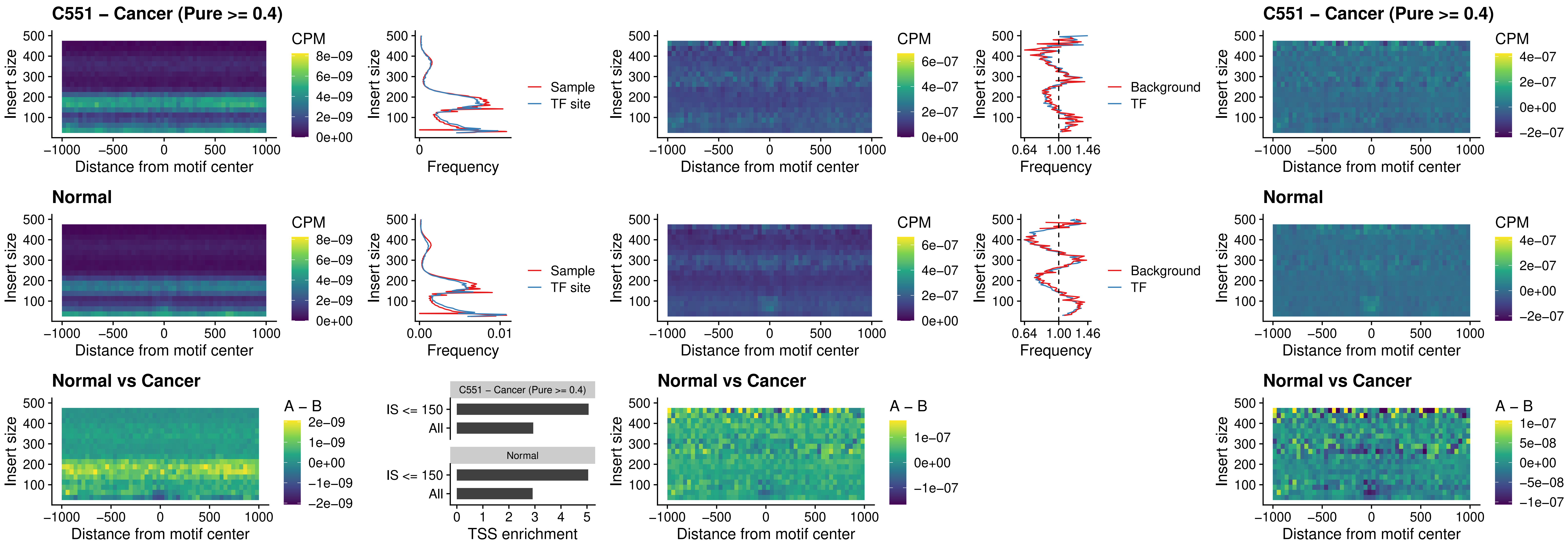

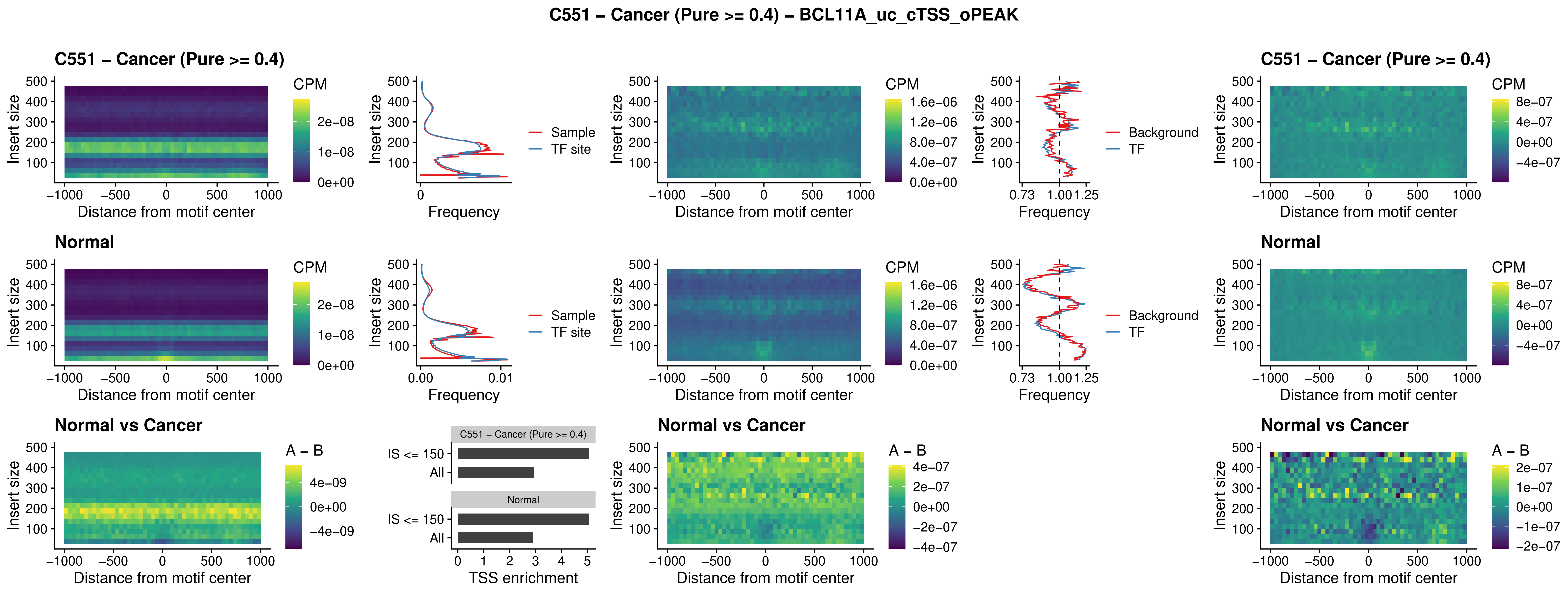
C551 – Cancer (Pure >= 0.4) – BCL11A_uc_cTSS_oPEAK
C551 – Cancer (Pure >= 0.4)
Normal
Normal vs Cancer
Insert size
Distance from motif center
Frequency
CPM
A – B
Sample
TF site
Background
TF
C551 – Cancer (Pure >= 0.4)
Normal
IS <= 150
All
TSS enrichment

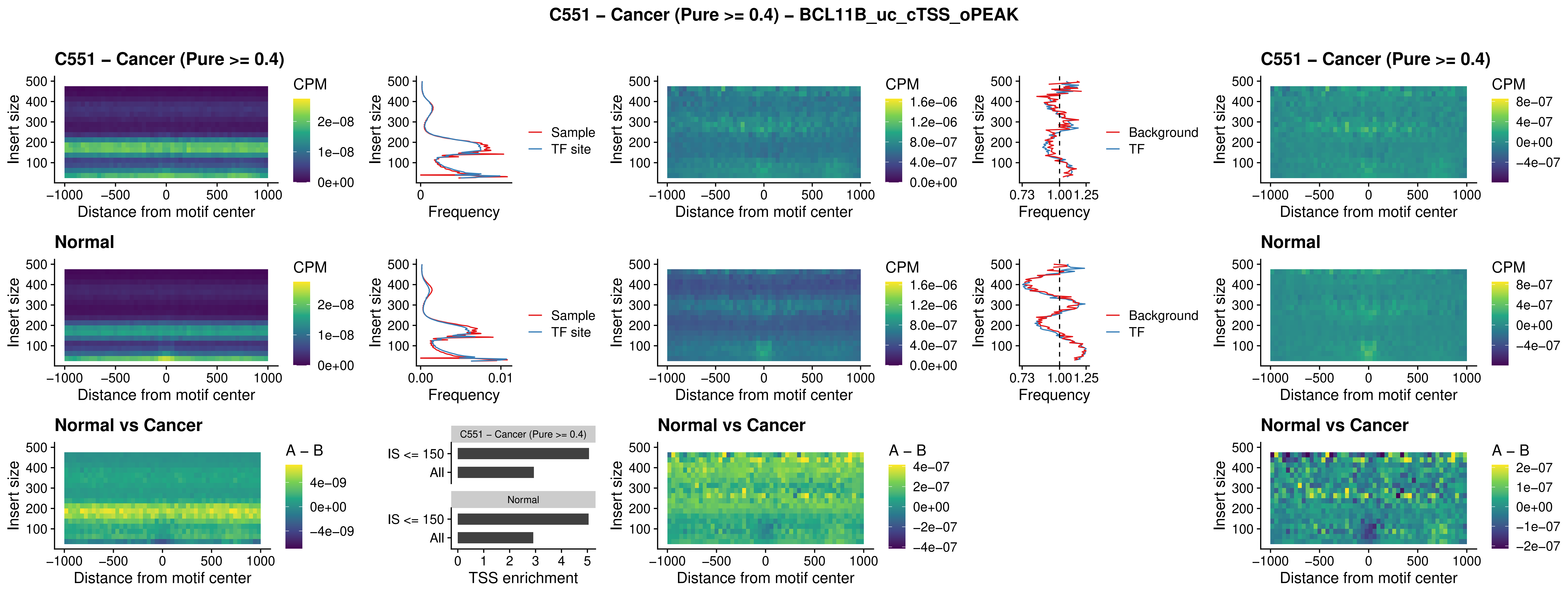
C551 – Cancer (Pure >= 0.4) – BCL11B_uc_cTSS_oPEAK
C551 – Cancer (Pure >= 0.4)
Normal
Normal vs Cancer
Insert size
Distance from motif center
CPM
A – B
Frequency
Sample
TF site
Background
TF
TSS enrichment
C551 – Cancer (Pure >= 0.4)
Normal
IS <= 150
All

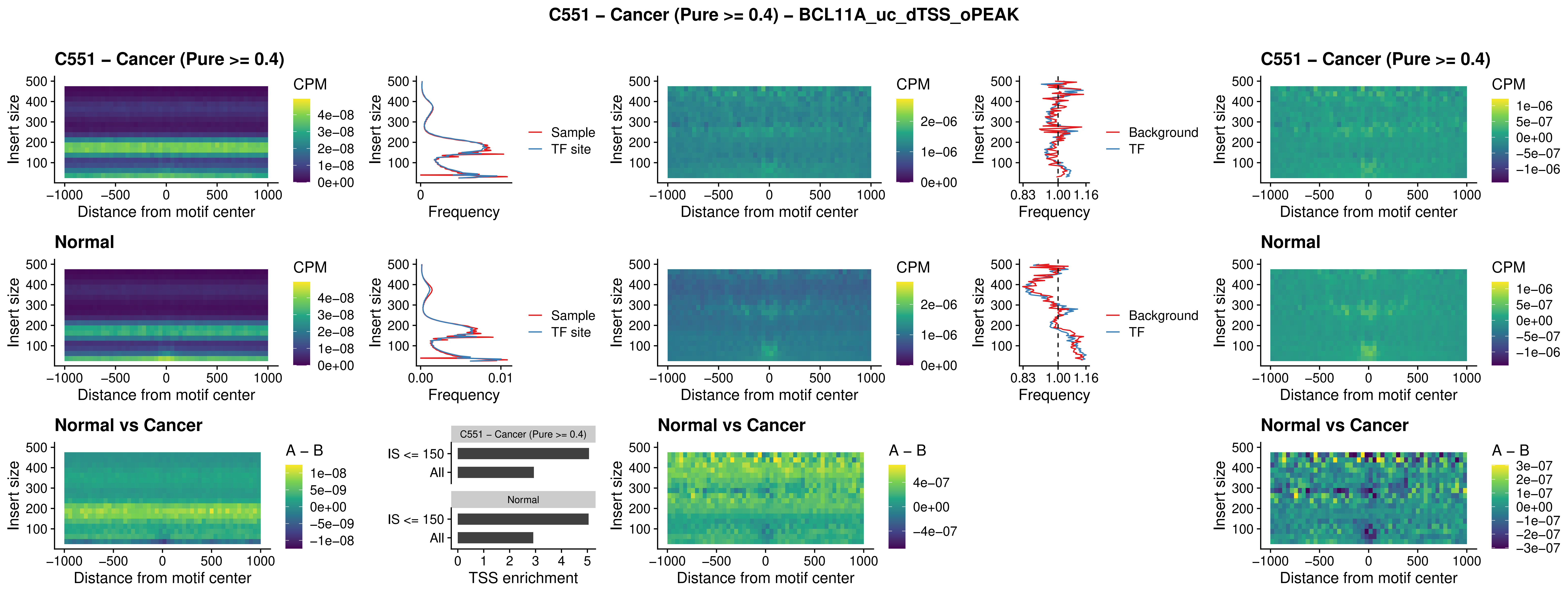
C551 – Cancer (Pure >= 0.4) – BCL11A_uc_dTSS_oPEAK
C551 – Cancer (Pure >= 0.4)
Normal
Normal vs Cancer
Insert size
Distance from motif center
Frequency
CPM
A – B
Sample
TF site
Background
TF
C551 – Cancer (Pure >= 0.4)
Normal
IS <= 150
All
TSS enrichment

#### C551 – Cancer (Pure >= 0.4) – BCL11B_uc_dTSS_oPEAK

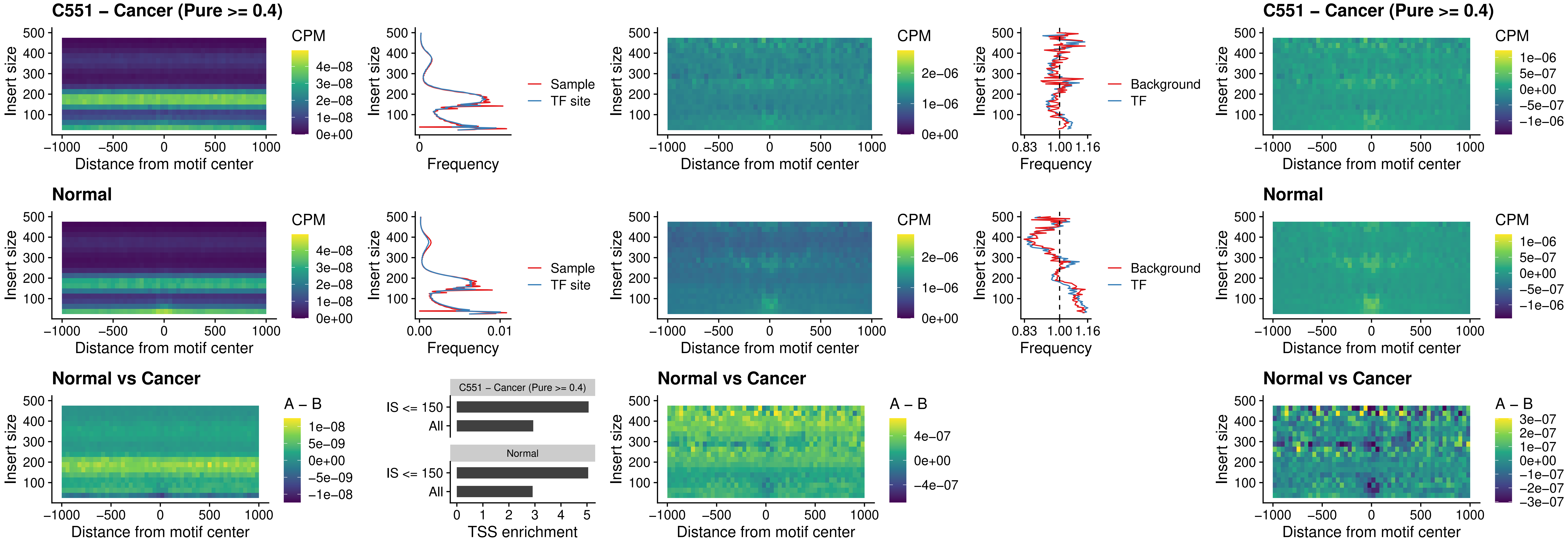

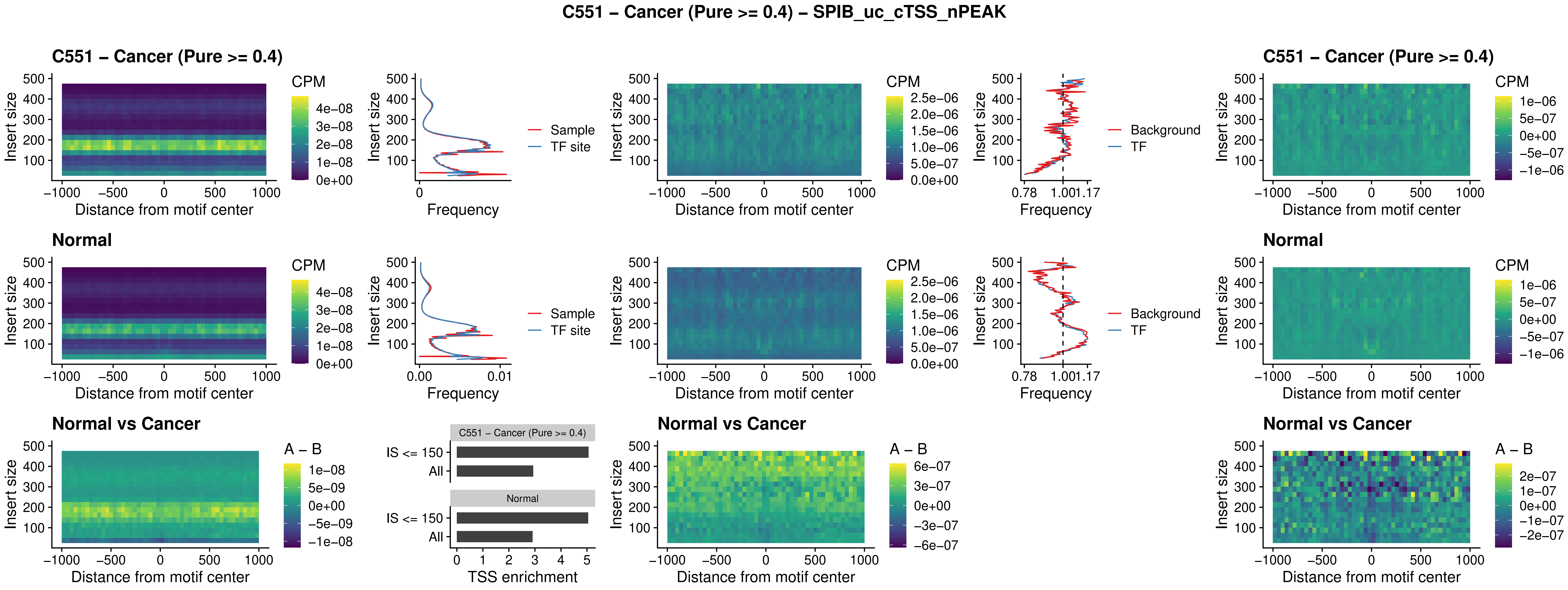
C551 – Cancer (Pure >= 0.4) – SPIB_uc_cTSS_nPEAK
C551 – Cancer (Pure >= 0.4)
Normal
Normal vs Cancer
Insert size
Distance from motif center
Frequency
CPM
A – B
Sample
TF site
Background
TF
C551 – Cancer (Pure >= 0.4)
Normal
IS <= 150
All
TSS enrichment

#### C551 – Cancer (Pure >= 0.4) – SPIC_uc_cTSS_nPEAK

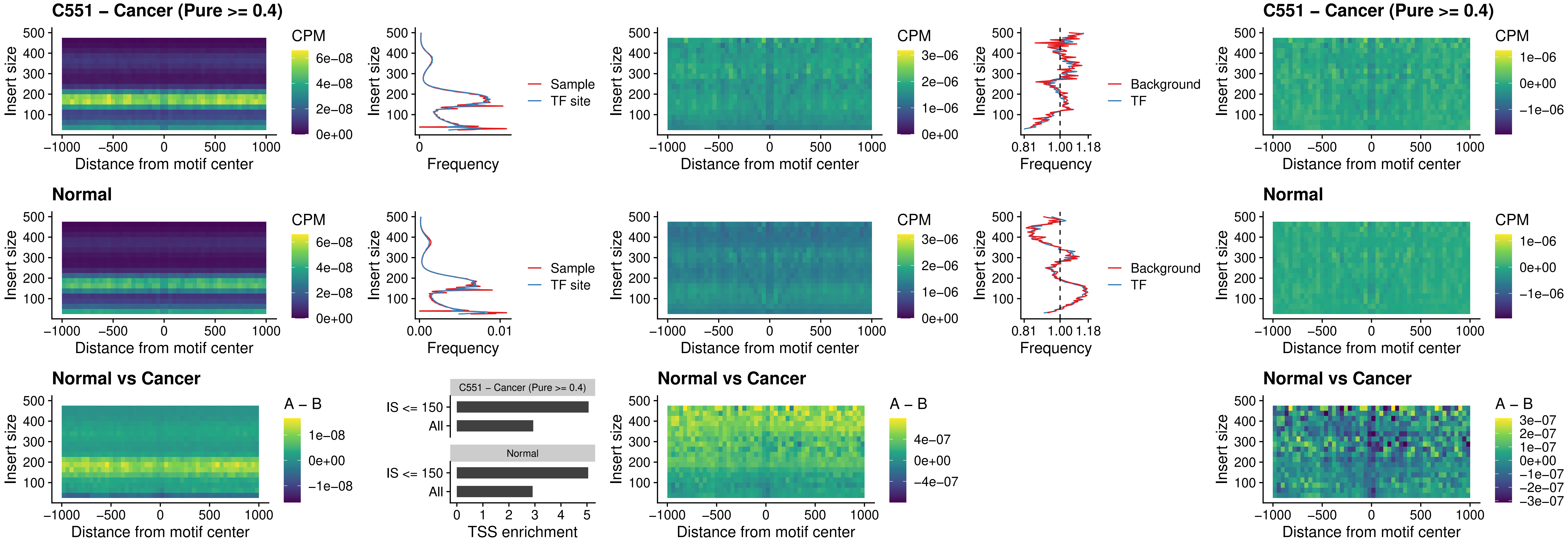

#### C551 – Cancer (Pure >= 0.4) – SPI1_uc_pTSS_oPEAK

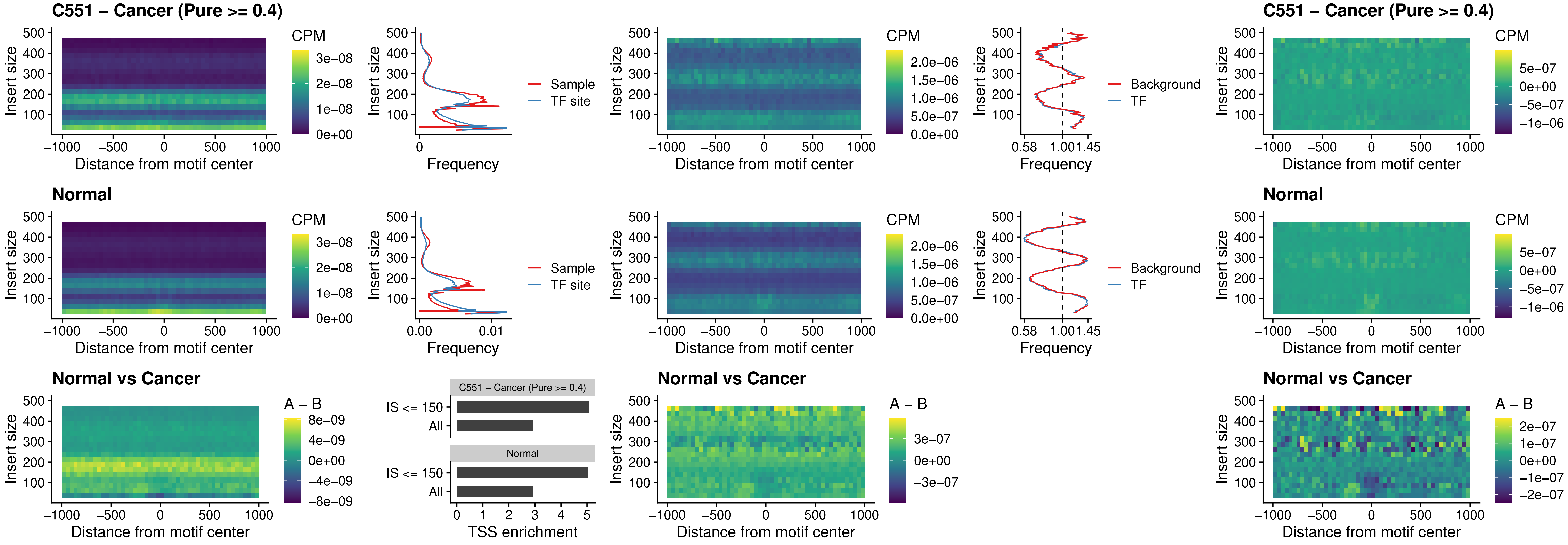

#### C551 – Cancer (Pure >= 0.4) – BCL11A_uc_pTSS_oPEAK

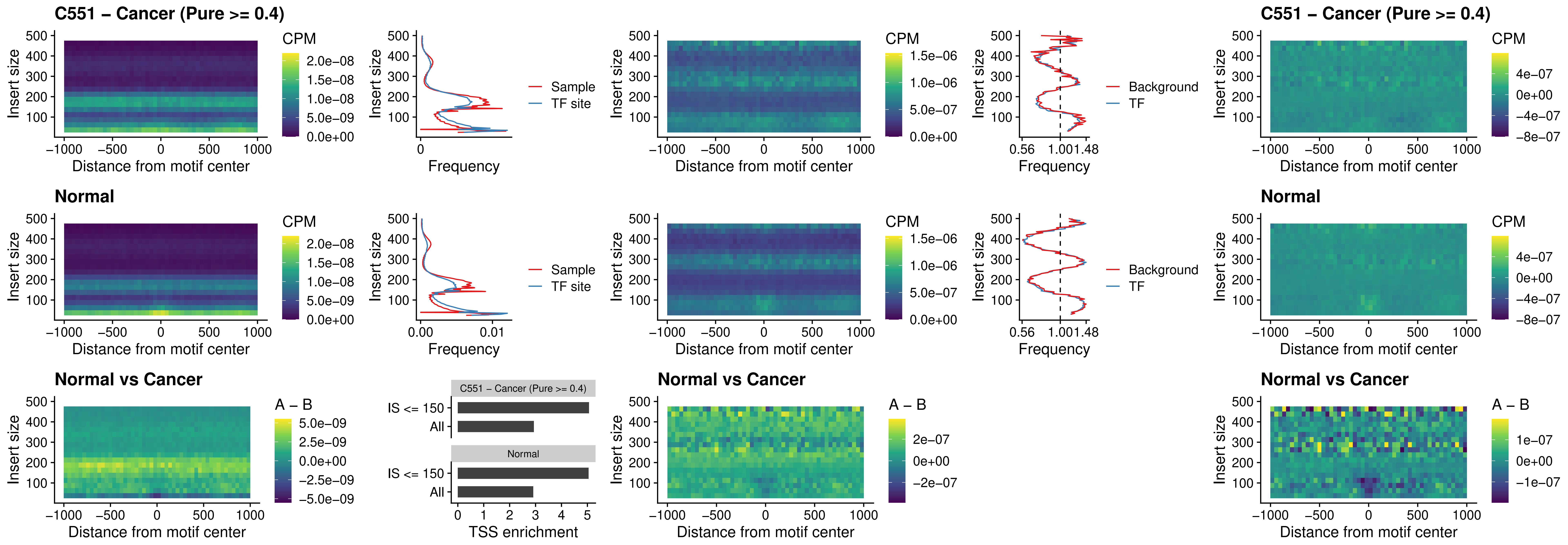

#### C551 – Cancer (Pure >= 0.4) – BCL11B_uc_pTSS_oPEAK

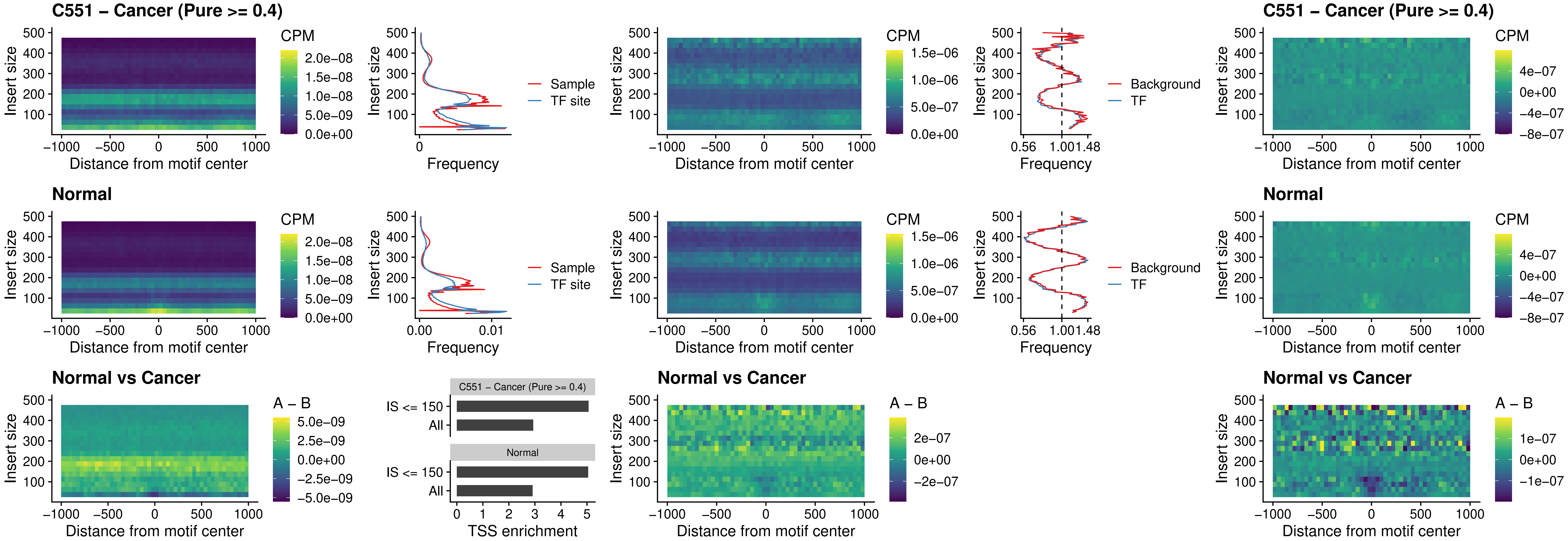

### C551 – Cancer (Pure >= 0.4) – SPI1_uc_cTSS_oPEAK

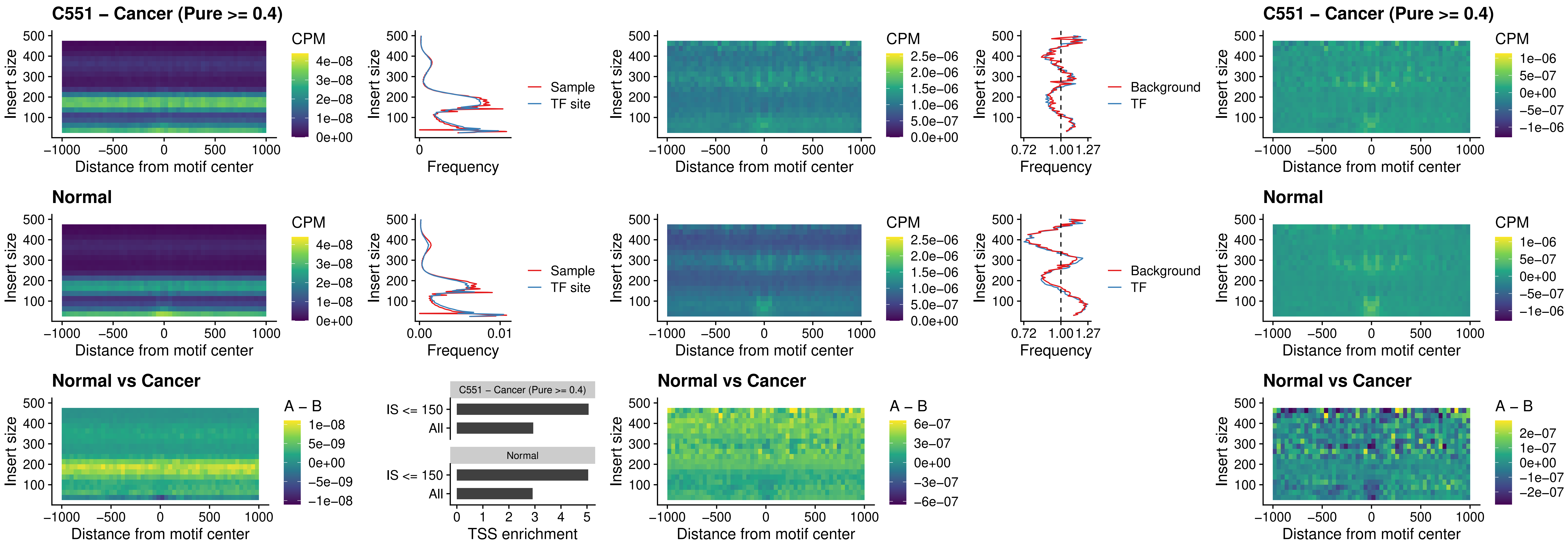

### C551 – Cancer (Pure >= 0.4) – SPIB_uc_cTSS_oPEAK

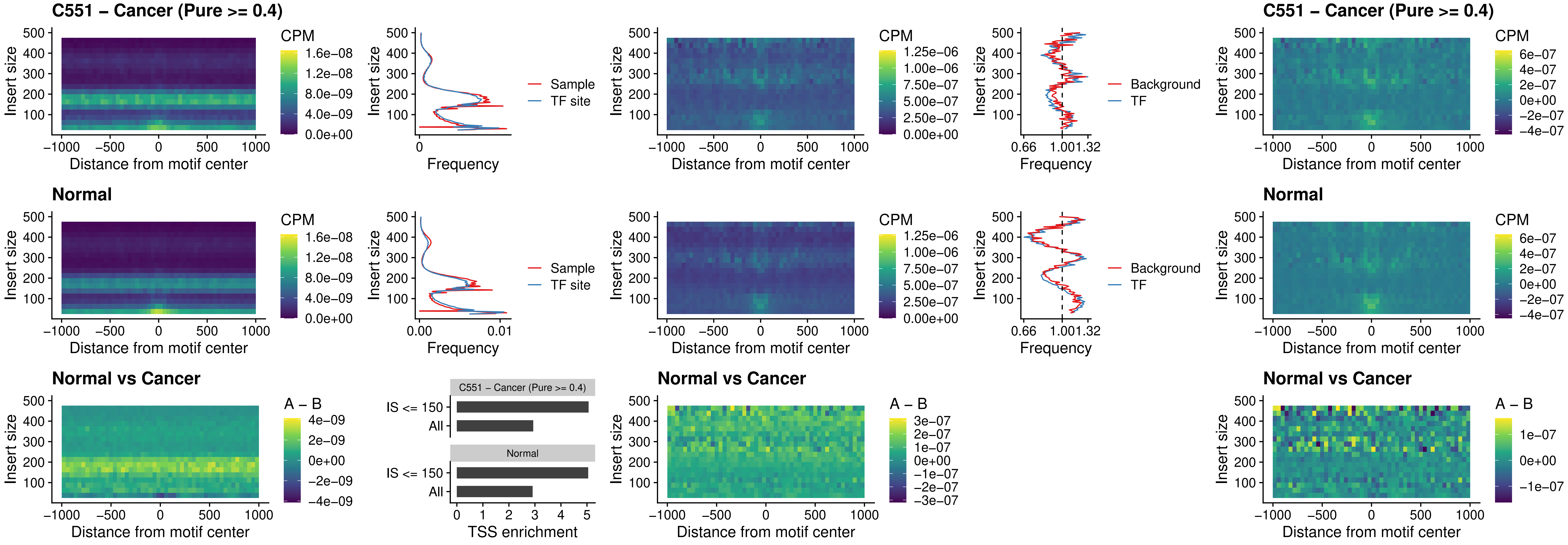

#### C551 – Cancer (Pure >= 0.4) – SPIB_uc_pTSS_oPEAK

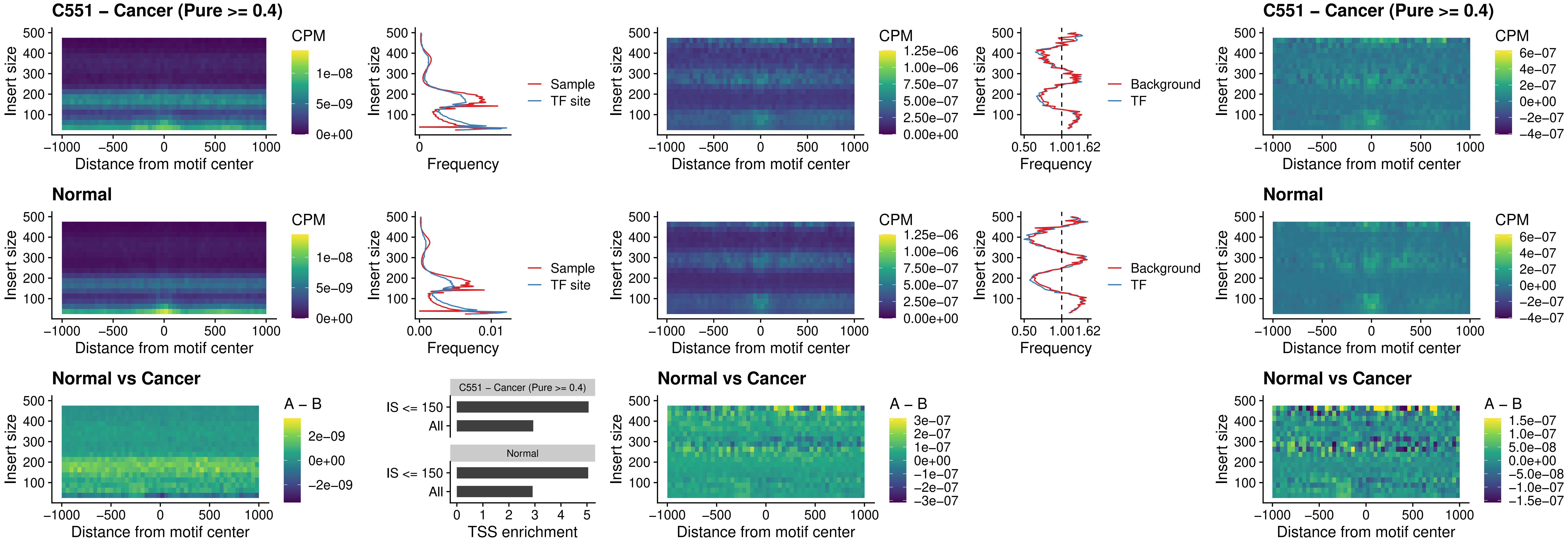

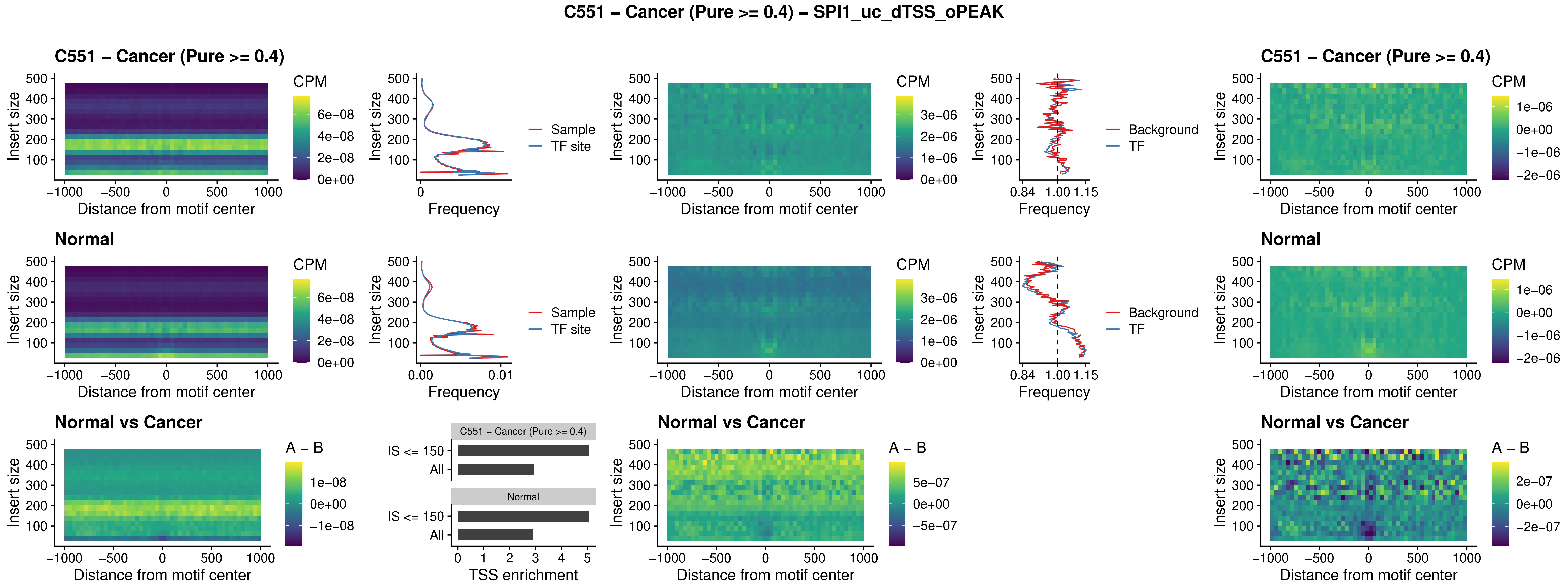
C551 – Cancer (Pure >= 0.4) – SPI1_uc_dTSS_oPEAK
C551 – Cancer (Pure >= 0.4)
Normal
Normal vs Cancer
Insert size
Distance from motif center
Frequency
CPM
A – B
Sample
TF site
Background
TF
C551 – Cancer (Pure >= 0.4)
Normal
IS <= 150
All
TSS enrichment

### C551 – Cancer (Pure >= 0.4) – SPIB_uc_dTSS_oPEAK

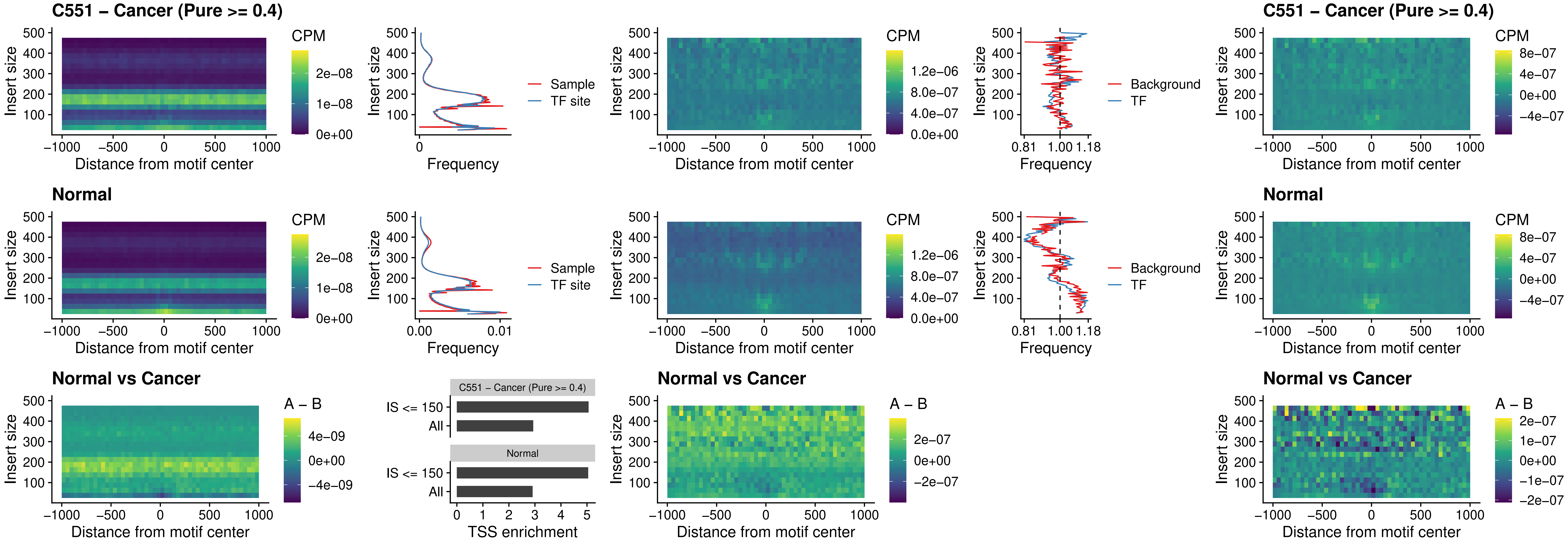

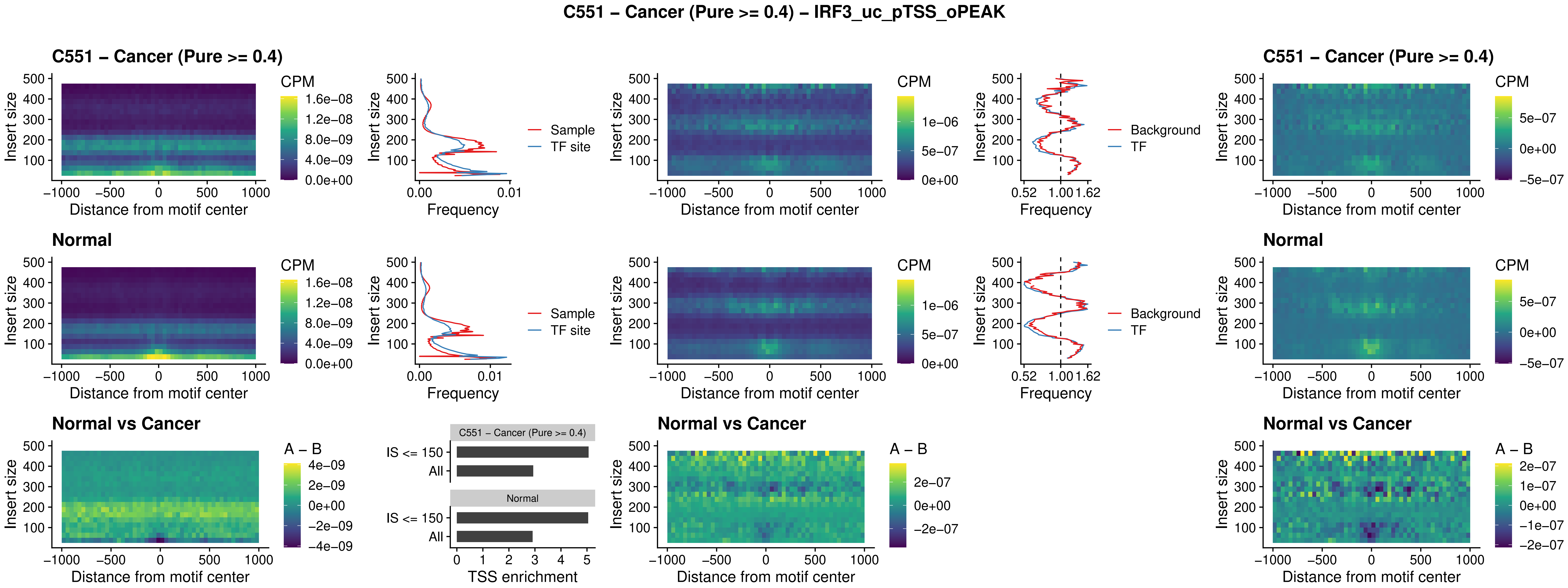
C551 – Cancer (Pure >= 0.4) – IRF3_uc_pTSS_oPEAK
C551 – Cancer (Pure >= 0.4)
Normal
Normal vs Cancer
Insert size
Distance from motif center
CPM
A – B
Frequency
Sample
TF site
Background
TF
C551 – Cancer (Pure >= 0.4)
Normal
IS <= 150
All
TSS enrichment

### C551 – Cancer (Pure >= 0.4) – IRF2_uc_dTSS_oPEAK

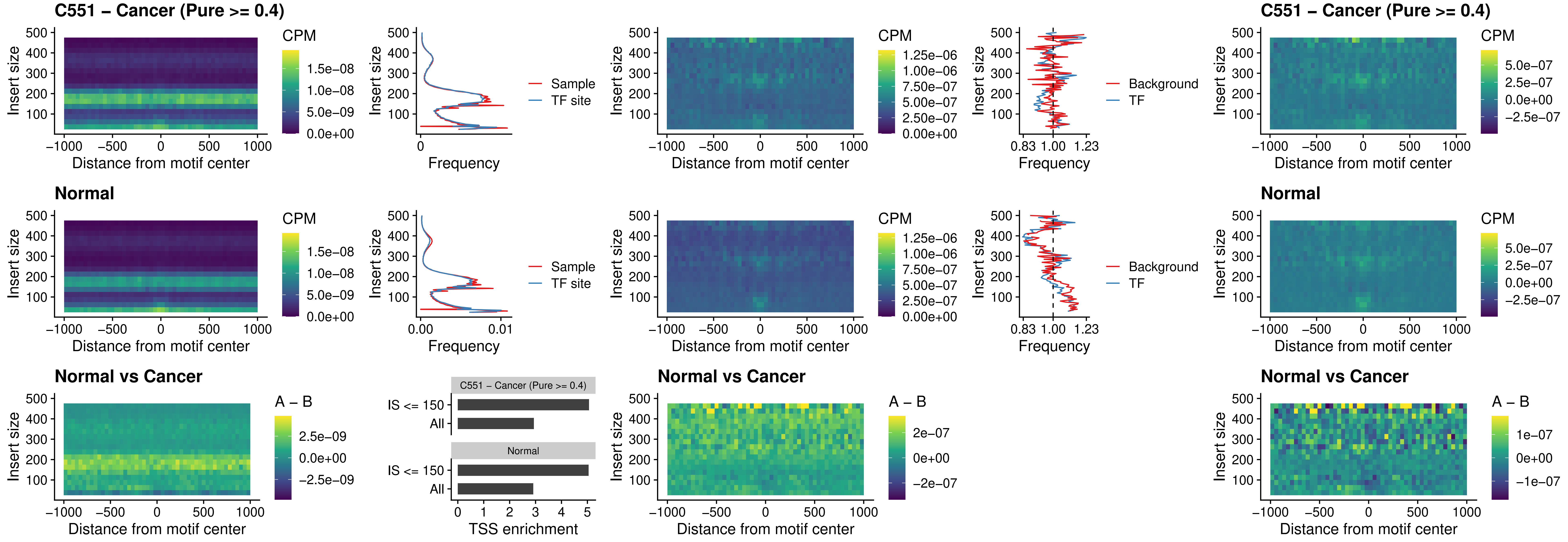

#### C551 – Cancer (Pure >= 0.4) – IRF2_uc_pTSS_oPEAK

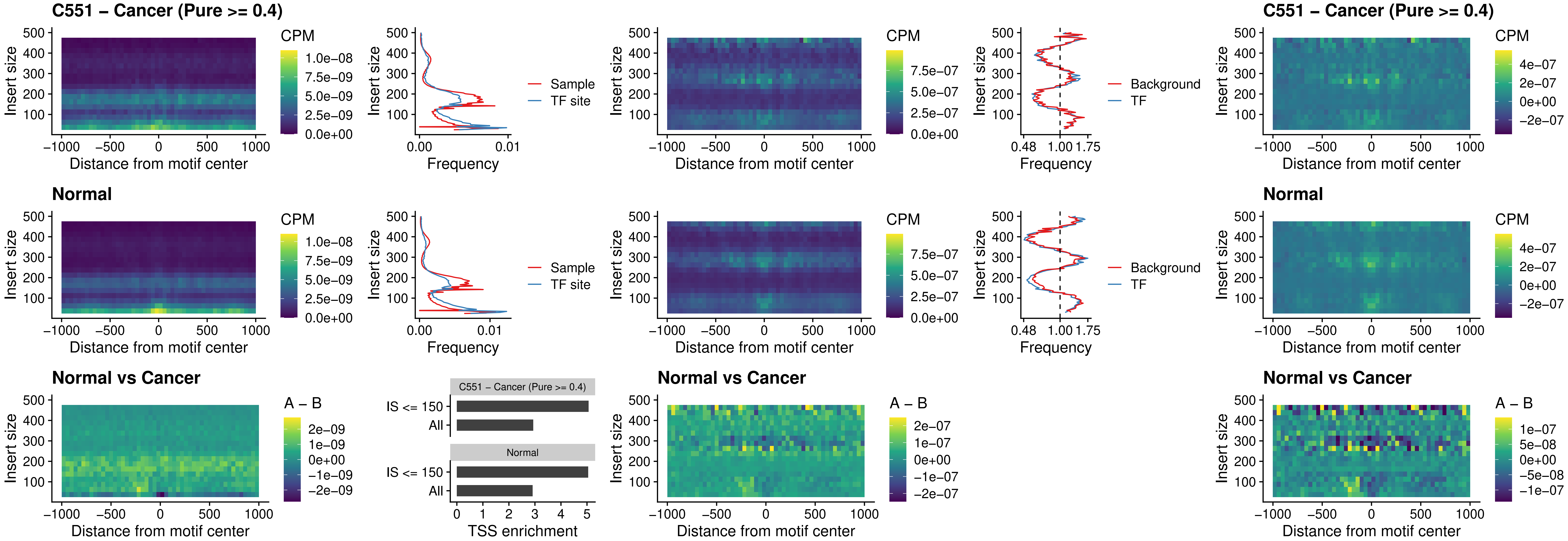

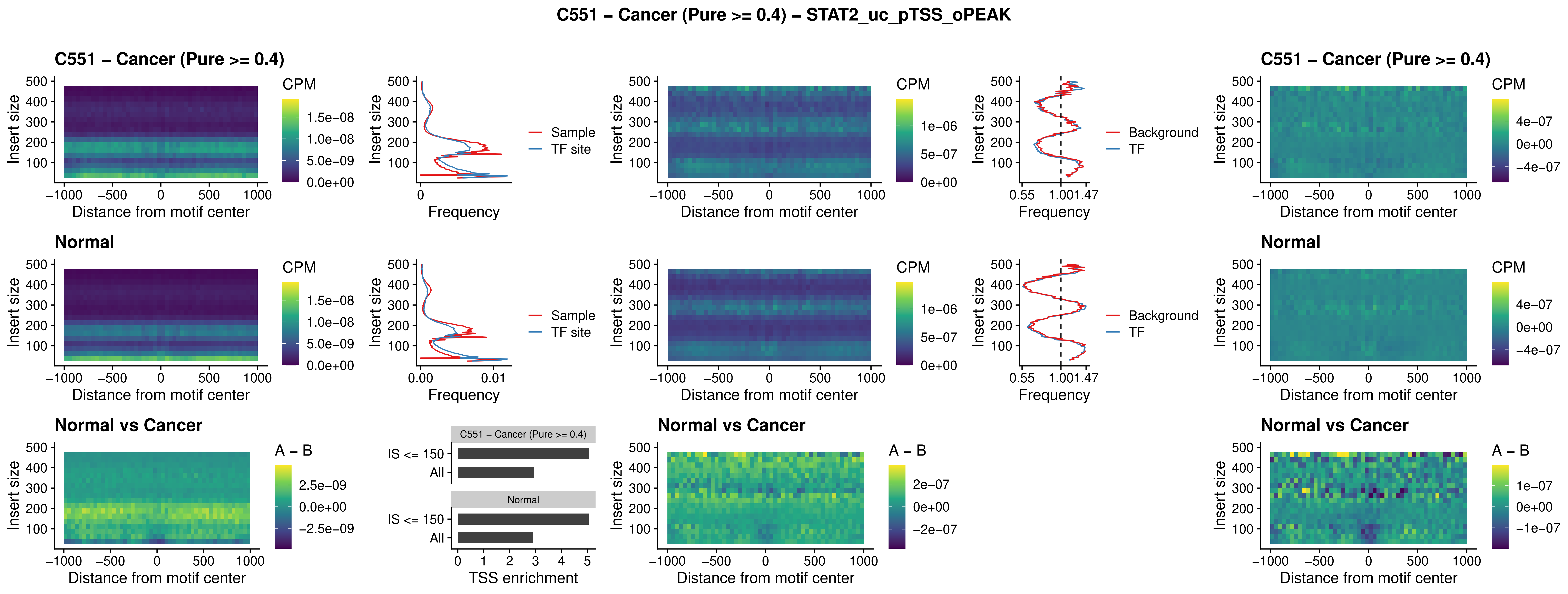
C551 – Cancer (Pure >= 0.4) – STAT2_uc_pTSS_oPEAK
C551 – Cancer (Pure >= 0.4)
Normal
Normal vs Cancer
Insert size
Distance from motif center
Frequency
CPM
A – B
Sample
TF site
Background
TF
C551 – Cancer (Pure >= 0.4)
Normal
IS <= 150
All
TSS enrichment

#### C551 – Cancer (Pure >= 0.4) – SPIB_uc_dTSS_nPEAK

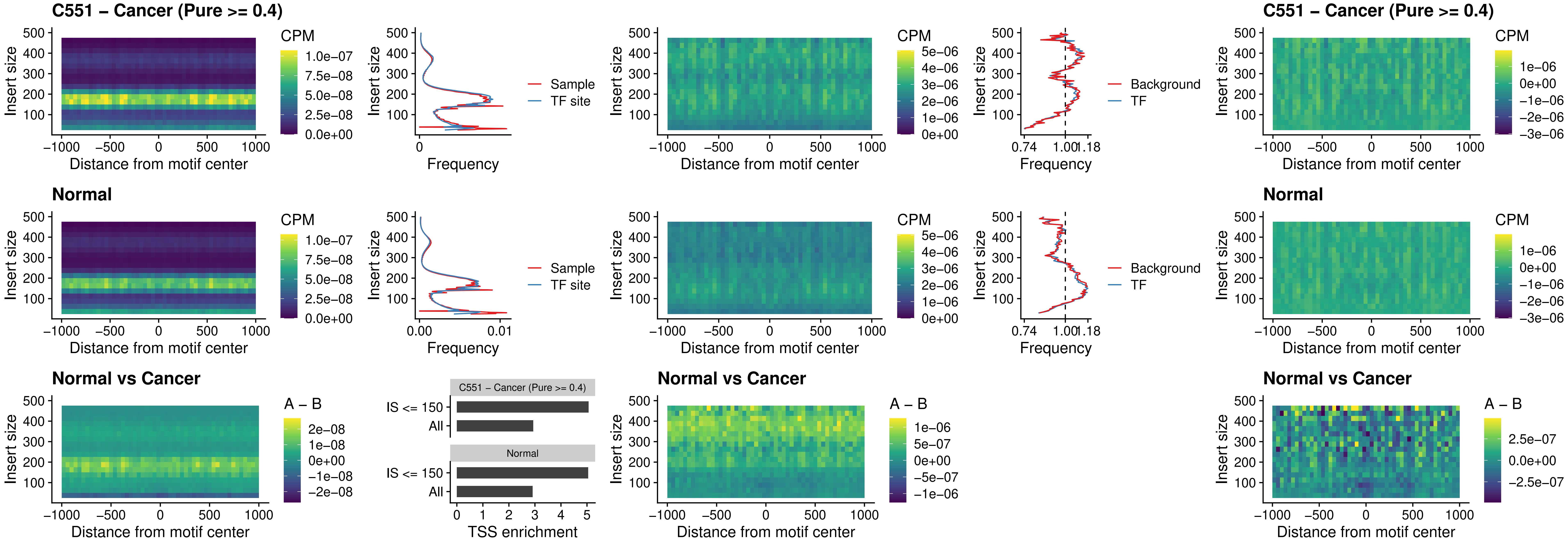

### C551 – Cancer (Pure >= 0.4) – SPIC_uc_cTSS_oPEAK

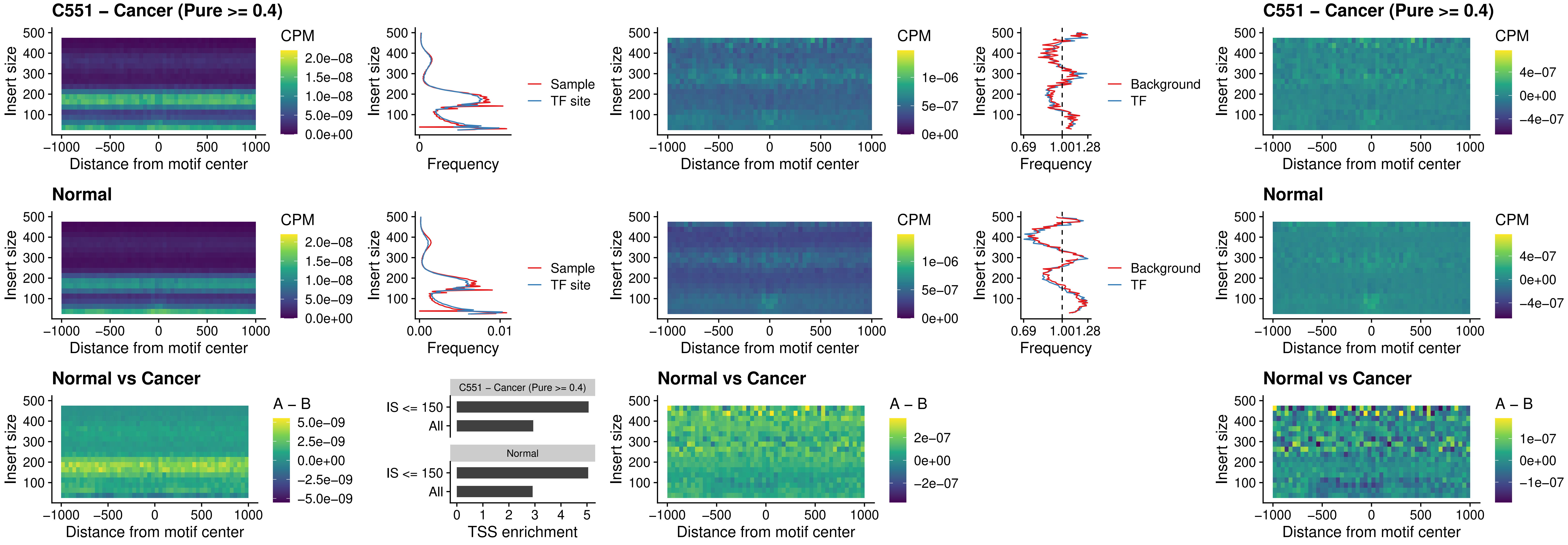

#### C551 – Cancer (Pure >= 0.4) – PRDM1_uc_cTSS_oPEAK

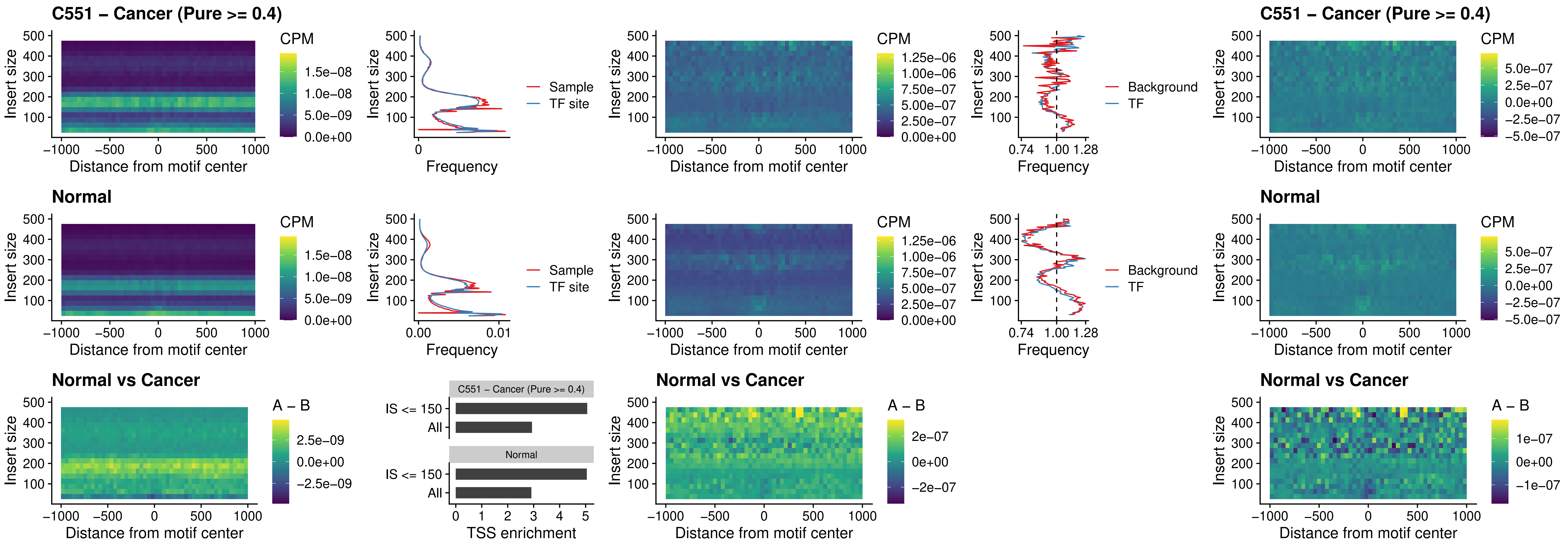

#### C551 – Cancer (Pure >= 0.4) – IRF3_uc_cTSS_oPEAK

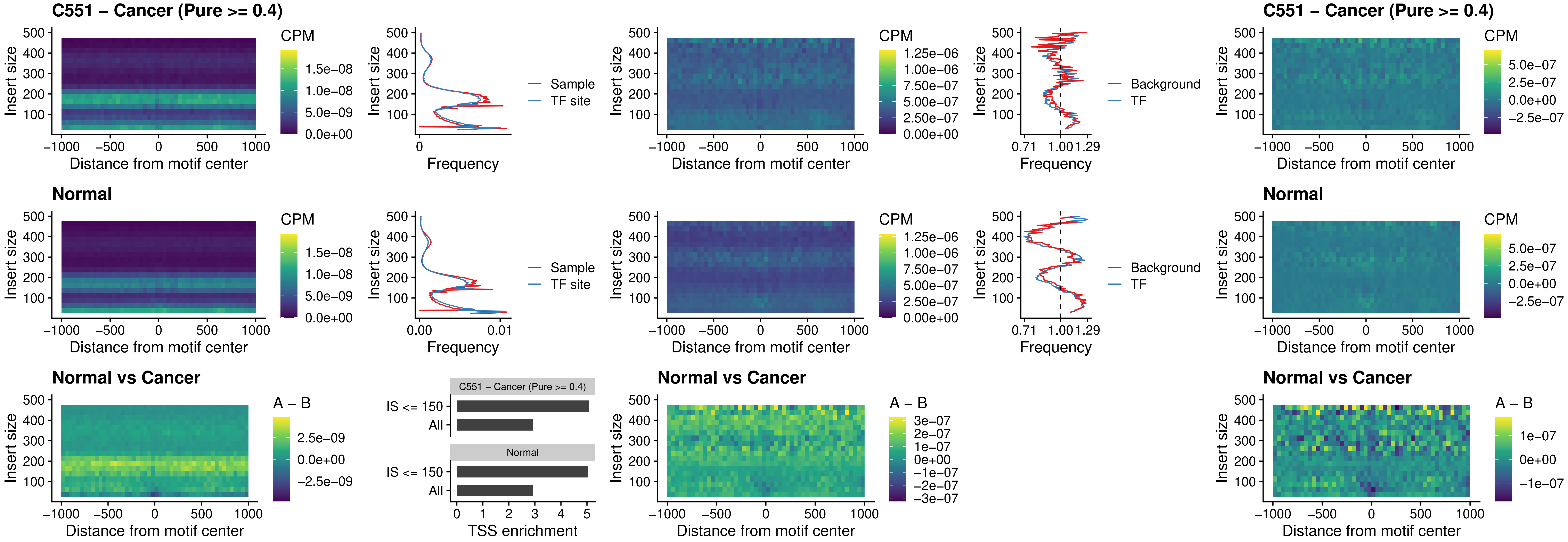

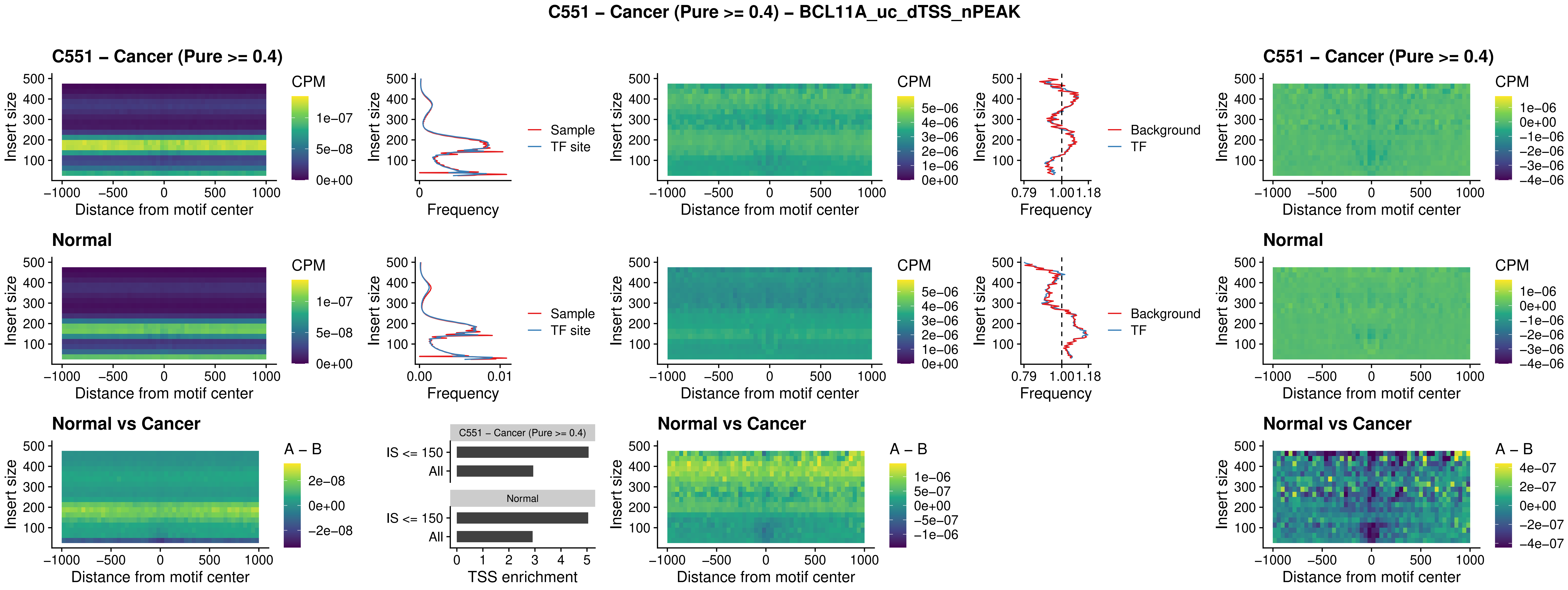
C551 – Cancer (Pure >= 0.4) – BCL11A_uc_dTSS_nPEAK
C551 – Cancer (Pure >= 0.4)
Normal
Normal vs Cancer
Insert size
Distance from motif center
Frequency
CPM
A – B
Sample
TF site
Background
TF
C551 – Cancer (Pure >= 0.4)
Normal
IS <= 150
All
TSS enrichment

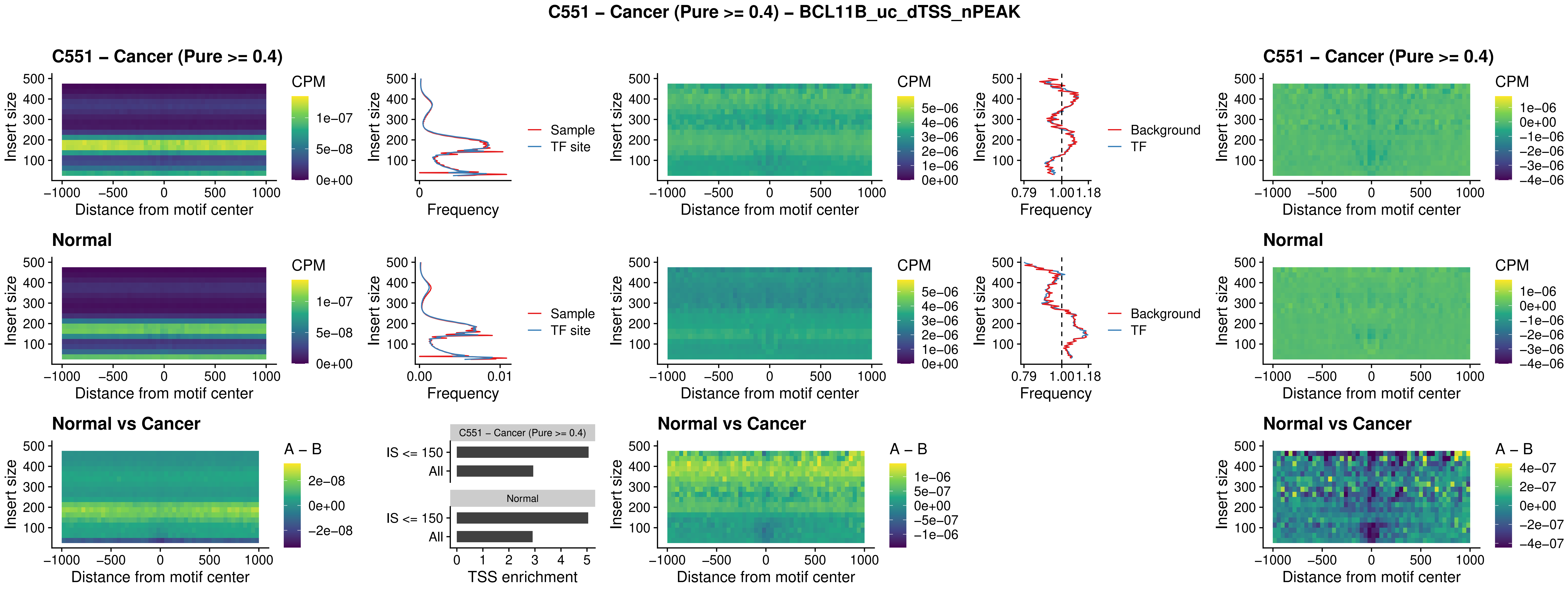
C551 – Cancer (Pure >= 0.4) – BCL11B_uc_dTSS_nPEAK
C551 – Cancer (Pure >= 0.4)
Normal
Normal vs Cancer
Insert size
Distance from motif center
Frequency
CPM
A – B
Sample
TF site
Background
TF
C551 – Cancer (Pure >= 0.4)
Normal
IS <= 150
All
TSS enrichment

#### C551 – Cancer (Pure >= 0.4) – IRF1_uc_cTSS_oPEAK

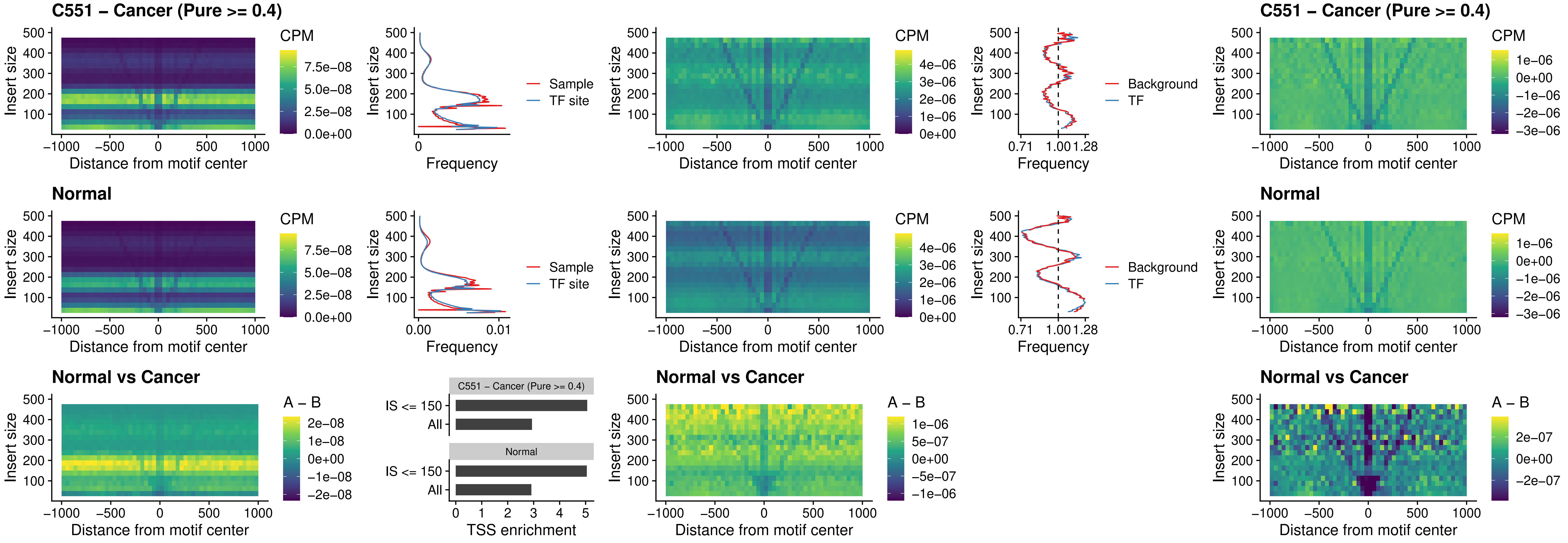

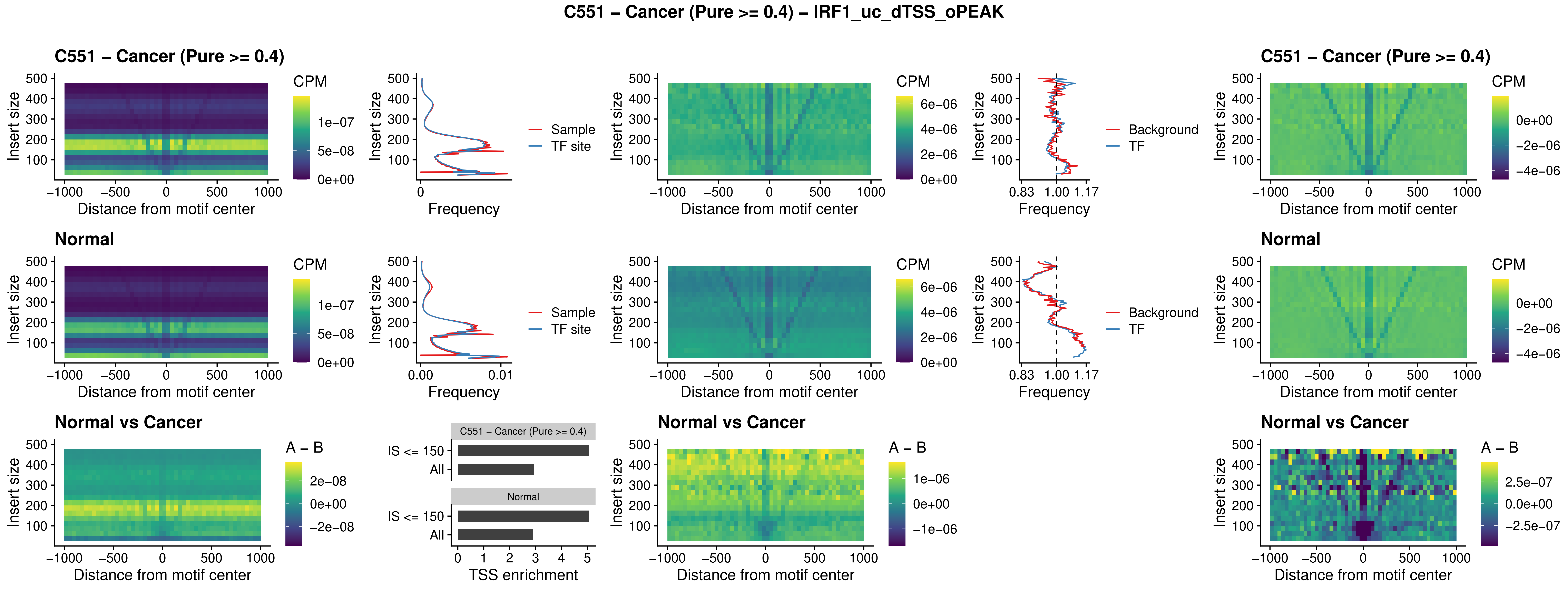
C551 – Cancer (Pure >= 0.4) – IRF1_uc_dTSS_oPEAK
C551 – Cancer (Pure >= 0.4)
Normal
Normal vs Cancer
Insert size
Distance from motif center
Frequency
CPM
A – B
Sample
TF site
Background
TF
C551 – Cancer (Pure >= 0.4)
Normal
IS <= 150
All
TSS enrichment

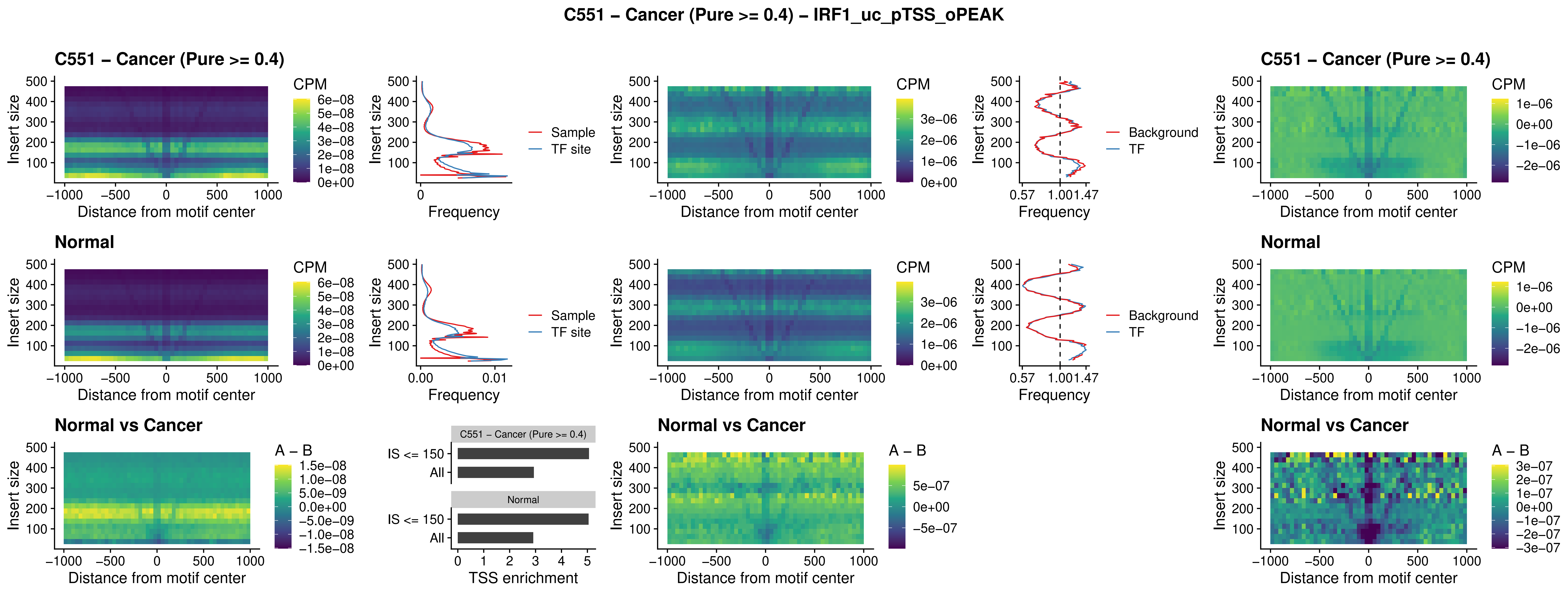
C551 – Cancer (Pure >= 0.4) – IRF1_uc_pTSS_oPEAK
C551 – Cancer (Pure >= 0.4)
Normal
Normal vs Cancer
Insert size
Distance from motif center
Frequency
CPM
A – B
Sample
TF site
Background
TF
C551 – Cancer (Pure >= 0.4)
Normal
IS <= 150
All
TSS enrichment

#### C551 – Cancer (Pure >= 0.4) – SPIC_uc_dTSS_nPEAK

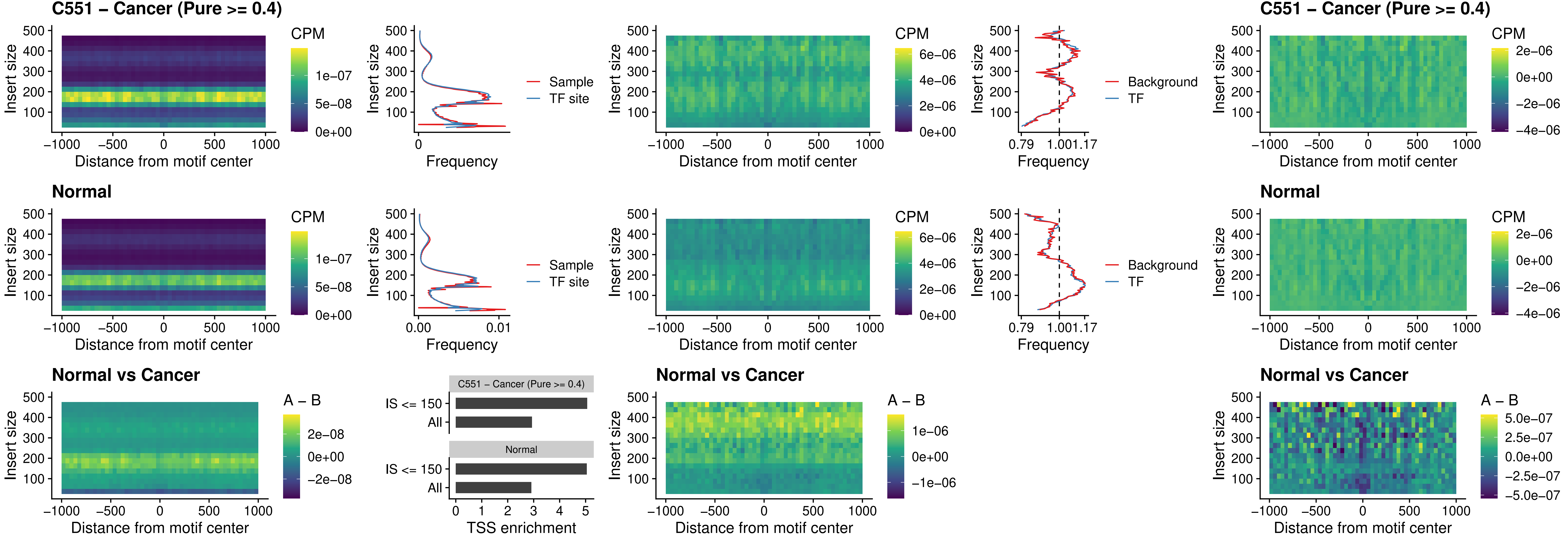

#### C551 – Cancer (Pure >= 0.4) – SPIC_uc_dTSS_oPEAK

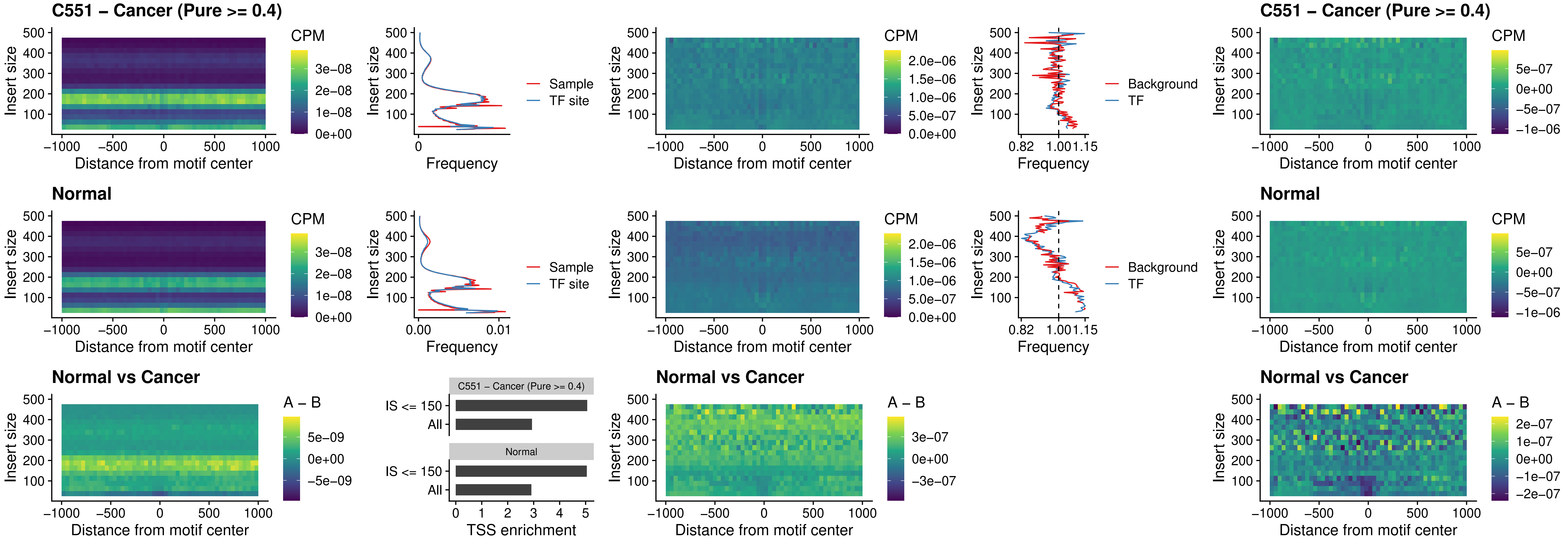

### C551 – Cancer (Pure >= 0.4) – SPIB_uc_pTSS_nPEAK

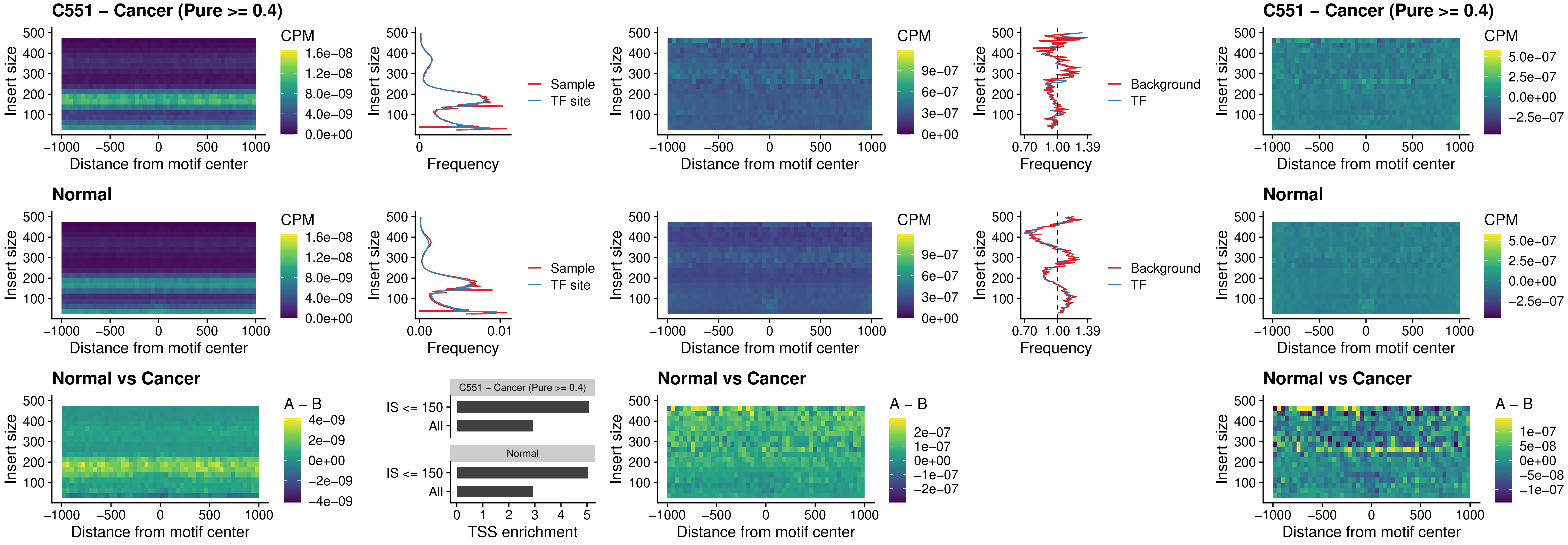

#### C551 – Cancer (Pure >= 0.4) – STAT2_uc_cTSS_oPEAK

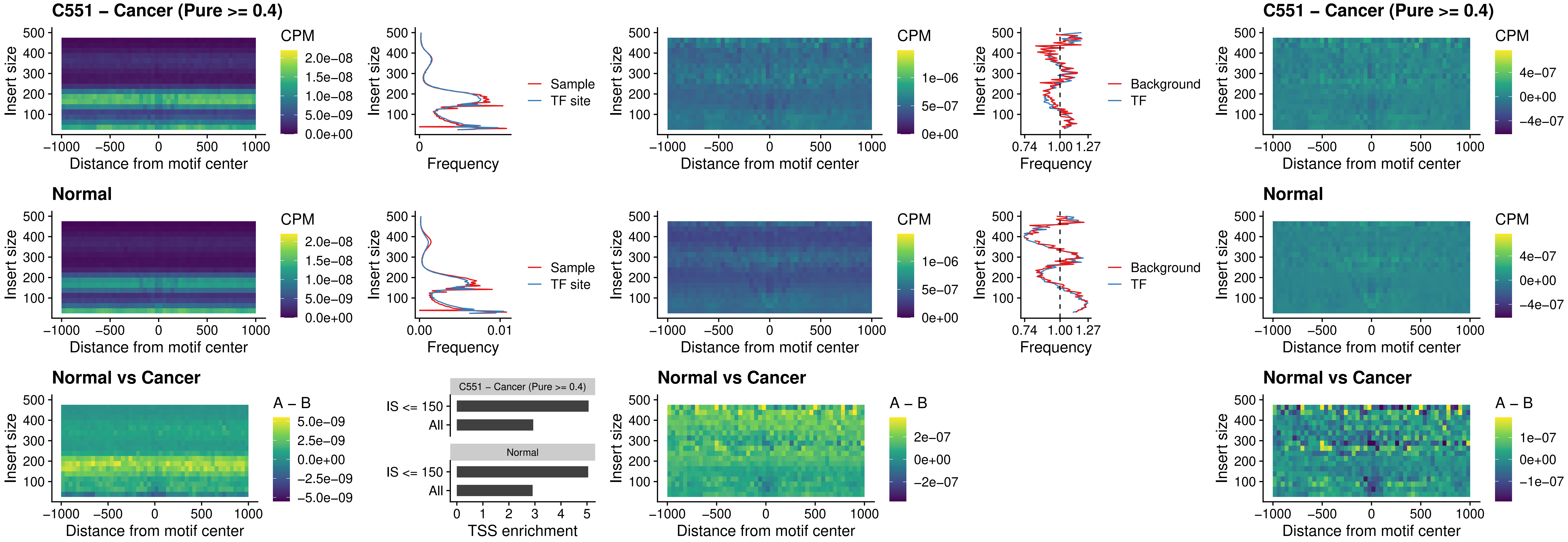

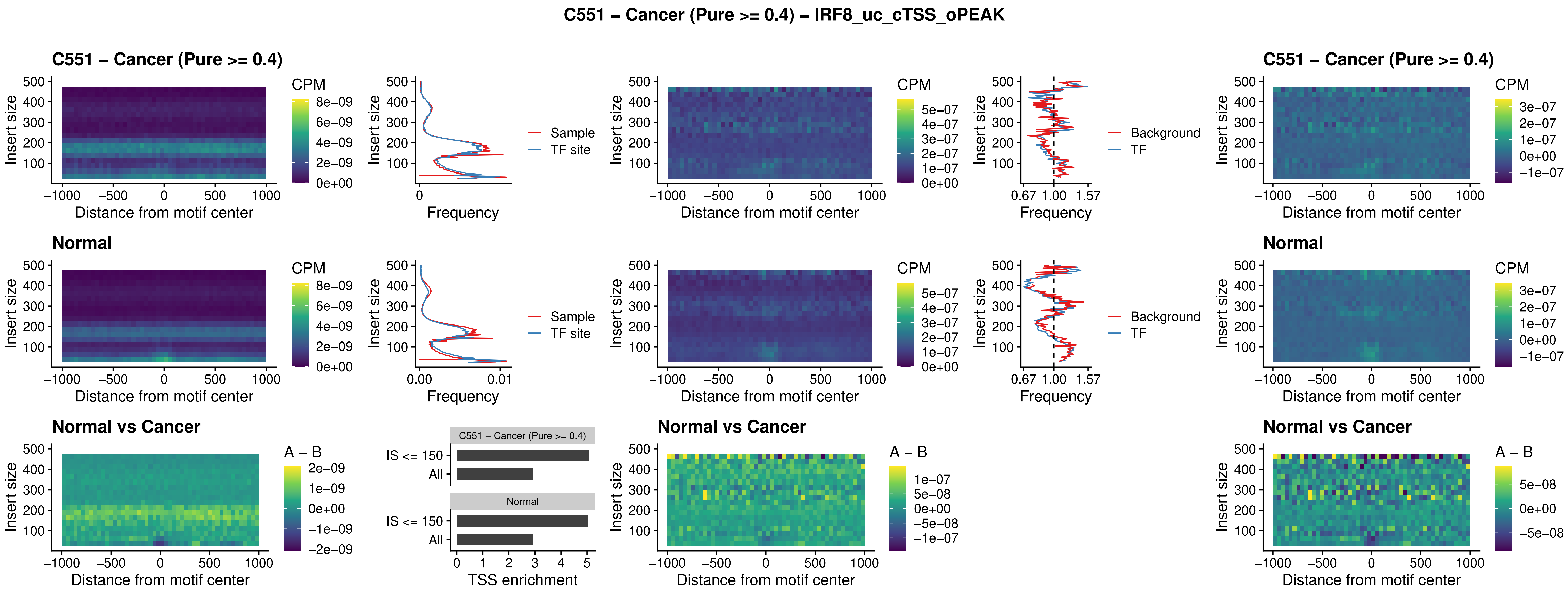
C551 – Cancer (Pure >= 0.4) – IRF8_uc_cTSS_oPEAK
C551 – Cancer (Pure >= 0.4)
Normal
Normal vs Cancer
Insert size
Distance from motif center
CPM
A – B
Frequency
Sample
TF site
Background
TF
C551 – Cancer (Pure >= 0.4)
Normal
IS <= 150
All
TSS enrichment

#### C551 – Cancer (Pure >= 0.4) – IRF3_uc_dTSS_oPEAK

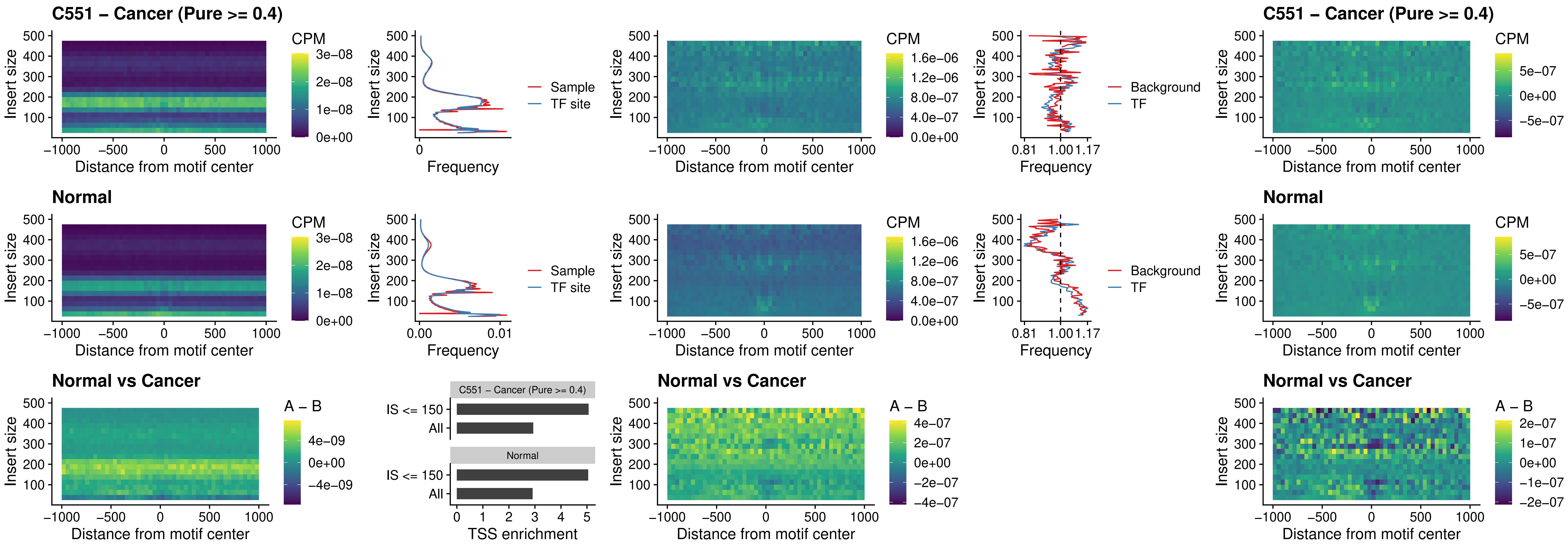

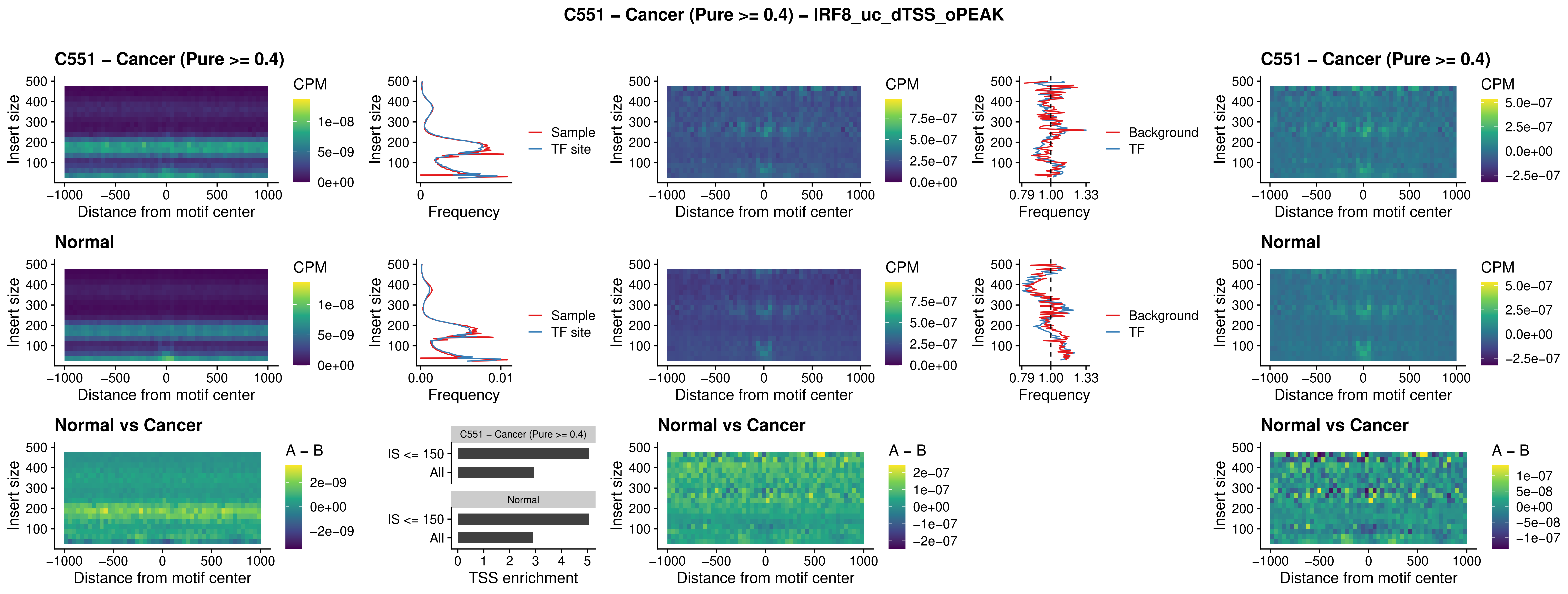
C551 – Cancer (Pure >= 0.4) – IRF8_uc_dTSS_oPEAK
C551 – Cancer (Pure >= 0.4)
Normal
Normal vs Cancer
Insert size
Distance from motif center
Frequency
CPM
A – B
Sample
TF site
Background
TF
C551 – Cancer (Pure >= 0.4)
Normal
IS <= 150
All
TSS enrichment

#### C551 – Cancer (Pure >= 0.4) – ETS2_uc_dTSS_oPEAK

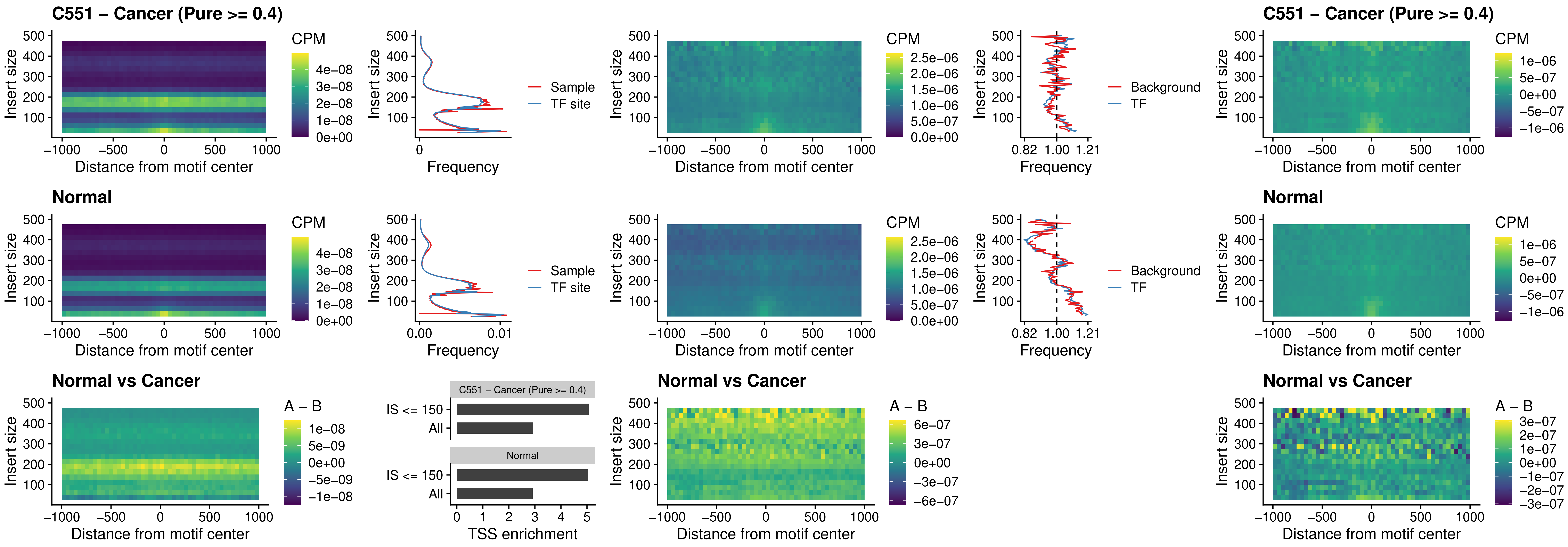

#### C551 – Cancer (Pure >= 0.4) – SMARCC2_uc_pTSS_oPEAK

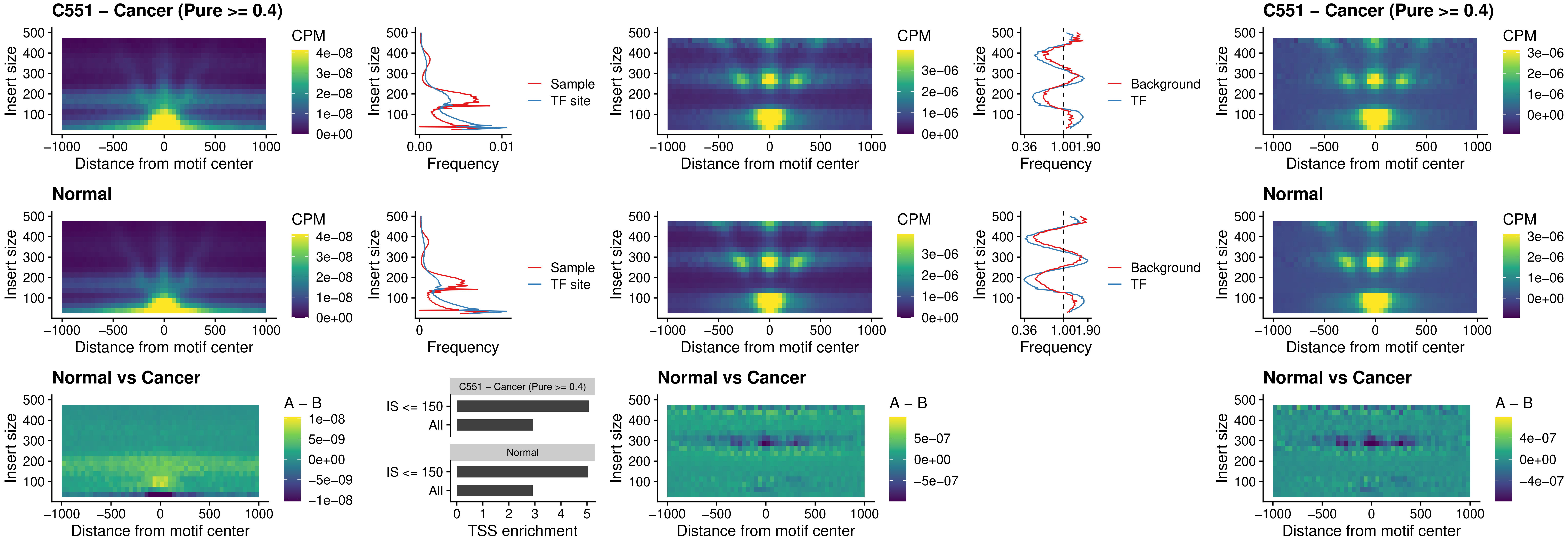

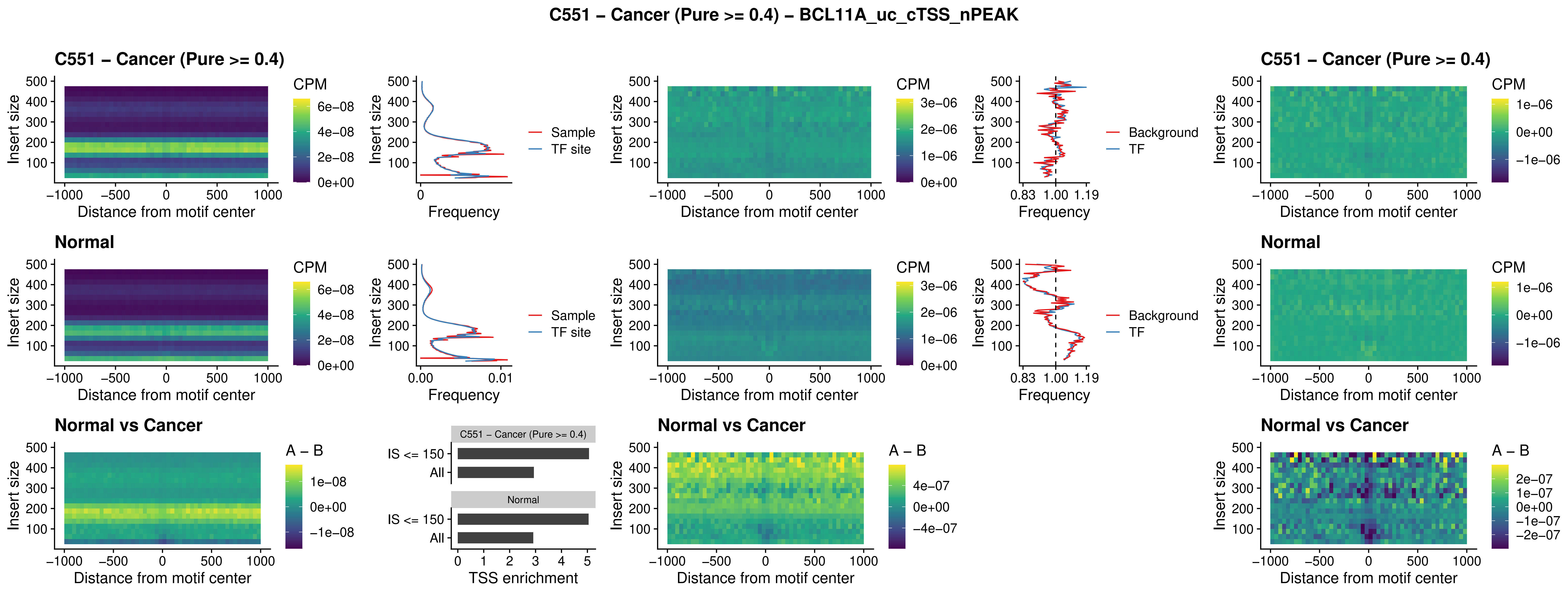
C551 – Cancer (Pure >= 0.4) – BCL11A_uc_cTSS_nPEAK
C551 – Cancer (Pure >= 0.4)
Normal
Normal vs Cancer
Insert size
Distance from motif center
CPM
A – B
Frequency
Sample
TF site
Background
TF
C551 – Cancer (Pure >= 0.4)
Normal
IS <= 150
All
TSS enrichment

#### C551 – Cancer (Pure >= 0.4) – BCL11B_uc_cTSS_nPEAK

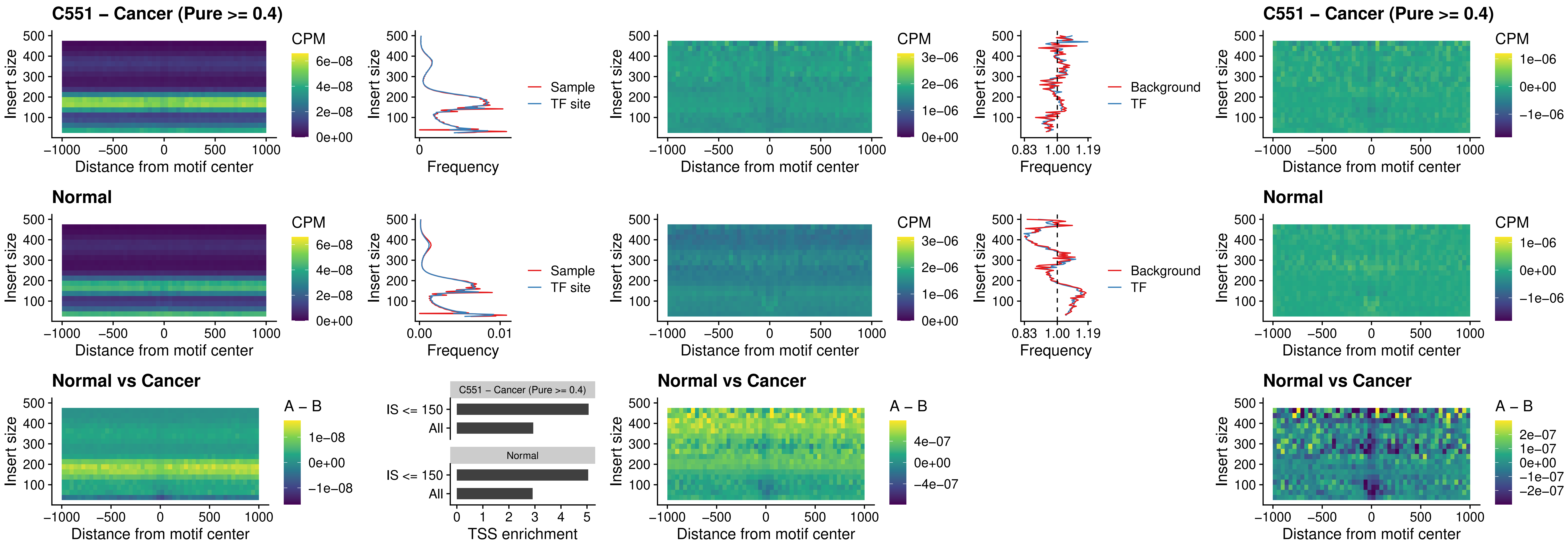

#### C551 – Cancer (Pure >= 0.4) – STAT2_uc_dTSS_oPEAK

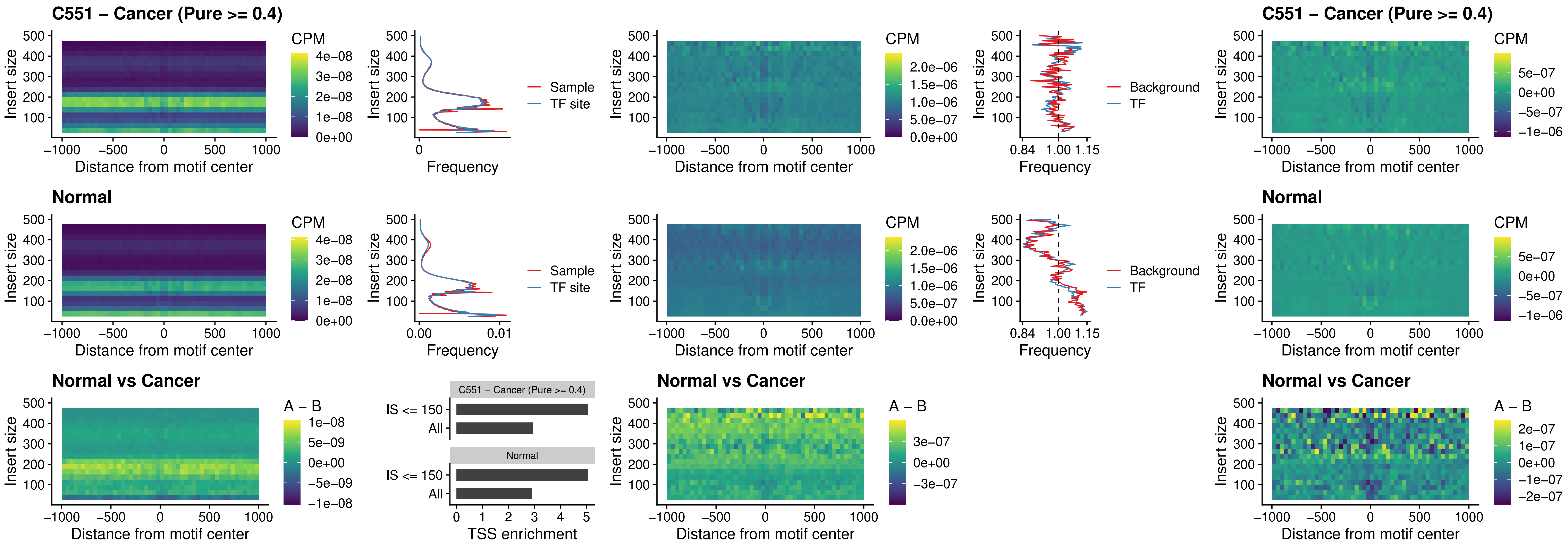

#### C551 – Cancer (Pure >= 0.4) – ZNF683_uc_cTSS_oPEAK

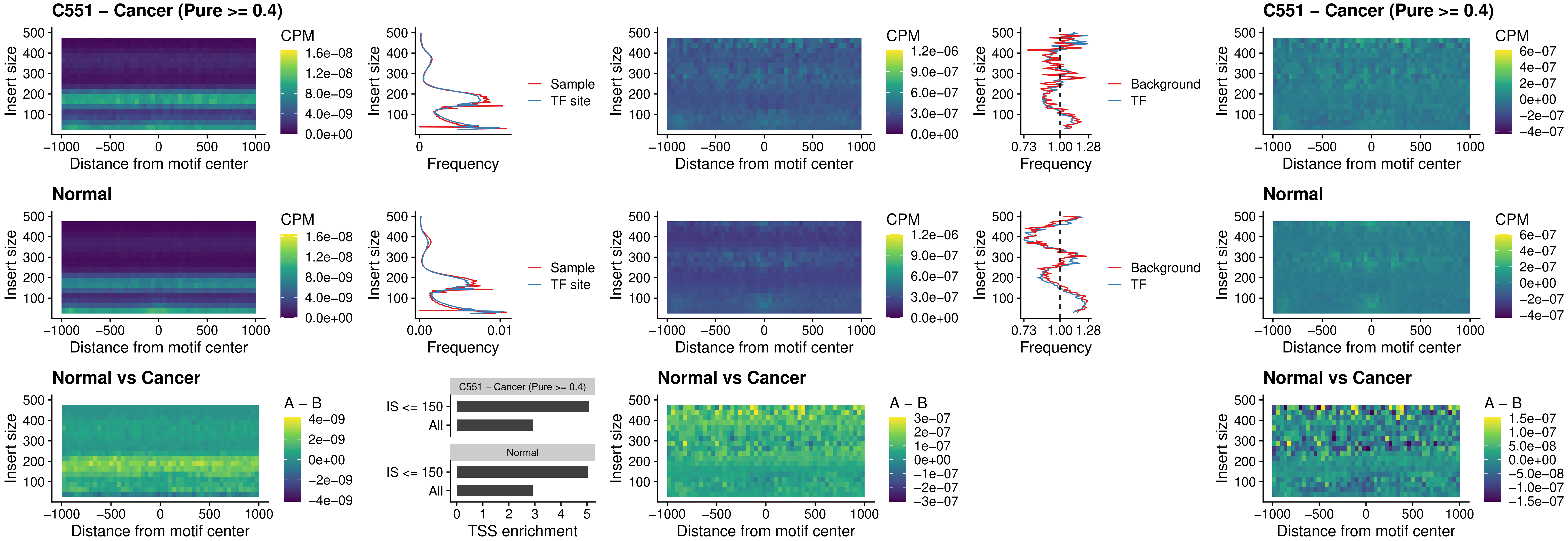

#### C551 – Cancer (Pure >= 0.4) – IRF8_uc_dTSS_nPEAK

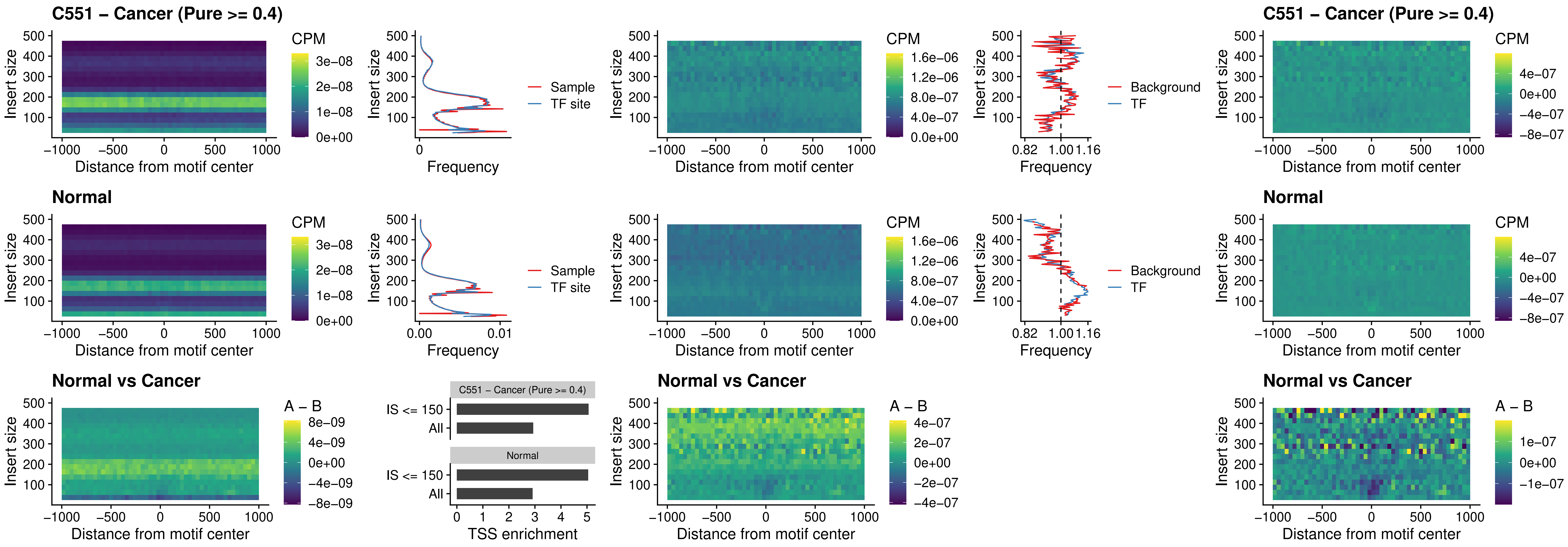

#### C551 – Cancer (Pure >= 0.4) – ETV7_uc_dTSS_oPEAK

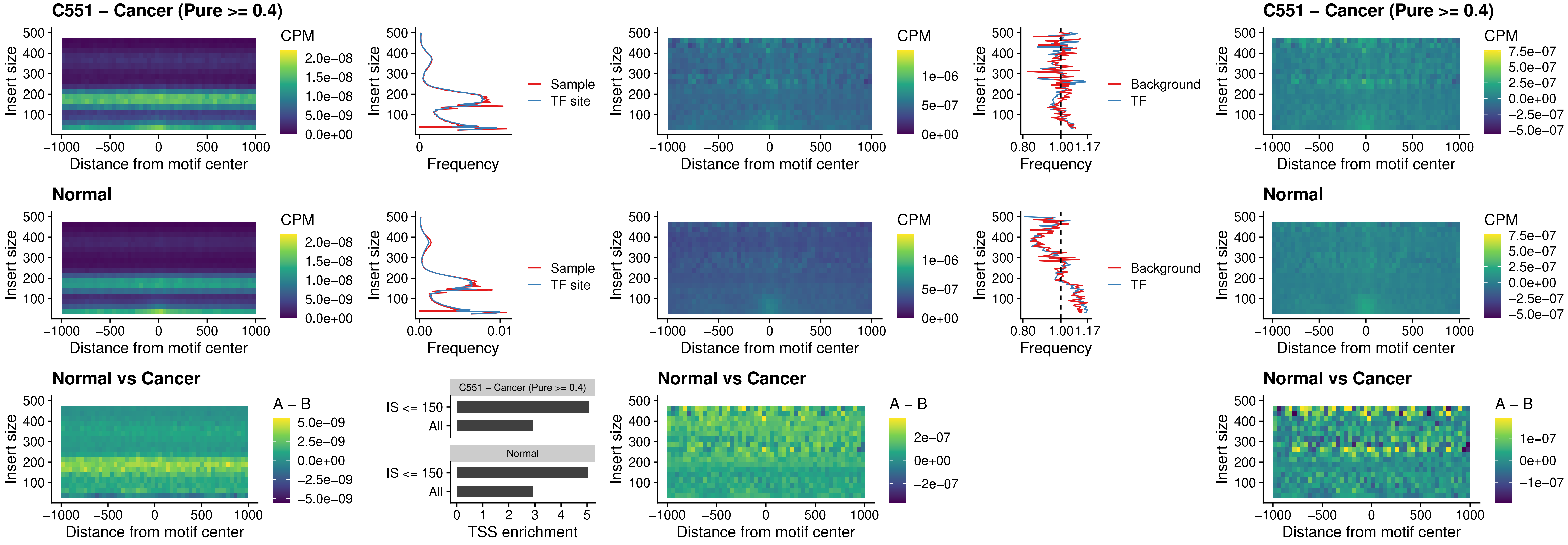

### FOXL1_uc_pTSS_oPEAK

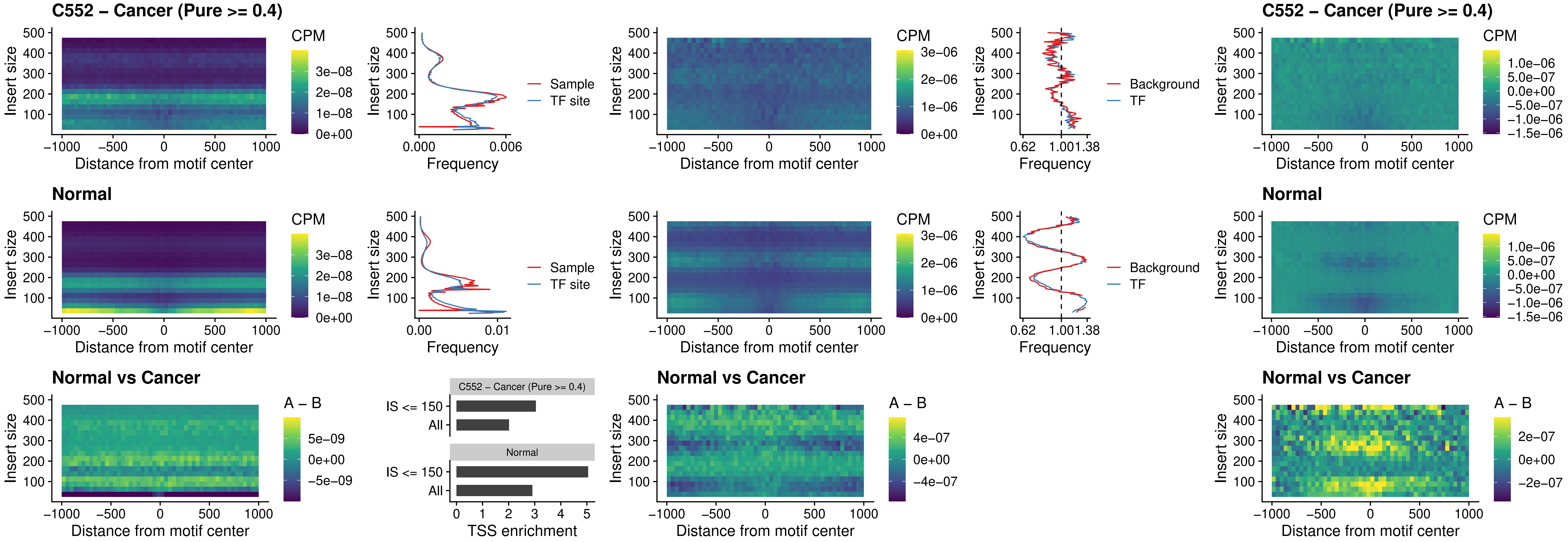

### CPEB1_uc_cTSS_oPEAK

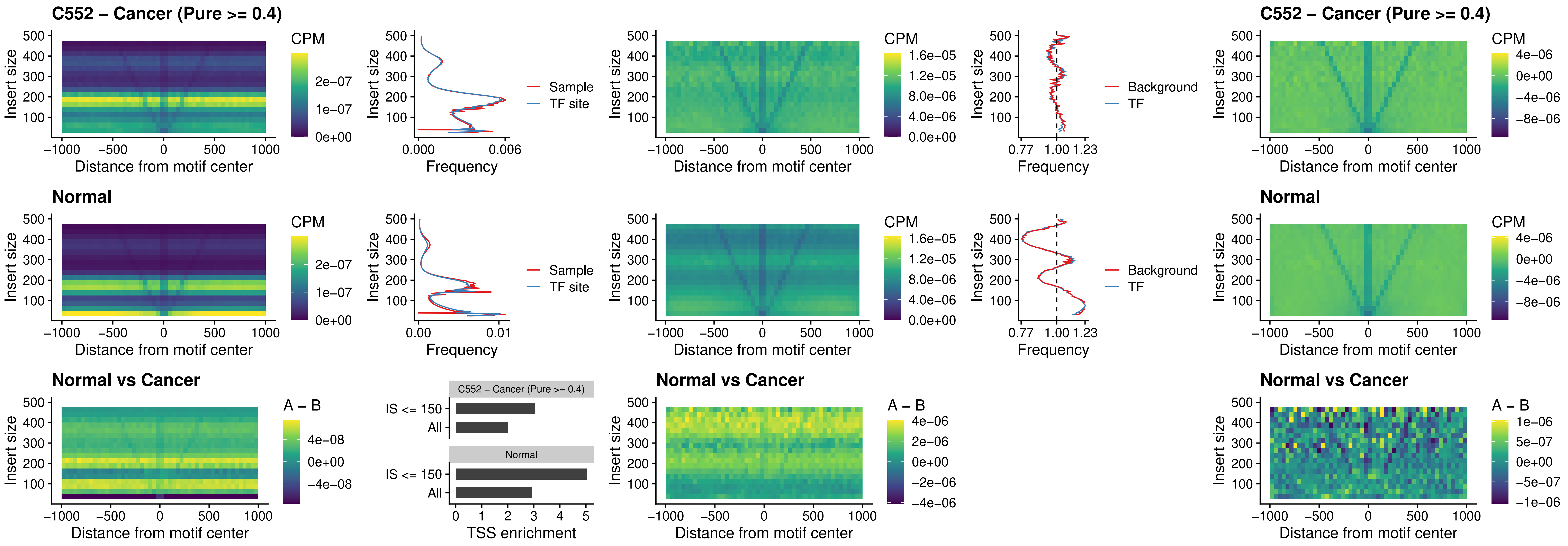

#### HMGA1_uc_pTSS_nPEAK

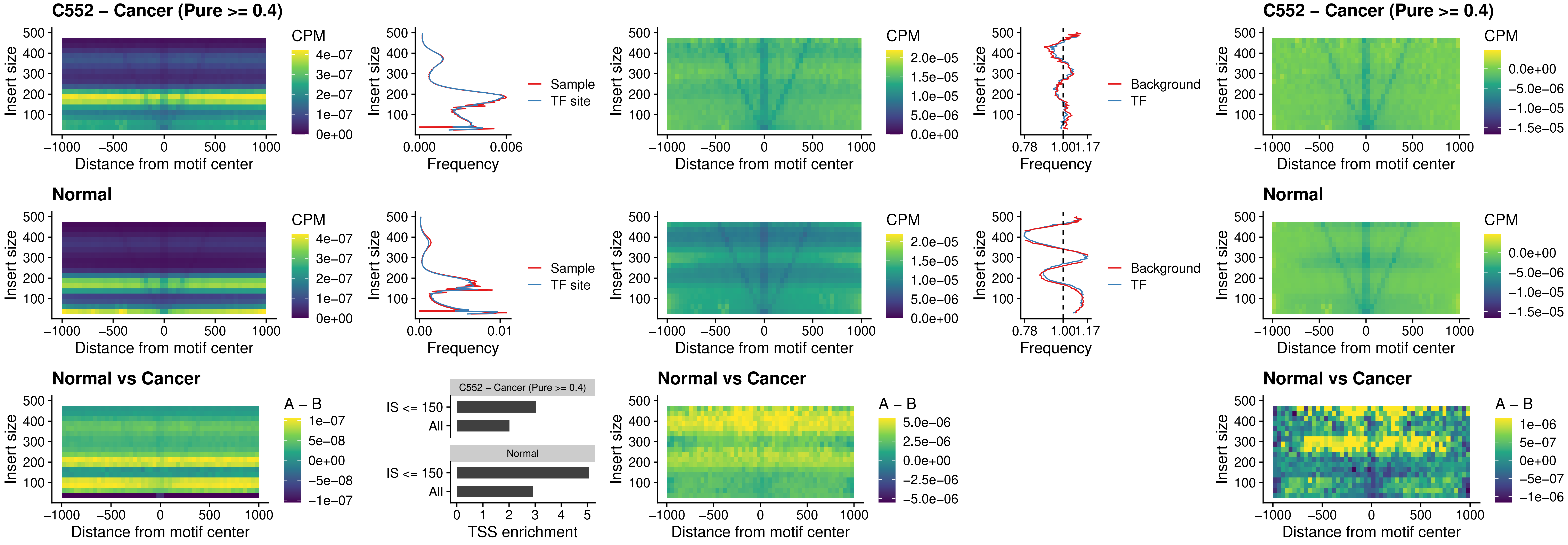

### SOX13_uc_cTSS_nPEAK

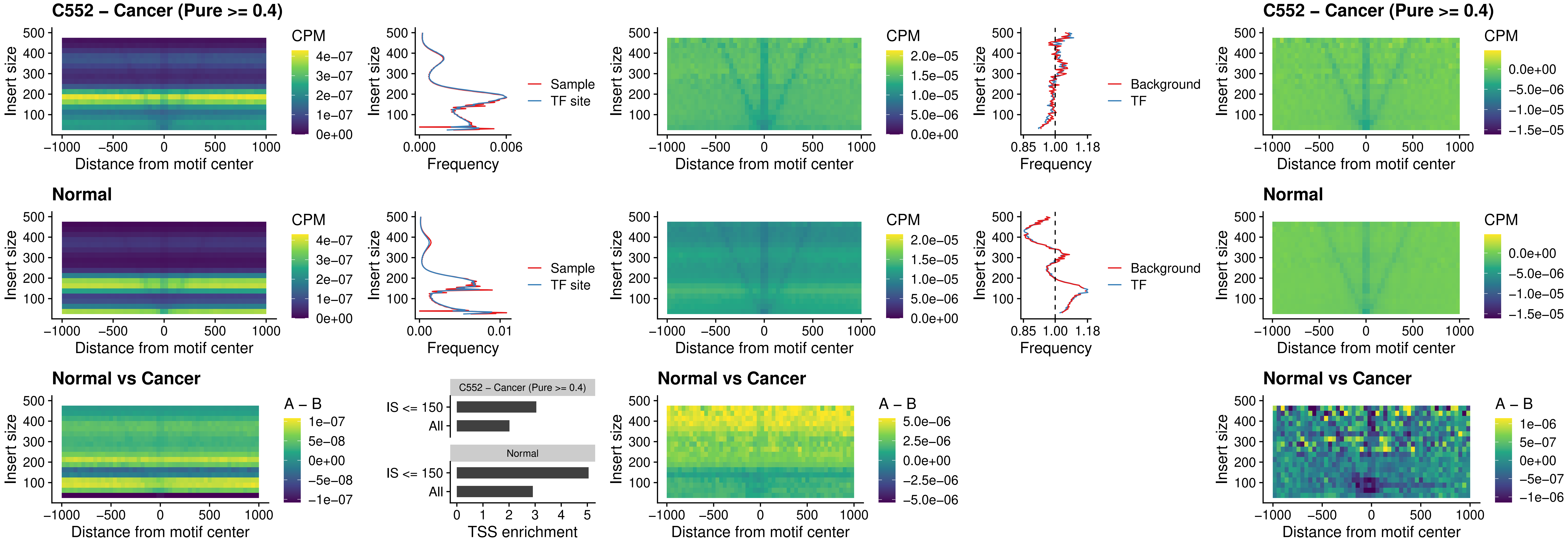

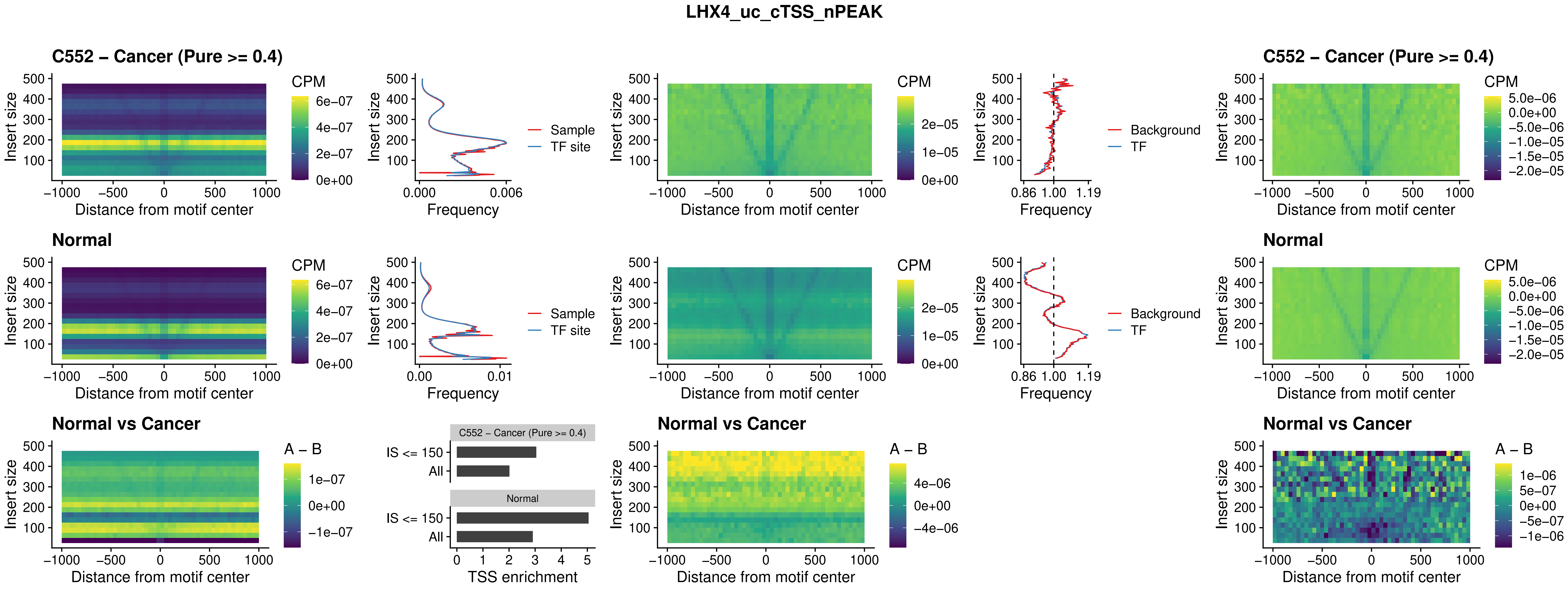

LHX4_uc_cTSS_nPEAK
C552 – Cancer (Pure >= 0.4)
Normal
Normal vs Cancer
Insert size
Distance from motif center
Frequency
CPM
A – B
Sample
TF site
Background
TF
C552 – Cancer (Pure >= 0.4)
Normal
IS <= 150
All
TSS enrichment

### HOXA4_uc_cTSS_nPEAK

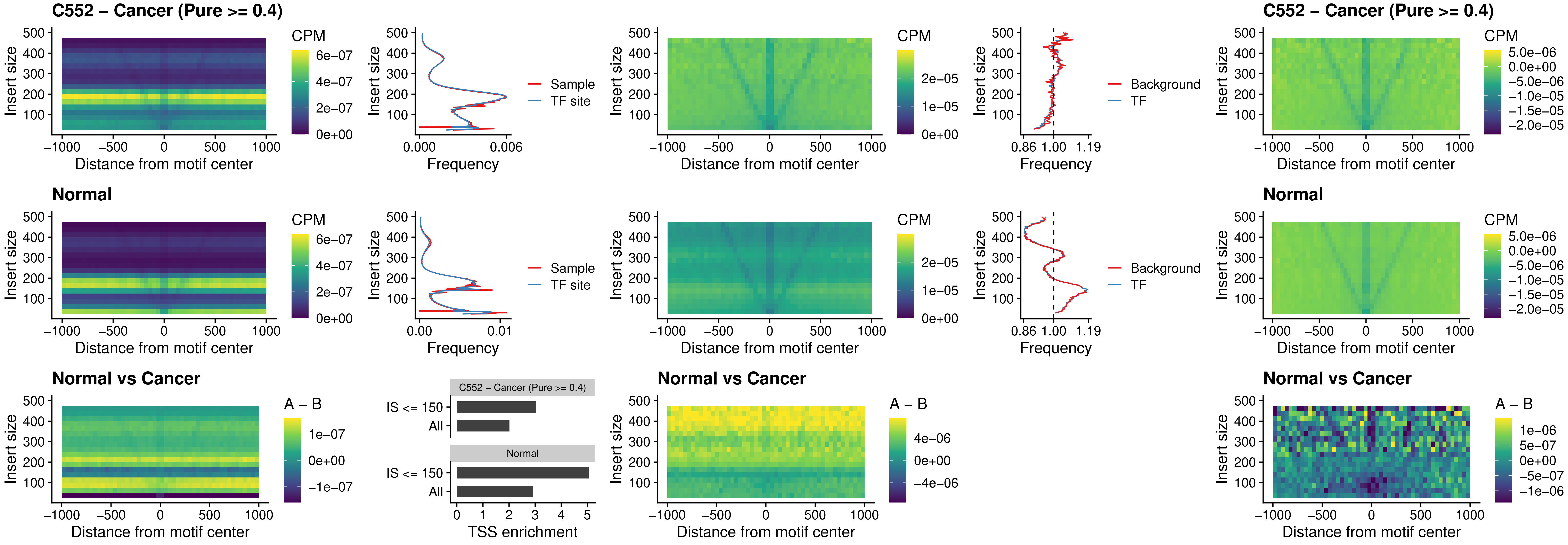

#### HLX_uc_cTSS_nPEAK

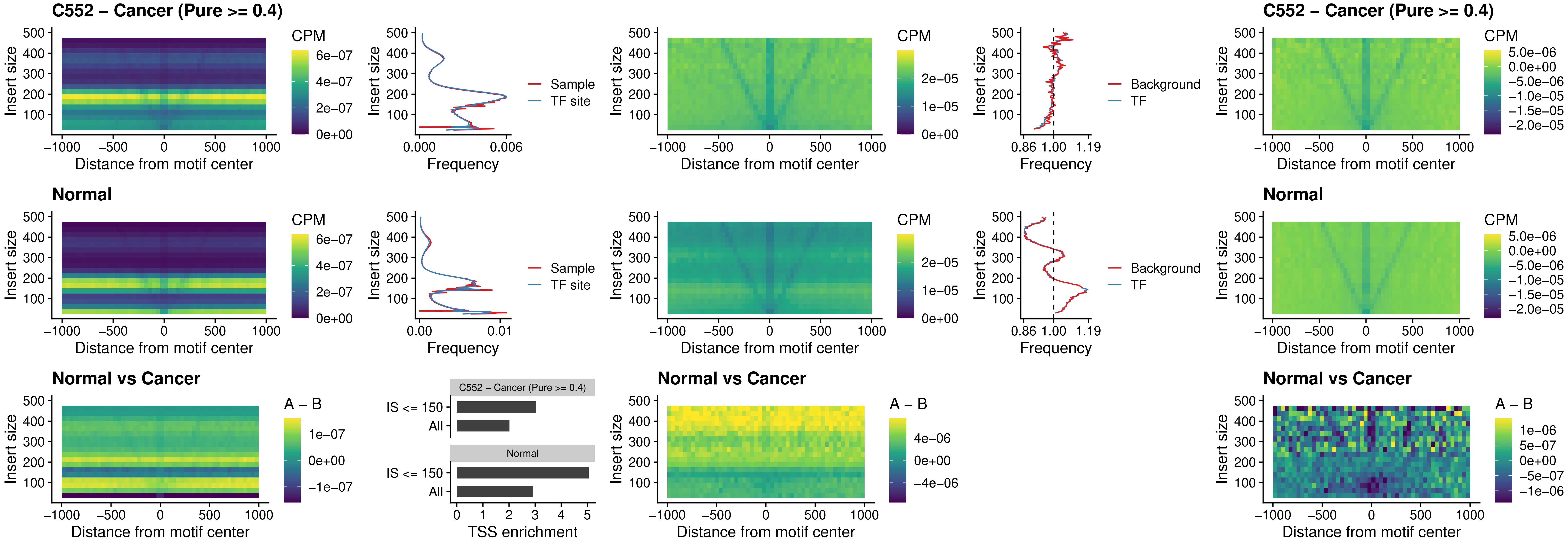

### AC0125311_uc_cTSS_nPEAK

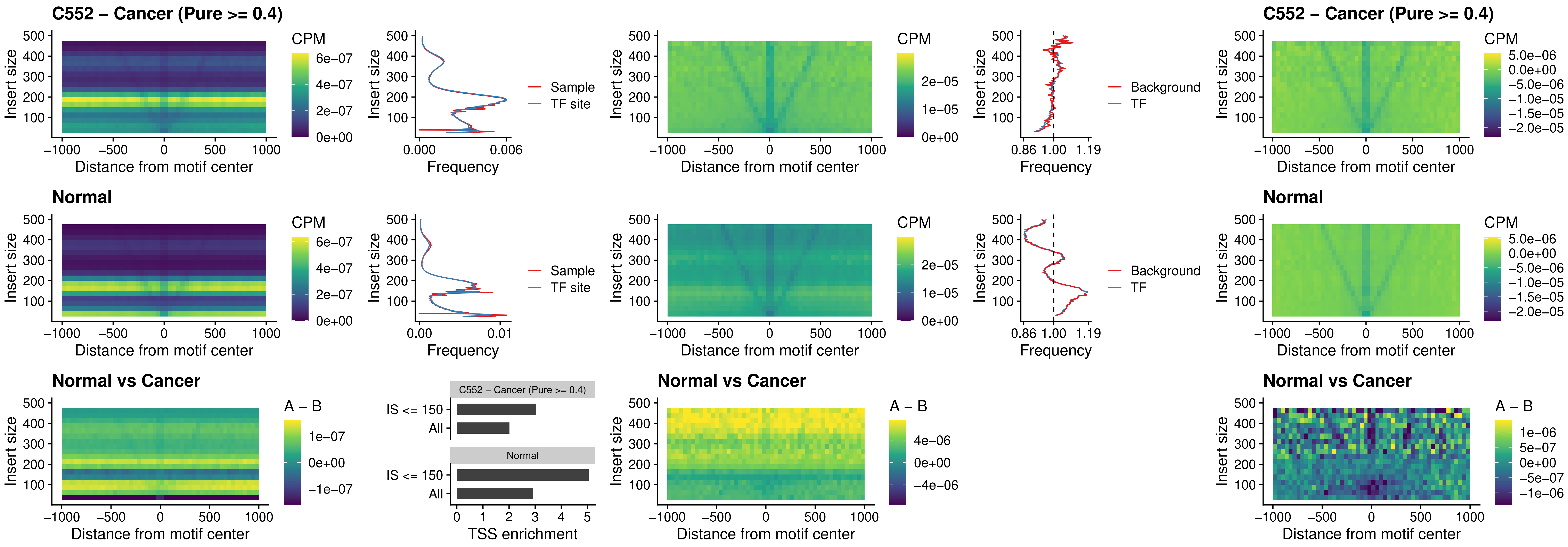

### DBX2_uc_cTSS_nPEAK

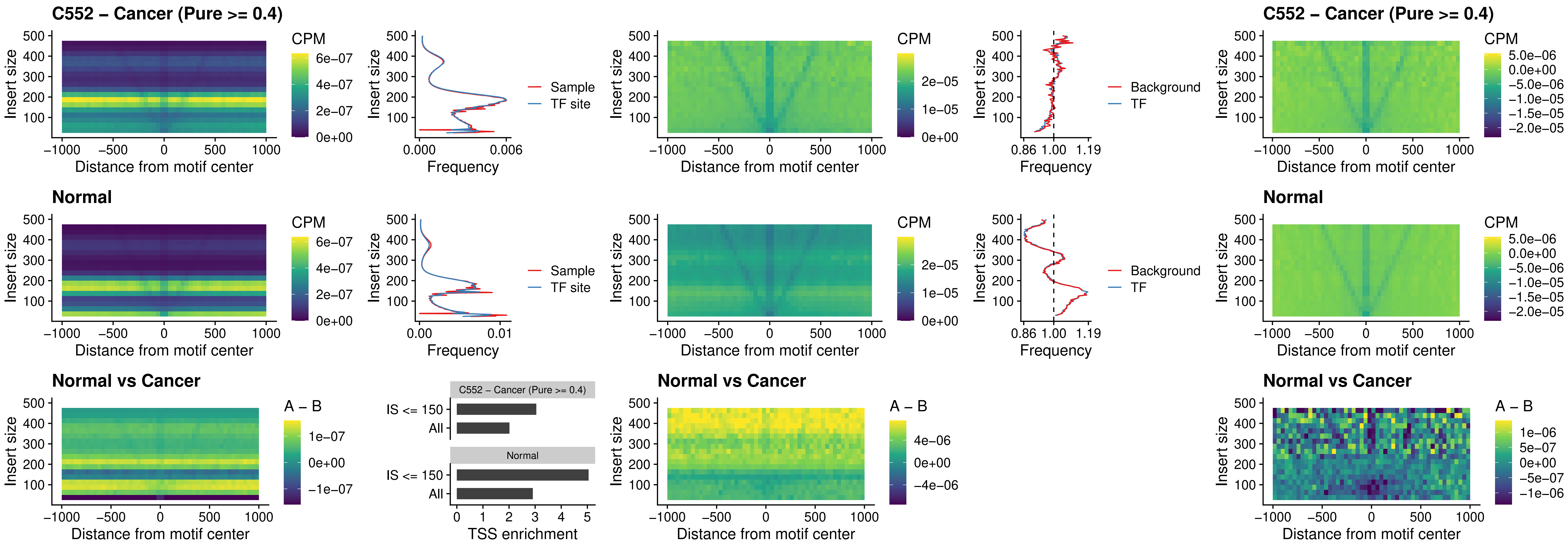

### LHX6_uc_dTSS_nPEAK

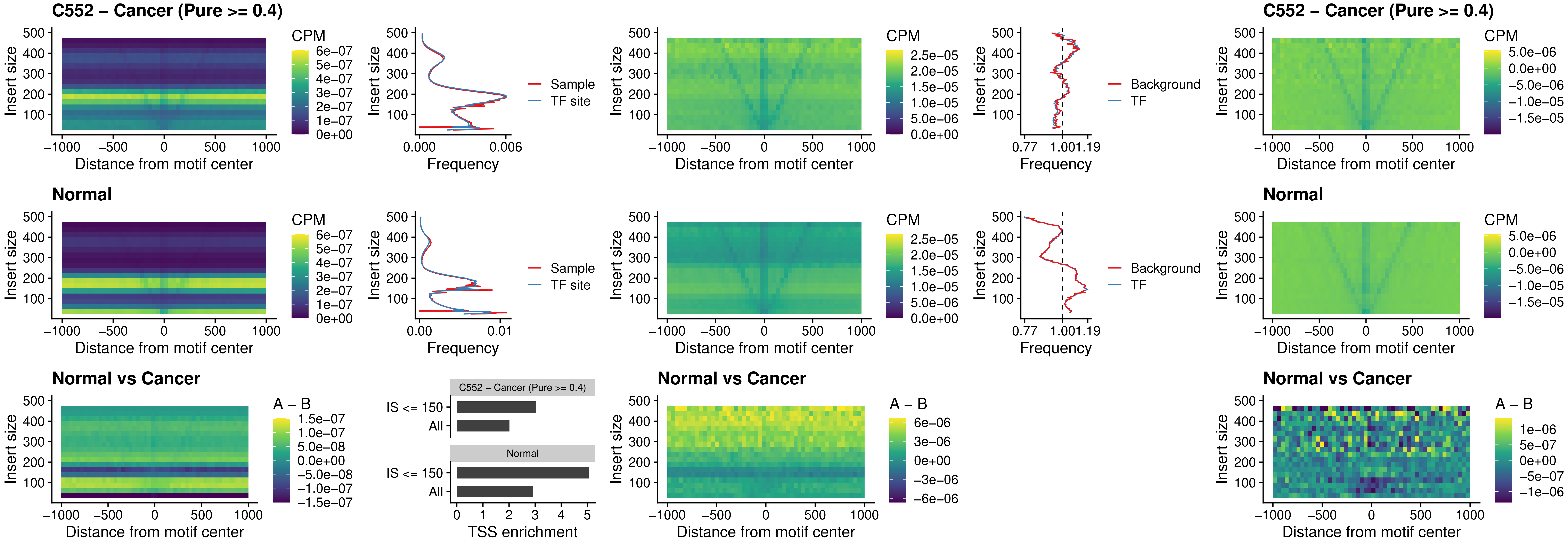

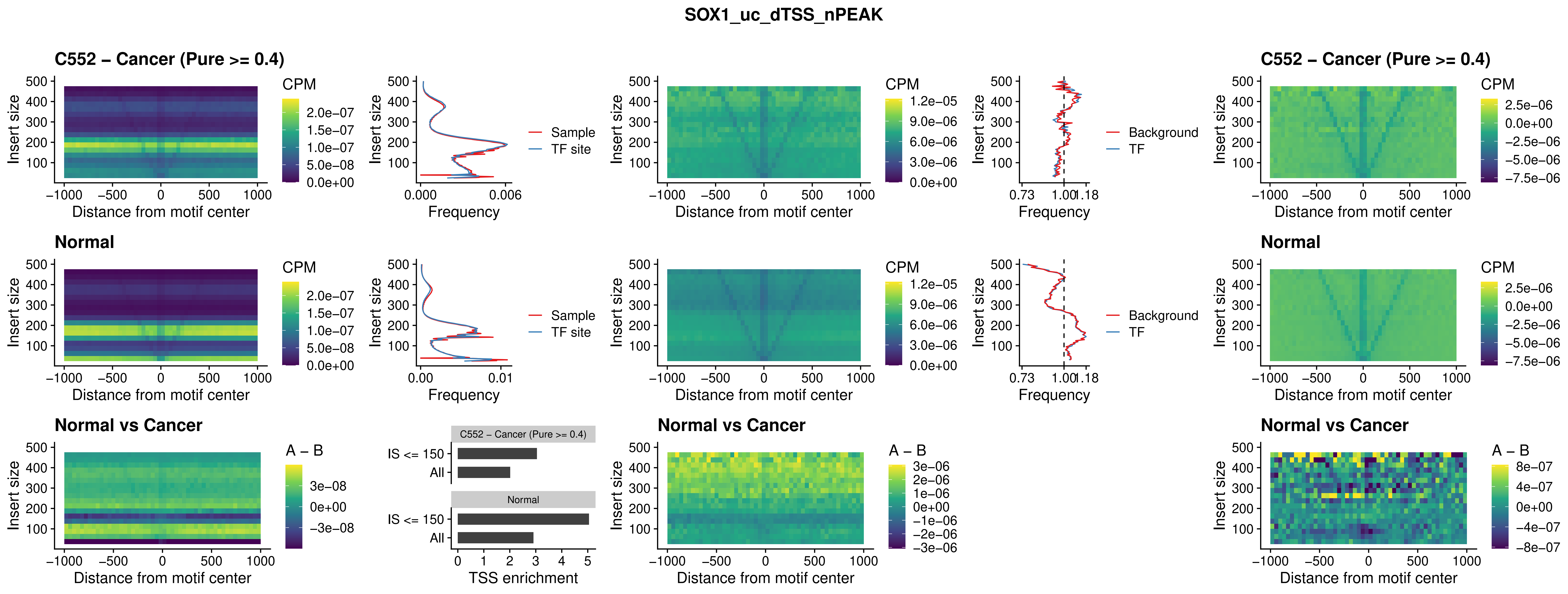
SOX1_uc_dTSS_nPEAK
C552 – Cancer (Pure >= 0.4)
Normal
Normal vs Cancer
Insert size
Distance from motif center
Frequency
CPM
A – B
Sample
TF site
Background
TF
C552 – Cancer (Pure >= 0.4)
Normal
IS <= 150
All
TSS enrichment

#### NKX24_uc_dTSS_nPEAK

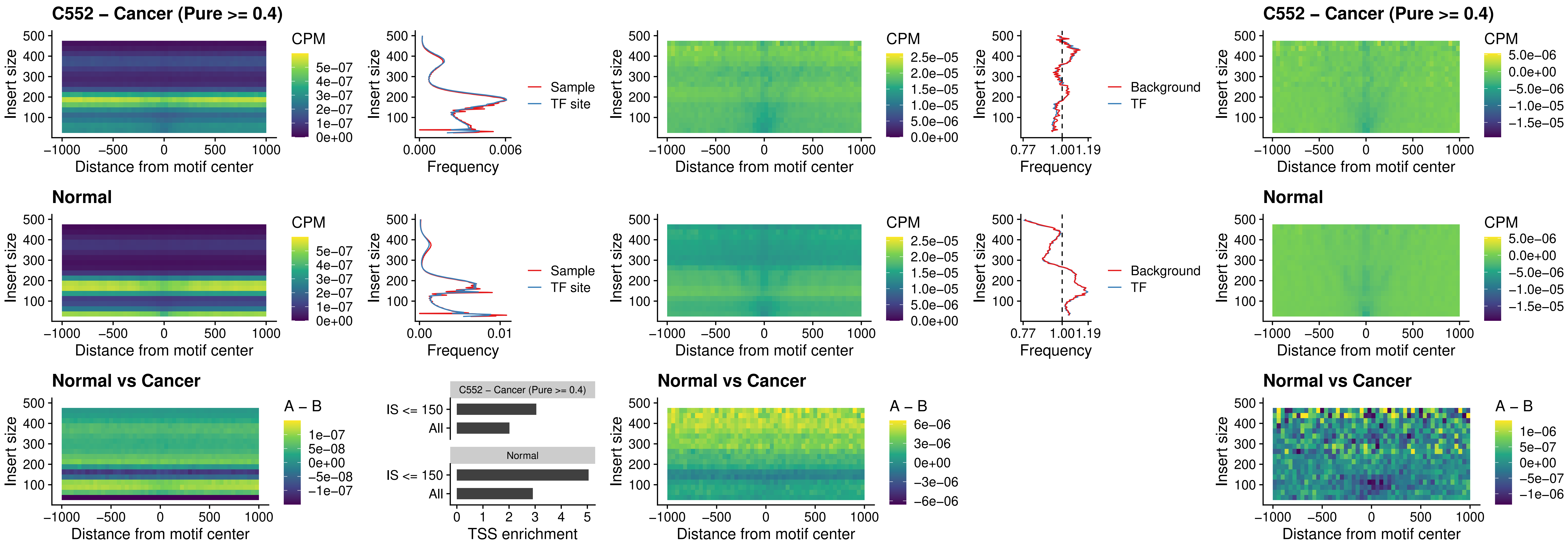

### DLX3_uc_cTSS_nPEAK

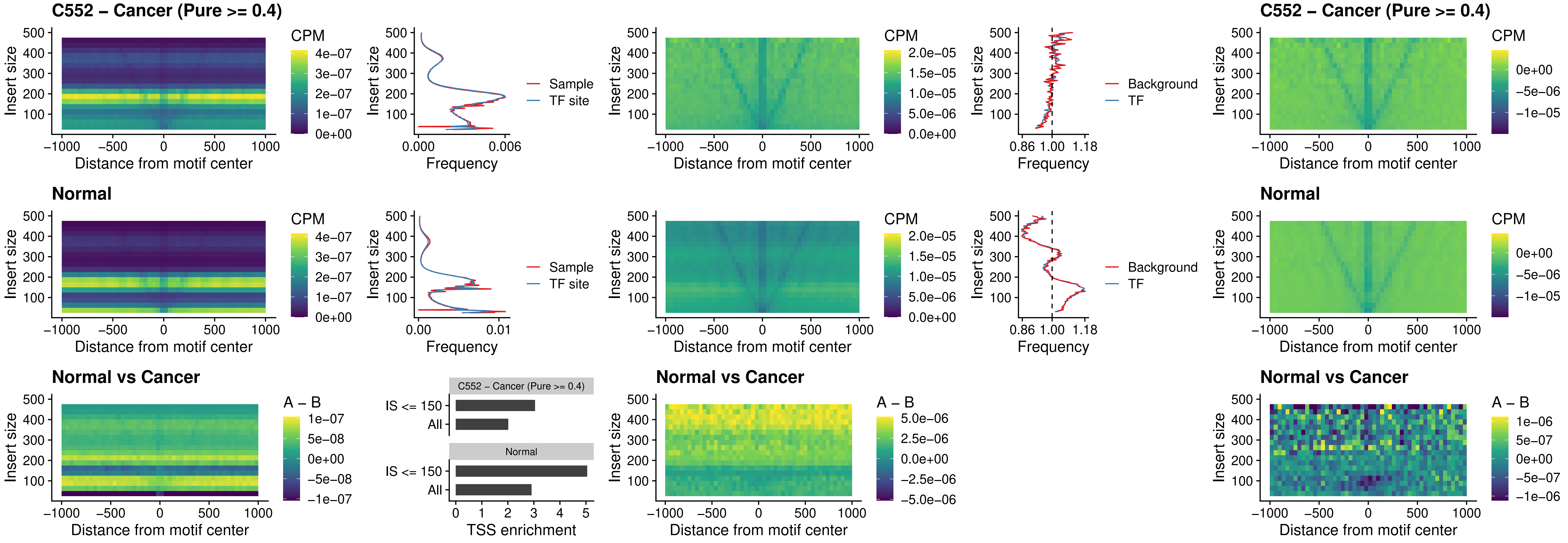

#### HOXD3_uc_cTSS_nPEAK

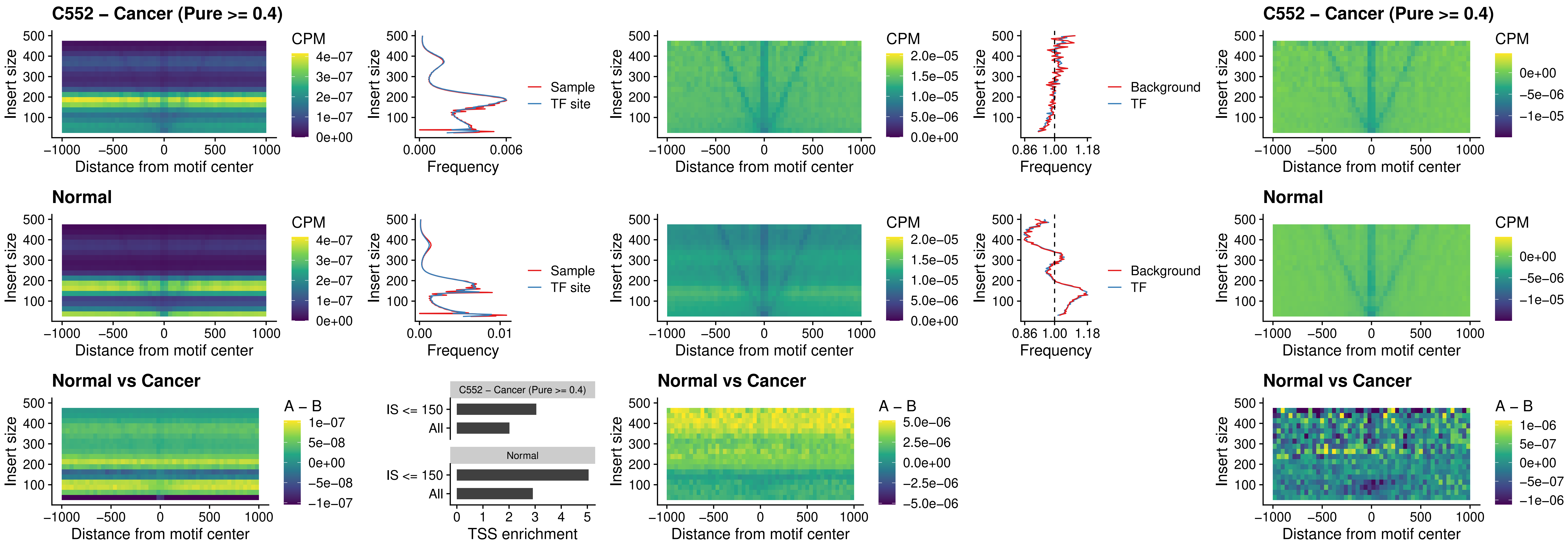

#### ISL1_uc_dTSS_oPEAK

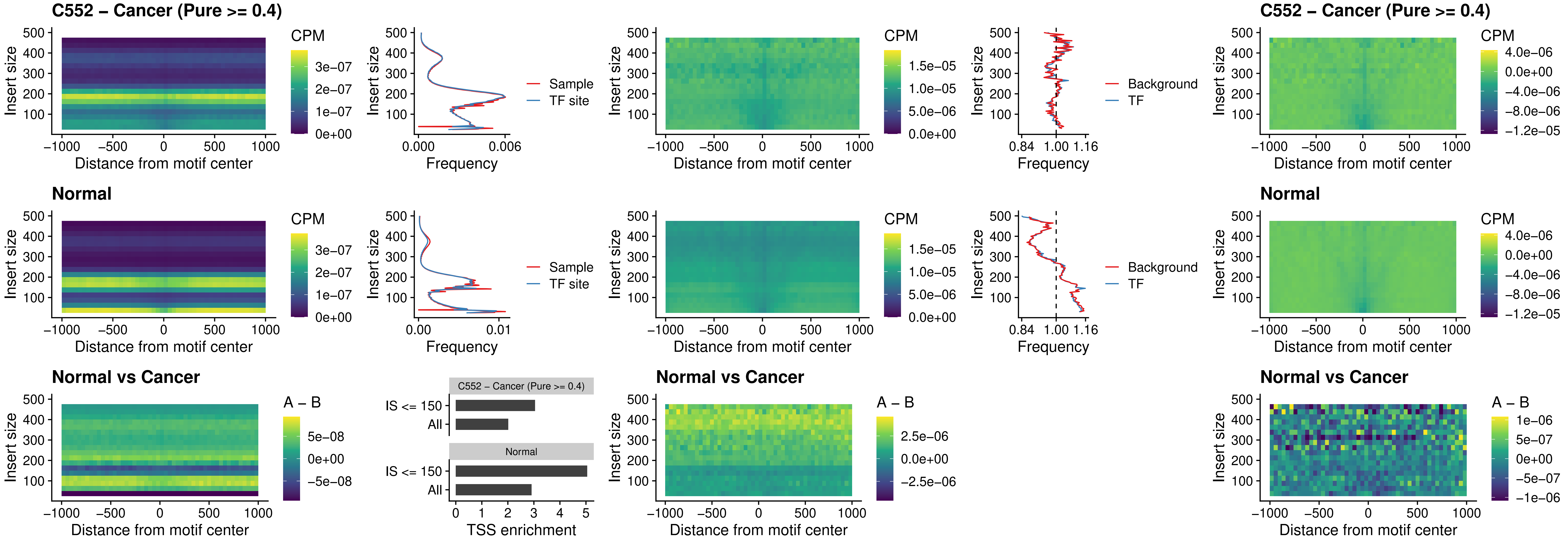

### HOXB4_uc_dTSS_oPEAK

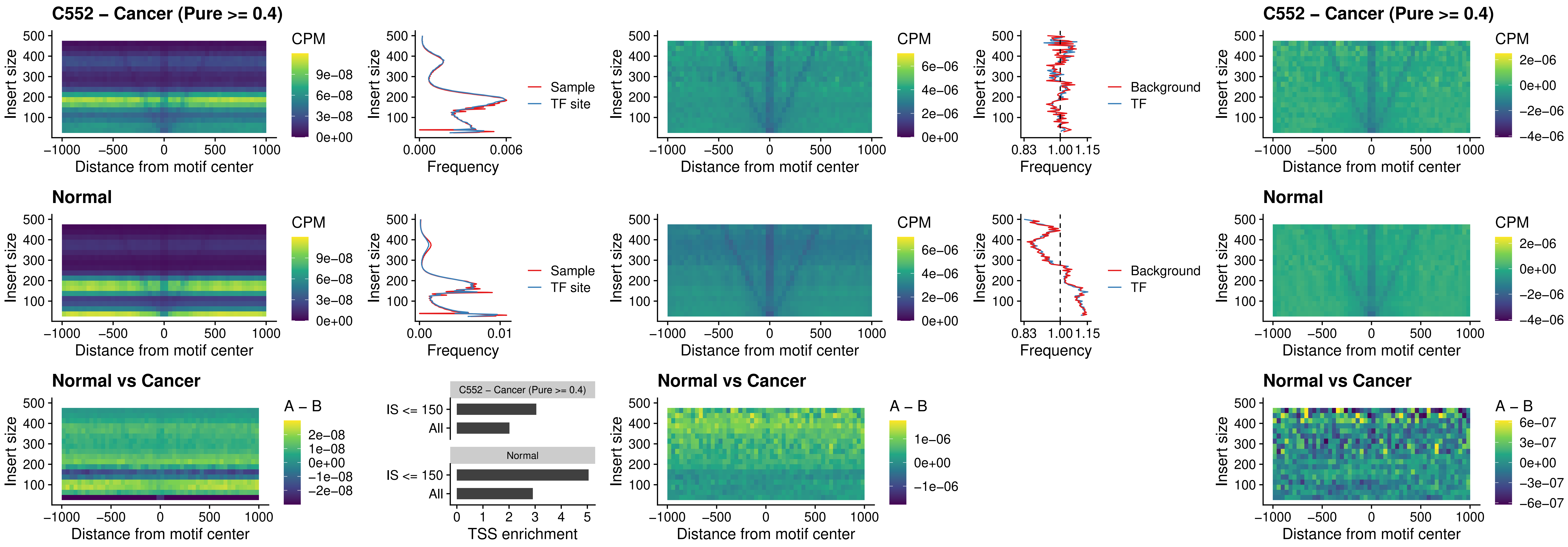

### VAX1_uc_dTSS_oPEAK

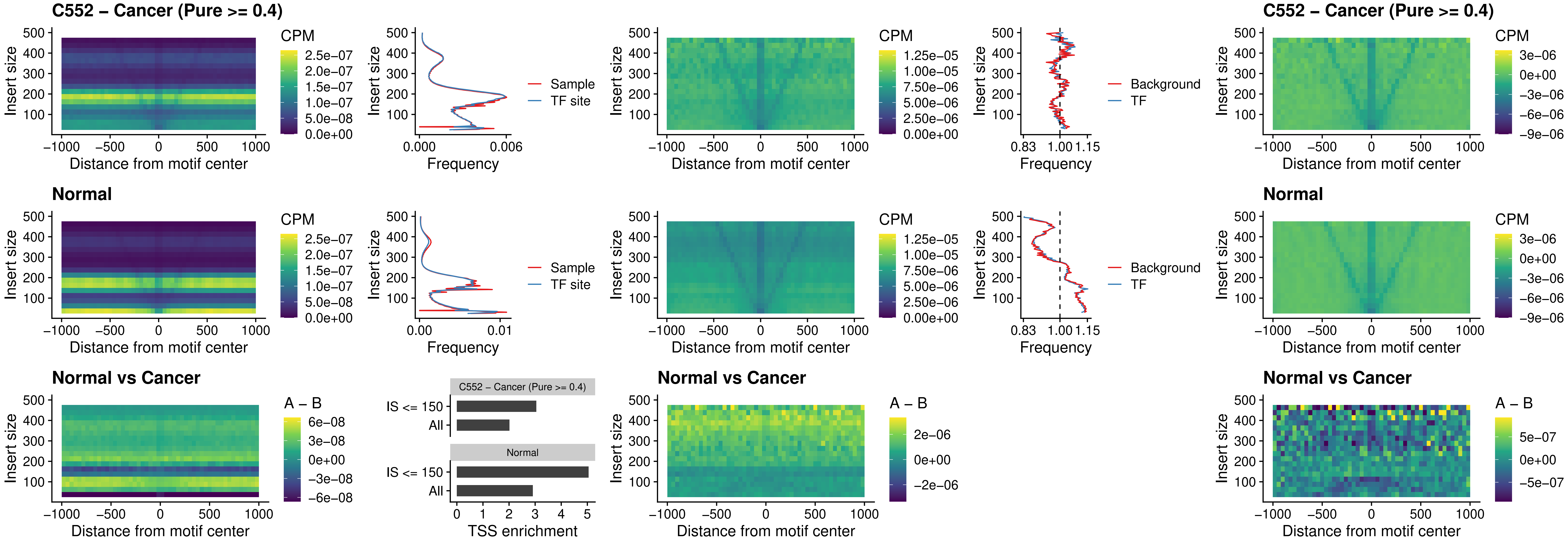

#### DLX3_uc_dTSS_oPEAK

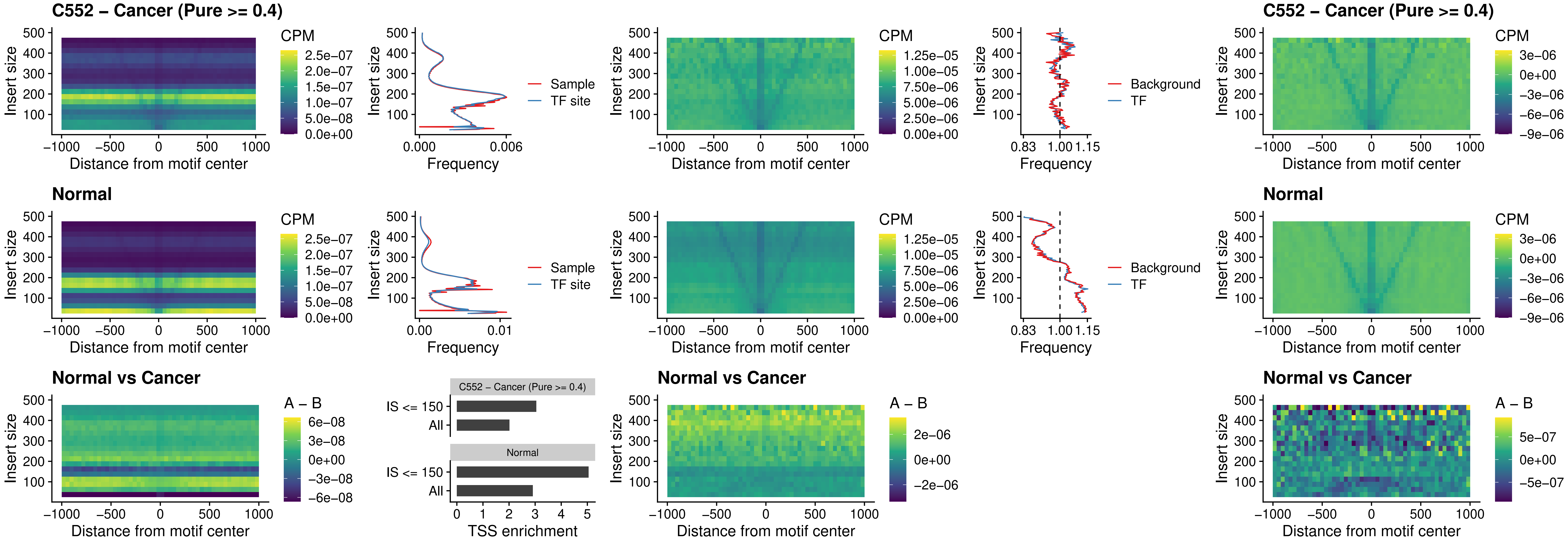

### HOXD3_uc_dTSS_oPEAK

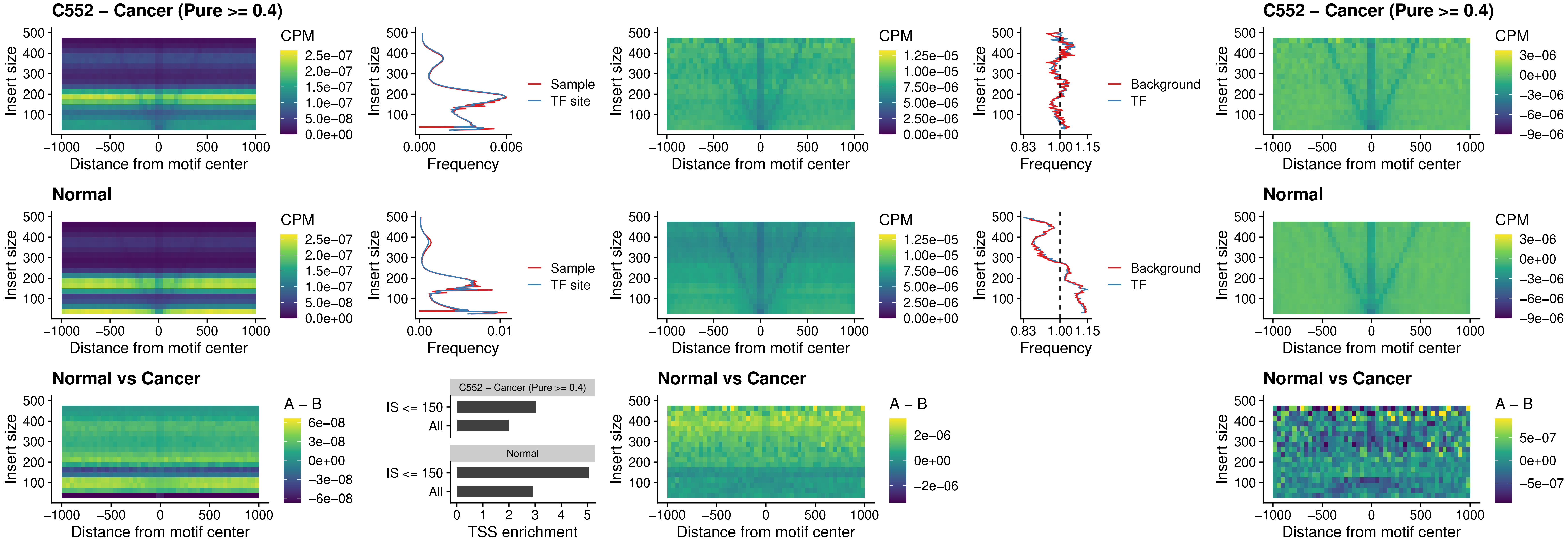

### BBX_uc_dTSS_nPEAK

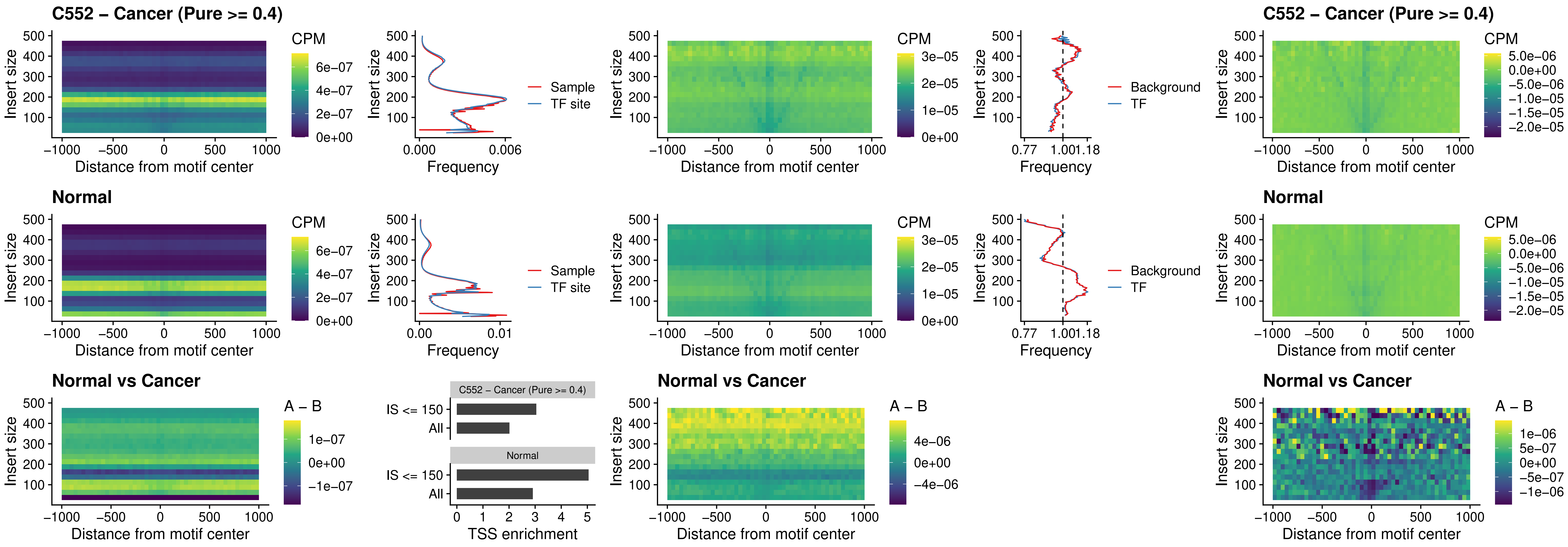

#### SOX3_uc_cTSS_nPEAK

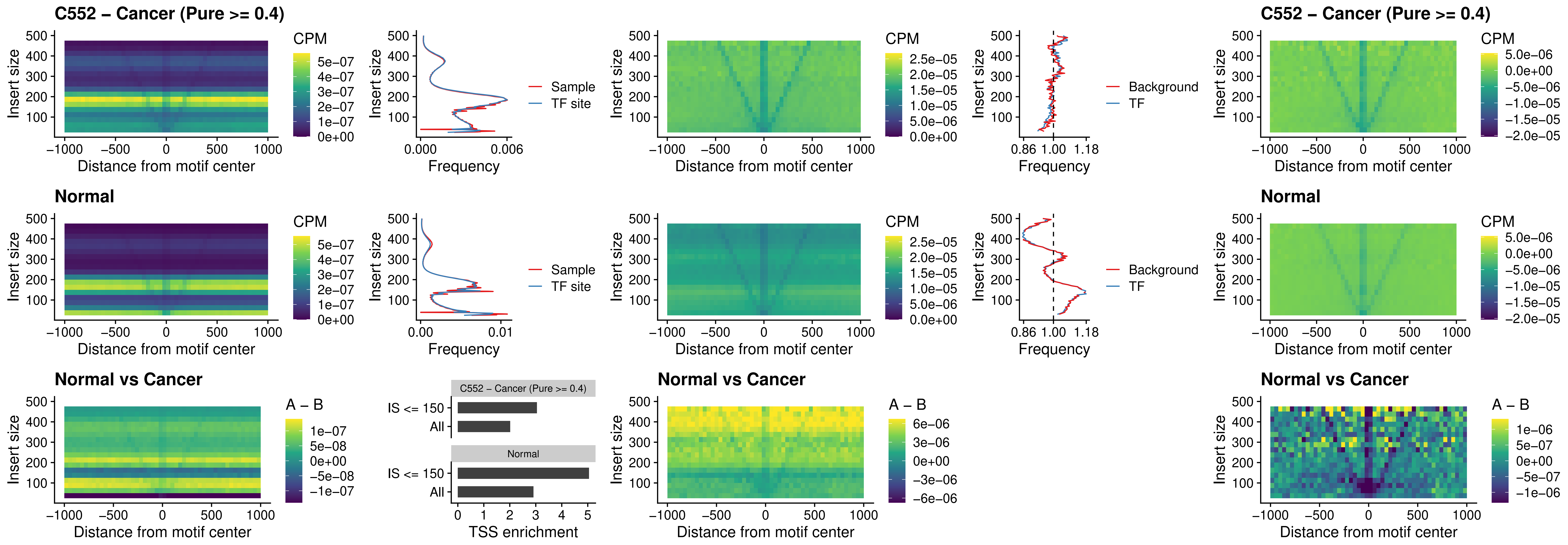

### TBPL2_uc_cTSS_nPEAK

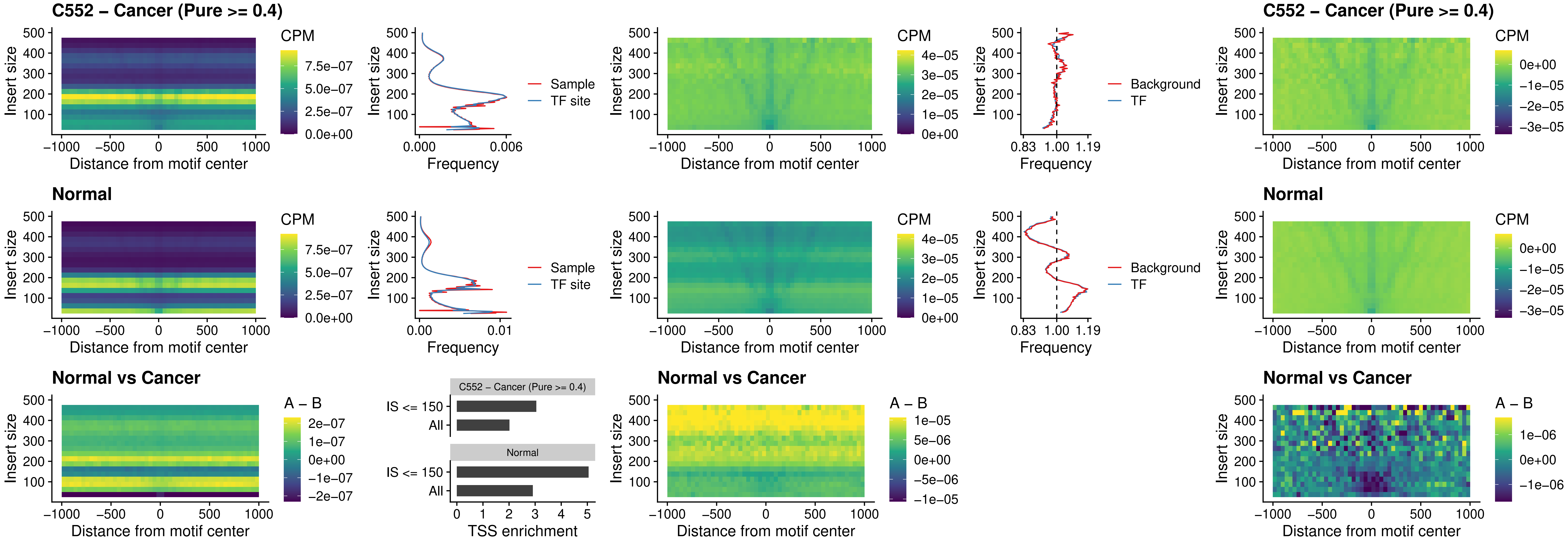

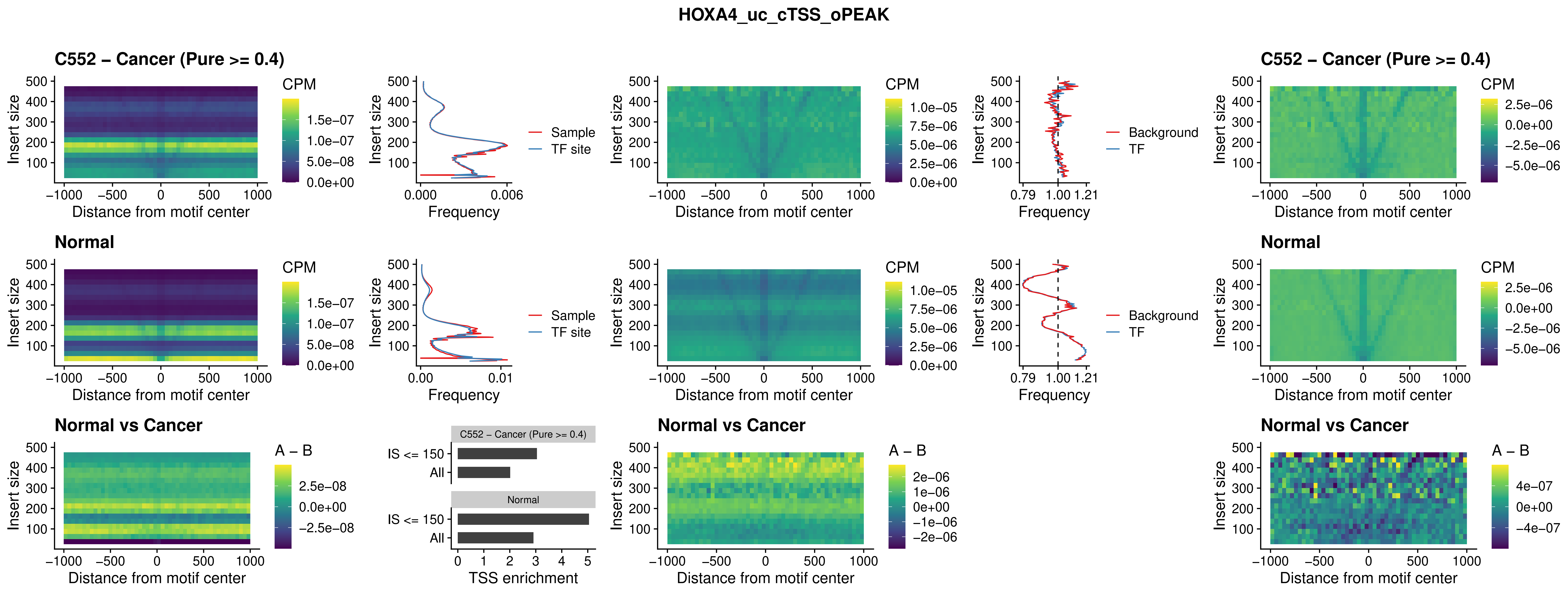
HOXA4_uc_cTSS_oPEAK
C552 – Cancer (Pure >= 0.4)
Normal
Normal vs Cancer
Insert size
Distance from motif center
Frequency
CPM
A – B
Sample
TF site
Background
TF
C552 – Cancer (Pure >= 0.4)
Normal
IS <= 150
All
TSS enrichment

### PDX1_uc_dTSS_oPEAK

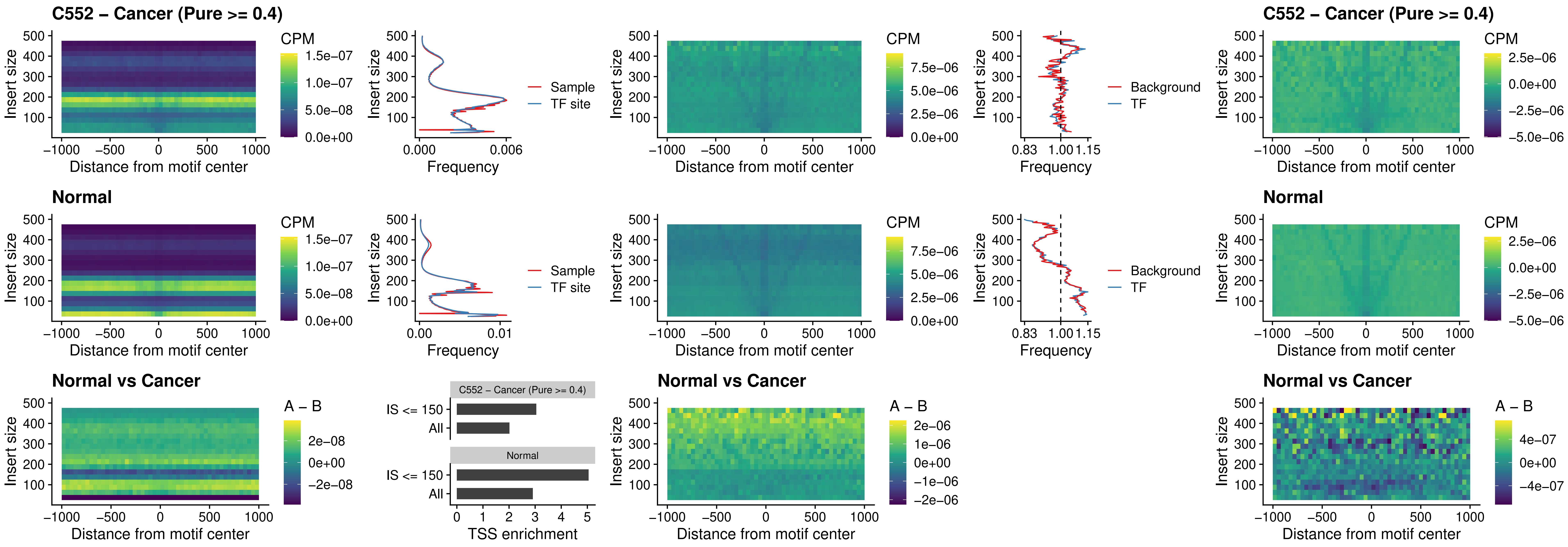

#### LHX4_uc_dTSS_oPEAK

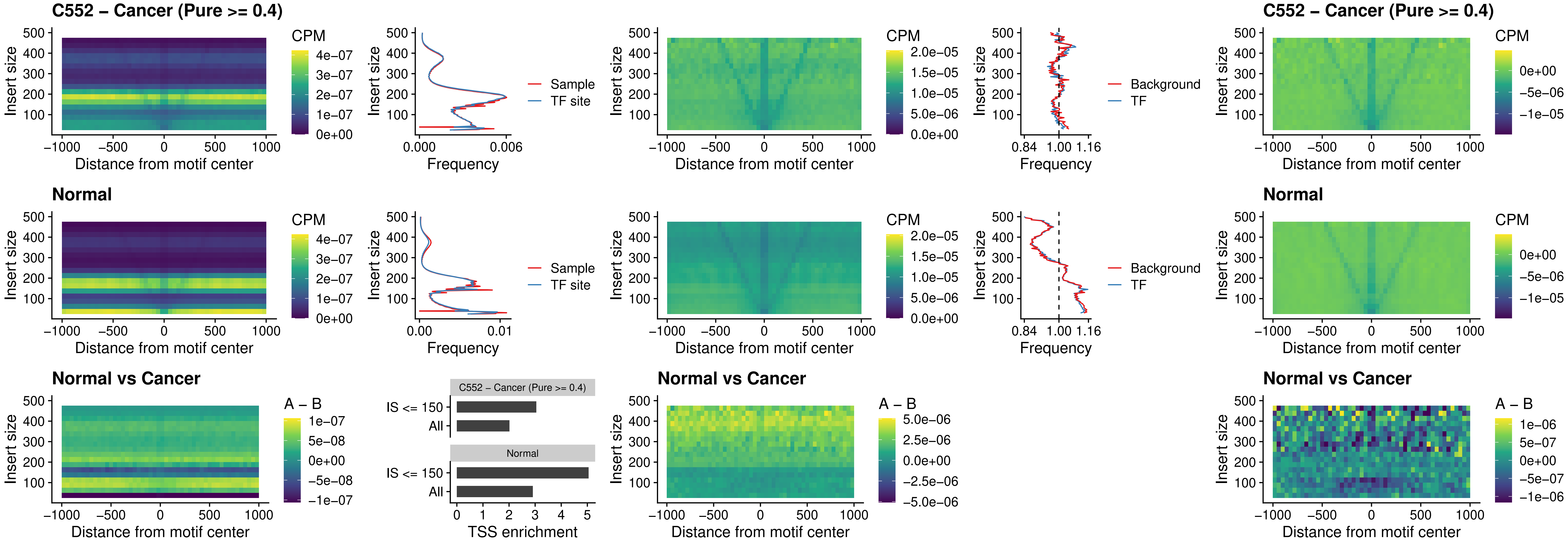

### HOXA4_uc_dTSS_oPEAK

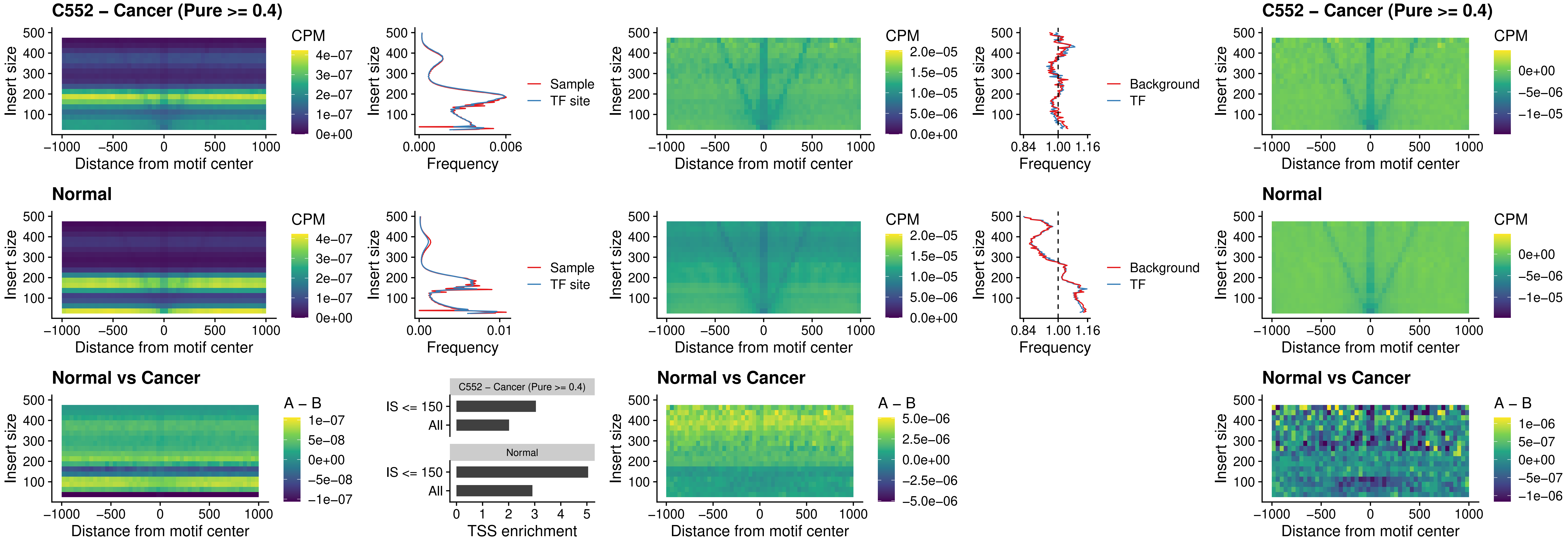

### HLX_uc_dTSS_oPEAK

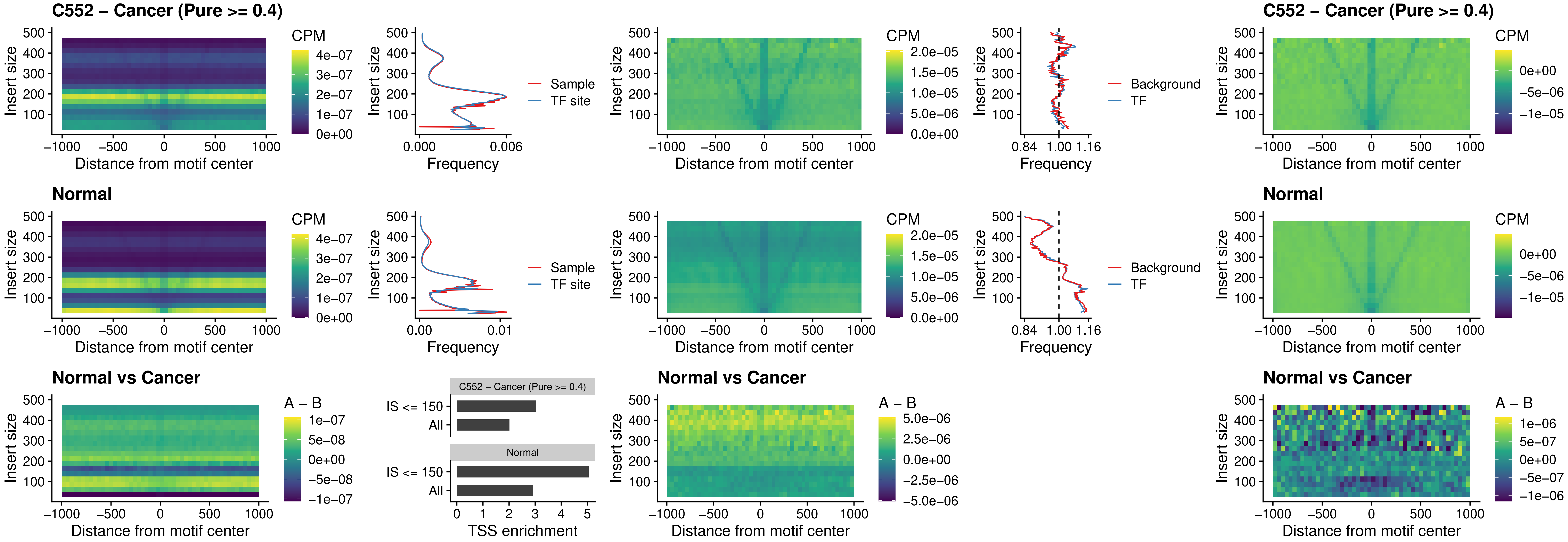

### AC0125311_uc_dTSS_oPEAK

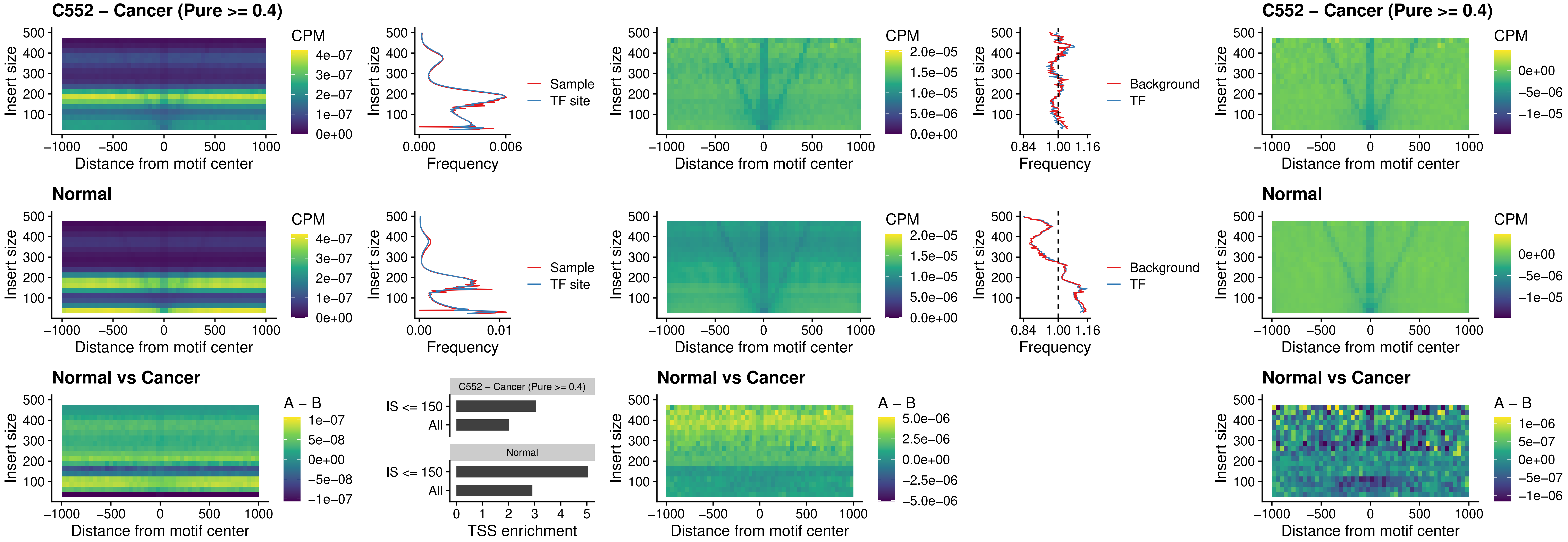

#### DBX2_uc_dTSS_oPEAK

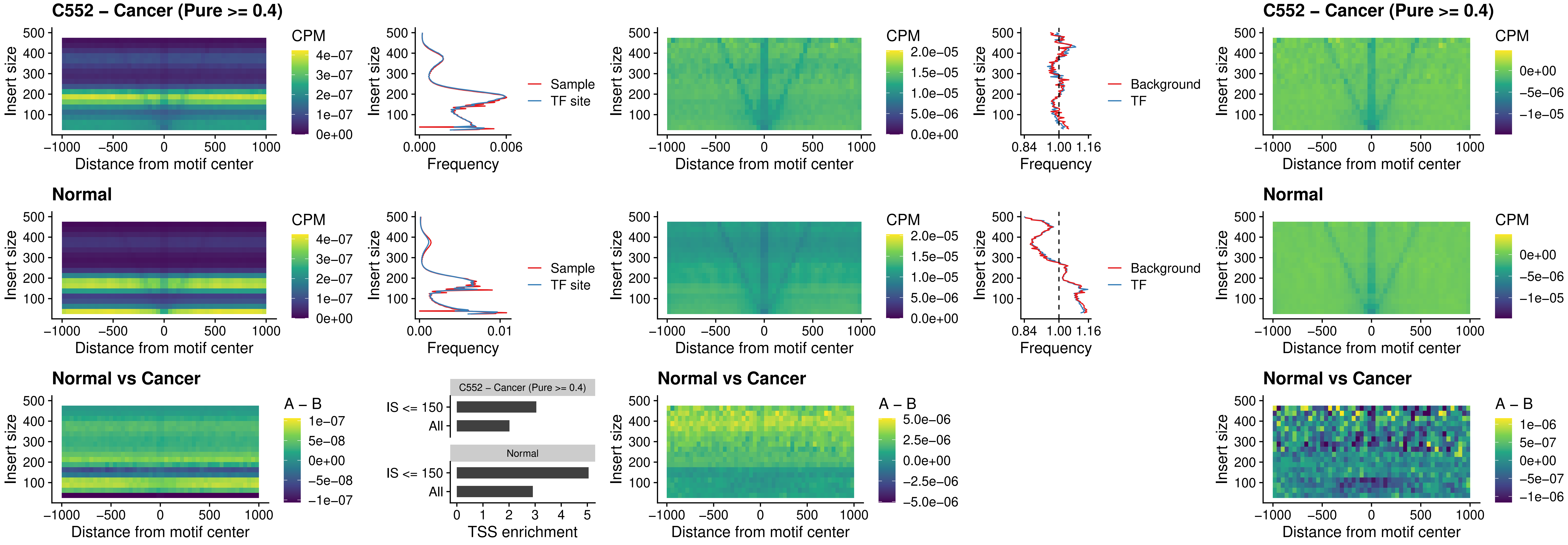

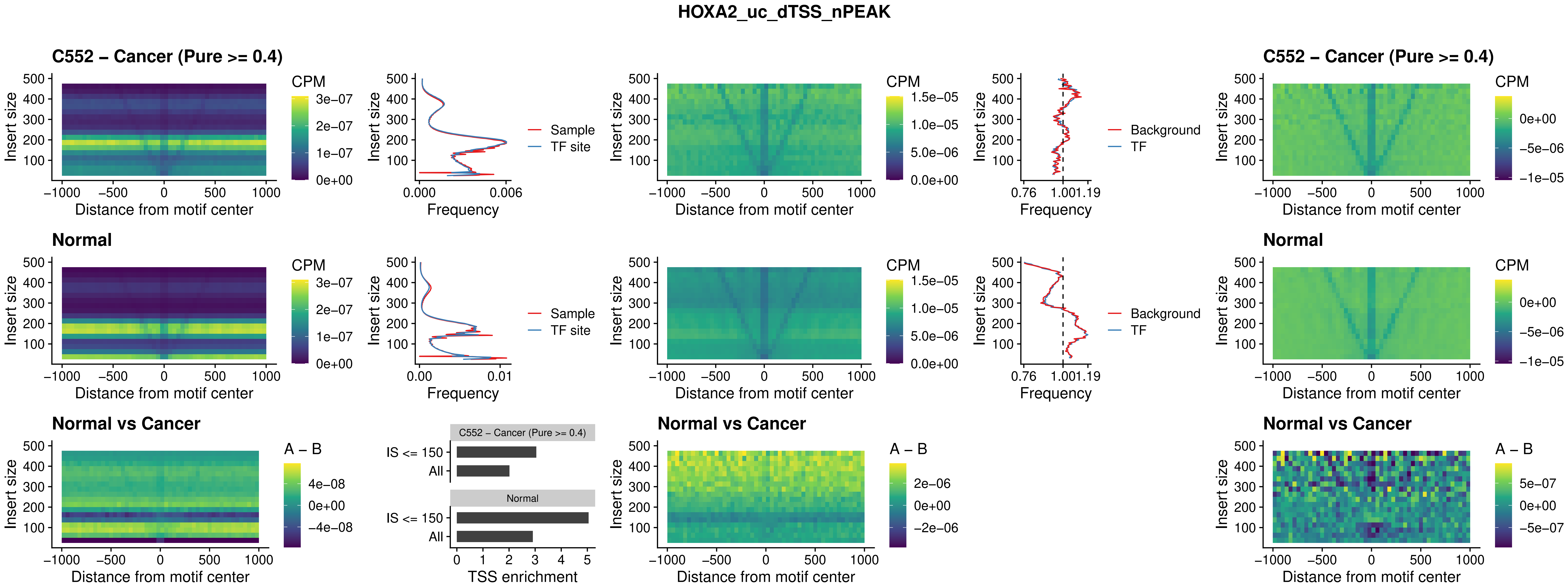
HOXA2_uc_dTSS_nPEAK
C552 – Cancer (Pure >= 0.4)
Normal
Normal vs Cancer
Insert size
Distance from motif center
Frequency
CPM
A – B
Sample
TF site
Background
TF
C552 – Cancer (Pure >= 0.4)
Normal
IS <= 150
All
TSS enrichment

### SHOX_uc_dTSS_nPEAK

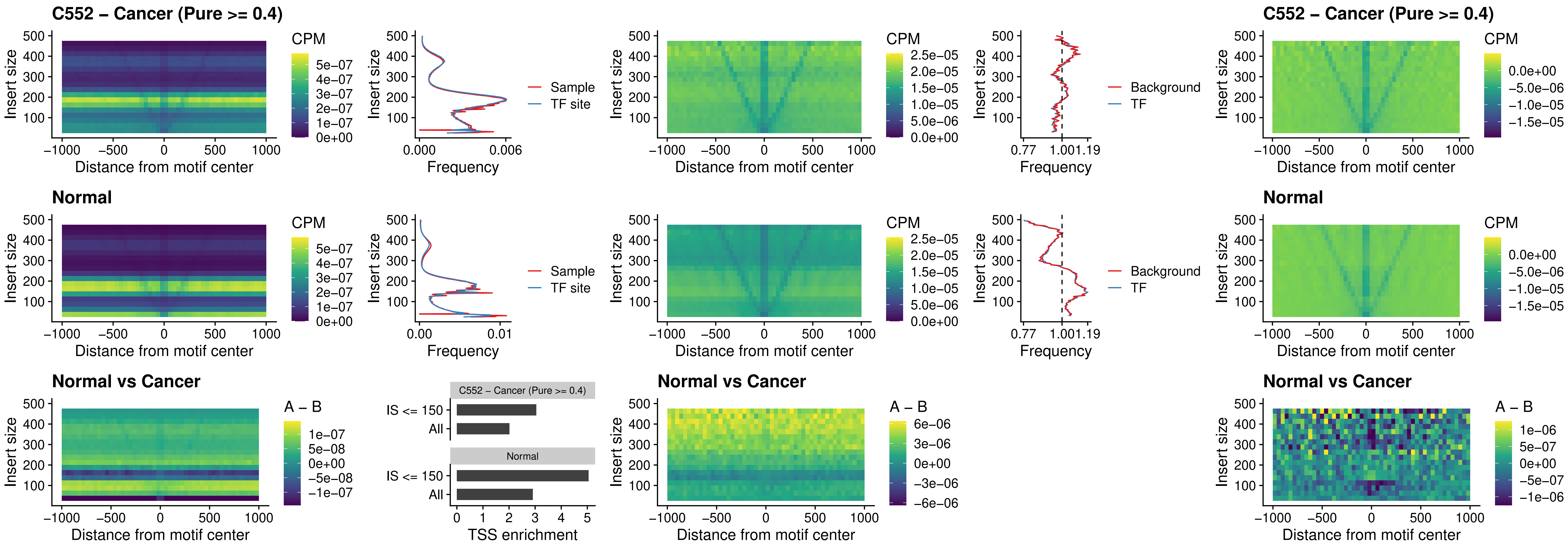

### PRRX1_uc_dTSS_nPEAK

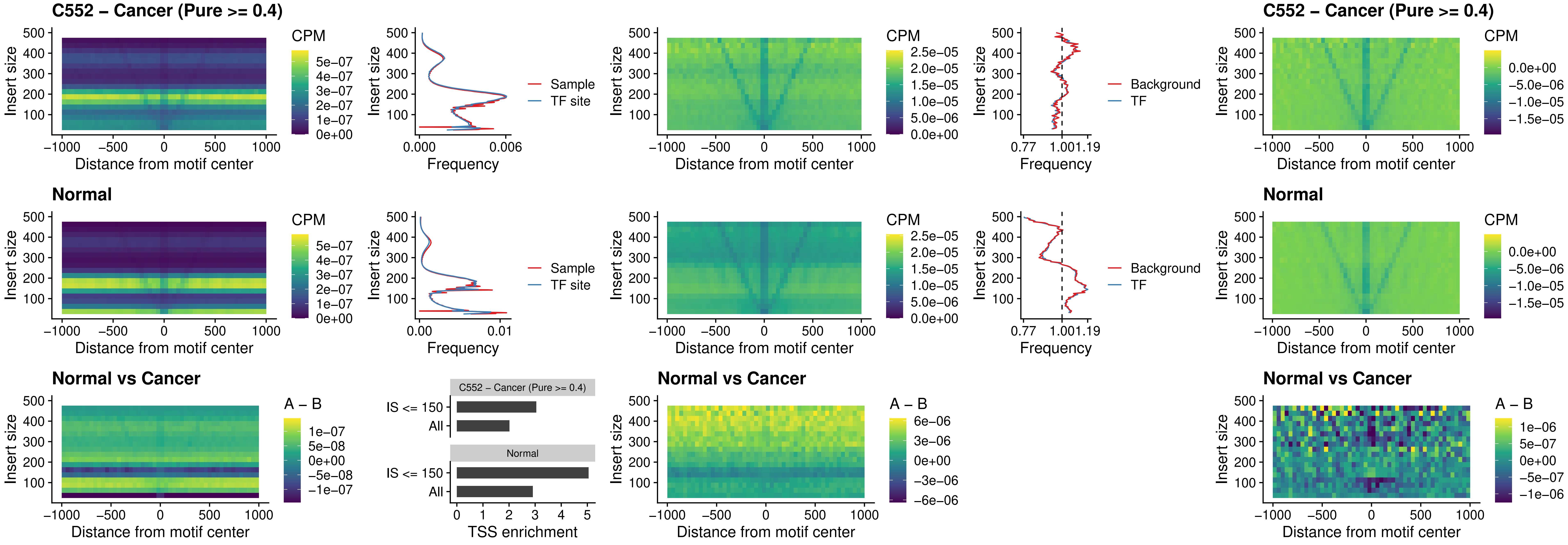

### LMX1A_uc_dTSS_nPEAK

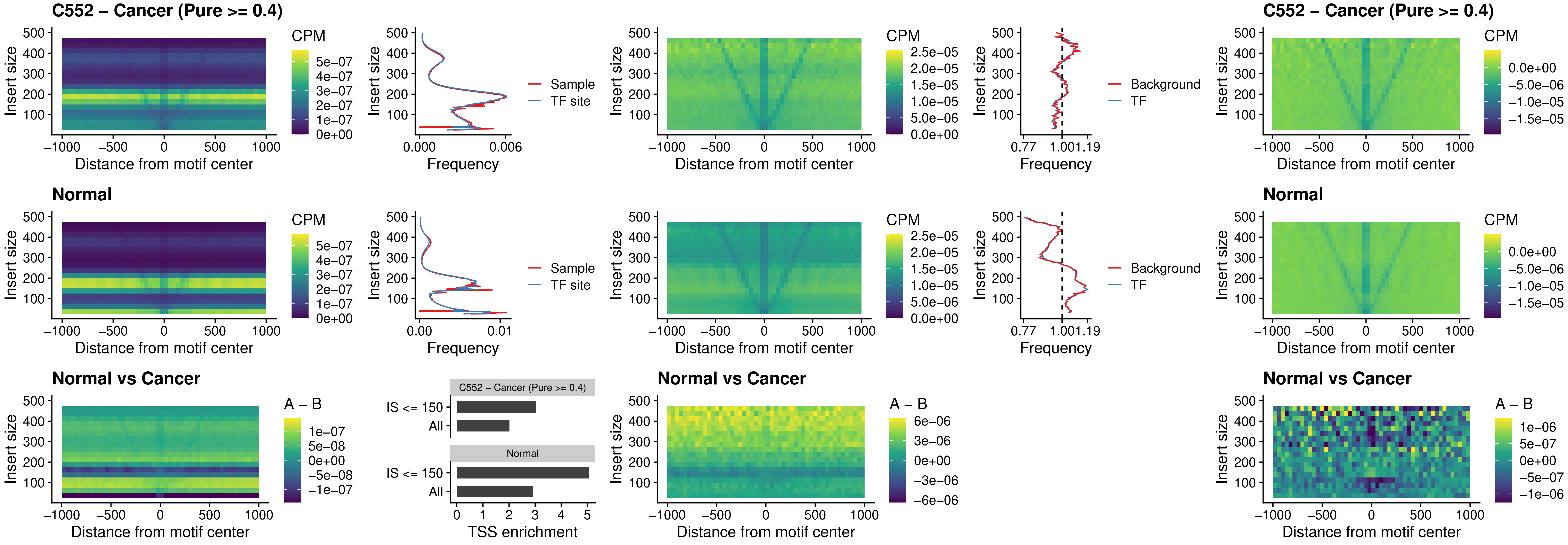

### LMX1B_uc_dTSS_nPEAK

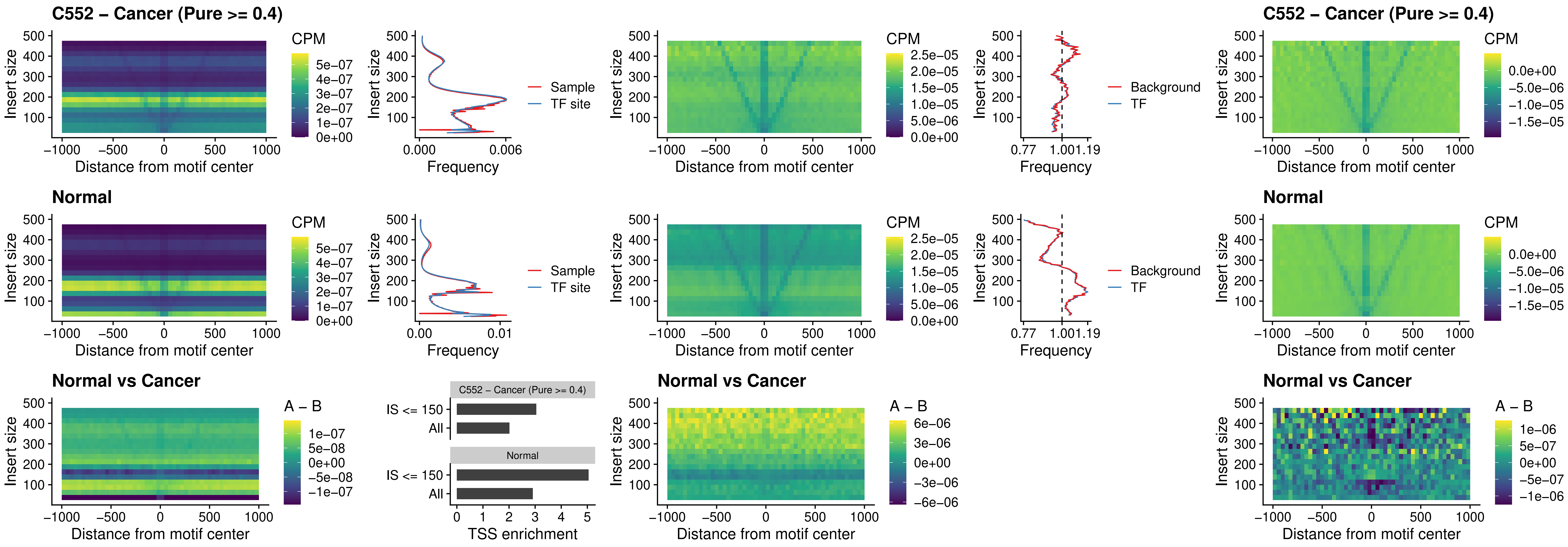

#### DLX5_uc_dTSS_oPEAK

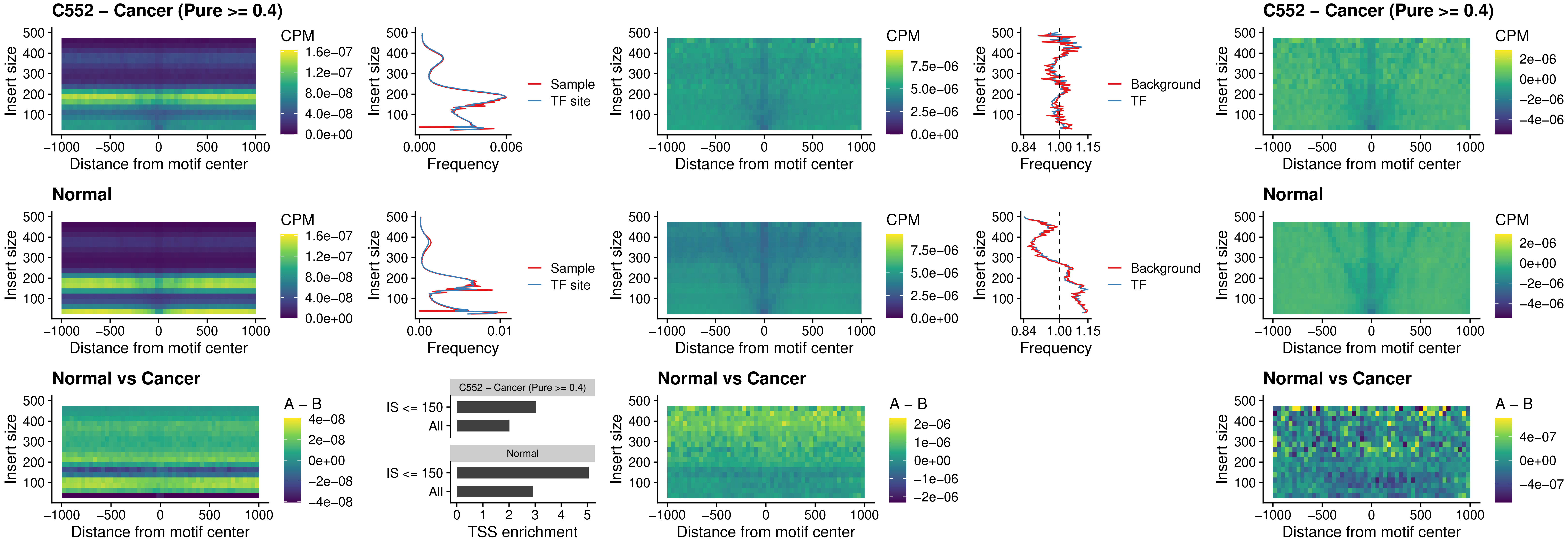

### ARID3C_uc_dTSS_nPEAK

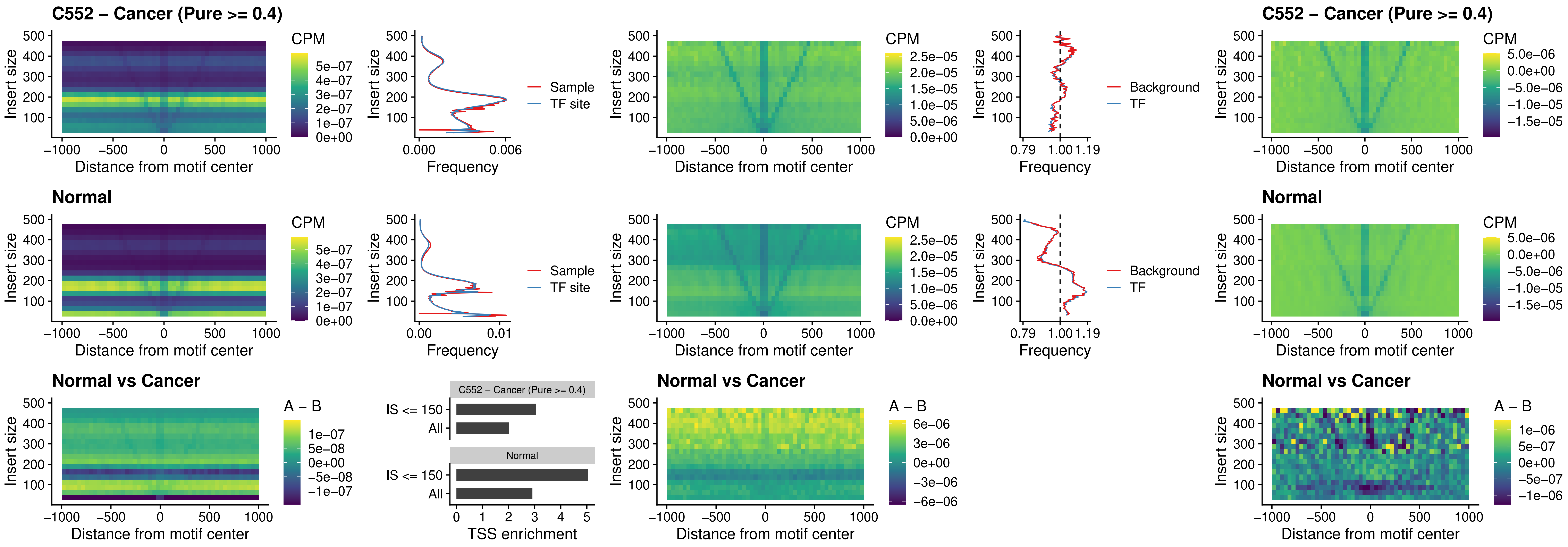

### HOXD12_uc_dTSS_nPEAK

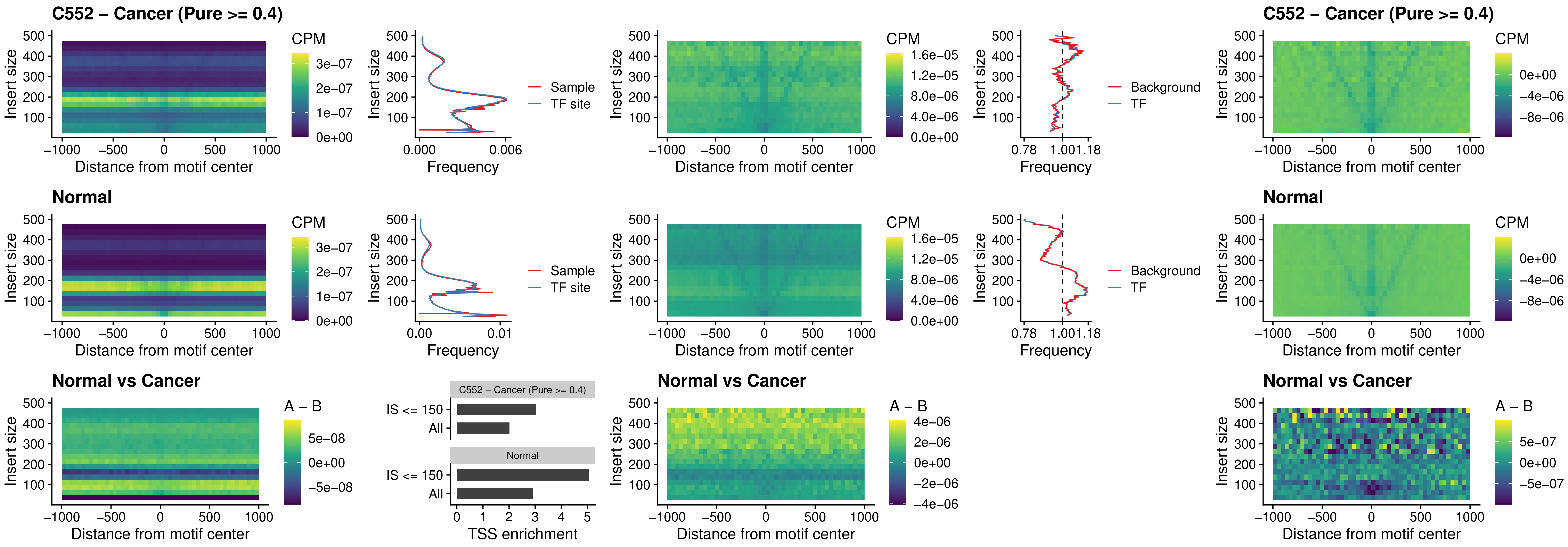

### CDX2_uc_dTSS_nPEAK

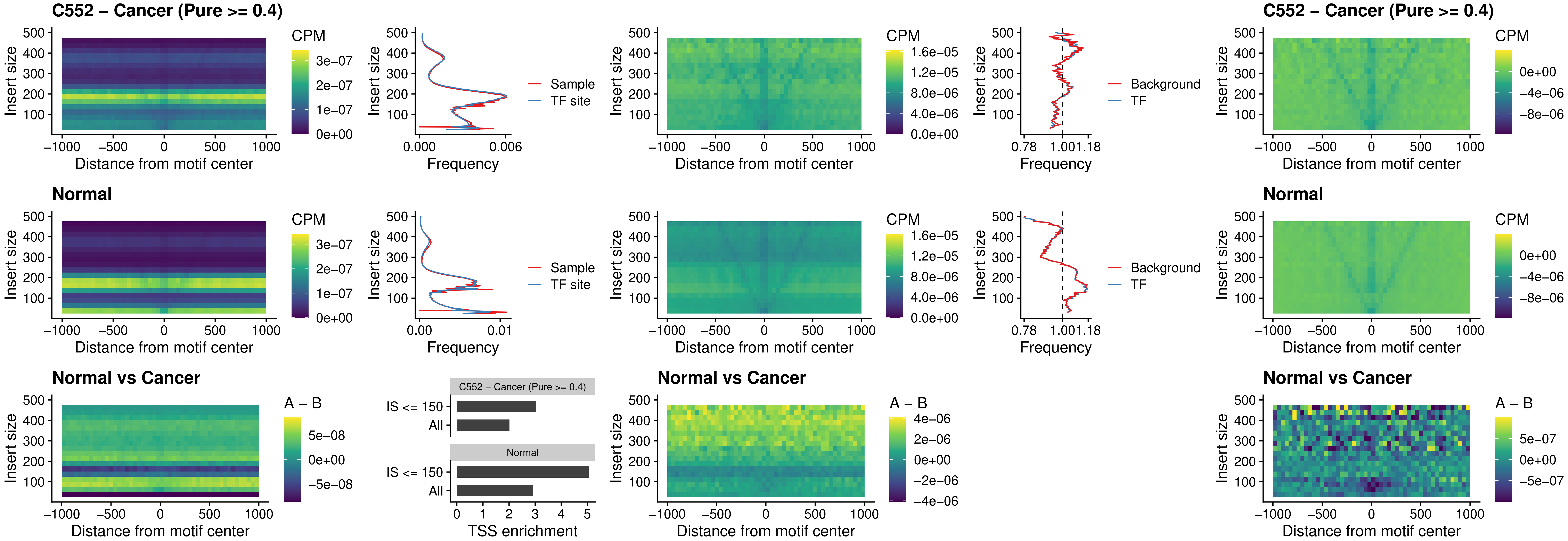

### HOXC12_uc_dTSS_nPEAK

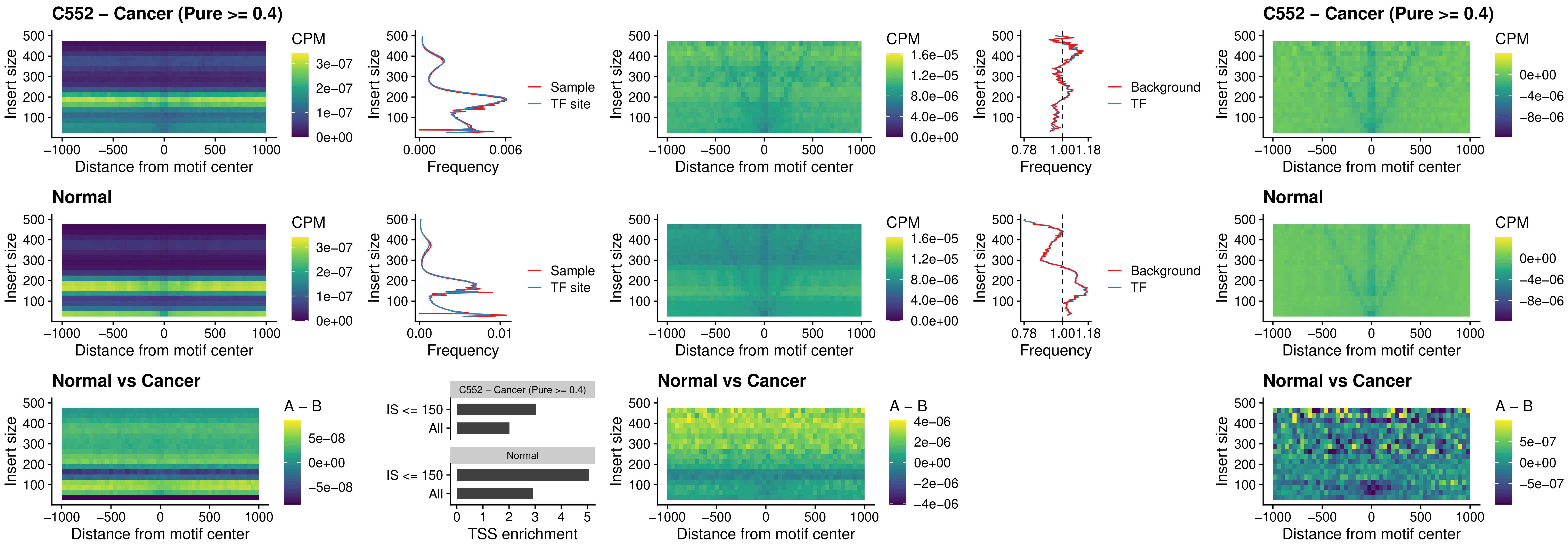

### HHEX_uc_dTSS_oPEAK

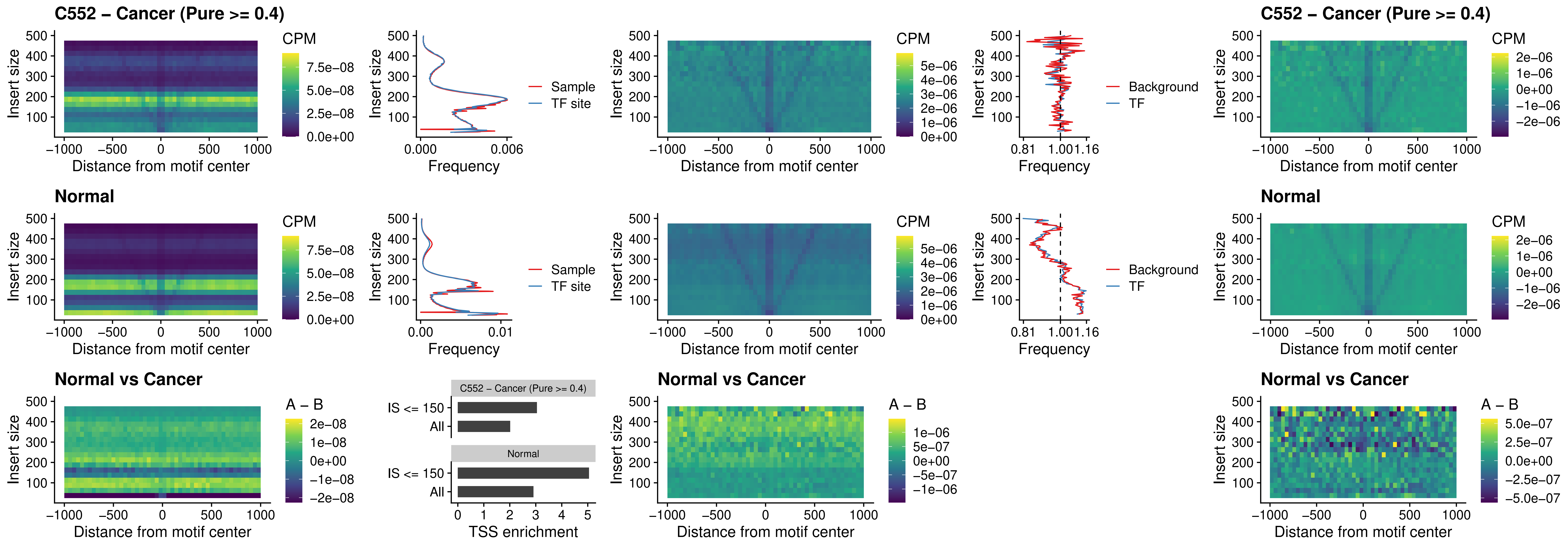

#### HOXA11_uc_dTSS_nPEAK

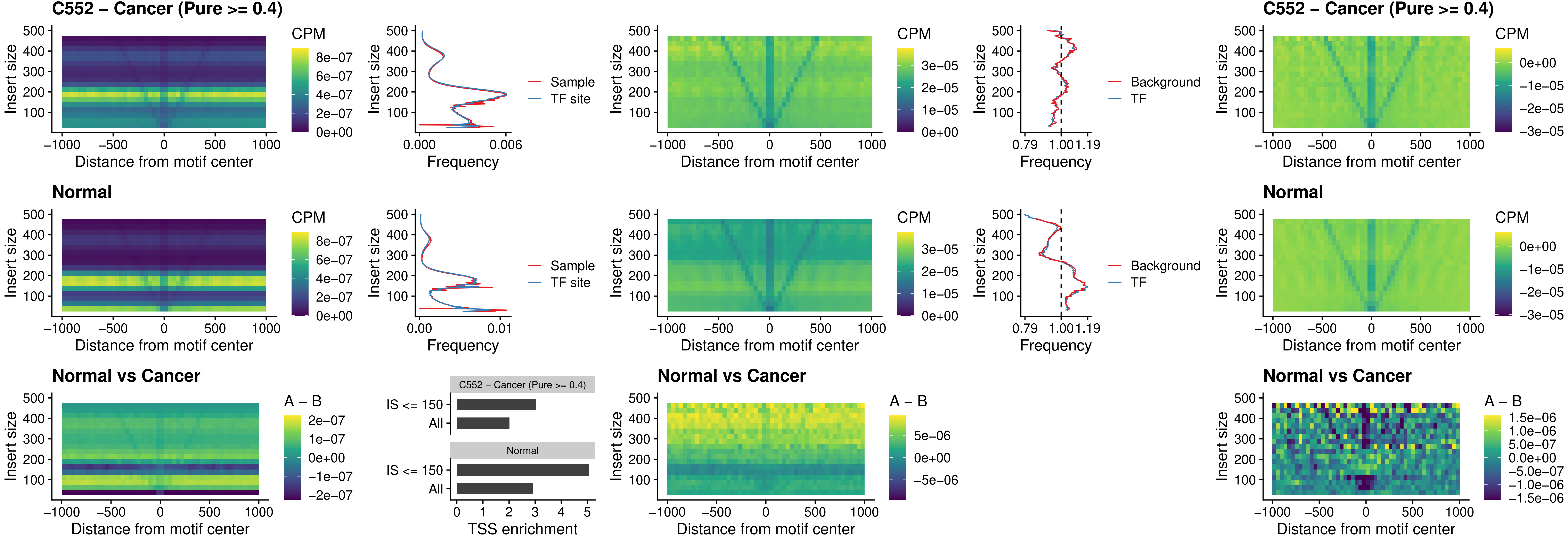

#### HOXA11_uc_cTSS_nPEAK

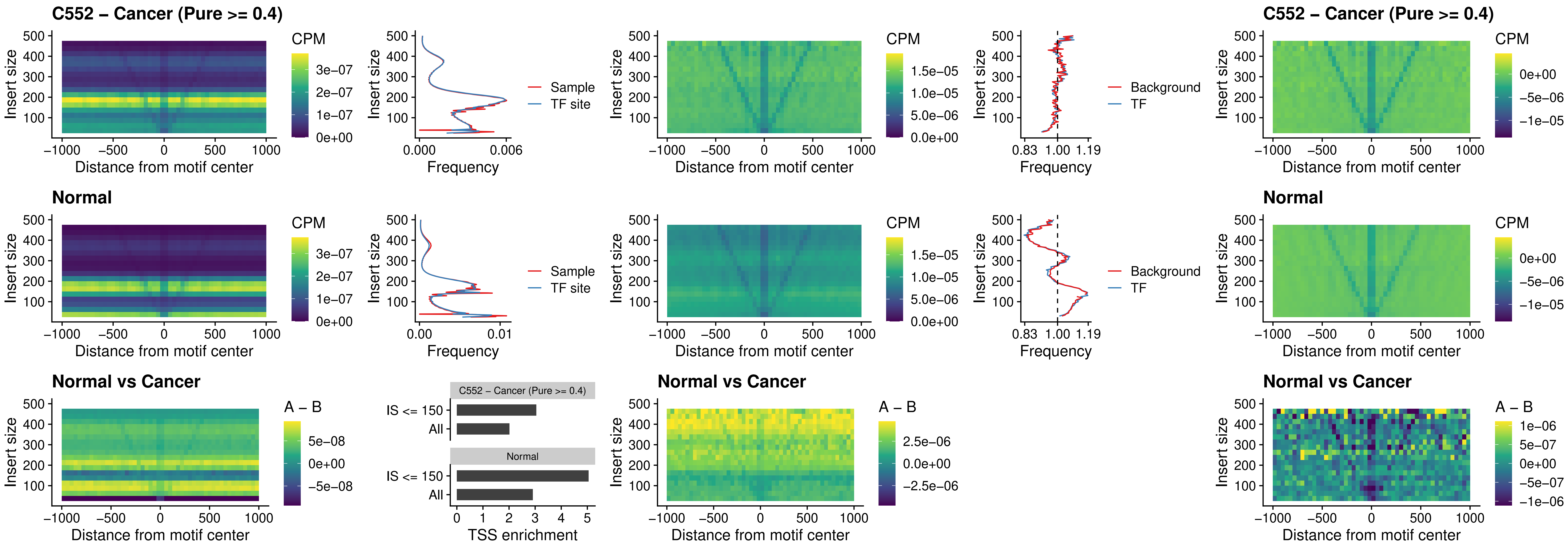

### ZFHX3_uc_dTSS_nPEAK

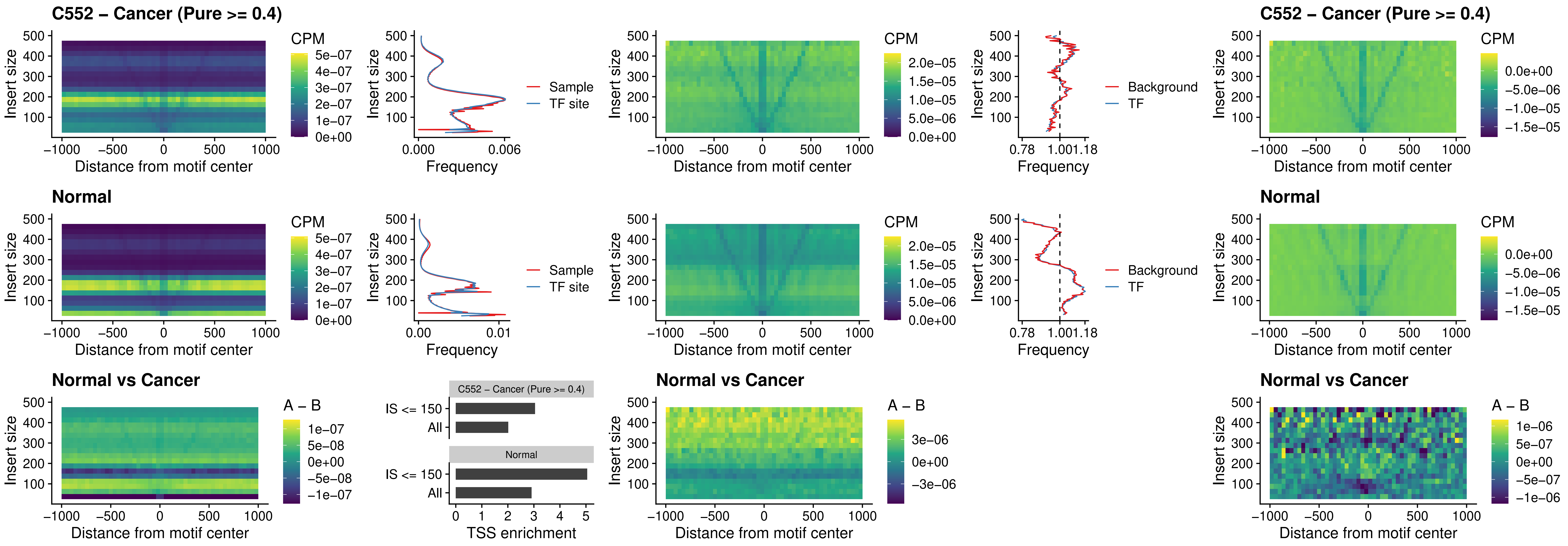

### HOXB4_uc_dTSS_nPEAK

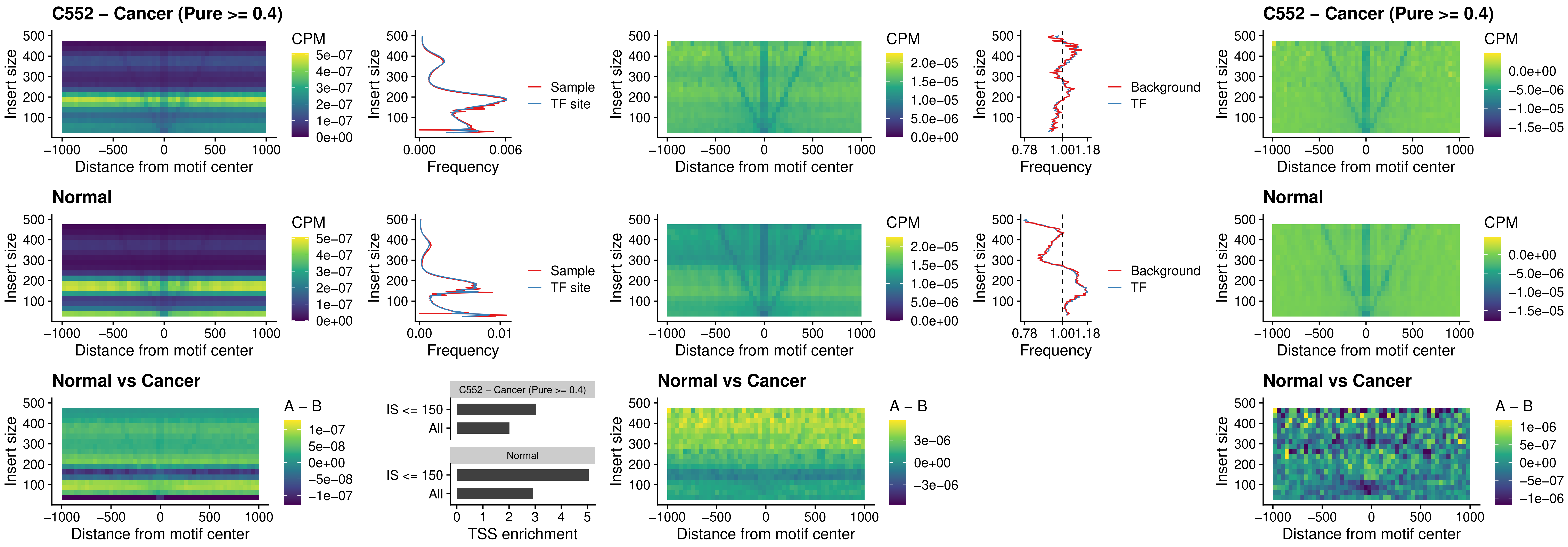

### HOXC5_uc_dTSS_nPEAK

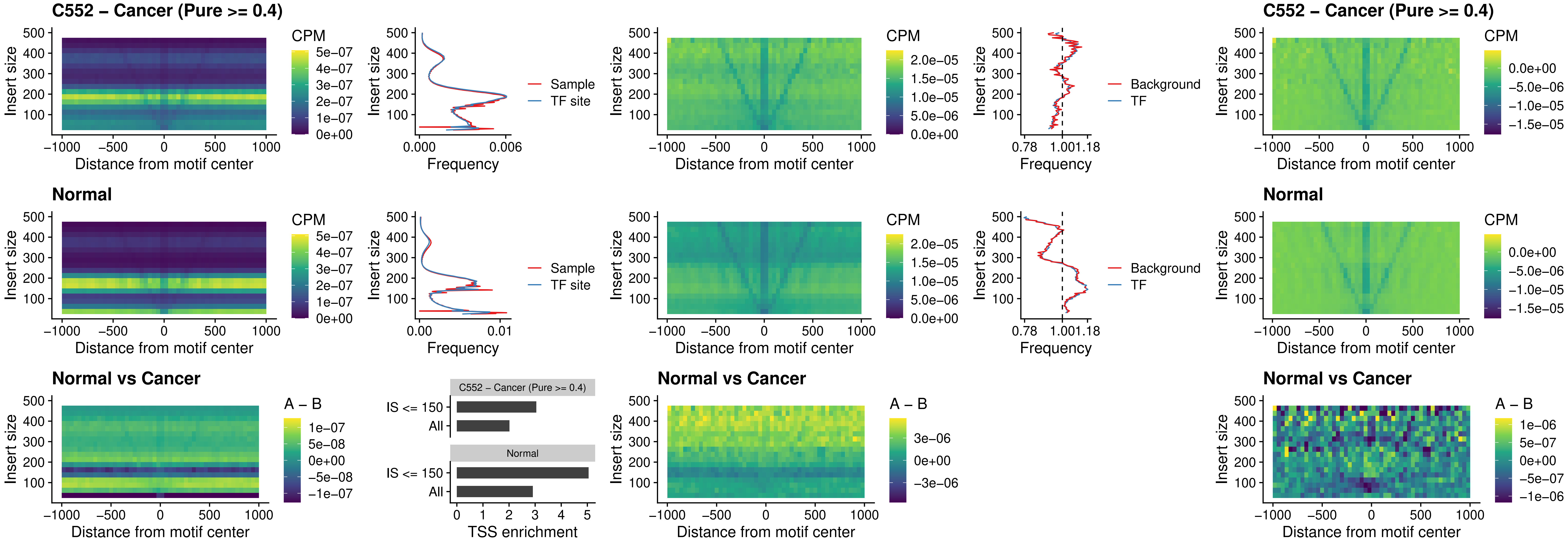

### HOXD1_uc_dTSS_nPEAK

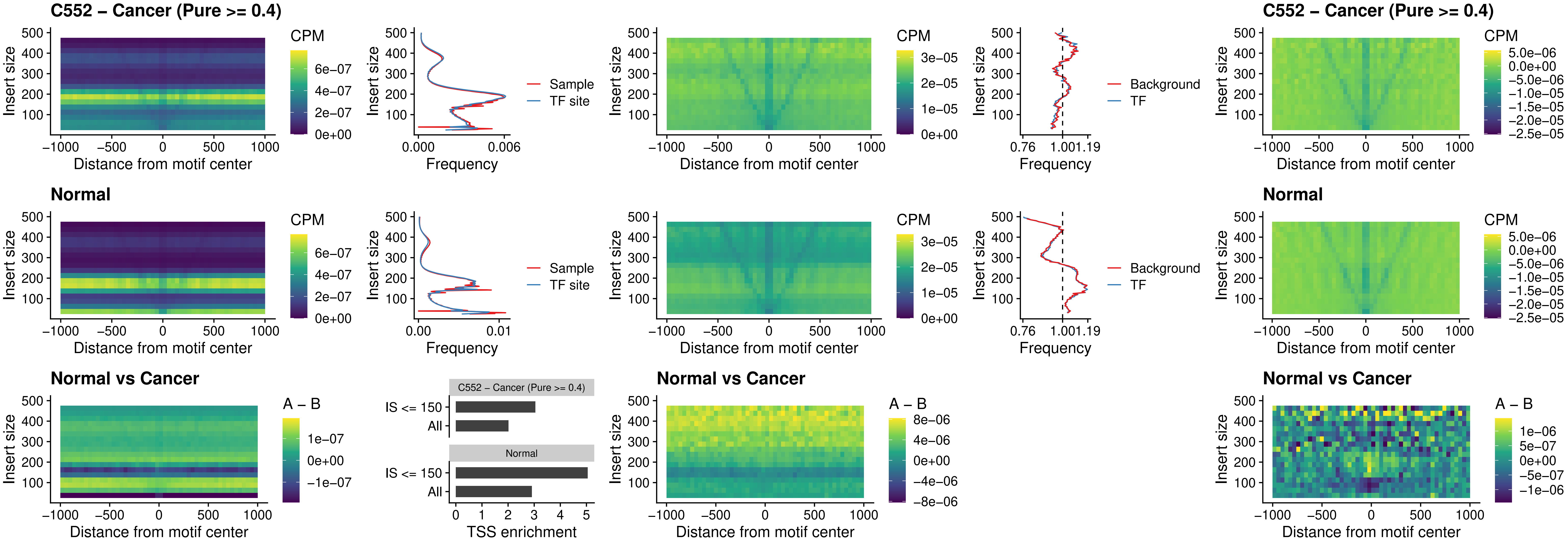

### OTP_uc_dTSS_nPEAK

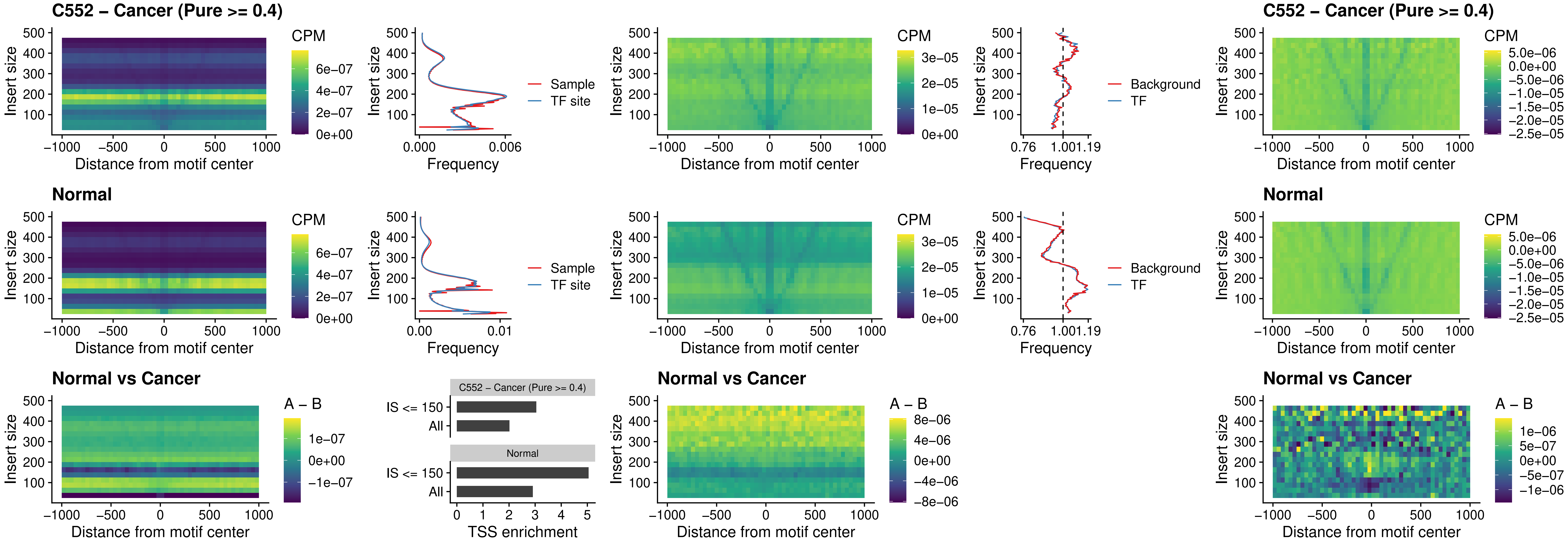

### FOXL1_uc_dTSS_oPEAK

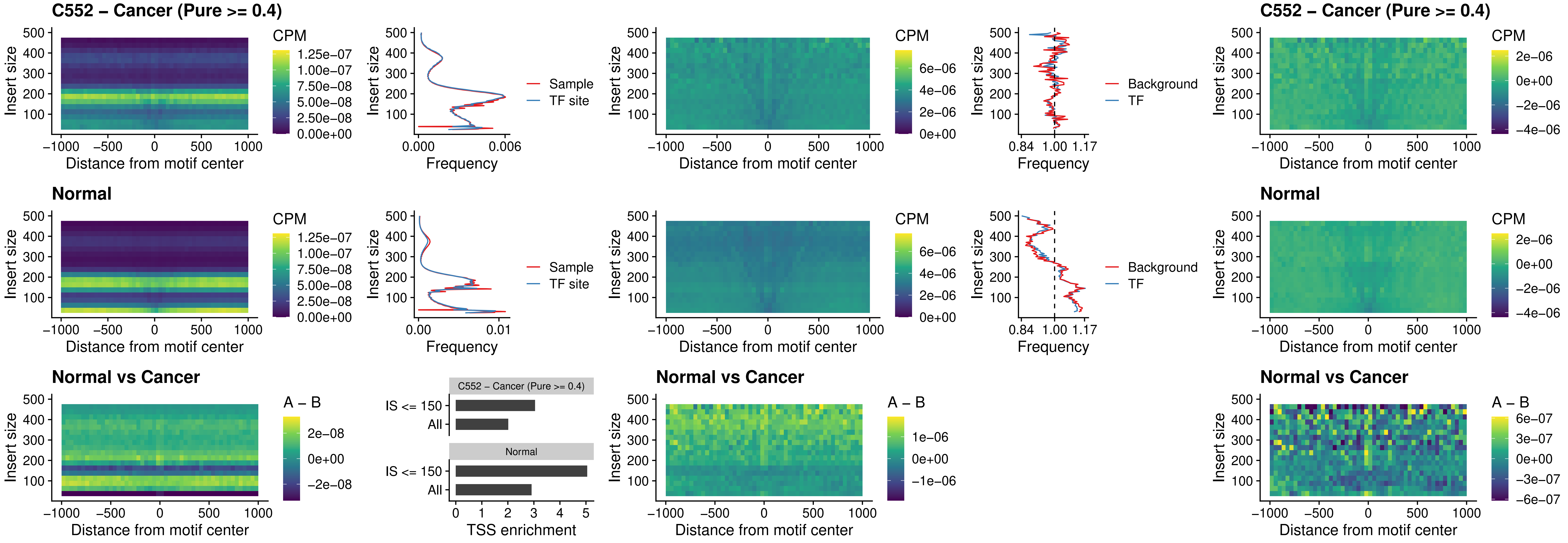

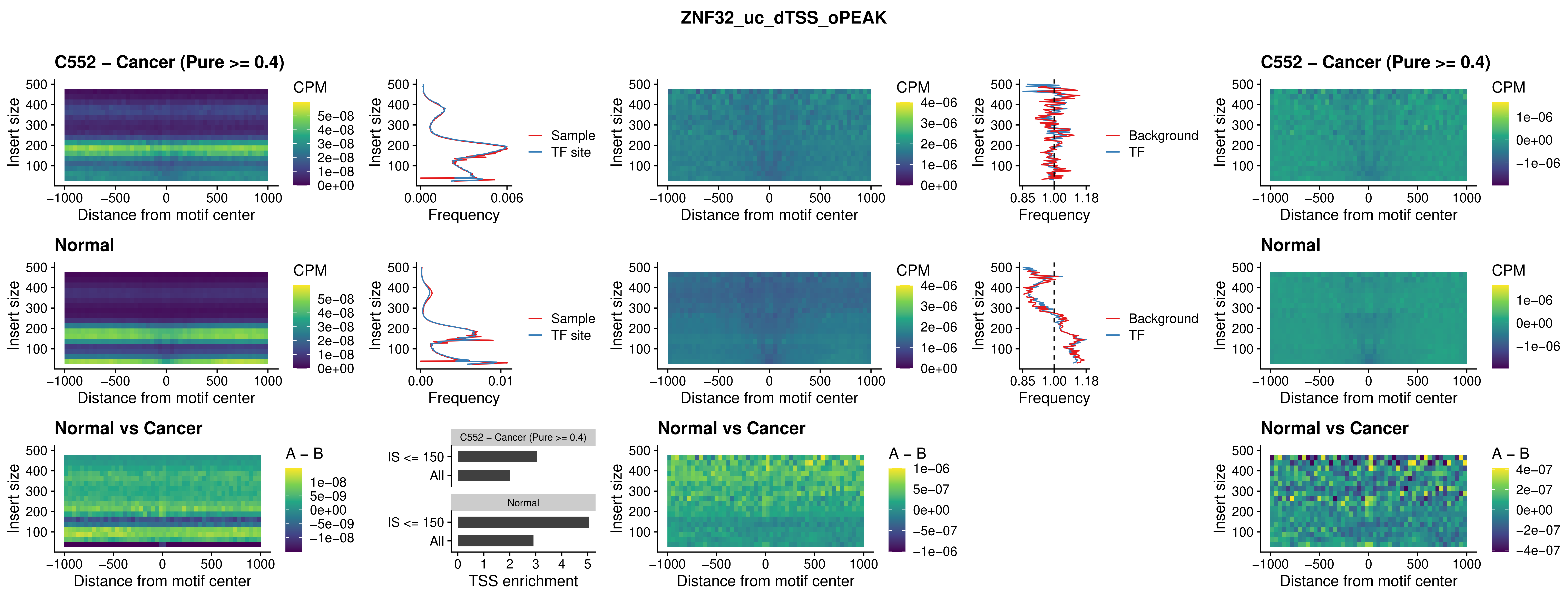
ZNF32_uc_dTSS_oPEAK
C552 – Cancer (Pure >= 0.4)
Normal
Normal vs Cancer
Insert size
Distance from motif center
Frequency
CPM
A – B
Sample
TF site
Background
TF
C552 – Cancer (Pure >= 0.4)
Normal
IS <= 150
All
TSS enrichment

#### FOXL1_uc_dTSS_nPEAK

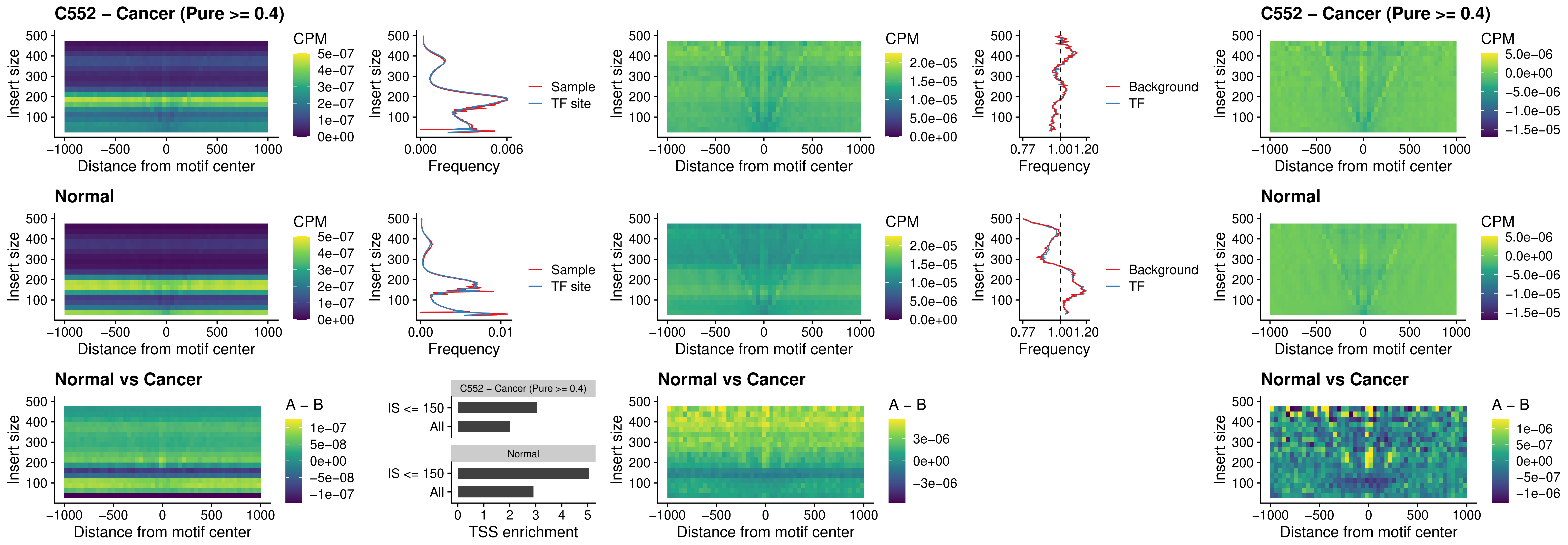

### ZNF32_uc_dTSS_nPEAK

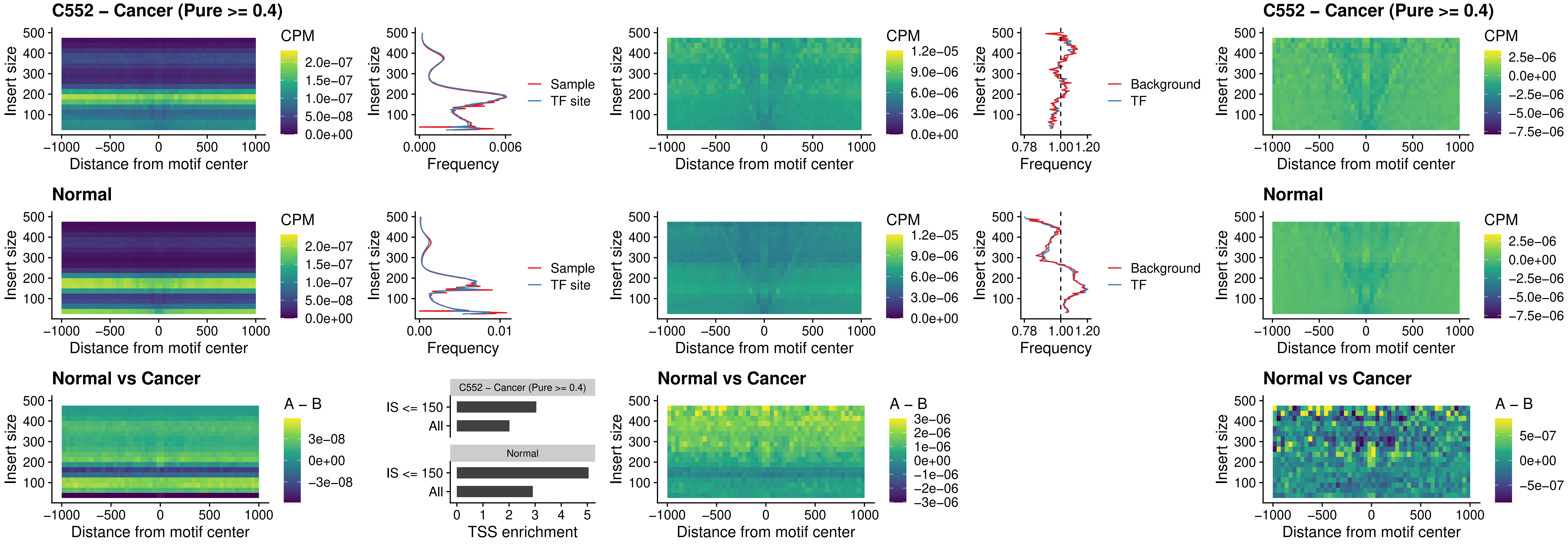

### DLX5_uc_dTSS_nPEAK

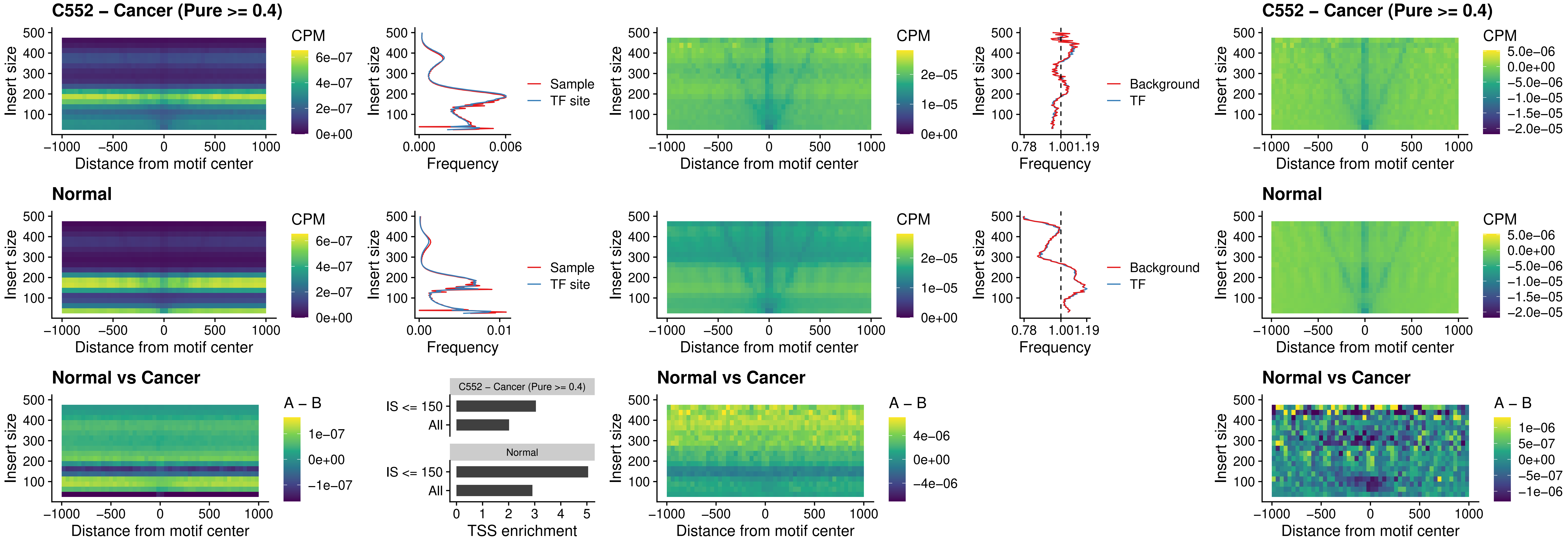

### DMRT2_uc_dTSS_nPEAK

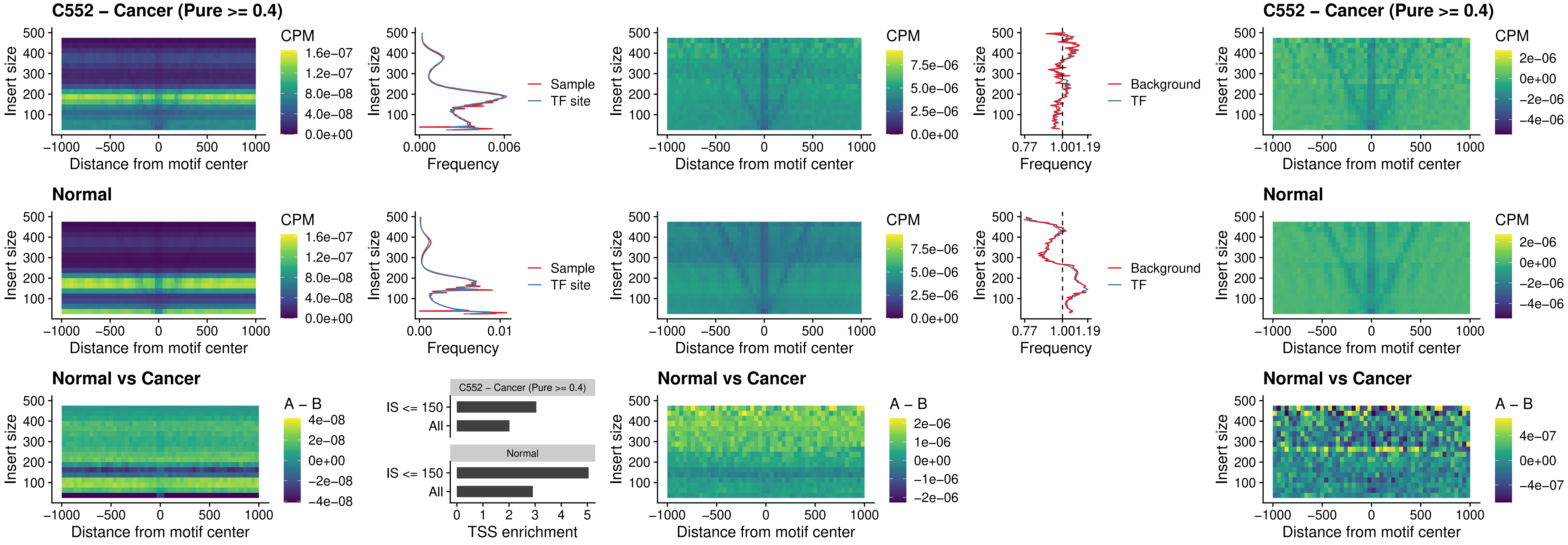

#### DMRTC2_uc_cTSS_nPEAK

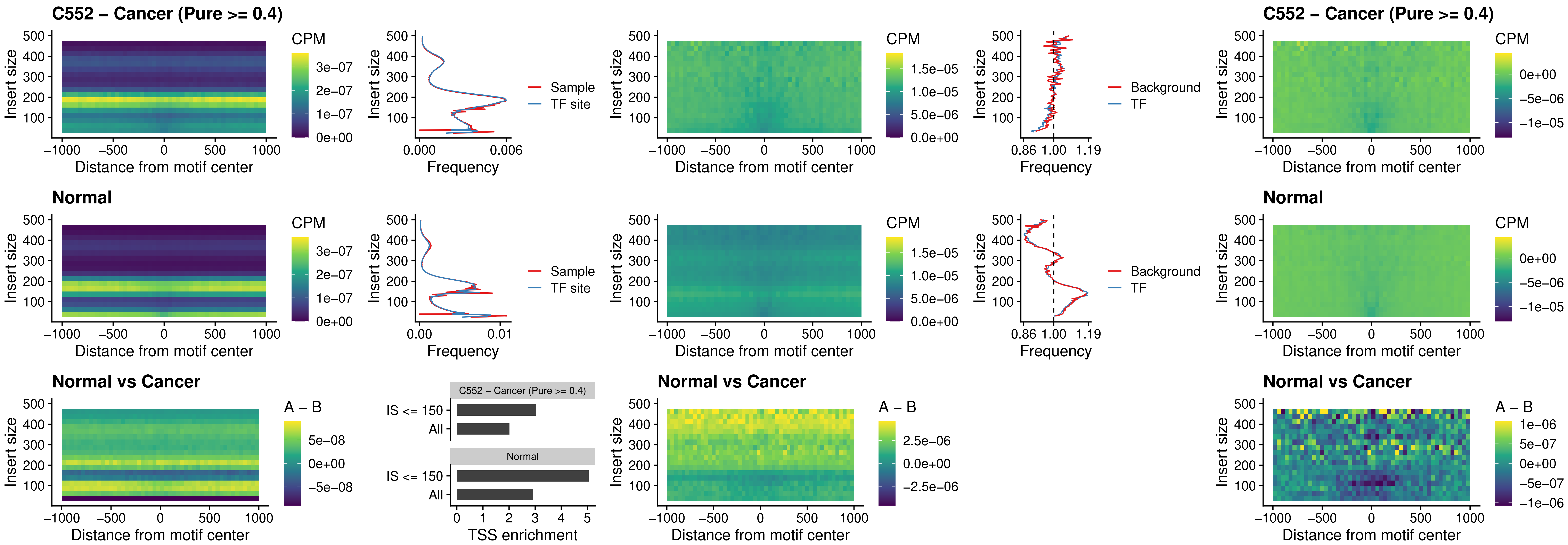

### HOXD9_uc_dTSS_nPEAK

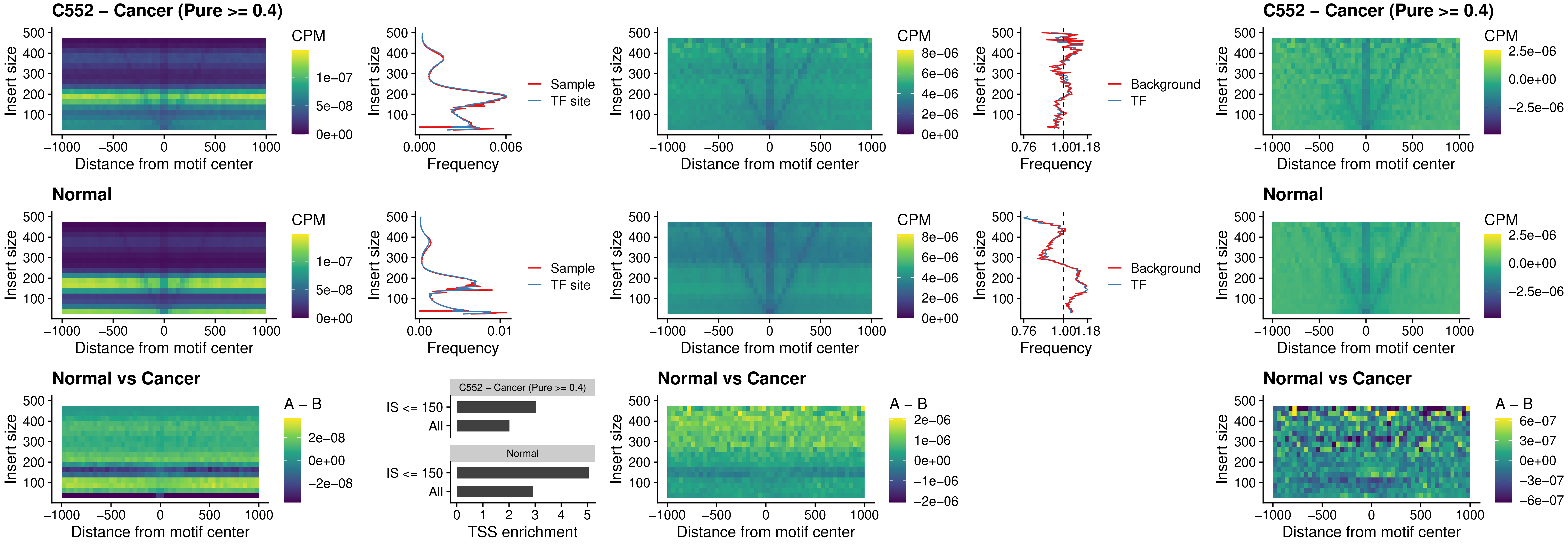

### HOXD4_uc_dTSS_nPEAK

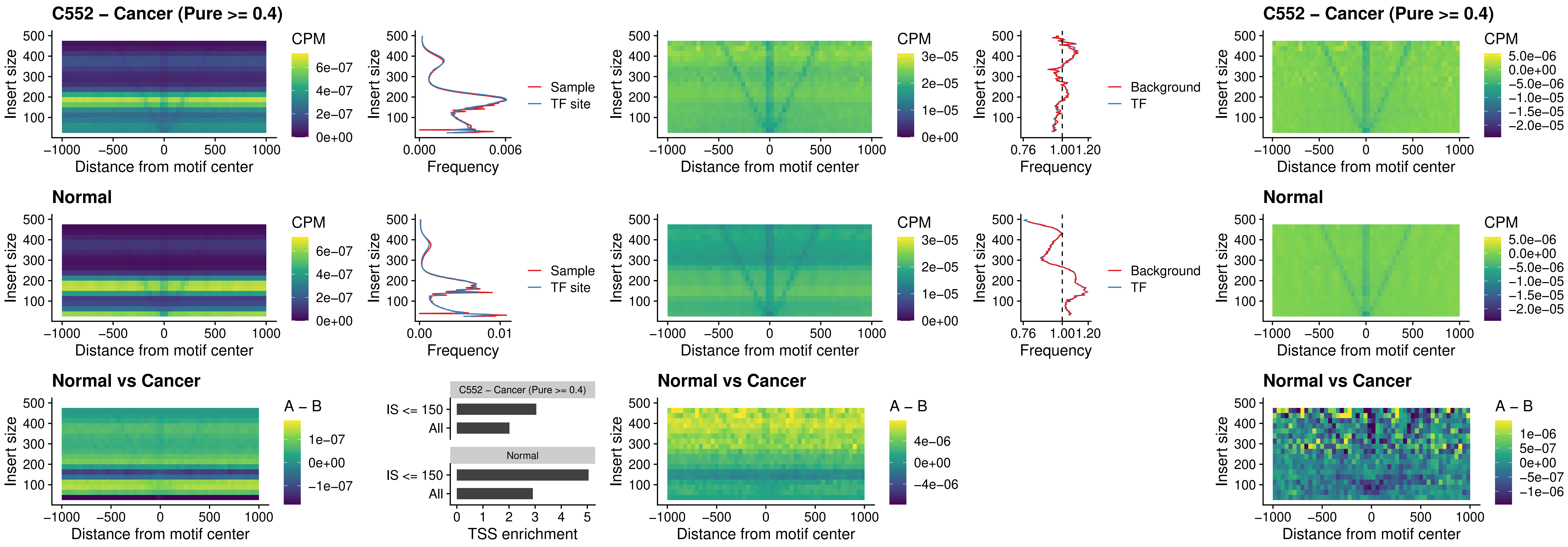

### FOXP3_uc_cTSS_nPEAK

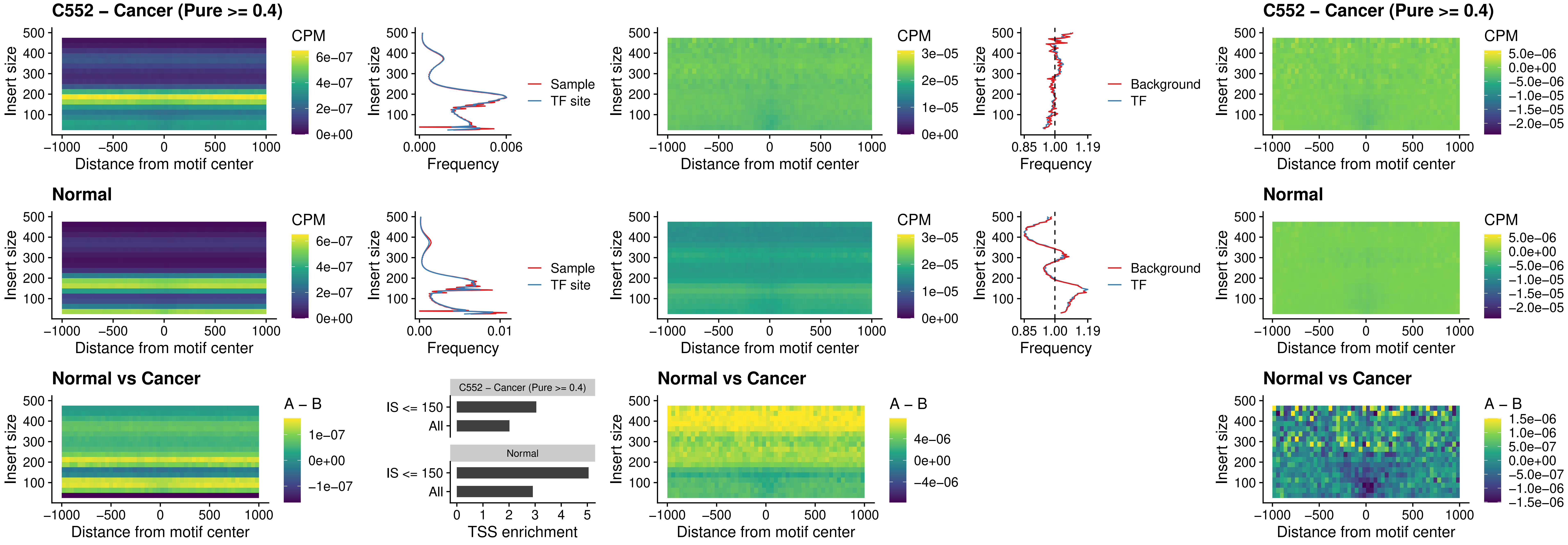

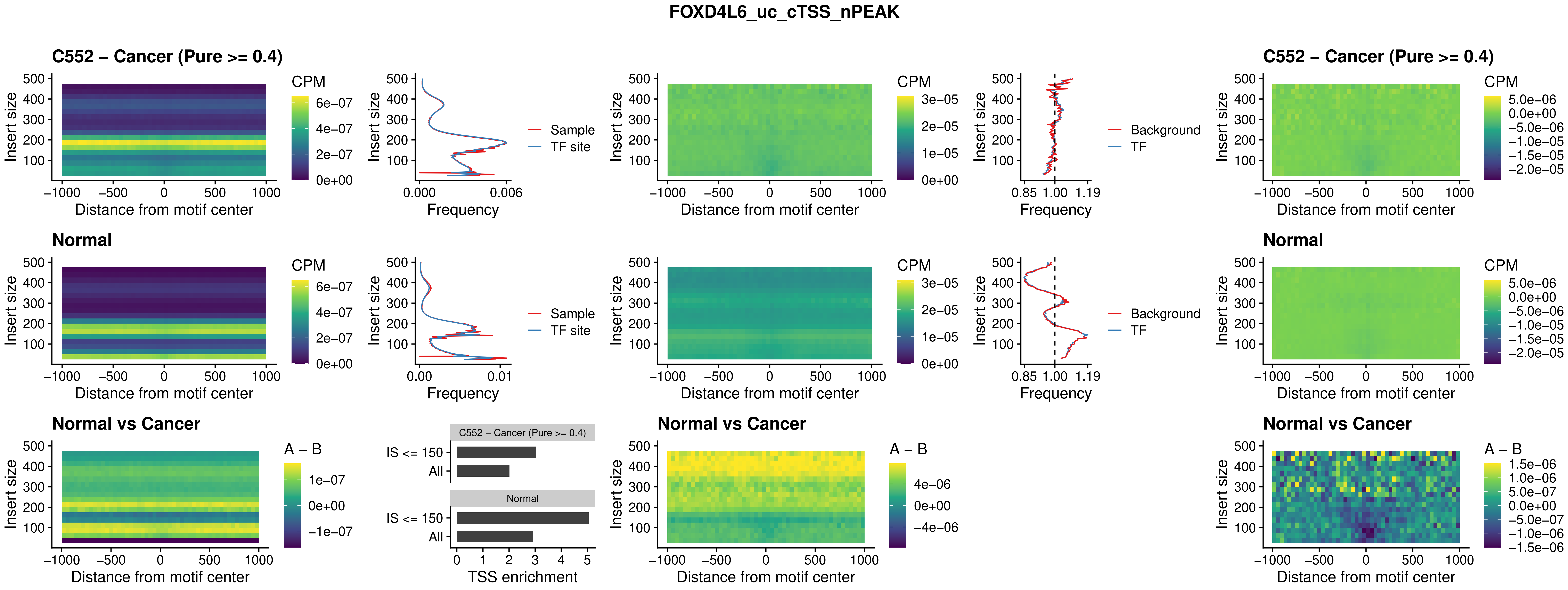
FOXD4L6_uc_cTSS_nPEAK
C552 – Cancer (Pure >= 0.4)
Normal
Normal vs Cancer
Insert size
Distance from motif center
Frequency
CPM
A – B
Sample
TF site
Background
TF
C552 – Cancer (Pure >= 0.4)
Normal
IS <= 150
All
TSS enrichment

#### FOXD4L5_uc_cTSS_nPEAK

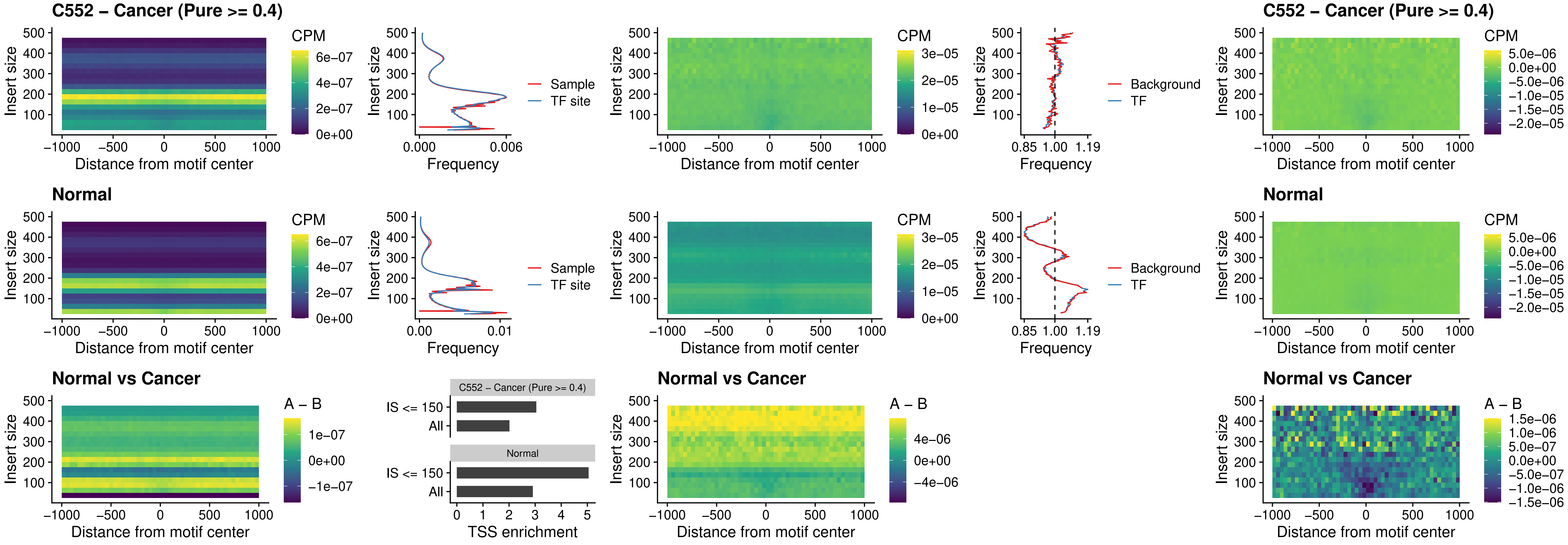

### FOXD4L4_uc_cTSS_nPEAK

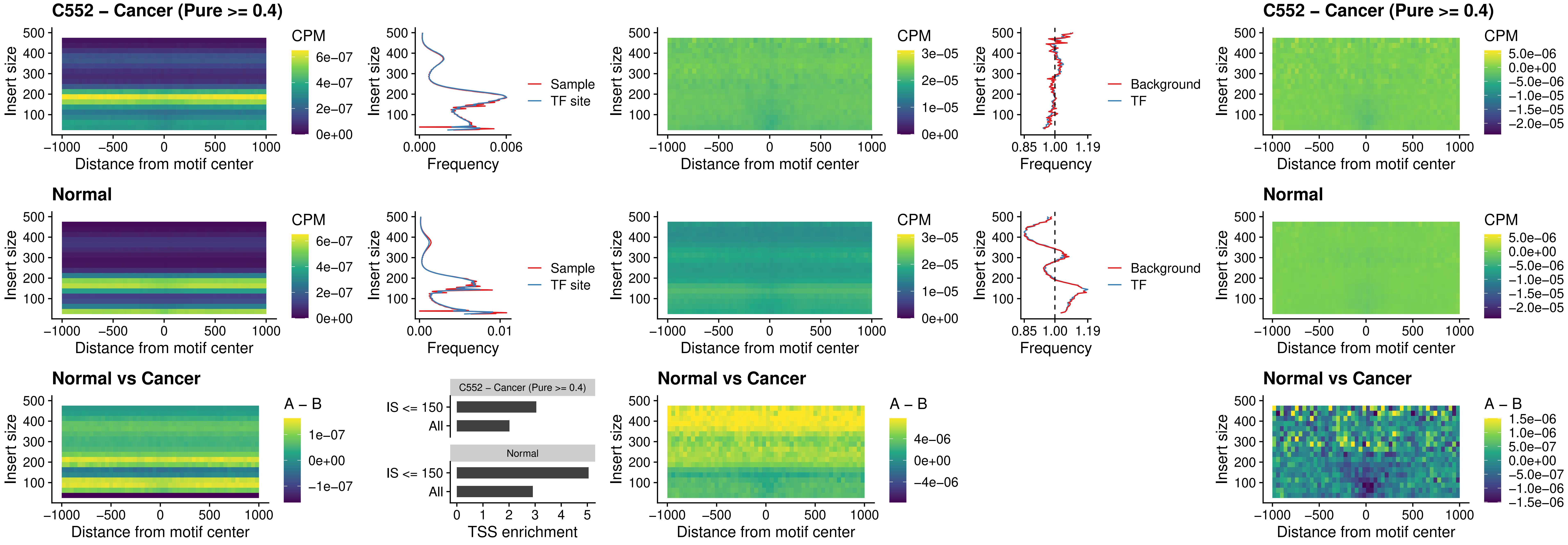

### FOXD4L2_uc_cTSS_nPEAK

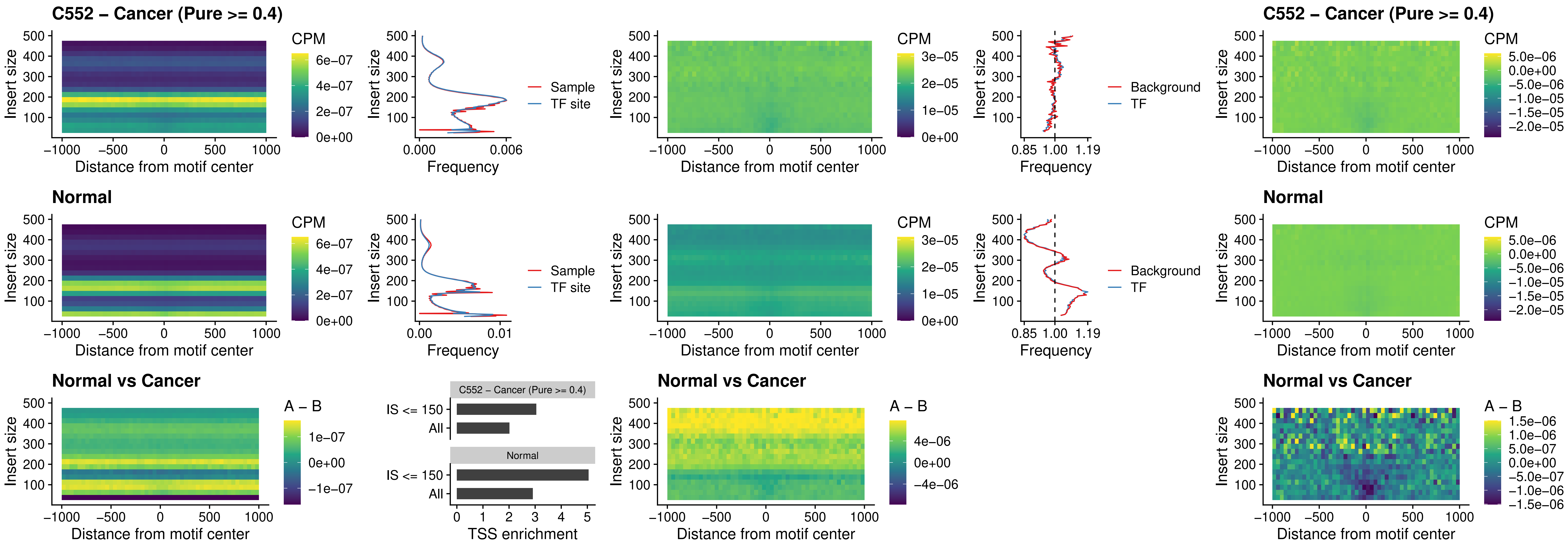

### FOXD4L3_uc_cTSS_nPEAK

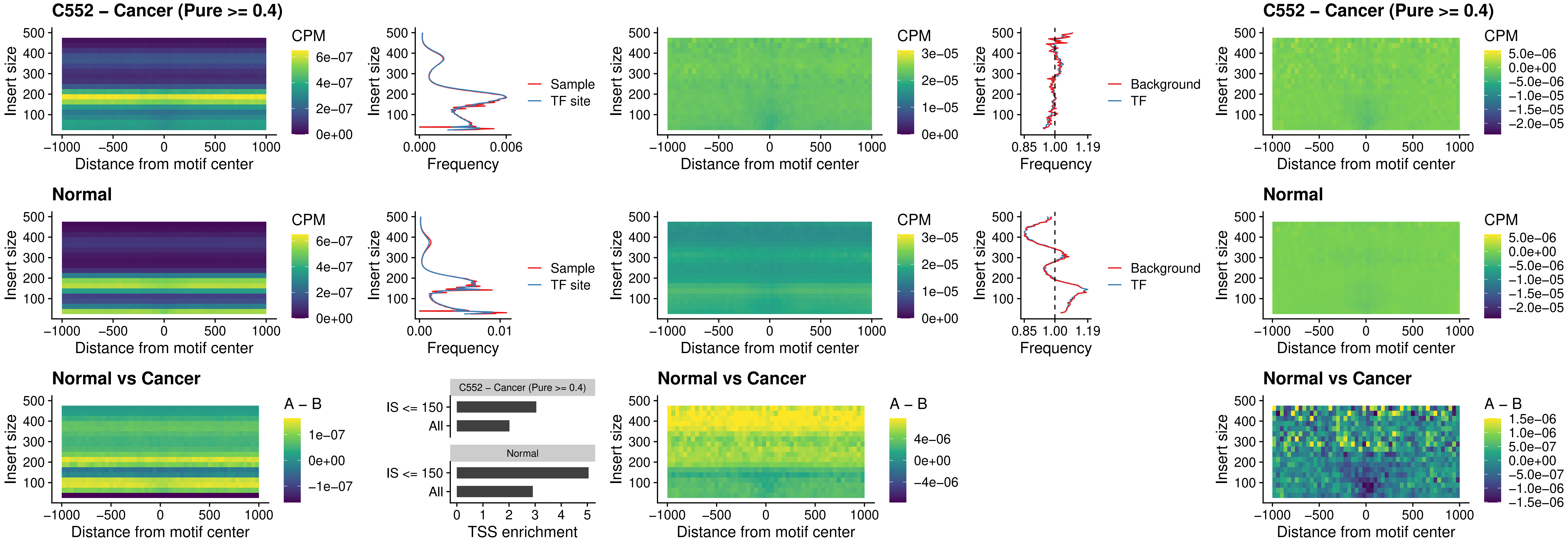

#### HOXD1_uc_dTSS_oPEAK

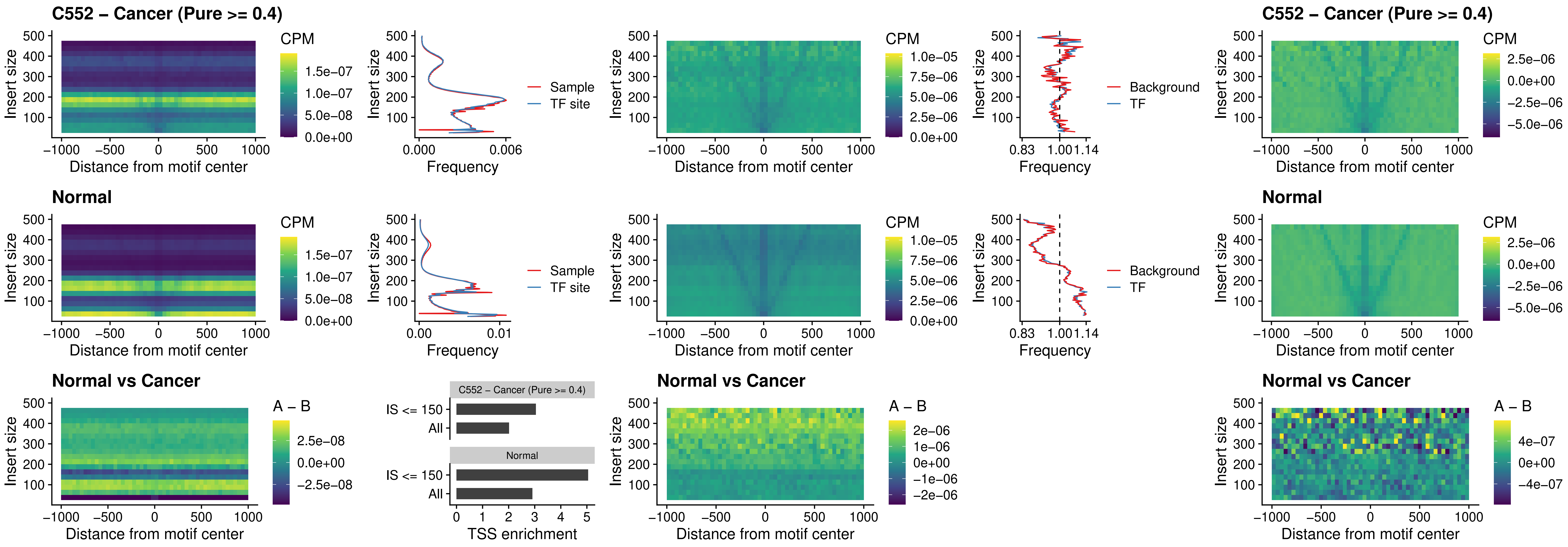

#### VAX1_uc_pTSS_nPEAK

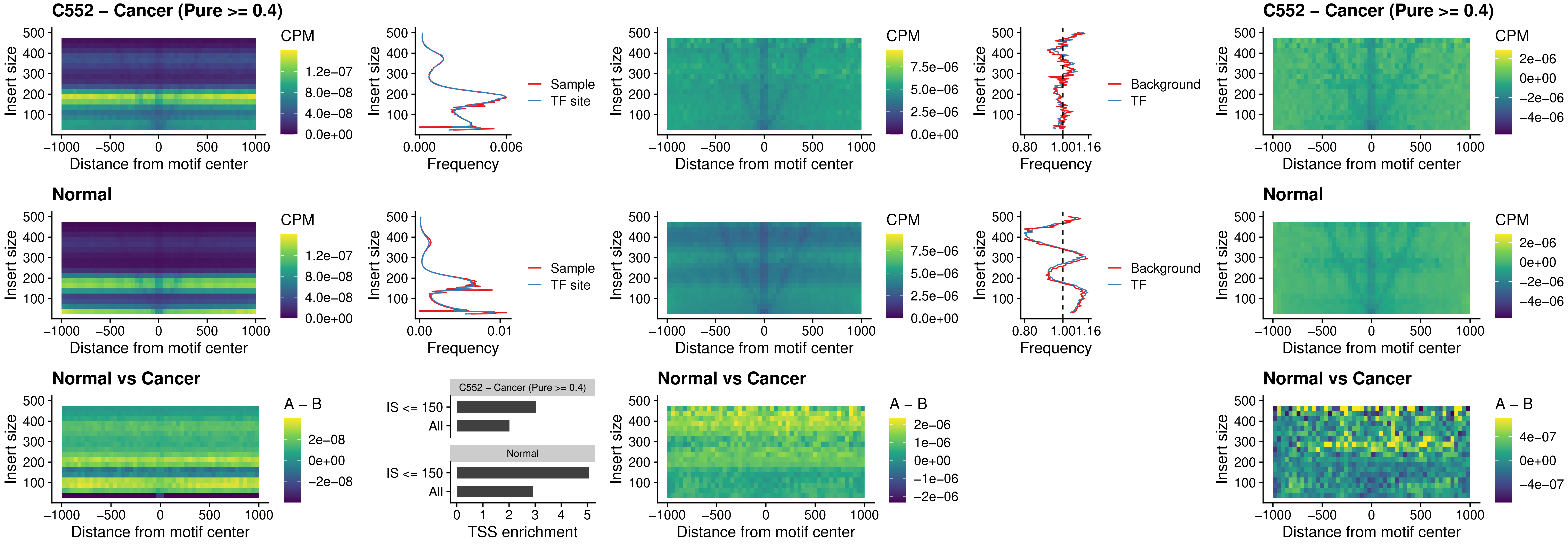

### DLX3_uc_pTSS_nPEAK

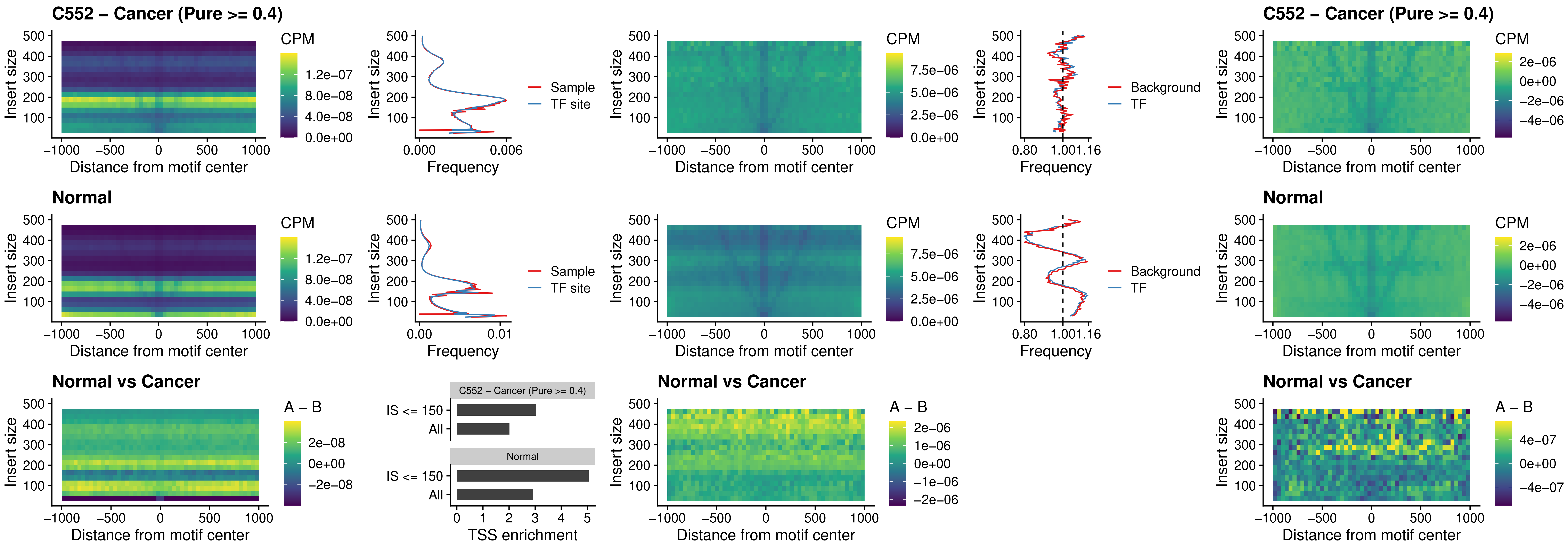

### HOXD3_uc_pTSS_nPEAK

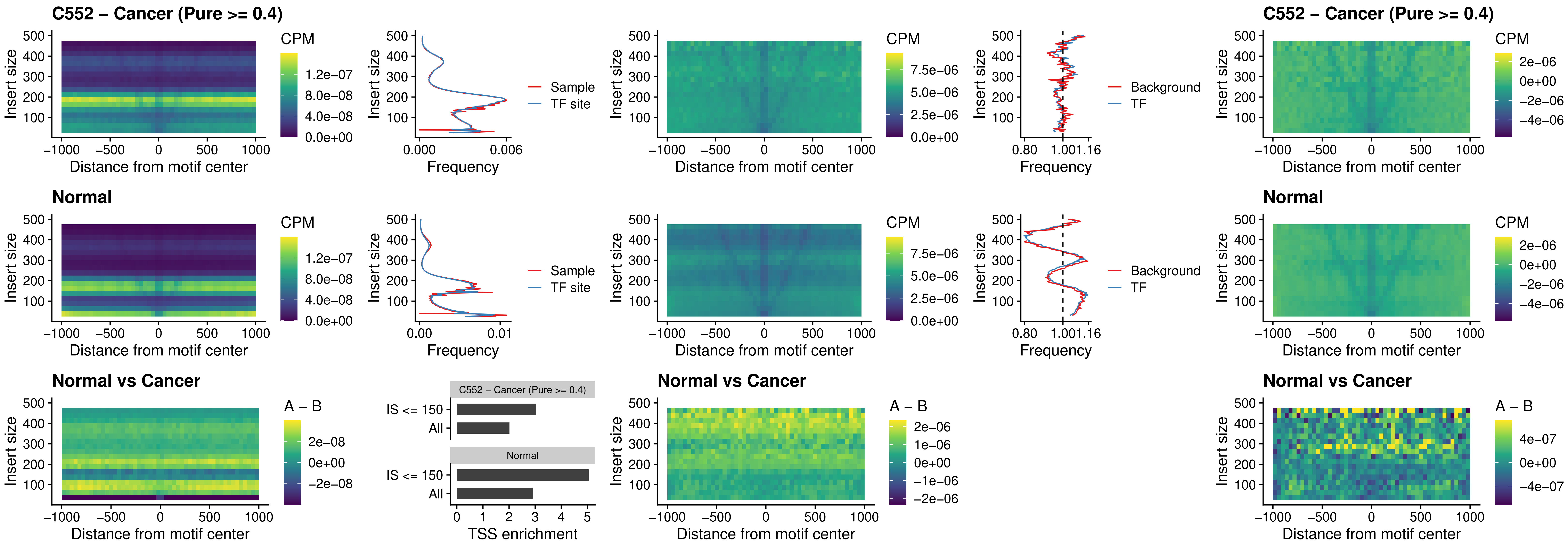

### PDX1_uc_dTSS_nPEAK

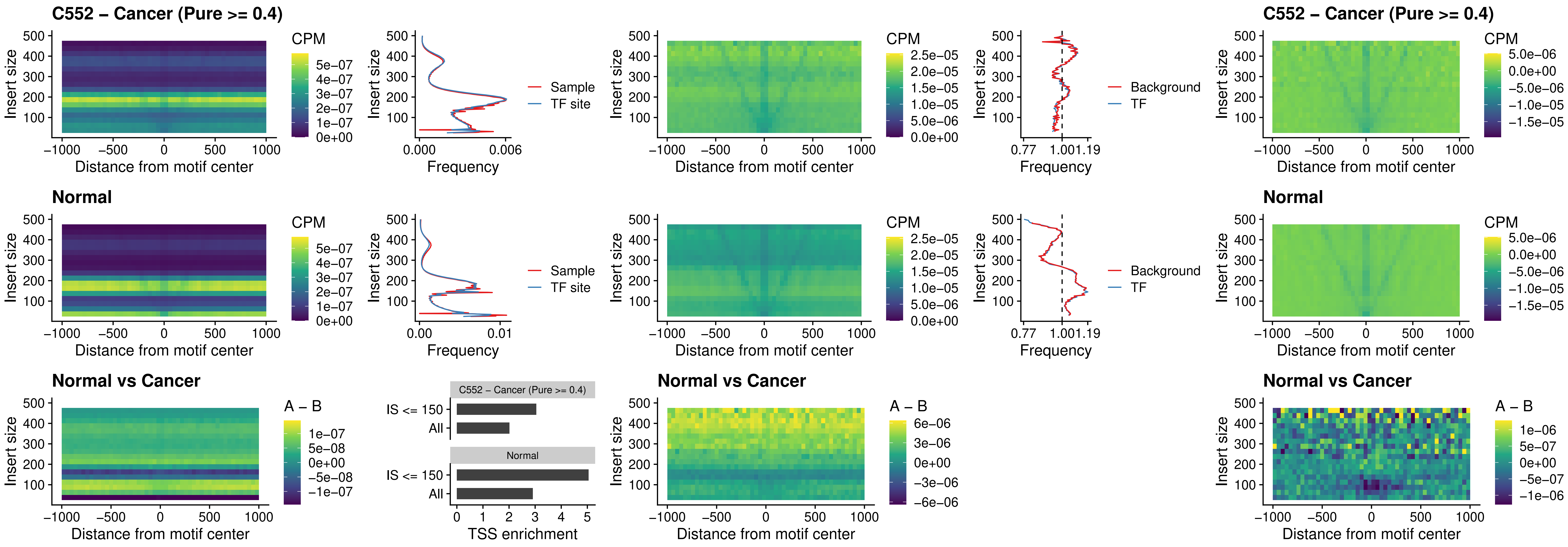

#### VAX2_uc_dTSS_nPEAK

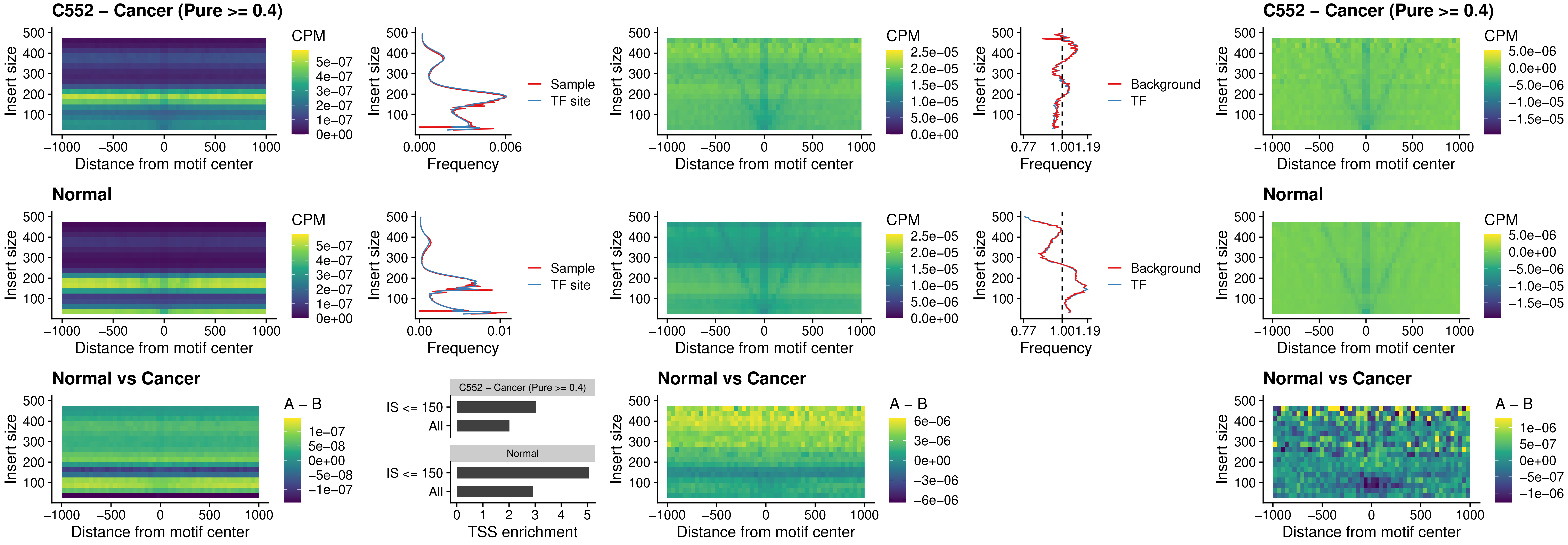

### SOX5_uc_dTSS_nPEAK

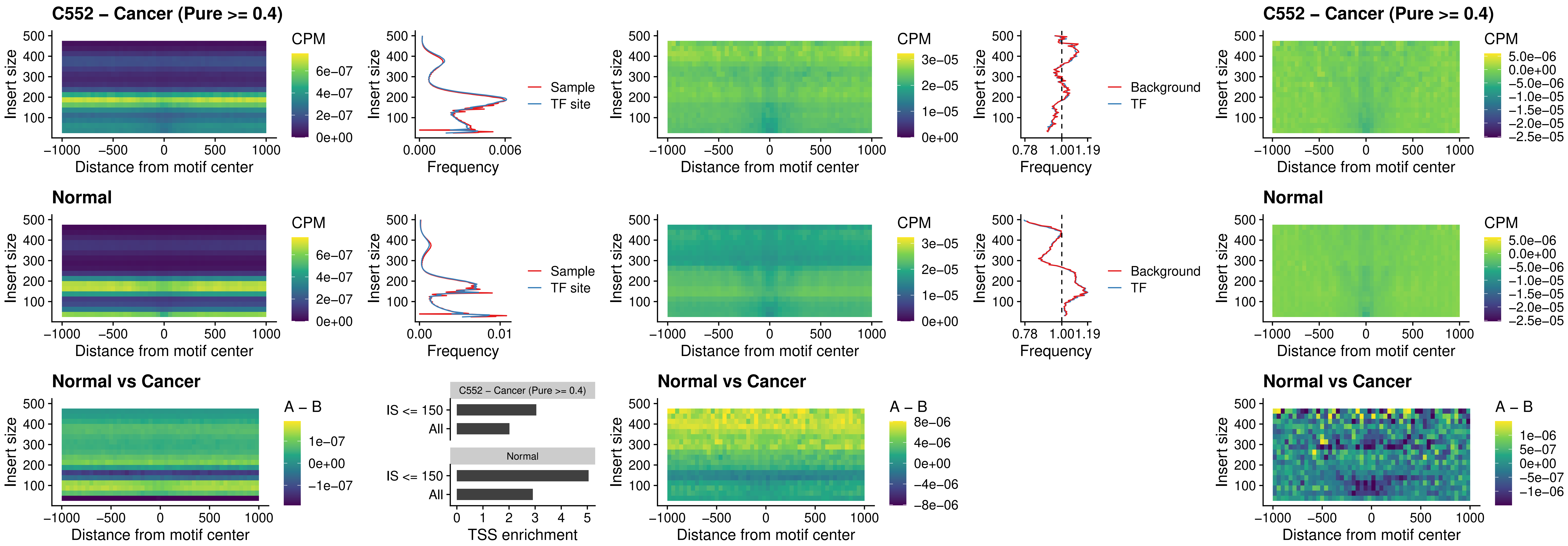

### SOX6_uc_dTSS_nPEAK

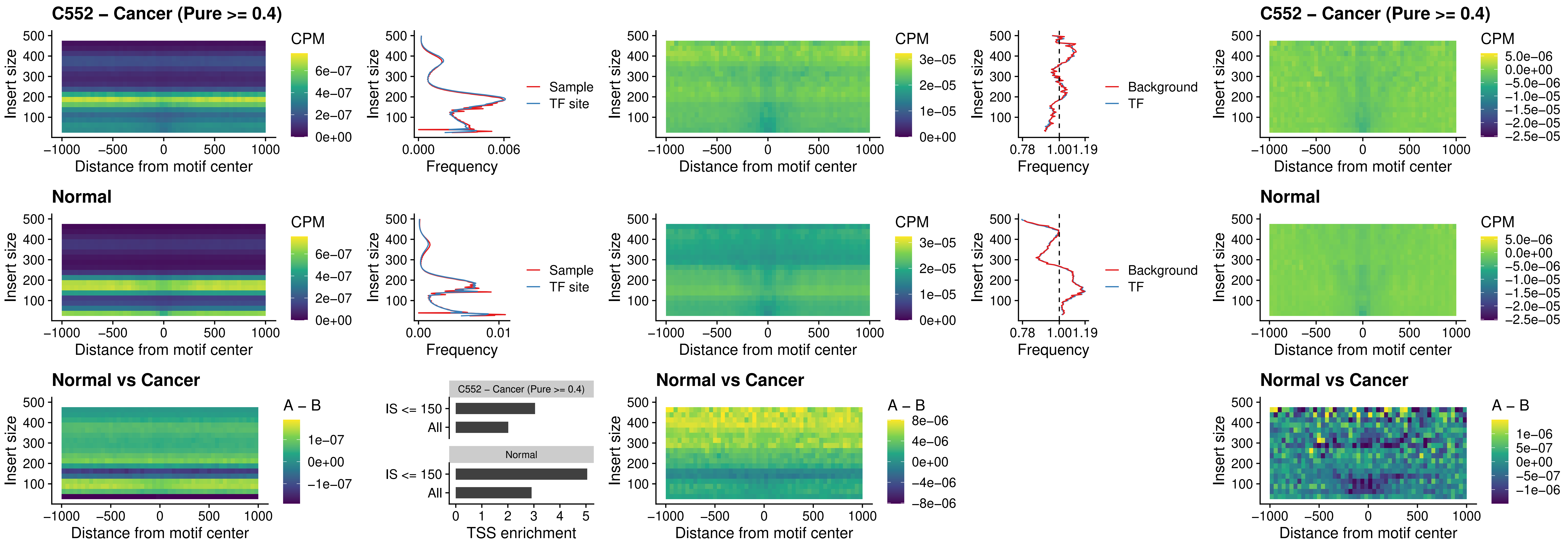

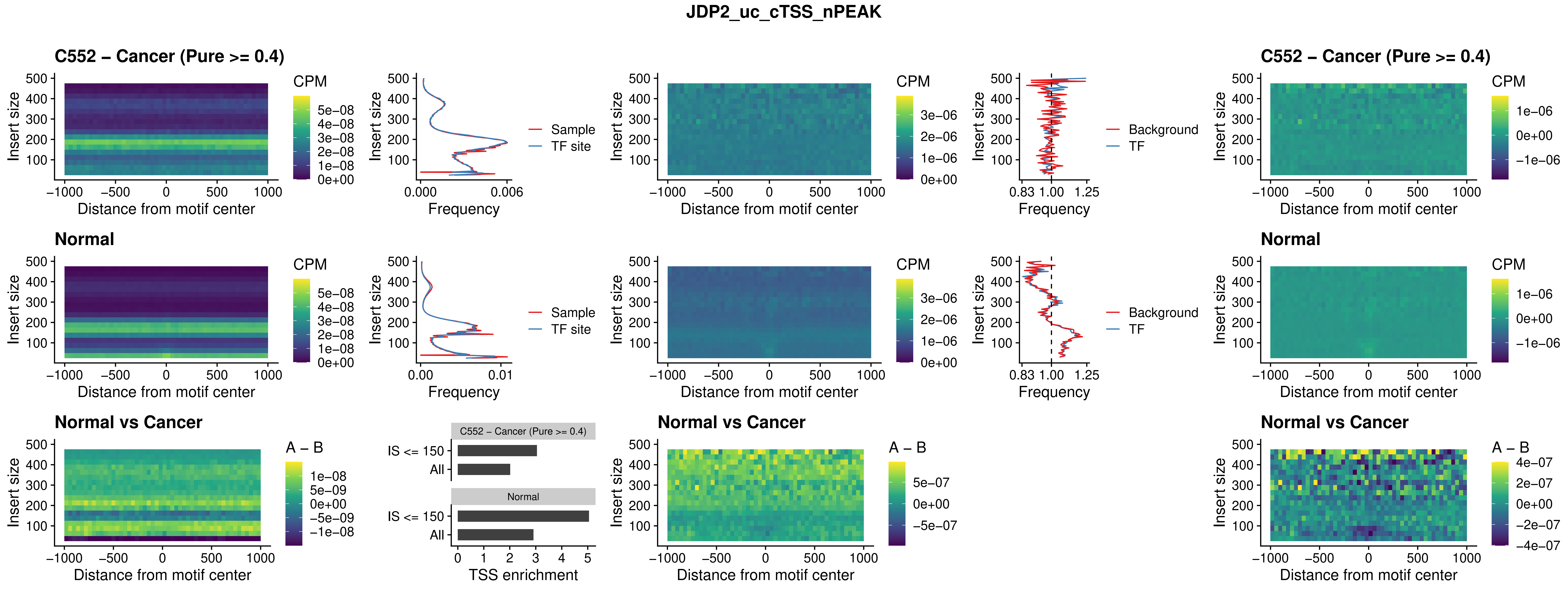
JDP2_uc_cTSS_nPEAK
C552 – Cancer (Pure >= 0.4)
Normal
Normal vs Cancer
Insert size
Distance from motif center
Frequency
CPM
A – B
Sample
TF site
Background
TF
C552 – Cancer (Pure >= 0.4)
Normal
IS <= 150
All
TSS enrichment

### PURA_uc_dTSS_oPEAK

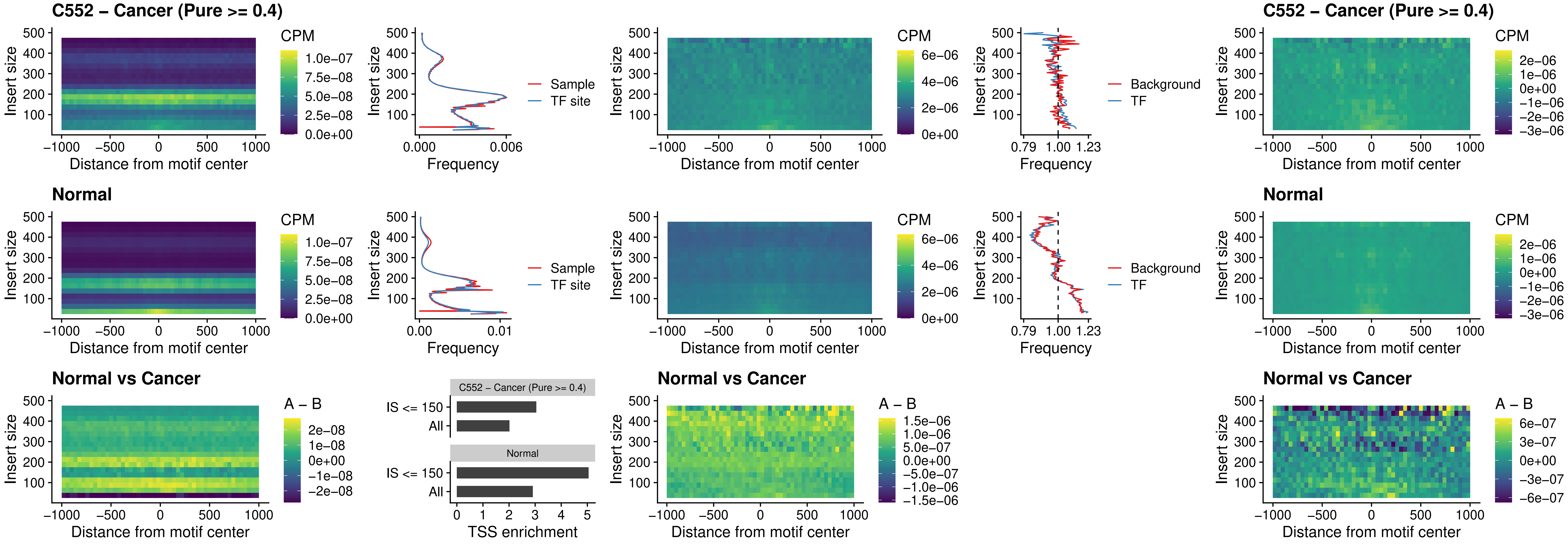

### HHEX_uc_dTSS_nPEAK

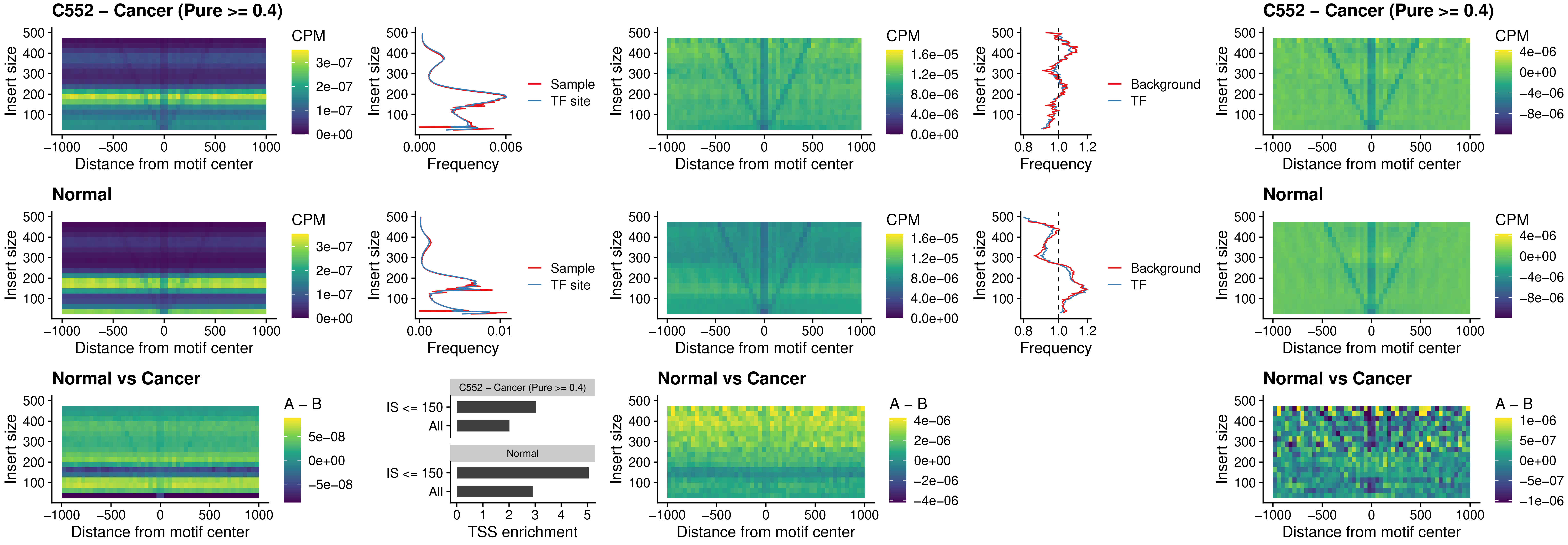

### HMG20B_uc_dTSS_nPEAK

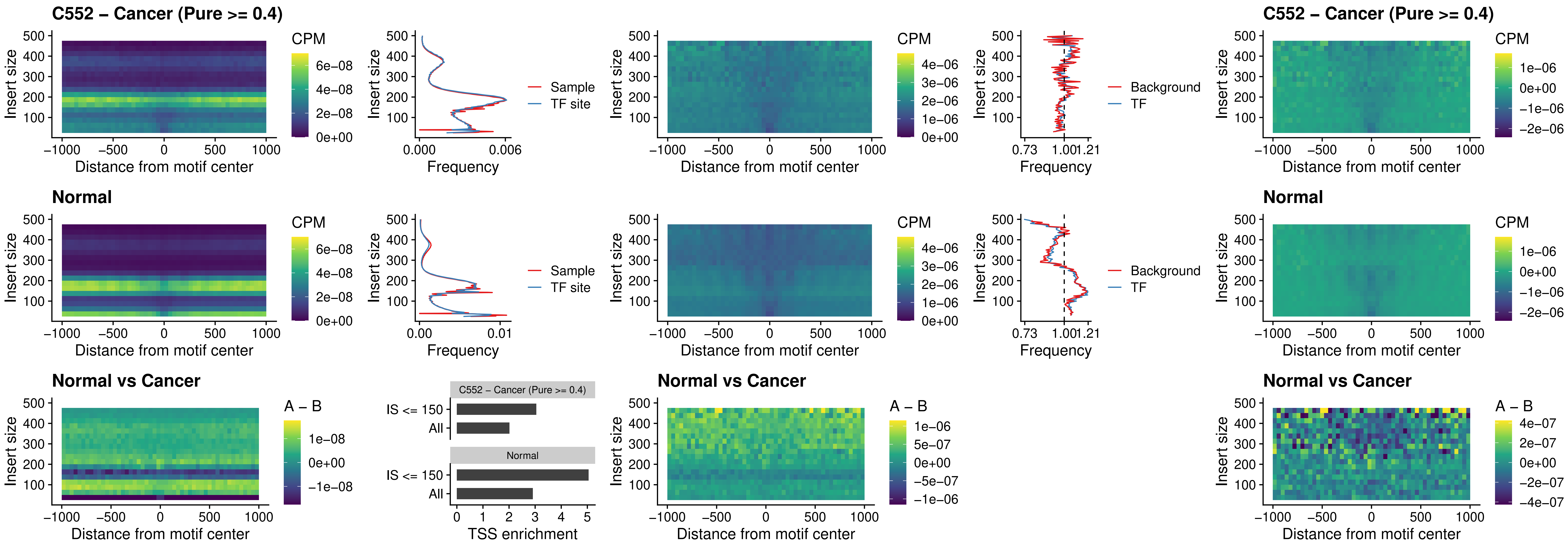

#### FOXB2_uc_dTSS_nPEAK

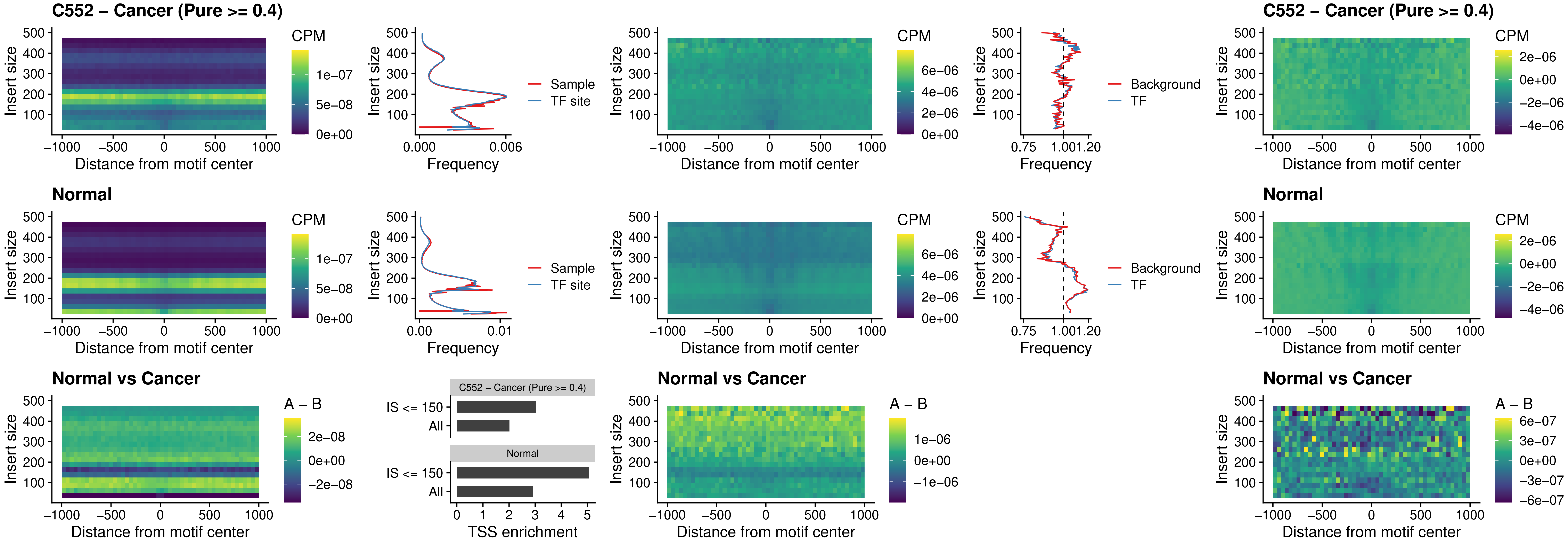

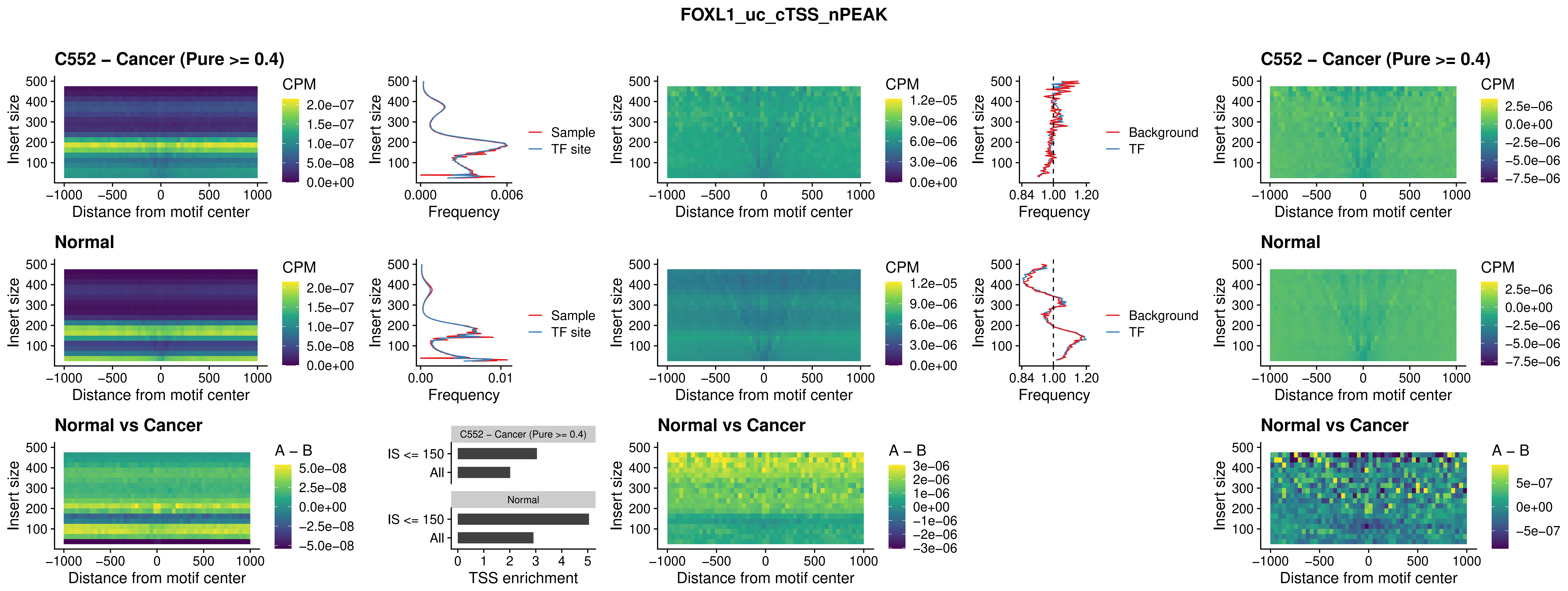
FOXL1_uc_cTSS_nPEAK
C552 – Cancer (Pure >= 0.4)
Normal
Normal vs Cancer
Insert size
Distance from motif center
Frequency
CPM
A – B
Sample
TF site
Background
TF
C552 – Cancer (Pure >= 0.4)
Normal
IS <= 150
All
TSS enrichment

### PBX3_uc_pTSS_oPEAK

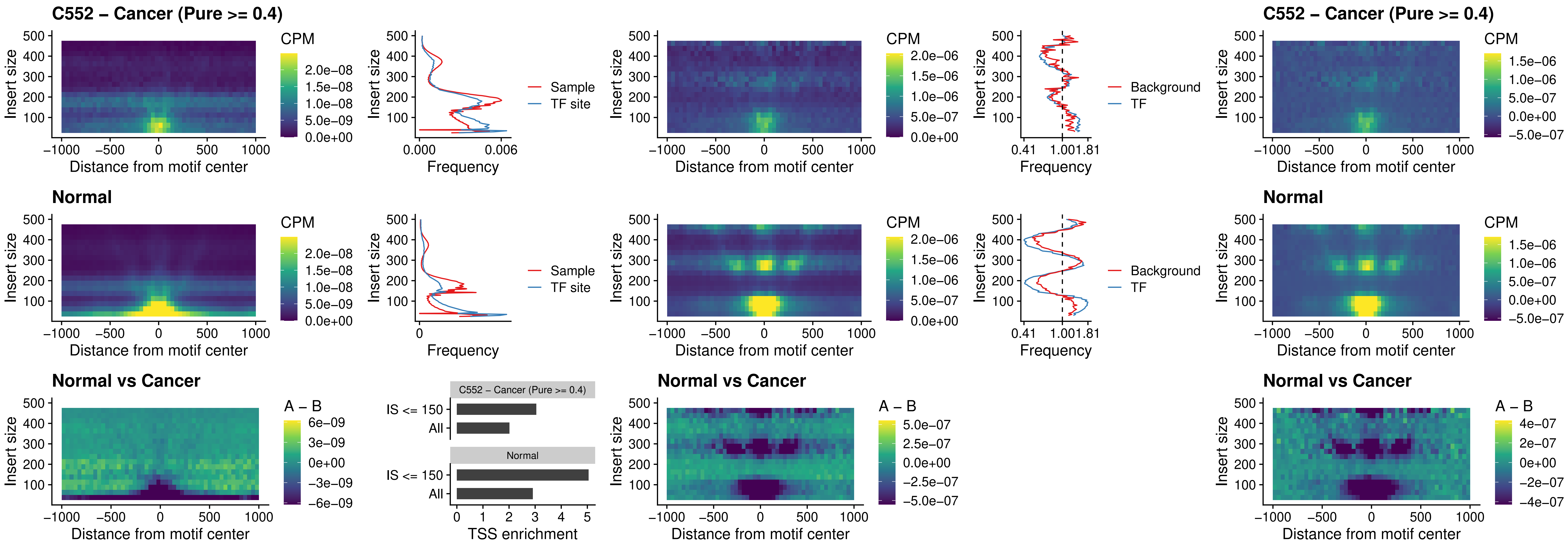

### SNAI2_uc_cTSS_nPEAK

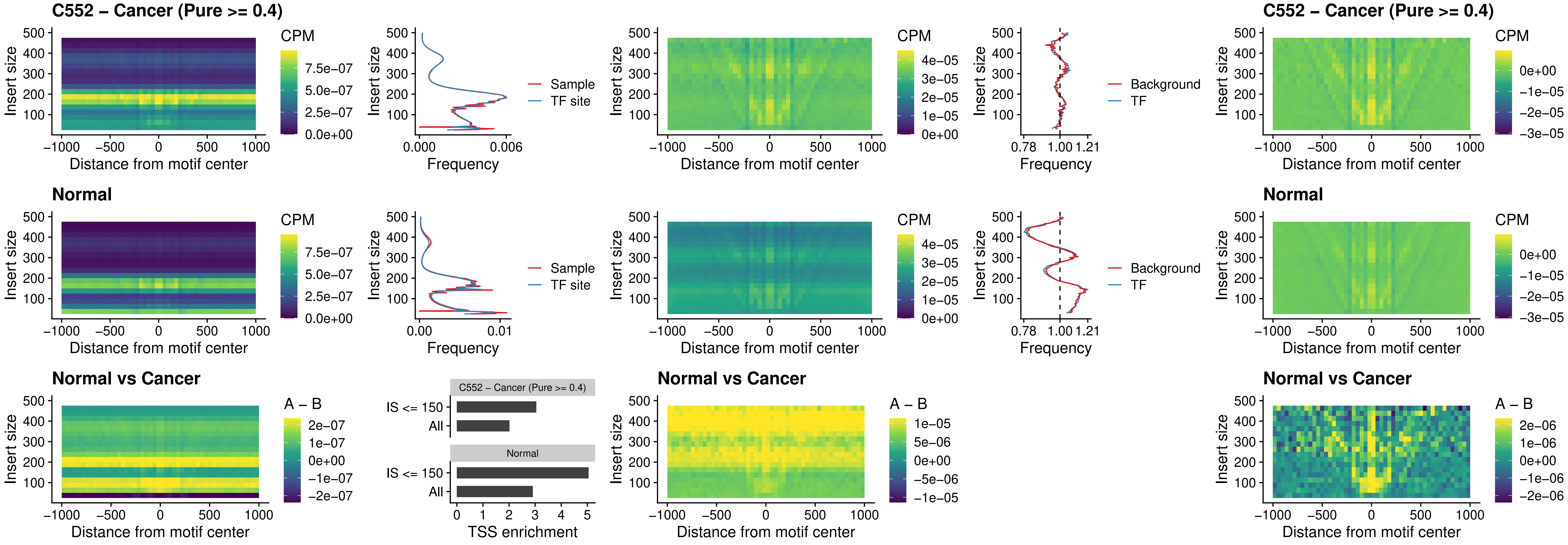

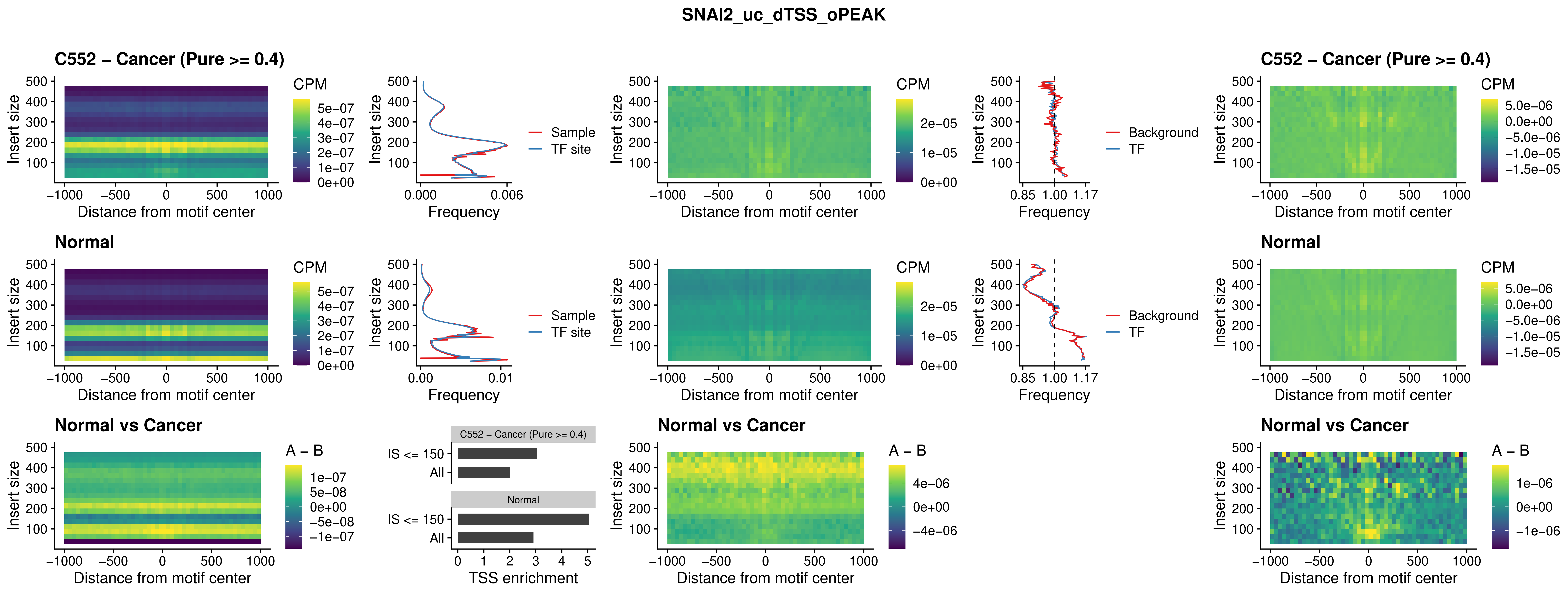
SNAI2_uc_dTSS_oPEAK
C552 – Cancer (Pure >= 0.4)
Normal
Normal vs Cancer
CPM
A – B
Insert size
Distance from motif center
Frequency
Sample
TF site
Background
TF
C552 – Cancer (Pure >= 0.4)
Normal
IS <= 150
All
TSS enrichment

#### SPI1_uc_pTSS_nPEAK

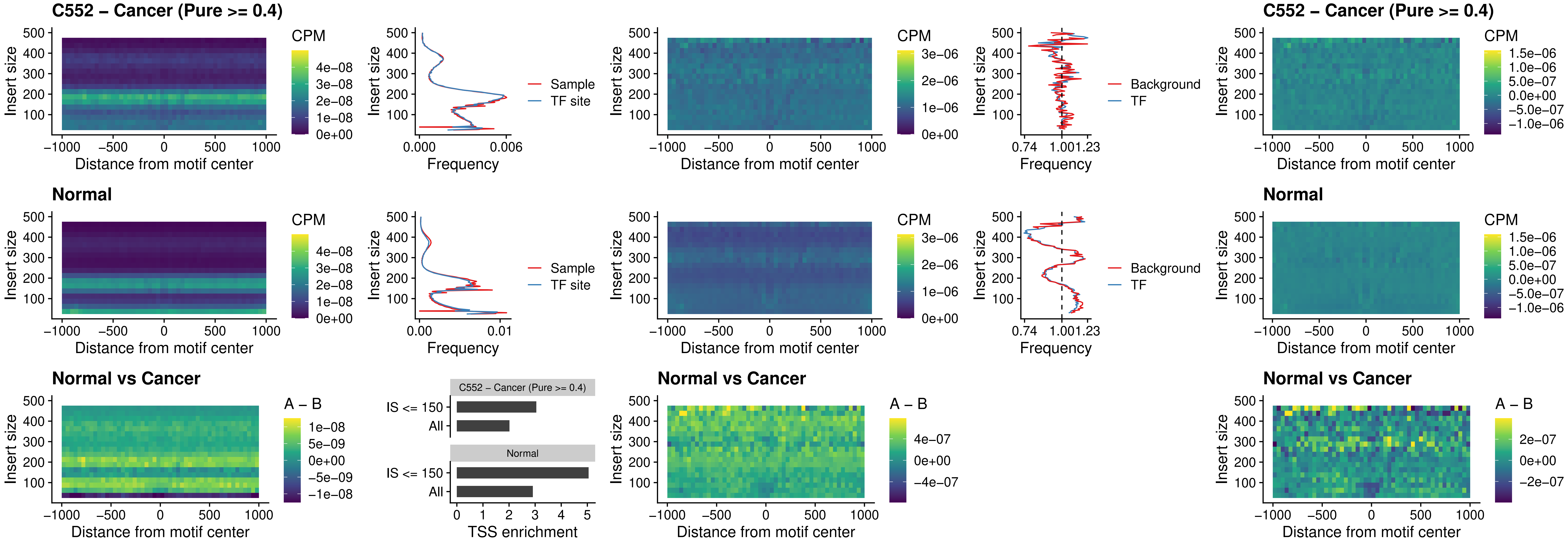

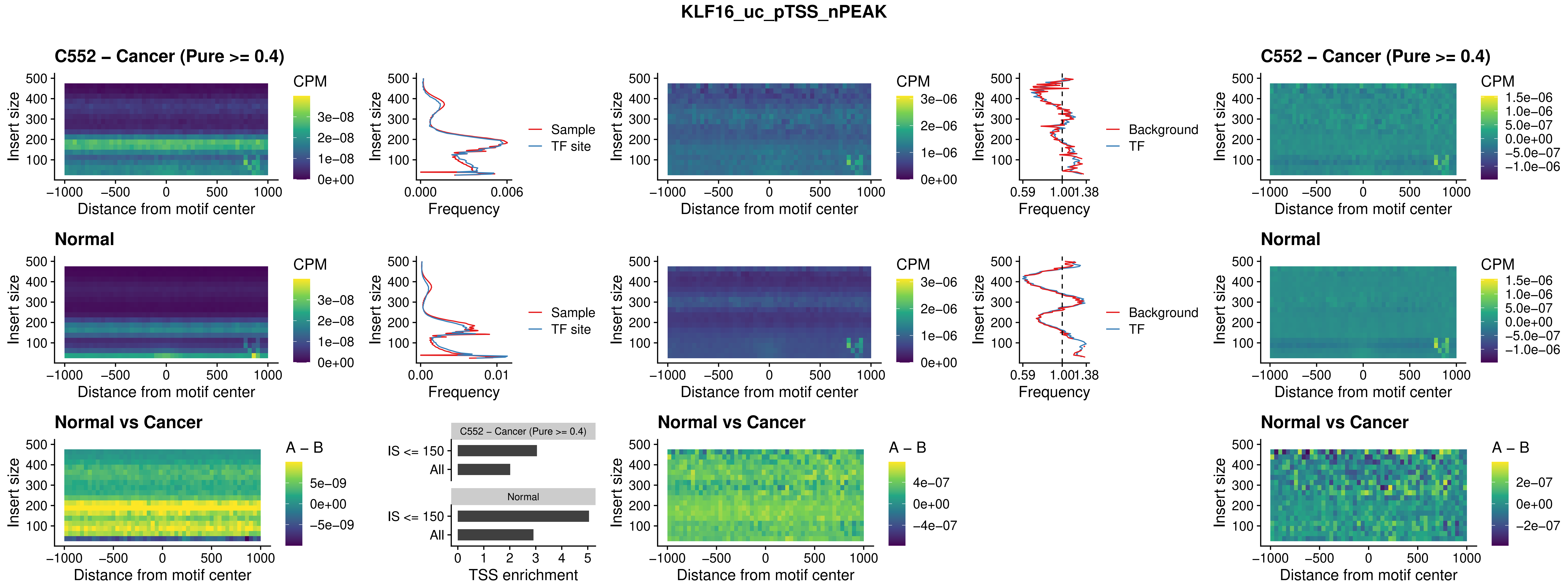
KLF16_uc_pTSS_nPEAK
C552 – Cancer (Pure >= 0.4)
Normal
Normal vs Cancer
Insert size
Distance from motif center
Frequency
CPM
A – B
Sample
TF site
Background
TF
C552 – Cancer (Pure >= 0.4)
Normal
IS <= 150
All
TSS enrichment

#### SP3_uc_pTSS_nPEAK

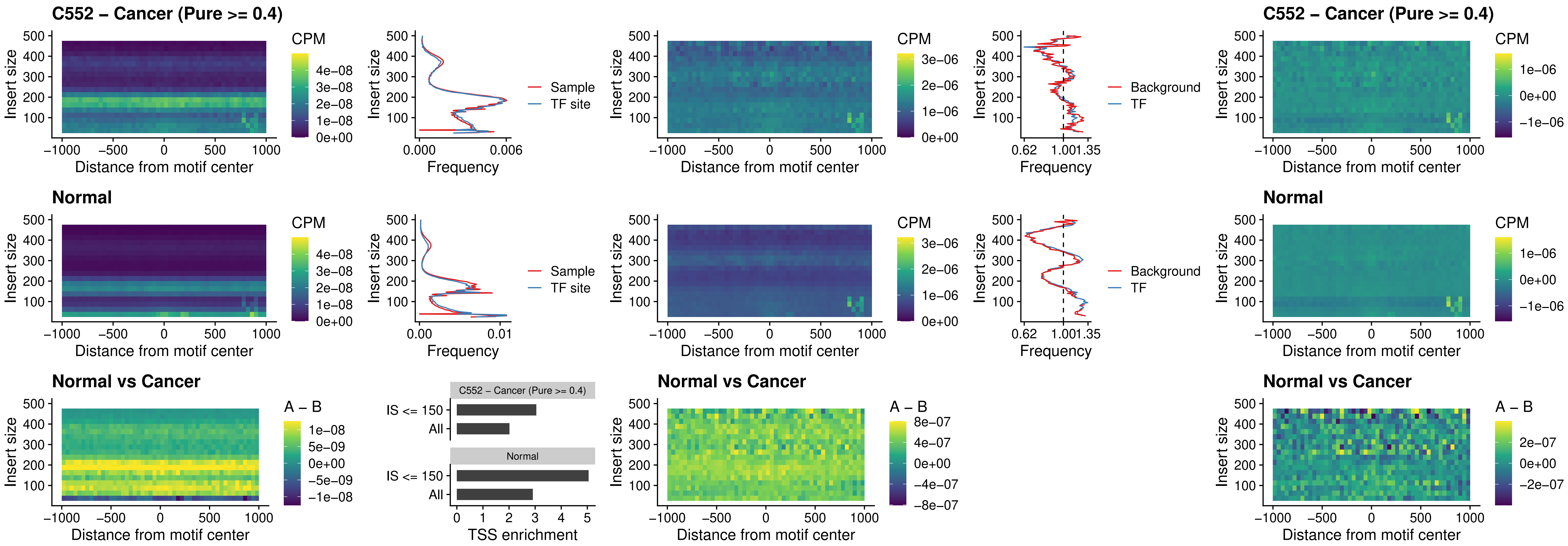

### NEURODD1_uc_dTSS_oPEAK

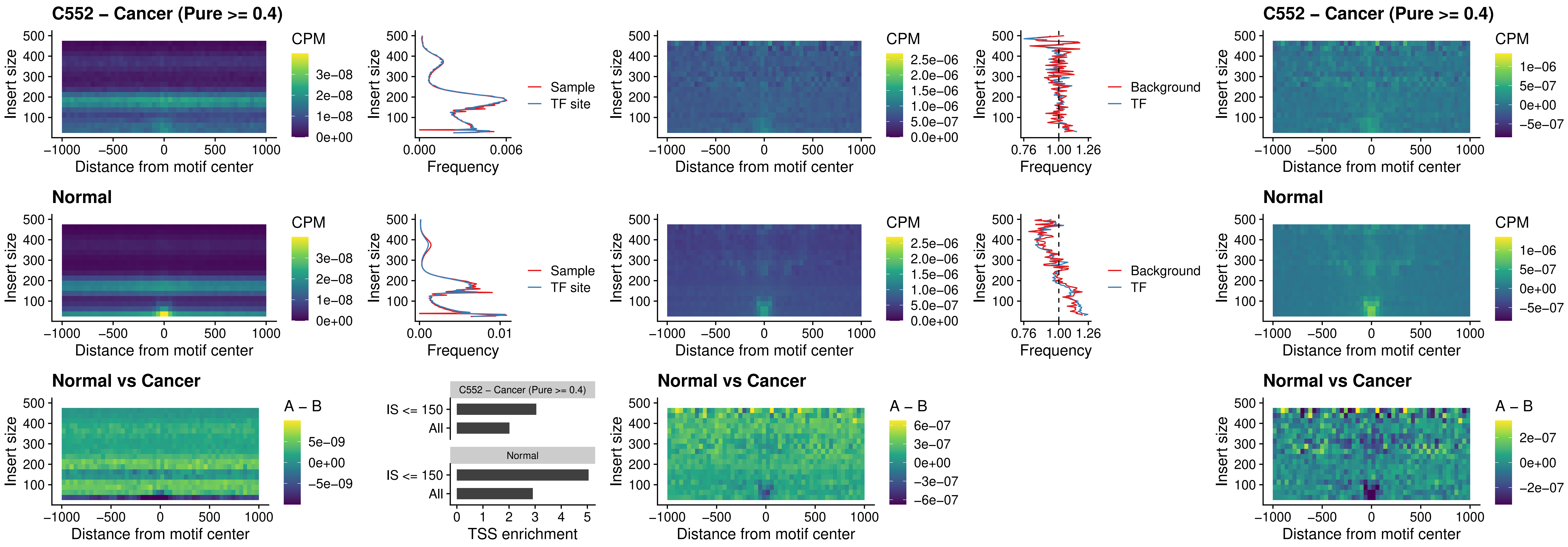

#### CTCF_uc_dTSS_oPEAK

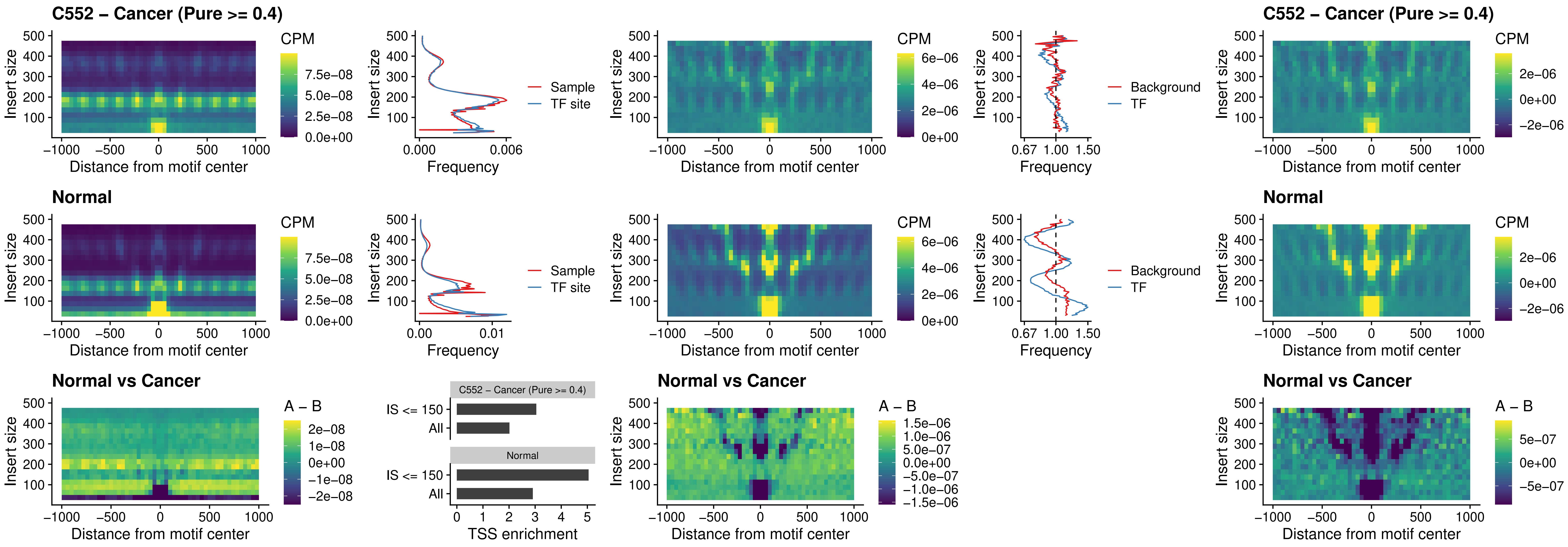

### CTCF_uc_cTSS_oPEAK

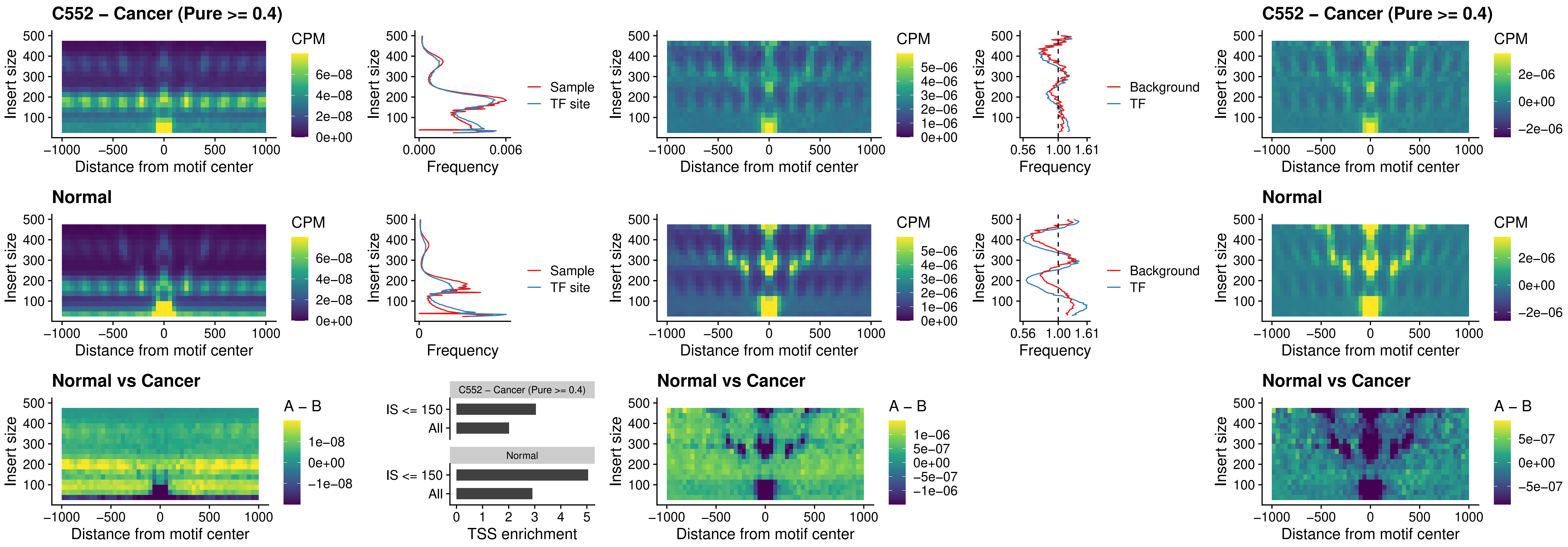

### CTCFL_uc_dTSS_oPEAK

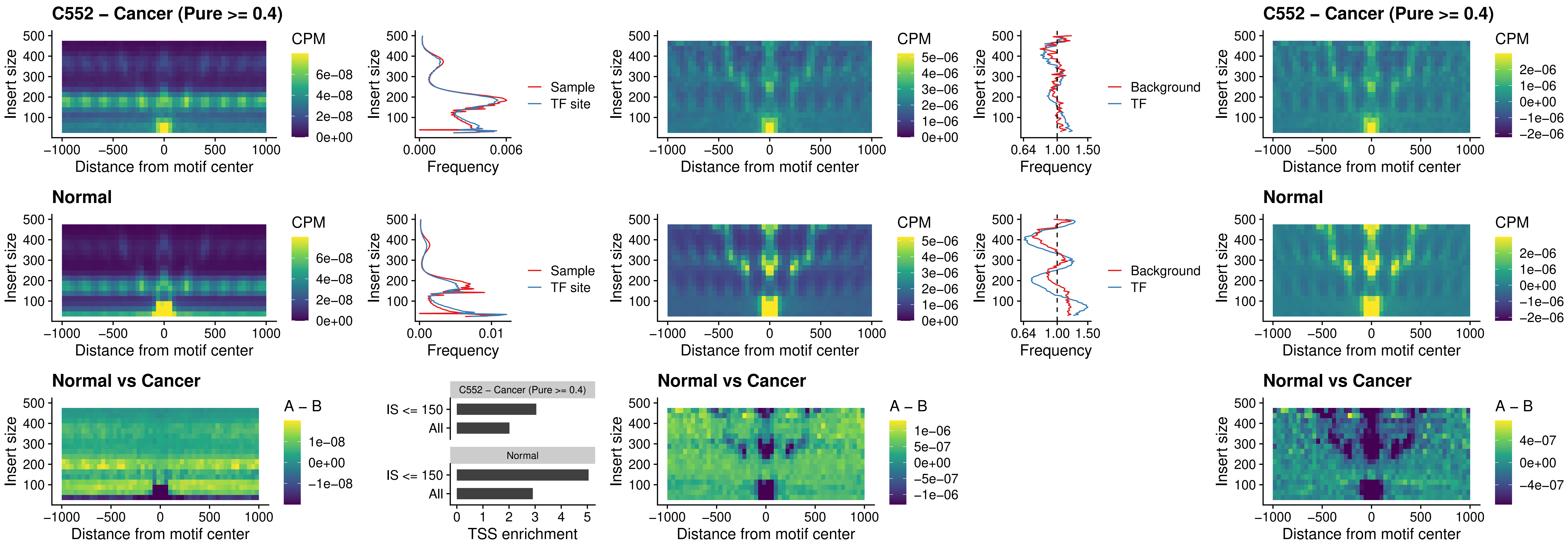

### ENCODE_CTCF_sites_COLON

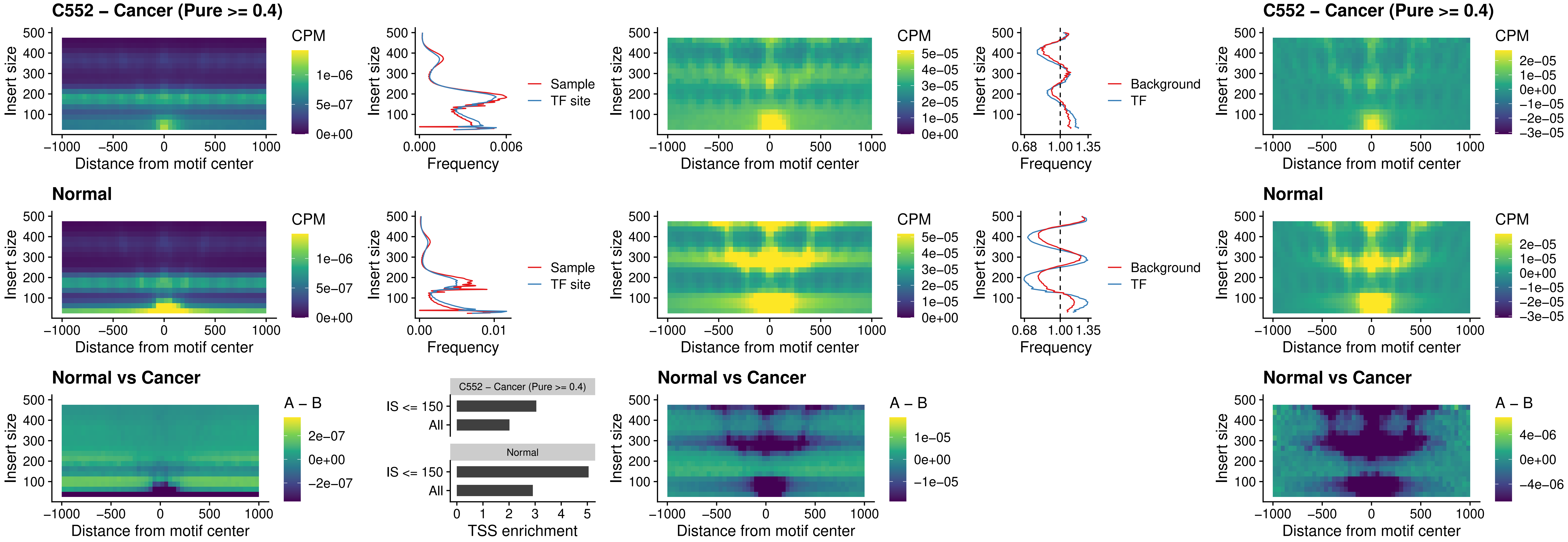

### ENCODE_CTCF_sites_COLON_loop_ctcf

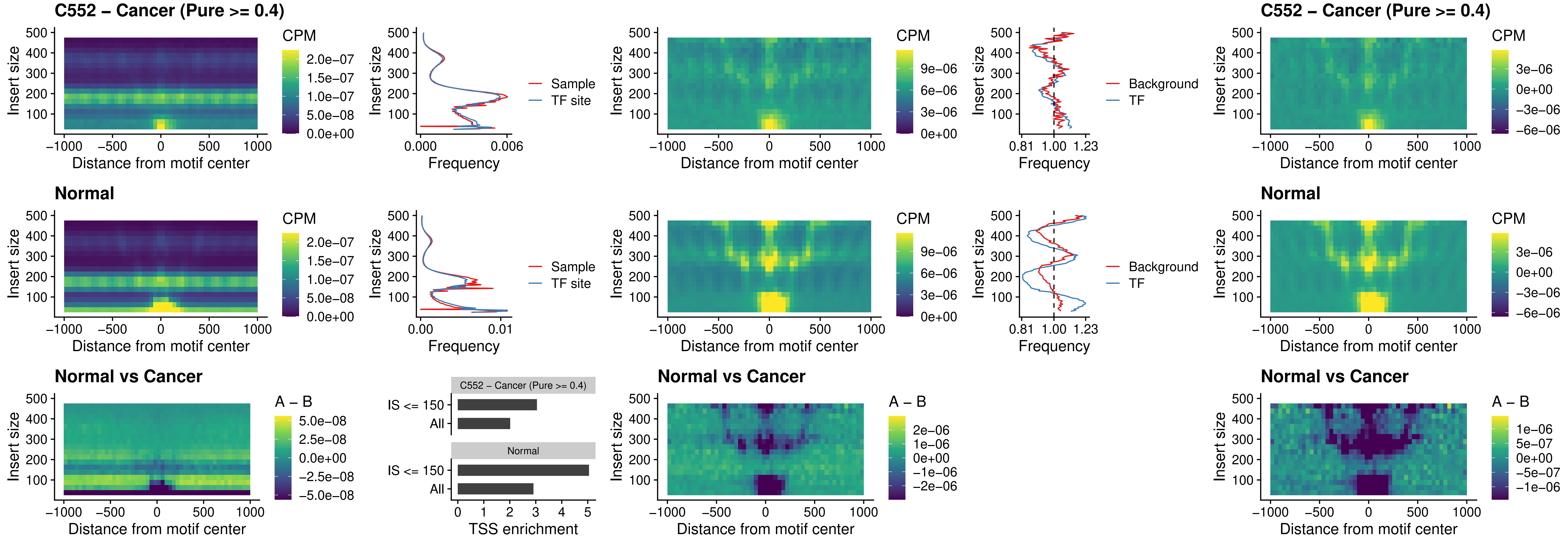

#### ENCODE_CTCF_sites_COLON_intergenic_ctcf

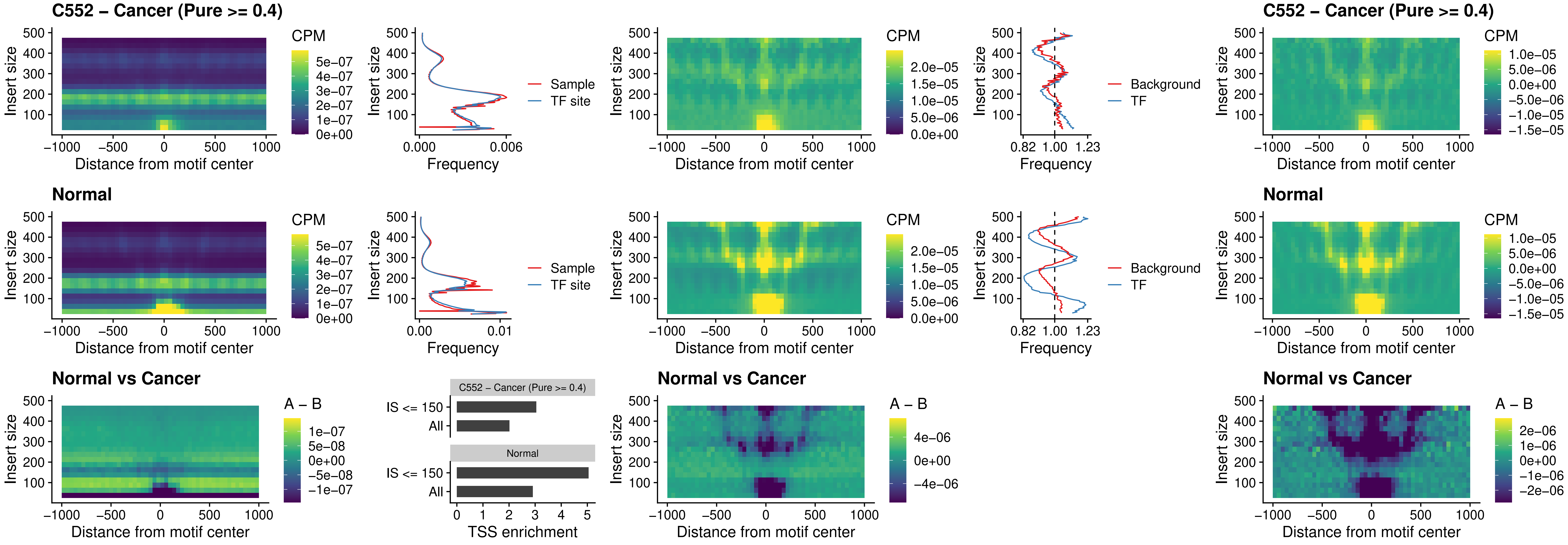

#### ENCODE_CTCF_sites_COLON_nonpromotor

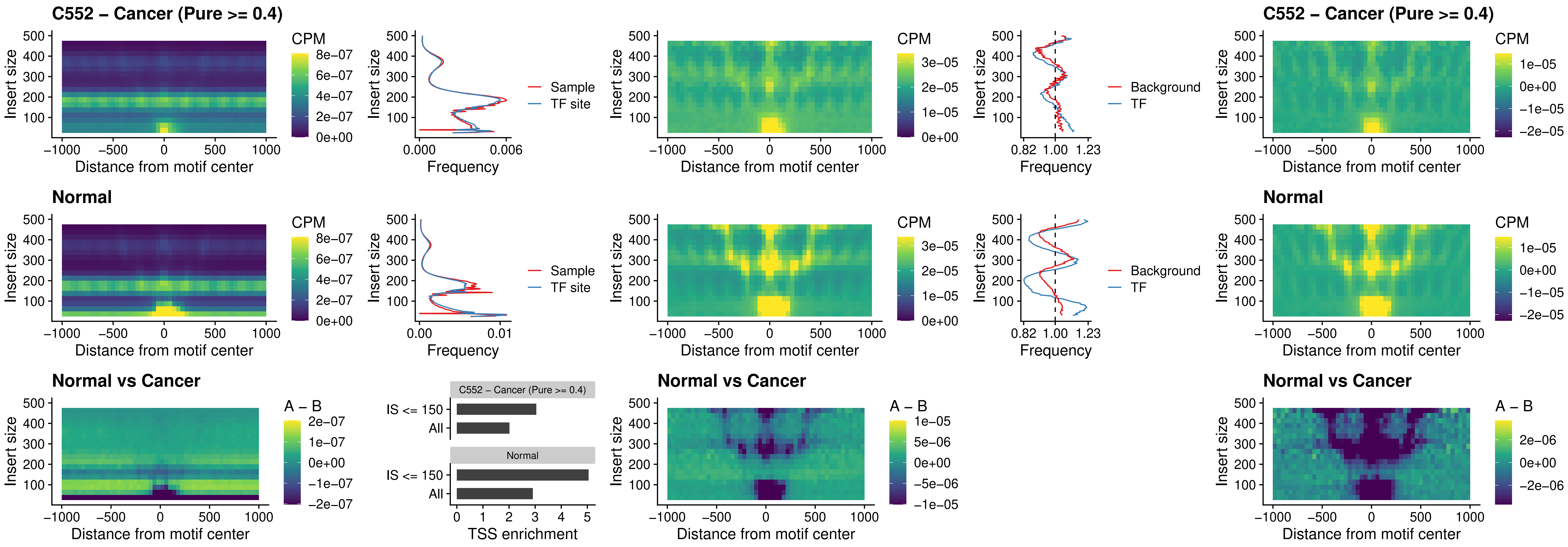

### CTCFL_uc_cTSS_oPEAK

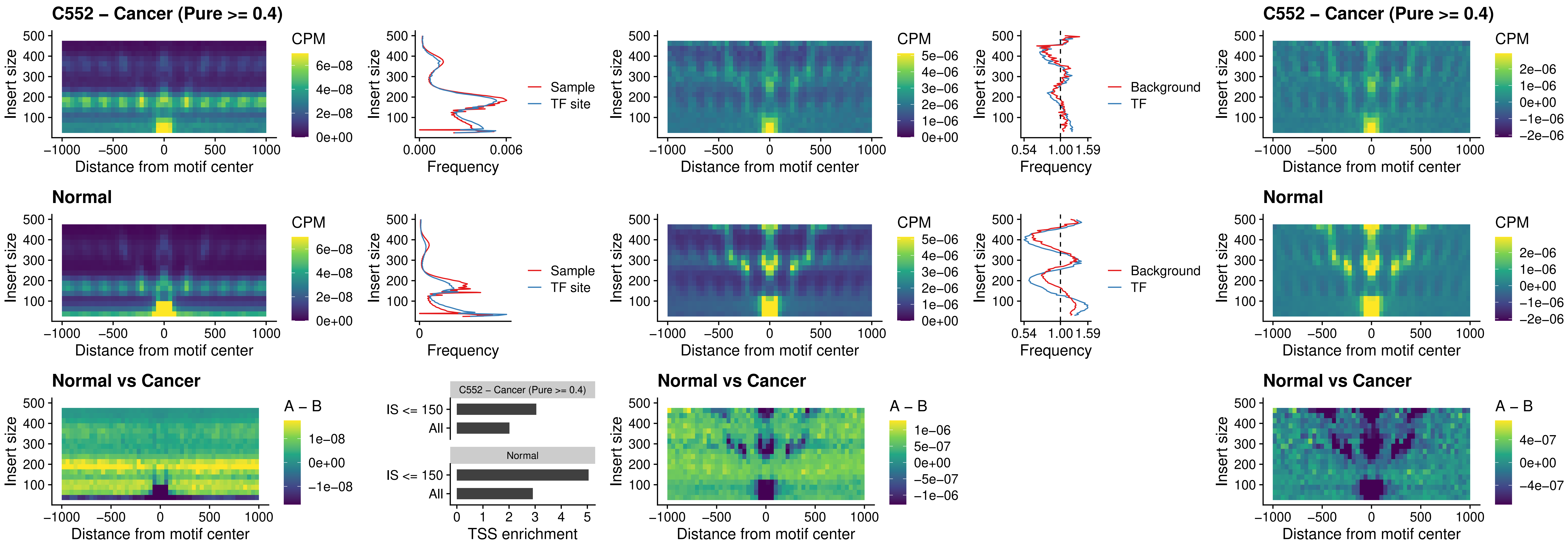

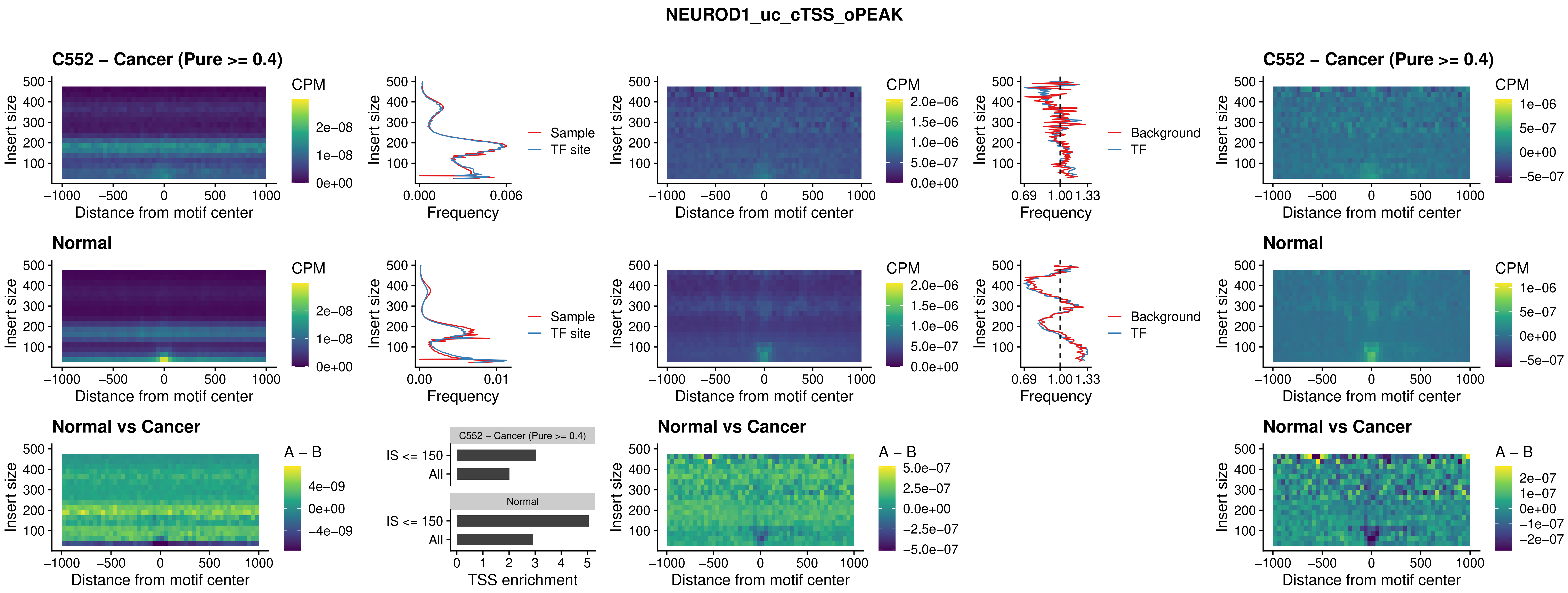
NEUROD1_uc_cTSS_oPEAK
C552 – Cancer (Pure >= 0.4)
Normal
Normal vs Cancer
Insert size
Distance from motif center
CPM
Frequency
Sample
TF site
Background
TF
A – B
C552 – Cancer (Pure >= 0.4)
Normal
IS <= 150
All
TSS enrichment

### ENCODE_YY1_sites_COLON_nonpromotor

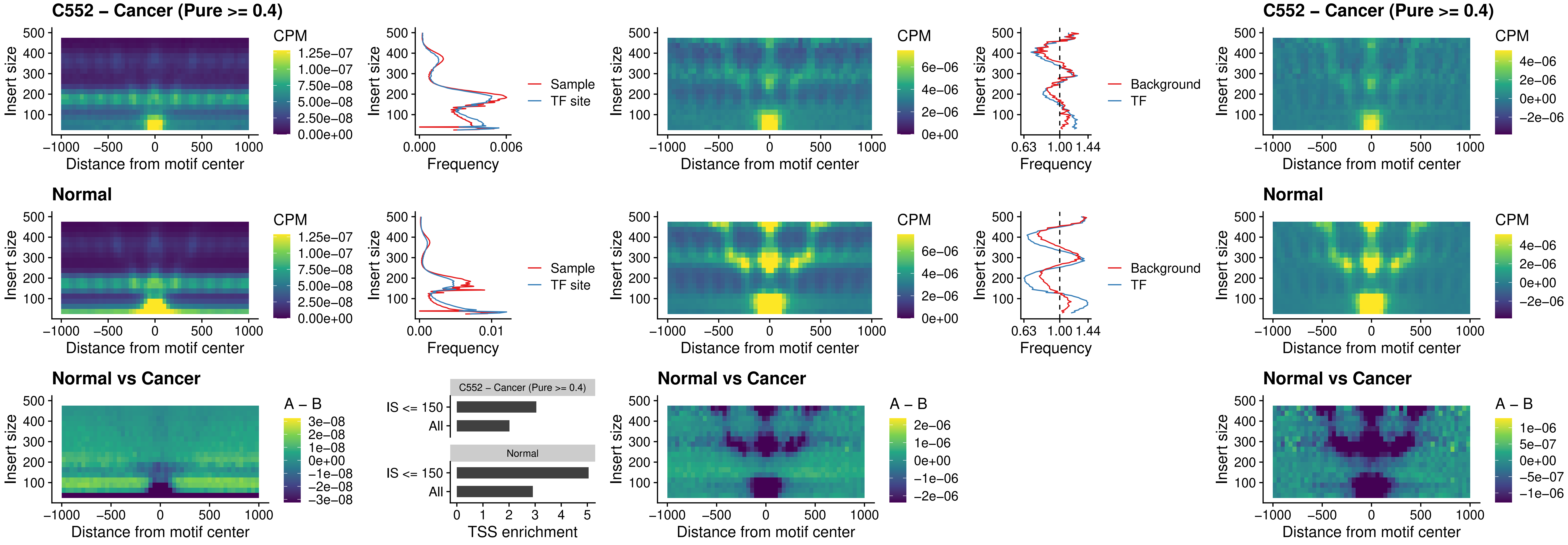

#### CTCF_uc_pTSS_oPEAK

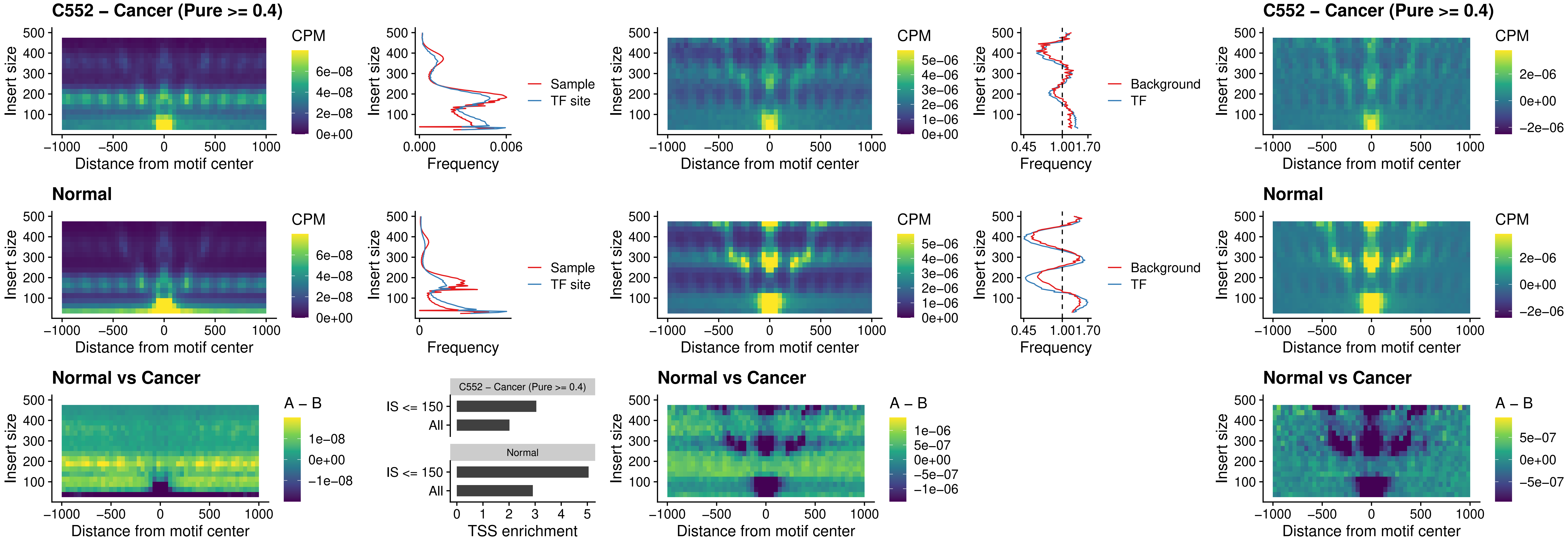

#### ENCODE_CTCF_sites_COLON_promotor

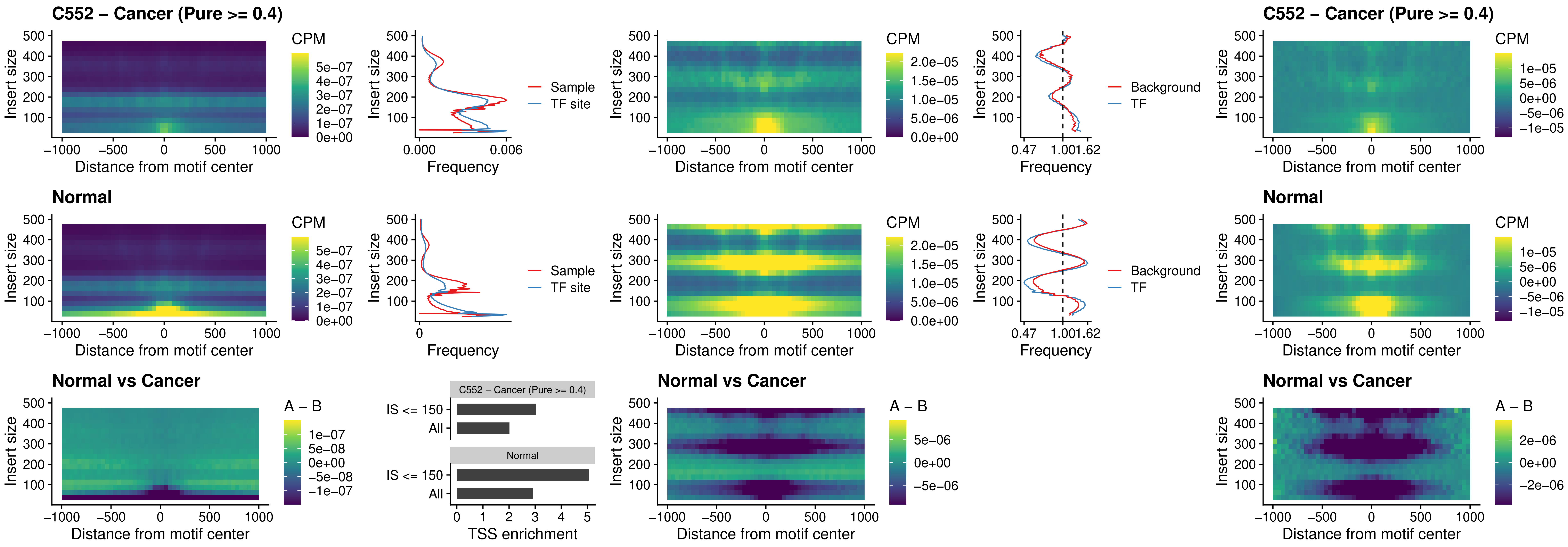

#### ENCODE_CTCF_sites_COLON_promotor_ctcf

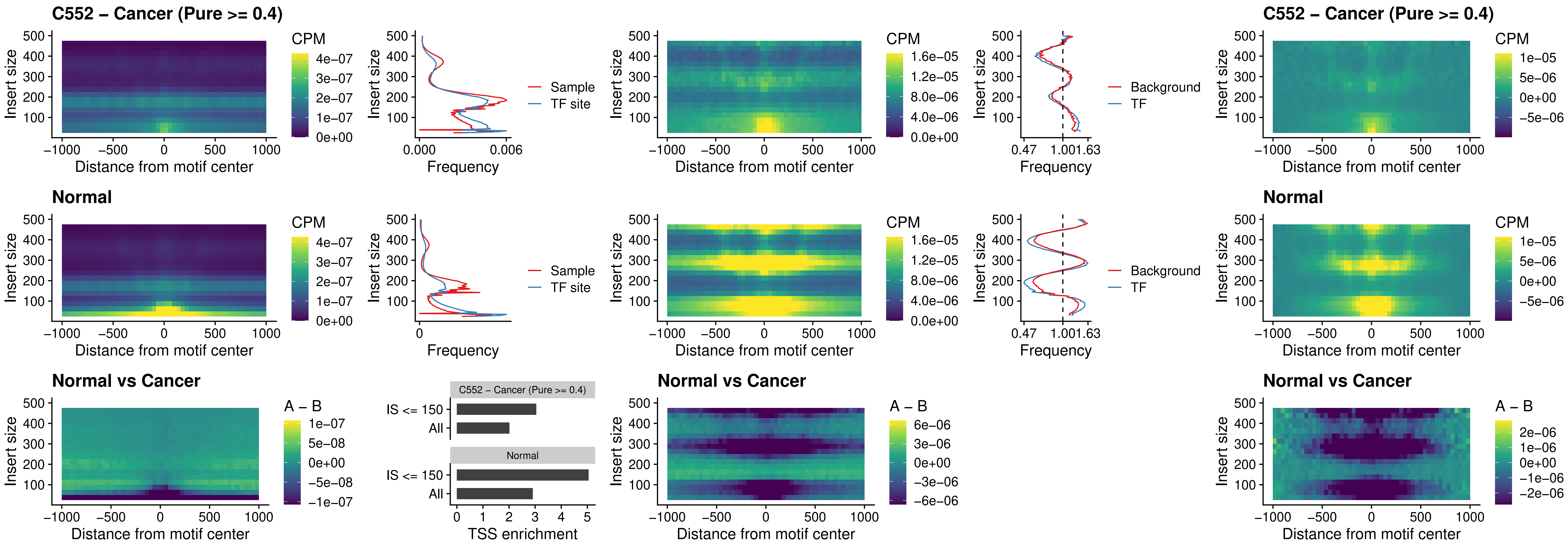

#### SPIC_uc_pTSS_oPEAK

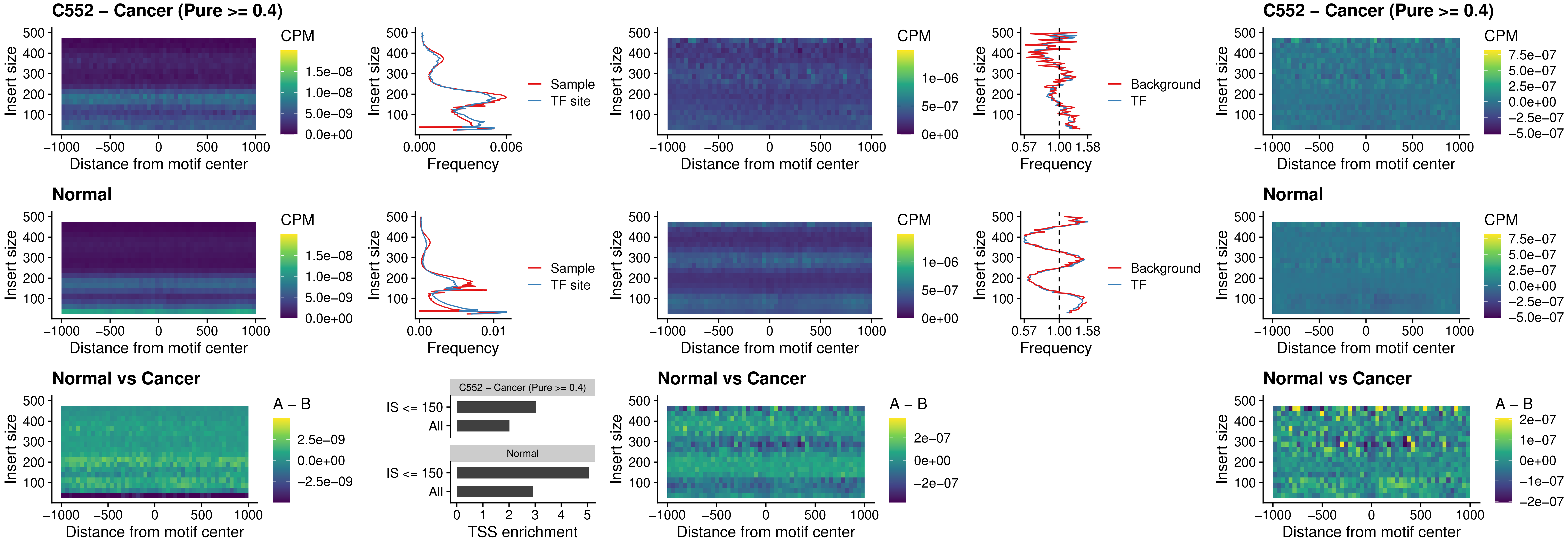

### SPI1_uc_dTSS_nPEAK

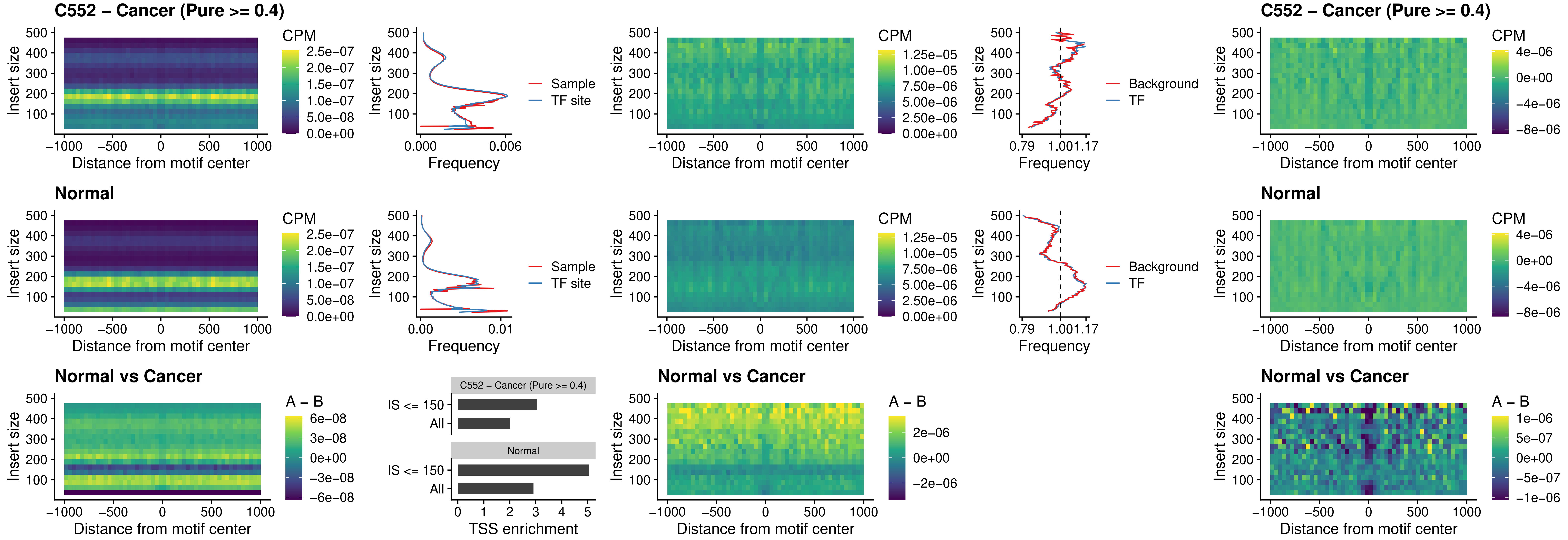

### SPI1_uc_cTSS_nPEAK

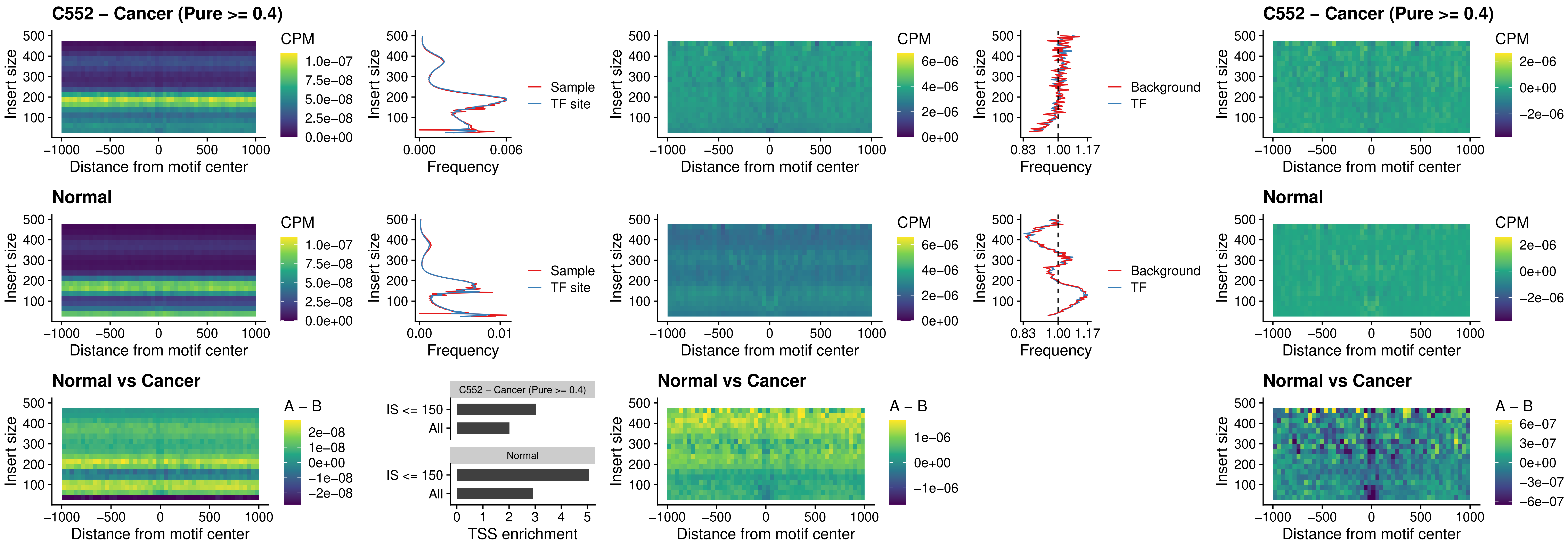

**BCL11A_uc_pTSS_nPEAK**

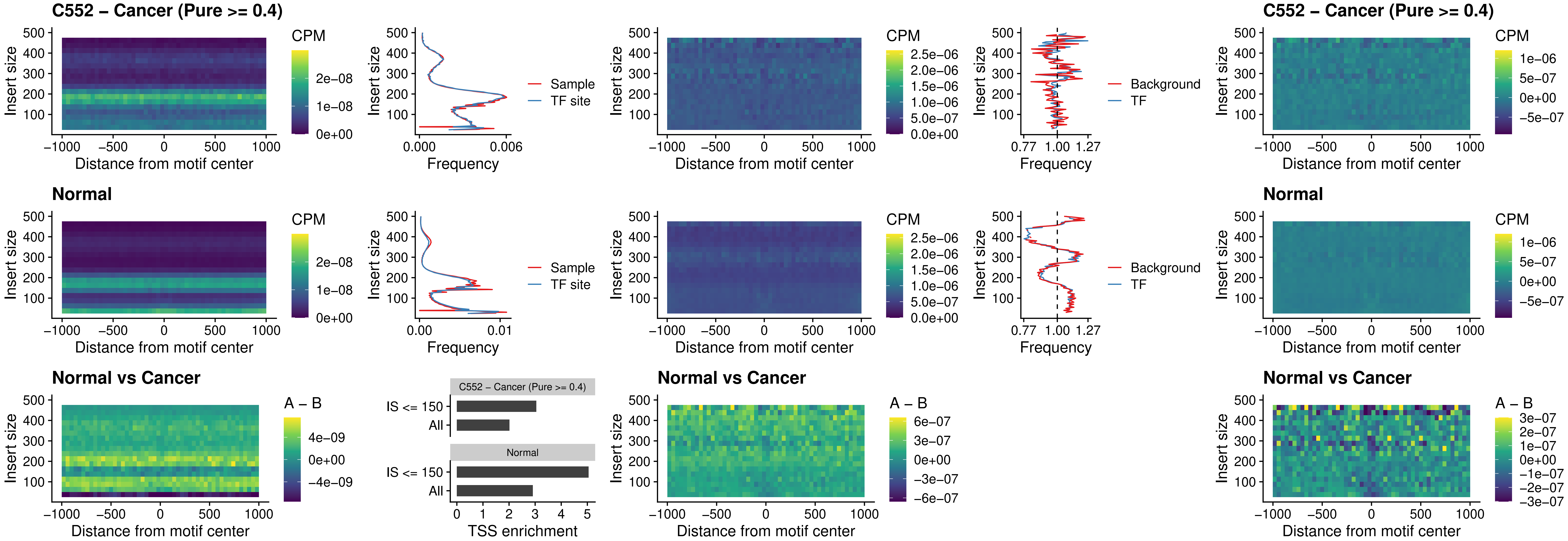

### BCL11B_uc_pTSS_nPEAK

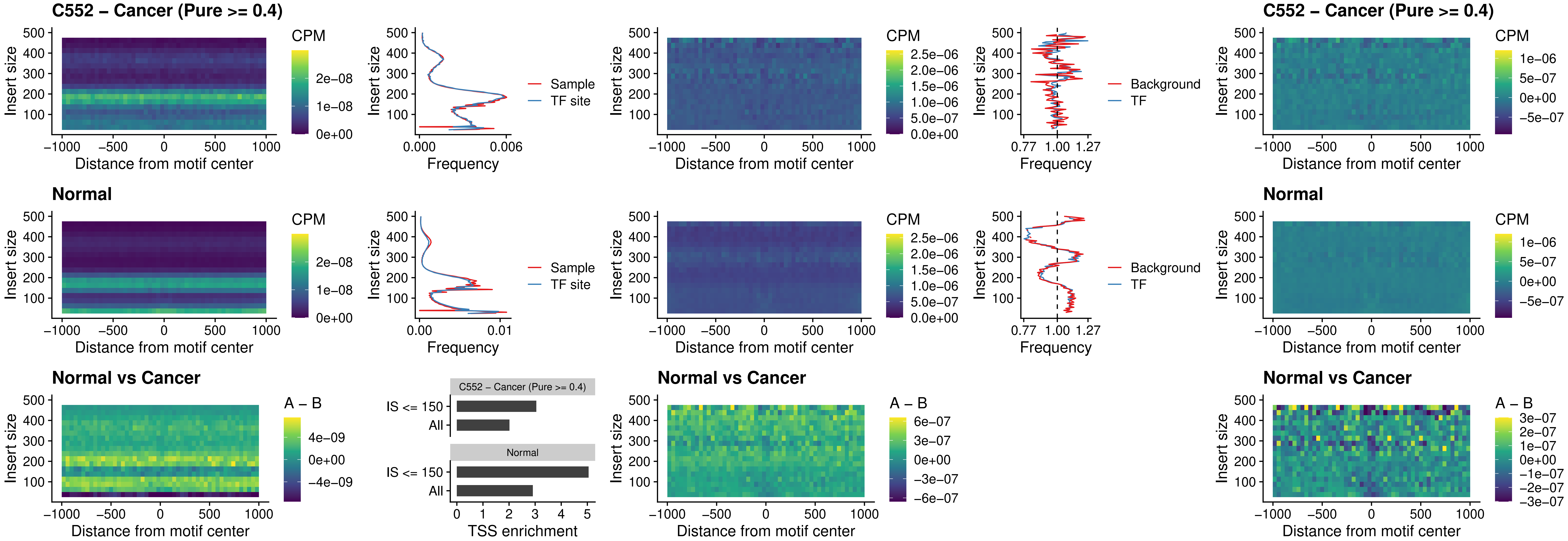

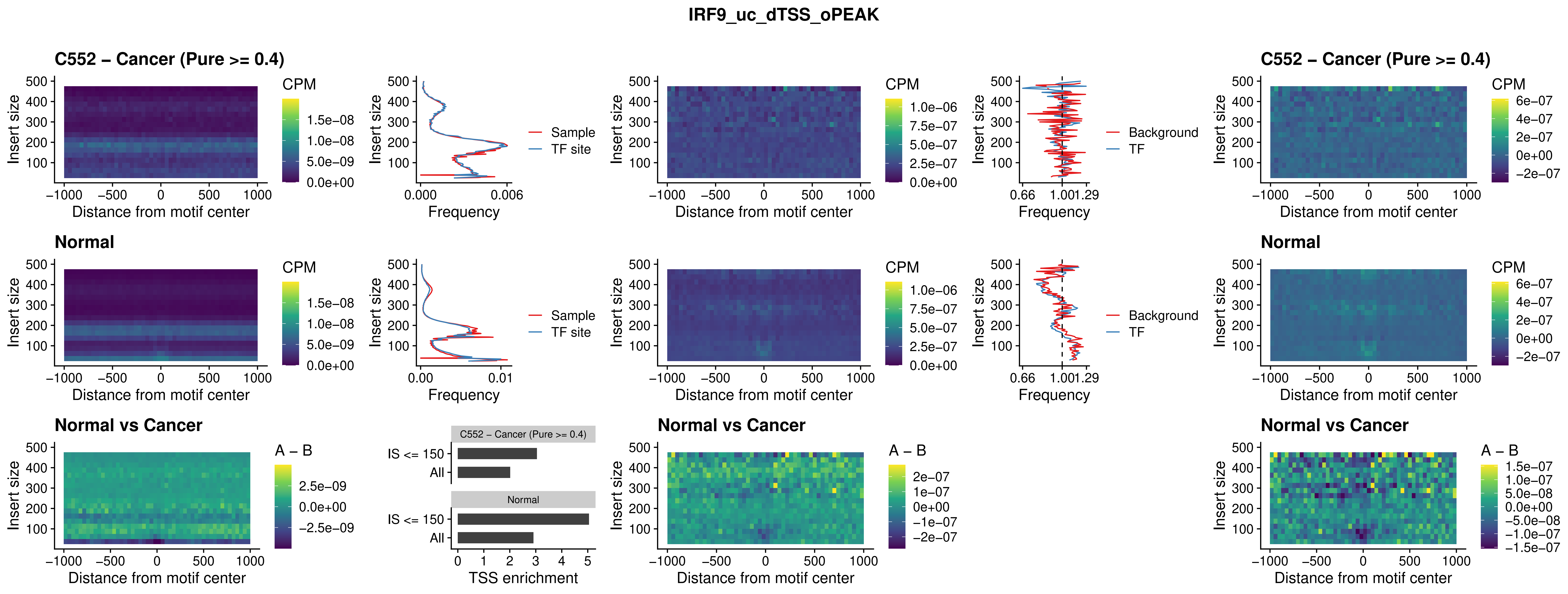
IRF9_uc_dTSS_oPEAK
C552 – Cancer (Pure >= 0.4)
Normal
Normal vs Cancer
Insert size
Distance from motif center
Frequency
CPM
A – B
Sample
TF site
Background
TF
C552 – Cancer (Pure >= 0.4)
Normal
IS <= 150
All
TSS enrichment

### IRF9_uc_cTSS_oPEAK

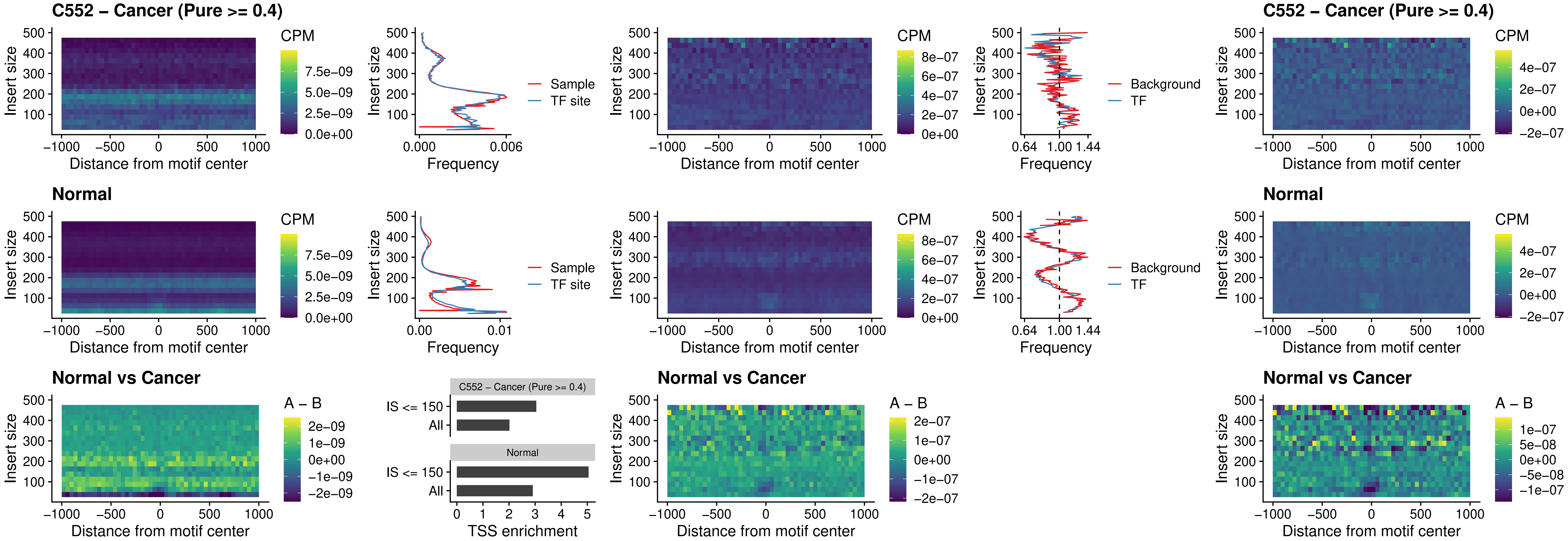

### BCL11A_uc_cTSS_oPEAK

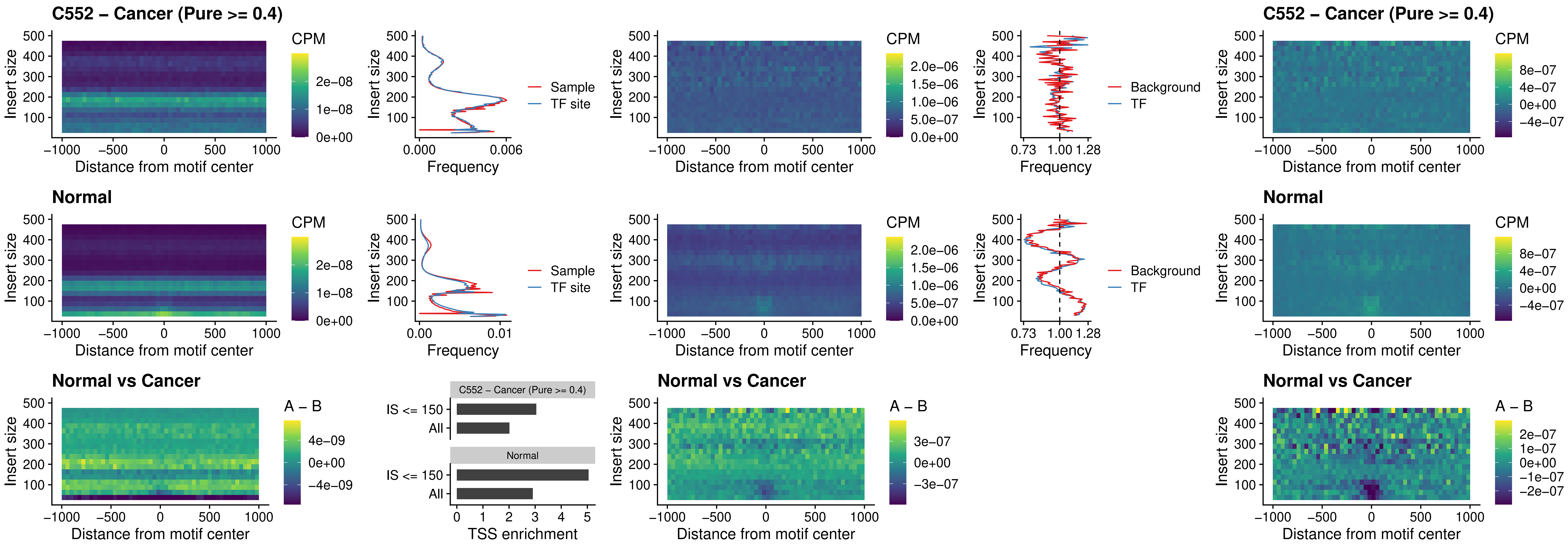

### BCL11B_uc_cTSS_oPEAK

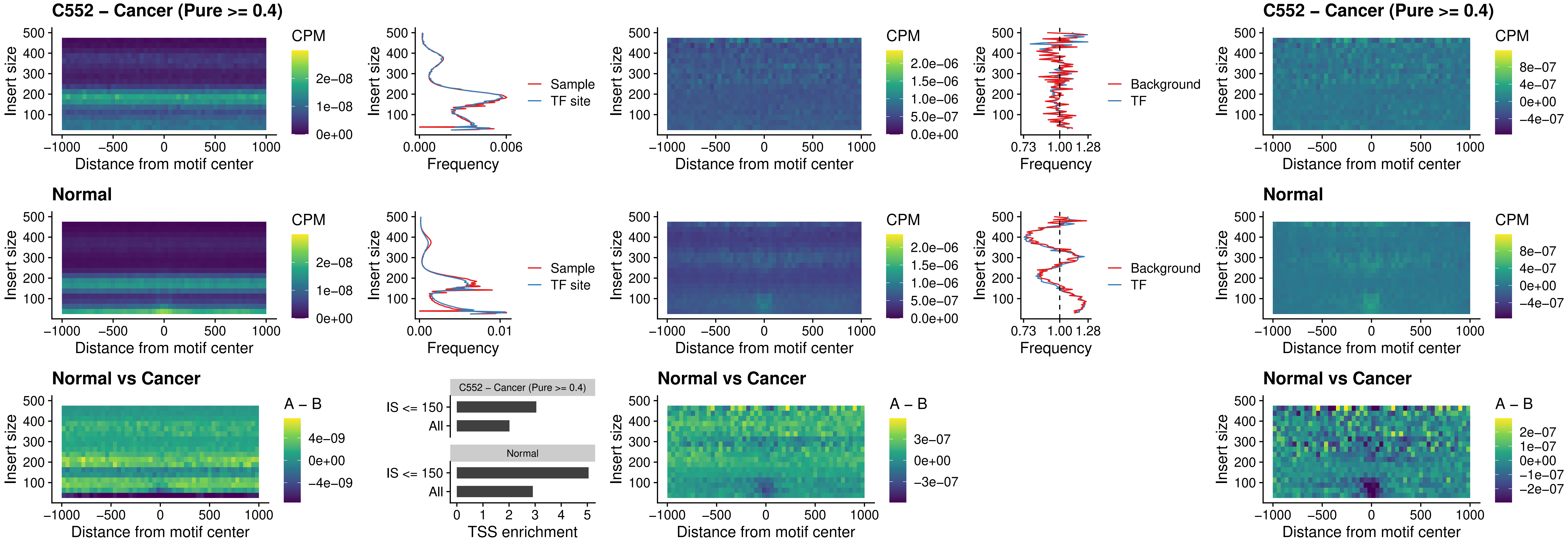

### BCL11A_uc_dTSS_oPEAK

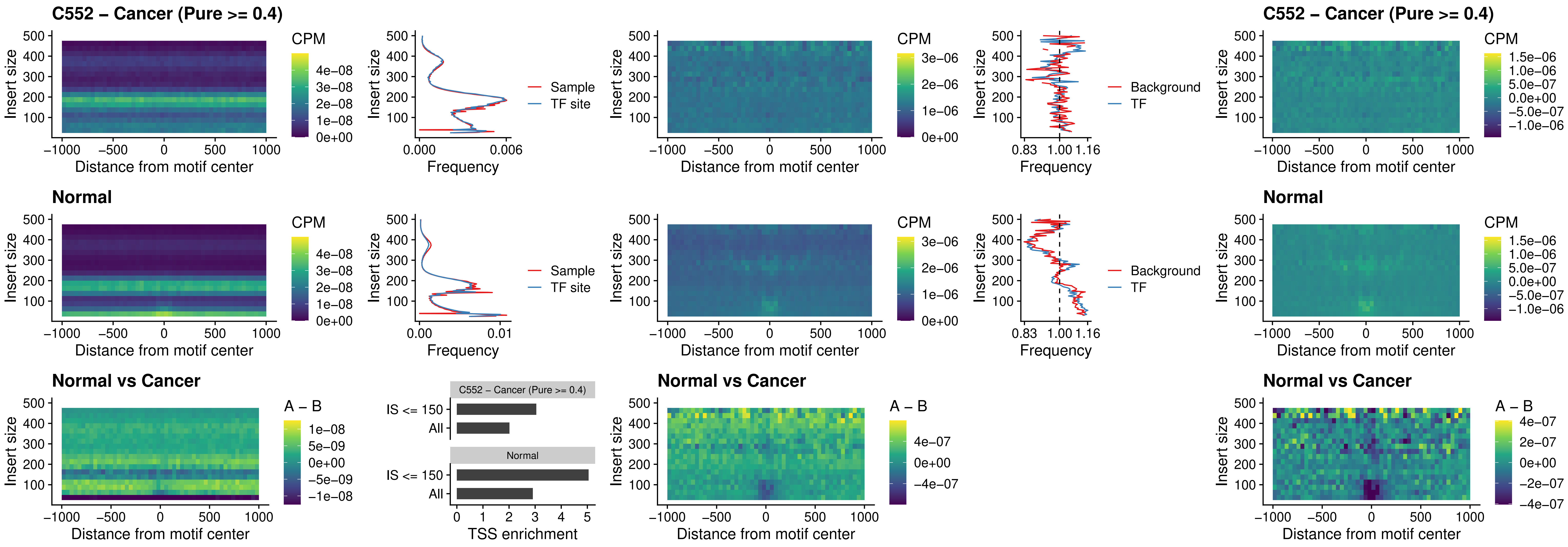

### BCL11B_uc_dTSS_oPEAK

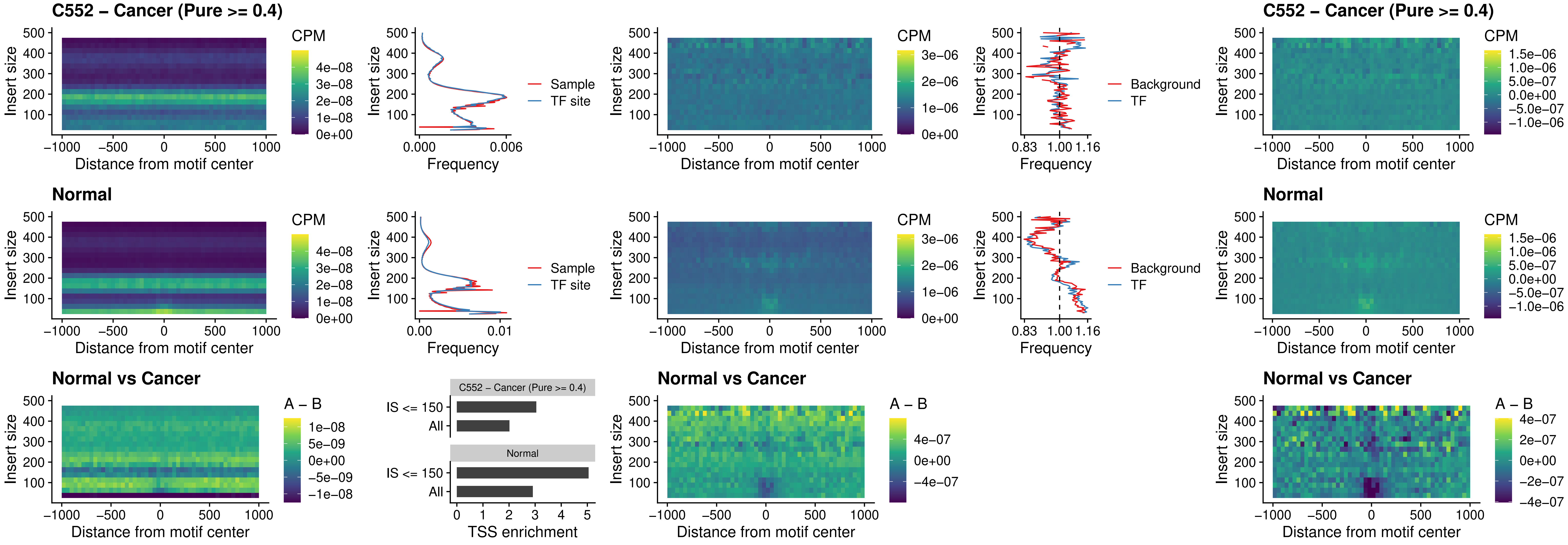

### SPIB_uc_cTSS_nPEAK

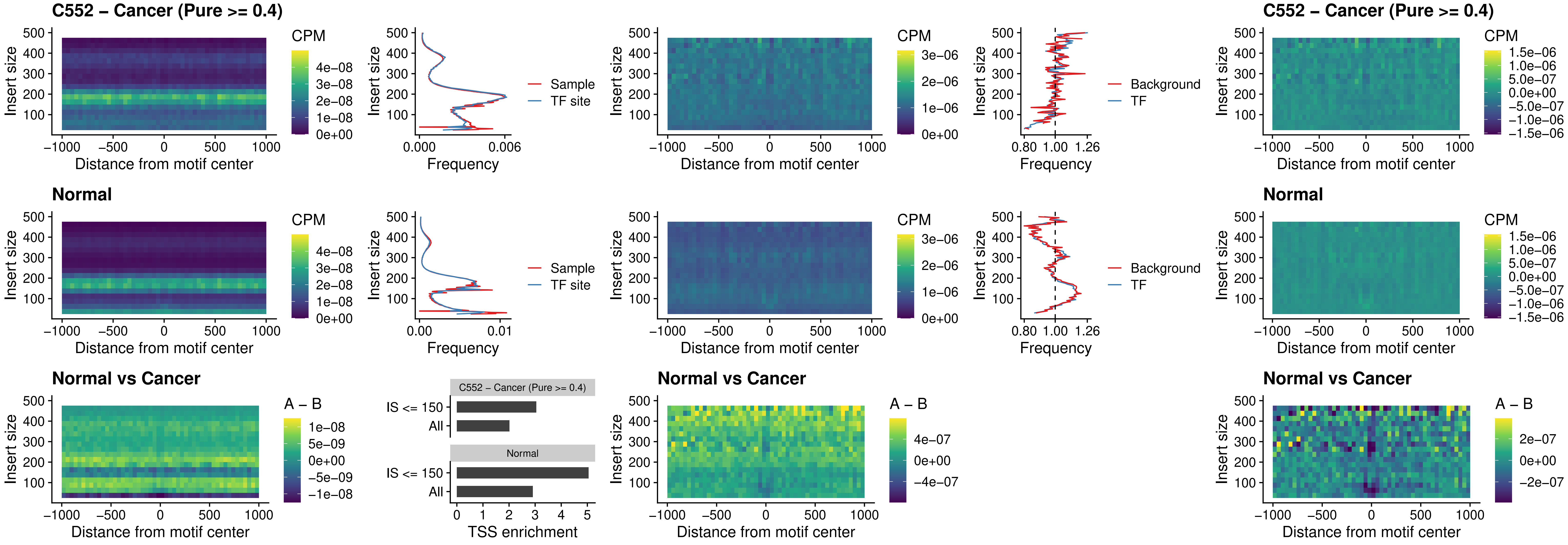

### SPIC_uc_cTSS_nPEAK

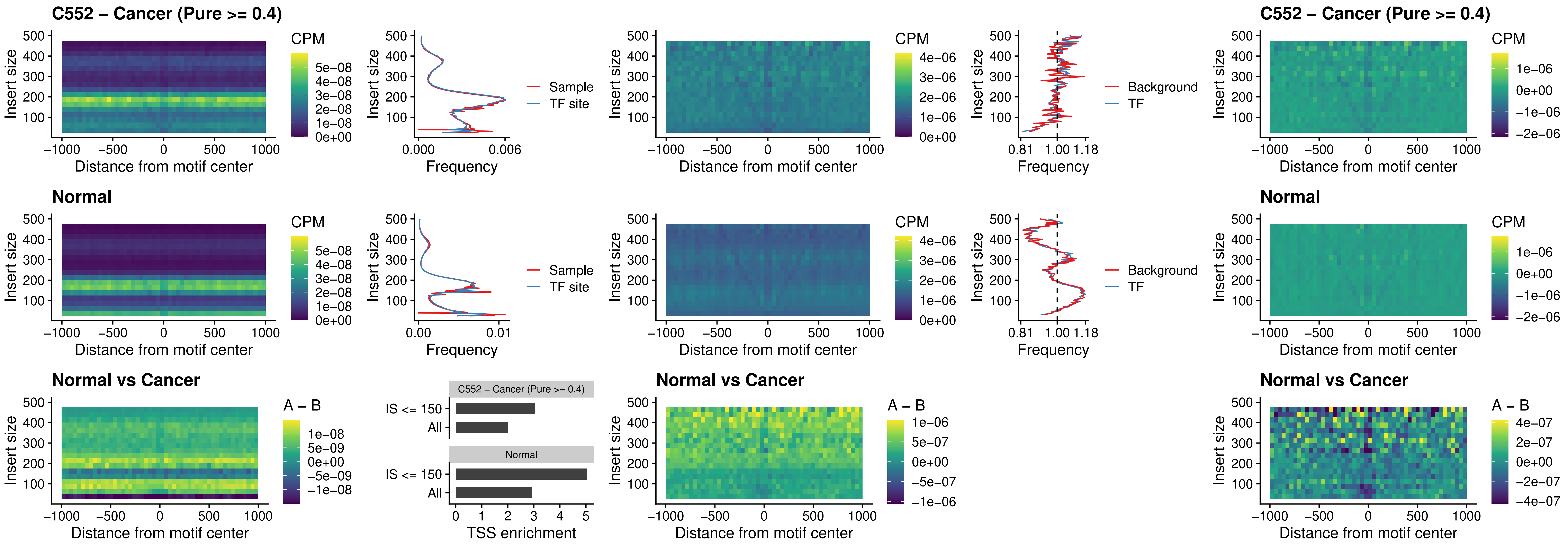

### SPI1_uc_pTSS_oPEAK

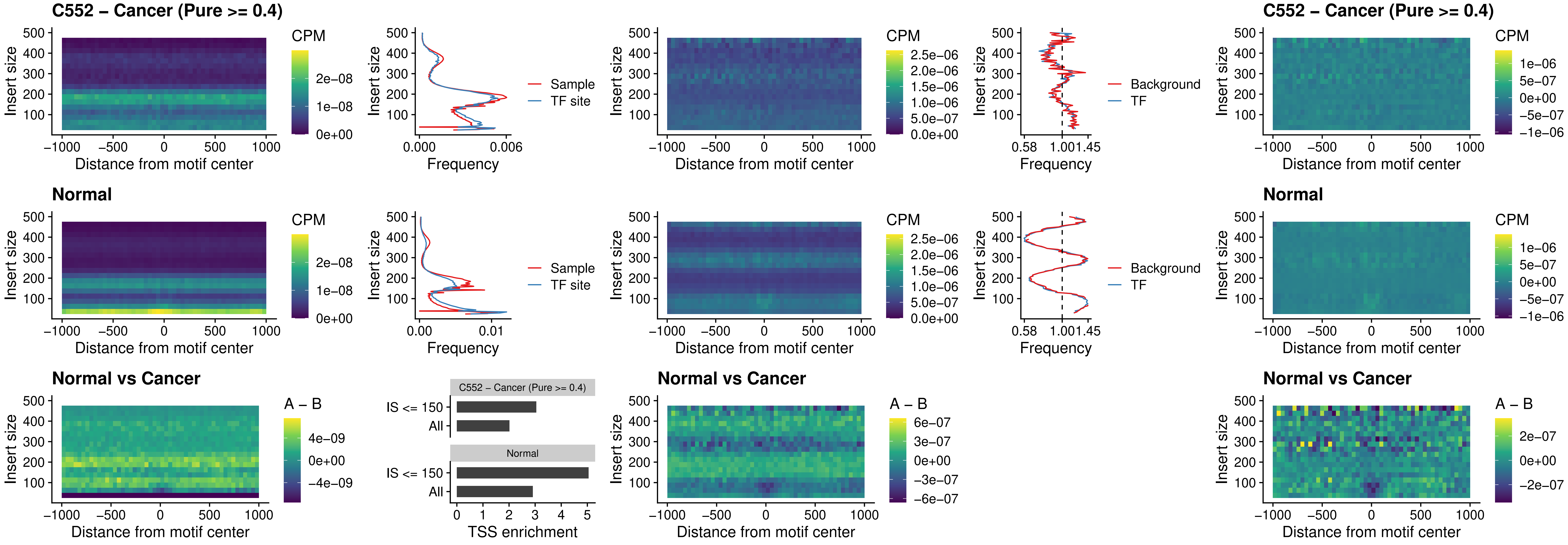

### BCL11A_uc_pTSS_oPEAK

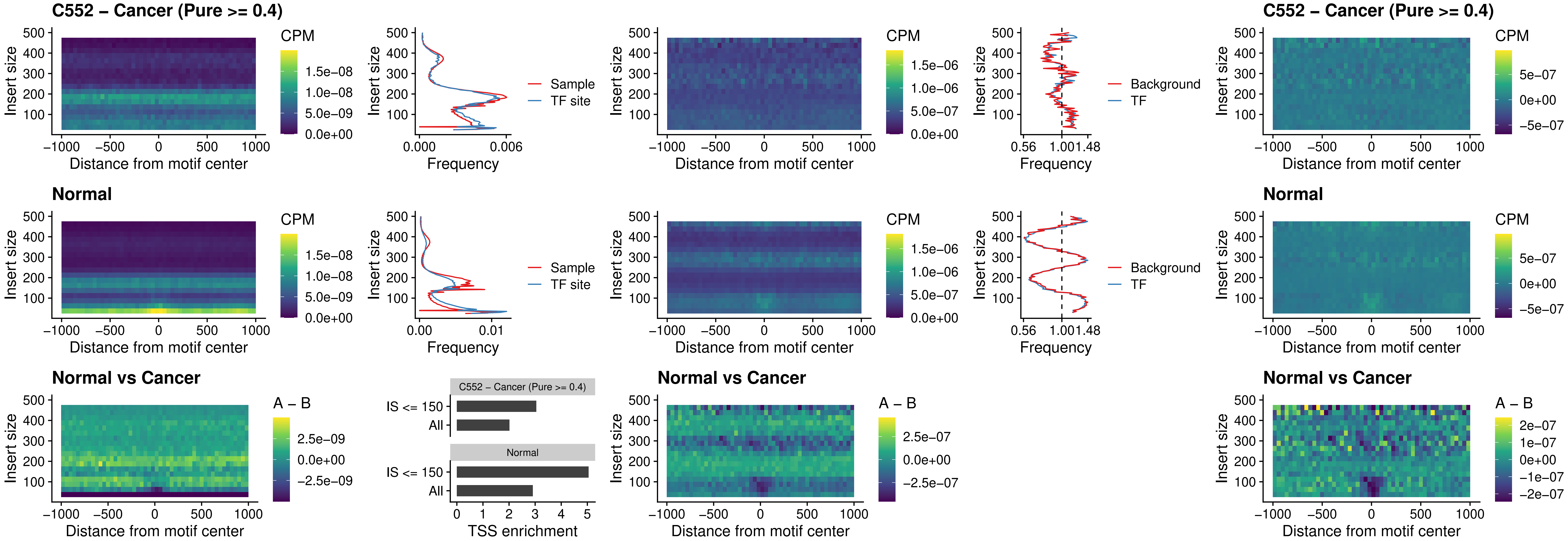

#### BCL11B_uc_pTSS_oPEAK

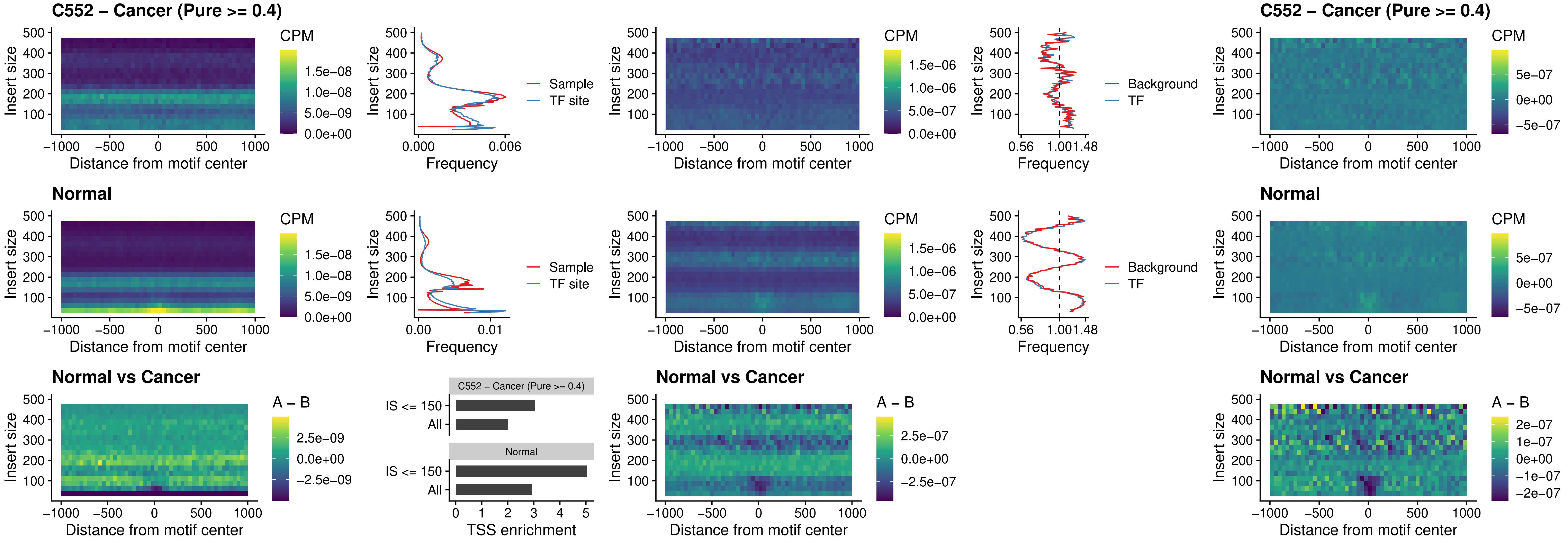

#### SPI1_uc_cTSS_oPEAK

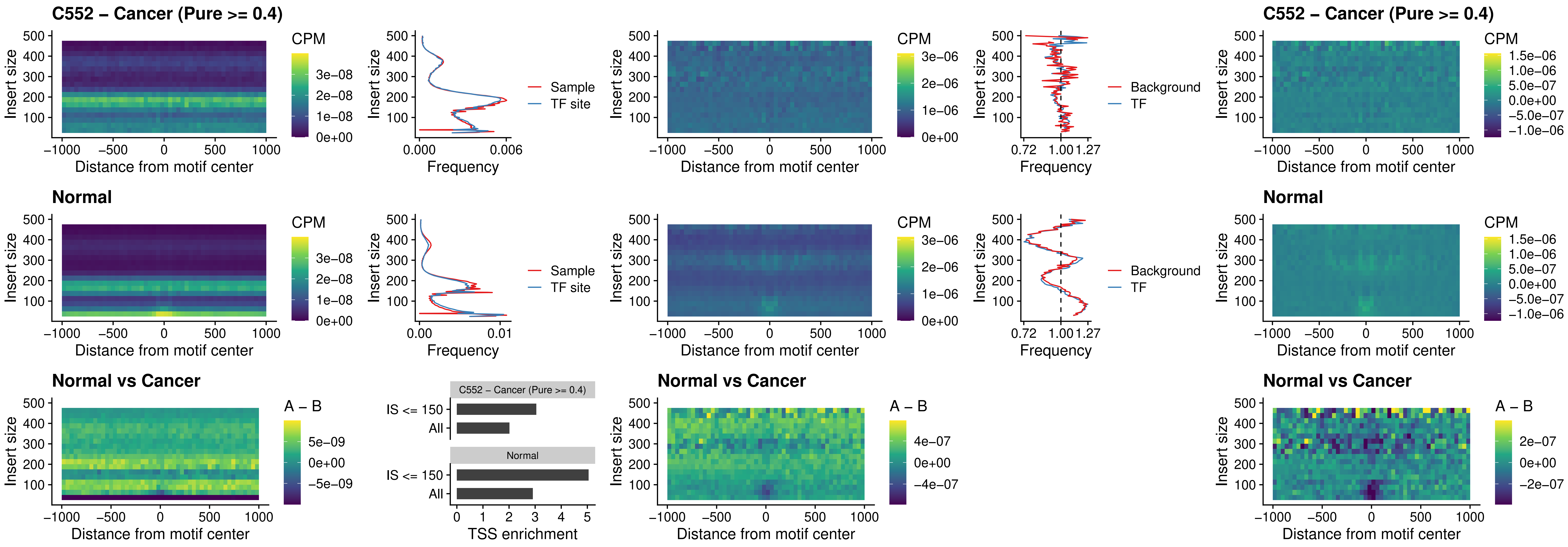

#### SPIB_uc_cTSS_oPEAK

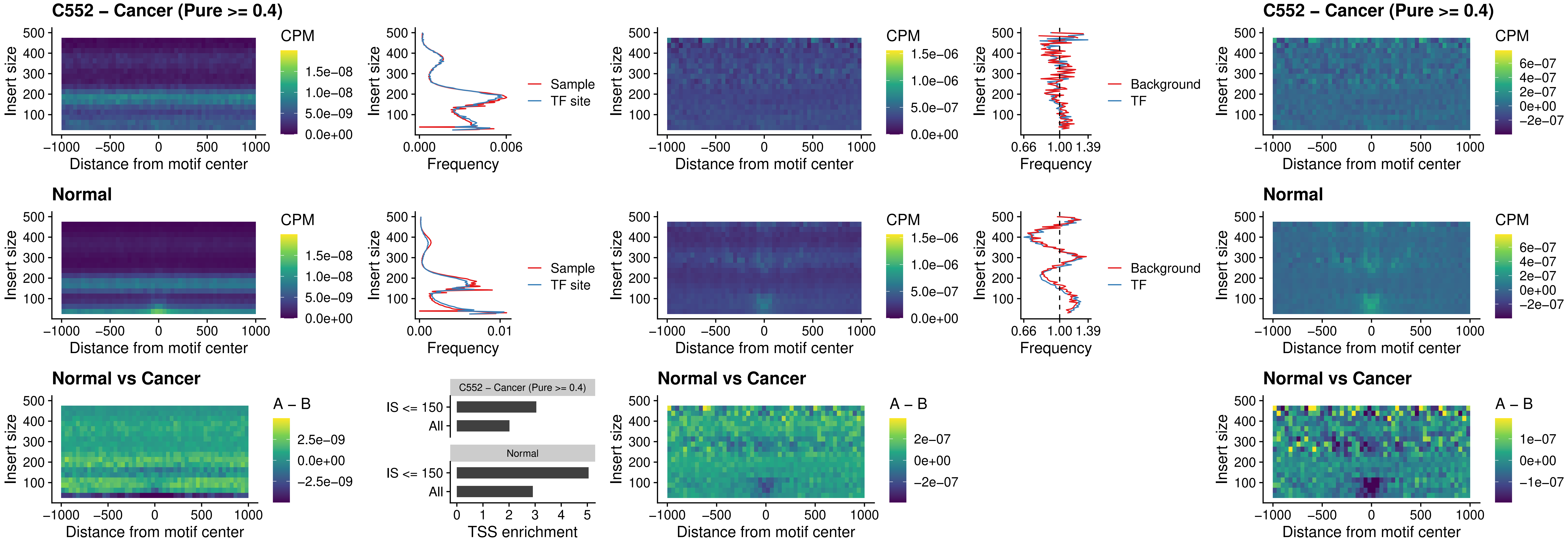

### SPIB_uc_pTSS_oPEAK

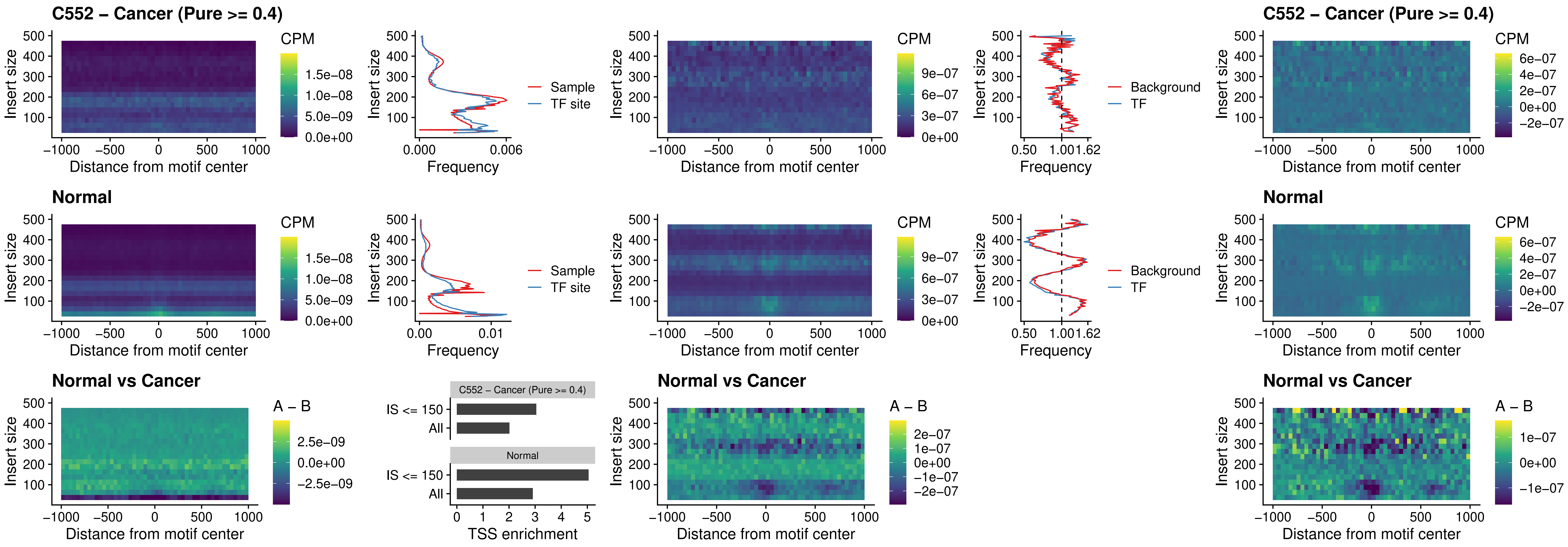

### SPI1_uc_dTSS_oPEAK

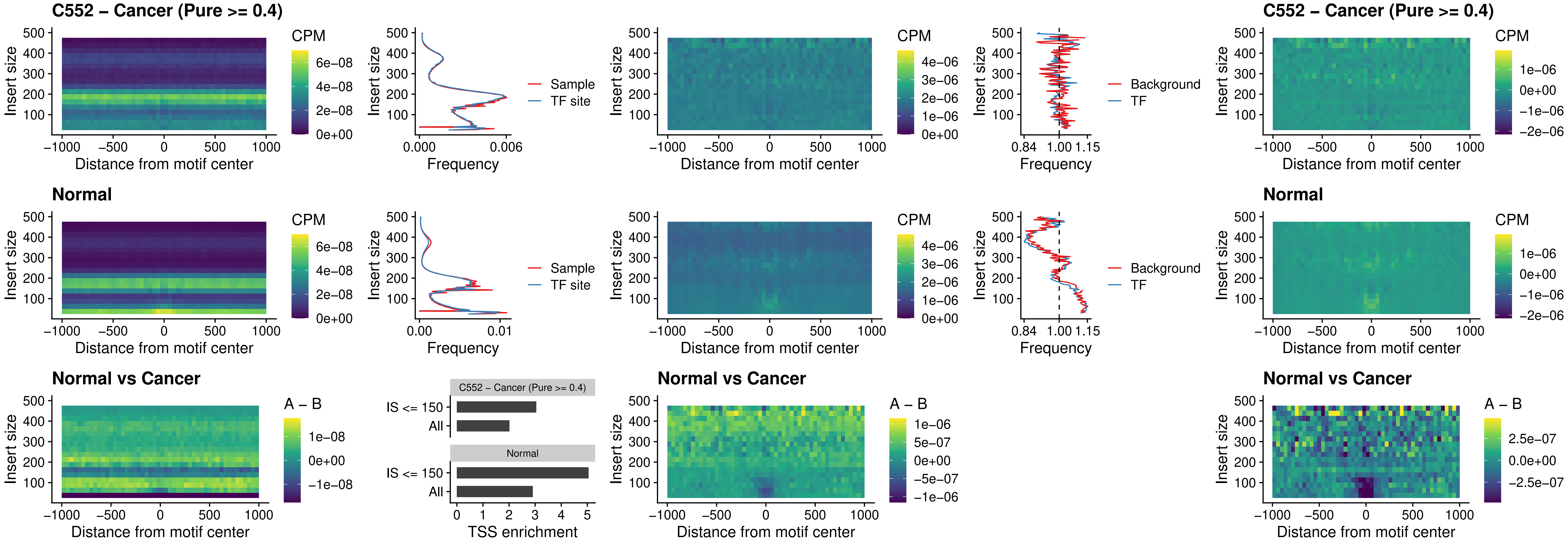

#### SPIB_uc_dTSS_oPEAK

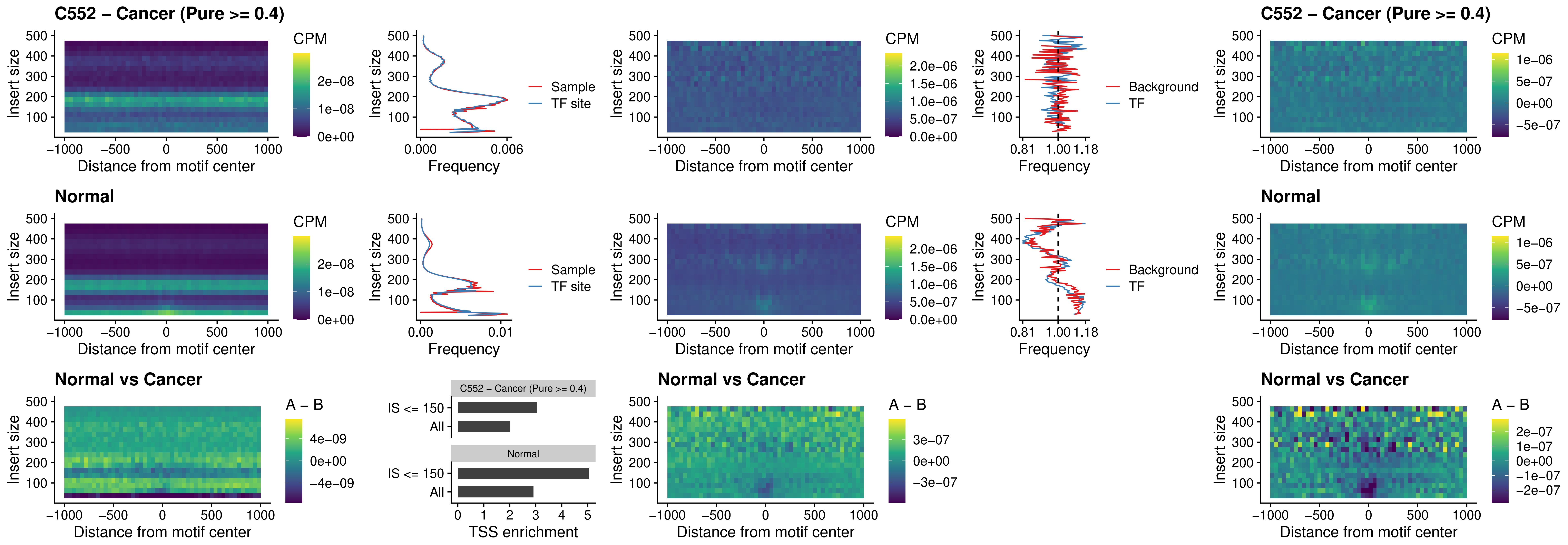

### IRF3_uc_pTSS_oPEAK

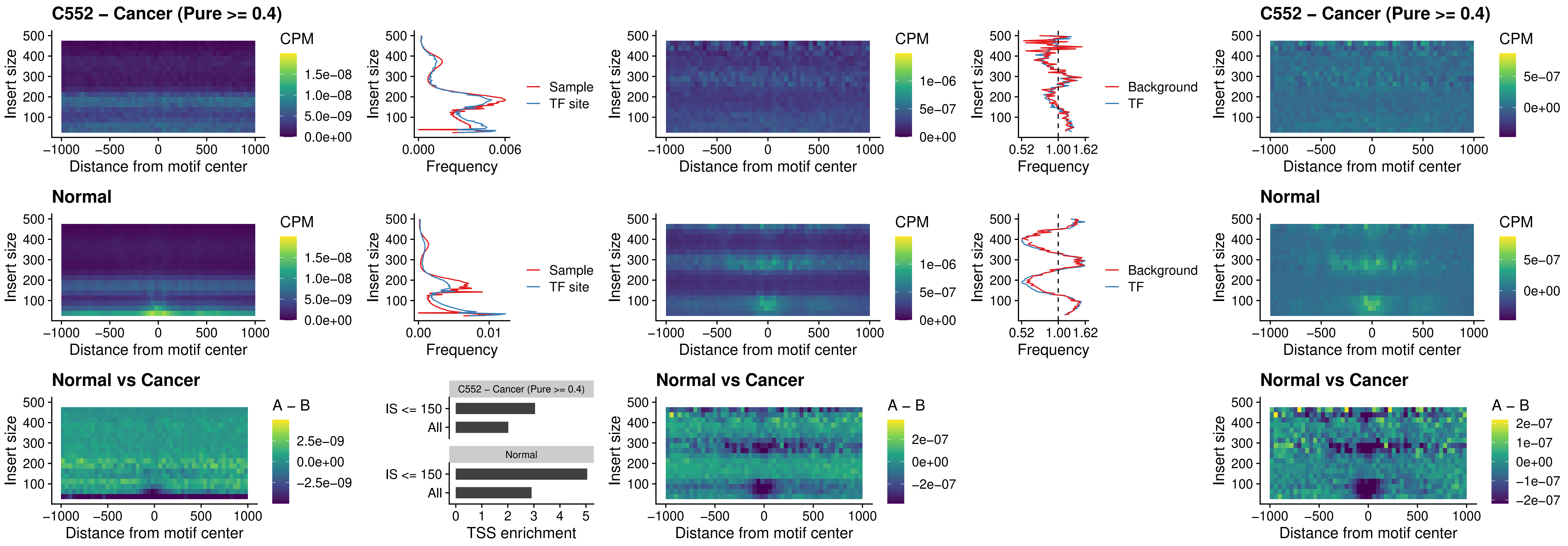

### IRF2_uc_dTSS_oPEAK

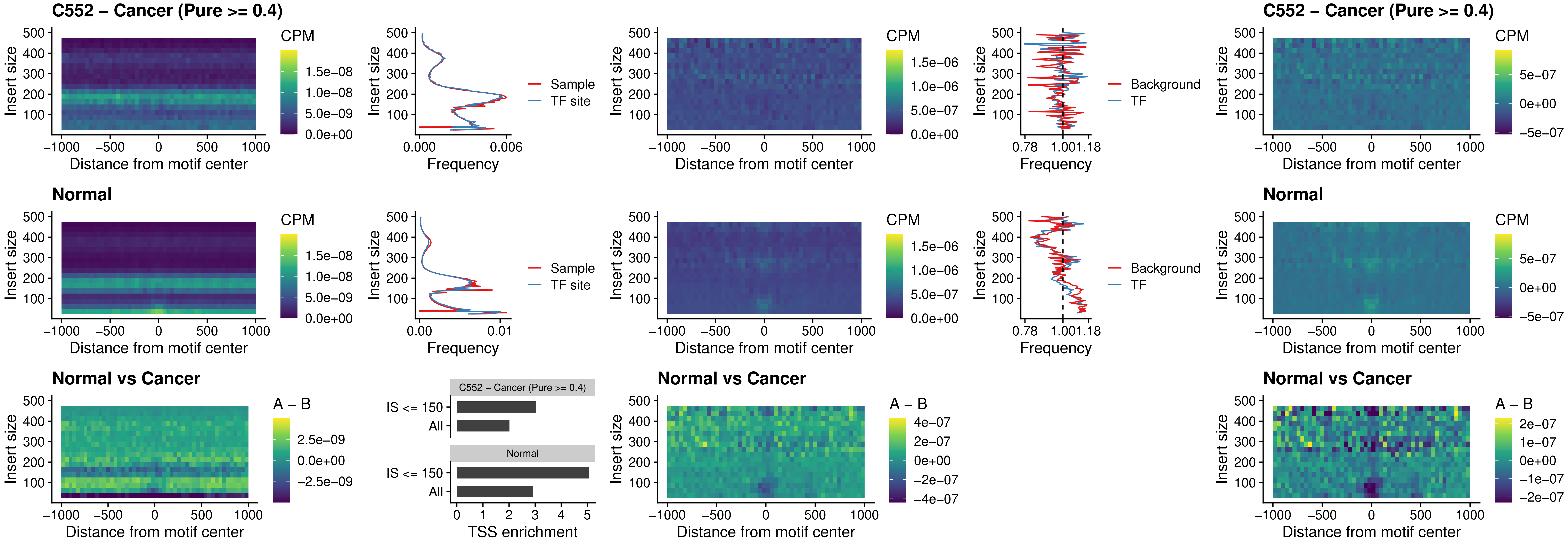

#### IRF2_uc_pTSS_oPEAK

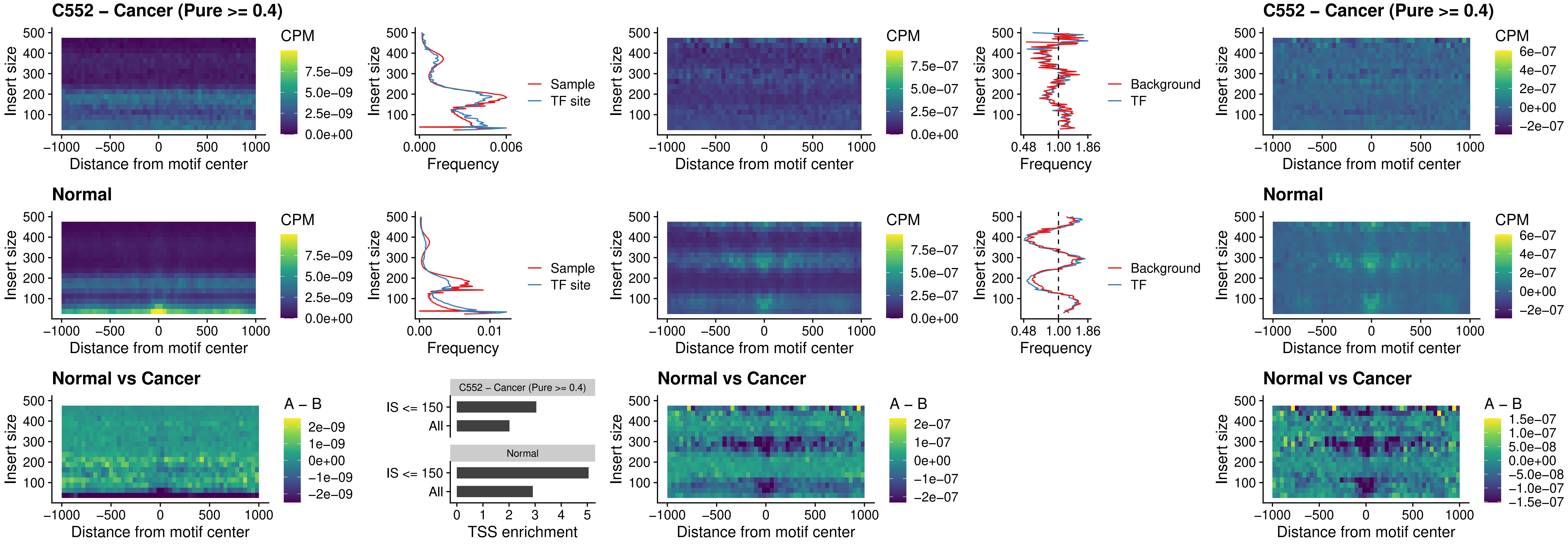

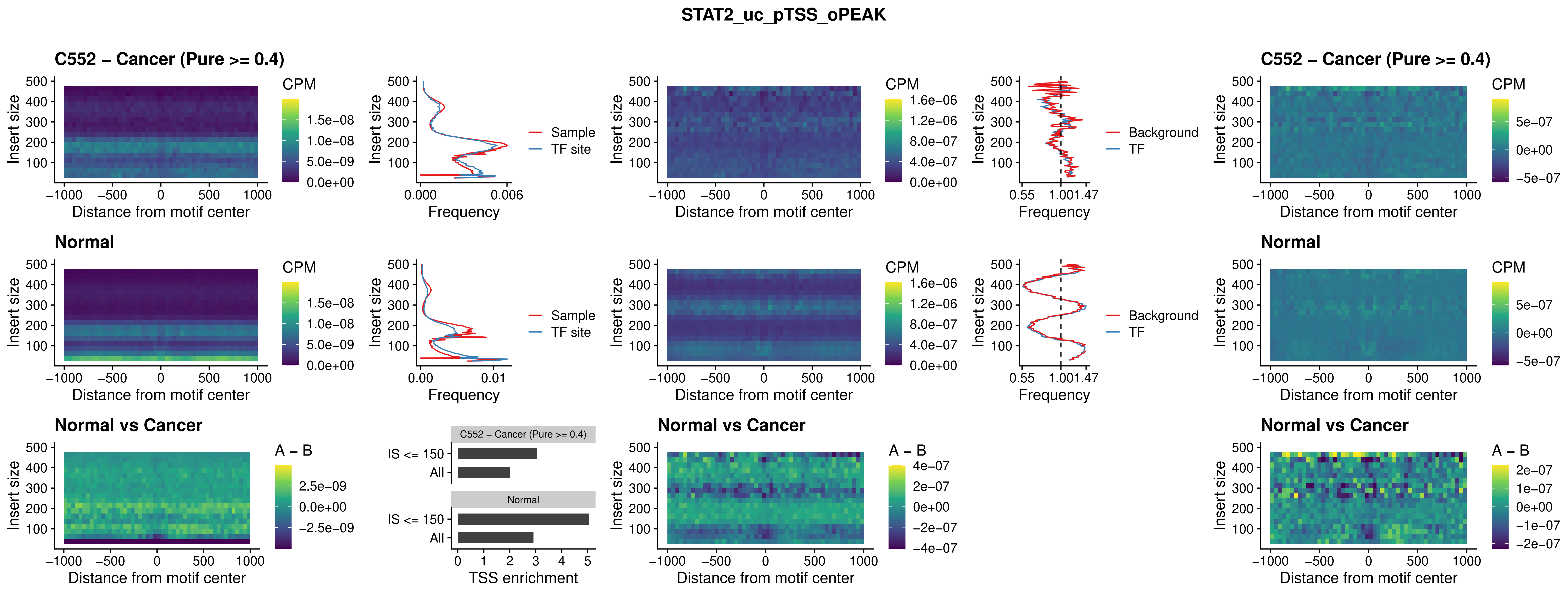
STAT2_uc_pTSS_oPEAK
C552 – Cancer (Pure >= 0.4)
Normal
Normal vs Cancer
Insert size
Distance from motif center
Frequency
CPM
A – B
Sample
TF site
Background
TF
C552 – Cancer (Pure >= 0.4)
Normal
IS <= 150
All
TSS enrichment

### SPIB_uc_dTSS_nPEAK

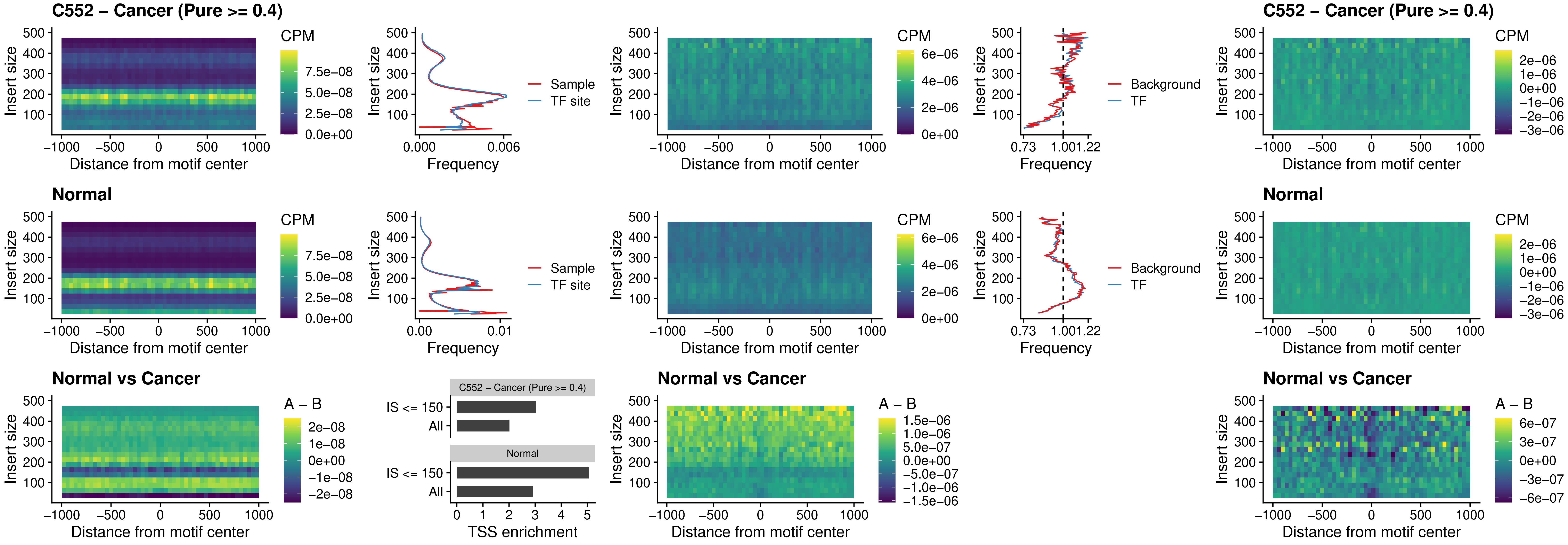

#### SPIC_uc_cTSS_oPEAK

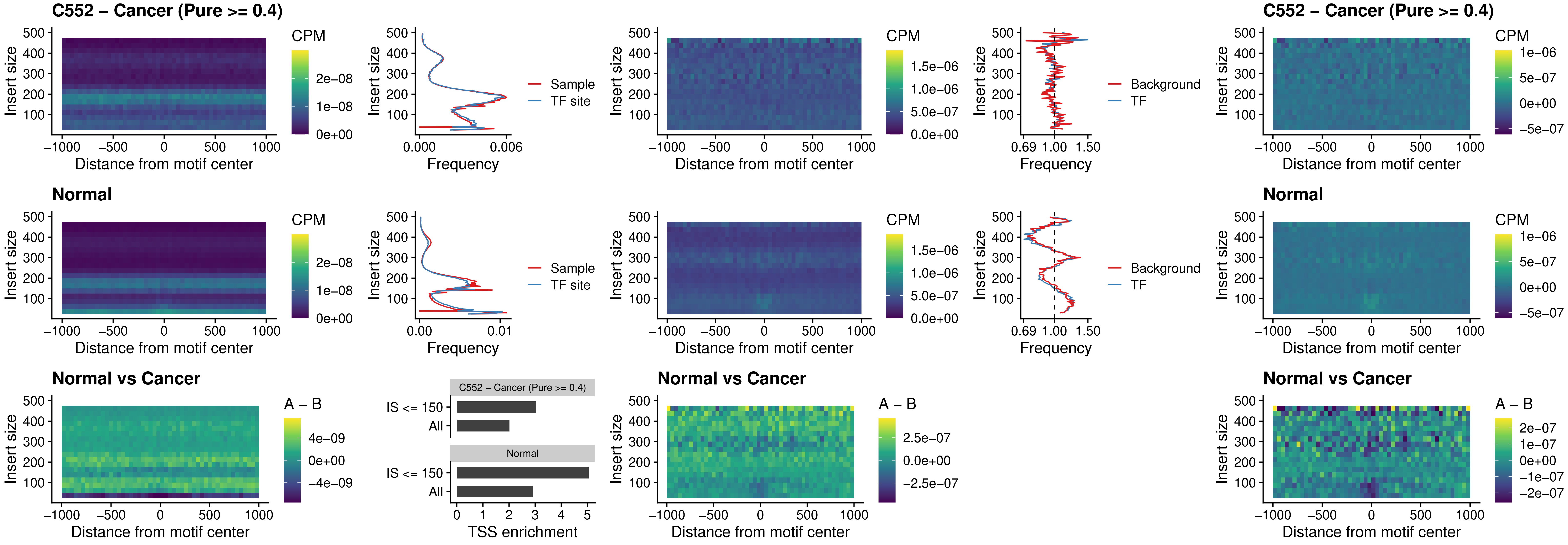

### PRDM1_uc_cTSS_oPEAK

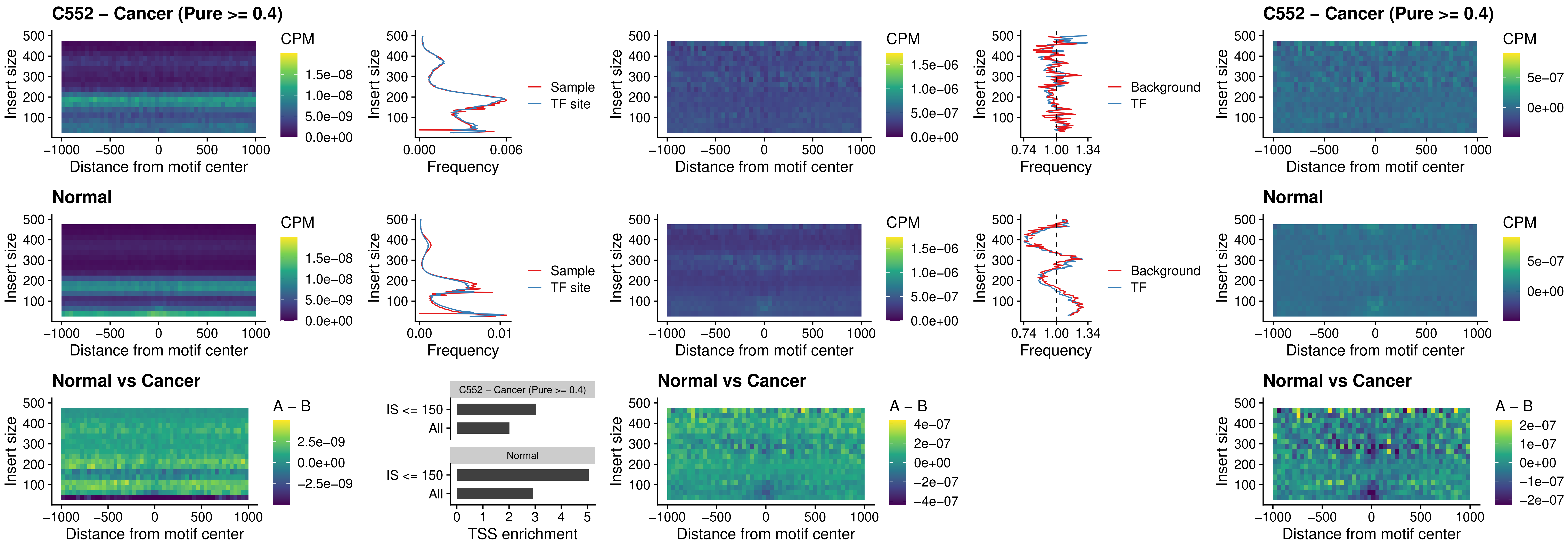

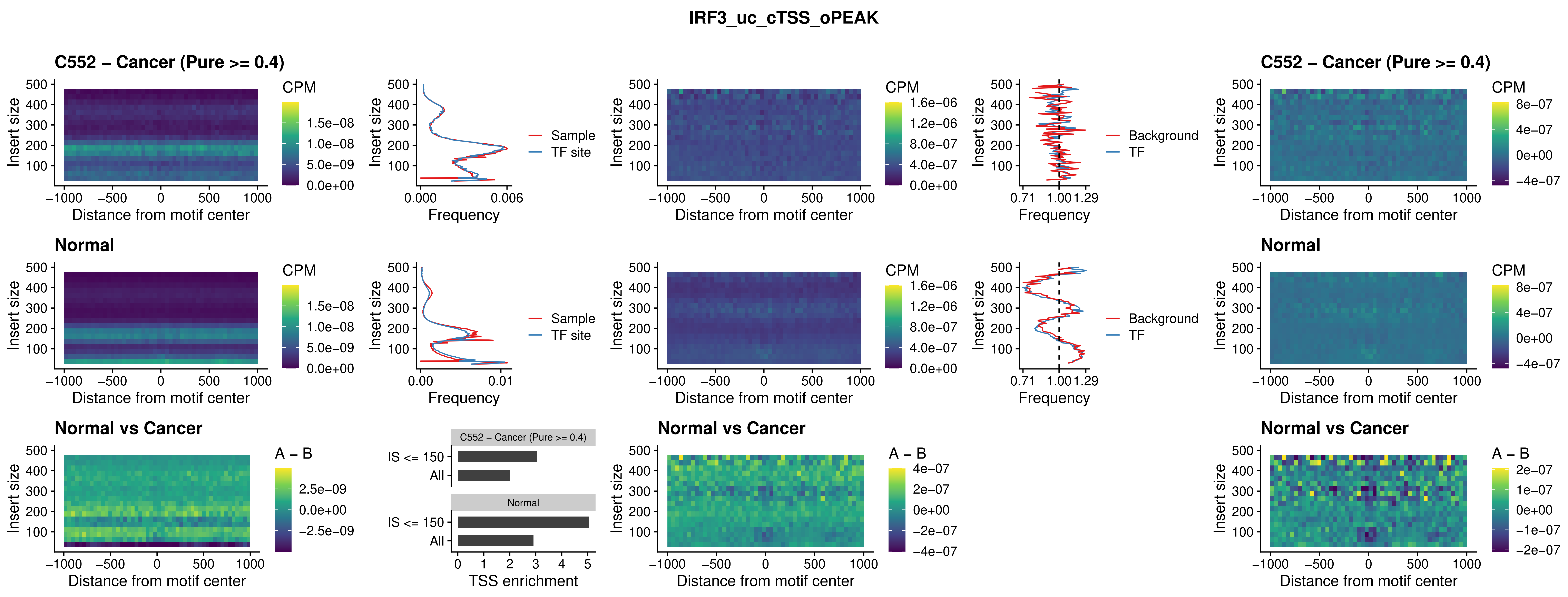
IRF3_uc_cTSS_oPEAK
C552 – Cancer (Pure >= 0.4)
Normal
Normal vs Cancer
Insert size
Distance from motif center
Frequency
CPM
A – B
Sample
TF site
Background
TF
IS <= 150
All
TSS enrichment

#### BCL11A_uc_dTSS_nPEAK

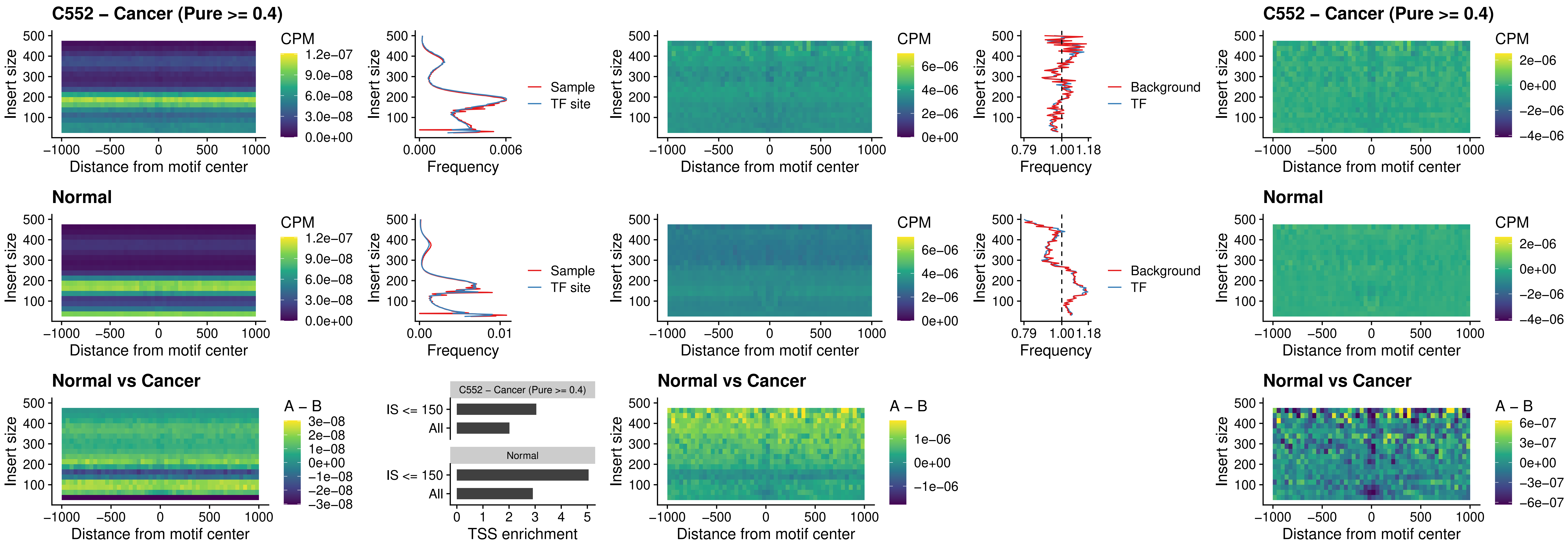

#### BCL11B_uc_dTSS_nPEAK

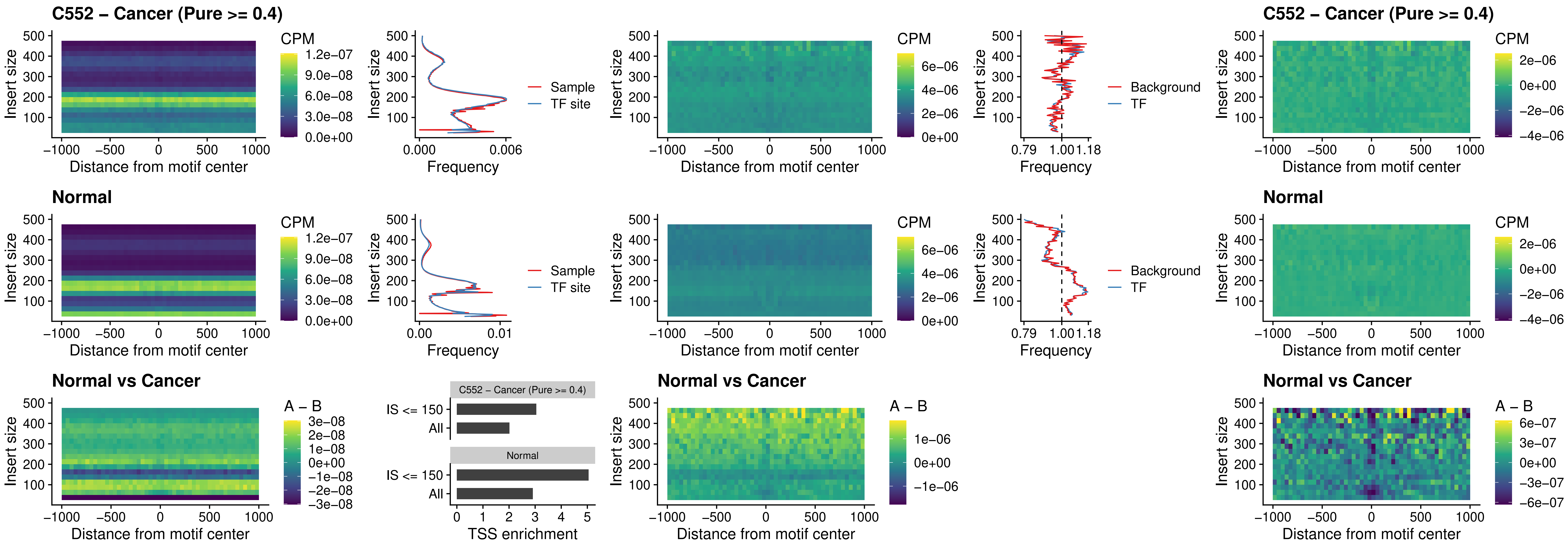

#### IRF1_uc_cTSS_oPEAK

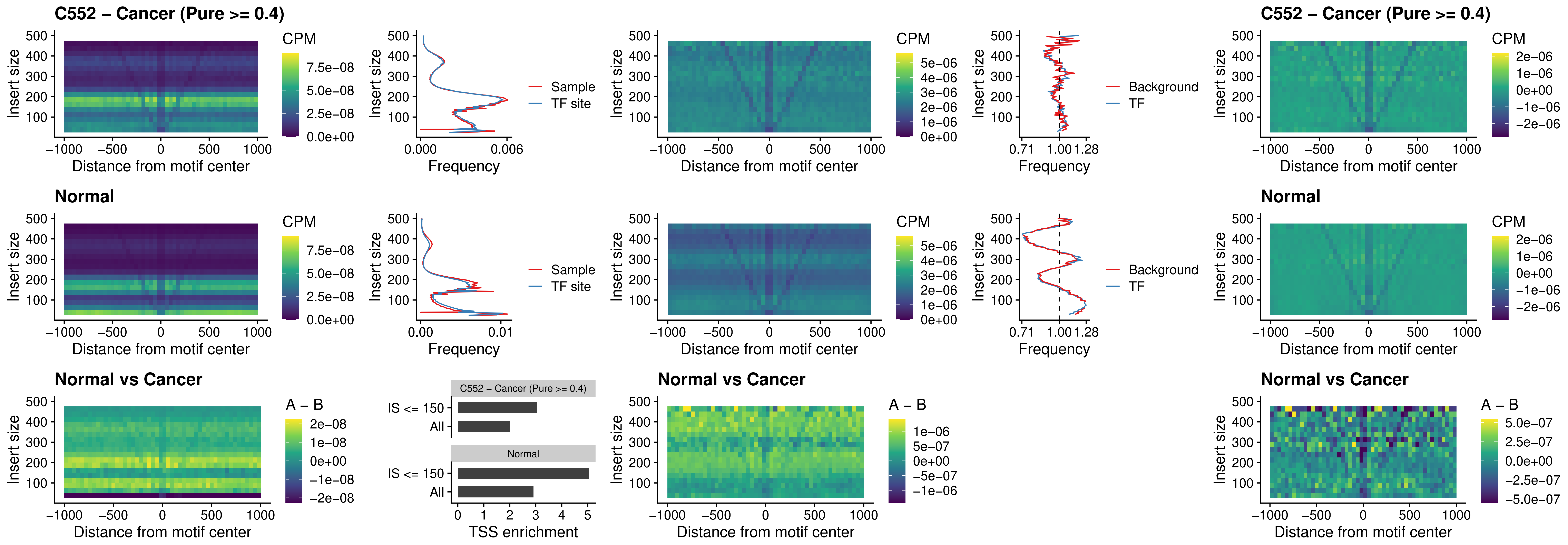

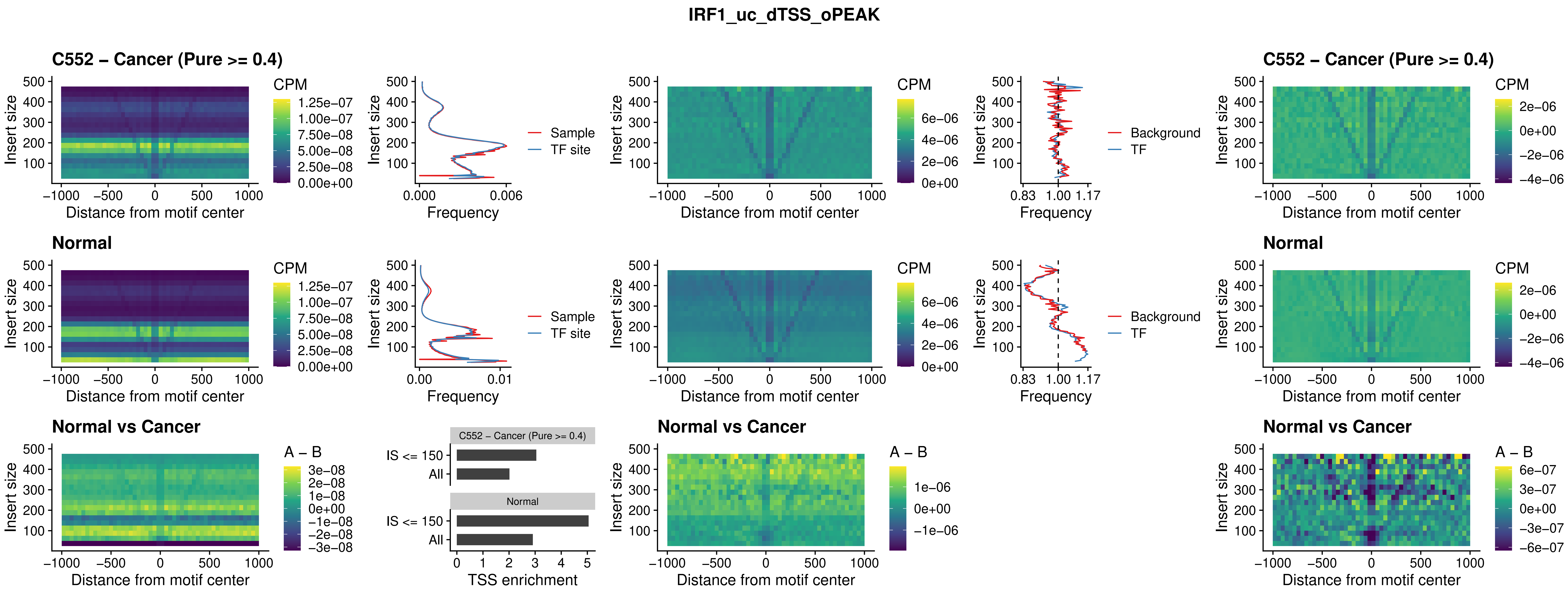

IRF1_uc_dTSS_oPEAK
C552 – Cancer (Pure >= 0.4)
Normal
Normal vs Cancer
Insert size
Distance from motif center
Frequency
CPM
A – B
Sample
TF site
Background
TF
C552 – Cancer (Pure >= 0.4)
Normal
IS <= 150
All
TSS enrichment

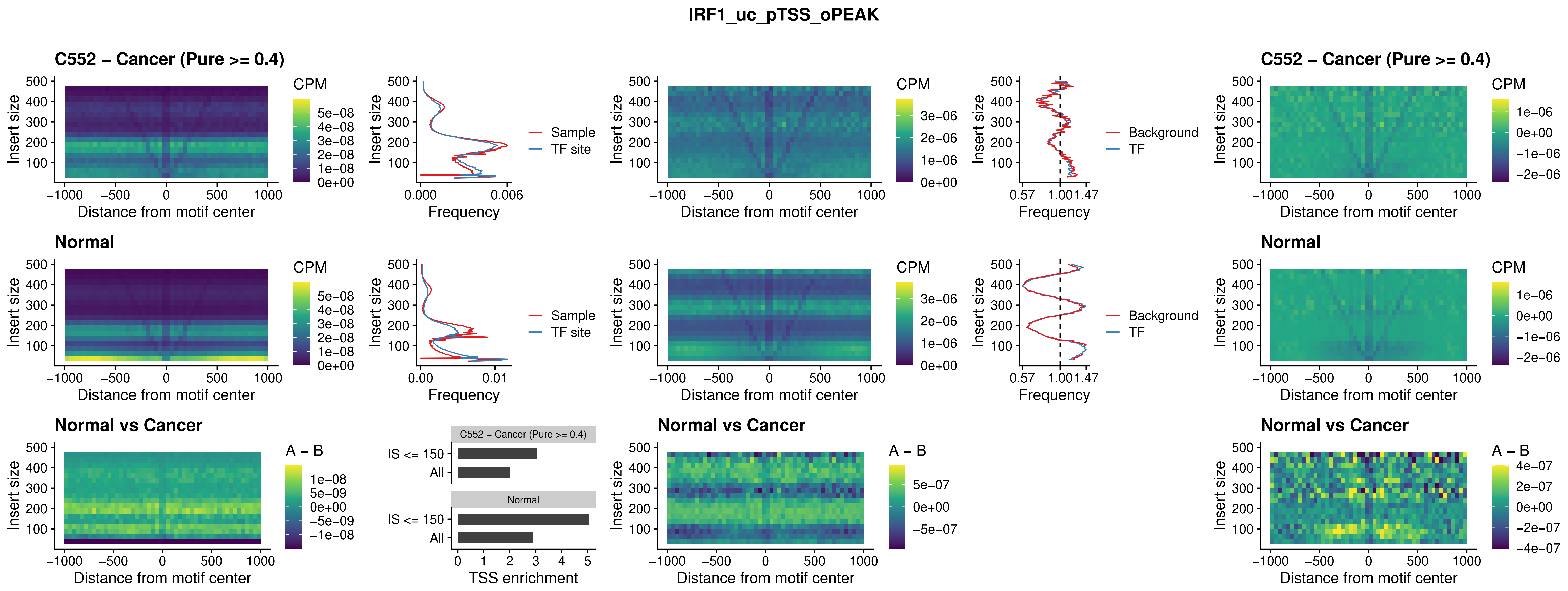
IRF1_uc_pTSS_oPEAK
C552 – Cancer (Pure >= 0.4)
Normal
Normal vs Cancer
Insert size
Distance from motif center
CPM
A – B
Frequency
Sample
TF site
Background
TF
TSS enrichment
C552 – Cancer (Pure >= 0.4)
Normal
IS <= 150
All

#### SPIC_uc_dTSS_nPEAK

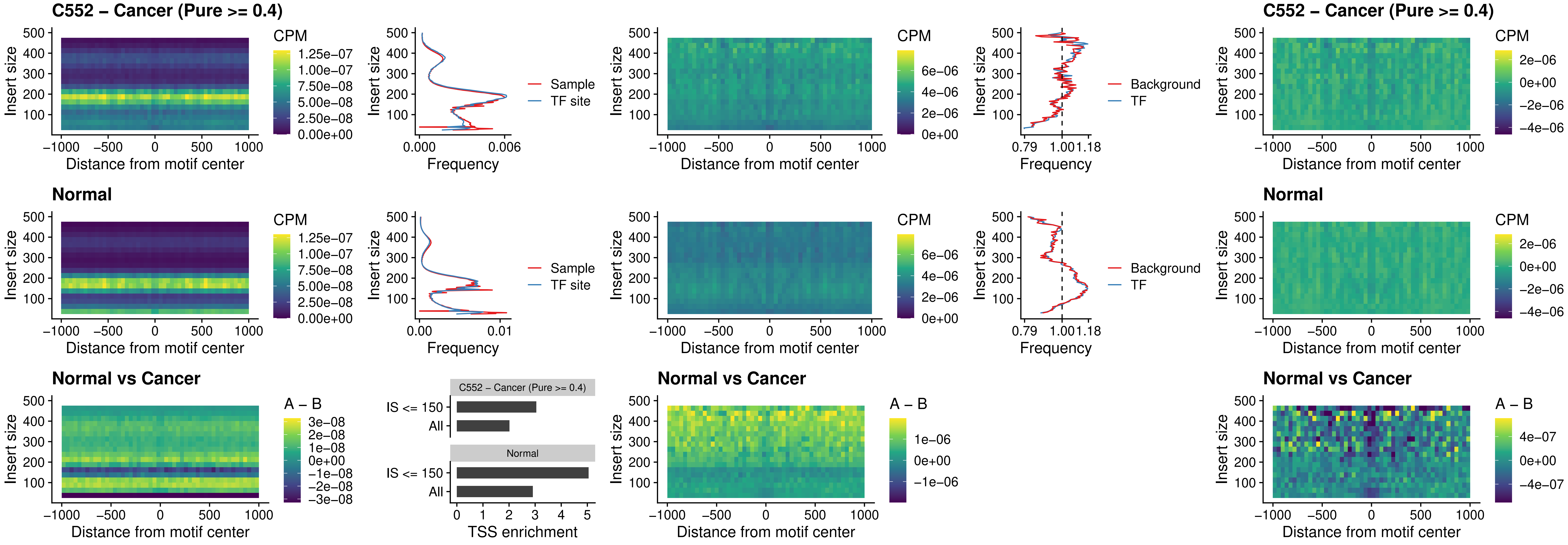

#### SPIC_uc_dTSS_oPEAK

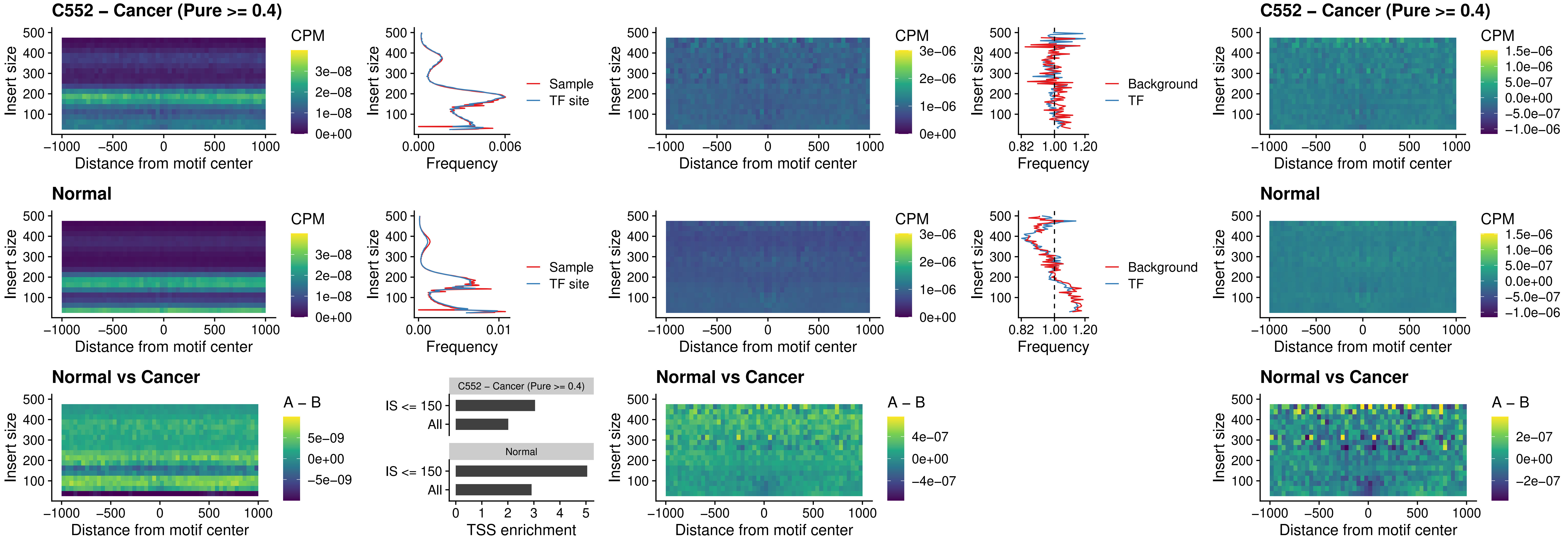

#### SPIB_uc_pTSS_nPEAK

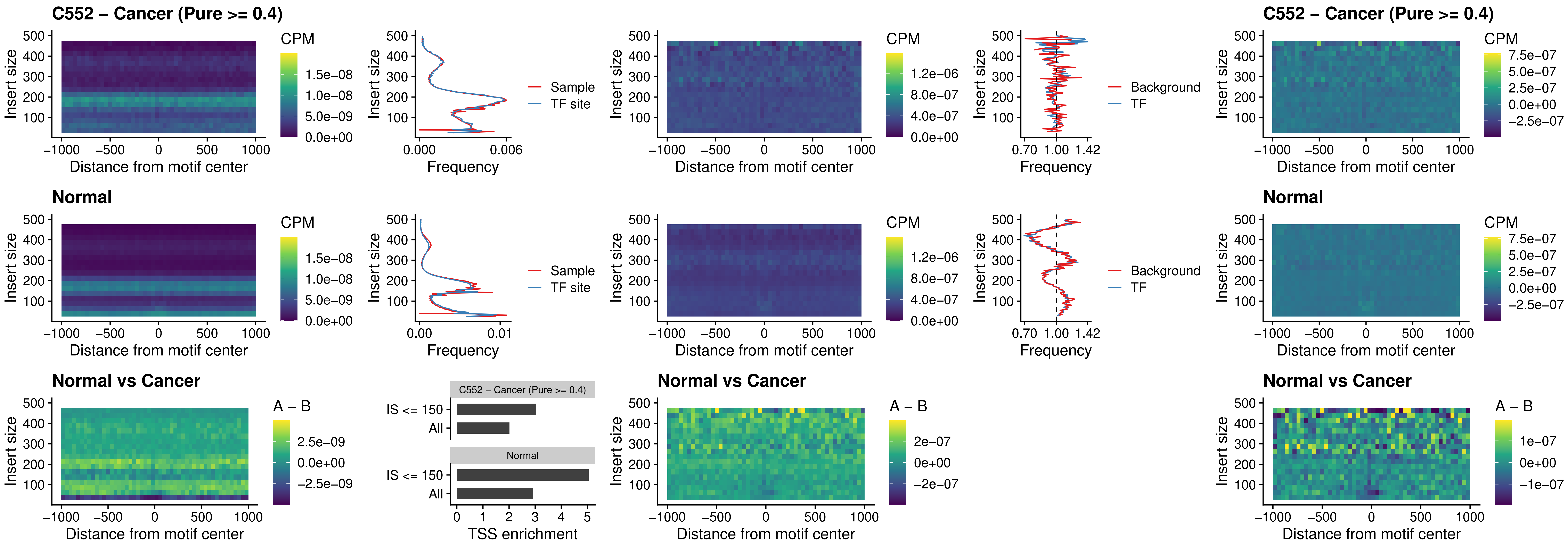

#### STAT2_uc_cTSS_oPEAK

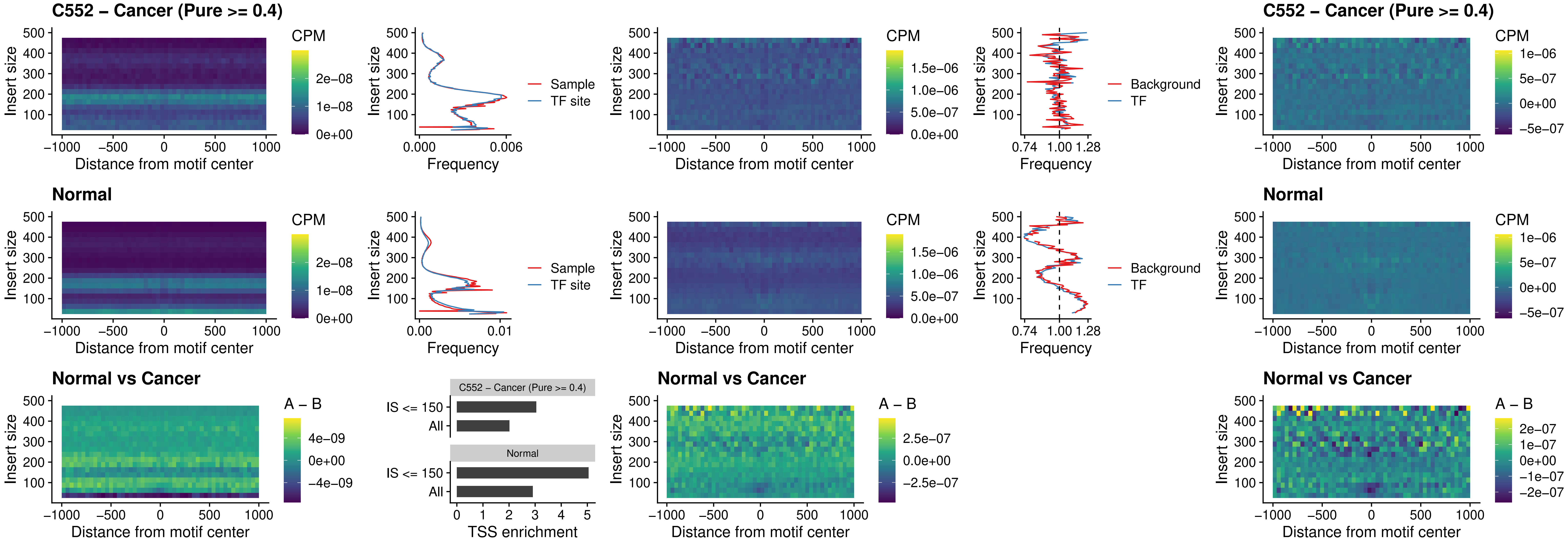

#### IRF8_uc_cTSS_oPEAK

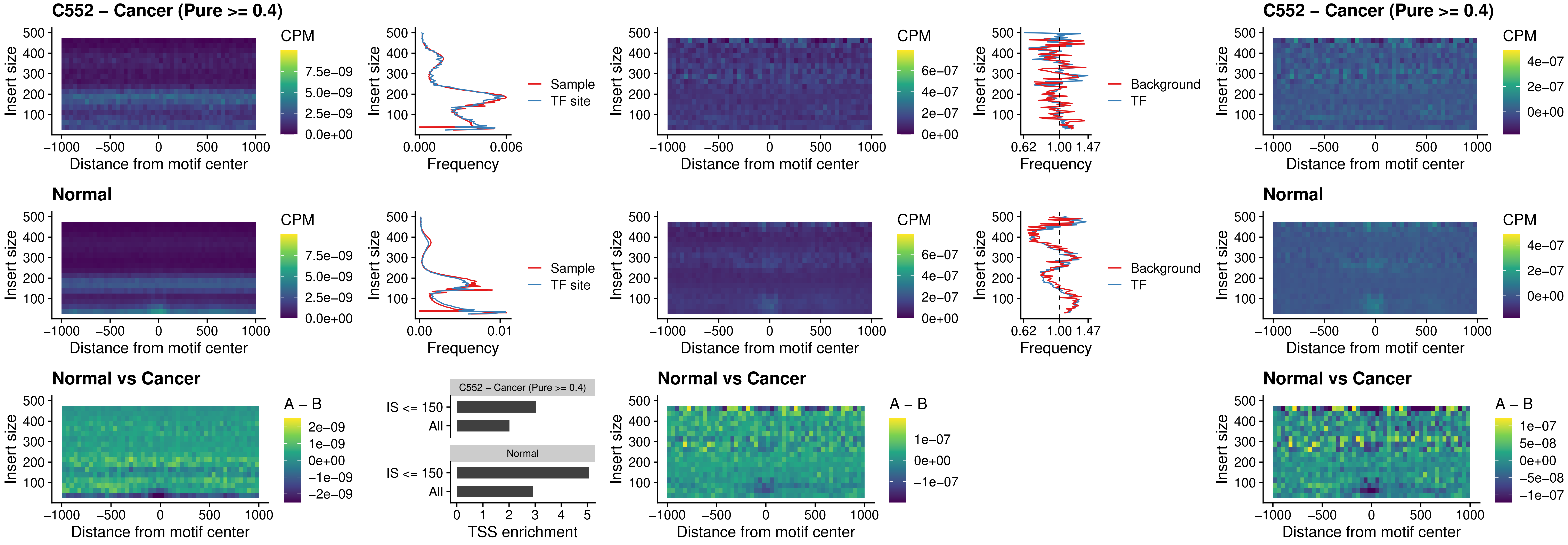

### IRF3_uc_dTSS_oPEAK

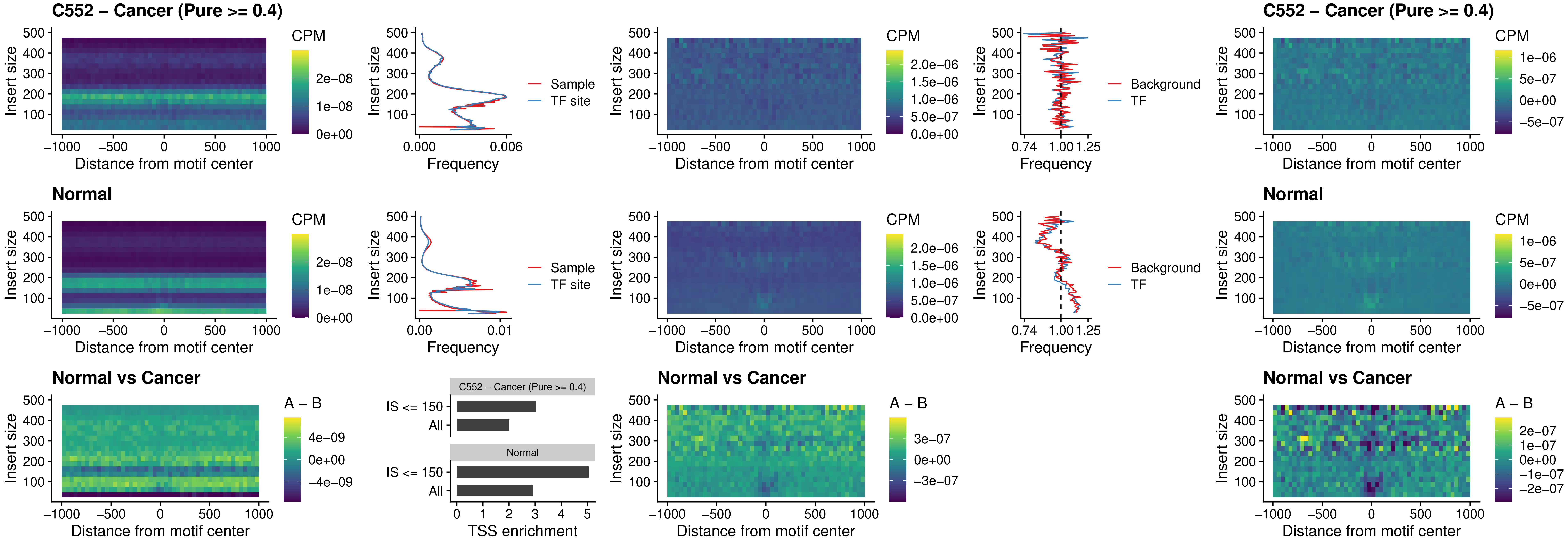

### IRF8_uc_dTSS_oPEAK

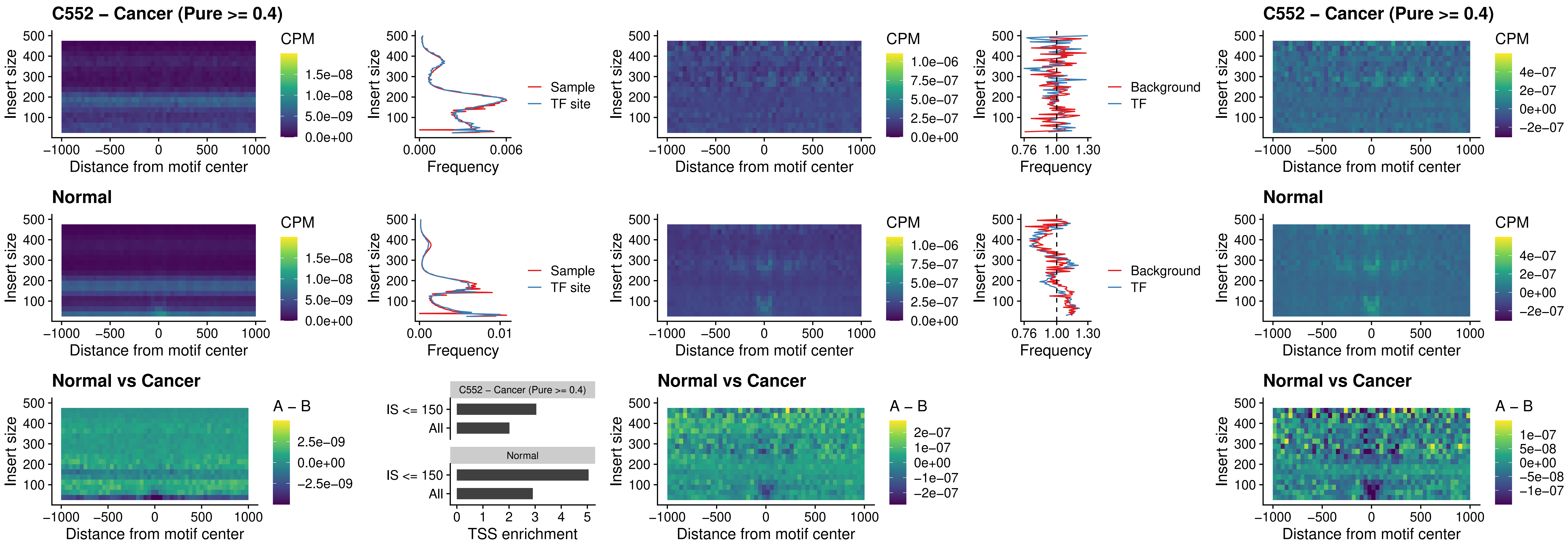

#### ETS2_uc_dTSS_oPEAK

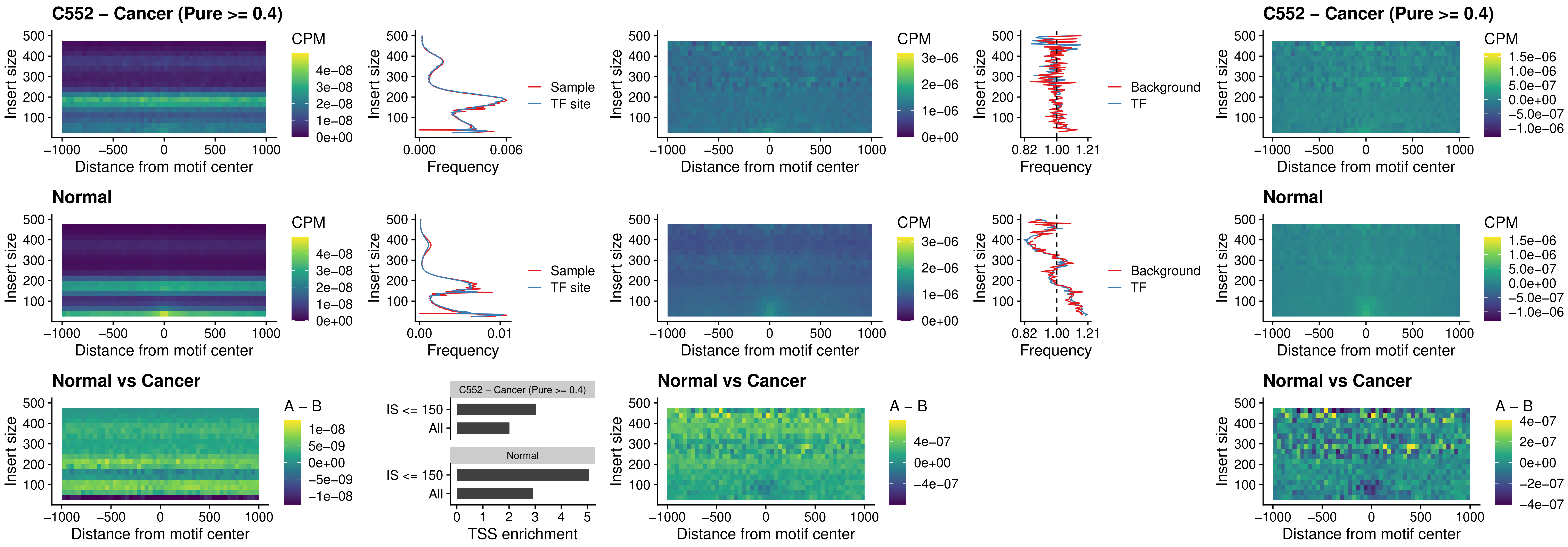

### SMARCC2_uc_pTSS_oPEAK

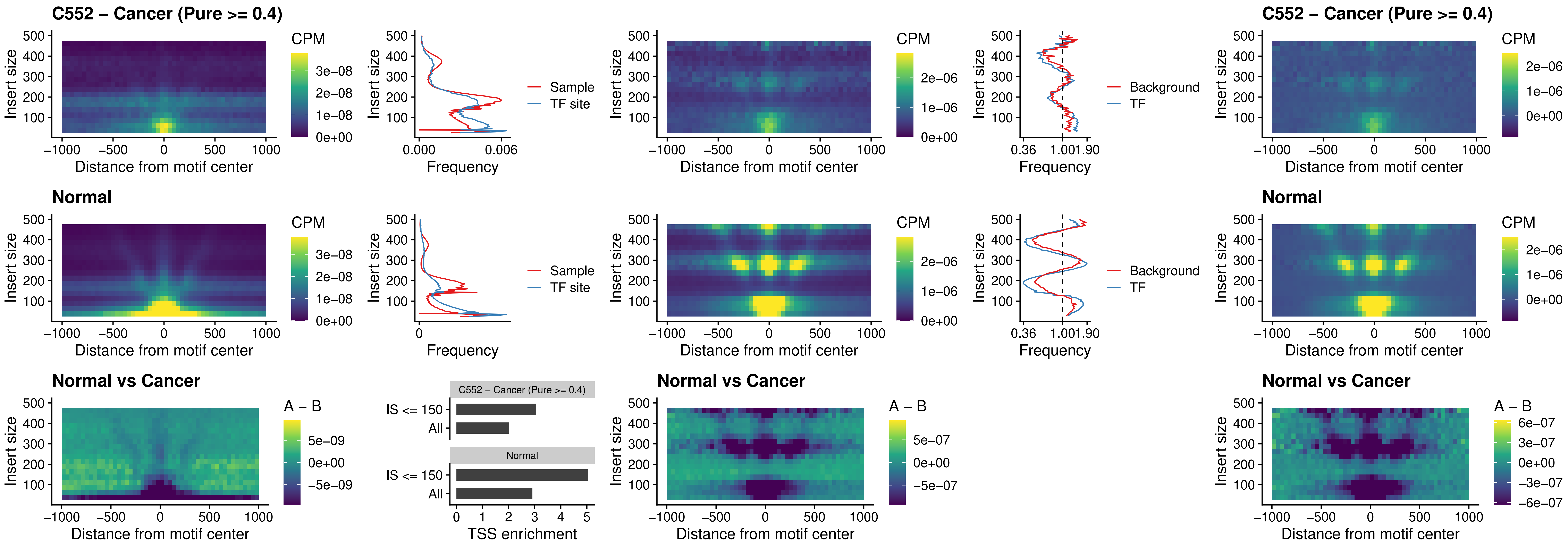

### BCL11A_uc_cTSS_nPEAK

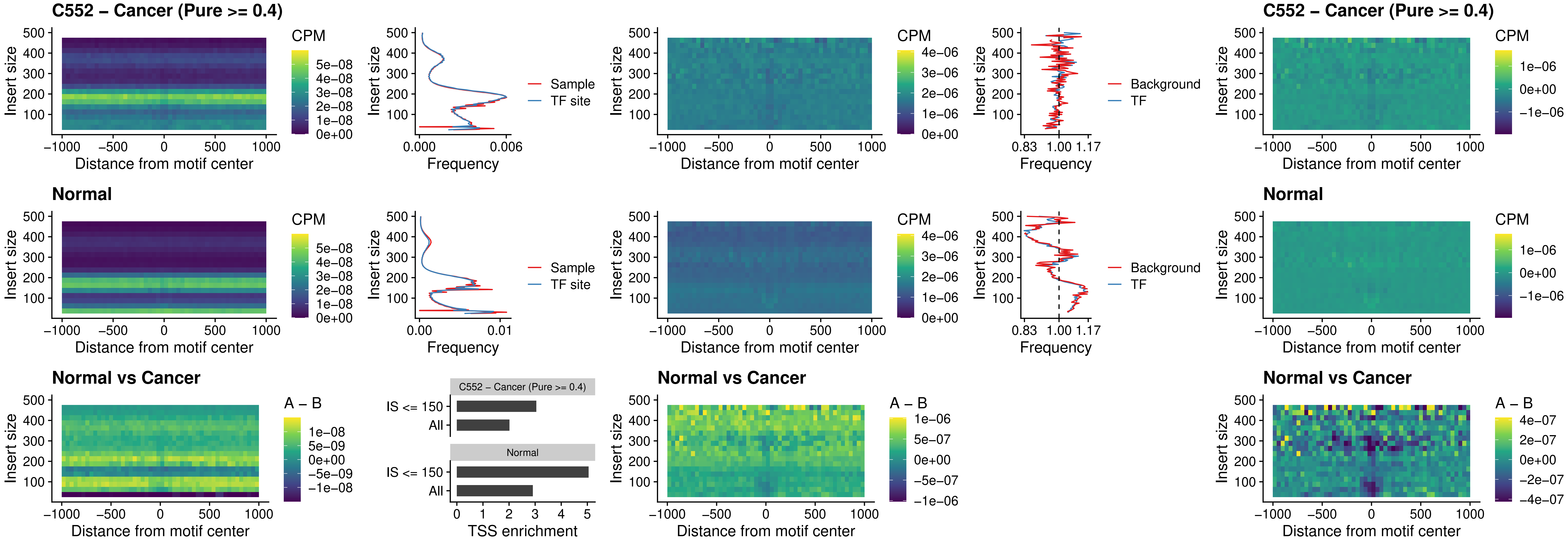

### BCL11B_uc_cTSS_nPEAK

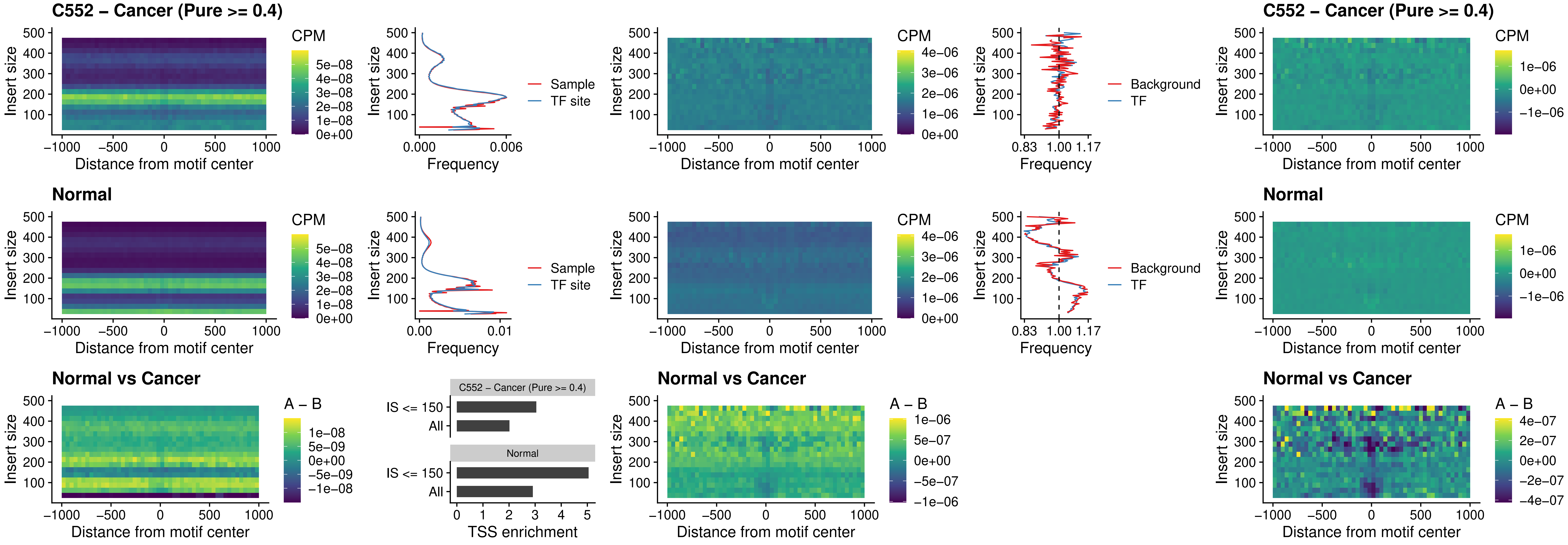

#### STAT2_uc_dTSS_oPEAK

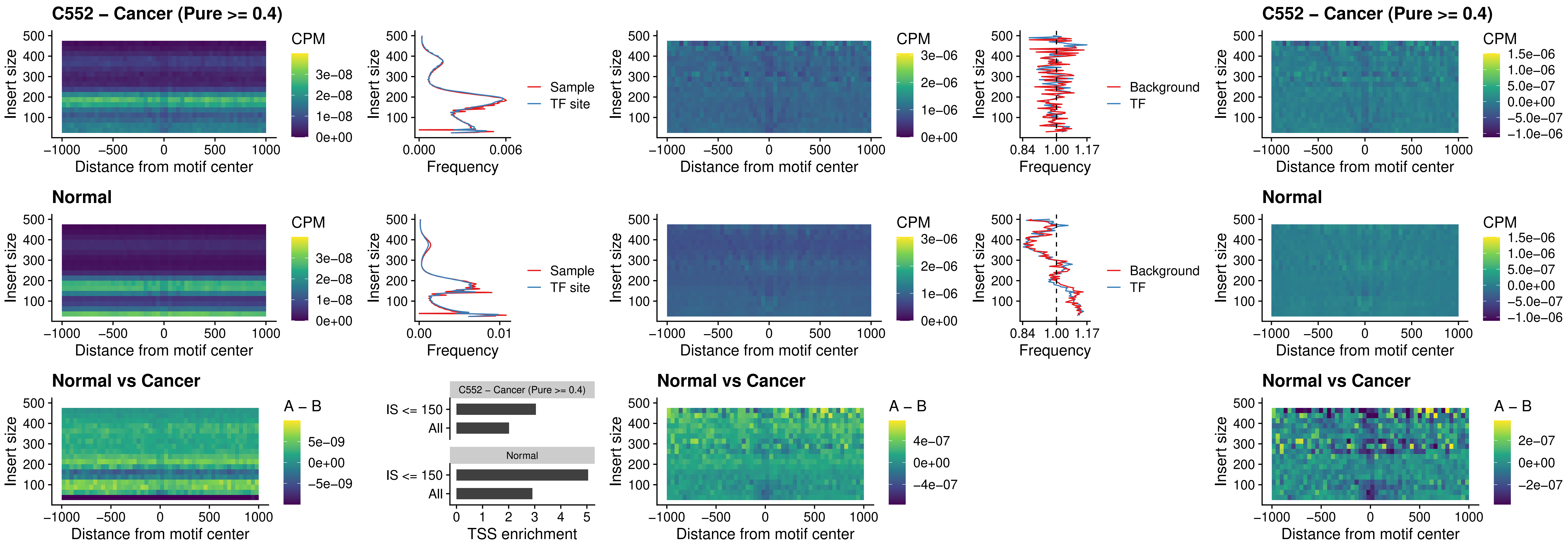

### ZNF683_uc_cTSS_oPEAK

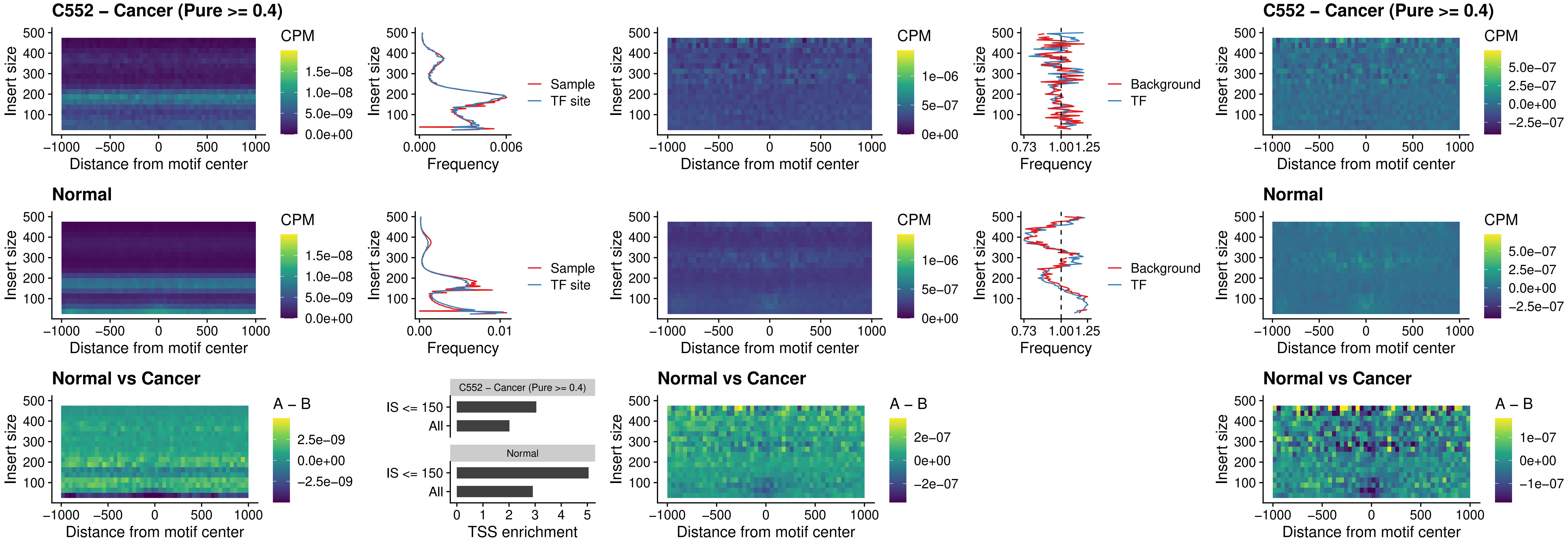

### IRF8_uc_dTSS_nPEAK

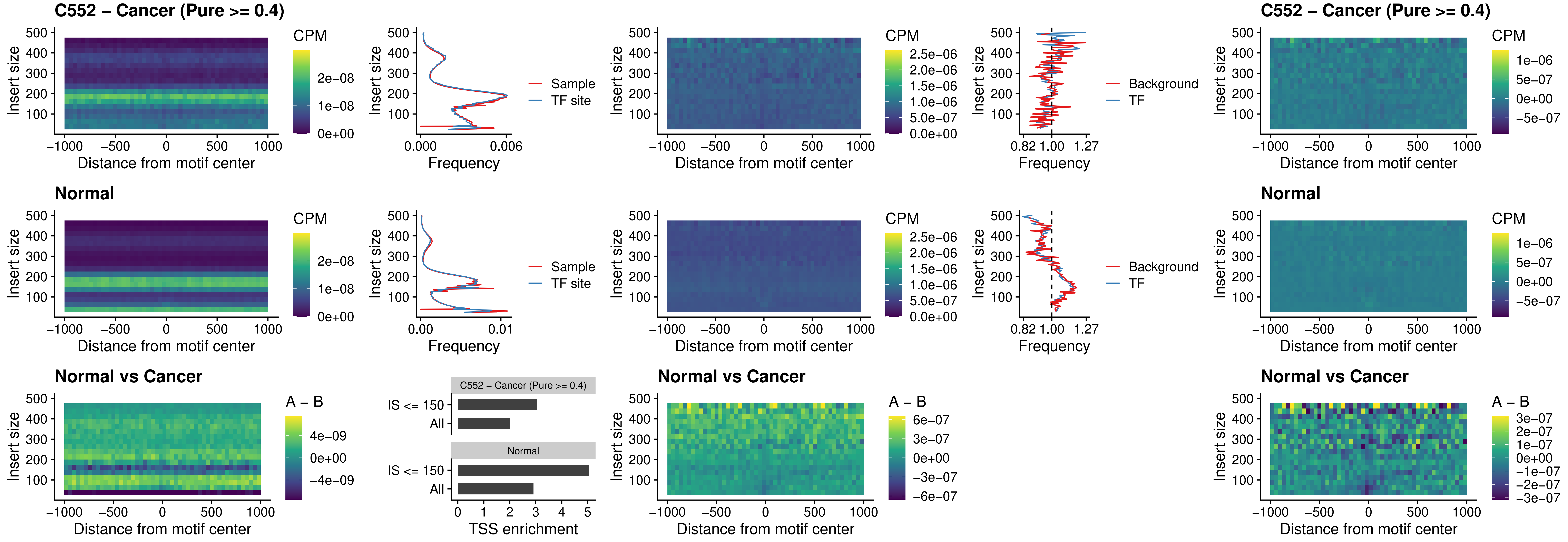

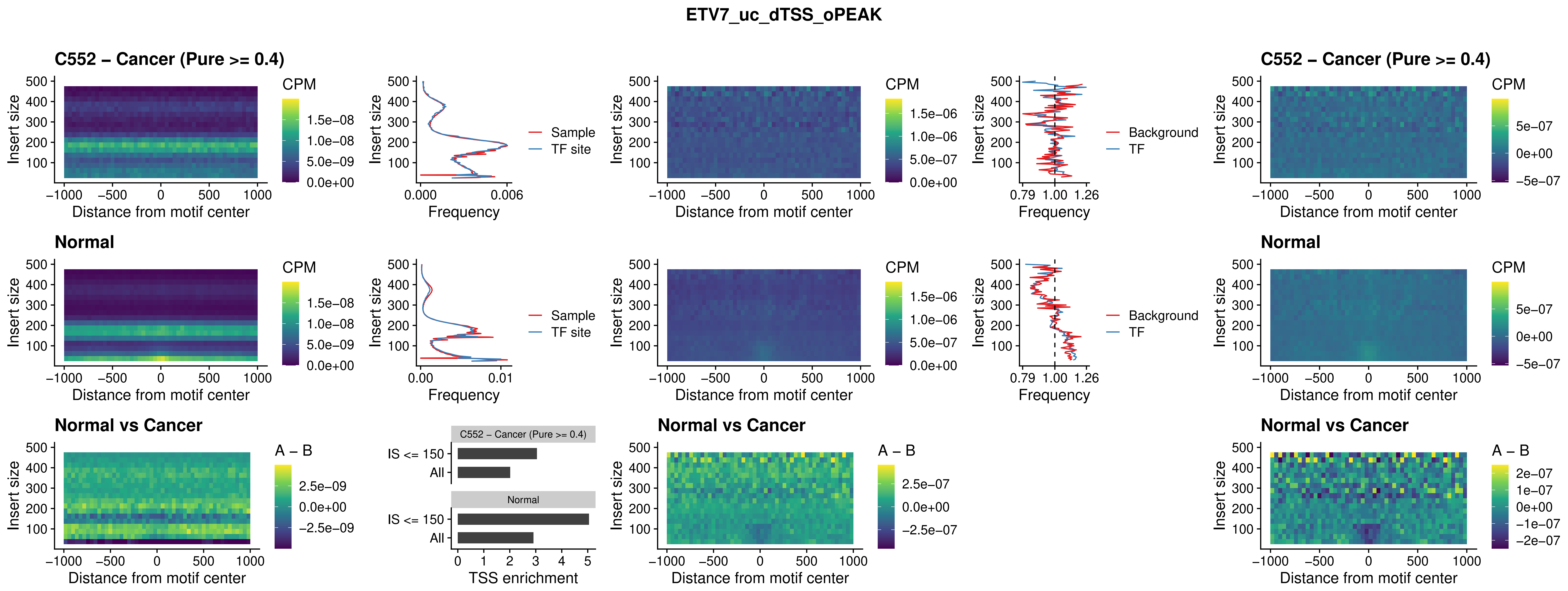
ETV7_uc_dTSS_oPEAK
C552 – Cancer (Pure >= 0.4)
Normal
Normal vs Cancer
Insert size
Distance from motif center
Frequency
CPM
A – B
Sample
TF site
Background
TF
C552 – Cancer (Pure >= 0.4)
Normal
IS <= 150
All
TSS enrichment

### FOXL1_uc_pTSS_oPEAK

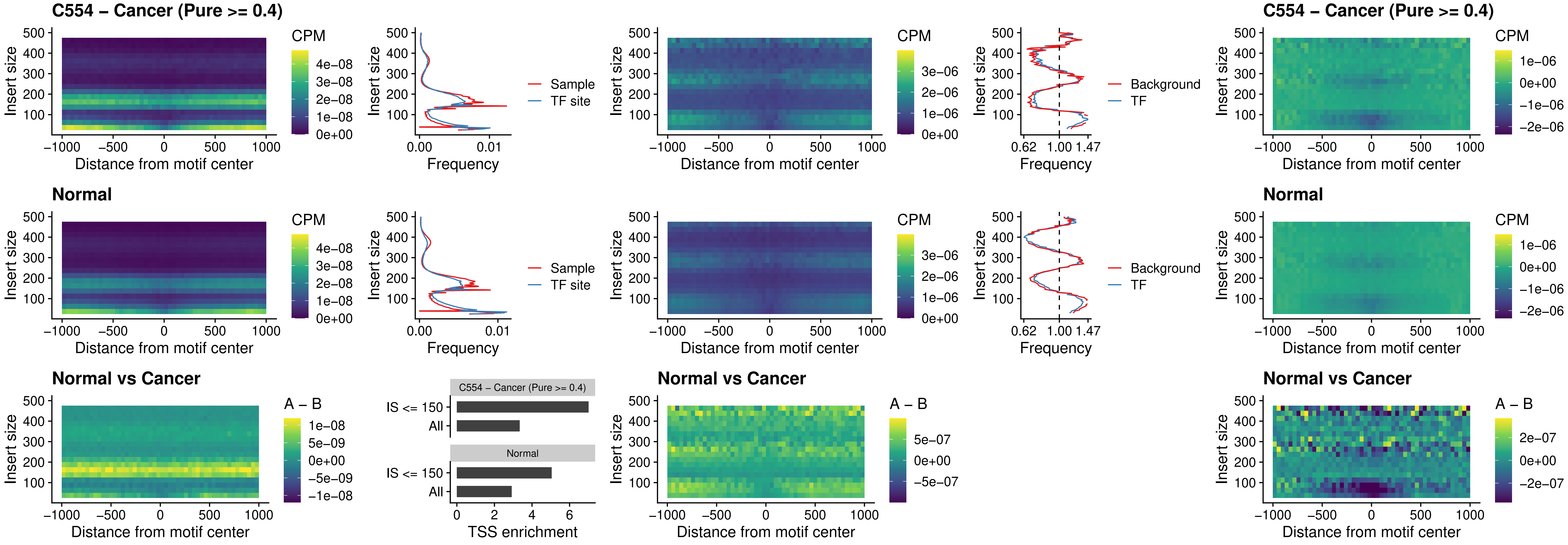

### CPEB1_uc_cTSS_oPEAK

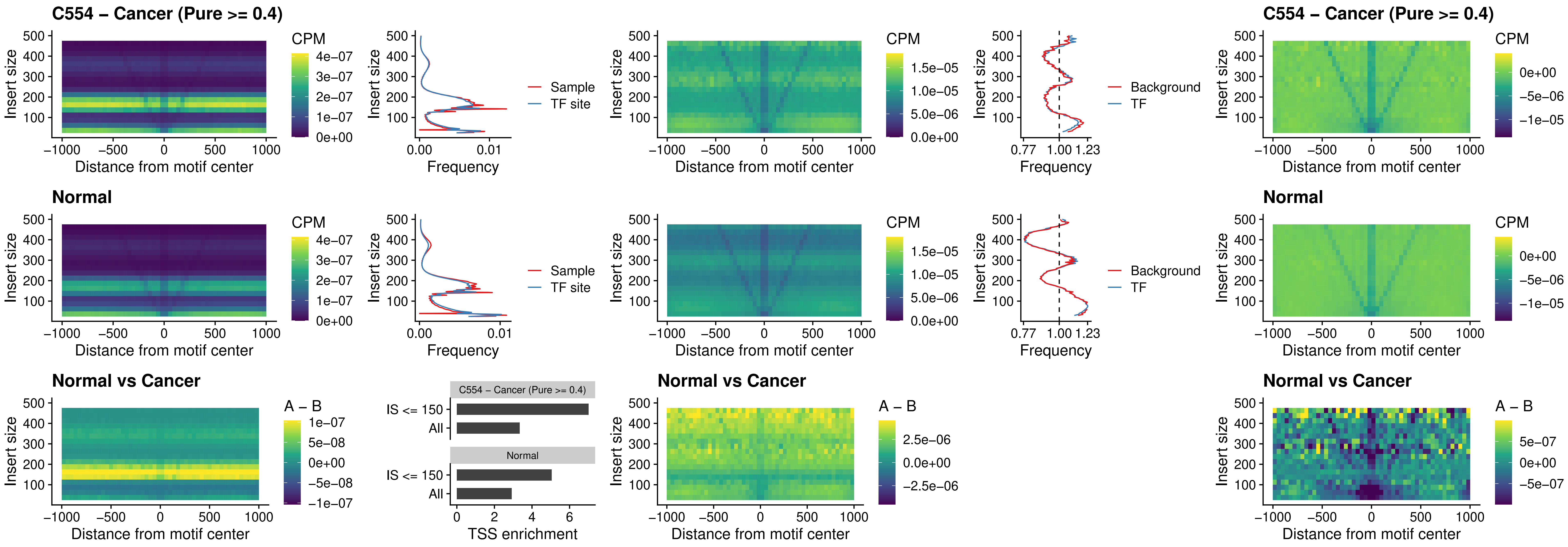

### HMGA1_uc_pTSS_nPEAK

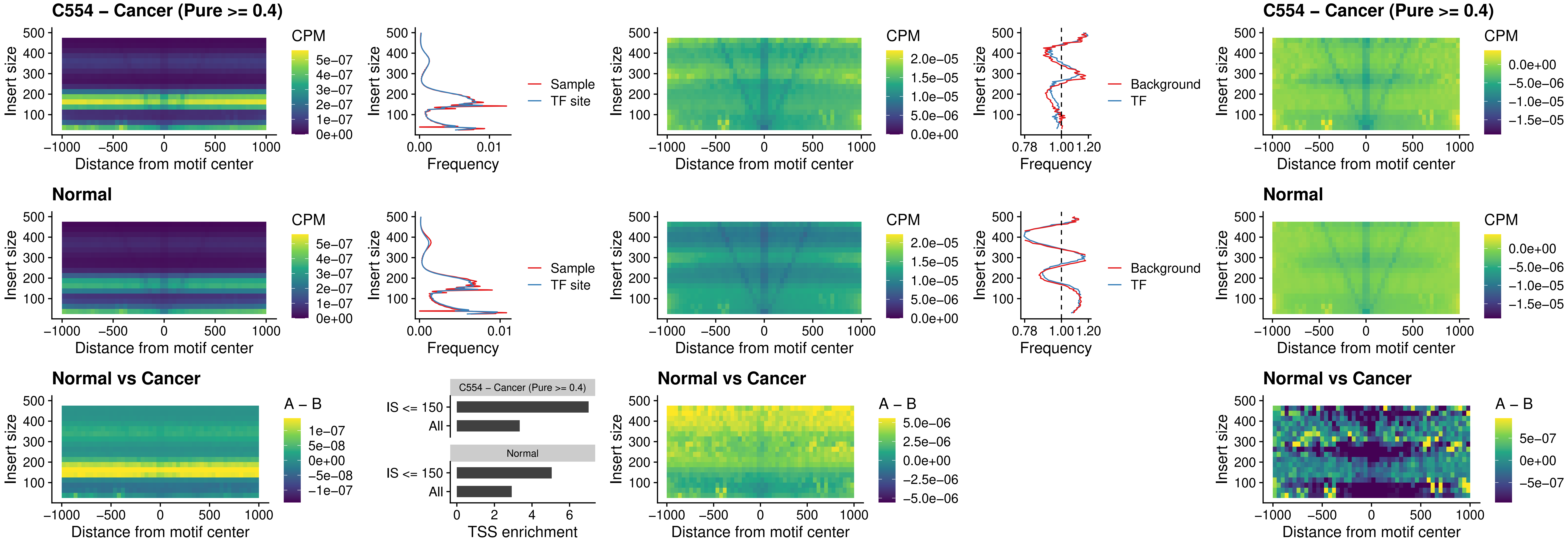

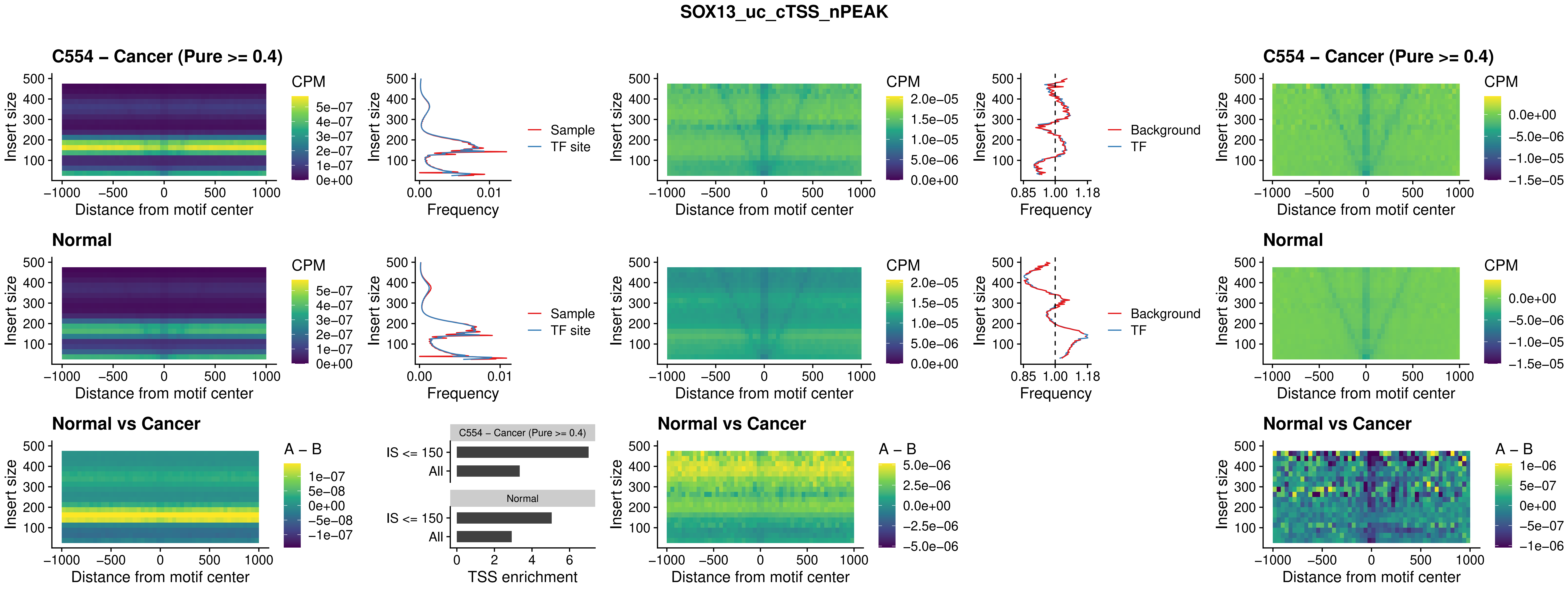
SOX13_uc_cTSS_nPEAK
C554 – Cancer (Pure >= 0.4)
Normal
Normal vs Cancer
Insert size
Distance from motif center
Frequency
CPM
A – B
Sample
TF site
Background
TF
C554 – Cancer (Pure >= 0.4)
Normal
IS <= 150
All
TSS enrichment

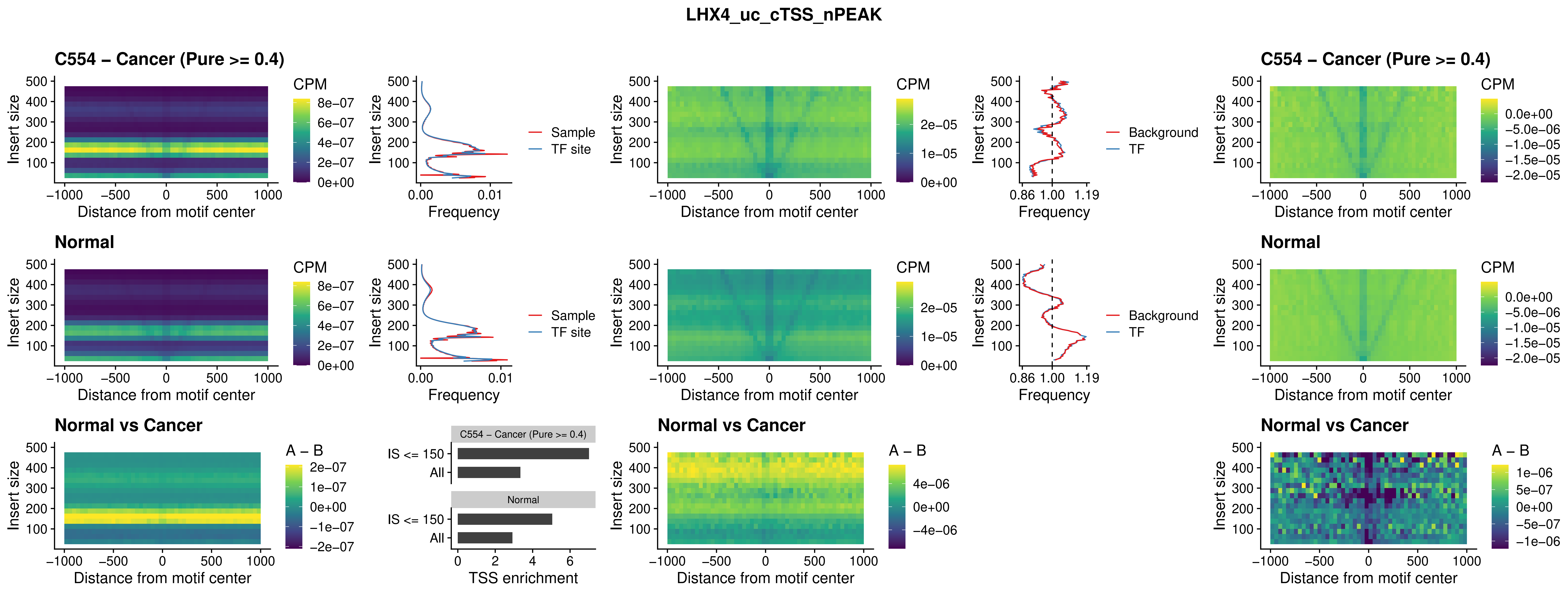

LHX4_uc_cTSS_nPEAK
C554 – Cancer (Pure >= 0.4)
Normal
Normal vs Cancer
Insert size
Distance from motif center
Frequency
CPM
A – B
Sample
TF site
Background
TF
C554 – Cancer (Pure >= 0.4)
Normal
IS <= 150
All
TSS enrichment

### HOXA4_uc_cTSS_nPEAK

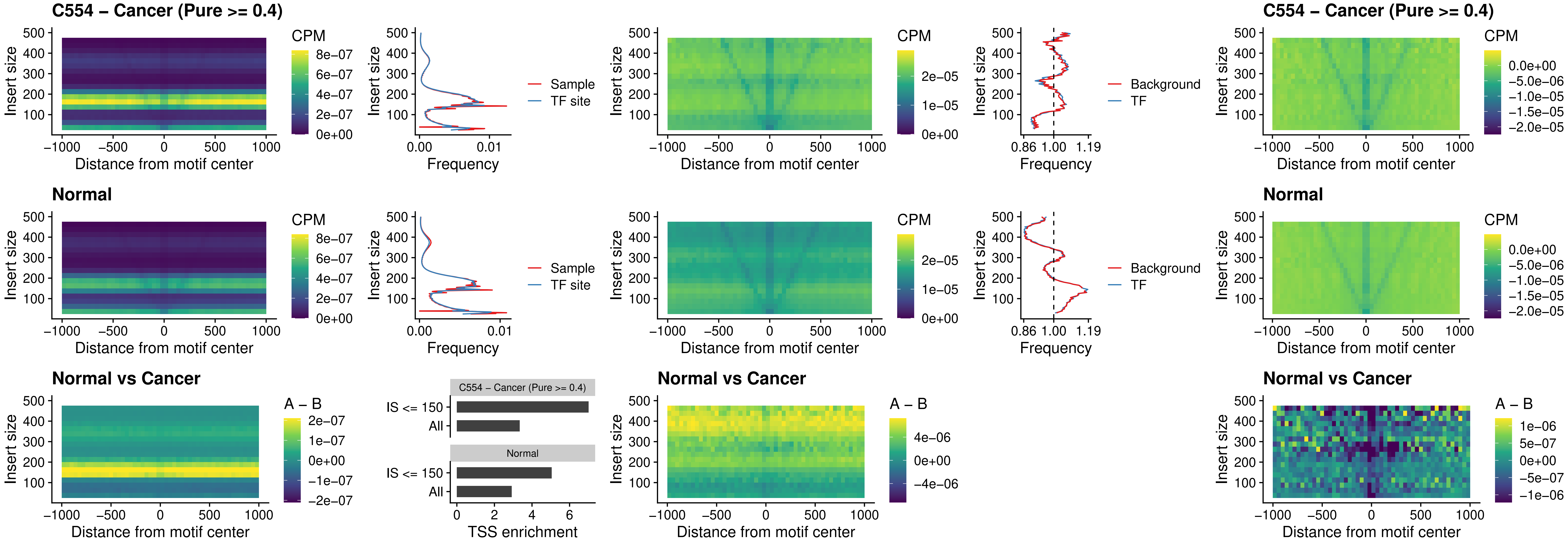

#### HLX_uc_cTSS_nPEAK

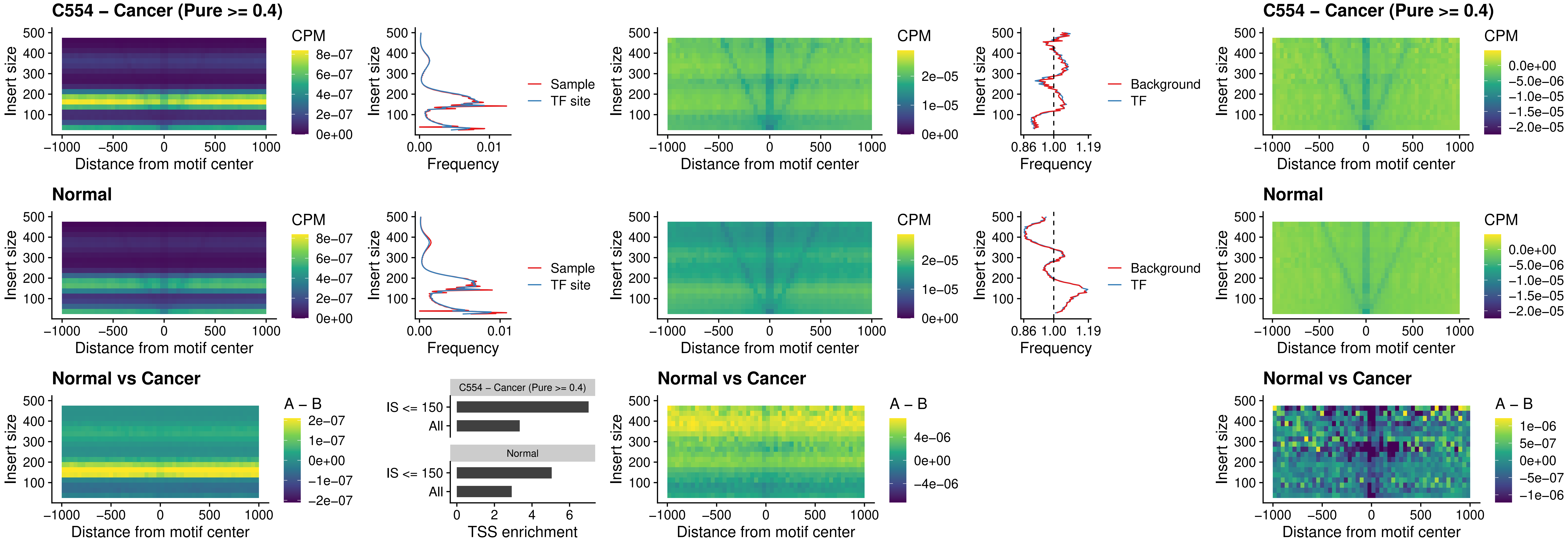

### AC0125311_uc_cTSS_nPEAK

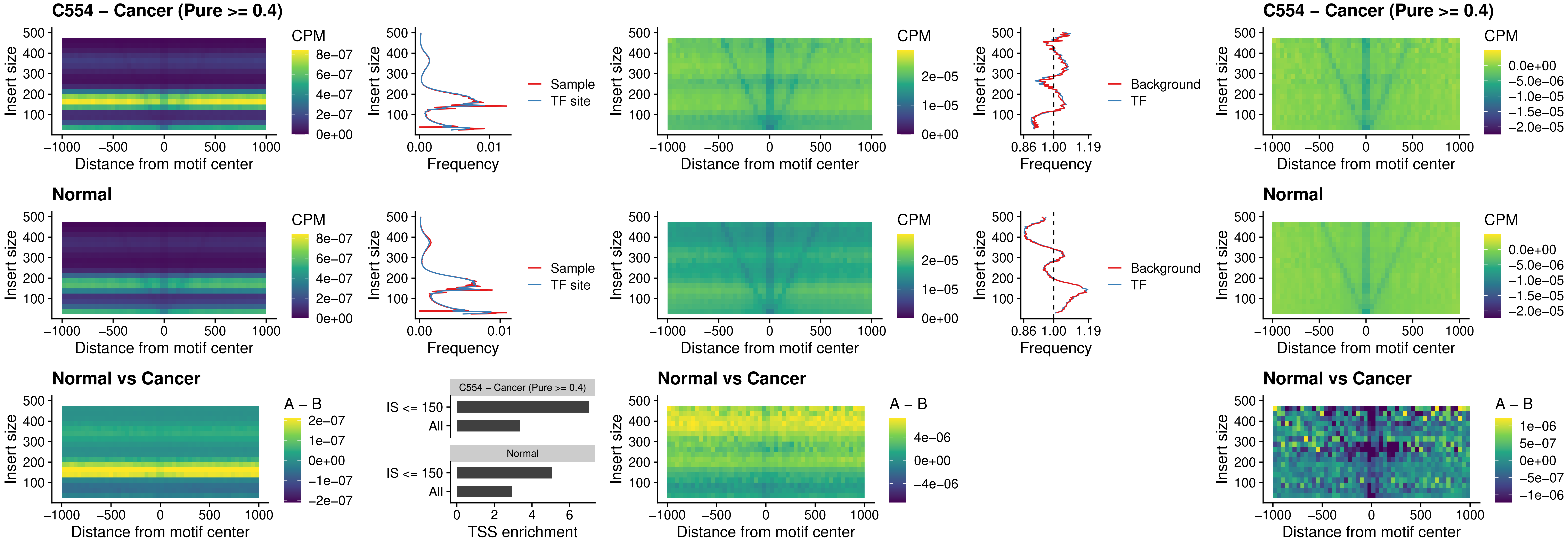

### DBX2_uc_cTSS_nPEAK

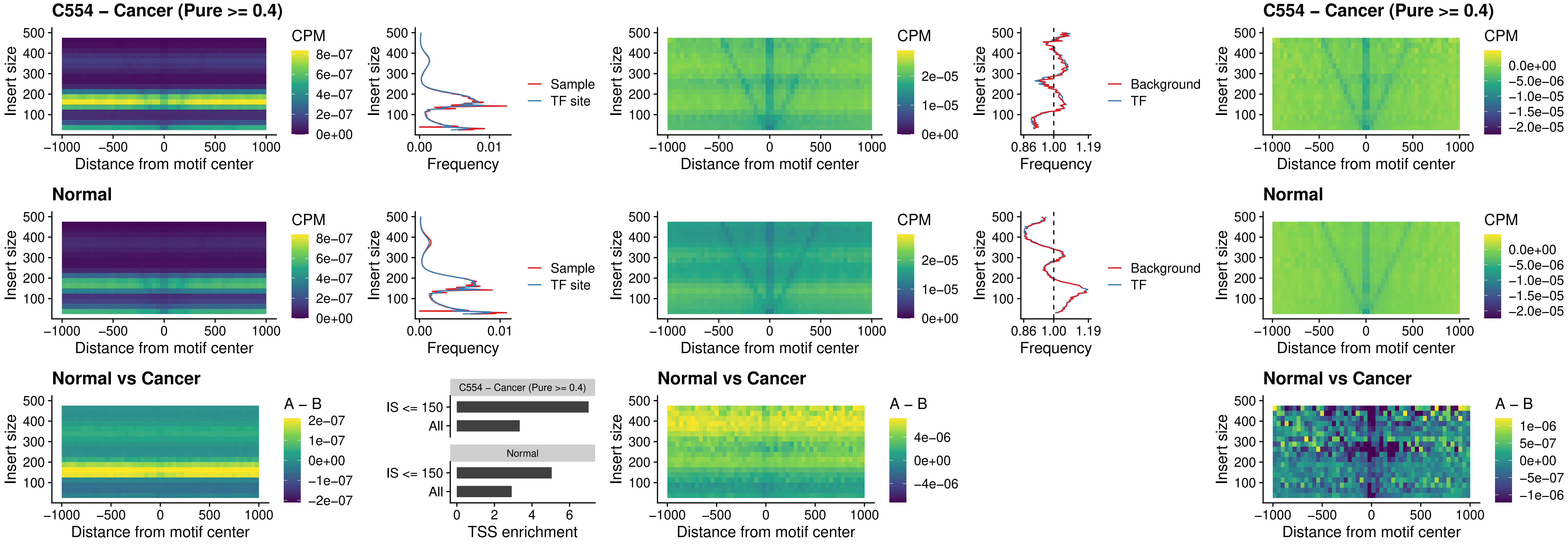

### LHX6_uc_dTSS_nPEAK

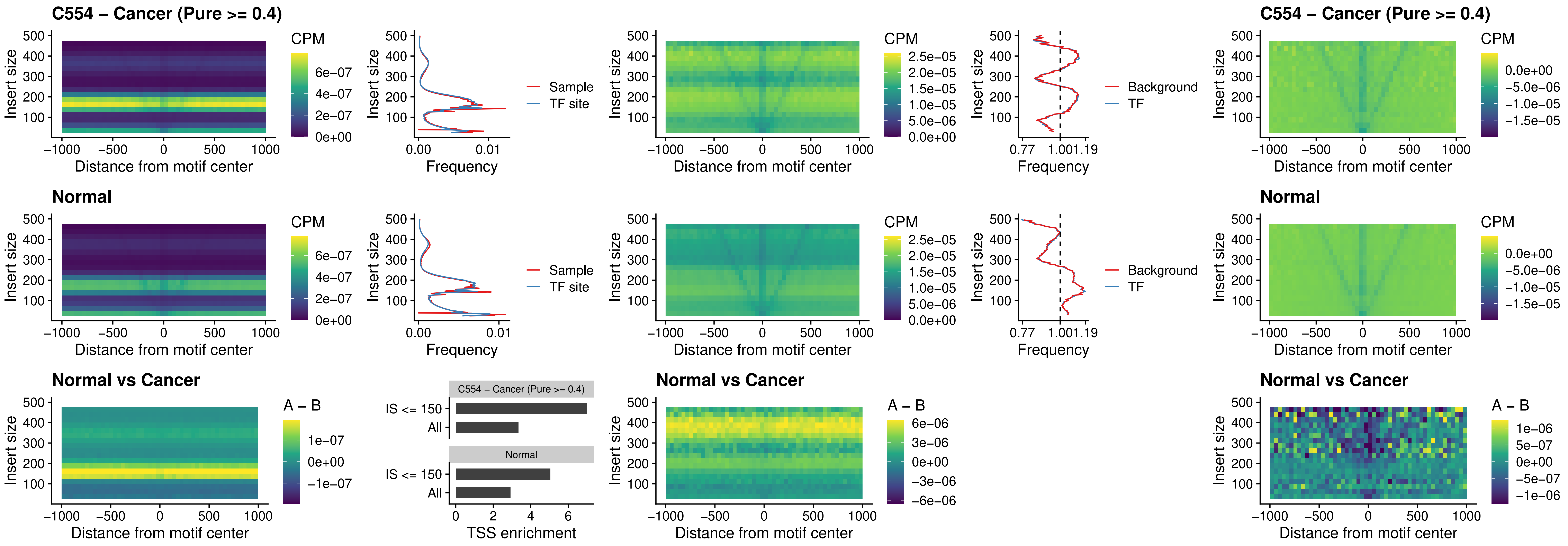

#### SOX1_uc_dTSS_nPEAK

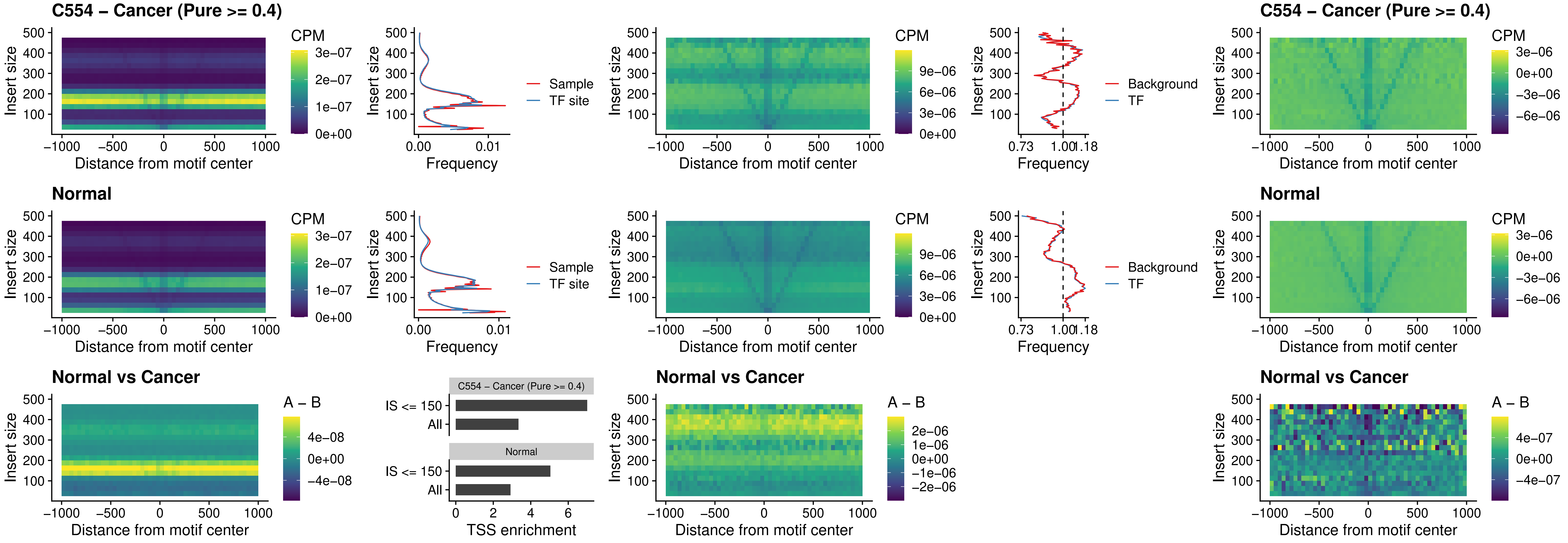

### NKX24_uc_dTSS_nPEAK

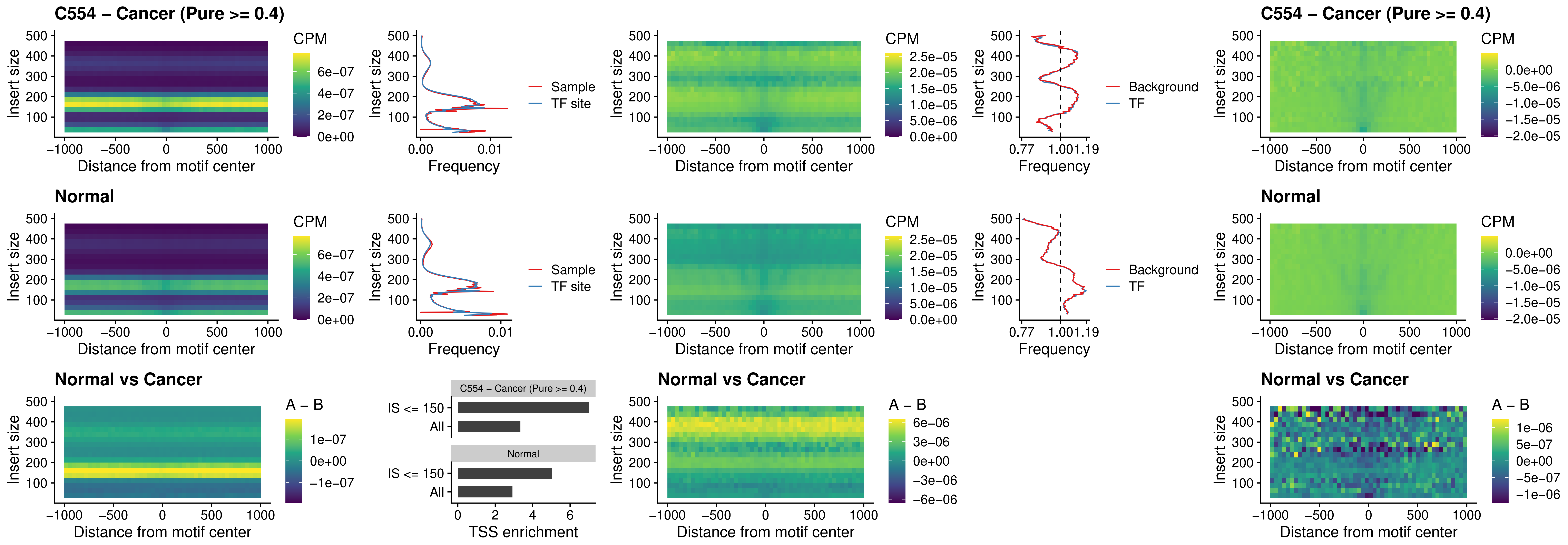

### DLX3_uc_cTSS_nPEAK

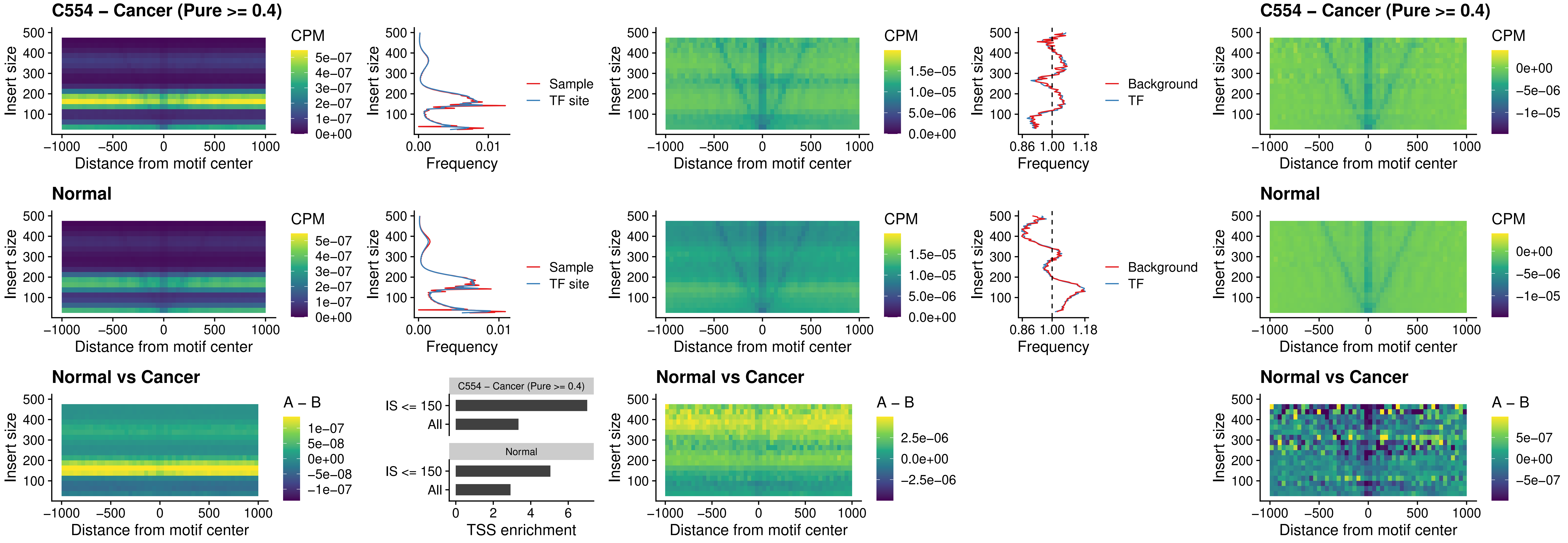

### HOXD3_uc_cTSS_nPEAK

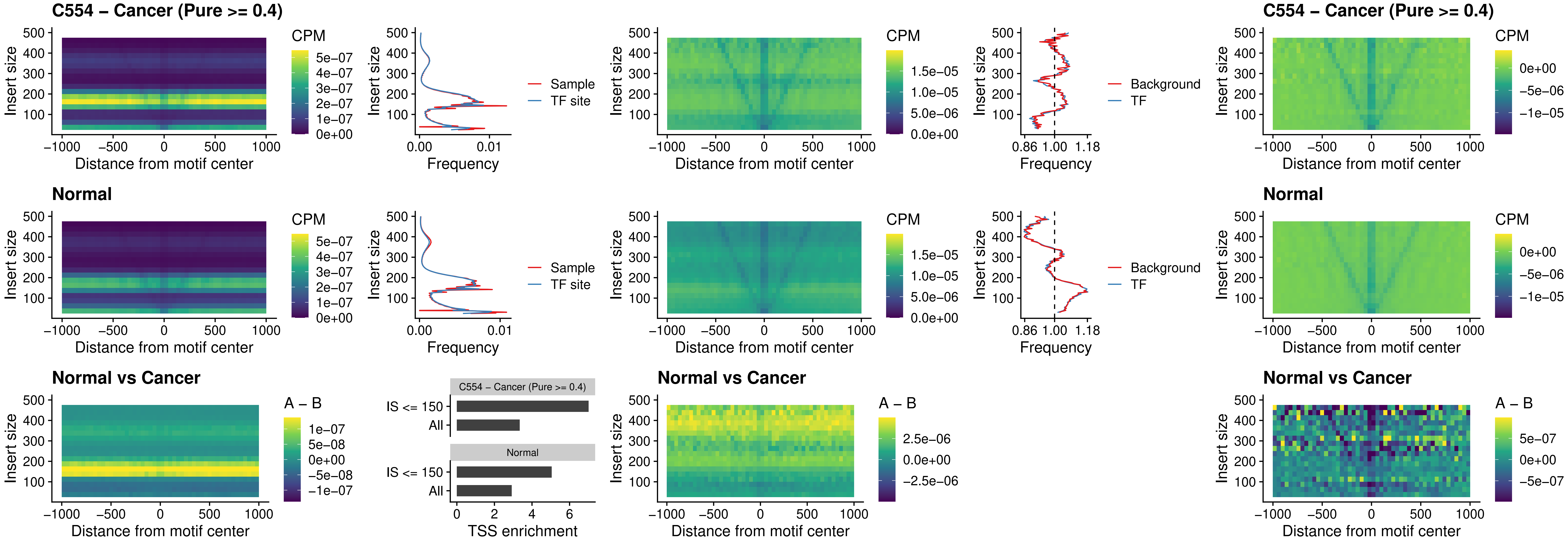

### ISL1_uc_dTSS_oPEAK

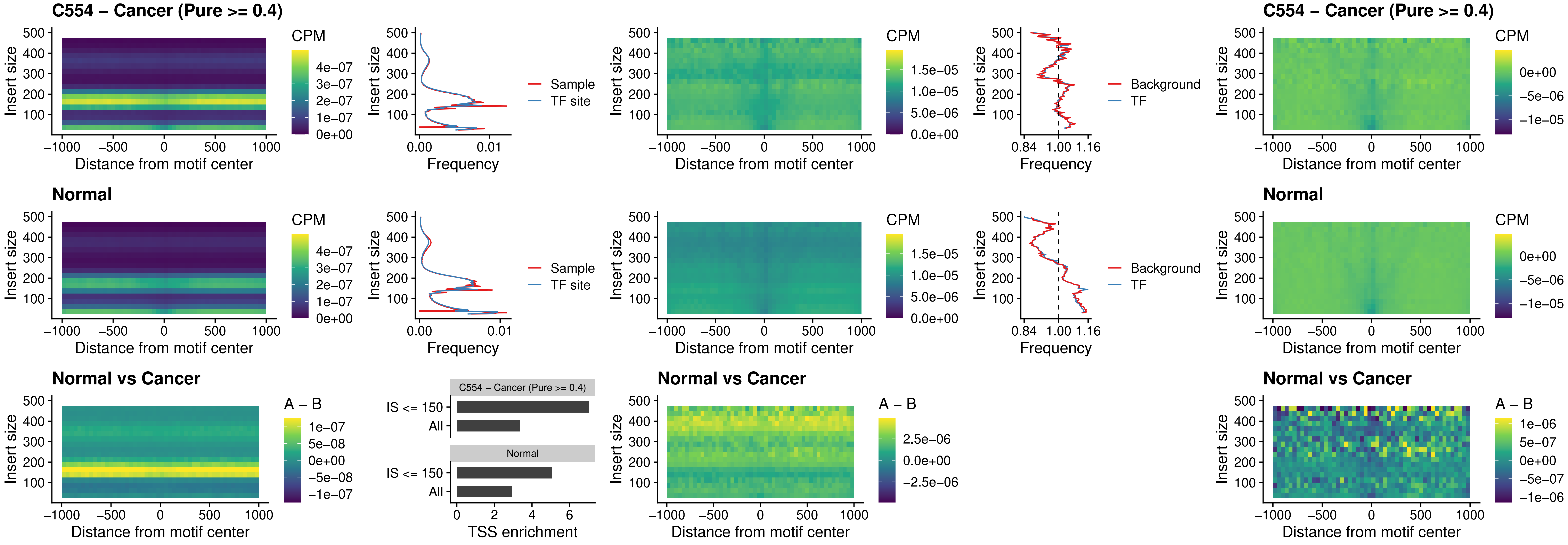

### HOXB4_uc_dTSS_oPEAK

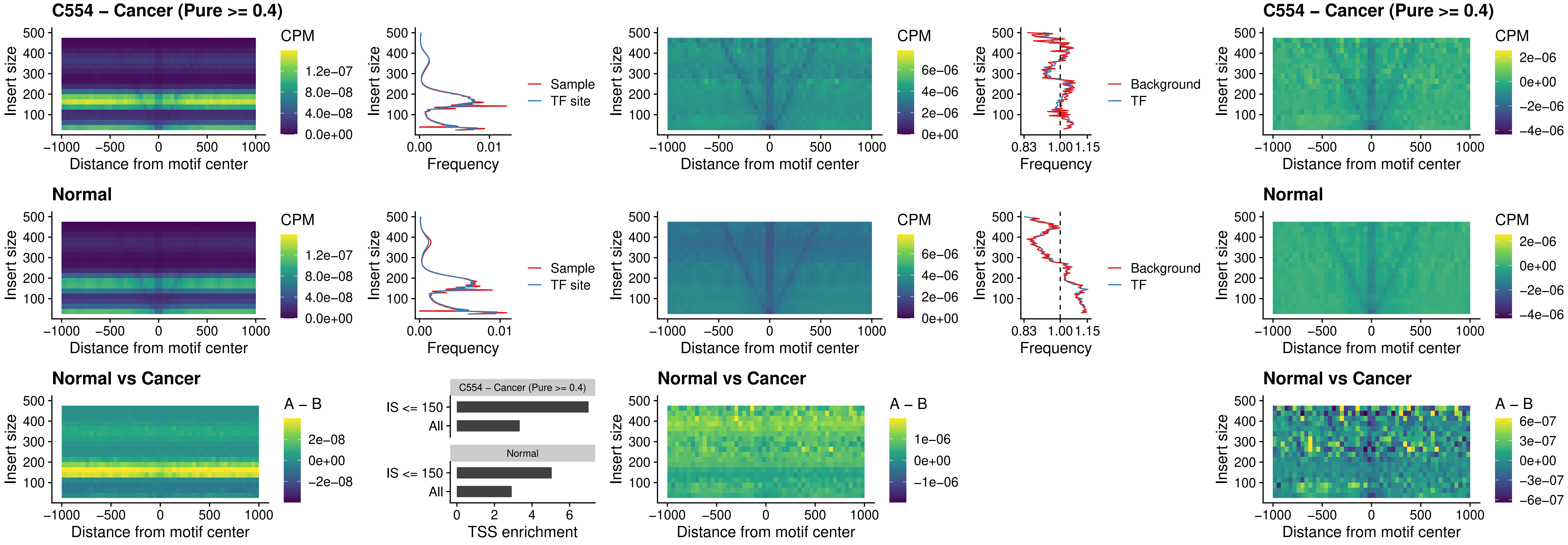

#### VAX1_uc_dTSS_oPEAK

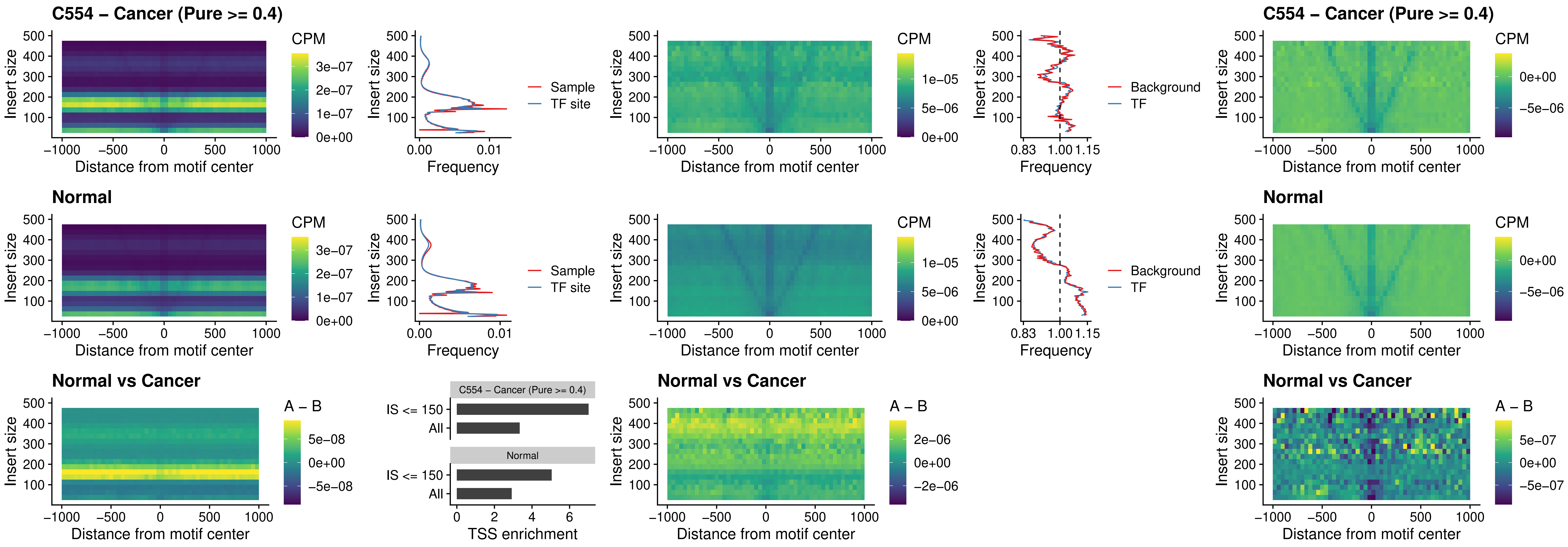

### DLX3_uc_dTSS_oPEAK

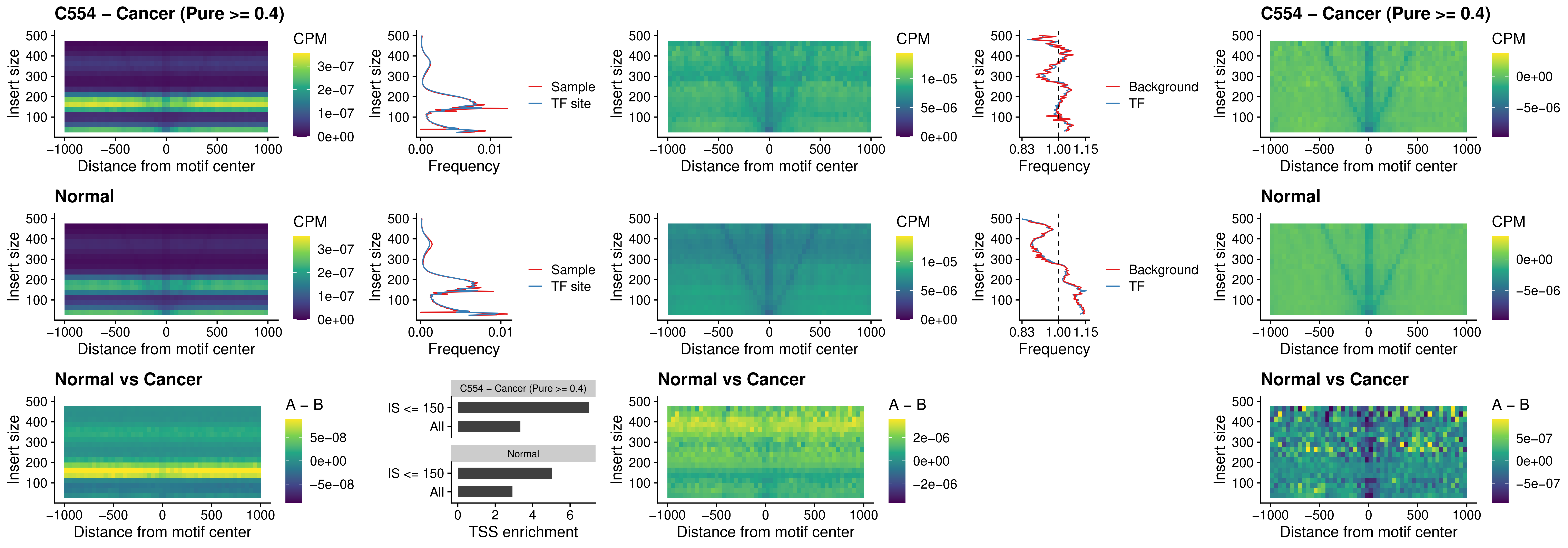

### HOXD3_uc_dTSS_oPEAK

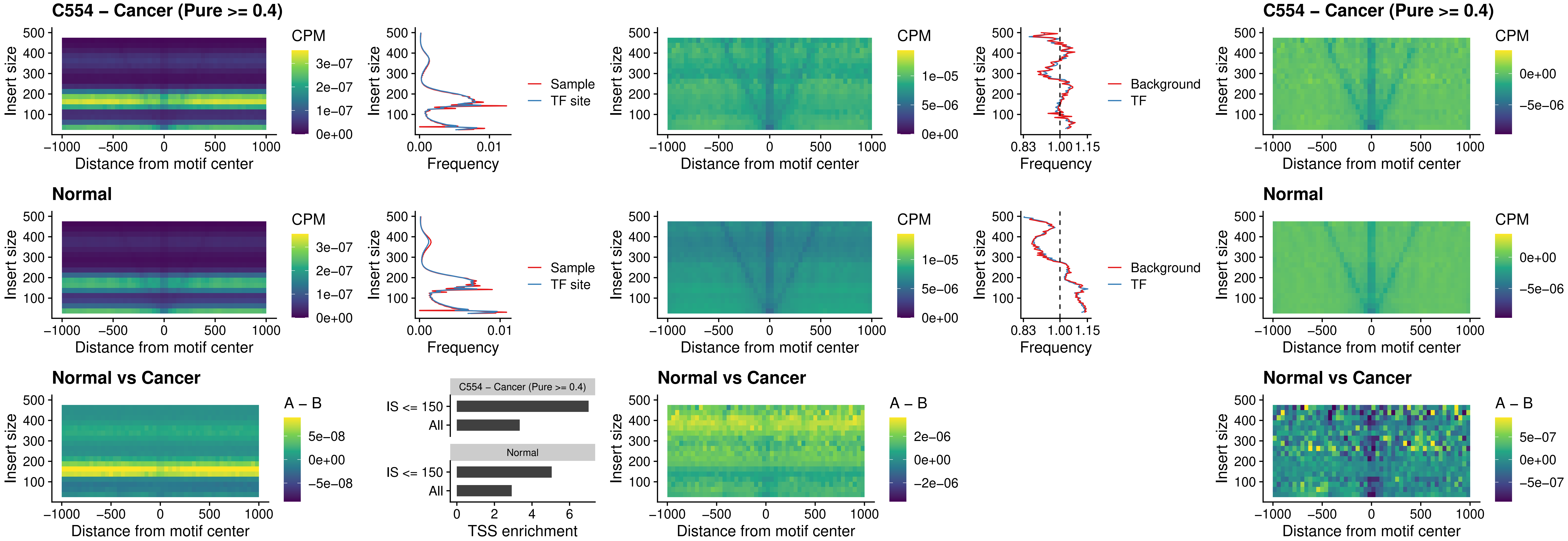

### BBX_uc_dTSS_nPEAK

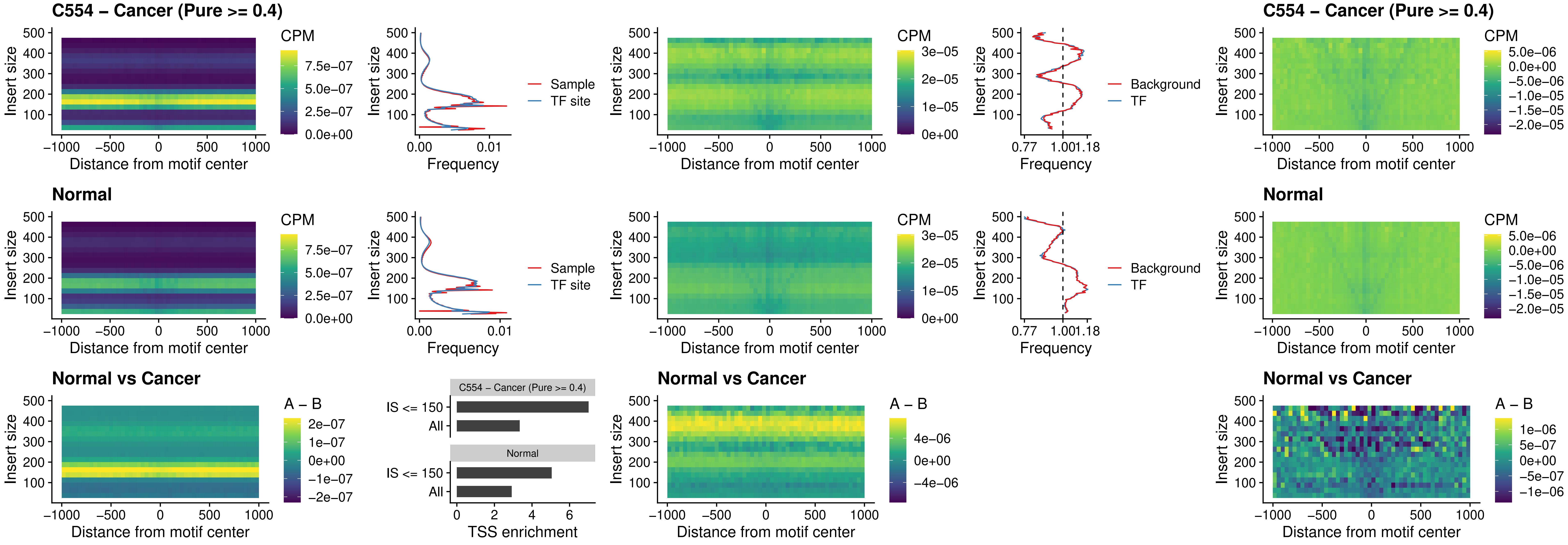

### SOX3_uc_cTSS_nPEAK

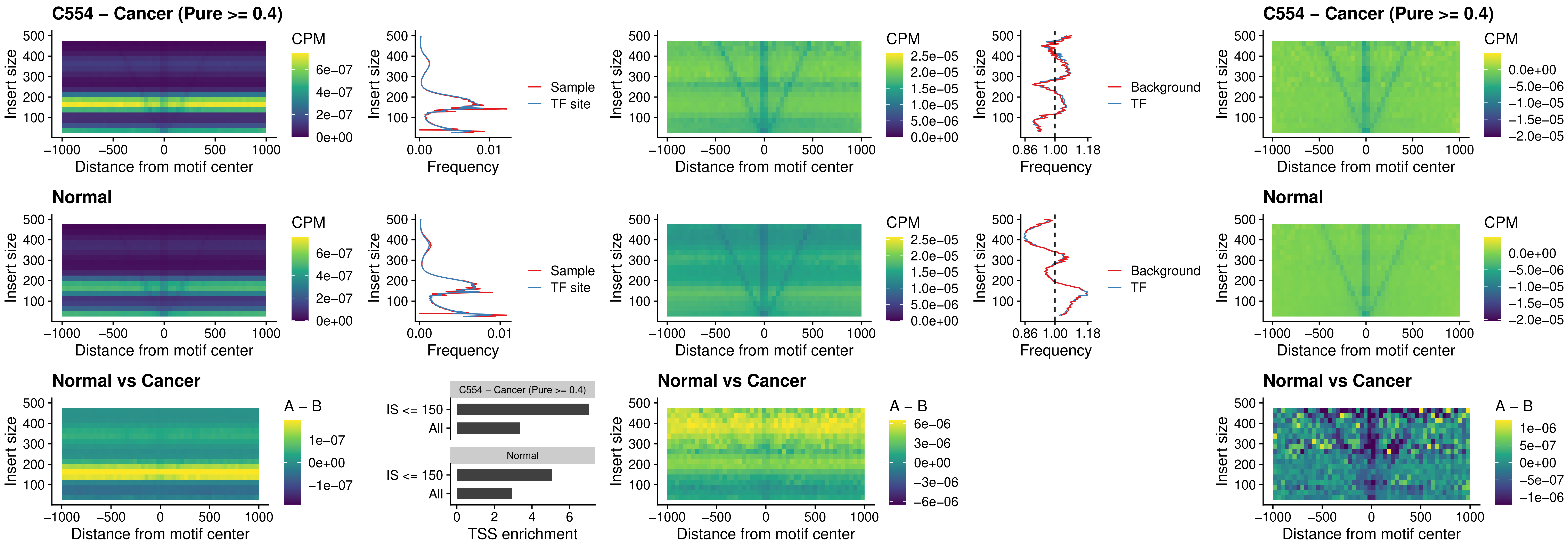

### TBPL2_uc_cTSS_nPEAK

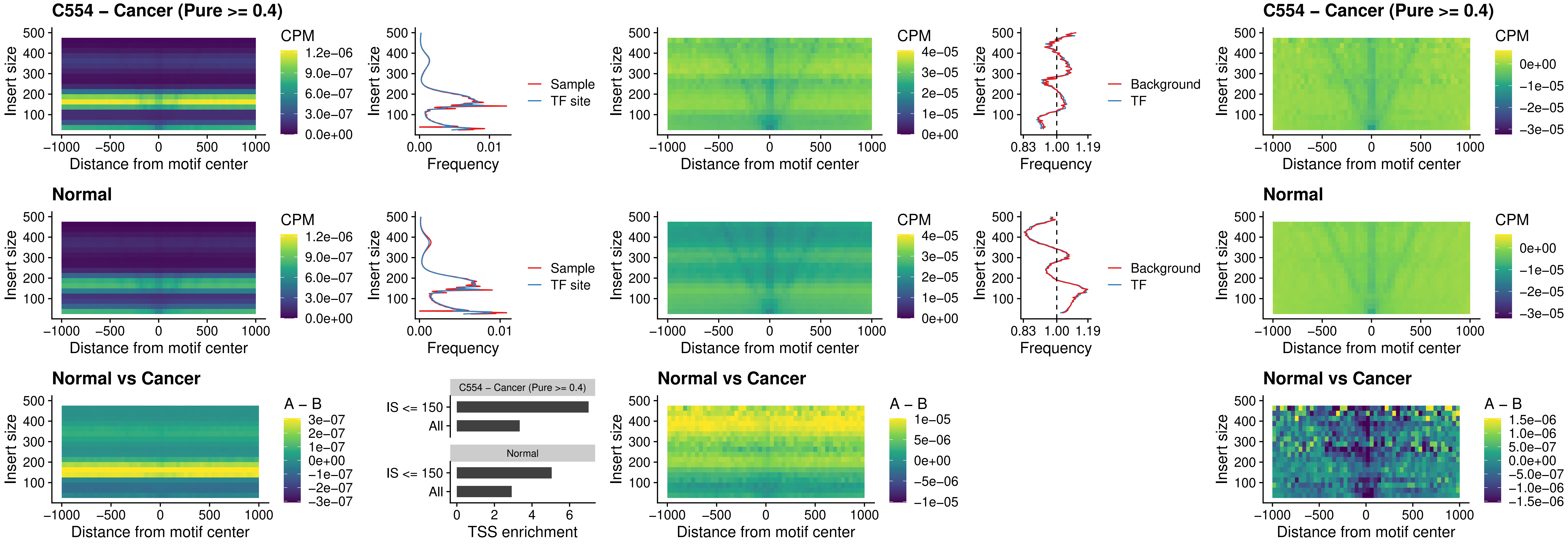

#### HOXA4_uc_cTSS_oPEAK

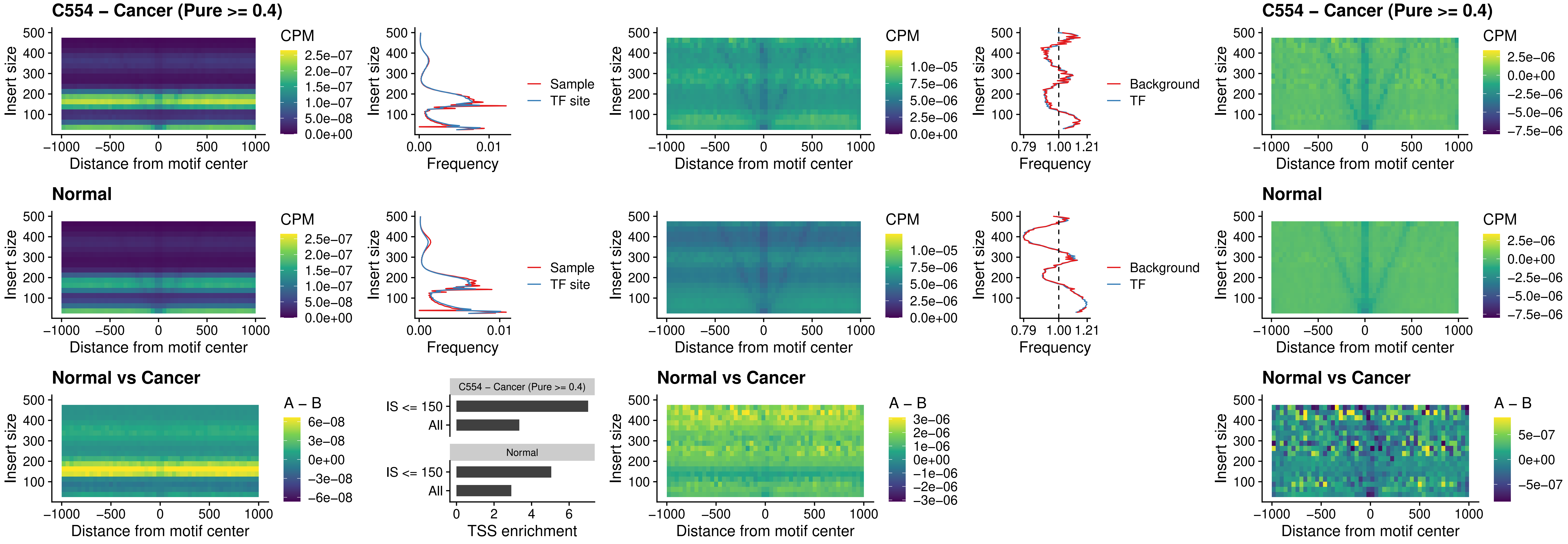

### PDX1_uc_dTSS_oPEAK

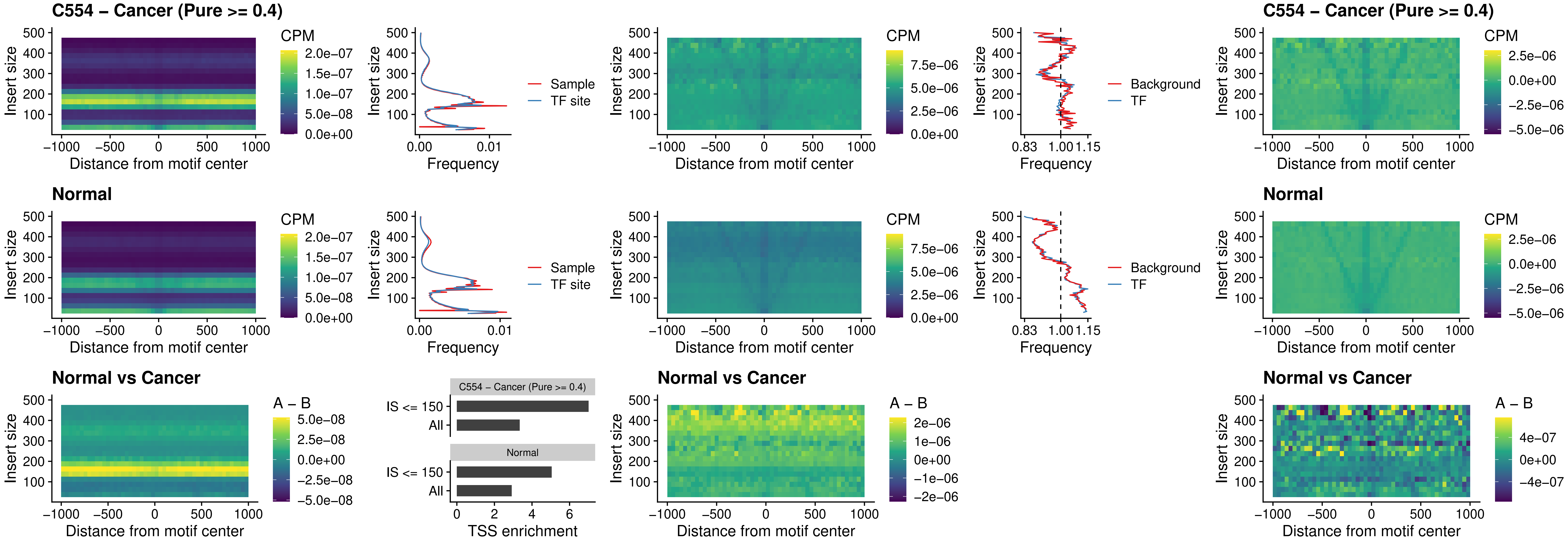

### LHX4_uc_dTSS_oPEAK

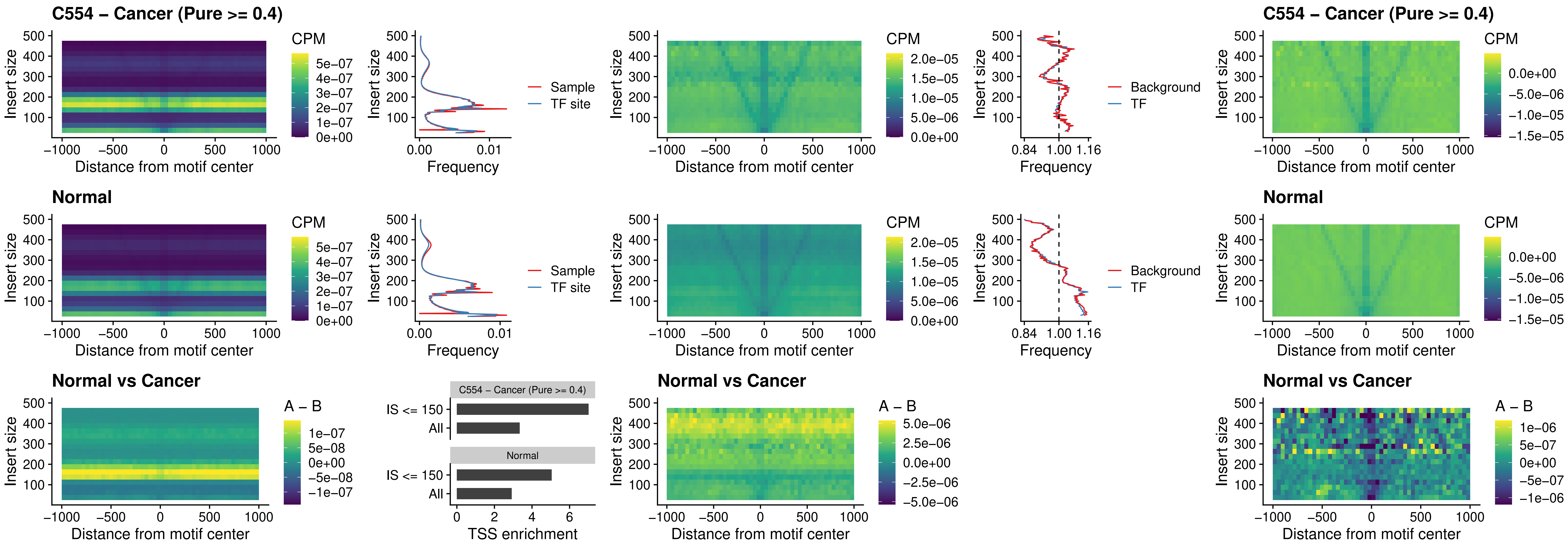

### HOXA4_uc_dTSS_oPEAK

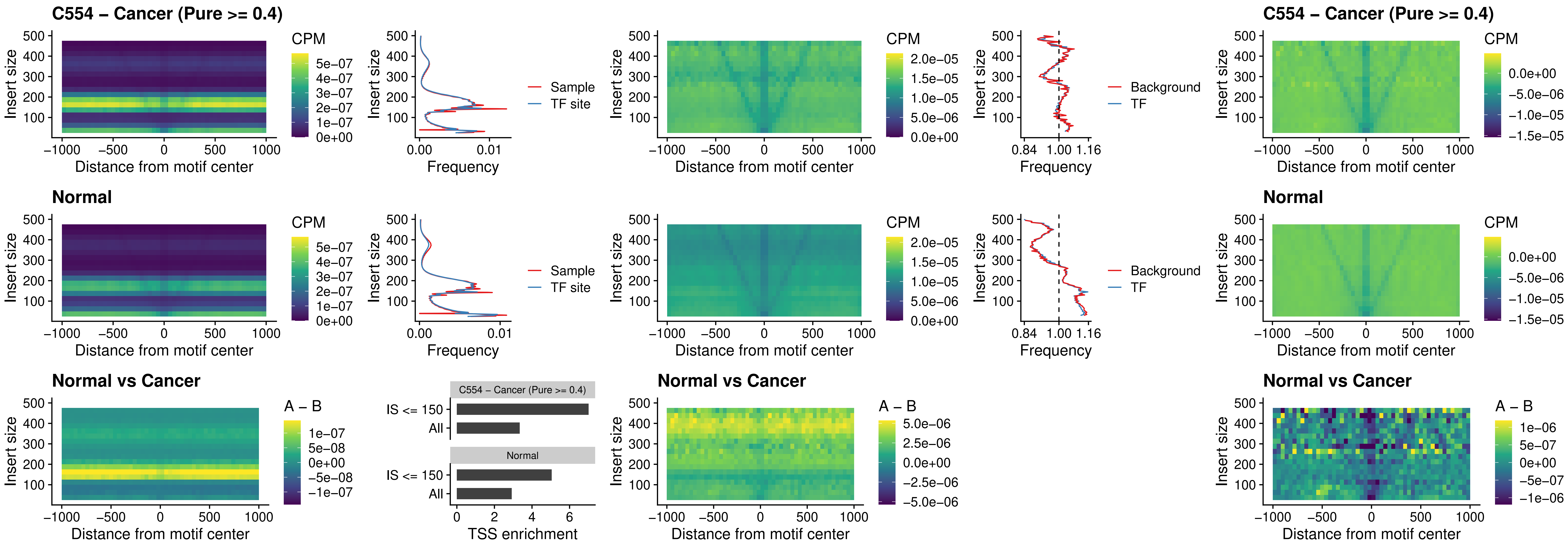

### HLX_uc_dTSS_oPEAK

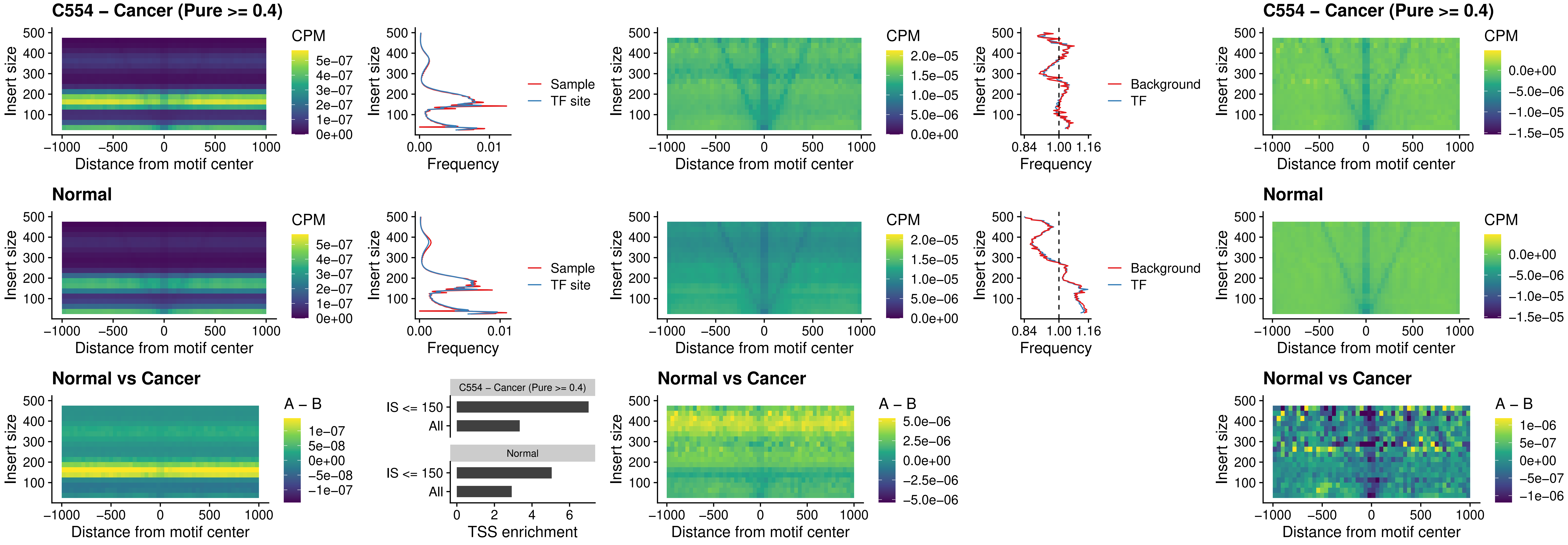

### AC0125311_uc_dTSS_oPEAK

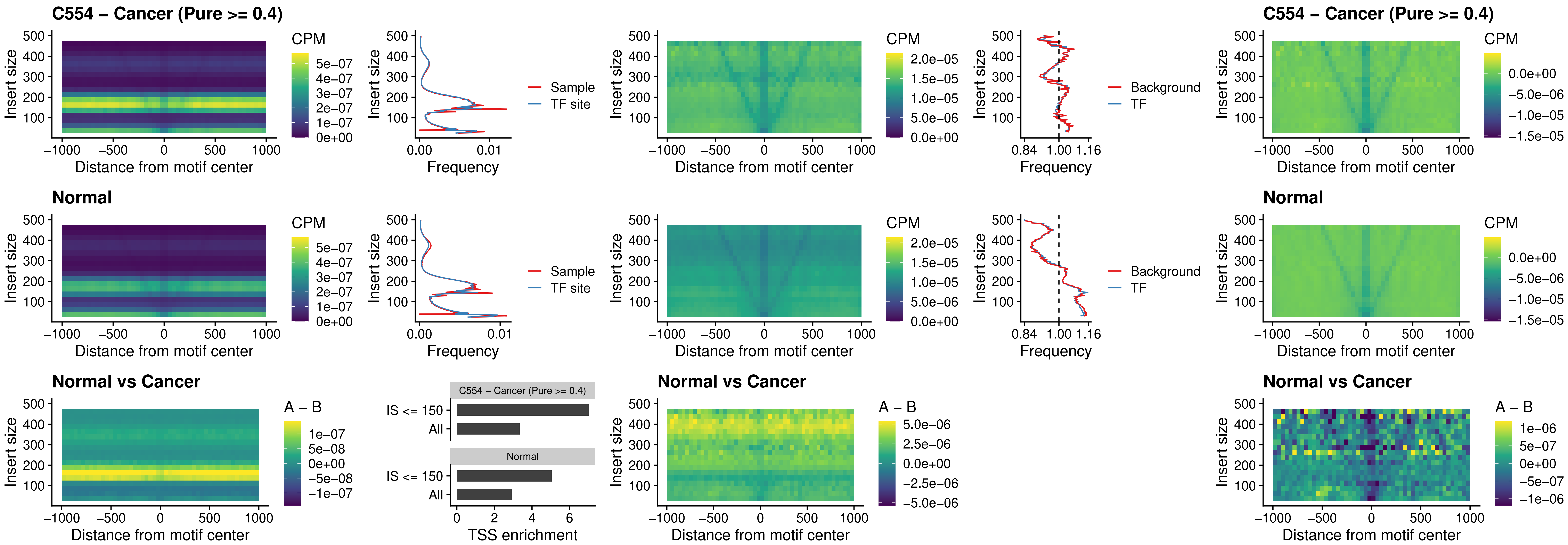

### DBX2_uc_dTSS_oPEAK

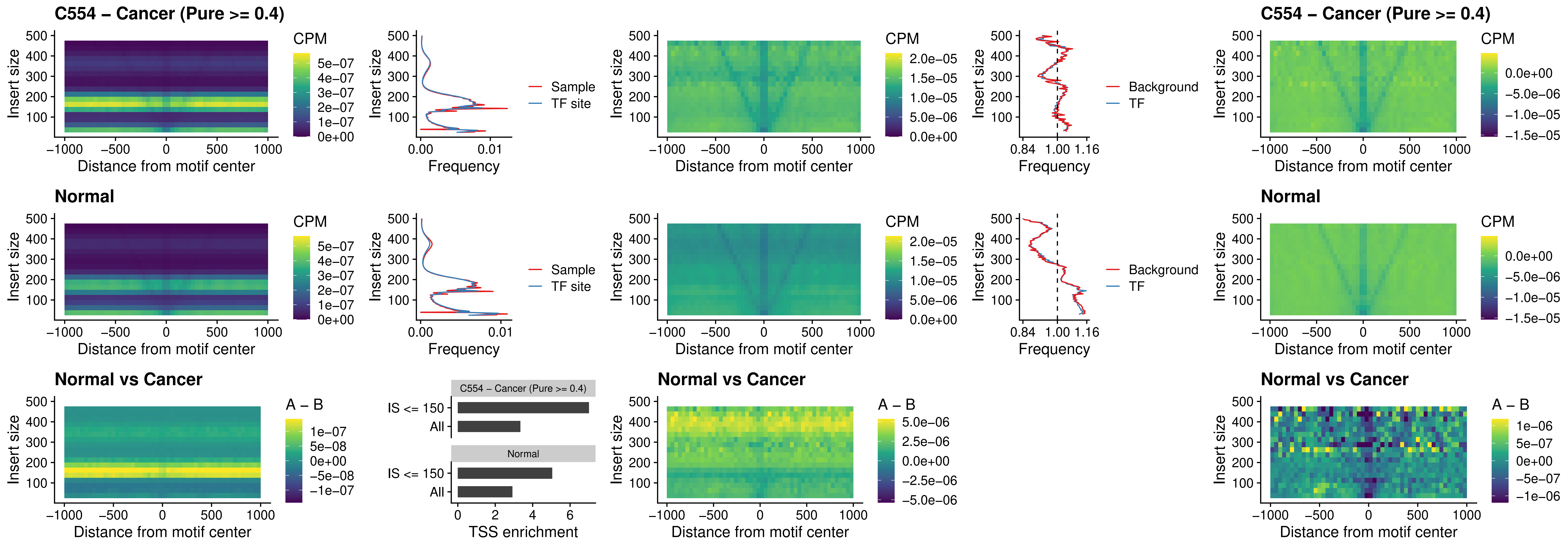

#### HOXA2_uc_dTSS_nPEAK

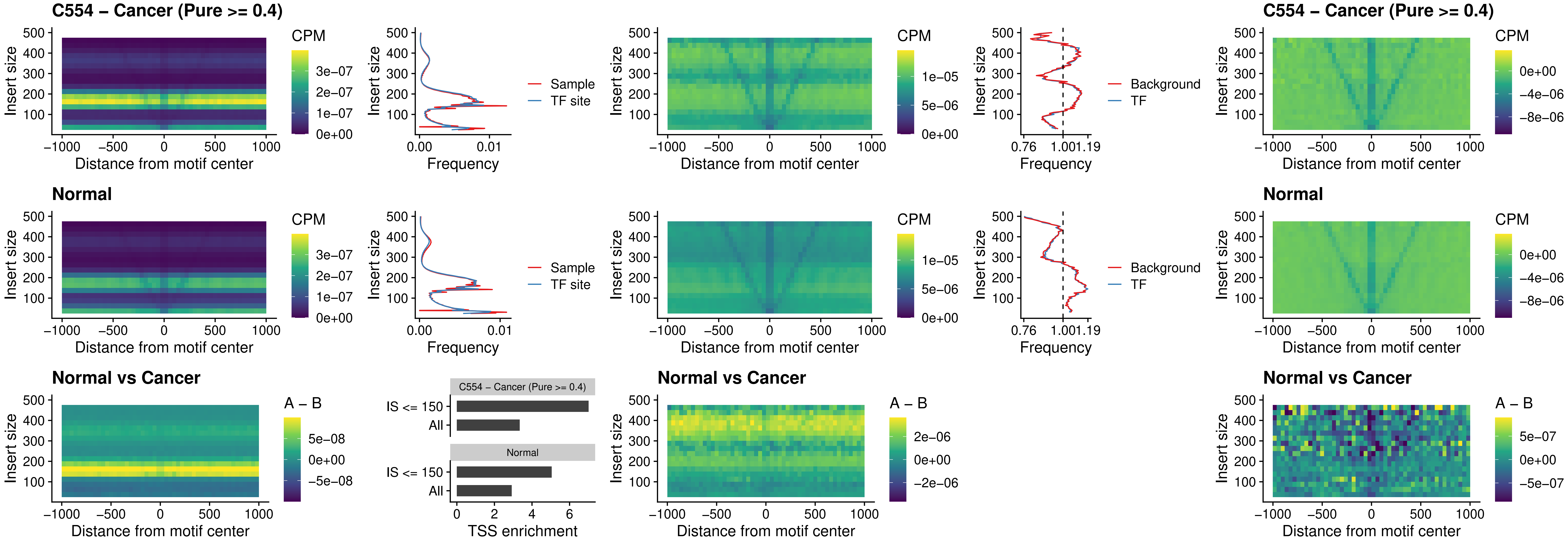

### SHOX_uc_dTSS_nPEAK

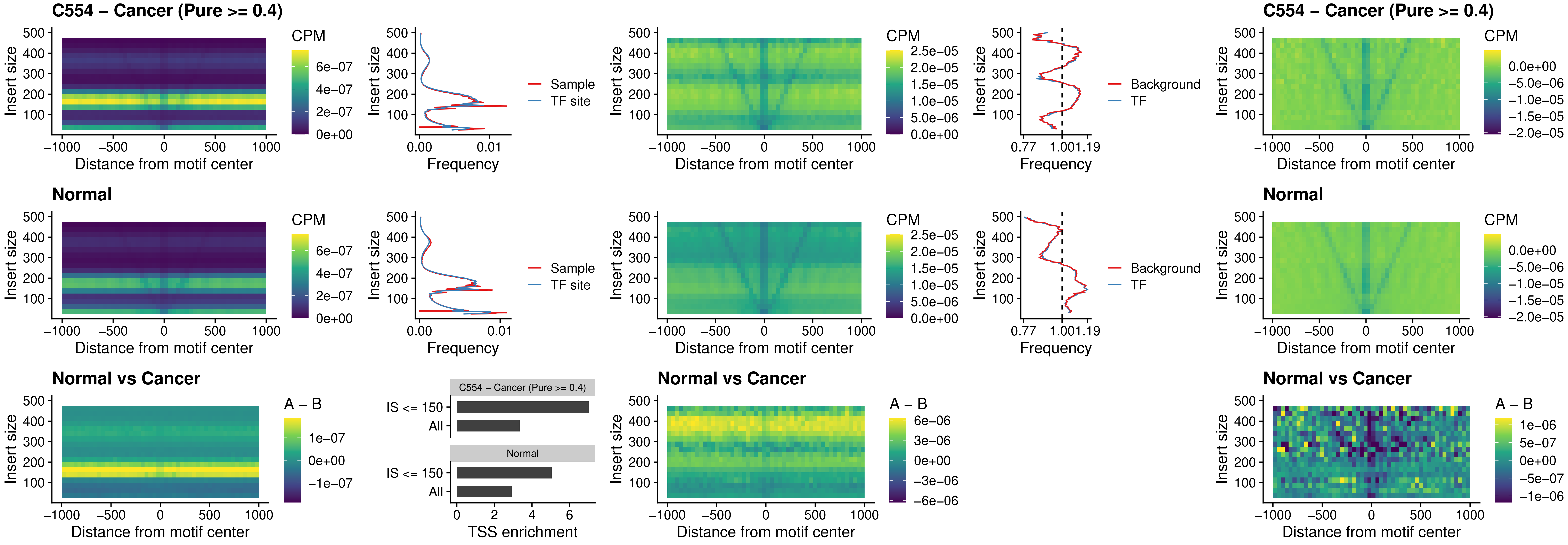

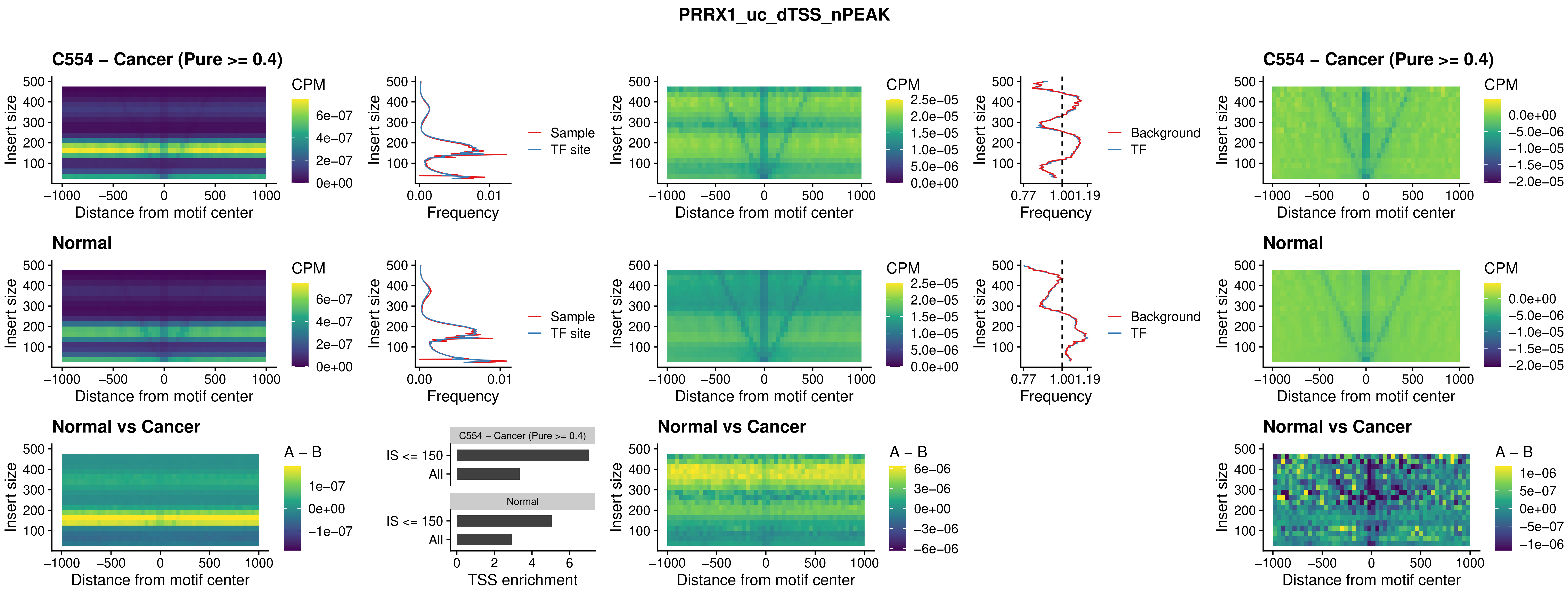
PRRX1_uc_dTSS_nPEAK
C554 – Cancer (Pure >= 0.4)
Normal
Normal vs Cancer
Insert size
Distance from motif center
Frequency
CPM
A – B
Sample
TF site
Background
TF
C554 – Cancer (Pure >= 0.4)
Normal
IS <= 150
All
TSS enrichment

### LMX1A_uc_dTSS_nPEAK

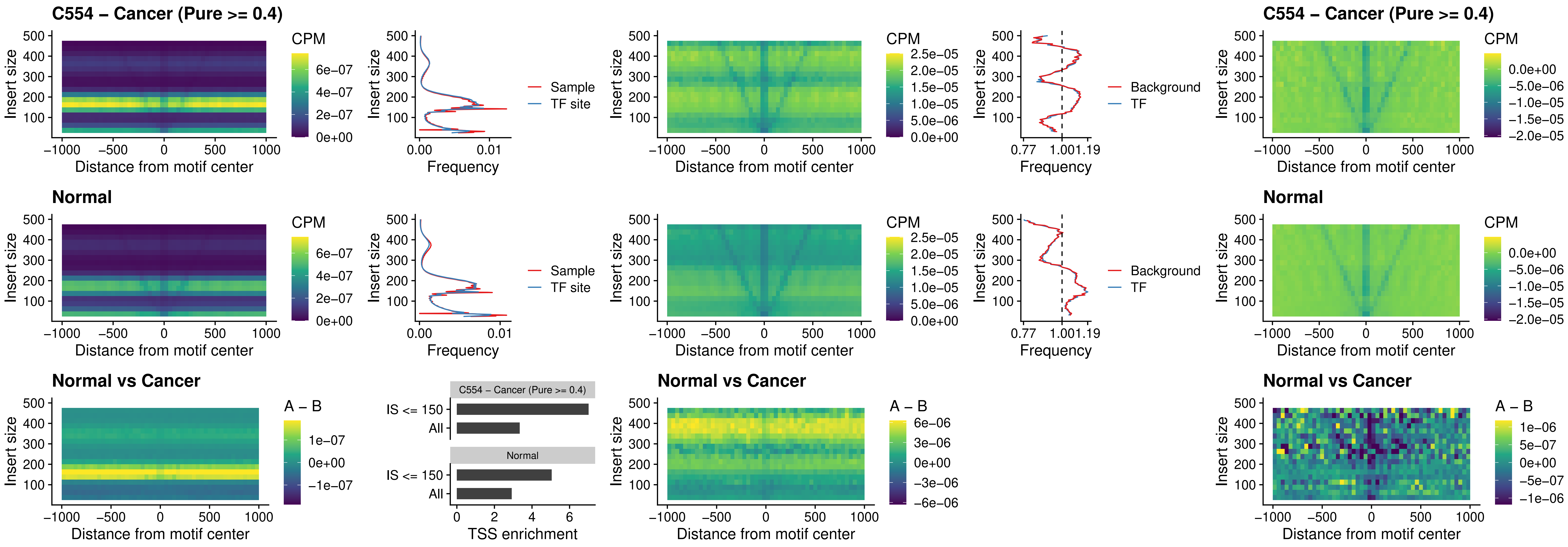

### LMX1B_uc_dTSS_nPEAK

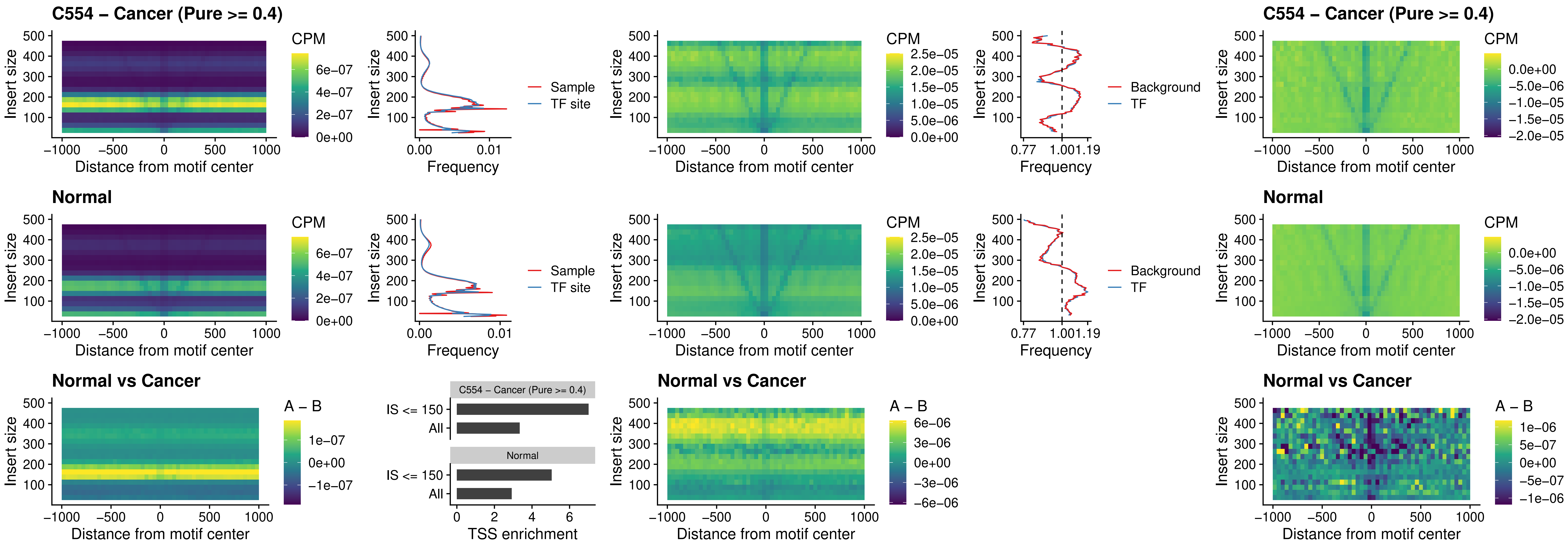

### DLX5_uc_dTSS_oPEAK

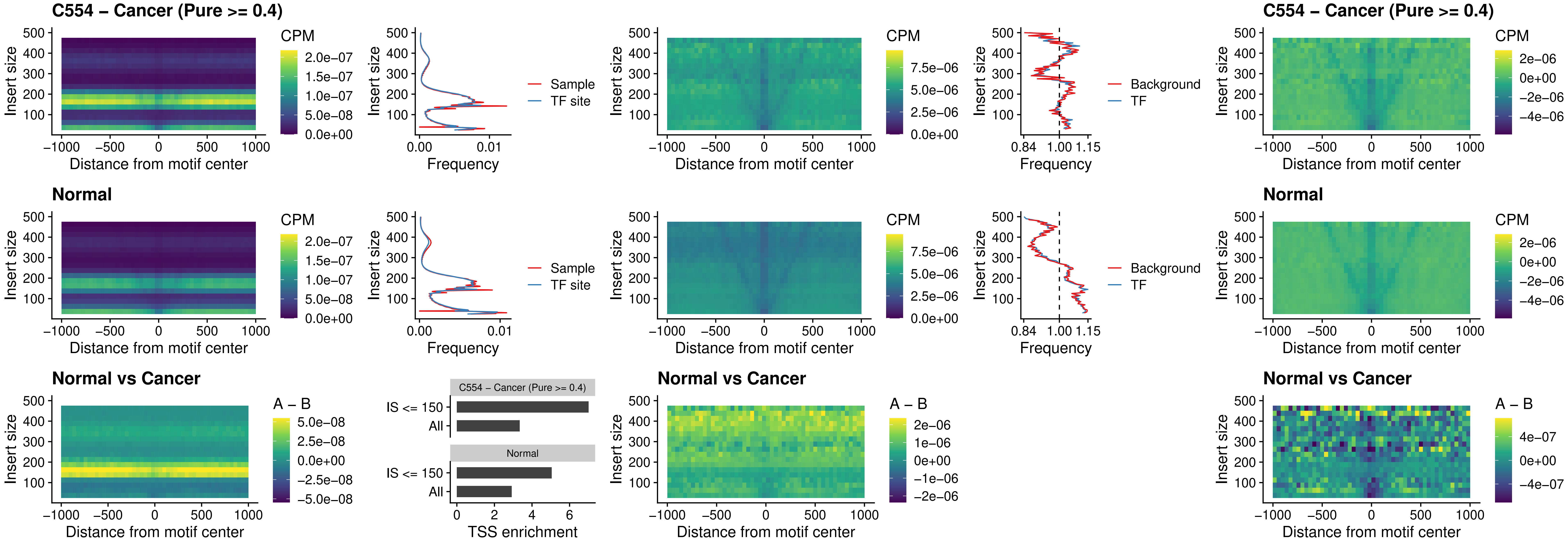

### ARID3C_uc_dTSS_nPEAK

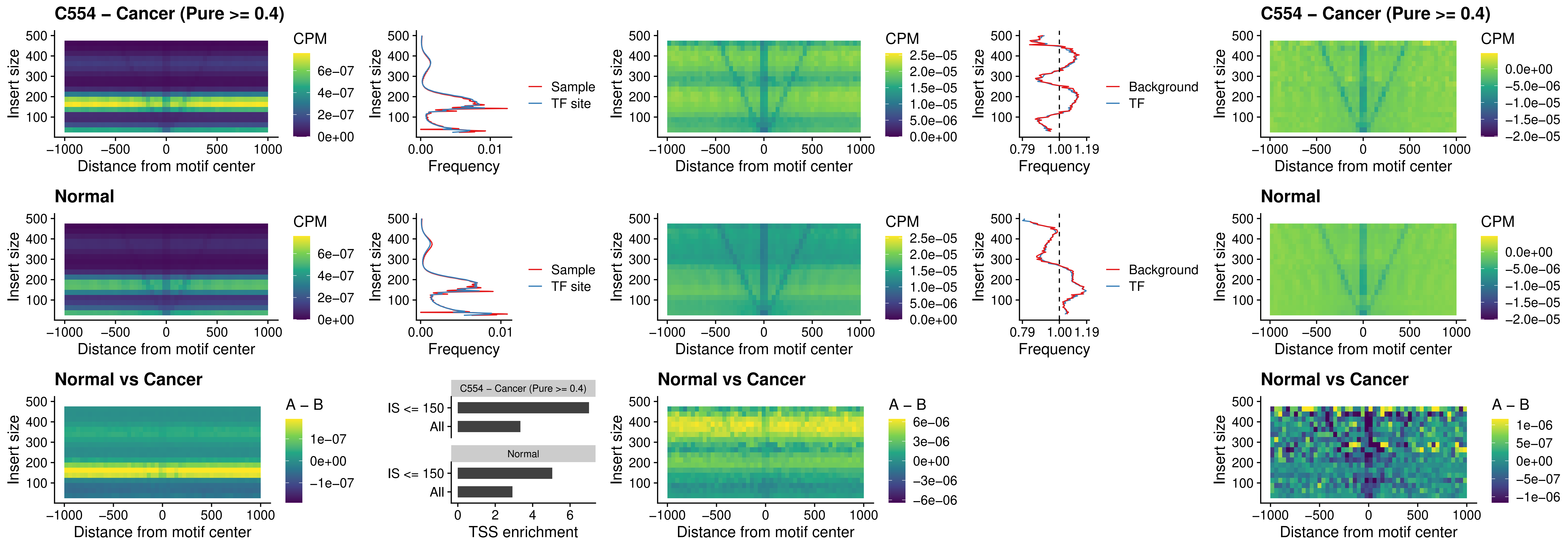

#### HOXD12_uc_dTSS_nPEAK

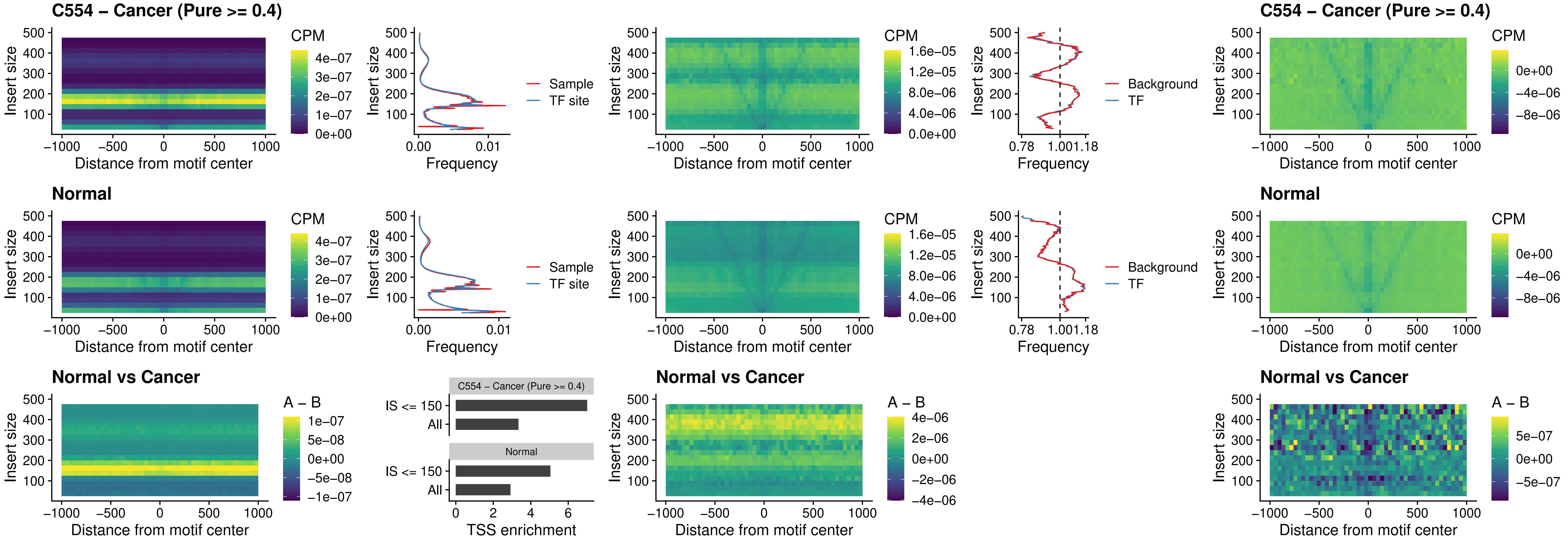

### CDX2_uc_dTSS_nPEAK

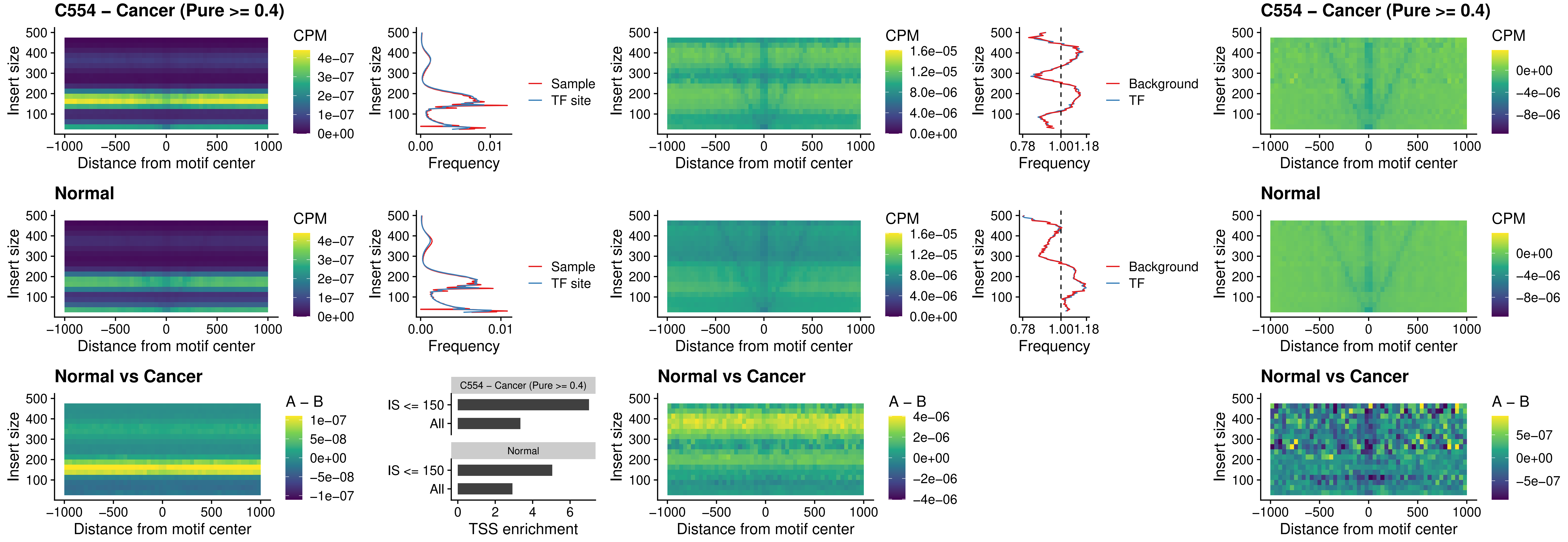

#### HOXC12_uc_dTSS_nPEAK

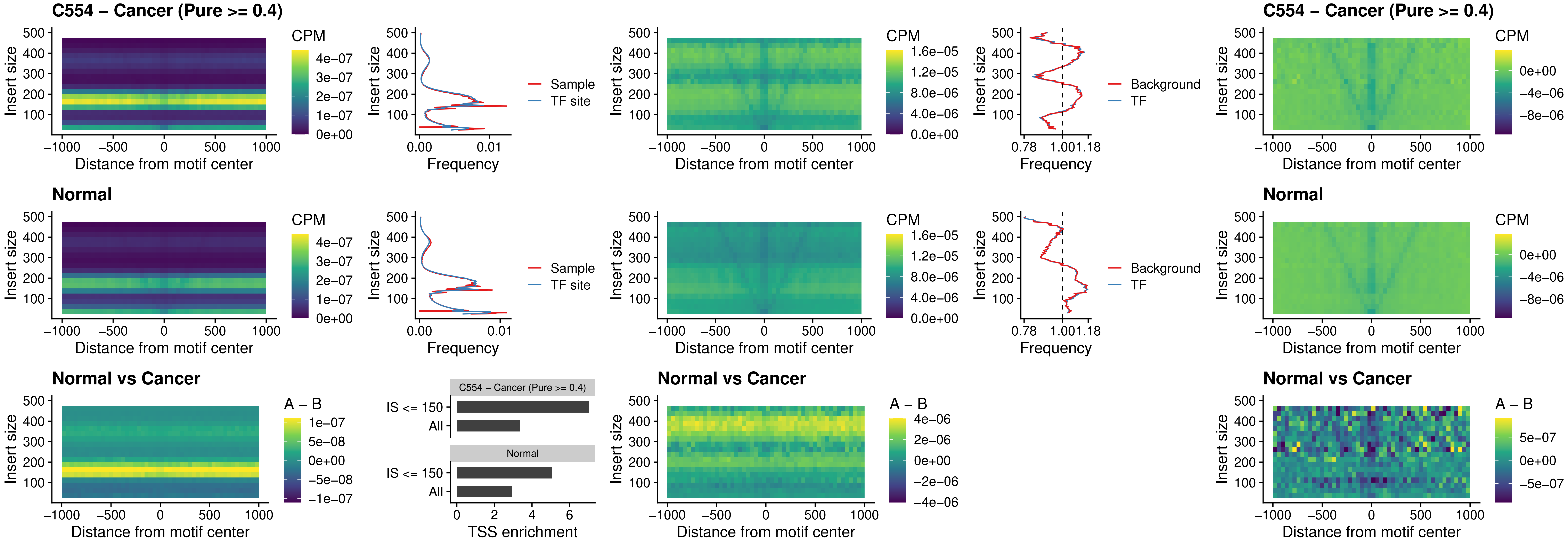

### HHEX_uc_dTSS_oPEAK

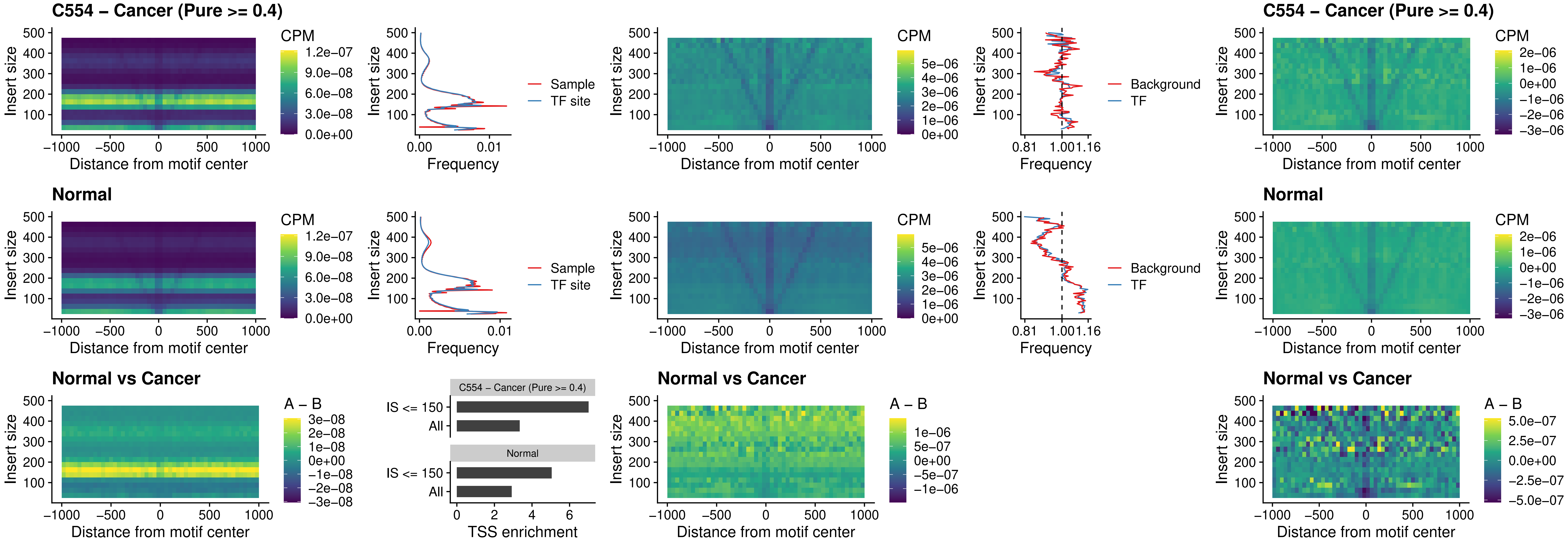

### HOXA11_uc_dTSS_nPEAK

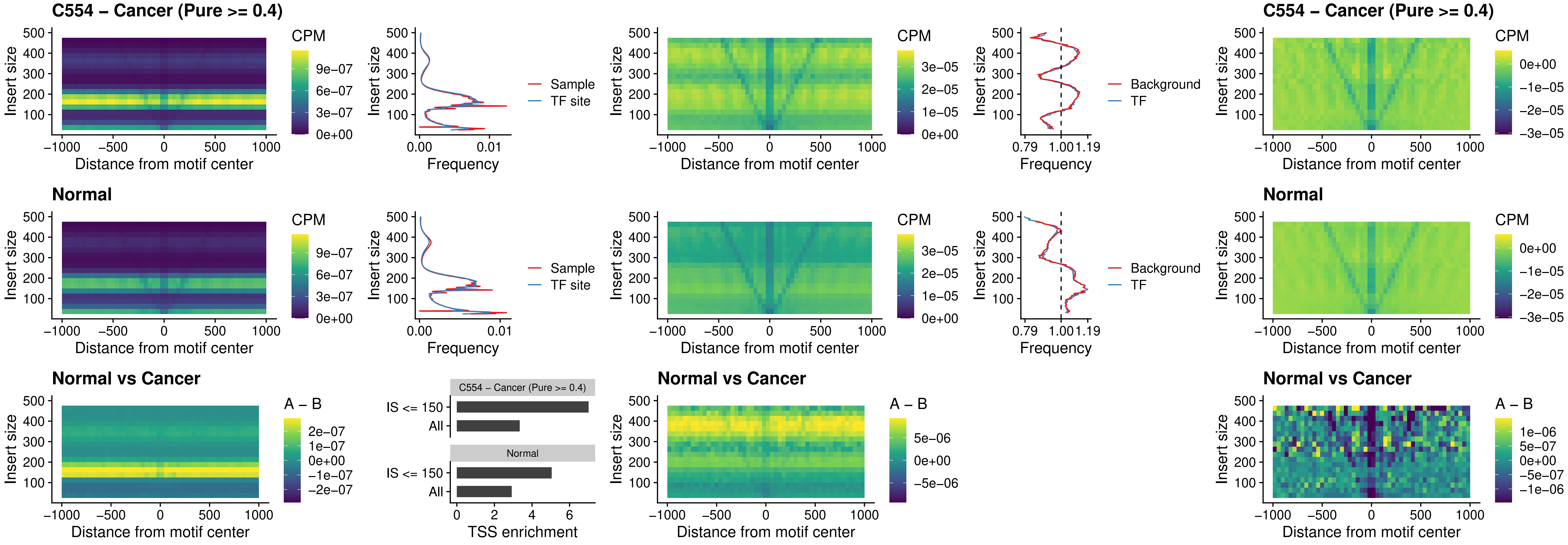

#### HOXA11_uc_cTSS_nPEAK

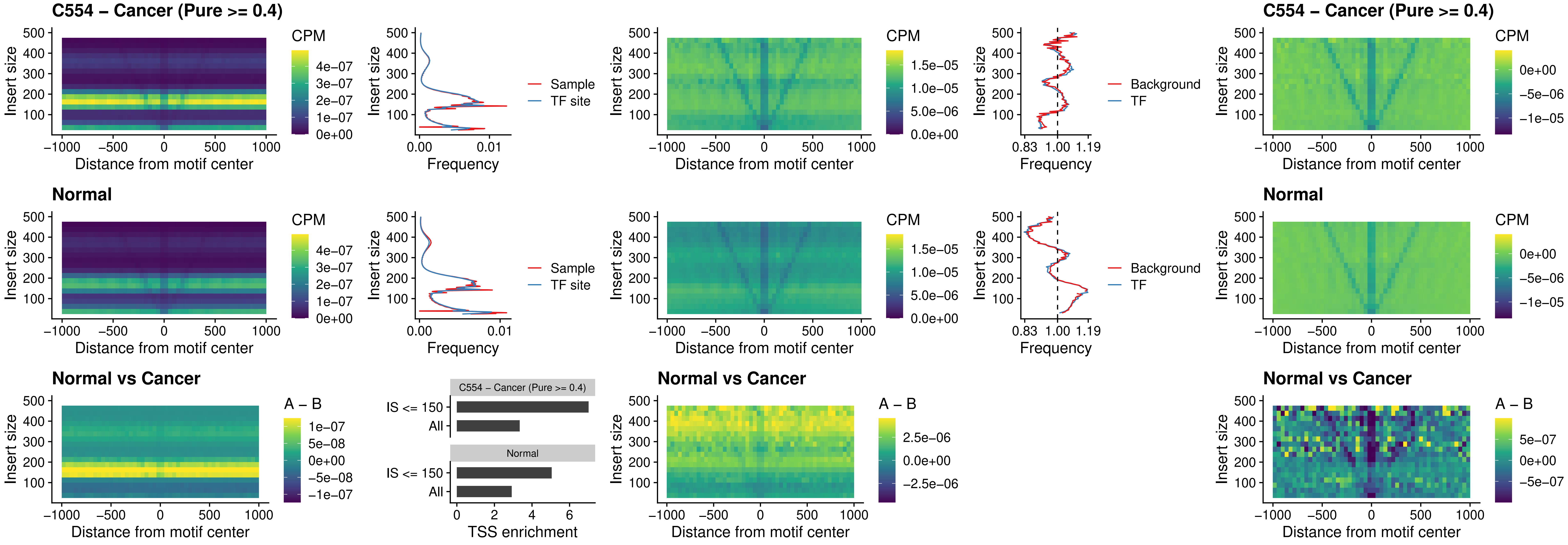

#### ZFHX3_uc_dTSS_nPEAK

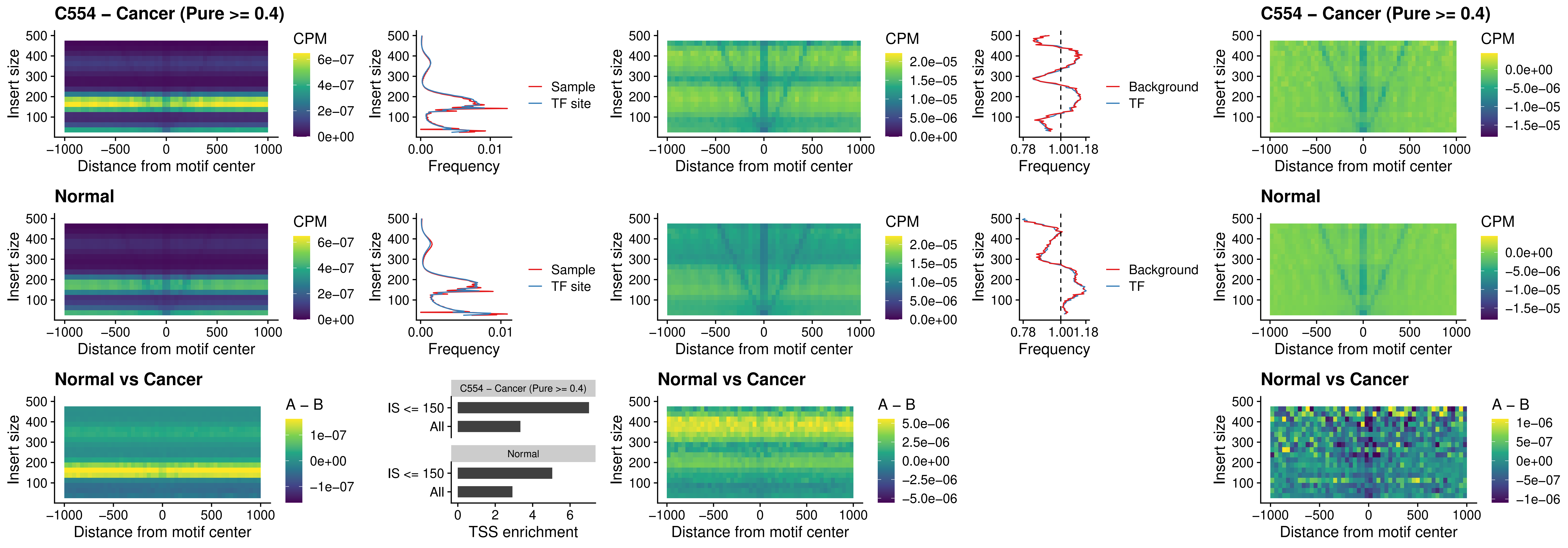

### HOXB4_uc_dTSS_nPEAK

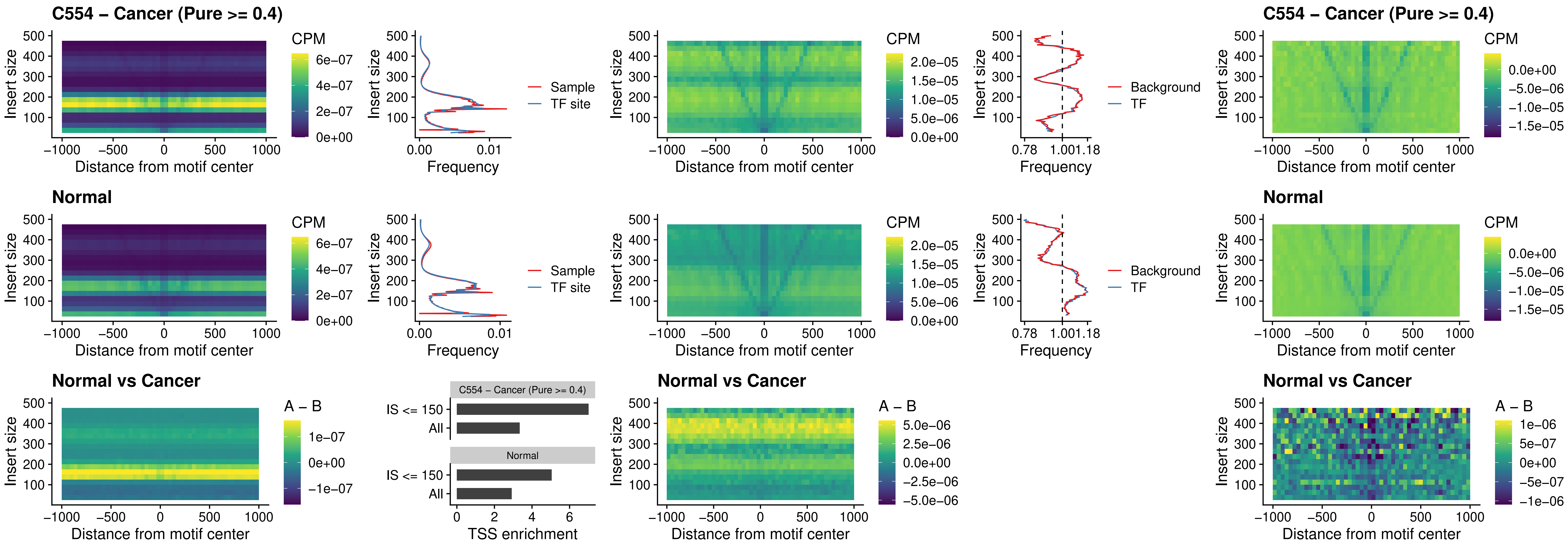

### HOXC5_uc_dTSS_nPEAK

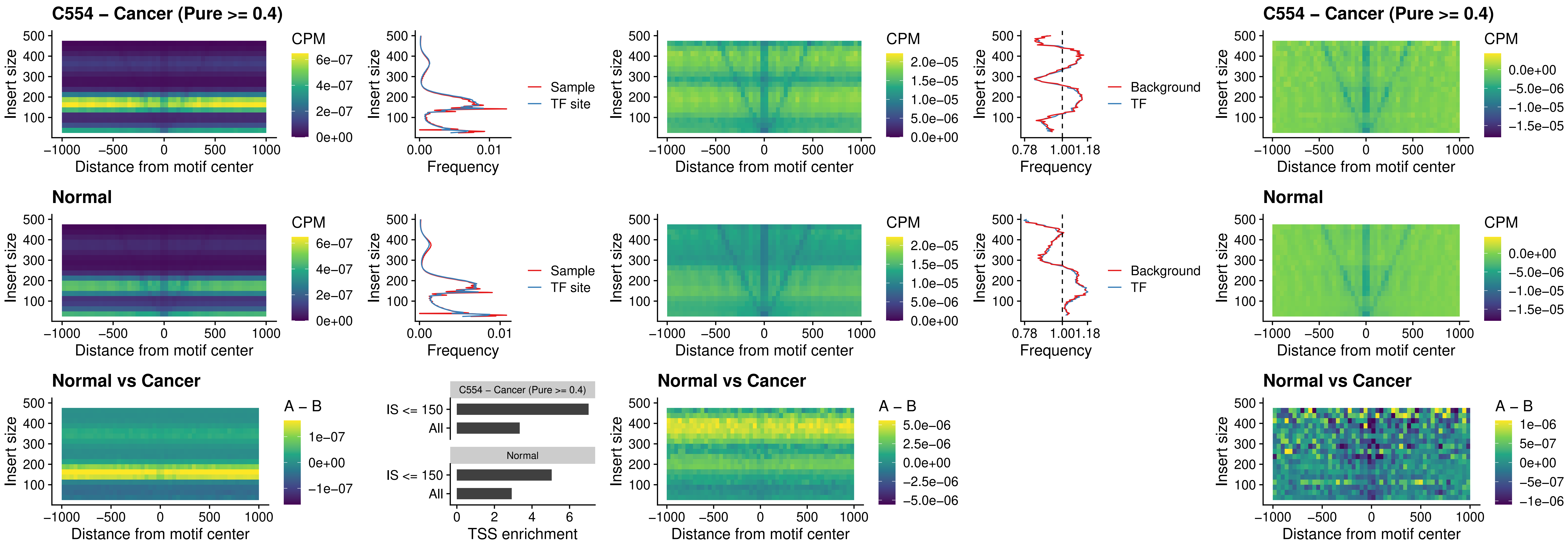

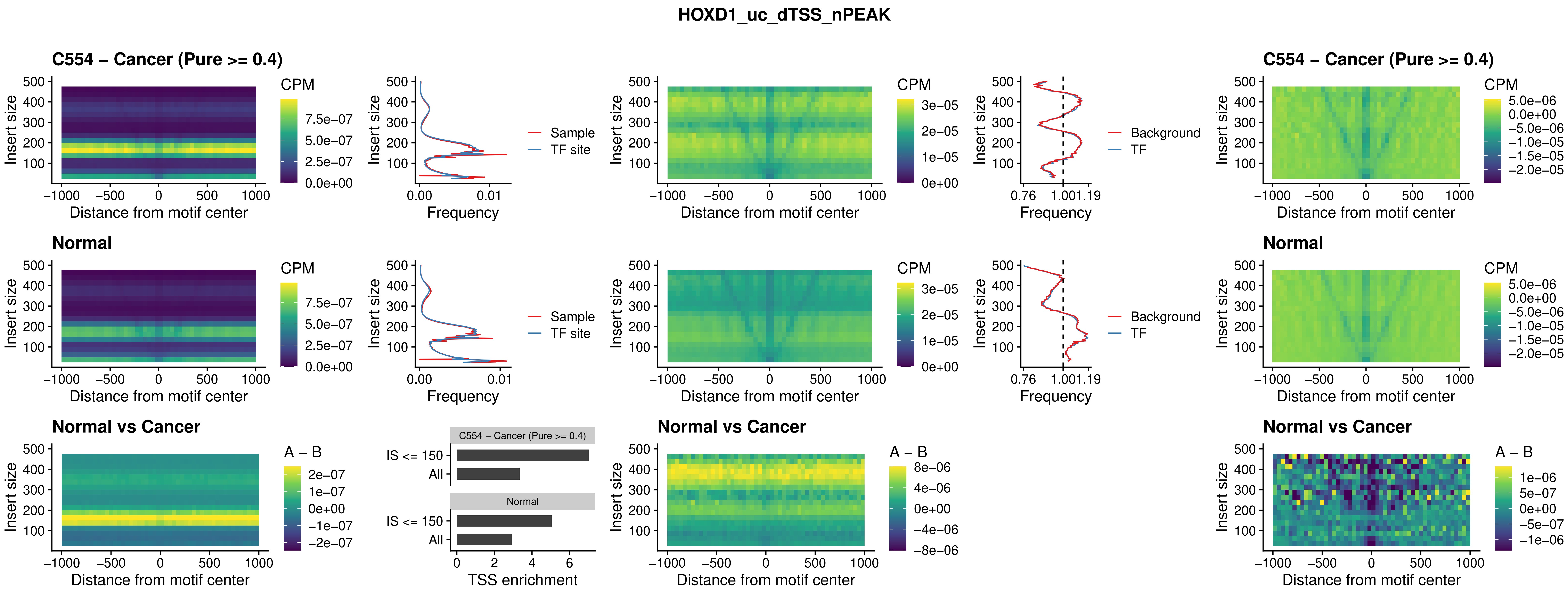
HOXD1_uc_dTSS_nPEAK
C554 – Cancer (Pure >= 0.4)
Normal
Normal vs Cancer
Insert size
Distance from motif center
Frequency
CPM
A – B
Sample
TF site
Background
TF
C554 – Cancer (Pure >= 0.4)
Normal
IS <= 150
All
TSS enrichment

### OTP_uc_dTSS_nPEAK

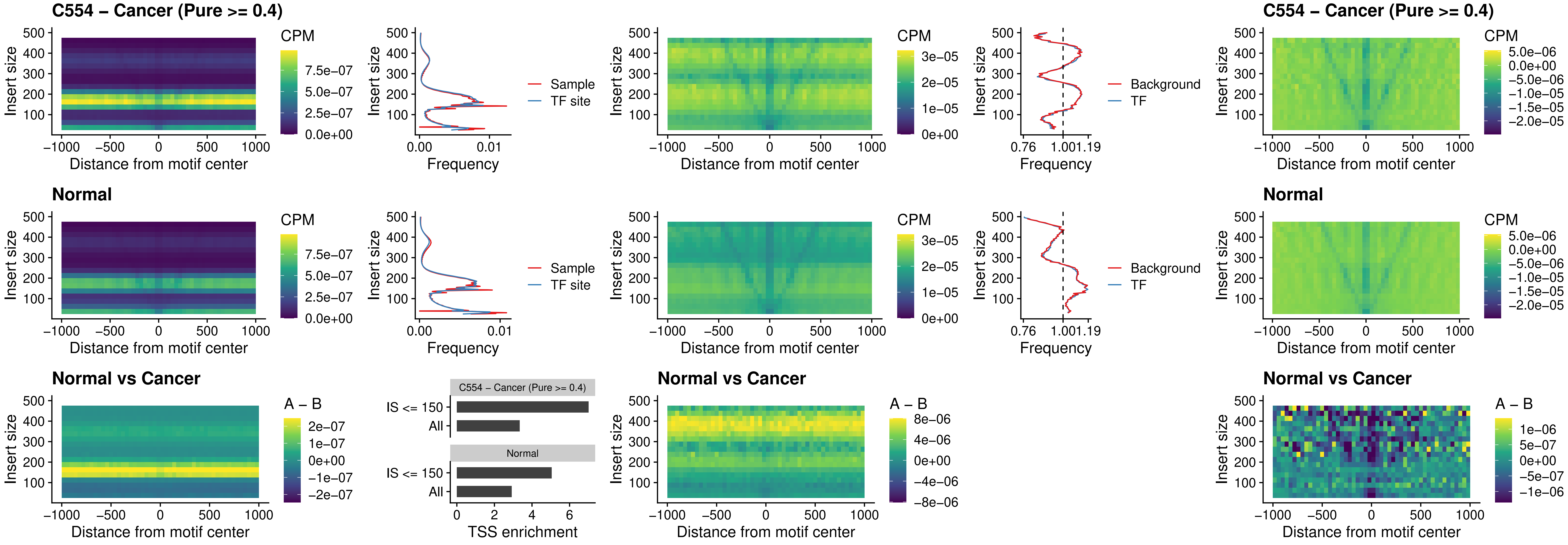

### FOXL1_uc_dTSS_oPEAK

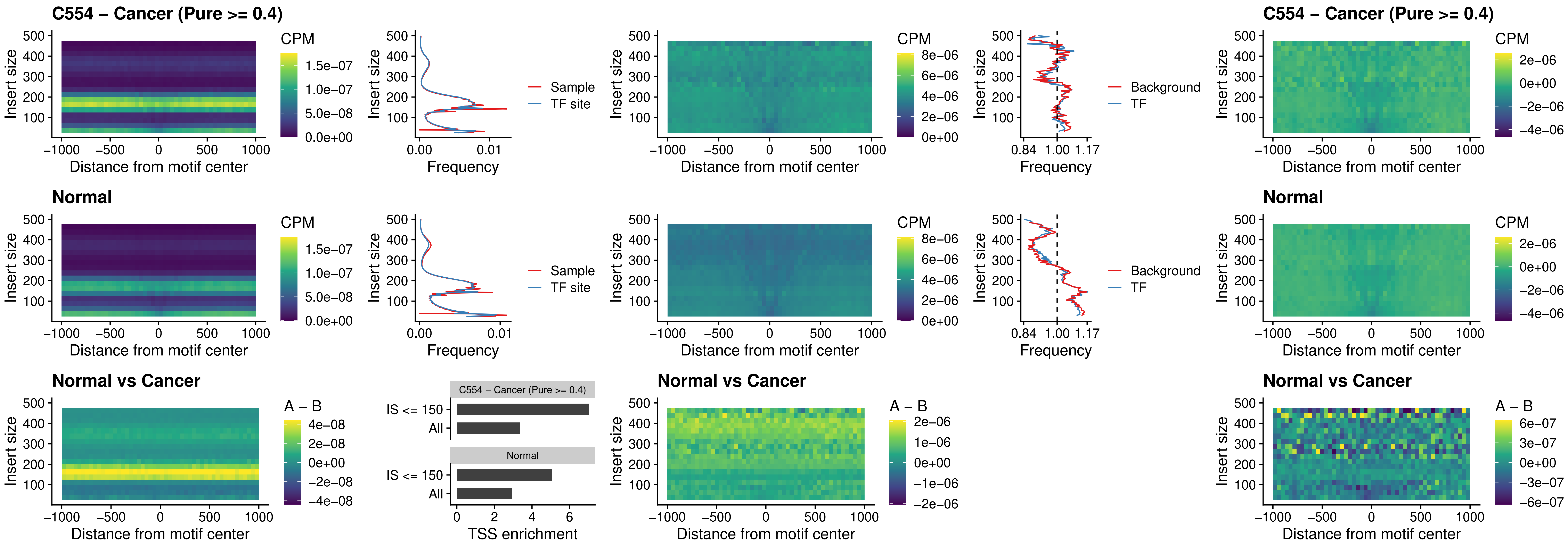

#### ZNF32_uc_dTSS_oPEAK

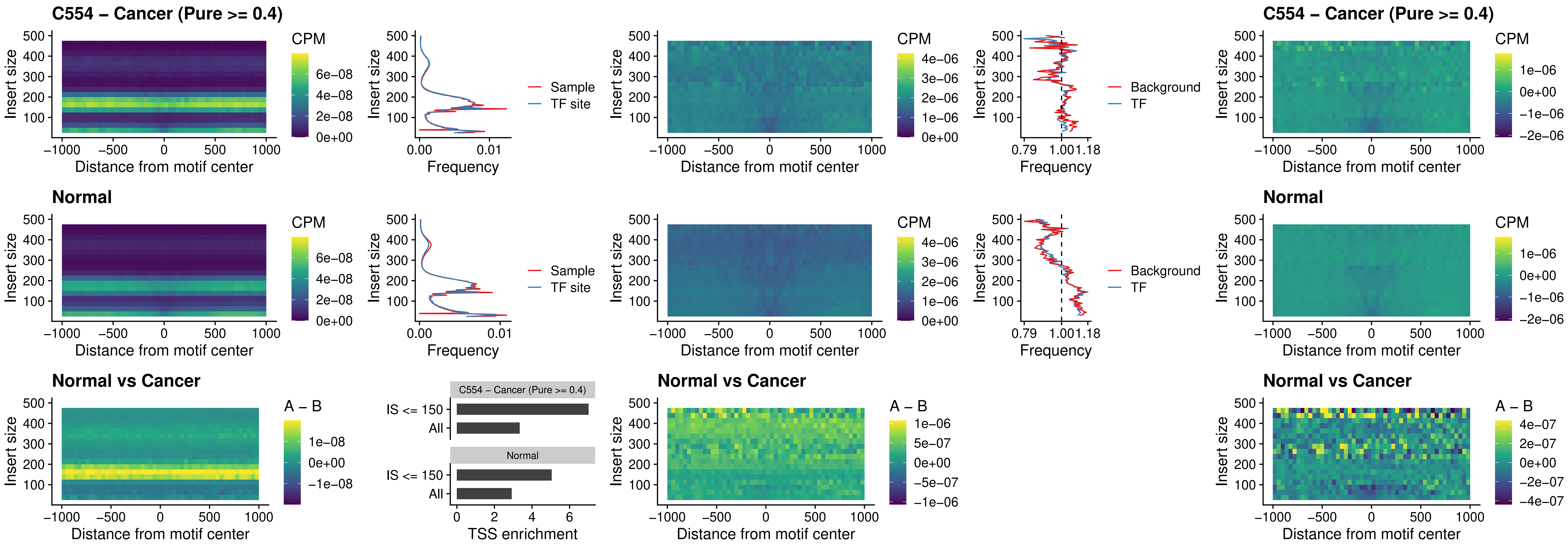

### FOXL1_uc_dTSS_nPEAK

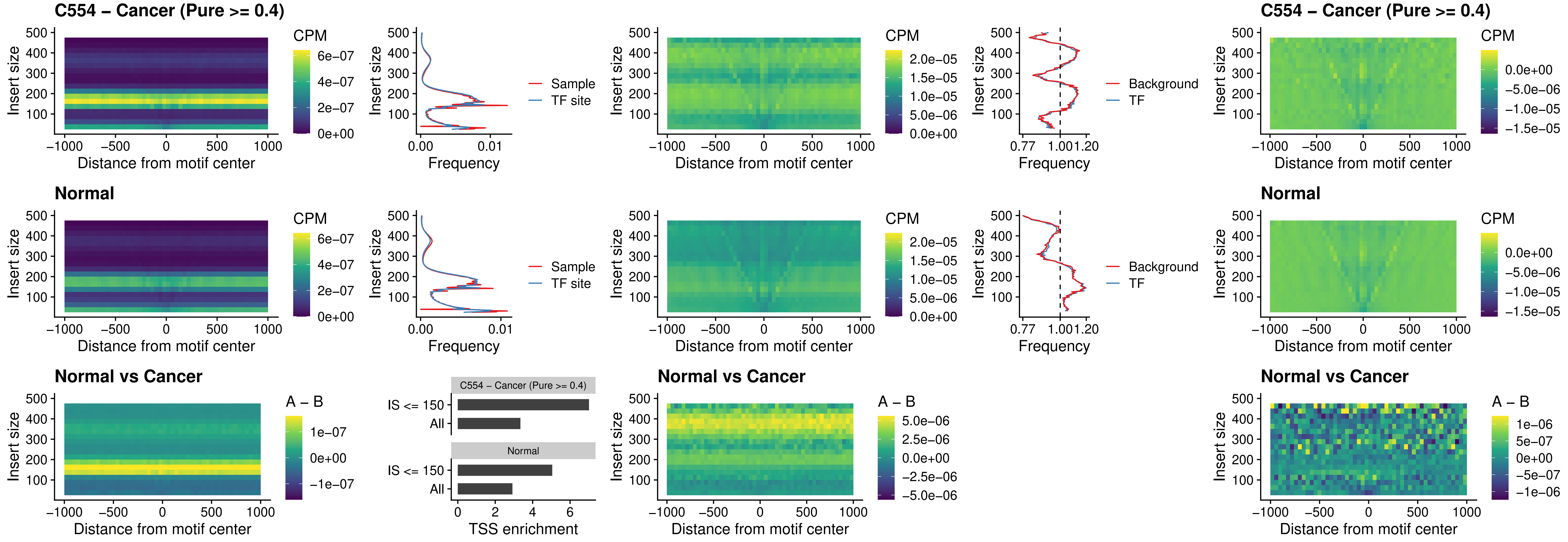

#### ZNF32_uc_dTSS_nPEAK

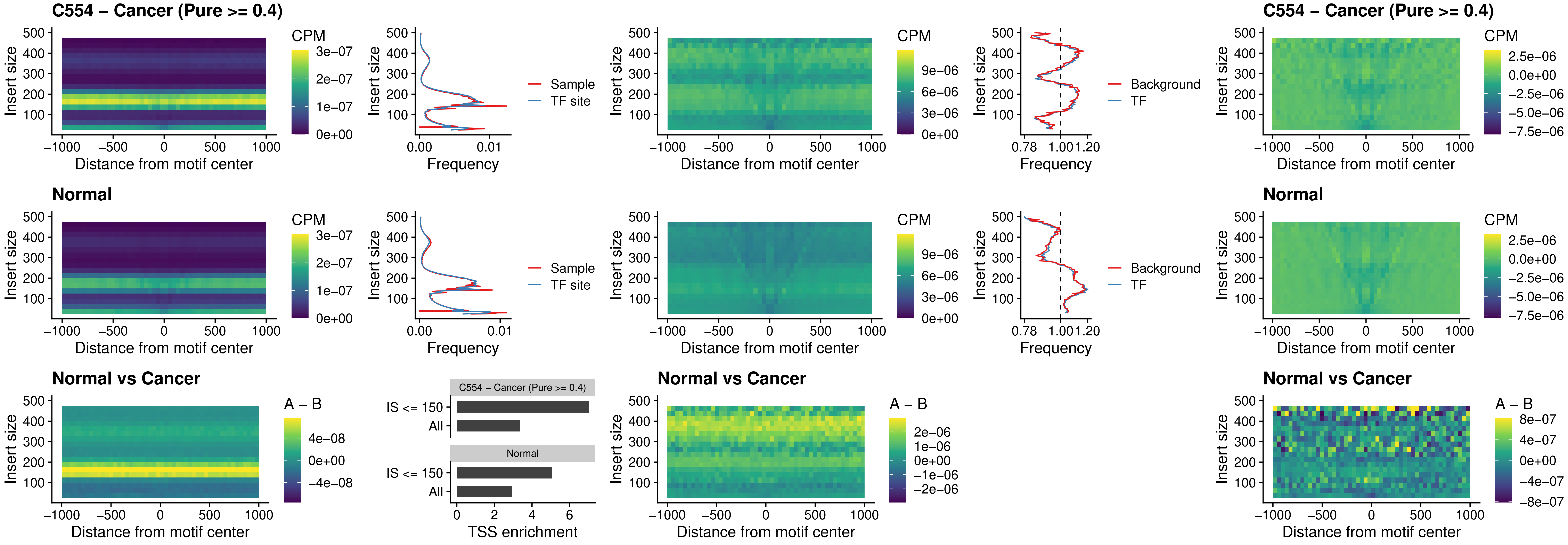

### DLX5_uc_dTSS_nPEAK

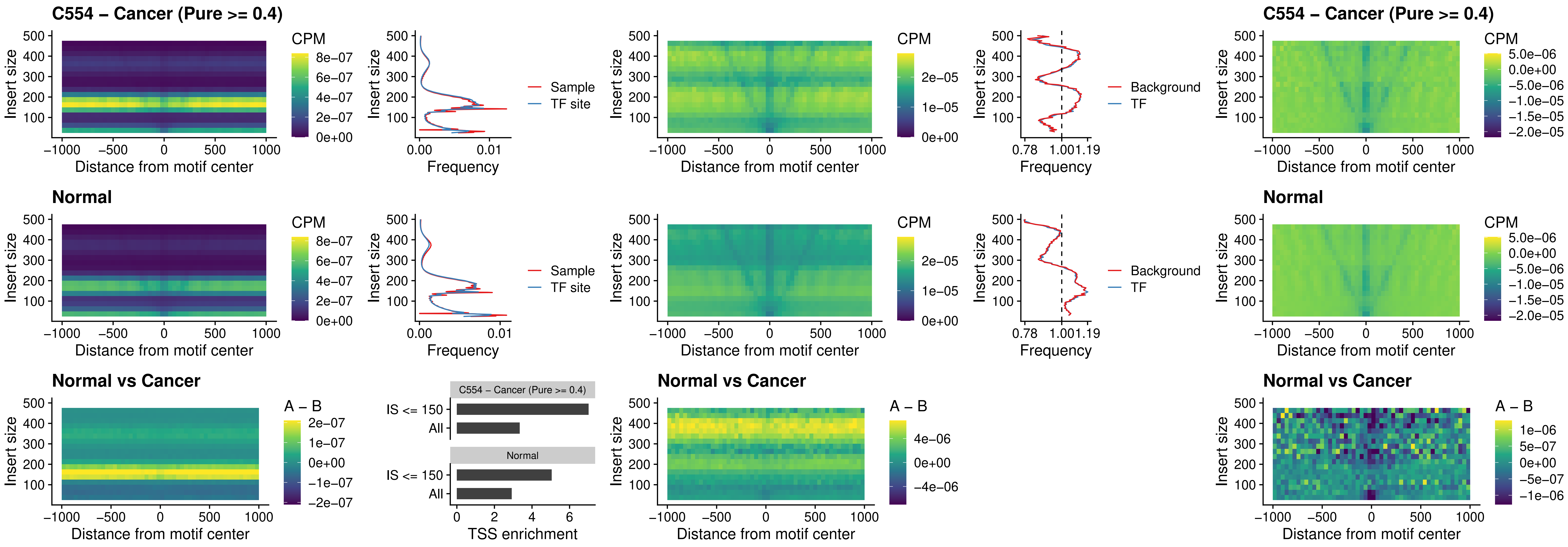

### DMRT2_uc_dTSS_nPEAK

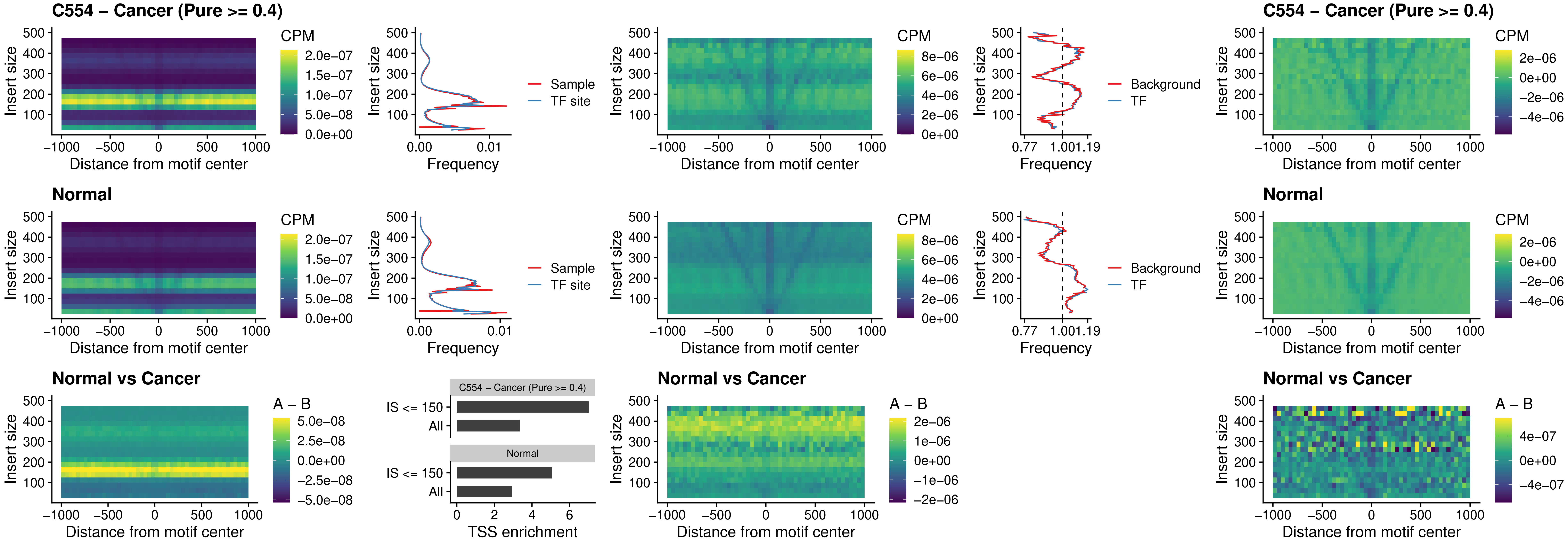

### DMRTC2_uc_cTSS_nPEAK

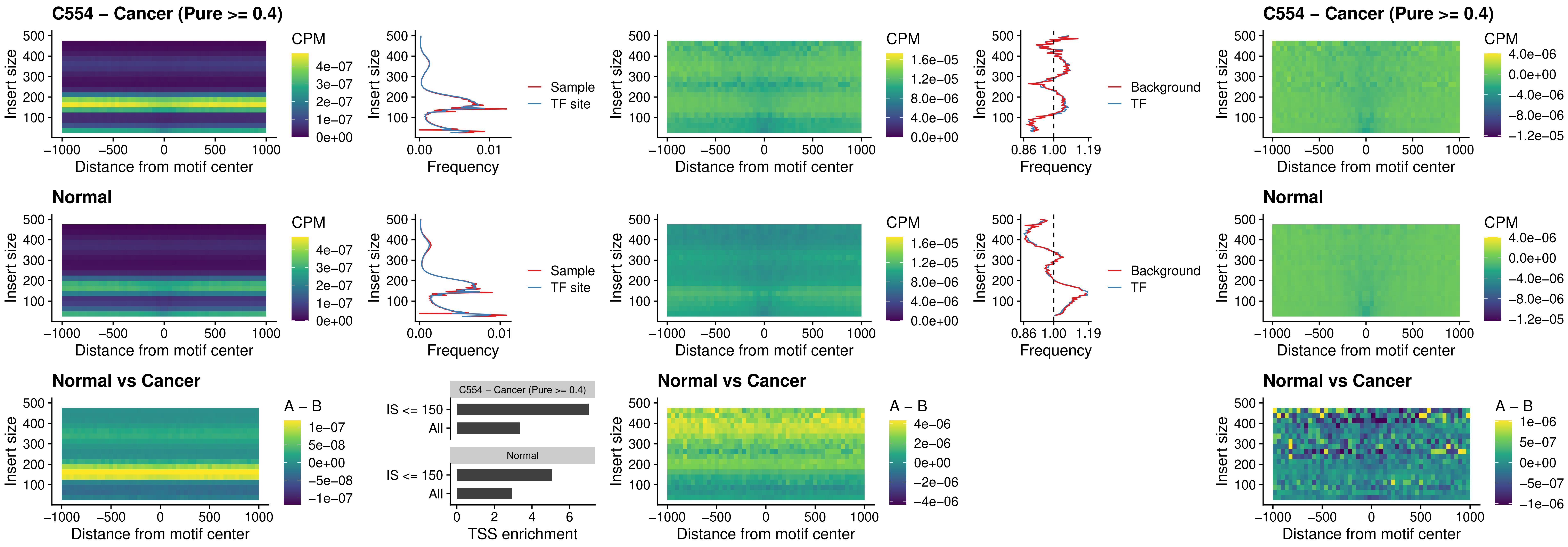

### HOXD9_uc_dTSS_nPEAK

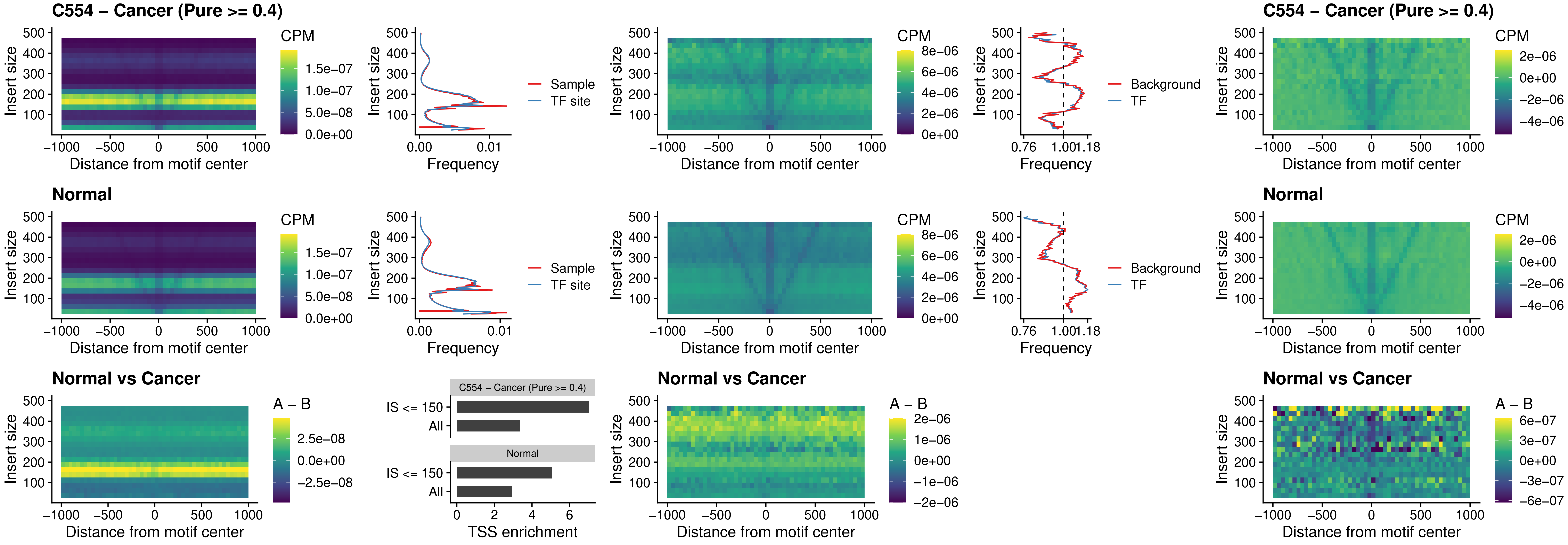

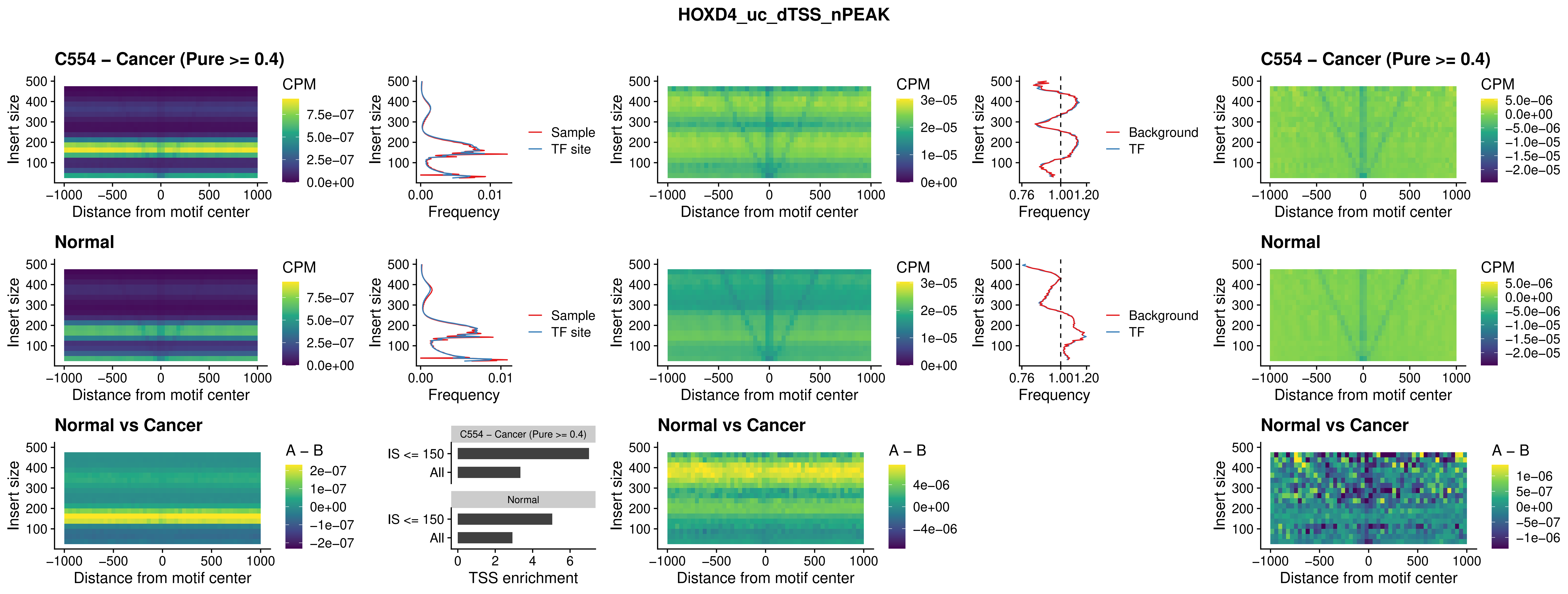
HOXD4_uc_dTSS_nPEAK
C554 – Cancer (Pure >= 0.4)
Normal
Normal vs Cancer
Insert size
Distance from motif center
Frequency
CPM
A – B
Sample
TF site
Background
TF
C554 – Cancer (Pure >= 0.4)
Normal
IS <= 150
All
TSS enrichment

#### FOXP3_uc_cTSS_nPEAK

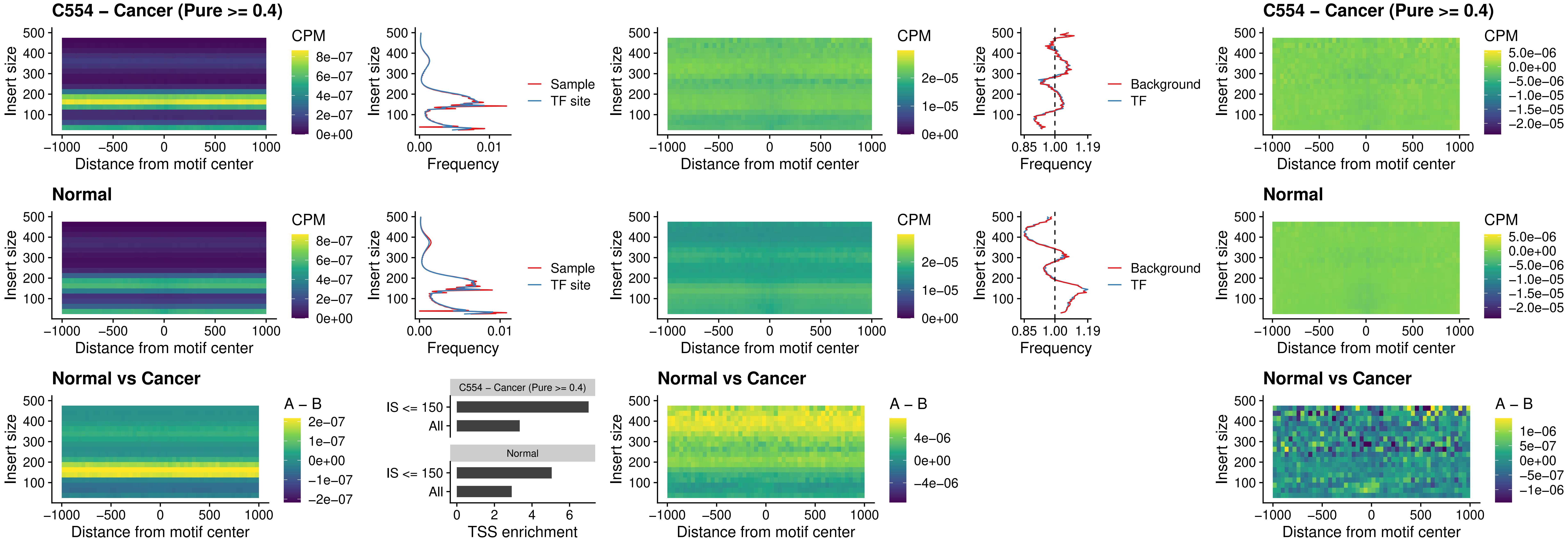

### FOXD4L6_uc_cTSS_nPEAK

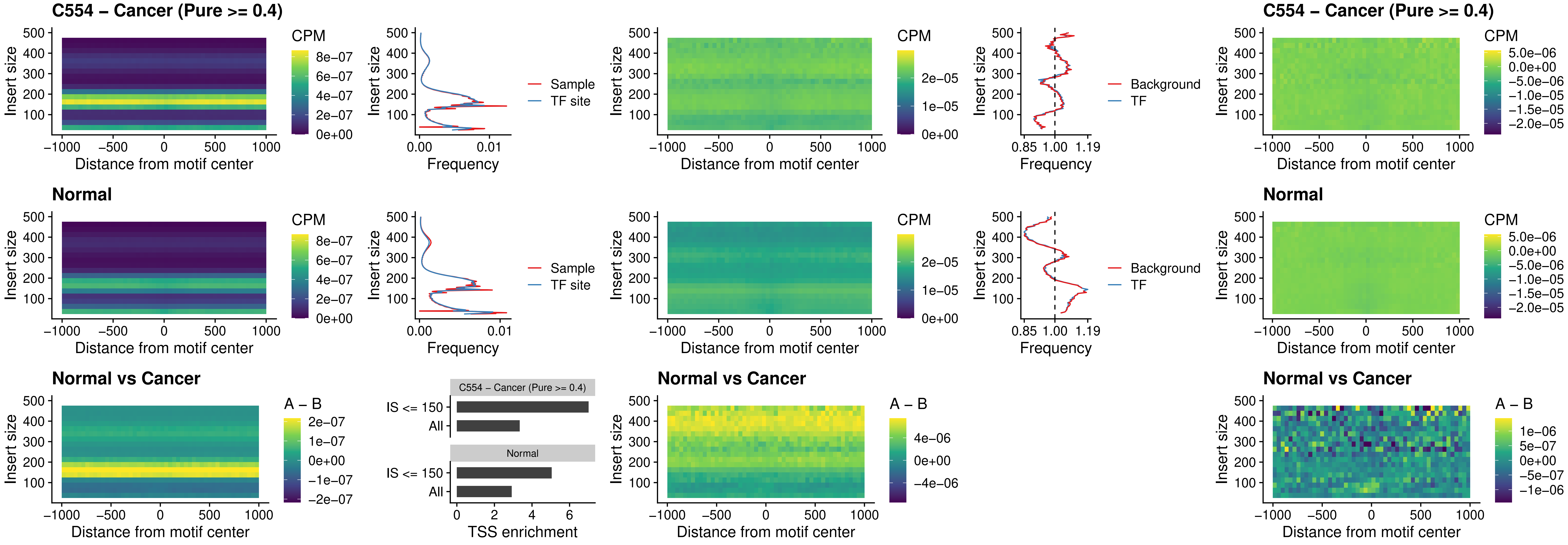

### FOXD4L5_uc_cTSS_nPEAK

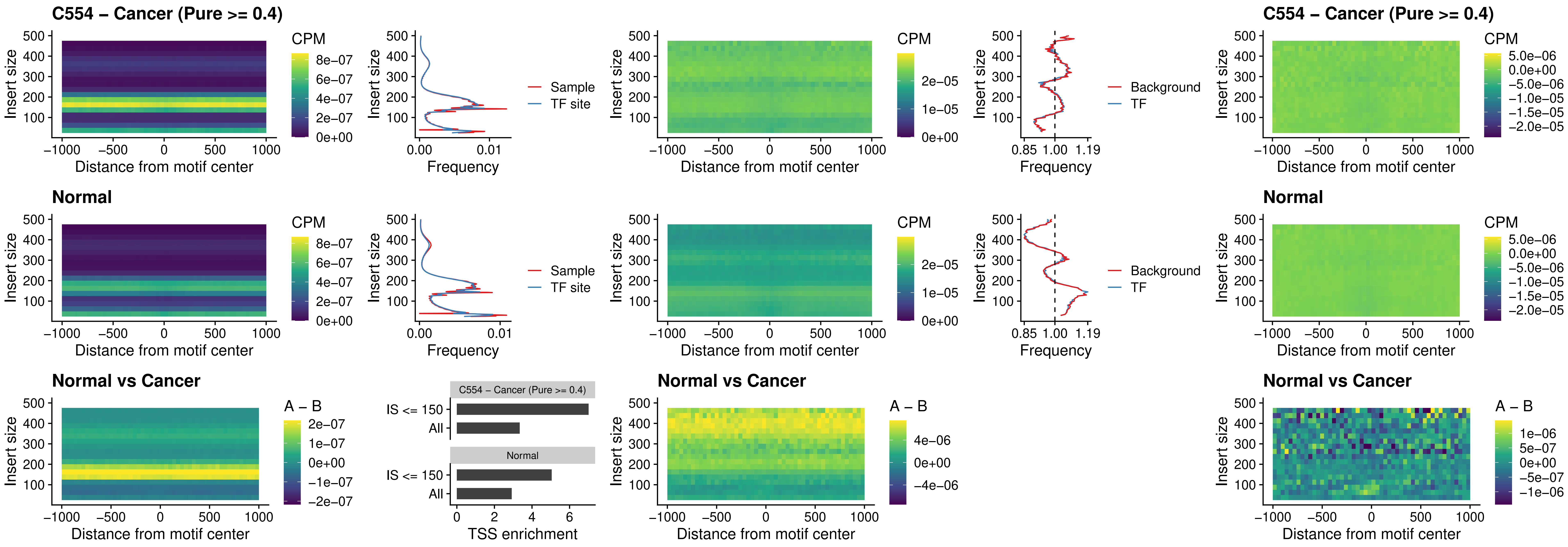

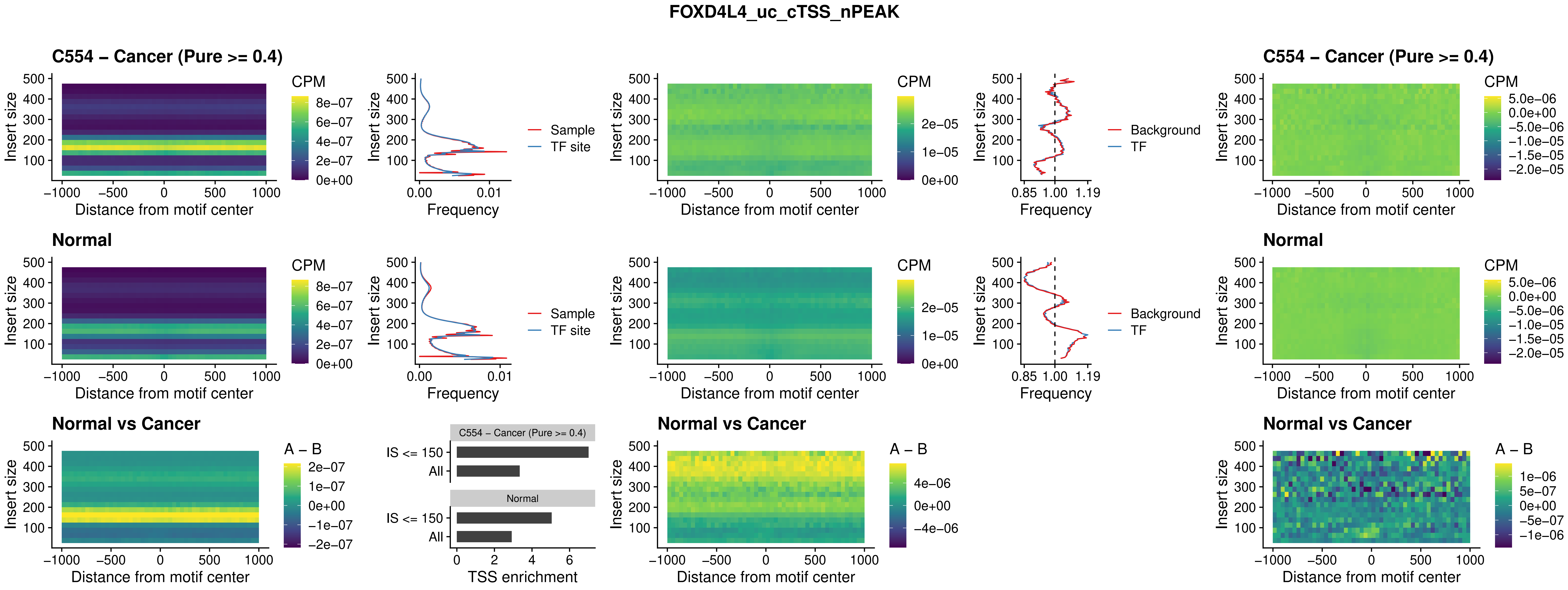

FOXD4L4_uc_cTSS_nPEAK
C554 – Cancer (Pure >= 0.4)
Normal
Normal vs Cancer
Insert size
Distance from motif center
Frequency
CPM
A – B
Sample
TF site
Background
TF
C554 – Cancer (Pure >= 0.4)
Normal
IS <= 150
All
TSS enrichment

### FOXD4L2_uc_cTSS_nPEAK

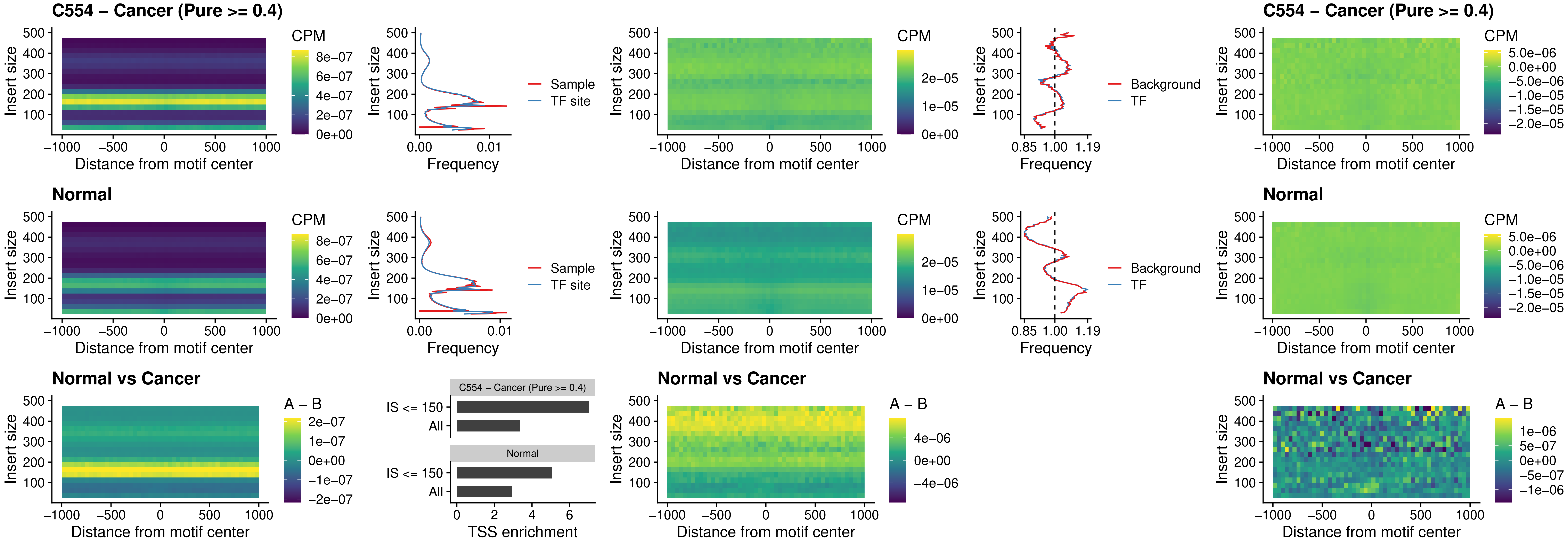

### FOXD4L3_uc_cTSS_nPEAK

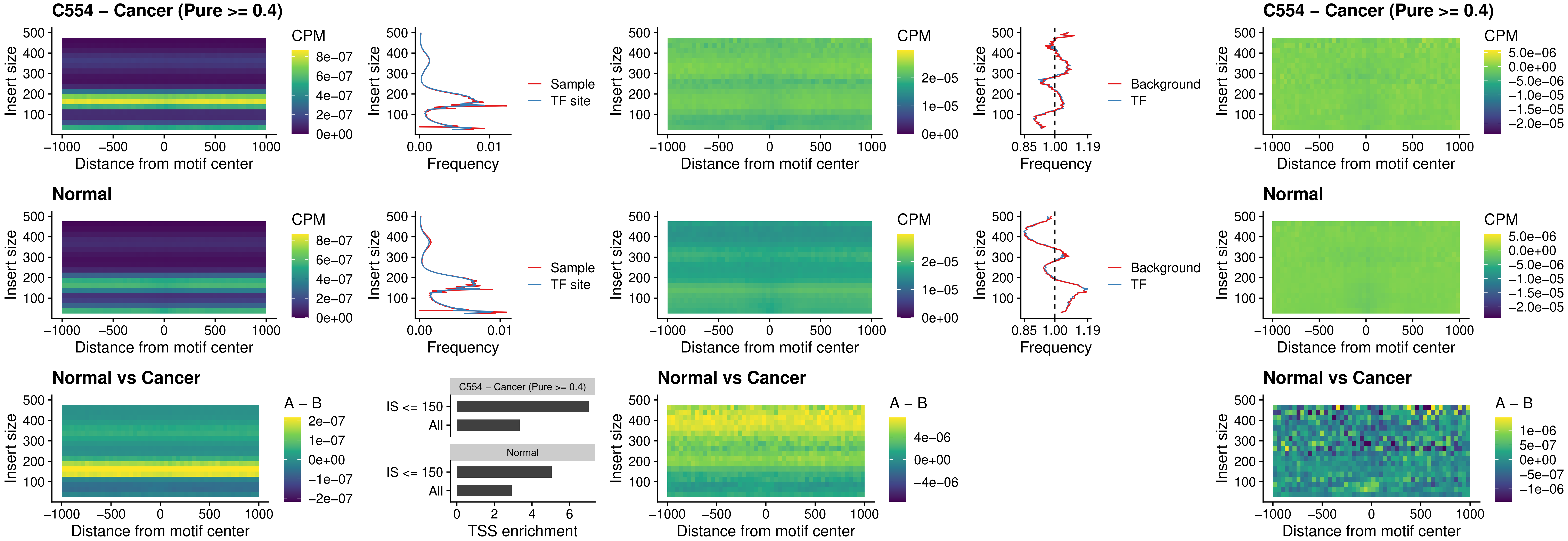

#### HOXD1_uc_dTSS_oPEAK

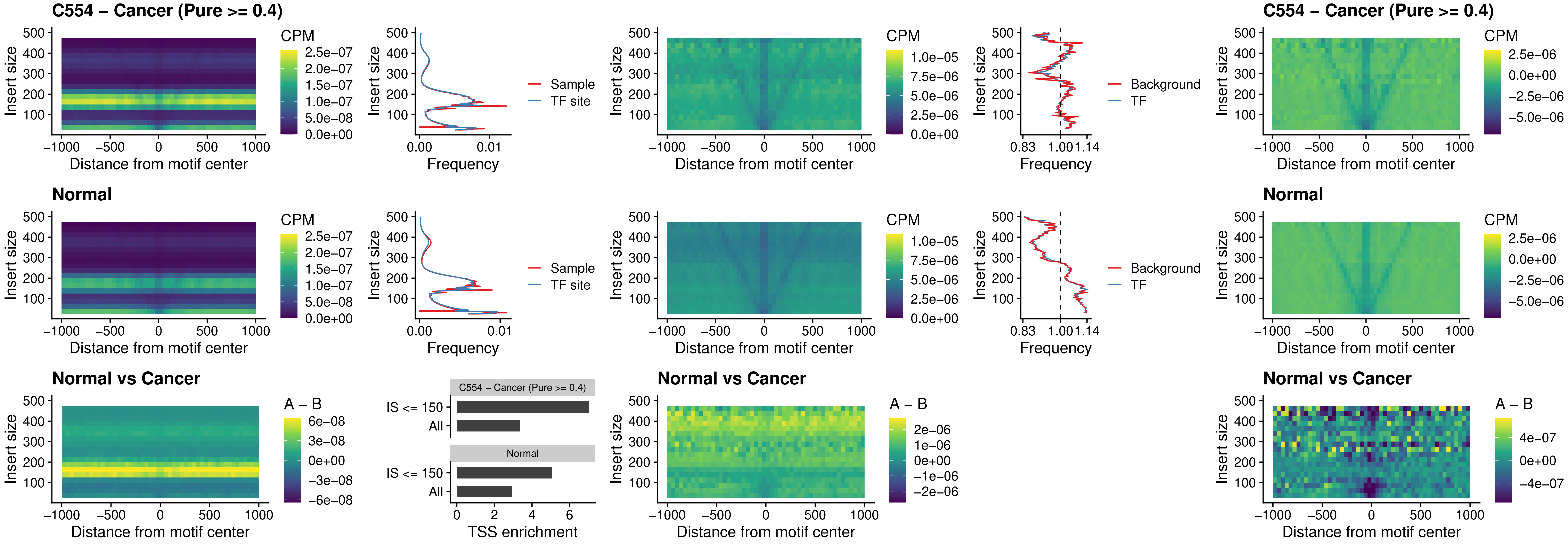

### VAX1_uc_pTSS_nPEAK

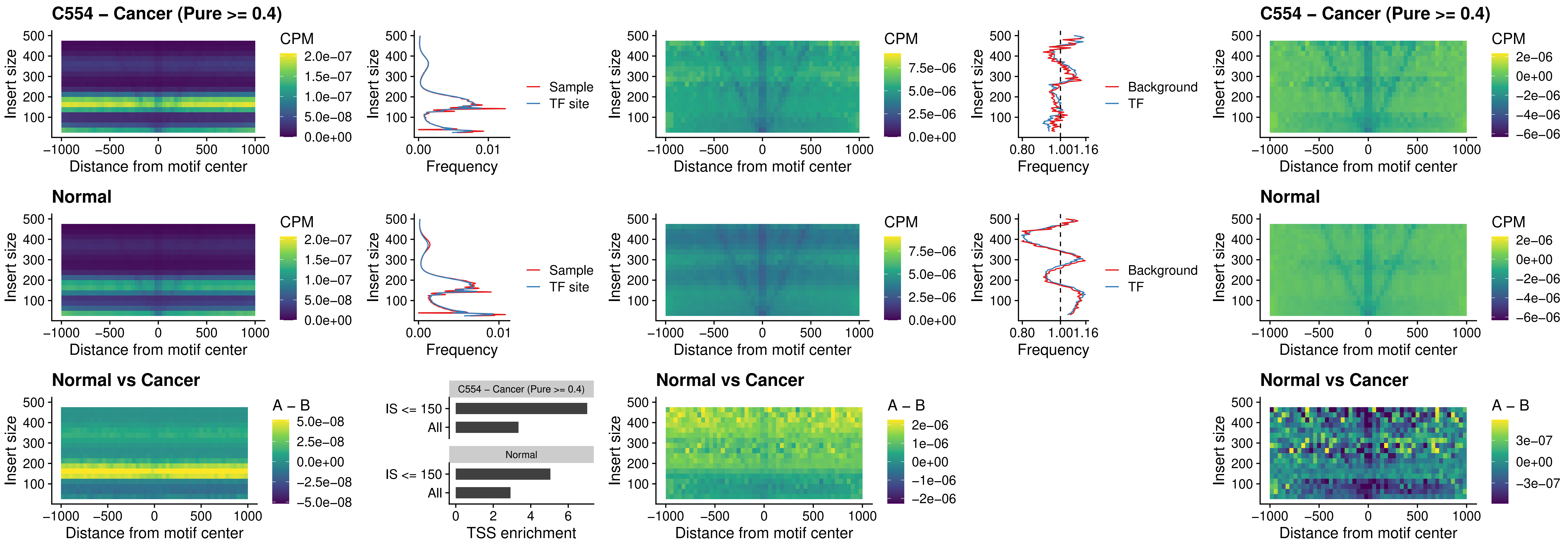

#### DLX3_uc_pTSS_nPEAK

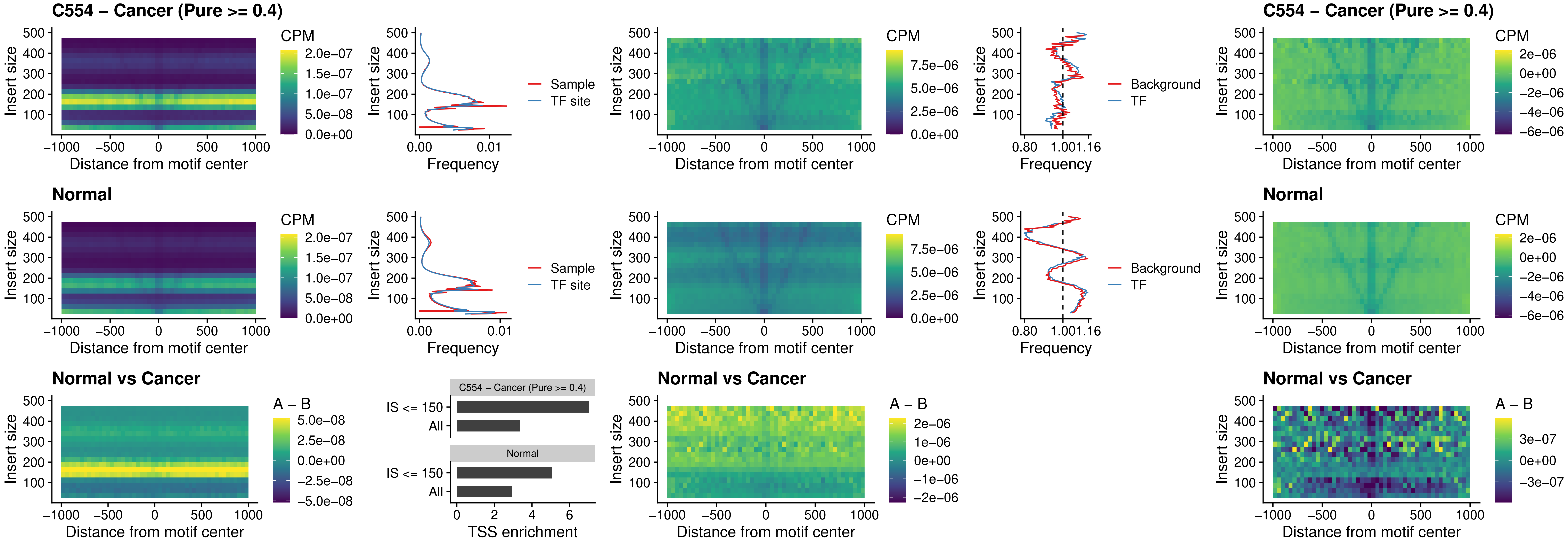

### HOXD3_uc_pTSS_nPEAK

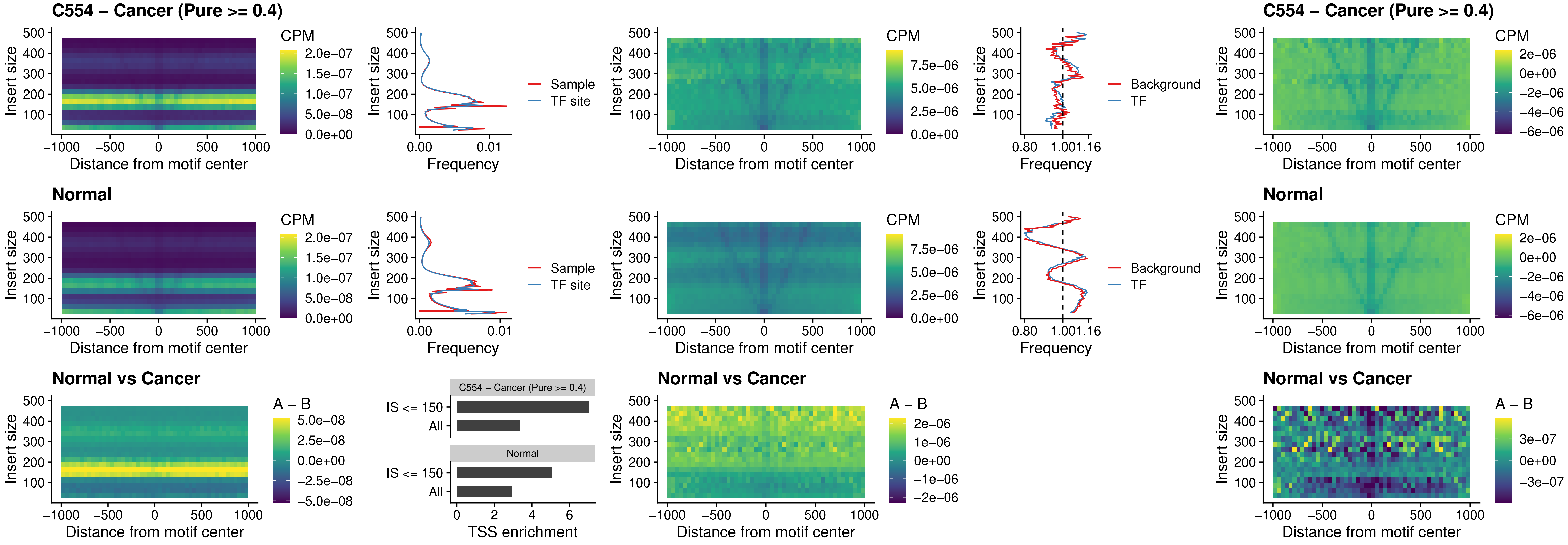

### PDX1_uc_dTSS_nPEAK

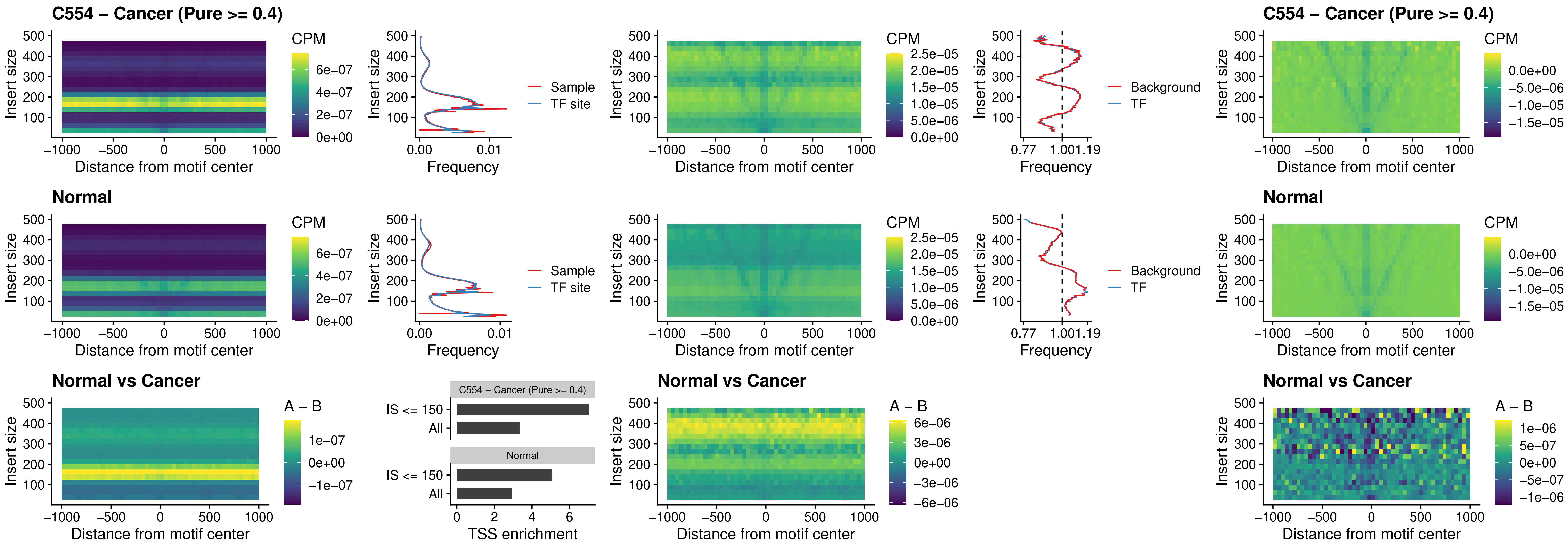

### VAX2_uc_dTSS_nPEAK

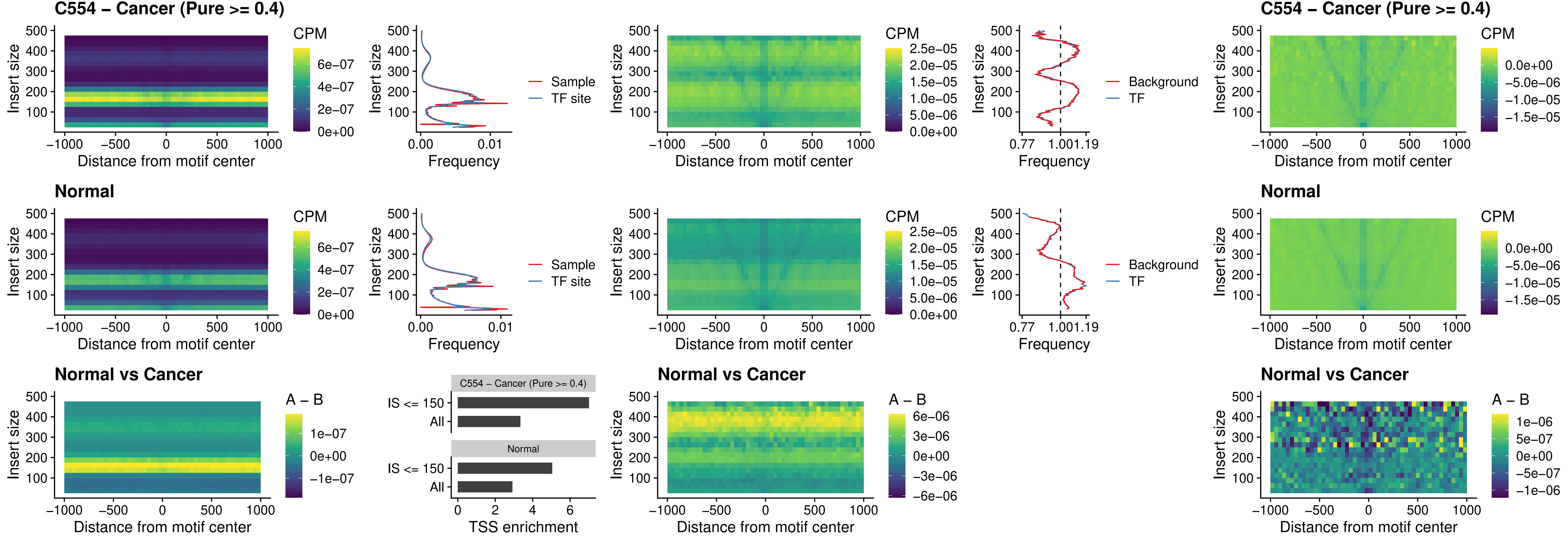

### SOX5_uc_dTSS_nPEAK

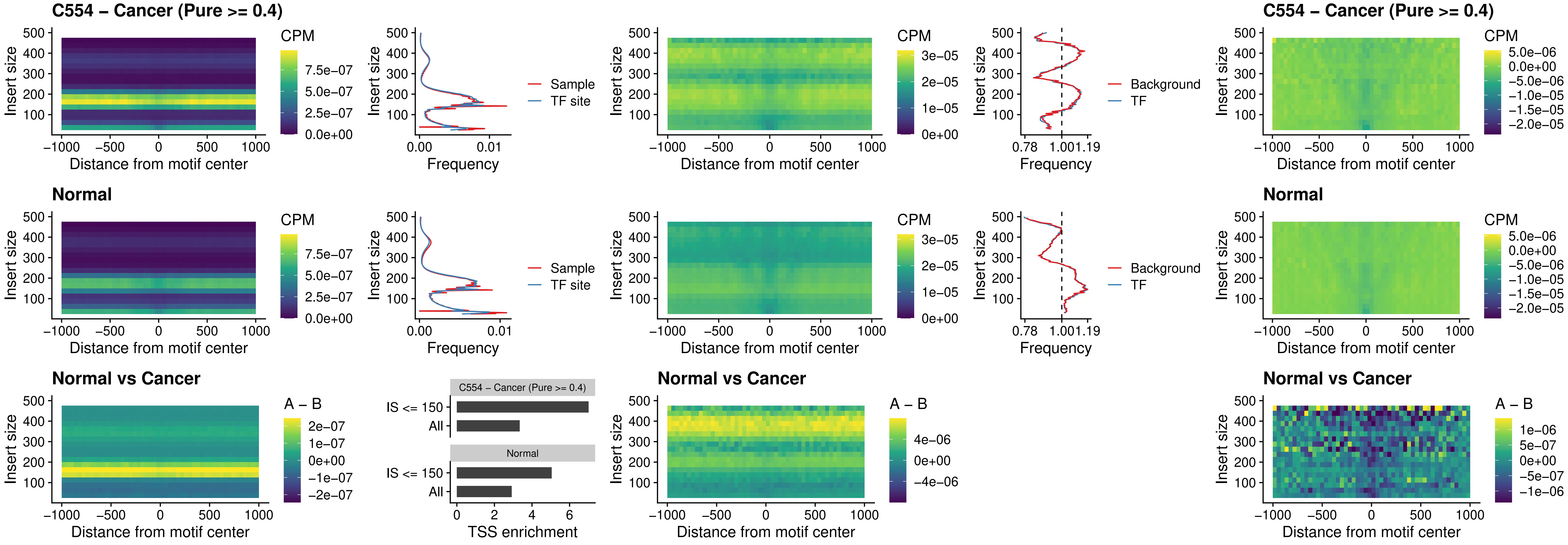

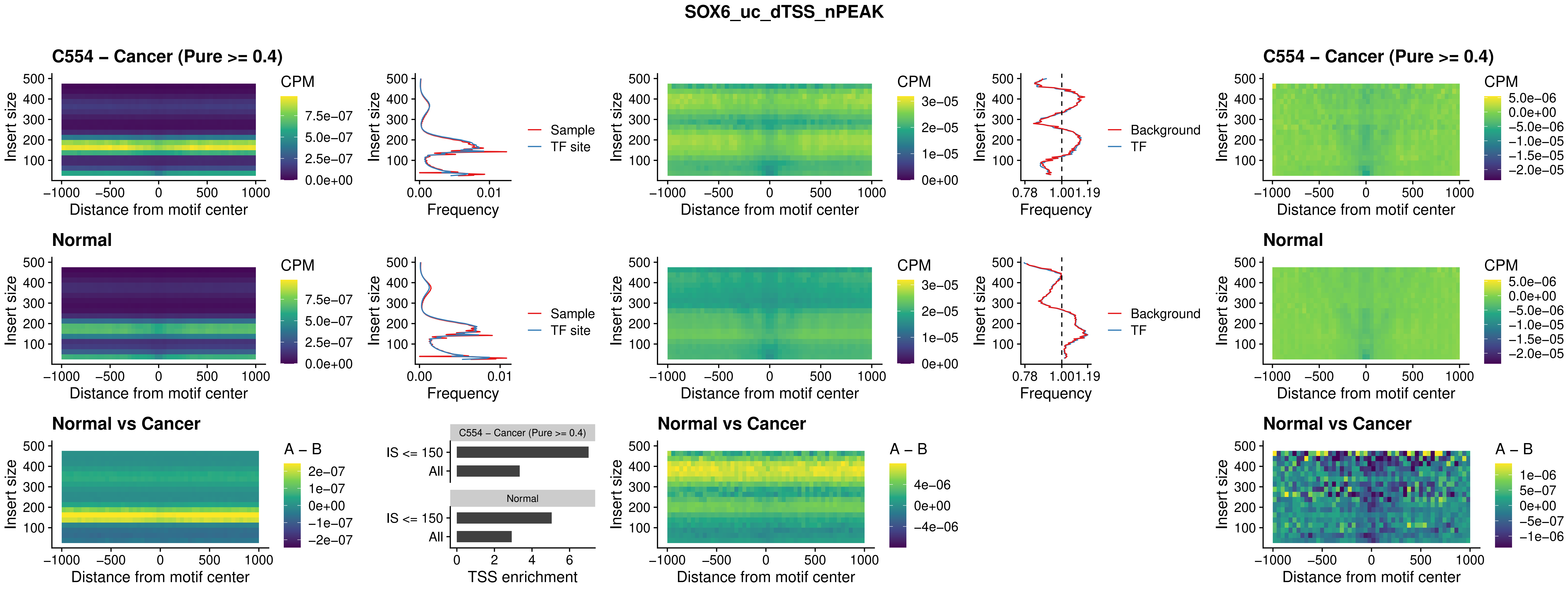
SOX6_uc_dTSS_nPEAK
C554 – Cancer (Pure >= 0.4)
Normal
Normal vs Cancer
Insert size
Distance from motif center
Frequency
CPM
A – B
Sample
TF site
Background
TF
TSS enrichment
IS <= 150
All
C554 – Cancer (Pure >= 0.4)
Normal

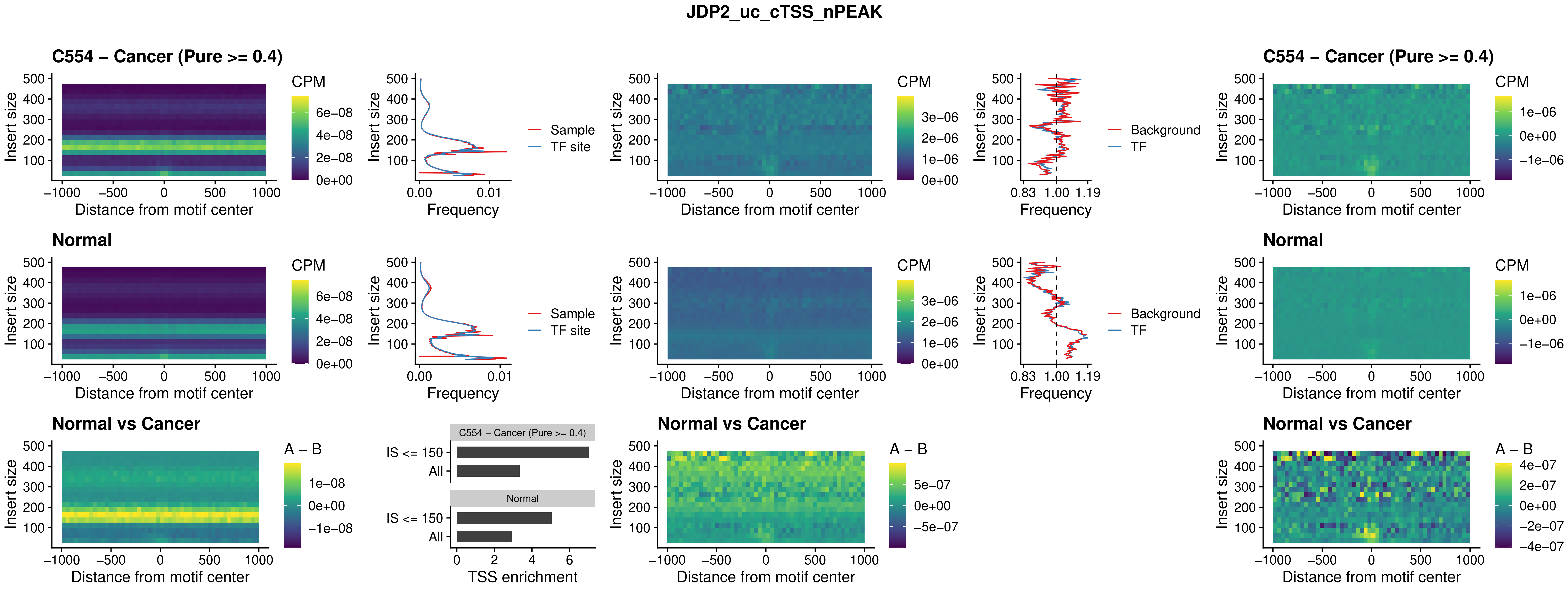
JDP2_uc_cTSS_nPEAK
C554 – Cancer (Pure >= 0.4)
Normal
Normal vs Cancer
Insert size
Distance from motif center
Frequency
CPM
A – B
Sample
TF site
Background
TF
C554 – Cancer (Pure >= 0.4)
Normal
IS <= 150
All
TSS enrichment

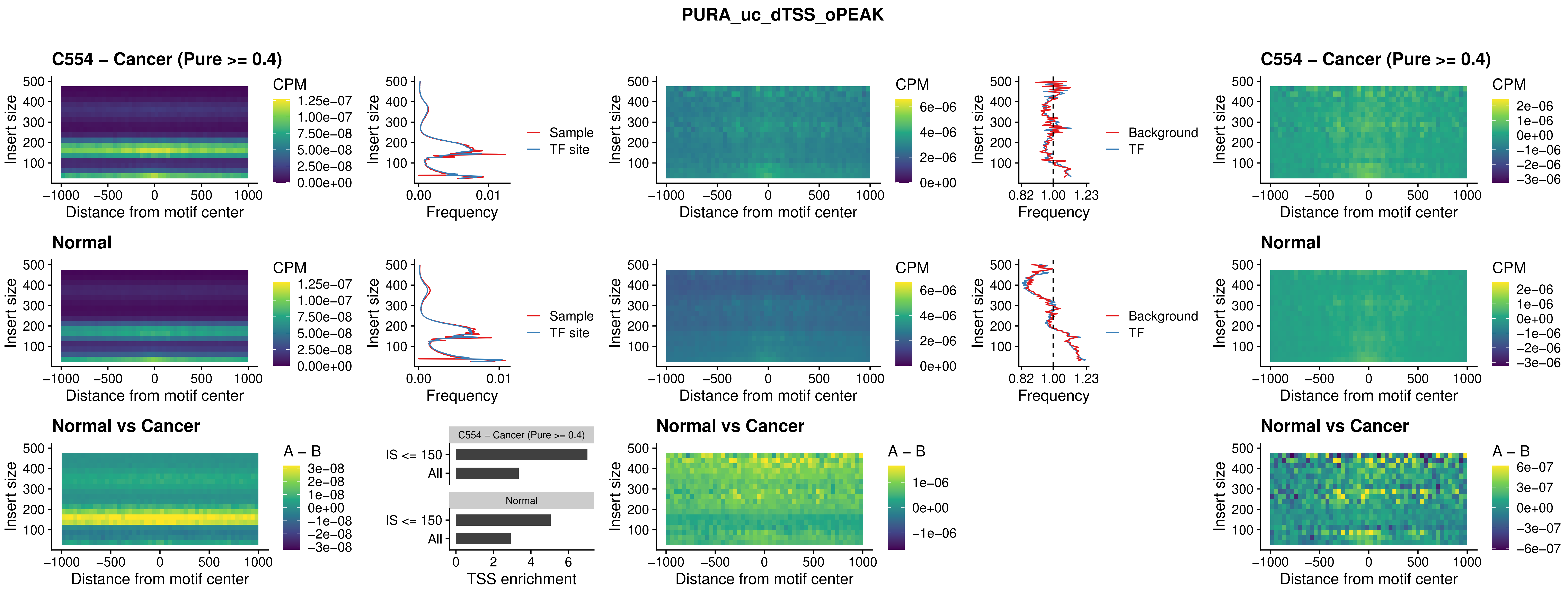

PURA_uc_dTSS_oPEAK
C554 – Cancer (Pure >= 0.4)
Insert size
Distance from motif center
CPM
Frequency
Sample
TF site
Background
TF
Normal
Normal vs Cancer
A – B
C554 – Cancer (Pure >= 0.4)
Normal
IS <= 150
All
TSS enrichment

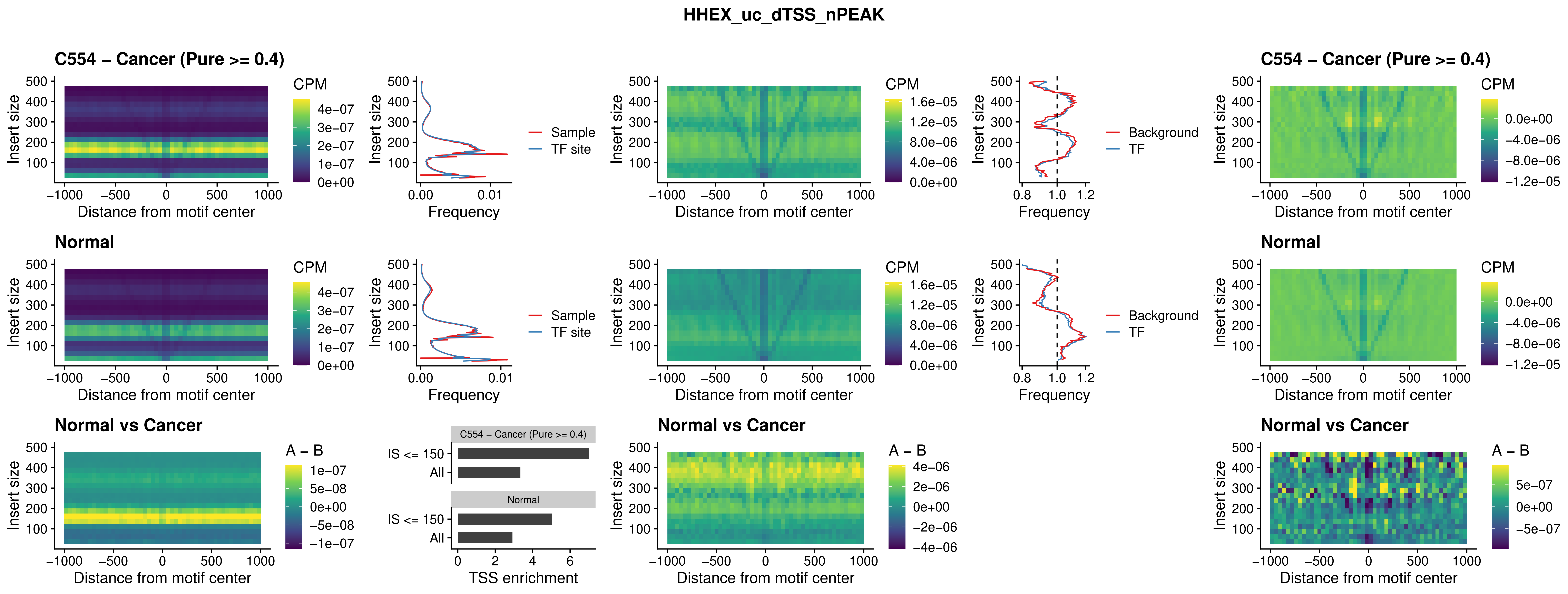
HHEX_uc_dTSS_nPEAK
C554 – Cancer (Pure >= 0.4)
Normal
Normal vs Cancer
Insert size
Distance from motif center
Frequency
CPM
A – B
Sample
TF site
Background
TF
C554 – Cancer (Pure >= 0.4)
Normal
IS <= 150
All
TSS enrichment

### HMG20B_uc_dTSS_nPEAK

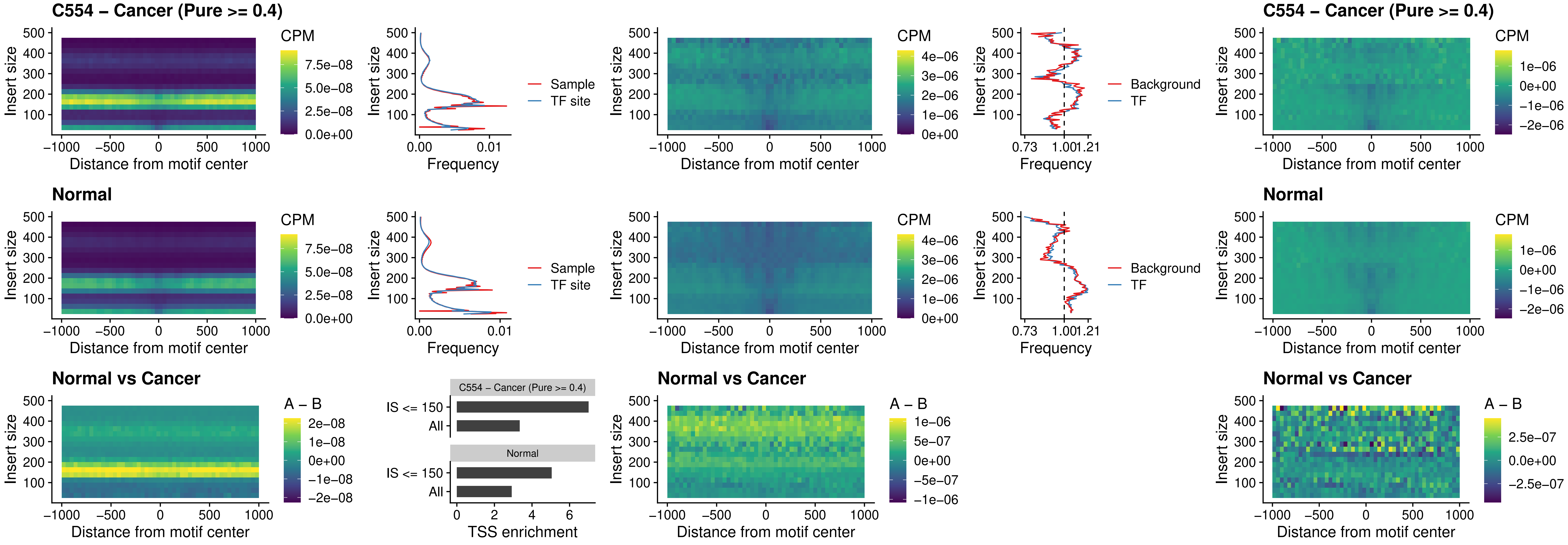

#### FOXB2_uc_dTSS_nPEAK

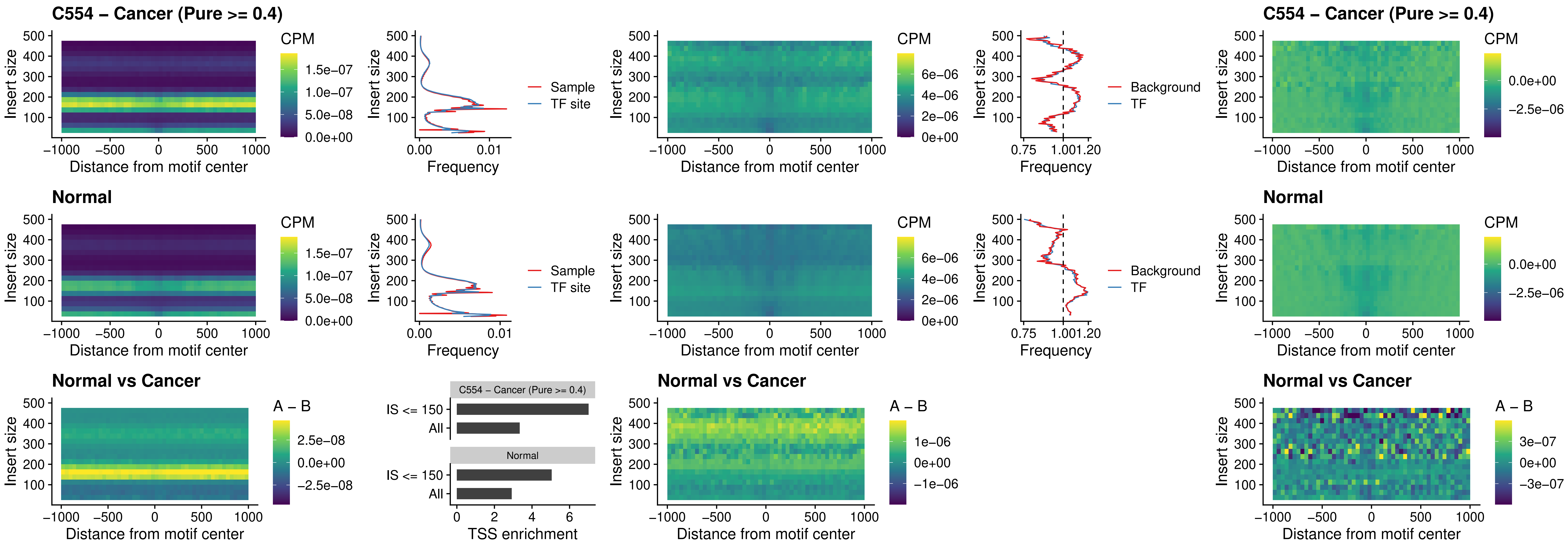

#### FOXL1_uc_cTSS_nPEAK

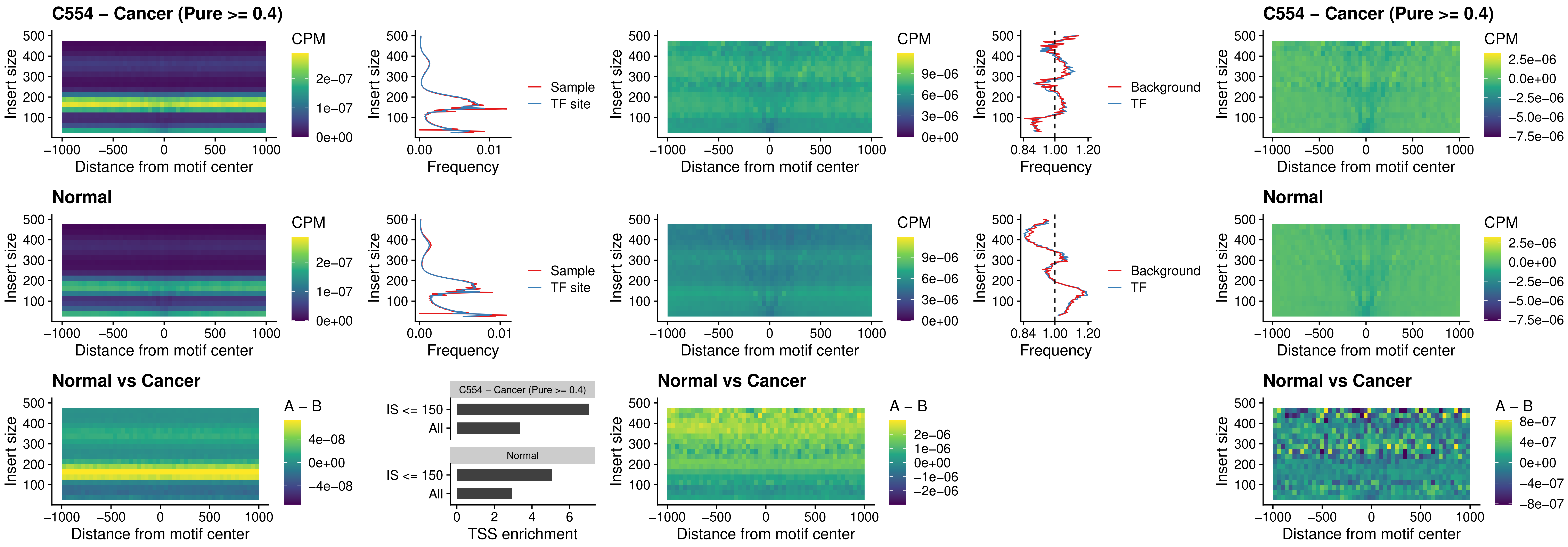

### PBX3_uc_pTSS_oPEAK

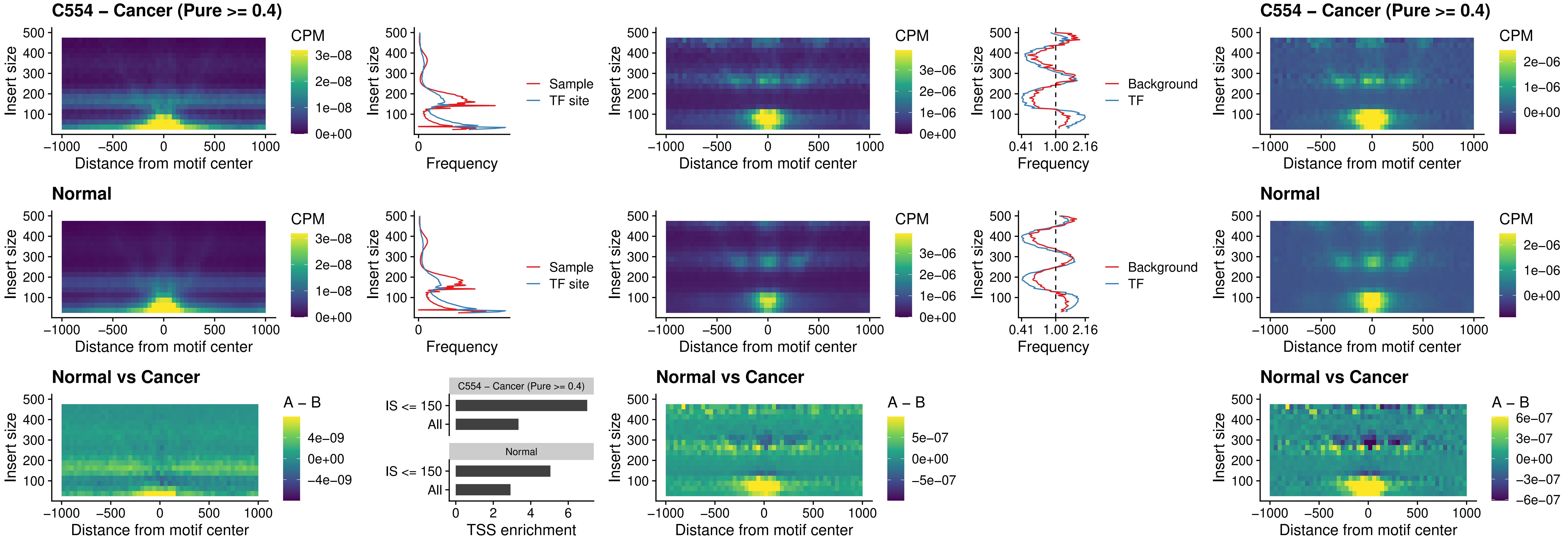

#### SNAI2_uc_cTSS_nPEAK

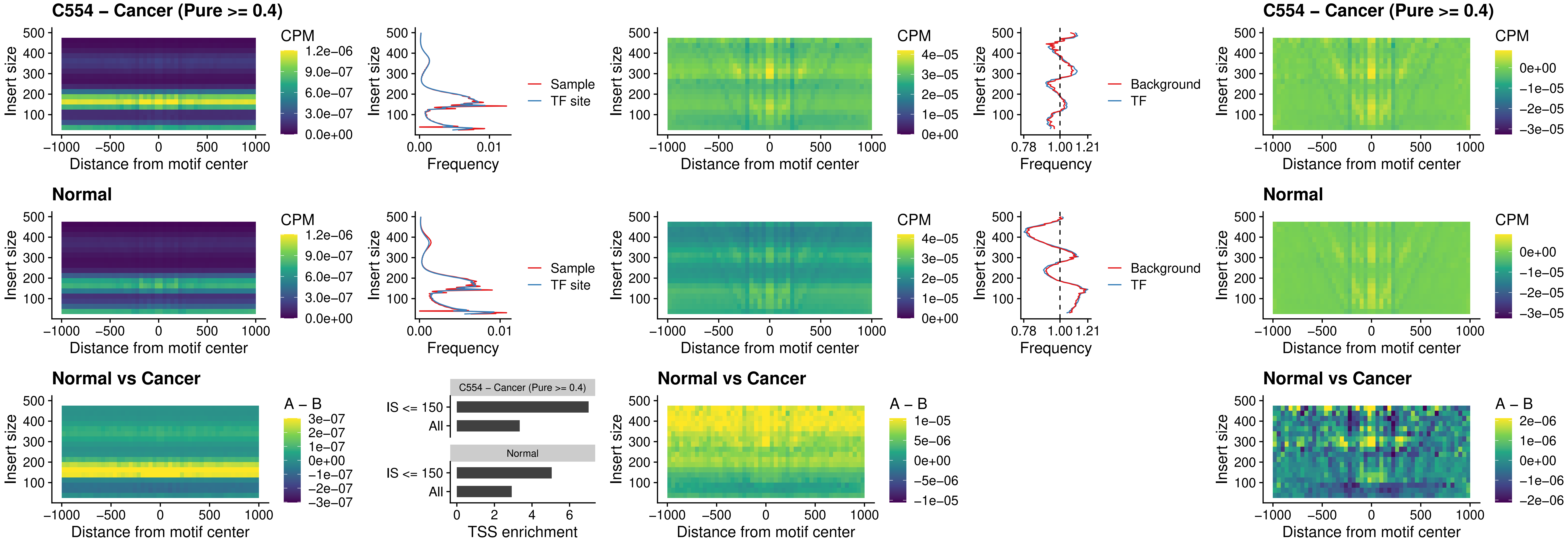

#### SNAI2_uc_dTSS_oPEAK

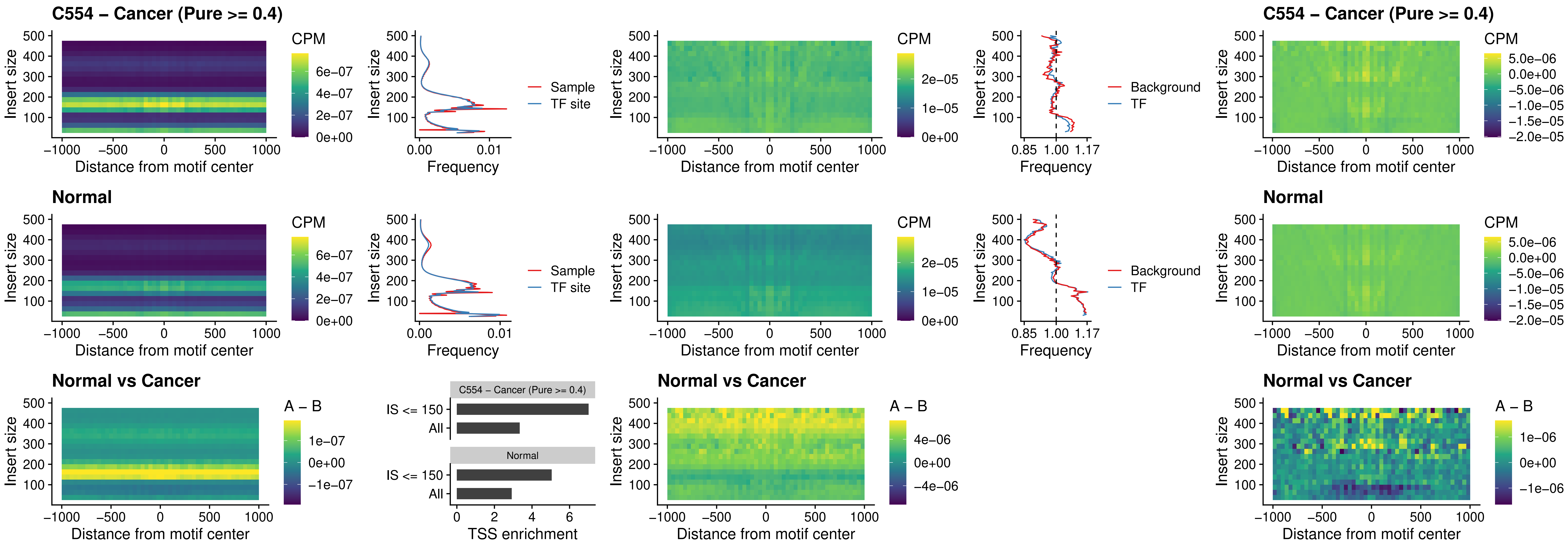

### SPI1_uc_pTSS_nPEAK

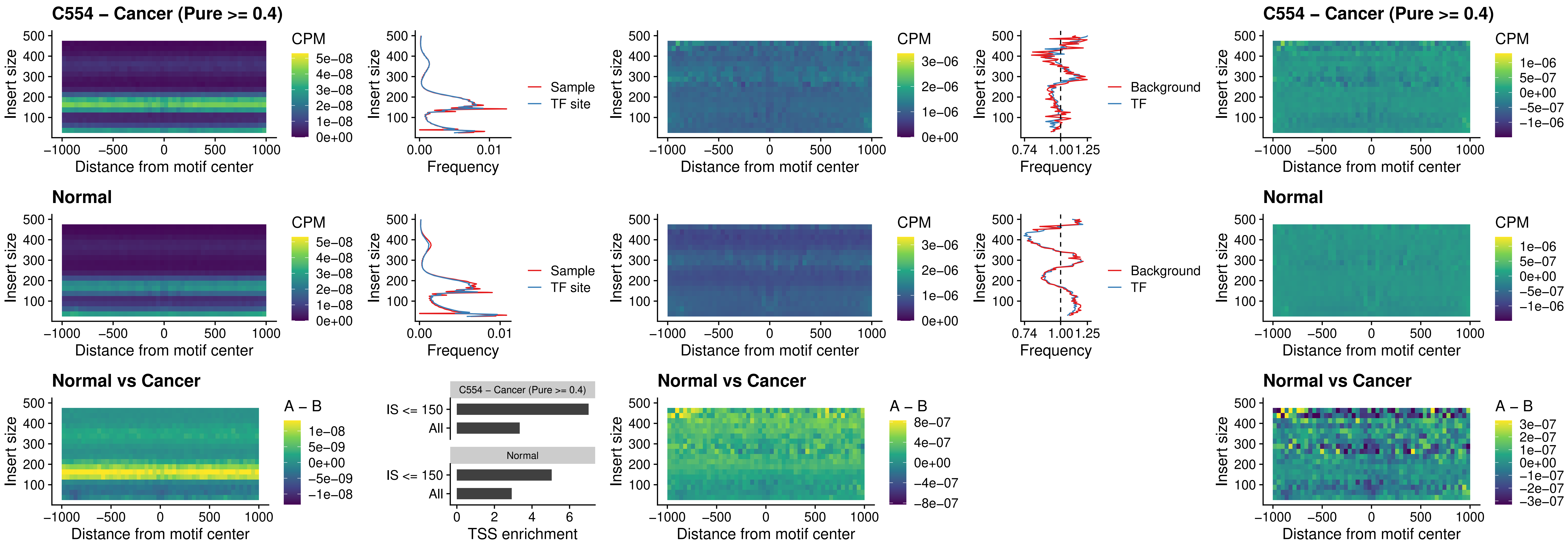

#### KLF16_uc_pTSS_nPEAK

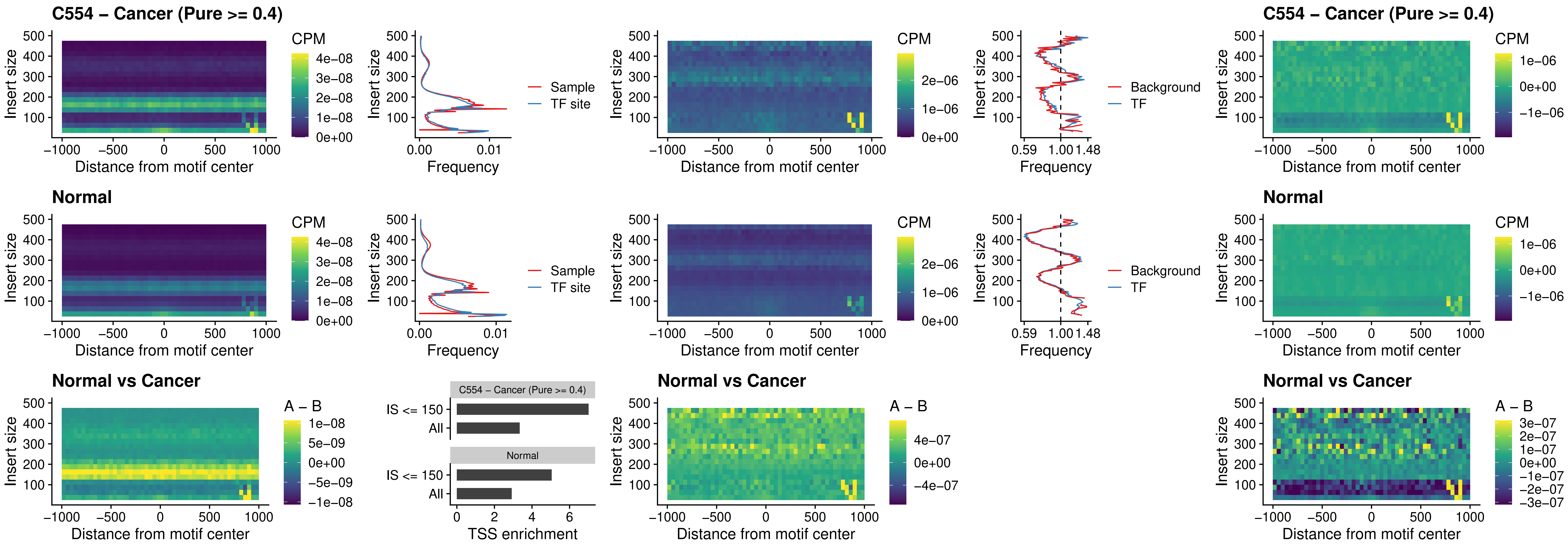

### SP3_uc_pTSS_nPEAK

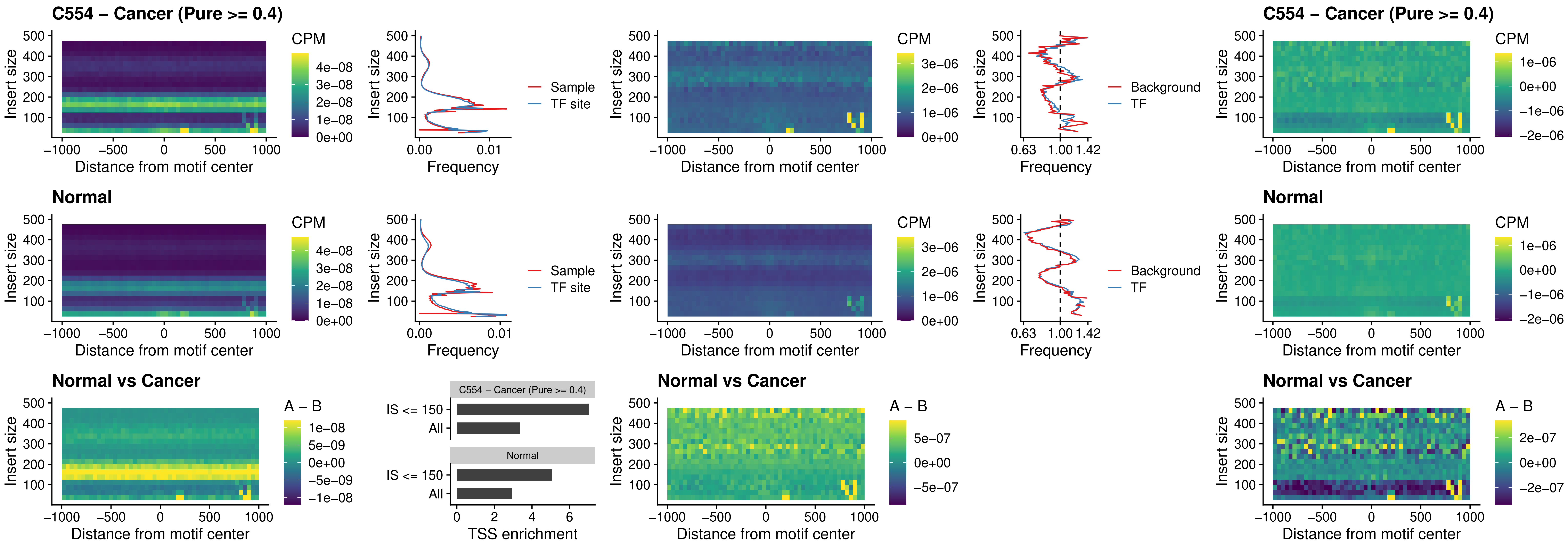

### NEUROD1_uc_dTSS_oPEAK

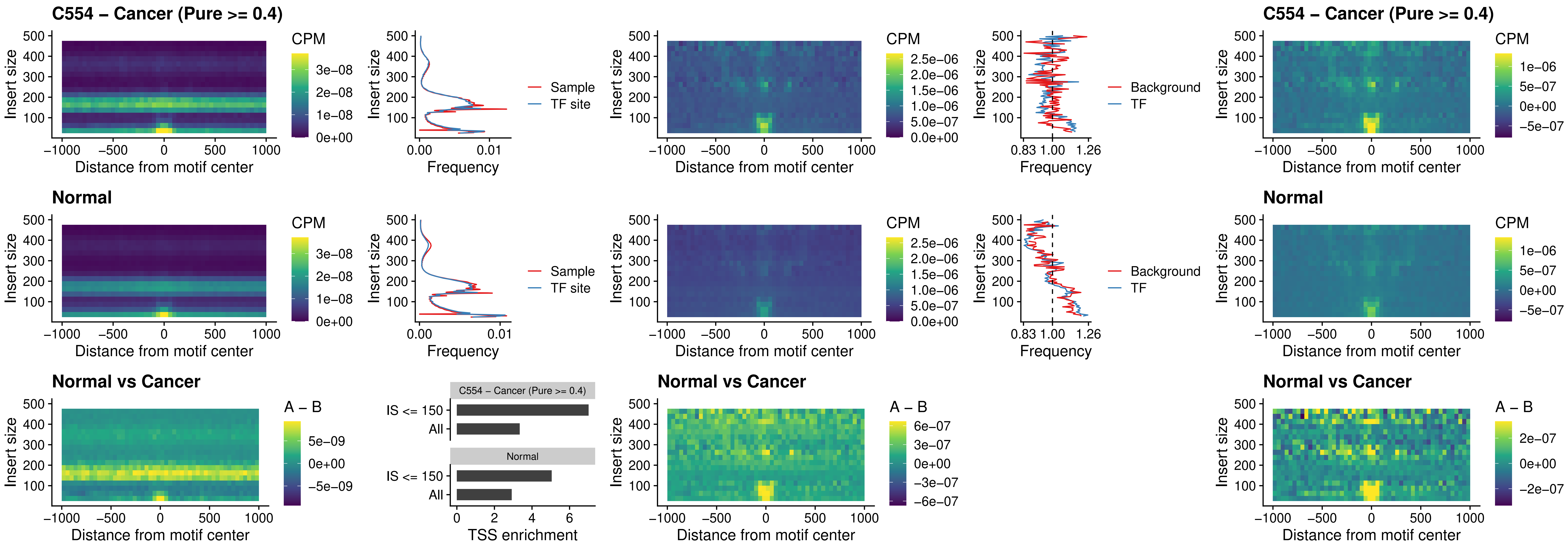

### CTCF_uc_dTSS_oPEAK

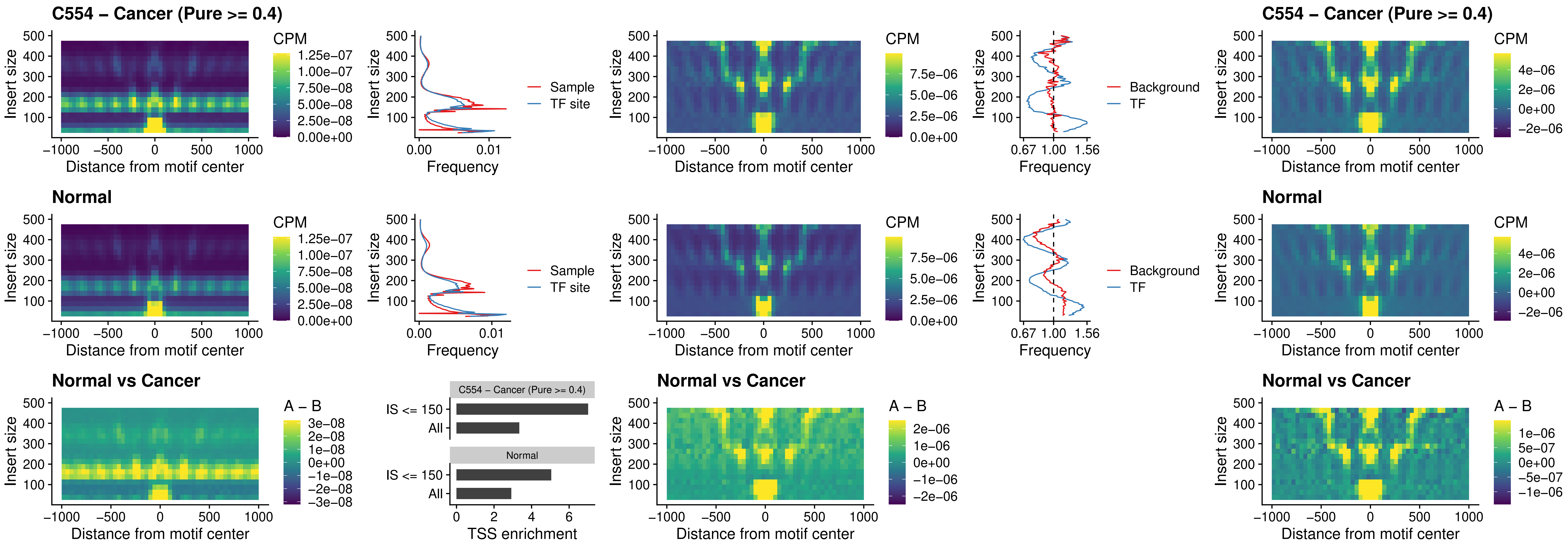

#### CTCF_uc_cTSS_oPEAK

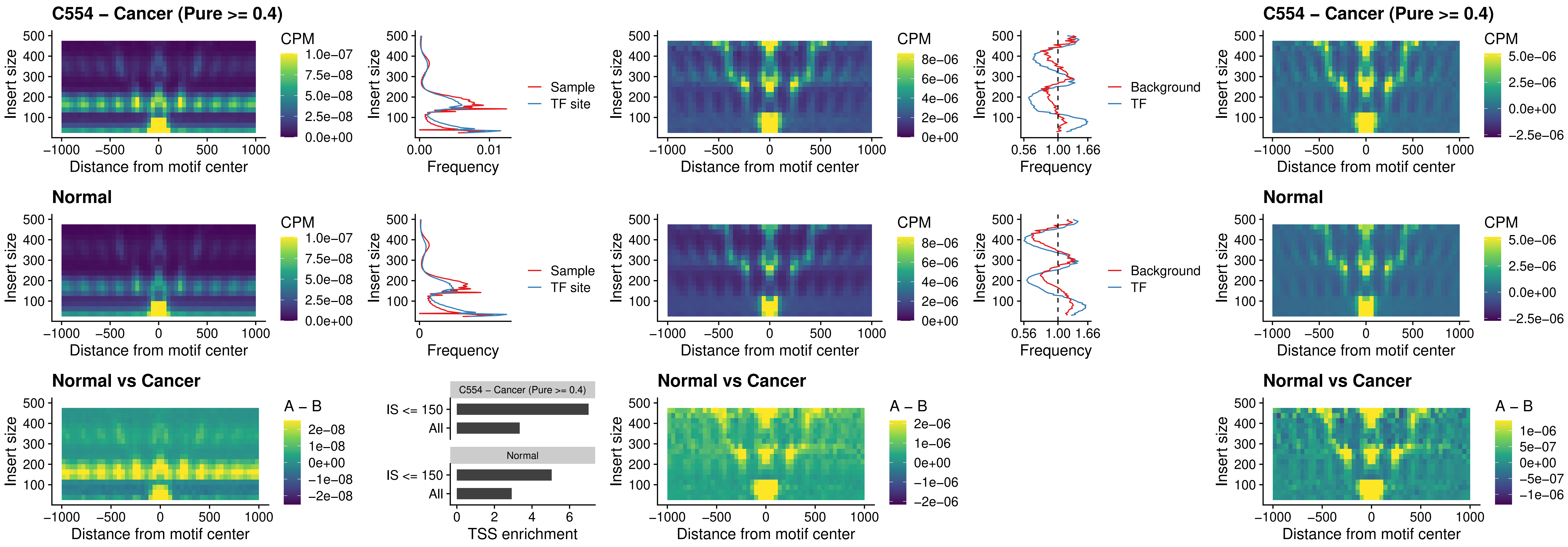

### CTCFL_uc_dTSS_oPEAK

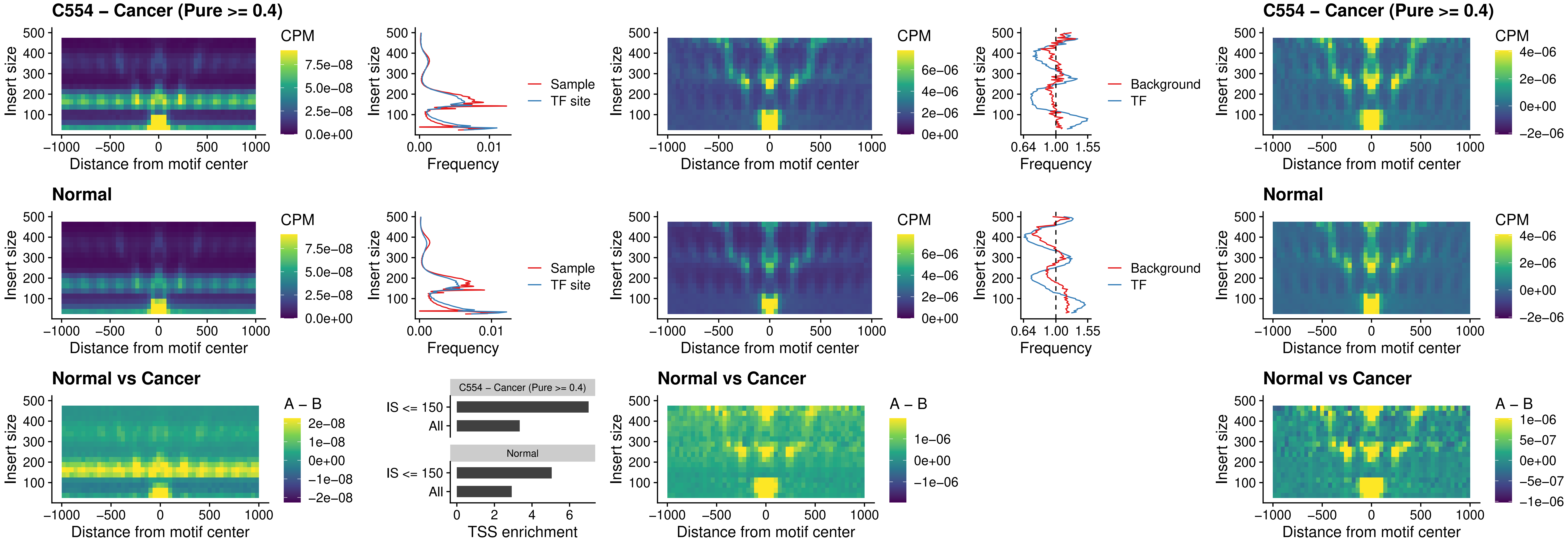

### ENCODE_CTCF_sites_COLON

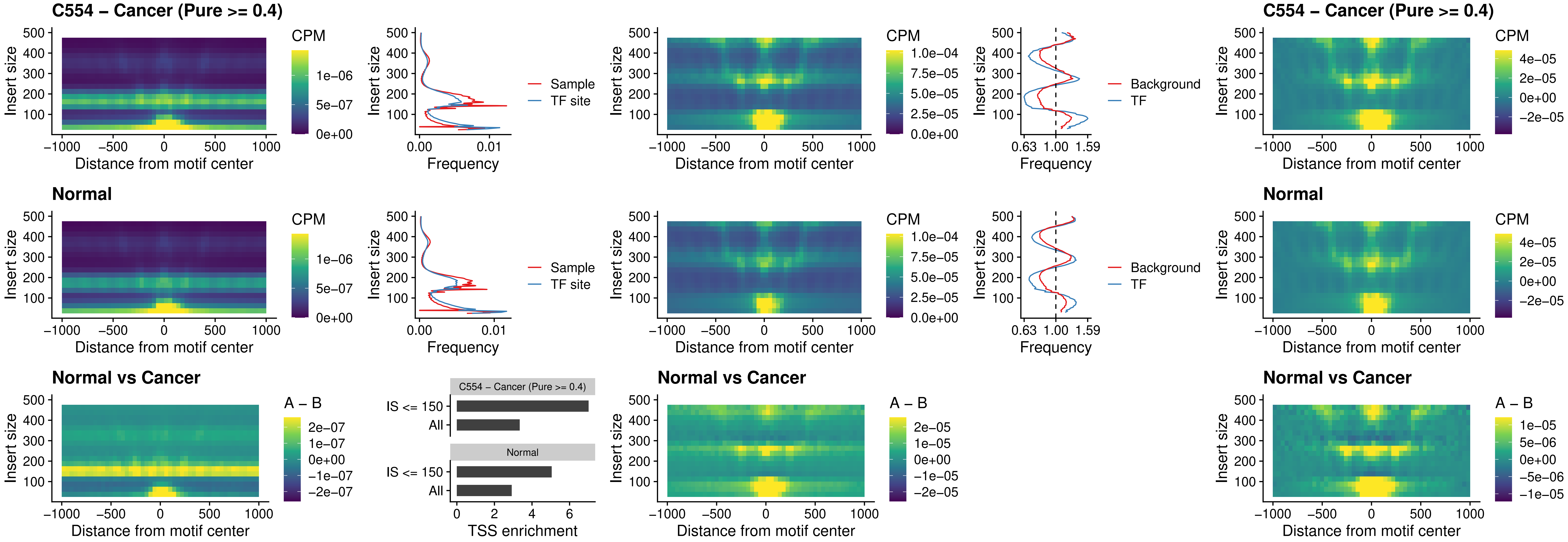

#### ENCODE_CTCF_sites_COLON_loop_ctcf

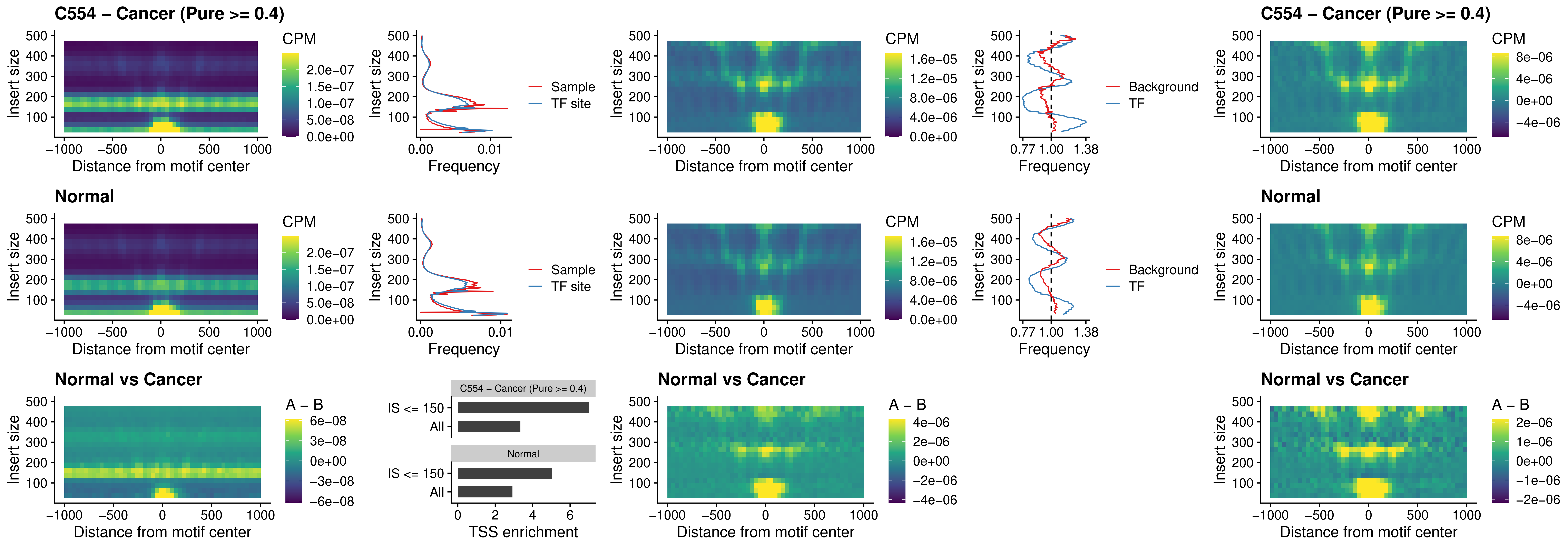

### ENCODE_CTCF_sites_COLON_intergenic_ctcf

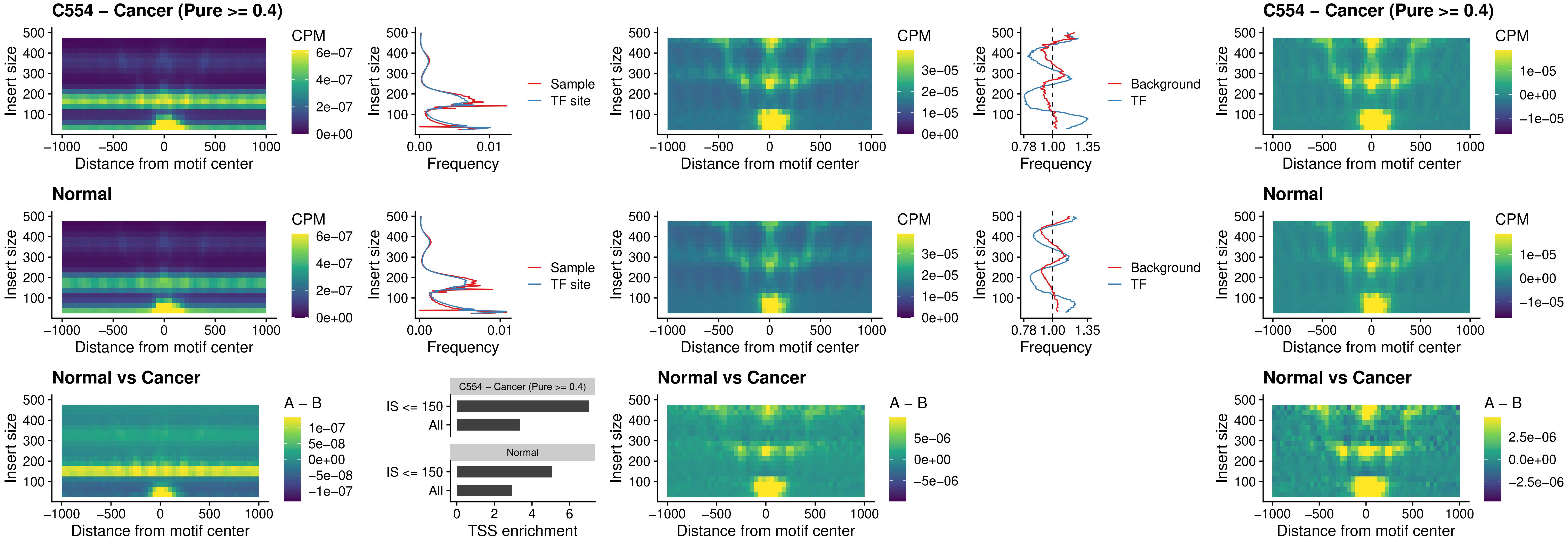

#### ENCODE_CTCF_sites_COLON_nonpromotor

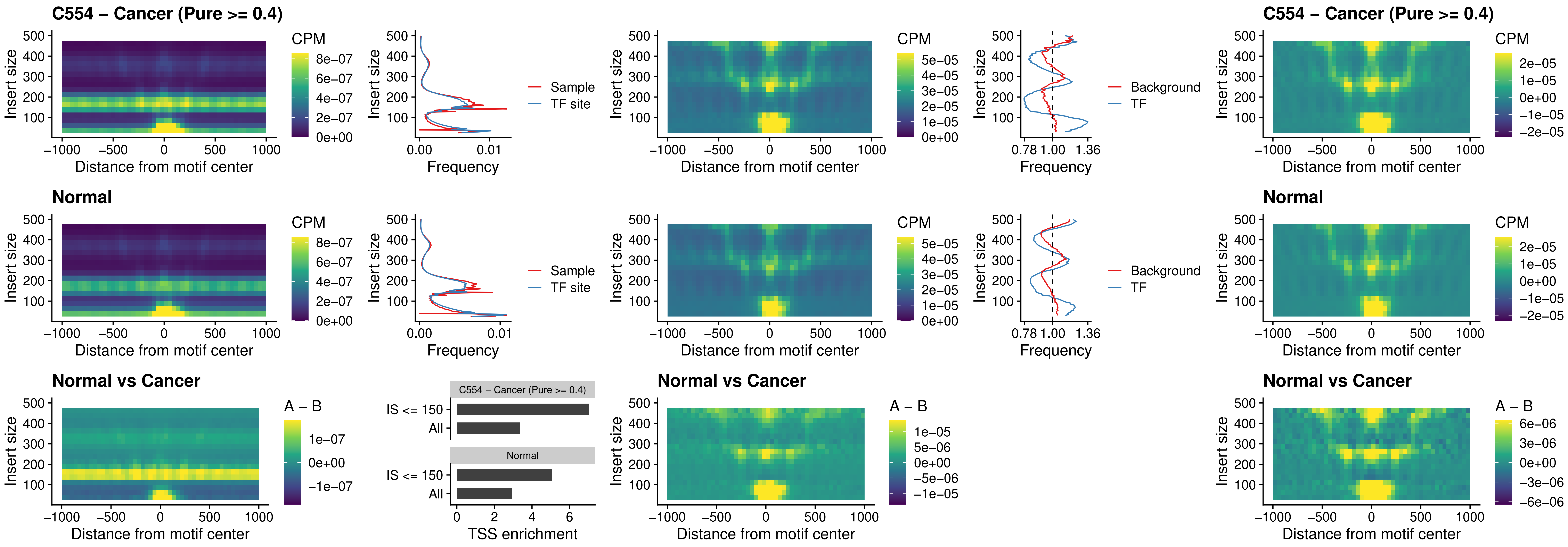

### CTCFL_uc_cTSS_oPEAK

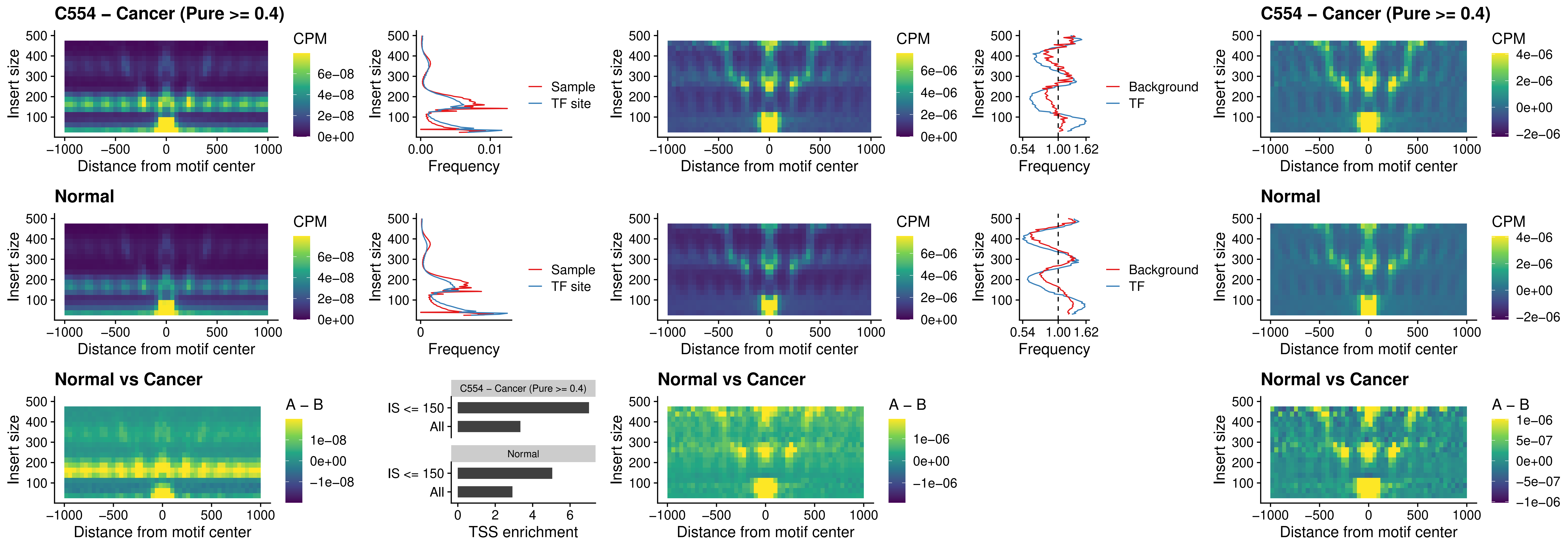

#### NEURODI_uc_cTSS_oPEAK

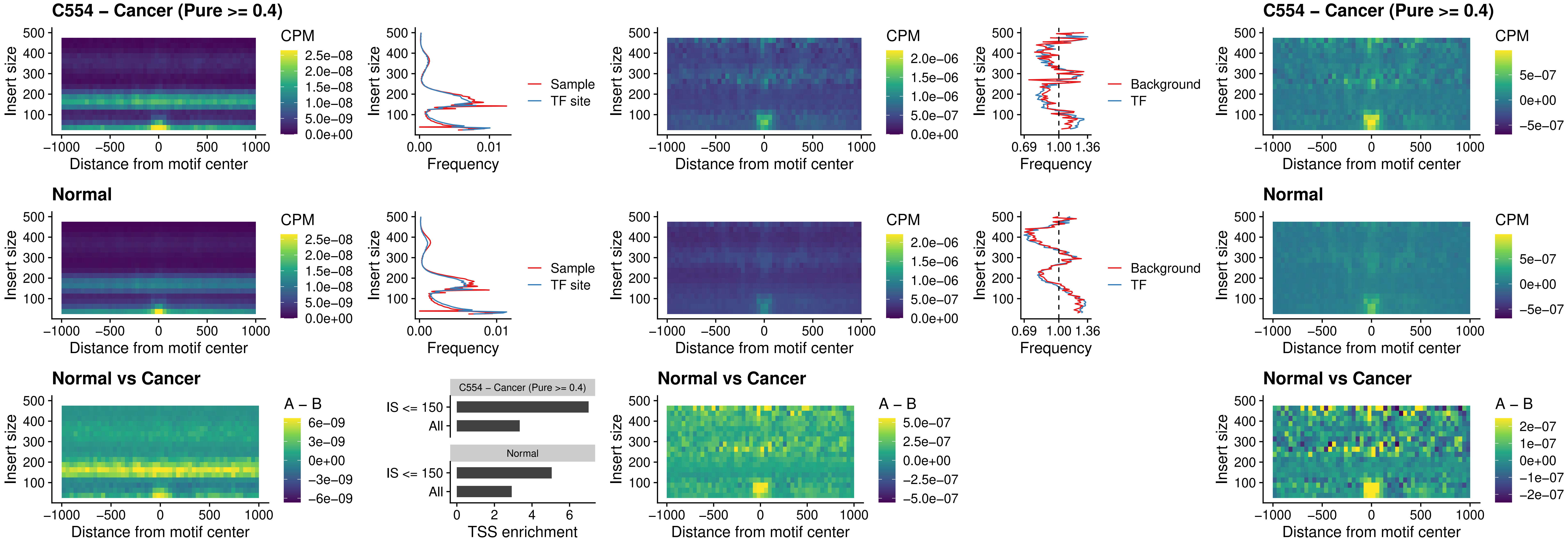

#### ENCODE_YY1_sites_COLON_nonpromotor

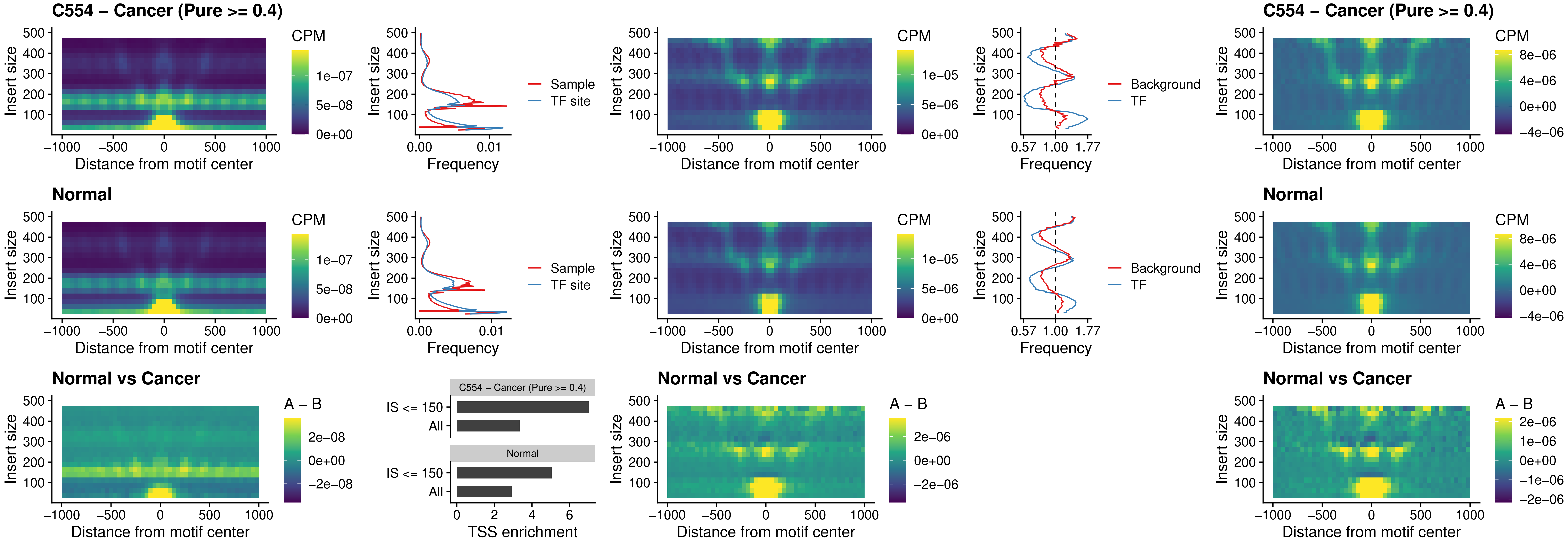

### CTCF_uc_pTSS_oPEAK

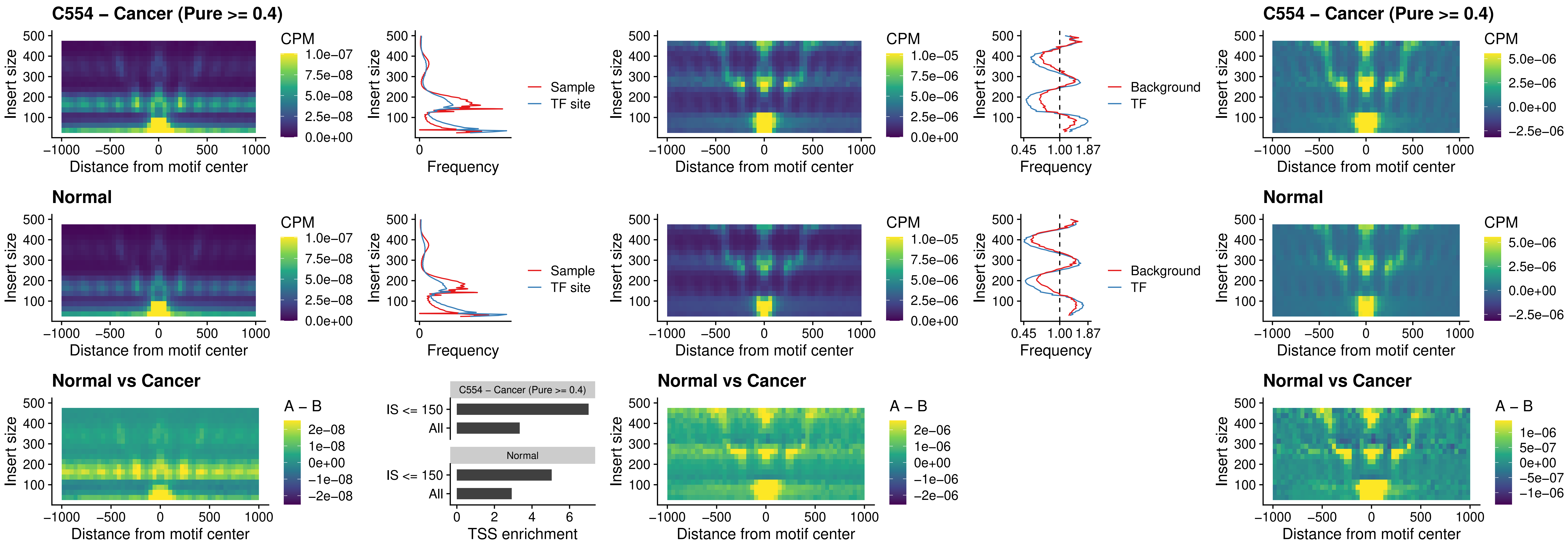

### ENCODE_CTCF_sites_COLON_promotor

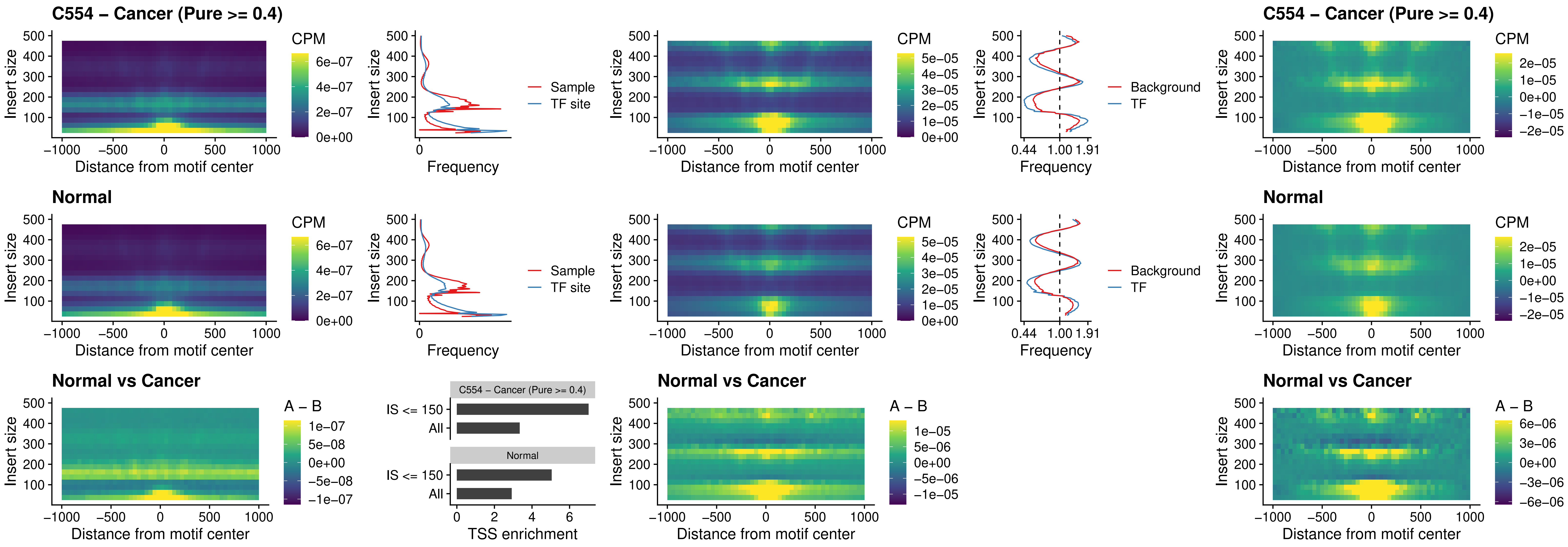

#### ENCODE_CTCF_sites_COLON_promotor_ctcf

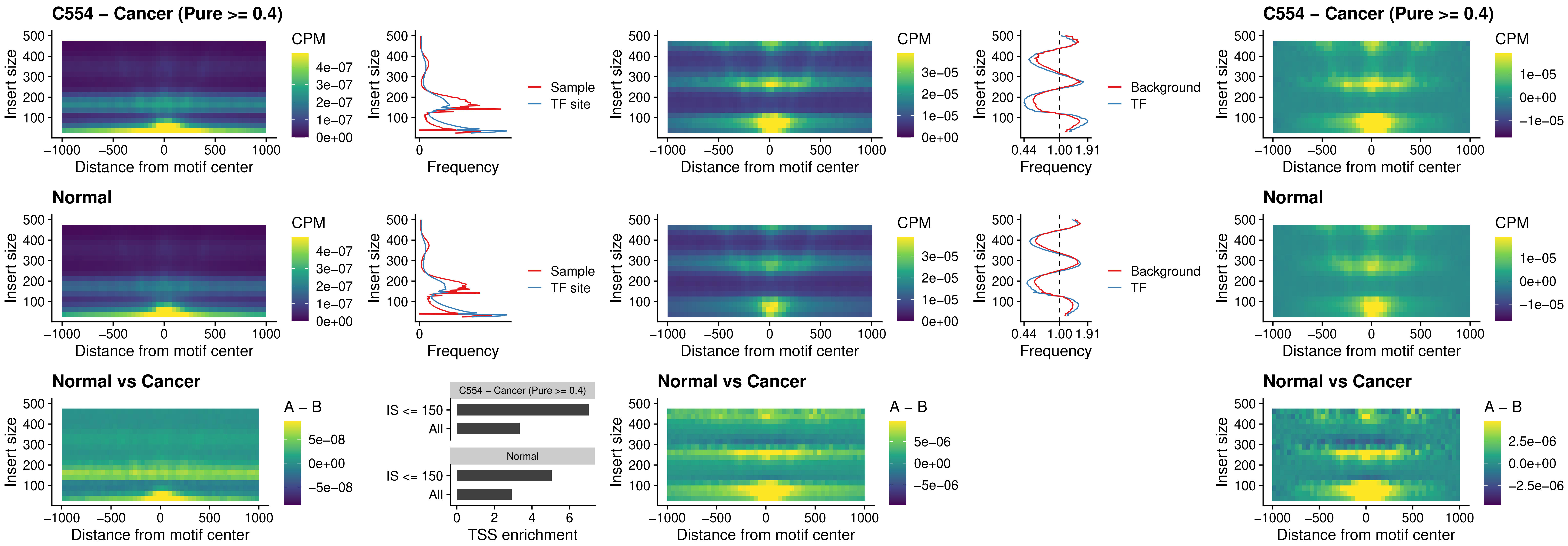

### SPIC_uc_pTSS_oPEAK

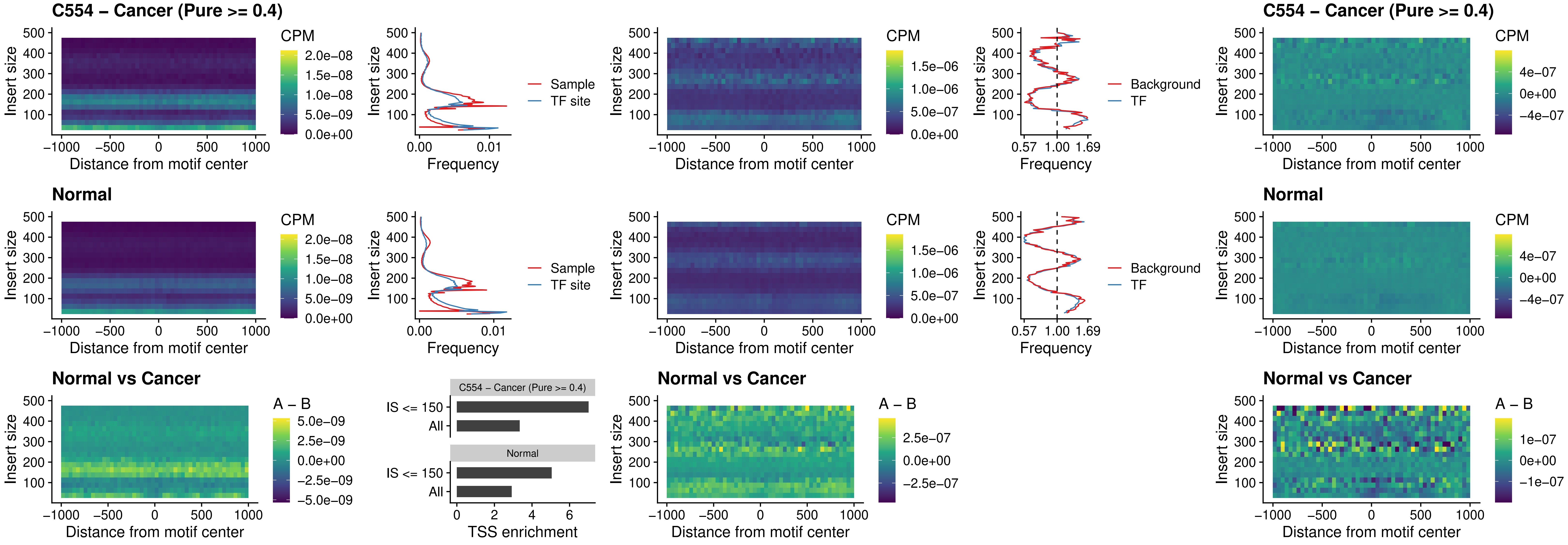

#### SPI1_uc_dTSS_nPEAK

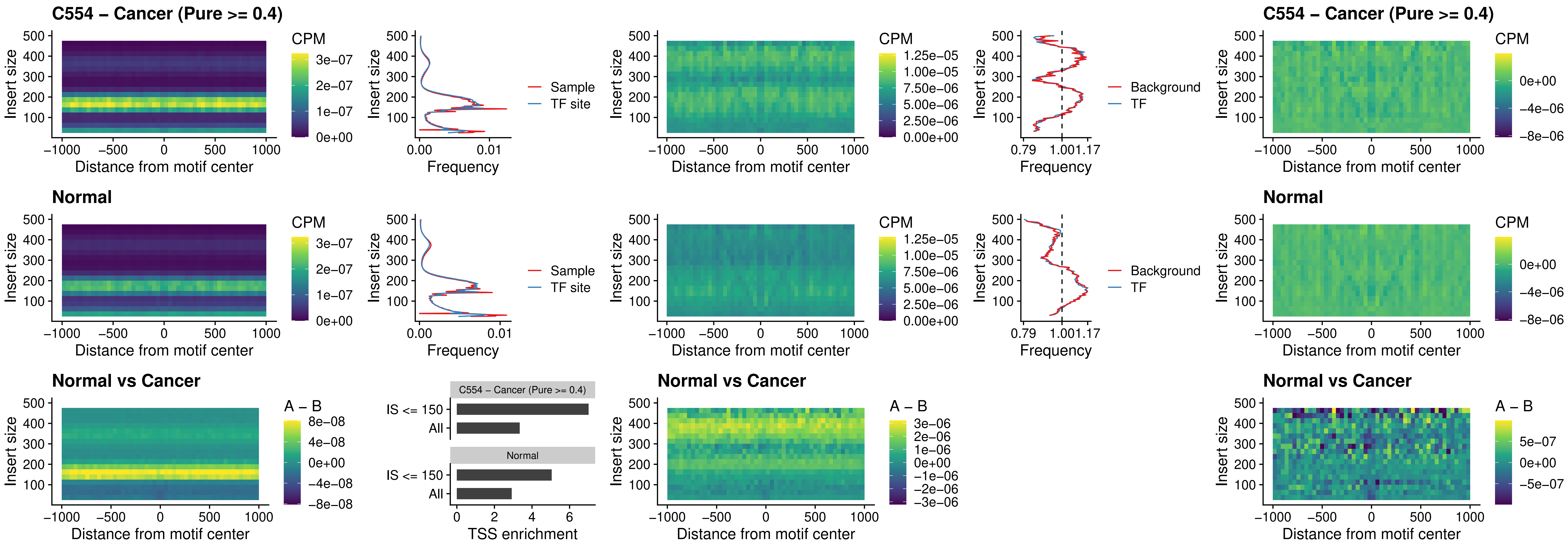

#### SPI1_uc_cTSS_nPEAK

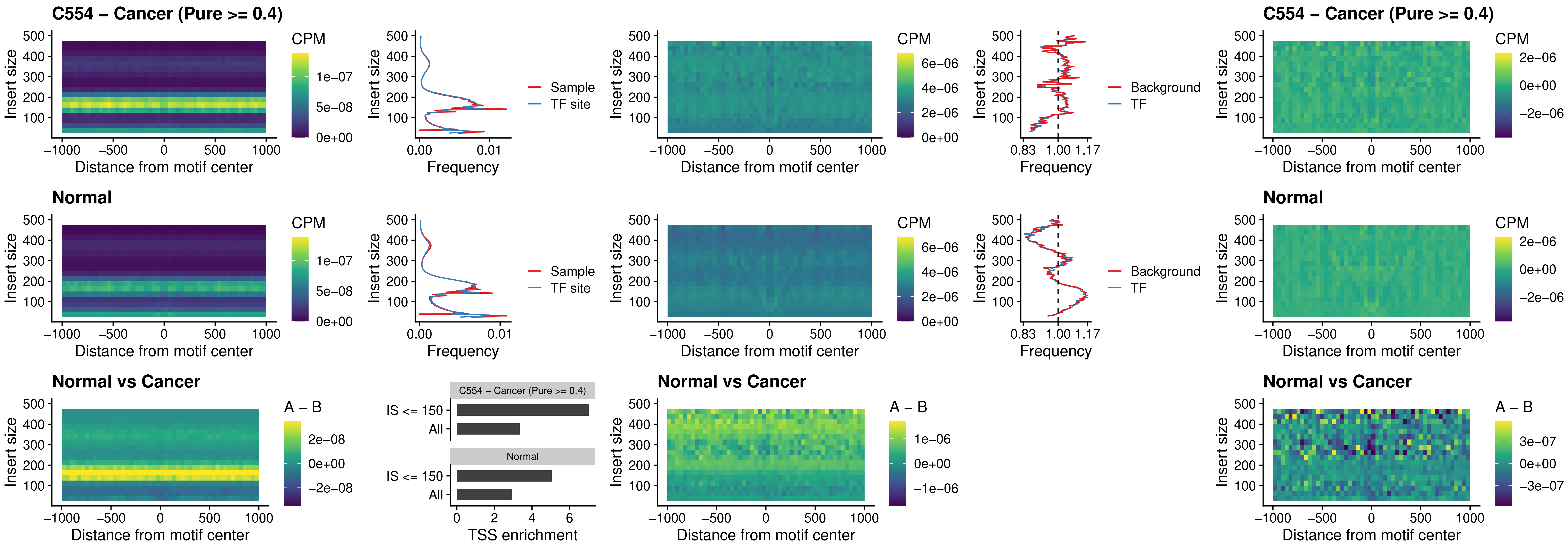

### BCL11A_uc_pTSS_nPEAK

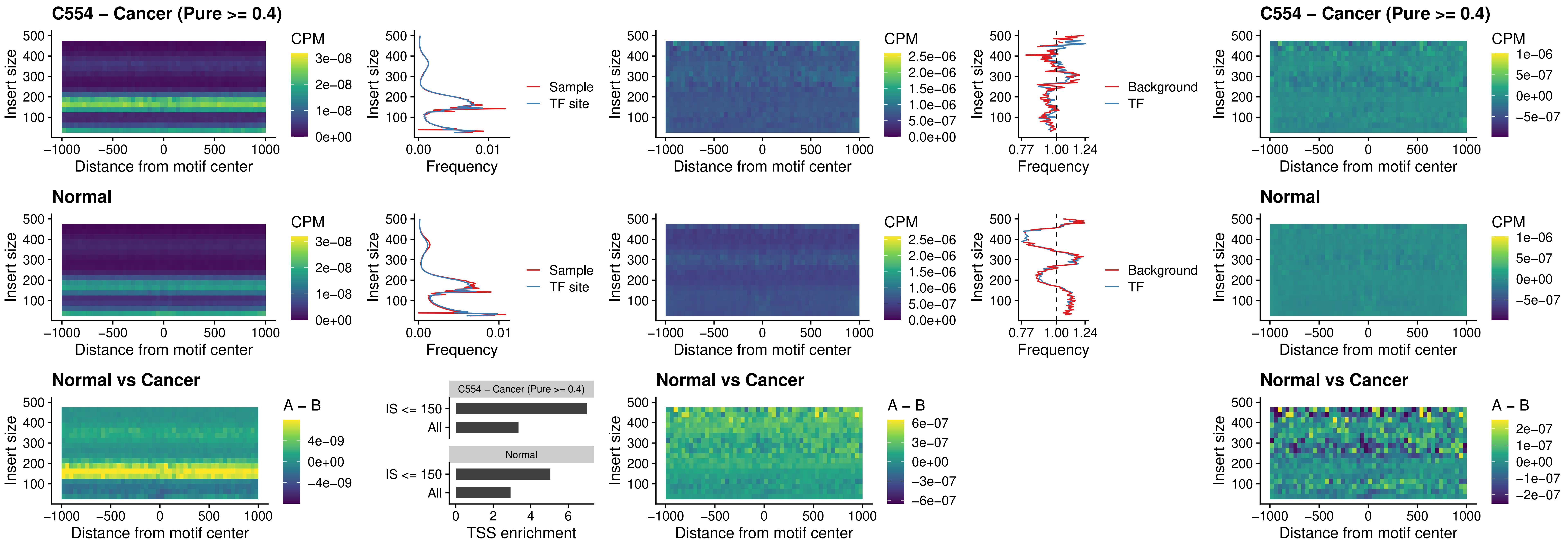

#### BCL11B_uc_pTSS_nPEAK

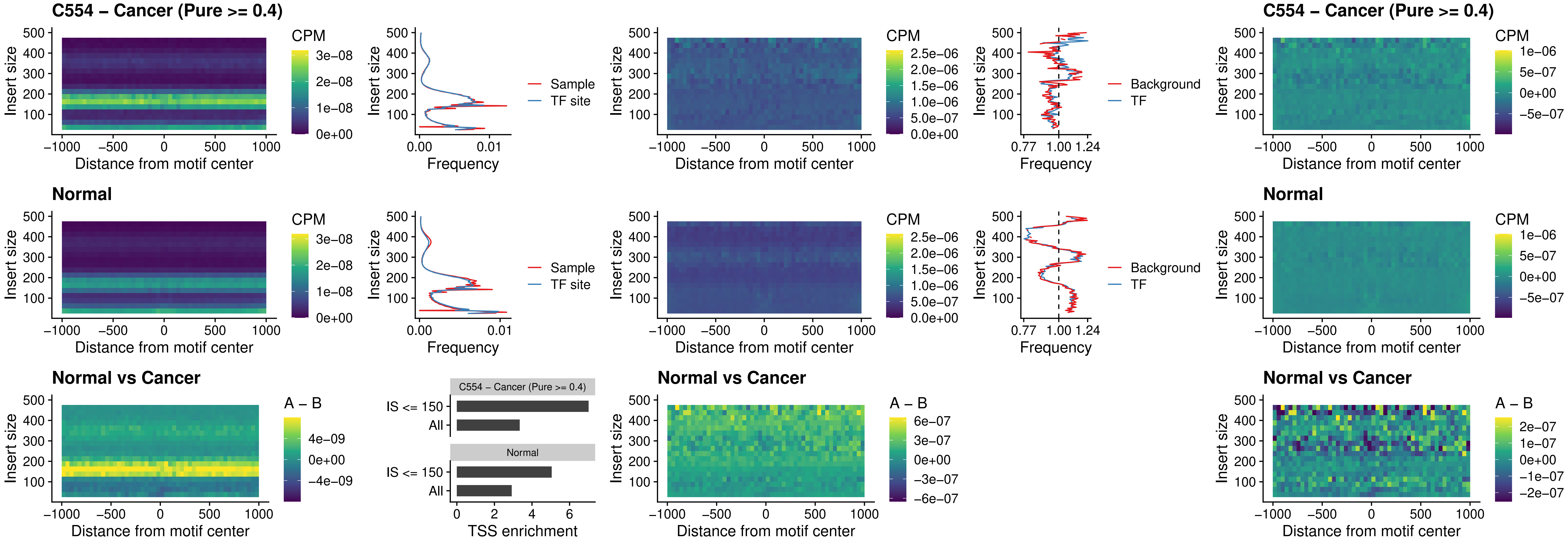

### IRF9_uc_dTSS_oPEAK

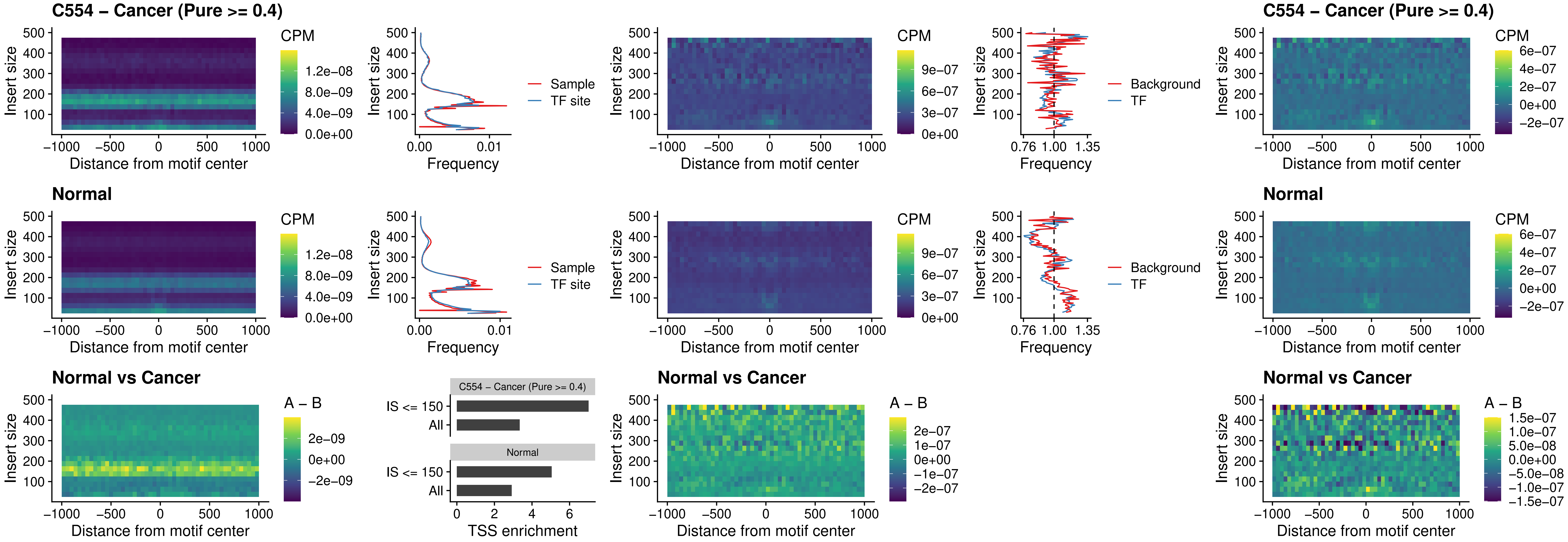

### IRF9_uc_cTSS_oPEAK

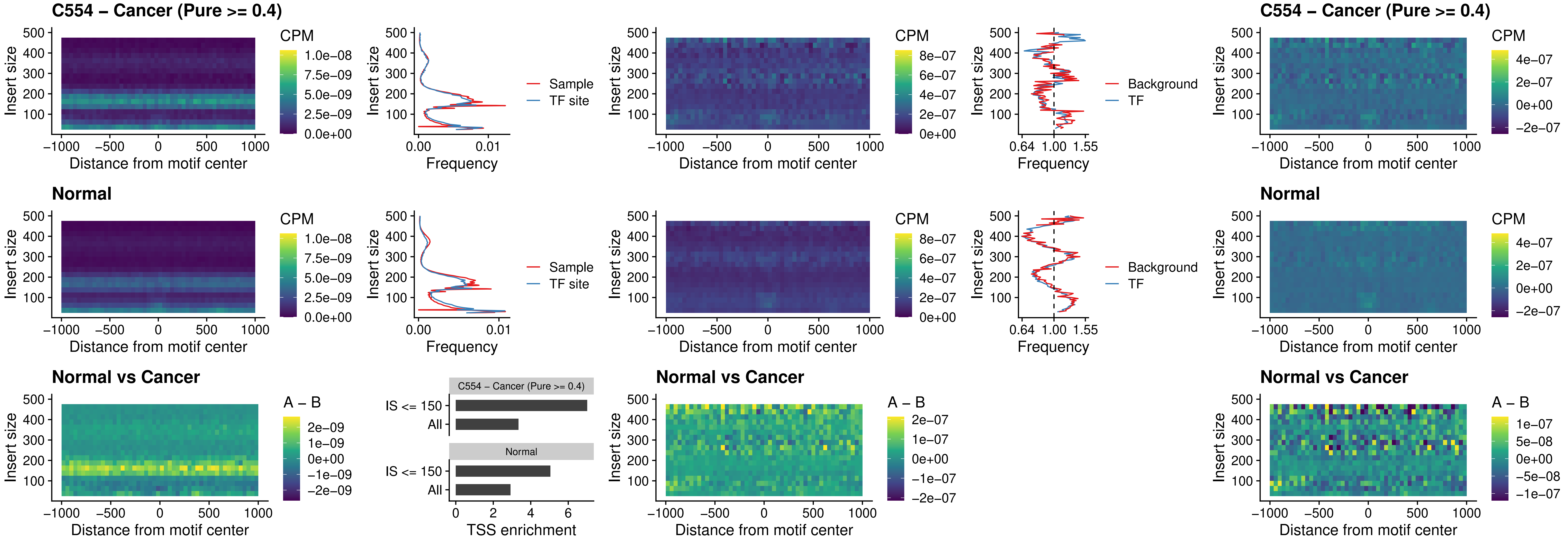

#### BCL11A_uc_cTSS_oPEAK

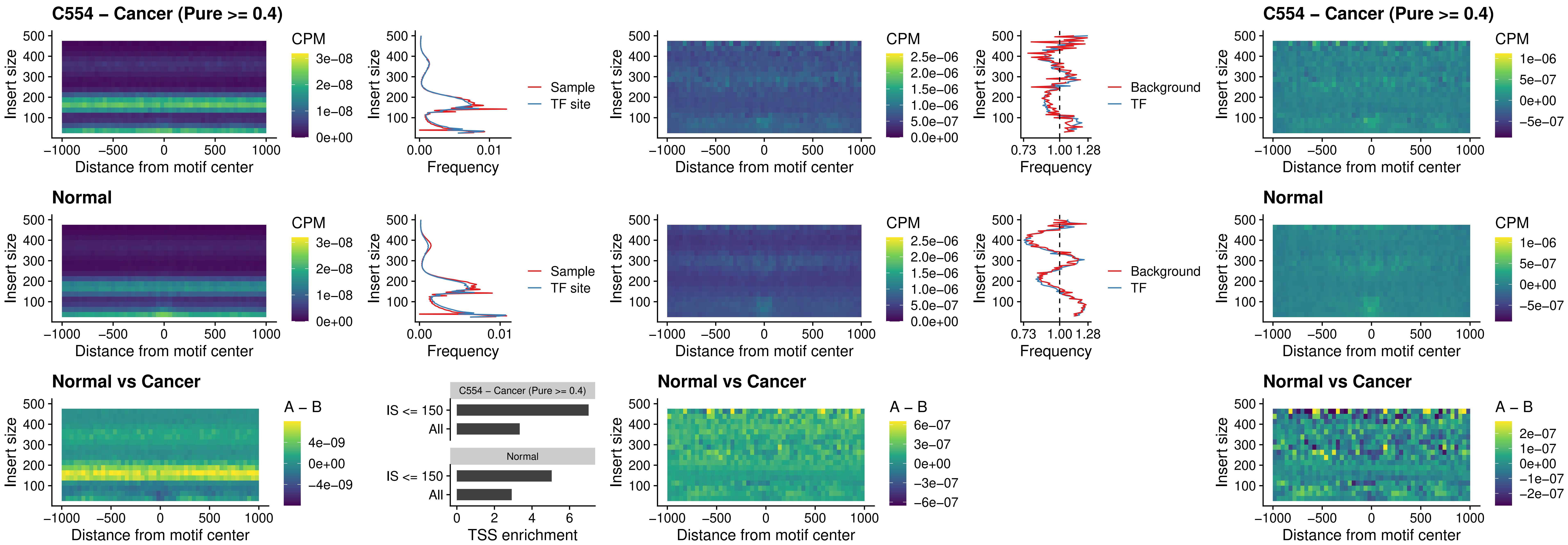

#### BCL11B_uc_cTSS_oPEAK

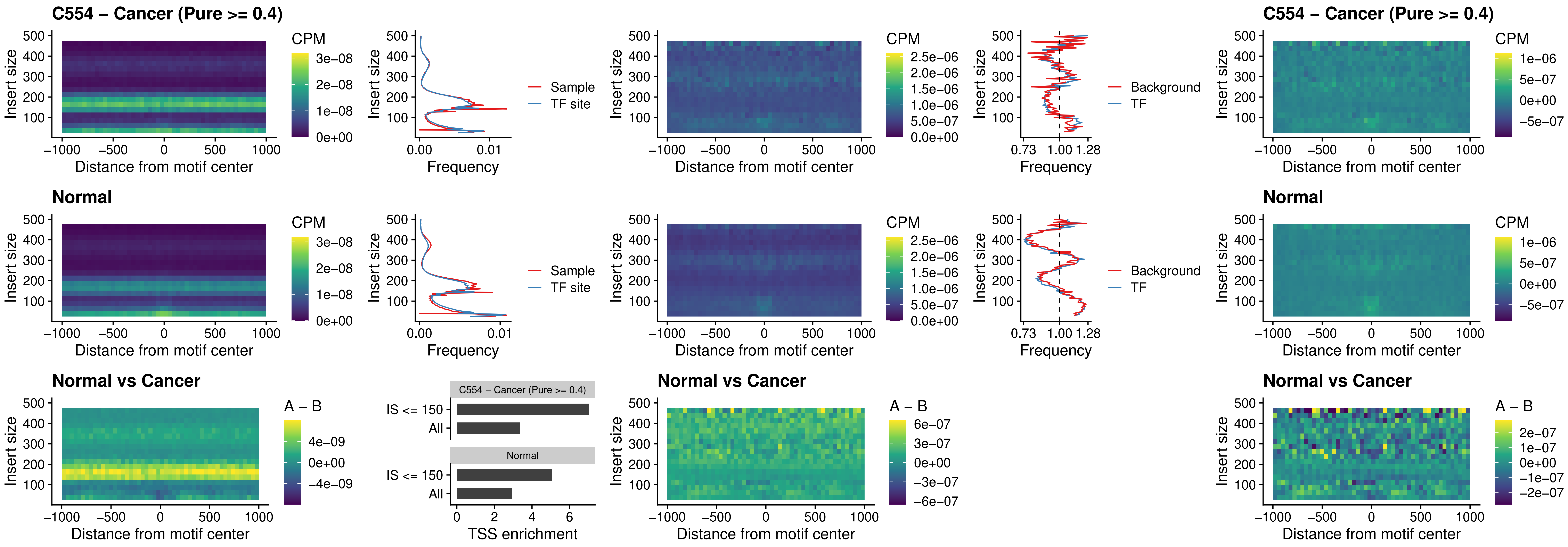

### BCL11A_uc_dTSS_oPEAK

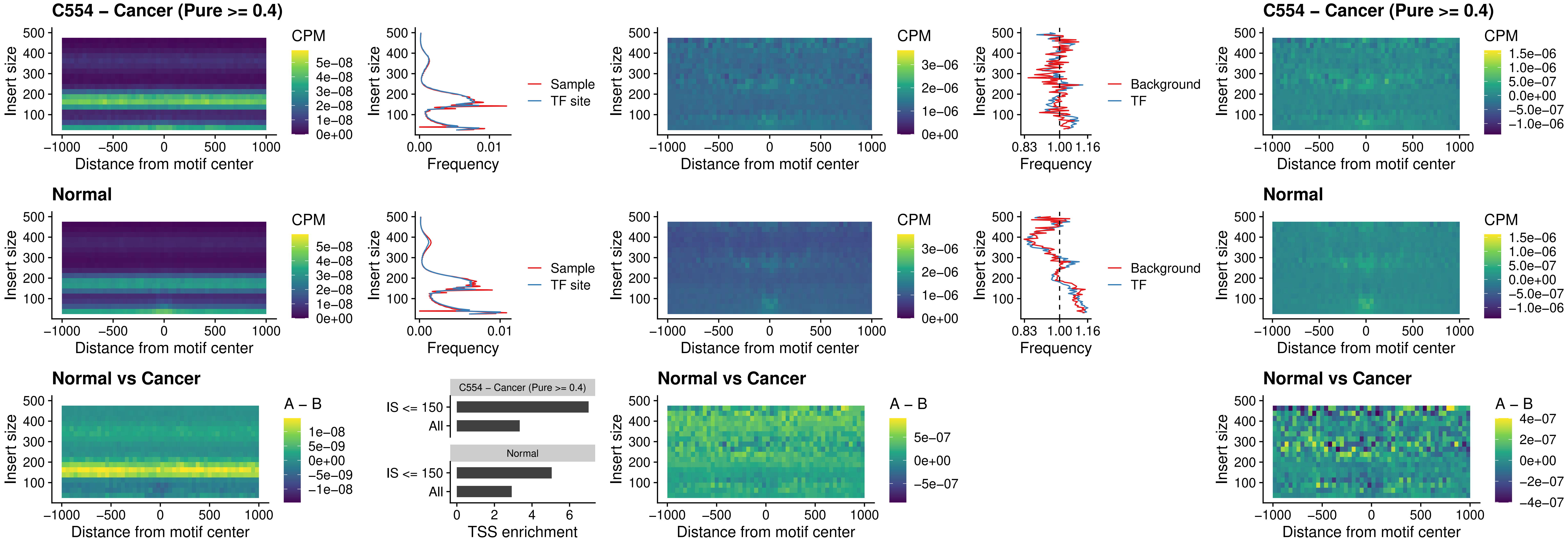

### BCL11B_uc_dTSS_oPEAK

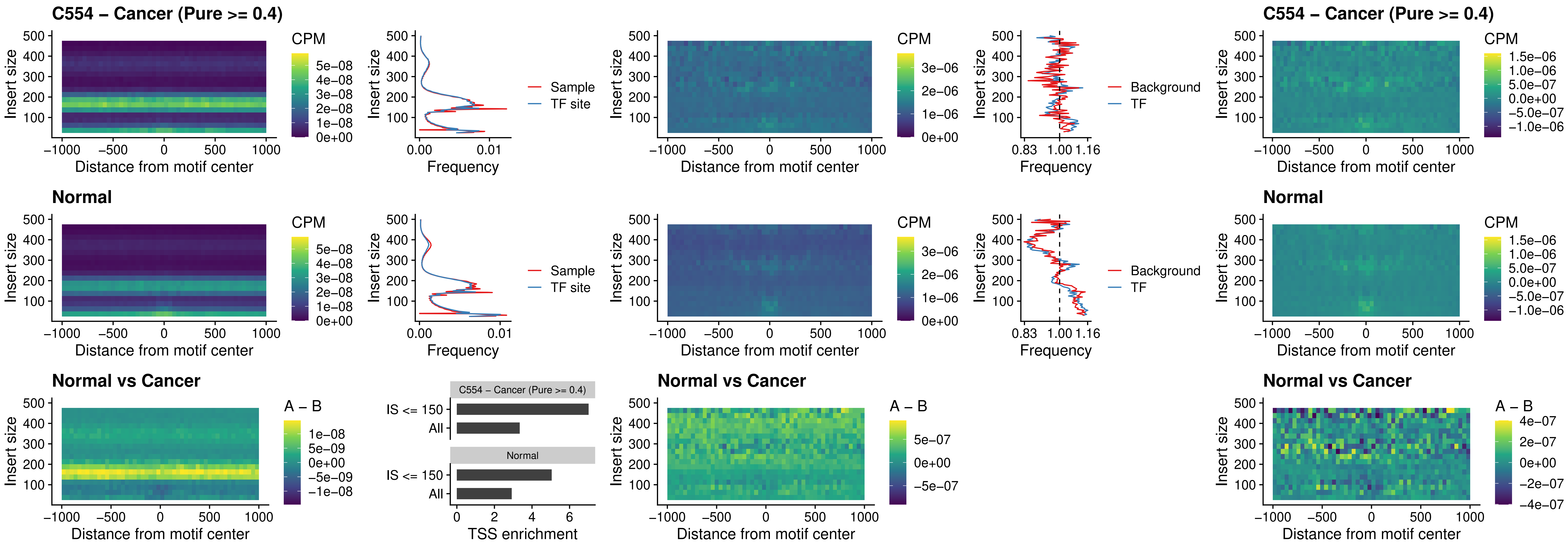

#### SPIB_uc_cTSS_nPEAK

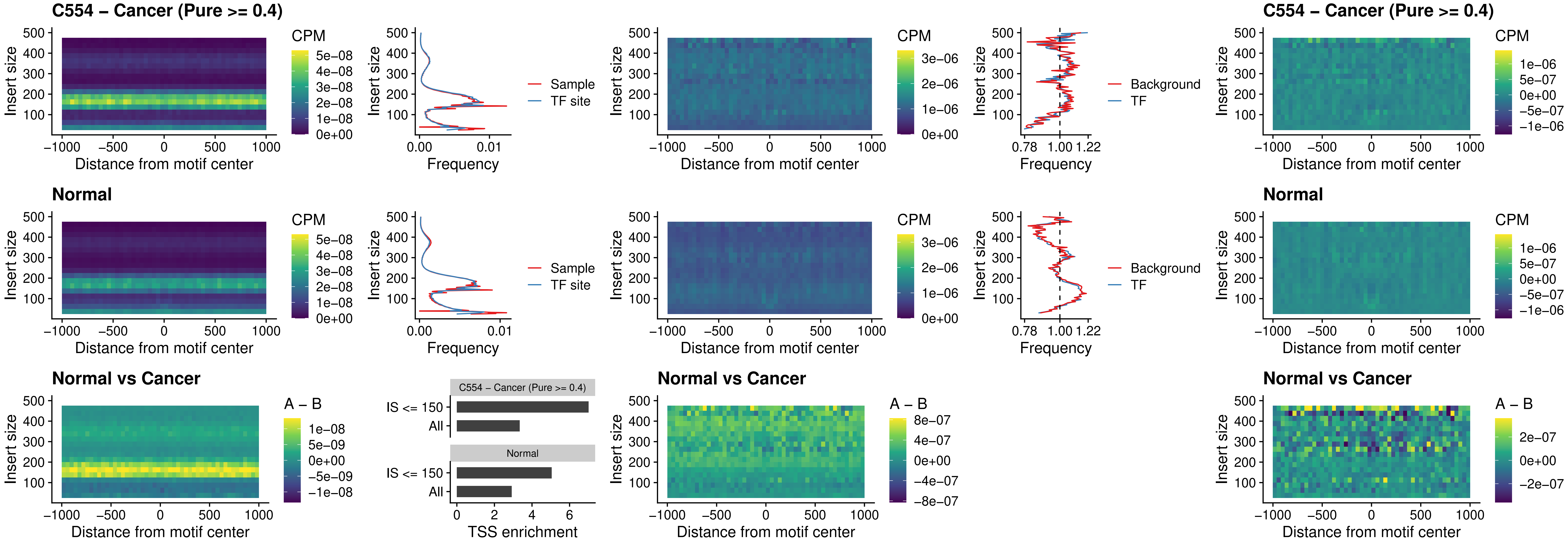

#### SPIC_uc_cTSS_nPEAK

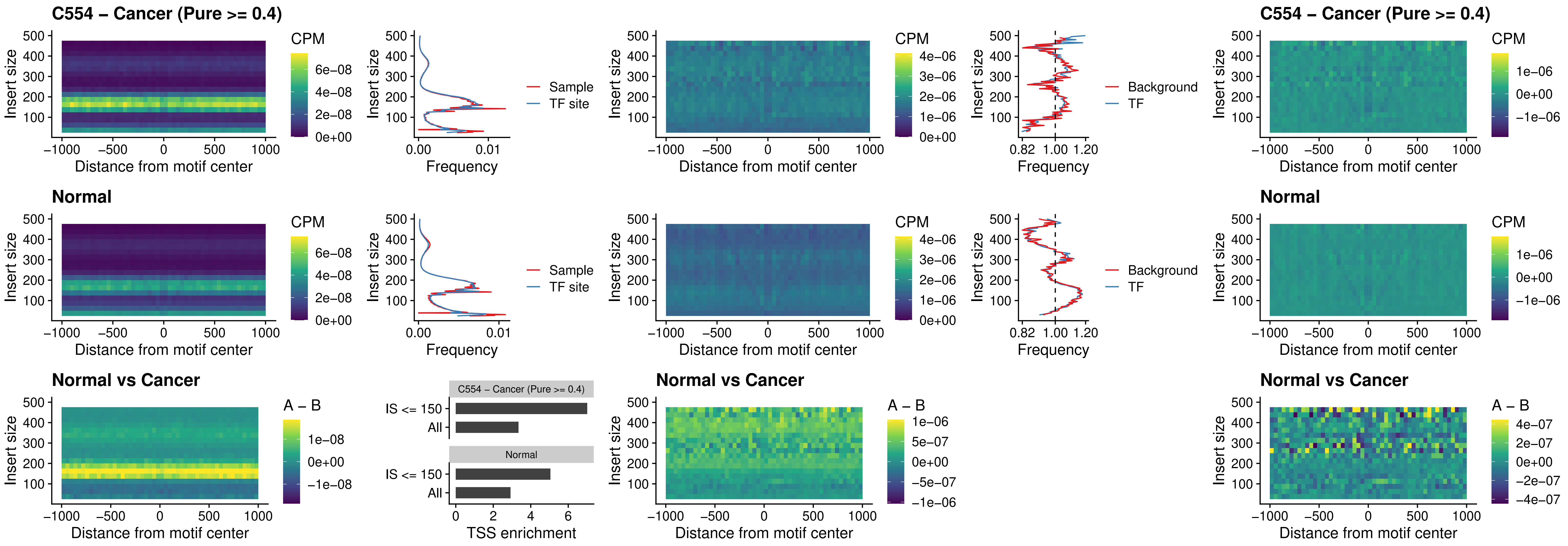

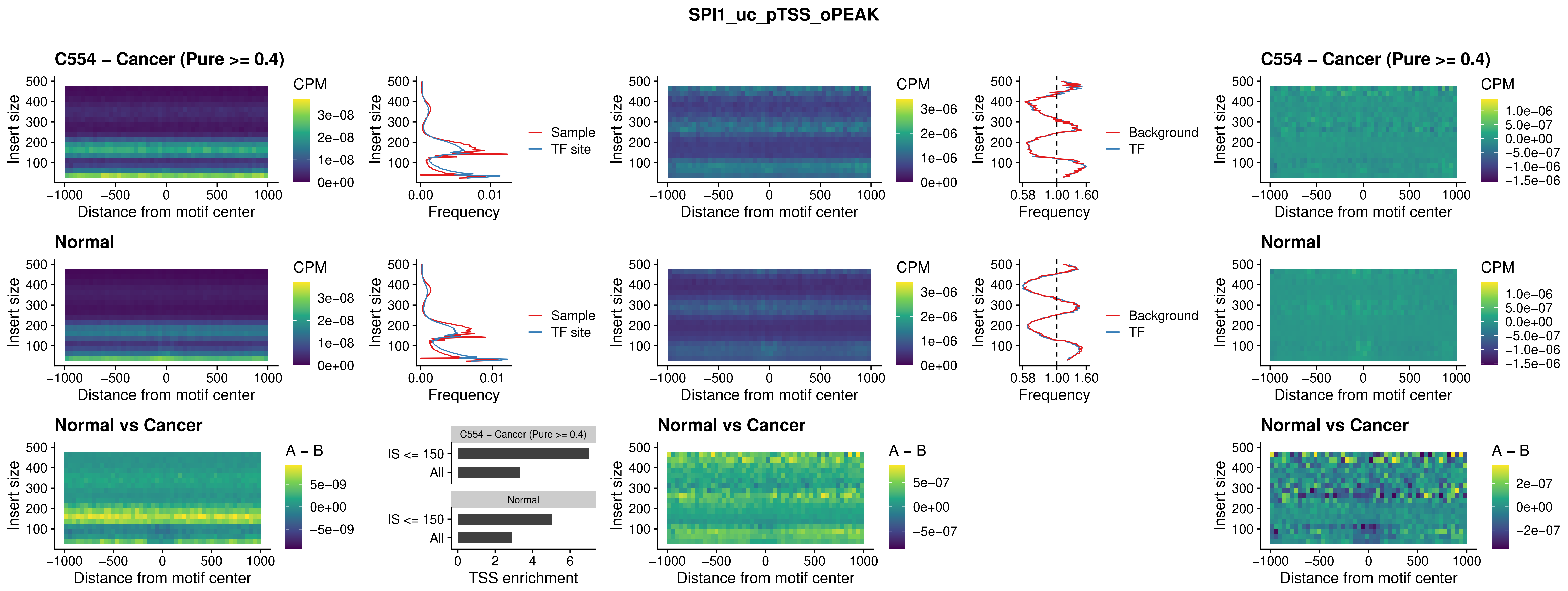
SPI1_uc_pTSS_oPEAK
C554 – Cancer (Pure >= 0.4)
Normal
Normal vs Cancer
Insert size
Distance from motif center
Frequency
CPM
A – B
Sample
TF site
Background
TF
C554 – Cancer (Pure >= 0.4)
Normal
IS <= 150
All
TSS enrichment

### BCL11A_uc_pTSS_oPEAK

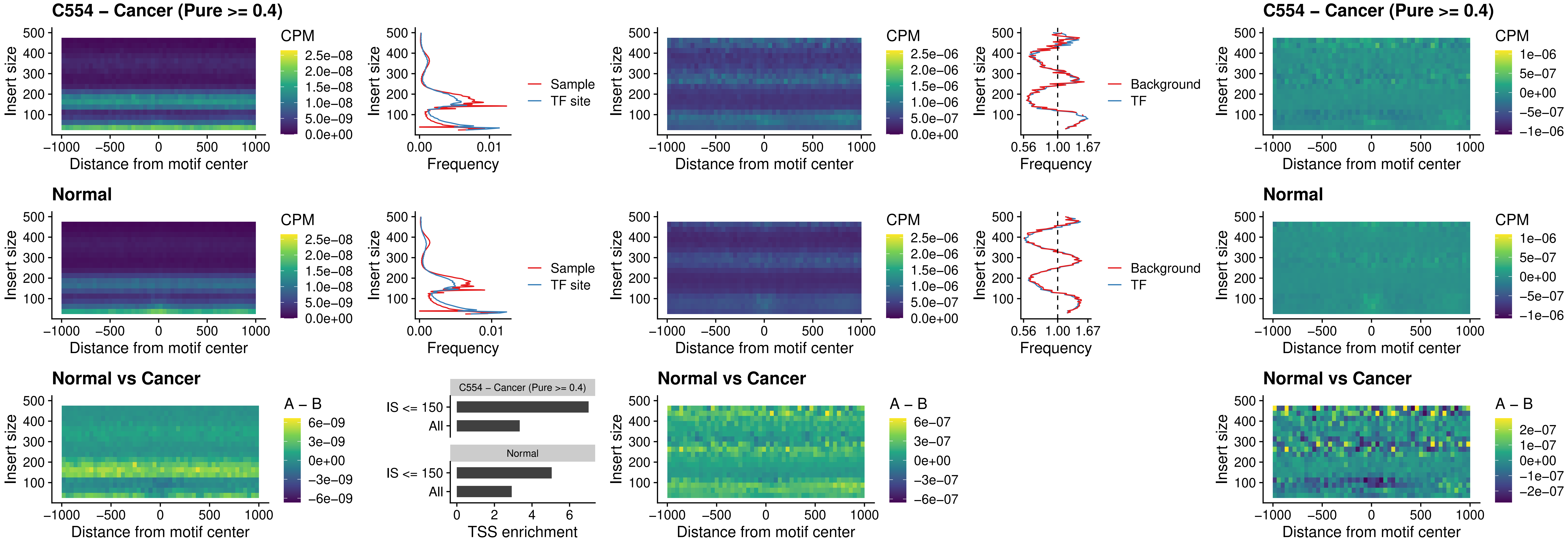

### BCL11B_uc_pTSS_oPEAK

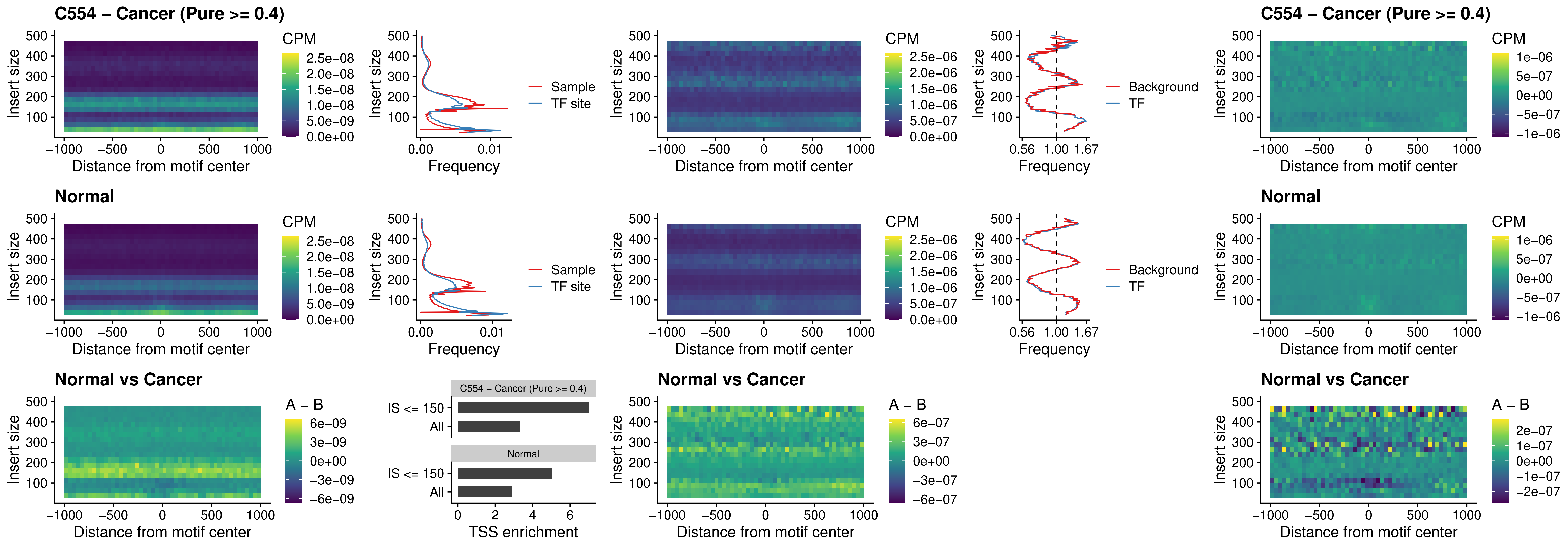

#### SPI1_uc_cTSS_oPEAK

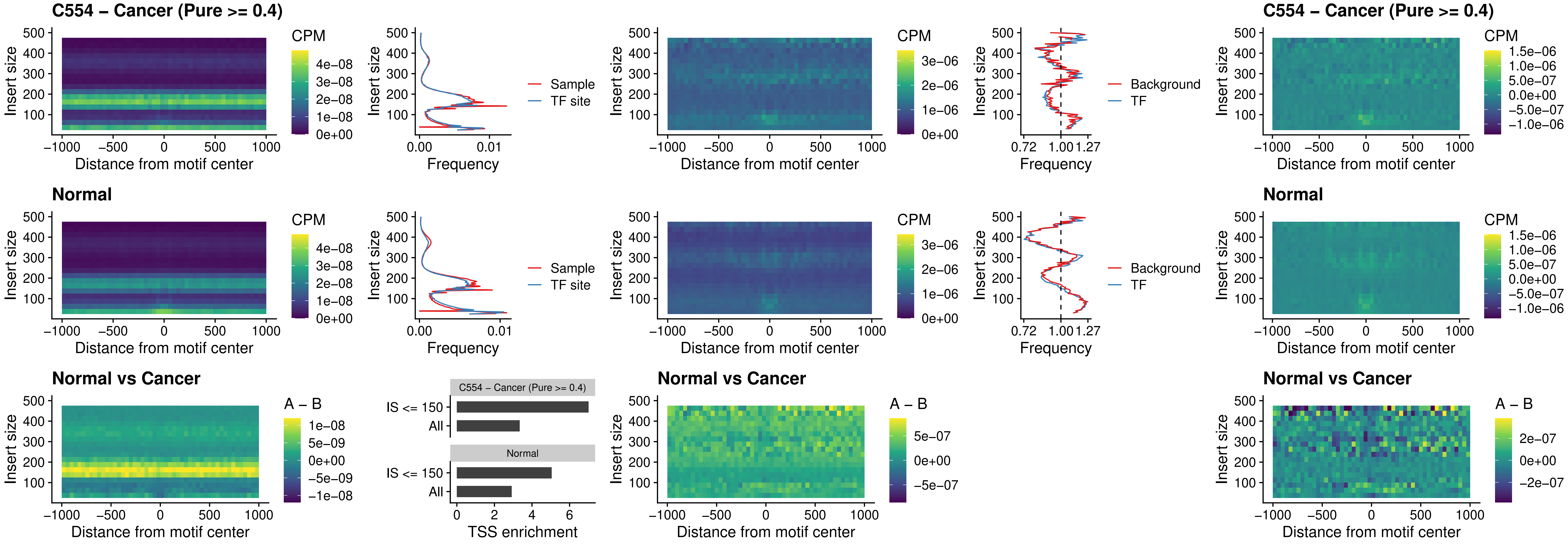

#### SPIB_uc_cTSS_oPEAK

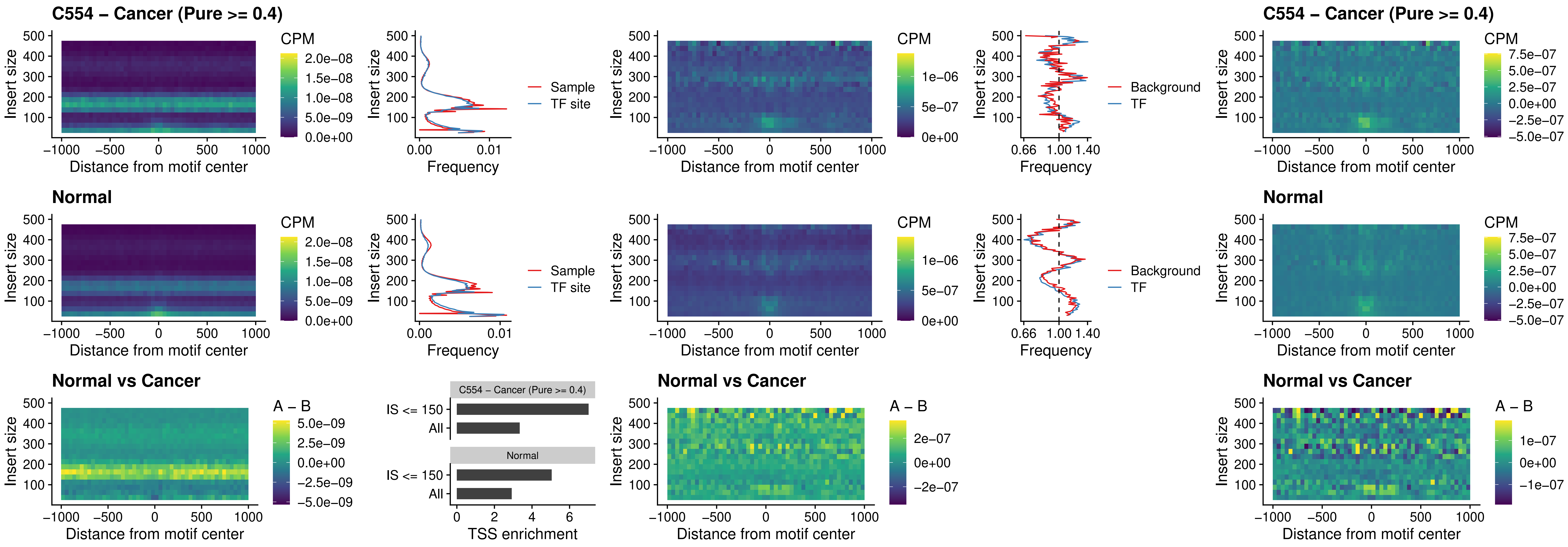

### SPIB_uc_pTSS_oPEAK

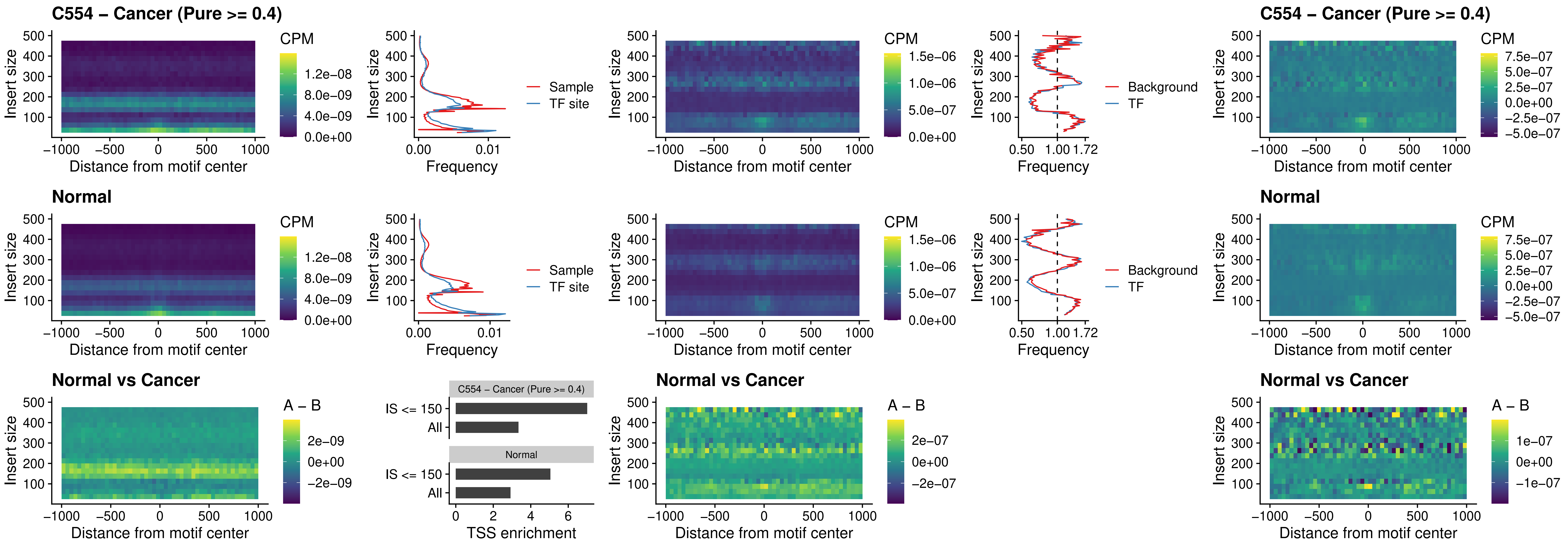

### SPI1_uc_dTSS_oPEAK

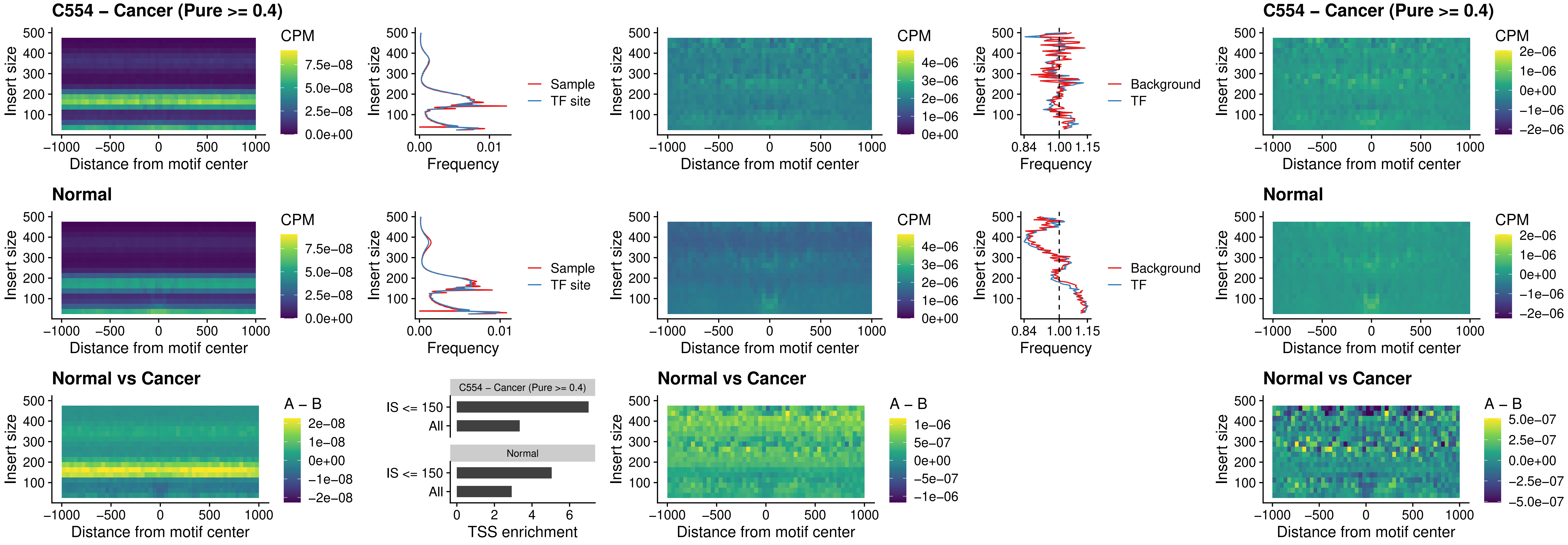

#### SPIB_uc_dTSS_oPEAK

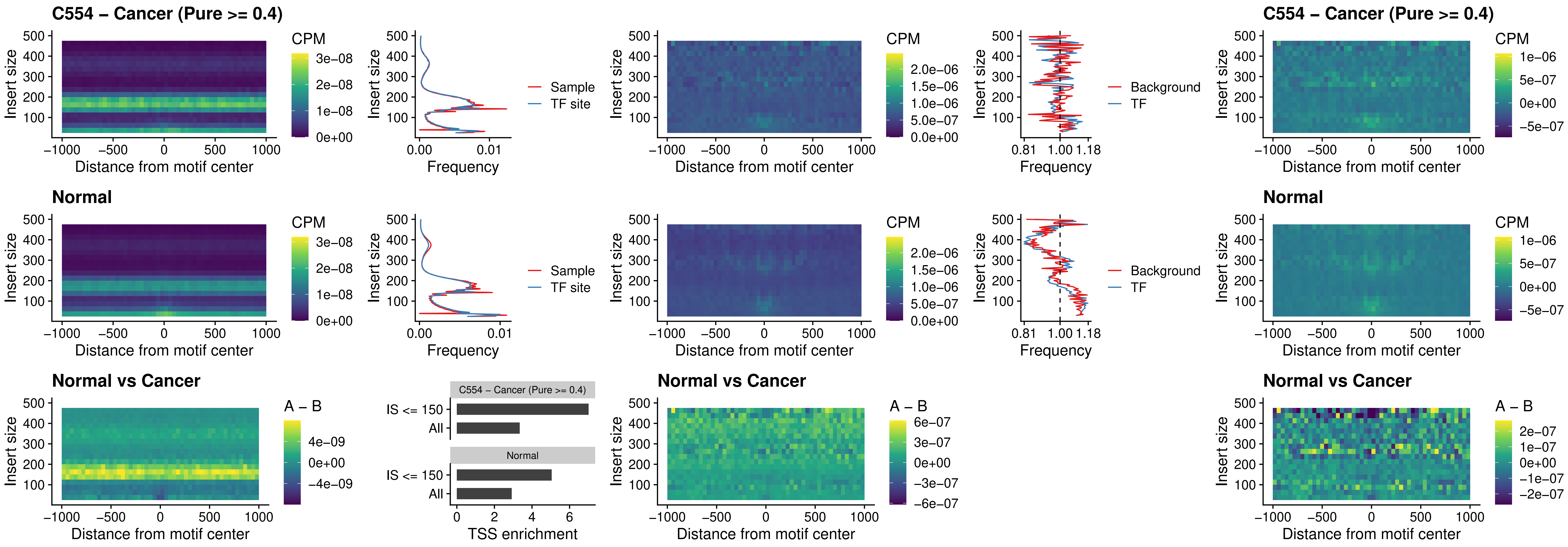

### IRF3_uc_pTSS_oPEAK

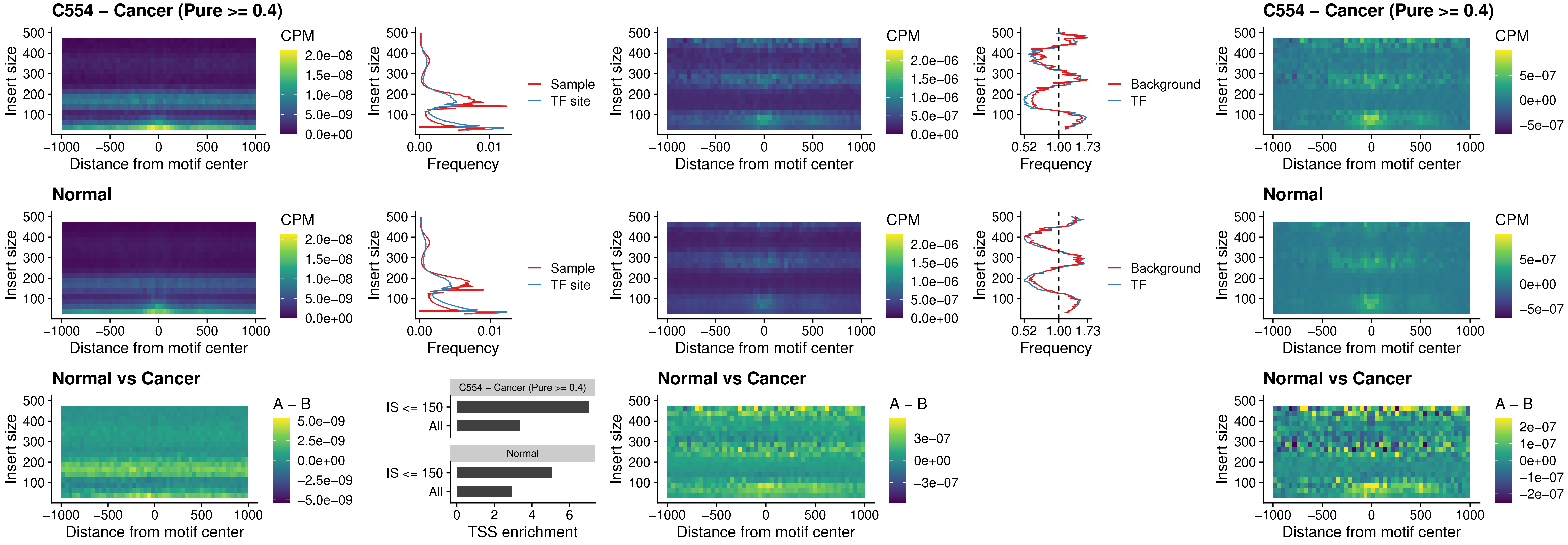

### IRF2_uc_dTSS_oPEAK

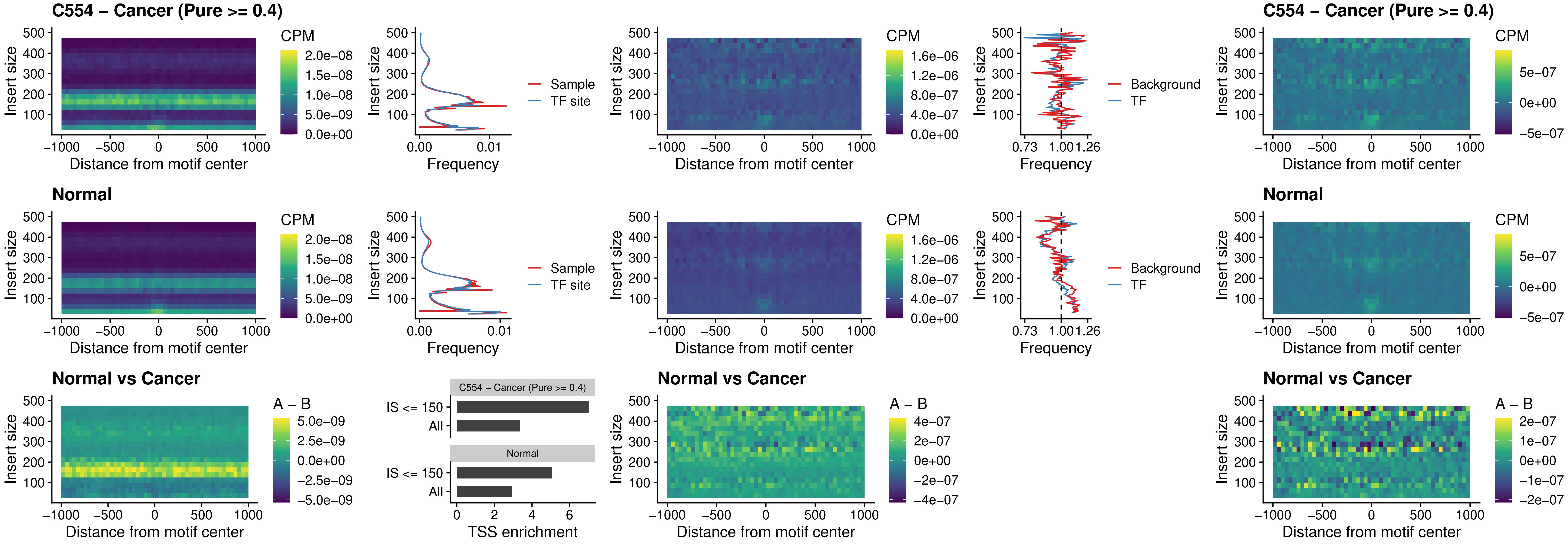

### IRF2_uc_pTSS_oPEAK

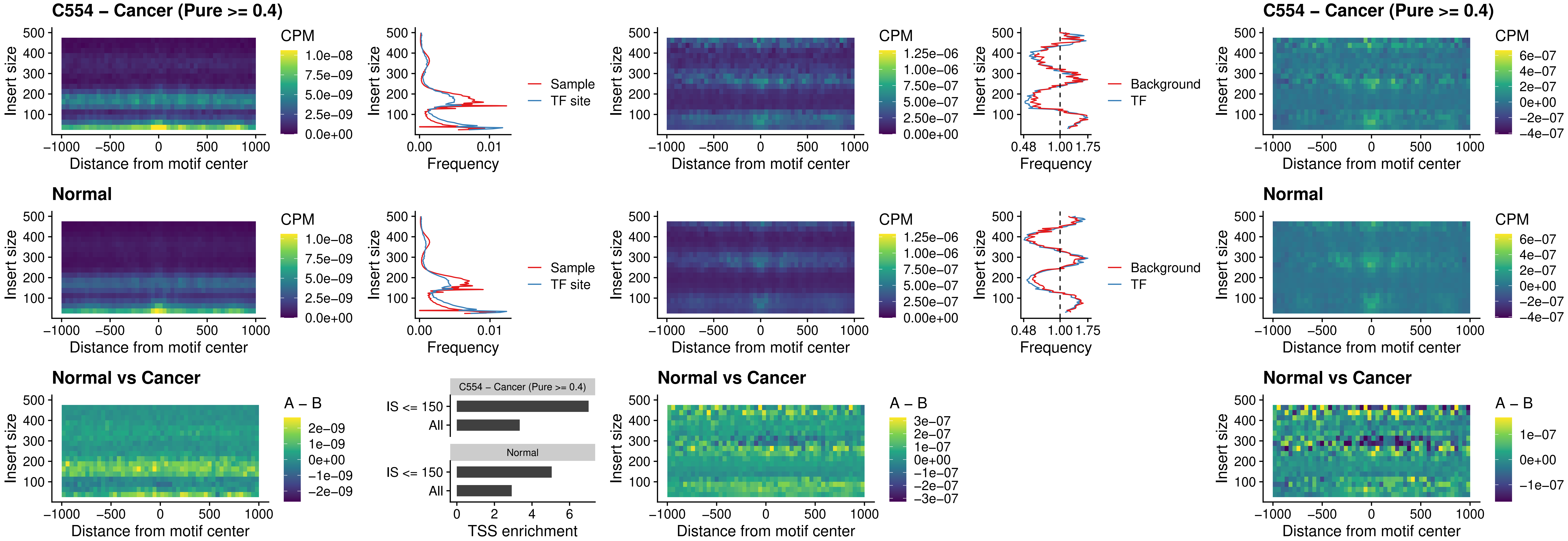

#### STAT2_uc_pTSS_oPEAK

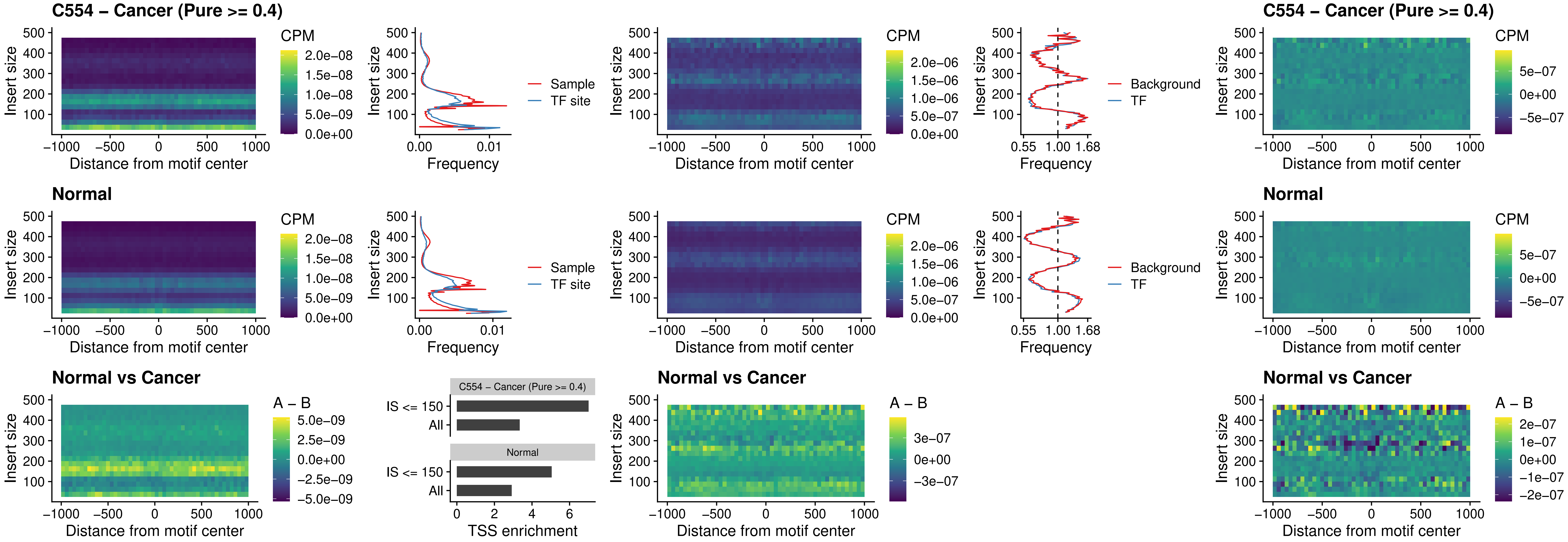

#### SPIB_uc_dTSS_nPEAK

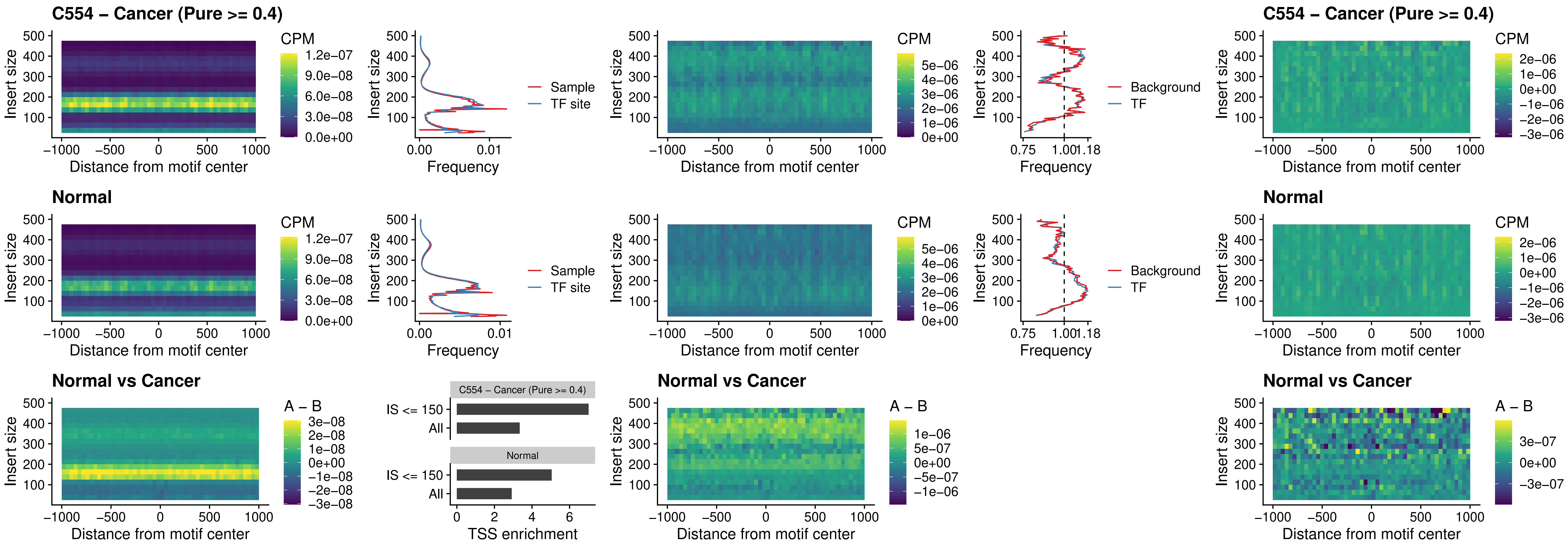

### SPIC_uc_cTSS_oPEAK

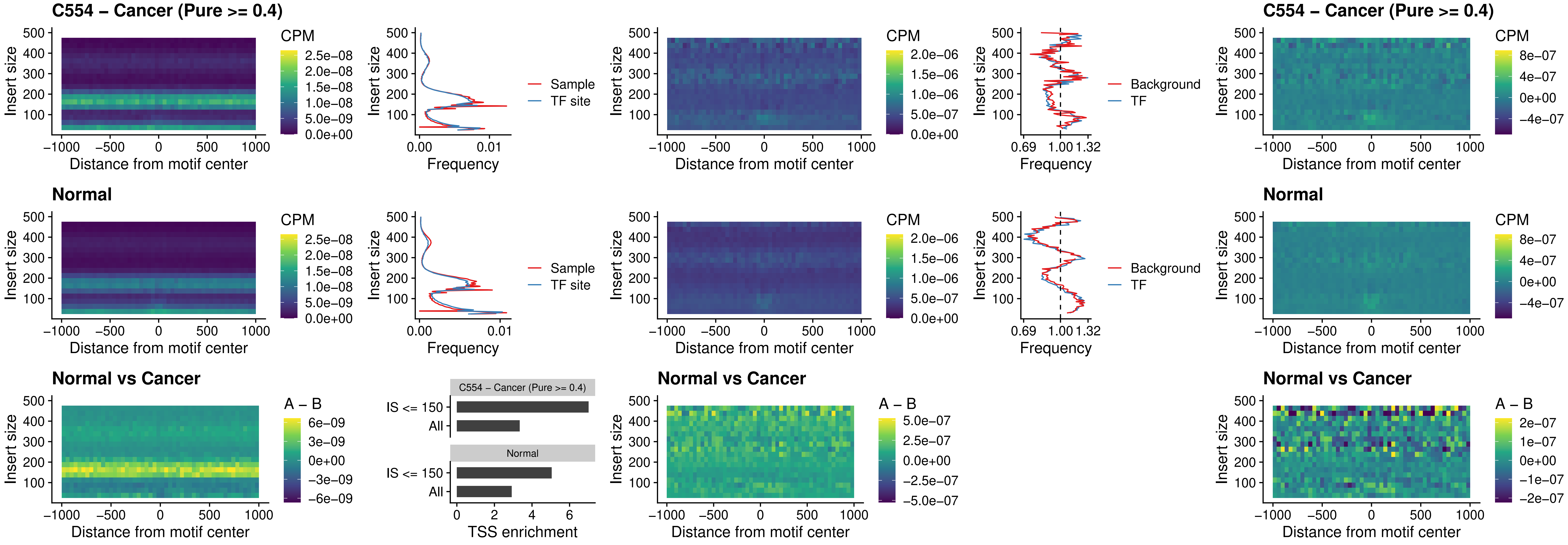

### PRDM1_uc_cTSS_oPEAK

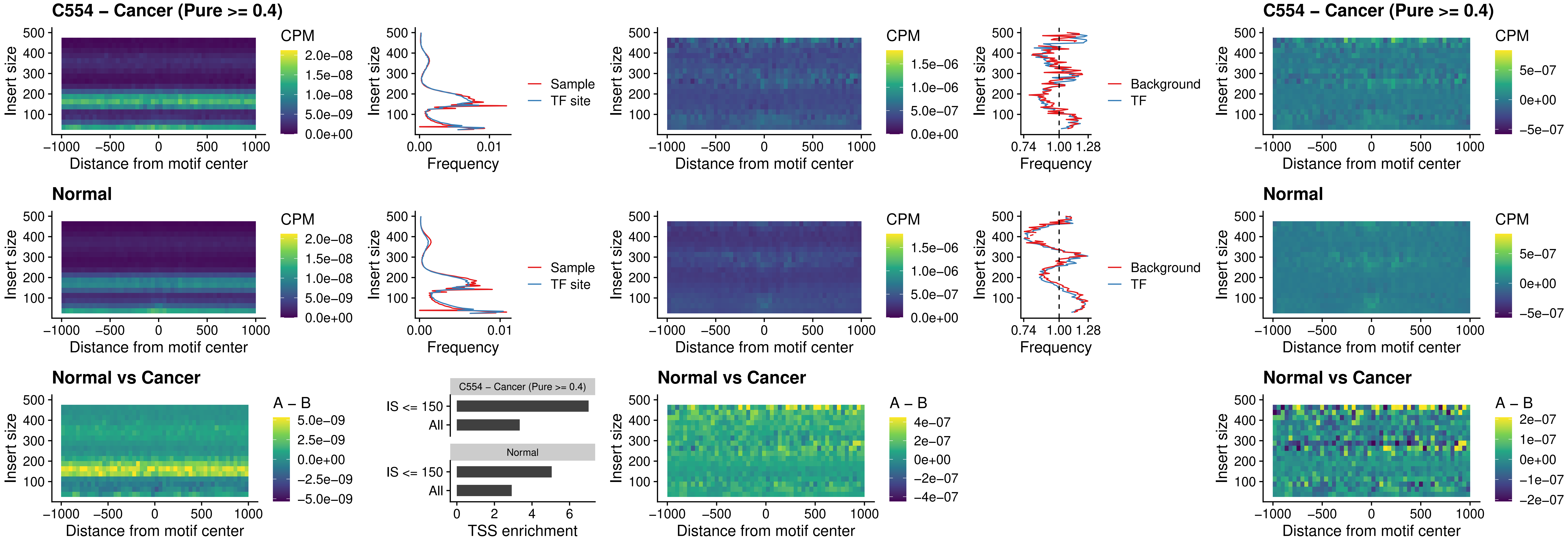

### IRF3_uc_cTSS_oPEAK

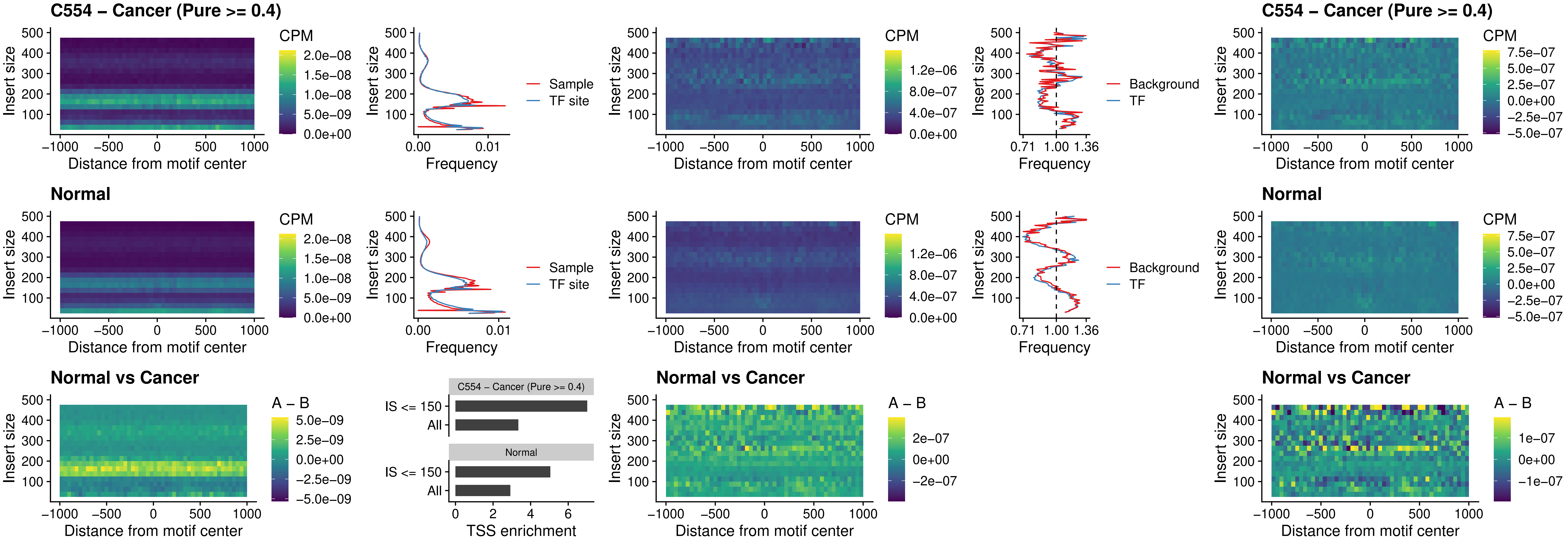

### BCL11A_uc_dTSS_nPEAK

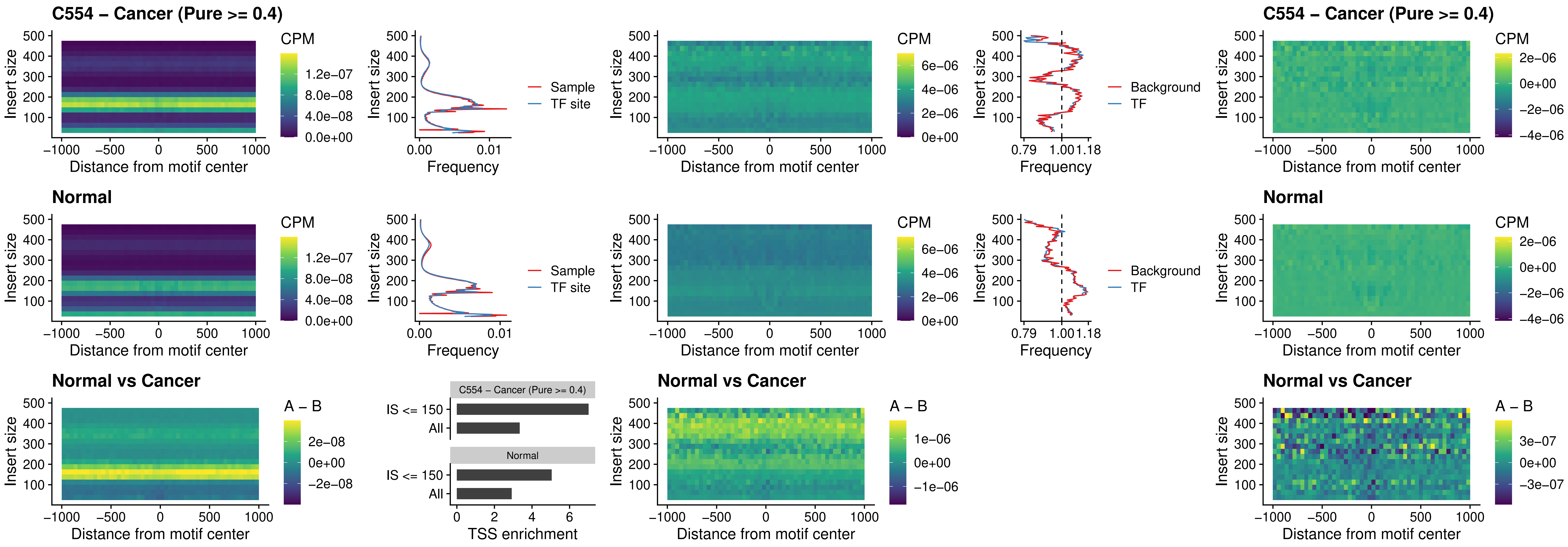

### BCL11B_uc_dTSS_nPEAK

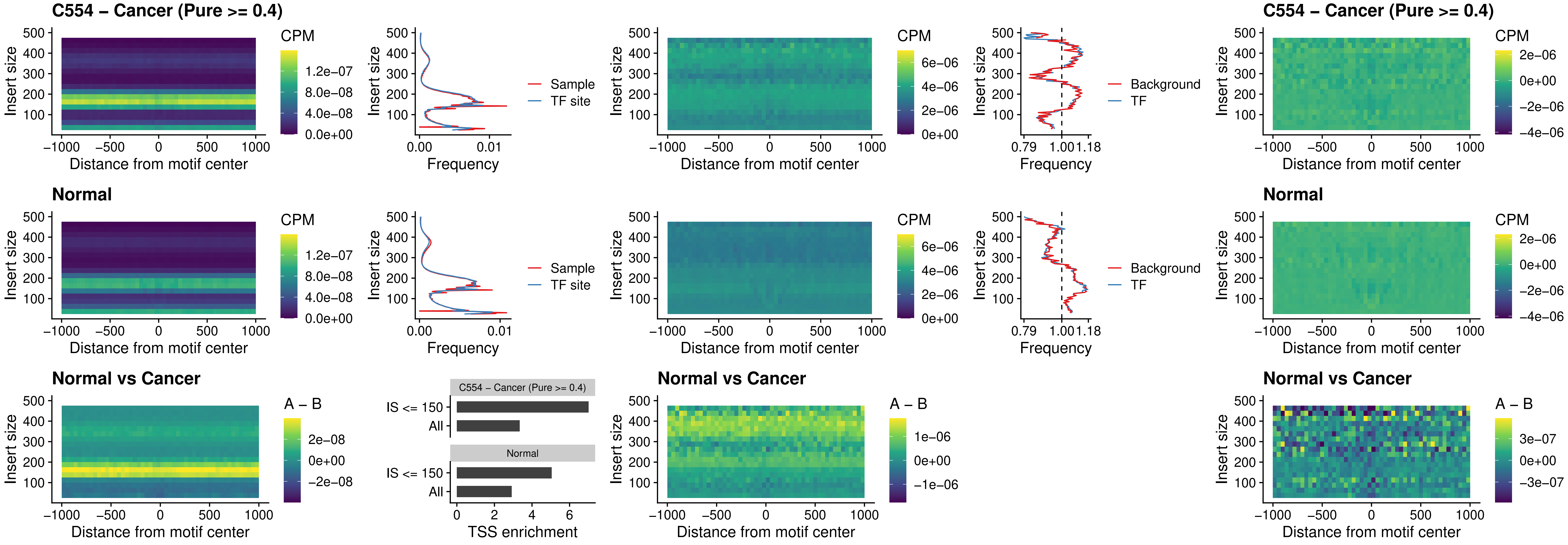

### IRF1_uc_cTSS_oPEAK

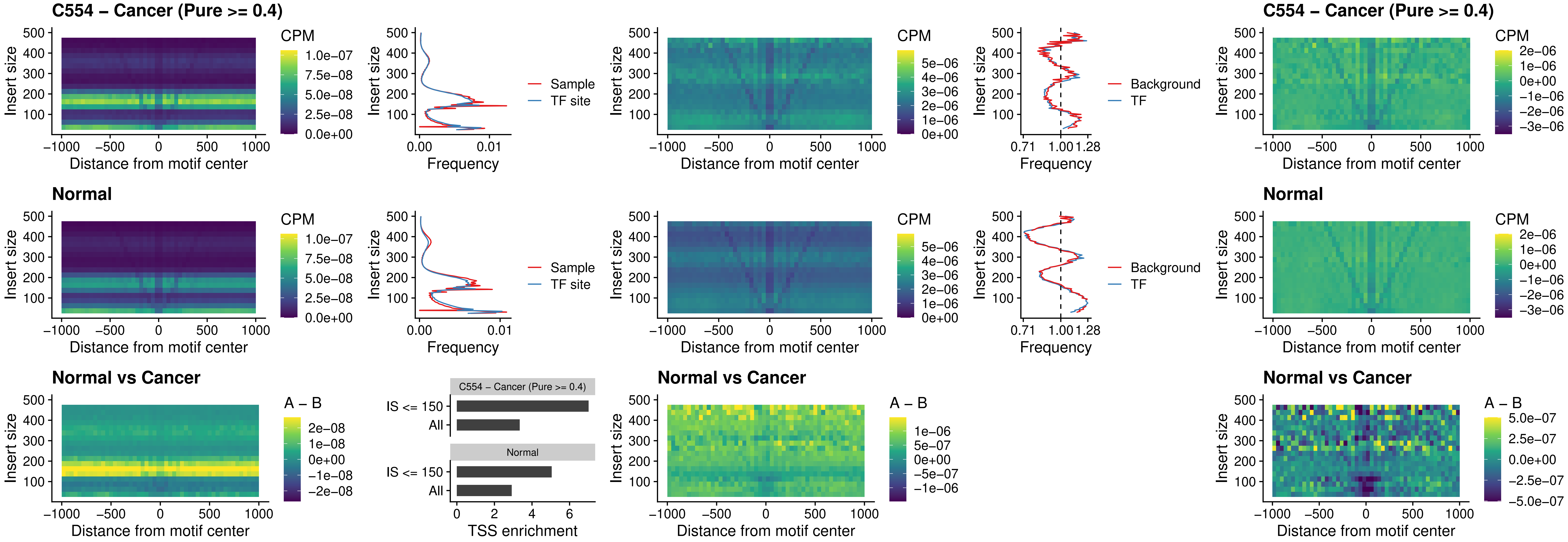

### IRF1_uc_dTSS_oPEAK

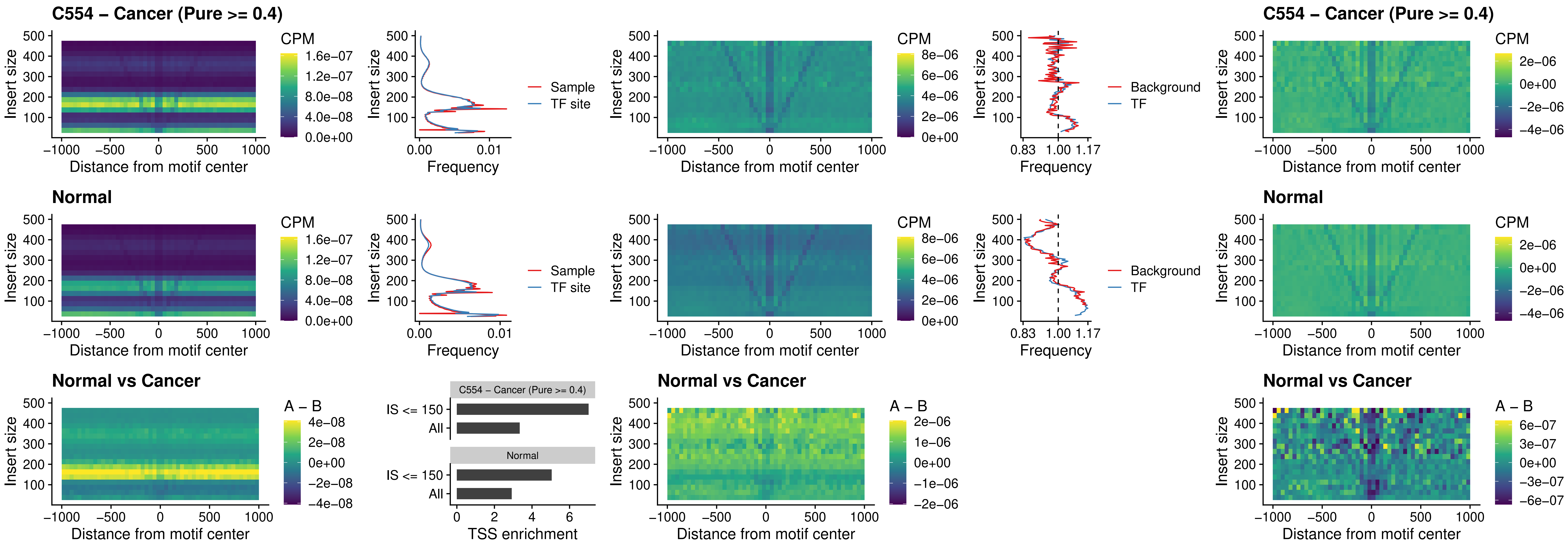

### IRF1_uc_pTSS_oPEAK

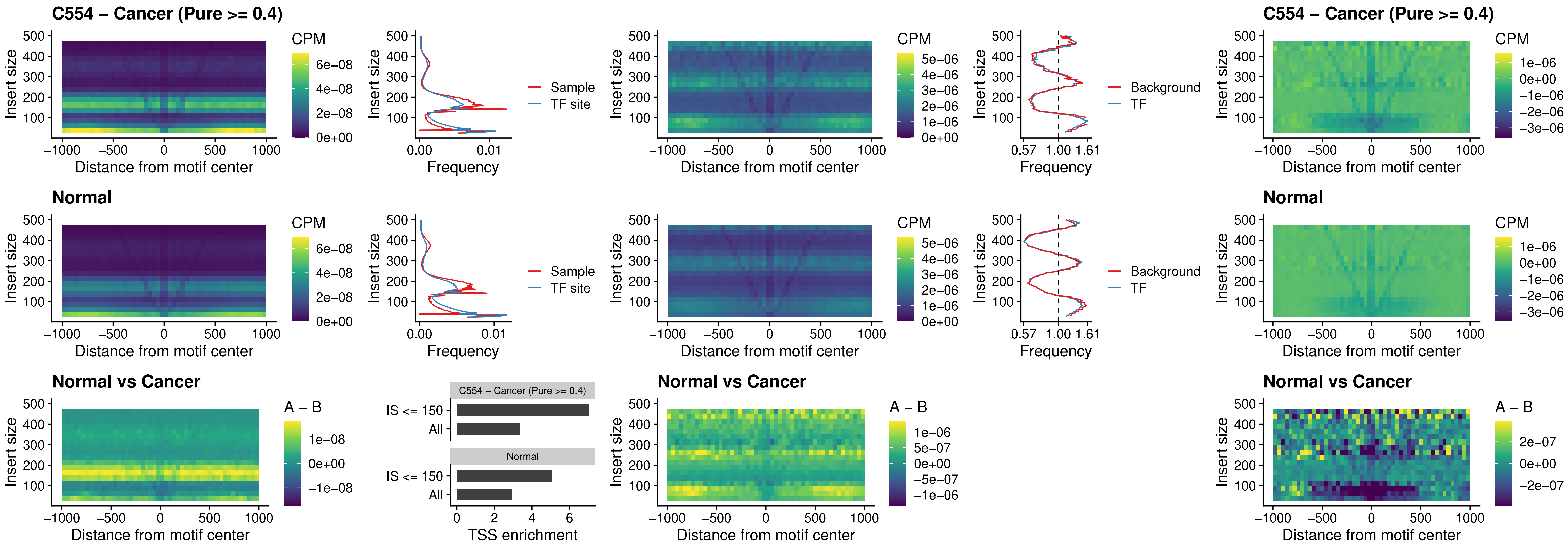

### SPIC_uc_dTSS_nPEAK

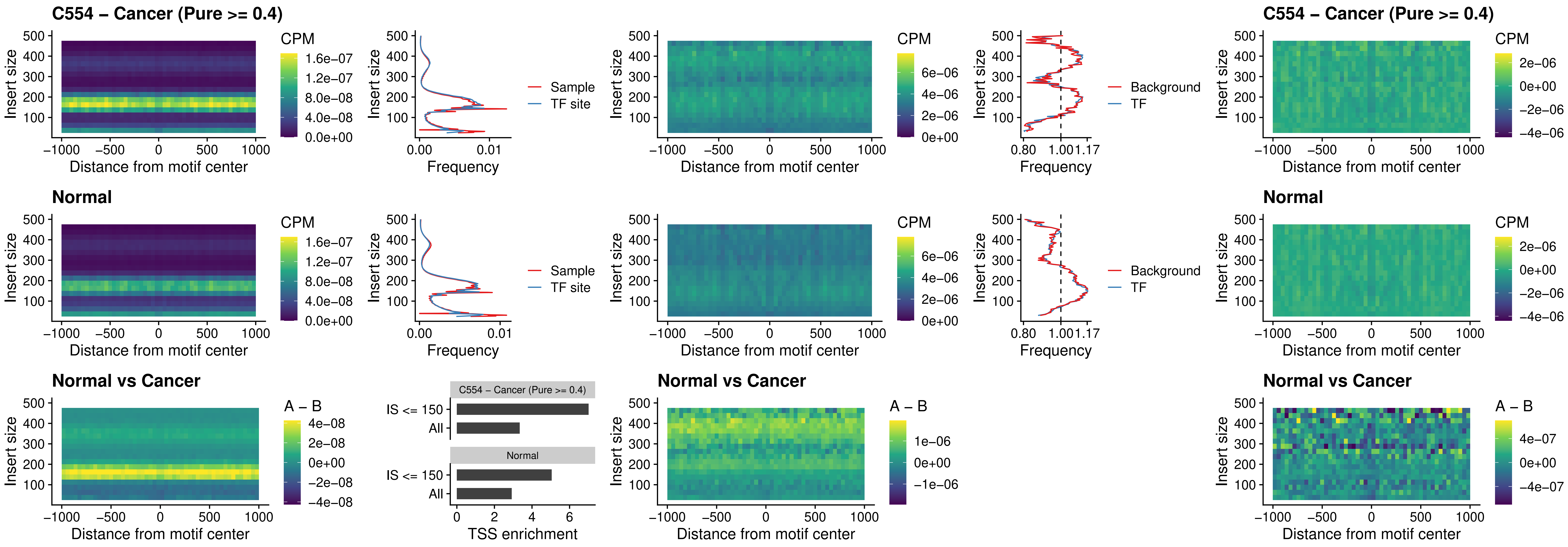

### SPIC_uc_dTSS_oPEAK

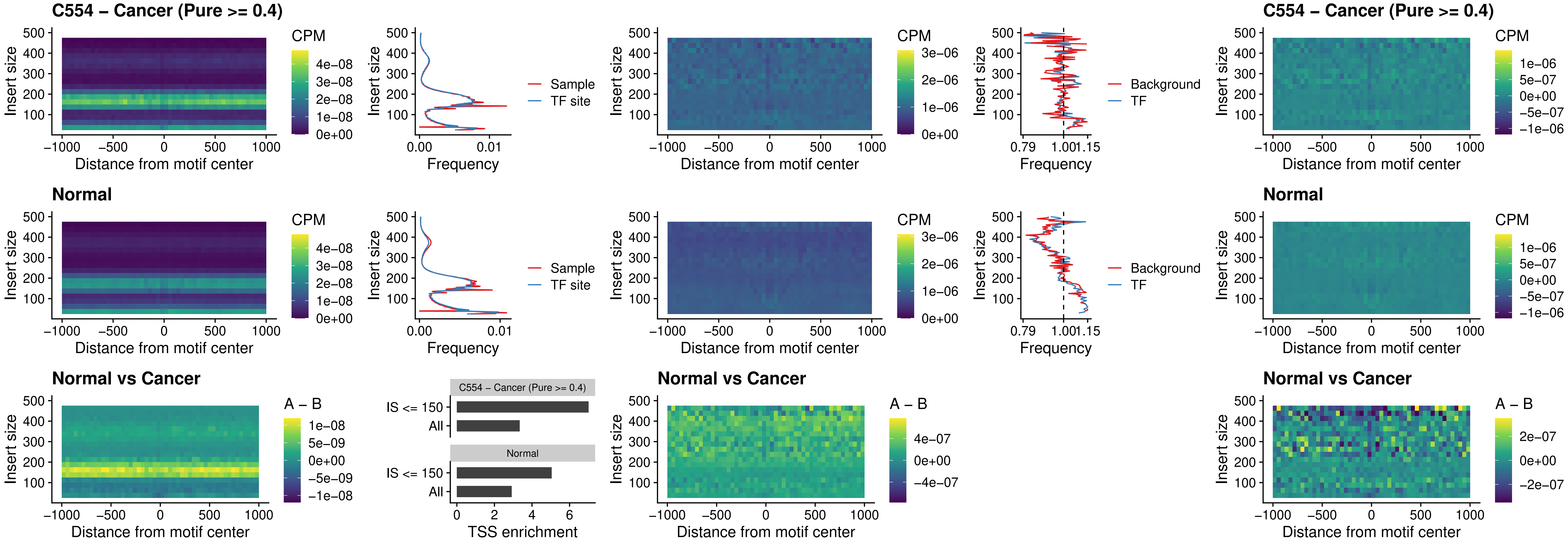

### SPIB_uc_pTSS_nPEAK

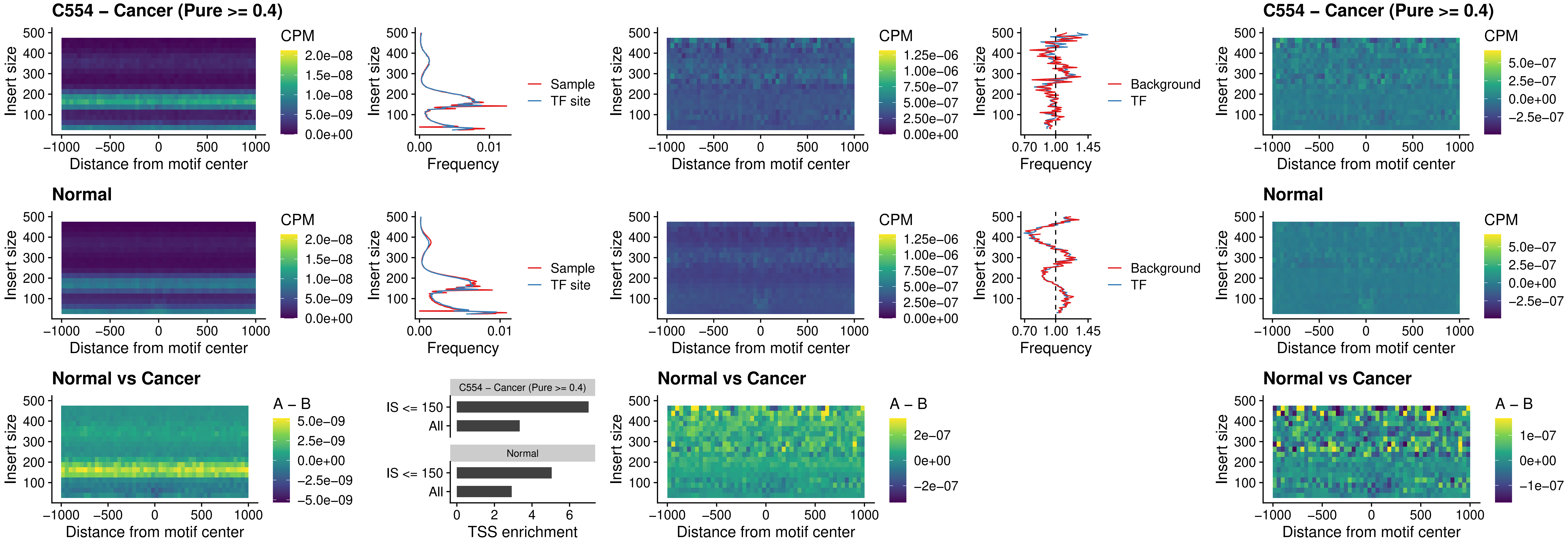

#### STAT2_uc_cTSS_oPEAK

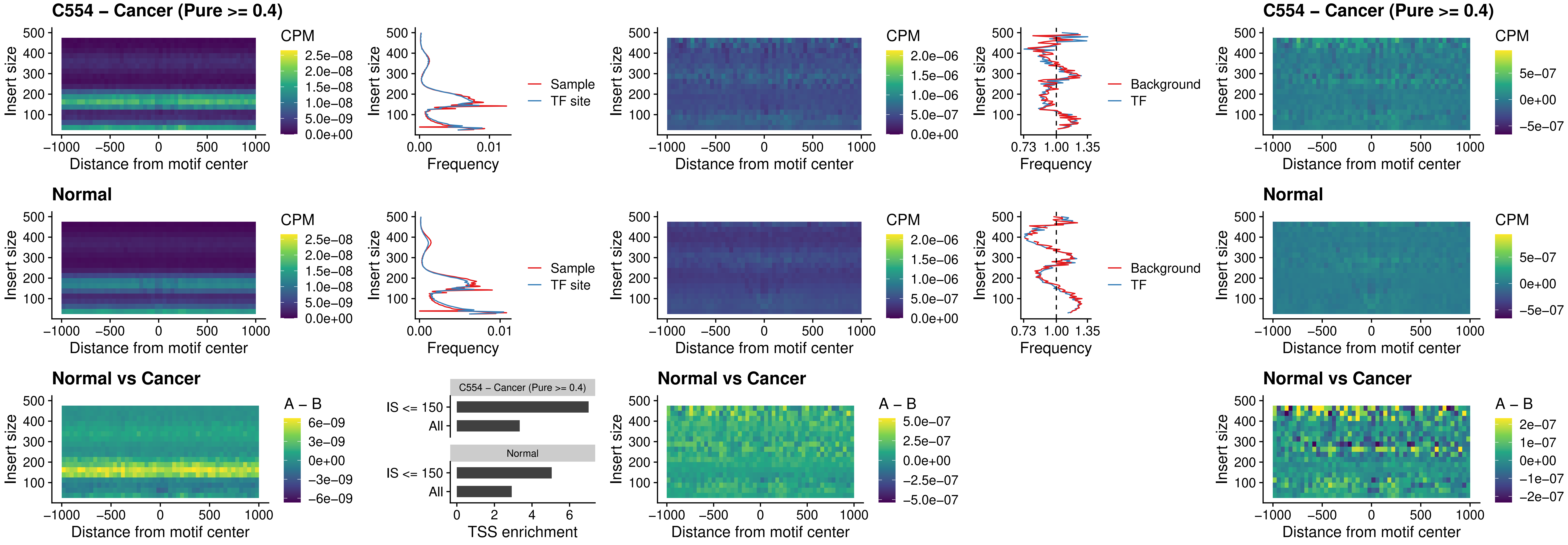

### IRF8_uc_cTSS_oPEAK

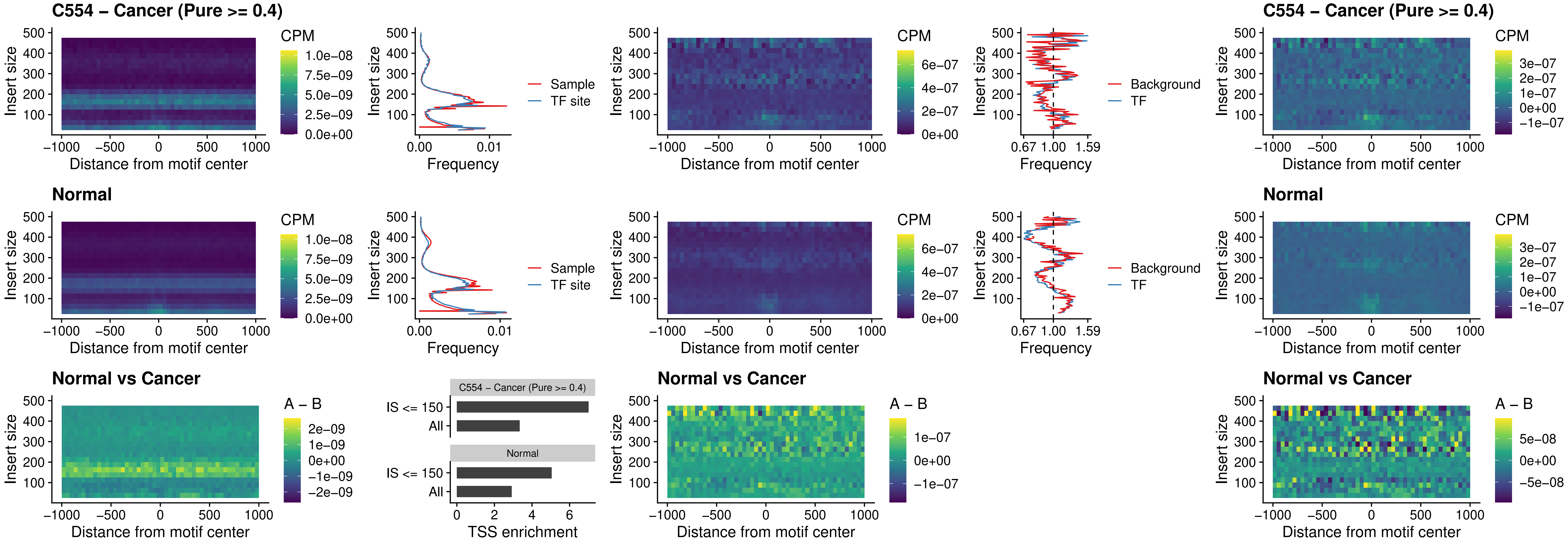

#### IRF3_uc_dTSS_oPEAK

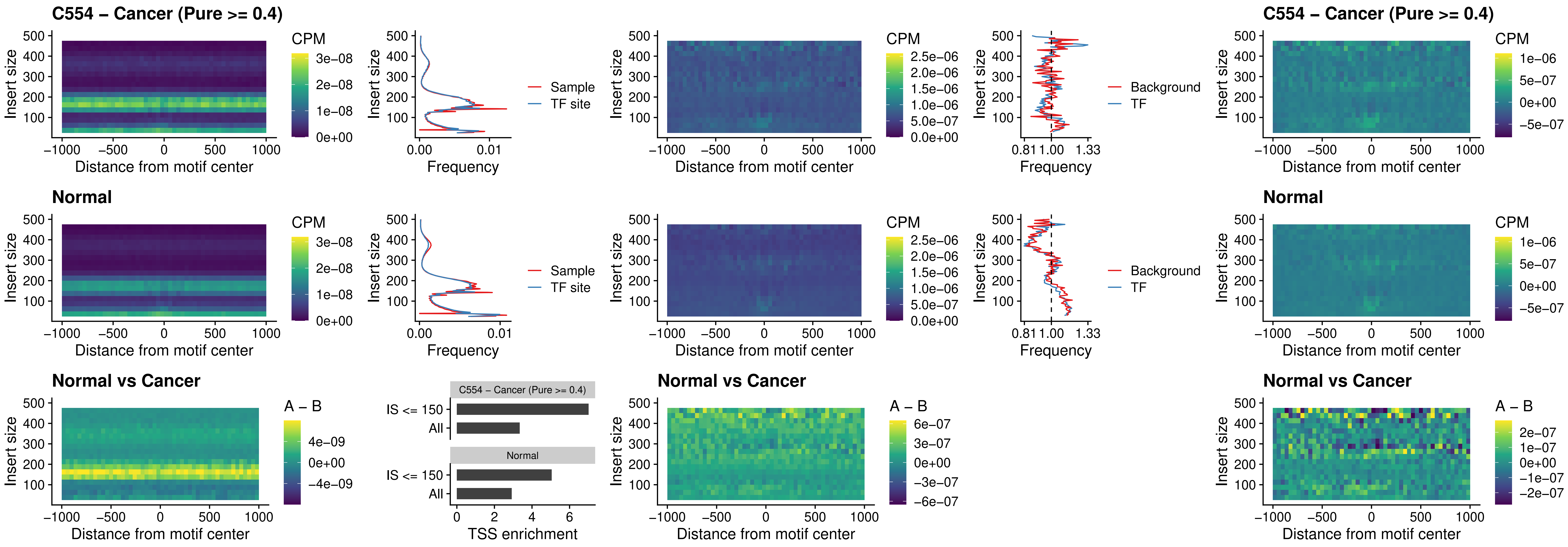

### IRF8_uc_dTSS_oPEAK

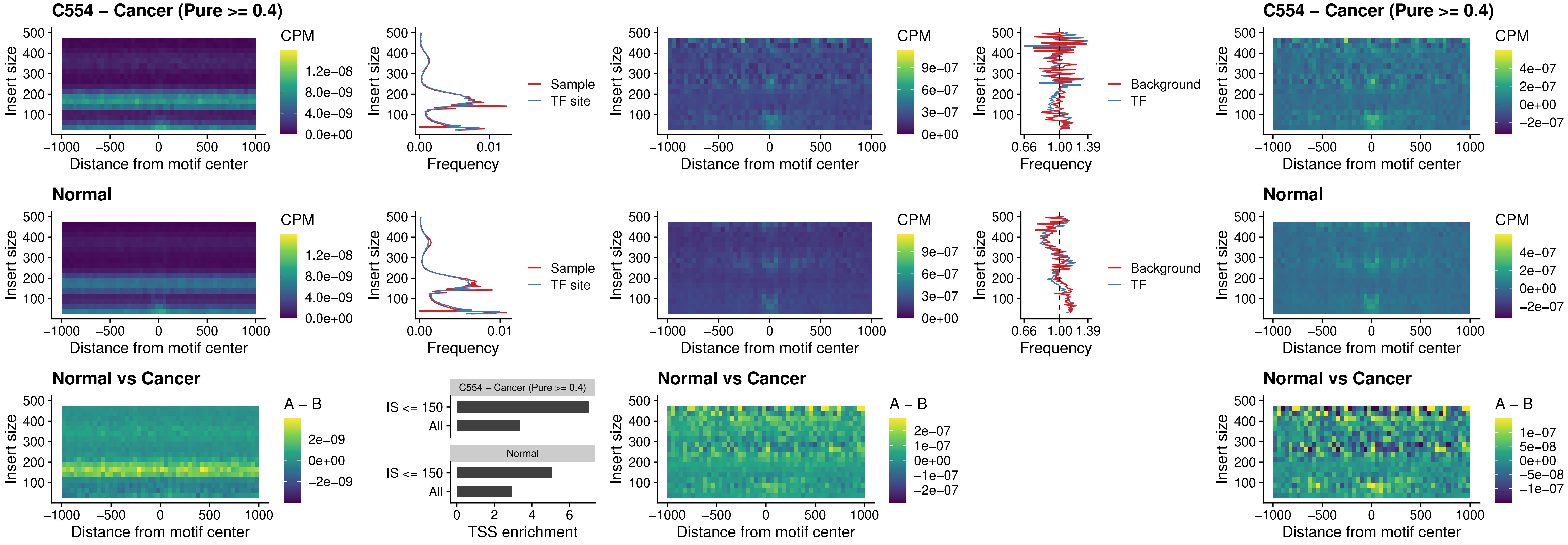

### ETS2_uc_dTSS_oPEAK

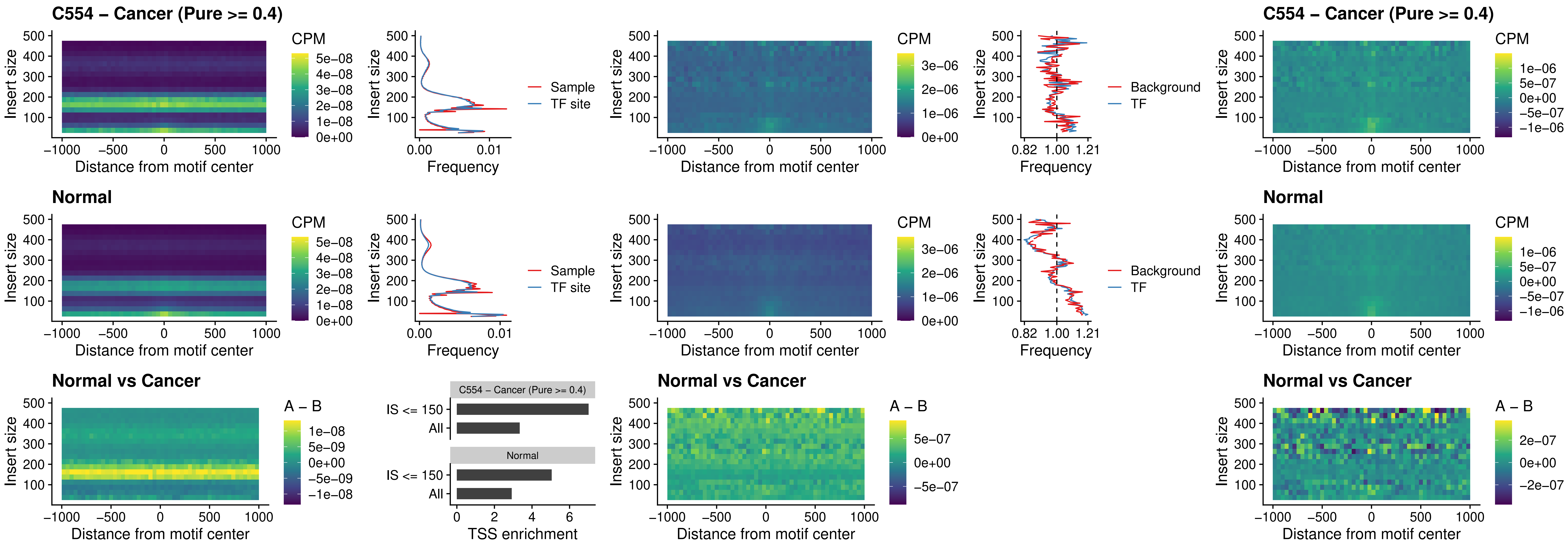

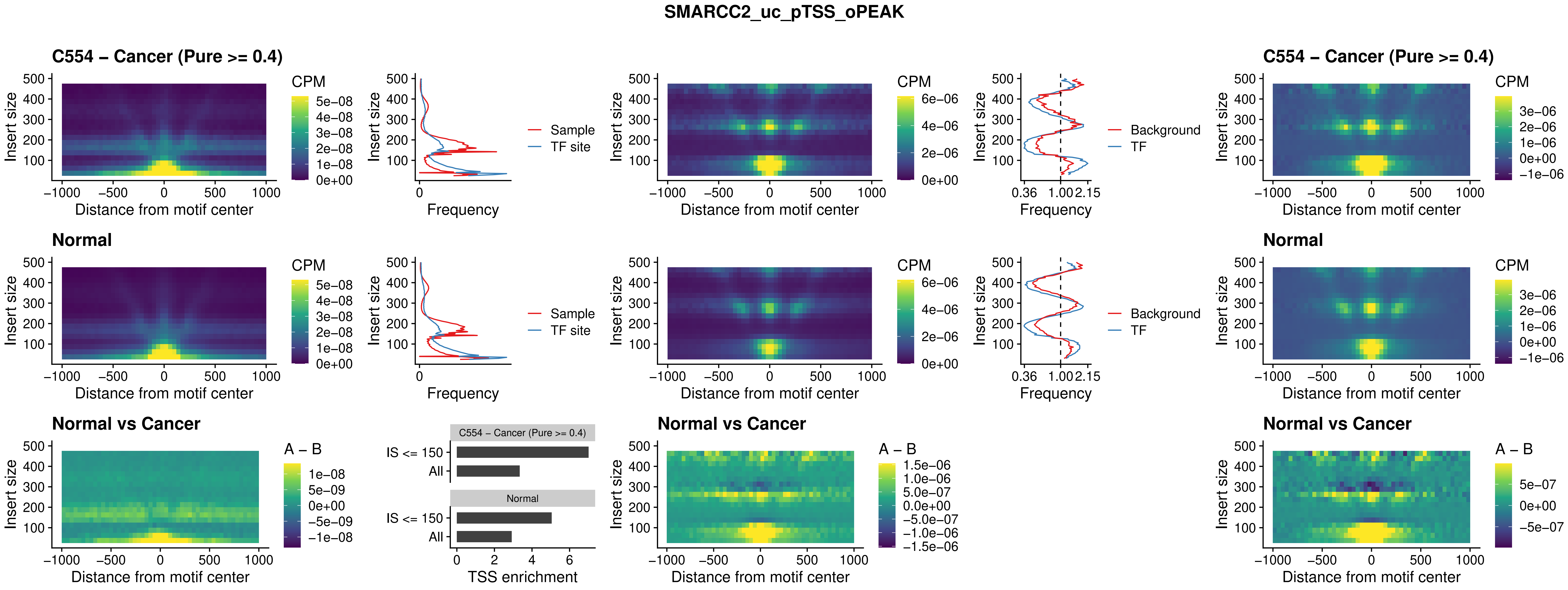
SMARCC2_uc_pTSS_oPEAK
C554 – Cancer (Pure >= 0.4)
Normal
Normal vs Cancer
Insert size
Distance from motif center
Frequency
CPM
A – B
Sample
TF site
Background
TF
C554 – Cancer (Pure >= 0.4)
Normal
IS <= 150
All
TSS enrichment

#### BCL11A_uc_cTSS_nPEAK

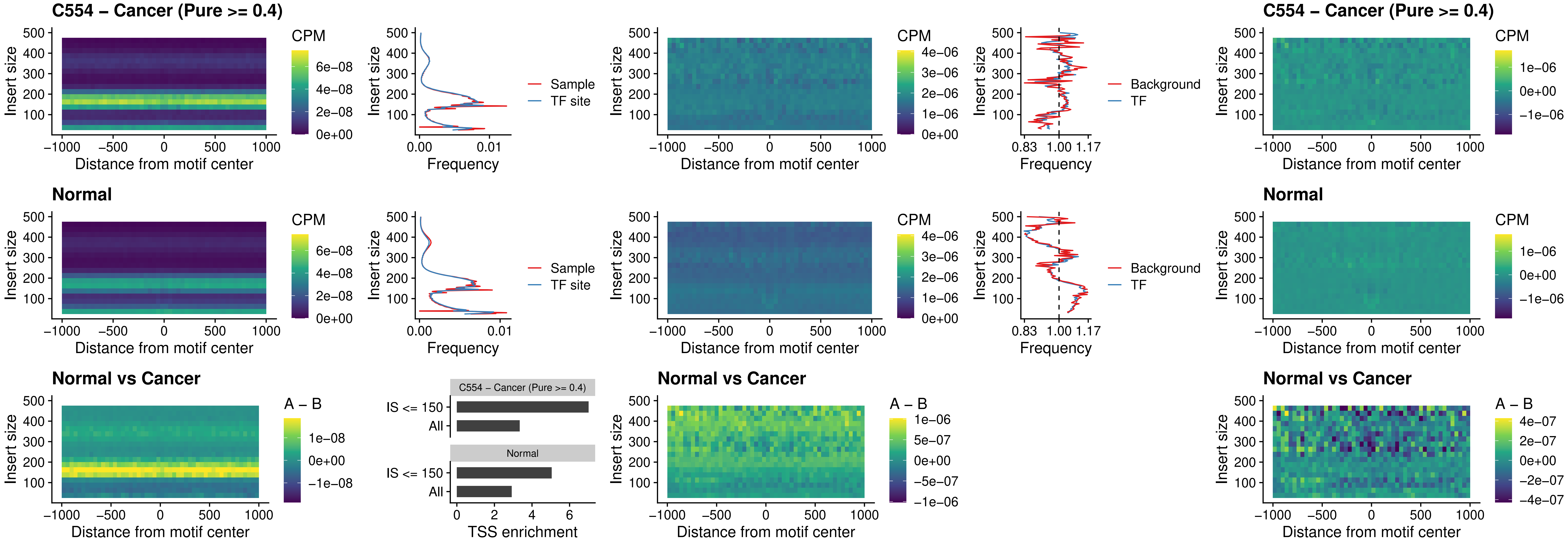

### BCL11B_uc_cTSS_nPEAK

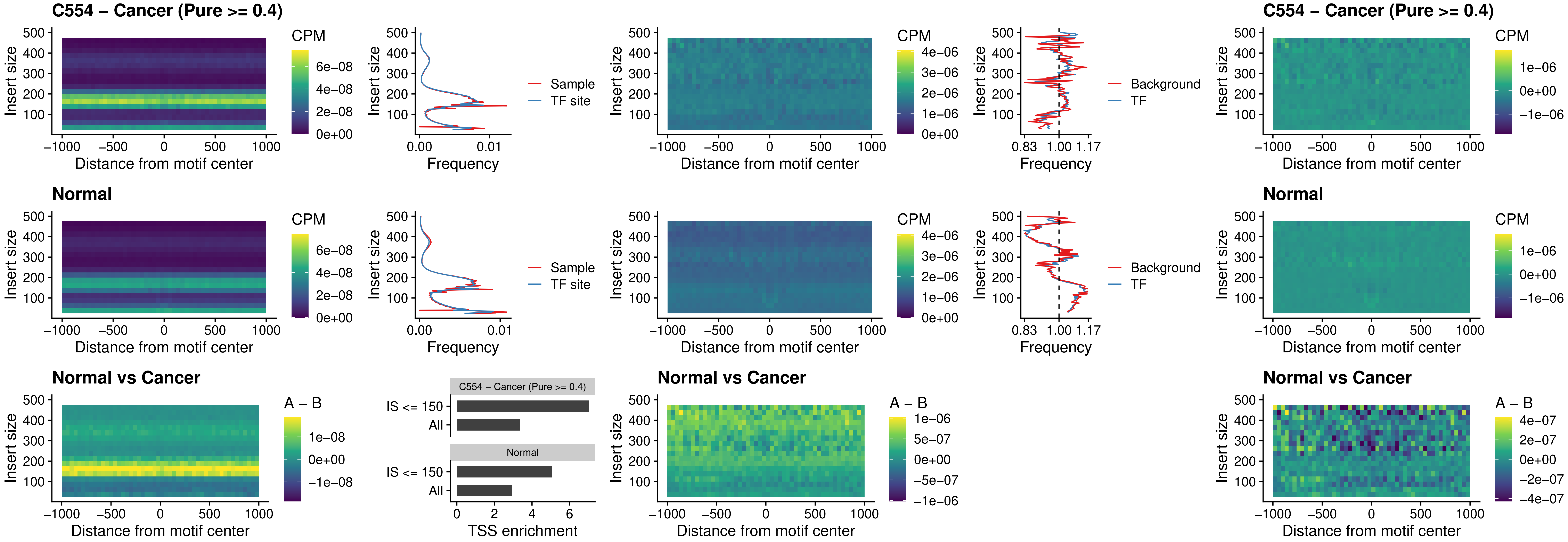

#### STAT2_uc_dTSS_oPEAK

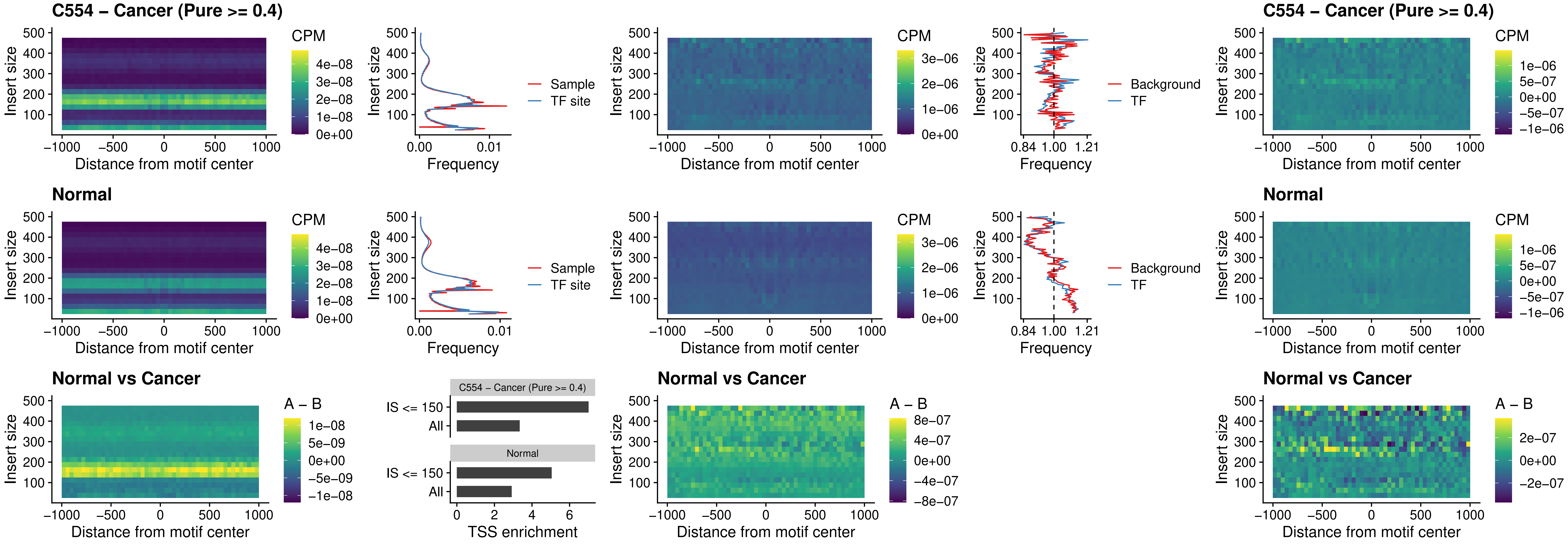

#### ZNF683_uc_cTSS_oPEAK

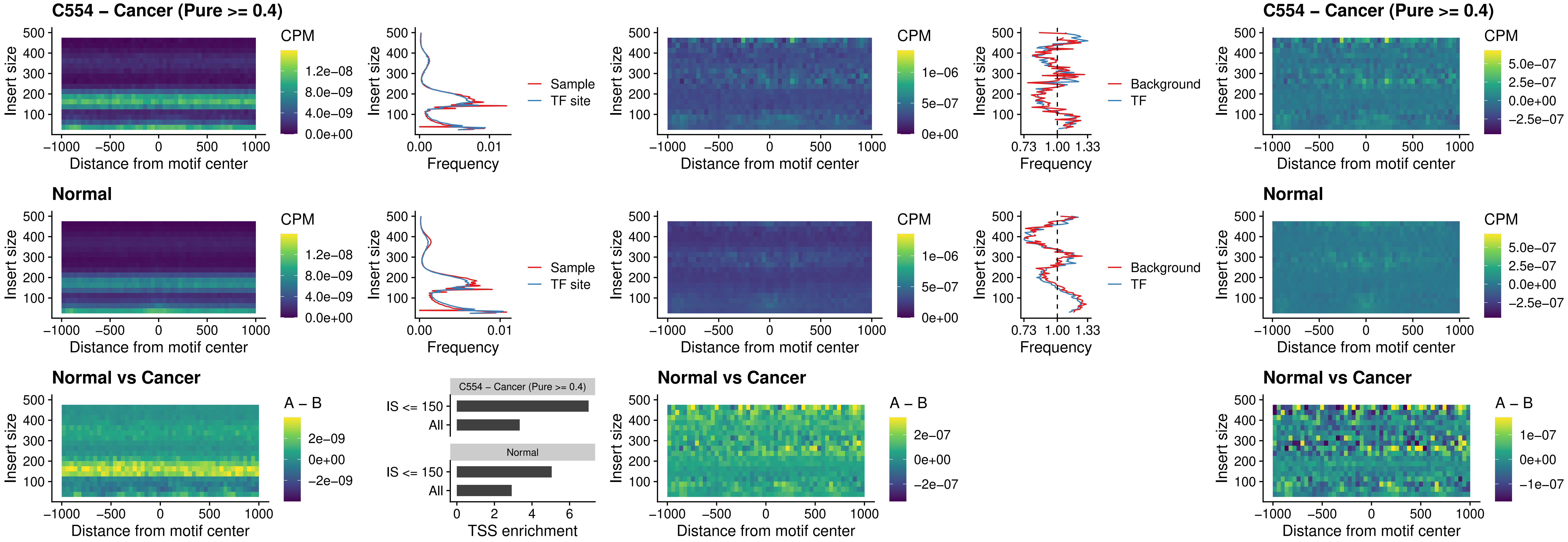

### IRF8_uc_dTSS_nPEAK

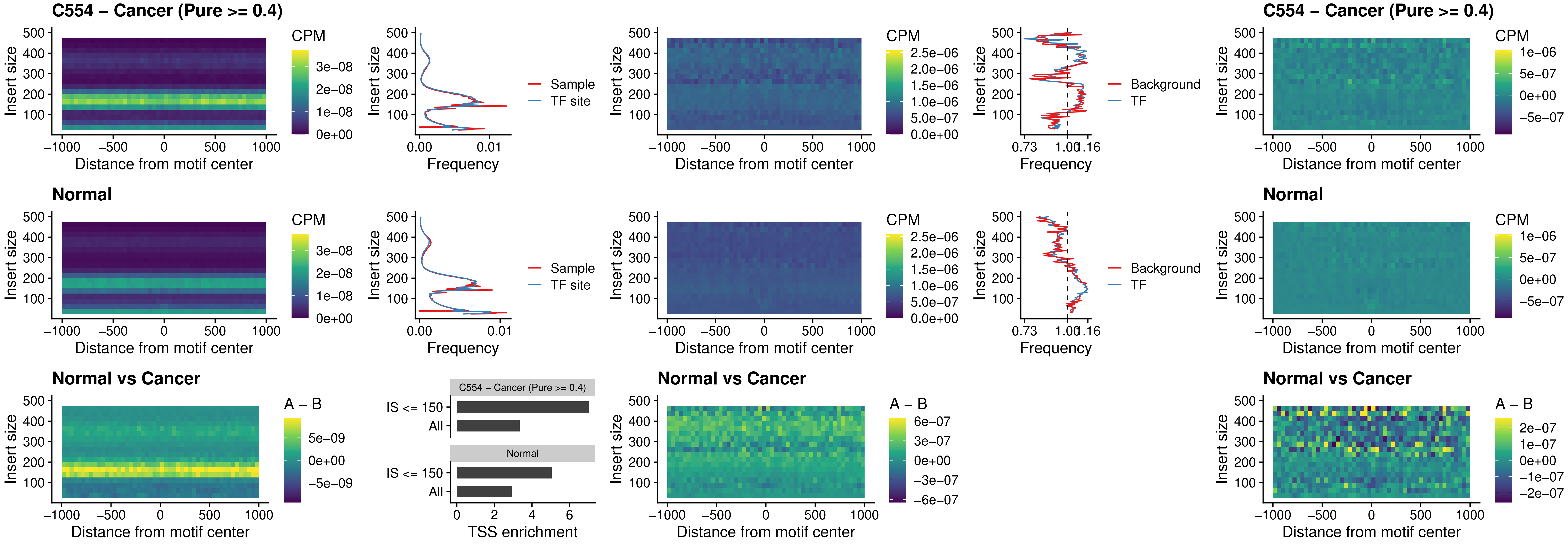

### ETV7_uc_dTSS_oPEAK

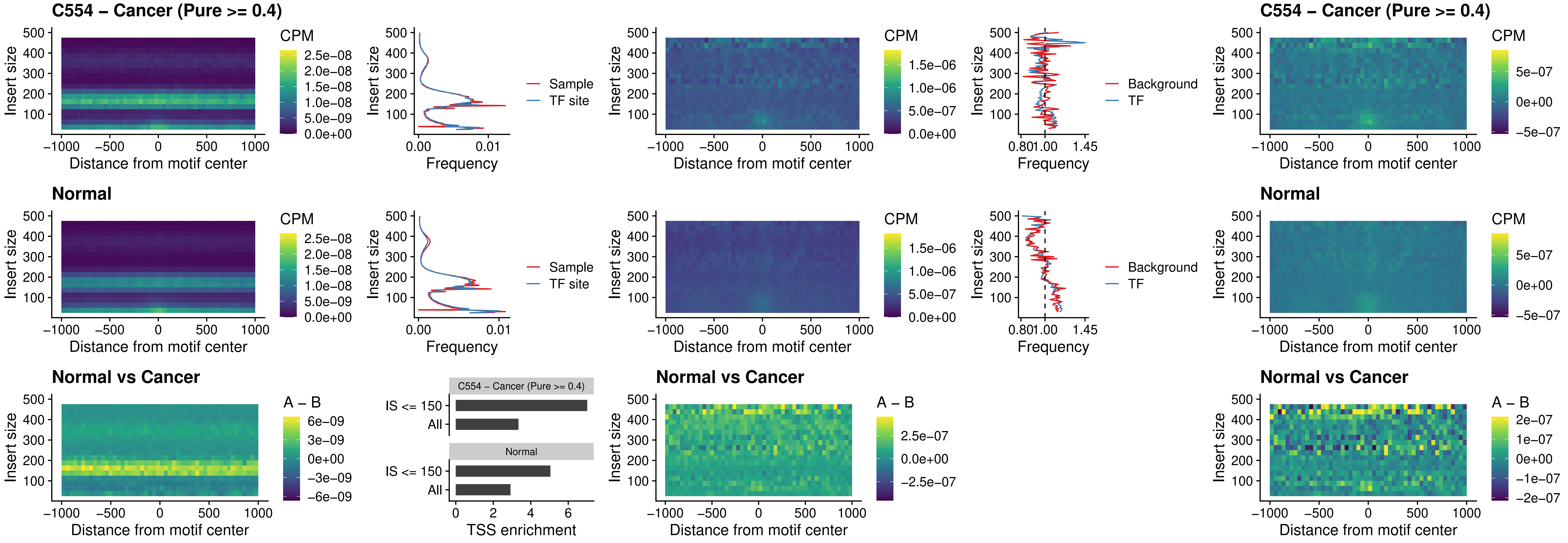

### FOXL1_uc_pTSS_oPEAK

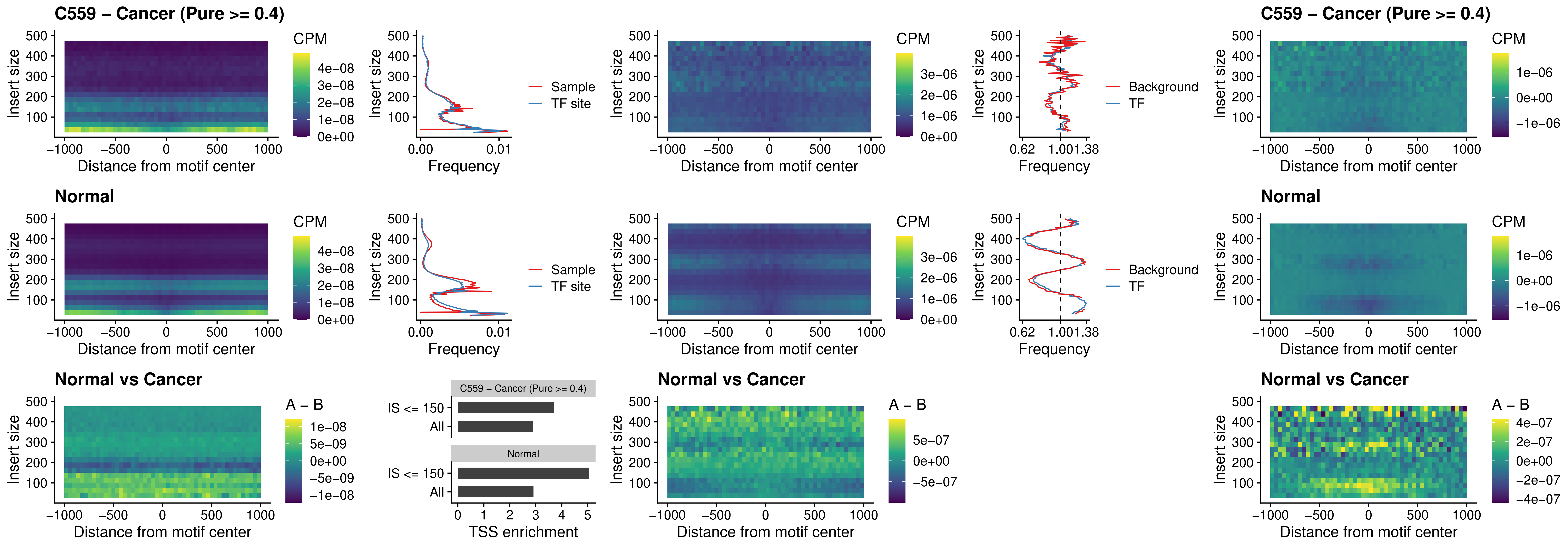

### CPEB1_uc_cTSS_oPEAK

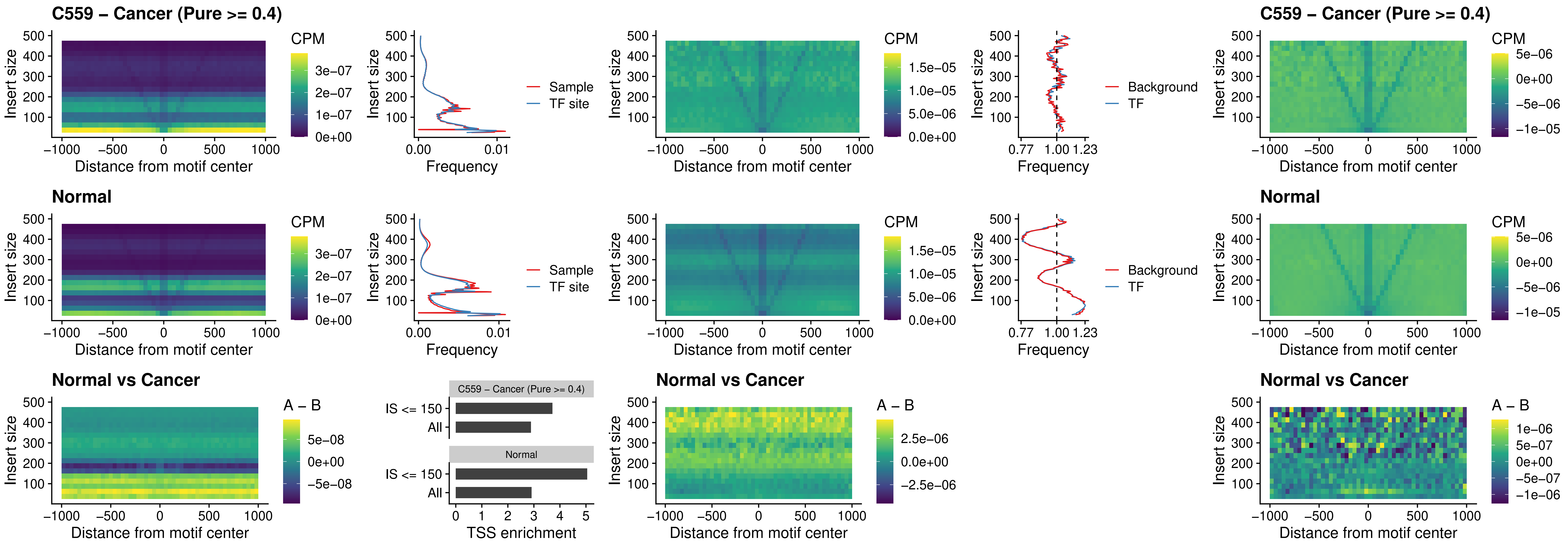

### HMGA1_uc_pTSS_nPEAK

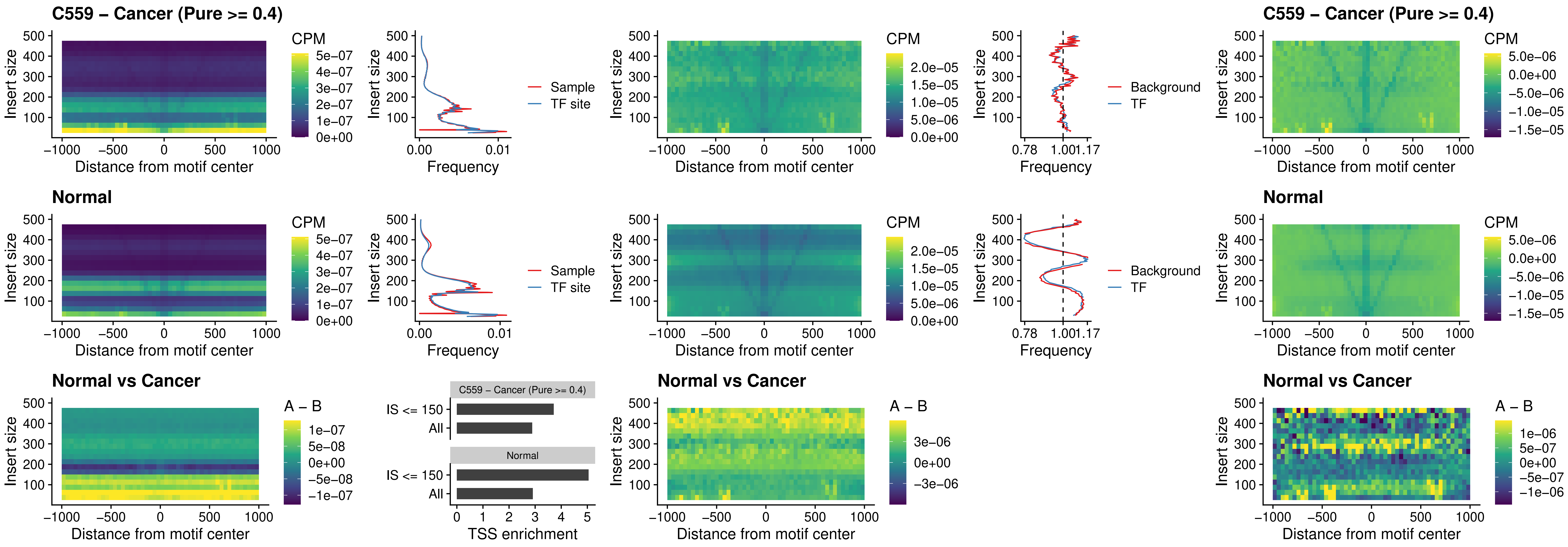

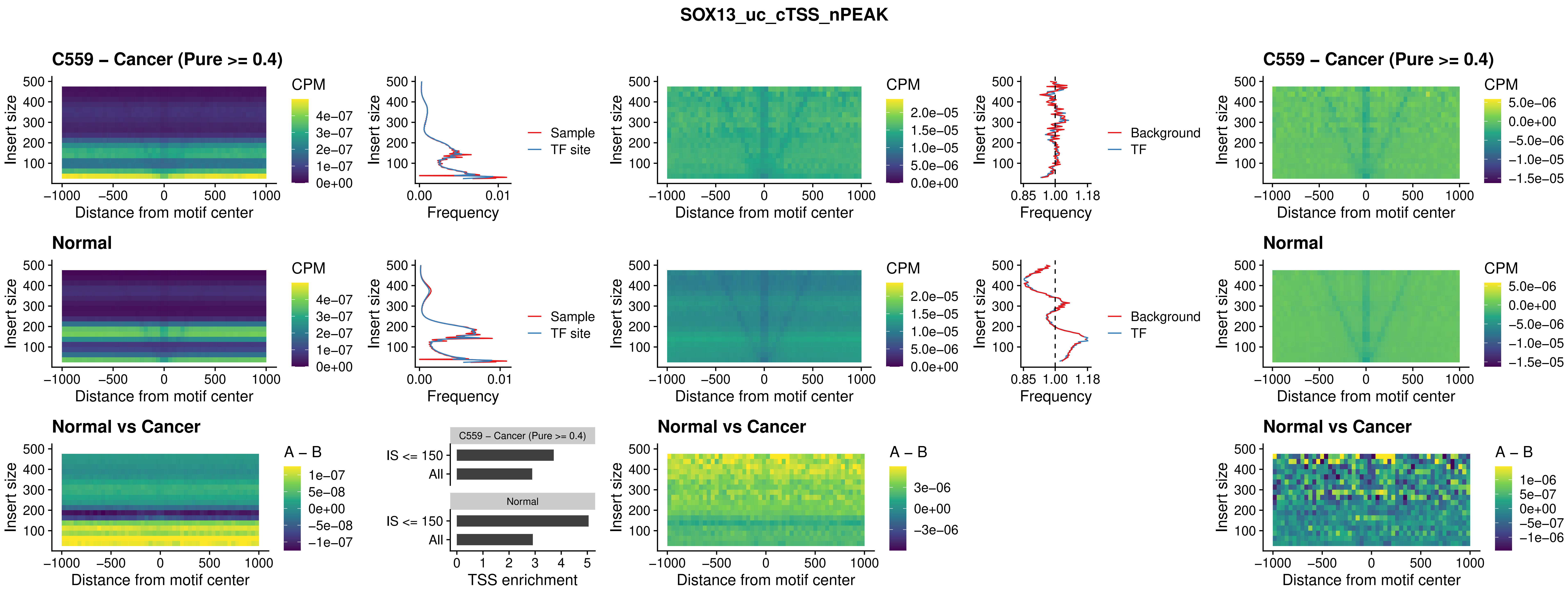
SOX13_uc_cTSS_nPEAK
C559 – Cancer (Pure >= 0.4)
Normal
Normal vs Cancer
Insert size
Distance from motif center
Frequency
CPM
A – B
Sample
TF site
Background
TF
C559 – Cancer (Pure >= 0.4)
Normal
IS <= 150
All
TSS enrichment

#### LHX4_uc_cTSS_nPEAK

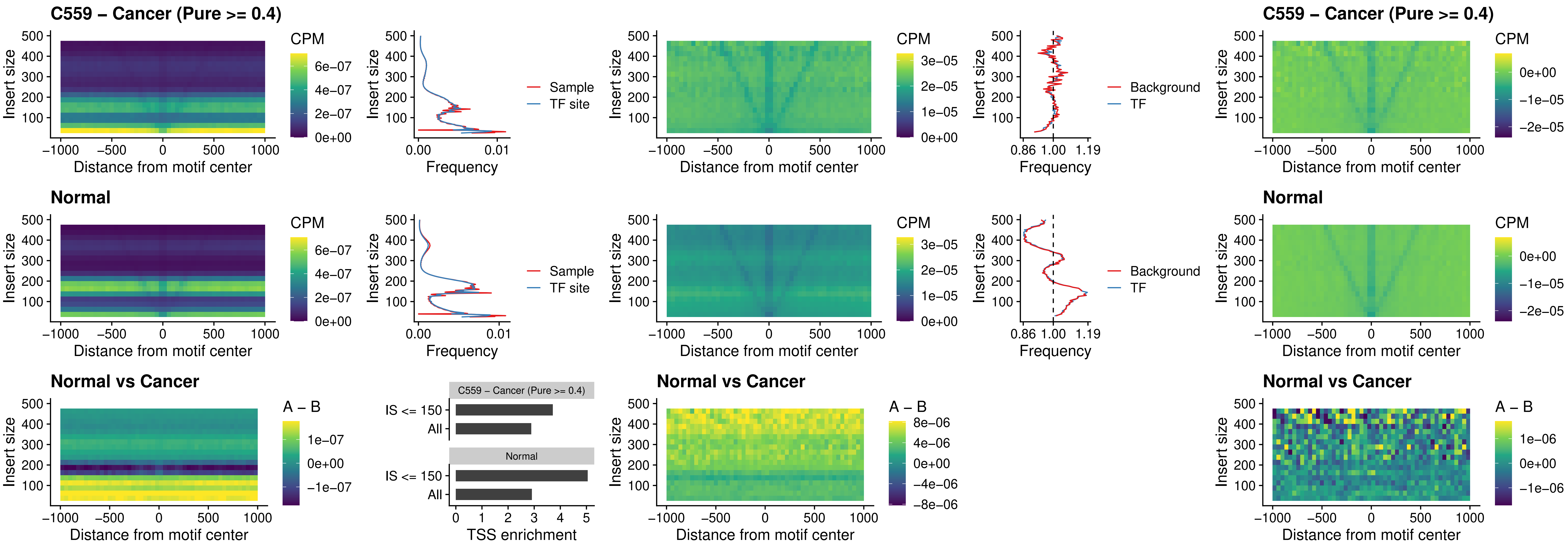

### HOXA4_uc_cTSS_nPEAK

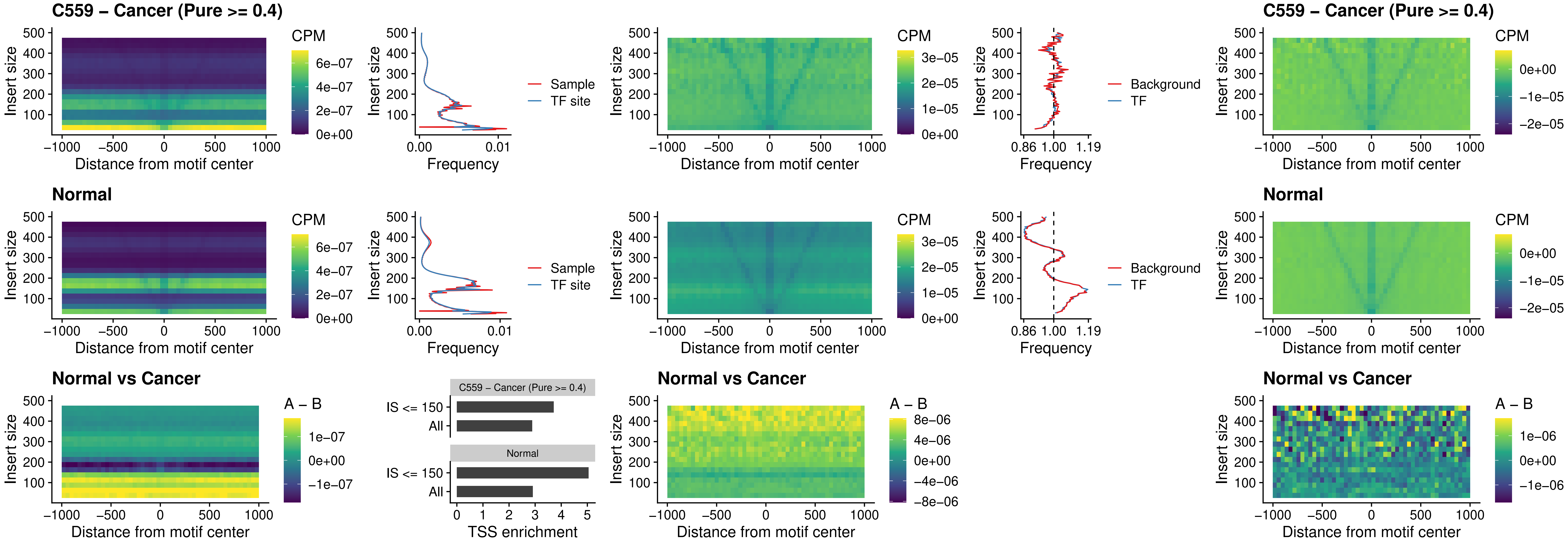

### HLX_uc_cTSS_nPEAK

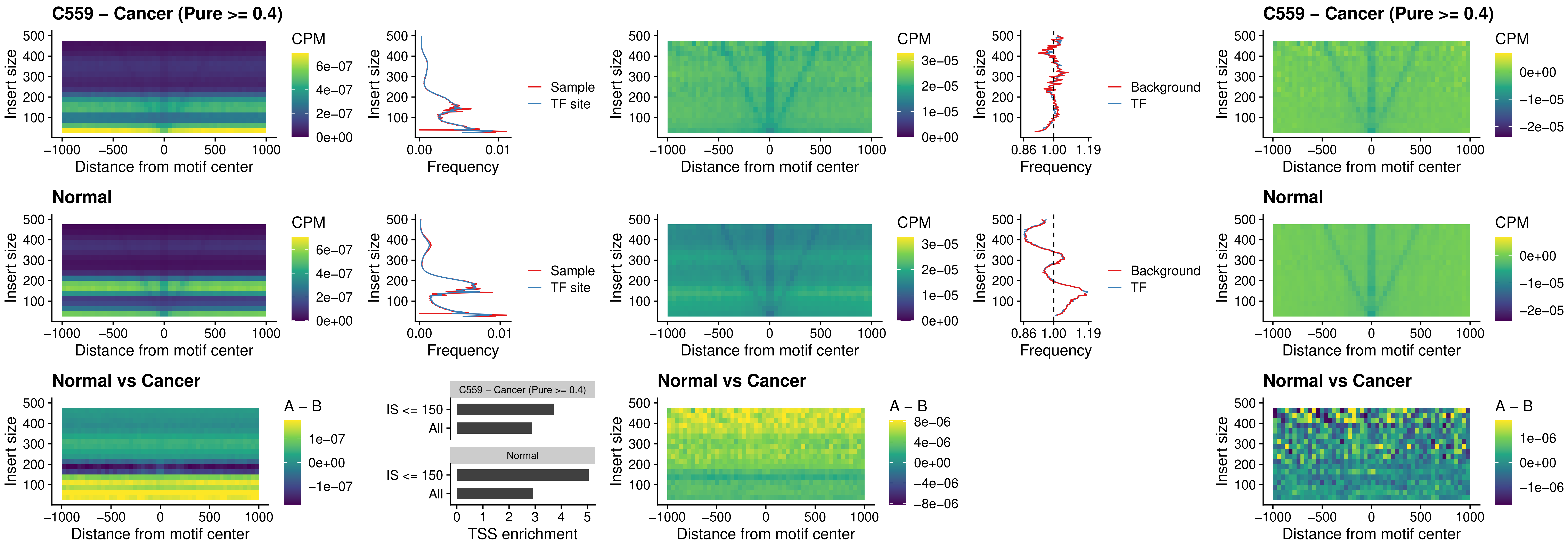

#### AC0125311_uc_cTSS_nPEAK

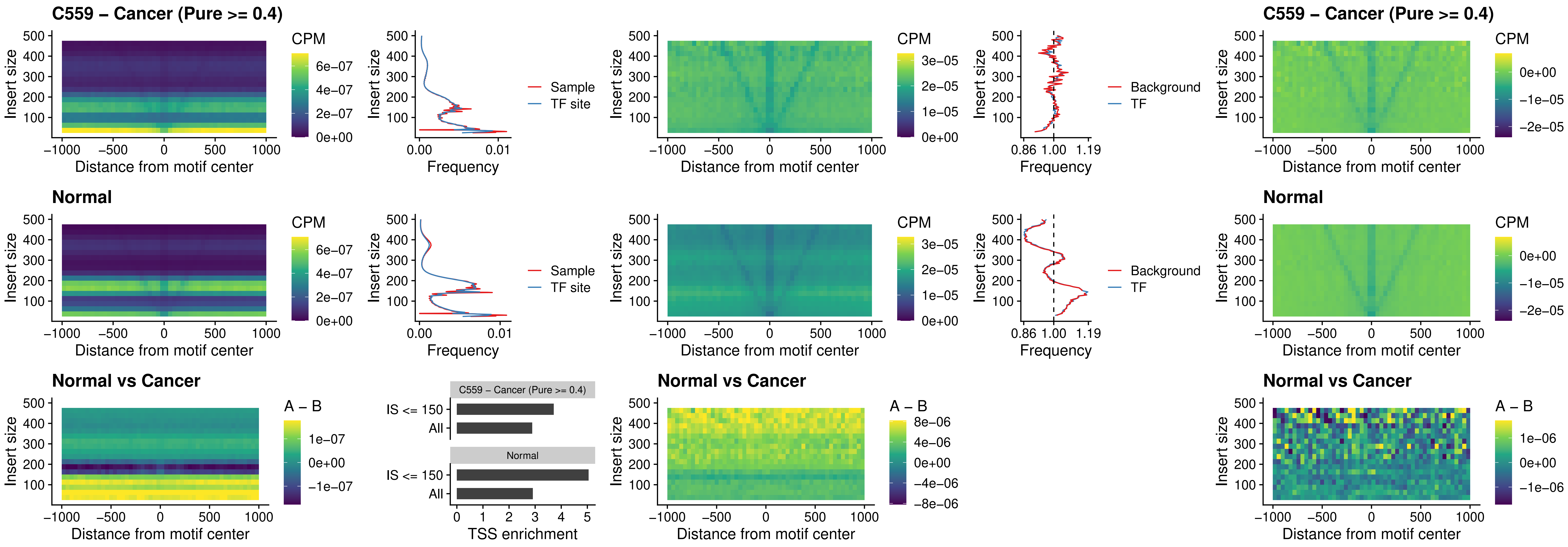

### DBX2_uc_cTSS_nPEAK

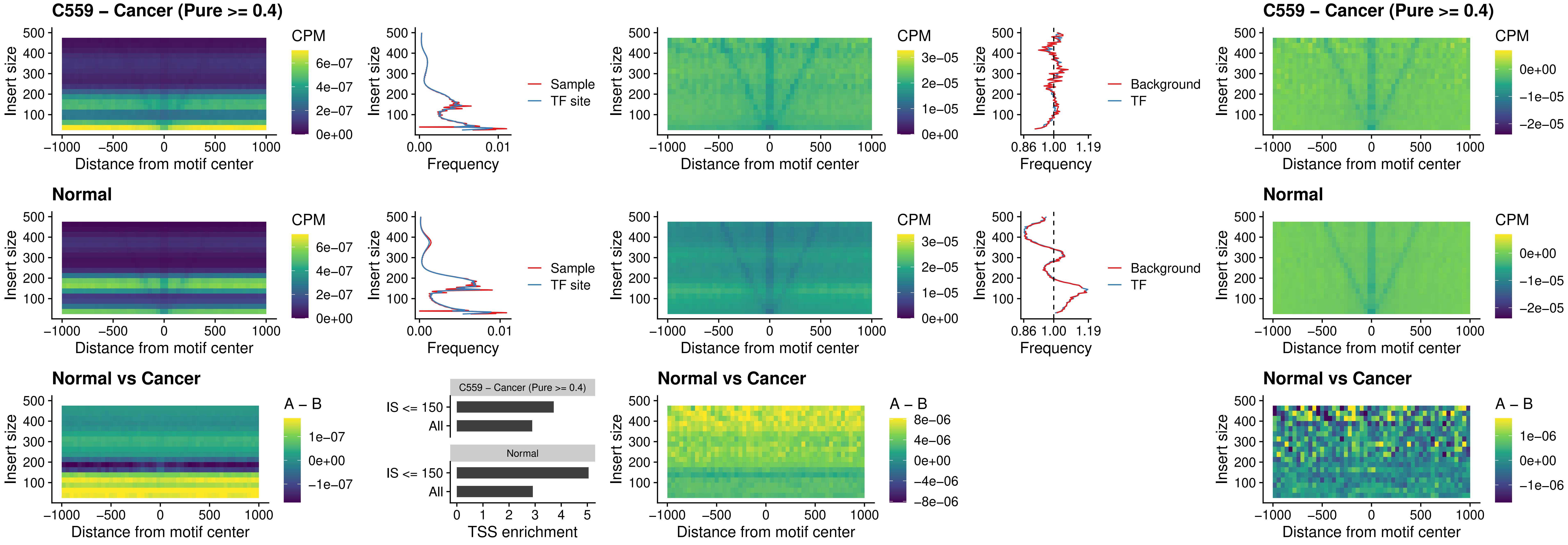

### LHX6_uc_dTSS_nPEAK

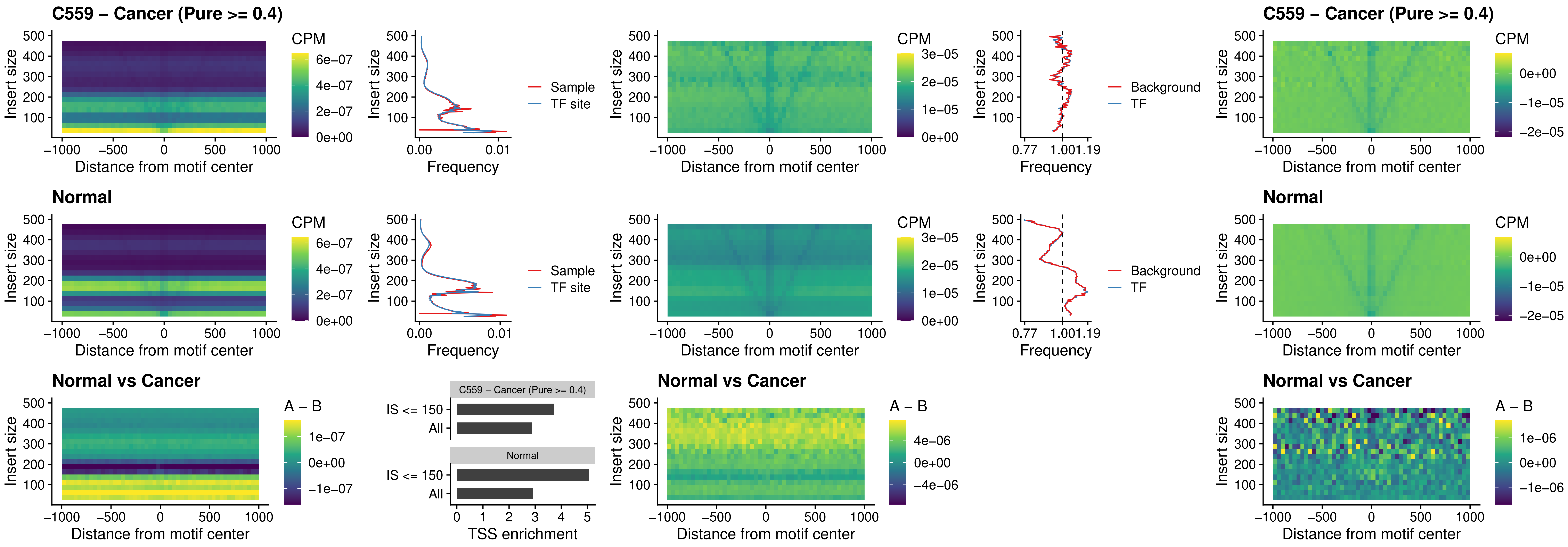

### SOX1_uc_dTSS_nPEAK

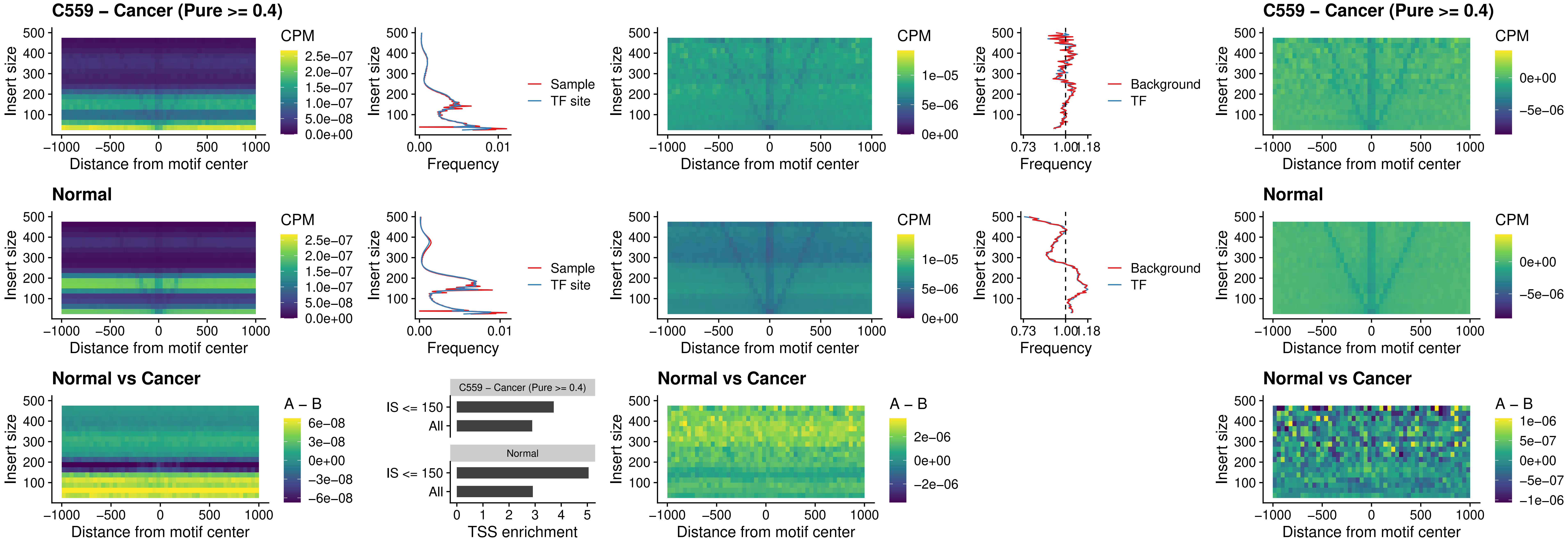

#### NKX24_uc_dTSS_nPEAK

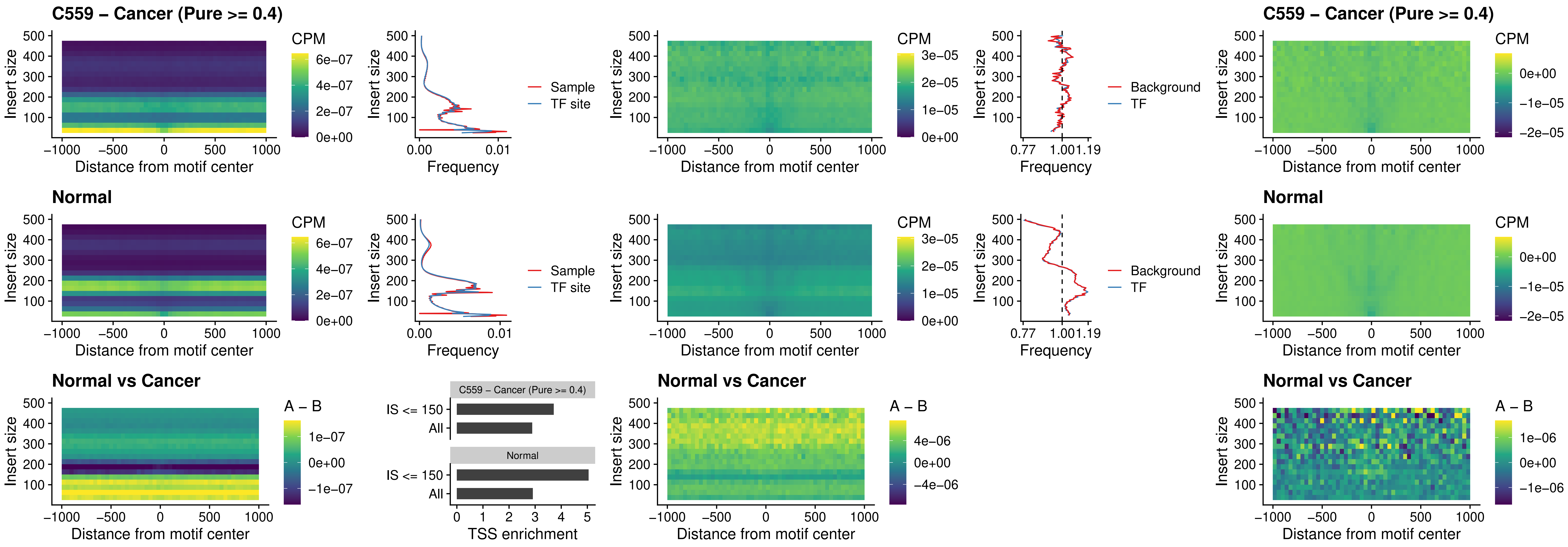

### DLX3_uc_cTSS_nPEAK

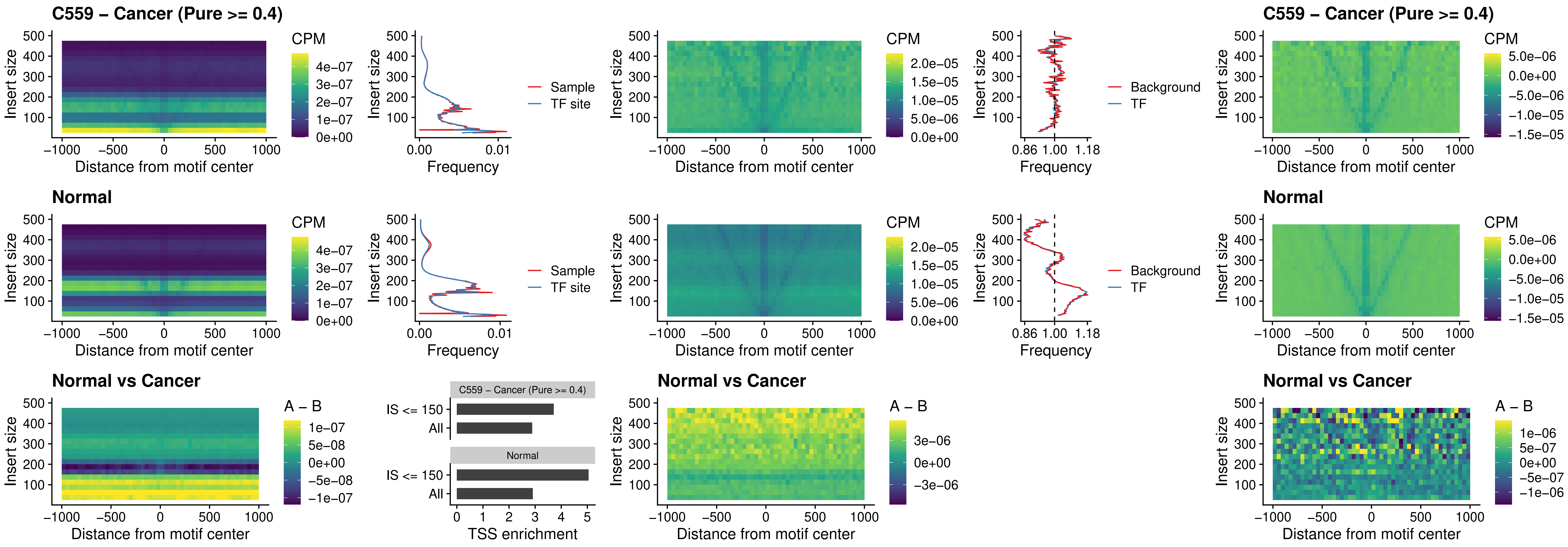

### HOXD3_uc_cTSS_nPEAK

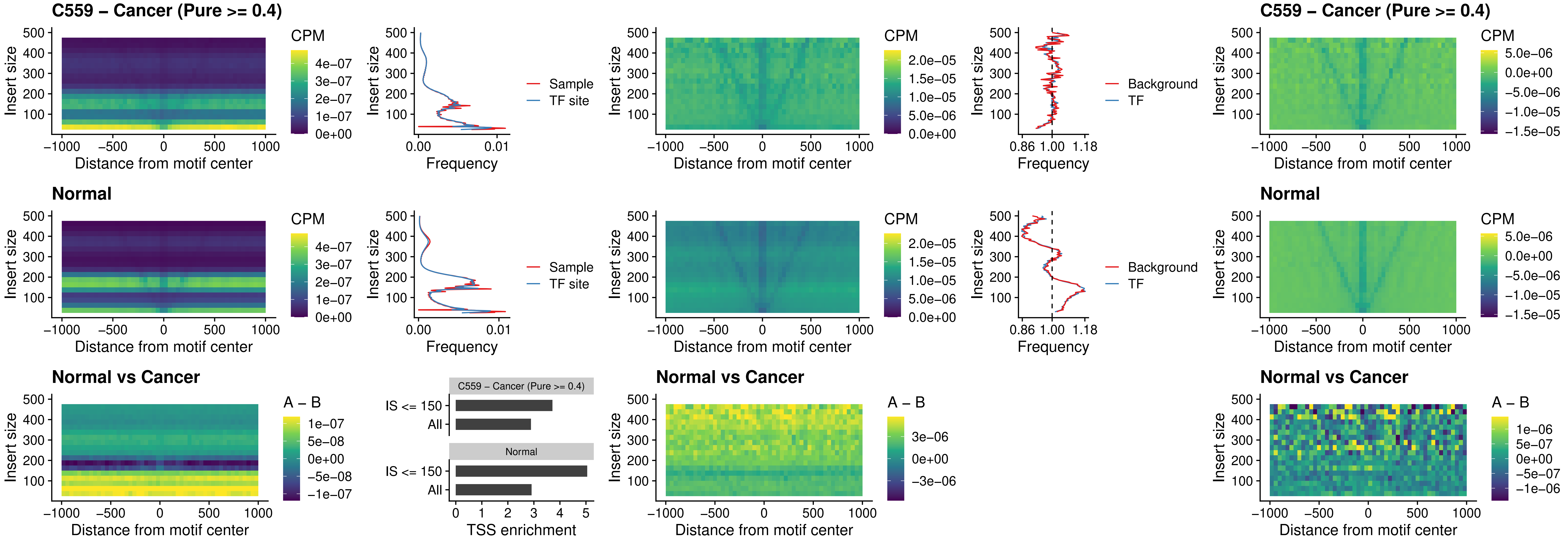

### ISL1_uc_dTSS_oPEAK

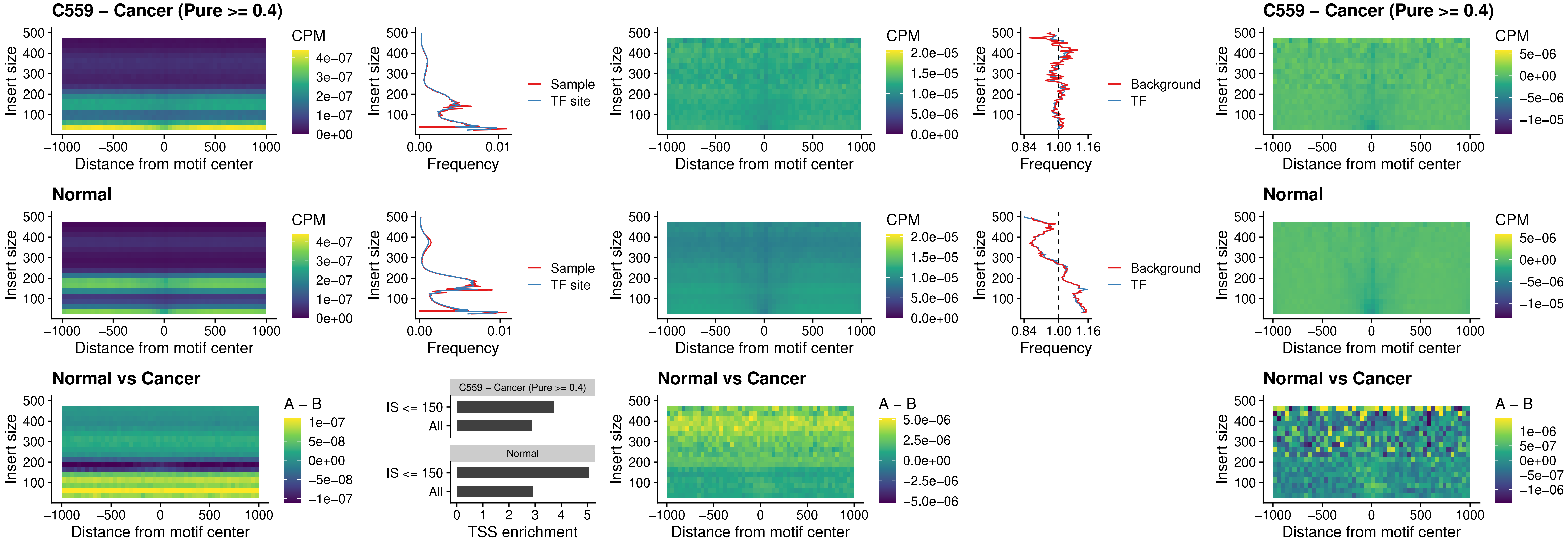

### HOXB4_uc_dTSS_oPEAK

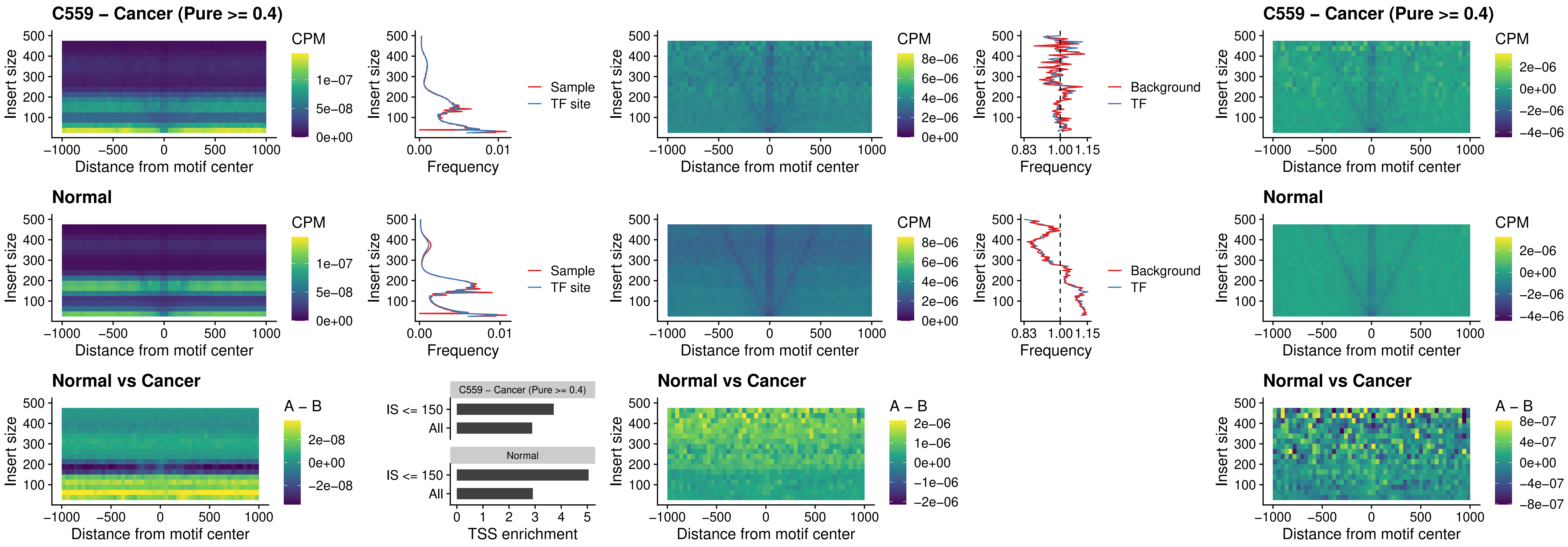

### VAX1_uc_dTSS_oPEAK

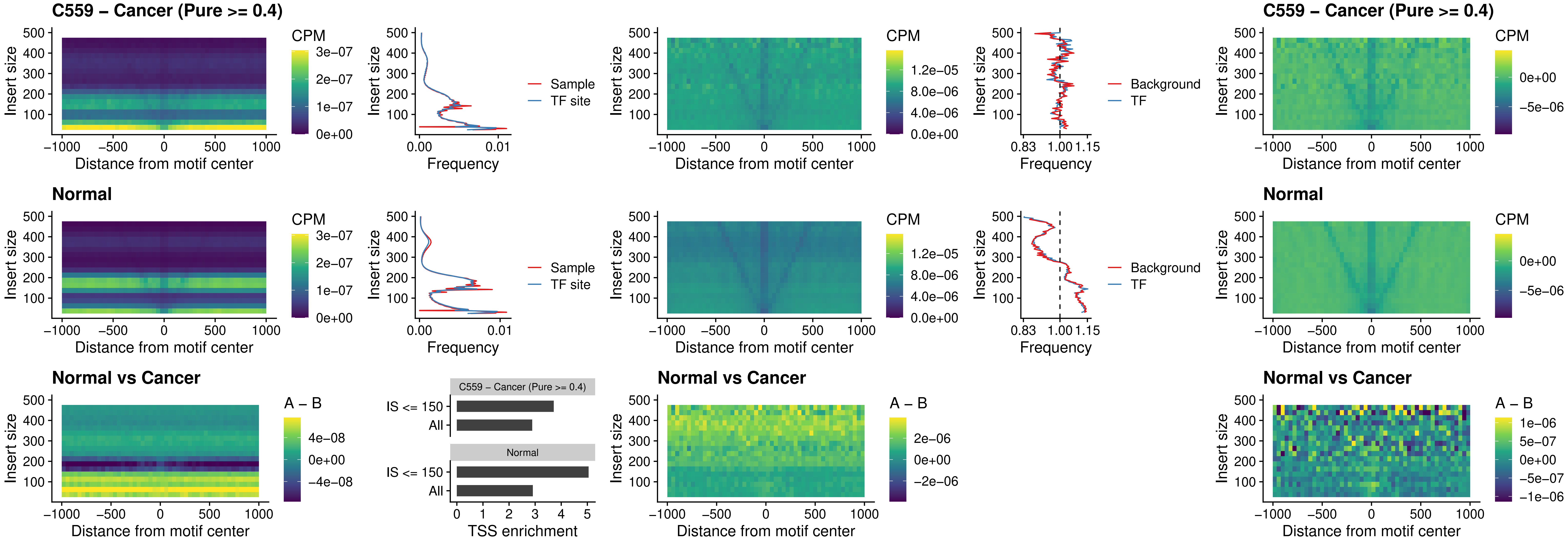

### DLX3_uc_dTSS_oPEAK

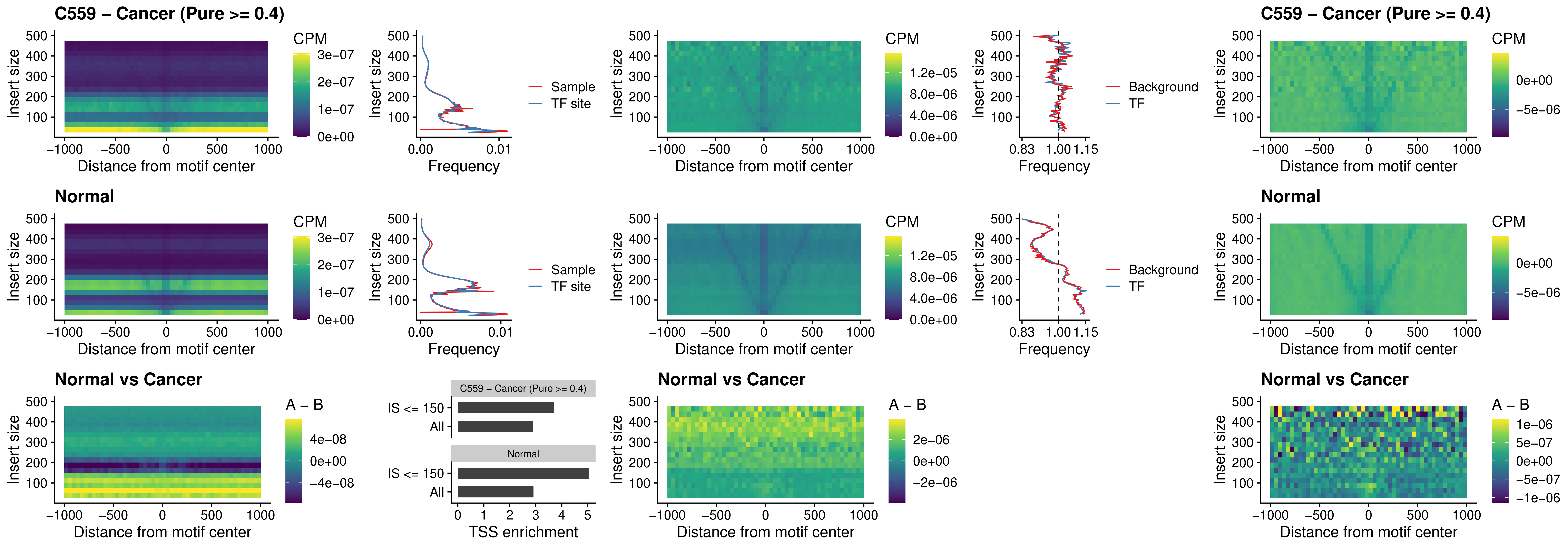

### HOXD3_uc_dTSS_oPEAK

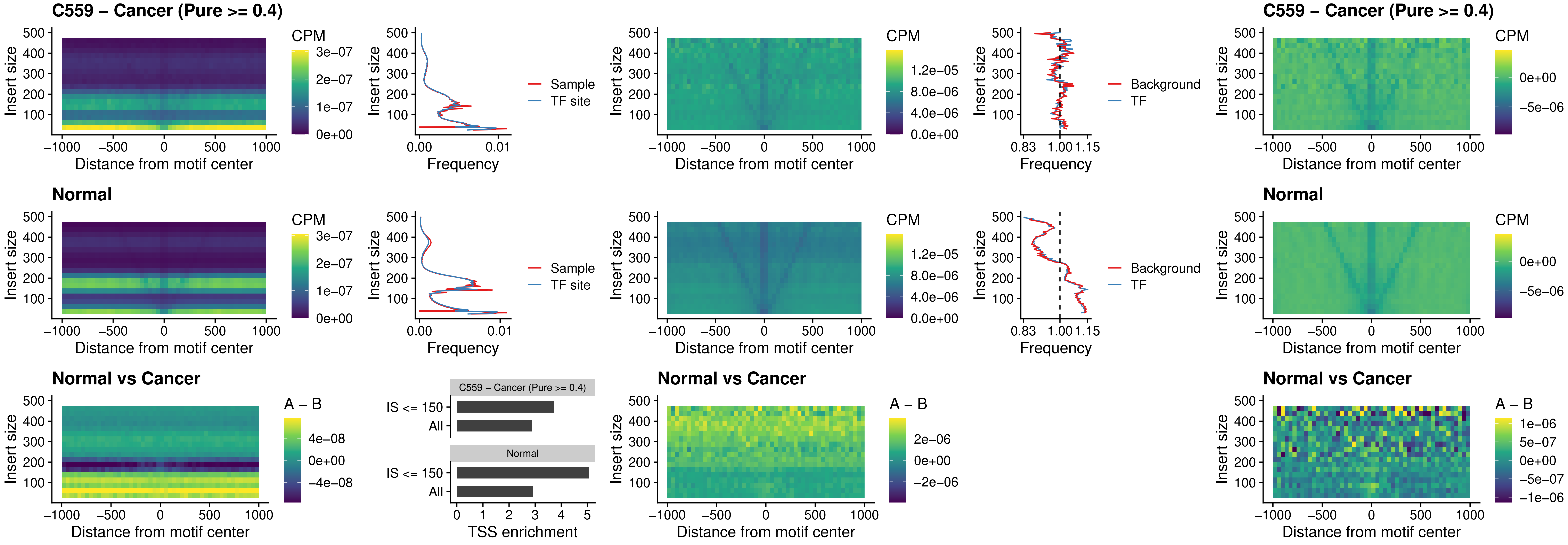

### BBX_uc_dTSS_nPEAK

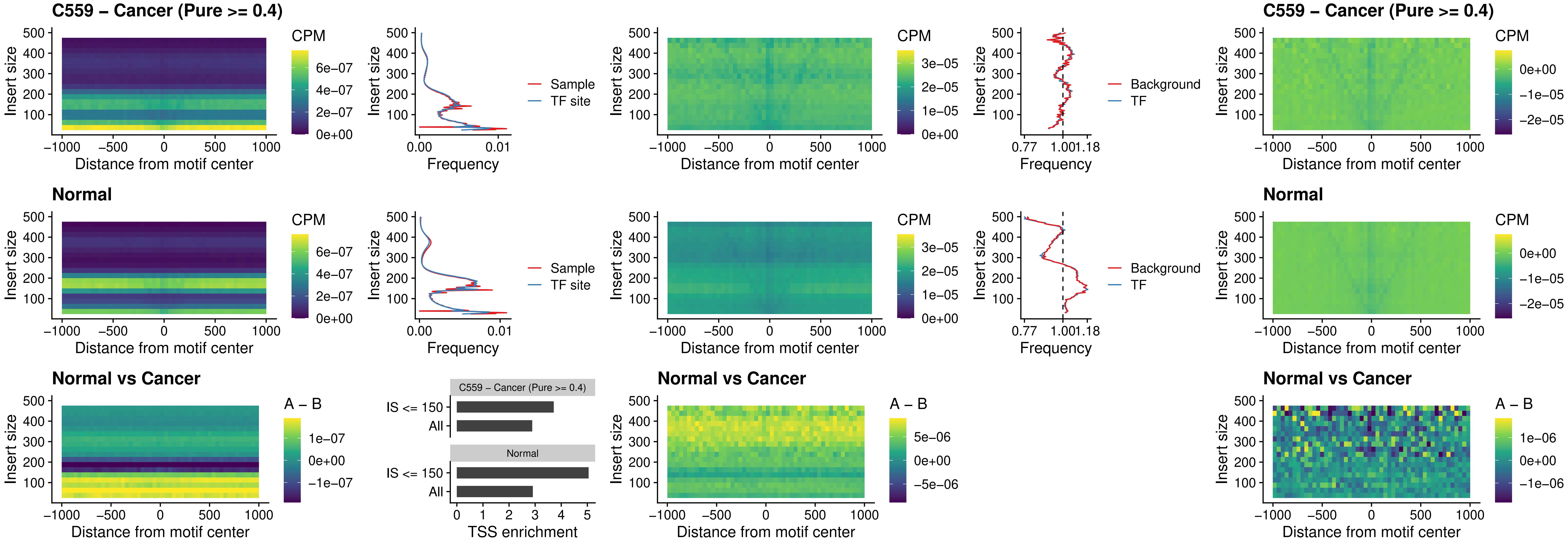

#### SOX3_uc_cTSS_nPEAK

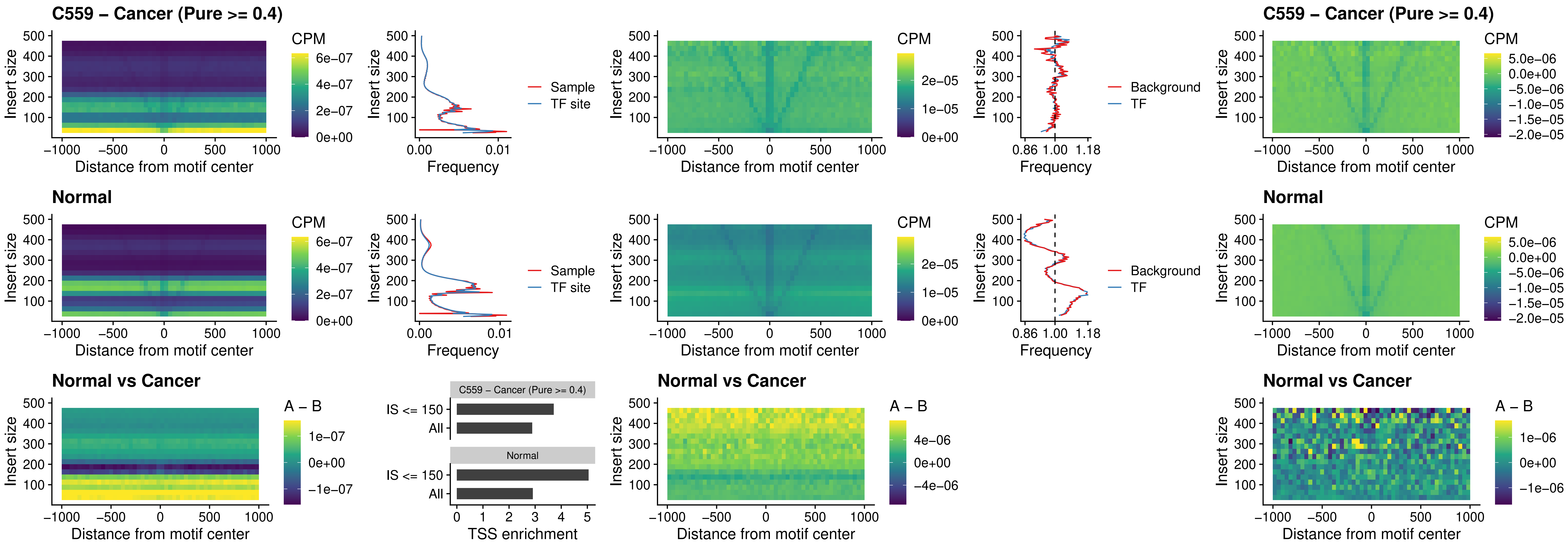

### TBPL2_uc_cTSS_nPEAK

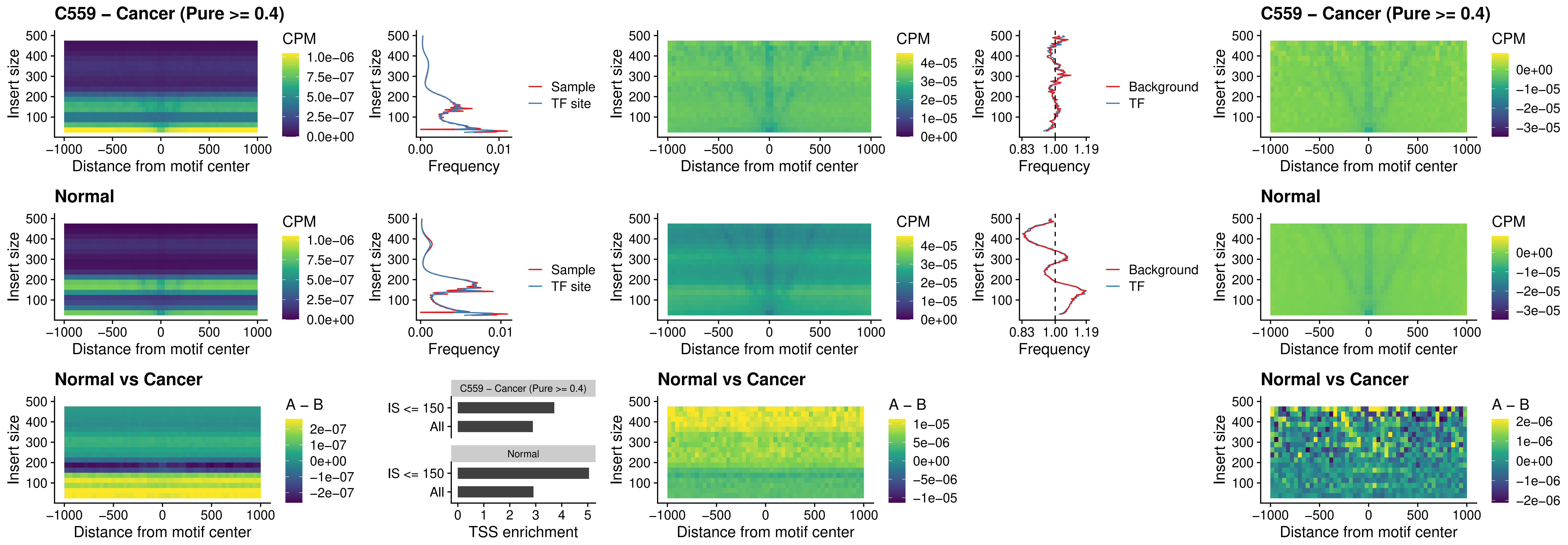

#### HOXA4_uc_cTSS_oPEAK

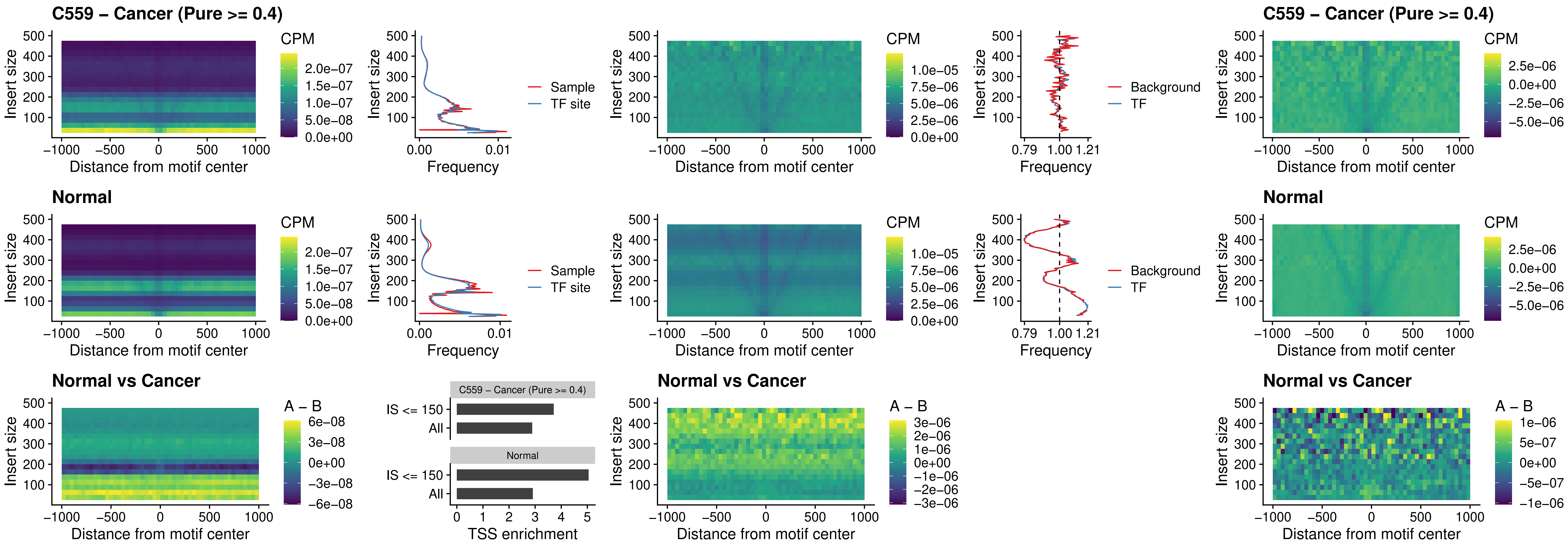

### PDX1_uc_dTSS_oPEAK

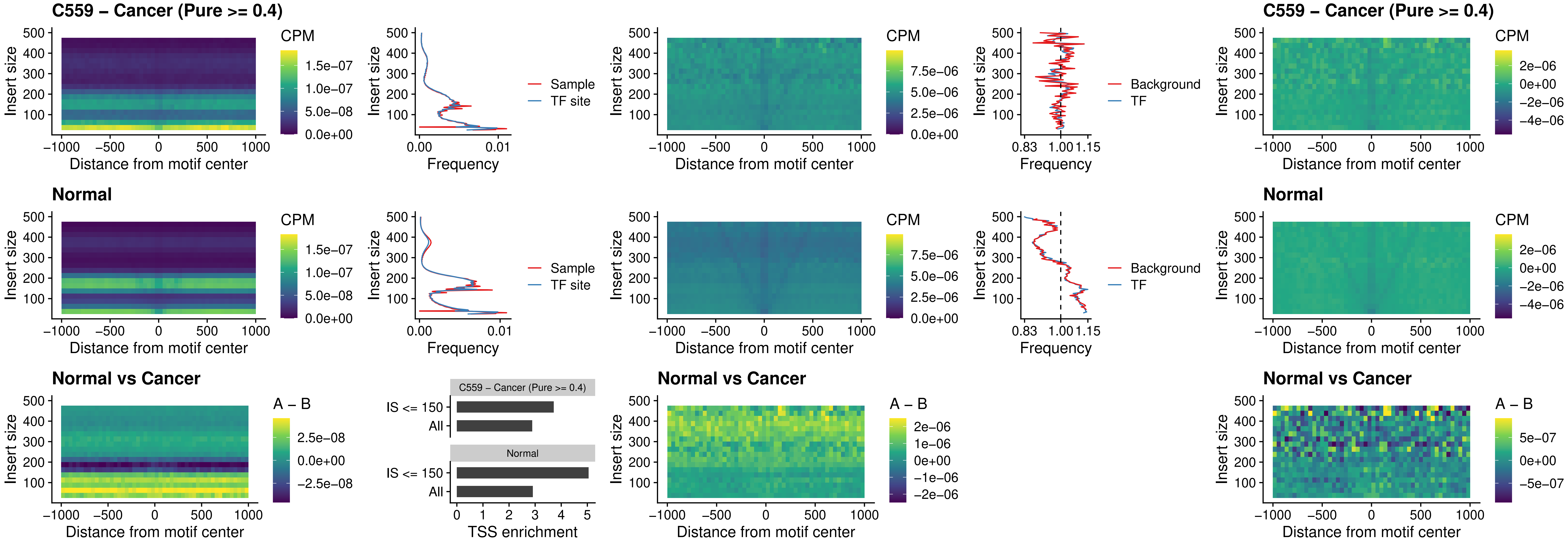

### LHX4_uc_dTSS_oPEAK

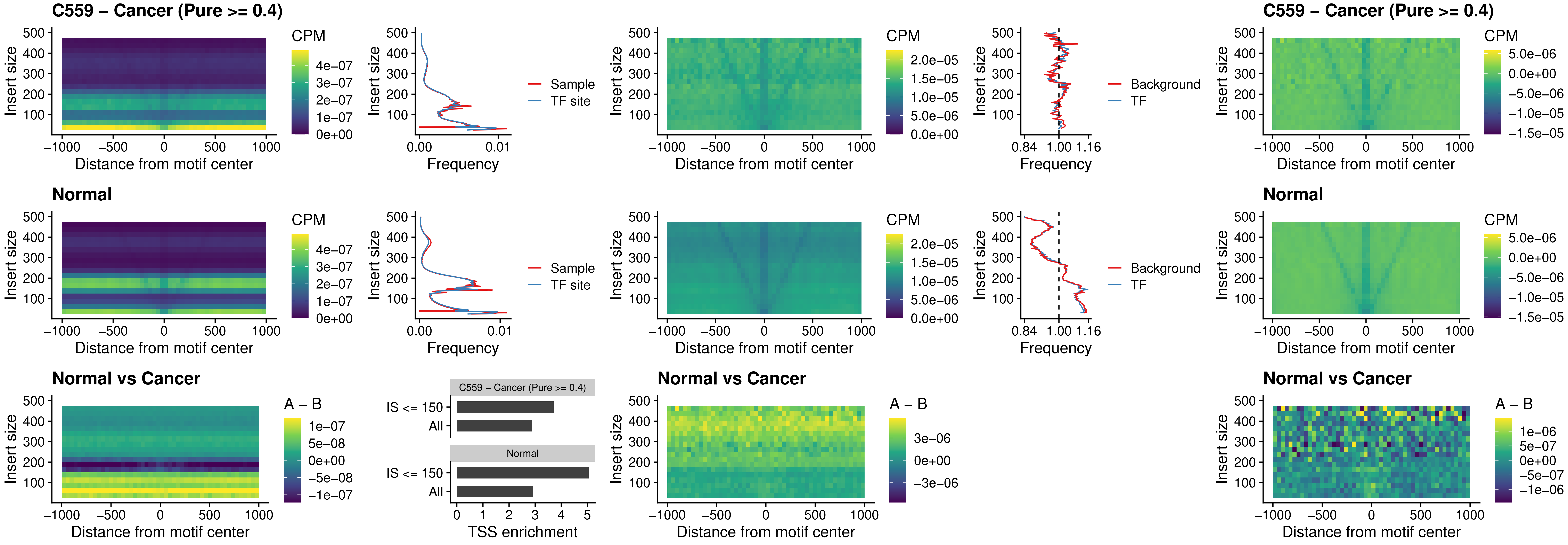

### HOXA4_uc_dTSS_oPEAK

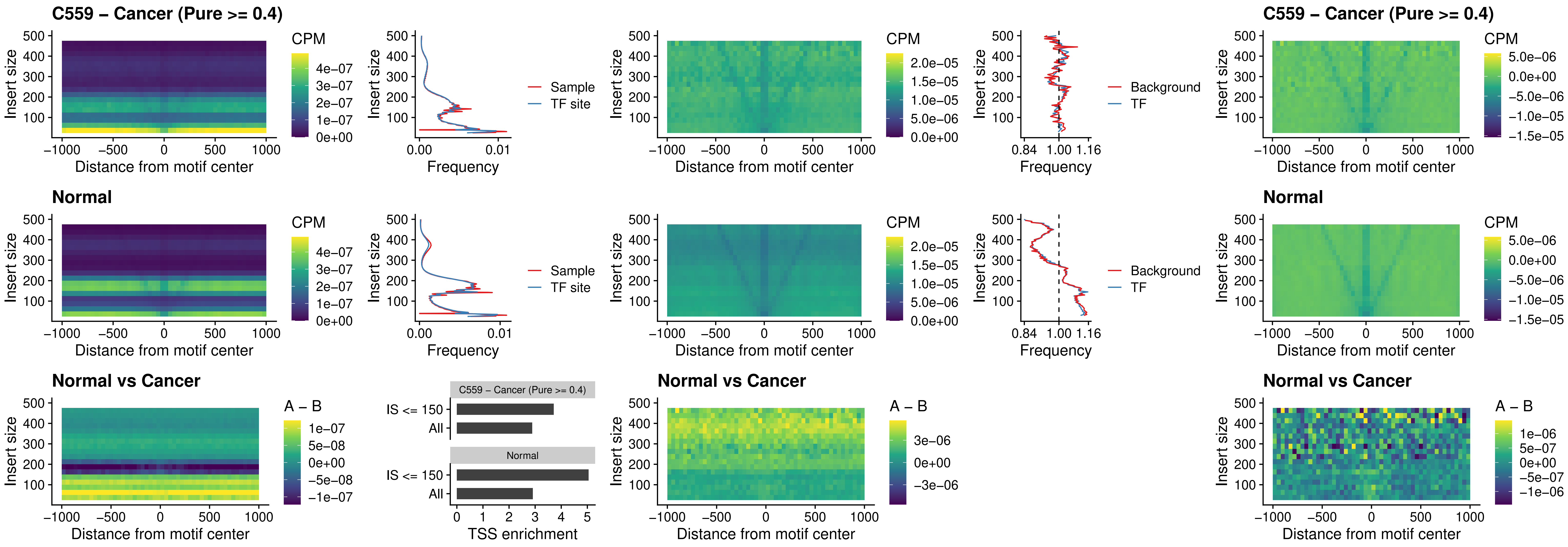

#### HLX_uc_dTSS_oPEAK

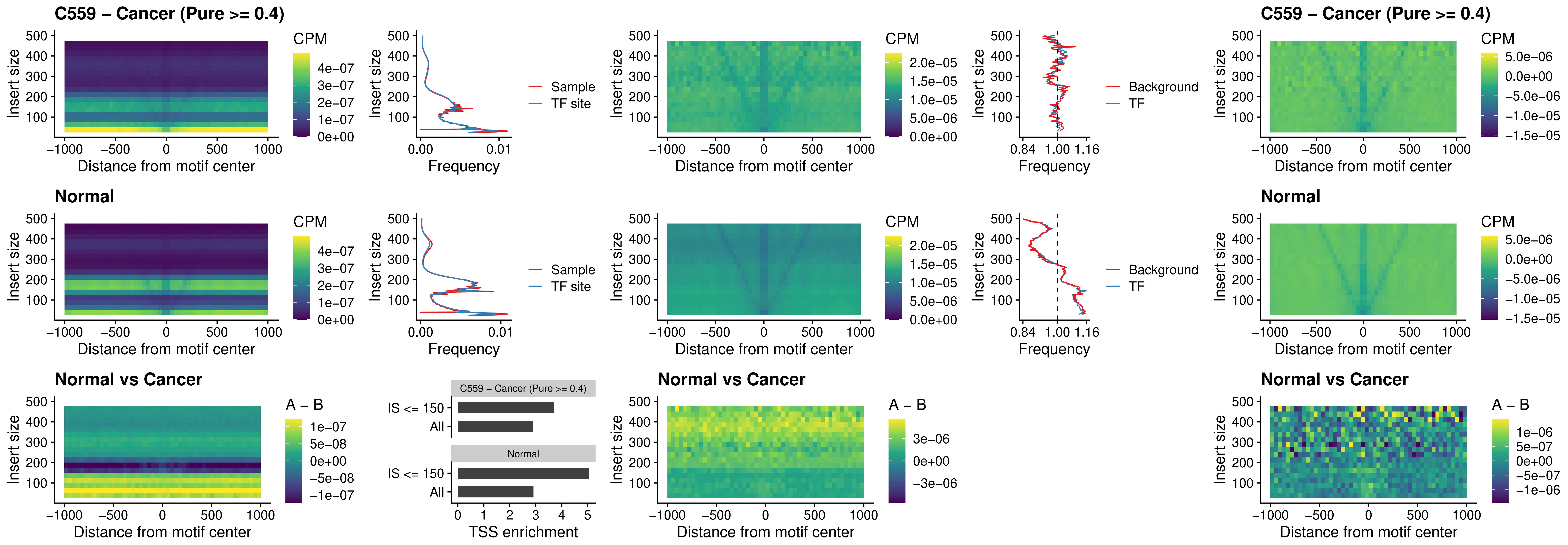

### AC0125311_uc_dTSS_oPEAK

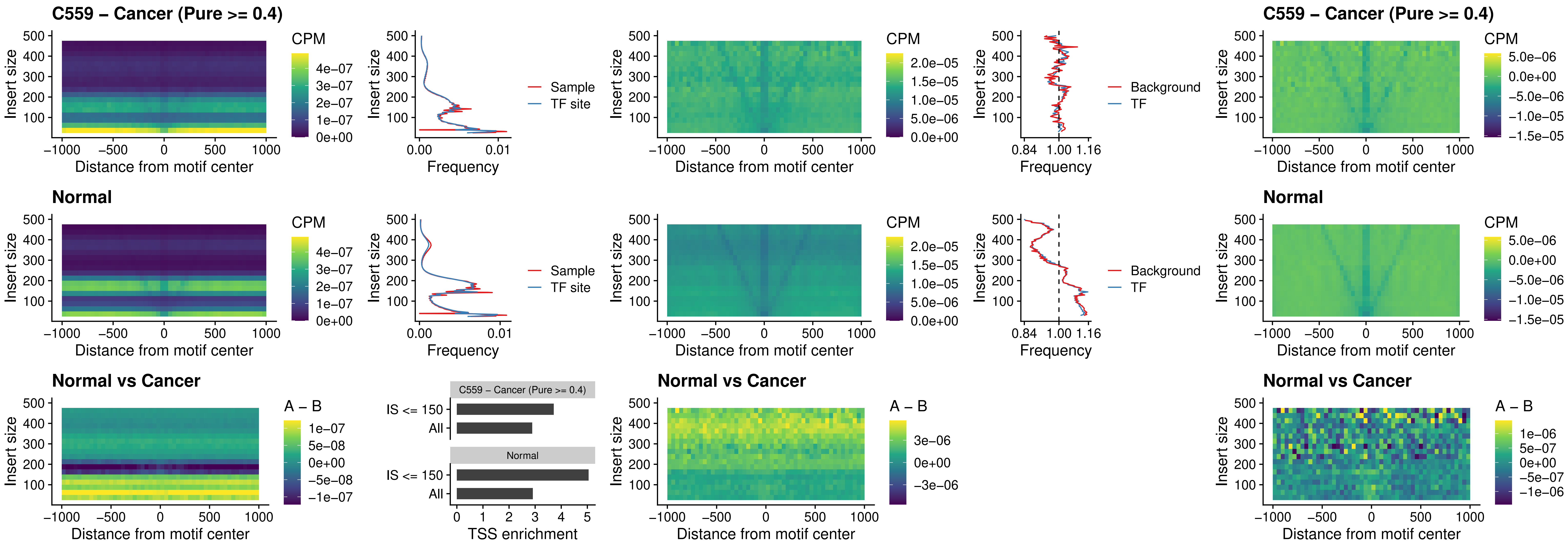

#### DBX2_uc_dTSS_oPEAK

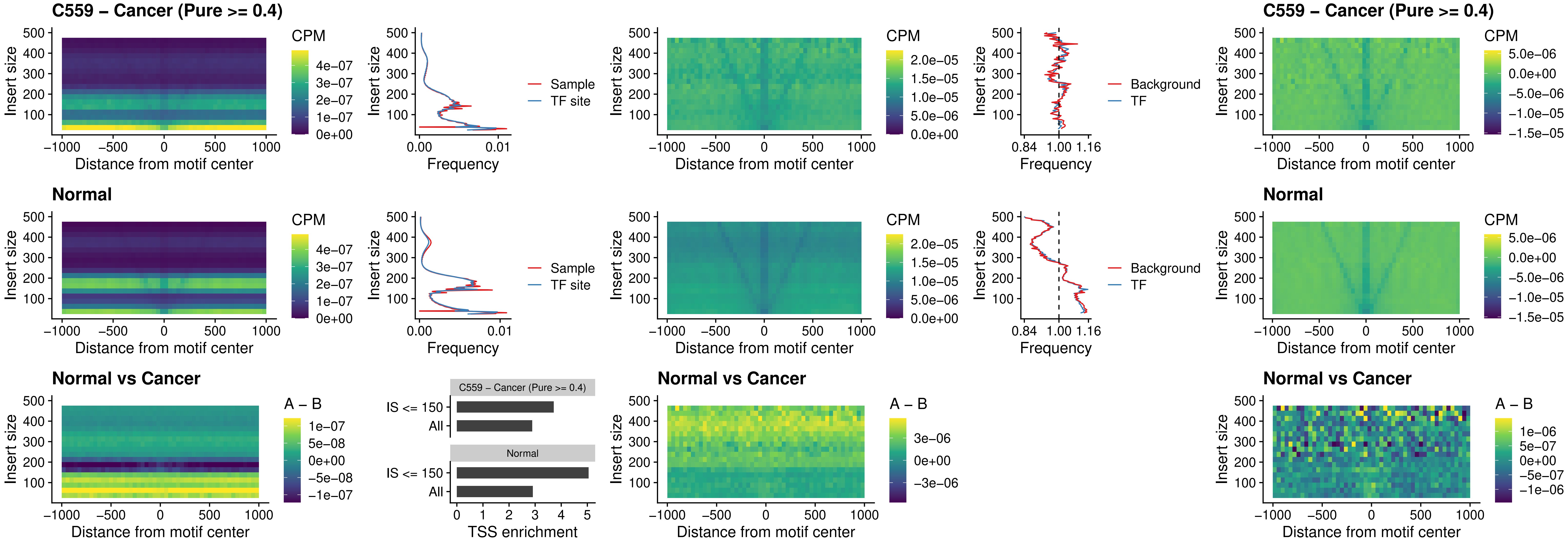

### HOXA2_uc_dTSS_nPEAK

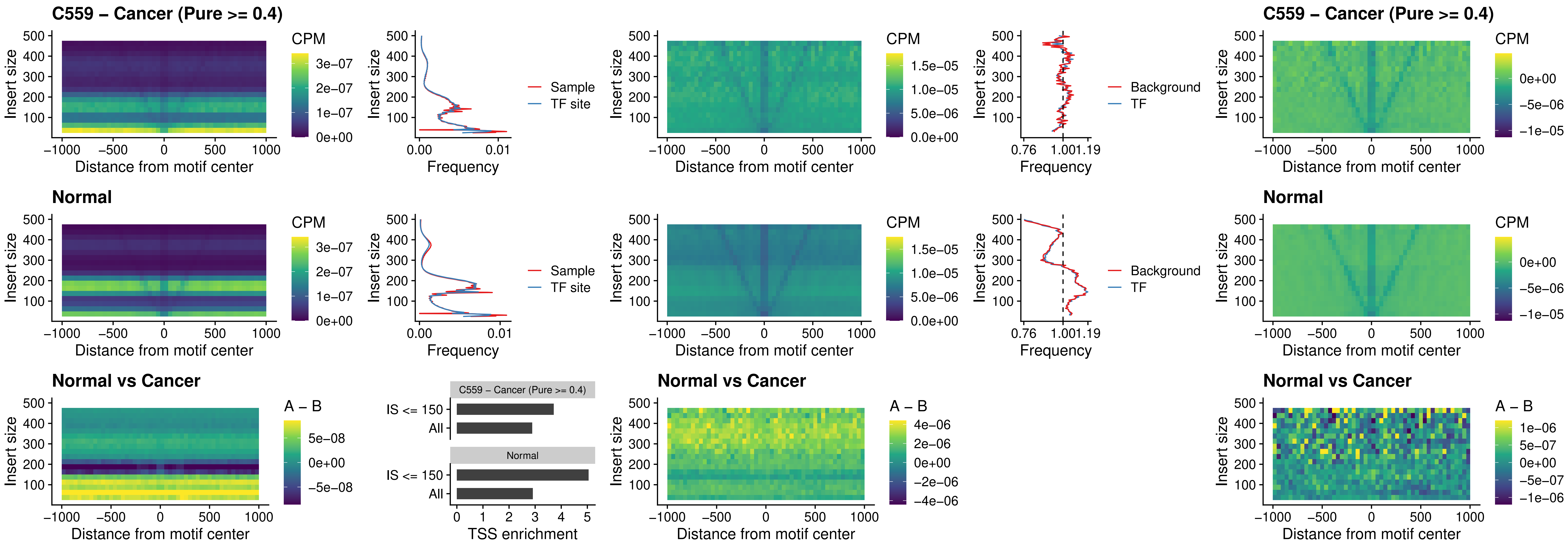

### SHOX_uc_dTSS_nPEAK

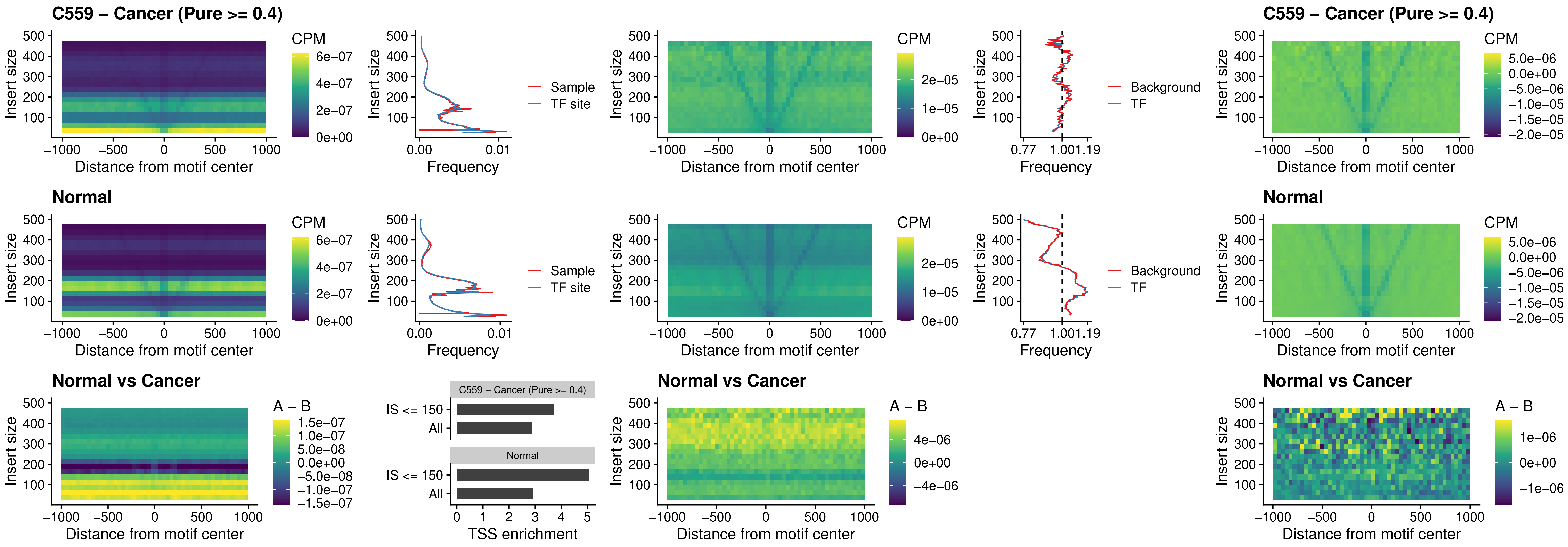

### PRRX1_uc_dTSS_nPEAK

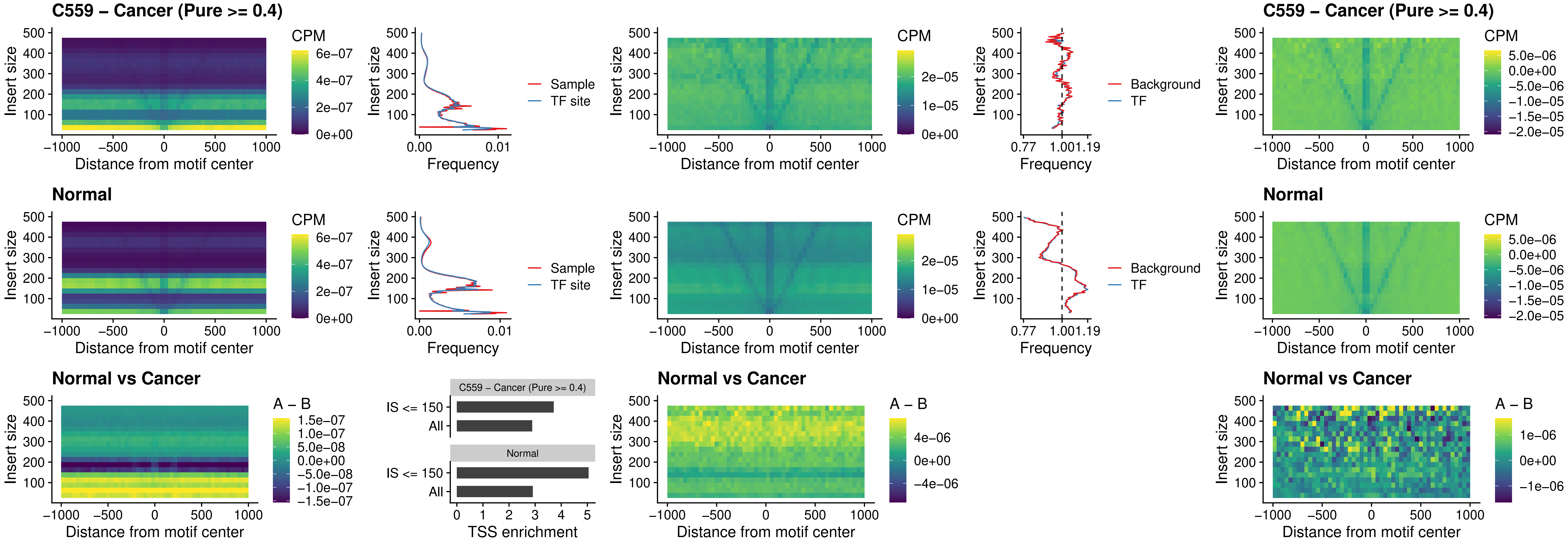

#### LMX1A_uc_dTSS_nPEAK

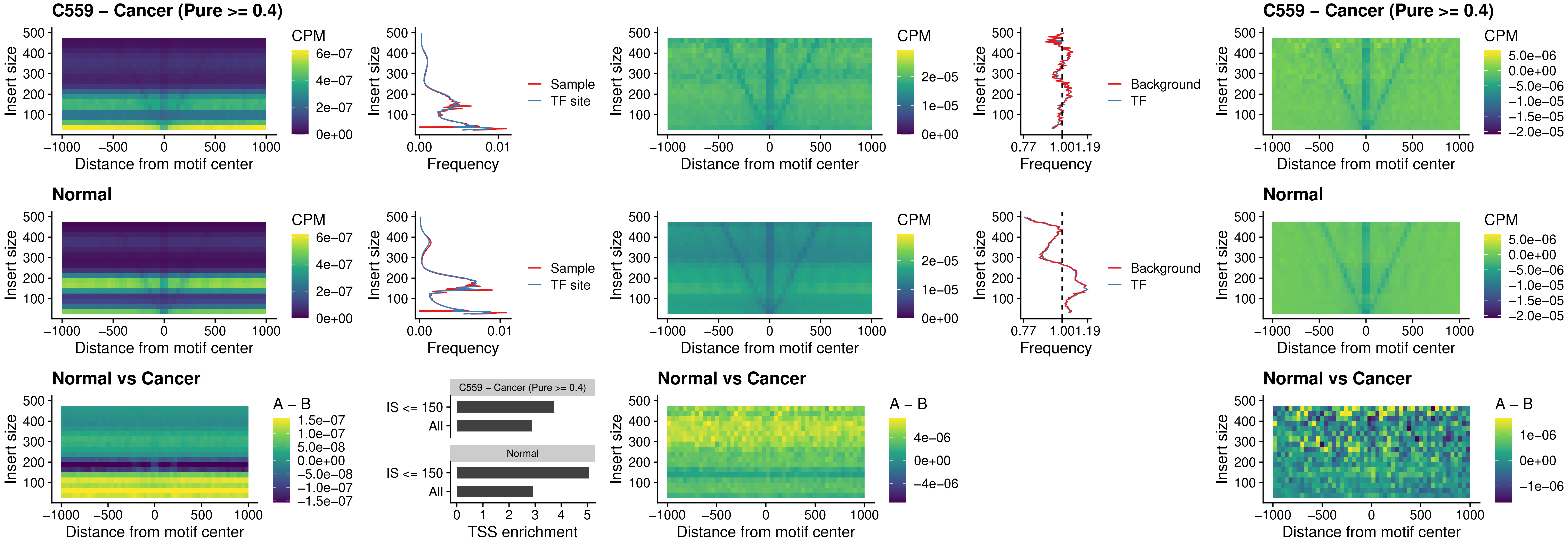

### LMX1B_uc_dTSS_nPEAK

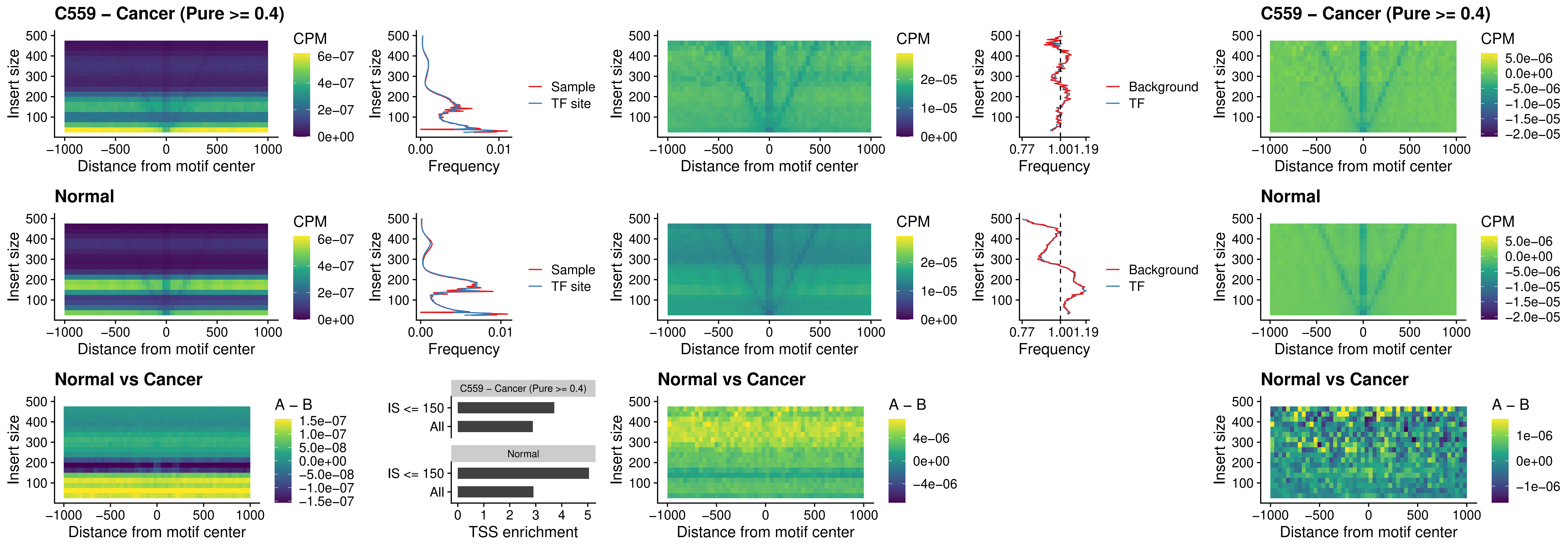

### DLX5_uc_dTSS_oPEAK

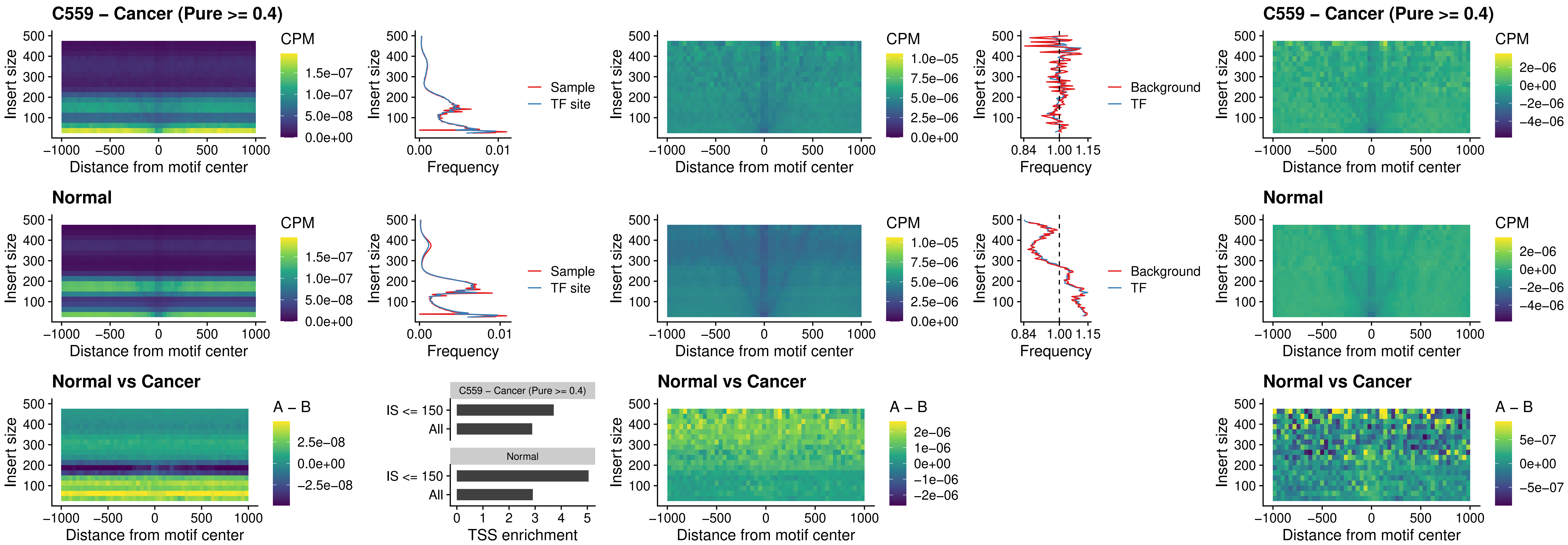

### ARID3C_uc_dTSS_nPEAK

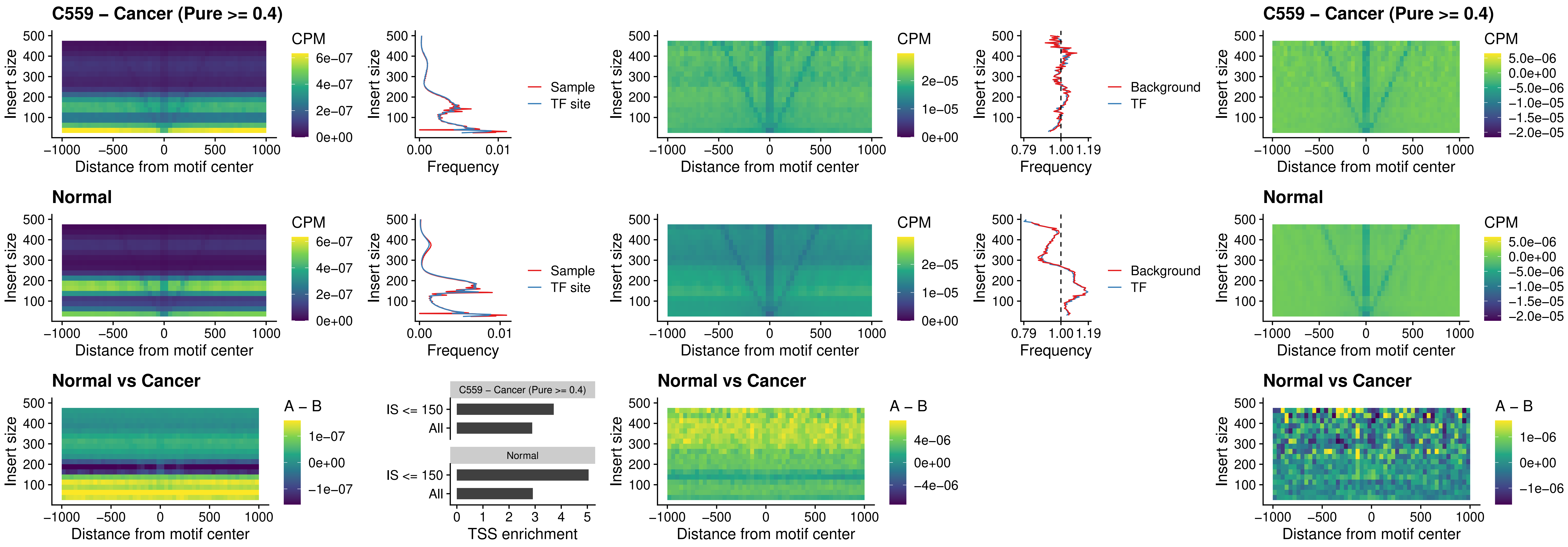

### HOXD12_uc_dTSS_nPEAK

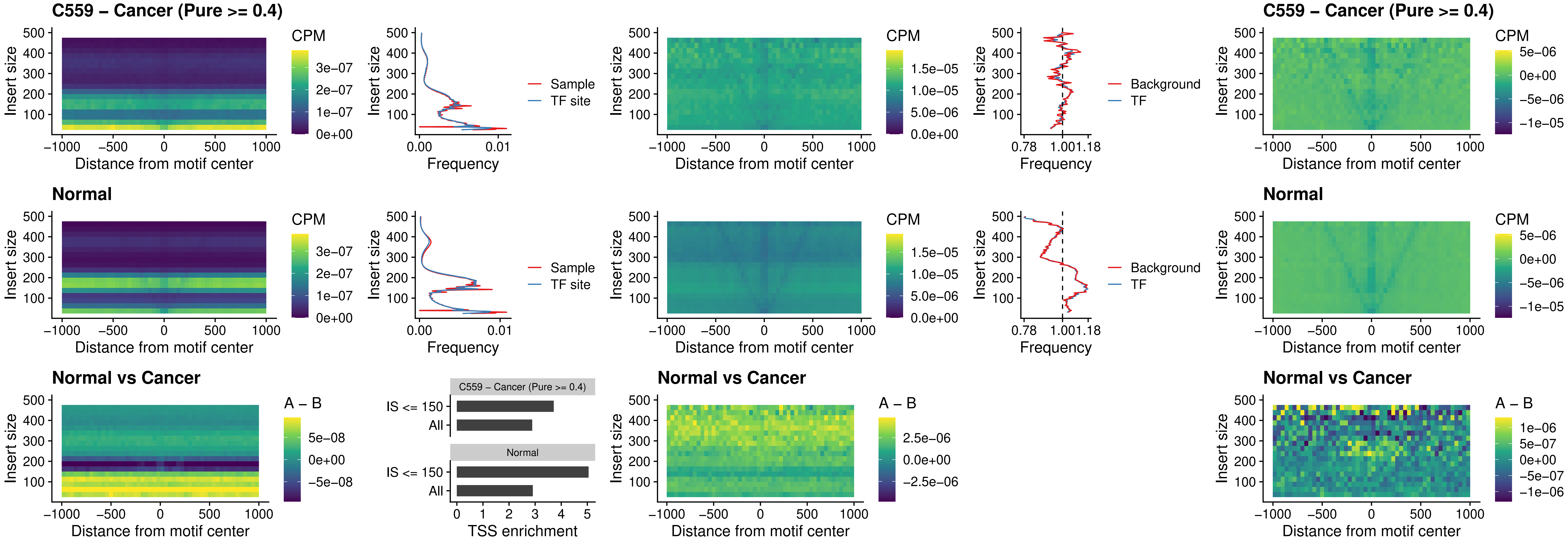

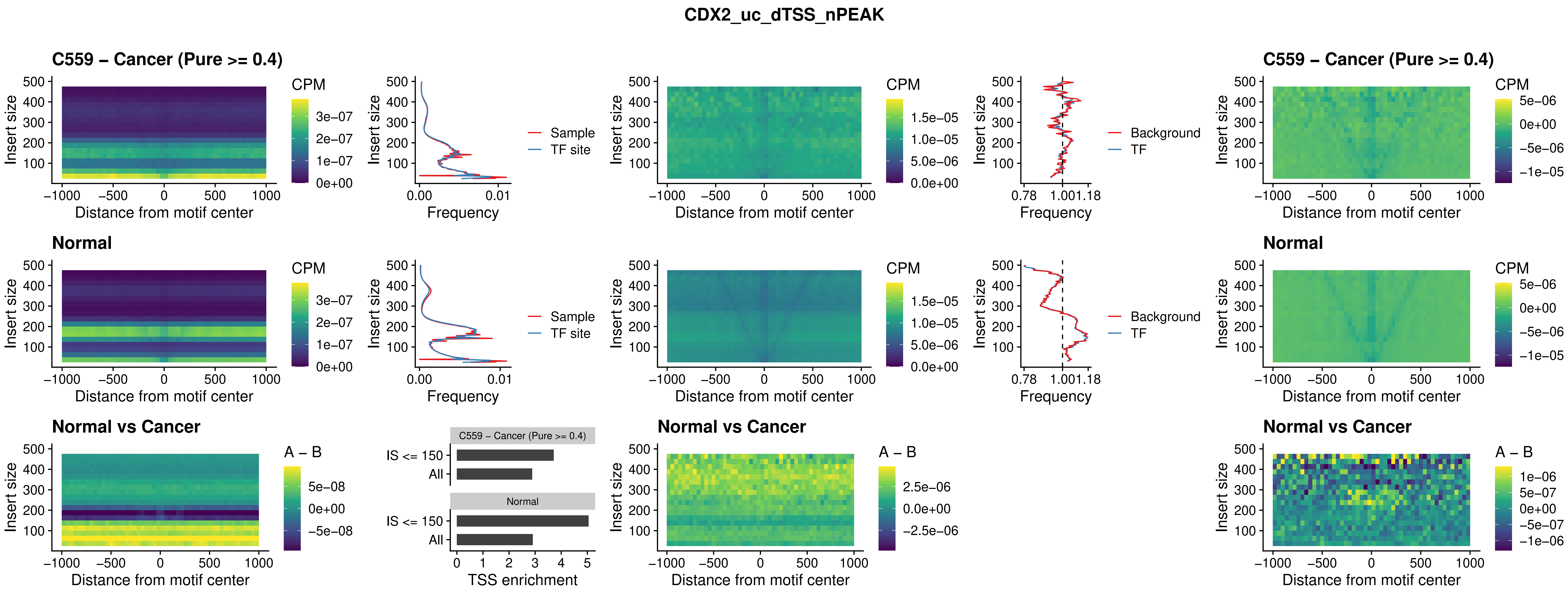
CDX2_uc_dTSS_nPEAK
C559 – Cancer (Pure >= 0.4)
Normal
Normal vs Cancer
Insert size
Distance from motif center
Frequency
CPM
A – B
Sample
TF site
Background
TF
C559 – Cancer (Pure >= 0.4)
Normal
IS <= 150
All
TSS enrichment

#### HOXC12_uc_dTSS_nPEAK

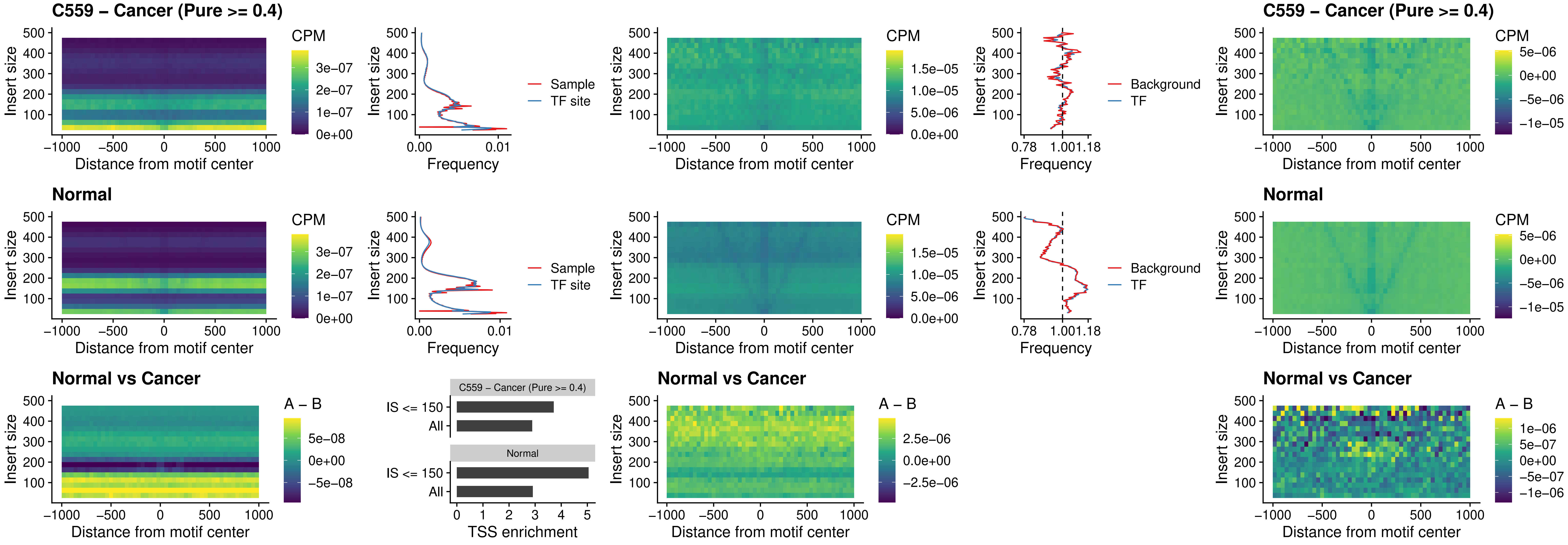

### HHEX_uc_dTSS_oPEAK

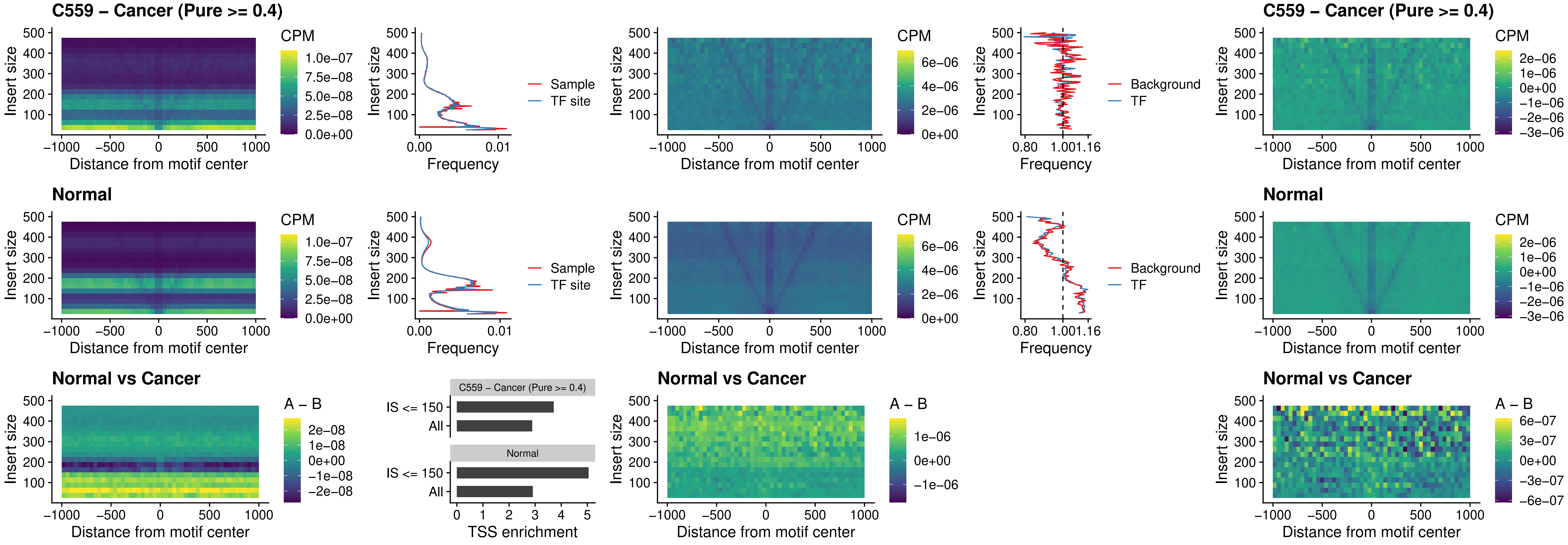

### HOXA11_uc_dTSS_nPEAK

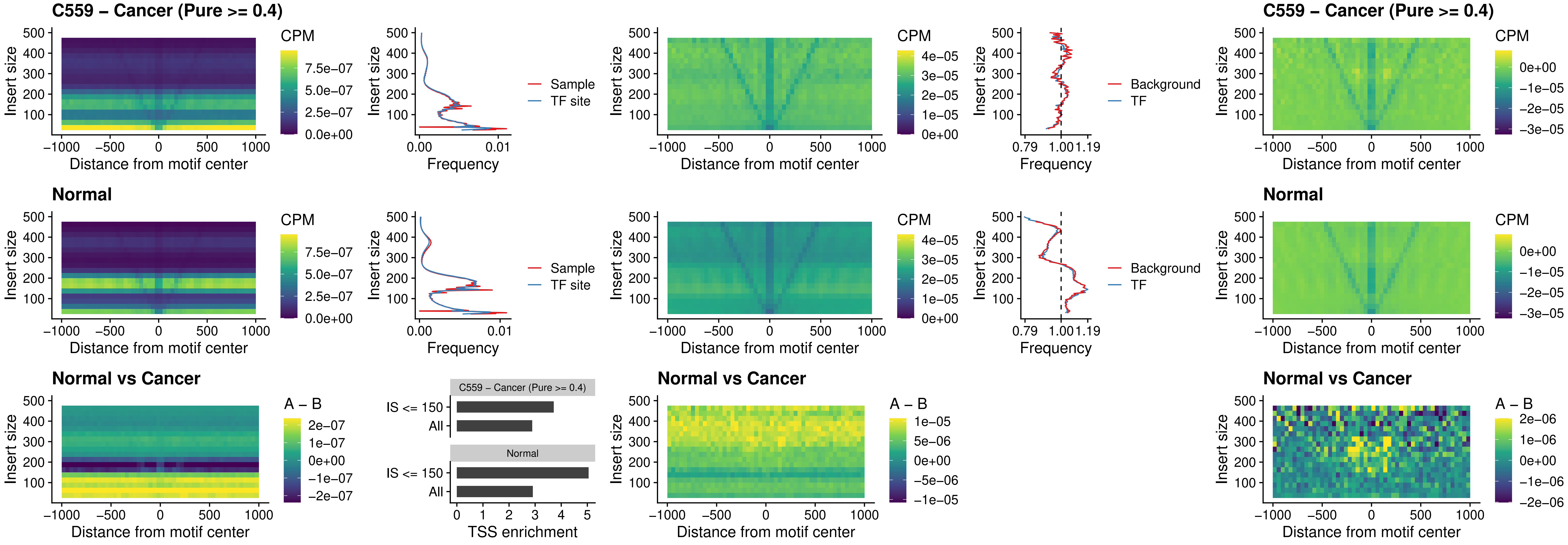

### HOXA11_uc_cTSS_nPEAK

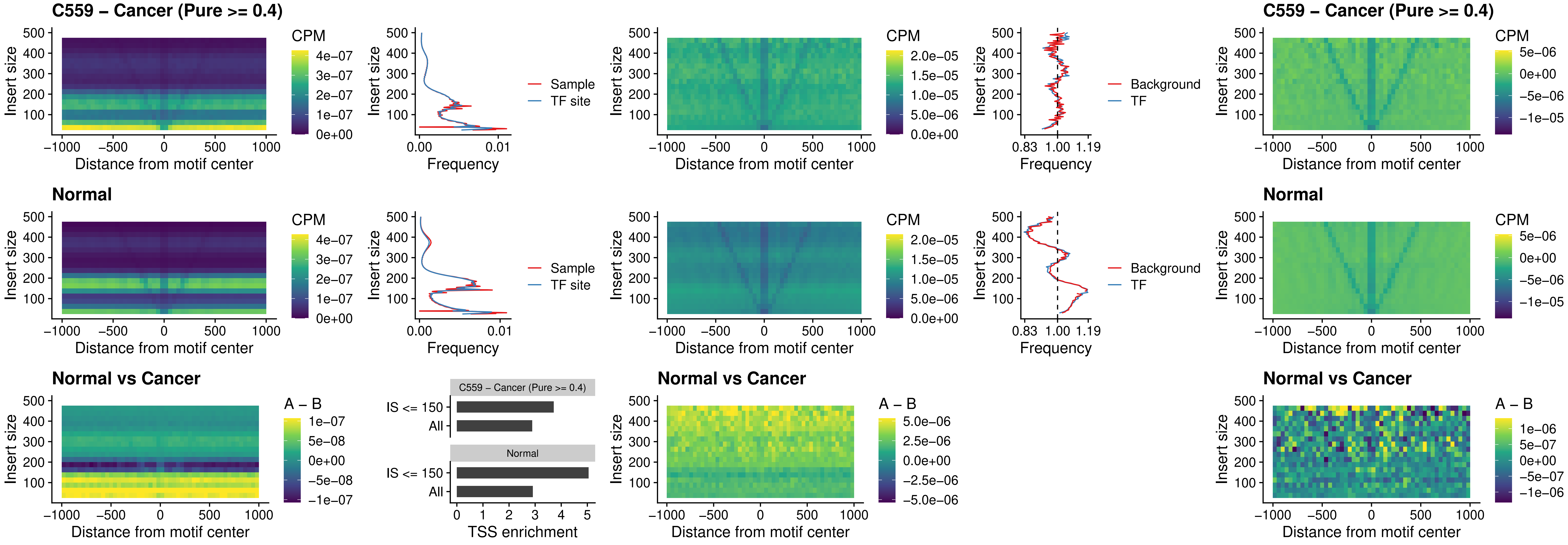

#### ZFHX3_uc_dTSS_nPEAK

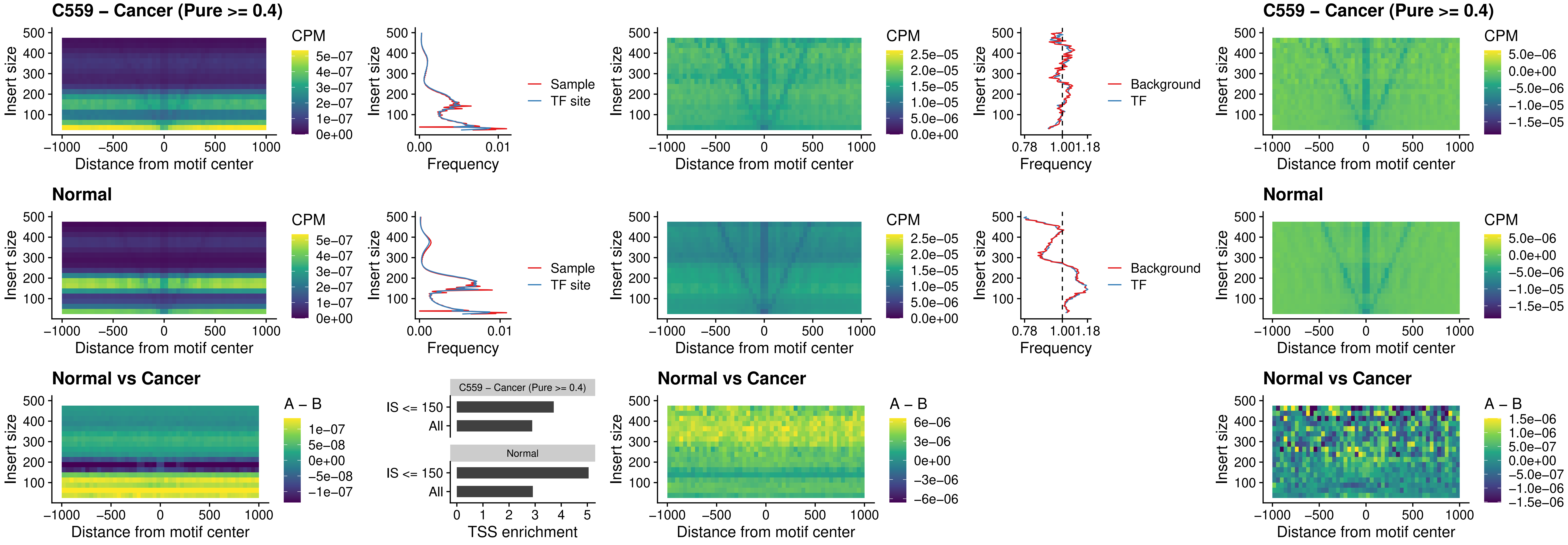

### HOXB4_uc_dTSS_nPEAK

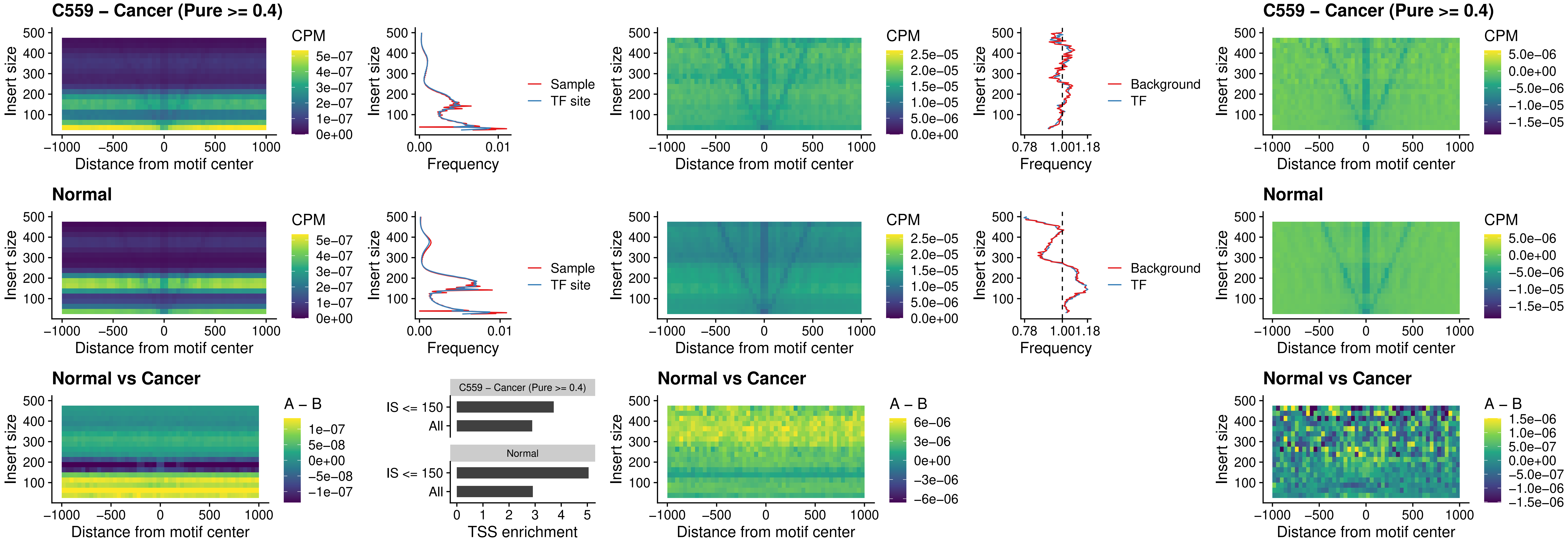

### HOXC5_uc_dTSS_nPEAK

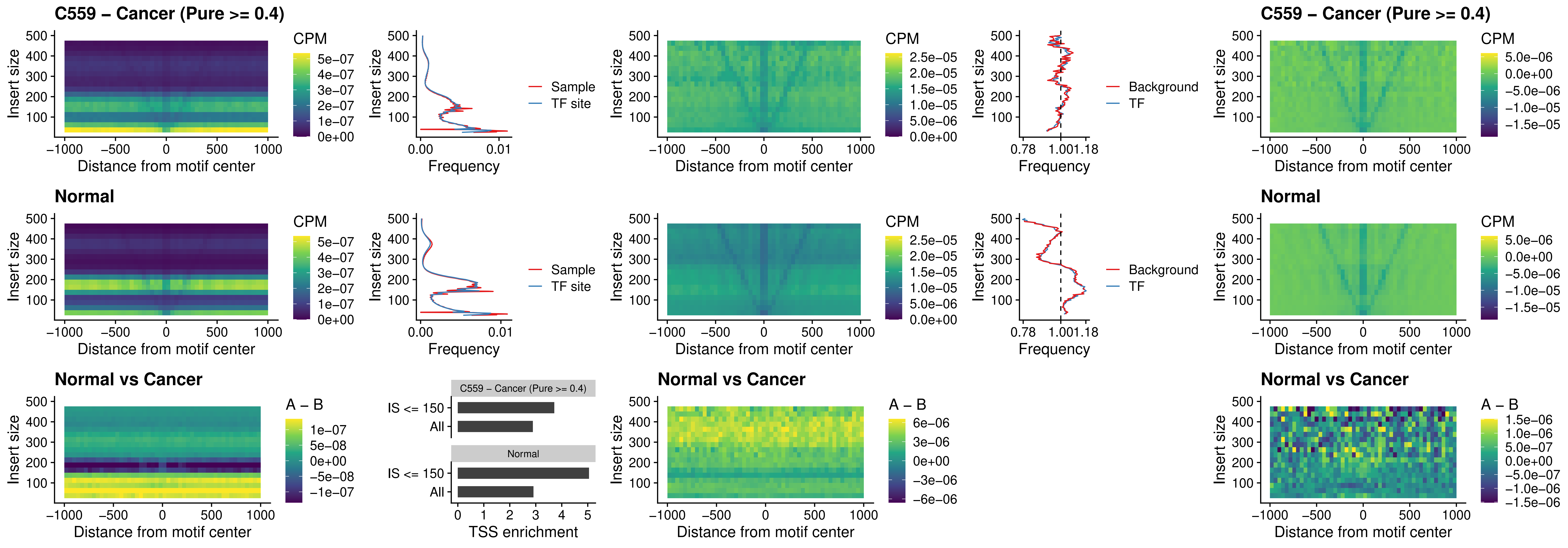

### HOXD1_uc_dTSS_nPEAK

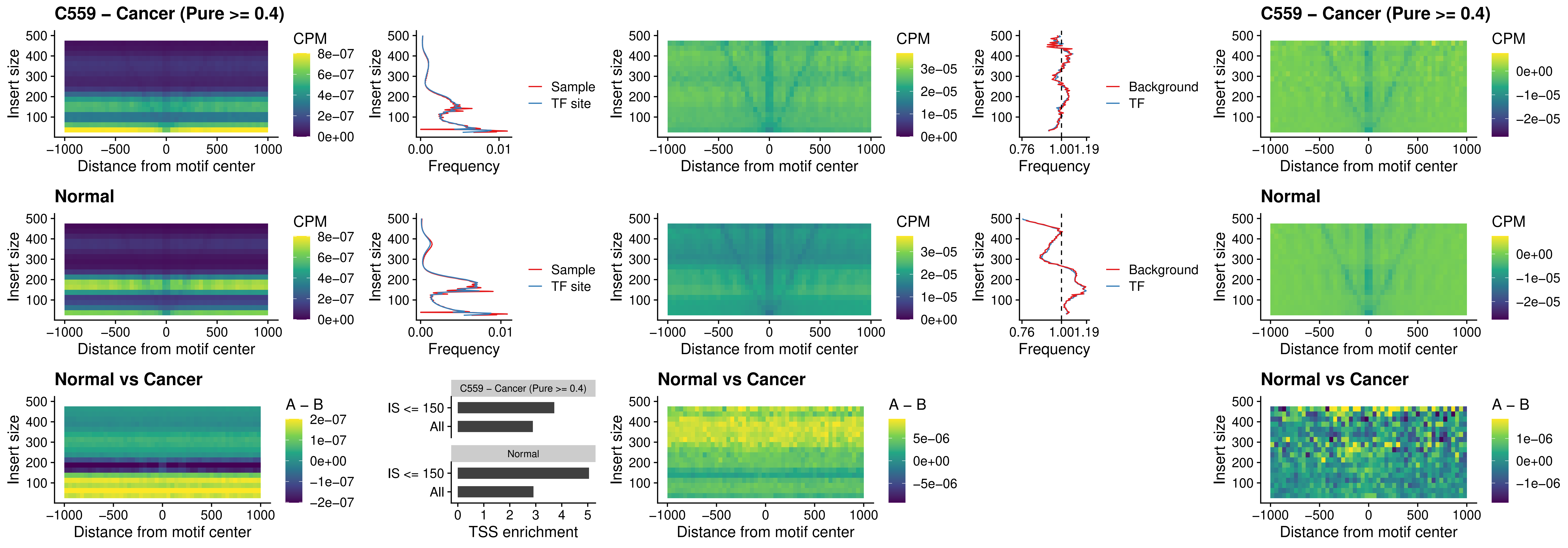

### OTP_uc_dTSS_nPEAK

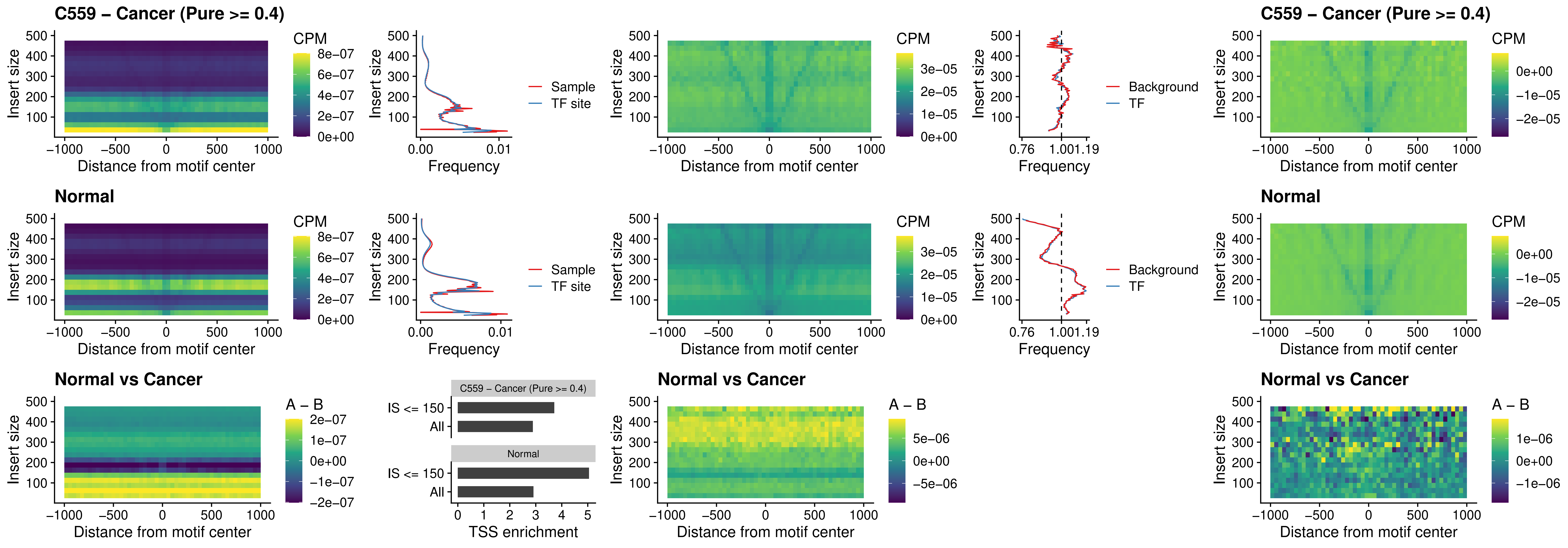

#### FOXL1_uc_dTSS_oPEAK

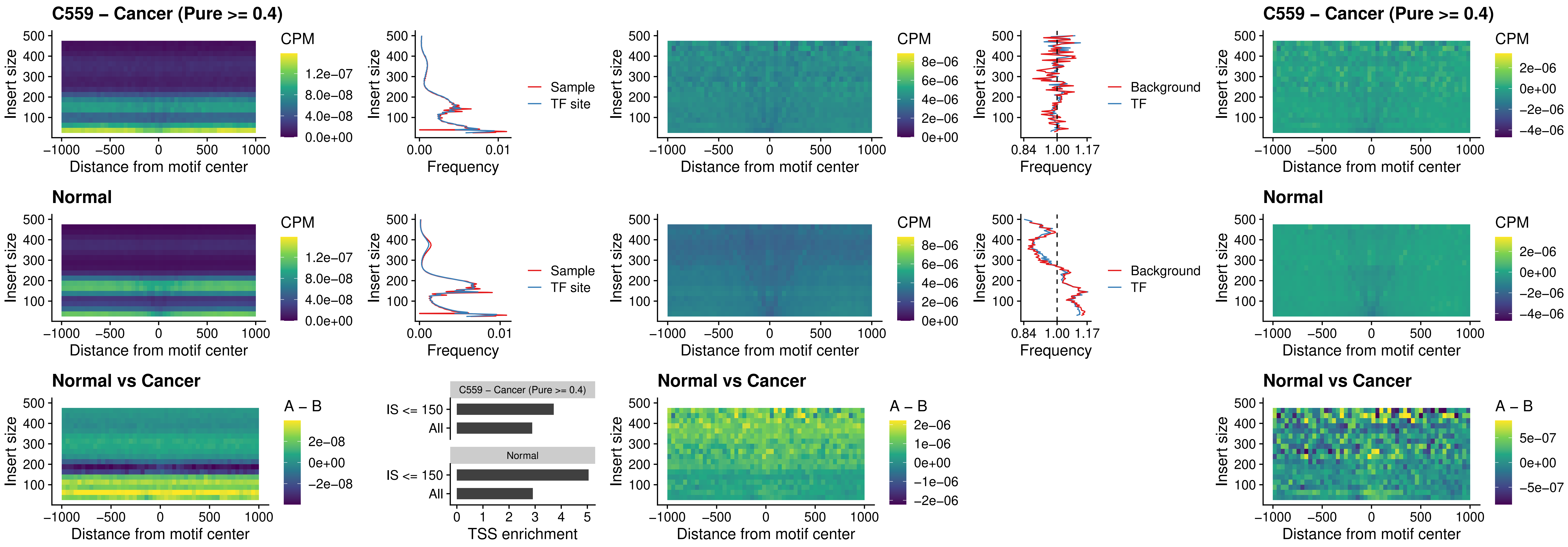

#### ZNF32_uc_dTSS_oPEAK

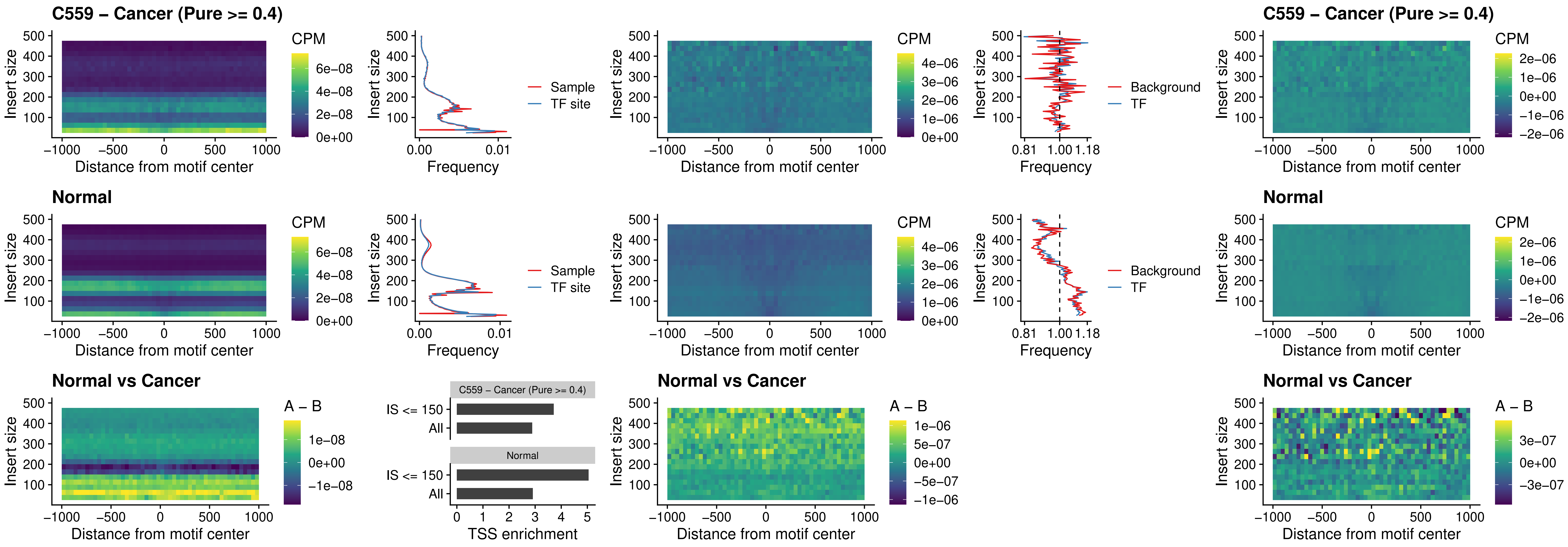

### FOXL1_uc_dTSS_nPEAK

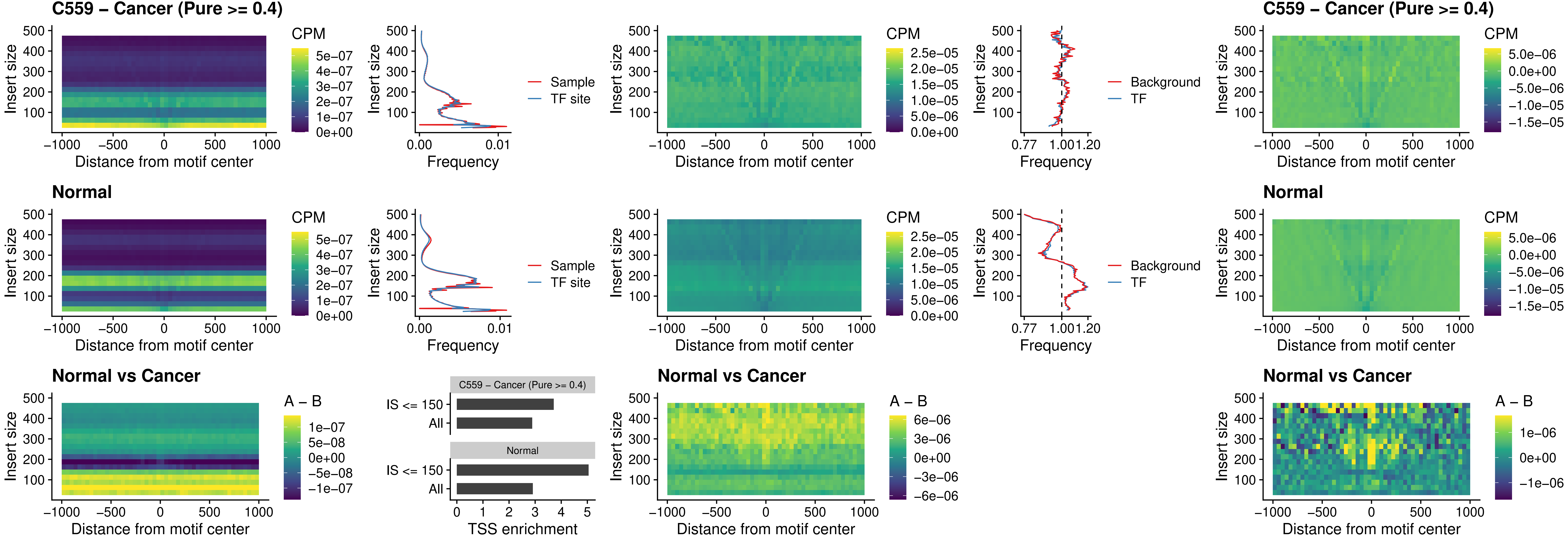

### ZNF32_uc_dTSS_nPEAK

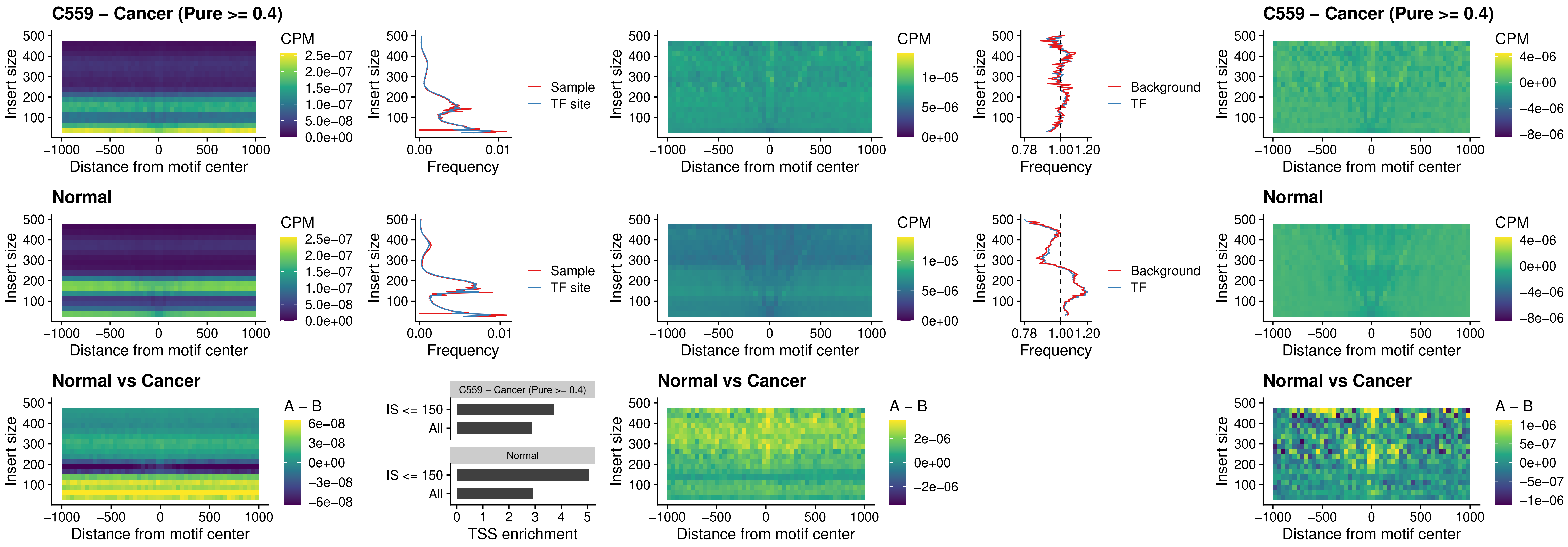

### DLX5_uc_dTSS_nPEAK

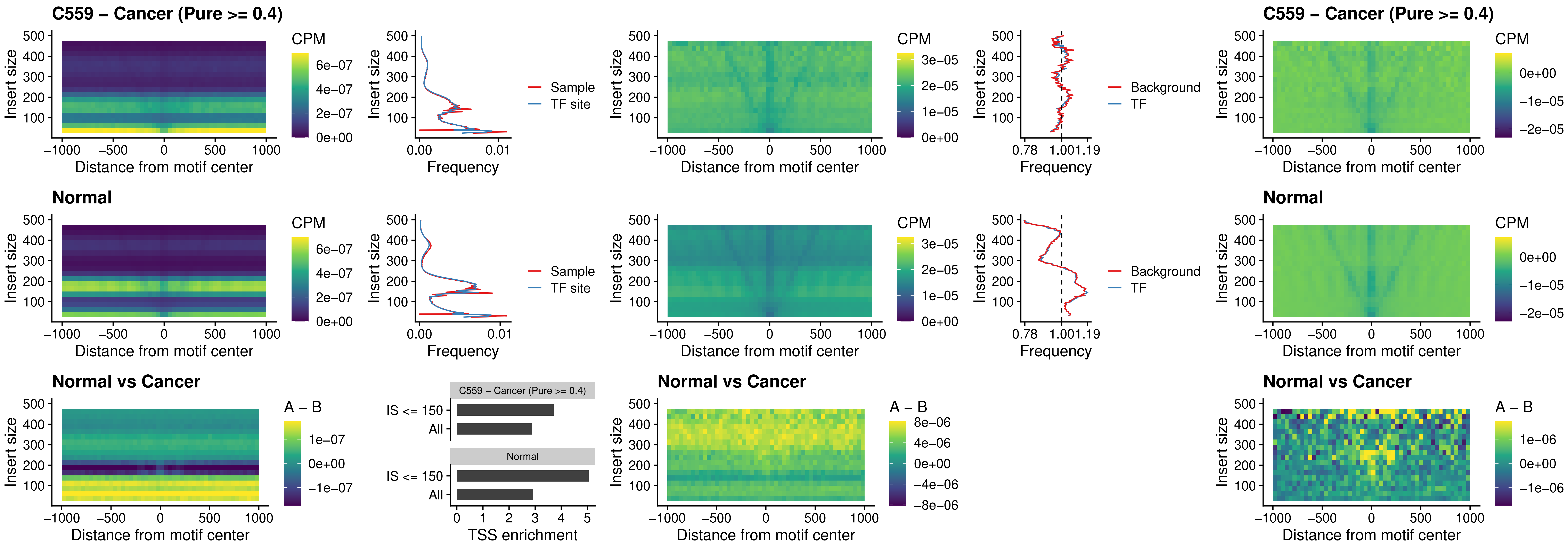

### DMRT2_uc_dTSS_nPEAK

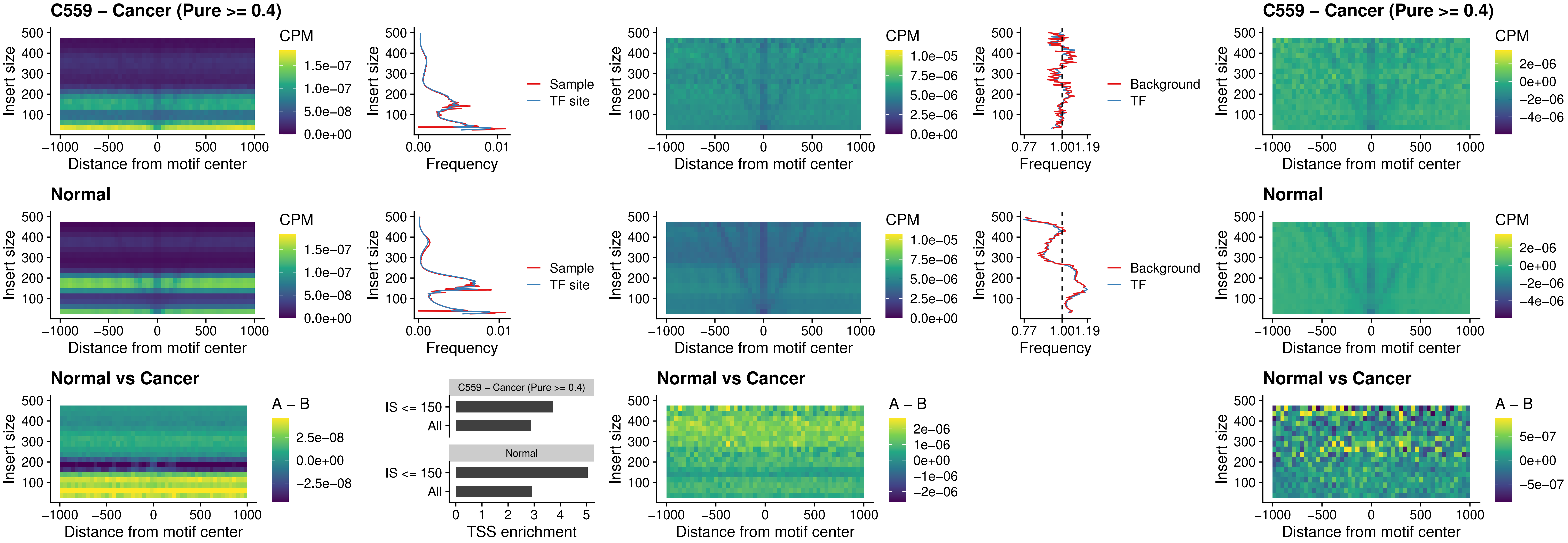

### DMRTC2_uc_cTSS_nPEAK

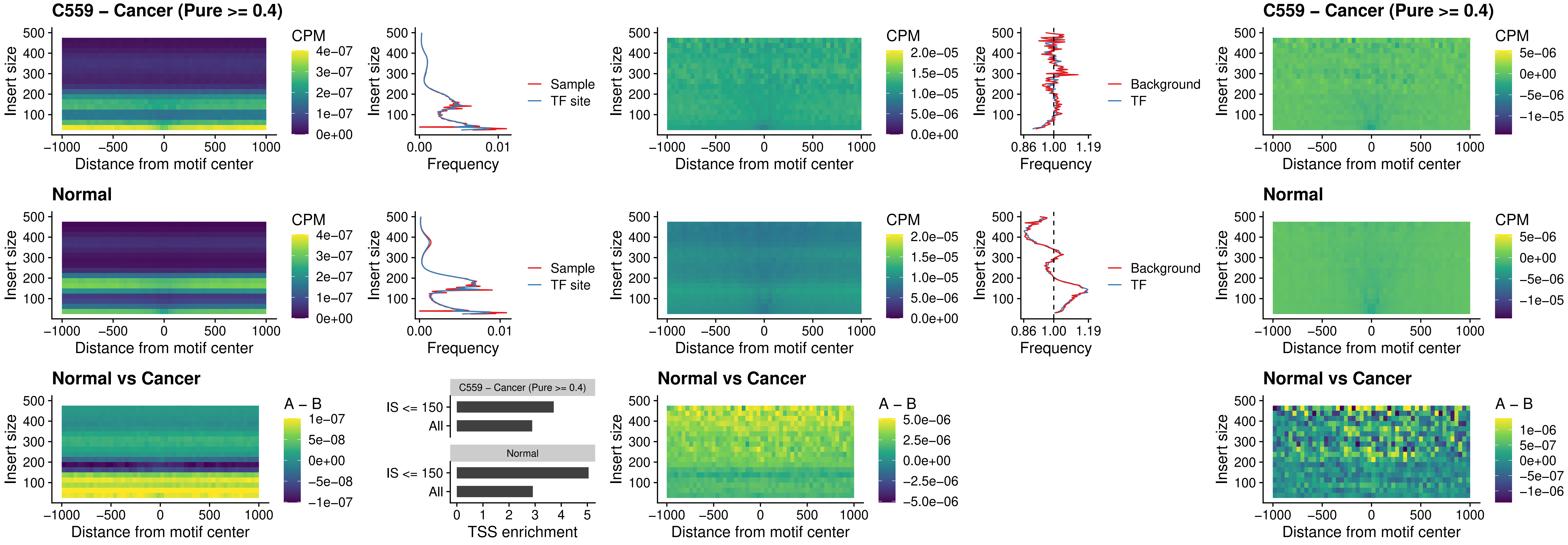

### HOXD9_uc_dTSS_nPEAK

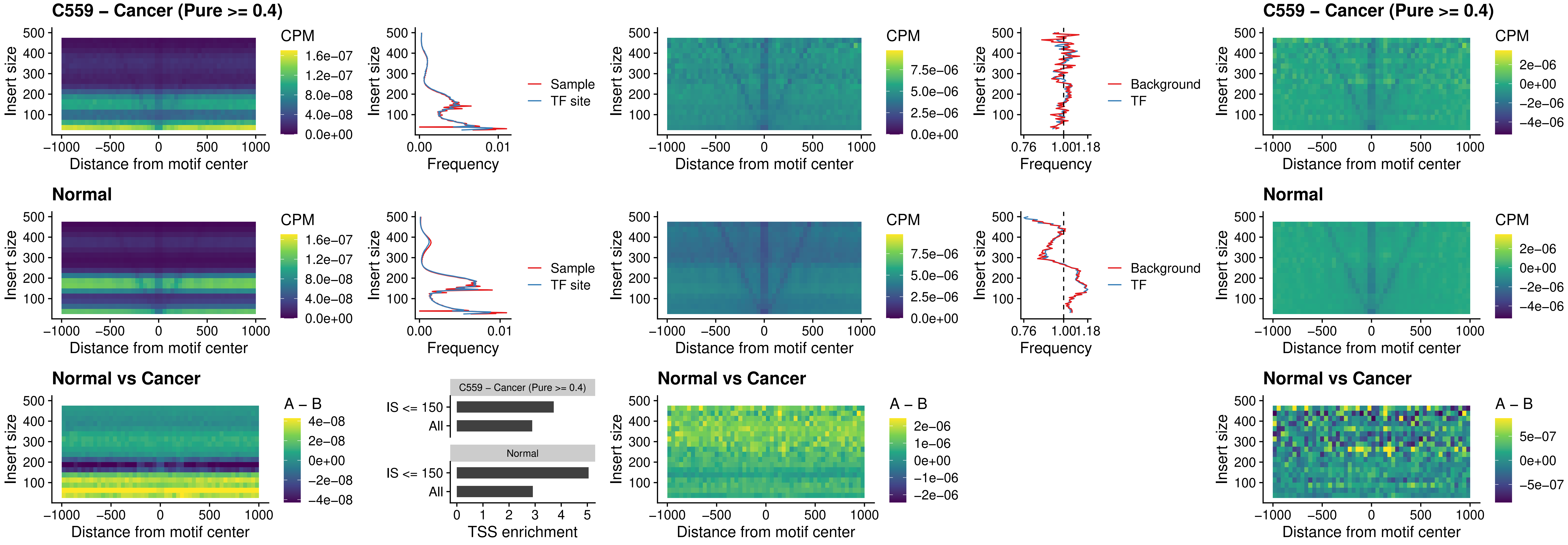

### HOXD4_uc_dTSS_nPEAK

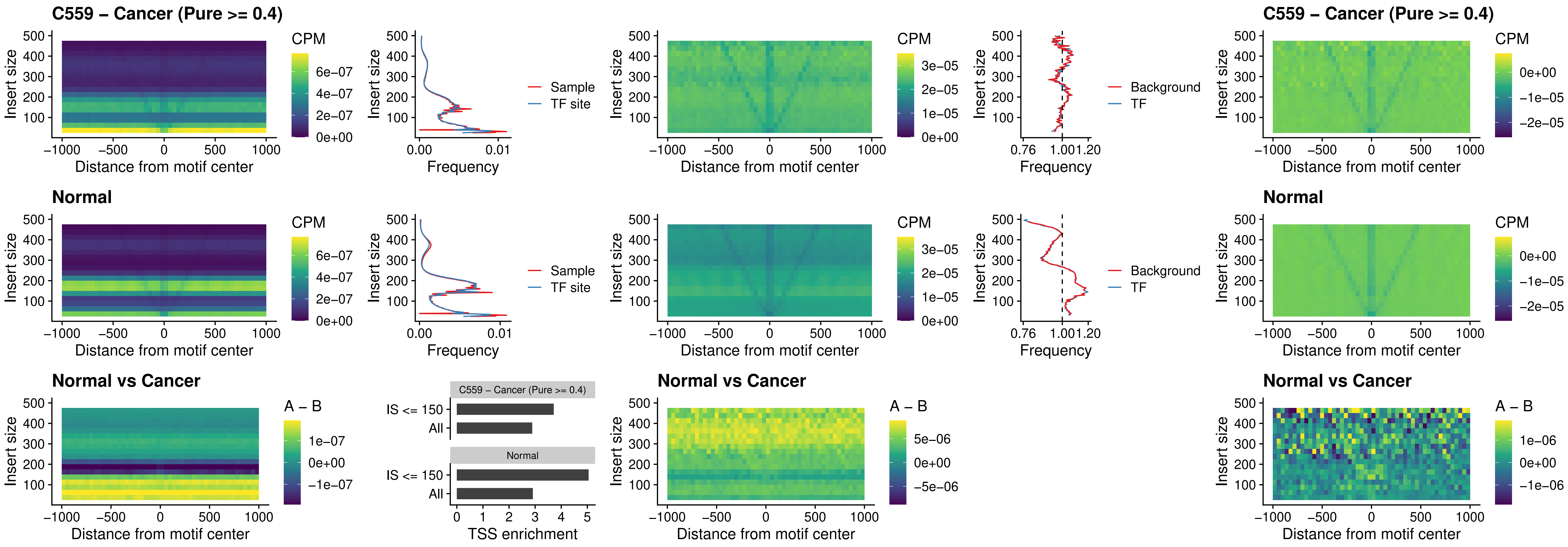

#### FOXP3_uc_cTSS_nPEAK

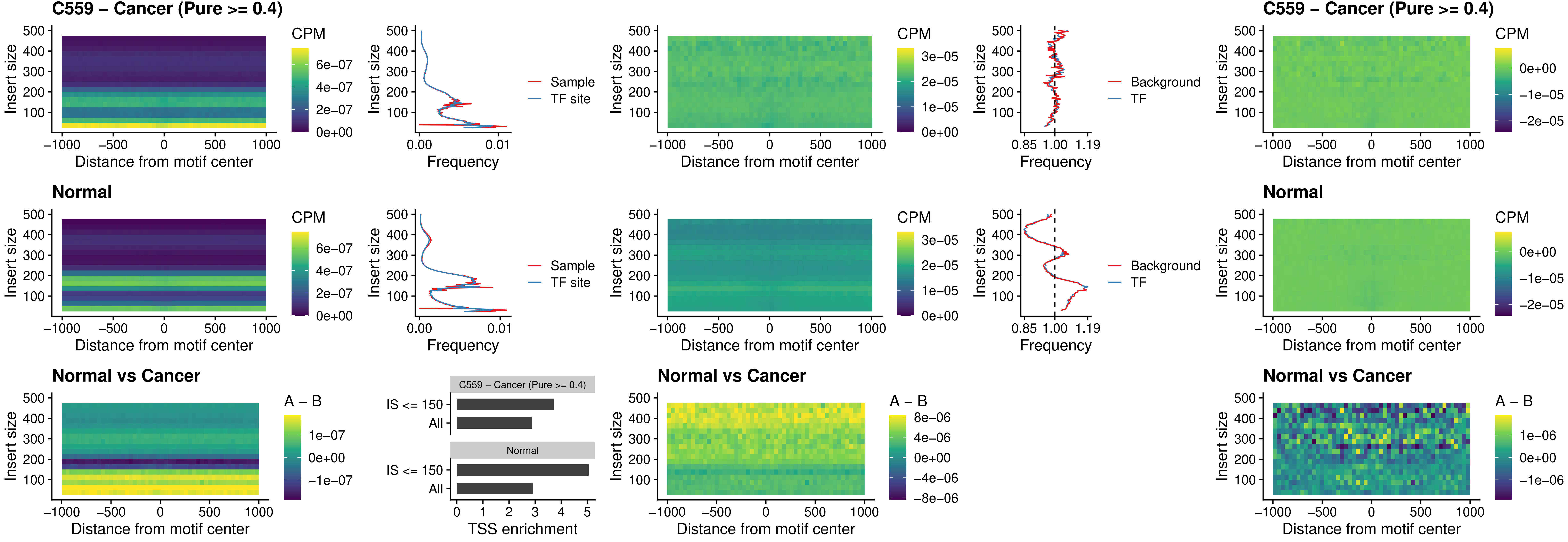

### FOXD4L6_uc_cTSS_nPEAK

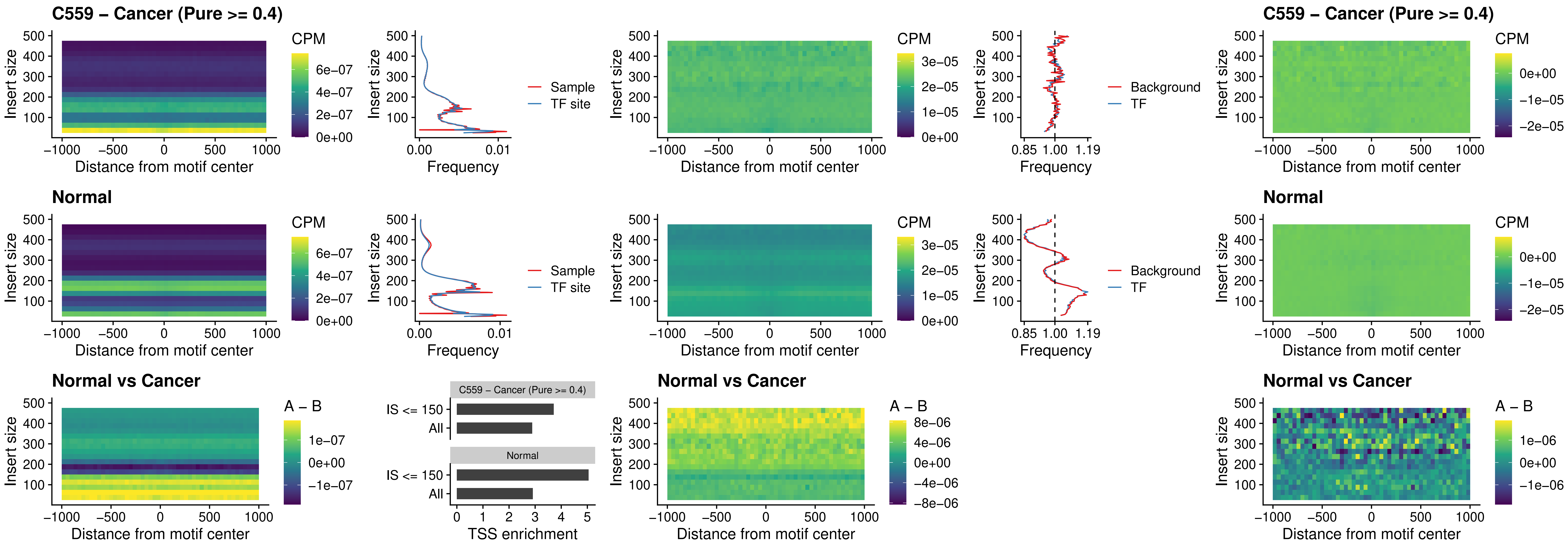

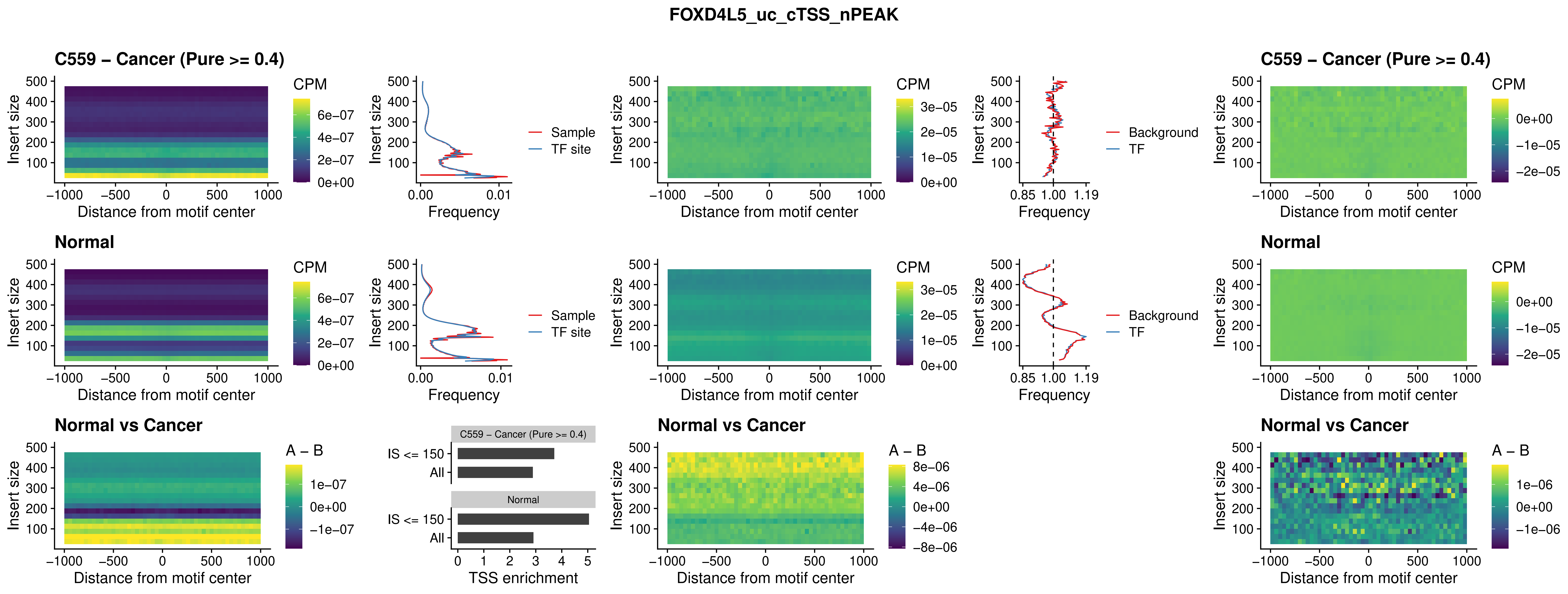
FOXD4L5_uc_cTSS_nPEAK
C559 – Cancer (Pure >= 0.4)
Normal
Normal vs Cancer
Insert size
Distance from motif center
CPM
A – B
Frequency
Sample
TF site
Background
TF
C559 – Cancer (Pure >= 0.4)
Normal
IS <= 150
All
TSS enrichment

#### FOXD4L4_uc_cTSS_nPEAK

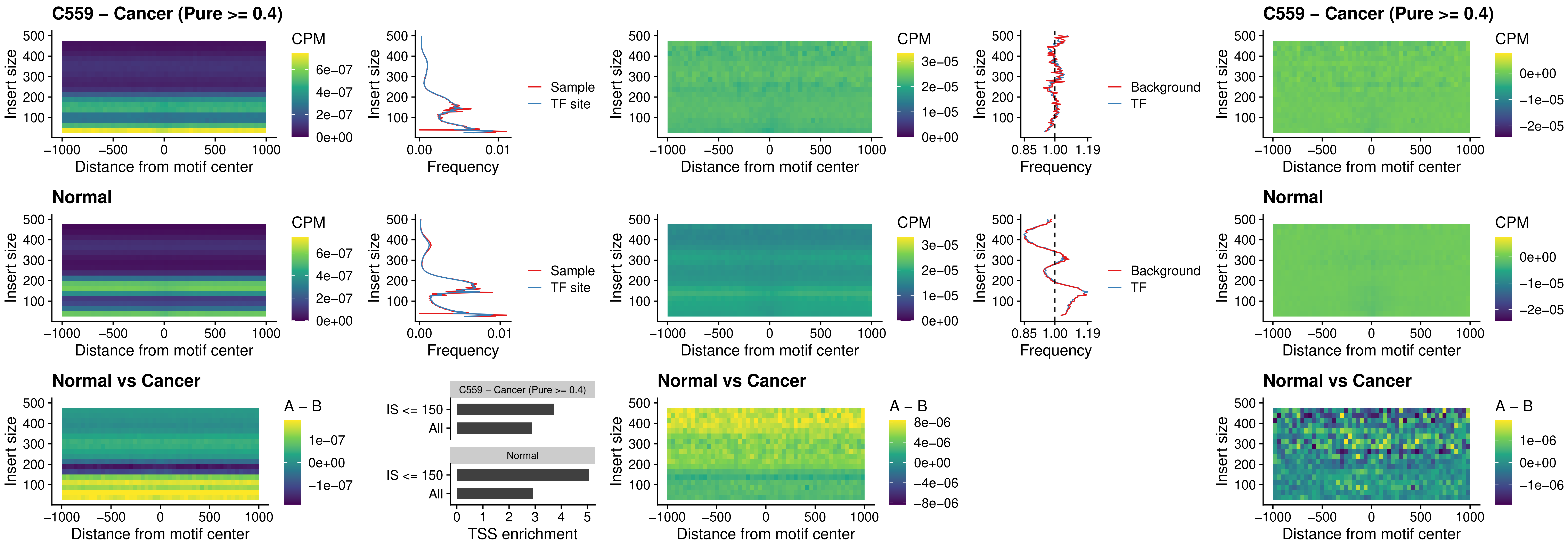

#### FOXD4L2_uc_cTSS_nPEAK

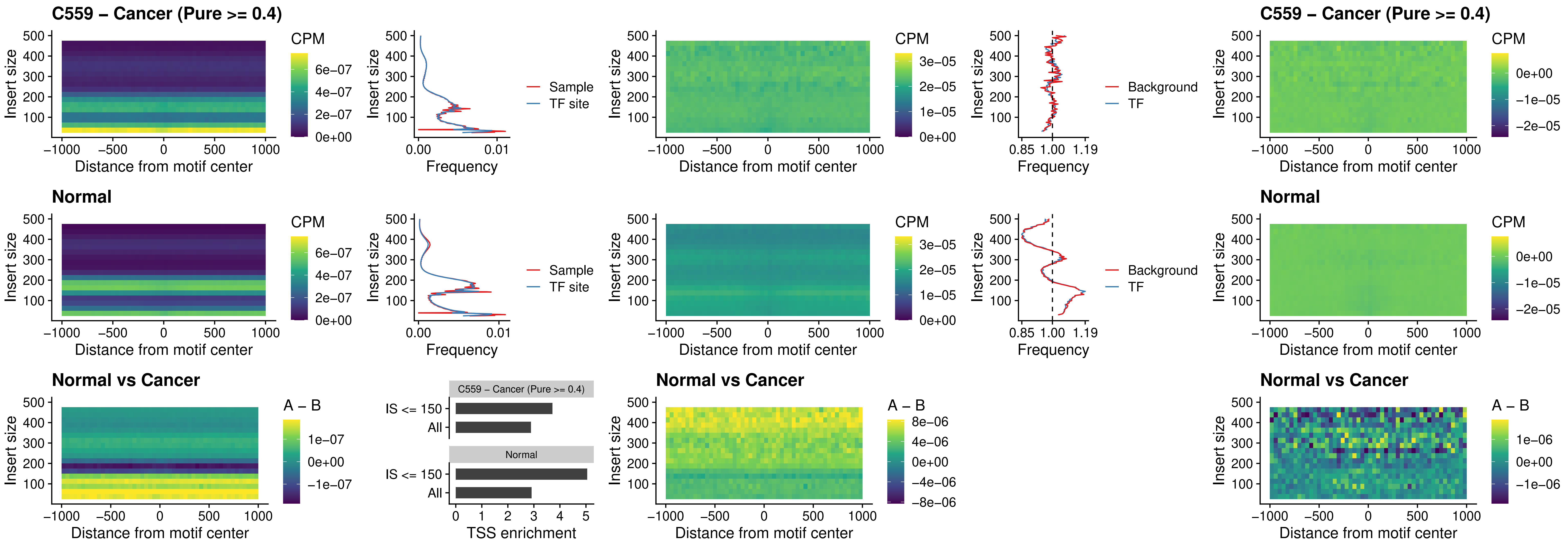

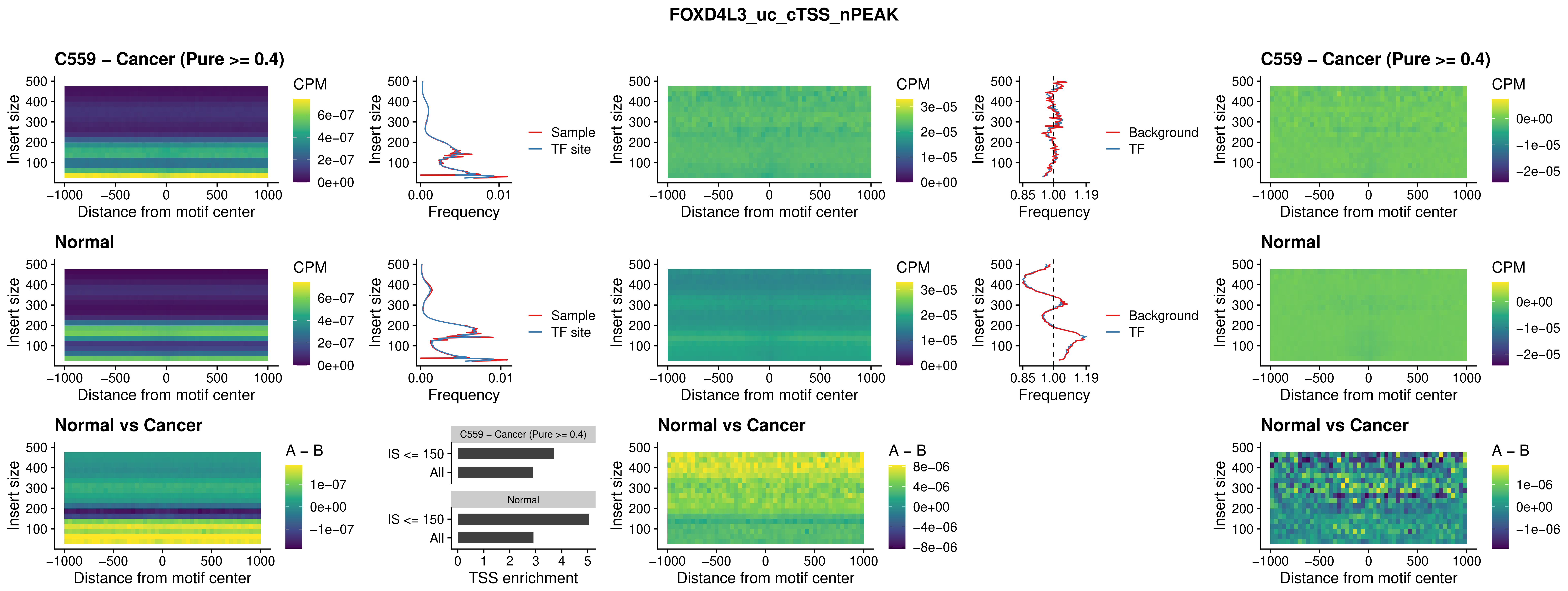
FOXD4L3_uc_cTSS_nPEAK
C559 – Cancer (Pure >= 0.4)
Normal
Normal vs Cancer
Insert size
Distance from motif center
CPM
A – B
Frequency
Sample
TF site
Background
TF
C559 – Cancer (Pure >= 0.4)
Normal
IS <= 150
All
TSS enrichment

### HOXD1_uc_dTSS_oPEAK

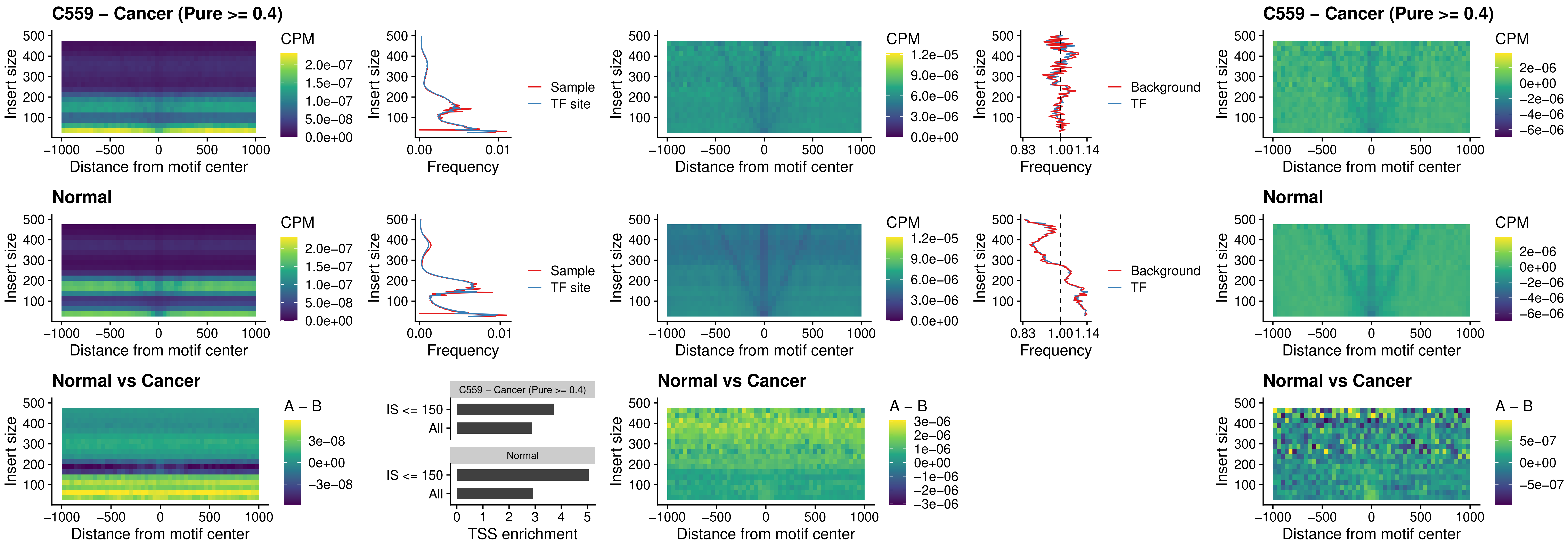

#### VAX1_uc_pTSS_nPEAK

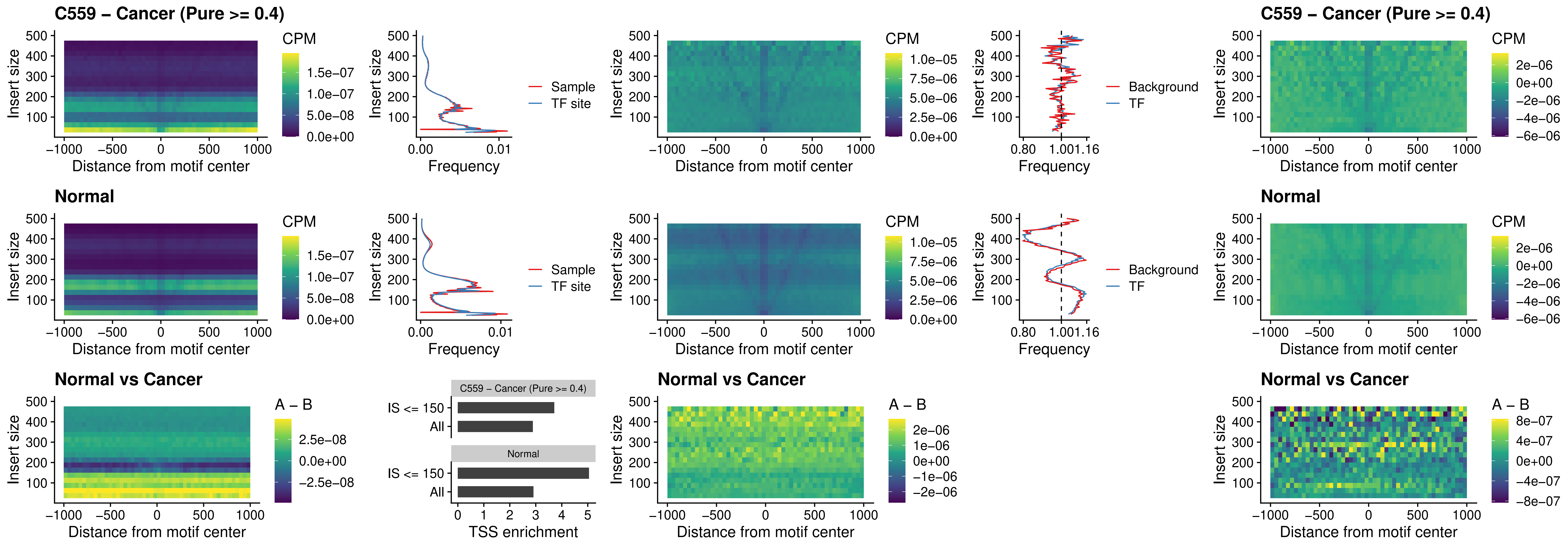

### DLX3_uc_pTSS_nPEAK

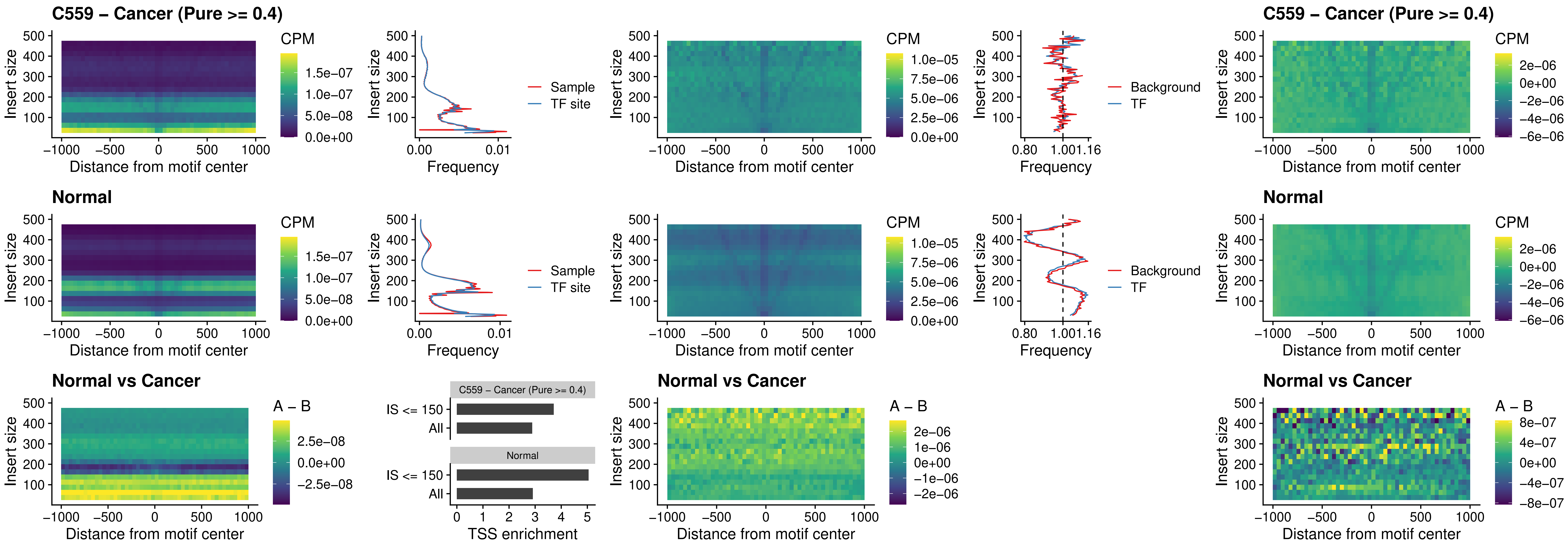

#### HOXD3_uc_pTSS_nPEAK

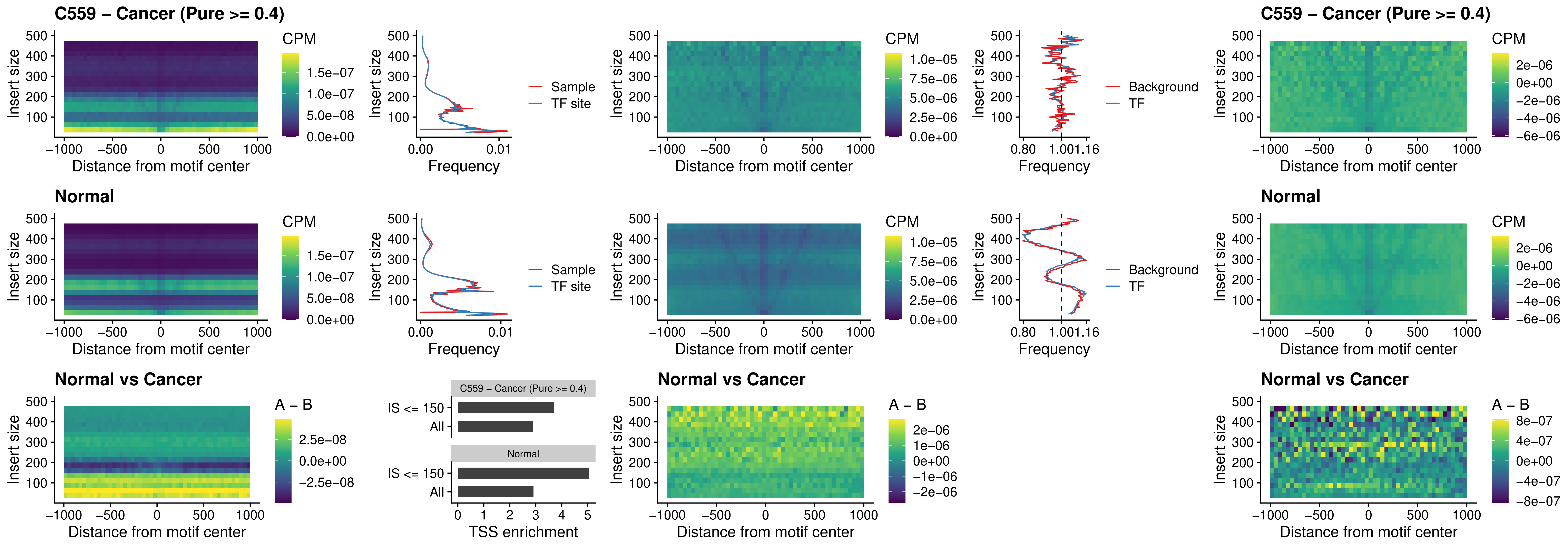

### PDX1_uc_dTSS_nPEAK

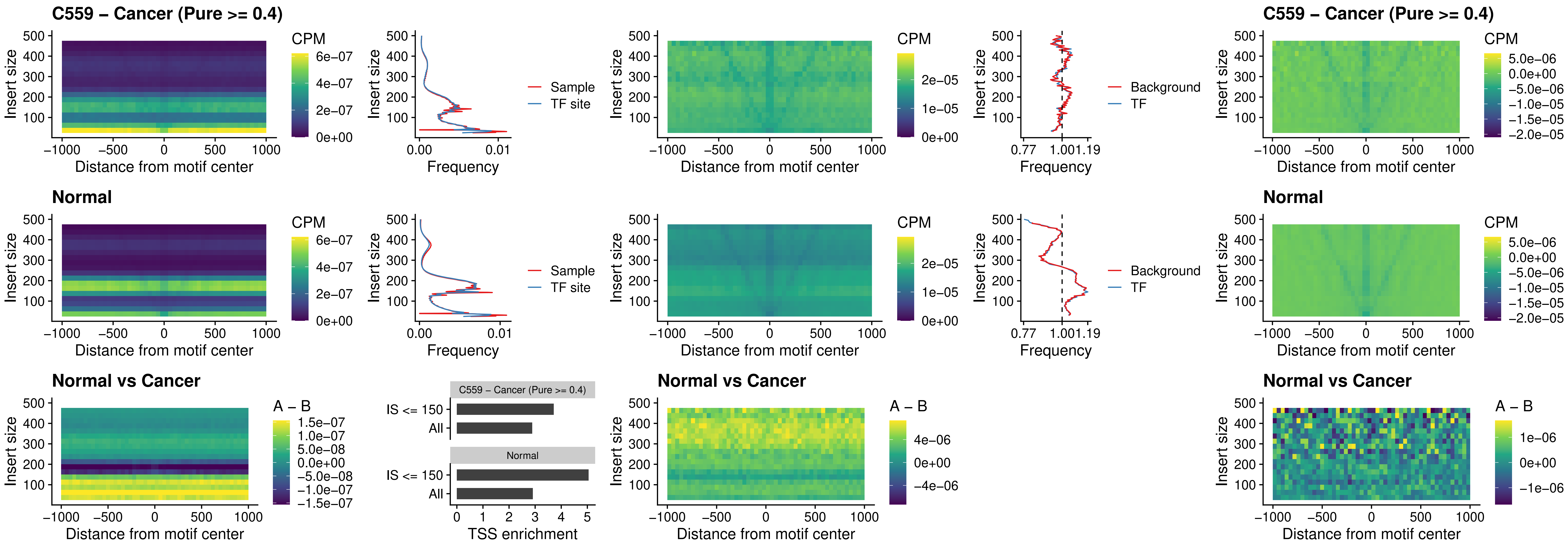

#### VAX2_uc_dTSS_nPEAK

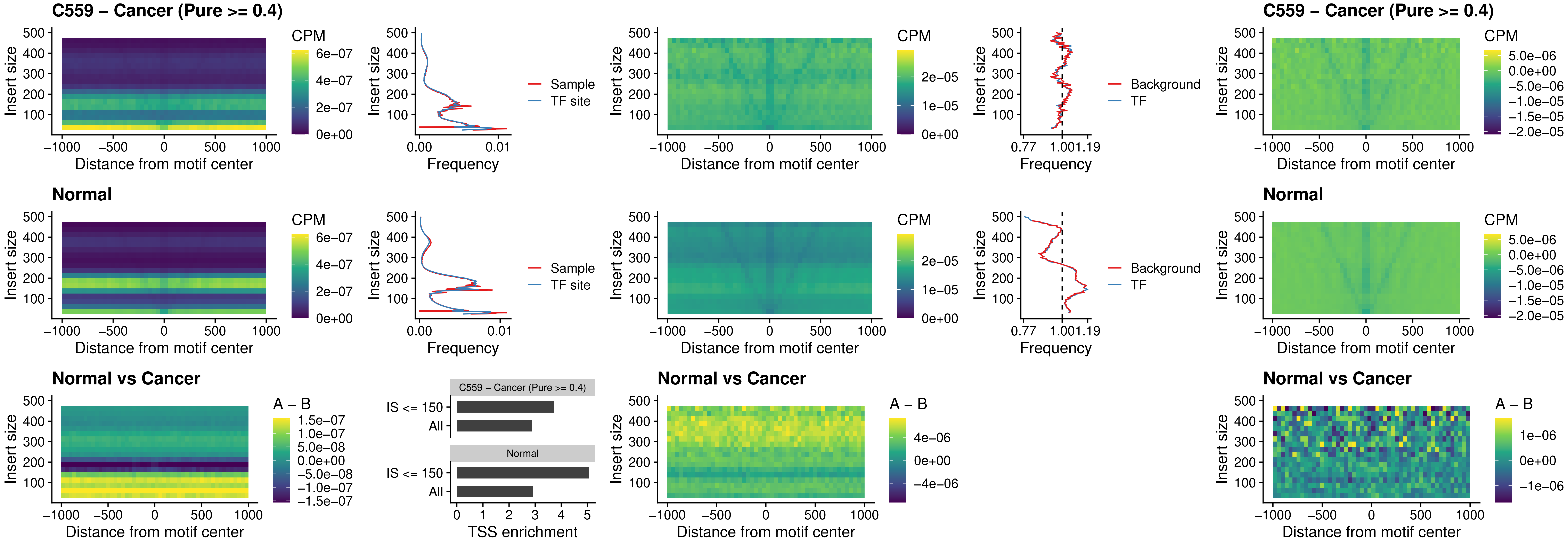

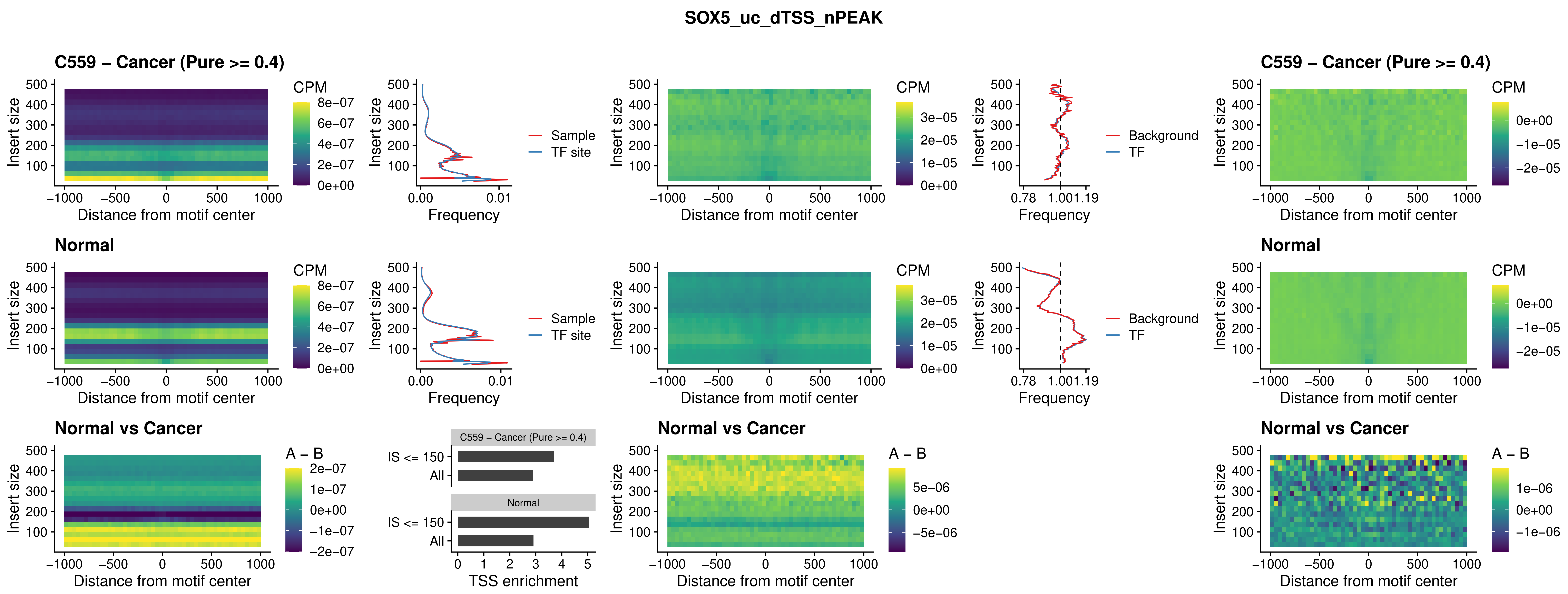

SOX5_uc_dTSS_nPEAK
C559 – Cancer (Pure >= 0.4)
Normal
Normal vs Cancer
Insert size
Distance from motif center
Frequency
CPM
A – B
Sample
TF site
Background
TF
C559 – Cancer (Pure >= 0.4)
Normal
IS <= 150
All
TSS enrichment

### SOX6_uc_dTSS_nPEAK

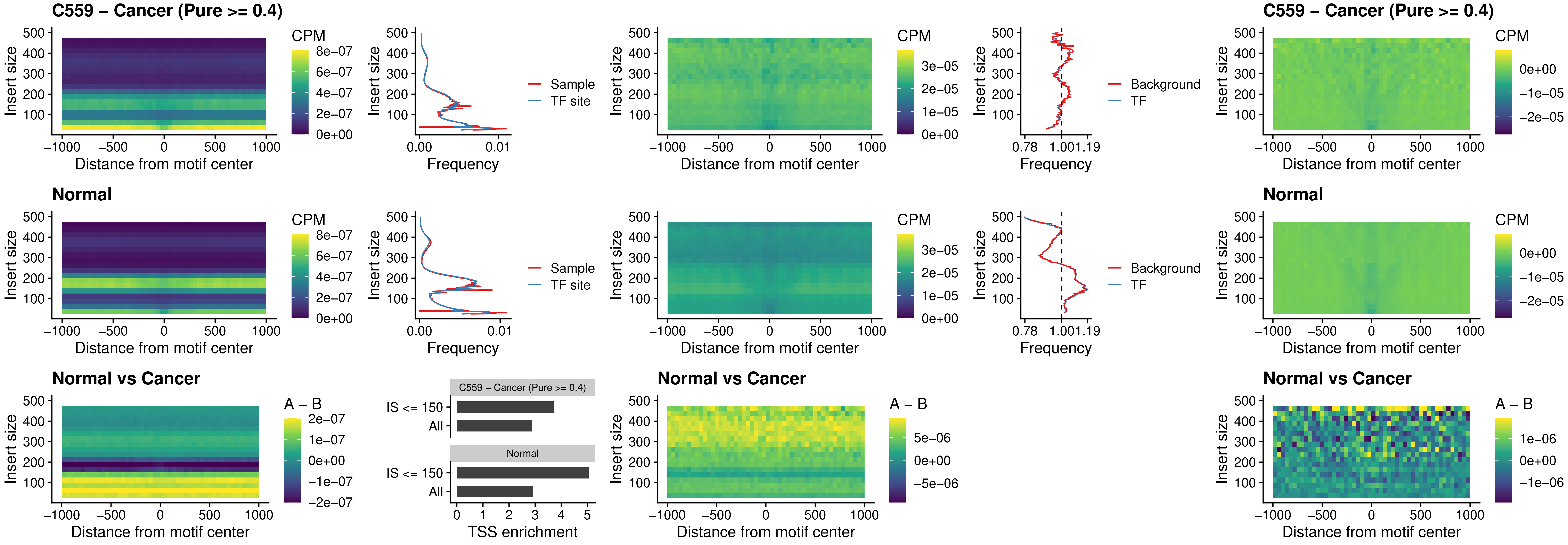

### JDP2_uc_cTSS_nPEAK

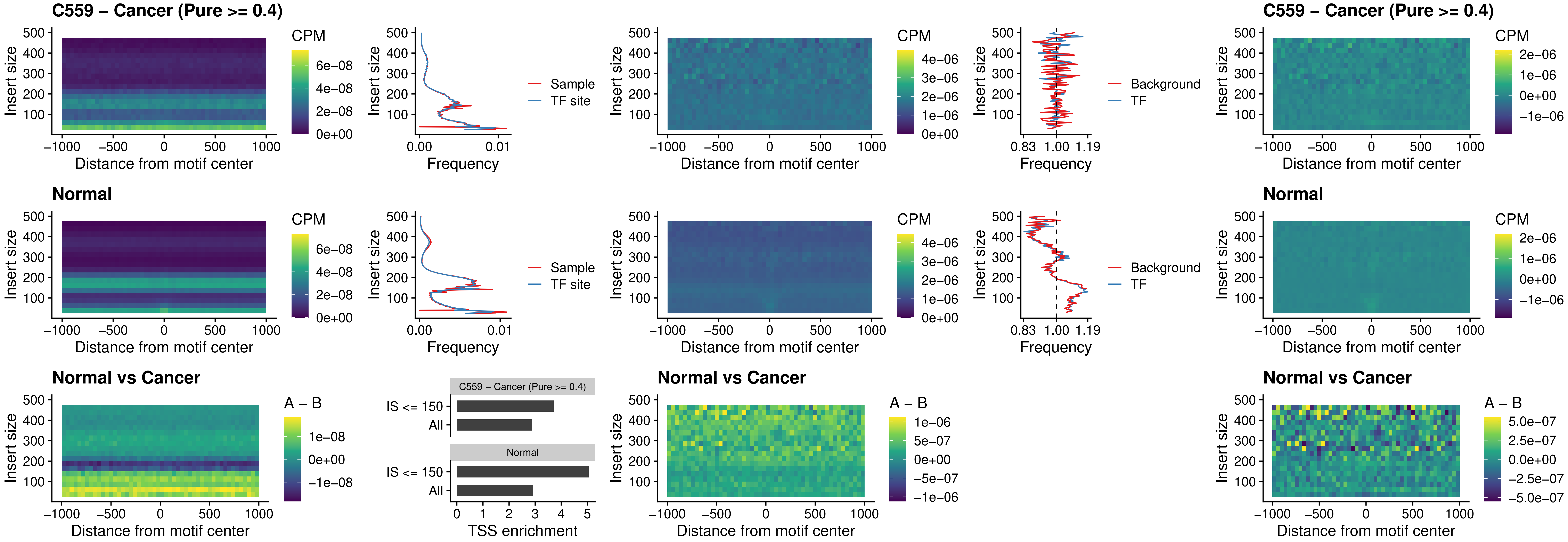

### PURA_uc_dTSS_oPEAK

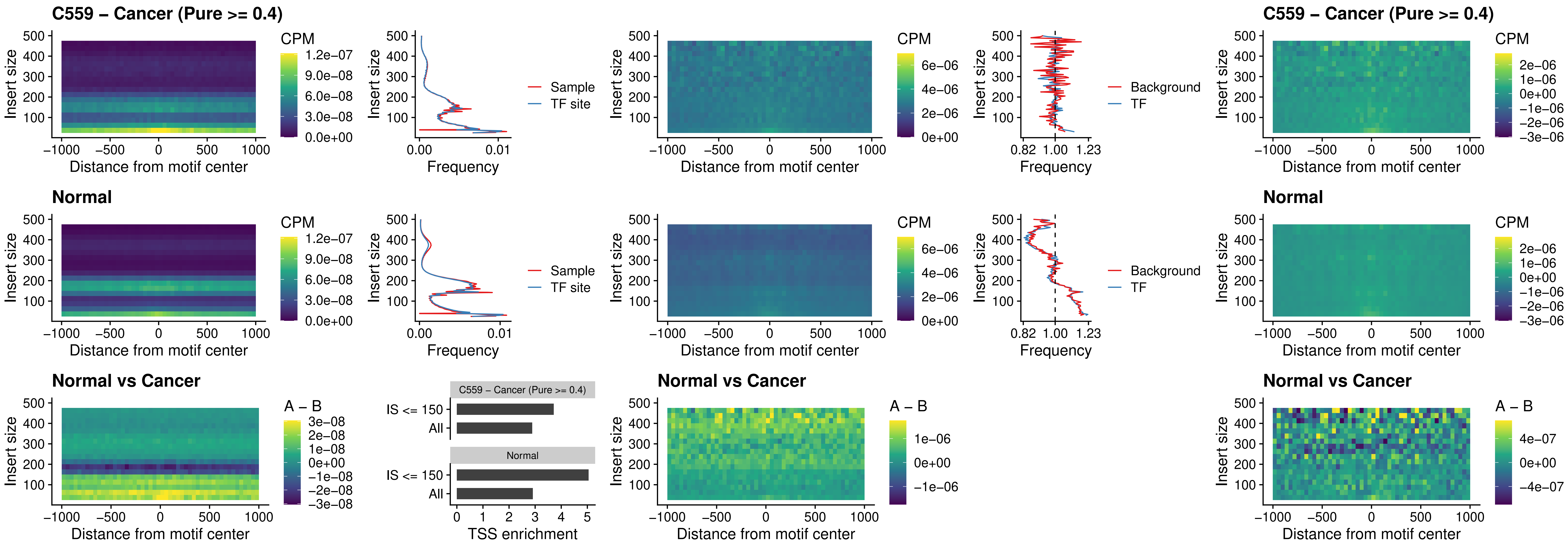

#### HHEX_uc_dTSS_nPEAK

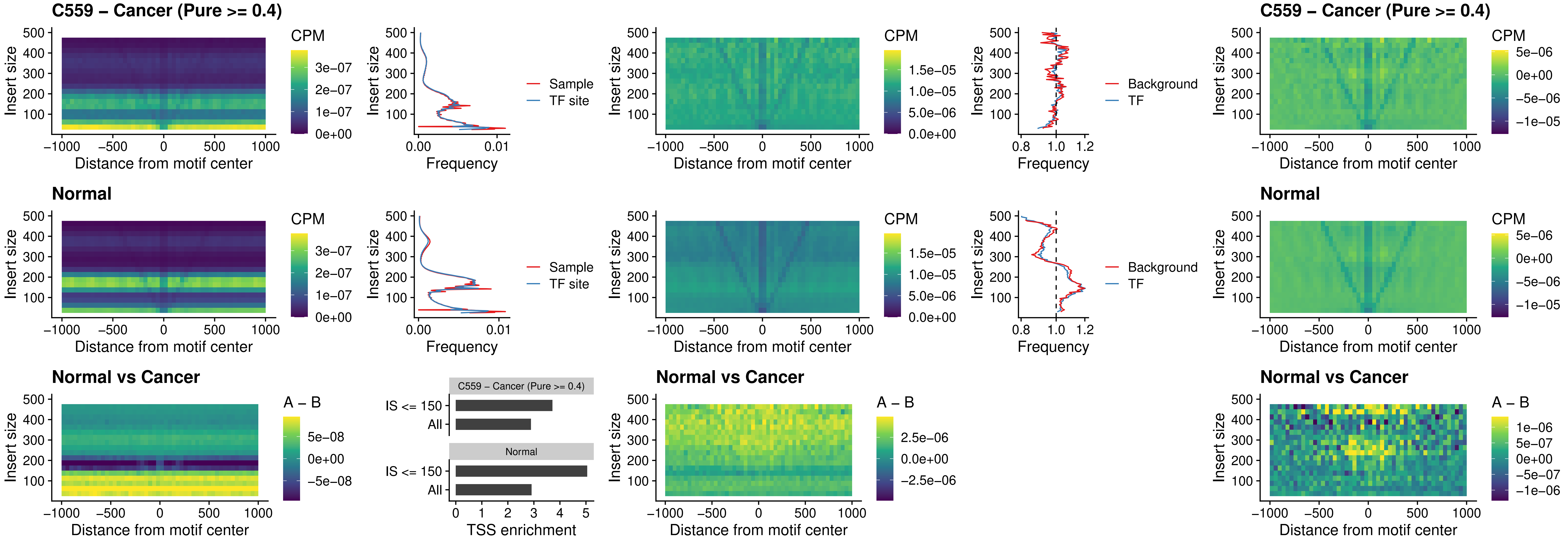

### HMG20B_uc_dTSS_nPEAK

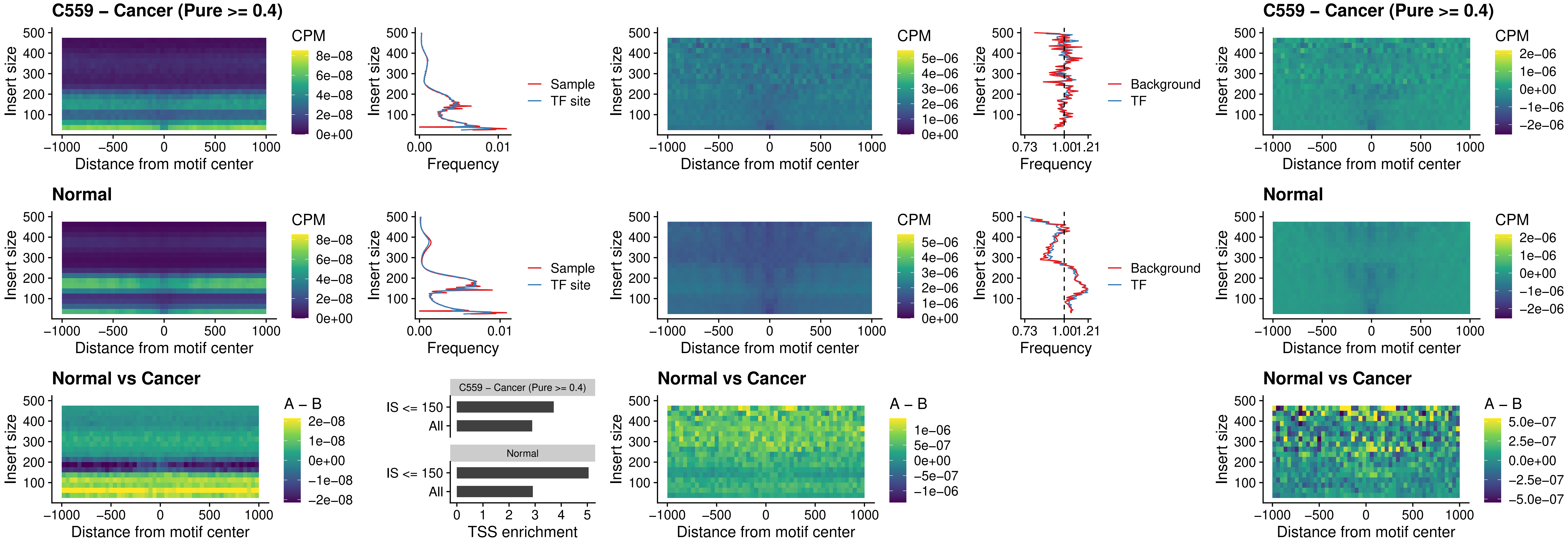

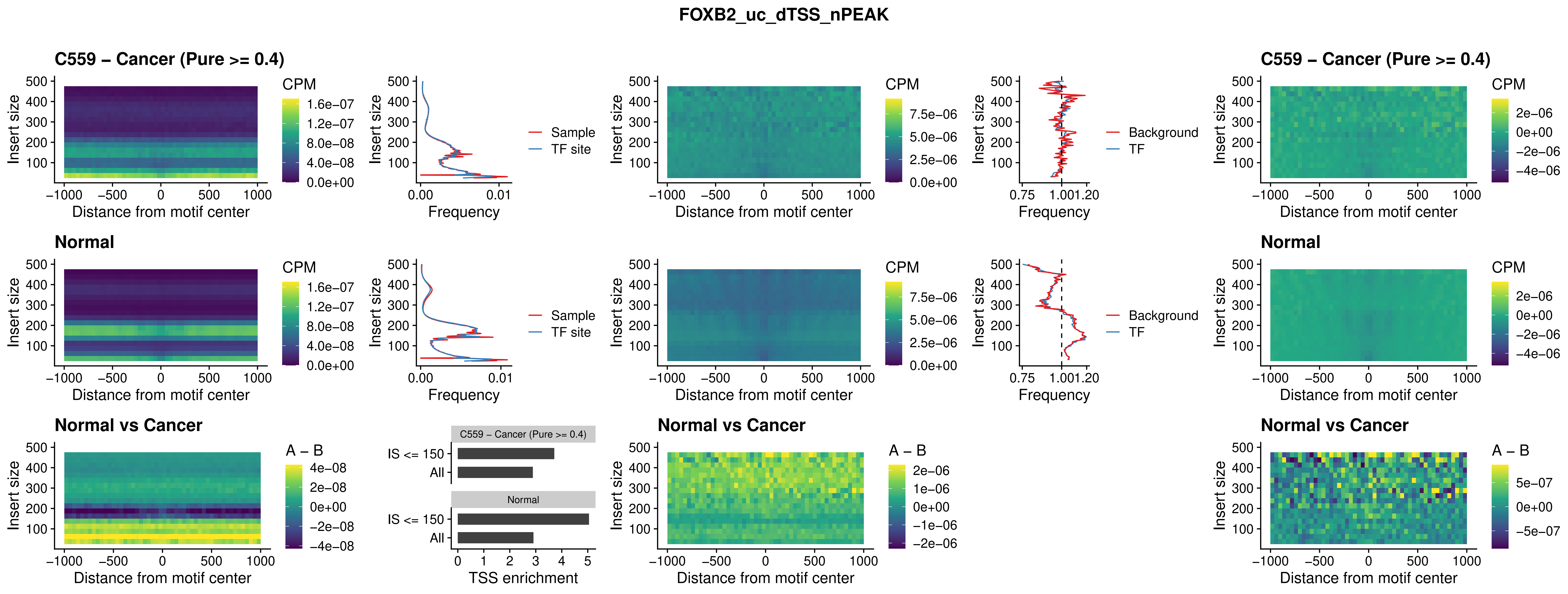
FOXB2_uc_dTSS_nPEAK
C559 – Cancer (Pure >= 0.4)
Normal
Normal vs Cancer
Insert size
Distance from motif center
Frequency
CPM
A – B
Sample
TF site
Background
TF
C559 – Cancer (Pure >= 0.4)
Normal
IS <= 150
All
TSS enrichment

### FOXL1_uc_cTSS_nPEAK

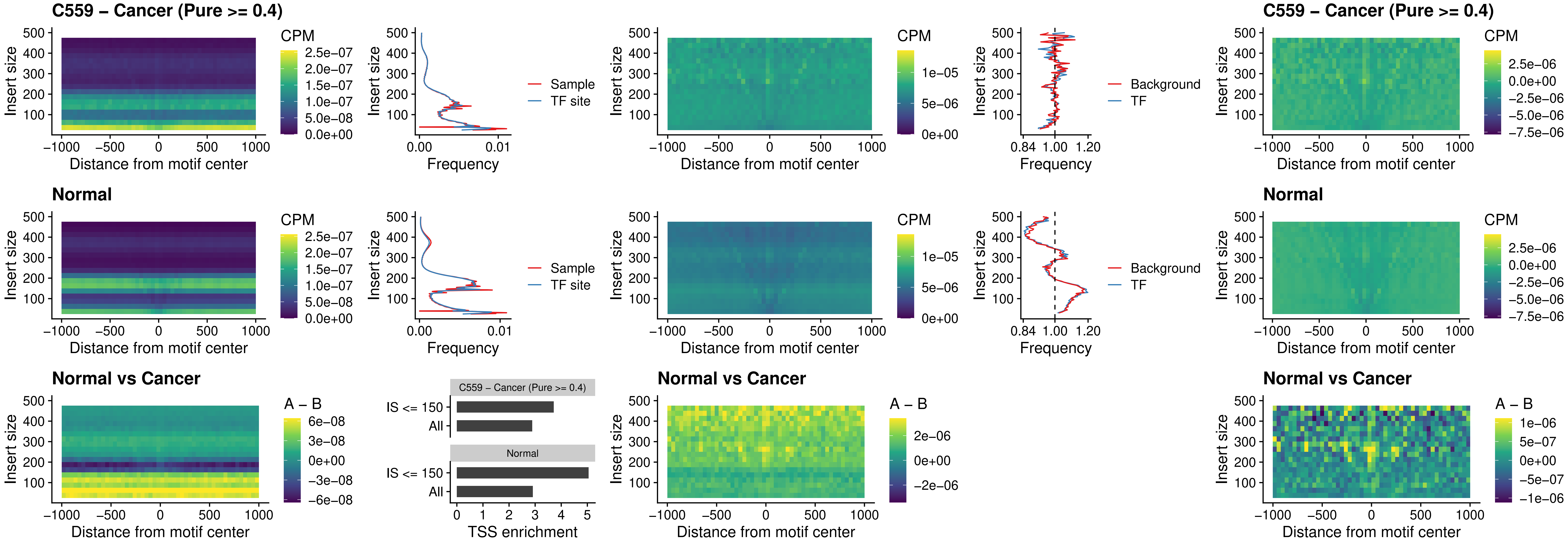

### PBX3_uc_pTSS_oPEAK

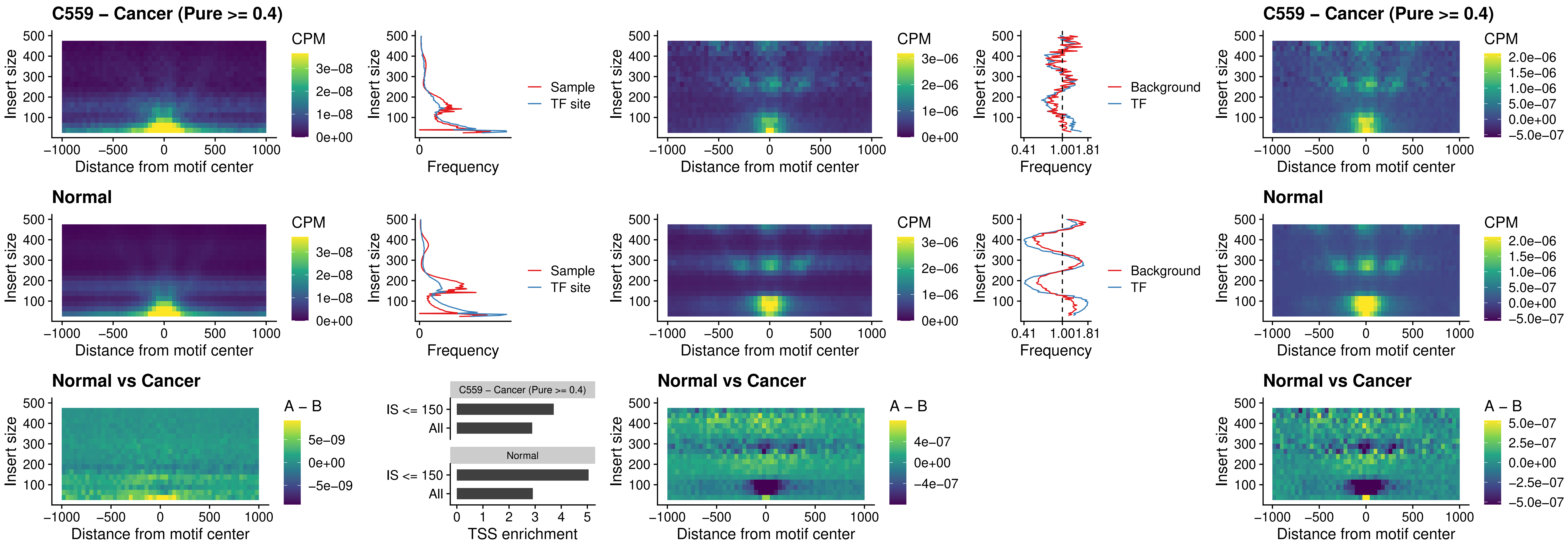

#### SNAI2_uc_cTSS_nPEAK

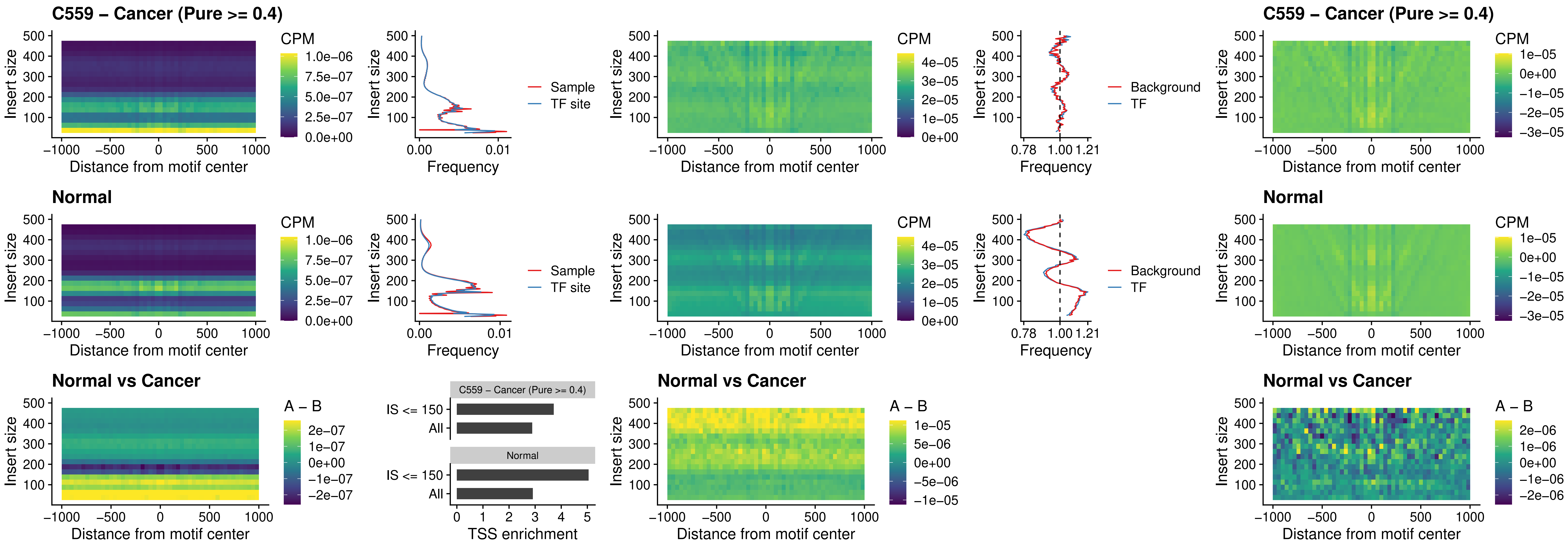

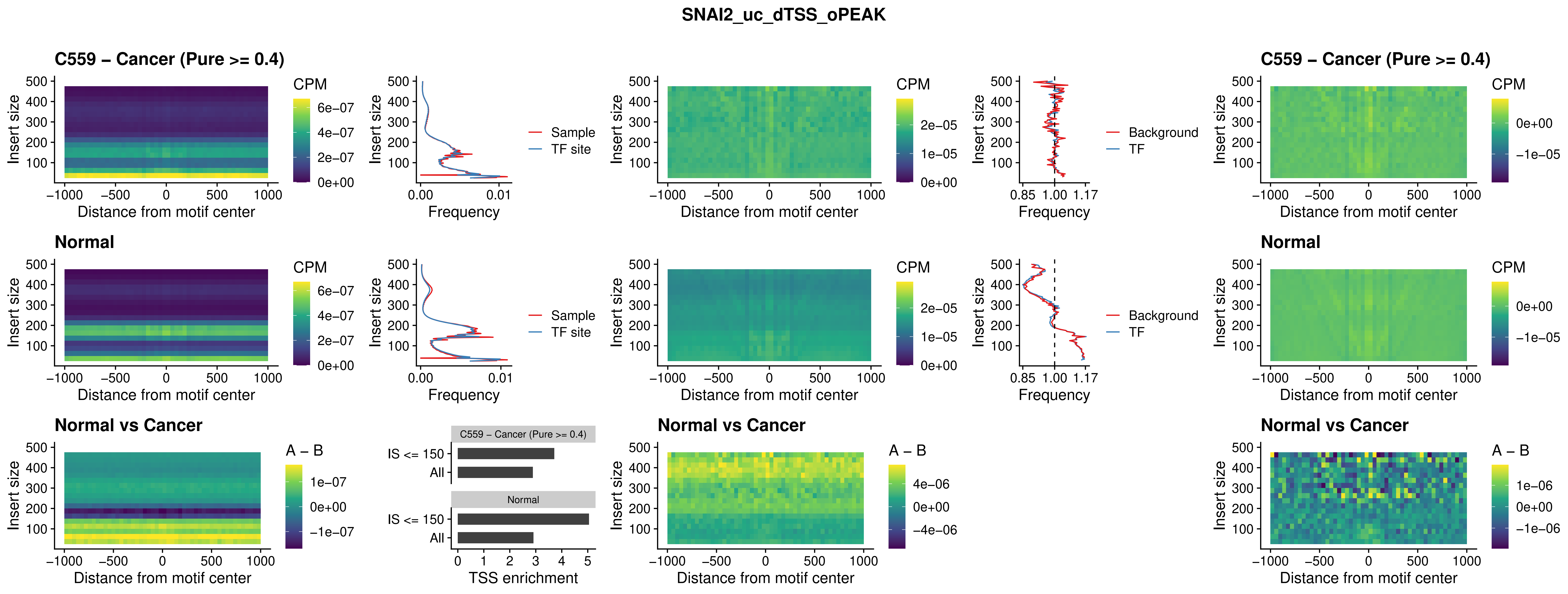
SNAI2_uc_dTSS_oPEAK
C559 – Cancer (Pure >= 0.4)
Normal
Normal vs Cancer
Insert size
Distance from motif center
Frequency
CPM
A – B
Sample
TF site
Background
TF
C559 – Cancer (Pure >= 0.4)
Normal
IS <= 150
All
TSS enrichment

### SPI1_uc_pTSS_nPEAK

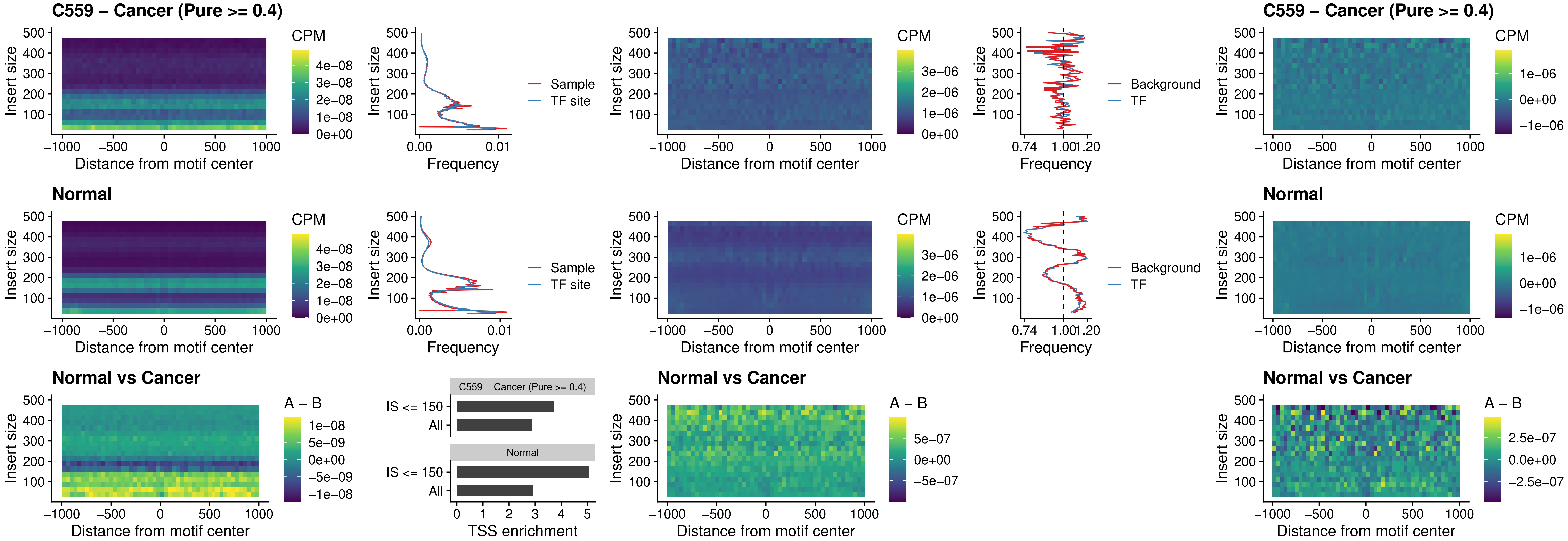

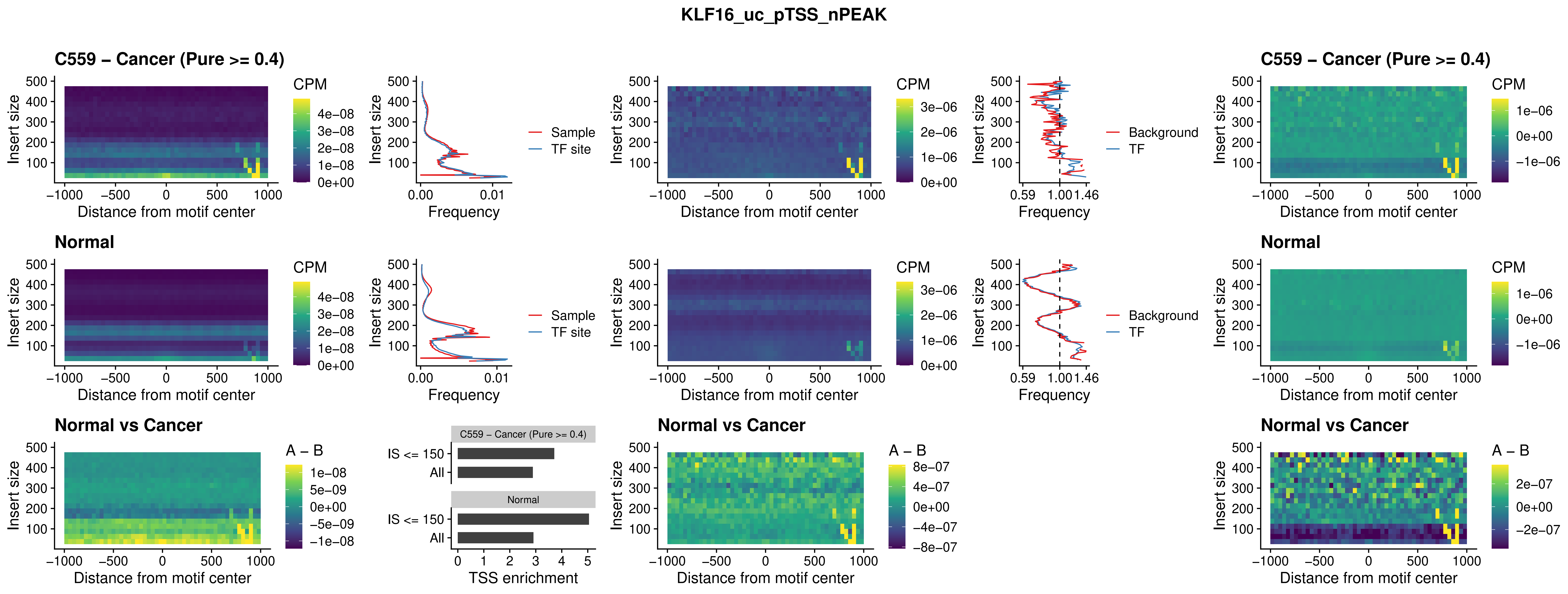
KLF16_uc_pTSS_nPEAK
C559 – Cancer (Pure >= 0.4)
Normal
Normal vs Cancer
Insert size
Distance from motif center
Frequency
CPM
A – B
Sample
TF site
Background
TF
C559 – Cancer (Pure >= 0.4)
Normal
IS <= 150
All
TSS enrichment

### SP3_uc_pTSS_nPEAK

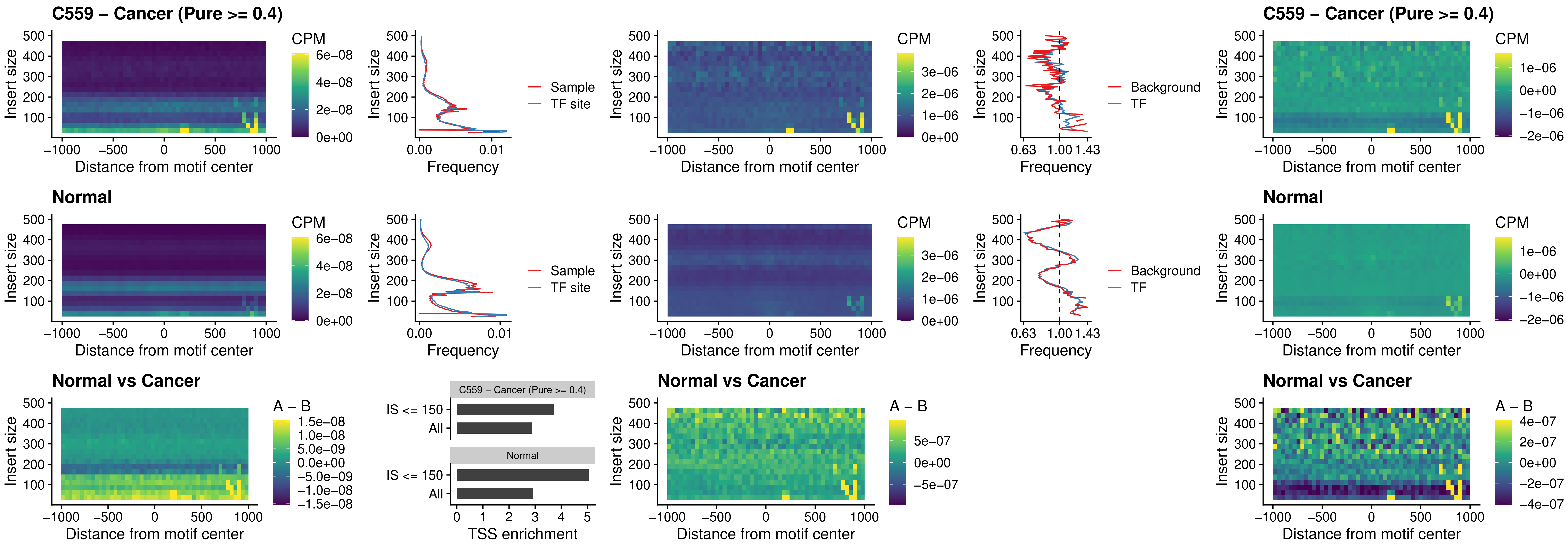

### NEUROD1_uc_dTSS_oPEAK

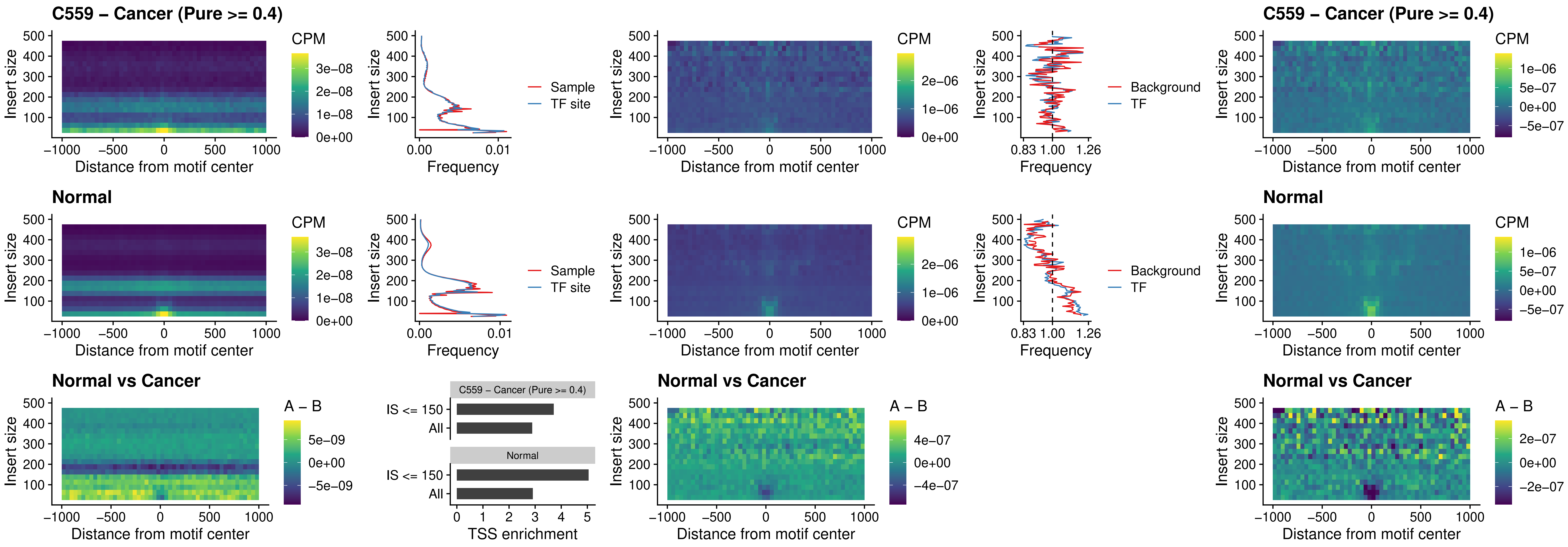

#### CTCF_uc_dTSS_oPEAK

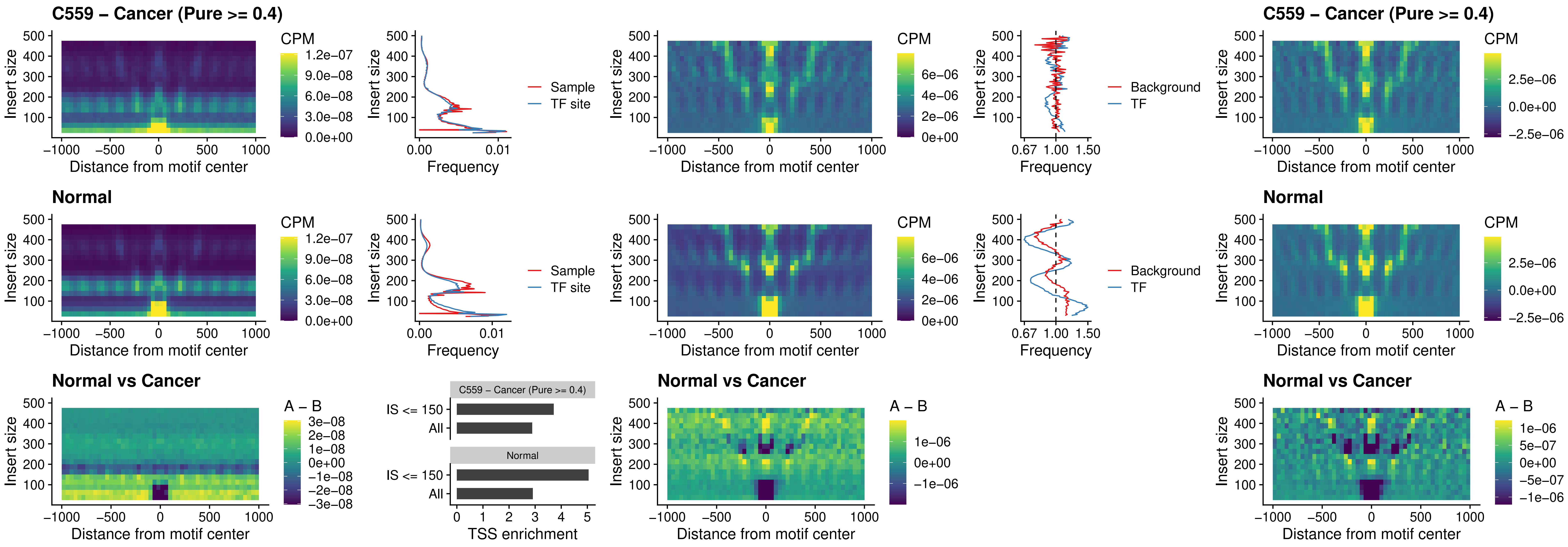

### CTCF_uc_cTSS_oPEAK

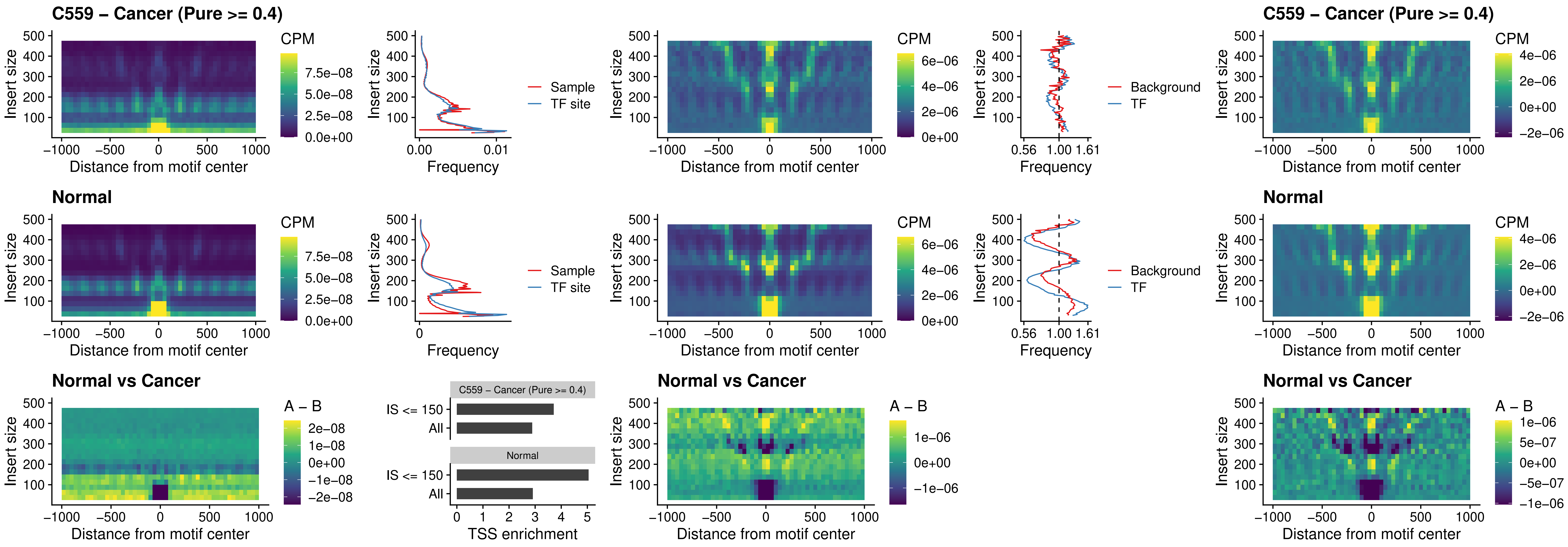

### CTCFL_uc_dTSS_oPEAK

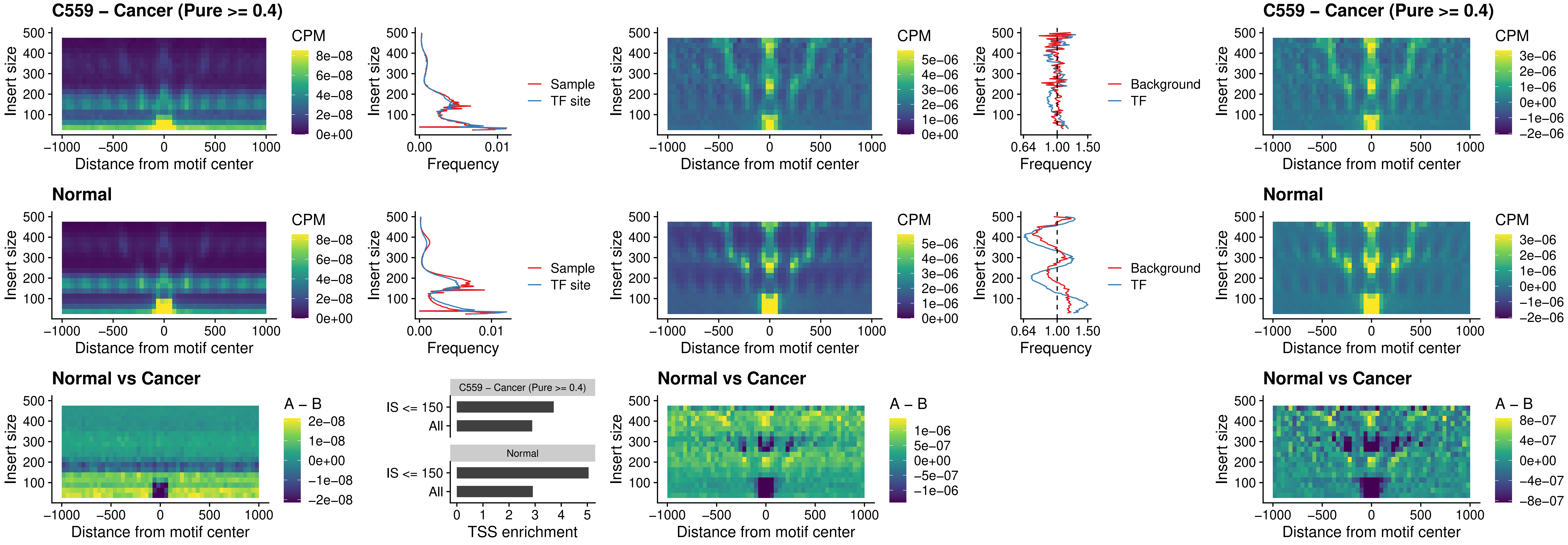

#### ENCODE_CTCF_sites_COLON

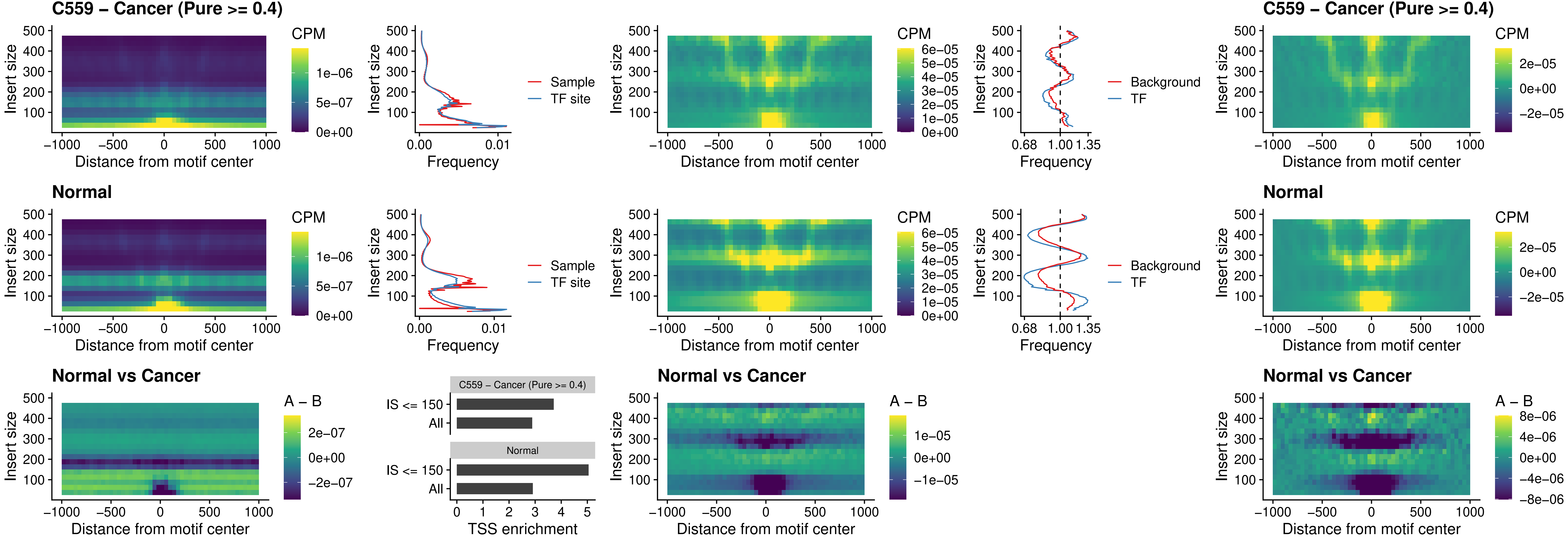

### ENCODE_CTCF_sites_COLON_loop_ctcf

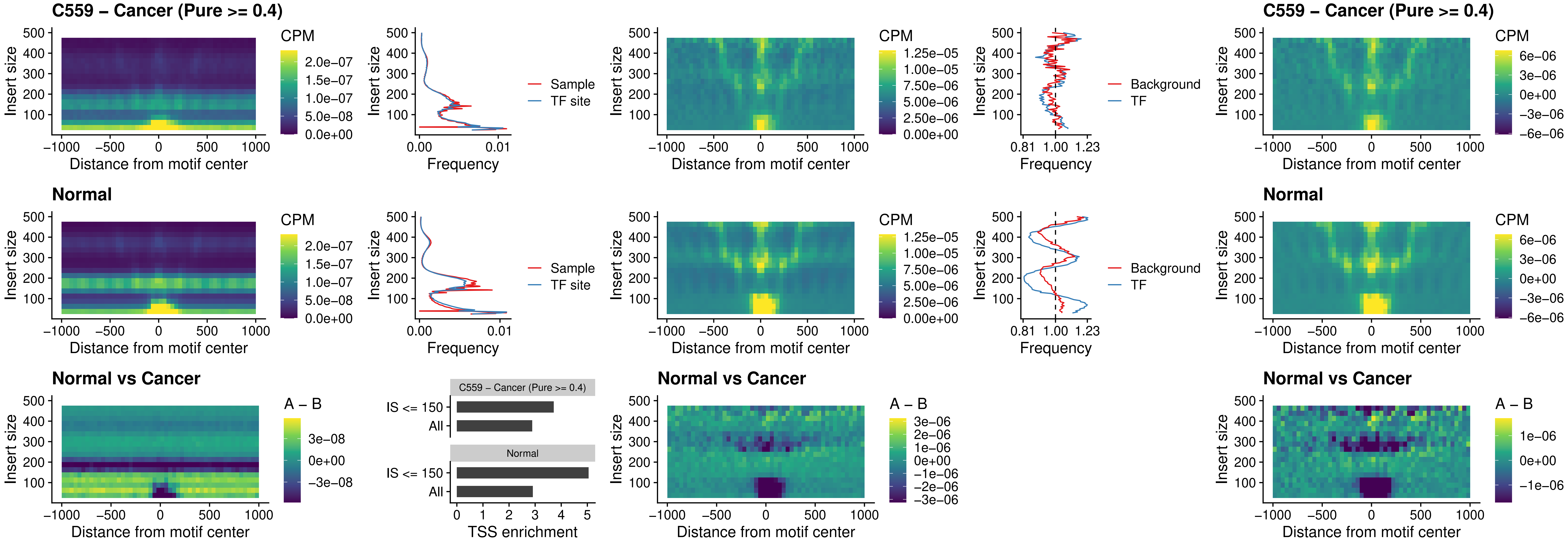

#### ENCODE_CTCF_sites_COLON_intergenic_ctcf

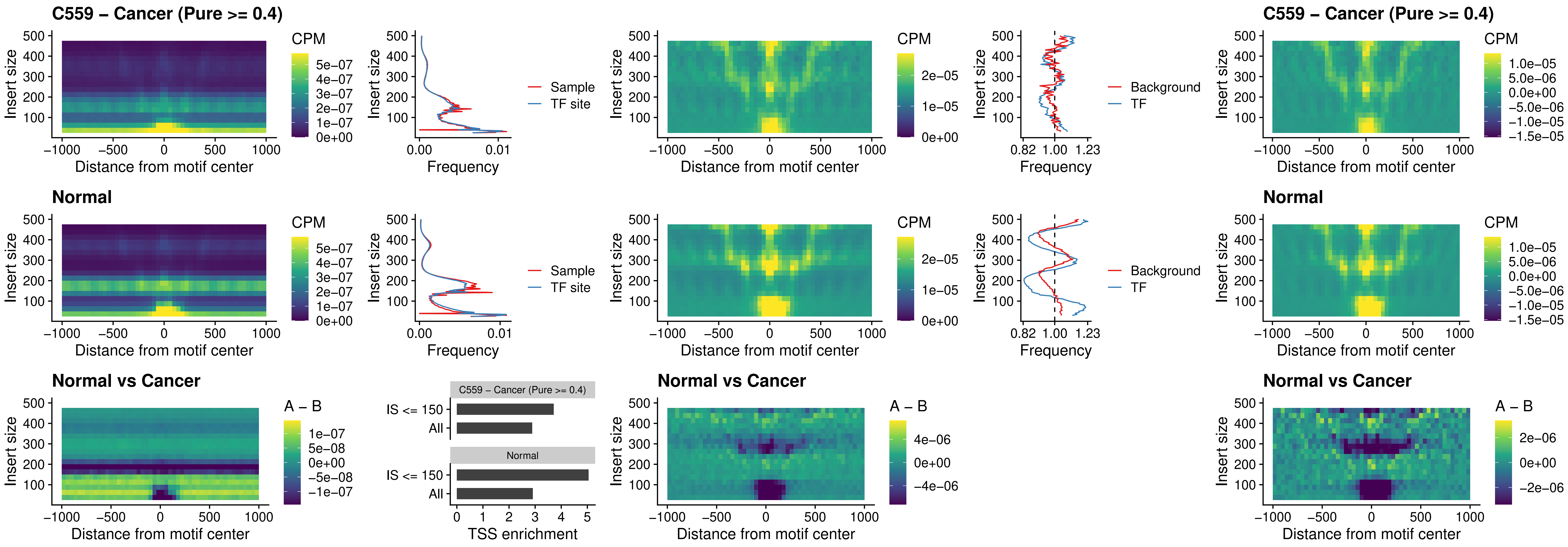

### ENCODE_CTCF_sites_COLON_nonpromotor

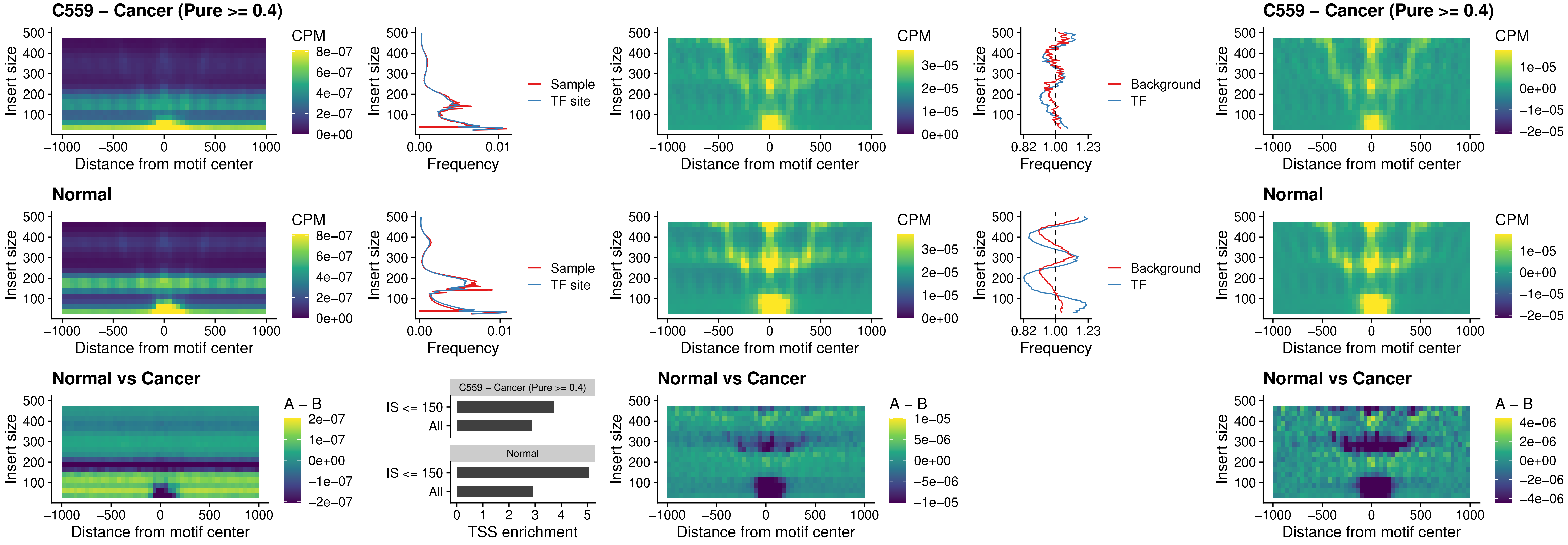

#### CTCFL_uc_cTSS_oPEAK

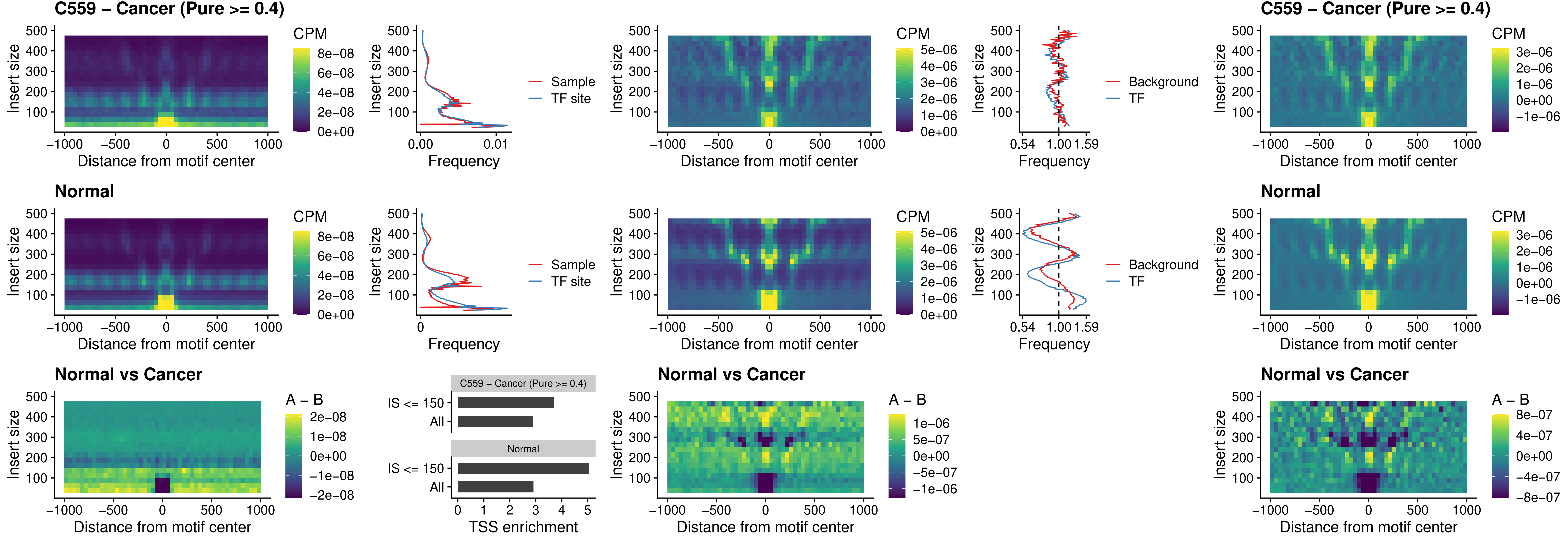

### NEUROD1_uc_cTSS_oPEAK

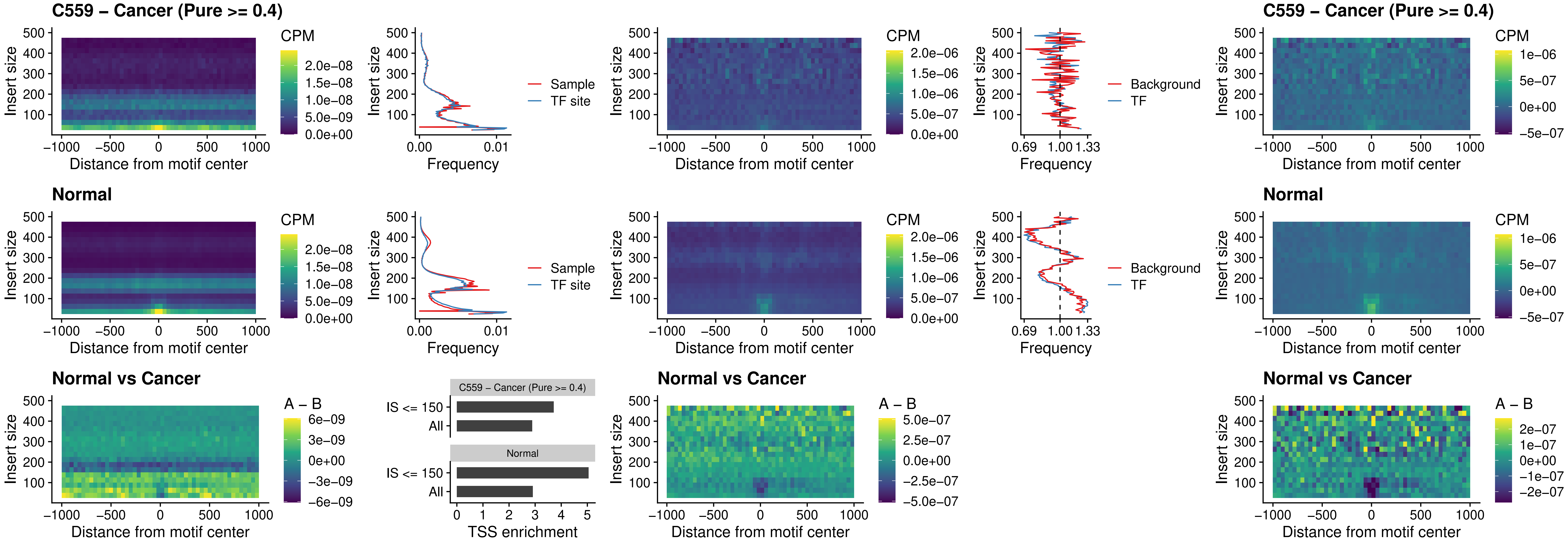

#### ENCODE_YY1_sites_COLON_nonpromotor

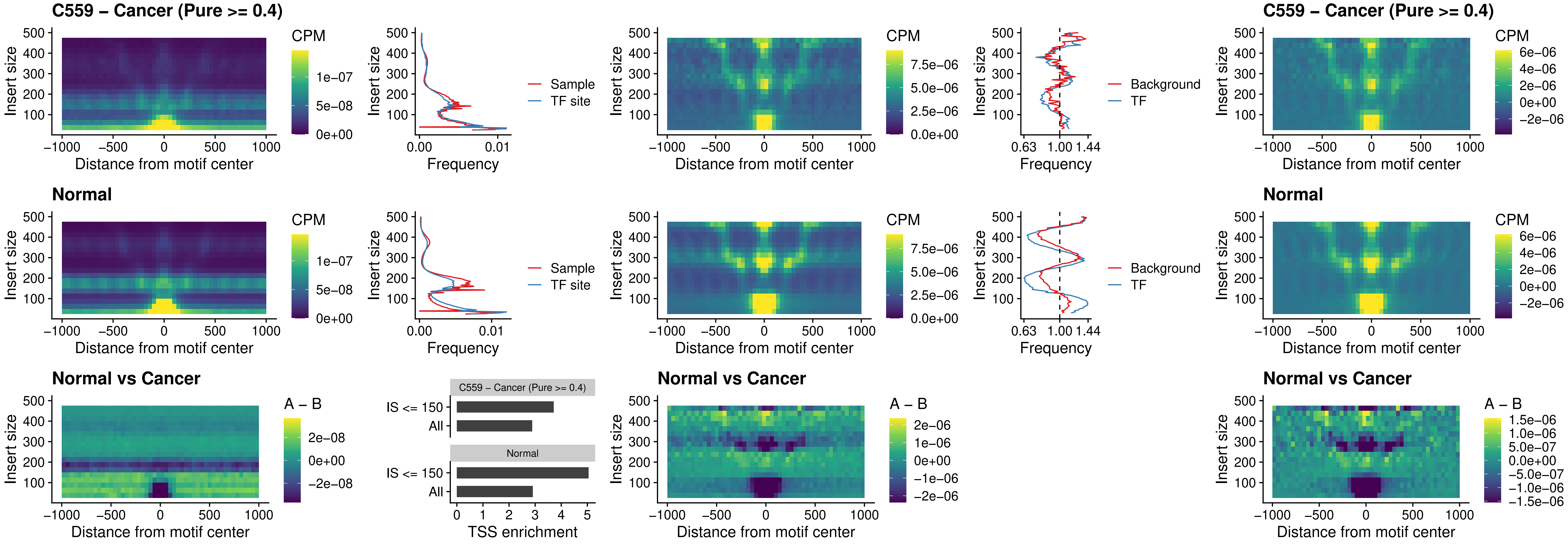

### CTCF_uc_pTSS_oPEAK

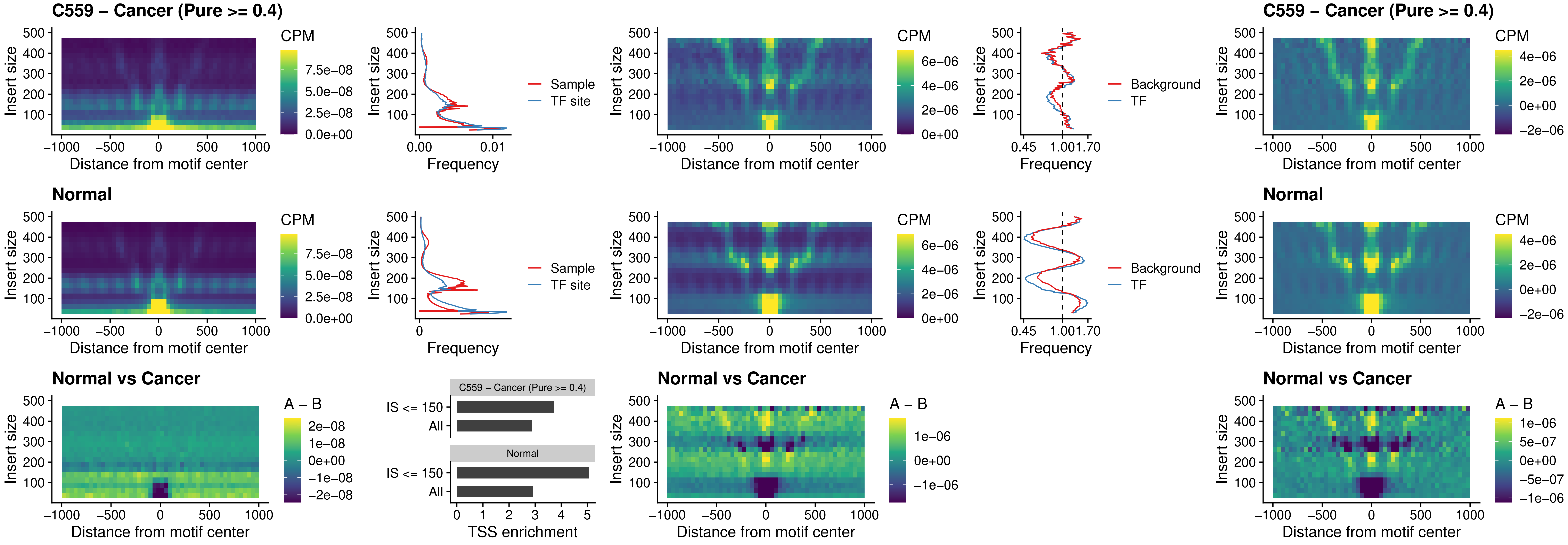

### ENCODE_CTCF_sites_COLON_promotor

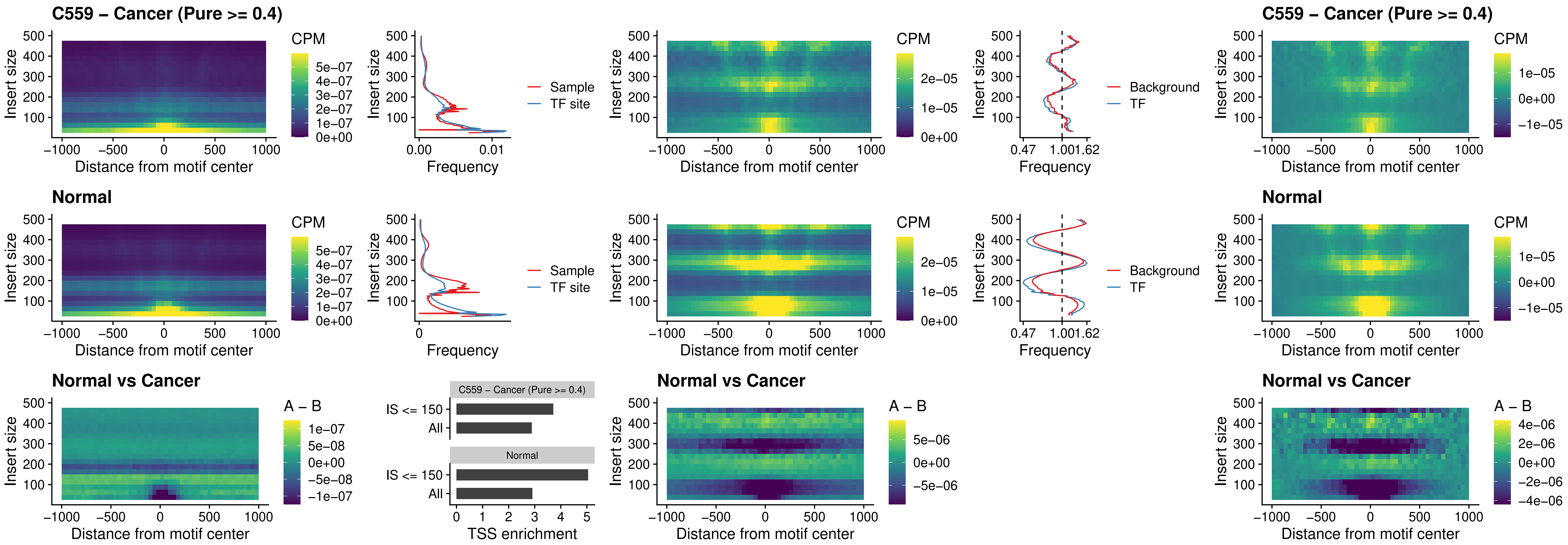

### ENCODE_CTCF_sites_COLON_promotor_ctcf

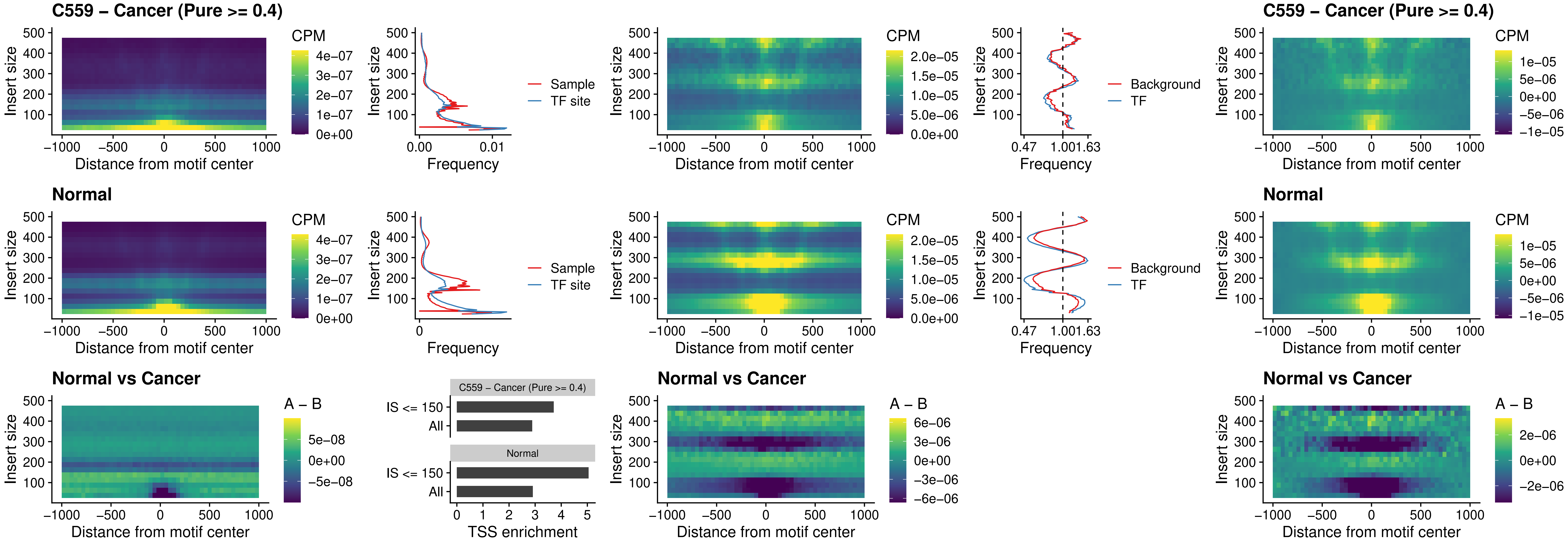

### SPIC_uc_pTSS_oPEAK

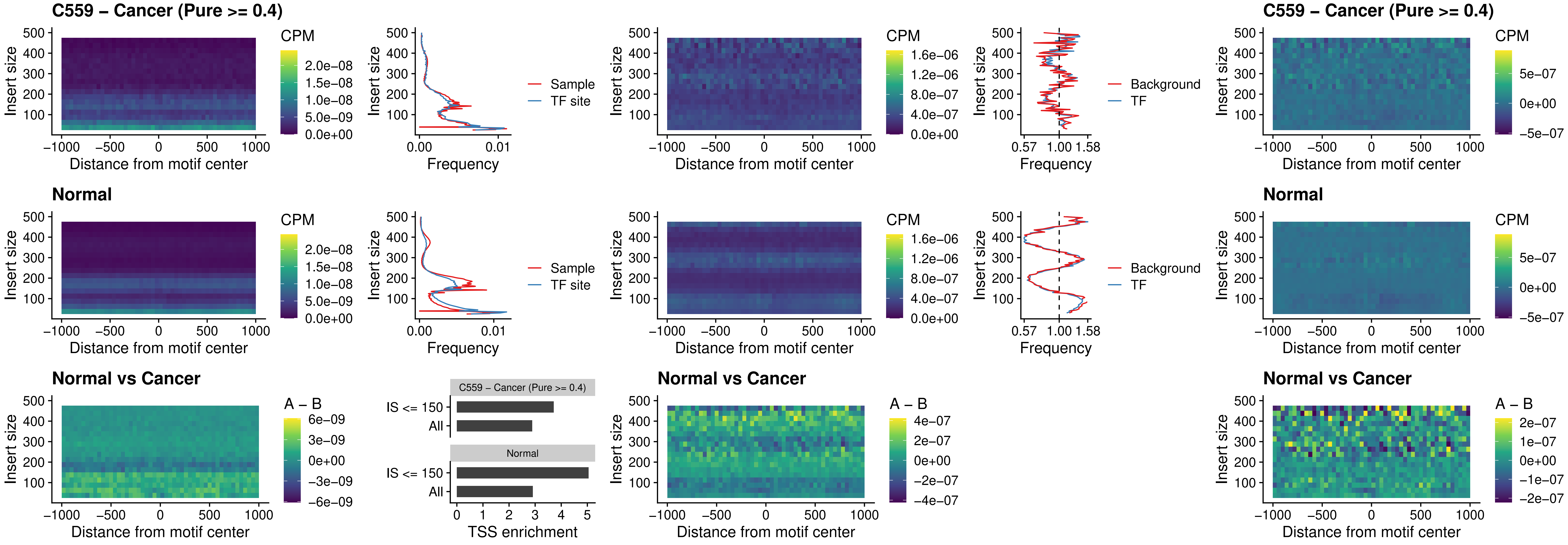

### SPI1_uc_dTSS_nPEAK

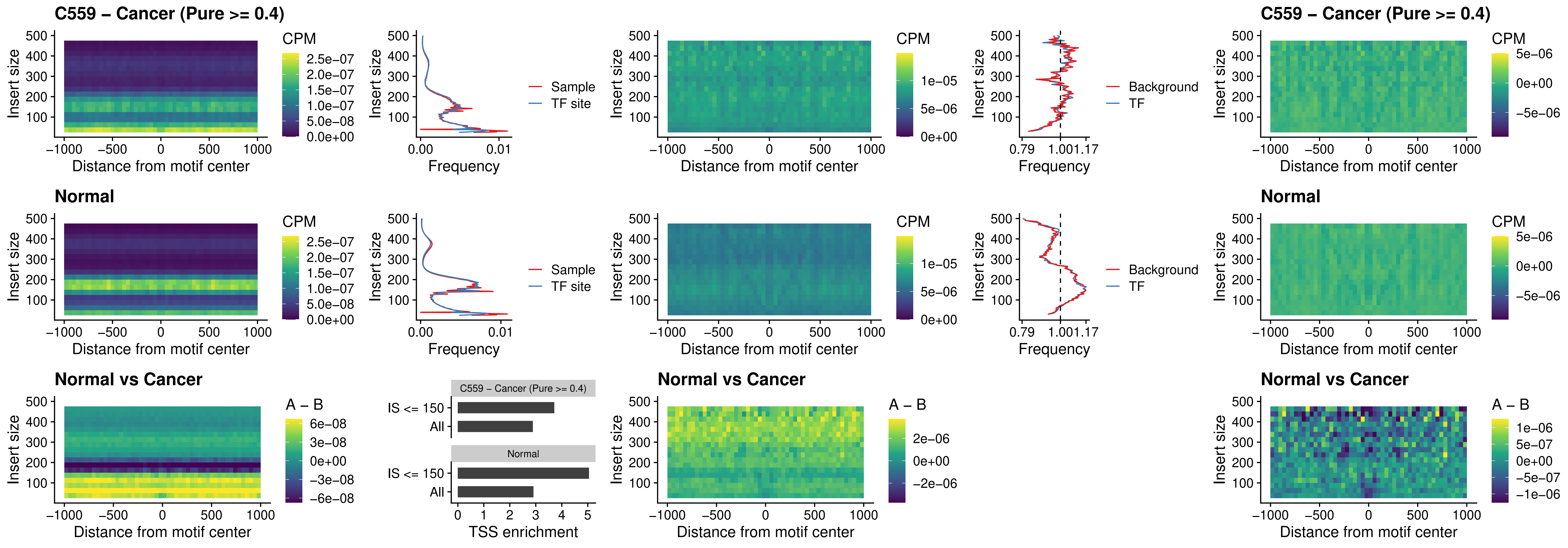

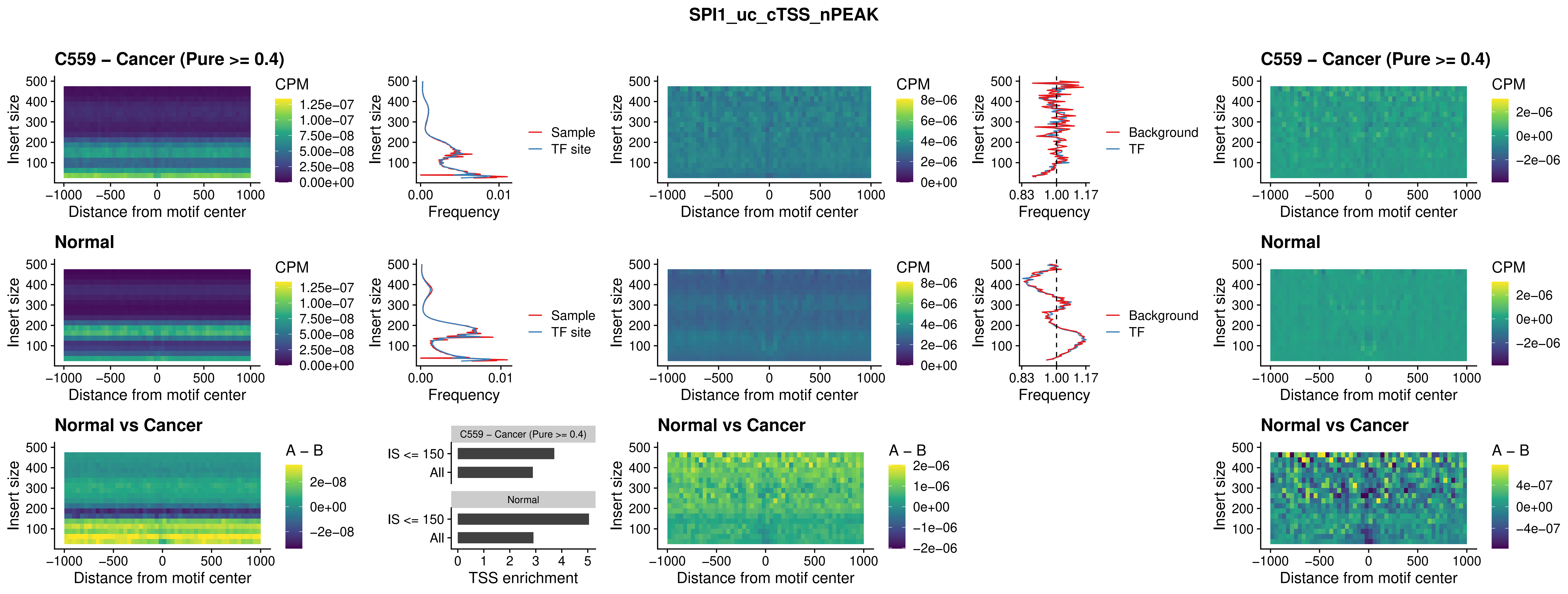
SPI1_uc_cTSS_nPEAK
C559 – Cancer (Pure >= 0.4)
Normal
Normal vs Cancer
Insert size
Distance from motif center
Frequency
CPM
A – B
Sample
TF site
Background
TF
C559 – Cancer (Pure >= 0.4)
Normal
IS <= 150
All
TSS enrichment

### BCL11A_uc_pTSS_nPEAK

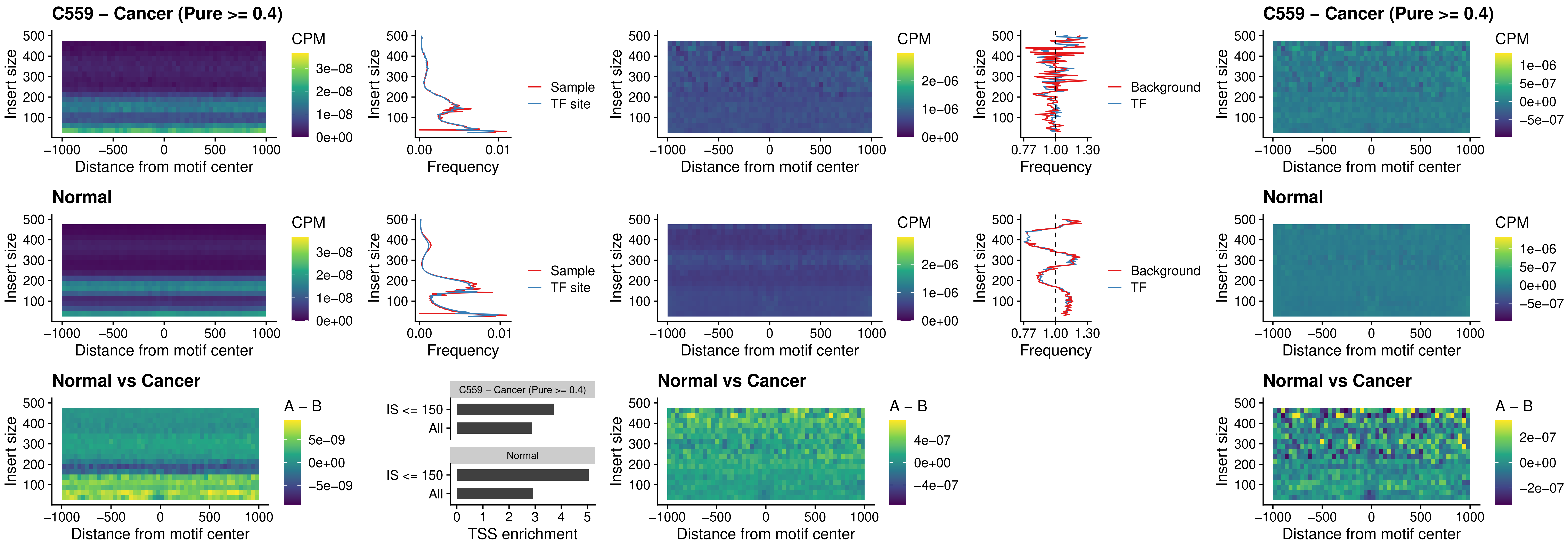

### BCL11B_uc_pTSS_nPEAK

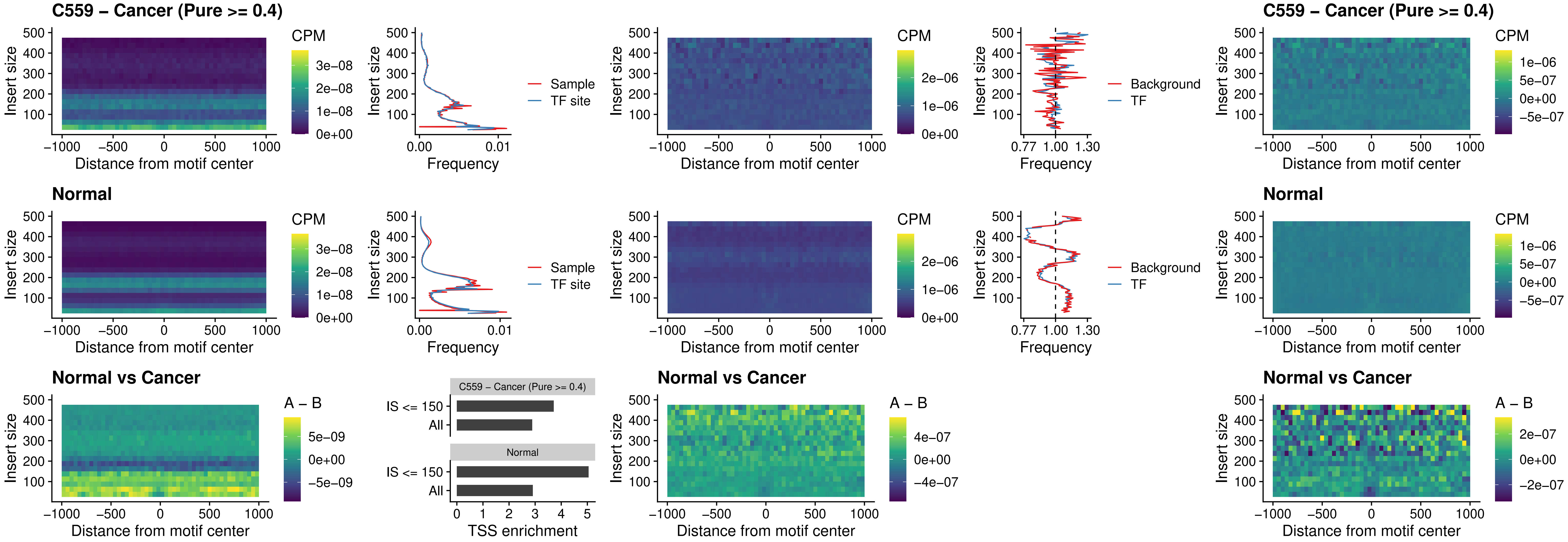

### IRF9_uc_dTSS_oPEAK

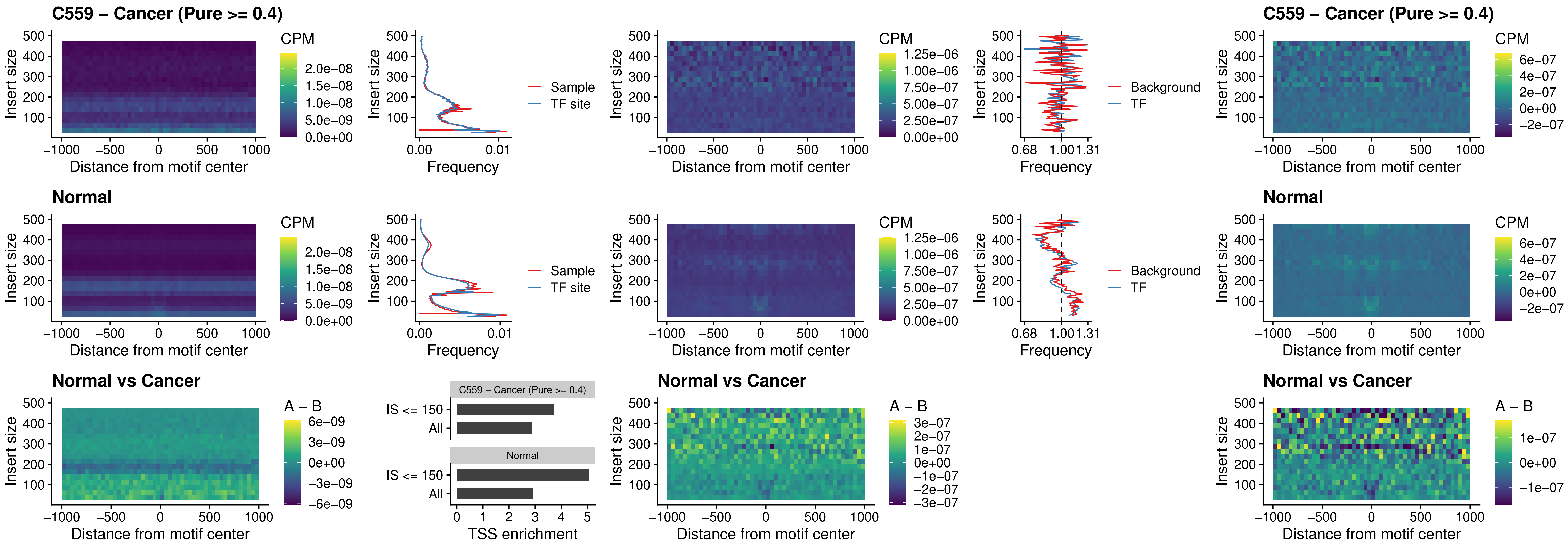

### IRF9_uc_cTSS_oPEAK

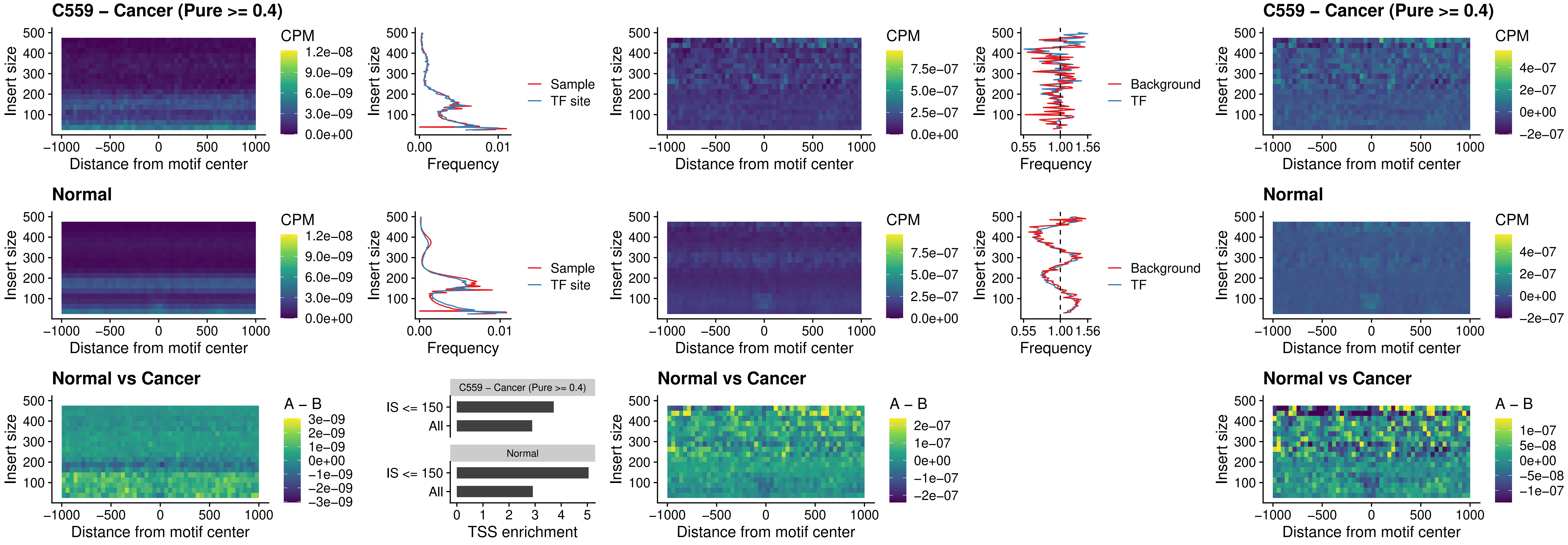

### BCL11A_uc_cTSS_oPEAK

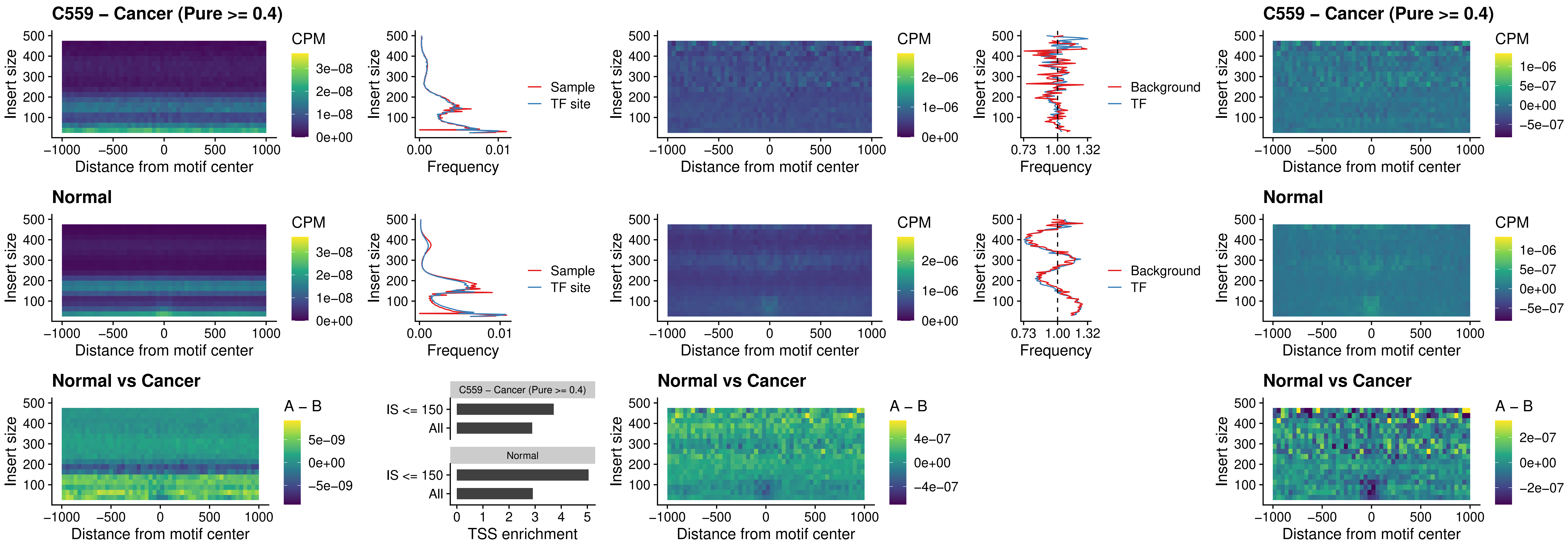

### BCL11B_uc_cTSS_oPEAK

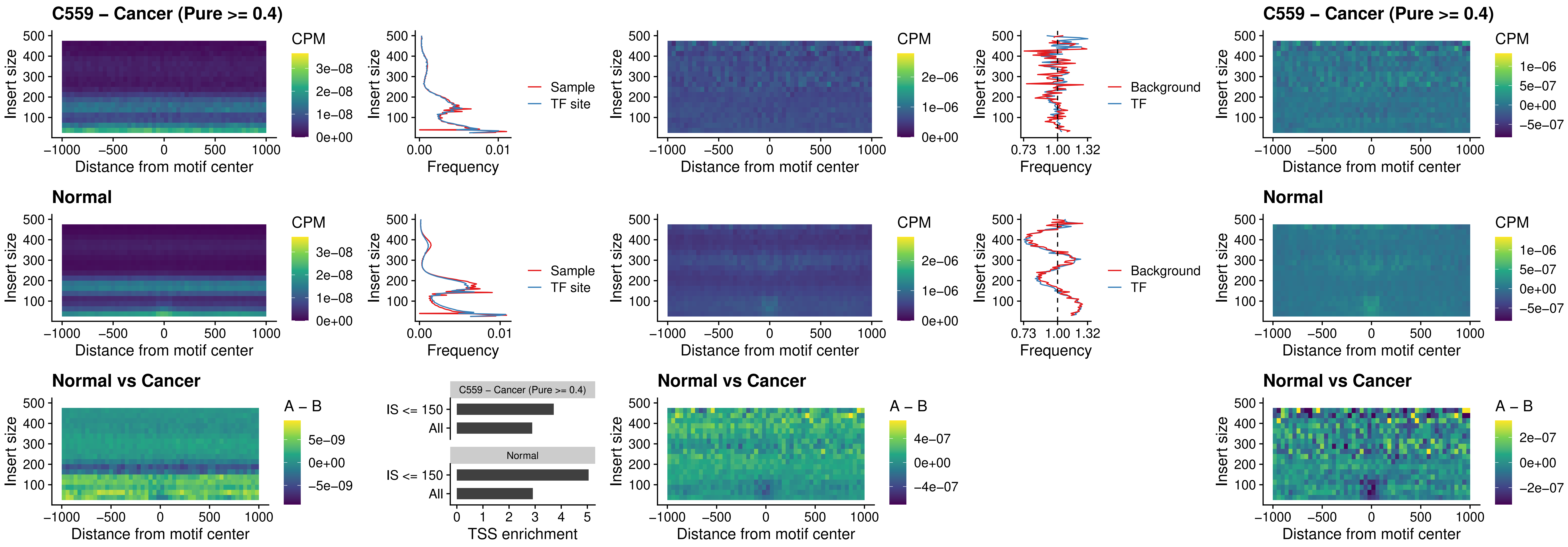

### BCL11A_uc_dTSS_oPEAK

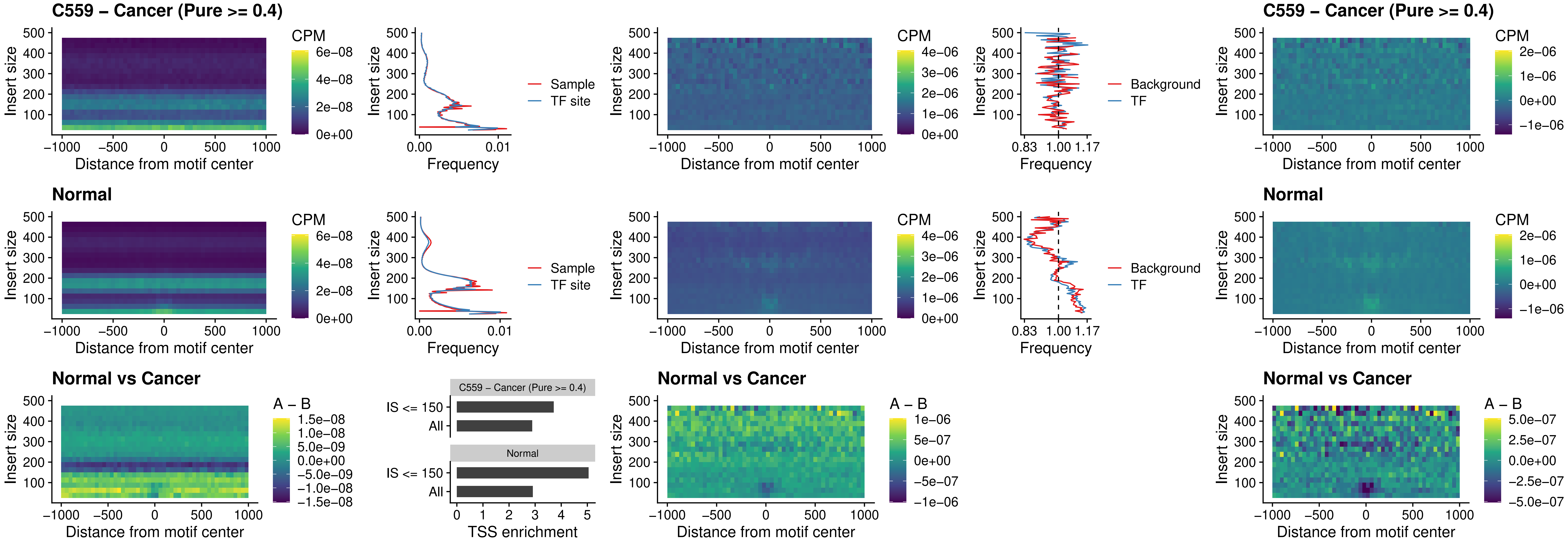

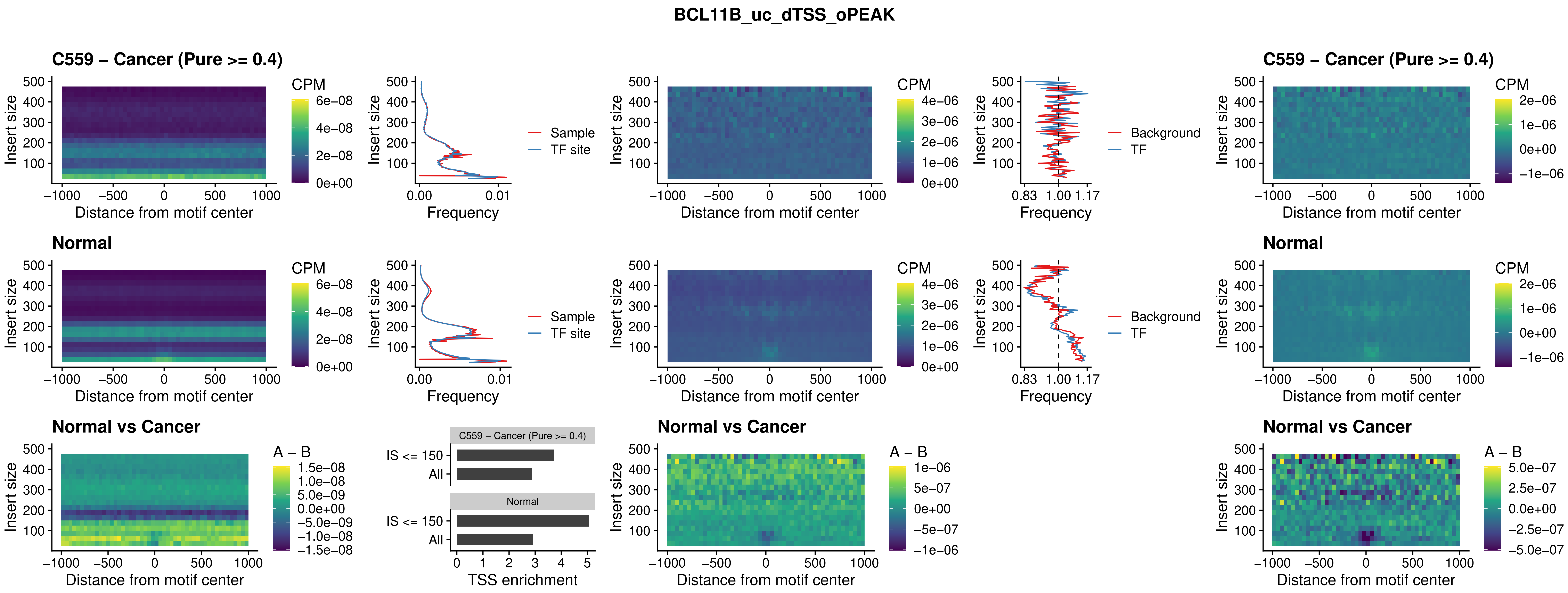
BCL11B_uc_dTSS_oPEAK
C559 – Cancer (Pure >= 0.4)
Normal
Normal vs Cancer
Insert size
Distance from motif center
Frequency
CPM
A – B
Sample
TF site
Background
TF
C559 – Cancer (Pure >= 0.4)
Normal
IS <= 150
All
TSS enrichment

### SPIB_uc_cTSS_nPEAK

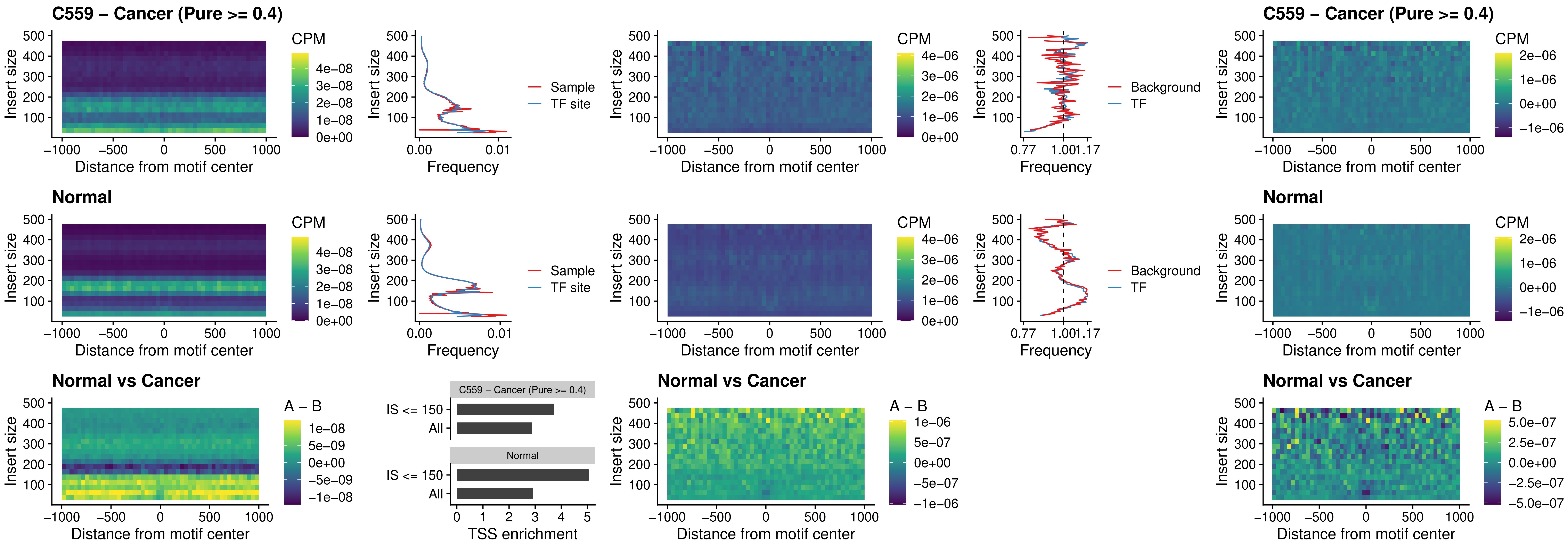

#### SPIC_uc_cTSS_nPEAK

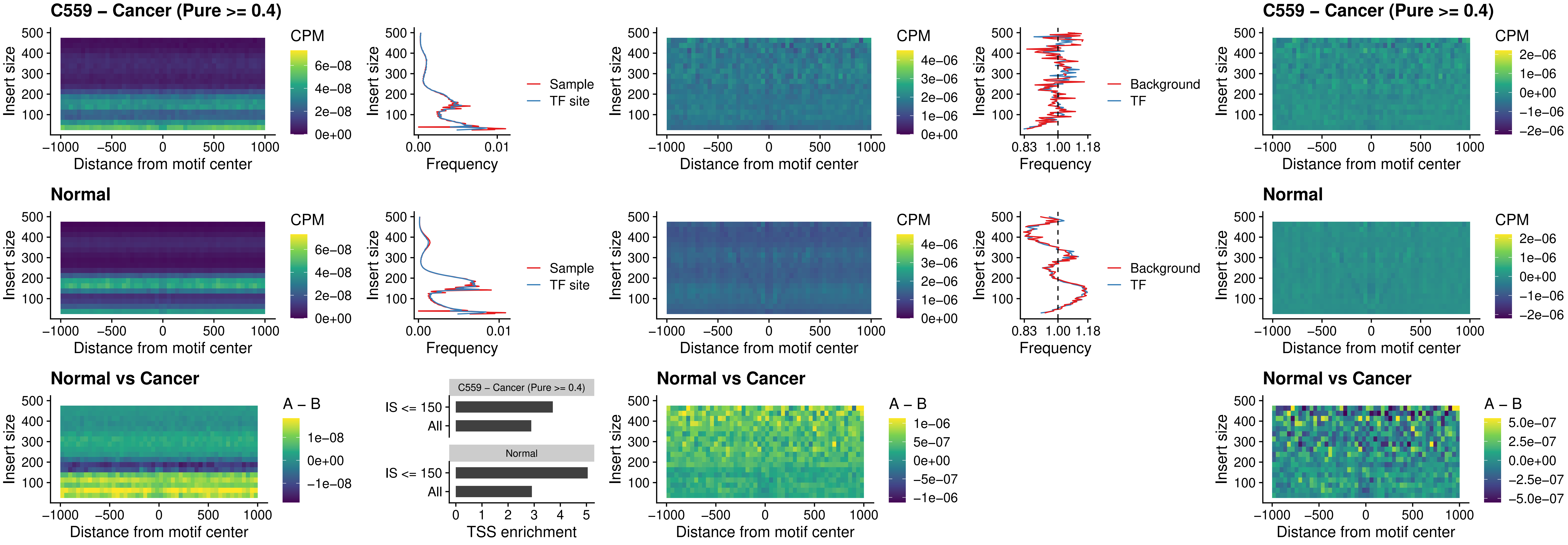

#### SPI1_uc_pTSS_oPEAK

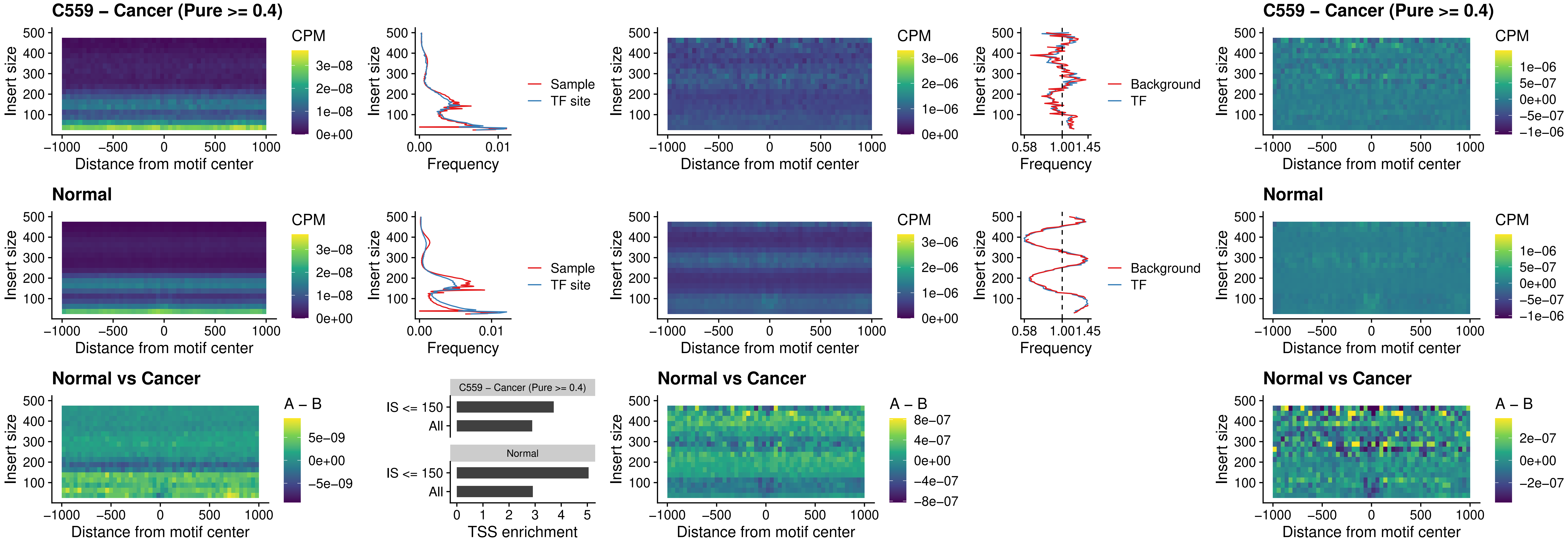

#### BCL11A_uc_pTSS_oPEAK

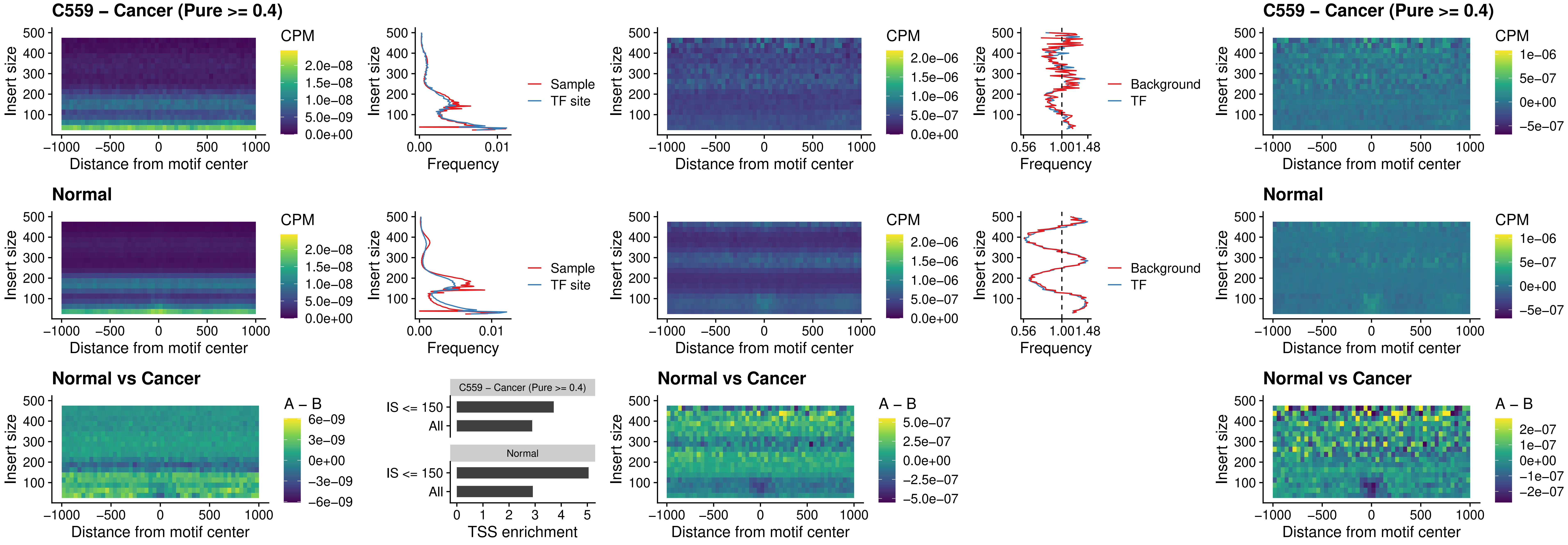

### BCL11B_uc_pTSS_oPEAK

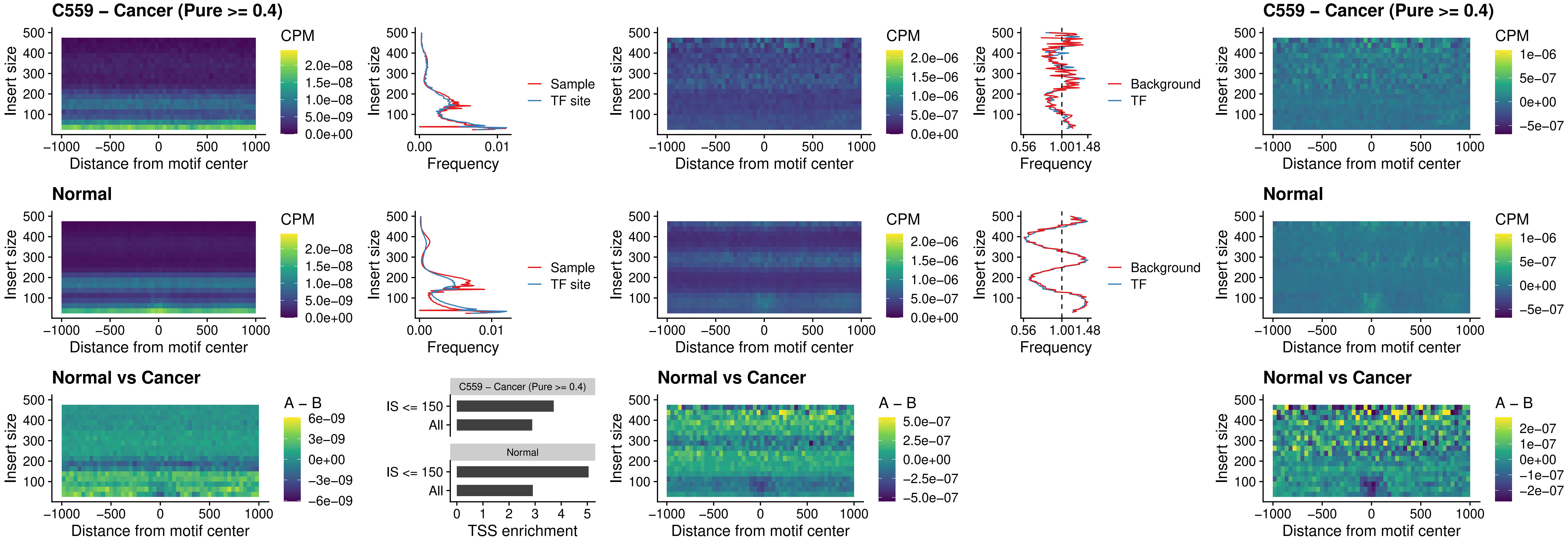

### SPI1_uc_cTSS_oPEAK

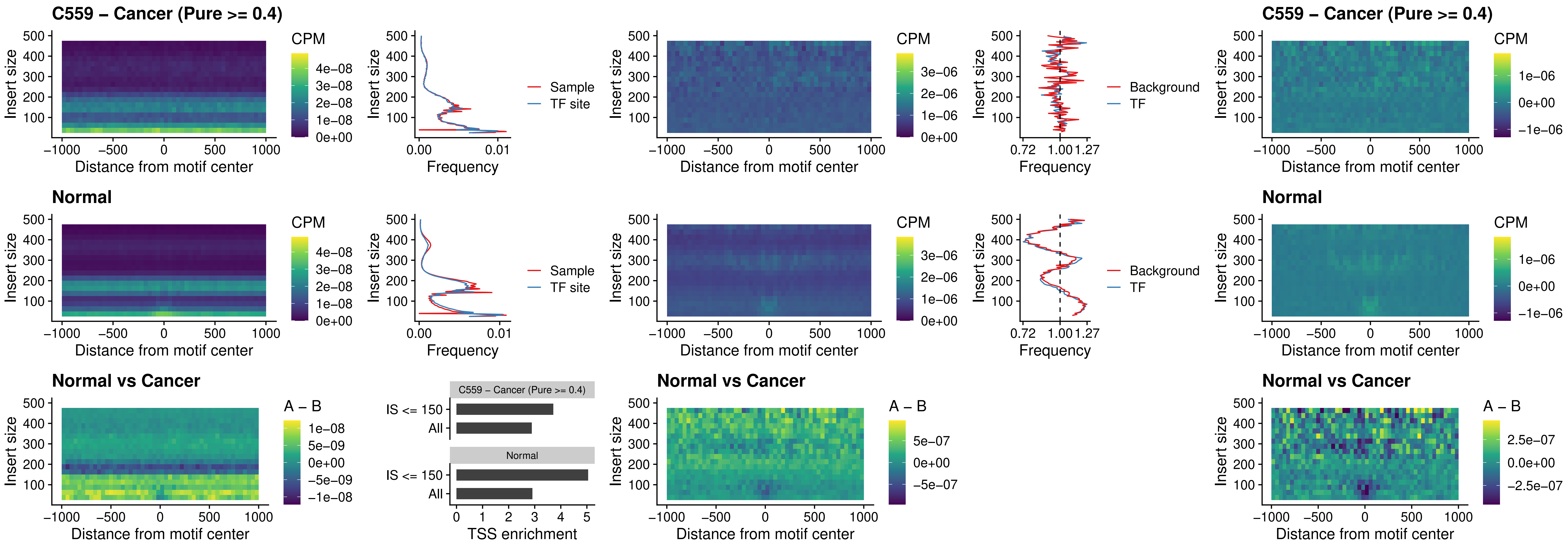

### SPIB_uc_cTSS_oPEAK

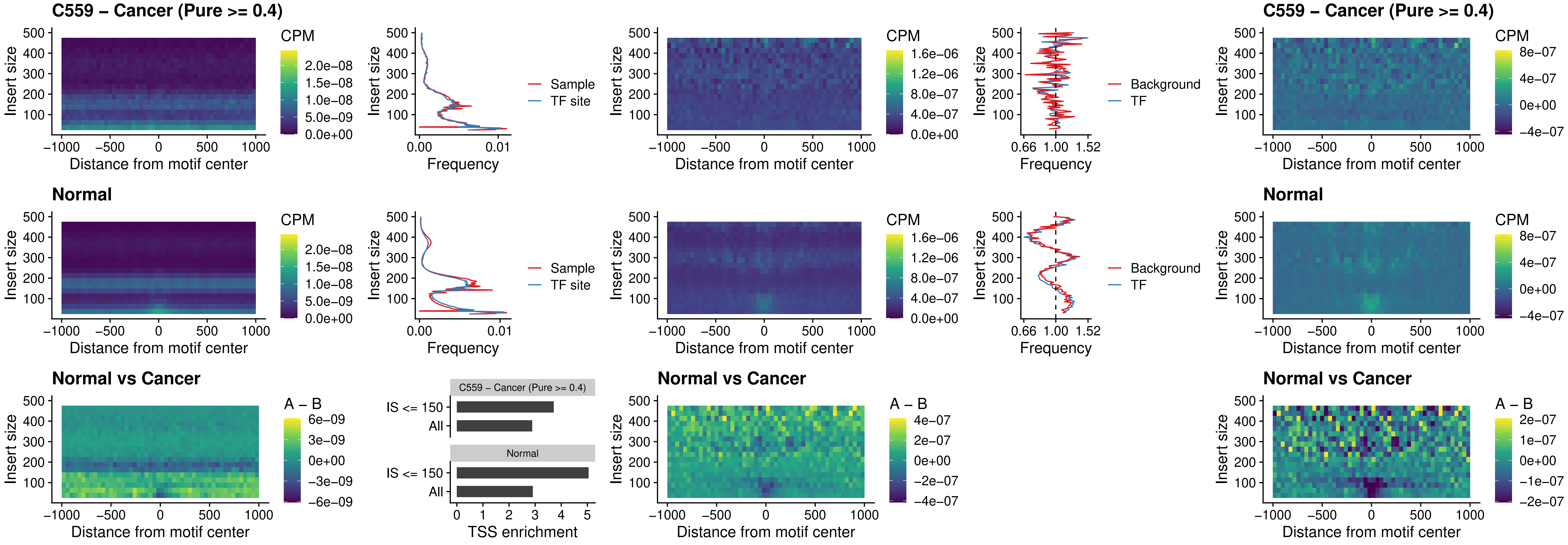

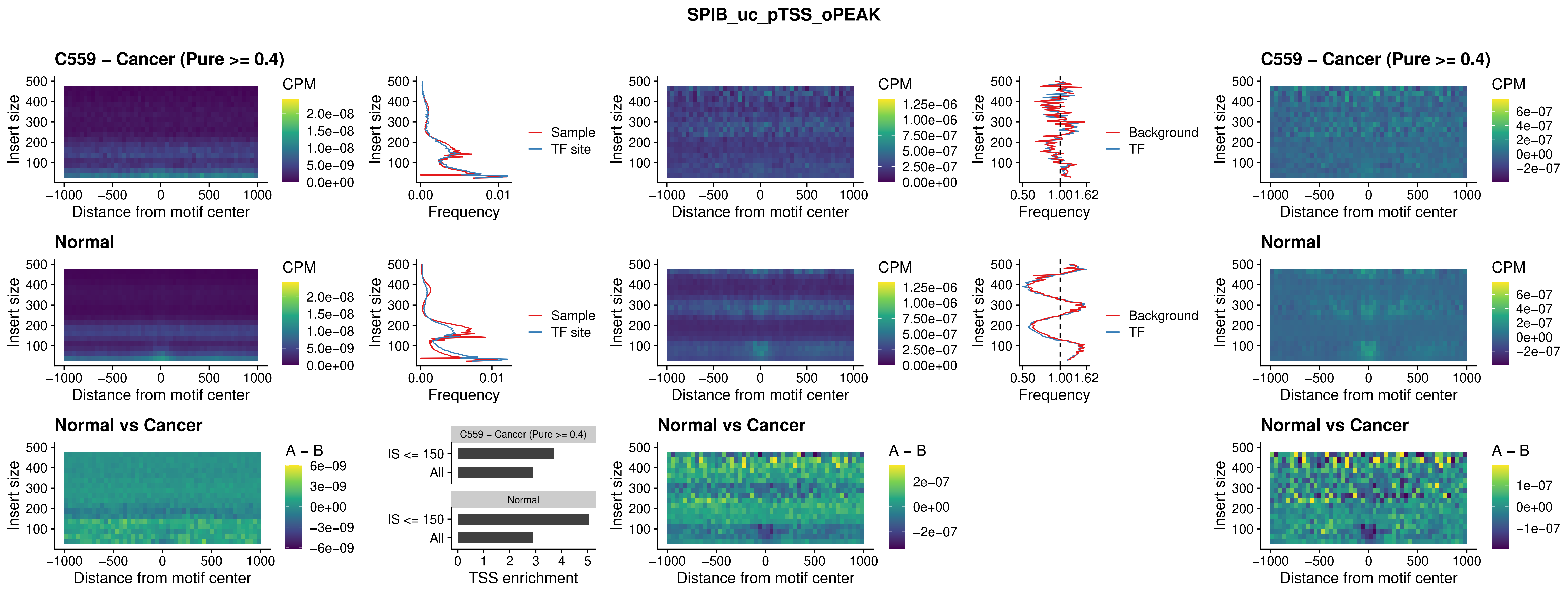
SPIB_uc_pTSS_oPEAK
C559 – Cancer (Pure >= 0.4)
Normal
Normal vs Cancer
Insert size
Distance from motif center
Frequency
CPM
A – B
Sample
TF site
Background
TF
C559 – Cancer (Pure >= 0.4)
Normal
IS <= 150
All
TSS enrichment

#### SPI1_uc_dTSS_oPEAK

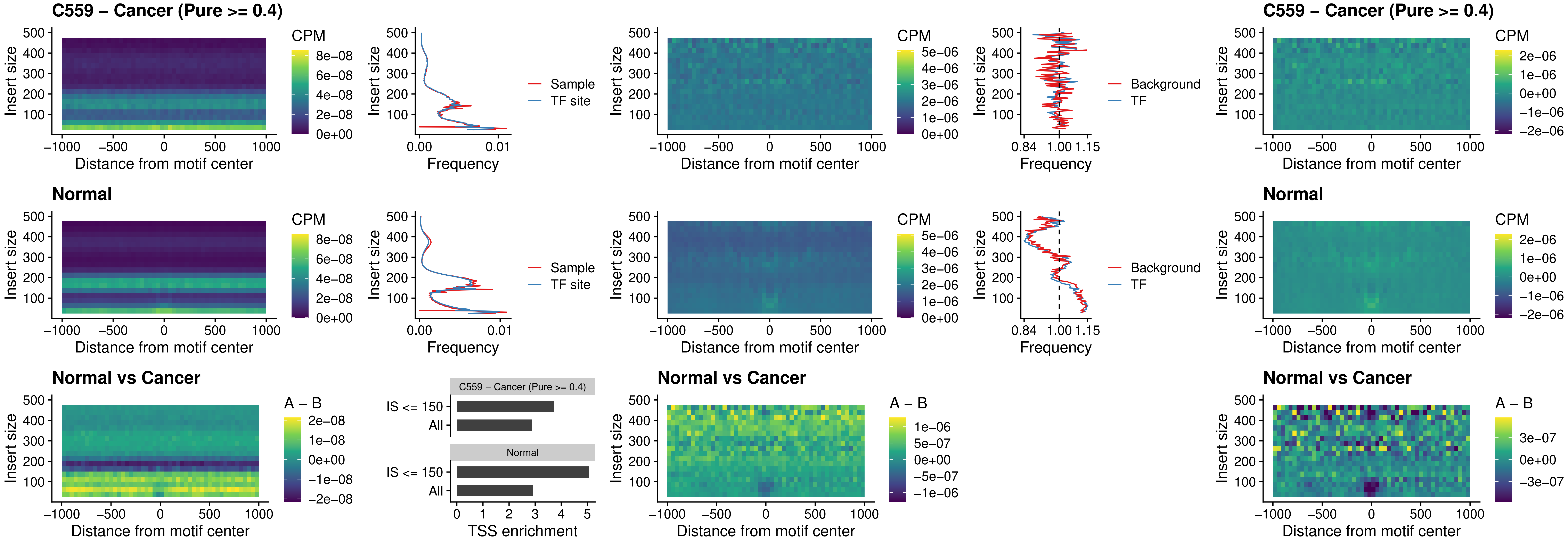

### SPIB_uc_dTSS_oPEAK

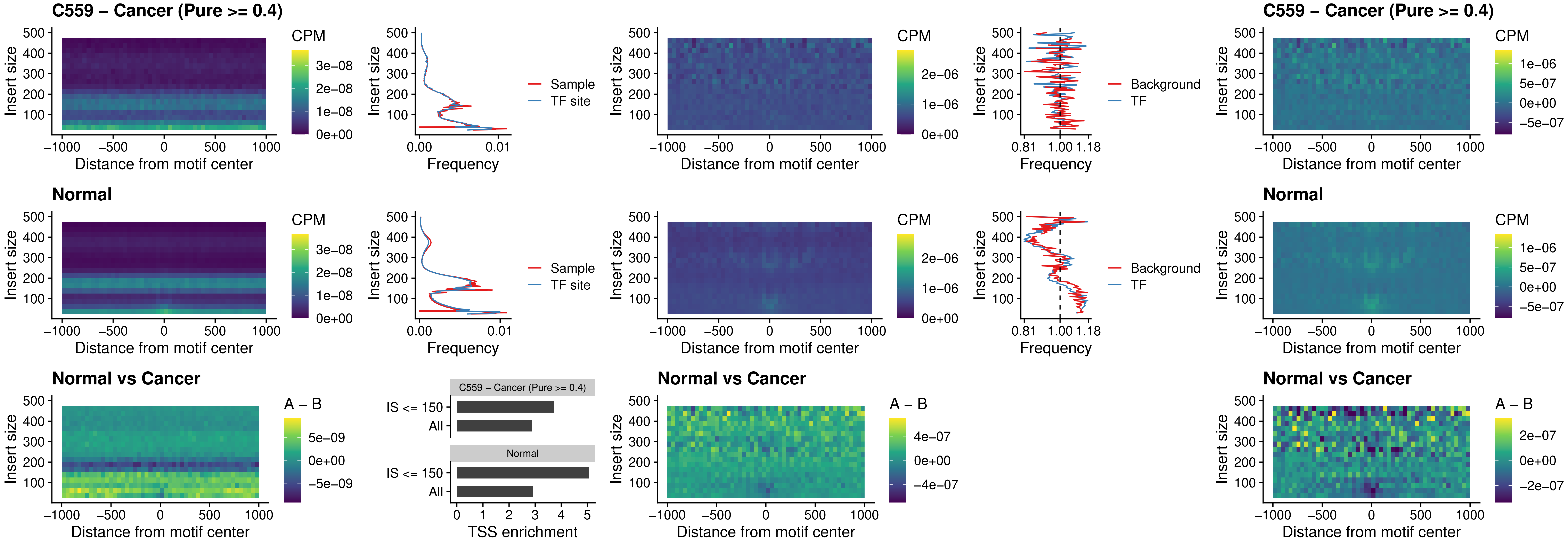

### IRF3_uc_pTSS_oPEAK

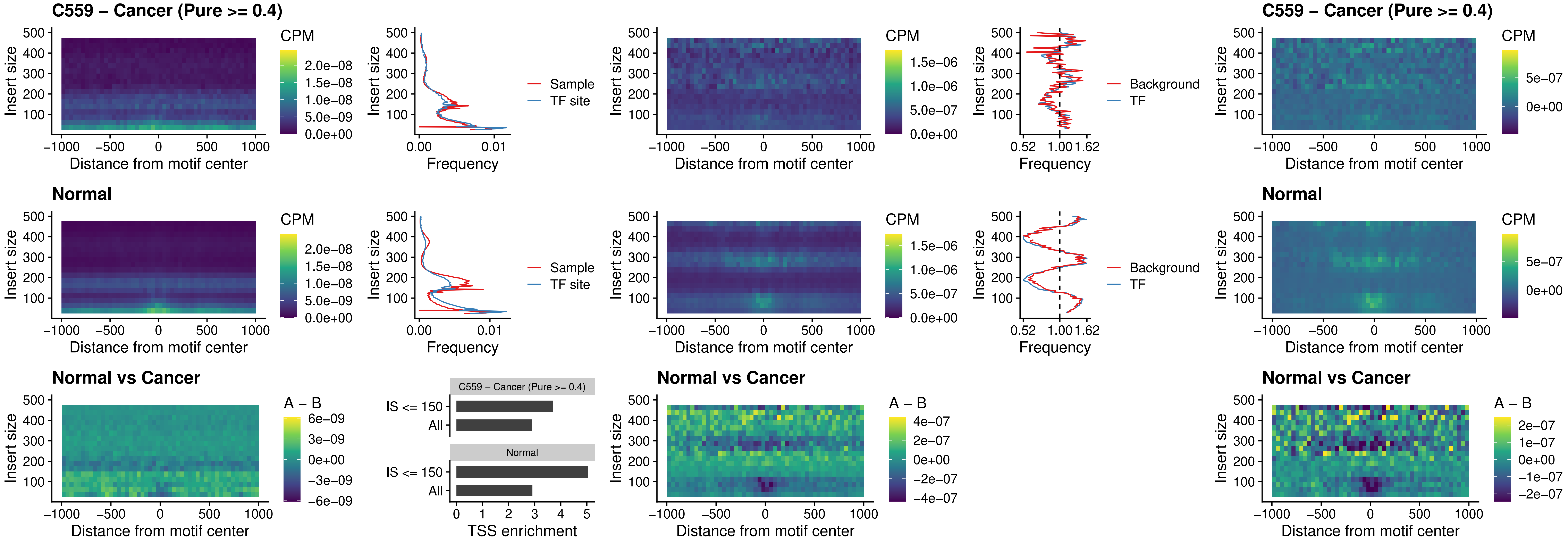

#### IRF2_uc_dTSS_oPEAK

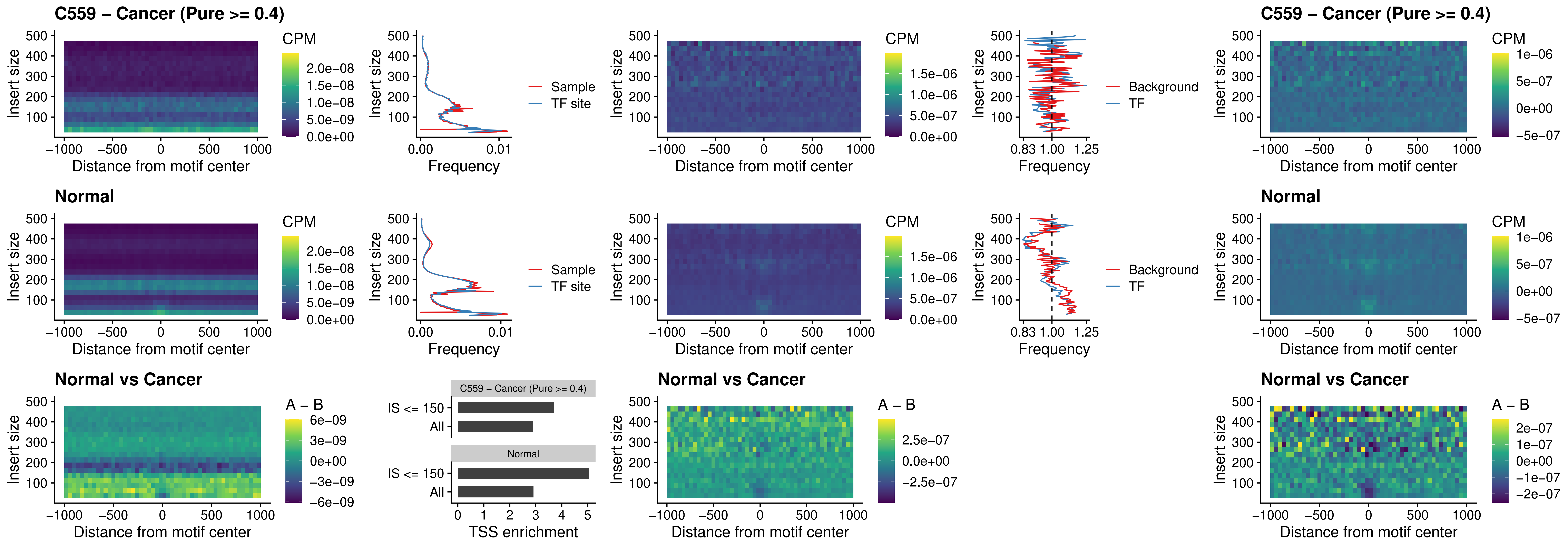

### IRF2_uc_pTSS_oPEAK

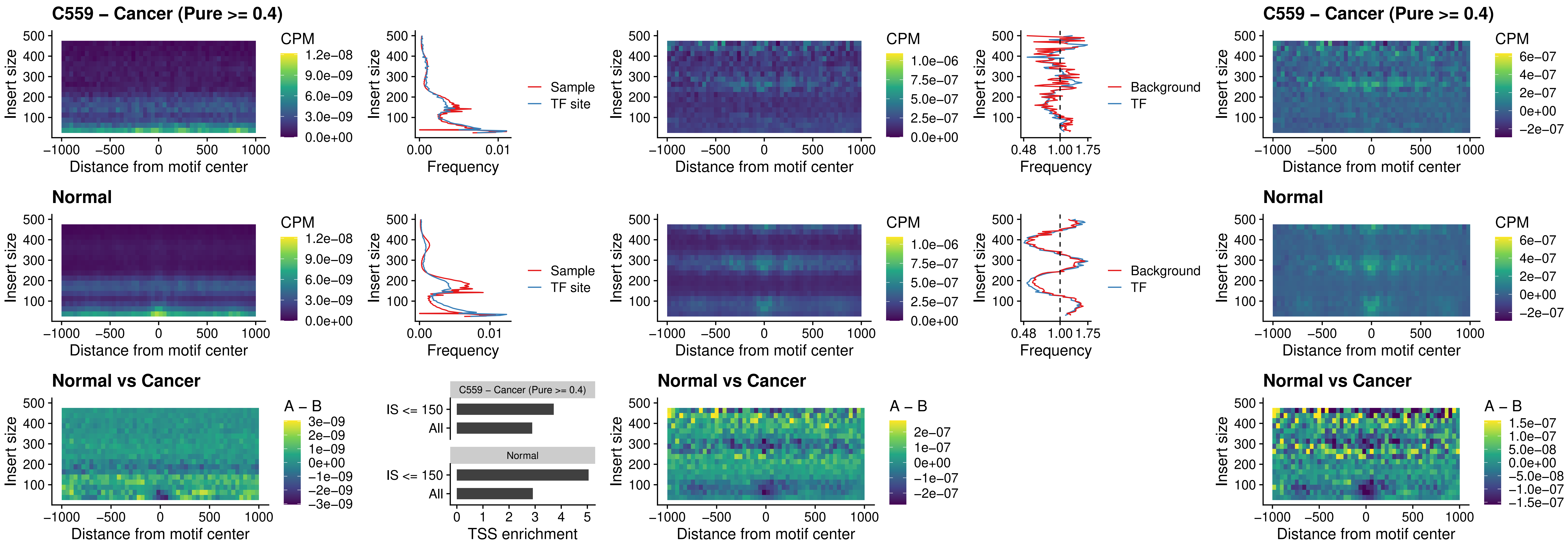

#### STAT2_uc_pTSS_oPEAK

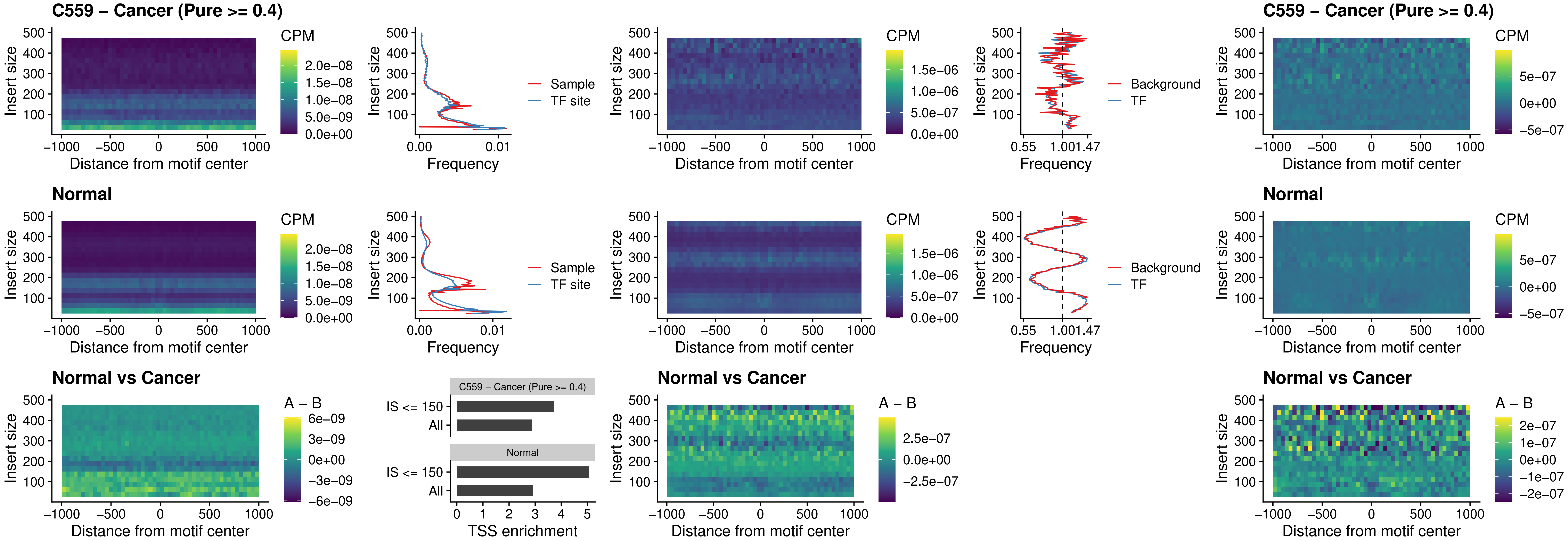

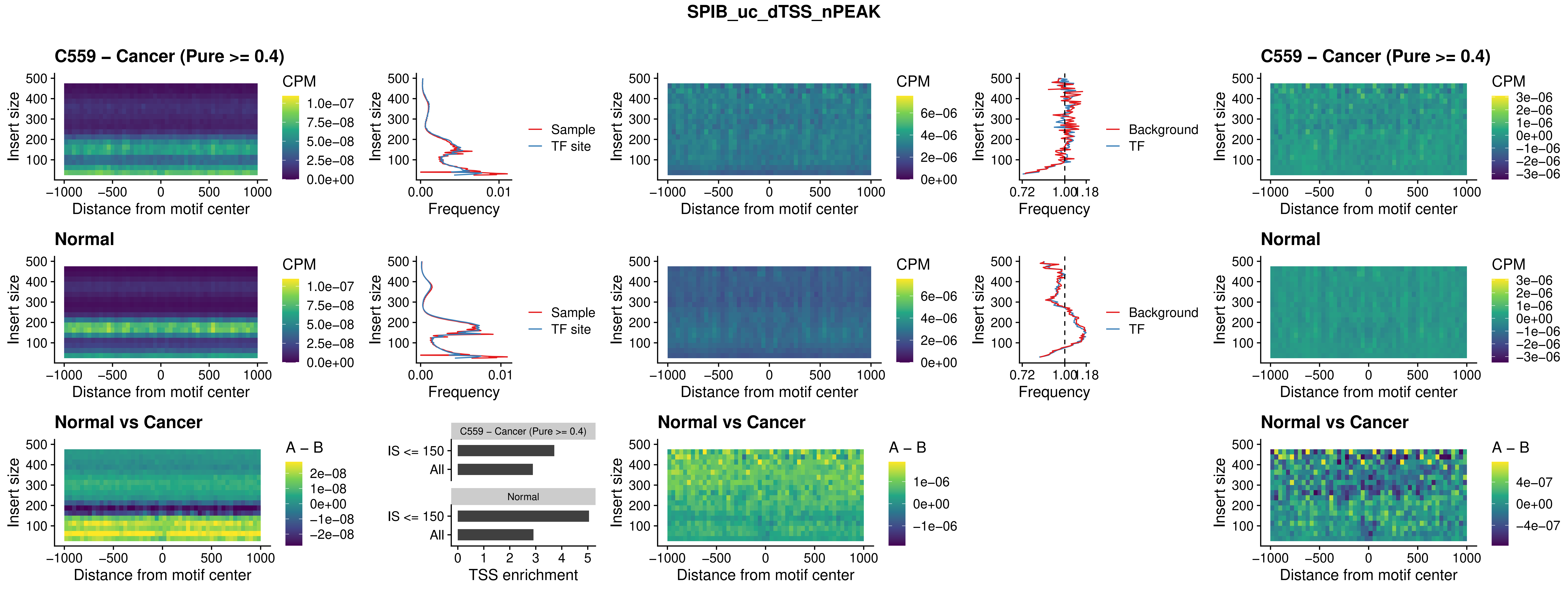
SPIB_uc_dTSS_nPEAK
C559 – Cancer (Pure >= 0.4)
Normal
Normal vs Cancer
Insert size
Distance from motif center
Frequency
CPM
A – B
Sample
TF site
Background
TF
C559 – Cancer (Pure >= 0.4)
Normal
IS <= 150
All
TSS enrichment

#### SPIC_uc_cTSS_oPEAK

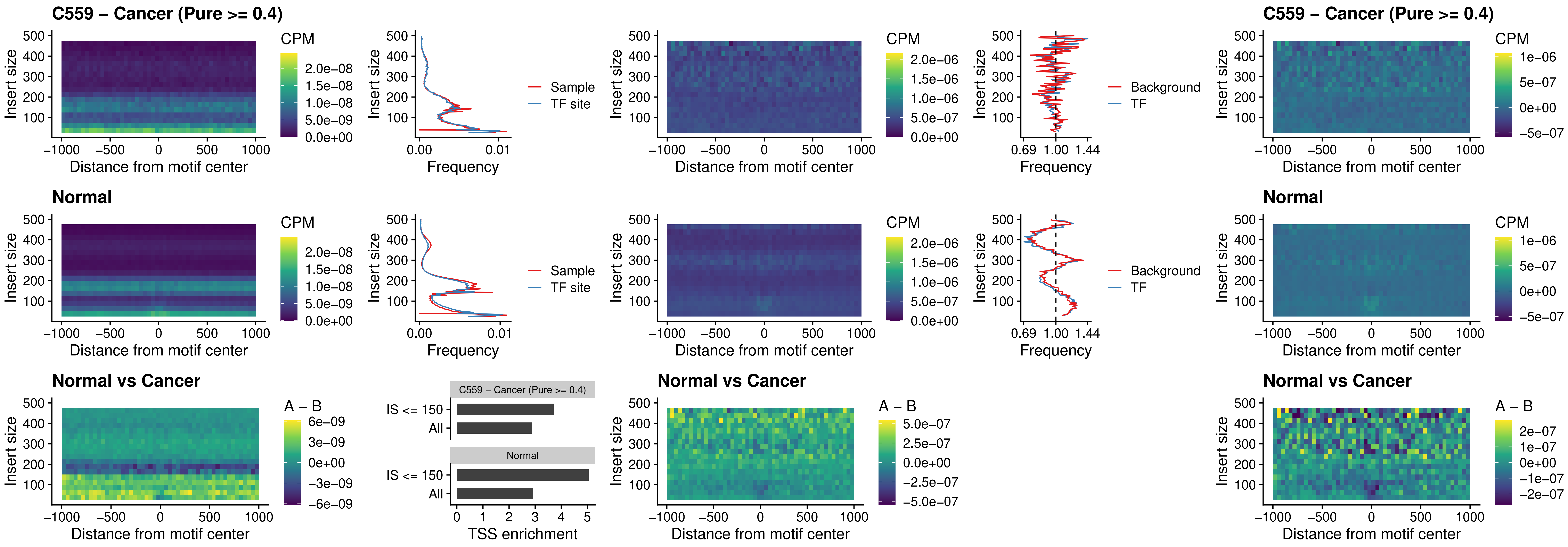

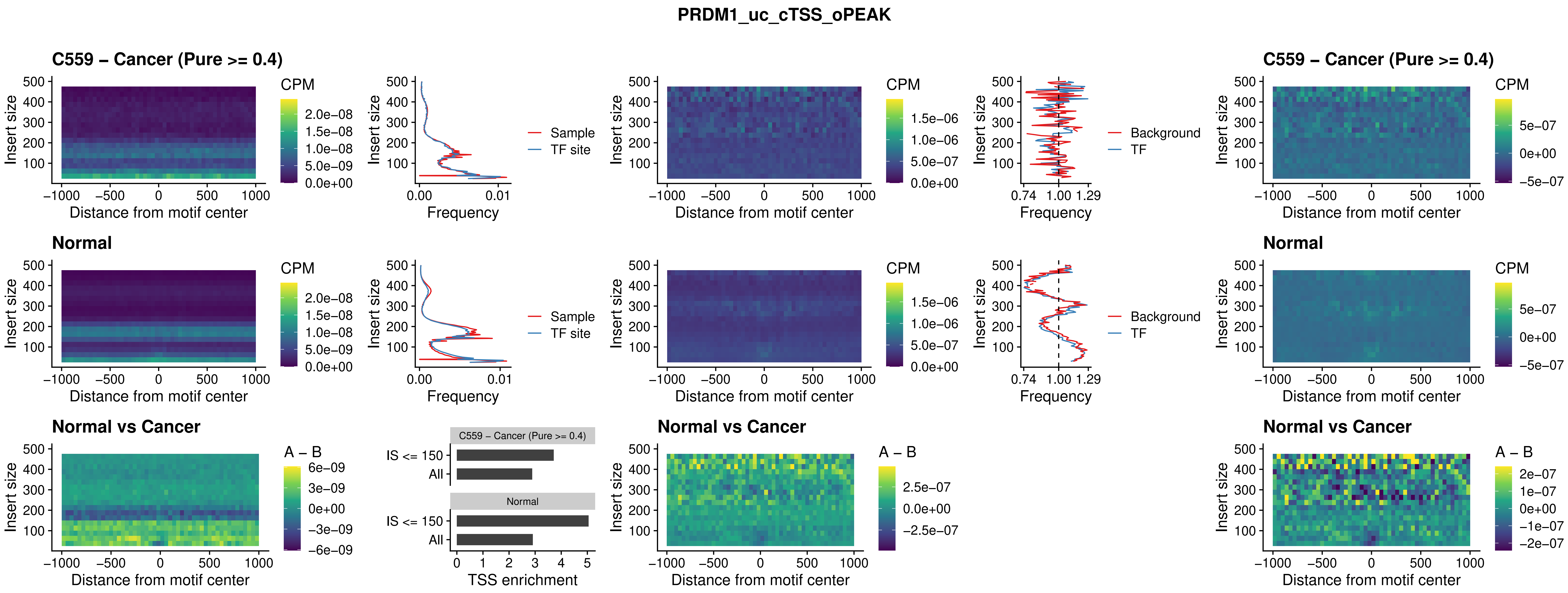
PRDM1_uc_cTSS_oPEAK
C559 – Cancer (Pure >= 0.4)
Normal
Normal vs Cancer
Insert size
Distance from motif center
Frequency
CPM
A – B
Sample
TF site
Background
TF
TSS enrichment
IS <= 150
All
C559 – Cancer (Pure >= 0.4)
Normal

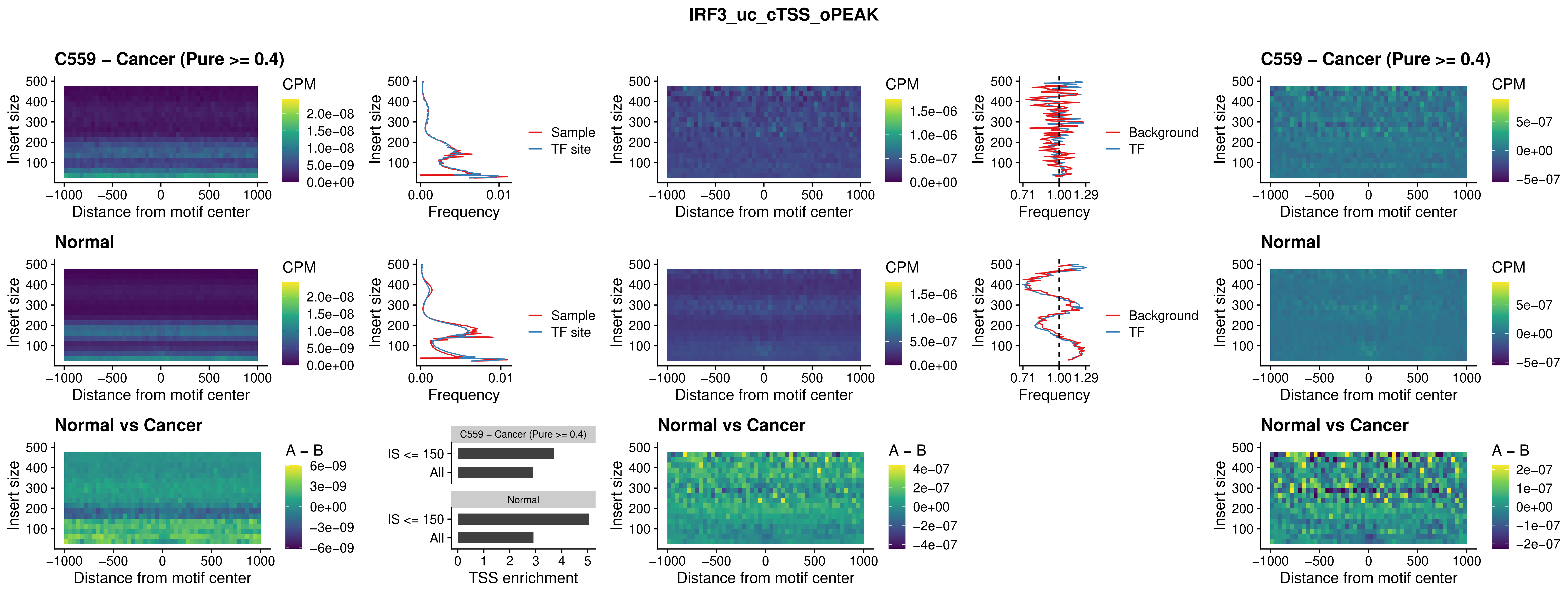

IRF3_uc_cTSS_oPEAK
C559 – Cancer (Pure >= 0.4)
Normal
Normal vs Cancer
CPM
A – B
Insert size
Distance from motif center
Frequency
Sample
TF site
Background
TF
C559 – Cancer (Pure >= 0.4)
Normal
IS <= 150
All
TSS enrichment

### BCL11A_uc_dTSS_nPEAK

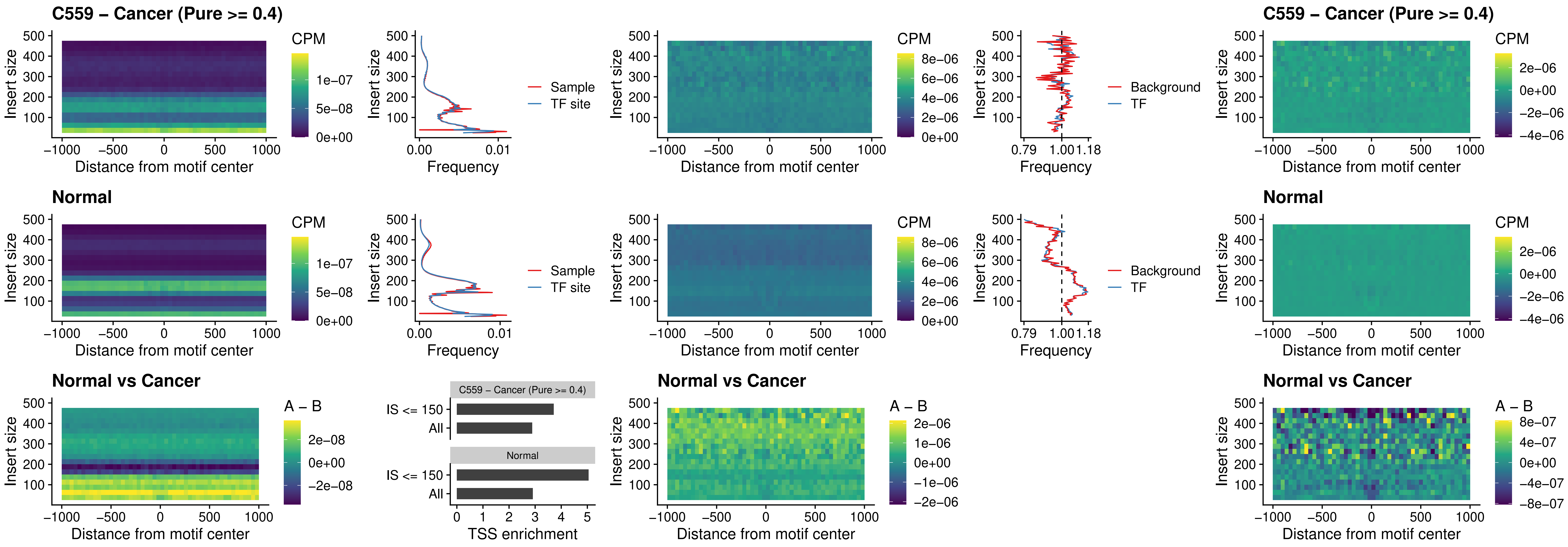

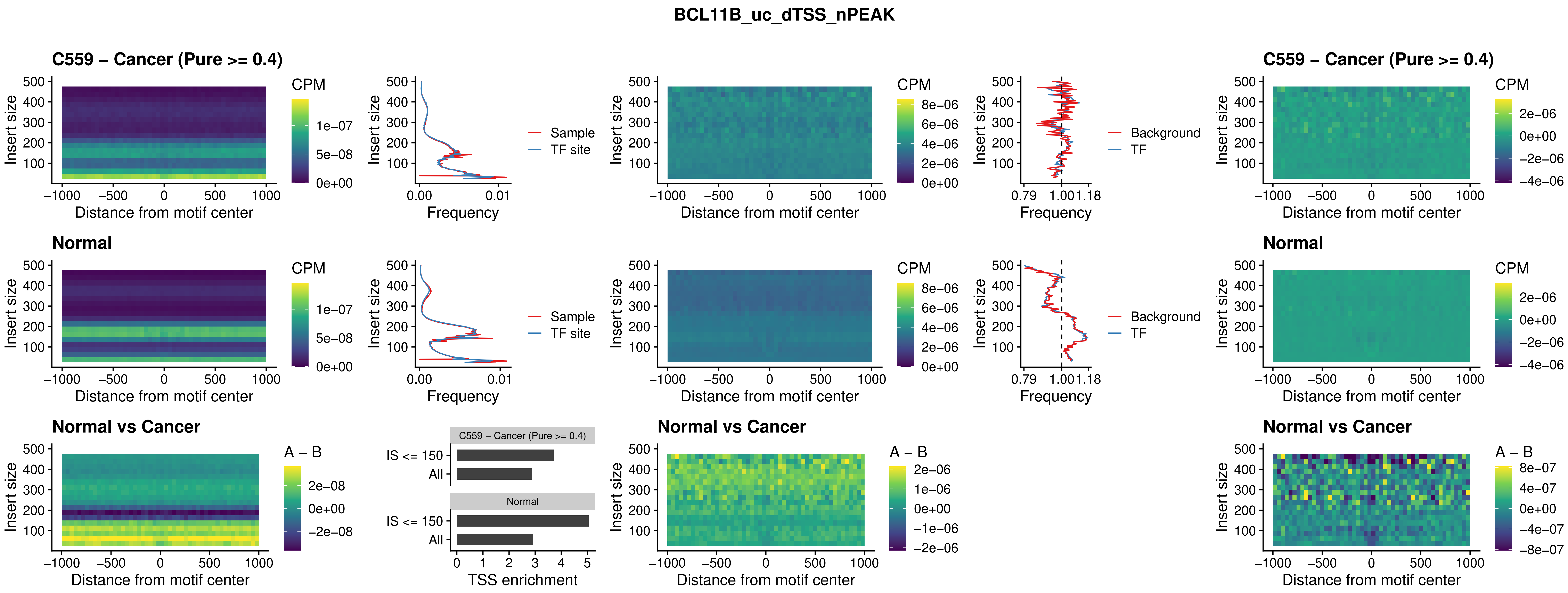

BCL11B_uc_dTSS_nPEAK
C559 – Cancer (Pure >= 0.4)
Normal
Normal vs Cancer
Insert size
Distance from motif center
Frequency
CPM
A – B
Sample
TF site
Background
TF
C559 – Cancer (Pure >= 0.4)
Normal
IS <= 150
All
TSS enrichment

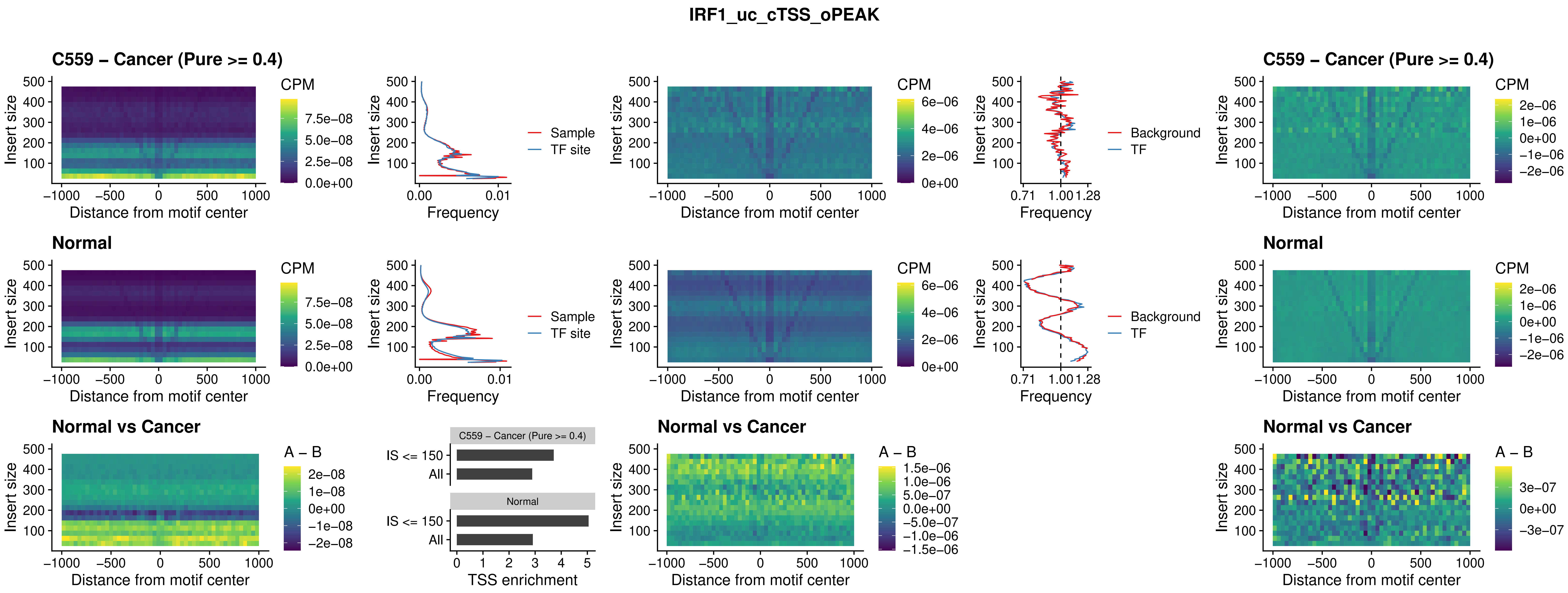
IRF1_uc_cTSS_oPEAK
C559 – Cancer (Pure >= 0.4)
Normal
Normal vs Cancer
Insert size
Distance from motif center
Frequency
CPM
A – B
Sample
TF site
Background
TF
C559 – Cancer (Pure >= 0.4)
Normal
IS <= 150
All
TSS enrichment

### IRF1_uc_dTSS_oPEAK

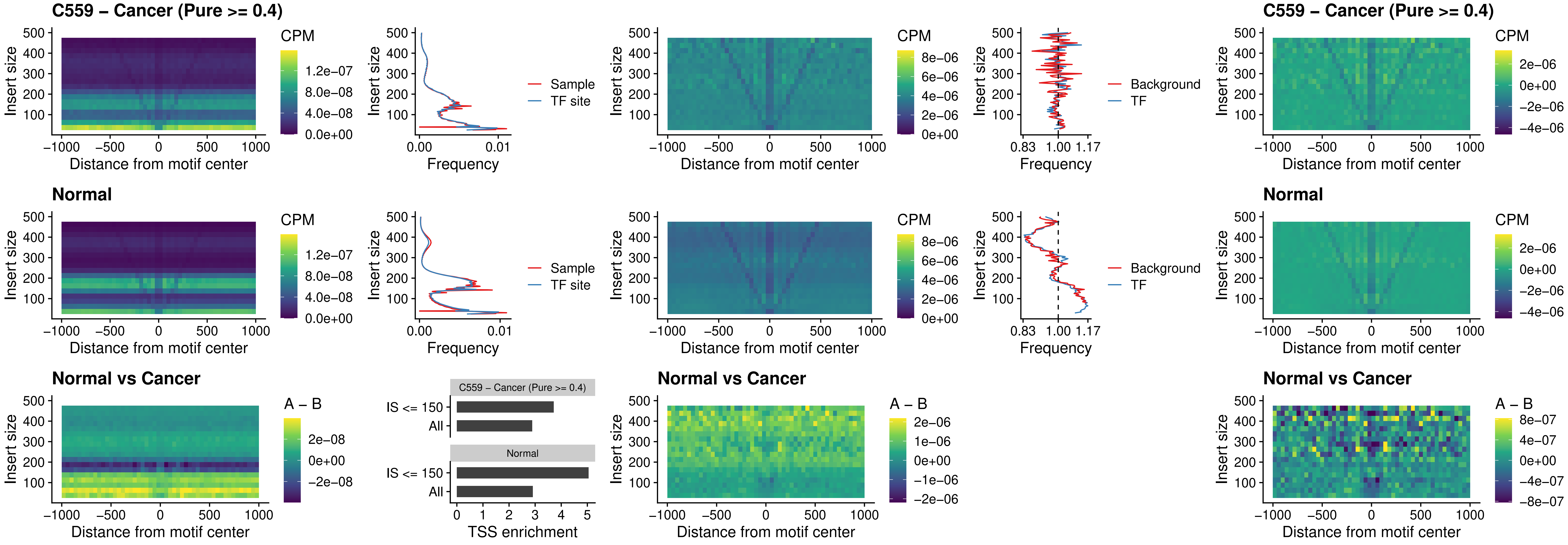

#### IRF1_uc_pTSS_oPEAK

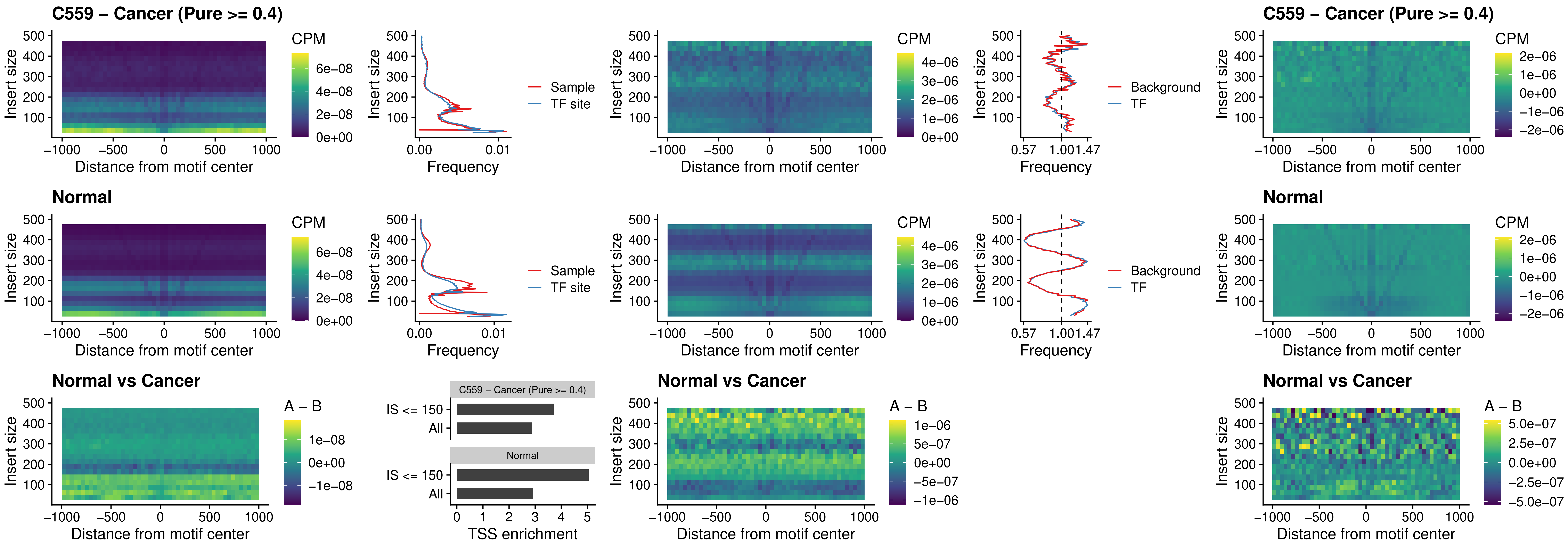

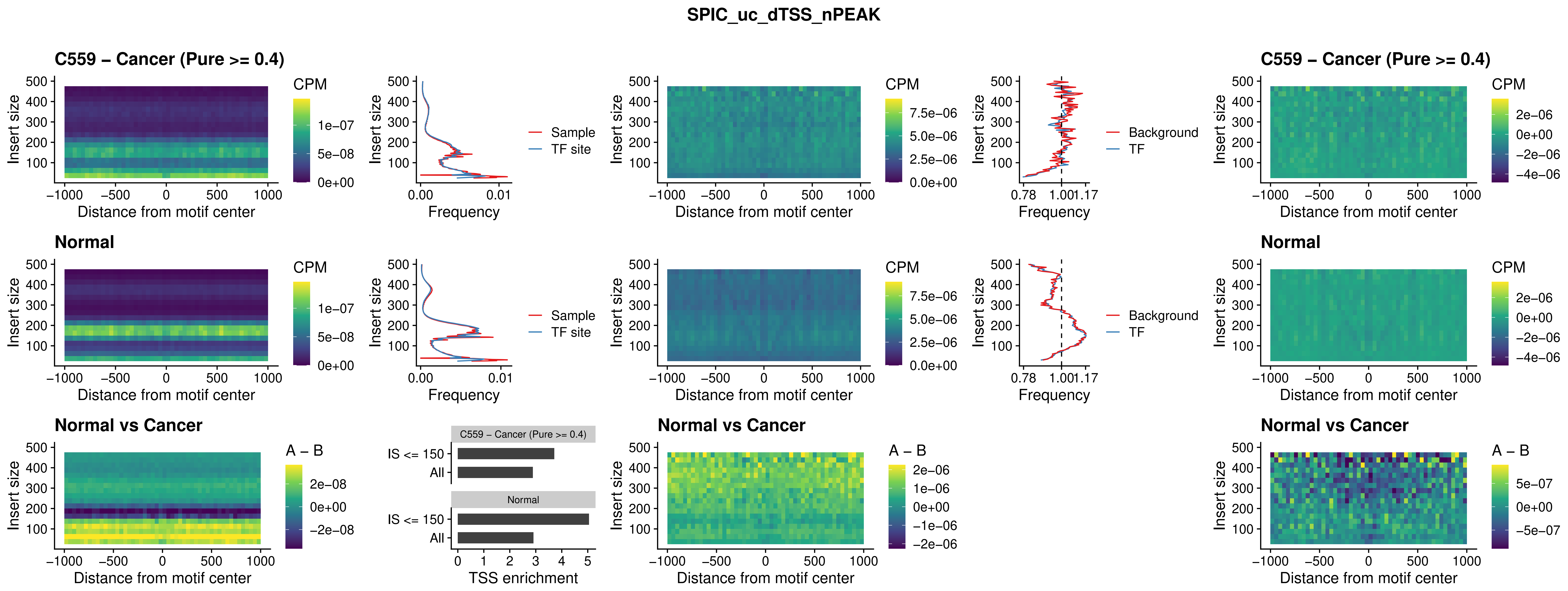
SPIC_uc_dTSS_nPEAK
C559 – Cancer (Pure >= 0.4)
Normal
Normal vs Cancer
Insert size
Distance from motif center
CPM
A – B
Frequency
Sample
TF site
Background
TF
TSS enrichment
C559 – Cancer (Pure >= 0.4)
Normal
IS <= 150
All

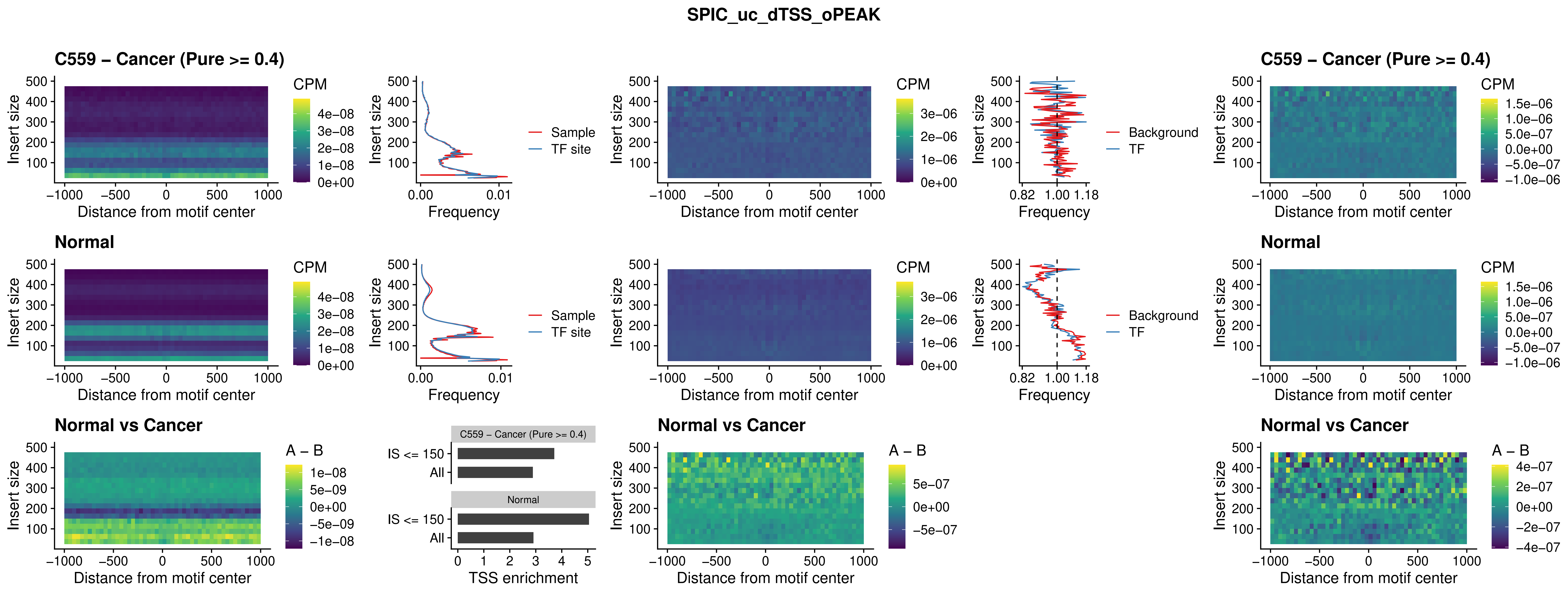
SPIC_uc_dTSS_oPEAK
C559 – Cancer (Pure >= 0.4)
Normal
Normal vs Cancer
Insert size
Distance from motif center
Frequency
CPM
A – B
Sample
TF site
Background
TF
C559 – Cancer (Pure >= 0.4)
Normal
IS <= 150
All
TSS enrichment

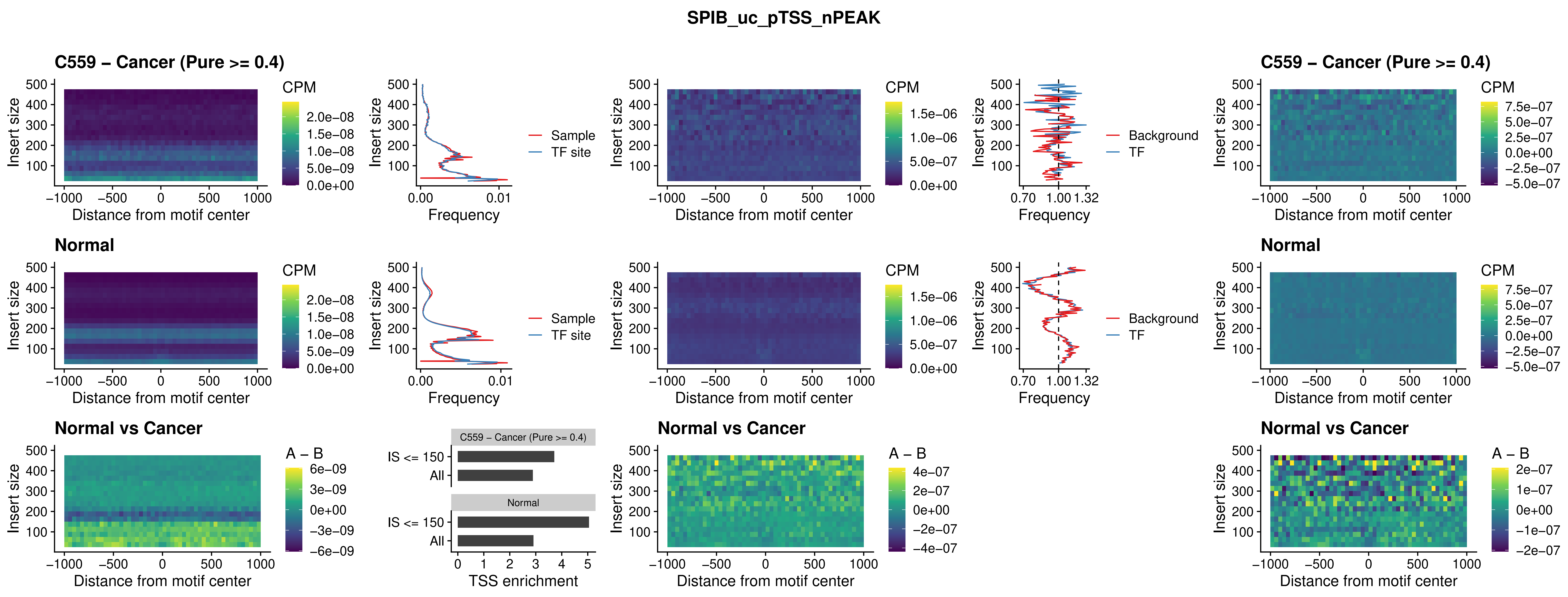
SPIB_uc_pTSS_nPEAK
C559 – Cancer (Pure >= 0.4)
Normal
Normal vs Cancer
Insert size
Distance from motif center
CPM
A – B
Frequency
Sample
TF site
Background
TF
TSS enrichment
IS <= 150
All
C559 – Cancer (Pure >= 0.4)
Normal

#### STAT2_uc_cTSS_oPEAK

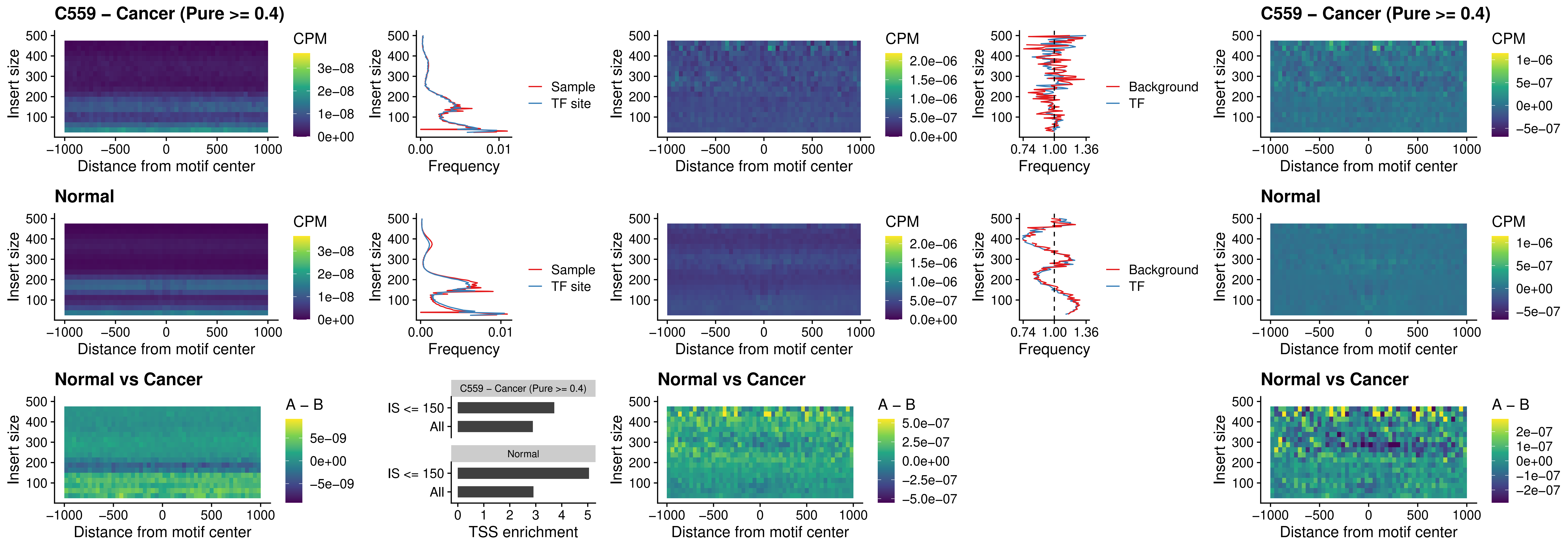

### IRF8_uc_cTSS_oPEAK

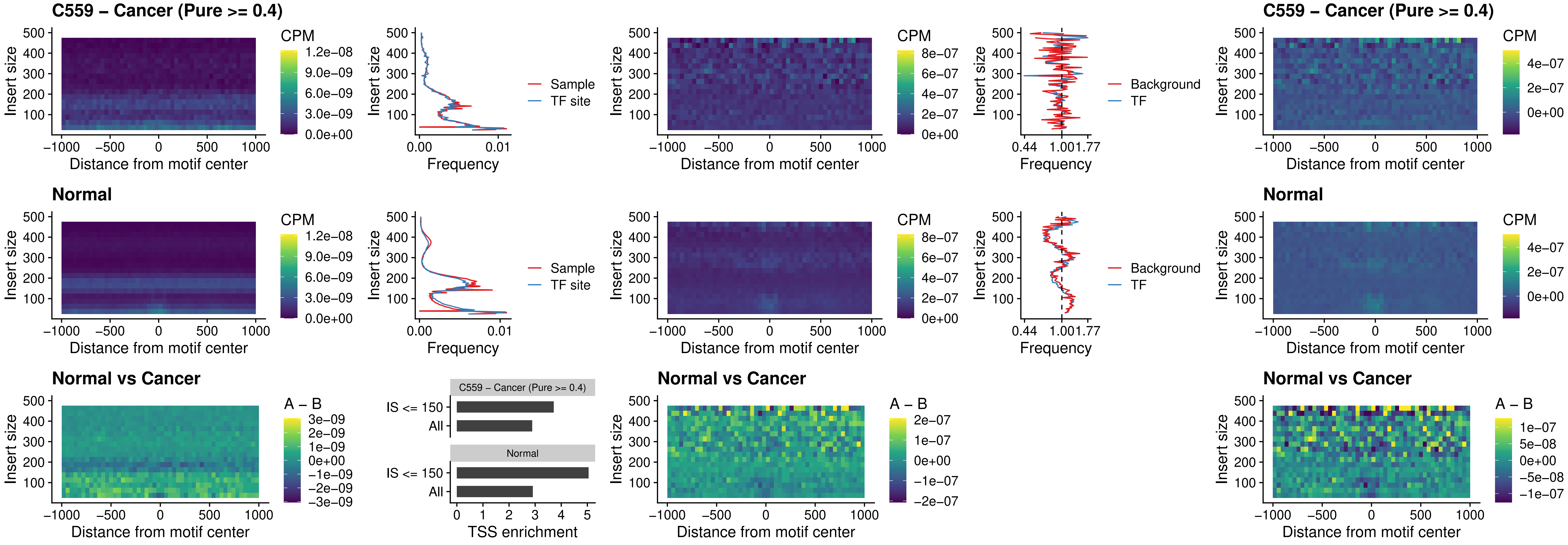

### IRF3_uc_dTSS_oPEAK

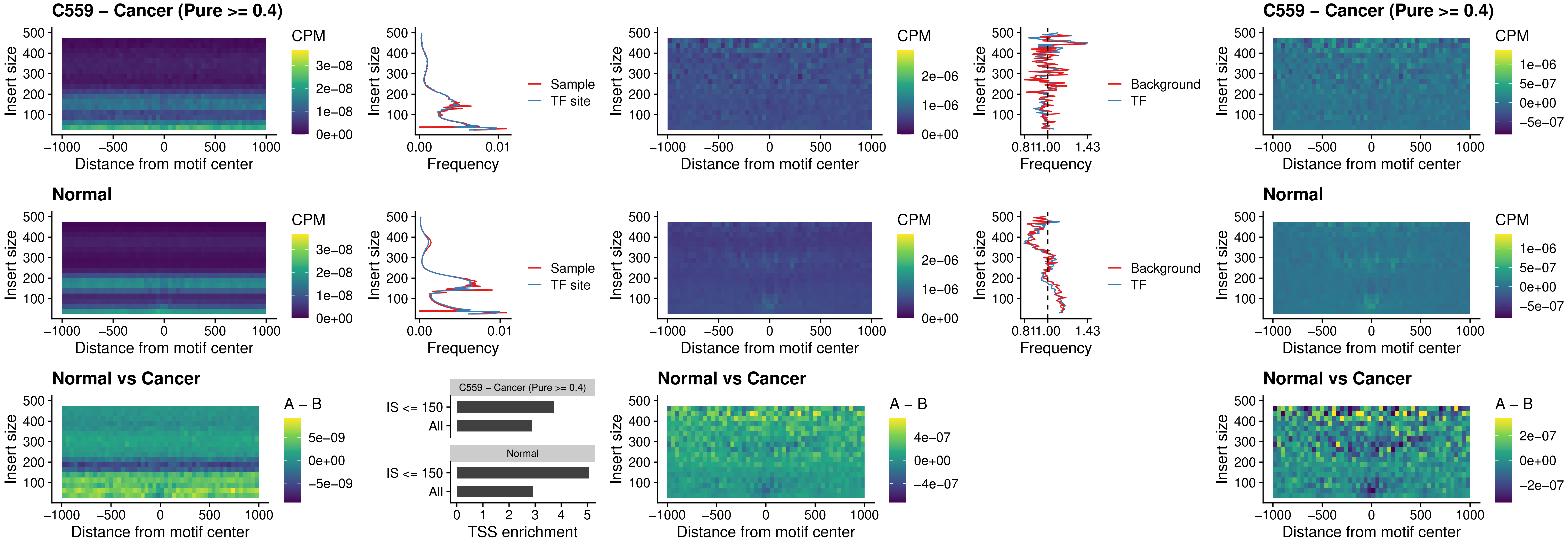

#### IRF8_uc_dTSS_oPEAK

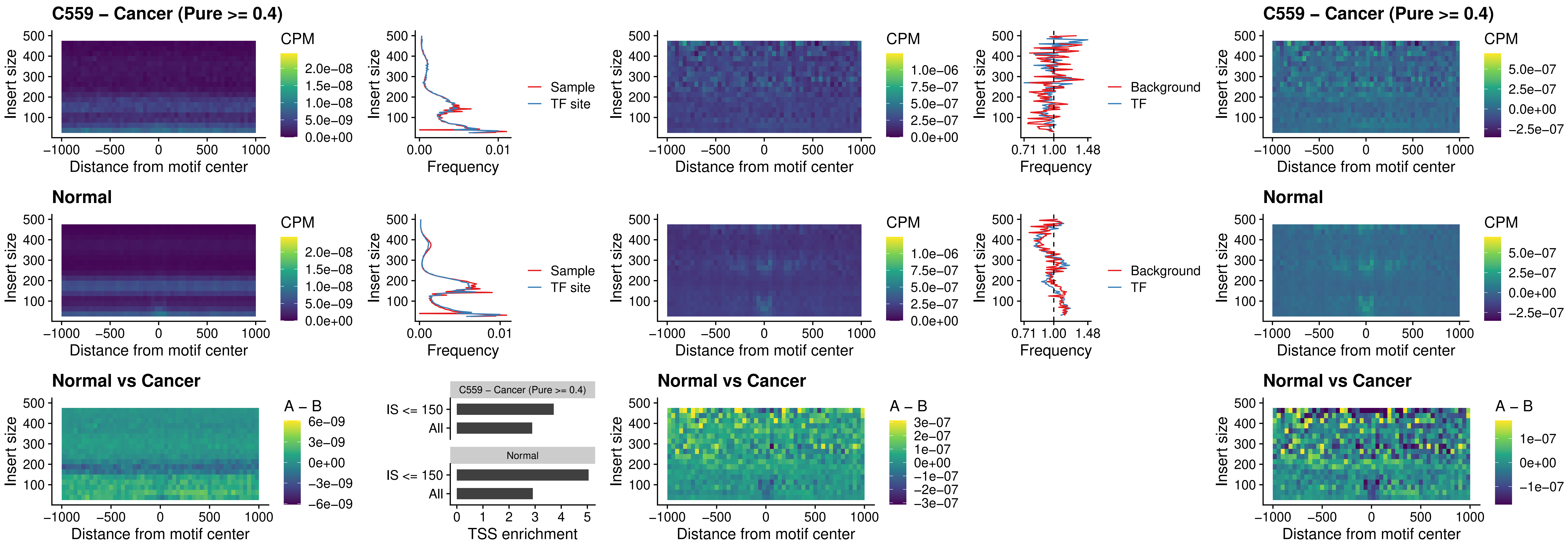

### ETS2_uc_dTSS_oPEAK

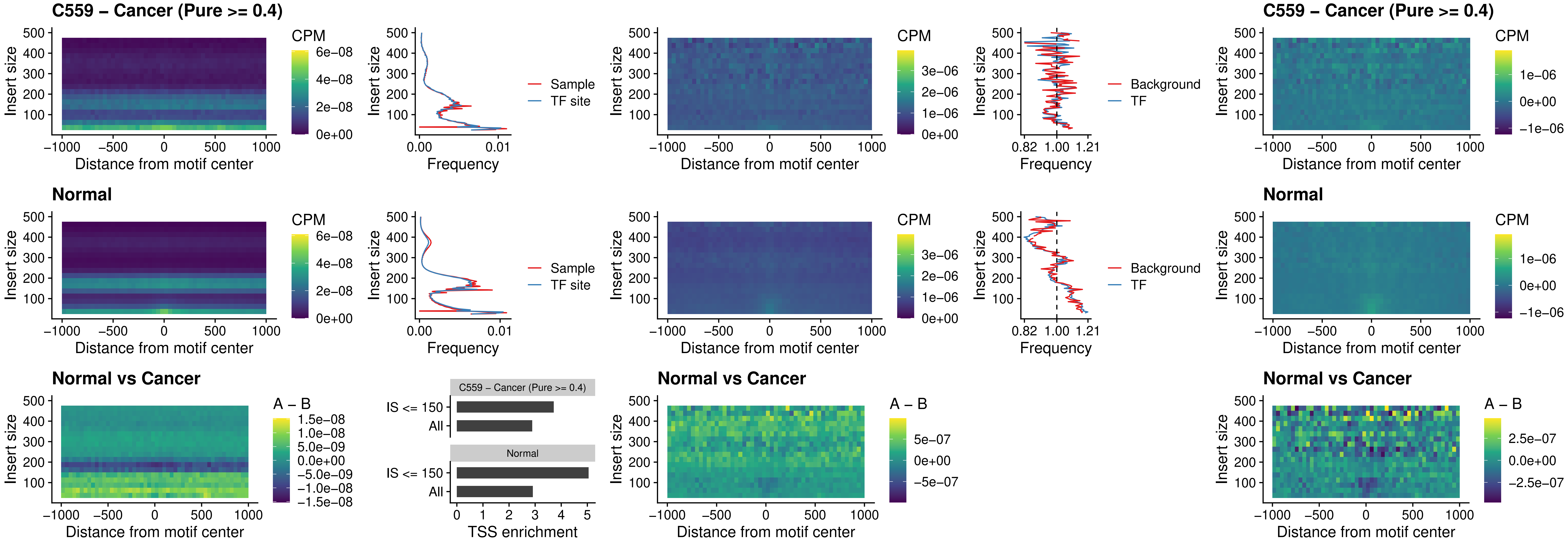

### SMARCC2_uc_pTSS_oPEAK

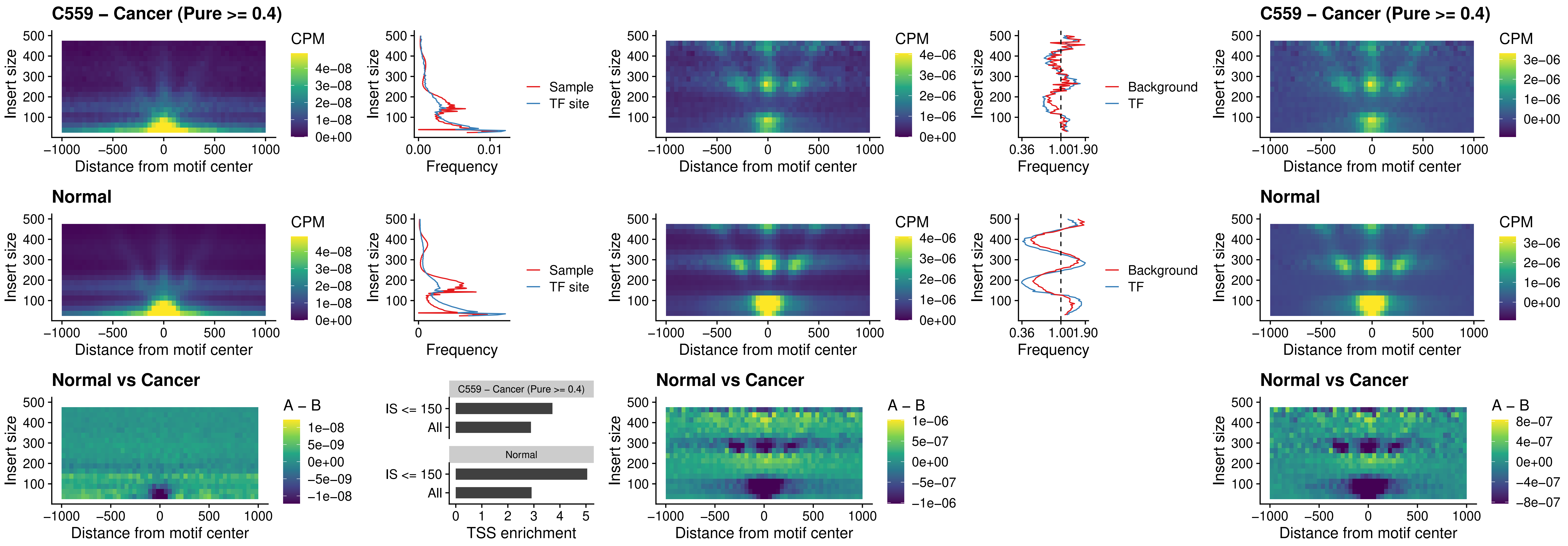

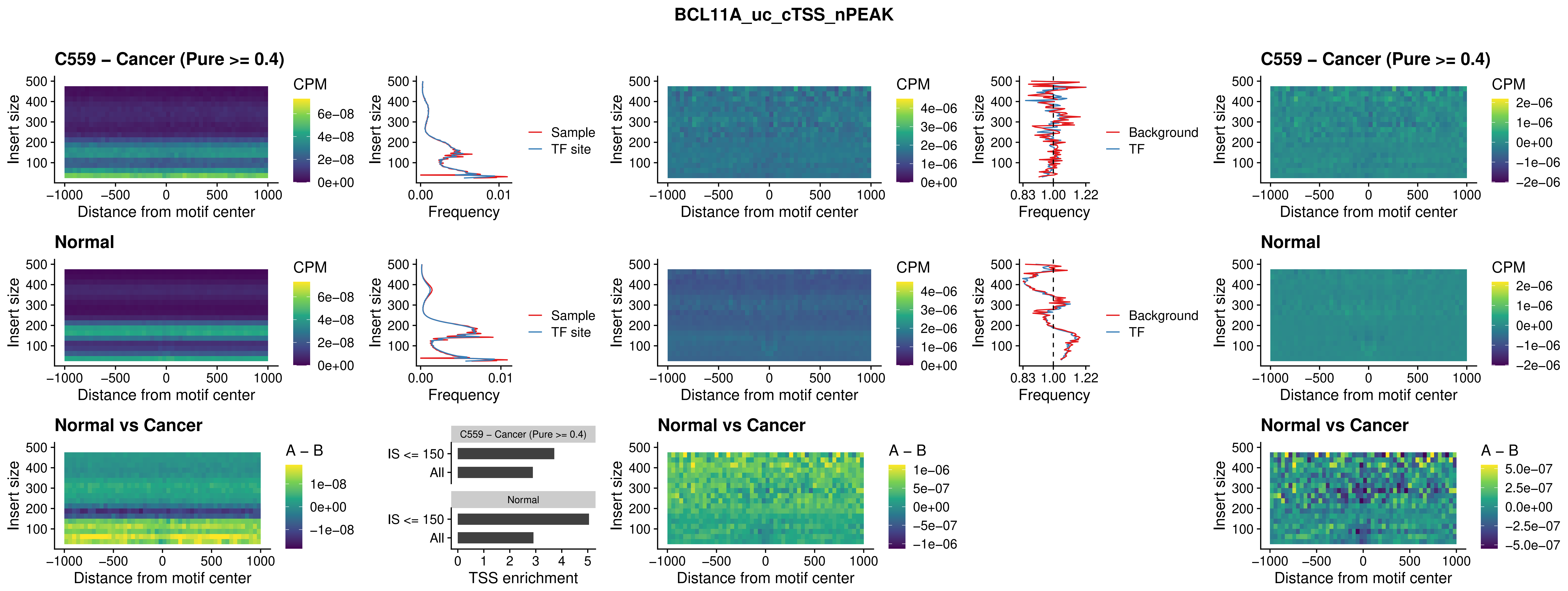
BCL11A_uc_cTSS_nPEAK
C559 – Cancer (Pure >= 0.4)
Normal
Normal vs Cancer
Insert size
Distance from motif center
Frequency
CPM
A – B
Sample
TF site
Background
TF
C559 – Cancer (Pure >= 0.4)
Normal
IS <= 150
All
TSS enrichment

#### BCL11B_uc_cTSS_nPEAK

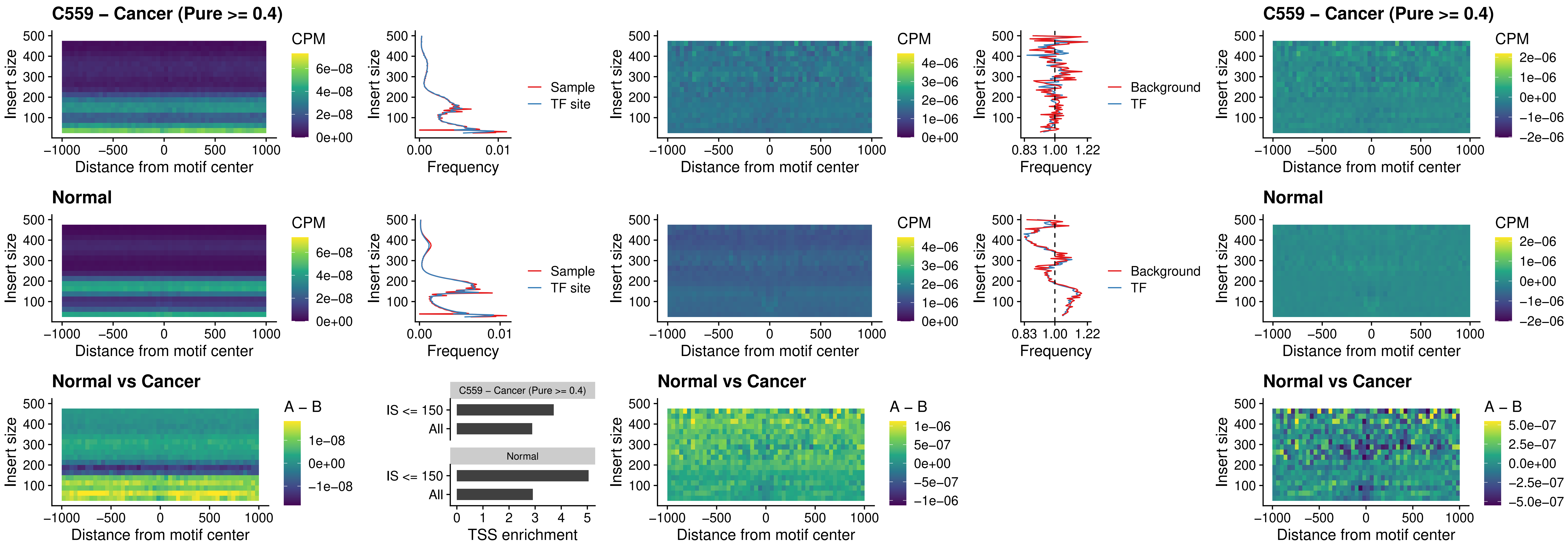

#### STAT2_uc_dTSS_oPEAK

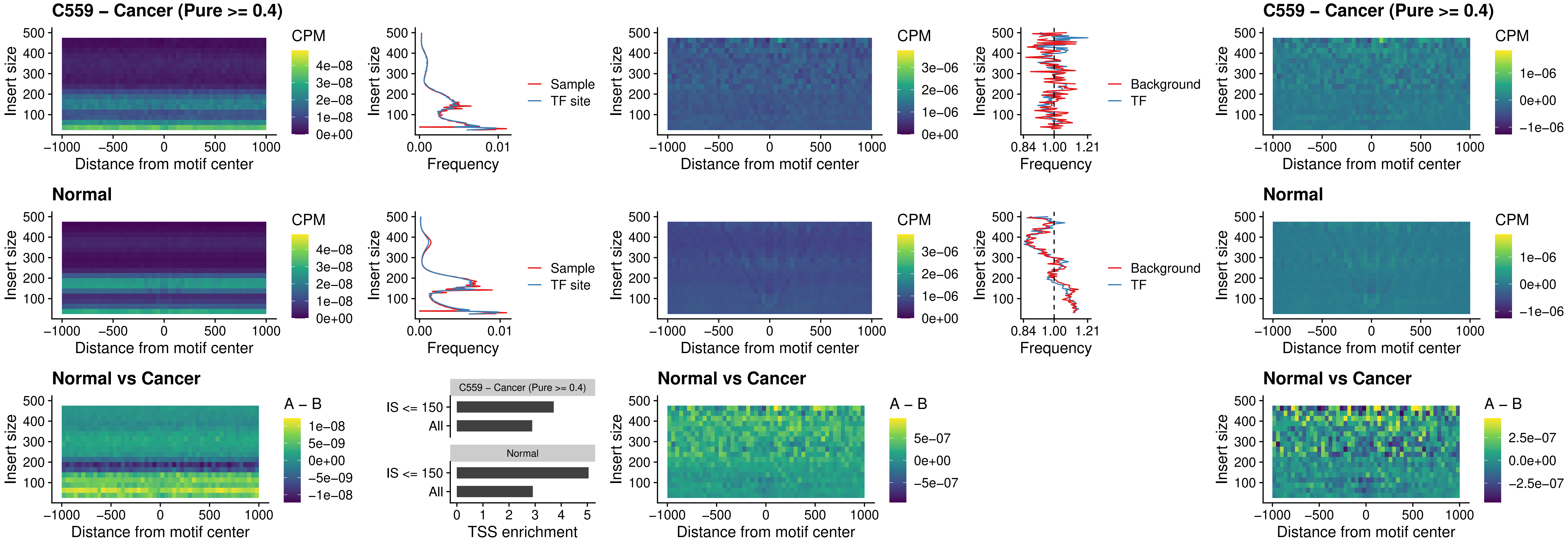

### ZNF683_uc_cTSS_oPEAK

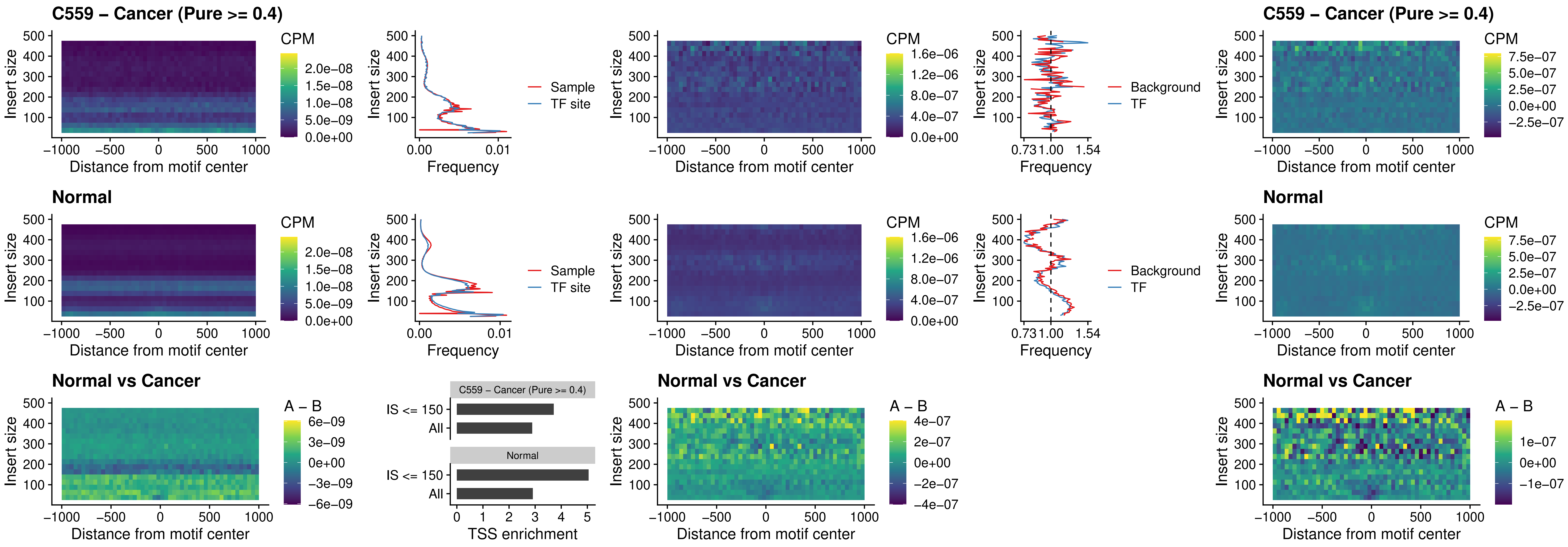

#### IRF8_uc_dTSS_nPEAK

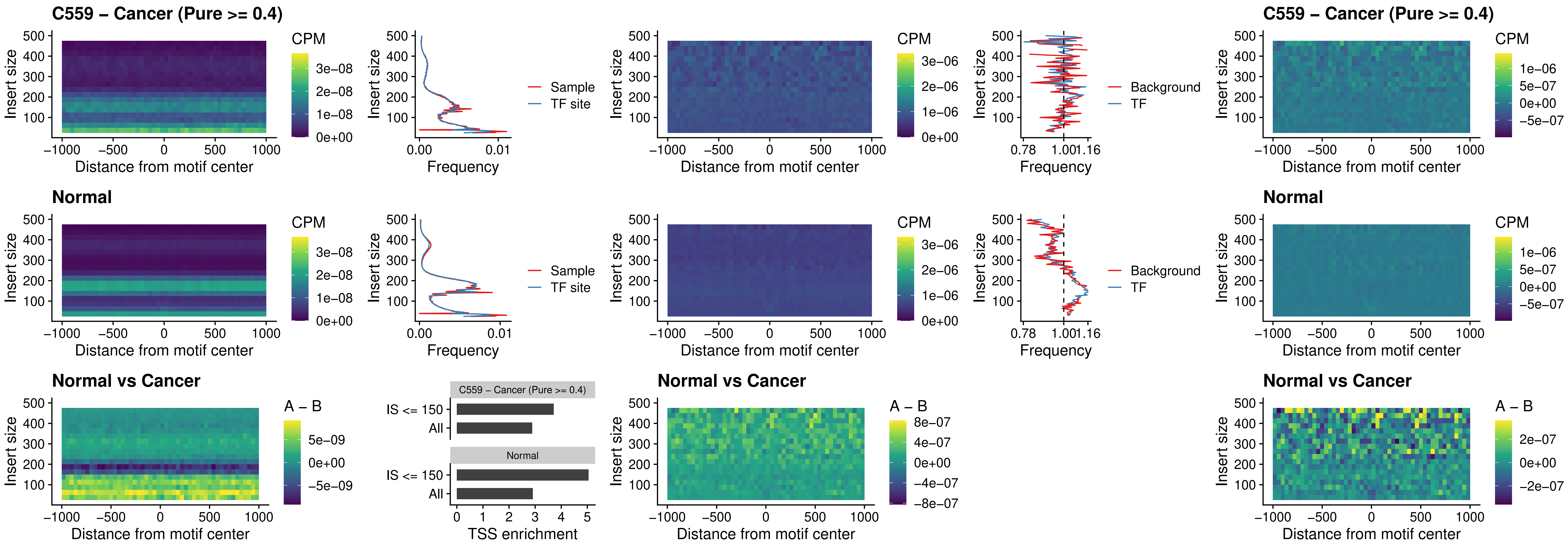

#### ETV7_uc_dTSS_oPEAK

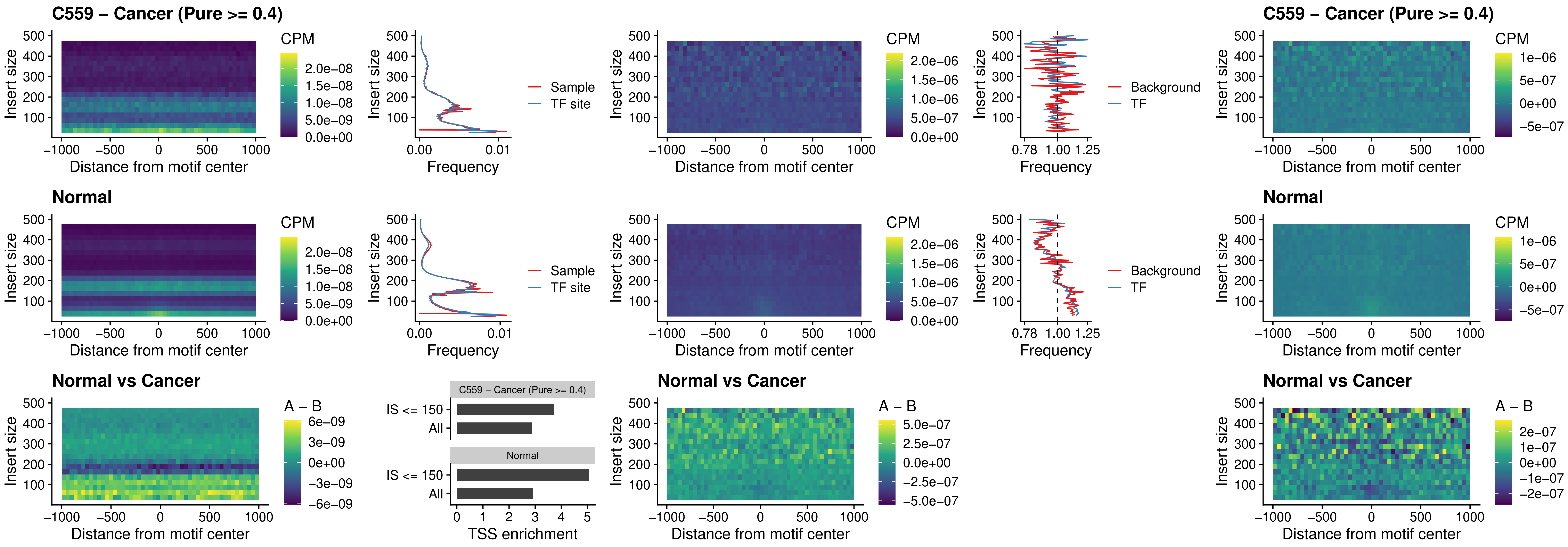

### FOXL1_uc_pTSS_oPEAK

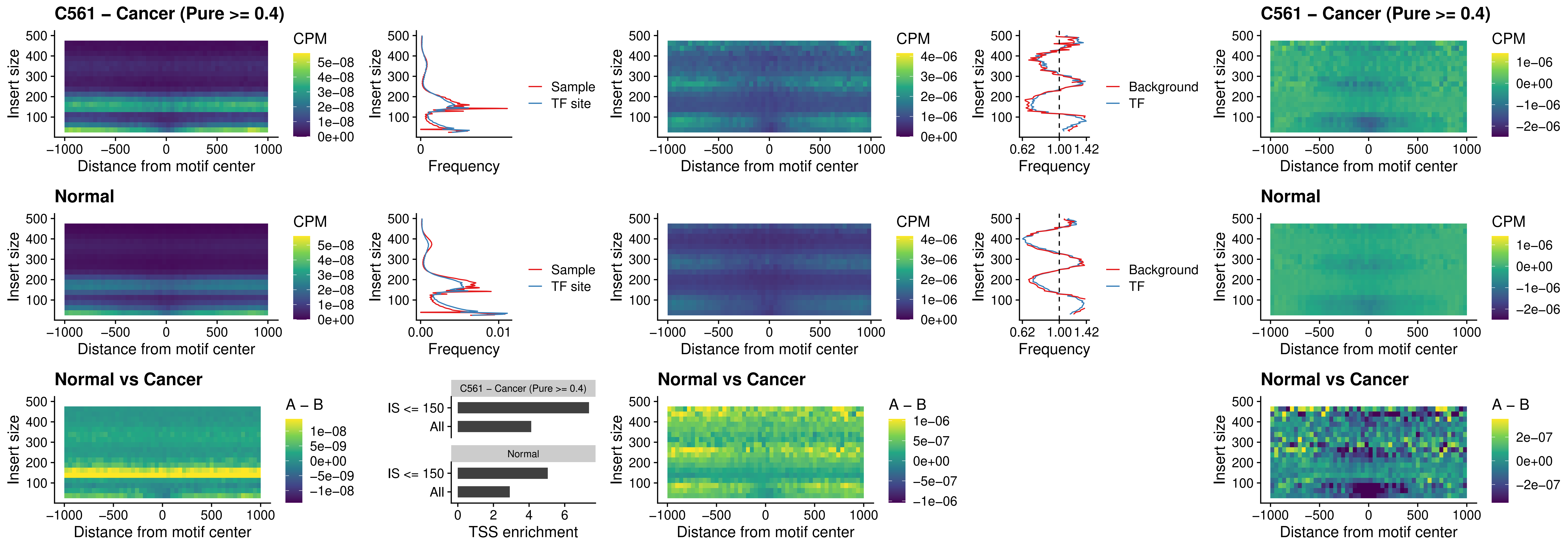

#### CPEB1_uc_cTSS_oPEAK

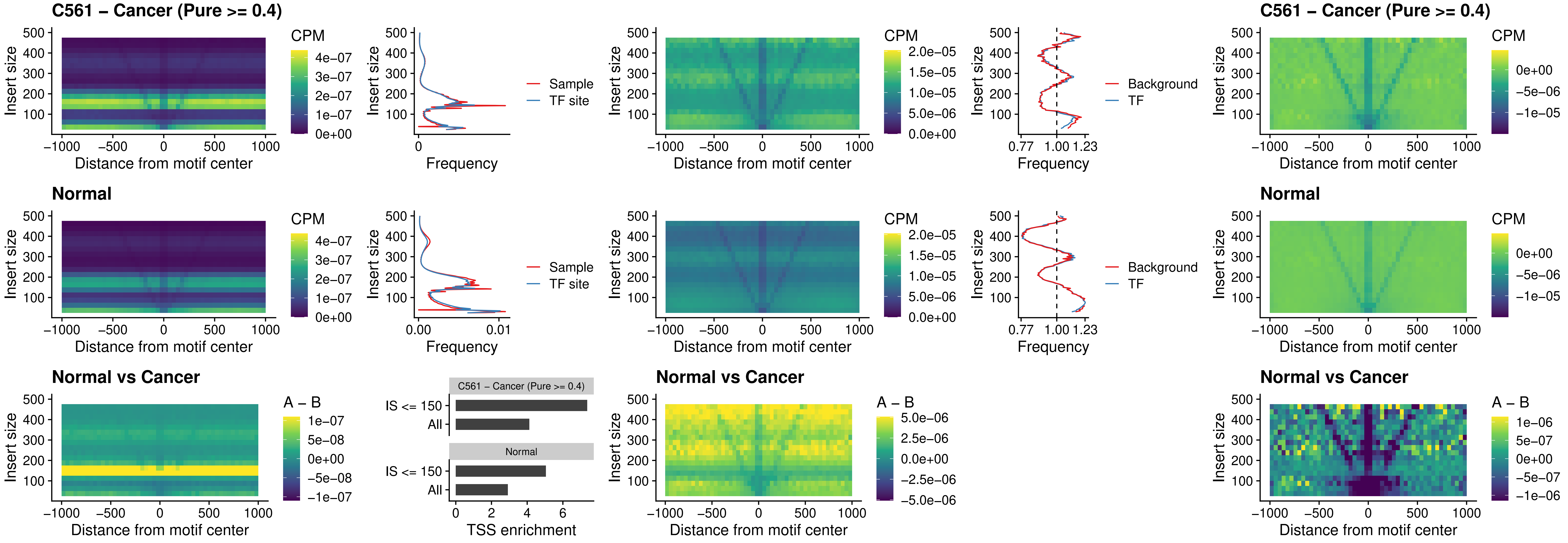

### HMGA1_uc_pTSS_nPEAK

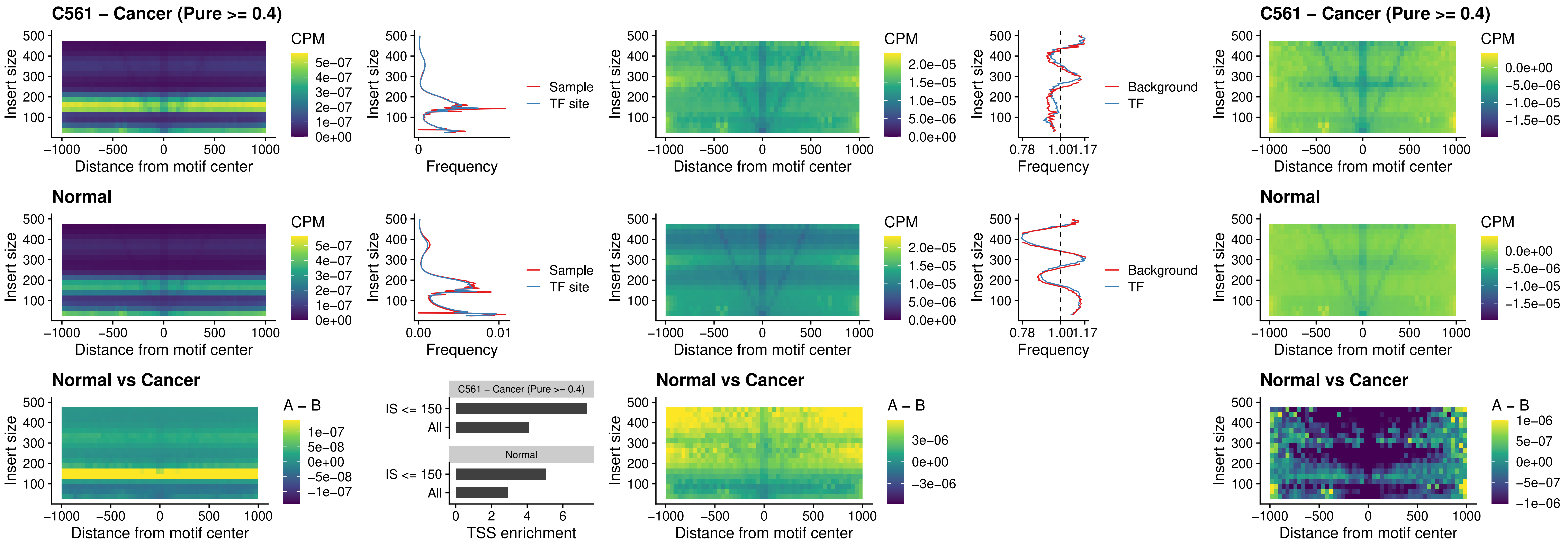

### SOX13_uc_cTSS_nPEAK

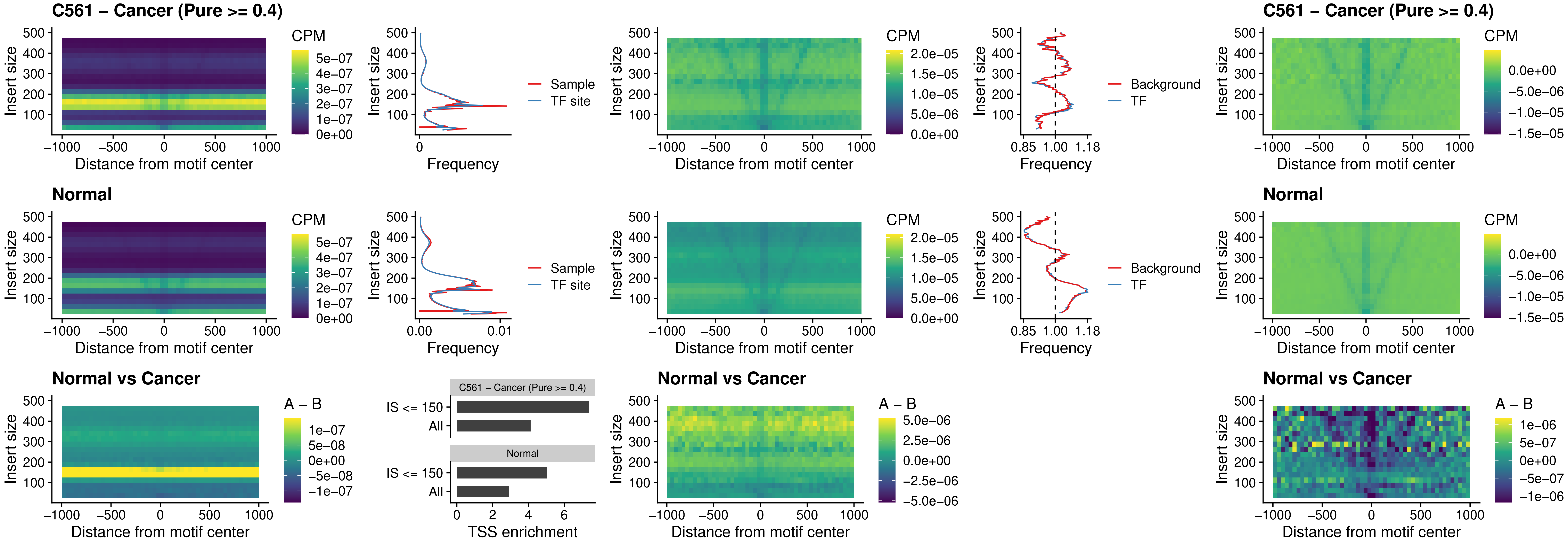

### LHX4_uc_cTSS_nPEAK

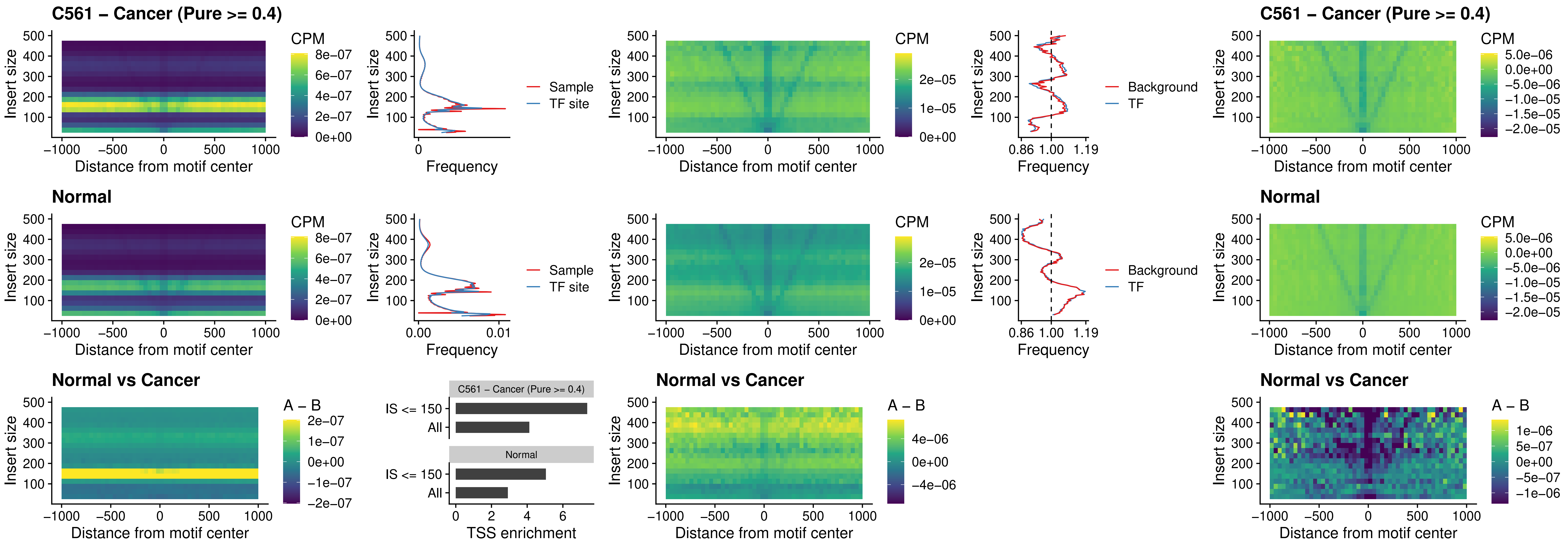

### HOXA4_uc_cTSS_nPEAK

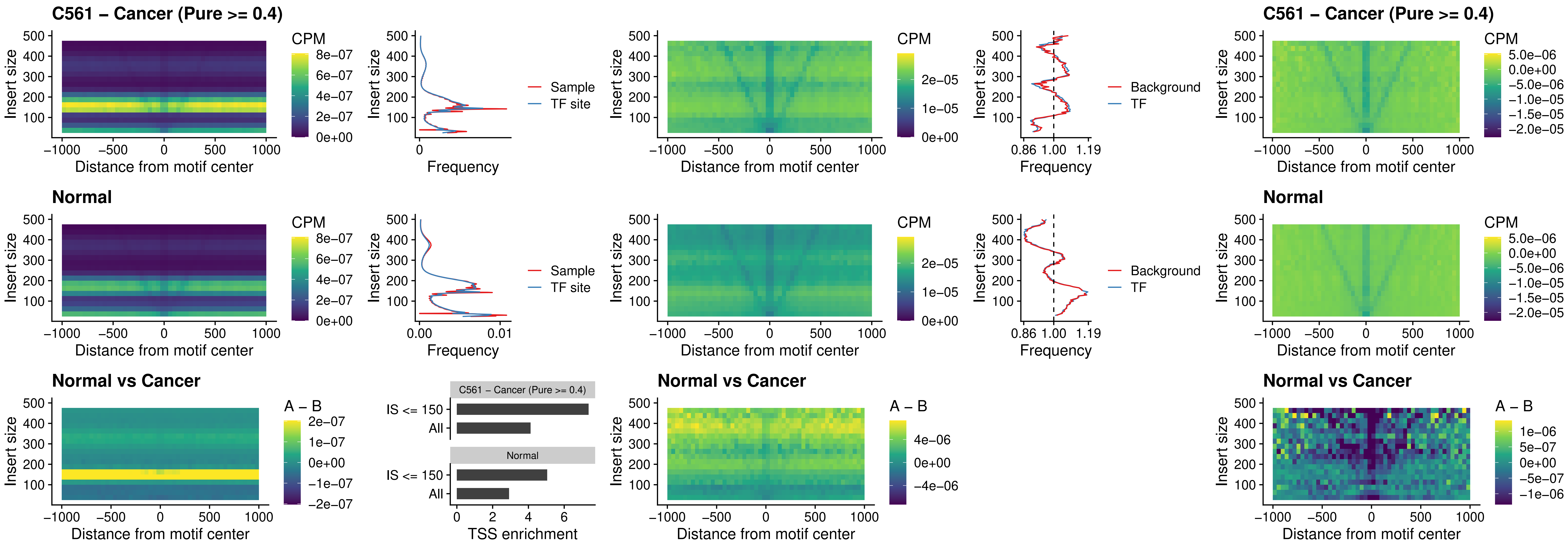

#### HLX_uc_cTSS_nPEAK

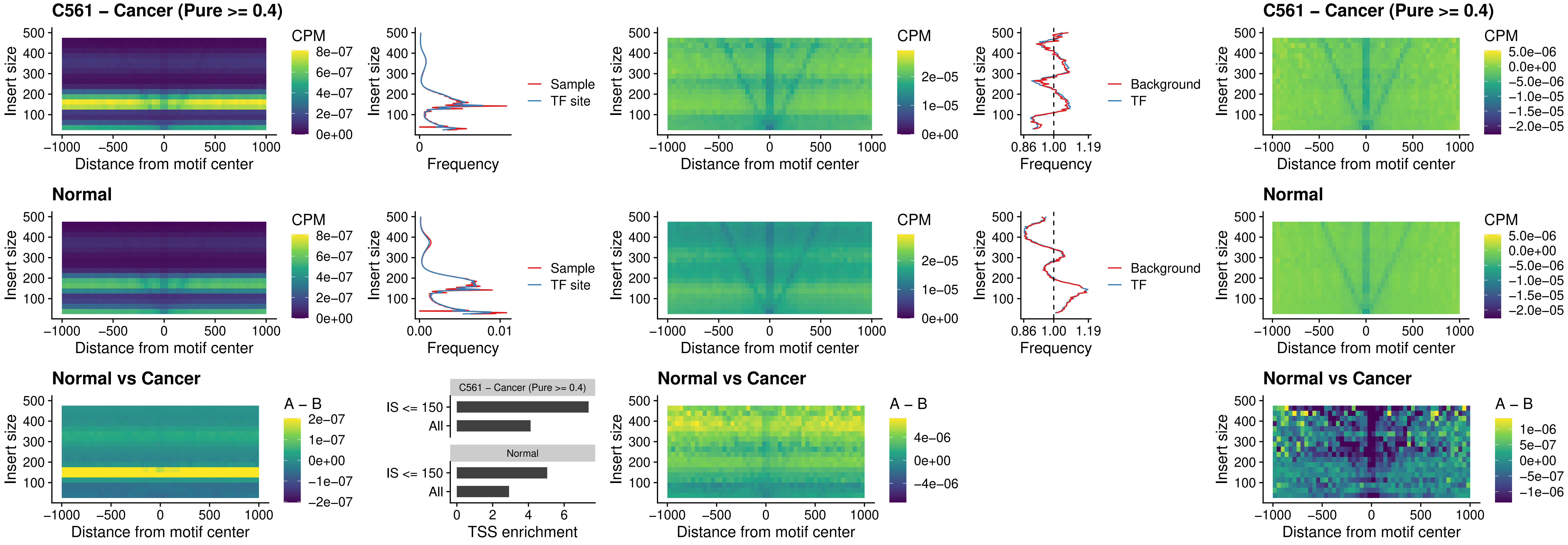

### AC0125311_uc_cTSS_nPEAK

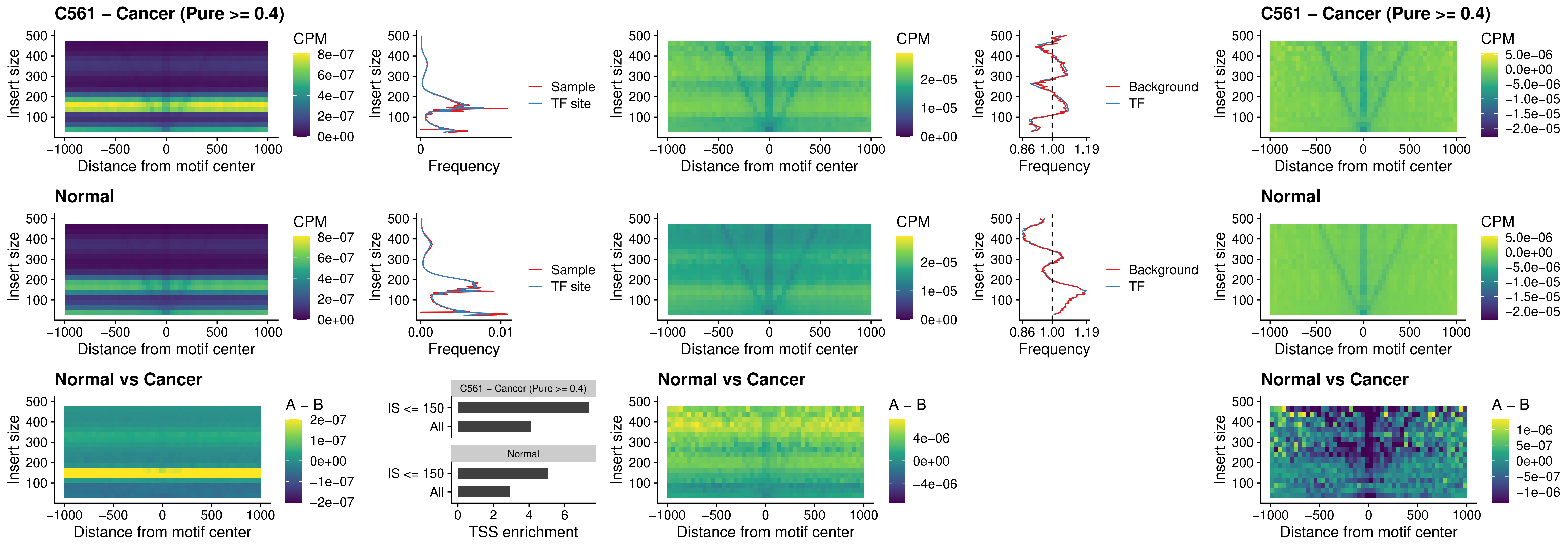

### DBX2_uc_cTSS_nPEAK

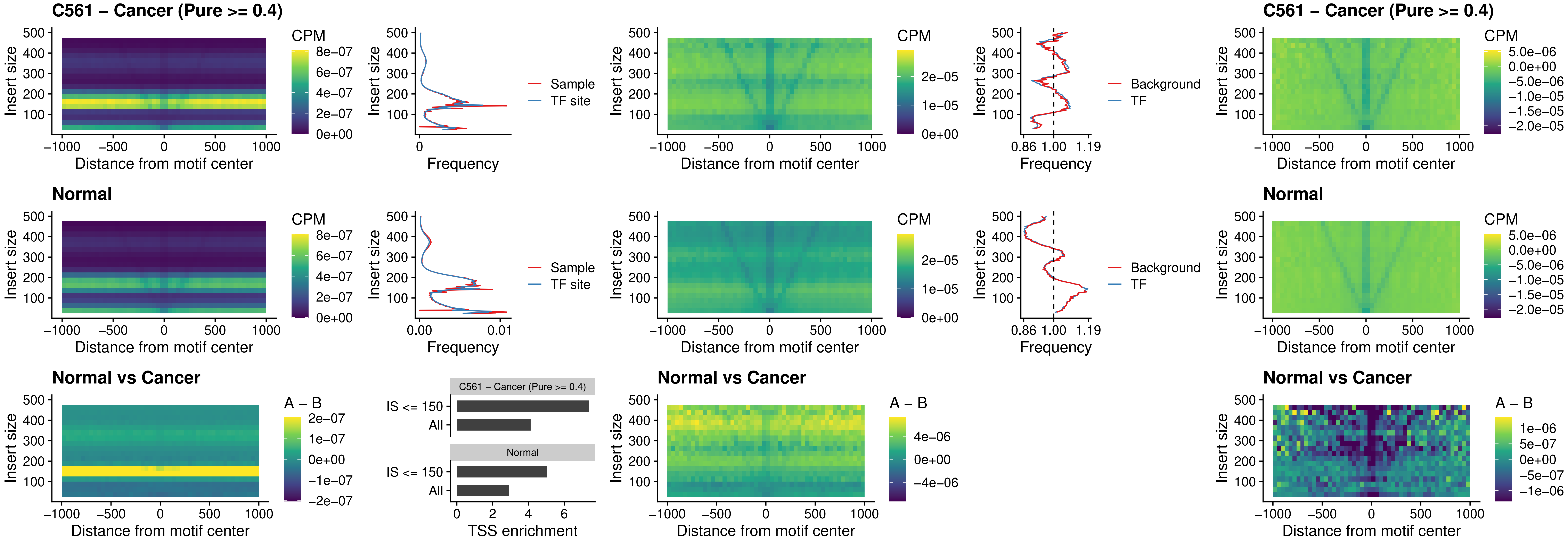

### LHX6_uc_dTSS_nPEAK

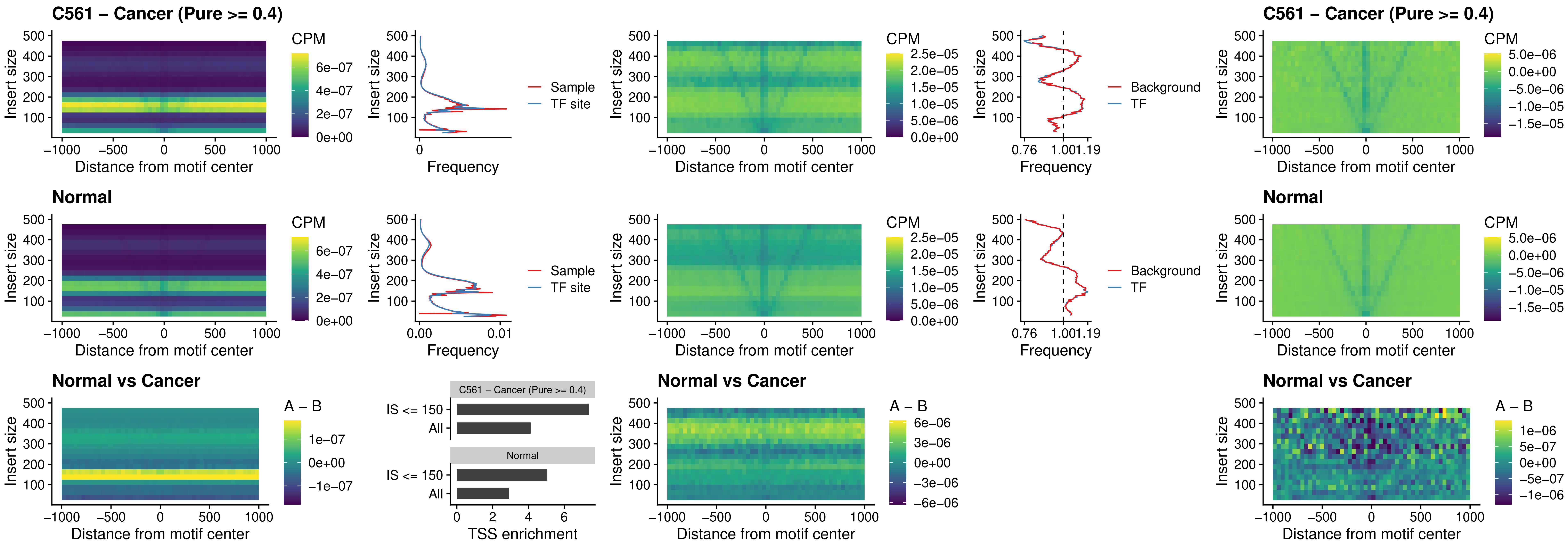

#### SOX1_uc_dTSS_nPEAK

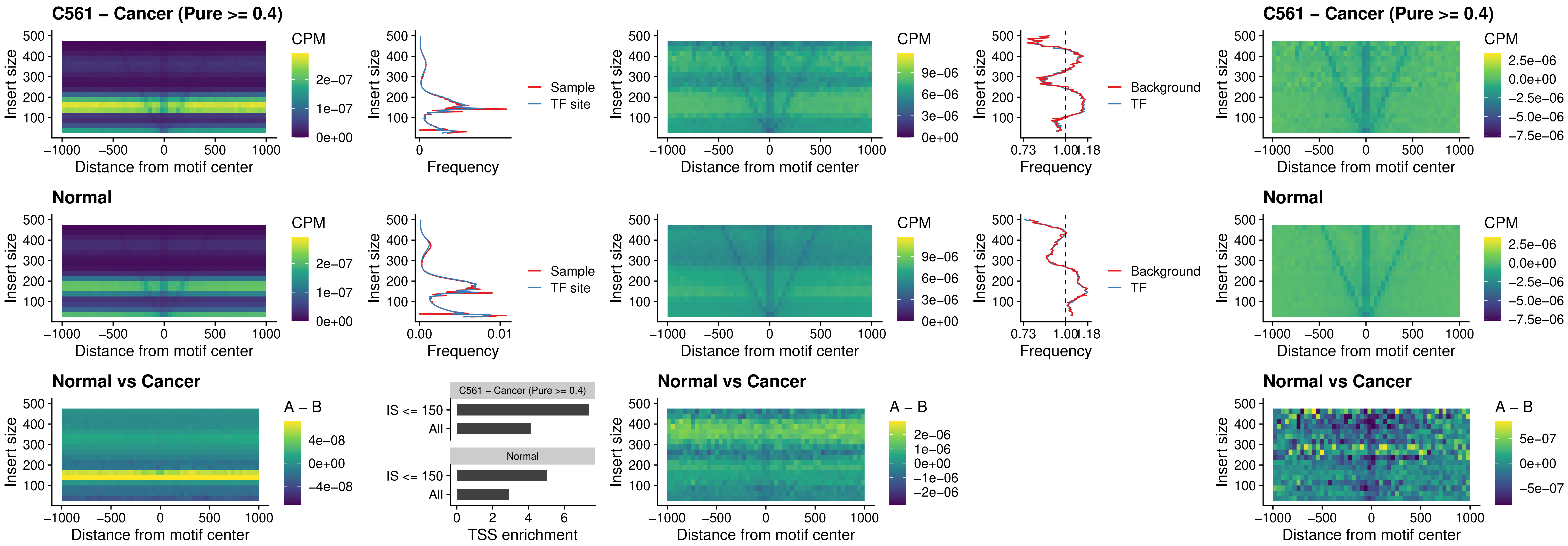

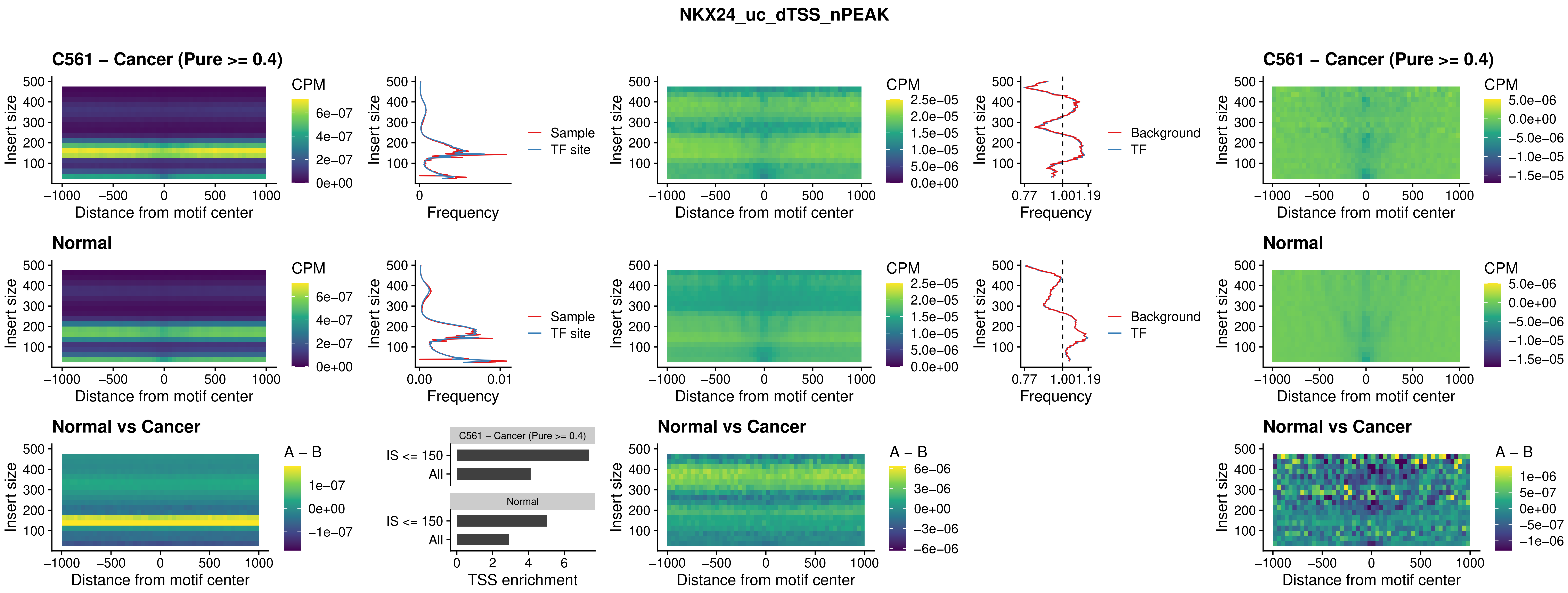

NKX24_uc_dTSS_nPEAK
C561 – Cancer (Pure >= 0.4)
Normal
Normal vs Cancer
Insert size
Distance from motif center
Frequency
CPM
A – B
Sample
TF site
Background
TF
C561 – Cancer (Pure >= 0.4)
Normal
IS <= 150
All
TSS enrichment

### DLX3_uc_cTSS_nPEAK

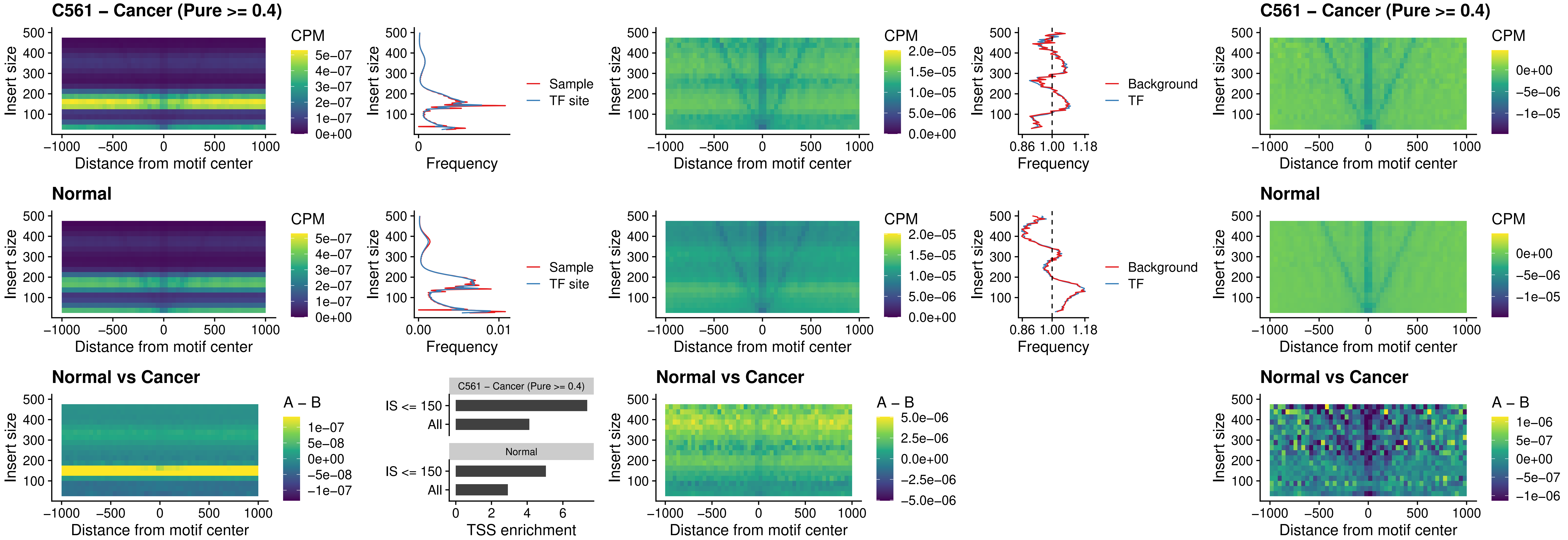

### HOXD3_uc_cTSS_nPEAK

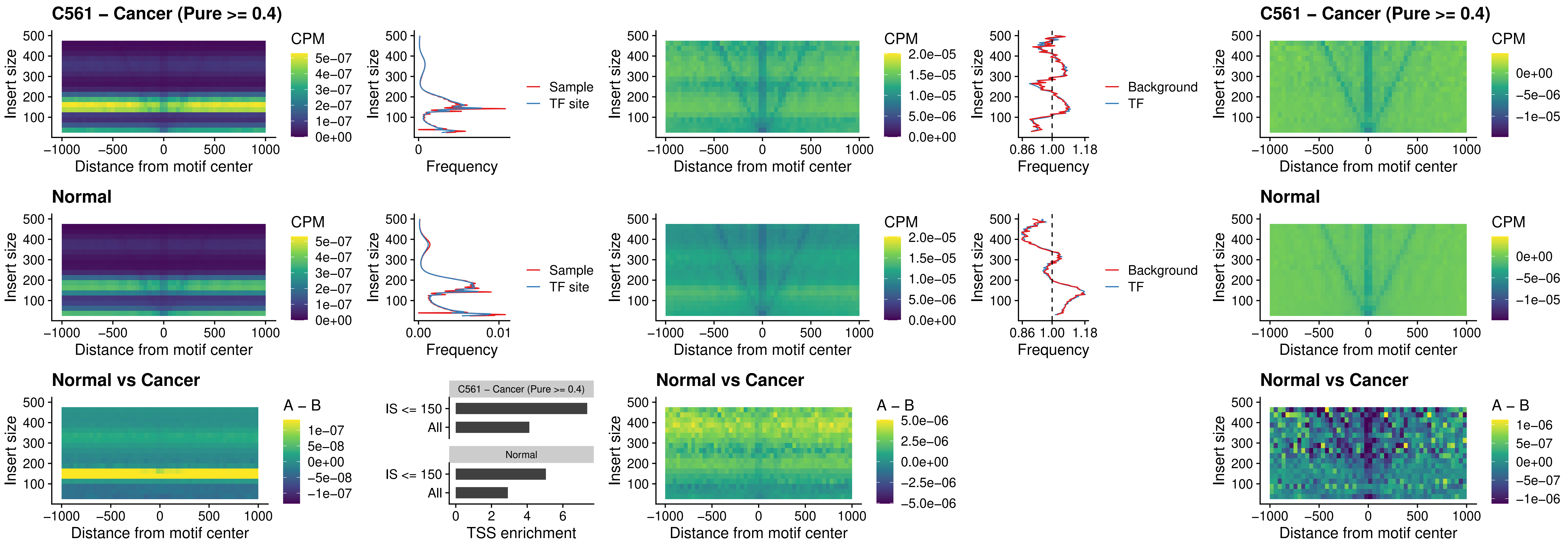

### ISL1_uc_dTSS_oPEAK

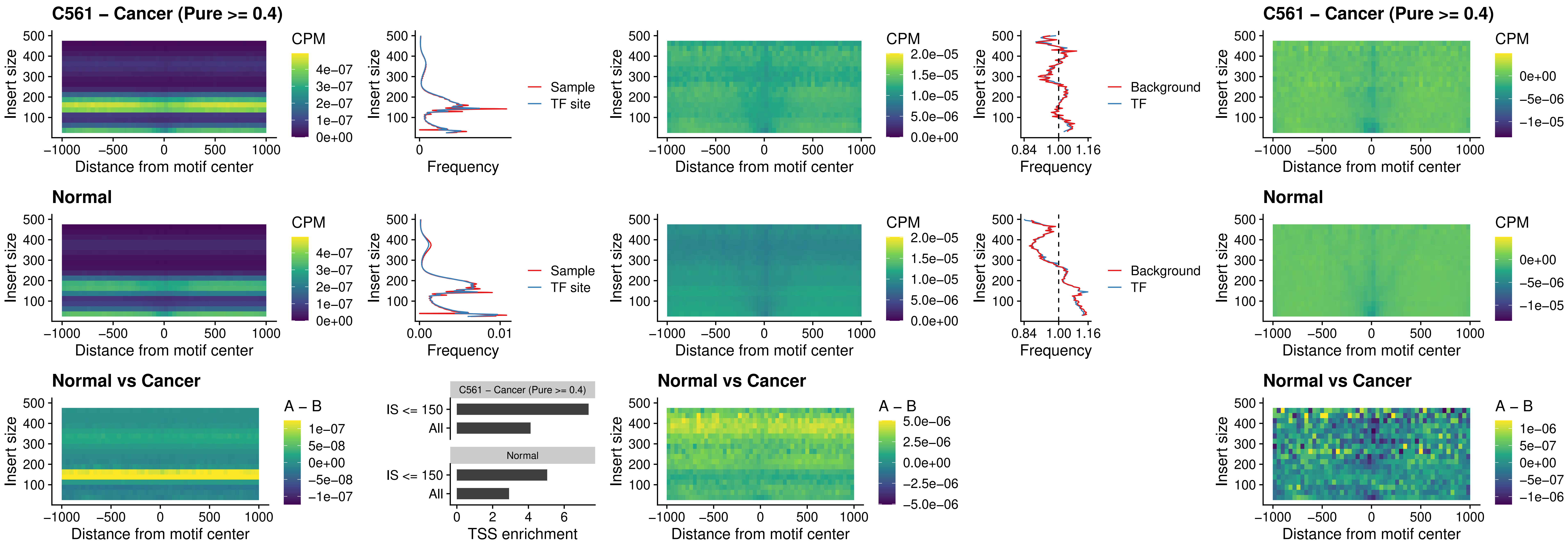

#### HOXB4_uc_dTSS_oPEAK

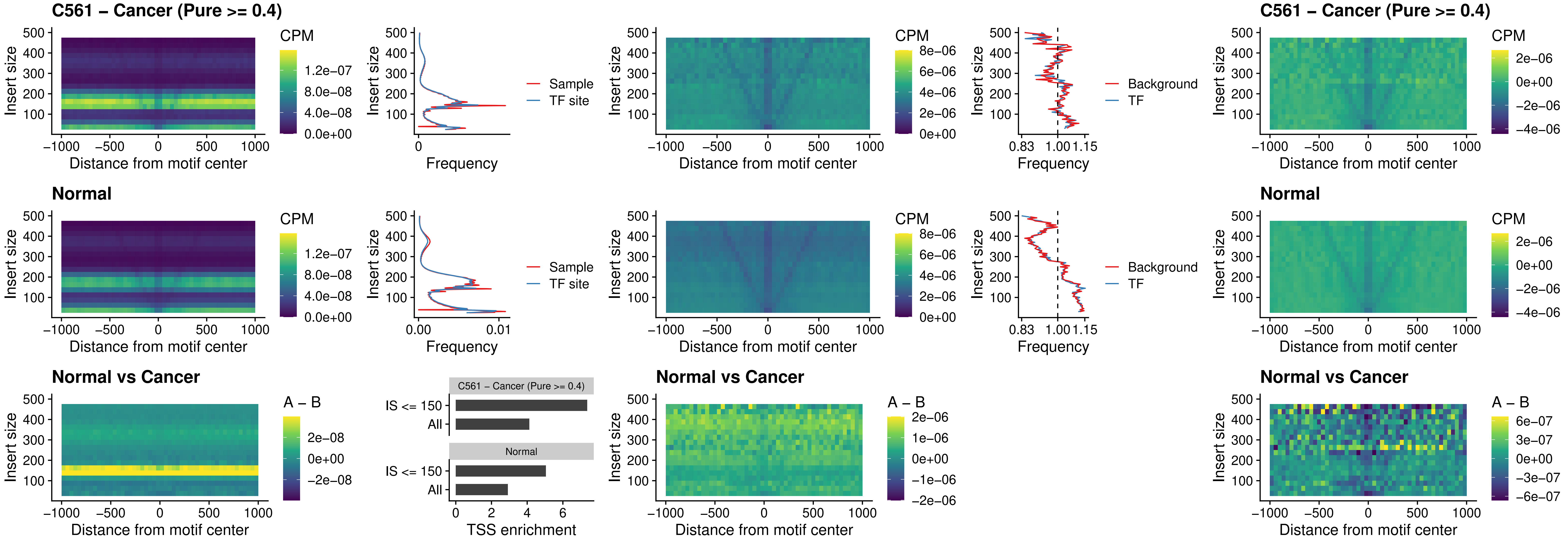

### VAX1_uc_dTSS_oPEAK

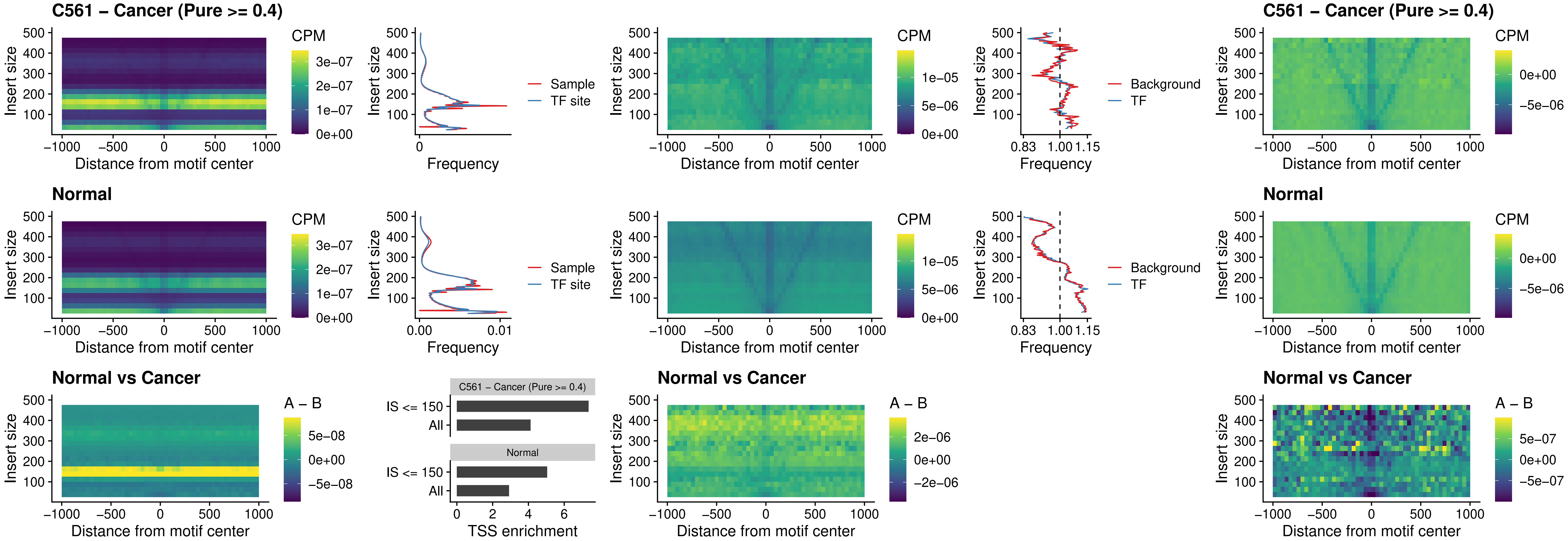

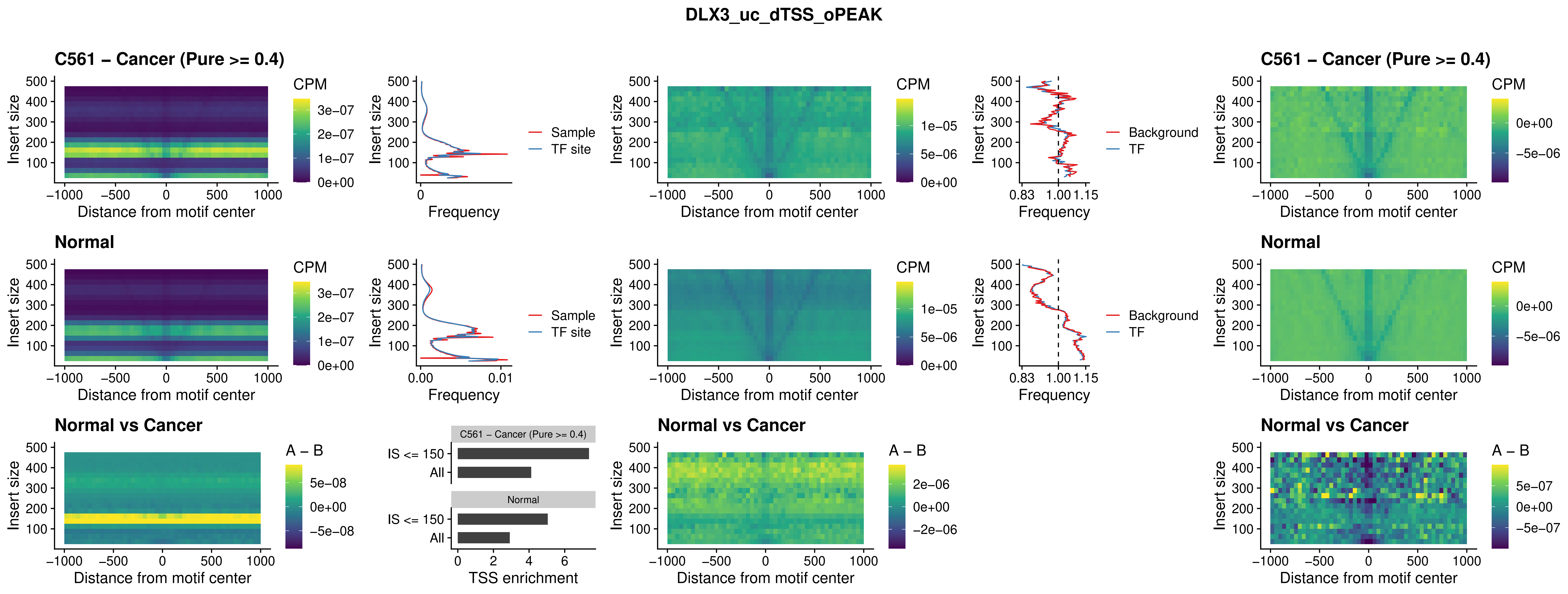
DLX3_uc_dTSS_oPEAK
C561 – Cancer (Pure >= 0.4)
Normal
Normal vs Cancer
Insert size
Distance from motif center
Frequency
CPM
A – B
Sample
TF site
Background
TF
C561 – Cancer (Pure >= 0.4)
Normal
IS <= 150
All
TSS enrichment

#### HOXD3_uc_dTSS_oPEAK

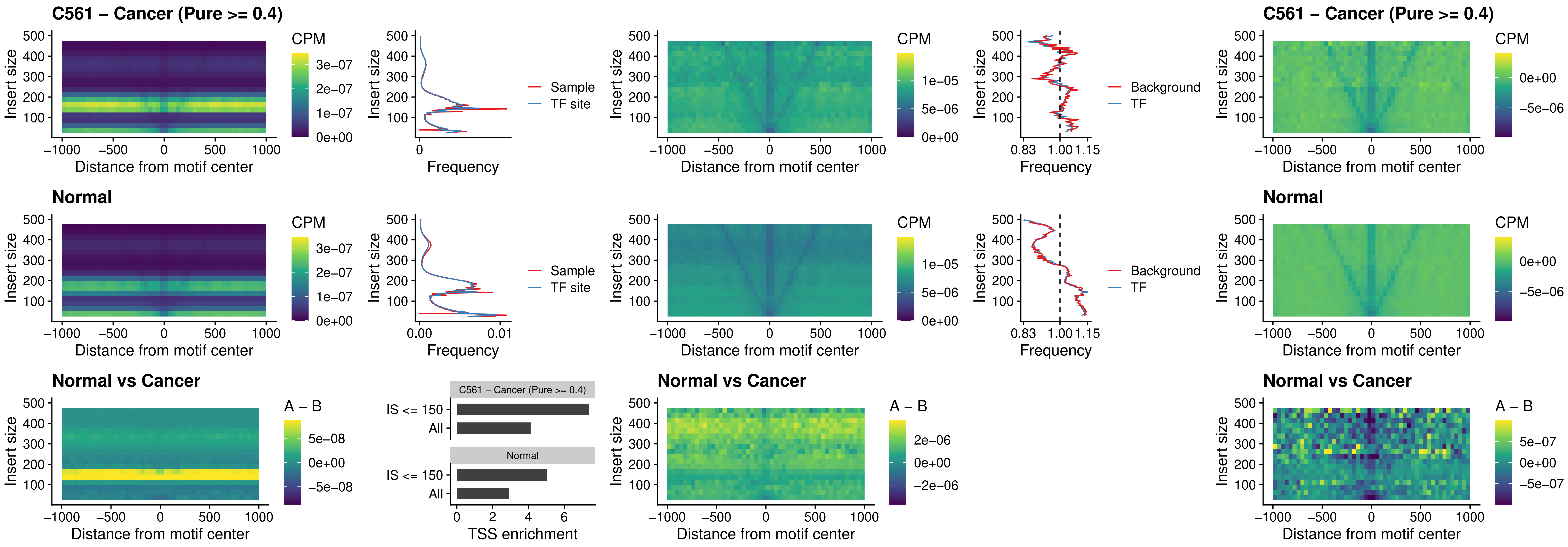

### BBX_uc_dTSS_nPEAK

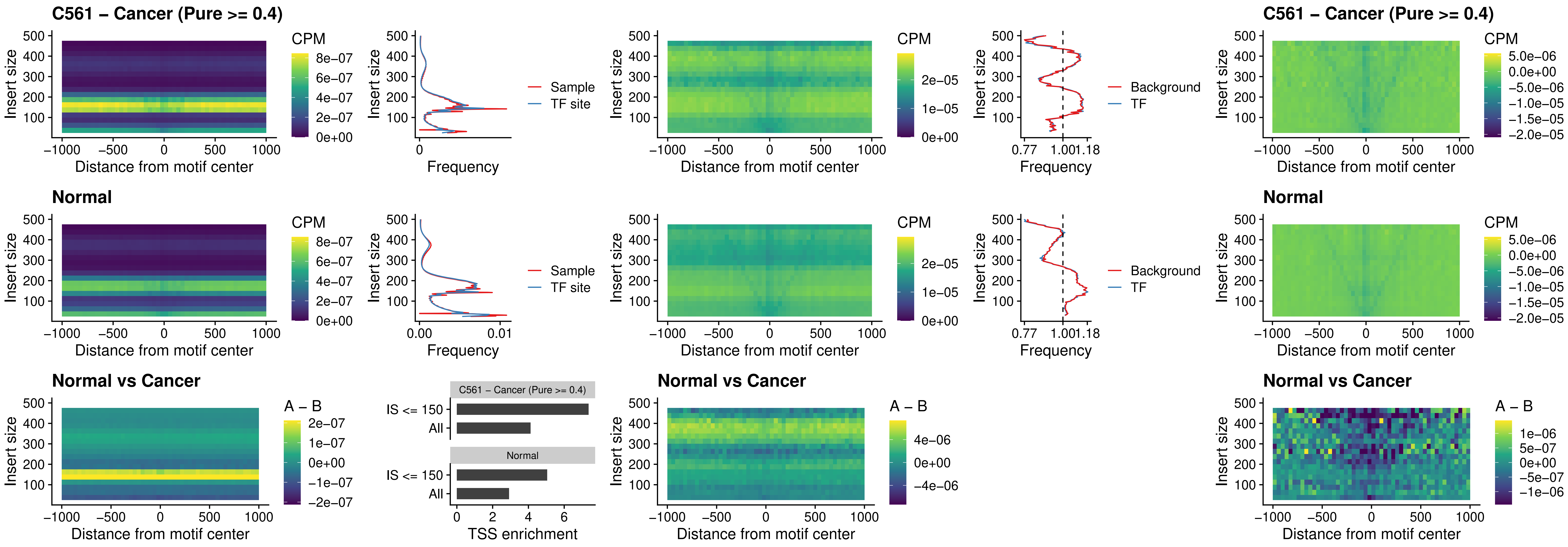

#### SOX3_uc_cTSS_nPEAK

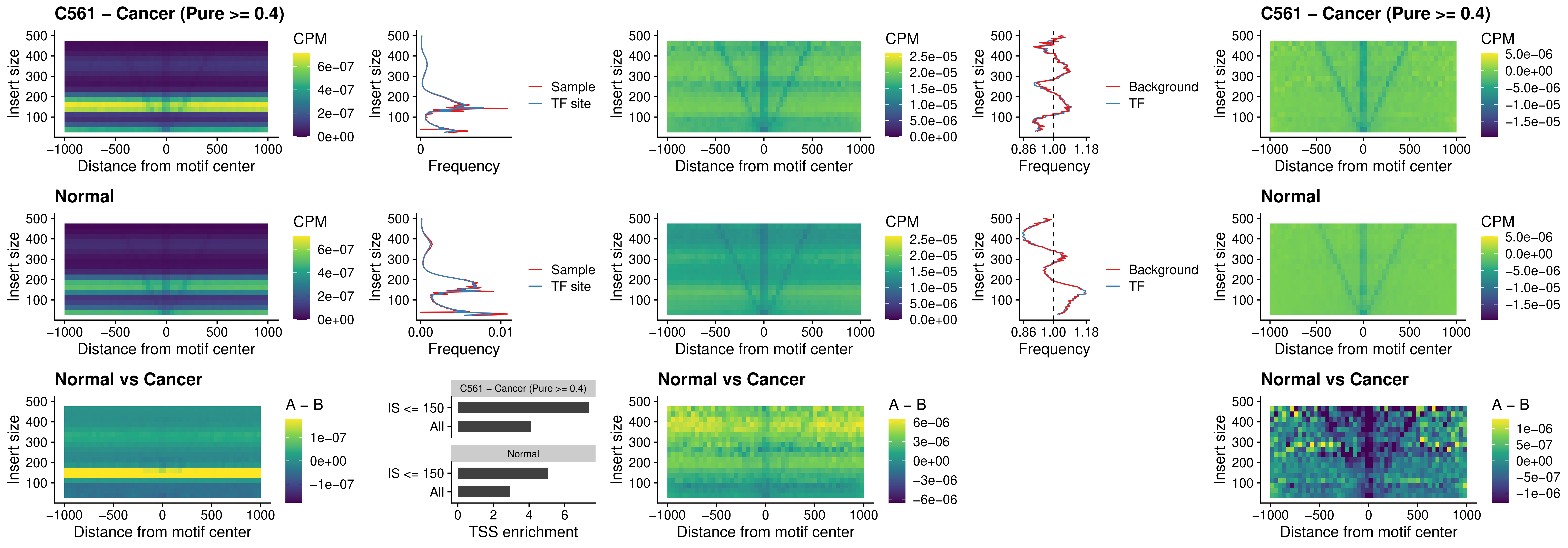

#### TBPL2_uc_cTSS_nPEAK

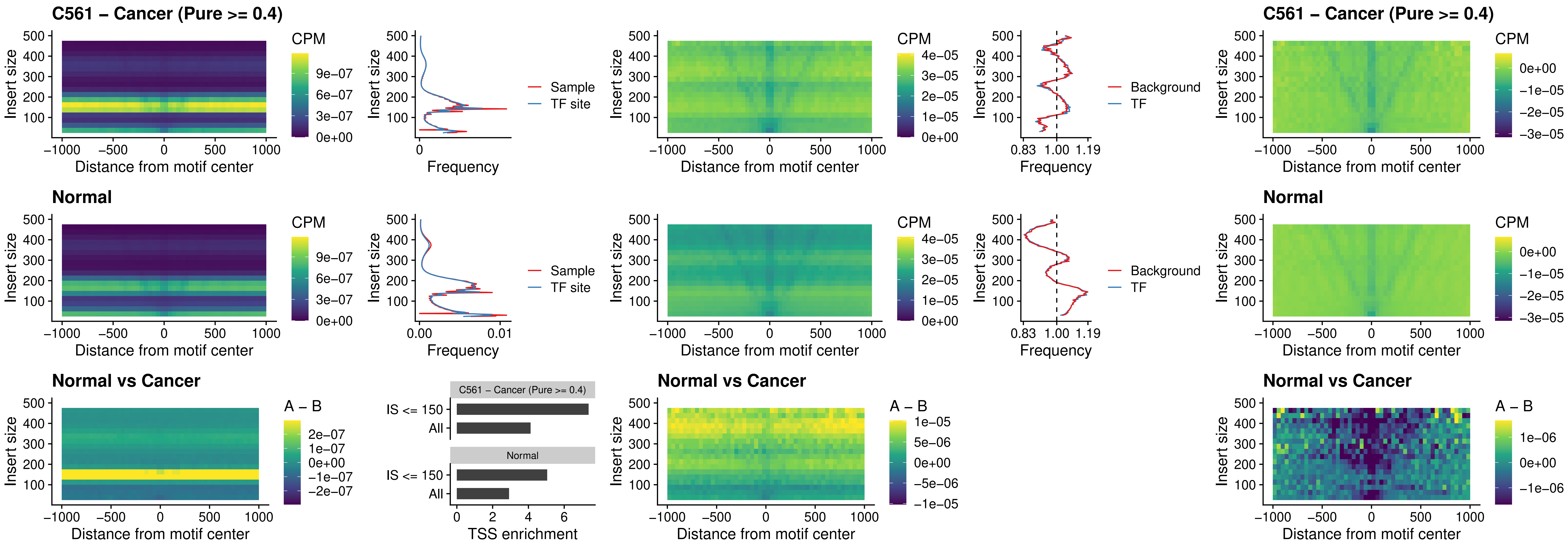

#### HOXA4_uc_cTSS_oPEAK

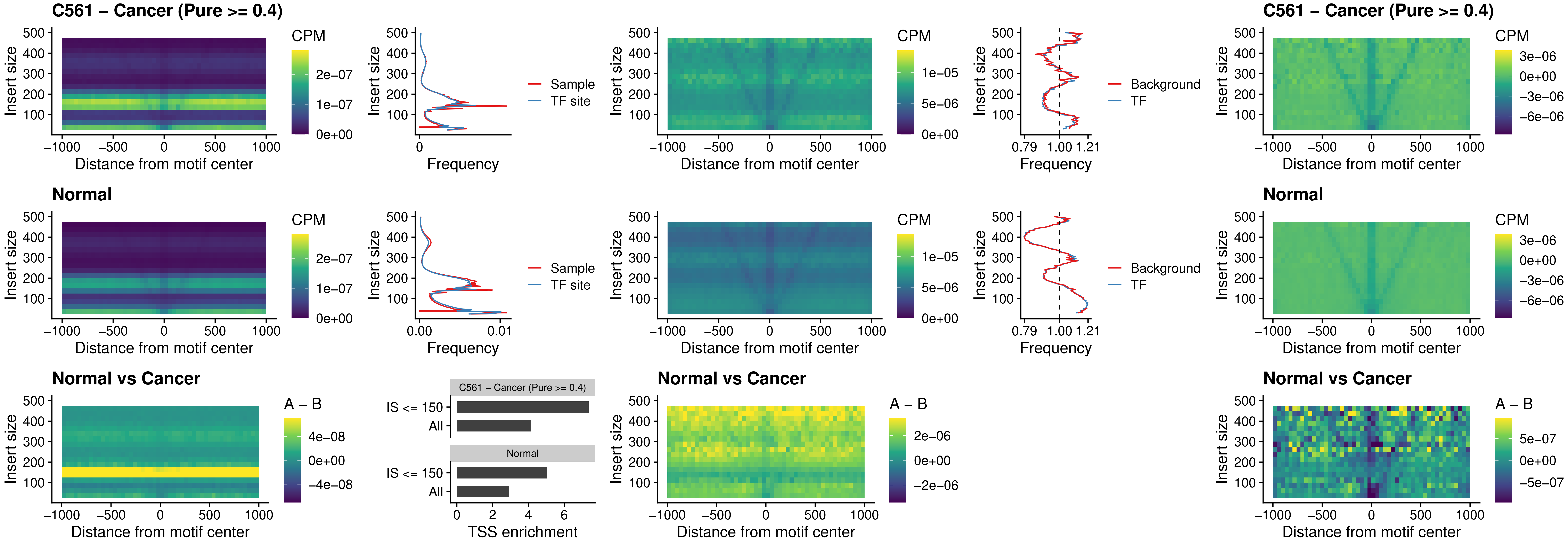

### PDX1_uc_dTSS_oPEAK

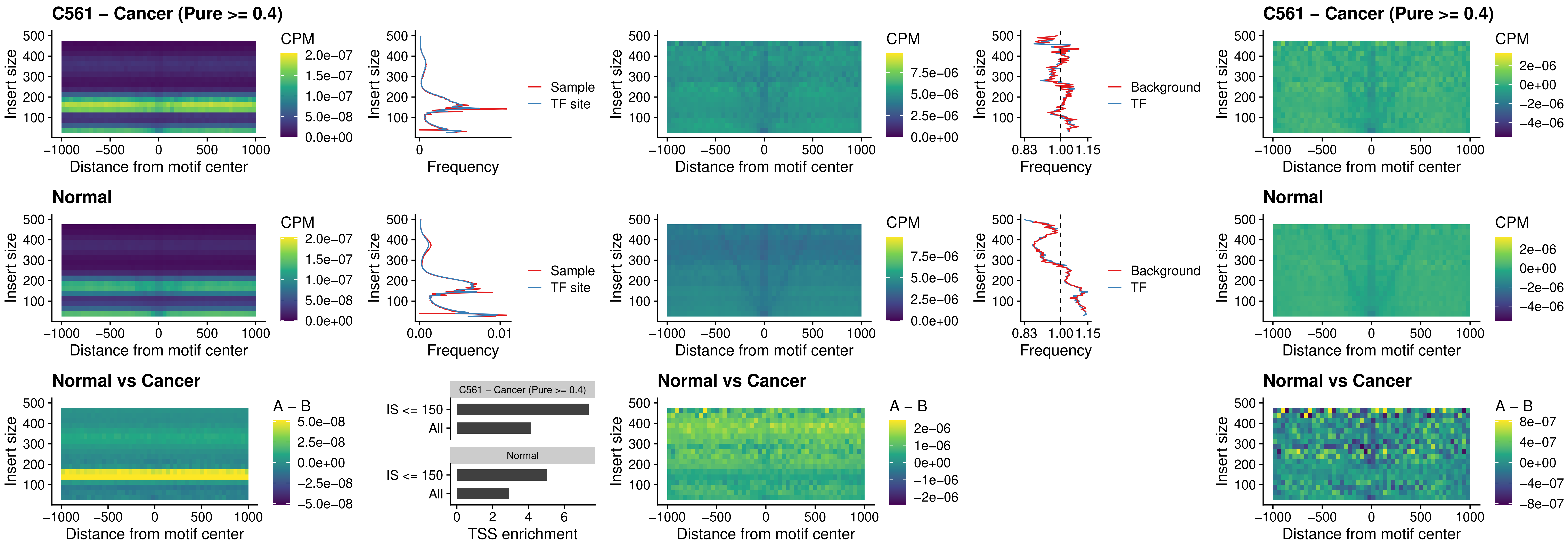

#### LHX4_uc_dTSS_oPEAK

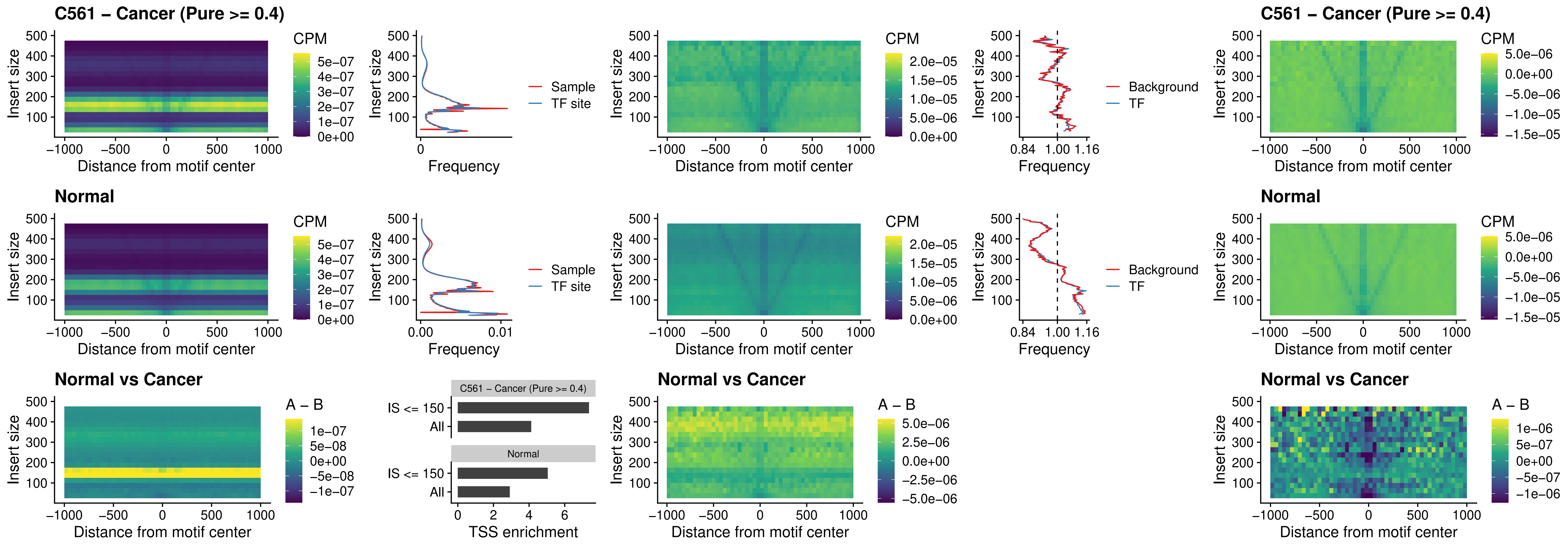

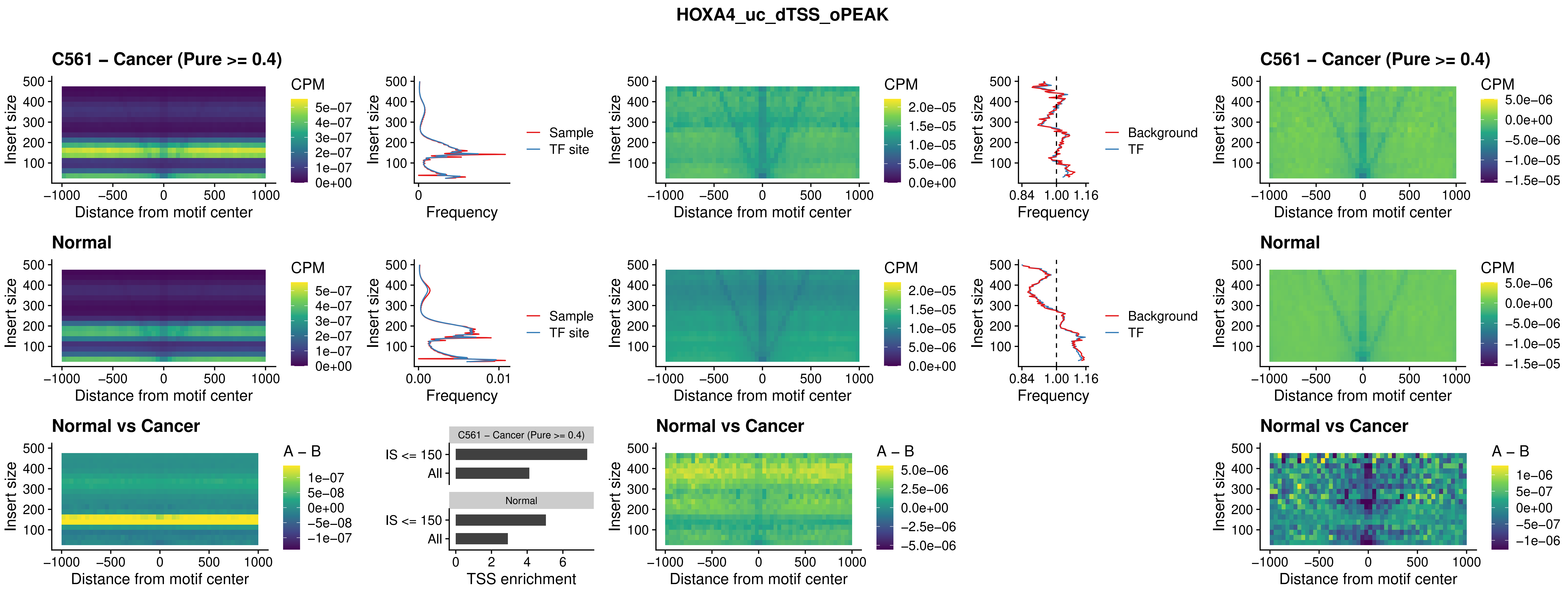
HOXA4_uc_dTSS_oPEAK
C561 – Cancer (Pure >= 0.4)
Normal
Normal vs Cancer
Insert size
Distance from motif center
Frequency
CPM
A – B
Sample
TF site
Background
TF
C561 – Cancer (Pure >= 0.4)
Normal
IS <= 150
All
TSS enrichment

#### HLX_uc_dTSS_oPEAK

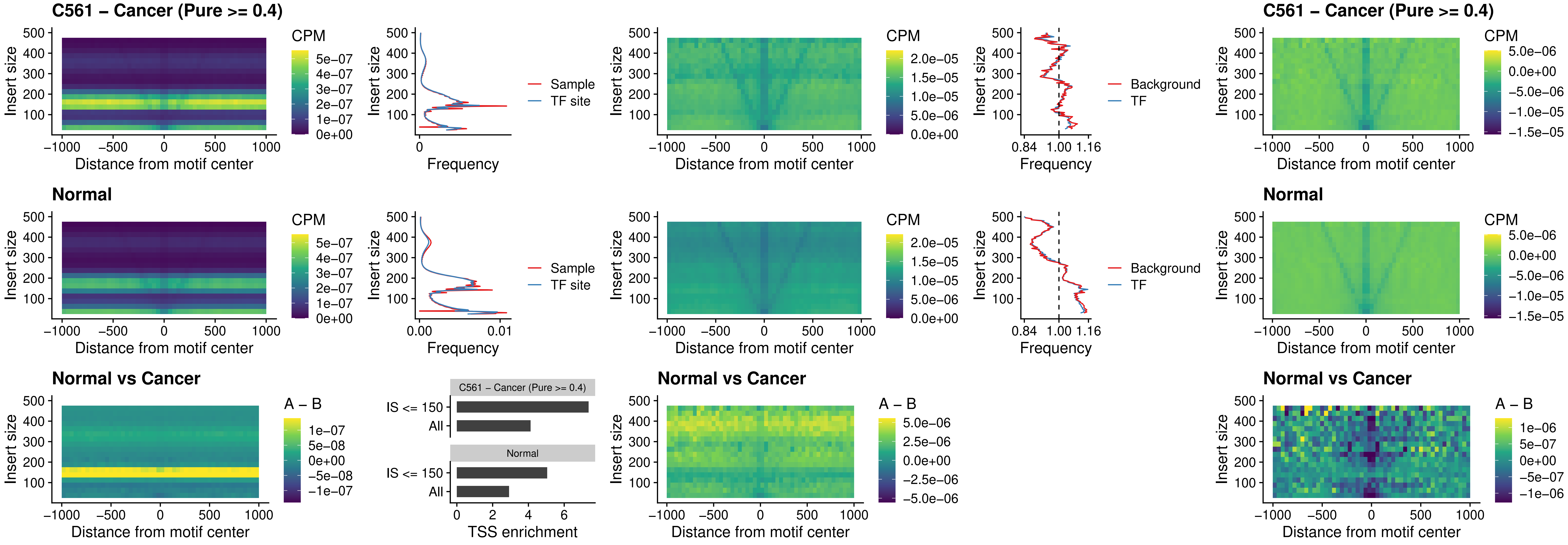

### AC0125311_uc_dTSS_oPEAK

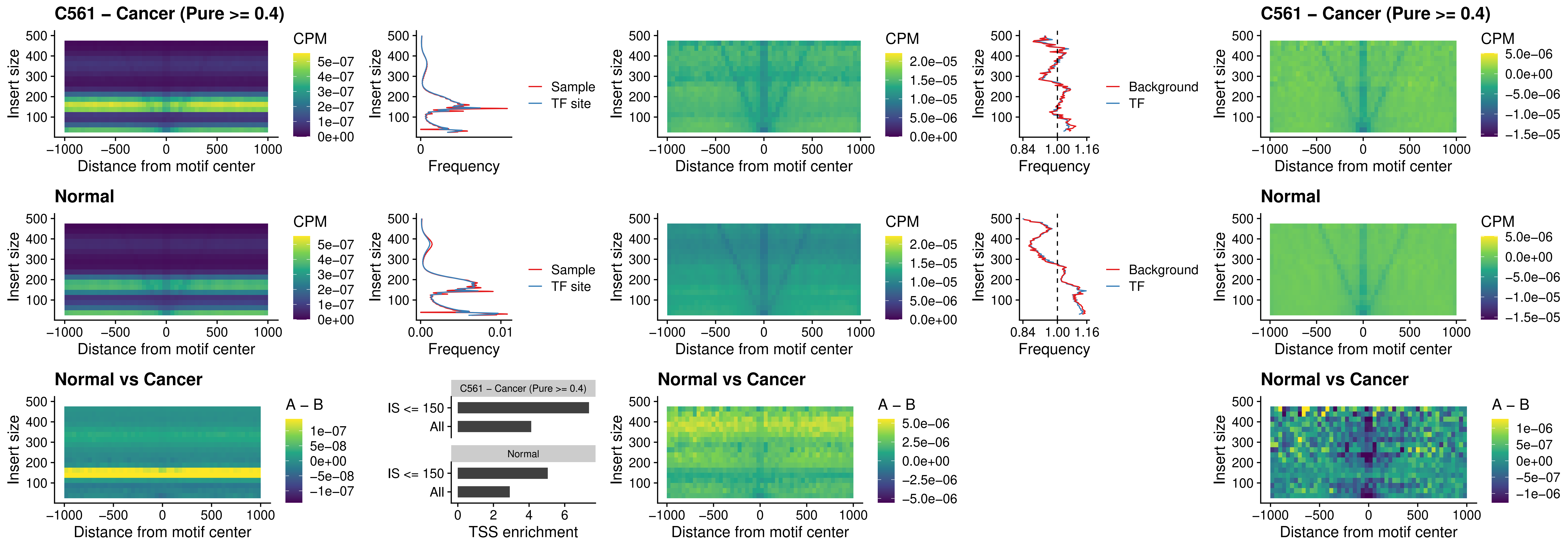

### DBX2_uc_dTSS_oPEAK

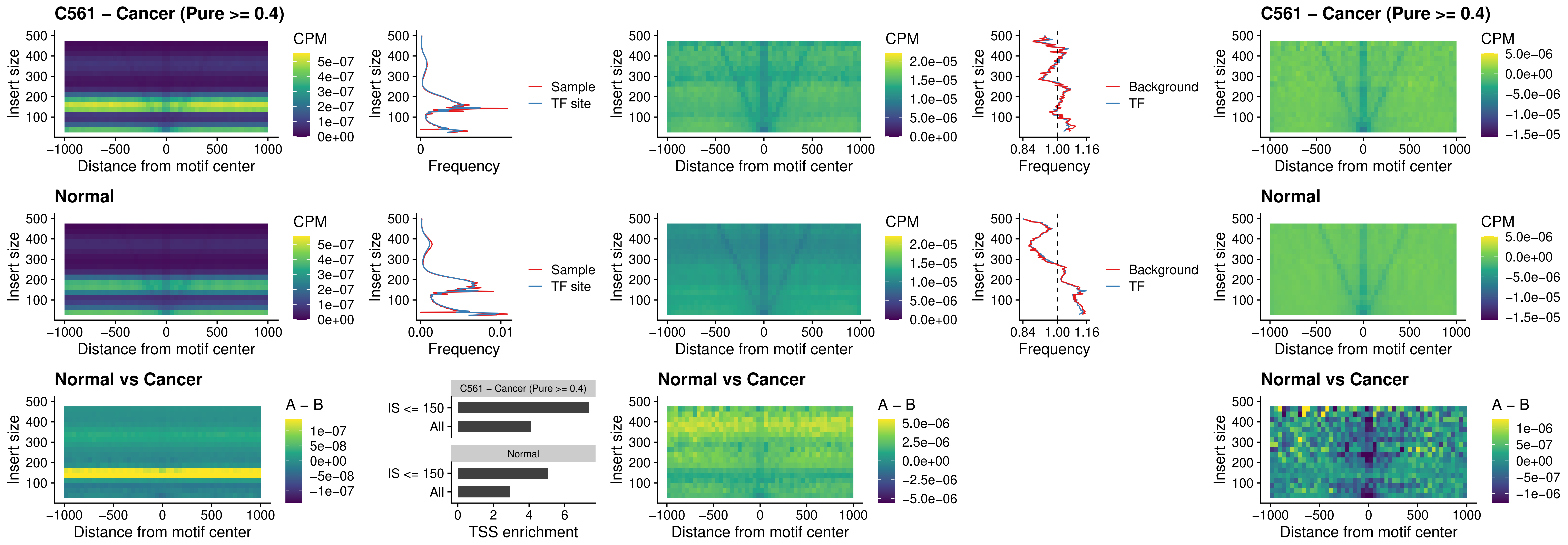

### HOXA2_uc_dTSS_nPEAK

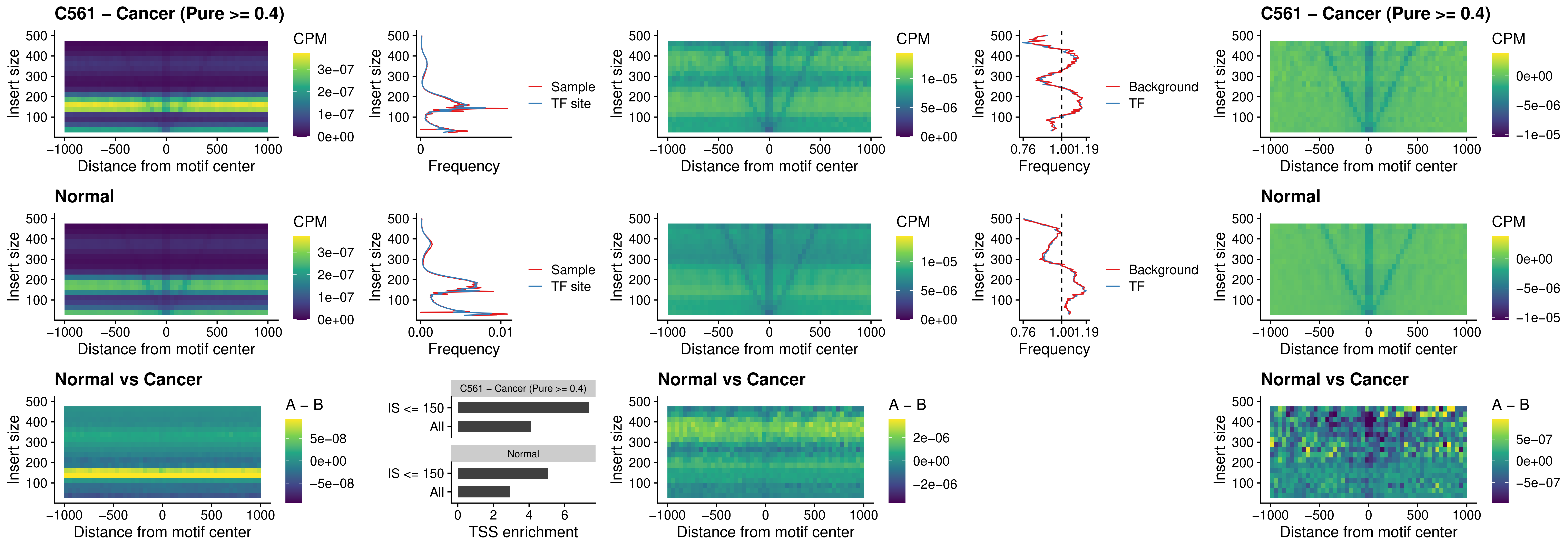

#### SHOX_uc_dTSS_nPEAK

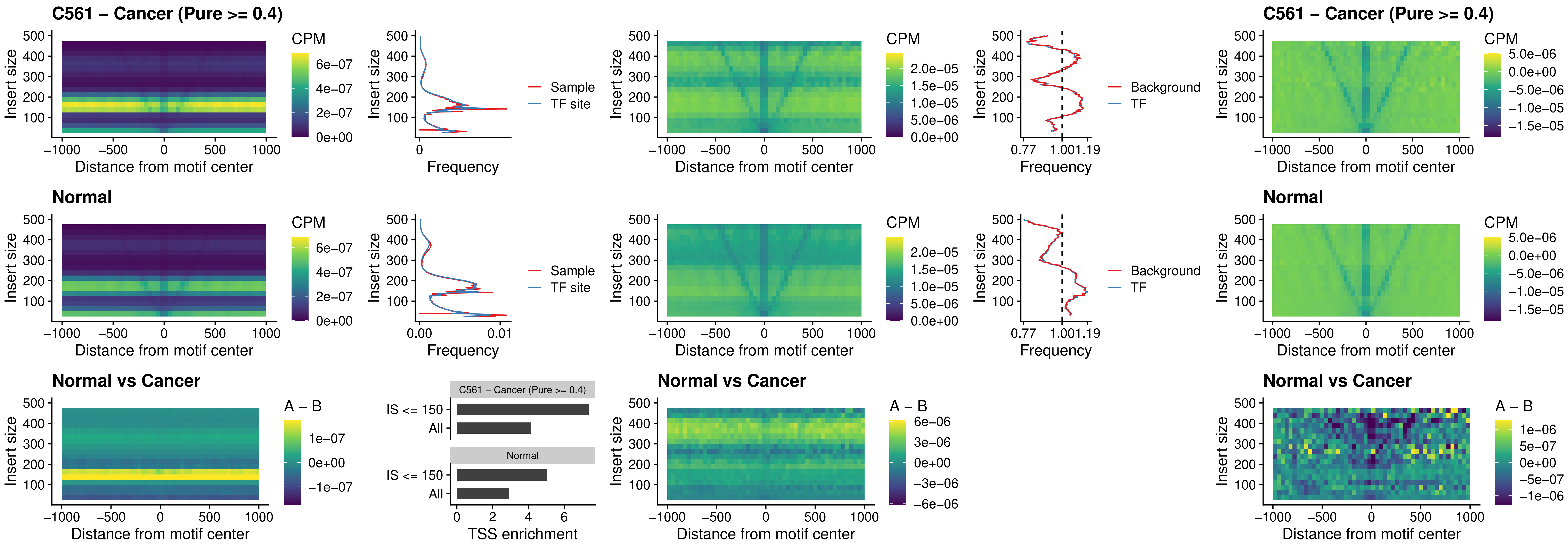

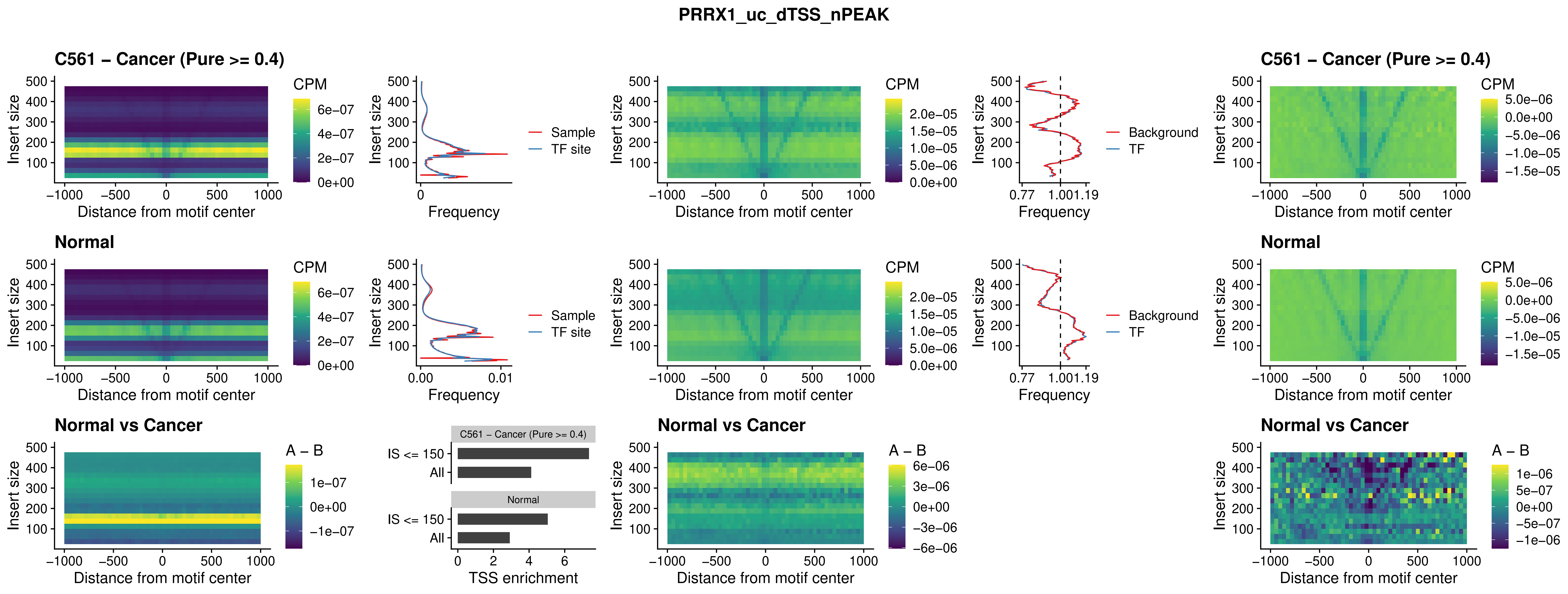
PRRX1_uc_dTSS_nPEAK
C561 – Cancer (Pure >= 0.4)
Normal
Normal vs Cancer
Insert size
Distance from motif center
Frequency
CPM
A – B
Sample
TF site
Background
TF
C561 – Cancer (Pure >= 0.4)
Normal
IS <= 150
All
TSS enrichment

#### LMX1A_uc_dTSS_nPEAK

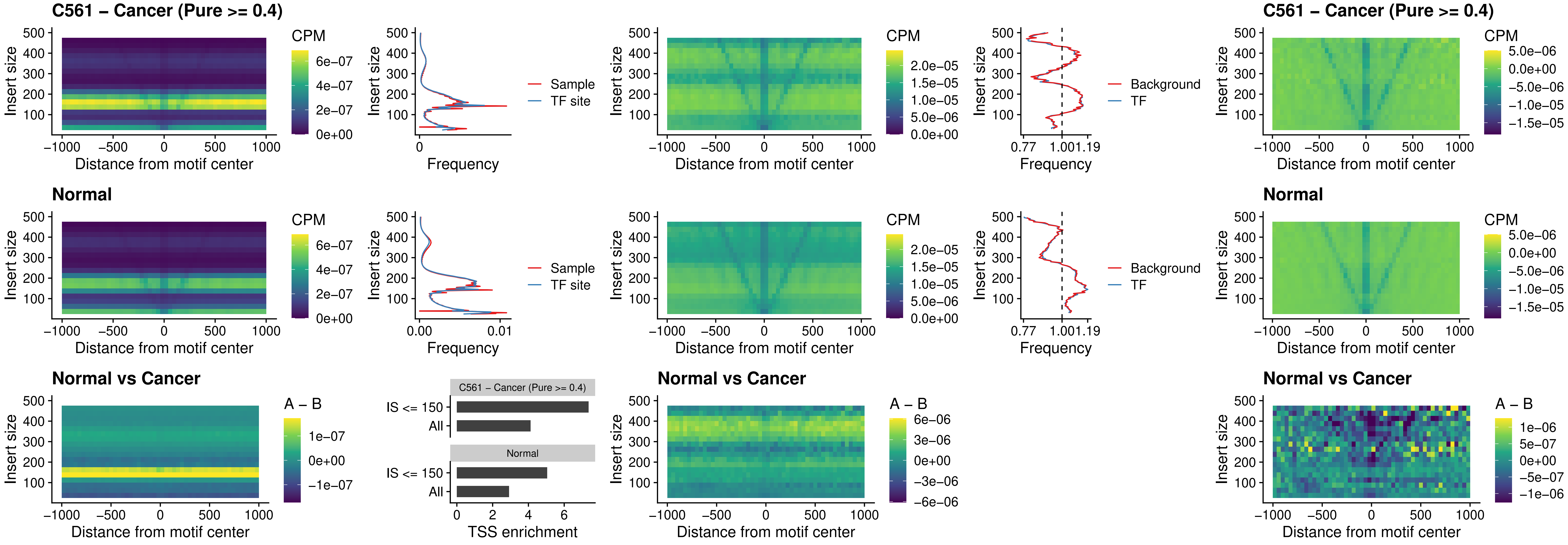

### LMX1B_uc_dTSS_nPEAK

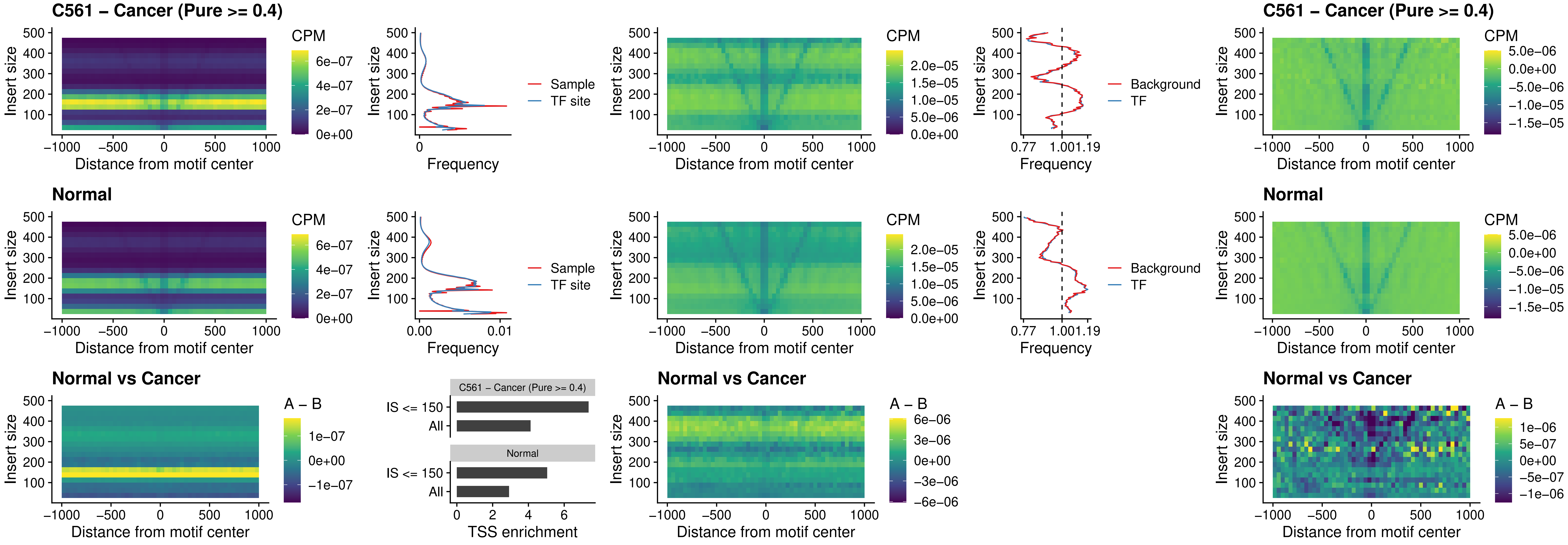

### DLX5_uc_dTSS_oPEAK

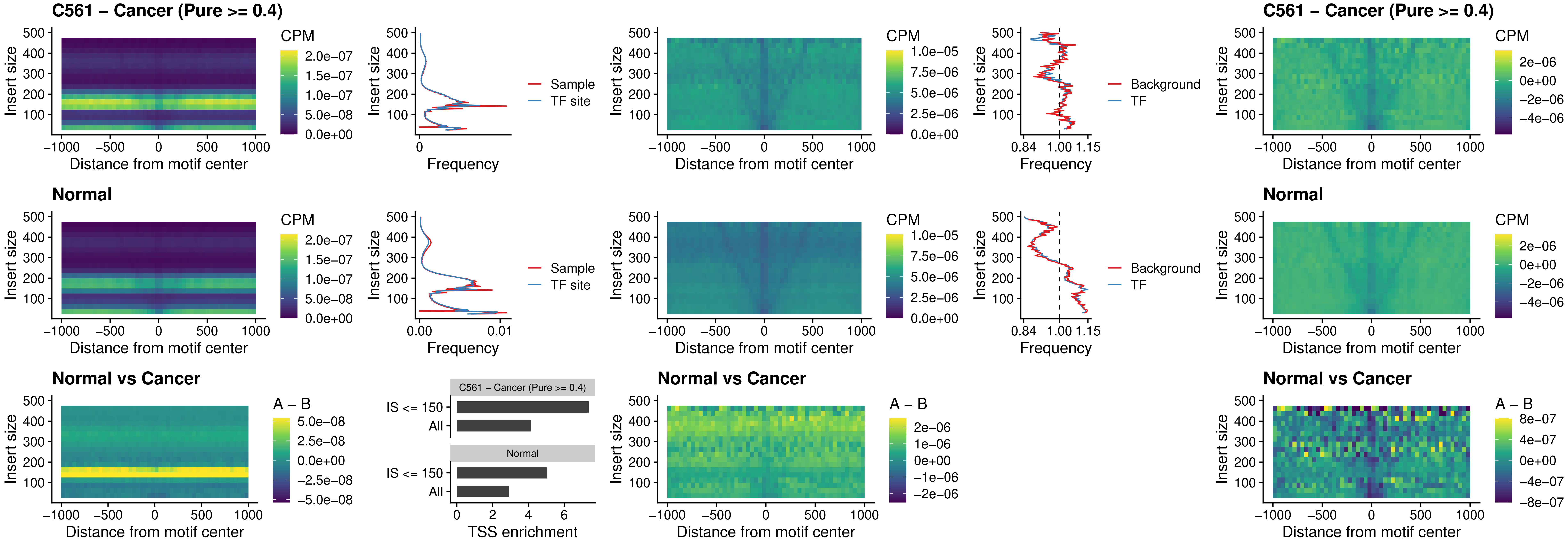

### ARID3C_uc_dTSS_nPEAK

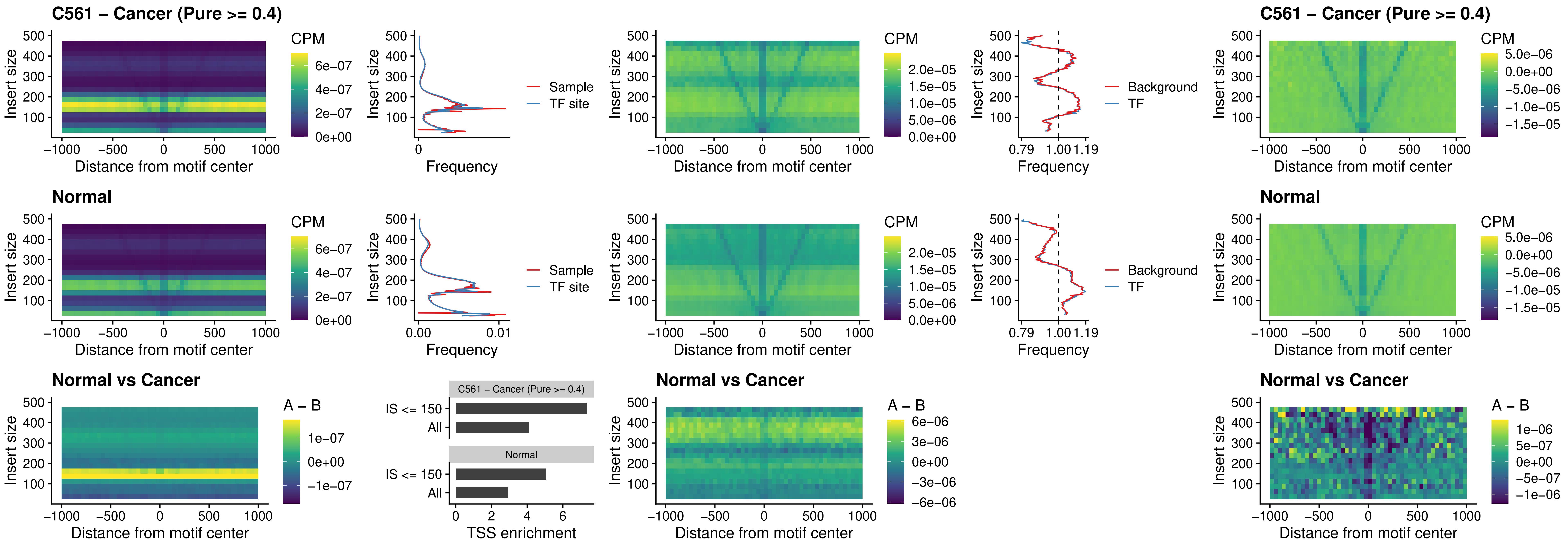

### HOXD12_uc_dTSS_nPEAK

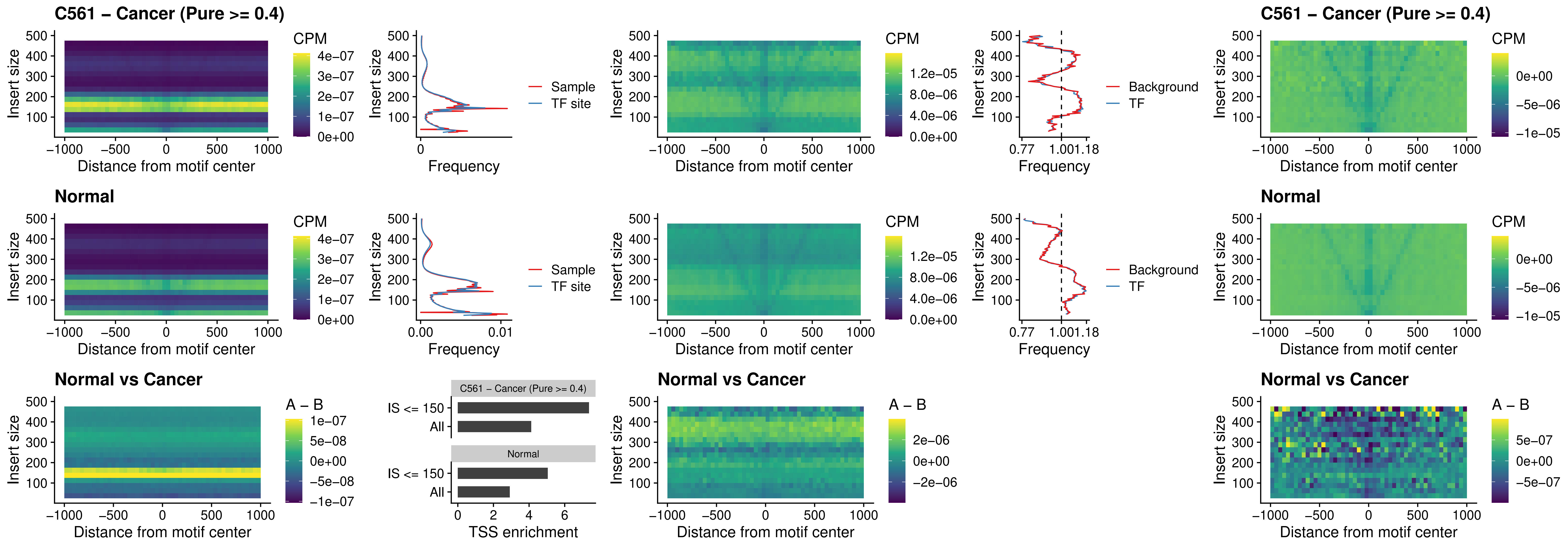

### CDX2_uc_dTSS_nPEAK

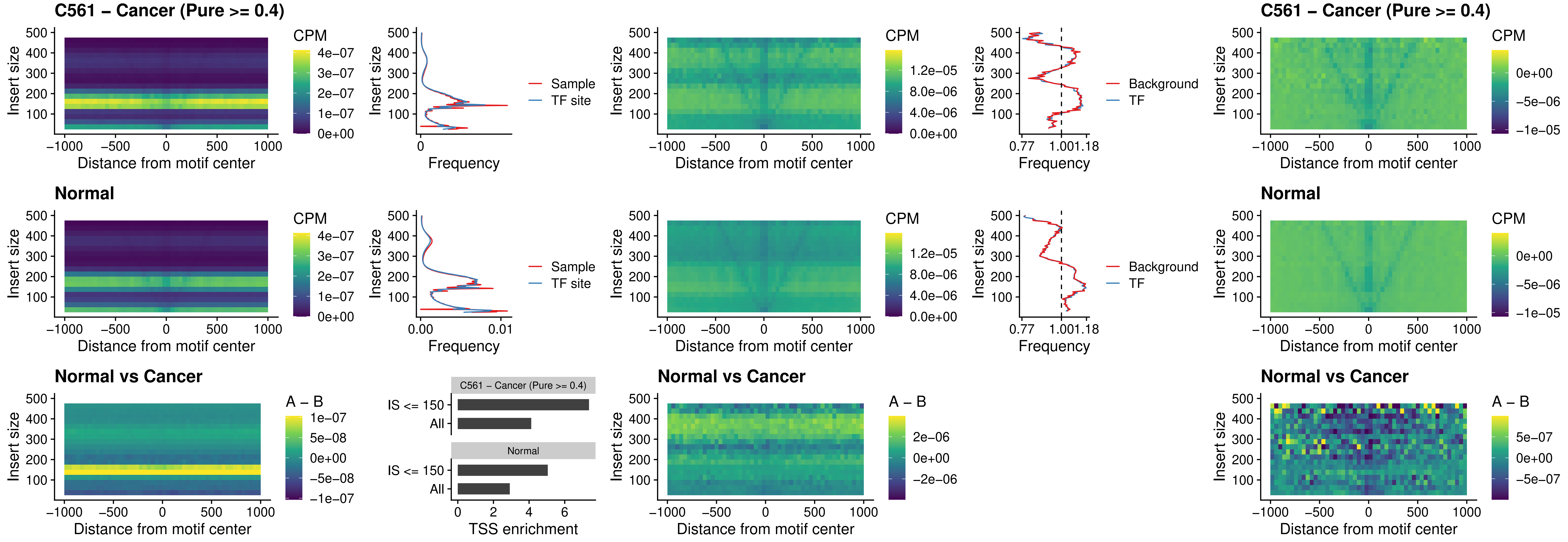

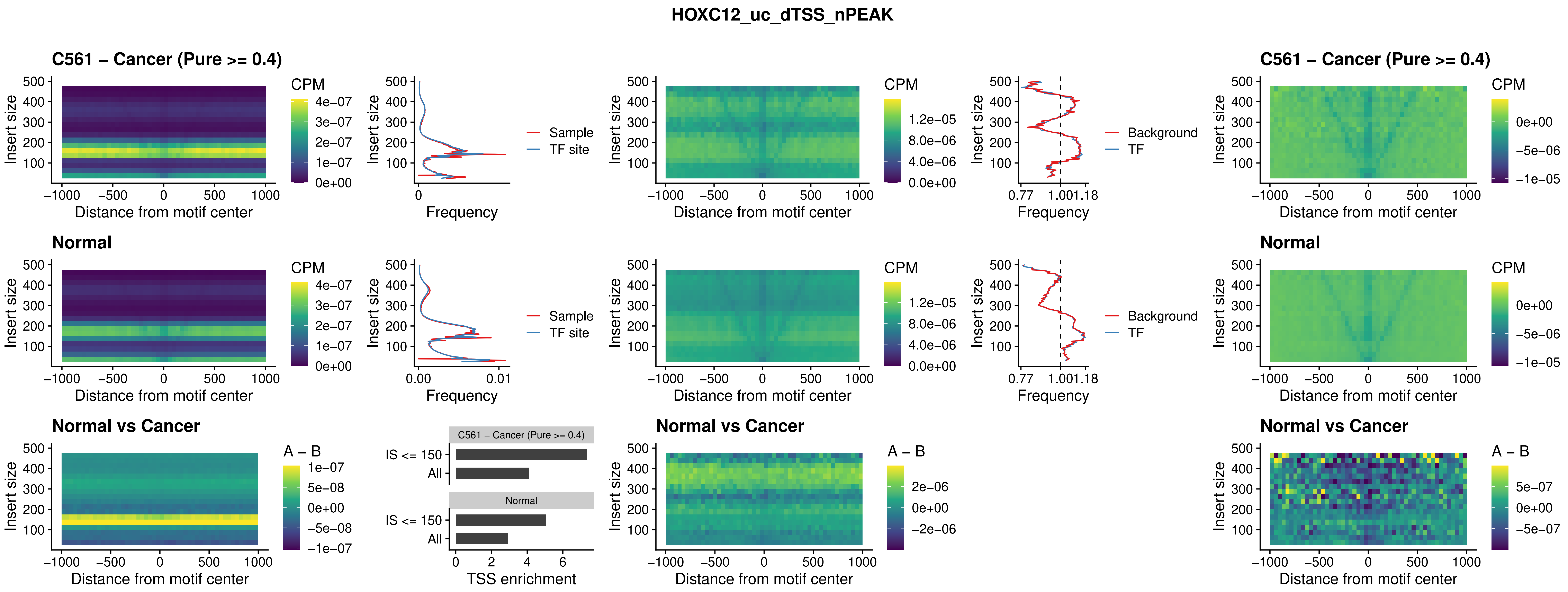
HOXC12_uc_dTSS_nPEAK
C561 – Cancer (Pure >= 0.4)
Normal
Normal vs Cancer
Insert size
Distance from motif center
Frequency
CPM
A – B
Sample
TF site
Background
TF
C561 – Cancer (Pure >= 0.4)
Normal
IS <= 150
All
TSS enrichment

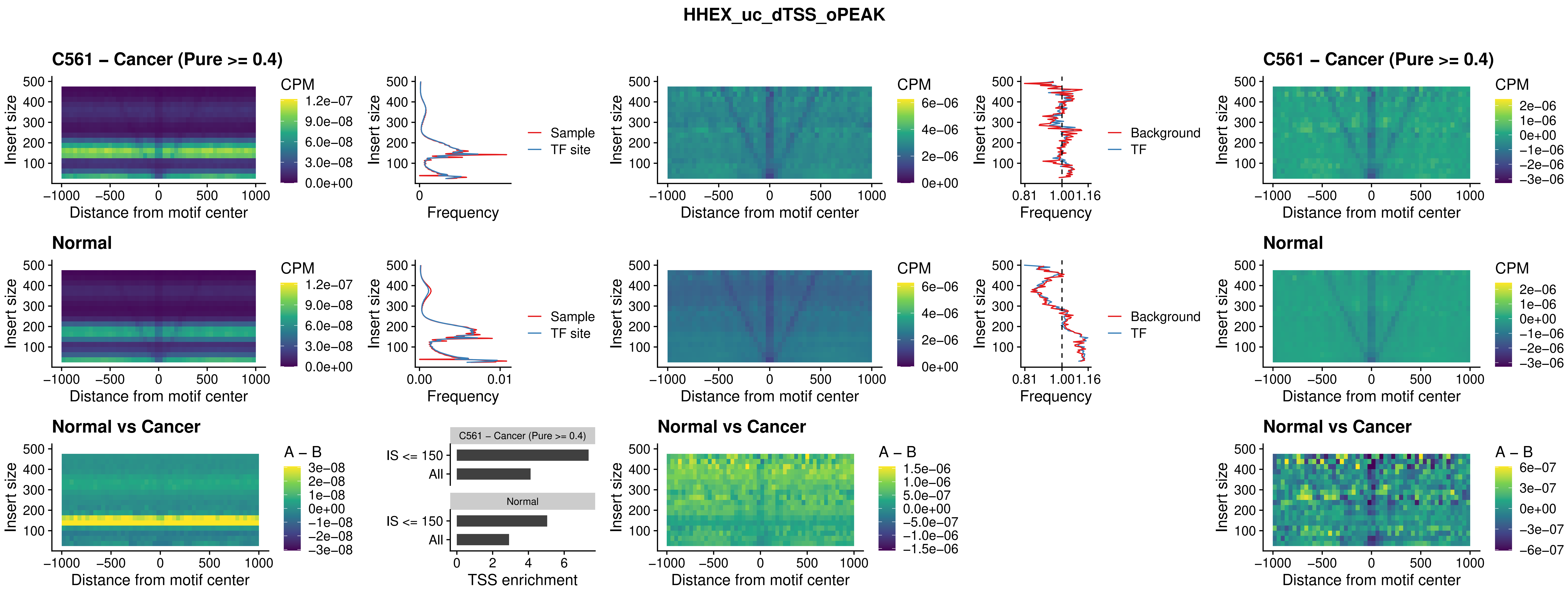

HHEX_uc_dTSS_oPEAK
C561 – Cancer (Pure >= 0.4)
Normal
Normal vs Cancer
Insert size
Distance from motif center
Frequency
CPM
A – B
Sample
TF site
Background
TF
TSS enrichment
IS <= 150
All

### HOXA11_uc_dTSS_nPEAK

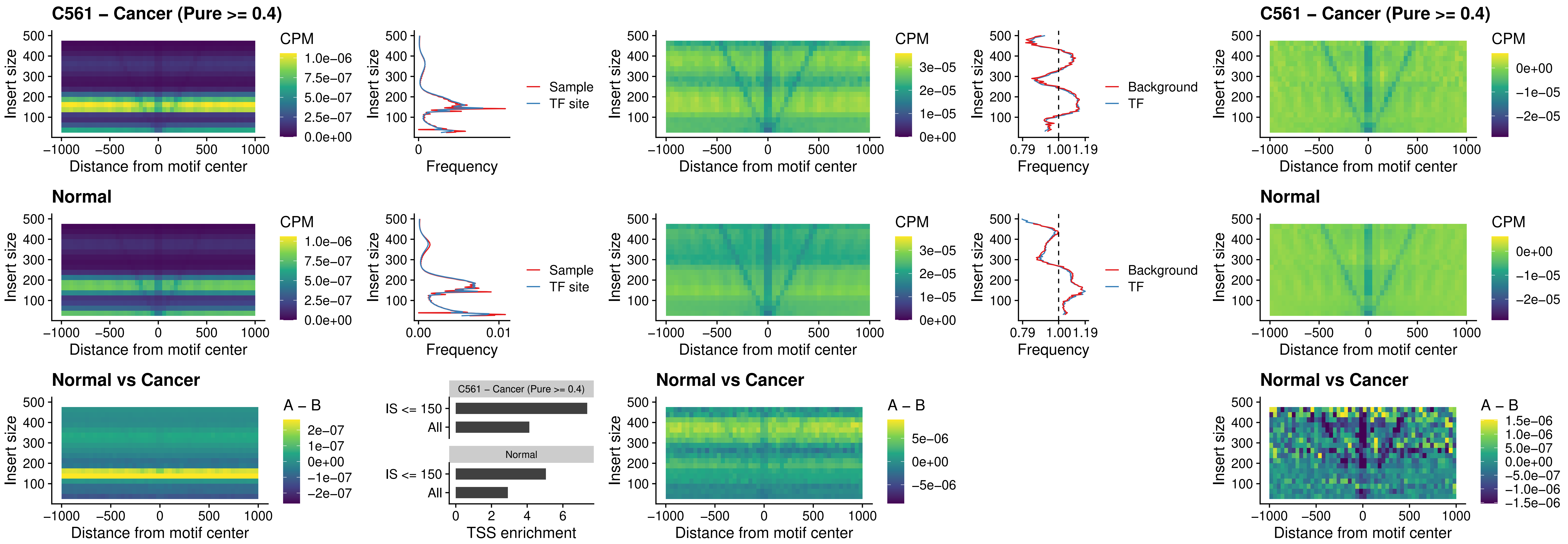

#### HOXA11_uc_cTSS_nPEAK

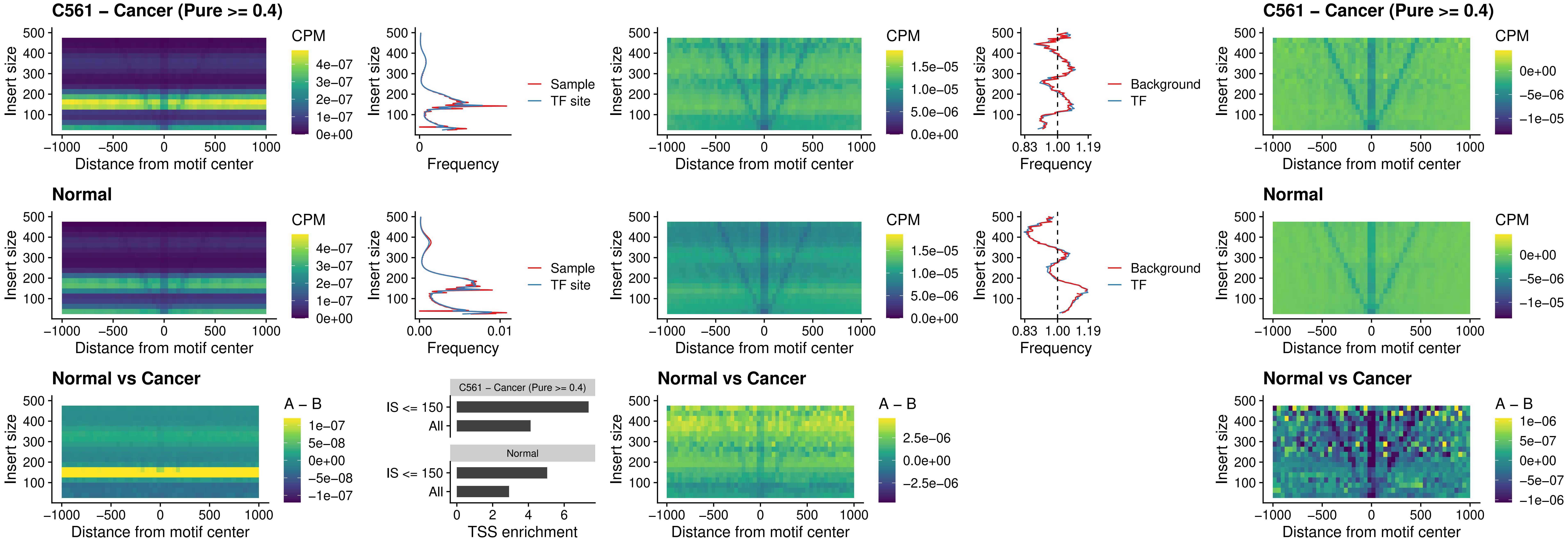

**ZFHX3_uc_dTSS_nPEAK**

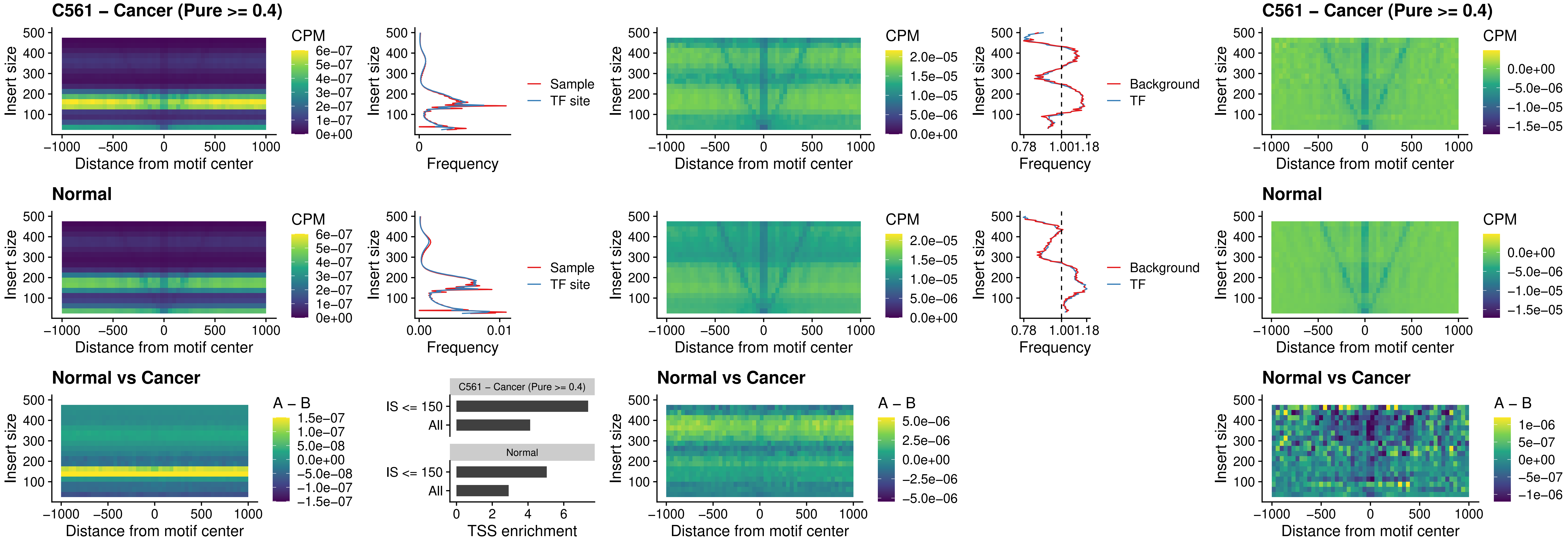

### HOXB4_uc_dTSS_nPEAK

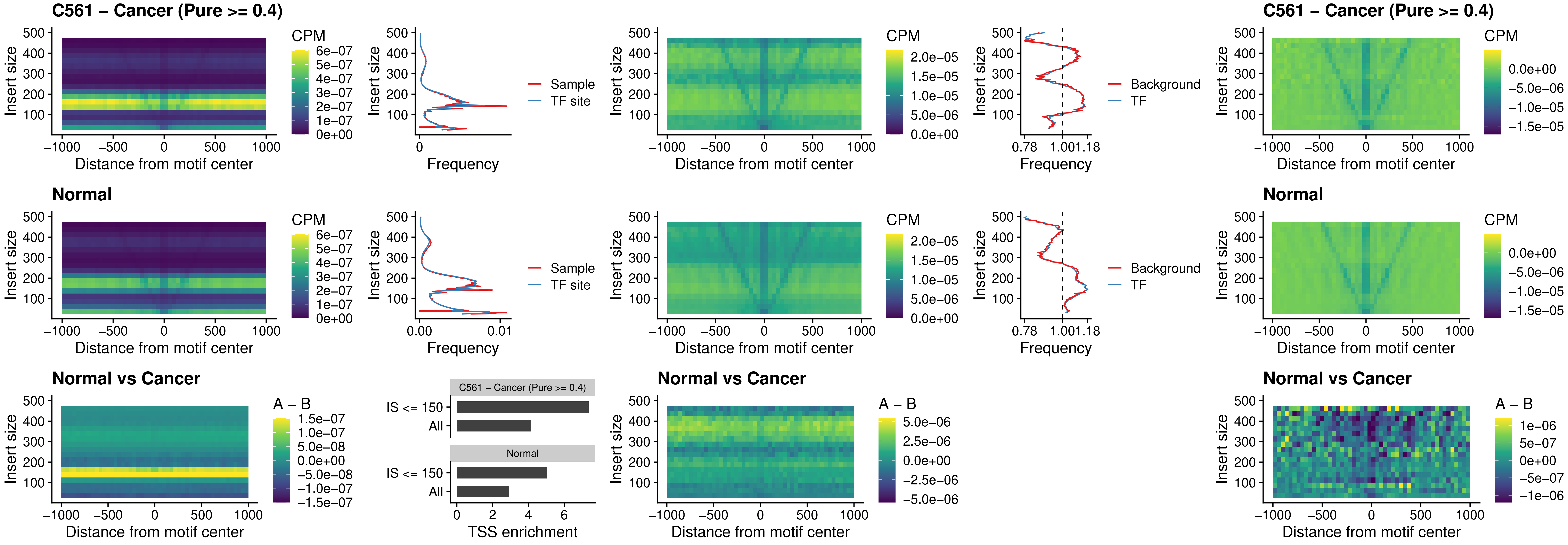

### HOXC5_uc_dTSS_nPEAK

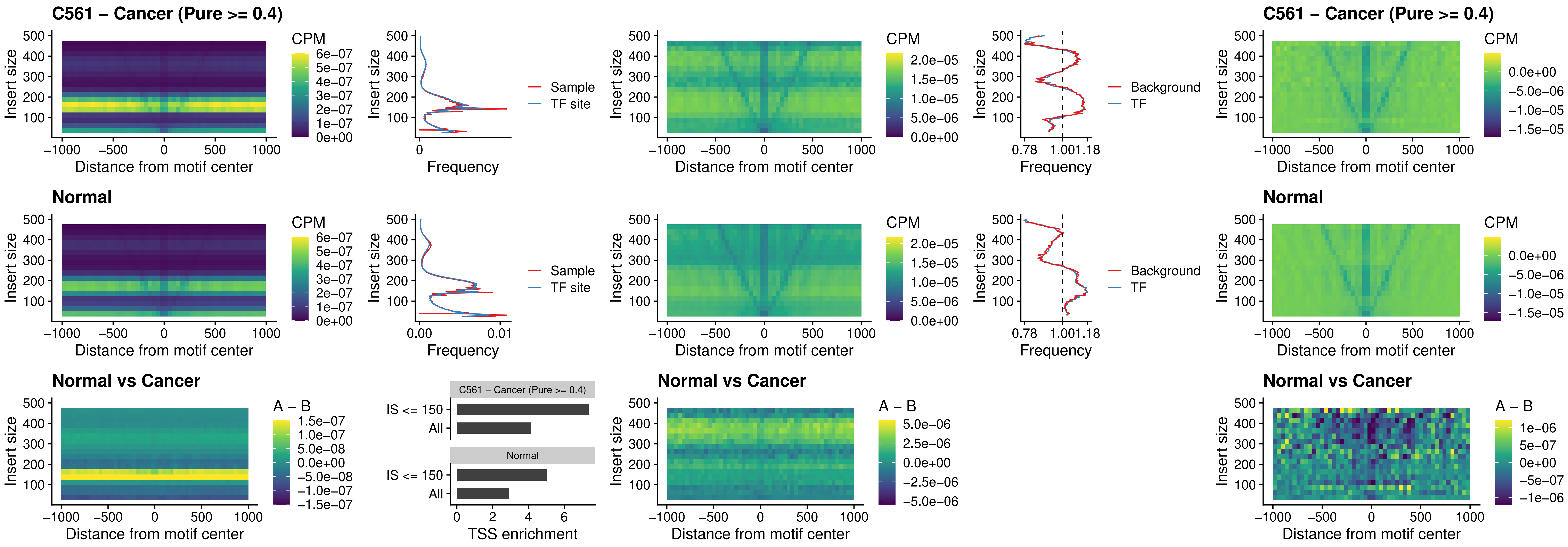

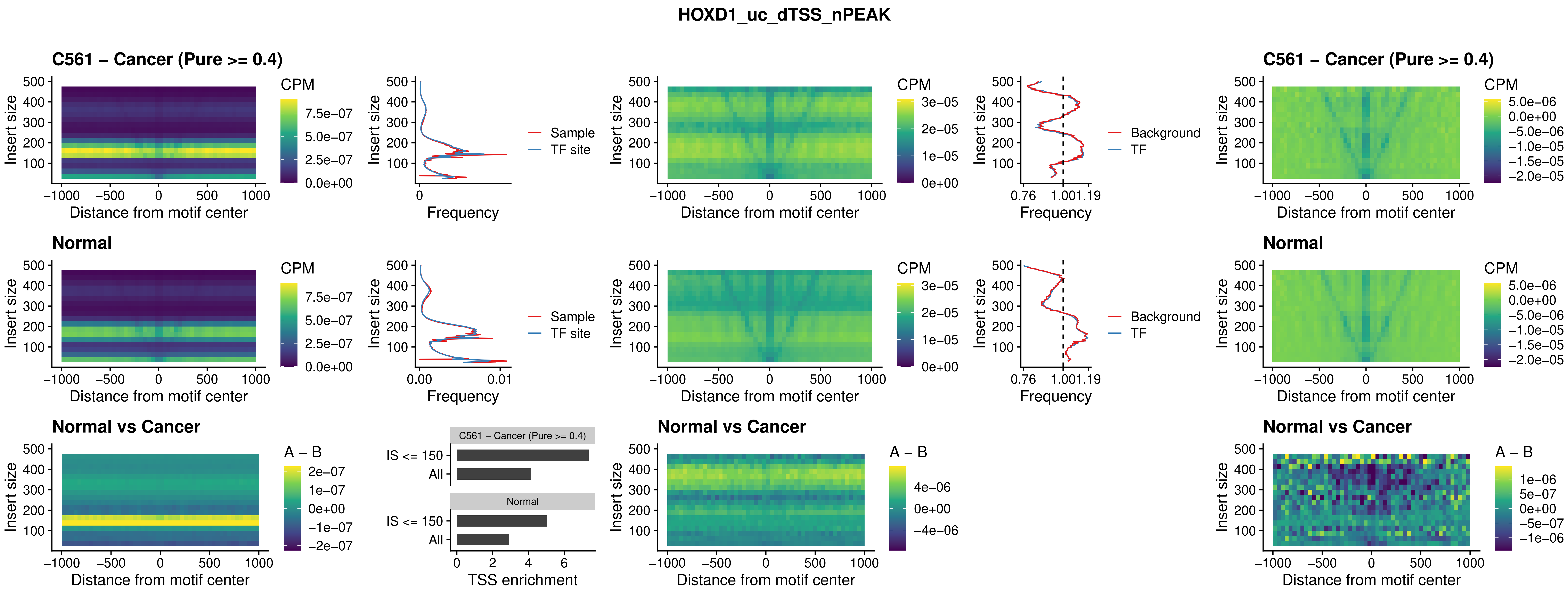
HOXD1_uc_dTSS_nPEAK
C561 – Cancer (Pure >= 0.4)
Normal
Normal vs Cancer
Insert size
Distance from motif center
Frequency
CPM
A – B
Sample
TF site
Background
TF
C561 – Cancer (Pure >= 0.4)
Normal
IS <= 150
All
TSS enrichment

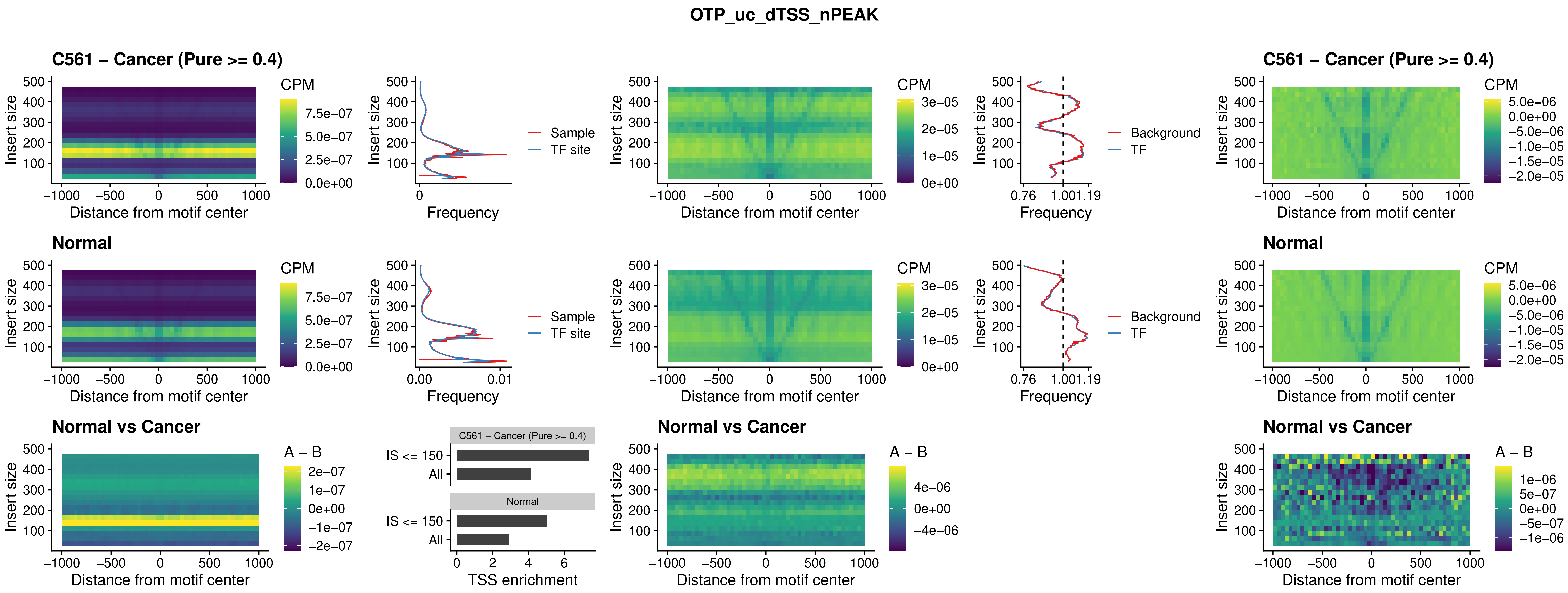
OTP_uc_dTSS_nPEAK
C561 – Cancer (Pure >= 0.4)
Normal
Normal vs Cancer
Insert size
Distance from motif center
Frequency
CPM
A – B
Sample
TF site
Background
TF
C561 – Cancer (Pure >= 0.4)
Normal
IS <= 150
All
TSS enrichment

### FOXL1_uc_dTSS_oPEAK

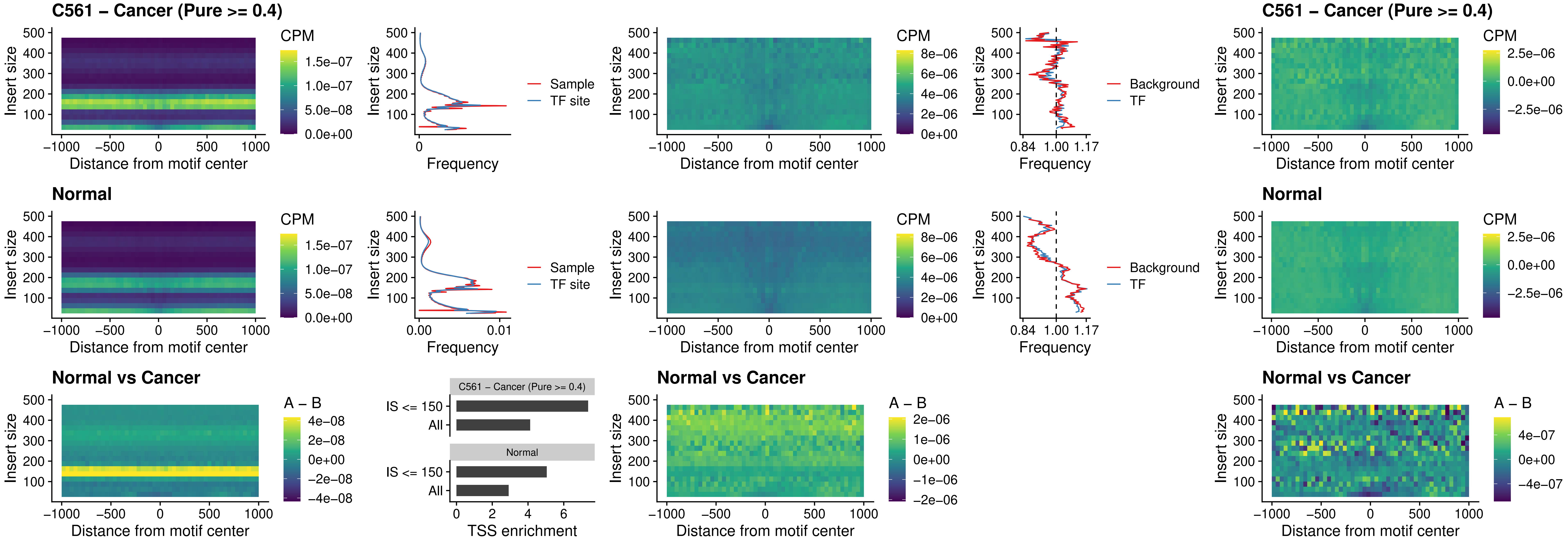

### ZNF32_uc_dTSS_oPEAK

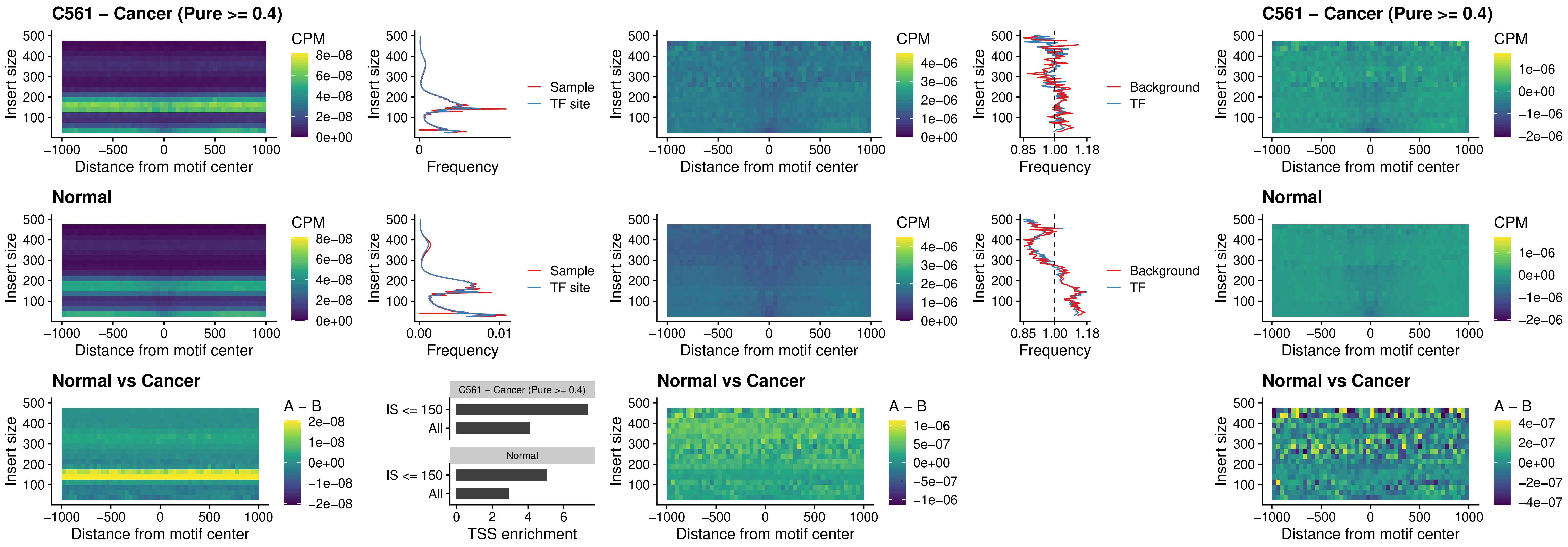

### FOXL1_uc_dTSS_nPEAK

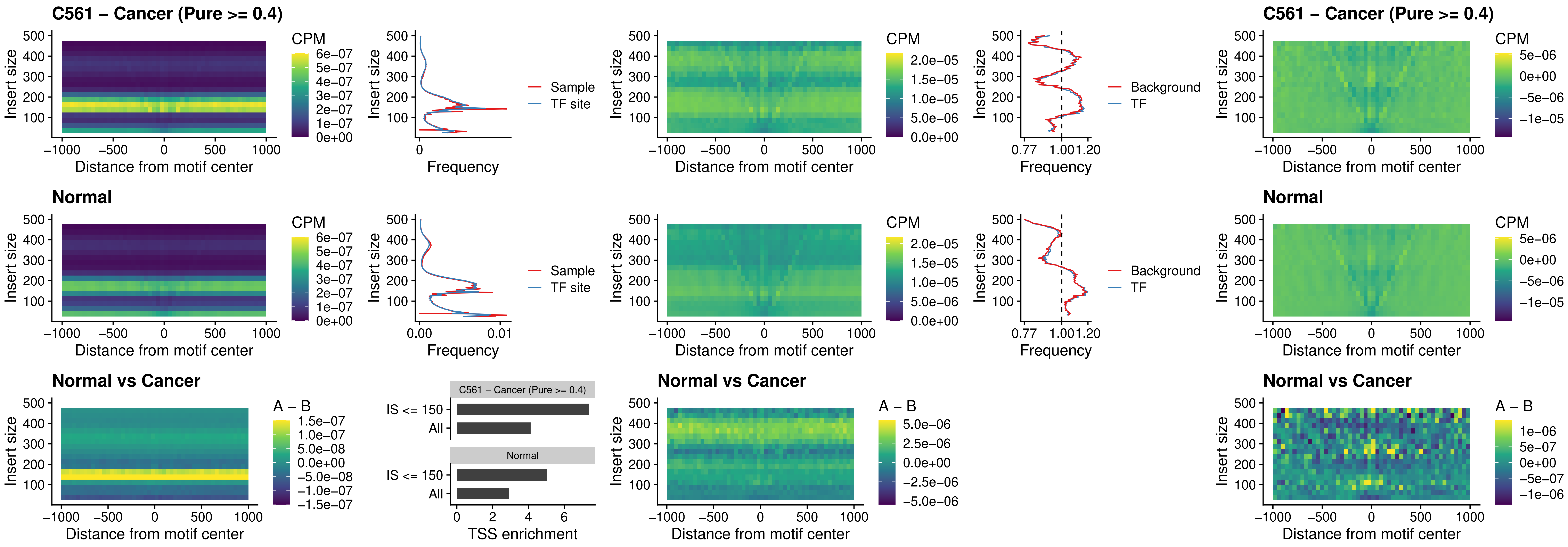

#### ZNF32_uc_dTSS_nPEAK

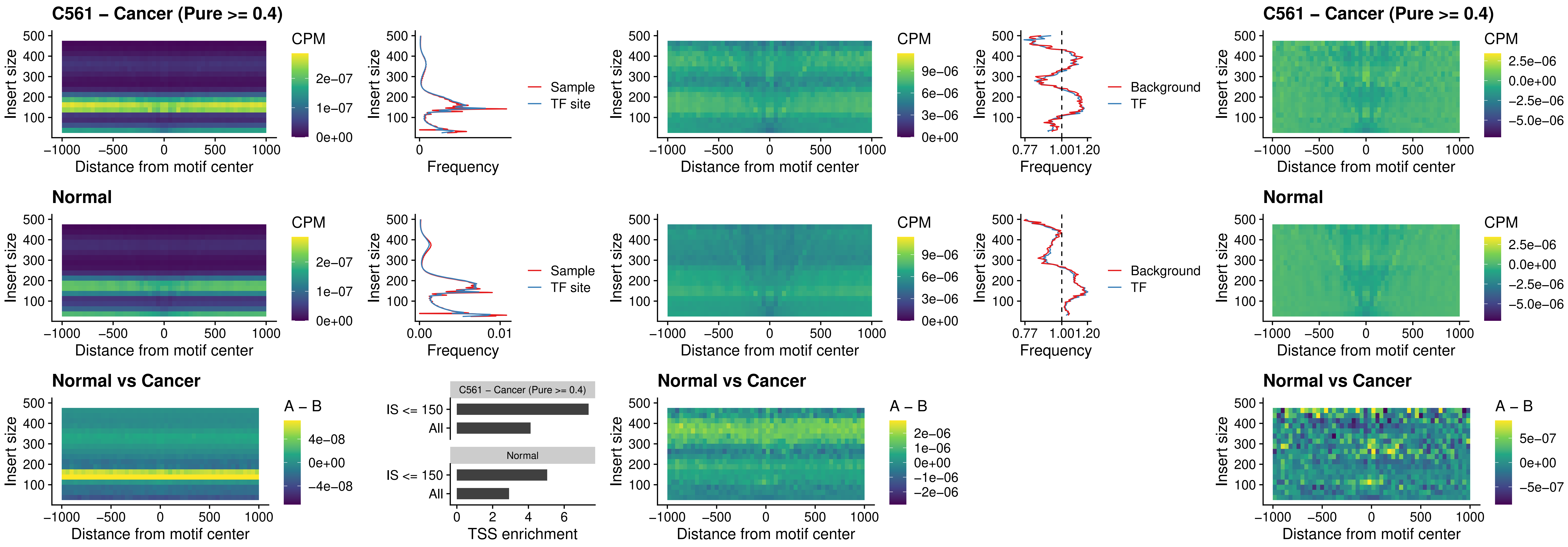

### DLX5_uc_dTSS_nPEAK

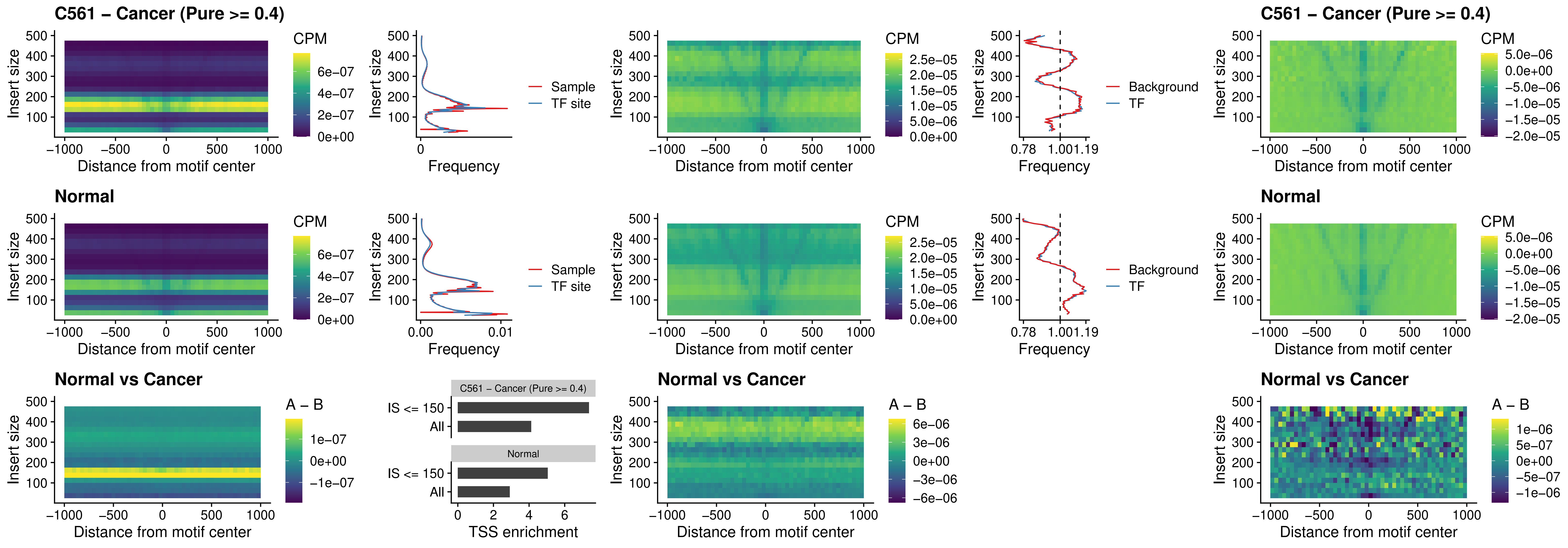

### DMRT2_uc_dTSS_nPEAK

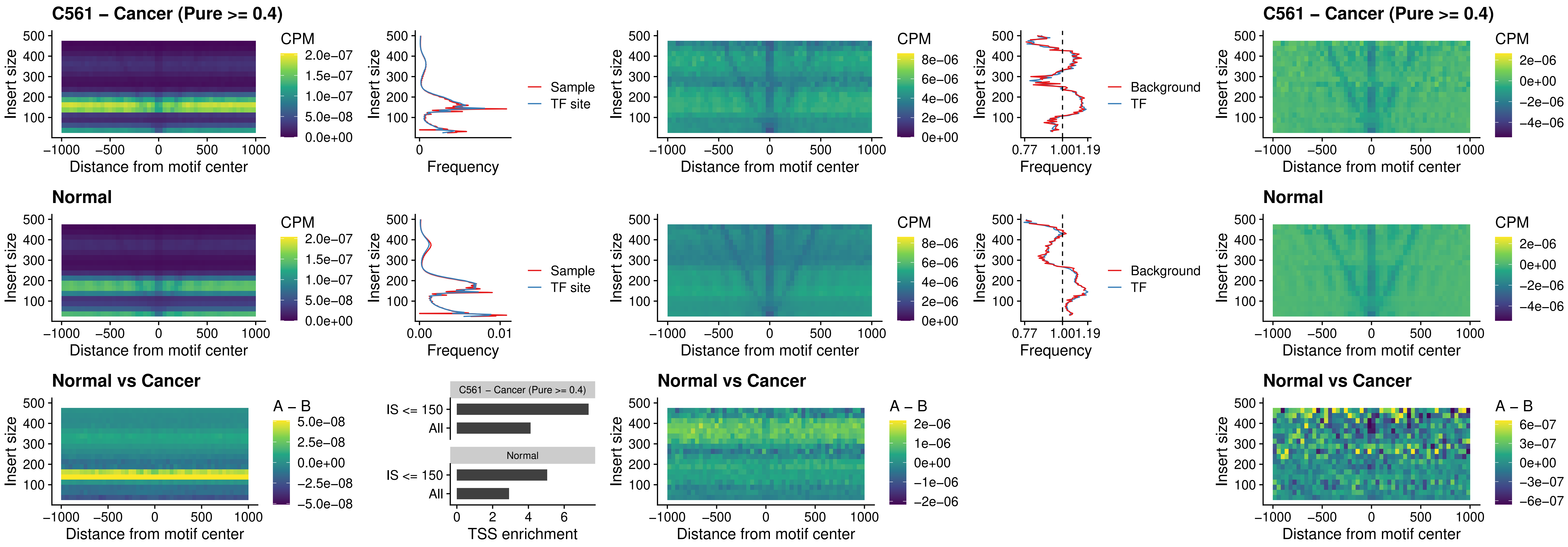

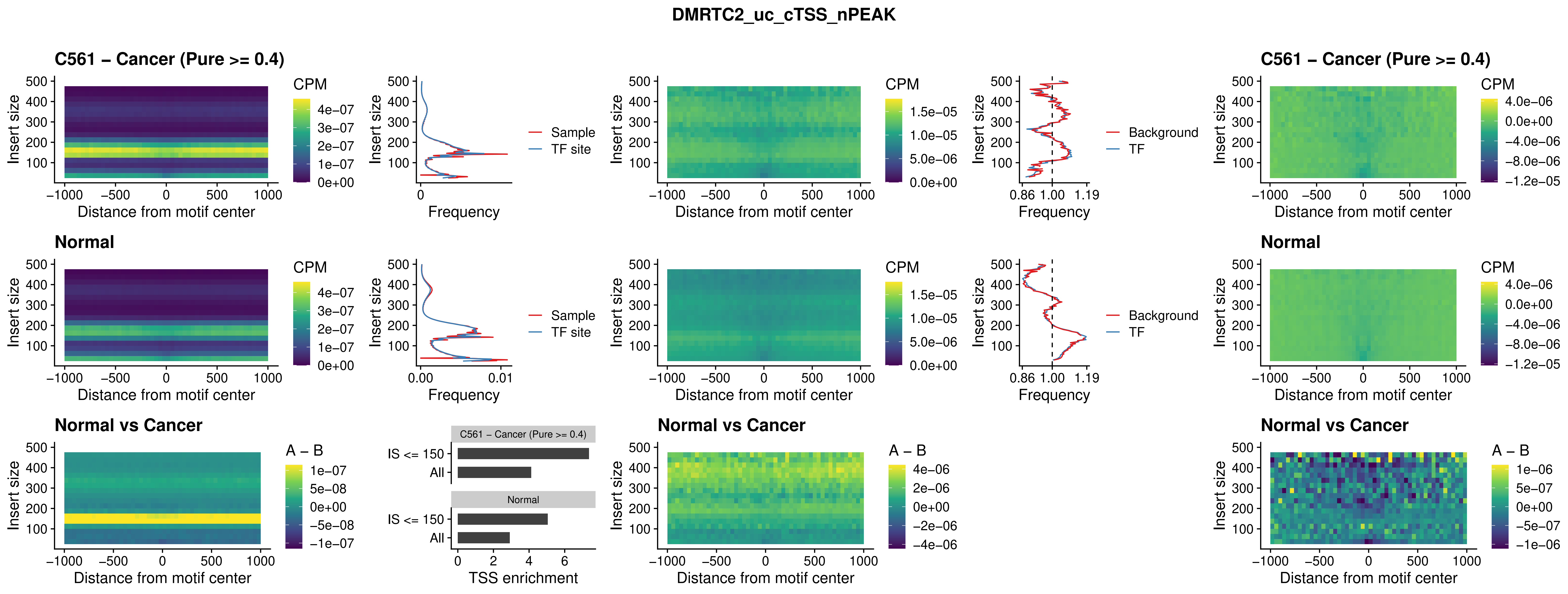
DMRTC2_uc_cTSS_nPEAK
C561 – Cancer (Pure >= 0.4)
Normal
Normal vs Cancer
Insert size
Distance from motif center
Frequency
CPM
A – B
Sample
TF site
Background
TF
C561 – Cancer (Pure >= 0.4)
Normal
IS <= 150
All
TSS enrichment

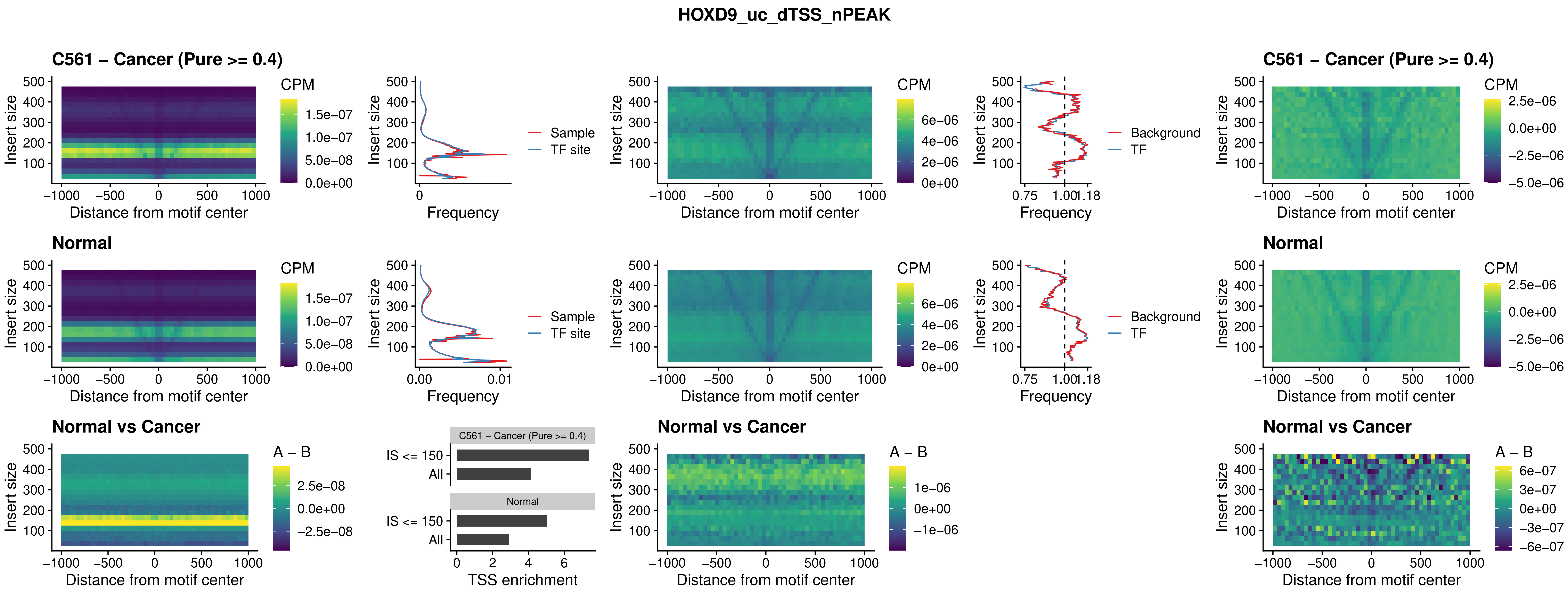
HOXD9_uc_dTSS_nPEAK
C561 – Cancer (Pure >= 0.4)
Normal
Normal vs Cancer
Insert size
Distance from motif center
Frequency
CPM
A – B
Sample
TF site
Background
TF
C561 – Cancer (Pure >= 0.4)
Normal
IS <= 150
All
TSS enrichment

### HOXD4_uc_dTSS_nPEAK

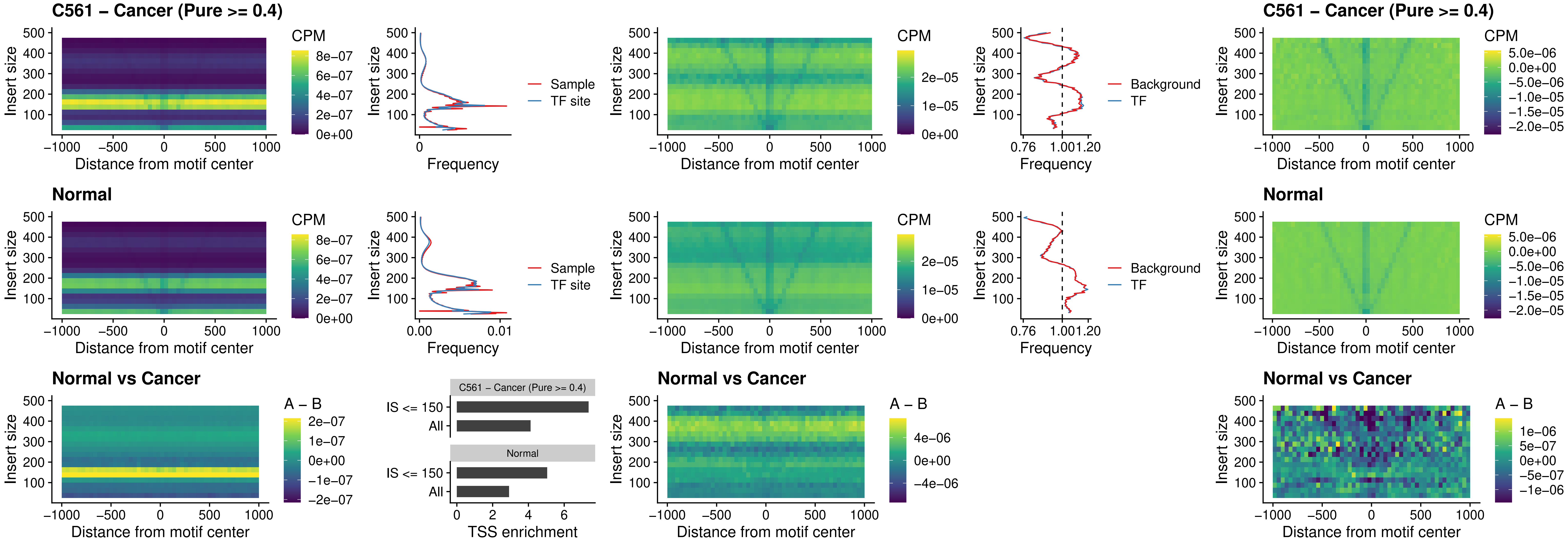

#### FOXP3_uc_cTSS_nPEAK

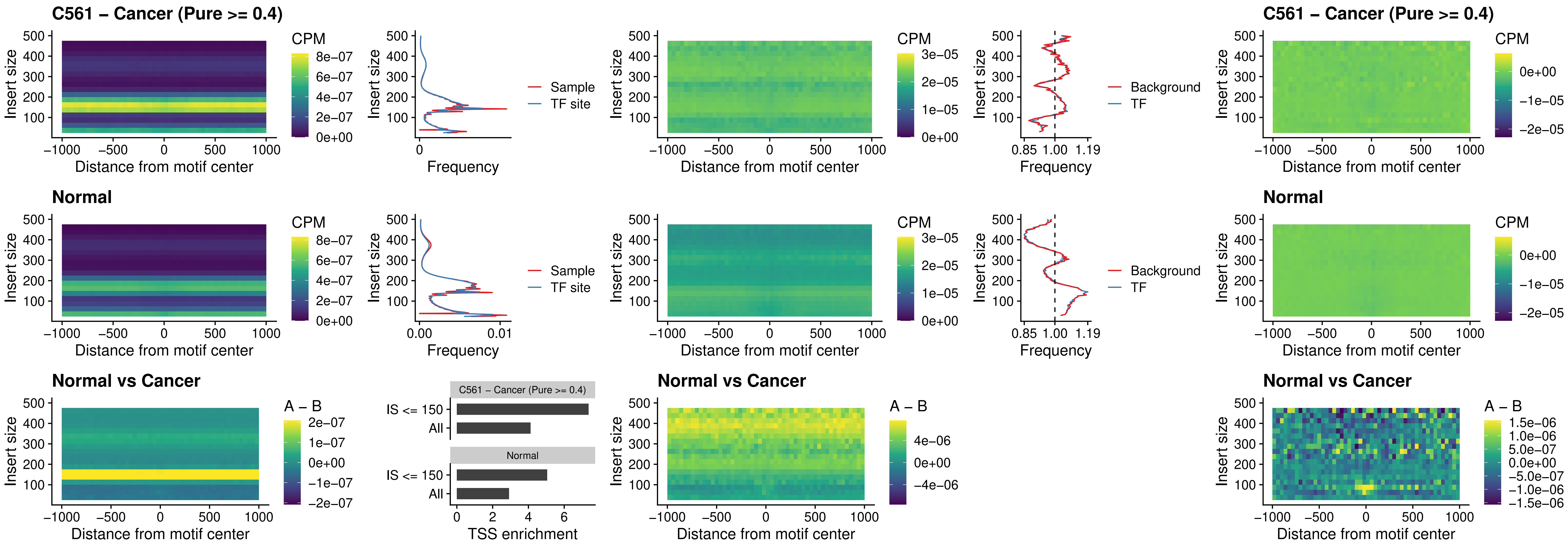

#### FOXD4L6_uc_cTSS_nPEAK

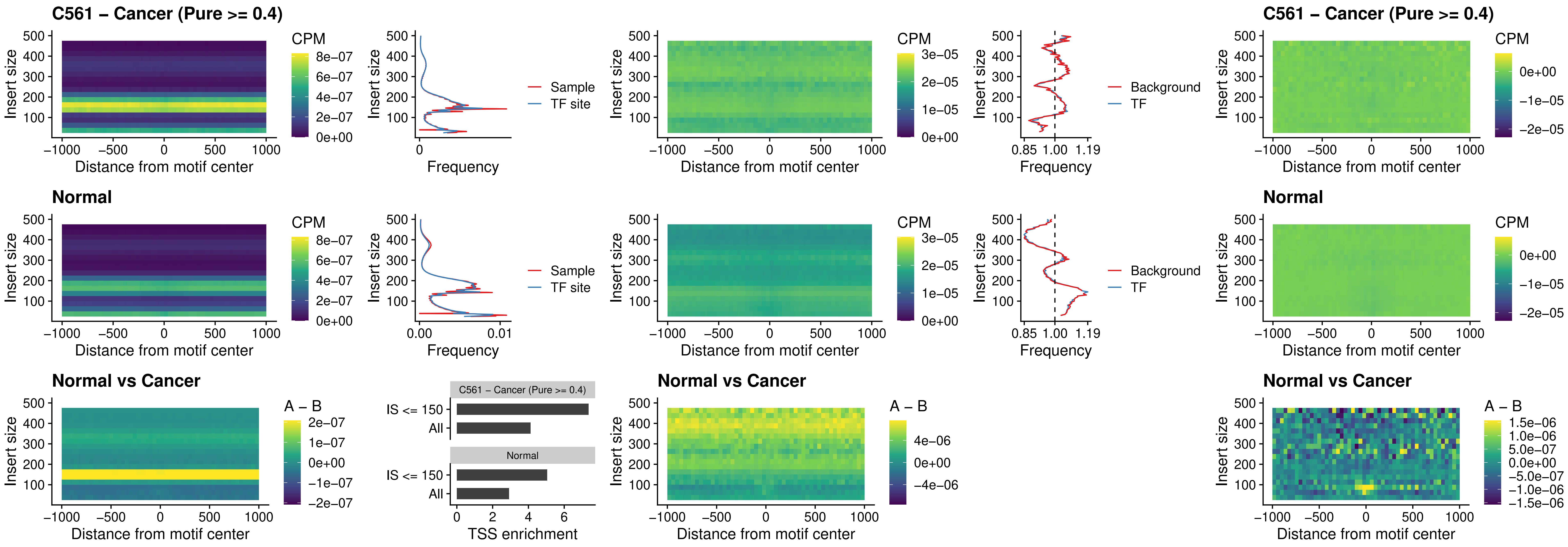

### FOXD4L5_uc_cTSS_nPEAK

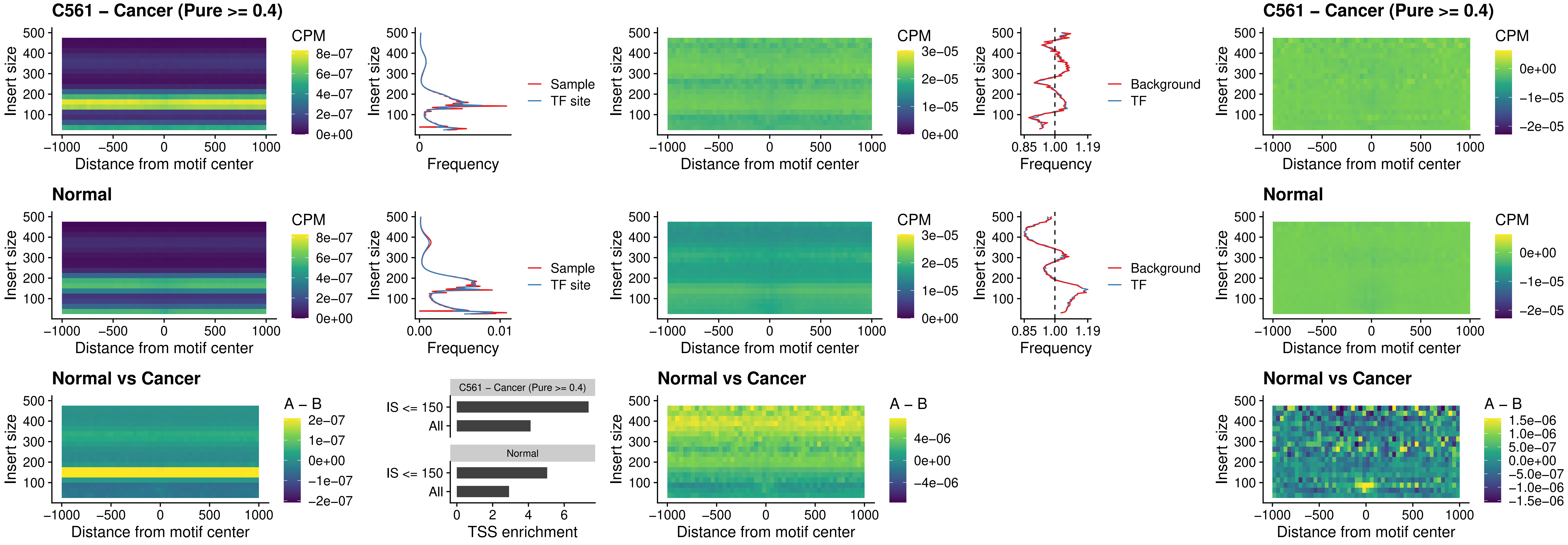

### FOXD4L4_uc_cTSS_nPEAK

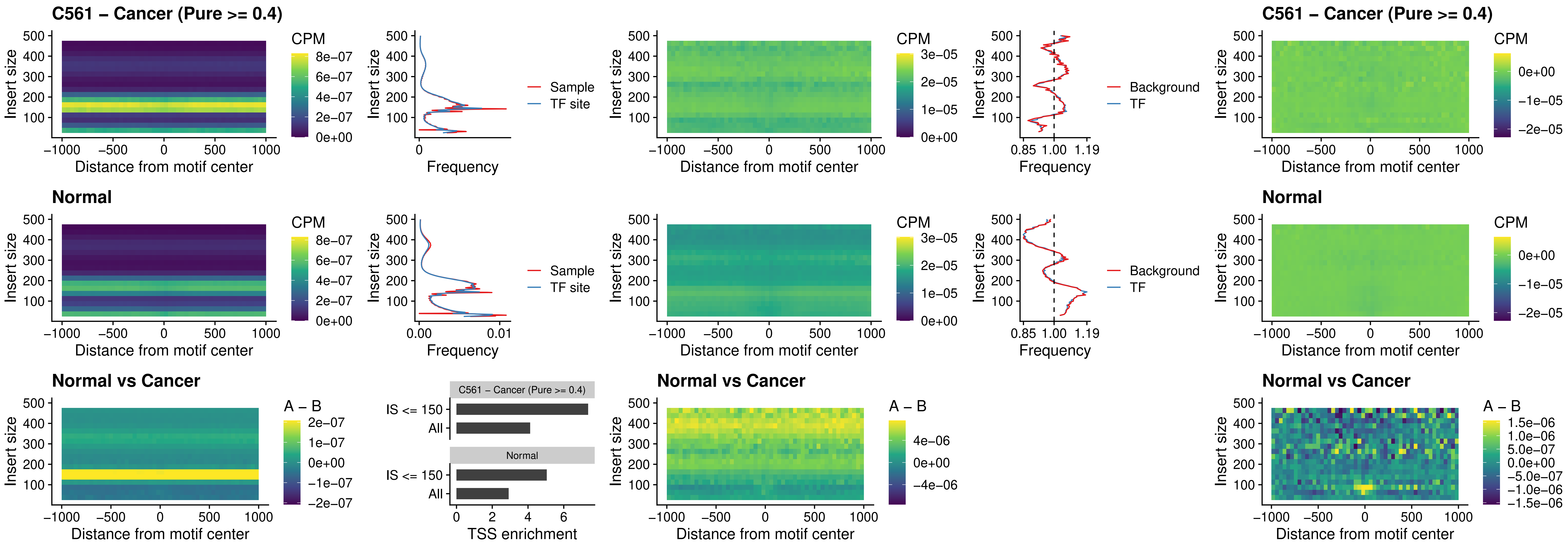

### FOXD4L2_uc_cTSS_nPEAK

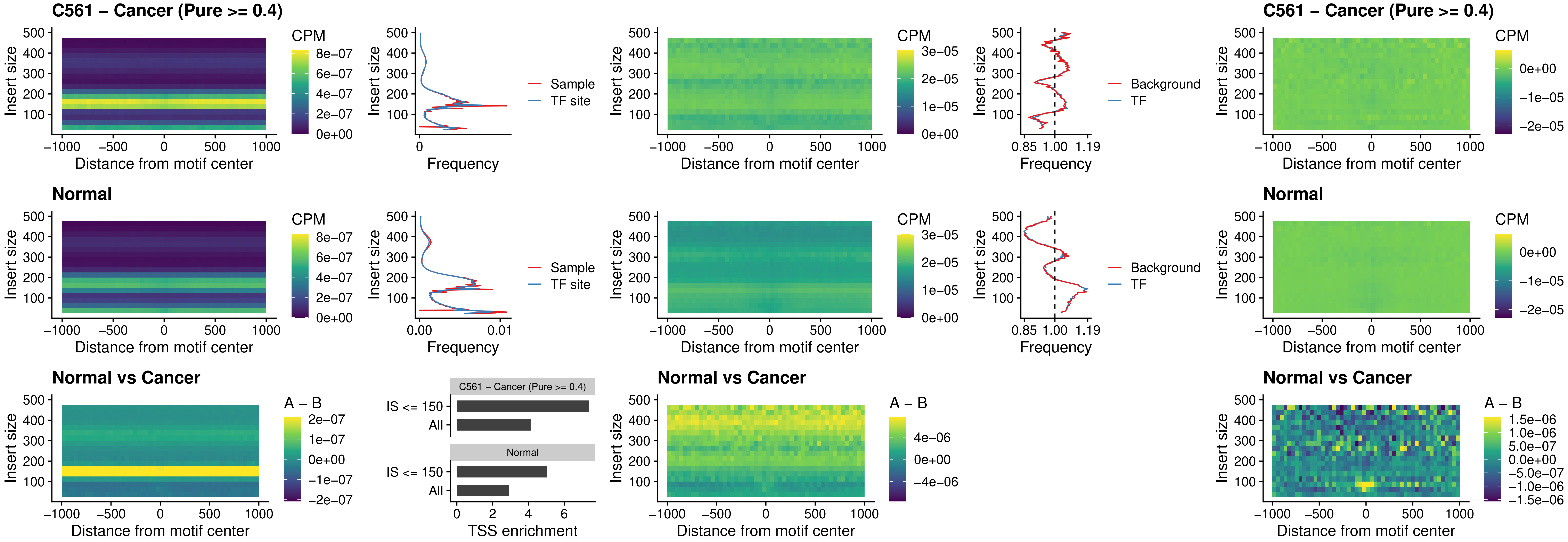

### FOXD4L3_uc_cTSS_nPEAK

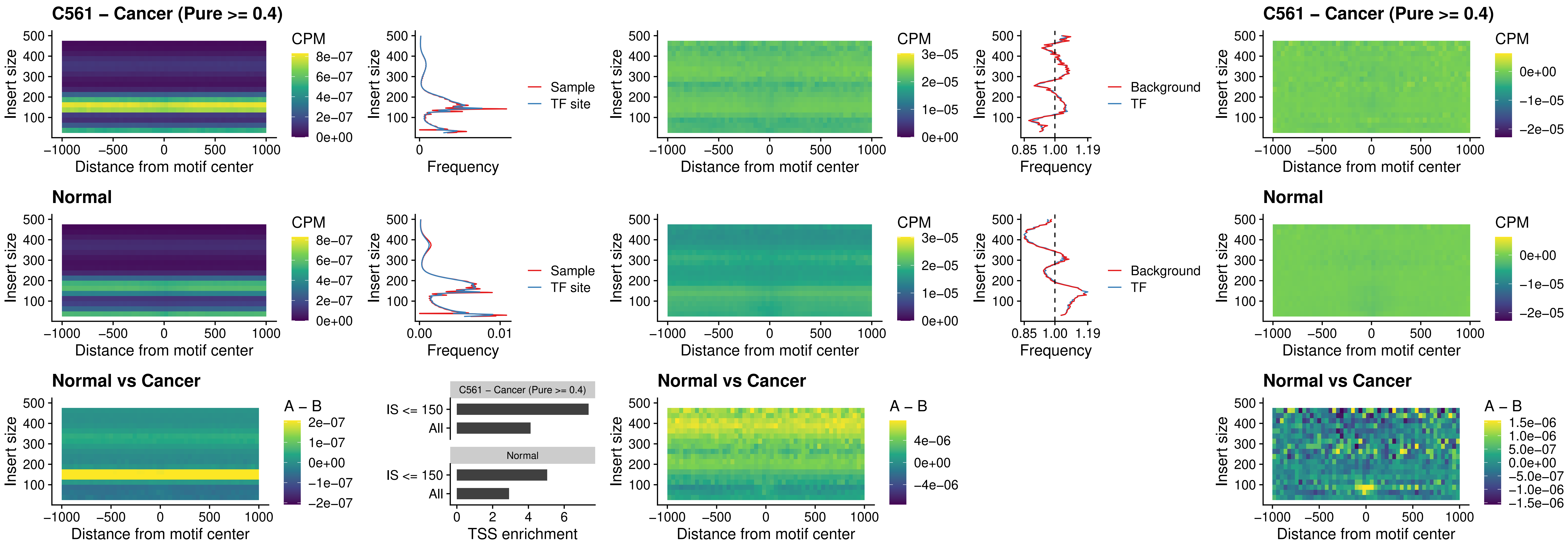

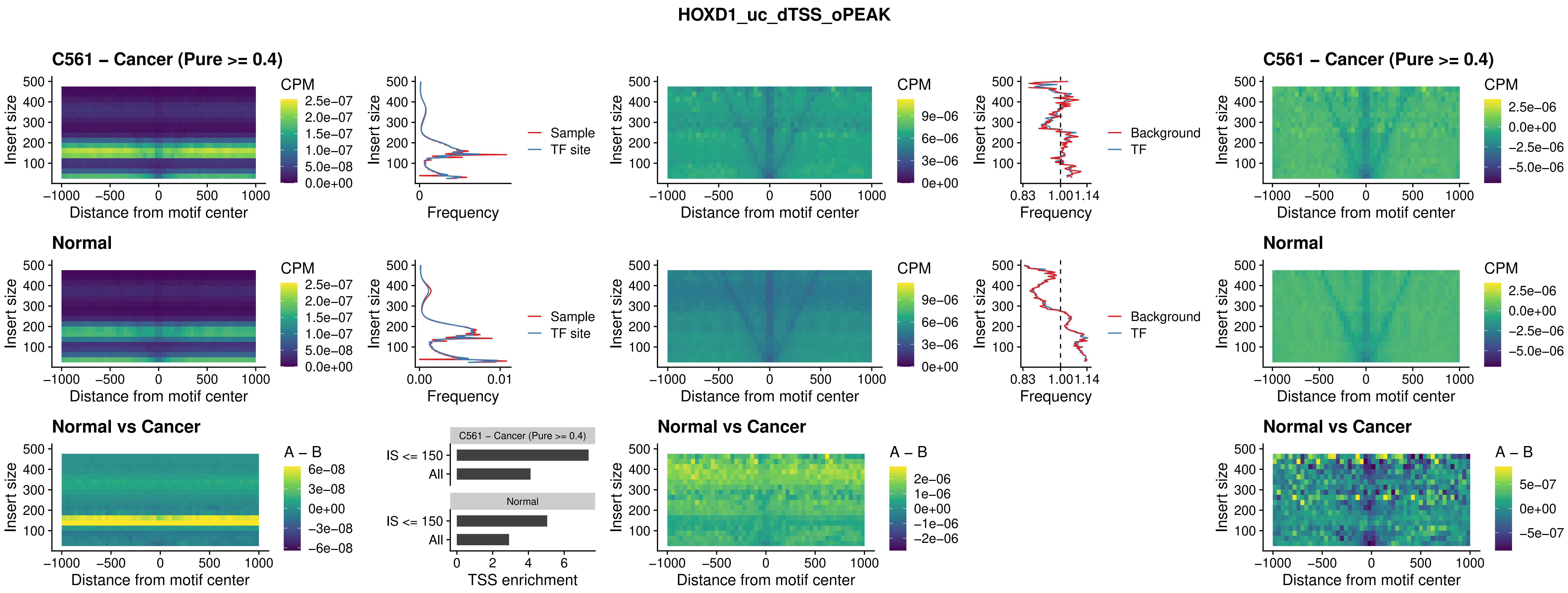
HOXD1_uc_dTSS_oPEAK
C561 – Cancer (Pure >= 0.4)
Normal
Normal vs Cancer
Insert size
Distance from motif center
Frequency
CPM
A – B
Sample
TF site
Background
TF
TSS enrichment
IS <= 150
All
C561 – Cancer (Pure >= 0.4)
Normal

### VAX1_uc_pTSS_nPEAK

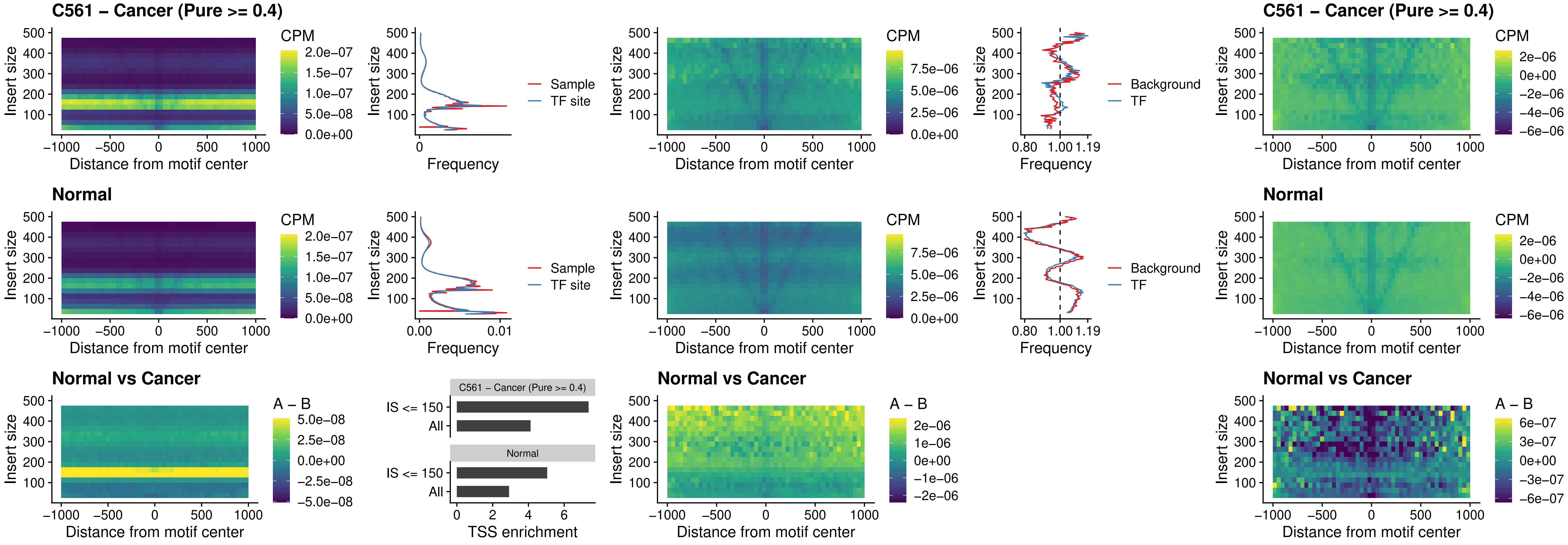

### DLX3_uc_pTSS_nPEAK

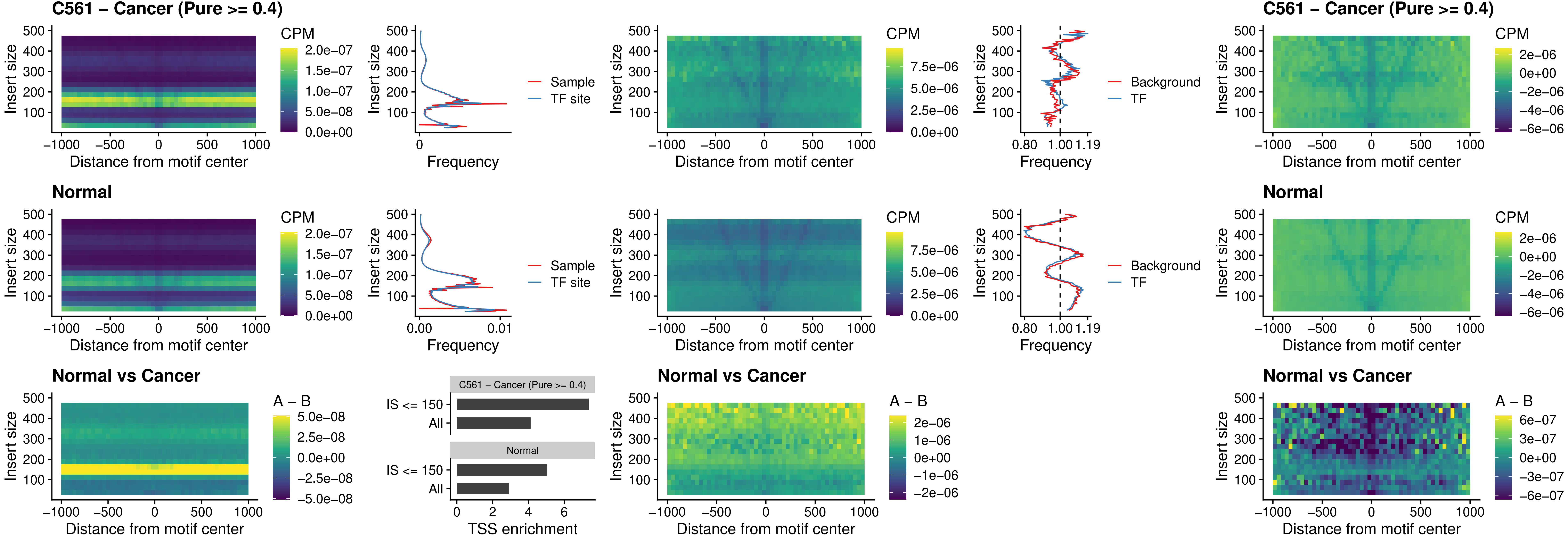

### HOXD3_uc_pTSS_nPEAK

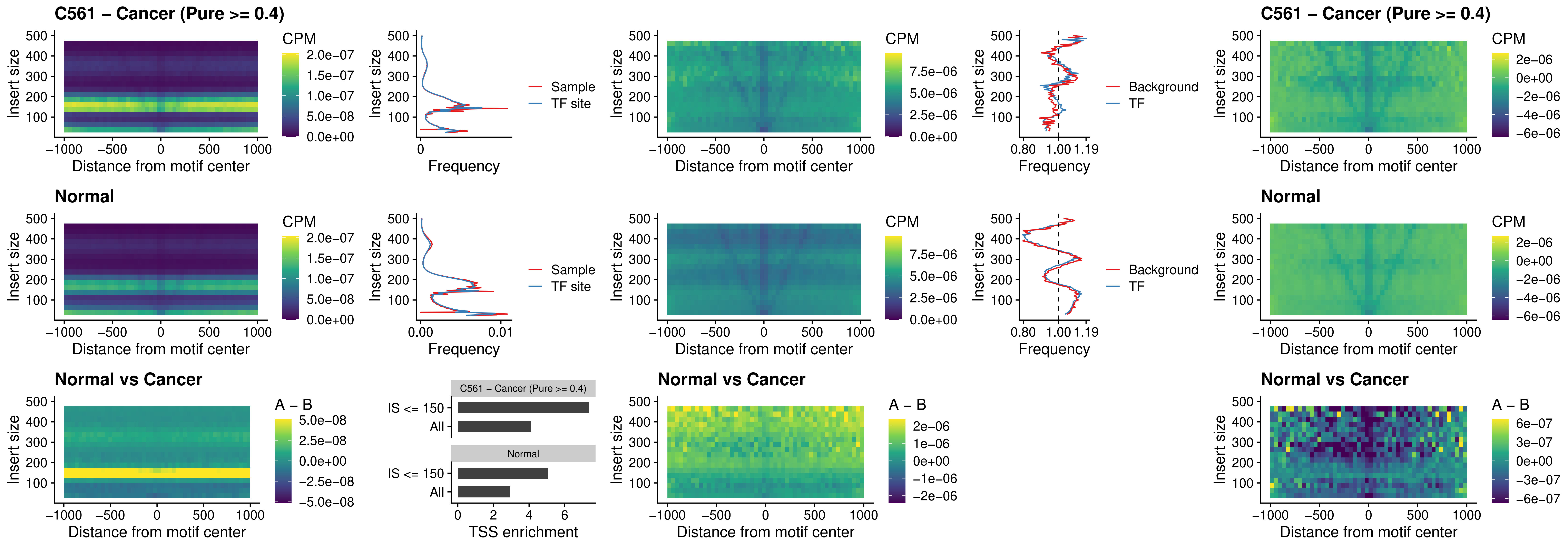

### PDX1_uc_dTSS_nPEAK

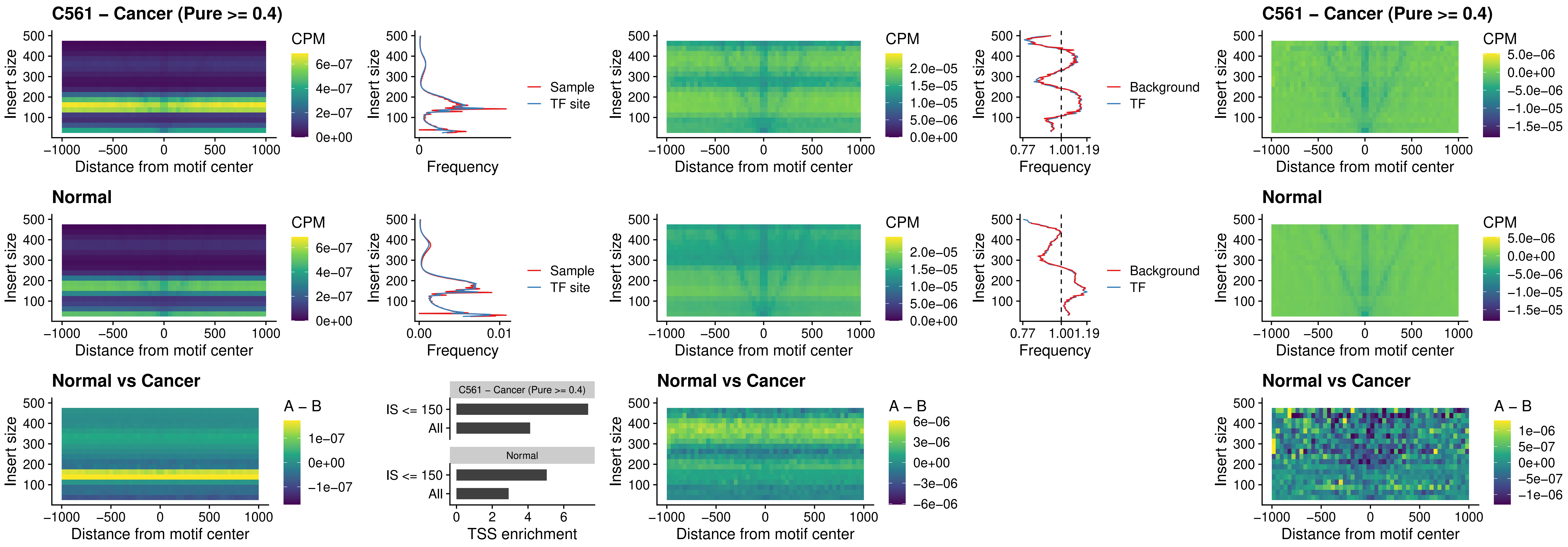

### VAX2_uc_dTSS_nPEAK

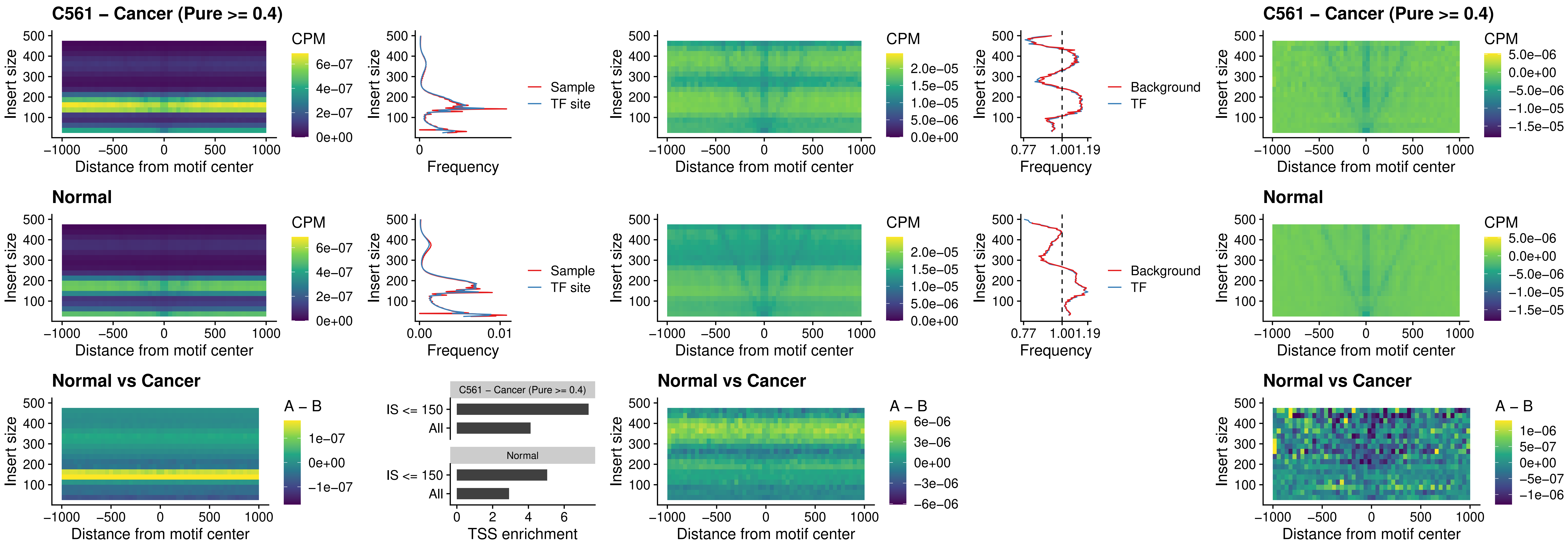

### SOX5_uc_dTSS_nPEAK

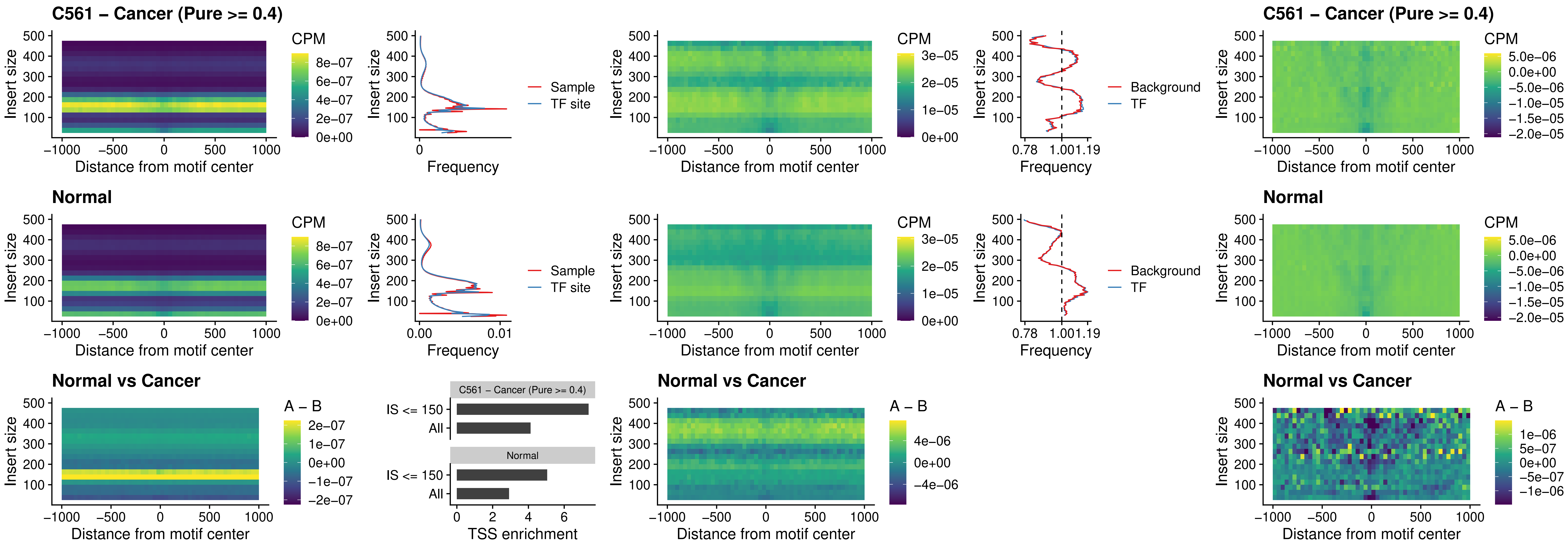

### SOX6_uc_dTSS_nPEAK

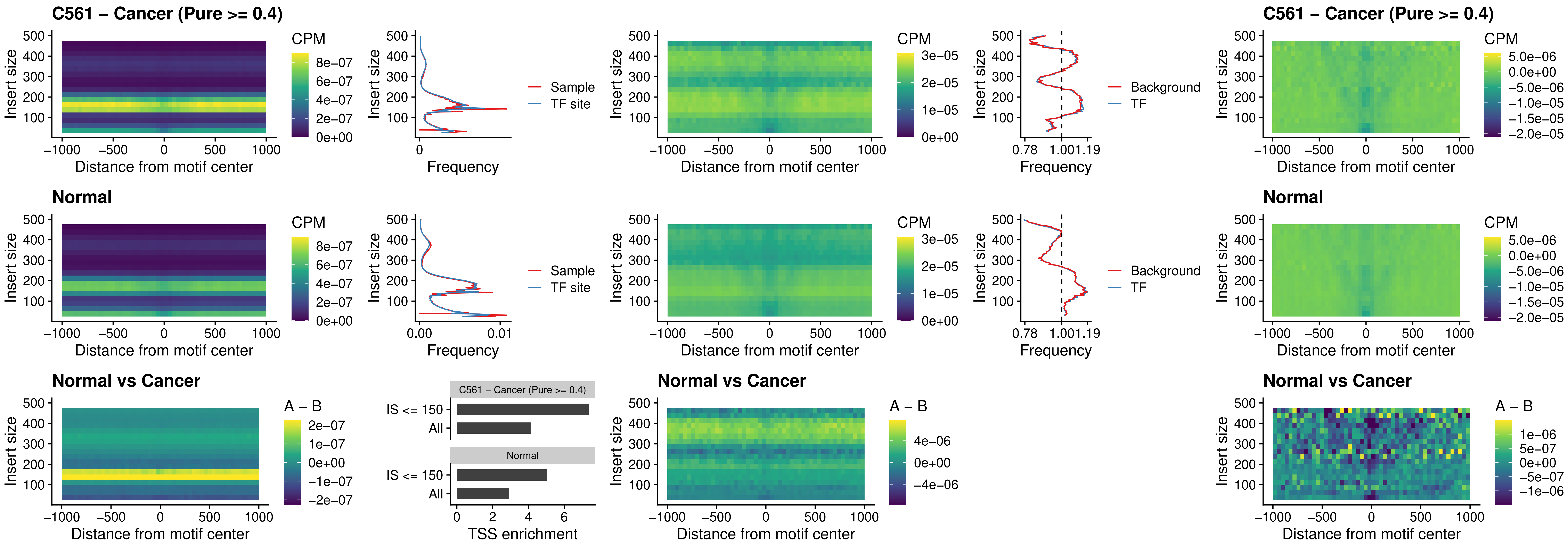

### JDP2_uc_cTSS_nPEAK

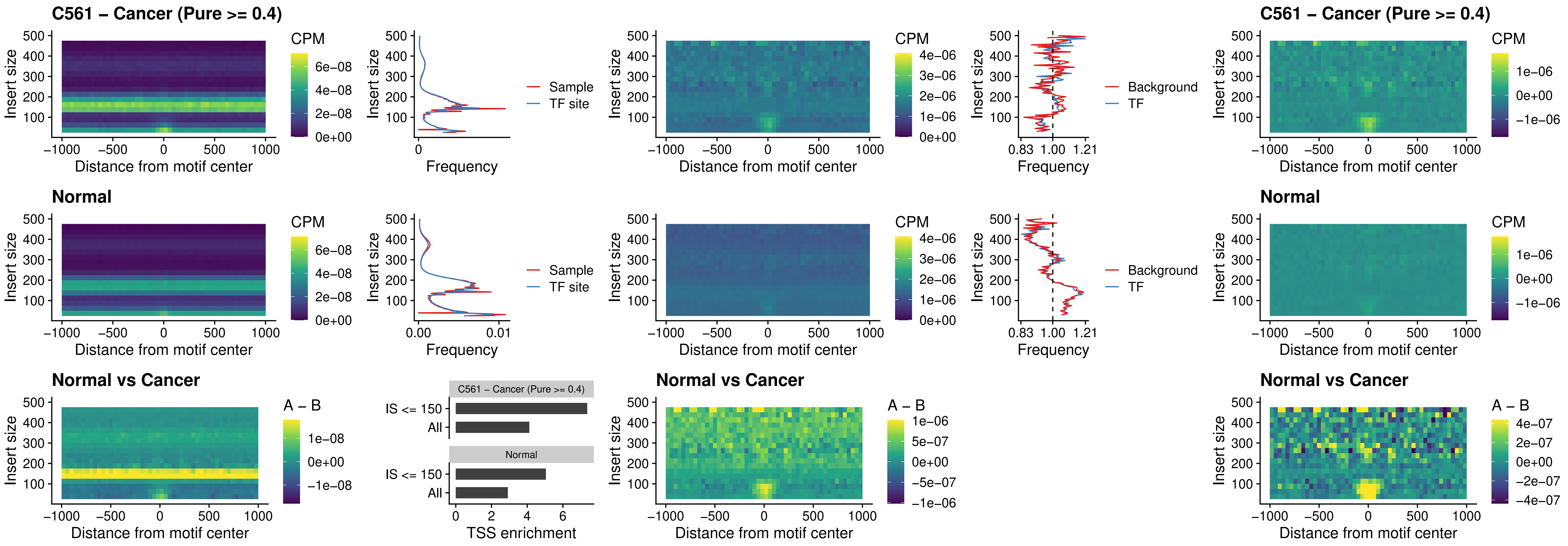

### PURA_uc_dTSS_oPEAK

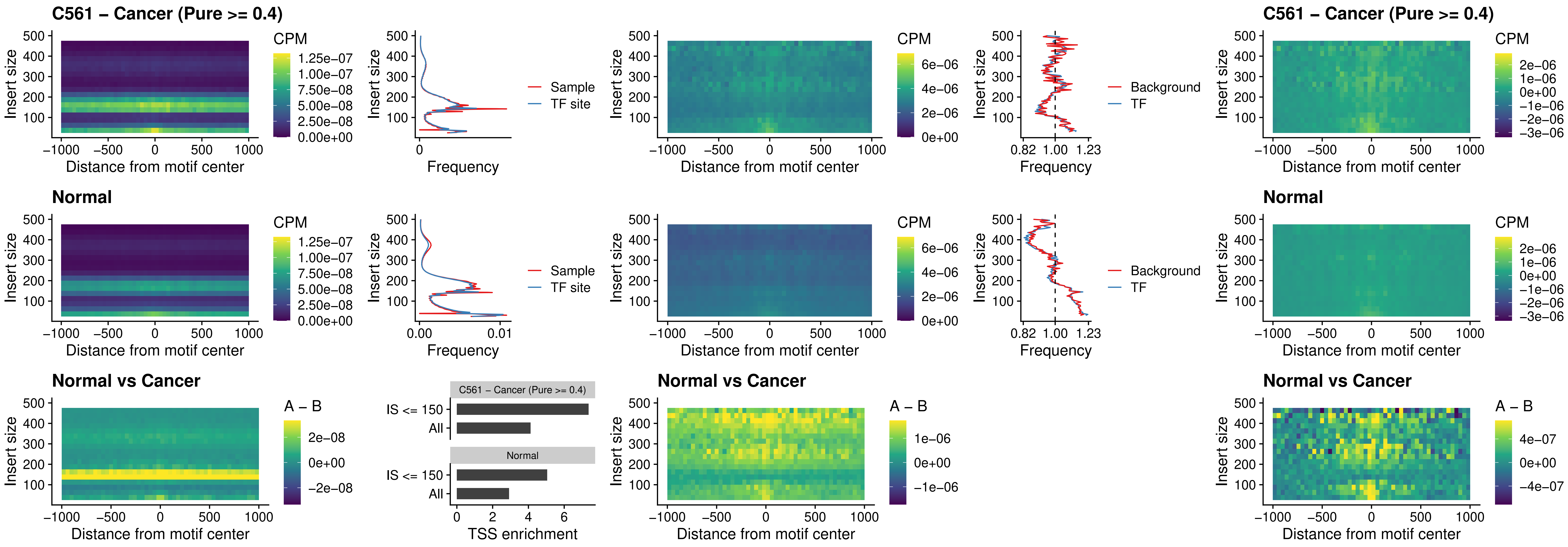

### HHEX_uc_dTSS_nPEAK

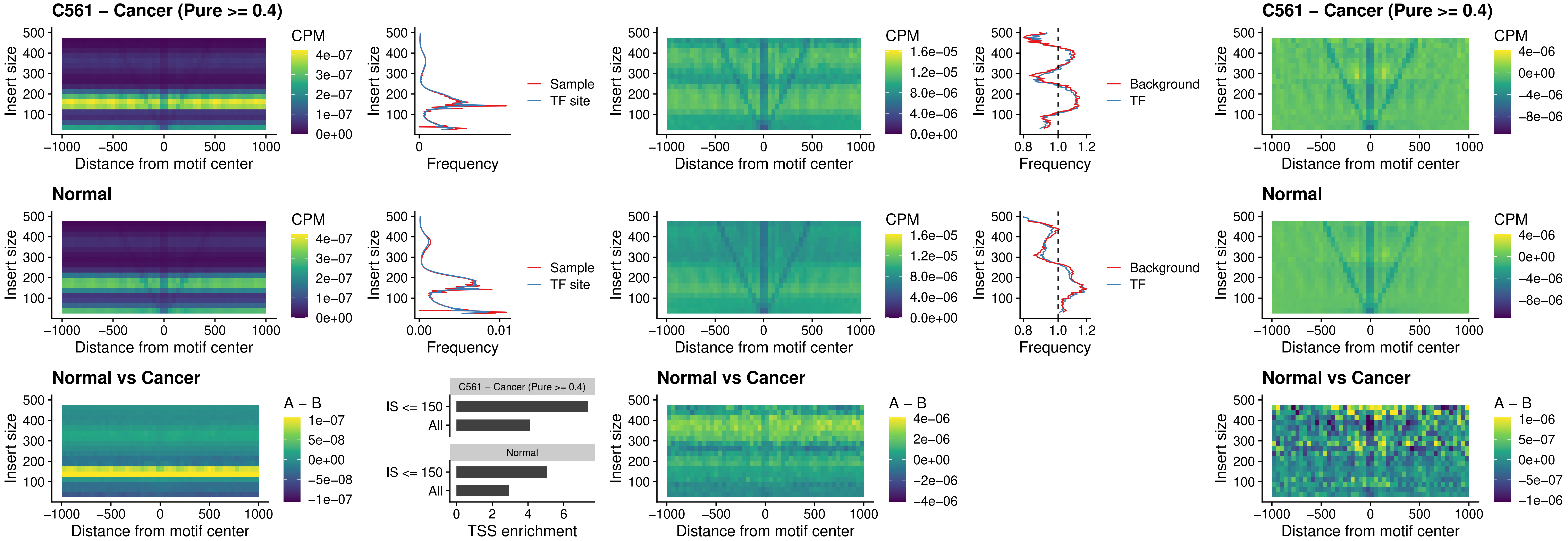

### HMG20B_uc_dTSS_nPEAK

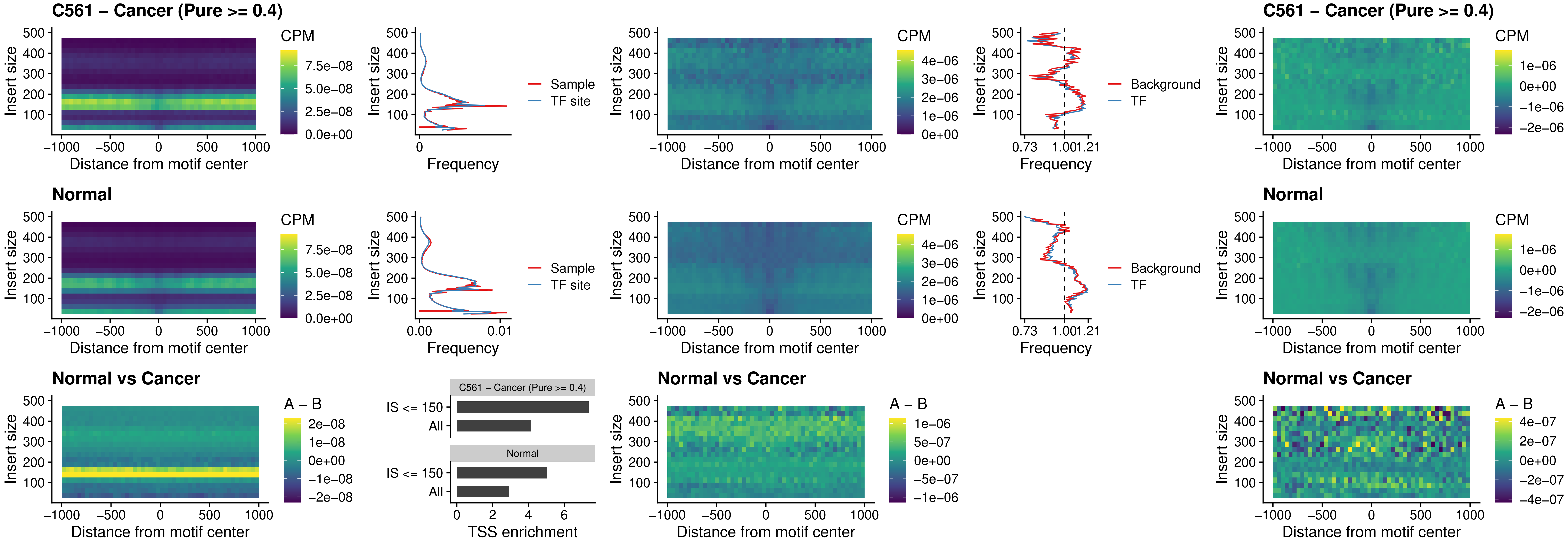

### FOXB2_uc_dTSS_nPEAK

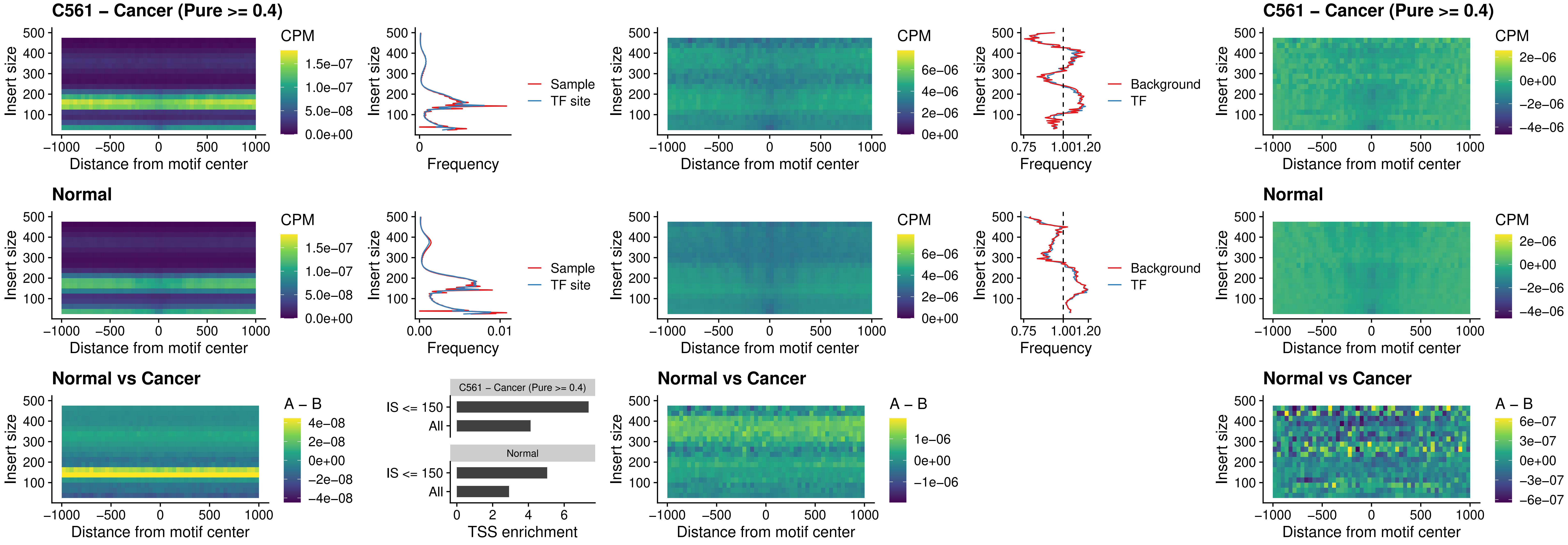

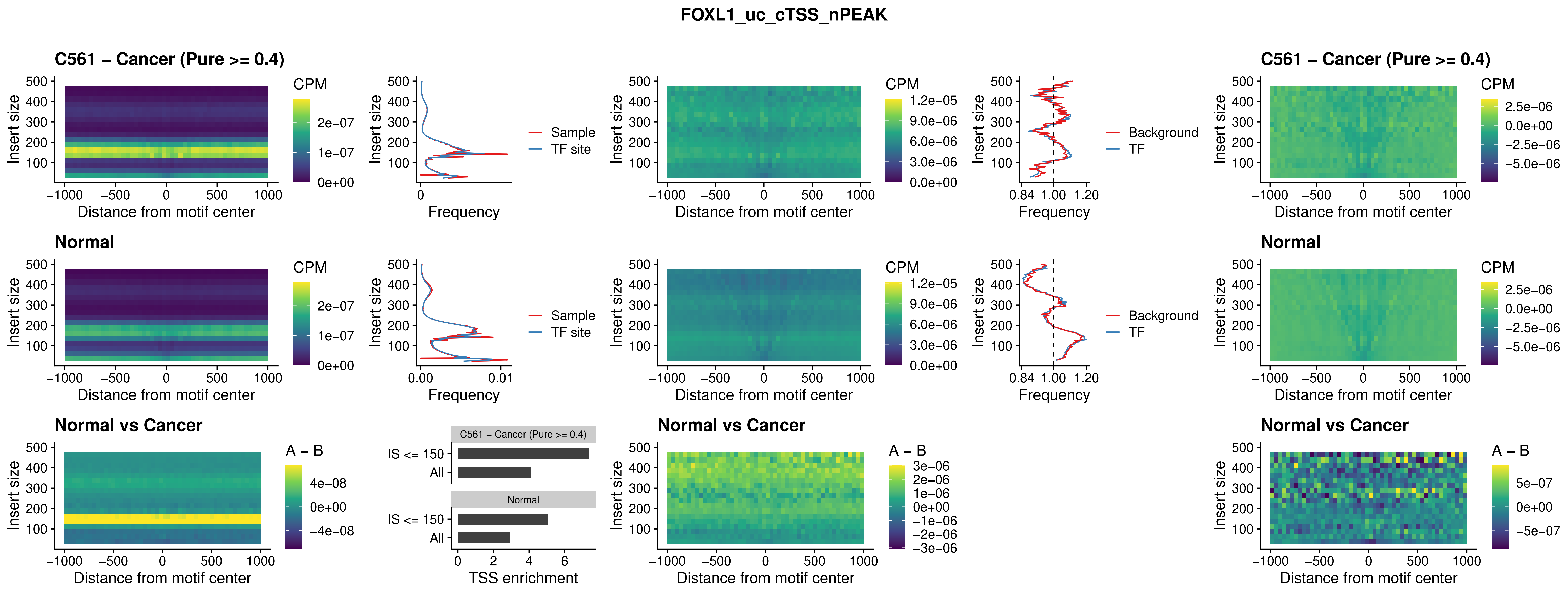
FOXL1_uc_cTSS_nPEAK
C561 – Cancer (Pure >= 0.4)
Normal
Normal vs Cancer
Insert size
Distance from motif center
Frequency
CPM
A – B
Sample
TF site
Background
TF
C561 – Cancer (Pure >= 0.4)
Normal
IS <= 150
All
TSS enrichment

### PBX3_uc_pTSS_oPEAK

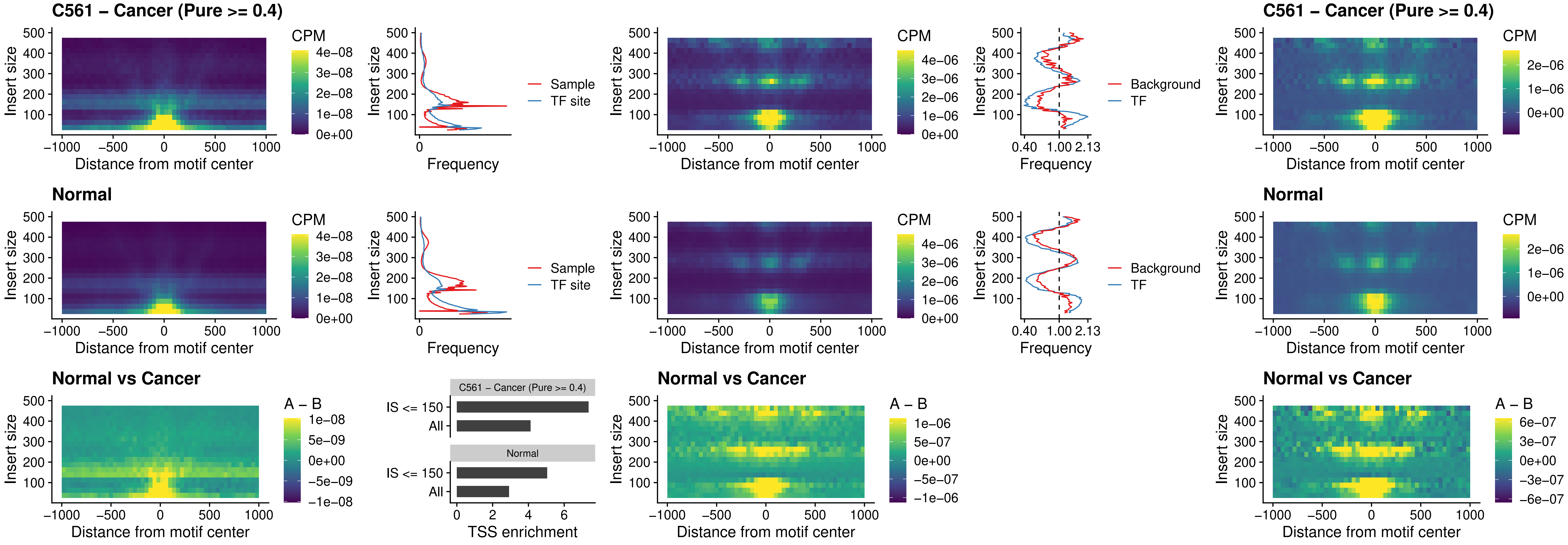

### SNAI2_uc_cTSS_nPEAK

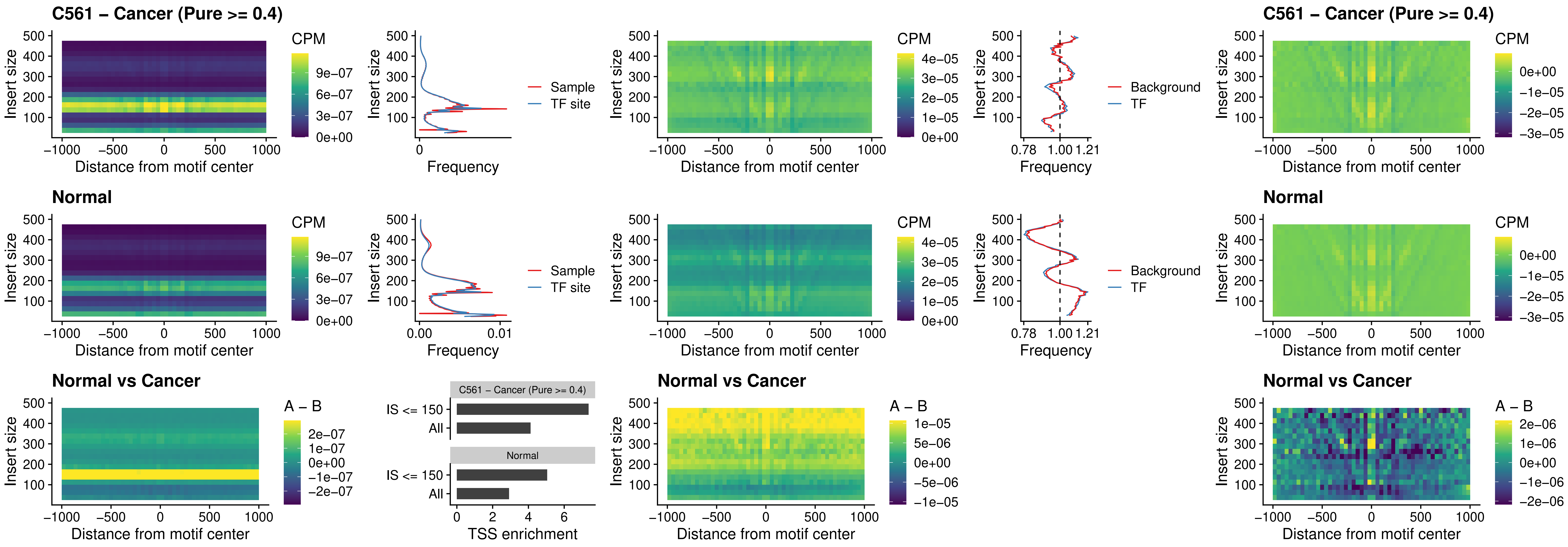

#### SNAI2_uc_dTSS_oPEAK

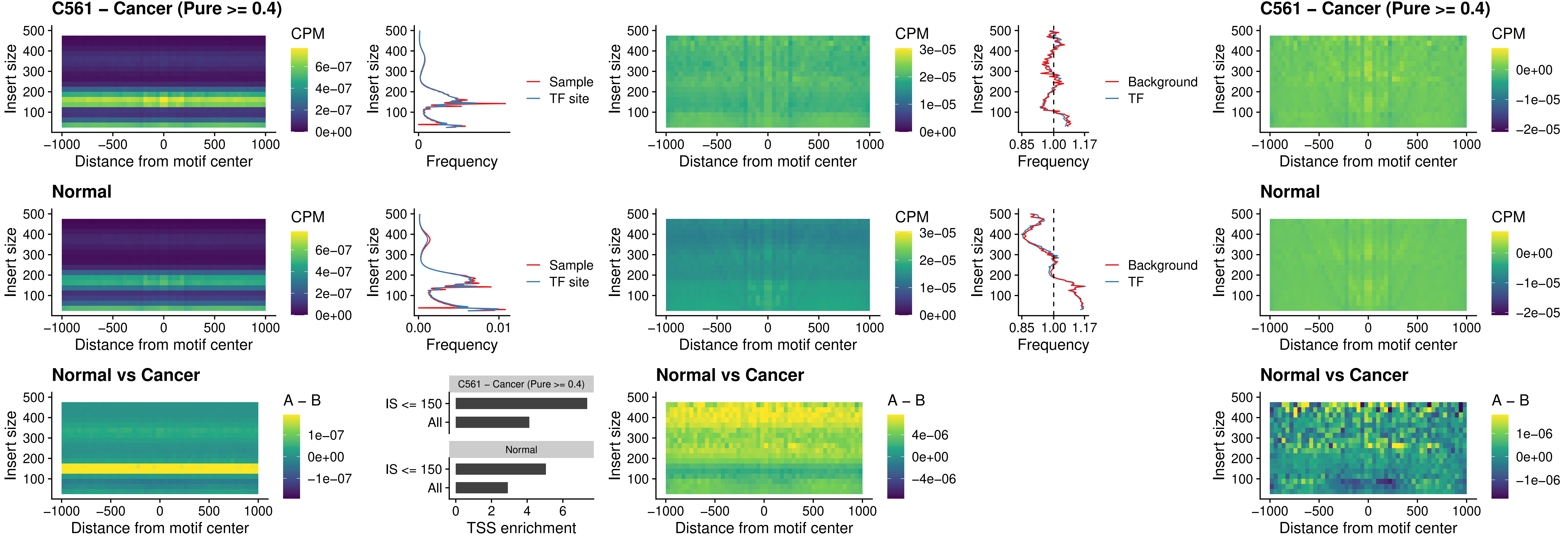

#### SPI1_uc_pTSS_nPEAK

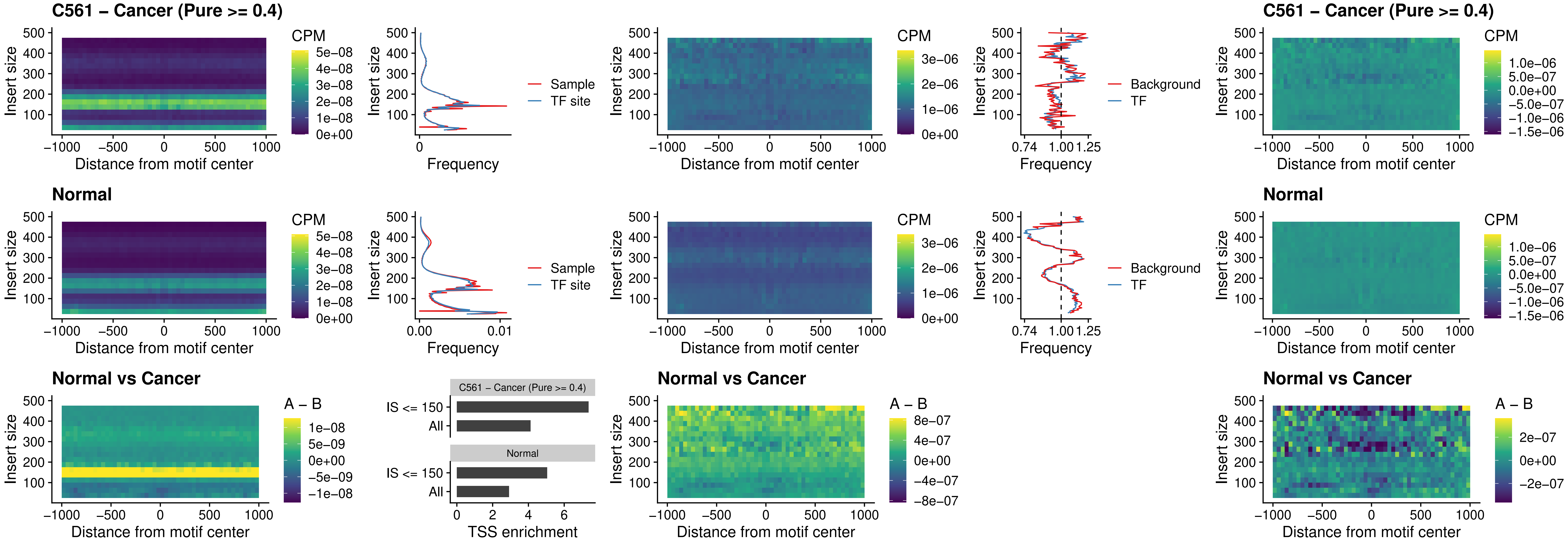

### KLF16_uc_pTSS_nPEAK

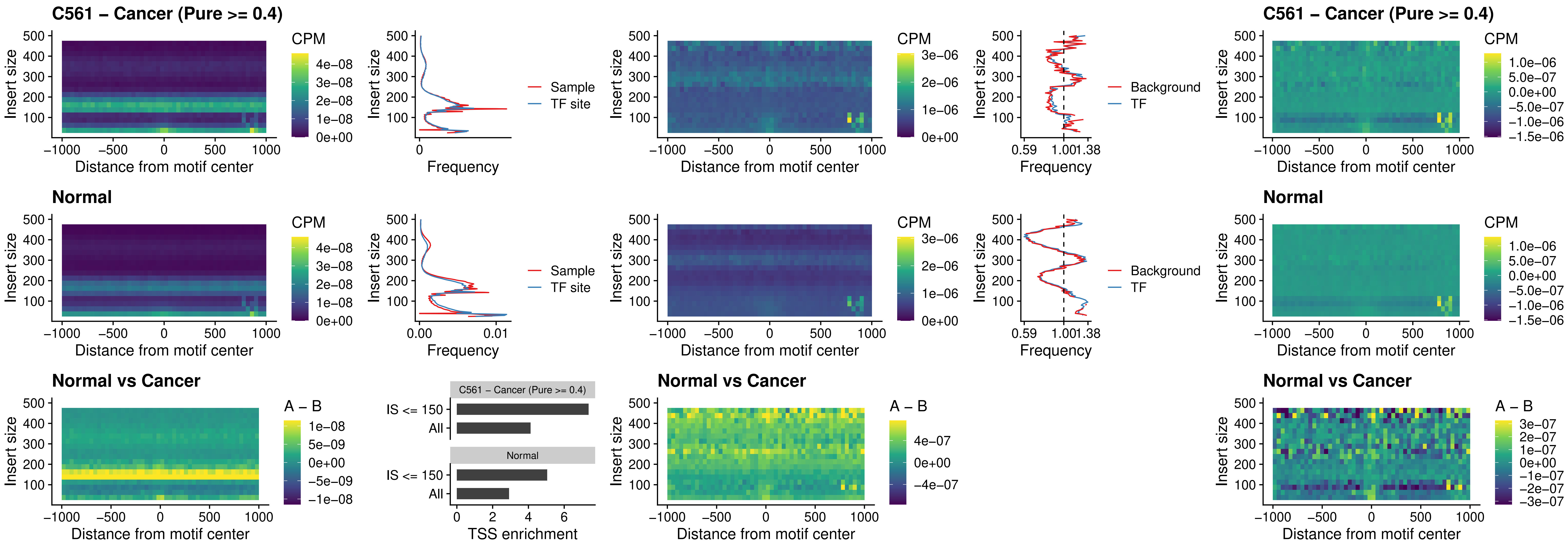

#### SP3_uc_pTSS_nPEAK

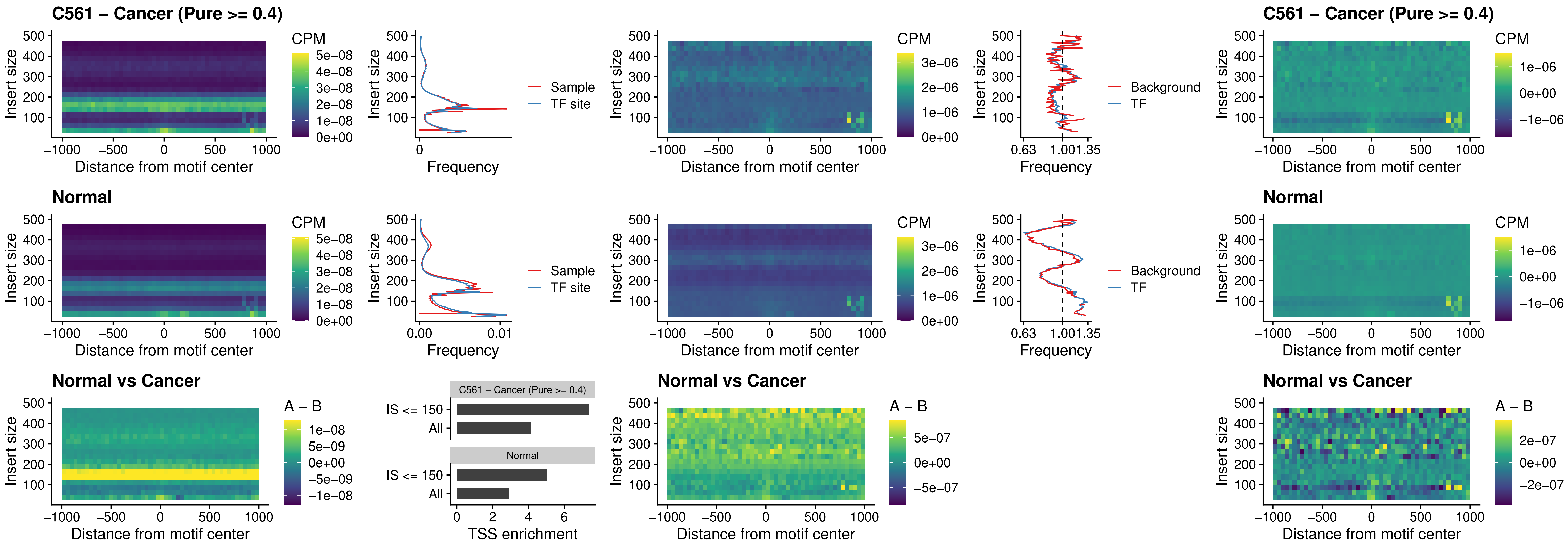

### NEUROD1_uc_dTSS_oPEAK

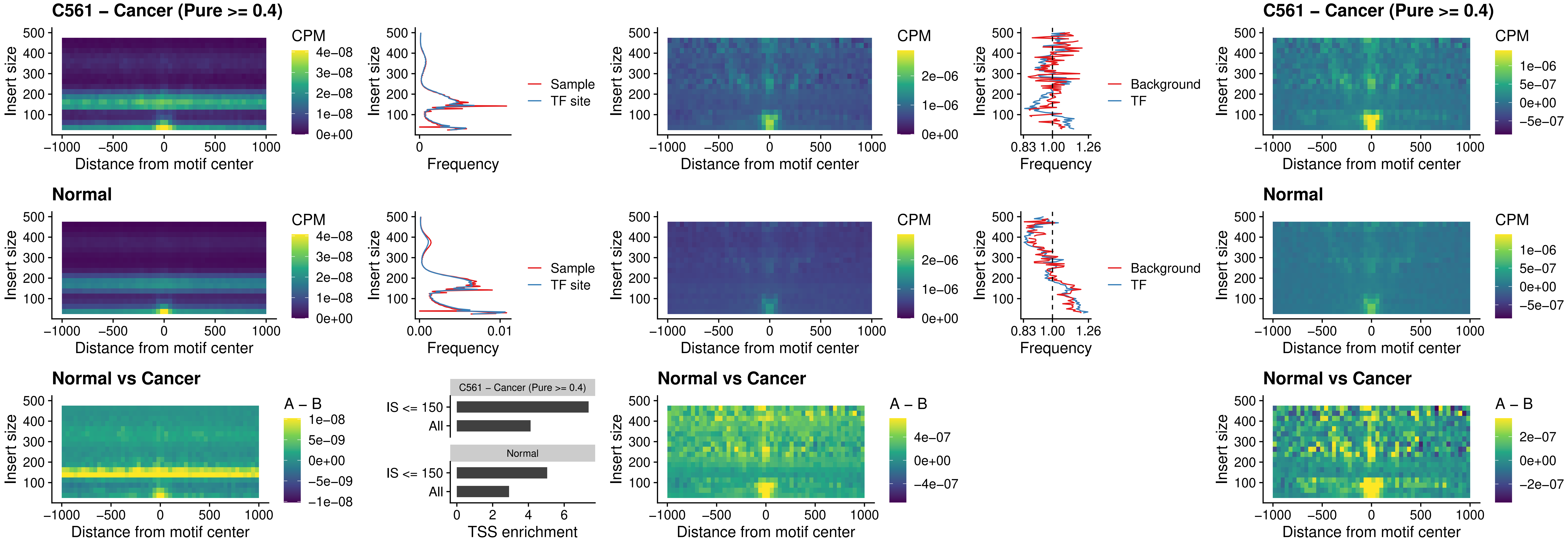

#### CTCF_uc_dTSS_oPEAK

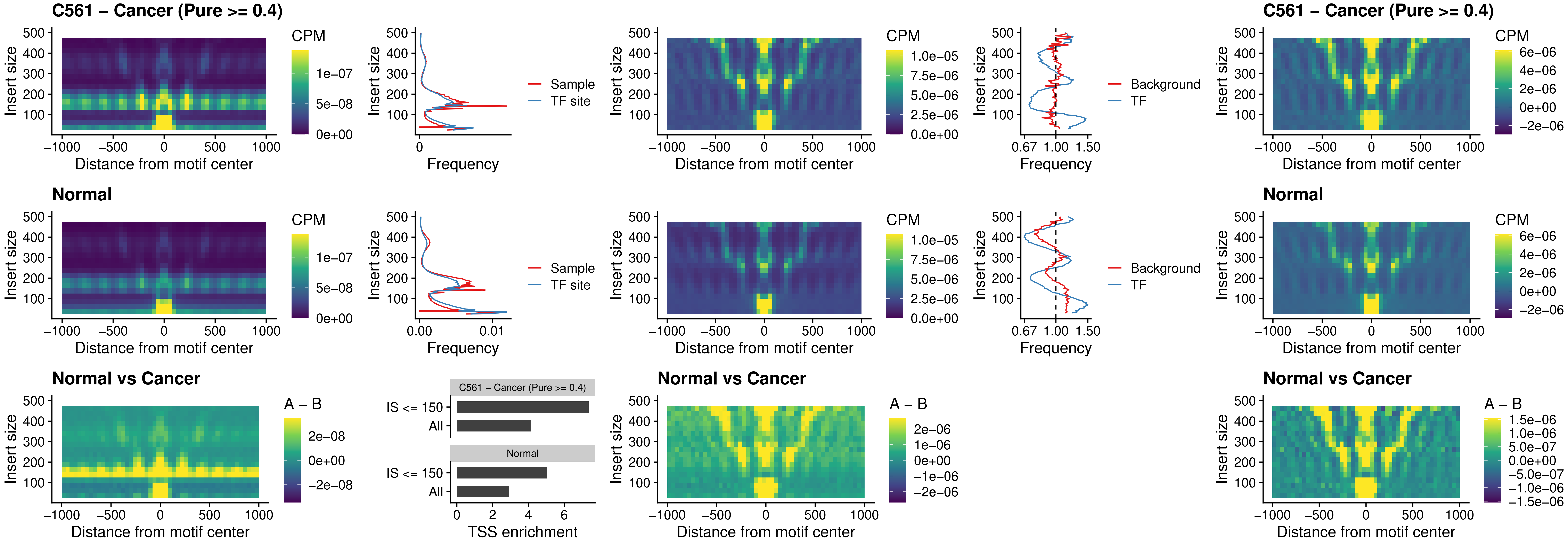

### CTCF_uc_cTSS_oPEAK

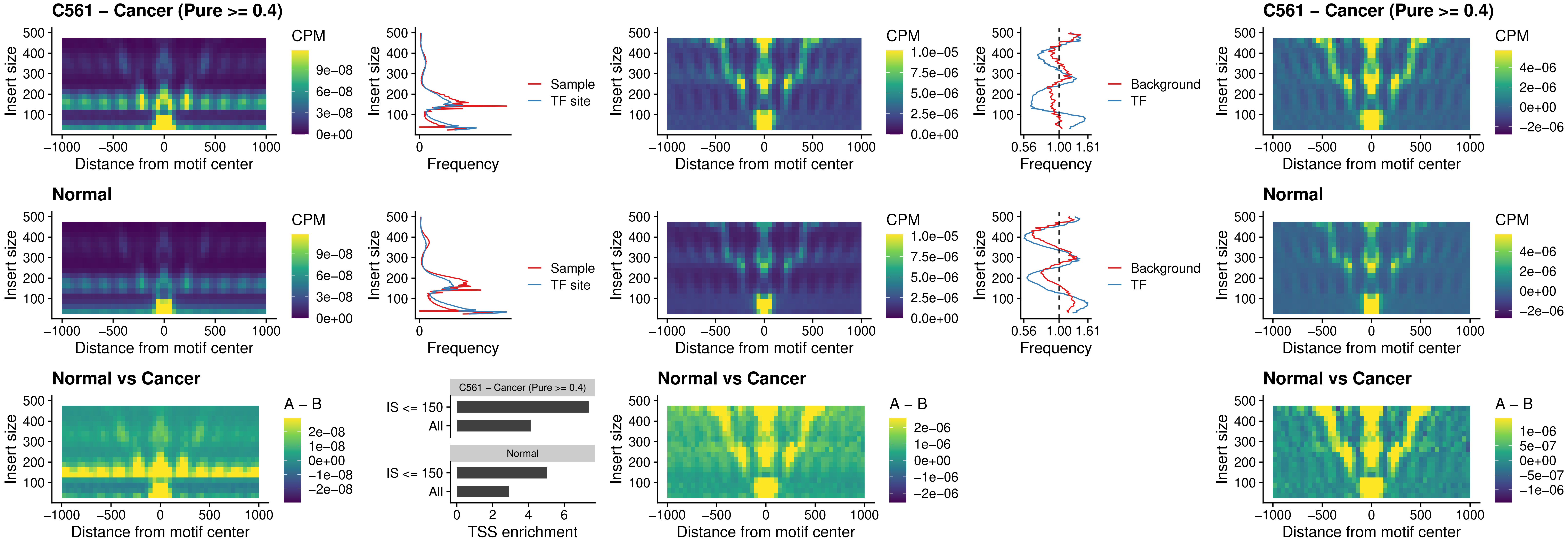

### CTCFL_uc_dTSS_oPEAK

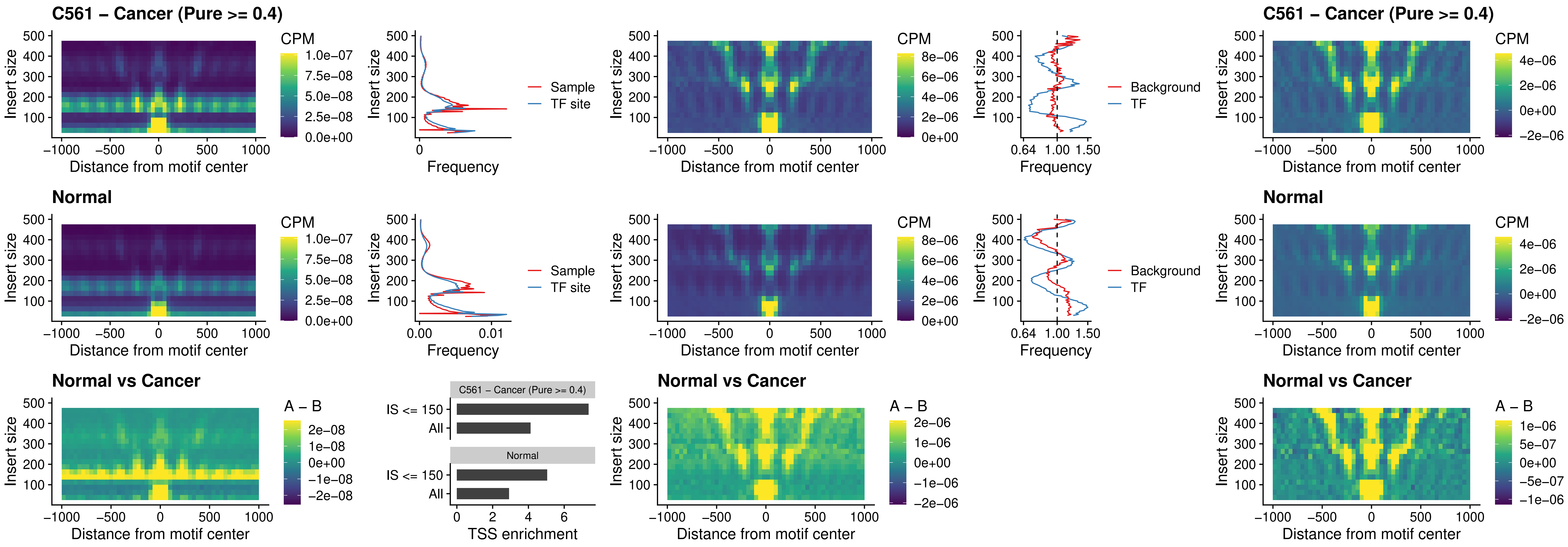

#### ENCODE_CTCF_sites_COLON

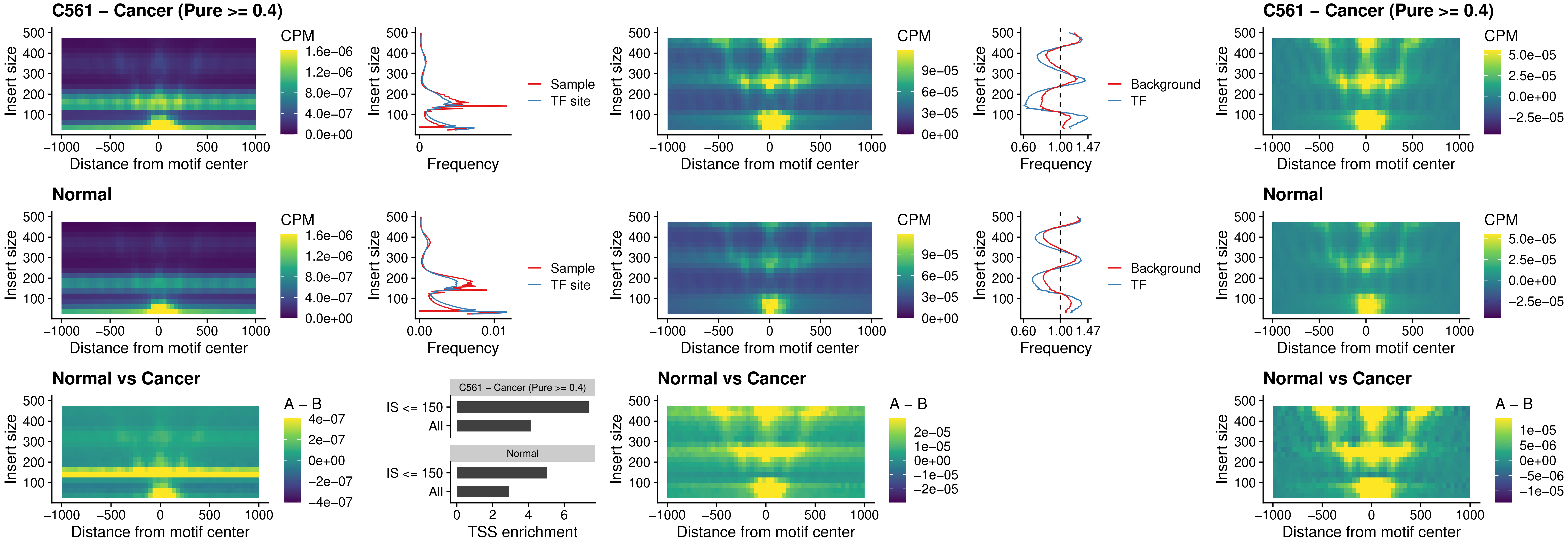

#### ENCODE_CTCF_sites_COLON_loop_ctcf

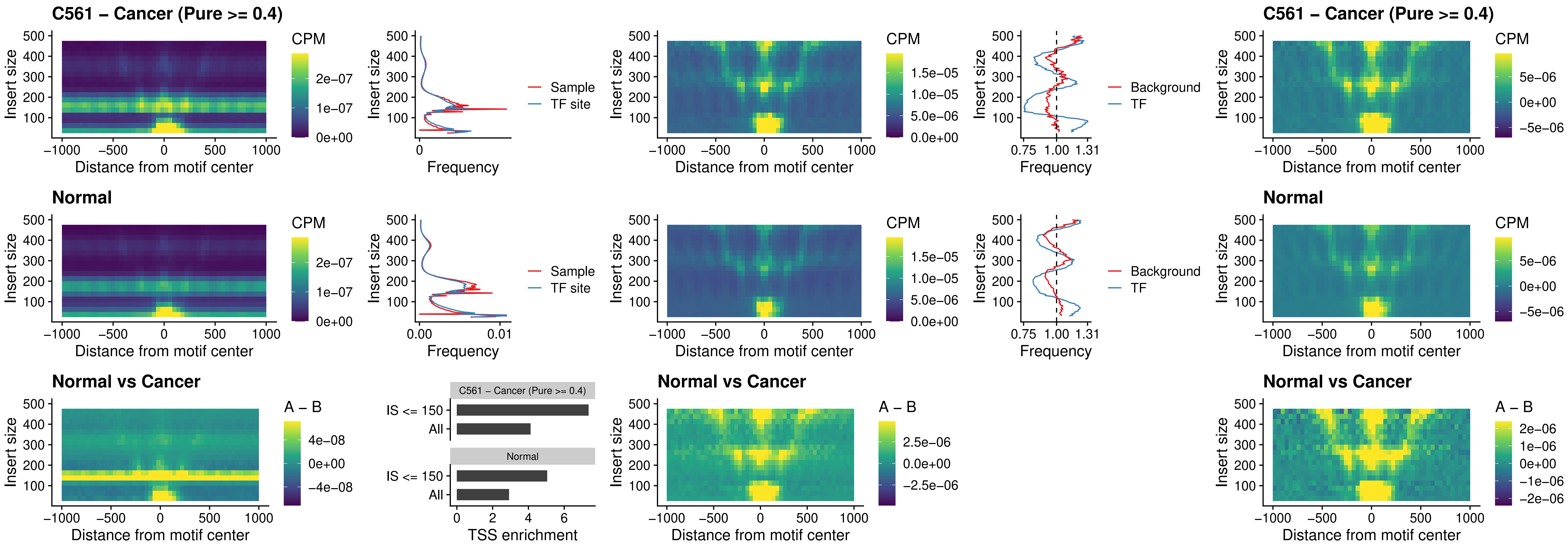

#### ENCODE_CTCF_sites_COLON_intergenic_ctcf

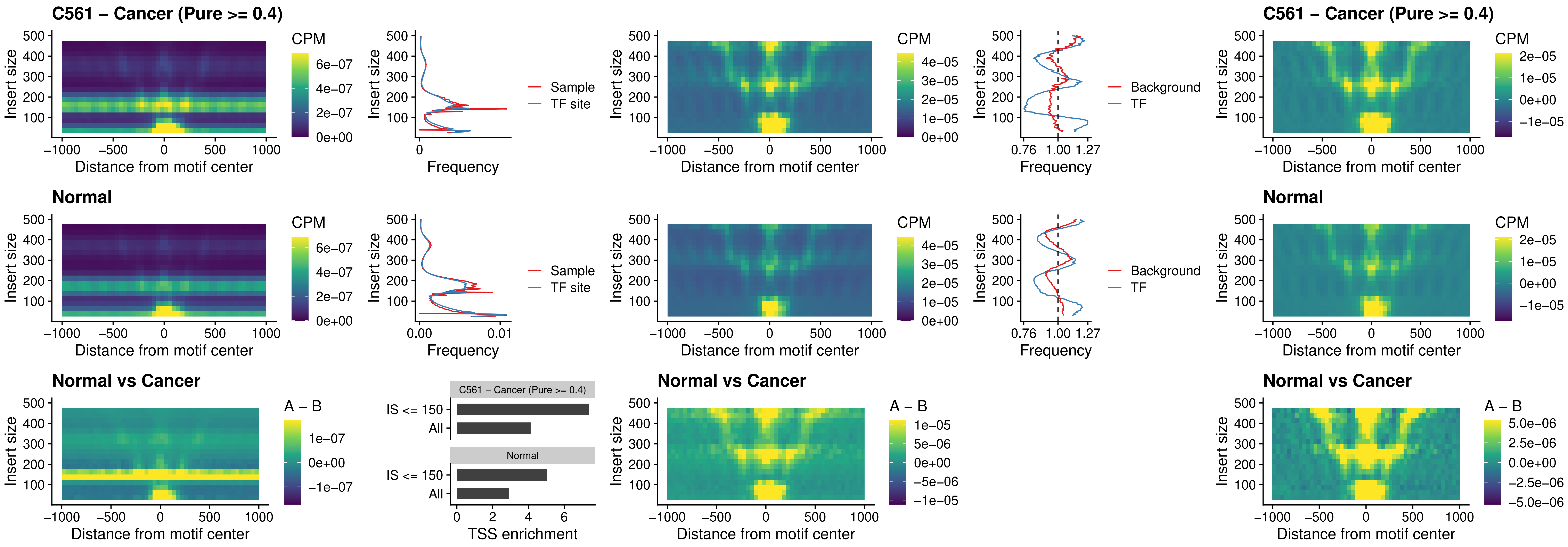

### ENCODE_CTCF_sites_COLON_nonpromotor

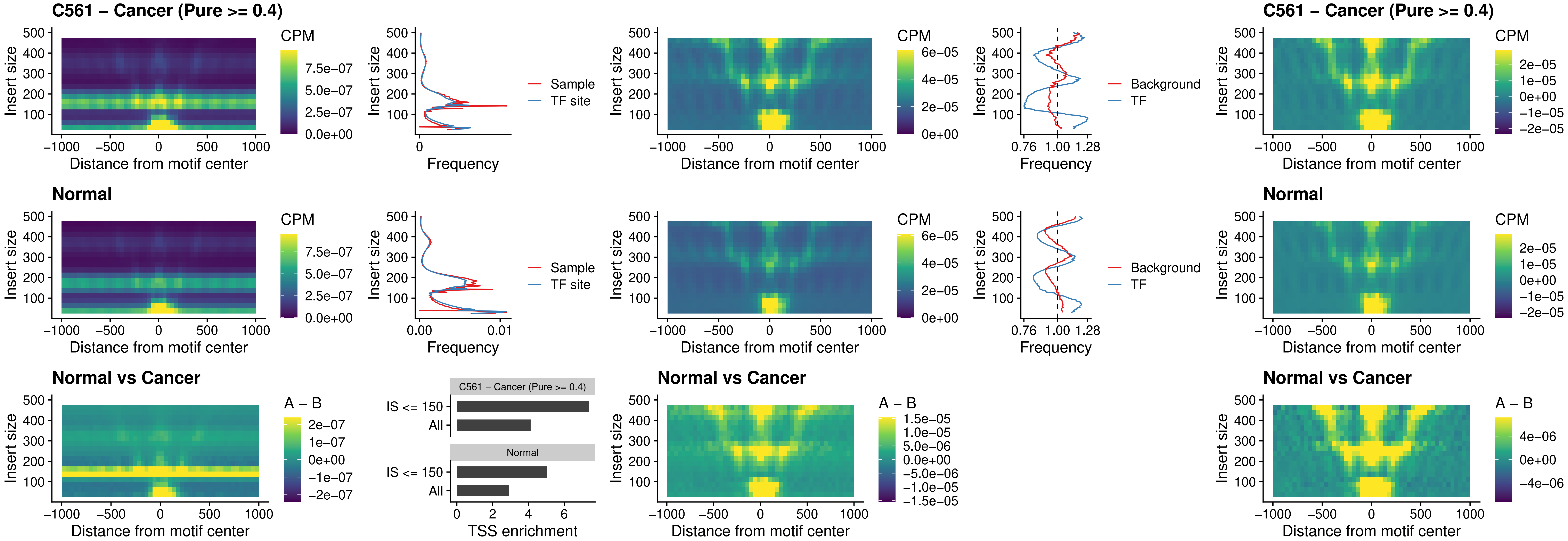

### CTCFL_uc_cTSS_oPEAK

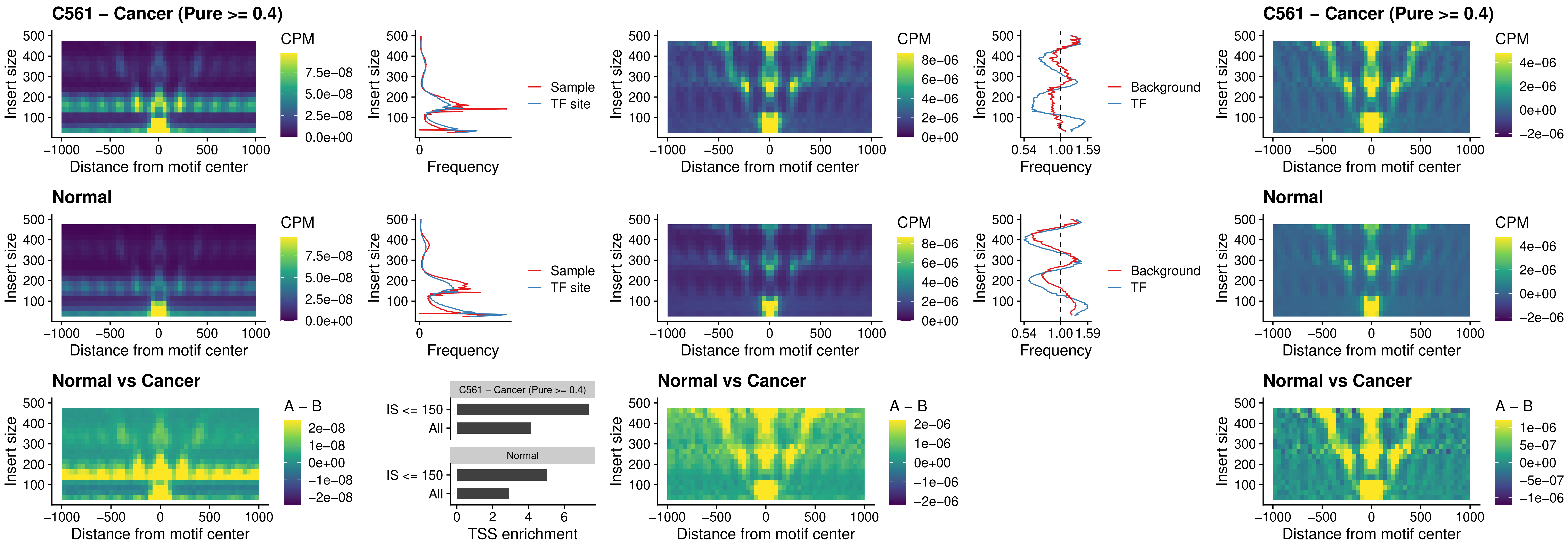

#### NEURОD1_uc_cTSS_oPEAK

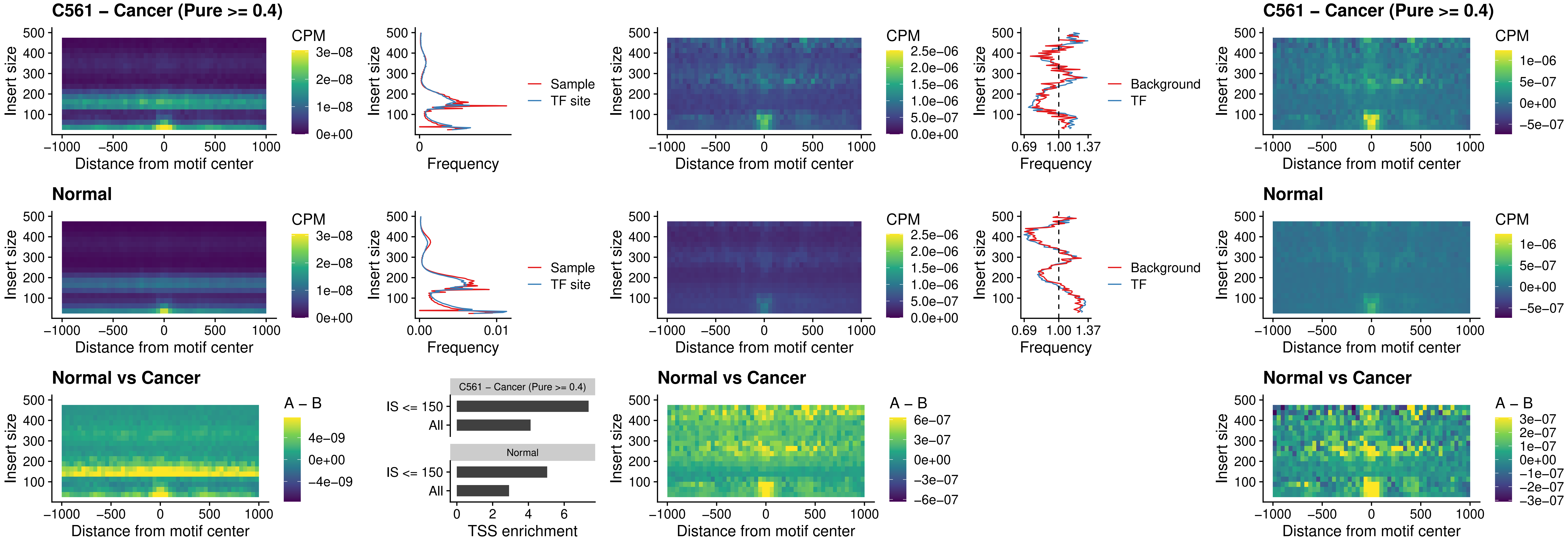

#### ENCODE_YY1_sites_COLON_nonpromotor

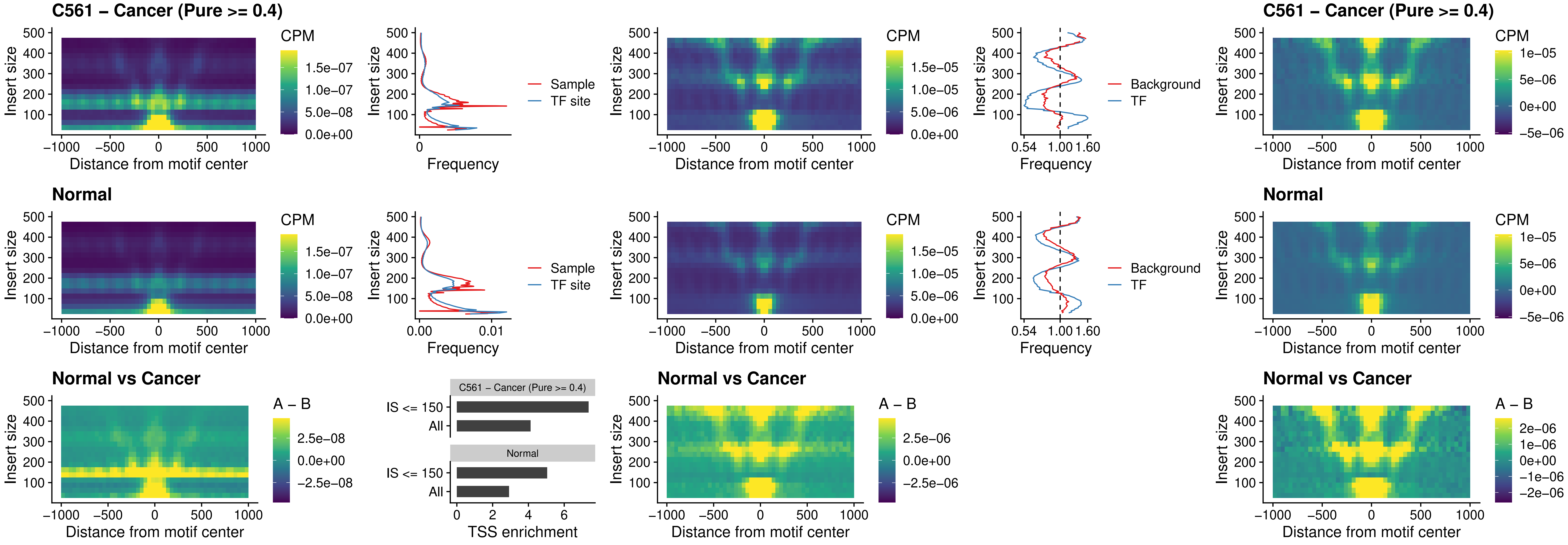

### CTCF_uc_pTSS_oPEAK

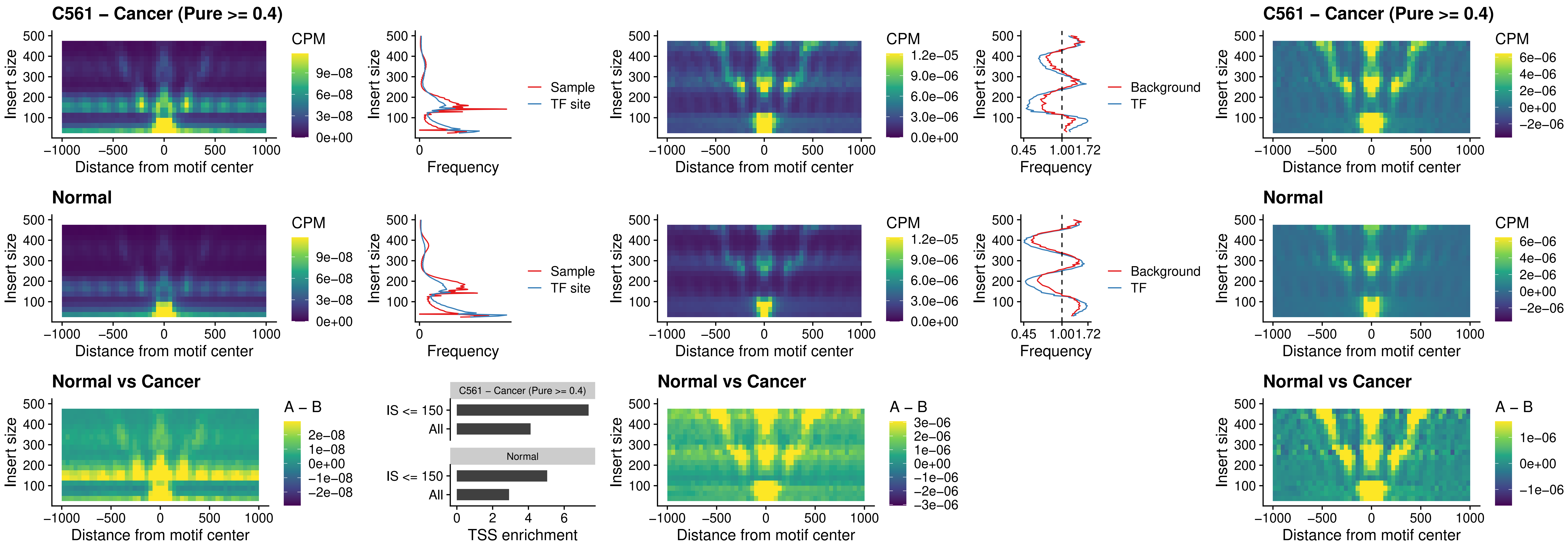

### ENCODE_CTCF_sites_COLON_promotor

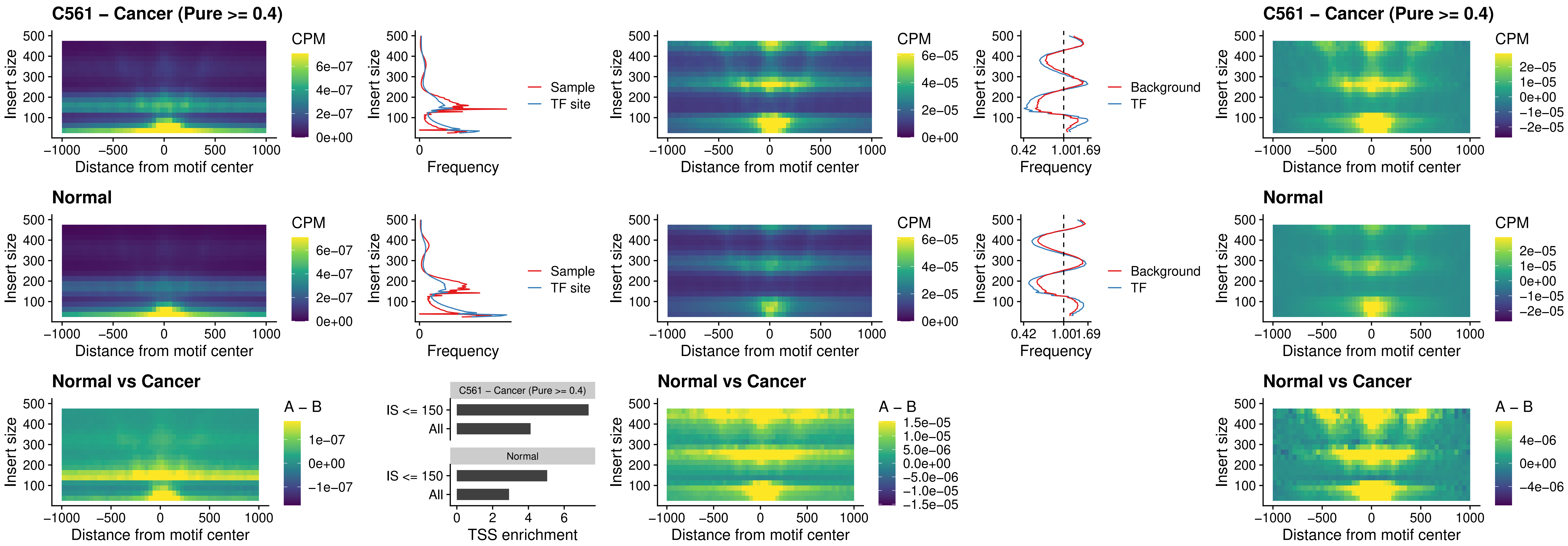

### ENCODE_CTCF_sites_COLON_promotor_ctcf

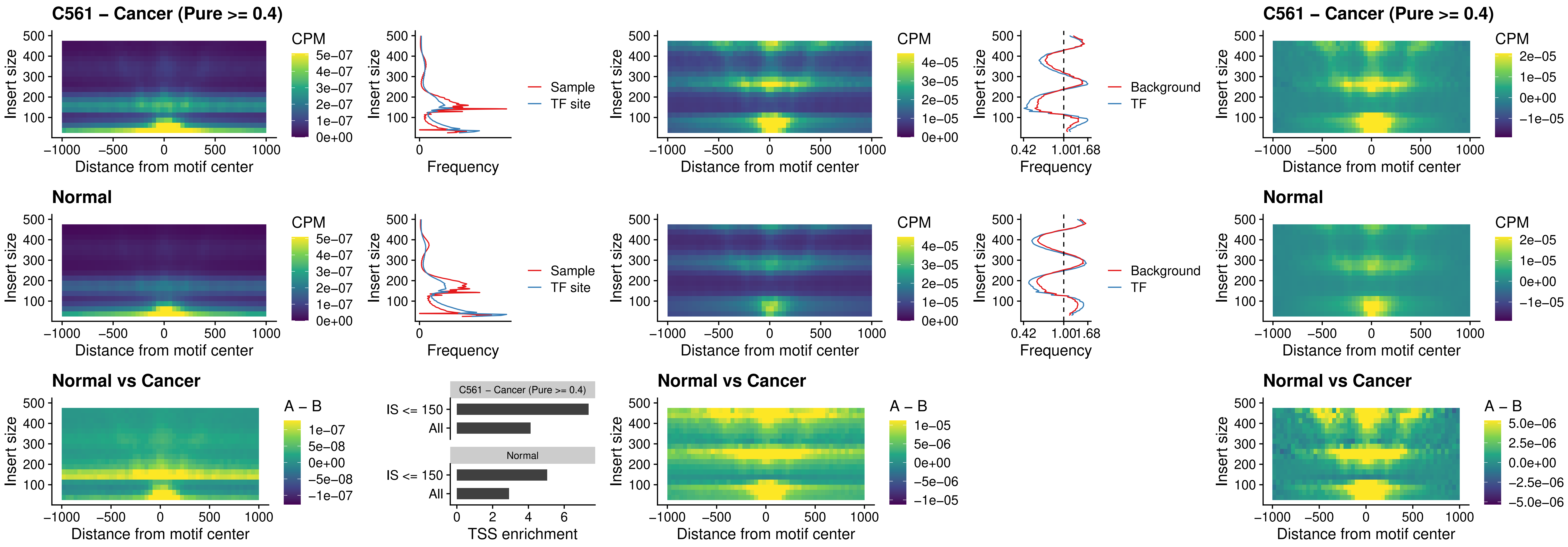

#### SPIC_uc_pTSS_oPEAK

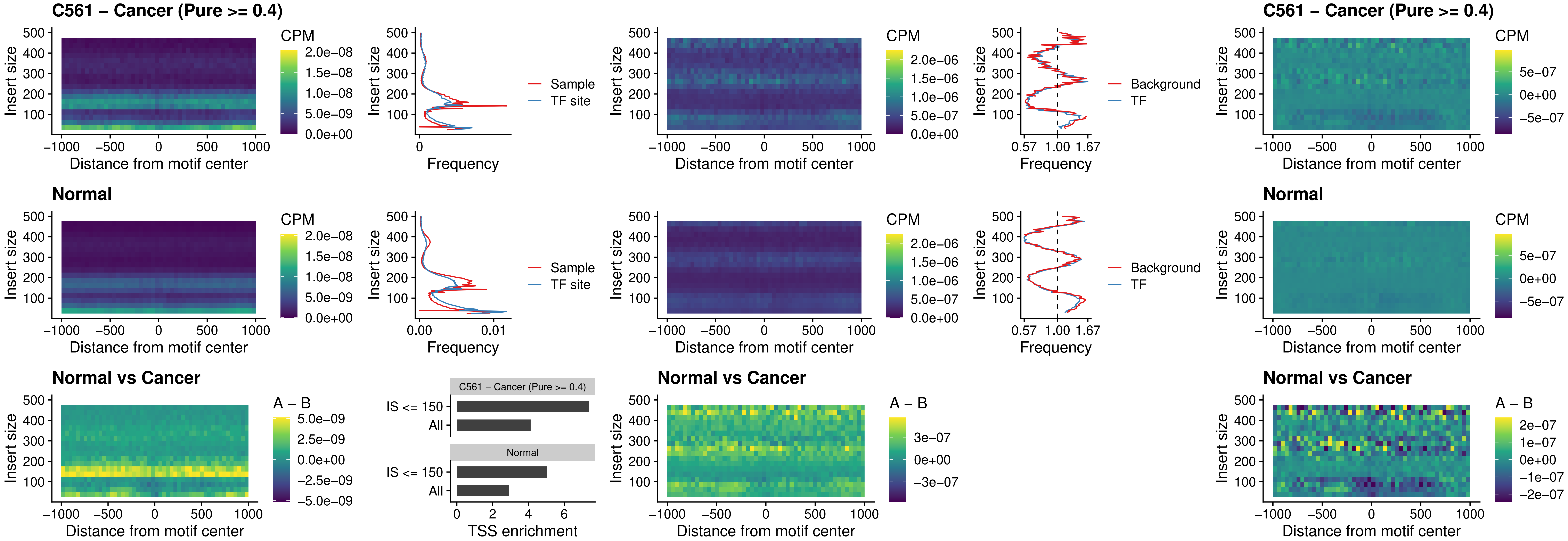

#### SPI1_uc_dTSS_nPEAK

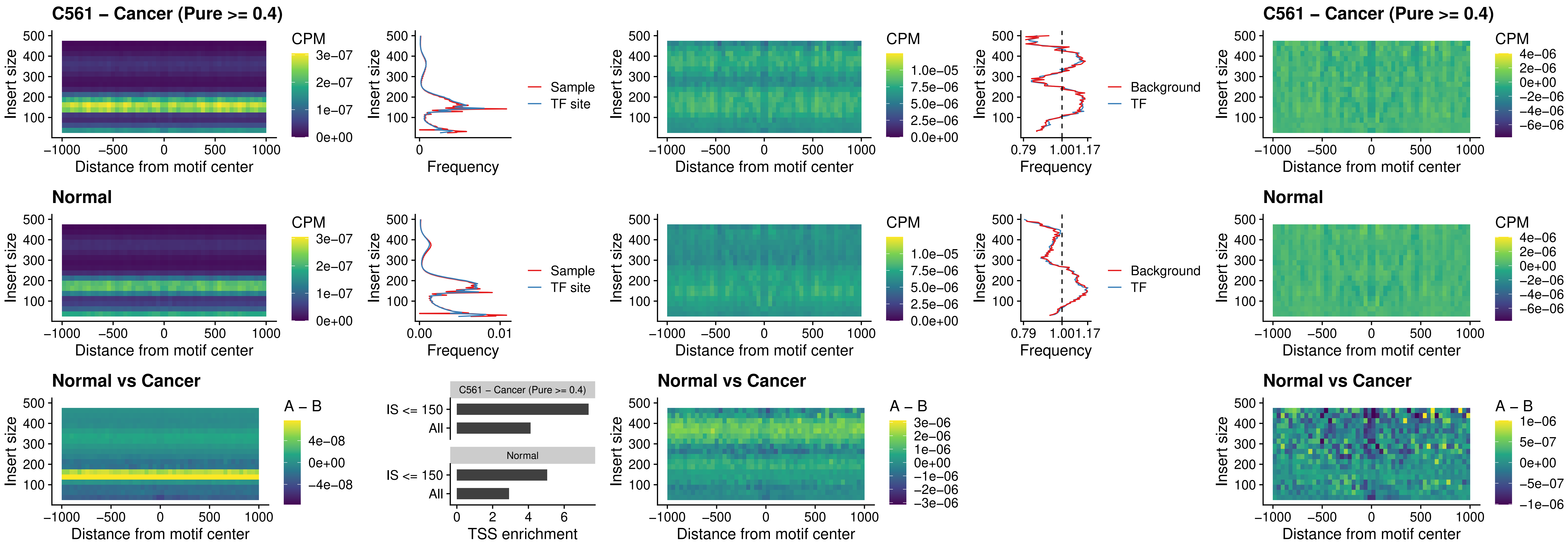

### SPI1_uc_cTSS_nPEAK

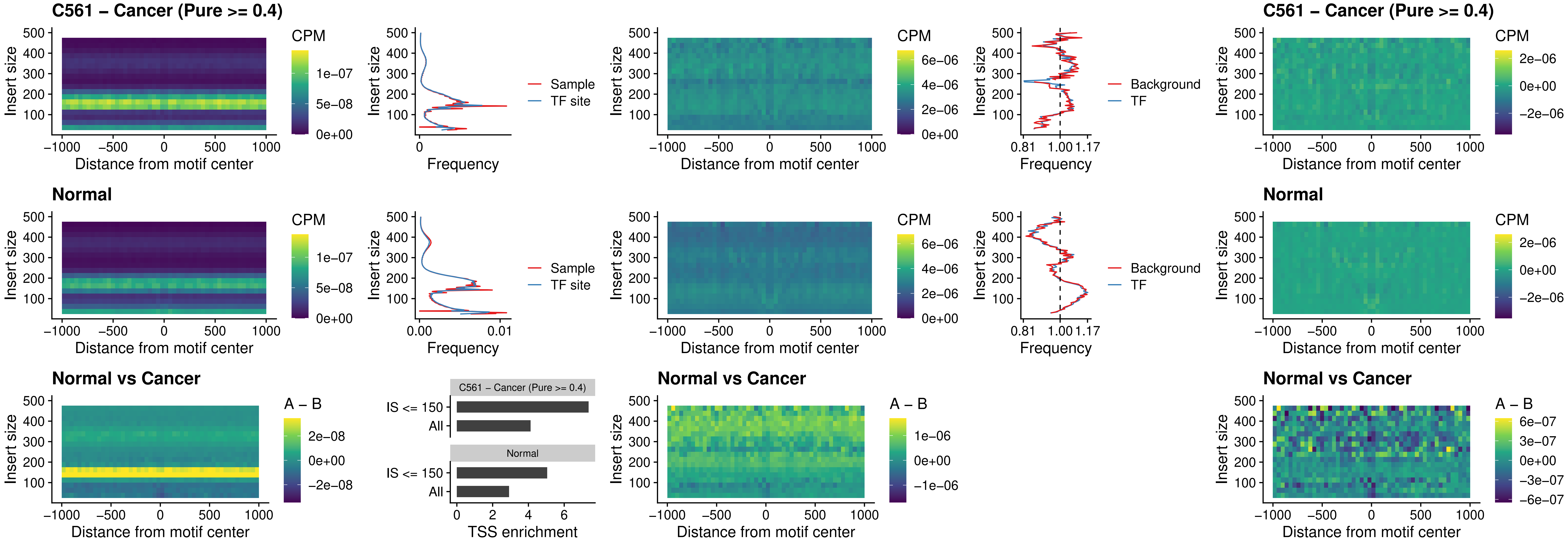

### BCL11A_uc_pTSS_nPEAK

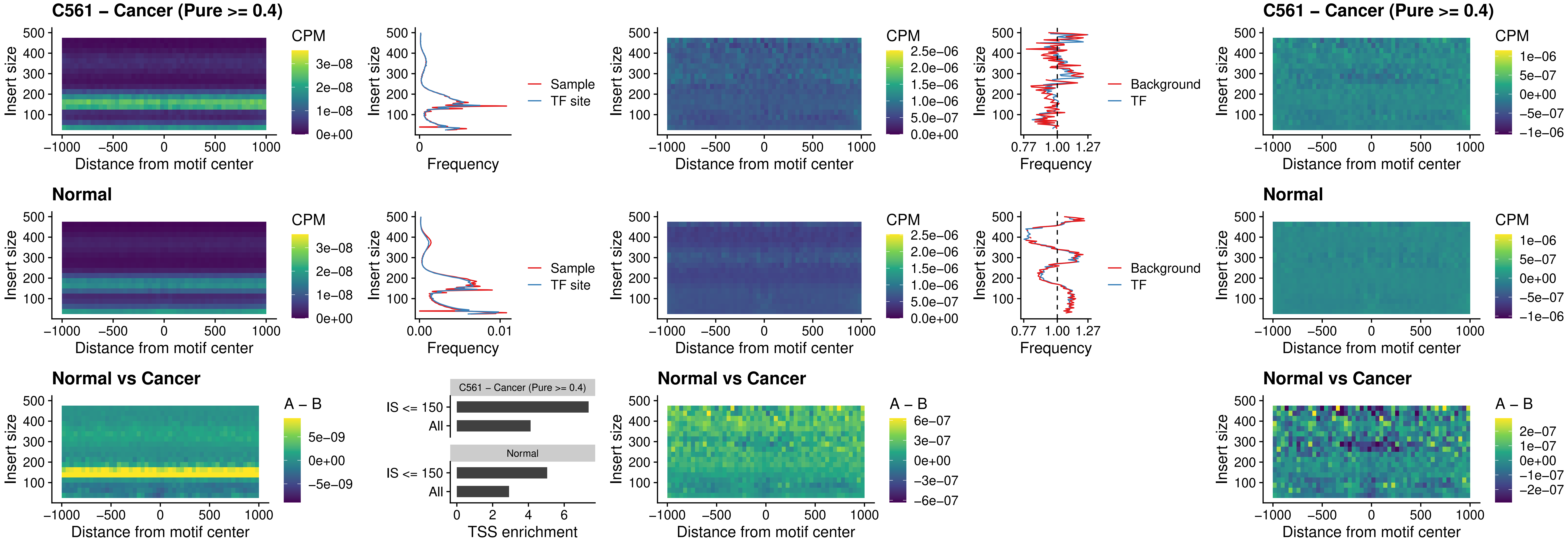

#### BCL11B_uc_pTSS_nPEAK

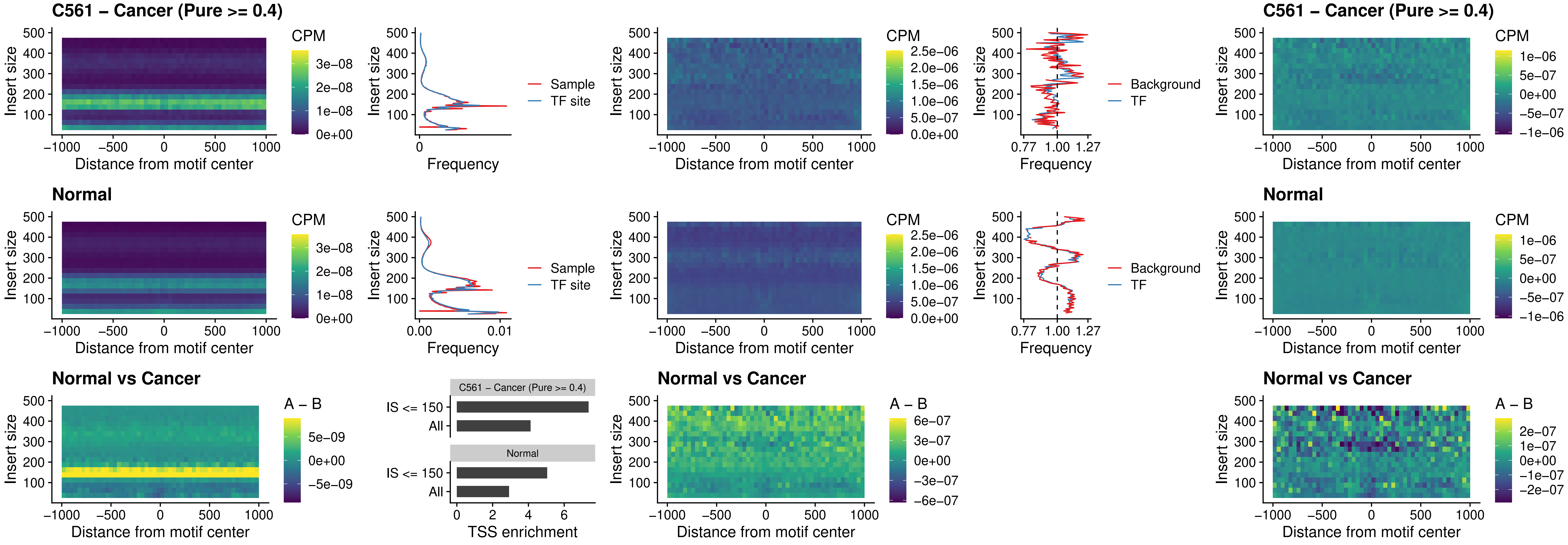

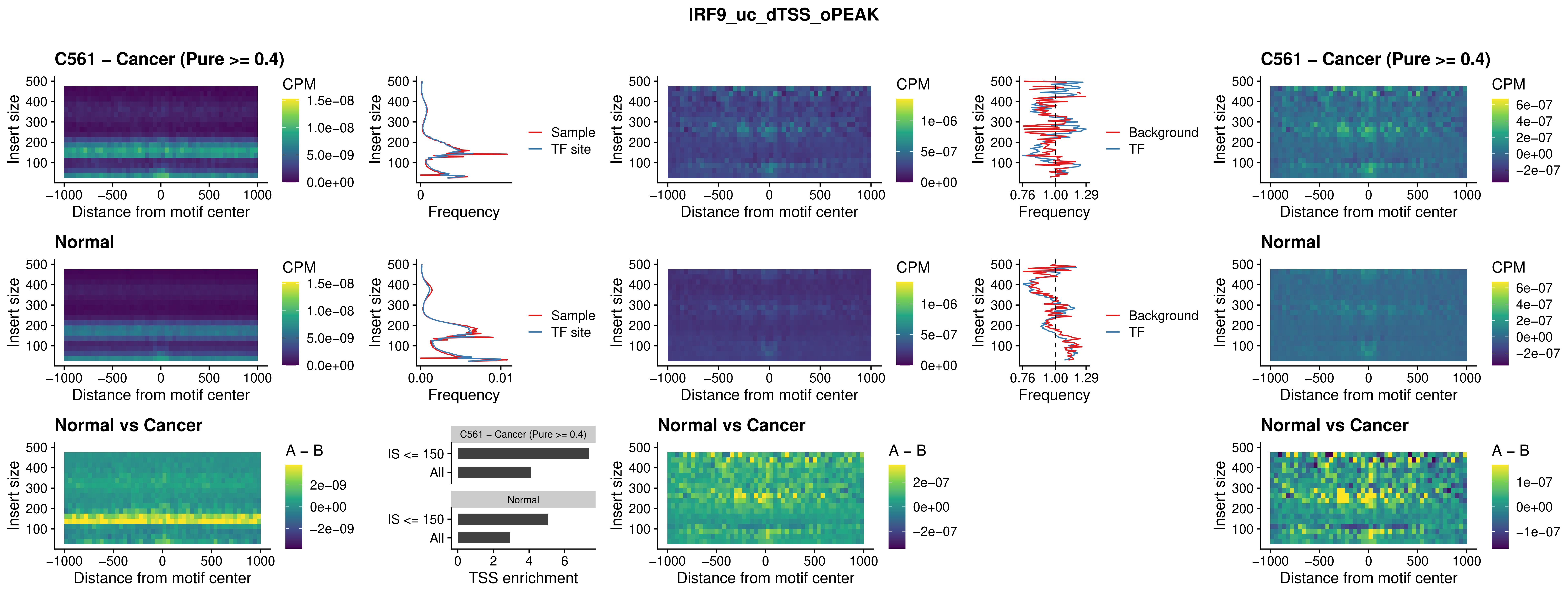
IRF9_uc_dTSS_oPEAK
C561 – Cancer (Pure >= 0.4)
Normal
Normal vs Cancer
Insert size
Distance from motif center
CPM
Frequency
Sample
TF site
Background
TF
A – B
C561 – Cancer (Pure >= 0.4)
Normal
IS <= 150
All
TSS enrichment

#### IRF9_uc_cTSS_oPEAK

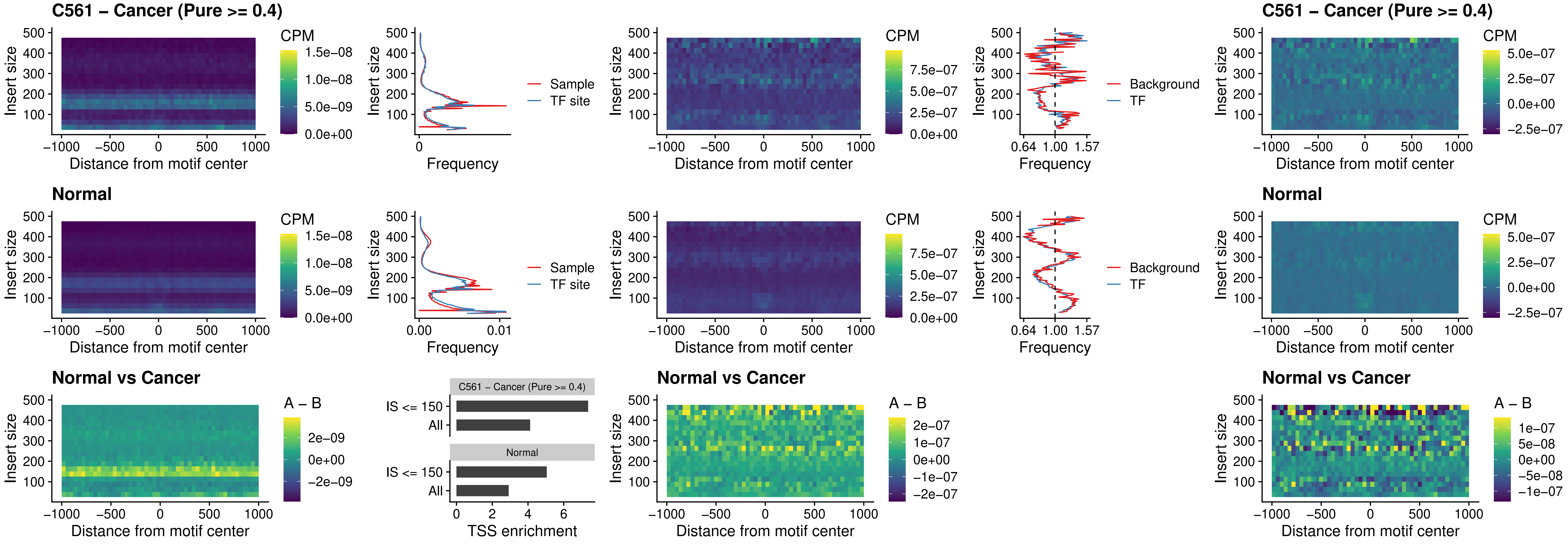

### BCL11A_uc_cTSS_oPEAK

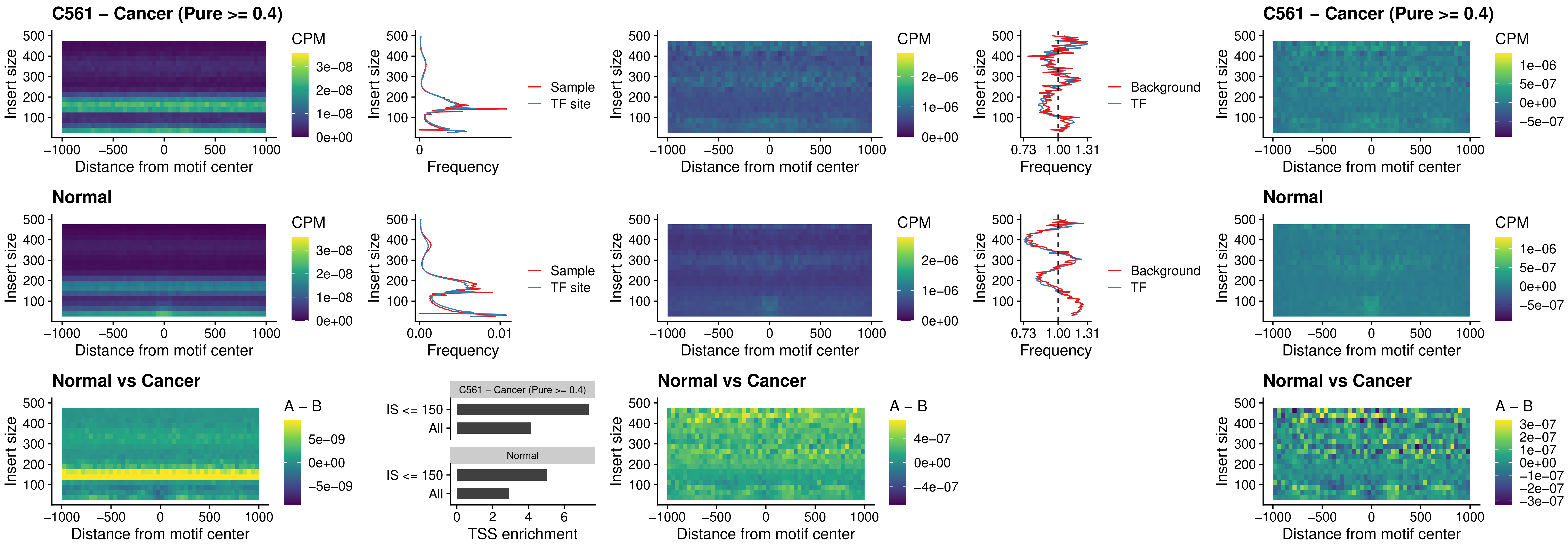

### BCL11B_uc_cTSS_oPEAK

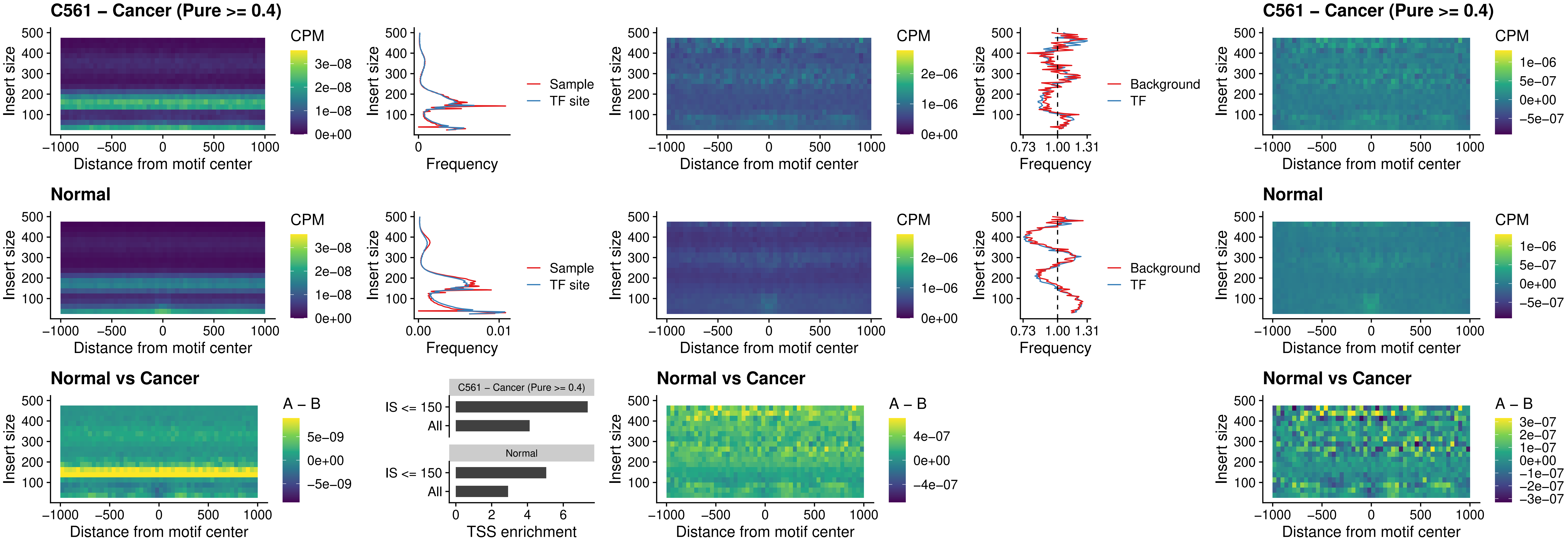

### BCL11A_uc_dTSS_oPEAK

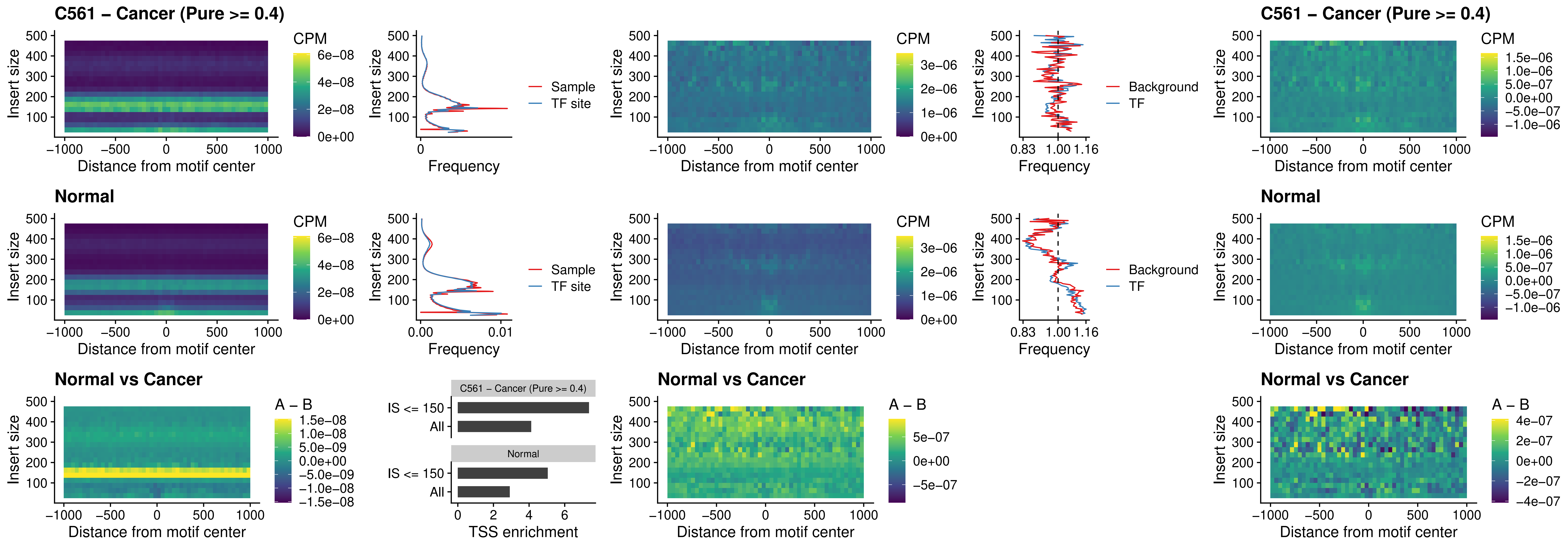

### BCL11B_uc_dTSS_oPEAK

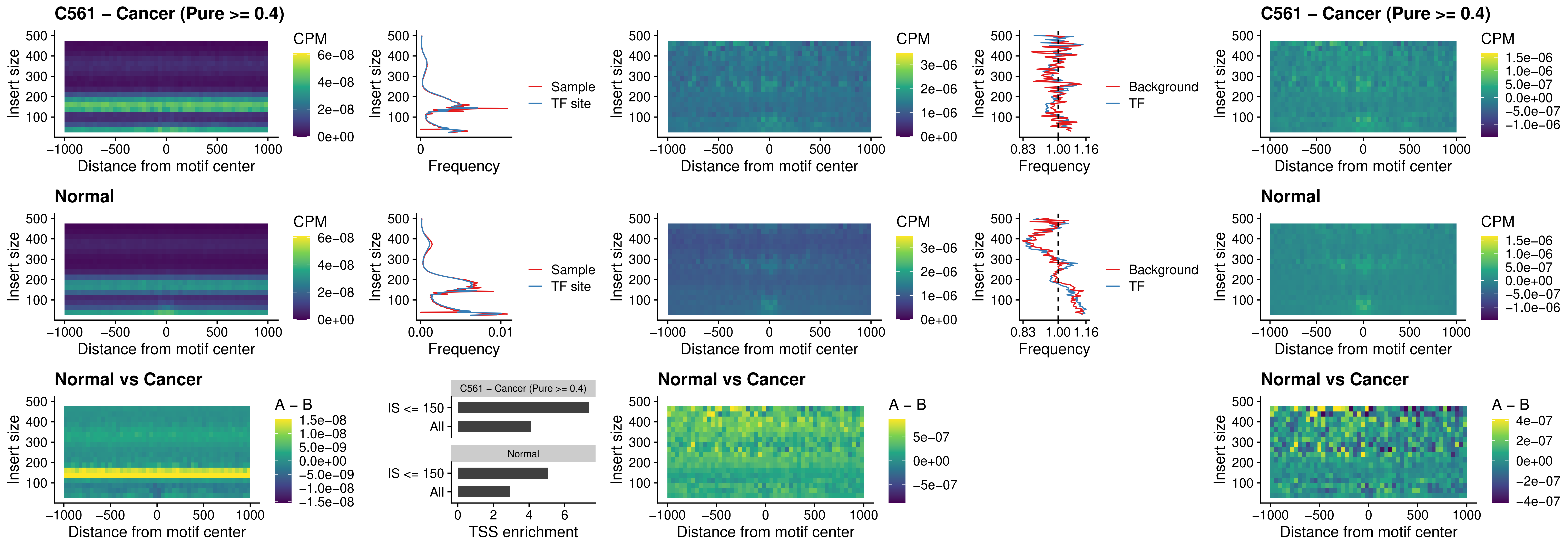

#### SPIB_uc_cTSS_nPEAK

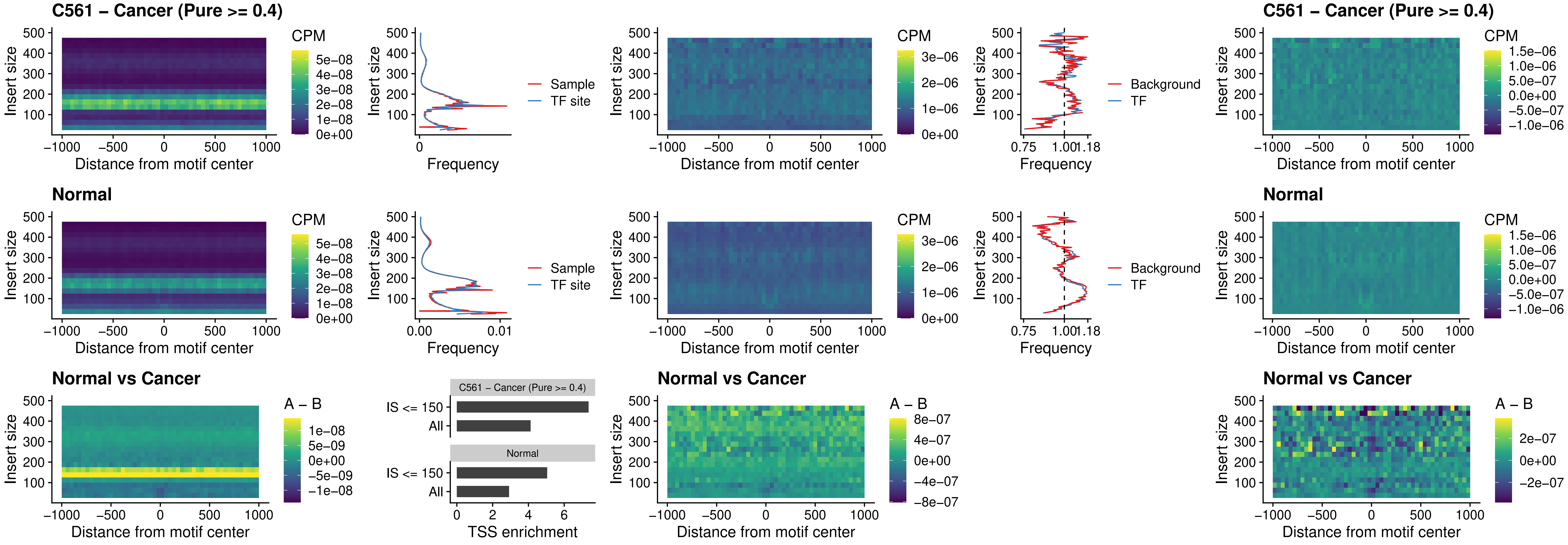

#### SPIC_uc_cTSS_nPEAK

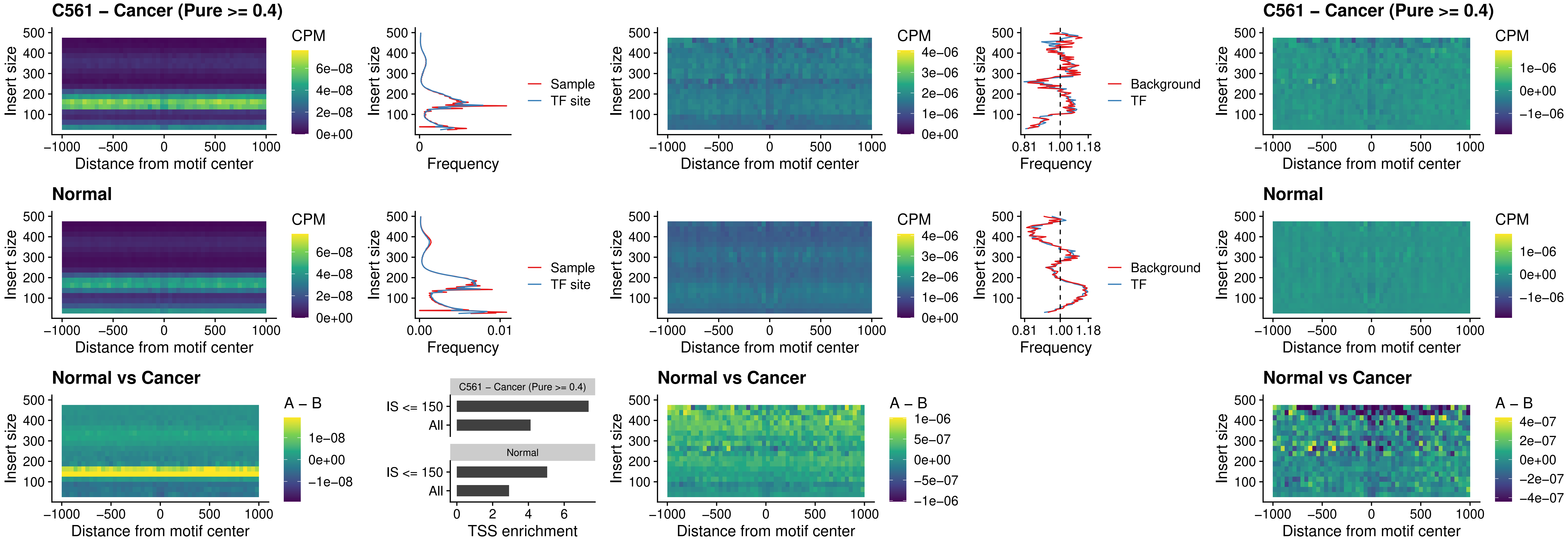

#### SPI1_uc_pTSS_oPEAK

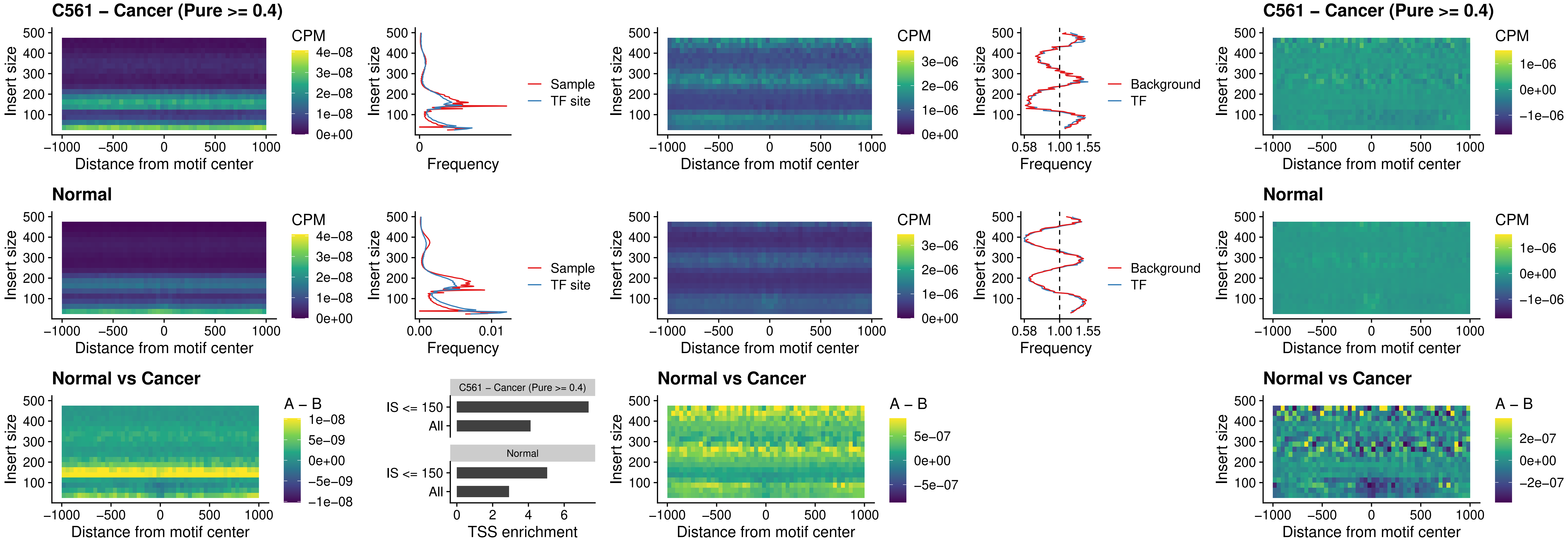

#### BCL11A_uc_pTSS_oPEAK

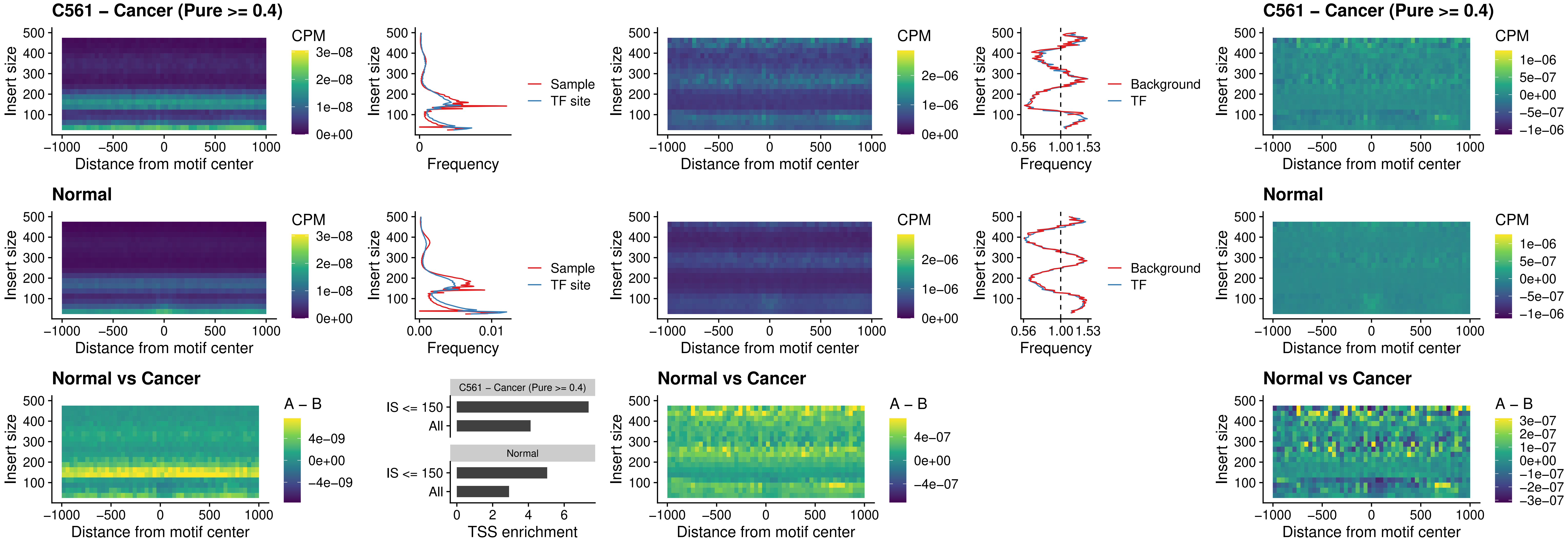

#### BCL11B_uc_pTSS_oPEAK

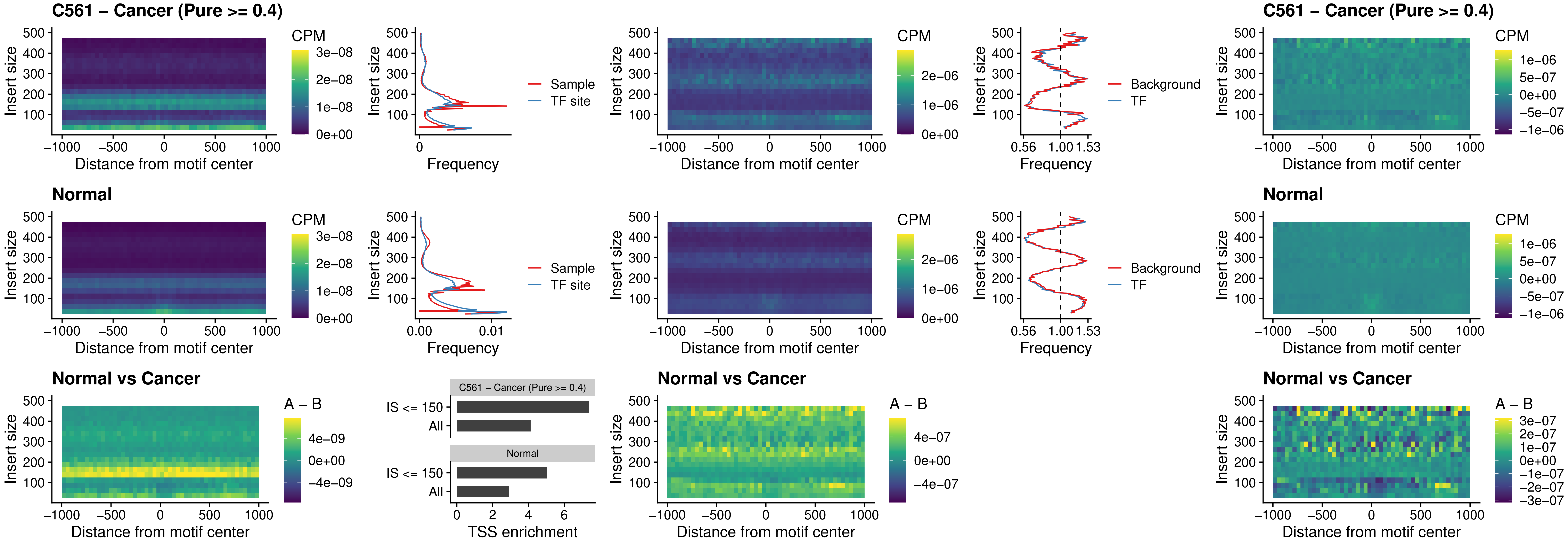

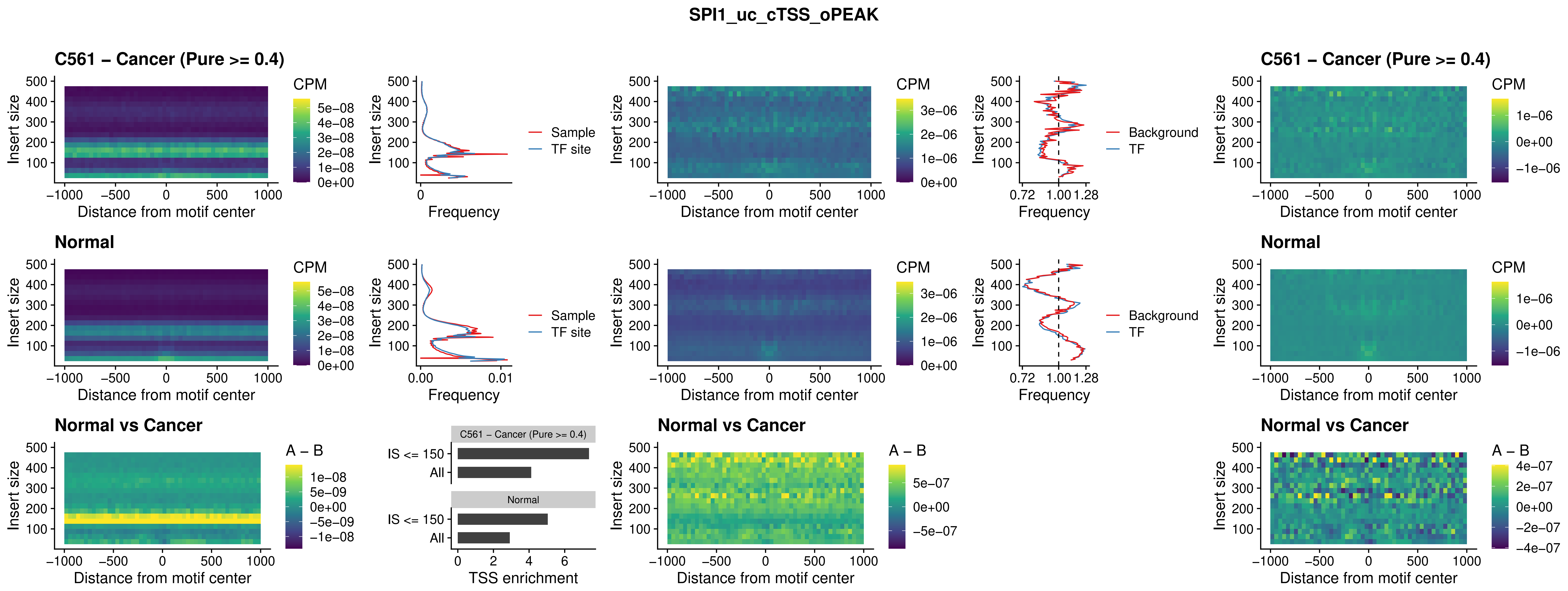
SPI1_uc_cTSS_oPEAK
C561 – Cancer (Pure >= 0.4)
Normal
Normal vs Cancer
Insert size
Distance from motif center
Frequency
CPM
A – B
Sample
TF site
Background
TF
TSS enrichment
IS <= 150
All

### SPIB_uc_cTSS_oPEAK

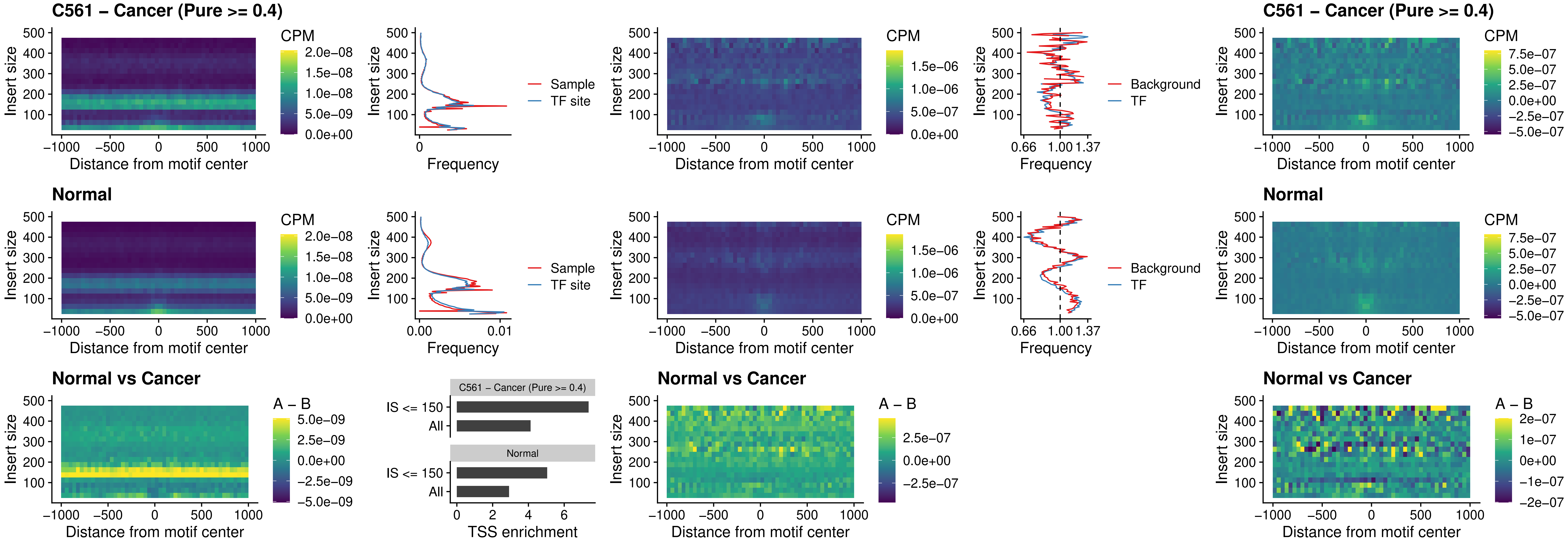

#### SPIB_uc_pTSS_oPEAK

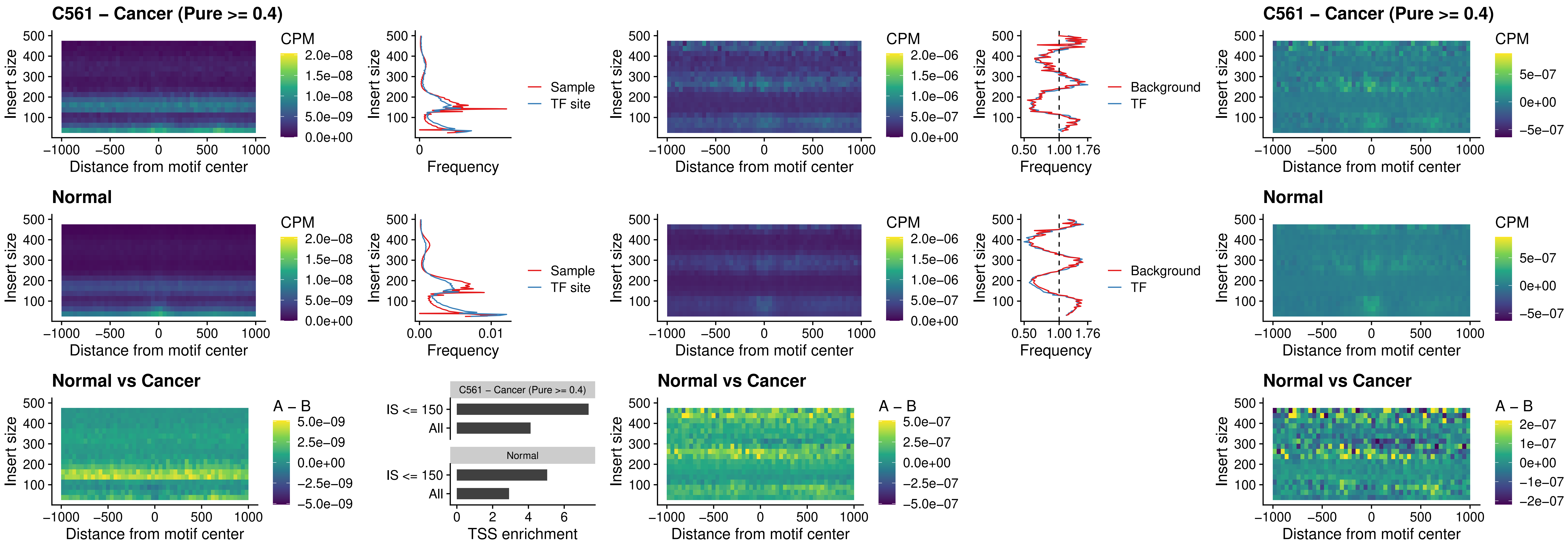

#### SPI1_uc_dTSS_oPEAK

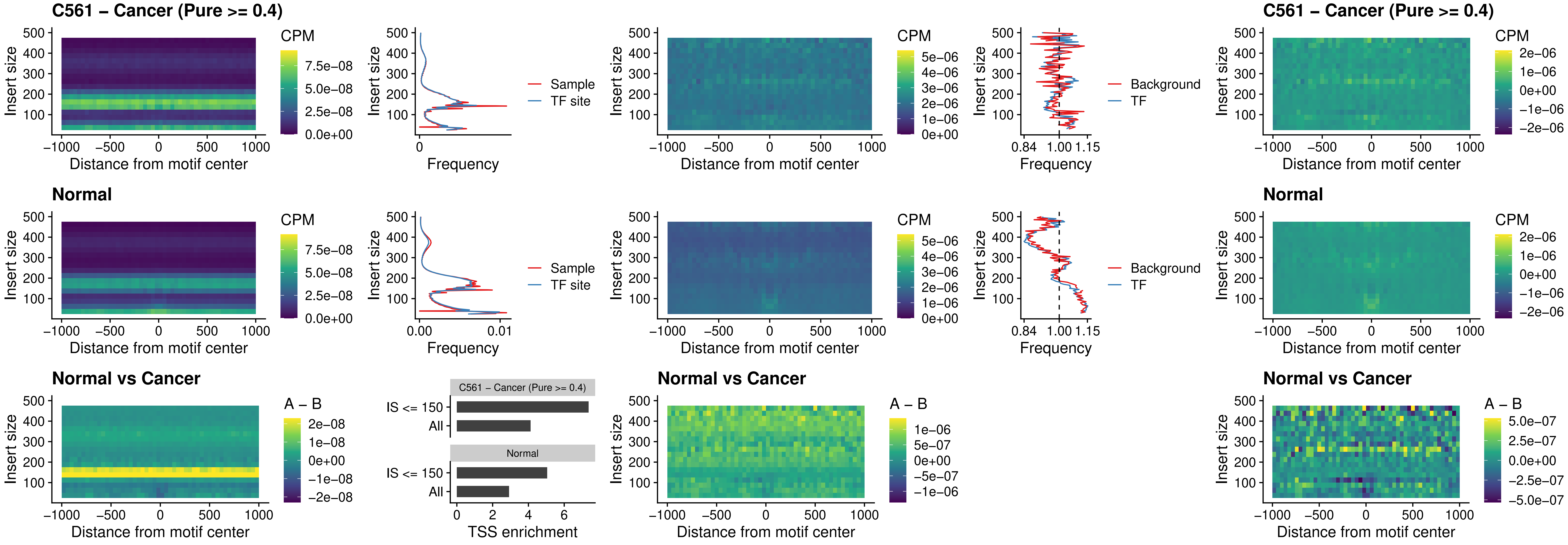

#### SPIB_uc_dTSS_oPEAK

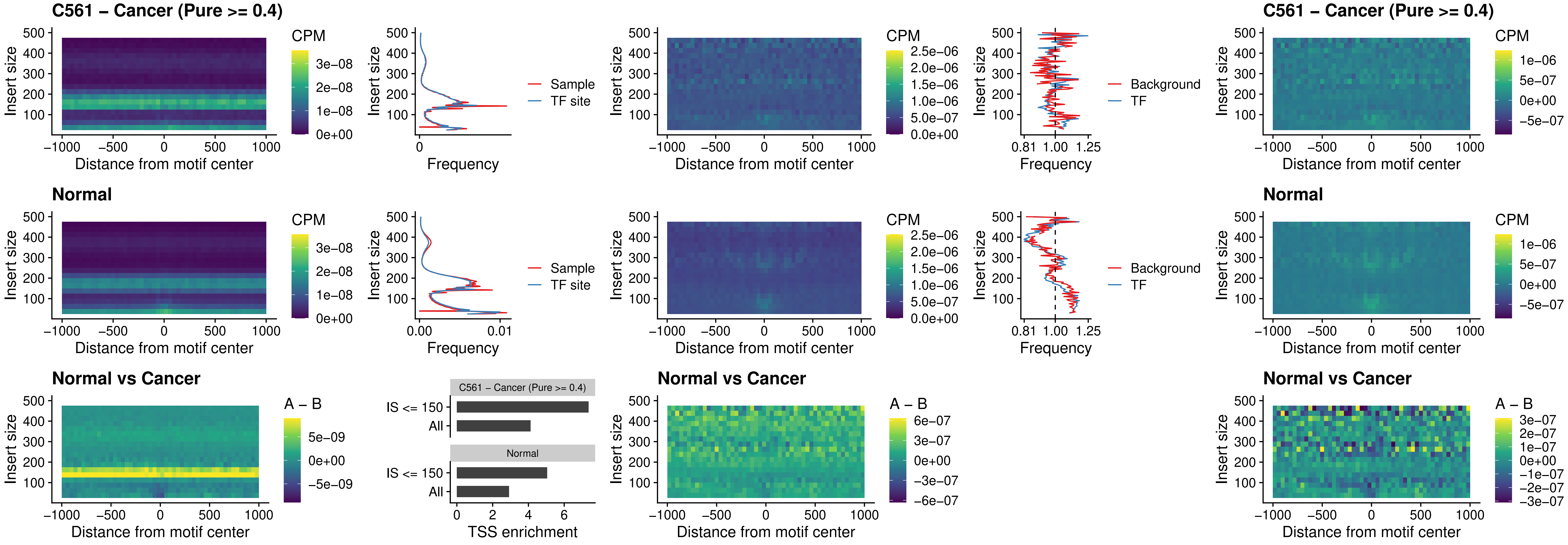

### IRF3_uc_pTSS_oPEAK

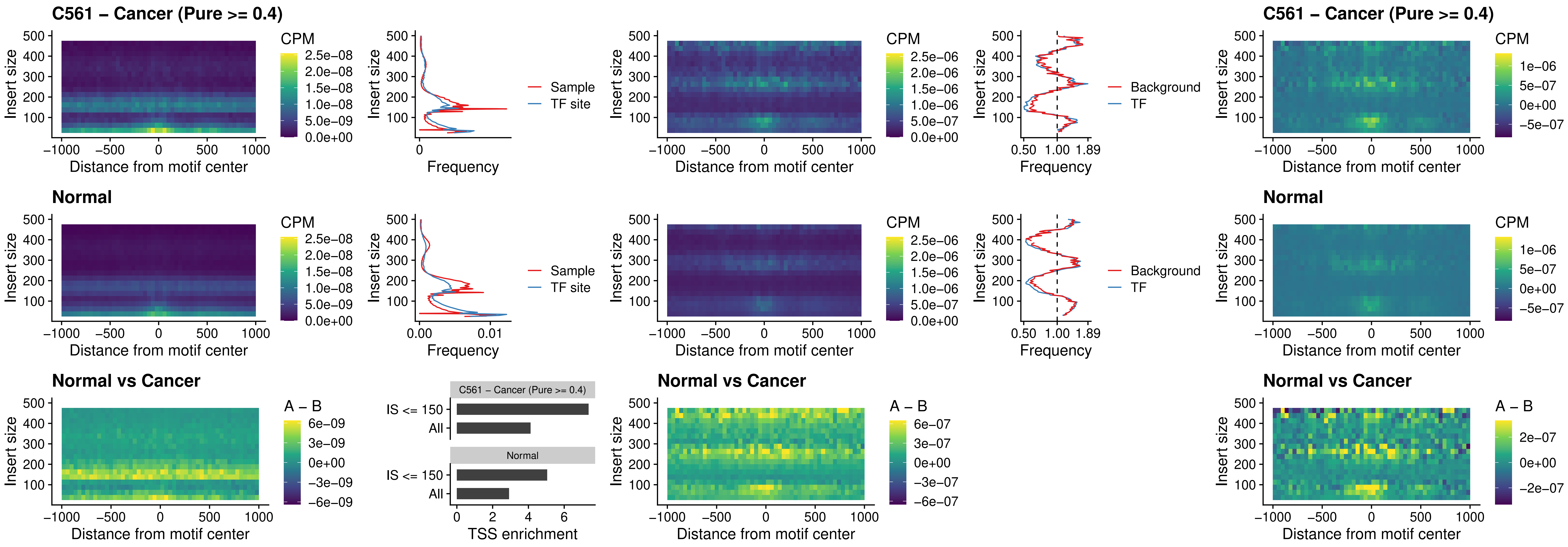

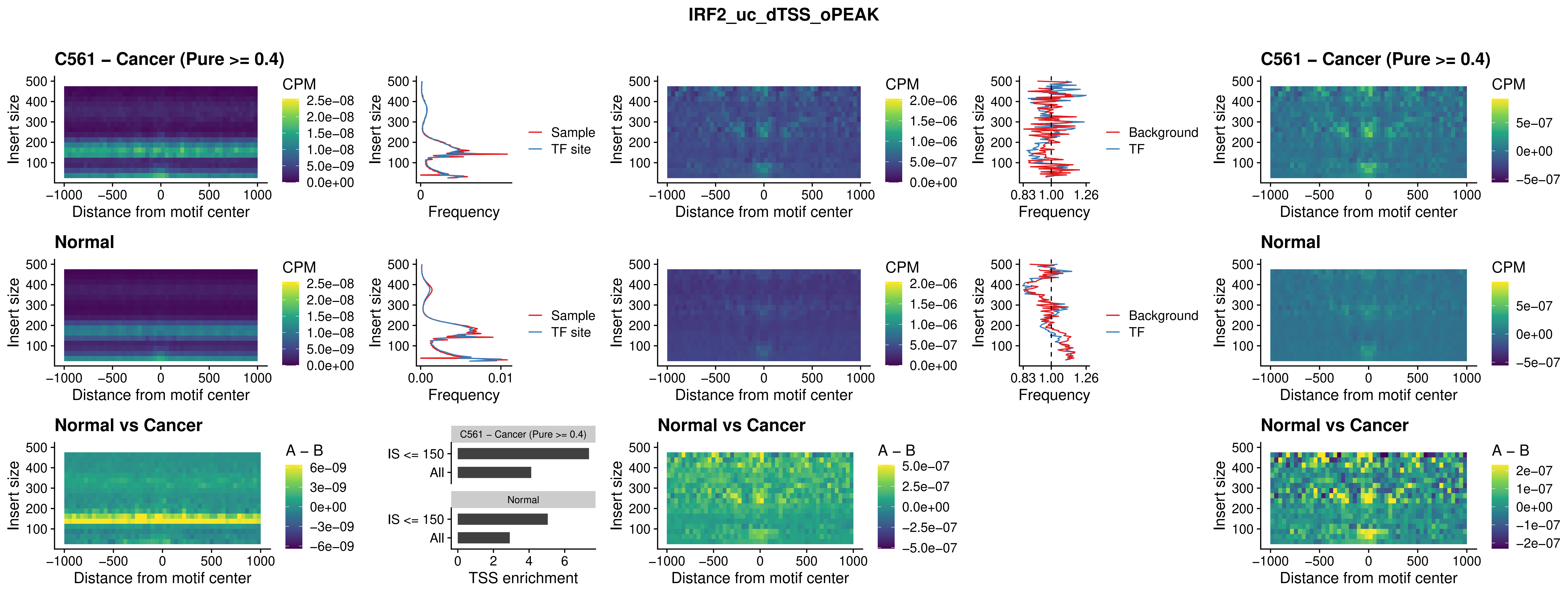
IRF2_uc_dTSS_oPEAK
C561 – Cancer (Pure >= 0.4)
Normal
Normal vs Cancer
Insert size
Distance from motif center
Frequency
CPM
A – B
Sample
TF site
Background
TF
C561 – Cancer (Pure >= 0.4)
Normal
IS <= 150
All
TSS enrichment

### IRF2_uc_pTSS_oPEAK

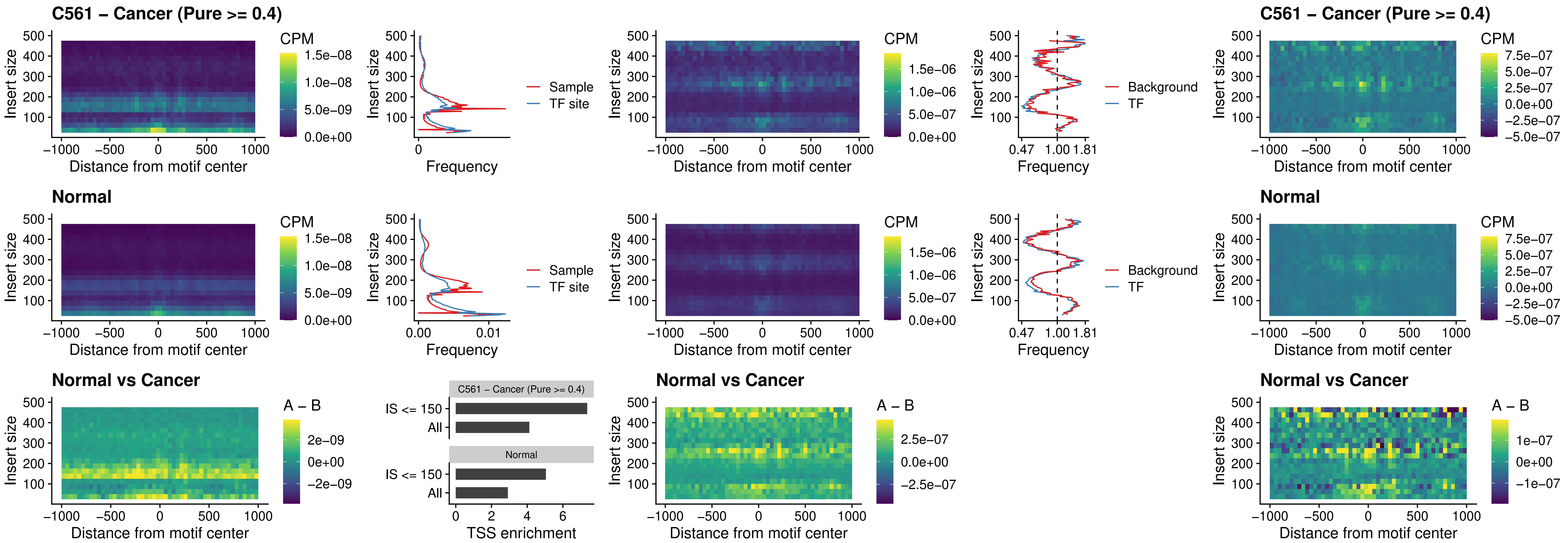

#### STAT2_uc_pTSS_oPEAK

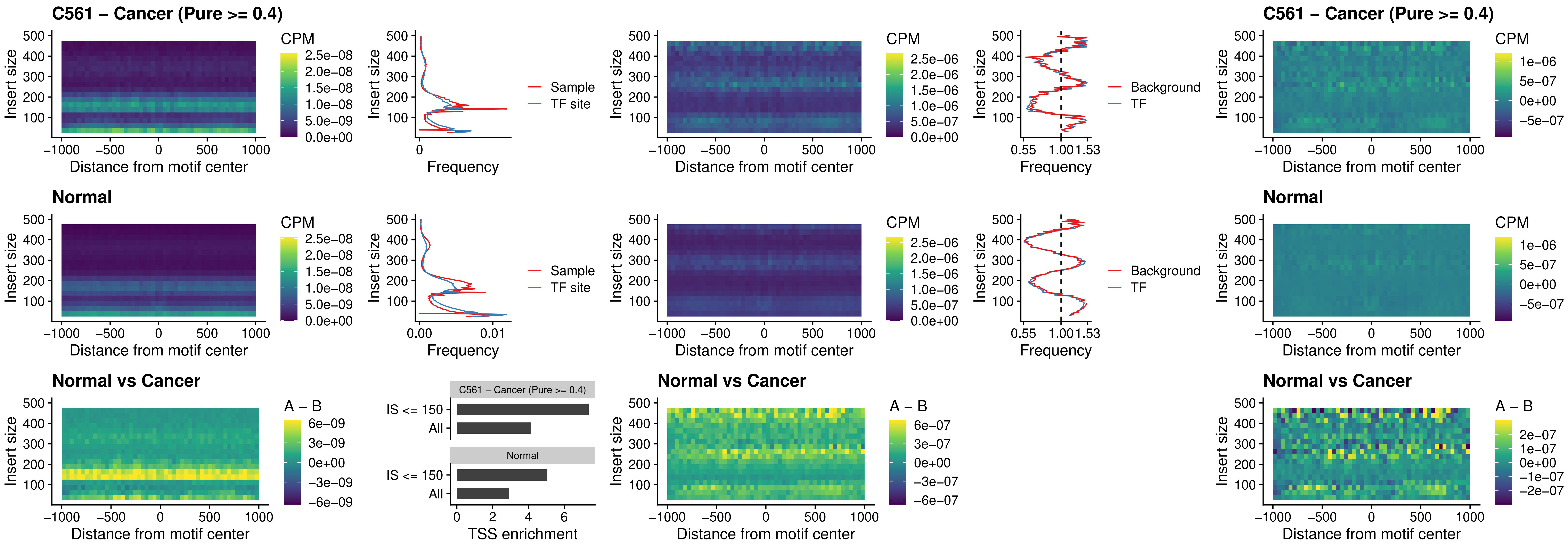

#### SPIB_uc_dTSS_nPEAK

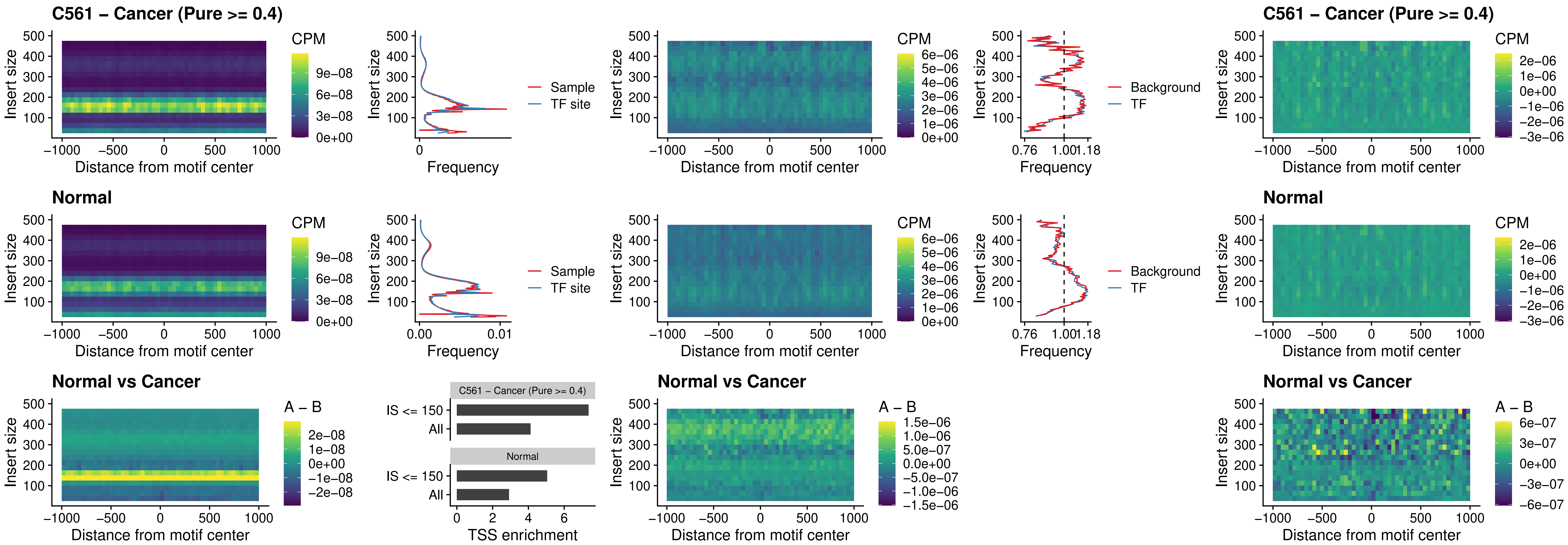

#### SPIC_uc_cTSS_oPEAK

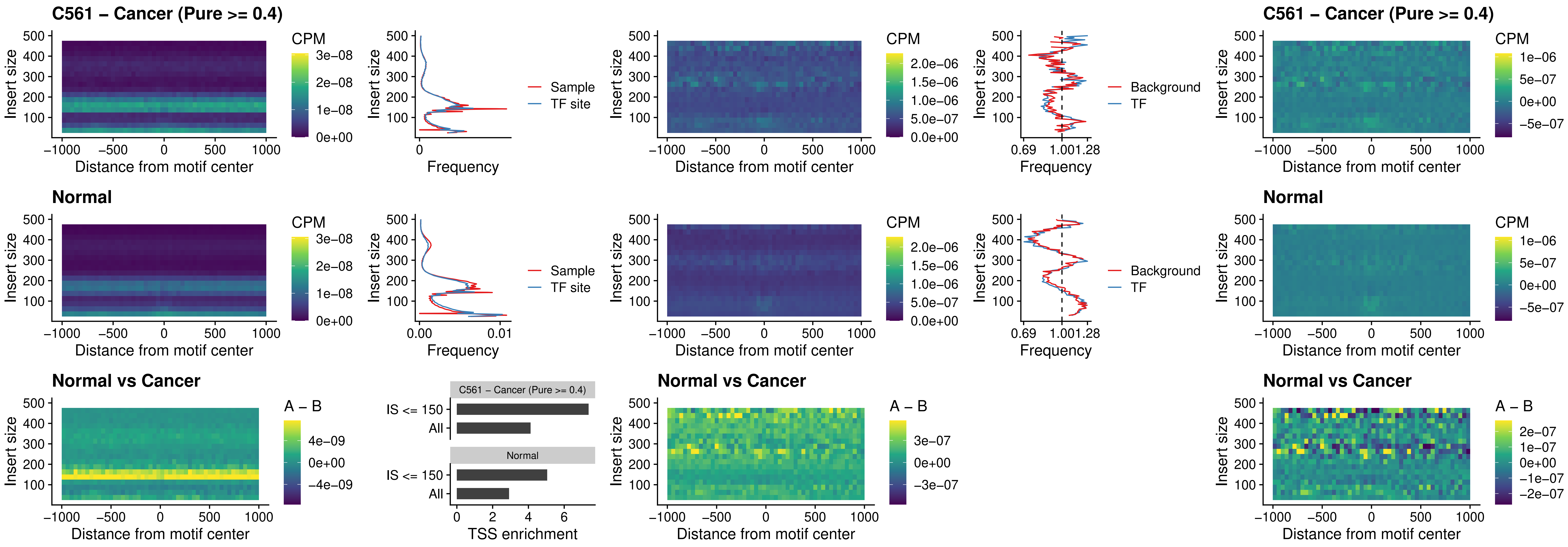

### PRDM1_uc_cTSS_oPEAK

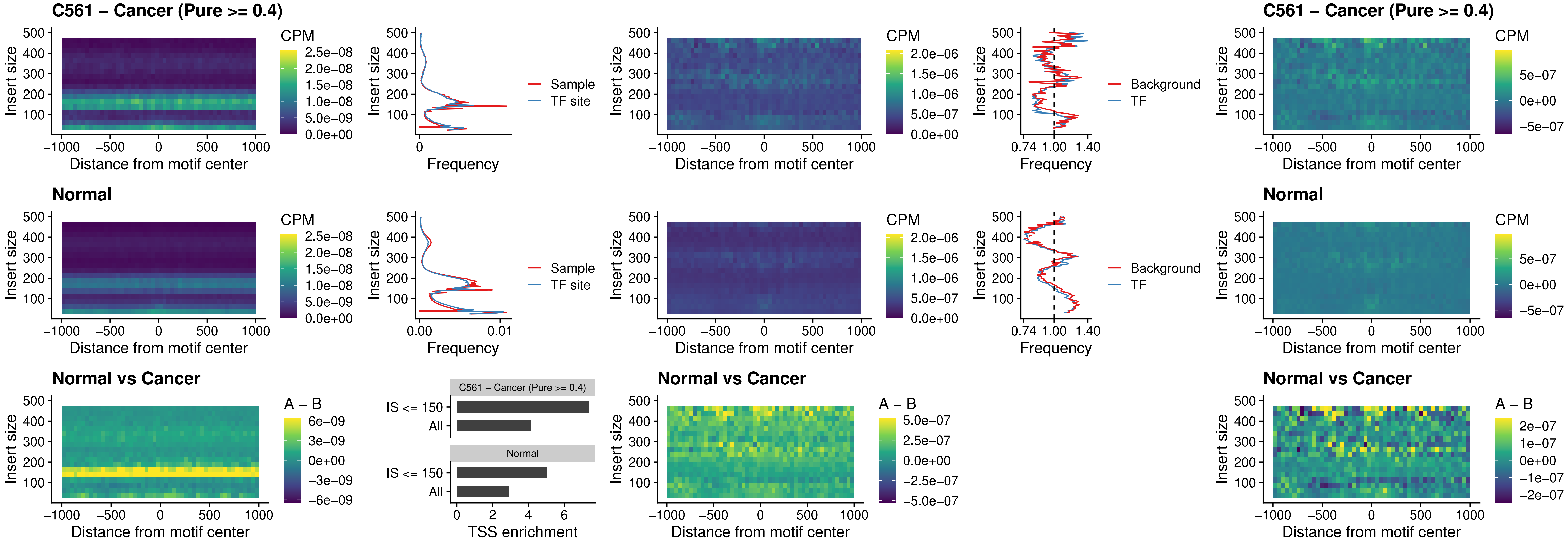

### IRF3_uc_cTSS_oPEAK

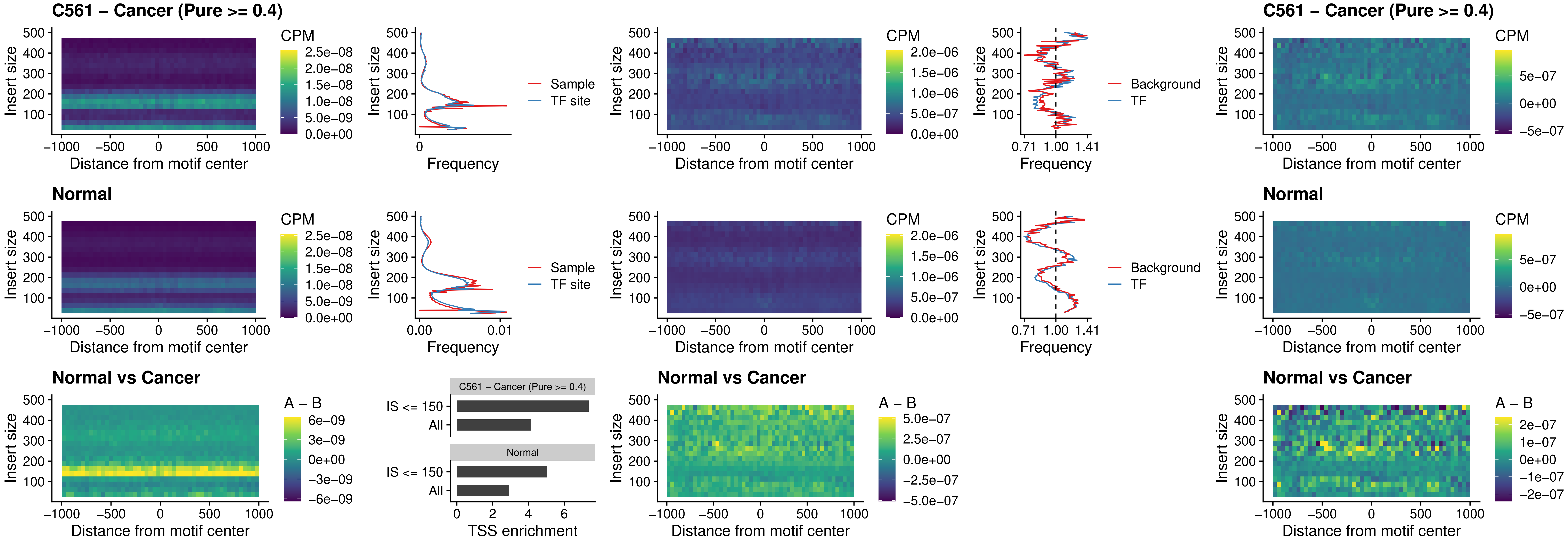

### BCL11A_uc_dTSS_nPEAK

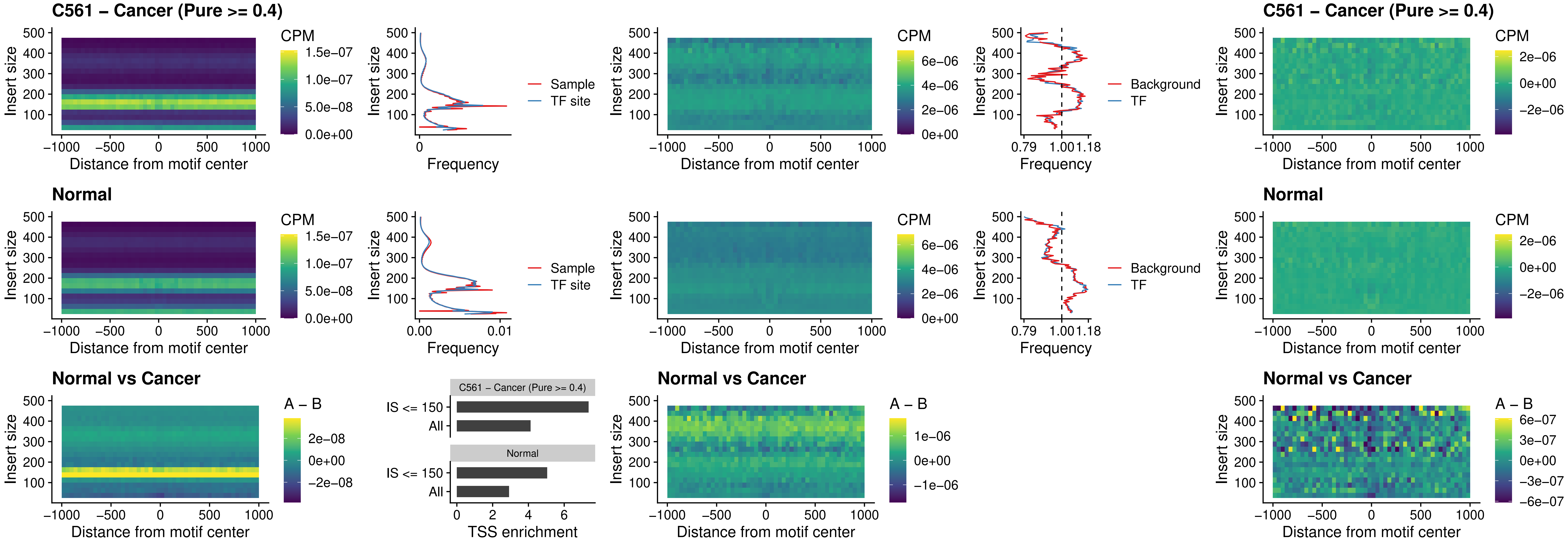

### BCL11B_uc_dTSS_nPEAK

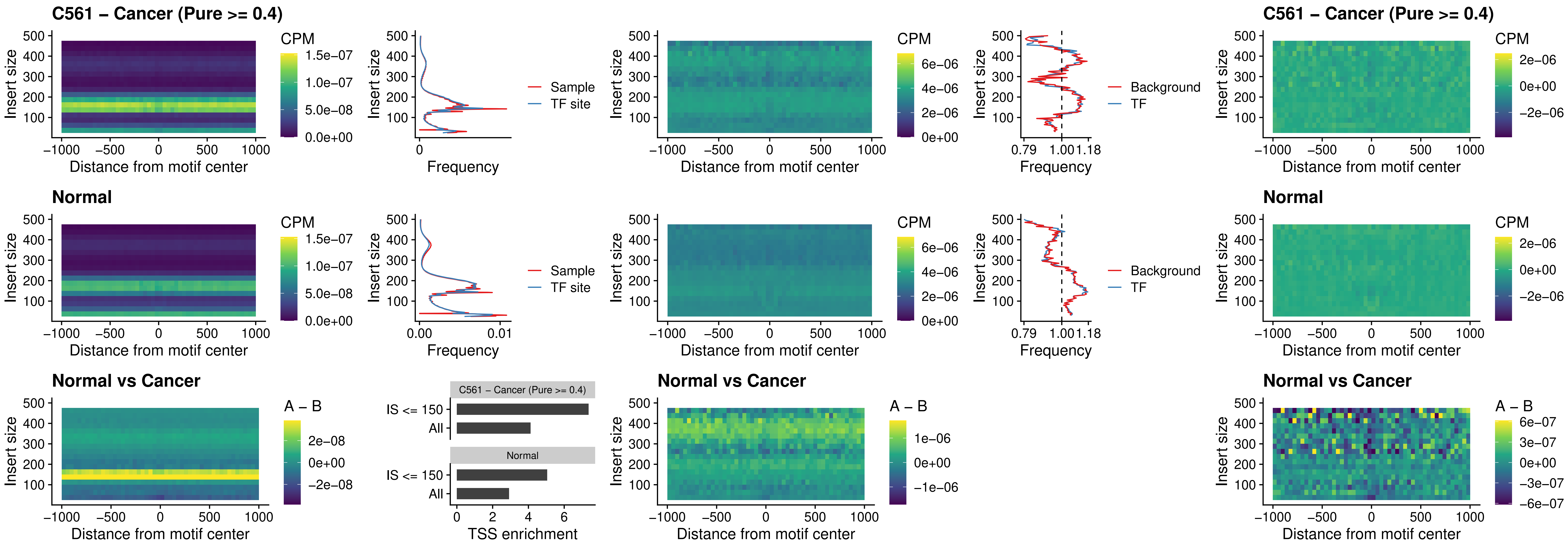

### IRF1_uc_cTSS_oPEAK

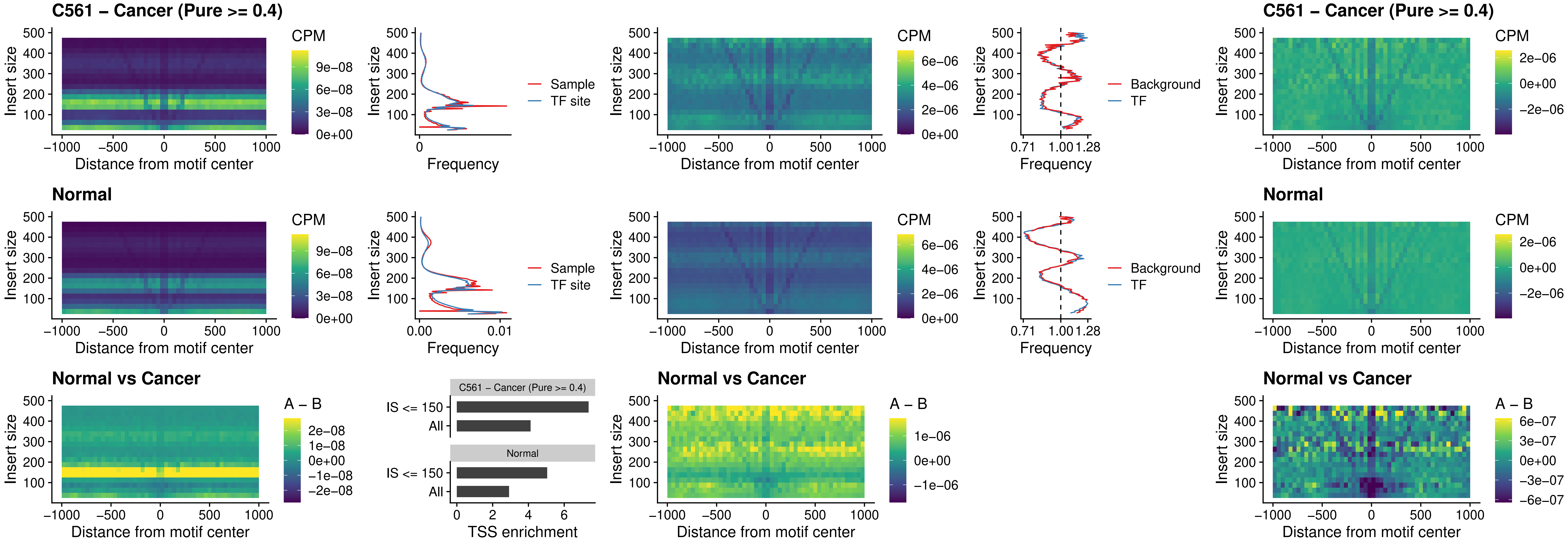

### IRF1_uc_dTSS_oPEAK

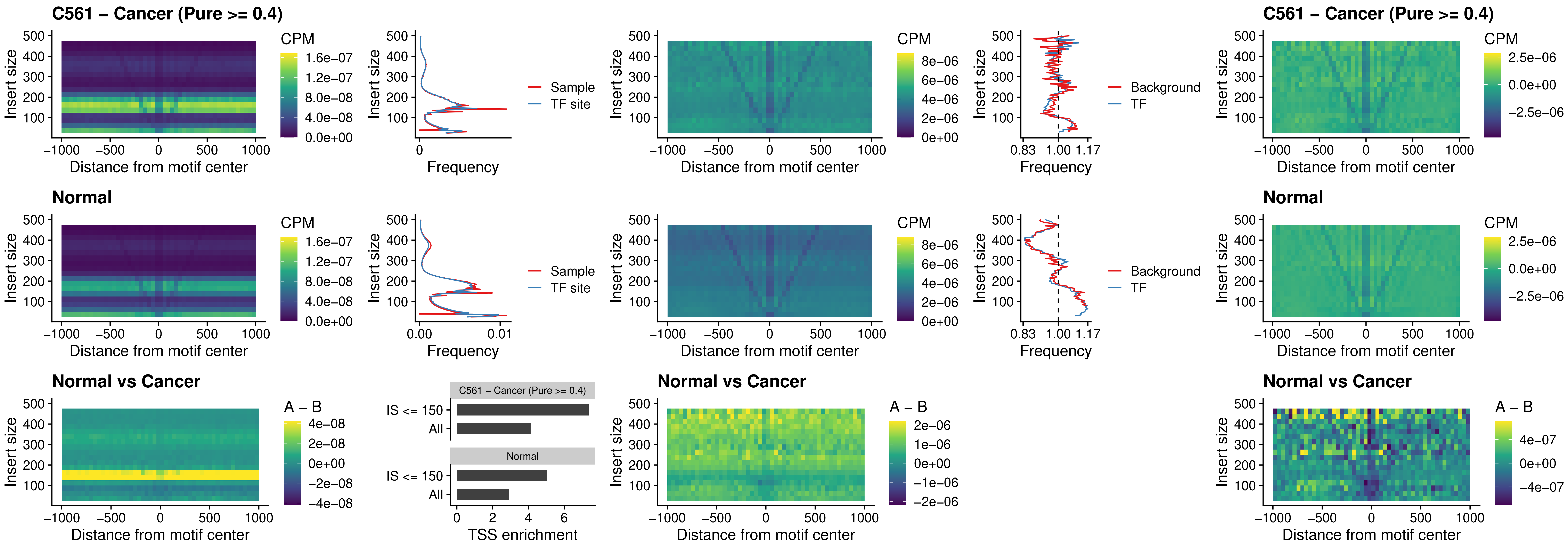

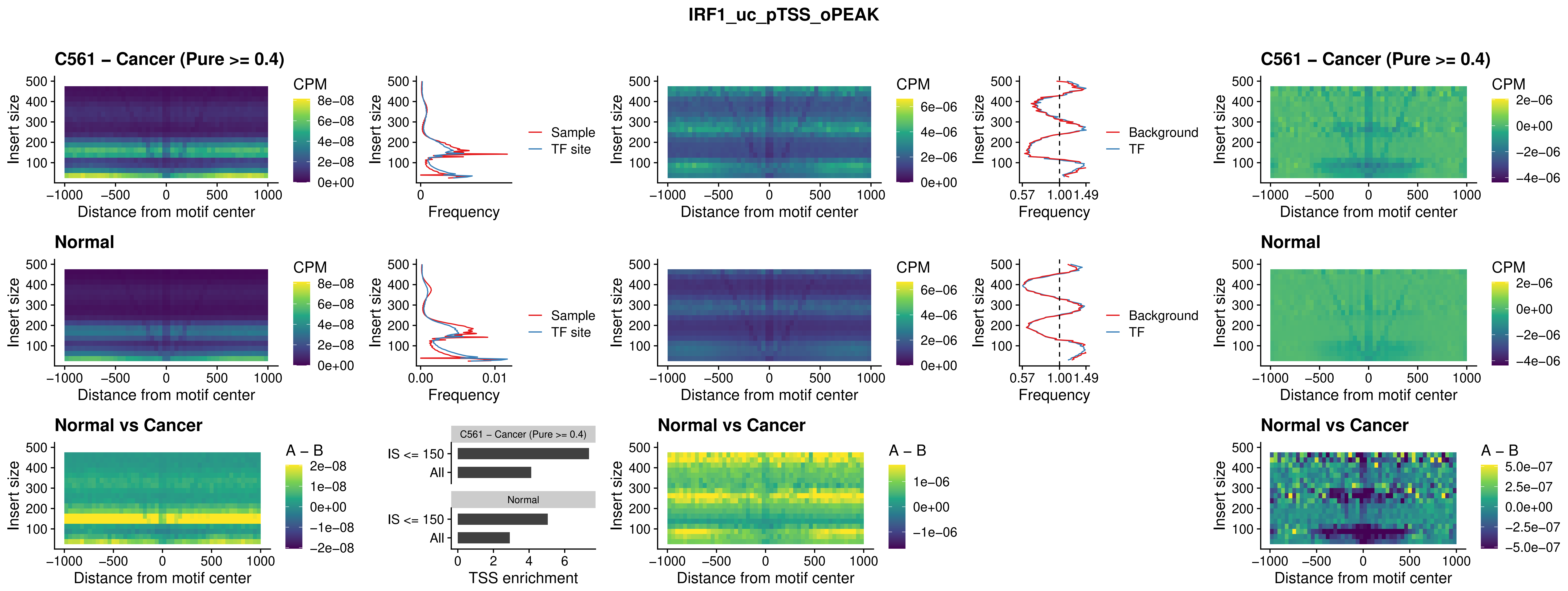
IRF1_uc_pTSS_oPEAK
C561 – Cancer (Pure >= 0.4)
Normal
Normal vs Cancer
Insert size
Distance from motif center
Frequency
CPM
A – B
Sample
TF site
Background
TF
C561 – Cancer (Pure >= 0.4)
Normal
IS <= 150
All
TSS enrichment

#### SPIC_uc_dTSS_nPEAK

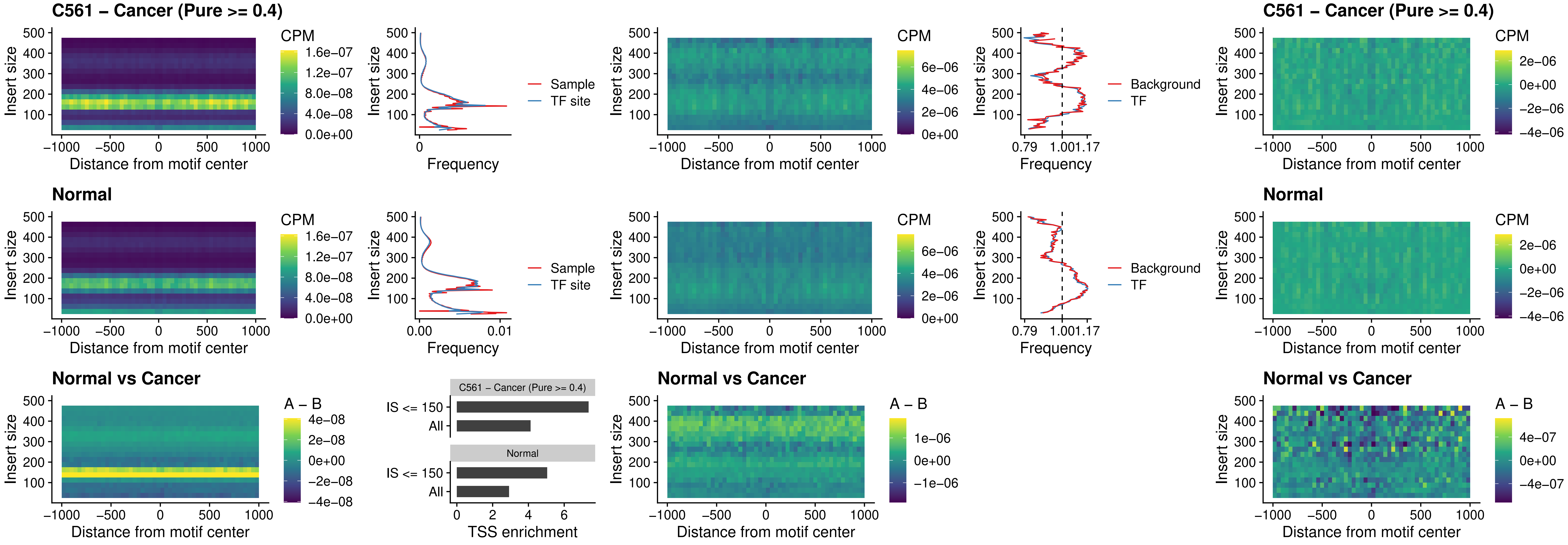

#### SPIC_uc_dTSS_oPEAK

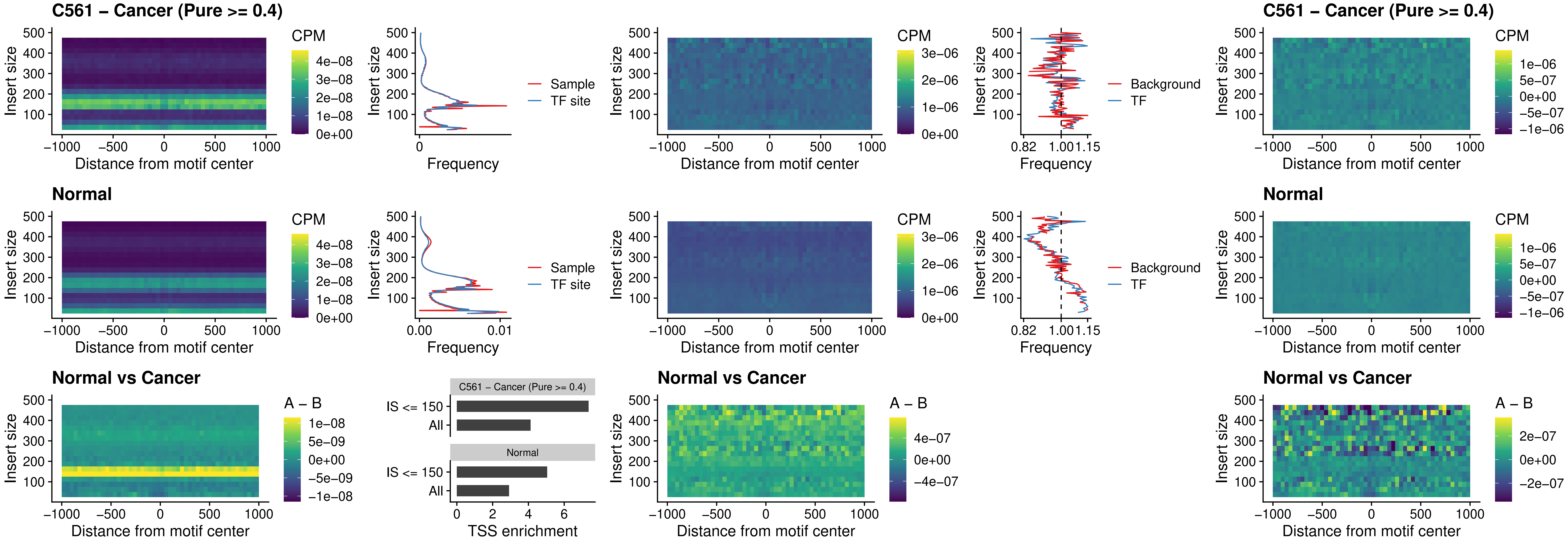

#### SPIB_uc_pTSS_nPEAK

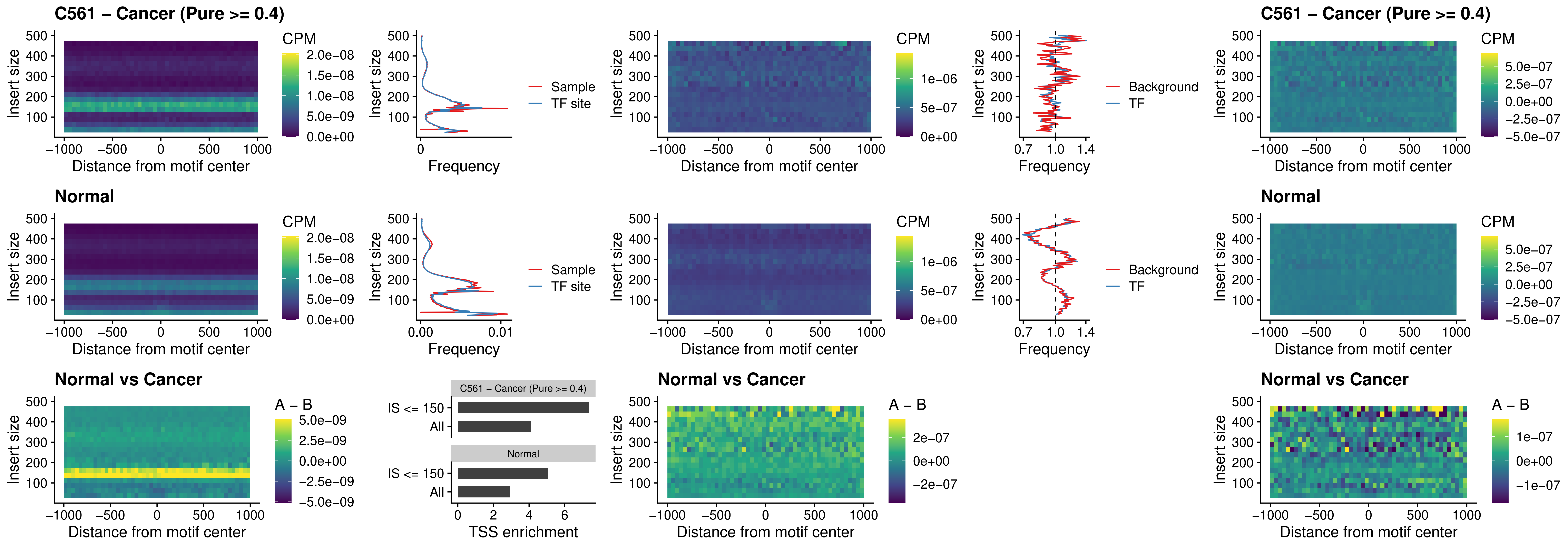

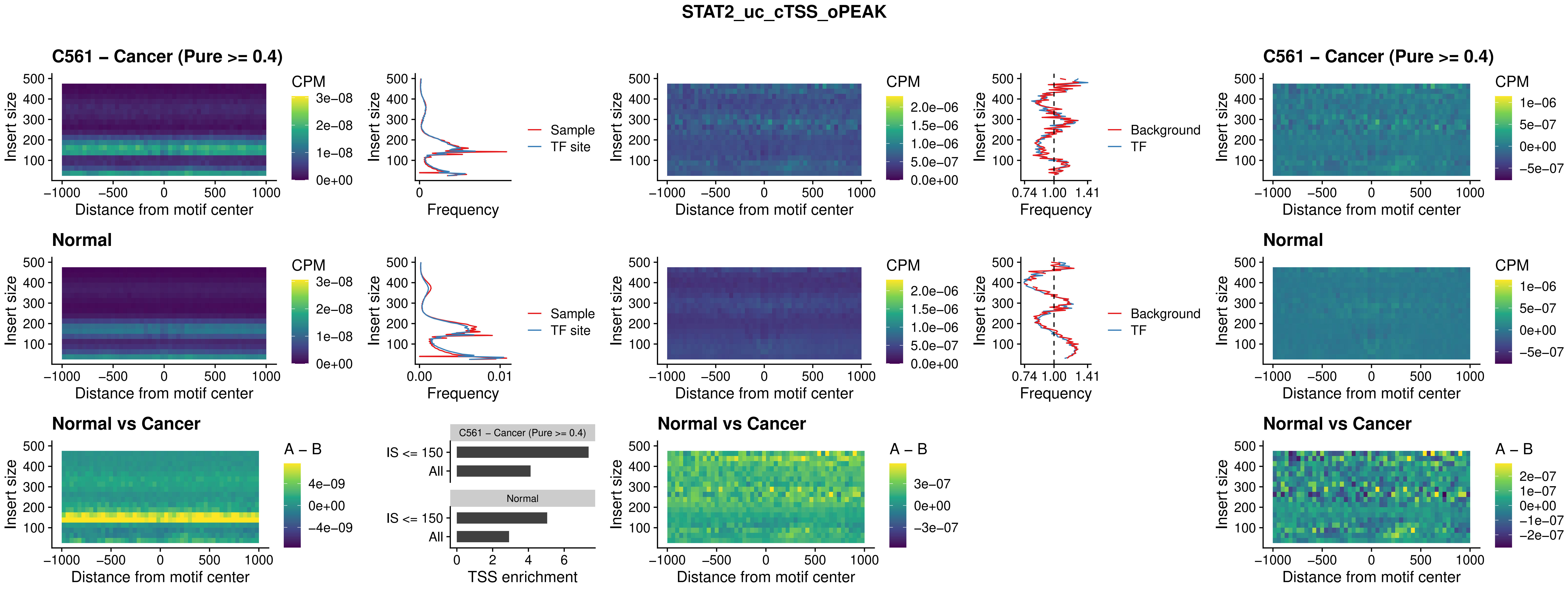
STAT2_uc_cTSS_oPEAK
C561 – Cancer (Pure >= 0.4)
Normal
Normal vs Cancer
Insert size
Distance from motif center
Frequency
CPM
A – B
Sample
TF site
Background
TF
C561 – Cancer (Pure >= 0.4)
Normal
IS <= 150
All
TSS enrichment

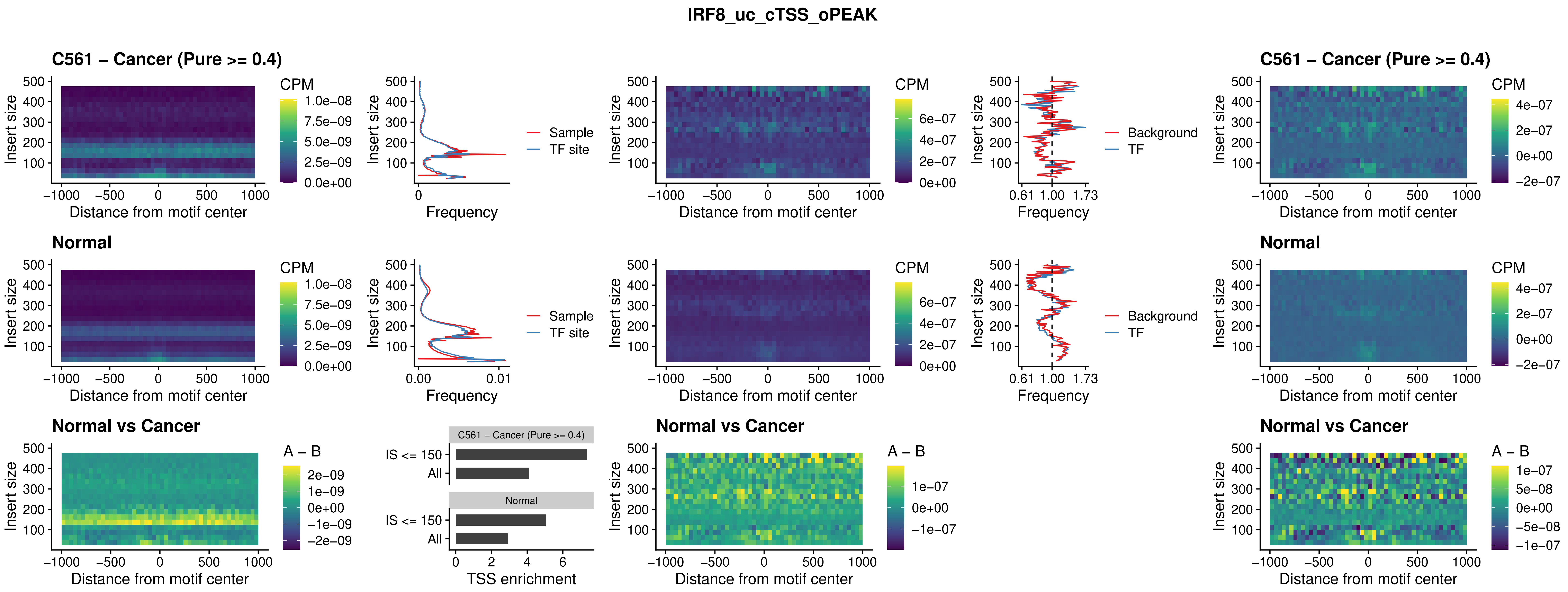
IRF8_uc_cTSS_oPEAK
C561 – Cancer (Pure >= 0.4)
Normal
Normal vs Cancer
Insert size
Distance from motif center
Frequency
CPM
A – B
Sample
TF site
Background
TF
TSS enrichment
IS <= 150
All

### IRF3_uc_dTSS_oPEAK

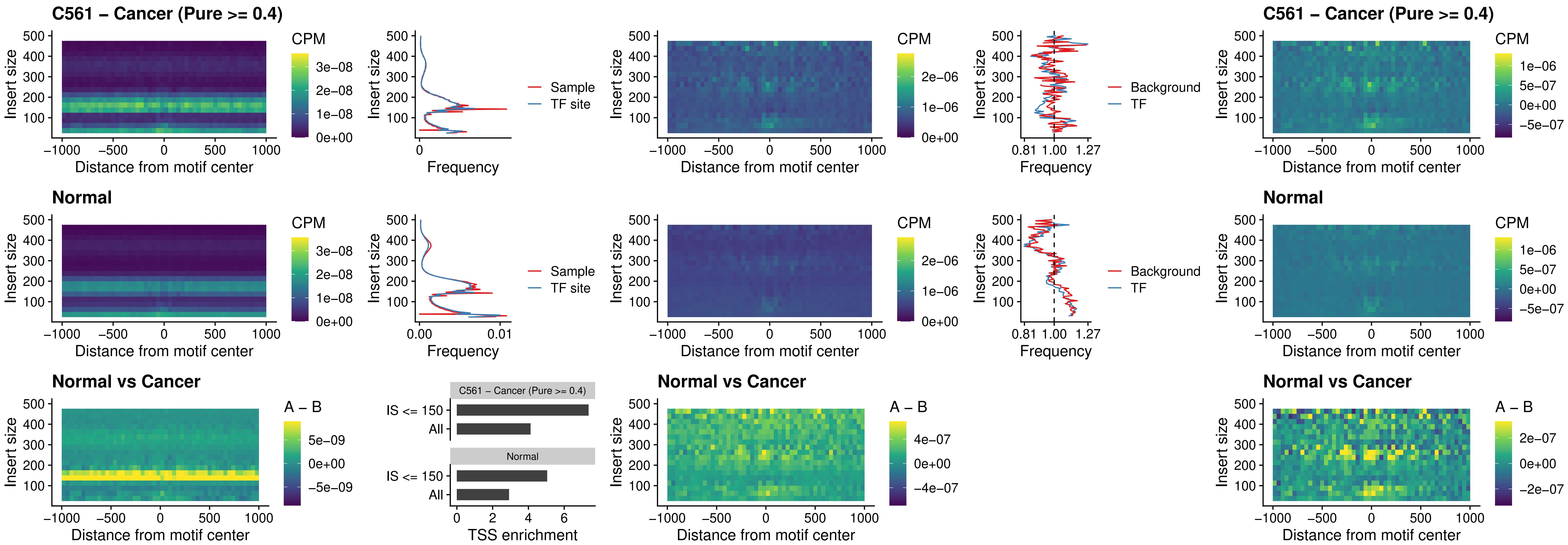

### IRF8_uc_dTSS_oPEAK

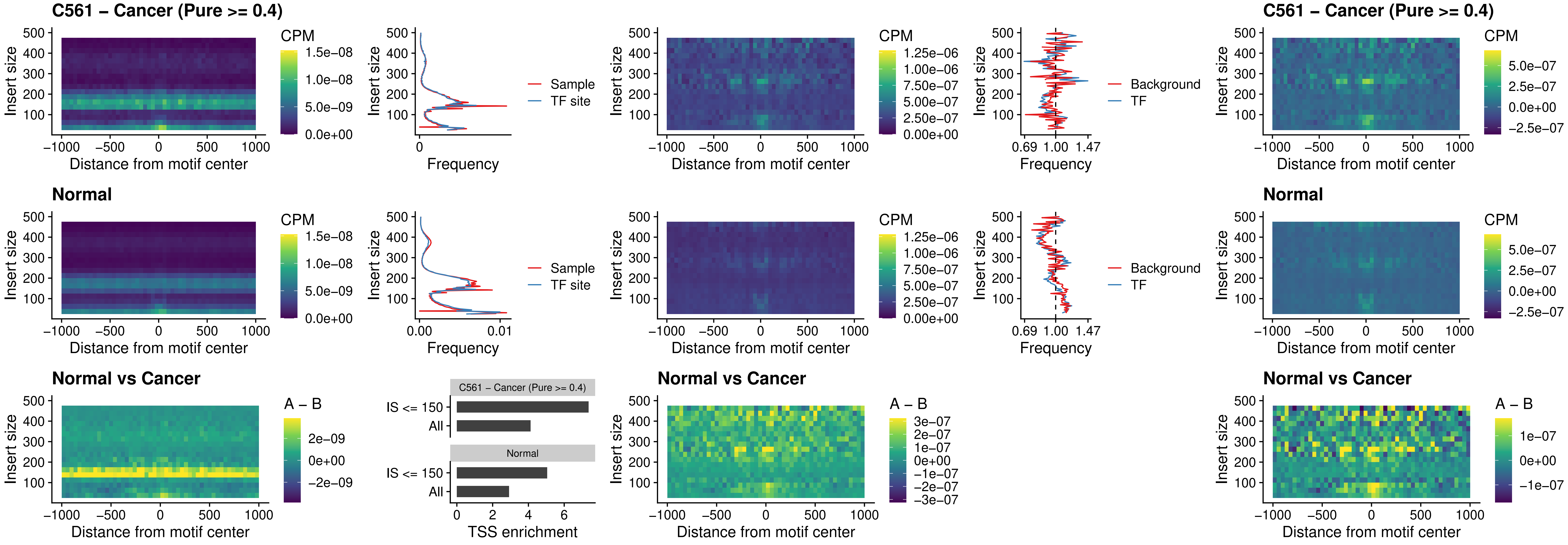

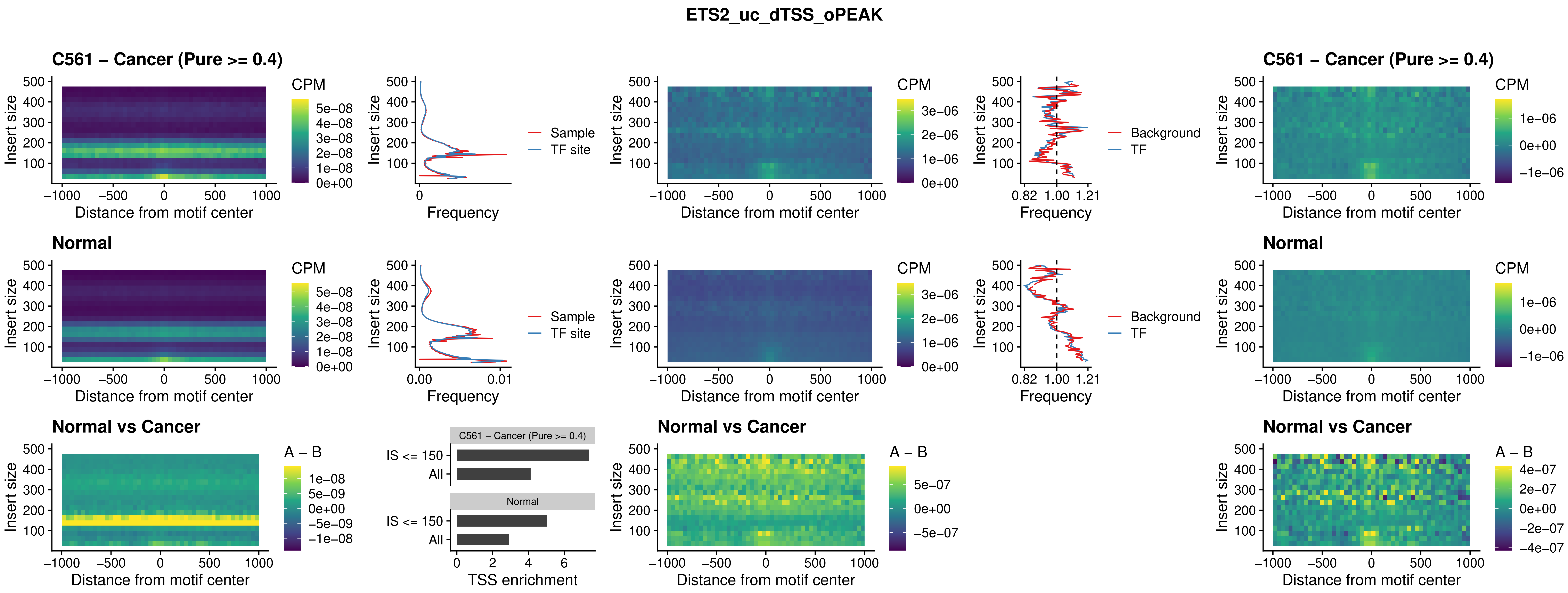
ETS2_uc_dTSS_oPEAK
C561 – Cancer (Pure >= 0.4)
Normal
Normal vs Cancer
Insert size
Distance from motif center
Frequency
CPM
A – B
Sample
TF site
Background
TF
C561 – Cancer (Pure >= 0.4)
Normal
IS <= 150
All
TSS enrichment

#### SMARCC2_uc_pTSS_oPEAK

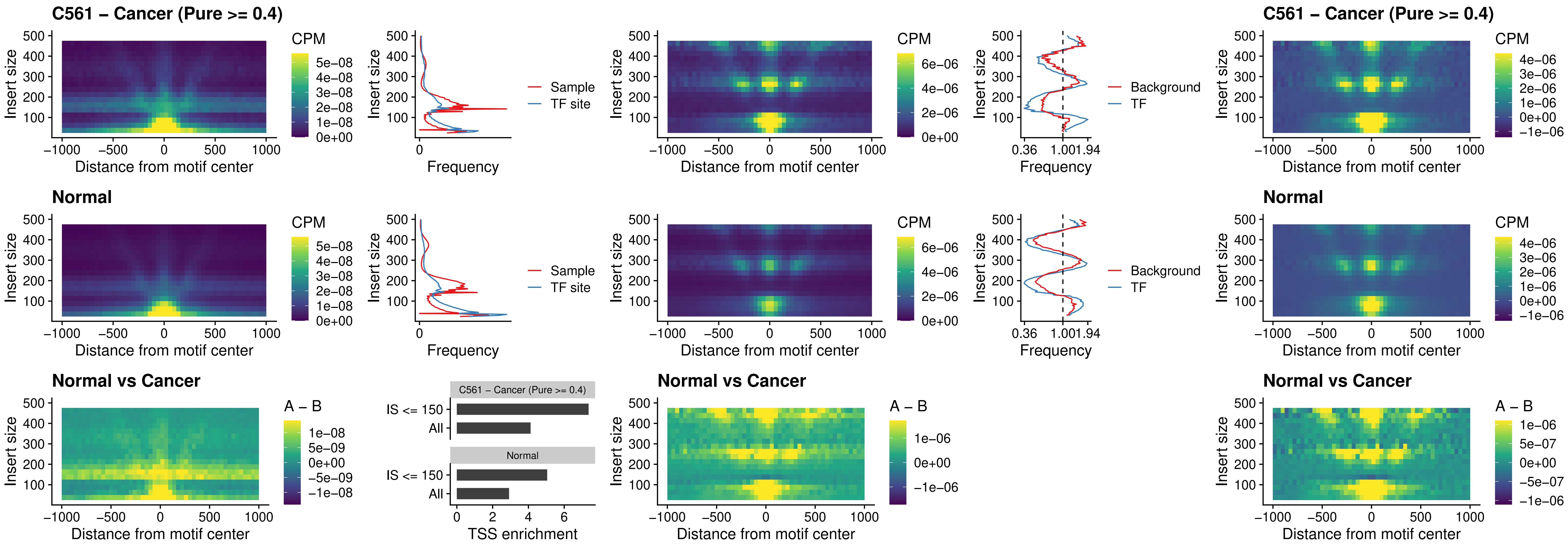

### BCL11A_uc_cTSS_nPEAK

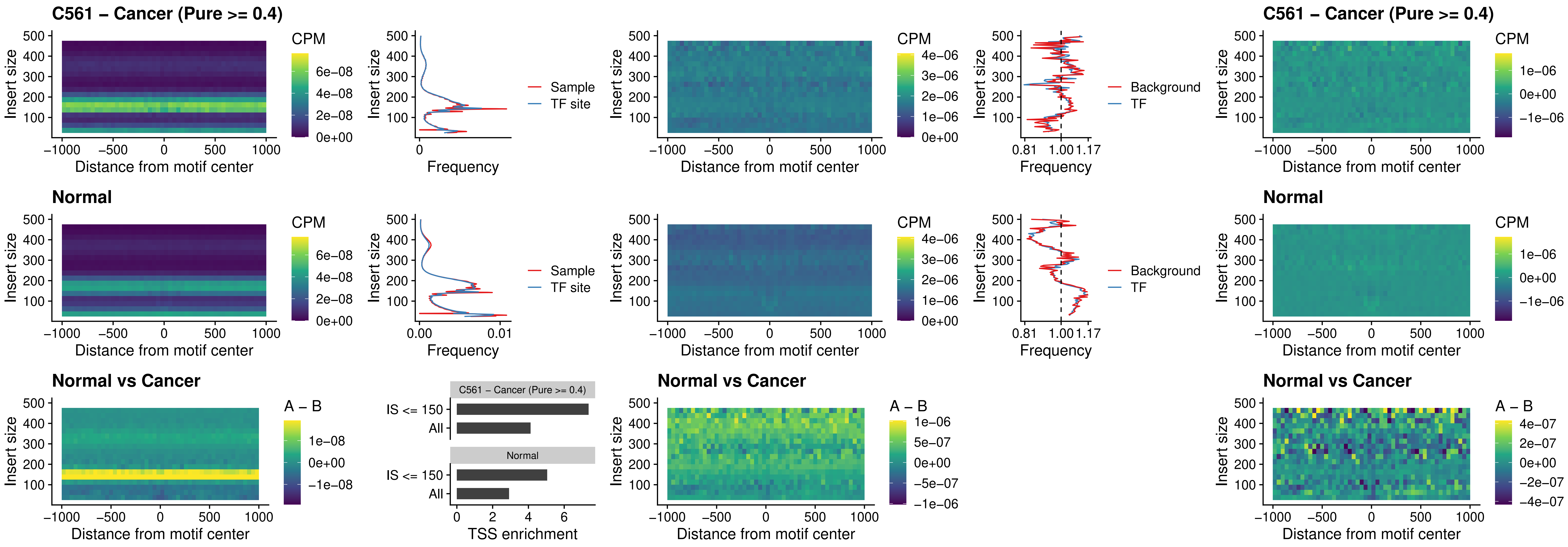

### BCL11B_uc_cTSS_nPEAK

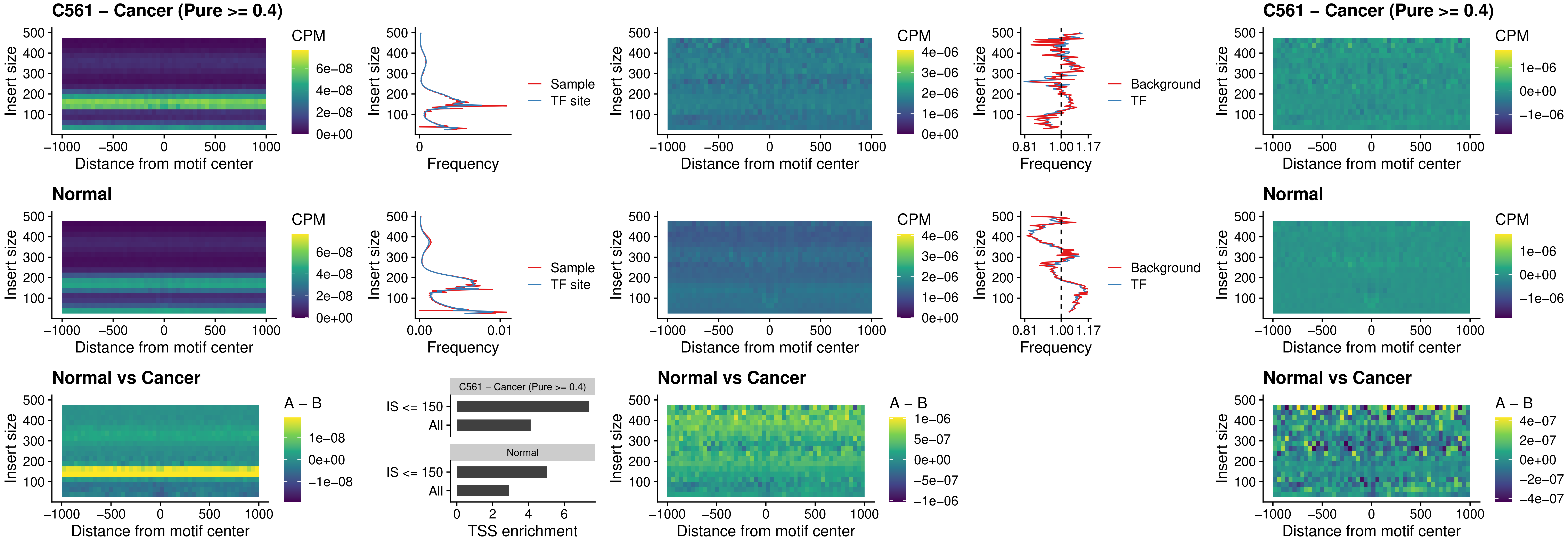

### STAT2_uc_dTSS_oPEAK

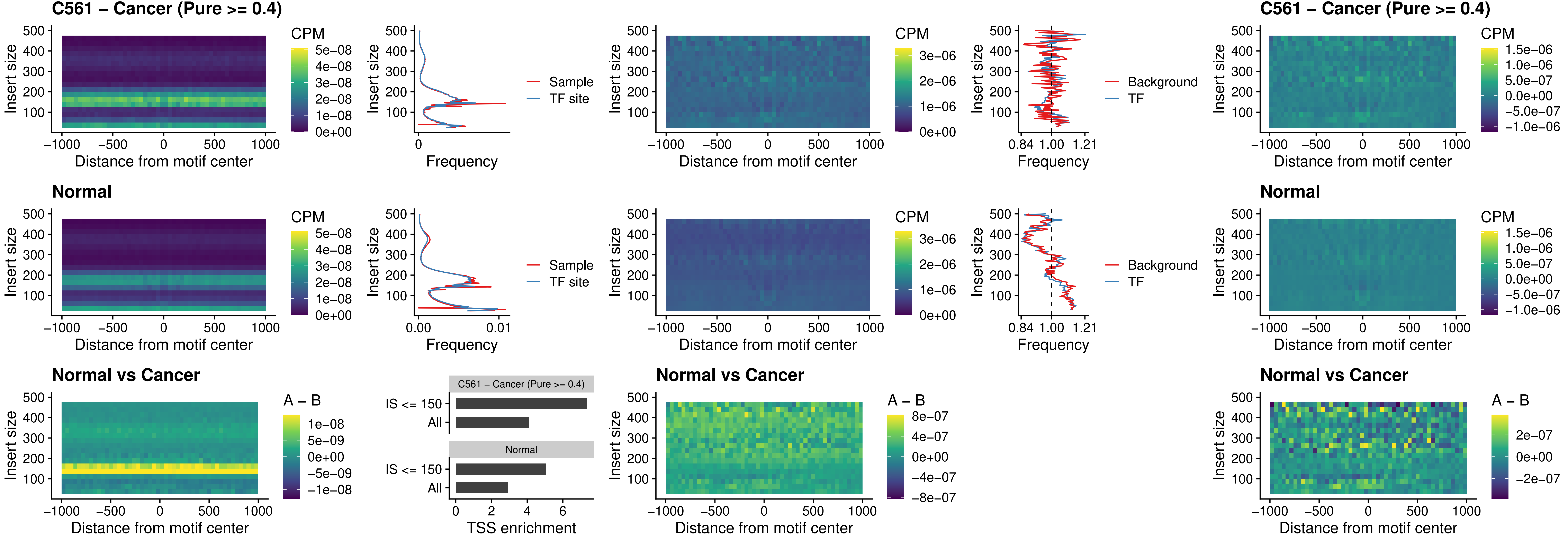

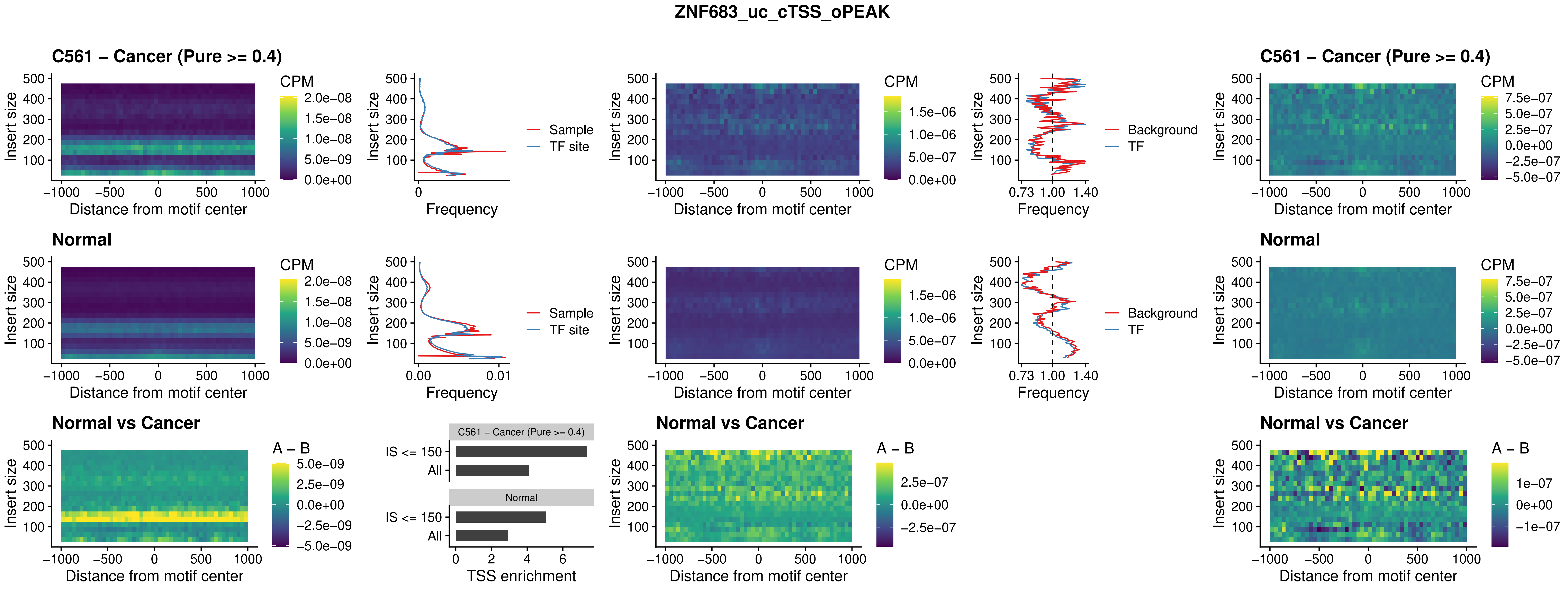
ZNF683_uc_cTSS_oPEAK
C561 – Cancer (Pure >= 0.4)
Normal
Normal vs Cancer
Insert size
Distance from motif center
Frequency
CPM
A – B
Sample
TF site
Background
TF
IS <= 150
All
TSS enrichment
C561 – Cancer (Pure >= 0.4)
Normal

#### IRF8_uc_dTSS_nPEAK

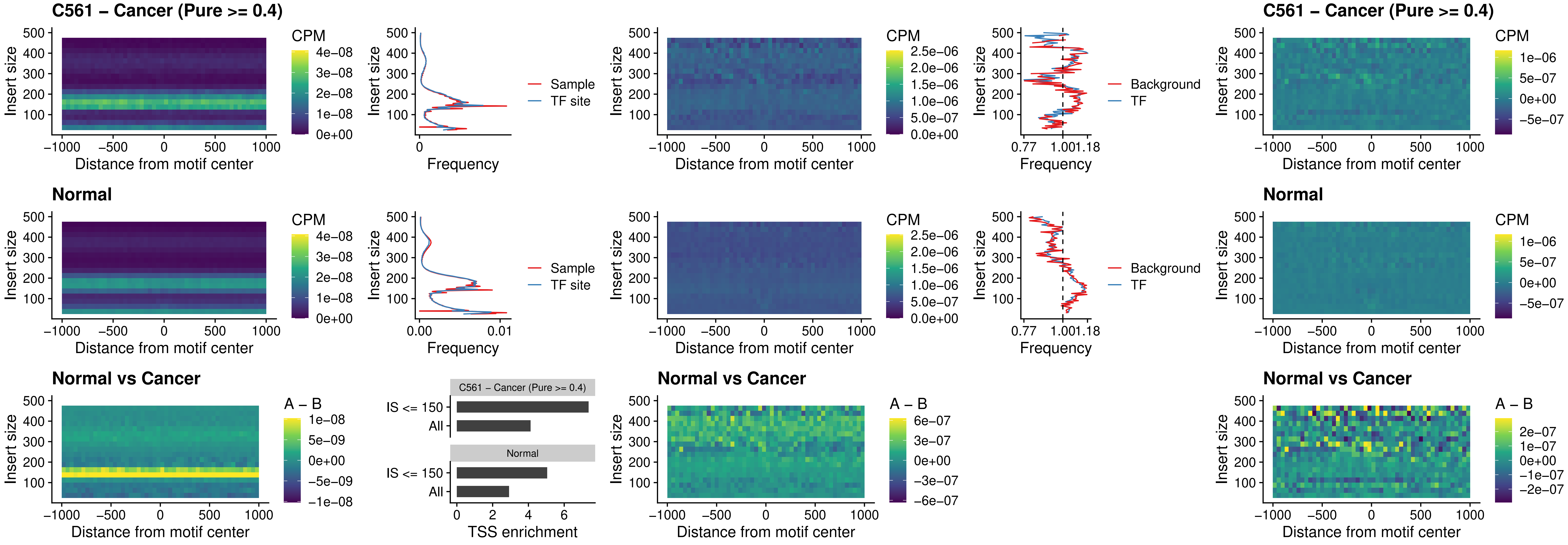

#### ETV7_uc_dTSS_oPEAK

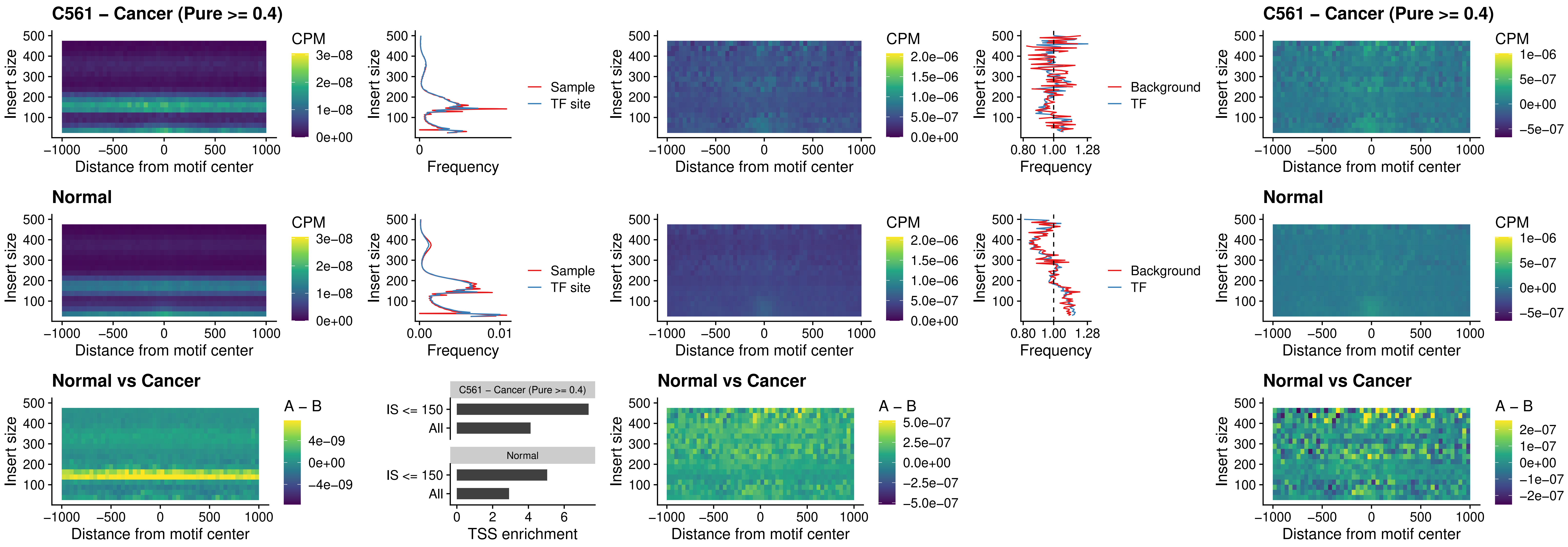

### FOXL1_uc_pTSS_oPEAK

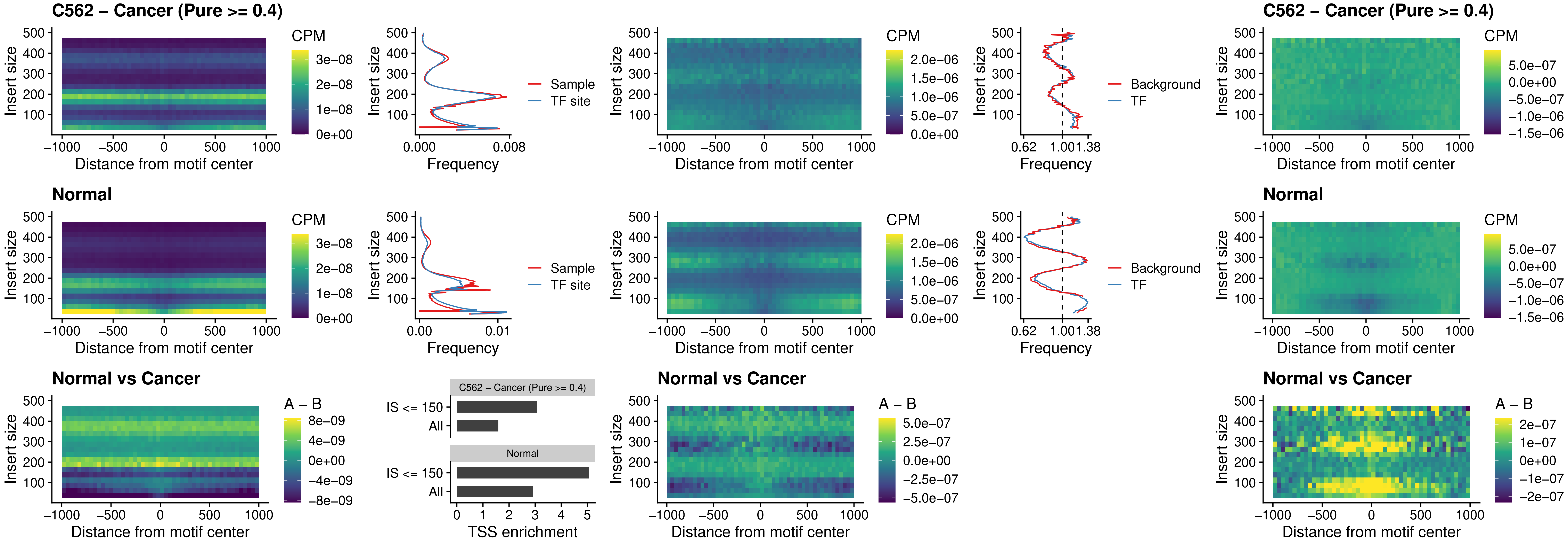

### CPEB1_uc_cTSS_oPEAK

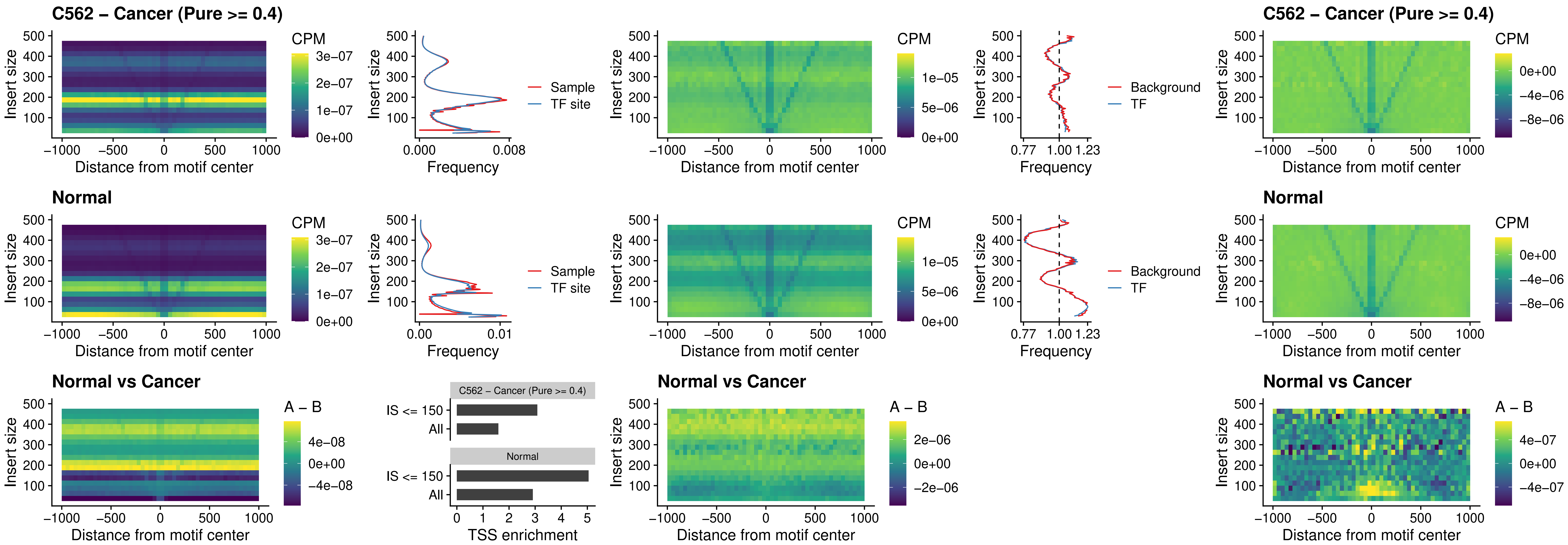

#### HMGA1_uc_pTSS_nPEAK

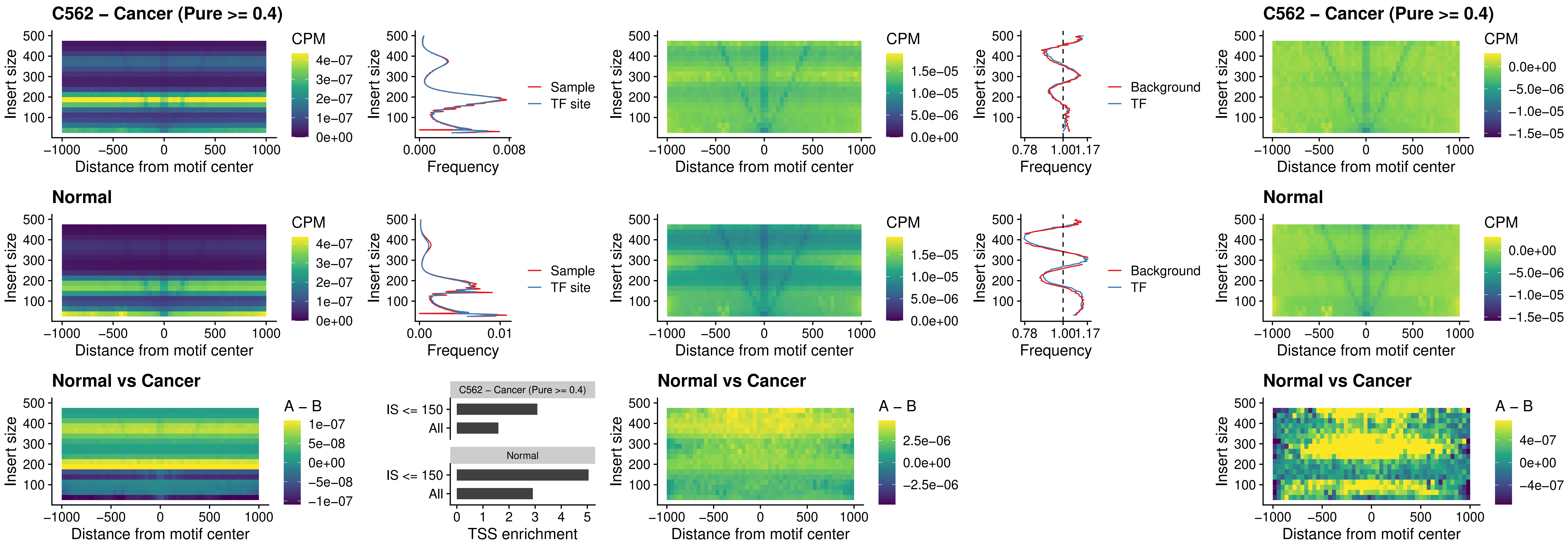

### SOX13_uc_cTSS_nPEAK

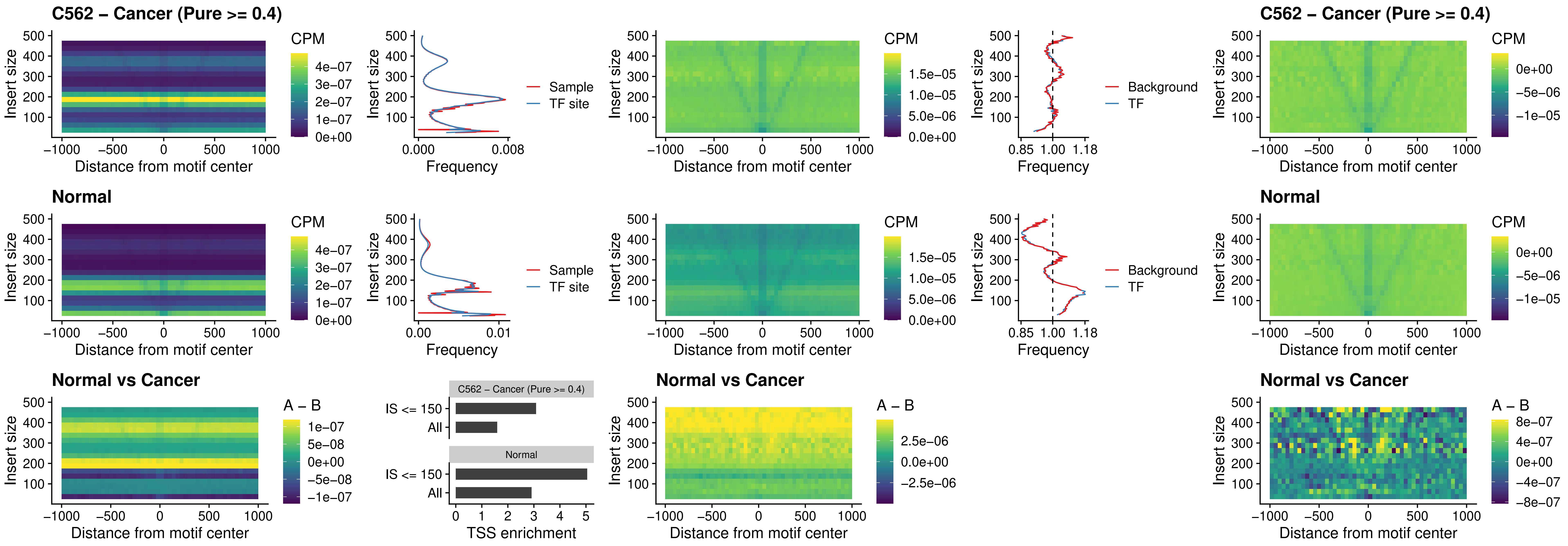

### LHX4_uc_cTSS_nPEAK

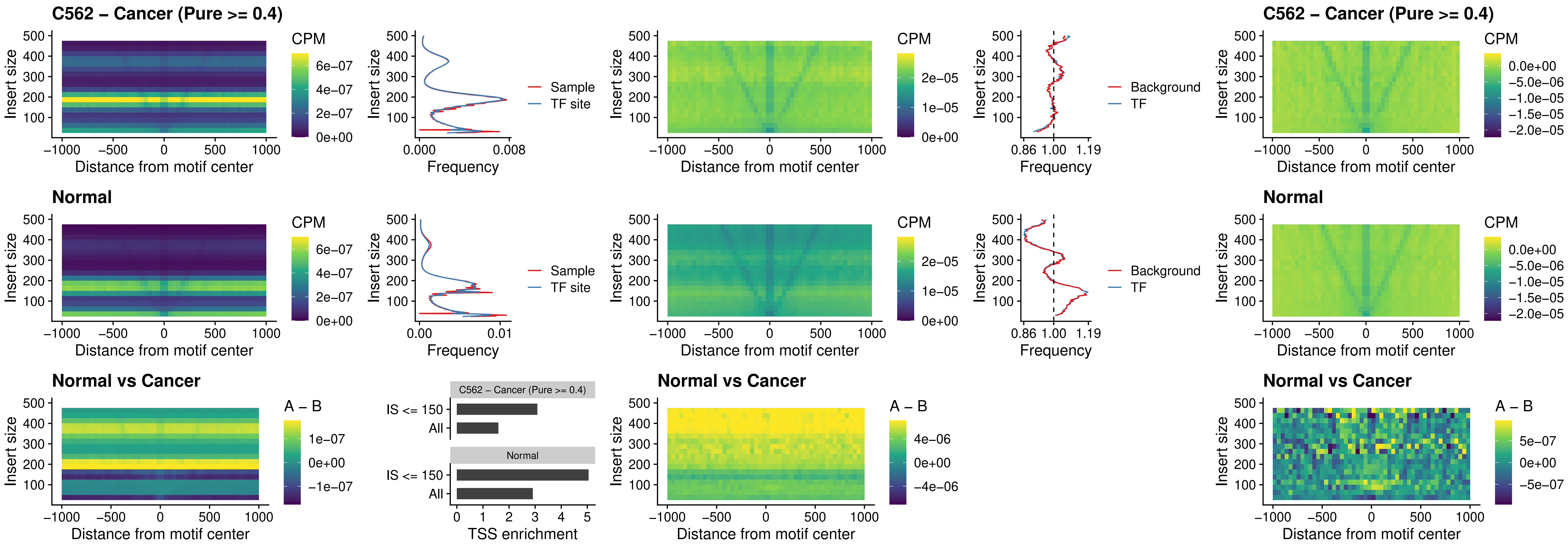

#### HOXA4_uc_cTSS_nPEAK

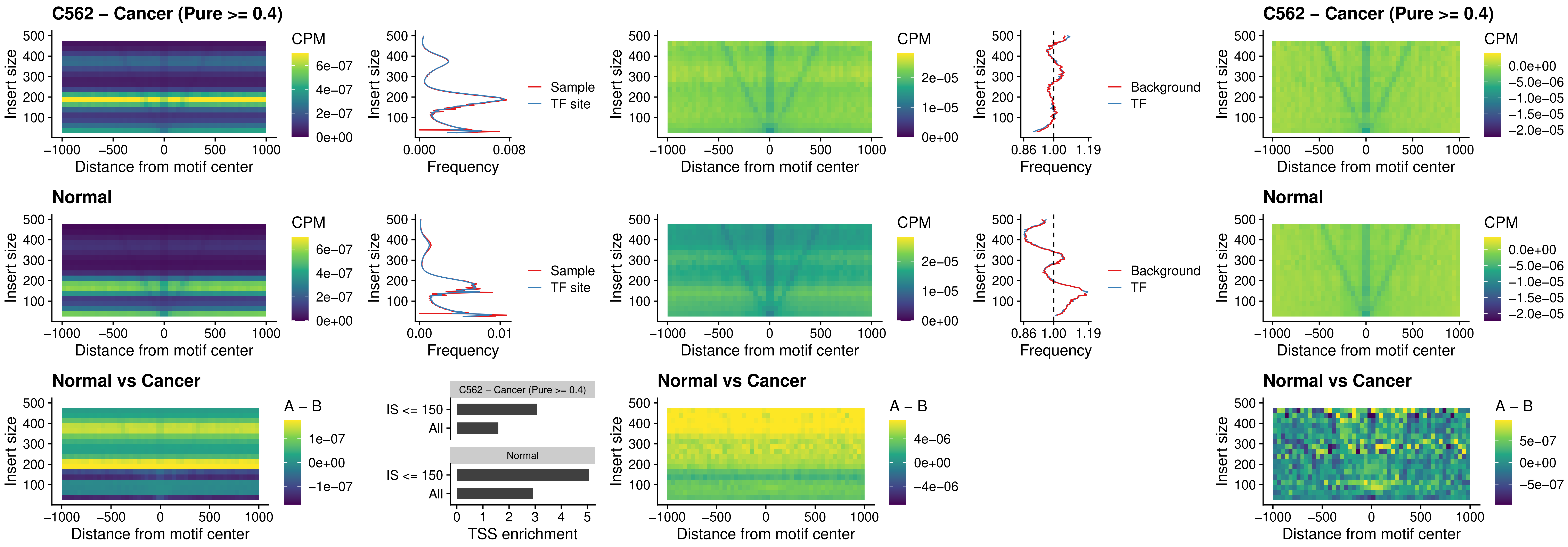

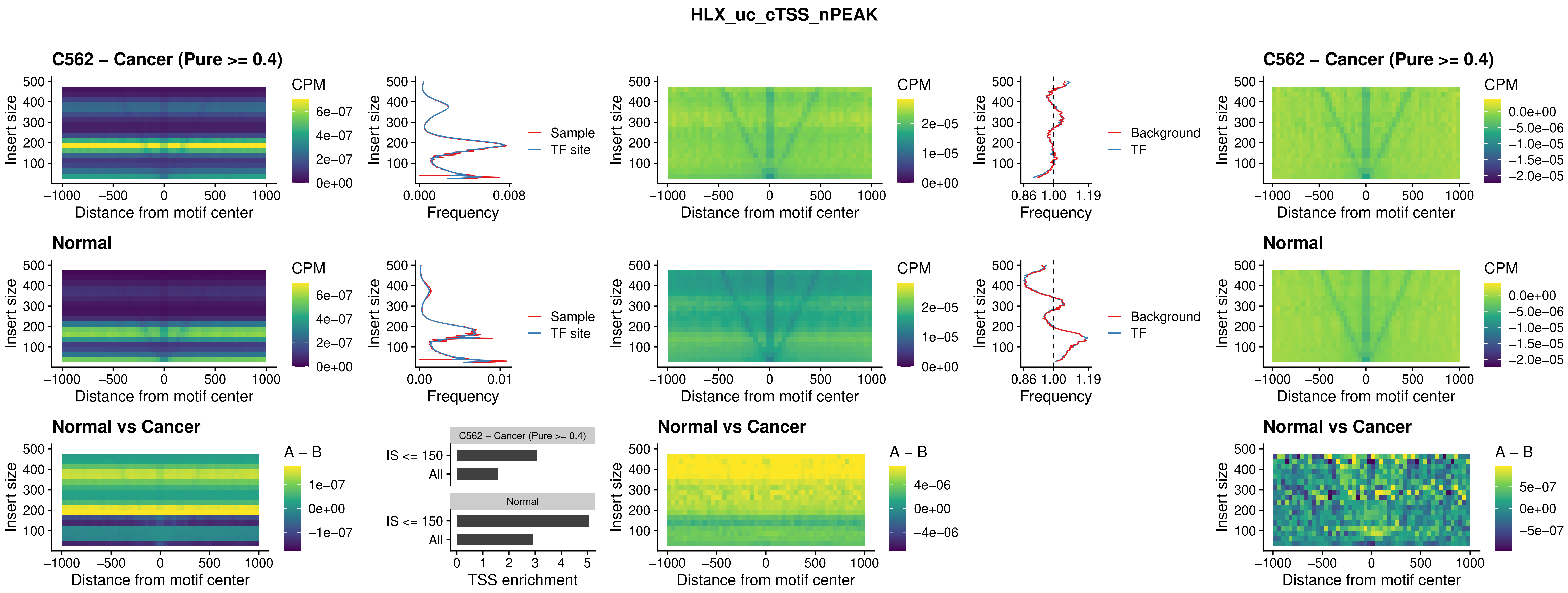
HLX_uc_cTSS_nPEAK
C562 – Cancer (Pure >= 0.4)
Normal
Normal vs Cancer
Insert size
Distance from motif center
Frequency
CPM
A – B
Sample
TF site
Background
TF
C562 – Cancer (Pure >= 0.4)
Normal
IS <= 150
All
TSS enrichment

### AC0125311_uc_cTSS_nPEAK

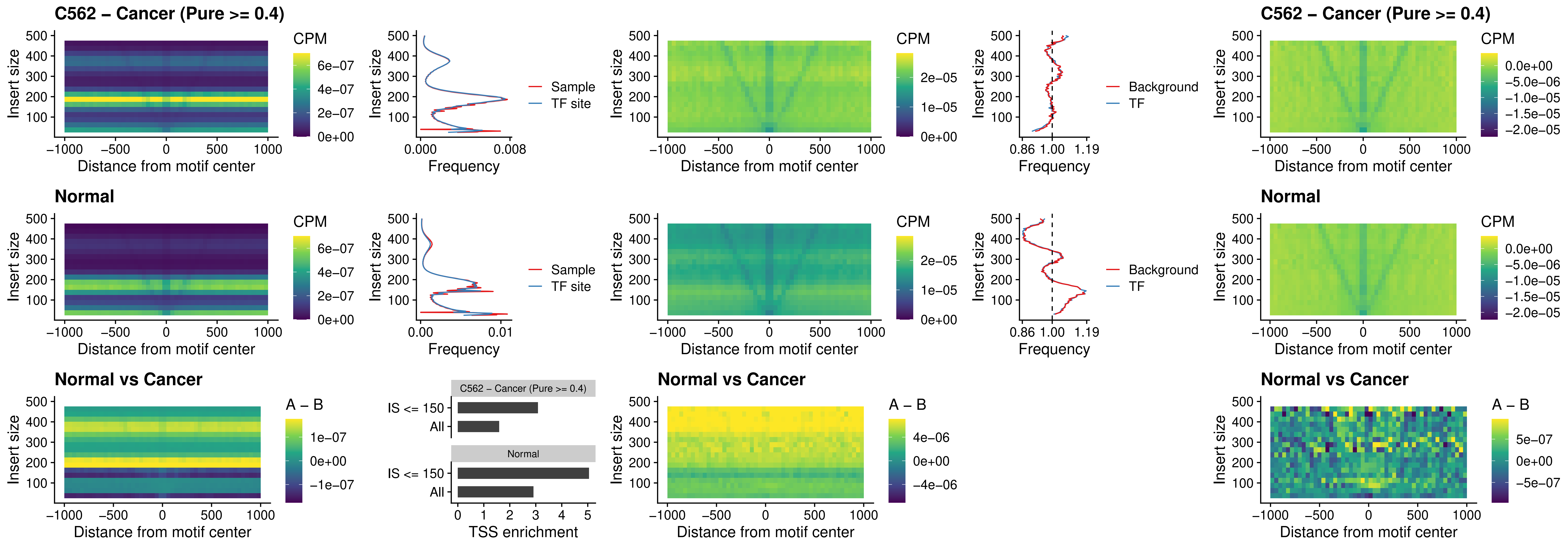

### DBX2_uc_cTSS_nPEAK

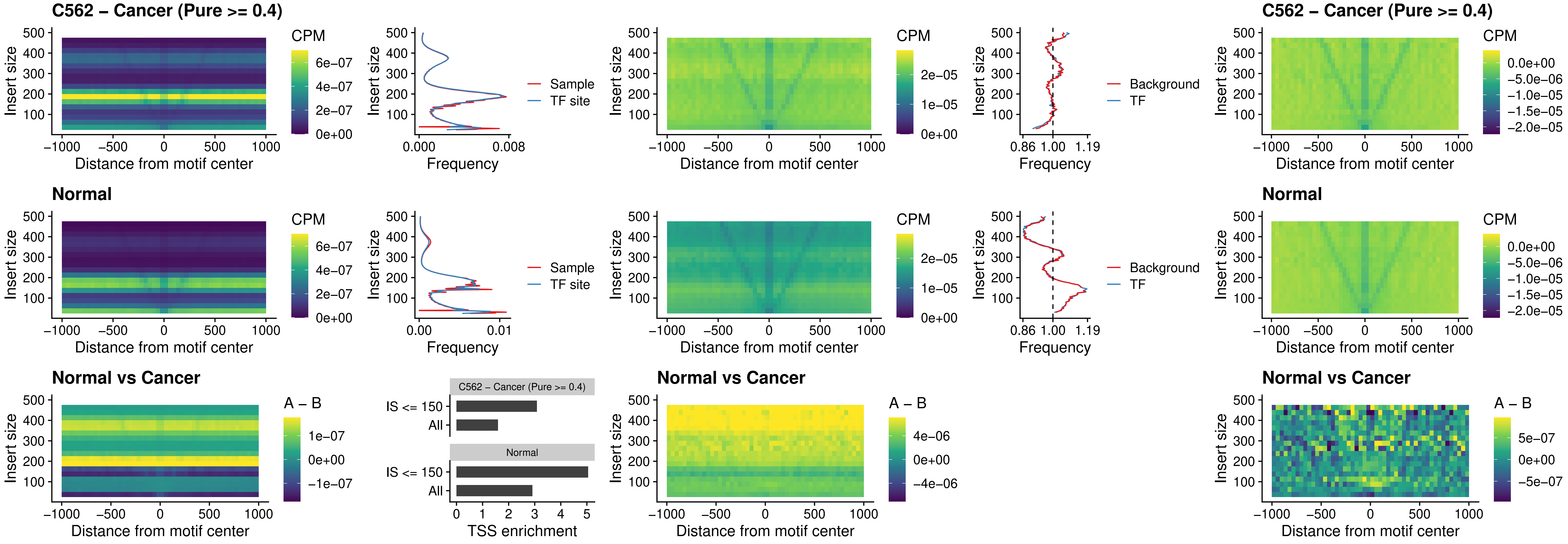

### LHX6_uc_dTSS_nPEAK

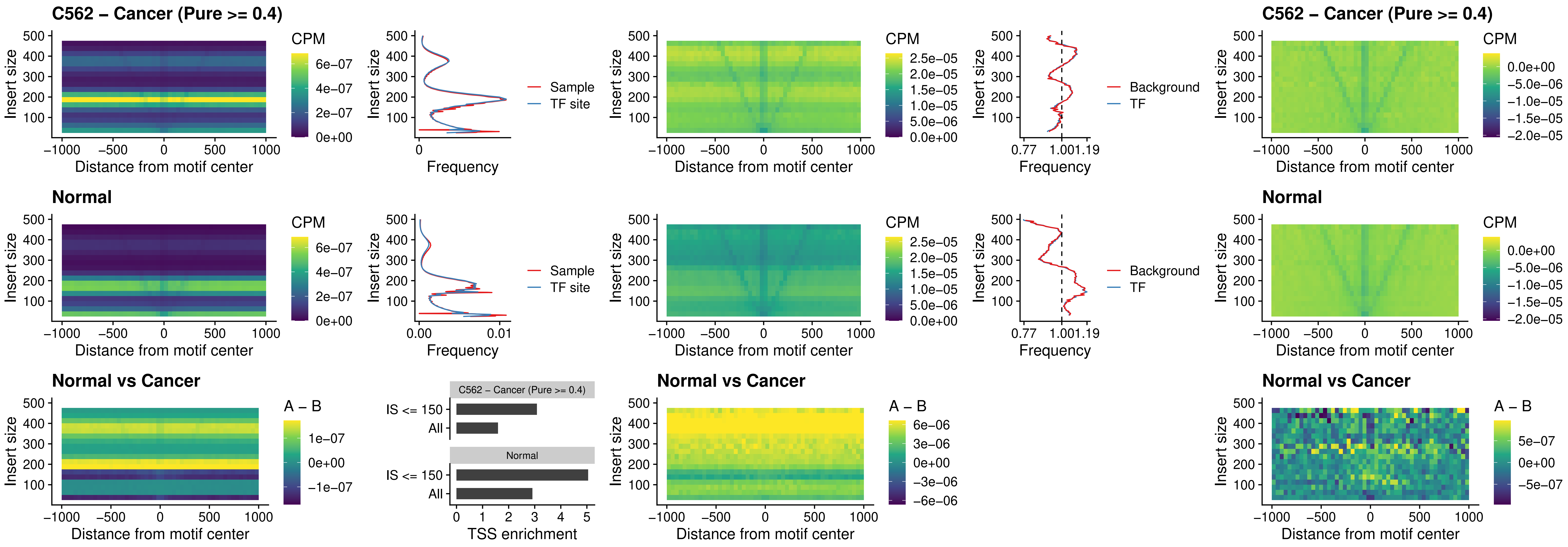

#### SOX1_uc_dTSS_nPEAK

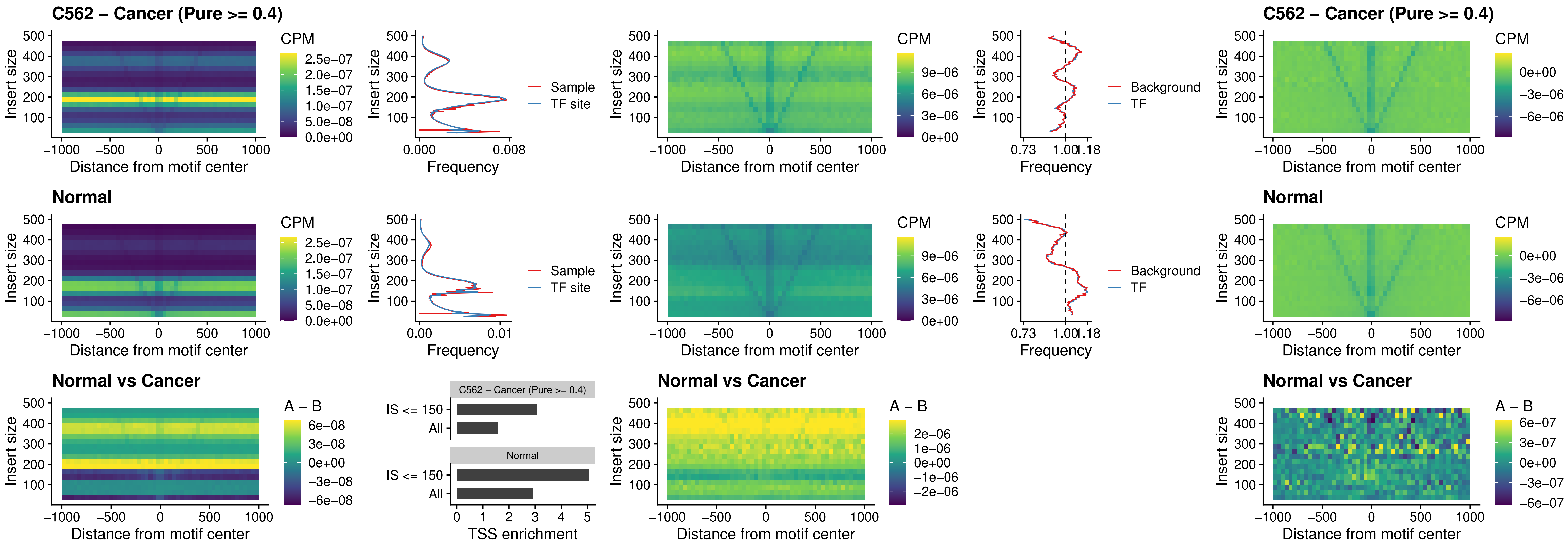

#### NKX24_uc_dTSS_nPEAK

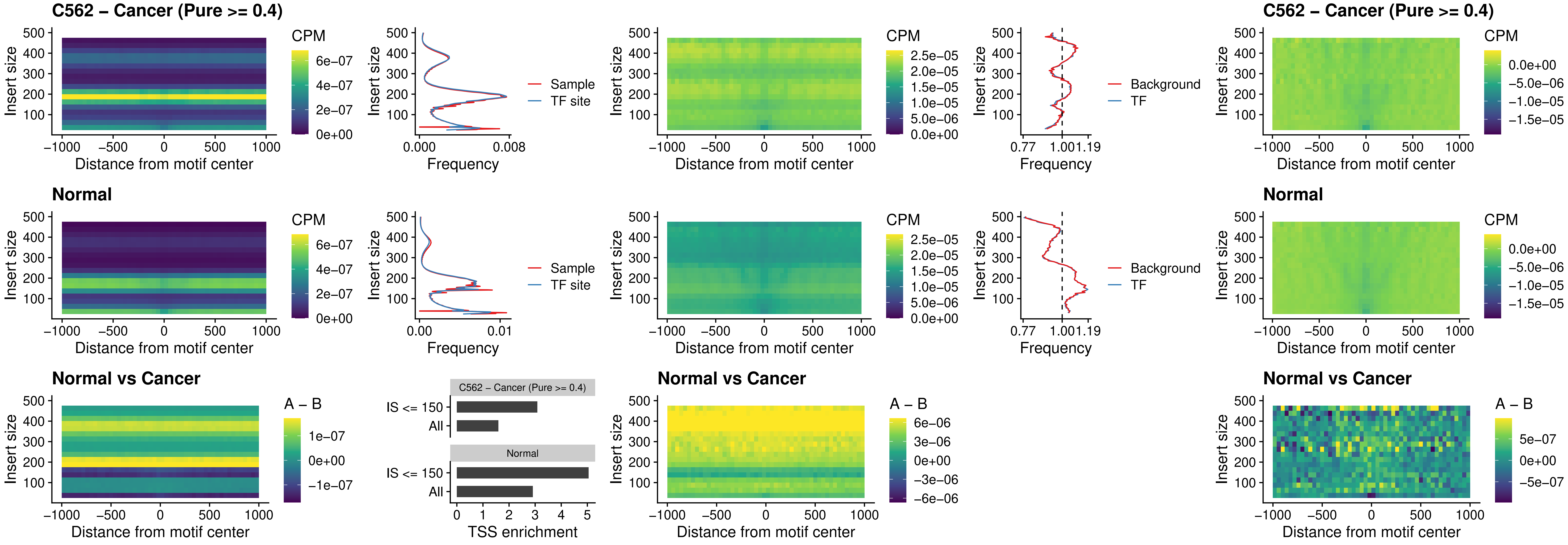

### DLX3_uc_cTSS_nPEAK

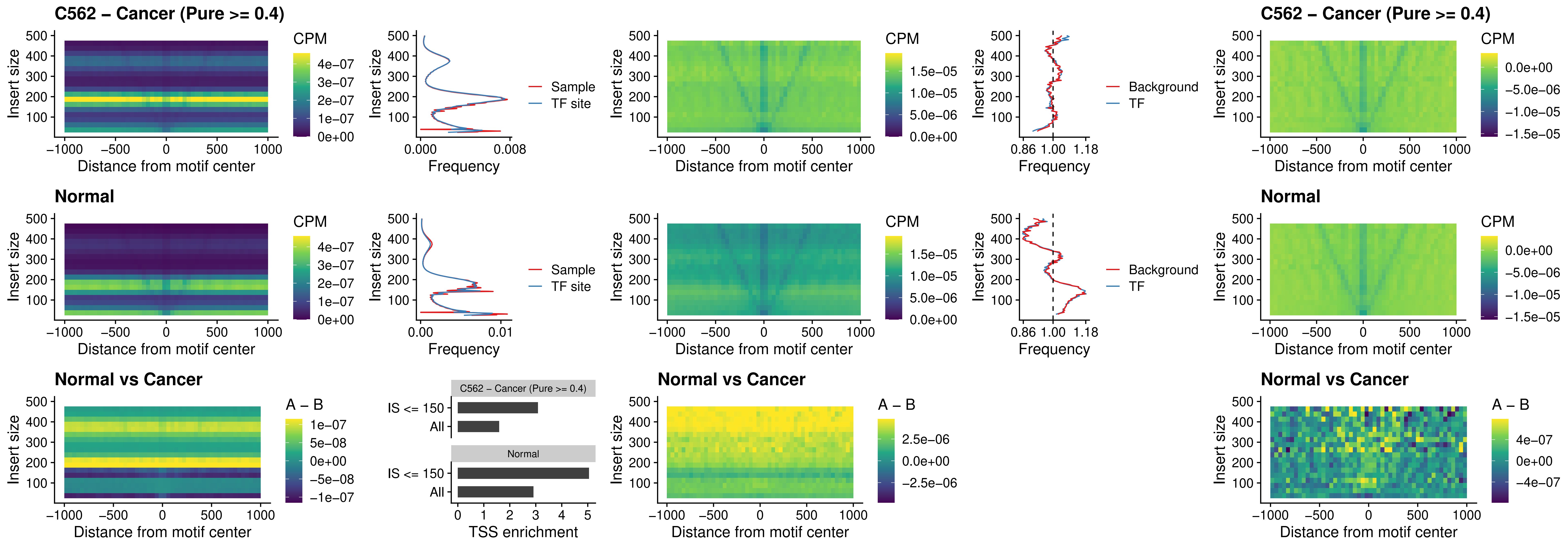

#### HOXD3_uc_cTSS_nPEAK

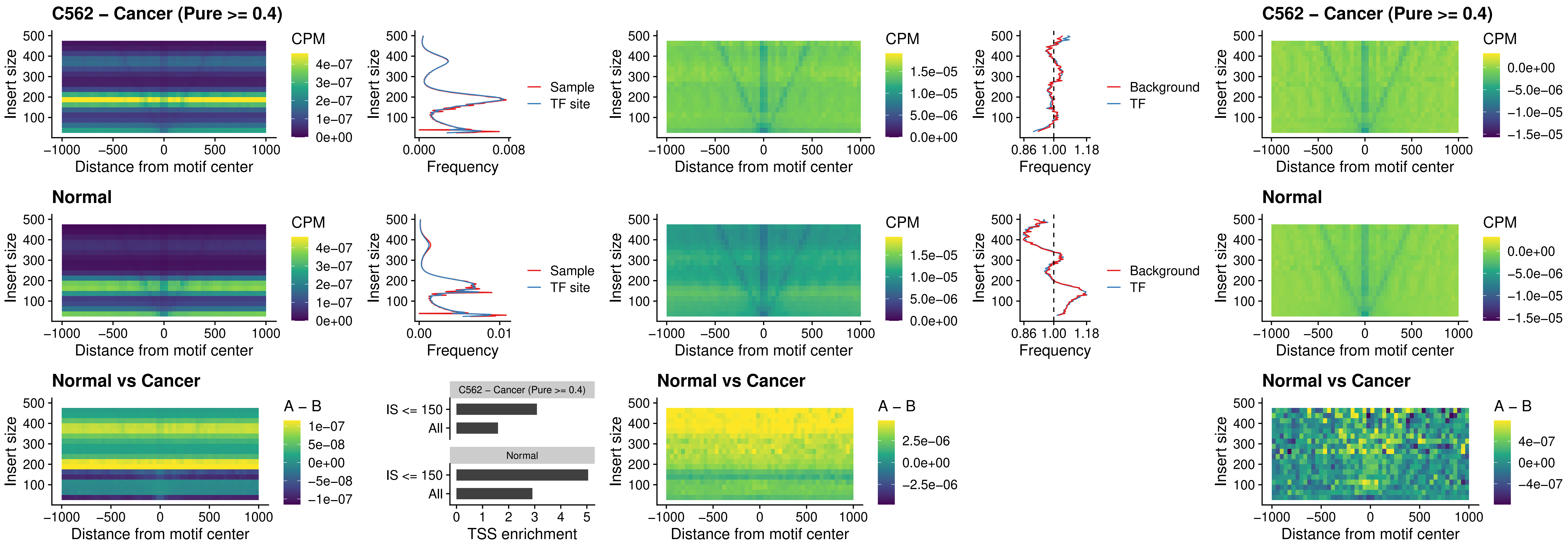

### ISL1_uc_dTSS_oPEAK

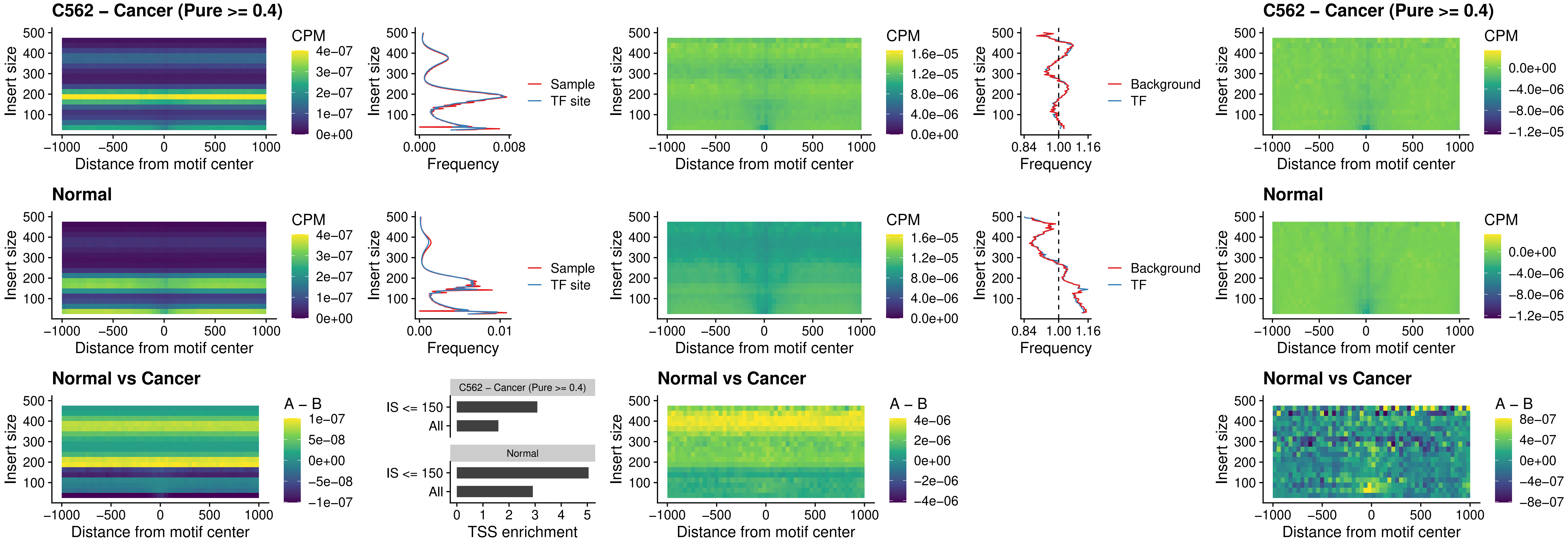

### HOXB4_uc_dTSS_oPEAK

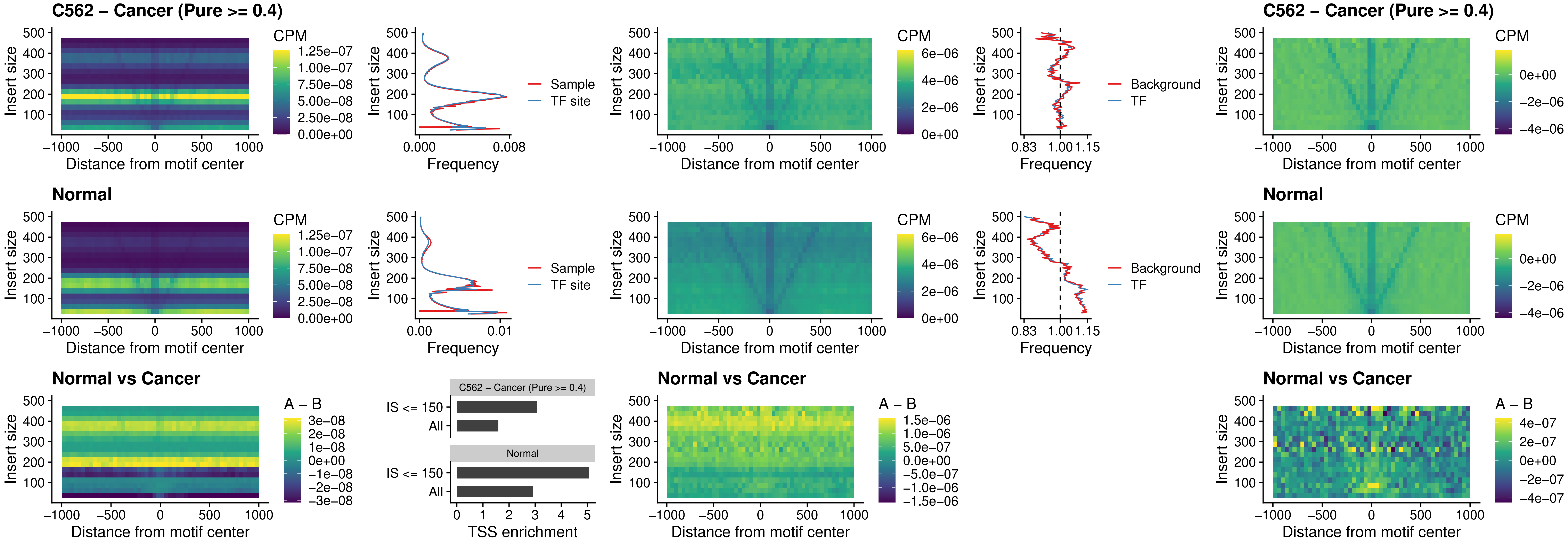

### VAX1_uc_dTSS_oPEAK

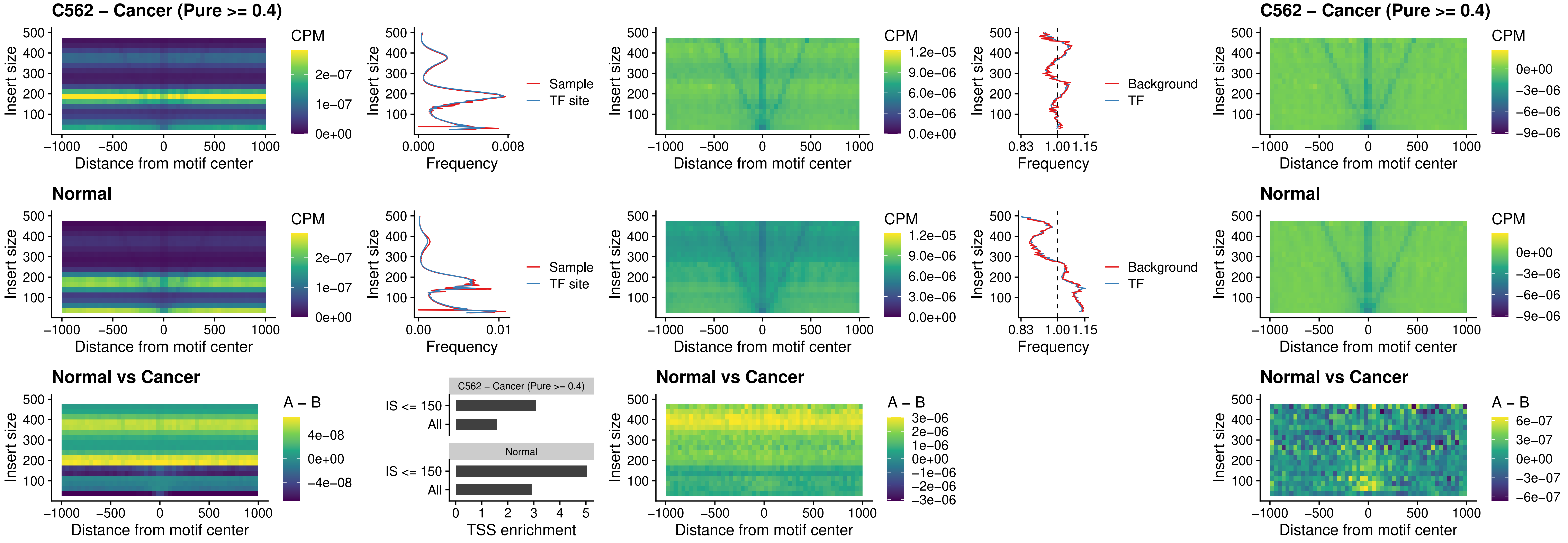

#### DLX3_uc_dTSS_oPEAK

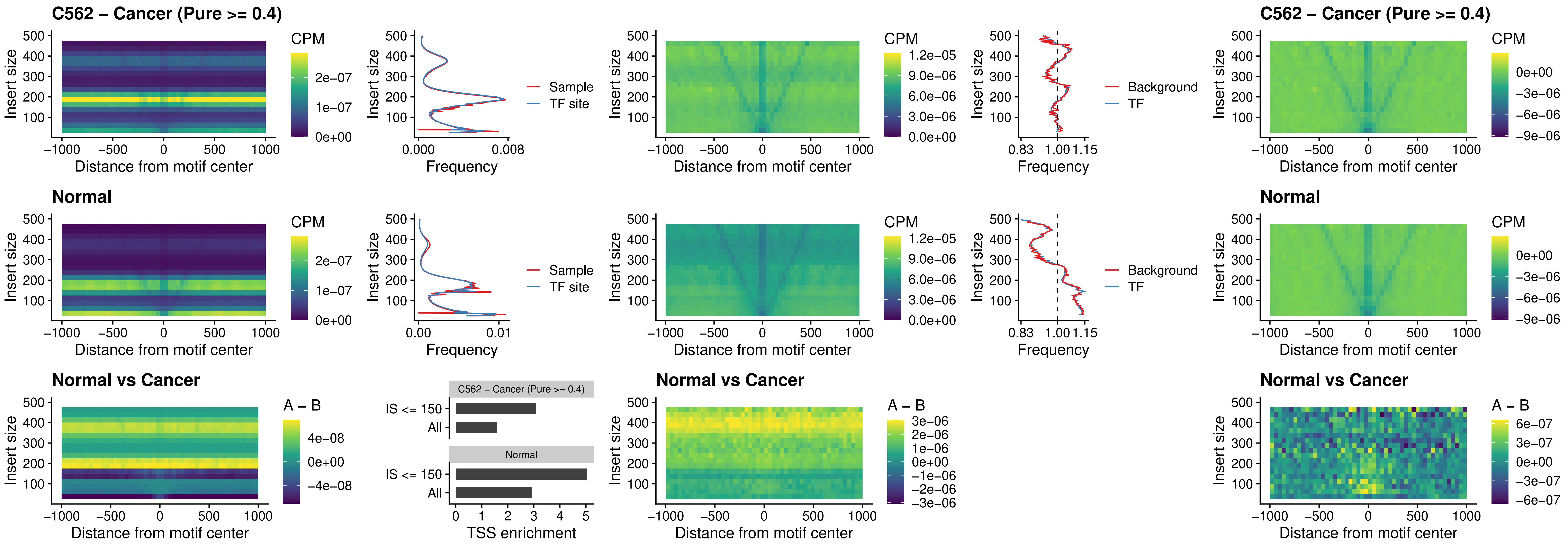

#### HOXD3_uc_dTSS_oPEAK

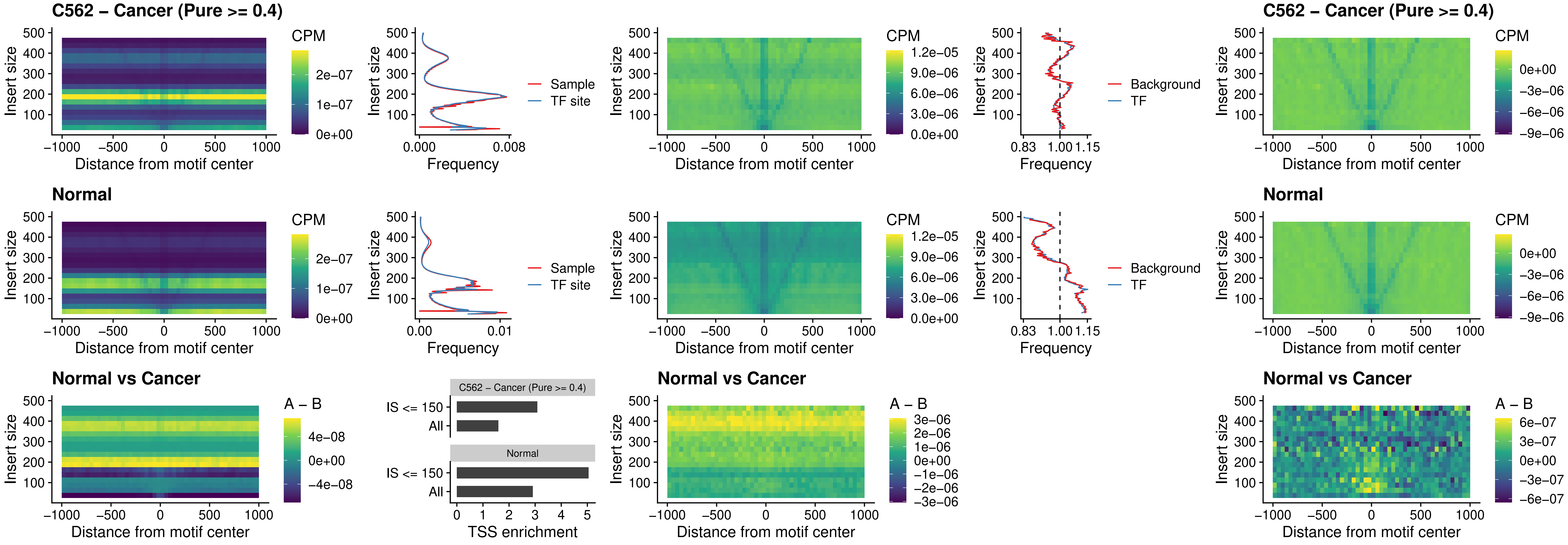

### BBX_uc_dTSS_nPEAK

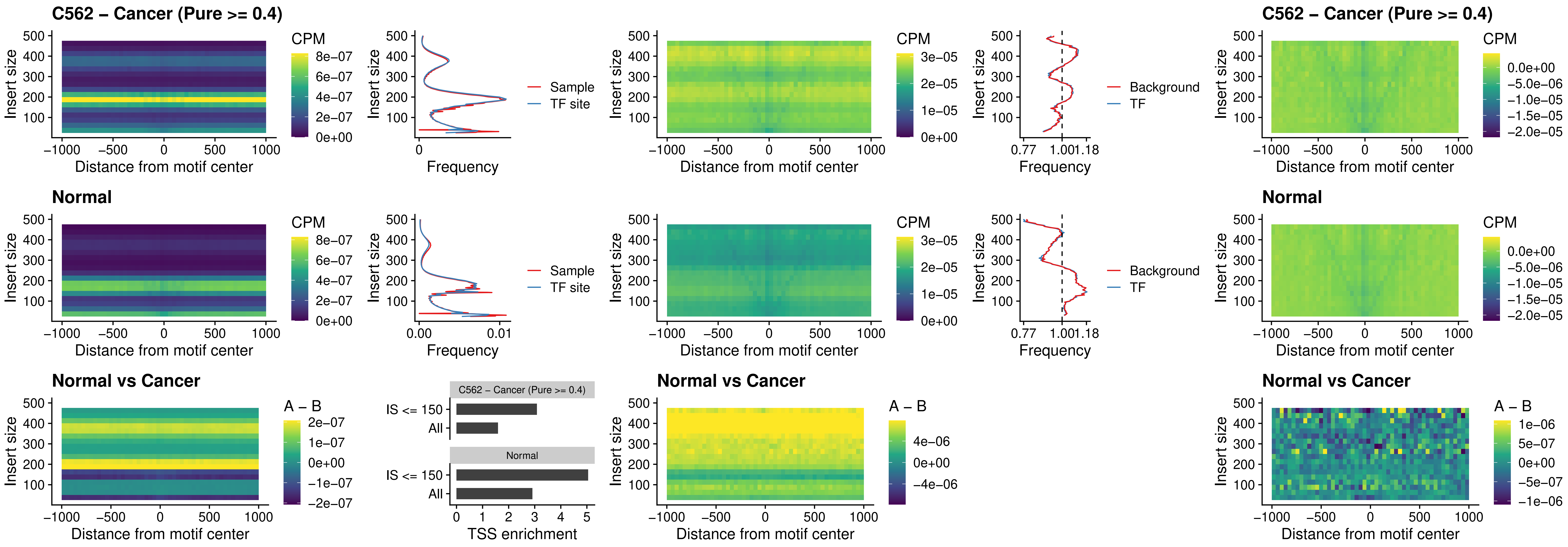

### SOX3_uc_cTSS_nPEAK

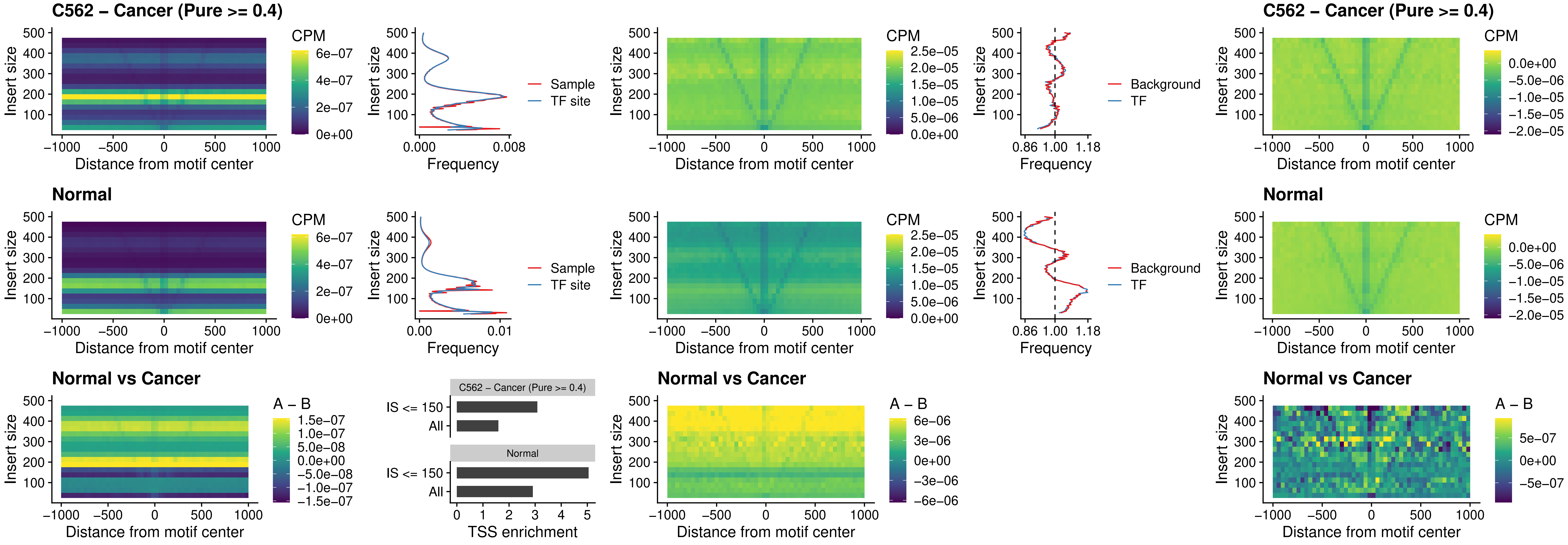

### TBPL2_uc_cTSS_nPEAK

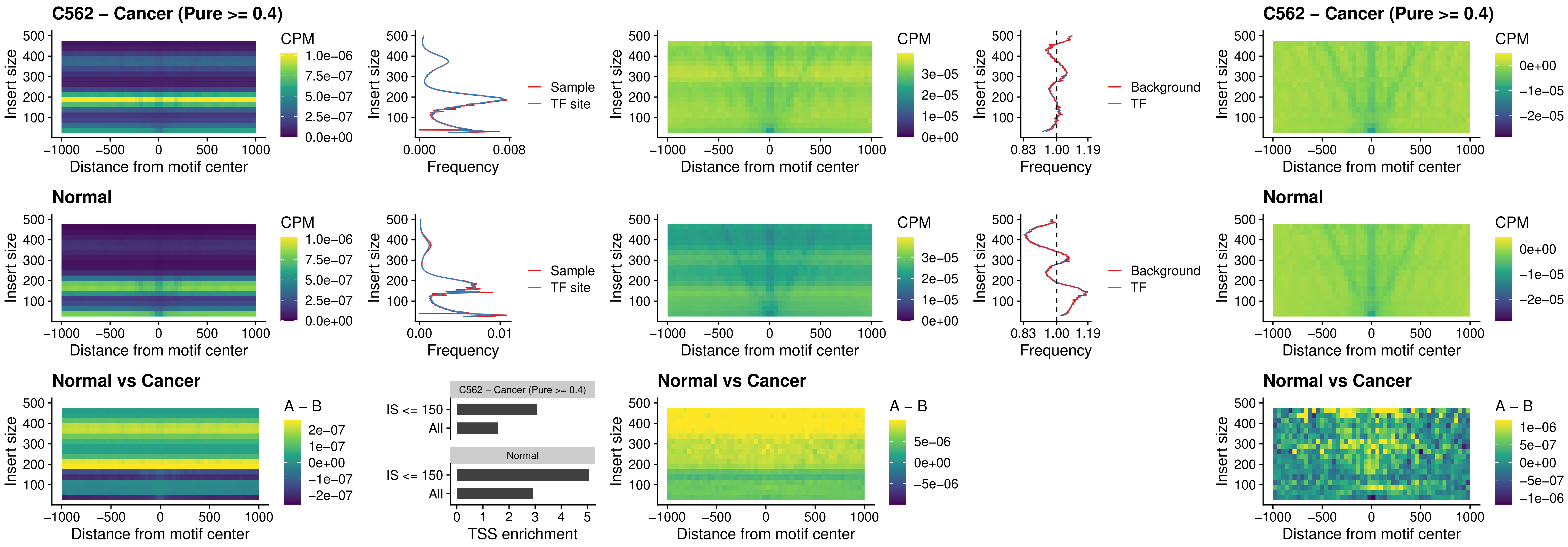

### HOXA4_uc_cTSS_oPEAK

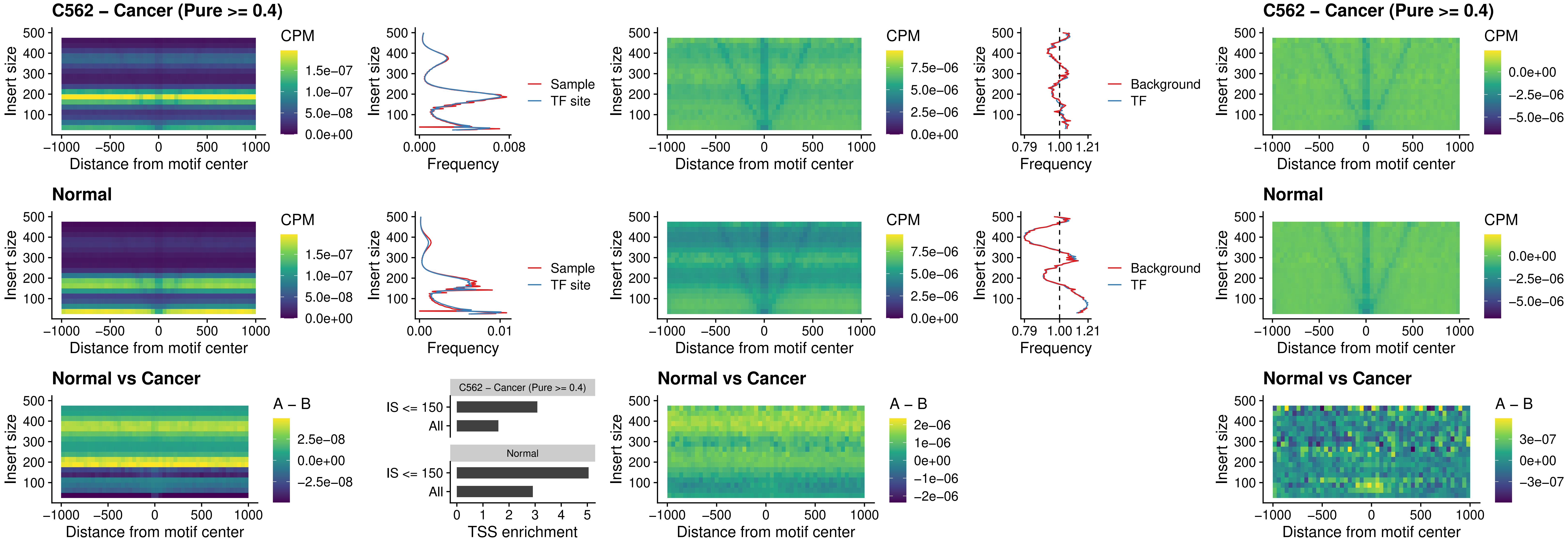

### PDX1_uc_dTSS_oPEAK

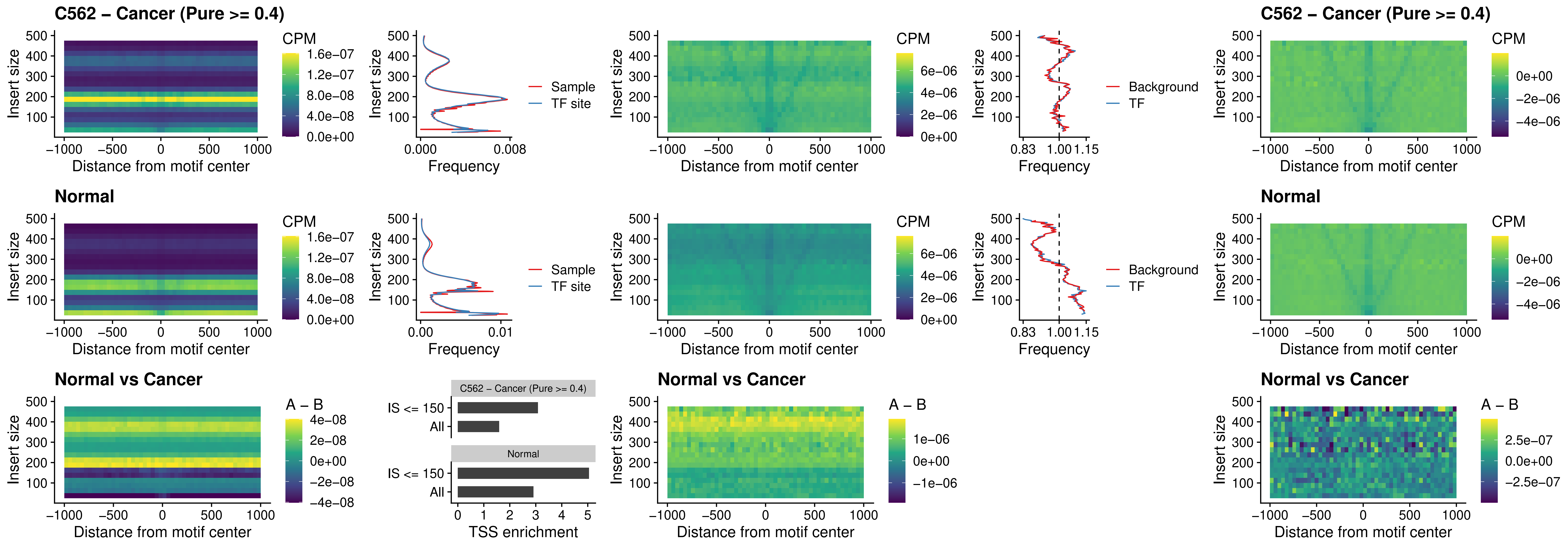

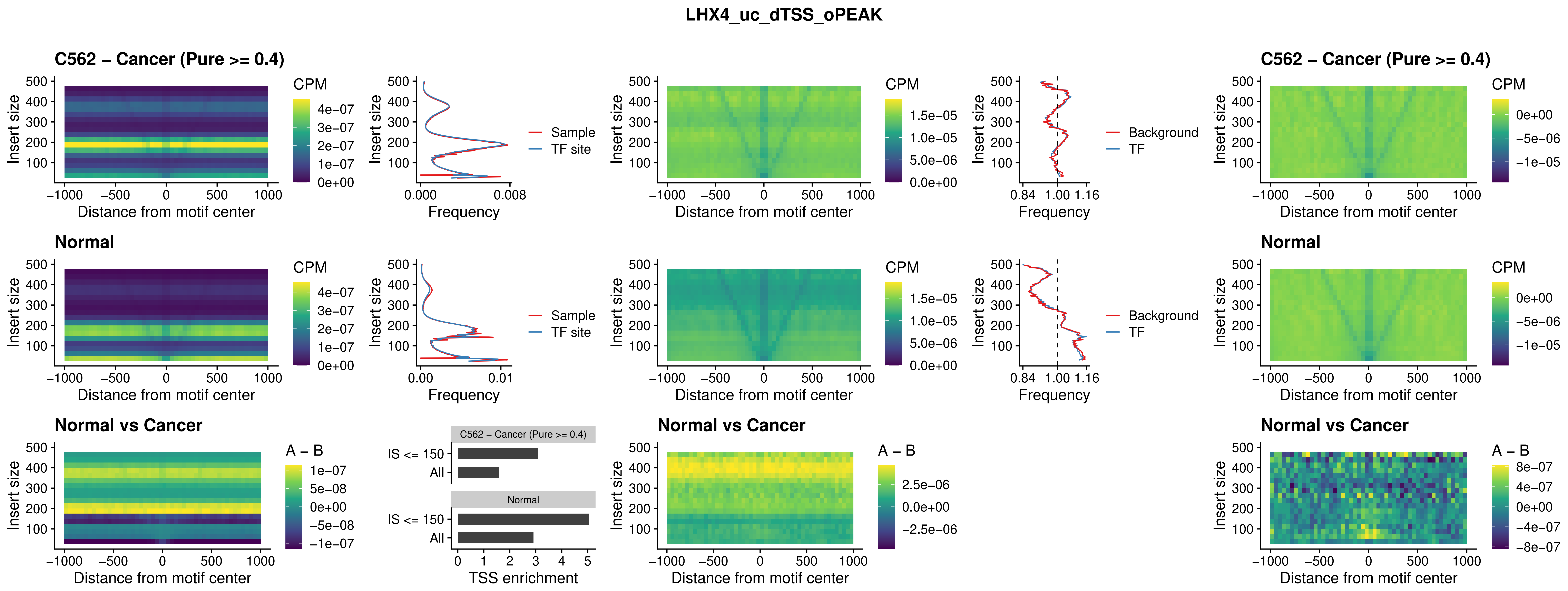
LHX4_uc_dTSS_oPEAK
C562 – Cancer (Pure >= 0.4)
Normal
Normal vs Cancer
Insert size
Distance from motif center
Frequency
CPM
A – B
Sample
TF site
Background
TF
C562 – Cancer (Pure >= 0.4)
Normal
IS <= 150
All
TSS enrichment

#### HOXA4_uc_dTSS_oPEAK

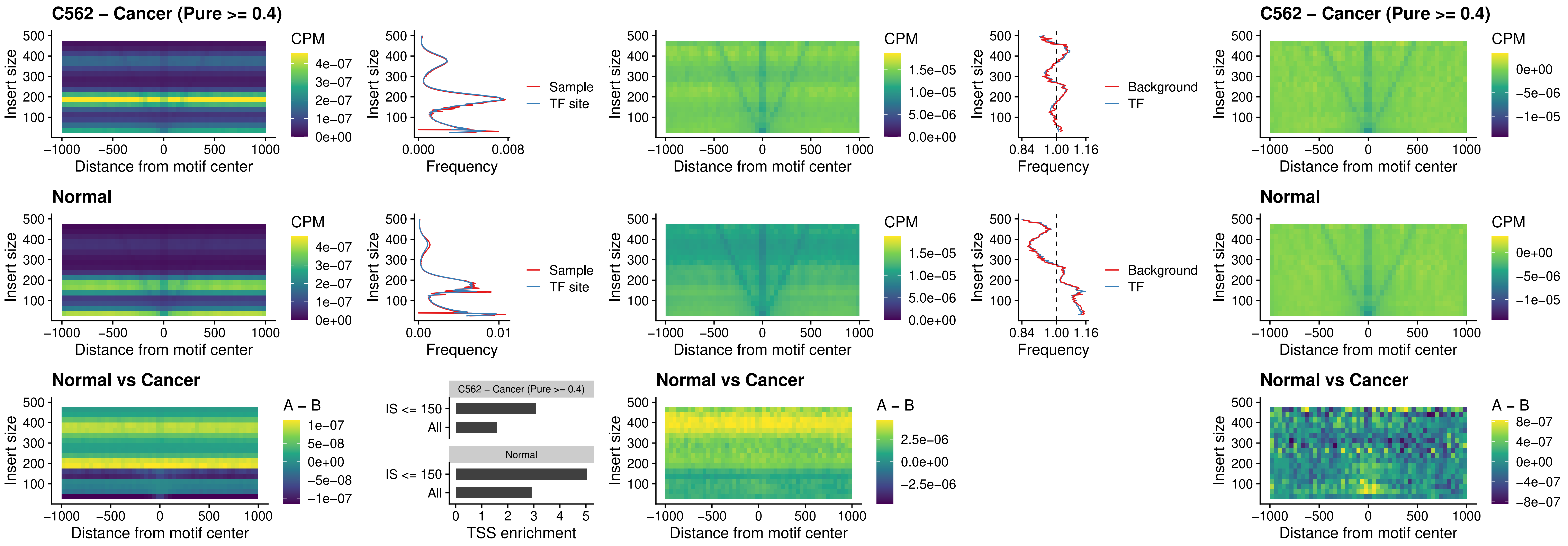

### HLX_uc_dTSS_oPEAK

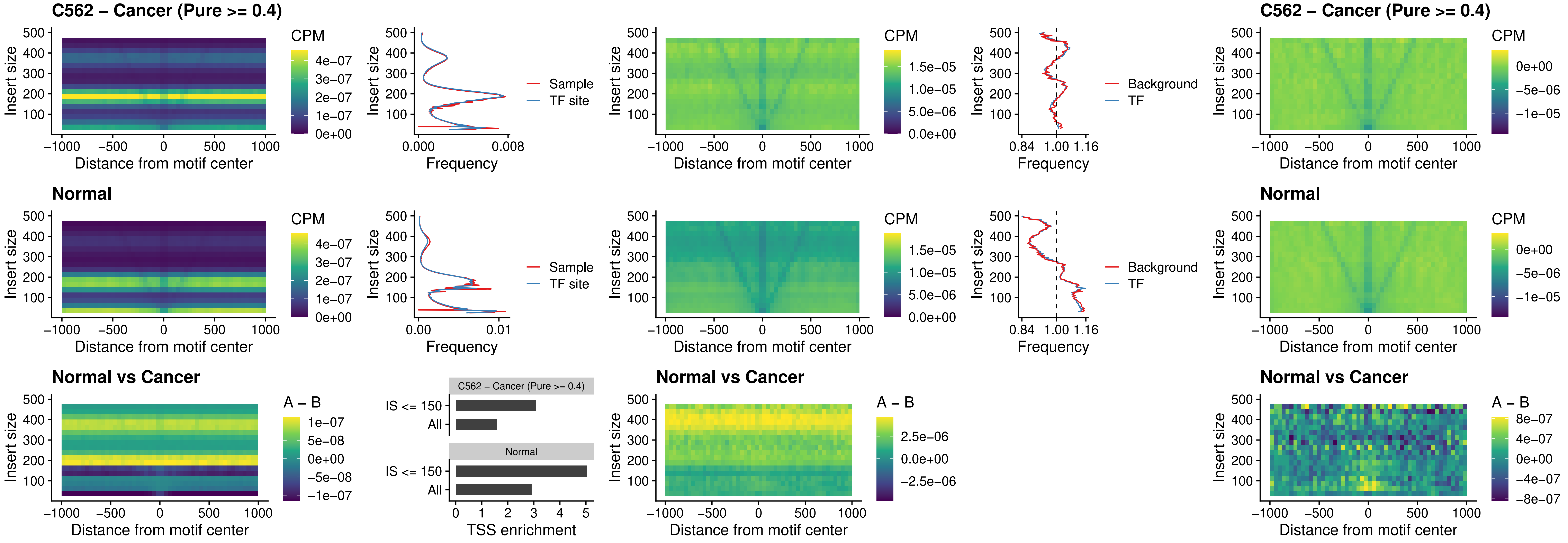

### AC0125311_uc_dTSS_oPEAK

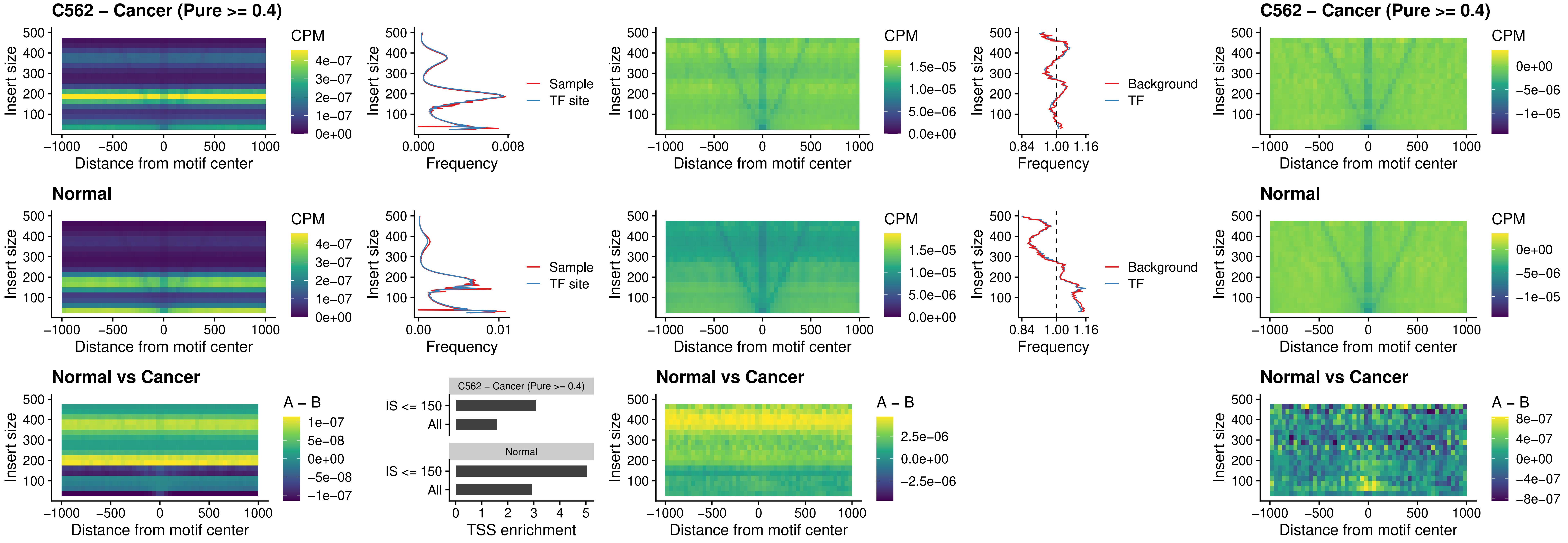

### DBX2_uc_dTSS_oPEAK

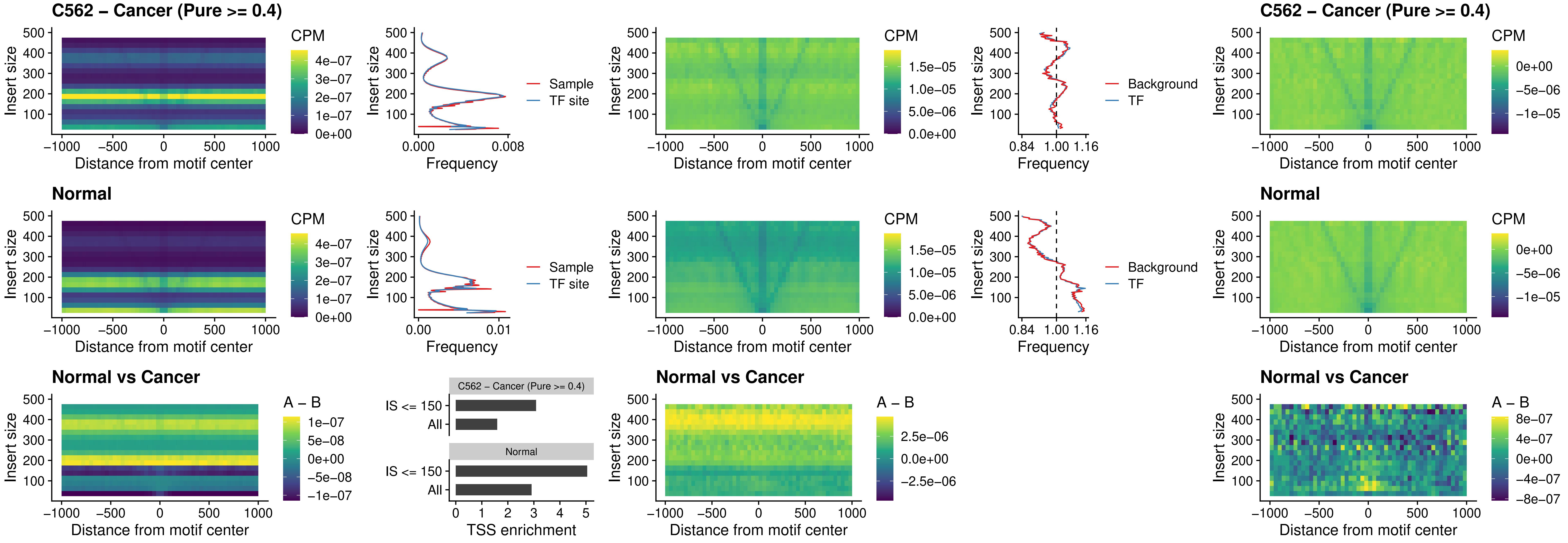

### HOXA2_uc_dTSS_nPEAK

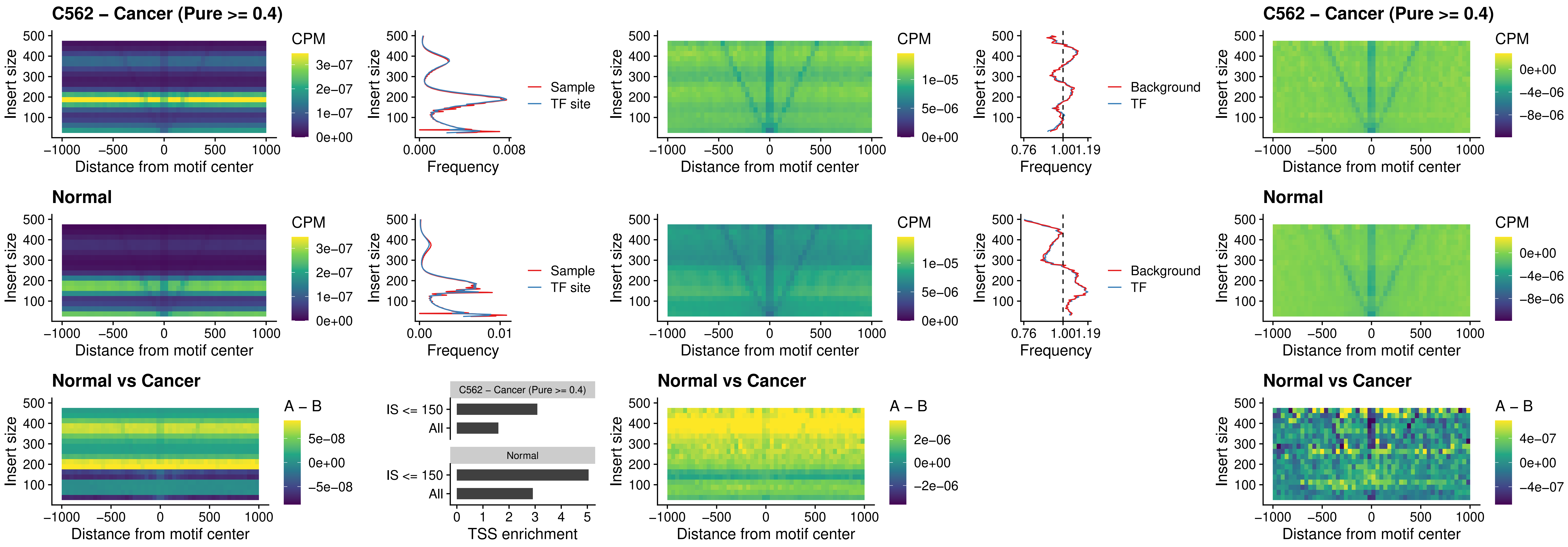

### SHOX_uc_dTSS_nPEAK

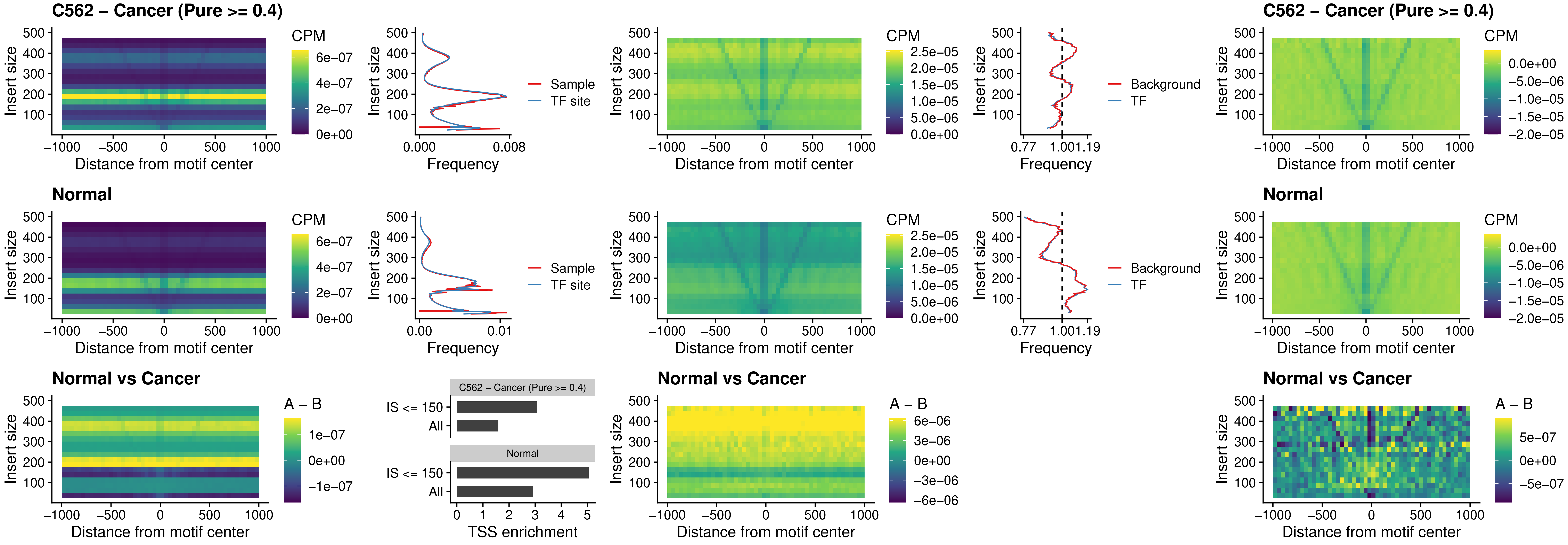

### PRRX1_uc_dTSS_nPEAK

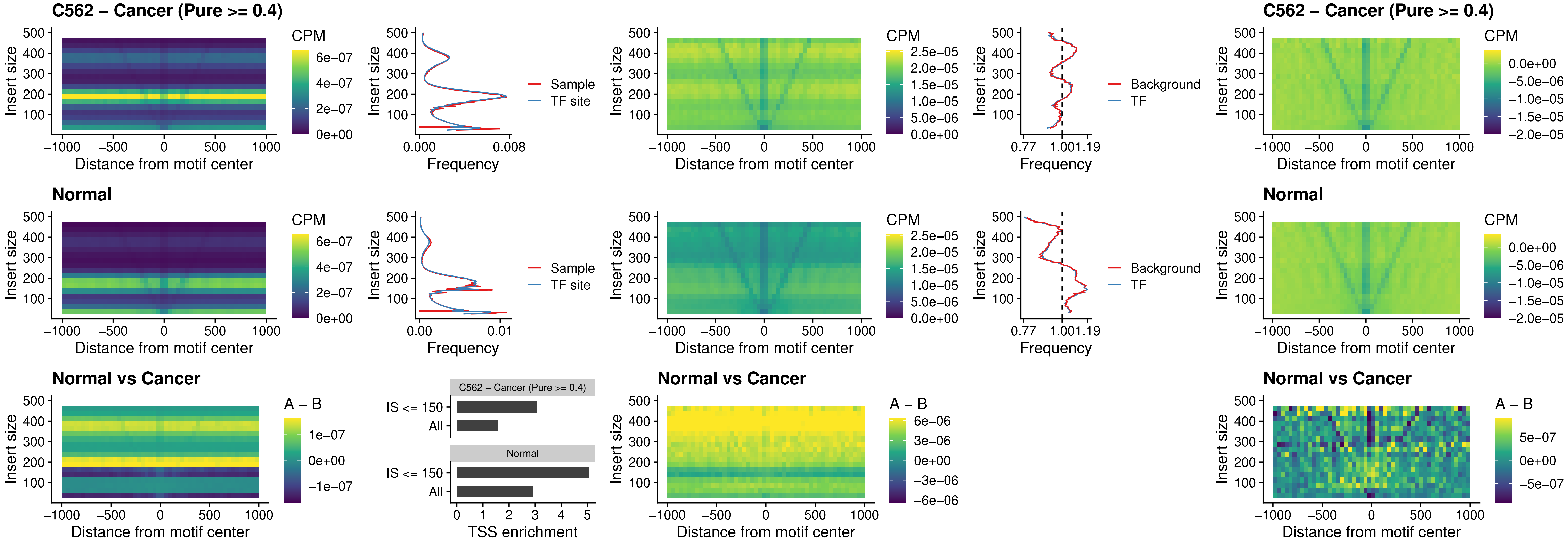

### LMX1A_uc_dTSS_nPEAK

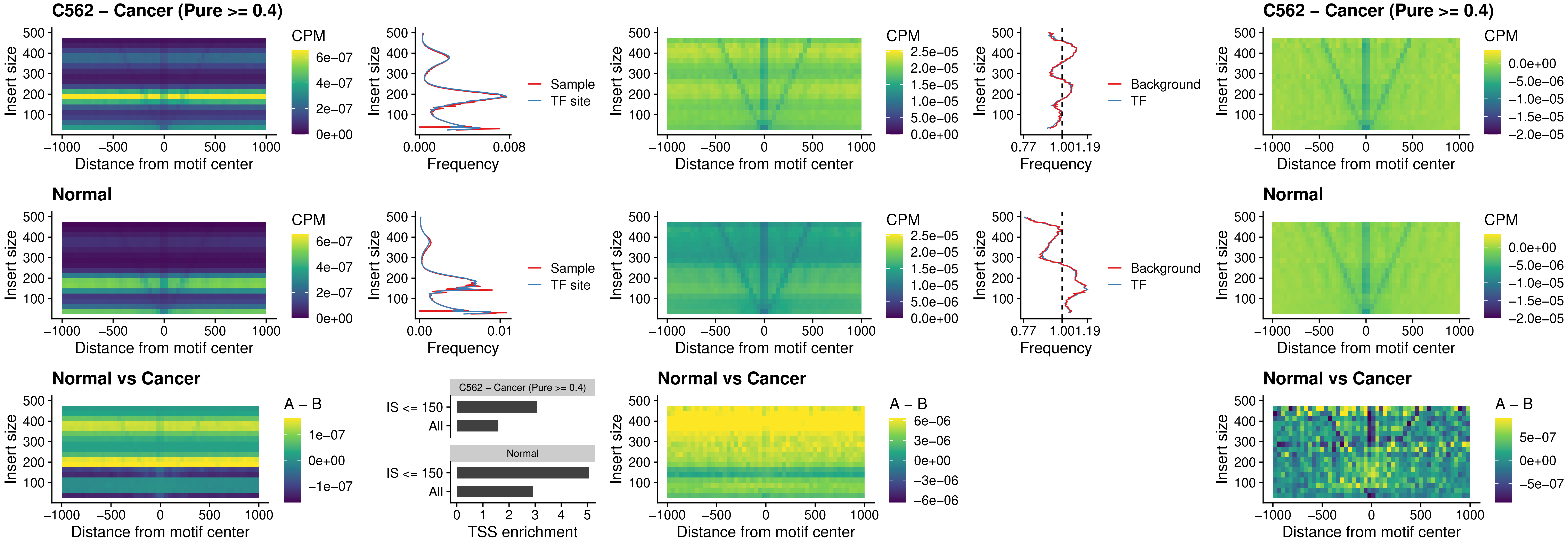

### LMX1B_uc_dTSS_nPEAK

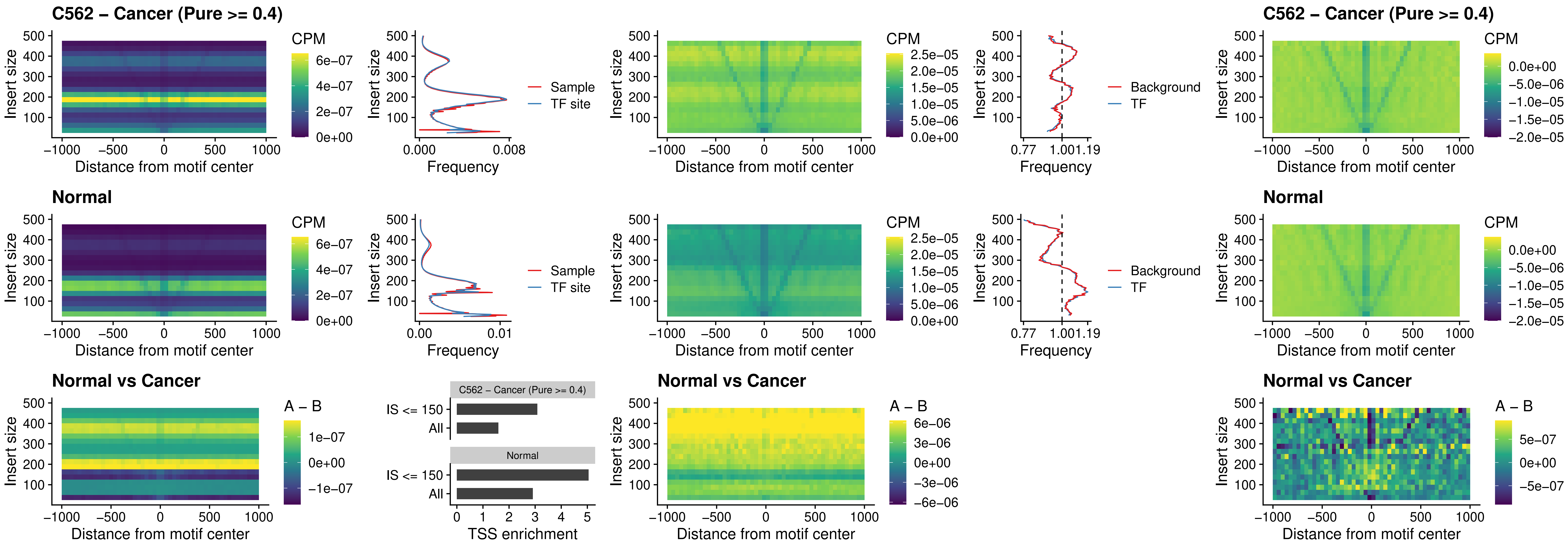

### DLX5_uc_dTSS_oPEAK

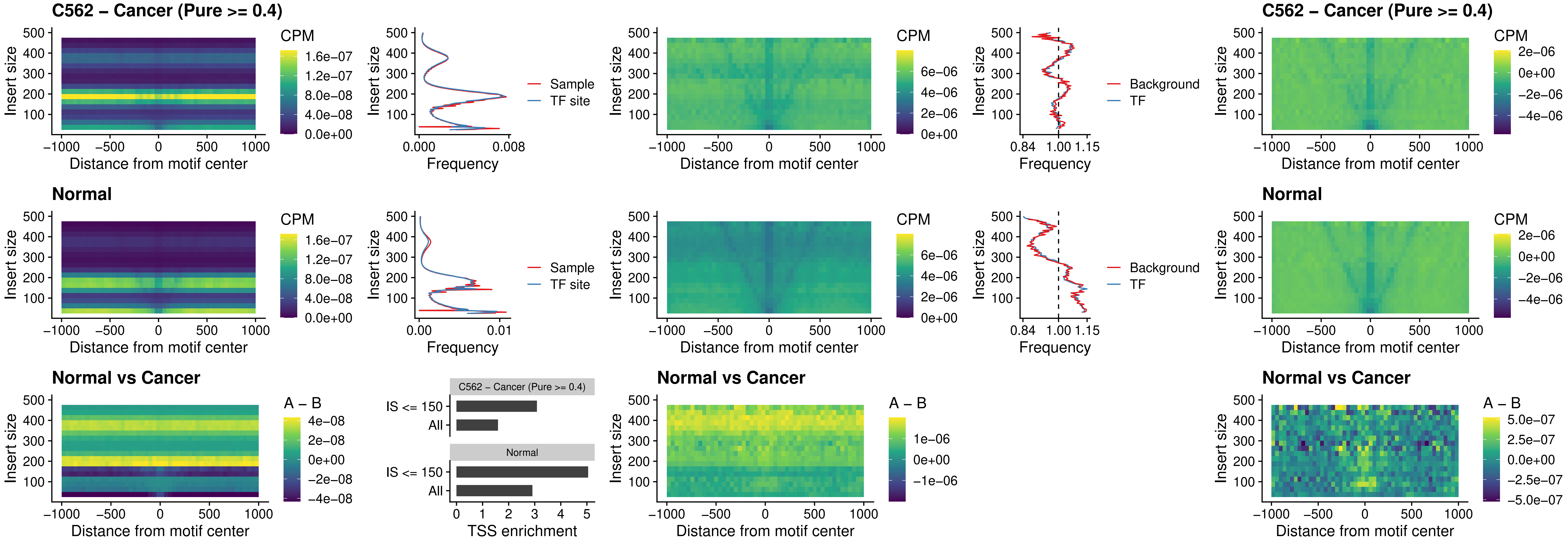

### ARID3C_uc_dTSS_nPEAK

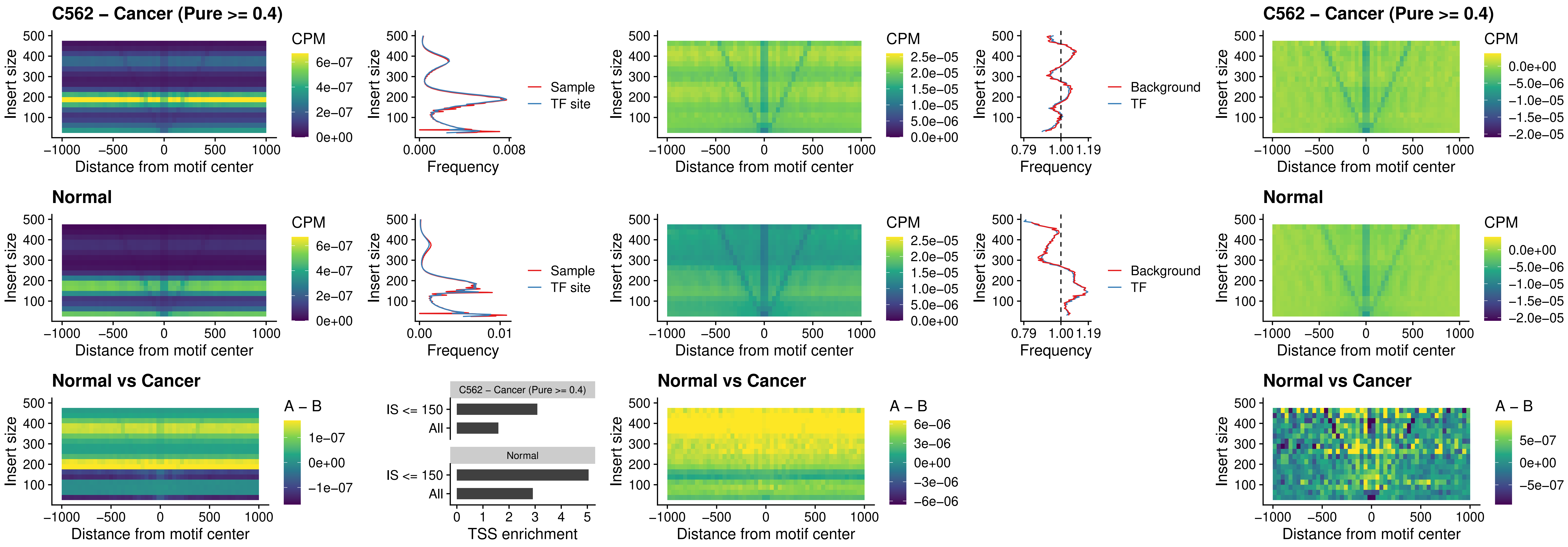

### HOXD12_uc_dTSS_nPEAK

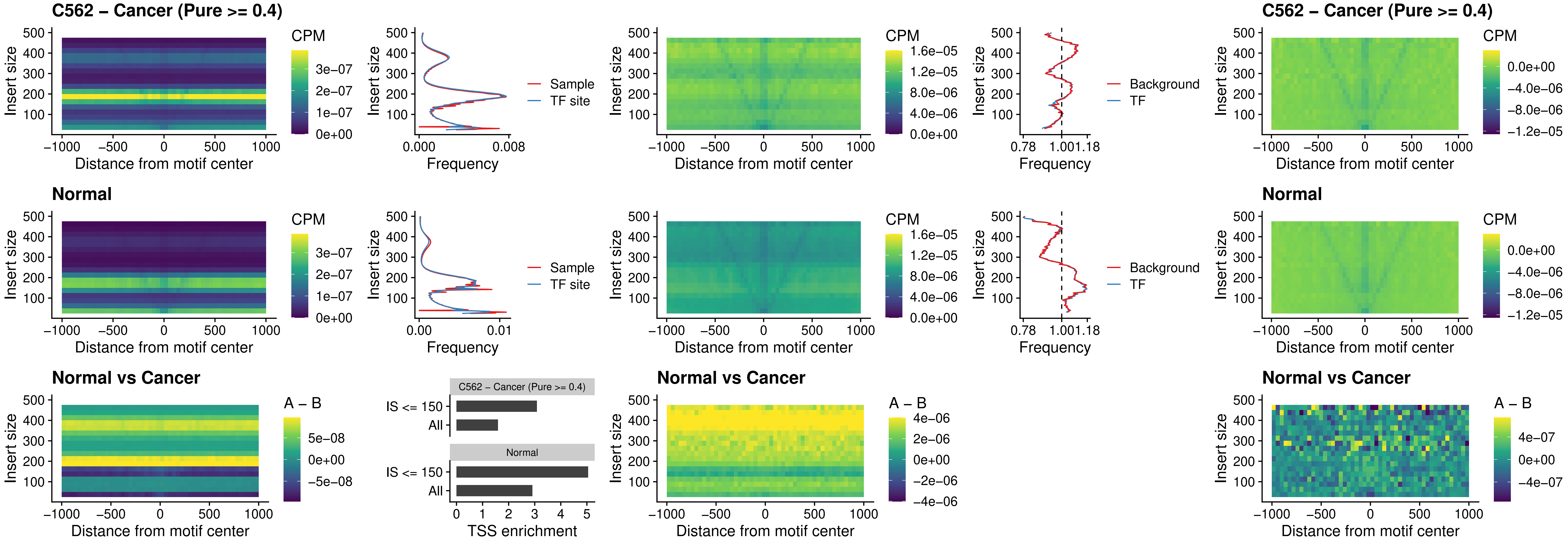

#### CDX2_uc_dTSS_nPEAK

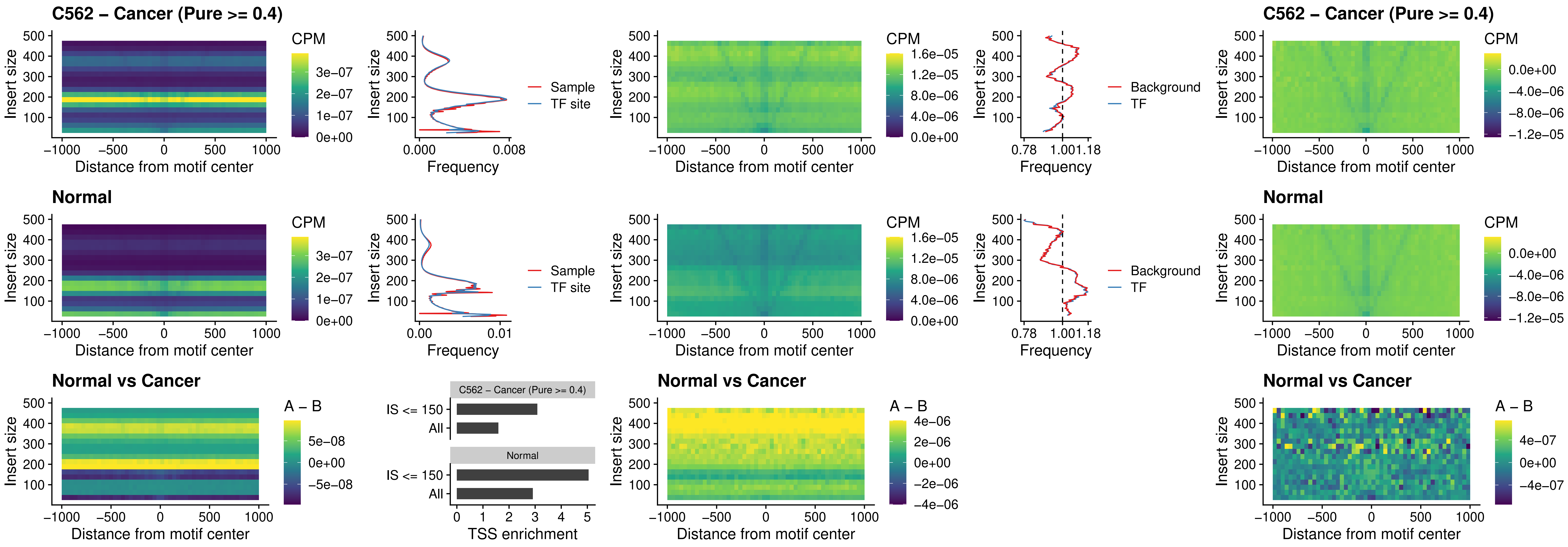

### HOXC12_uc_dTSS_nPEAK

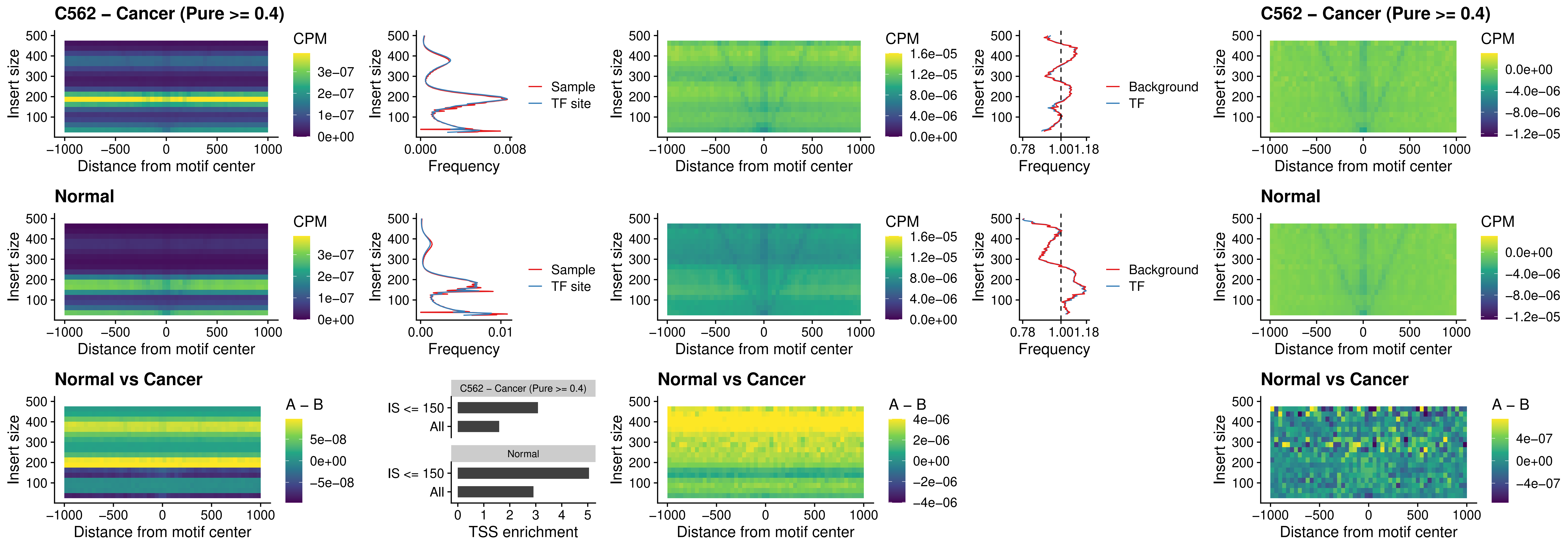

### HHEX_uc_dTSS_oPEAK

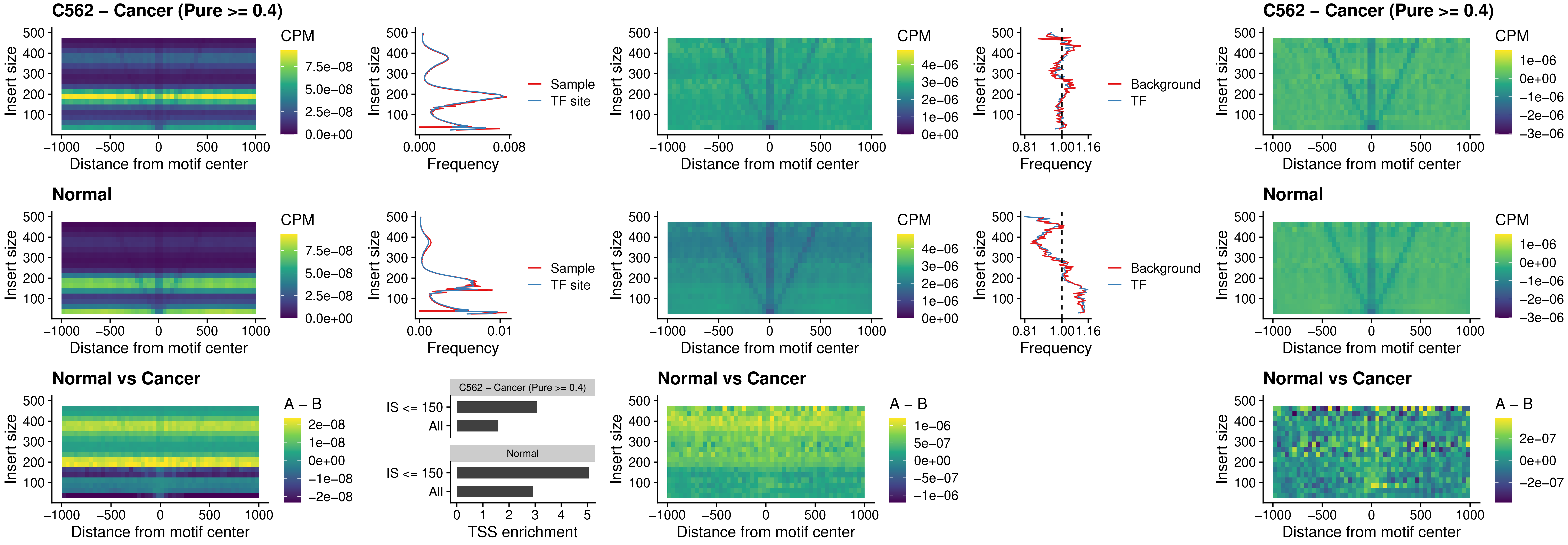

### HOXA11_uc_dTSS_nPEAK

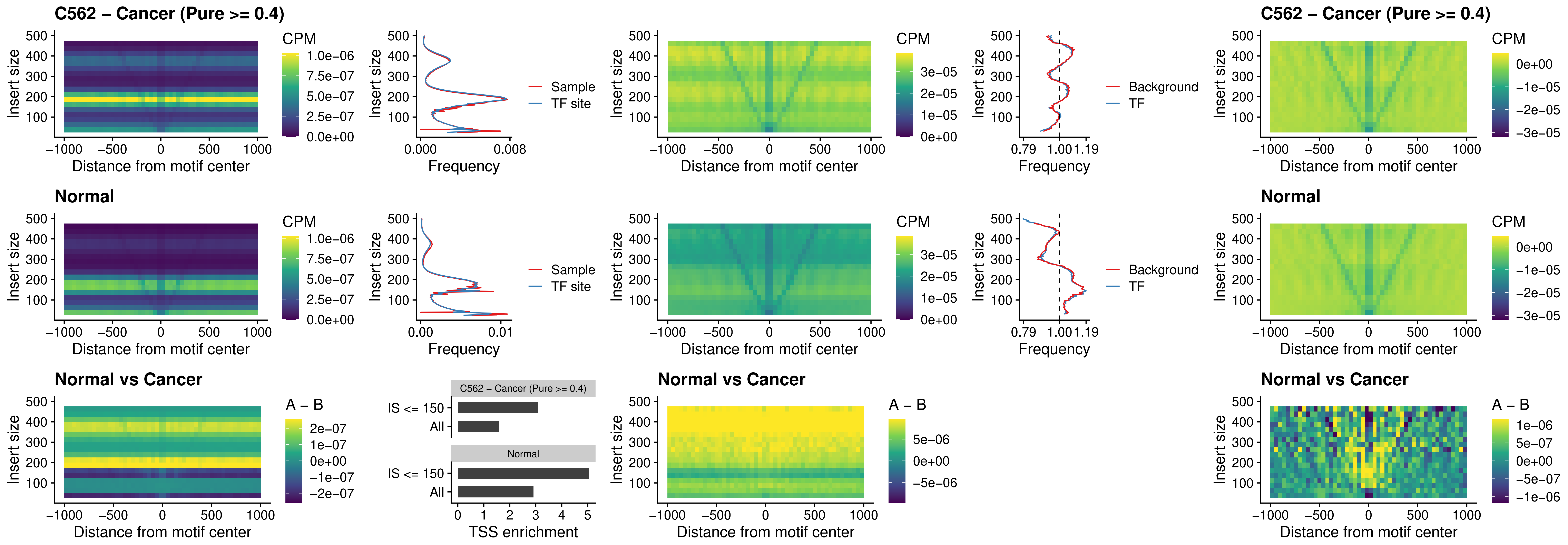

### HOXA11_uc_cTSS_nPEAK

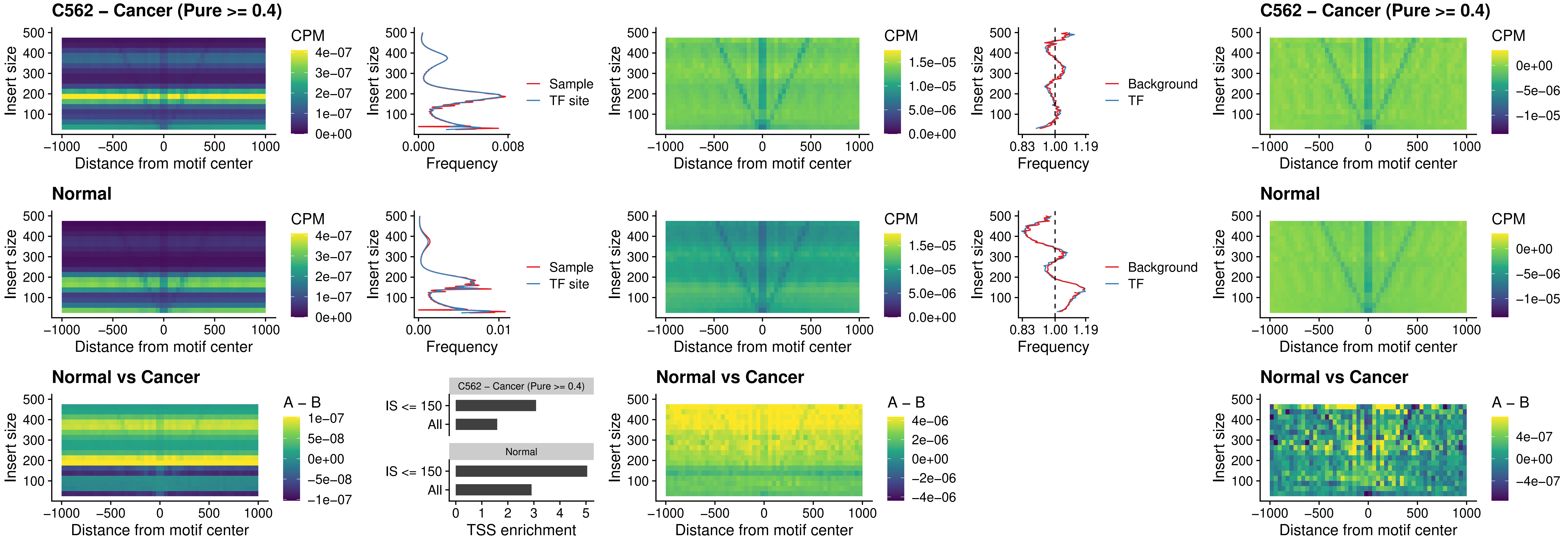

### ZFHX3_uc_dTSS_nPEAK

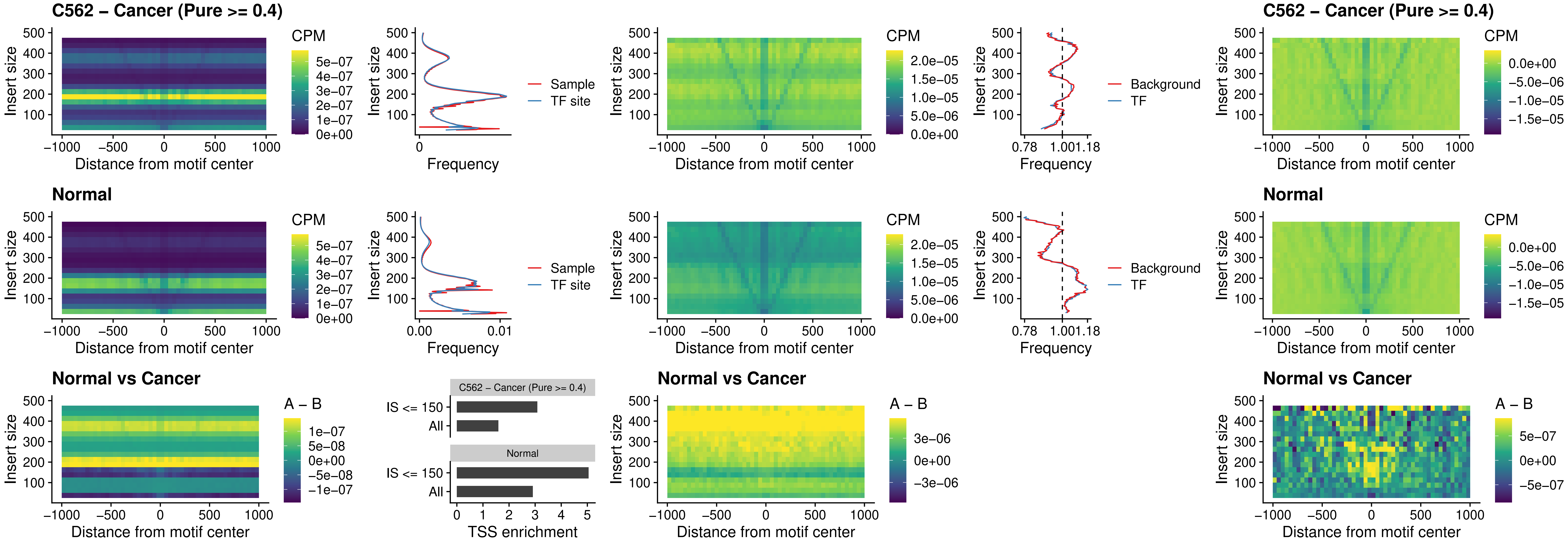

### HOXB4_uc_dTSS_nPEAK

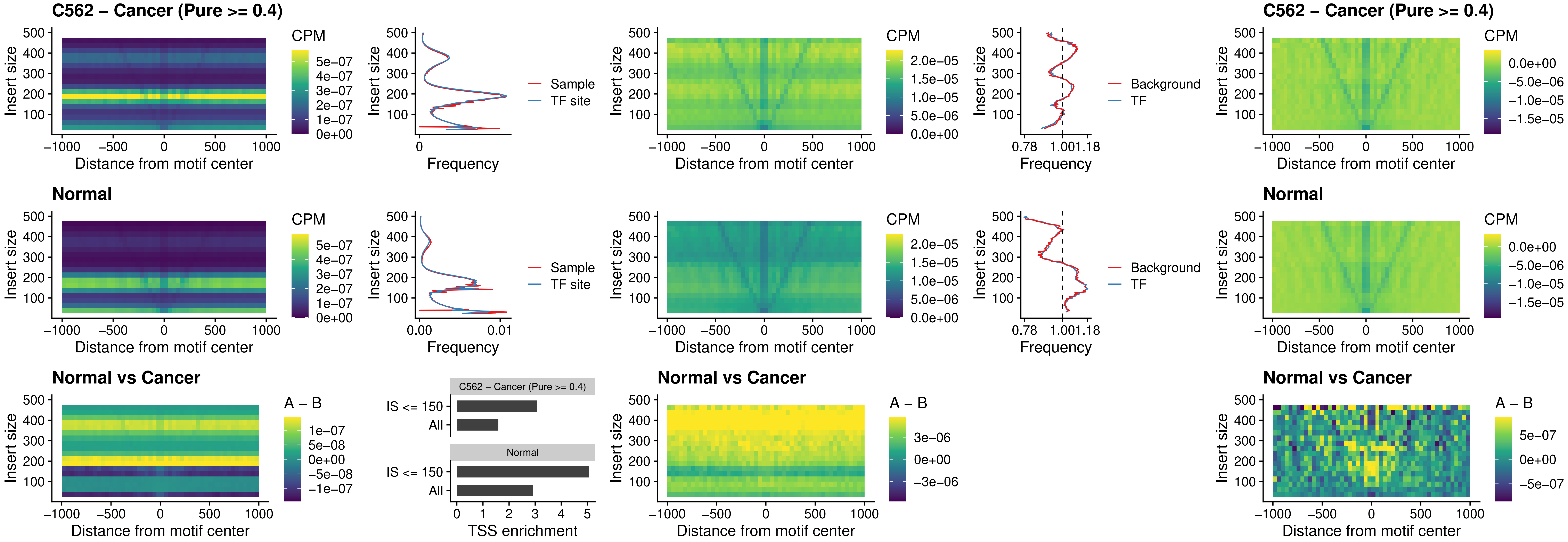

### HOXC5_uc_dTSS_nPEAK

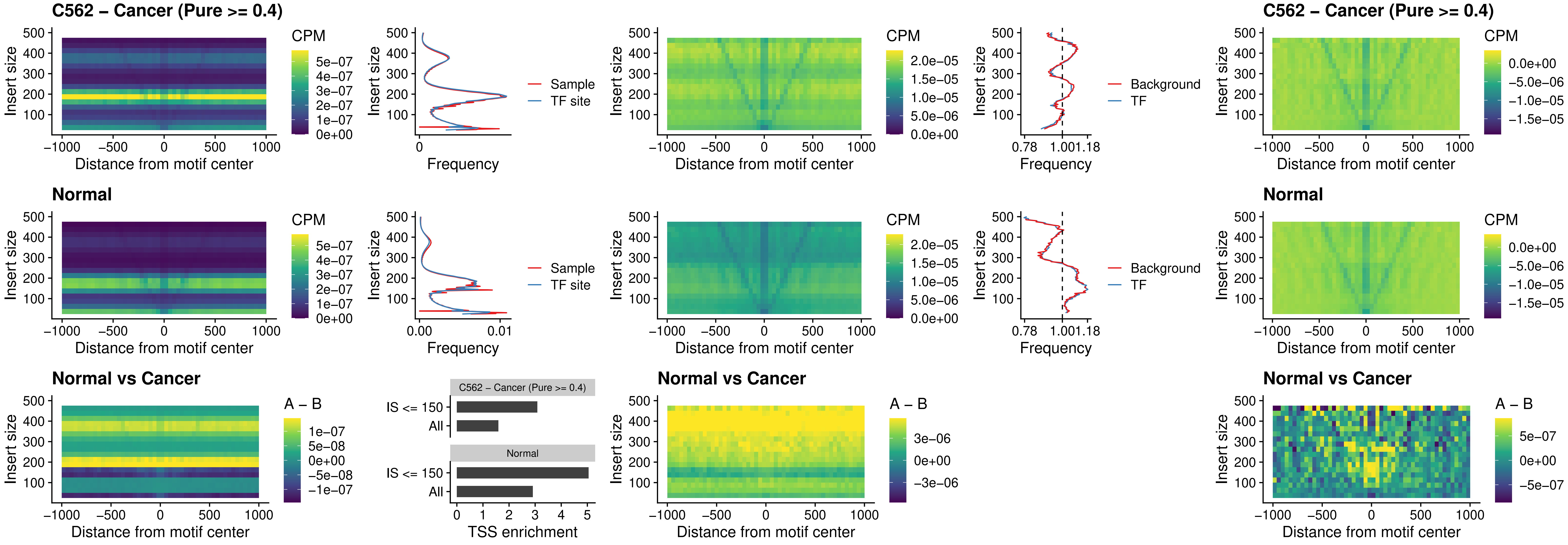

### HOXD1_uc_dTSS_nPEAK

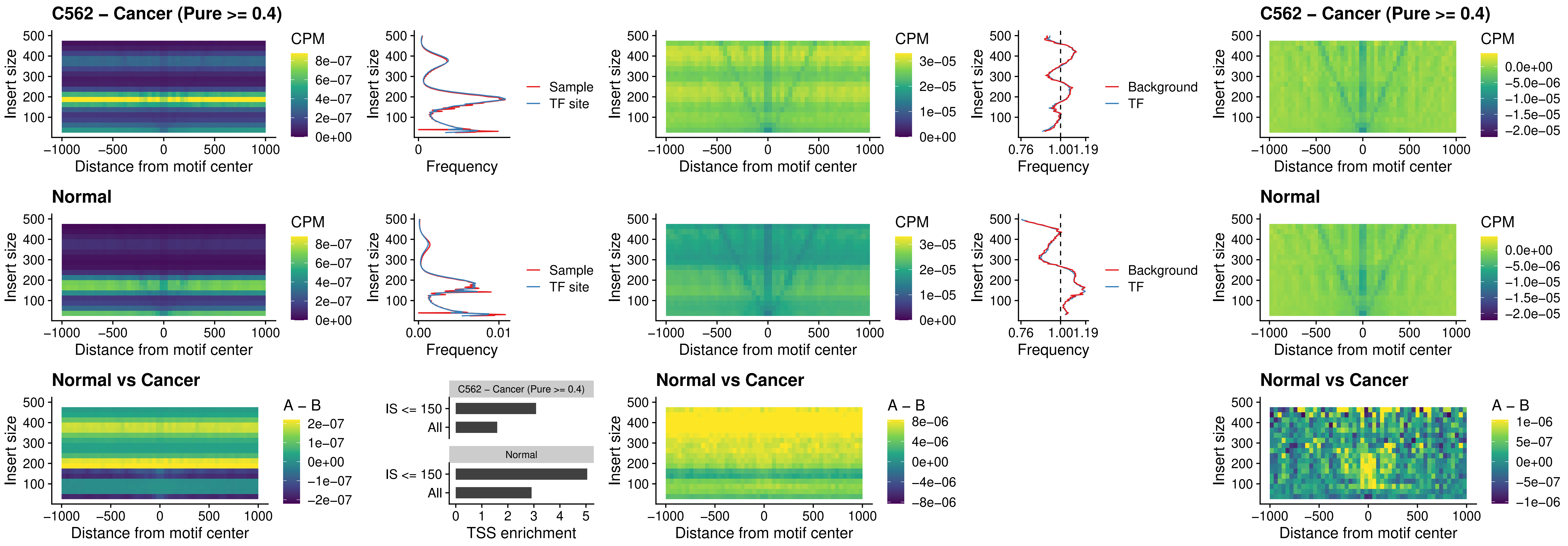

#### OTP_uc_dTSS_nPEAK

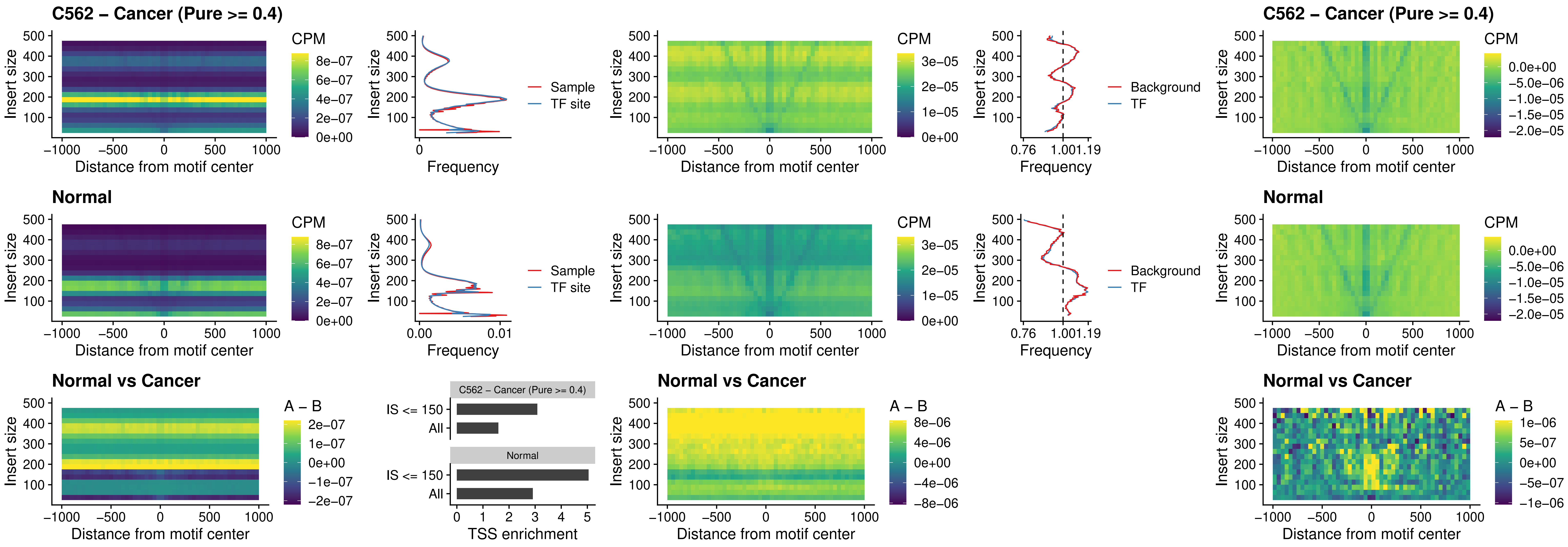

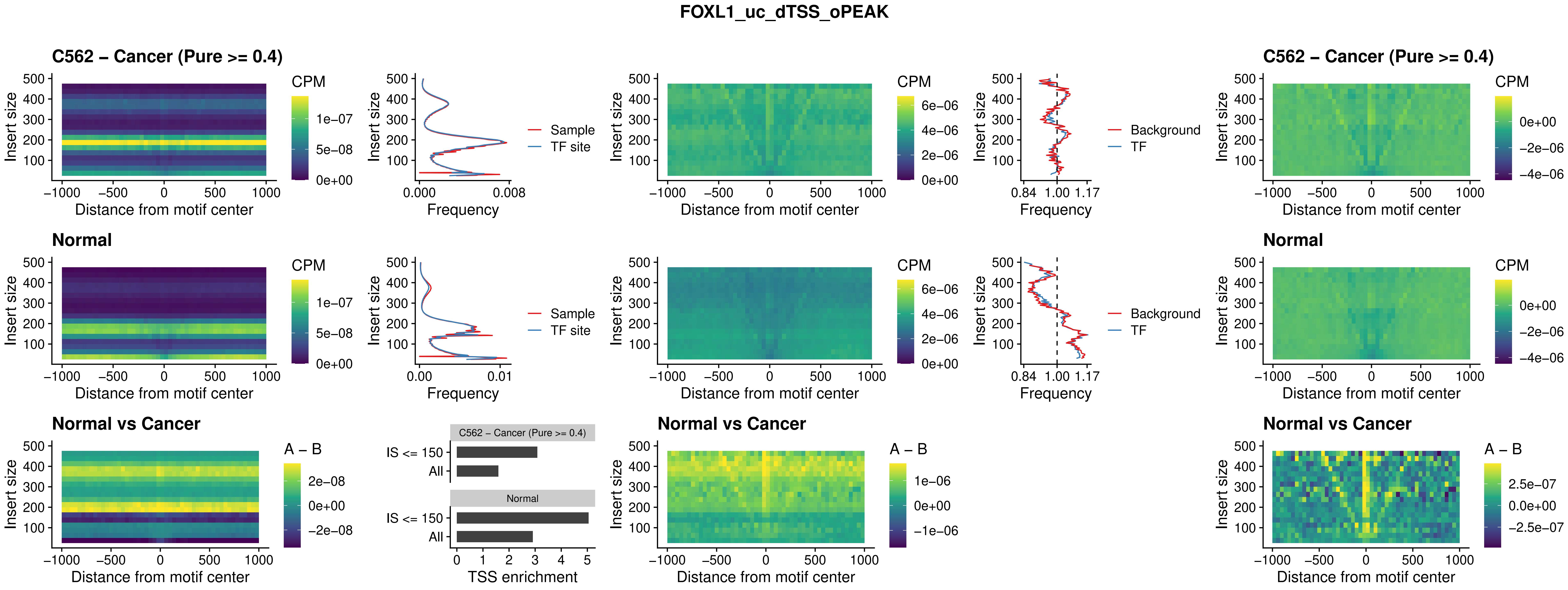

FOXL1_uc_dTSS_oPEAK
C562 – Cancer (Pure >= 0.4)
Normal
Normal vs Cancer
Insert size
Distance from motif center
Frequency
CPM
A – B
Sample
TF site
Background
TF
TSS enrichment
IS <= 150
All
C562 – Cancer (Pure >= 0.4)
Normal

#### ZNF32_uc_dTSS_oPEAK

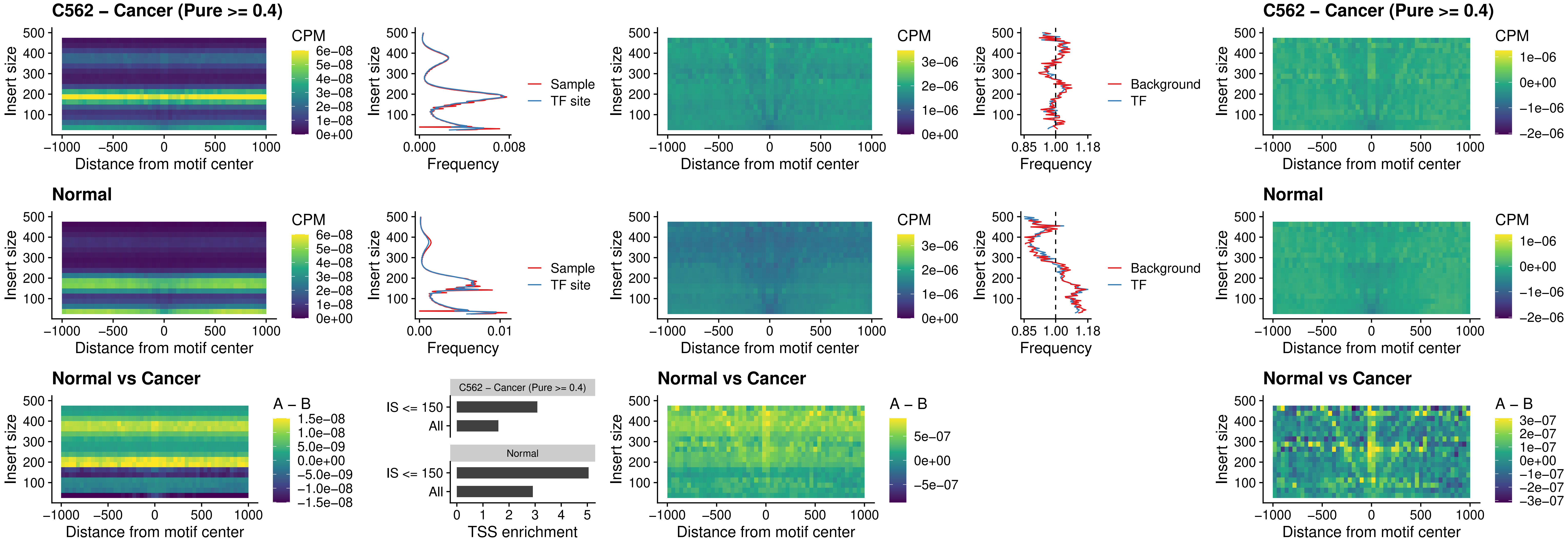

### FOXL1_uc_dTSS_nPEAK

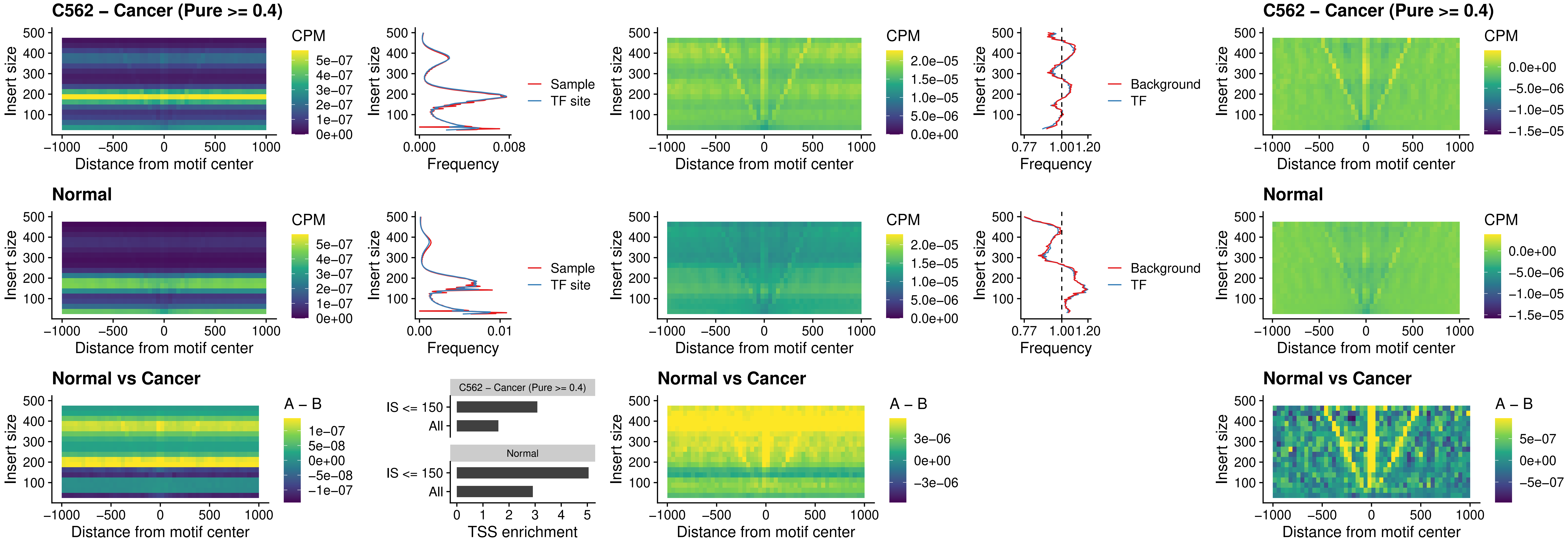

### ZNF32_uc_dTSS_nPEAK

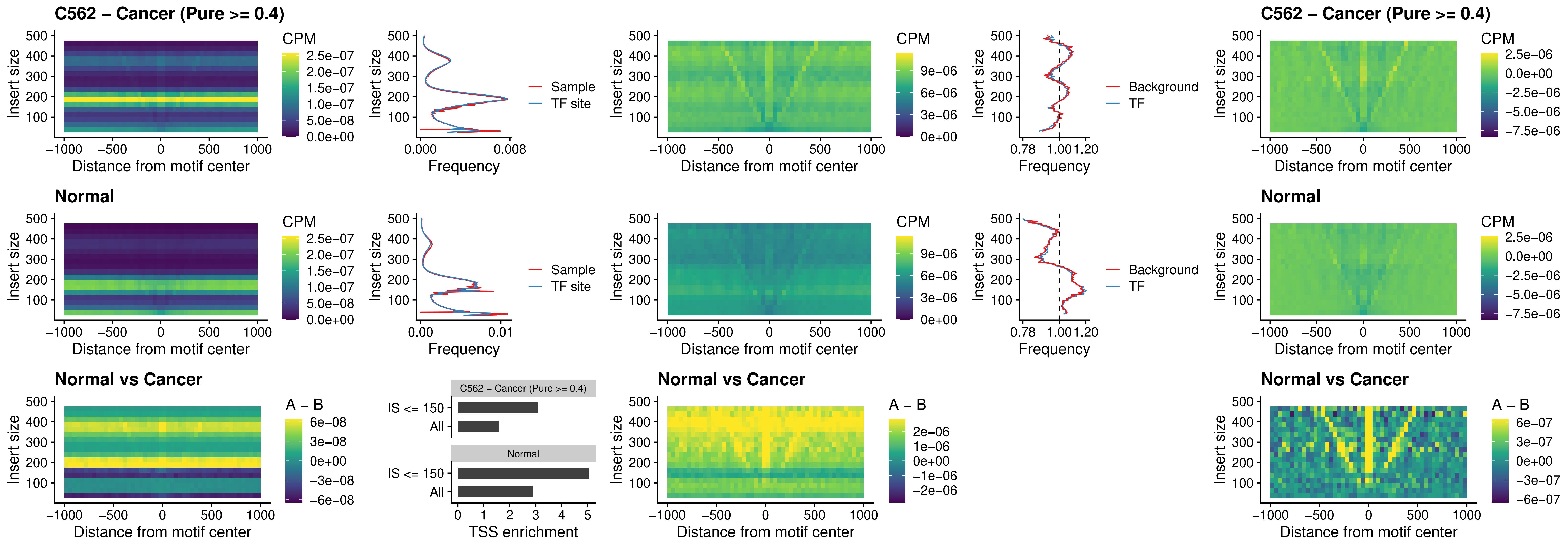

### DLX5_uc_dTSS_nPEAK

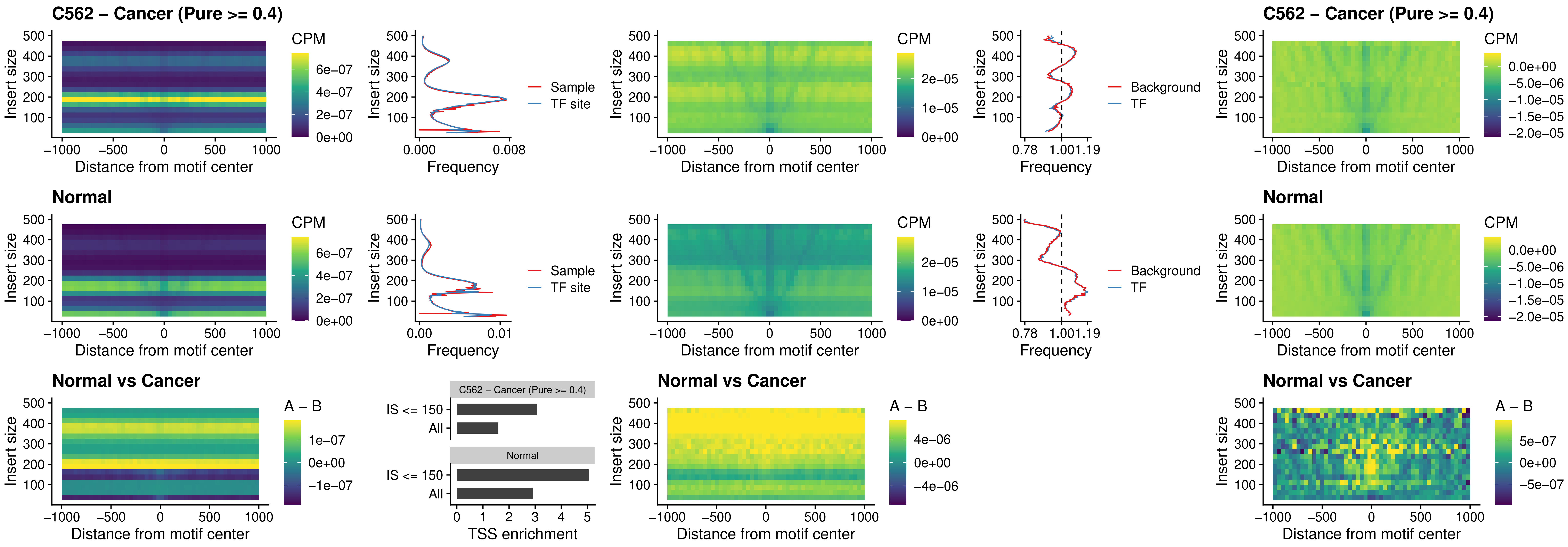

#### DMRT2_uc_dTSS_nPEAK

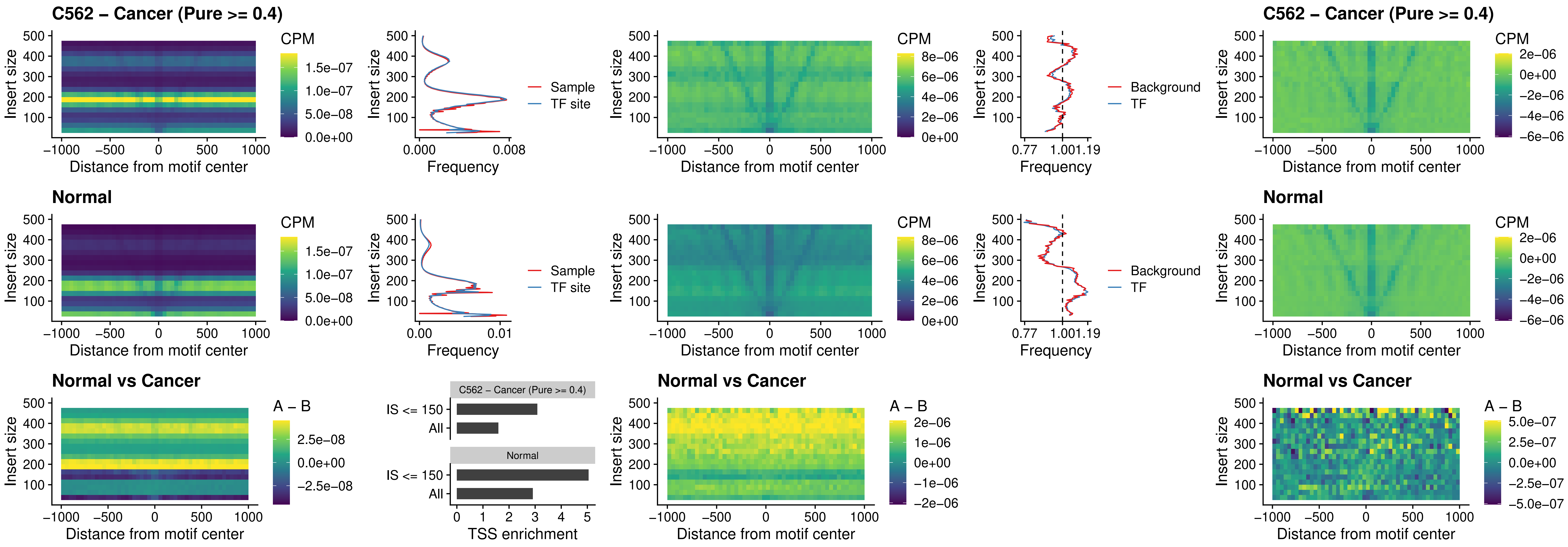

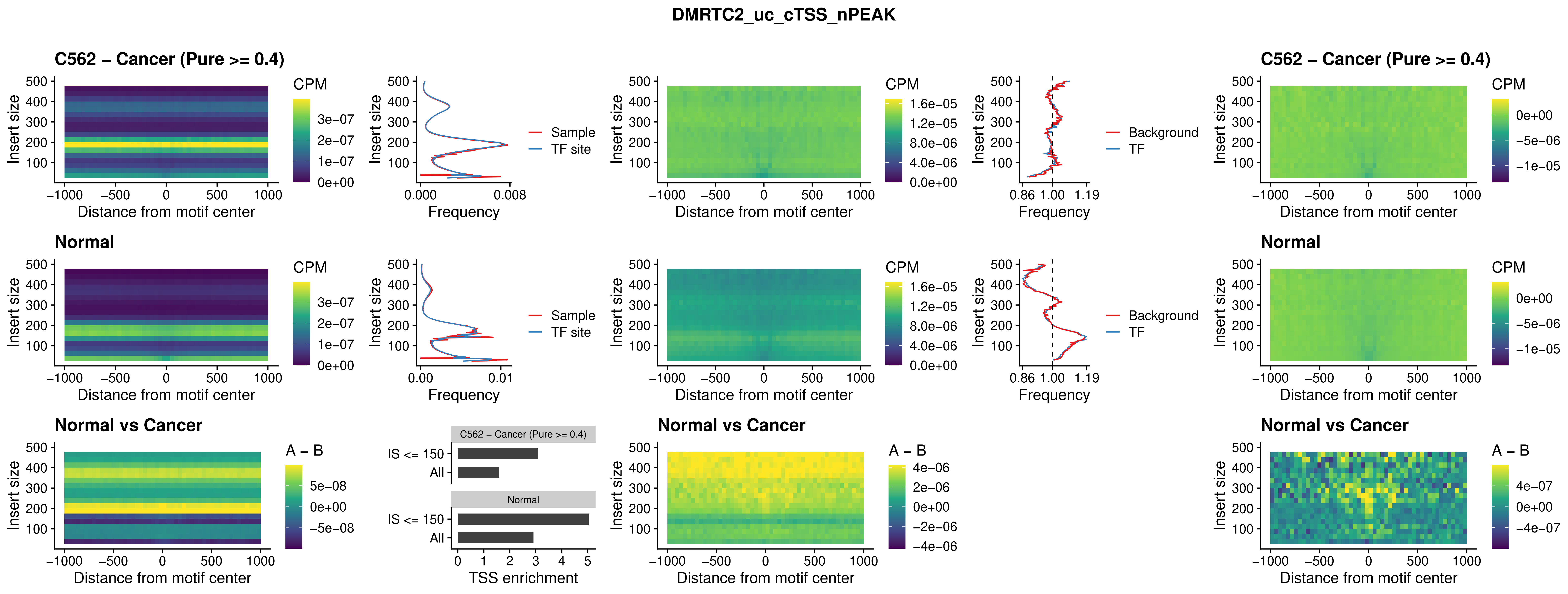
DMRTC2_uc_cTSS_nPEAK
C562 – Cancer (Pure >= 0.4)
Normal
Normal vs Cancer
Insert size
Distance from motif center
Frequency
CPM
A – B
Sample
TF site
Background
TF
C562 – Cancer (Pure >= 0.4)
Normal
IS <= 150
All
TSS enrichment

#### HOXD9_uc_dTSS_nPEAK

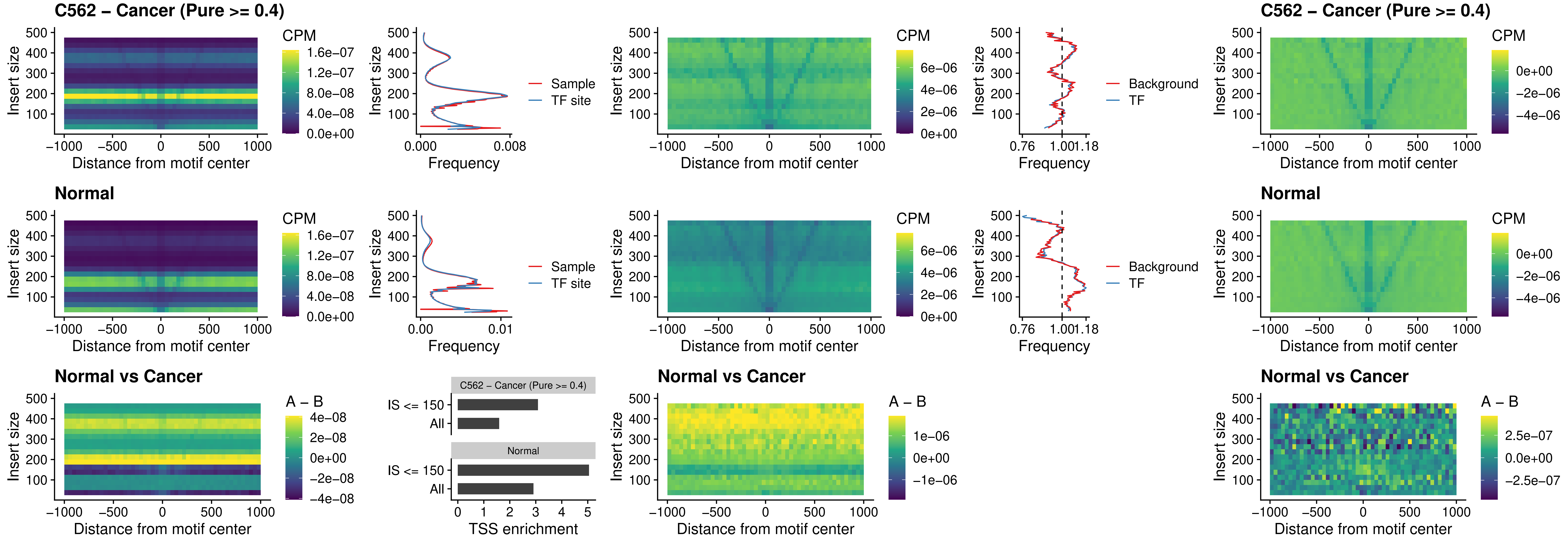

### HOXD4_uc_dTSS_nPEAK

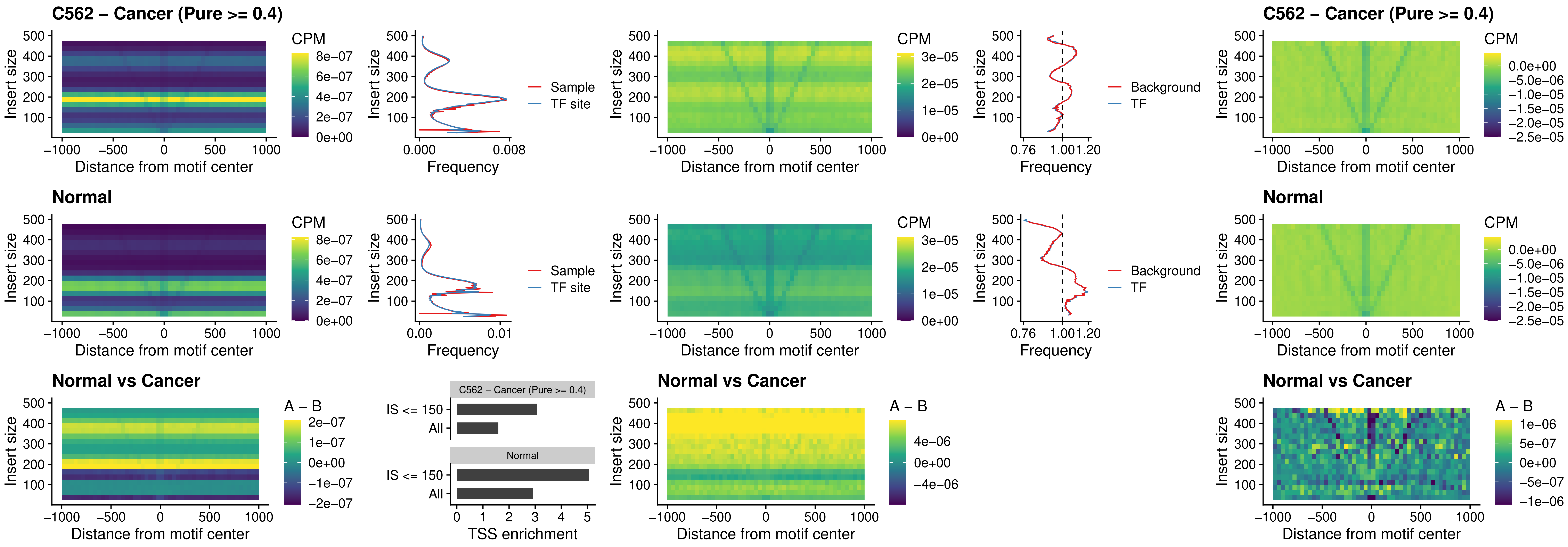

### FOXP3_uc_cTSS_nPEAK

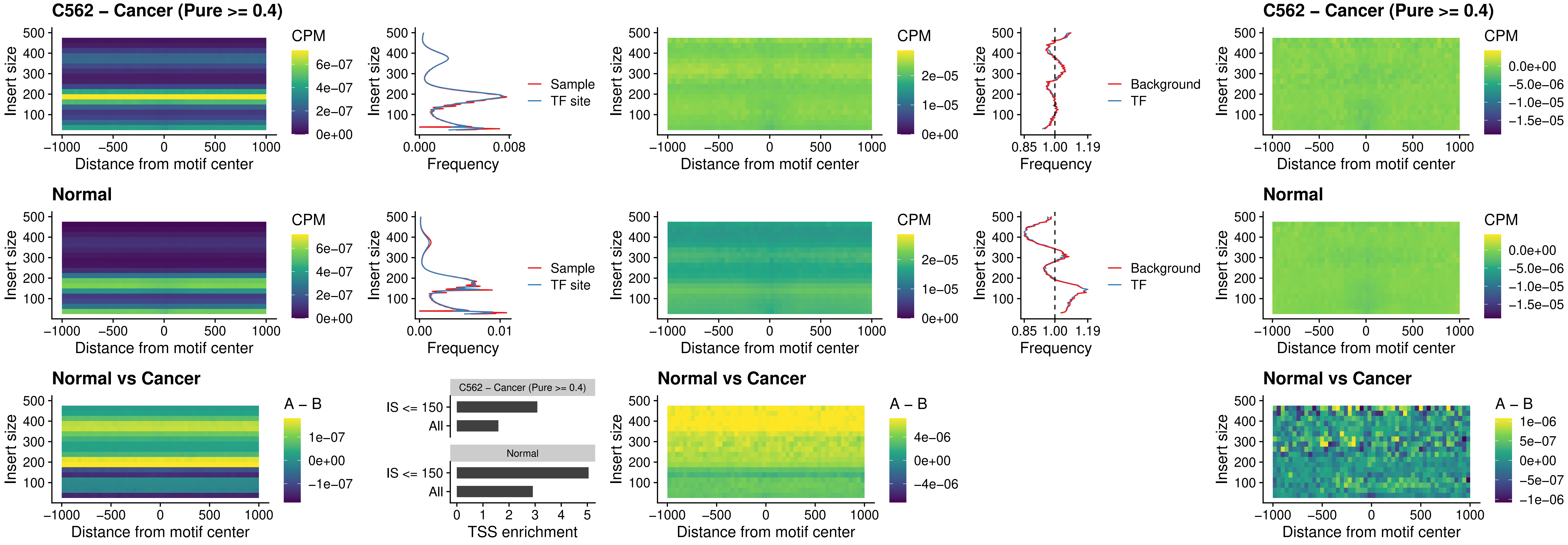

### FOXD4L6_uc_cTSS_nPEAK

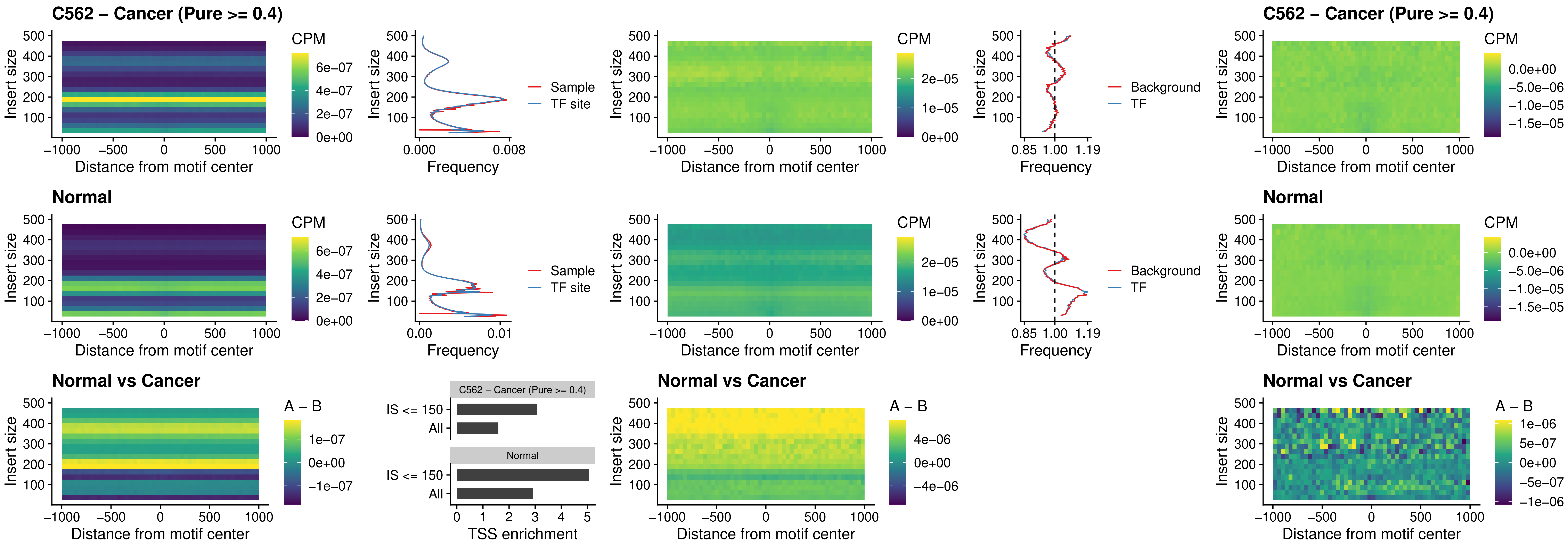

#### FOXD4L5_uc_cTSS_nPEAK

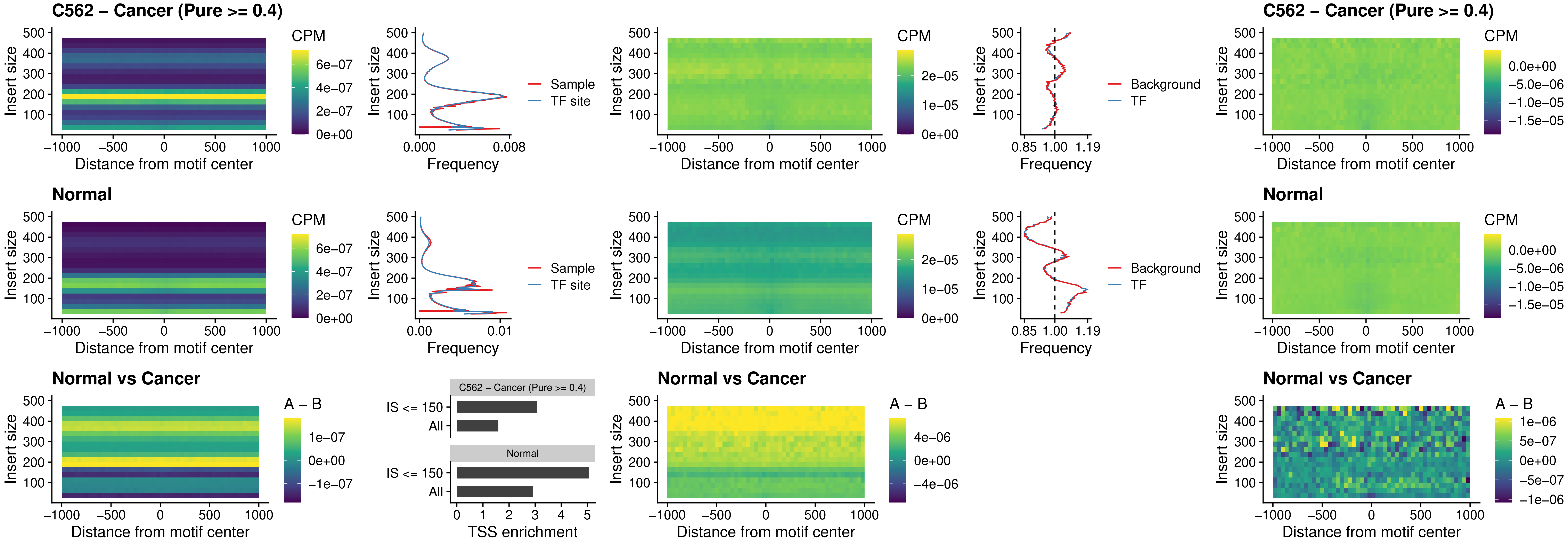

### FOXD4L4_uc_cTSS_nPEAK

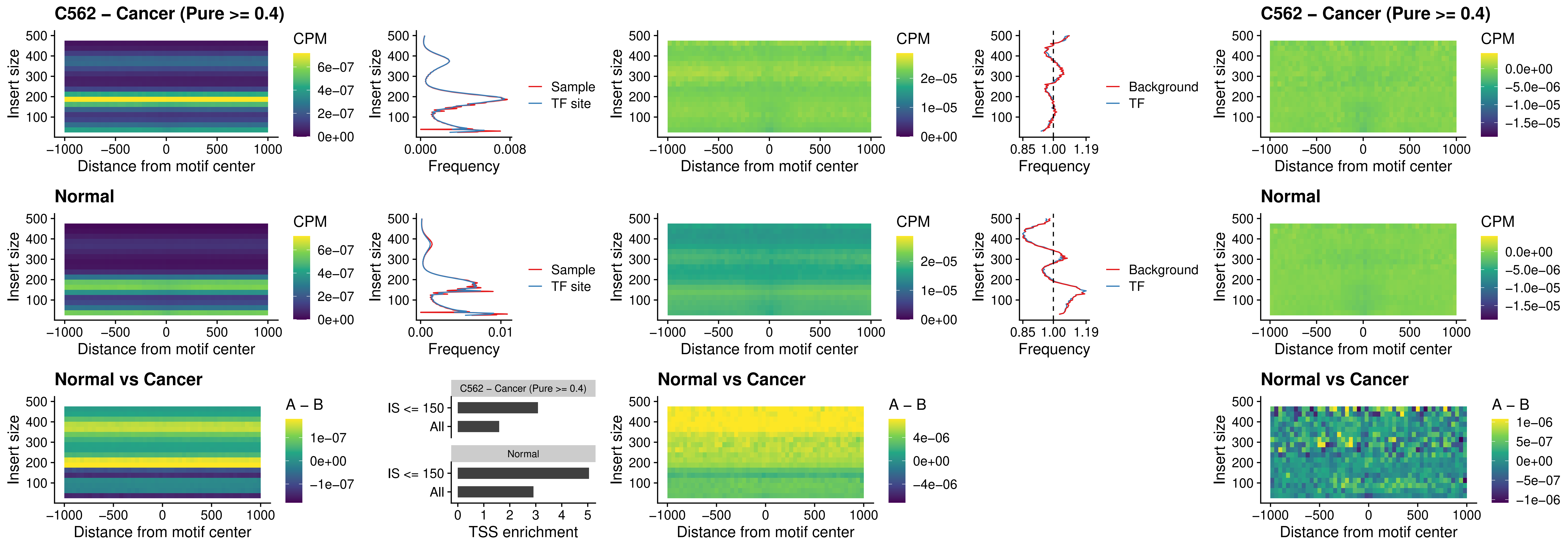

### FOXD4L2_uc_cTSS_nPEAK

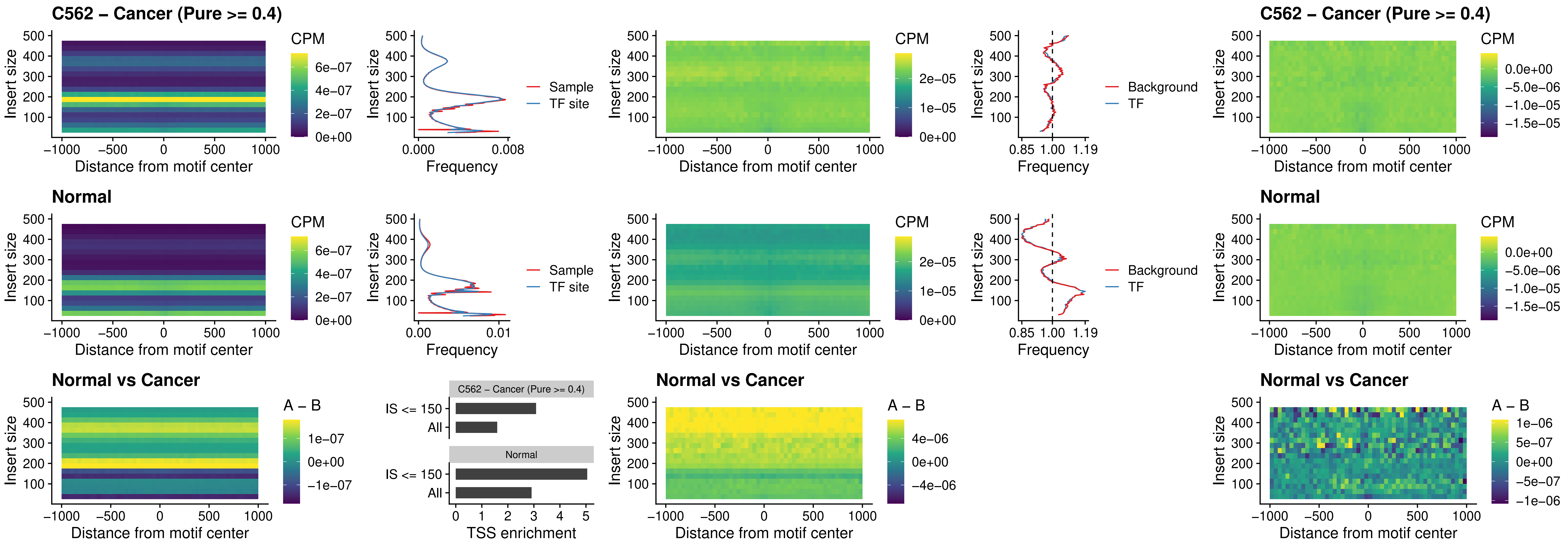

### FOXD4L3_uc_cTSS_nPEAK

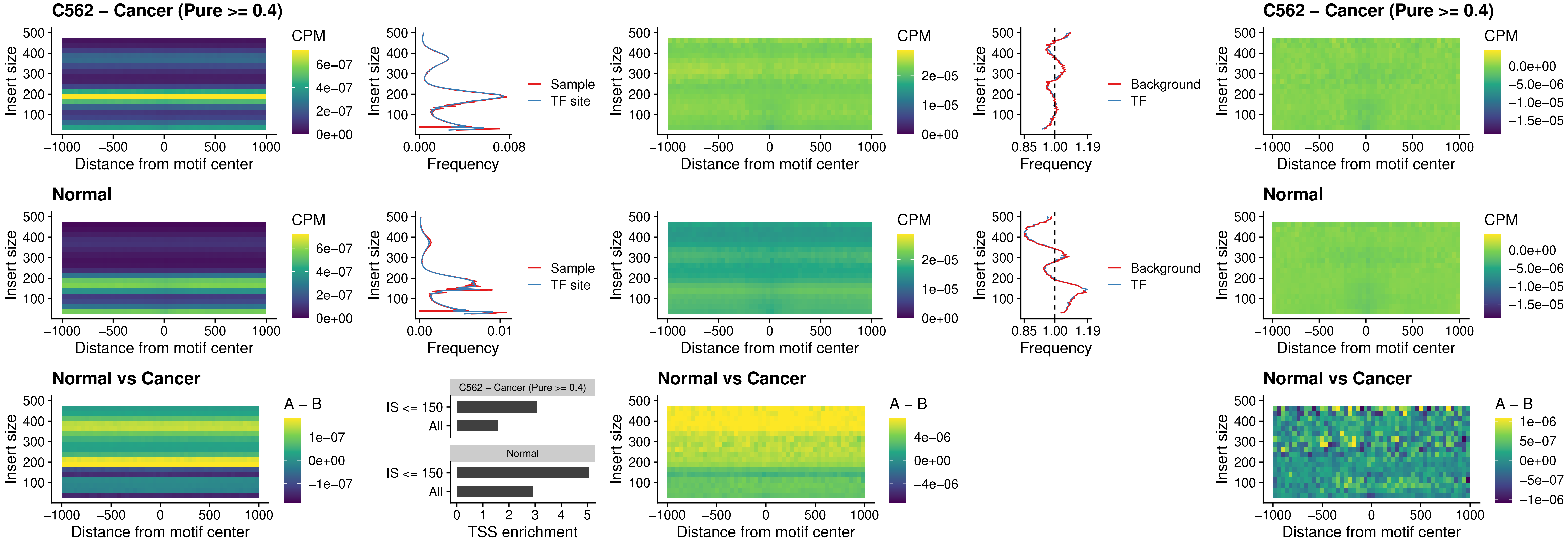

### HOXD1_uc_dTSS_oPEAK

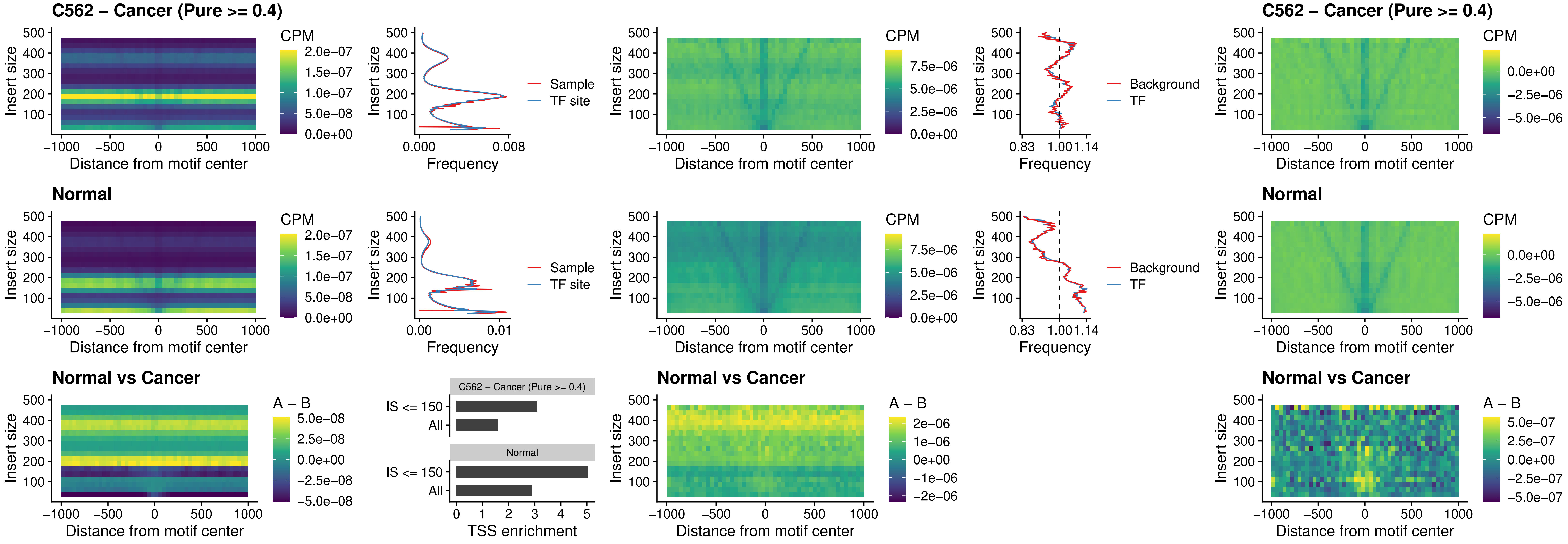

### VAX1_uc_pTSS_nPEAK

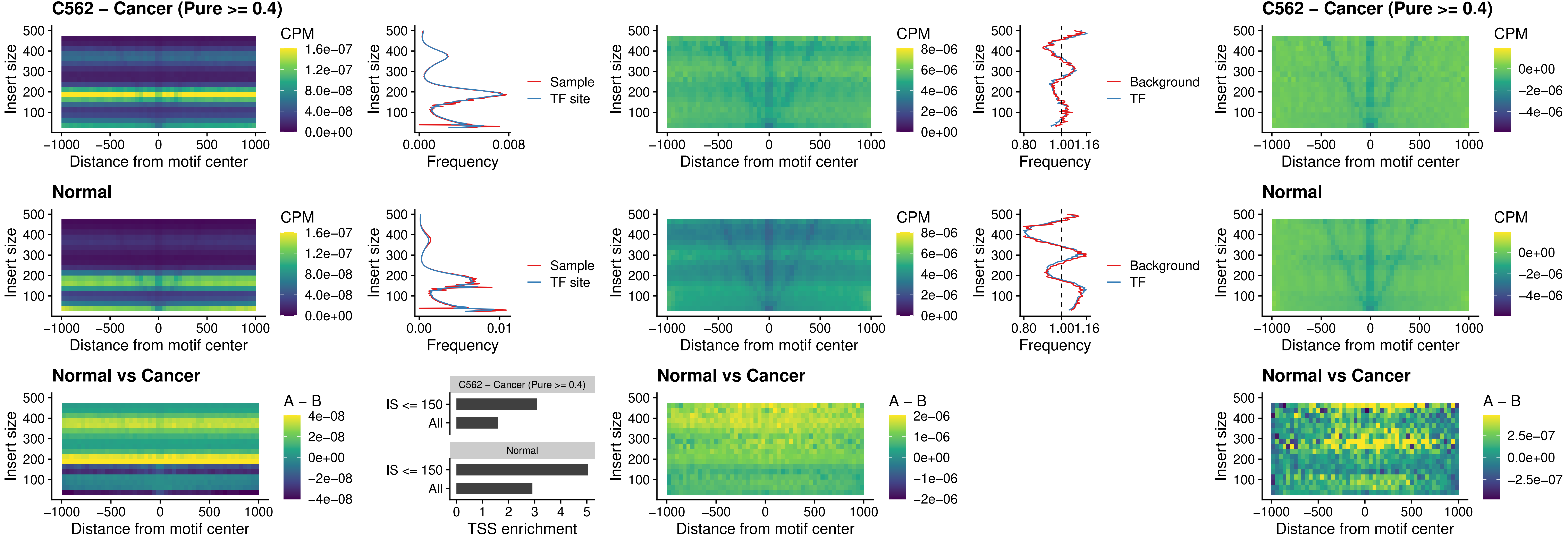

### DLX3_uc_pTSS_nPEAK

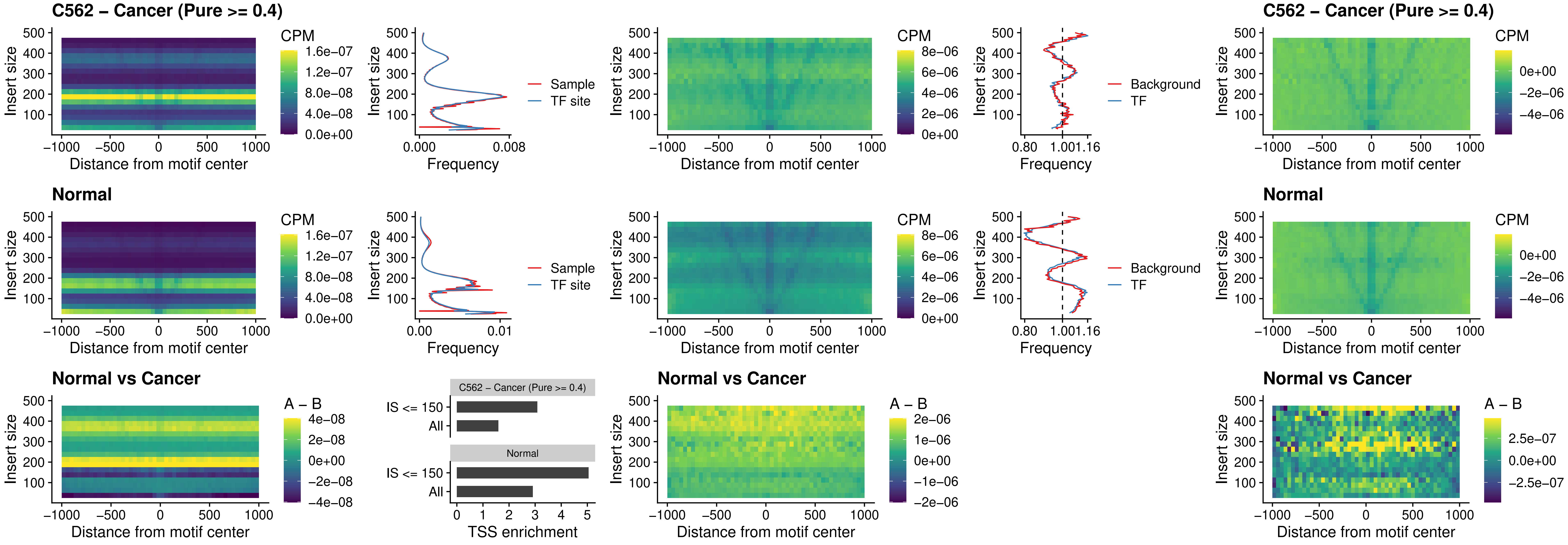

### HOXD3_uc_pTSS_nPEAK

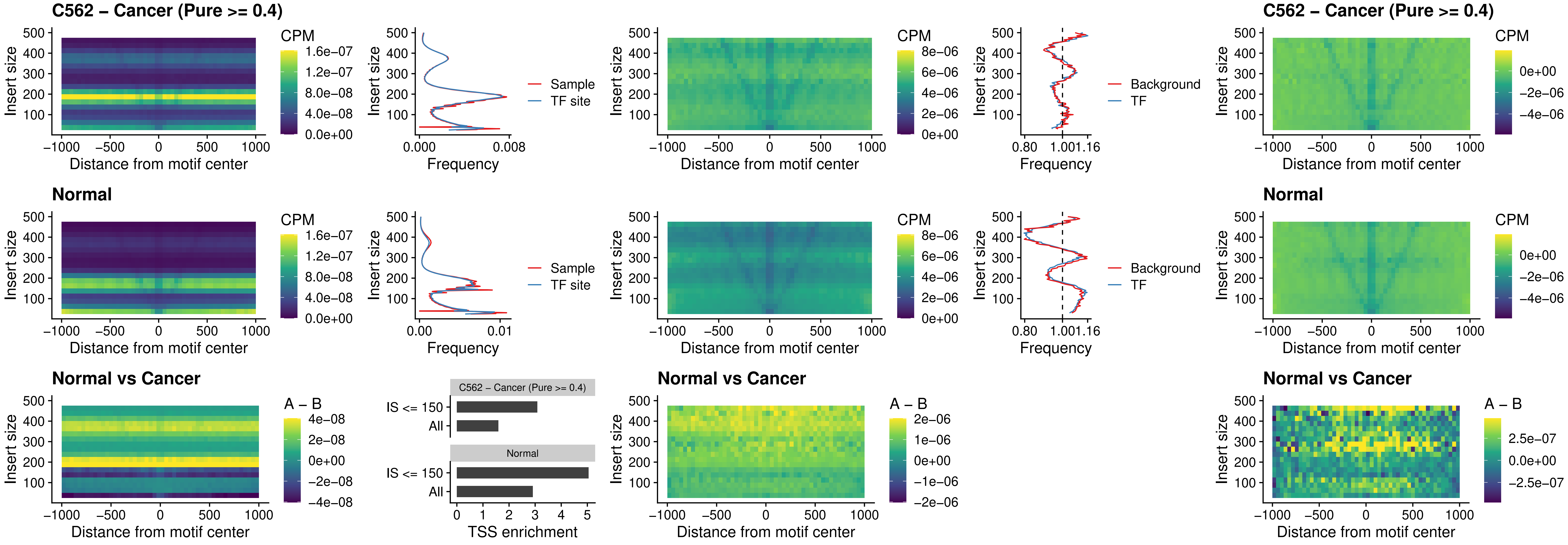

#### PDX1_uc_dTSS_nPEAK

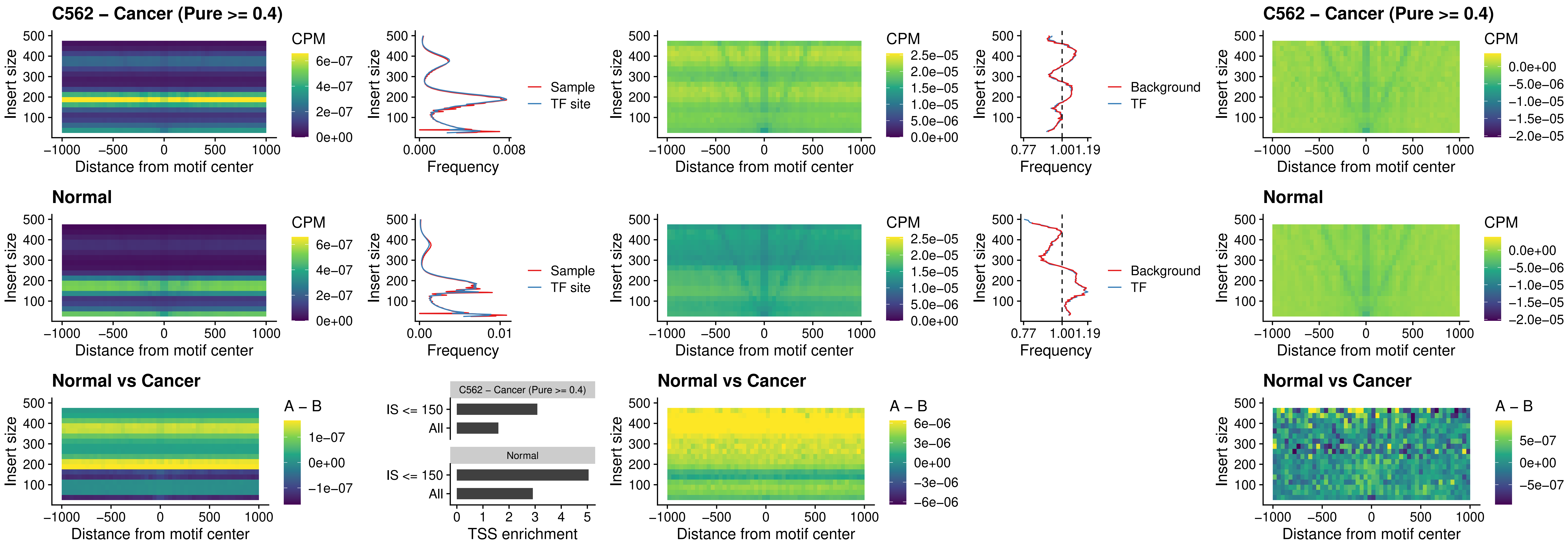

### VAX2_uc_dTSS_nPEAK

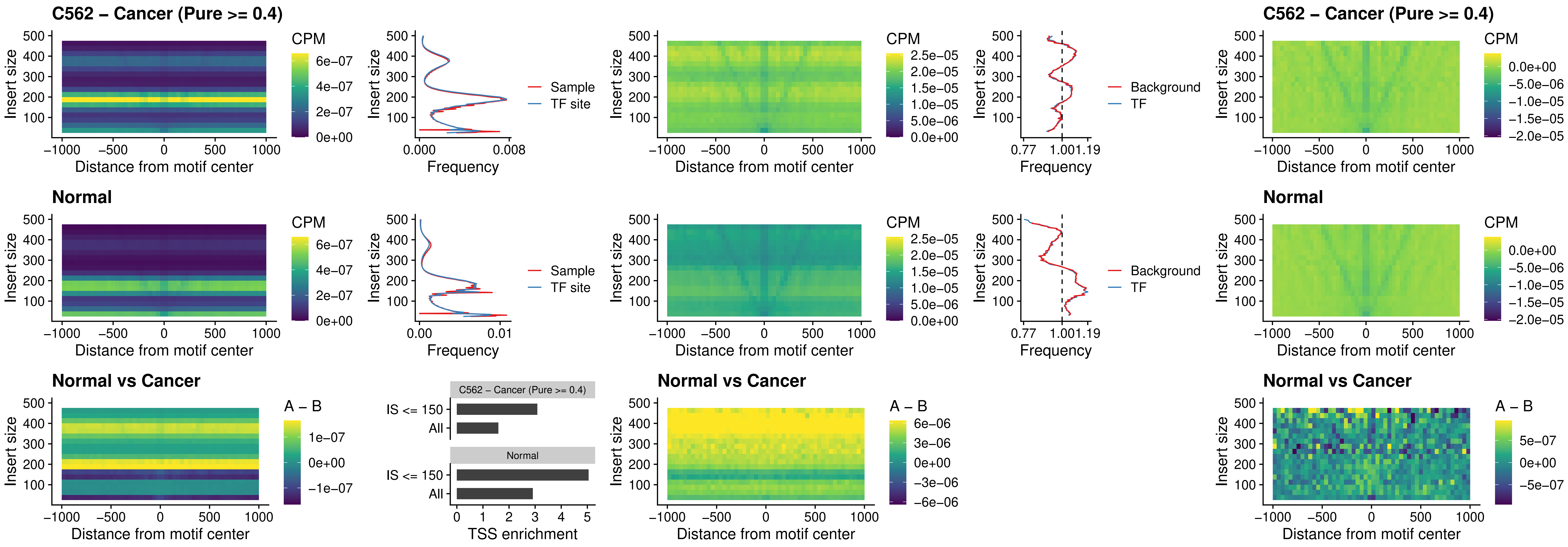

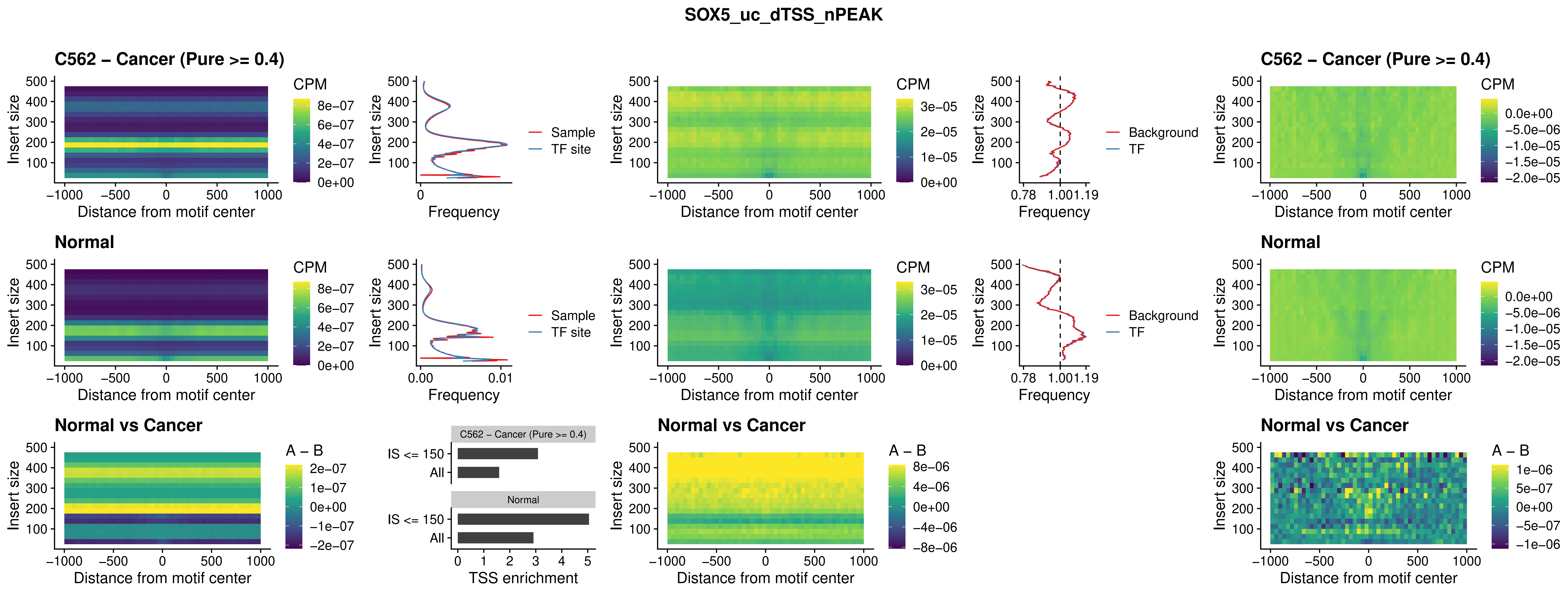
SOX5_uc_dTSS_nPEAK
C562 – Cancer (Pure >= 0.4)
Normal
Normal vs Cancer
Insert size
Distance from motif center
CPM
A – B
Frequency
Sample
TF site
Background
TF
C562 – Cancer (Pure >= 0.4)
Normal
IS <= 150
All
TSS enrichment

#### SOX6_uc_dTSS_nPEAK

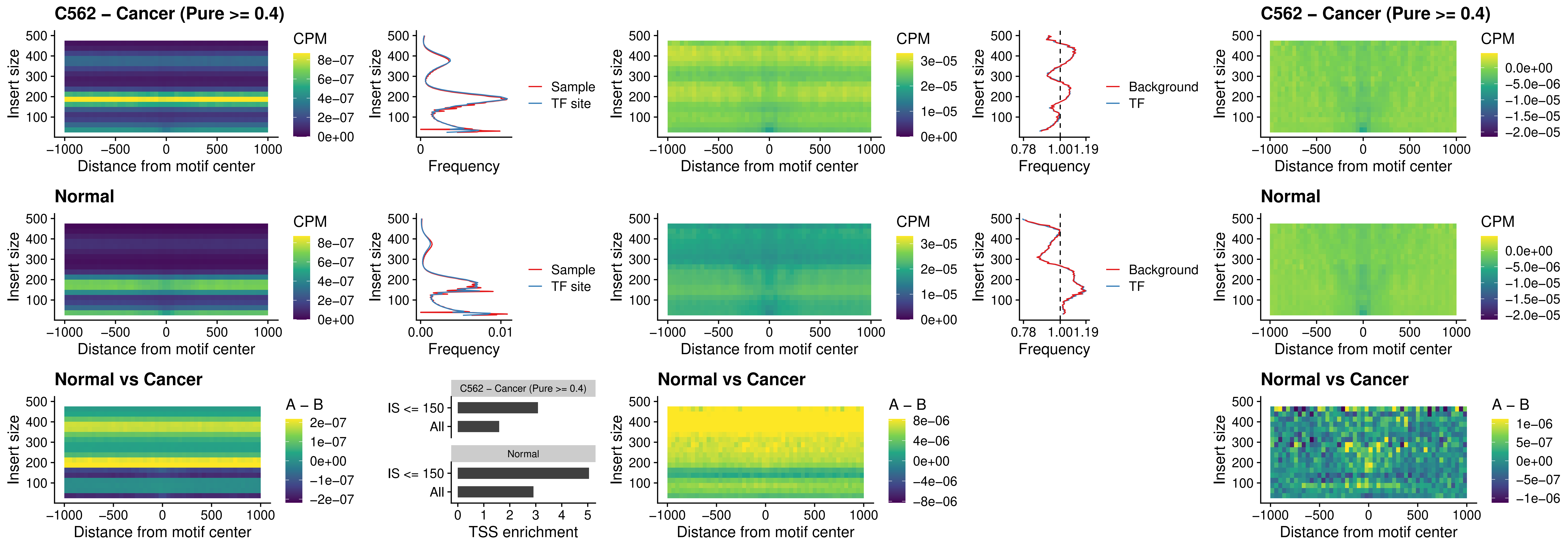

### JDP2_uc_cTSS_nPEAK

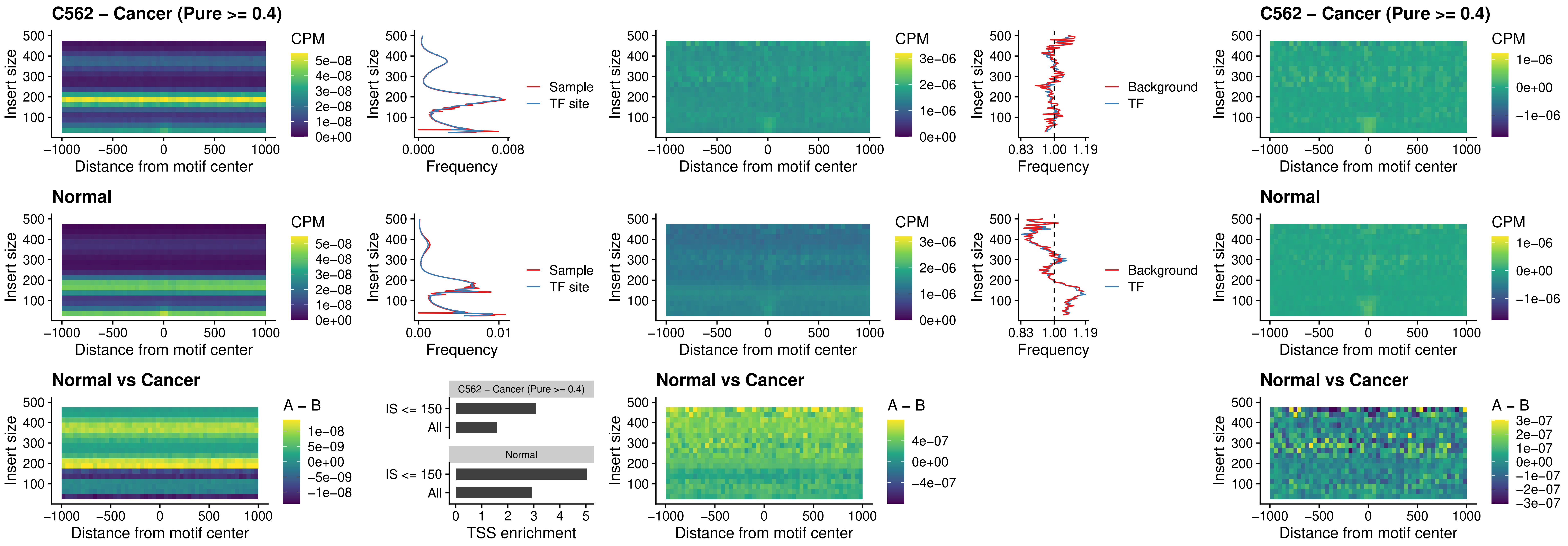

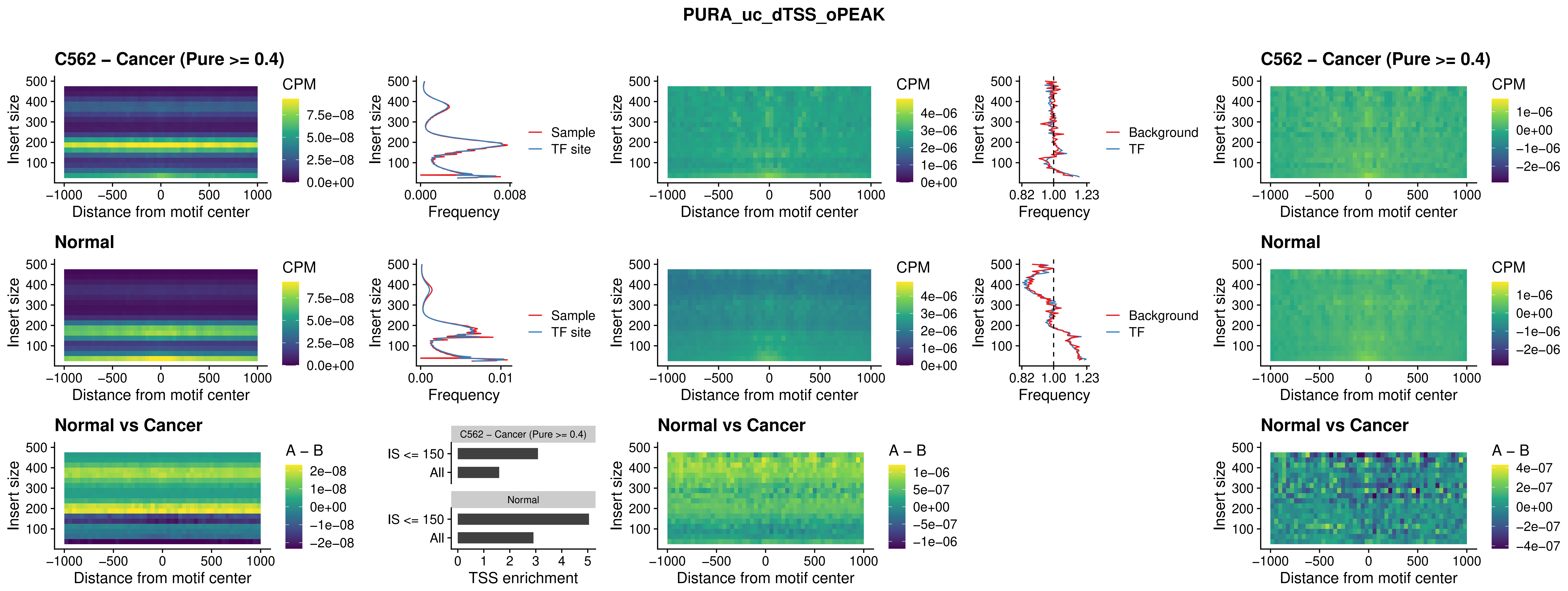
PURA_uc_dTSS_oPEAK
C562 – Cancer (Pure >= 0.4)
Normal
Normal vs Cancer
Insert size
Distance from motif center
Frequency
CPM
A – B
Sample
TF site
Background
TF
C562 – Cancer (Pure >= 0.4)
Normal
IS <= 150
All
TSS enrichment

### HHEX_uc_dTSS_nPEAK

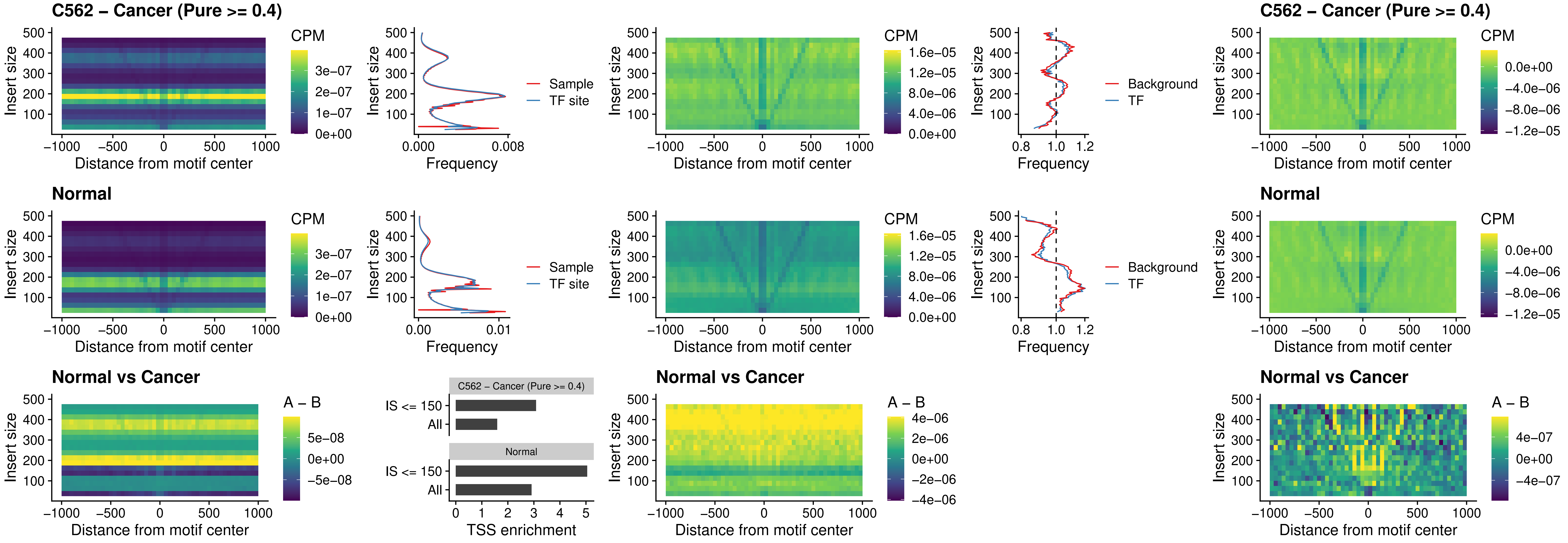

### HMG20B_uc_dTSS_nPEAK

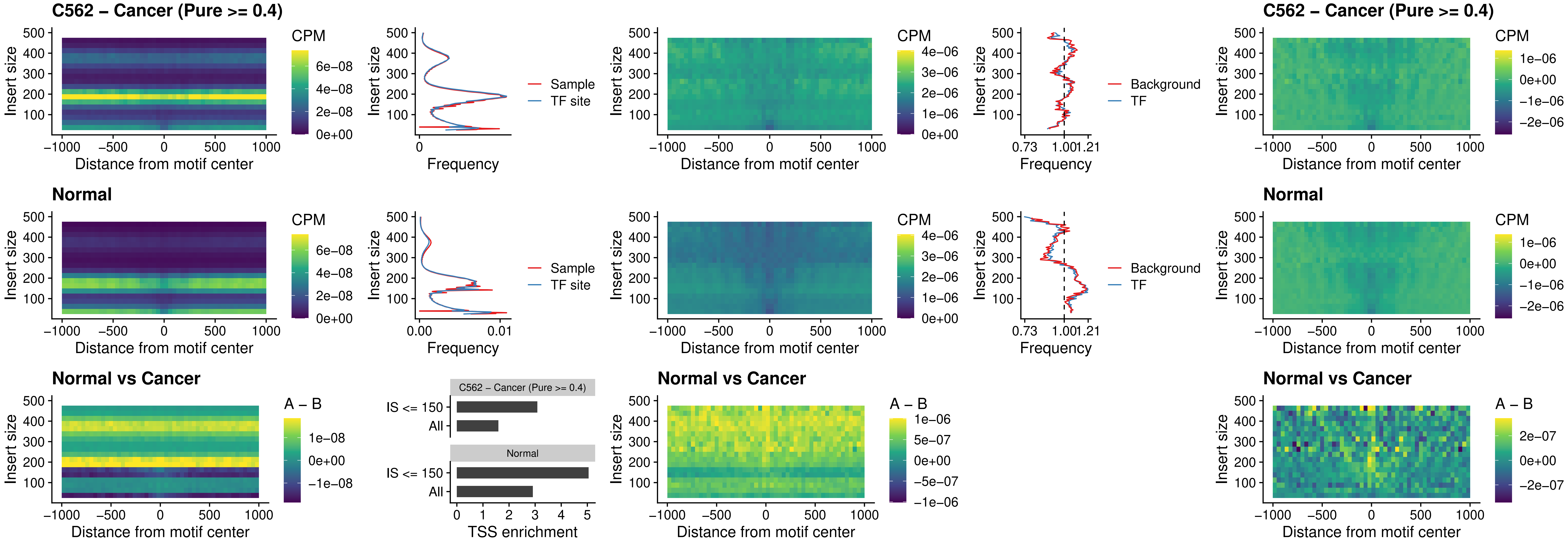

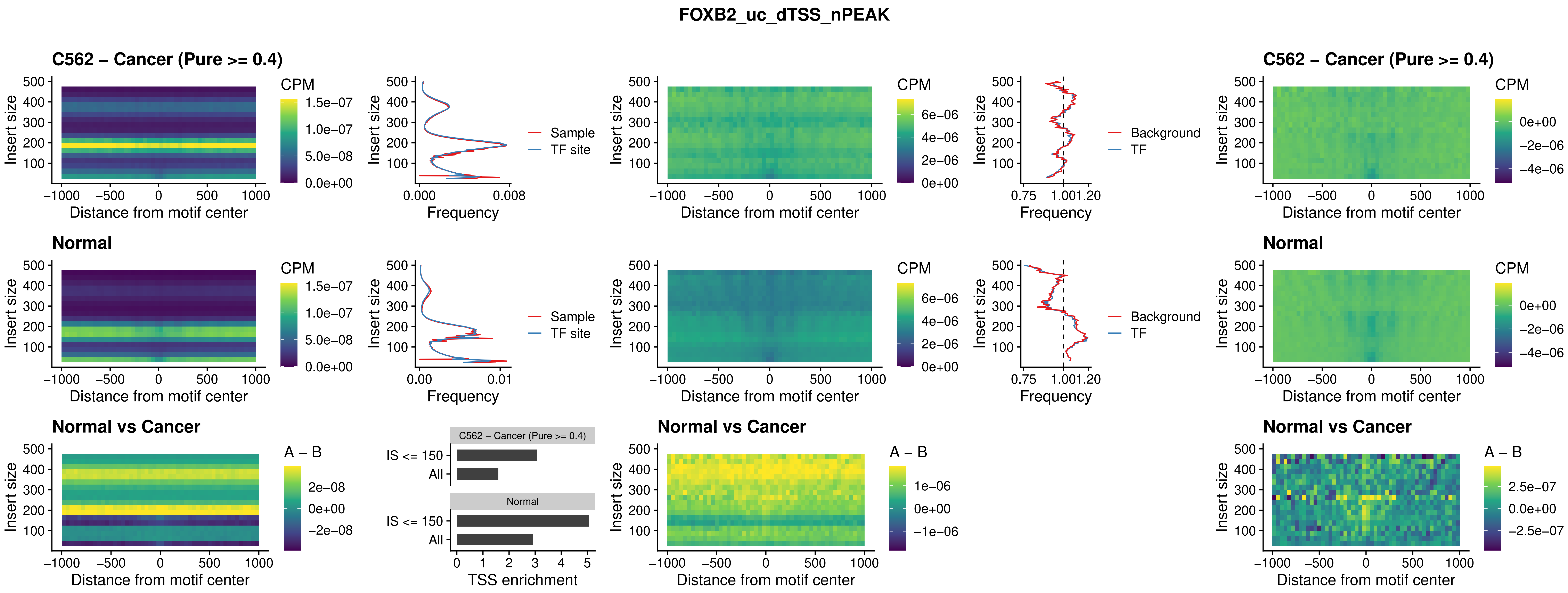
FOXB2_uc_dTSS_nPEAK
C562 – Cancer (Pure >= 0.4)
Normal
Normal vs Cancer
Insert size
Distance from motif center
Frequency
CPM
A – B
Sample
TF site
Background
TF
C562 – Cancer (Pure >= 0.4)
Normal
IS <= 150
All
TSS enrichment

### FOXL1_uc_cTSS_nPEAK

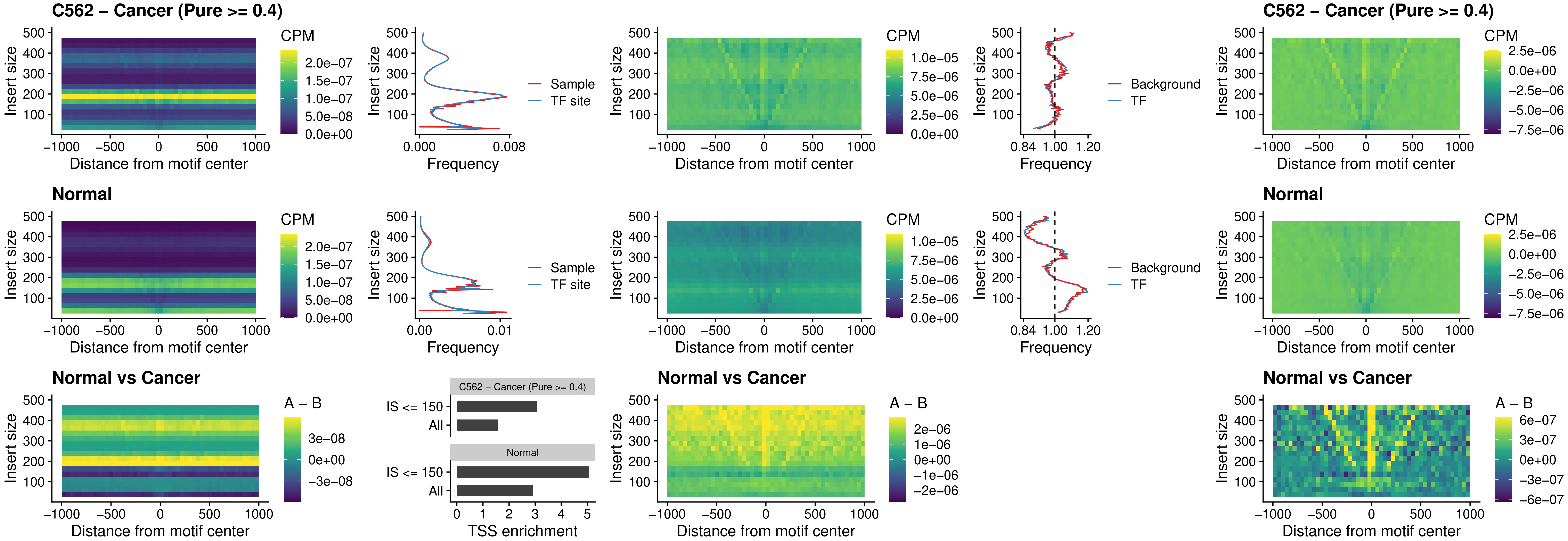

#### PBX3_uc_pTSS_oPEAK

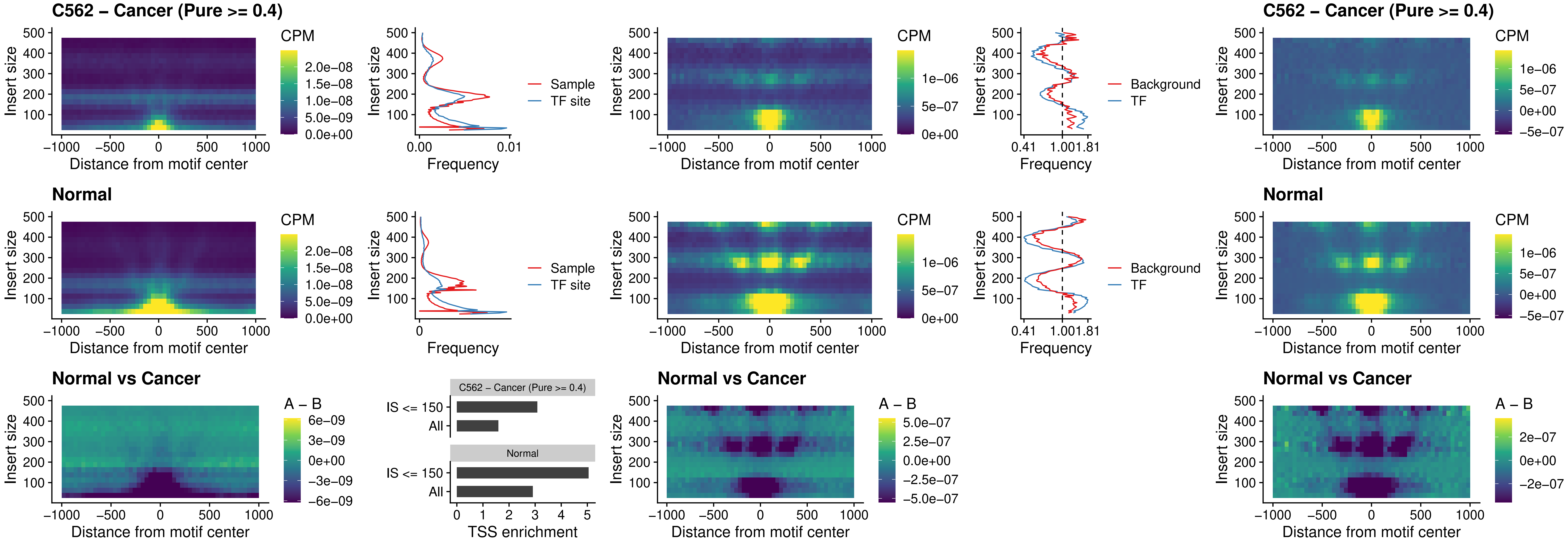

### SNAI2_uc_cTSS_nPEAK

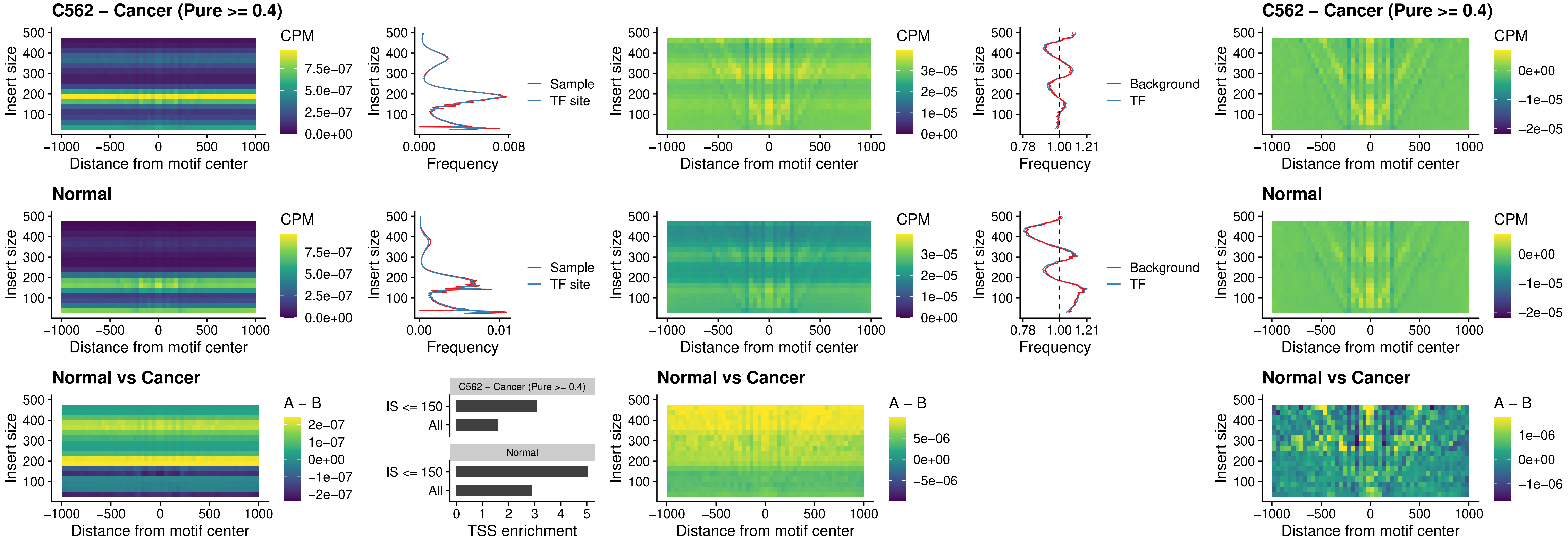

#### SNAI2_uc_dTSS_oPEAK

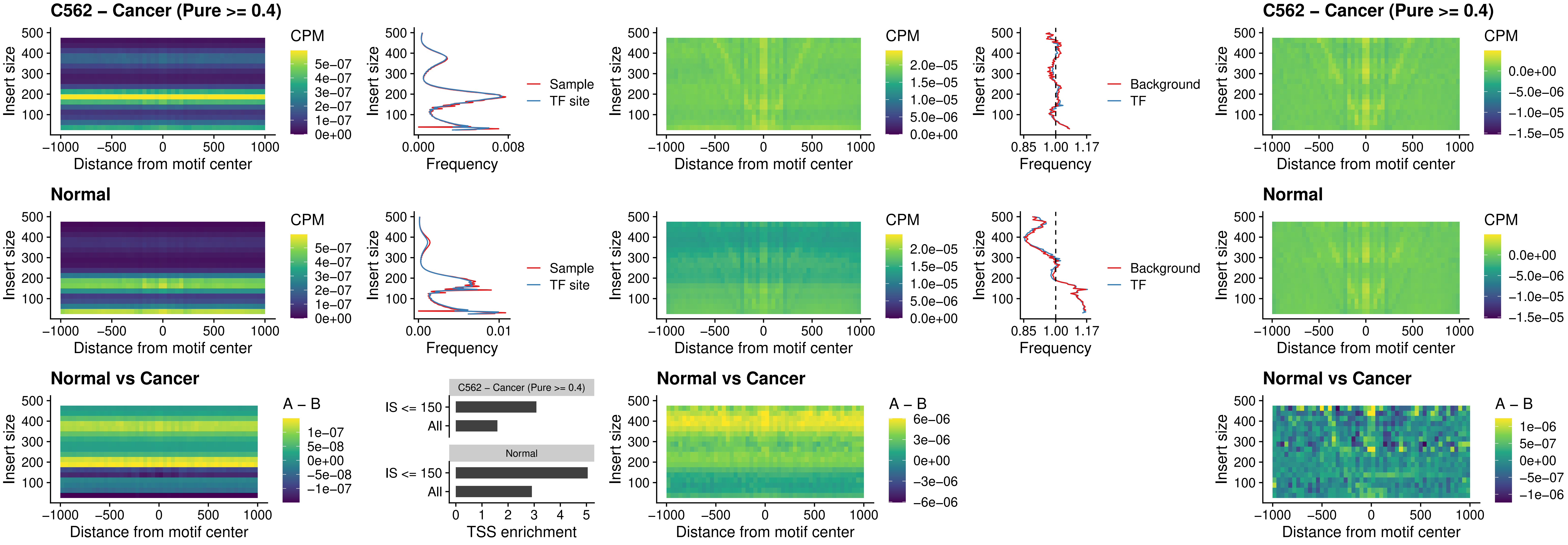

### SPI1_uc_pTSS_nPEAK

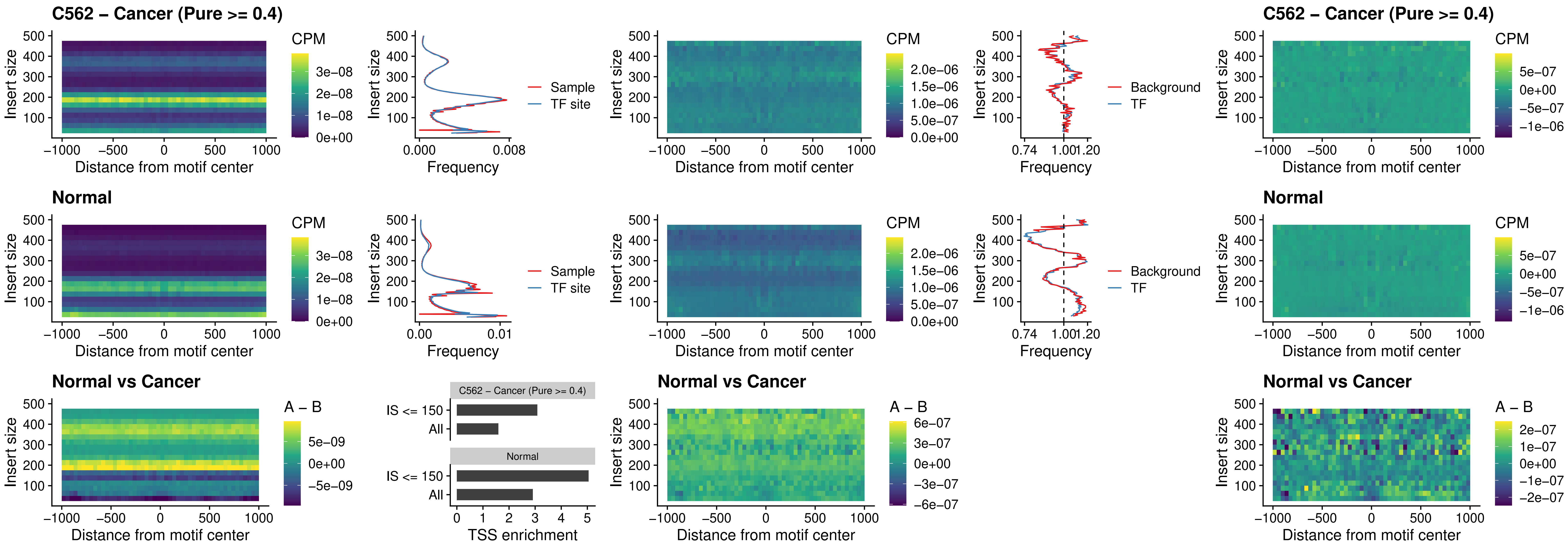

### KLF16_uc_pTSS_nPEAK

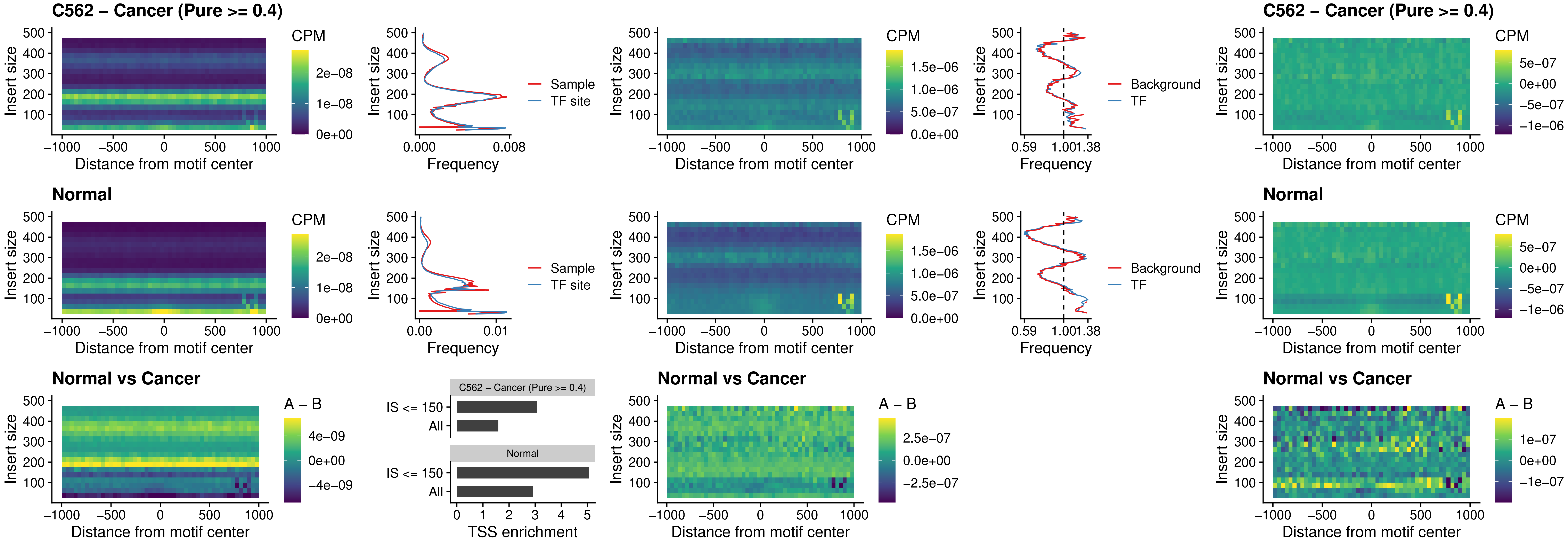

### SP3_uc_pTSS_nPEAK

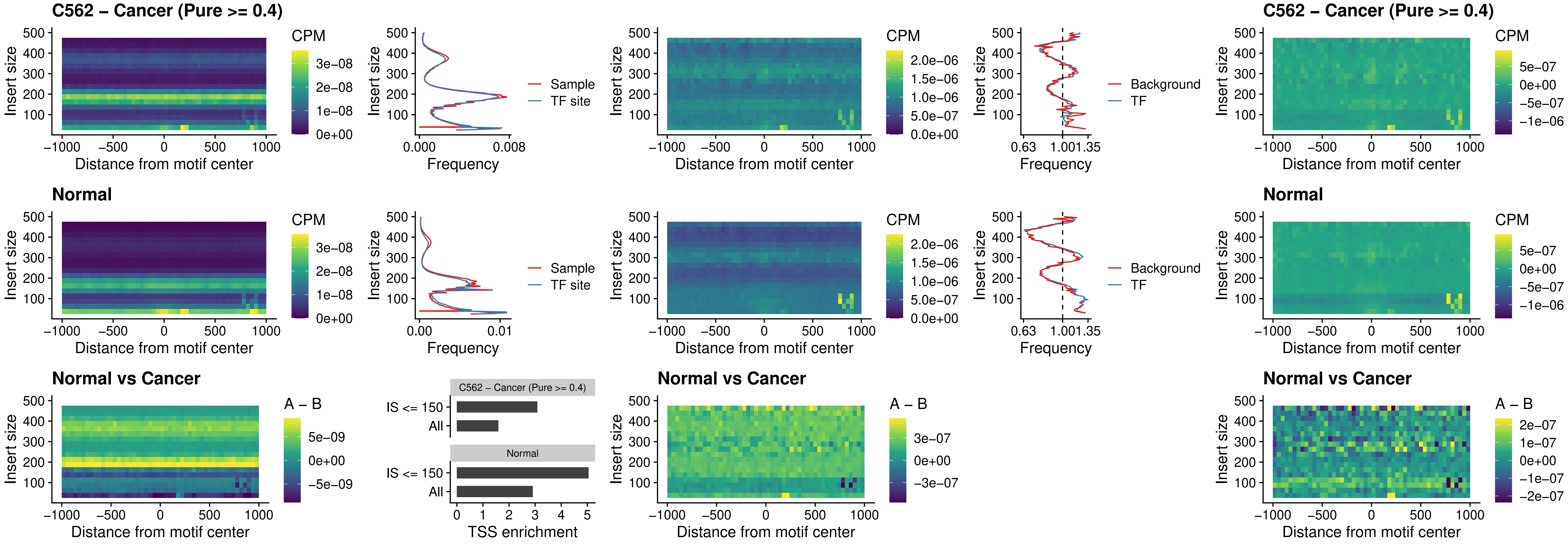

#### NEURODD1_uc_dTSS_oPEAK

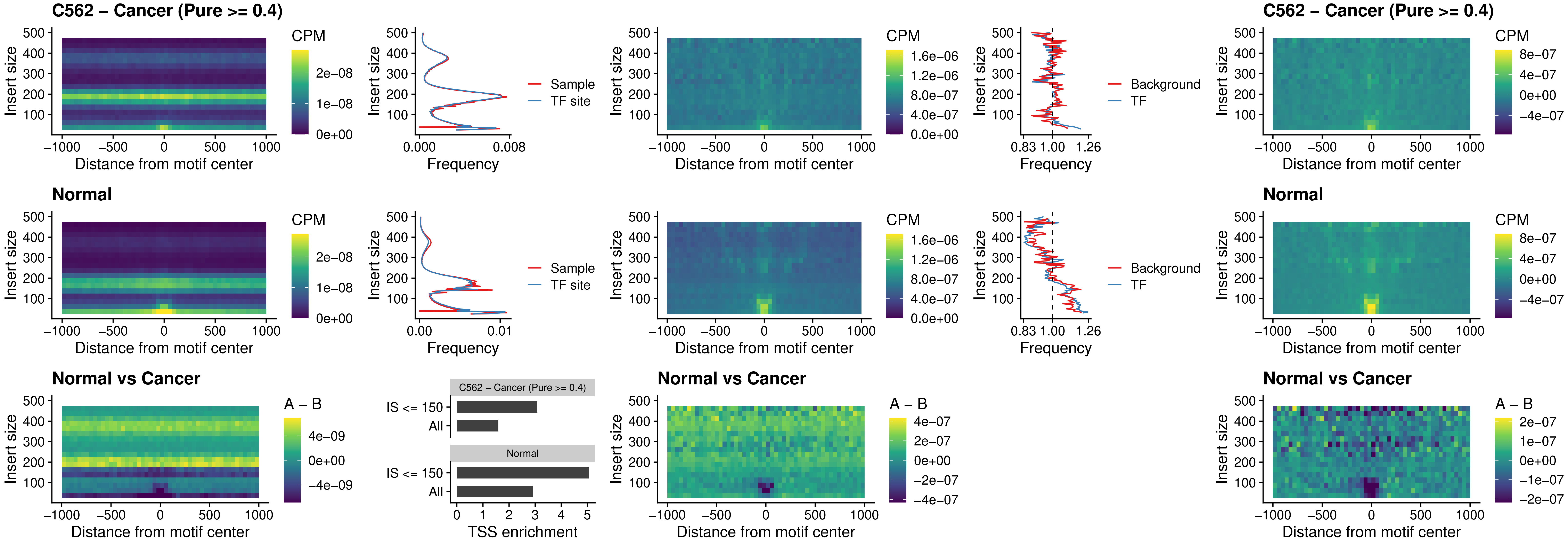

#### CTCF_uc_dTSS_oPEAK

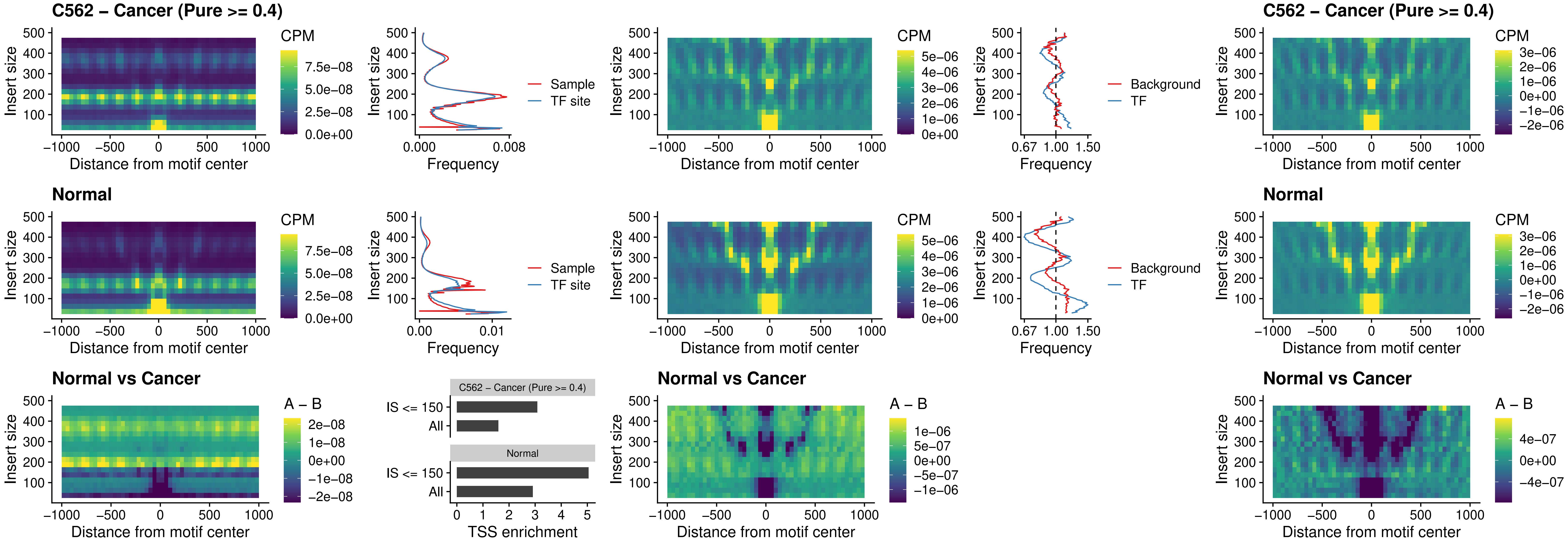

#### CTCF_uc_cTSS_oPEAK

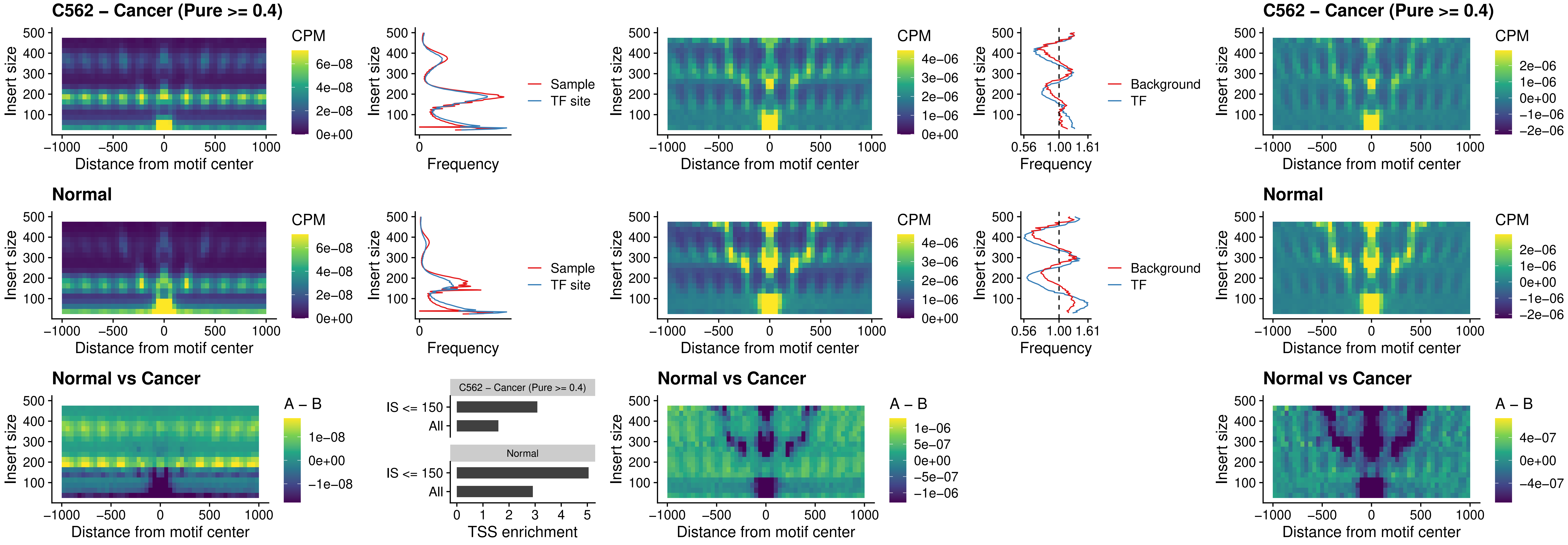

### CTCFL_uc_dTSS_oPEAK

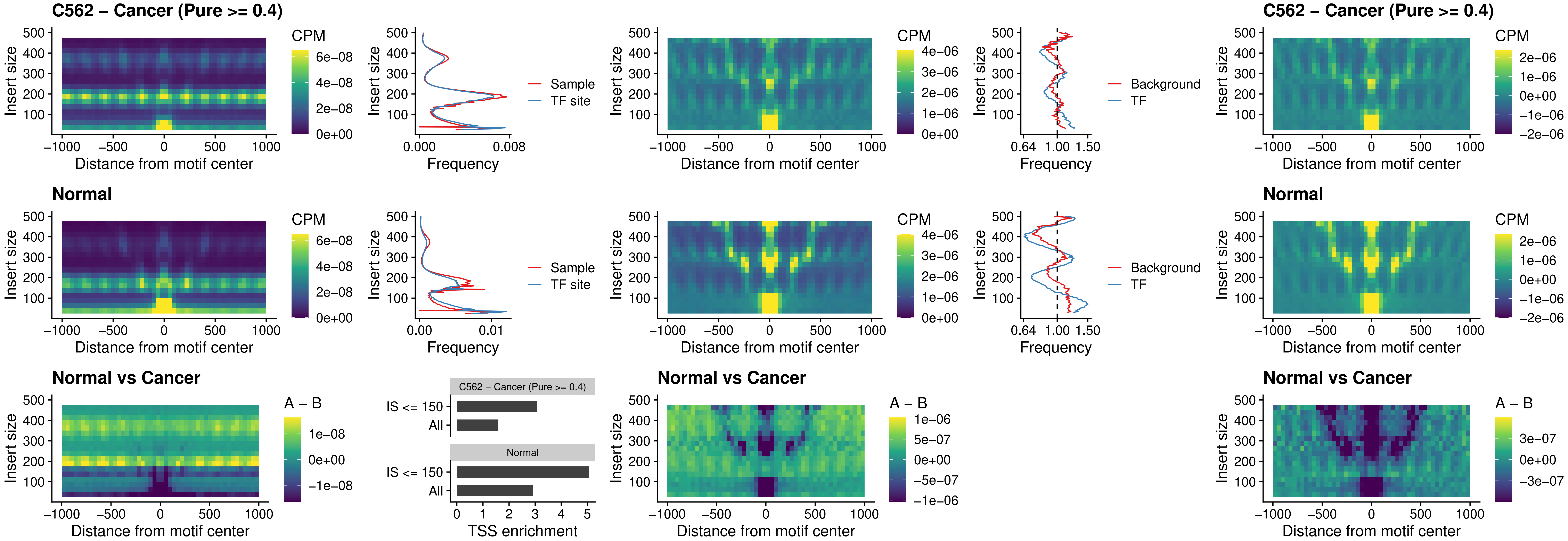

### ENCODE_CTCF_sites_COLON

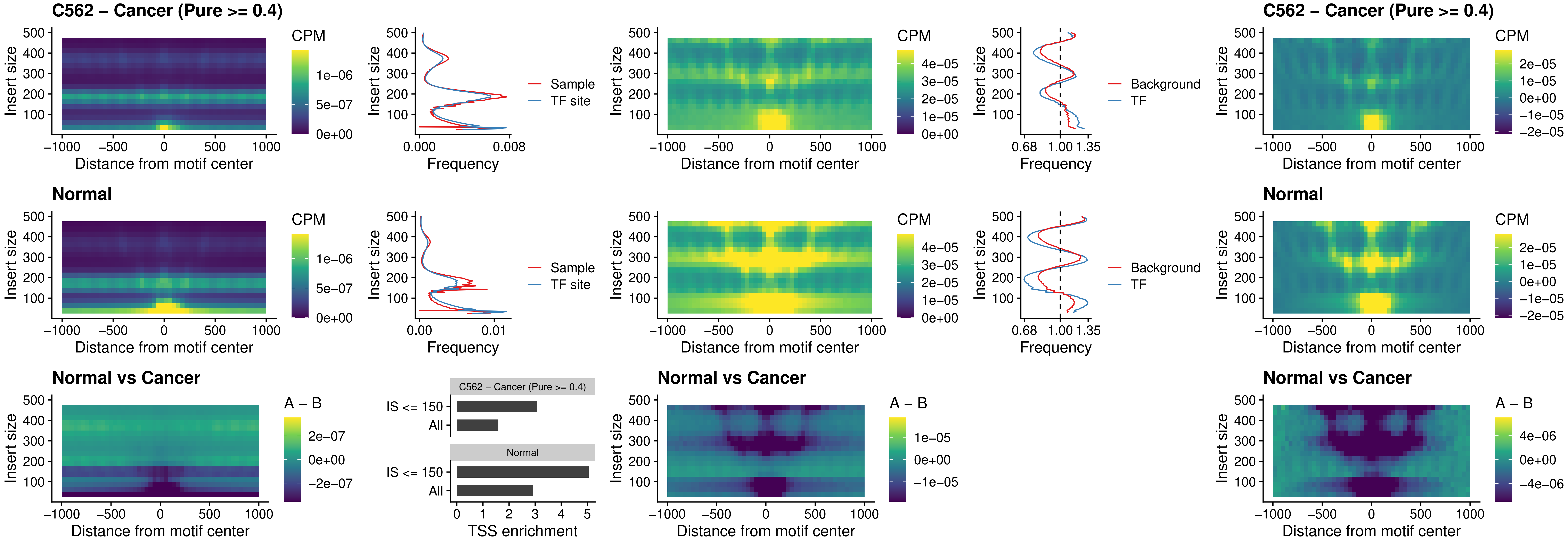

#### ENCODE_CTCF_sites_COLON_loop_ctcf

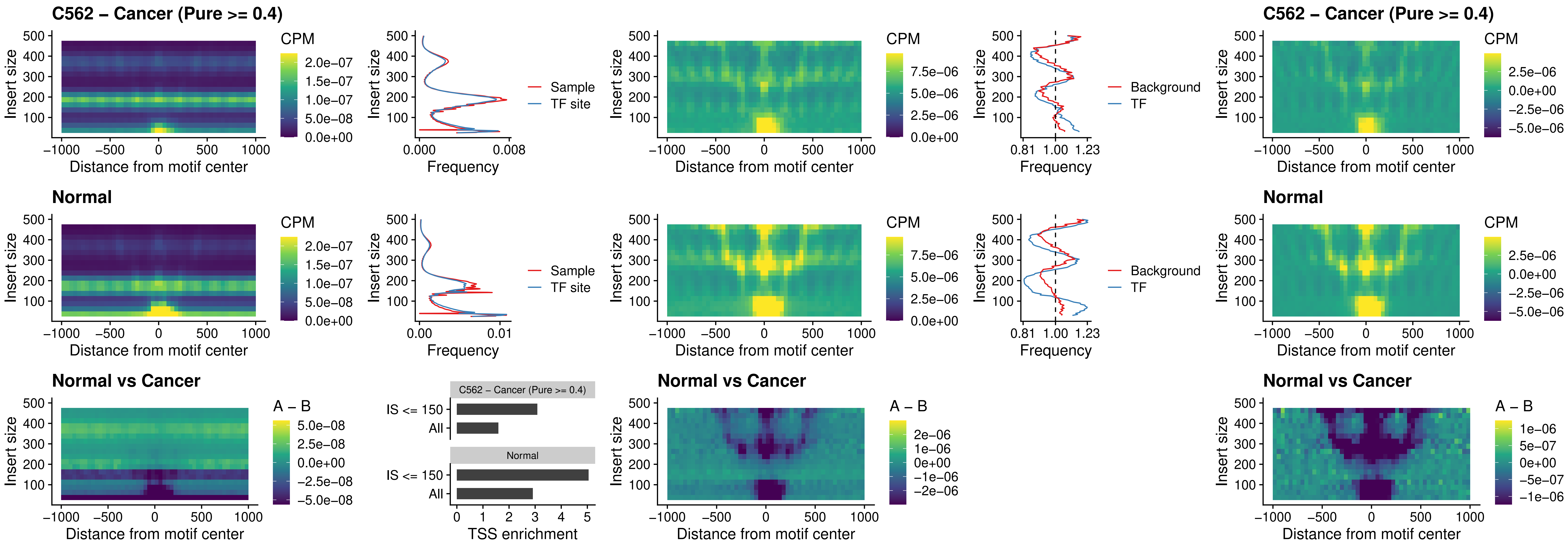

### ENCODE_CTCF_sites_COLON_intergenic_ctcf

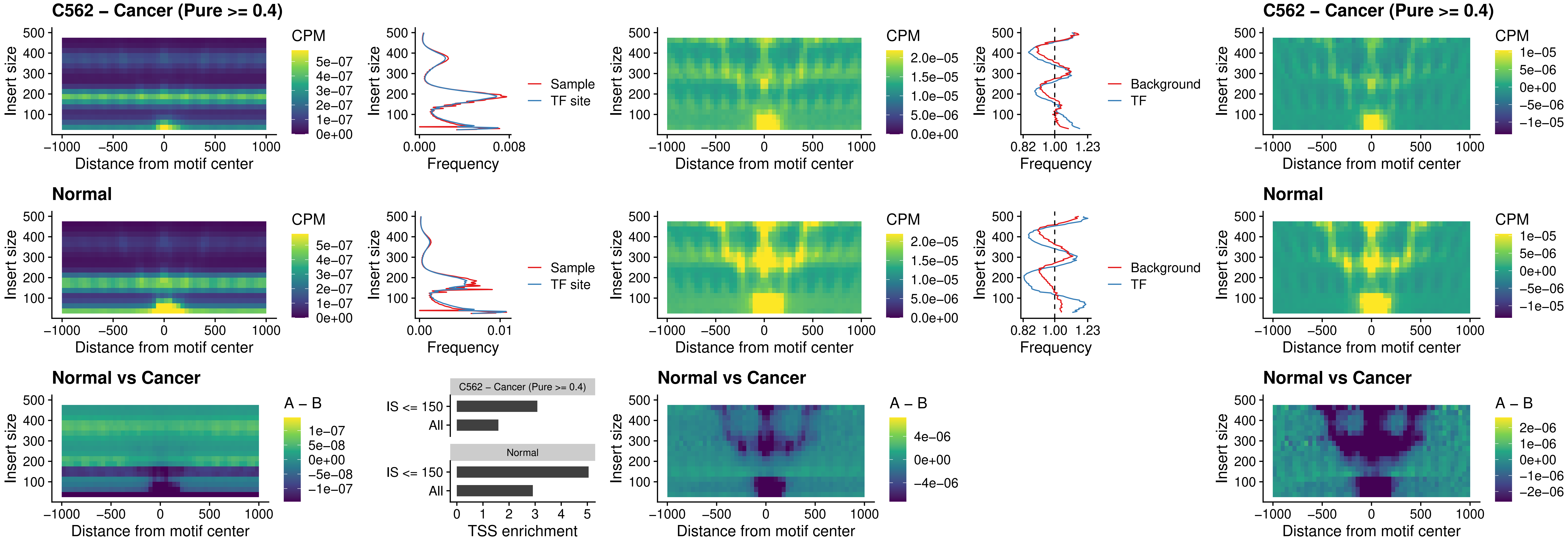

### ENCODE_CTCF_sites_COLON_nonpromotor

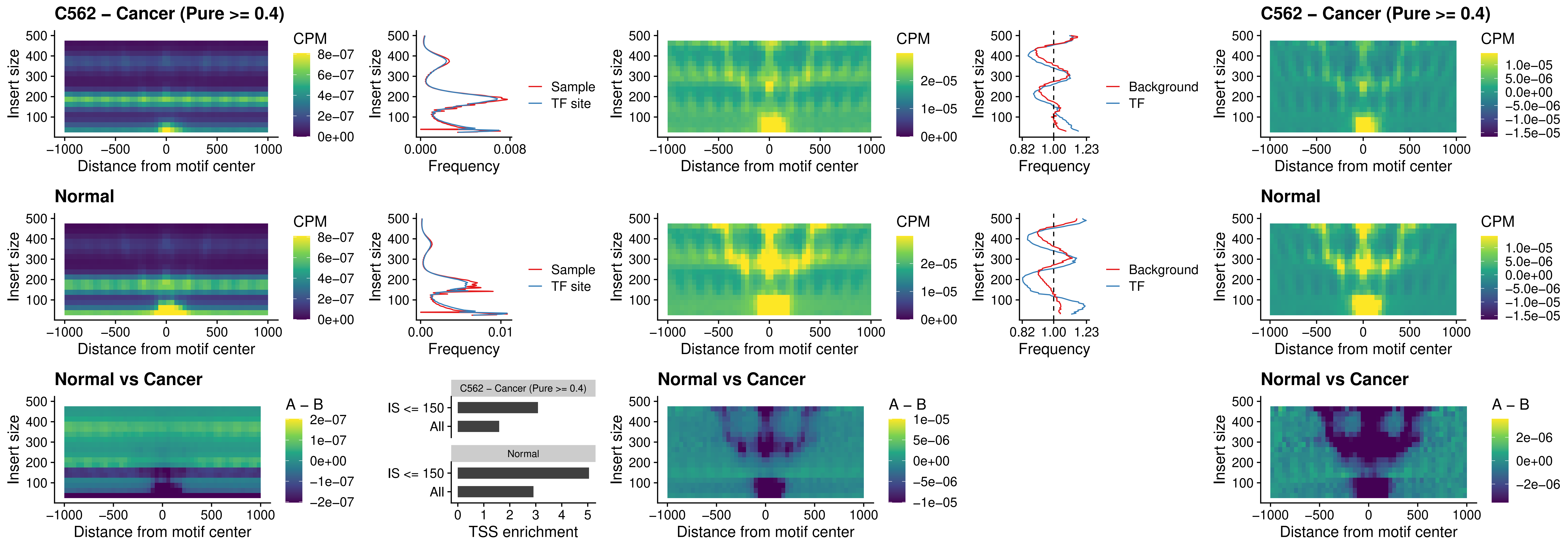

### CTCFL_uc_cTSS_oPEAK

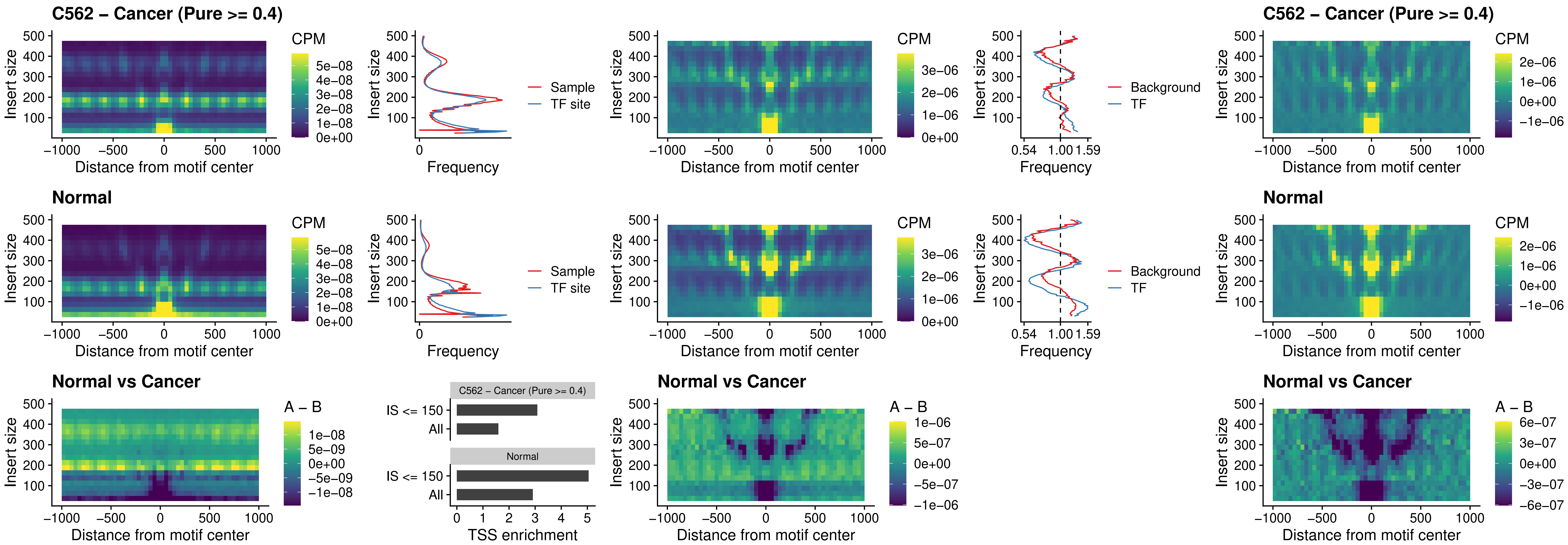

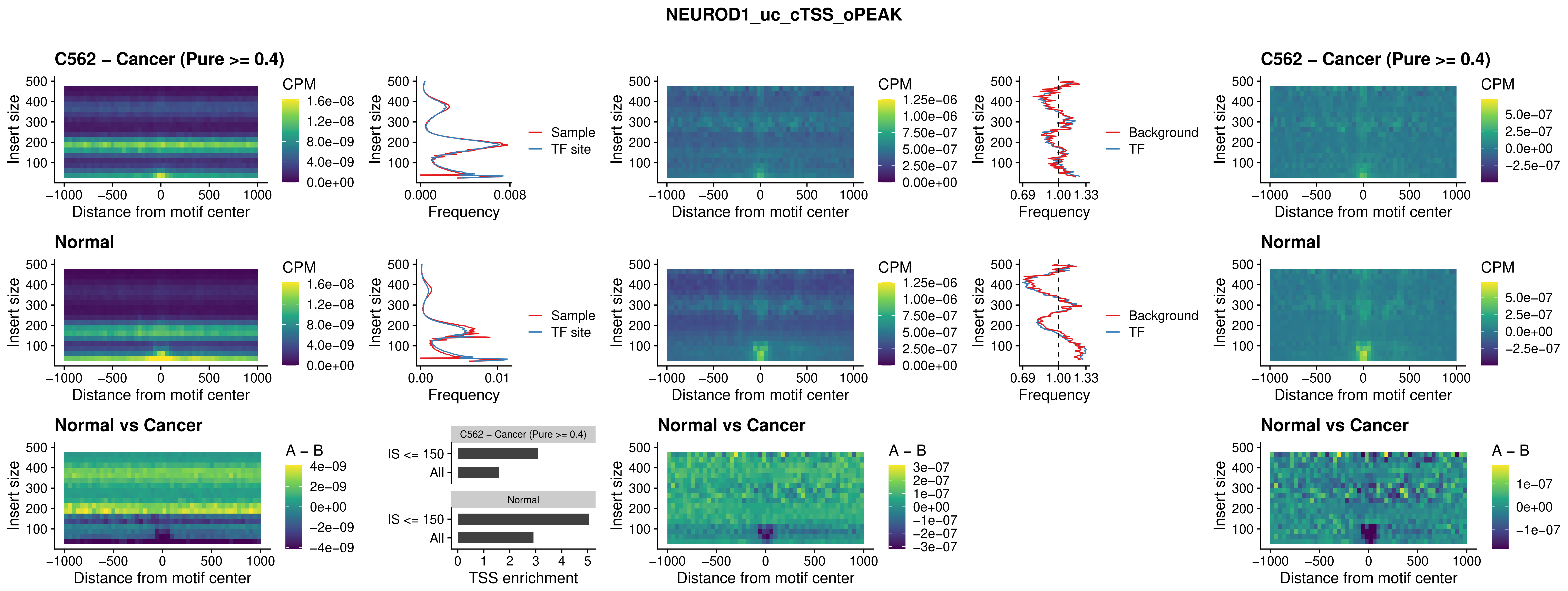
NEUROD1_uc_cTSS_oPEAK
C562 – Cancer (Pure >= 0.4)
Normal
Normal vs Cancer
Insert size
Distance from motif center
Frequency
CPM
A – B
Sample
TF site
Background
TF
C562 – Cancer (Pure >= 0.4)
Normal
IS <= 150
All
TSS enrichment

### ENCODE_YY1_sites_COLON_nonpromotor

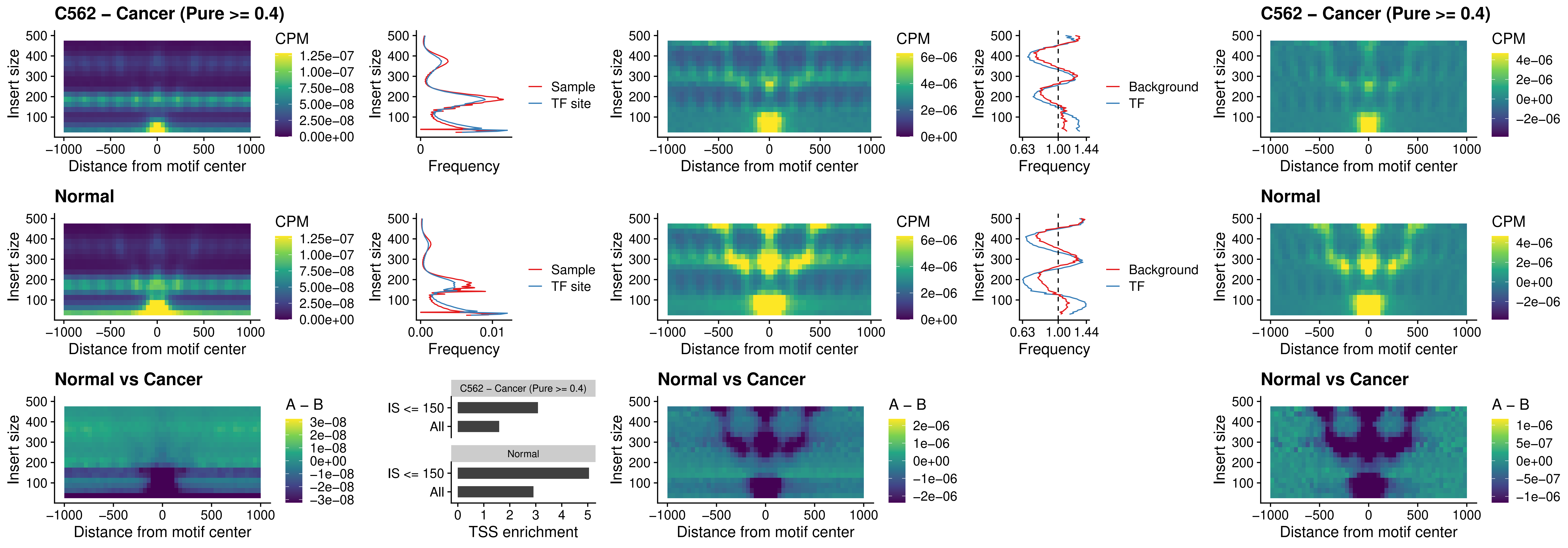

#### CTCF_uc_pTSS_oPEAK

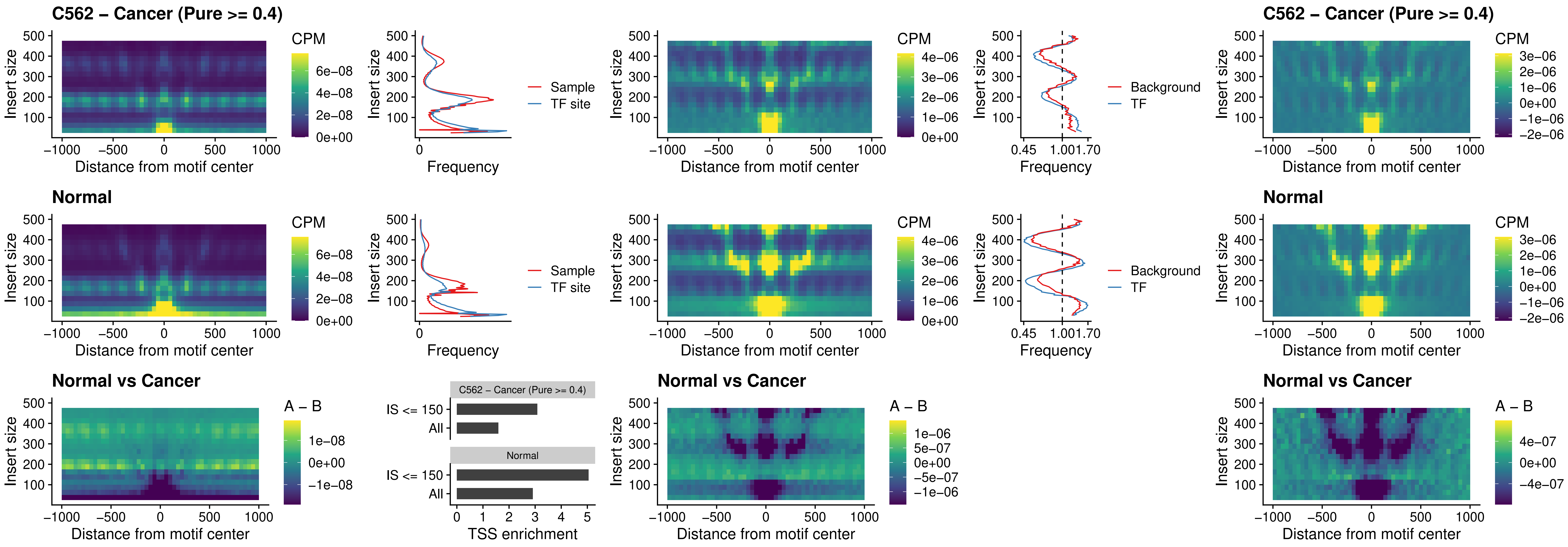

### ENCODE_CTCF_sites_COLON_promotor

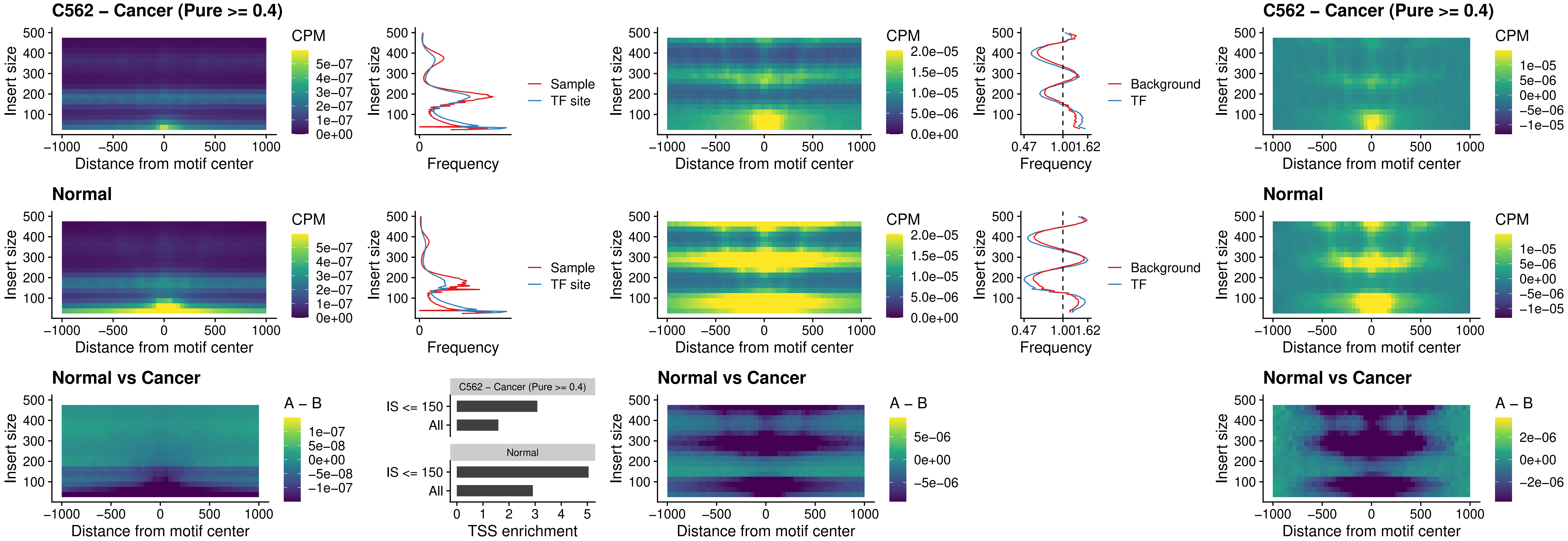

### ENCODE_CTCF_sites_COLON_promotor_ctcf

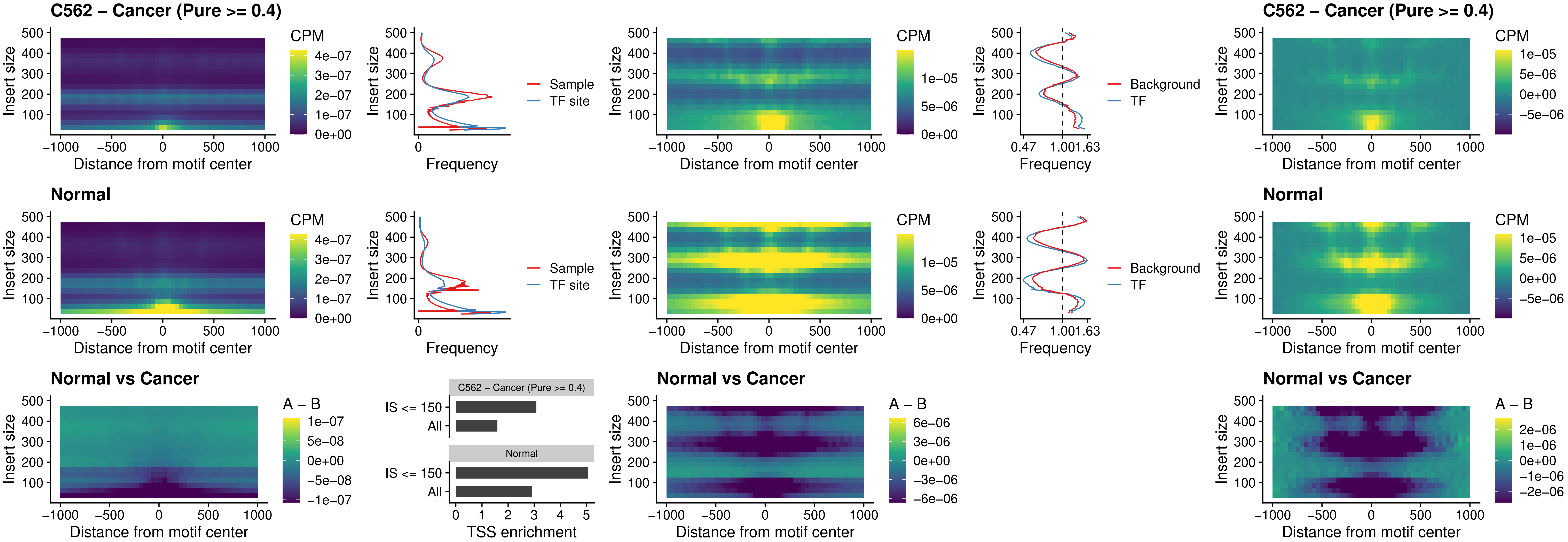

### SPIC_uc_pTSS_oPEAK

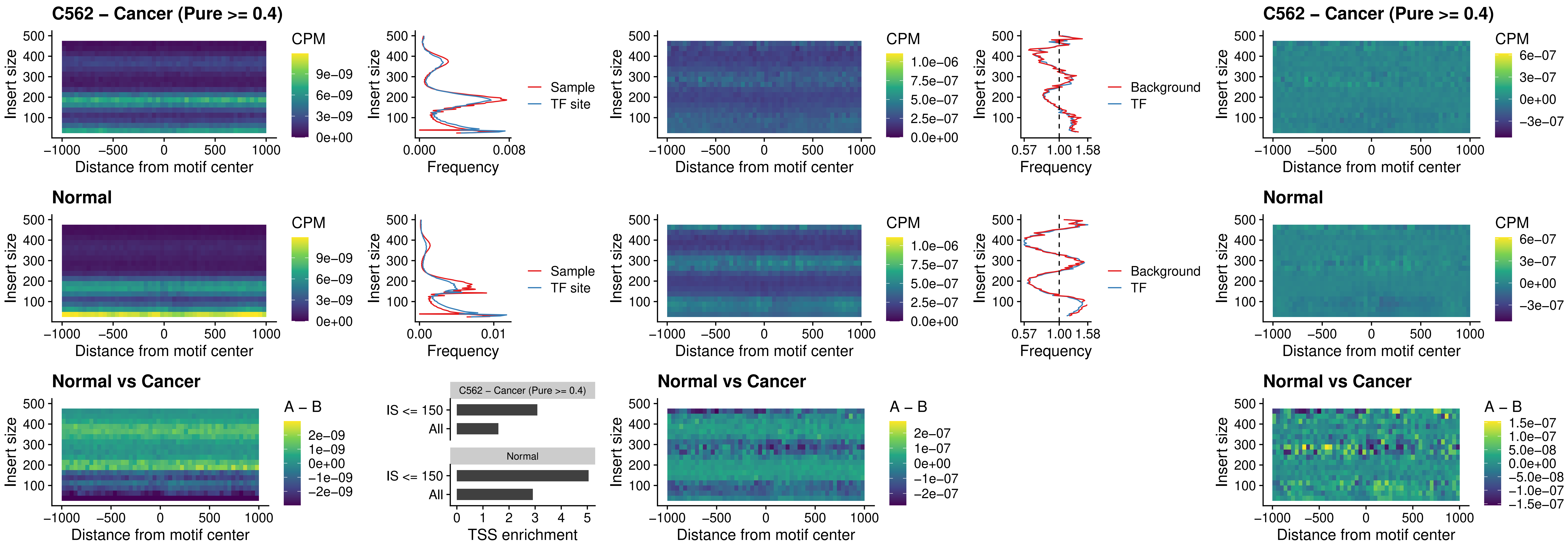

#### SPI1_uc_dTSS_nPEAK

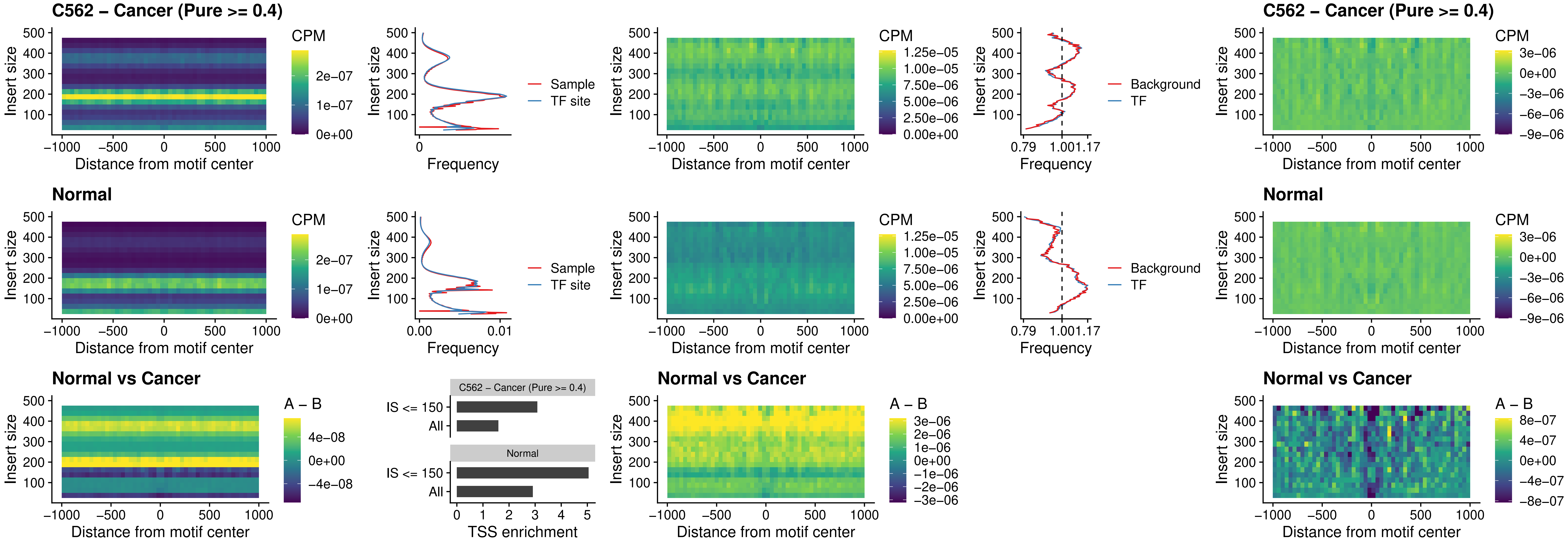

#### SPI1_uc_cTSS_nPEAK

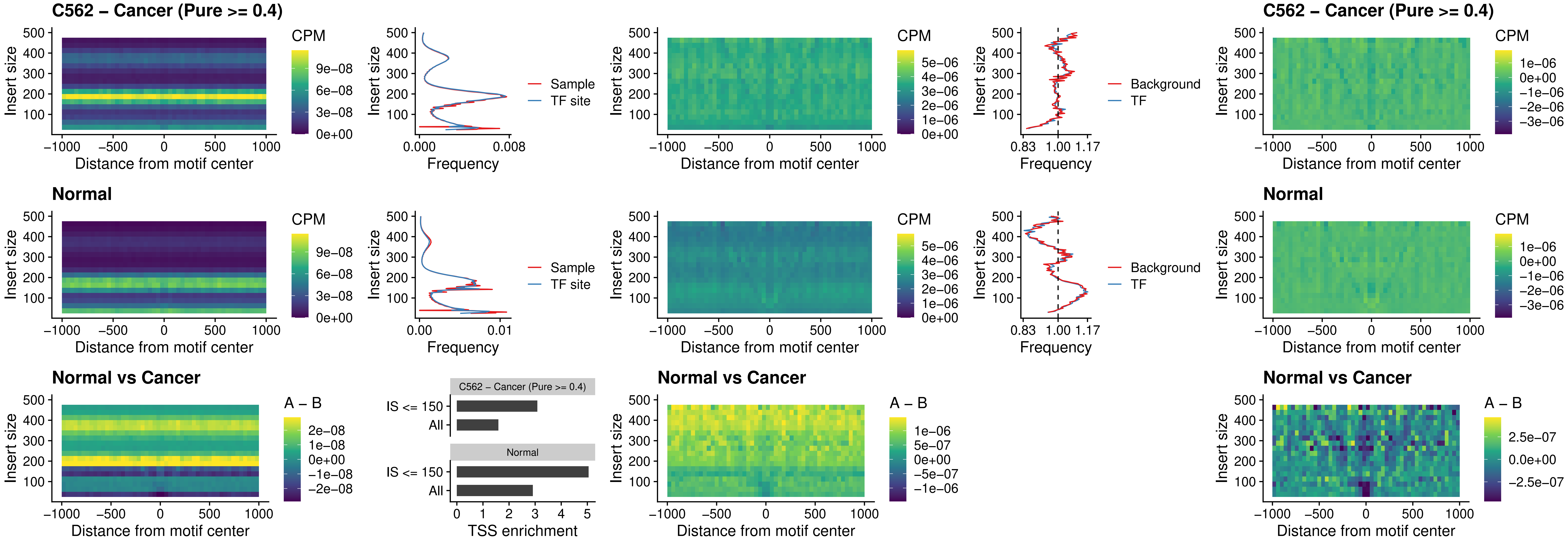

### BCL11A_uc_pTSS_nPEAK

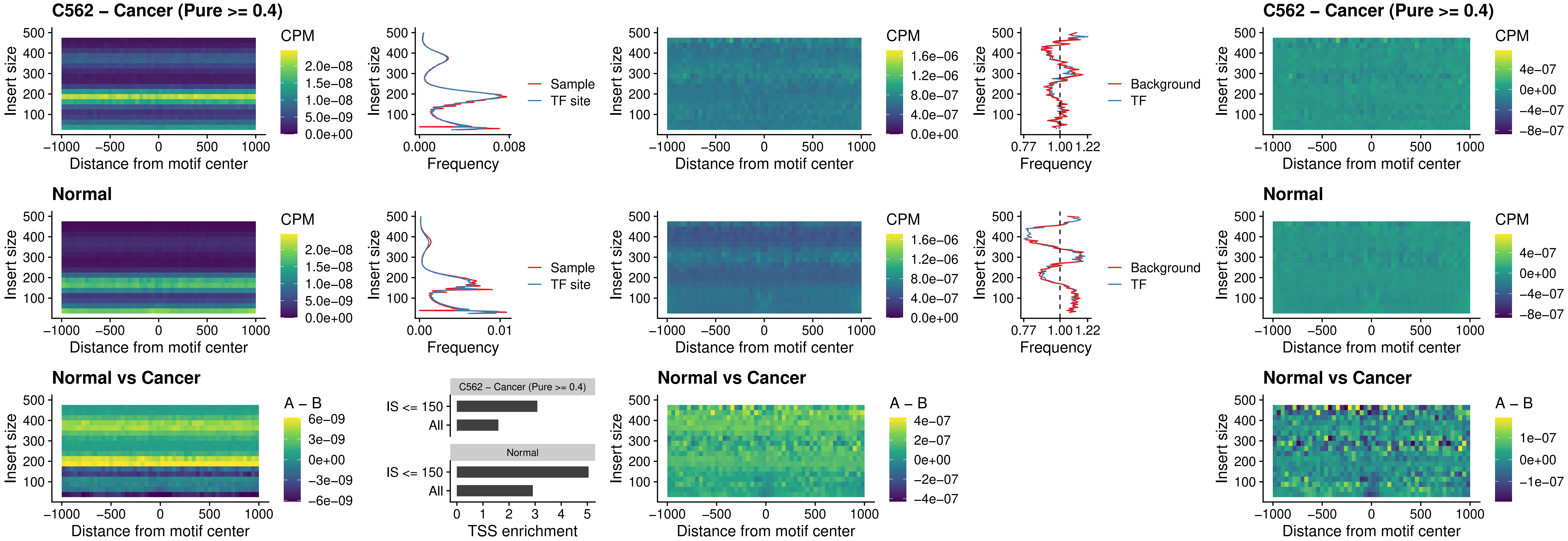

#### BCL11B_uc_pTSS_nPEAK

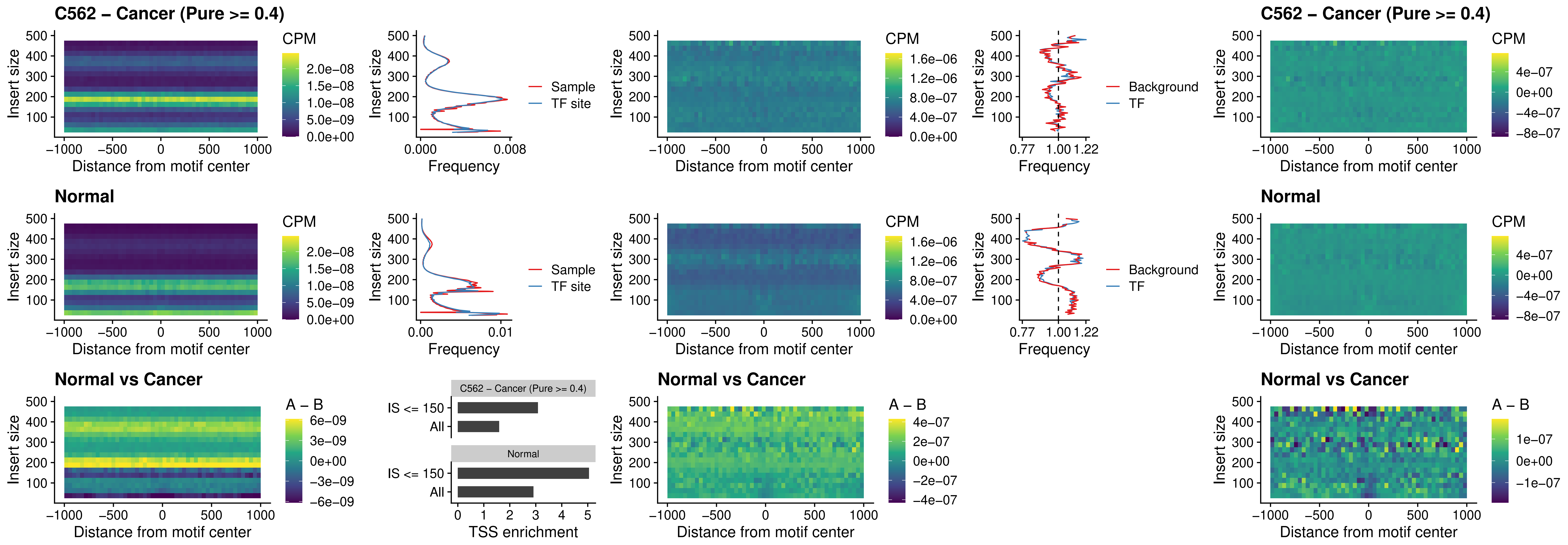

### IRF9_uc_dTSS_oPEAK

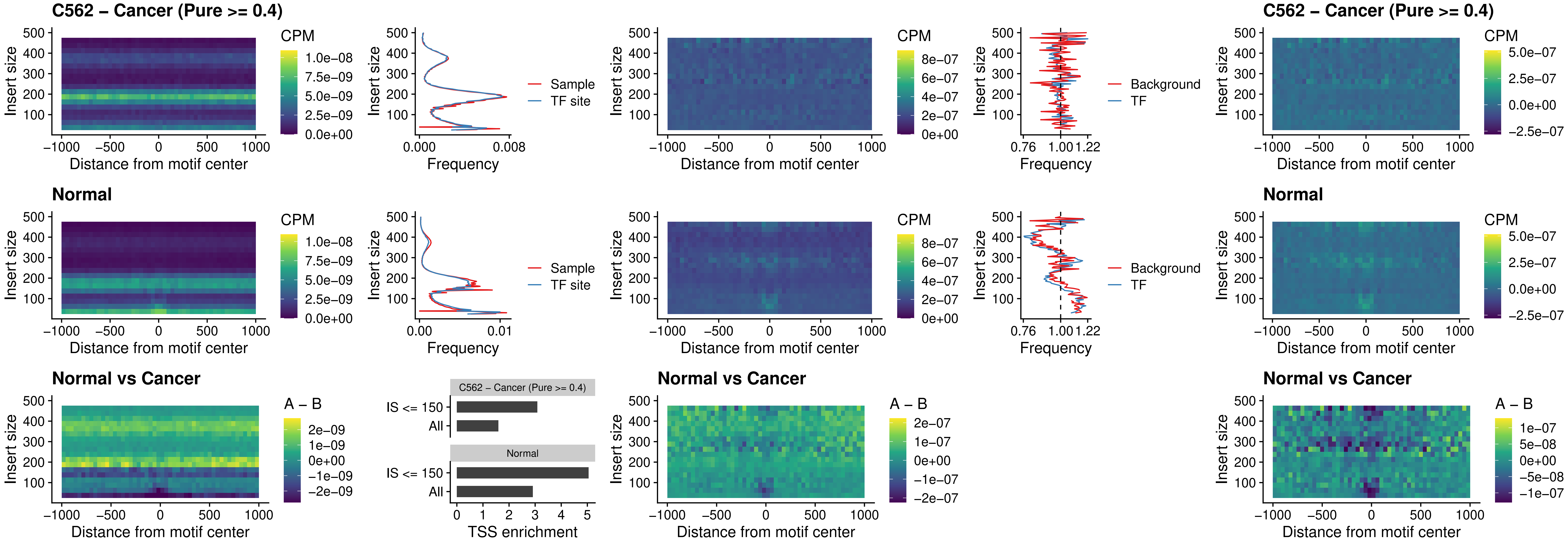

### IRF9_uc_cTSS_oPEAK

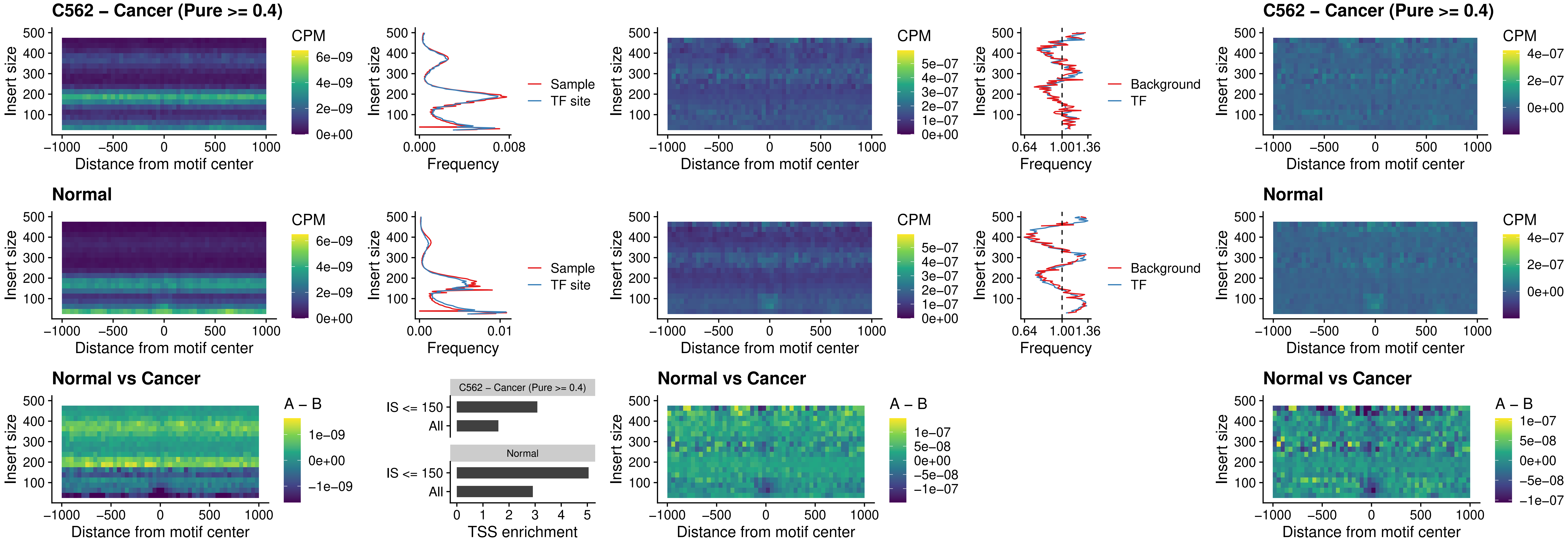

### BCL11A_uc_cTSS_oPEAK

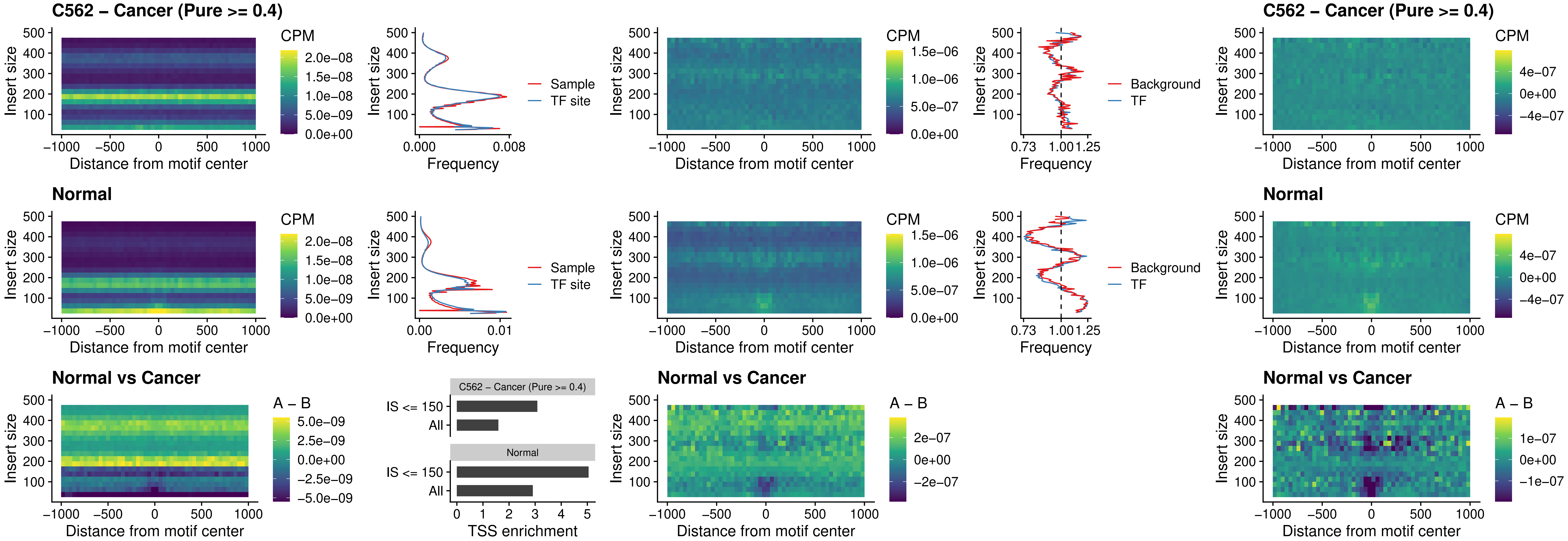

### BCL11B_uc_cTSS_oPEAK

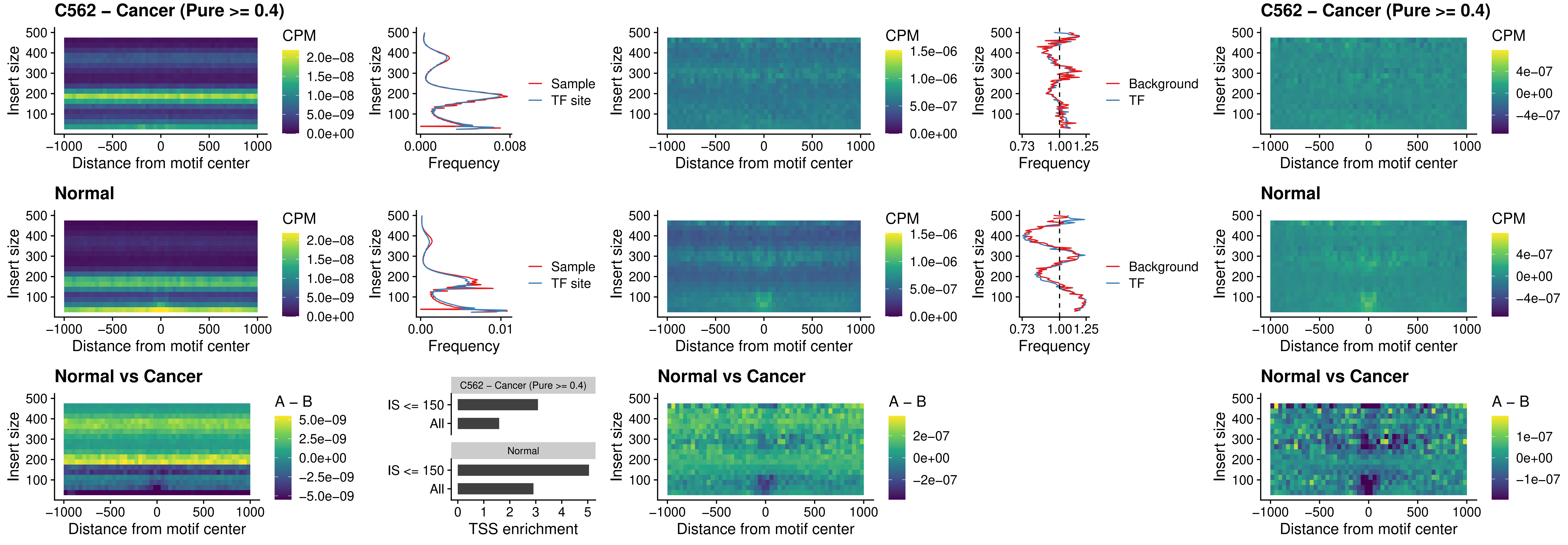

### BCL11A_uc_dTSS_oPEAK

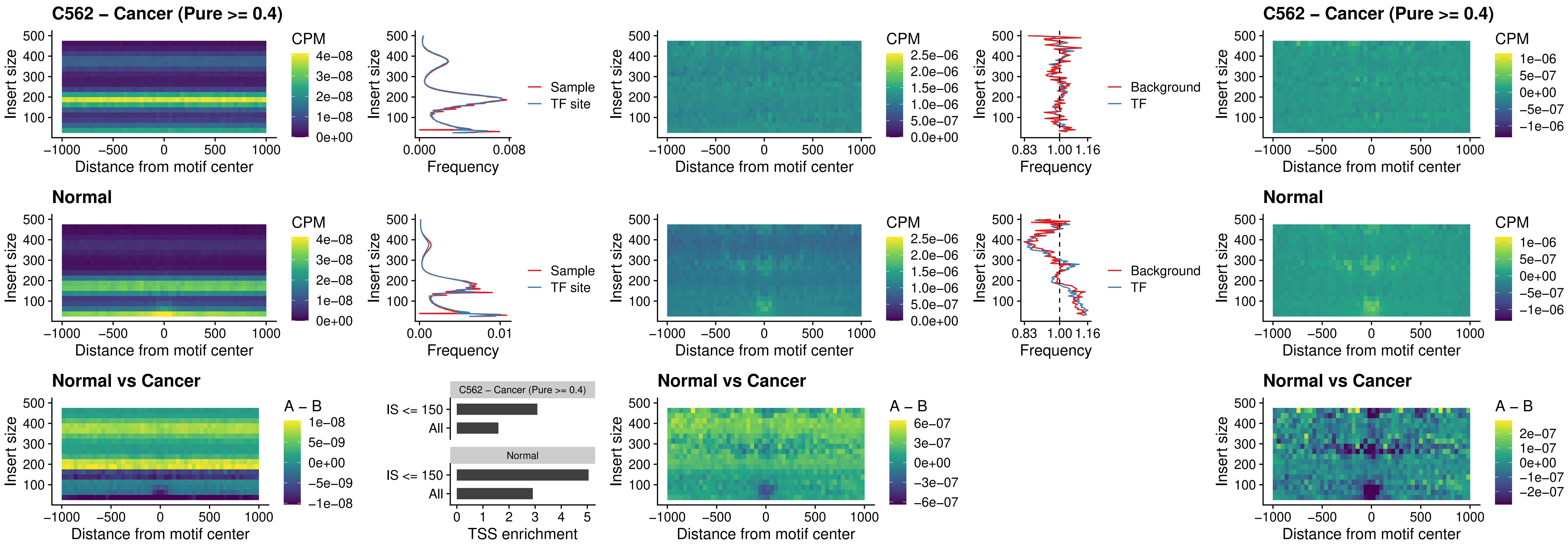

### BCL11B_uc_dTSS_oPEAK

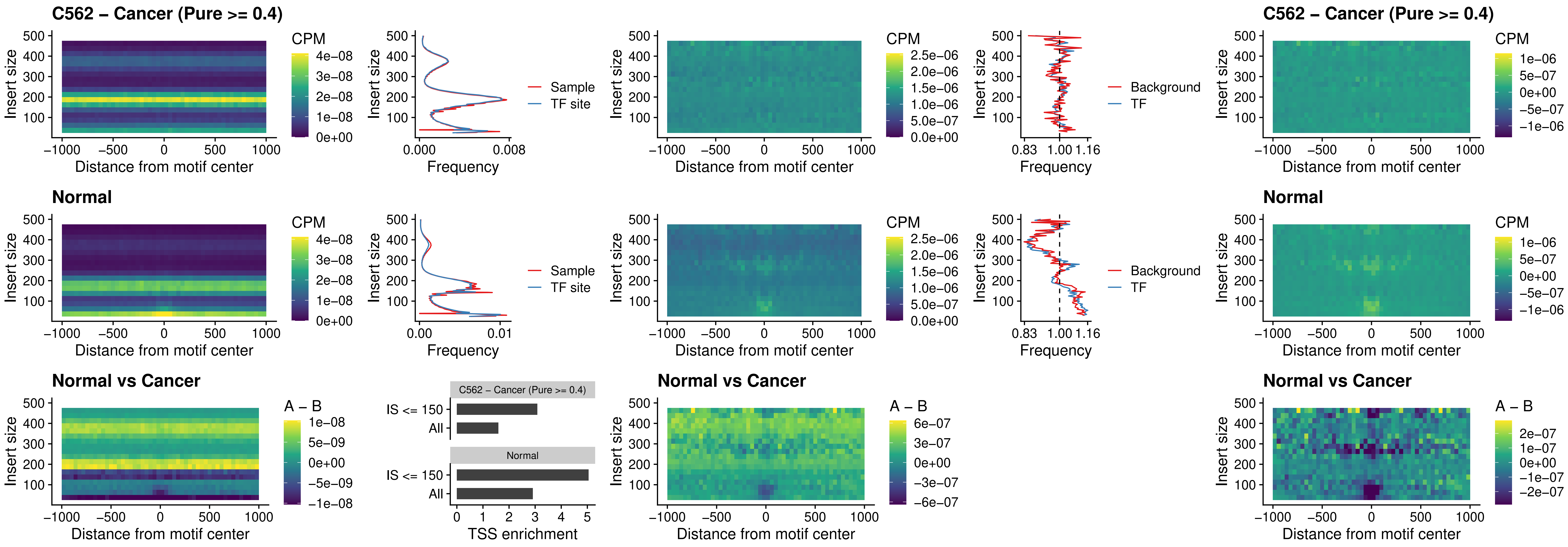

#### SPIB_uc_cTSS_nPEAK

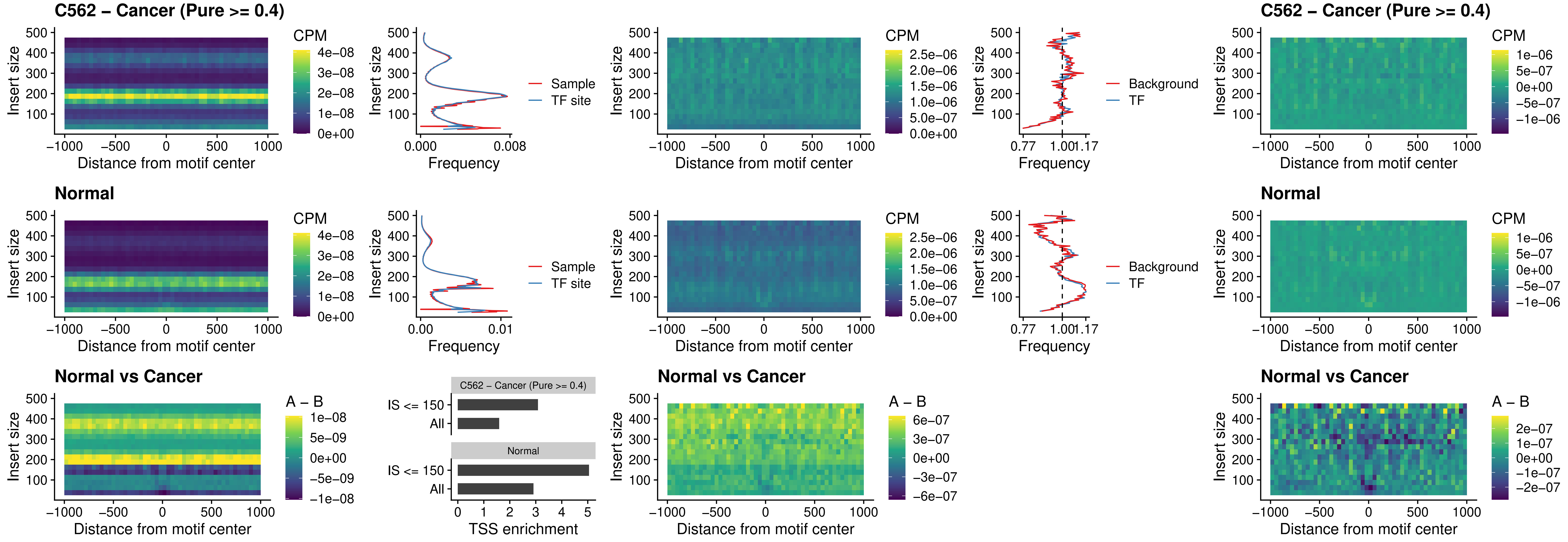

### SPIC_uc_cTSS_nPEAK

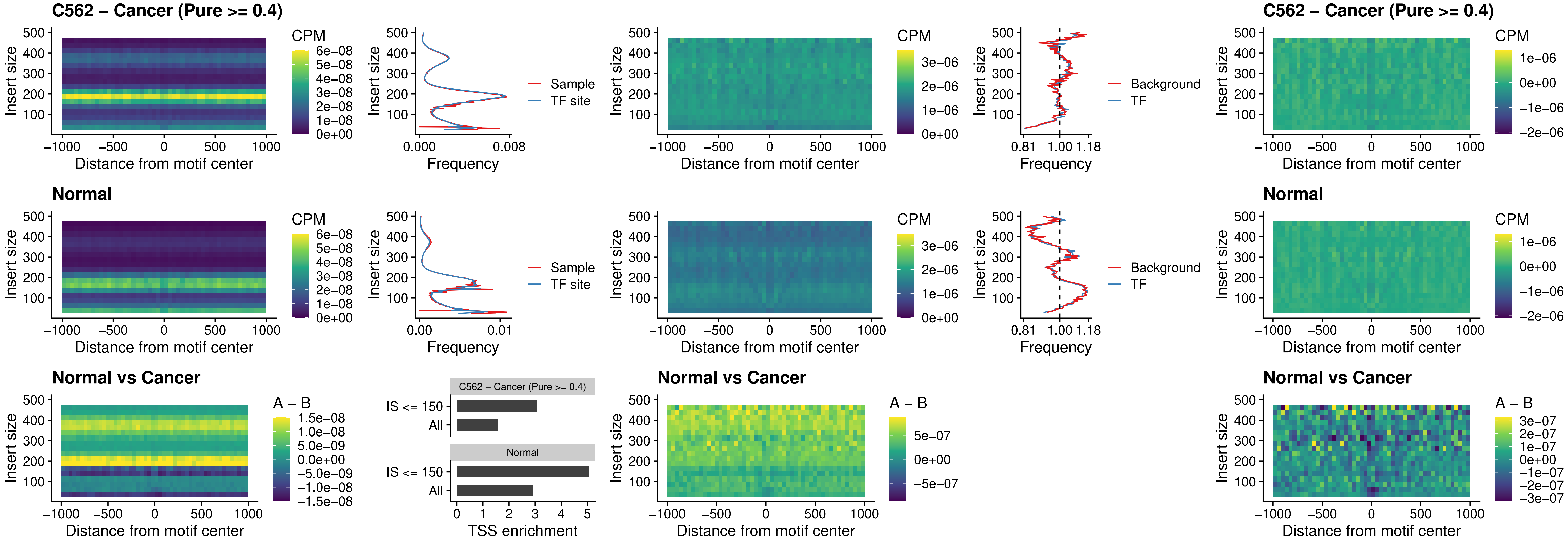

#### SPI1_uc_pTSS_oPEAK

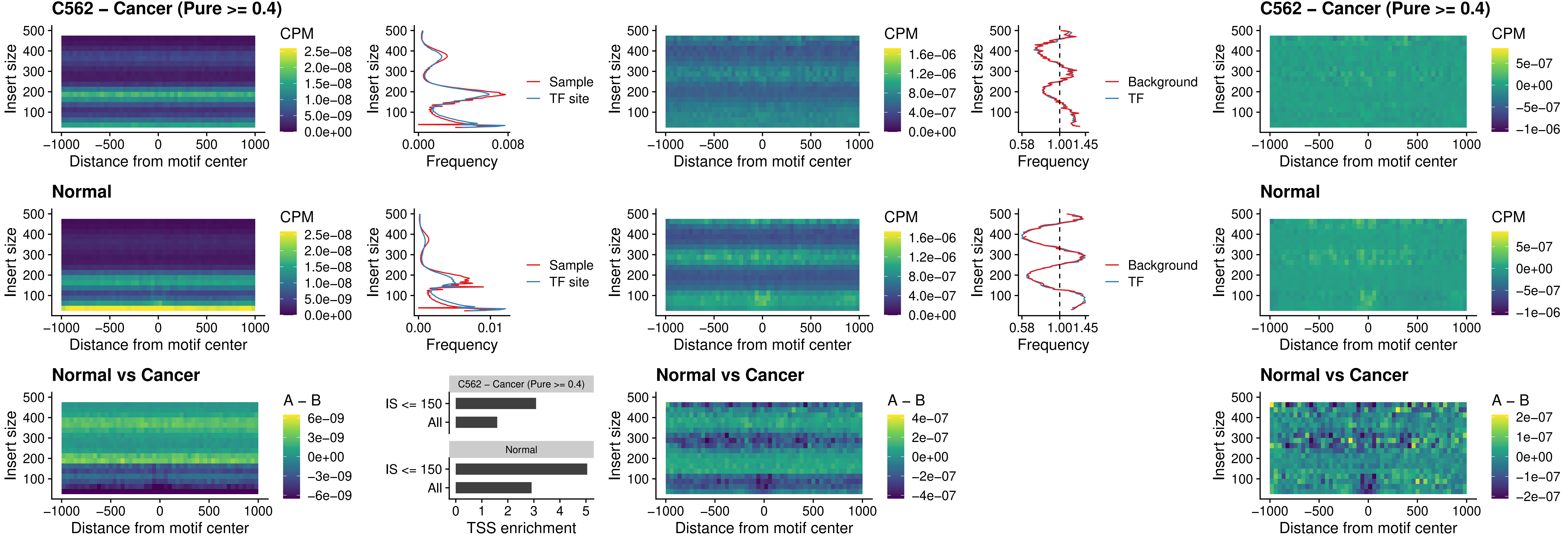

### BCL11A_uc_pTSS_oPEAK

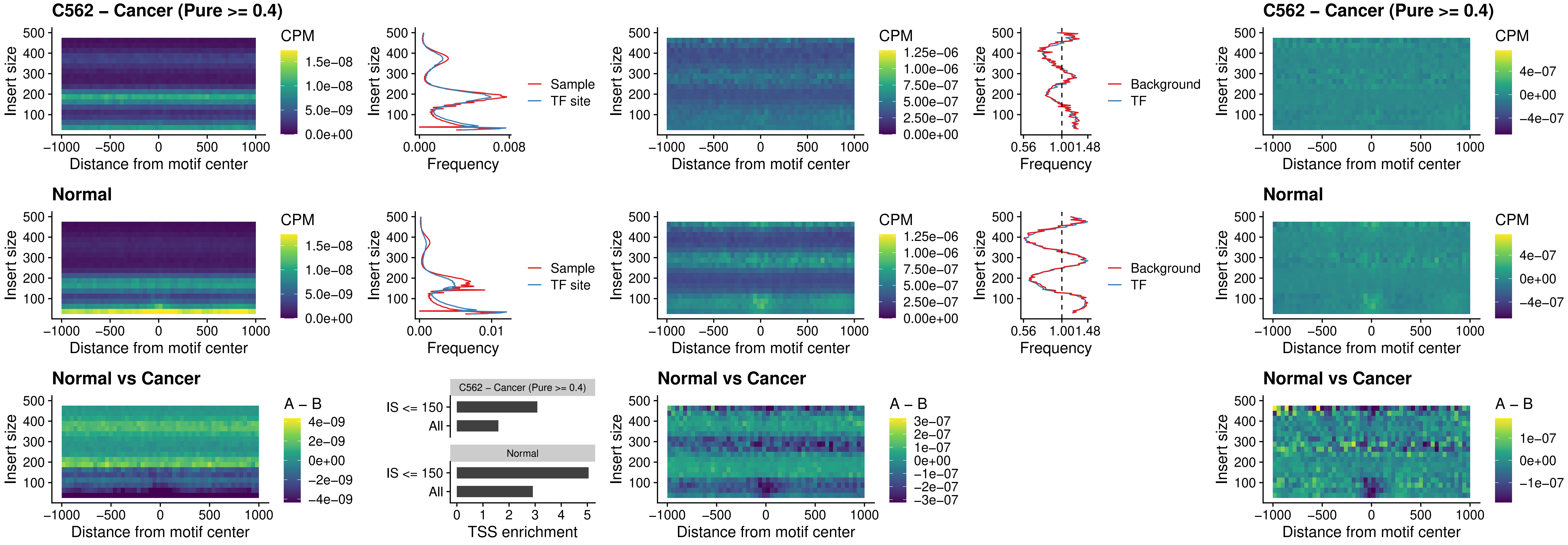

### BCL11B_uc_pTSS_oPEAK

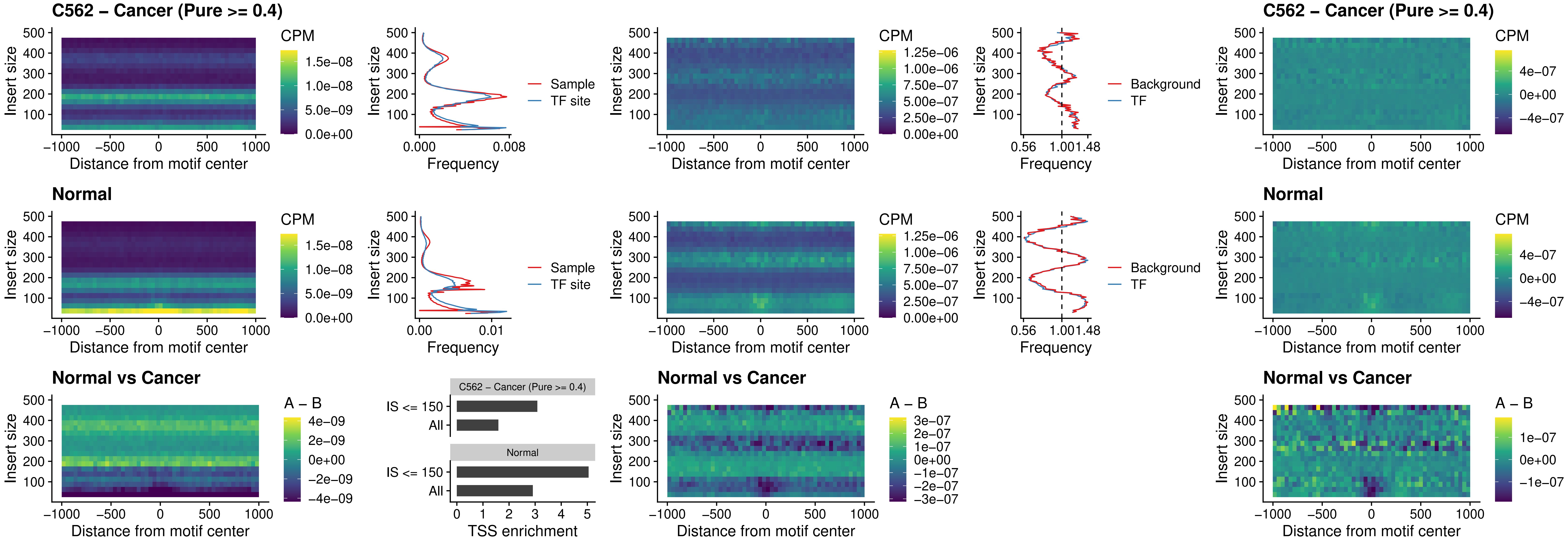

### SPI1_uc_cTSS_oPEAK

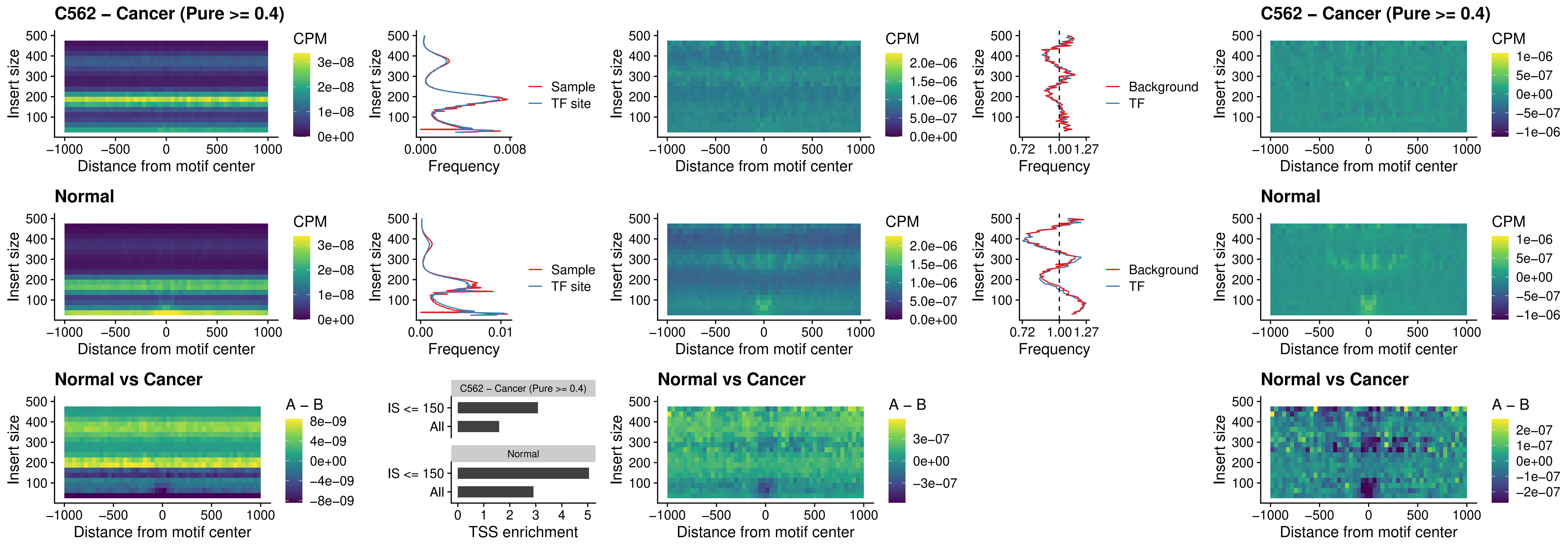

#### SPIB_uc_cTSS_oPEAK

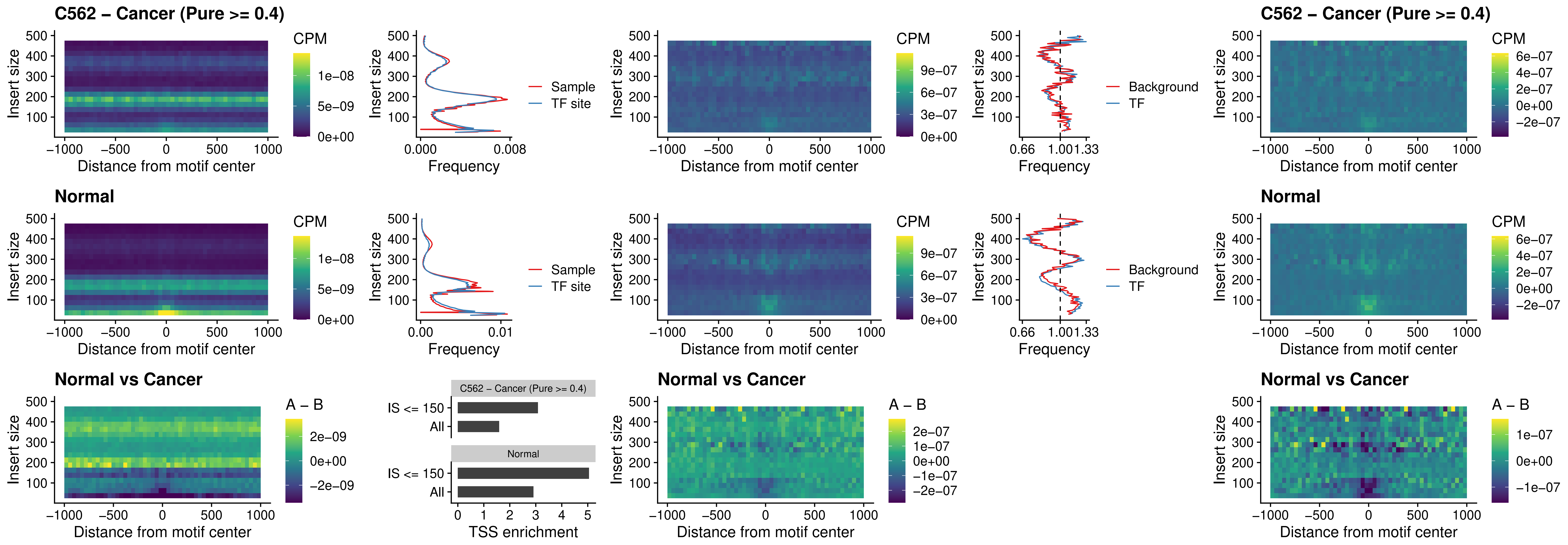

#### SPIB_uc_pTSS_oPEAK

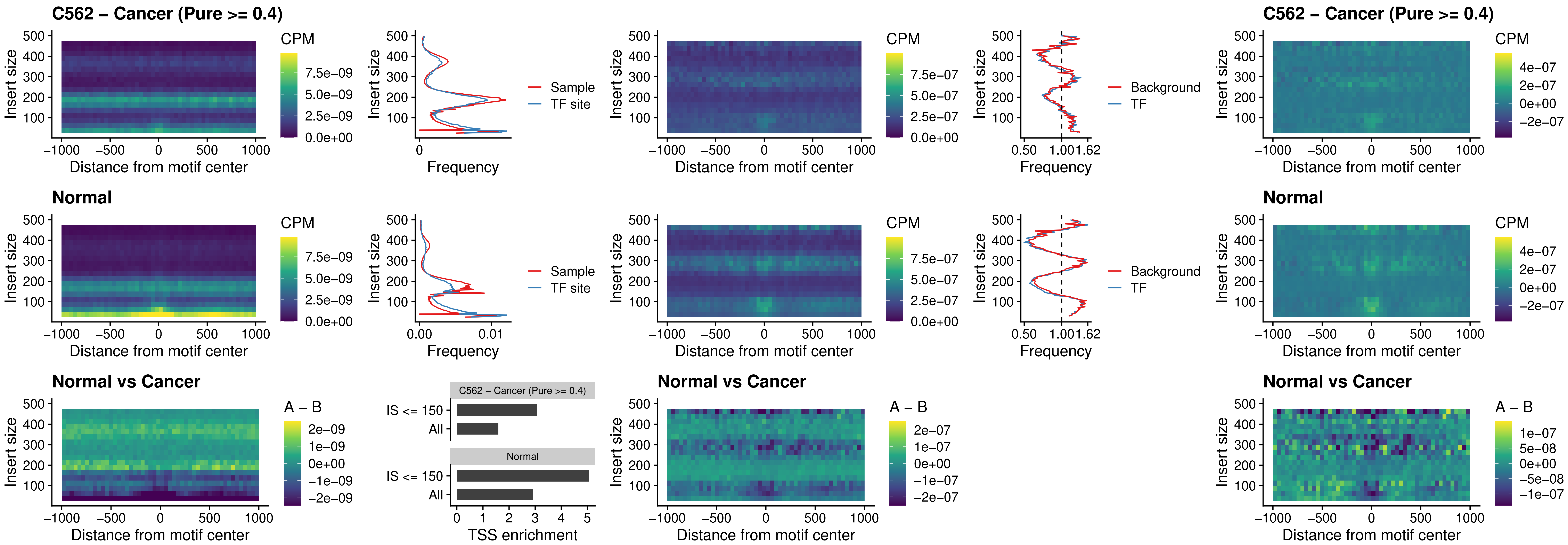

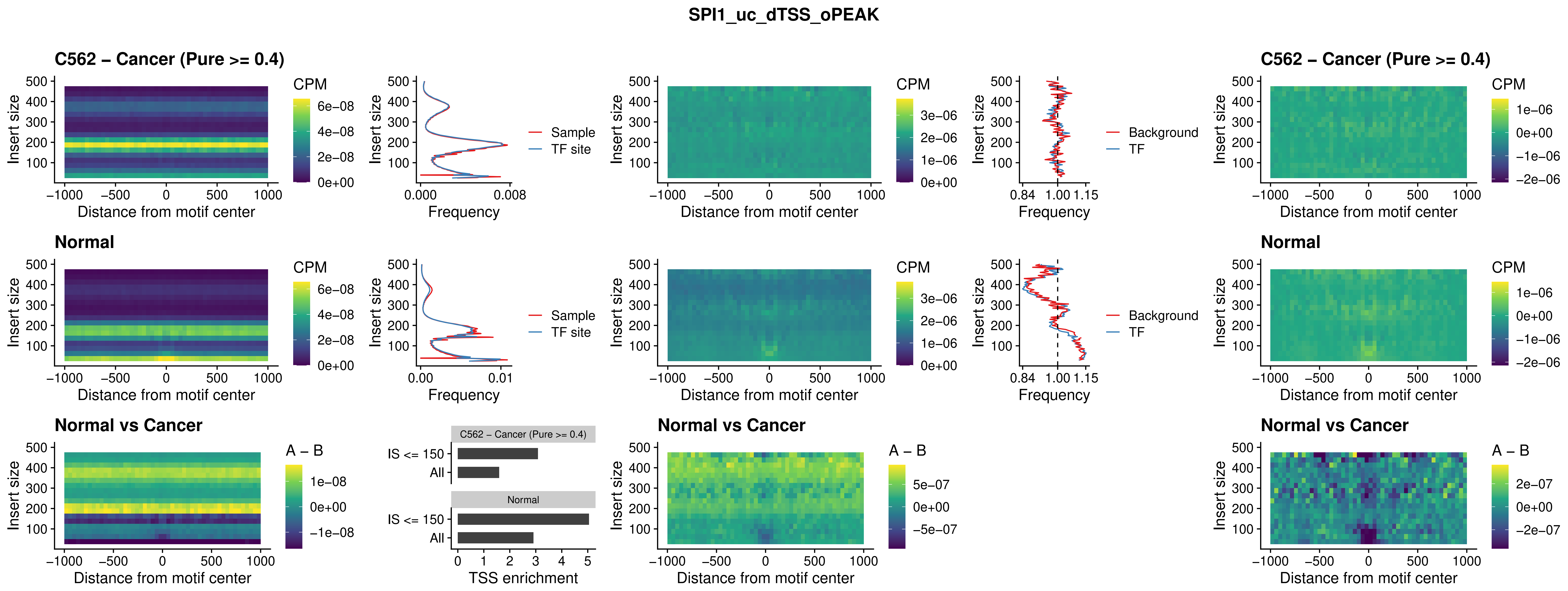
SPI1_uc_dTSS_oPEAK
C562 – Cancer (Pure >= 0.4)
Normal
Normal vs Cancer
Insert size
Distance from motif center
Frequency
CPM
A – B
Sample
TF site
Background
TF
C562 – Cancer (Pure >= 0.4)
Normal
IS <= 150
All
TSS enrichment

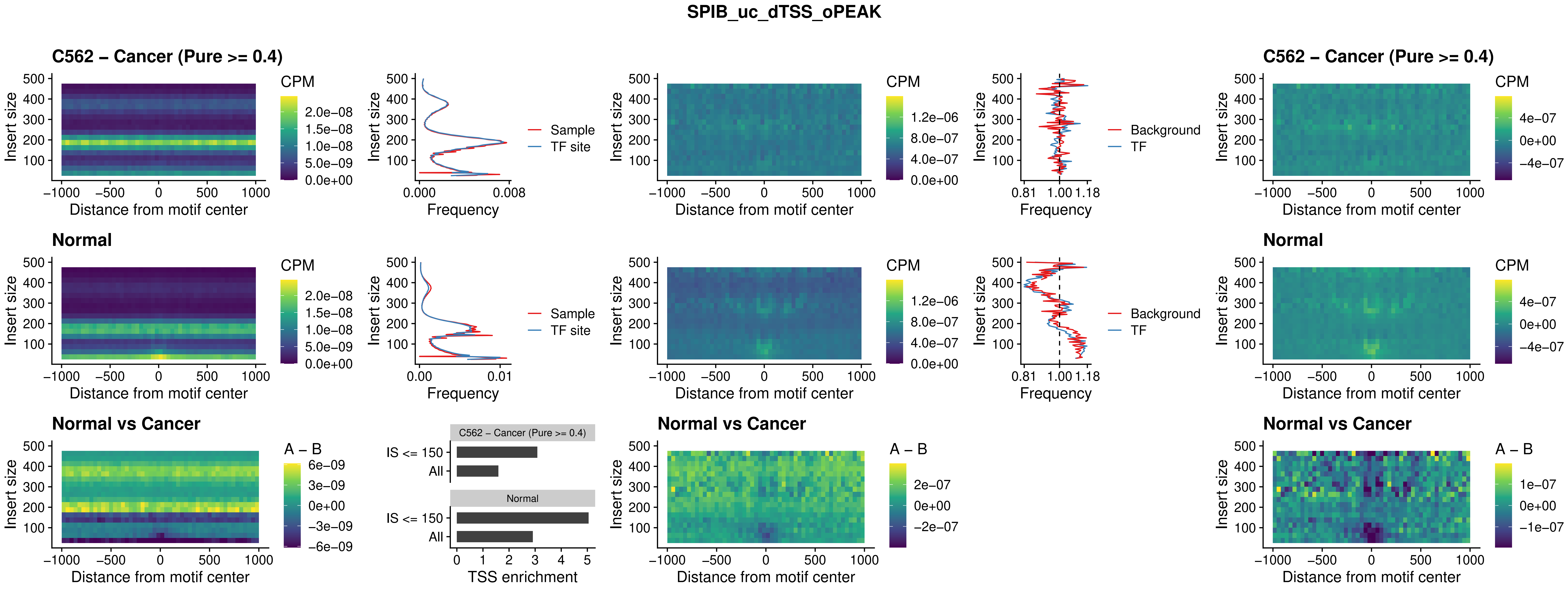
SPIB_uc_dTSS_oPEAK
C562 – Cancer (Pure >= 0.4)
Normal
Normal vs Cancer
Insert size
Distance from motif center
Frequency
CPM
A – B
Sample
TF site
Background
TF
C562 – Cancer (Pure >= 0.4)
Normal
IS <= 150
All
TSS enrichment

### IRF3_uc_pTSS_oPEAK

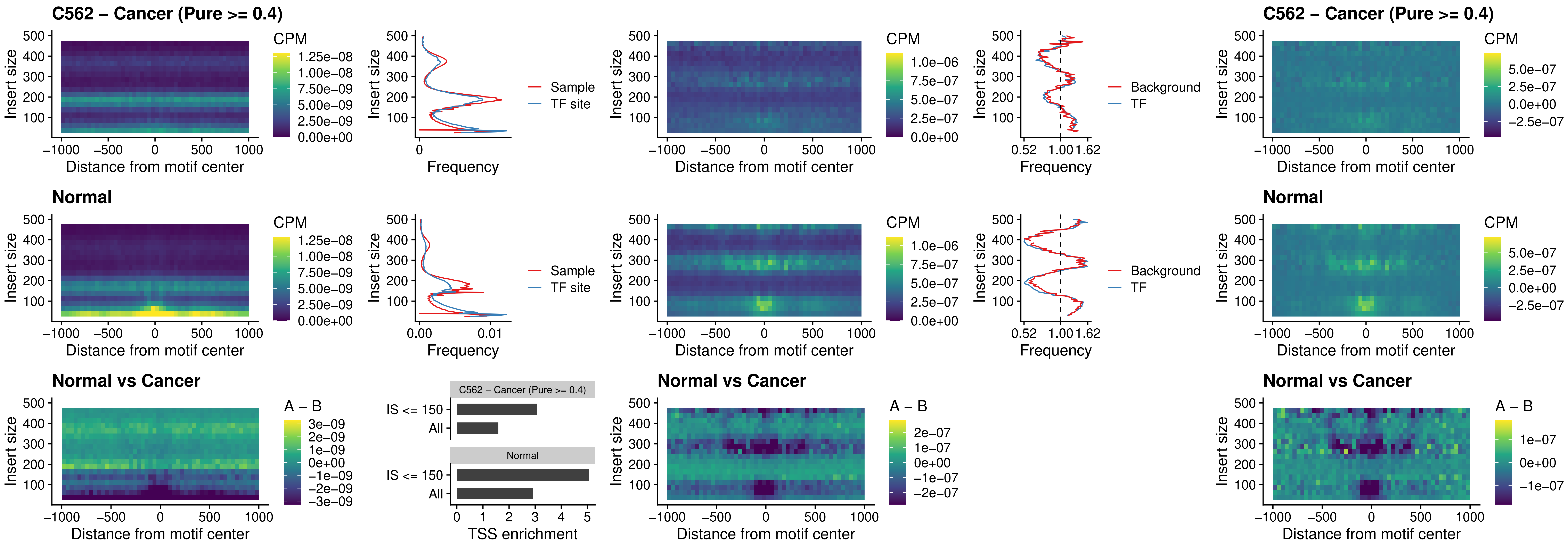

### IRF2_uc_dTSS_oPEAK

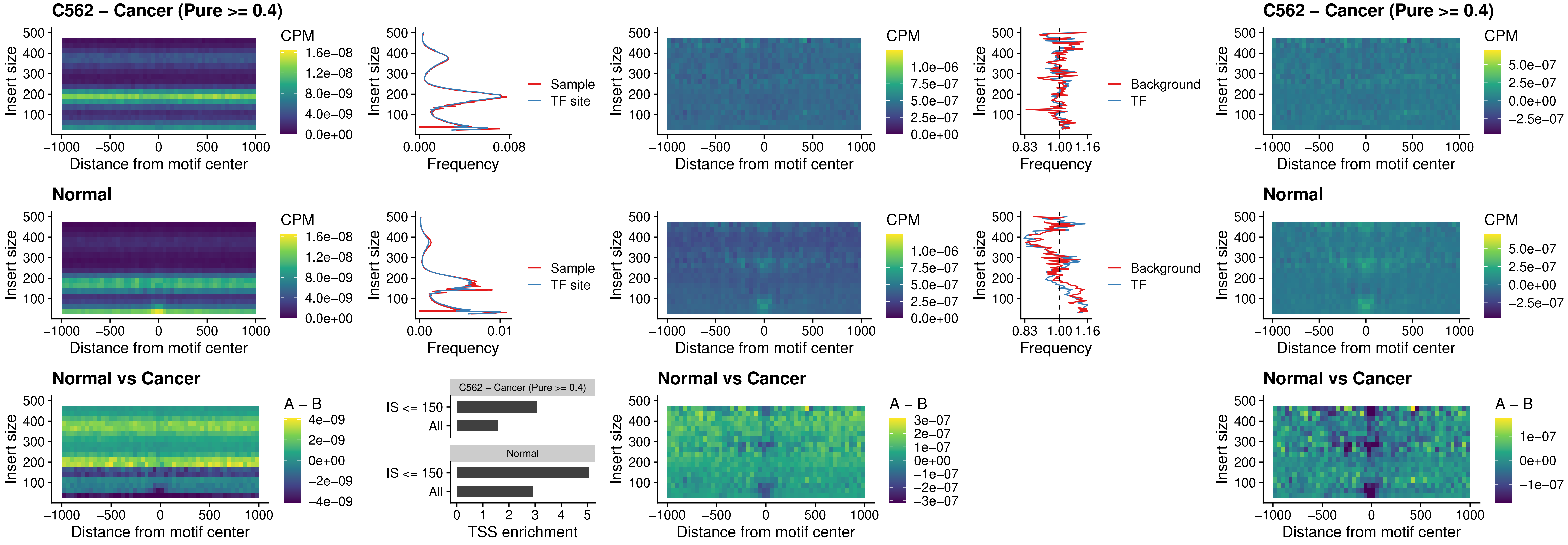

### IRF2_uc_pTSS_oPEAK

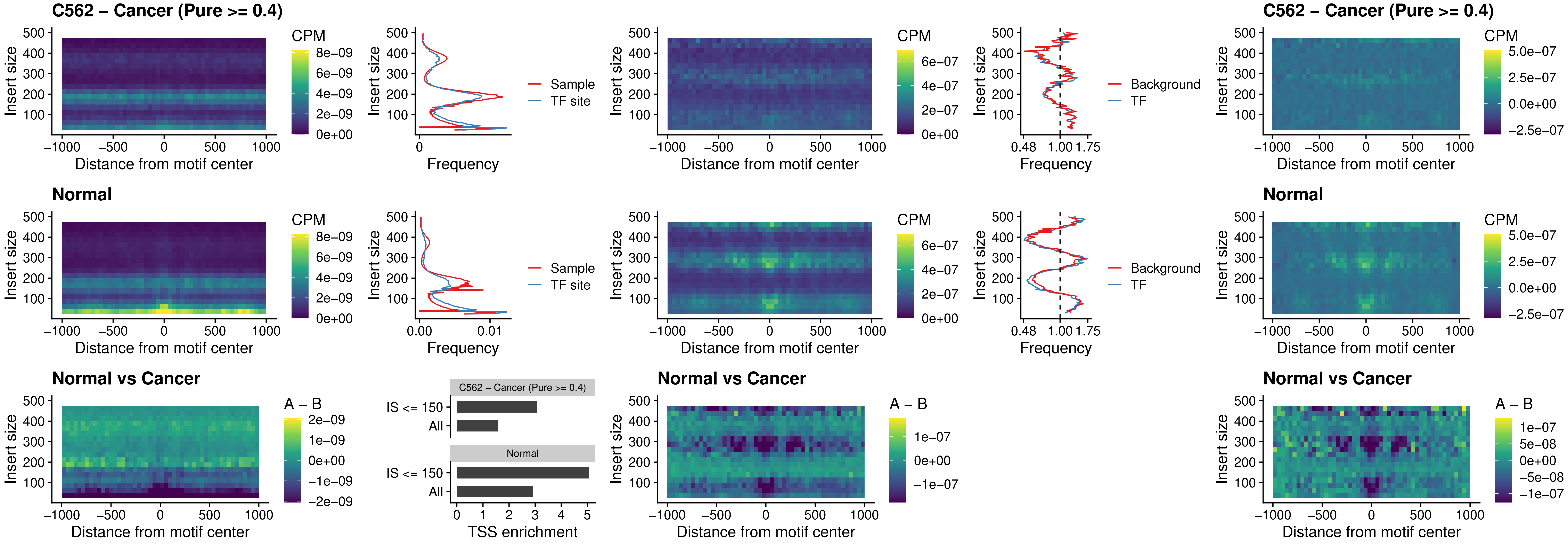

#### STAT2_uc_pTSS_oPEAK

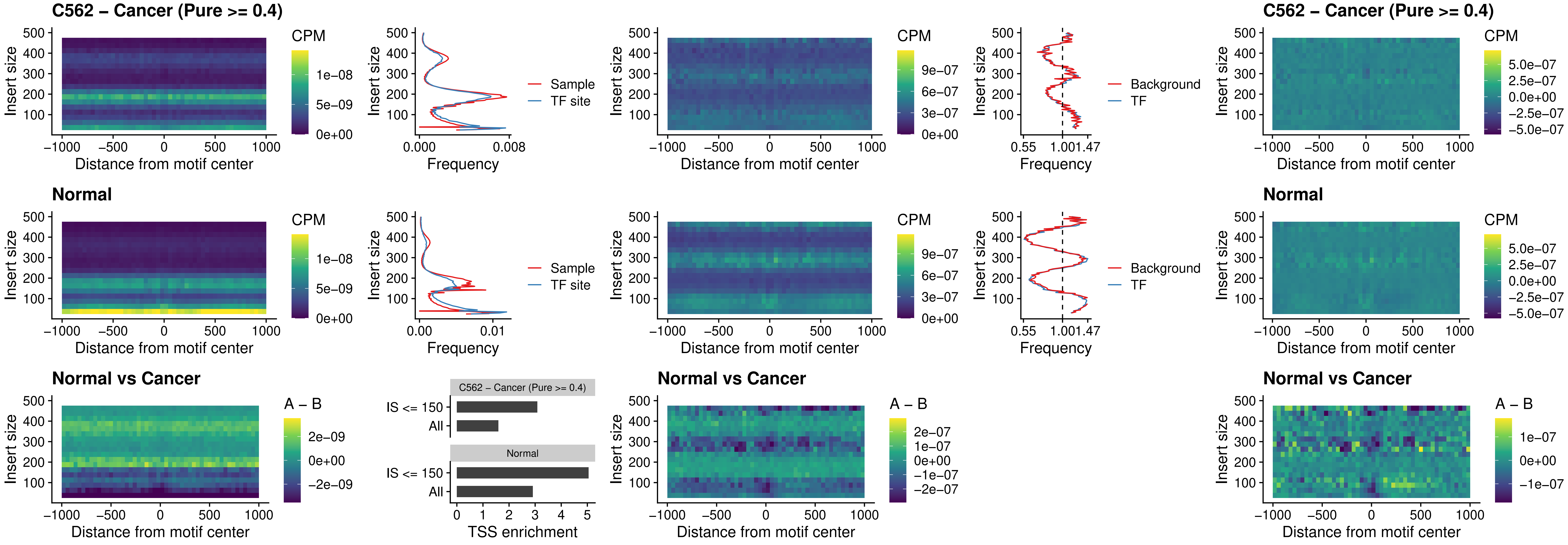

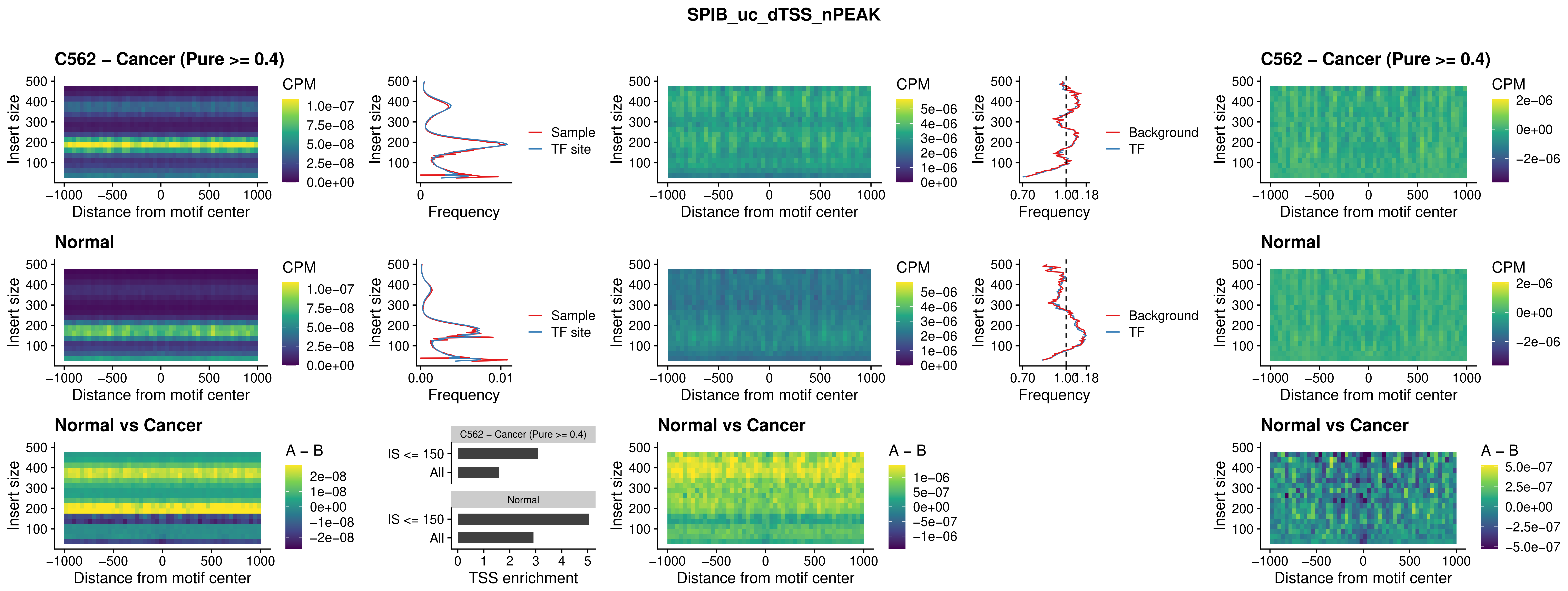
SPIB_uc_dTSS_nPEAK
C562 – Cancer (Pure >= 0.4)
Normal
Normal vs Cancer
Insert size
Distance from motif center
Frequency
CPM
A – B
Sample
TF site
Background
TF
C562 – Cancer (Pure >= 0.4)
Normal
IS <= 150
All
TSS enrichment

#### SPIC_uc_cTSS_oPEAK

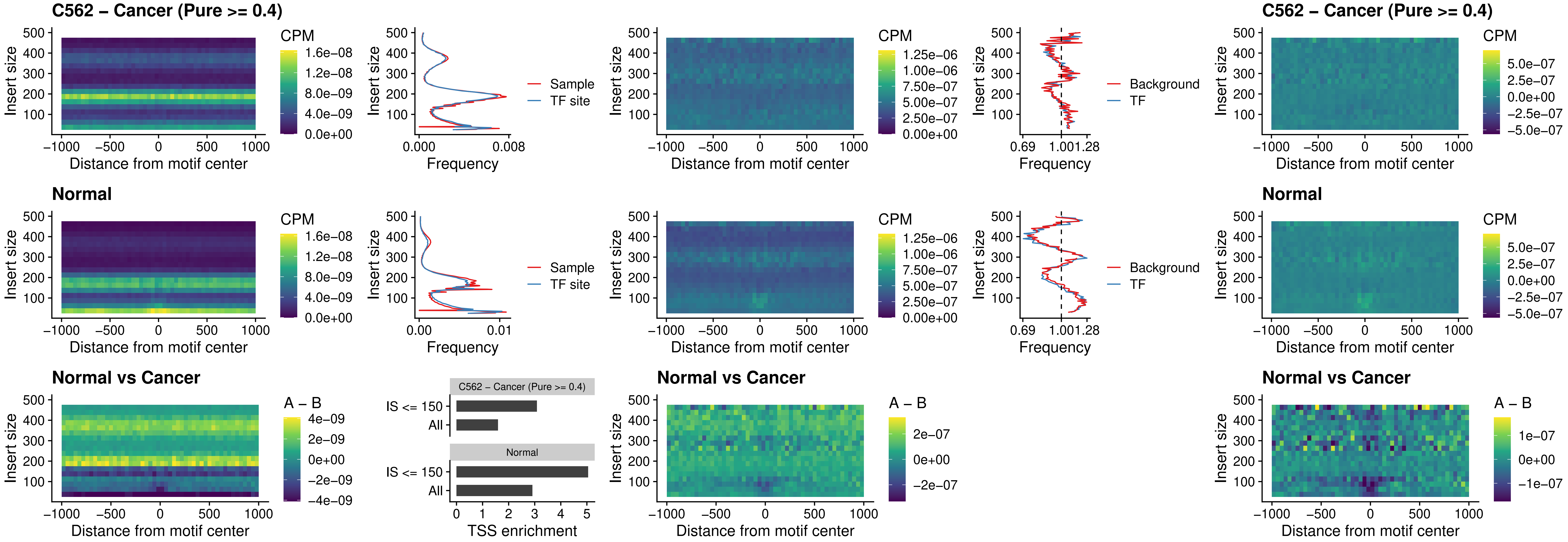

### PRDM1_uc_cTSS_oPEAK

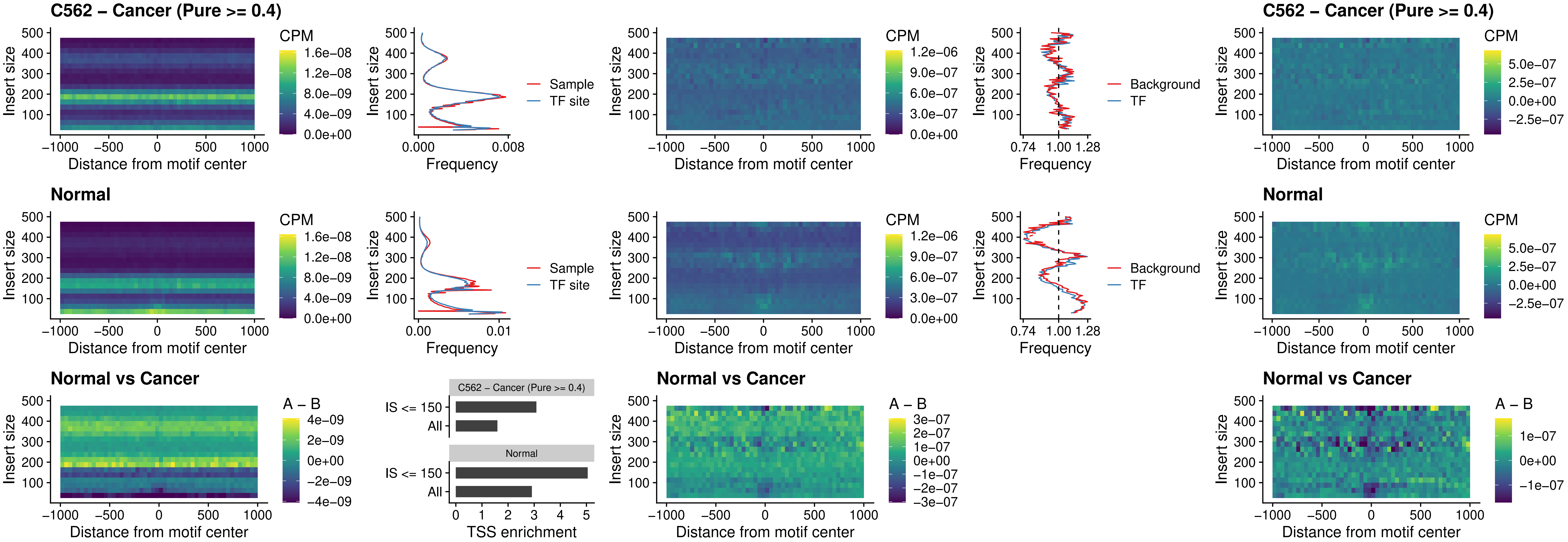

#### IRF3_uc_cTSS_oPEAK

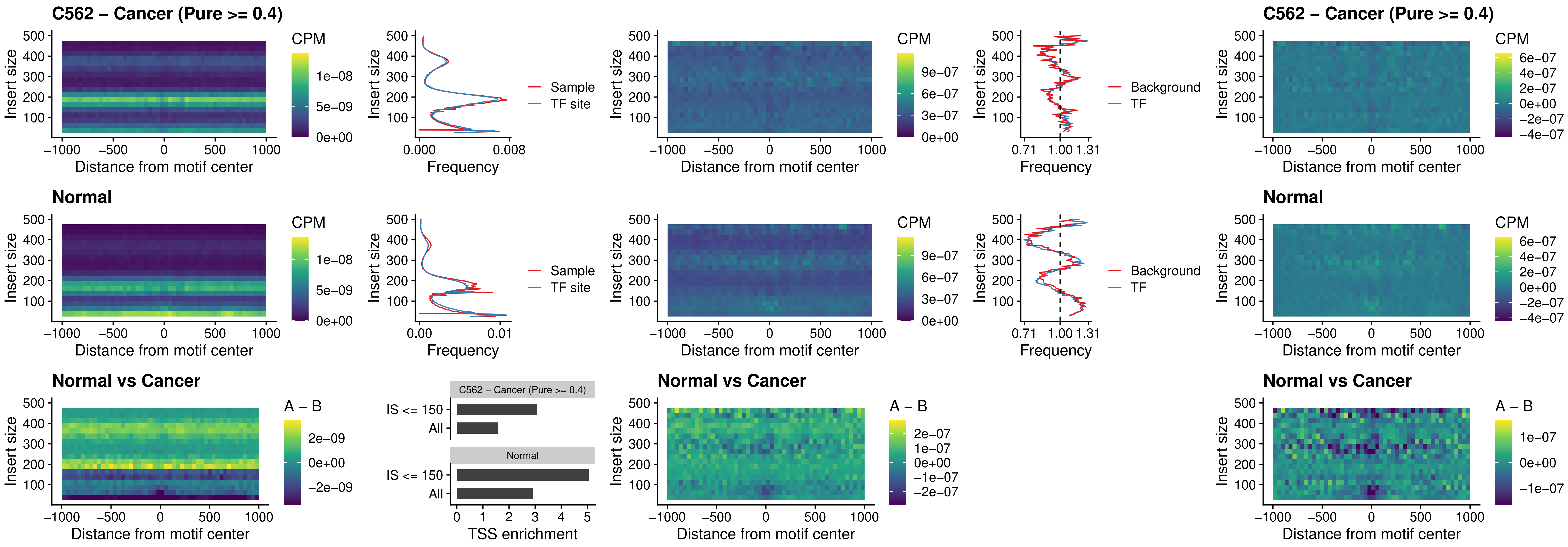

### BCL11A_uc_dTSS_nPEAK

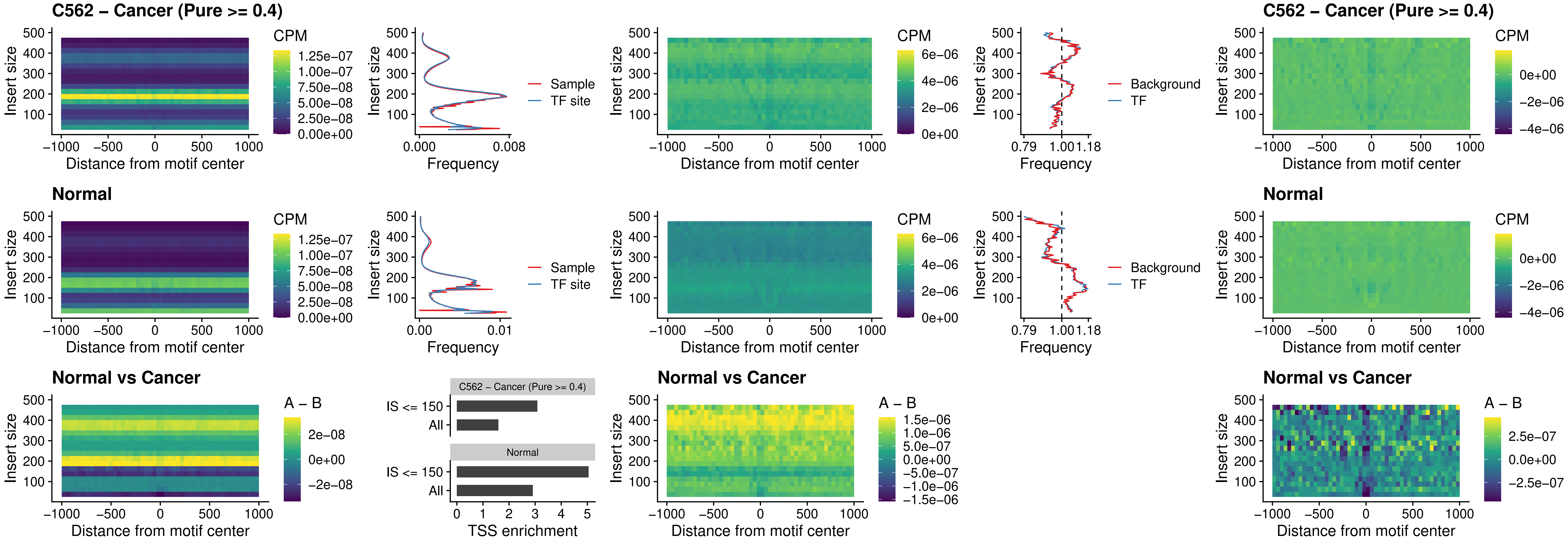

### BCL11B_uc_dTSS_nPEAK

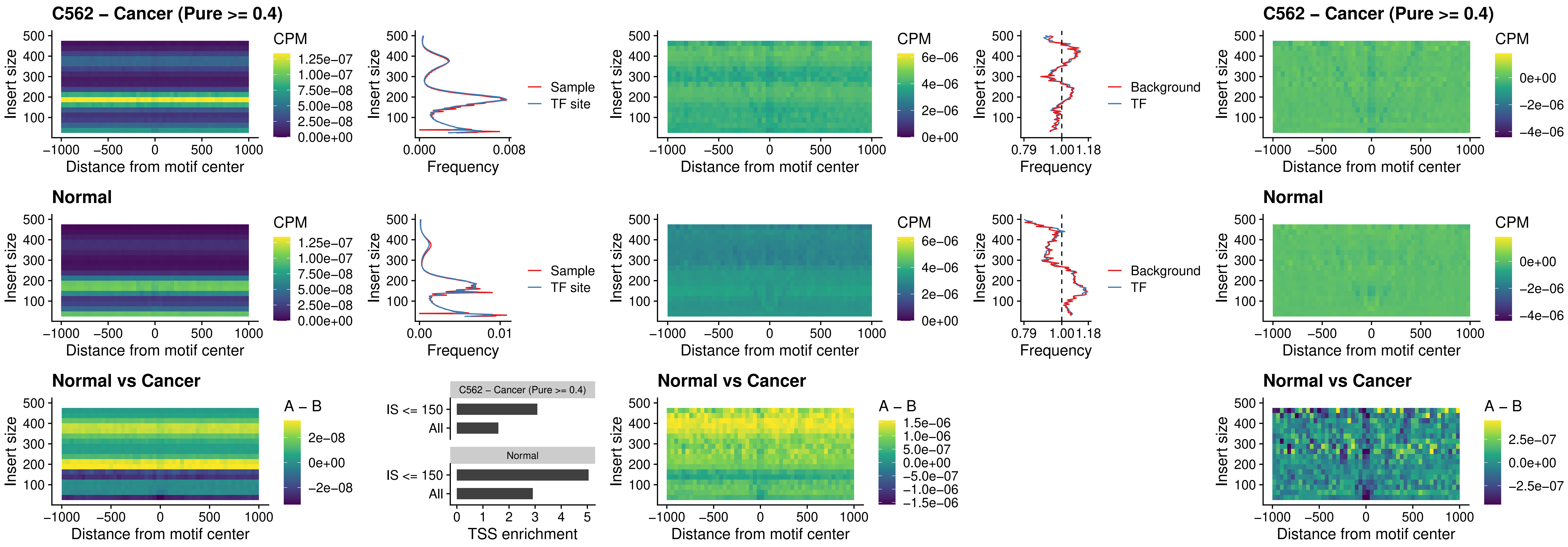

### IRF1_uc_cTSS_oPEAK

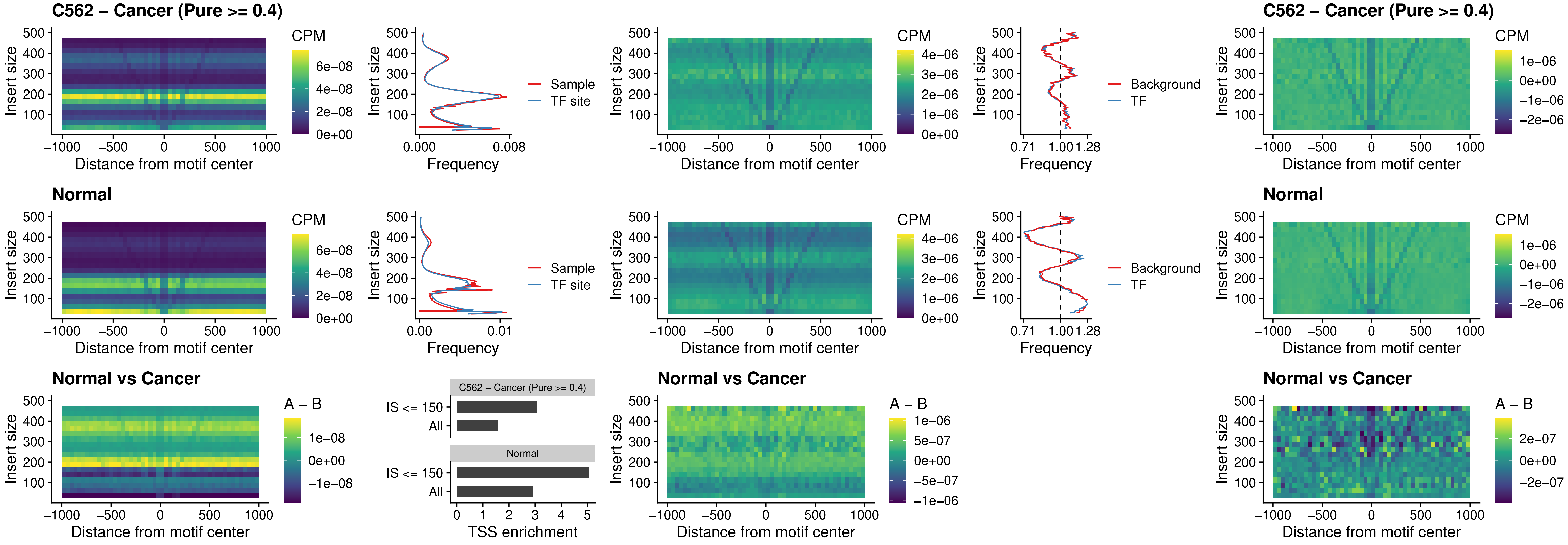

### IRF1_uc_dTSS_oPEAK

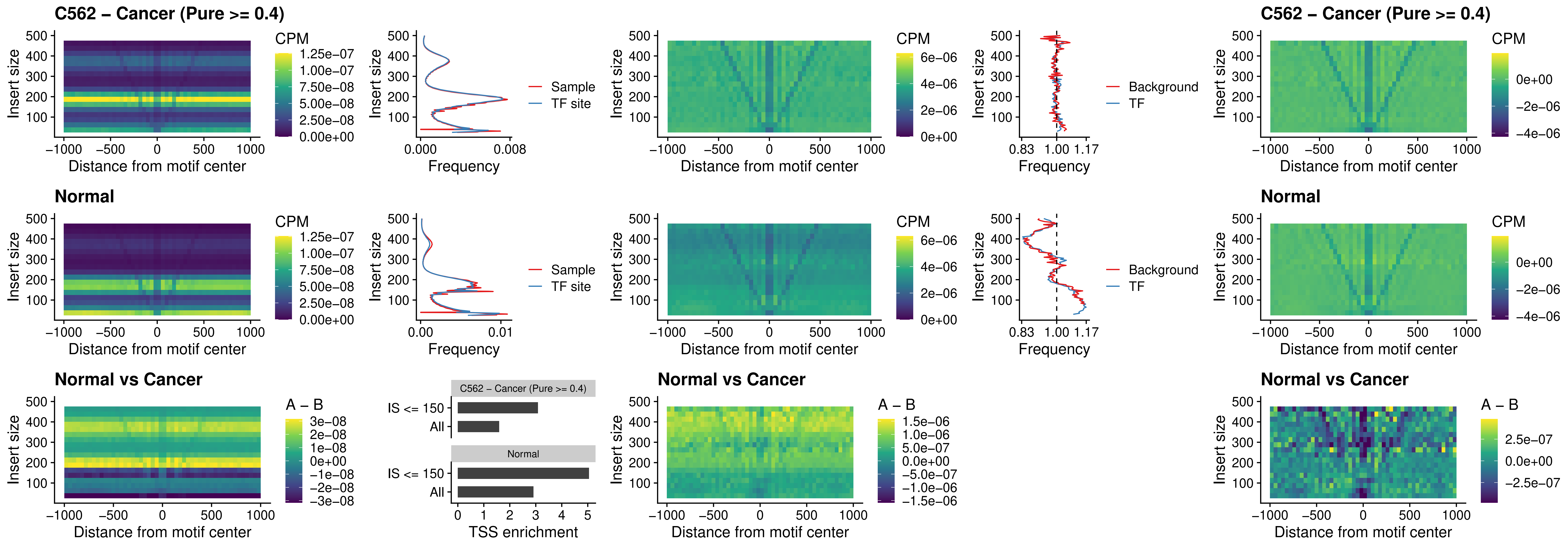

### IRF1_uc_pTSS_oPEAK

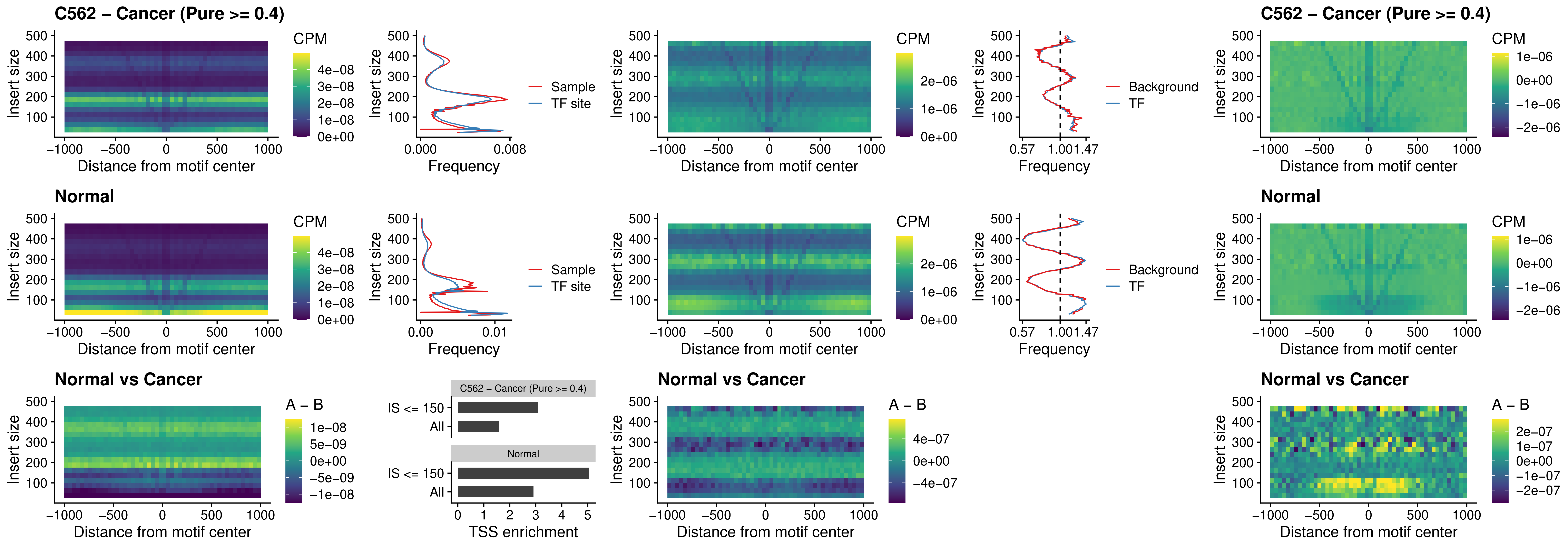

#### SPIC_uc_dTSS_nPEAK

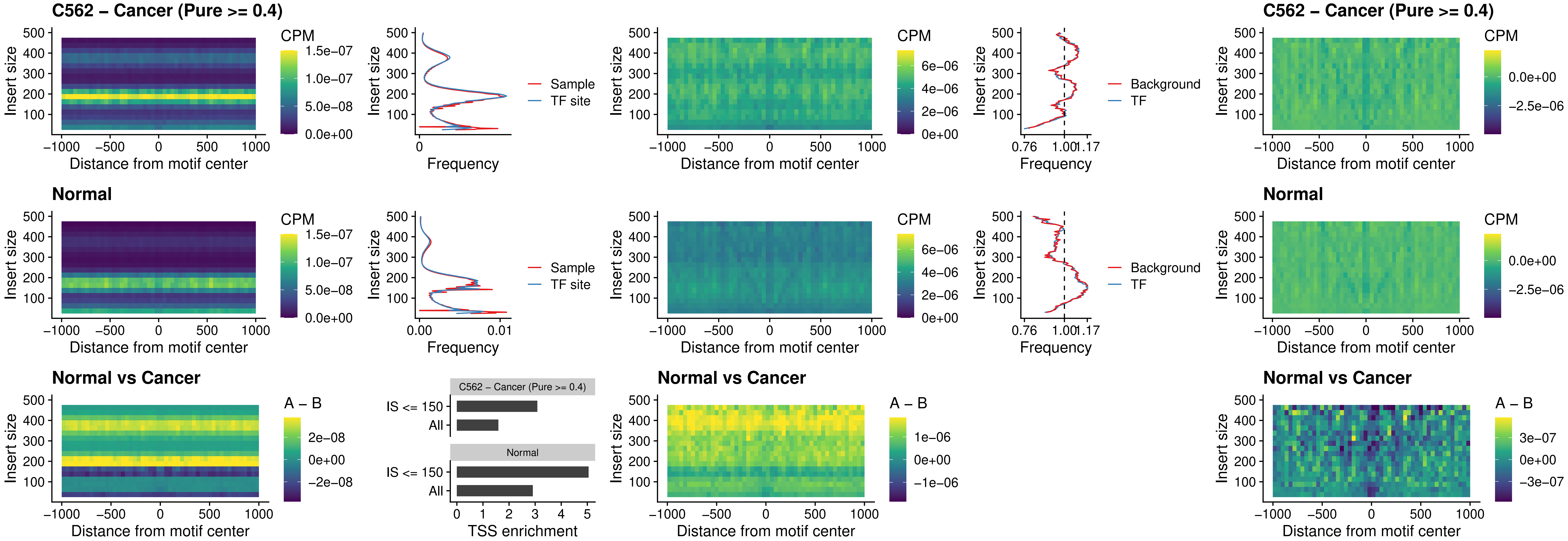

#### SPIC_uc_dTSS_oPEAK

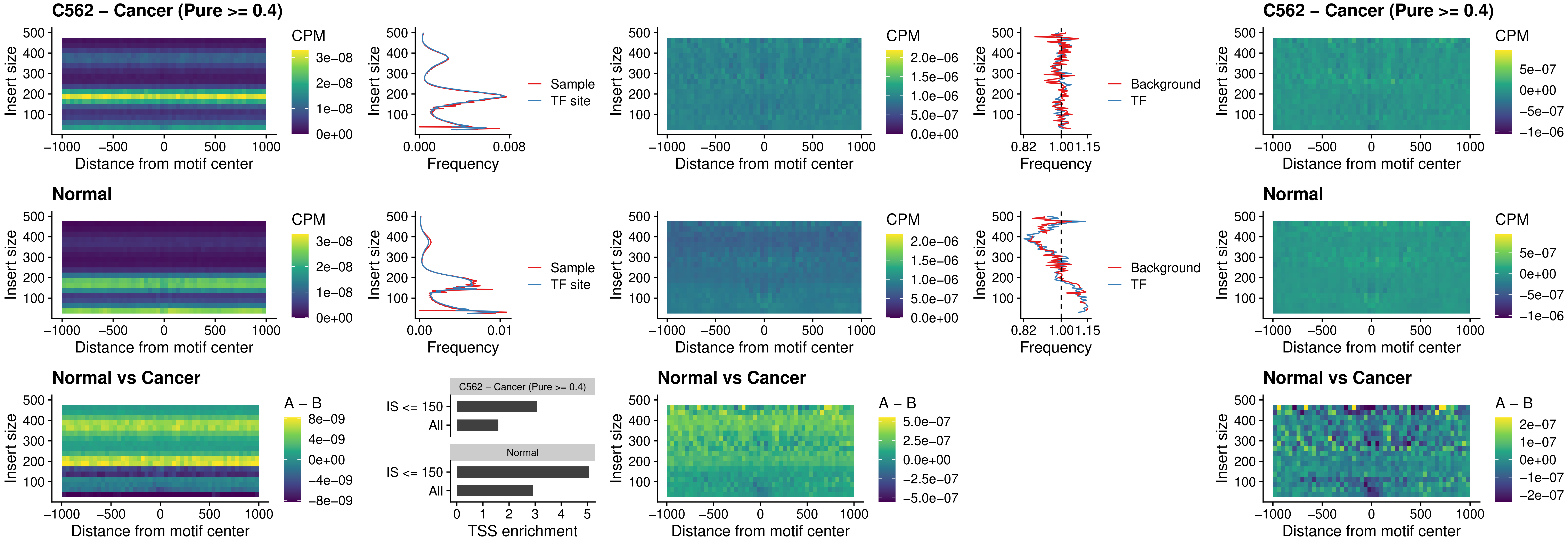

### SPIB_uc_pTSS_nPEAK

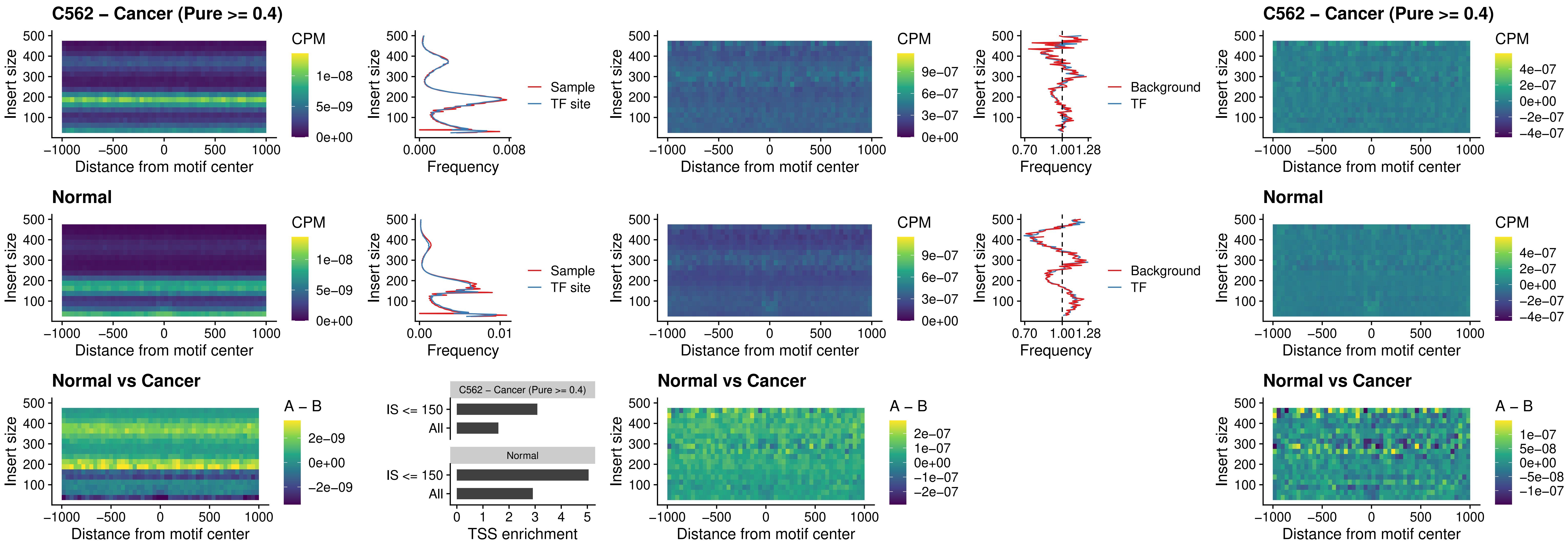

#### STAT2_uc_cTSS_oPEAK

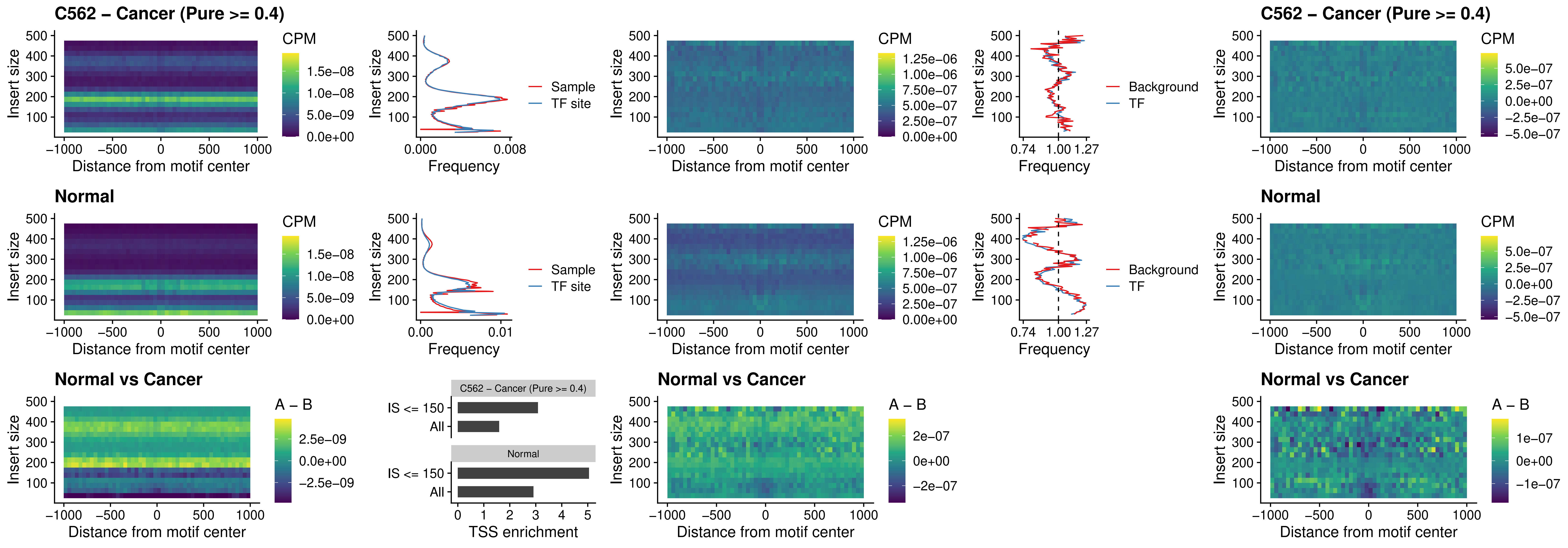

### IRF8_uc_cTSS_oPEAK

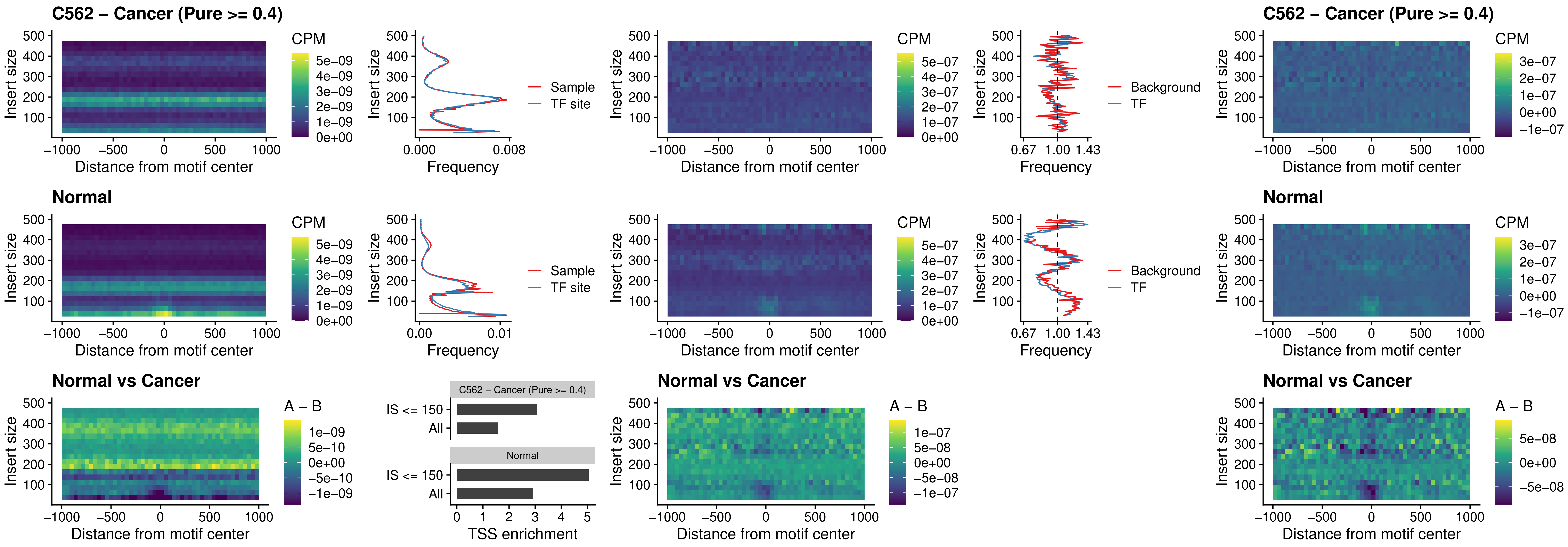

### IRF3_uc_dTSS_oPEAK

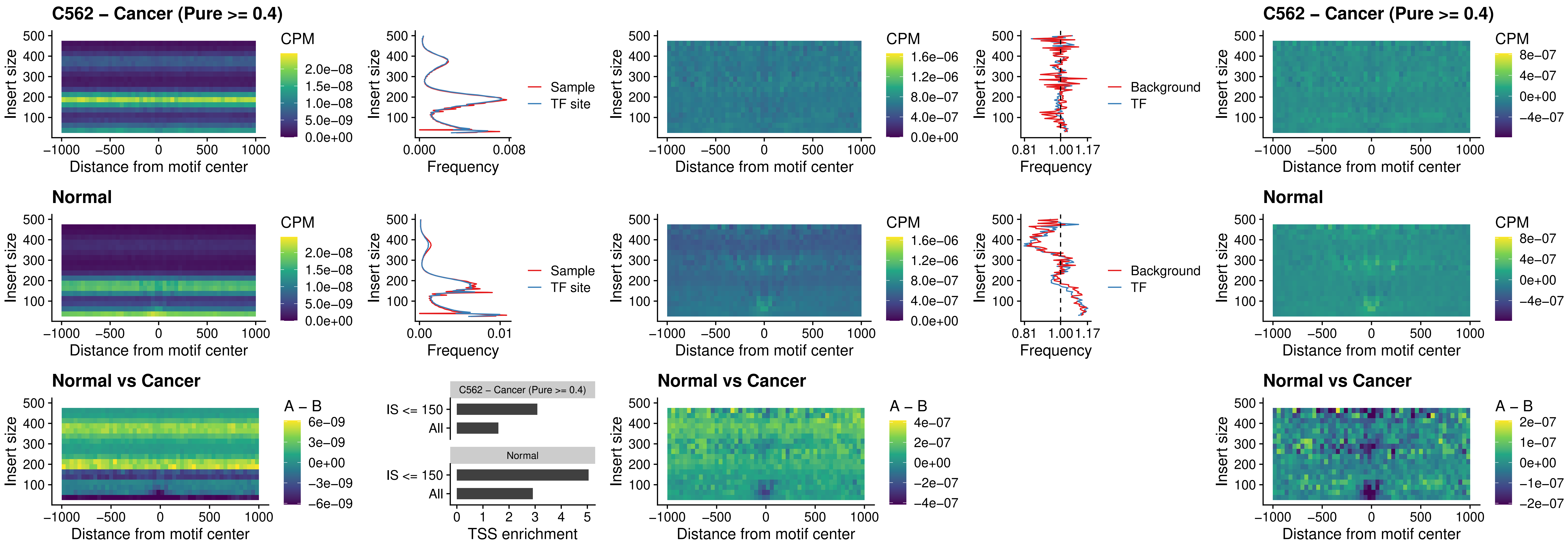

### IRF8_uc_dTSS_oPEAK

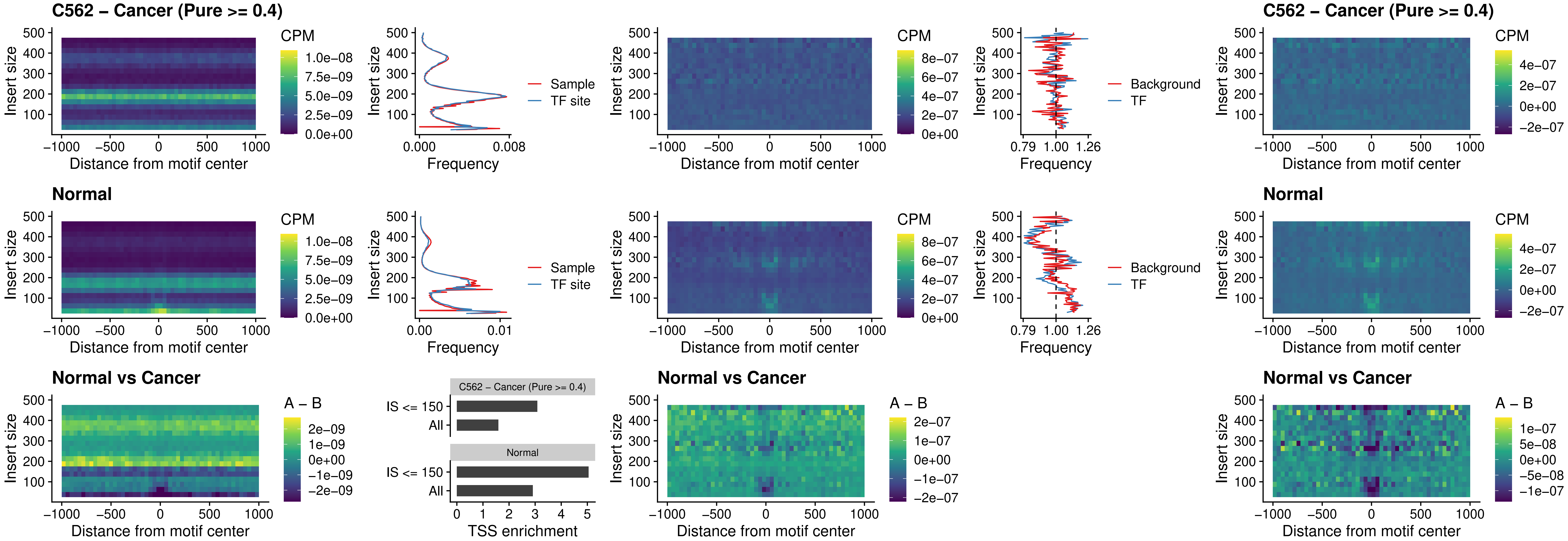

### ETS2_uc_dTSS_oPEAK

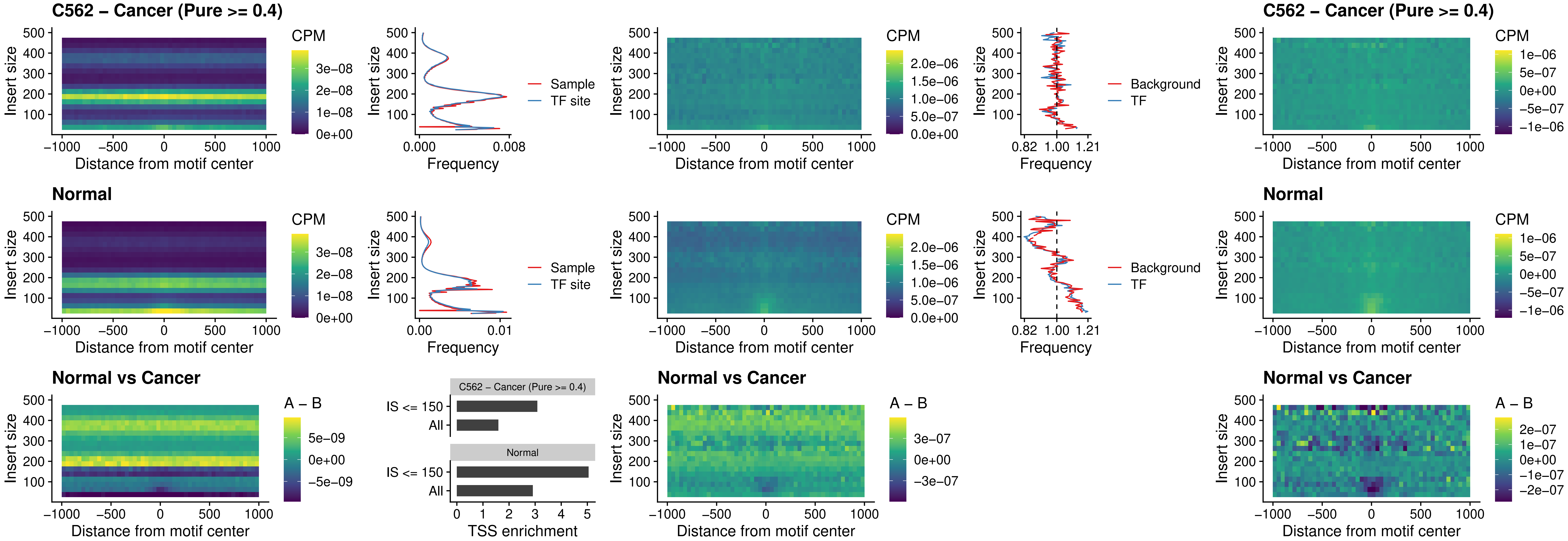

### SMARCC2_uc_pTSS_oPEAK

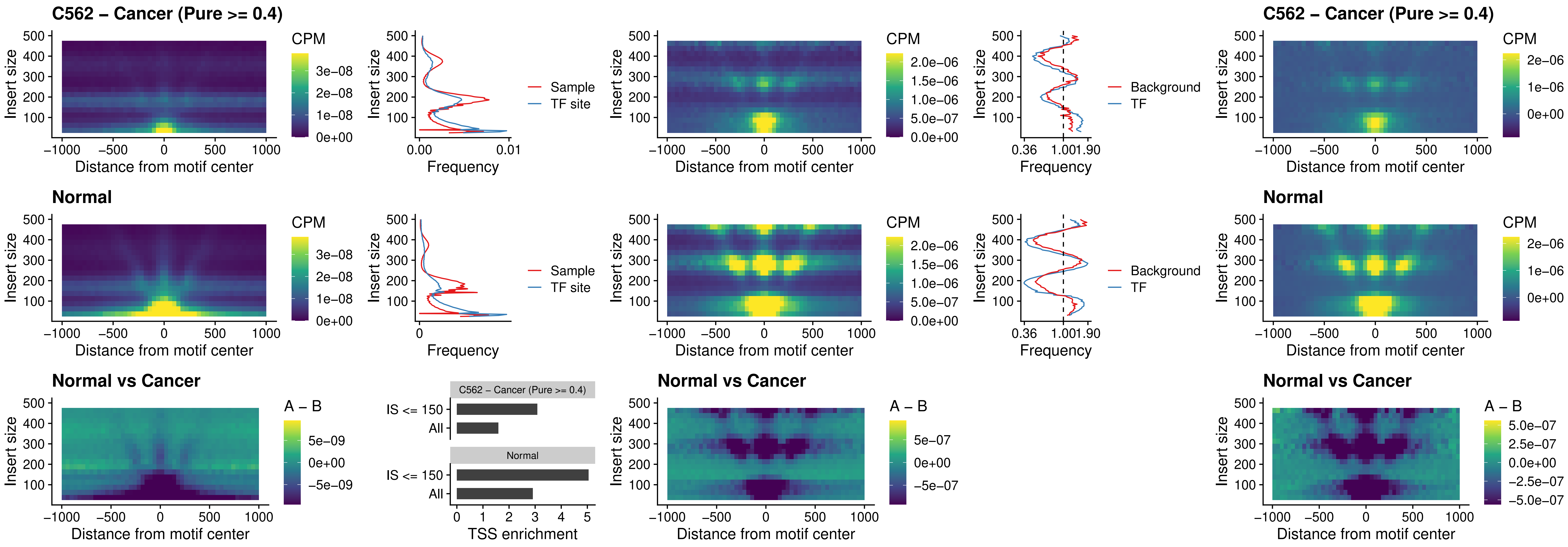

### BCL11A_uc_cTSS_nPEAK

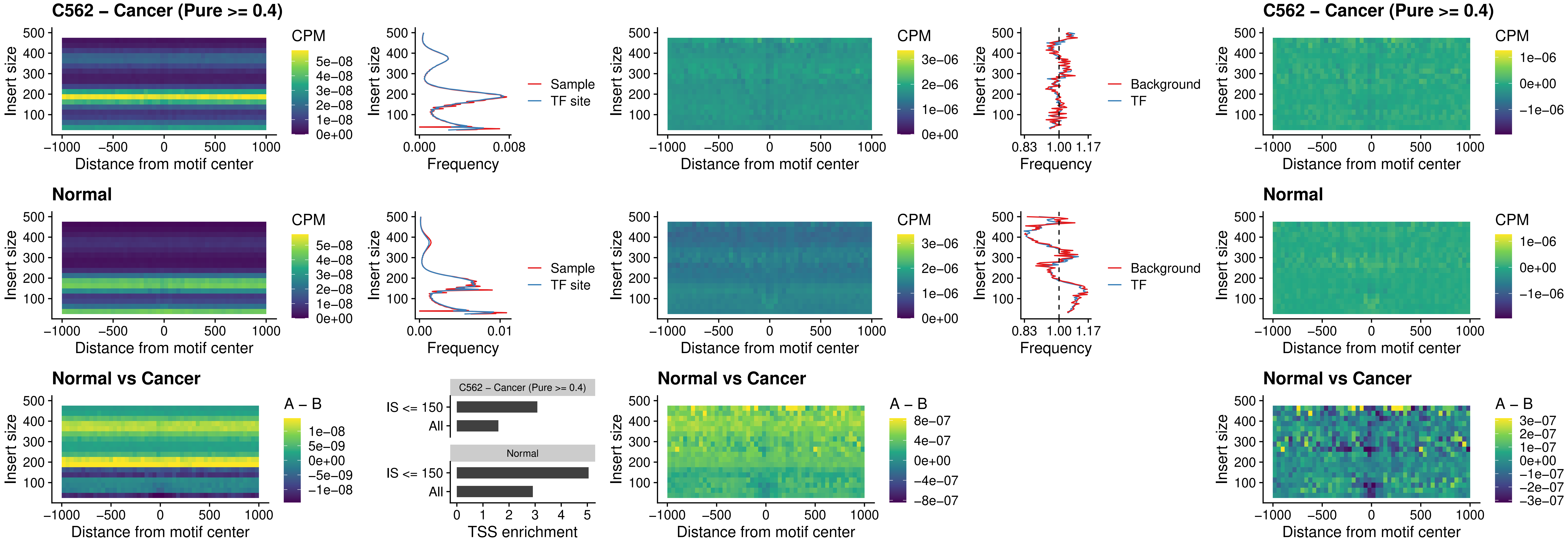

#### BCL11B_uc_cTSS_nPEAK

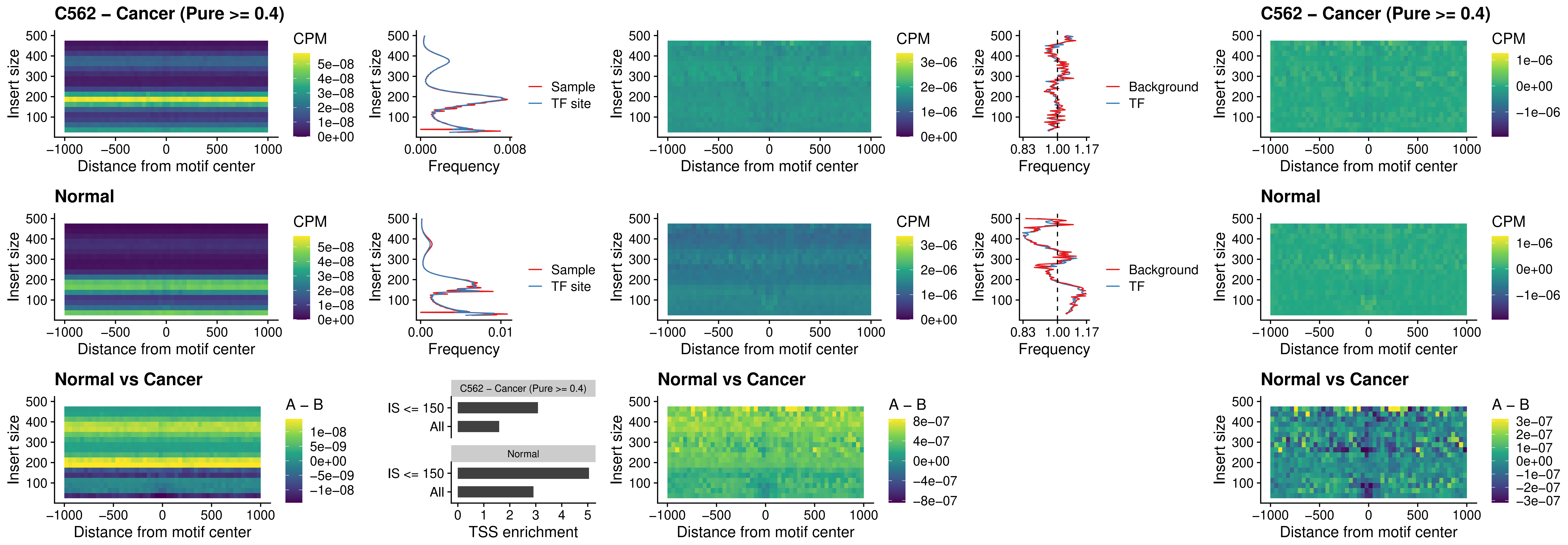

#### STAT2_uc_dTSS_oPEAK

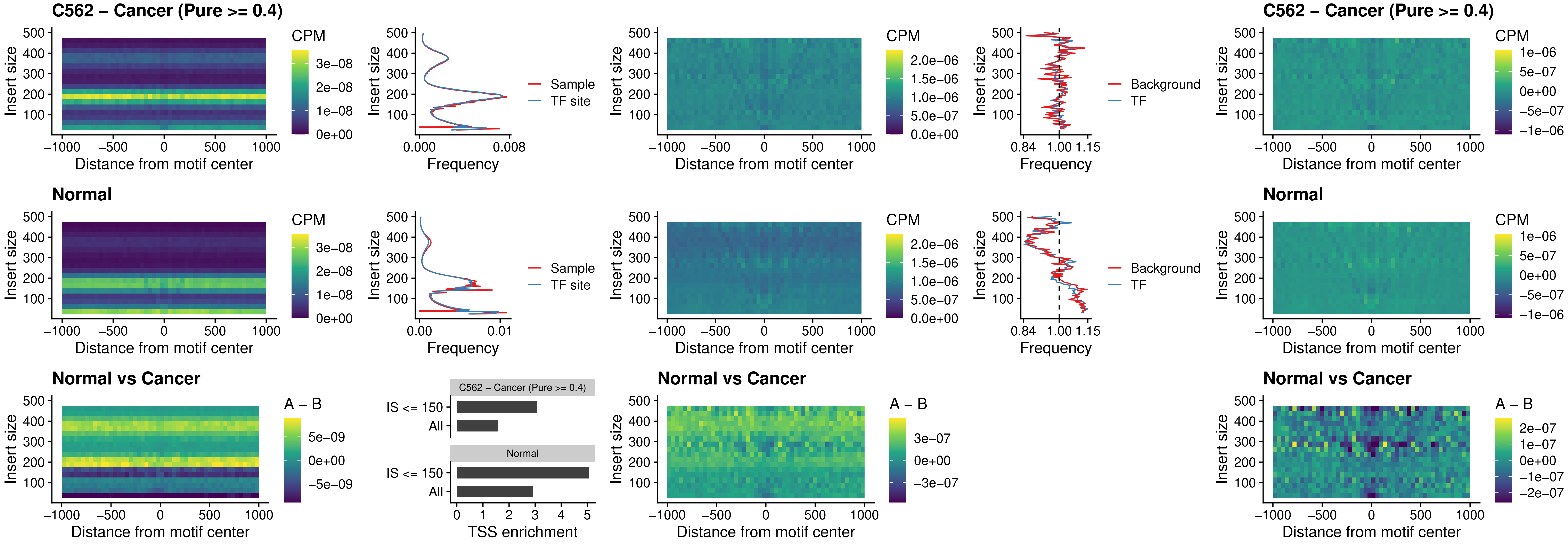

### ZNF683_uc_cTSS_oPEAK

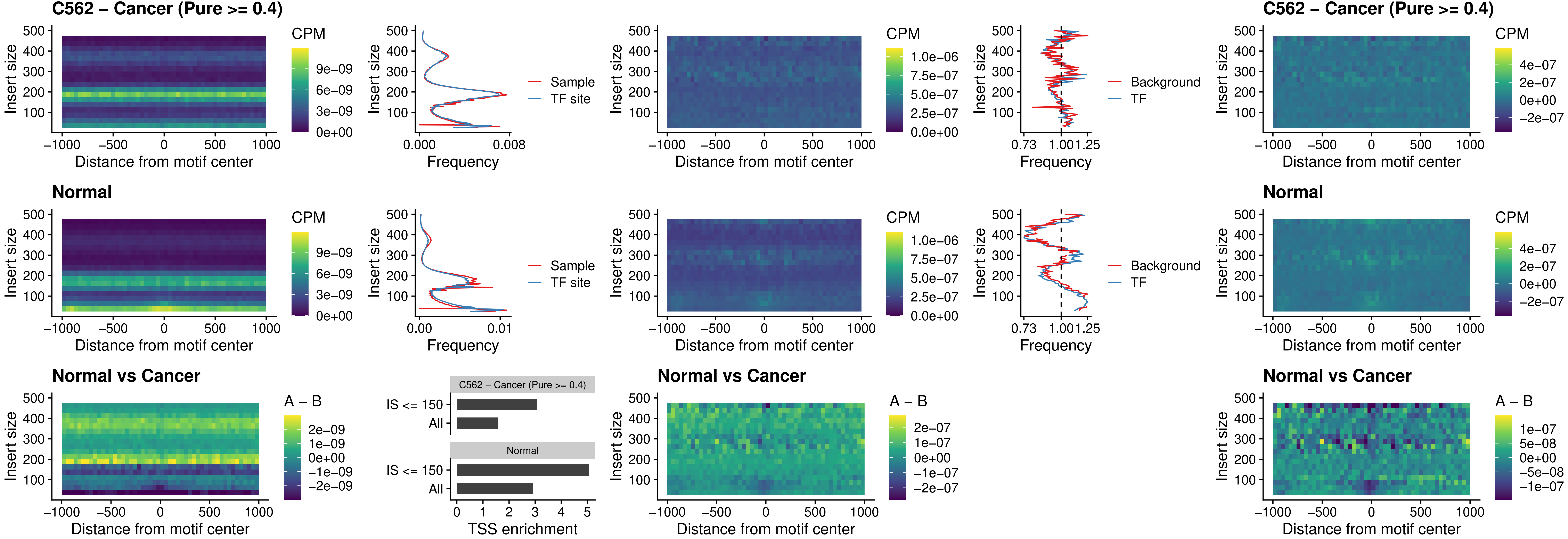

#### IRF8_uc_dTSS_nPEAK

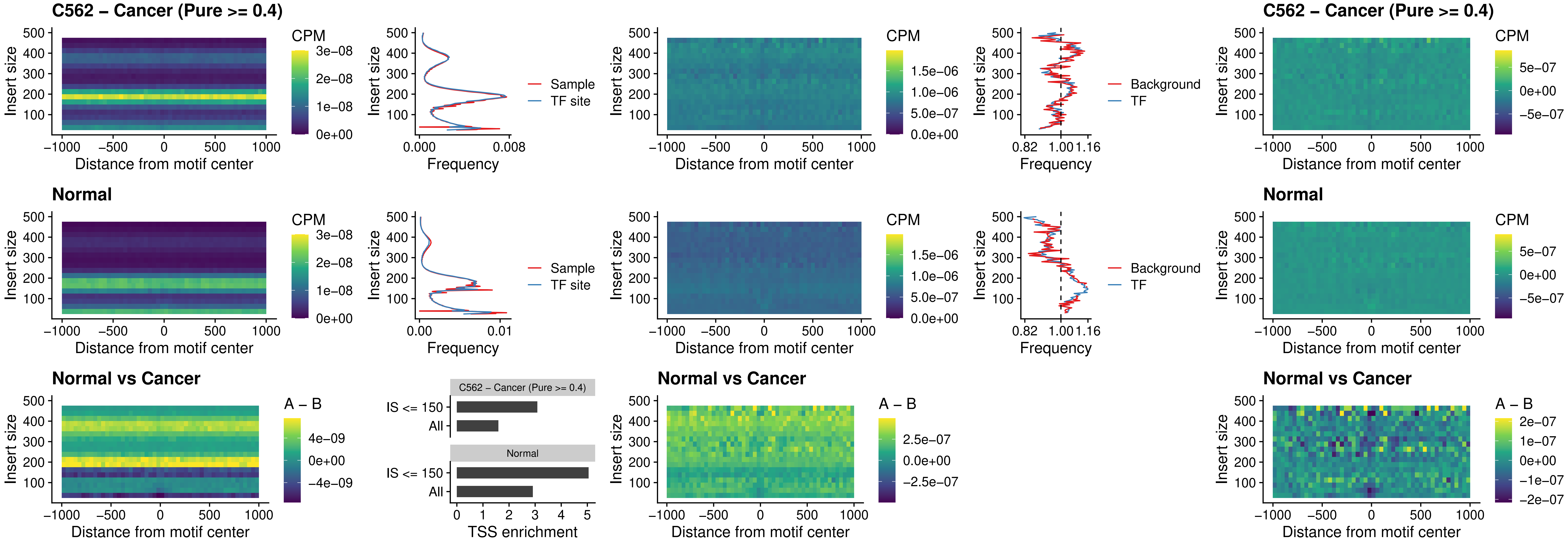

### ETV7_uc_dTSS_oPEAK

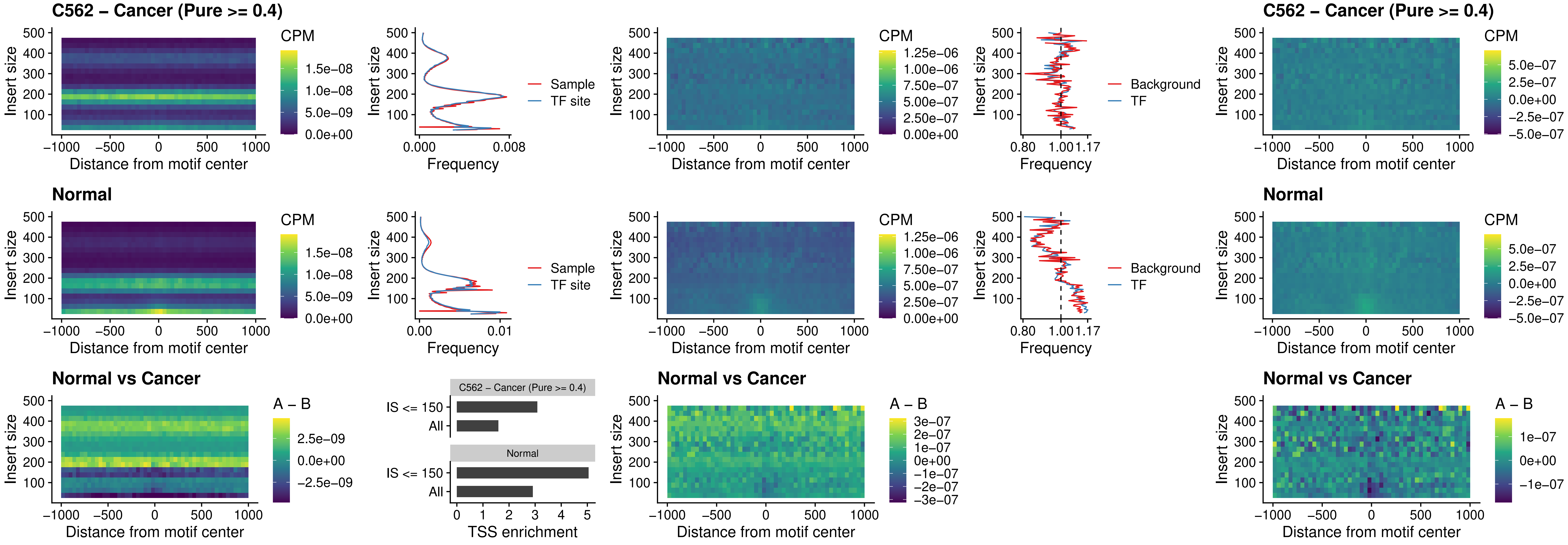
