## Supplementary Figures for "The co-evolution of the genome and epigenome in colorectal cancer": figS21.pdf

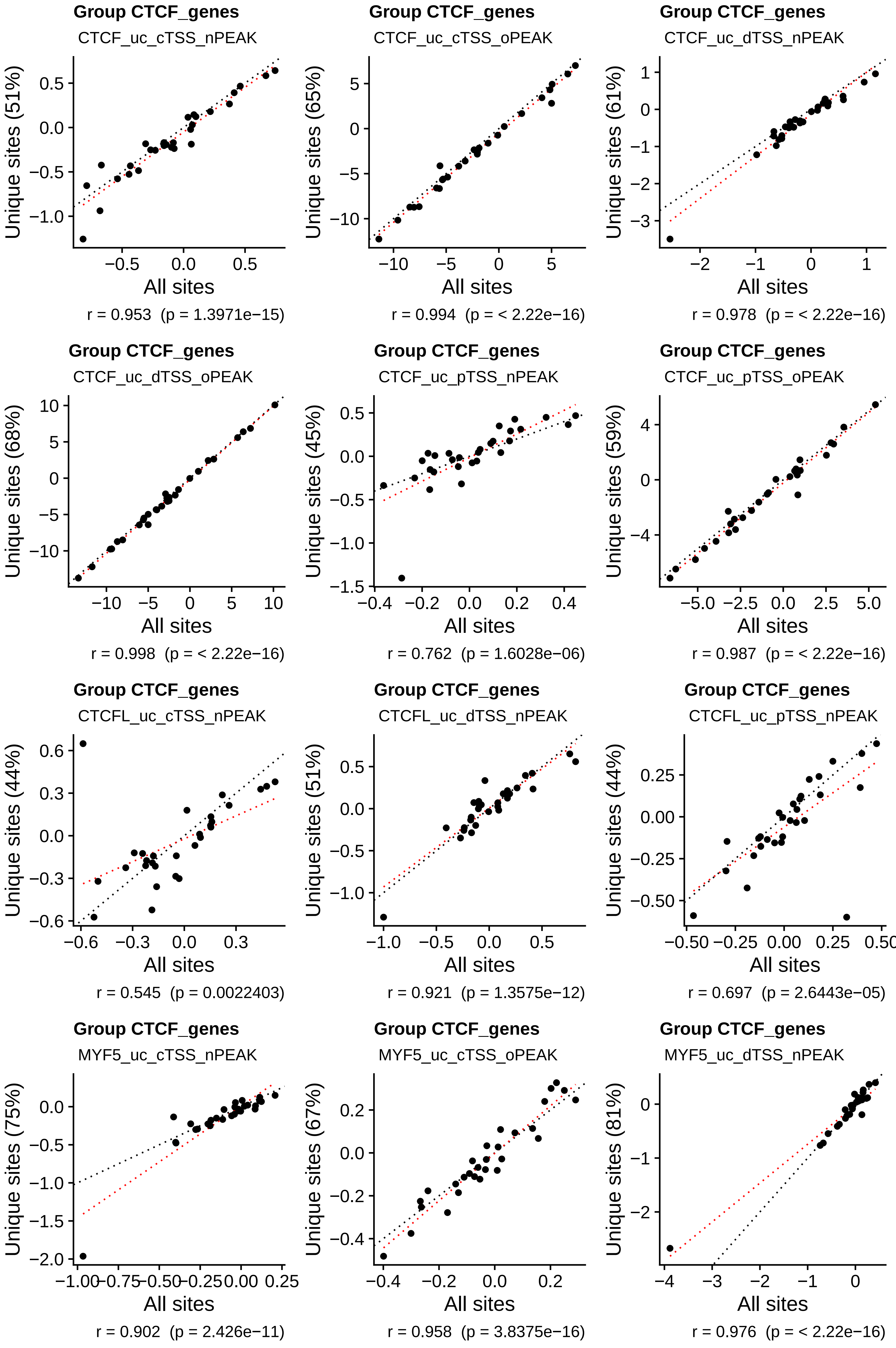
Group CTCF_genes
CTCF_uc_cTSS_nPEAK
Unique sites (51%)
All sites
r = 0.953  (p = 1.3971e−15)
Group CTCF_genes
CTCF_uc_cTSS_oPEAK
Unique sites (65%)
All sites
r = 0.994  (p = < 2.22e−16)
Group CTCF_genes
CTCF_uc_dTSS_nPEAK
Unique sites (61%)
All sites
r = 0.978  (p = < 2.22e−16)
Group CTCF_genes
CTCF_uc_dTSS_oPEAK
Unique sites (68%)
All sites
r = 0.998  (p = < 2.22e−16)
Group CTCF_genes
CTCF_uc_pTSS_nPEAK
Unique sites (45%)
All sites
r = 0.762  (p = 1.6028e−06)
Group CTCF_genes
CTCF_uc_pTSS_oPEAK
Unique sites (59%)
All sites
r = 0.987  (p = < 2.22e−16)
Group CTCF_genes
CTCFL_uc_cTSS_nPEAK
Unique sites (44%)
All sites
r = 0.545  (p = 0.0022403)
Group CTCF_genes
CTCFL_uc_dTSS_nPEAK
Unique sites (51%)
All sites
r = 0.921  (p = 1.3575e−12)
Group CTCF_genes
CTCFL_uc_pTSS_nPEAK
Unique sites (44%)
All sites
r = 0.697  (p = 2.6443e−05)
Group CTCF_genes
MYF5_uc_cTSS_nPEAK
Unique sites (75%)
All sites
r = 0.902  (p = 2.426e−11)
Group CTCF_genes
MYF5_uc_cTSS_oPEAK
Unique sites (67%)
All sites
r = 0.958  (p = 3.8375e−16)
Group CTCF_genes
MYF5_uc_dTSS_nPEAK
Unique sites (81%)
All sites
r = 0.976  (p = < 2.22e−16)

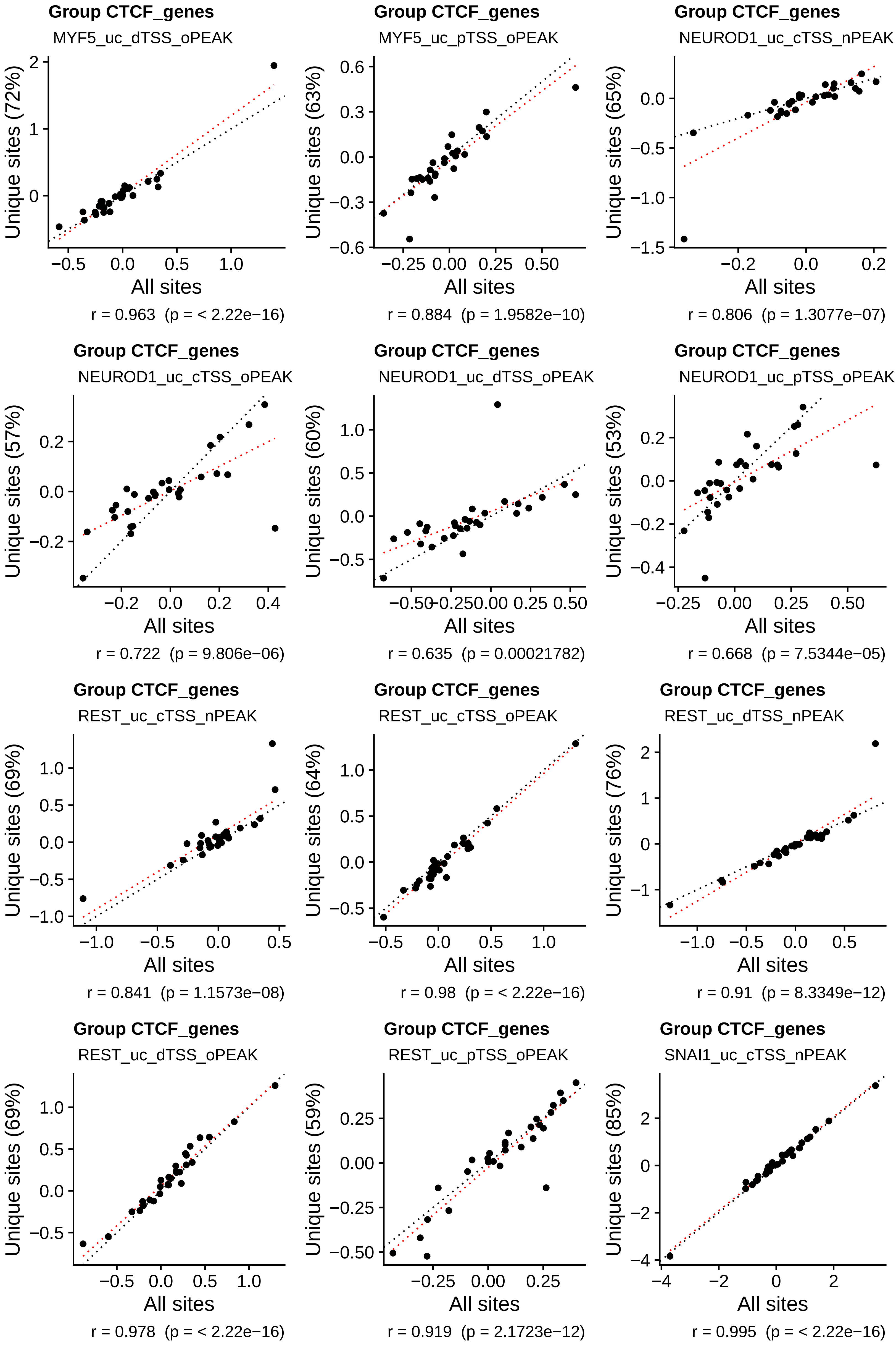
Group CTCF_genes
MYF5_uc_dTSS_oPEAK
Unique sites (72%)
All sites
r = 0.963 (p = < 2.22e−16)
Group CTCF_genes
MYF5_uc_pTSS_oPEAK
Unique sites (63%)
All sites
r = 0.884 (p = 1.9582e−10)
Group CTCF_genes
NEUROD1_uc_cTSS_nPEAK
Unique sites (65%)
All sites
r = 0.806 (p = 1.3077e−07)
Group CTCF_genes
NEUROD1_uc_cTSS_oPEAK
Unique sites (57%)
All sites
r = 0.722 (p = 9.806e−06)
Group CTCF_genes
NEUROD1_uc_dTSS_oPEAK
Unique sites (60%)
All sites
r = 0.635 (p = 0.00021782)
Group CTCF_genes
NEUROD1_uc_pTSS_oPEAK
Unique sites (53%)
All sites
r = 0.668 (p = 7.5344e−05)
Group CTCF_genes
REST_uc_cTSS_nPEAK
Unique sites (69%)
All sites
r = 0.841 (p = 1.1573e−08)
Group CTCF_genes
REST_uc_cTSS_oPEAK
Unique sites (64%)
All sites
r = 0.98 (p = < 2.22e−16)
Group CTCF_genes
REST_uc_dTSS_nPEAK
Unique sites (76%)
All sites
r = 0.91 (p = 8.3349e−12)
Group CTCF_genes
REST_uc_dTSS_oPEAK
Unique sites (69%)
All sites
r = 0.978 (p = < 2.22e−16)
Group CTCF_genes
REST_uc_pTSS_oPEAK
Unique sites (59%)
All sites
r = 0.919 (p = 2.1723e−12)
Group CTCF_genes
SNAI1_uc_cTSS_nPEAK
Unique sites (85%)
All sites
r = 0.995 (p = < 2.22e−16)

**Group CTCF_genes**

SNAI1_uc_cTSS_oPEAK

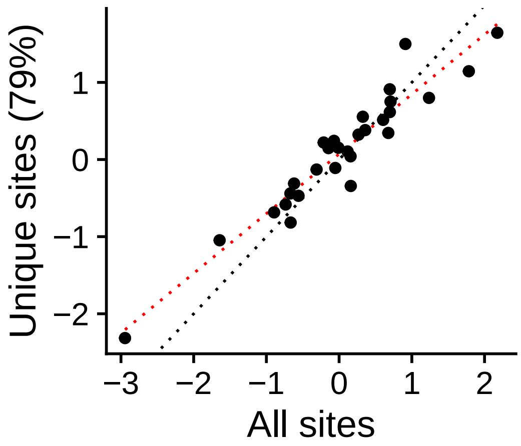


r = 0.953 (p = 1.4579e−15)

**Group CTCF_genes**

SNAI1_uc_dTSS_nPEAK

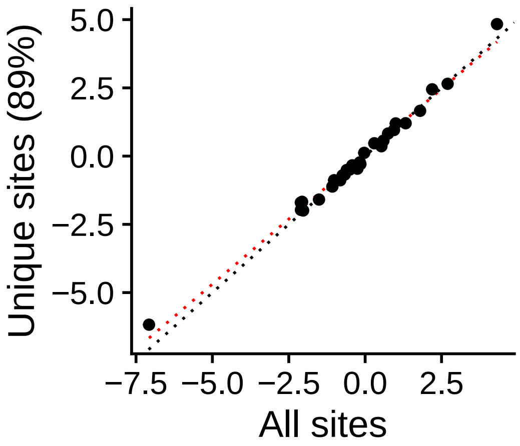


r = 0.994 (p = < 2.22e−16)

**Group CTCF_genes**

SNAI1_uc_dTSS_oPEAK

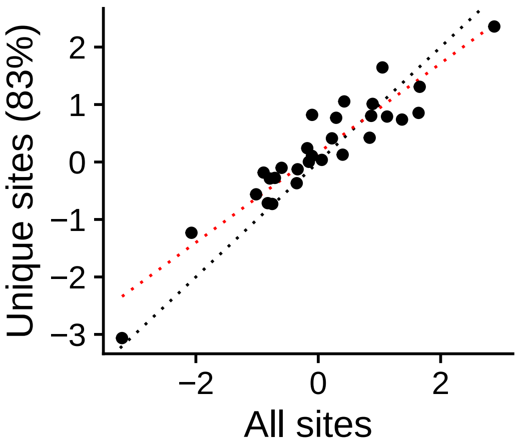


r = 0.935 (p = 1.1336e−13)

**Group CTCF_genes**

SNAI1_uc_pTSS_nPEAK

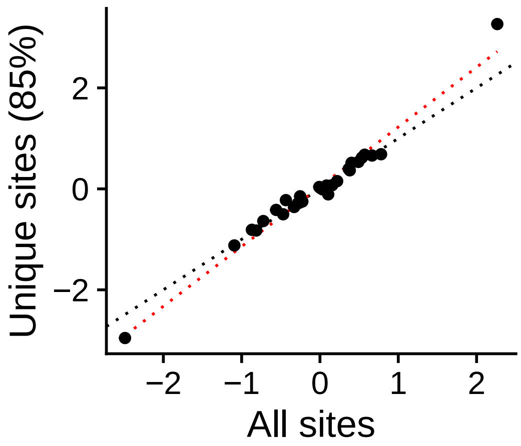


r = 0.985 (p = < 2.22e−16)

**Group CTCF_genes**

SNAI1_uc_pTSS_oPEAK

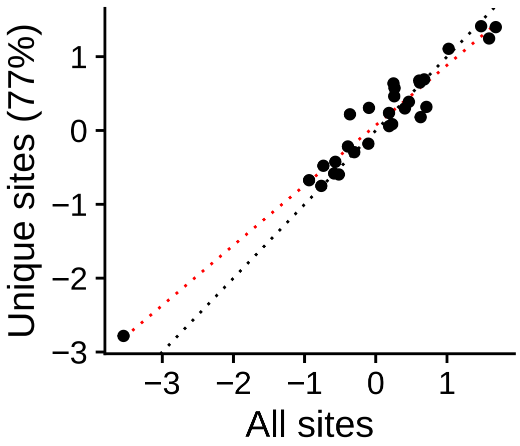


r = 0.969 (p = < 2.22e−16)

**Group CTCF_genes**

YY1_uc_pTSS_oPEAK

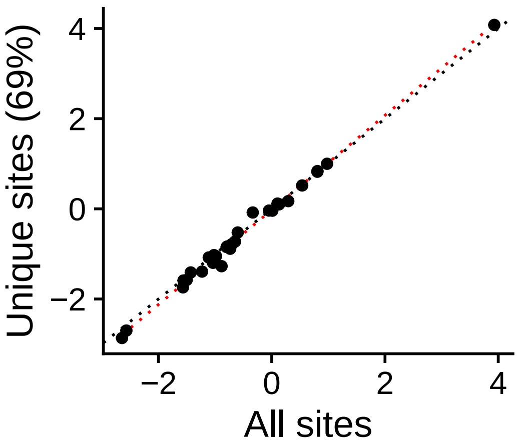


r = 0.997 (p = < 2.22e−16)

**Group IMUNO_**

BCL11A_uc_cTSS_nPEAK

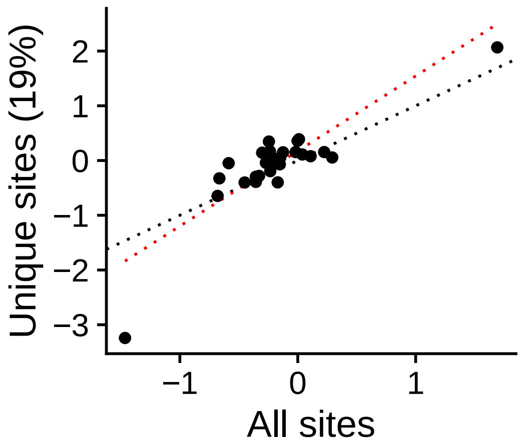


r = 0.872 (p = 7.4034e−10)

**Group IMUNO_**

BCL11A_uc_cTSS_oPEAK

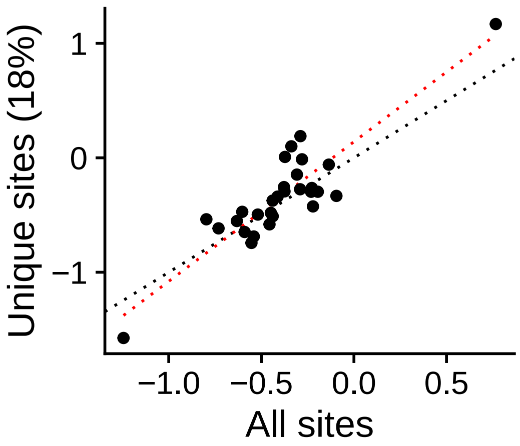


r = 0.896 (p = 4.8603e−11)

**Group IMUNO_**

BCL11A_uc_dTSS_nPEAK

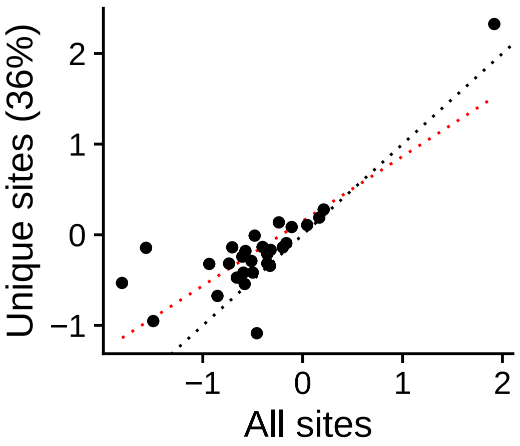


r = 0.814 (p = 7.8603e−08)

**Group IMUNO_**

BCL11A_uc_dTSS_oPEAK

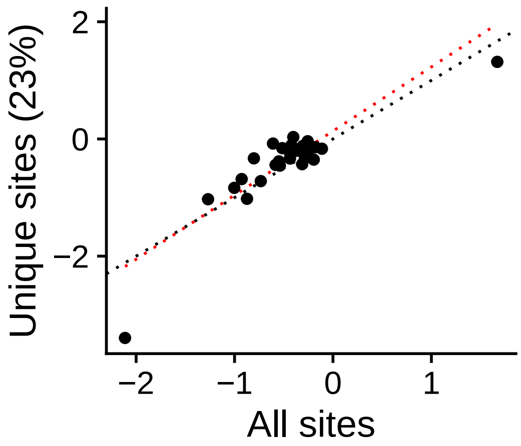


r = 0.889 (p = 1.139e−10)

**Group IMUNO_**

BCL11A_uc_pTSS_nPEAK

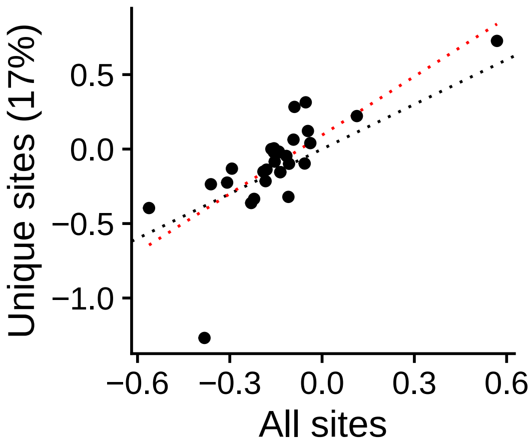


r = 0.755 (p = 2.2342e−06)

**Group IMUNO_**

BCL11A_uc_pTSS_oPEAK

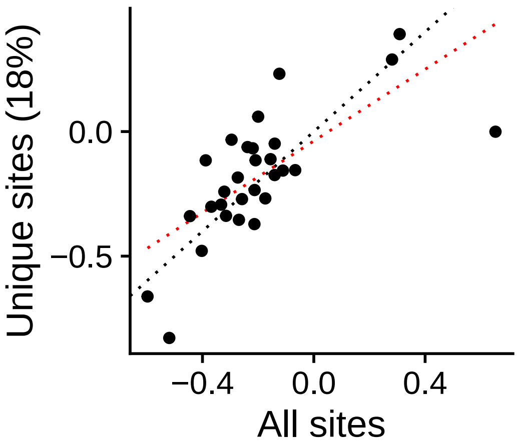


r = 0.711 (p = 1.5349e−05)

**Group IMUNO_**
ETS2_uc_cTSS_nPEAK

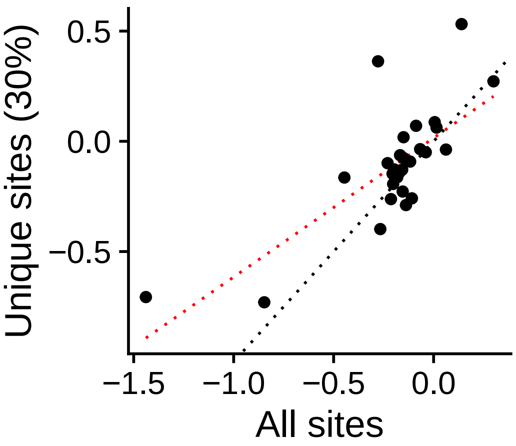


r = 0.753  (p = 2.4943e−06)

**Group IMUNO_**
ETS2_uc_cTSS_oPEAK

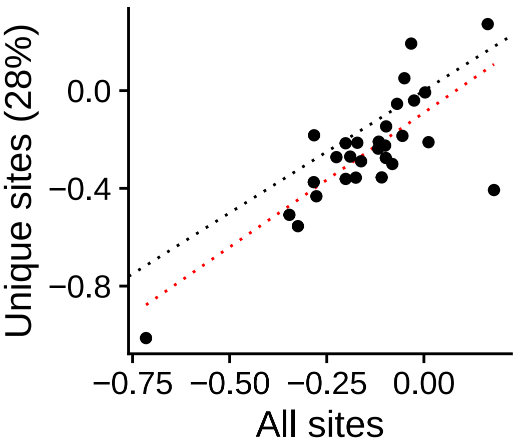


r = 0.773  (p = 8.8374e−07)

**Group IMUNO_**
ETS2_uc_dTSS_nPEAK

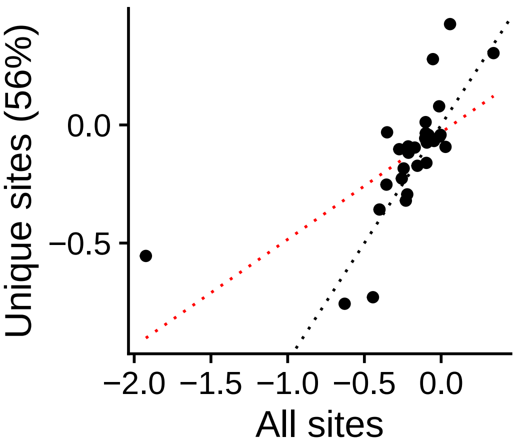


r = 0.657  (p = 0.00010719)

**Group IMUNO_**
ETS2_uc_dTSS_oPEAK

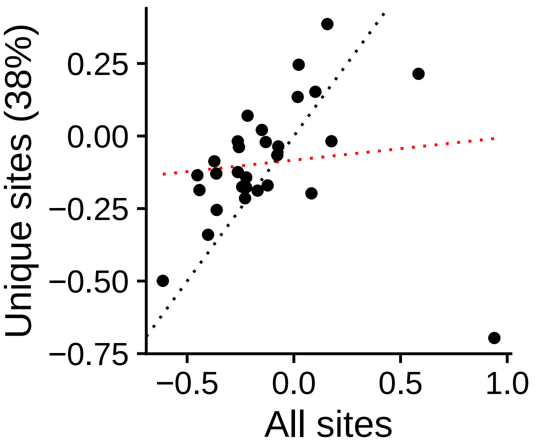


r = 0.115  (p = 0.55151)

**Group IMUNO_**
ETS2_uc_pTSS_oPEAK

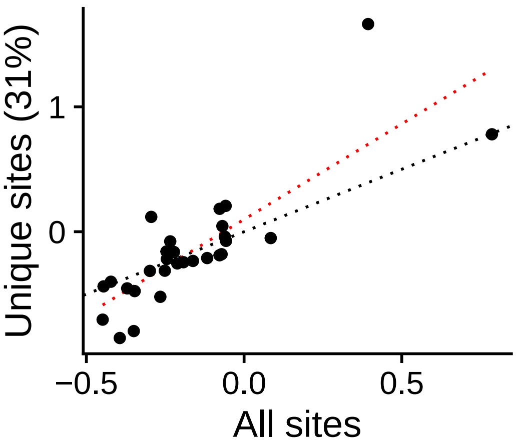


r = 0.818  (p = 5.8119e−08)

**Group IMUNO_**
ETV6_uc_cTSS_nPEAK

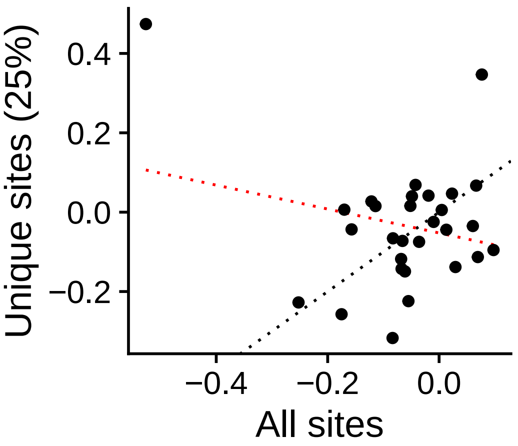


r = −0.231  (p = 0.2288)

**Group IMUNO_**
ETV6_uc_cTSS_oPEAK

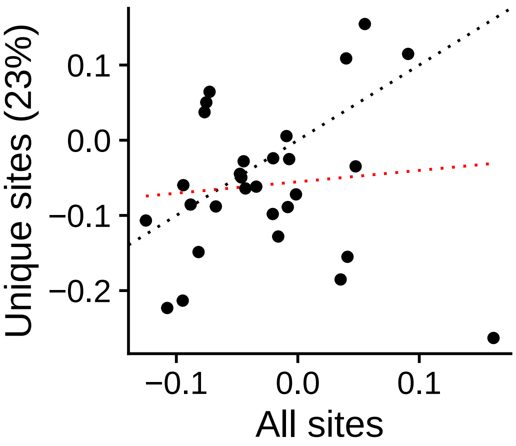


r = 0.0976  (p = 0.61453)

**Group IMUNO_**
ETV6_uc_dTSS_oPEAK

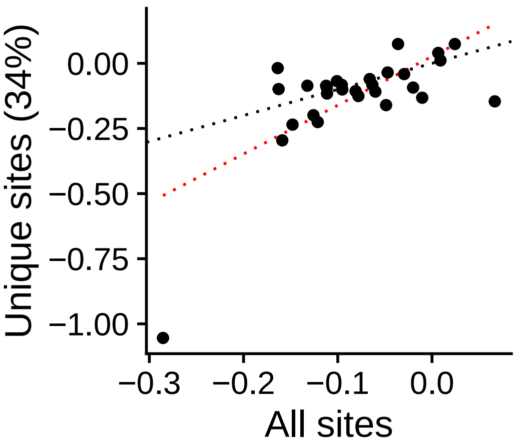


r = 0.672  (p = 6.5068e−05)

**Group IMUNO_**
ETV6_uc_pTSS_nPEAK

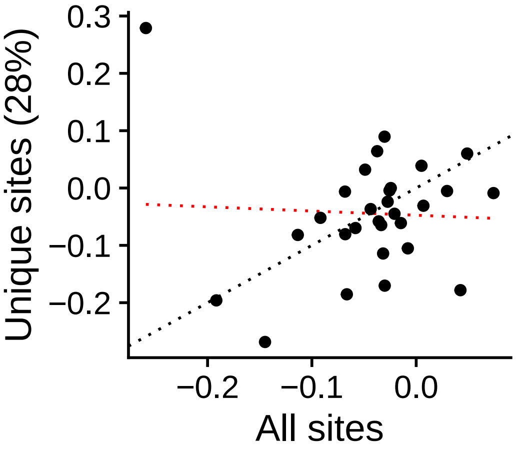


r = −0.048  (p = 0.80464)

**Group IMUNO_**
ETV6_uc_pTSS_oPEAK

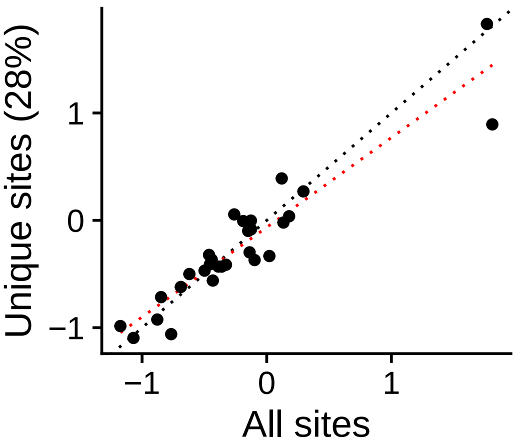


r = 0.937  (p = 7.866e−14)

**Group IMUNO_**
IRF1_uc_cTSS_nPEAK

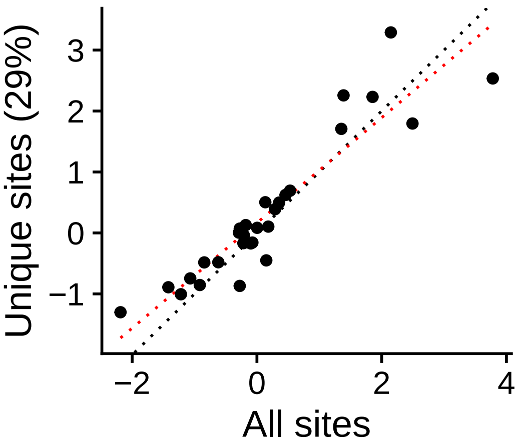


r = 0.924  (p = 9.4087e−13)

**Group IMUNO_**
IRF1_uc_cTSS_oPEAK

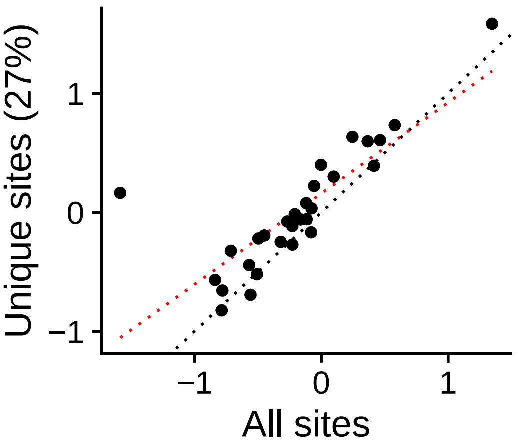


r = 0.81  (p = 1.0501e−07)

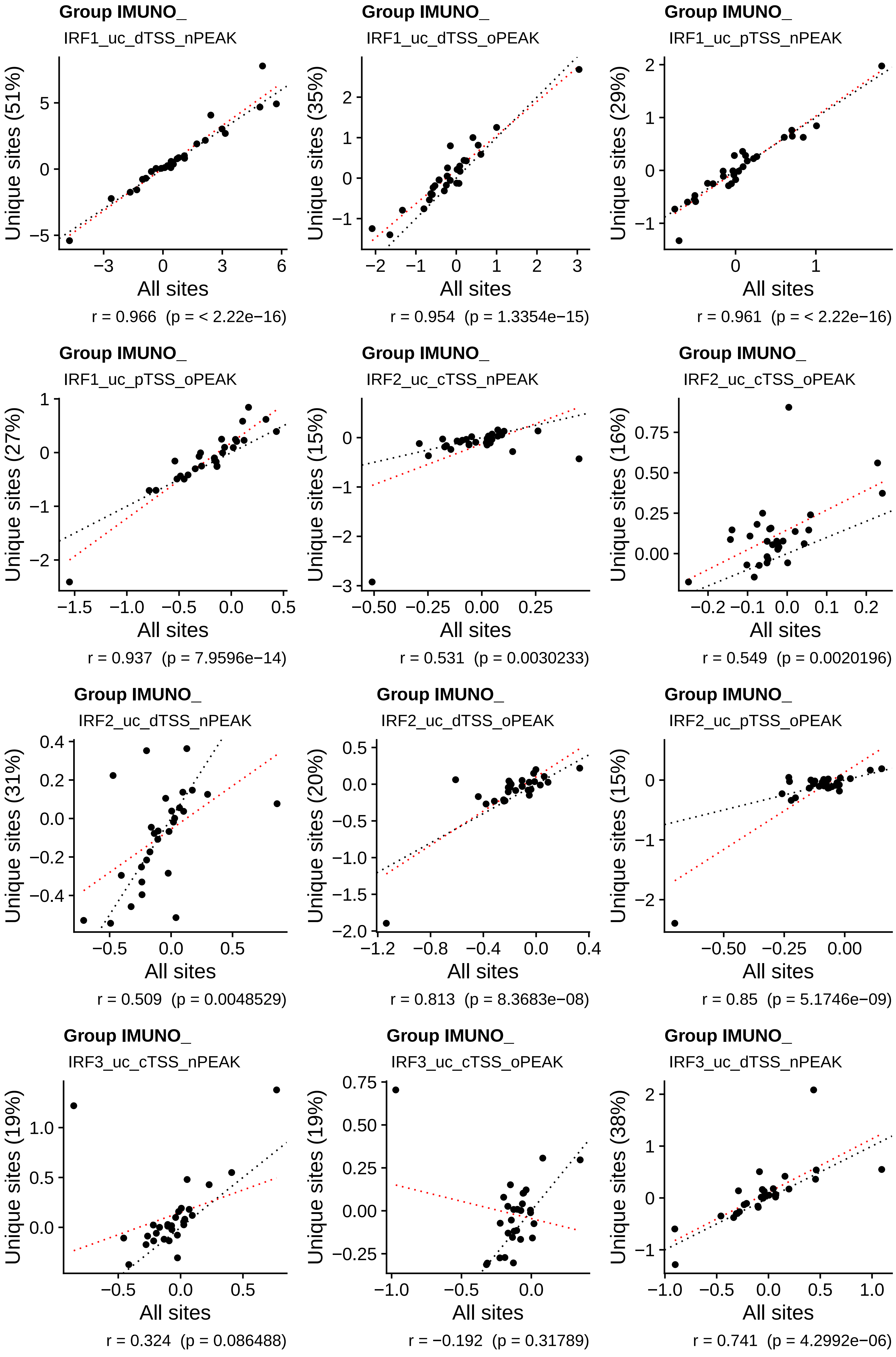
Group IMUNO_
IRF1_uc_dTSS_nPEAK
Unique sites (51%)
All sites
r = 0.966  (p = < 2.22e−16)
Group IMUNO_
IRF1_uc_dTSS_oPEAK
Unique sites (35%)
All sites
r = 0.954  (p = 1.3354e−15)
Group IMUNO_
IRF1_uc_pTSS_nPEAK
Unique sites (29%)
All sites
r = 0.961  (p = < 2.22e−16)
Group IMUNO_
IRF1_uc_pTSS_oPEAK
Unique sites (27%)
All sites
r = 0.937  (p = 7.9596e−14)
Group IMUNO_
IRF2_uc_cTSS_nPEAK
Unique sites (15%)
All sites
r = 0.531  (p = 0.0030233)
Group IMUNO_
IRF2_uc_cTSS_oPEAK
Unique sites (16%)
All sites
r = 0.549  (p = 0.0020196)
Group IMUNO_
IRF2_uc_dTSS_nPEAK
Unique sites (31%)
All sites
r = 0.509  (p = 0.0048529)
Group IMUNO_
IRF2_uc_dTSS_oPEAK
Unique sites (20%)
All sites
r = 0.813  (p = 8.3683e−08)
Group IMUNO_
IRF2_uc_pTSS_oPEAK
Unique sites (15%)
All sites
r = 0.85  (p = 5.1746e−09)
Group IMUNO_
IRF3_uc_cTSS_nPEAK
Unique sites (19%)
All sites
r = 0.324  (p = 0.086488)
Group IMUNO_
IRF3_uc_cTSS_oPEAK
Unique sites (19%)
All sites
r = −0.192  (p = 0.31789)
Group IMUNO_
IRF3_uc_dTSS_nPEAK
Unique sites (38%)
All sites
r = 0.741  (p = 4.2992e−06)

**Group IMUNO_**
IRF3_uc_dTSS_oPEAK

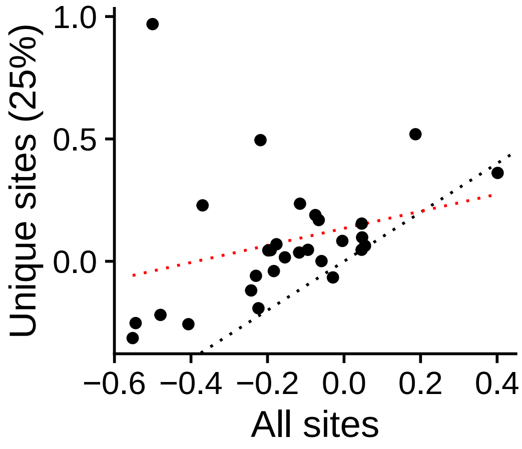


r = 0.287 (p = 0.1309)

**Group IMUNO_**
IRF3_uc_pTSS_nPEAK

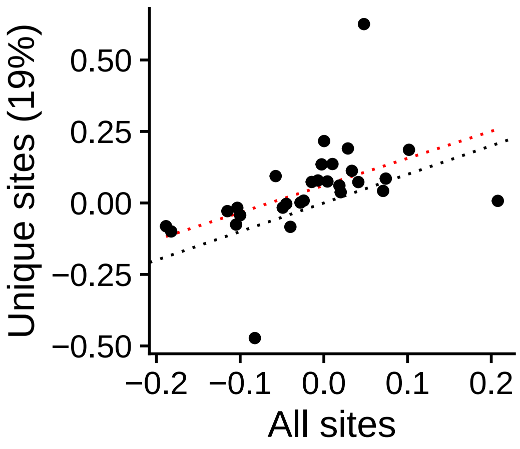


r = 0.467 (p = 0.010673)

**Group IMUNO_**
IRF3_uc_pTSS_oPEAK

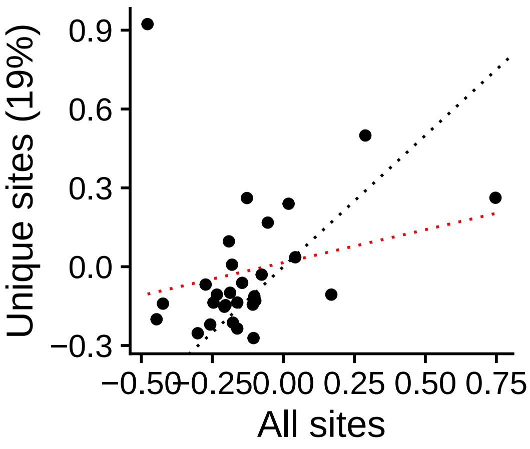


r = 0.227 (p = 0.23582)

**Group IMUNO_**
IRF8_uc_cTSS_nPEAK

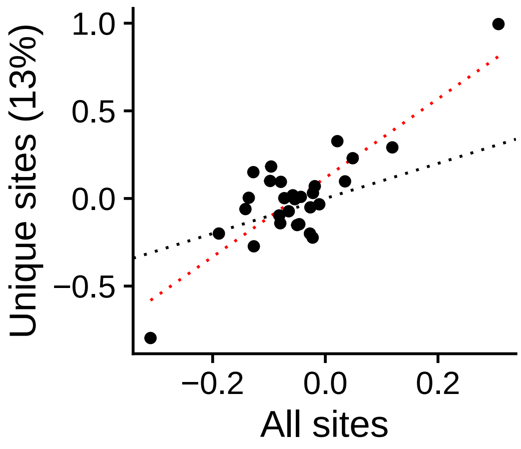


r = 0.832 (p = 2.284e−08)

**Group IMUNO_**
IRF8_uc_cTSS_oPEAK

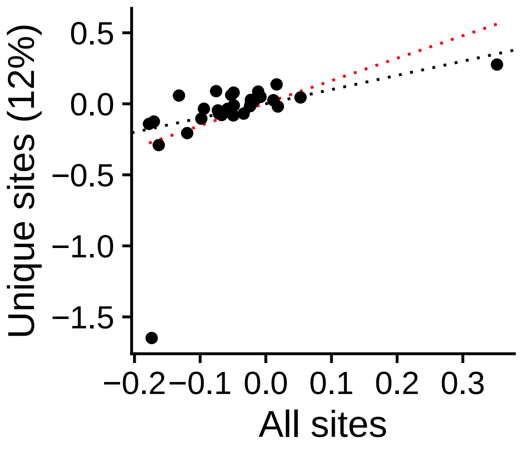


r = 0.48 (p = 0.0083471)

**Group IMUNO_**
IRF8_uc_dTSS_nPEAK

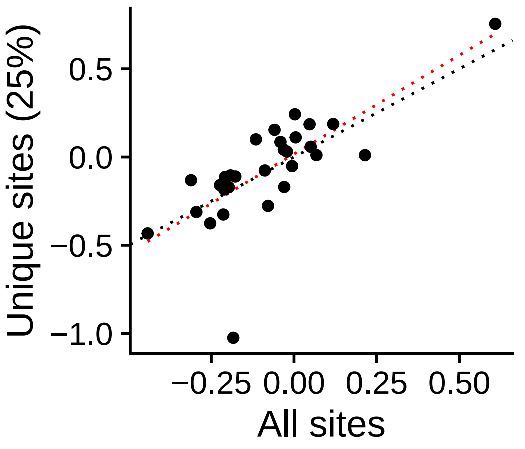


r = 0.739 (p = 4.7174e−06)

**Group IMUNO_**
IRF8_uc_dTSS_oPEAK

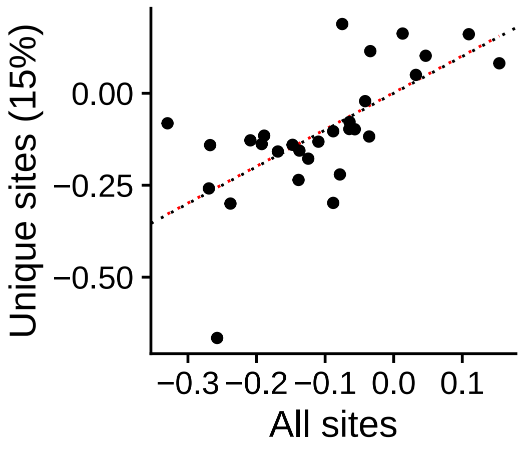


r = 0.661 (p = 9.3506e−05)

**Group IMUNO_**
IRF8_uc_pTSS_nPEAK

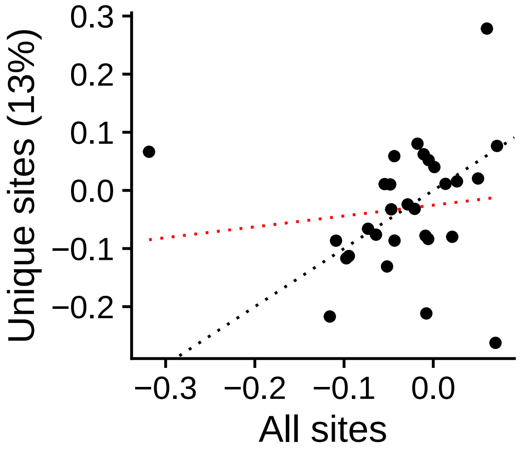


r = 0.129 (p = 0.50555)

**Group IMUNO_**
IRF9_uc_cTSS_nPEAK

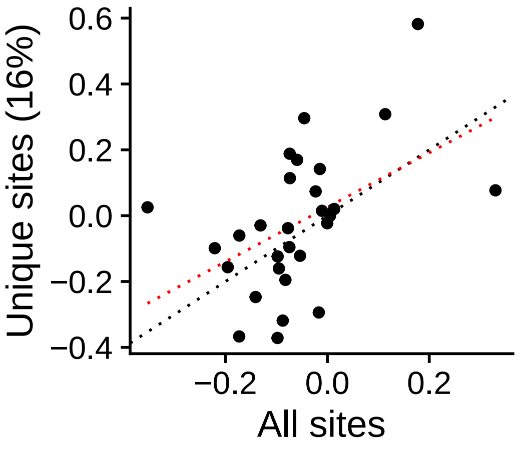


r = 0.481 (p = 0.0082344)

**Group IMUNO_**
IRF9_uc_cTSS_oPEAK

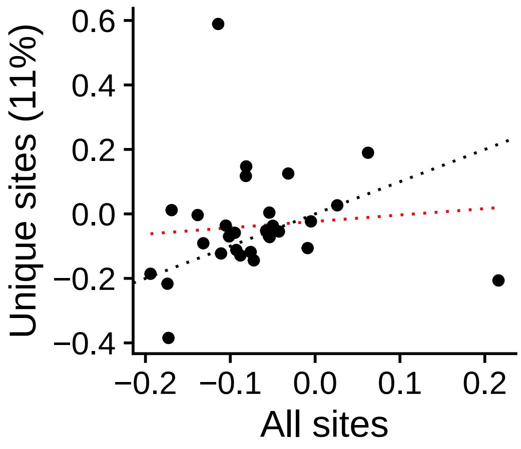


r = 0.095 (p = 0.62398)

**Group IMUNO_**
IRF9_uc_dTSS_nPEAK

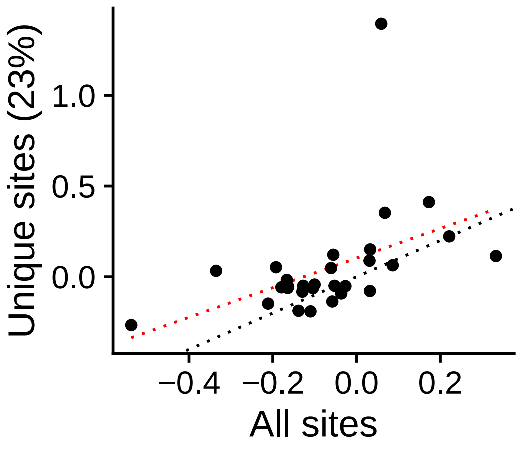


r = 0.454 (p = 0.013401)

**Group IMUNO_**
IRF9_uc_dTSS_oPEAK

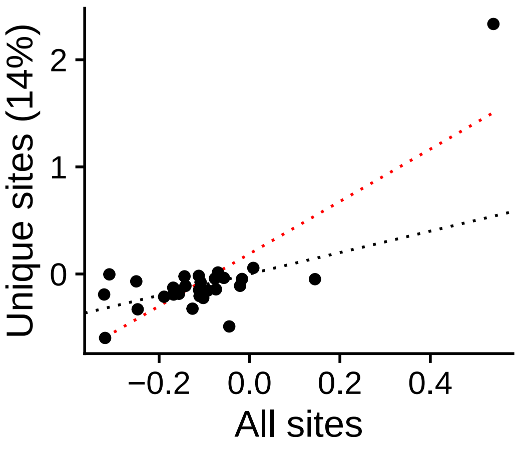


r = 0.808 (p = 1.181e−07)

**Group IMUNO_**
IRF9_uc_pTSS_nPEAK

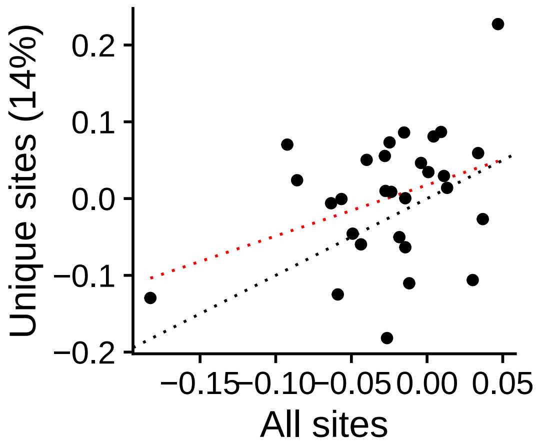


r = 0.364 (p = 0.052301)

**Group IMUNO_**
PRDM1_uc_cTSS_nPEAK

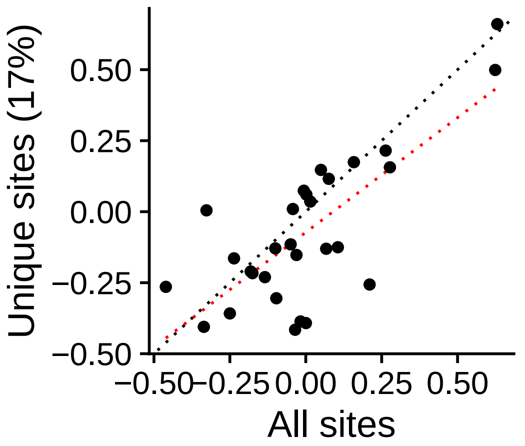


r = 0.754 (p = 2.2871e−06)

**Group IMUNO_**
PRDM1_uc_cTSS_oPEAK

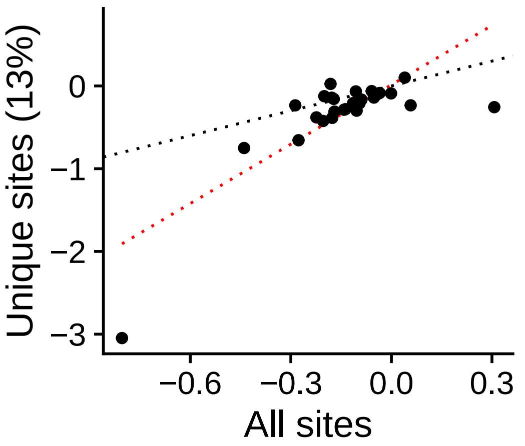


r = 0.787 (p = 4.149e−07)

**Group IMUNO_**
PRDM1_uc_dTSS_nPEAK

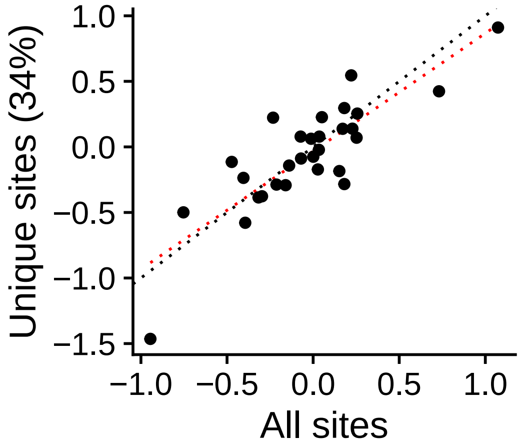


r = 0.845 (p = 8.3078e−09)

**Group IMUNO_**
PRDM1_uc_dTSS_oPEAK

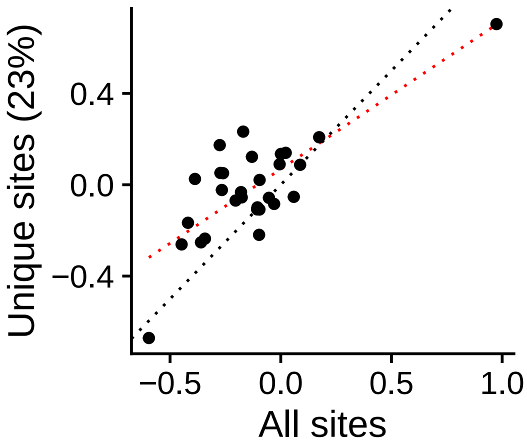


r = 0.784 (p = 4.8787e−07)

**Group IMUNO_**
PRDM1_uc_pTSS_nPEAK

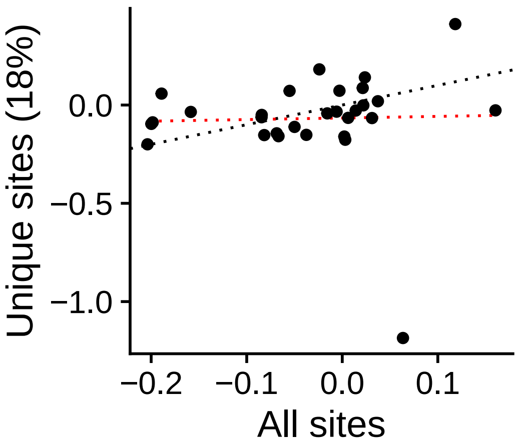


r = 0.0295 (p = 0.87932)

**Group IMUNO_**
PRDM1_uc_pTSS_oPEAK

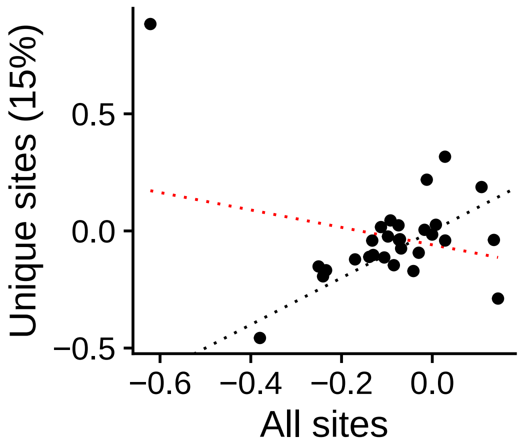


r = −0.25 (p = 0.19063)

**Group IMUNO_**
SPI1_uc_cTSS_nPEAK

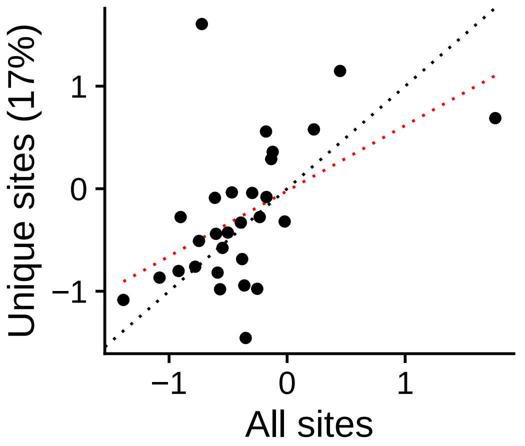


r = 0.507 (p = 0.0049836)

**Group IMUNO_**
SPI1_uc_cTSS_oPEAK

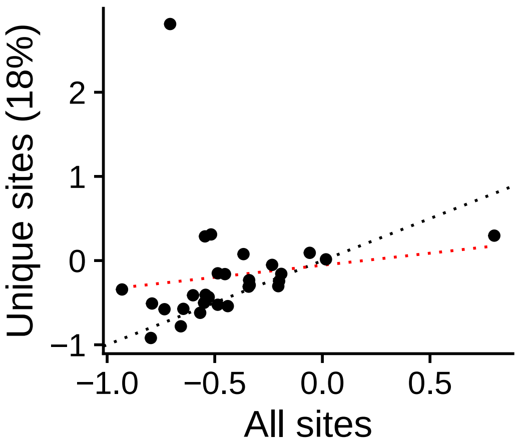


r = 0.143 (p = 0.46044)

**Group IMUNO_**
SPI1_uc_dTSS_nPEAK

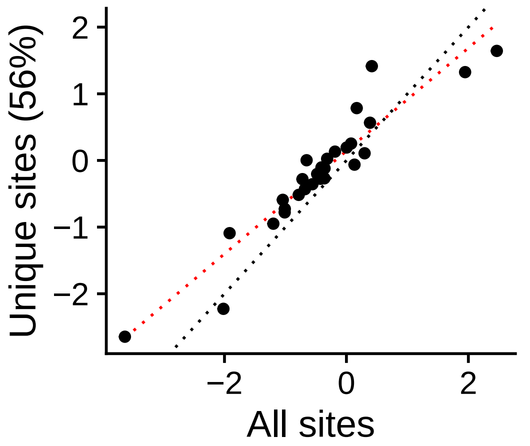


r = 0.941 (p = 3.0355e−14)

**Group IMUNO_**
SPI1_uc_dTSS_oPEAK

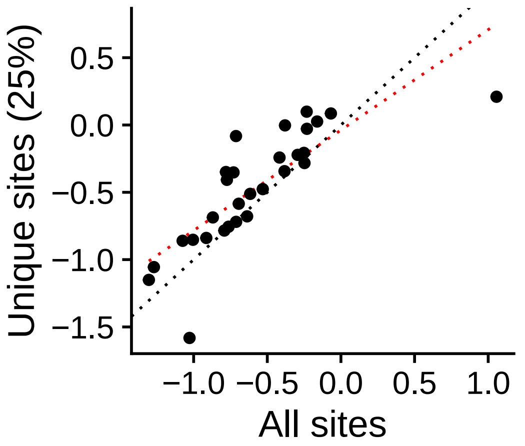


r = 0.805 (p = 1.3925e−07)

**Group IMUNO_**
SPI1_uc_pTSS_nPEAK

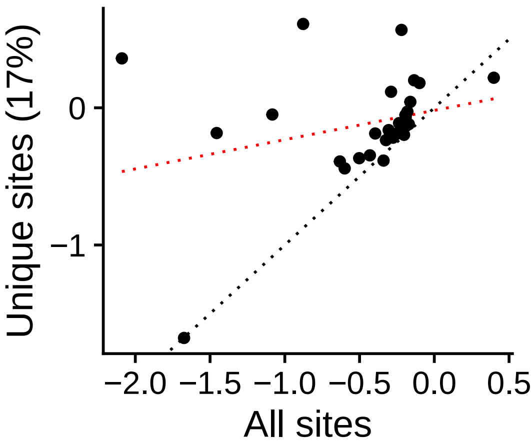


r = 0.277 (p = 0.14547)

**Group IMUNO_**
SPI1_uc_pTSS_oPEAK

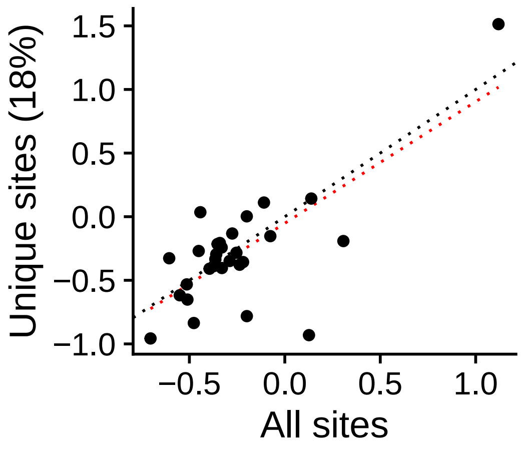


r = 0.735 (p = 5.6173e−06)

**Group IMUNO_**
SPIB_uc_cTSS_nPEAK

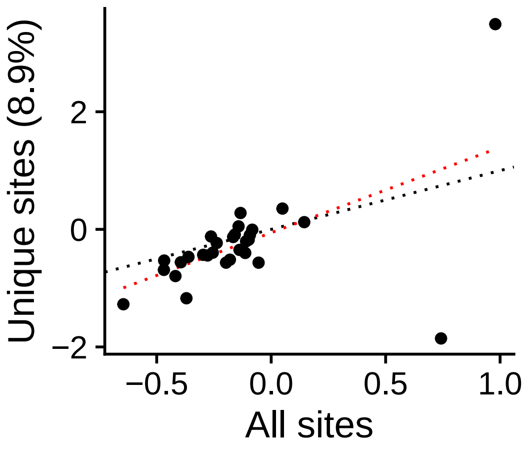


r = 0.551 (p = 0.0019398)

**Group IMUNO_**
SPIB_uc_cTSS_oPEAK

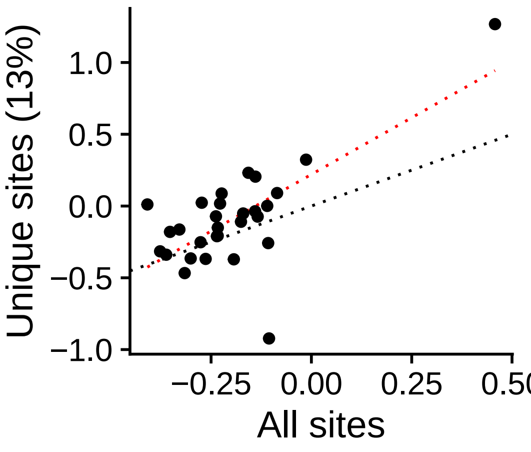


r = 0.701 (p = 2.2945e−05)

**Group IMUNO_**
SPIB_uc_dTSS_oPEAK

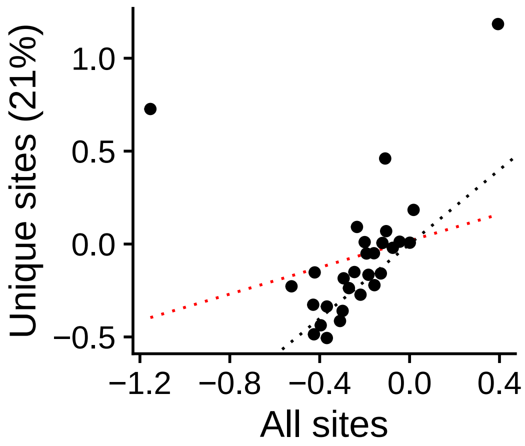


r = 0.25 (p = 0.19084)

**Group IMUNO_**
SPIB_uc_pTSS_nPEAK

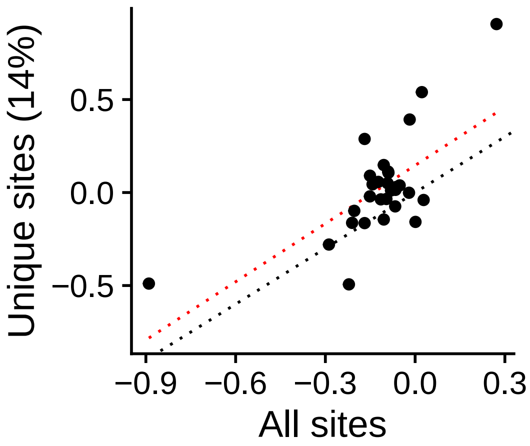


r = 0.687 (p = 3.8158e−05)

**Group IMUNO_**
SPIB_uc_pTSS_oPEAK

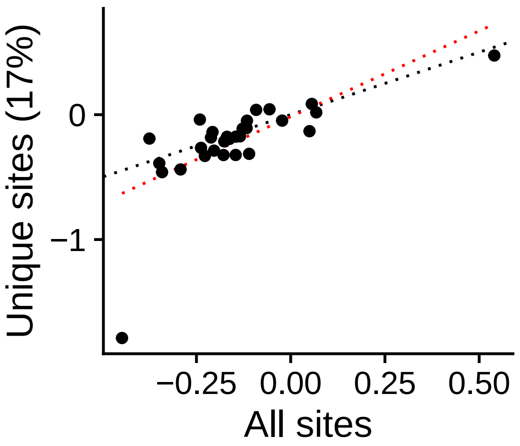


r = 0.694 (p = 2.9417e−05)

**Group IMUNO_**
STAT2_uc_cTSS_nPEAK

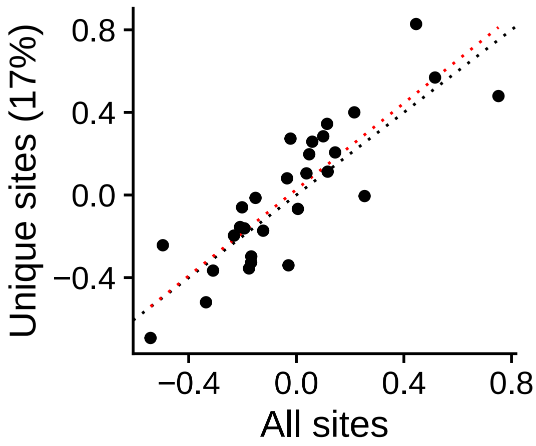


r = 0.855 (p = 3.6127e−09)

**Group IMUNO_**
STAT2_uc_cTSS_oPEAK

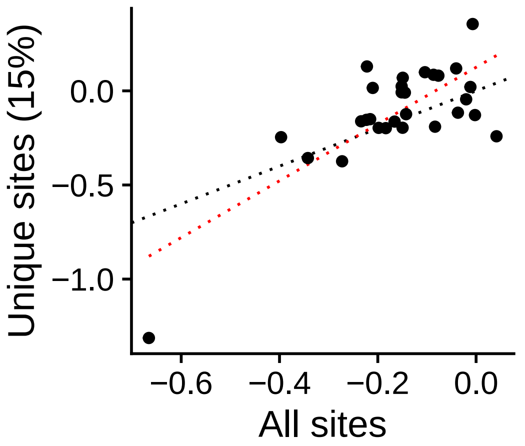


r = 0.761 (p = 1.6654e−06)

**Group IMUNO_**
STAT2_uc_dTSS_nPEAK

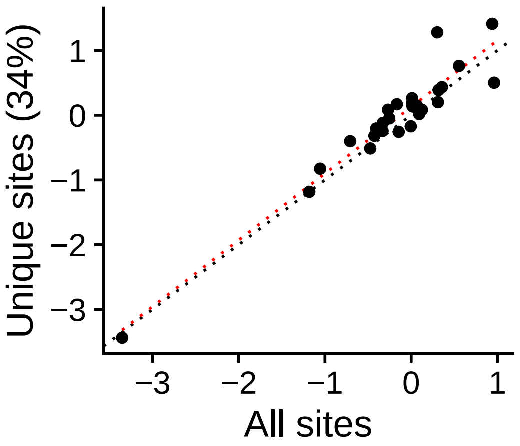


r = 0.955 (p = 8.733e−16)

**Group IMUNO_**
STAT2_uc_dTSS_oPEAK

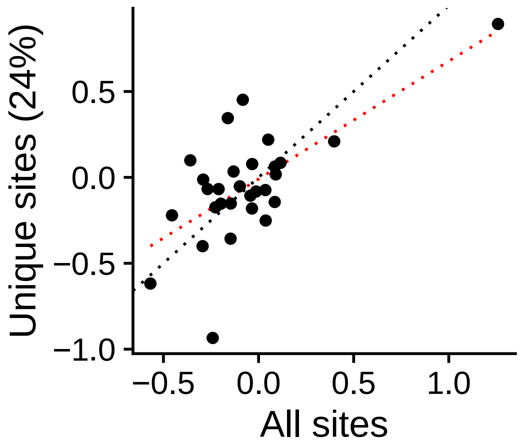


r = 0.674 (p = 6.2109e−05)

**Group IMUNO_**
STAT2_uc_pTSS_nPEAK

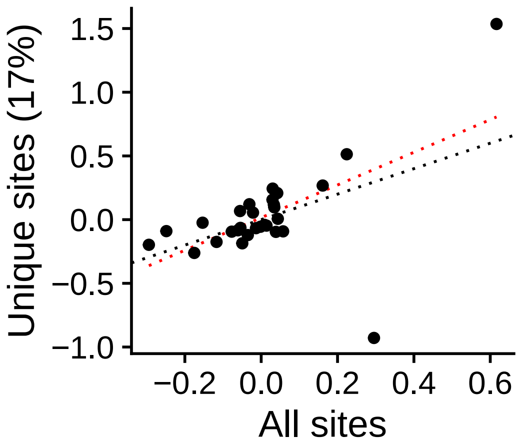


r = 0.574 (p = 0.0011417)

**Group IMUNO_**
STAT2_uc_pTSS_oPEAK

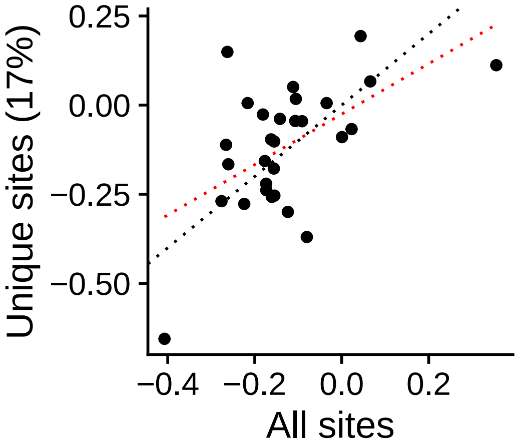


r = 0.558 (p = 0.0016501)

**Group IMUNO_**
THAP1_uc_cTSS_oPEAK

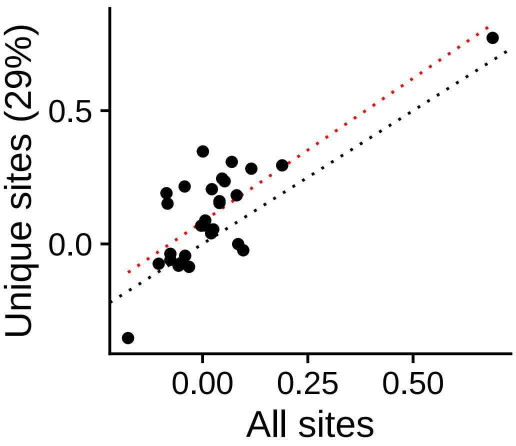


r = 0.792 (p = 3.0206e−07)

**Group IMUNO_**
THAP1_uc_dTSS_nPEAK

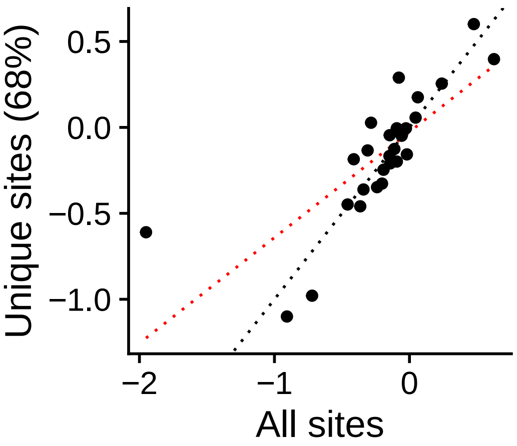


r = 0.77 (p = 1.054e−06)

**Group IMUNO_**
FUBP1_uc_cTSS_nPEAK

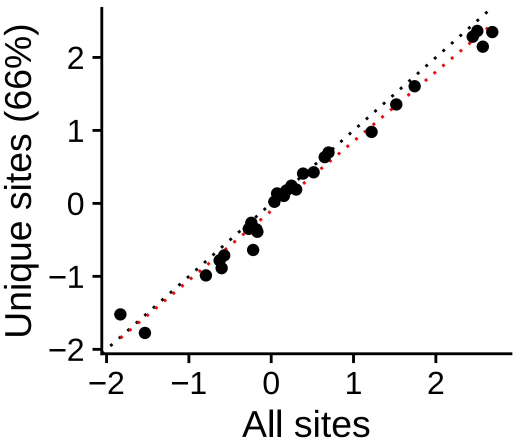


r = 0.992 (p = < 2.22e−16)

**Group IMUNO_**
FUBP1_uc_cTSS_oPEAK

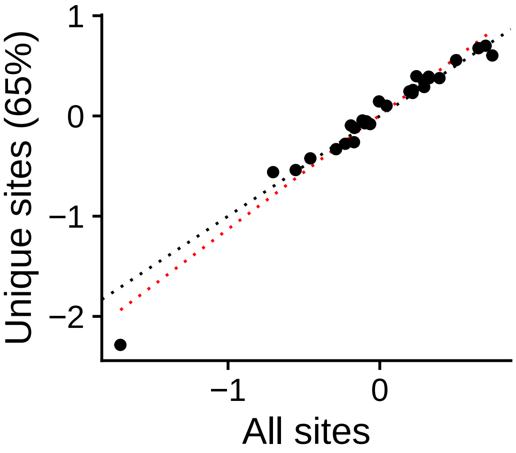


r = 0.981 (p = < 2.22e−16)

**Group IMUNO_**
FUBP1_uc_dTSS_nPEAK

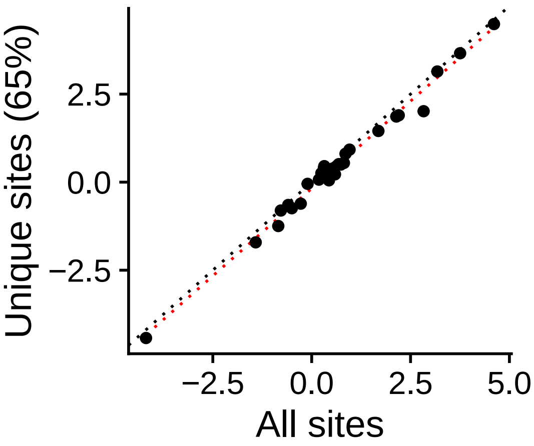


r = 0.994 (p = < 2.22e−16)

**Group IMUNO_**
FUBP1_uc_dTSS_oPEAK

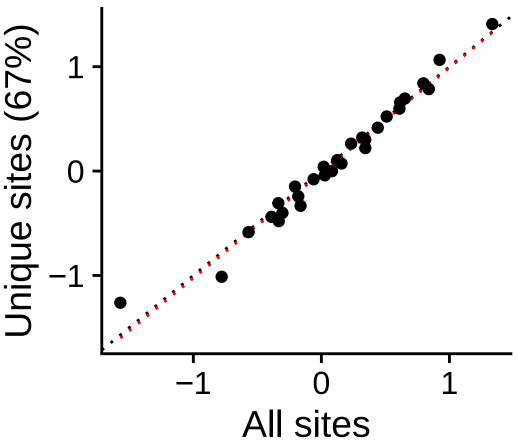


r = 0.986 (p = < 2.22e−16)

**Group IMUNO_**
FUBP1_uc_pTSS_nPEAK

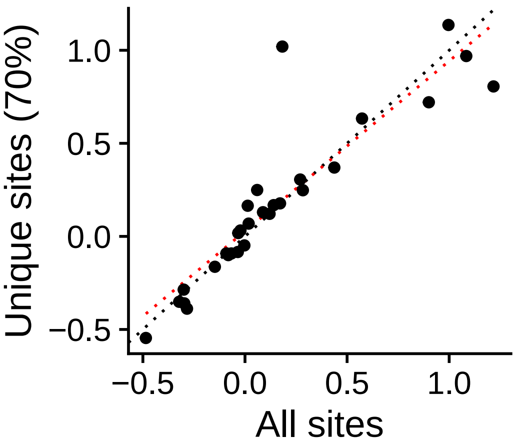


r = 0.902 (p = 2.2391e−11)

**Group IMUNO_**
FUBP1_uc_pTSS_oPEAK

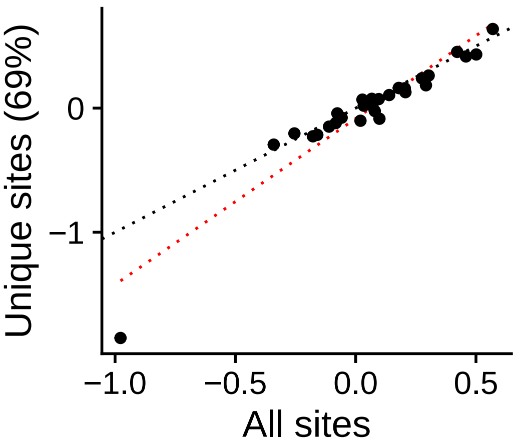


r = 0.95 (p = 3.7129e−15)

**Group IMUNO_**
UBP1_uc_cTSS_nPEAK

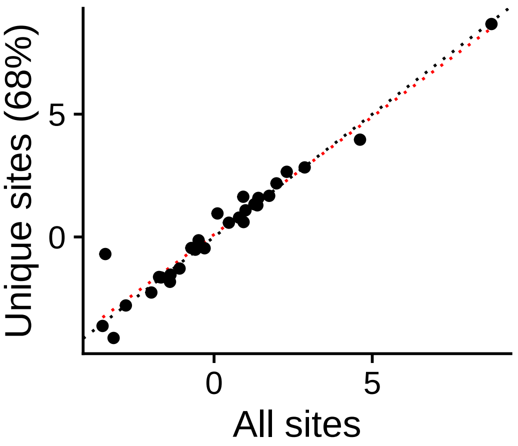


r = 0.971 (p = < 2.22e−16)

**Group IMUNO_**
UBP1_uc_cTSS_oPEAK

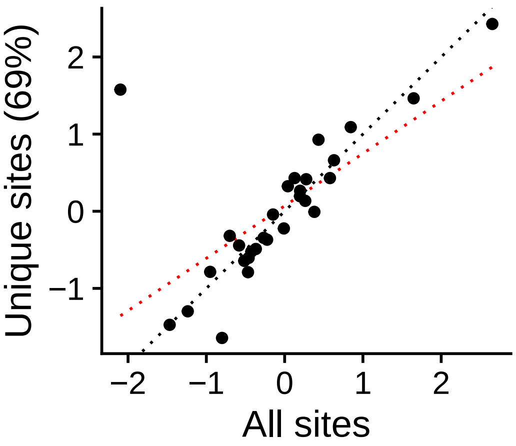


r = 0.675 (p = 5.9116e−05)

**Group IMUNO_**
UBP1_uc_dTSS_oPEAK

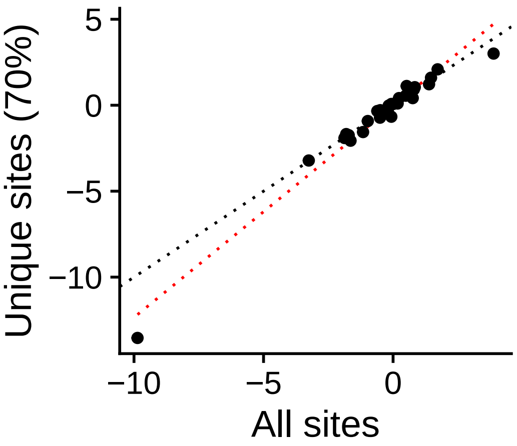


r = 0.982 (p = < 2.22e−16)

**Group IMUNO_**
UBP1_uc_pTSS_nPEAK

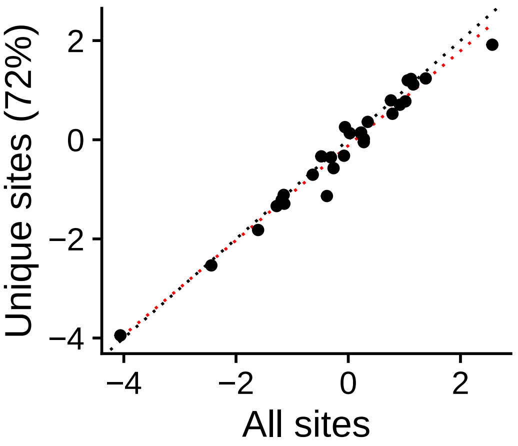


r = 0.985 (p = < 2.22e−16)

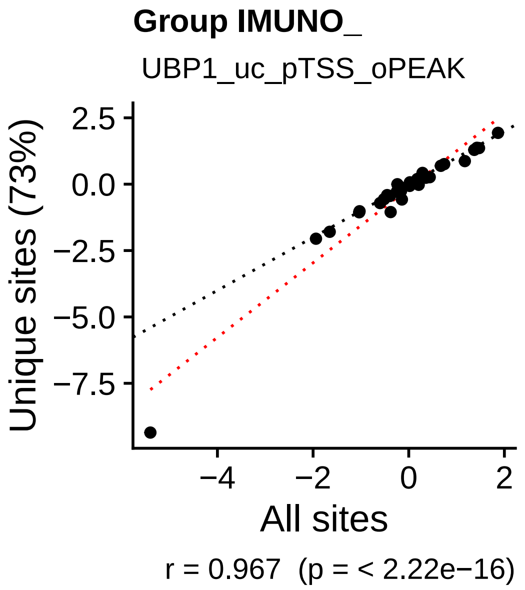
Group IMUNO_
UBP1_uc_pTSS_oPEAK
Unique sites (73%)
All sites
r = 0.967  (p = < 2.22e−16)

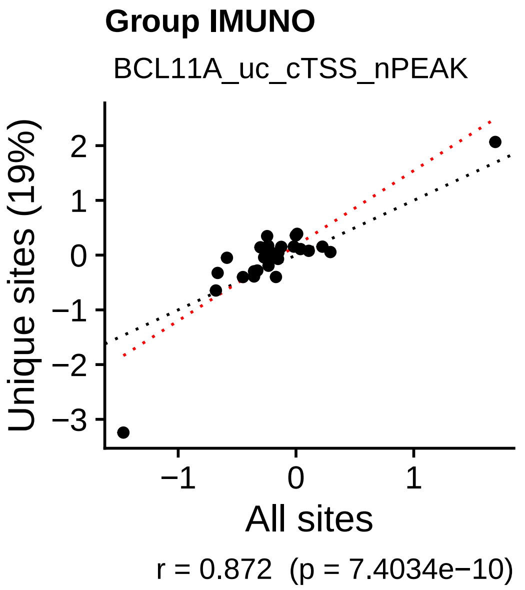
Group IMUNO
BCL11A_uc_cTSS_nPEAK
Unique sites (19%)
All sites
r = 0.872  (p = 7.4034e−10)

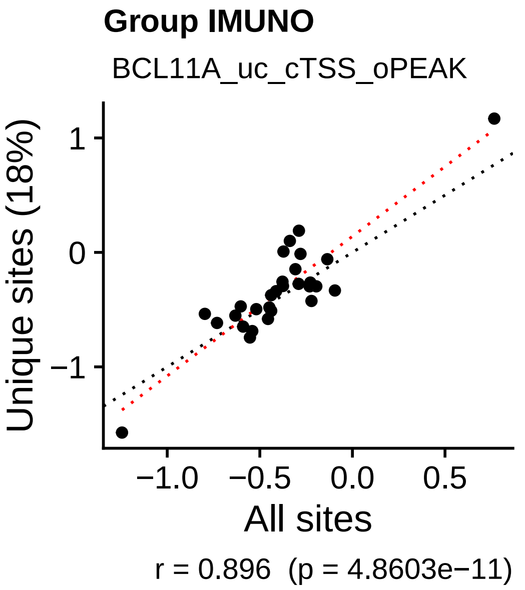
Group IMUNO
BCL11A_uc_cTSS_oPEAK
Unique sites (18%)
All sites
r = 0.896  (p = 4.8603e−11)

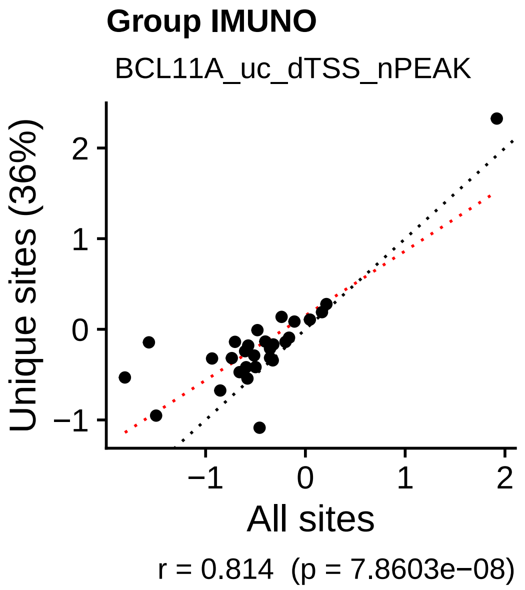
Group IMUNO
BCL11A_uc_dTSS_nPEAK
Unique sites (36%)
All sites
r = 0.814  (p = 7.8603e−08)

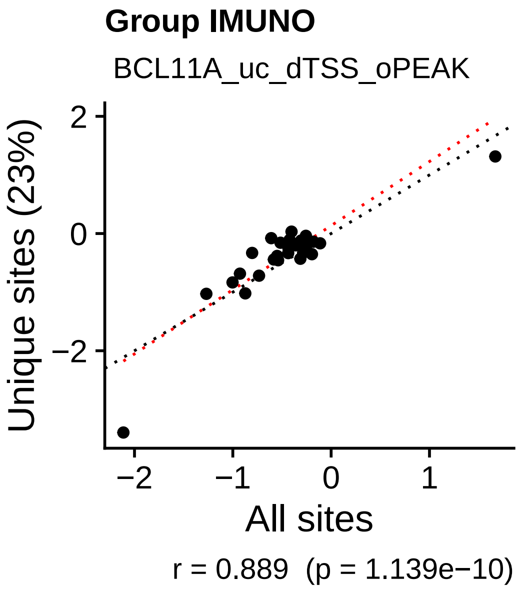
Group IMUNO
BCL11A_uc_dTSS_oPEAK
Unique sites (23%)
All sites
r = 0.889  (p = 1.139e−10)

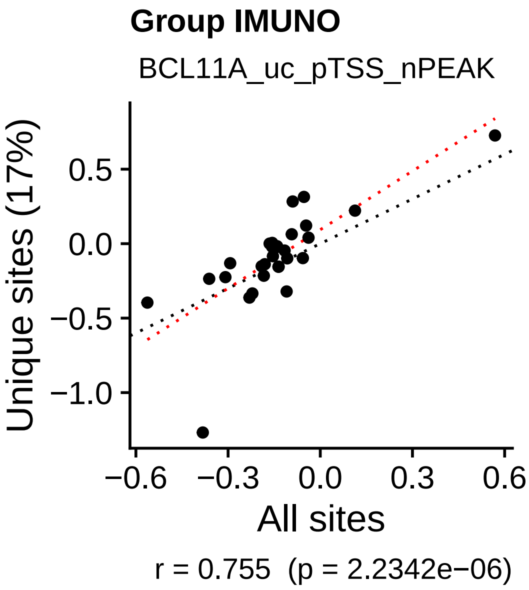
Group IMUNO
BCL11A_uc_pTSS_nPEAK
Unique sites (17%)
All sites
r = 0.755  (p = 2.2342e−06)

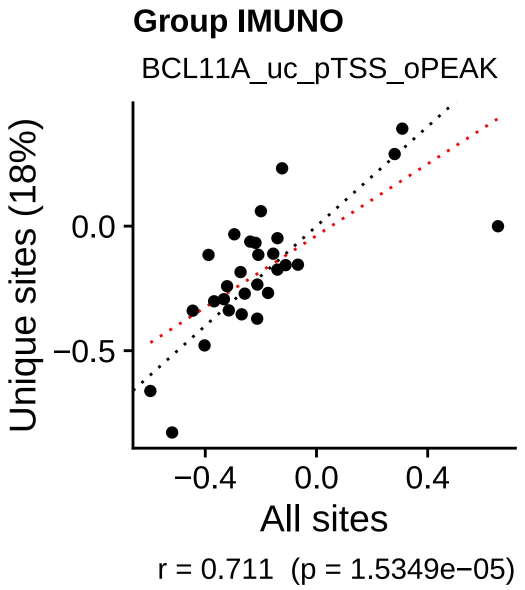
Group IMUNO
BCL11A_uc_pTSS_oPEAK
Unique sites (18%)
All sites
r = 0.711  (p = 1.5349e−05)

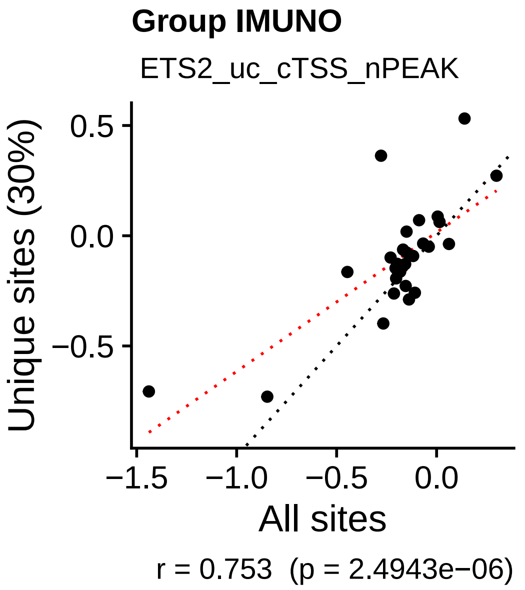
Group IMUNO
ETS2_uc_cTSS_nPEAK
Unique sites (30%)
All sites
r = 0.753  (p = 2.4943e−06)

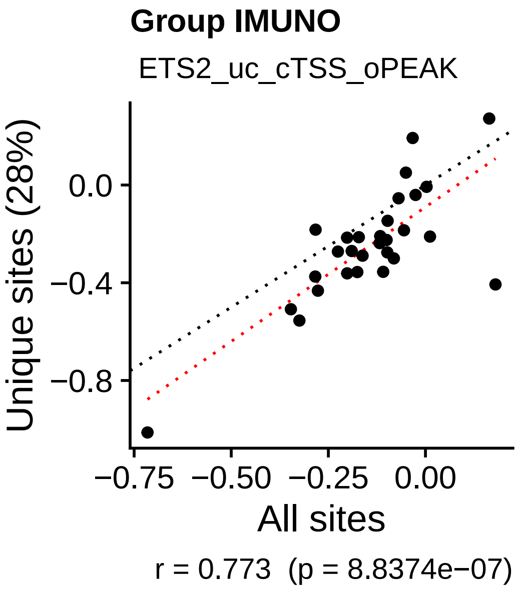
Group IMUNO
ETS2_uc_cTSS_oPEAK
Unique sites (28%)
All sites
r = 0.773  (p = 8.8374e−07)

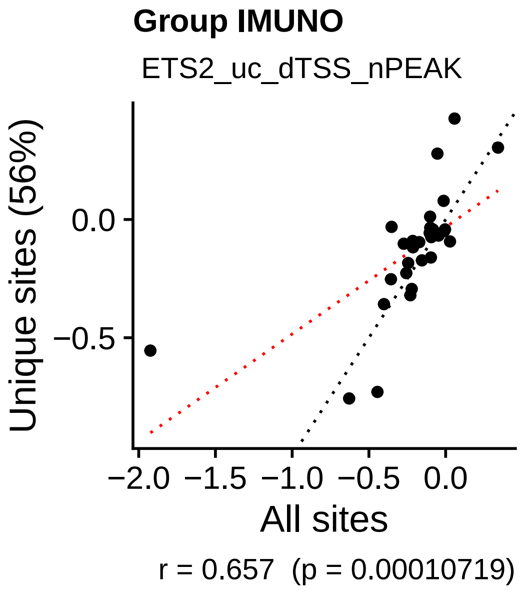
Group IMUNO
ETS2_uc_dTSS_nPEAK
Unique sites (56%)
All sites
r = 0.657  (p = 0.00010719)

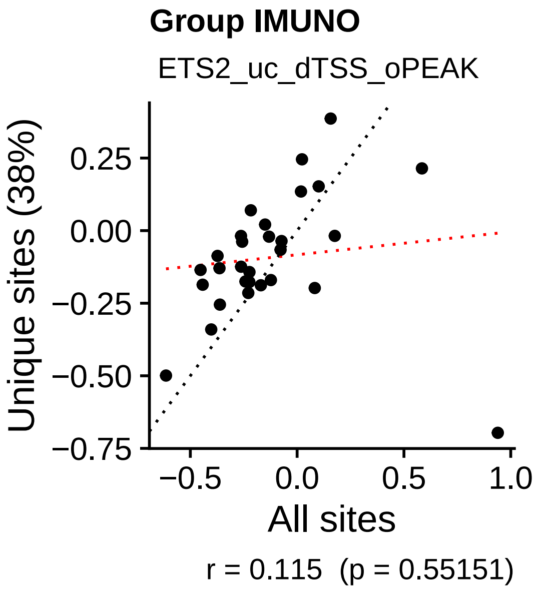
Group IMUNO
ETS2_uc_dTSS_oPEAK
Unique sites (38%)
All sites
r = 0.115  (p = 0.55151)

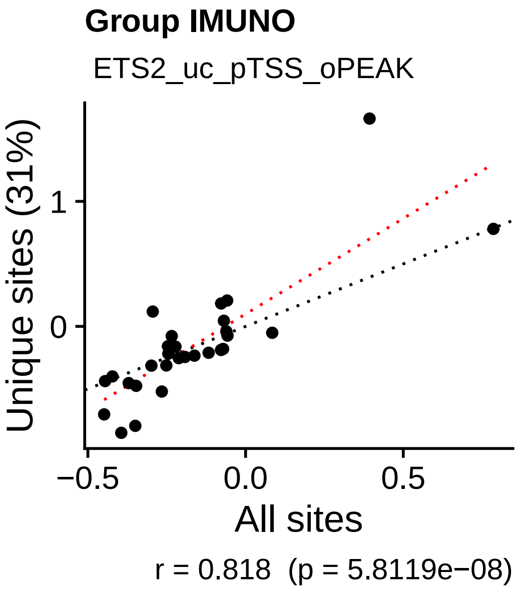
Group IMUNO
ETS2_uc_pTSS_oPEAK
Unique sites (31%)
All sites
r = 0.818  (p = 5.8119e−08)

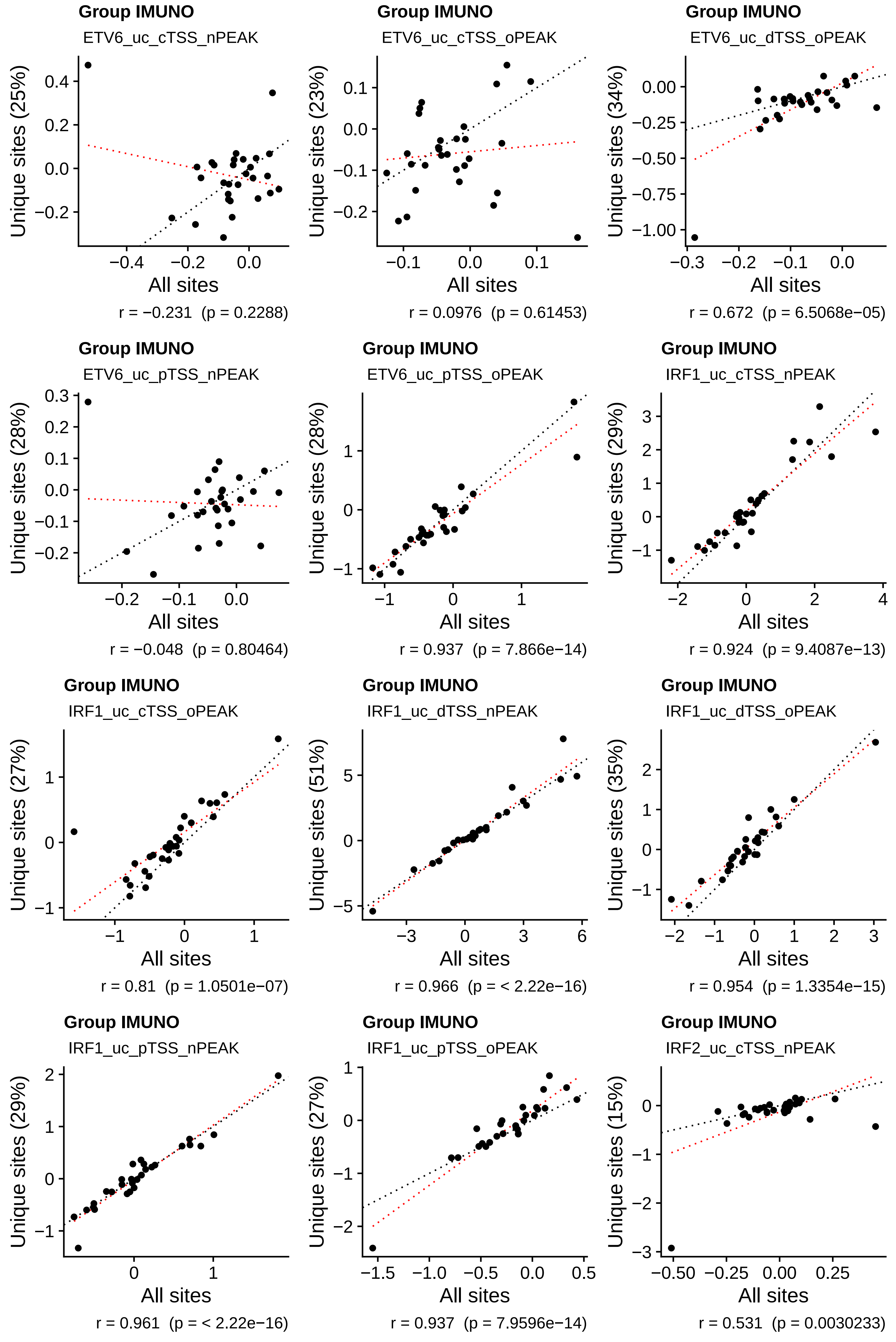

Group IMUNO
ETV6_uc_cTSS_nPEAK
Unique sites (25%)
All sites
r = −0.231 (p = 0.2288)
Group IMUNO
ETV6_uc_cTSS_oPEAK
Unique sites (23%)
All sites
r = 0.0976 (p = 0.61453)
Group IMUNO
ETV6_uc_dTSS_oPEAK
Unique sites (34%)
All sites
r = 0.672 (p = 6.5068e−05)
Group IMUNO
ETV6_uc_pTSS_nPEAK
Unique sites (28%)
All sites
r = −0.048 (p = 0.80464)
Group IMUNO
ETV6_uc_pTSS_oPEAK
Unique sites (28%)
All sites
r = 0.937 (p = 7.866e−14)
Group IMUNO
IRF1_uc_cTSS_nPEAK
Unique sites (29%)
All sites
r = 0.924 (p = 9.4087e−13)
Group IMUNO
IRF1_uc_cTSS_oPEAK
Unique sites (27%)
All sites
r = 0.81 (p = 1.0501e−07)
Group IMUNO
IRF1_uc_dTSS_nPEAK
Unique sites (51%)
All sites
r = 0.966 (p = < 2.22e−16)
Group IMUNO
IRF1_uc_dTSS_oPEAK
Unique sites (35%)
All sites
r = 0.954 (p = 1.3354e−15)
Group IMUNO
IRF1_uc_pTSS_nPEAK
Unique sites (29%)
All sites
r = 0.961 (p = < 2.22e−16)
Group IMUNO
IRF1_uc_pTSS_oPEAK
Unique sites (27%)
All sites
r = 0.937 (p = 7.9596e−14)
Group IMUNO
IRF2_uc_cTSS_nPEAK
Unique sites (15%)
All sites
r = 0.531 (p = 0.0030233)

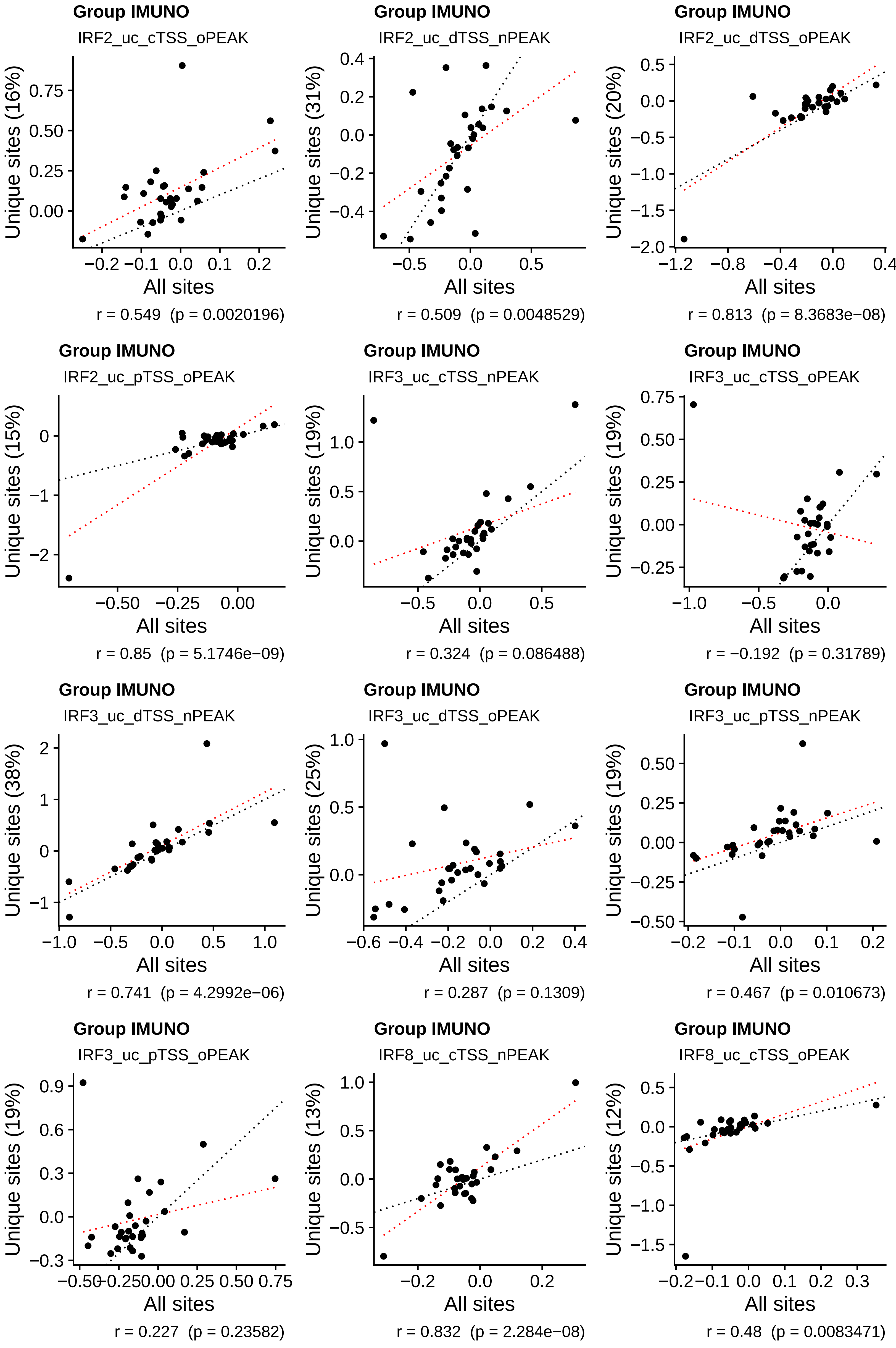
Group IMUNO
IRF2_uc_cTSS_oPEAK
Unique sites (16%)
All sites
r = 0.549 (p = 0.0020196)
Group IMUNO
IRF2_uc_dTSS_nPEAK
Unique sites (31%)
All sites
r = 0.509 (p = 0.0048529)
Group IMUNO
IRF2_uc_dTSS_oPEAK
Unique sites (20%)
All sites
r = 0.813 (p = 8.3683e−08)
Group IMUNO
IRF2_uc_pTSS_oPEAK
Unique sites (15%)
All sites
r = 0.85 (p = 5.1746e−09)
Group IMUNO
IRF3_uc_cTSS_nPEAK
Unique sites (19%)
All sites
r = 0.324 (p = 0.086488)
Group IMUNO
IRF3_uc_cTSS_oPEAK
Unique sites (19%)
All sites
r = −0.192 (p = 0.31789)
Group IMUNO
IRF3_uc_dTSS_nPEAK
Unique sites (38%)
All sites
r = 0.741 (p = 4.2992e−06)
Group IMUNO
IRF3_uc_dTSS_oPEAK
Unique sites (25%)
All sites
r = 0.287 (p = 0.1309)
Group IMUNO
IRF3_uc_pTSS_nPEAK
Unique sites (19%)
All sites
r = 0.467 (p = 0.010673)
Group IMUNO
IRF3_uc_pTSS_oPEAK
Unique sites (19%)
All sites
r = 0.227 (p = 0.23582)
Group IMUNO
IRF8_uc_cTSS_nPEAK
Unique sites (13%)
All sites
r = 0.832 (p = 2.284e−08)
Group IMUNO
IRF8_uc_cTSS_oPEAK
Unique sites (12%)
All sites
r = 0.48 (p = 0.0083471)

**Group IMUNO**
IRF8_uc_dTSS_nPEAK

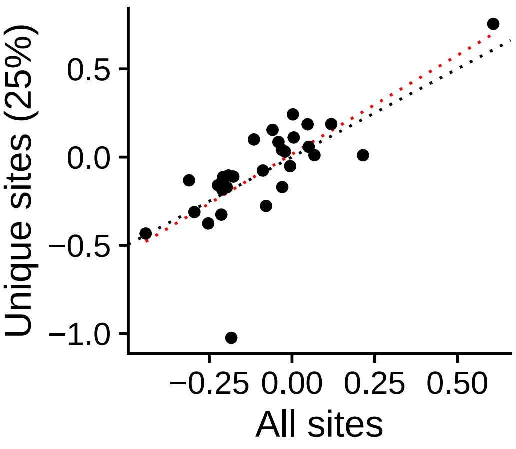


r = 0.739 (p = 4.7174e−06)

**Group IMUNO**
IRF8_uc_dTSS_oPEAK

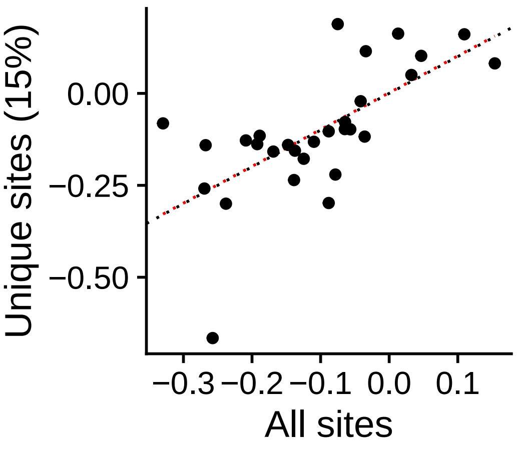


r = 0.661 (p = 9.3506e−05)

**Group IMUNO**
IRF8_uc_pTSS_nPEAK

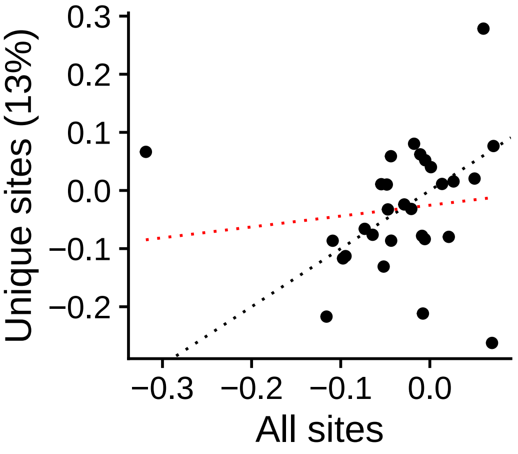


r = 0.129 (p = 0.50555)

**Group IMUNO**
IRF9_uc_cTSS_nPEAK

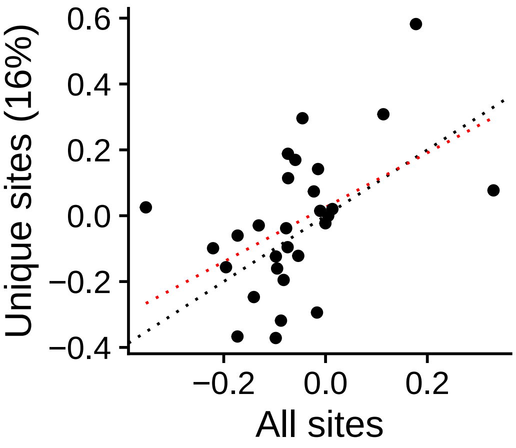


r = 0.481 (p = 0.0082344)

**Group IMUNO**
IRF9_uc_cTSS_oPEAK

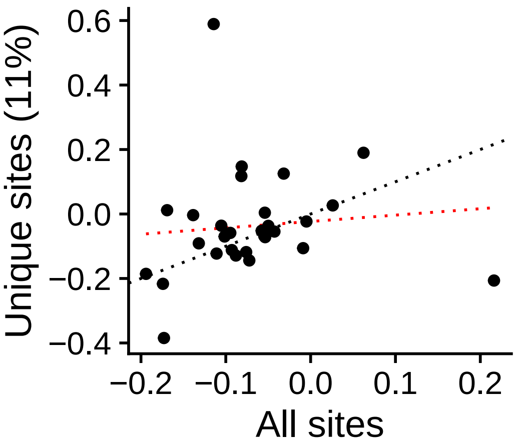


r = 0.095 (p = 0.62398)

**Group IMUNO**
IRF9_uc_dTSS_nPEAK

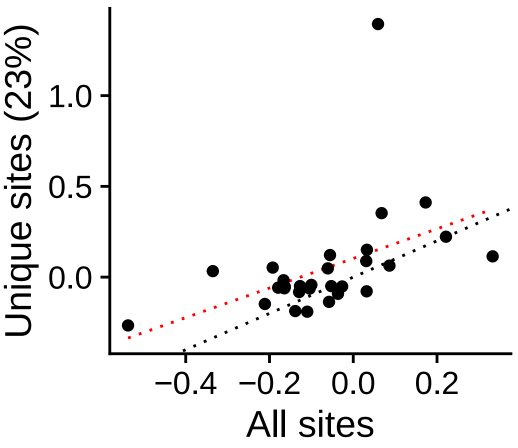


r = 0.454 (p = 0.013401)

**Group IMUNO**
IRF9_uc_dTSS_oPEAK

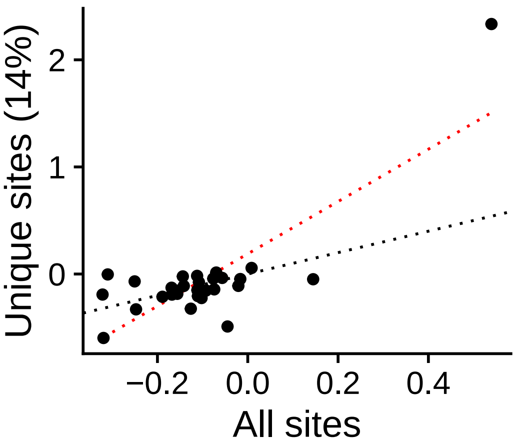


r = 0.808 (p = 1.181e−07)

**Group IMUNO**
IRF9_uc_pTSS_nPEAK

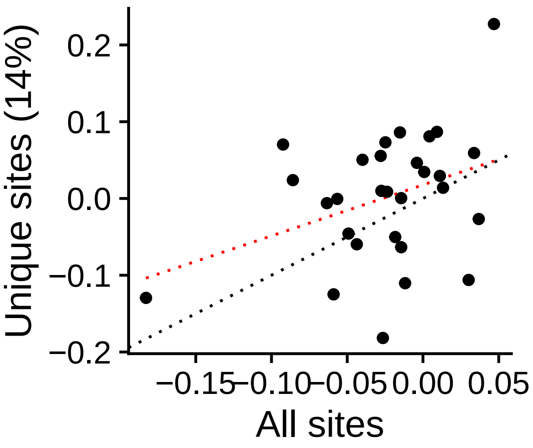


r = 0.364 (p = 0.052301)

**Group IMUNO**
STAT2_uc_cTSS_nPEAK

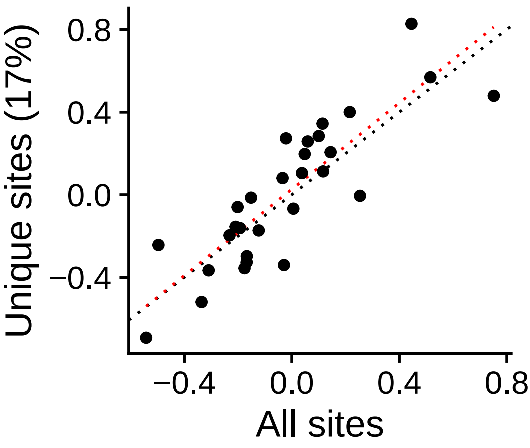


r = 0.855 (p = 3.6127e−09)

**Group IMUNO**
STAT2_uc_cTSS_oPEAK

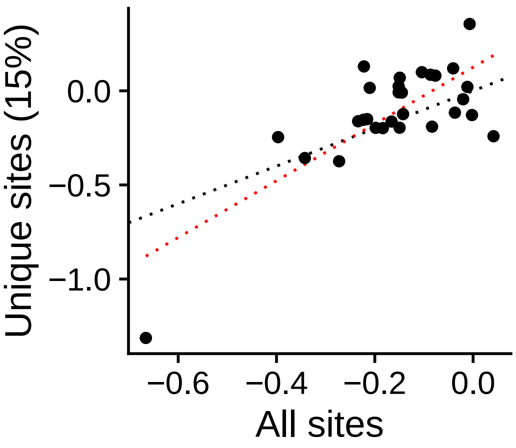


r = 0.761 (p = 1.6654e−06)

**Group IMUNO**
STAT2_uc_dTSS_nPEAK

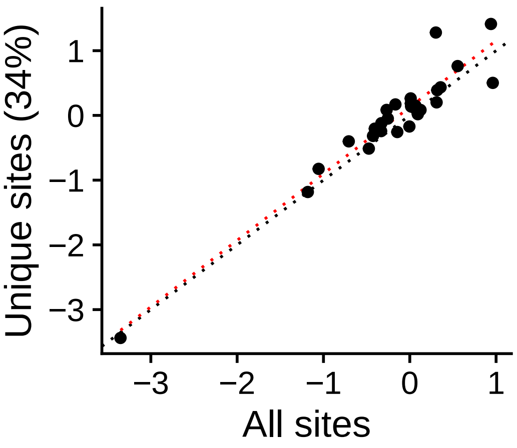


r = 0.955 (p = 8.733e−16)

**Group IMUNO**
STAT2_uc_dTSS_oPEAK

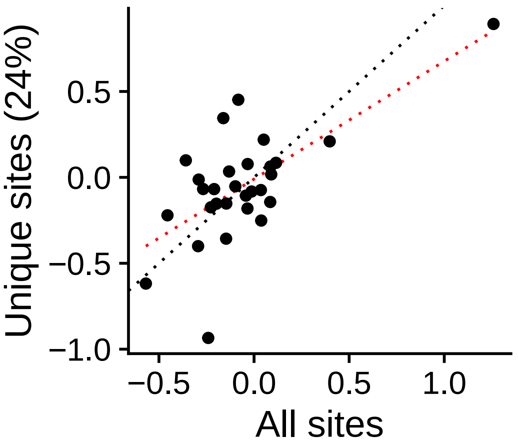


r = 0.674 (p = 6.2109e−05)

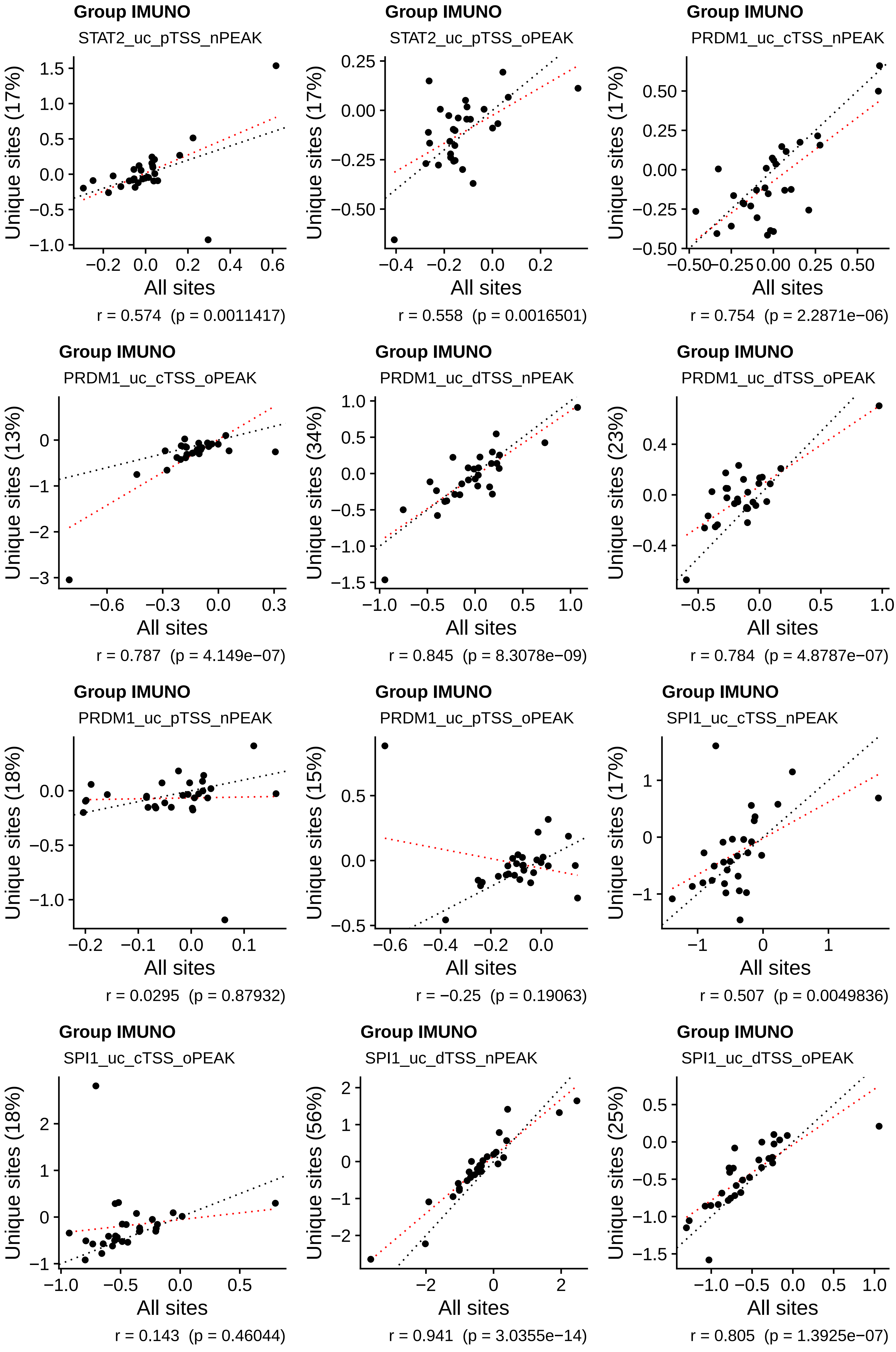
Group IMUNO
STAT2_uc_pTSS_nPEAK
Unique sites (17%)
All sites
r = 0.574 (p = 0.0011417)
Group IMUNO
STAT2_uc_pTSS_oPEAK
Unique sites (17%)
All sites
r = 0.558 (p = 0.0016501)
Group IMUNO
PRDM1_uc_cTSS_nPEAK
Unique sites (17%)
All sites
r = 0.754 (p = 2.2871e−06)
Group IMUNO
PRDM1_uc_cTSS_oPEAK
Unique sites (13%)
All sites
r = 0.787 (p = 4.149e−07)
Group IMUNO
PRDM1_uc_dTSS_nPEAK
Unique sites (34%)
All sites
r = 0.845 (p = 8.3078e−09)
Group IMUNO
PRDM1_uc_dTSS_oPEAK
Unique sites (23%)
All sites
r = 0.784 (p = 4.8787e−07)
Group IMUNO
PRDM1_uc_pTSS_nPEAK
Unique sites (18%)
All sites
r = 0.0295 (p = 0.87932)
Group IMUNO
PRDM1_uc_pTSS_oPEAK
Unique sites (15%)
All sites
r = −0.25 (p = 0.19063)
Group IMUNO
SPI1_uc_cTSS_nPEAK
Unique sites (17%)
All sites
r = 0.507 (p = 0.0049836)
Group IMUNO
SPI1_uc_cTSS_oPEAK
Unique sites (18%)
All sites
r = 0.143 (p = 0.46044)
Group IMUNO
SPI1_uc_dTSS_nPEAK
Unique sites (56%)
All sites
r = 0.941 (p = 3.0355e−14)
Group IMUNO
SPI1_uc_dTSS_oPEAK
Unique sites (25%)
All sites
r = 0.805 (p = 1.3925e−07)

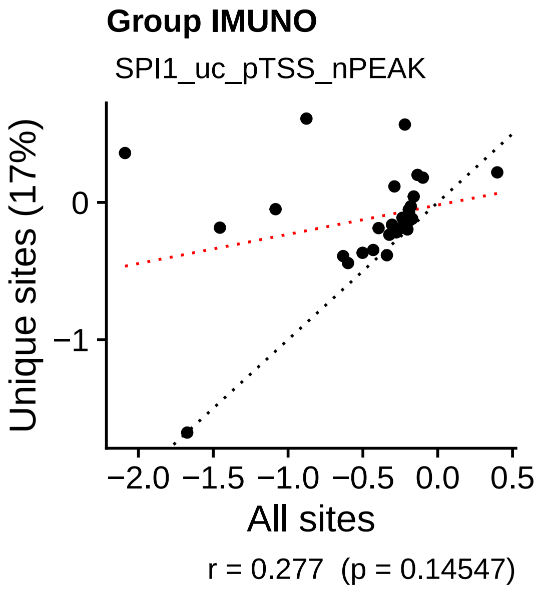
Group IMUNO
SPI1_uc_pTSS_nPEAK
Unique sites (17%)
All sites
r = 0.277 (p = 0.14547)

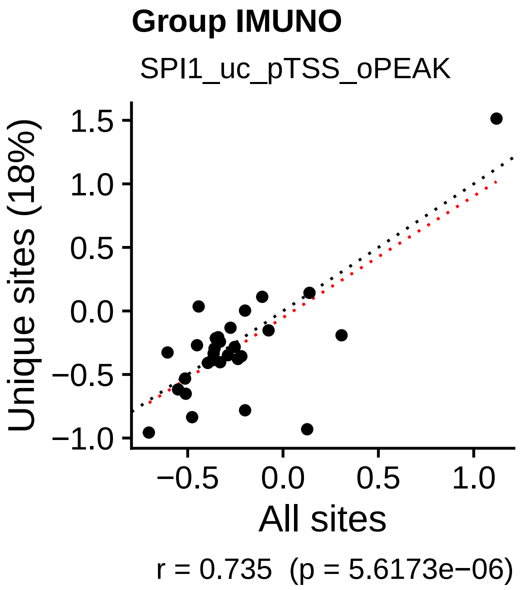
Group IMUNO
SPI1_uc_pTSS_oPEAK
Unique sites (18%)
All sites
r = 0.735 (p = 5.6173e−06)

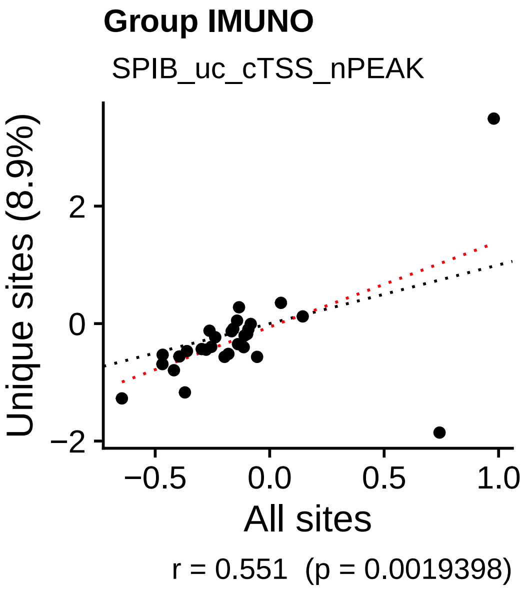
Group IMUNO
SPIB_uc_cTSS_nPEAK
Unique sites (8.9%)
All sites
r = 0.551 (p = 0.0019398)

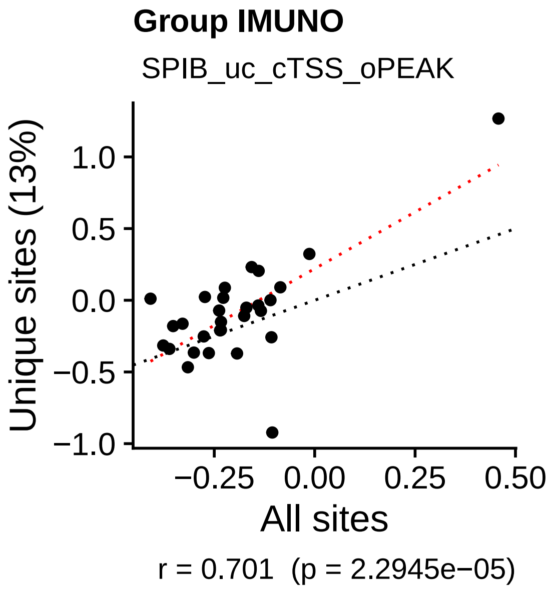
Group IMUNO
SPIB_uc_cTSS_oPEAK
Unique sites (13%)
All sites
r = 0.701 (p = 2.2945e−05)

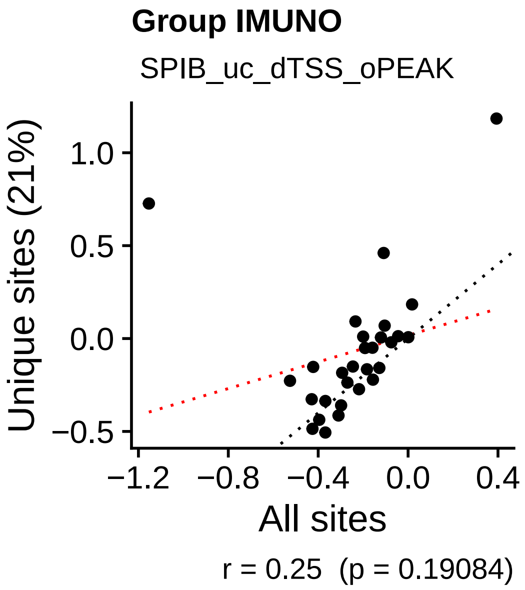
Group IMUNO
SPIB_uc_dTSS_oPEAK
Unique sites (21%)
All sites
r = 0.25 (p = 0.19084)

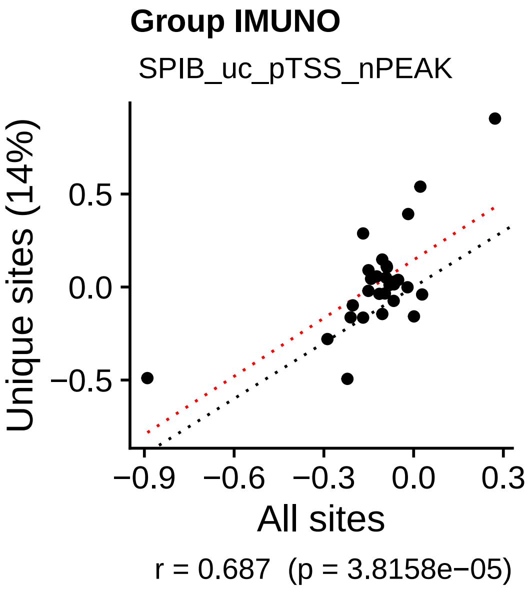
Group IMUNO
SPIB_uc_pTSS_nPEAK
Unique sites (14%)
All sites
r = 0.687 (p = 3.8158e−05)

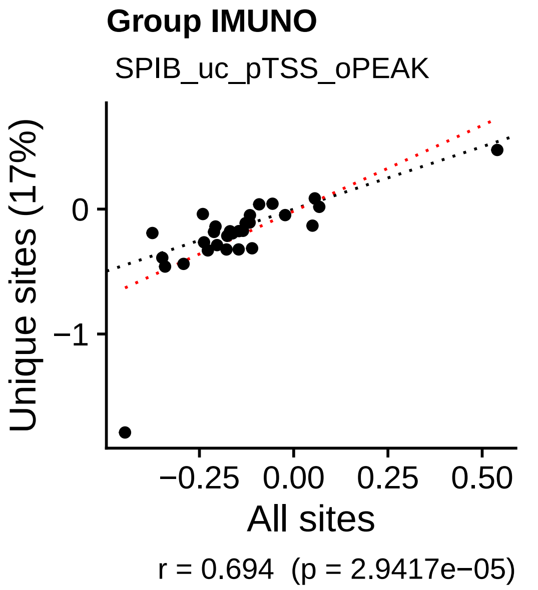
Group IMUNO
SPIB_uc_pTSS_oPEAK
Unique sites (17%)
All sites
r = 0.694 (p = 2.9417e−05)

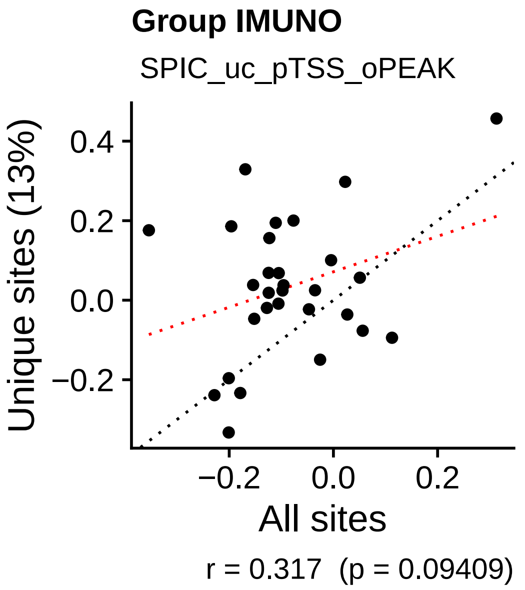
Group IMUNO
SPIC_uc_pTSS_oPEAK
Unique sites (13%)
All sites
r = 0.317 (p = 0.09409)

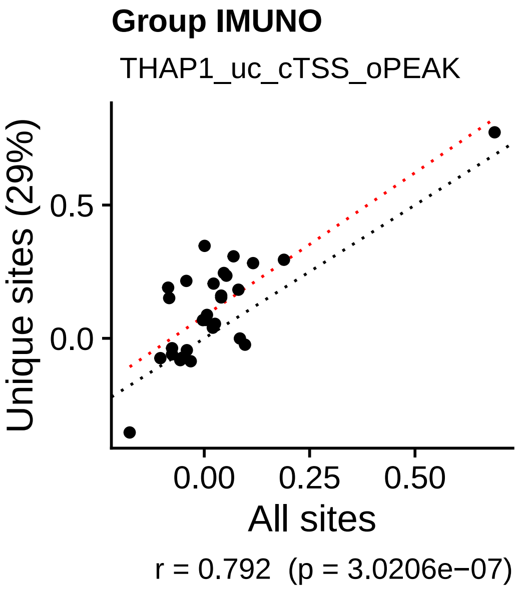
Group IMUNO
THAP1_uc_cTSS_oPEAK
Unique sites (29%)
All sites
r = 0.792 (p = 3.0206e−07)

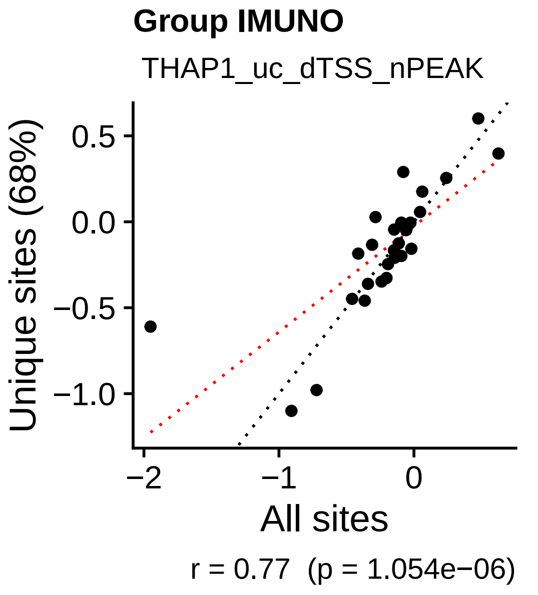
Group IMUNO
THAP1_uc_dTSS_nPEAK
Unique sites (68%)
All sites
r = 0.77 (p = 1.054e−06)

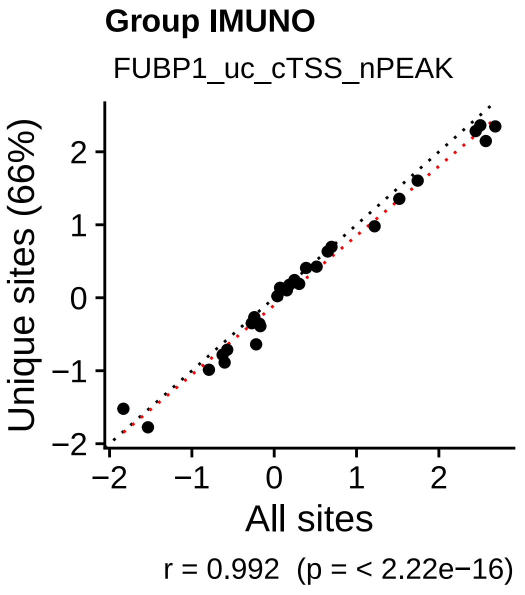
Group IMUNO
FUBP1_uc_cTSS_nPEAK
Unique sites (66%)
All sites
r = 0.992 (p = < 2.22e−16)

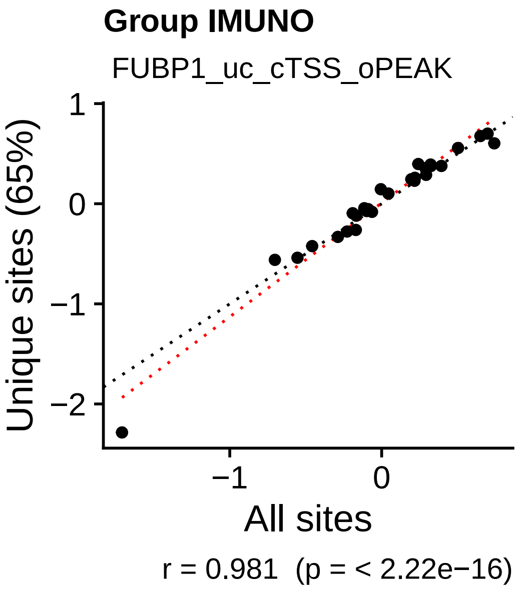
Group IMUNO
FUBP1_uc_cTSS_oPEAK
Unique sites (65%)
All sites
r = 0.981 (p = < 2.22e−16)

**Group IMUNO**

FUBP1_uc_dTSS_nPEAK

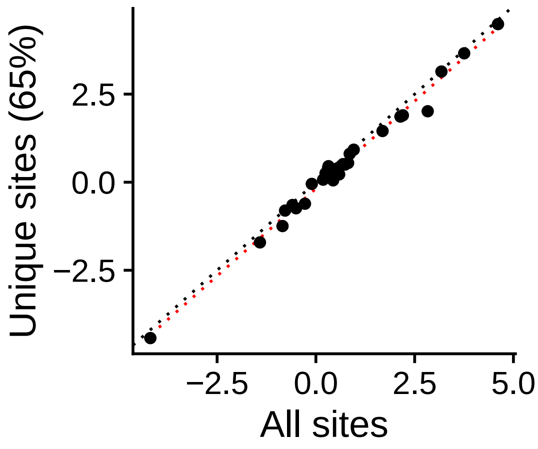


r = 0.994 (p = < 2.22e−16)

**Group IMUNO**

FUBP1_uc_dTSS_oPEAK

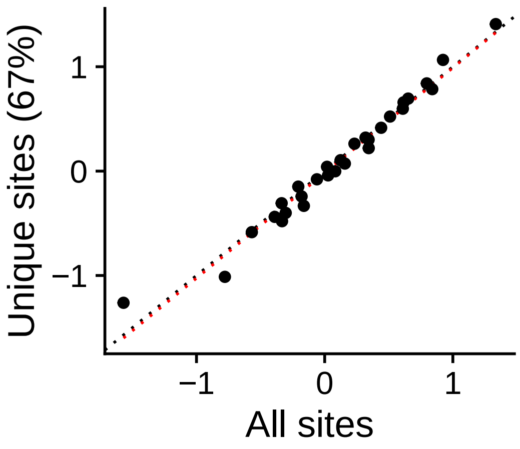


r = 0.986 (p = < 2.22e−16)

**Group IMUNO**

FUBP1_uc_pTSS_nPEAK

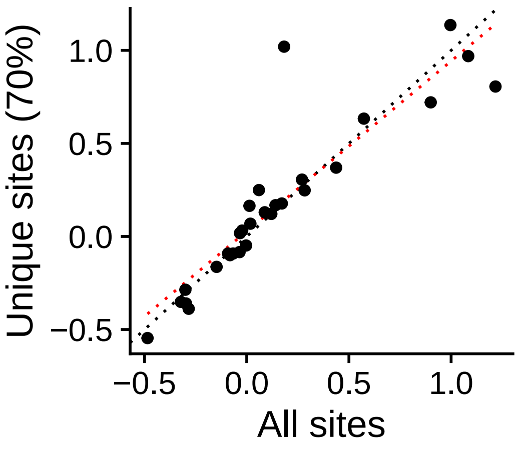


r = 0.902 (p = 2.2391e−11)

**Group IMUNO**

FUBP1_uc_pTSS_oPEAK

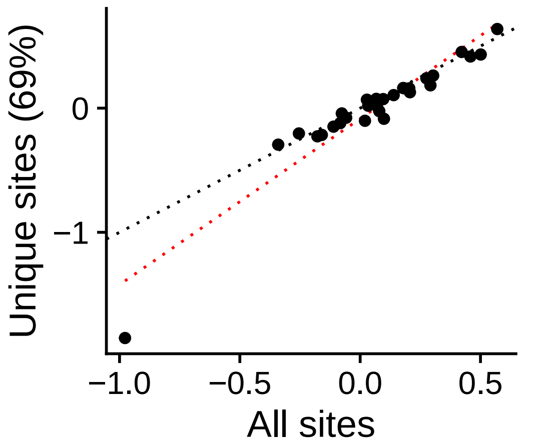


r = 0.95 (p = 3.7129e−15)

**Group IMUNO**

UBP1_uc_cTSS_nPEAK

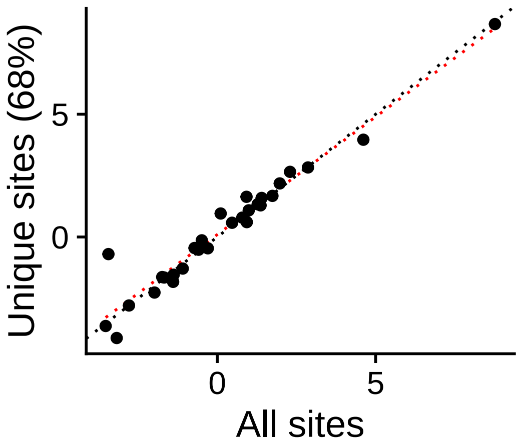


r = 0.971 (p = < 2.22e−16)

**Group IMUNO**

UBP1_uc_cTSS_oPEAK

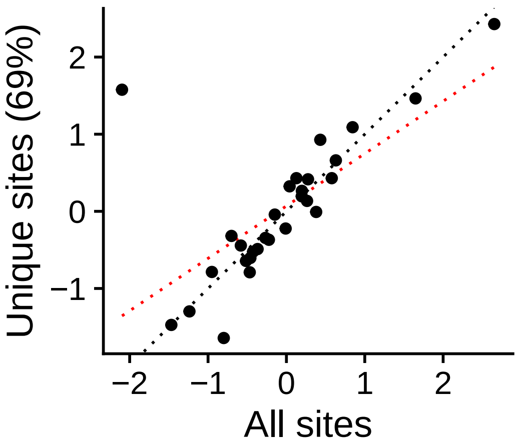


r = 0.675 (p = 5.9116e−05)

**Group IMUNO**

UBP1_uc_dTSS_oPEAK

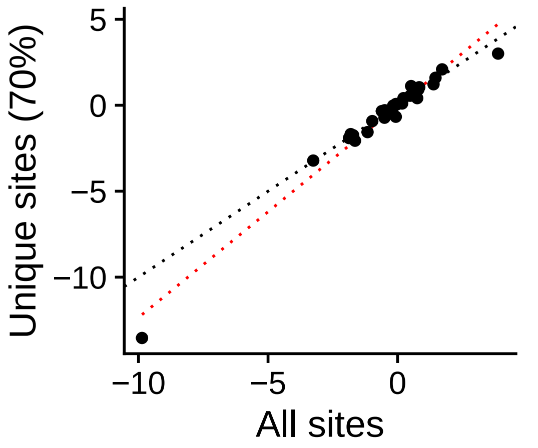


r = 0.982 (p = < 2.22e−16)

**Group IMUNO**

UBP1_uc_pTSS_nPEAK

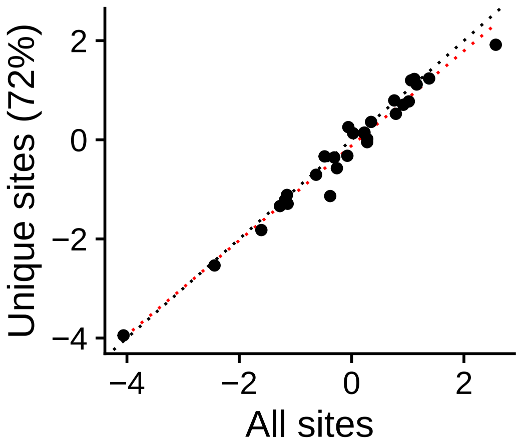


r = 0.985 (p = < 2.22e−16)

**Group IMUNO**

UBP1_uc_pTSS_oPEAK

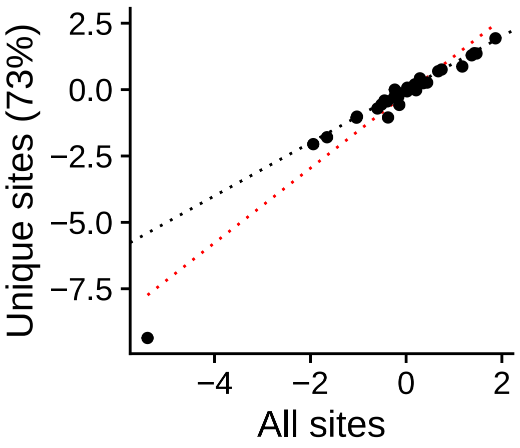


r = 0.967 (p = < 2.22e−16)

**Group STEM**

ARID3A_uc_cTSS_nPEAK

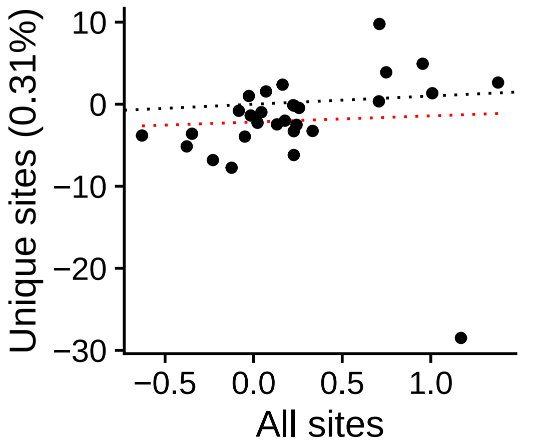


r = 0.0565 (p = 0.77106)

**Group STEM**

ARID3A_uc_cTSS_oPEAK

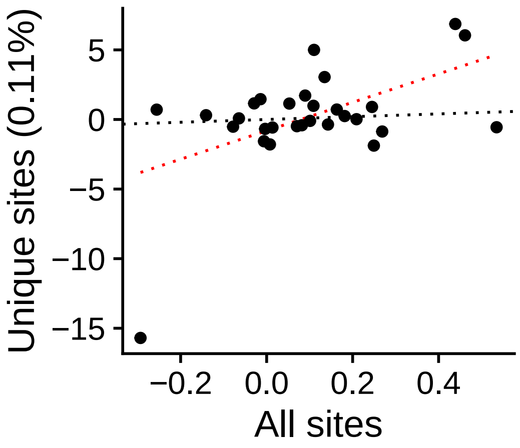


r = 0.52 (p = 0.0038244)

**Group STEM**

ARID3A_uc_dTSS_nPEAK

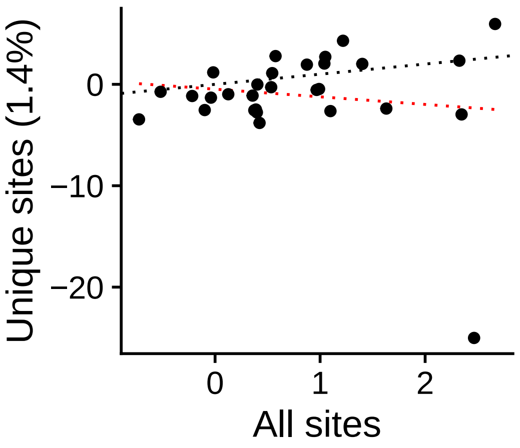


r = −0.126 (p = 0.51576)

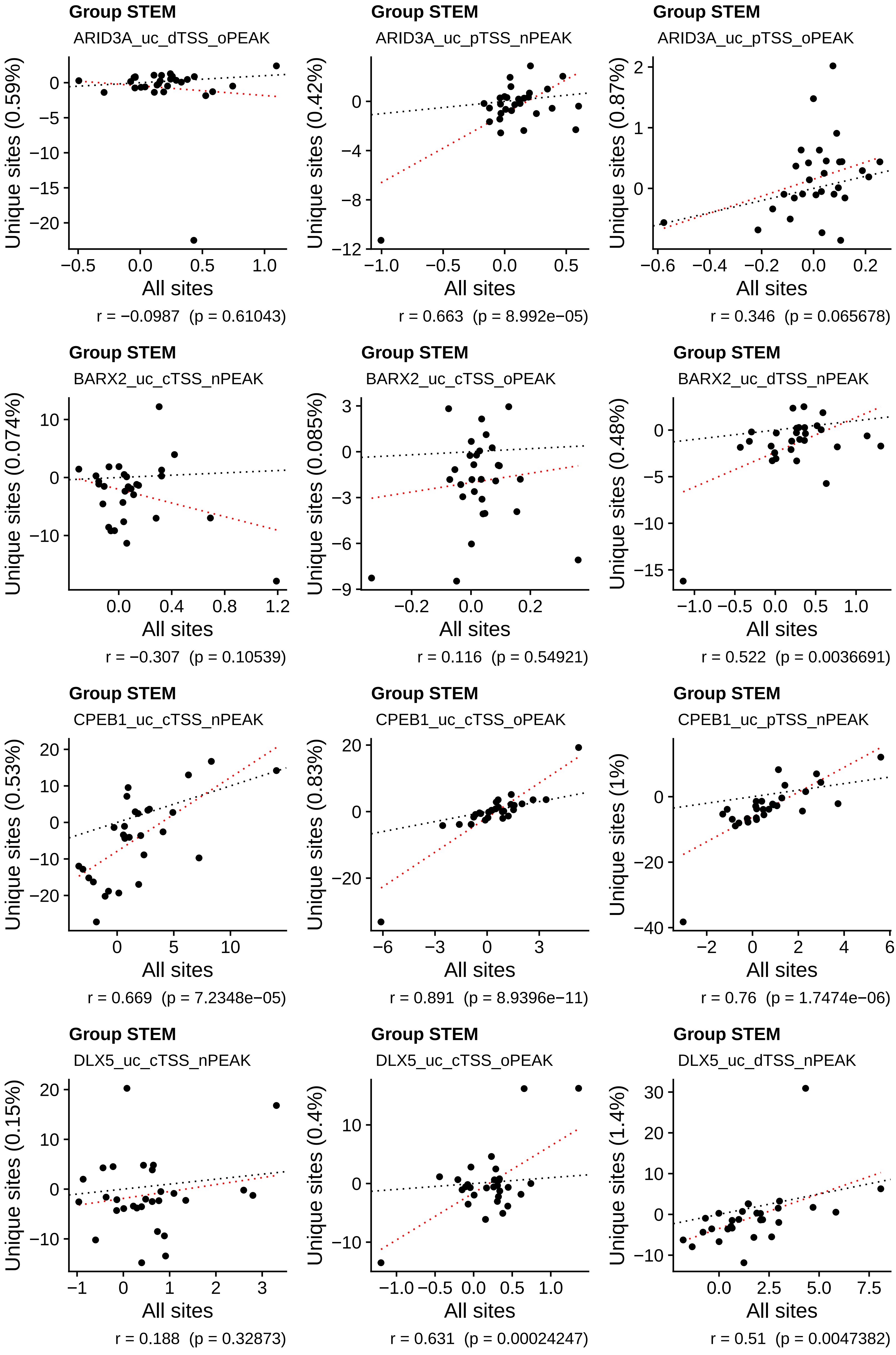
Group STEM
ARID3A_uc_dTSS_oPEAK
Unique sites (0.59%)
All sites
r = −0.0987 (p = 0.61043)
Group STEM
ARID3A_uc_pTSS_nPEAK
Unique sites (0.42%)
All sites
r = 0.663 (p = 8.992e−05)
Group STEM
ARID3A_uc_pTSS_oPEAK
Unique sites (0.87%)
All sites
r = 0.346 (p = 0.065678)
Group STEM
BARX2_uc_cTSS_nPEAK
Unique sites (0.074%)
All sites
r = −0.307 (p = 0.10539)
Group STEM
BARX2_uc_cTSS_oPEAK
Unique sites (0.085%)
All sites
r = 0.116 (p = 0.54921)
Group STEM
BARX2_uc_dTSS_nPEAK
Unique sites (0.48%)
All sites
r = 0.522 (p = 0.0036691)
Group STEM
CPEB1_uc_cTSS_nPEAK
Unique sites (0.53%)
All sites
r = 0.669 (p = 7.2348e−05)
Group STEM
CPEB1_uc_cTSS_oPEAK
Unique sites (0.83%)
All sites
r = 0.891 (p = 8.9396e−11)
Group STEM
CPEB1_uc_pTSS_nPEAK
Unique sites (1%)
All sites
r = 0.76 (p = 1.7474e−06)
Group STEM
DLX5_uc_cTSS_nPEAK
Unique sites (0.15%)
All sites
r = 0.188 (p = 0.32873)
Group STEM
DLX5_uc_cTSS_oPEAK
Unique sites (0.4%)
All sites
r = 0.631 (p = 0.00024247)
Group STEM
DLX5_uc_dTSS_nPEAK
Unique sites (1.4%)
All sites
r = 0.51 (p = 0.0047382)

**Group STEM**

DLX5_uc_dTSS_oPEAK

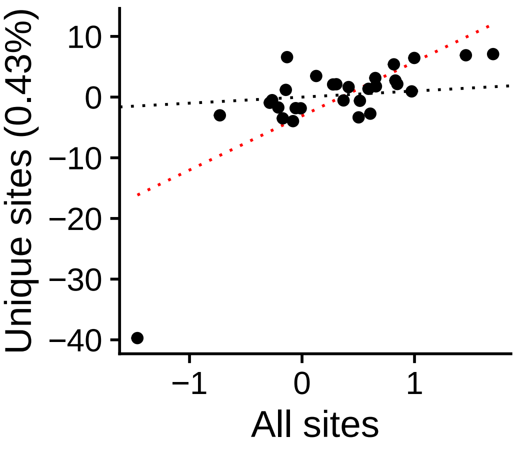


r = 0.699  (p = 2.5044e−05)

**Group STEM**

DLX5_uc_pTSS_nPEAK

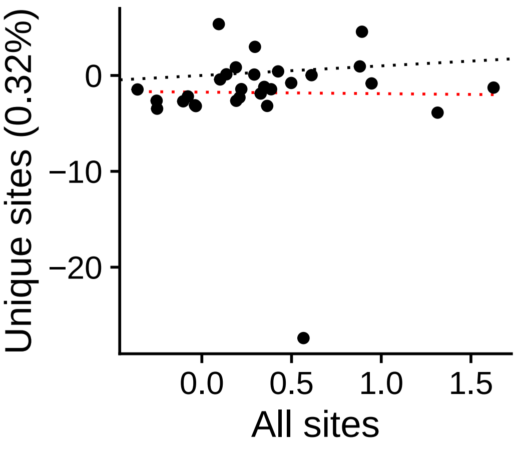


r = −0.0135  (p = 0.94456)

**Group STEM**

DLX5_uc_pTSS_oPEAK

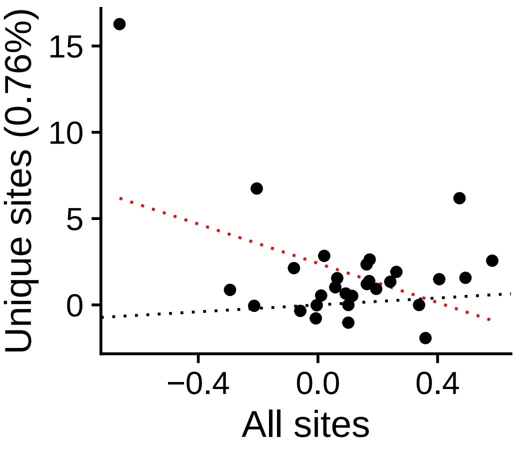


r = −0.439  (p = 0.01731)

**Group STEM**

DMRT2_uc_cTSS_nPEAK

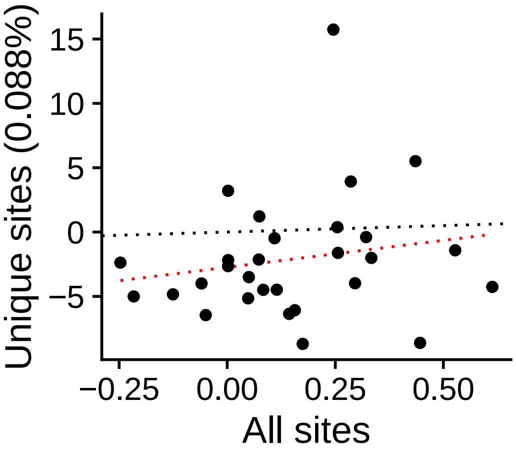


r = 0.181  (p = 0.34853)

**Group STEM**

DMRT2_uc_cTSS_oPEAK

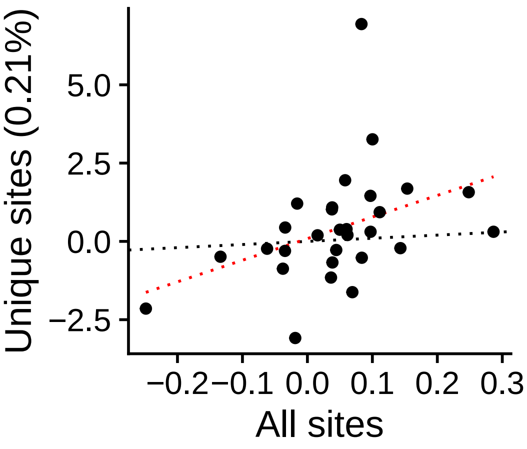


r = 0.394  (p = 0.034288)

**Group STEM**

DMRT2_uc_dTSS_nPEAK

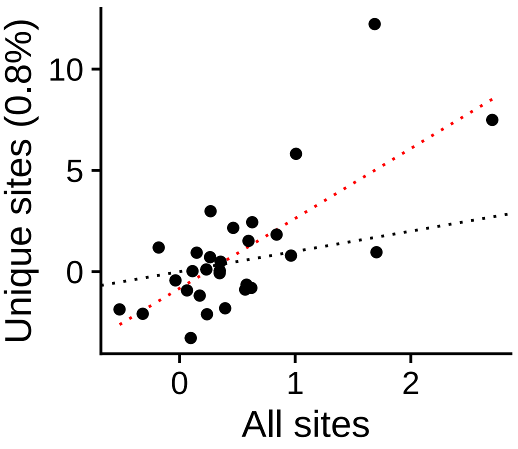


r = 0.715  (p = 1.3112e−05)

**Group STEM**

DMRT2_uc_dTSS_oPEAK

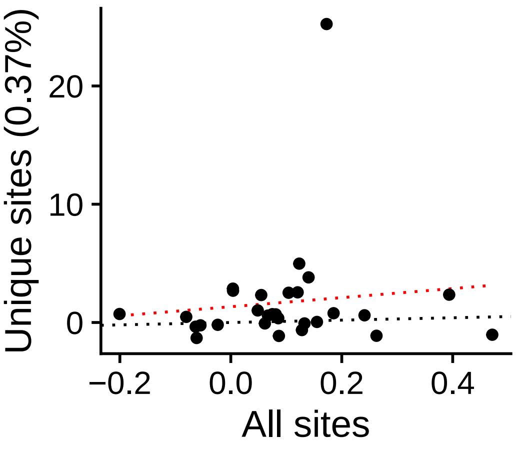


r = 0.11  (p = 0.56838)

**Group STEM**

DMRT2_uc_pTSS_nPEAK

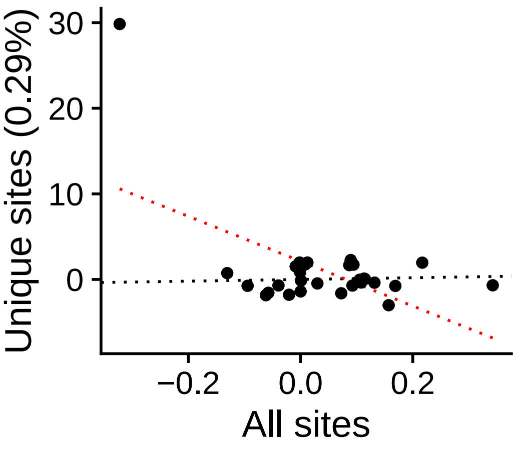


r = −0.556  (p = 0.0017356)

**Group STEM**

DMRTC2_uc_cTSS_nPEAK

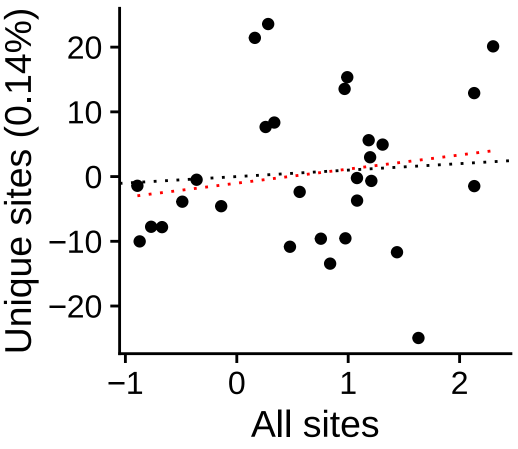


r = 0.171  (p = 0.37581)

**Group STEM**

DMRTC2_uc_cTSS_oPEAK

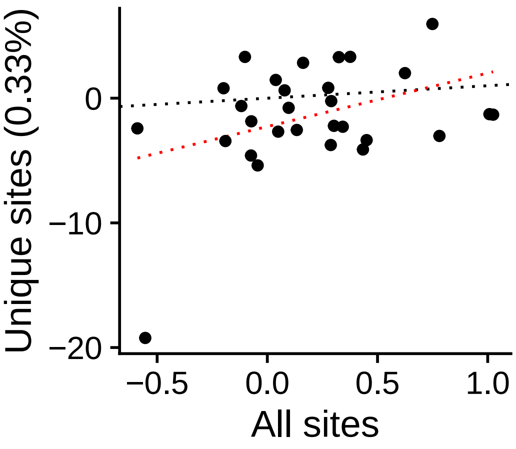


r = 0.382  (p = 0.040801)

**Group STEM**

DMRTC2_uc_dTSS_oPEAK

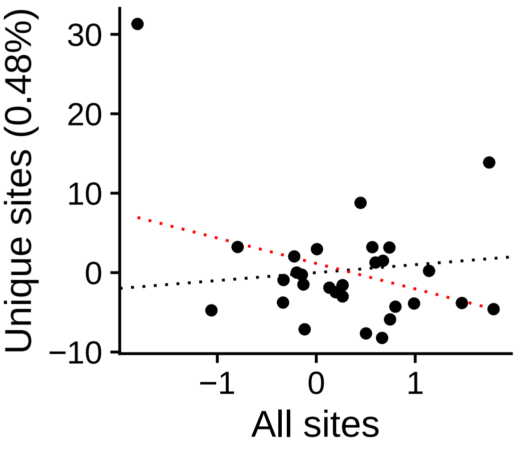


r = −0.331  (p = 0.07917)

**Group STEM**

DMRTC2_uc_pTSS_nPEAK

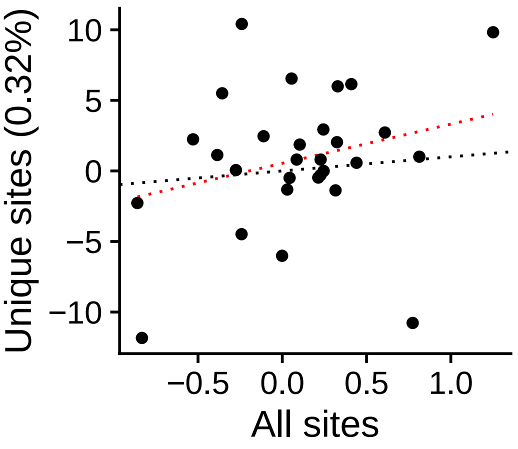


r = 0.26  (p = 0.17338)

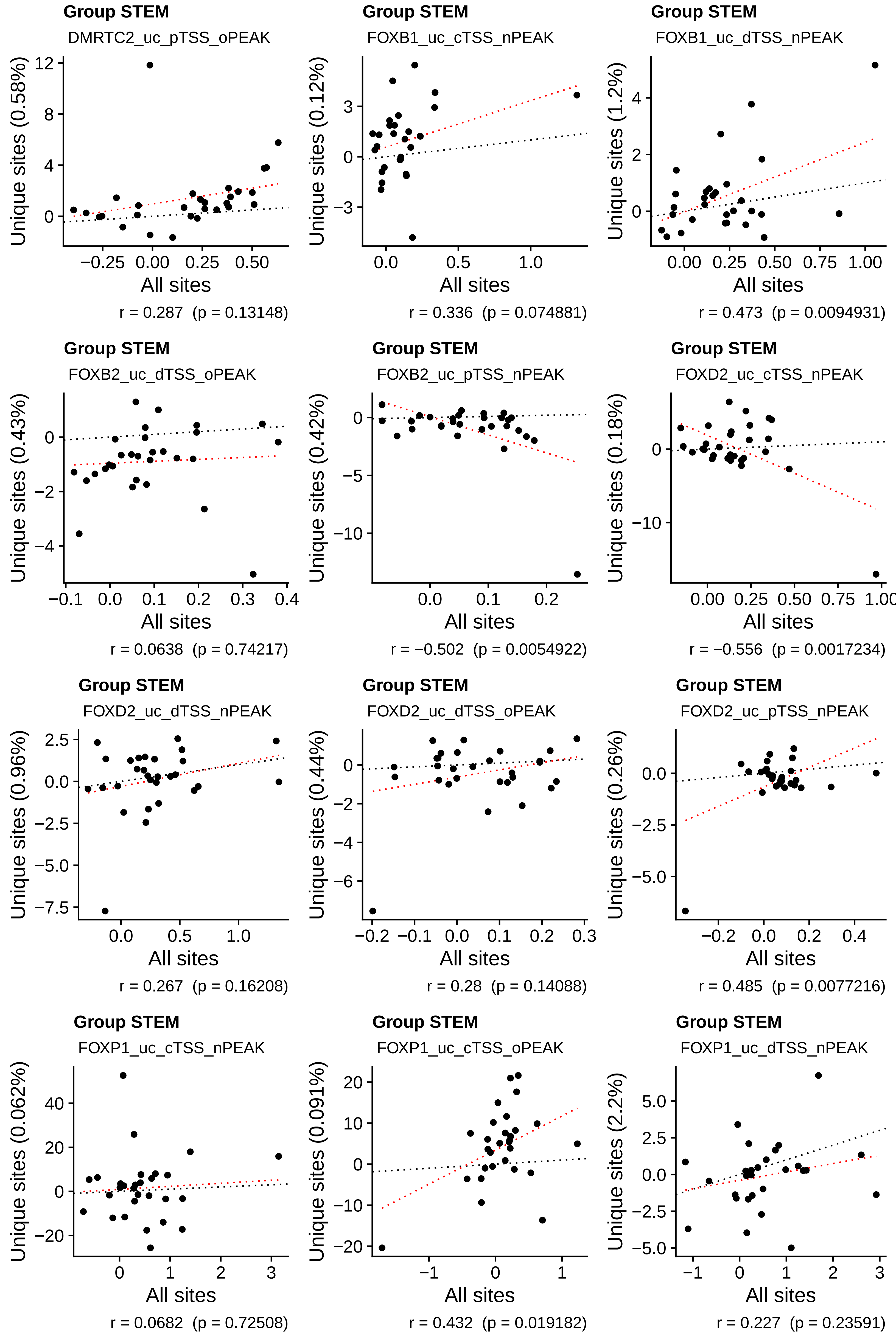
Group STEM
DMRTC2_uc_pTSS_oPEAK
Unique sites (0.58%)
All sites
r = 0.287 (p = 0.13148)
Group STEM
FOXB1_uc_cTSS_nPEAK
Unique sites (0.12%)
All sites
r = 0.336 (p = 0.074881)
Group STEM
FOXB1_uc_dTSS_nPEAK
Unique sites (1.2%)
All sites
r = 0.473 (p = 0.0094931)
Group STEM
FOXB2_uc_dTSS_oPEAK
Unique sites (0.43%)
All sites
r = 0.0638 (p = 0.74217)
Group STEM
FOXB2_uc_pTSS_nPEAK
Unique sites (0.42%)
All sites
r = −0.502 (p = 0.0054922)
Group STEM
FOXD2_uc_cTSS_nPEAK
Unique sites (0.18%)
All sites
r = −0.556 (p = 0.0017234)
Group STEM
FOXD2_uc_dTSS_nPEAK
Unique sites (0.96%)
All sites
r = 0.267 (p = 0.16208)
Group STEM
FOXD2_uc_dTSS_oPEAK
Unique sites (0.44%)
All sites
r = 0.28 (p = 0.14088)
Group STEM
FOXD2_uc_pTSS_nPEAK
Unique sites (0.26%)
All sites
r = 0.485 (p = 0.0077216)
Group STEM
FOXP1_uc_cTSS_nPEAK
Unique sites (0.062%)
All sites
r = 0.0682 (p = 0.72508)
Group STEM
FOXP1_uc_cTSS_oPEAK
Unique sites (0.091%)
All sites
r = 0.432 (p = 0.019182)
Group STEM
FOXP1_uc_dTSS_nPEAK
Unique sites (2.2%)
All sites
r = 0.227 (p = 0.23591)

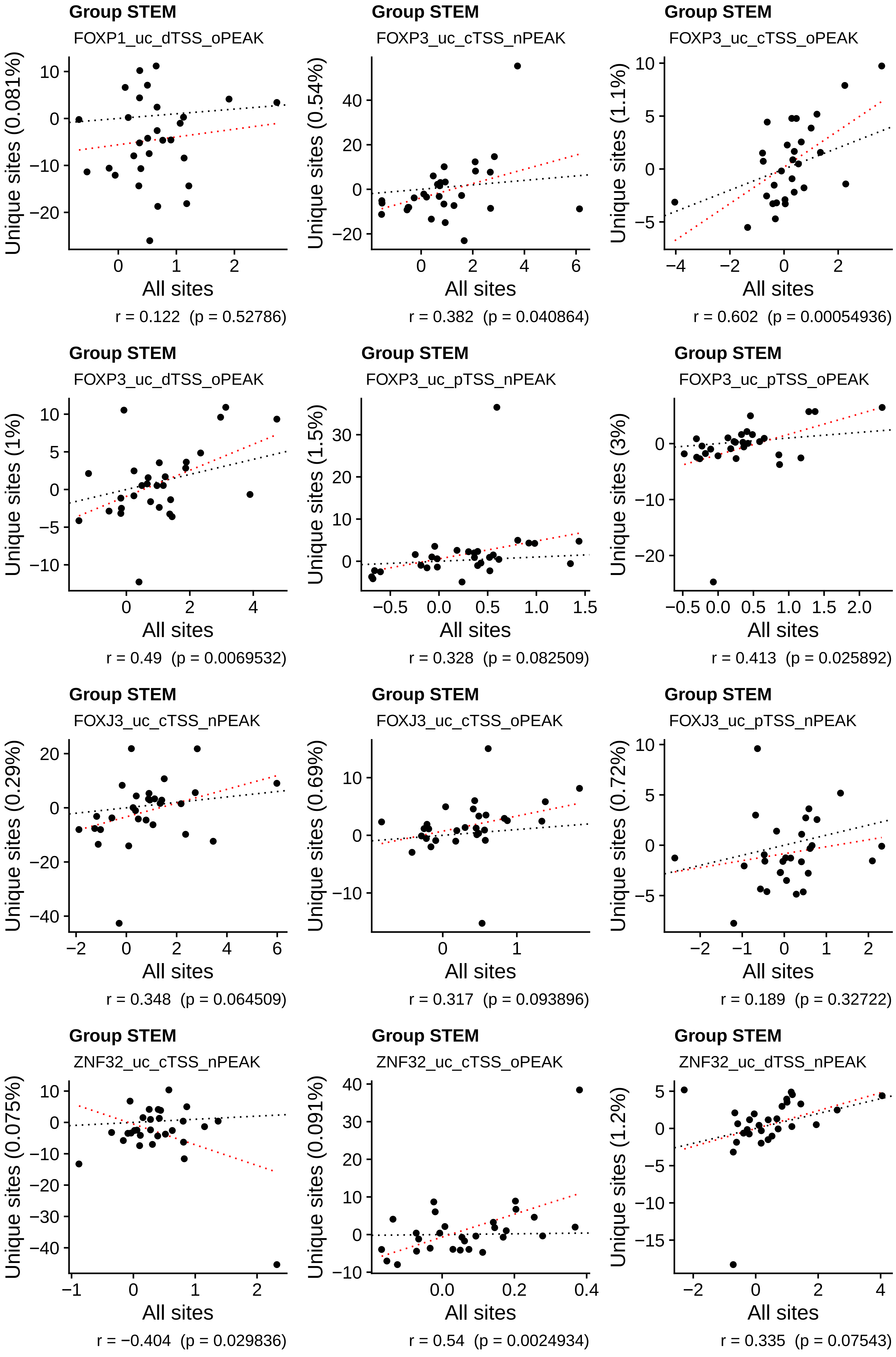
Group STEM
FOXP1_uc_dTSS_oPEAK
Unique sites (0.081%)
All sites
r = 0.122  (p = 0.52786)
Group STEM
FOXP3_uc_cTSS_nPEAK
Unique sites (0.54%)
All sites
r = 0.382  (p = 0.040864)
Group STEM
FOXP3_uc_cTSS_oPEAK
Unique sites (1.1%)
All sites
r = 0.602  (p = 0.00054936)
Group STEM
FOXP3_uc_dTSS_oPEAK
Unique sites (1%)
All sites
r = 0.49  (p = 0.0069532)
Group STEM
FOXP3_uc_pTSS_nPEAK
Unique sites (1.5%)
All sites
r = 0.328  (p = 0.082509)
Group STEM
FOXP3_uc_pTSS_oPEAK
Unique sites (3%)
All sites
r = 0.413  (p = 0.025892)
Group STEM
FOXJ3_uc_cTSS_nPEAK
Unique sites (0.29%)
All sites
r = 0.348  (p = 0.064509)
Group STEM
FOXJ3_uc_cTSS_oPEAK
Unique sites (0.69%)
All sites
r = 0.317  (p = 0.093896)
Group STEM
FOXJ3_uc_pTSS_nPEAK
Unique sites (0.72%)
All sites
r = 0.189  (p = 0.32722)
Group STEM
ZNF32_uc_cTSS_nPEAK
Unique sites (0.075%)
All sites
r = −0.404  (p = 0.029836)
Group STEM
ZNF32_uc_cTSS_oPEAK
Unique sites (0.091%)
All sites
r = 0.54  (p = 0.0024934)
Group STEM
ZNF32_uc_dTSS_nPEAK
Unique sites (1.2%)
All sites
r = 0.335  (p = 0.07543)

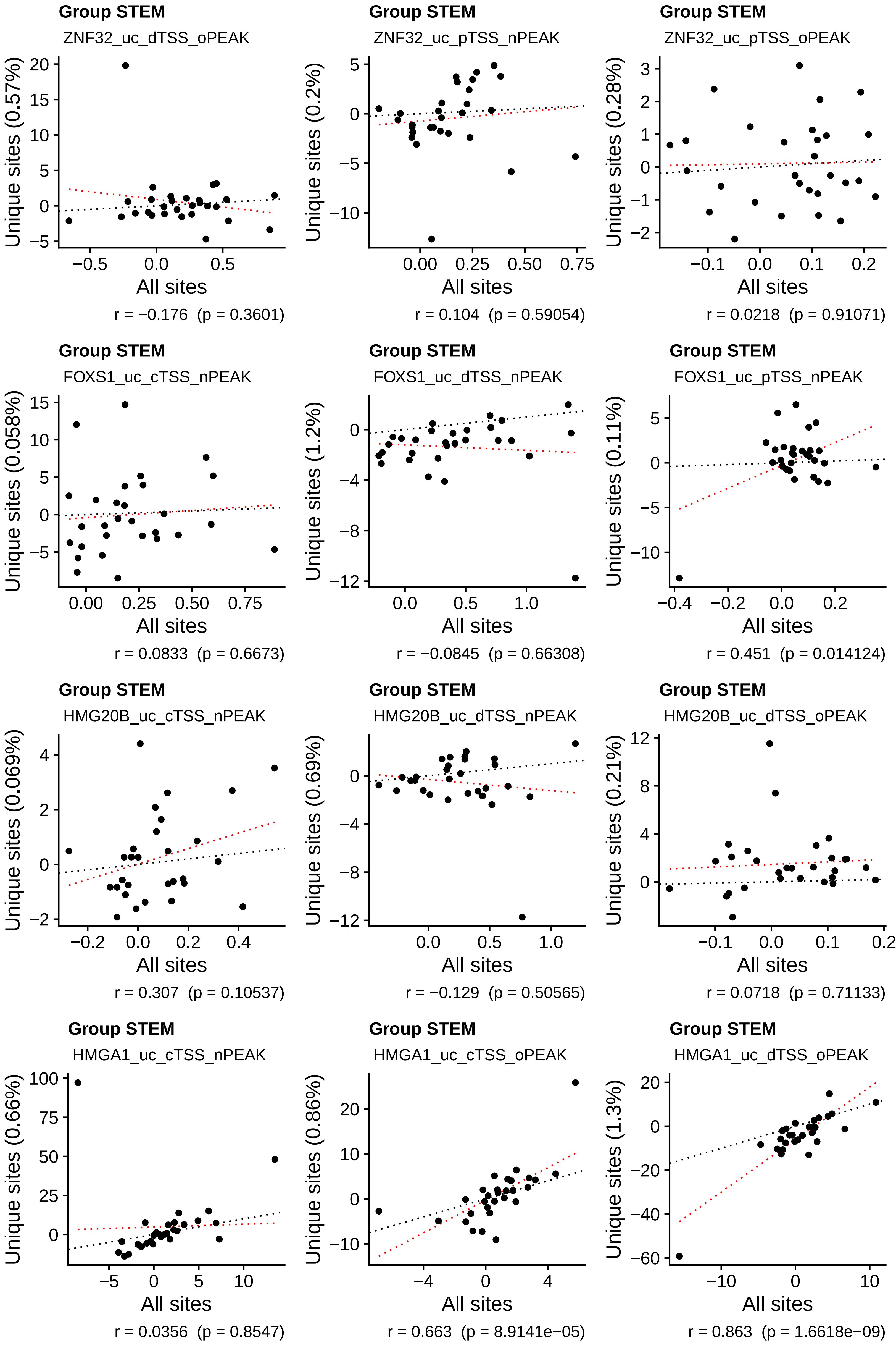
Group STEM
ZNF32_uc_dTSS_oPEAK
Unique sites (0.57%)
All sites
r = −0.176 (p = 0.3601)
Group STEM
ZNF32_uc_pTSS_nPEAK
Unique sites (0.2%)
All sites
r = 0.104 (p = 0.59054)
Group STEM
ZNF32_uc_pTSS_oPEAK
Unique sites (0.28%)
All sites
r = 0.0218 (p = 0.91071)
Group STEM
FOXS1_uc_cTSS_nPEAK
Unique sites (0.058%)
All sites
r = 0.0833 (p = 0.6673)
Group STEM
FOXS1_uc_dTSS_nPEAK
Unique sites (1.2%)
All sites
r = −0.0845 (p = 0.66308)
Group STEM
FOXS1_uc_pTSS_nPEAK
Unique sites (0.11%)
All sites
r = 0.451 (p = 0.014124)
Group STEM
HMG20B_uc_cTSS_nPEAK
Unique sites (0.069%)
All sites
r = 0.307 (p = 0.10537)
Group STEM
HMG20B_uc_dTSS_nPEAK
Unique sites (0.69%)
All sites
r = −0.129 (p = 0.50565)
Group STEM
HMG20B_uc_dTSS_oPEAK
Unique sites (0.21%)
All sites
r = 0.0718 (p = 0.71133)
Group STEM
HMGA1_uc_cTSS_nPEAK
Unique sites (0.66%)
All sites
r = 0.0356 (p = 0.8547)
Group STEM
HMGA1_uc_cTSS_oPEAK
Unique sites (0.86%)
All sites
r = 0.663 (p = 8.9141e−05)
Group STEM
HMGA1_uc_dTSS_oPEAK
Unique sites (1.3%)
All sites
r = 0.863 (p = 1.6618e−09)

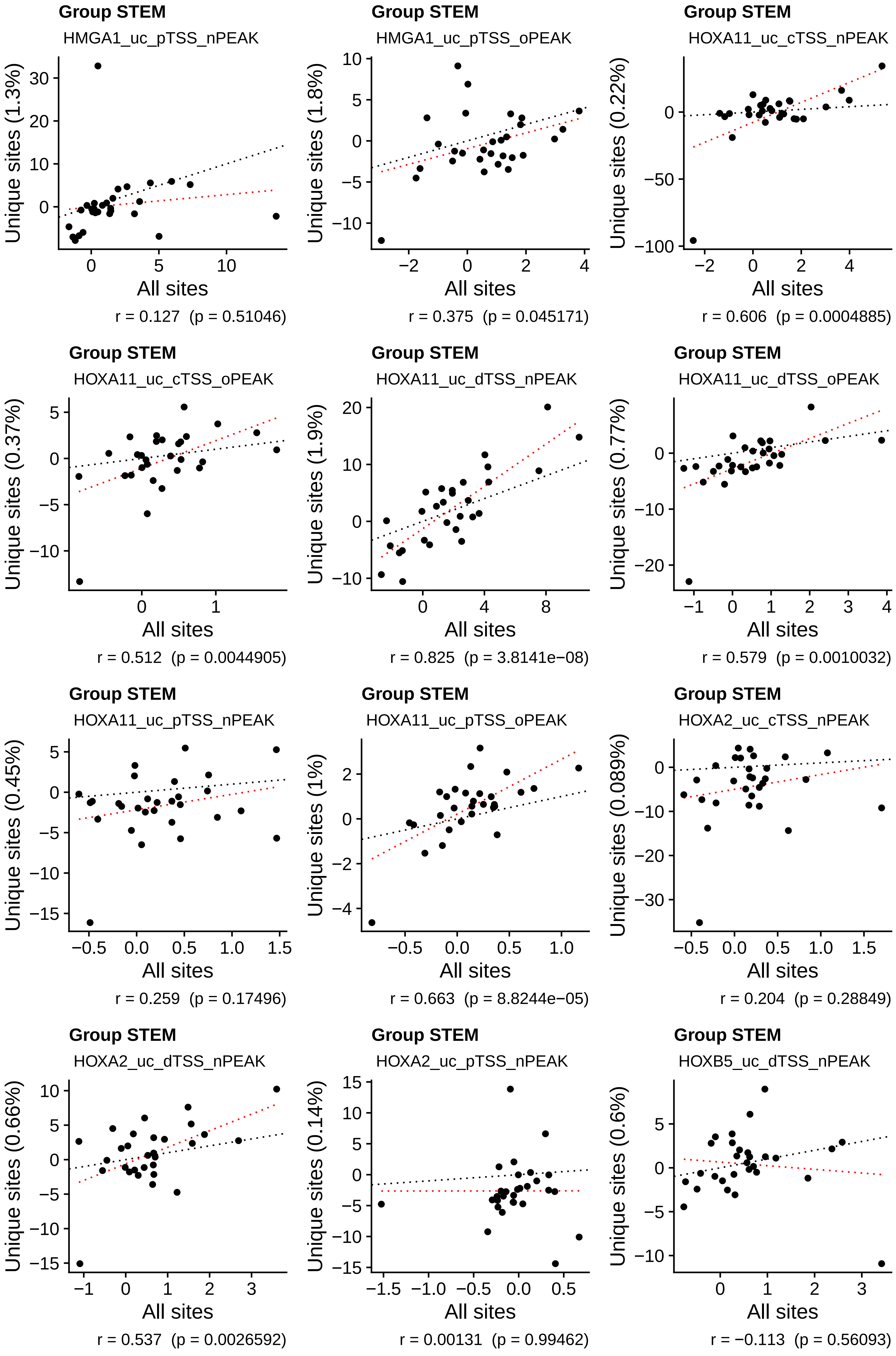
Group STEM
HMGA1_uc_pTSS_nPEAK
Unique sites (1.3%)
All sites
r = 0.127 (p = 0.51046)
Group STEM
HMGA1_uc_pTSS_oPEAK
Unique sites (1.8%)
All sites
r = 0.375 (p = 0.045171)
Group STEM
HOXA11_uc_cTSS_nPEAK
Unique sites (0.22%)
All sites
r = 0.606 (p = 0.0004885)
Group STEM
HOXA11_uc_cTSS_oPEAK
Unique sites (0.37%)
All sites
r = 0.512 (p = 0.0044905)
Group STEM
HOXA11_uc_dTSS_nPEAK
Unique sites (1.9%)
All sites
r = 0.825 (p = 3.8141e−08)
Group STEM
HOXA11_uc_dTSS_oPEAK
Unique sites (0.77%)
All sites
r = 0.579 (p = 0.0010032)
Group STEM
HOXA11_uc_pTSS_nPEAK
Unique sites (0.45%)
All sites
r = 0.259 (p = 0.17496)
Group STEM
HOXA11_uc_pTSS_oPEAK
Unique sites (1%)
All sites
r = 0.663 (p = 8.8244e−05)
Group STEM
HOXA2_uc_cTSS_nPEAK
Unique sites (0.089%)
All sites
r = 0.204 (p = 0.28849)
Group STEM
HOXA2_uc_dTSS_nPEAK
Unique sites (0.66%)
All sites
r = 0.537 (p = 0.0026592)
Group STEM
HOXA2_uc_pTSS_nPEAK
Unique sites (0.14%)
All sites
r = 0.00131 (p = 0.99462)
Group STEM
HOXB5_uc_dTSS_nPEAK
Unique sites (0.6%)
All sites
r = −0.113 (p = 0.56093)

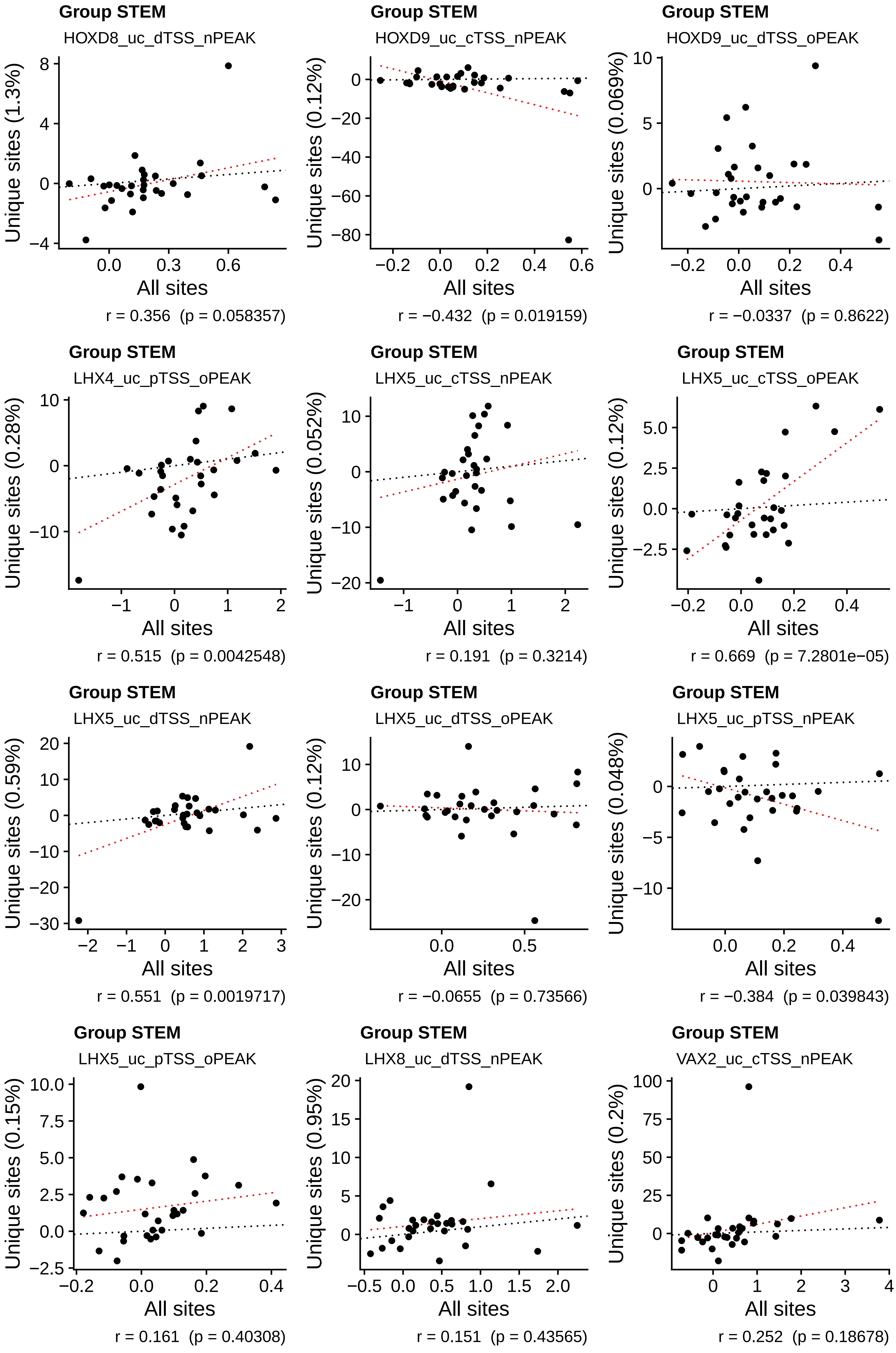
Group STEM
HOXD8_uc_dTSS_nPEAK
Unique sites (1.3%)
All sites
r = 0.356 (p = 0.058357)
Group STEM
HOXD9_uc_cTSS_nPEAK
Unique sites (0.12%)
All sites
r = −0.432 (p = 0.019159)
Group STEM
HOXD9_uc_dTSS_oPEAK
Unique sites (0.069%)
All sites
r = −0.0337 (p = 0.8622)
Group STEM
LHX4_uc_pTSS_oPEAK
Unique sites (0.28%)
All sites
r = 0.515 (p = 0.0042548)
Group STEM
LHX5_uc_cTSS_nPEAK
Unique sites (0.052%)
All sites
r = 0.191 (p = 0.3214)
Group STEM
LHX5_uc_cTSS_oPEAK
Unique sites (0.12%)
All sites
r = 0.669 (p = 7.2801e−05)
Group STEM
LHX5_uc_dTSS_nPEAK
Unique sites (0.59%)
All sites
r = 0.551 (p = 0.0019717)
Group STEM
LHX5_uc_dTSS_oPEAK
Unique sites (0.12%)
All sites
r = −0.0655 (p = 0.73566)
Group STEM
LHX5_uc_pTSS_nPEAK
Unique sites (0.048%)
All sites
r = −0.384 (p = 0.039843)
Group STEM
LHX5_uc_pTSS_oPEAK
Unique sites (0.15%)
All sites
r = 0.161 (p = 0.40308)
Group STEM
LHX8_uc_dTSS_nPEAK
Unique sites (0.95%)
All sites
r = 0.151 (p = 0.43565)
Group STEM
VAX2_uc_cTSS_nPEAK
Unique sites (0.2%)
All sites
r = 0.252 (p = 0.18678)

**Group STEM**

VAX2_uc_cTSS_oPEAK

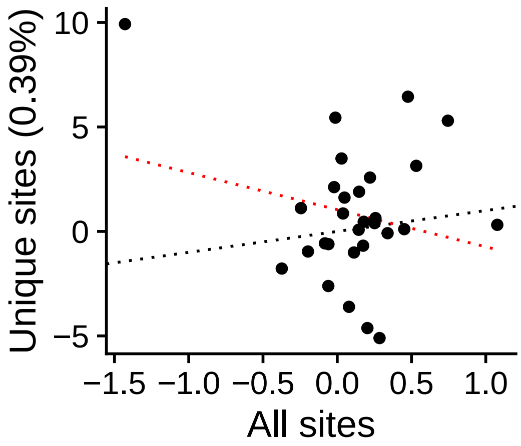


r = −0.229 (p = 0.23118)

**Group STEM**

VAX2_uc_dTSS_nPEAK

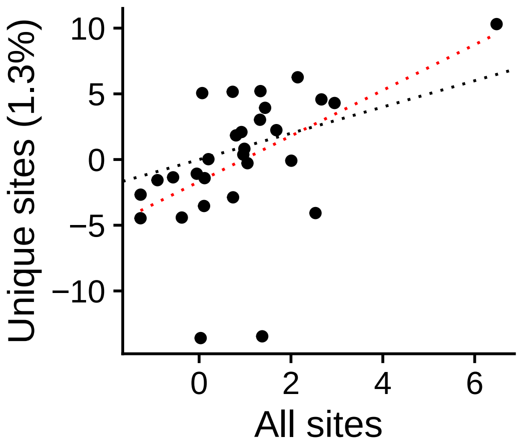


r = 0.512 (p = 0.0045657)

**Group STEM**

VAX2_uc_pTSS_nPEAK

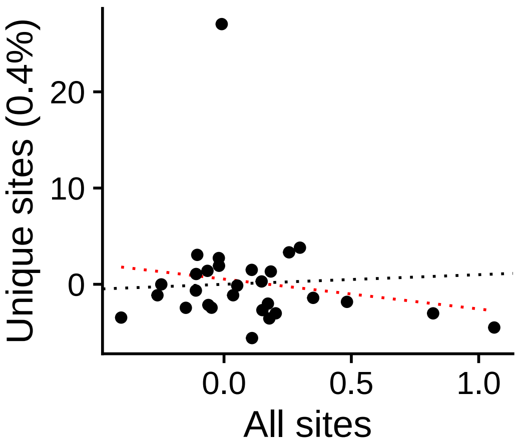


r = −0.165 (p = 0.39329)

**Group STEM**

VAX2_uc_pTSS_oPEAK

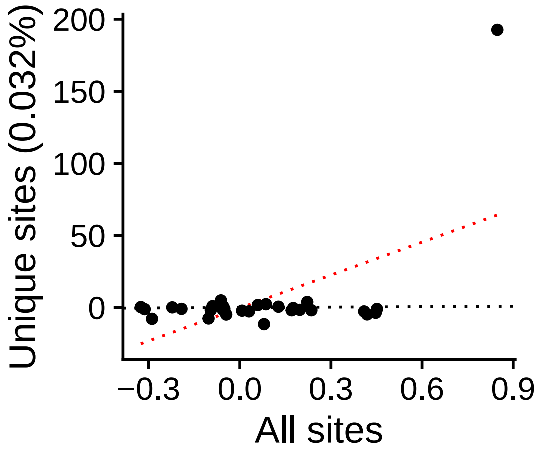


r = 0.558 (p = 0.0016712)

**Group STEM**

DLX3_uc_cTSS_nPEAK

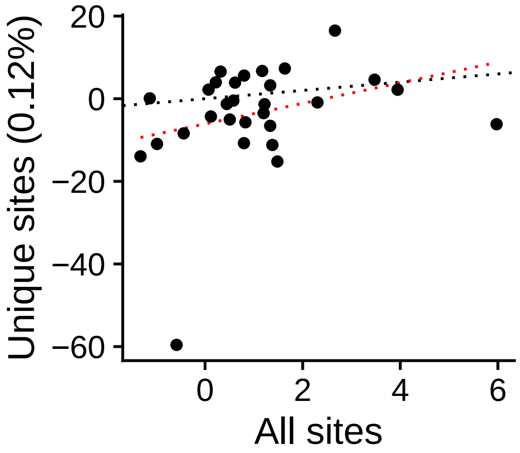


r = 0.296 (p = 0.11847)

**Group STEM**

DLX3_uc_cTSS_oPEAK

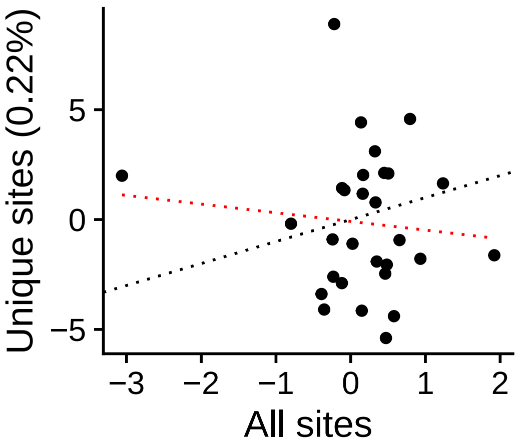


r = −0.101 (p = 0.60096)

**Group STEM**

DLX3_uc_dTSS_oPEAK

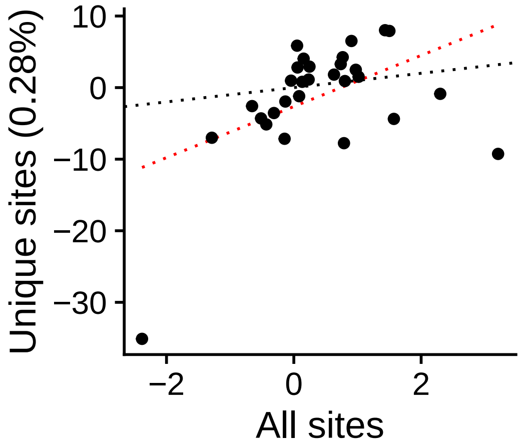


r = 0.474 (p = 0.0094585)

**Group STEM**

DLX3_uc_pTSS_nPEAK

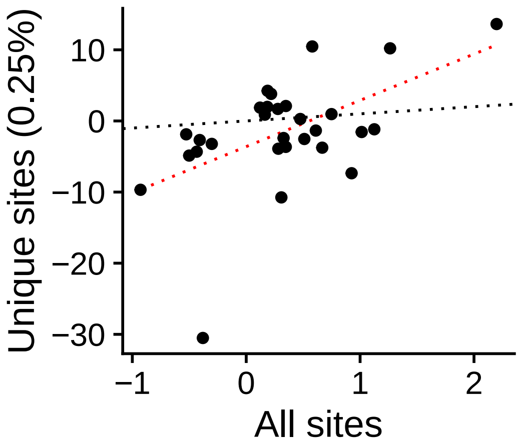


r = 0.534 (p = 0.0028392)

**Group STEM**

DLX3_uc_pTSS_oPEAK

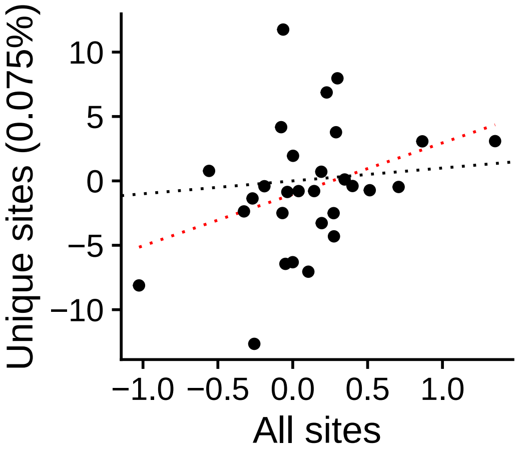


r = 0.35 (p = 0.062886)

**Group STEM**

NKX63_uc_dTSS_oPEAK

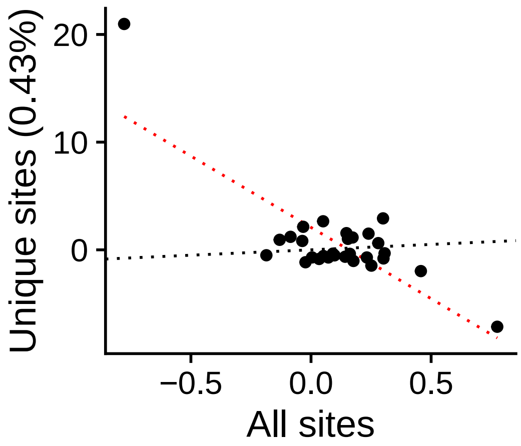


r = −0.782 (p = 5.3812e−07)

**Group STEM**

NKX62_uc_cTSS_oPEAK

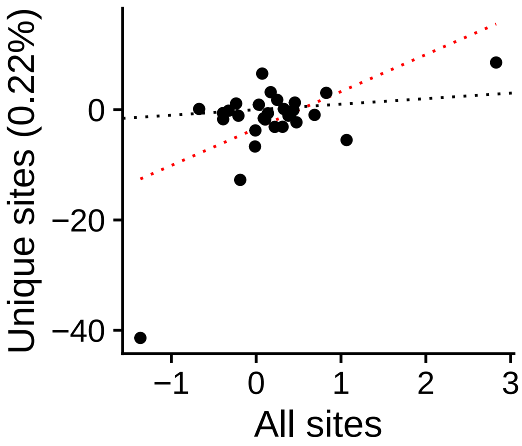


r = 0.548 (p = 0.0020909)

**Group STEM**

NKX62_uc_dTSS_oPEAK

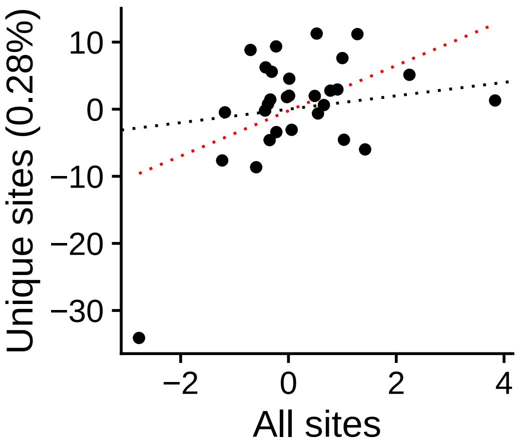


r = 0.472 (p = 0.0097054)

**Group STEM**
NKX62_uc_pTSS_oPEAK

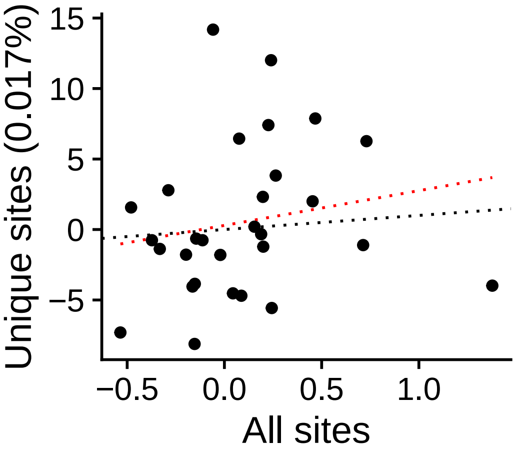


r = 0.183 (p = 0.34303)

**Group STEM**
ARID3C_uc_cTSS_nPEAK

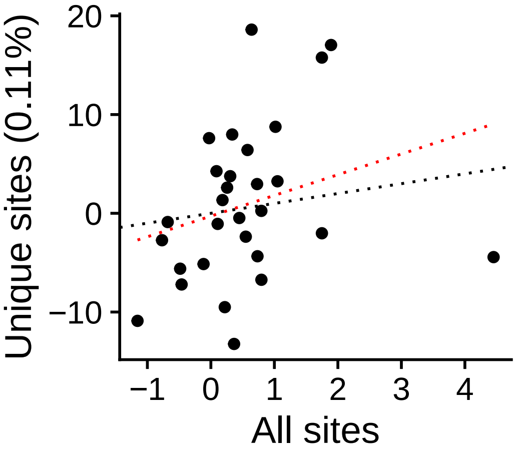


r = 0.276 (p = 0.14782)

**Group STEM**
ARID3C_uc_dTSS_nPEAK

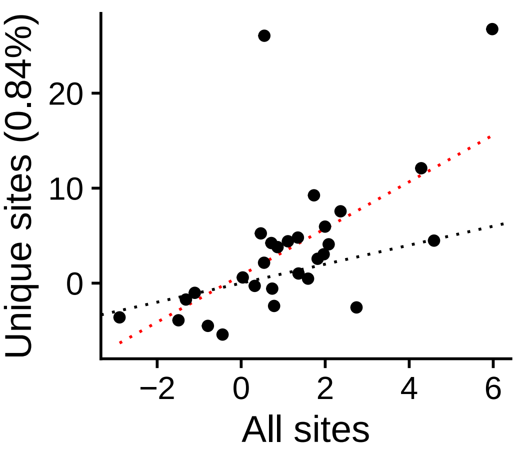


r = 0.603 (p = 0.00053005)

**Group STEM**
ARID3C_uc_pTSS_nPEAK

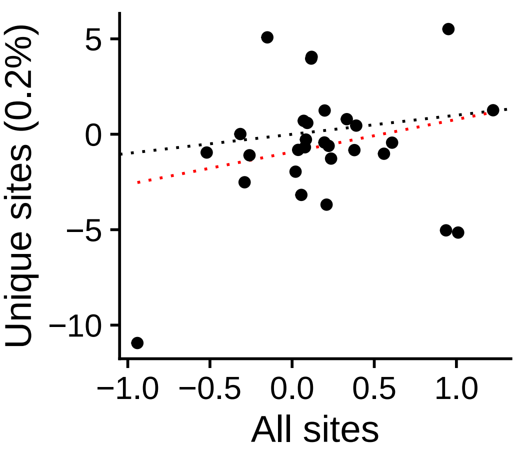


r = 0.243 (p = 0.20471)

**Group STEM**
CDX1_uc_cTSS_nPEAK

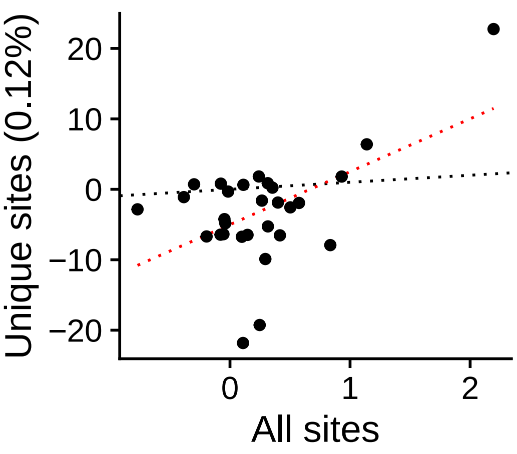


r = 0.528 (p = 0.00324)

**Group STEM**
CDX1_uc_cTSS_oPEAK

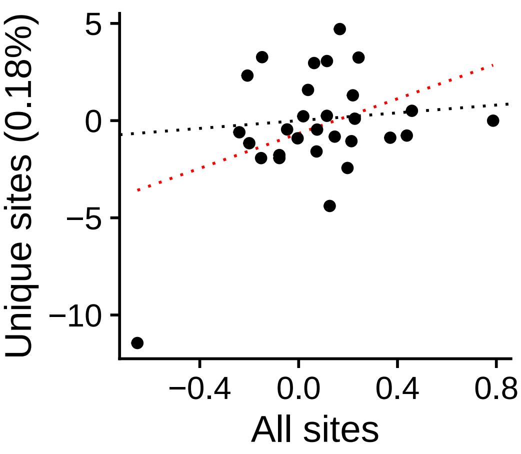


r = 0.4 (p = 0.031356)

**Group STEM**
CDX1_uc_dTSS_nPEAK

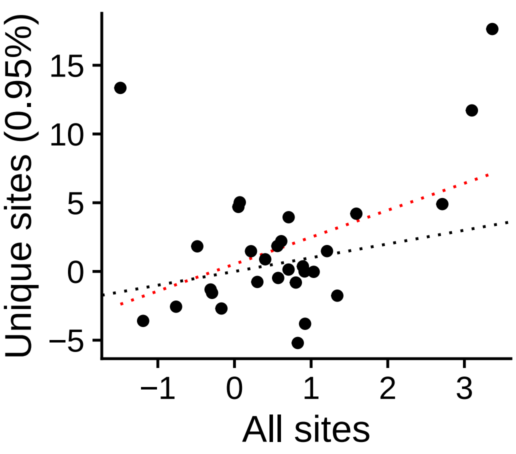


r = 0.425 (p = 0.021637)

**Group STEM**
CDX1_uc_dTSS_oPEAK

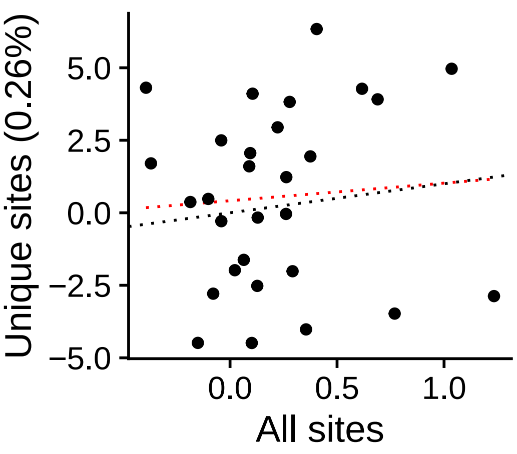


r = 0.0732 (p = 0.70597)

**Group STEM**
CDX1_uc_pTSS_nPEAK

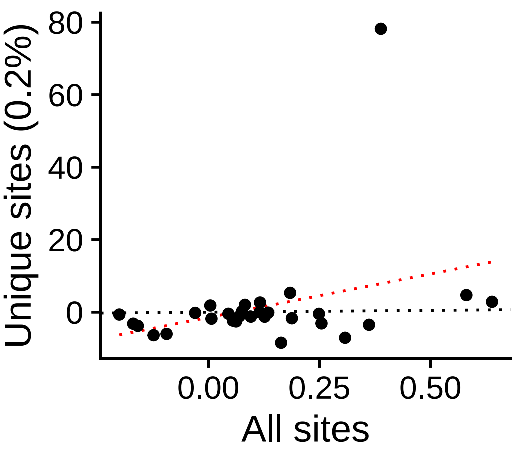


r = 0.319 (p = 0.091573)

**Group STEM**
CDX1_uc_pTSS_oPEAK

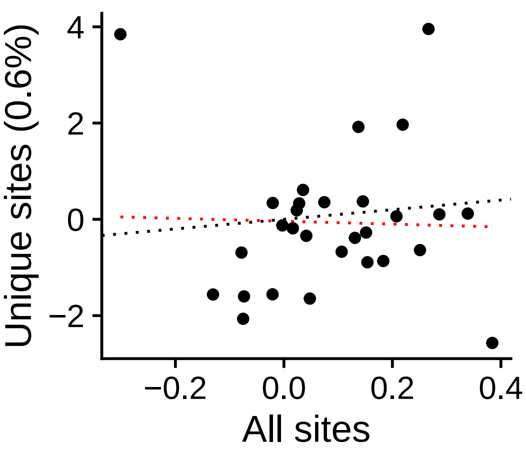


r = −0.0301 (p = 0.877)

**Group STEM**
CDX4_uc_cTSS_nPEAK

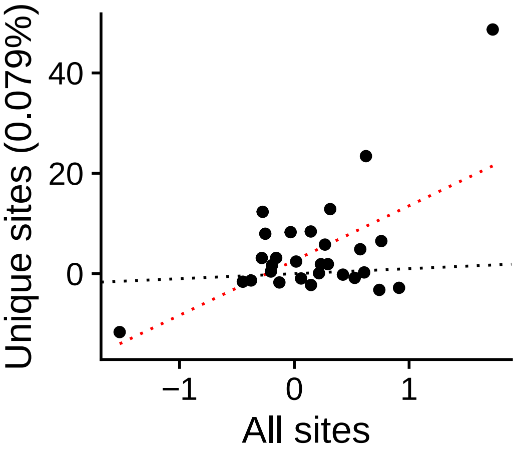


r = 0.585 (p = 0.00086834)

**Group STEM**
CDX4_uc_cTSS_oPEAK

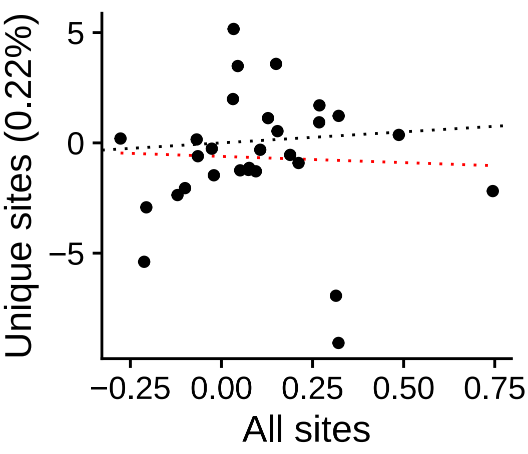


r = −0.0413 (p = 0.83168)

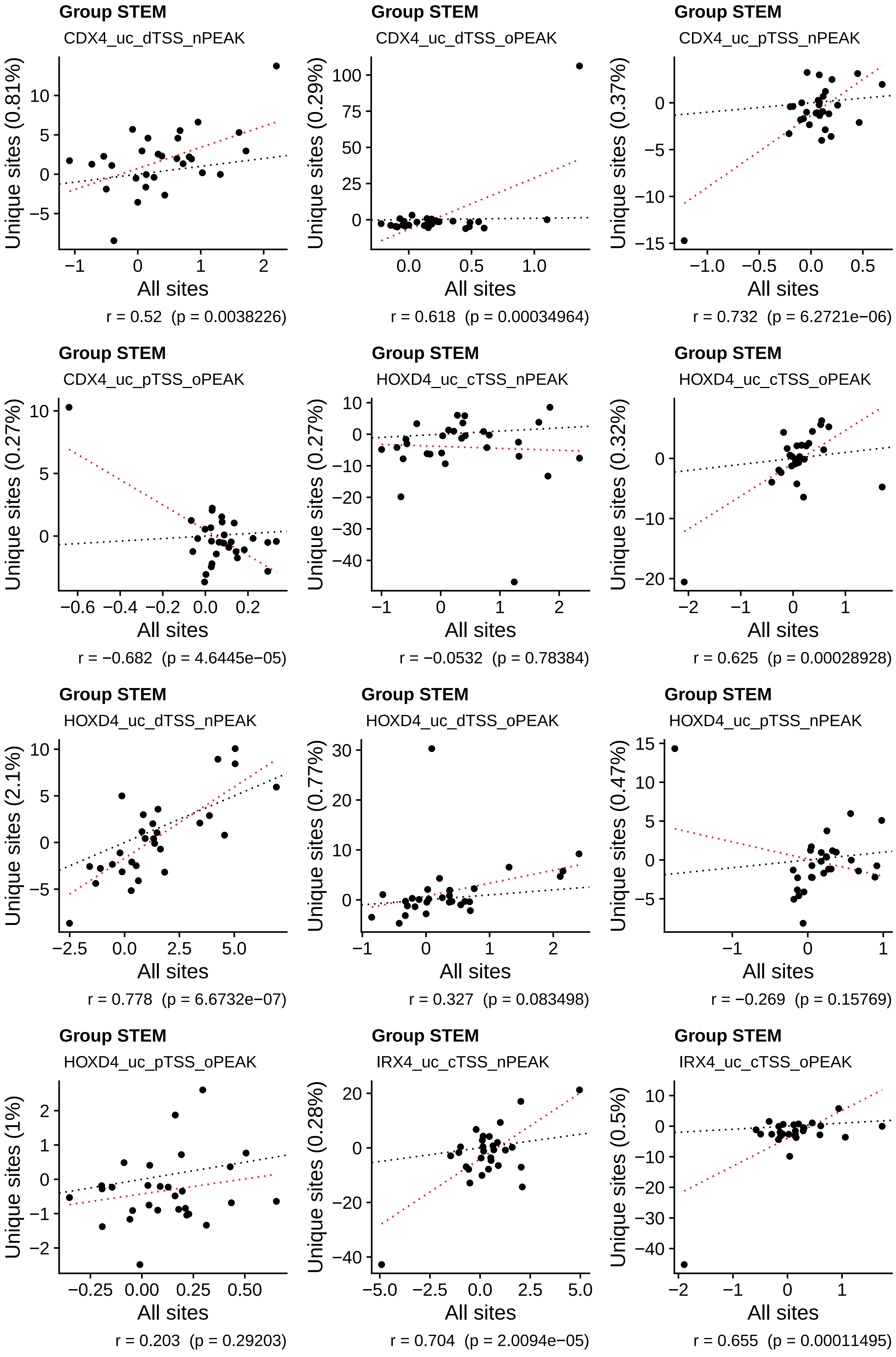
Group STEM
CDX4_uc_dTSS_nPEAK
Unique sites (0.81%)
All sites
r = 0.52  (p = 0.0038226)
Group STEM
CDX4_uc_dTSS_oPEAK
Unique sites (0.29%)
All sites
r = 0.618  (p = 0.00034964)
Group STEM
CDX4_uc_pTSS_nPEAK
Unique sites (0.37%)
All sites
r = 0.732  (p = 6.2721e−06)
Group STEM
CDX4_uc_pTSS_oPEAK
Unique sites (0.27%)
All sites
r = −0.682  (p = 4.6445e−05)
Group STEM
HOXD4_uc_cTSS_nPEAK
Unique sites (0.27%)
All sites
r = −0.0532  (p = 0.78384)
Group STEM
HOXD4_uc_cTSS_oPEAK
Unique sites (0.32%)
All sites
r = 0.625  (p = 0.00028928)
Group STEM
HOXD4_uc_dTSS_nPEAK
Unique sites (2.1%)
All sites
r = 0.778  (p = 6.6732e−07)
Group STEM
HOXD4_uc_dTSS_oPEAK
Unique sites (0.77%)
All sites
r = 0.327  (p = 0.083498)
Group STEM
HOXD4_uc_pTSS_nPEAK
Unique sites (0.47%)
All sites
r = −0.269  (p = 0.15769)
Group STEM
HOXD4_uc_pTSS_oPEAK
Unique sites (1%)
All sites
r = 0.203  (p = 0.29203)
Group STEM
IRX4_uc_cTSS_nPEAK
Unique sites (0.28%)
All sites
r = 0.704  (p = 2.0094e−05)
Group STEM
IRX4_uc_cTSS_oPEAK
Unique sites (0.5%)
All sites
r = 0.655  (p = 0.00011495)

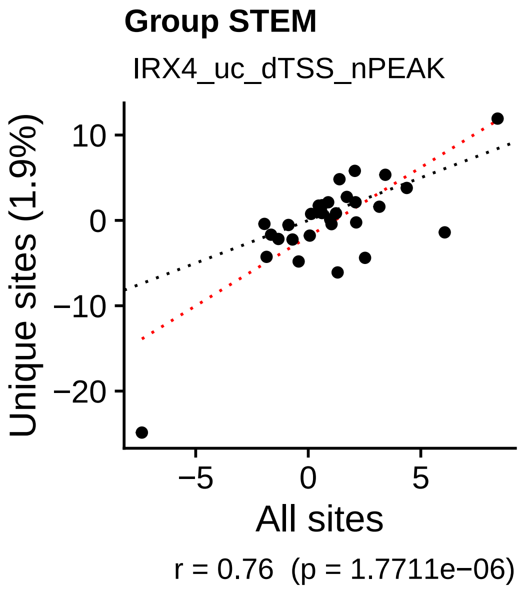
Group STEM
IRX4_uc_dTSS_nPEAK
Unique sites (1.9%)
All sites
r = 0.76 (p = 1.7711e−06)

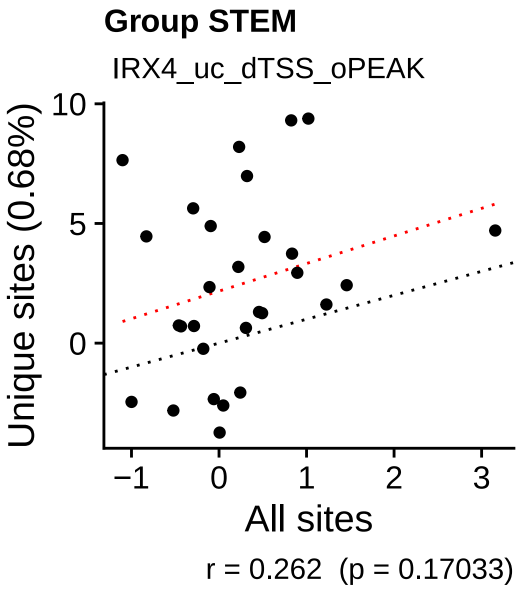
Group STEM
IRX4_uc_dTSS_oPEAK
Unique sites (0.68%)
All sites
r = 0.262 (p = 0.17033)

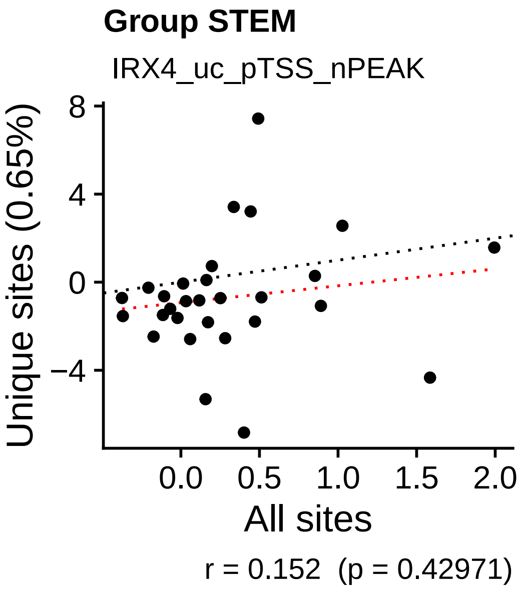
Group STEM
IRX4_uc_pTSS_nPEAK
Unique sites (0.65%)
All sites
r = 0.152 (p = 0.42971)

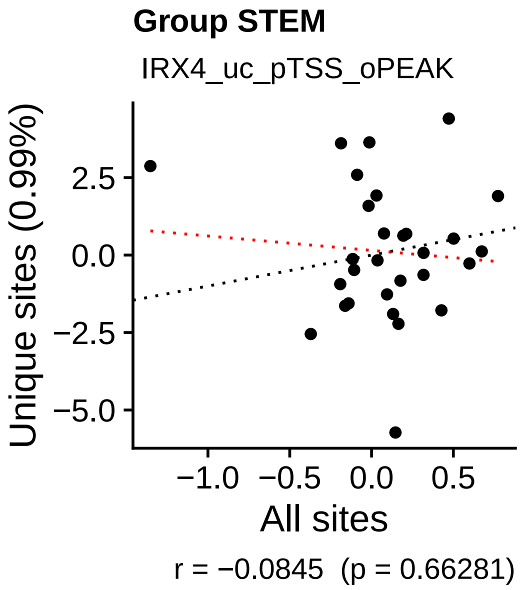
Group STEM
IRX4_uc_pTSS_oPEAK
Unique sites (0.99%)
All sites
r = −0.0845 (p = 0.66281)

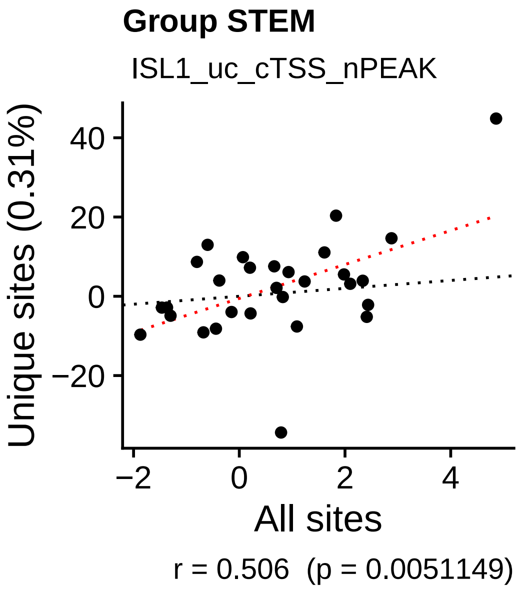
Group STEM
ISL1_uc_cTSS_nPEAK
Unique sites (0.31%)
All sites
r = 0.506 (p = 0.0051149)

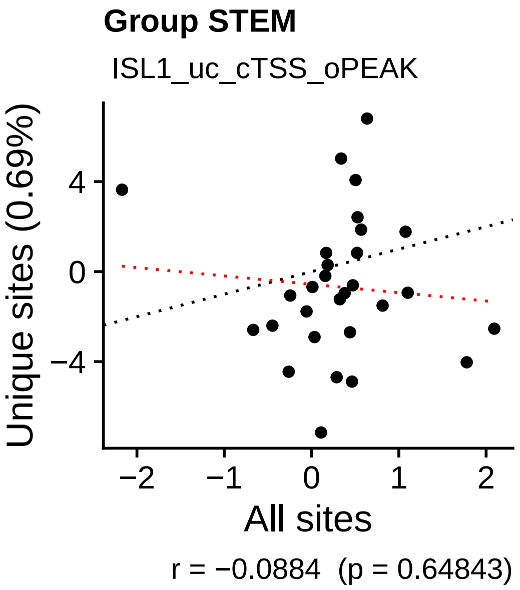
Group STEM
ISL1_uc_cTSS_oPEAK
Unique sites (0.69%)
All sites
r = −0.0884 (p = 0.64843)

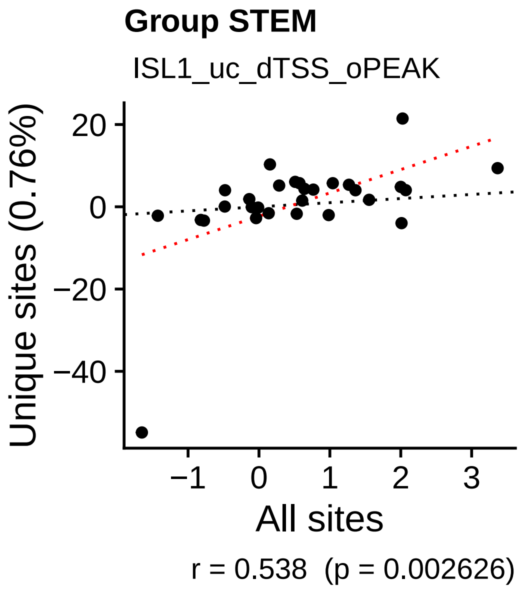
Group STEM
ISL1_uc_dTSS_oPEAK
Unique sites (0.76%)
All sites
r = 0.538 (p = 0.002626)

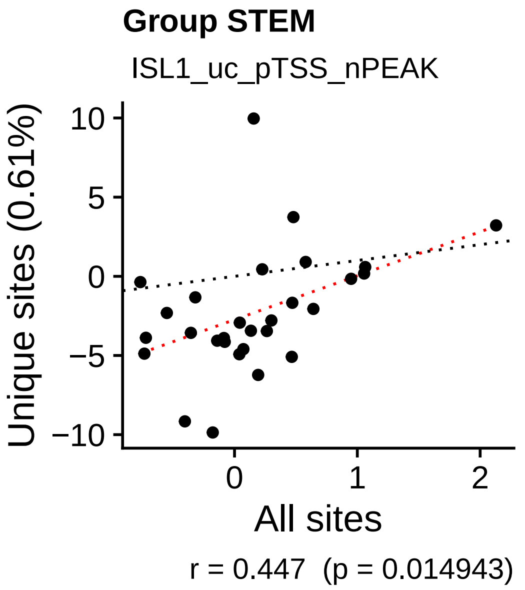
Group STEM
ISL1_uc_pTSS_nPEAK
Unique sites (0.61%)
All sites
r = 0.447 (p = 0.014943)

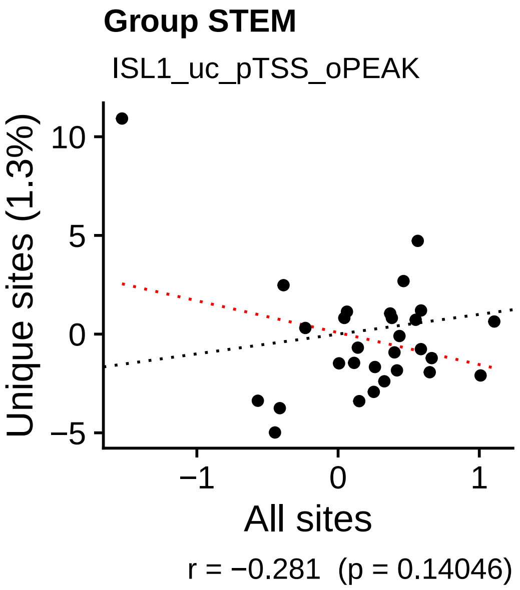
Group STEM
ISL1_uc_pTSS_oPEAK
Unique sites (1.3%)
All sites
r = −0.281 (p = 0.14046)

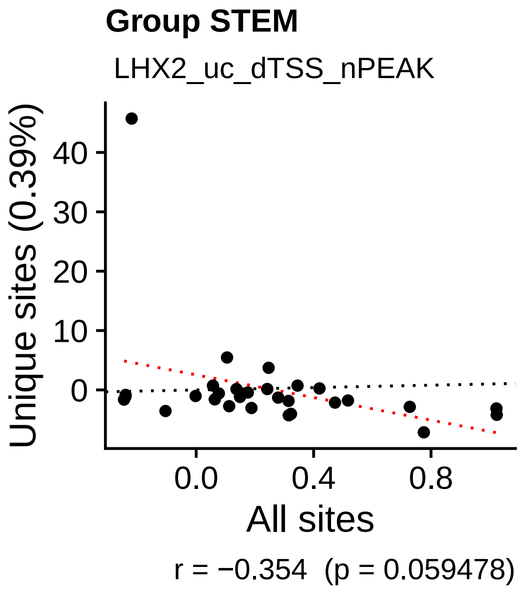
Group STEM
LHX2_uc_dTSS_nPEAK
Unique sites (0.39%)
All sites
r = −0.354 (p = 0.059478)

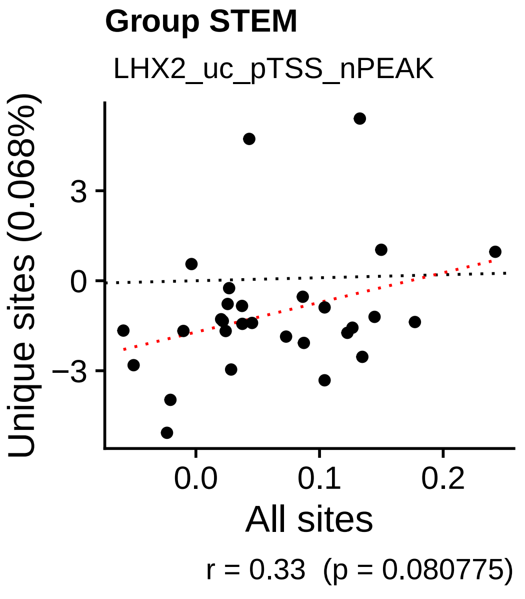
Group STEM
LHX2_uc_pTSS_nPEAK
Unique sites (0.068%)
All sites
r = 0.33 (p = 0.080775)

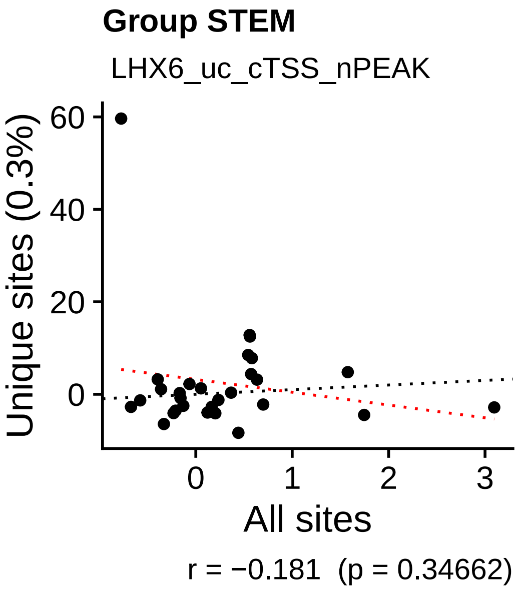
Group STEM
LHX6_uc_cTSS_nPEAK
Unique sites (0.3%)
All sites
r = −0.181 (p = 0.34662)

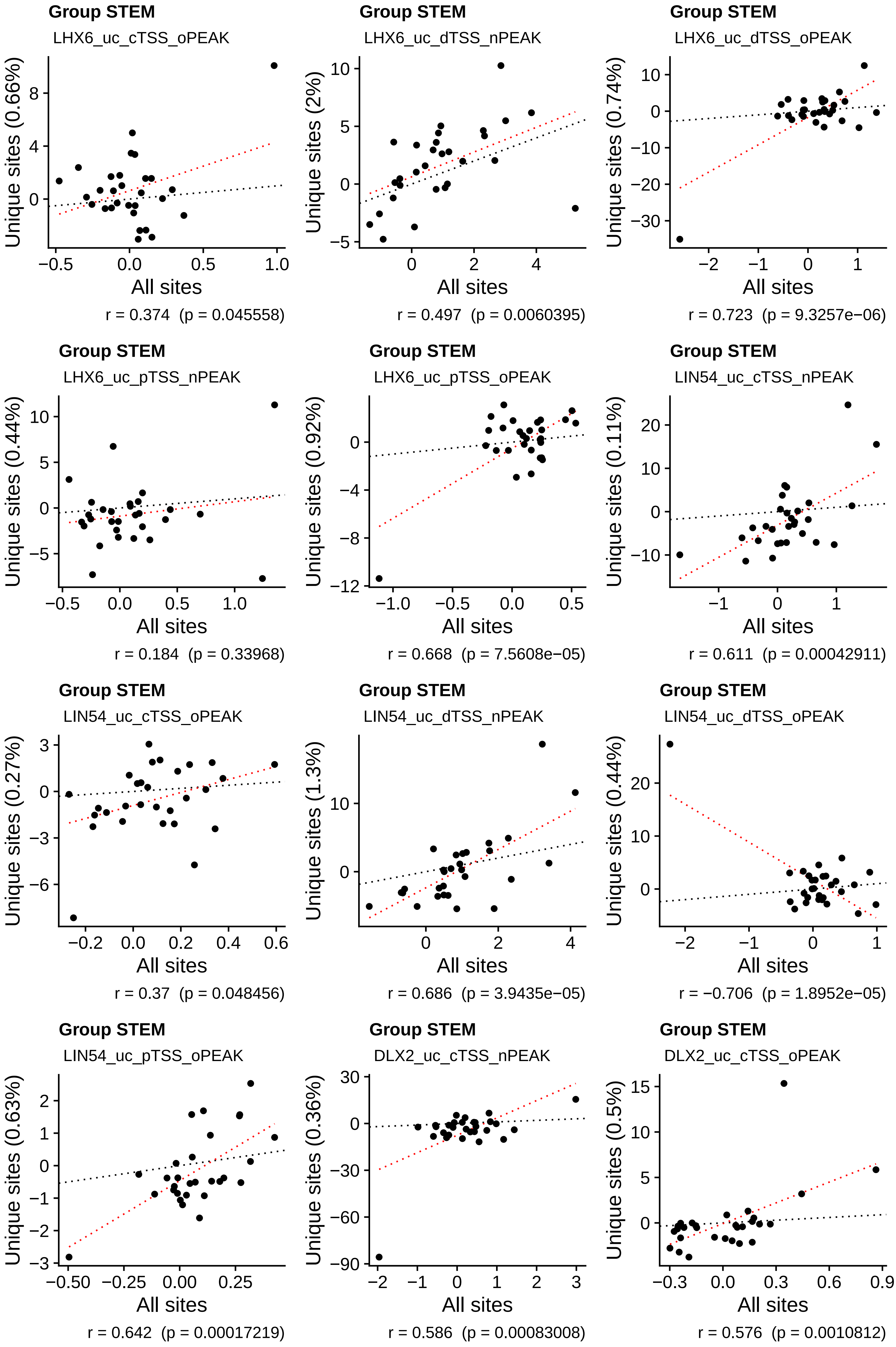
Group STEM
LHX6_uc_cTSS_oPEAK
Unique sites (0.66%)
All sites
r = 0.374  (p = 0.045558)
Group STEM
LHX6_uc_dTSS_nPEAK
Unique sites (2%)
All sites
r = 0.497  (p = 0.0060395)
Group STEM
LHX6_uc_dTSS_oPEAK
Unique sites (0.74%)
All sites
r = 0.723  (p = 9.3257e−06)
Group STEM
LHX6_uc_pTSS_nPEAK
Unique sites (0.44%)
All sites
r = 0.184  (p = 0.33968)
Group STEM
LHX6_uc_pTSS_oPEAK
Unique sites (0.92%)
All sites
r = 0.668  (p = 7.5608e−05)
Group STEM
LIN54_uc_cTSS_nPEAK
Unique sites (0.11%)
All sites
r = 0.611  (p = 0.00042911)
Group STEM
LIN54_uc_cTSS_oPEAK
Unique sites (0.27%)
All sites
r = 0.37  (p = 0.048456)
Group STEM
LIN54_uc_dTSS_nPEAK
Unique sites (1.3%)
All sites
r = 0.686  (p = 3.9435e−05)
Group STEM
LIN54_uc_dTSS_oPEAK
Unique sites (0.44%)
All sites
r = −0.706  (p = 1.8952e−05)
Group STEM
LIN54_uc_pTSS_oPEAK
Unique sites (0.63%)
All sites
r = 0.642  (p = 0.00017219)
Group STEM
DLX2_uc_cTSS_nPEAK
Unique sites (0.36%)
All sites
r = 0.586  (p = 0.00083008)
Group STEM
DLX2_uc_cTSS_oPEAK
Unique sites (0.5%)
All sites
r = 0.576  (p = 0.0010812)

**Group STEM**
DLX2_uc_pTSS_nPEAK

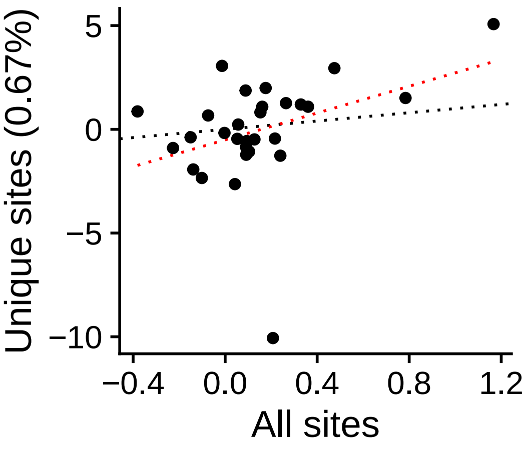


r = 0.372 (p = 0.047048)

**Group STEM**
DLX2_uc_pTSS_oPEAK

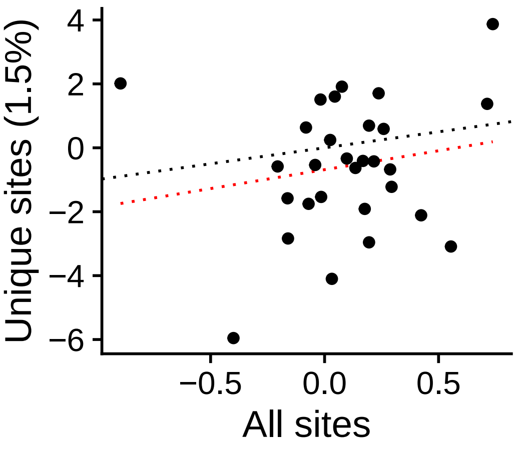


r = 0.181 (p = 0.34865)

**Group STEM**
NANOG_uc_cTSS_nPEAK

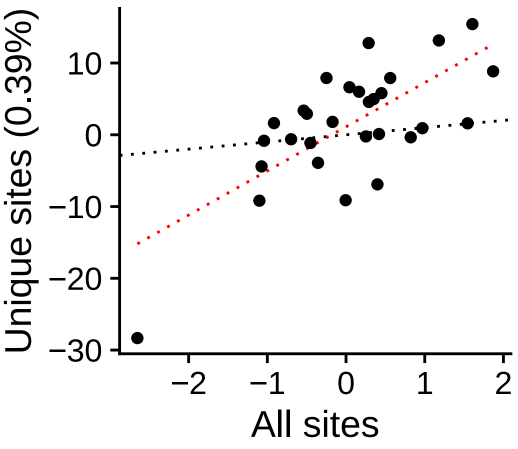


r = 0.7 (p = 2.3645e−05)

**Group STEM**
NANOG_uc_cTSS_oPEAK

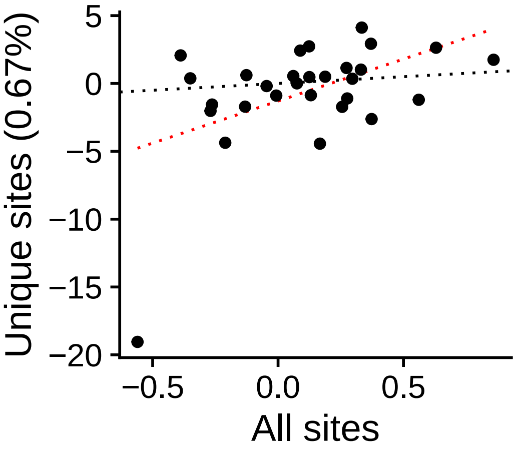


r = 0.481 (p = 0.0082141)

**Group STEM**
NANOG_uc_dTSS_nPEAK

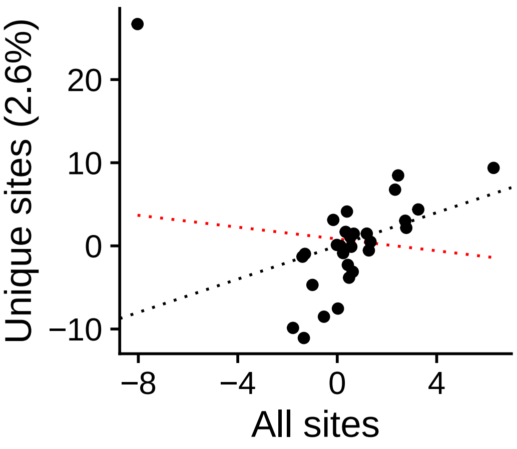


r = −0.118 (p = 0.54081)

**Group STEM**
NANOG_uc_dTSS_oPEAK

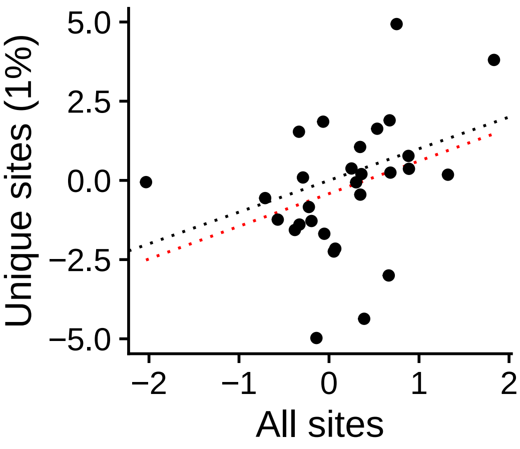


r = 0.347 (p = 0.06519)

**Group STEM**
NANOG_uc_pTSS_nPEAK

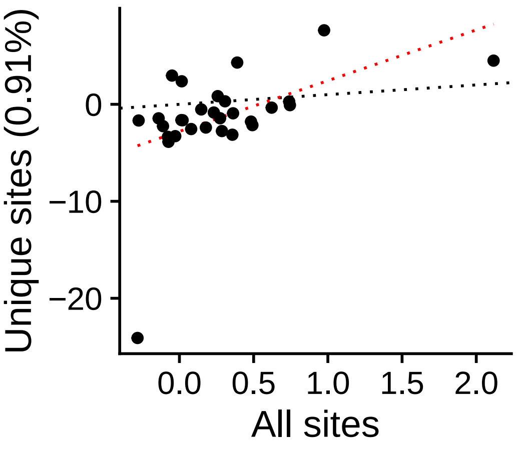


r = 0.481 (p = 0.0082928)

**Group STEM**
NANOG_uc_pTSS_oPEAK

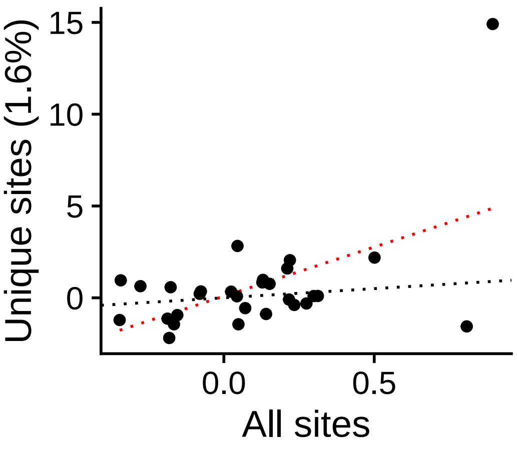


r = 0.53 (p = 0.0031346)

**Group STEM**
NKX24_uc_cTSS_nPEAK

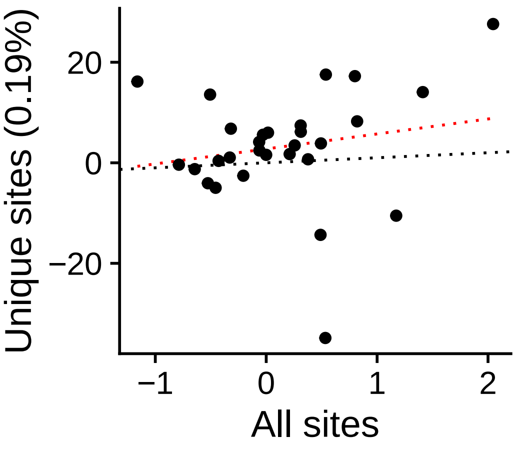


r = 0.181 (p = 0.34874)

**Group STEM**
NKX24_uc_cTSS_oPEAK

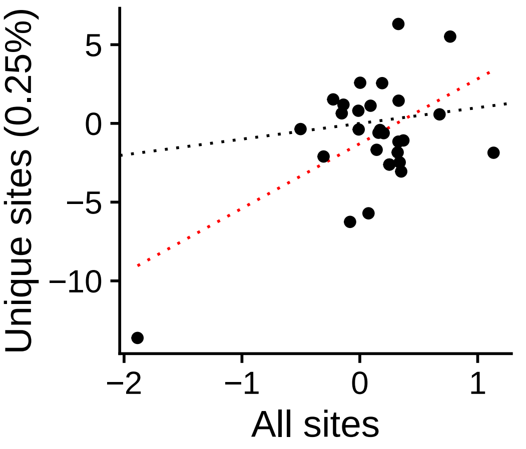


r = 0.565 (p = 0.0014129)

**Group STEM**
NKX24_uc_dTSS_nPEAK

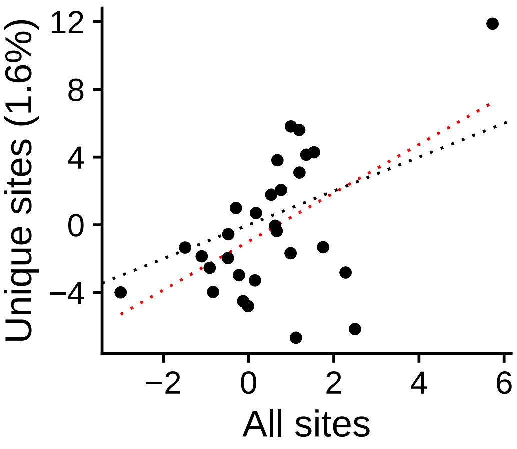


r = 0.533 (p = 0.0028907)

**Group STEM**
NKX24_uc_dTSS_oPEAK

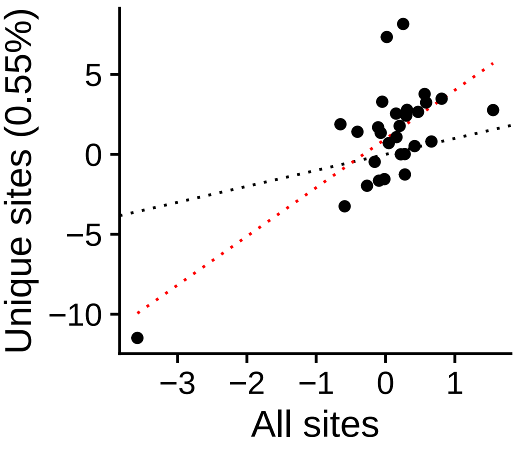


r = 0.723 (p = 9.5107e−06)

**Group STEM**
NKX24_uc_pTSS_nPEAK

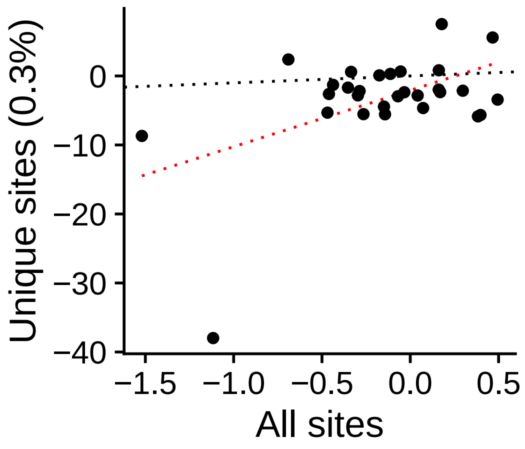


r = 0.487 (p = 0.0073767)

**Group STEM**
NKX24_uc_pTSS_oPEAK

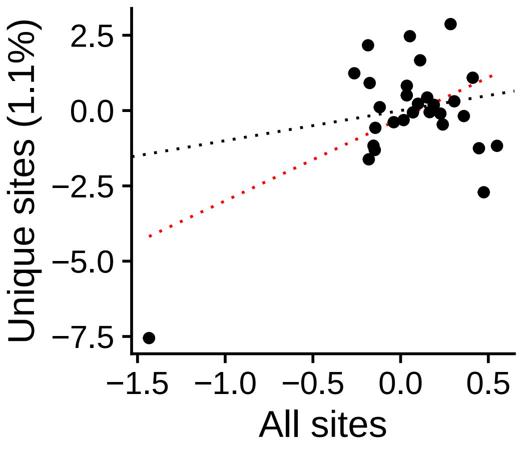


r = 0.523 (p = 0.0035959)

**Group STEM**
OTX2_uc_cTSS_nPEAK

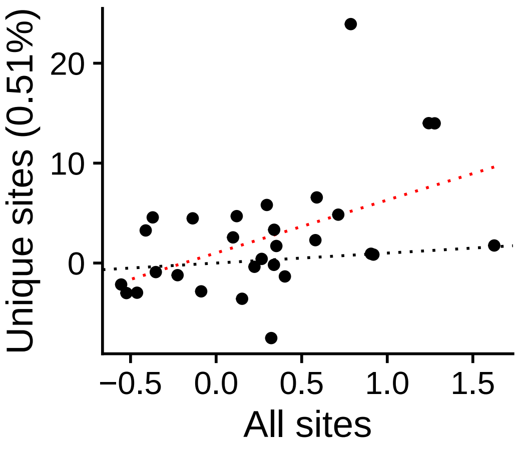


r = 0.483 (p = 0.0079075)

**Group STEM**
OTX2_uc_dTSS_nPEAK

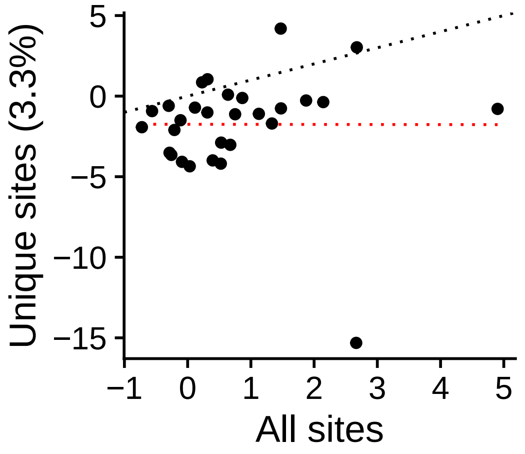


r = −0.00154 (p = 0.99369)

**Group STEM**
OTX2_uc_dTSS_oPEAK

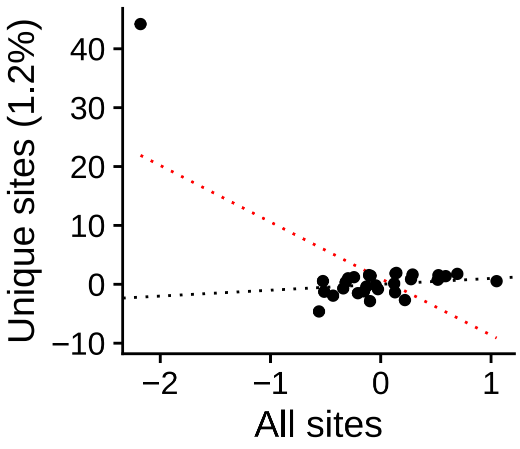


r = −0.648 (p = 0.00014542)

**Group STEM**
OTX2_uc_pTSS_nPEAK

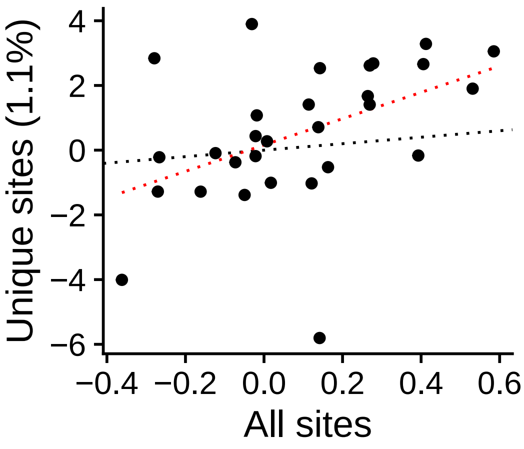


r = 0.459 (p = 0.012158)

**Group STEM**
OTX2_uc_pTSS_oPEAK

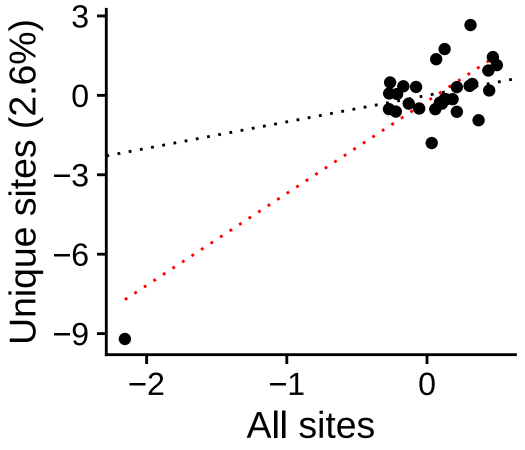


r = 0.855 (p = 3.6517e−09)

**Group STEM**
PBX4_uc_cTSS_nPEAK

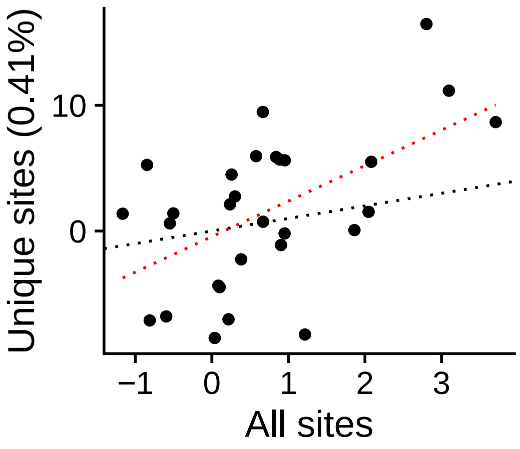


r = 0.548 (p = 0.0020874)

**Group STEM**
PBX4_uc_cTSS_oPEAK

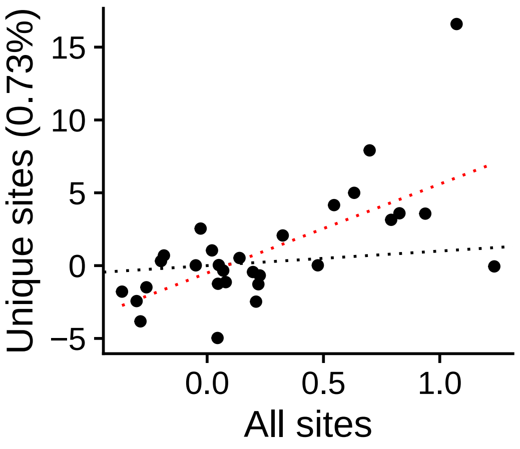


r = 0.653 (p = 0.00012133)

**Group STEM**
PBX4_uc_dTSS_nPEAK

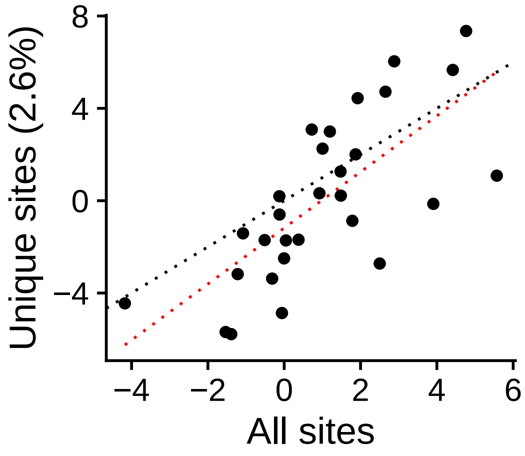


r = 0.723 (p = 9.3546e−06)

**Group STEM**
PBX4_uc_dTSS_oPEAK

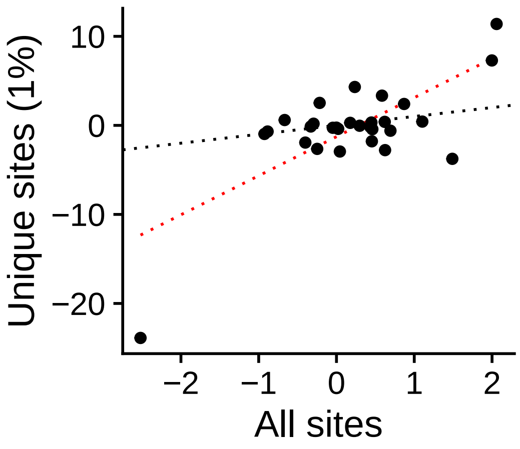


r = 0.72 (p = 1.0469e−05)

**Group STEM**
PBX4_uc_pTSS_nPEAK

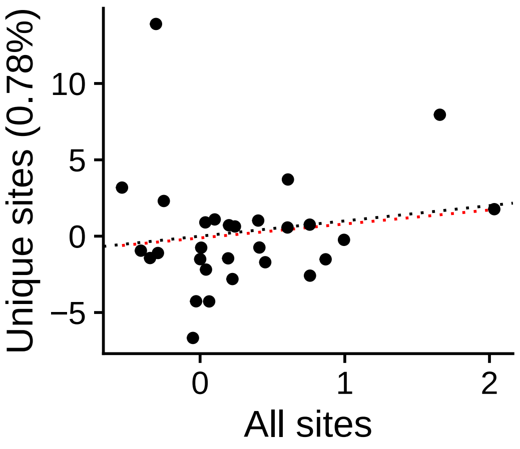


r = 0.14 (p = 0.46762)

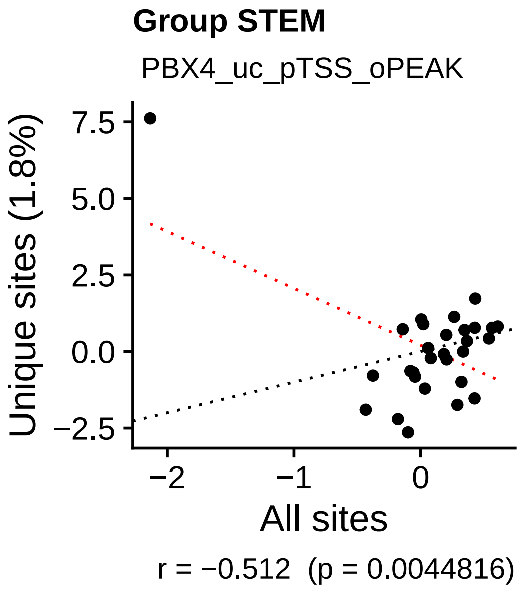
Group STEM
PBX4_uc_pTSS_oPEAK
Unique sites (1.8%)
All sites
r = −0.512 (p = 0.0044816)

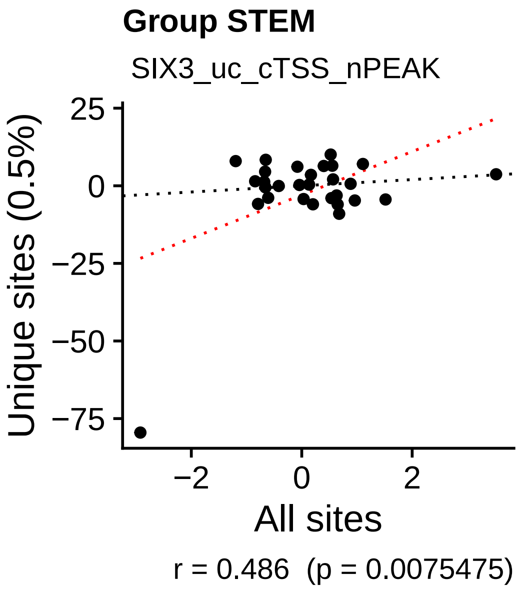
Group STEM
SIX3_uc_cTSS_nPEAK
Unique sites (0.5%)
All sites
r = 0.486 (p = 0.0075475)

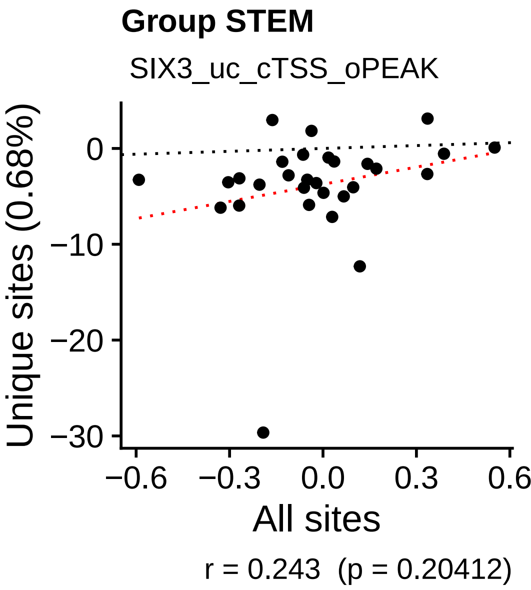
Group STEM
SIX3_uc_cTSS_oPEAK
Unique sites (0.68%)
All sites
r = 0.243 (p = 0.20412)

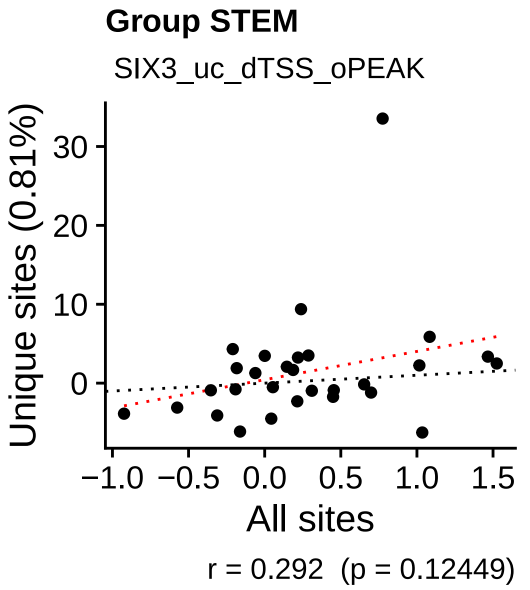
Group STEM
SIX3_uc_dTSS_oPEAK
Unique sites (0.81%)
All sites
r = 0.292 (p = 0.12449)

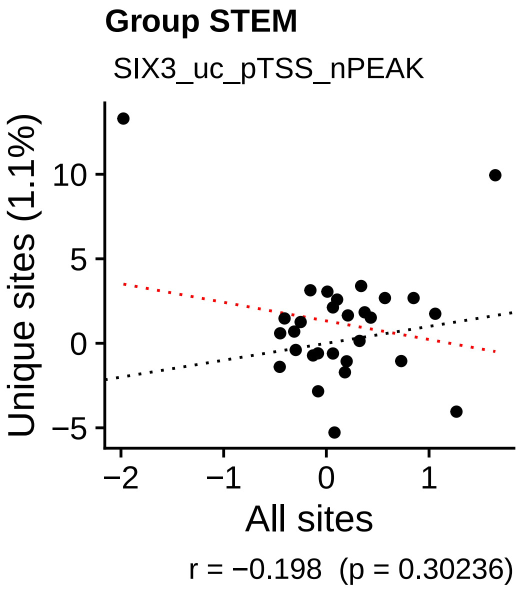
Group STEM
SIX3_uc_pTSS_nPEAK
Unique sites (1.1%)
All sites
r = −0.198 (p = 0.30236)

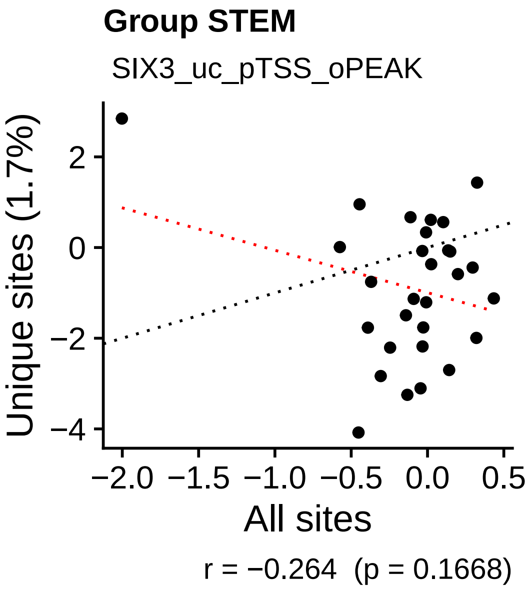
Group STEM
SIX3_uc_pTSS_oPEAK
Unique sites (1.7%)
All sites
r = −0.264 (p = 0.1668)

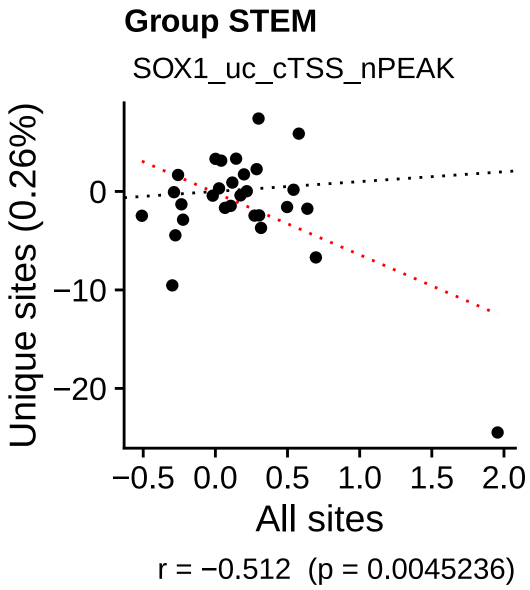
Group STEM
SOX1_uc_cTSS_nPEAK
Unique sites (0.26%)
All sites
r = −0.512 (p = 0.0045236)

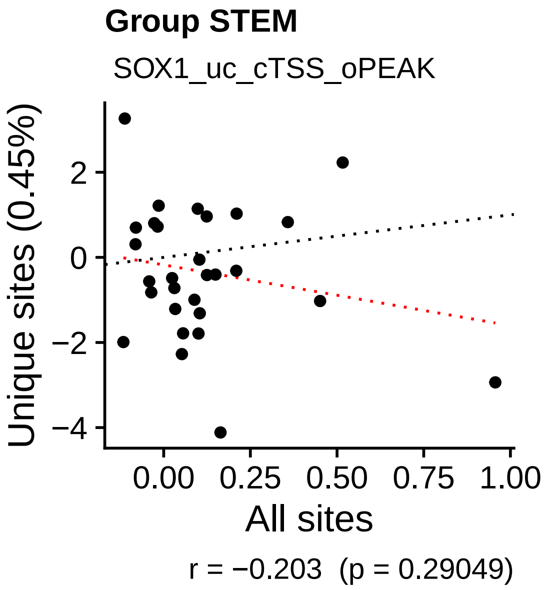
Group STEM
SOX1_uc_cTSS_oPEAK
Unique sites (0.45%)
All sites
r = −0.203 (p = 0.29049)

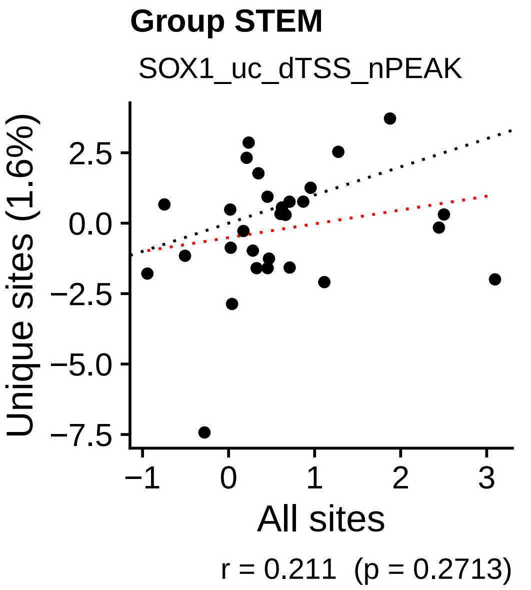
Group STEM
SOX1_uc_dTSS_nPEAK
Unique sites (1.6%)
All sites
r = 0.211 (p = 0.2713)

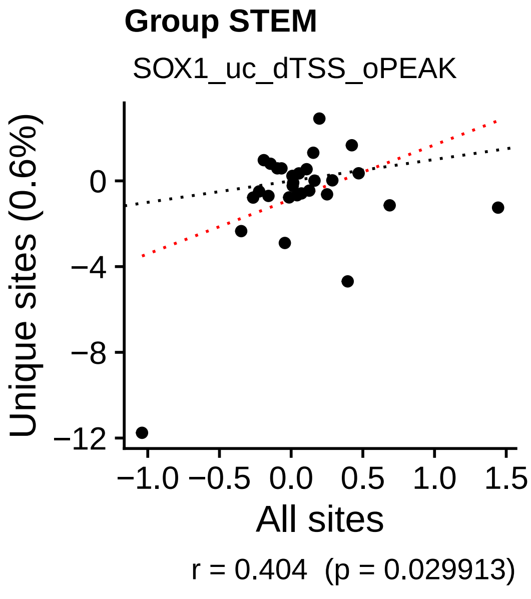
Group STEM
SOX1_uc_dTSS_oPEAK
Unique sites (0.6%)
All sites
r = 0.404 (p = 0.029913)

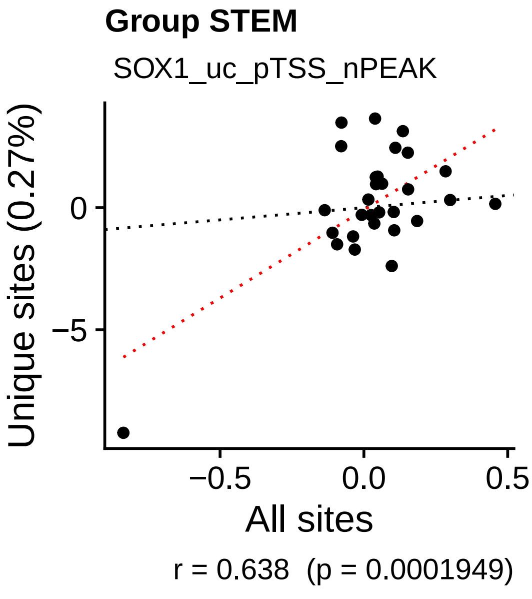
Group STEM
SOX1_uc_pTSS_nPEAK
Unique sites (0.27%)
All sites
r = 0.638 (p = 0.0001949)

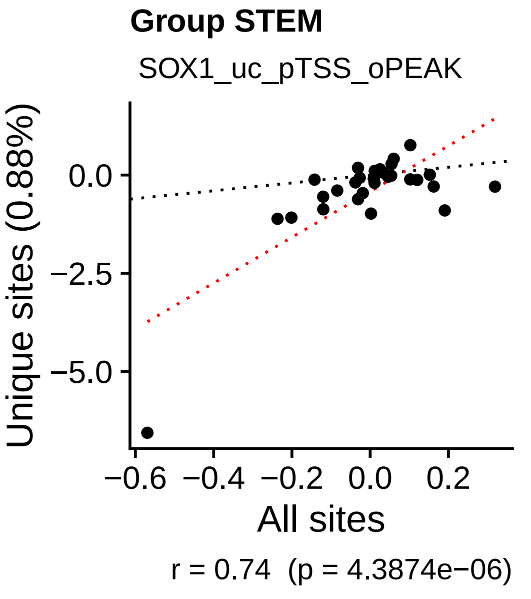
Group STEM
SOX1_uc_pTSS_oPEAK
Unique sites (0.88%)
All sites
r = 0.74 (p = 4.3874e−06)

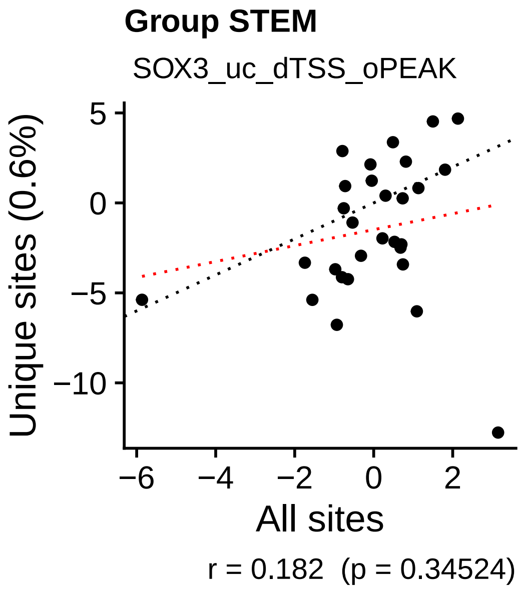
Group STEM
SOX3_uc_dTSS_oPEAK
Unique sites (0.6%)
All sites
r = 0.182 (p = 0.34524)

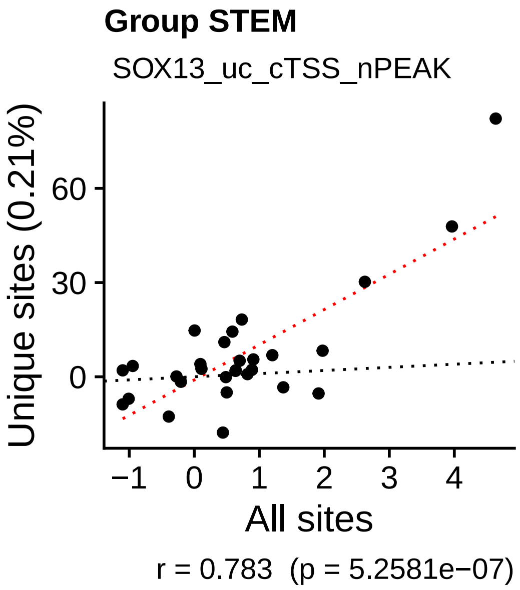
Group STEM
SOX13_uc_cTSS_nPEAK
Unique sites (0.21%)
All sites
r = 0.783 (p = 5.2581e−07)

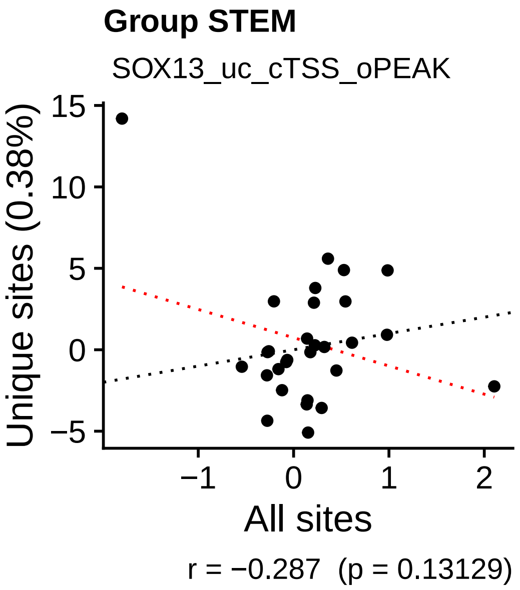
Group STEM
SOX13_uc_cTSS_oPEAK
Unique sites (0.38%)
All sites
r = −0.287 (p = 0.13129)

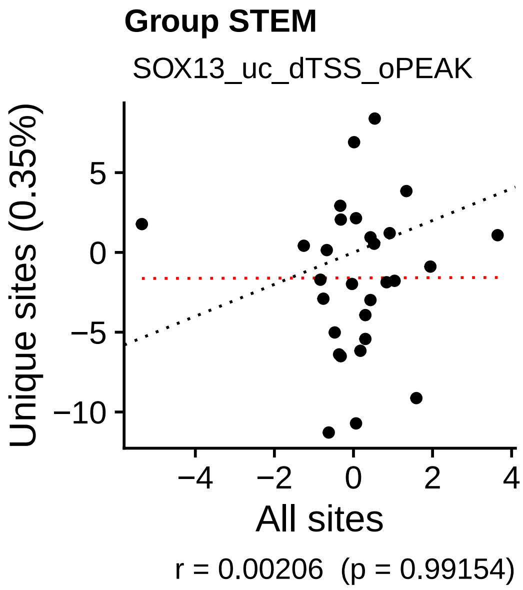
Group STEM
SOX13_uc_dTSS_oPEAK
Unique sites (0.35%)
All sites
r = 0.00206 (p = 0.99154)

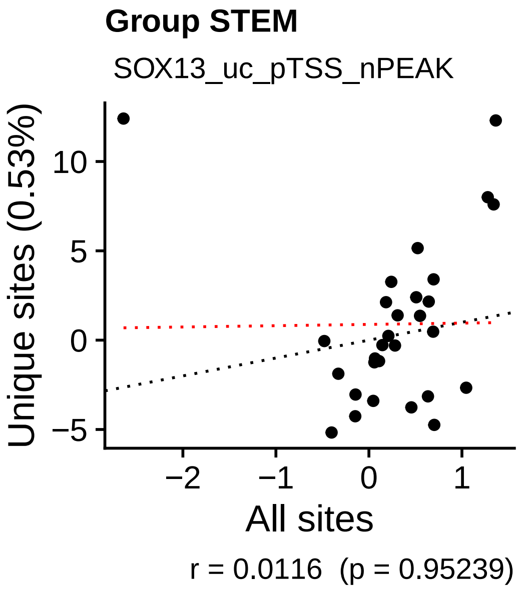
Group STEM
SOX13_uc_pTSS_nPEAK
Unique sites (0.53%)
All sites
r = 0.0116 (p = 0.95239)

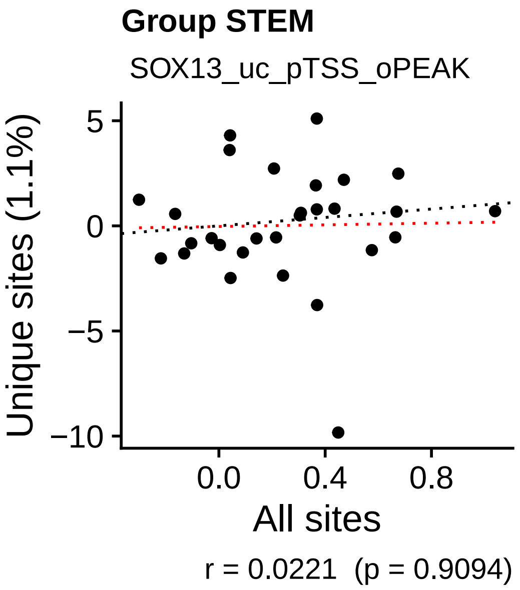
Group STEM
SOX13_uc_pTSS_oPEAK
Unique sites (1.1%)
All sites
r = 0.0221 (p = 0.9094)

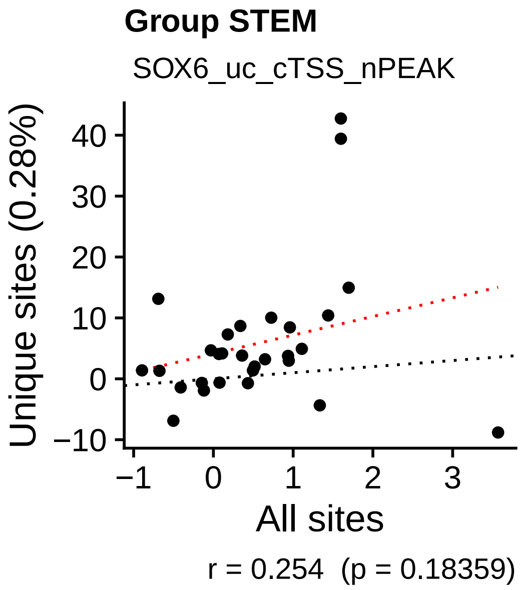
Group STEM
SOX6_uc_cTSS_nPEAK
Unique sites (0.28%)
All sites
r = 0.254 (p = 0.18359)

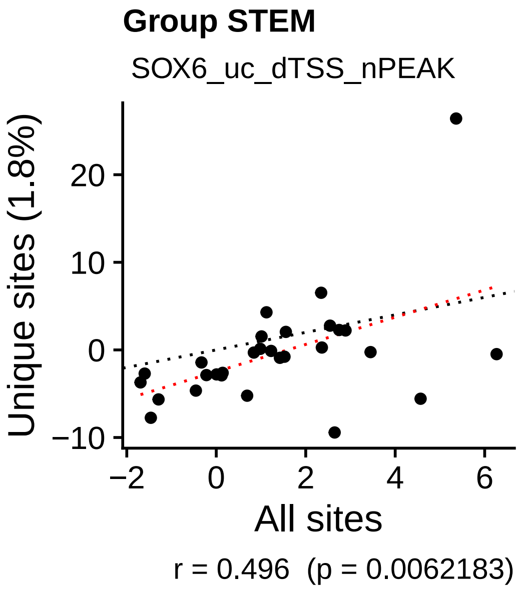
Group STEM
SOX6_uc_dTSS_nPEAK
Unique sites (1.8%)
All sites
r = 0.496 (p = 0.0062183)

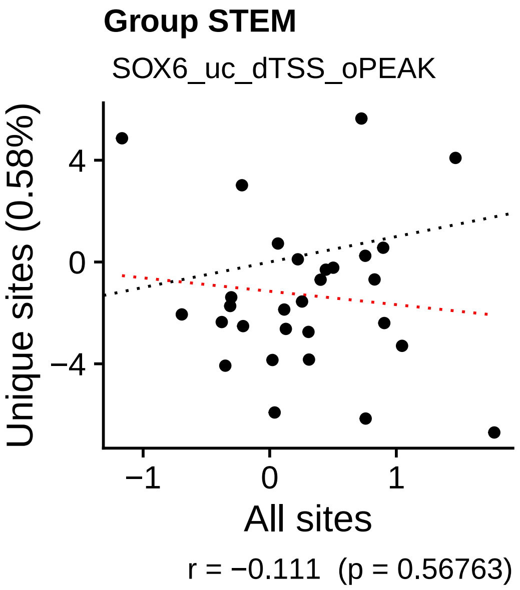
Group STEM
SOX6_uc_dTSS_oPEAK
Unique sites (0.58%)
All sites
r = −0.111 (p = 0.56763)

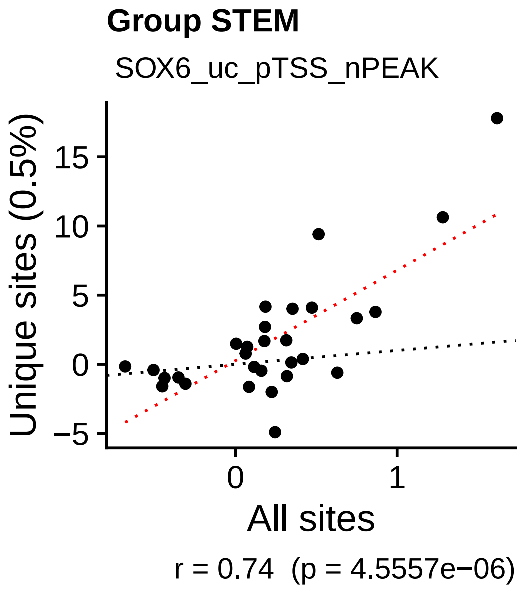
Group STEM
SOX6_uc_pTSS_nPEAK
Unique sites (0.5%)
All sites
r = 0.74 (p = 4.5557e−06)

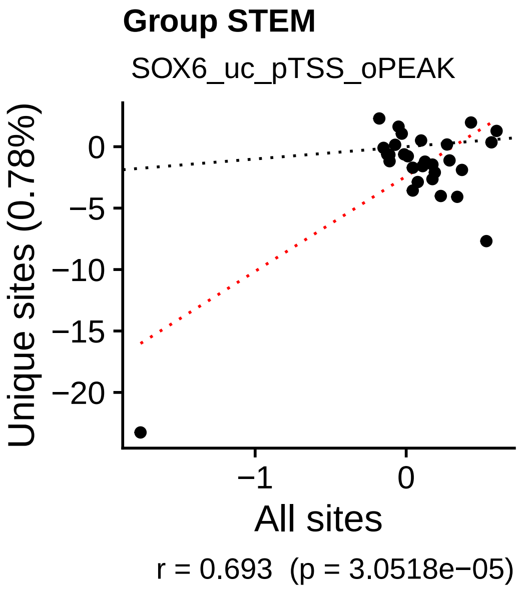
Group STEM
SOX6_uc_pTSS_oPEAK
Unique sites (0.78%)
All sites
r = 0.693 (p = 3.0518e−05)

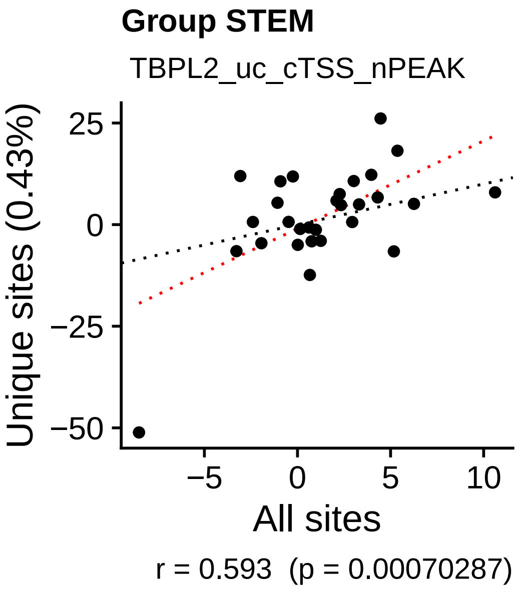
Group STEM
TBPL2_uc_cTSS_nPEAK
Unique sites (0.43%)
All sites
r = 0.593 (p = 0.00070287)

**Group STEM**

TBPL2_uc_cTSS_oPEAK

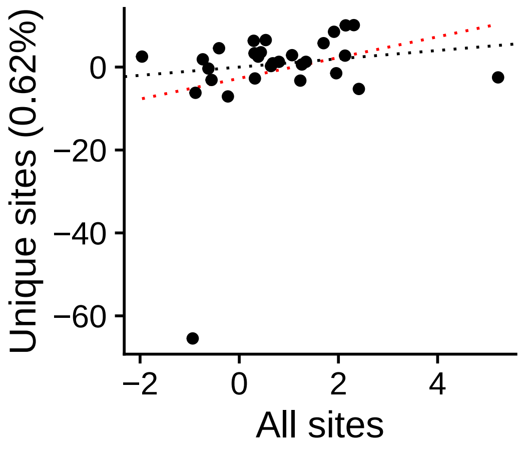


r = 0.267  (p = 0.16177)

**Group STEM**

TBPL2_uc_dTSS_oPEAK

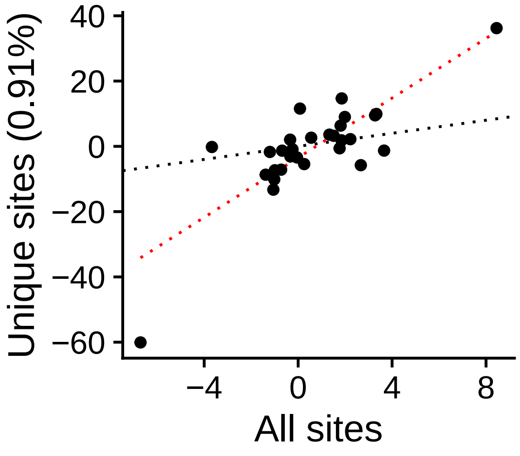


r = 0.813  (p = 8.2463e−08)

**Group STEM**

TBPL2_uc_pTSS_nPEAK

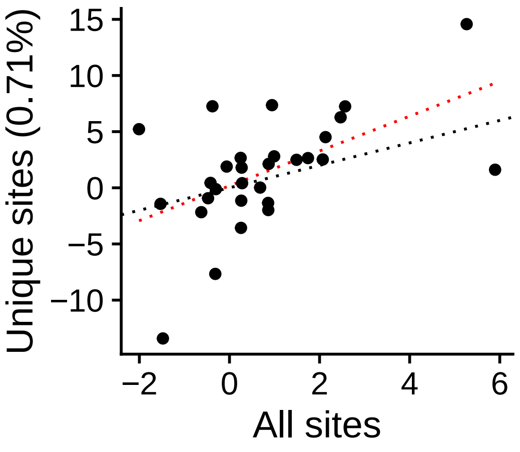


r = 0.537  (p = 0.0026451)

**Group STEM**

TBPL2_uc_pTSS_oPEAK

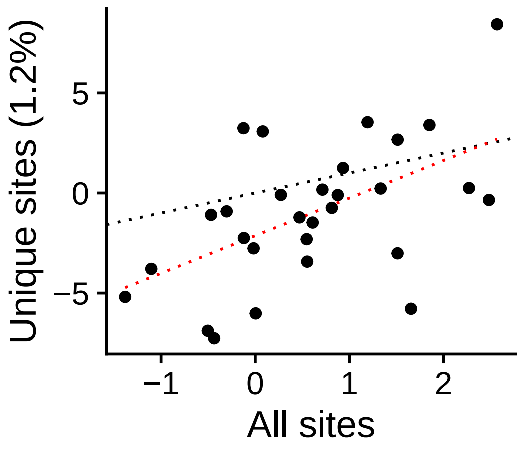


r = 0.538  (p = 0.0025876)

**Group STEM**

TCF7L2_uc_cTSS_nPEAK

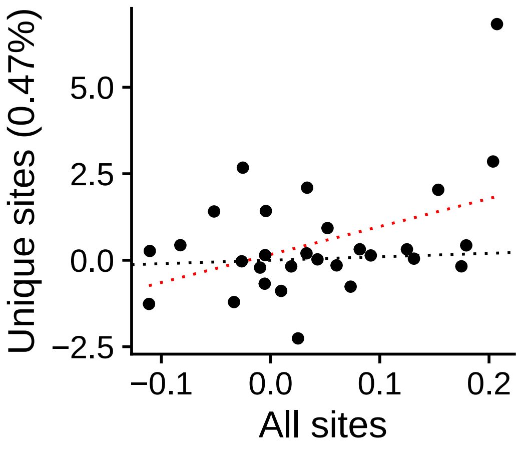


r = 0.428  (p = 0.0206)

**Group STEM**

TCF7L2_uc_cTSS_oPEAK

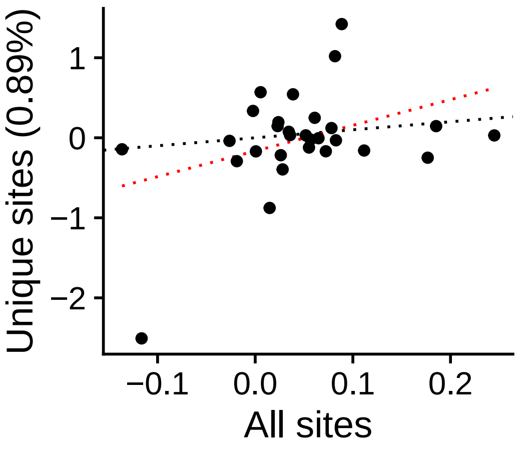


r = 0.385  (p = 0.039112)

**Group STEM**

TCF7L2_uc_dTSS_nPEAK

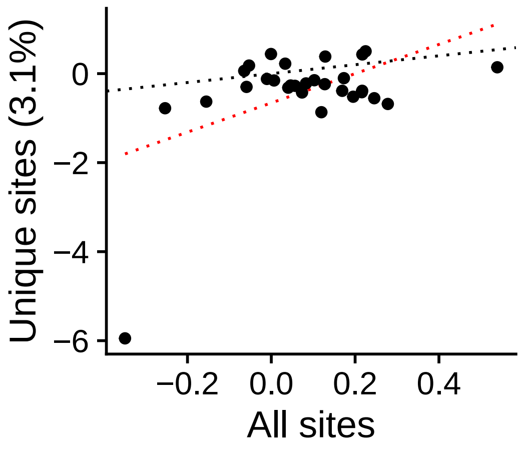


r = 0.499  (p = 0.0058515)

**Group STEM**

TCF7L2_uc_dTSS_oPEAK

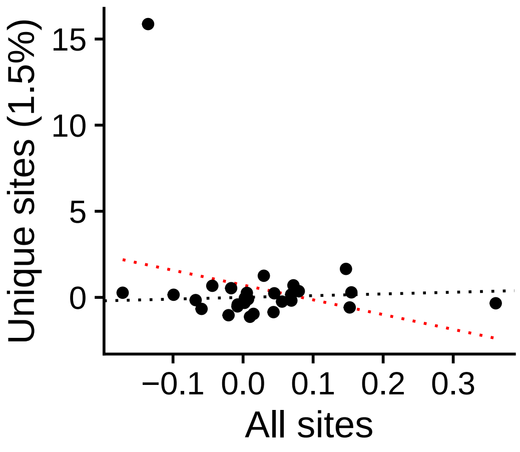


r = −0.284  (p = 0.13571)

**Group STEM**

ZKSCAN1_uc_dTSS_nPEAK

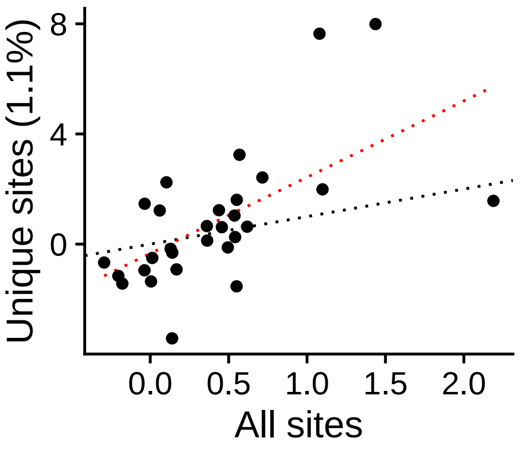


r = 0.606  (p = 0.00048715)

**Group STEM**

ZKSCAN1_uc_pTSS_oPEAK

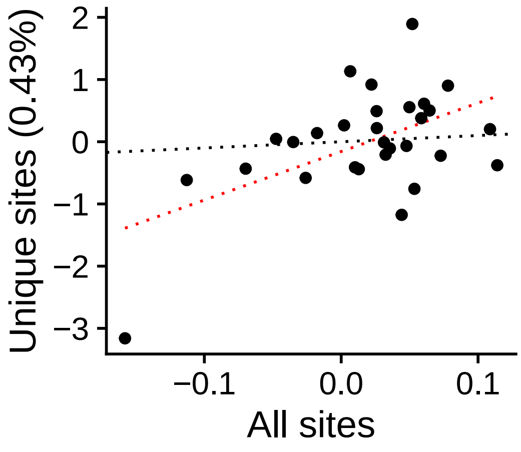


r = 0.538  (p = 0.0026146)

**Group STEM**

ZNF35_uc_cTSS_nPEAK

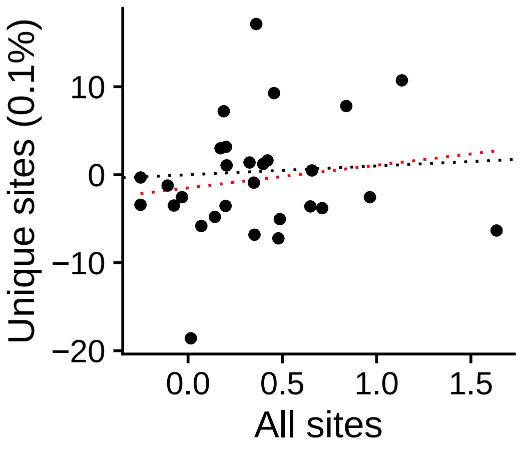


r = 0.159  (p = 0.40879)

**Group STEM**

ZNF35_uc_cTSS_oPEAK

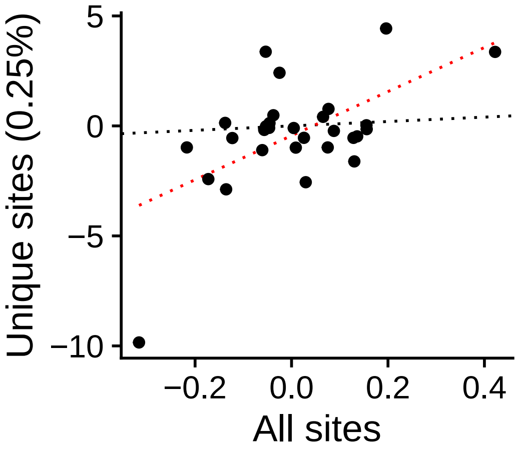


r = 0.588  (p = 0.00078628)

**Group STEM**

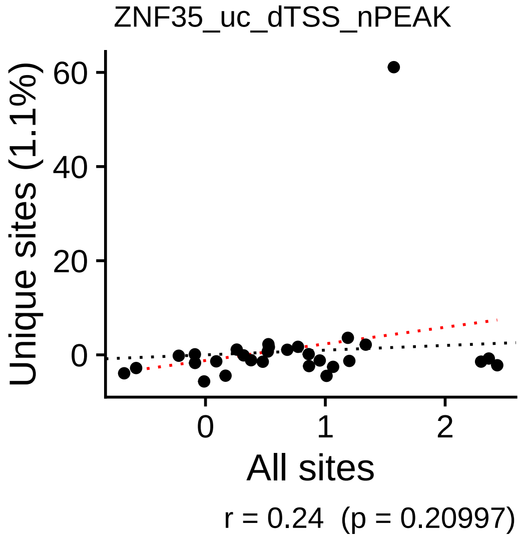


**Group STEM**

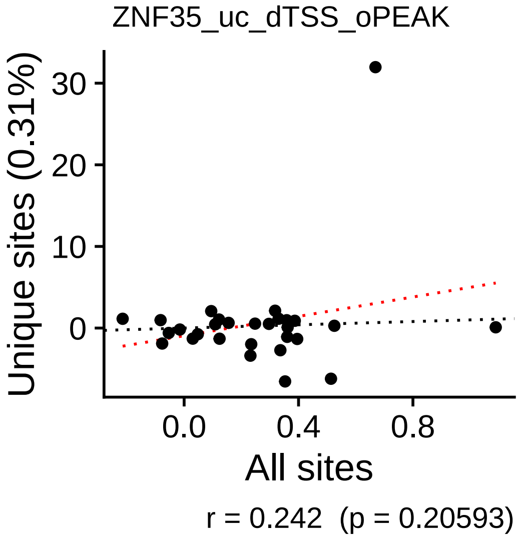


**Group STEM**

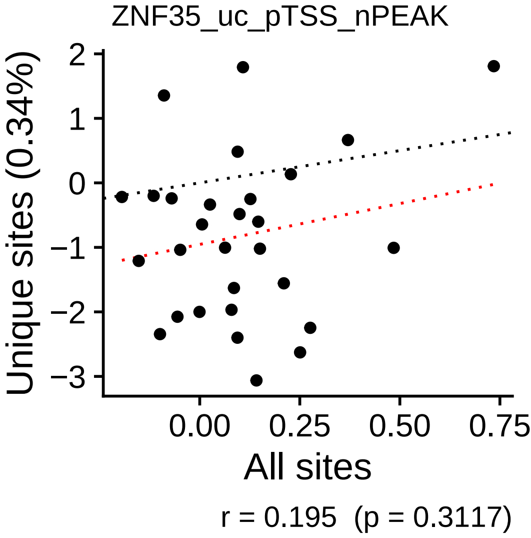


**Group STEM**

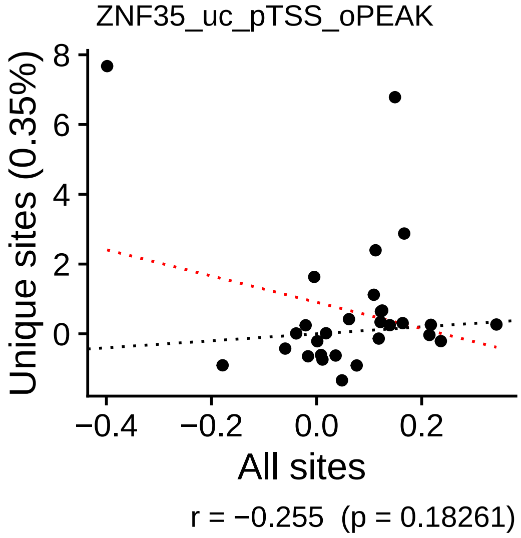
