## Supplementary Figures for "The co-evolution of the genome and epigenome in colorectal cancer": figS24.pdf

### AC0125311_uc_cTSS_nPEAK

+ 10/33 (1/116), – 4/33 (39/116)

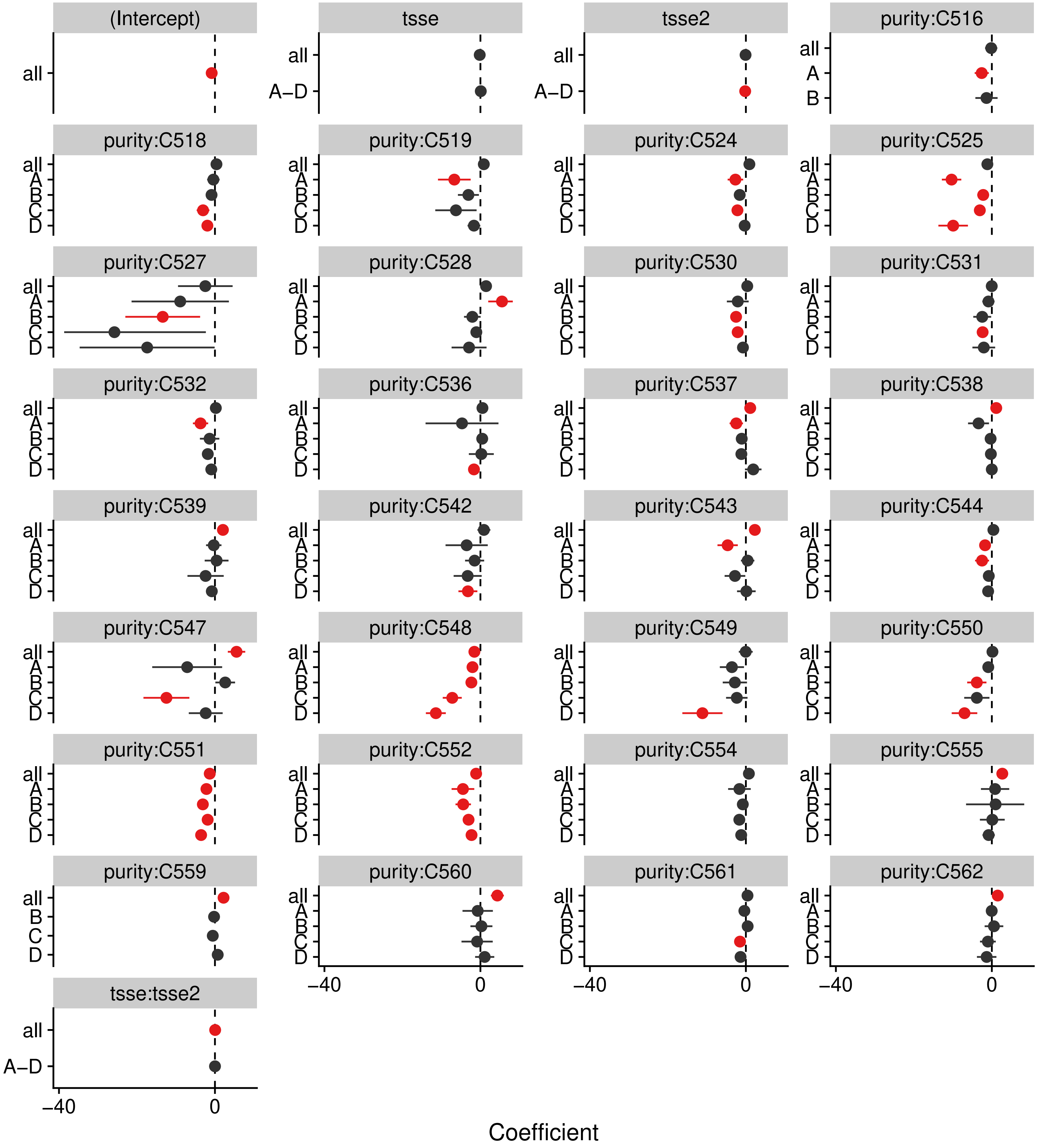


p_c = < 2.22e–16, p_sc = 0.021685

#### AC0125311_uc_dTSS_oPEAK

+ 12/33 (0/116), – 4/33 (42/116)

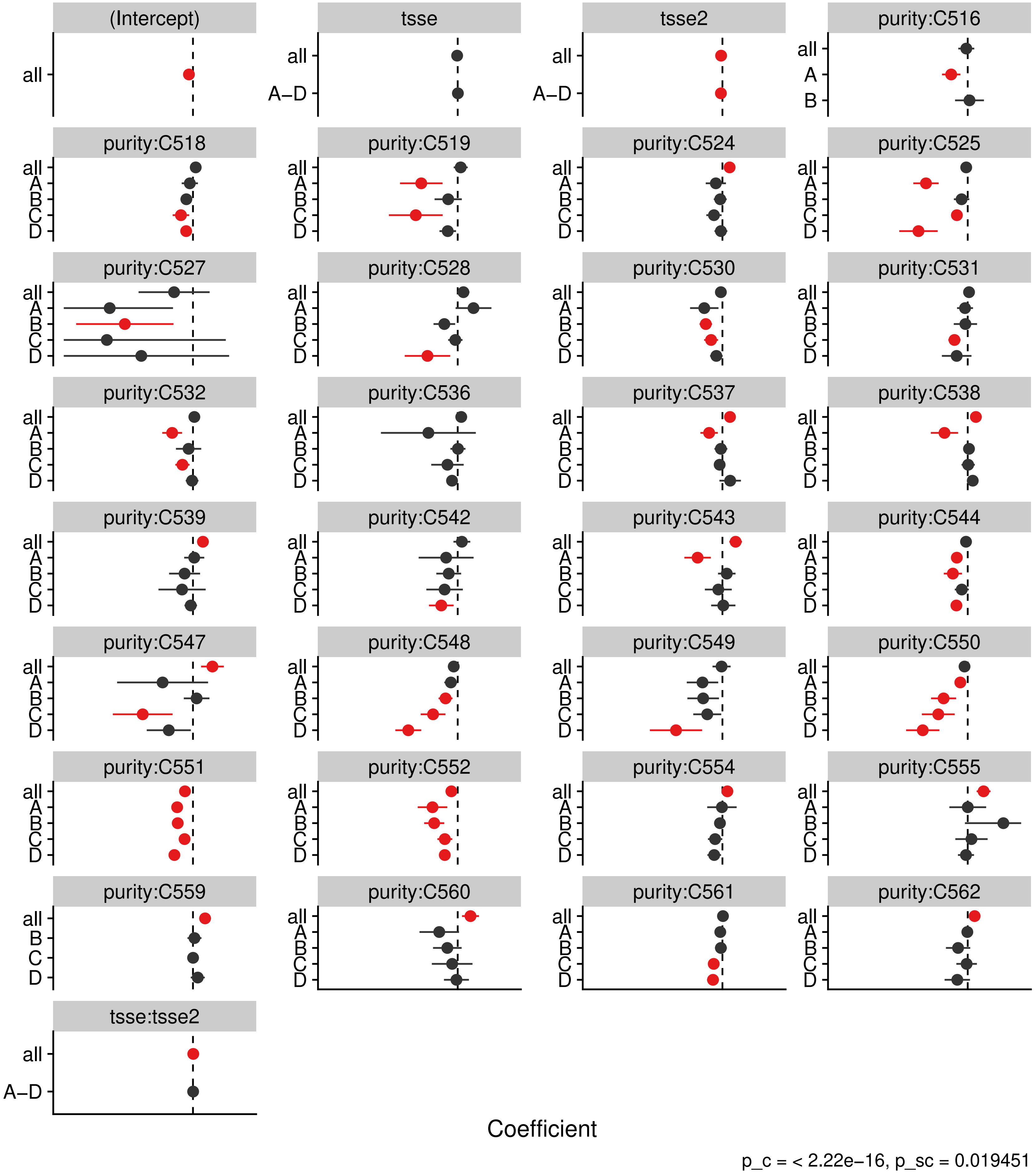

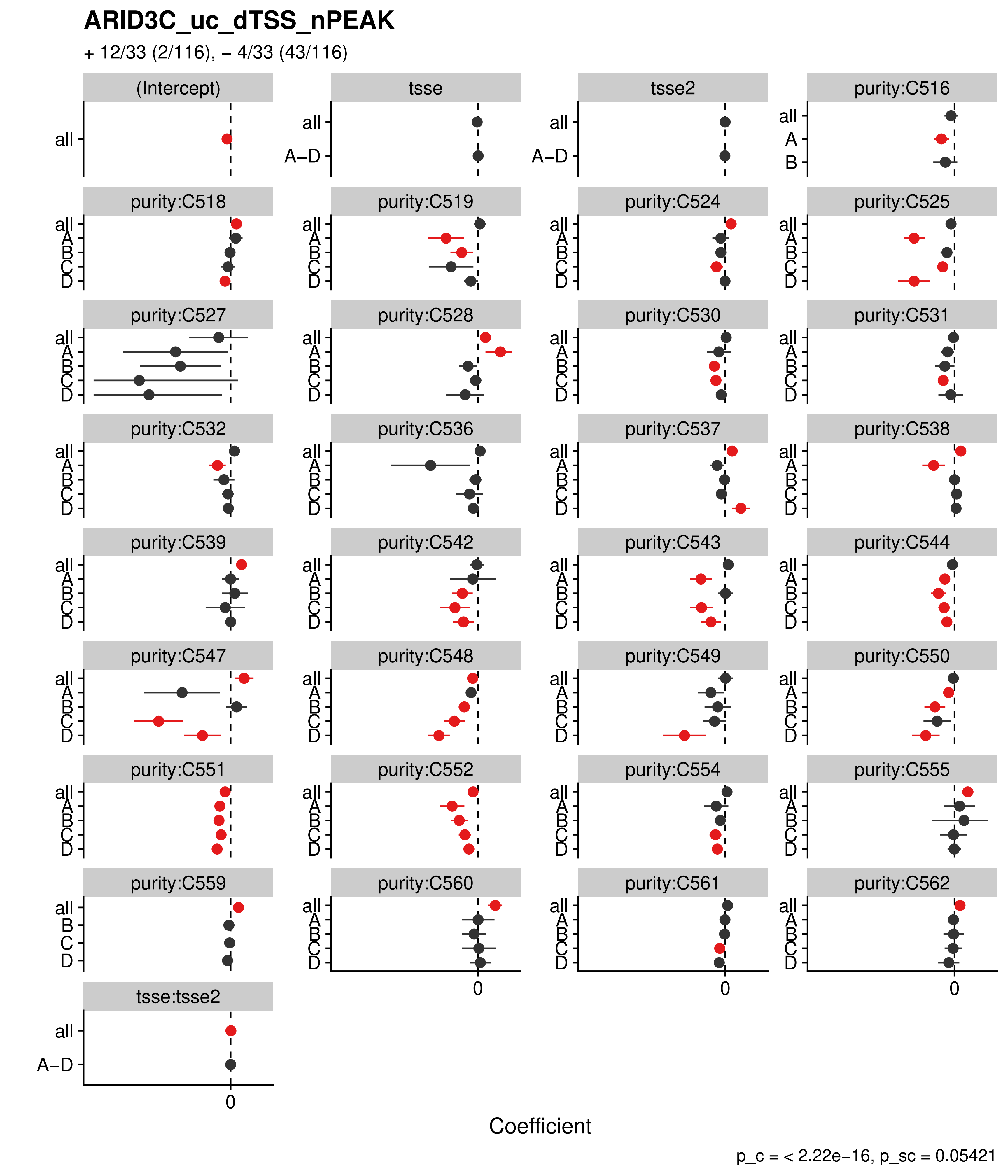
ARID3C_uc_dTSS_nPEAK
+ 12/33 (2/116), – 4/33 (43/116)
(Intercept)
tsse
tsse2
purity:C516
purity:C518
purity:C519
purity:C524
purity:C525
purity:C527
purity:C528
purity:C530
purity:C531
purity:C532
purity:C536
purity:C537
purity:C538
purity:C539
purity:C542
purity:C543
purity:C544
purity:C547
purity:C548
purity:C549
purity:C550
purity:C551
purity:C552
purity:C554
purity:C555
purity:C559
purity:C560
purity:C561
purity:C562
tsse:tsse2
all
A–D
A
B
C
D
0
Coefficient
p_c = < 2.22e–16, p_sc = 0.05421

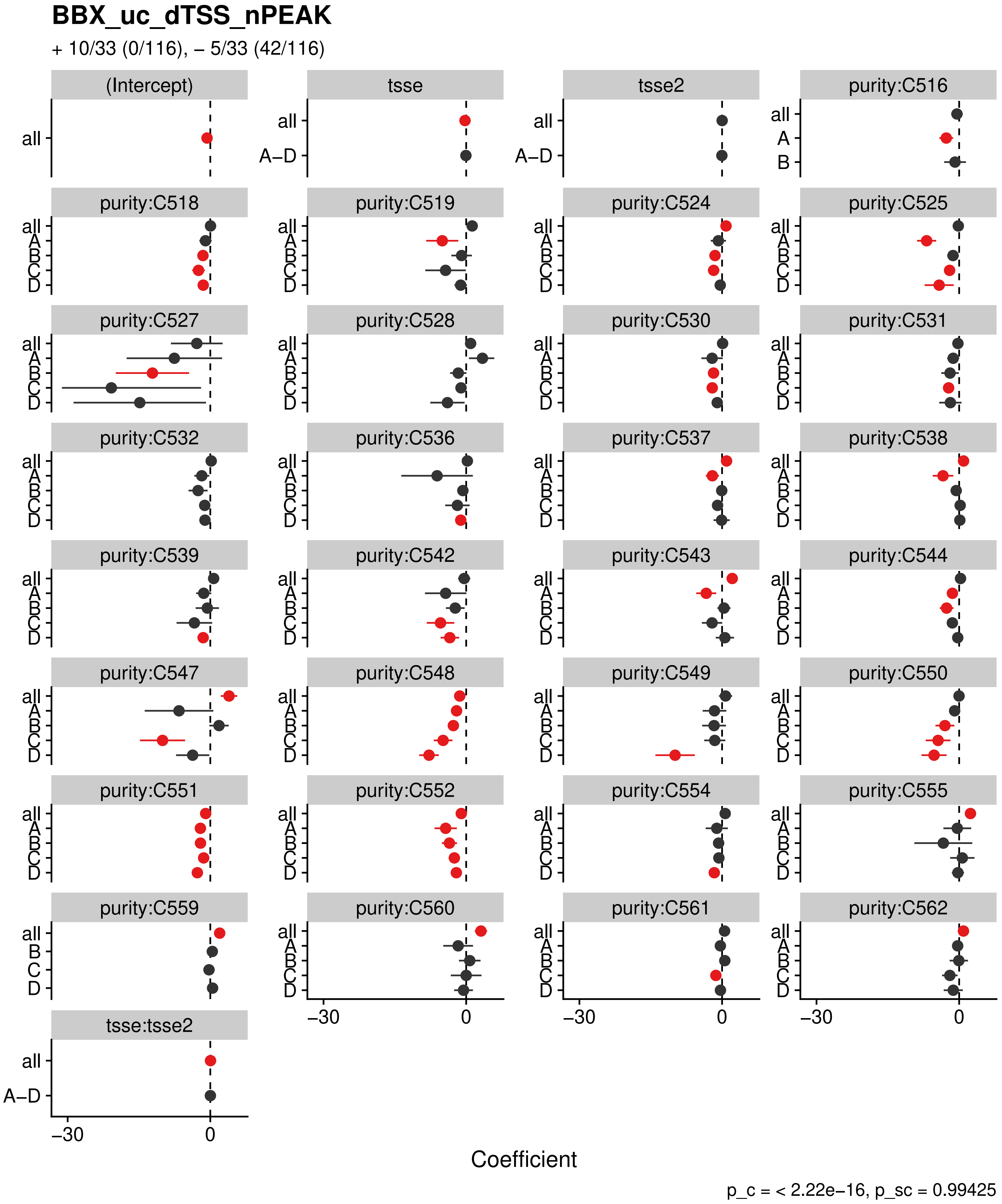

BBX_uc_dTSS_nPEAK
+ 10/33 (0/116), – 5/33 (42/116)
(Intercept)
tsse
tsse2
purity:C516
purity:C518
purity:C519
purity:C524
purity:C525
purity:C527
purity:C528
purity:C530
purity:C531
purity:C532
purity:C536
purity:C537
purity:C538
purity:C539
purity:C542
purity:C543
purity:C544
purity:C547
purity:C548
purity:C549
purity:C550
purity:C551
purity:C552
purity:C554
purity:C555
purity:C559
purity:C560
purity:C561
purity:C562
tsse:tsse2
all
A–D
A
B
C
D
–30
0
Coefficient
p_c = < 2.22e–16, p_sc = 0.99425

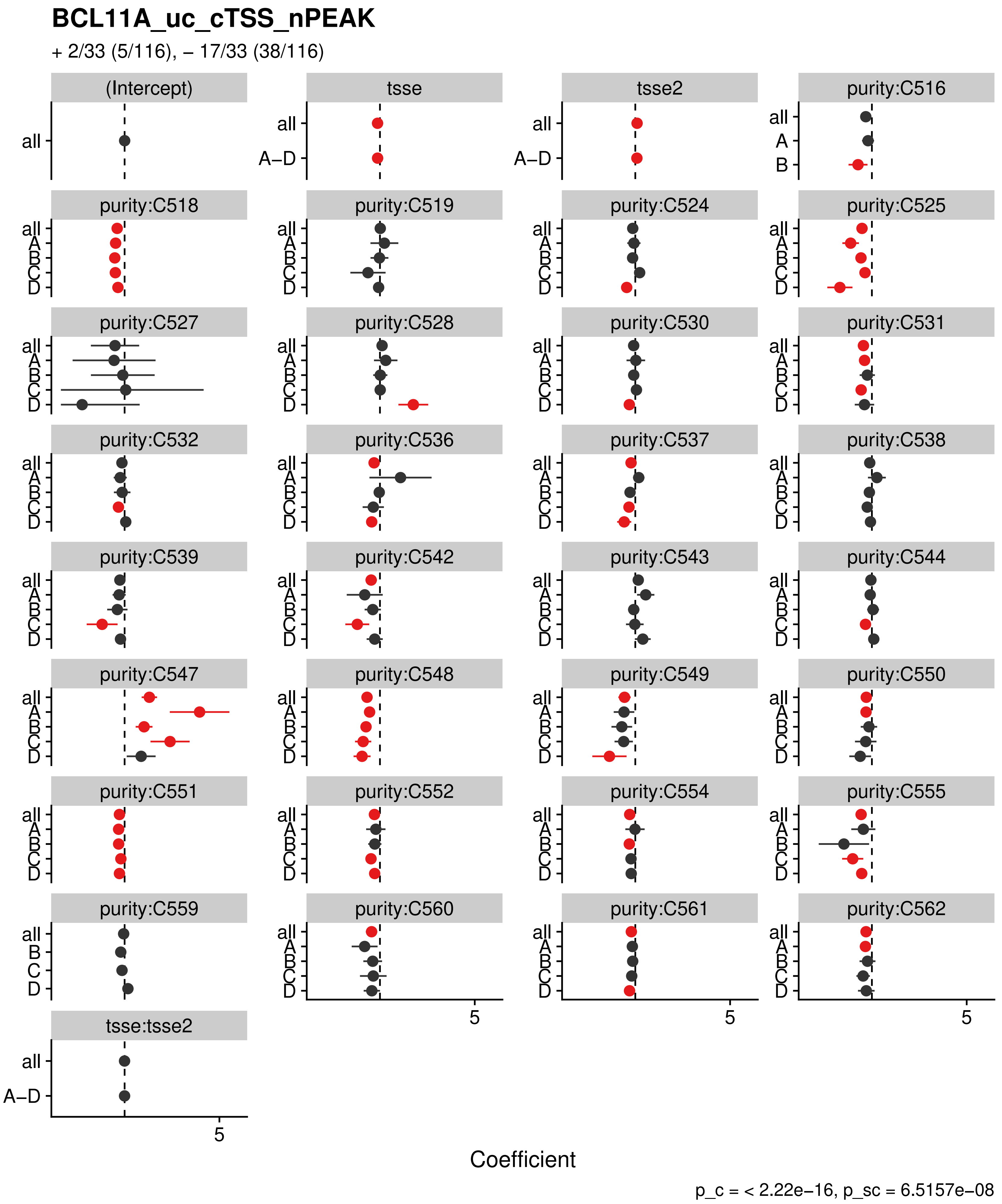

BCL11A_uc_cTSS_nPEAK
+ 2/33 (5/116), – 17/33 (38/116)
(Intercept)
tsse
tsse2
purity:C516
purity:C518
purity:C519
purity:C524
purity:C525
purity:C527
purity:C528
purity:C530
purity:C531
purity:C532
purity:C536
purity:C537
purity:C538
purity:C539
purity:C542
purity:C543
purity:C544
purity:C547
purity:C548
purity:C549
purity:C550
purity:C551
purity:C552
purity:C554
purity:C555
purity:C559
purity:C560
purity:C561
purity:C562
tsse:tsse2
all
A–D
A
B
C
D
5
Coefficient
p_c = < 2.22e–16, p_sc = 6.5157e–08

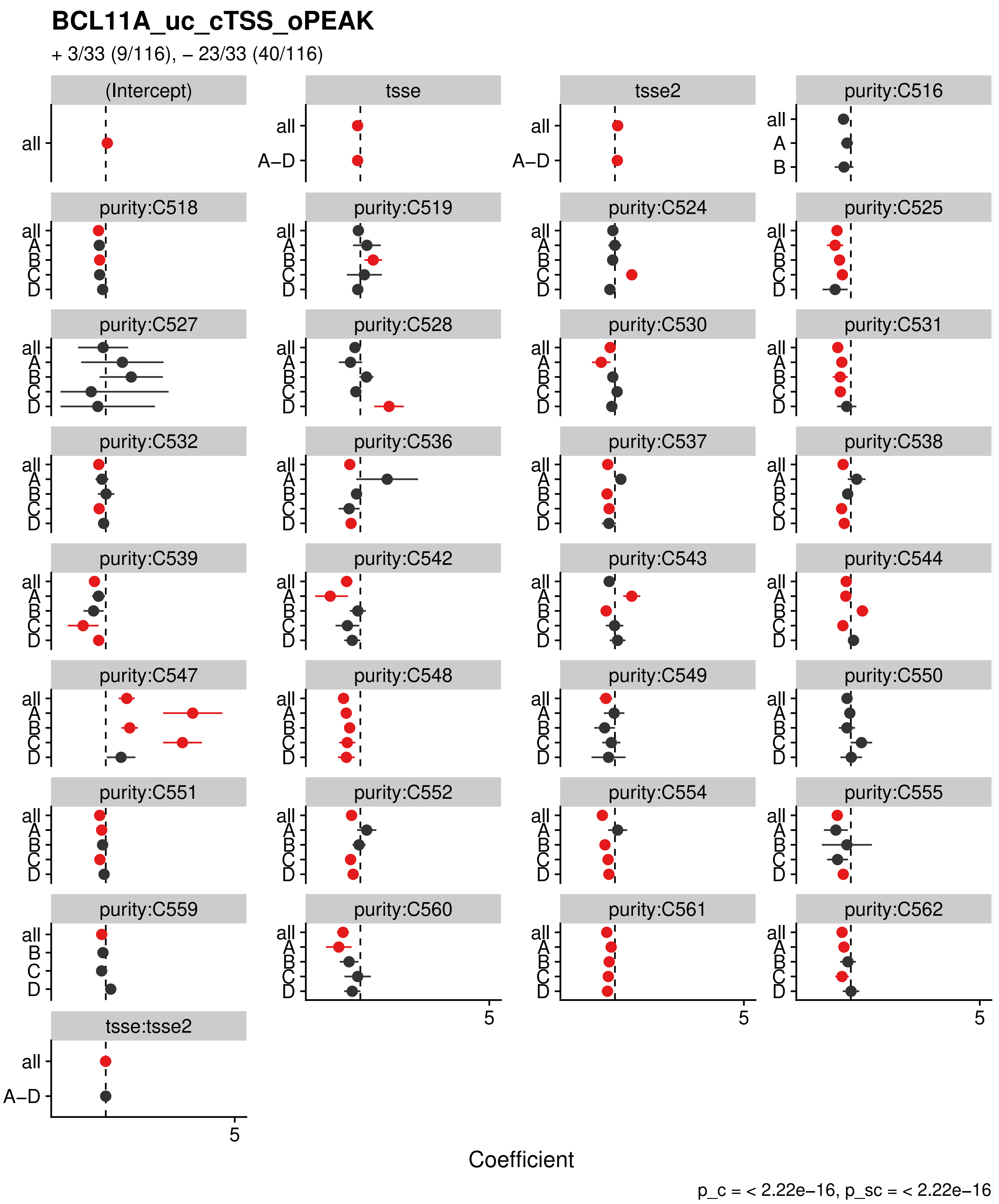
BCL11A_uc_cTSS_oPEAK
+ 3/33 (9/116), − 23/33 (40/116)
(Intercept)
tsse
tsse2
purity:C516
purity:C518
purity:C519
purity:C524
purity:C525
purity:C527
purity:C528
purity:C530
purity:C531
purity:C532
purity:C536
purity:C537
purity:C538
purity:C539
purity:C542
purity:C543
purity:C544
purity:C547
purity:C548
purity:C549
purity:C550
purity:C551
purity:C552
purity:C554
purity:C555
purity:C559
purity:C560
purity:C561
purity:C562
tsse:tsse2
all
A−D
A
B
C
D
5
Coefficient
p_c = < 2.22e−16, p_sc = < 2.22e−16

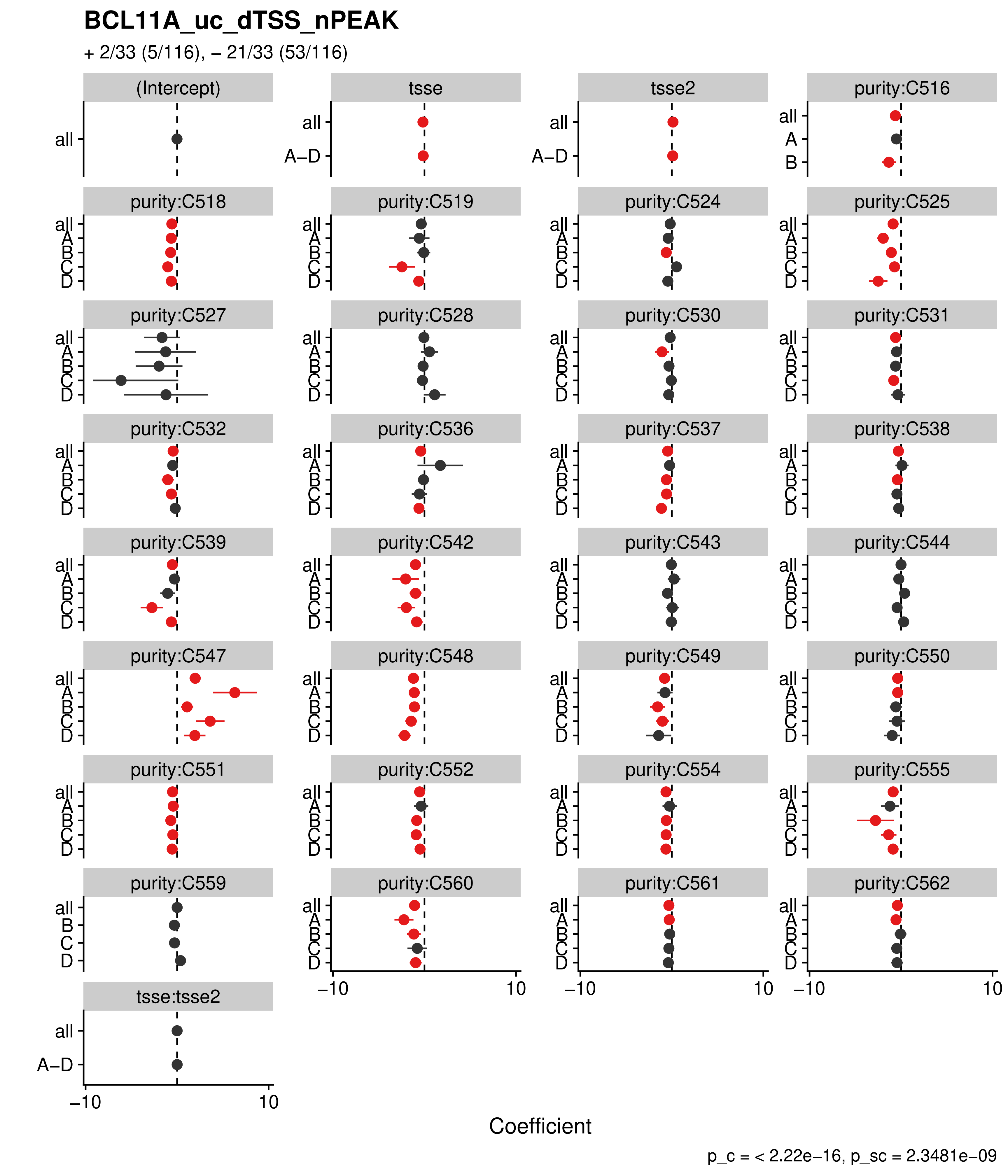
BCL11A_uc_dTSS_nPEAK
+ 2/33 (5/116), – 21/33 (53/116)
(Intercept)
tsse
tsse2
purity:C516
purity:C518
purity:C519
purity:C524
purity:C525
purity:C527
purity:C528
purity:C530
purity:C531
purity:C532
purity:C536
purity:C537
purity:C538
purity:C539
purity:C542
purity:C543
purity:C544
purity:C547
purity:C548
purity:C549
purity:C550
purity:C551
purity:C552
purity:C554
purity:C555
purity:C559
purity:C560
purity:C561
purity:C562
tsse:tsse2
all
A–D
A
B
C
D
–10
10
Coefficient
p_c = < 2.22e–16, p_sc = 2.3481e–09

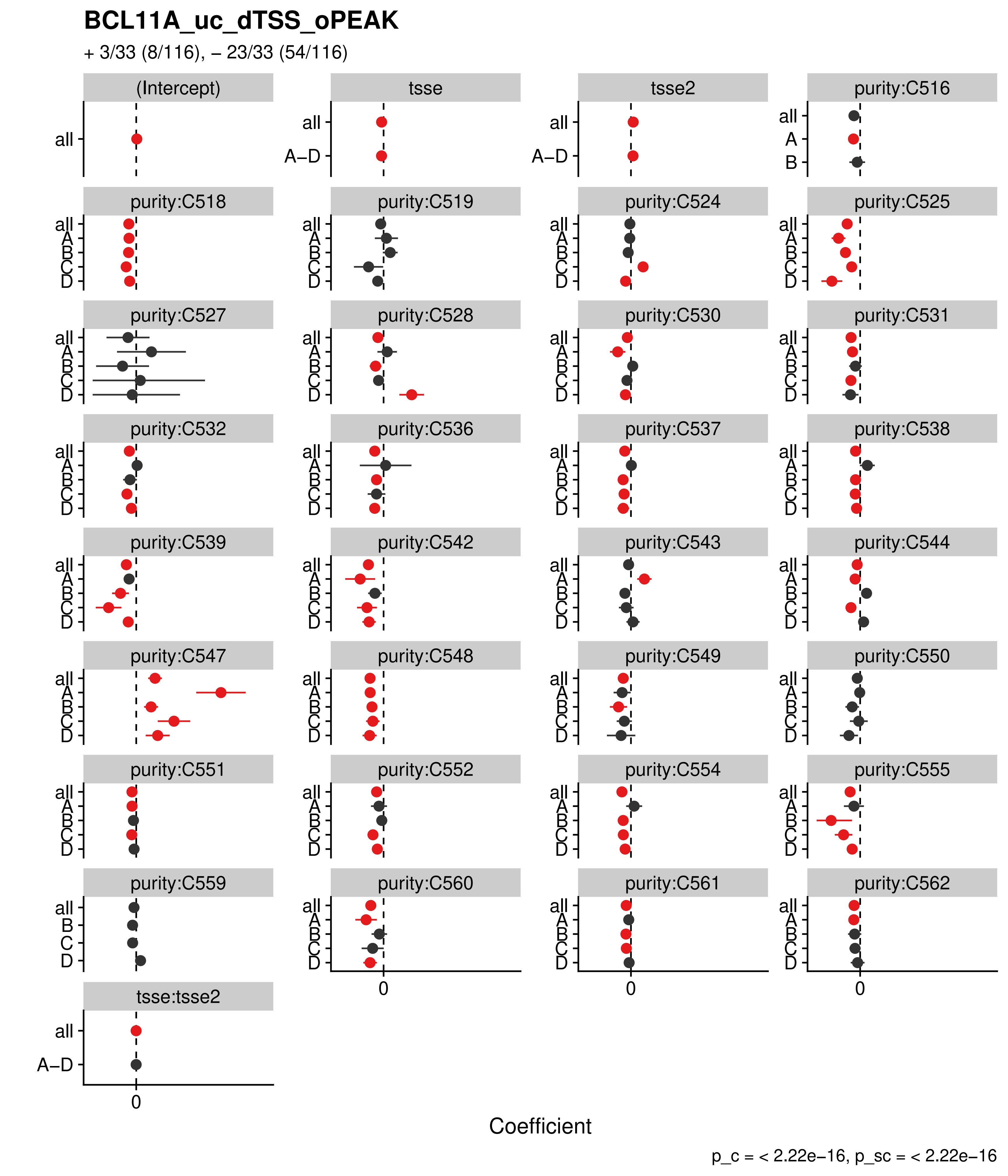

BCL11A_uc_dTSS_oPEAK
+ 3/33 (8/116), – 23/33 (54/116)
(Intercept)
tsse
tsse2
purity:C516
purity:C518
purity:C519
purity:C524
purity:C525
purity:C527
purity:C528
purity:C530
purity:C531
purity:C532
purity:C536
purity:C537
purity:C538
purity:C539
purity:C542
purity:C543
purity:C544
purity:C547
purity:C548
purity:C549
purity:C550
purity:C551
purity:C552
purity:C554
purity:C555
purity:C559
purity:C560
purity:C561
purity:C562
tsse:tsse2
all
A–D
A
B
C
D
0
Coefficient
p_c = < 2.22e–16, p_sc = < 2.22e–16

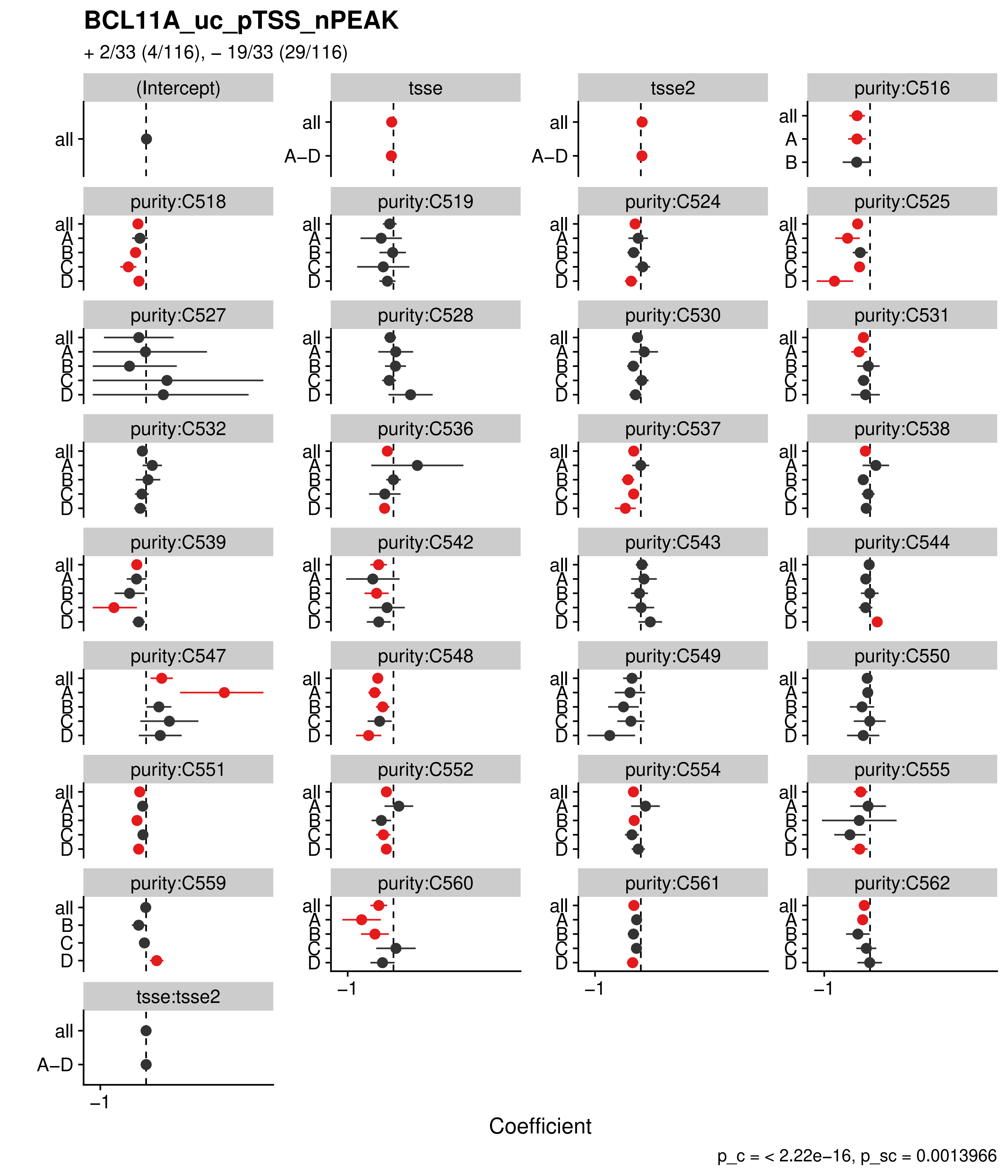
BCL11A_uc_pTSS_nPEAK
+ 2/33 (4/116), – 19/33 (29/116)
(Intercept)
tsse
tsse2
purity:C516
purity:C518
purity:C519
purity:C524
purity:C525
purity:C527
purity:C528
purity:C530
purity:C531
purity:C532
purity:C536
purity:C537
purity:C538
purity:C539
purity:C542
purity:C543
purity:C544
purity:C547
purity:C548
purity:C549
purity:C550
purity:C551
purity:C552
purity:C554
purity:C555
purity:C559
purity:C560
purity:C561
purity:C562
tsse:tsse2
all
A–D
A
B
C
D
–1
Coefficient
p_c = < 2.22e–16, p_sc = 0.0013966

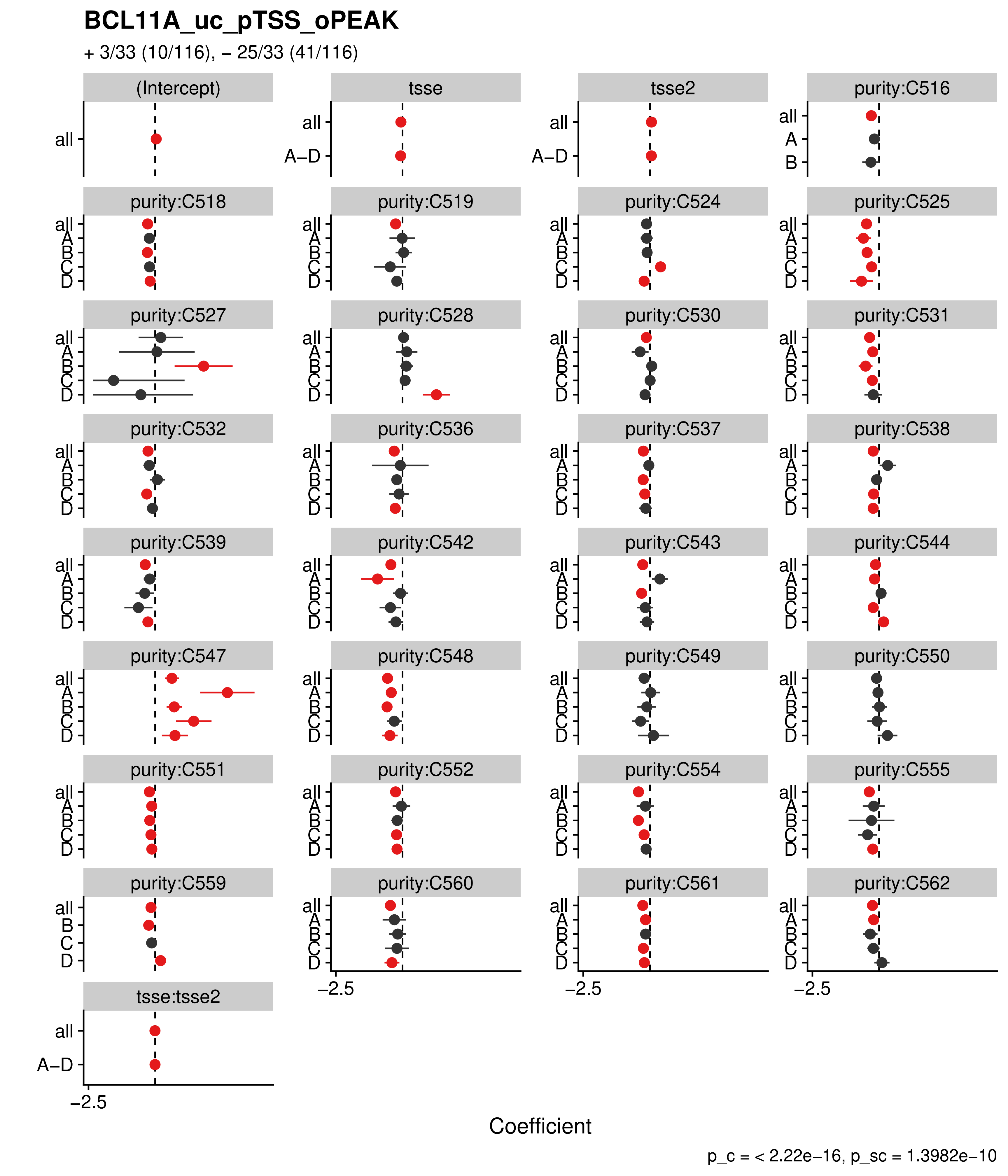
BCL11A_uc_pTSS_oPEAK
+ 3/33 (10/116), – 25/33 (41/116)
(Intercept)
tsse
tsse2
purity:C516
purity:C518
purity:C519
purity:C524
purity:C525
purity:C527
purity:C528
purity:C530
purity:C531
purity:C532
purity:C536
purity:C537
purity:C538
purity:C539
purity:C542
purity:C543
purity:C544
purity:C547
purity:C548
purity:C549
purity:C550
purity:C551
purity:C552
purity:C554
purity:C555
purity:C559
purity:C560
purity:C561
purity:C562
tsse:tsse2
all
A–D
A
B
C
D
−2.5
Coefficient
p_c = < 2.22e−16, p_sc = 1.3982e−10

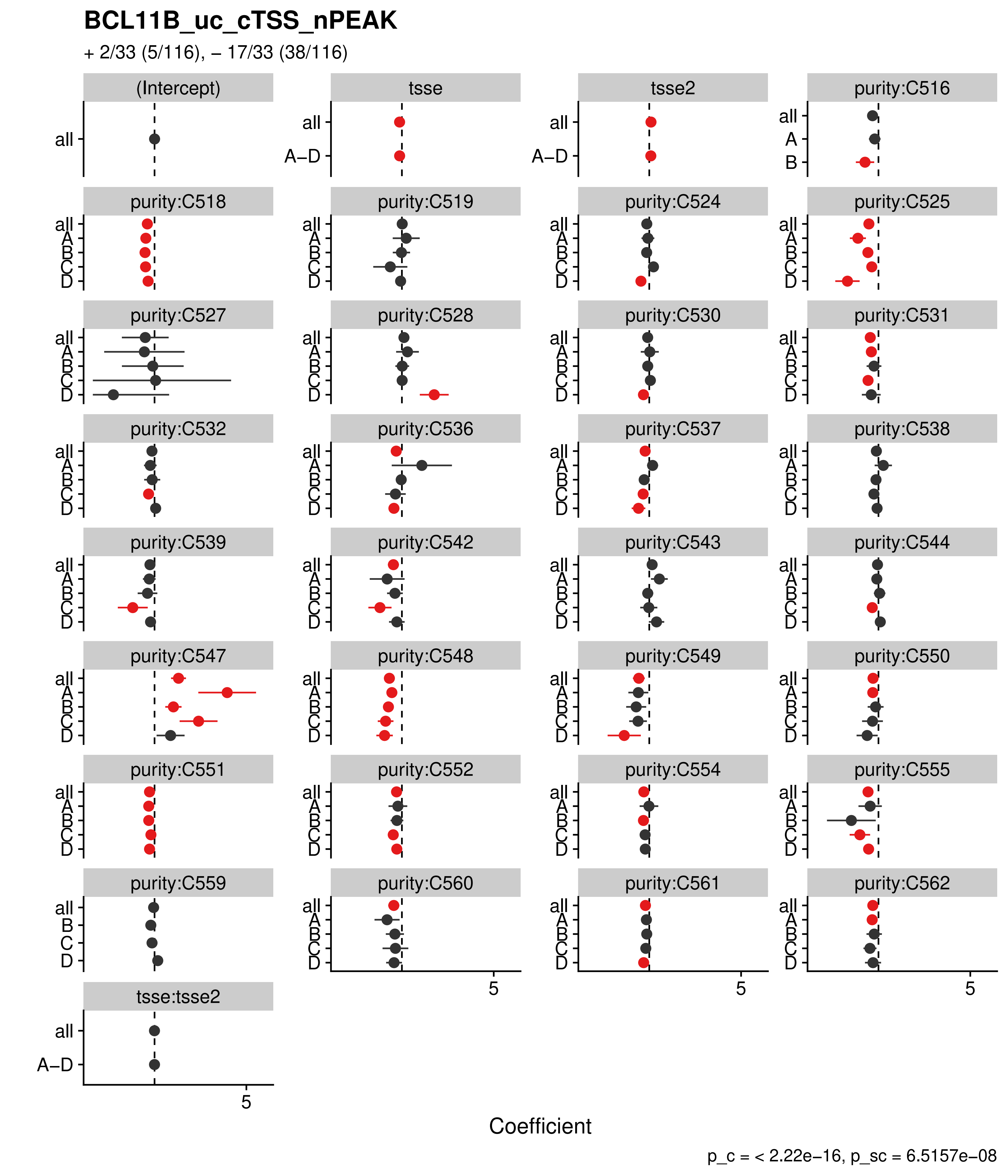

BCL11B_uc_cTSS_nPEAK
+ 2/33 (5/116), – 17/33 (38/116)
(Intercept)
tsse
tsse2
purity:C516
purity:C518
purity:C519
purity:C524
purity:C525
purity:C527
purity:C528
purity:C530
purity:C531
purity:C532
purity:C536
purity:C537
purity:C538
purity:C539
purity:C542
purity:C543
purity:C544
purity:C547
purity:C548
purity:C549
purity:C550
purity:C551
purity:C552
purity:C554
purity:C555
purity:C559
purity:C560
purity:C561
purity:C562
tsse:tsse2
all
A–D
A
B
C
D
5
Coefficient
p_c = < 2.22e–16, p_sc = 6.5157e–08

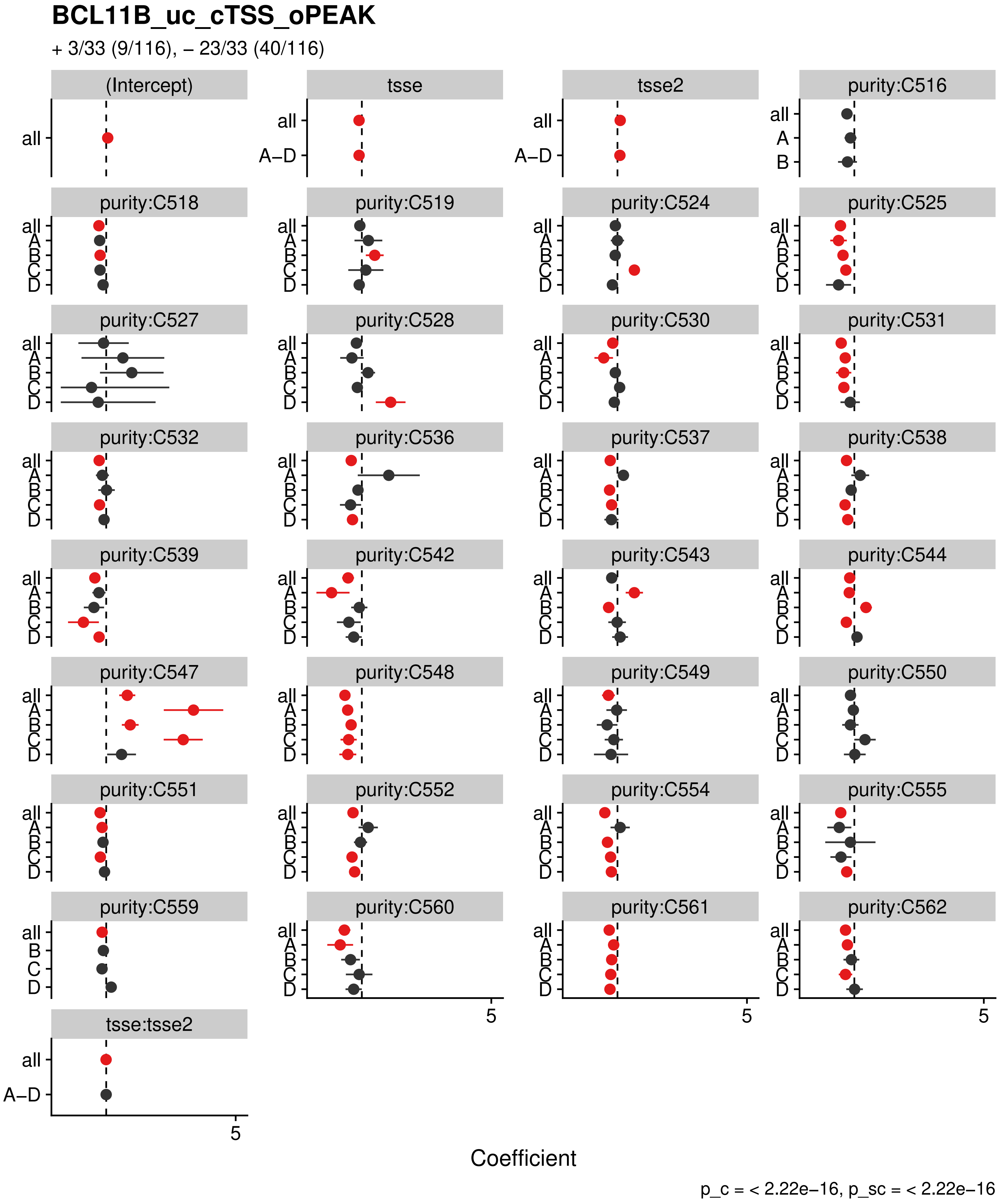

BCL11B_uc_cTSS_oPEAK
+ 3/33 (9/116), − 23/33 (40/116)
(Intercept)
tsse
tsse2
purity:C516
purity:C518
purity:C519
purity:C524
purity:C525
purity:C527
purity:C528
purity:C530
purity:C531
purity:C532
purity:C536
purity:C537
purity:C538
purity:C539
purity:C542
purity:C543
purity:C544
purity:C547
purity:C548
purity:C549
purity:C550
purity:C551
purity:C552
purity:C554
purity:C555
purity:C559
purity:C560
purity:C561
purity:C562
tsse:tsse2
all
A–D
A
B
C
D
5
Coefficient
p_c = < 2.22e−16, p_sc = < 2.22e−16

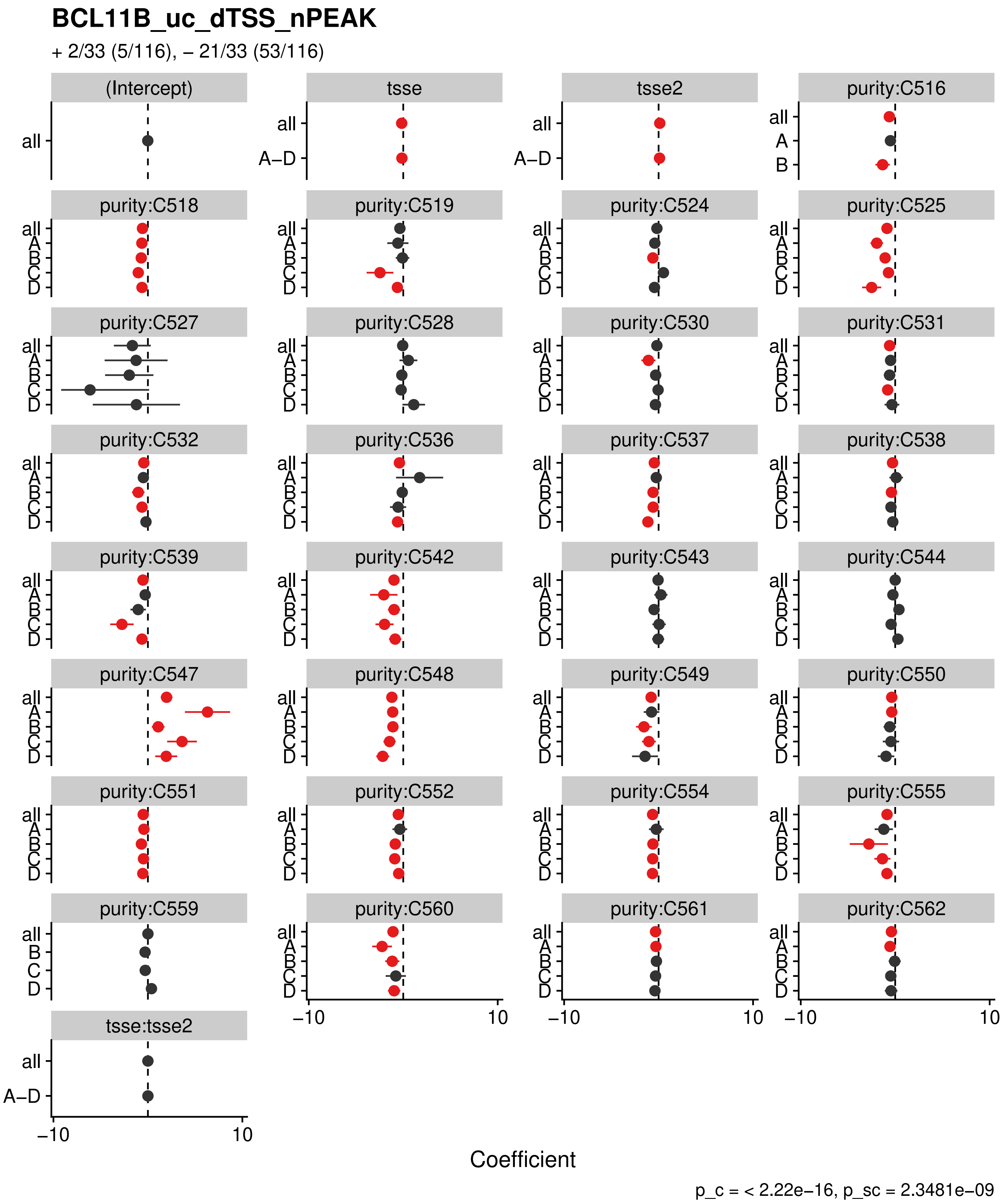

BCL11B_uc_dTSS_nPEAK
+ 2/33 (5/116), – 21/33 (53/116)
(Intercept)
tsse
tsse2
purity:C516
purity:C518
purity:C519
purity:C524
purity:C525
purity:C527
purity:C528
purity:C530
purity:C531
purity:C532
purity:C536
purity:C537
purity:C538
purity:C539
purity:C542
purity:C543
purity:C544
purity:C547
purity:C548
purity:C549
purity:C550
purity:C551
purity:C552
purity:C554
purity:C555
purity:C559
purity:C560
purity:C561
purity:C562
tsse:tsse2
all
A–D
A
B
C
D
–10
10
Coefficient
p_c = < 2.22e–16, p_sc = 2.3481e–09

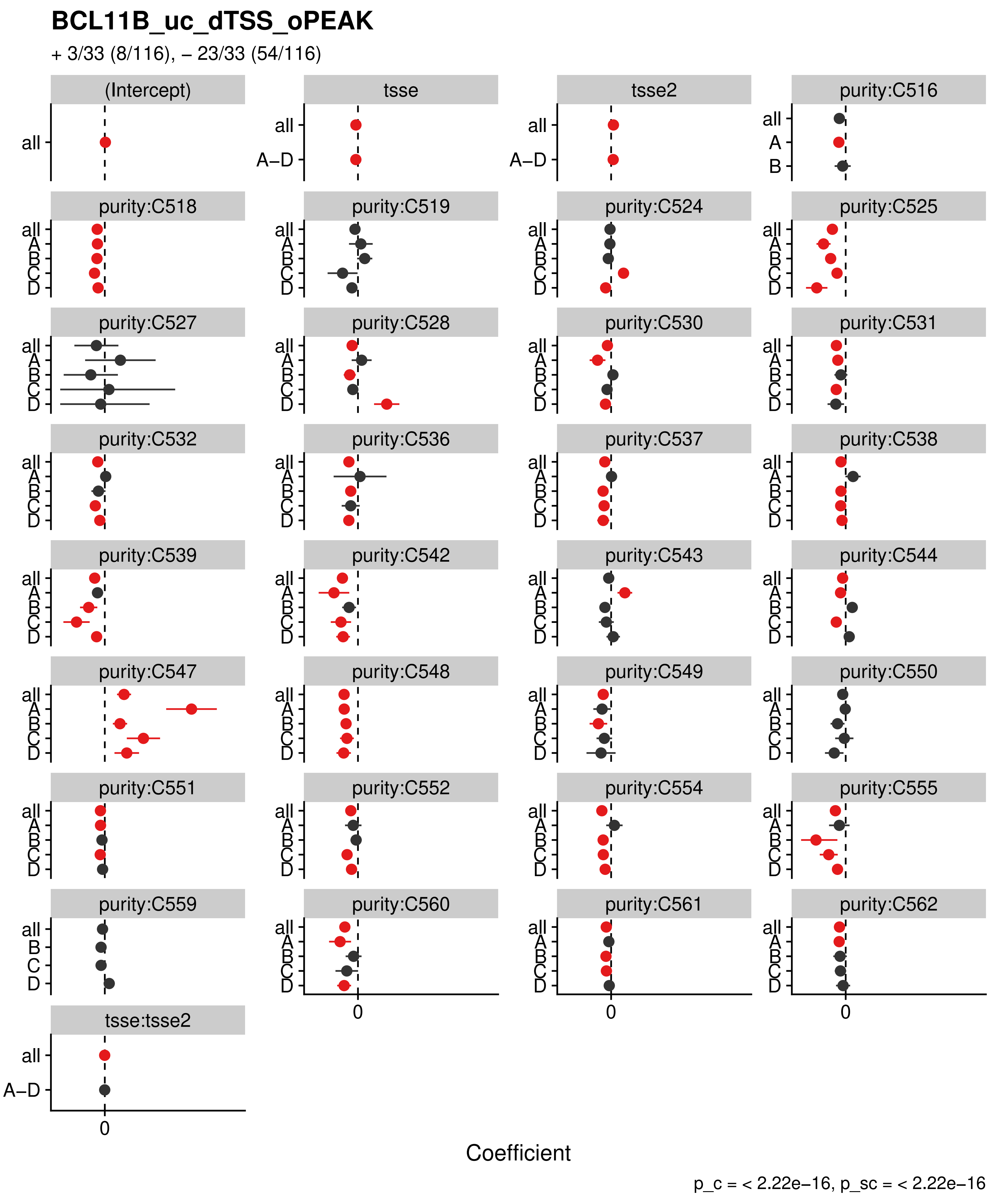

BCL11B_uc_dTSS_oPEAK
+ 3/33 (8/116), – 23/33 (54/116)
(Intercept)
tsse
tsse2
purity:C516
purity:C518
purity:C519
purity:C524
purity:C525
purity:C527
purity:C528
purity:C530
purity:C531
purity:C532
purity:C536
purity:C537
purity:C538
purity:C539
purity:C542
purity:C543
purity:C544
purity:C547
purity:C548
purity:C549
purity:C550
purity:C551
purity:C552
purity:C554
purity:C555
purity:C559
purity:C560
purity:C561
purity:C562
tsse:tsse2
all
A–D
A
B
C
D
0
Coefficient
p_c = < 2.22e–16, p_sc = < 2.22e–16

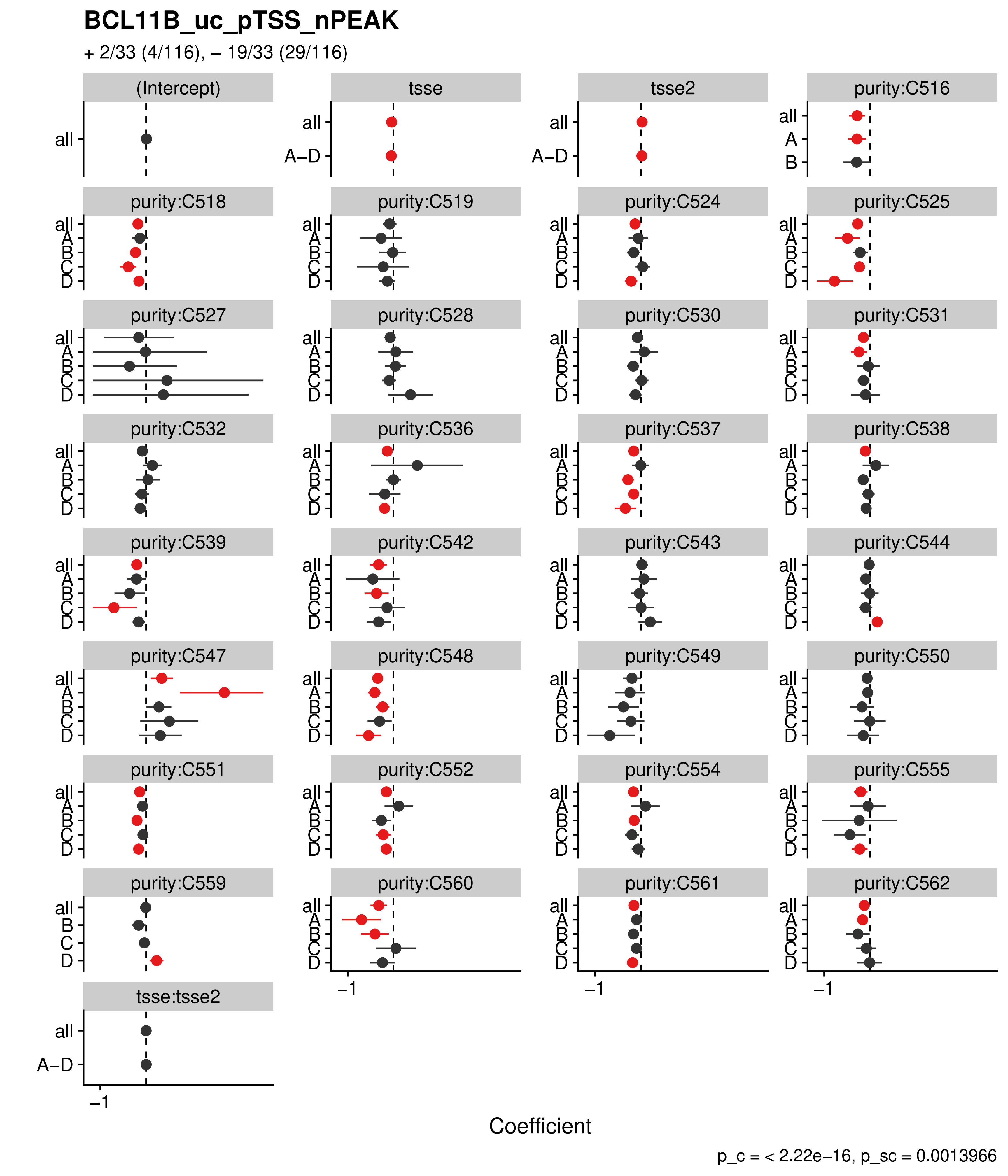

BCL11B_uc_pTSS_nPEAK
+ 2/33 (4/116), – 19/33 (29/116)
(Intercept)
tsse
tsse2
purity:C516
purity:C518
purity:C519
purity:C524
purity:C525
purity:C527
purity:C528
purity:C530
purity:C531
purity:C532
purity:C536
purity:C537
purity:C538
purity:C539
purity:C542
purity:C543
purity:C544
purity:C547
purity:C548
purity:C549
purity:C550
purity:C551
purity:C552
purity:C554
purity:C555
purity:C559
purity:C560
purity:C561
purity:C562
tsse:tsse2
all
A–D
A
B
C
D
–1
Coefficient
p_c = < 2.22e–16, p_sc = 0.0013966

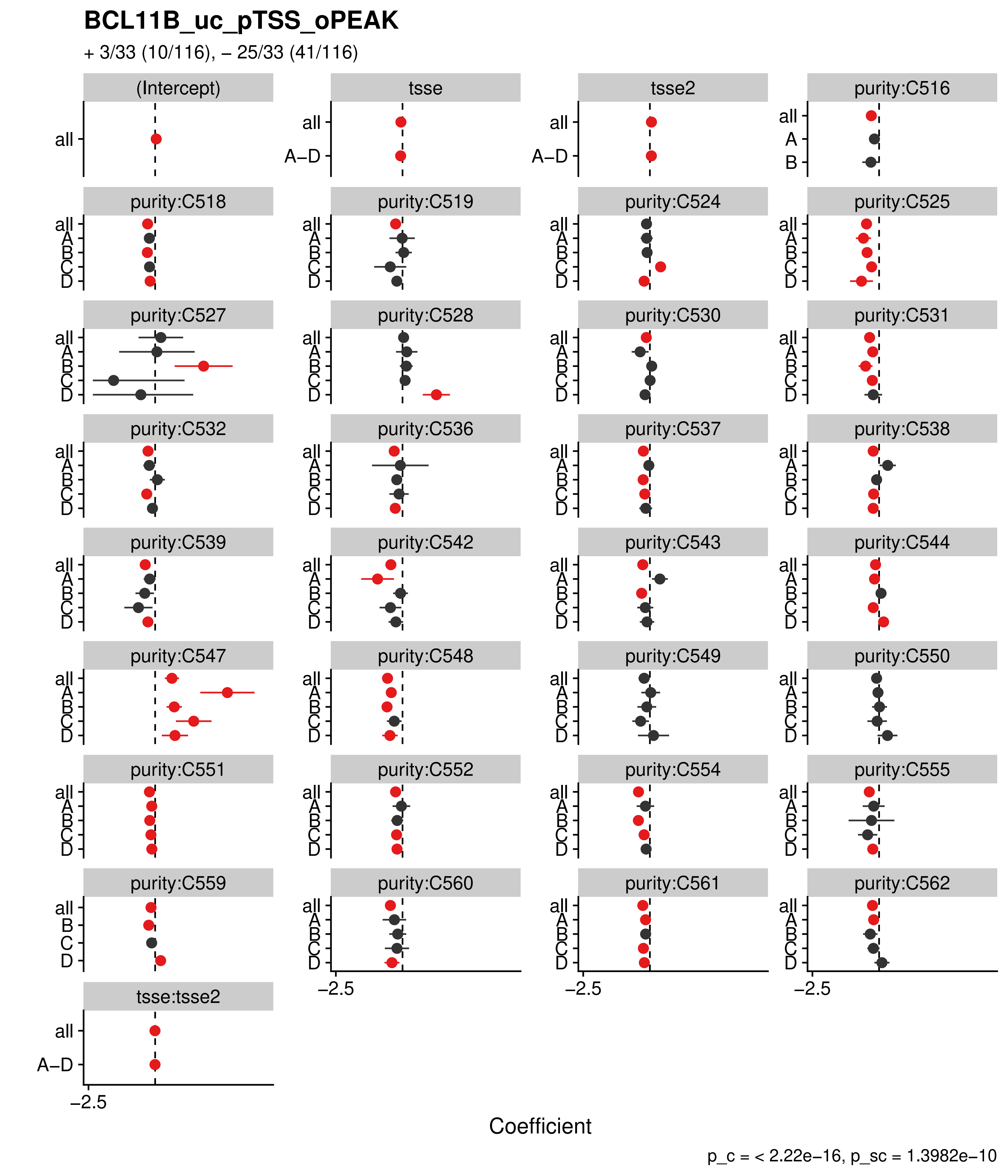

BCL11B_uc_pTSS_oPEAK
+ 3/33 (10/116), – 25/33 (41/116)
(Intercept)
tsse
tsse2
purity:C516
purity:C518
purity:C519
purity:C524
purity:C525
purity:C527
purity:C528
purity:C530
purity:C531
purity:C532
purity:C536
purity:C537
purity:C538
purity:C539
purity:C542
purity:C543
purity:C544
purity:C547
purity:C548
purity:C549
purity:C550
purity:C551
purity:C552
purity:C554
purity:C555
purity:C559
purity:C560
purity:C561
purity:C562
tsse:tsse2
all
A–D
A
B
C
D
−2.5
Coefficient
p_c = < 2.22e−16, p_sc = 1.3982e−10

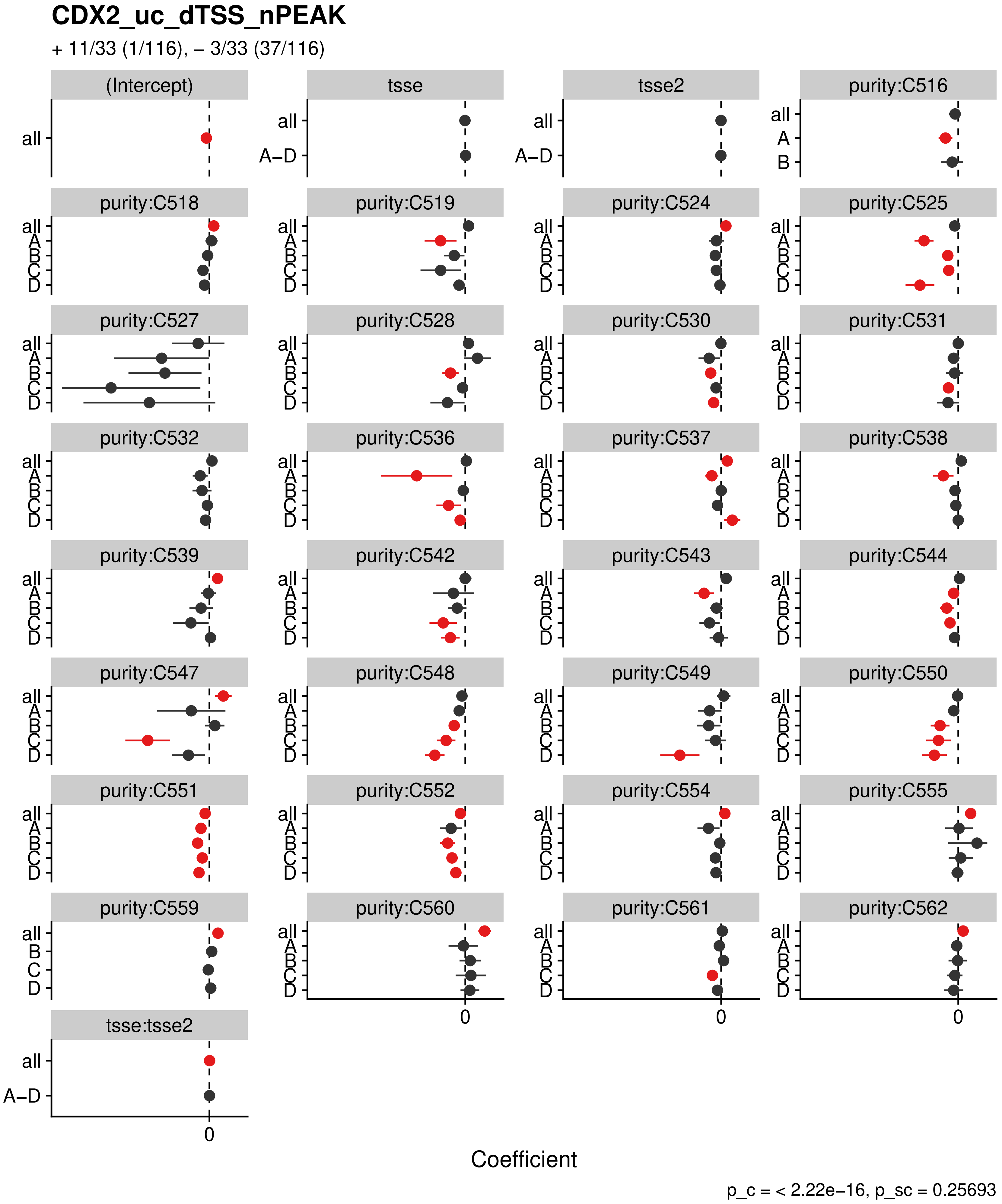
CDX2_uc_dTSS_nPEAK
+ 11/33 (1/116), – 3/33 (37/116)
(Intercept)
tsse
tsse2
purity:C516
purity:C518
purity:C519
purity:C524
purity:C525
purity:C527
purity:C528
purity:C530
purity:C531
purity:C532
purity:C536
purity:C537
purity:C538
purity:C539
purity:C542
purity:C543
purity:C544
purity:C547
purity:C548
purity:C549
purity:C550
purity:C551
purity:C552
purity:C554
purity:C555
purity:C559
purity:C560
purity:C561
purity:C562
tsse:tsse2
all
A–D
A
B
C
D
0
Coefficient
p_c = < 2.22e–16, p_sc = 0.25693

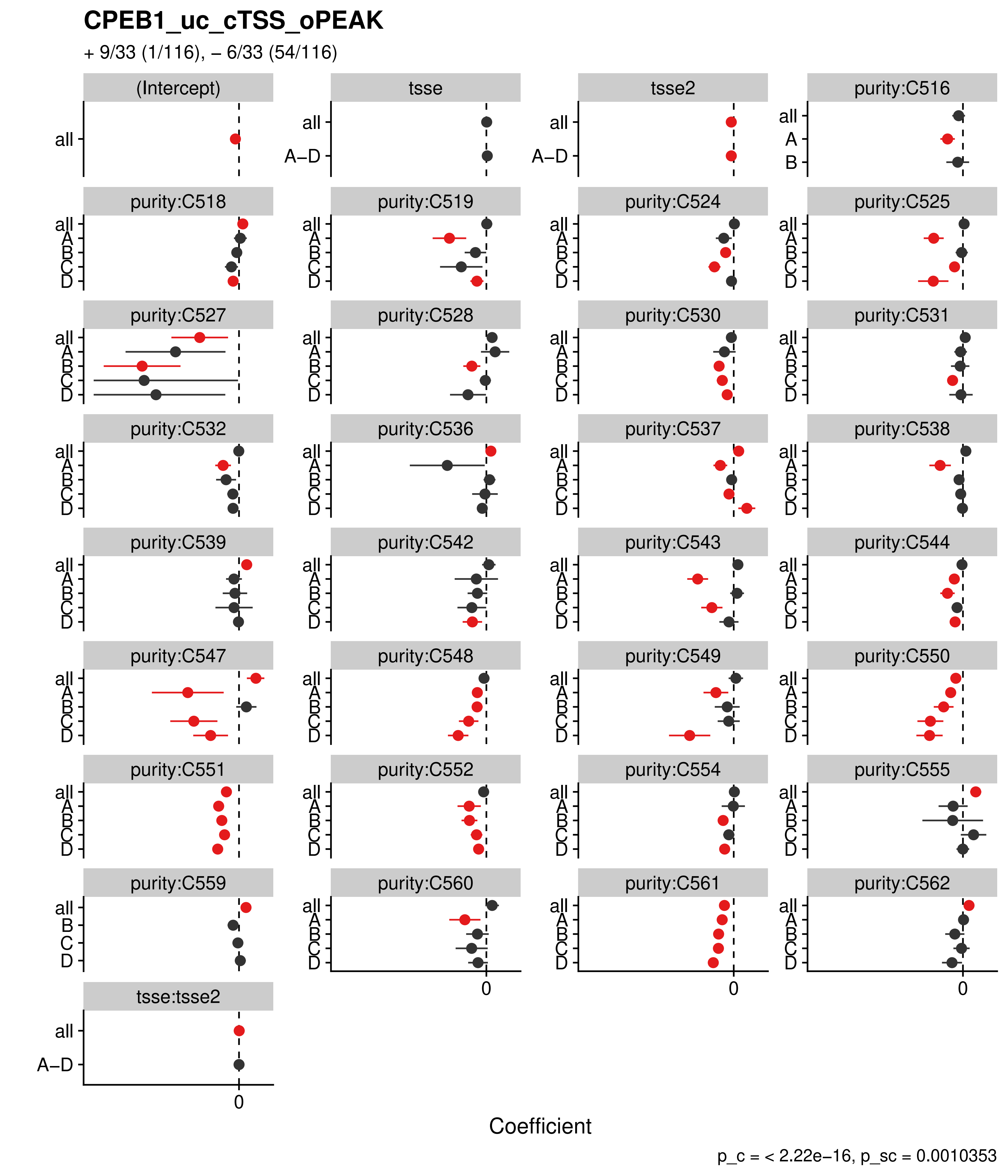
CPEB1_uc_cTSS_oPEAK
+ 9/33 (1/116), – 6/33 (54/116)
(Intercept)
tsse
tsse2
purity:C516
purity:C518
purity:C519
purity:C524
purity:C525
purity:C527
purity:C528
purity:C530
purity:C531
purity:C532
purity:C536
purity:C537
purity:C538
purity:C539
purity:C542
purity:C543
purity:C544
purity:C547
purity:C548
purity:C549
purity:C550
purity:C551
purity:C552
purity:C554
purity:C555
purity:C559
purity:C560
purity:C561
purity:C562
tsse:tsse2
all
A–D
A
B
C
D
0
Coefficient
p_c = < 2.22e–16, p_sc = 0.0010353

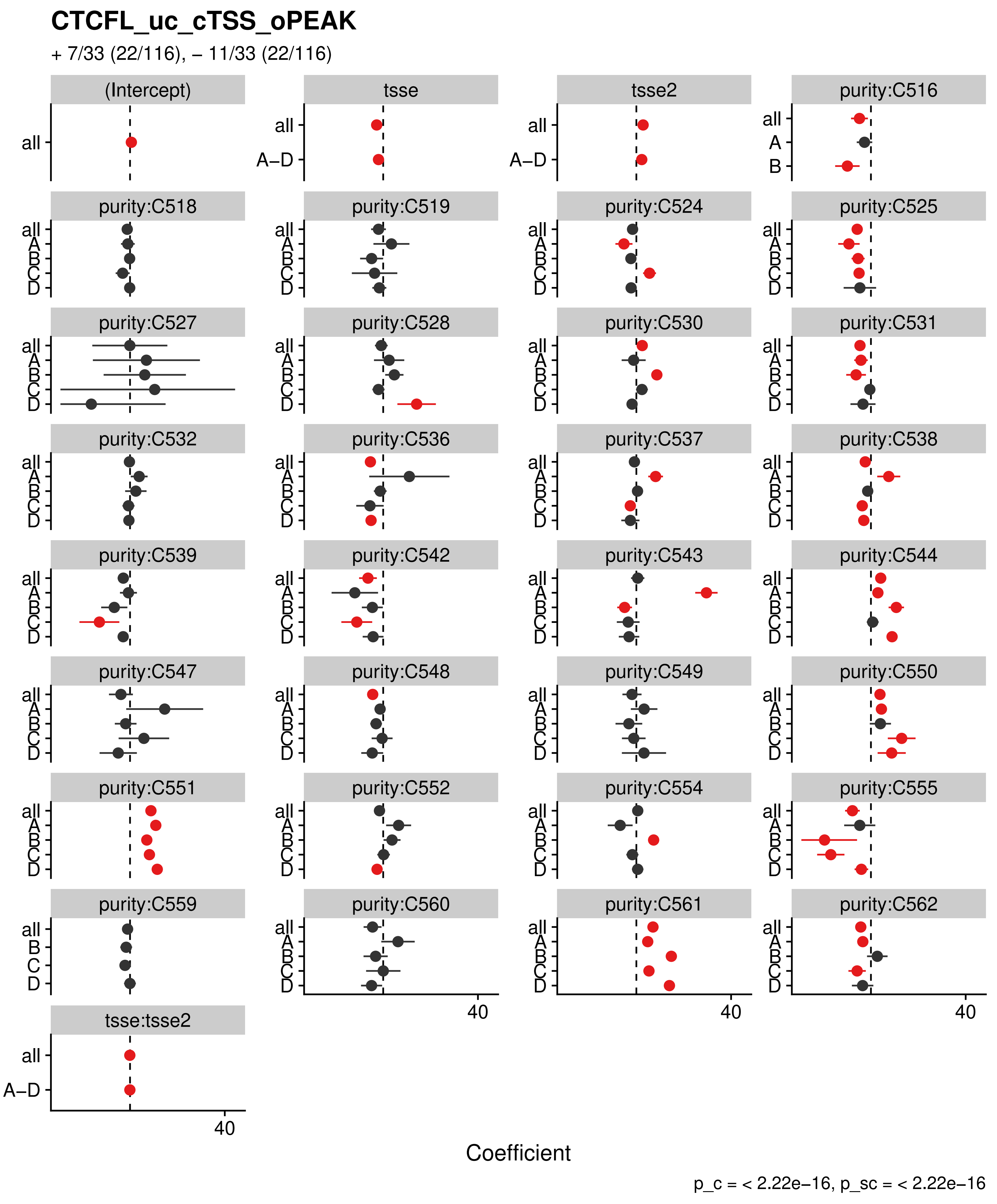

CTCFL_uc_cTSS_oPEAK
+ 7/33 (22/116), – 11/33 (22/116)
(Intercept)
tsse
tsse2
purity:C516
purity:C518
purity:C519
purity:C524
purity:C525
purity:C527
purity:C528
purity:C530
purity:C531
purity:C532
purity:C536
purity:C537
purity:C538
purity:C539
purity:C542
purity:C543
purity:C544
purity:C547
purity:C548
purity:C549
purity:C550
purity:C551
purity:C552
purity:C554
purity:C555
purity:C559
purity:C560
purity:C561
purity:C562
tsse:tsse2
all
A–D
A
B
C
D
40
Coefficient
p_c = < 2.22e–16, p_sc = < 2.22e–16

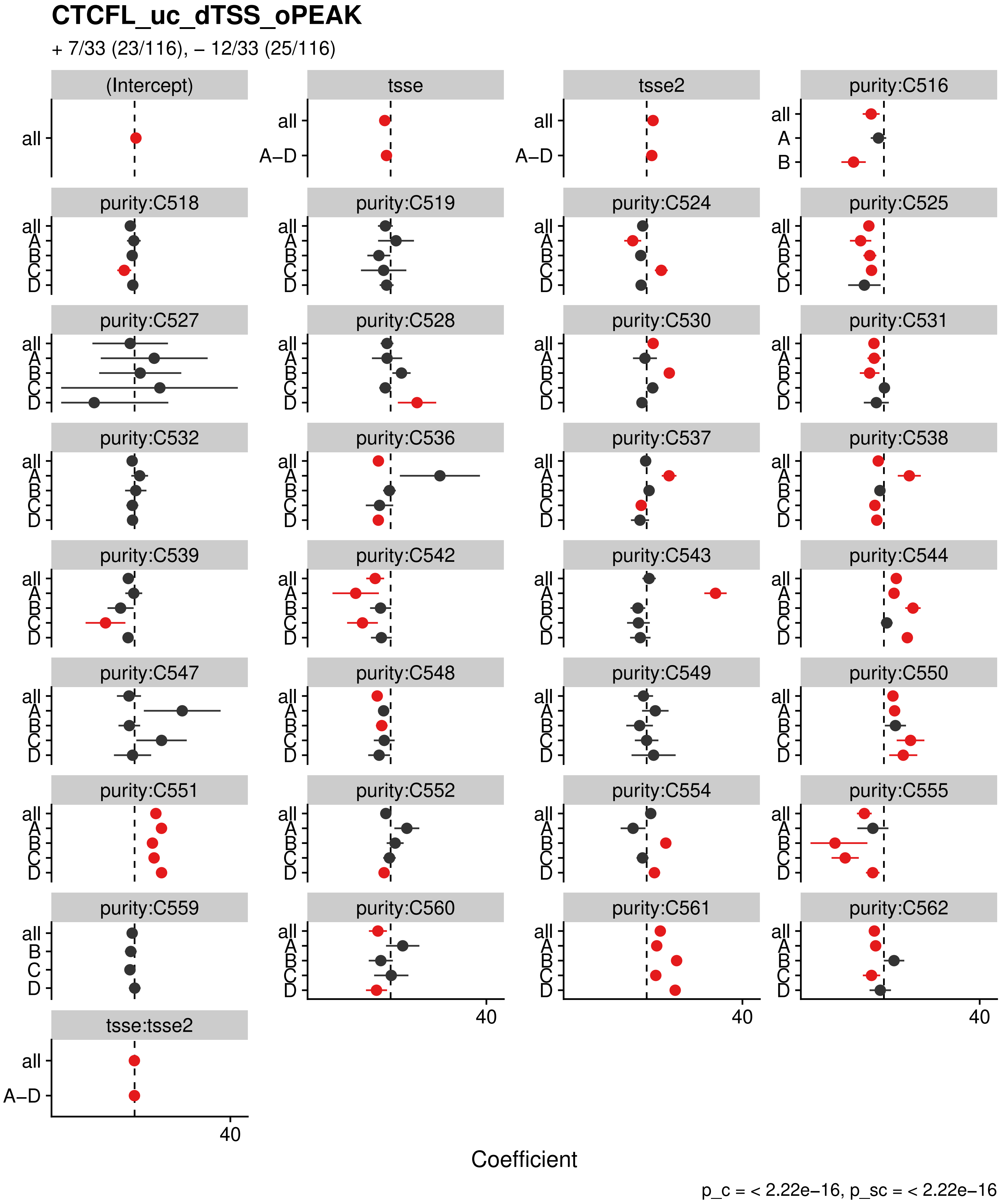
CTCFL_uc_dTSS_oPEAK
+ 7/33 (23/116), – 12/33 (25/116)
(Intercept)
tsse
tsse2
purity:C516
purity:C518
purity:C519
purity:C524
purity:C525
purity:C527
purity:C528
purity:C530
purity:C531
purity:C532
purity:C536
purity:C537
purity:C538
purity:C539
purity:C542
purity:C543
purity:C544
purity:C547
purity:C548
purity:C549
purity:C550
purity:C551
purity:C552
purity:C554
purity:C555
purity:C559
purity:C560
purity:C561
purity:C562
tsse:tsse2
all
A–D
A
B
C
D
40
Coefficient
p_c = < 2.22e−16, p_sc = < 2.22e−16

#### CTCF_uc_cTSS_oPEAK

+ 7/33 (21/116), – 14/33 (29/116)

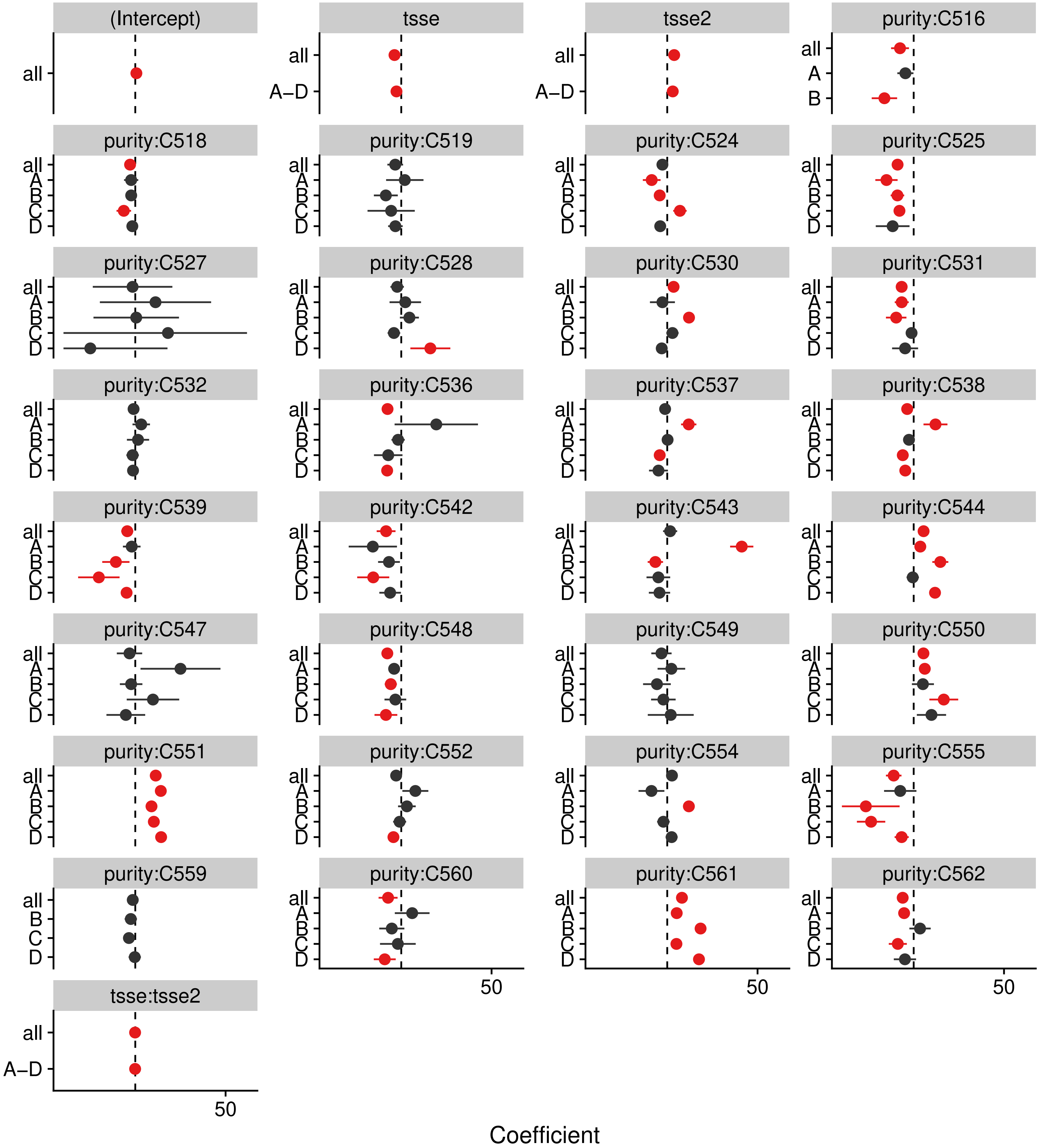


p_c = < 2.22e−16, p_sc = < 2.22e−16

#### CTCF_uc_dTSS_oPEAK

+ 8/33 (23/116), – 14/33 (31/116)

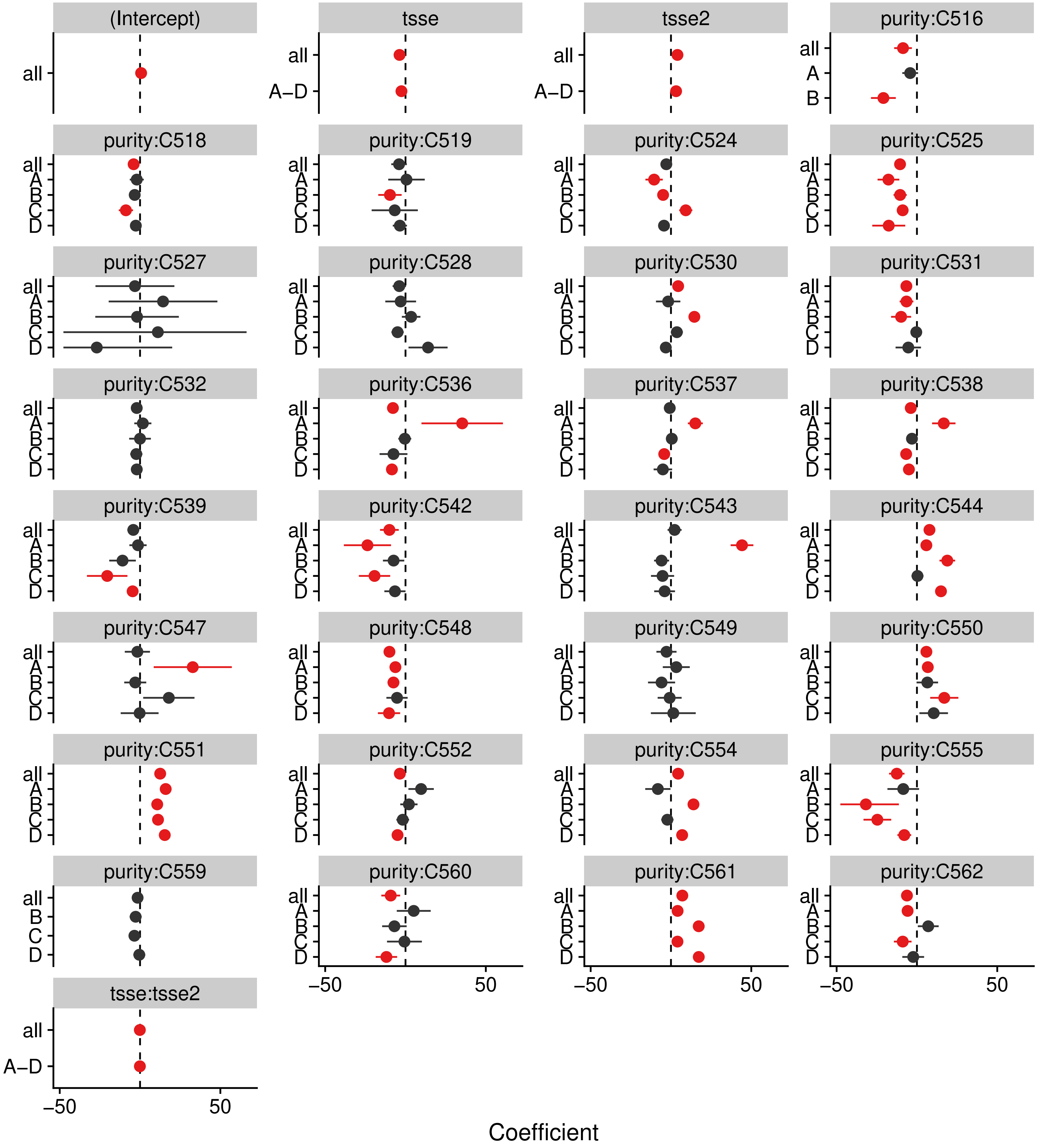


p_c = < 2.22e–16, p_sc = < 2.22e–16

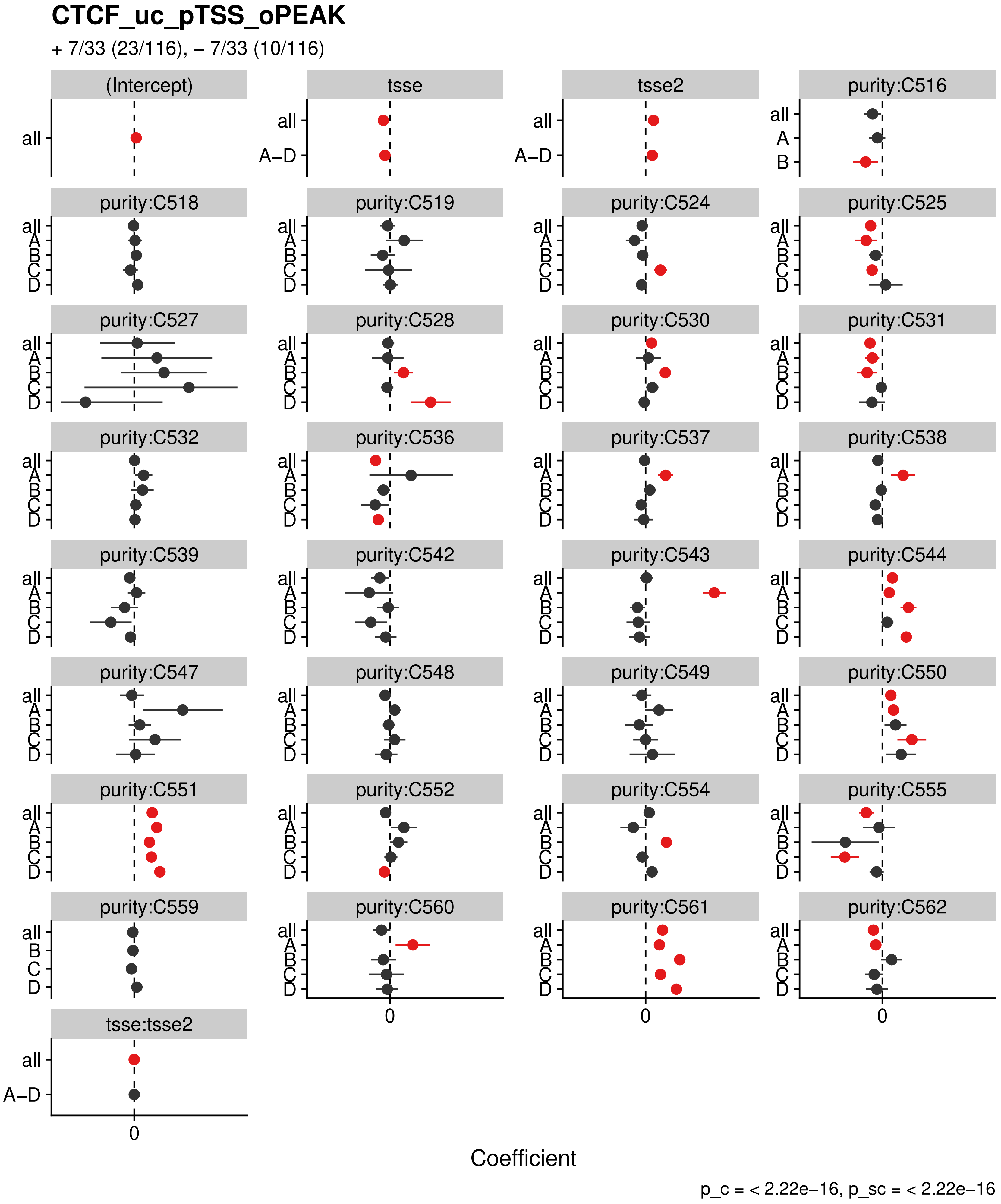
CTCF_uc_pTSS_oPEAK
+ 7/33 (23/116), – 7/33 (10/116)
(Intercept)
tsse
tsse2
purity:C516
purity:C518
purity:C519
purity:C524
purity:C525
purity:C527
purity:C528
purity:C530
purity:C531
purity:C532
purity:C536
purity:C537
purity:C538
purity:C539
purity:C542
purity:C543
purity:C544
purity:C547
purity:C548
purity:C549
purity:C550
purity:C551
purity:C552
purity:C554
purity:C555
purity:C559
purity:C560
purity:C561
purity:C562
tsse:tsse2
all
A–D
A
B
C
D
0
Coefficient
p_c = < 2.22e–16, p_sc = < 2.22e–16

**DBX2_uc_cTSS_nPEAK**

+ 10/33 (1/116), – 4/33 (39/116)

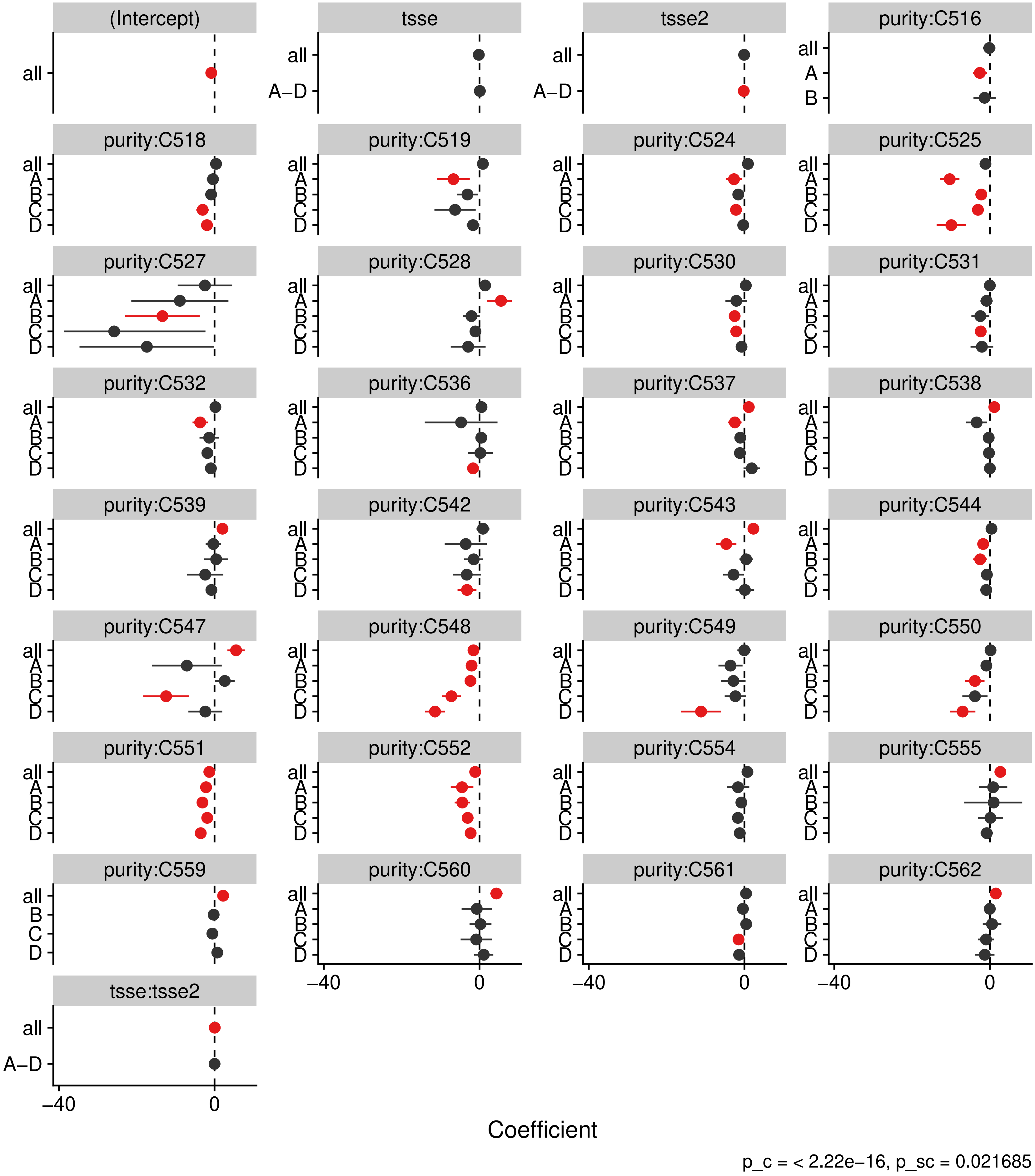

### DBX2_uc_dTSS_oPEAK

+ 12/33 (0/116), – 4/33 (42/116)

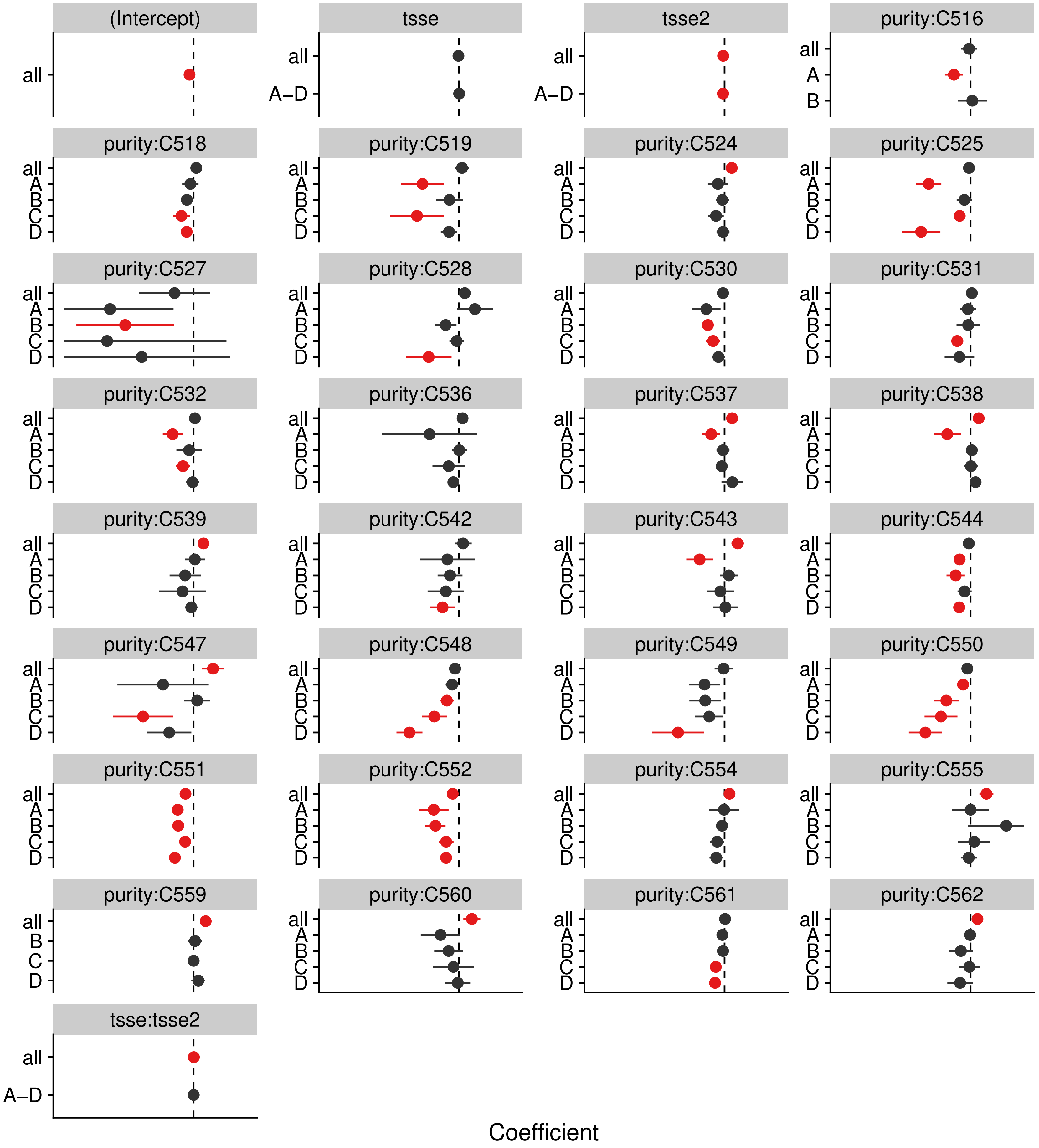


p_c = < 2.22e−16, p_sc = 0.019451

### DLX3_uc_cTSS_nPEAK

+ 11/33 (1/116), – 2/33 (33/116)

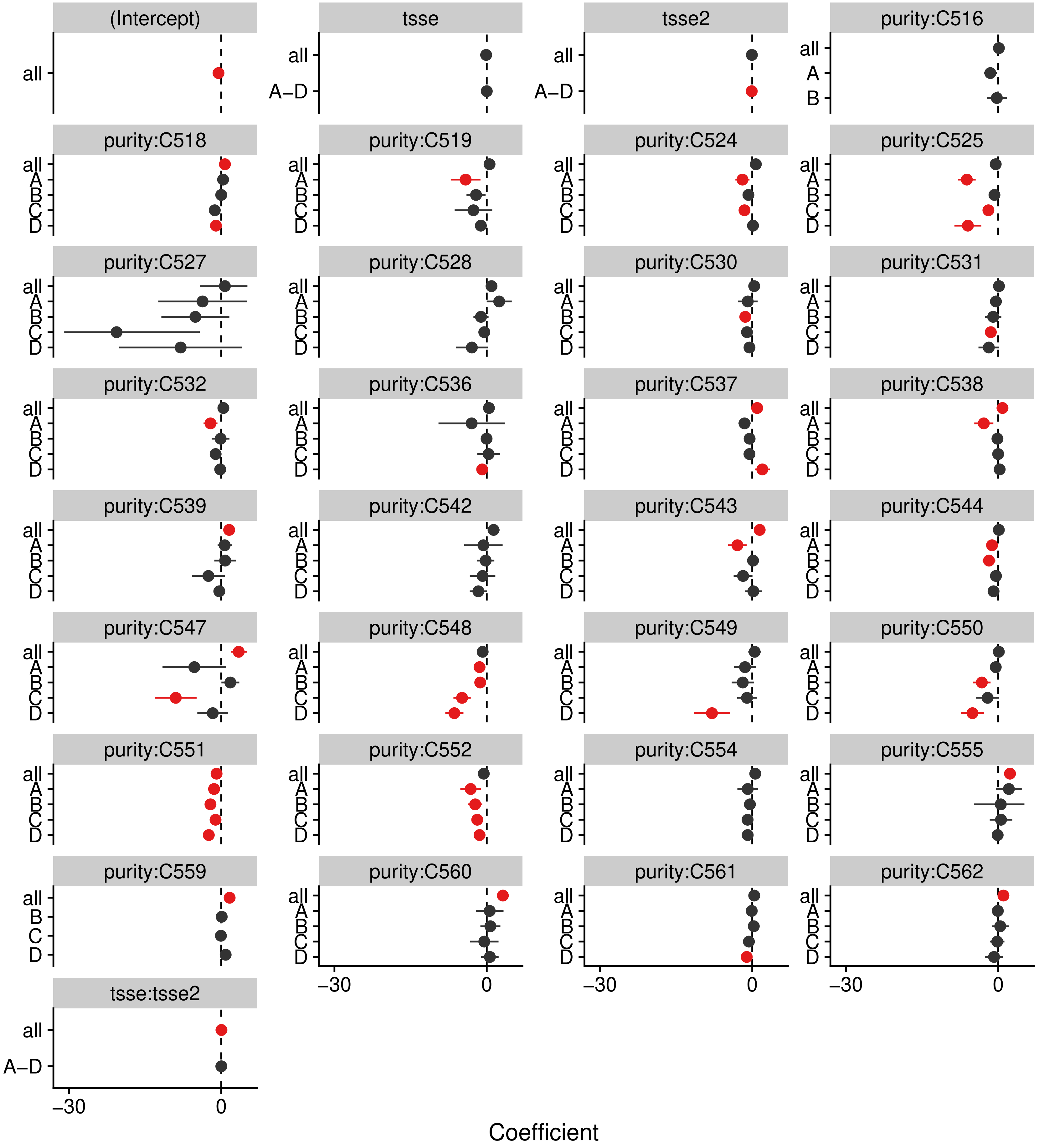


p_c = < 2.22e–16, p_sc = 0.0056747

**DLX3_uc_dTSS_oPEAK**

+ 13/33 (1/116), – 3/33 (41/116)

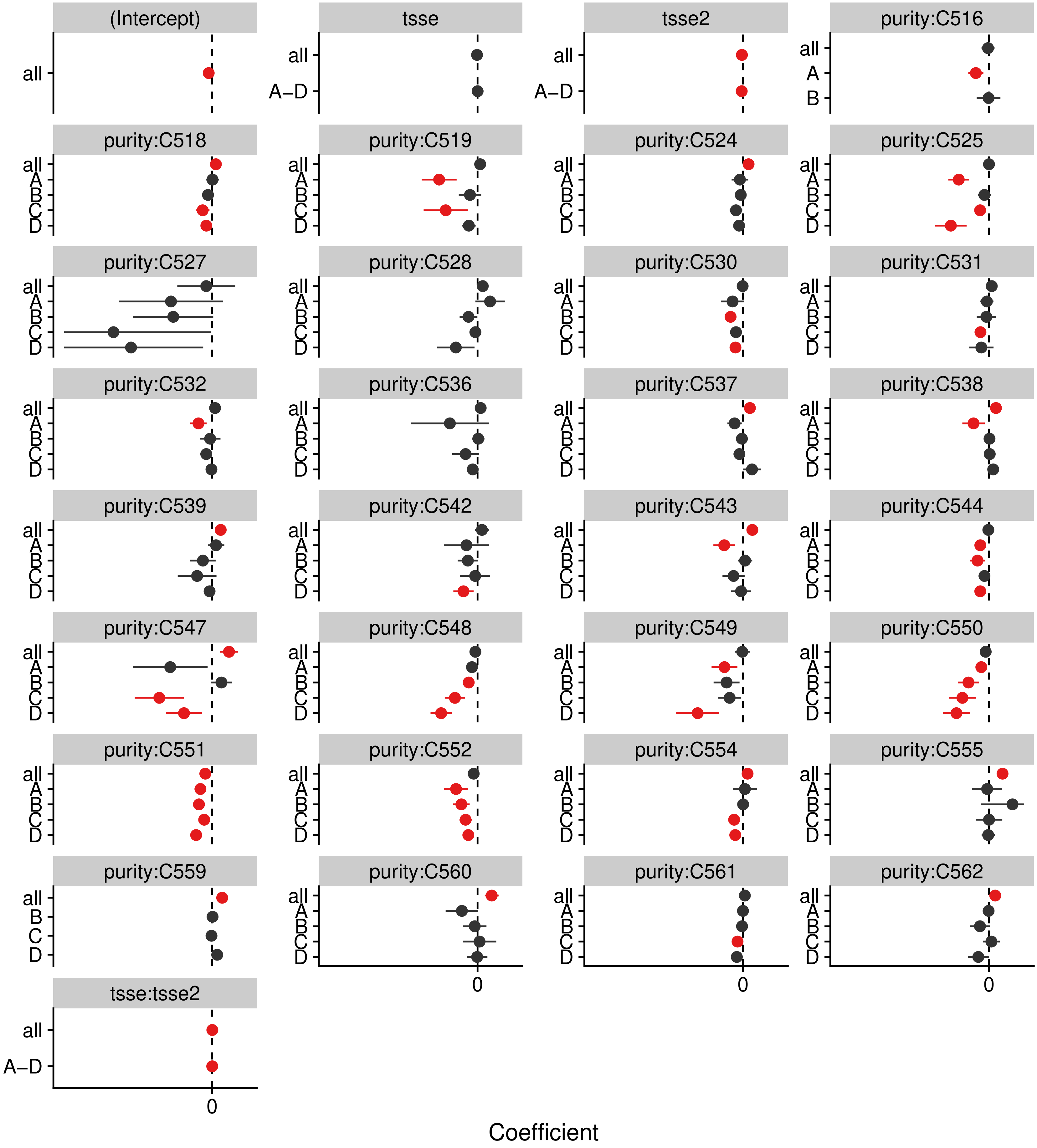


p_c = < 2.22e–16, p_sc = 0.1284

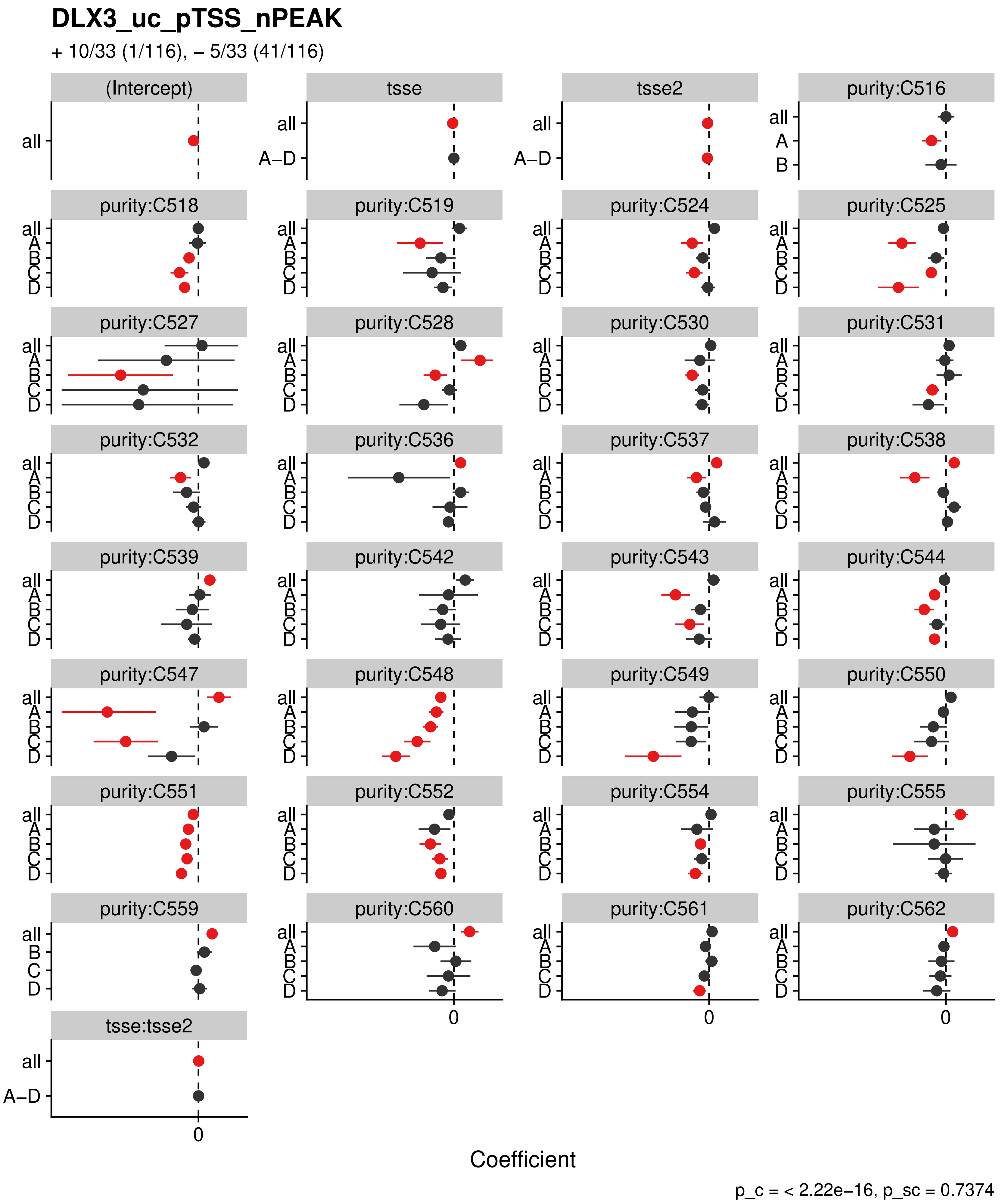
DLX3_uc_pTSS_nPEAK
+ 10/33 (1/116), – 5/33 (41/116)
(Intercept)
tsse
tsse2
purity:C516
purity:C518
purity:C519
purity:C524
purity:C525
purity:C527
purity:C528
purity:C530
purity:C531
purity:C532
purity:C536
purity:C537
purity:C538
purity:C539
purity:C542
purity:C543
purity:C544
purity:C547
purity:C548
purity:C549
purity:C550
purity:C551
purity:C552
purity:C554
purity:C555
purity:C559
purity:C560
purity:C561
purity:C562
tsse:tsse2
all
A–D
A
B
C
D
0
Coefficient
p_c = < 2.22e–16, p_sc = 0.7374

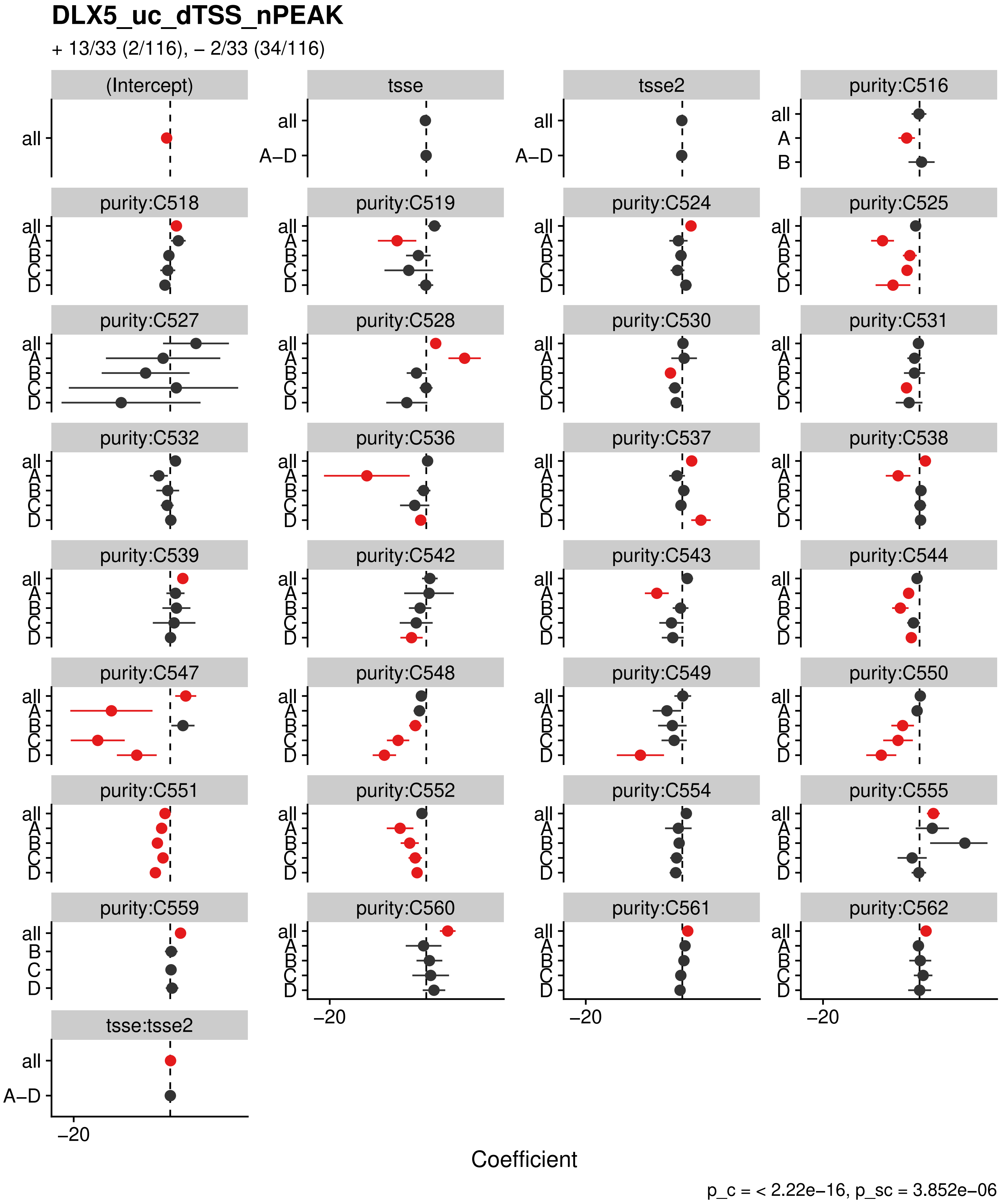

DLX5_uc_dTSS_nPEAK
+ 13/33 (2/116), − 2/33 (34/116)
(Intercept)
tsse
tsse2
purity:C516
purity:C518
purity:C519
purity:C524
purity:C525
purity:C527
purity:C528
purity:C530
purity:C531
purity:C532
purity:C536
purity:C537
purity:C538
purity:C539
purity:C542
purity:C543
purity:C544
purity:C547
purity:C548
purity:C549
purity:C550
purity:C551
purity:C552
purity:C554
purity:C555
purity:C559
purity:C560
purity:C561
purity:C562
tsse:tsse2
all
A−D
A
B
C
D
−20
Coefficient
p_c = < 2.22e−16, p_sc = 3.852e−06

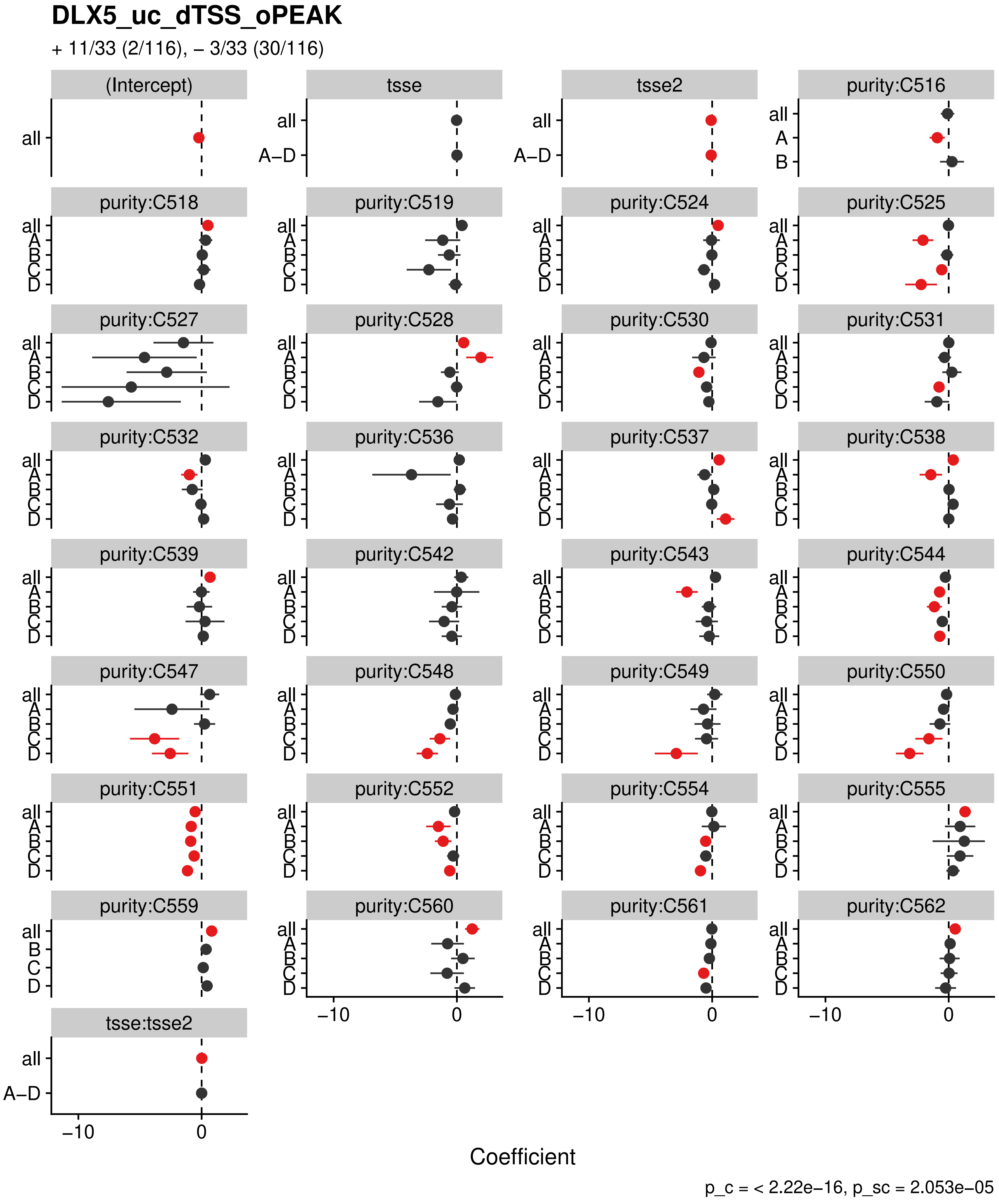
DLX5_uc_dTSS_oPEAK
+ 11/33 (2/116), – 3/33 (30/116)
(Intercept)
tsse
tsse2
purity:C516
purity:C518
purity:C519
purity:C524
purity:C525
purity:C527
purity:C528
purity:C530
purity:C531
purity:C532
purity:C536
purity:C537
purity:C538
purity:C539
purity:C542
purity:C543
purity:C544
purity:C547
purity:C548
purity:C549
purity:C550
purity:C551
purity:C552
purity:C554
purity:C555
purity:C559
purity:C560
purity:C561
purity:C562
tsse:tsse2
all
A–D
A
B
C
D
−10
0
Coefficient
p_c = < 2.22e−16, p_sc = 2.053e−05

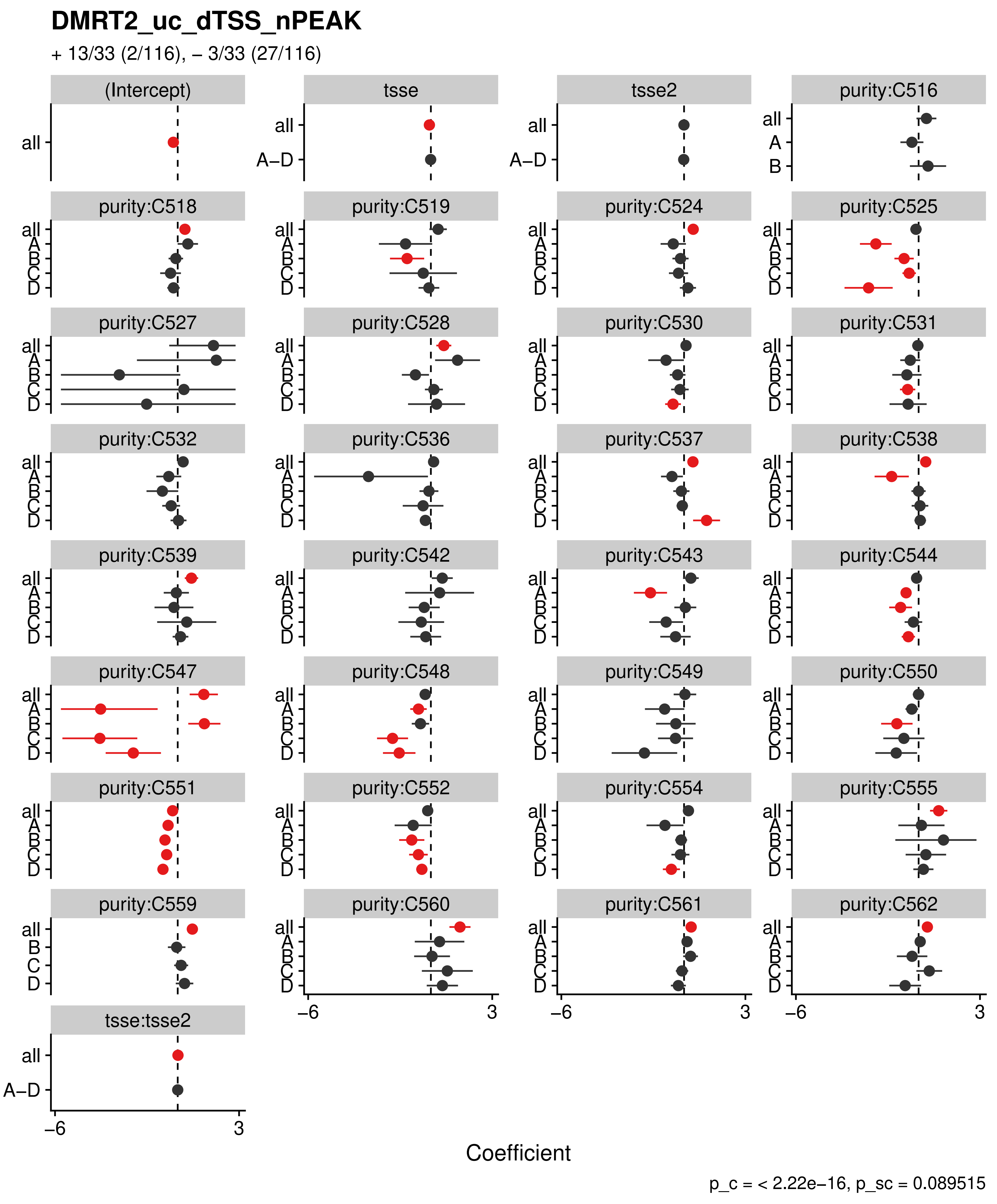
DMRT2_uc_dTSS_nPEAK
+ 13/33 (2/116), – 3/33 (27/116)
(Intercept)
tsse
tsse2
purity:C516
purity:C518
purity:C519
purity:C524
purity:C525
purity:C527
purity:C528
purity:C530
purity:C531
purity:C532
purity:C536
purity:C537
purity:C538
purity:C539
purity:C542
purity:C543
purity:C544
purity:C547
purity:C548
purity:C549
purity:C550
purity:C551
purity:C552
purity:C554
purity:C555
purity:C559
purity:C560
purity:C561
purity:C562
tsse:tsse2
all
A–D
A
B
C
D
–6
3
Coefficient
p_c = < 2.22e–16, p_sc = 0.089515

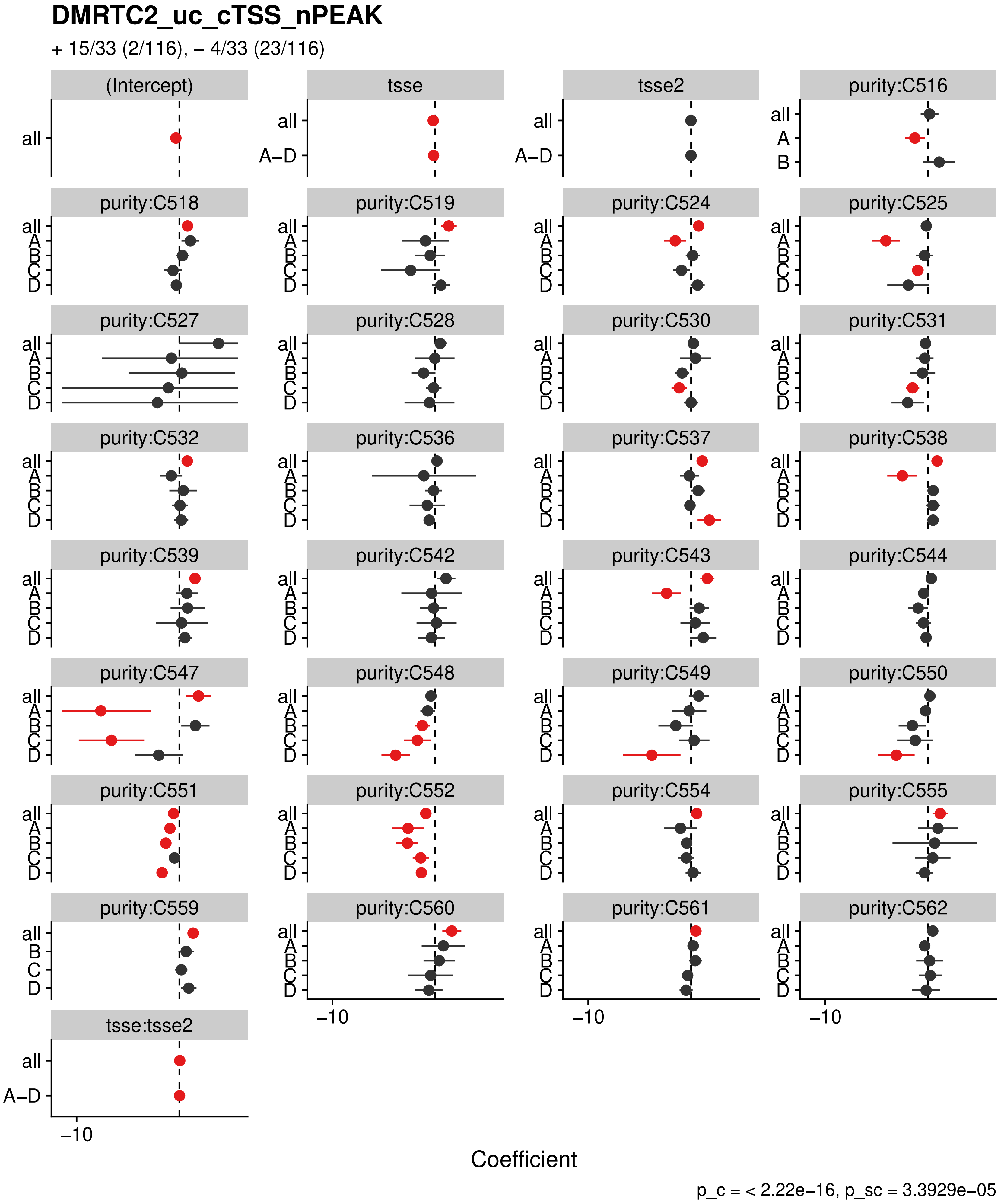
DMRTC2_uc_cTSS_nPEAK
+ 15/33 (2/116), – 4/33 (23/116)
(Intercept)
tsse
tsse2
purity:C516
purity:C518
purity:C519
purity:C524
purity:C525
purity:C527
purity:C528
purity:C530
purity:C531
purity:C532
purity:C536
purity:C537
purity:C538
purity:C539
purity:C542
purity:C543
purity:C544
purity:C547
purity:C548
purity:C549
purity:C550
purity:C551
purity:C552
purity:C554
purity:C555
purity:C559
purity:C560
purity:C561
purity:C562
tsse:tsse2
all
A–D
A
B
C
D
−10
Coefficient
p_c = < 2.22e−16, p_sc = 3.3929e−05

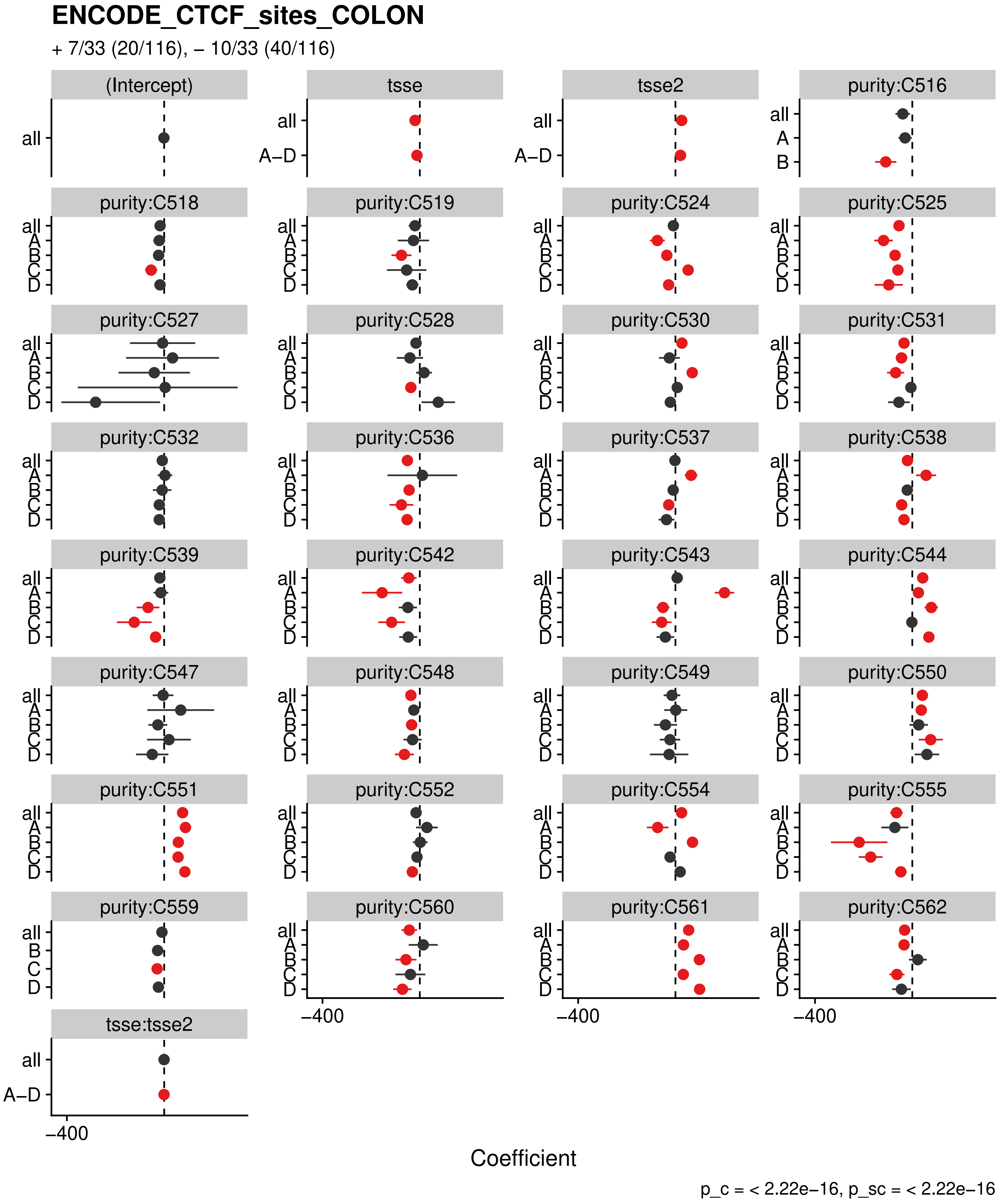
ENCODE_CTCF_sites_COLON
+ 7/33 (20/116), – 10/33 (40/116)
(Intercept)
tsse
tsse2
purity:C516
purity:C518
purity:C519
purity:C524
purity:C525
purity:C527
purity:C528
purity:C530
purity:C531
purity:C532
purity:C536
purity:C537
purity:C538
purity:C539
purity:C542
purity:C543
purity:C544
purity:C547
purity:C548
purity:C549
purity:C550
purity:C551
purity:C552
purity:C554
purity:C555
purity:C559
purity:C560
purity:C561
purity:C562
tsse:tsse2
all
A–D
A
B
C
D
–400
Coefficient
p_c = < 2.22e–16, p_sc = < 2.22e–16

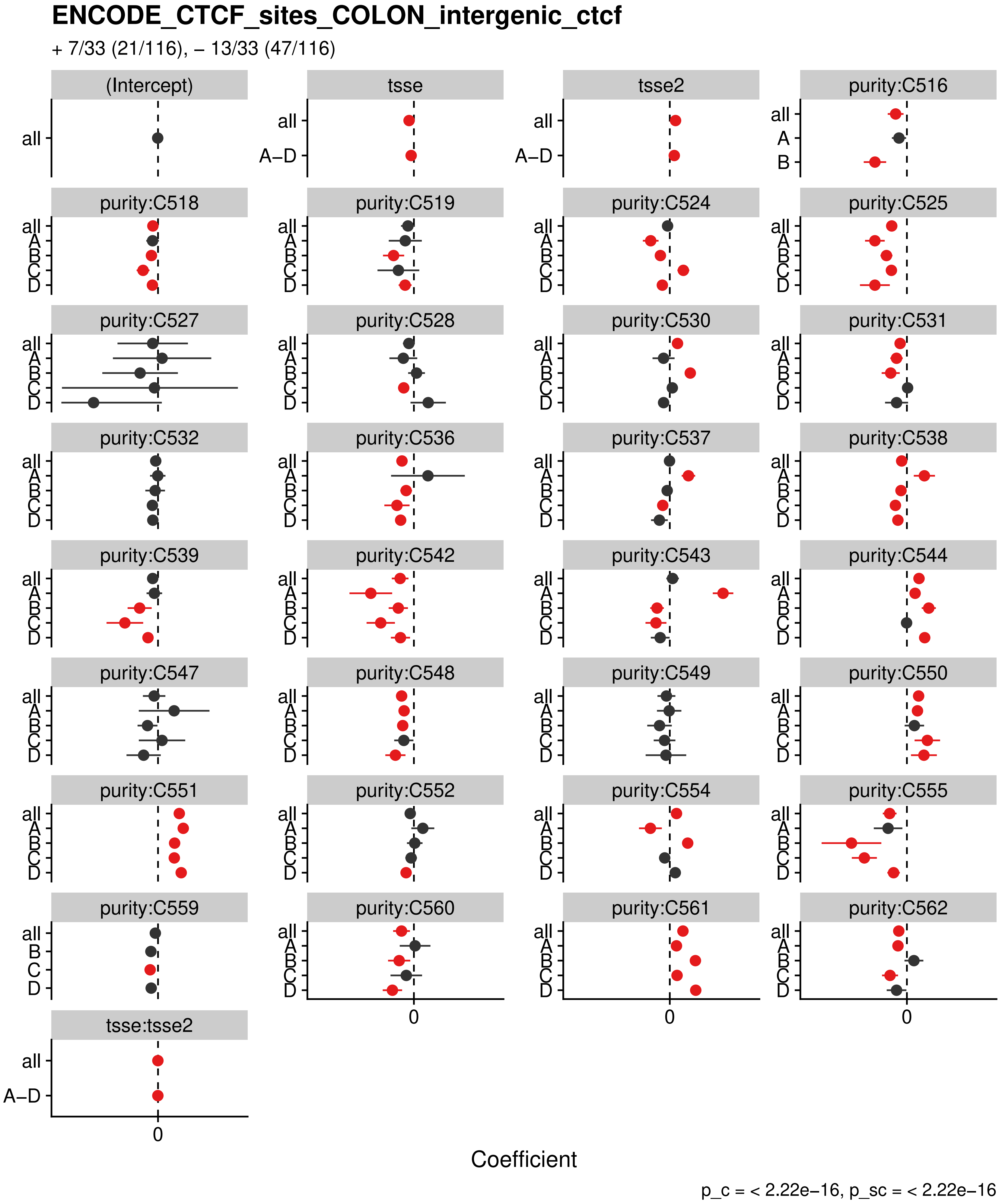

ENCODE_CTCF_sites_COLON_intergenic_ctcf
+ 7/33 (21/116), – 13/33 (47/116)
(Intercept)
tsse
tsse2
purity:C516
purity:C518
purity:C519
purity:C524
purity:C525
purity:C527
purity:C528
purity:C530
purity:C531
purity:C532
purity:C536
purity:C537
purity:C538
purity:C539
purity:C542
purity:C543
purity:C544
purity:C547
purity:C548
purity:C549
purity:C550
purity:C551
purity:C552
purity:C554
purity:C555
purity:C559
purity:C560
purity:C561
purity:C562
tsse:tsse2
all
A–D
A
B
C
D
0
Coefficient
p_c = < 2.22e−16, p_sc = < 2.22e−16

**ENCODE_CTCF_sites_COLON_loop_ctcf**

+ 7/33 (20/116), – 13/33 (48/116)

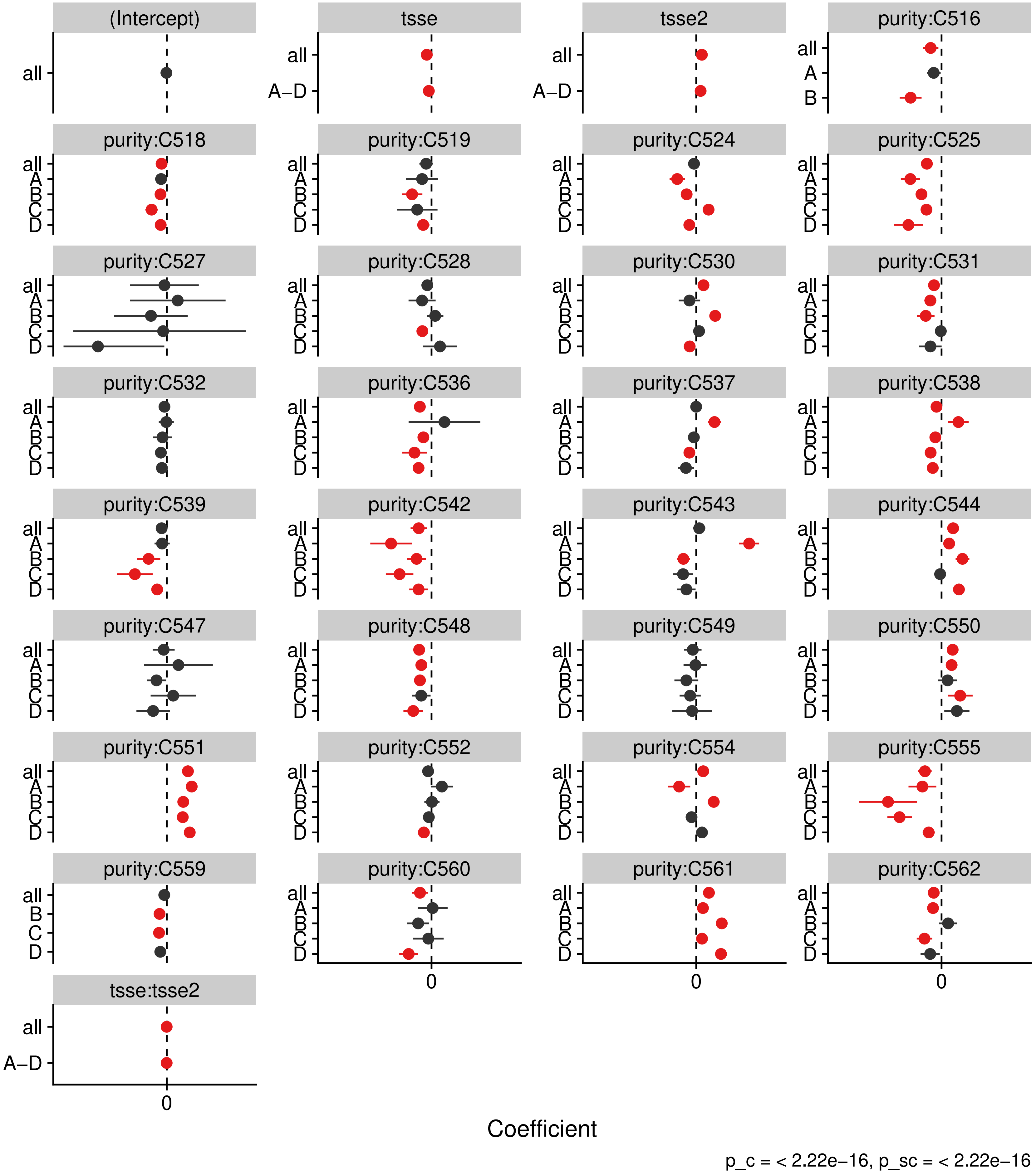

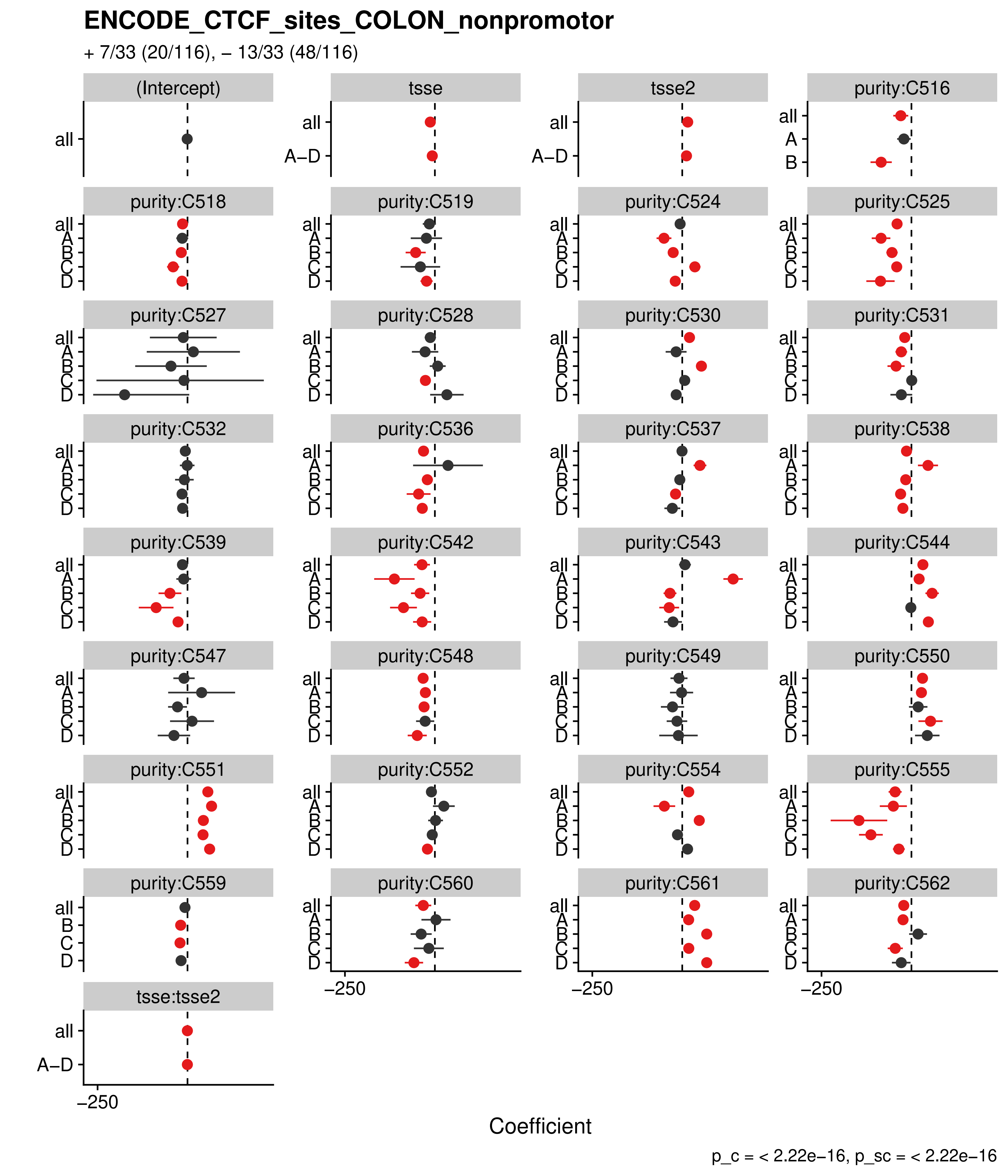
ENCODE_CTCF_sites_COLON_nonpromotor
+ 7/33 (20/116), – 13/33 (48/116)
(Intercept)
tsse
tsse2
purity:C516
purity:C518
purity:C519
purity:C524
purity:C525
purity:C527
purity:C528
purity:C530
purity:C531
purity:C532
purity:C536
purity:C537
purity:C538
purity:C539
purity:C542
purity:C543
purity:C544
purity:C547
purity:C548
purity:C549
purity:C550
purity:C551
purity:C552
purity:C554
purity:C555
purity:C559
purity:C560
purity:C561
purity:C562
tsse:tsse2
all
A–D
A
B
C
D
–250
Coefficient
p_c = < 2.22e–16, p_sc = < 2.22e–16

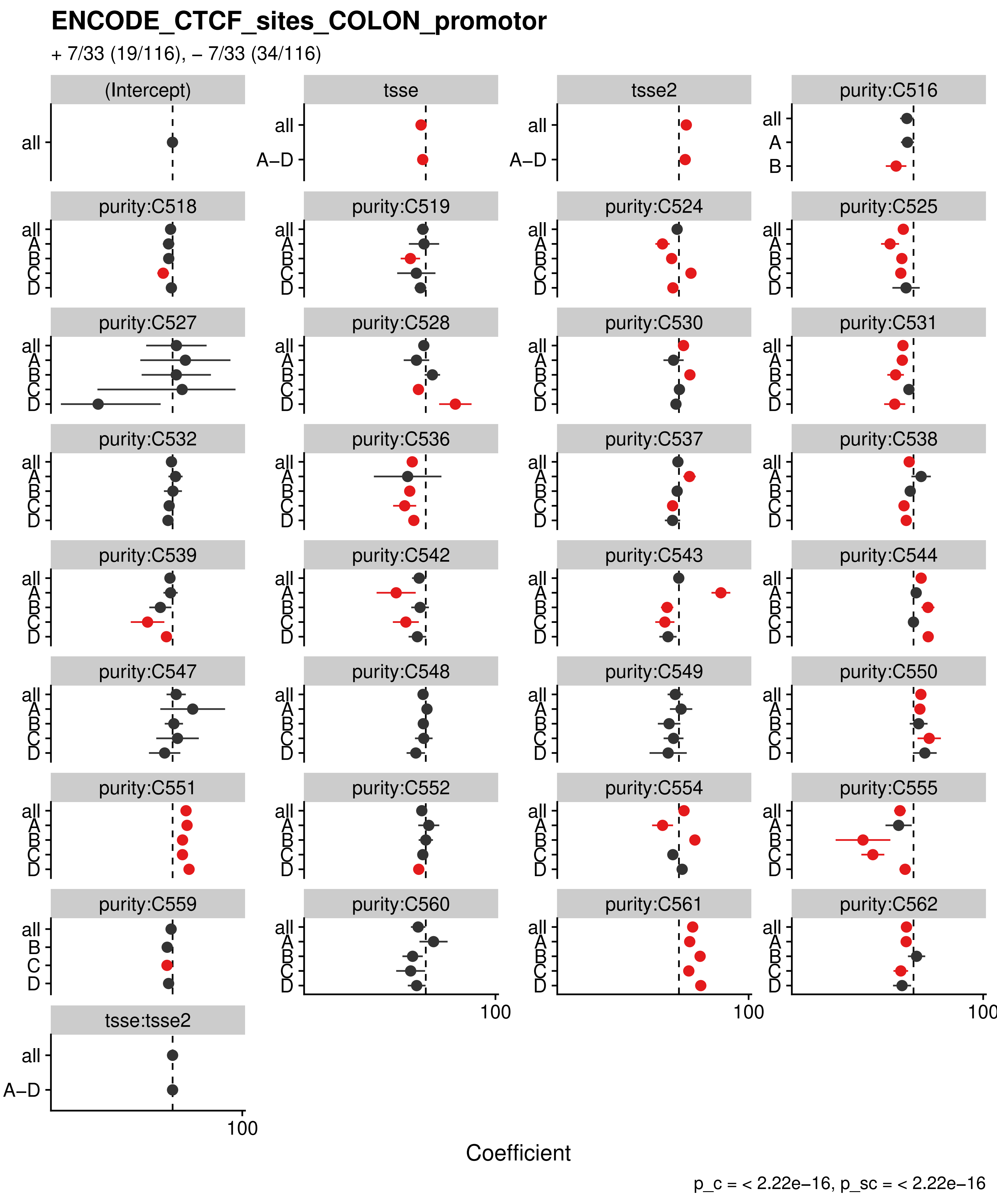
ENCODE_CTCF_sites_COLON_promotor
+ 7/33 (19/116), – 7/33 (34/116)
(Intercept)
tsse
tsse2
purity:C516
purity:C518
purity:C519
purity:C524
purity:C525
purity:C527
purity:C528
purity:C530
purity:C531
purity:C532
purity:C536
purity:C537
purity:C538
purity:C539
purity:C542
purity:C543
purity:C544
purity:C547
purity:C548
purity:C549
purity:C550
purity:C551
purity:C552
purity:C554
purity:C555
purity:C559
purity:C560
purity:C561
purity:C562
tsse:tsse2
all
A–D
A
B
C
D
100
Coefficient
p_c = < 2.22e–16, p_sc = < 2.22e–16

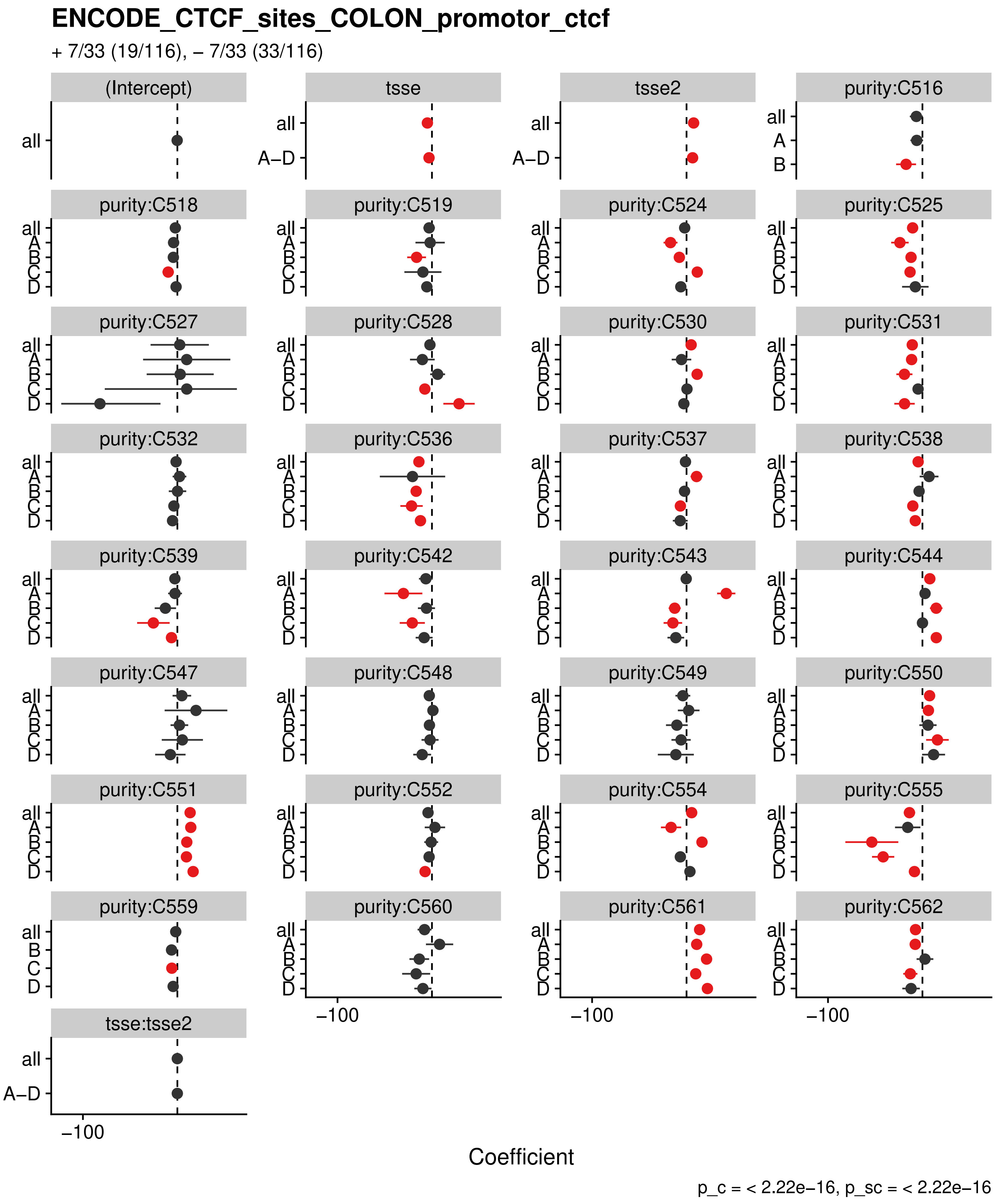
ENCODE_CTCF_sites_COLON_promotor_ctcf
+ 7/33 (19/116), – 7/33 (33/116)
(Intercept)
tsse
tsse2
purity:C516
purity:C518
purity:C519
purity:C524
purity:C525
purity:C527
purity:C528
purity:C530
purity:C531
purity:C532
purity:C536
purity:C537
purity:C538
purity:C539
purity:C542
purity:C543
purity:C544
purity:C547
purity:C548
purity:C549
purity:C550
purity:C551
purity:C552
purity:C554
purity:C555
purity:C559
purity:C560
purity:C561
purity:C562
tsse:tsse2
all
A–D
A
B
C
D
−100
Coefficient
p_c = < 2.22e−16, p_sc = < 2.22e−16

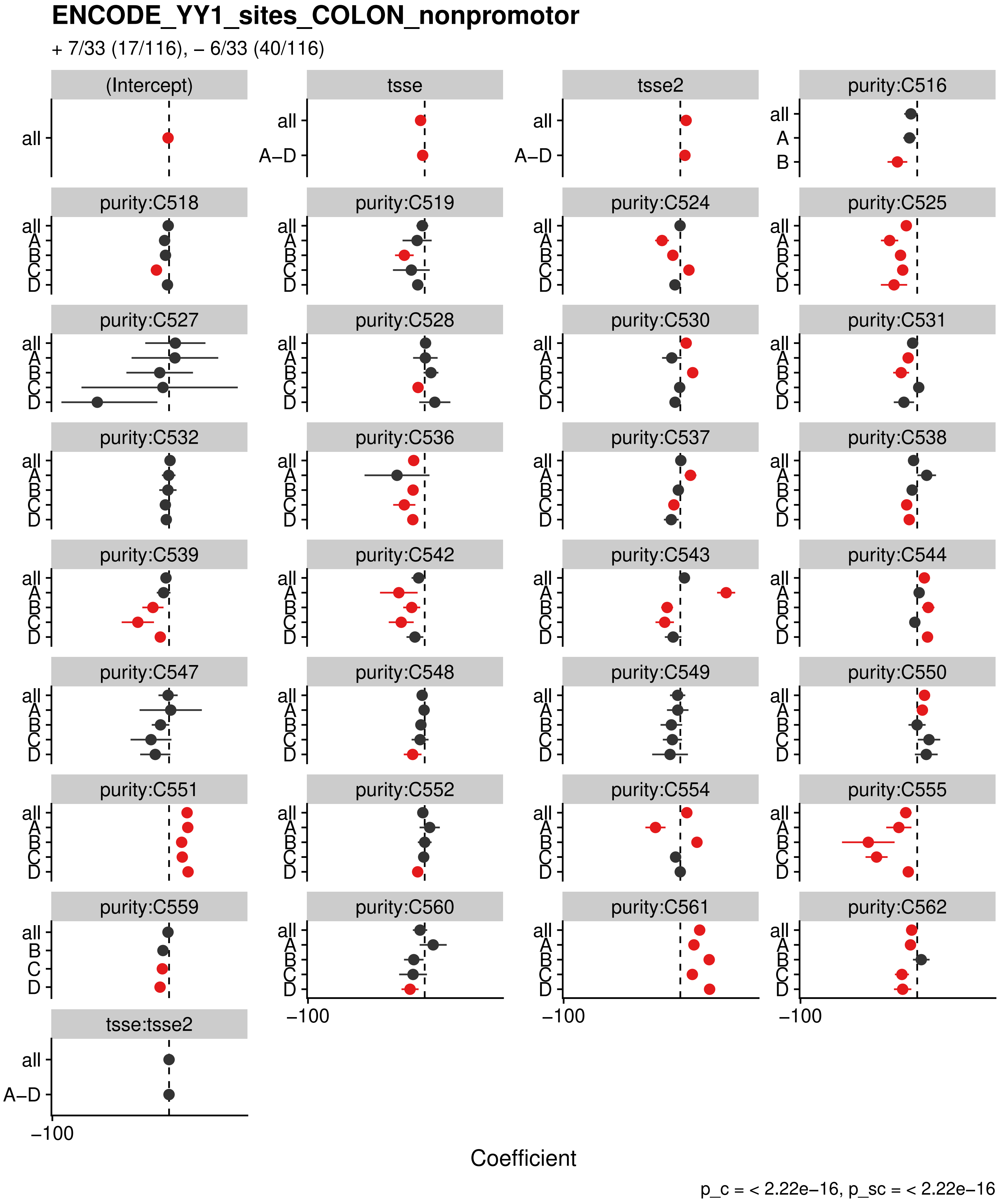
ENCODE_YY1_sites_COLON_nonpromotor
+ 7/33 (17/116), – 6/33 (40/116)
(Intercept)
tsse
tsse2
purity:C516
purity:C518
purity:C519
purity:C524
purity:C525
purity:C527
purity:C528
purity:C530
purity:C531
purity:C532
purity:C536
purity:C537
purity:C538
purity:C539
purity:C542
purity:C543
purity:C544
purity:C547
purity:C548
purity:C549
purity:C550
purity:C551
purity:C552
purity:C554
purity:C555
purity:C559
purity:C560
purity:C561
purity:C562
tsse:tsse2
all
A–D
A
B
C
D
−100
Coefficient
p_c = < 2.22e−16, p_sc = < 2.22e−16

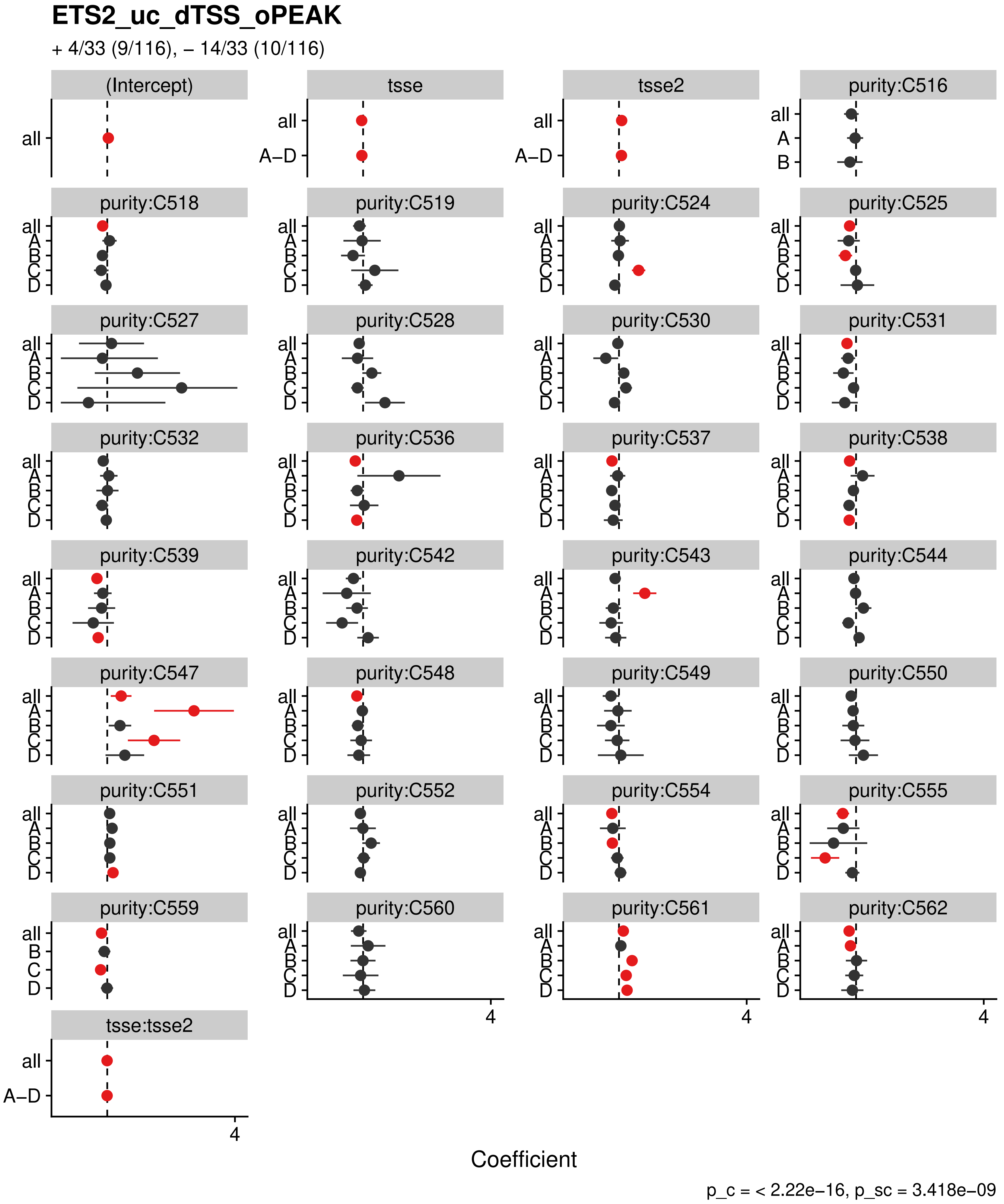
ETS2_uc_dTSS_oPEAK
+ 4/33 (9/116), – 14/33 (10/116)
(Intercept)
tsse
tsse2
purity:C516
purity:C518
purity:C519
purity:C524
purity:C525
purity:C527
purity:C528
purity:C530
purity:C531
purity:C532
purity:C536
purity:C537
purity:C538
purity:C539
purity:C542
purity:C543
purity:C544
purity:C547
purity:C548
purity:C549
purity:C550
purity:C551
purity:C552
purity:C554
purity:C555
purity:C559
purity:C560
purity:C561
purity:C562
tsse:tsse2
all
A–D
A
B
C
D
4
Coefficient
p_c = < 2.22e−16, p_sc = 3.418e−09

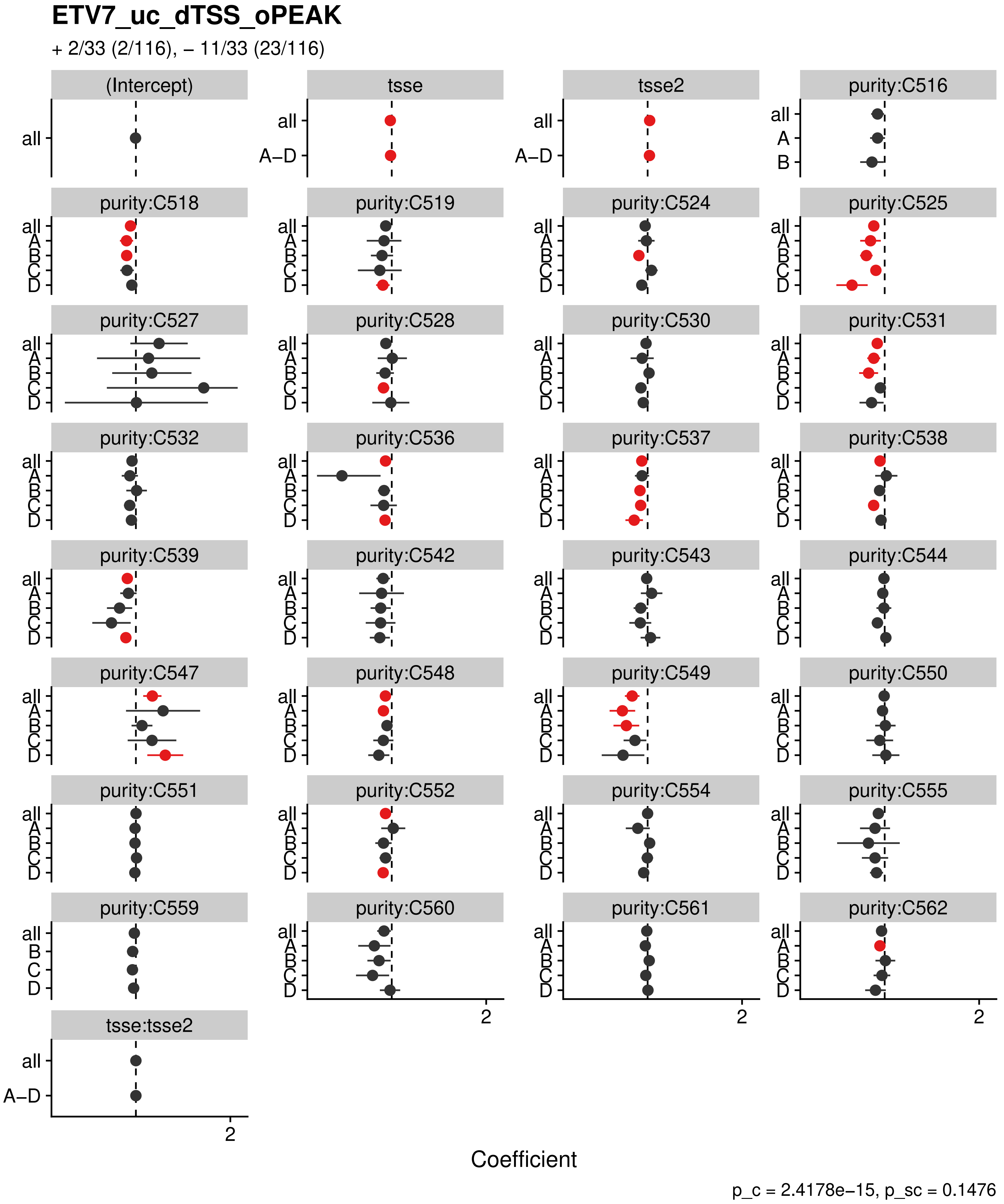

ETV7_uc_dTSS_oPEAK
+ 2/33 (2/116), – 11/33 (23/116)
(Intercept)
tsse
tsse2
purity:C516
purity:C518
purity:C519
purity:C524
purity:C525
purity:C527
purity:C528
purity:C530
purity:C531
purity:C532
purity:C536
purity:C537
purity:C538
purity:C539
purity:C542
purity:C543
purity:C544
purity:C547
purity:C548
purity:C549
purity:C550
purity:C551
purity:C552
purity:C554
purity:C555
purity:C559
purity:C560
purity:C561
purity:C562
tsse:tsse2
all
A–D
A
B
C
D
2
Coefficient
p_c = 2.4178e–15, p_sc = 0.1476

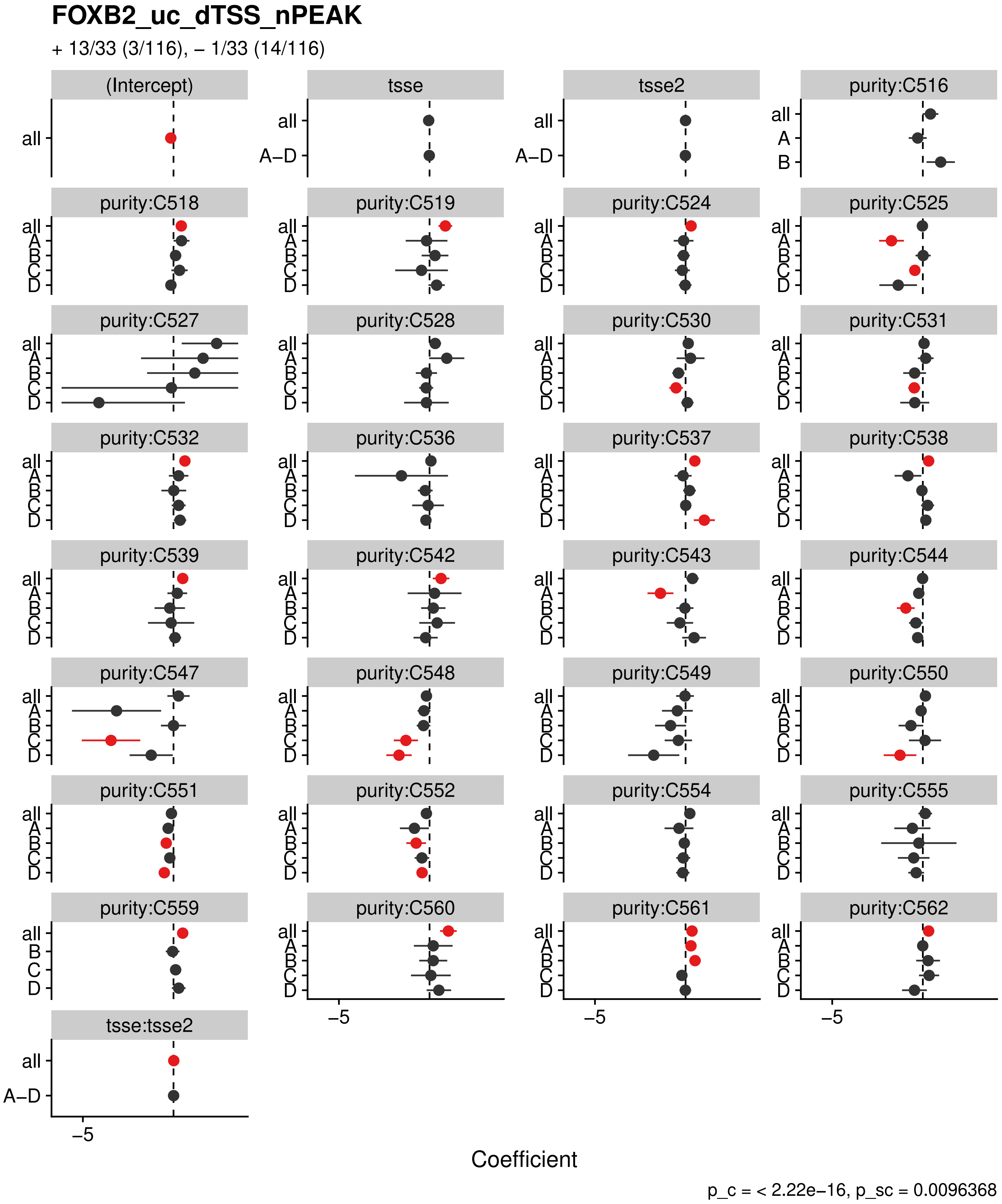
FOXB2_uc_dTSS_nPEAK
+ 13/33 (3/116), – 1/33 (14/116)
(Intercept)
tsse
tsse2
purity:C516
purity:C518
purity:C519
purity:C524
purity:C525
purity:C527
purity:C528
purity:C530
purity:C531
purity:C532
purity:C536
purity:C537
purity:C538
purity:C539
purity:C542
purity:C543
purity:C544
purity:C547
purity:C548
purity:C549
purity:C550
purity:C551
purity:C552
purity:C554
purity:C555
purity:C559
purity:C560
purity:C561
purity:C562
tsse:tsse2
all
A–D
A
B
C
D
–5
Coefficient
p_c = < 2.22e–16, p_sc = 0.0096368

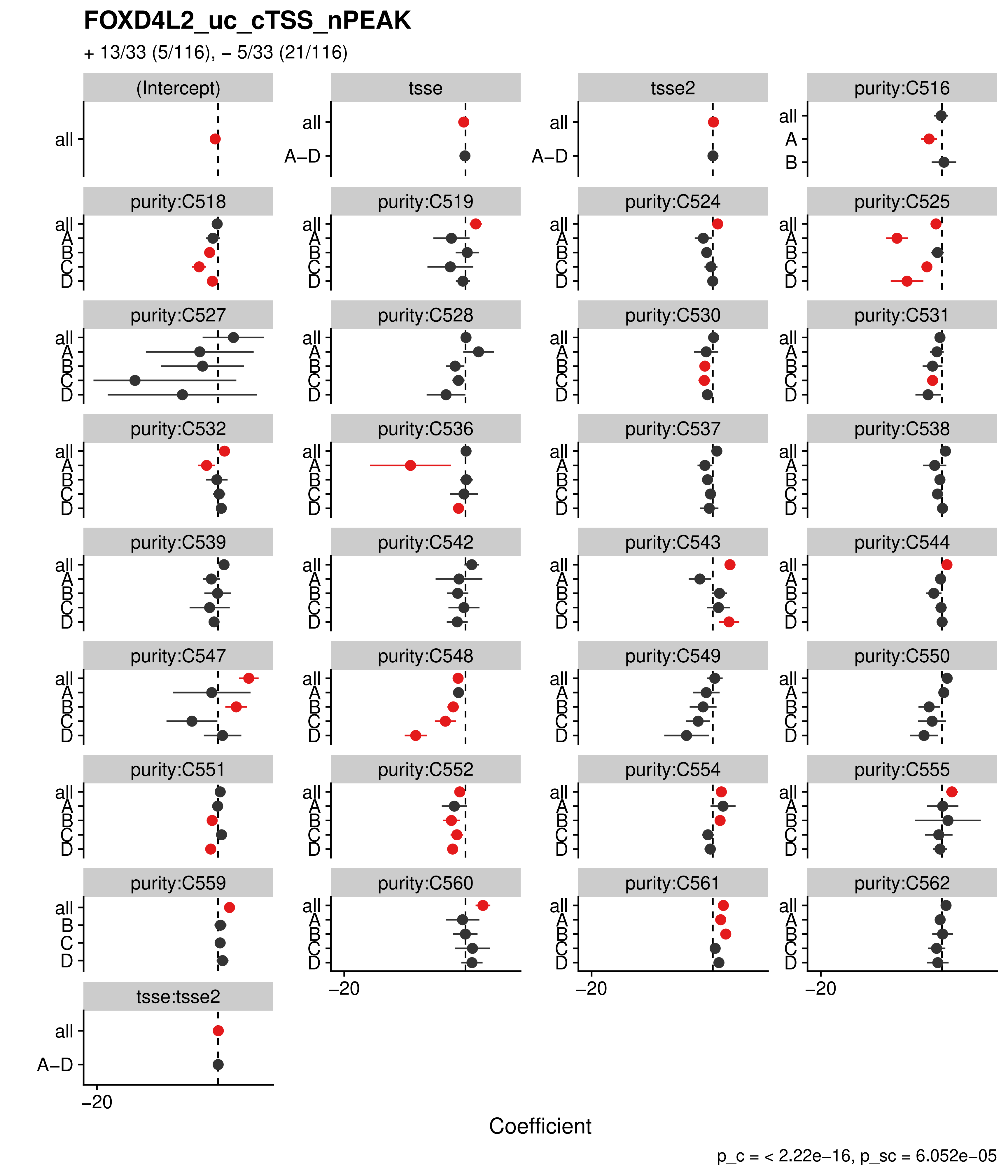
FOXD4L2_uc_cTSS_nPEAK
+ 13/33 (5/116), – 5/33 (21/116)
(Intercept)
tsse
tsse2
purity:C516
purity:C518
purity:C519
purity:C524
purity:C525
purity:C527
purity:C528
purity:C530
purity:C531
purity:C532
purity:C536
purity:C537
purity:C538
purity:C539
purity:C542
purity:C543
purity:C544
purity:C547
purity:C548
purity:C549
purity:C550
purity:C551
purity:C552
purity:C554
purity:C555
purity:C559
purity:C560
purity:C561
purity:C562
tsse:tsse2
all
A–D
A
B
C
D
–20
Coefficient
p_c = < 2.22e–16, p_sc = 6.052e–05

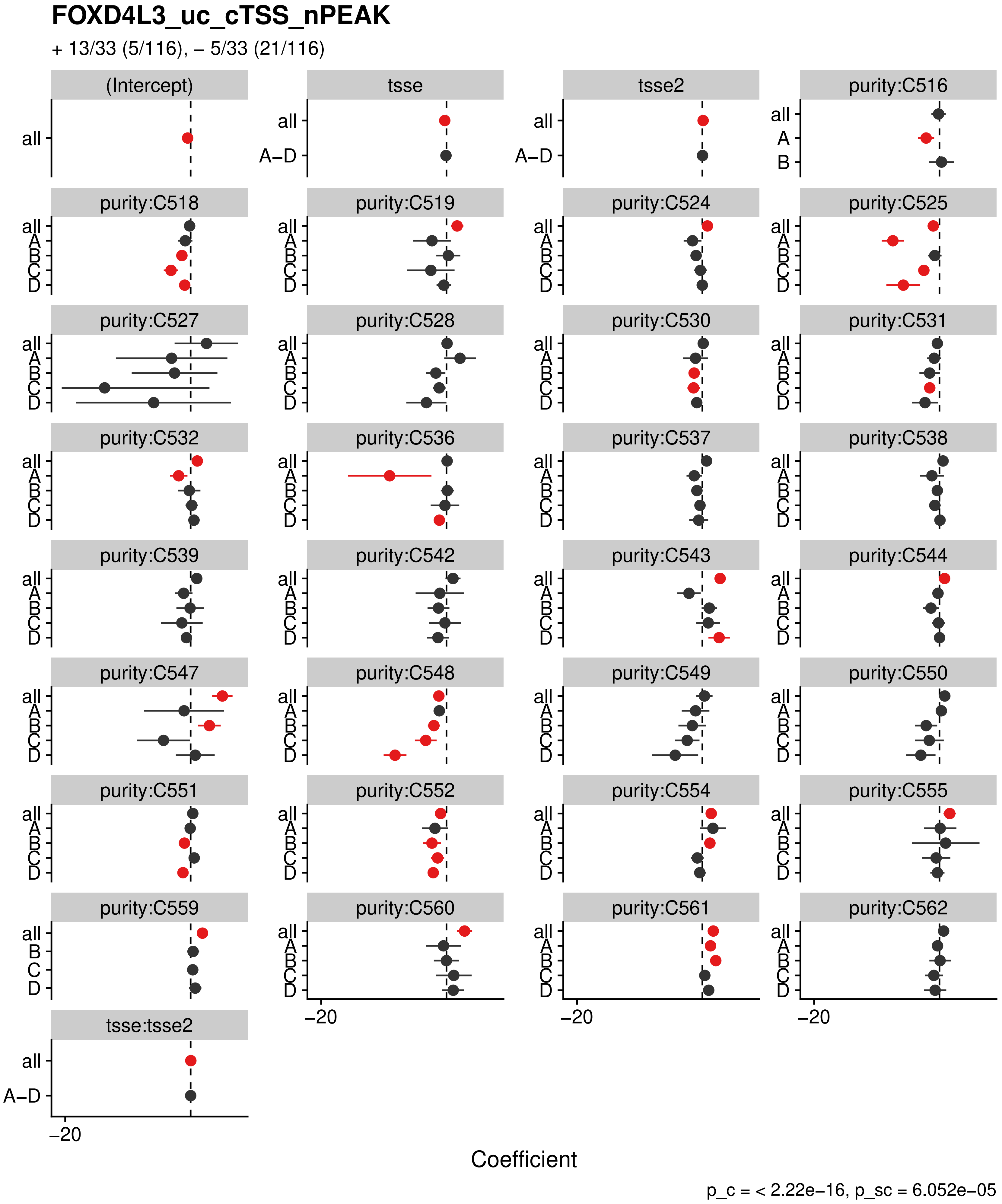
FOXD4L3_uc_cTSS_nPEAK
+ 13/33 (5/116), – 5/33 (21/116)
(Intercept)
tsse
tsse2
purity:C516
purity:C518
purity:C519
purity:C524
purity:C525
purity:C527
purity:C528
purity:C530
purity:C531
purity:C532
purity:C536
purity:C537
purity:C538
purity:C539
purity:C542
purity:C543
purity:C544
purity:C547
purity:C548
purity:C549
purity:C550
purity:C551
purity:C552
purity:C554
purity:C555
purity:C559
purity:C560
purity:C561
purity:C562
tsse:tsse2
all
A–D
A
B
C
D
–20
Coefficient
p_c = < 2.22e–16, p_sc = 6.052e–05

**FOXD4L4_uc_cTSS_nPEAK**

+ 13/33 (5/116), – 5/33 (21/116)

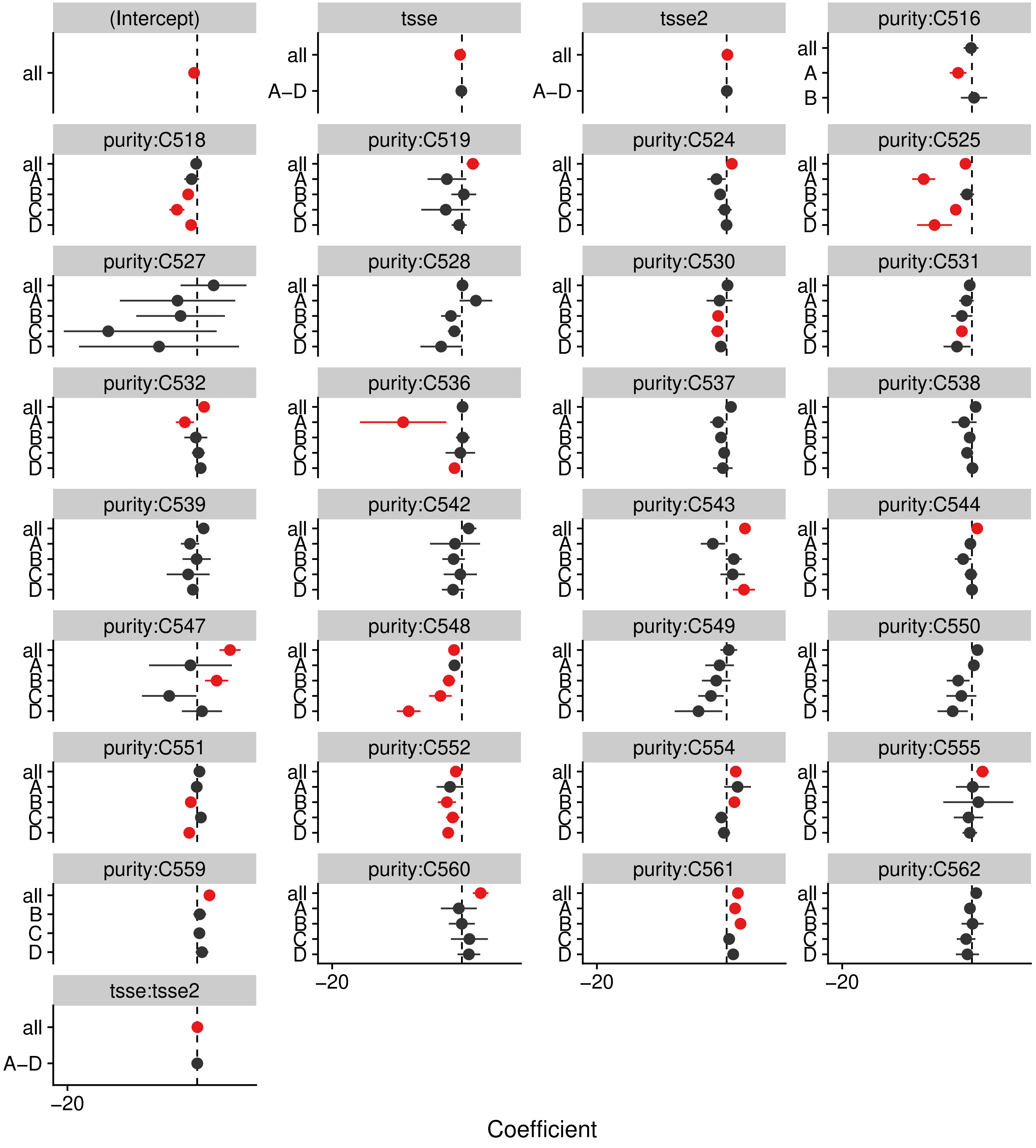


p_c = < 2.22e–16, p_sc = 6.052e–05

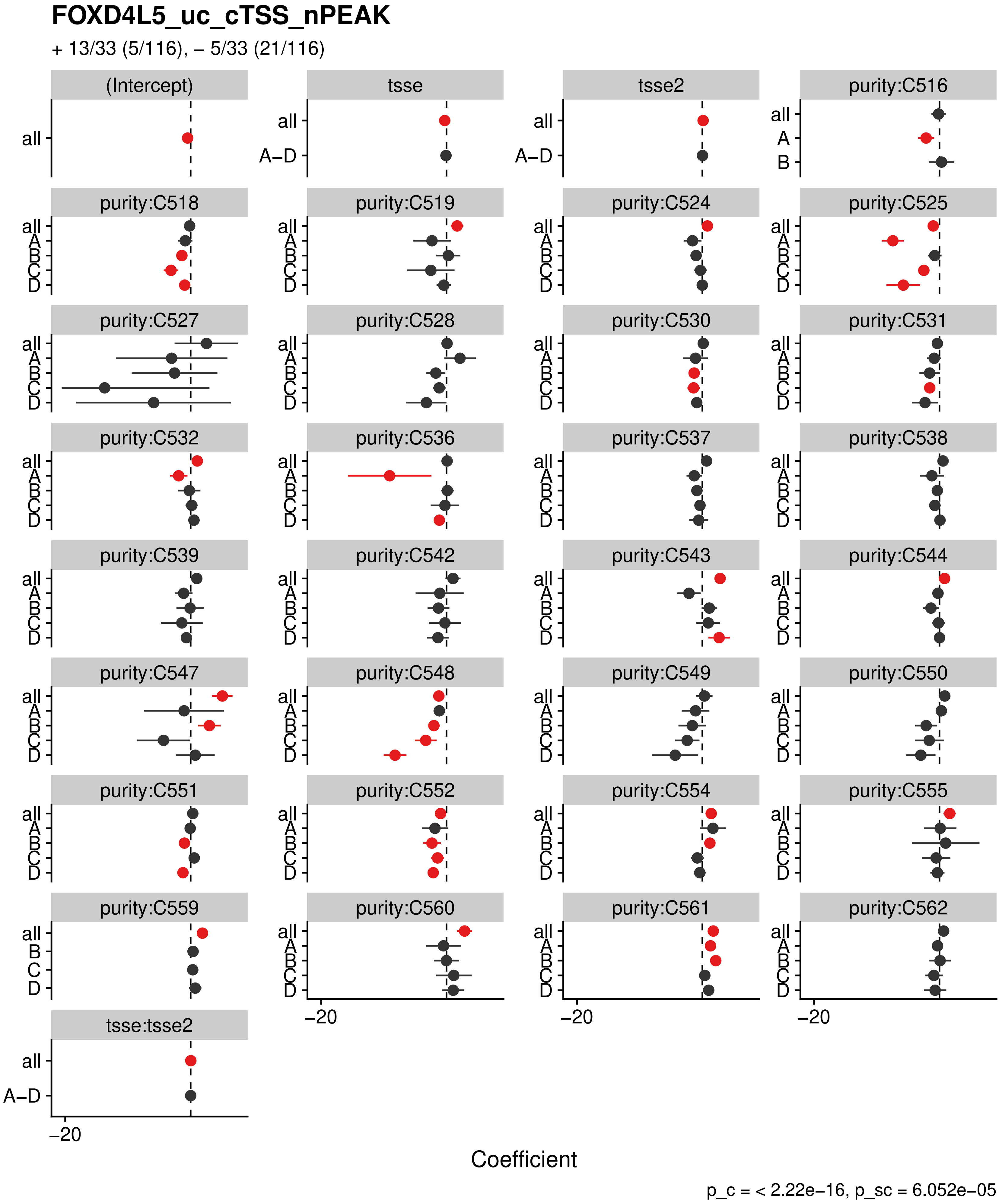
FOXD4L5_uc_cTSS_nPEAK
+ 13/33 (5/116), – 5/33 (21/116)
(Intercept)
tsse
tsse2
purity:C516
purity:C518
purity:C519
purity:C524
purity:C525
purity:C527
purity:C528
purity:C530
purity:C531
purity:C532
purity:C536
purity:C537
purity:C538
purity:C539
purity:C542
purity:C543
purity:C544
purity:C547
purity:C548
purity:C549
purity:C550
purity:C551
purity:C552
purity:C554
purity:C555
purity:C559
purity:C560
purity:C561
purity:C562
tsse:tsse2
all
A–D
A
B
C
D
–20
Coefficient
p_c = < 2.22e–16, p_sc = 6.052e–05

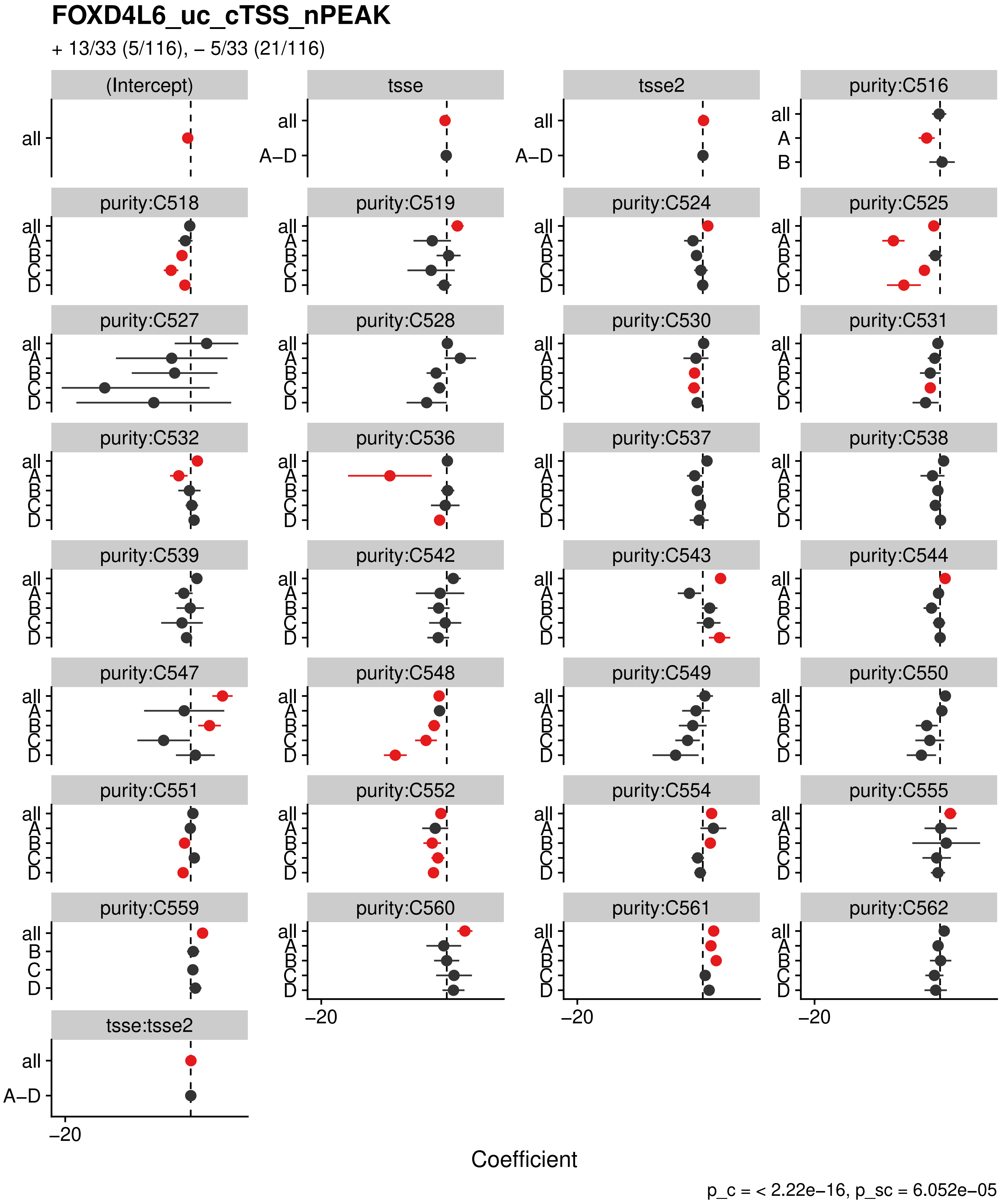

FOXD4L6_uc_cTSS_nPEAK
+ 13/33 (5/116), – 5/33 (21/116)
(Intercept)
tsse
tsse2
purity:C516
purity:C518
purity:C519
purity:C524
purity:C525
purity:C527
purity:C528
purity:C530
purity:C531
purity:C532
purity:C536
purity:C537
purity:C538
purity:C539
purity:C542
purity:C543
purity:C544
purity:C547
purity:C548
purity:C549
purity:C550
purity:C551
purity:C552
purity:C554
purity:C555
purity:C559
purity:C560
purity:C561
purity:C562
tsse:tsse2
all
A–D
A
B
C
D
–20
Coefficient
p_c = < 2.22e–16, p_sc = 6.052e–05

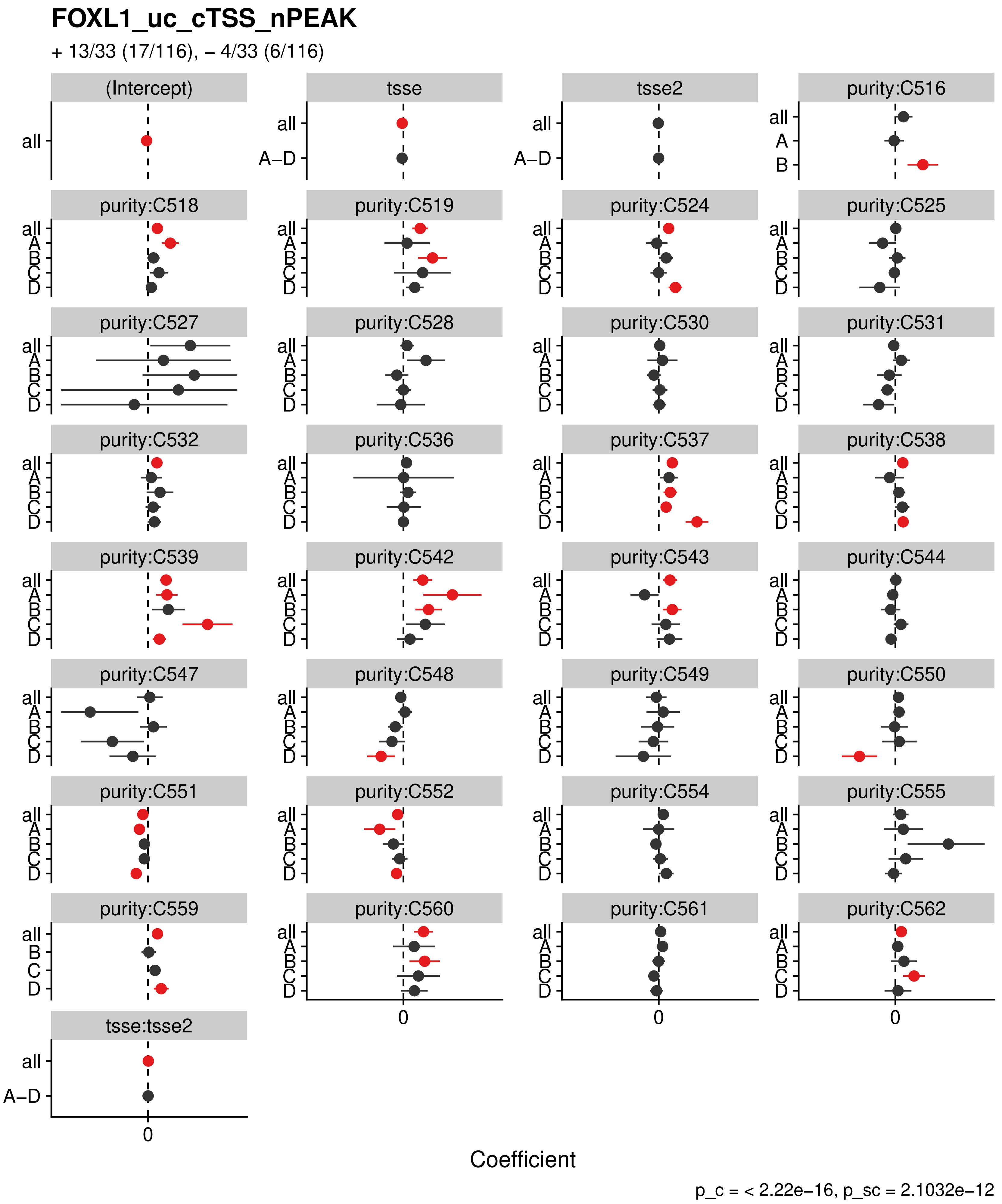
FOXL1_uc_cTSS_nPEAK
+ 13/33 (17/116), – 4/33 (6/116)
(Intercept)
tsse
tsse2
purity:C516
purity:C518
purity:C519
purity:C524
purity:C525
purity:C527
purity:C528
purity:C530
purity:C531
purity:C532
purity:C536
purity:C537
purity:C538
purity:C539
purity:C542
purity:C543
purity:C544
purity:C547
purity:C548
purity:C549
purity:C550
purity:C551
purity:C552
purity:C554
purity:C555
purity:C559
purity:C560
purity:C561
purity:C562
tsse:tsse2
all
A–D
A
B
C
D
0
Coefficient
p_c = < 2.22e–16, p_sc = 2.1032e–12

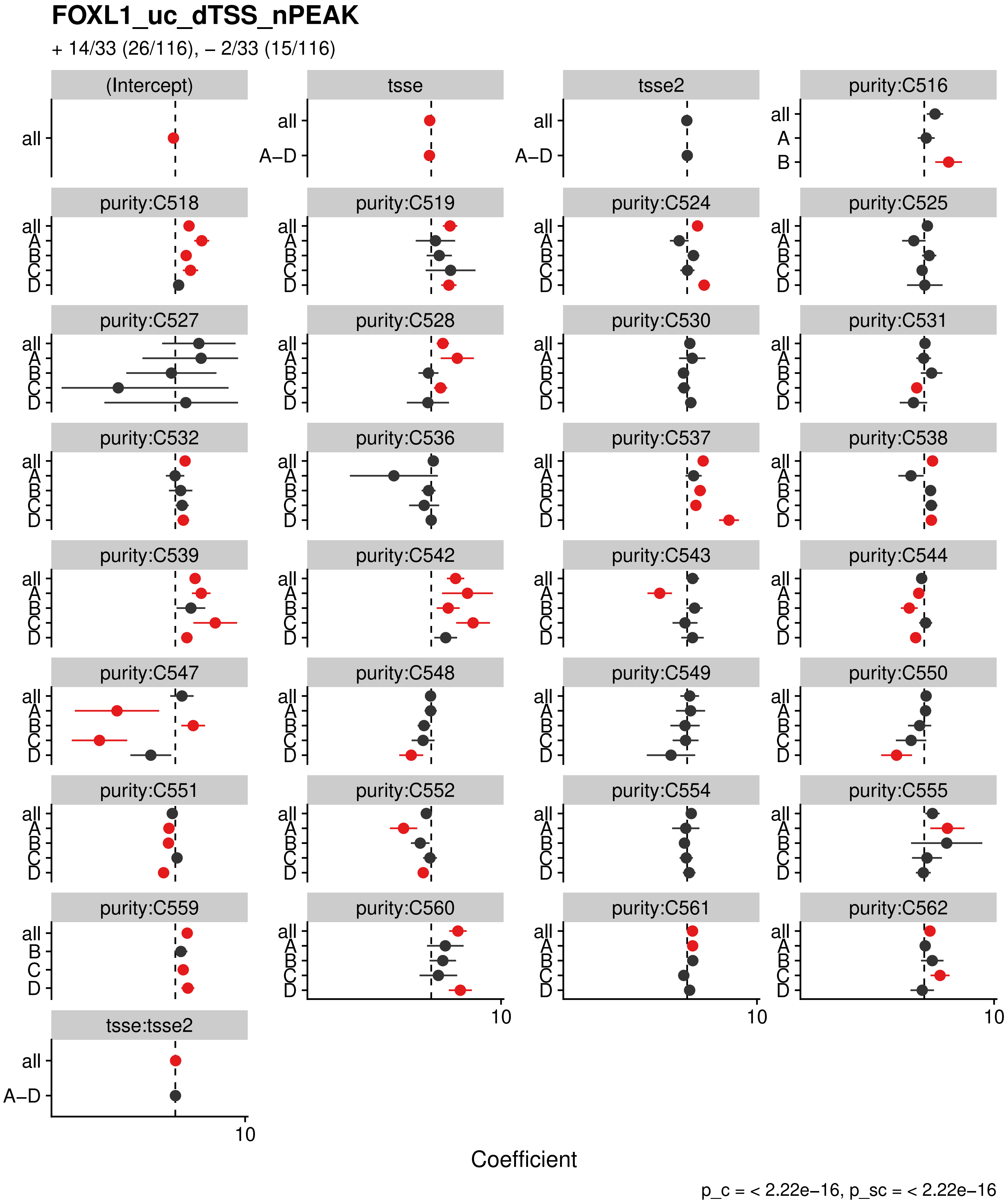
FOXL1_uc_dTSS_nPEAK
+ 14/33 (26/116), – 2/33 (15/116)
(Intercept)
tsse
tsse2
purity:C516
purity:C518
purity:C519
purity:C524
purity:C525
purity:C527
purity:C528
purity:C530
purity:C531
purity:C532
purity:C536
purity:C537
purity:C538
purity:C539
purity:C542
purity:C543
purity:C544
purity:C547
purity:C548
purity:C549
purity:C550
purity:C551
purity:C552
purity:C554
purity:C555
purity:C559
purity:C560
purity:C561
purity:C562
tsse:tsse2
all
A–D
A
B
C
D
10
Coefficient
p_c = < 2.22e–16, p_sc = < 2.22e–16

#### FOXL1_uc_dTSS_oPEAK

+ 16/33 (15/116), – 3/33 (15/116)

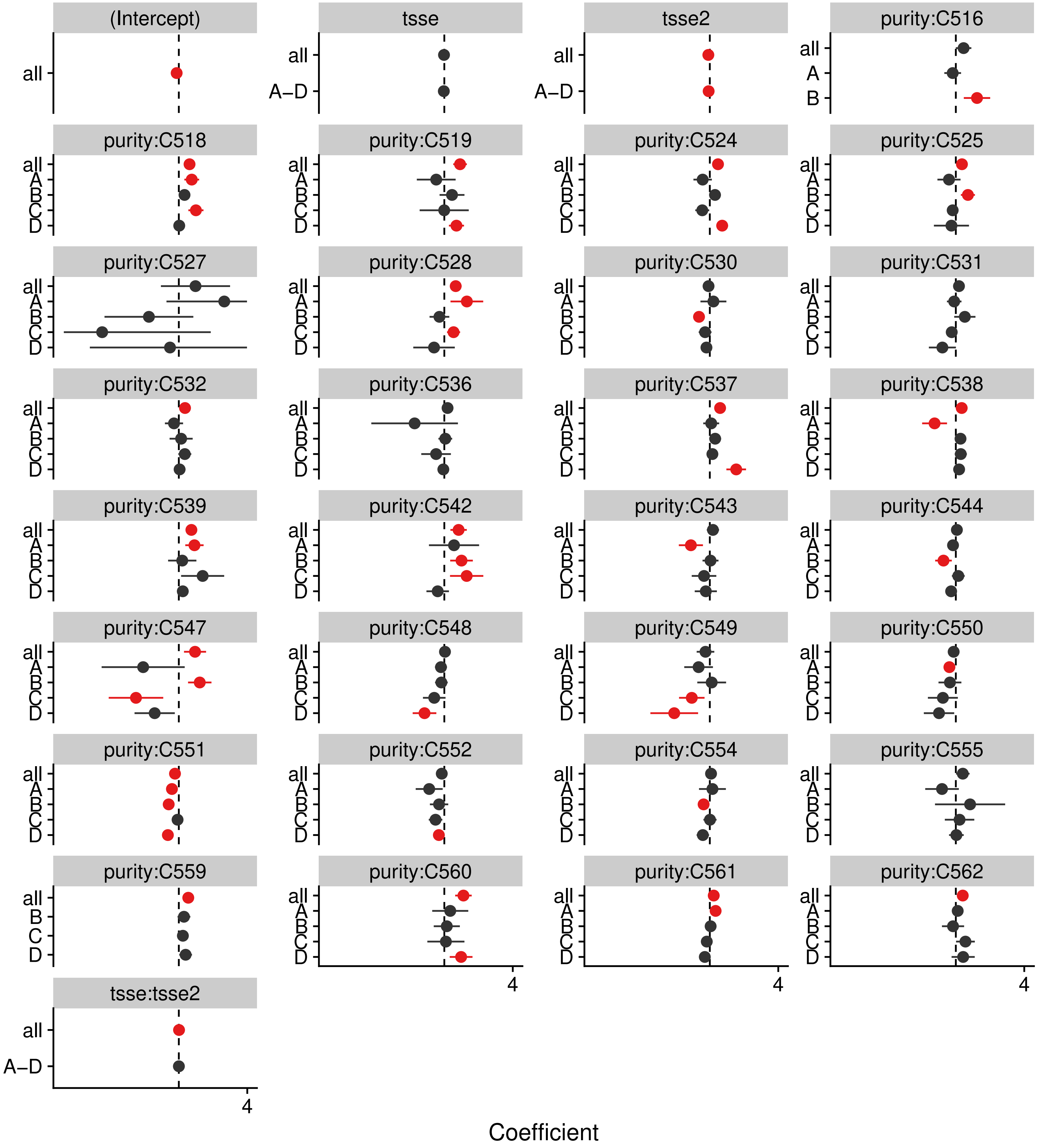


p_c = < 2.22e–16, p_sc = 1.9192e–13

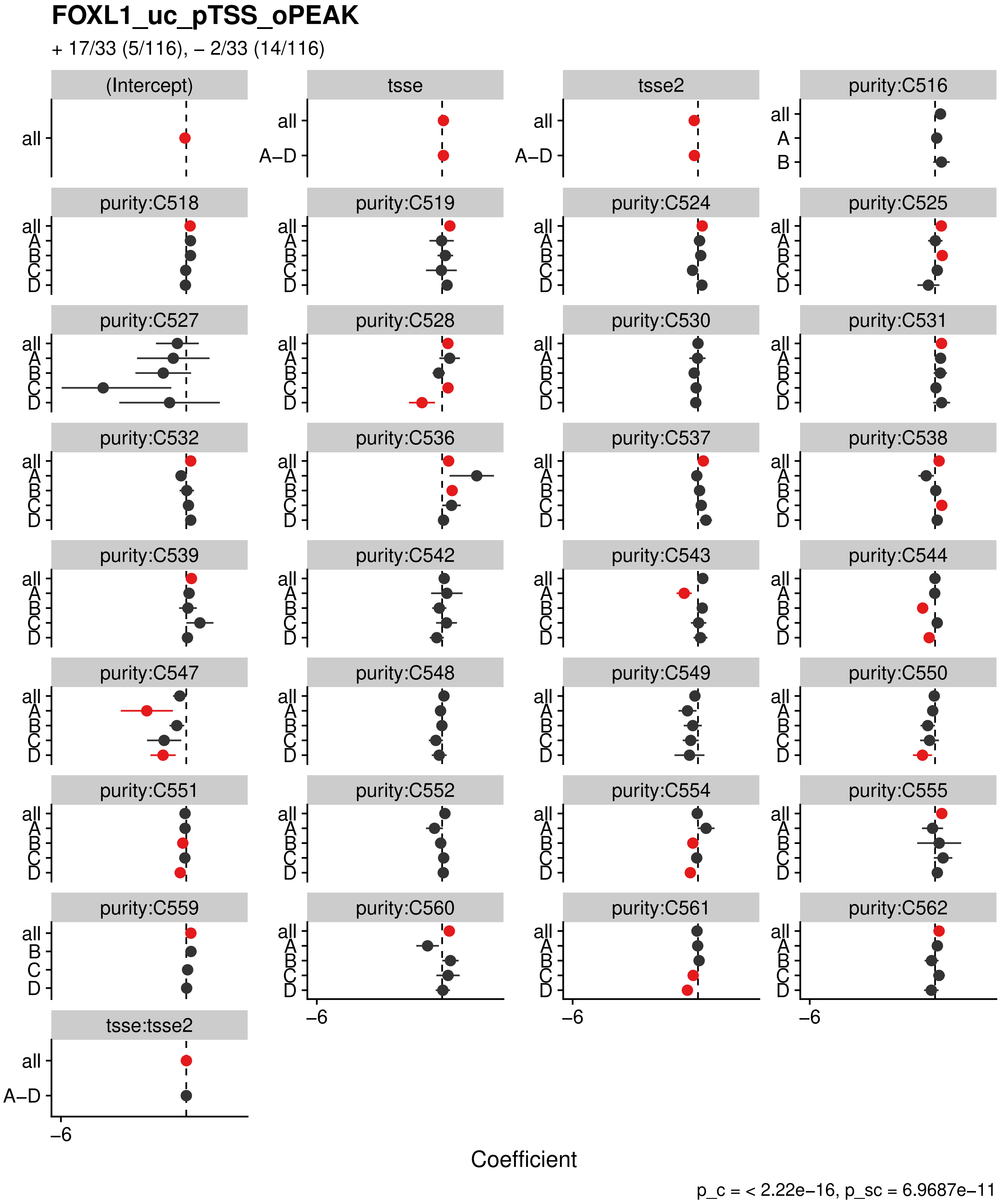
FOXL1_uc_pTSS_oPEAK
+ 17/33 (5/116), – 2/33 (14/116)
(Intercept)
tsse
tsse2
purity:C516
purity:C518
purity:C519
purity:C524
purity:C525
purity:C527
purity:C528
purity:C530
purity:C531
purity:C532
purity:C536
purity:C537
purity:C538
purity:C539
purity:C542
purity:C543
purity:C544
purity:C547
purity:C548
purity:C549
purity:C550
purity:C551
purity:C552
purity:C554
purity:C555
purity:C559
purity:C560
purity:C561
purity:C562
tsse:tsse2
all
A
B
C
D
A–D
–6
Coefficient
p_c = < 2.22e–16, p_sc = 6.9687e–11

#### FOXP3_uc_cTSS_nPEAK

+ 13/33 (5/116), – 5/33 (21/116)

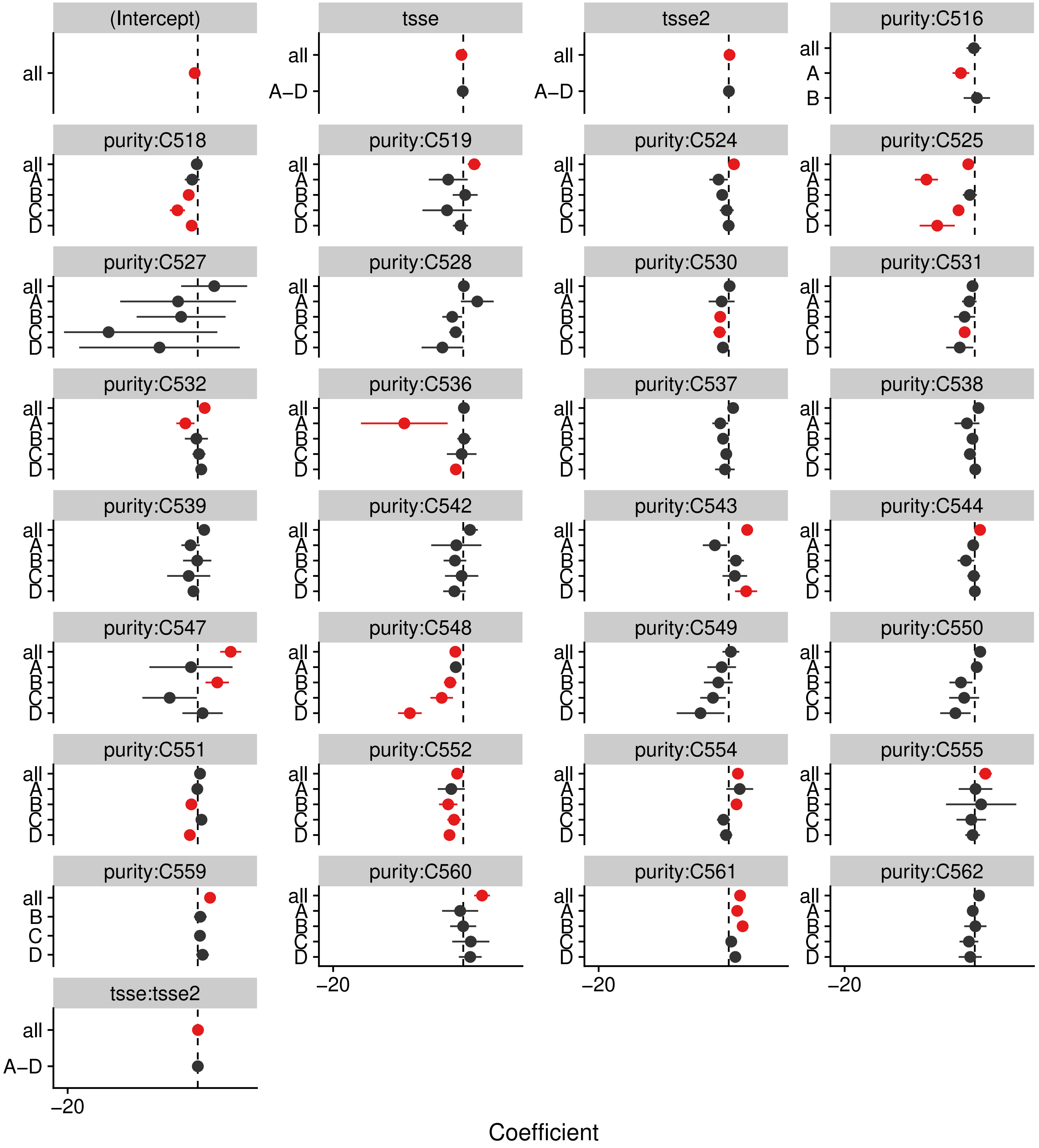


p_c = < 2.22e–16, p_sc = 6.052e–05

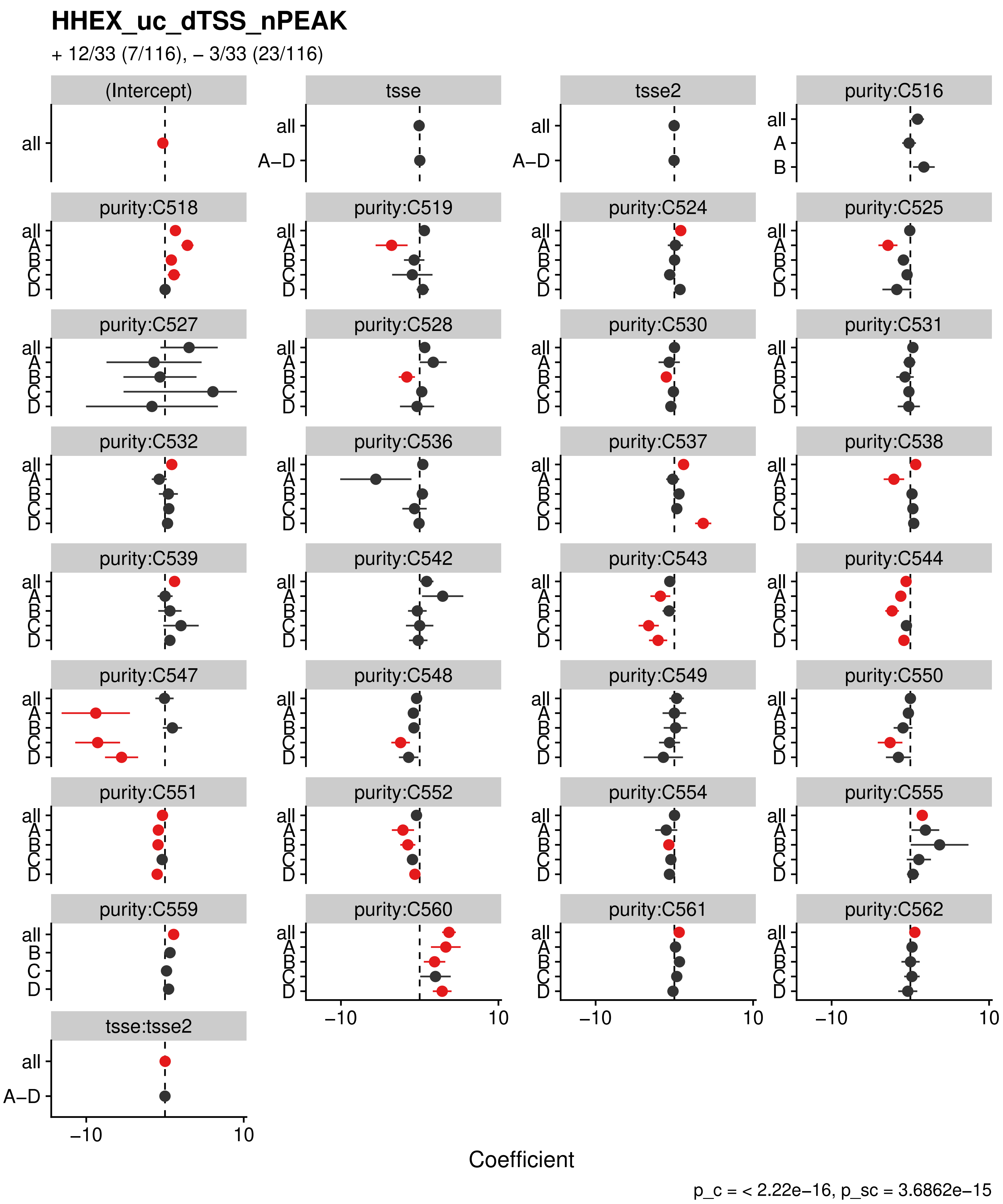

HHEX_uc_dTSS_nPEAK
+ 12/33 (7/116), – 3/33 (23/116)
(Intercept)
tsse
tsse2
purity:C516
purity:C518
purity:C519
purity:C524
purity:C525
purity:C527
purity:C528
purity:C530
purity:C531
purity:C532
purity:C536
purity:C537
purity:C538
purity:C539
purity:C542
purity:C543
purity:C544
purity:C547
purity:C548
purity:C549
purity:C550
purity:C551
purity:C552
purity:C554
purity:C555
purity:C559
purity:C560
purity:C561
purity:C562
tsse:tsse2
all
A–D
A
B
C
D
−10
10
Coefficient
p_c = < 2.22e−16, p_sc = 3.6862e−15

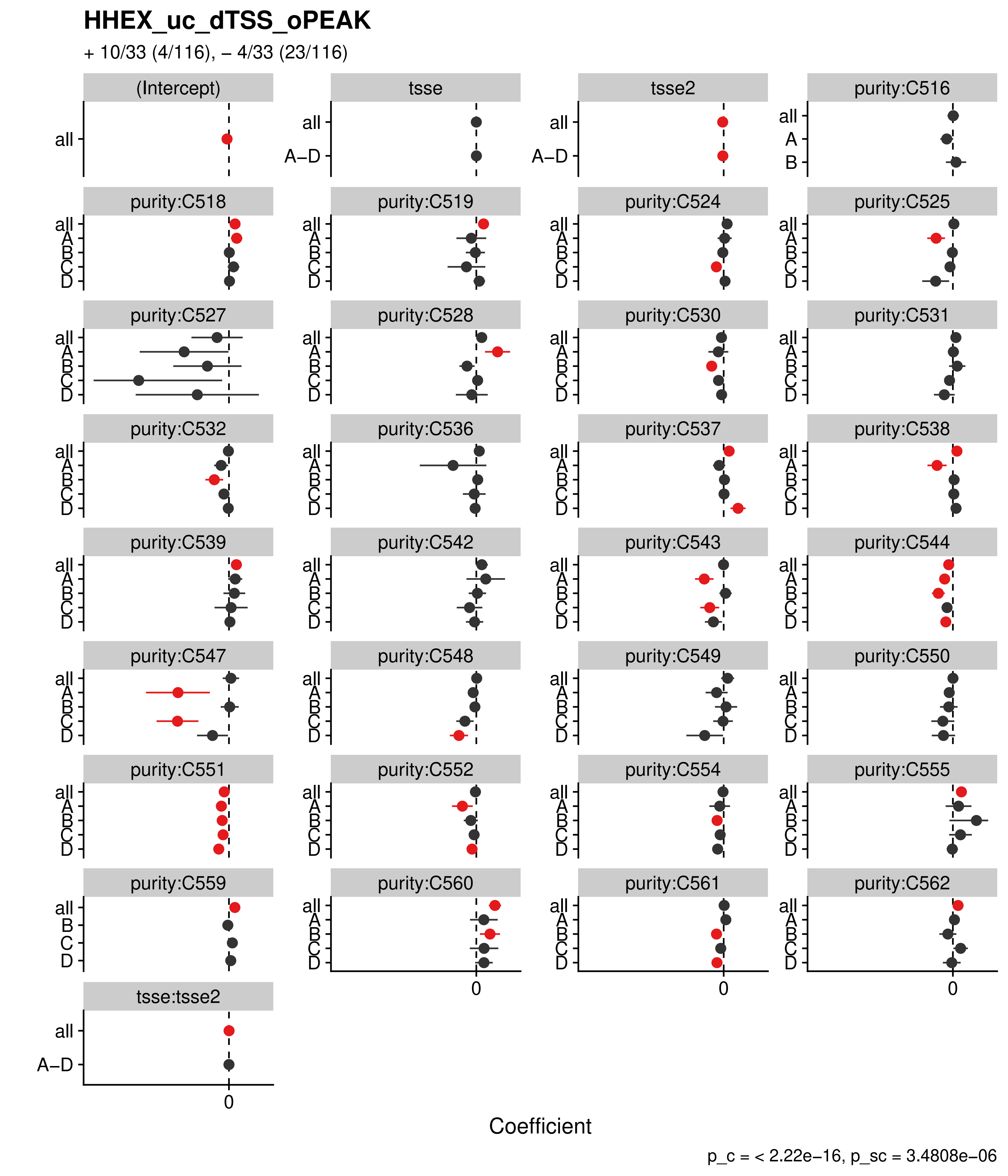
HHEX_uc_dTSS_oPEAK
+ 10/33 (4/116), – 4/33 (23/116)
(Intercept)
tsse
tsse2
purity:C516
purity:C518
purity:C519
purity:C524
purity:C525
purity:C527
purity:C528
purity:C530
purity:C531
purity:C532
purity:C536
purity:C537
purity:C538
purity:C539
purity:C542
purity:C543
purity:C544
purity:C547
purity:C548
purity:C549
purity:C550
purity:C551
purity:C552
purity:C554
purity:C555
purity:C559
purity:C560
purity:C561
purity:C562
tsse:tsse2
all
A–D
A
B
C
D
0
Coefficient
p_c = < 2.22e−16, p_sc = 3.4808e−06

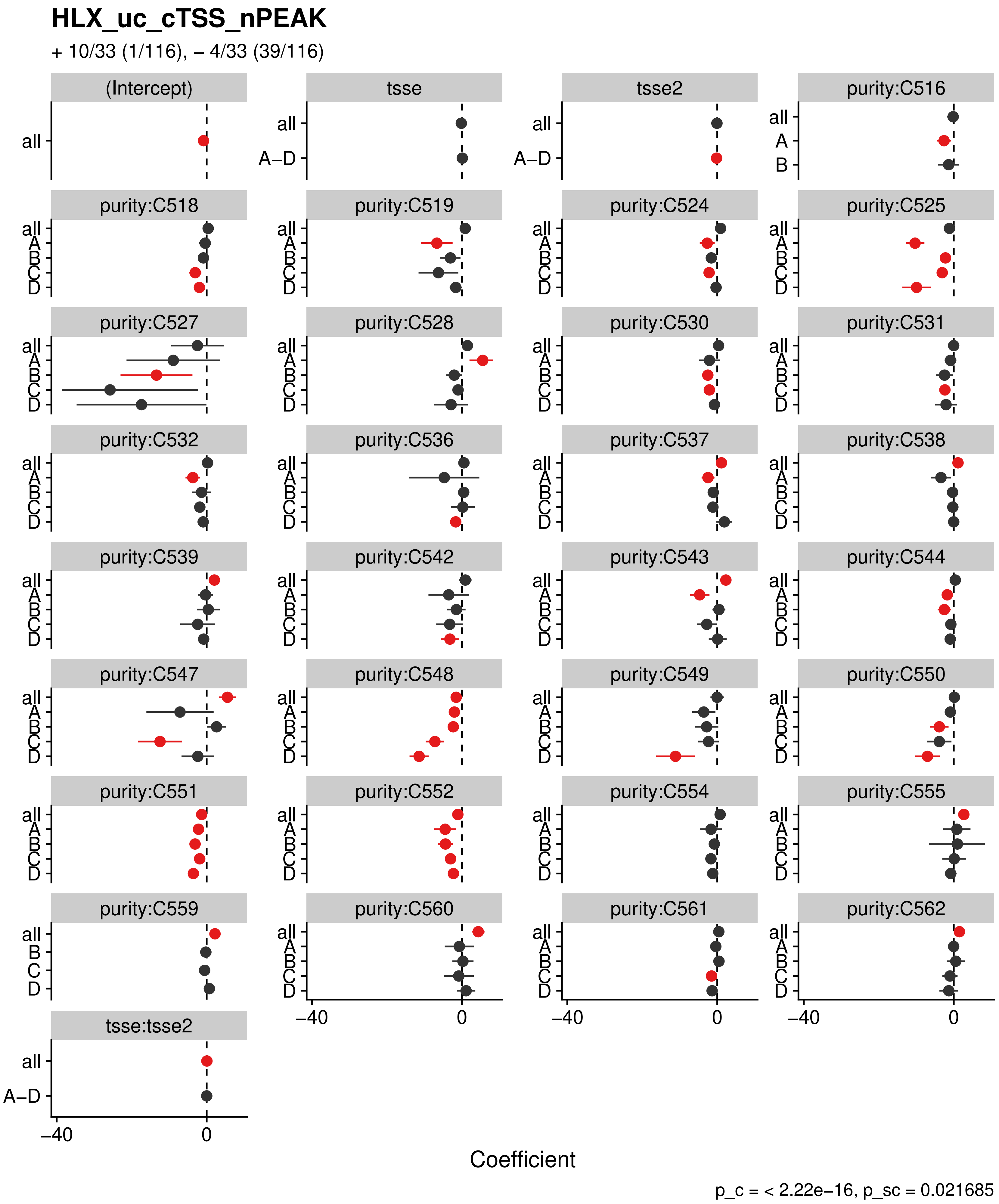
HLX_uc_cTSS_nPEAK
+ 10/33 (1/116), – 4/33 (39/116)
(Intercept)
tsse
tsse2
purity:C516
purity:C518
purity:C519
purity:C524
purity:C525
purity:C527
purity:C528
purity:C530
purity:C531
purity:C532
purity:C536
purity:C537
purity:C538
purity:C539
purity:C542
purity:C543
purity:C544
purity:C547
purity:C548
purity:C549
purity:C550
purity:C551
purity:C552
purity:C554
purity:C555
purity:C559
purity:C560
purity:C561
purity:C562
tsse:tsse2
all
A–D
A
B
C
D
–40
0
Coefficient
p_c = < 2.22e–16, p_sc = 0.021685

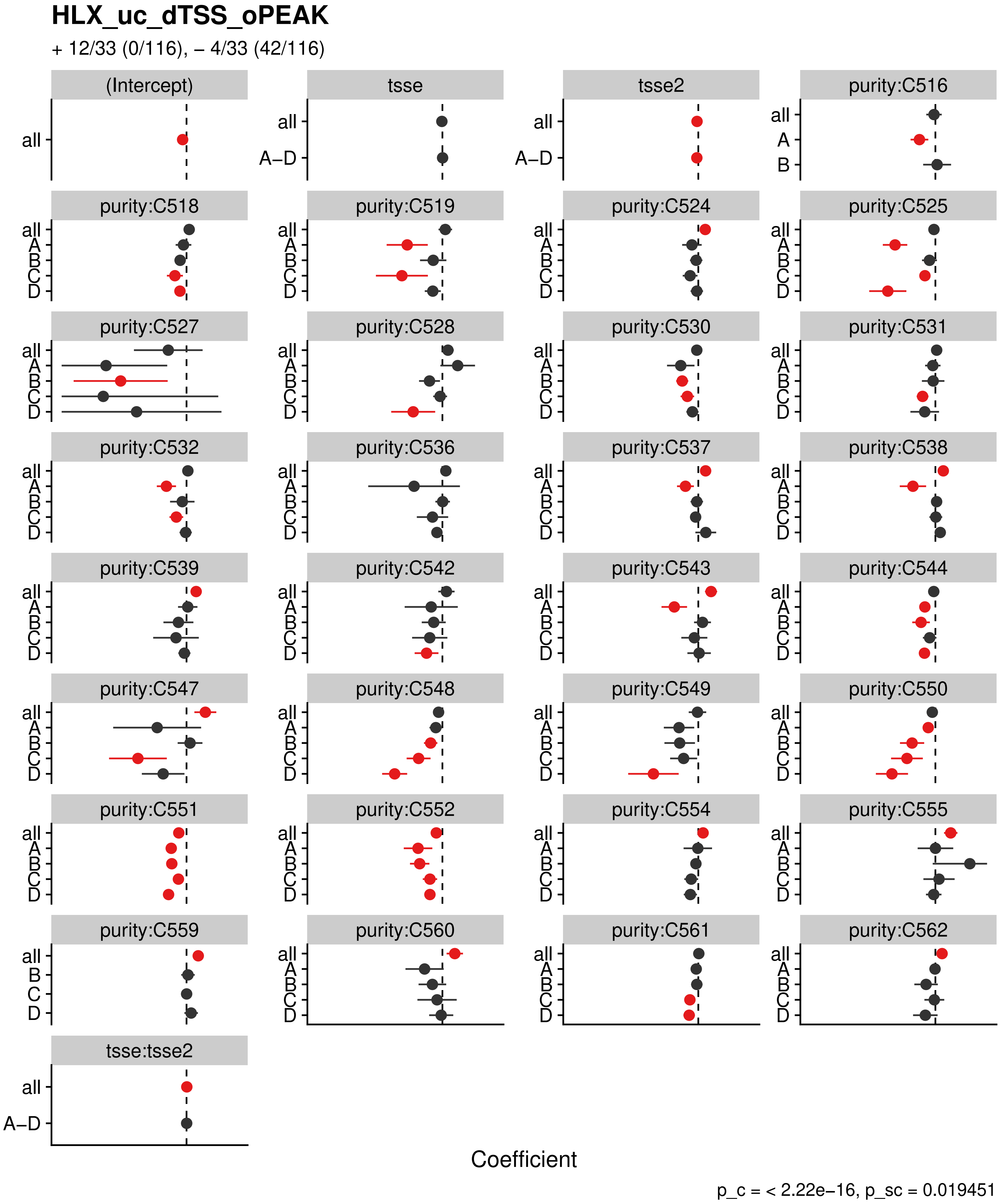
HLX_uc_dTSS_oPEAK
+ 12/33 (0/116), – 4/33 (42/116)
(Intercept)
tsse
tsse2
purity:C516
purity:C518
purity:C519
purity:C524
purity:C525
purity:C527
purity:C528
purity:C530
purity:C531
purity:C532
purity:C536
purity:C537
purity:C538
purity:C539
purity:C542
purity:C543
purity:C544
purity:C547
purity:C548
purity:C549
purity:C550
purity:C551
purity:C552
purity:C554
purity:C555
purity:C559
purity:C560
purity:C561
purity:C562
tsse:tsse2
all
A–D
A
B
C
D
Coefficient
p_c = < 2.22e–16, p_sc = 0.019451

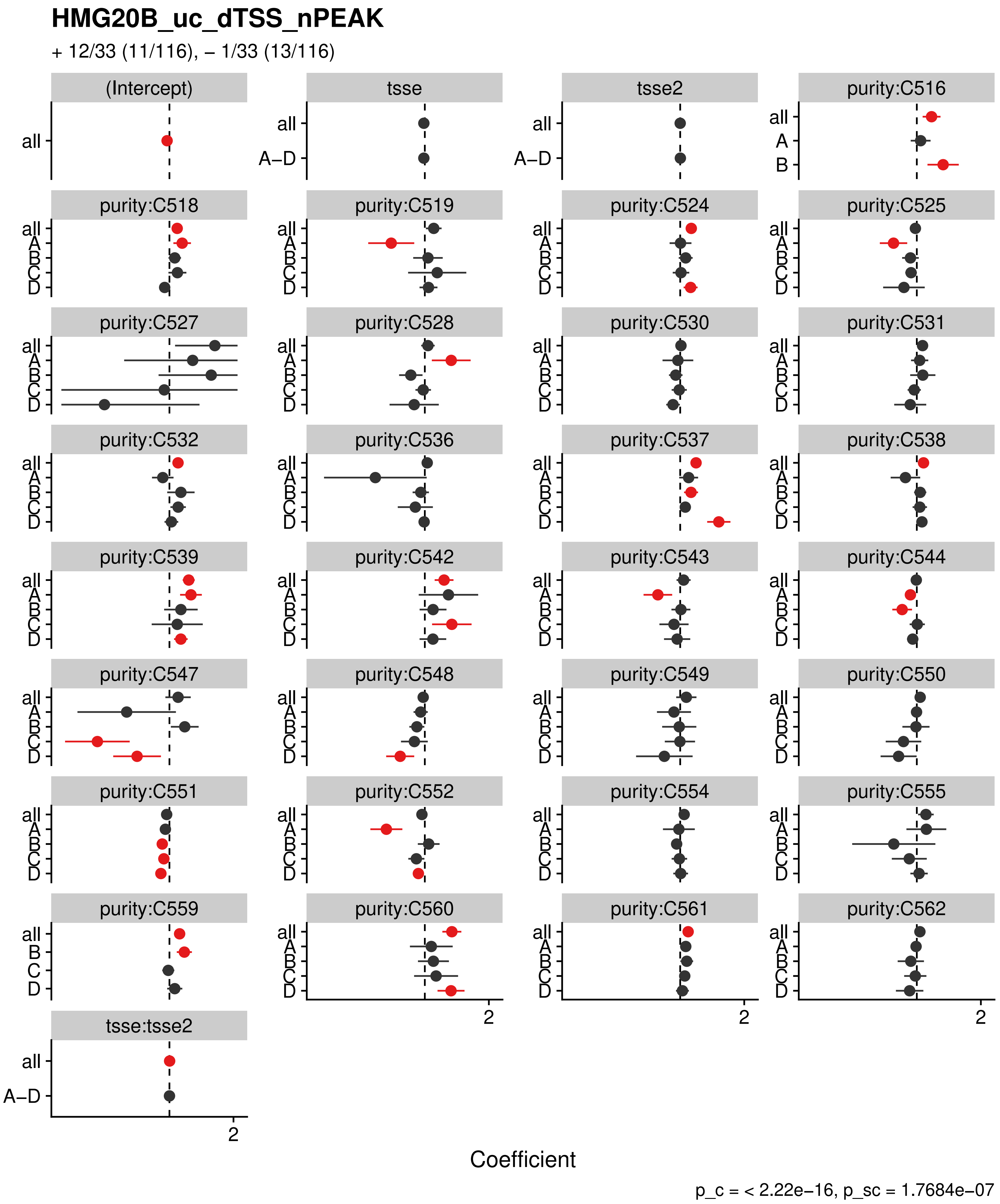
HMG20B_uc_dTSS_nPEAK
+ 12/33 (11/116), – 1/33 (13/116)
(Intercept)
tsse
tsse2
purity:C516
purity:C518
purity:C519
purity:C524
purity:C525
purity:C527
purity:C528
purity:C530
purity:C531
purity:C532
purity:C536
purity:C537
purity:C538
purity:C539
purity:C542
purity:C543
purity:C544
purity:C547
purity:C548
purity:C549
purity:C550
purity:C551
purity:C552
purity:C554
purity:C555
purity:C559
purity:C560
purity:C561
purity:C562
tsse:tsse2
all
A–D
A
B
C
D
2
Coefficient
p_c = < 2.22e–16, p_sc = 1.7684e–07

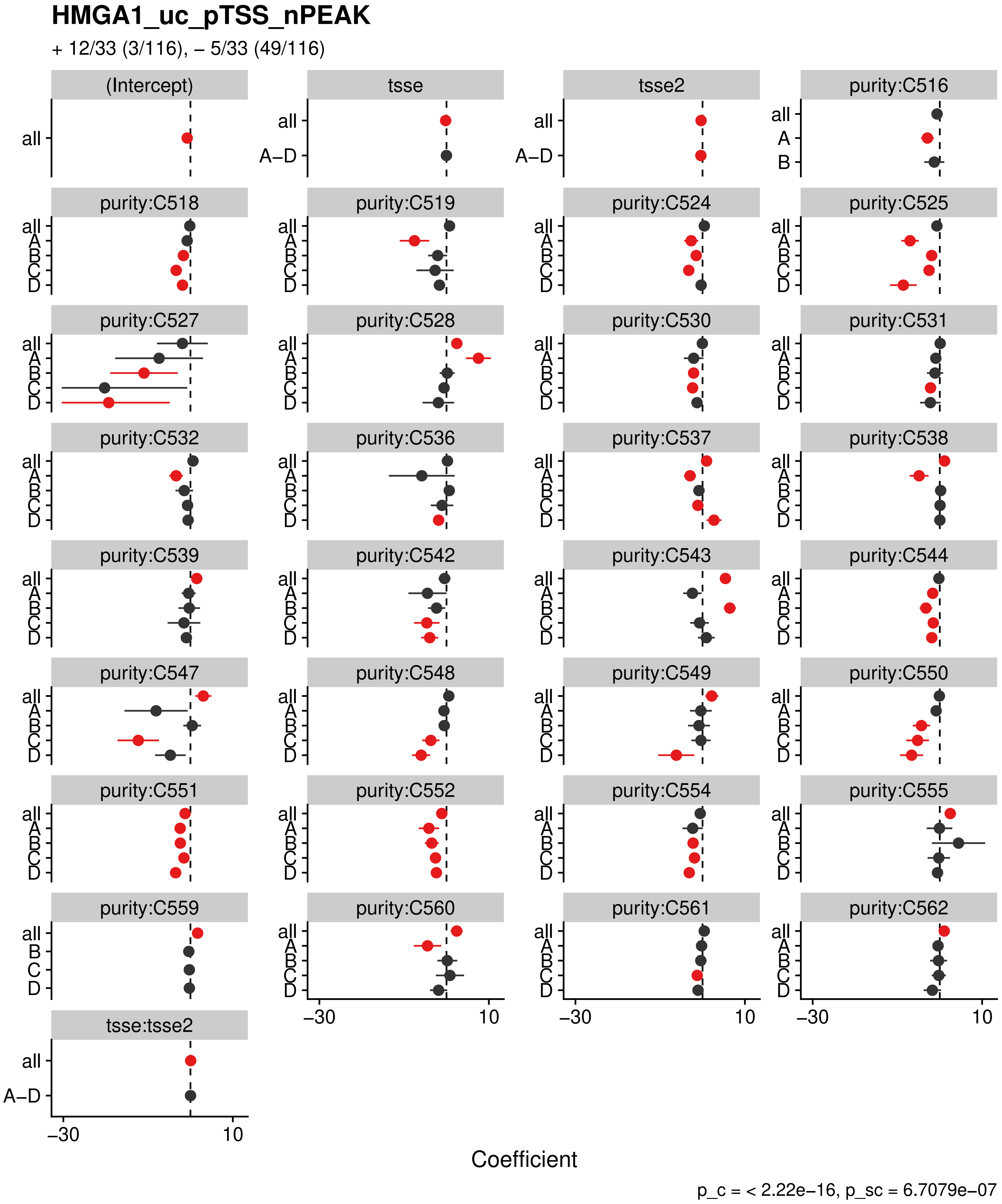
HMGA1_uc_pTSS_nPEAK
+ 12/33 (3/116), – 5/33 (49/116)
(Intercept)
tsse
tsse2
purity:C516
purity:C518
purity:C519
purity:C524
purity:C525
purity:C527
purity:C528
purity:C530
purity:C531
purity:C532
purity:C536
purity:C537
purity:C538
purity:C539
purity:C542
purity:C543
purity:C544
purity:C547
purity:C548
purity:C549
purity:C550
purity:C551
purity:C552
purity:C554
purity:C555
purity:C559
purity:C560
purity:C561
purity:C562
tsse:tsse2
all
A–D
A
B
C
D
–30
10
Coefficient
p_c = < 2.22e–16, p_sc = 6.7079e–07

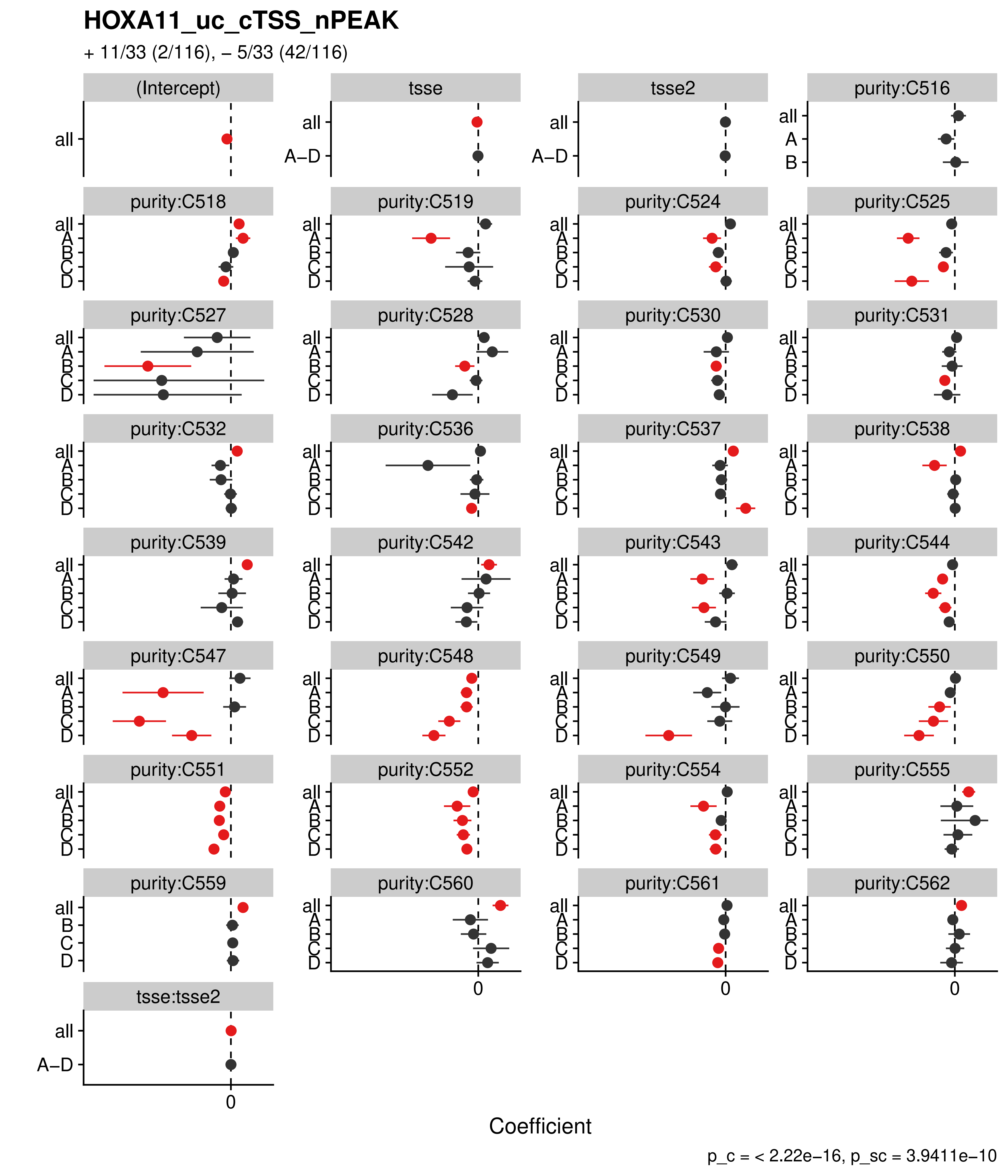

HOXA11_uc_cTSS_nPEAK
+ 11/33 (2/116), – 5/33 (42/116)
(Intercept)
tsse
tsse2
purity:C516
purity:C518
purity:C519
purity:C524
purity:C525
purity:C527
purity:C528
purity:C530
purity:C531
purity:C532
purity:C536
purity:C537
purity:C538
purity:C539
purity:C542
purity:C543
purity:C544
purity:C547
purity:C548
purity:C549
purity:C550
purity:C551
purity:C552
purity:C554
purity:C555
purity:C559
purity:C560
purity:C561
purity:C562
tsse:tsse2
all
A–D
A
B
C
D
0
Coefficient
p_c = < 2.22e−16, p_sc = 3.9411e−10

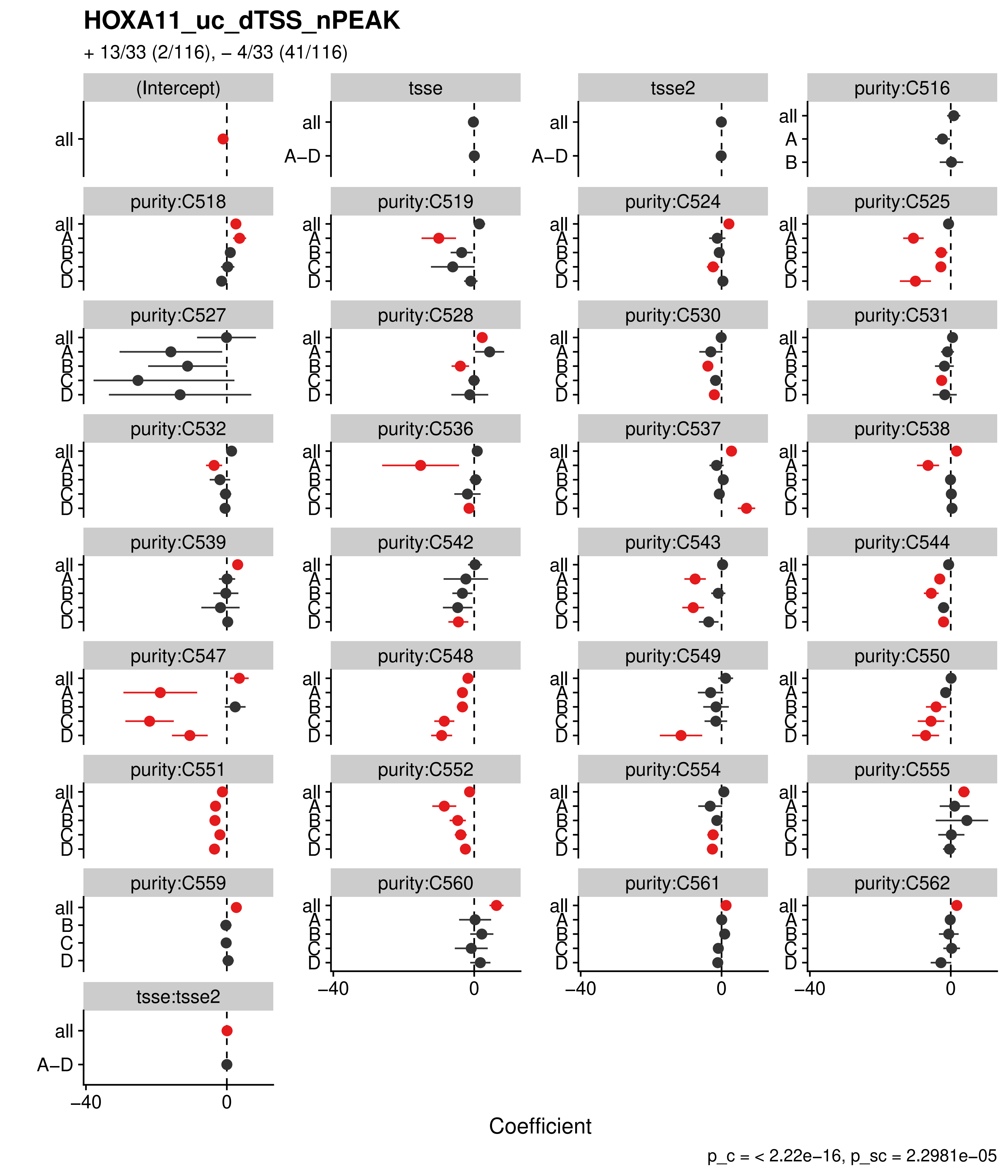

HOXA11_uc_dTSS_nPEAK
+ 13/33 (2/116), – 4/33 (41/116)
(Intercept)
tsse
tsse2
purity:C516
purity:C518
purity:C519
purity:C524
purity:C525
purity:C527
purity:C528
purity:C530
purity:C531
purity:C532
purity:C536
purity:C537
purity:C538
purity:C539
purity:C542
purity:C543
purity:C544
purity:C547
purity:C548
purity:C549
purity:C550
purity:C551
purity:C552
purity:C554
purity:C555
purity:C559
purity:C560
purity:C561
purity:C562
tsse:tsse2
all
A–D
A
B
C
D
–40
0
Coefficient
p_c = < 2.22e–16, p_sc = 2.2981e–05

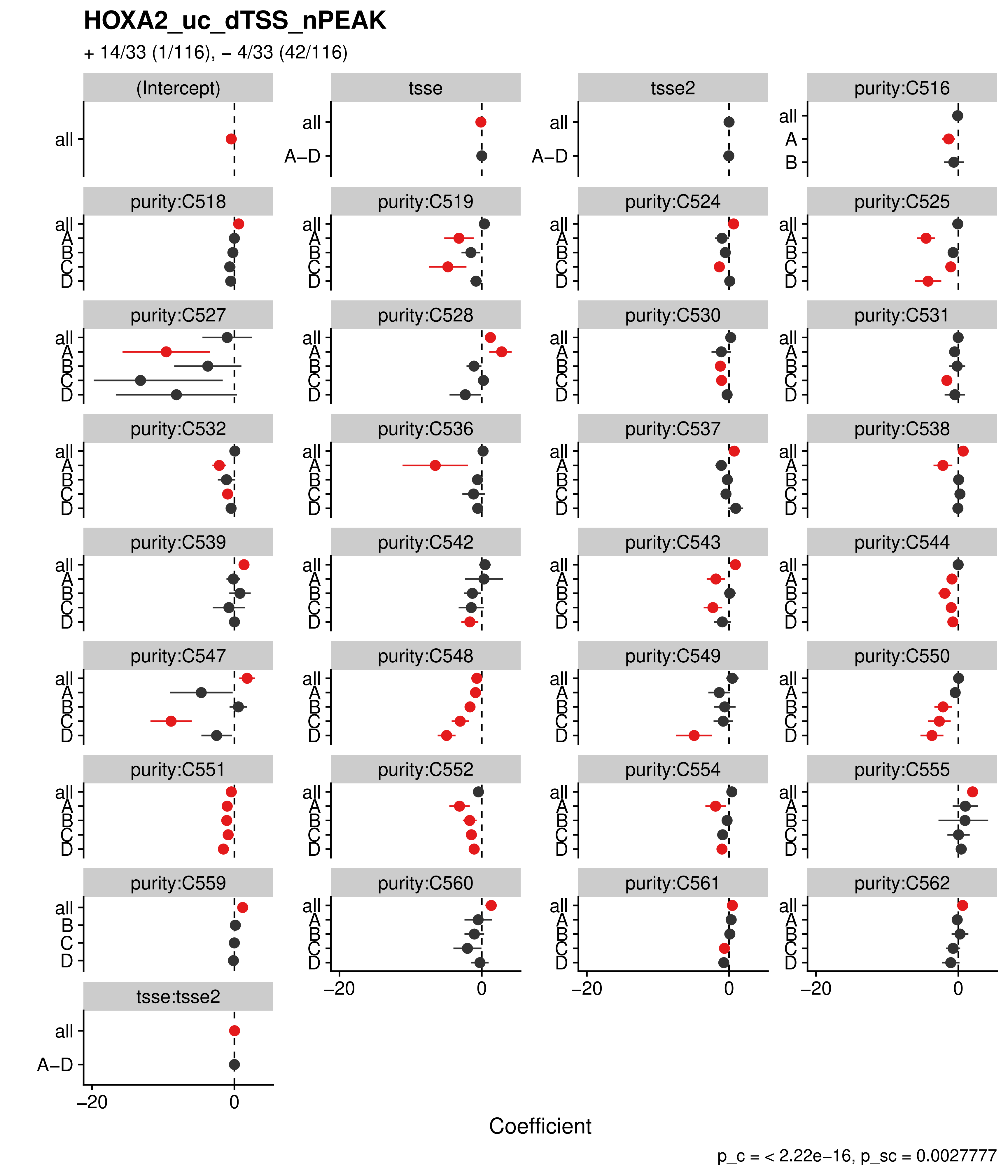

HOXA2_uc_dTSS_nPEAK
+ 14/33 (1/116), – 4/33 (42/116)
(Intercept)
tsse
tsse2
purity:C516
purity:C518
purity:C519
purity:C524
purity:C525
purity:C527
purity:C528
purity:C530
purity:C531
purity:C532
purity:C536
purity:C537
purity:C538
purity:C539
purity:C542
purity:C543
purity:C544
purity:C547
purity:C548
purity:C549
purity:C550
purity:C551
purity:C552
purity:C554
purity:C555
purity:C559
purity:C560
purity:C561
purity:C562
tsse:tsse2
all
A–D
A
B
C
D
–20
0
Coefficient
p_c = < 2.22e–16, p_sc = 0.0027777

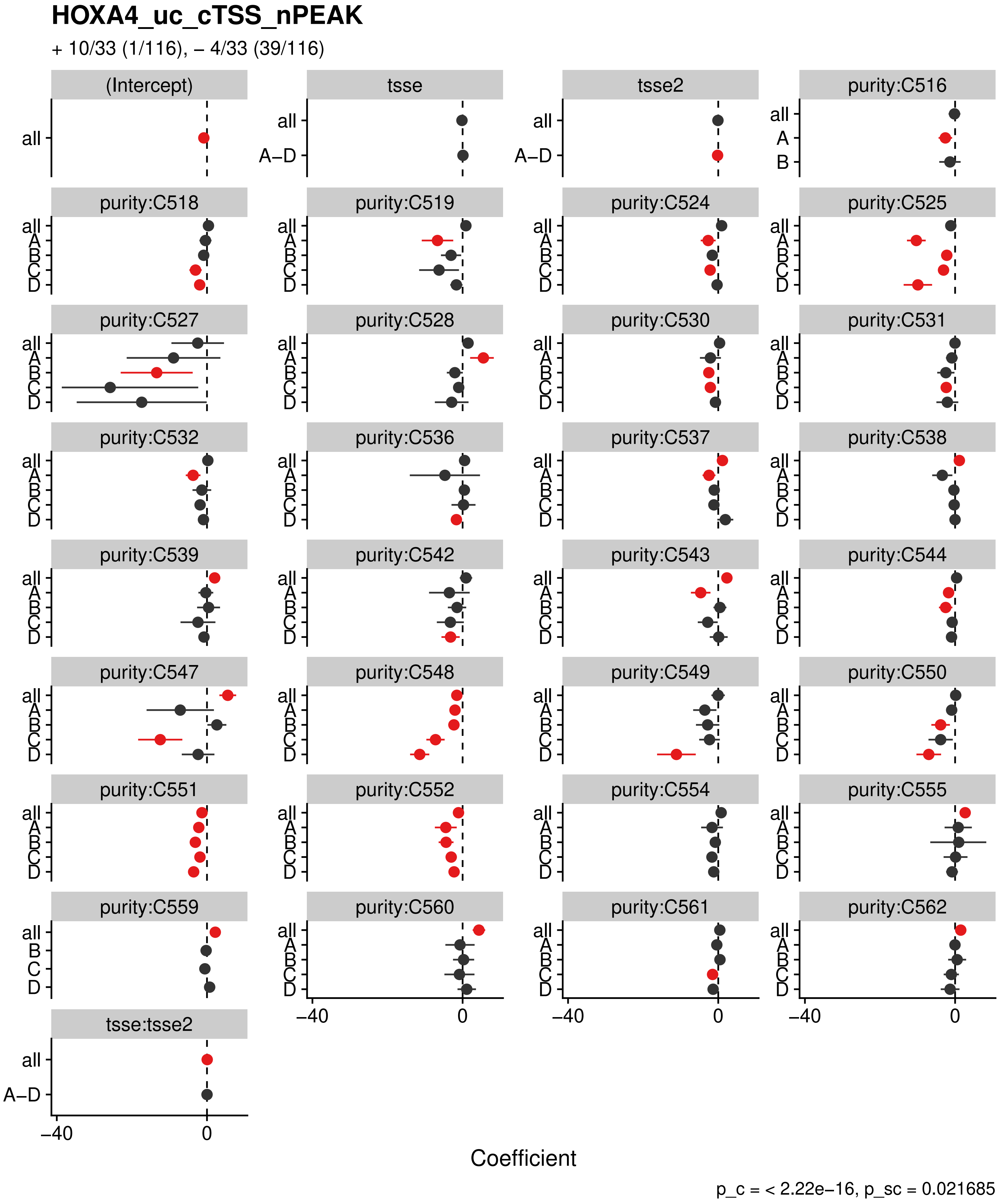
HOXA4_uc_cTSS_nPEAK
+ 10/33 (1/116), – 4/33 (39/116)
(Intercept)
tsse
tsse2
purity:C516
purity:C518
purity:C519
purity:C524
purity:C525
purity:C527
purity:C528
purity:C530
purity:C531
purity:C532
purity:C536
purity:C537
purity:C538
purity:C539
purity:C542
purity:C543
purity:C544
purity:C547
purity:C548
purity:C549
purity:C550
purity:C551
purity:C552
purity:C554
purity:C555
purity:C559
purity:C560
purity:C561
purity:C562
tsse:tsse2
all
A–D
A
B
C
D
–40
0
Coefficient
p_c = < 2.22e–16, p_sc = 0.021685

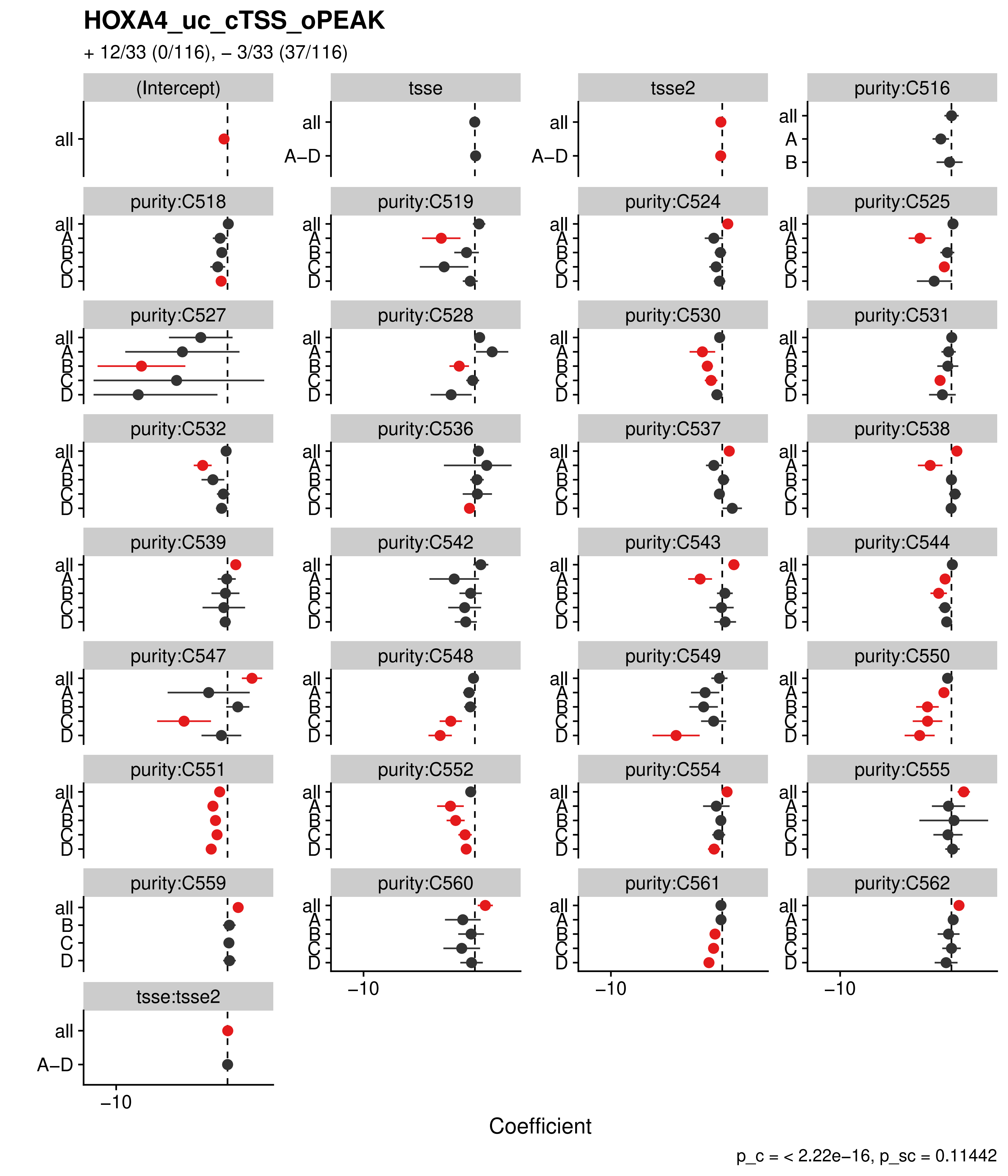
HOXA4_uc_cTSS_oPEAK
+ 12/33 (0/116), – 3/33 (37/116)
(Intercept)
tsse
tsse2
purity:C516
purity:C518
purity:C519
purity:C524
purity:C525
purity:C527
purity:C528
purity:C530
purity:C531
purity:C532
purity:C536
purity:C537
purity:C538
purity:C539
purity:C542
purity:C543
purity:C544
purity:C547
purity:C548
purity:C549
purity:C550
purity:C551
purity:C552
purity:C554
purity:C555
purity:C559
purity:C560
purity:C561
purity:C562
tsse:tsse2
all
A–D
A
B
C
D
−10
Coefficient
p_c = < 2.22e−16, p_sc = 0.11442

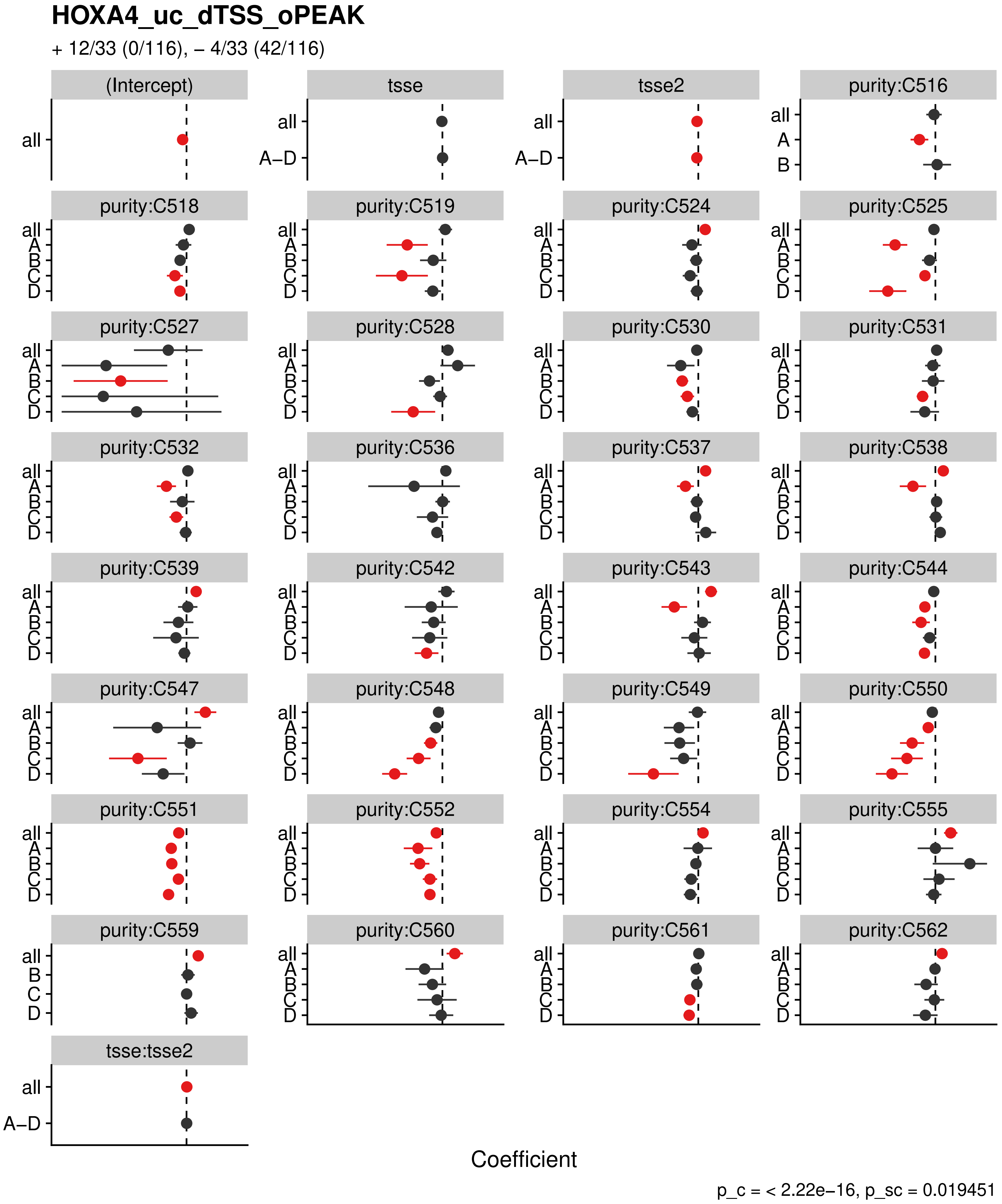
HOXA4_uc_dTSS_oPEAK
+ 12/33 (0/116), – 4/33 (42/116)
(Intercept)
tsse
tsse2
purity:C516
purity:C518
purity:C519
purity:C524
purity:C525
purity:C527
purity:C528
purity:C530
purity:C531
purity:C532
purity:C536
purity:C537
purity:C538
purity:C539
purity:C542
purity:C543
purity:C544
purity:C547
purity:C548
purity:C549
purity:C550
purity:C551
purity:C552
purity:C554
purity:C555
purity:C559
purity:C560
purity:C561
purity:C562
tsse:tsse2
all
A–D
A
B
C
D
Coefficient
p_c = < 2.22e−16, p_sc = 0.019451

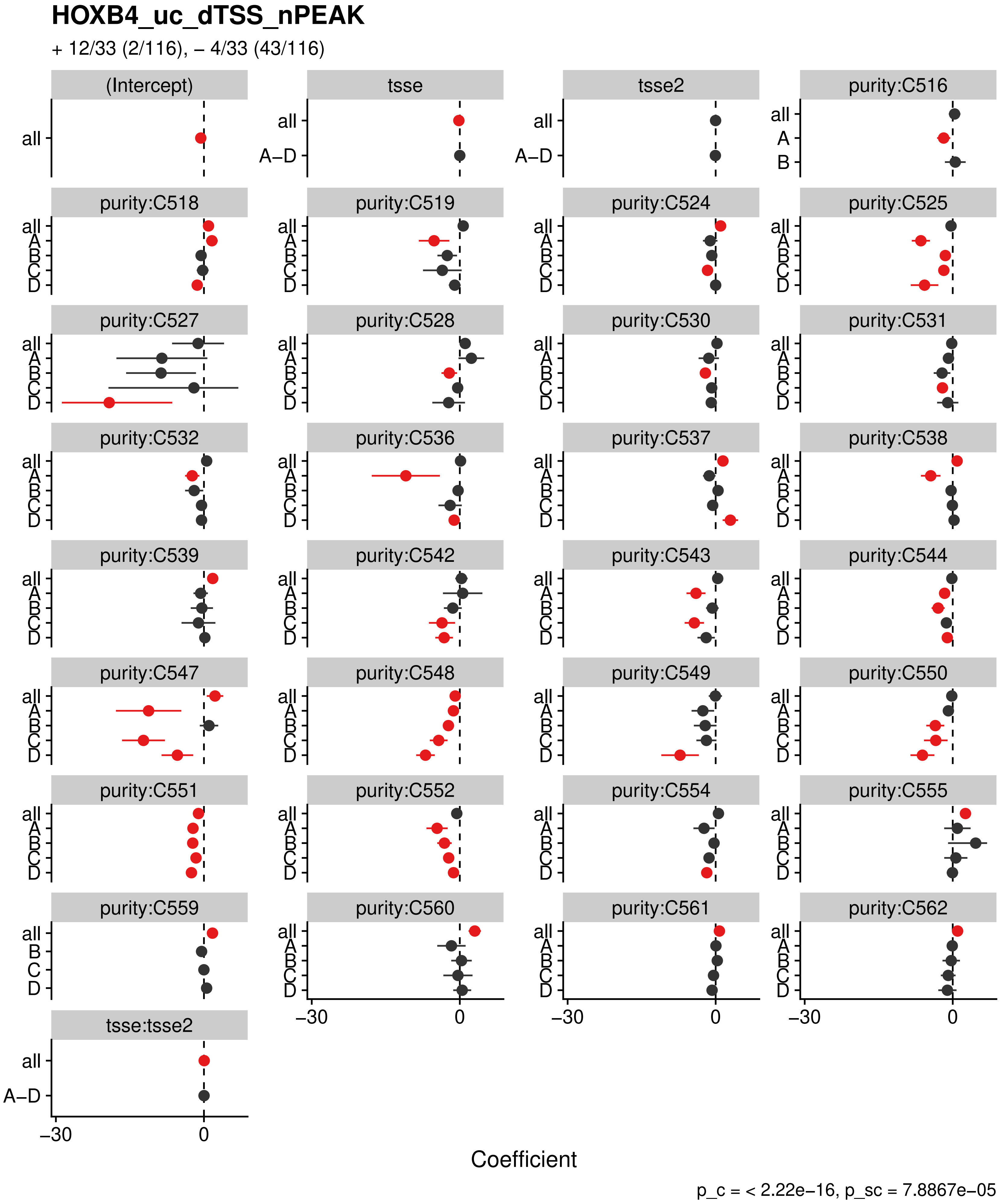
HOXB4_uc_dTSS_nPEAK
+ 12/33 (2/116), – 4/33 (43/116)
(Intercept)
tsse
tsse2
purity:C516
purity:C518
purity:C519
purity:C524
purity:C525
purity:C527
purity:C528
purity:C530
purity:C531
purity:C532
purity:C536
purity:C537
purity:C538
purity:C539
purity:C542
purity:C543
purity:C544
purity:C547
purity:C548
purity:C549
purity:C550
purity:C551
purity:C552
purity:C554
purity:C555
purity:C559
purity:C560
purity:C561
purity:C562
tsse:tsse2
all
A–D
A
B
C
D
–30
0
Coefficient
p_c = < 2.22e–16, p_sc = 7.8867e–05

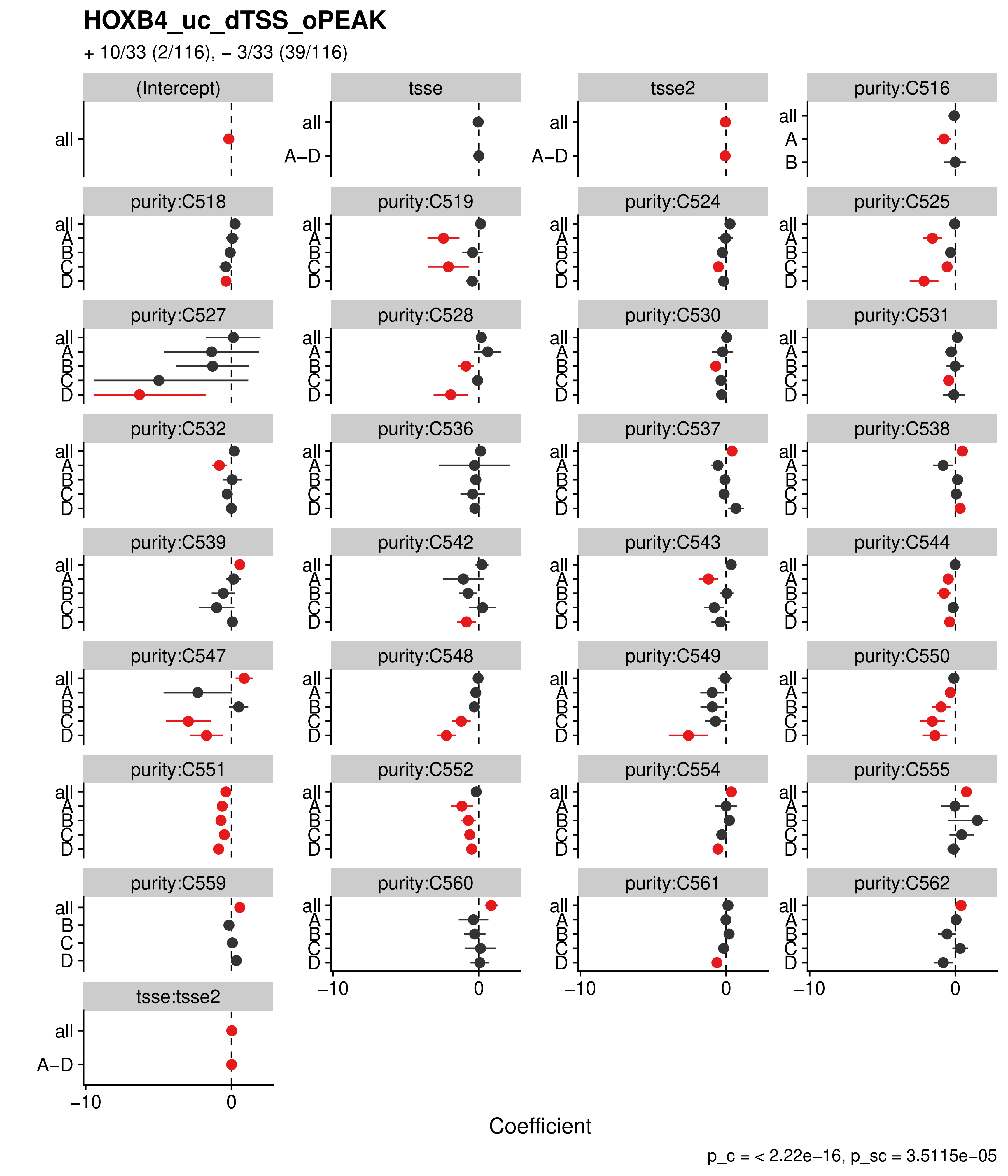
HOXB4_uc_dTSS_oPEAK
+ 10/33 (2/116), − 3/33 (39/116)
(Intercept)
tsse
tsse2
purity:C516
purity:C518
purity:C519
purity:C524
purity:C525
purity:C527
purity:C528
purity:C530
purity:C531
purity:C532
purity:C536
purity:C537
purity:C538
purity:C539
purity:C542
purity:C543
purity:C544
purity:C547
purity:C548
purity:C549
purity:C550
purity:C551
purity:C552
purity:C554
purity:C555
purity:C559
purity:C560
purity:C561
purity:C562
tsse:tsse2
all
A−D
A
B
C
D
−10
0
Coefficient
p_c = < 2.22e−16, p_sc = 3.5115e−05

**HOXC12_uc_dTSS_nPEAK**

+ 11/33 (1/116), – 3/33 (37/116)

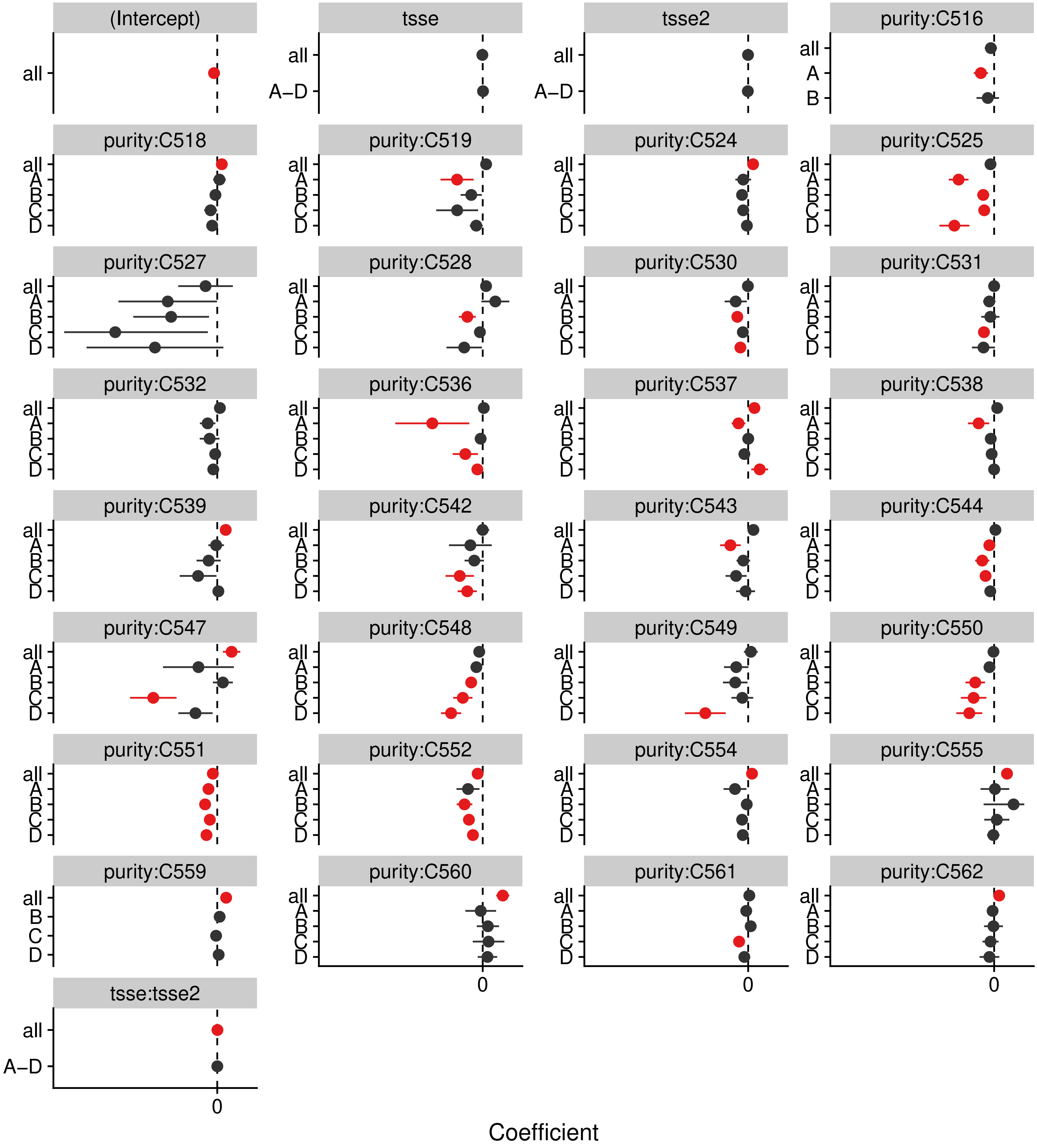


p_c = < 2.22e–16, p_sc = 0.25693

#### HOXC5_uc_dTSS_nPEAK

+ 12/33 (2/116), – 4/33 (43/116)

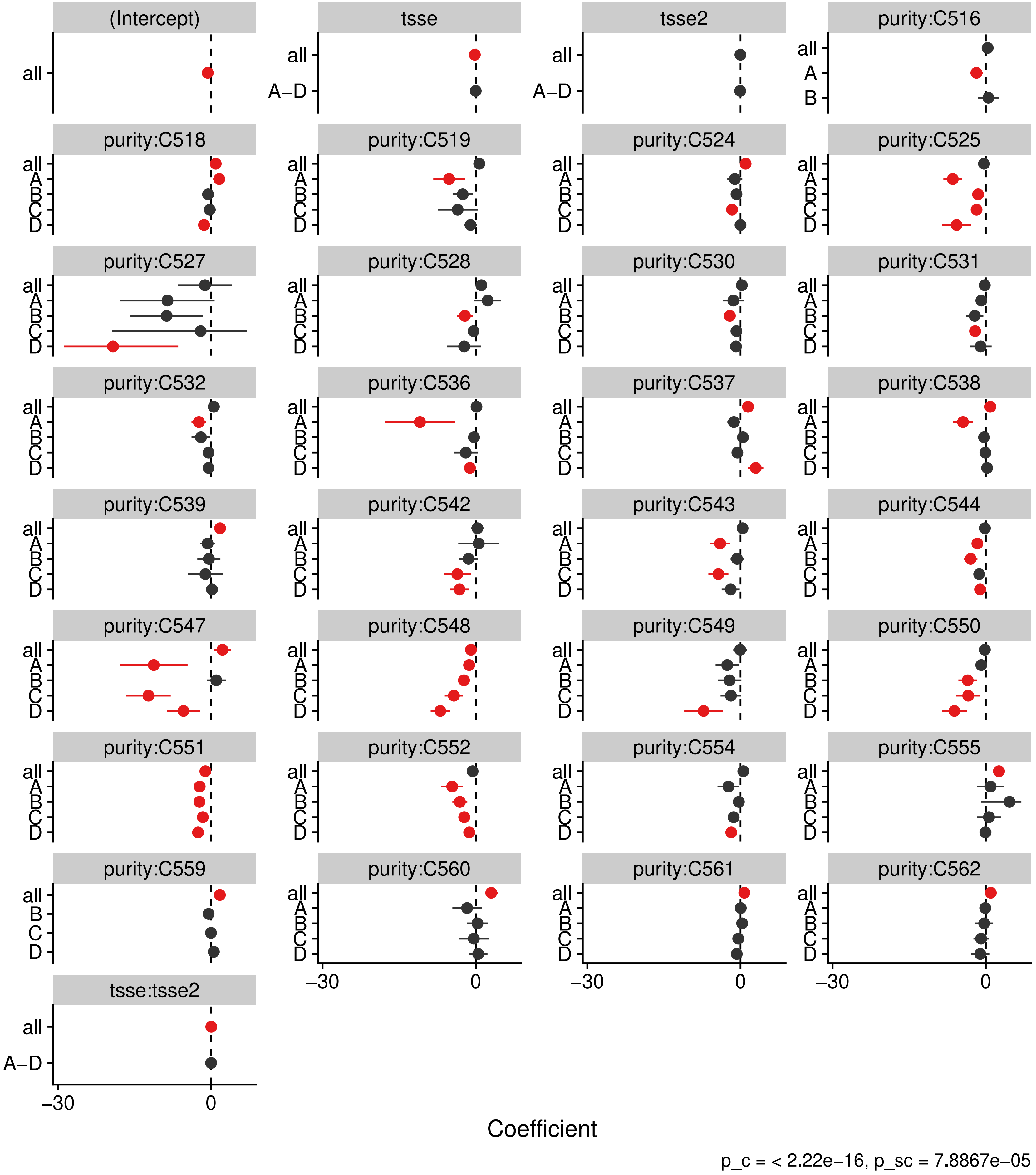

#### HOXD12_uc_dTSS_nPEAK

+ 11/33 (1/116), – 3/33 (37/116)

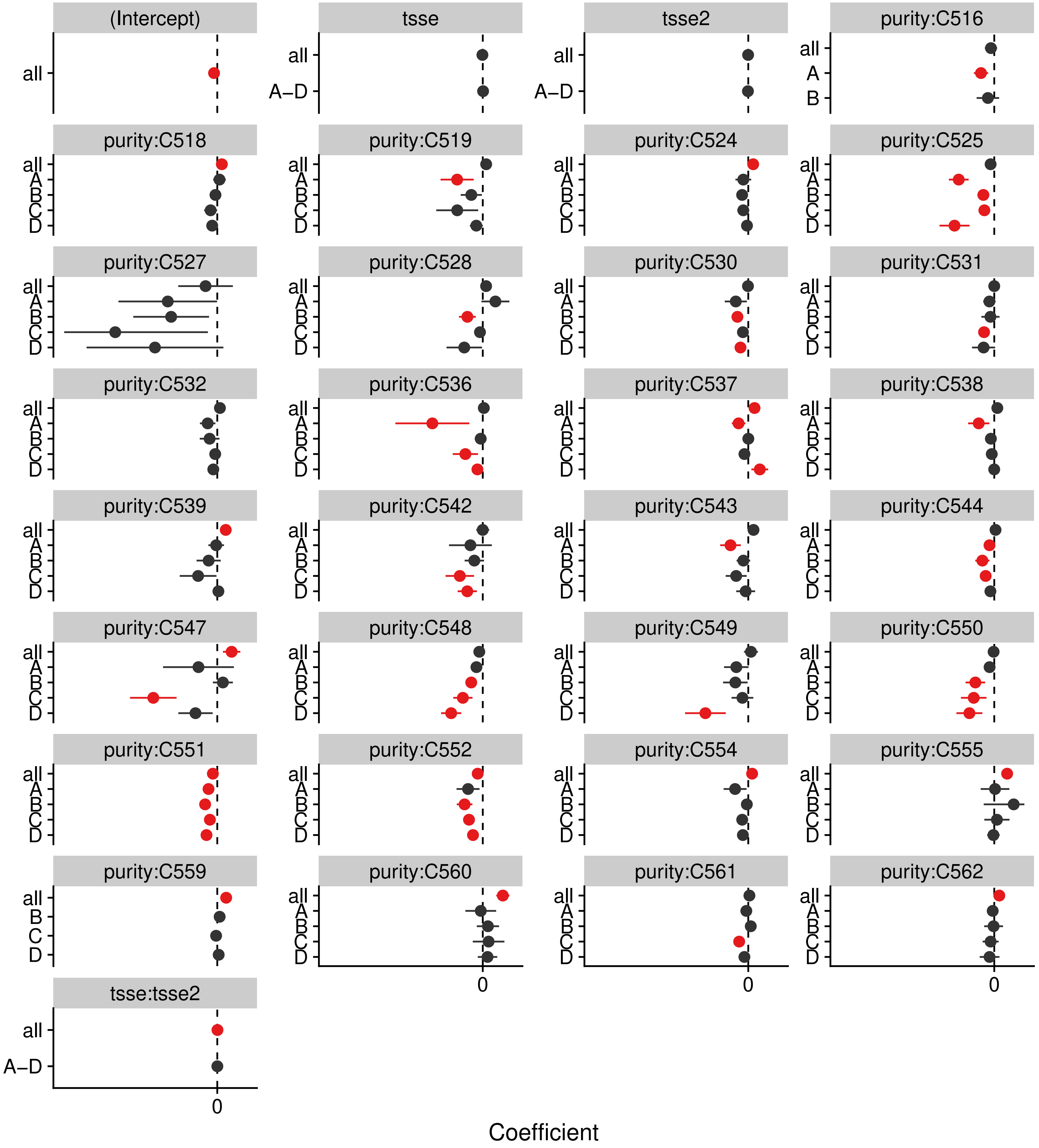


p_c = < 2.22e−16, p_sc = 0.25693

**HOXD1_uc_dTSS_nPEAK**

+ 14/33 (2/116), – 3/33 (34/116)

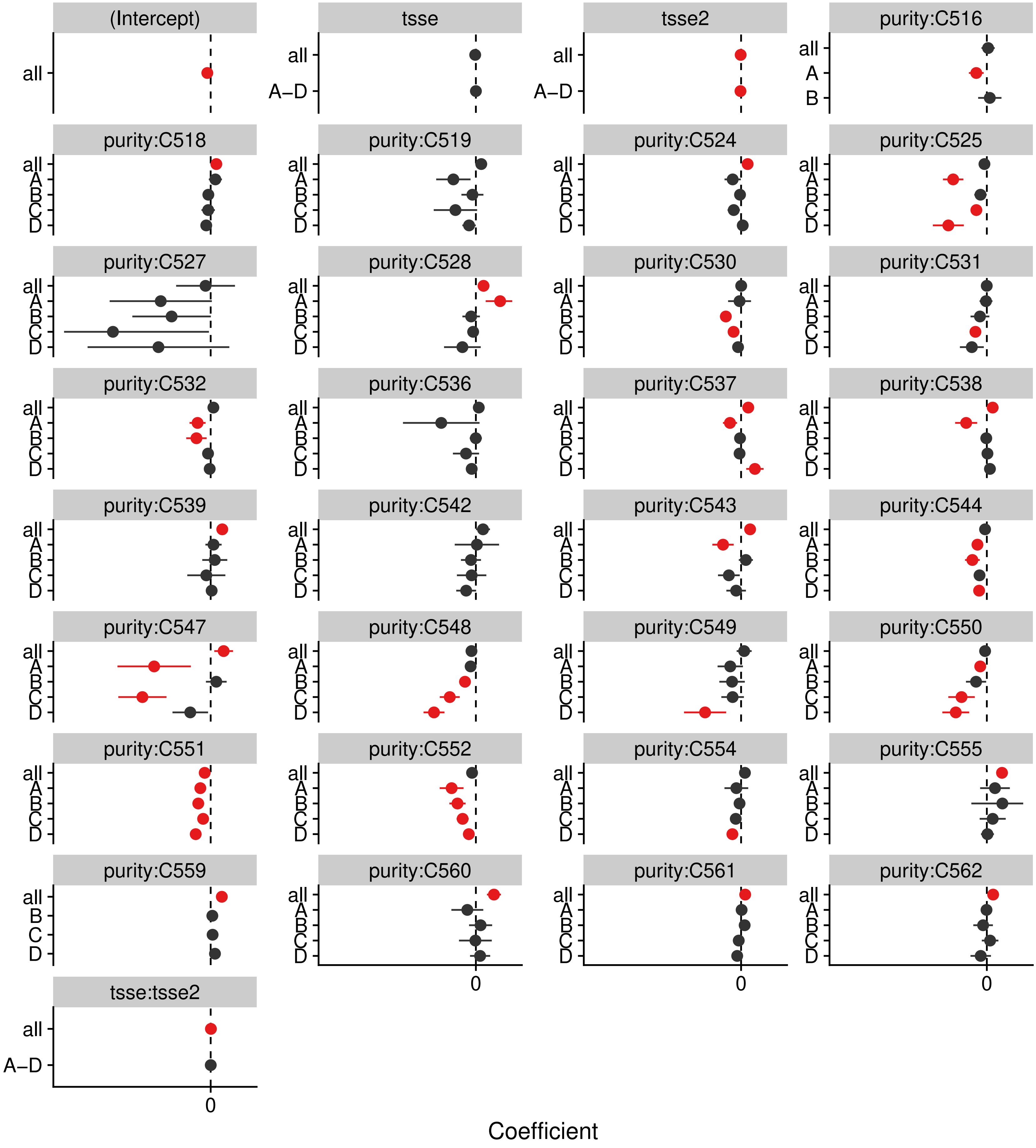


p_c = < 2.22e–16, p_sc = 4.9549e–06

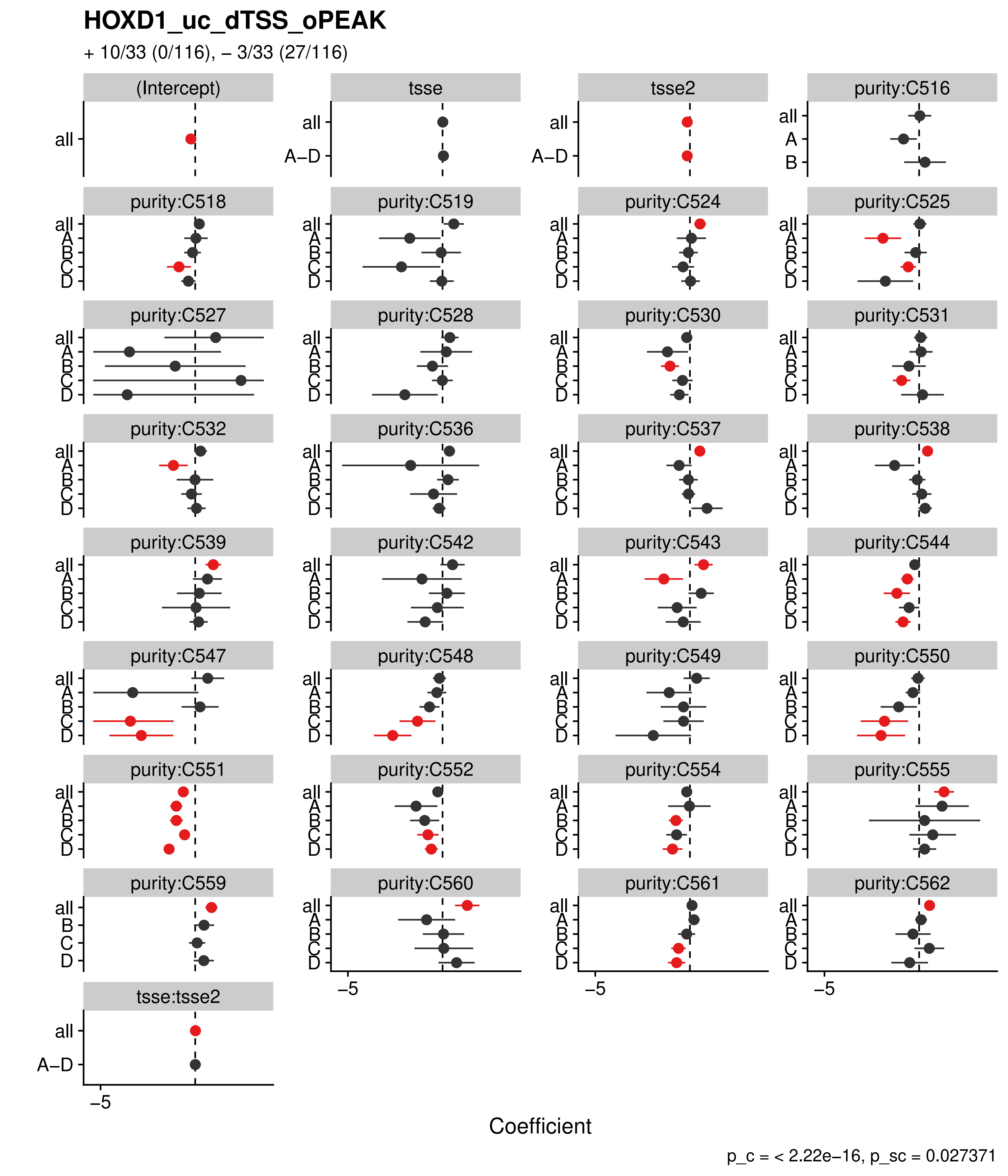

HOXD1_uc_dTSS_oPEAK
+ 10/33 (0/116), – 3/33 (27/116)
(Intercept)
tsse
tsse2
purity:C516
purity:C518
purity:C519
purity:C524
purity:C525
purity:C527
purity:C528
purity:C530
purity:C531
purity:C532
purity:C536
purity:C537
purity:C538
purity:C539
purity:C542
purity:C543
purity:C544
purity:C547
purity:C548
purity:C549
purity:C550
purity:C551
purity:C552
purity:C554
purity:C555
purity:C559
purity:C560
purity:C561
purity:C562
tsse:tsse2
all
A–D
A
B
C
D
–5
Coefficient
p_c = < 2.22e–16, p_sc = 0.027371

### HOXD3_uc_cTSS_nPEAK

+ 11/33 (1/116), – 2/33 (33/116)

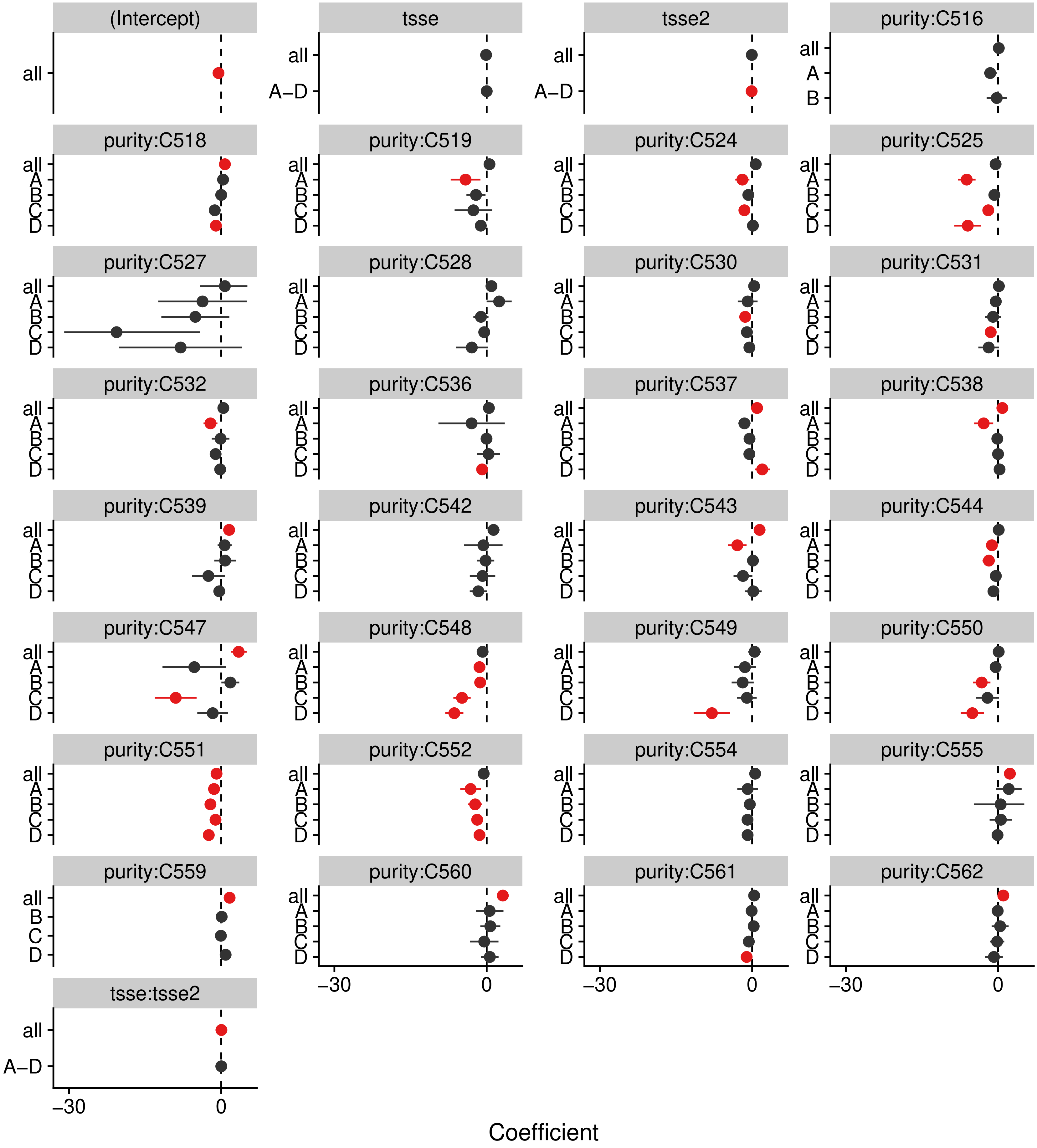


p_c = < 2.22e–16, p_sc = 0.0056747

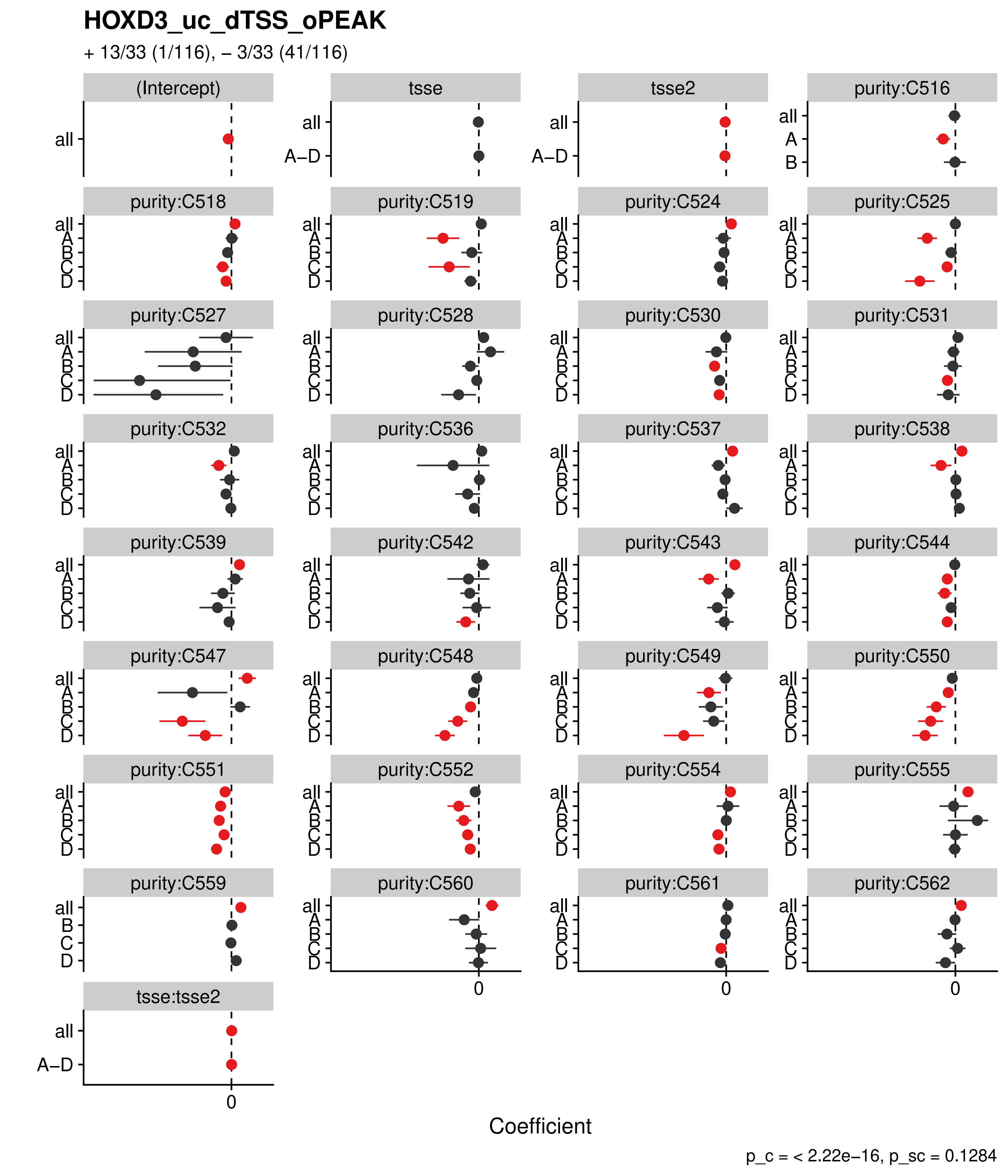

HOXD3_uc_dTSS_oPEAK
+ 13/33 (1/116), – 3/33 (41/116)
(Intercept)
tsse
tsse2
purity:C516
purity:C518
purity:C519
purity:C524
purity:C525
purity:C527
purity:C528
purity:C530
purity:C531
purity:C532
purity:C536
purity:C537
purity:C538
purity:C539
purity:C542
purity:C543
purity:C544
purity:C547
purity:C548
purity:C549
purity:C550
purity:C551
purity:C552
purity:C554
purity:C555
purity:C559
purity:C560
purity:C561
purity:C562
tsse:tsse2
all
A–D
A
B
C
D
0
Coefficient
p_c = < 2.22e–16, p_sc = 0.1284

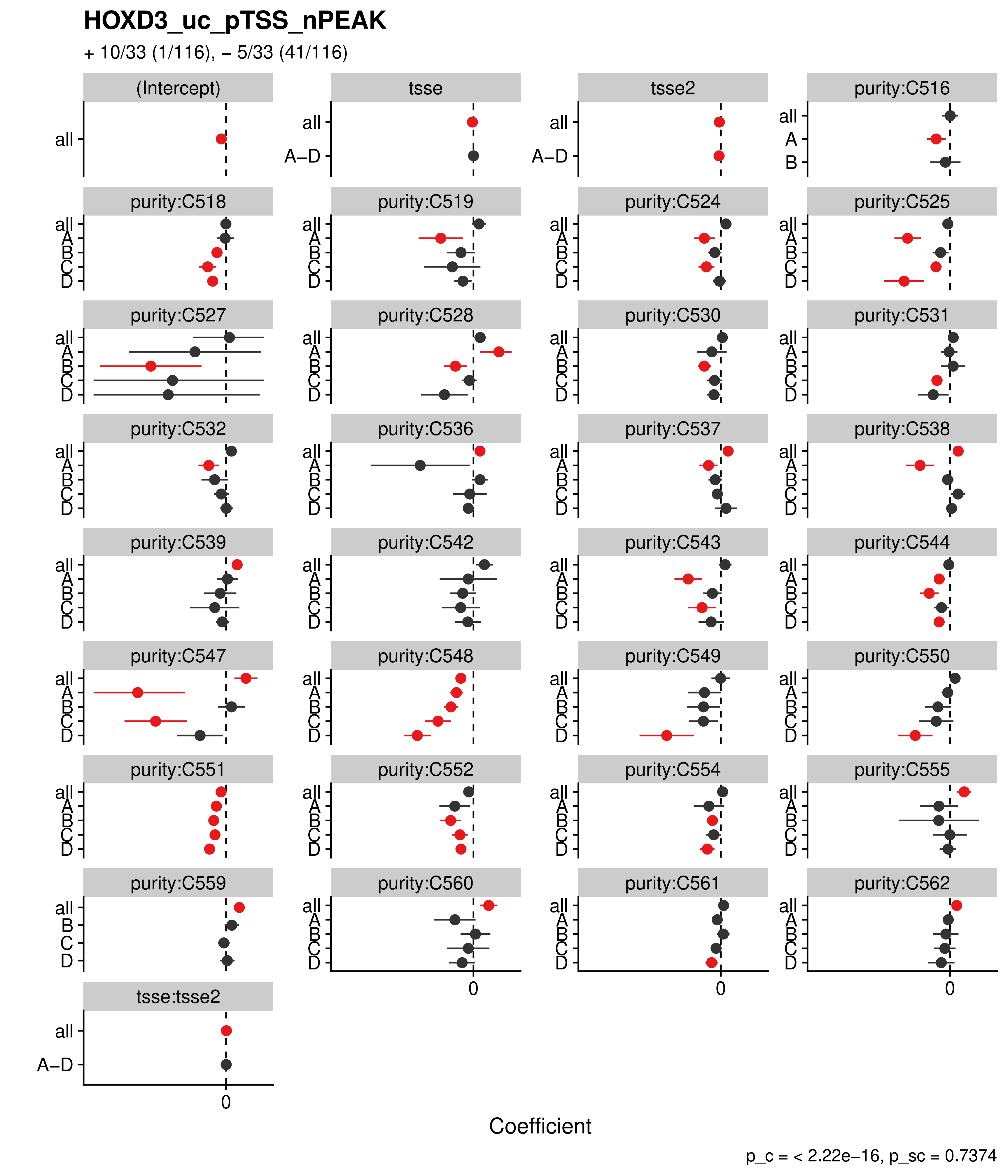

HOXD3_uc_pTSS_nPEAK
+ 10/33 (1/116), – 5/33 (41/116)
(Intercept)
tsse
tsse2
purity:C516
purity:C518
purity:C519
purity:C524
purity:C525
purity:C527
purity:C528
purity:C530
purity:C531
purity:C532
purity:C536
purity:C537
purity:C538
purity:C539
purity:C542
purity:C543
purity:C544
purity:C547
purity:C548
purity:C549
purity:C550
purity:C551
purity:C552
purity:C554
purity:C555
purity:C559
purity:C560
purity:C561
purity:C562
tsse:tsse2
all
A–D
A
B
C
D
0
Coefficient
p_c = < 2.22e–16, p_sc = 0.7374

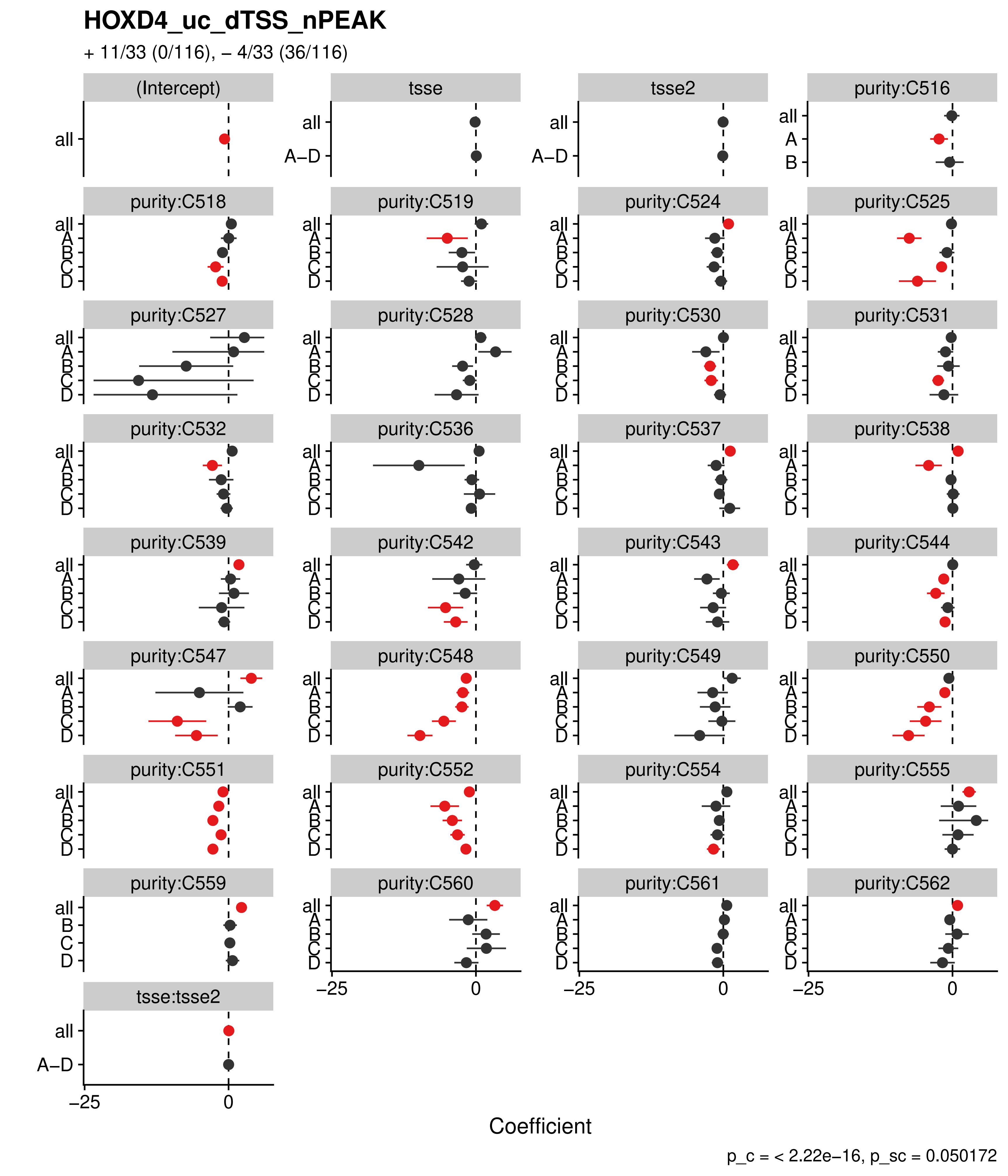

HOXD4_uc_dTSS_nPEAK
+ 11/33 (0/116), – 4/33 (36/116)
(Intercept)
tsse
tsse2
purity:C516
purity:C518
purity:C519
purity:C524
purity:C525
purity:C527
purity:C528
purity:C530
purity:C531
purity:C532
purity:C536
purity:C537
purity:C538
purity:C539
purity:C542
purity:C543
purity:C544
purity:C547
purity:C548
purity:C549
purity:C550
purity:C551
purity:C552
purity:C554
purity:C555
purity:C559
purity:C560
purity:C561
purity:C562
tsse:tsse2
all
A–D
A
B
C
D
–25
0
Coefficient
p_c = < 2.22e–16, p_sc = 0.050172

**HOXD9_uc_dTSS_nPEAK**

+ 11/33 (0/116), – 2/33 (24/116)

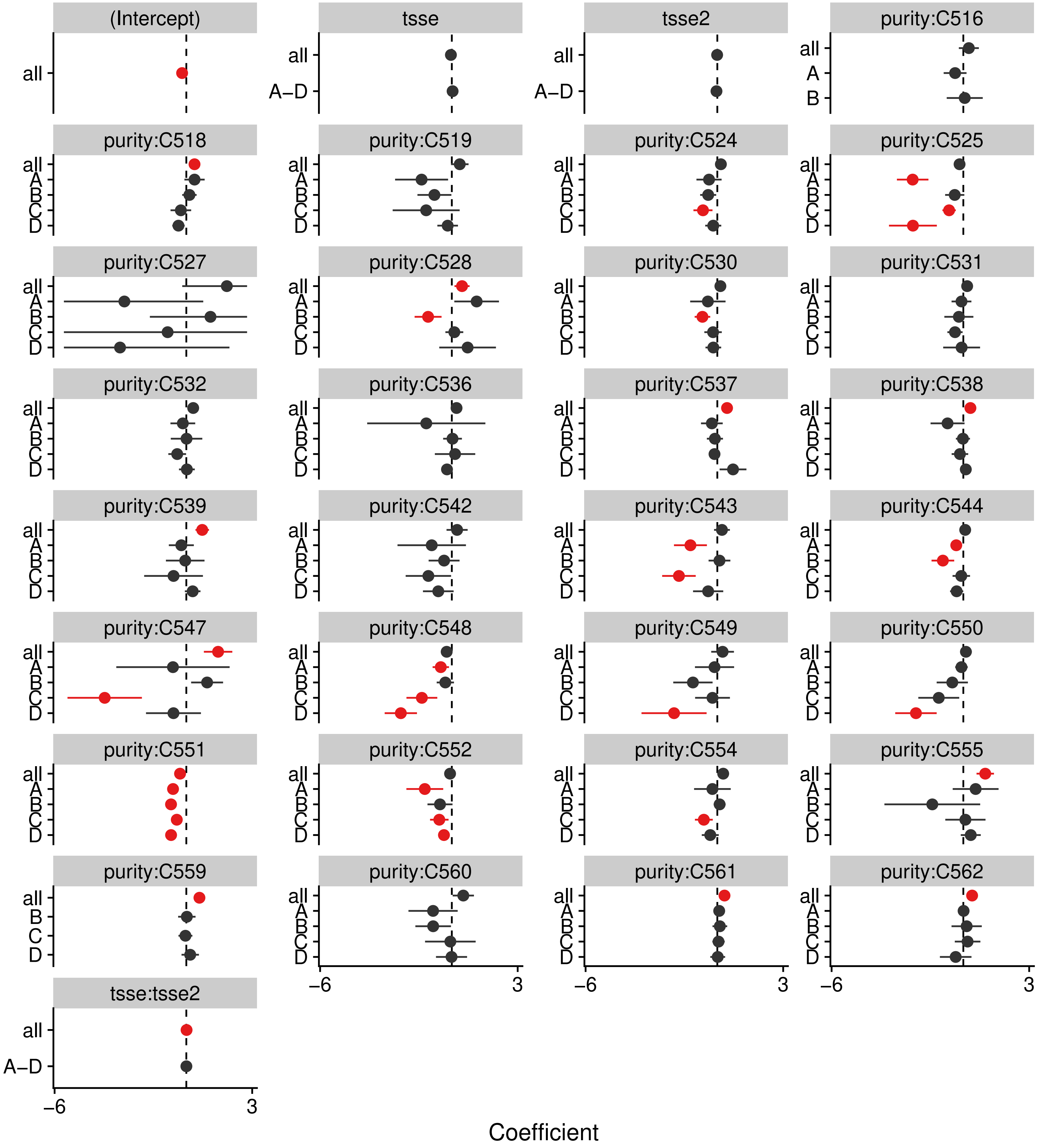


p_c = < 2.22e–16, p_sc = 0.0027069

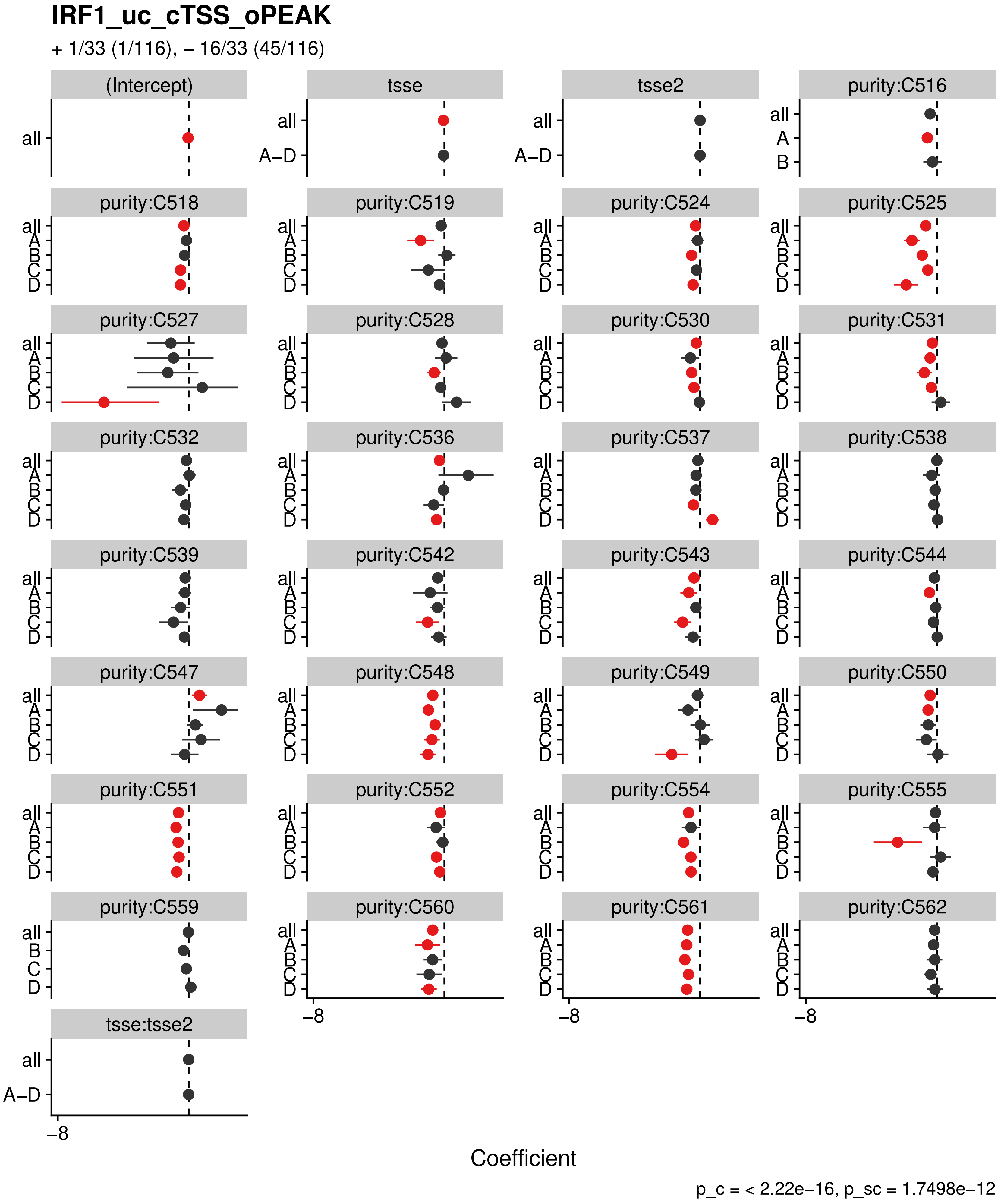

IRF1_uc_cTSS_oPEAK
+ 1/33 (1/116), – 16/33 (45/116)
(Intercept)
tsse
tsse2
purity:C516
purity:C518
purity:C519
purity:C524
purity:C525
purity:C527
purity:C528
purity:C530
purity:C531
purity:C532
purity:C536
purity:C537
purity:C538
purity:C539
purity:C542
purity:C543
purity:C544
purity:C547
purity:C548
purity:C549
purity:C550
purity:C551
purity:C552
purity:C554
purity:C555
purity:C559
purity:C560
purity:C561
purity:C562
tsse:tsse2
all
A–D
A
B
C
D
−8
Coefficient
p_c = < 2.22e−16, p_sc = 1.7498e−12

**IRF1_uc_dTSS_oPEAK**

+ 2/33 (2/116), – 14/33 (64/116)

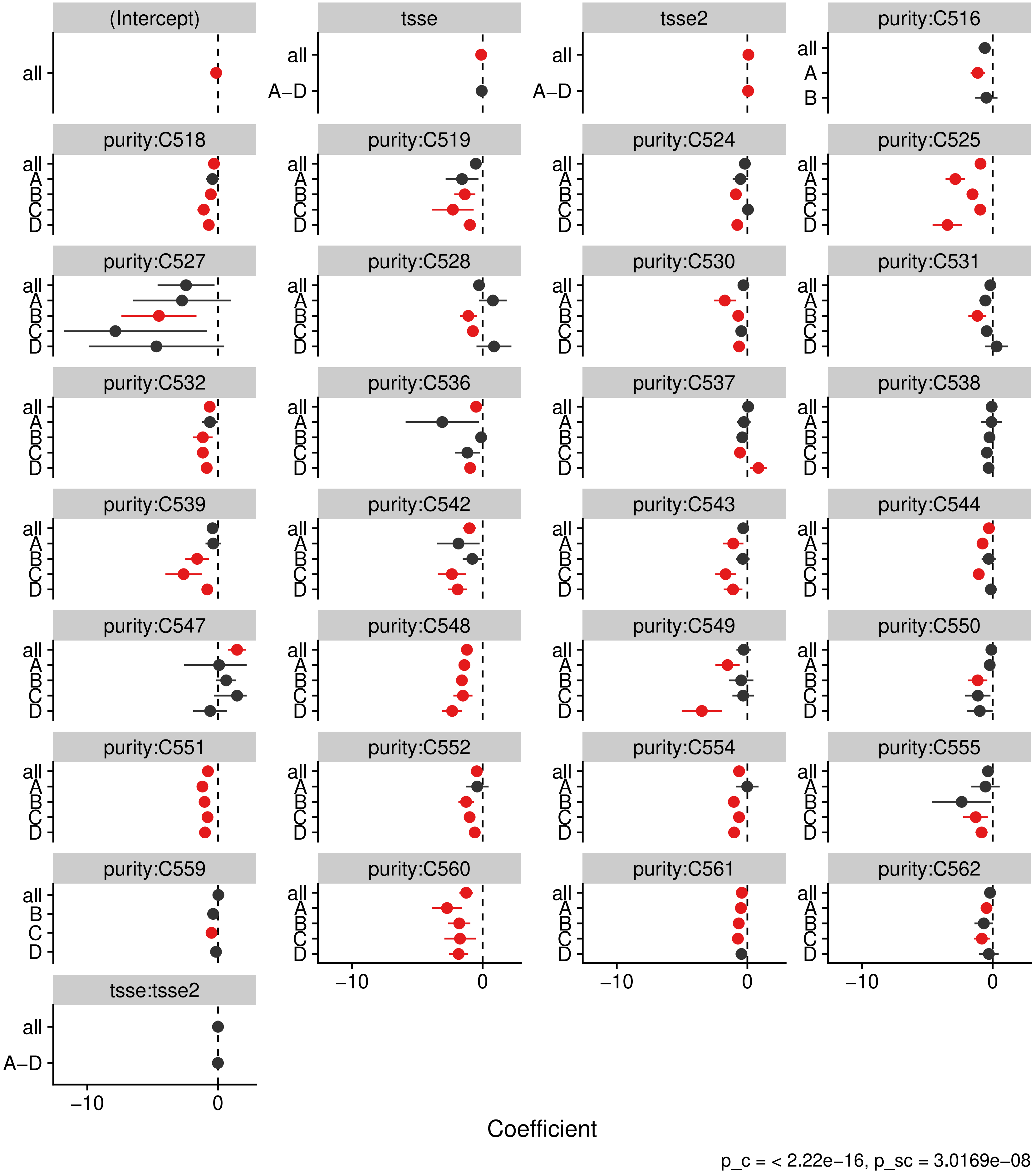

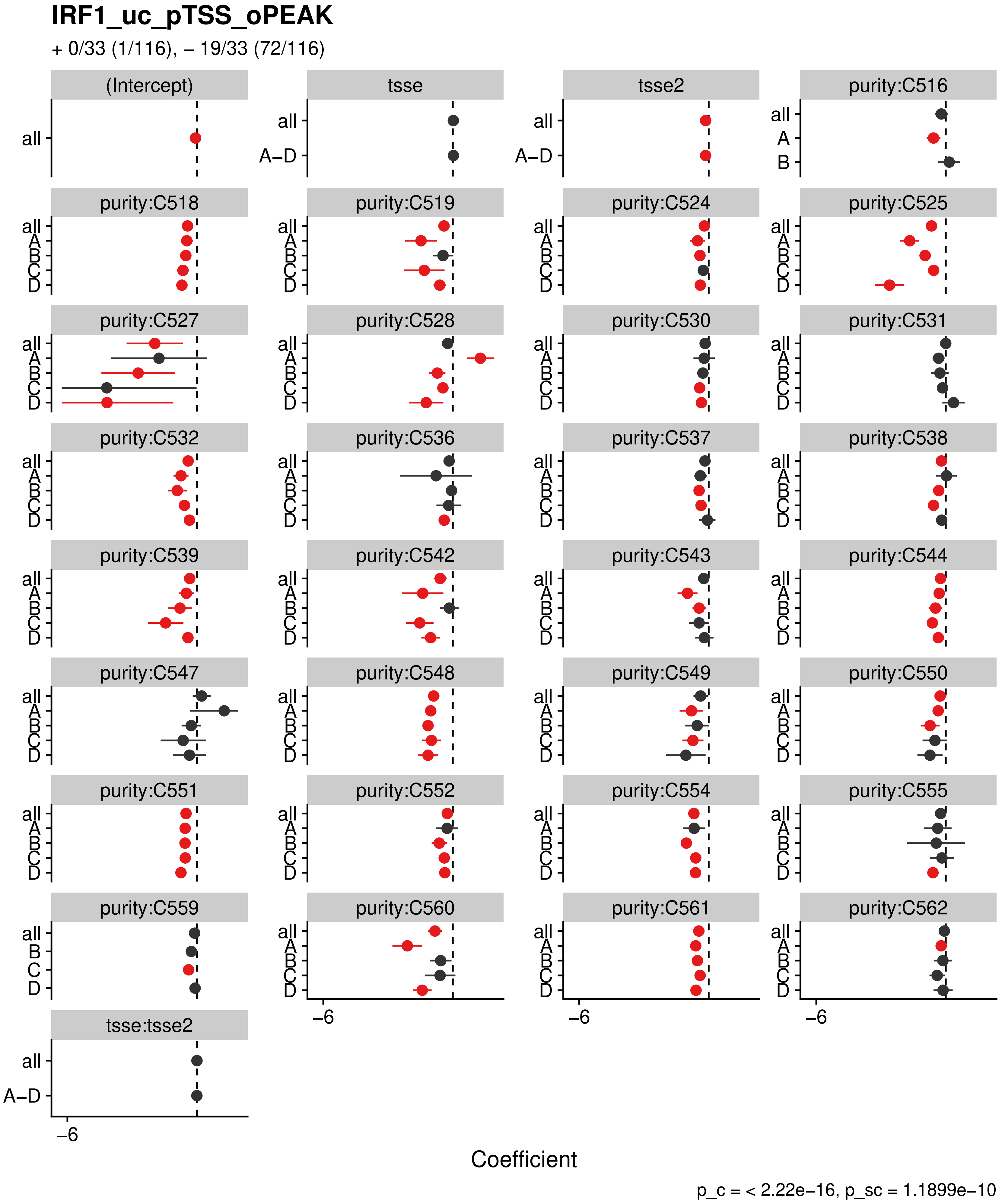

IRF1_uc_pTSS_oPEAK
+ 0/33 (1/116), – 19/33 (72/116)
(Intercept)
tsse
tsse2
purity:C516
purity:C518
purity:C519
purity:C524
purity:C525
purity:C527
purity:C528
purity:C530
purity:C531
purity:C532
purity:C536
purity:C537
purity:C538
purity:C539
purity:C542
purity:C543
purity:C544
purity:C547
purity:C548
purity:C549
purity:C550
purity:C551
purity:C552
purity:C554
purity:C555
purity:C559
purity:C560
purity:C561
purity:C562
tsse:tsse2
all
A–D
A
B
C
D
–6
Coefficient
p_c = < 2.22e–16, p_sc = 1.1899e–10

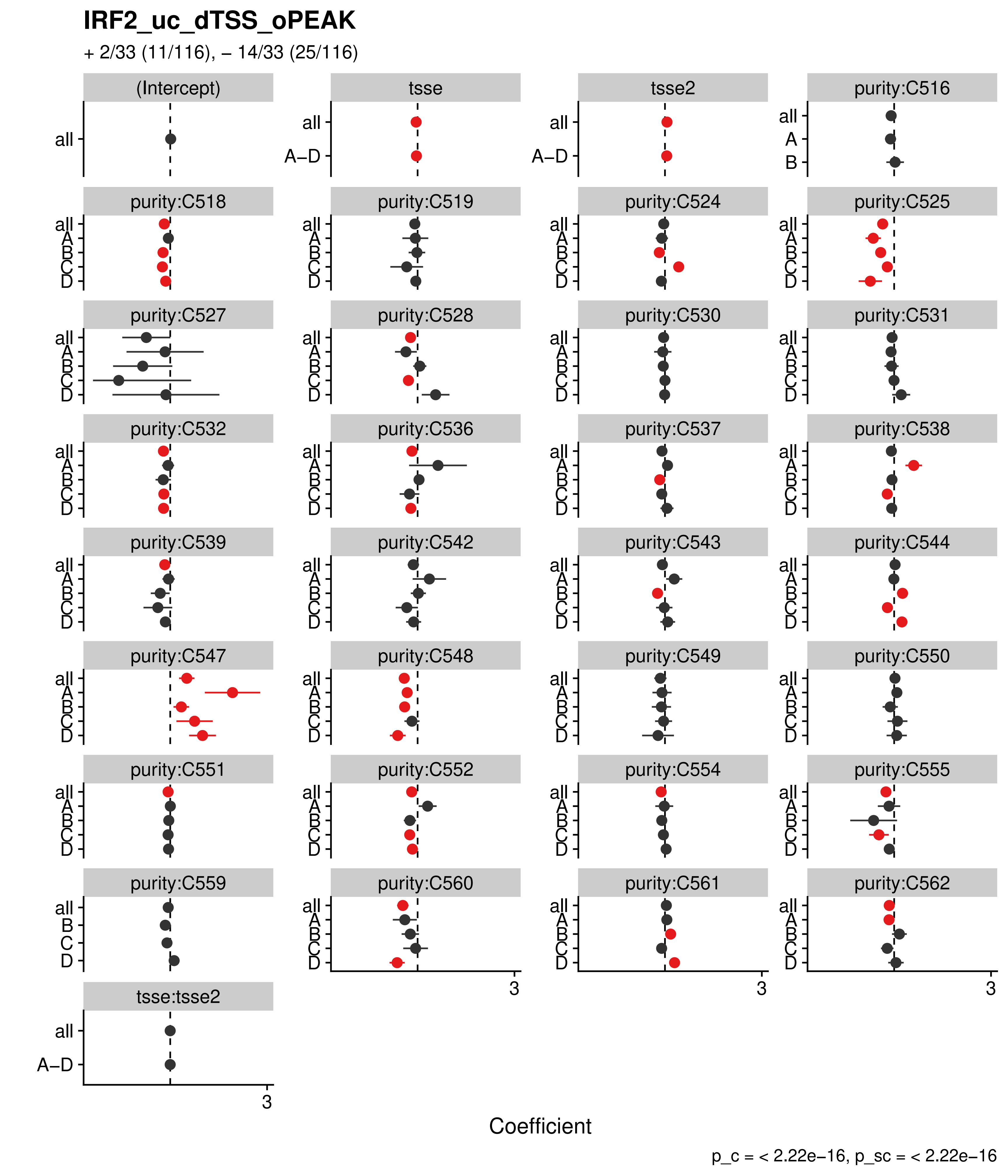

IRF2_uc_dTSS_oPEAK
+ 2/33 (11/116), – 14/33 (25/116)
(Intercept)
tsse
tsse2
purity:C516
purity:C518
purity:C519
purity:C524
purity:C525
purity:C527
purity:C528
purity:C530
purity:C531
purity:C532
purity:C536
purity:C537
purity:C538
purity:C539
purity:C542
purity:C543
purity:C544
purity:C547
purity:C548
purity:C549
purity:C550
purity:C551
purity:C552
purity:C554
purity:C555
purity:C559
purity:C560
purity:C561
purity:C562
tsse:tsse2
all
A–D
A
B
C
D
3
Coefficient
p_c = < 2.22e–16, p_sc = < 2.22e–16

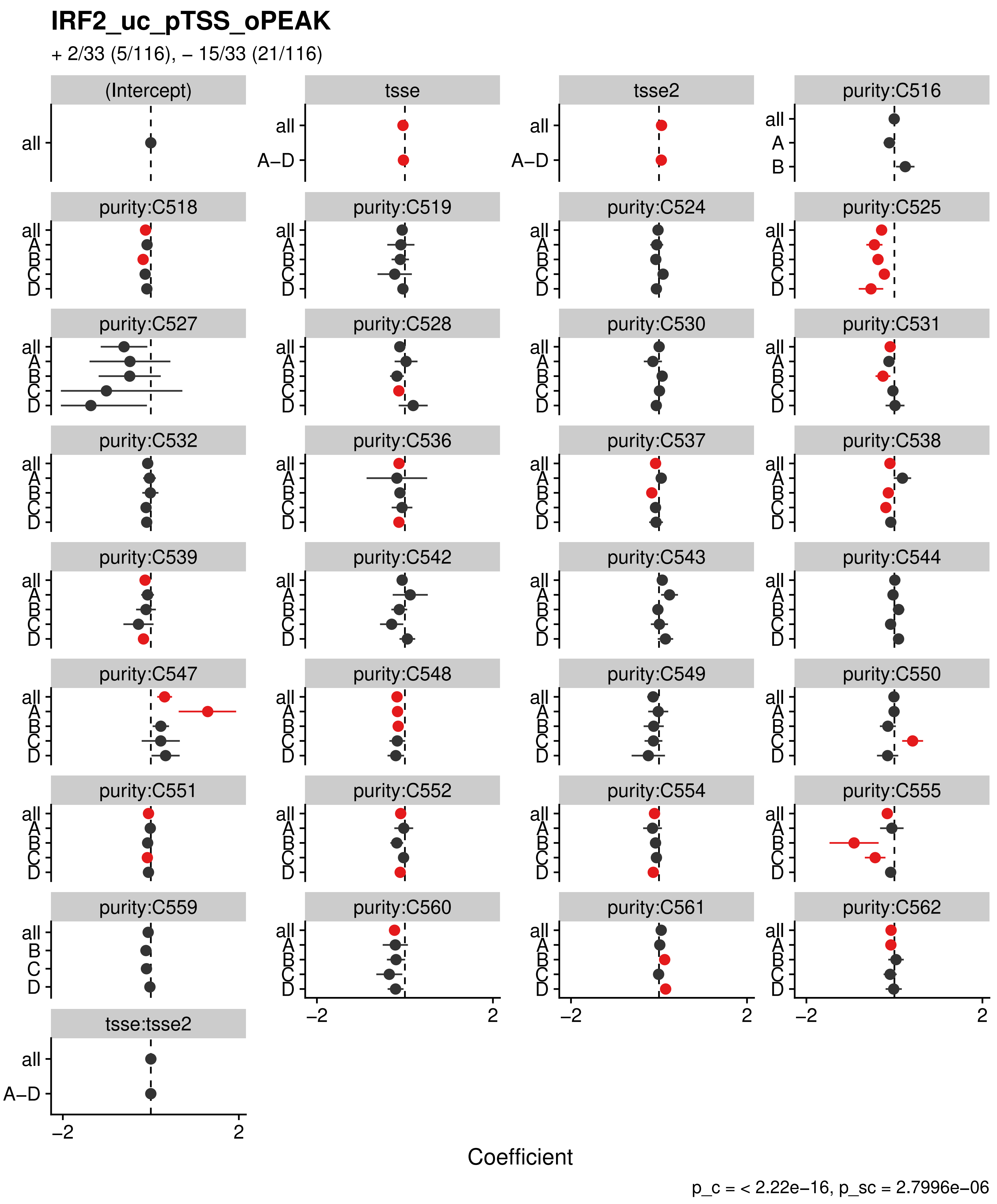

IRF2_uc_pTSS_oPEAK
+ 2/33 (5/116), – 15/33 (21/116)
(Intercept)
tsse
tsse2
purity:C516
purity:C518
purity:C519
purity:C524
purity:C525
purity:C527
purity:C528
purity:C530
purity:C531
purity:C532
purity:C536
purity:C537
purity:C538
purity:C539
purity:C542
purity:C543
purity:C544
purity:C547
purity:C548
purity:C549
purity:C550
purity:C551
purity:C552
purity:C554
purity:C555
purity:C559
purity:C560
purity:C561
purity:C562
tsse:tsse2
all
A–D
A
B
C
D
–2
2
Coefficient
p_c = < 2.22e–16, p_sc = 2.7996e–06

**IRF3_uc_cTSS_oPEAK**

+ 3/33 (8/116), – 17/33 (20/116)

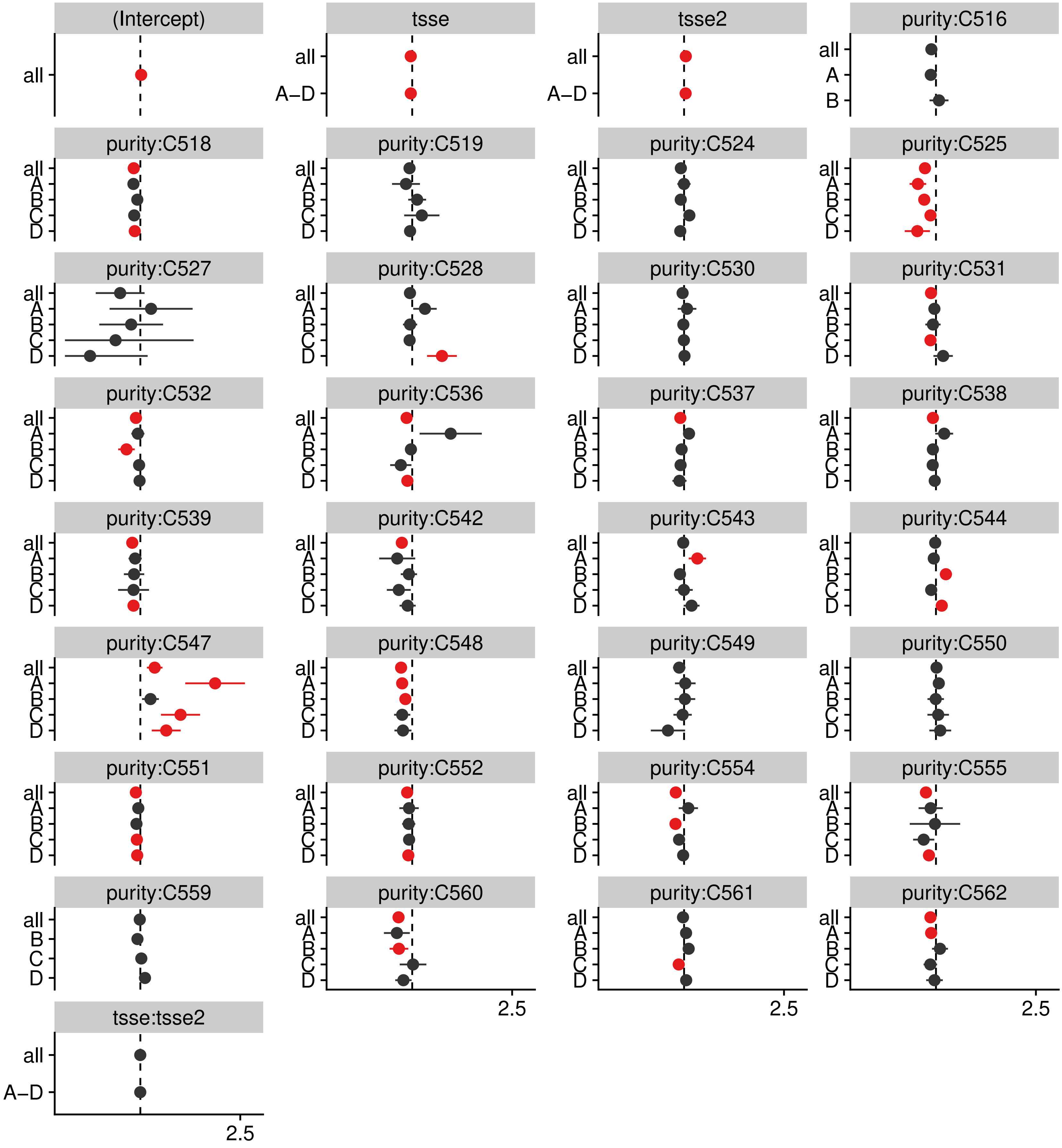


p_c = < 2.22e–16, p_sc = 8.6561e–10

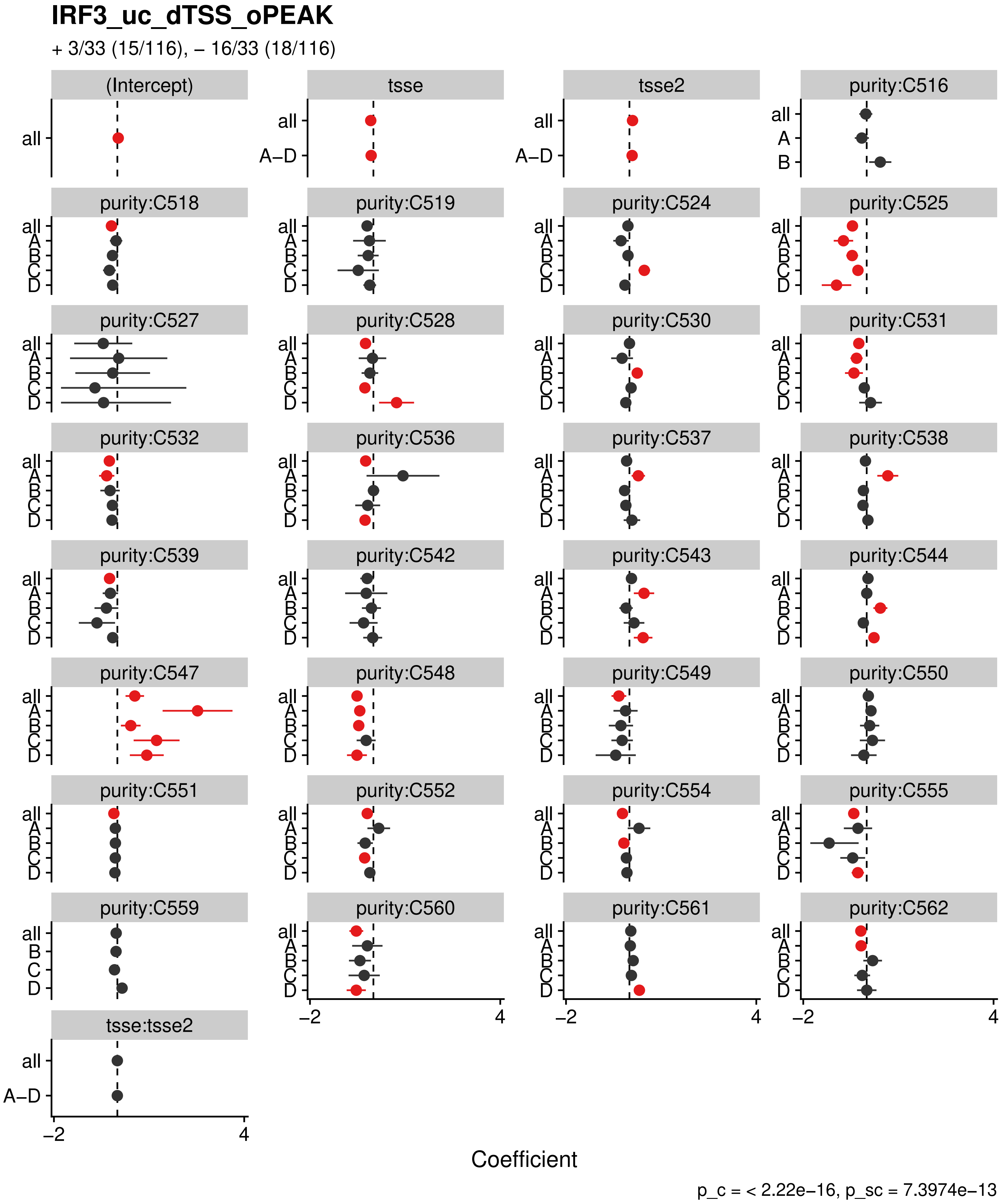
IRF3_uc_dTSS_oPEAK
+ 3/33 (15/116), − 16/33 (18/116)
(Intercept)
tsse
tsse2
purity:C516
purity:C518
purity:C519
purity:C524
purity:C525
purity:C527
purity:C528
purity:C530
purity:C531
purity:C532
purity:C536
purity:C537
purity:C538
purity:C539
purity:C542
purity:C543
purity:C544
purity:C547
purity:C548
purity:C549
purity:C550
purity:C551
purity:C552
purity:C554
purity:C555
purity:C559
purity:C560
purity:C561
purity:C562
tsse:tsse2
all
A
B
C
D
A−D
−2
4
Coefficient
p_c = < 2.22e−16, p_sc = 7.3974e−13

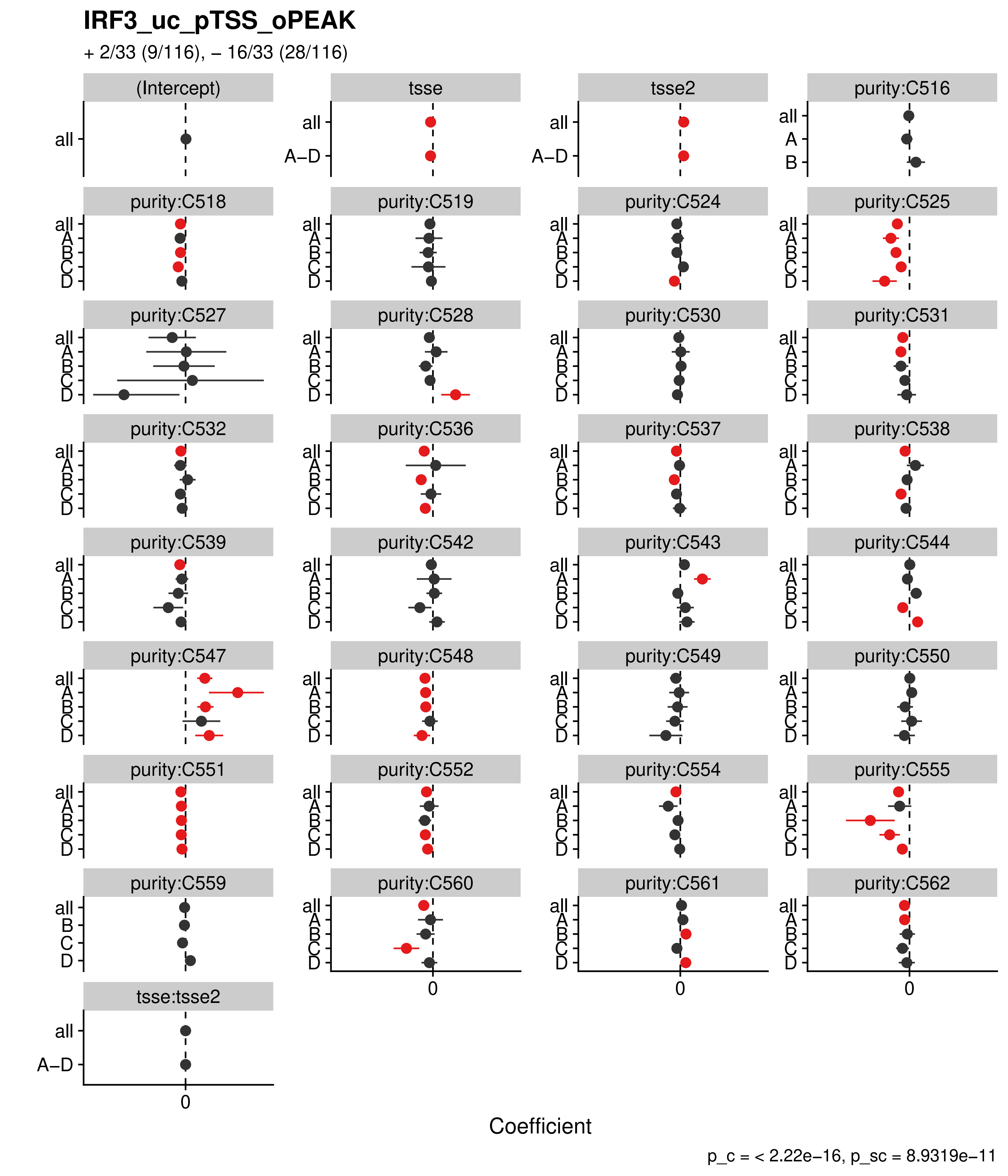

IRF3_uc_pTSS_oPEAK
+ 2/33 (9/116), – 16/33 (28/116)
(Intercept)
tsse
tsse2
purity:C516
purity:C518
purity:C519
purity:C524
purity:C525
purity:C527
purity:C528
purity:C530
purity:C531
purity:C532
purity:C536
purity:C537
purity:C538
purity:C539
purity:C542
purity:C543
purity:C544
purity:C547
purity:C548
purity:C549
purity:C550
purity:C551
purity:C552
purity:C554
purity:C555
purity:C559
purity:C560
purity:C561
purity:C562
tsse:tsse2
all
A–D
A
B
C
D
0
Coefficient
p_c = < 2.22e−16, p_sc = 8.9319e−11

### IRF8_uc_cTSS_oPEAK

+ 3/33 (7/116), – 15/33 (15/116)

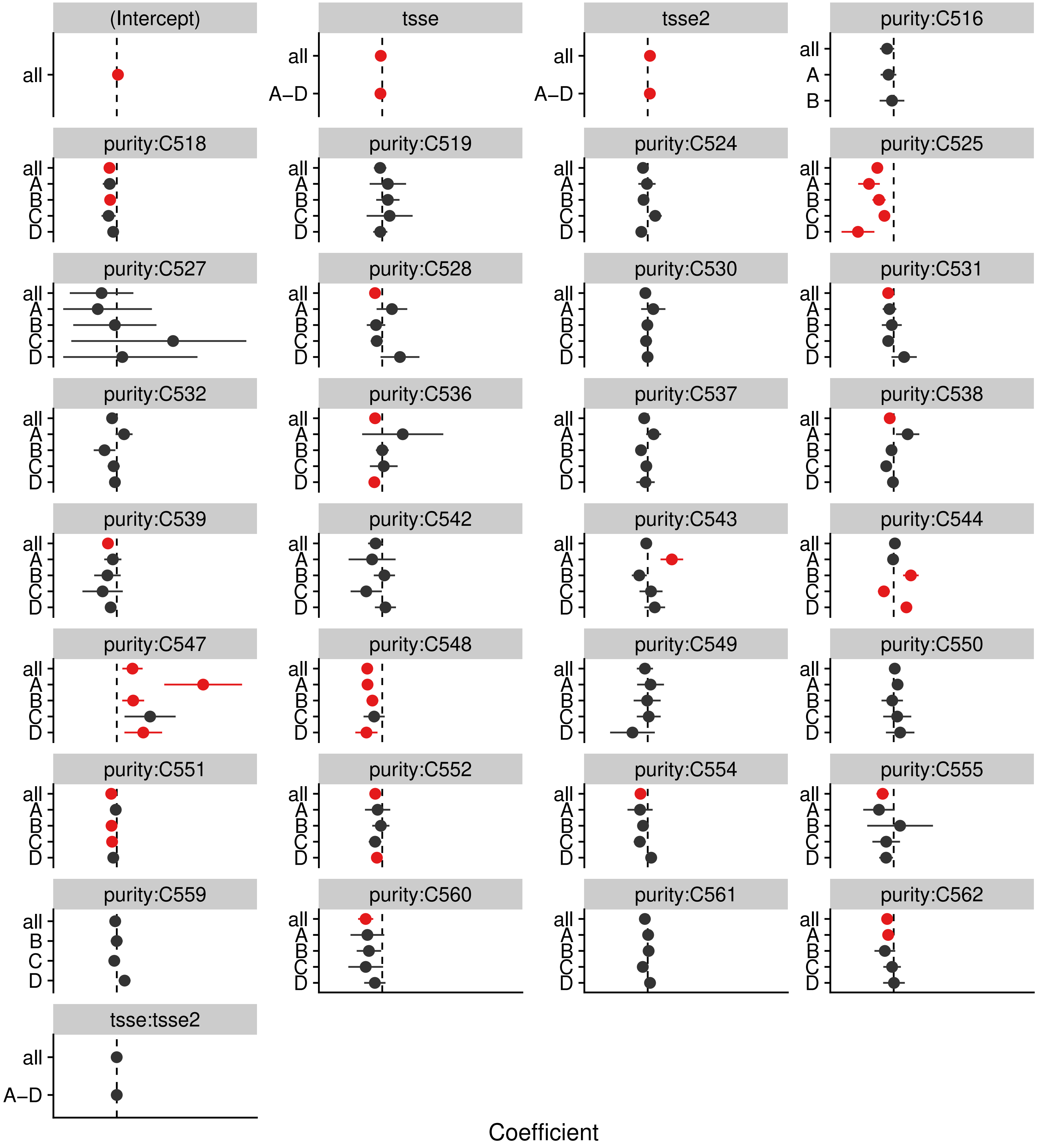


p_c = < 2.22e–16, p_sc = 1.5528e–09

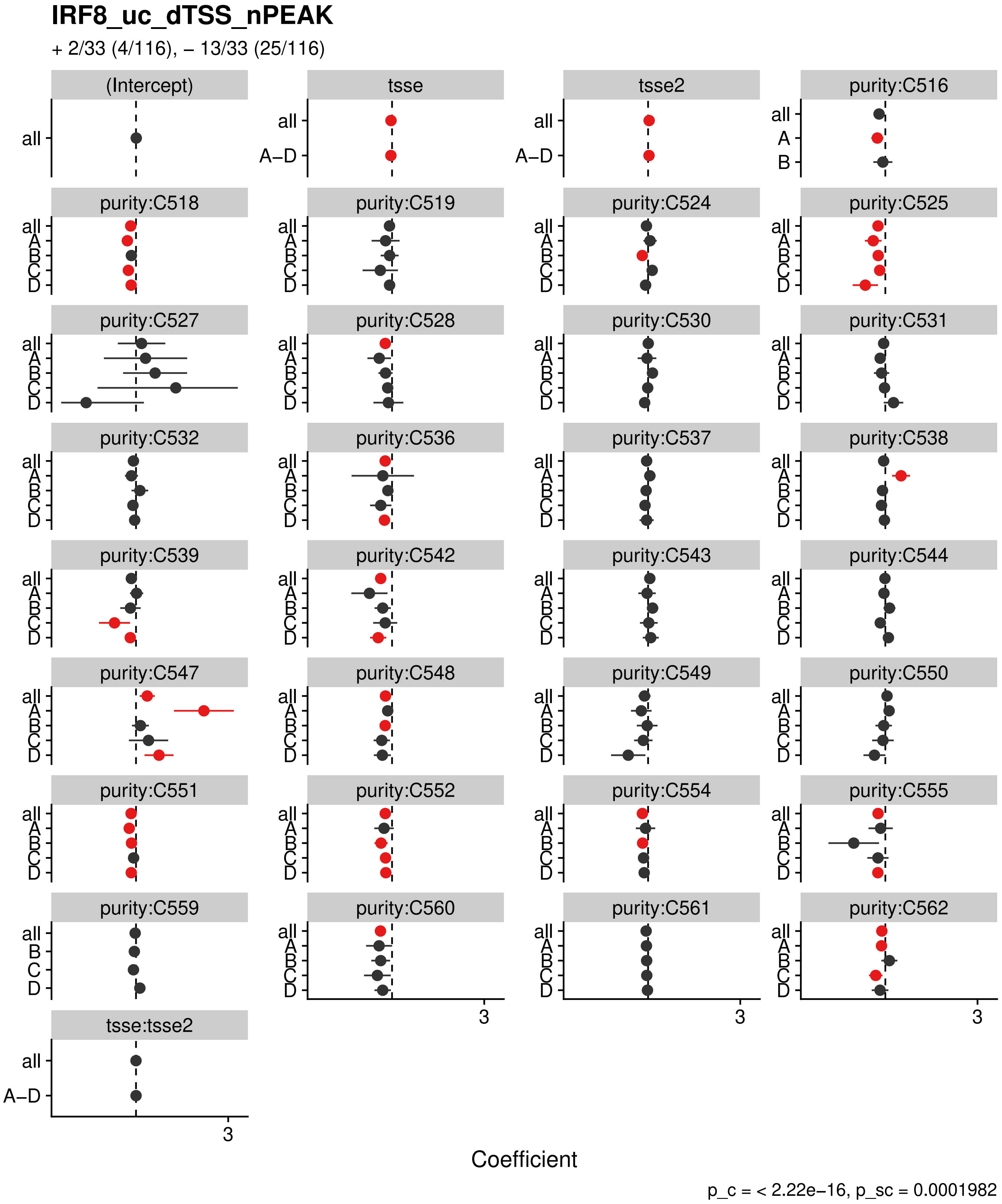
IRF8_uc_dTSS_nPEAK
+ 2/33 (4/116), – 13/33 (25/116)
(Intercept)
tsse
tsse2
purity:C516
purity:C518
purity:C519
purity:C524
purity:C525
purity:C527
purity:C528
purity:C530
purity:C531
purity:C532
purity:C536
purity:C537
purity:C538
purity:C539
purity:C542
purity:C543
purity:C544
purity:C547
purity:C548
purity:C549
purity:C550
purity:C551
purity:C552
purity:C554
purity:C555
purity:C559
purity:C560
purity:C561
purity:C562
tsse:tsse2
all
A–D
A
B
C
D
3
Coefficient
p_c = < 2.22e–16, p_sc = 0.0001982

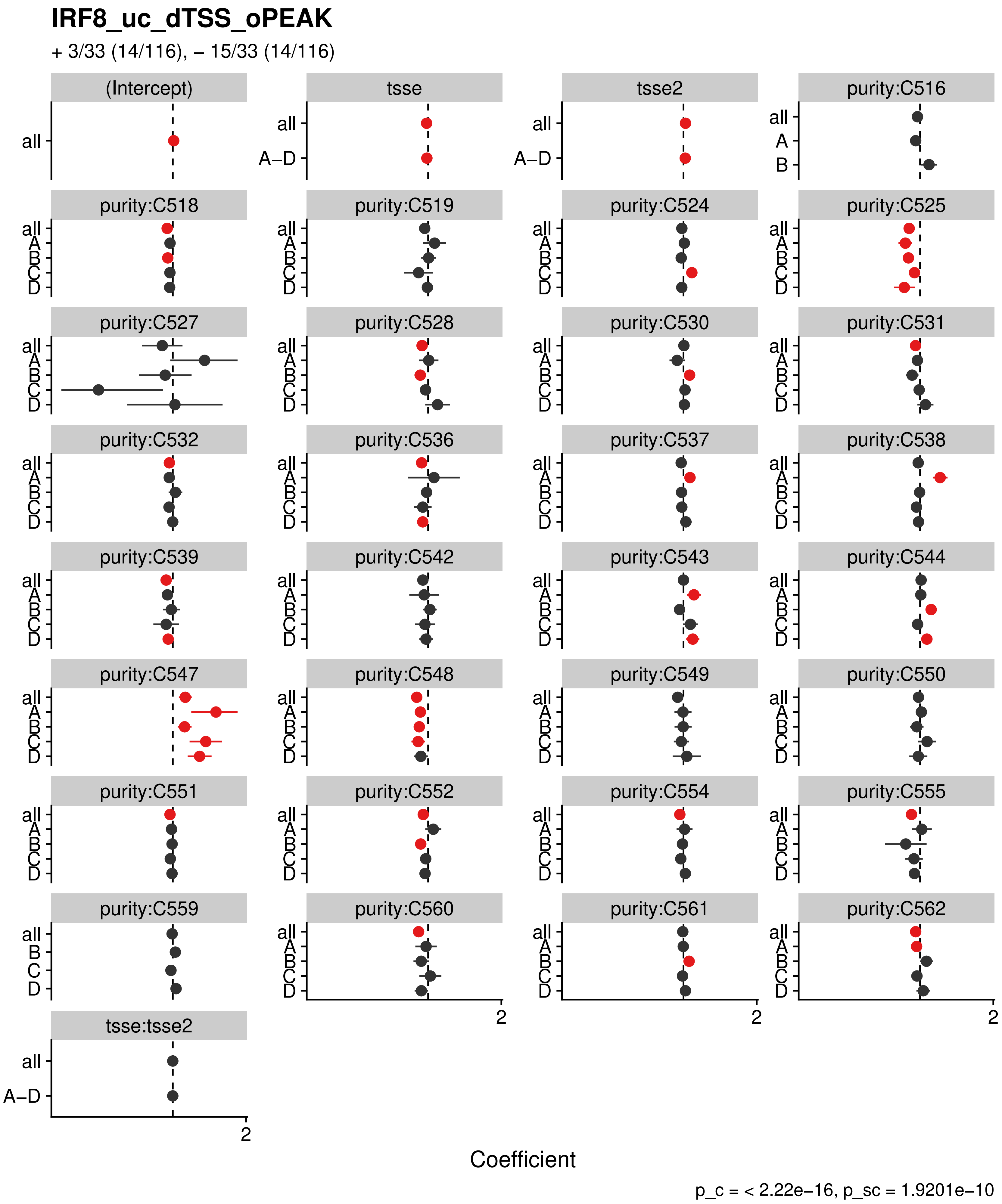
IRF8_uc_dTSS_oPEAK
+ 3/33 (14/116), – 15/33 (14/116)
(Intercept)
tsse
tsse2
purity:C516
purity:C518
purity:C519
purity:C524
purity:C525
purity:C527
purity:C528
purity:C530
purity:C531
purity:C532
purity:C536
purity:C537
purity:C538
purity:C539
purity:C542
purity:C543
purity:C544
purity:C547
purity:C548
purity:C549
purity:C550
purity:C551
purity:C552
purity:C554
purity:C555
purity:C559
purity:C560
purity:C561
purity:C562
tsse:tsse2
all
A–D
A
B
C
D
2
Coefficient
p_c = < 2.22e–16, p_sc = 1.9201e–10

### IRF9_uc_cTSS_oPEAK

+ 2/33 (7/116), – 19/33 (33/116)

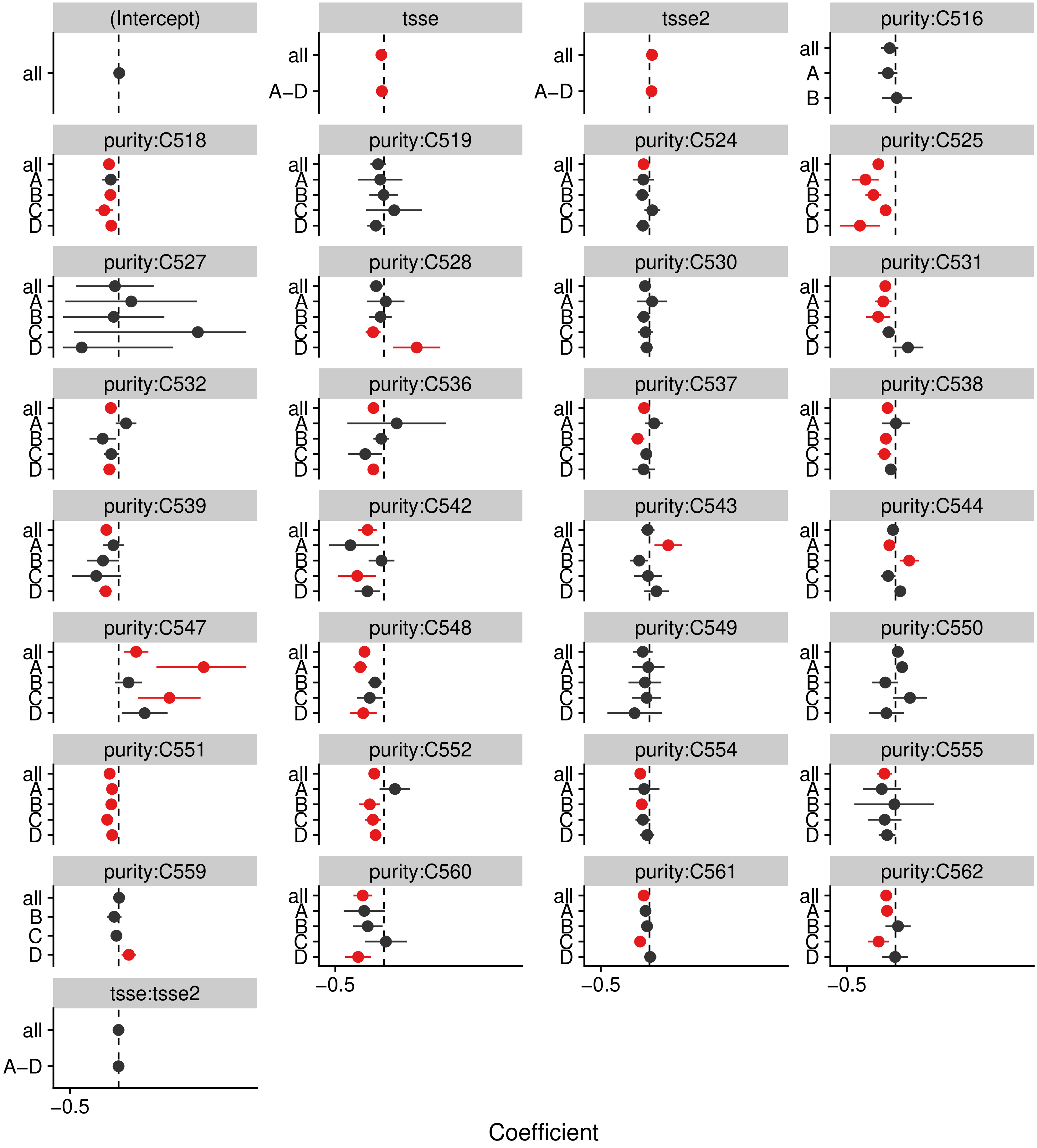


p_c = < 2.22e–16, p_sc = 1.9339e–09

#### IRF9_uc_dTSS_oPEAK

+ 3/33 (14/116), – 17/33 (15/116)

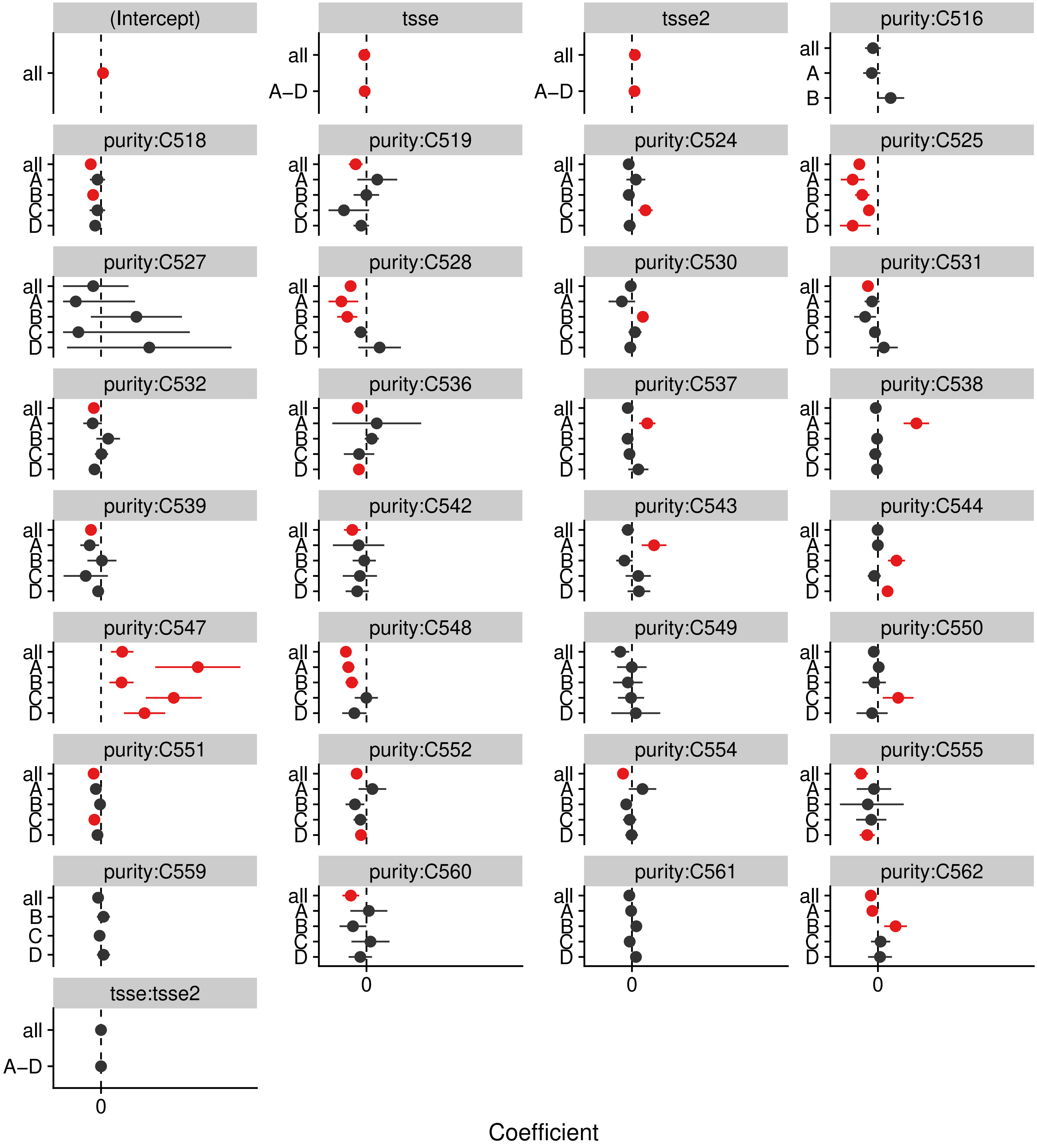


p_c = < 2.22e−16, p_sc = 1.7335e−12

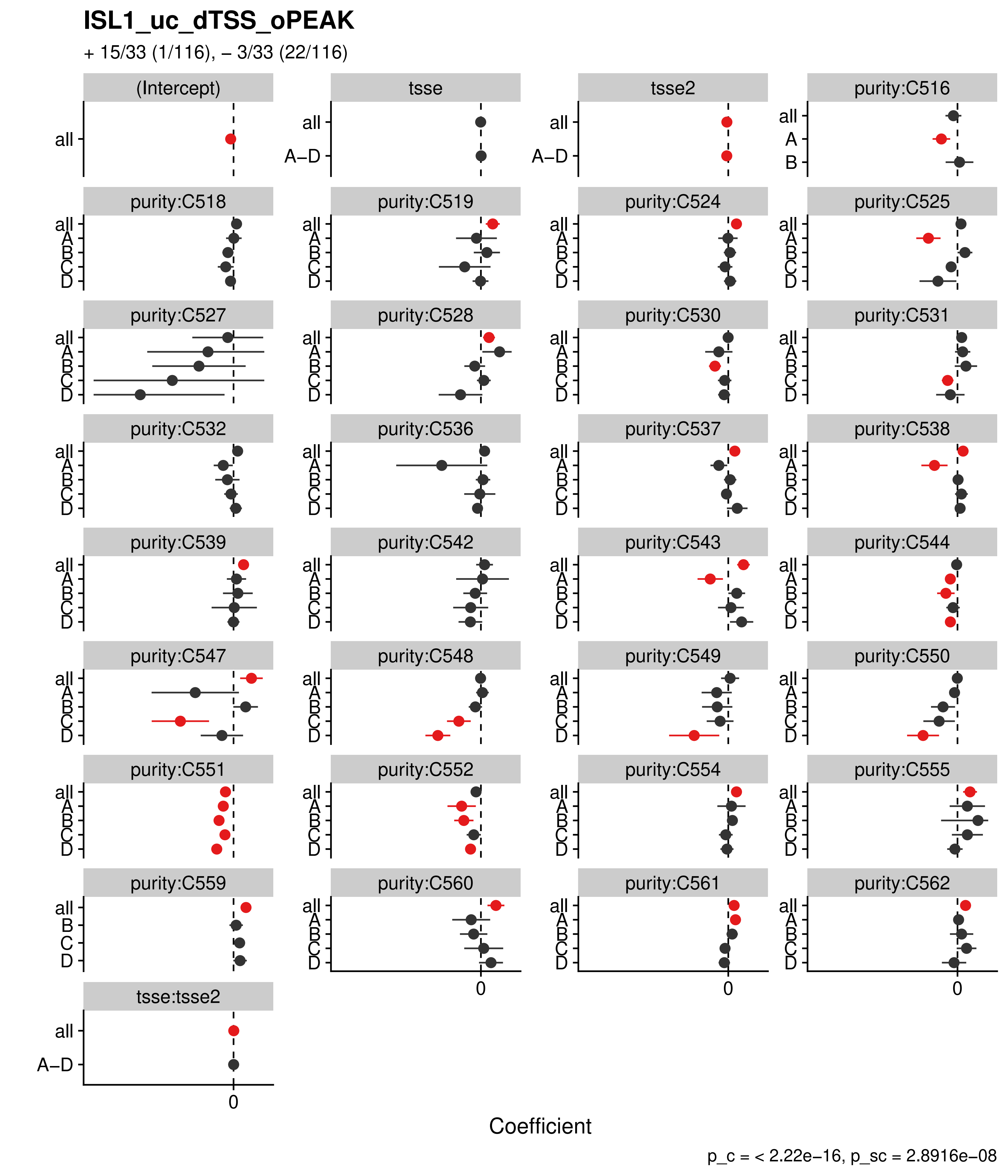
ISL1_uc_dTSS_oPEAK
+ 15/33 (1/116), – 3/33 (22/116)
(Intercept)
tsse
tsse2
purity:C516
purity:C518
purity:C519
purity:C524
purity:C525
purity:C527
purity:C528
purity:C530
purity:C531
purity:C532
purity:C536
purity:C537
purity:C538
purity:C539
purity:C542
purity:C543
purity:C544
purity:C547
purity:C548
purity:C549
purity:C550
purity:C551
purity:C552
purity:C554
purity:C555
purity:C559
purity:C560
purity:C561
purity:C562
tsse:tsse2
all
A–D
A
B
C
D
0
Coefficient
p_c = < 2.22e–16, p_sc = 2.8916e–08

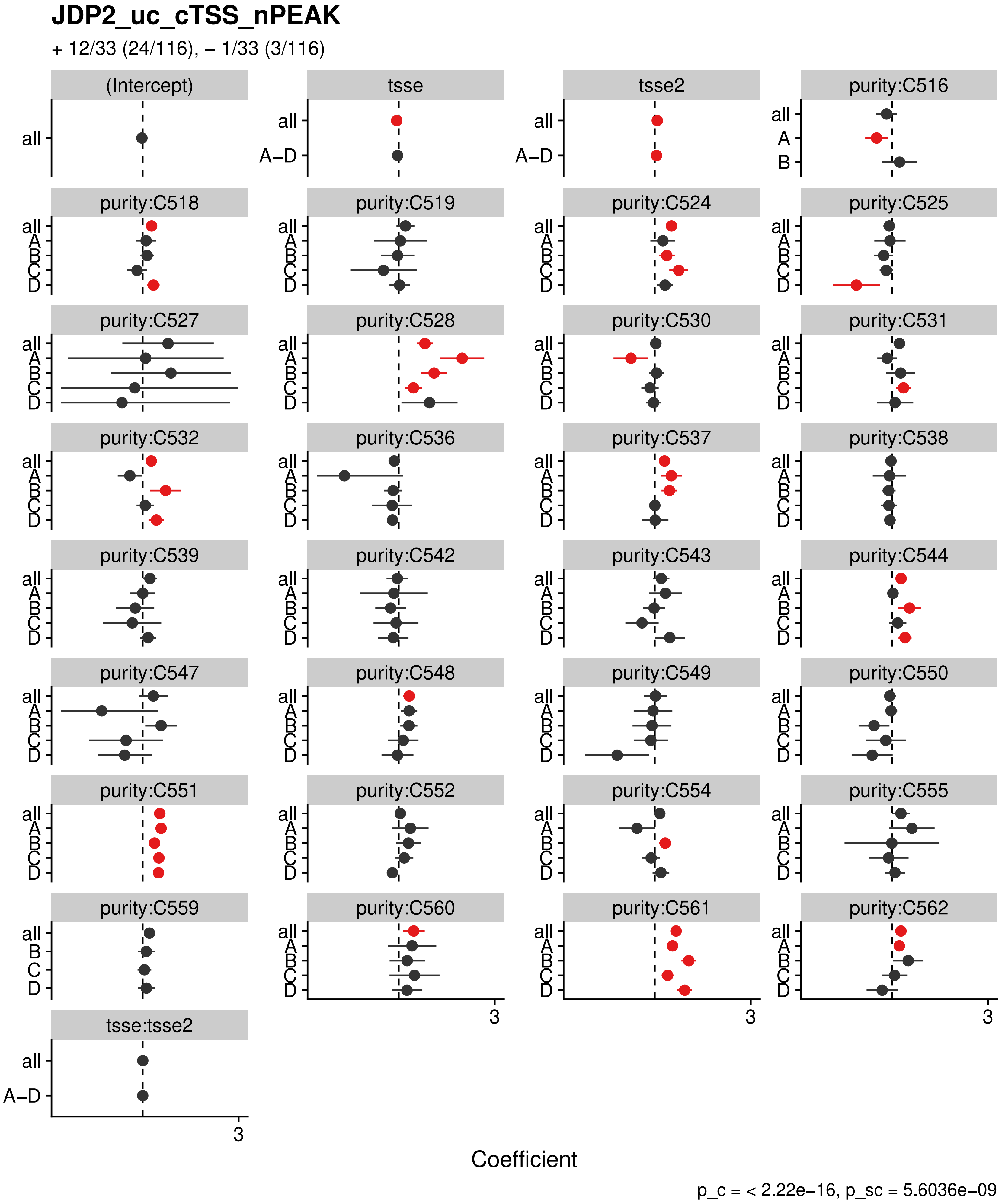
JDP2_uc_cTSS_nPEAK
+ 12/33 (24/116), – 1/33 (3/116)
(Intercept)
tsse
tsse2
purity:C516
purity:C518
purity:C519
purity:C524
purity:C525
purity:C527
purity:C528
purity:C530
purity:C531
purity:C532
purity:C536
purity:C537
purity:C538
purity:C539
purity:C542
purity:C543
purity:C544
purity:C547
purity:C548
purity:C549
purity:C550
purity:C551
purity:C552
purity:C554
purity:C555
purity:C559
purity:C560
purity:C561
purity:C562
tsse:tsse2
all
A–D
A
B
C
D
3
Coefficient
p_c = < 2.22e–16, p_sc = 5.6036e–09

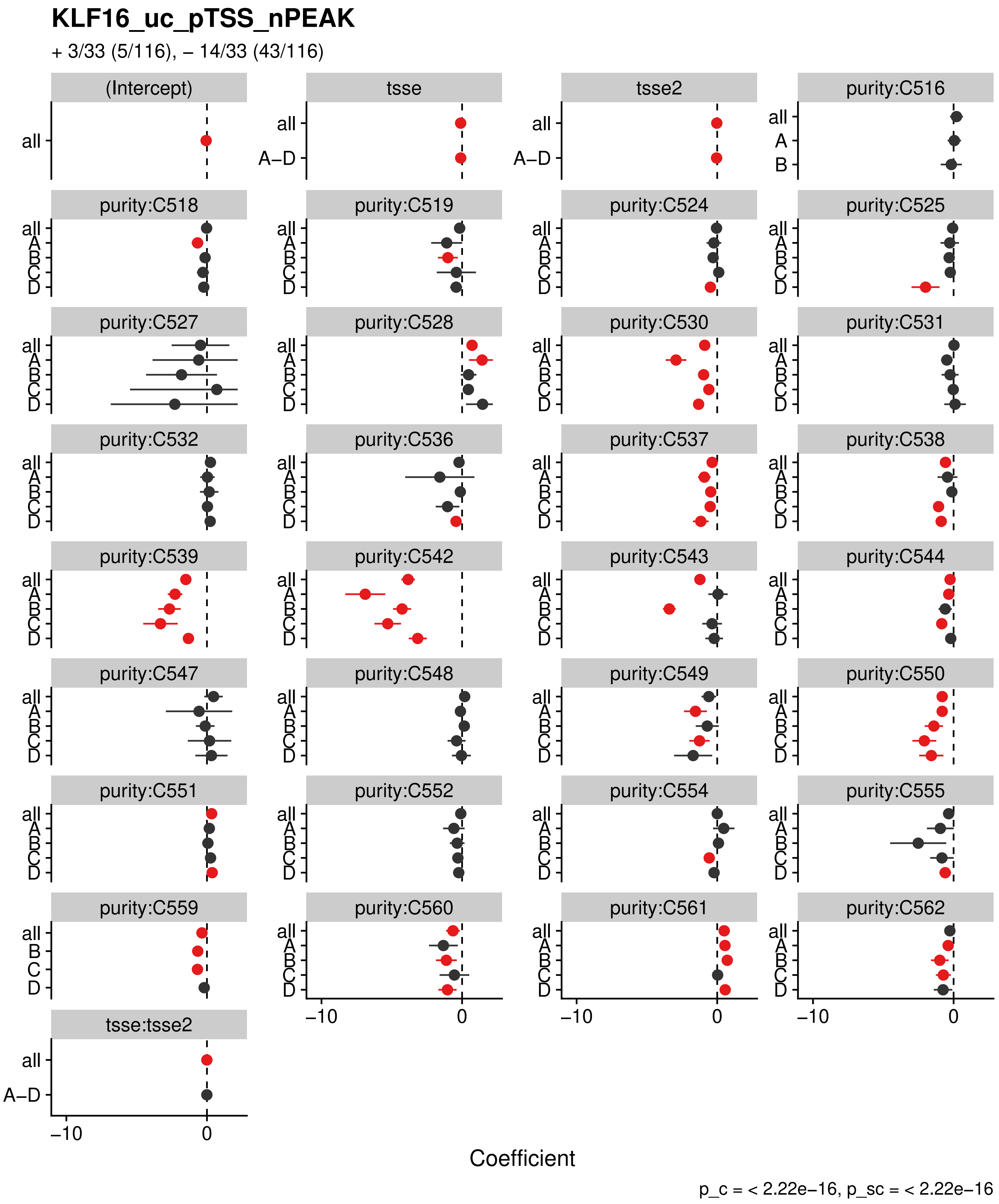

KLF16_uc_pTSS_nPEAK
+ 3/33 (5/116), – 14/33 (43/116)
(Intercept)
tsse
tsse2
purity:C516
purity:C518
purity:C519
purity:C524
purity:C525
purity:C527
purity:C528
purity:C530
purity:C531
purity:C532
purity:C536
purity:C537
purity:C538
purity:C539
purity:C542
purity:C543
purity:C544
purity:C547
purity:C548
purity:C549
purity:C550
purity:C551
purity:C552
purity:C554
purity:C555
purity:C559
purity:C560
purity:C561
purity:C562
tsse:tsse2
all
A–D
A
B
C
D
–10
0
Coefficient
p_c = < 2.22e–16, p_sc = < 2.22e–16

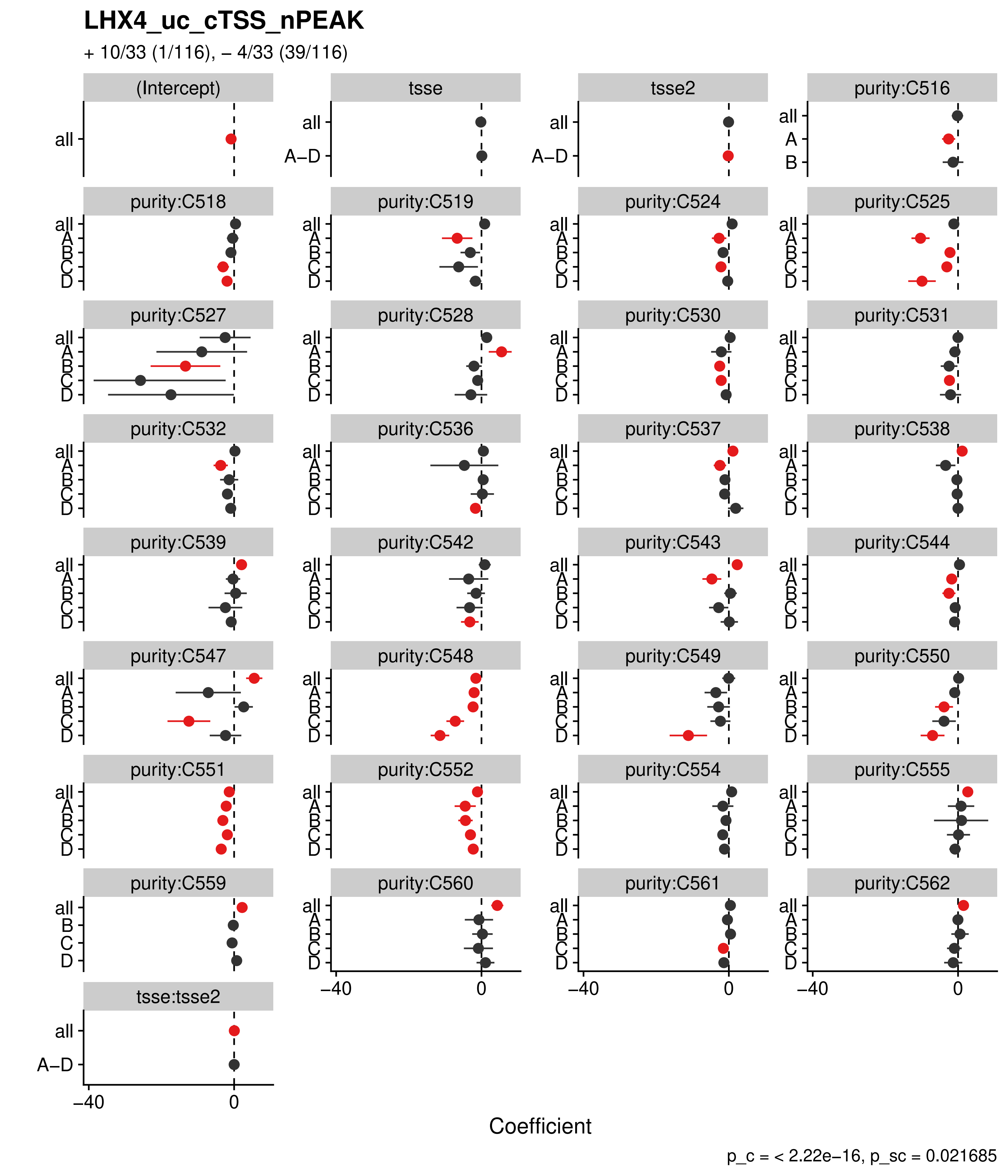
LHX4_uc_cTSS_nPEAK
+ 10/33 (1/116), – 4/33 (39/116)
(Intercept)
tsse
tsse2
purity:C516
purity:C518
purity:C519
purity:C524
purity:C525
purity:C527
purity:C528
purity:C530
purity:C531
purity:C532
purity:C536
purity:C537
purity:C538
purity:C539
purity:C542
purity:C543
purity:C544
purity:C547
purity:C548
purity:C549
purity:C550
purity:C551
purity:C552
purity:C554
purity:C555
purity:C559
purity:C560
purity:C561
purity:C562
tsse:tsse2
Coefficient
p_c = < 2.22e–16, p_sc = 0.021685

#### LHX4_uc_dTSS_oPEAK

+ 12/33 (0/116), – 4/33 (42/116)

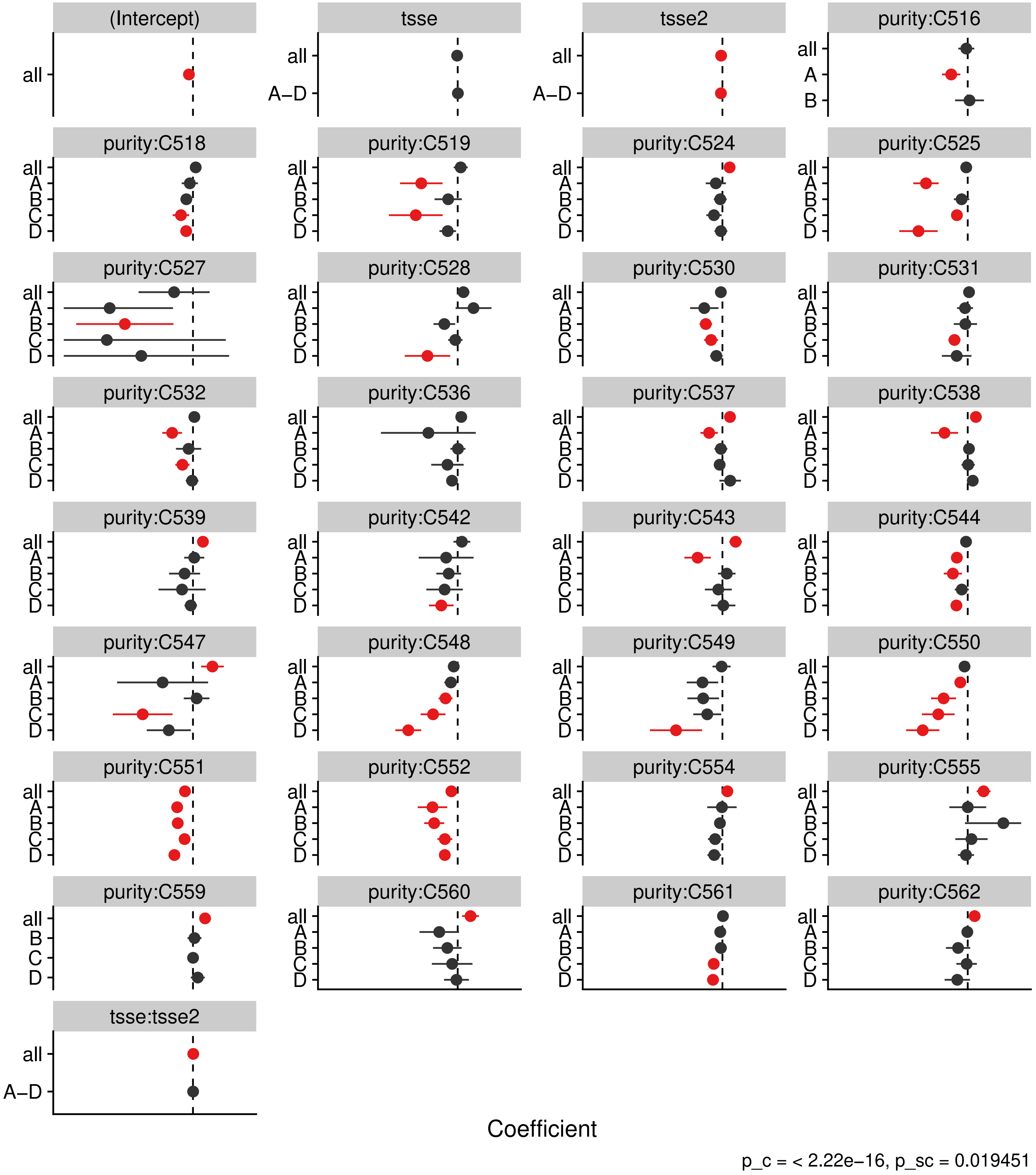

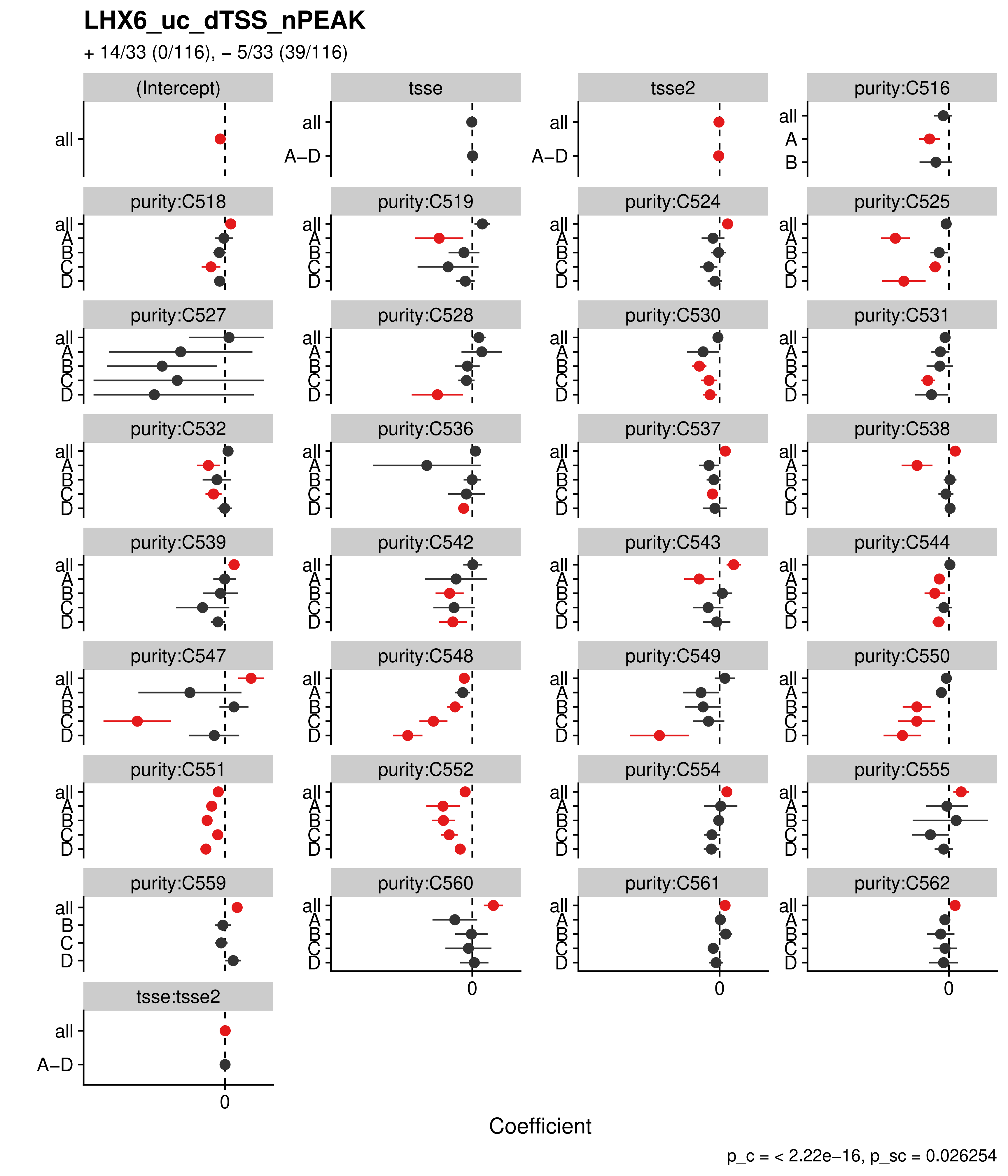

LHX6_uc_dTSS_nPEAK
+ 14/33 (0/116), – 5/33 (39/116)
(Intercept)
tsse
tsse2
purity:C516
purity:C518
purity:C519
purity:C524
purity:C525
purity:C527
purity:C528
purity:C530
purity:C531
purity:C532
purity:C536
purity:C537
purity:C538
purity:C539
purity:C542
purity:C543
purity:C544
purity:C547
purity:C548
purity:C549
purity:C550
purity:C551
purity:C552
purity:C554
purity:C555
purity:C559
purity:C560
purity:C561
purity:C562
tsse:tsse2
all
A–D
A
B
C
D
0
Coefficient
p_c = < 2.22e–16, p_sc = 0.026254

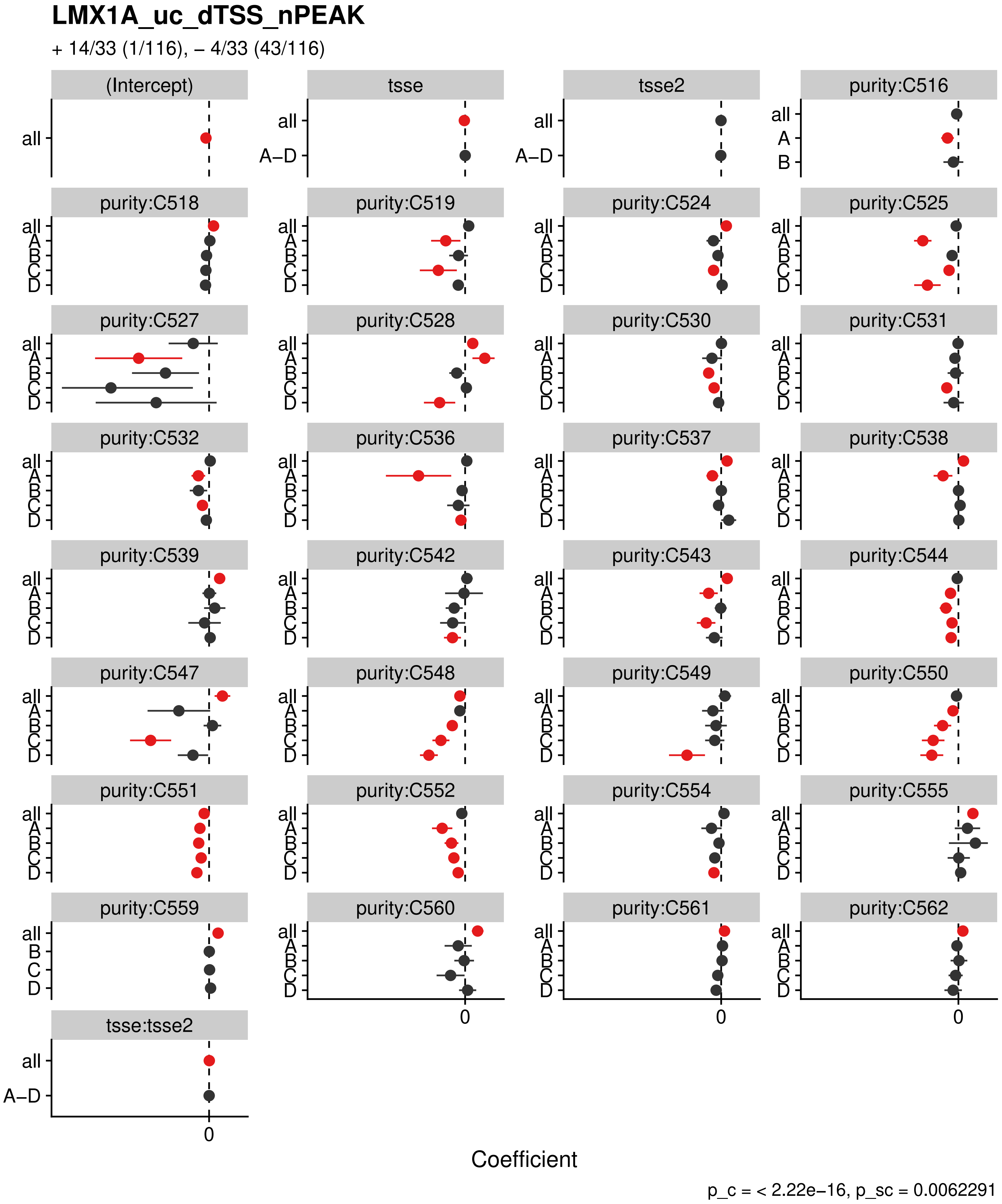
LMX1A_uc_dTSS_nPEAK
+ 14/33 (1/116), – 4/33 (43/116)
(Intercept)
tsse
tsse2
purity:C516
purity:C518
purity:C519
purity:C524
purity:C525
purity:C527
purity:C528
purity:C530
purity:C531
purity:C532
purity:C536
purity:C537
purity:C538
purity:C539
purity:C542
purity:C543
purity:C544
purity:C547
purity:C548
purity:C549
purity:C550
purity:C551
purity:C552
purity:C554
purity:C555
purity:C559
purity:C560
purity:C561
purity:C562
tsse:tsse2
all
A–D
A
B
C
D
0
Coefficient
p_c = < 2.22e–16, p_sc = 0.0062291

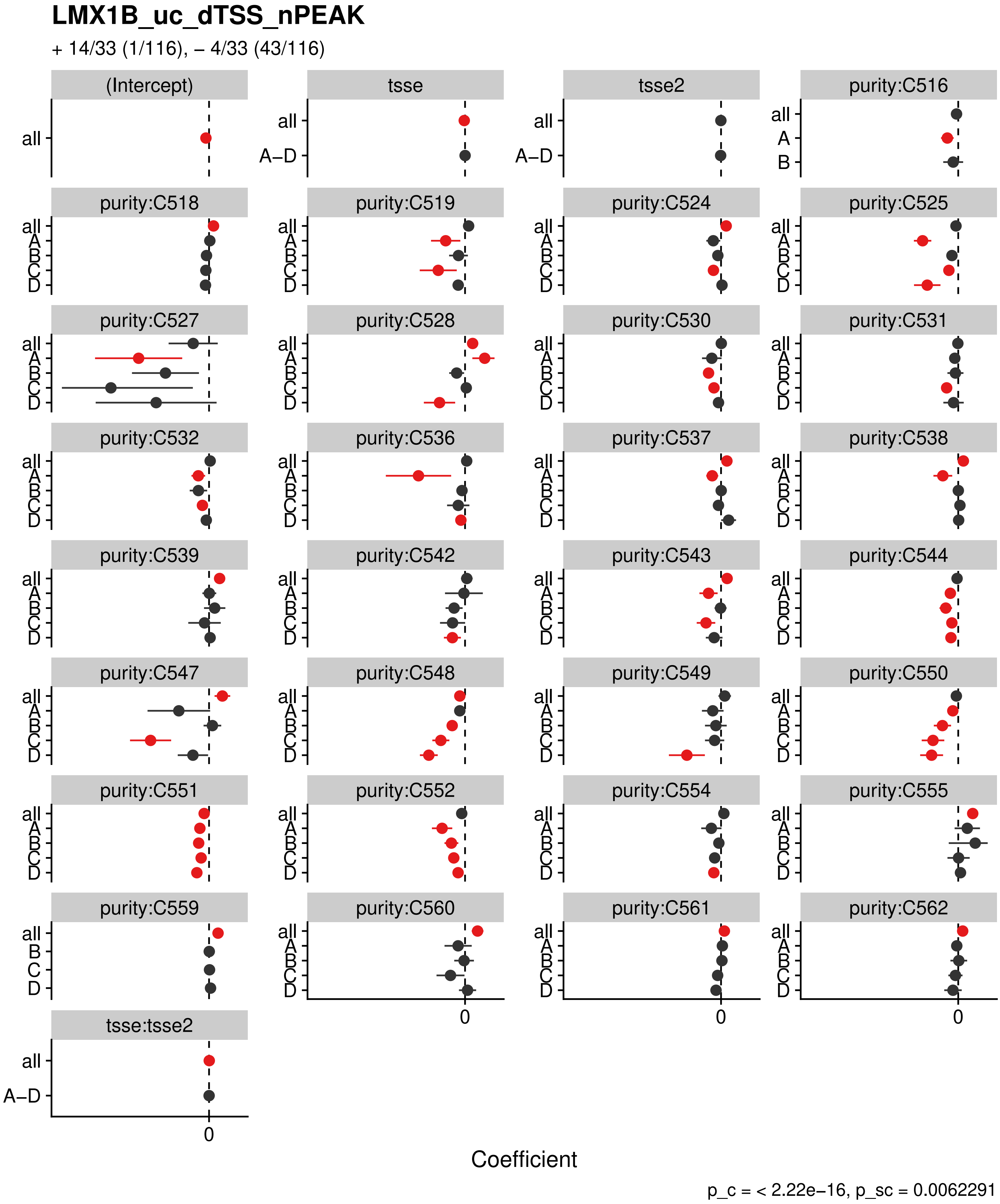

LMX1B_uc_dTSS_nPEAK
+ 14/33 (1/116), – 4/33 (43/116)
(Intercept)
tsse
tsse2
purity:C516
purity:C518
purity:C519
purity:C524
purity:C525
purity:C527
purity:C528
purity:C530
purity:C531
purity:C532
purity:C536
purity:C537
purity:C538
purity:C539
purity:C542
purity:C543
purity:C544
purity:C547
purity:C548
purity:C549
purity:C550
purity:C551
purity:C552
purity:C554
purity:C555
purity:C559
purity:C560
purity:C561
purity:C562
tsse:tsse2
all
A–D
A
B
C
D
0
Coefficient
p_c = < 2.22e–16, p_sc = 0.0062291

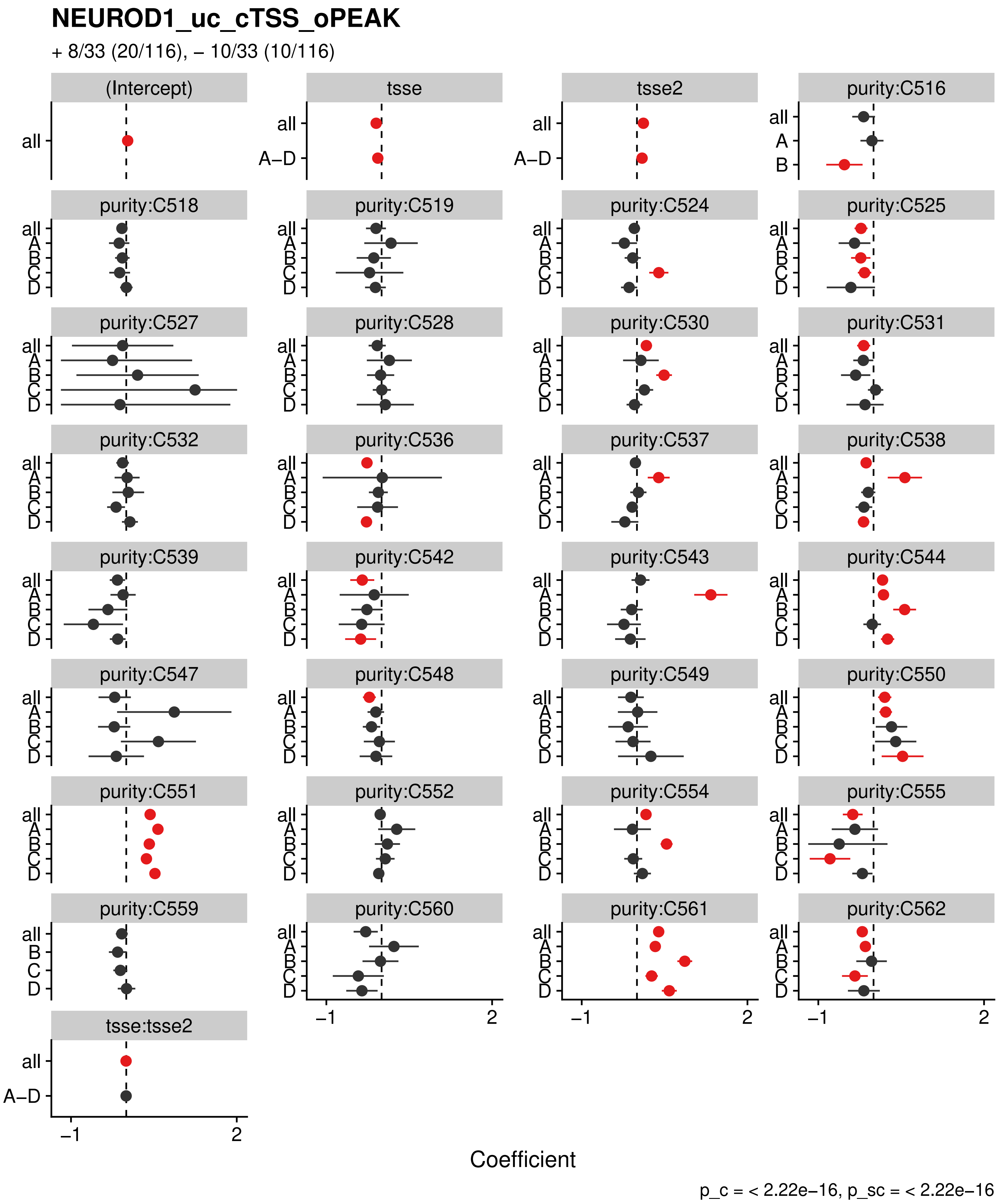
NEUROD1_uc_cTSS_oPEAK
+ 8/33 (20/116), – 10/33 (10/116)
(Intercept)
tsse
tsse2
purity:C516
purity:C518
purity:C519
purity:C524
purity:C525
purity:C527
purity:C528
purity:C530
purity:C531
purity:C532
purity:C536
purity:C537
purity:C538
purity:C539
purity:C542
purity:C543
purity:C544
purity:C547
purity:C548
purity:C549
purity:C550
purity:C551
purity:C552
purity:C554
purity:C555
purity:C559
purity:C560
purity:C561
purity:C562
tsse:tsse2
all
A–D
A
B
C
D
−1
2
Coefficient
p_c = < 2.22e−16, p_sc = < 2.22e−16

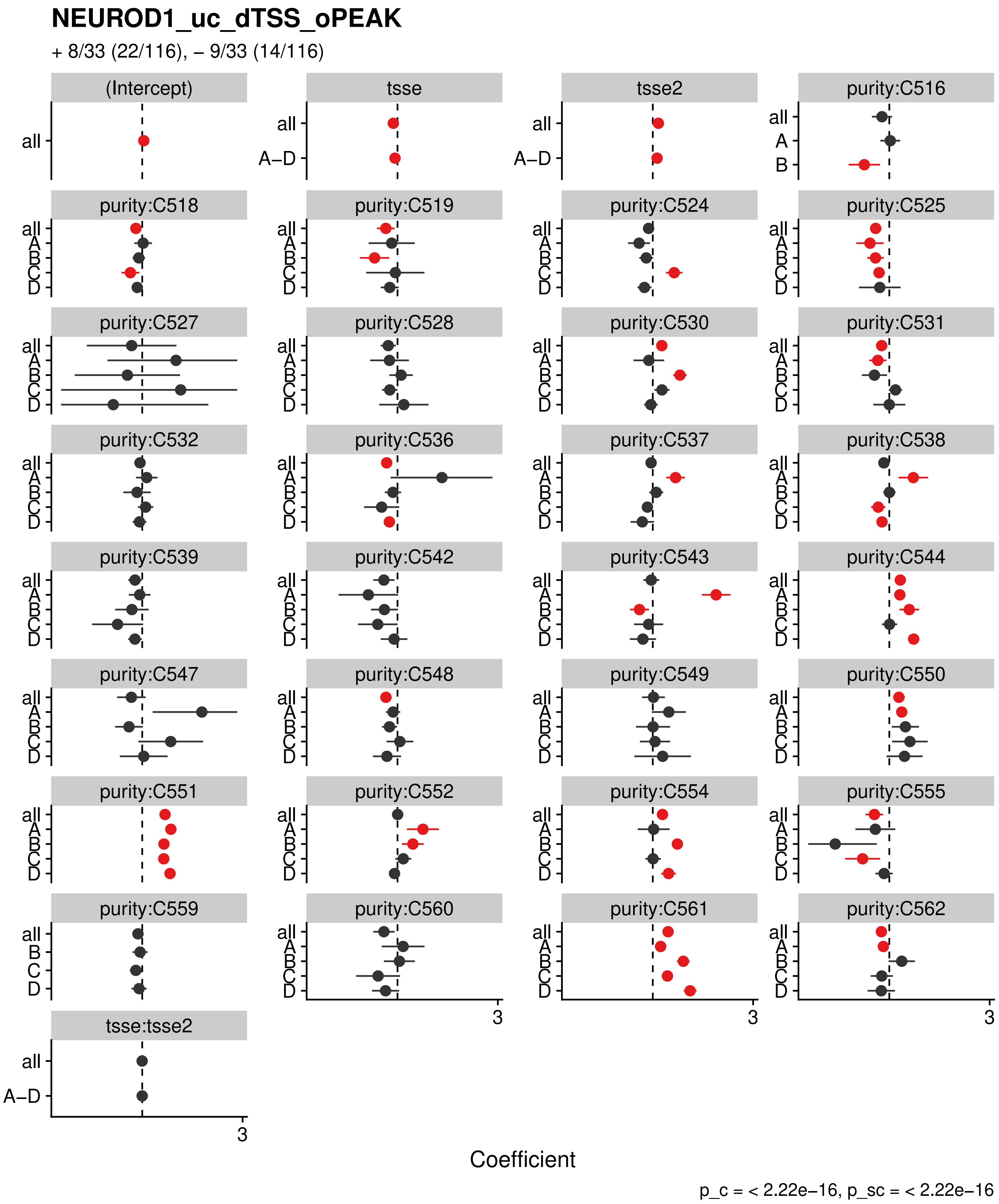

NEUROD1_uc_dTSS_oPEAK
+ 8/33 (22/116), − 9/33 (14/116)
(Intercept)
tsse
tsse2
purity:C516
purity:C518
purity:C519
purity:C524
purity:C525
purity:C527
purity:C528
purity:C530
purity:C531
purity:C532
purity:C536
purity:C537
purity:C538
purity:C539
purity:C542
purity:C543
purity:C544
purity:C547
purity:C548
purity:C549
purity:C550
purity:C551
purity:C552
purity:C554
purity:C555
purity:C559
purity:C560
purity:C561
purity:C562
tsse:tsse2
all
A–D
A
B
C
D
3
Coefficient
p_c = < 2.22e−16, p_sc = < 2.22e−16

### NKX24_uc_dTSS_nPEAK

+ 14/33 (1/116), – 5/33 (33/116)

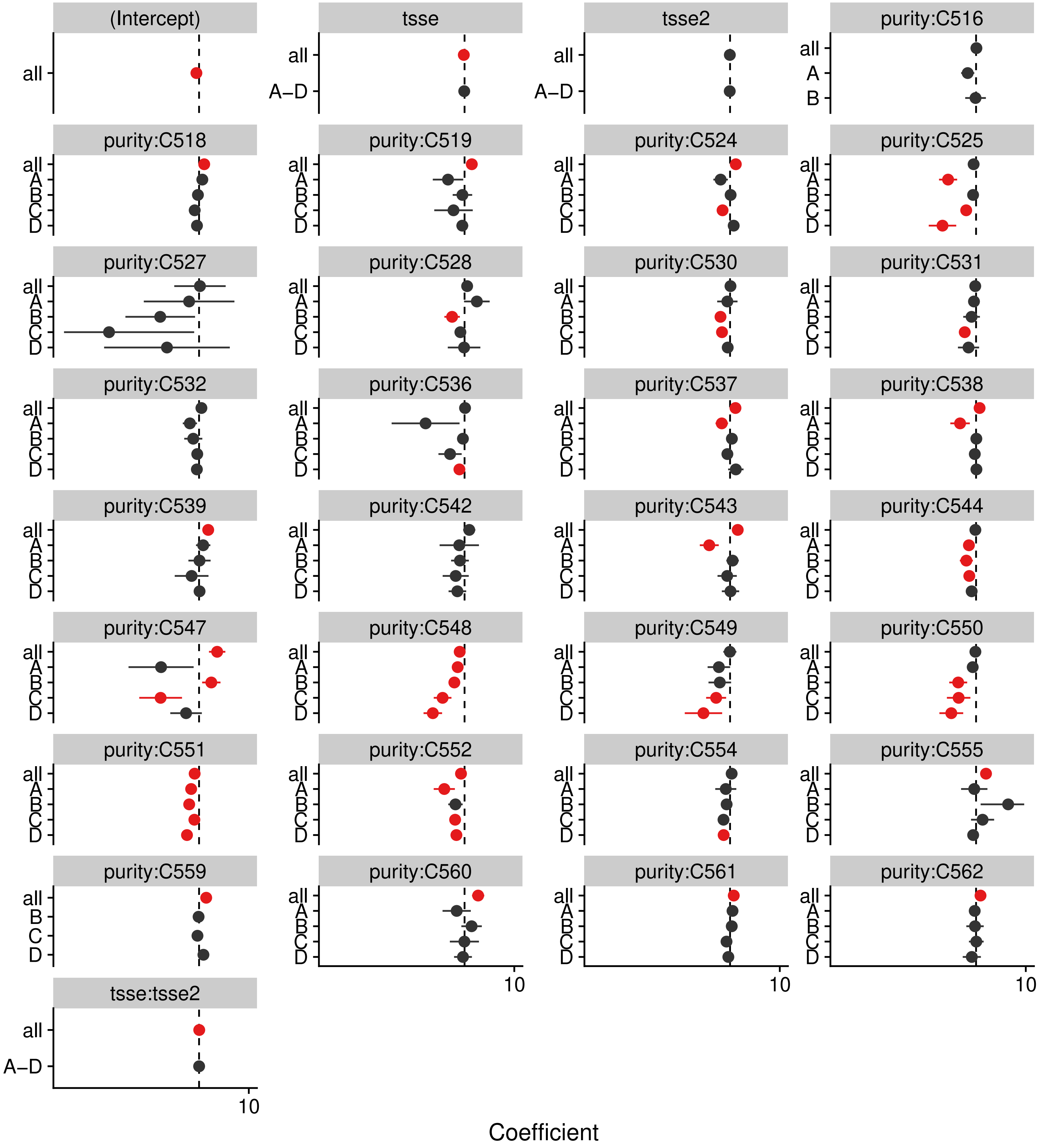


p_c = < 2.22e–16, p_sc = 0.00092284

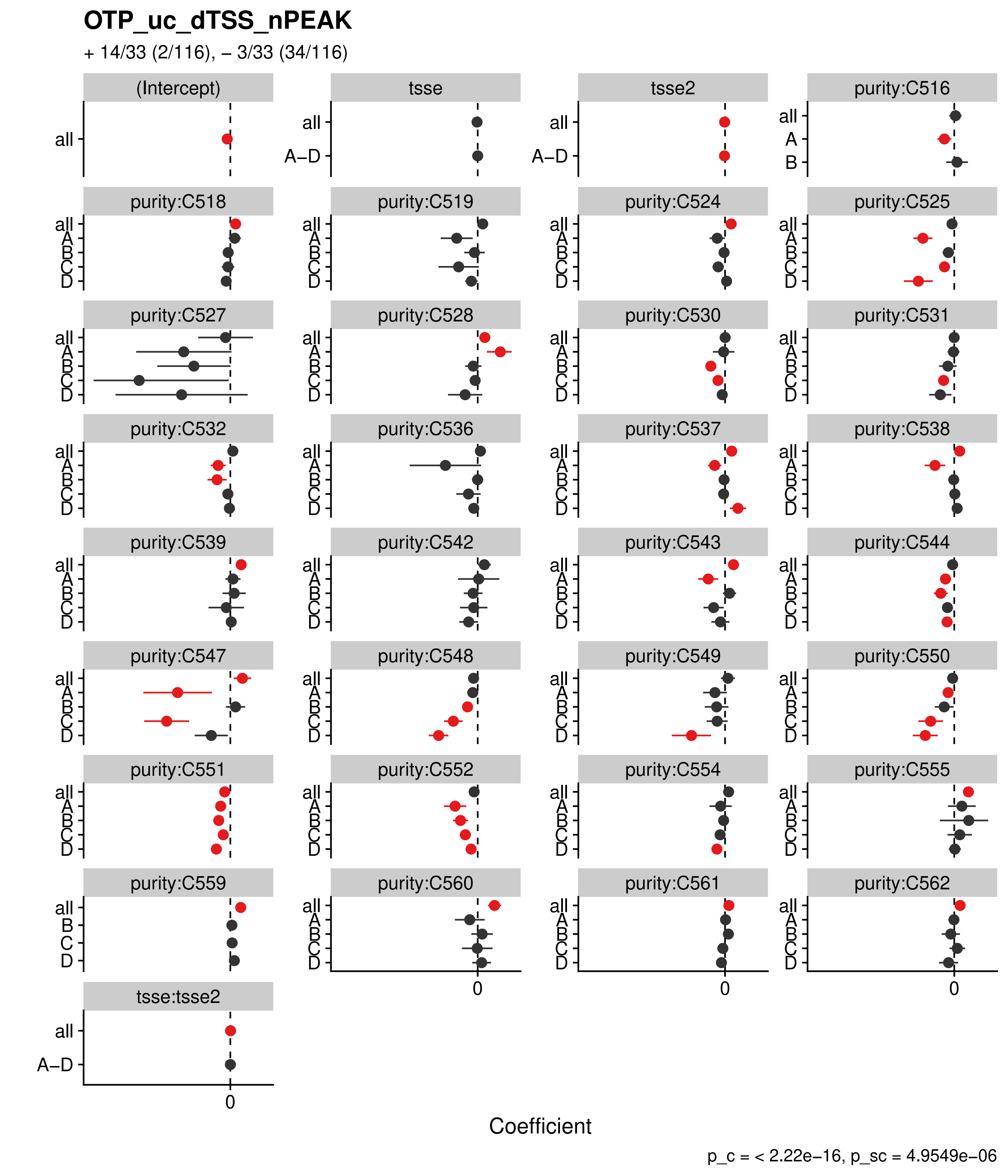

OTP_uc_dTSS_nPEAK
+ 14/33 (2/116), – 3/33 (34/116)
(Intercept)
tsse
tsse2
purity:C516
purity:C518
purity:C519
purity:C524
purity:C525
purity:C527
purity:C528
purity:C530
purity:C531
purity:C532
purity:C536
purity:C537
purity:C538
purity:C539
purity:C542
purity:C543
purity:C544
purity:C547
purity:C548
purity:C549
purity:C550
purity:C551
purity:C552
purity:C554
purity:C555
purity:C559
purity:C560
purity:C561
purity:C562
tsse:tsse2
all
A–D
A
B
C
D
0
Coefficient
p_c = < 2.22e–16, p_sc = 4.9549e–06

#### PBX3_uc_pTSS_oPEAK

+ 8/33 (58/116), – 6/33 (8/116)

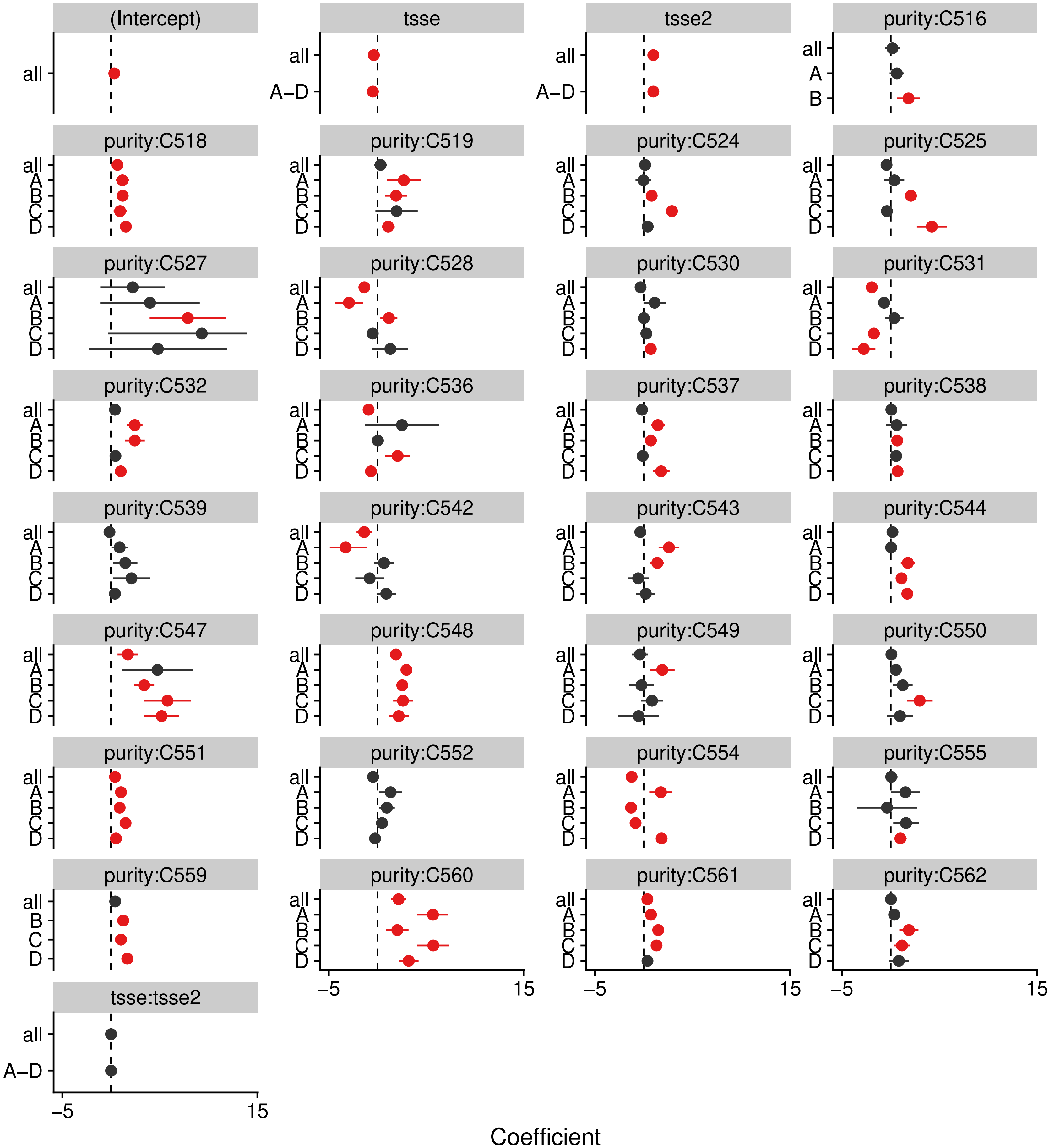


p_c = < 2.22e–16, p_sc = < 2.22e–16

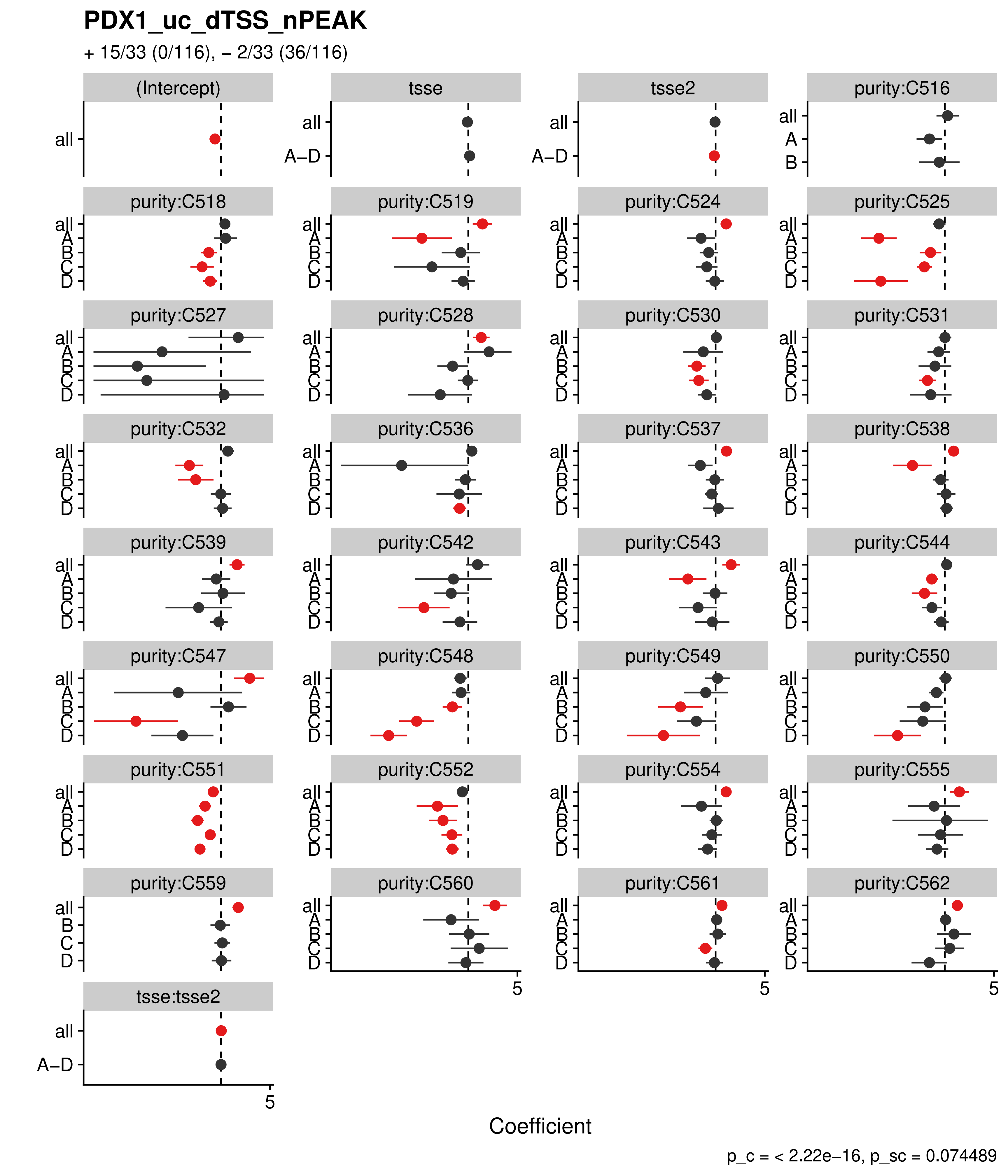

PDX1_uc_dTSS_nPEAK
+ 15/33 (0/116), – 2/33 (36/116)
(Intercept)
tsse
tsse2
purity:C516
purity:C518
purity:C519
purity:C524
purity:C525
purity:C527
purity:C528
purity:C530
purity:C531
purity:C532
purity:C536
purity:C537
purity:C538
purity:C539
purity:C542
purity:C543
purity:C544
purity:C547
purity:C548
purity:C549
purity:C550
purity:C551
purity:C552
purity:C554
purity:C555
purity:C559
purity:C560
purity:C561
purity:C562
tsse:tsse2
all
A–D
A
B
C
D
5
Coefficient
p_c = < 2.22e–16, p_sc = 0.074489

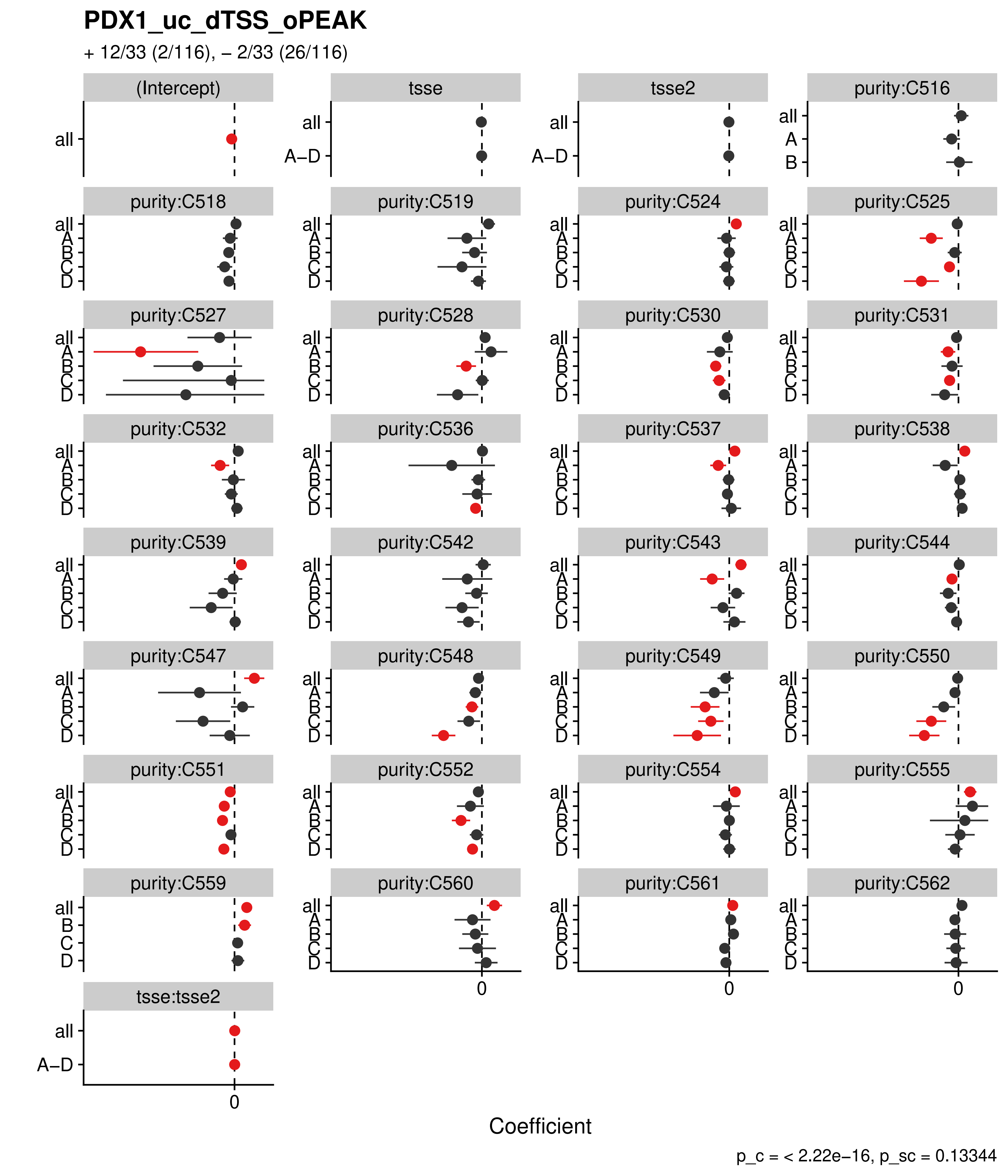
PDX1_uc_dTSS_oPEAK
+ 12/33 (2/116), – 2/33 (26/116)
(Intercept)
tsse
tsse2
purity:C516
purity:C518
purity:C519
purity:C524
purity:C525
purity:C527
purity:C528
purity:C530
purity:C531
purity:C532
purity:C536
purity:C537
purity:C538
purity:C539
purity:C542
purity:C543
purity:C544
purity:C547
purity:C548
purity:C549
purity:C550
purity:C551
purity:C552
purity:C554
purity:C555
purity:C559
purity:C560
purity:C561
purity:C562
tsse:tsse2
all
A–D
A
B
C
D
0
Coefficient
p_c = < 2.22e–16, p_sc = 0.13344

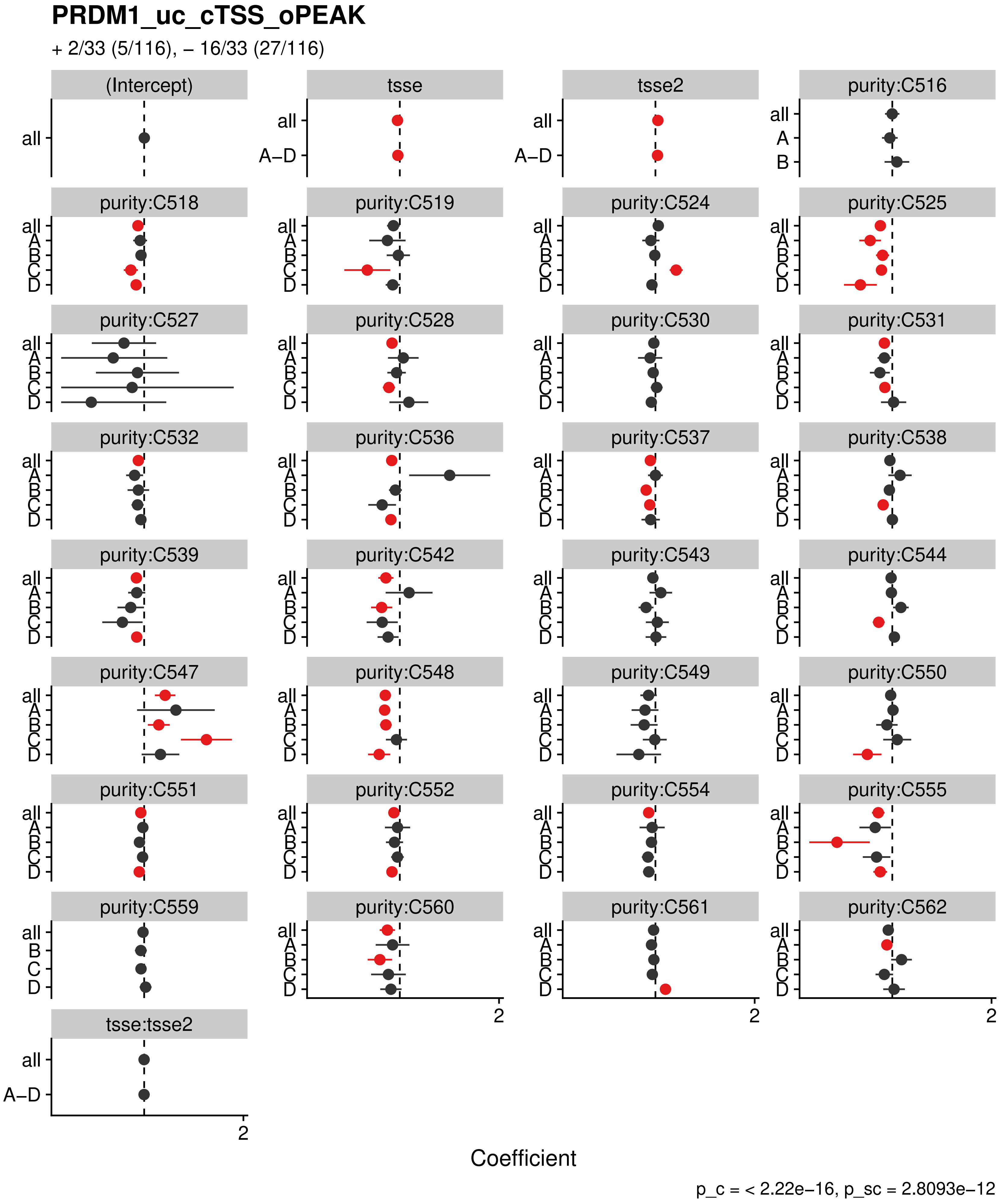

PRDM1_uc_cTSS_oPEAK
+ 2/33 (5/116), – 16/33 (27/116)
(Intercept)
tsse
tsse2
purity:C516
purity:C518
purity:C519
purity:C524
purity:C525
purity:C527
purity:C528
purity:C530
purity:C531
purity:C532
purity:C536
purity:C537
purity:C538
purity:C539
purity:C542
purity:C543
purity:C544
purity:C547
purity:C548
purity:C549
purity:C550
purity:C551
purity:C552
purity:C554
purity:C555
purity:C559
purity:C560
purity:C561
purity:C562
tsse:tsse2
all
A–D
A
B
C
D
2
Coefficient
p_c = < 2.22e–16, p_sc = 2.8093e–12

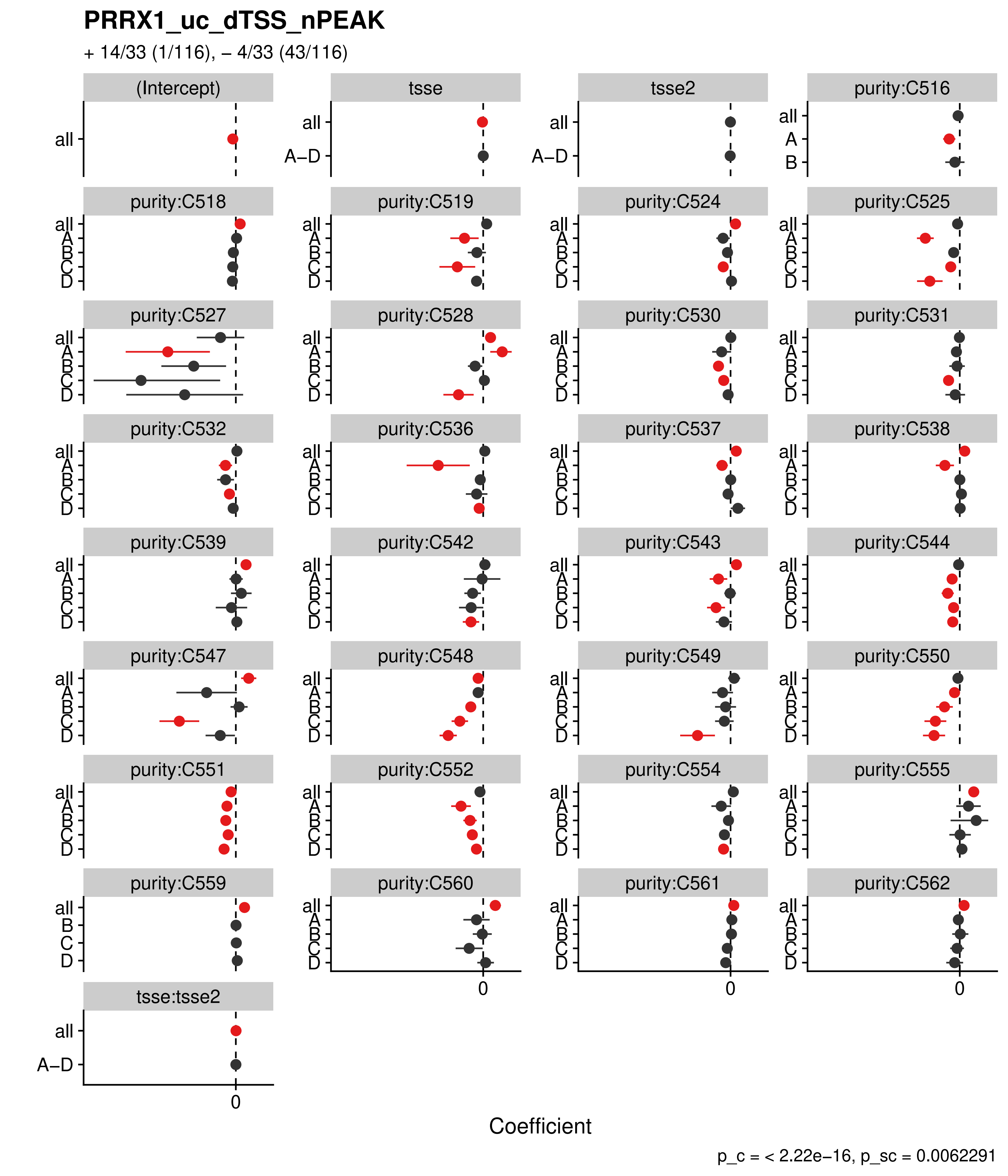
PRRX1_uc_dTSS_nPEAK
+ 14/33 (1/116), – 4/33 (43/116)
(Intercept)
tsse
tsse2
purity:C516
purity:C518
purity:C519
purity:C524
purity:C525
purity:C527
purity:C528
purity:C530
purity:C531
purity:C532
purity:C536
purity:C537
purity:C538
purity:C539
purity:C542
purity:C543
purity:C544
purity:C547
purity:C548
purity:C549
purity:C550
purity:C551
purity:C552
purity:C554
purity:C555
purity:C559
purity:C560
purity:C561
purity:C562
tsse:tsse2
all
A–D
A
B
C
D
0
Coefficient
p_c = < 2.22e–16, p_sc = 0.0062291

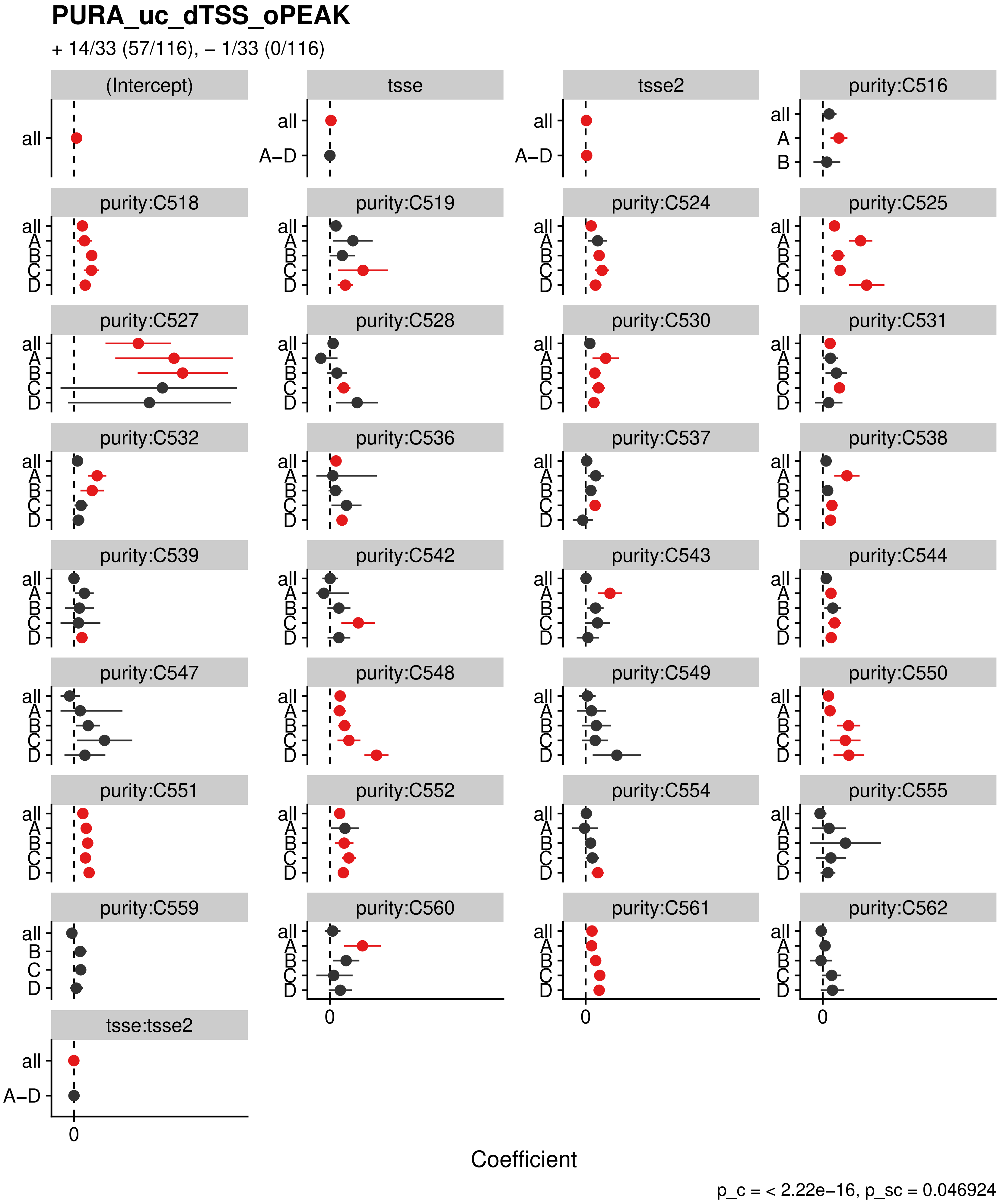
PURA_uc_dTSS_oPEAK
+ 14/33 (57/116), – 1/33 (0/116)
(Intercept)
tsse
tsse2
purity:C516
purity:C518
purity:C519
purity:C524
purity:C525
purity:C527
purity:C528
purity:C530
purity:C531
purity:C532
purity:C536
purity:C537
purity:C538
purity:C539
purity:C542
purity:C543
purity:C544
purity:C547
purity:C548
purity:C549
purity:C550
purity:C551
purity:C552
purity:C554
purity:C555
purity:C559
purity:C560
purity:C561
purity:C562
tsse:tsse2
all
A–D
A
B
C
D
0
Coefficient
p_c = < 2.22e–16, p_sc = 0.046924

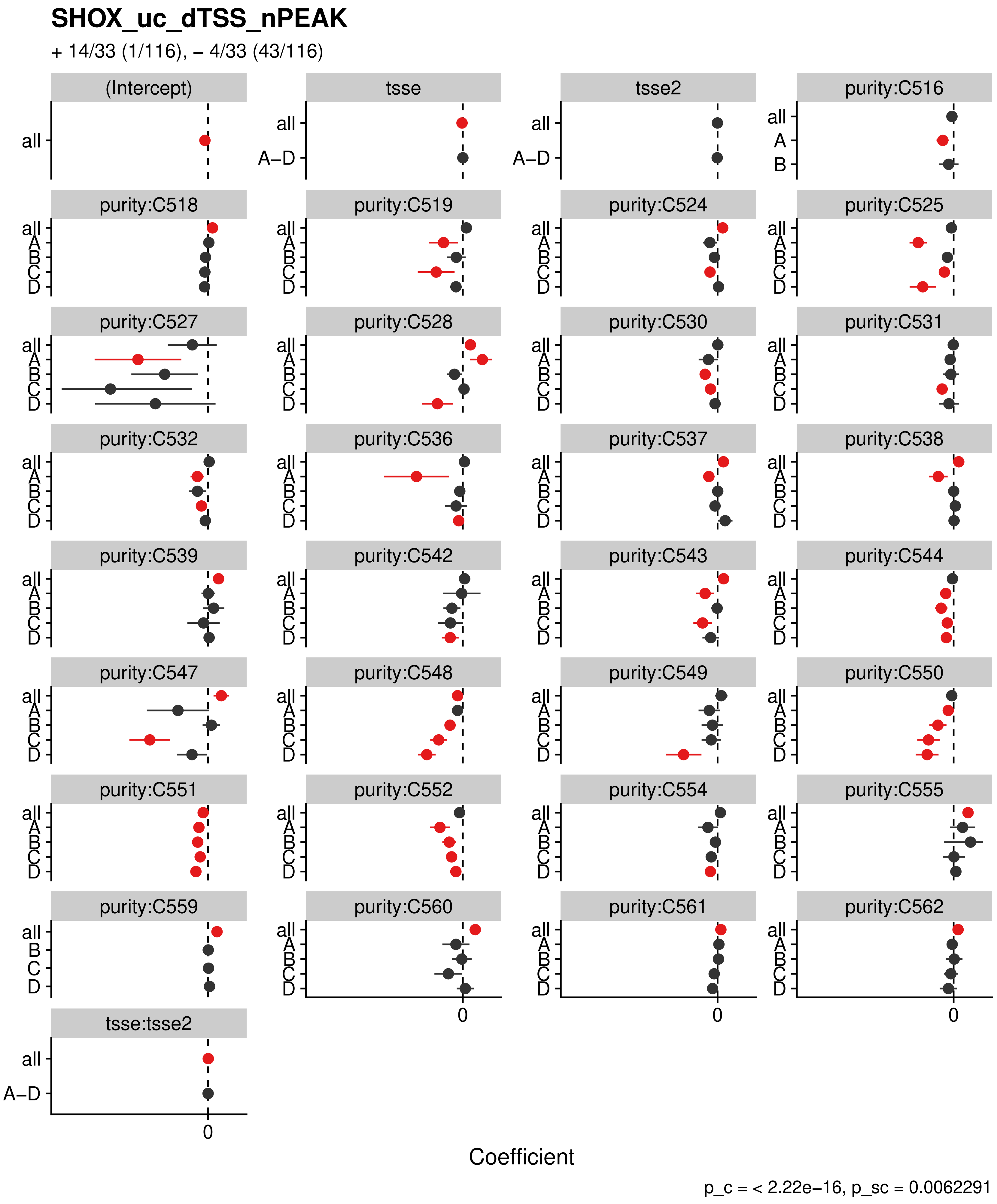

SHOX_uc_dTSS_nPEAK
+ 14/33 (1/116), – 4/33 (43/116)
(Intercept)
tsse
tsse2
purity:C516
purity:C518
purity:C519
purity:C524
purity:C525
purity:C527
purity:C528
purity:C530
purity:C531
purity:C532
purity:C536
purity:C537
purity:C538
purity:C539
purity:C542
purity:C543
purity:C544
purity:C547
purity:C548
purity:C549
purity:C550
purity:C551
purity:C552
purity:C554
purity:C555
purity:C559
purity:C560
purity:C561
purity:C562
tsse:tsse2
all
A–D
A
B
C
D
0
Coefficient
p_c = < 2.22e–16, p_sc = 0.0062291

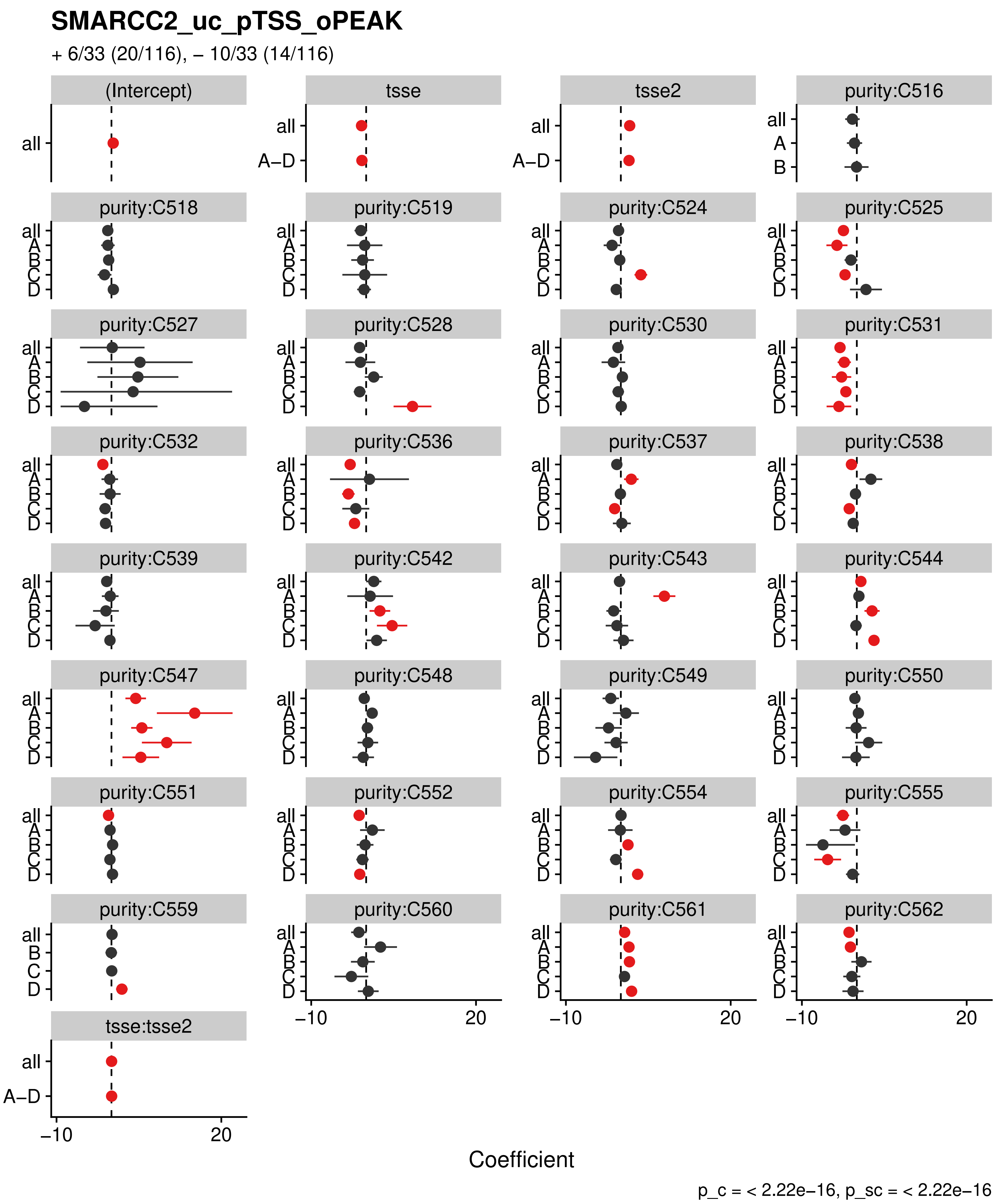

SMARCC2_uc_pTSS_oPEAK
+ 6/33 (20/116), – 10/33 (14/116)
(Intercept)
tsse
tsse2
purity:C516
purity:C518
purity:C519
purity:C524
purity:C525
purity:C527
purity:C528
purity:C530
purity:C531
purity:C532
purity:C536
purity:C537
purity:C538
purity:C539
purity:C542
purity:C543
purity:C544
purity:C547
purity:C548
purity:C549
purity:C550
purity:C551
purity:C552
purity:C554
purity:C555
purity:C559
purity:C560
purity:C561
purity:C562
tsse:tsse2
all
A–D
A
B
C
D
–10
20
Coefficient
p_c = < 2.22e–16, p_sc = < 2.22e–16

#### SNAI2_uc_cTSS_nPEAK

+ 5/33 (48/116), – 12/33 (2/116)

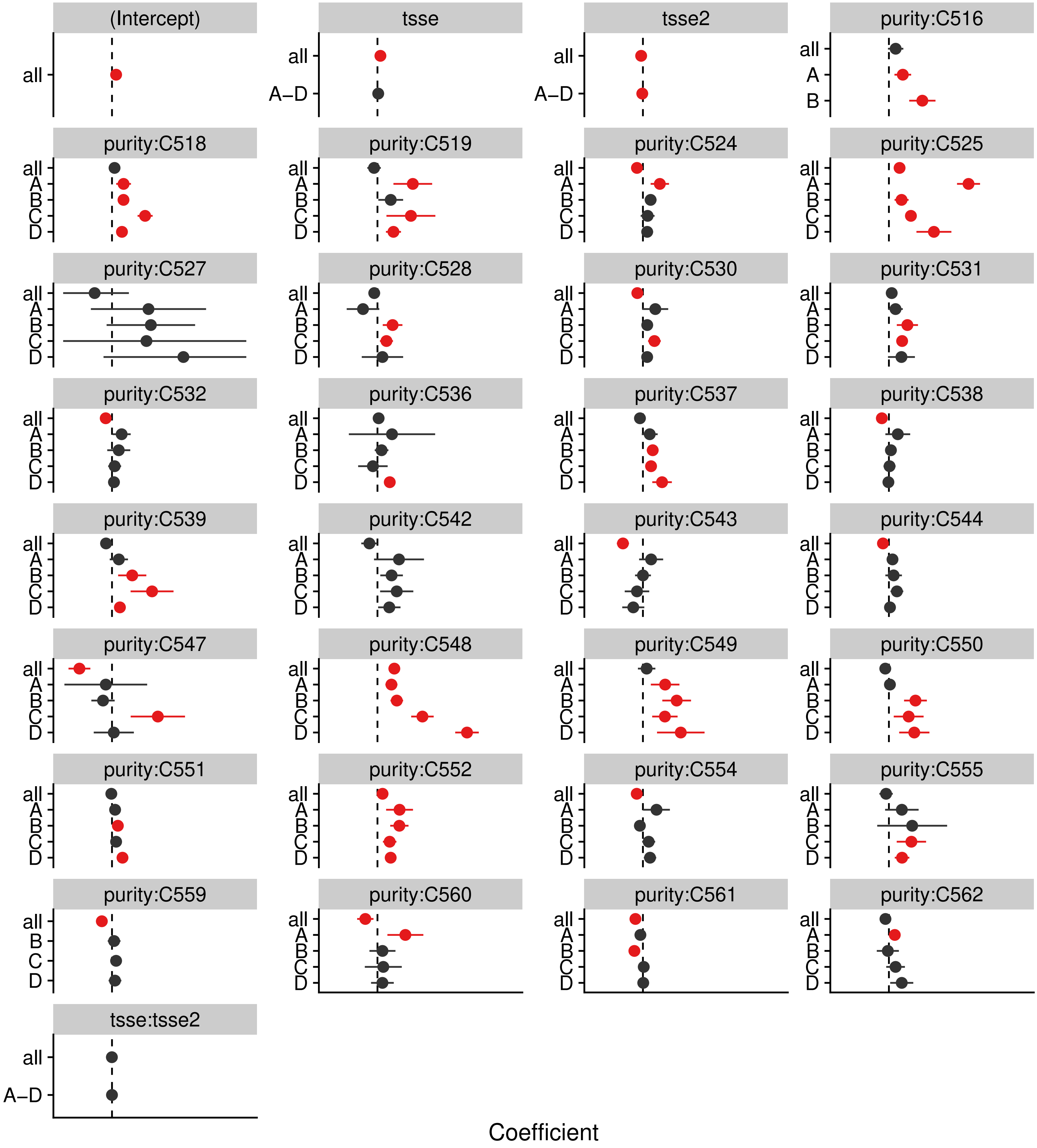


p_c = < 2.22e−16, p_sc = 3.045e−12

#### SNAI2_uc_dTSS_oPEAK

+ 6/33 (40/116), – 8/33 (2/116)

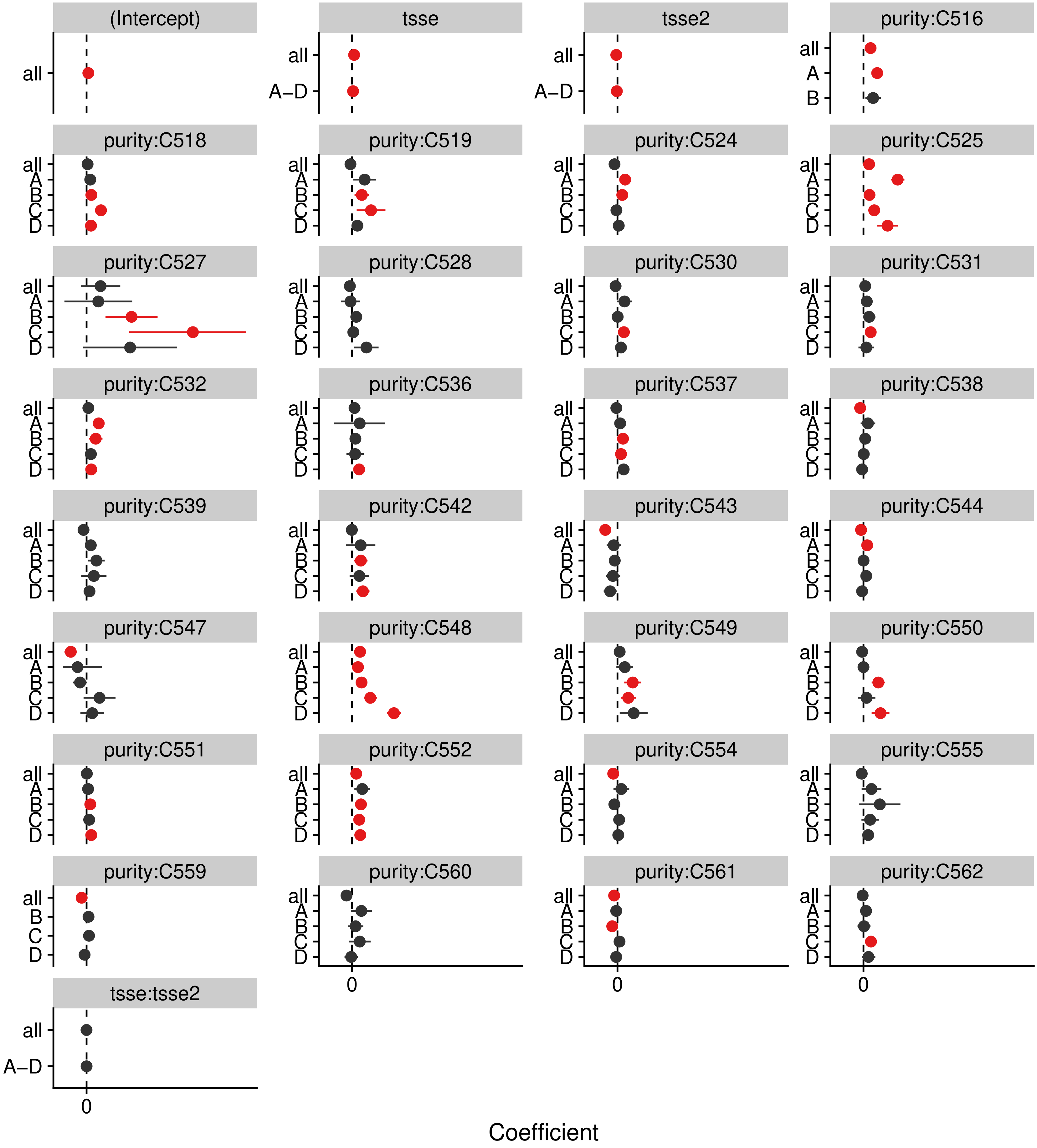


p_c = < 2.22e–16, p_sc = 4.9881e–08

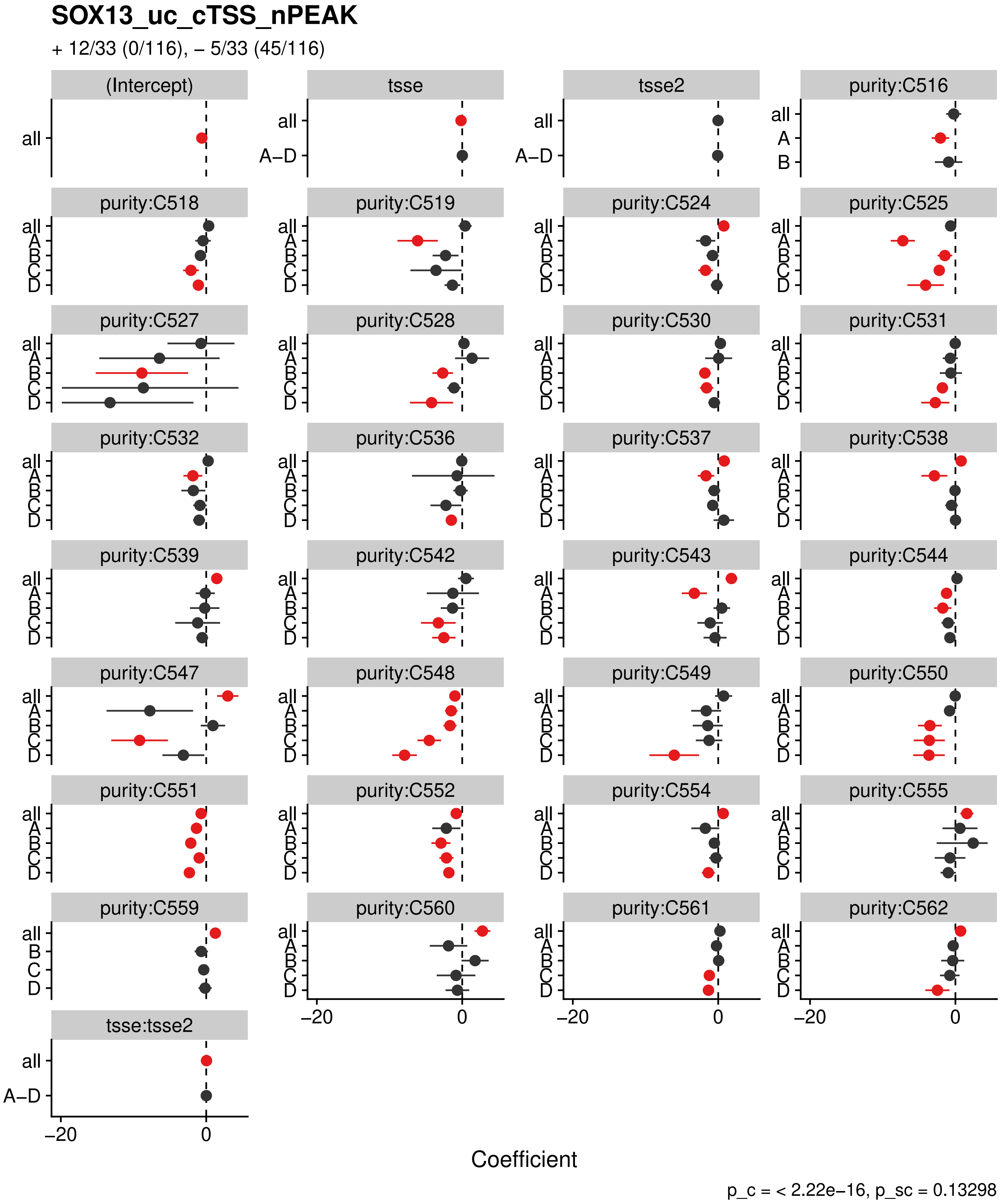

SOX13_uc_cTSS_nPEAK
+ 12/33 (0/116), – 5/33 (45/116)
(Intercept)
tsse
tsse2
purity:C516
purity:C518
purity:C519
purity:C524
purity:C525
purity:C527
purity:C528
purity:C530
purity:C531
purity:C532
purity:C536
purity:C537
purity:C538
purity:C539
purity:C542
purity:C543
purity:C544
purity:C547
purity:C548
purity:C549
purity:C550
purity:C551
purity:C552
purity:C554
purity:C555
purity:C559
purity:C560
purity:C561
purity:C562
tsse:tsse2
all
A–D
A
B
C
D
–20
0
Coefficient
p_c = < 2.22e–16, p_sc = 0.13298

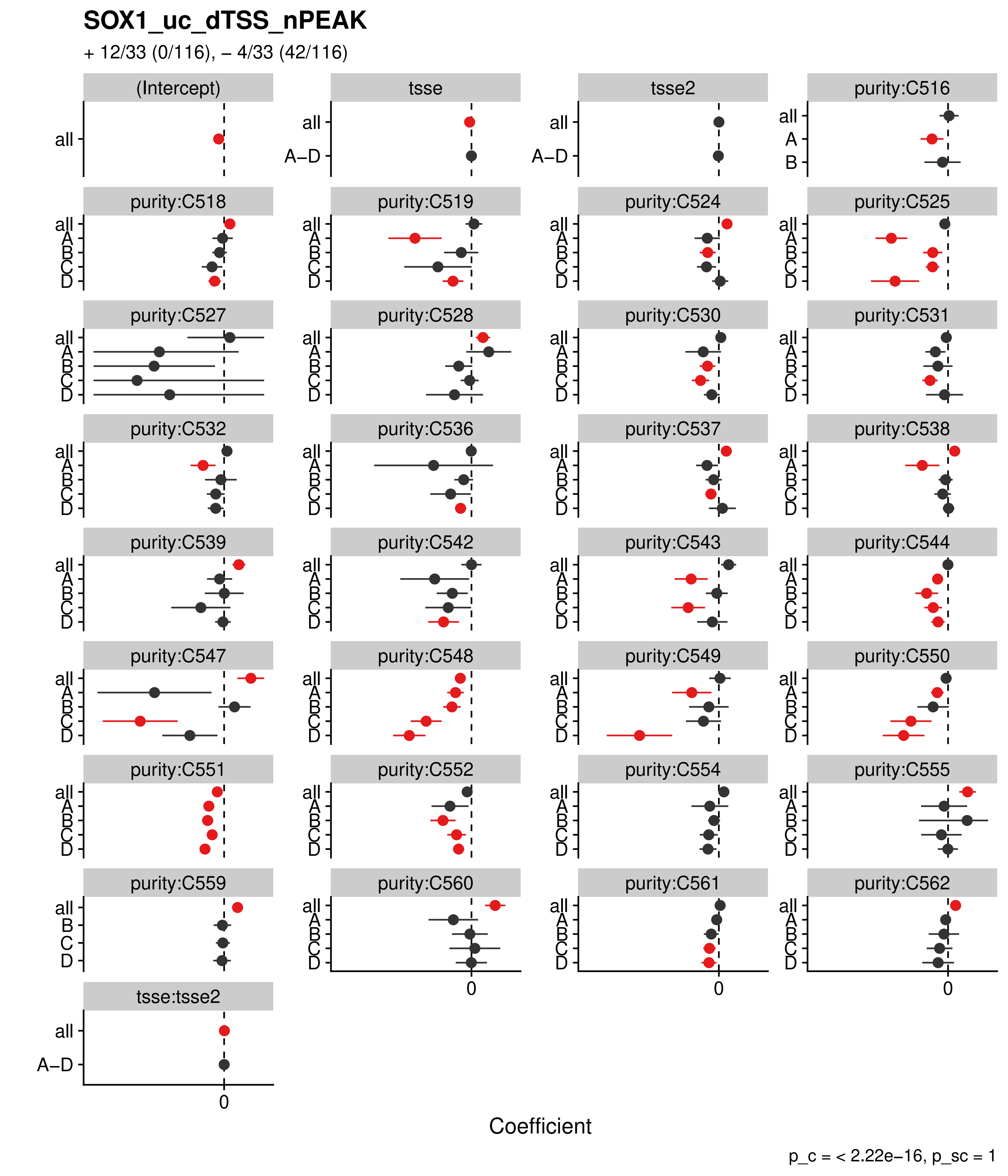
SOX1_uc_dTSS_nPEAK
+ 12/33 (0/116), – 4/33 (42/116)
(Intercept)
tsse
tsse2
purity:C516
purity:C518
purity:C519
purity:C524
purity:C525
purity:C527
purity:C528
purity:C530
purity:C531
purity:C532
purity:C536
purity:C537
purity:C538
purity:C539
purity:C542
purity:C543
purity:C544
purity:C547
purity:C548
purity:C549
purity:C550
purity:C551
purity:C552
purity:C554
purity:C555
purity:C559
purity:C560
purity:C561
purity:C562
tsse:tsse2
all
A–D
A
B
C
D
0
Coefficient
p_c = < 2.22e–16, p_sc = 1

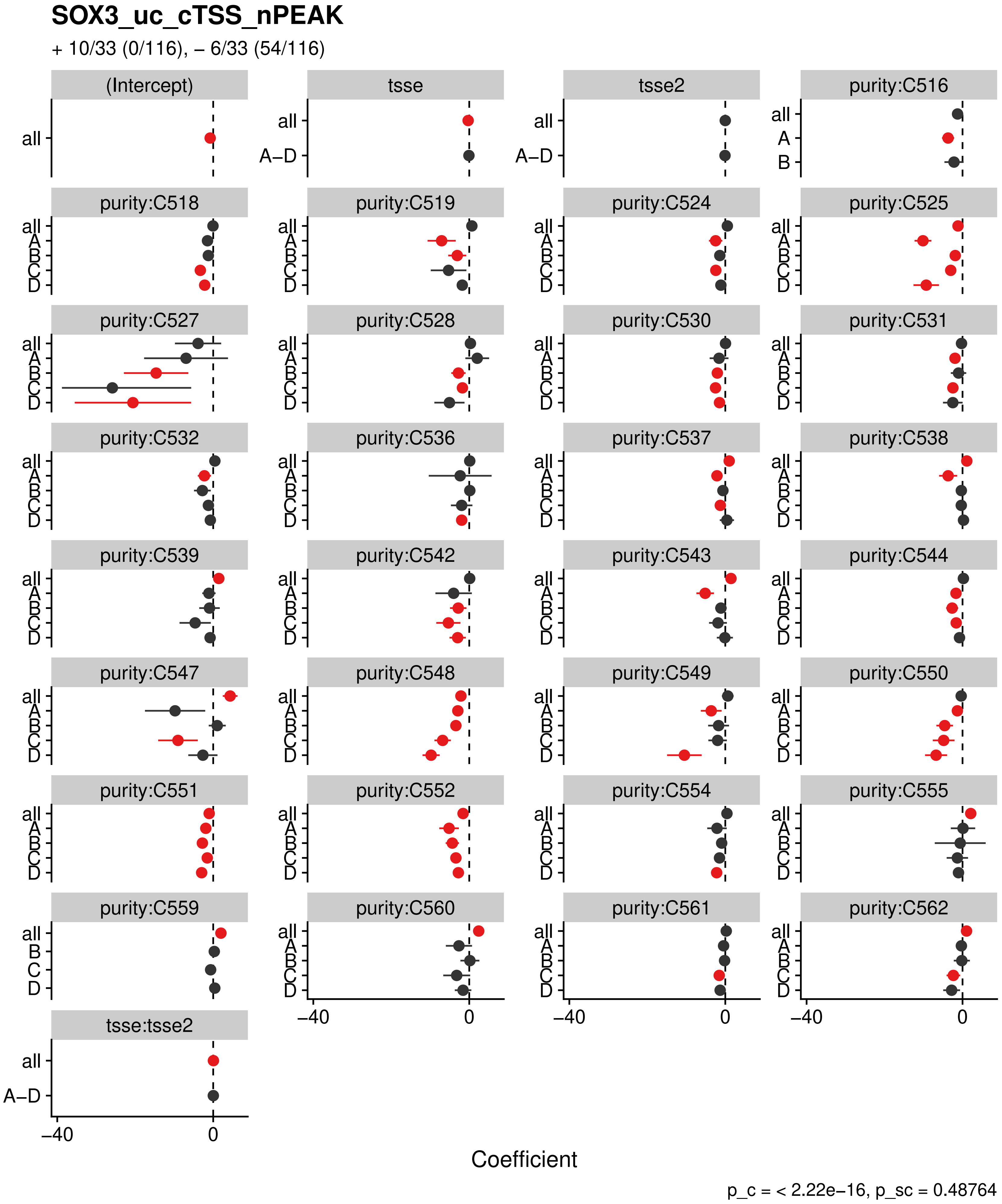

SOX3_uc_cTSS_nPEAK
+ 10/33 (0/116), – 6/33 (54/116)
(Intercept)
tsse
tsse2
purity:C516
purity:C518
purity:C519
purity:C524
purity:C525
purity:C527
purity:C528
purity:C530
purity:C531
purity:C532
purity:C536
purity:C537
purity:C538
purity:C539
purity:C542
purity:C543
purity:C544
purity:C547
purity:C548
purity:C549
purity:C550
purity:C551
purity:C552
purity:C554
purity:C555
purity:C559
purity:C560
purity:C561
purity:C562
tsse:tsse2
all
A–D
A
B
C
D
–40
0
Coefficient
p_c = < 2.22e–16, p_sc = 0.48764

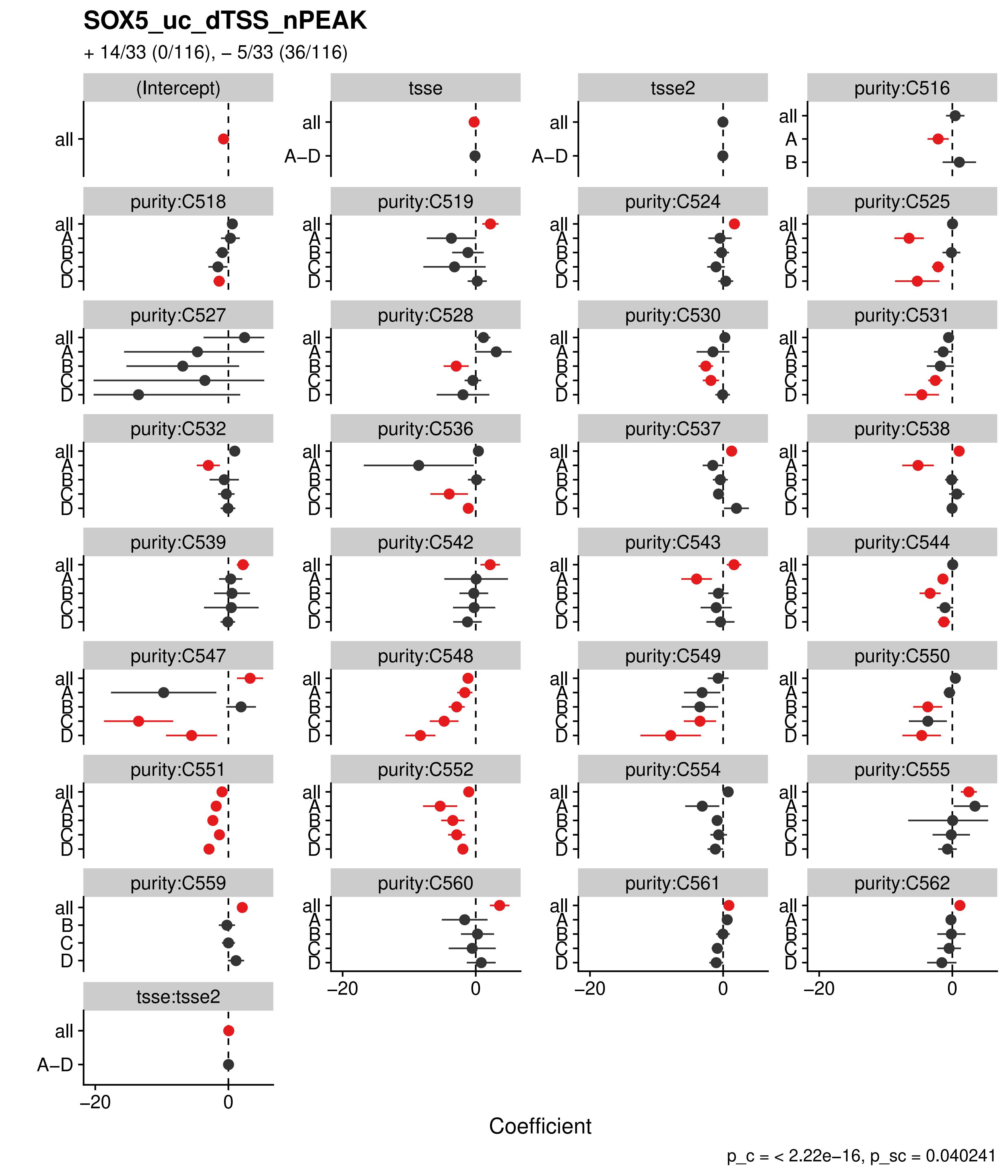
SOX5_uc_dTSS_nPEAK
+ 14/33 (0/116), – 5/33 (36/116)
(Intercept)
tsse
tsse2
purity:C516
purity:C518
purity:C519
purity:C524
purity:C525
purity:C527
purity:C528
purity:C530
purity:C531
purity:C532
purity:C536
purity:C537
purity:C538
purity:C539
purity:C542
purity:C543
purity:C544
purity:C547
purity:C548
purity:C549
purity:C550
purity:C551
purity:C552
purity:C554
purity:C555
purity:C559
purity:C560
purity:C561
purity:C562
tsse:tsse2
all
A–D
A
B
C
D
–20
0
Coefficient
p_c = < 2.22e–16, p_sc = 0.040241

#### SOX6_uc_dTSS_nPEAK

+ 14/33 (0/116), – 5/33 (36/116)

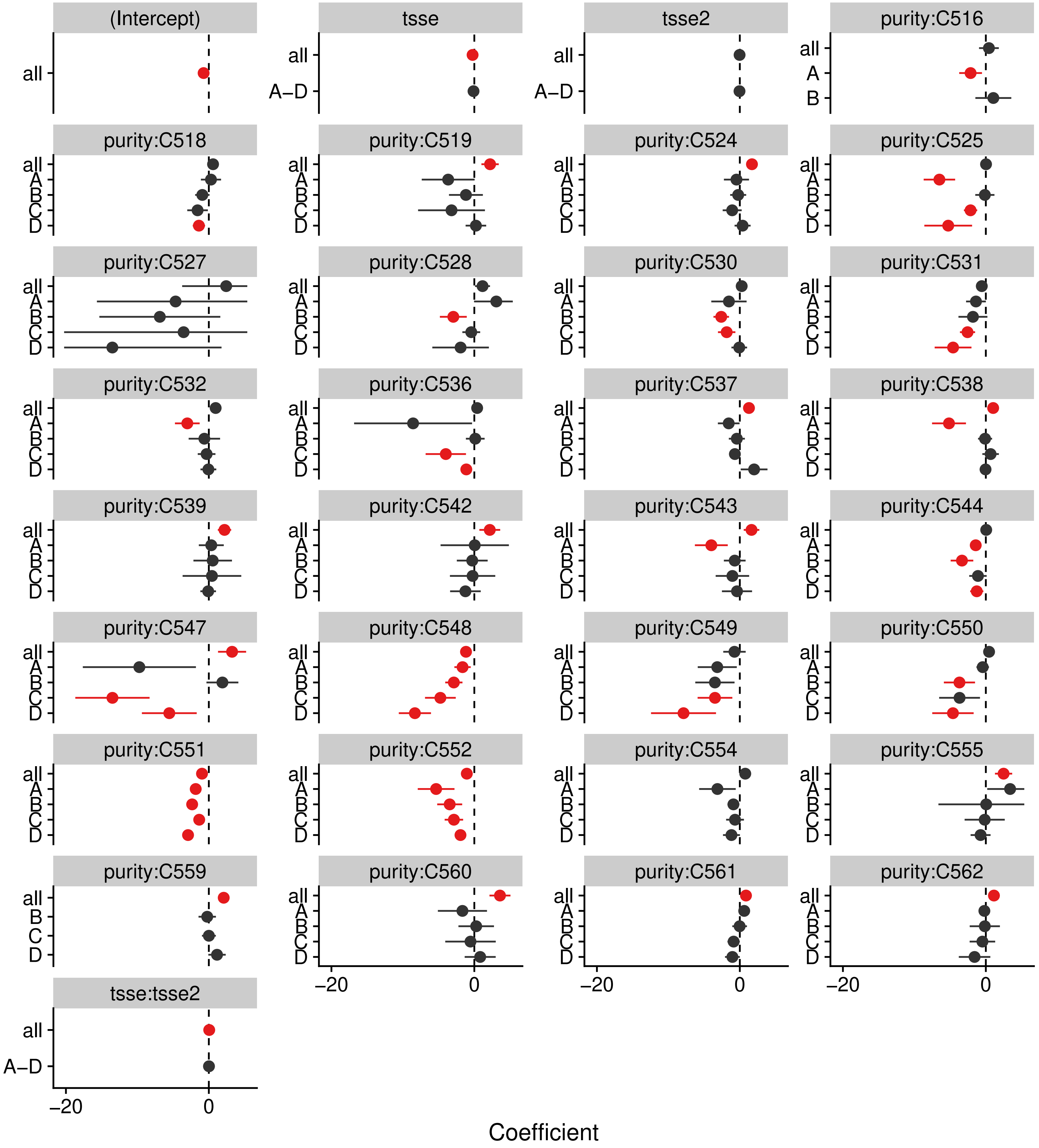


p_c = < 2.22e–16, p_sc = 0.040241

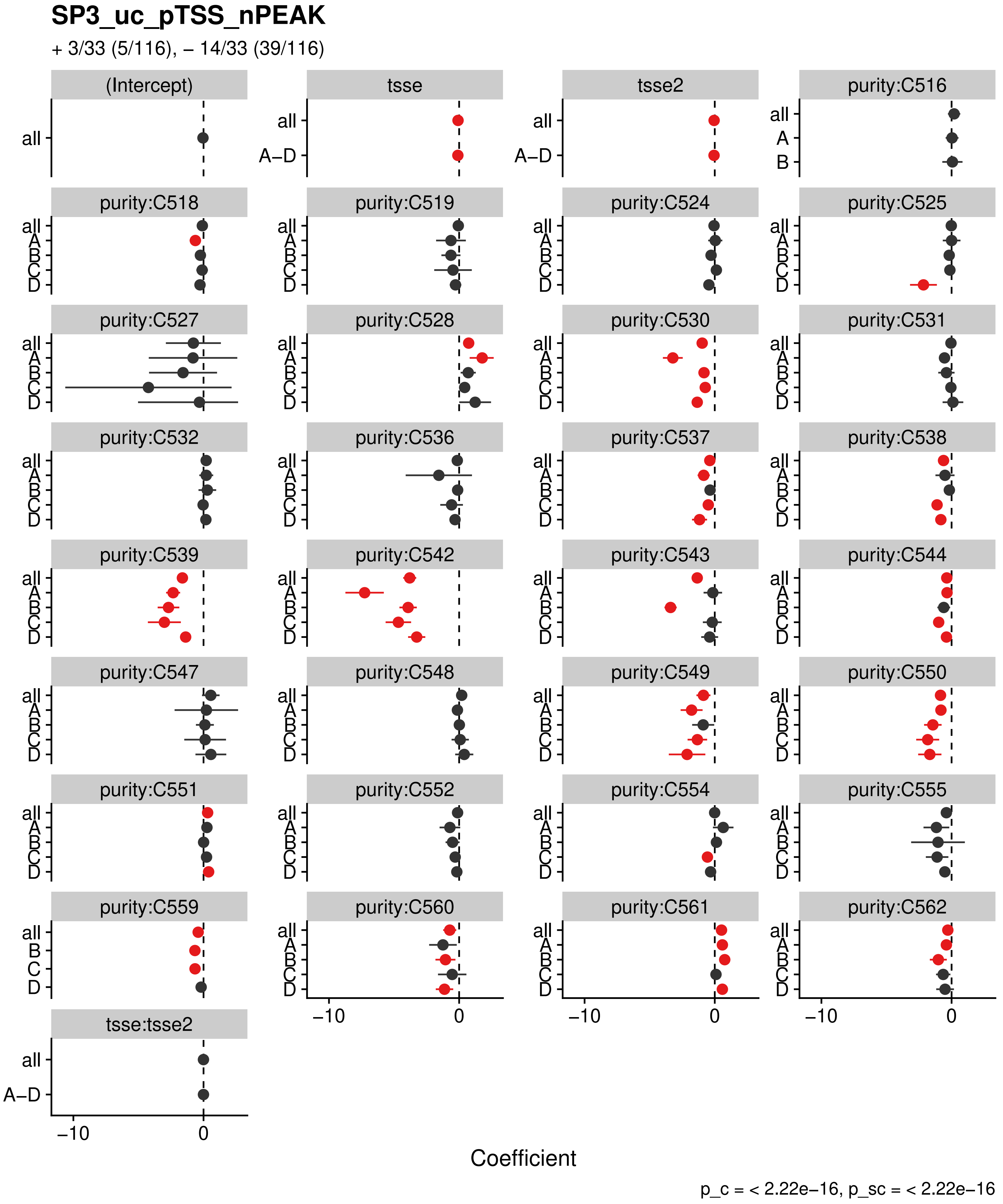
SP3_uc_pTSS_nPEAK
+ 3/33 (5/116), – 14/33 (39/116)
(Intercept)
tsse
tsse2
purity:C516
purity:C518
purity:C519
purity:C524
purity:C525
purity:C527
purity:C528
purity:C530
purity:C531
purity:C532
purity:C536
purity:C537
purity:C538
purity:C539
purity:C542
purity:C543
purity:C544
purity:C547
purity:C548
purity:C549
purity:C550
purity:C551
purity:C552
purity:C554
purity:C555
purity:C559
purity:C560
purity:C561
purity:C562
tsse:tsse2
all
A–D
A
B
C
D
–10
0
Coefficient
p_c = < 2.22e–16, p_sc = < 2.22e–16

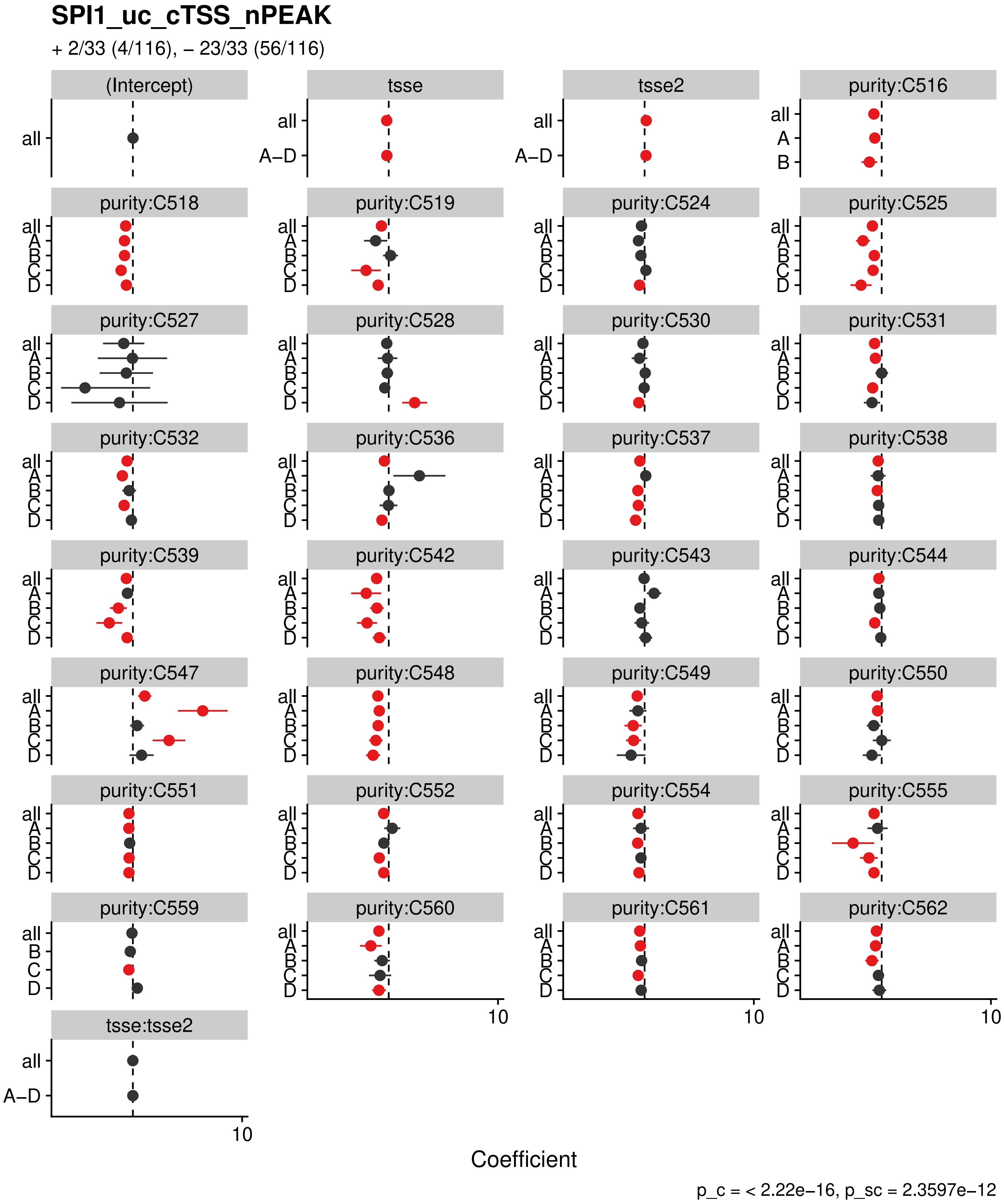
SPI1_uc_cTSS_nPEAK
+ 2/33 (4/116), – 23/33 (56/116)
(Intercept)
tsse
tsse2
purity:C516
purity:C518
purity:C519
purity:C524
purity:C525
purity:C527
purity:C528
purity:C530
purity:C531
purity:C532
purity:C536
purity:C537
purity:C538
purity:C539
purity:C542
purity:C543
purity:C544
purity:C547
purity:C548
purity:C549
purity:C550
purity:C551
purity:C552
purity:C554
purity:C555
purity:C559
purity:C560
purity:C561
purity:C562
tsse:tsse2
all
A–D
A
B
C
D
10
Coefficient
p_c = < 2.22e–16, p_sc = 2.3597e–12

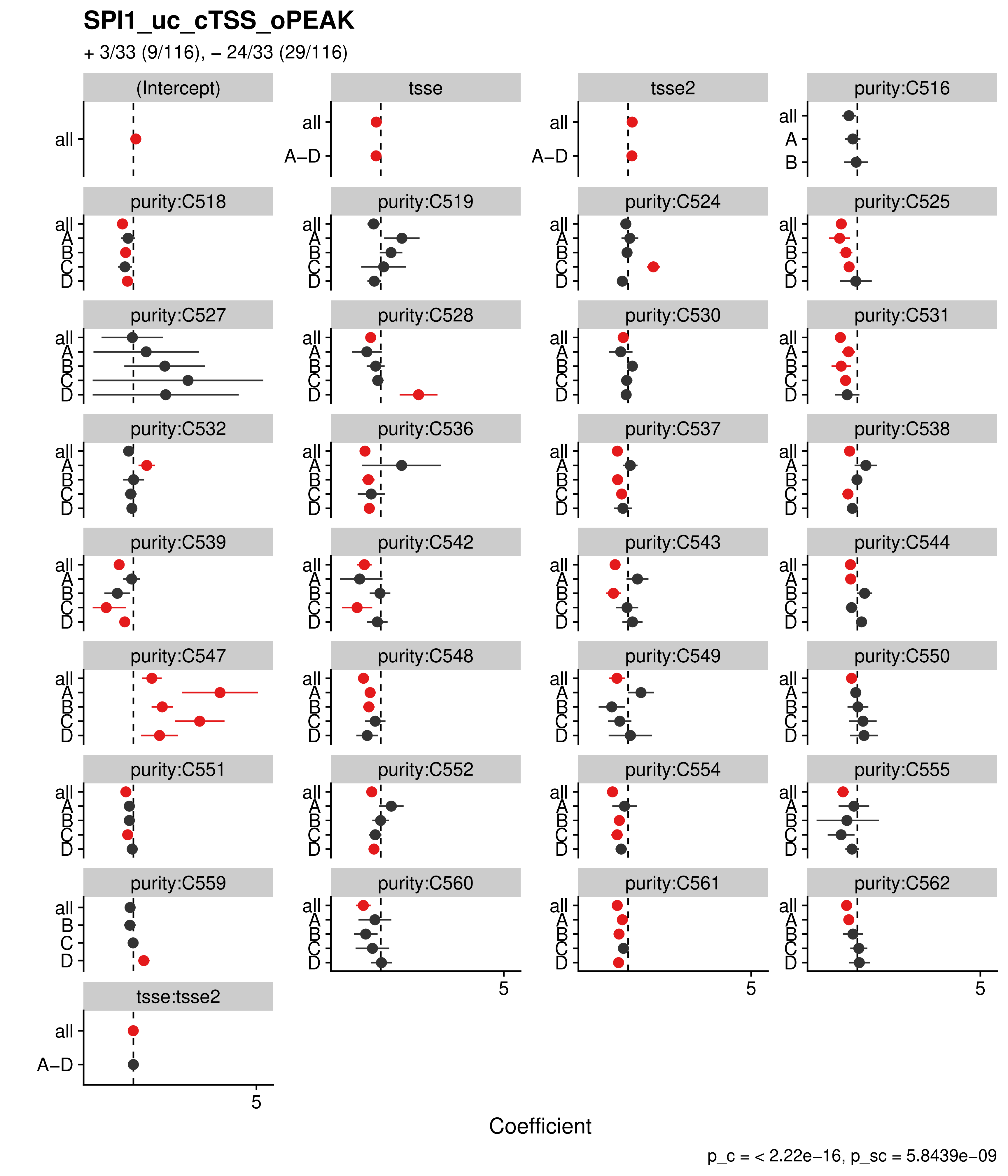
SPI1_uc_cTSS_oPEAK
+ 3/33 (9/116), – 24/33 (29/116)
(Intercept)
tsse
tsse2
purity:C516
purity:C518
purity:C519
purity:C524
purity:C525
purity:C527
purity:C528
purity:C530
purity:C531
purity:C532
purity:C536
purity:C537
purity:C538
purity:C539
purity:C542
purity:C543
purity:C544
purity:C547
purity:C548
purity:C549
purity:C550
purity:C551
purity:C552
purity:C554
purity:C555
purity:C559
purity:C560
purity:C561
purity:C562
tsse:tsse2
all
A–D
A
B
C
D
5
Coefficient
p_c = < 2.22e–16, p_sc = 5.8439e–09

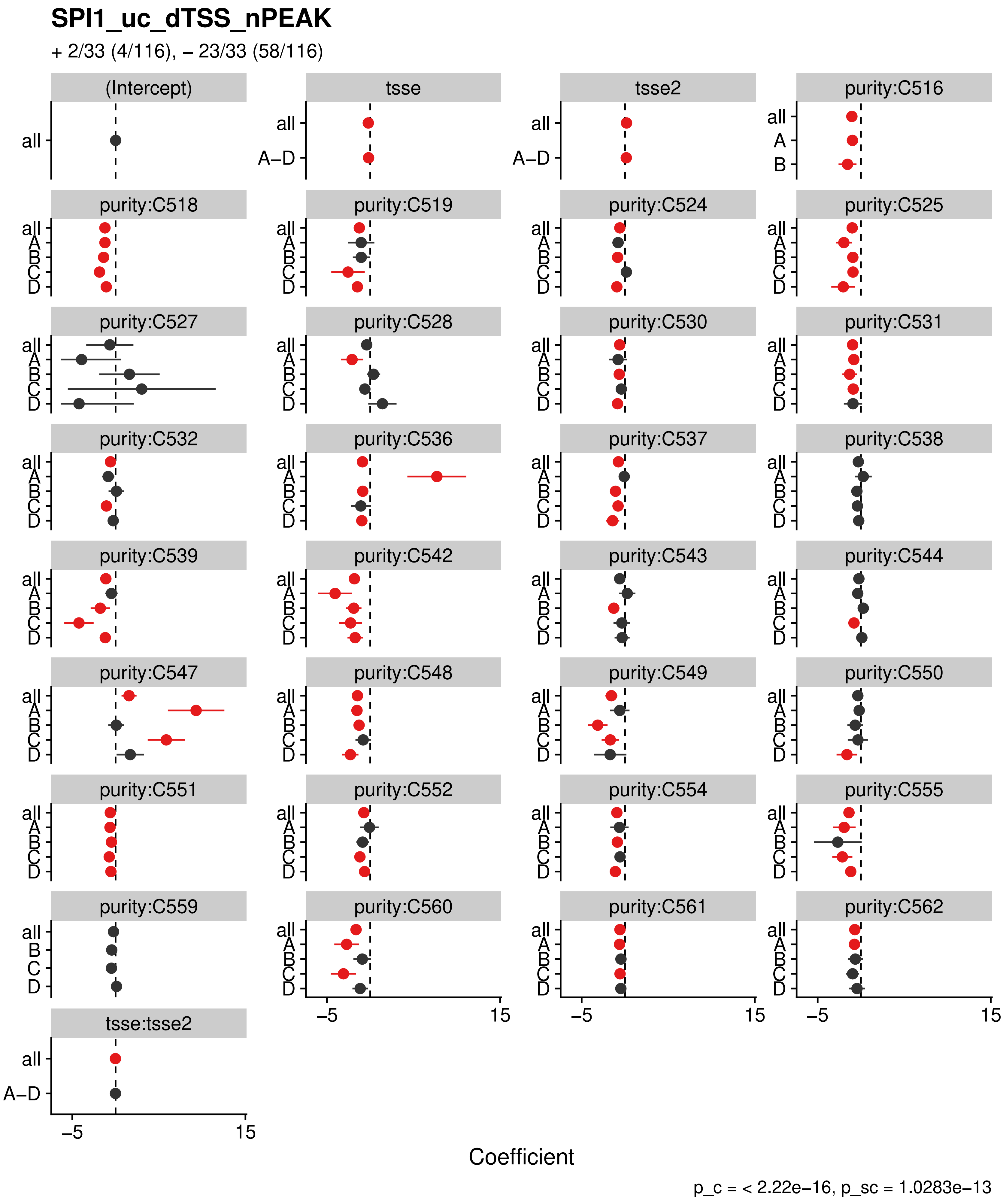
SPI1_uc_dTSS_nPEAK
+ 2/33 (4/116), – 23/33 (58/116)
(Intercept)
tsse
tsse2
purity:C516
purity:C518
purity:C519
purity:C524
purity:C525
purity:C527
purity:C528
purity:C530
purity:C531
purity:C532
purity:C536
purity:C537
purity:C538
purity:C539
purity:C542
purity:C543
purity:C544
purity:C547
purity:C548
purity:C549
purity:C550
purity:C551
purity:C552
purity:C554
purity:C555
purity:C559
purity:C560
purity:C561
purity:C562
tsse:tsse2
all
A–D
A
B
C
D
–5
15
Coefficient
p_c = < 2.22e–16, p_sc = 1.0283e–13

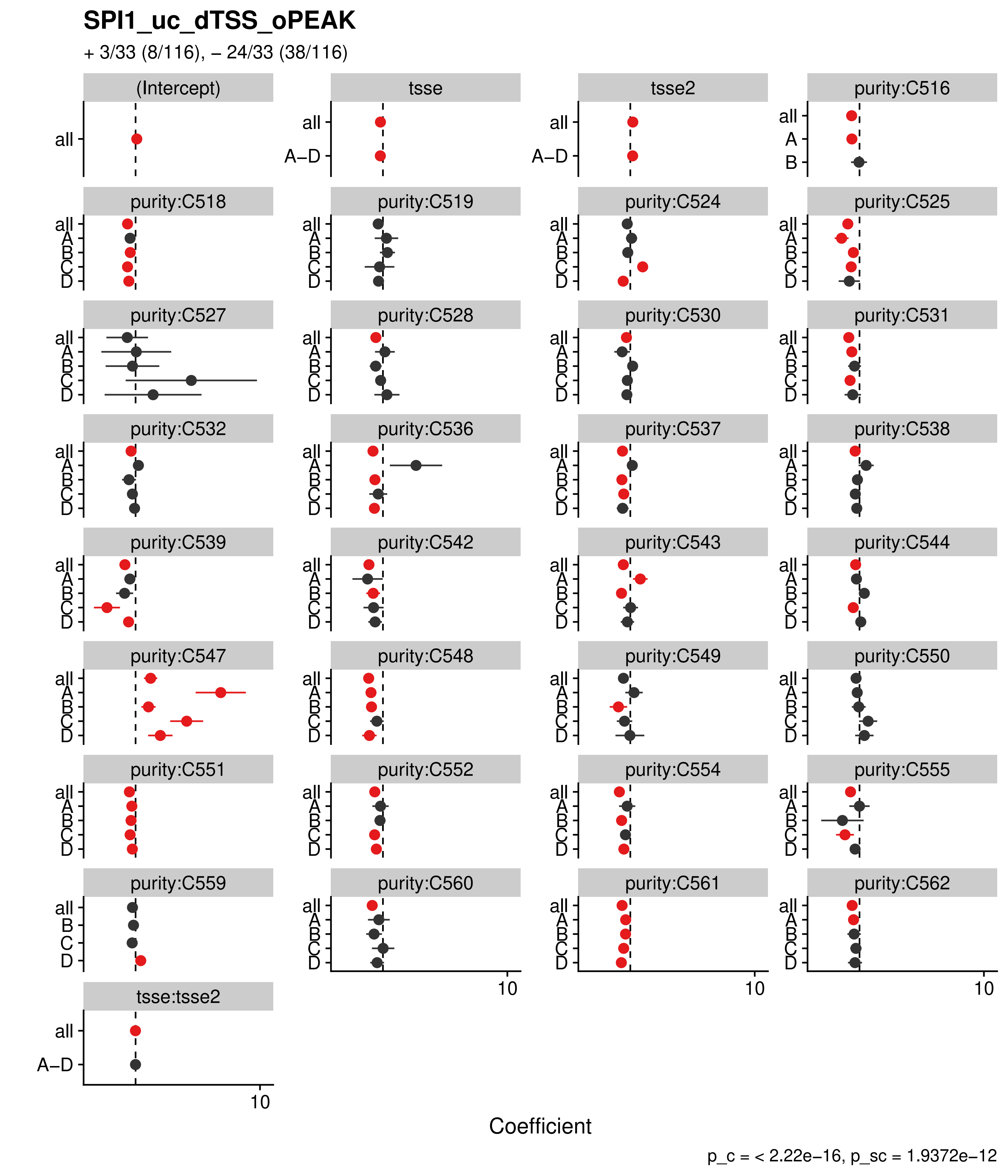
SPI1_uc_dTSS_oPEAK
+ 3/33 (8/116), – 24/33 (38/116)
(Intercept)
tsse
tsse2
purity:C516
purity:C518
purity:C519
purity:C524
purity:C525
purity:C527
purity:C528
purity:C530
purity:C531
purity:C532
purity:C536
purity:C537
purity:C538
purity:C539
purity:C542
purity:C543
purity:C544
purity:C547
purity:C548
purity:C549
purity:C550
purity:C551
purity:C552
purity:C554
purity:C555
purity:C559
purity:C560
purity:C561
purity:C562
tsse:tsse2
all
A–D
A
B
C
D
10
Coefficient
p_c = < 2.22e–16, p_sc = 1.9372e–12

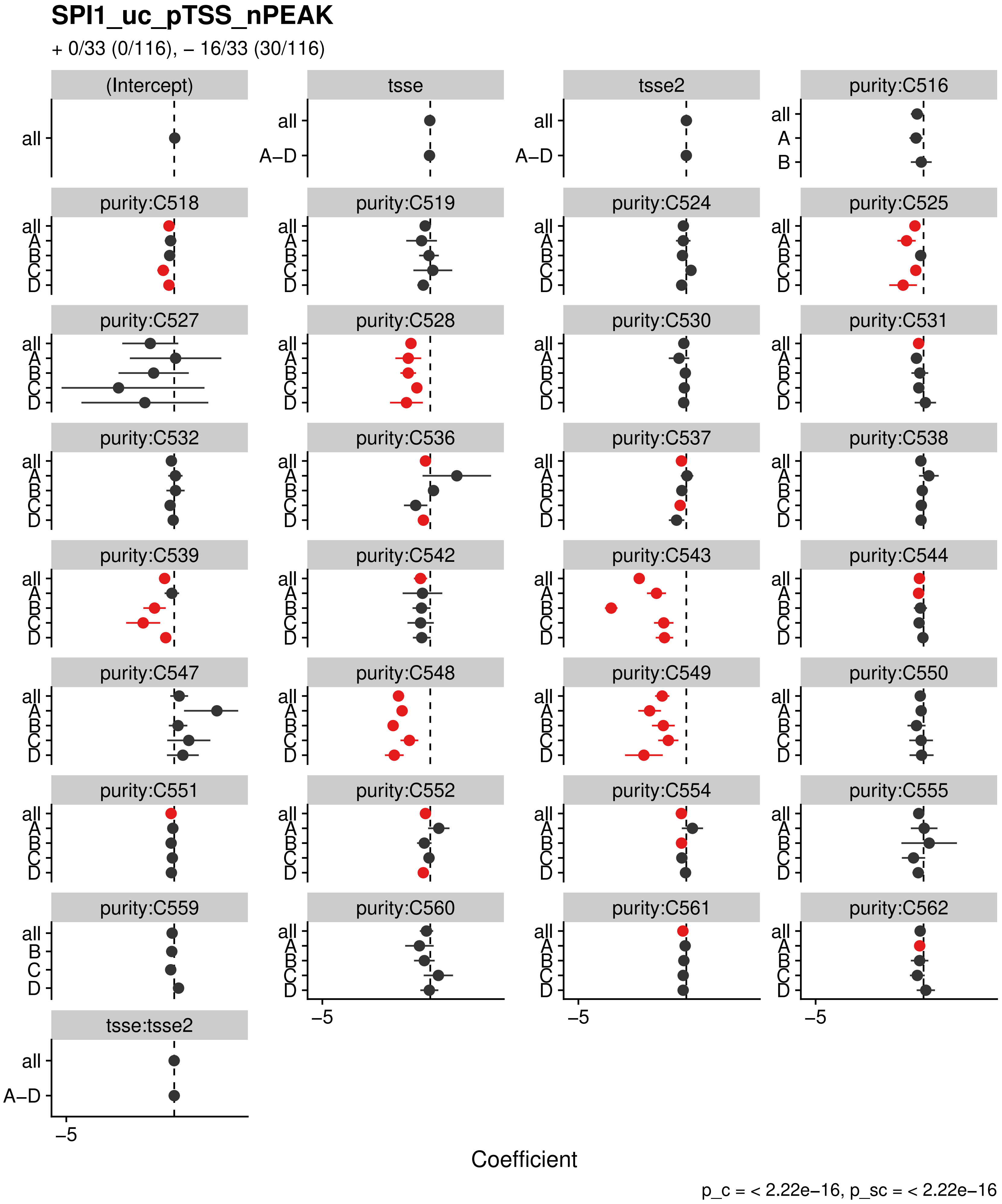
SPI1_uc_pTSS_nPEAK
+ 0/33 (0/116), – 16/33 (30/116)
(Intercept)
tsse
tsse2
tsse:tsse2
purity:C516
purity:C518
purity:C519
purity:C524
purity:C525
purity:C527
purity:C528
purity:C530
purity:C531
purity:C532
purity:C536
purity:C537
purity:C538
purity:C539
purity:C542
purity:C543
purity:C544
purity:C547
purity:C548
purity:C549
purity:C550
purity:C551
purity:C552
purity:C554
purity:C555
purity:C559
purity:C560
purity:C561
purity:C562
all
A–D
A
B
C
D
–5
Coefficient
p_c = < 2.22e–16, p_sc = < 2.22e–16

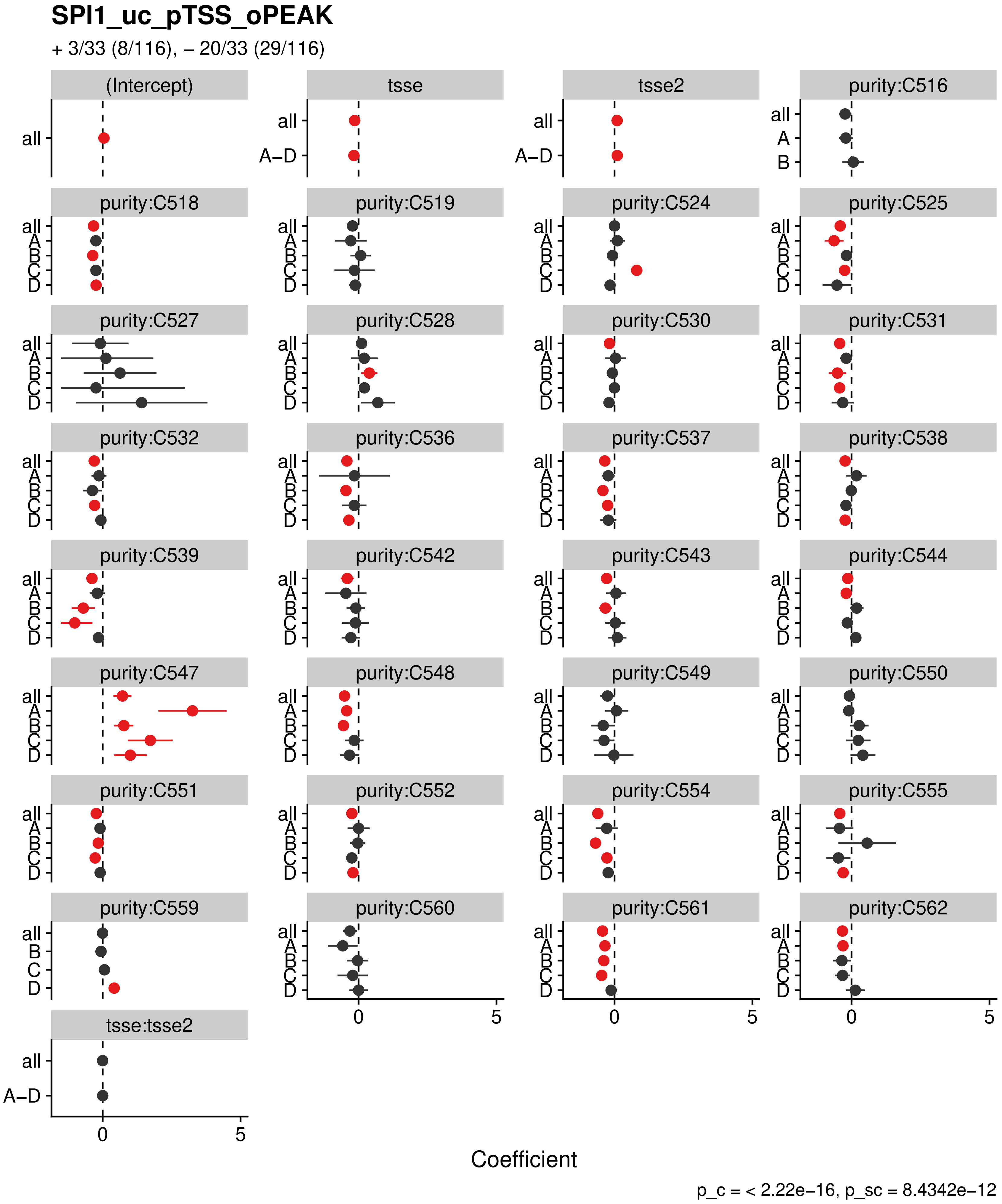

SPI1_uc_pTSS_oPEAK
+ 3/33 (8/116), – 20/33 (29/116)
(Intercept)
tsse
tsse2
purity:C516
purity:C518
purity:C519
purity:C524
purity:C525
purity:C527
purity:C528
purity:C530
purity:C531
purity:C532
purity:C536
purity:C537
purity:C538
purity:C539
purity:C542
purity:C543
purity:C544
purity:C547
purity:C548
purity:C549
purity:C550
purity:C551
purity:C552
purity:C554
purity:C555
purity:C559
purity:C560
purity:C561
purity:C562
tsse:tsse2
all
A–D
A
B
C
D
0
5
Coefficient
p_c = < 2.22e−16, p_sc = 8.4342e−12

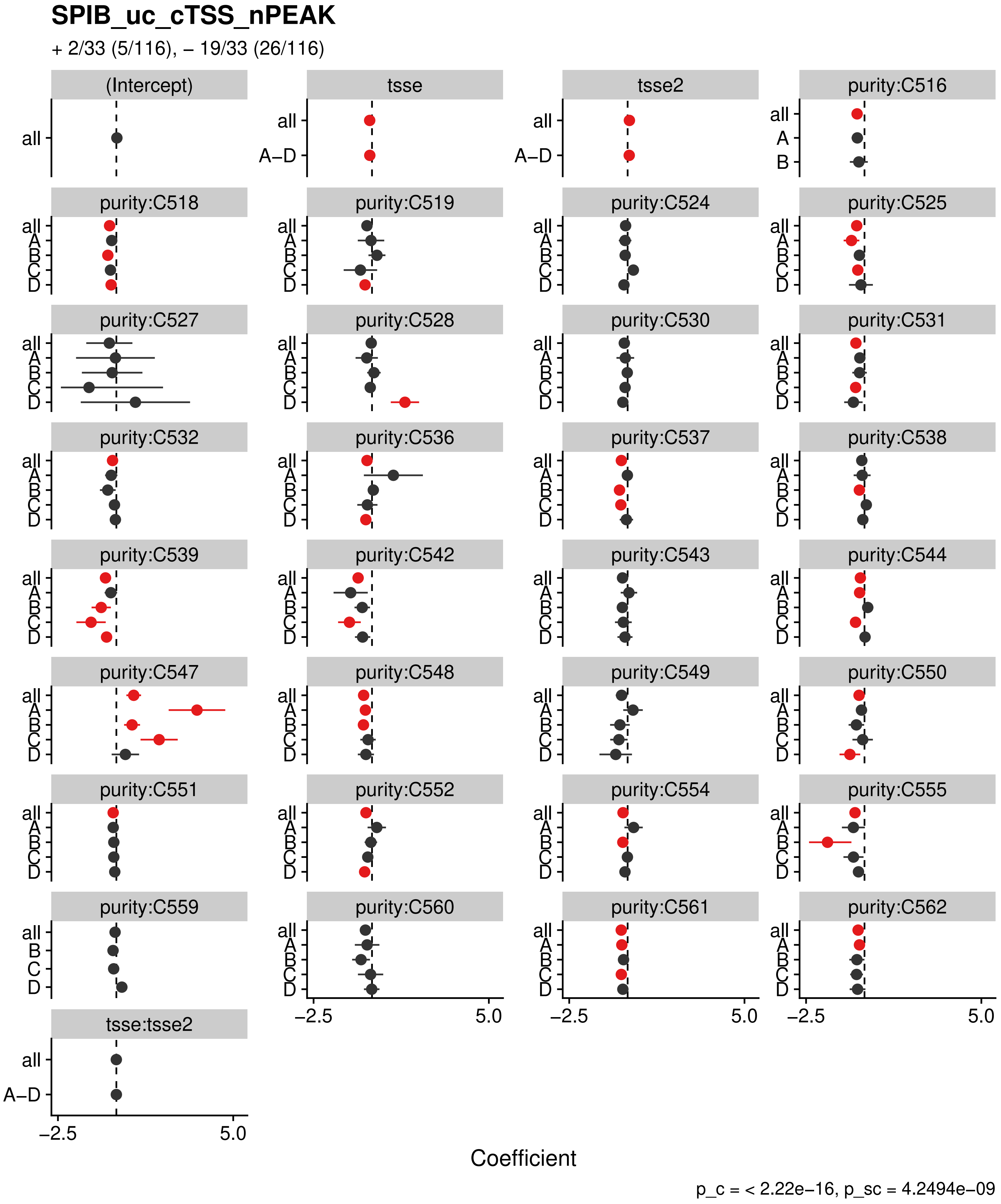

SPIB_uc_cTSS_nPEAK
+ 2/33 (5/116), – 19/33 (26/116)
(Intercept)
tsse
tsse2
purity:C516
purity:C518
purity:C519
purity:C524
purity:C525
purity:C527
purity:C528
purity:C530
purity:C531
purity:C532
purity:C536
purity:C537
purity:C538
purity:C539
purity:C542
purity:C543
purity:C544
purity:C547
purity:C548
purity:C549
purity:C550
purity:C551
purity:C552
purity:C554
purity:C555
purity:C559
purity:C560
purity:C561
purity:C562
tsse:tsse2
all
A–D
A
B
C
D
–2.5
5.0
Coefficient
p_c = < 2.22e–16, p_sc = 4.2494e–09

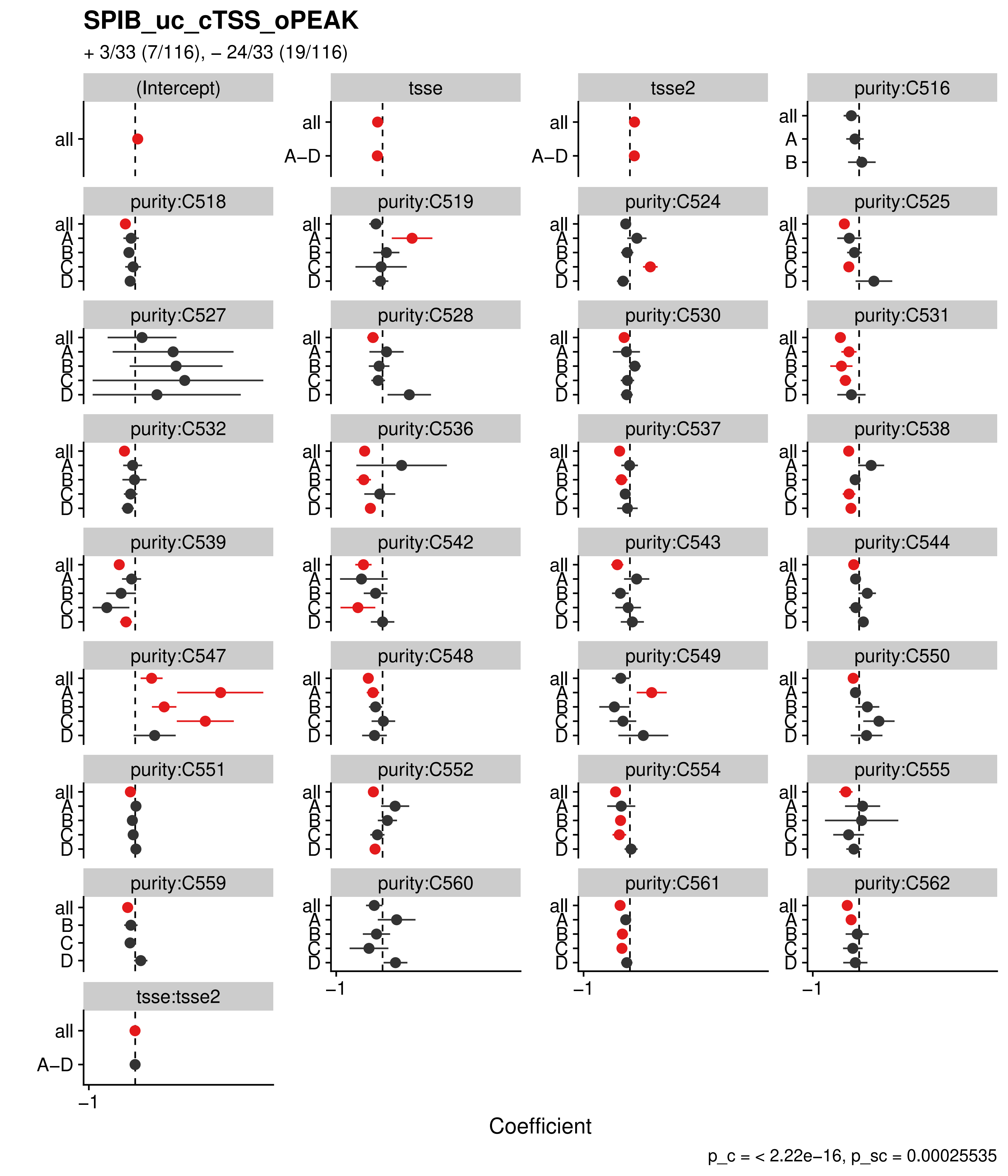

SPIB_uc_cTSS_oPEAK
+ 3/33 (7/116), – 24/33 (19/116)
(Intercept)
tsse
tsse2
purity:C516
purity:C518
purity:C519
purity:C524
purity:C525
purity:C527
purity:C528
purity:C530
purity:C531
purity:C532
purity:C536
purity:C537
purity:C538
purity:C539
purity:C542
purity:C543
purity:C544
purity:C547
purity:C548
purity:C549
purity:C550
purity:C551
purity:C552
purity:C554
purity:C555
purity:C559
purity:C560
purity:C561
purity:C562
tsse:tsse2
all
A–D
A
B
C
D
–1
Coefficient
p_c = < 2.22e–16, p_sc = 0.00025535

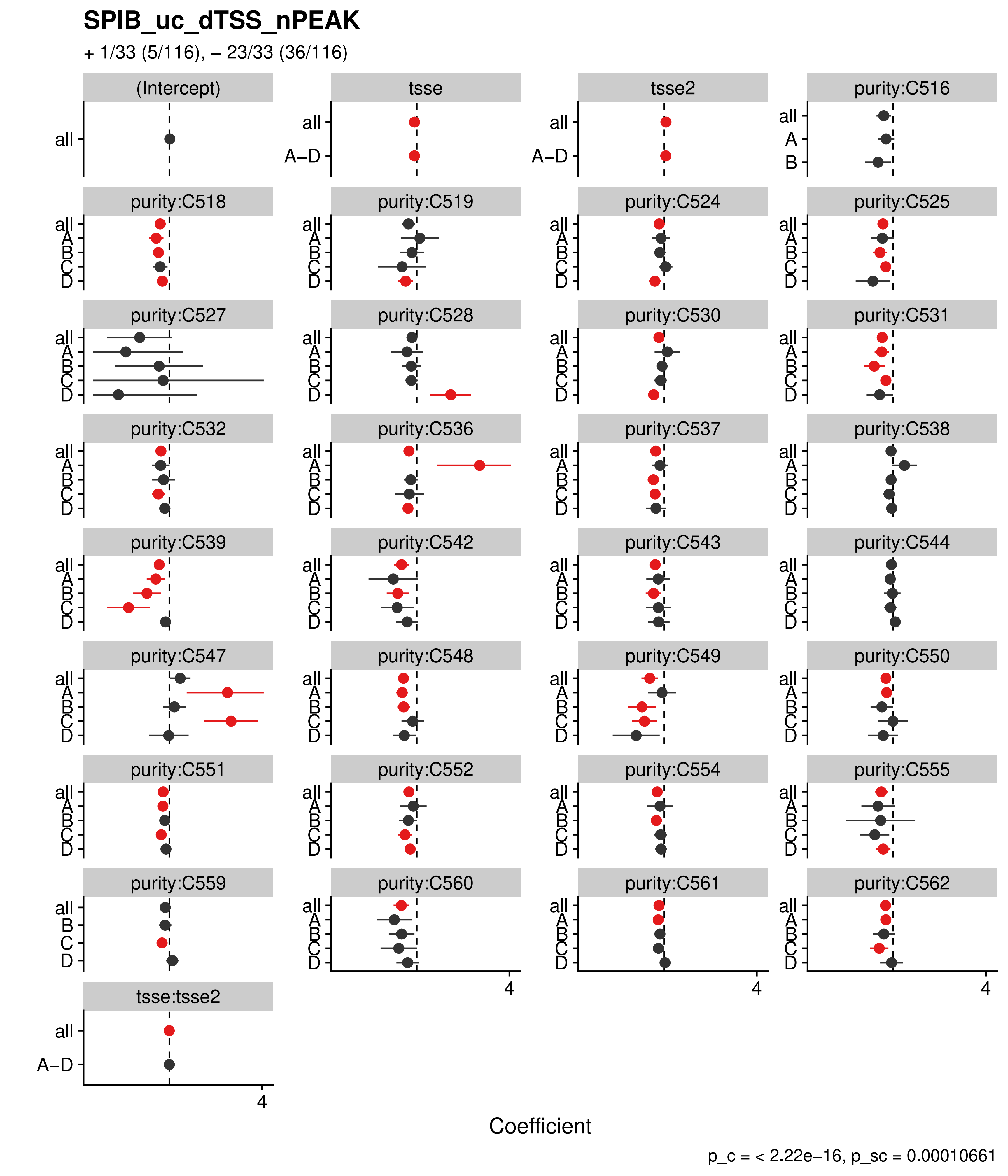
SPIB_uc_dTSS_nPEAK
+ 1/33 (5/116), – 23/33 (36/116)
(Intercept)
tsse
tsse2
purity:C516
purity:C518
purity:C519
purity:C524
purity:C525
purity:C527
purity:C528
purity:C530
purity:C531
purity:C532
purity:C536
purity:C537
purity:C538
purity:C539
purity:C542
purity:C543
purity:C544
purity:C547
purity:C548
purity:C549
purity:C550
purity:C551
purity:C552
purity:C554
purity:C555
purity:C559
purity:C560
purity:C561
purity:C562
tsse:tsse2
all
A–D
A
B
C
D
4
Coefficient
p_c = < 2.22e–16, p_sc = 0.00010661

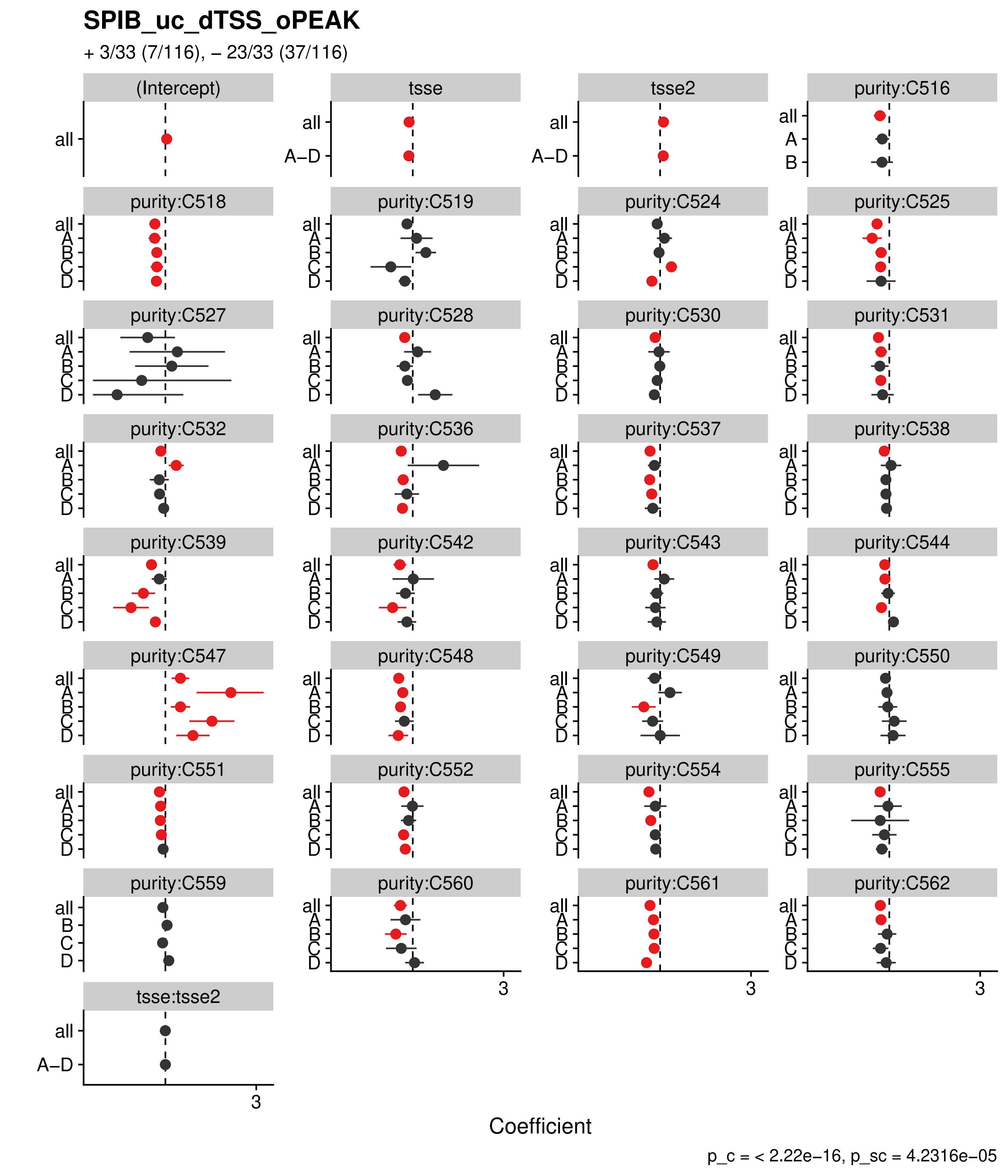

SPIB_uc_dTSS_oPEAK
+ 3/33 (7/116), – 23/33 (37/116)
(Intercept)
tsse
tsse2
purity:C516
purity:C518
purity:C519
purity:C524
purity:C525
purity:C527
purity:C528
purity:C530
purity:C531
purity:C532
purity:C536
purity:C537
purity:C538
purity:C539
purity:C542
purity:C543
purity:C544
purity:C547
purity:C548
purity:C549
purity:C550
purity:C551
purity:C552
purity:C554
purity:C555
purity:C559
purity:C560
purity:C561
purity:C562
tsse:tsse2
all
A–D
A
B
C
D
3
Coefficient
p_c = < 2.22e–16, p_sc = 4.2316e–05

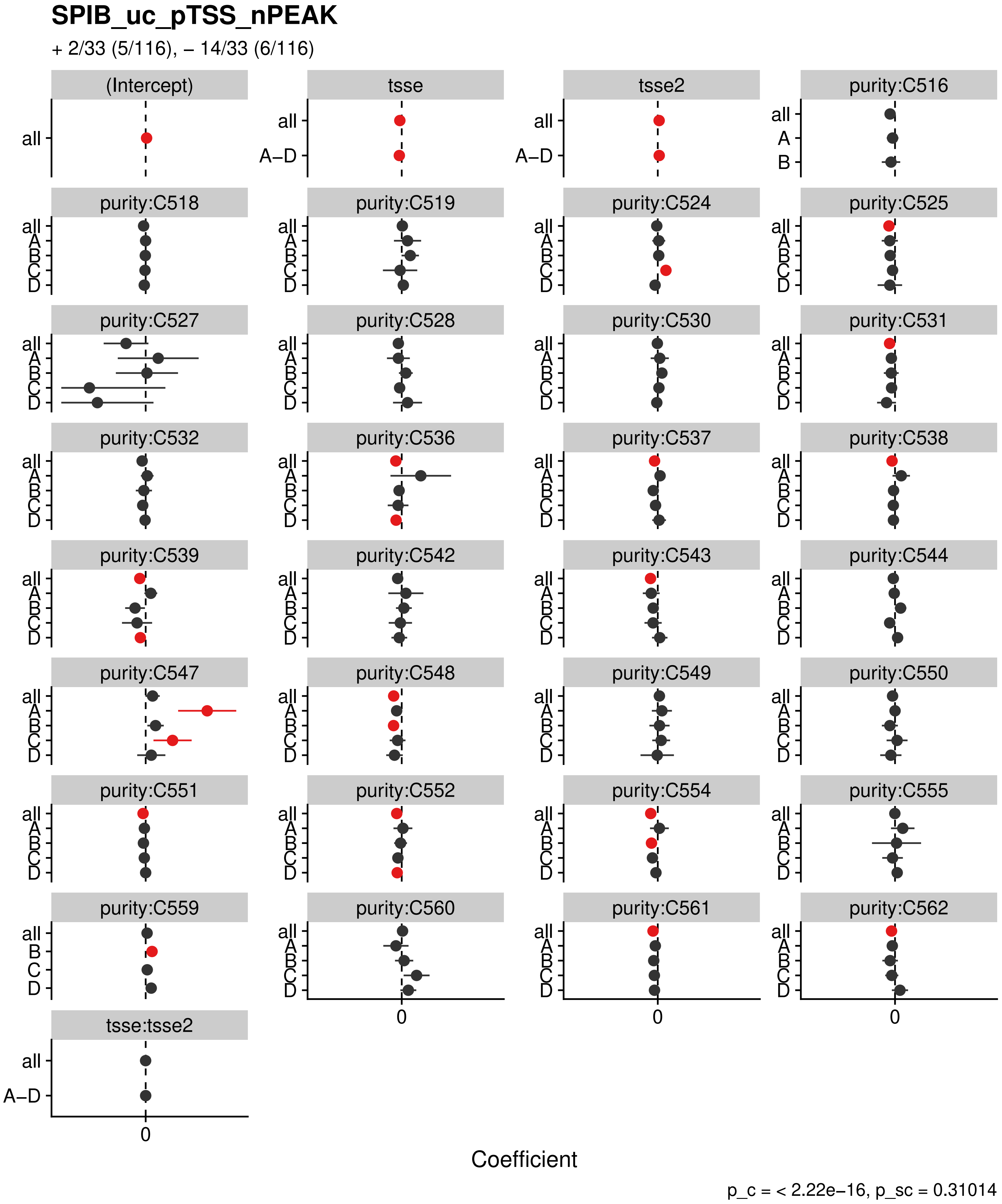

SPIB_uc_pTSS_nPEAK
+ 2/33 (5/116), – 14/33 (6/116)
(Intercept)
tsse
tsse2
purity:C516
purity:C518
purity:C519
purity:C524
purity:C525
purity:C527
purity:C528
purity:C530
purity:C531
purity:C532
purity:C536
purity:C537
purity:C538
purity:C539
purity:C542
purity:C543
purity:C544
purity:C547
purity:C548
purity:C549
purity:C550
purity:C551
purity:C552
purity:C554
purity:C555
purity:C559
purity:C560
purity:C561
purity:C562
tsse:tsse2
all
A–D
A
B
C
D
0
Coefficient
p_c = < 2.22e–16, p_sc = 0.31014

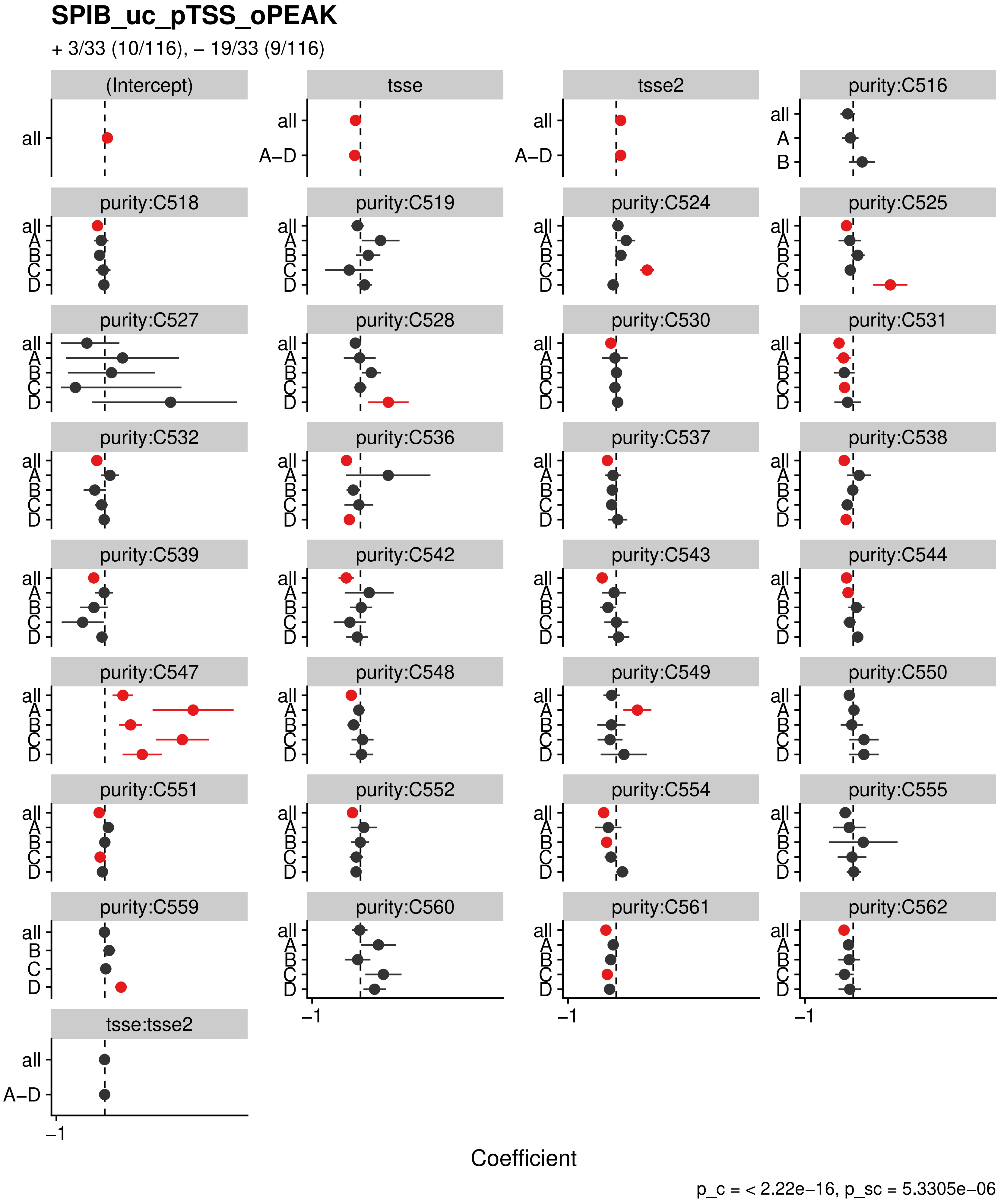

SPIB_uc_pTSS_oPEAK
+ 3/33 (10/116), – 19/33 (9/116)
(Intercept)
tsse
tsse2
purity:C516
purity:C518
purity:C519
purity:C524
purity:C525
purity:C527
purity:C528
purity:C530
purity:C531
purity:C532
purity:C536
purity:C537
purity:C538
purity:C539
purity:C542
purity:C543
purity:C544
purity:C547
purity:C548
purity:C549
purity:C550
purity:C551
purity:C552
purity:C554
purity:C555
purity:C559
purity:C560
purity:C561
purity:C562
tsse:tsse2
all
A–D
A
B
C
D
–1
Coefficient
p_c = < 2.22e–16, p_sc = 5.3305e–06

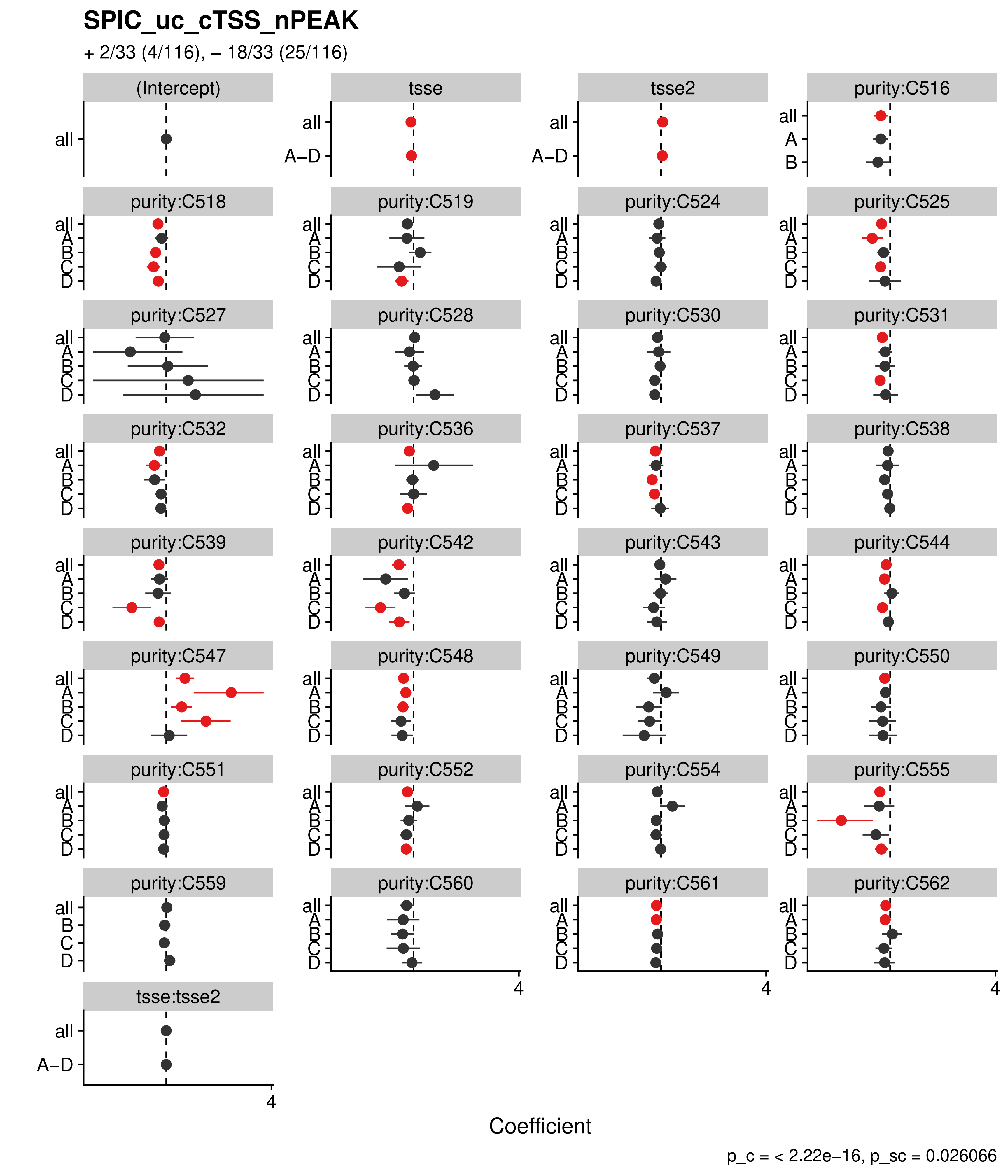

SPIC_uc_cTSS_nPEAK
+ 2/33 (4/116), − 18/33 (25/116)
(Intercept)
tsse
tsse2
purity:C516
purity:C518
purity:C519
purity:C524
purity:C525
purity:C527
purity:C528
purity:C530
purity:C531
purity:C532
purity:C536
purity:C537
purity:C538
purity:C539
purity:C542
purity:C543
purity:C544
purity:C547
purity:C548
purity:C549
purity:C550
purity:C551
purity:C552
purity:C554
purity:C555
purity:C559
purity:C560
purity:C561
purity:C562
tsse:tsse2
all
A–D
A
B
C
D
4
Coefficient
p_c = < 2.22e−16, p_sc = 0.026066

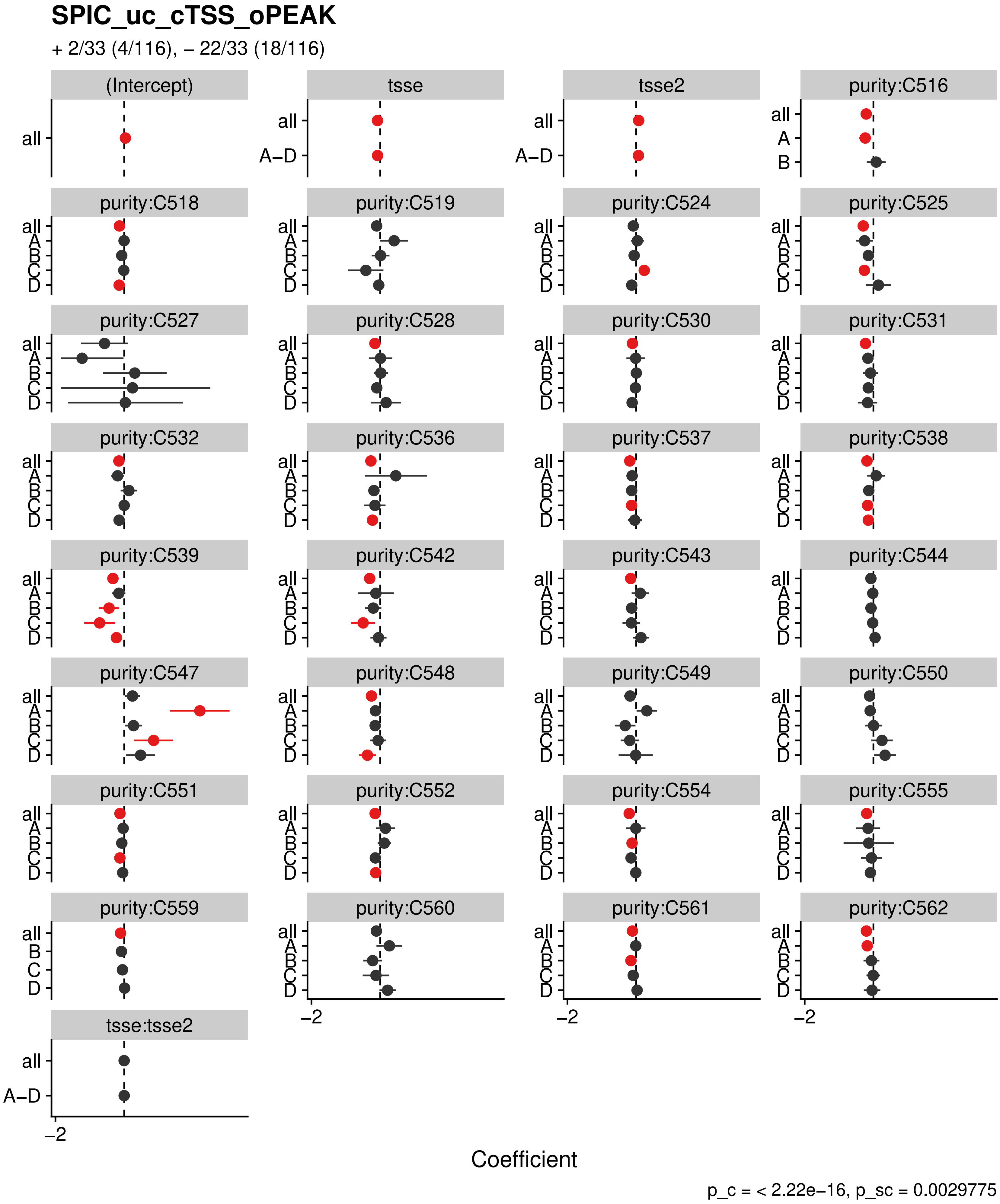

SPIC_uc_cTSS_oPEAK
+ 2/33 (4/116), – 22/33 (18/116)
(Intercept)
tsse
tsse2
purity:C516
purity:C518
purity:C519
purity:C524
purity:C525
purity:C527
purity:C528
purity:C530
purity:C531
purity:C532
purity:C536
purity:C537
purity:C538
purity:C539
purity:C542
purity:C543
purity:C544
purity:C547
purity:C548
purity:C549
purity:C550
purity:C551
purity:C552
purity:C554
purity:C555
purity:C559
purity:C560
purity:C561
purity:C562
tsse:tsse2
all
A–D
A
B
C
D
–2
Coefficient
p_c = < 2.22e–16, p_sc = 0.0029775

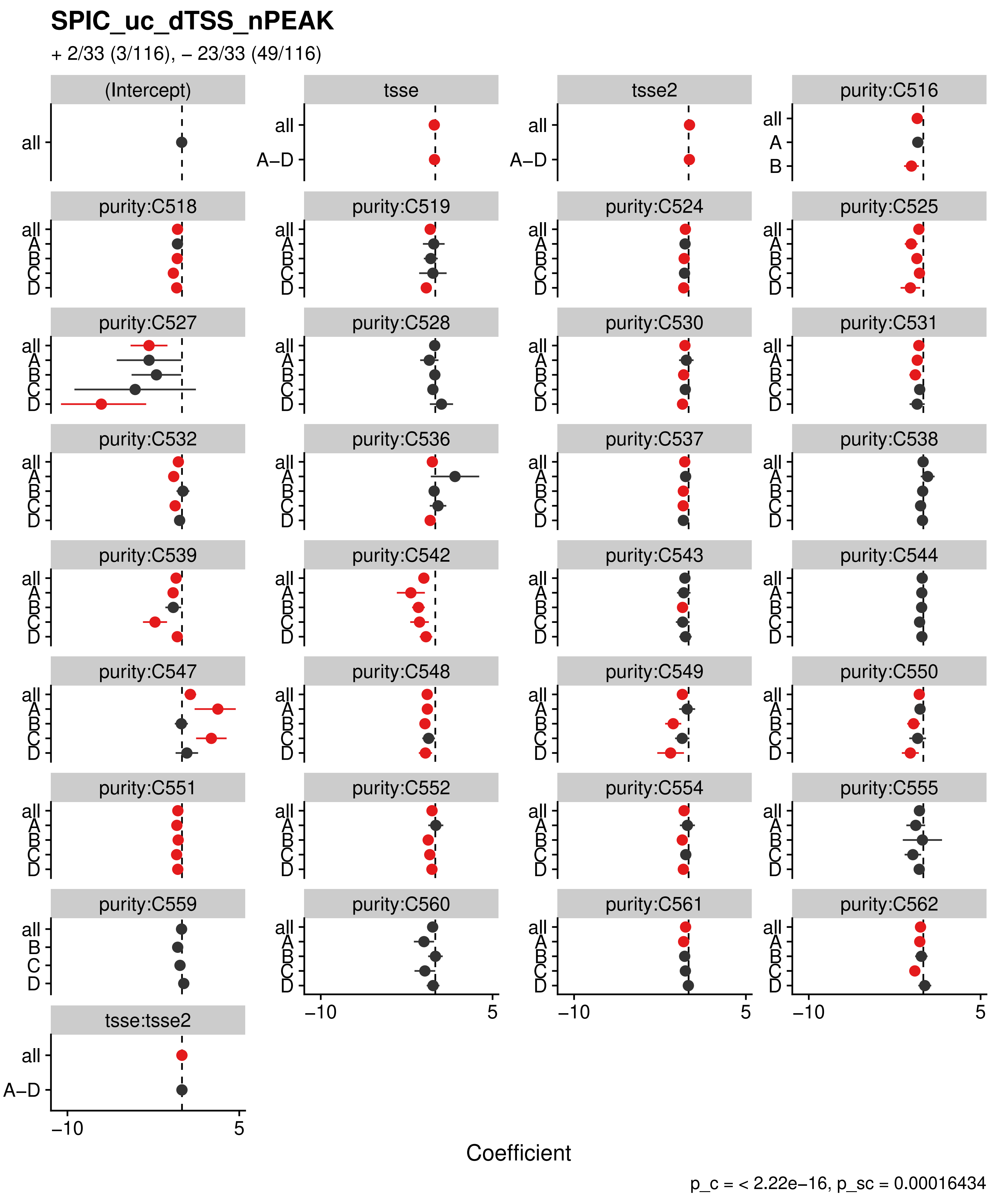

SPIC_uc_dTSS_nPEAK
+ 2/33 (3/116), – 23/33 (49/116)
(Intercept)
tsse
tsse2
purity:C516
purity:C518
purity:C519
purity:C524
purity:C525
purity:C527
purity:C528
purity:C530
purity:C531
purity:C532
purity:C536
purity:C537
purity:C538
purity:C539
purity:C542
purity:C543
purity:C544
purity:C547
purity:C548
purity:C549
purity:C550
purity:C551
purity:C552
purity:C554
purity:C555
purity:C559
purity:C560
purity:C561
purity:C562
tsse:tsse2
all
A–D
A
B
C
D
–10
5
Coefficient
p_c = < 2.22e–16, p_sc = 0.00016434

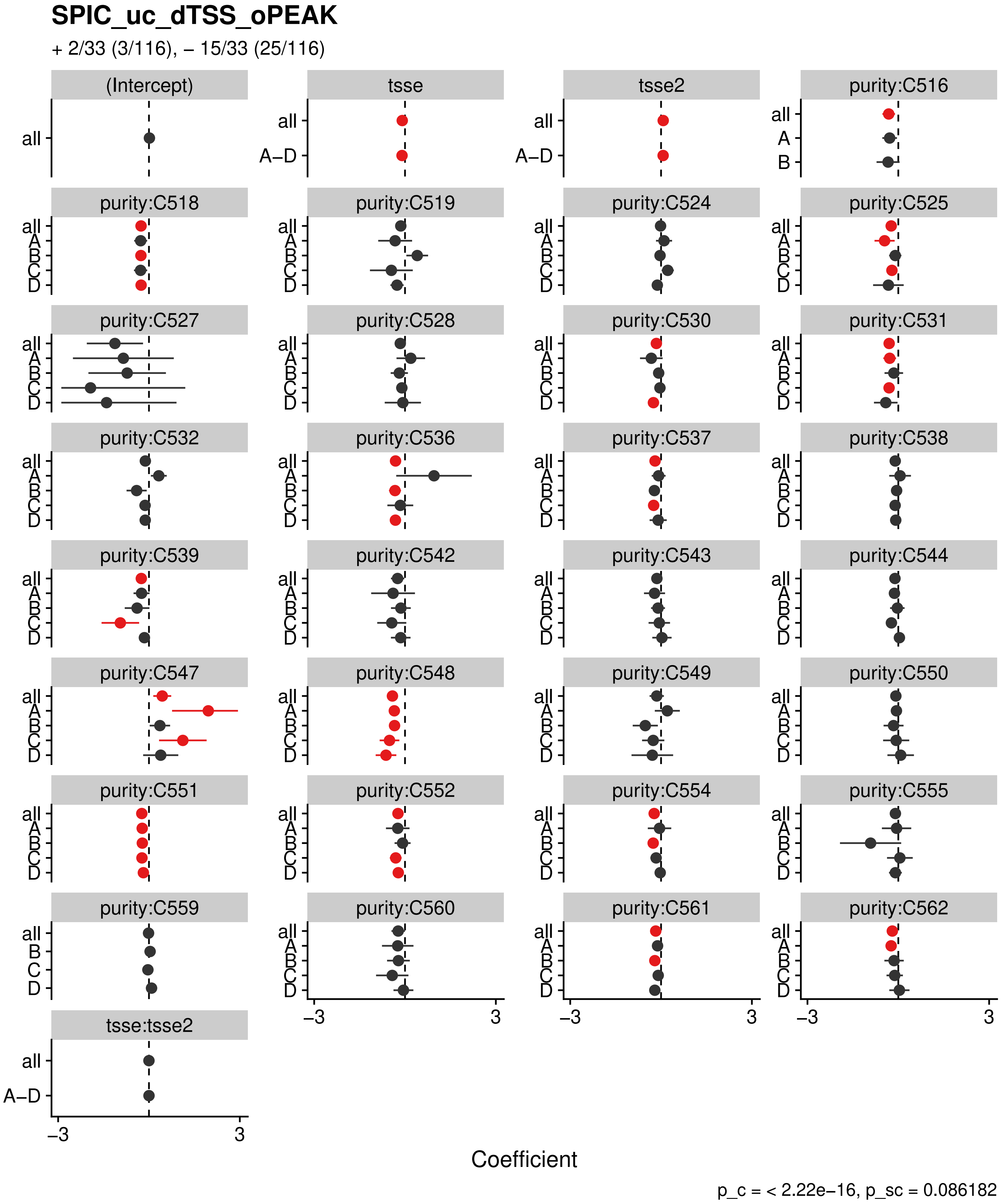
SPIC_uc_dTSS_oPEAK
+ 2/33 (3/116), – 15/33 (25/116)
(Intercept)
tsse
tsse2
purity:C516
purity:C518
purity:C519
purity:C524
purity:C525
purity:C527
purity:C528
purity:C530
purity:C531
purity:C532
purity:C536
purity:C537
purity:C538
purity:C539
purity:C542
purity:C543
purity:C544
purity:C547
purity:C548
purity:C549
purity:C550
purity:C551
purity:C552
purity:C554
purity:C555
purity:C559
purity:C560
purity:C561
purity:C562
tsse:tsse2
all
A–D
A
B
C
D
–3
3
Coefficient
p_c = < 2.22e–16, p_sc = 0.086182

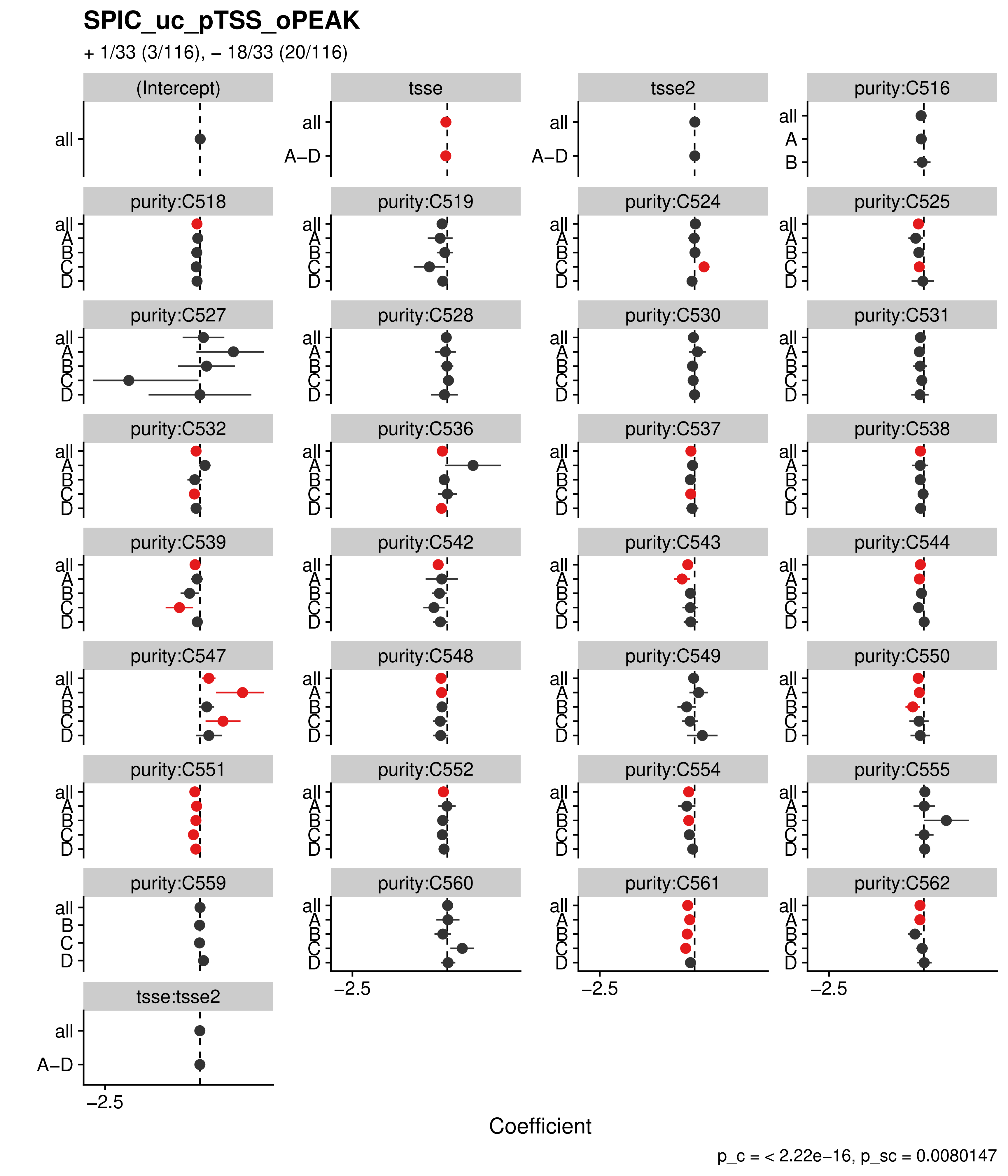

SPIC_uc_pTSS_oPEAK
+ 1/33 (3/116), – 18/33 (20/116)
(Intercept)
tsse
tsse2
purity:C516
purity:C518
purity:C519
purity:C524
purity:C525
purity:C527
purity:C528
purity:C530
purity:C531
purity:C532
purity:C536
purity:C537
purity:C538
purity:C539
purity:C542
purity:C543
purity:C544
purity:C547
purity:C548
purity:C549
purity:C550
purity:C551
purity:C552
purity:C554
purity:C555
purity:C559
purity:C560
purity:C561
purity:C562
tsse:tsse2
all
A–D
A
B
C
D
−2.5
Coefficient
p_c = < 2.22e−16, p_sc = 0.0080147

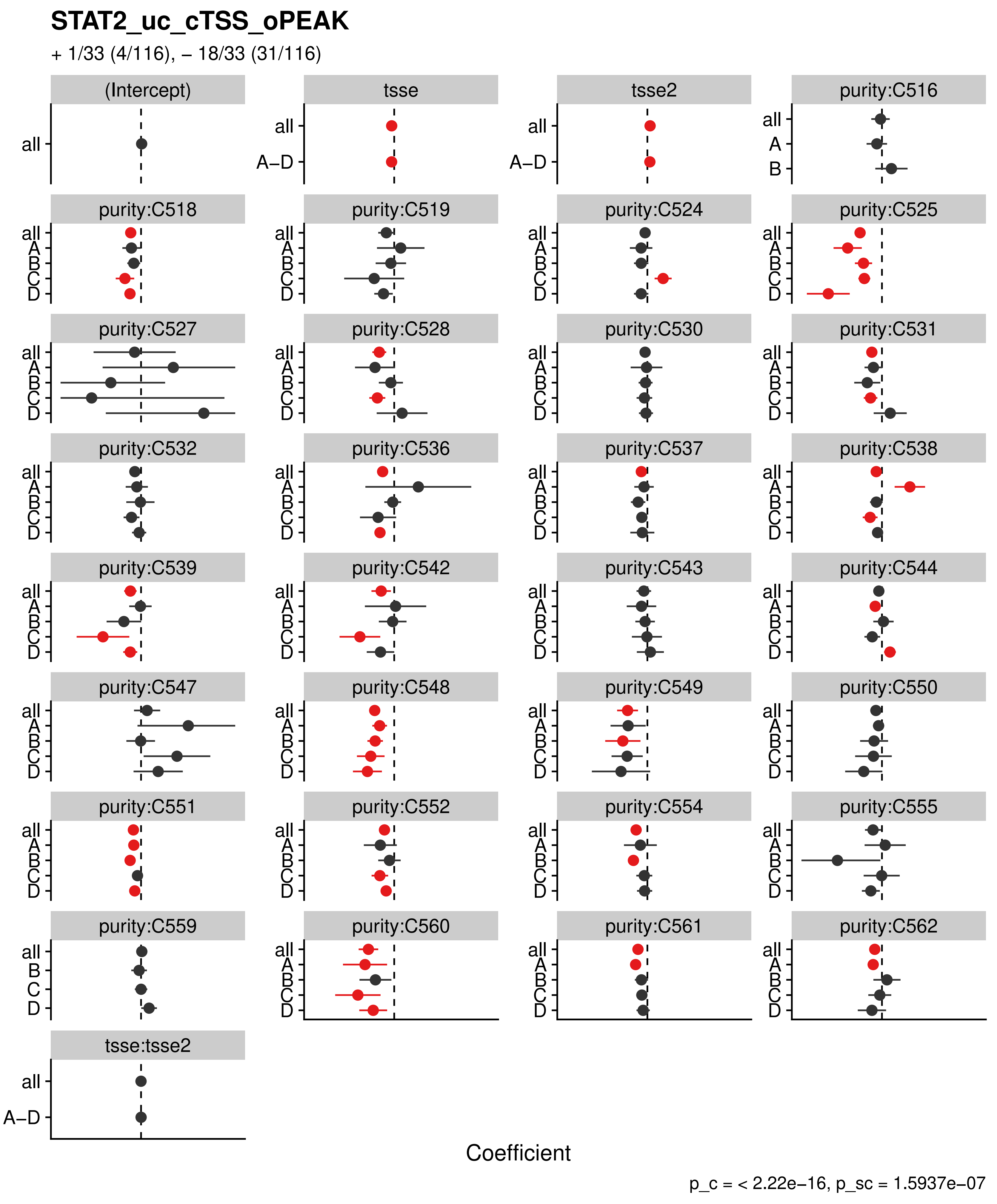

STAT2_uc_cTSS_oPEAK
+ 1/33 (4/116), – 18/33 (31/116)
(Intercept)
tsse
tsse2
purity:C516
purity:C518
purity:C519
purity:C524
purity:C525
purity:C527
purity:C528
purity:C530
purity:C531
purity:C532
purity:C536
purity:C537
purity:C538
purity:C539
purity:C542
purity:C543
purity:C544
purity:C547
purity:C548
purity:C549
purity:C550
purity:C551
purity:C552
purity:C554
purity:C555
purity:C559
purity:C560
purity:C561
purity:C562
tsse:tsse2
all
A–D
A
B
C
D
Coefficient
p_c = < 2.22e–16, p_sc = 1.5937e–07

### STAT2_uc_dTSS_oPEAK

+ 2/33 (6/116), – 14/33 (43/116)

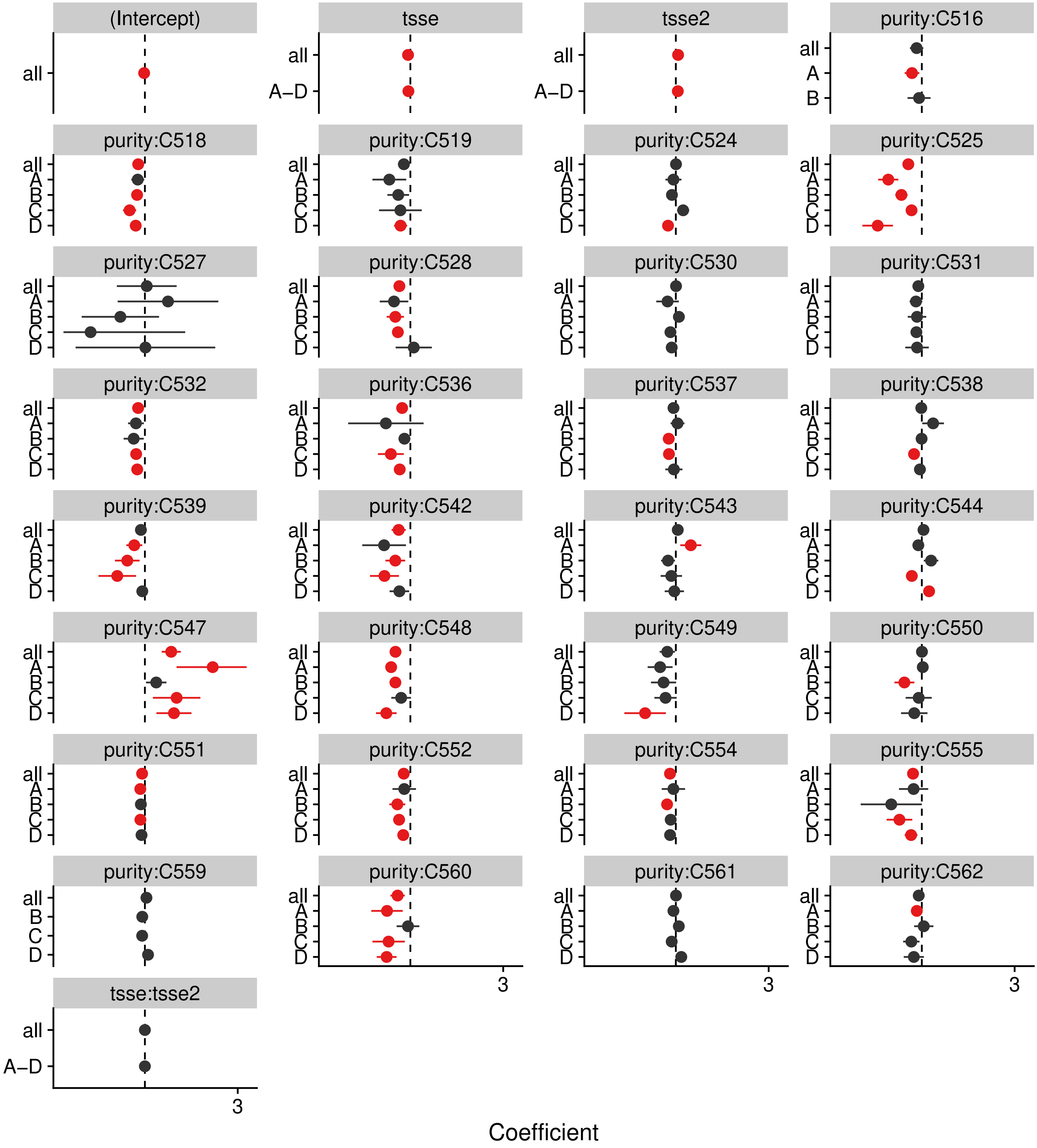


p_c = < 2.22e–16, p_sc = 4.8064e–12

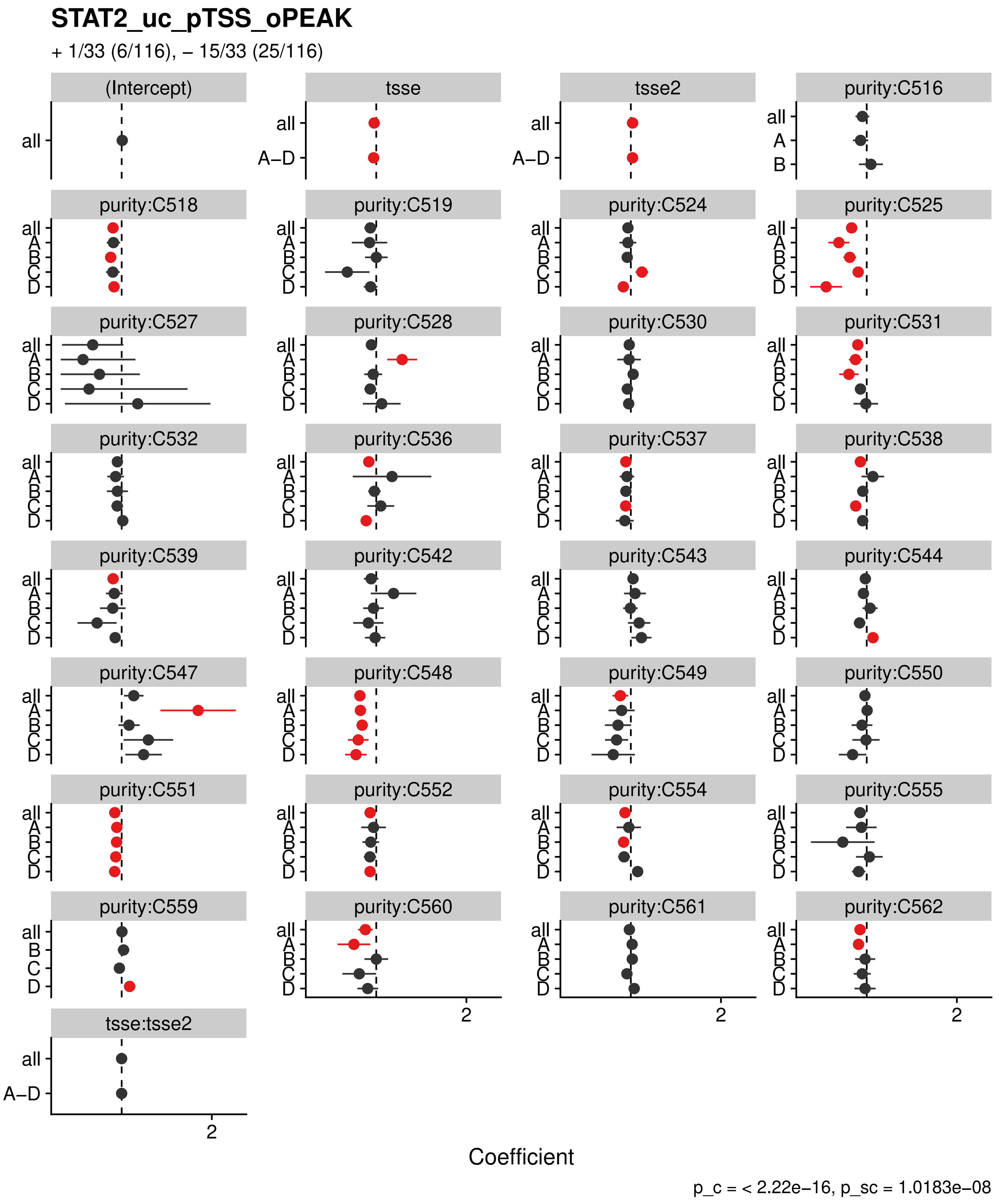
STAT2_uc_pTSS_oPEAK
+ 1/33 (6/116), – 15/33 (25/116)
(Intercept)
tsse
tsse2
purity:C516
purity:C518
purity:C519
purity:C524
purity:C525
purity:C527
purity:C528
purity:C530
purity:C531
purity:C532
purity:C536
purity:C537
purity:C538
purity:C539
purity:C542
purity:C543
purity:C544
purity:C547
purity:C548
purity:C549
purity:C550
purity:C551
purity:C552
purity:C554
purity:C555
purity:C559
purity:C560
purity:C561
purity:C562
tsse:tsse2
all
A–D
A
B
C
D
2
Coefficient
p_c = < 2.22e–16, p_sc = 1.0183e–08

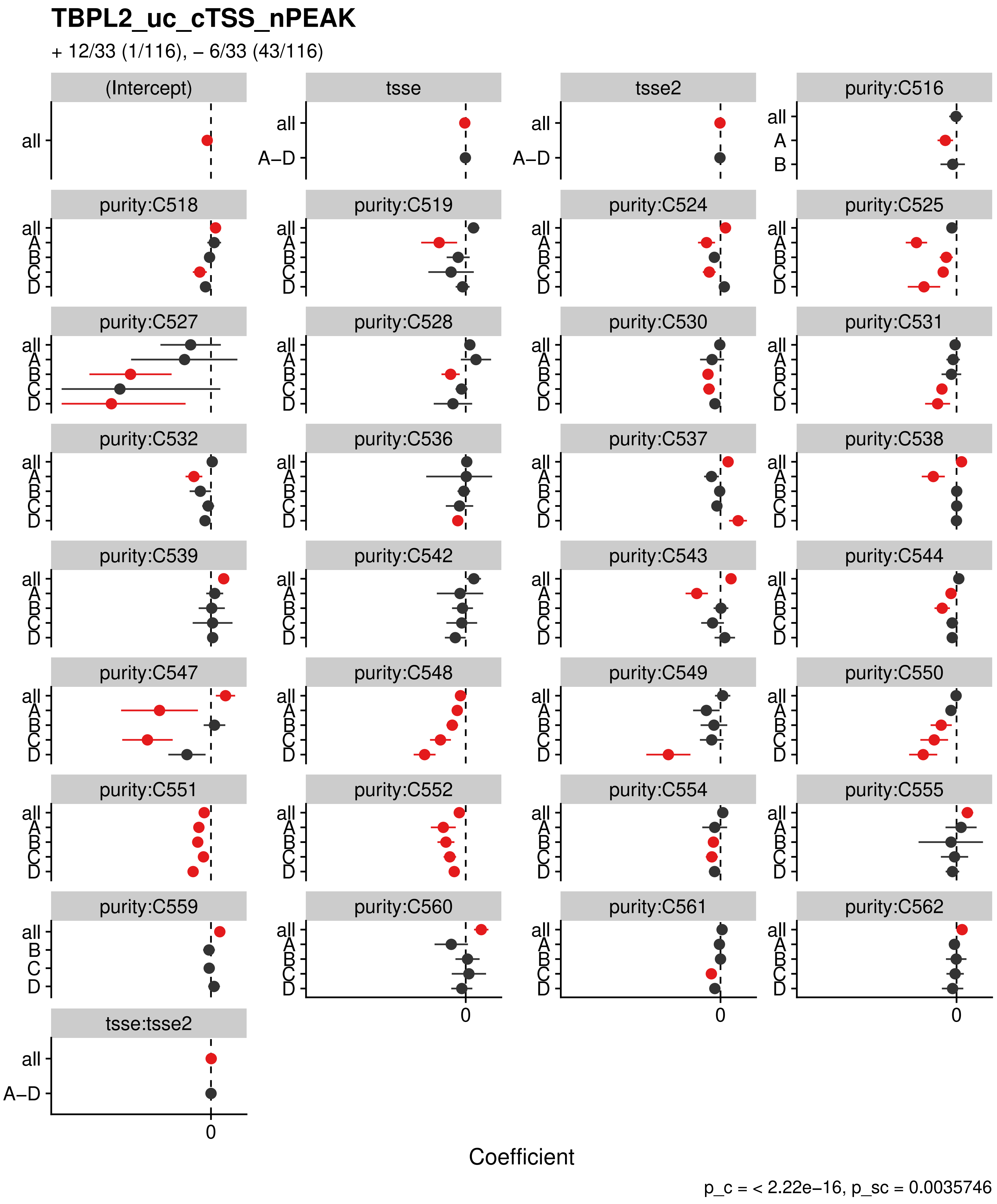
TBPL2_uc_cTSS_nPEAK
+ 12/33 (1/116), – 6/33 (43/116)
(Intercept)
tsse
tsse2
purity:C516
purity:C518
purity:C519
purity:C524
purity:C525
purity:C527
purity:C528
purity:C530
purity:C531
purity:C532
purity:C536
purity:C537
purity:C538
purity:C539
purity:C542
purity:C543
purity:C544
purity:C547
purity:C548
purity:C549
purity:C550
purity:C551
purity:C552
purity:C554
purity:C555
purity:C559
purity:C560
purity:C561
purity:C562
tsse:tsse2
all
A–D
A
B
C
D
0
Coefficient
p_c = < 2.22e–16, p_sc = 0.0035746

### VAX1_uc_dTSS_oPEAK

+ 13/33 (1/116), – 3/33 (41/116)

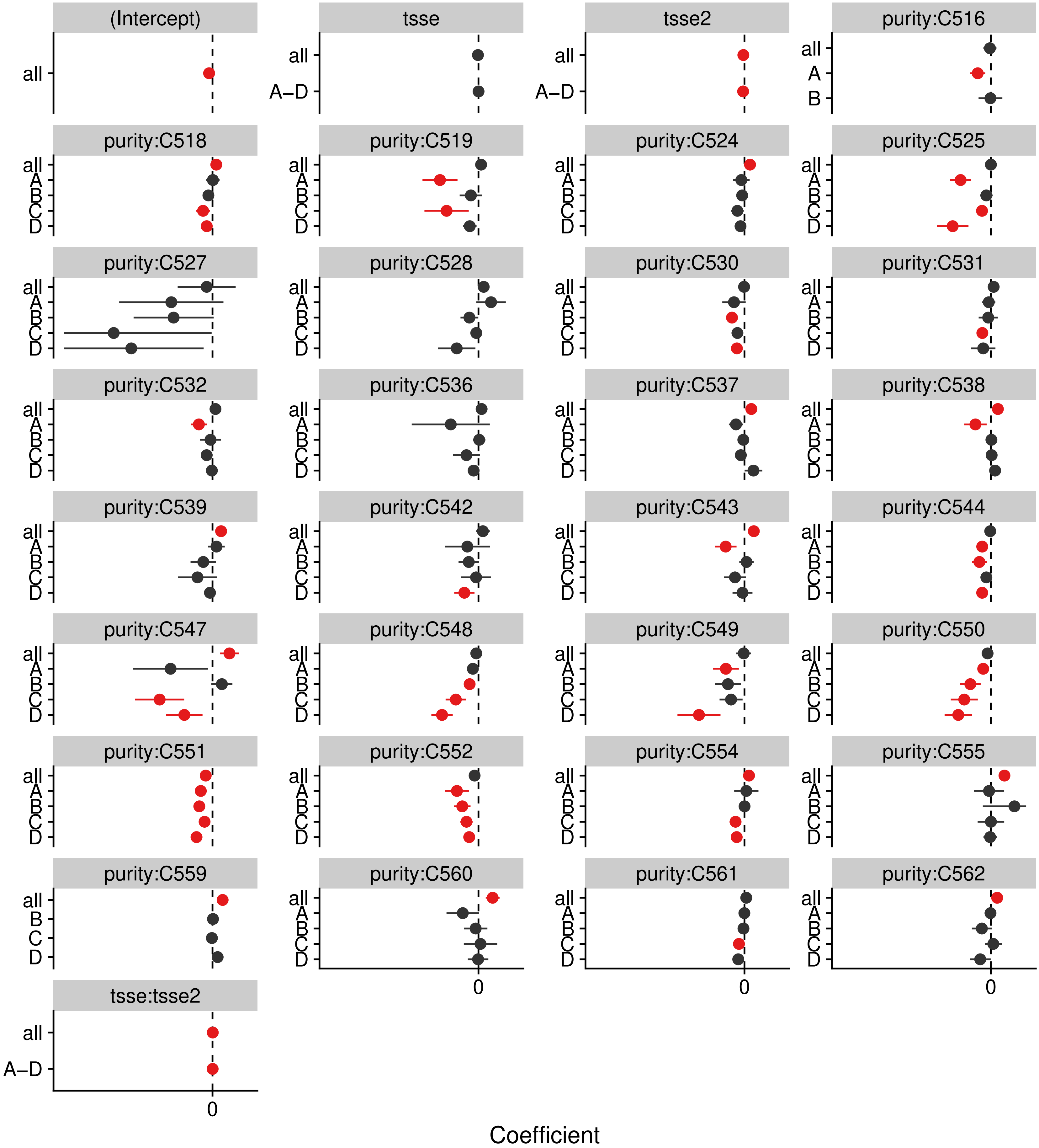


p_c = < 2.22e–16, p_sc = 0.1284

#### VAX1_uc_pTSS_nPEAK

+ 10/33 (1/116), – 5/33 (41/116)

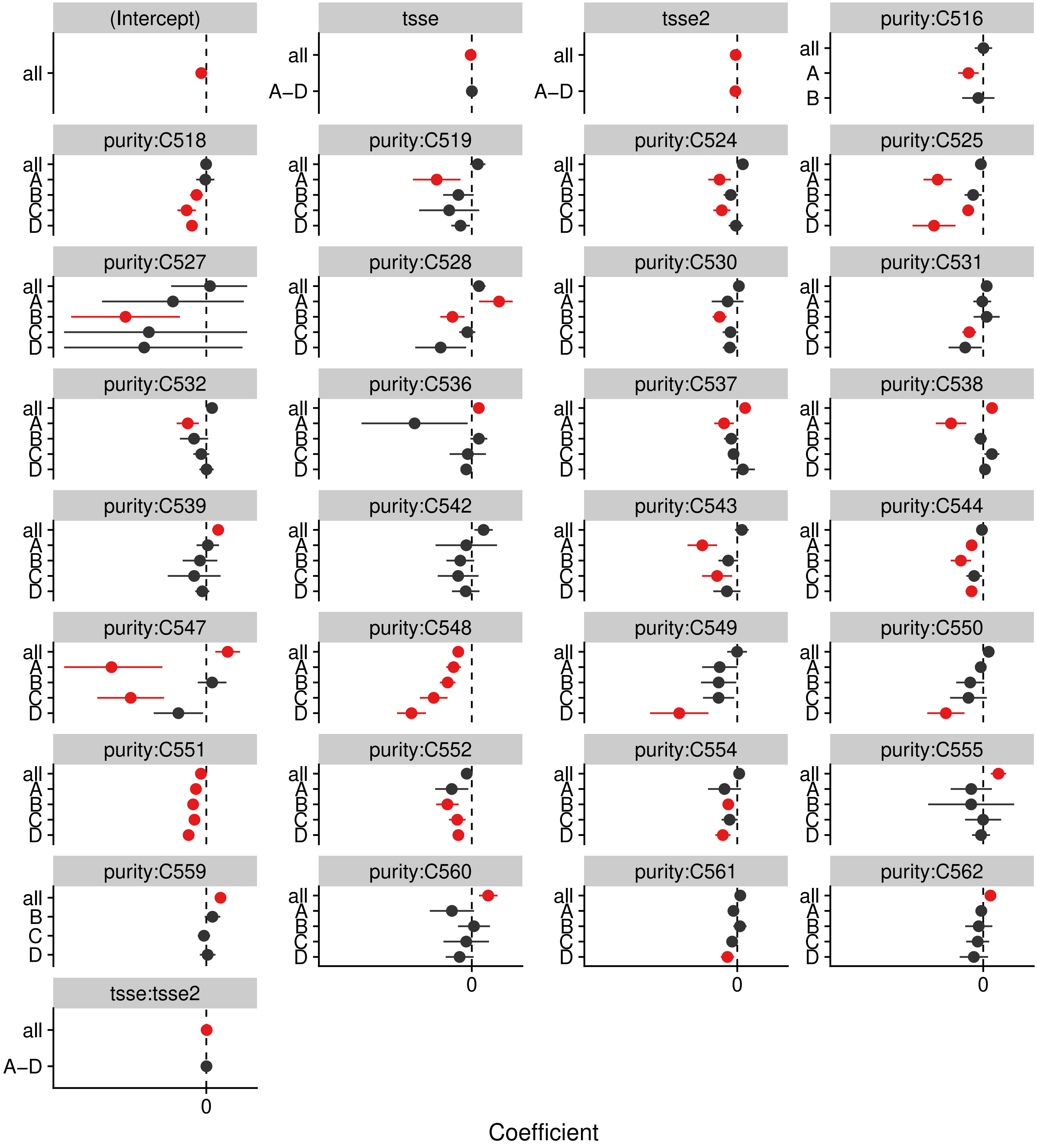


p_c = < 2.22e−16, p_sc = 0.7374

### VAX2_uc_dTSS_nPEAK

+ 15/33 (0/116), – 2/33 (36/116)

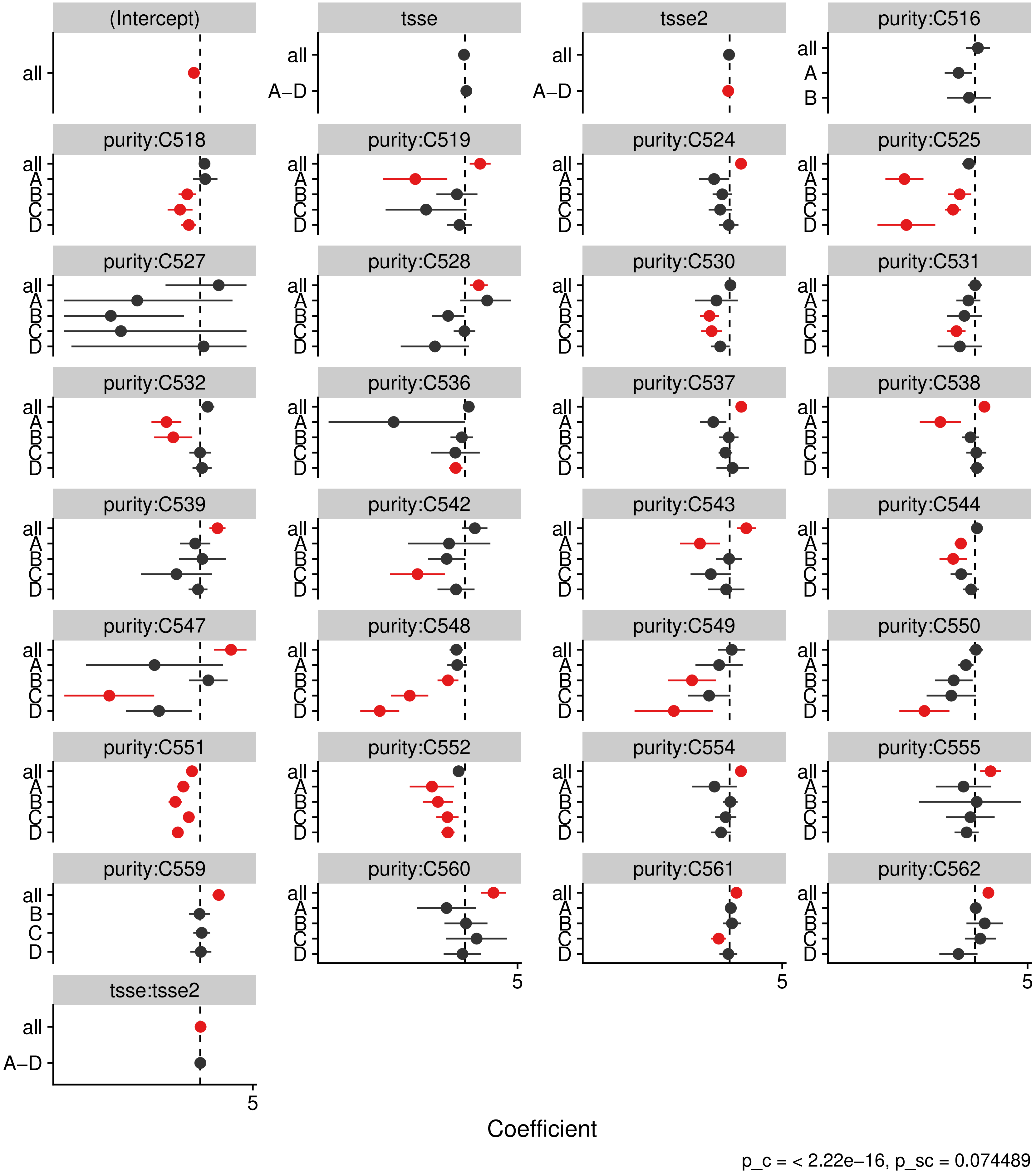

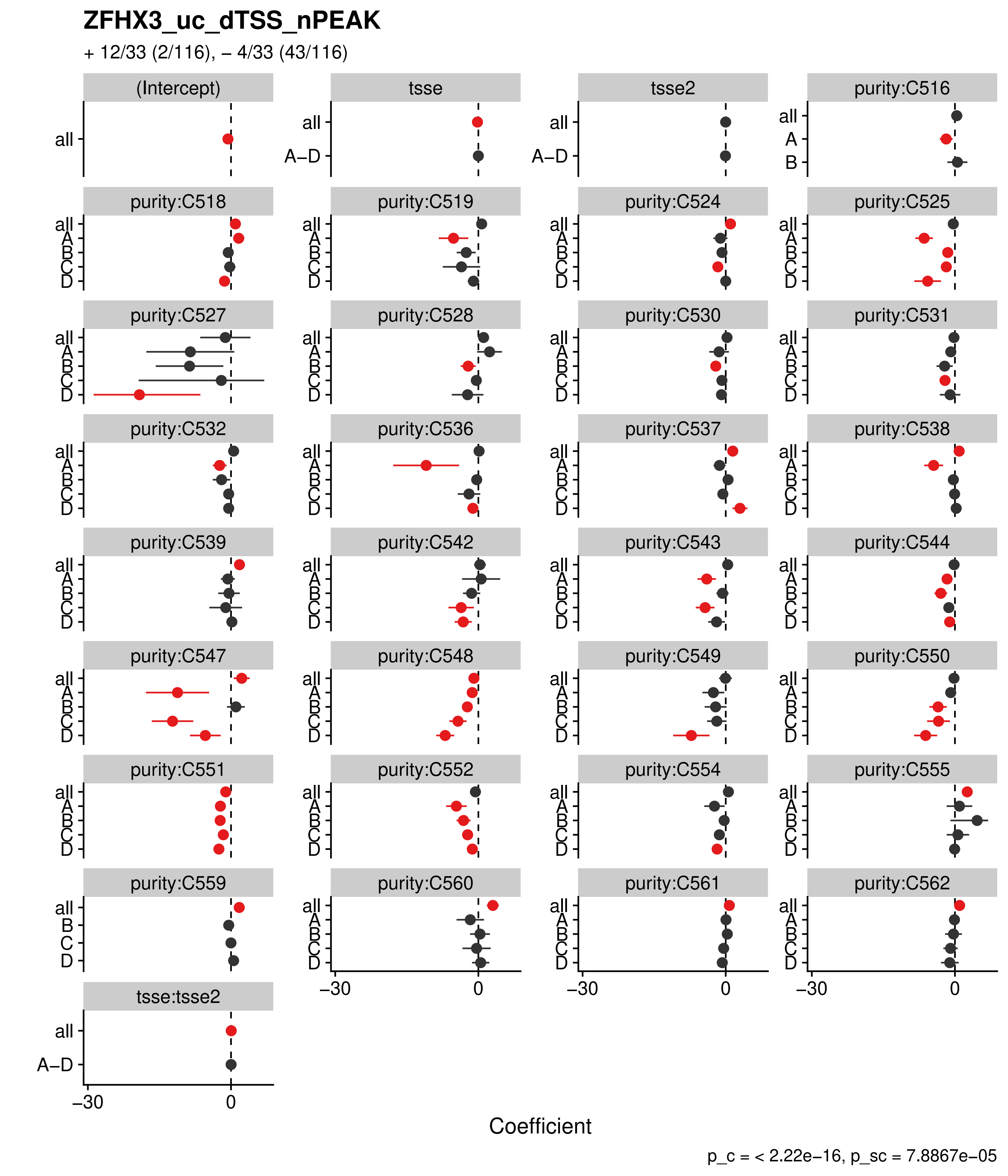
ZFHX3_uc_dTSS_nPEAK
+ 12/33 (2/116), – 4/33 (43/116)
(Intercept)
tsse
tsse2
purity:C516
purity:C518
purity:C519
purity:C524
purity:C525
purity:C527
purity:C528
purity:C530
purity:C531
purity:C532
purity:C536
purity:C537
purity:C538
purity:C539
purity:C542
purity:C543
purity:C544
purity:C547
purity:C548
purity:C549
purity:C550
purity:C551
purity:C552
purity:C554
purity:C555
purity:C559
purity:C560
purity:C561
purity:C562
tsse:tsse2
all
A–D
A
B
C
D
–30
0
Coefficient
p_c = < 2.22e–16, p_sc = 7.8867e–05

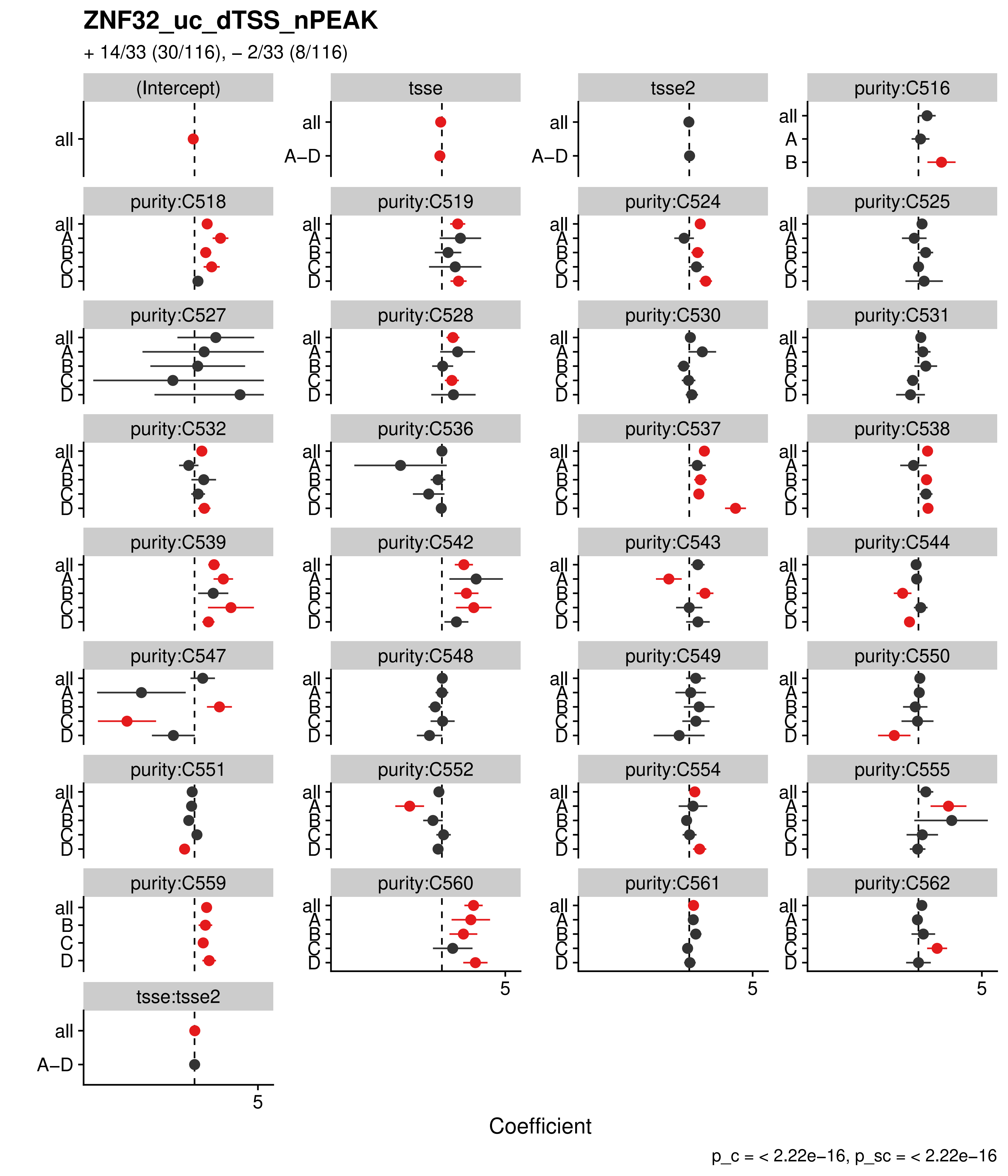
ZNF32_uc_dTSS_nPEAK
+ 14/33 (30/116), – 2/33 (8/116)
(Intercept)
tsse
tsse2
purity:C516
purity:C518
purity:C519
purity:C524
purity:C525
purity:C527
purity:C528
purity:C530
purity:C531
purity:C532
purity:C536
purity:C537
purity:C538
purity:C539
purity:C542
purity:C543
purity:C544
purity:C547
purity:C548
purity:C549
purity:C550
purity:C551
purity:C552
purity:C554
purity:C555
purity:C559
purity:C560
purity:C561
purity:C562
tsse:tsse2
all
A–D
A
B
C
D
5
Coefficient
p_c = < 2.22e–16, p_sc = < 2.22e–16

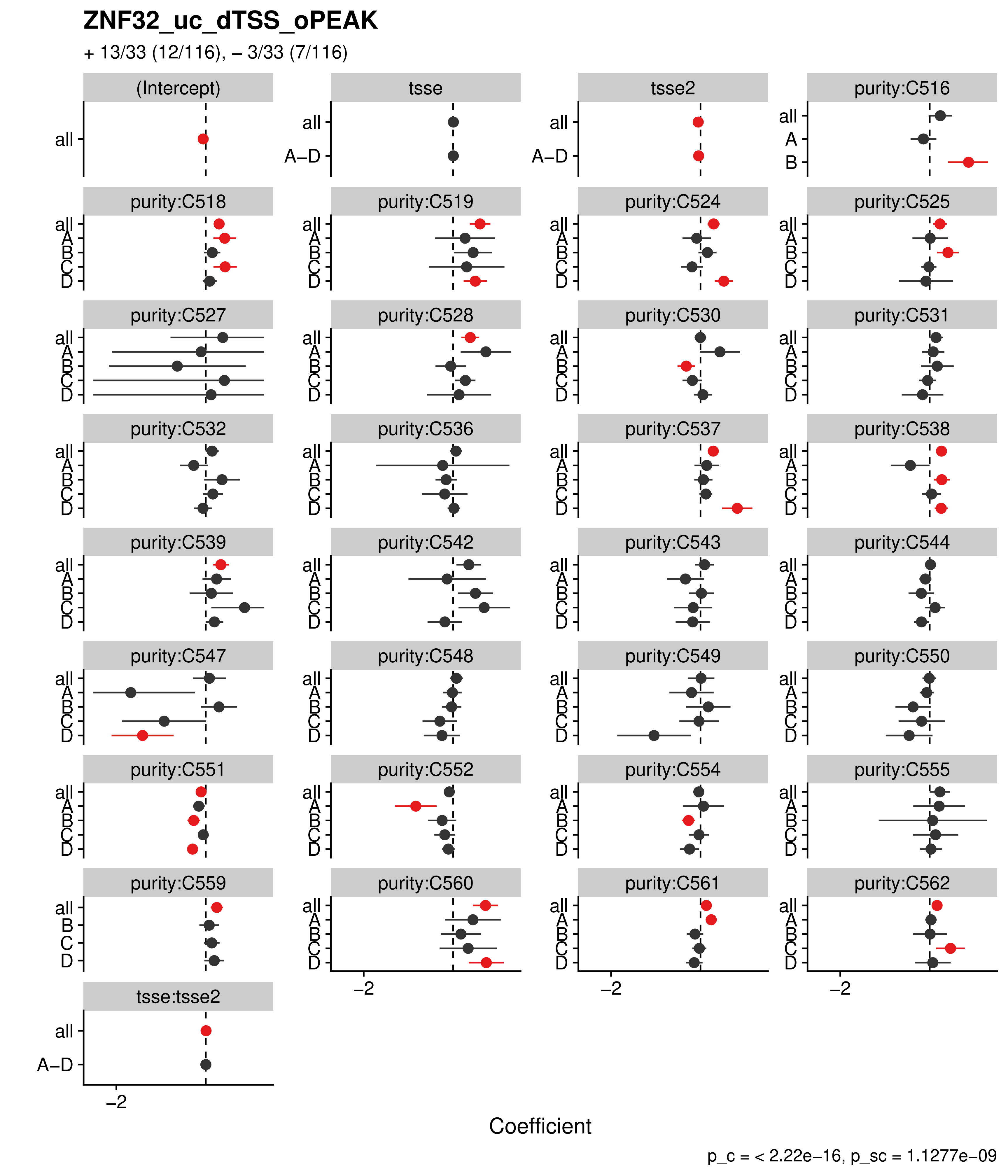

ZNF32_uc_dTSS_oPEAK
+ 13/33 (12/116), – 3/33 (7/116)
(Intercept)
tsse
tsse2
purity:C516
purity:C518
purity:C519
purity:C524
purity:C525
purity:C527
purity:C528
purity:C530
purity:C531
purity:C532
purity:C536
purity:C537
purity:C538
purity:C539
purity:C542
purity:C543
purity:C544
purity:C547
purity:C548
purity:C549
purity:C550
purity:C551
purity:C552
purity:C554
purity:C555
purity:C559
purity:C560
purity:C561
purity:C562
tsse:tsse2
all
A–D
A
B
C
D
–2
Coefficient
p_c = < 2.22e–16, p_sc = 1.1277e–09

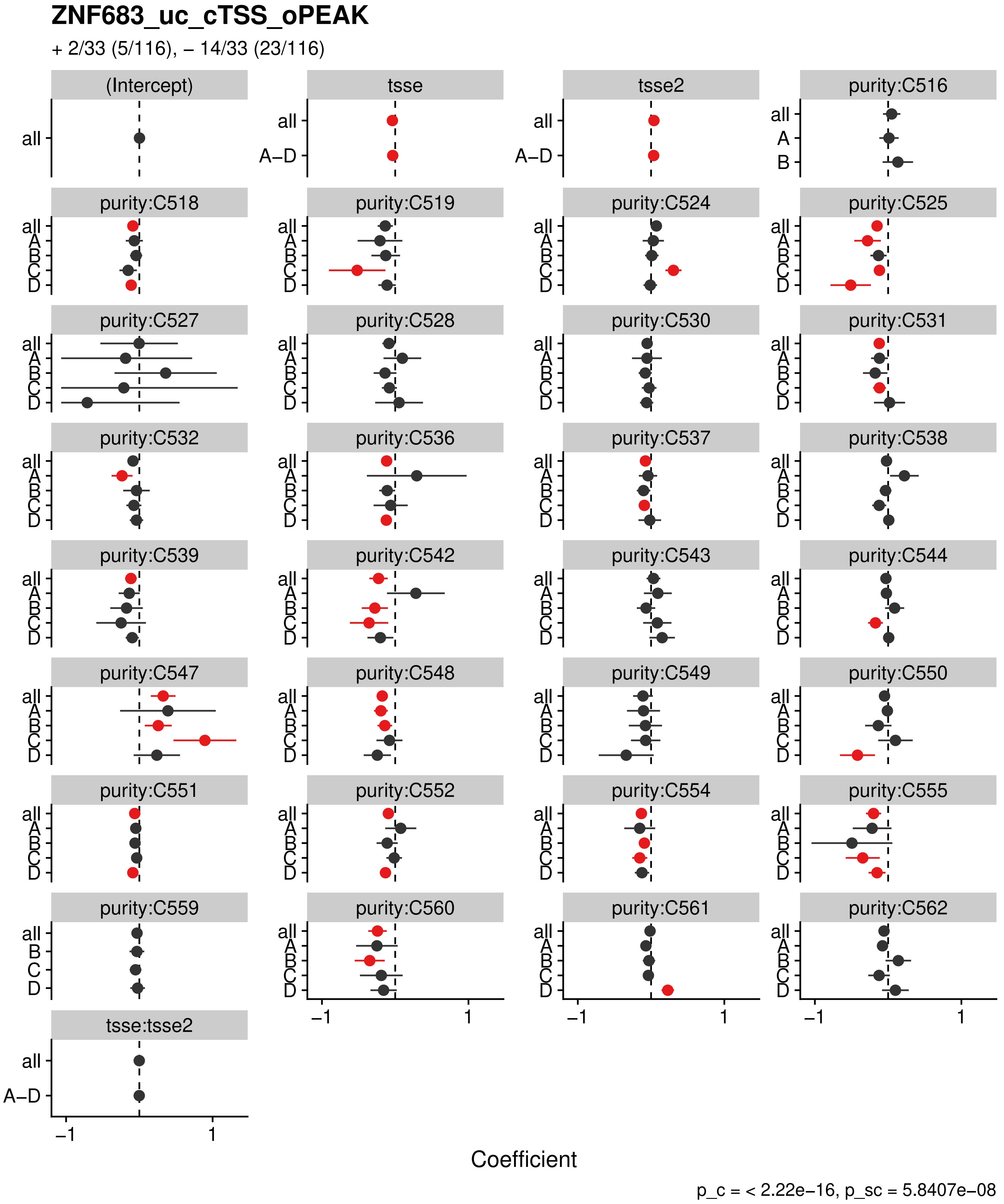
ZNF683_uc_cTSS_oPEAK
+ 2/33 (5/116), – 14/33 (23/116)
(Intercept)
tsse
tsse2
purity:C516
purity:C518
purity:C519
purity:C524
purity:C525
purity:C527
purity:C528
purity:C530
purity:C531
purity:C532
purity:C536
purity:C537
purity:C538
purity:C539
purity:C542
purity:C543
purity:C544
purity:C547
purity:C548
purity:C549
purity:C550
purity:C551
purity:C552
purity:C554
purity:C555
purity:C559
purity:C560
purity:C561
purity:C562
tsse:tsse2
all
A–D
A
B
C
D
–1
1
Coefficient
p_c = < 2.22e–16, p_sc = 5.8407e–08
