## Supplementary figures and images for "The co-evolution of the genome and epigenome in colorectal cancer"

### figS1_reduced.png

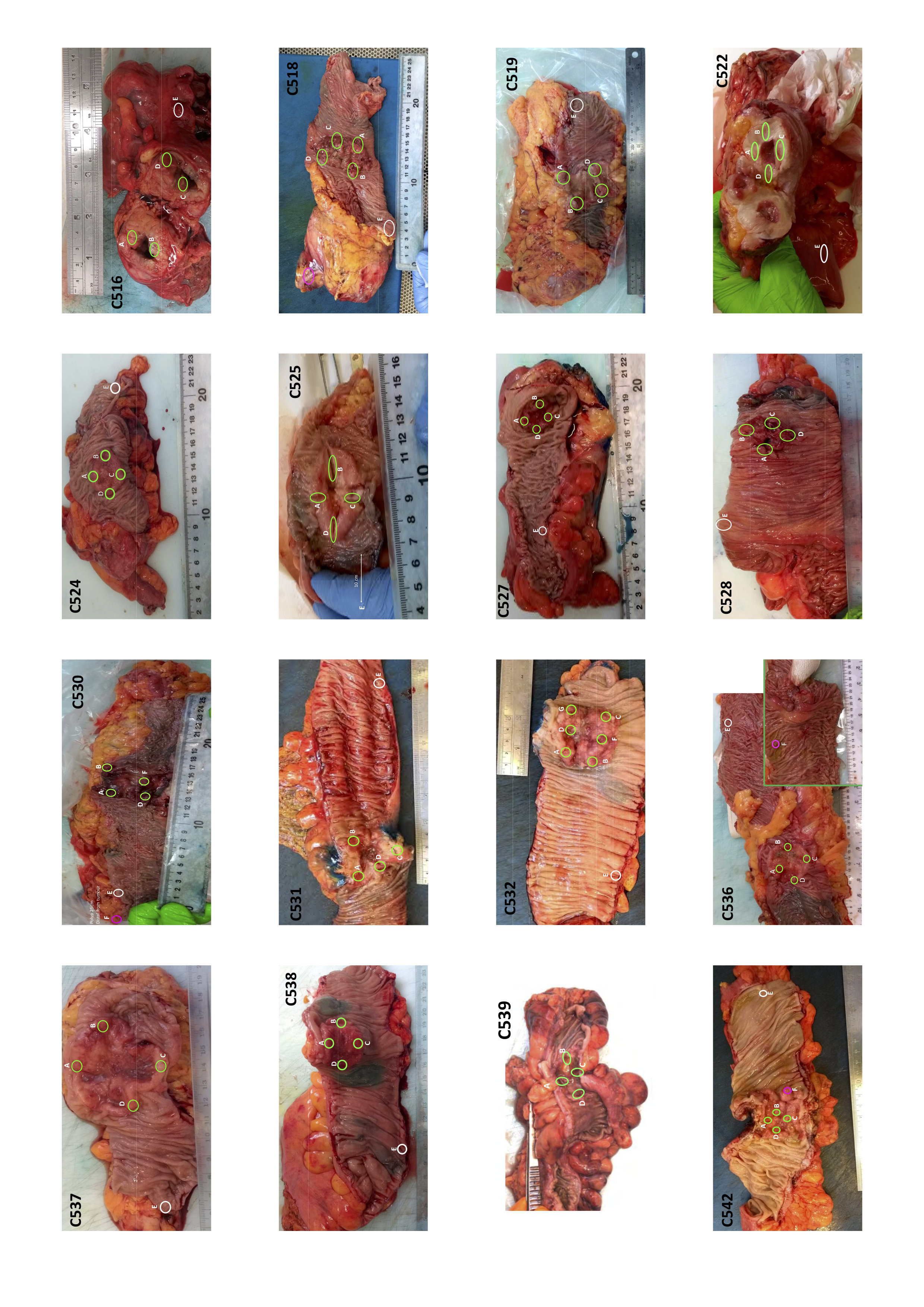

### figS3.pdf

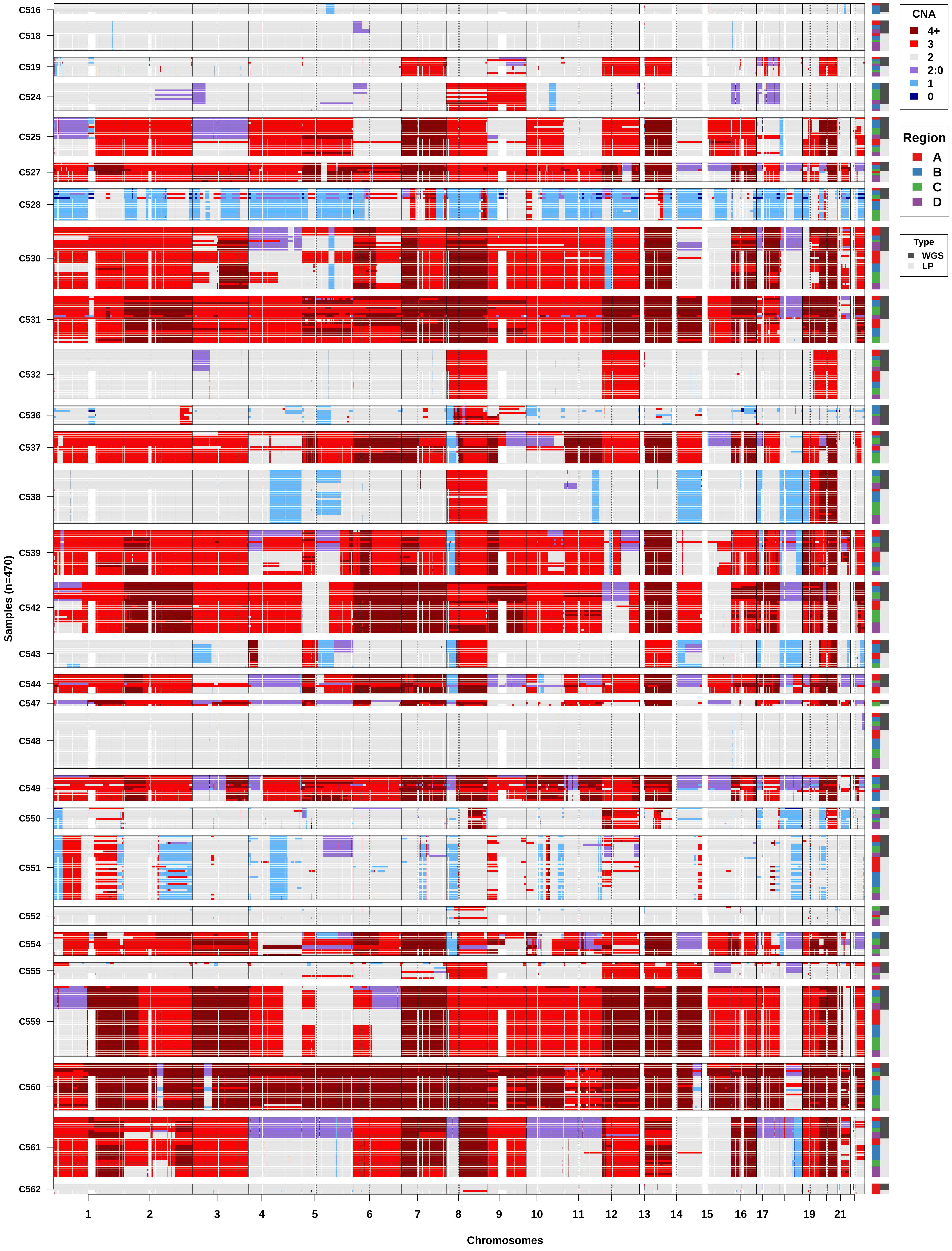

### figS5.pdf

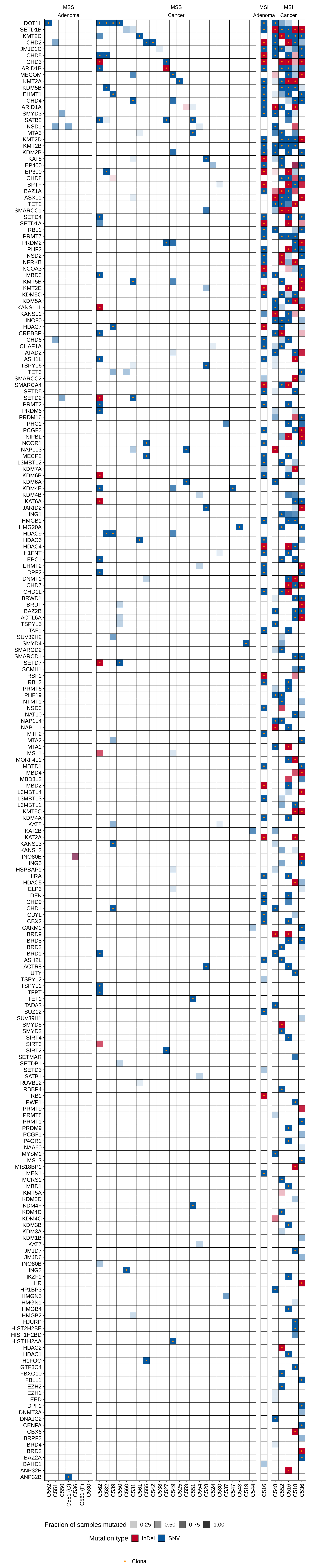

### figS6.pdf

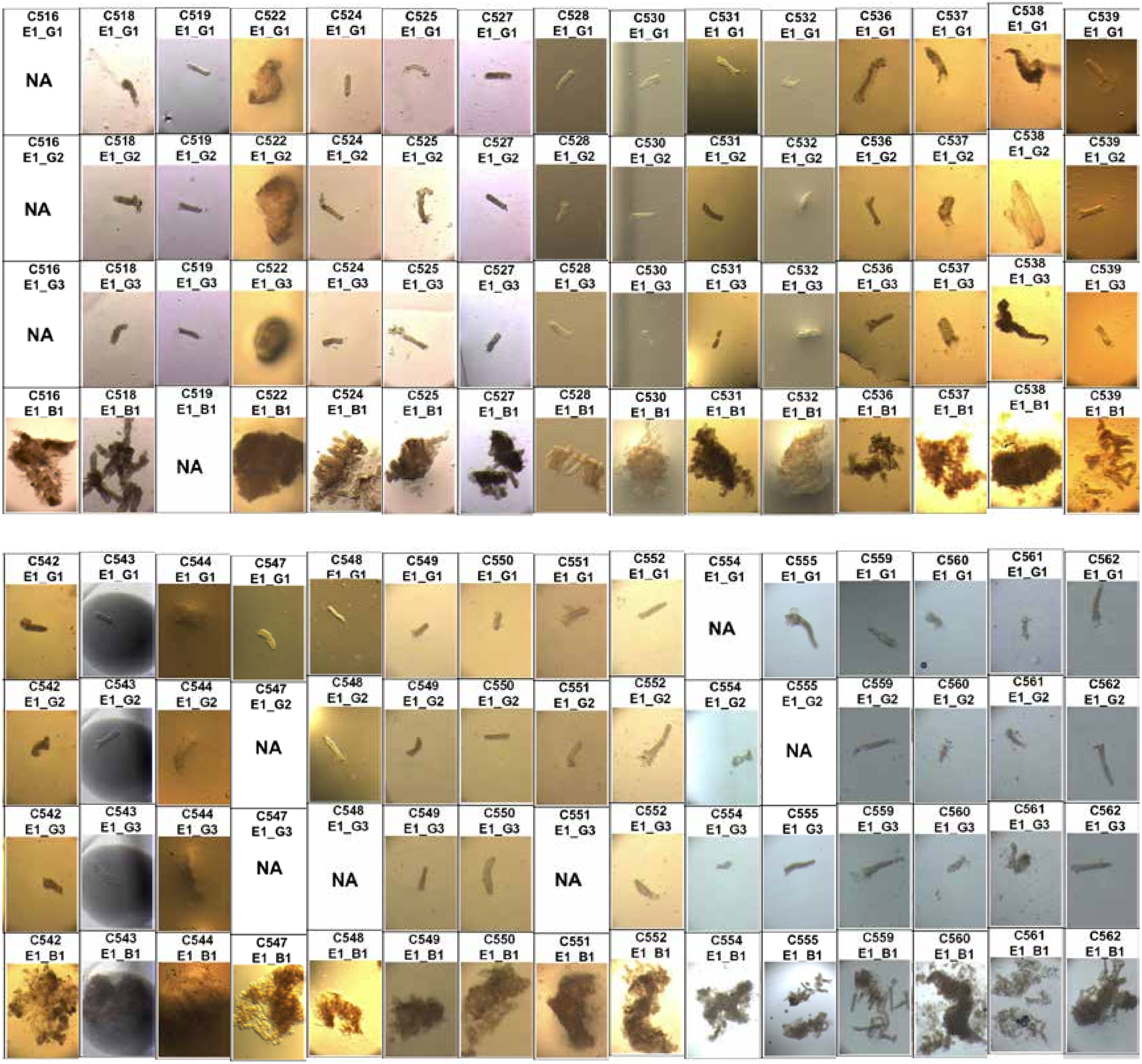

### figS7.pdf

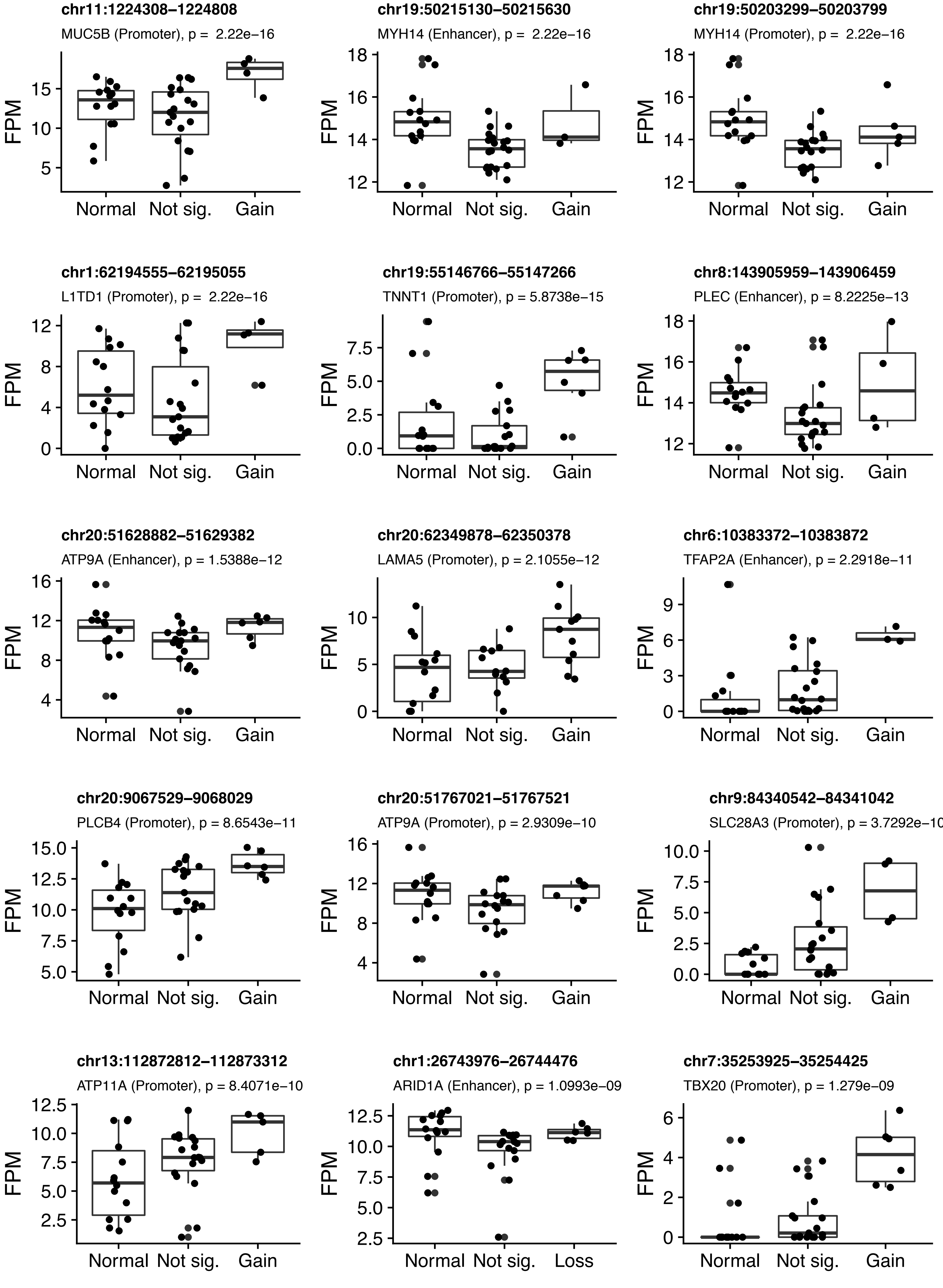

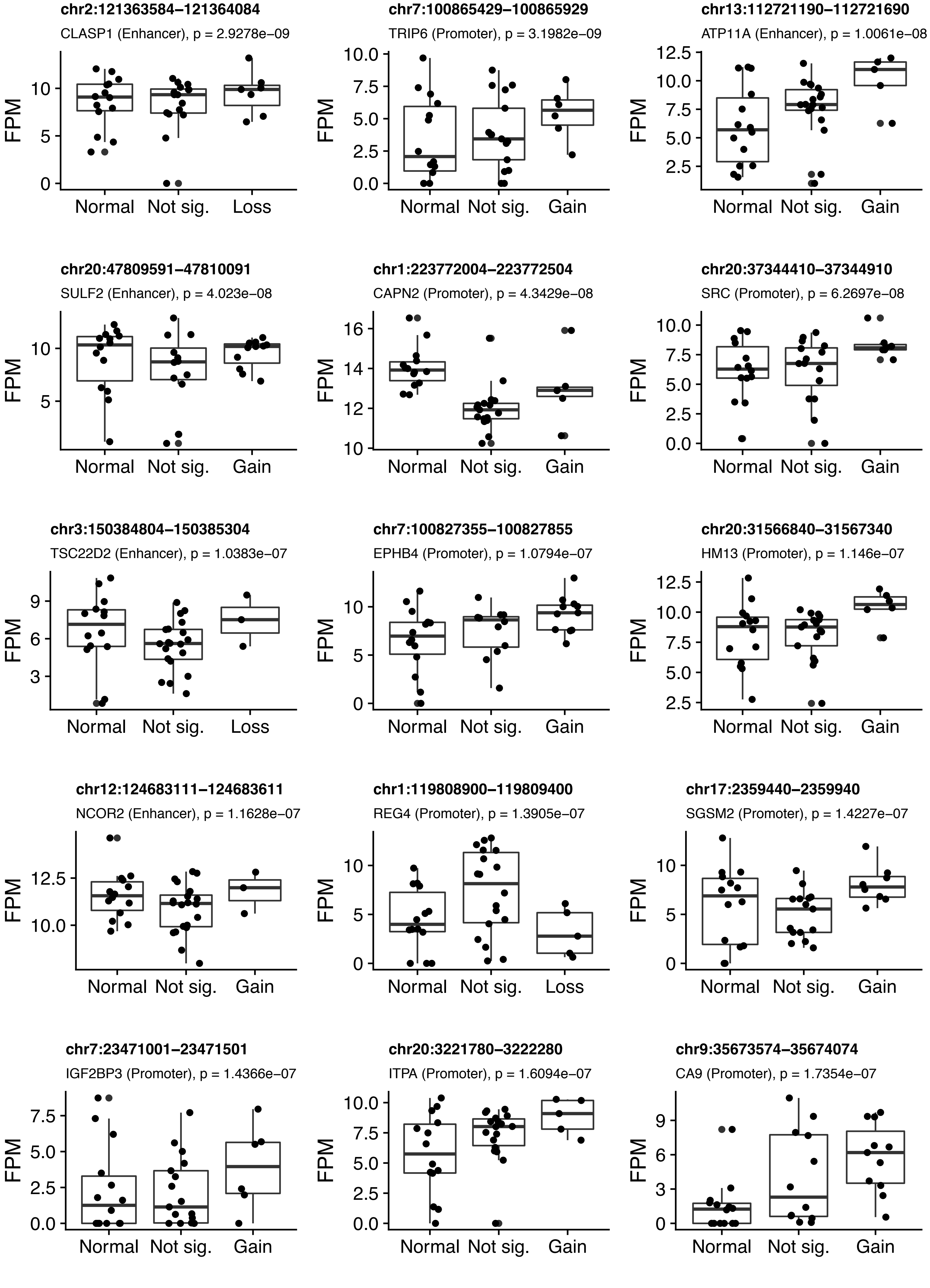

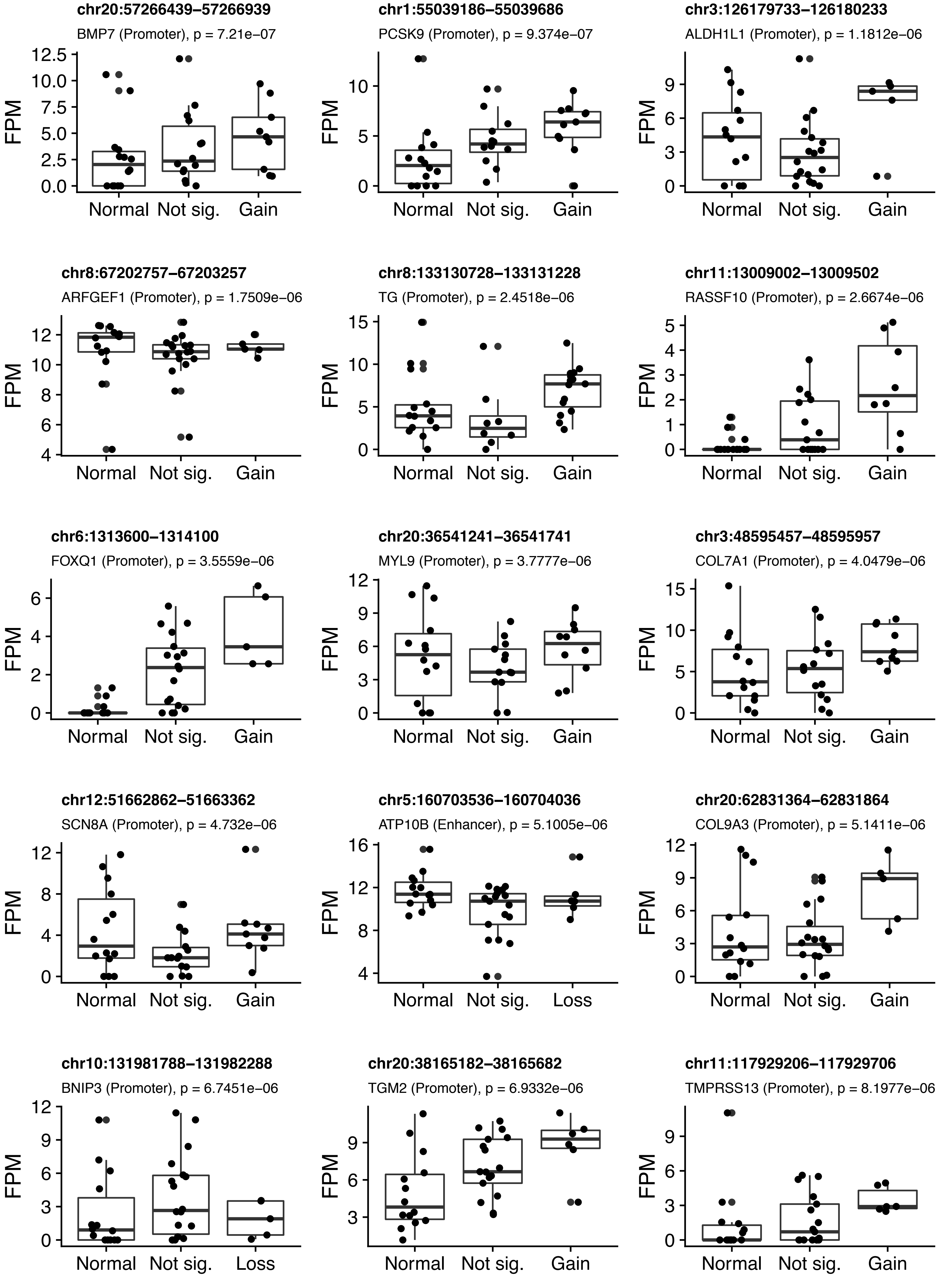

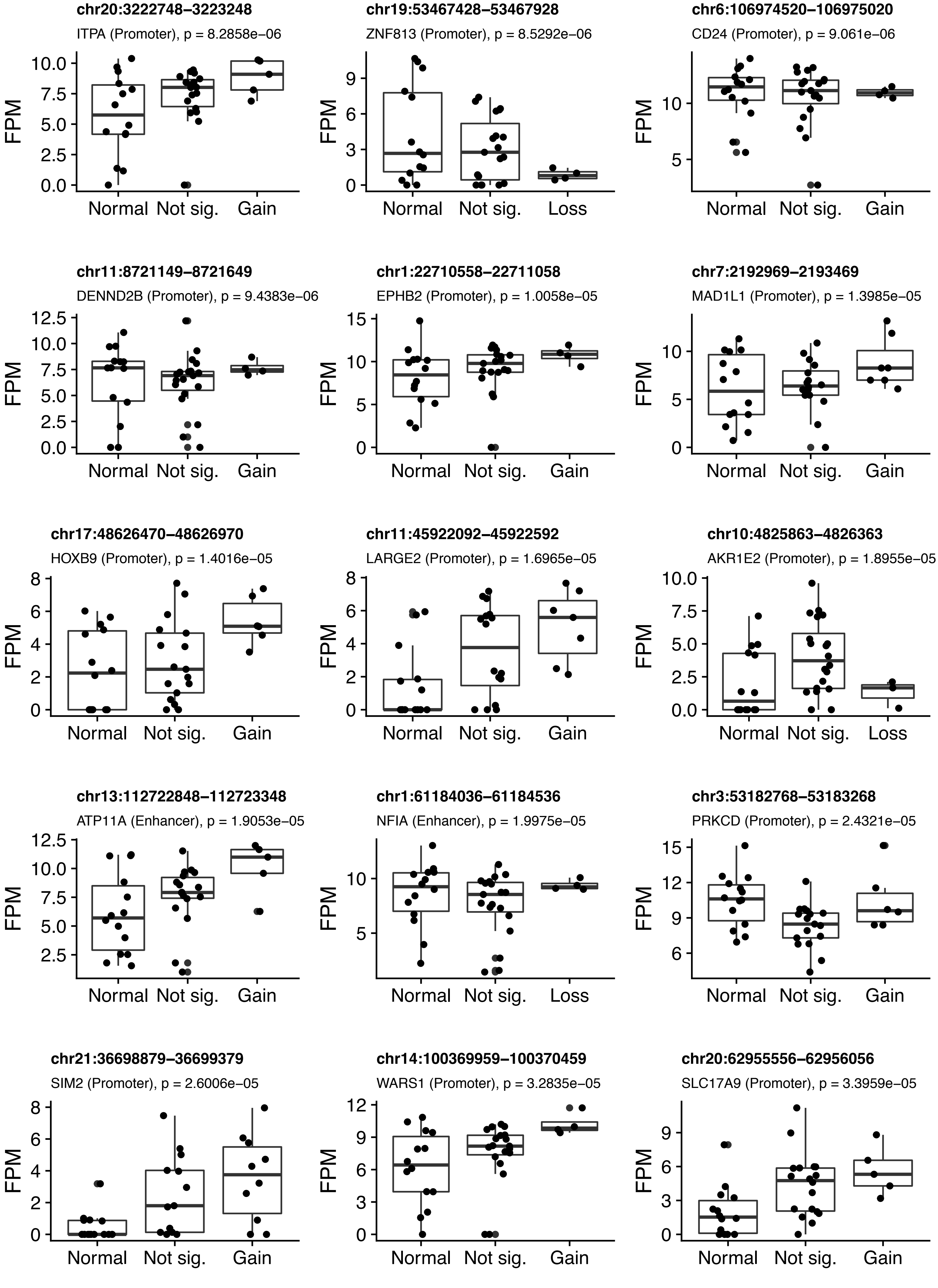

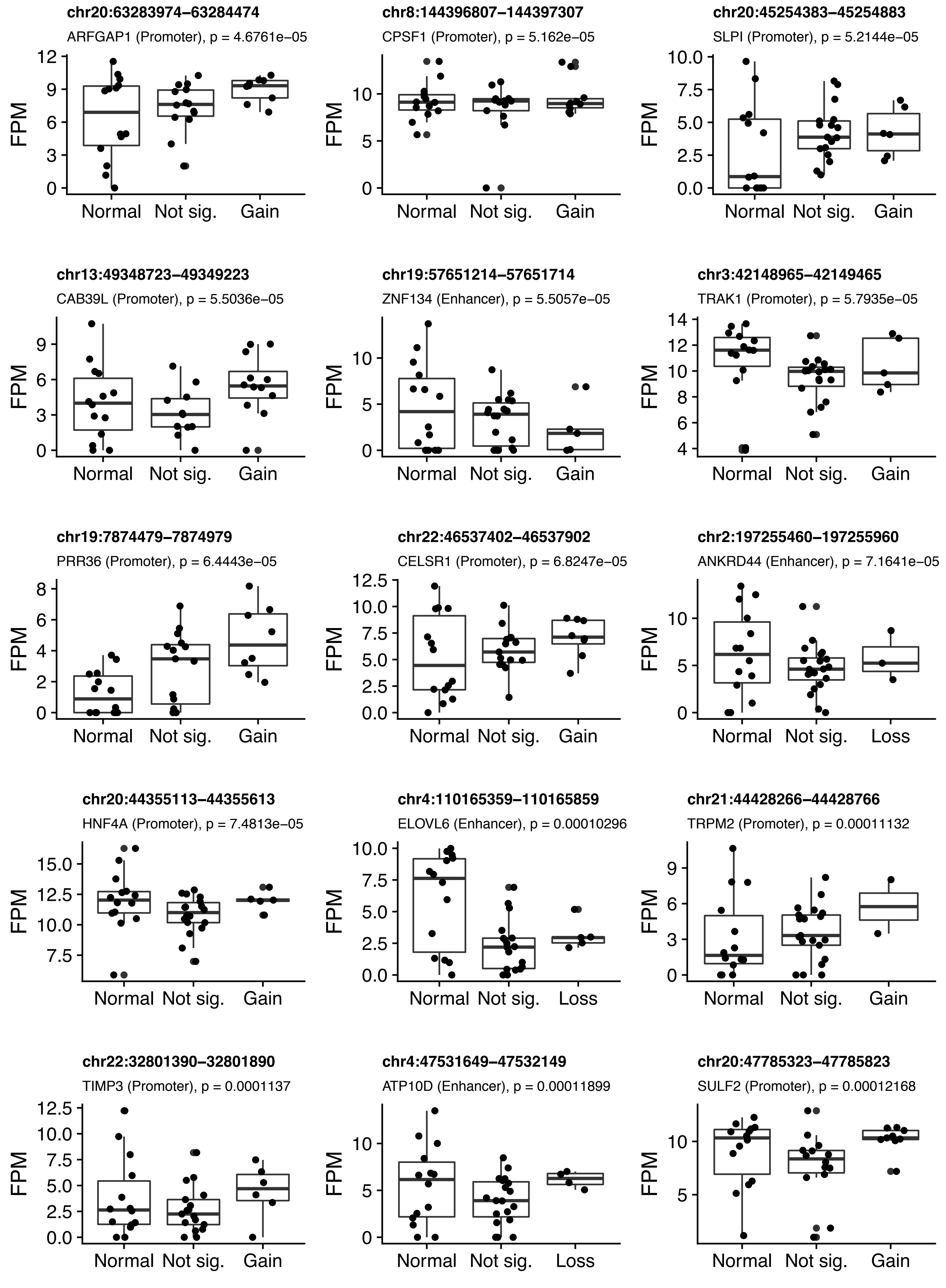

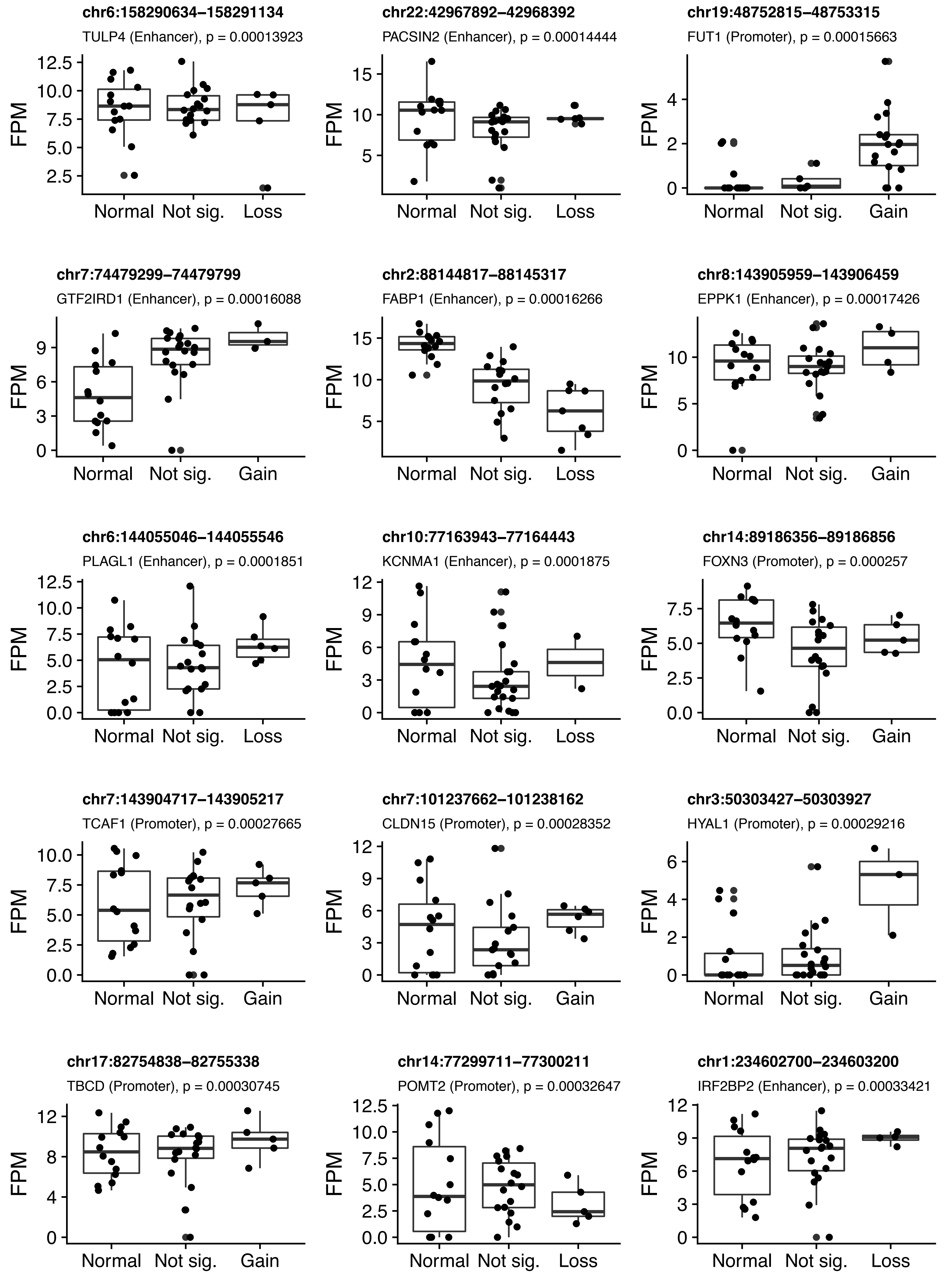

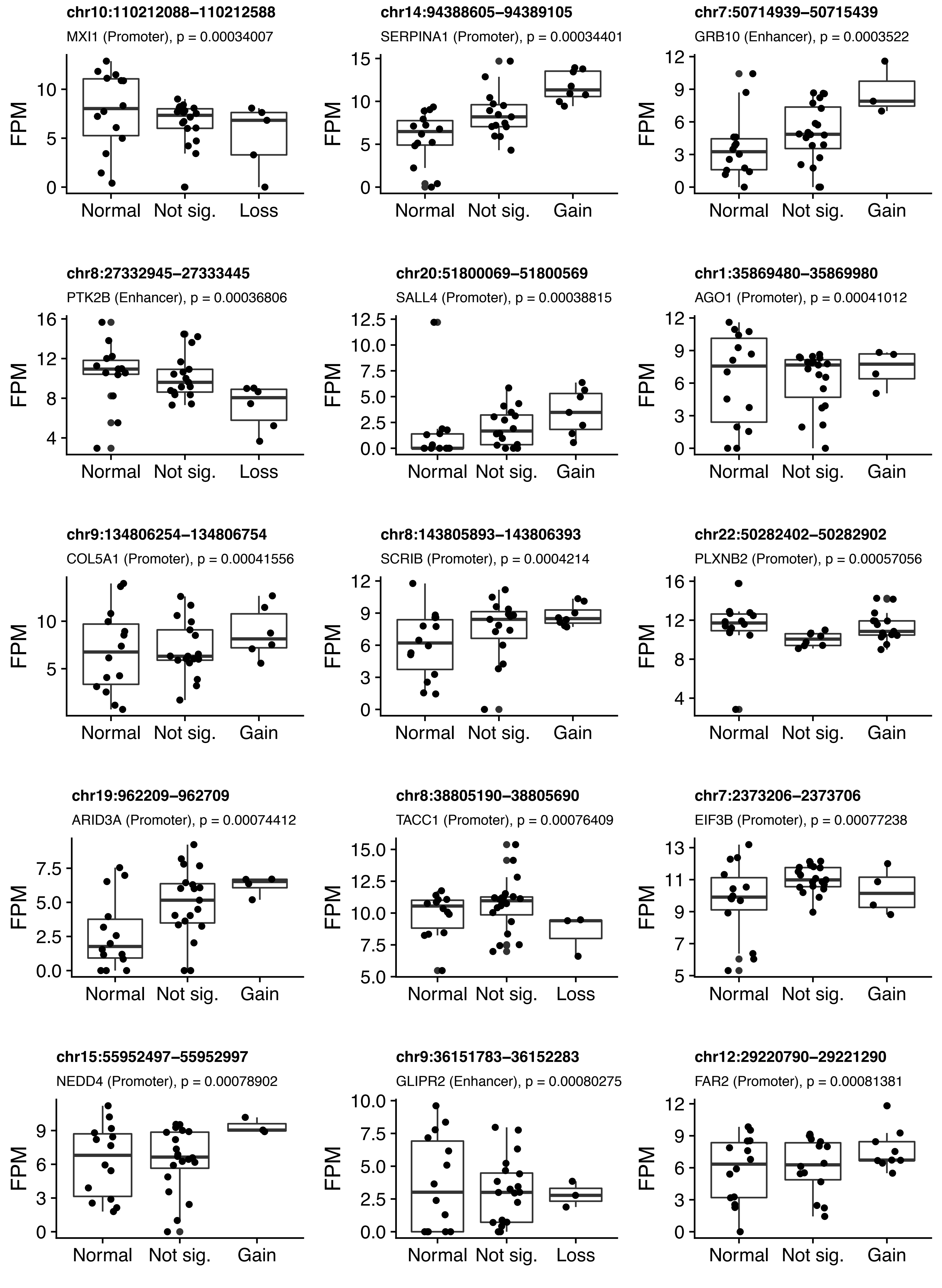

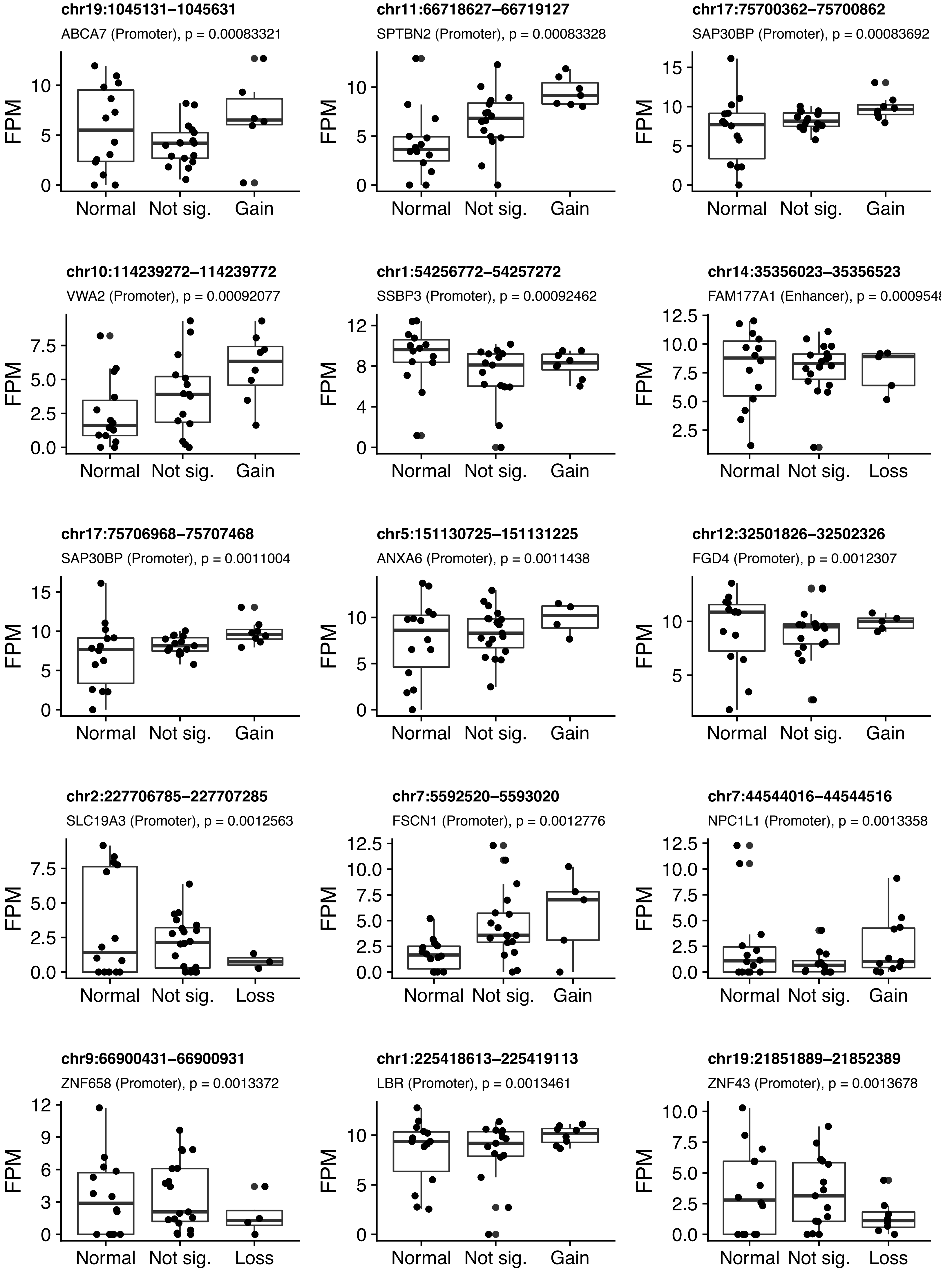

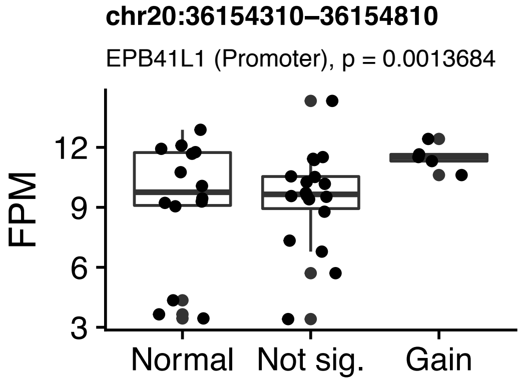

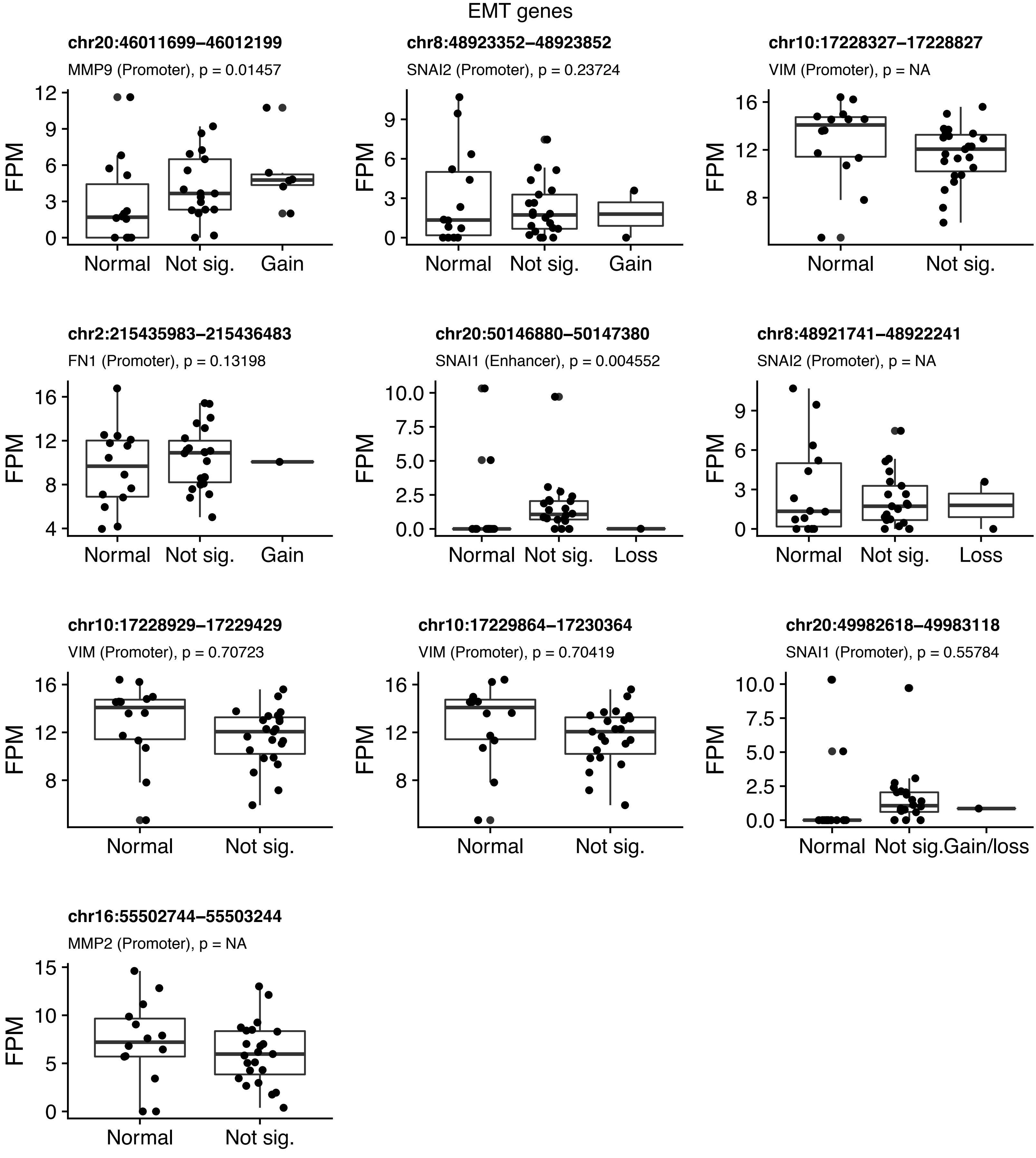

### figS8.png

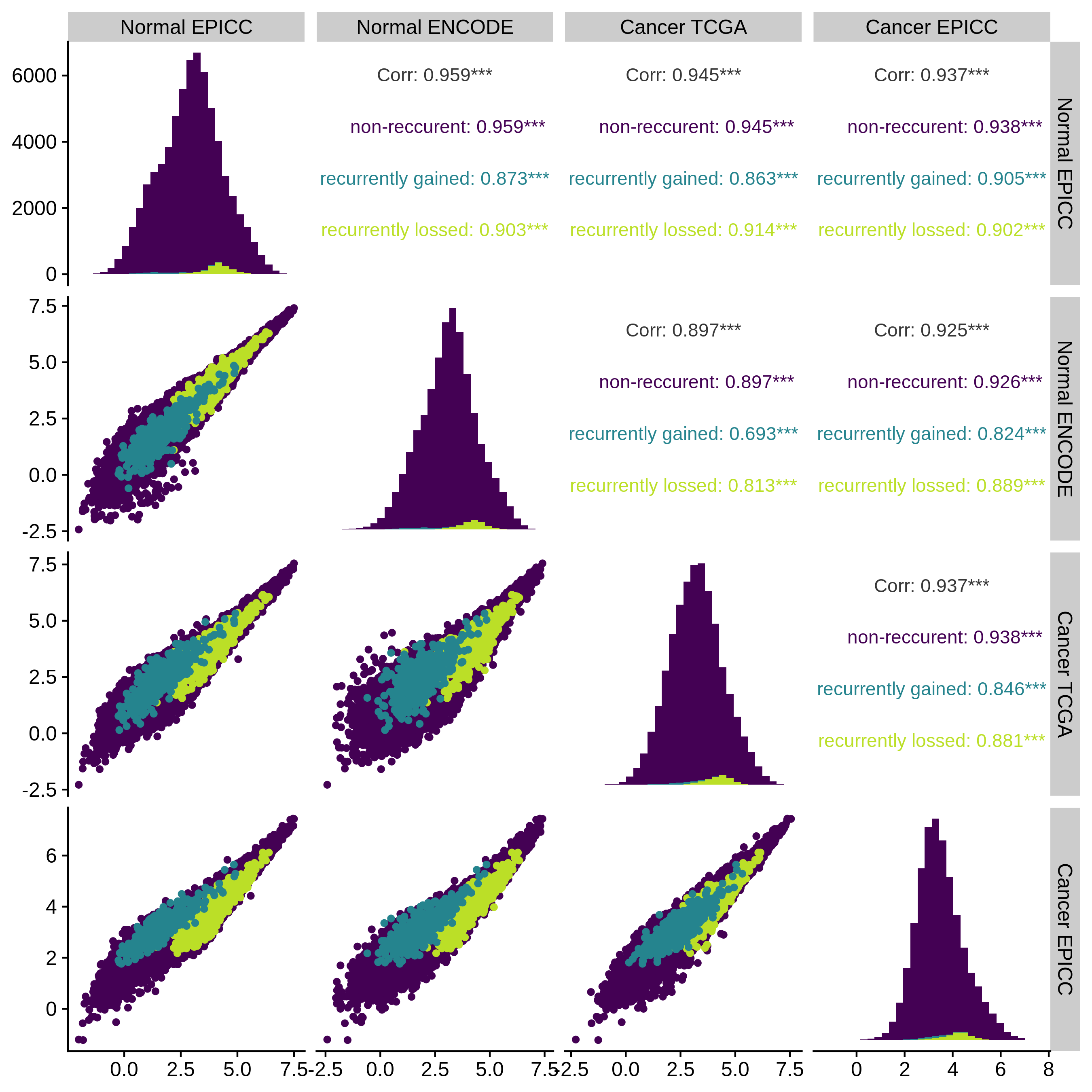

### figS20.pdf

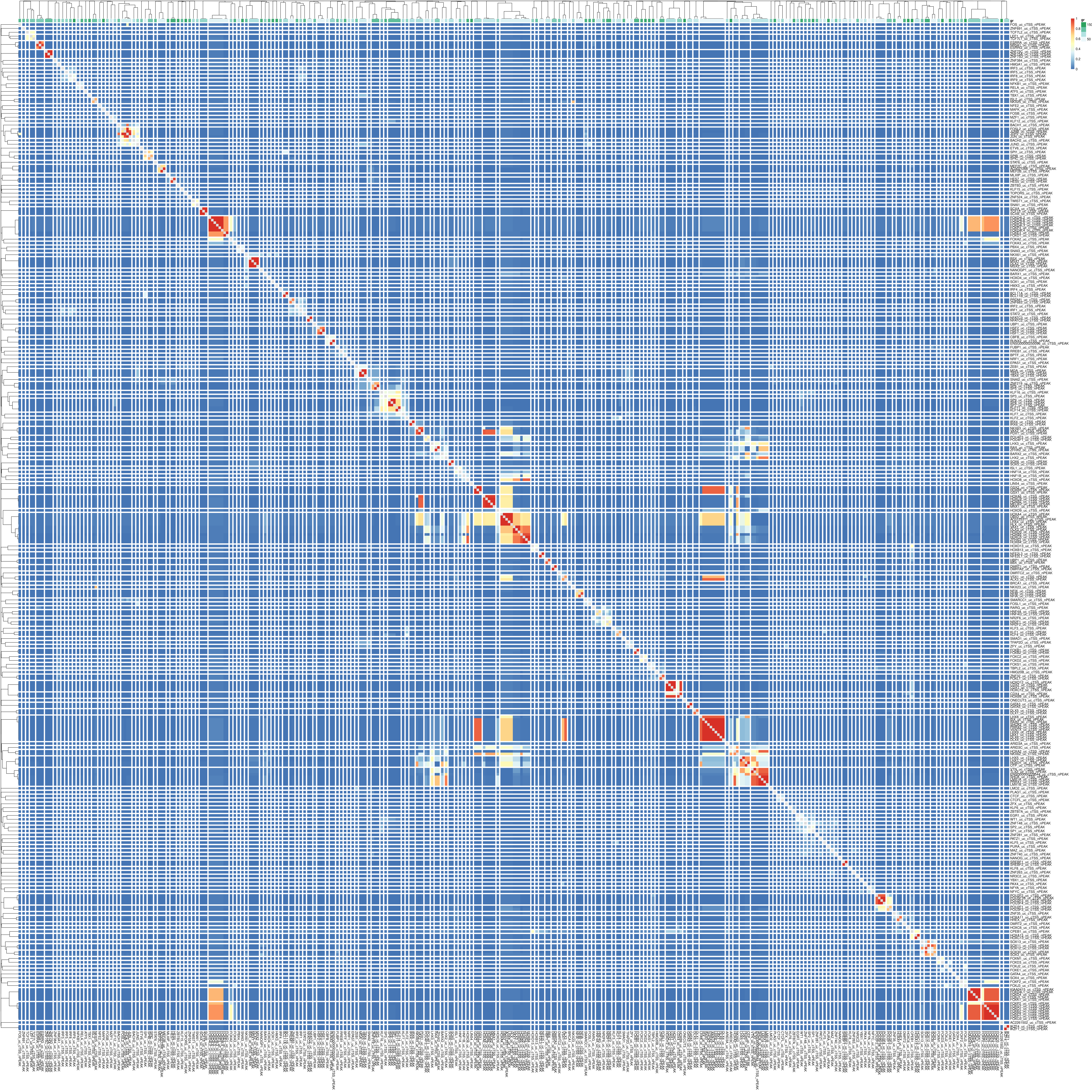

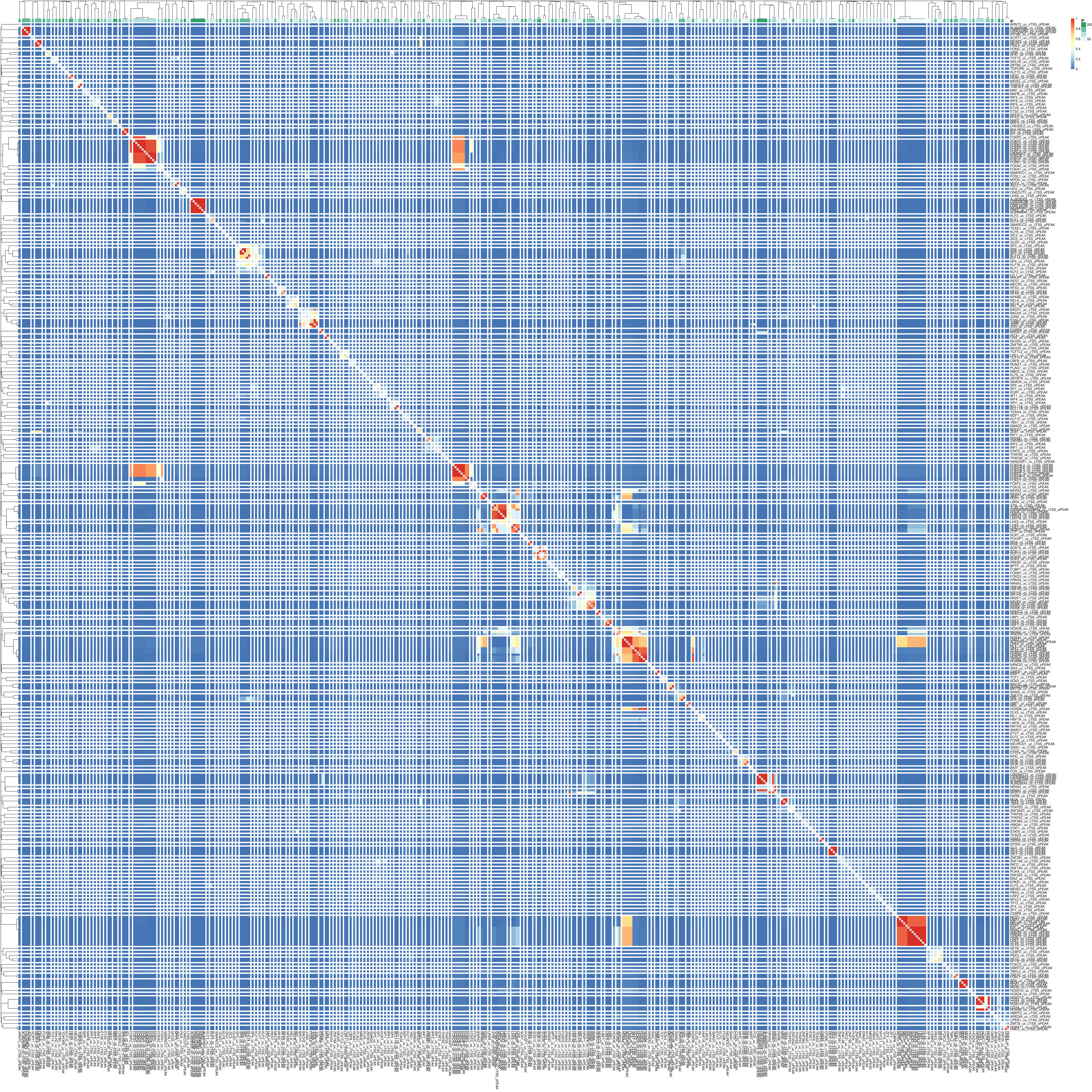

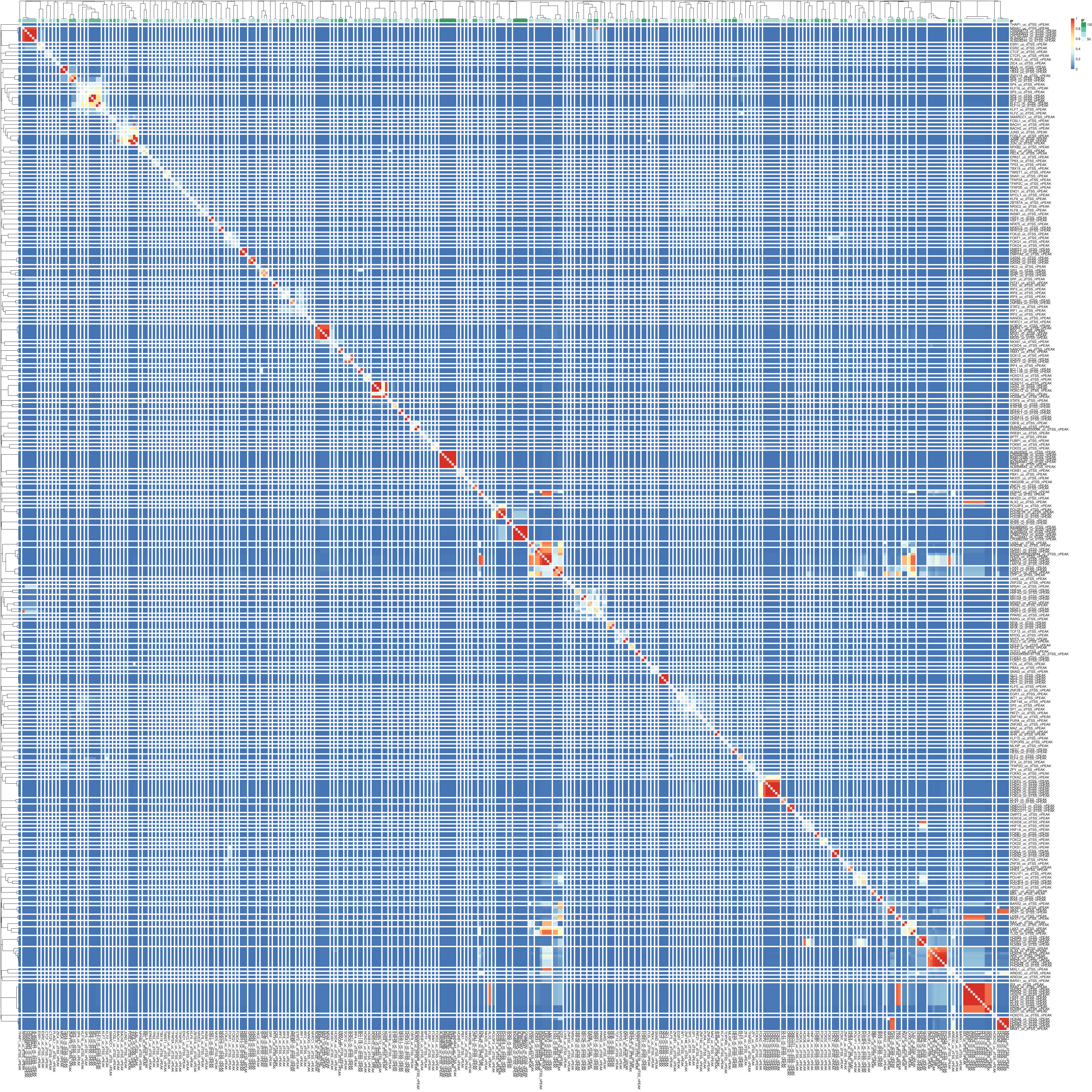

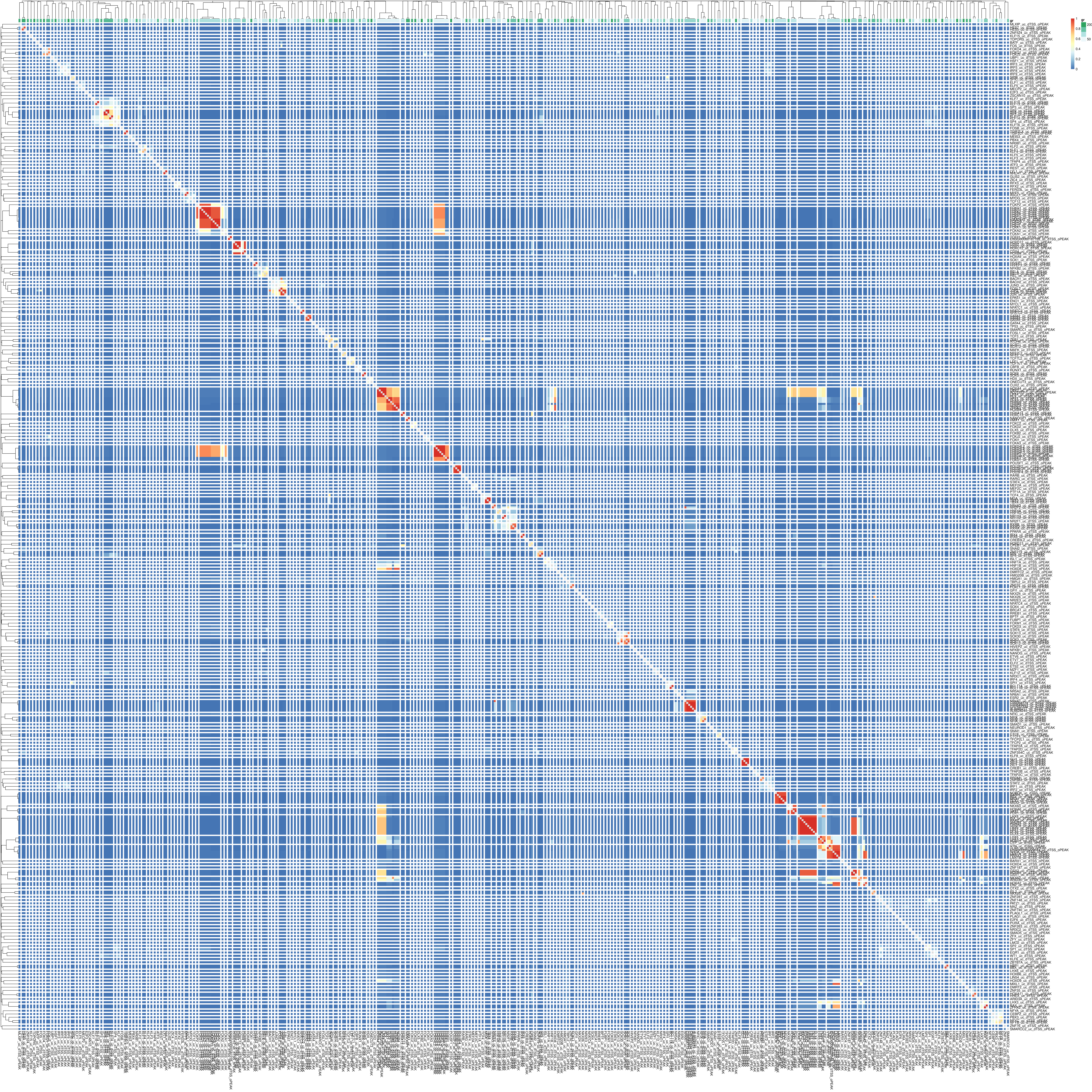

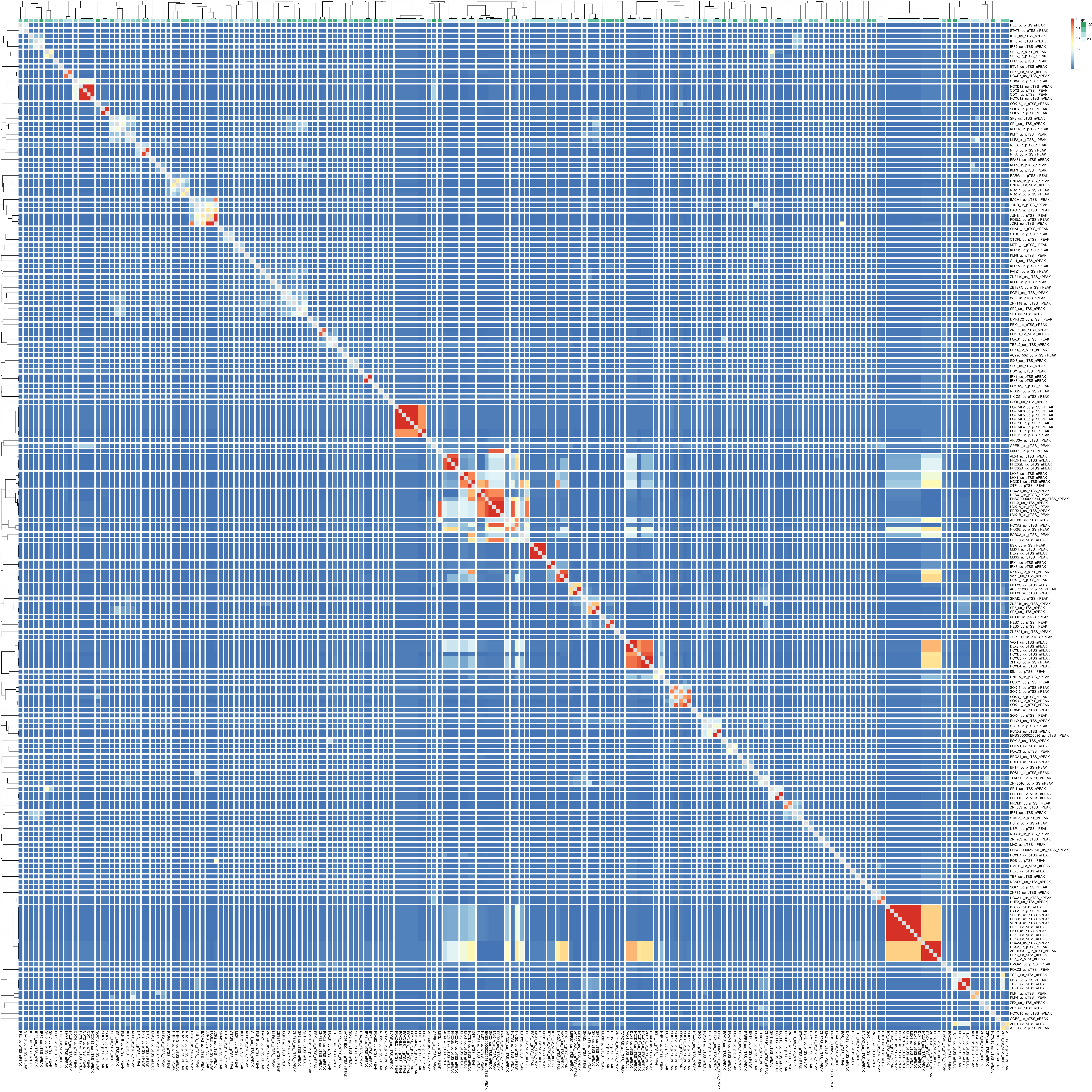

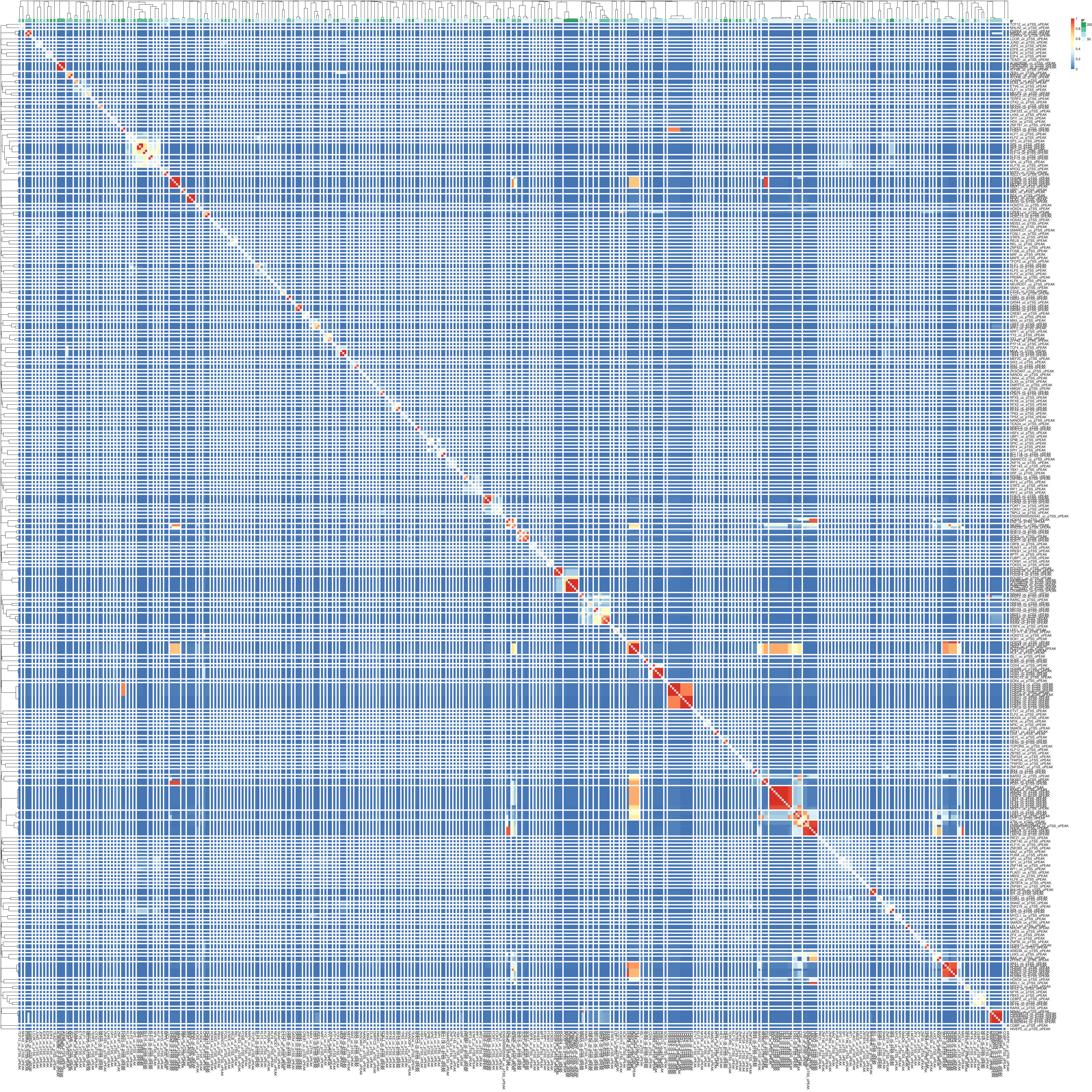

### figS22.pdf

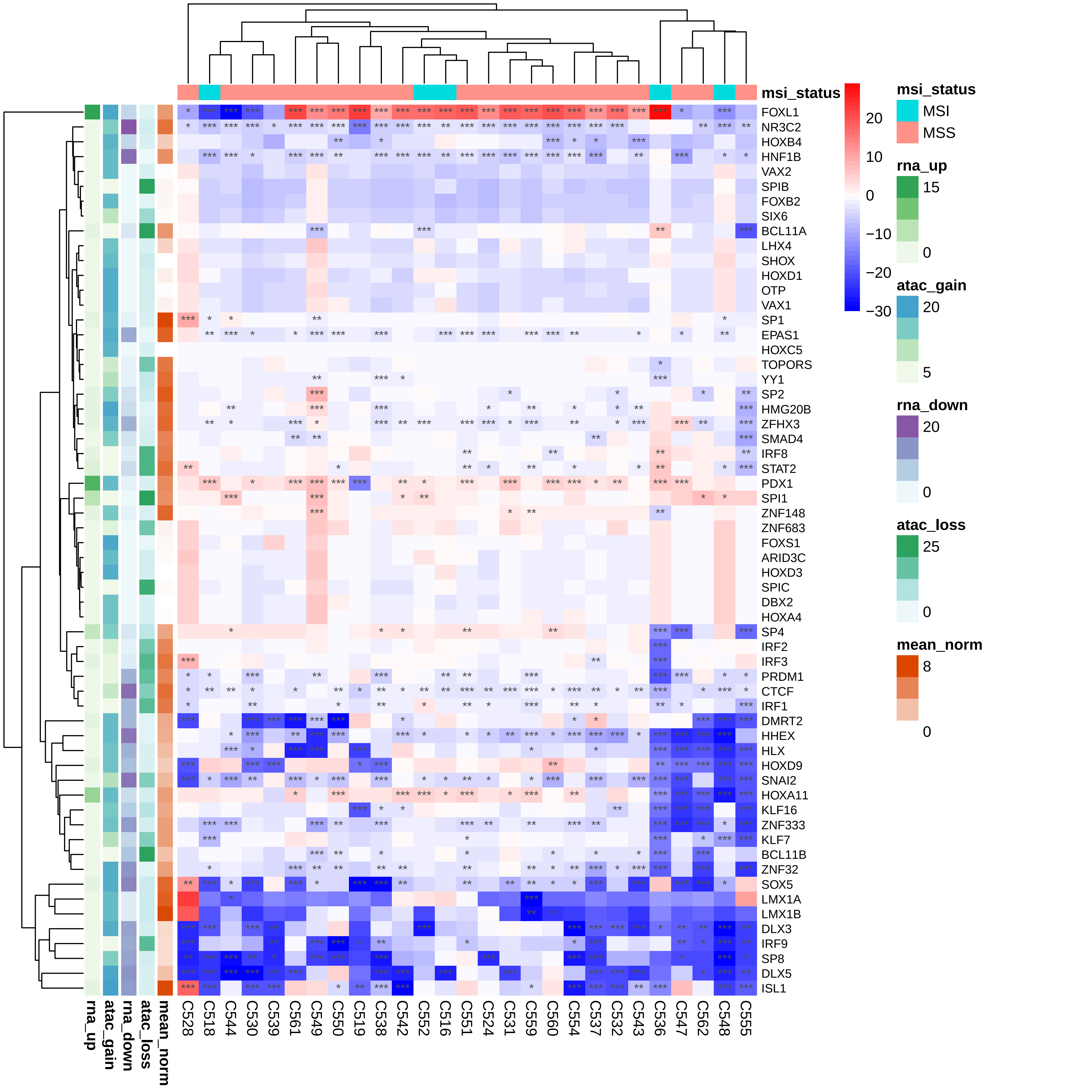

### figS23_coefs_anova_within_between_regions_per_region.pdf

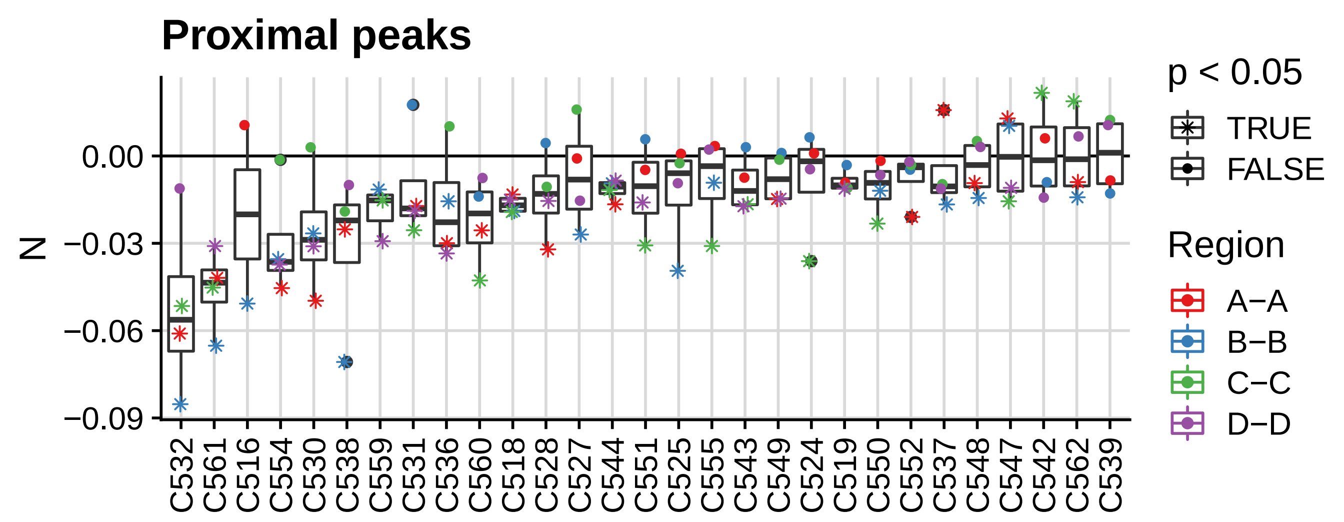

### figS25_SigProfiler.pdf

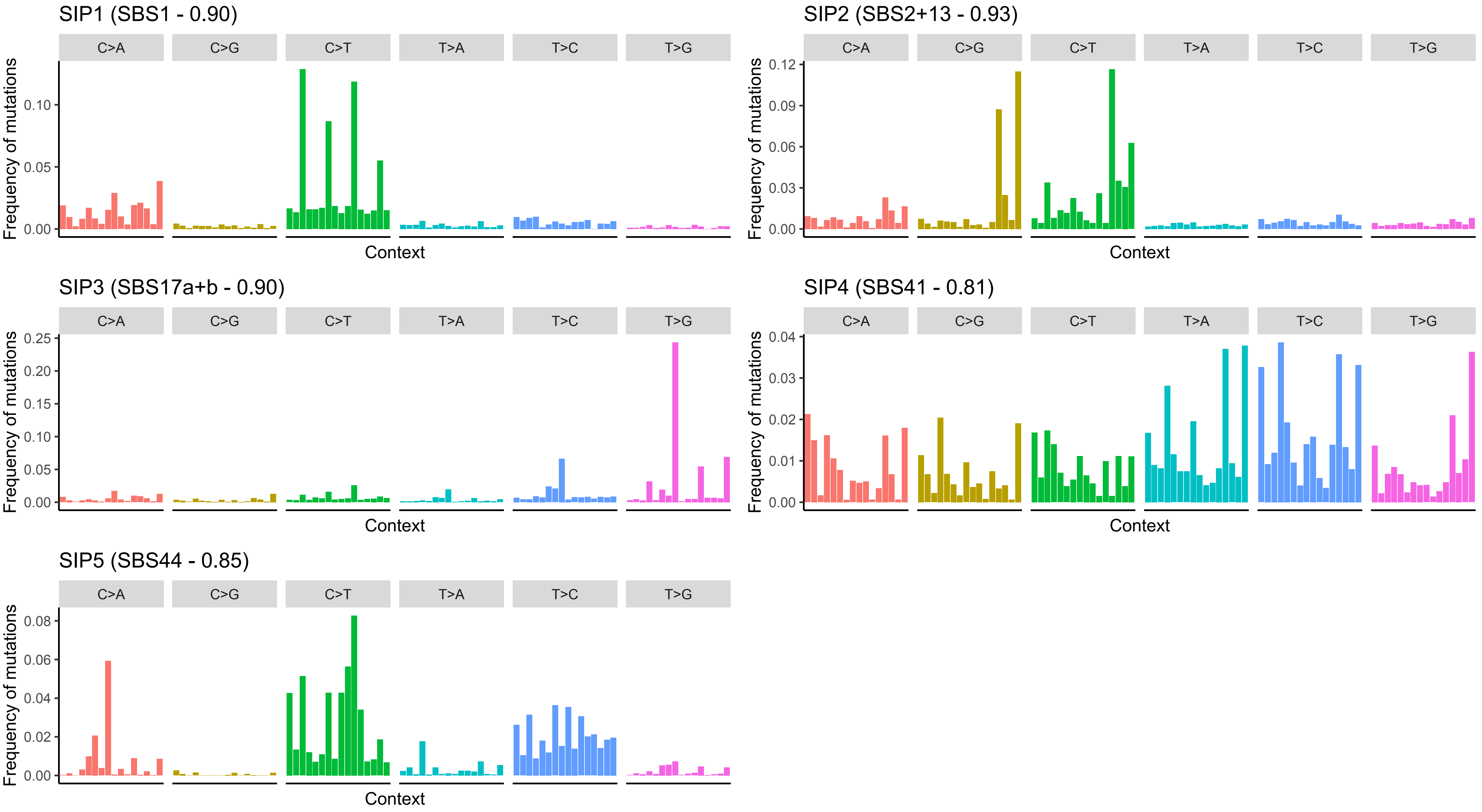

### figS26.pdf

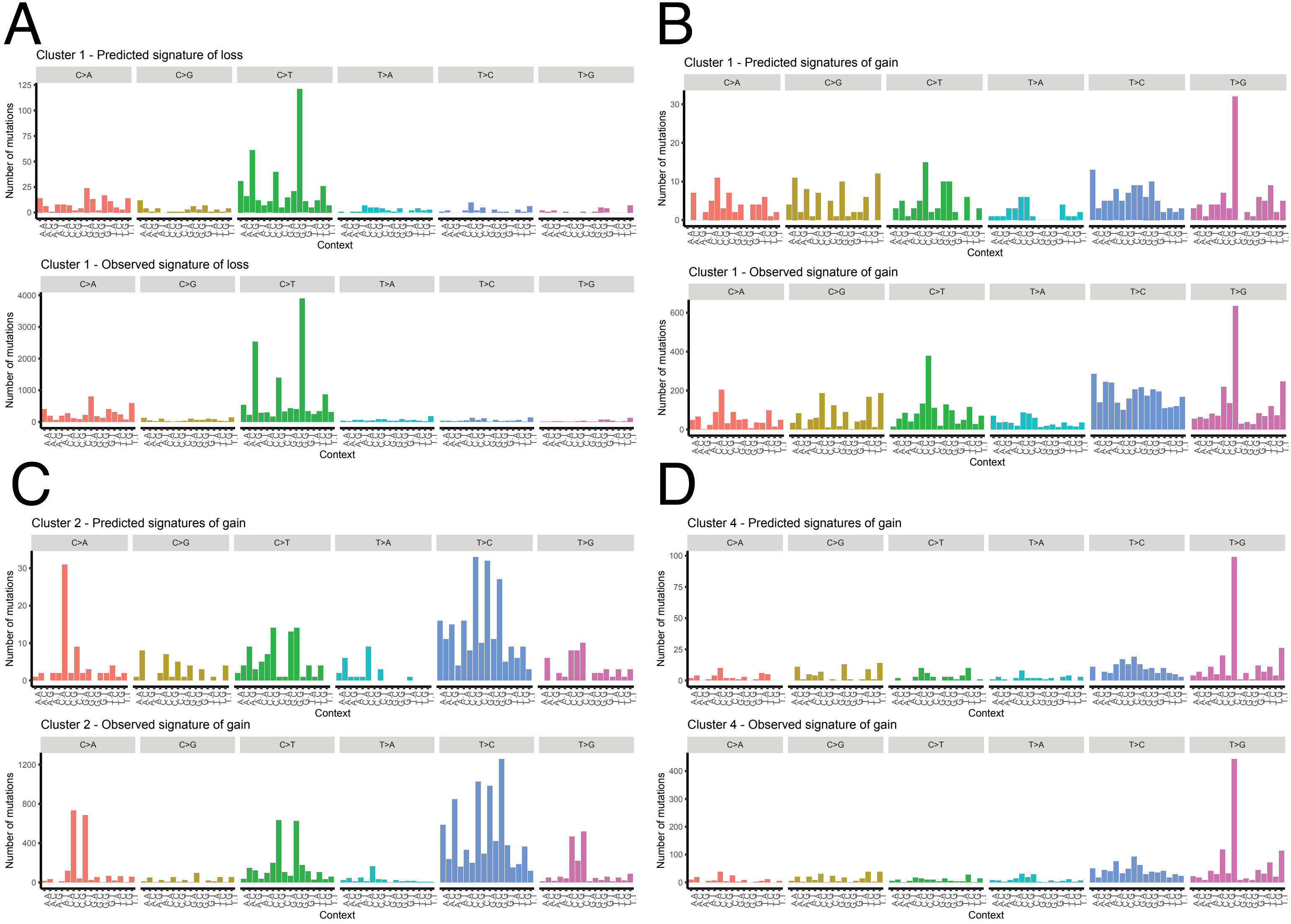

### figS27.pdf

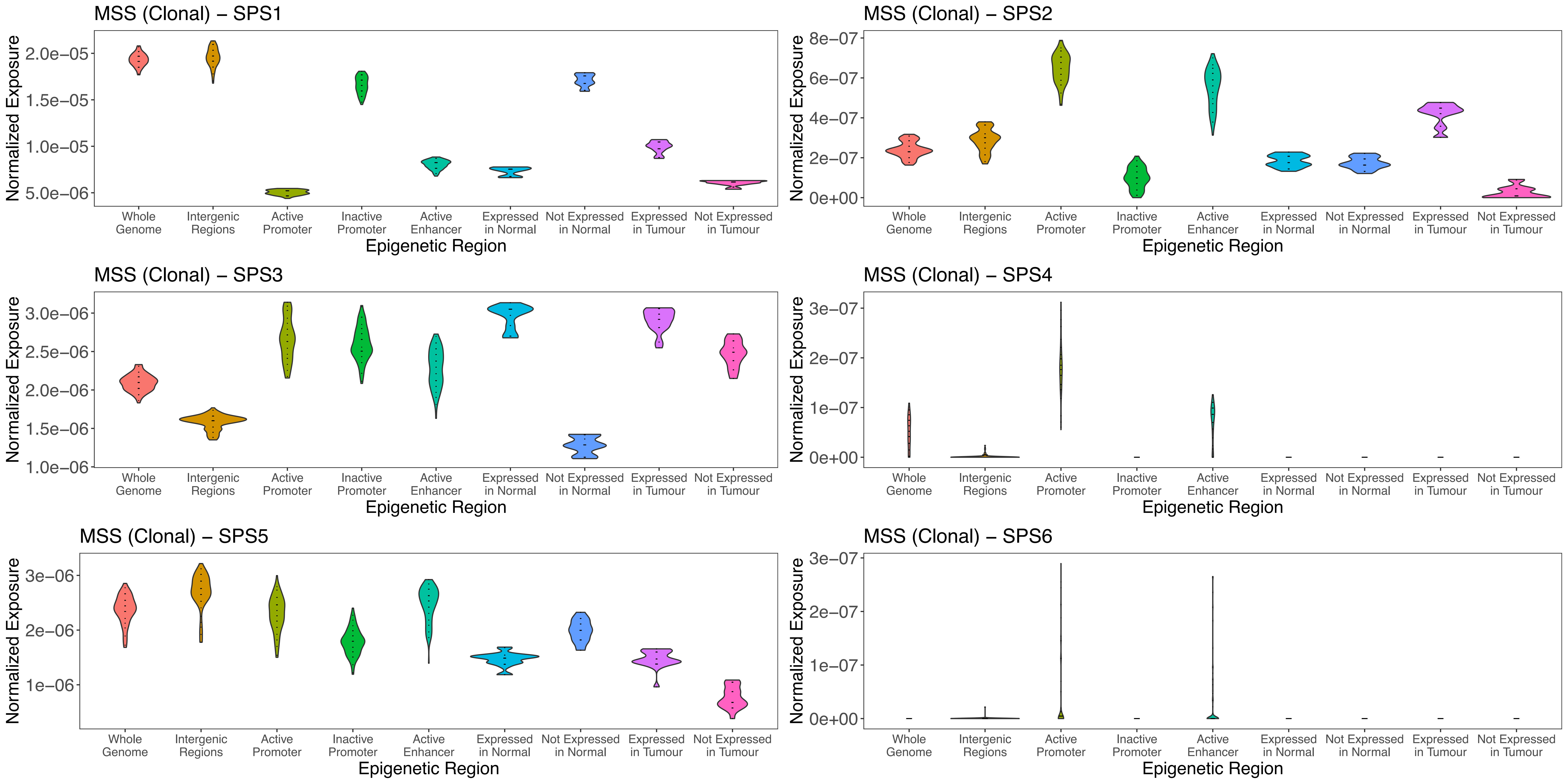

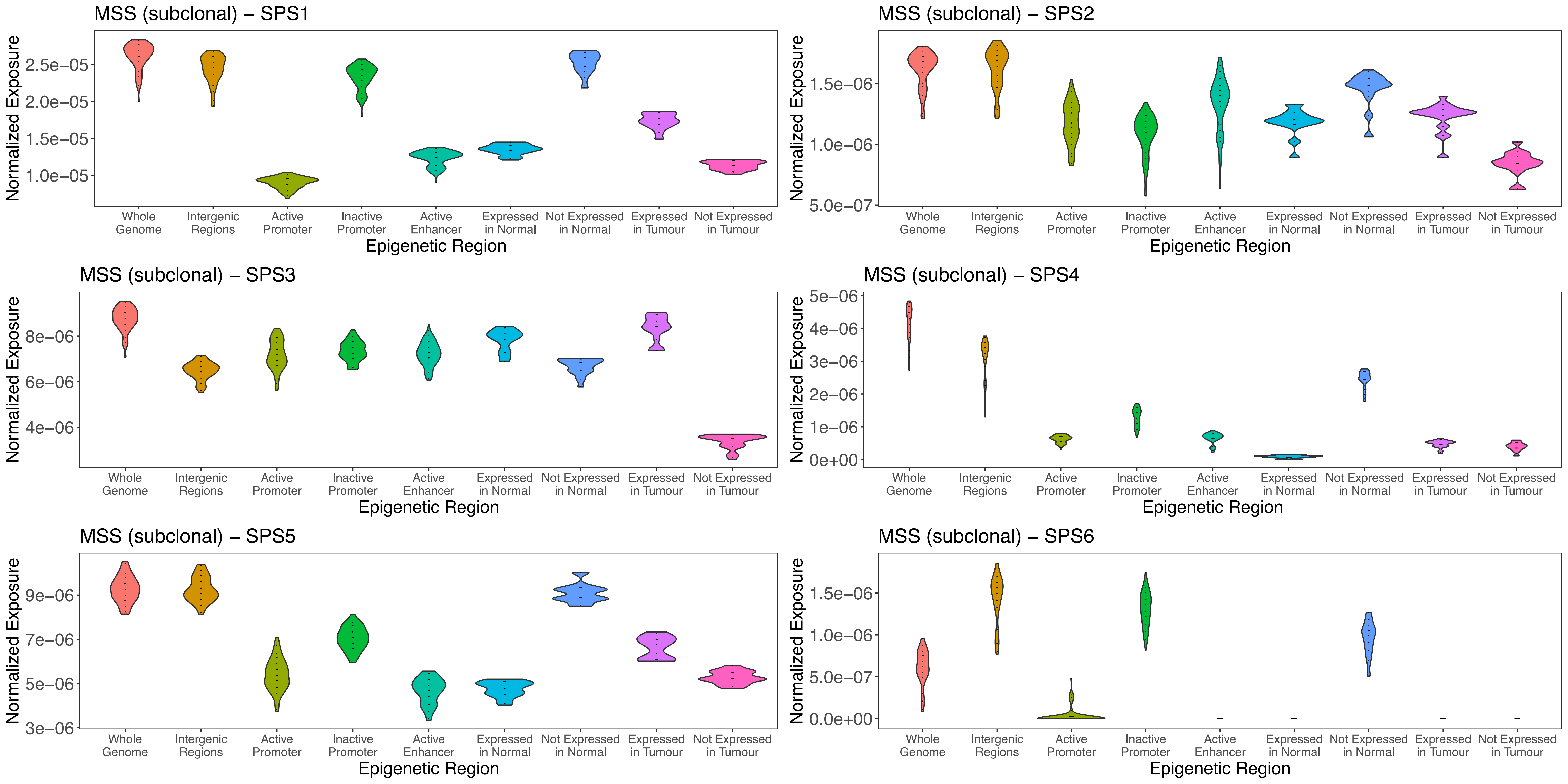

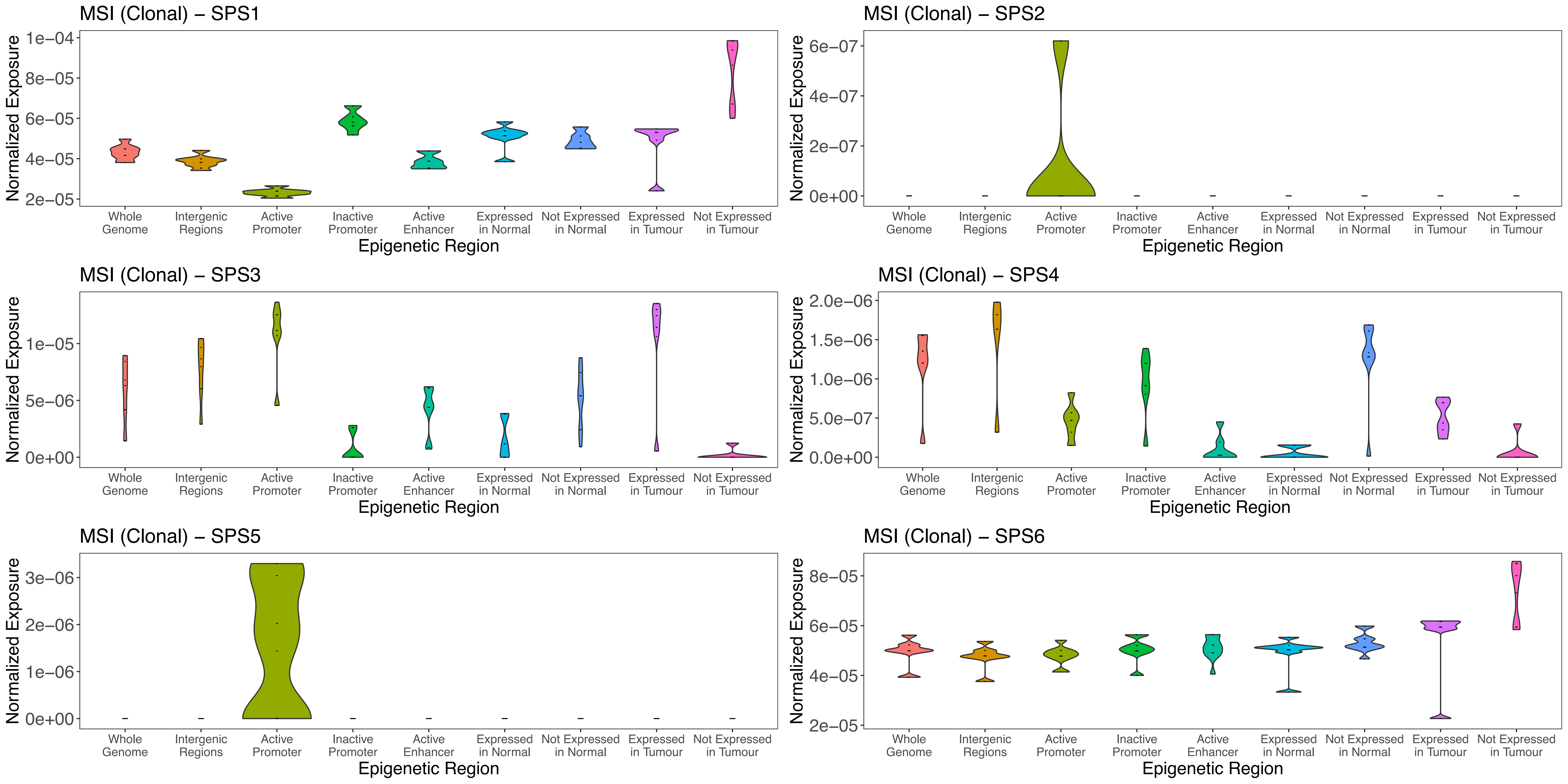

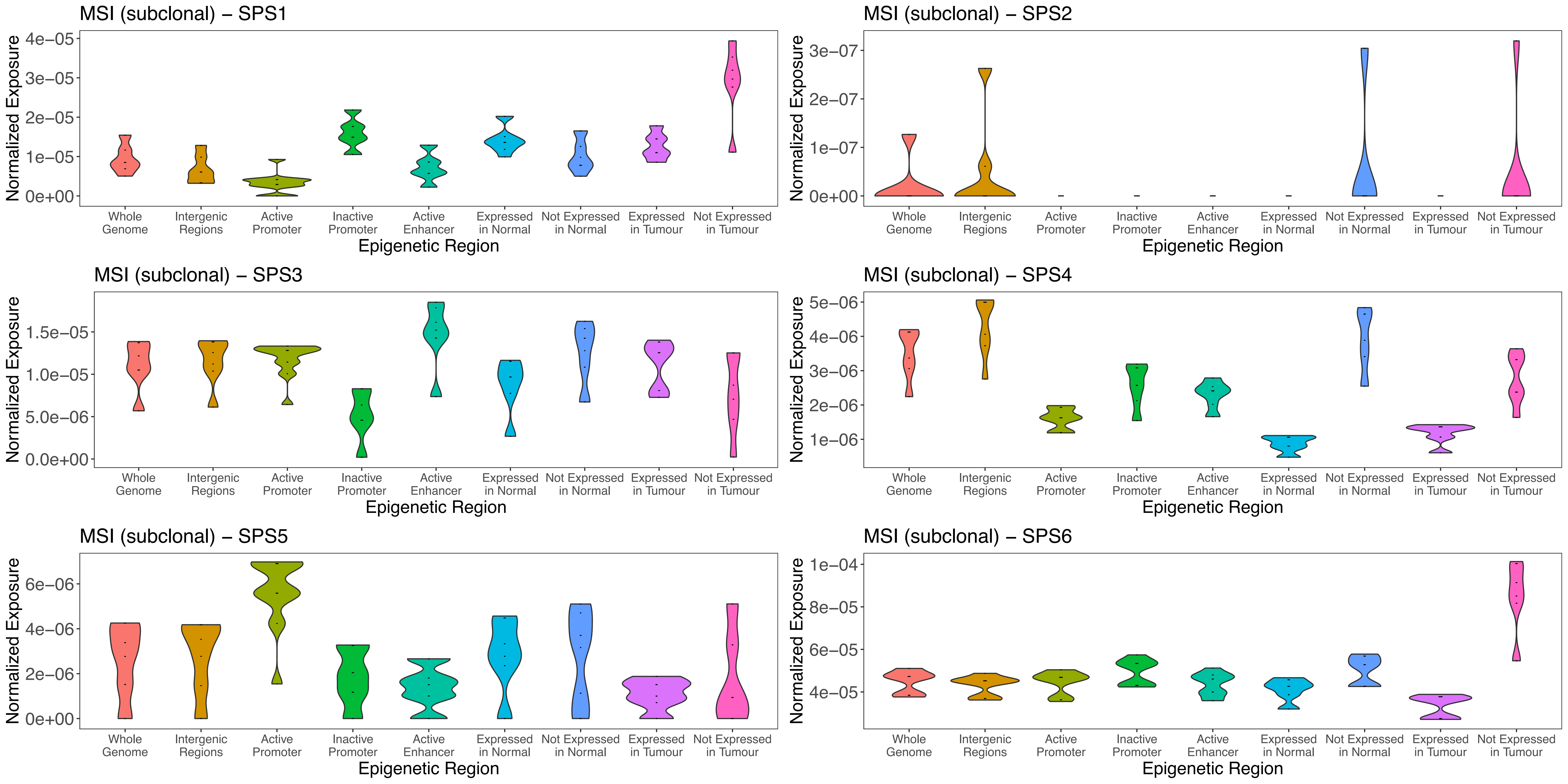
